# Chitinase-3-like 1 regulates T_H_2 cells, T_FH_ cells and IgE following helminth infection

**DOI:** 10.1101/2022.09.26.507843

**Authors:** Miranda L. Curtiss, Alexander F. Rosenberg, Christopher D. Scharer, Betty Mousseau, Natalia A. Ballesteros Benavides, John E. Bradley, Beatriz León, Chad Steele, Troy D. Randall, Frances E. Lund

## Abstract

Data from patient cohorts and mouse models of atopic dermatitis, food allergy and asthma strongly support a role for the chitinase-like protein ChI3L1 in allergic disease. To address whether CHI3L1 also contributes to T_H_2 responses following nematode infection, we infected *Chi3l1*^-/-^ mice with *Heligmosomoides polygyrus* (*Hp*) and analyzed T cell responses. Not surprisingly, we observed impaired T_H_2 responses in *Hp*-infected *Chi3l1*^-/-^ mice. However, we also found that T cell intrinsic expression of *Chi3l1* was required for ICOS upregulation following activation of naïve CD4 T cells and was necessary for the development of the IL-4^+^ T_FH_ subset, which supports germinal center (GC) B cell reactions and IgE responses. The requirement for *Chi3l1* in T_FH_ and IgE responses was also seen following alum-adjuvanted vaccination. While Chi3l1 was critical for IgE humoral responses it was not required for vaccine or infection induced IgG1 responses. These results suggest that *Chi3l1* specifically modulates IgE responses that are highly dependent on help from IL-4-producing T_FH_ cells.

## Introduction

Studies of human allergic diseases reveal that expression of CH3L1 protein and transcripts is increased in the lungs and serum of some asthma cohorts^1, 2^. Increased expression of CH3L1/*CHI3L1* is positively associated with pathogenesis in allergic rhinitis^3, 4^, atopic dermatitis^5–7^ and food allergy^8^ and *CHI3L1* SNPs confer risk of asthma development and airway remodeling^2, 9–11^ as well as increased serum IgE and atopy^12^. Similarly, experiments using *Chi3l1*^-/-^ mice show that Chi3L1 regulates type 2 cytokines and IgE in models of asthma, atopic dermatitis and food allergy^7, 8, 13–15^. Thus, both mouse and human data support a role for Chi3L1 in promoting atopic disease.

Parasitic infections are recognized as the likely impetus for the evolution of T_H_2 immunity in mammals, and studies using these infection models show that mediators regulating protective immunity to parasite infections often contribute to pathologic responses to allergens^16–19^. Thus, it was somewhat surprising that *Chi3l1*^-/-^ mice, which make attenuated allergic responses^7, 8, 13–15^, cleared an infection with the nematode *Nippostrongylus brasiliensis* (*Nb*)^20^. However, other types of immunity can contribute to parasite elimination and the *Chi3l1*^-/-^ *Nb* study did not measure T_H_2 or IgE responses^20^. Therefore, we used the *Heligmosomoides polygyrus bakerii* (*Hp*) nematode model, which elicits strong T_H_2^18, 21^, T_FH22-25_ and polyclonal IgE responses^25, 26^, to assess the role for Chi3L1 in parasite-driven T and B cell responses. Here, we show that Chi3L1 regulates the establishment of the IL-4 producing T_FH_ response to *Hp* infection and controls T_FH_-dependent B cell IgE responses to infection and immunization.

## Results

### *Chi3l1* regulates CD4 T_H_2 effector responses to *Hp* infection

Experiments using allergic airway models revealed that type 2 cytokine responses are blunted in *Chi3l1*^-/-^ animals^13^ and *Chi3l1*^-/-^ CD4 T cells primed *in vitro* with antibodies to CD3 and CD28 in the presence of T_H_2-polarizing conditions are impaired in IL-4 production ^27^. To test whether *Chi3l1* regulates T_H_2 responses to helminth infection, we measured mesenteric LN (msLN) CD4 T cell responses in wild-type (WT) BALB/c and *Chi3l1*^-/-^ BALB/c mice that were orally infected with *Hp*. Uninfected *Chi3l1*^-/-^ mice did not exhibit changes in the numbers of total msLN cells, CD19 B cells, or CD4 T cells (Fig. S1A-D). However, the msLN CD4 and CD44^hi^CD62L^lo^ CD4 T cell responses were attenuated in *Hp*-infected *Chi3l1*^-/-^ mice (Fig. S1A-E). To determine whether the remaining *Chi3l1*^-/-^ CD4 T cells produce T_H_2 cytokines after *Hp* infection, we analyzed intracellular IL-4 and IL-13 levels in anti-CD3 restimulated msLN CD44^hi^ CD4 T cells from D8 *Hp*-infected mice. We observed that the percentage and number of CD44^hi^ *Chi3l1*^-/-^ CD4 T_H_2 cells making IL-4 and IL-13 were significantly decreased relative to the restimulated WT T cells (Fig. S1F-J). Moreover, this impairment could not be rescued with PMA+calcimycin stimulation (Fig. S1F, K-L), indicating that the effector function of parasite-elicited CD4 T cells was compromised in *Chi3l1*^-/-^ mice. Consistent with the dominant T_H_2 response to *Hp* infection^28^, few CD4 T cells from BALB/c or *Chi3l1*^-/-^ mice produced IFNγ following anti-CD3 stimulation (Fig. S1M-O).

### Chi3L1 regulates T_FH_ responses to *Hp* infection

Helminth infections also induce robust T follicular helper (T_FH_) responses^23, 29^. Since T_FH_ responses have not been reported in *Chi3l1*^-/-^ mice, we enumerated T_FH_ (CXCR5^+^PD1^hi^) cells in the msLN of uninfected and *Hp*-infected animals. T_FH_ cells (Fig. 1A) were detected in the msLN of uninfected BALB/c mice and increased in both frequency and number (Fig. 1B-C) over the first two weeks following *Hp* infection. Although the kinetics of the T_FH_ responses were similar between the *Hp*-infected BALB/c and *Chi3l1*^-/-^ mice, the magnitude of the *Chi3l1*^-/-^ T_FH_ response was significantly decreased (Fig. 1B-C). Moreover, while the T_FH_ master regulator^30^ Bcl-6 was expressed at similar levels in the msLN BALB/c and *Chi3l1*^-/-^ T_FH_ cells (Fig. 1D-E), expression of ICOS, a key co-stimulatory molecule for T_FH_ cells^30–34^, was significantly decreased on the *Chi3l1*^-/-^ CXCR5^+^PD-1^hi^ T_FH_ cells throughout infection (Fig. 2F-G). To assess whether the ICOS reduction was due to an intrinsic defect in *Chi3l1*^-/-^ T cells, we activated splenic CD4 T cells from uninfected WT and *Chi3l1*^-/-^ mice under early T_FH_ differentiation conditions^35^ with platebound anti-CD3 + anti-CD28 and IL-6 plus IL-2 blocking antibodies. We found decreased ICOS upregulation in the *in vitro* activated *Chi3l1*^-/-^ CD4 T cells (Fig. 1H). Therefore, Chi3L1 does not control T cell commitment to the T_FH_ lineage but does play a cell intrinsic role in regulating ICOS expression by CD4 T cells.

**Figure 1.**
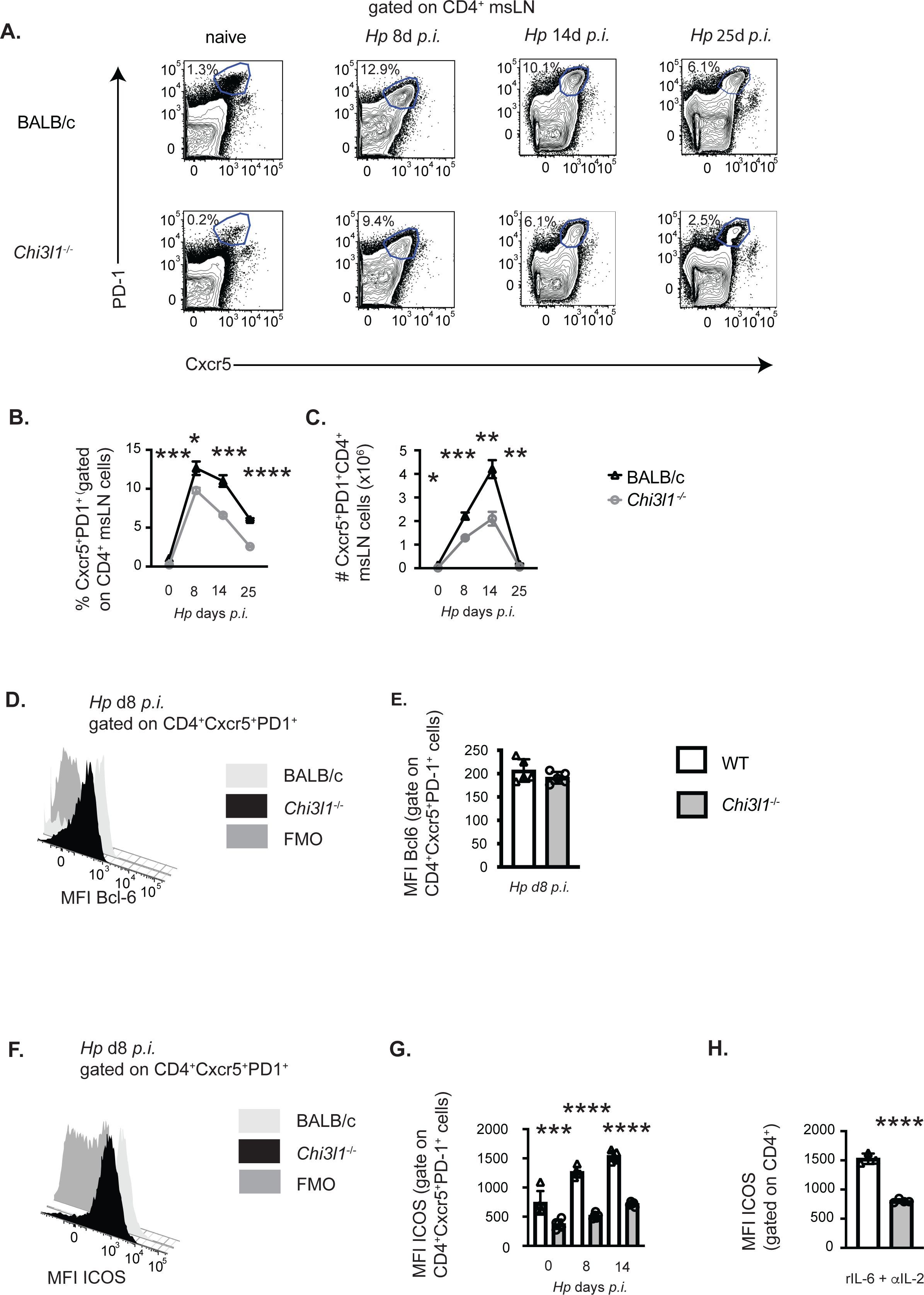
*Chi3l1* regulates T_FH_ responses to *Hp* infection. (**A-G**) Enumeration and characterization of msLN T_FH_ cells from uninfected (n=5/group) or *Hp*-infected BALB/c (white bars) and *Chi3l1*^-/-^ (grey bars) mice (n=5/group). (**A-C**) T_FH_ response to *Hp* infection with flow plots (**A**) showing CXCR5^+^PD-1^+^ T_FH_ cells between D0 (uninfected) and D25 post-*Hp* with frequency (**B**) and number (**C**) of T_FH_ cells. Kinetics of the msLN response (total, CD19 B cells, CD4 T cell and activated CD44^hi^CD62L^lo^) shown in Fig. S1A-E. Cytokine production by CD44^hi^ CD4 T cells shown in Fig. S1F-N. (**D-G**) T_FH_ phenotype in msLN of uninfected (D0) and *Hp*-infected mice. Bcl-6 (**D-E**) and ICOS (**F-G**) expression by T_FH_ on D8 (**D-F**) or between D0-D25 (**G**). Flow cytometry plots for Bcl-6 (**D**) and ICOS (**F**) are shown. Expression levels of Bcl-6 (**E**) or ICOS (**F**) in T_FH_ cells presented as the geometric Mean Fluorescence Intensity (gMFI). **(A)** (**H**) gMFI of ICOS expression by purified splenic CD4^+^ T cells from uninfected BALB/c (white bars) and *Chi3l1*^-/-^ (grey bars) mice (n=4/group) that were stimulated *in vitro* for 48 hours with anti-CD3 plus anti-CD28 in the presence of rIL-6 plus anti-IL-2. Data representative of 2 (**D-E**) or ≥ 3 independent experiments (all others). Data displayed as the mean±SEM (**B-C**) or mean±SD (**E, G, H**) of each group with individual animals depicted as circles or triangles. Statistical significance determined using unpaired 2-tailed student’s t test. *p≤0.05, **p≤0.01, ***p≤0.001, ****p≤0.0001.

**Figure 2.**
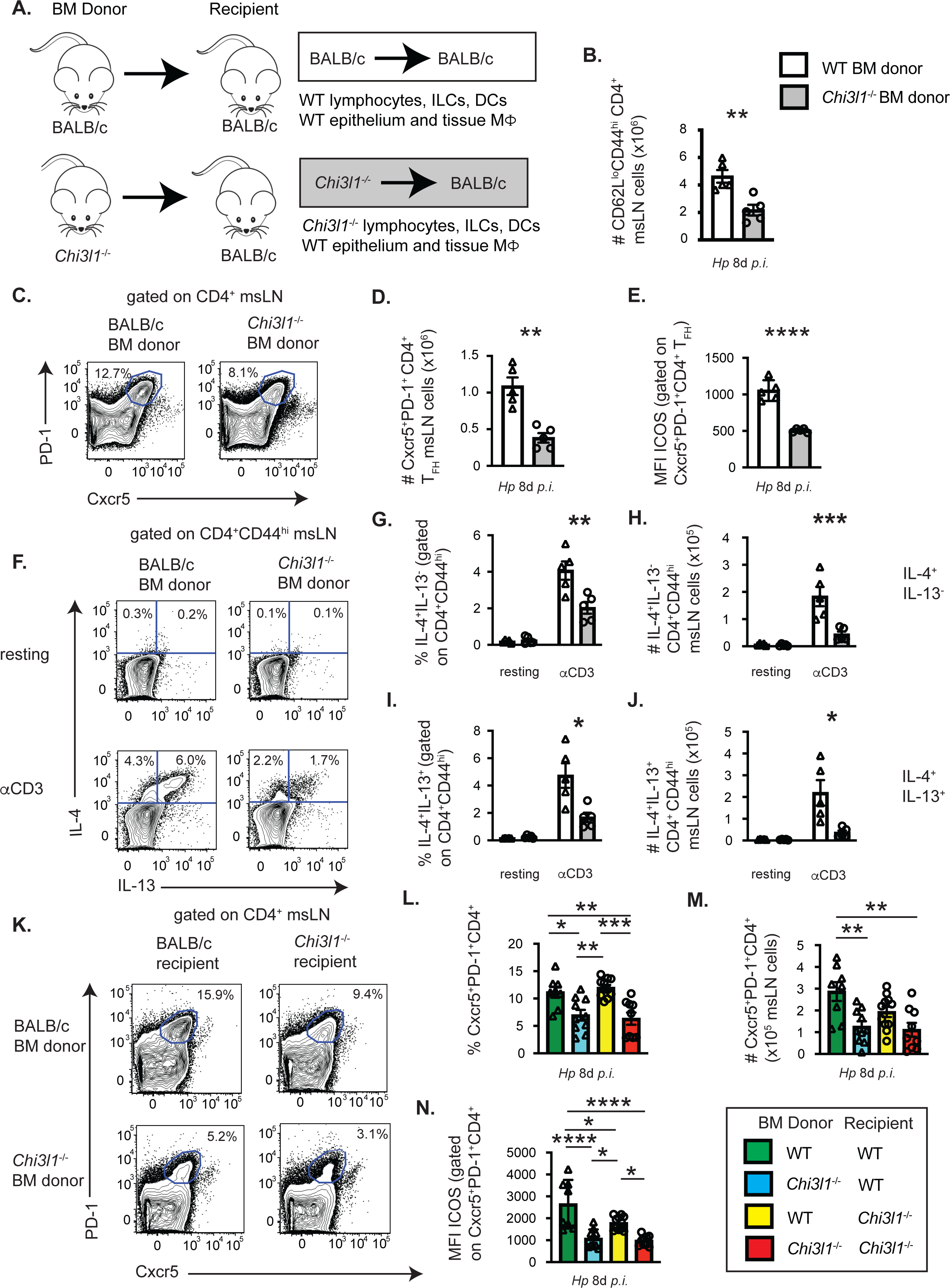
*Chi3l1* expressing hematopoietic cells are necessary and sufficient for T_FH_ and T_H_2 responses to *Hp*. (**A-J**) Enumeration of T cell responses in msLNs of D8 *Hp*-infected BM chimeric mice (n=5 mice/group) that were generated (**A**) by reconstituting lethally-irradiated BALB/c recipient mice with BALB/c (white bars) or *Chi3l1*^-/-^ (grey bars) BM. msLN CD4 cells from D8 *Hp*-infected chimeric mice were analyzed directly *ex vivo* (**B-E**) or following restimulation for 4 hours with anti-CD3 in the presence of BFA (**F-J**). (**B-E**) Representative flow cytometry plot (**C**) showing CXCR5^+^PD-1^+^ T_FH_ cells in D8 msLNs with the numbers (**D**) of T_FH_ cells and ICOS expression levels (**E**) by T_FH_ cells represented as gMFI. (**F-J**) IL-4 and IL-13 production by resting and anti-CD3 stimulated msLN cells. Representative flow plot **(A)** (**F**) showing IL-4 and IL-13 expression by CD44^hi^ CD4 cells from D8 *Hp*-infected chimeras with the percentage and number of IL-4^+^IL-13^neg^ (**G-H**) and IL-4^+^IL-13^+^ (**I-J**) producers. (**K-N**) T_FH_ responses in msLNs from *Hp*-infected BM chimeric mice that were generated by reconstituting lethally-irradiated: (i) BALB/c recipients with BALB/c BM (green bars), (ii) BALB/c recipients with *Chi3l1*^-/-^ BM (blue bars), (iii) *Chi3l1*^-/-^ recipients with BALB/c BM (yellow bars) and (iv) *Chi3l1*^-/-^ recipients with *Chi3l1*^-/-^ BM (red bars). msLN T_FH_ cells from reciprocal BM chimeras (n=5 mice/group) were analyzed on D8 post-*Hp* infection. Representative flow plots (**K**) showing CXCR5^+^PD-1^hi^ T_FH_ cells with the frequency (**L**) and number (**M**) of msLN T_FH_ cells in each group. gMFI of ICOS expression (**N**) by T_FH_ cells in each of the 4 groups of chimeras. Data representative of ≥ 3 independent experiments (**A-J**) or cumulation of 2 experiments repeated twice (**K-N**). Data displayed as the mean±SD (**E, N**) or mean±SEM (**all others**) of each group with cells from individual animals depicted as circles or triangles. Unpaired 2-tailed student’s t-test (**B, E-J**) or one-way ANOVA (**L-N**) was used to assess statistical significance. *p≤≤0.05, **p≤0.01, ***p≤0.001, ****p≤0.0001.

### Hematopoietic cell expression of *Chi3l1* regulates T_H_2 and T_FH_ responses to *Hp*

Transgene-directed *Chi3l1* expression by epithelial cells is reported to restore lung T_H_2 cytokine levels in allergen-exposed *Chi3l1*^-/-^ mice^13^. To assess whether the attenuated T_H_2 and T_FH_ responses to *Hp* were due to *Chi3l1* expression by radiation resistant cells, like epithelial cells, or to *Chi3l1* expression by radiation sensitive bone marrow (BM)-derived immune cells, we analyzed *Hp*-elicited T cell responses in BALB/c mice that were lethally irradiated and reconstituted with either BALB/c BM or *Chi3l1*^-/-^ BM (Fig. 2A). We observed a significant reduction in the numbers of total activated CD62L^lo^CD44^hi^ CD4 cells (Fig. 2B) and CXCR5^hi^PD1^hi^ T_FH_ cells (Fig. 2C-D) in the *Hp*-infected BALB/c recipients reconstituted with *Chi3l1*^-/-^ BM relative to animals reconstituted with BALB/c BM. Expression of ICOS on CXCR5^+^PD1^hi^ T_FH_ cells was also reduced in mice reconstituted with *Chi3l1*^-/-^ BM (Fig. 2E). Moreover, the effector function of CD4 T cells from BALB/c mice reconstituted with *Chi3l1*^-/-^ BM chimeras was impaired, as the frequencies and numbers of IL-4^+^ single producers (Fig. 2F-H) and IL-4^+^IL-13^+^ double producers (Fig. 2F, 2I-J) were decreased following anti-CD3 stimulation.

Next, we asked whether *Chi3l1* expression by the non-hematopoietic radiation resistant cells might also contribute to the development of robust T_FH_ responses to *Hp*. We therefore enumerated T_FH_ cells in D8 *Hp*-infected BM chimeras that were competent to express *Chi3l1* in all cell types (WT donor/WT recipient), lacked *Chi3l1* in all cell types (KO donor/KO recipient), lacked expression of *Chi3l1* specifically in the hematopoietic compartment (KO donor/WT recipient) or lacked expression of *Chi3l1* in the radiation resistant compartment (WT donor/KO recipient). Loss of *Chi3l1* expression specifically in radiation sensitive hematopoietic cells was sufficient to account for the reduction in percentage and number of msLN T_FH_ cells (Fig. 2K-M) and the attenuated ICOS expression by the T_FH_ cells (Fig. 2N). Thus, *Chi3l1* expression selectively within radiation sensitive immune cells is required for optimal T_FH_ responses to *Hp* infection.

### *Chi3l1* regulates B cell responses to *Hp* infection and immunization

Following *Hp* infection, Chi3L1 regulates T_FH_ ICOS expression and the size of the T_FH_ response. ICOS is a key co-stimulatory molecule that modulates T_FH_-B cell interactions, particularly during the germinal center B cell (GCB) response^33, 36^. To test whether *Chi3l1*^-/-^ mice mount impaired B cell responses to *Hp*, we analyzed B cell subsets in mLNs of D14 *Hp*-infected BALB/c and *Chi3l1*^-/-^ mice (Fig. 3A, Fig. S2A-C). The numbers of total msLN cells (Fig. 3B) and B220^+^CD138^-^ B cells (Fig. 3C) in D14 *Hp*-infected mice were decreased in the *Chi3l1*^-/-^ mice relative to WT mice. This reduction was not due to changes in the number of naïve B cells (Fig. 3D). Instead, antigen-experienced isotype-switched B cells (Fig. 3E), isotype-switched GCB cells (Fig. 3F), and antibody secreting cells (ASCs) (Fig. 3G) were decreased in the D14 *Hp*-infected *Chi3l1*^-/-^ msLNs. Thus, *Chi3l1* regulates both T cell and B cell responses to *Hp*.

**Figure 3.**
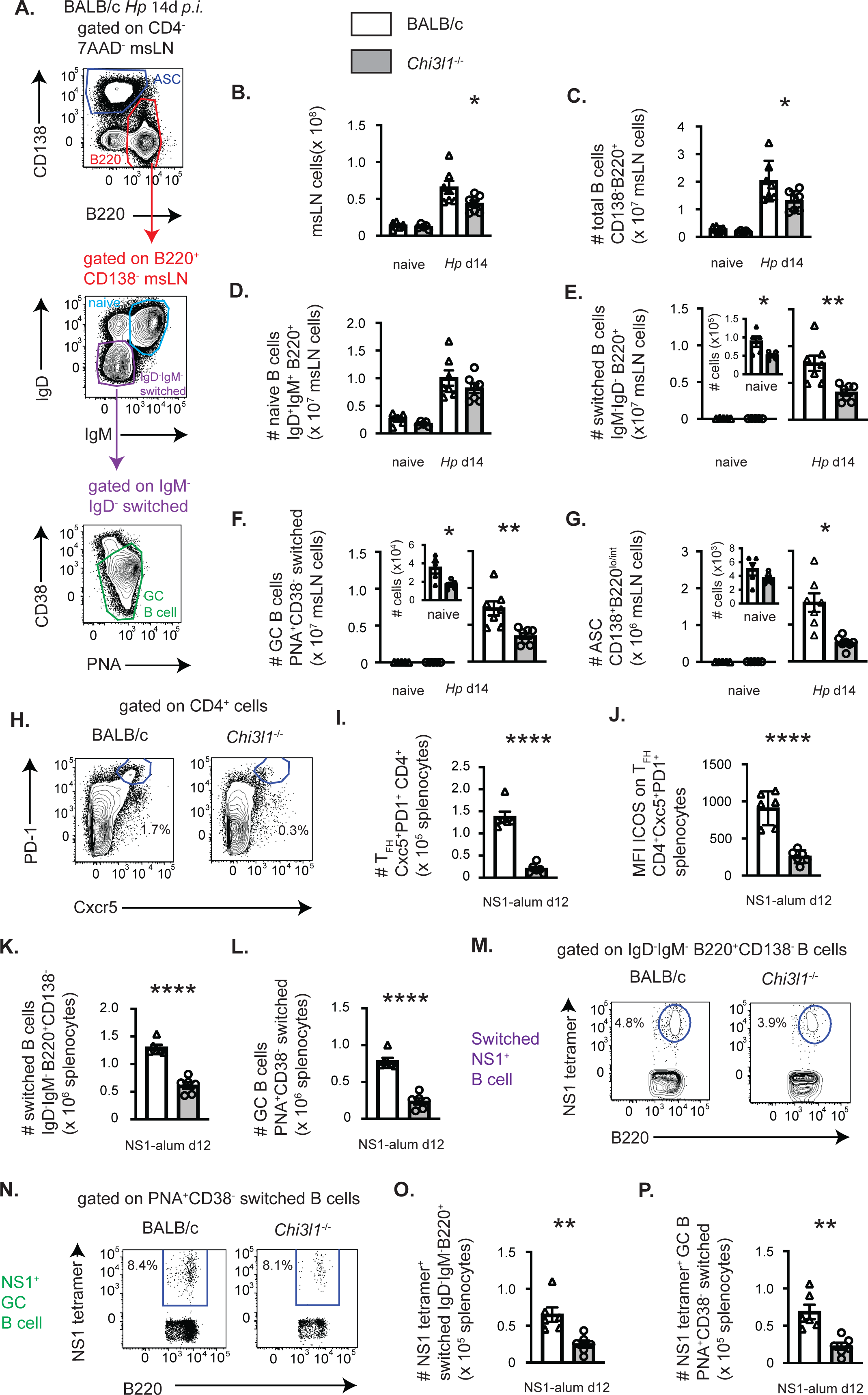
*Chi3l1* regulates B cell responses to *Hp* infection and alum-adjuvanted protein immunization. Enumeration of B cell responses in BALB/c (white bars) and *Chi3l1*^-/-^ (grey bars) mice following infection with *Hp* (**A-G**) or vaccination with NS1 protein in alum (**H-P**). (**A-G**) Characterization of B cell responses in msLNs of uninfected (n=5/group) and D14 *Hp*-infected (n=7/group) BALB/c and *Chi3l1*^-/-^ mice. Flow plots (**A**) showing the gating strategy to identify B220^lo^CD138^hi^ ASCs (blue gate), total B cells (red gate), IgD^neg^IgM^neg^ isotype-switched B cells (purple gate), naïve B cells (cerulean gate) and PNA^hi^CD38^lo^ GCB cells (green gate). Gating strategies to identify B cell subsets with representative flow plots from BALB/c and *Chi31l*^-/-^ mice shown in Fig. S2A-C. Numbers of total msLN cells (**B**), B cells (**C**), naïve B cells (**D**), isotype-switched B cells (**E**), GCB cells (**F**) and ASCs (**G**). (**H-P**) BALB/c and *Chi3l1*^-/-^ mice (n=6/group) on D12 post-immunization with NS1 protein adsorbed to alum. Enumeration of vaccine-elicited splenic T_FH_ (**H-J**) and B cells (**K-P**). (**H-J**) Representative flow plots showing the gating strategy (**H**) to identify and enumerate (**I**) splenic CXCR5^+^PD1^+^ T_FH_ cells. ICOS expression (**J**) by T_FH_ cells reported as gMFI. Numbers of total splenocytes and CD4 T cells shown in Fig. S2D-E. (**K-P**) Numbers of vaccine-elicited splenic isotype-switched B cells (**K**) and GCB cells (**L**). Representative flow plots with B cell subsets gating strategies in vaccinated BALB/c and *Chi3l1*^-/-^ mice shown in Fig. S2F. Numbers of naïve B cells and ASCs shown in Fig. S2G-H. Representative flow plots to identify NS1-specific (NS1^+^) isotype-switched B cells (**M**), NS1^+^ isotype-switched GCB (**N**) and NS1^+^ ASCs (Fig. S2I). Numbers of NS1^+^ isotype-switched cells (**O**), NS1^+^ GCB cells (**P**) and NS1^+^ ASCs (Fig. S2J). Data is representative of 3 (**A-G**) or 2 (**H-P**) independent experiments. Data displayed as the mean±SD (**J**) or mean±SEM (all others) of each group with individual animals depicted as circles or triangles. Statistical analysis was performed with unpaired 2-tailed student’s t-test. *p≤0.05, **p≤0.01, ***p≤0.001, ****p≤0.0001.

To determine whether the *Chi3l1*-dependent attenuated B cell responses was limited to *Hp* infections, we immunized mice with recombinant influenza NS1 protein that was adsorbed to the T_H_2-biasing adjuvant alum and measured splenic responses on D12. The numbers of total splenocytes (Fig. S2D), CD4 T cells (Fig. S2E) and naïve B cells (Fig. S2F-G) did not change following vaccination of either group. However, the number of splenic T_FH_ cells in immunized *Chi3l1*^-/-^ mice was significantly reduced (Fig. 3H-I) and the remaining *Ch3l1*^-/-^ T_FH_ cells expressed decreased levels of ICOS (Fig. 3J). Similarly, the numbers of splenic isotype-switched B cells (Fig. 3K), GCB cells (Fig. 3L) and ASCs (Fig. S2H) were decreased in vaccinated *Chi3l1*^-/-^ mice. Next, we assessed NS1-specific B cells using fluorochrome-labeled NS1 protein tetramers (Fig. 3M-N, Fig. S2I-J). We found very few NS1-specific splenic ASCs at this timepoint post-vaccination in either group (Fig. S2I-J). However, the numbers of NS1-specific isotype-switched B cells (Fig. 3O) and NS1-specific GC B cells (Fig. 3P) were significantly decreased in vaccinated *Chi3l1*^-/-^ mice. Thus, *Chi3l1* regulates the magnitude of T_FH_ and B cell responses to both infection and immunization.

### *Chi3l1* regulates IgE responses

Given our data and published reports showing that *Hp*-induced IgE responses depend on T_FH_ cells and IL-4^25^, we next analyzed IgE responses in the *Hp*-infected *Chi3l1*^-/-^ mice. We observed a significant reduction in the frequency and number of IgE^+^ ASCs in the msLNs from *Hp*-infected *Chi3l1*^-/-^ mice (Fig. 4A-C) compared to the WT animals. Moreover, the impaired IgE^+^ ASC response was accompanied by a significant reduction in total IgE levels in the serum of D21 *Hp*-infected *Chi3l1*^-/-^ mice (Fig. 4D). In contrast, *Hp-*specific IgG1 serum antibody titers were equivalent between *Chi3l1*^-/-^ and WT mice (Fig. 4E). Thus, Chi3L1 selectively regulates the polyclonal IgE ASC response following *Hp* infection.

**Figure 4.**
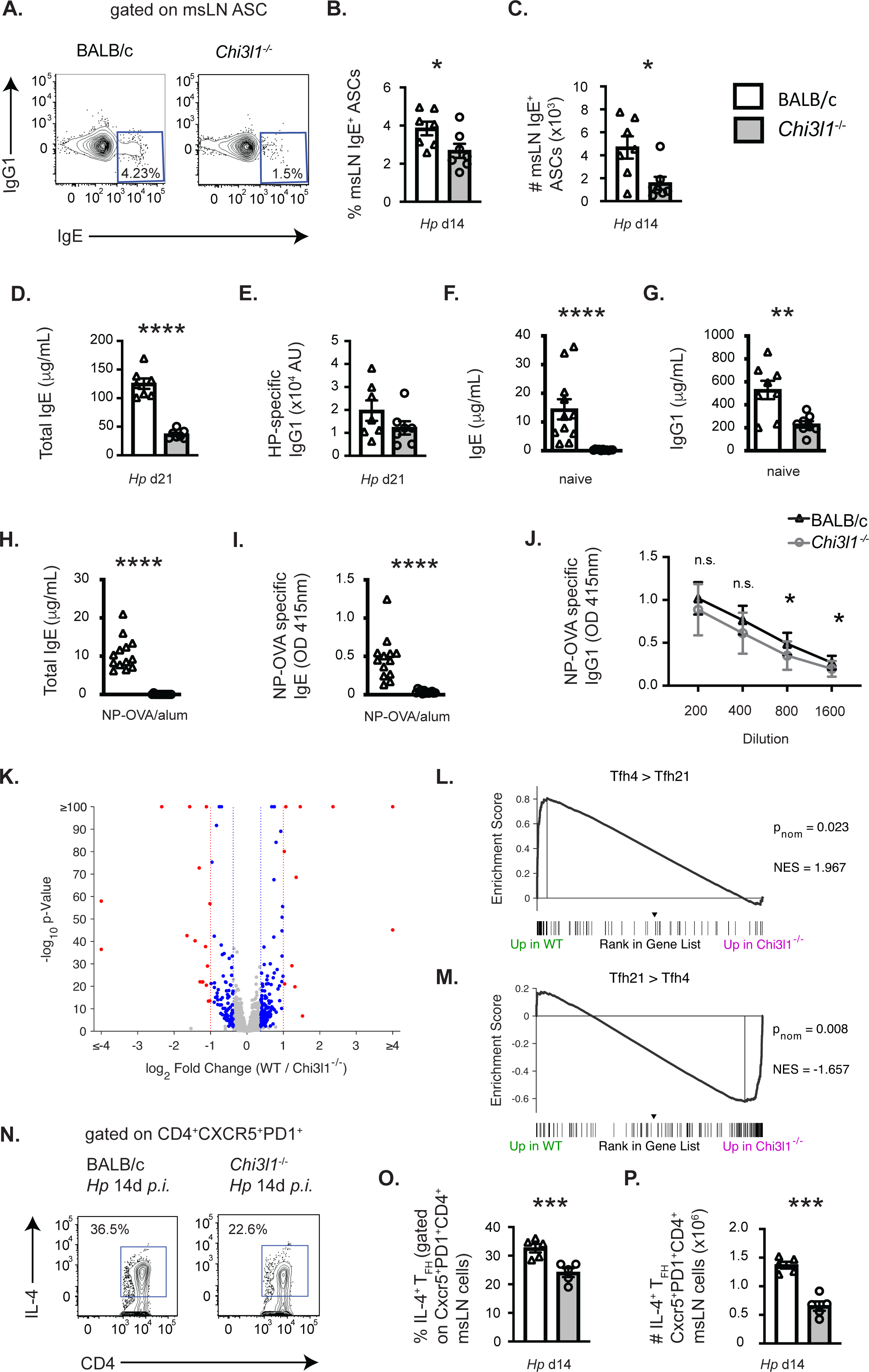
*Chi3l1* regulates T_FH4_ programming and IgE responses to infection and immunization. (**A-J**) Evaluation of IgE and antibody responses in *Hp*-infected (**A-E**), naïve (**F-G**) and NP-OVA immunized (**H-J**) BALB/c (white bars) and *Chi3l1*^-/-^ (grey bars) mice. (**A-E**) Quantitation of D14 IgE-expressing ASCs (**A-C**) and D21 serum IgE antibody titers (**D-E**) from *Hp*-infected mice (n=7 mice/group). Representative flow plots showing msLN IgE-expressing ASCs (**A**) with the frequency (**B**) and number (**C**) of IgE-expressing ASCs. Gating strategy to identify *Hp*-induced ASCs shown in Fig. S2C. ELISA quantitation of D21 total serum IgE (**D**) and *Hp*-specific IgG1 (**E**). (**F-G**) Serum antibody levels in naïve mice (n=8/group). Data is reported in μg/ml for total IgE (**F**), and IgG1 (**G**). (**H-J**) NP-specific antibody responses measured by ELISA on D14 post-immunization with NP-OVA adsorbed to alum (n=13/group). Total serum IgE (**H**, reported in μg/ml), NP-specific IgE (**I**) and NP-specific IgG1 (**J**) are shown. NP-specific IgE and IgG1 reported as OD values for serum (1:10 dilution for **I** and 1:200-1:1600 for **J**). (**K-M**) RNA-seq analysis of sorted-purified CXCR5^+^PD1^+^ msLN T_FH_ isolated from D14 *Hp*-infected WT and *Chi3l1*^-/-^ mice (n=9 pooled samples/group). Volcano plot (**K**) showing 9853 expressed genes. 1465 genes were expressed at significantly different levels (FDR q<0.05) between WT and *Chi3l1*^-/-^ T_FH_ cells. Of those, genes meeting a log_2_FC of ±0.3785 (252 genes) are indicated with blue or red symbols. Red symbols indicate the 28 genes meeting a threshold of abs(log_2_FC) of ±1, blue indicate the 224 genes with abs(log2FC) between 0.3785 and 1.0. Gene set enrichment analysis (GSEA) using the ranked gene list of WT and *Chi3l1*^-/-^ T_FH_ cells and DEG identified as up in IL4^+^IL-21^neg^ T_FH_ cells (T_FH4_, panel **L**) or up in IL-4^neg^IL-21^+^ T_FH_ cells (T_FH21_, panel **M**) isolated from *Nippostrongylus brasiliensis*-infected mice^40^. p_nom_ and NES are provided. Principal component analysis and absolute expression levels of the 28 genes meeting log_2_FC of ±1 shown in Fig. S3A-B. IPA analysis of the 252 genes meeting the log_2_FC of ±0.3785 are shown in Fig. S3C-E. Complete gene list used in all analyses provided in Table S1. (**N-P**) Cytokine production by D14 *Hp*-infected WT and *Chi3l1*^-/-^ T_FH_ cells (N=5 mice/group) following *in vitro* anti-CD3 restimulation. Flow plots (**N**) measuring IL-4 production and the percentage (**O**) and number (**P**) of T_FH_4 in each culture. Data representative of 3 (**A-E, K-O**), 5 (**F**), 1 (**G**) or 2 (**H-J**) independent experiments. Data displayed as the mean±SEM of each group with individual animals depicted as circles or triangles. Statistical analysis was performed with unpaired 2-tailed student’s t-test. *p≤0.05, **p≤0.01, ***p≤0.001, ****p≤0.0001.

To assess whether the defective IgE response in *Chi3l1*^-/-^ mice was restricted to *Hp* infection, we first examined IgE levels in non-infected animals. Although serum IgE levels were low in WT naïve mice, IgE was detected (Fig. 4F). In contrast, serum IgE levels were undetectable (at least 100-fold lower) in the *Chi3l1*^-/-^ serum (Fig. 4F). However, total IgG1 in serum from uninfected *Chi3l1*^-/-^ mice was only decreased ∼2.5 fold relative to the WT animals (Fig. 4G), again suggesting a more profound deficit in IgE production by *Chi3l1*^-/-^ mice. To address whether antigen-specific IgE responses are also dependent on *Chi3l1*, we vaccinated mice with nitrophenyl haptenated ovalbumin (NP-OVA) adsorbed to alum and measured total and NP-specific IgE on D14. Again, total IgE levels were significantly lower in the vaccinated *Chi3l1*^-/-^ mice (Fig. 4H). Moreover, NP-specific IgE, which was easily measured in vaccinated WT mice, was undetectable in *Chi3l1*^-/-^ serum (Fig. 4I). In contrast, the IgG1 NP-specific Ab response was reduced no more than 2-fold in *Chi3l1*^-/-^ mice relative to the WT controls. (Fig. 4J). Thus, *Chi3l1* plays a central role in IgE responses but is dispensable for IgG1 responses.

### *Chi3l1* regulates IL-4^+^ T_FH_ programming

IL-4 producing T_FH_ cells are required for the polyclonal IgE response to *Hp*^25^. Since T_FH_ cells were decreased but not ablated in the *Hp*-infected *Chi3l1*^-/-^ mice, we suspected that the large reduction in IgE responses seen in these mice might reflect additional functional impairments in the *Chi3l1*^-/-^ T_FH_ cells. To assess this in an unbiased fashion, we compared the transcriptome of T_FH_ cells from msLNs of WT and *Chi3l1*^-/-^ mice on D14 post *Hp* infection. We identified 9853 expressed genes (Fig. 4K) and 1465 differentially expressed genes (DEG) between the two T_FH_ populations that met an FDR q<0.05 cutoff (Table S1) with the major principal component differentiating the samples being genotype (Fig. S3A). 252 DEG exhibited at least a ±0.3785 log_2_FC (Fig. 4K) and 28 of those DEG met a ± 1 log_2_FC threshold (Fig. 4K, Fig. S3B). Since the number of DEG was small, we performed an unsupervised gene set enrichment analysis (GSEA) comparing the rank-ordered WT and *Chi3l1*^-/-^ T_FH_ gene set to the 5219 C7 Immunologic Signature Gene Sets from MSigDB. None of the C7 gene sets were preferentially enriched (FDR q<0.05) in the WT or *Chi3l1*^-/-^ T_FH_ transcriptomes, including the 54 C7 gene sets derived from T_FH_ cells (*data not shown*). This result, which was consistent with normal expression of the T_FH_ lineage master regulator Bcl-6 by *Chi3l1*^-/-^ T_FH_ cells (Fig. 1D-E), indicated that *Chi3l1* is not required for establishment of the core T_FH_ transcriptional program^37^.

We next performed Ingenuity Pathway Analysis (IPA) with the 252 genes meeting an FDR q<0.05 and ± 0.3785 log_2_FC threshold to assess whether changes in expression of suites of genes might reflect alterations in *Chi3l1*^-/-^ T_FH_ cell function or signaling. IPA-defined pathways that were predicted to be most significantly different (B-H p < 0.05) between the WT and *Chi3l1*^-/-^ T_FH_ cells included T_H_2 and T_H_1/T_H_2 activation (Fig. S3C). IPA functions that were predicted to be most different between the WT and *Chi3l1*^-/-^ T_FH_ cells (B-H p value < 0.001) included those related to cell movement, leukocyte quantity, glucose metabolism and inflammation/immune mediated disease (Fig. S3D). 94 of the 119 (79%) unique DEGs associated with these 15 functions were associated with cell movement (72 DEG) or cell quantity (62 DEG) and several overlapped into both functional categories (40 DEG) (Fig. S3E). Thus, transcriptional networks associated with cell migration and cell quantity are predicted to be altered in *Chi3l1*^-/-^ T_FH_.

IL-4 produced by T_FH_ cells is required for B cell class switching to IgE following *Hp* infection^25, 38^ and even modest reductions in the amount of IL-4 present can prevent B cell switching to IgE while having no impact on the IgG1 response^39^. Prior studies examining the T_FH_ response after infection with the nematode *Nb* showed that T_FH_ cells change over time following infection and proceed from producing IL-21 alone (T_FH21_ cells) to producing primarily IL-4 (T_FH4_ cells)^40^. This change in the T_FH_ cytokine profile from IL-21 to IL-4 is associated with migration of T_FH_ cells into the B cell follicle^41^, formation of ICOS-regulated T_FH_/B cell conjugates that support T_FH_ cell survival and expansion^42^ and acquisition of a transcriptionally distinct T_FH4_ program^40^. Given our data, we hypothesized that Chi3L1 regulates acquisition of the T_FH4_ transcriptional program. To assess this, we performed GSEA using published lists of genes that are differentially expressed in T_FH4_ and T_FH21_ cells from *Nb-*infected mice^40^. We found that DEG that are normally upregulated in *Nb* T_FH4_ cells compared to *Nb* T_FH21_ cells^40^ were significantly enriched in *Hp* WT T_FH_ cells compared to *Hp Chi3l1*^-/-^ T_FH_ (Fig. 4L). In contrast, we observed significant enrichment in the transcriptome of the *Hp*-induced *Chi3l1*^-/-^ T_FH_ cells for genes that are increased in *Nb* T_FH21_ cells relative to *Nb* T_FH4_ cells. To address whether loss of *Chi3l1* also altered IL-4 production specifically by T_FH_ cells, we measured IL-4 in anti-CD3 restimulated CXCR5^+^PD1^hi^ T_FH_ cells from D14 *Hp*-infected WT and *Chi3l1*^-/-^ mice. Consistent with the GSEA data, we observed a significant decrease in frequency and number (Fig. 4O-P) of *Chi3l1*^-/-^ T_FH4_ cells. Together, the data support the conclusion that *Chi3l1* contributes to the development or maintenance of the ICOS-expressing T_FH4_ subset that facilitates B cell IgE responses.

Here, we provide the first evidence of a role for *Chi3l1* in T_FH_ responses. Our data reveal new functions for Chi3L1 in regulating: expression of ICOS by T_FH_ cells, expansion of the T_FH_ compartment, acquisition of the T_FH4_ program, establishment of T_FH_-dependent B cell responses and production of T_FH_ and B cell-dependent IgE. We show that these effects are due to Chi3L1 expressed by hematopoietic cells and that deletion of *Chi3l1* selectively ablates IgE responses to immunization without affecting IgG1 responses to the same immunogen. Given the known roles for ICOS/ICOSL interactions in the maintenance and expansion of T_FH_ cells, in T_FH_/B cell responses, and in IgE responses to allergens^42–45^, we postulate that some of the *Chi3l1*^-/-^ T_FH_ functional deficits can be explained by a *Chi3l1*-dependent T cell intrinsic defect that prevents normal ICOS upregulation following TCR and CD28 engagement. In the future, it will be important to assess how Chi3L1 regulates ICOS expression and whether blocking Chi3L1 activity can attenuate T_H_2 responses and selectively block T_FH_-driven IgE responses in pathologic settings such as atopy and allergic airway disease.

## Methods

### Mice

Animals were bred and maintained in the UAB animal facilities. All procedures were approved by the UAB IACUC and were conducted in accordance with the principles outlined by the National Research Council. BALB/cByJ mice were purchased from The Jackson Laboratory. BALB/c BRP-39 (*Chi3l1*^-/-^) mice^13^ were provided by Dr. Allison Humbles (MedImmune). Bone marrow (BM) chimeras were generated by irradiating recipients with 850 Rads from a high-energy X-ray source (split dose 5 hours apart), and then reconstituting the recipients with 5x10^6^ total BM cells delivered iv. Chimeras were used in experiments 8-12 weeks post-reconstitution. Both male and female mice were used and animals were matched for age and sex within an experiment. No gender-specific differences were observed.

### Heligmosomoides polygyrus stock

*H polygyrus* (*Hp*) stocks were maintained as described previously^22, 23^. Mice were gavaged with 200 *Hp* L3 larvae.

### Flow cytometry

Single cell suspensions from msLN or spleens were preincubated with FcR blocking antibody (2.4G2) and then stained with cocktails of labeled antibodies. Antibodies from BD: CD3 (17A2), CD4 (GK1.5), CD25 (PC61 BD), CD44 (IM7), B220 (RA3-6B2), CD62L (MEL-14), CD138 (281-2). Cxcr5 (2G8) and IgE (R35-72). Antibodies from eBioscience: CD19 (1D3), CD38 (90), CD279/PD-1 (J43), CD278/ICOS (7E.17G9 and C398.4A), IgD (11-26), IgM (II/41) and B220 (RA3-6B2). Other reagents included: CD4 (GK1.5, Biolegend), AF488-labeled PNA (Life Technologies) or PNA (Sigma) labeled with Pacific Blue (Invitrogen), 7AAD (Invitrogen^TM^), and LIVE/DEAD® aqua or red (Life Technologies).

Intracellular Bcl-6 (K112-91, BD) was detected using eBioscience^TM^ Foxp3/Transcription Factor Staining Buffer set (Invitrogen^TM^). To detect intracellular cytokines, cells were restimulated with 2.5 μg/mL plate-bound anti-CD3 (145-2C11 BioXcell) or 5 ng/mL PMA (Sigma) with 1.25 μM calcimycin (CalBiochem) plus Brefeldin A (BFA, 12.5 μg/mL, Sigma) for 4 hours. Cells were washed, incubated with FcBlock, surface stained and washed. Cells were fixed in formalin, permeabilized with 0.1% NP-40, washed, incubated with anti-cytokine antibodies (IL-4 (11B11, BD and Invitrogen), IL-13 (eBio13A, eBioscience), IFNγ (XMG1.2 eBioscience) in permeabilization buffer and washed. All incubations prior to fixation were performed with BFA. All flow analysis was performed using the BD Canto.

### In vitro activation of CD4 T cells

CD4^+^ T cells were purified from spleens of uninfected BALB/c and *Chi3l1*^-/-^ mice using CD4 MACS L3T4 beads (Miltenyi Biotech) and stimulated for 48 hours as previously described^35^ with 2.5 μg/mL plate-bound anti-CD3 (145-2C11 BioXcell) and 2.5μg/ml plate-bound anti-CD28 (37.51) in the presence of 10μg/ml rIL-6 (R&D) and neutralizing antibodies to IL-2 (JES6-1A12 BioXcell, 5μg/ml and S4B6-1 BioXcell, 5μg/mL).

### Cell sorting

Single cell suspensions of msLN cells from D14 *Hp*-infected BALB/c ByJ and *Chi3l1*^-/-^ mice were incubated with FcBlock (BD), enriched with CD4 microbeads (Miltenyi), and then stained with fluorochrome-conjugated antibodies. CD4^+^Cxcr5^+^PD-1^neg^Lin^neg^ cells were sorted (BD Aria, UAB Flow Cytometry Core), pelleted, lysed in TRIzol (ThermoFisher), then stored at -80°C.

### Production of recombinant Influenza NS1 Antigen and NS1 tetramers

The influenza NS1 gene, modified to contain a 3’ in frame addition of the BirA enzymatic biotinylation site and the 6X-His purification tag (GeneArt), was mutated at R38A and K41A (to prevent aggregation of NS1 at high concentrations^46^), then cloned into the pTRC–His2c expression vector (Invitrogen) and expressed in the BirA-enzyme containing *E. coli* strain CVB101 (Avidity). Biotinylated recombinant NS1 was purified by FPLC and then tetramerized to fluorochrome-conjugated streptavidin (Prozyme). To detect NS1-specific B cells, PE-labeled NS1 tetramers (1:100) were incubated with cells for 30 minutes at 4°C.

### NP(15)-OVA or NS1 Immunization

50μg biotinylated NS1 protein or 50μg (4-hydroxy-3-nitrophenyl)-acetyl(15)-OVA (NP-OVA) was adsorbed to 100μg alhydrogel alum (InVivoGen) in a total of 200μL per mouse for 30 min at room temperature, then injected *i.p.* Animals were analyzed on D12 (splenic T and B cell responses) or D14 (serum antibody).

### ELISAs

#### IgG1 detection

Serum from uninfected mice was analyzed for total IgG1 using a mouse clonotyping kit and standards (Southern Biotech) according to manufacturer’s recommendations. *Hp-*specific IgG1 was detected as previously described^47^ using plates coated with *Hp* extract and detected using rat anti-mouse IgG1-HRP antibody (Southern Biotech). NP-OVA specific IgG1 was detected as previously described^48^ using plates were coated with TNP(5)-BSA and detected using goat anti-mouse IgG1-HRP (Southern Biotech).

#### IgE detection

For analysis of total IgE in serum from uninfected, D21 *Hp*-infected mice, or NS1 immunized mice, ELISAs were performed using paired rat anti-mouse IgE antibodies (capture with 1 μg/mL purified R35-72, BD Biosciences, detect with biotinylated R35-118, BD Biosciences) and streptavidin-HRP (BD Biosciences). Purified mouse IgE Κ was used as a standard (C38-2, BD). To detect NP-OVA specific IgE, plates were coated with 1 μg/mL purified anti-IgE (R35-72, BD). Samples were applied by 2-fold dilutions to the coated plates. Specific IgE was detected by biotinylated TNP(5)-BSA followed by streptavidin-HRP.

### RNA-seq analysis

RNA-seq libraries were prepared (see Fig. S3 for detailed description) from sort-purified msLN T_FH_ cells derived from D14 *Hp*-infected WT and *Chi3l1*^-/-^ mice. Expressed genes were defined as genes with at least 3 RPM in all samples of either the WT and/or *Chi3l1*^-/-^ T_FH_. Expressed genes were ranked by multiplying the -log_10_ of the P-value from DESeq2 by the sign of the FC and then used as input in the GSEA^49^ PreRanked analysis program (http://software.broadinstitute.org/gsea/index.jsp). RNA-seq data sets were deposited in the NCBI Gene Expression Omnibus (GEO) under accession GSE203113. All RNA-seq processing code can be found at https://github.com/cdschar/Curtiss_Tfh_RNAseq/.

### Statistical Analysis

Statistical details of all experiments including tests used, n, and number of experimental repeats are provided in figure legends. FlowJo (version 9, Tree Star) was used for flow cytometric analyses. Prism Graphpad (version 9) was used for statistical analyses of flow cytometry experiments. Statistical analysis of RNA-seq experiments is summarized within the text of the RNA-seq experimental design in Supplemental Figure 3.

## Supplemental Figures and Table Legends

**Supplemental Figure S1.**
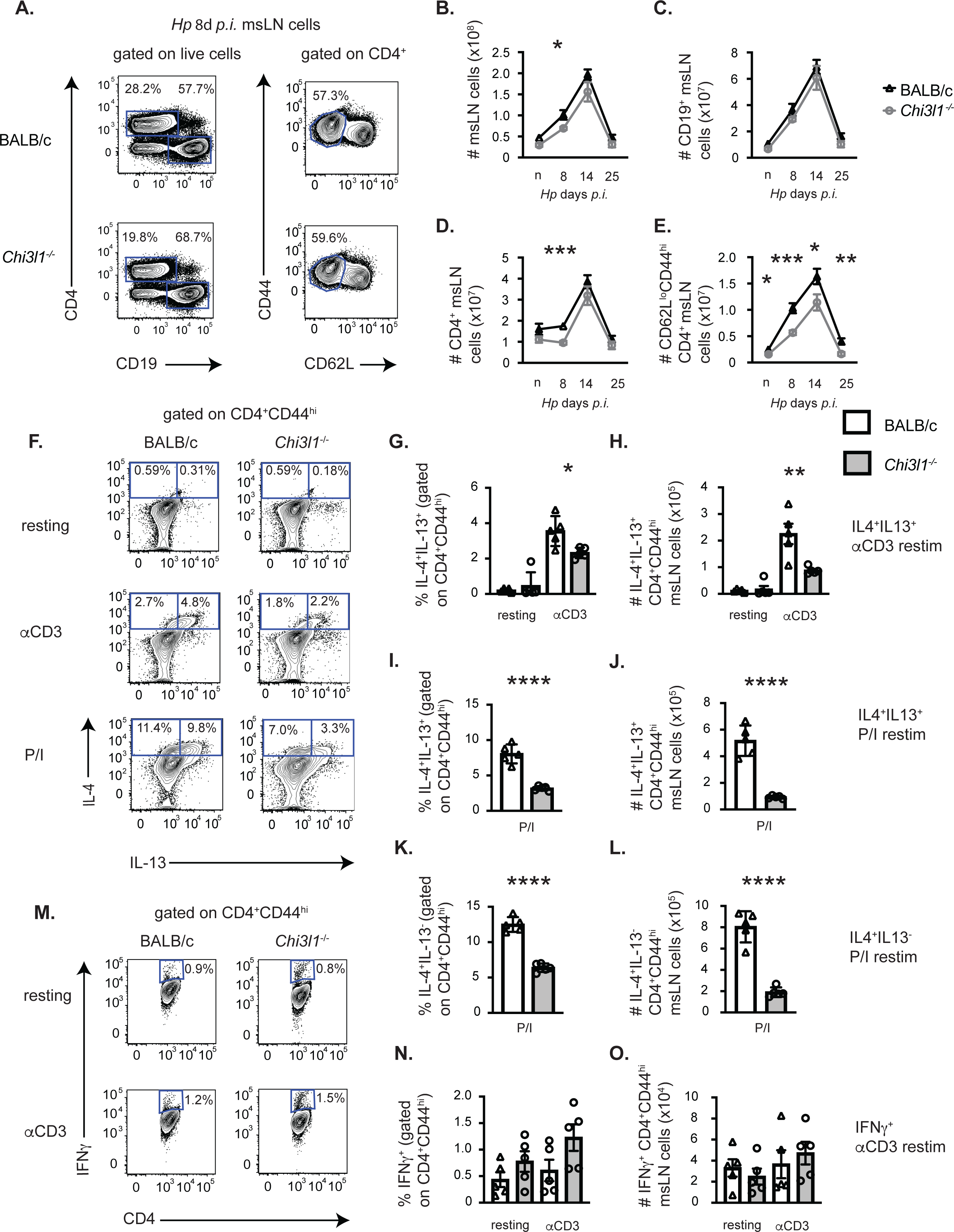
showing supporting information for Figure 1. CD4 effector type 2 cytokine responses are impaired in *Hp*-infected *Chi3l1*^-/-^ mice. (**A-D**) Enumeration and characterization of CD4 T cell subsets in msLNs of uninfected (n=5/group) and D8 *Hp*-infected (n=5/group/timepoint) BALB/c (white bars) and *Chi3l1*^-/-^ (grey bars) mice. Kinetic analysis of total msLN cells and lymphocytes between D0 (uninfected) and D25. gated as shown in (**A**). Data reported as number of total msLN cells (**B**), CD19^+^ B cells (**C**), CD4^+^ T cells (**D**), and CD62L^lo^CD44^hi^ CD4 cells (**E**) at each timepoint. (**F-O**) Cytokine production by CD4 cells from the msLN of D8 *Hp*-infected (n=5/group) BALB/c (white bars) and *Chi3l1*^-/-^ (grey bars) mice cultured for 4 hours with Brefeldin A (BFA). Intracellular IL-4, IL-13 and IFNγ were measured in CD44^hi^ gated CD4 T cells that were rested or restimulated for 4 hours with plate-bound anti-CD3 (**G-H, N-O**) or PMA and ionomycin (**I-L**). (**F-L**) IL-4 and IL-13 production by resting and anti-CD3 stimulated msLN CD44^hi^ CD4 cells (**F**) with the percentage and number of IL-4^+^IL-13^+^ (**G-J**) producers. Percentage and number of IL-4^+^IL-13^neg^ cells restimulated with PMA and ionomycin (**K-L**). **M-O**) IFNγ production by resting and anti-CD3 stimulated msLN CD44^hi^ CD4 cells (**M**) with the percentage and number of IFNγ^+^ (**N-O**) producers. Data representative of ≥ 3 independent experiments and displayed as the mean±SEM of each group with individual animals depicted as circles or triangles. Statistical analysis was performed with unpaired 2-tailed student’s t-test. *p≤0.05, **p≤0.01, ***p≤0.001, ****p≤0.0001.

**Supplemental Figure S2.**
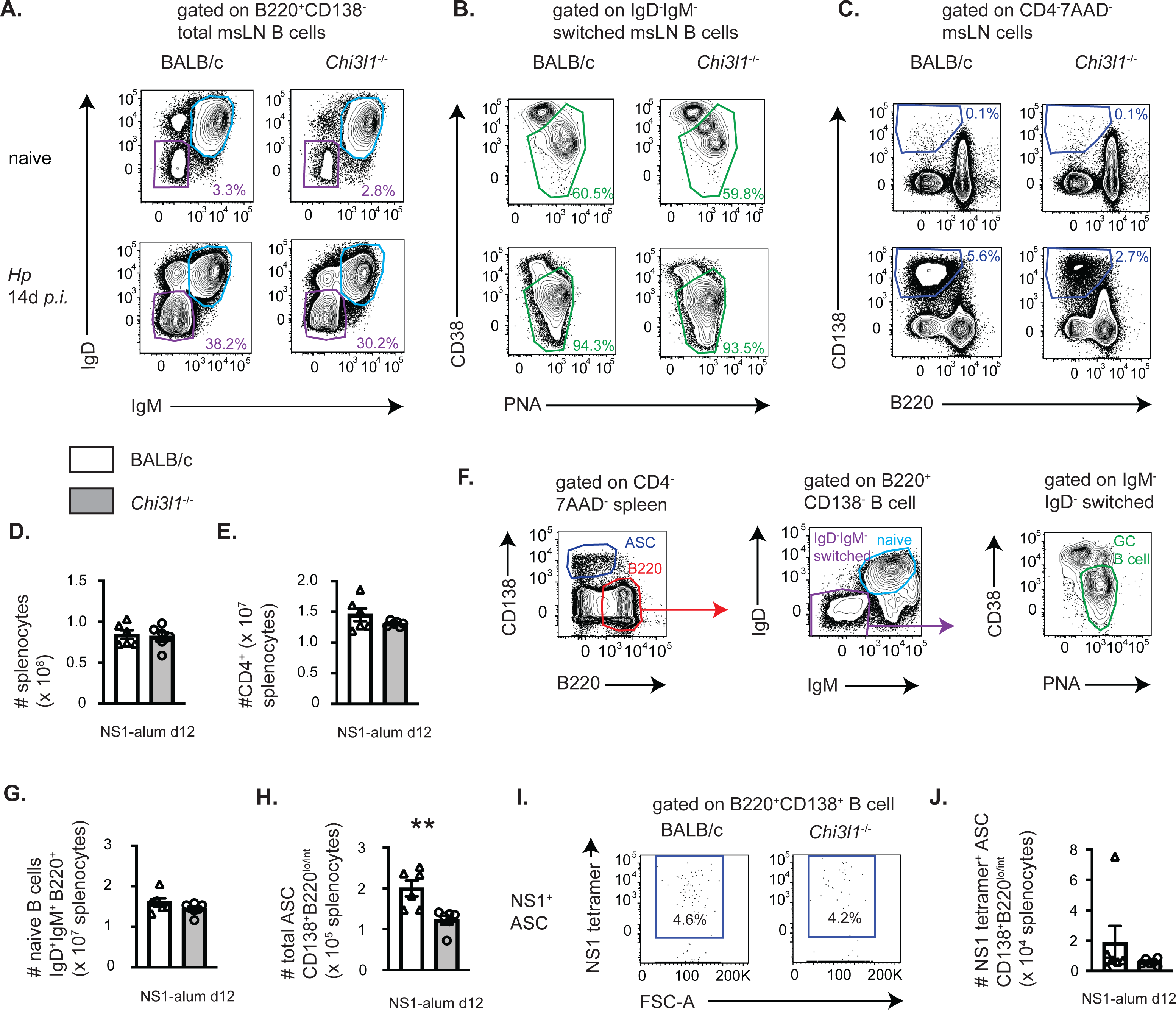
showing supporting information for Figure 3. Gating strategies to identify B cells subsets in *Hp*-infected and immunized mice. (**A-C**) Gating strategies to identify B cell subsets in msLNs from naïve and D14 *Hp*-infected WT and *Chi3l1*^-/-^ mice. Isotype-switched IgD^neg^IgM^neg^ B cells (purple gate, panel **A**), naïve IgD^+^IgM^+^ B220^+^ B cells (cerulean gate, panel **A**), isotype-switched PNA^hi^CD38^lo^ GCB cells (green gate, panel **B**) and CD138^+^B220^lo^ ASCs (blue gate, panel **C**) are shown. (**D-J**) Evaluation of vaccine-elicited splenic B and T cells at D12 after NS1-alum immunization. Number of total spleen cells (**D**) and splenic CD4^+^ T cells (**E**). Representative flow plots showing the gating strategy (**F**) to identify ASCs (blue gate), total B (red gate), naïve B (cyan gate), IgD^neg^IgM^neg^ isotype-switched B gate (purple gate) and isotype-switched GC B cells (green gate). Numbers of naïve B cells (**G**) and ASCs (**H**). Representative flow plots (**I**) showing splenic NS1-specific ASCs. Number of NS1-specific ASCs (**K**). Data displayed as mean±SEM of each group with individual animals depicted as circles or triangles. Statistical analysis performed with unpaired 2-tailed student’s t-test. *p≤0.05, **p≤0.01, ***p≤0.001, ****p≤0.0001

**Supplemental Figure 3.**
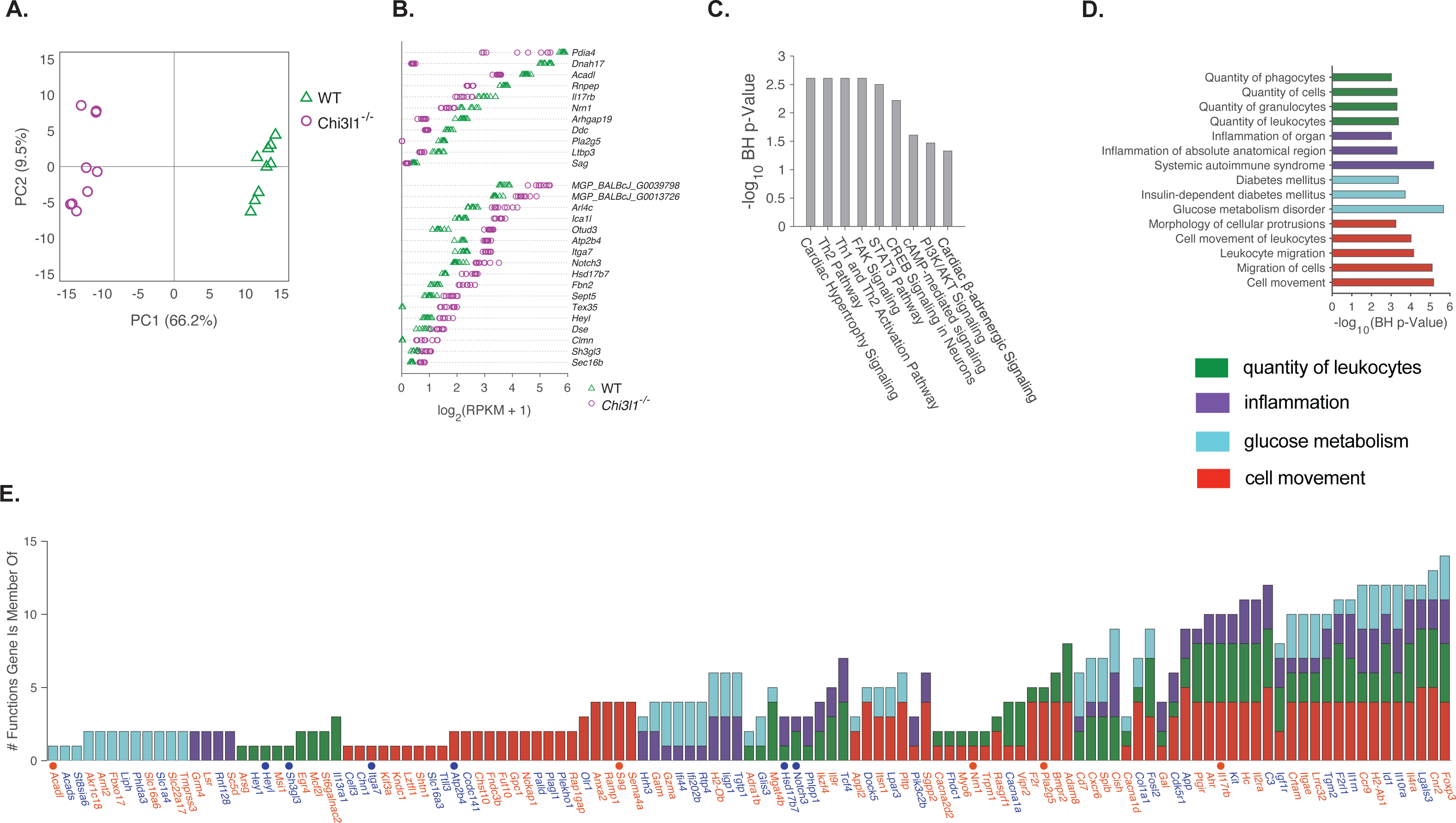
showing supporting information for Figure 4. RNAseq methodology and bioinformatic analyses. RNAseq analysis of T_FH_ cells sort-purified from msLN of D14 *Hp*-infected WT and *Chi3l1*^-/-^ mice (samples derived from 3 independent experiments with 2-3 mice/group/experiment). 9853 expressed genes (defined as genes with at least 3 RPM in all samples of either the WT and/or *Chi3l1*^-/-^ T_FH_) were identified. Using an FDR cutoff of q<0.05, 1465 genes were identified as significantly different between WT and *Chi3l1*^-/-^ T_FH_ cells. Of these, 252 genes met a threshold ±0.3785 log_2_FC and 28 genes met a ±1.0 log_2_FC. See Table S1 for gene expression levels and detailed methods. **(A)** Principal component analysis of 252 genes with ±0.3785 log_2_FC and FDR q<0.05. **(B)** Expression levels of 28 genes with ±1.0 log_2_FC and FDR q<0.05. (**C-D**) The gene list containing 252 genes meeting an FDR q< 0.05 and ± 0.3785 log_2_FC cutoff were imported into Ingenuity Pathway Analysis (IPA, QIAGEN Digital Insights) to identify significant signaling pathways (**C**) and functional categories (**D**). Pathways (**C**) with a log transformed Benjamini-Hochberg (B-H) corrected overlap p< 0.05 are shown. Functional categories (**D**) with a BH p-value <0.001 are shown and grouped according to shared functions (cell movement/migration (red, n=5 function gene sets), quantity of leukocytes (green, n=4 function gene sets), inflammation (violet, n=3 function gene sets), glucose metabolism (cyan, n=3 function gene sets)). (**E**) Analysis of the 119 unique genes associated with the 15 IPA predicted function gene sets. Number of instances (**E**) a gene was identified within each functional category. Genes upregulated in WT T_FH_ (orange) or *Chi3l1*^-/-^ T_FH_ (indigo) are demarcated with genes meeting the ±1.0log_2_ FC and FDR q<0.05 threshold indicated with a filled circle.

**Table S1.**
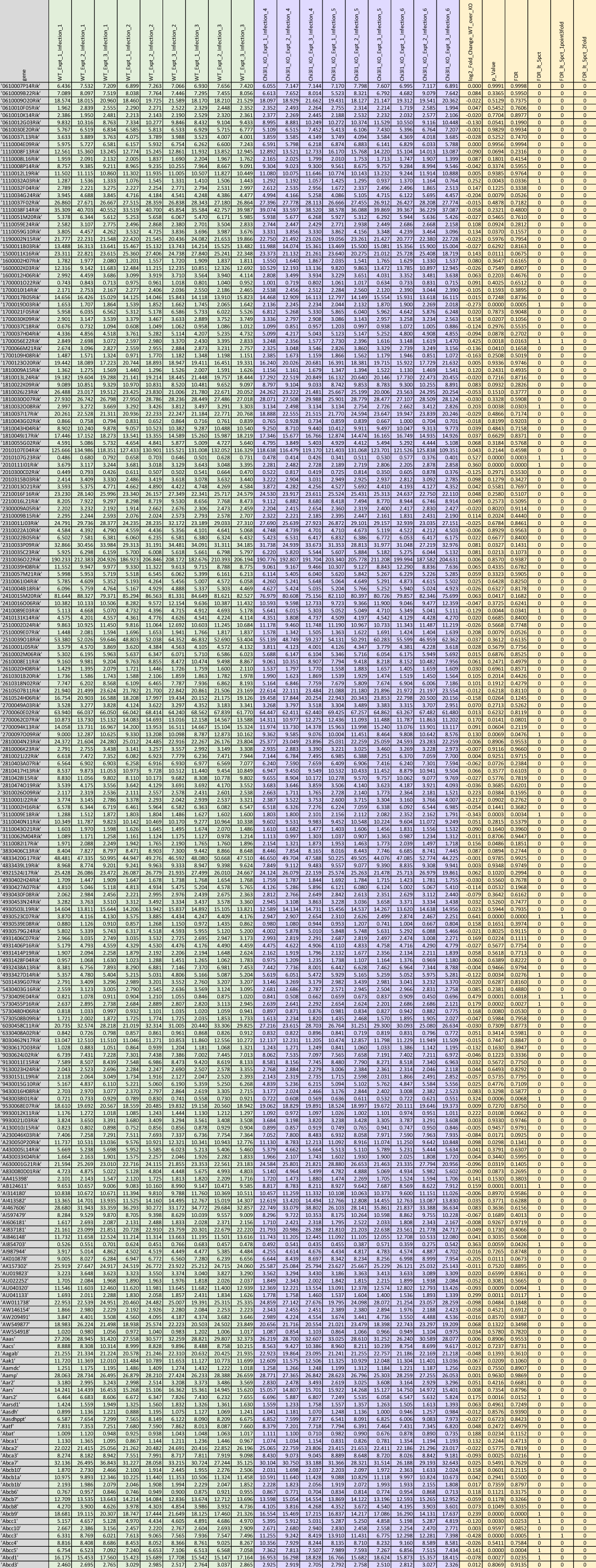
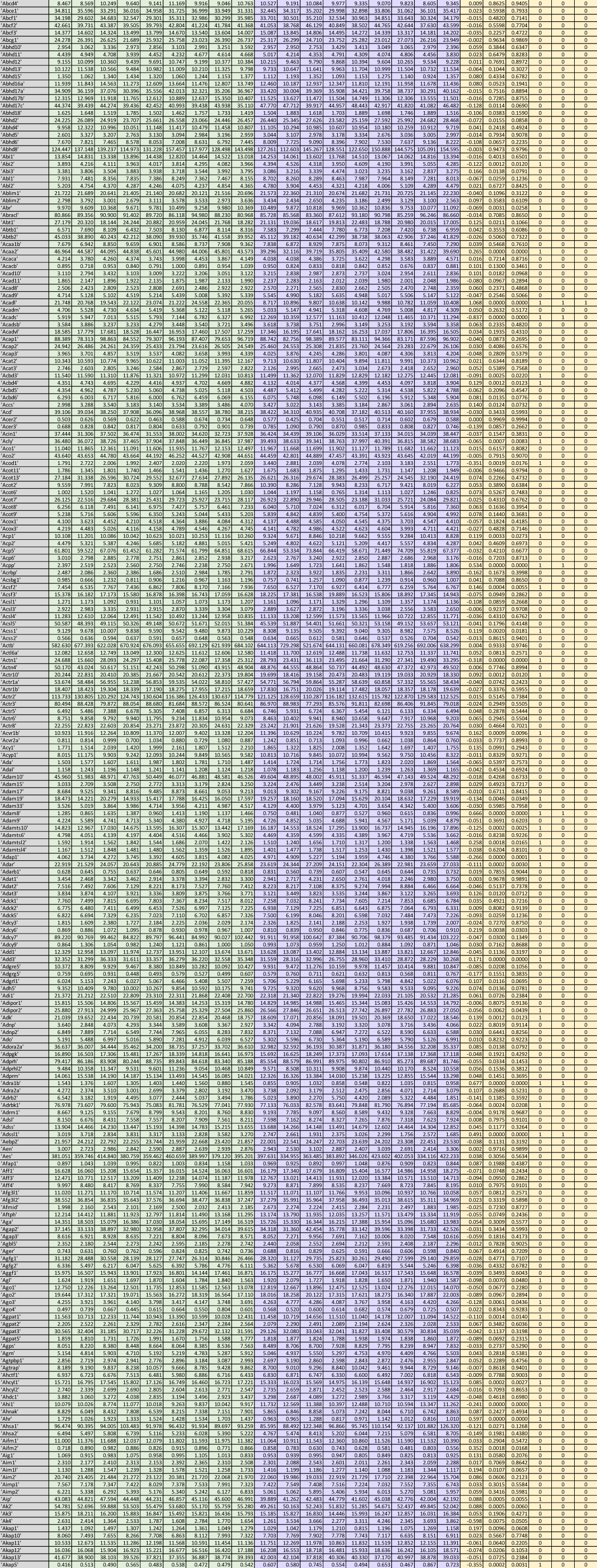
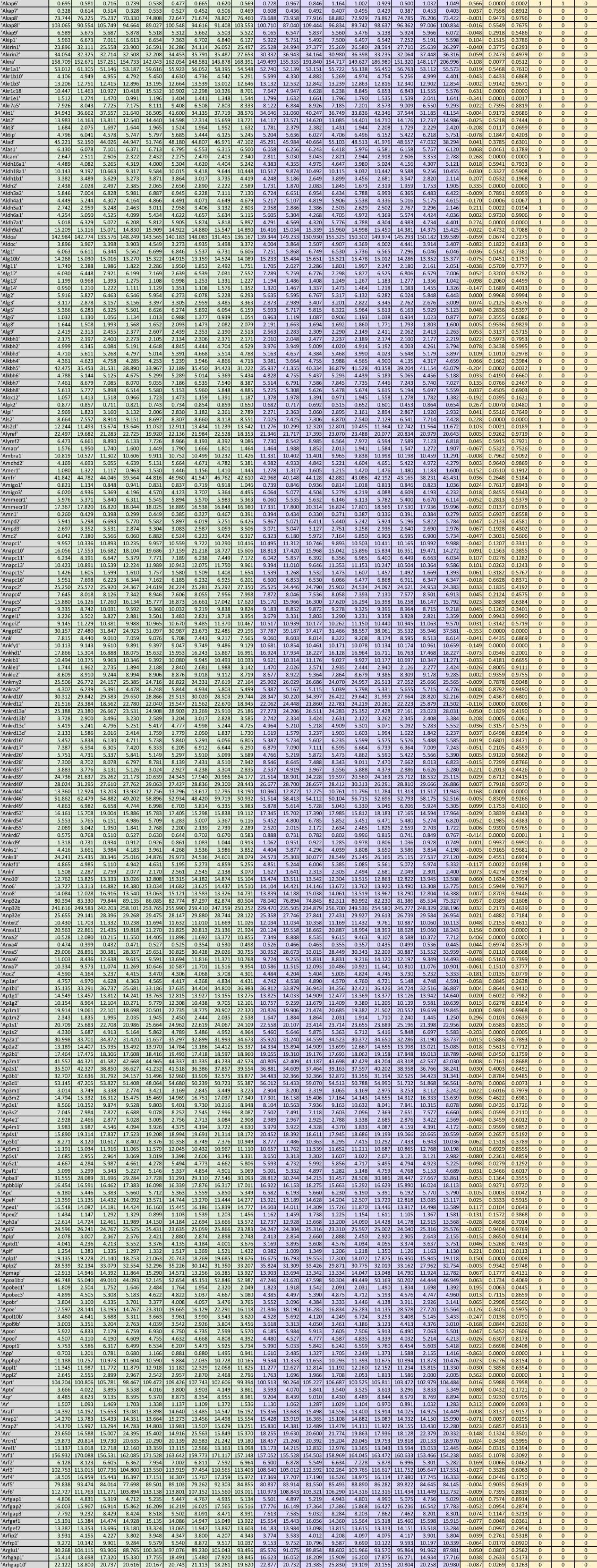
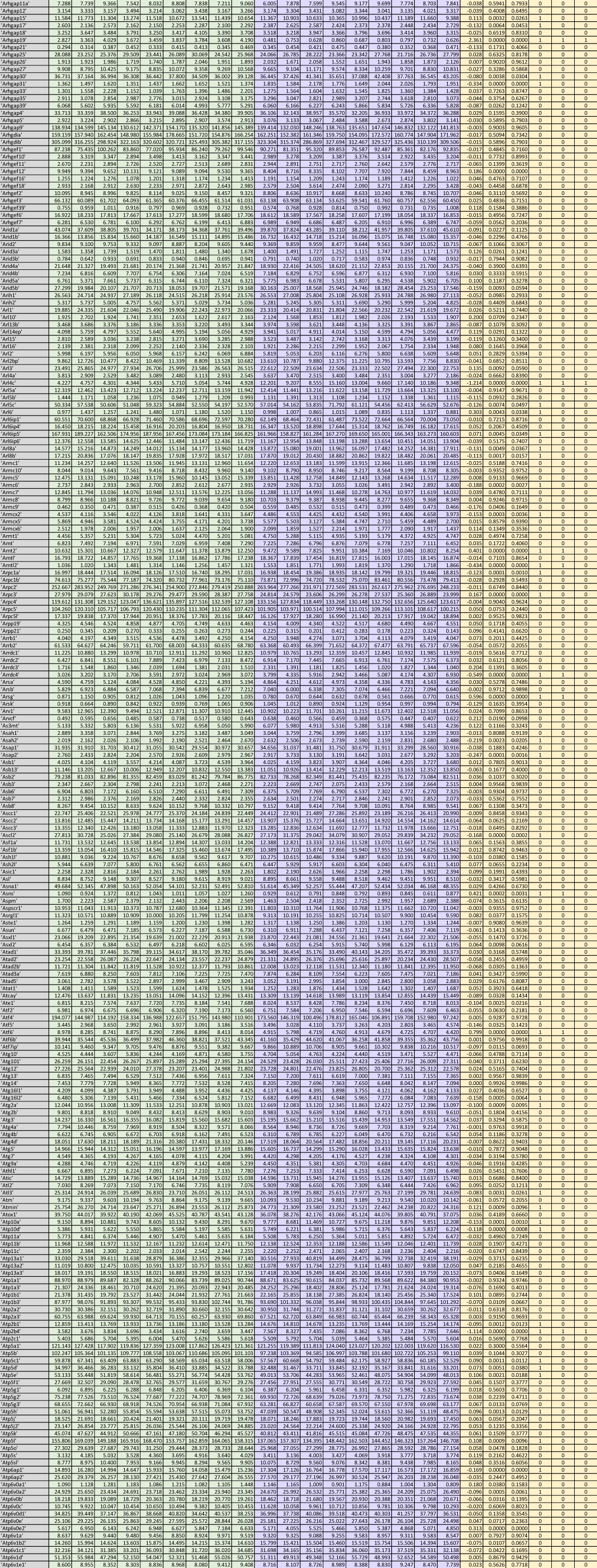
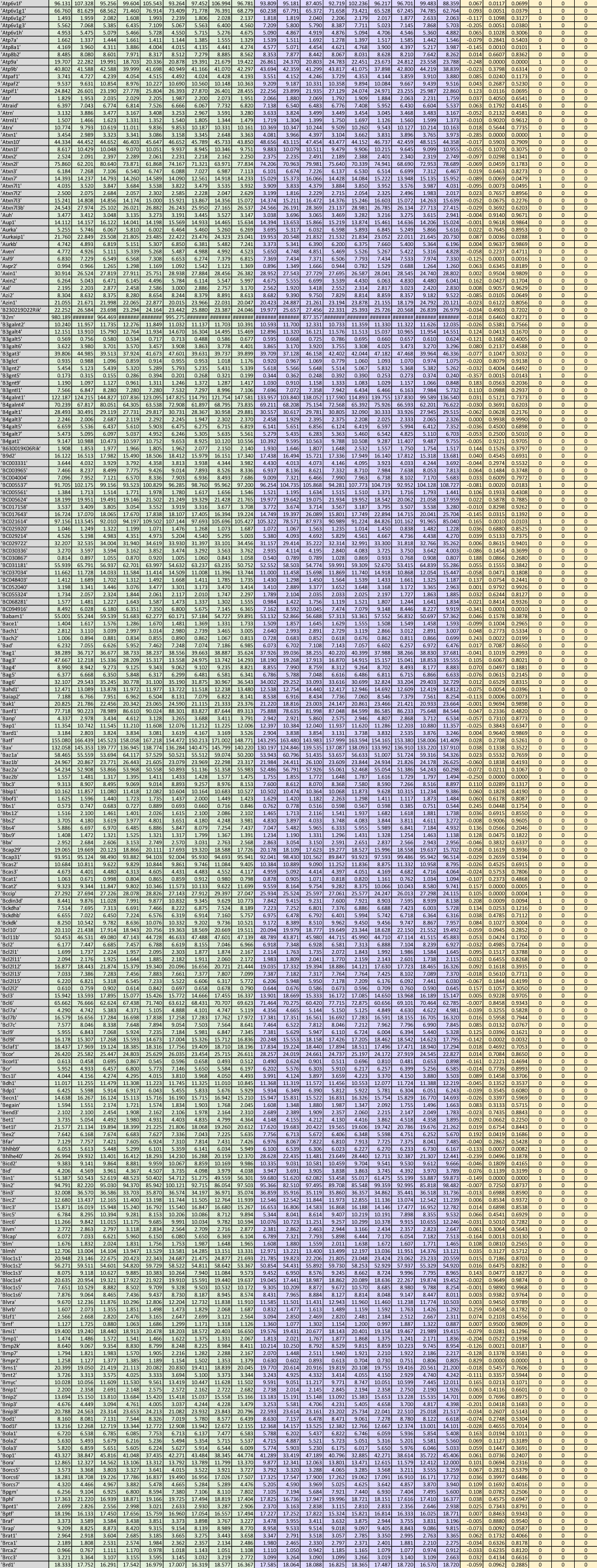
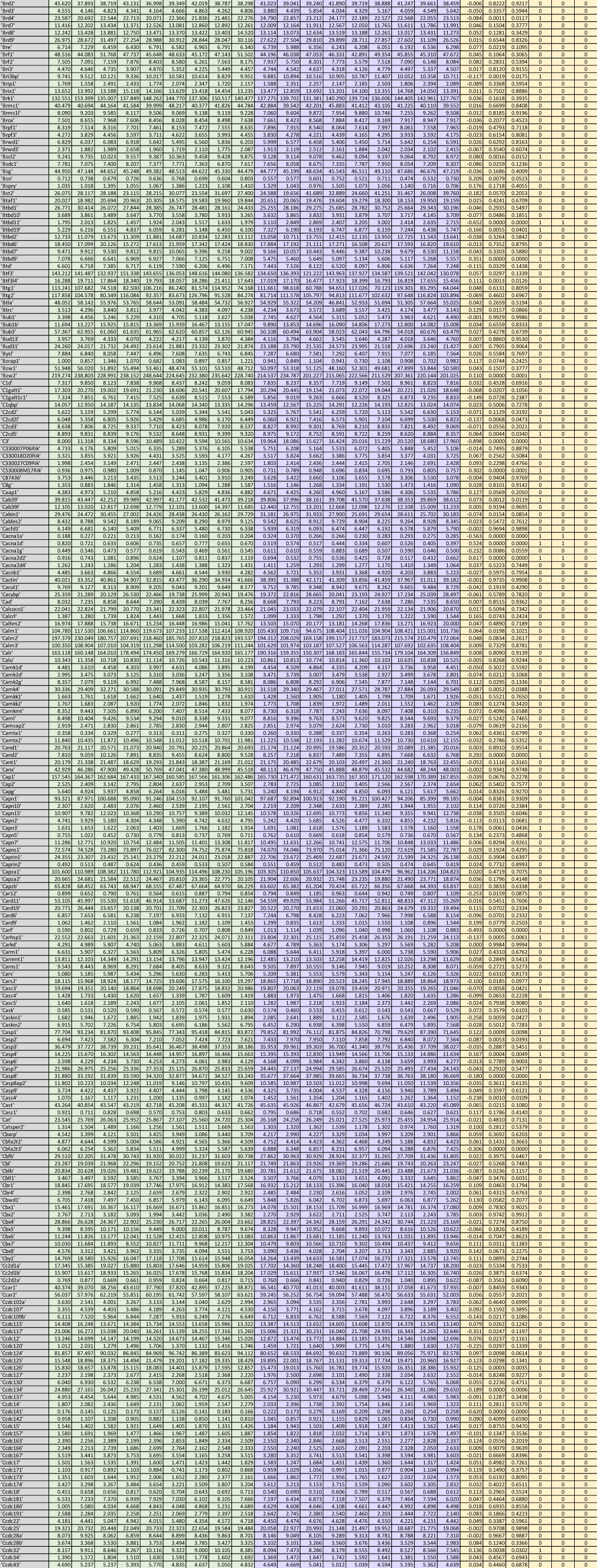
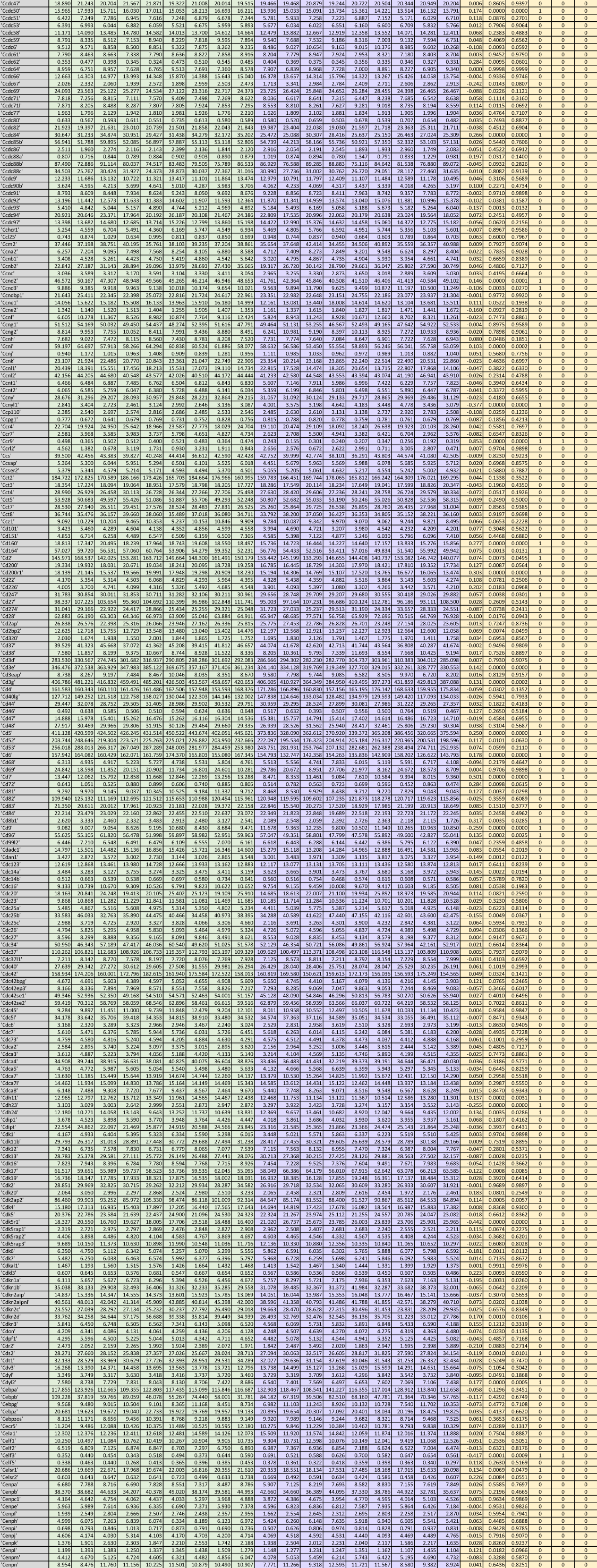
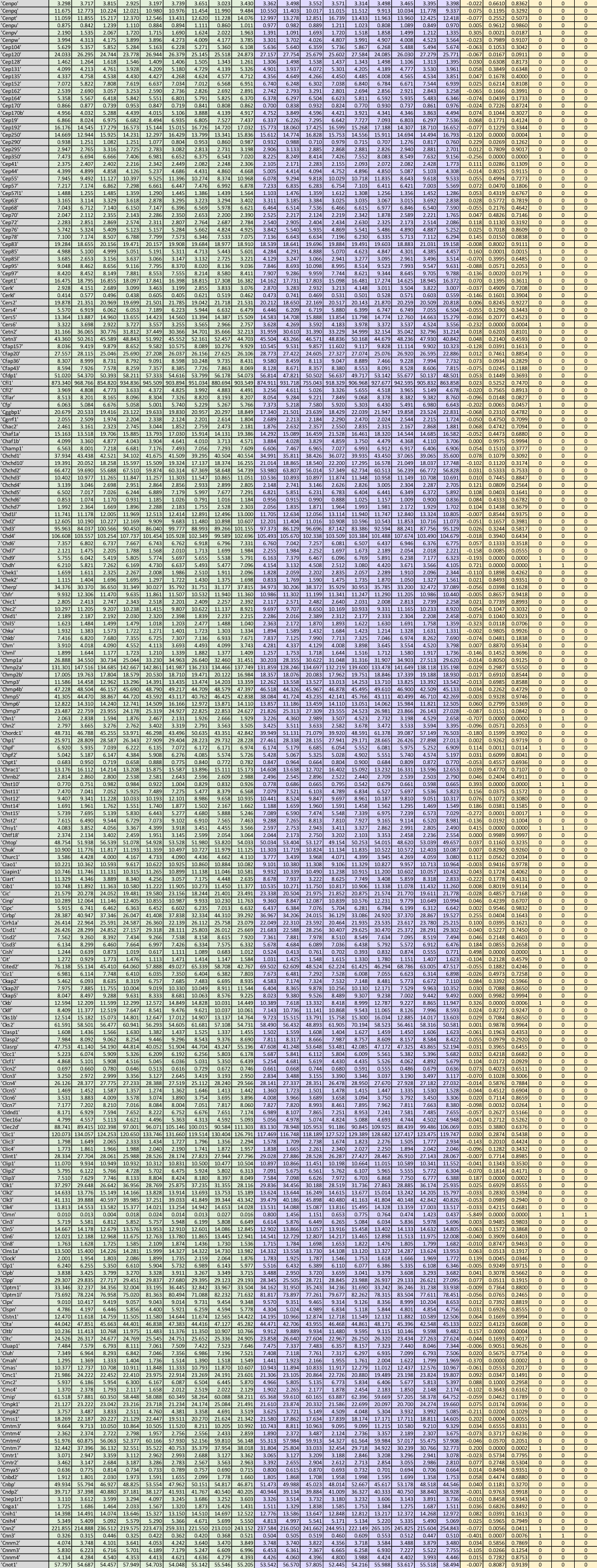
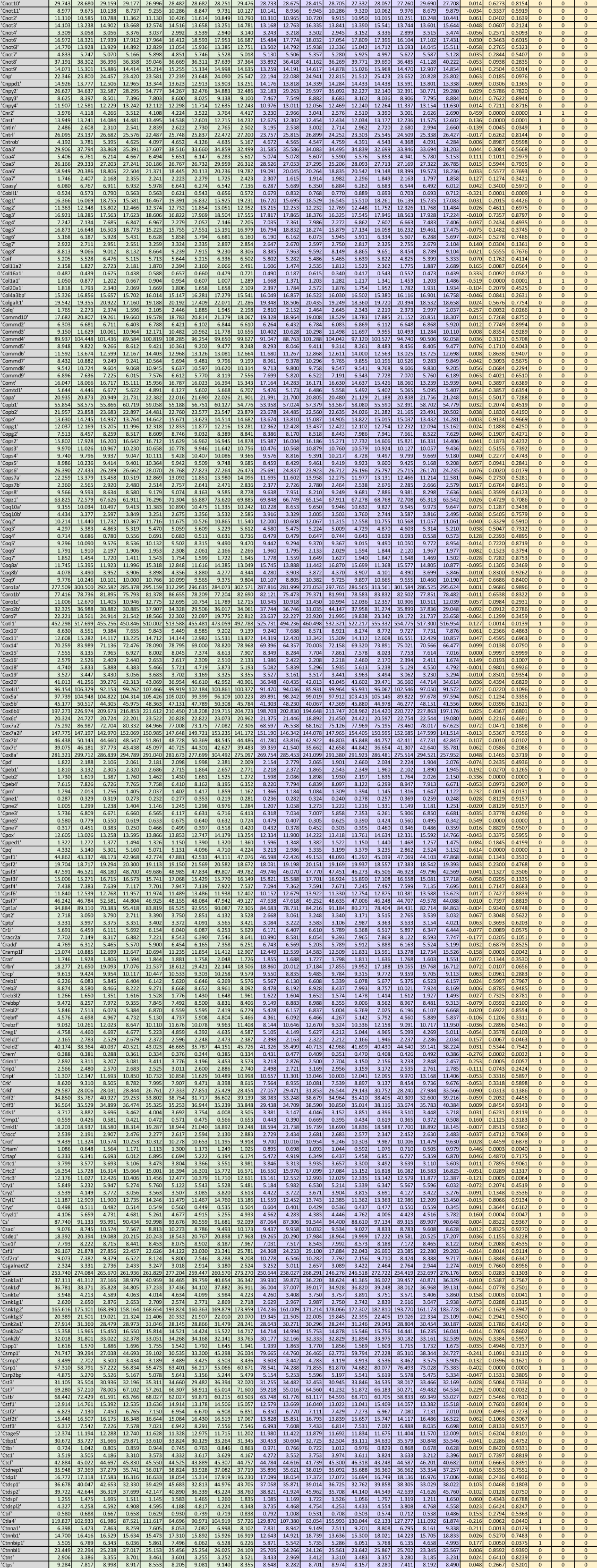
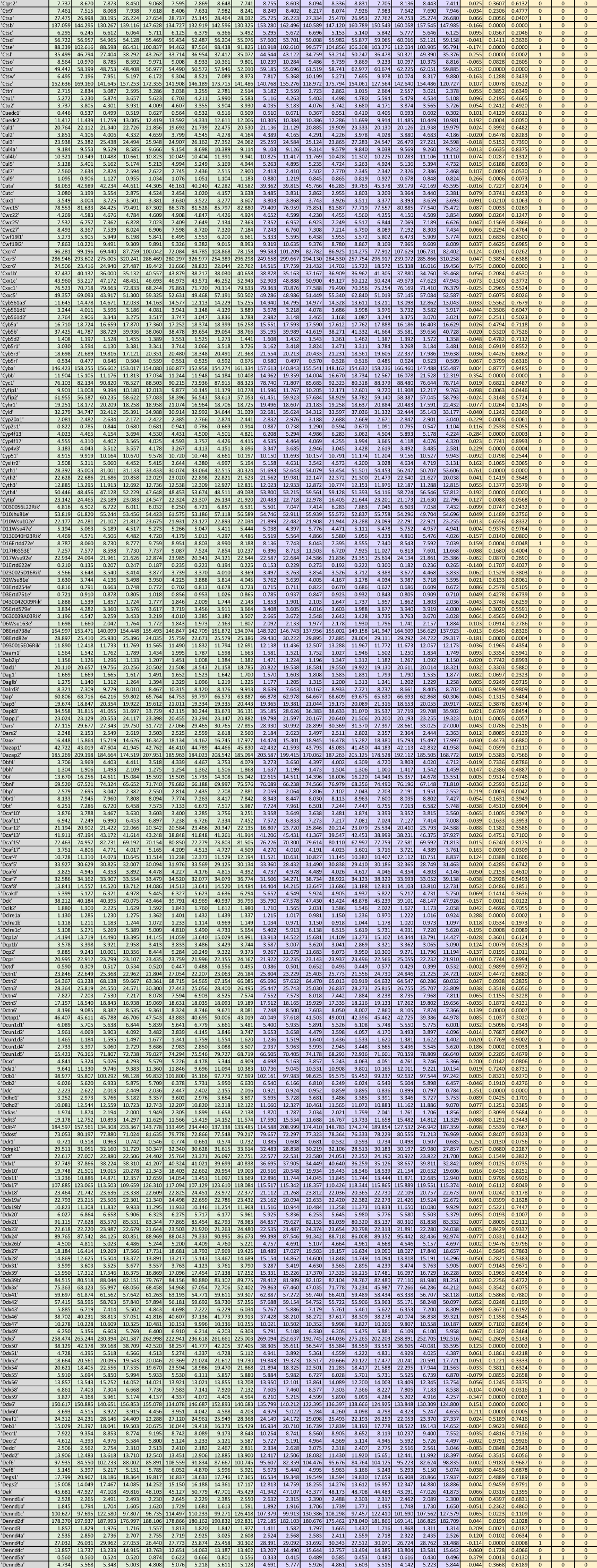
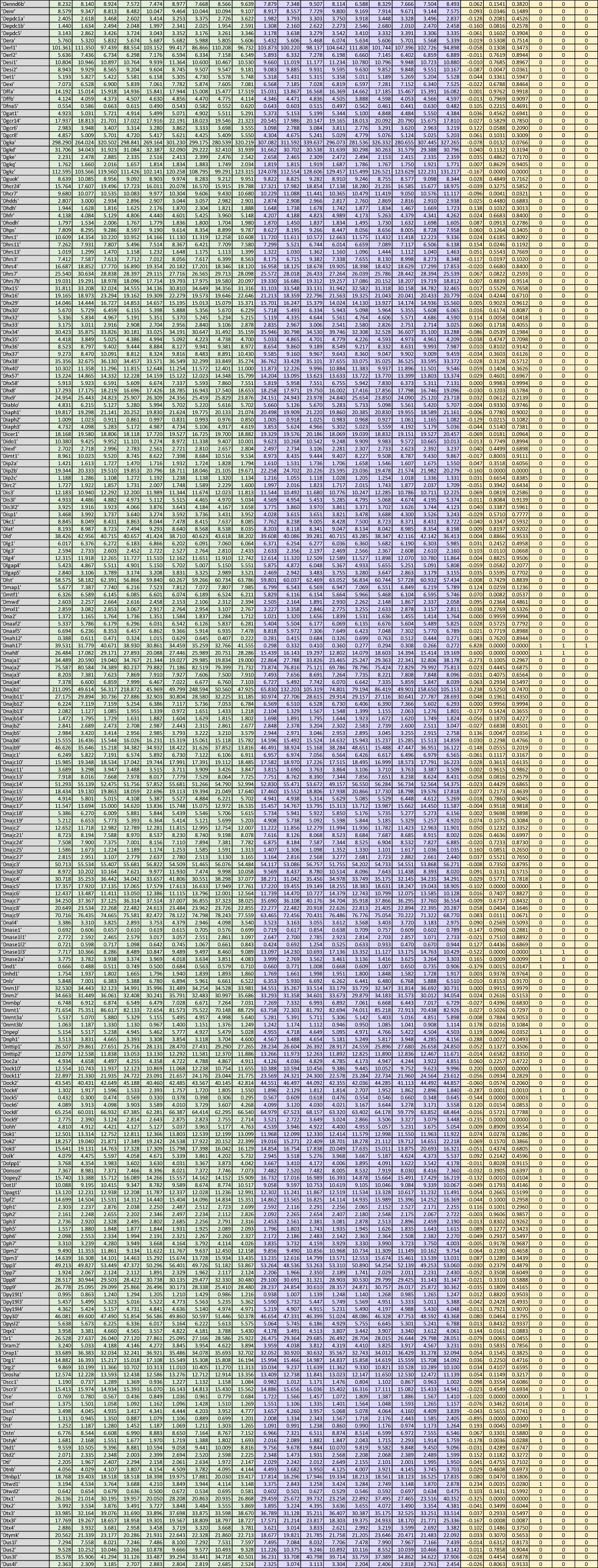
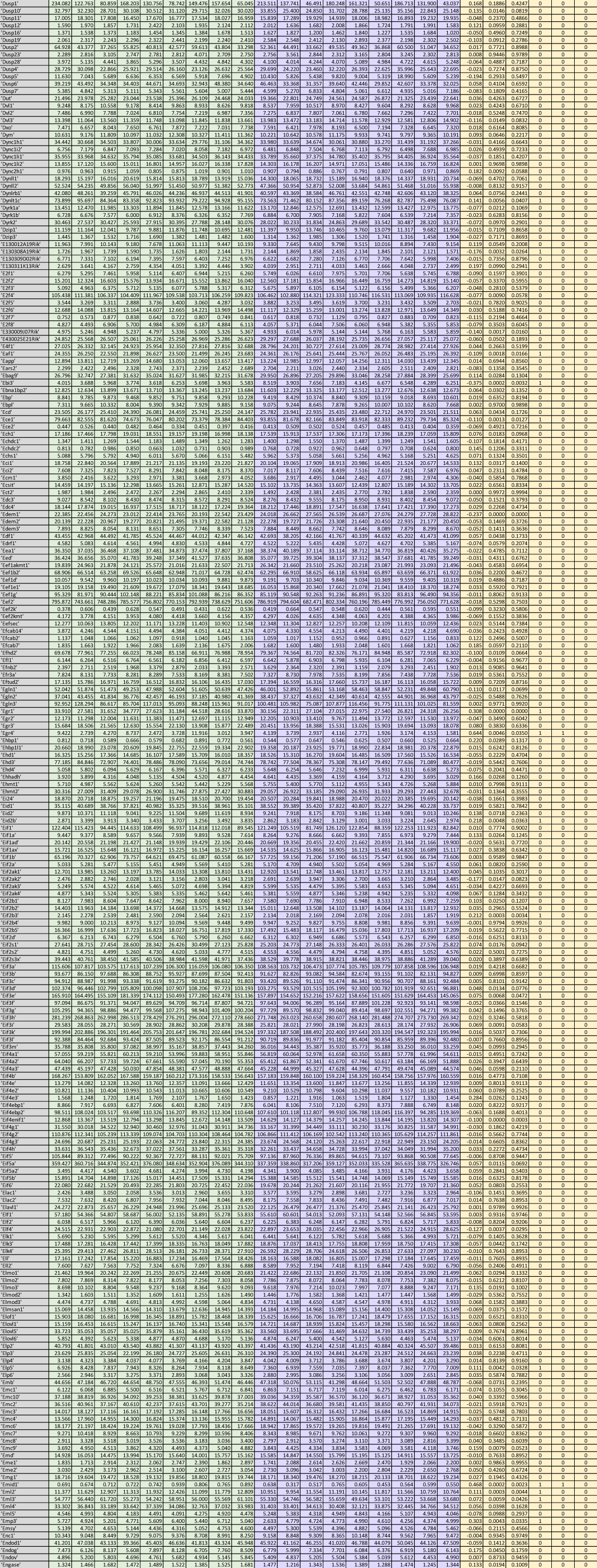
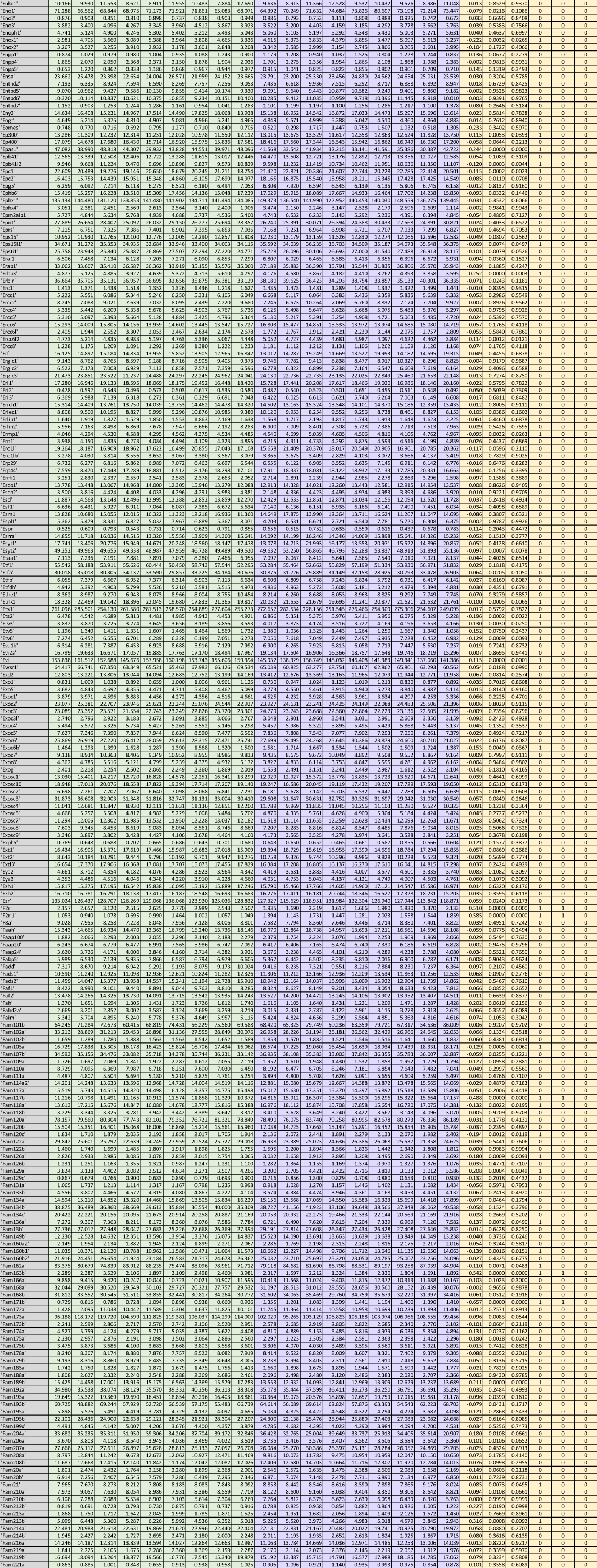
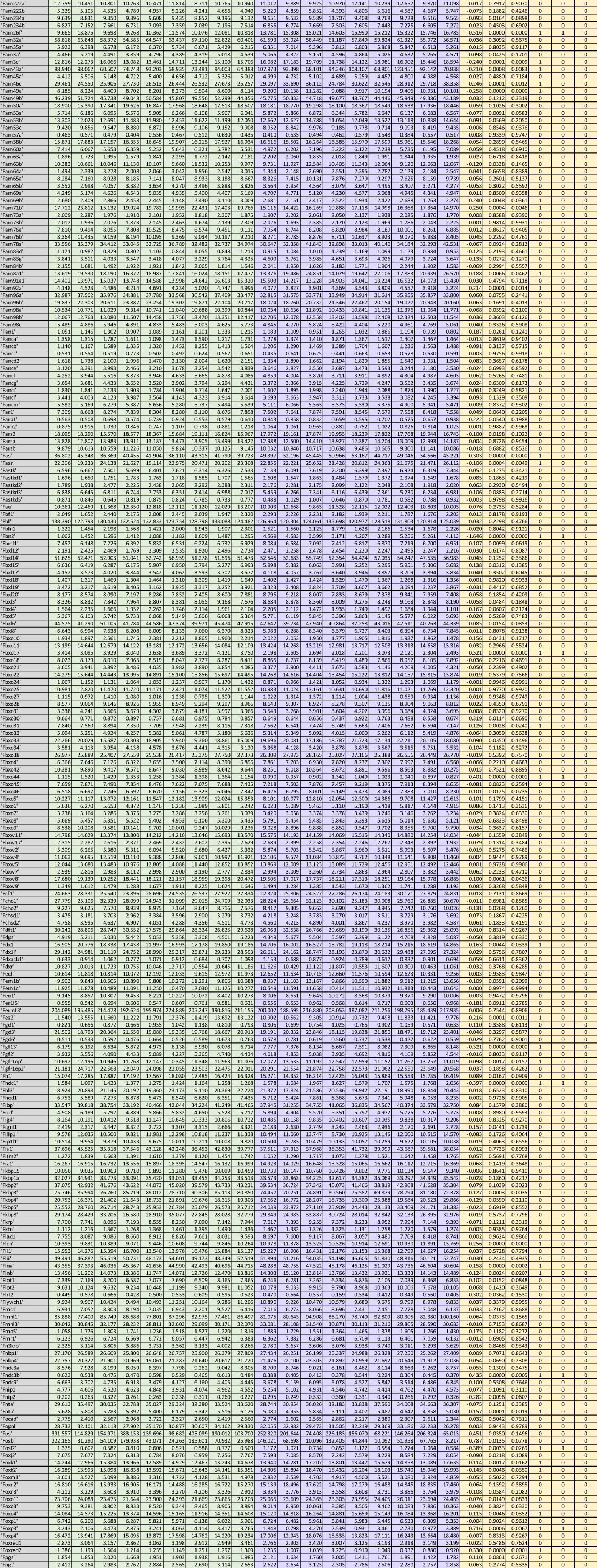
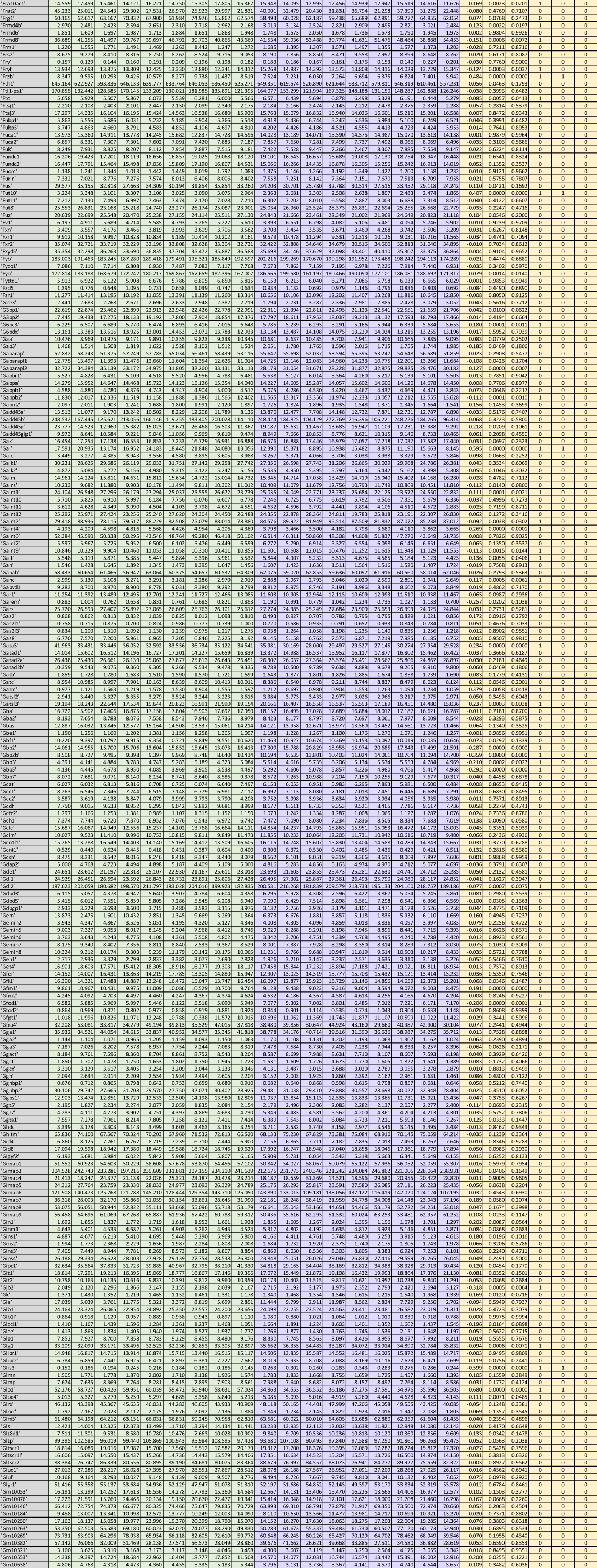
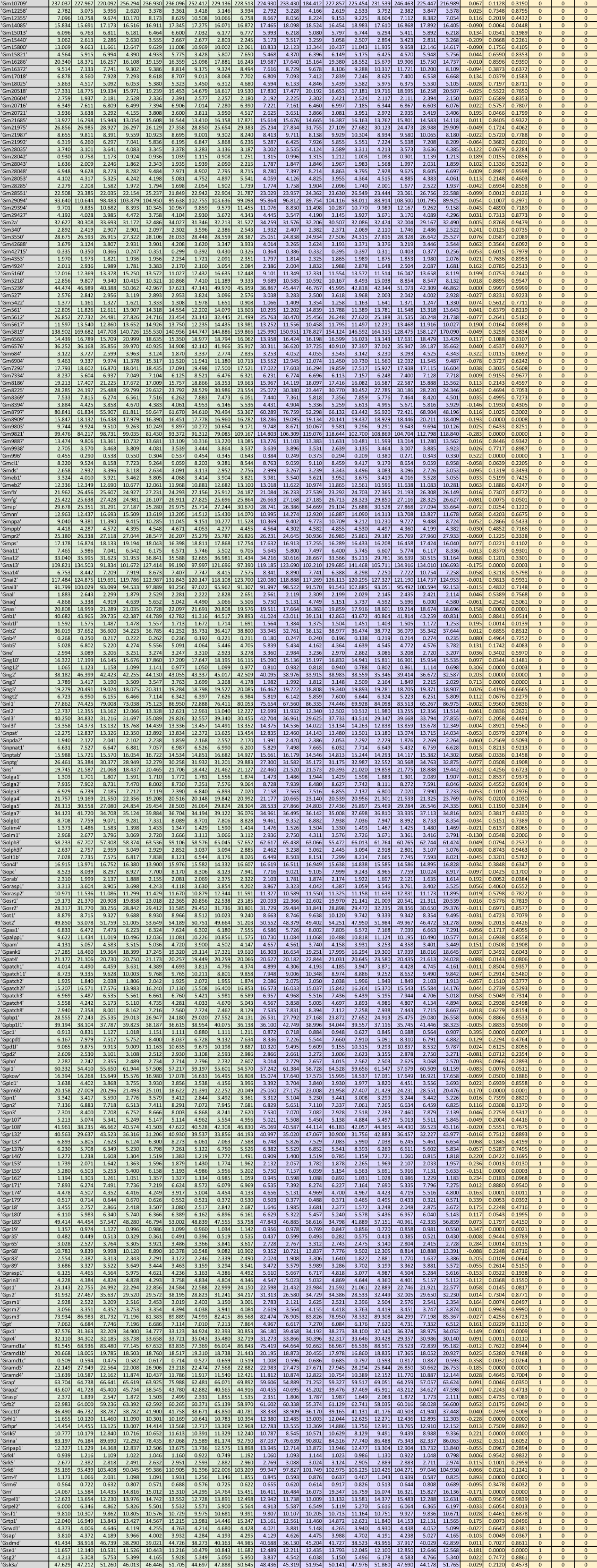
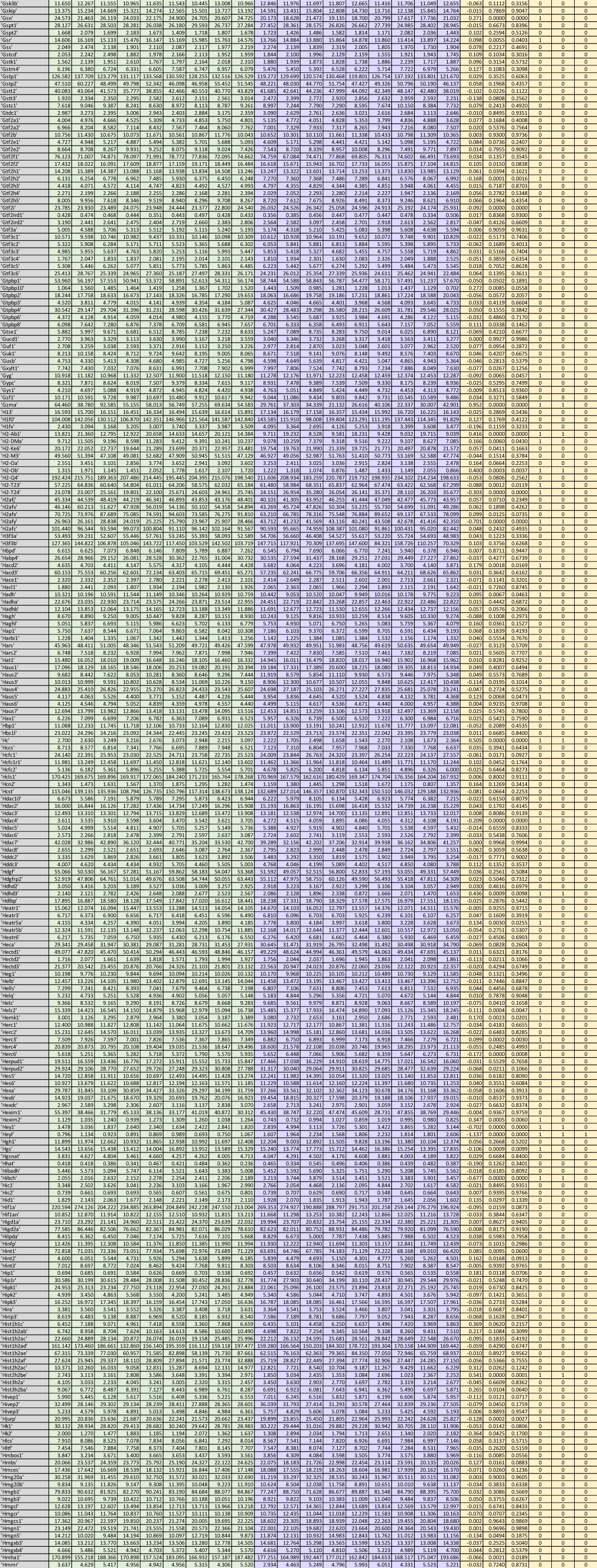
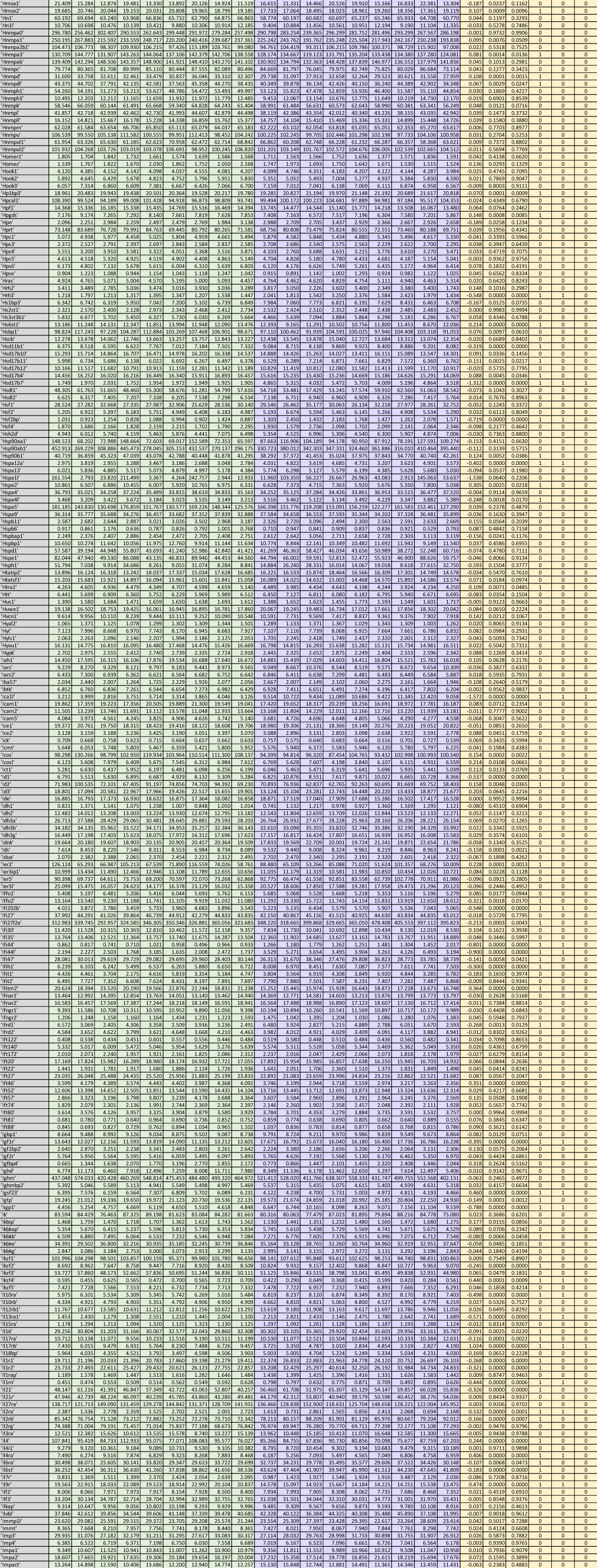
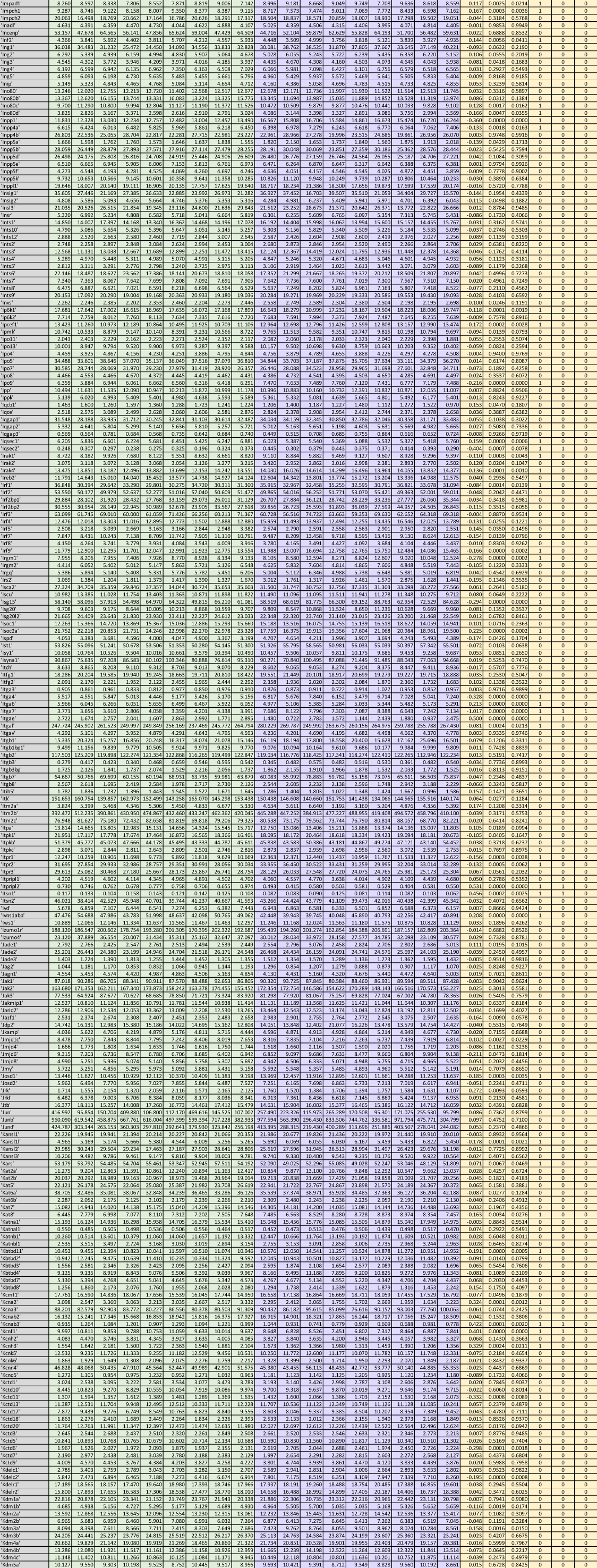
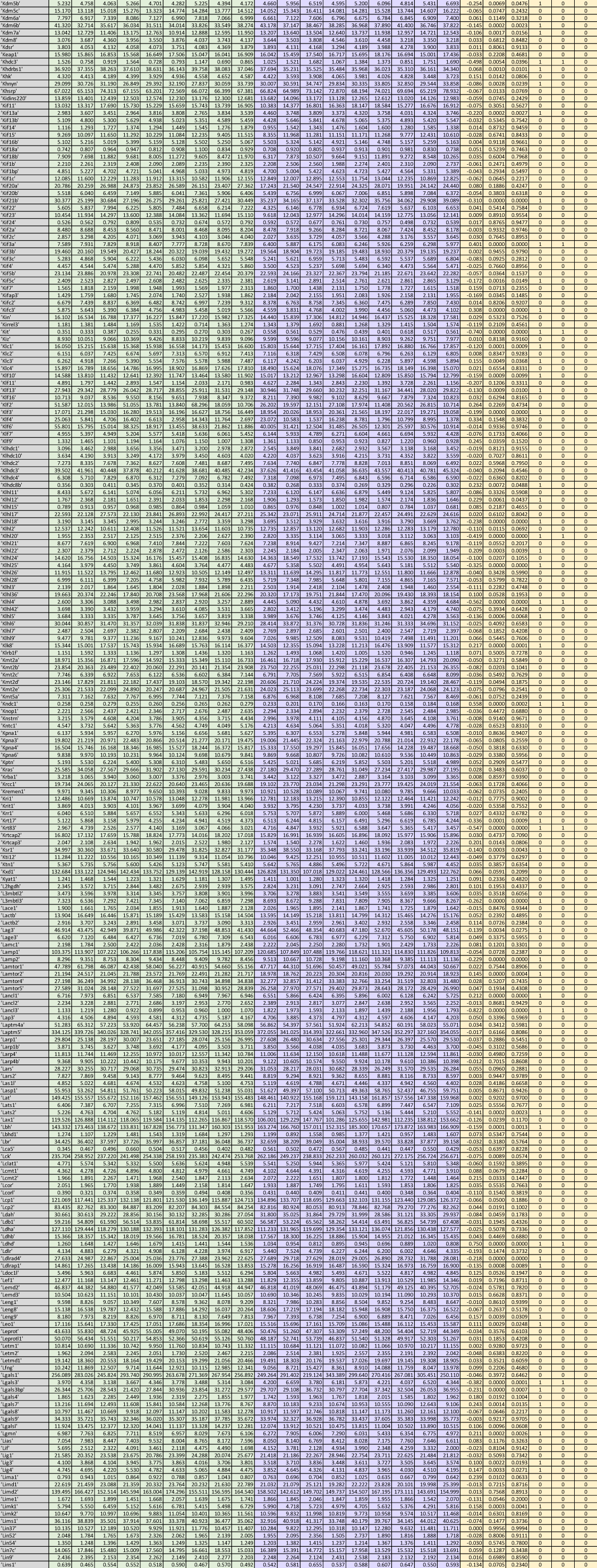
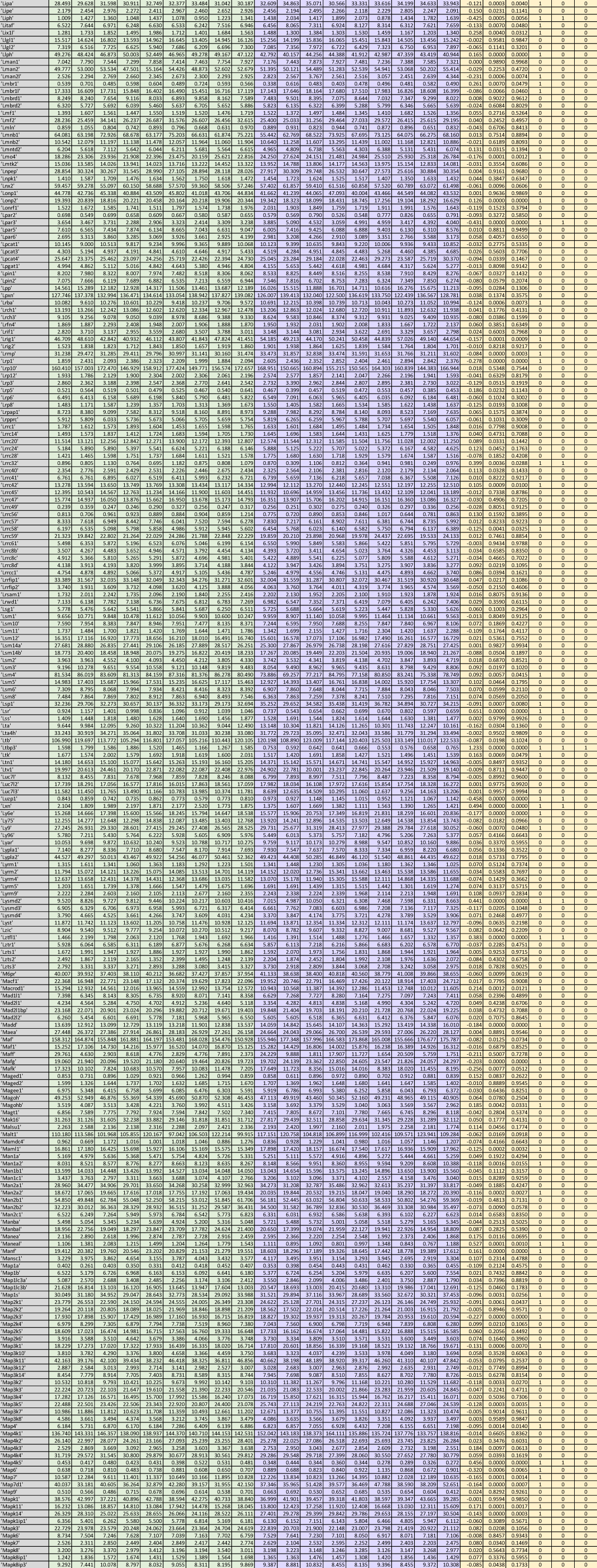
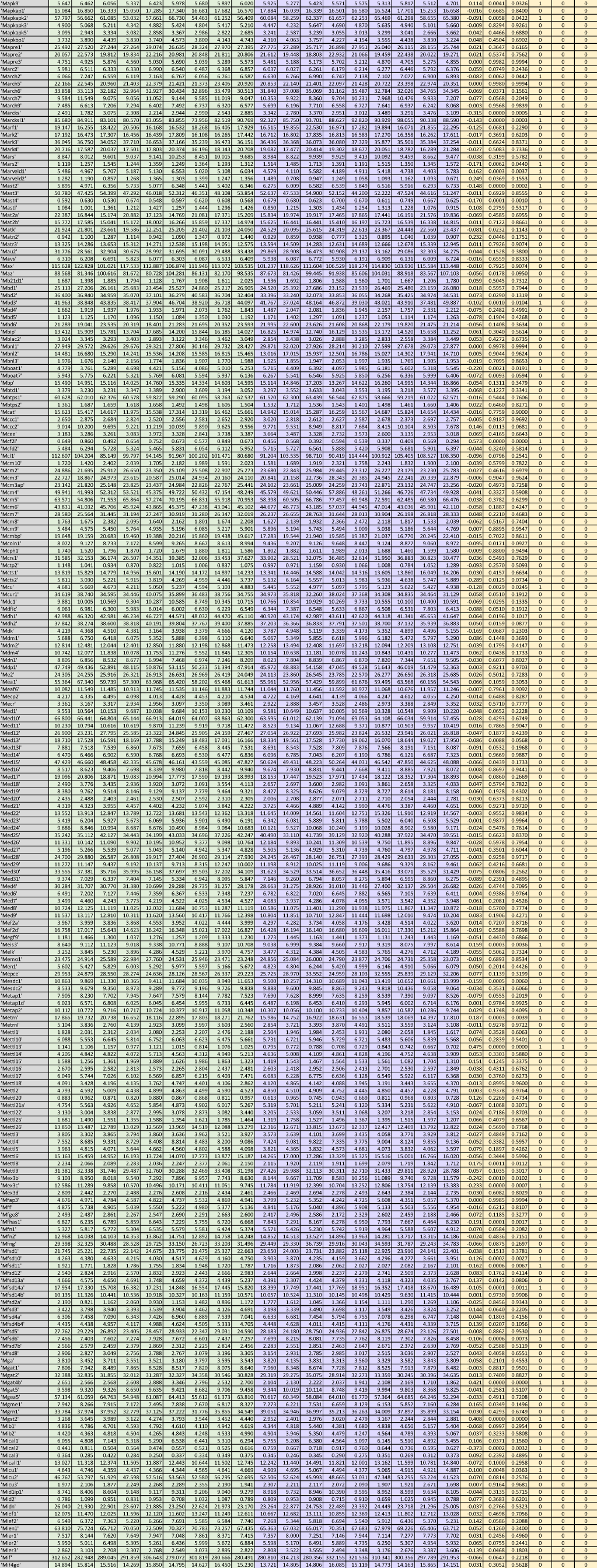
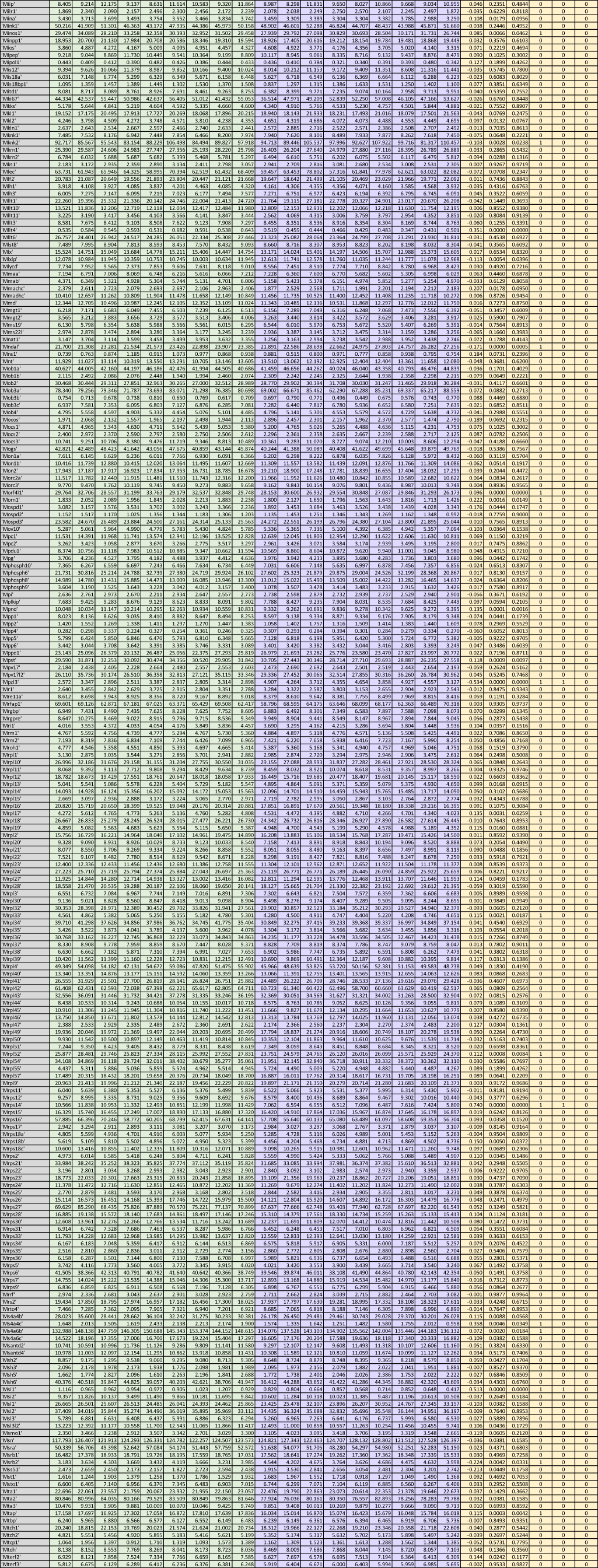
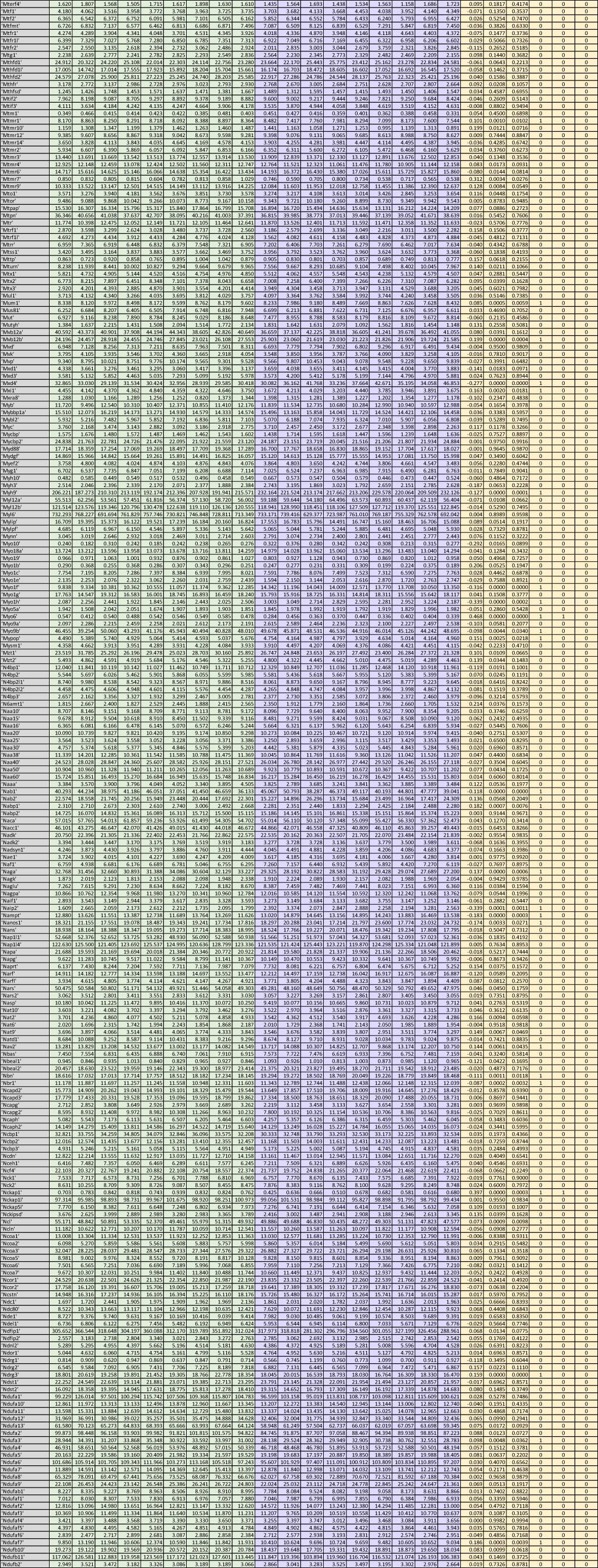
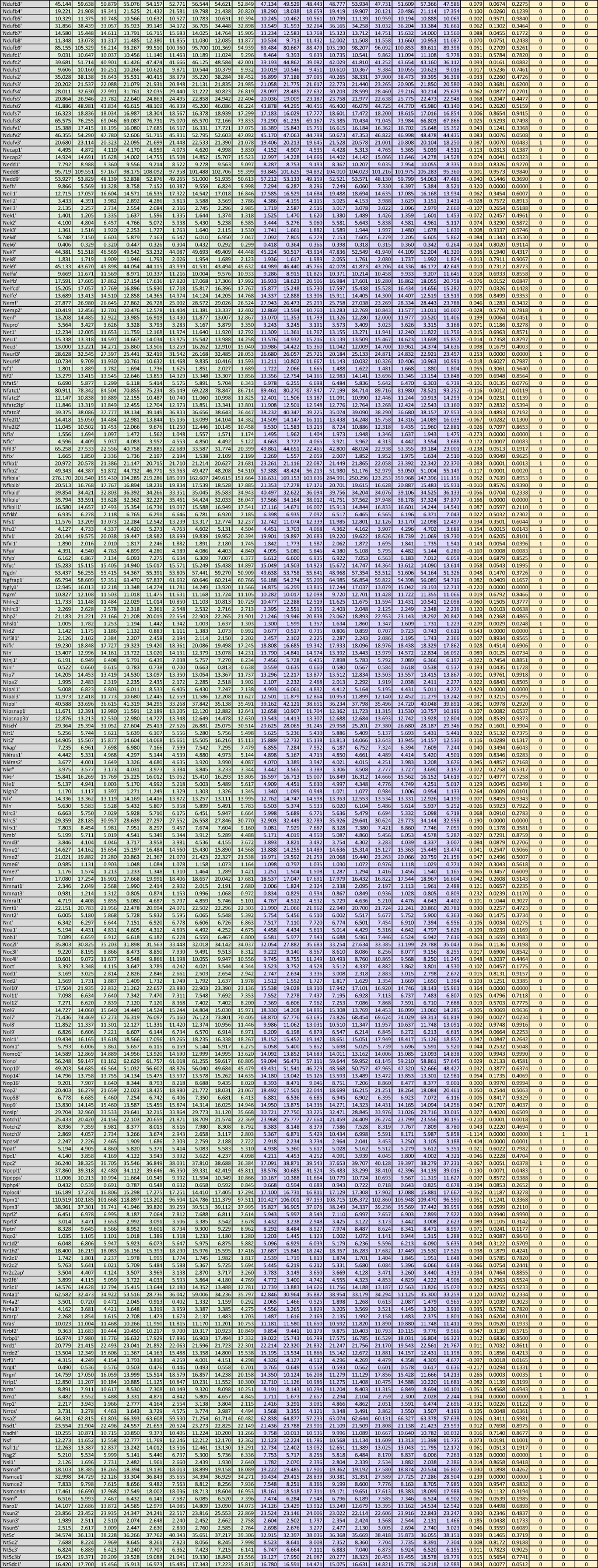
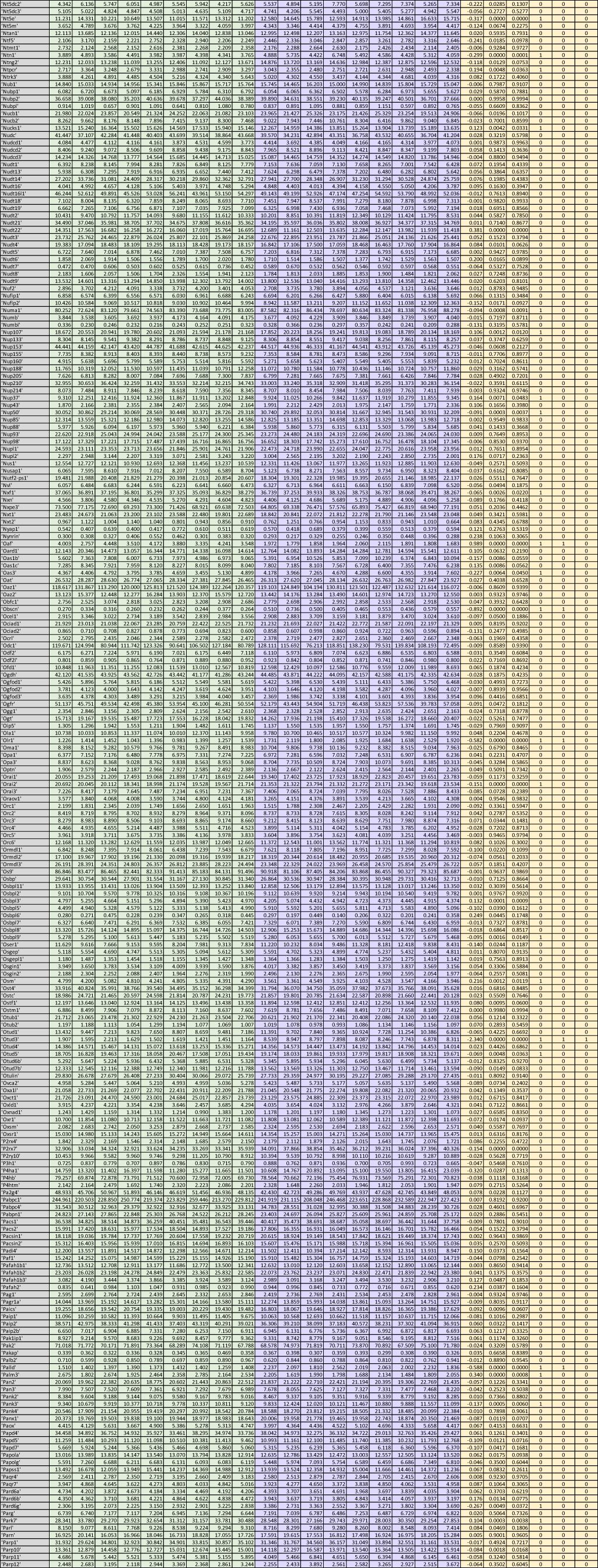
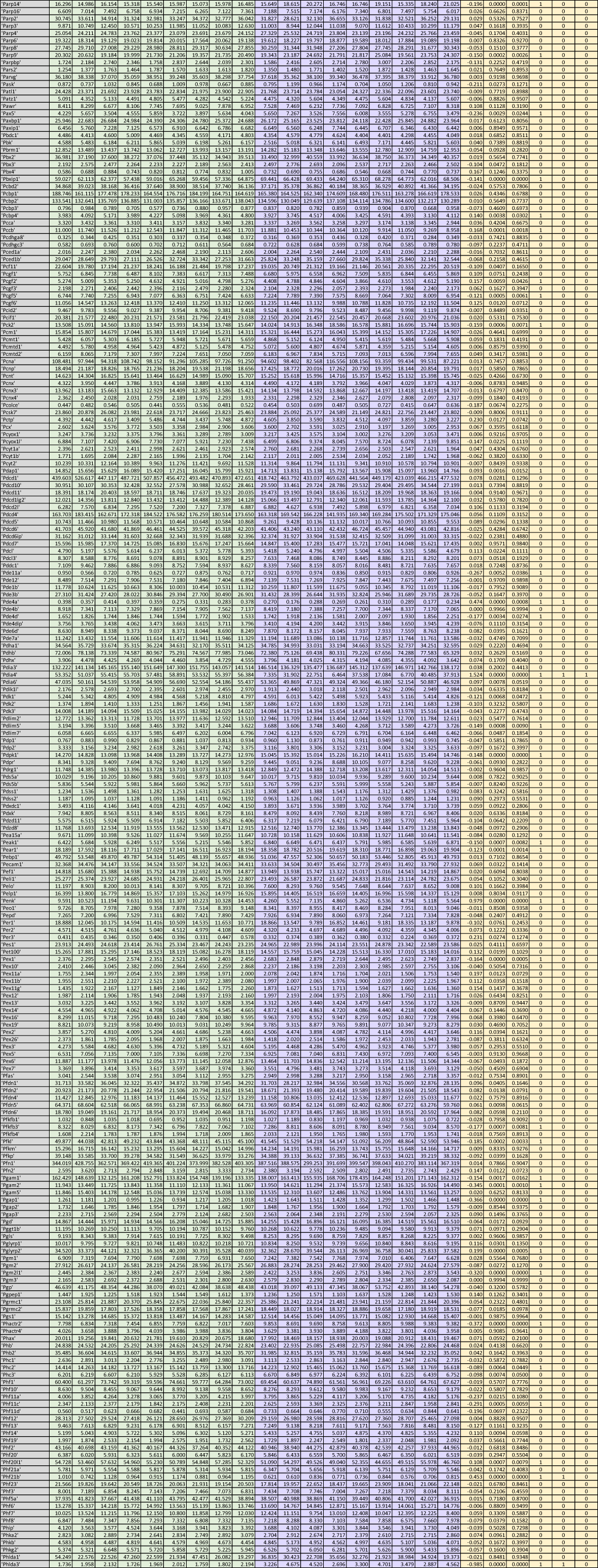
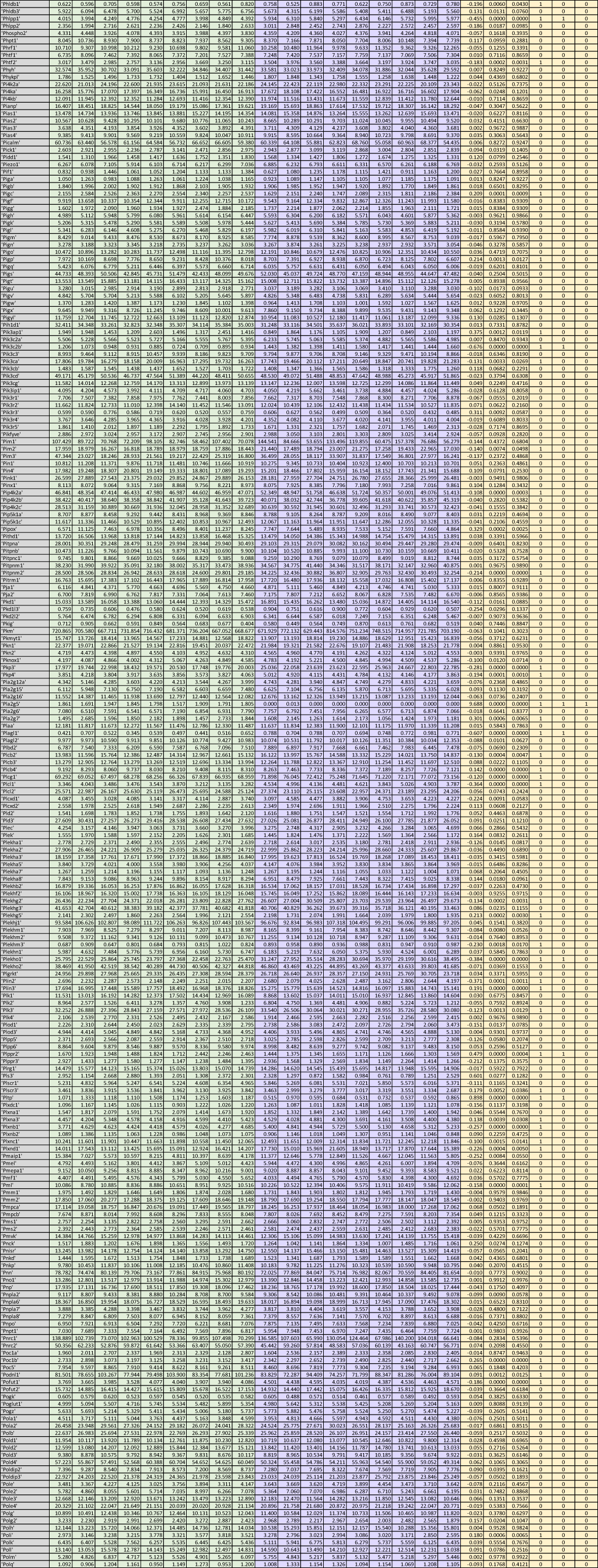
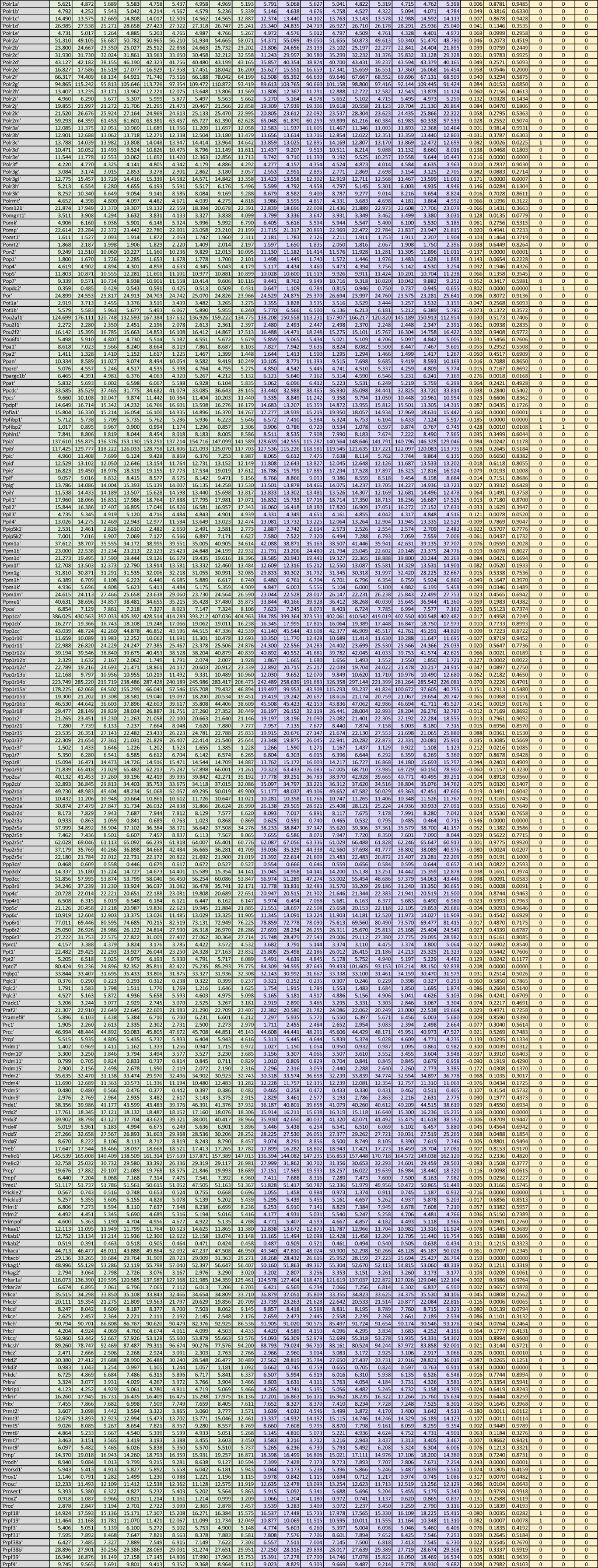
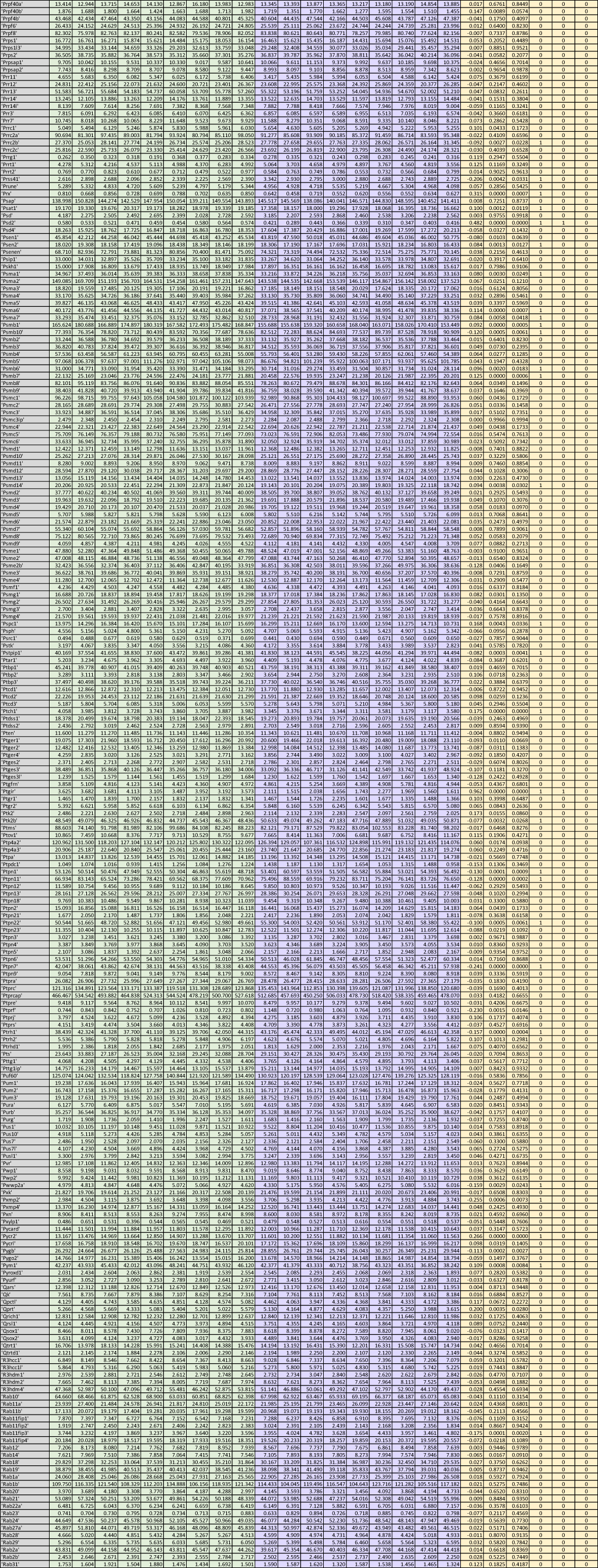
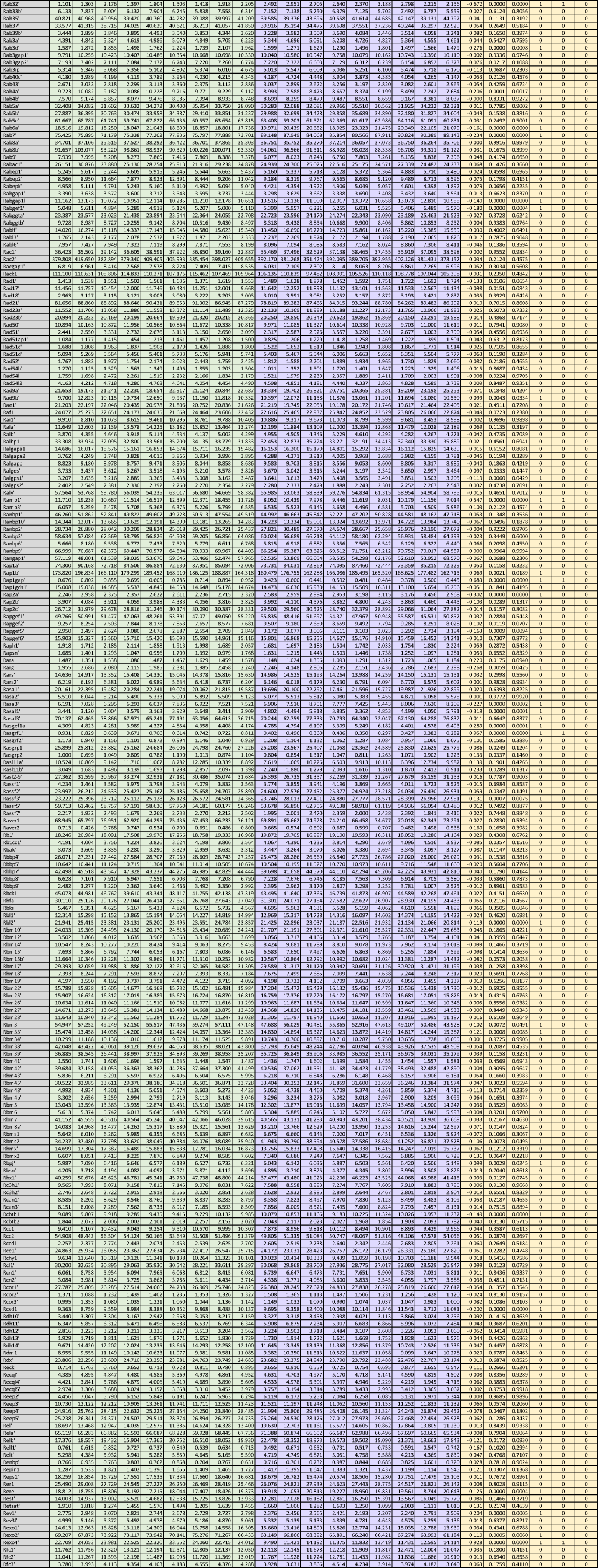
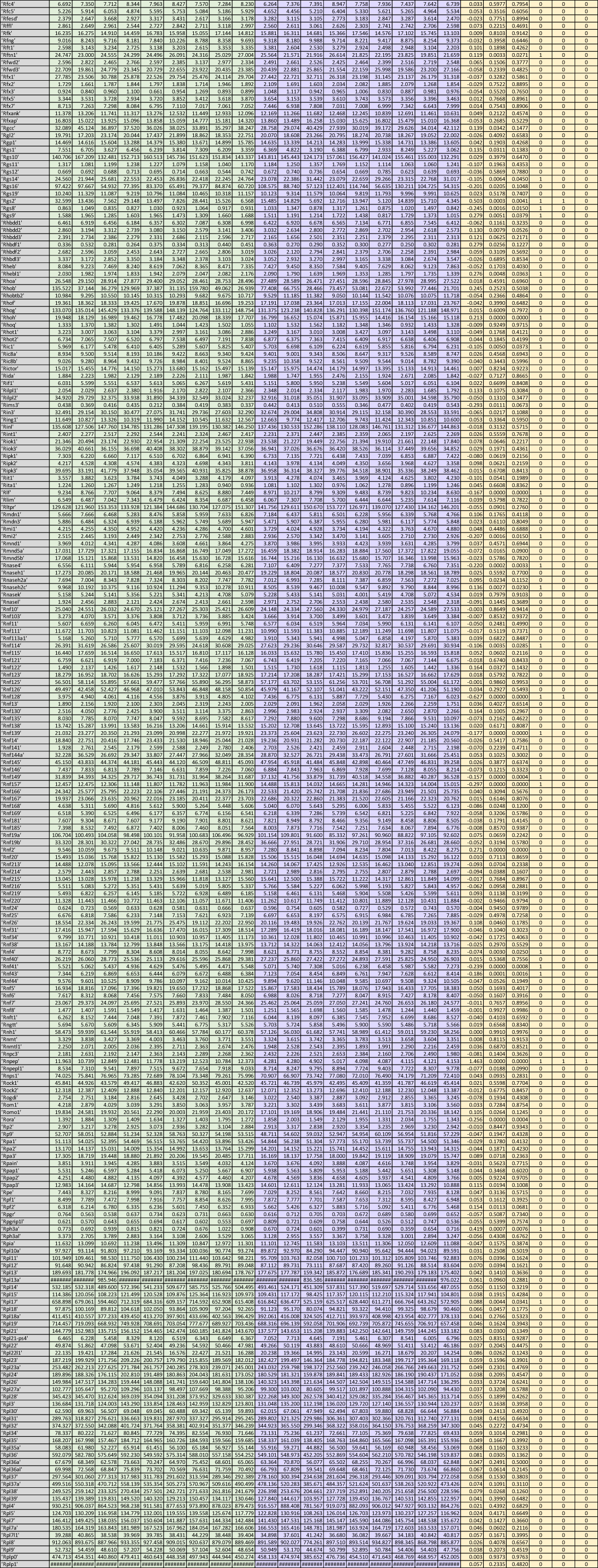
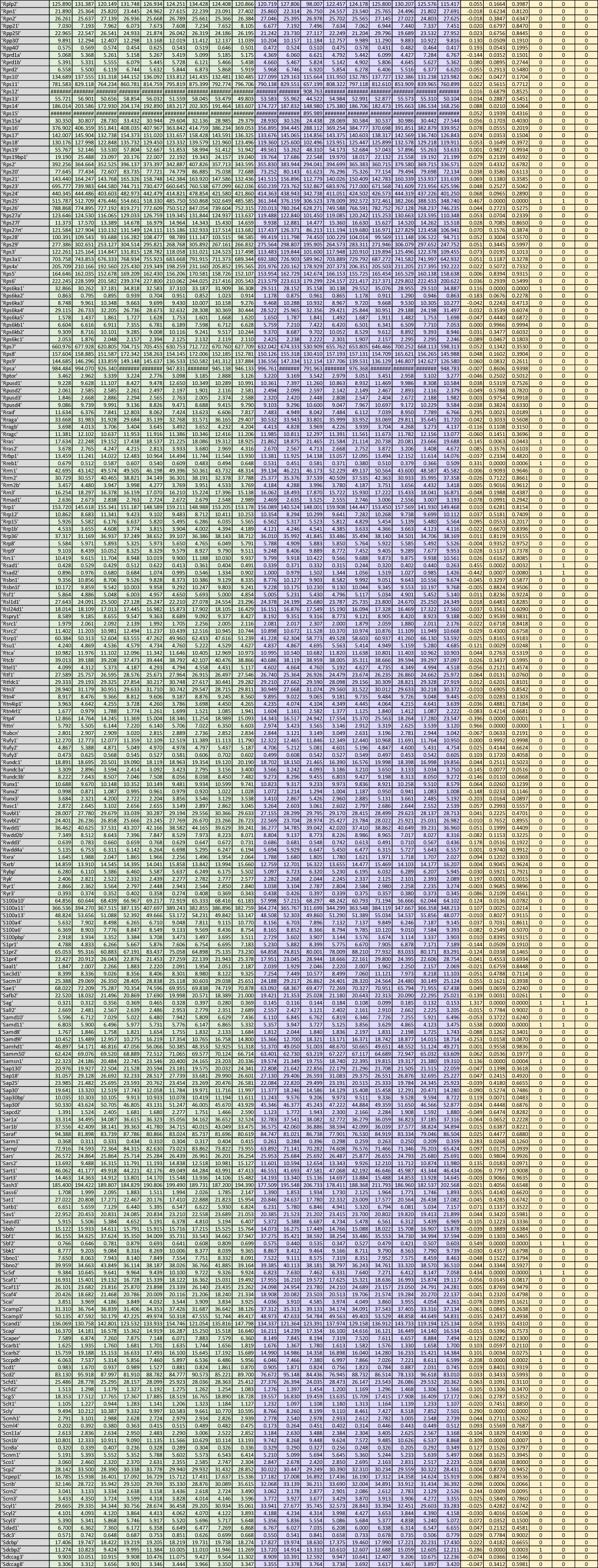
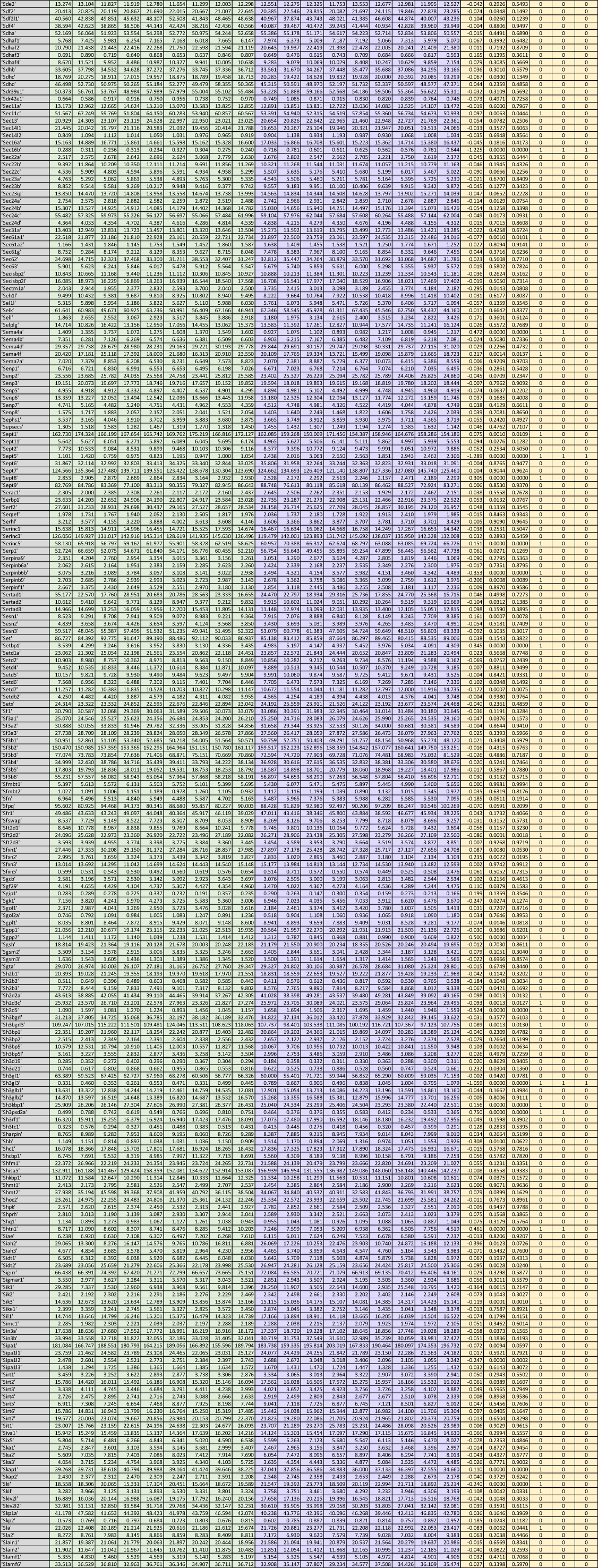
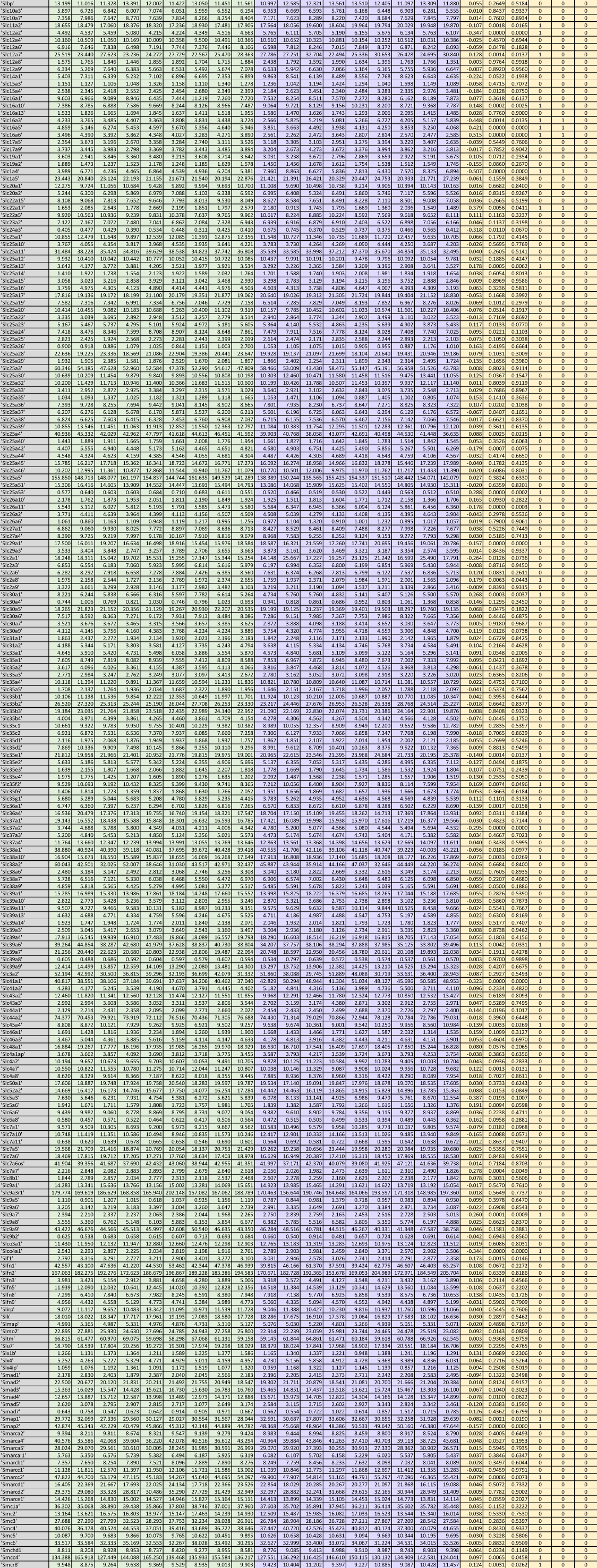
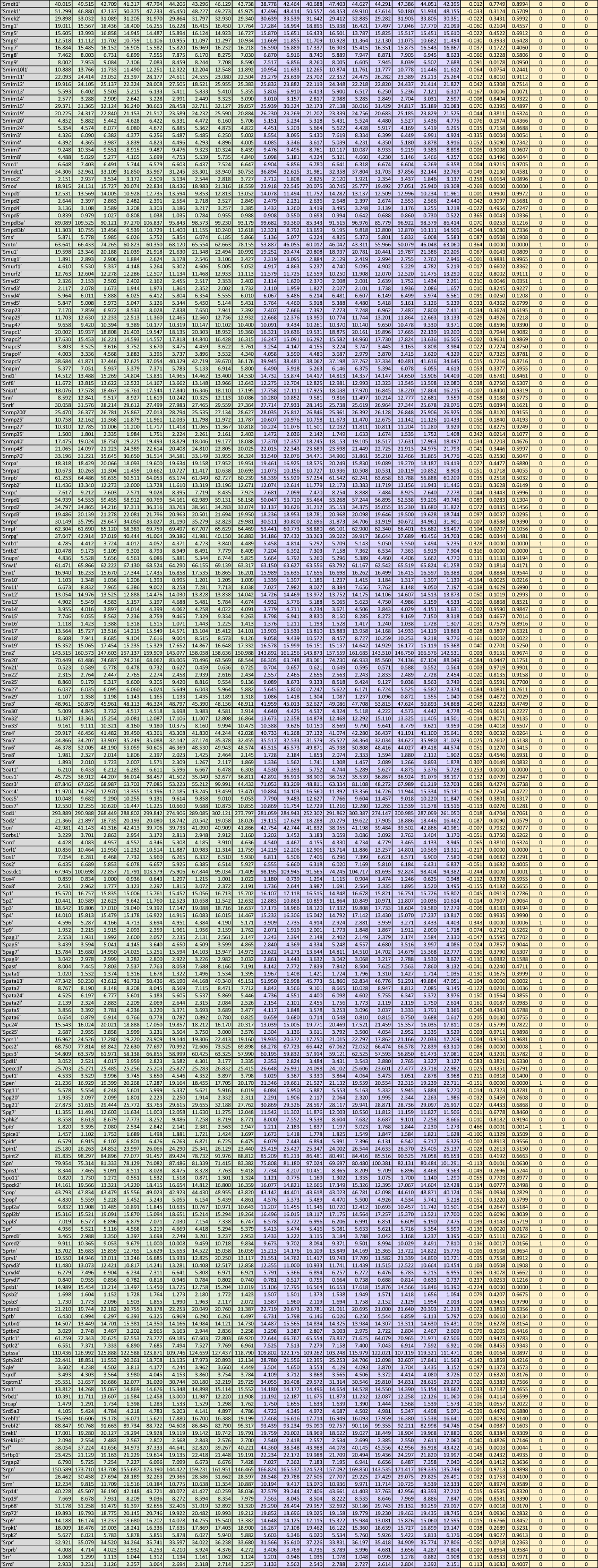
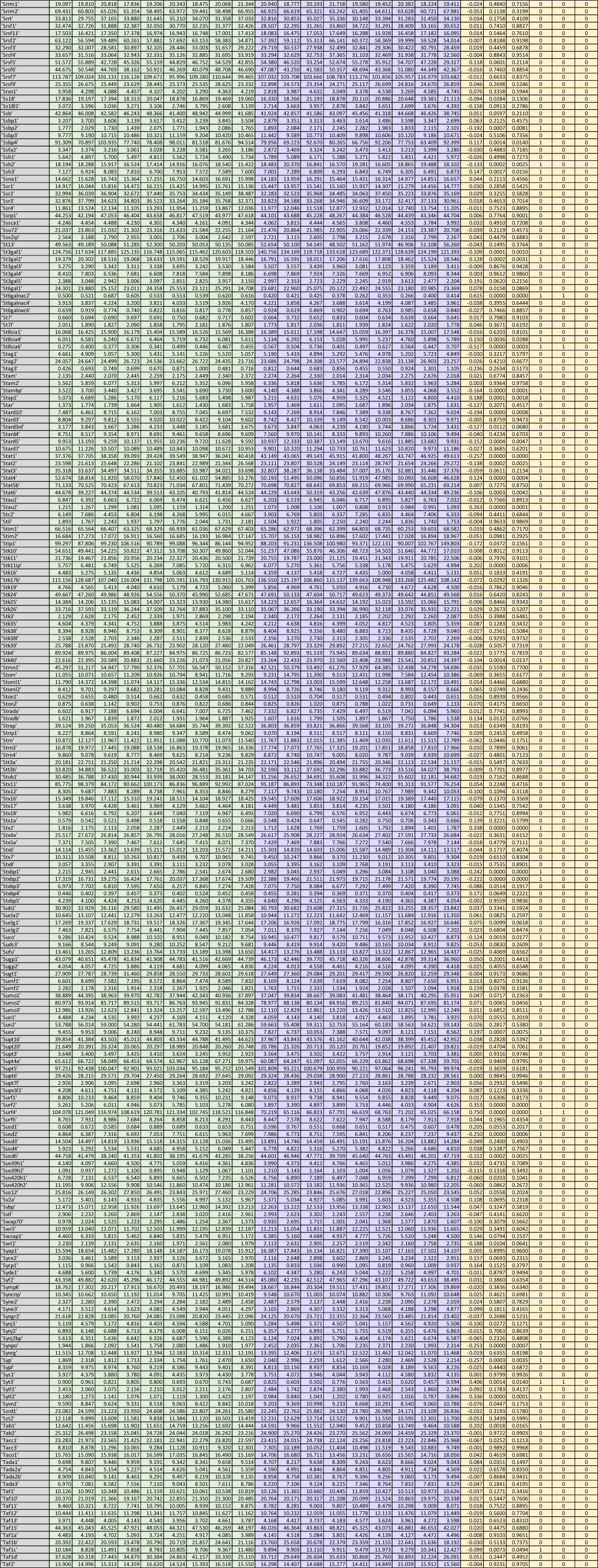
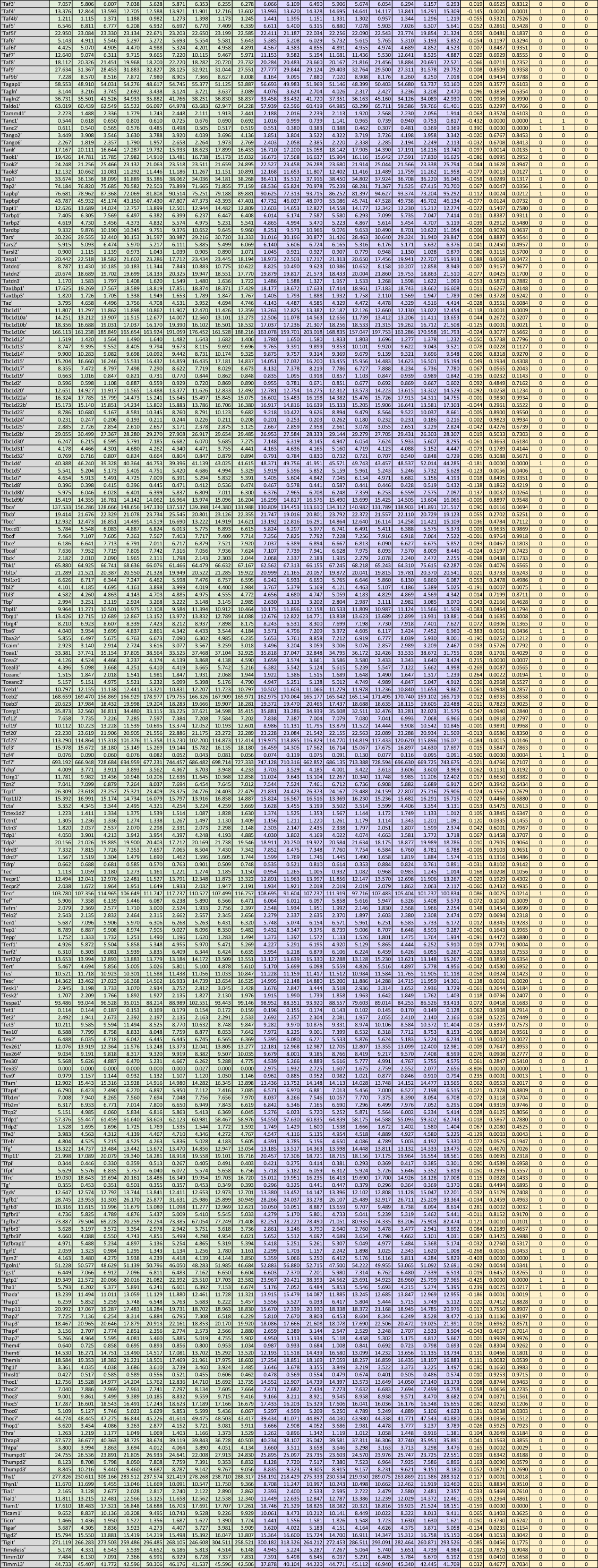
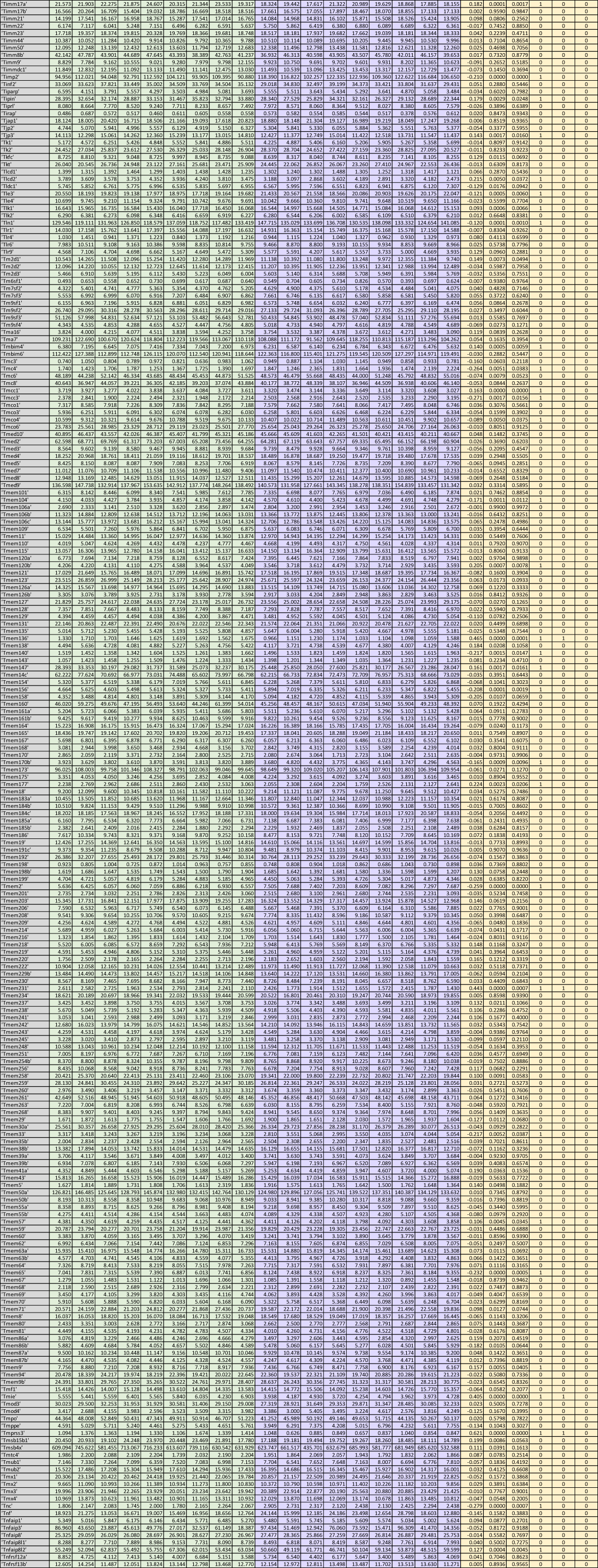
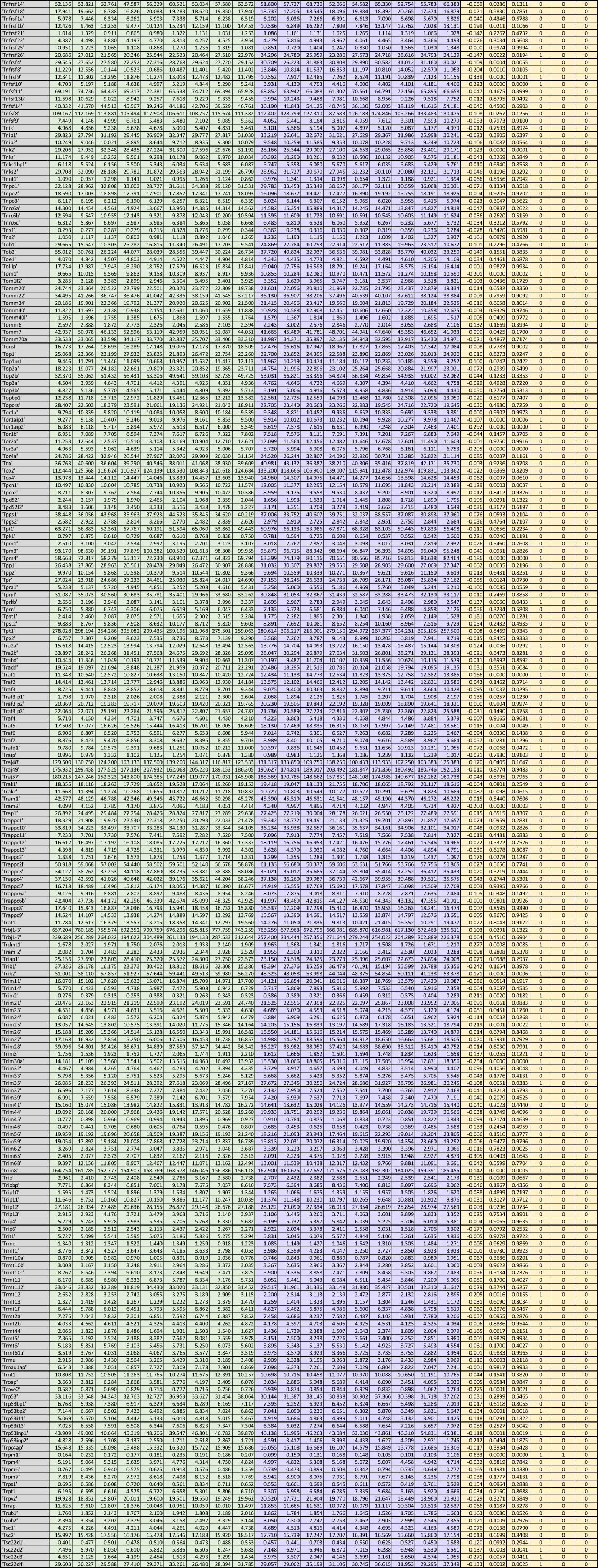
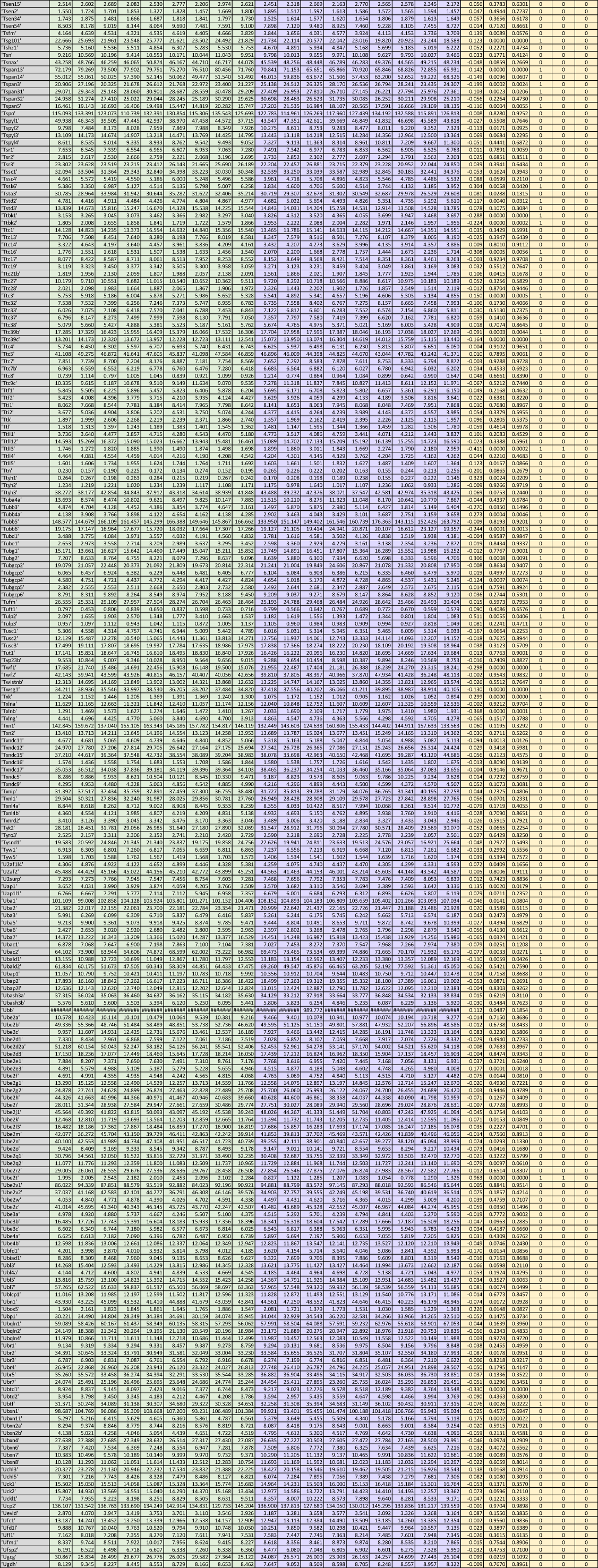
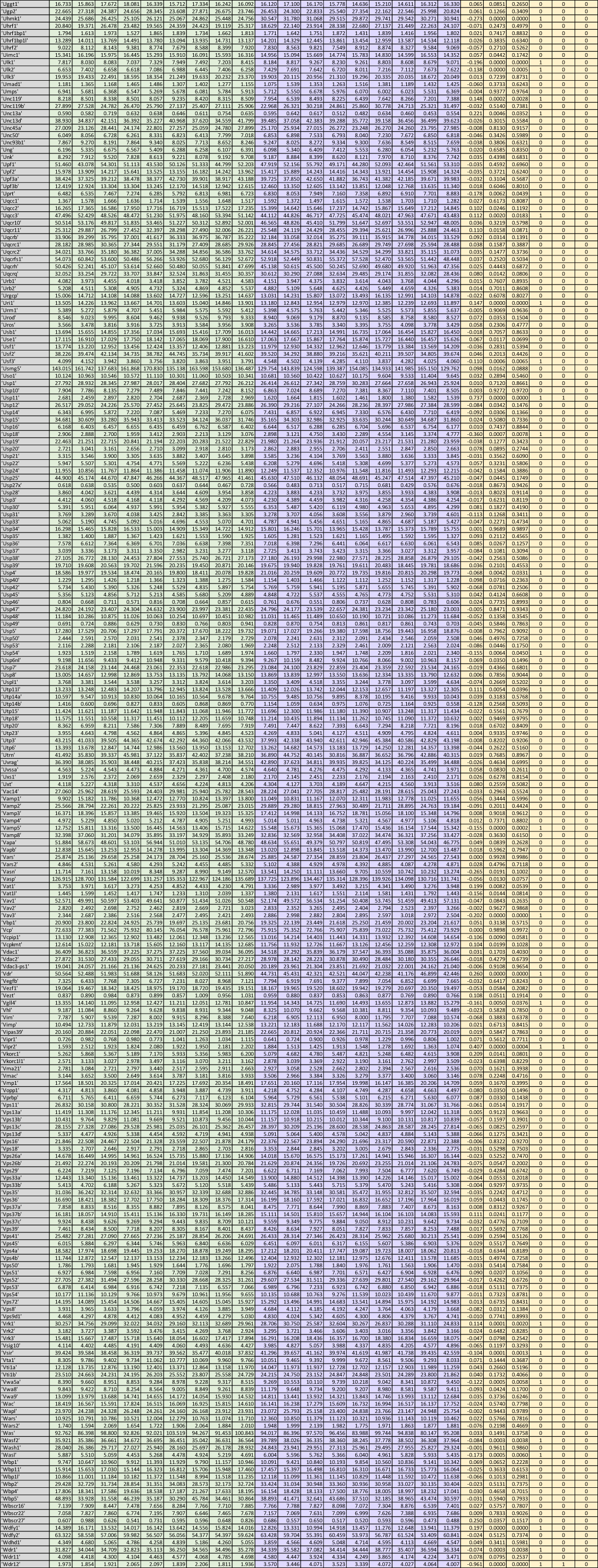
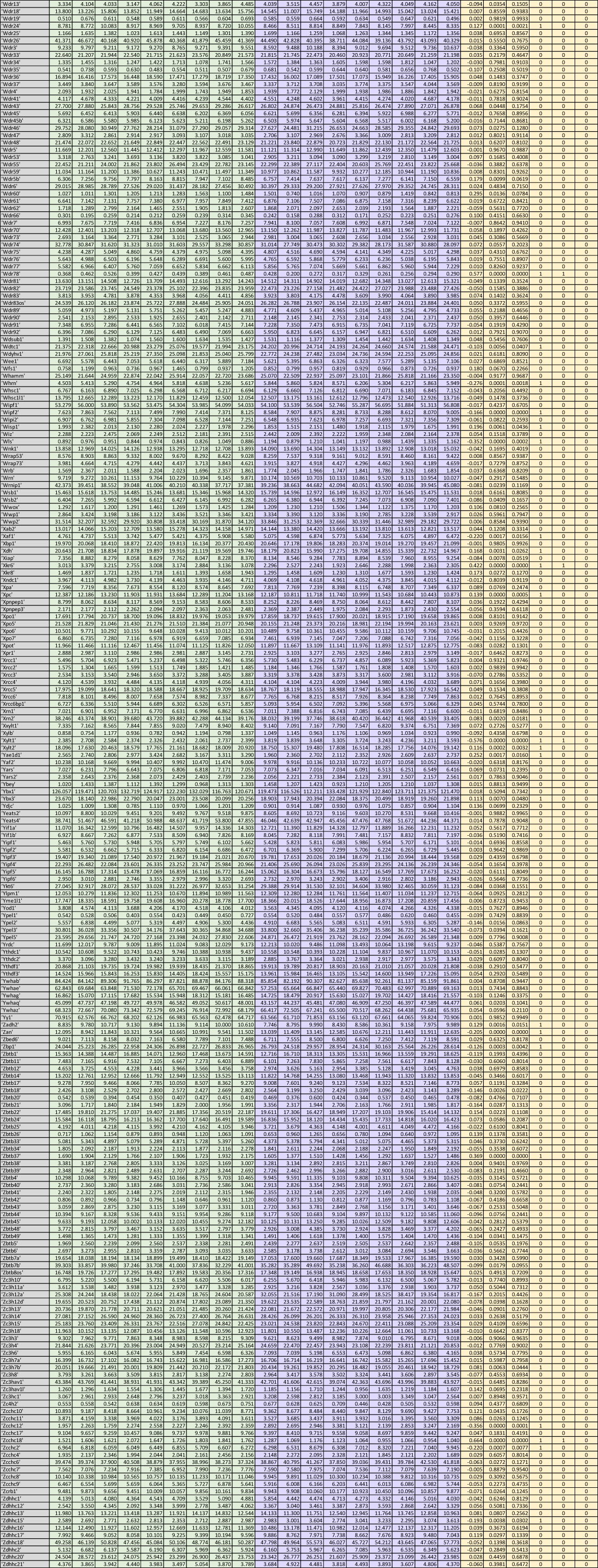
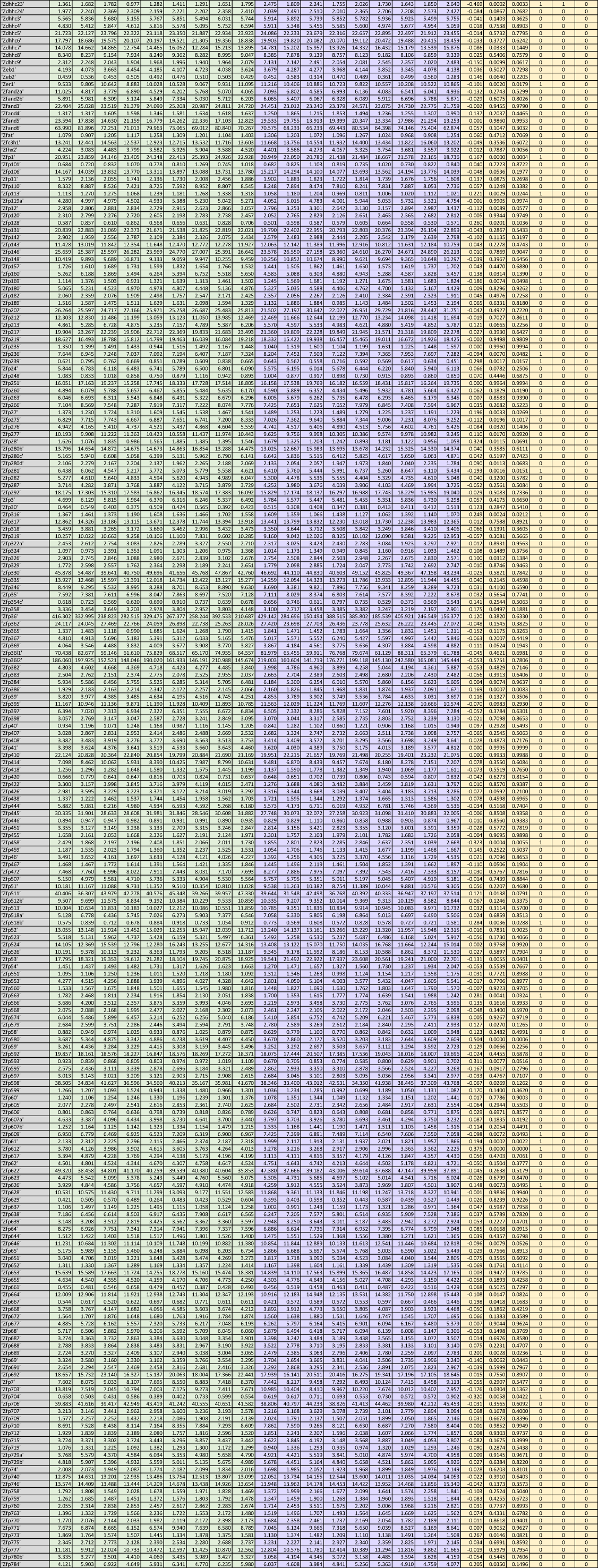
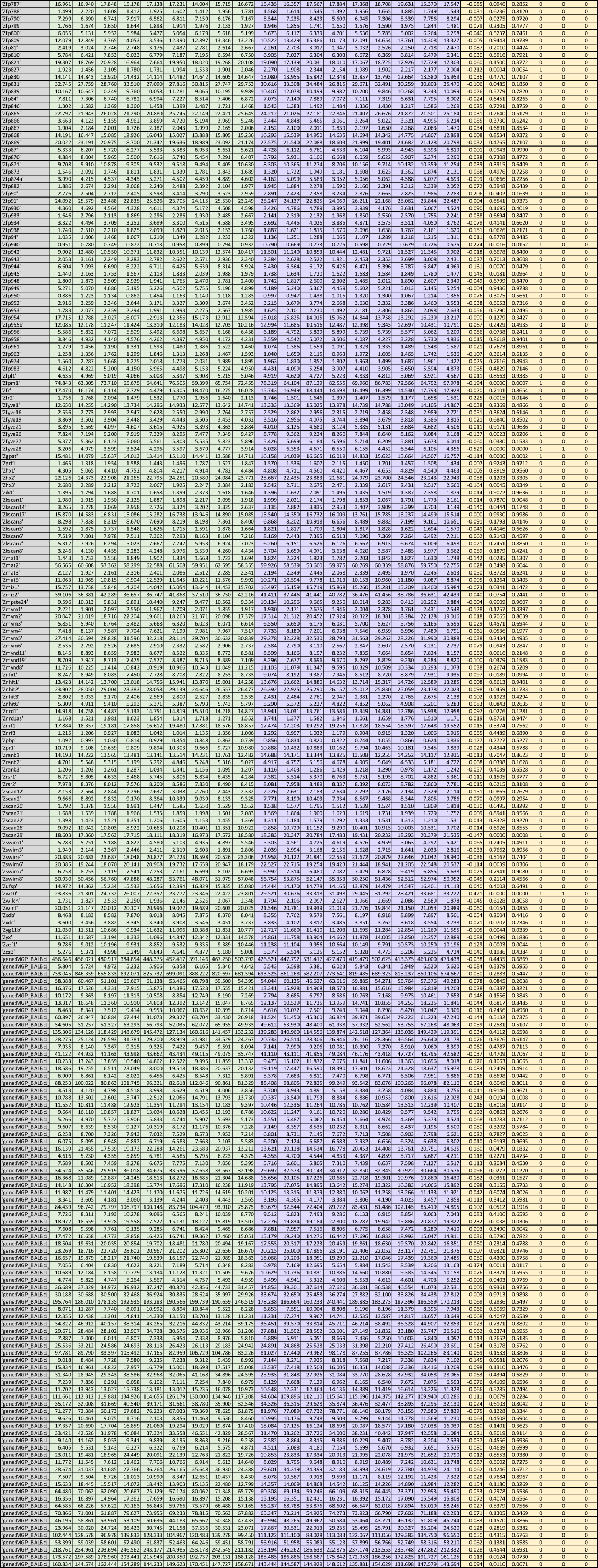
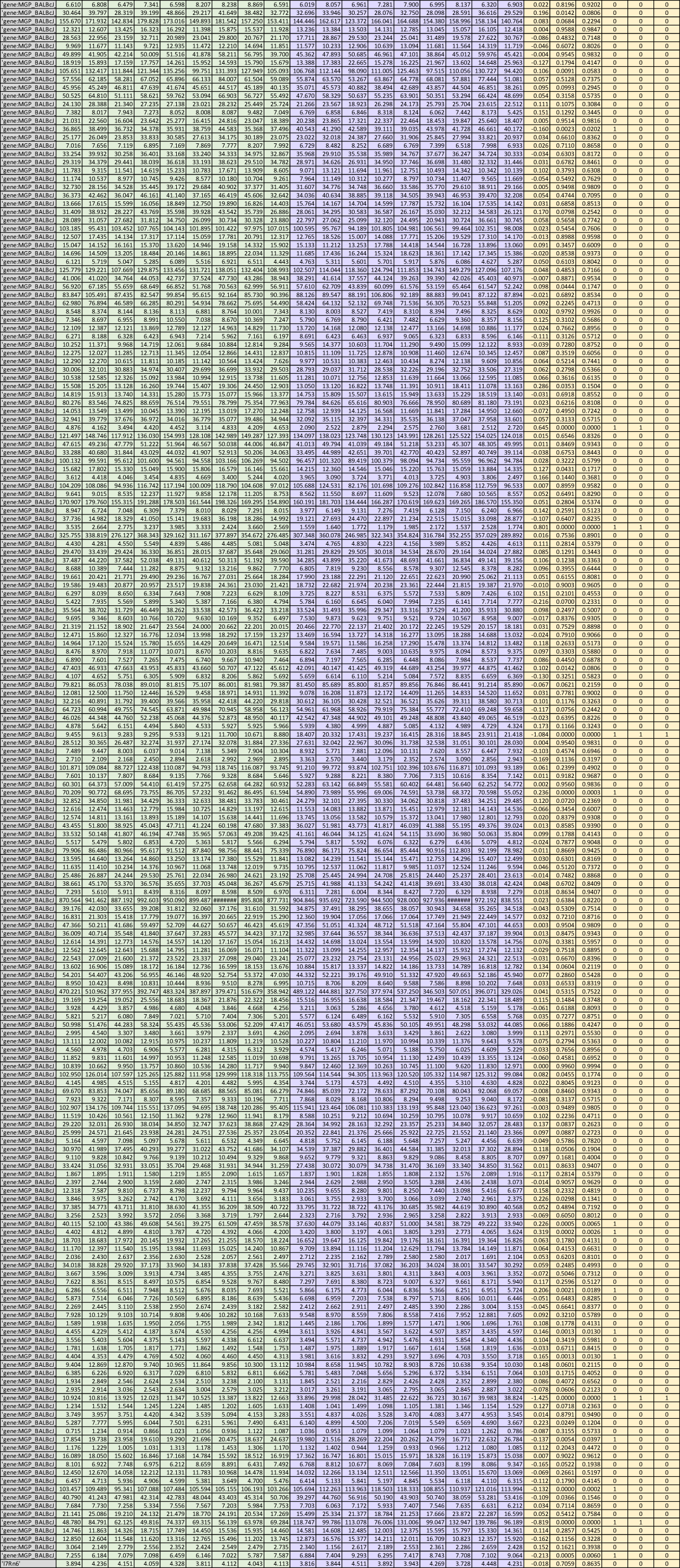
Supporting Figure 4. RNA-seq methods and analysis of msLN T_FH_ cells isolated from D14 *Hp*-infected WT and *Chi3l1*^-/-^ mice. RNA was isolated (RNeasy micro column (Qiagen)) from T_FH_ cells sorted into TRIzol as previously described^50^ and then enriched with Oligo(dT) beads. Sequencing libraries (NEBNext Ultra II RNA Library Prep Kit for Illumina (NEB, Ipswitch, MA, USA)) were prepared by GeneWiz LLC (South Plainfield, NJ, USA) and sequenced with a 2x150bp Paired End configuration on an Illumina HiSeq 4000. Image analysis and base calling were conducted using Hiseq Control Software (HCS). Raw sequence data (.bcl files) generated from Illumina HiSeq was converted into fastq files and de-multiplexed using Illumina’s bcl2fastq 2.17 software. One mismatch was allowed for index sequence identification. Samples were aligned to the ENSEMBLE BALB/c/j GCA_001632525 reference mouse genome and transcriptome to map data and annotate genes. After confirming that the first 8 exons of the *Chi3l1* gene, which were deleted in the *Chi3l1*^-/-^ mouse strain, were not expressed in the *Chi3l1*^-/-^ T_FH_ samples, we removed the *Chi3l1* gene (also called *Chil1*) from the RNA-seq analysis as exons 8-10 of the *Chi3l1* gene, which are directly downstream from the inserted promoter + neomycin cassette^13^, were expressed, presumably due to the insertion of the neomycin deletion cassette. Data shown are derived from 3 independent experiments with 3 mice/group/experiment. FDR, p value and log2 FC values are provided for all expressed genes, which are defined as genes with at least 3 reads per million in all samples of at least one group.

## Acknowledgements

We thank Scott Simpler, Rebecca Burnham, and Uma Mudunuru for mouse colony support, Sithy Allie and Amber Papillion for thoughtful comments, and Jessica Peel and Chris Risley for advice designing B cell panels. Allison Humbles (MEDIMMUNE) and Astra Zeneca provided mice. We thank Vidya Sigar Hanumanthu and the UAB Flow Cytometry Core Facility for sorting. Financial support provided by R01 AI104725 (FEL), R01 AI50740 (FEL), R01 AI153413 (TDR), R01 AI52476 (TDR), T32 2T32HL105346 (VT), UAB Frommeyer Investigative Fellowship (MLC), UAB Dixon Award (MLC), and AAAAI/ALA Respiratory Disease Research Award (MLC). The authors have no conflicting financial interests.

## Author Contributions

Conceptualization, MLC and FEL, Methodology, MLC, JEB, Investigation and Analysis, MLC, NBB, AFR, CDS, JEB and BM, Writing – Original Draft: MLC and FEL, Writing – Review and Editing: MLC, FEL, CDS, and AFR, Funding Acquisition: FEL, TDR and MLC, Resources, CDS, BL, TDR and AFR, Supervision, MLC and FEL.

## Notes

### Competing Interest Statement

The authors have declared no competing interest.

