## Supplementary figures and images for "Chitinase-3-like 1 regulates T_H_2 cells, T_FH_ cells and IgE following helminth infection"

### Supplemental Figure 1

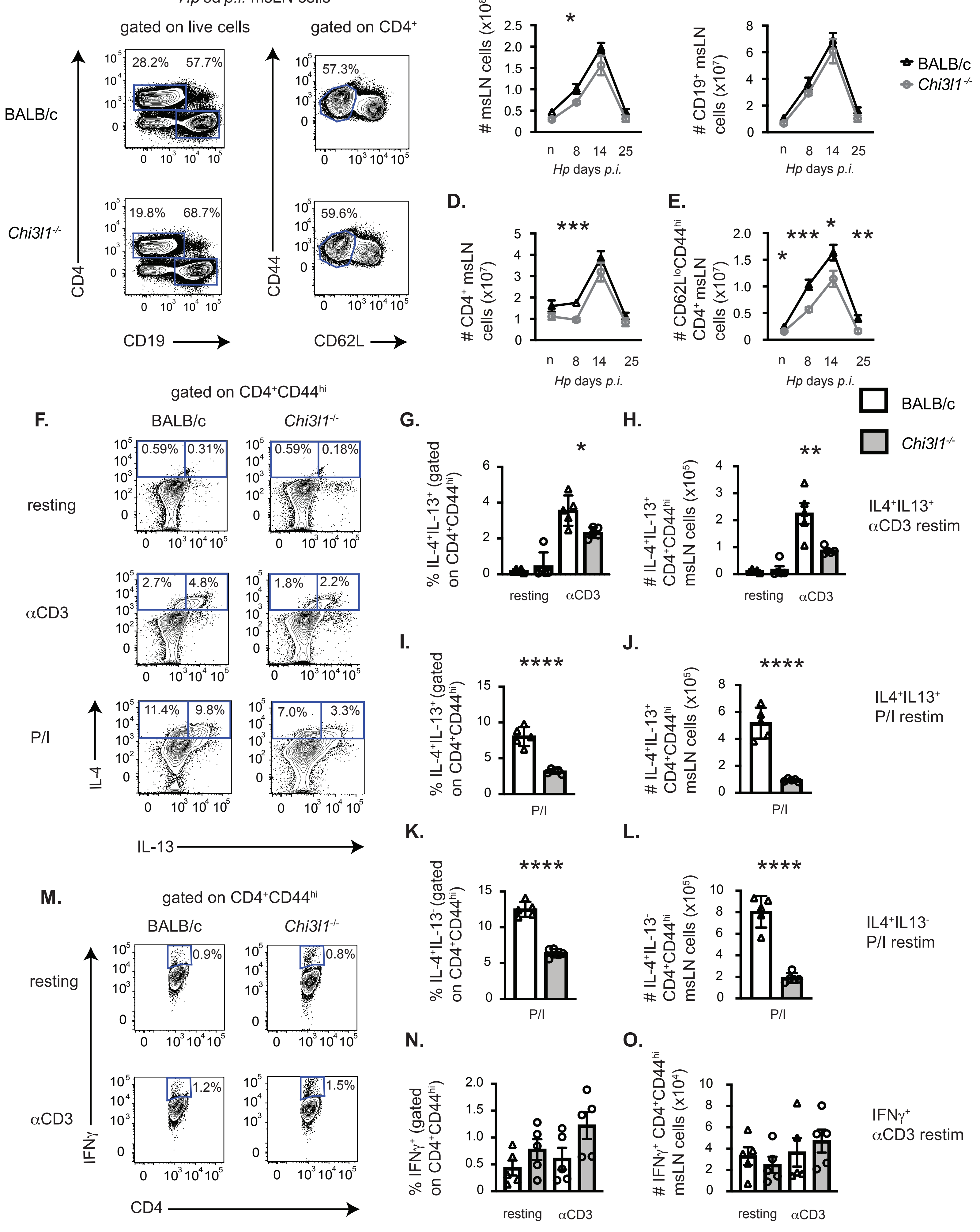

### Supplemental Figure 2

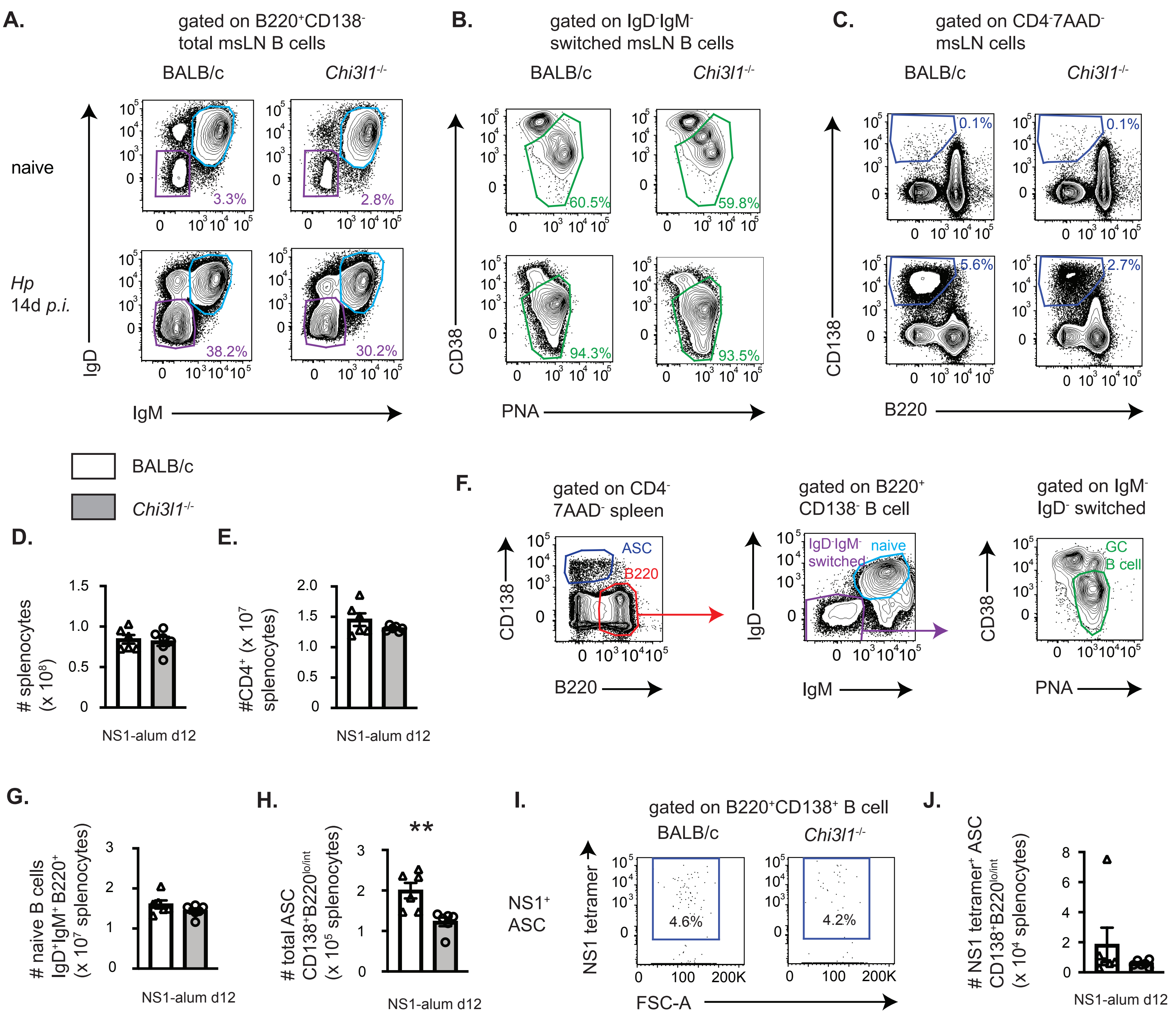

### Supplemental Figure 3

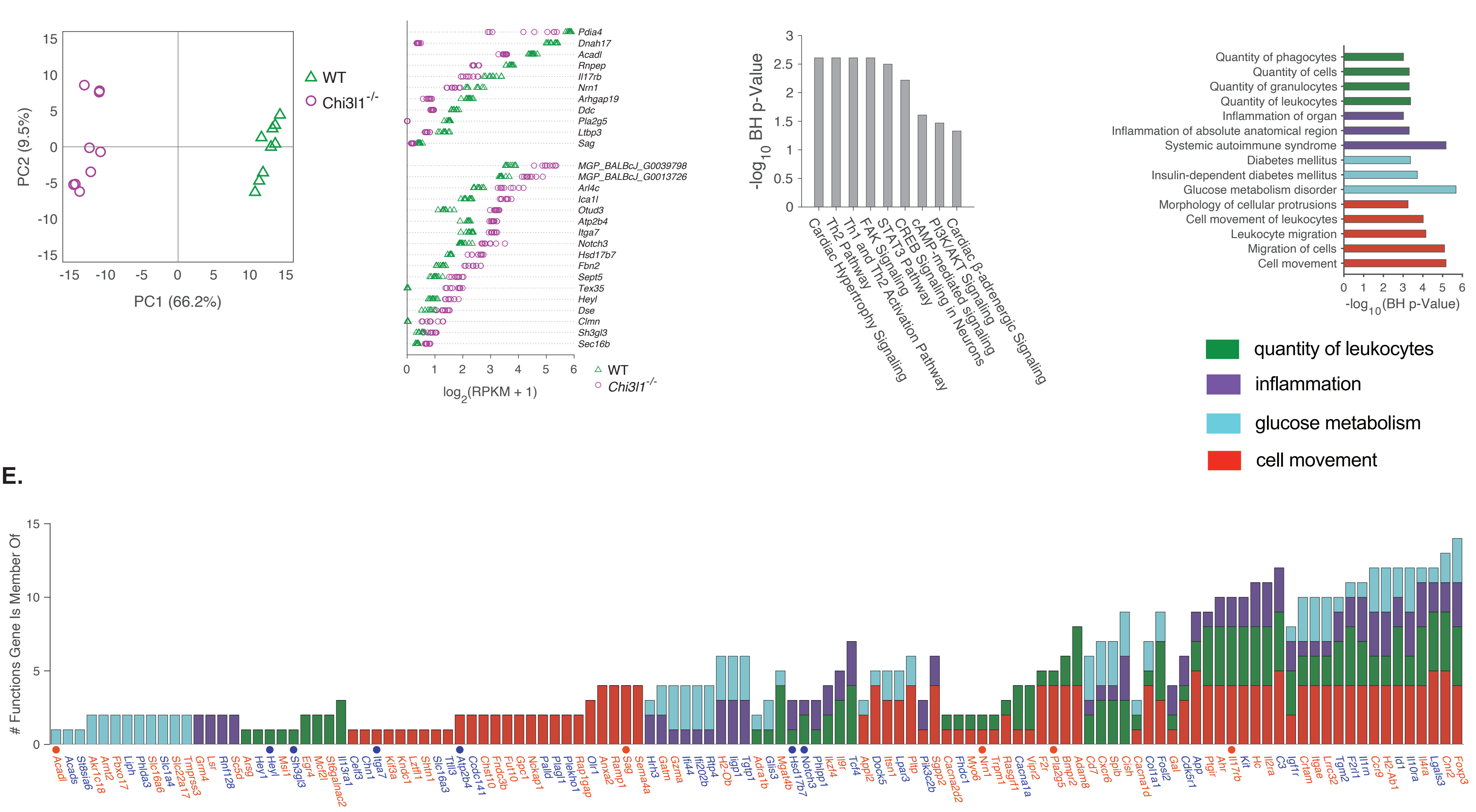
