## Supplemental Table 1 for "Chitinase-3-like 1 regulates T_H_2 cells, T_FH_ cells and IgE following helminth infection"

| gene | WT_Expt_1_Infection_1 | WT_Expt_2_Infection_1 | WT_Expt_3_Infection_1 | WT_Expt_1_Infection_2 | WT_Expt_2_Infection_2 | WT_Expt_3_Infection_2 | WT_Expt_1_Infection_3 | WT_Expt_2_Infection_3 | WT_Expt_3_Infection_3 | Chi3l1_KO_Expt_1_Infection_4 | Chi3l1_KO_Expt_2_Infection_4 | Chi3l1_KO_Expt_3_Infection_4 | Chi3l1_KO_Expt_1_Infection_5 | Chi3l1_KO_Expt_2_Infection_5 | Chi3l1_KO_Expt_3_Infection_5 | Chi3l1_KO_Expt_1_Infection_6 | Chi3l1_KO_Expt_2_Infection_6 | Chi3l1_KO_Expt_3_Infection_6 | log2_Fold_Change_WT_over_KO | p_Value | FDR | FDR_lt_5pct | FDR_lt_5pct_1point3fold | FDR_lt_5pct_2fold |
|---|---|---|---|---|---|---|---|---|---|---|---|---|---|---|---|---|---|---|---|---|---|---|---|---|
| '0610007P14Rik' | 6.436 | 7.532 | 7.209 | 6.899 | 7.263 | 7.066 | 6.930 | 7.656 | 7.420 | 6.055 | 7.147 | 7.144 | 7.170 | 7.798 | 7.607 | 6.995 | 7.117 | 6.891 | 0.000 | 0.9991 | 0.9998 | 0 | 0 | 0 |
| '0610009B22Rik' | 7.089 | 8.097 | 7.519 | 8.038 | 7.764 | 7.446 | 7.295 | 7.455 | 8.056 | 6.613 | 7.652 | 8.014 | 5.523 | 8.321 | 6.792 | 4.682 | 9.079 | 7.642 | 0.084 | 0.3365 | 0.5950 | 0 | 0 | 0 |
| '0610009O20Rik' | 18.574 | 18.015 | 20.960 | 18.460 | 19.725 | 21.589 | 18.170 | 18.210 | 21.529 | 18.097 | 18.929 | 21.662 | 19.431 | 18.127 | 21.147 | 19.312 | 19.541 | 20.362 | -0.022 | 0.5129 | 0.7375 | 0 | 0 | 0 |
| '0610010F05Rik' | 1.962 | 2.839 | 2.555 | 2.290 | 2.271 | 2.522 | 2.329 | 2.448 | 2.352 | 2.352 | 2.493 | 2.264 | 2.755 | 2.314 | 2.214 | 1.719 | 2.585 | 1.994 | 0.047 | 0.5452 | 0.7606 | 0 | 0 | 0 |
| '0610010K14Rik' | 2.386 | 1.950 | 2.481 | 2.213 | 2.143 | 2.190 | 2.529 | 2.320 | 2.361 | 2.377 | 2.269 | 2.445 | 2.188 | 2.532 | 2.232 | 2.032 | 2.577 | 2.106 | -0.020 | 0.7704 | 0.8977 | 0 | 0 | 0 |
| '0610012G03Rik' | 9.832 | 10.316 | 8.763 | 7.334 | 10.277 | 9.846 | 8.432 | 9.104 | 9.433 | 8.995 | 8.881 | 10.249 | 10.272 | 10.374 | 11.529 | 10.550 | 9.116 | 10.448 | -0.130 | 0.0541 | 0.1990 | 0 | 0 | 0 |
| '0610030E20Rik' | 5.767 | 6.519 | 6.834 | 6.585 | 5.813 | 6.533 | 6.929 | 5.715 | 6.777 | 5.109 | 6.515 | 7.452 | 5.413 | 6.106 | 7.430 | 5.396 | 6.764 | 7.207 | -0.001 | 0.9829 | 0.9934 | 0 | 0 | 0 |
| '0610037L13Rik' | 3.633 | 3.889 | 3.763 | 4.075 | 3.789 | 3.988 | 3.523 | 4.007 | 4.001 | 3.859 | 3.585 | 4.149 | 3.749 | 4.094 | 3.584 | 4.369 | 4.018 | 3.685 | -0.028 | 0.5252 | 0.7470 | 0 | 0 | 0 |
| '1110004E09Rik' | 5.975 | 5.727 | 6.581 | 6.157 | 5.932 | 6.754 | 6.262 | 6.600 | 7.243 | 6.591 | 5.798 | 6.218 | 6.874 | 6.883 | 6.141 | 6.829 | 6.033 | 5.788 | 0.000 | 0.9956 | 0.9994 | 0 | 0 | 0 |
| '1110008F13Rik' | 12.561 | 15.360 | 13.245 | 12.774 | 15.245 | 12.861 | 11.932 | 13.852 | 12.945 | 12.892 | 13.521 | 12.733 | 16.170 | 15.768 | 14.220 | 15.104 | 14.013 | 13.087 | -0.090 | 0.0694 | 0.2316 | 0 | 0 | 0 |
| '1110008L16Rik' | 1.959 | 2.091 | 2.132 | 2.005 | 1.837 | 1.690 | 2.204 | 1.967 | 1.762 | 2.165 | 2.025 | 1.799 | 2.010 | 1.753 | 1.713 | 1.747 | 1.907 | 1.399 | 0.087 | 0.1801 | 0.4154 | 0 | 0 | 0 |
| '1110008P14Rik' | 8.757 | 9.385 | 9.211 | 8.965 | 9.235 | 10.255 | 7.964 | 8.667 | 9.091 | 9.304 | 9.023 | 9.300 | 9.561 | 8.675 | 9.757 | 9.284 | 8.994 | 9.546 | -0.042 | 0.3374 | 0.5955 | 0 | 0 | 0 |
| '1110012L19Rik' | 11.502 | 11.115 | 10.860 | 11.302 | 11.935 | 11.005 | 10.507 | 11.827 | 10.449 | 11.080 | 10.075 | 11.646 | 10.774 | 10.143 | 13.232 | 9.244 | 11.914 | 10.888 | 0.005 | 0.9385 | 0.9764 | 0 | 0 | 0 |
| '1110032A03Rik' | 1.287 | 1.536 | 1.333 | 1.076 | 1.545 | 1.331 | 1.410 | 1.506 | 1.443 | 1.292 | 1.192 | 1.057 | 1.425 | 1.295 | 0.937 | 1.370 | 1.164 | 0.764 | 0.252 | 0.0043 | 0.0336 | 1 | 0 | 0 |
| '1110032F04Rik' | 2.789 | 2.221 | 3.275 | 2.227 | 2.254 | 2.771 | 2.794 | 2.531 | 2.997 | 2.612 | 2.535 | 2.956 | 1.672 | 2.337 | 2.496 | 2.496 | 1.865 | 2.513 | 0.147 | 0.1225 | 0.3338 | 0 | 0 | 0 |
| '1110034G24Rik' | 3.945 | 4.688 | 3.845 | 4.716 | 4.184 | 4.541 | 4.248 | 4.386 | 4.477 | 4.994 | 4.166 | 5.258 | 4.086 | 5.105 | 4.715 | 6.122 | 5.695 | 4.457 | -0.204 | 0.0079 | 0.0526 | 0 | 0 | 0 |
| '1110037F02Rik' | 26.860 | 27.671 | 26.667 | 27.515 | 28.359 | 26.838 | 28.343 | 27.180 | 26.864 | 27.396 | 27.778 | 28.113 | 26.666 | 27.455 | 26.912 | 26.427 | 28.208 | 27.774 | -0.015 | 0.4878 | 0.7182 | 0 | 0 | 0 |
| '1110038F14Rik' | 35.309 | 40.703 | 40.552 | 33.519 | 40.700 | 45.854 | 35.584 | 42.757 | 39.987 | 39.074 | 33.597 | 38.520 | 38.578 | 36.088 | 39.869 | 39.367 | 36.229 | 37.087 | 0.058 | 0.2321 | 0.4800 | 0 | 0 | 0 |
| '1110051M20Rik' | 5.378 | 6.344 | 5.612 | 5.253 | 5.658 | 6.067 | 5.470 | 6.171 | 5.985 | 5.938 | 5.677 | 6.268 | 5.927 | 5.312 | 6.292 | 5.944 | 5.636 | 5.426 | -0.027 | 0.5465 | 0.7610 | 0 | 0 | 0 |
| '1110059E24Rik' | 2.582 | 3.107 | 2.775 | 2.496 | 2.868 | 2.380 | 2.701 | 3.504 | 2.833 | 2.744 | 2.447 | 2.429 | 2.771 | 2.938 | 2.449 | 2.686 | 2.668 | 2.158 | 0.108 | 0.0924 | 0.2812 | 0 | 0 | 0 |
| '1110059G10Rik' | 3.805 | 4.457 | 4.262 | 3.532 | 4.725 | 3.836 | 3.696 | 3.987 | 3.676 | 3.331 | 3.856 | 3.330 | 3.862 | 4.156 | 3.348 | 4.239 | 3.464 | 3.096 | 0.134 | 0.0370 | 0.1557 | 0 | 0 | 0 |
| '1190002N15Rik' | 21.777 | 22.231 | 21.548 | 22.420 | 21.545 | 20.436 | 24.082 | 21.653 | 19.866 | 22.750 | 21.492 | 23.026 | 19.056 | 23.261 | 21.427 | 20.777 | 22.380 | 22.728 | -0.023 | 0.5976 | 0.7954 | 0 | 0 | 0 |
| '1500011B03Rik' | 13.488 | 16.313 | 13.641 | 15.467 | 15.132 | 13.743 | 14.214 | 15.525 | 13.482 | 11.988 | 14.074 | 15.361 | 13.469 | 15.500 | 15.081 | 15.356 | 15.900 | 15.004 | -0.027 | 0.6292 | 0.8163 | 0 | 0 | 0 |
| '1500011K16Rik' | 23.311 | 22.821 | 23.615 | 25.360 | 27.406 | 24.738 | 27.840 | 25.241 | 22.348 | 23.373 | 21.132 | 21.261 | 23.640 | 20.275 | 21.012 | 25.728 | 25.408 | 18.719 | 0.143 | 0.0111 | 0.0675 | 0 | 0 | 0 |
| '1600002H07Rik' | 1.782 | 1.977 | 2.080 | 1.201 | 1.557 | 1.720 | 1.909 | 1.837 | 1.811 | 1.550 | 1.640 | 1.867 | 2.035 | 1.541 | 1.765 | 1.629 | 1.330 | 1.537 | 0.080 | 0.3647 | 0.6165 | 0 | 0 | 0 |
| '1600002K03Rik' | 12.316 | 9.142 | 11.683 | 12.484 | 11.215 | 12.235 | 10.851 | 12.326 | 12.692 | 10.529 | 12.193 | 13.136 | 9.820 | 9.863 | 13.472 | 13.785 | 10.897 | 12.945 | -0.026 | 0.7549 | 0.8907 | 0 | 0 | 0 |
| '1600012H06Rik' | 2.992 | 4.459 | 3.686 | 3.099 | 3.919 | 3.710 | 3.564 | 3.940 | 4.114 | 2.808 | 3.499 | 3.934 | 3.229 | 3.651 | 4.031 | 3.352 | 3.481 | 3.638 | 0.063 | 0.2203 | 0.4676 | 0 | 0 | 0 |
| '1700001O22Rik' | 0.743 | 0.843 | 0.713 | 0.975 | 0.961 | 1.018 | 0.801 | 1.040 | 0.952 | 1.001 | 0.719 | 0.802 | 1.061 | 1.017 | 0.634 | 0.733 | 0.831 | 0.715 | 0.091 | 0.4025 | 0.6512 | 0 | 0 | 0 |
| '1700010I14Rik' | 2.171 | 2.753 | 2.167 | 2.277 | 2.406 | 2.036 | 2.550 | 2.186 | 2.465 | 2.538 | 2.456 | 2.512 | 2.284 | 2.560 | 2.120 | 2.390 | 3.044 | 2.390 | -0.105 | 0.1593 | 0.3895 | 0 | 0 | 0 |
| '1700017B05Rik' | 14.656 | 16.426 | 15.029 | 14.125 | 14.046 | 15.843 | 14.118 | 13.910 | 15.823 | 14.468 | 12.909 | 16.113 | 12.797 | 14.149 | 15.554 | 15.931 | 13.618 | 16.115 | 0.015 | 0.7248 | 0.8736 | 0 | 0 | 0 |
| '1700019D03Rik' | 1.653 | 1.707 | 1.864 | 1.539 | 1.852 | 1.662 | 1.745 | 2.065 | 1.642 | 2.136 | 2.245 | 2.234 | 2.044 | 2.132 | 1.870 | 1.900 | 2.269 | 2.018 | -0.273 | 0.0000 | 0.0005 | 1 | 0 | 0 |
| '1700021F05Rik' | 5.958 | 6.035 | 6.562 | 5.312 | 5.178 | 6.586 | 5.733 | 6.022 | 5.526 | 6.812 | 5.268 | 5.330 | 5.865 | 6.040 | 5.962 | 4.642 | 5.876 | 6.248 | 0.020 | 0.7873 | 0.9048 | 0 | 0 | 0 |
| '1700030K09Rik' | 2.901 | 3.147 | 3.539 | 3.379 | 3.467 | 3.633 | 2.889 | 3.752 | 3.749 | 3.336 | 2.797 | 2.908 | 3.086 | 3.143 | 2.957 | 3.258 | 3.234 | 2.563 | 0.158 | 0.0207 | 0.1056 | 0 | 0 | 0 |
| '1700037C18Rik' | 0.676 | 0.732 | 1.094 | 0.608 | 1.049 | 1.062 | 0.958 | 1.086 | 1.012 | 1.099 | 0.851 | 0.957 | 1.203 | 0.997 | 0.938 | 1.072 | 1.005 | 0.886 | -0.124 | 0.2976 | 0.5535 | 0 | 0 | 0 |
| '1700037H04Rik' | 4.336 | 4.856 | 4.518 | 3.761 | 5.282 | 5.114 | 4.207 | 5.235 | 4.732 | 5.099 | 4.217 | 5.043 | 5.123 | 5.147 | 5.252 | 4.800 | 4.908 | 4.855 | -0.094 | 0.0878 | 0.2702 | 0 | 0 | 0 |
| '1700056E22Rik' | 2.849 | 2.698 | 3.072 | 2.597 | 2.980 | 3.370 | 2.430 | 3.395 | 2.833 | 3.248 | 2.356 | 1.577 | 2.730 | 2.396 | 1.616 | 3.148 | 1.619 | 1.470 | 0.425 | 0.0018 | 0.0163 | 1 | 1 | 0 |
| '1700066M21Rik' | 2.674 | 3.096 | 2.827 | 2.559 | 2.955 | 2.884 | 2.873 | 3.231 | 2.757 | 2.325 | 3.048 | 3.546 | 2.826 | 3.860 | 3.329 | 2.739 | 3.249 | 3.156 | -0.136 | 0.0410 | 0.1658 | 0 | 0 | 0 |
| '1700109H08Rik' | 1.487 | 1.571 | 1.324 | 0.971 | 1.770 | 1.182 | 1.348 | 1.198 | 1.151 | 2.385 | 1.673 | 1.159 | 1.866 | 1.562 | 1.179 | 1.946 | 0.851 | 1.072 | -0.163 | 0.2508 | 0.5019 | 0 | 0 | 0 |
| '1700123O20Rik' | 19.442 | 18.089 | 17.223 | 20.744 | 18.893 | 18.947 | 19.411 | 16.451 | 19.331 | 16.240 | 20.026 | 20.681 | 16.391 | 18.381 | 19.715 | 15.922 | 17.729 | 21.632 | 0.005 | 0.9336 | 0.9746 | 0 | 0 | 0 |
| '1810009A15Rik' | 1.362 | 1.275 | 1.569 | 1.440 | 1.296 | 1.526 | 2.007 | 1.591 | 1.626 | 1.156 | 1.161 | 1.679 | 1.347 | 1.394 | 1.522 | 1.130 | 1.469 | 1.541 | 0.120 | 0.2431 | 0.4935 | 0 | 0 | 0 |
| '1810013L24Rik' | 19.182 | 19.604 | 19.288 | 21.141 | 19.214 | 18.445 | 21.448 | 19.757 | 18.444 | 17.292 | 22.519 | 20.849 | 16.132 | 20.640 | 20.146 | 17.730 | 22.473 | 20.455 | -0.020 | 0.7216 | 0.8716 | 0 | 0 | 0 |
| '1810022K09Rik' | 9.089 | 10.851 | 9.329 | 10.970 | 10.831 | 8.520 | 10.481 | 9.652 | 9.097 | 8.797 | 9.104 | 9.033 | 8.742 | 9.853 | 8.783 | 9.300 | 10.255 | 8.891 | 0.083 | 0.0932 | 0.2826 | 0 | 0 | 0 |
| '1810026J23Rik' | 26.488 | 23.017 | 19.512 | 23.425 | 23.830 | 21.006 | 21.780 | 22.671 | 20.052 | 24.262 | 23.222 | 21.481 | 25.667 | 25.199 | 20.006 | 23.563 | 24.295 | 20.254 | -0.053 | 0.1510 | 0.3777 | 0 | 0 | 0 |
| '1810030O07Rik' | 27.930 | 26.742 | 26.798 | 27.950 | 28.786 | 28.236 | 28.449 | 27.486 | 27.018 | 28.071 | 27.508 | 29.988 | 25.901 | 28.779 | 28.477 | 27.107 | 28.509 | 28.124 | -0.030 | 0.3328 | 0.5908 | 0 | 0 | 0 |
| '1810032O08Rik' | 2.997 | 3.272 | 3.669 | 3.292 | 3.426 | 3.812 | 3.497 | 3.291 | 3.303 | 3.134 | 2.498 | 3.134 | 3.134 | 2.754 | 2.726 | 2.662 | 3.412 | 2.826 | 0.203 | 0.0038 | 0.0303 | 1 | 0 | 0 |
| '1810037I17Rik' | 20.261 | 22.528 | 21.311 | 20.936 | 22.233 | 22.247 | 21.184 | 22.771 | 20.768 | 18.888 | 22.555 | 21.515 | 21.770 | 24.594 | 23.647 | 19.947 | 23.839 | 20.246 | -0.029 | 0.4866 | 0.7174 | 0 | 0 | 0 |
| '1810043G02Rik' | 0.866 | 0.758 | 0.794 | 0.831 | 0.652 | 0.864 | 0.716 | 0.761 | 0.839 | 0.765 | 0.928 | 0.734 | 0.859 | 0.839 | 0.667 | 1.000 | 0.704 | 0.701 | -0.018 | 0.8199 | 0.9203 | 0 | 0 | 0 |
| '1810043H04Rik' | 8.902 | 10.240 | 9.878 | 9.057 | 10.523 | 10.382 | 9.287 | 10.488 | 10.540 | 9.250 | 8.710 | 9.440 | 10.412 | 9.911 | 9.497 | 10.047 | 9.313 | 9.773 | 0.039 | 0.4843 | 0.7158 | 0 | 0 | 0 |
| '1810049J17Rik' | 17.446 | 17.152 | 18.273 | 13.541 | 13.355 | 14.589 | 15.260 | 15.987 | 18.219 | 17.346 | 15.677 | 16.766 | 12.874 | 14.474 | 16.165 | 16.749 | 14.935 | 14.926 | 0.037 | 0.6629 | 0.8371 | 0 | 0 | 0 |
| '1810055G02Rik' | 4.591 | 5.086 | 5.732 | 4.654 | 4.841 | 5.877 | 5.009 | 4.727 | 5.640 | 4.795 | 3.849 | 5.403 | 4.929 | 4.412 | 5.494 | 5.292 | 4.444 | 5.108 | 0.068 | 0.3184 | 0.5768 | 0 | 0 | 0 |
| '2010107E04Rik' | 125.666 | 134.986 | 118.351 | 127.433 | 130.901 | 115.521 | 131.008 | 132.052 | 116.329 | 118.638 | 116.479 | 119.170 | 121.403 | 131.068 | 123.701 | 121.526 | 125.838 | 109.351 | 0.043 | 0.2144 | 0.4598 | 0 | 0 | 0 |
| '2010107G23Rik' | 0.486 | 0.680 | 0.792 | 0.658 | 0.703 | 0.646 | 0.501 | 0.628 | 0.731 | 0.478 | 0.414 | 0.426 | 0.341 | 0.511 | 0.530 | 0.577 | 0.376 | 0.401 | 0.527 | 0.0000 | 0.0003 | 1 | 1 | 0 |
| '2010111I01Rik' | 3.679 | 3.117 | 3.244 | 3.681 | 3.018 | 3.129 | 3.643 | 3.048 | 3.395 | 2.281 | 2.482 | 2.728 | 2.189 | 2.719 | 2.806 | 2.205 | 2.878 | 2.858 | 0.360 | 0.0000 | 0.0000 | 1 | 0 | 0 |
| '2010300C02Rik' | 0.449 | 0.793 | 0.426 | 0.611 | 0.507 | 0.502 | 0.541 | 0.664 | 0.470 | 0.522 | 0.817 | 0.419 | 0.725 | 0.814 | 0.350 | 0.605 | 0.878 | 0.376 | -0.125 | 0.2971 | 0.5530 | 0 | 0 | 0 |
| '2010315B03Rik' | 2.414 | 3.409 | 3.330 | 2.486 | 3.419 | 3.618 | 3.078 | 3.632 | 3.440 | 3.222 | 2.904 | 3.031 | 2.949 | 2.925 | 2.937 | 2.812 | 3.092 | 2.785 | 0.098 | 0.1279 | 0.3427 | 0 | 0 | 0 |
| '2210013O21Rik' | 3.593 | 5.375 | 4.771 | 4.662 | 4.890 | 4.422 | 4.748 | 4.269 | 4.584 | 3.872 | 4.282 | 4.256 | 4.527 | 5.692 | 4.410 | 4.193 | 4.127 | 4.352 | 0.042 | 0.5581 | 0.7697 | 0 | 0 | 0 |
| '2210016F16Rik' | 23.230 | 28.140 | 25.996 | 23.340 | 26.157 | 27.349 | 22.341 | 25.717 | 24.579 | 24.530 | 23.917 | 23.611 | 25.524 | 25.431 | 25.313 | 24.637 | 22.750 | 22.110 | 0.048 | 0.2580 | 0.5107 | 0 | 0 | 0 |
| '2210016L21Rik' | 8.205 | 7.922 | 9.297 | 8.298 | 8.719 | 9.530 | 8.656 | 7.768 | 8.473 | 9.112 | 6.882 | 8.680 | 8.418 | 7.494 | 8.770 | 8.944 | 6.746 | 8.914 | 0.049 | 0.2573 | 0.5095 | 0 | 0 | 0 |
| '2300009A05Rik' | 2.202 | 3.232 | 2.192 | 1.914 | 2.662 | 2.676 | 2.306 | 2.473 | 2.459 | 2.204 | 2.415 | 2.654 | 2.360 | 2.319 | 2.400 | 2.417 | 2.830 | 2.427 | -0.020 | 0.8020 | 0.9114 | 0 | 0 | 0 |
| '2310009B15Rik' | 2.295 | 2.244 | 2.593 | 2.076 | 2.024 | 2.573 | 2.793 | 2.578 | 2.707 | 2.322 | 2.221 | 2.185 | 2.395 | 2.447 | 2.161 | 1.831 | 2.431 | 2.190 | 0.114 | 0.2024 | 0.4440 | 0 | 0 | 0 |
| '2310011J03Rik' | 24.791 | 29.736 | 28.377 | 24.235 | 28.235 | 32.172 | 23.189 | 29.033 | 27.310 | 27.690 | 25.639 | 27.923 | 26.872 | 29.101 | 29.157 | 32.939 | 23.035 | 27.151 | -0.025 | 0.6784 | 0.8461 | 0 | 0 | 0 |
| '2310022A10Rik' | 4.584 | 4.392 | 4.790 | 4.559 | 4.436 | 5.356 | 4.101 | 4.641 | 5.068 | 4.748 | 4.739 | 4.701 | 4.710 | 4.673 | 5.119 | 4.522 | 4.212 | 4.503 | -0.006 | 0.8929 | 0.9563 | 0 | 0 | 0 |
| '2310022B05Rik' | 6.502 | 7.581 | 6.381 | 6.060 | 6.235 | 6.581 | 6.380 | 6.324 | 6.432 | 5.423 | 6.531 | 6.417 | 6.832 | 6.386 | 6.772 | 6.053 | 6.417 | 6.175 | 0.022 | 0.6677 | 0.8400 | 0 | 0 | 0 |
| '2310033P09Rik' | 32.866 | 30.456 | 33.984 | 29.313 | 31.191 | 34.481 | 34.091 | 31.311 | 34.185 | 31.738 | 24.939 | 33.673 | 31.353 | 28.813 | 31.977 | 31.048 | 27.219 | 32.976 | 0.081 | 0.0327 | 0.1431 | 0 | 0 | 0 |
| '2310035C23Rik' | 5.925 | 6.298 | 6.159 | 5.700 | 6.008 | 5.618 | 5.661 | 6.798 | 5.797 | 6.220 | 5.820 | 5.544 | 5.607 | 5.884 | 5.182 | 5.275 | 6.044 | 5.132 | 0.081 | 0.0213 | 0.1073 | 0 | 0 | 0 |
| '2310036O22Rik' | 190.233 | 212.383 | 204.926 | 186.923 | 206.846 | 208.172 | 182.676 | 210.393 | 206.194 | 190.776 | 192.807 | 191.704 | 203.340 | 205.778 | 211.208 | 199.994 | 187.582 | 204.631 | 0.006 | 0.8570 | 0.9387 | 0 | 0 | 0 |
| '2310039H08Rik' | 11.552 | 9.947 | 9.977 | 9.330 | 11.322 | 9.613 | 9.715 | 8.788 | 8.775 | 9.061 | 9.167 | 9.466 | 10.307 | 9.127 | 8.843 | 12.290 | 8.836 | 7.636 | 0.065 | 0.4335 | 0.6782 | 0 | 0 | 0 |
| '2310057M21Rik' | 5.998 | 5.953 | 5.719 | 5.518 | 6.545 | 6.062 | 5.399 | 6.161 | 6.213 | 6.114 | 5.405 | 6.040 | 5.620 | 5.842 | 5.267 | 6.229 | 5.226 | 5.285 | 0.059 | 0.3323 | 0.5905 | 0 | 0 | 0 |
| '2310061I04Rik' | 5.785 | 4.609 | 5.352 | 5.193 | 4.264 | 5.456 | 5.007 | 4.572 | 6.058 | 4.260 | 5.241 | 5.648 | 5.064 | 4.649 | 5.291 | 4.873 | 4.615 | 5.502 | 0.025 | 0.6428 | 0.8250 | 0 | 0 | 0 |
| '2410004B18Rik' | 6.096 | 5.759 | 4.764 | 5.167 | 4.929 | 4.888 | 5.337 | 5.303 | 4.469 | 4.627 | 5.424 | 5.035 | 5.204 | 5.766 | 5.252 | 5.940 | 5.024 | 4.923 | -0.026 | 0.6327 | 0.8178 | 0 | 0 | 0 |
| '2410015M20Rik' | 81.644 | 88.327 | 79.371 | 85.294 | 86.563 | 81.331 | 84.649 | 81.621 | 82.527 | 76.979 | 80.608 | 75.156 | 82.110 | 80.397 | 80.726 | 79.857 | 82.346 | 75.699 | 0.063 | 0.0417 | 0.1682 | 0 | 0 | 0 |
| '2410016O06Rik' | 10.382 | 10.133 | 10.506 | 8.282 | 9.572 | 12.154 | 9.636 | 10.387 | 11.432 | 10.593 | 9.598 | 12.733 | 9.723 | 9.366 | 11.900 | 9.046 | 9.477 | 12.359 | -0.047 | 0.3725 | 0.6241 | 0 | 0 | 0 |
| '2410089E03Rik' | 5.113 | 4.668 | 5.070 | 4.732 | 4.396 | 4.715 | 4.912 | 4.693 | 5.178 | 5.641 | 6.015 | 5.303 | 5.052 | 5.049 | 4.710 | 5.349 | 5.041 | 5.111 | -0.129 | 0.0044 | 0.0341 | 1 | 0 | 0 |
| '2410131K14Rik' | 4.575 | 4.201 | 4.557 | 4.361 | 4.776 | 4.626 | 4.541 | 4.224 | 4.114 | 4.351 | 3.808 | 4.737 | 4.509 | 4.197 | 4.542 | 4.129 | 4.428 | 4.270 | 0.020 | 0.6685 | 0.8400 | 0 | 0 | 0 |
| '2510002D24Rik' | 9.863 | 10.925 | 11.450 | 9.816 | 11.064 | 12.692 | 10.603 | 11.245 | 10.684 | 11.178 | 9.460 | 11.748 | 11.190 | 10.967 | 10.733 | 11.343 | 11.487 | 11.219 | -0.026 | 0.5668 | 0.7748 | 0 | 0 | 0 |
| '2510009E07Rik' | 1.448 | 2.081 | 1.594 | 1.696 | 1.653 | 1.941 | 1.766 | 1.817 | 1.837 | 1.578 | 1.342 | 1.505 | 1.363 | 1.622 | 1.691 | 1.424 | 1.404 | 1.639 | 0.208 | 0.0079 | 0.0526 | 0 | 0 | 0 |
| '2510039O18Rik' | 53.380 | 52.026 | 59.646 | 48.803 | 52.038 | 64.352 | 46.832 | 52.690 | 53.404 | 55.139 | 48.749 | 59.237 | 54.131 | 50.291 | 60.283 | 55.599 | 46.959 | 62.362 | -0.037 | 0.3612 | 0.6135 | 0 | 0 | 0 |
| '2610001J05Rik' | 3.579 | 4.570 | 3.869 | 3.620 | 4.384 | 4.563 | 4.105 | 4.572 | 4.132 | 3.811 | 4.123 | 4.001 | 4.126 | 4.347 | 3.779 | 4.381 | 4.228 | 3.618 | 0.028 | 0.5679 | 0.7756 | 0 | 0 | 0 |
| '2610002M06Rik' | 5.302 | 6.195 | 5.963 | 5.637 | 6.347 | 6.071 | 5.710 | 6.586 | 6.023 | 5.688 | 6.147 | 6.104 | 5.346 | 5.716 | 6.054 | 6.175 | 5.949 | 5.692 | 0.015 | 0.6876 | 0.8525 | 0 | 0 | 0 |
| '2610008E11Rik' | 9.160 | 9.981 | 9.204 | 9.763 | 8.855 | 8.472 | 10.474 | 9.498 | 8.867 | 9.061 | 10.351 | 8.907 | 7.794 | 9.418 | 8.218 | 8.152 | 10.482 | 7.956 | 0.061 | 0.2471 | 0.4979 | 0 | 0 | 0 |
| '2610020H08Rik' | 1.429 | 1.395 | 2.079 | 1.721 | 1.446 | 1.726 | 1.759 | 1.600 | 2.110 | 1.537 | 1.797 | 1.770 | 1.558 | 1.883 | 1.657 | 1.405 | 1.659 | 1.609 | 0.030 | 0.6961 | 0.8571 | 0 | 0 | 0 |
| '2610301B20Rik' | 1.736 | 1.586 | 1.743 | 1.588 | 2.106 | 1.859 | 1.863 | 1.782 | 1.978 | 1.990 | 1.623 | 1.869 | 1.539 | 1.929 | 1.474 | 1.519 | 1.450 | 1.564 | 0.105 | 0.2014 | 0.4426 | 0 | 0 | 0 |
| '2610318N02Rik' | 7.747 | 6.202 | 8.568 | 6.109 | 6.465 | 7.787 | 7.936 | 6.862 | 8.193 | 5.164 | 6.846 | 7.759 | 7.679 | 5.809 | 7.674 | 6.904 | 6.006 | 7.186 | 0.101 | 0.1912 | 0.4279 | 0 | 0 | 0 |
| '2610507B11Rik' | 21.940 | 21.499 | 23.624 | 21.782 | 21.700 | 22.842 | 20.861 | 21.506 | 23.169 | 22.614 | 22.111 | 23.484 | 21.088 | 21.180 | 21.896 | 21.972 | 21.197 | 23.554 | -0.012 | 0.6218 | 0.8110 | 0 | 0 | 0 |
| '2610524H06Rik' | 16.754 | 20.903 | 16.588 | 18.208 | 17.997 | 19.434 | 20.152 | 21.175 | 19.126 | 19.458 | 17.844 | 20.254 | 22.943 | 20.343 | 23.853 | 22.798 | 20.500 | 20.156 | -0.158 | 0.0264 | 0.1245 | 0 | 0 | 0 |
| '2700049A03Rik' | 3.528 | 3.277 | 3.828 | 4.124 | 3.622 | 3.297 | 4.352 | 3.183 | 3.341 | 3.268 | 3.797 | 3.518 | 3.304 | 3.489 | 3.383 | 3.315 | 3.707 | 2.951 | 0.070 | 0.2713 | 0.5262 | 0 | 0 | 0 |
| '2700060E02Rik' | 63.940 | 66.037 | 66.050 | 66.042 | 68.414 | 64.240 | 68.562 | 67.839 | 61.770 | 64.447 | 62.411 | 62.440 | 69.425 | 67.257 | 64.862 | 63.267 | 67.482 | 61.480 | 0.013 | 0.6232 | 0.8119 | 0 | 0 | 0 |
| '2700062C07Rik' | 10.873 | 13.730 | 15.132 | 14.083 | 14.693 | 13.016 | 12.158 | 14.567 | 13.588 | 14.311 | 10.977 | 12.275 | 12.436 | 11.093 | 11.488 | 11.787 | 11.863 | 11.202 | 0.170 | 0.0141 | 0.0801 | 0 | 0 | 0 |
| '2700094K13Rik' | 14.058 | 13.731 | 16.967 | 14.200 | 13.953 | 16.511 | 14.667 | 15.104 | 15.324 | 11.974 | 13.730 | 14.378 | 15.963 | 13.998 | 15.240 | 13.076 | 13.901 | 13.117 | 0.091 | 0.0604 | 0.2119 | 0 | 0 | 0 |
| '2700097O09Rik' | 9.000 | 12.287 | 10.625 | 9.330 | 13.208 | 10.098 | 8.787 | 12.873 | 10.162 | 9.362 | 9.585 | 9.076 | 10.004 | 11.451 | 8.464 | 9.808 | 10.642 | 8.576 | 0.130 | 0.0069 | 0.0476 | 1 | 0 | 0 |
| '2810004N23Rik' | 24.372 | 21.604 | 24.280 | 25.012 | 24.485 | 22.916 | 22.267 | 26.176 | 23.804 | 25.377 | 23.049 | 23.896 | 25.031 | 22.259 | 25.059 | 24.593 | 23.283 | 22.259 | -0.006 | 0.8906 | 0.9553 | 0 | 0 | 0 |
| '2810006K23Rik' | 2.791 | 2.755 | 3.438 | 3.141 | 3.257 | 3.557 | 2.992 | 3.149 | 3.308 | 2.935 | 2.883 | 3.390 | 3.221 | 3.025 | 3.460 | 3.269 | 3.228 | 2.973 | -0.007 | 0.9116 | 0.9660 | 0 | 0 | 0 |
| '2810021J22Rik' | 6.618 | 7.472 | 7.352 | 6.082 | 6.923 | 7.779 | 6.236 | 7.471 | 7.944 | 7.144 | 6.784 | 7.495 | 6.985 | 6.388 | 7.251 | 6.370 | 7.059 | 7.700 | 0.004 | 0.9251 | 0.9715 | 0 | 0 | 0 |
| '2810403A07Rik' | 6.564 | 6.902 | 6.903 | 6.258 | 6.916 | 6.930 | 6.977 | 6.569 | 7.077 | 6.240 | 7.590 | 7.659 | 6.409 | 6.906 | 7.416 | 6.240 | 7.301 | 7.594 | -0.062 | 0.0726 | 0.2384 | 0 | 0 | 0 |
| '2810417H13Rik' | 8.537 | 9.873 | 11.053 | 10.973 | 9.728 | 10.512 | 11.440 | 9.454 | 10.849 | 6.947 | 9.450 | 9.549 | 10.532 | 10.433 | 11.452 | 8.879 | 10.941 | 9.504 | 0.066 | 0.3577 | 0.6103 | 0 | 0 | 0 |
| '2810428I15Rik' | 8.830 | 11.056 | 9.802 | 8.110 | 10.173 | 9.682 | 8.308 | 10.778 | 9.802 | 9.655 | 8.904 | 10.172 | 10.278 | 9.570 | 9.757 | 10.062 | 9.077 | 9.769 | -0.027 | 0.5776 | 0.7819 | 0 | 0 | 0 |
| '2810474O19Rik' | 3.539 | 4.175 | 3.556 | 3.642 | 4.129 | 3.691 | 3.692 | 4.170 | 3.552 | 3.683 | 3.646 | 3.859 | 3.506 | 4.140 | 3.623 | 4.187 | 3.921 | 4.093 | -0.036 | 0.3685 | 0.6201 | 0 | 0 | 0 |
| '3010026O09Rik' | 2.117 | 2.319 | 2.536 | 2.111 | 2.557 | 2.578 | 2.431 | 2.601 | 2.538 | 2.663 | 1.711 | 1.765 | 2.728 | 2.140 | 1.773 | 2.364 | 2.181 | 1.521 | 0.223 | 0.0384 | 0.1595 | 0 | 0 | 0 |
| '3110001I22Rik' | 3.774 | 3.145 | 2.786 | 3.378 | 2.293 | 2.042 | 2.939 | 2.537 | 3.321 | 2.387 | 3.522 | 3.753 | 2.600 | 3.715 | 3.304 | 3.160 | 3.766 | 4.007 | -0.217 | 0.0902 | 0.2762 | 0 | 0 | 0 |
| '3110002H16Rik' | 6.578 | 6.344 | 6.719 | 6.461 | 5.964 | 6.582 | 6.363 | 6.082 | 6.547 | 6.518 | 6.326 | 7.276 | 6.224 | 7.059 | 6.338 | 6.092 | 6.544 | 6.985 | -0.054 | 0.1441 | 0.3682 | 0 | 0 | 0 |
| '3110009E18Rik' | 1.288 | 1.512 | 1.872 | 1.803 | 1.804 | 1.486 | 1.627 | 1.602 | 1.600 | 1.803 | 1.800 | 2.101 | 2.156 | 2.112 | 2.082 | 2.352 | 2.162 | 1.791 | -0.343 | 0.0003 | 0.0034 | 1 | 0 | 0 |
| '3110040N11Rik' | 10.349 | 11.787 | 9.823 | 10.142 | 10.469 | 10.170 | 9.277 | 10.964 | 10.338 | 9.602 | 9.531 | 9.983 | 9.452 | 10.548 | 10.224 | 9.604 | 11.072 | 9.249 | 0.051 | 0.2815 | 0.5379 | 0 | 0 | 0 |
| '3110043O21Rik' | 1.603 | 1.970 | 1.598 | 1.626 | 1.645 | 1.495 | 1.674 | 2.070 | 1.486 | 1.610 | 1.682 | 1.477 | 1.403 | 1.606 | 1.456 | 1.831 | 1.556 | 1.532 | 0.090 | 0.1640 | 0.3960 | 0 | 0 | 0 |
| '3110062M04Rik' | 1.089 | 1.171 | 1.258 | 1.161 | 1.124 | 1.175 | 1.127 | 0.978 | 1.214 | 1.113 | 0.997 | 1.303 | 1.037 | 0.907 | 1.363 | 0.987 | 1.234 | 1.312 | -0.011 | 0.8706 | 0.9447 | 0 | 0 | 0 |
| '3110082I17Rik' | 1.971 | 2.088 | 2.249 | 1.942 | 1.765 | 2.190 | 1.765 | 1.760 | 1.896 | 2.154 | 1.321 | 1.873 | 1.953 | 1.463 | 1.773 | 2.039 | 1.497 | 1.718 | 0.156 | 0.0486 | 0.1851 | 0 | 0 | 0 |
| '3830406C13Rik' | 8.404 | 7.827 | 8.797 | 8.471 | 8.903 | 7.300 | 9.442 | 8.866 | 8.648 | 8.446 | 7.854 | 8.165 | 8.016 | 8.443 | 7.746 | 6.685 | 8.741 | 7.445 | 0.087 | 0.0894 | 0.2744 | 0 | 0 | 0 |
| '4833420G17Rik' | 48.481 | 47.335 | 50.995 | 44.947 | 49.276 | 46.592 | 48.080 | 50.668 | 47.510 | 46.650 | 49.704 | 47.588 | 50.225 | 49.505 | 44.076 | 47.085 | 52.774 | 44.225 | -0.001 | 0.9785 | 0.9925 | 0 | 0 | 0 |
| '4833439L19Rik' | 8.968 | 8.774 | 9.201 | 9.241 | 9.963 | 9.333 | 8.947 | 9.398 | 9.624 | 7.849 | 9.112 | 9.483 | 9.557 | 9.077 | 9.390 | 8.835 | 9.308 | 9.941 | 0.003 | 0.9348 | 0.9749 | 0 | 0 | 0 |
| '4921524J17Rik' | 25.428 | 26.086 | 23.472 | 26.087 | 26.779 | 21.935 | 27.499 | 26.010 | 24.667 | 24.124 | 26.079 | 22.159 | 25.574 | 25.263 | 21.478 | 25.713 | 26.979 | 19.861 | 0.062 | 0.1020 | 0.2994 | 0 | 0 | 0 |
| '4930402H24Rik' | 1.709 | 1.447 | 1.909 | 1.647 | 1.678 | 1.738 | 1.768 | 1.654 | 1.768 | 1.759 | 1.787 | 1.844 | 1.692 | 1.784 | 1.715 | 1.423 | 1.781 | 1.755 | -0.030 | 0.5560 | 0.7678 | 0 | 0 | 0 |
| '4930427A07Rik' | 4.810 | 5.046 | 5.118 | 4.813 | 4.934 | 5.475 | 5.204 | 4.578 | 5.765 | 4.126 | 5.286 | 5.896 | 6.121 | 6.080 | 6.124 | 5.002 | 5.067 | 5.410 | -0.114 | 0.0532 | 0.1968 | 0 | 0 | 0 |
| '4930430F08Rik' | 2.062 | 2.984 | 2.456 | 2.221 | 2.995 | 2.976 | 2.439 | 2.675 | 2.363 | 2.812 | 2.766 | 2.649 | 2.842 | 2.613 | 2.351 | 2.629 | 3.112 | 2.440 | -0.079 | 0.3642 | 0.6162 | 0 | 0 | 0 |
| '4930453N24Rik' | 3.282 | 3.763 | 3.510 | 3.312 | 3.492 | 3.334 | 3.437 | 3.578 | 3.360 | 2.945 | 3.108 | 3.863 | 3.228 | 3.036 | 3.658 | 3.371 | 3.334 | 3.438 | 0.032 | 0.5260 | 0.7477 | 0 | 0 | 0 |
| '4930503L19Rik' | 14.604 | 13.811 | 15.644 | 14.206 | 13.942 | 15.837 | 14.892 | 15.105 | 13.821 | 12.589 | 14.134 | 14.731 | 15.456 | 14.537 | 14.267 | 13.620 | 14.638 | 14.956 | 0.023 | 0.5944 | 0.7935 | 0 | 0 | 0 |
| '4930523C07Rik' | 3.870 | 4.116 | 4.130 | 3.575 | 3.885 | 4.434 | 4.247 | 4.409 | 4.176 | 2.947 | 2.907 | 2.654 | 2.310 | 2.626 | 2.499 | 2.874 | 2.467 | 2.251 | 0.641 | 0.0000 | 0.0000 | 1 | 1 | 0 |
| '4930539E08Rik' | 0.880 | 1.126 | 0.910 | 0.857 | 1.268 | 1.150 | 0.972 | 1.435 | 0.862 | 0.980 | 1.080 | 0.944 | 0.953 | 1.207 | 0.741 | 1.004 | 0.667 | 0.804 | 0.158 | 0.1651 | 0.3974 | 0 | 0 | 0 |
| '4930579G24Rik' | 5.802 | 5.339 | 5.743 | 6.317 | 4.518 | 4.593 | 5.955 | 5.120 | 5.200 | 4.002 | 5.878 | 5.010 | 5.848 | 5.748 | 5.631 | 5.292 | 6.088 | 5.466 | -0.021 | 0.8025 | 0.9115 | 0 | 0 | 0 |
| '4931406C07Rik' | 2.966 | 3.035 | 2.749 | 3.035 | 3.532 | 2.725 | 2.695 | 2.947 | 3.173 | 2.993 | 2.819 | 2.291 | 2.687 | 2.819 | 2.497 | 2.474 | 3.008 | 2.271 | 0.169 | 0.0224 | 0.1111 | 0 | 0 | 0 |
| '4931406P16Rik' | 5.179 | 4.793 | 4.559 | 4.329 | 4.530 | 4.476 | 4.176 | 4.490 | 4.459 | 4.475 | 4.622 | 4.906 | 4.110 | 4.833 | 4.758 | 4.716 | 4.290 | 4.779 | -0.027 | 0.5677 | 0.7754 | 0 | 0 | 0 |
| '4931414P19Rik' | 1.907 | 2.094 | 2.258 | 1.879 | 2.192 | 2.206 | 2.194 | 1.648 | 2.624 | 2.162 | 1.919 | 1.796 | 2.132 | 1.677 | 2.356 | 2.134 | 2.211 | 1.839 | 0.058 | 0.5618 | 0.7713 | 0 | 0 | 0 |
| '4931428F04Rik' | 0.957 | 1.068 | 1.630 | 1.023 | 1.288 | 1.451 | 1.265 | 1.062 | 1.783 | 0.975 | 1.209 | 1.235 | 1.738 | 1.107 | 1.164 | 1.376 | 0.969 | 1.180 | 0.060 | 0.6389 | 0.8222 | 0 | 0 | 0 |
| '4932438A13Rik' | 8.381 | 6.756 | 7.893 | 8.290 | 6.881 | 7.146 | 7.370 | 6.981 | 7.453 | 7.442 | 7.736 | 8.001 | 6.442 | 6.628 | 7.462 | 6.964 | 7.344 | 8.788 | -0.004 | 0.9466 | 0.9794 | 0 | 0 | 0 |
| '4933427D14Rik' | 5.357 | 4.780 | 5.404 | 5.215 | 5.031 | 4.806 | 5.166 | 5.087 | 5.204 | 5.619 | 6.051 | 5.472 | 5.929 | 5.165 | 5.259 | 5.052 | 5.975 | 5.281 | -0.122 | 0.0034 | 0.0276 | 1 | 0 | 0 |
| '5031439G07Rik' | 2.791 | 3.409 | 3.296 | 2.989 | 3.201 | 3.552 | 2.760 | 3.207 | 3.207 | 3.146 | 3.269 | 3.179 | 2.982 | 3.439 | 2.981 | 3.041 | 3.232 | 3.370 | -0.020 | 0.6287 | 0.8160 | 0 | 0 | 0 |
| '5430403G16Rik' | 2.559 | 3.123 | 3.005 | 2.790 | 2.545 | 2.636 | 3.569 | 3.124 | 3.095 | 2.681 | 2.686 | 2.787 | 2.571 | 2.945 | 2.504 | 2.966 | 2.831 | 2.758 | 0.085 | 0.2381 | 0.4880 | 0 | 0 | 0 |
| '5730409E04Rik' | 0.821 | 1.078 | 0.911 | 0.904 | 1.210 | 1.055 | 0.846 | 0.875 | 1.020 | 0.841 | 0.508 | 0.662 | 0.659 | 0.673 | 0.837 | 0.909 | 0.450 | 0.696 | 0.479 | 0.0001 | 0.0018 | 1 | 1 | 0 |
| '5730455P16Rik' | 2.637 | 2.895 | 2.738 | 2.684 | 2.889 | 2.807 | 2.820 | 3.113 | 2.945 | 2.639 | 2.641 | 2.292 | 2.654 | 2.624 | 2.201 | 2.686 | 2.686 | 2.121 | 0.179 | 0.0002 | 0.0027 | 1 | 0 | 0 |
| '5730480H06Rik' | 0.818 | 1.033 | 0.997 | 0.932 | 1.101 | 1.035 | 1.020 | 1.059 | 0.941 | 0.897 | 0.871 | 0.876 | 0.981 | 0.834 | 0.827 | 0.942 | 0.882 | 0.775 | 0.168 | 0.0080 | 0.0530 | 0 | 0 | 0 |
| '5730508B09Rik' | 1.721 | 2.002 | 1.872 | 1.725 | 1.774 | 1.725 | 2.035 | 1.853 | 1.733 | 1.613 | 2.234 | 1.820 | 1.435 | 2.468 | 1.570 | 1.895 | 1.905 | 2.027 | -0.047 | 0.5984 | 0.7958 | 0 | 0 | 0 |
| '6030458C11Rik' | 20.735 | 32.574 | 28.218 | 21.019 | 32.314 | 31.005 | 20.440 | 33.306 | 29.825 | 27.216 | 23.615 | 28.703 | 26.764 | 31.251 | 29.300 | 30.093 | 25.080 | 26.634 | -0.030 | 0.7309 | 0.8773 | 0 | 0 | 0 |
| '6330408A02Rik' | 0.842 | 0.726 | 0.798 | 0.857 | 0.861 | 0.961 | 0.868 | 0.826 | 0.912 | 0.832 | 0.822 | 0.896 | 0.841 | 0.719 | 0.819 | 0.831 | 0.796 | 0.772 | 0.051 | 0.3414 | 0.5981 | 0 | 0 | 0 |
| '8030462N17Rik' | 13.047 | 12.510 | 11.510 | 11.046 | 11.271 | 10.853 | 11.860 | 12.556 | 10.272 | 12.137 | 12.231 | 11.205 | 10.474 | 12.857 | 11.798 | 11.229 | 11.949 | 11.509 | -0.015 | 0.7447 | 0.8847 | 0 | 0 | 0 |
| '9030617O03Rik' | 1.028 | 0.883 | 1.051 | 0.864 | 0.939 | 1.204 | 1.181 | 1.068 | 1.321 | 1.243 | 1.271 | 1.249 | 0.841 | 1.060 | 1.033 | 1.386 | 1.142 | 1.195 | -0.132 | 0.1630 | 0.3947 | 0 | 0 | 0 |
| '9030624J02Rik' | 6.739 | 7.431 | 7.228 | 7.301 | 7.438 | 7.386 | 7.002 | 7.445 | 7.013 | 8.062 | 7.535 | 7.097 | 7.565 | 7.658 | 7.191 | 7.402 | 7.211 | 6.972 | -0.046 | 0.1223 | 0.3336 | 0 | 0 | 0 |
| '9130011E15Rik' | 7.589 | 8.507 | 8.439 | 7.548 | 6.986 | 8.473 | 9.420 | 8.619 | 8.133 | 8.581 | 8.156 | 7.745 | 8.480 | 7.790 | 8.271 | 8.518 | 7.340 | 6.963 | 0.032 | 0.5672 | 0.7750 | 0 | 0 | 0 |
| '9130023H24Rik' | 2.043 | 2.523 | 2.696 | 2.284 | 2.247 | 2.690 | 2.507 | 2.578 | 3.355 | 2.768 | 2.884 | 2.279 | 3.006 | 2.384 | 2.361 | 2.314 | 2.046 | 2.118 | 0.044 | 0.6493 | 0.8292 | 0 | 0 | 0 |
| '9330151L19Rik' | 2.118 | 2.064 | 3.049 | 1.734 | 1.916 | 2.127 | 2.047 | 2.520 | 2.393 | 2.143 | 2.319 | 2.735 | 1.715 | 2.598 | 2.031 | 1.866 | 2.491 | 2.852 | -0.057 | 0.5735 | 0.7795 | 0 | 0 | 0 |
| '9430015G10Rik' | 5.167 | 4.837 | 6.110 | 5.221 | 5.060 | 6.190 | 5.359 | 5.250 | 6.268 | 4.839 | 5.236 | 6.215 | 5.094 | 5.102 | 5.762 | 4.847 | 5.584 | 5.556 | 0.025 | 0.4776 | 0.7109 | 0 | 0 | 0 |
| '9430016H08Rik' | 2.703 | 2.970 | 3.077 | 2.370 | 2.797 | 2.864 | 2.619 | 3.305 | 2.715 | 3.177 | 2.024 | 2.466 | 3.176 | 2.844 | 2.402 | 3.008 | 2.382 | 2.523 | 0.083 | 0.3298 | 0.5877 | 0 | 0 | 0 |
| '9430038I01Rik' | 0.721 | 0.733 | 0.929 | 0.789 | 0.830 | 0.741 | 0.558 | 0.730 | 0.921 | 0.722 | 0.608 | 0.569 | 0.636 | 0.611 | 0.532 | 0.722 | 0.621 | 0.551 | 0.324 | 0.0006 | 0.0068 | 1 | 0 | 0 |
| '9530068E07Rik' | 18.610 | 19.692 | 20.567 | 18.559 | 20.485 | 19.832 | 19.158 | 20.560 | 18.942 | 19.062 | 18.829 | 19.891 | 18.524 | 18.997 | 19.672 | 20.111 | 19.646 | 19.373 | 0.009 | 0.7270 | 0.8750 | 0 | 0 | 0 |
| '9930012K11Rik' | 1.176 | 1.272 | 1.018 | 1.085 | 1.243 | 1.444 | 1.130 | 1.212 | 1.297 | 1.092 | 0.972 | 1.097 | 1.026 | 1.002 | 1.101 | 0.974 | 0.951 | 1.011 | 0.222 | 0.0108 | 0.0662 | 0 | 0 | 0 |
| '9930021J03Rik' | 3.824 | 3.650 | 3.391 | 3.680 | 3.409 | 3.294 | 3.561 | 3.408 | 3.508 | 3.684 | 3.198 | 3.820 | 3.238 | 3.428 | 3.305 | 3.787 | 3.291 | 3.608 | 0.003 | 0.9330 | 0.9746 | 0 | 0 | 0 |
| 'A130010J15Rik' | 0.823 | 0.802 | 0.898 | 0.752 | 0.856 | 0.856 | 0.878 | 0.929 | 0.904 | 0.899 | 0.857 | 0.919 | 0.749 | 0.765 | 0.941 | 0.747 | 0.950 | 0.846 | -0.005 | 0.9457 | 0.9791 | 0 | 0 | 0 |
| 'A230046K03Rik' | 7.406 | 7.258 | 7.291 | 7.511 | 7.693 | 7.337 | 6.736 | 7.754 | 7.364 | 7.052 | 7.800 | 8.483 | 6.932 | 8.058 | 7.971 | 7.590 | 7.963 | 7.935 | -0.084 | 0.0171 | 0.0925 | 0 | 0 | 0 |
| 'A230050P20Rik' | 11.737 | 10.531 | 13.036 | 9.576 | 10.921 | 12.321 | 10.341 | 10.943 | 12.776 | 11.130 | 8.783 | 12.213 | 11.092 | 8.916 | 11.074 | 11.250 | 9.642 | 10.848 | 0.098 | 0.0298 | 0.1341 | 0 | 0 | 0 |
| 'A430005L14Rik' | 5.669 | 5.238 | 5.698 | 5.952 | 5.585 | 6.023 | 5.213 | 5.406 | 5.460 | 5.379 | 4.662 | 5.664 | 5.513 | 5.110 | 5.789 | 5.231 | 5.444 | 5.634 | 0.041 | 0.3791 | 0.6307 | 0 | 0 | 0 |
| 'A430033K04Rik' | 1.664 | 2.163 | 1.901 | 1.575 | 2.257 | 2.046 | 1.926 | 2.282 | 1.833 | 1.966 | 2.107 | 1.743 | 1.602 | 1.930 | 1.900 | 2.025 | 1.808 | 1.720 | 0.064 | 0.3440 | 0.5995 | 0 | 0 | 0 |
| 'A630001G21Rik' | 21.594 | 25.269 | 23.010 | 22.716 | 24.115 | 21.855 | 23.353 | 22.561 | 23.183 | 24.584 | 25.801 | 21.821 | 28.880 | 26.653 | 21.463 | 23.335 | 27.794 | 20.956 | -0.096 | 0.0319 | 0.1405 | 0 | 0 | 0 |
| 'A830080D01Rik' | 4.723 | 4.875 | 5.022 | 5.128 | 4.804 | 4.448 | 5.675 | 4.993 | 4.803 | 5.140 | 4.964 | 5.499 | 4.782 | 4.888 | 5.069 | 4.934 | 5.982 | 5.602 | -0.090 | 0.0873 | 0.2695 | 0 | 0 | 0 |
| 'AA415398' | 2.101 | 2.143 | 1.547 | 2.120 | 1.725 | 1.813 | 1.820 | 2.209 | 1.716 | 1.720 | 1.473 | 1.880 | 1.474 | 2.269 | 1.705 | 1.524 | 1.594 | 1.706 | 0.141 | 0.1530 | 0.3803 | 0 | 0 | 0 |
| 'AB124611' | 9.653 | 10.657 | 9.006 | 9.083 | 10.160 | 8.990 | 9.147 | 10.471 | 9.585 | 8.817 | 8.783 | 8.211 | 8.927 | 9.642 | 7.687 | 8.569 | 8.622 | 7.912 | 0.159 | 0.0001 | 0.0011 | 1 | 0 | 0 |
| 'AI314180' | 10.838 | 10.672 | 10.671 | 11.394 | 9.810 | 9.788 | 11.760 | 10.369 | 10.511 | 10.457 | 11.259 | 11.332 | 10.108 | 10.063 | 10.373 | 9.600 | 11.151 | 11.026 | -0.006 | 0.8970 | 0.9586 | 0 | 0 | 0 |
| 'AI413582' | 13.365 | 14.701 | 13.935 | 11.525 | 14.160 | 14.495 | 12.767 | 15.019 | 14.307 | 12.619 | 13.420 | 14.494 | 12.766 | 12.808 | 14.455 | 12.763 | 13.087 | 13.830 | 0.035 | 0.3771 | 0.6288 | 0 | 0 | 0 |
| 'AI467606' | 28.680 | 31.943 | 33.359 | 36.293 | 30.272 | 33.172 | 34.772 | 29.684 | 32.857 | 22.749 | 33.079 | 38.802 | 26.103 | 28.141 | 35.861 | 21.837 | 33.388 | 36.634 | 0.083 | 0.3636 | 0.6156 | 0 | 0 | 0 |
| 'AI597479' | 8.284 | 9.529 | 9.870 | 8.705 | 9.398 | 8.629 | 10.039 | 9.557 | 9.009 | 8.296 | 9.722 | 10.353 | 8.175 | 10.264 | 10.598 | 8.862 | 9.755 | 10.228 | -0.067 | 0.1689 | 0.4013 | 0 | 0 | 0 |
| 'AI606181' | 1.617 | 2.693 | 2.087 | 2.131 | 2.488 | 1.833 | 2.028 | 2.371 | 2.156 | 1.710 | 2.421 | 2.318 | 1.795 | 2.522 | 2.033 | 1.808 | 2.343 | 2.167 | -0.008 | 0.9267 | 0.9719 | 0 | 0 | 0 |
| 'AI837181' | 21.161 | 23.099 | 21.851 | 20.728 | 22.910 | 23.759 | 20.301 | 22.679 | 22.220 | 21.793 | 20.986 | 25.288 | 21.810 | 21.203 | 22.638 | 23.561 | 21.778 | 24.717 | -0.049 | 0.1730 | 0.4066 | 0 | 0 | 0 |
| 'AI846148' | 11.732 | 11.658 | 12.524 | 11.214 | 11.314 | 13.663 | 11.195 | 11.501 | 13.616 | 11.743 | 11.205 | 12.445 | 11.092 | 11.105 | 12.055 | 12.708 | 10.533 | 12.080 | 0.041 | 0.3035 | 0.5608 | 0 | 0 | 0 |
| 'AI854703' | 0.526 | 0.551 | 0.701 | 0.624 | 0.451 | 0.766 | 0.683 | 0.457 | 0.478 | 0.492 | 0.541 | 0.435 | 0.455 | 0.387 | 0.571 | 0.359 | 0.275 | 0.542 | 0.363 | 0.0059 | 0.0426 | 1 | 0 | 0 |
| 'AI987944' | 3.917 | 5.014 | 4.862 | 4.502 | 4.519 | 4.449 | 4.477 | 5.385 | 4.484 | 4.255 | 4.614 | 4.676 | 4.434 | 4.817 | 4.783 | 4.574 | 4.887 | 4.702 | -0.016 | 0.7265 | 0.8748 | 0 | 0 | 0 |
| 'AK010878' | 9.005 | 8.027 | 6.284 | 6.947 | 6.772 | 6.560 | 7.280 | 6.239 | 6.656 | 6.644 | 8.439 | 8.697 | 8.342 | 8.234 | 8.256 | 6.998 | 8.999 | 7.954 | -0.205 | 0.0111 | 0.0673 | 0 | 0 | 0 |
| 'AK157302' | 25.919 | 27.647 | 24.917 | 24.519 | 26.772 | 23.922 | 25.212 | 24.715 | 24.060 | 25.587 | 25.084 | 25.794 | 23.627 | 25.667 | 25.229 | 26.121 | 25.032 | 25.143 | -0.011 | 0.7520 | 0.8895 | 0 | 0 | 0 |
| 'AU019823' | 3.223 | 3.648 | 3.623 | 3.323 | 3.550 | 3.374 | 3.040 | 3.827 | 3.290 | 3.562 | 3.294 | 3.430 | 3.186 | 3.363 | 3.413 | 3.633 | 3.089 | 3.309 | 0.020 | 0.6599 | 0.8361 | 0 | 0 | 0 |
| 'AU022252' | 1.705 | 2.084 | 1.968 | 1.890 | 1.963 | 1.976 | 1.818 | 2.026 | 2.037 | 1.849 | 2.343 | 2.002 | 1.842 | 1.815 | 2.215 | 1.899 | 1.938 | 2.084 | -0.052 | 0.3081 | 0.5665 | 0 | 0 | 0 |
| 'AU040320' | 11.546 | 11.603 | 12.460 | 11.620 | 11.981 | 13.645 | 11.682 | 11.400 | 12.939 | 12.369 | 12.221 | 13.554 | 13.091 | 12.378 | 12.574 | 12.802 | 12.793 | 13.426 | -0.093 | 0.0009 | 0.0094 | 1 | 0 | 0 |
| 'AU041133' | 1.693 | 2.011 | 2.288 | 1.830 | 2.058 | 1.857 | 2.431 | 1.834 | 1.626 | 1.778 | 1.758 | 1.460 | 1.537 | 1.604 | 1.400 | 1.536 | 1.893 | 1.339 | 0.299 | 0.0011 | 0.0117 | 1 | 0 | 0 |
| 'AW011738' | 22.953 | 22.539 | 24.951 | 20.460 | 24.482 | 25.007 | 19.391 | 25.315 | 25.335 | 24.859 | 27.142 | 27.676 | 19.795 | 24.098 | 28.072 | 21.254 | 23.057 | 28.259 | -0.098 | 0.0484 | 0.1848 | 0 | 0 | 0 |
| 'AW146154' | 1.866 | 2.980 | 2.229 | 2.192 | 2.926 | 2.280 | 2.084 | 2.253 | 2.223 | 2.343 | 2.455 | 2.451 | 2.389 | 2.380 | 2.894 | 1.976 | 2.188 | 2.423 | -0.058 | 0.4521 | 0.6912 | 0 | 0 | 0 |
| 'AW209491' | 3.847 | 4.401 | 3.508 | 4.560 | 4.095 | 4.187 | 4.374 | 3.682 | 3.646 | 2.989 | 4.224 | 4.554 | 3.674 | 3.441 | 4.736 | 3.550 | 4.488 | 4.536 | -0.016 | 0.8570 | 0.9387 | 0 | 0 | 0 |
| 'AW549877' | 18.983 | 26.224 | 21.498 | 18.938 | 25.574 | 22.223 | 20.503 | 24.502 | 23.849 | 20.656 | 21.716 | 20.554 | 21.021 | 23.479 | 18.398 | 22.743 | 23.297 | 19.209 | 0.068 | 0.1322 | 0.3498 | 0 | 0 | 0 |
| 'AW554918' | 1.020 | 0.980 | 1.056 | 0.972 | 1.040 | 0.983 | 1.202 | 1.006 | 1.017 | 1.087 | 0.854 | 1.103 | 0.864 | 1.066 | 0.966 | 0.949 | 1.104 | 0.975 | 0.034 | 0.5780 | 0.7820 | 0 | 0 | 0 |
| 'Aaas' | 27.206 | 28.945 | 31.420 | 27.558 | 30.577 | 32.259 | 28.821 | 29.807 | 32.373 | 26.219 | 28.700 | 32.607 | 33.025 | 28.610 | 31.252 | 26.240 | 30.589 | 28.077 | 0.006 | 0.8906 | 0.9553 | 0 | 0 | 0 |
| 'Aacs' | 8.888 | 8.308 | 10.314 | 8.999 | 8.828 | 9.896 | 8.488 | 8.758 | 10.215 | 8.563 | 9.427 | 10.386 | 8.960 | 8.211 | 10.239 | 8.754 | 8.699 | 9.617 | -0.012 | 0.7237 | 0.8731 | 0 | 0 | 0 |
| 'Aagab' | 21.255 | 21.334 | 21.224 | 20.578 | 21.246 | 22.310 | 20.632 | 20.425 | 21.935 | 22.923 | 19.864 | 23.095 | 21.241 | 21.255 | 22.757 | 21.186 | 22.169 | 21.218 | -0.048 | 0.1393 | 0.3610 | 0 | 0 | 0 |
| 'Aak1' | 11.720 | 11.369 | 12.010 | 11.484 | 10.789 | 11.653 | 11.127 | 10.773 | 11.699 | 12.609 | 11.575 | 12.506 | 11.325 | 10.929 | 12.048 | 11.304 | 11.401 | 13.036 | -0.067 | 0.0209 | 0.1060 | 0 | 0 | 0 |
| 'Aamdc' | 1.251 | 1.175 | 1.195 | 1.486 | 1.409 | 1.274 | 1.432 | 1.222 | 1.018 | 1.258 | 1.266 | 1.248 | 1.199 | 1.312 | 1.184 | 1.221 | 1.187 | 1.256 | 0.023 | 0.7550 | 0.8907 | 0 | 0 | 0 |
| 'Aamp' | 28.063 | 28.734 | 26.495 | 26.879 | 28.210 | 27.424 | 26.233 | 28.388 | 26.659 | 28.771 | 27.365 | 26.842 | 28.623 | 26.796 | 25.303 | 28.259 | 27.255 | 26.053 | 0.001 | 0.9630 | 0.9869 | 0 | 0 | 0 |
| 'Aar2' | 3.180 | 2.995 | 3.243 | 2.998 | 2.514 | 3.208 | 3.373 | 3.486 | 3.569 | 2.830 | 2.478 | 3.493 | 2.619 | 3.025 | 3.608 | 3.164 | 2.929 | 3.296 | 0.051 | 0.4216 | 0.6681 | 0 | 0 | 0 |
| 'Aars' | 14.241 | 14.439 | 16.453 | 15.268 | 15.106 | 16.362 | 15.361 | 14.945 | 15.620 | 15.057 | 14.807 | 15.701 | 15.922 | 14.268 | 15.127 | 14.750 | 14.972 | 15.401 | 0.008 | 0.7354 | 0.8796 | 0 | 0 | 0 |
| 'Aars2' | 6.464 | 6.683 | 8.606 | 6.672 | 6.347 | 7.826 | 7.430 | 6.232 | 7.655 | 6.696 | 5.887 | 6.807 | 7.249 | 5.535 | 6.058 | 6.547 | 5.632 | 5.824 | 0.175 | 0.0016 | 0.0152 | 1 | 0 | 0 |
| 'Aarsd1' | 1.424 | 1.559 | 1.949 | 1.325 | 1.560 | 1.832 | 1.326 | 1.361 | 1.630 | 1.559 | 1.233 | 1.758 | 1.557 | 1.357 | 1.263 | 1.505 | 1.613 | 1.393 | 0.063 | 0.4961 | 0.7249 | 0 | 0 | 0 |
| 'Aasdh' | 0.899 | 1.136 | 1.221 | 0.888 | 1.195 | 1.075 | 1.127 | 1.069 | 1.241 | 1.041 | 1.181 | 1.070 | 1.248 | 1.136 | 1.000 | 0.946 | 1.257 | 0.984 | -0.012 | 0.8576 | 0.9390 | 0 | 0 | 0 |
| 'Aasdhppt' | 6.587 | 7.654 | 7.299 | 7.565 | 8.149 | 6.122 | 8.090 | 8.209 | 6.675 | 6.852 | 7.599 | 7.877 | 6.541 | 8.091 | 6.825 | 6.006 | 9.083 | 7.973 | -0.027 | 0.6723 | 0.8423 | 0 | 0 | 0 |
| 'Aatf' | 7.831 | 7.353 | 7.251 | 7.680 | 7.590 | 7.862 | 8.013 | 8.087 | 7.660 | 8.379 | 7.201 | 7.718 | 7.794 | 6.391 | 7.464 | 7.431 | 7.345 | 6.820 | 0.048 | 0.2472 | 0.4979 | 0 | 0 | 0 |
| 'Abat' | 1.009 | 1.120 | 0.948 | 0.925 | 0.938 | 1.043 | 1.048 | 1.063 | 1.017 | 1.111 | 1.100 | 0.710 | 0.982 | 0.990 | 0.676 | 0.878 | 0.890 | 0.735 | 0.188 | 0.0234 | 0.1152 | 0 | 0 | 0 |
| 'Abca1' | 1.130 | 1.365 | 1.095 | 0.867 | 1.144 | 1.211 | 1.236 | 1.446 | 0.967 | 1.074 | 1.034 | 1.154 | 0.831 | 0.826 | 0.781 | 1.354 | 1.194 | 1.193 | 0.132 | 0.2244 | 0.4713 | 0 | 0 | 0 |
| 'Abca2' | 22.022 | 21.415 | 25.056 | 21.262 | 20.482 | 24.691 | 20.416 | 22.852 | 26.196 | 25.065 | 22.759 | 23.806 | 23.415 | 21.653 | 22.411 | 22.186 | 21.296 | 23.017 | -0.022 | 0.5775 | 0.7819 | 0 | 0 | 0 |
| 'Abca3' | 8.274 | 8.182 | 8.942 | 7.551 | 7.991 | 8.717 | 7.811 | 7.919 | 9.098 | 8.430 | 9.073 | 9.045 | 8.889 | 8.648 | 8.720 | 8.026 | 8.842 | 9.181 | -0.093 | 0.0025 | 0.0216 | 1 | 0 | 0 |
| 'Abca7' | 32.136 | 26.495 | 36.843 | 31.227 | 28.058 | 33.215 | 30.724 | 27.244 | 35.125 | 30.104 | 30.750 | 33.188 | 31.366 | 28.321 | 31.514 | 26.188 | 29.193 | 32.643 | 0.025 | 0.5491 | 0.7629 | 0 | 0 | 0 |
| 'Abcb10' | 1.870 | 2.730 | 2.466 | 2.100 | 1.914 | 2.445 | 1.955 | 2.276 | 2.506 | 2.031 | 1.698 | 2.037 | 2.203 | 2.097 | 1.972 | 2.363 | 1.633 | 2.024 | 0.158 | 0.0601 | 0.2115 | 0 | 0 | 0 |
| 'Abcb1a' | 10.975 | 9.893 | 12.346 | 10.225 | 11.440 | 11.353 | 10.506 | 11.324 | 11.458 | 10.591 | 11.640 | 11.428 | 9.088 | 10.829 | 11.118 | 9.997 | 10.824 | 10.673 | 0.042 | 0.2941 | 0.5500 | 0 | 0 | 0 |
| 'Abcb1b' | 2.193 | 1.986 | 2.079 | 2.046 | 1.908 | 1.994 | 2.229 | 2.047 | 1.852 | 2.228 | 1.823 | 2.056 | 1.919 | 2.072 | 1.993 | 1.933 | 2.155 | 1.808 | 0.017 | 0.7359 | 0.8797 | 0 | 0 | 0 |
| 'Abcb6' | 0.767 | 0.957 | 0.846 | 0.746 | 0.949 | 0.900 | 0.875 | 0.921 | 0.955 | 0.867 | 0.771 | 0.704 | 0.834 | 0.814 | 0.774 | 0.954 | 0.868 | 0.713 | 0.118 | 0.1121 | 0.3175 | 0 | 0 | 0 |
| 'Abcb7' | 12.709 | 13.535 | 13.643 | 14.214 | 14.084 | 12.836 | 13.674 | 12.712 | 13.696 | 13.598 | 15.054 | 14.554 | 13.869 | 14.122 | 13.196 | 12.593 | 15.265 | 12.952 | -0.059 | 0.1178 | 0.3266 | 0 | 0 | 0 |
| 'Abcb8' | 4.270 | 3.900 | 4.626 | 3.978 | 4.303 | 4.854 | 3.986 | 3.932 | 4.736 | 4.105 | 3.816 | 4.268 | 4.352 | 3.672 | 4.540 | 4.195 | 3.903 | 3.601 | 0.073 | 0.1049 | 0.3035 | 0 | 0 | 0 |
| 'Abcb9' | 18.681 | 19.115 | 20.307 | 18.747 | 17.444 | 21.649 | 18.125 | 17.460 | 21.326 | 16.554 | 15.469 | 17.215 | 16.837 | 14.217 | 17.086 | 16.290 | 14.131 | 17.637 | 0.239 | 0.0000 | 0.0000 | 1 | 0 | 0 |
| 'Abcc1' | 5.157 | 4.657 | 5.128 | 4.970 | 4.434 | 4.605 | 4.891 | 4.686 | 4.970 | 5.395 | 5.912 | 5.031 | 5.287 | 5.250 | 4.858 | 5.198 | 5.287 | 4.819 | -0.120 | 0.0030 | 0.0253 | 1 | 0 | 0 |
| 'Abcc10' | 2.667 | 2.386 | 3.156 | 2.457 | 2.220 | 2.767 | 2.604 | 2.693 | 2.909 | 2.671 | 2.680 | 2.940 | 2.830 | 2.458 | 2.558 | 2.254 | 2.470 | 2.771 | 0.003 | 0.9597 | 0.9852 | 0 | 0 | 0 |
| 'Abcc3' | 6.331 | 8.769 | 6.021 | 7.613 | 9.065 | 7.565 | 7.936 | 7.547 | 7.496 | 11.255 | 9.242 | 8.419 | 13.910 | 11.431 | 6.775 | 12.298 | 12.281 | 7.398 | -0.428 | 0.0000 | 0.0005 | 1 | 1 | 0 |
| 'Abcc4' | 8.816 | 8.408 | 8.686 | 8.453 | 8.052 | 8.366 | 8.761 | 9.025 | 8.267 | 10.356 | 7.929 | 8.244 | 8.135 | 8.710 | 8.232 | 9.160 | 8.589 | 8.581 | -0.026 | 0.5411 | 0.7584 | 0 | 0 | 0 |
| 'Abcc5' | 6.754 | 6.523 | 7.092 | 7.240 | 6.653 | 7.106 | 6.513 | 6.568 | 7.058 | 7.362 | 7.813 | 7.507 | 7.989 | 7.593 | 7.267 | 6.856 | 7.515 | 7.434 | -0.141 | 0.0000 | 0.0004 | 1 | 0 | 0 |
| 'Abcd1' | 16.175 | 15.453 | 17.560 | 15.423 | 15.689 | 17.708 | 15.542 | 15.147 | 17.164 | 16.953 | 16.298 | 18.828 | 16.766 | 15.682 | 18.624 | 15.873 | 15.357 | 18.415 | -0.078 | 0.0027 | 0.0235 | 1 | 0 | 0 |
| 'Abcd3' | 2.460 | 2.695 | 2.765 | 3.029 | 2.985 | 2.517 | 2.764 | 3.037 | 2.865 | 2.925 | 2.919 | 2.705 | 2.792 | 2.758 | 2.510 | 2.812 | 3.027 | 2.326 | 0.012 | 0.8069 | 0.9135 | 0 | 0 | 0 |

| | | | | | | | | | | | | | | | | | | | | | | | | |
|---|---|---|---|---|---|---|---|---|---|---|---|---|---|---|---|---|---|---|---|---|---|---|---|---|
| 'Abcd4' | 8.467 | 8.569 | 10.249 | 9.640 | 9.141 | 11.169 | 9.916 | 9.046 | 10.763 | 10.527 | 9.191 | 10.084 | 9.977 | 9.335 | 9.070 | 9.823 | 8.605 | 9.345 | 0.009 | 0.8625 | 0.9405 | 0 | 0 | 0 |
| 'Abce1' | 34.811 | 35.596 | 33.291 | 36.016 | 34.958 | 31.725 | 36.999 | 33.949 | 31.331 | 32.445 | 34.317 | 35.202 | 29.998 | 32.898 | 33.806 | 31.062 | 36.101 | 35.417 | 0.023 | 0.5938 | 0.7933 | 0 | 0 | 0 |
| 'Abcf1' | 34.198 | 29.602 | 34.683 | 32.547 | 29.301 | 35.311 | 32.986 | 30.299 | 35.985 | 33.701 | 30.501 | 35.210 | 32.534 | 30.963 | 34.851 | 33.643 | 30.324 | 34.179 | -0.015 | 0.4820 | 0.7141 | 0 | 0 | 0 |
| 'Abcf2' | 42.661 | 39.731 | 43.387 | 39.505 | 39.793 | 42.804 | 41.224 | 41.784 | 41.368 | 41.053 | 38.768 | 46.129 | 40.849 | 38.502 | 44.765 | 42.644 | 37.630 | 43.599 | -0.016 | 0.5598 | 0.7704 | 0 | 0 | 0 |
| 'Abcf3' | 14.377 | 14.602 | 14.324 | 13.499 | 13.799 | 14.670 | 13.540 | 13.604 | 14.007 | 15.087 | 13.845 | 14.806 | 14.495 | 14.272 | 14.339 | 13.317 | 14.181 | 14.202 | -0.035 | 0.2257 | 0.4722 | 0 | 0 | 0 |
| 'Abcg1' | 24.278 | 26.391 | 26.625 | 21.689 | 25.932 | 25.758 | 23.023 | 26.390 | 26.737 | 25.317 | 26.299 | 24.710 | 23.752 | 25.282 | 23.012 | 27.073 | 26.216 | 23.949 | -0.002 | 0.9634 | 0.9869 | 0 | 0 | 0 |
| 'Abhd10' | 2.954 | 3.062 | 3.336 | 2.973 | 2.856 | 3.103 | 2.991 | 3.251 | 3.592 | 2.957 | 2.950 | 2.753 | 3.429 | 3.413 | 3.049 | 3.065 | 2.979 | 2.396 | 0.059 | 0.3844 | 0.6347 | 0 | 0 | 0 |
| 'Abhd11' | 4.439 | 4.949 | 4.708 | 3.939 | 4.452 | 4.232 | 4.677 | 4.614 | 4.668 | 5.017 | 4.214 | 4.353 | 4.791 | 4.309 | 4.074 | 4.806 | 4.456 | 3.830 | 0.023 | 0.6479 | 0.8283 | 0 | 0 | 0 |
| 'Abhd12' | 9.155 | 10.099 | 10.360 | 9.439 | 9.691 | 10.747 | 9.199 | 10.377 | 10.384 | 10.215 | 9.463 | 9.790 | 9.868 | 10.394 | 9.604 | 10.265 | 9.534 | 9.228 | 0.011 | 0.7691 | 0.8972 | 0 | 0 | 0 |
| 'Abhd13' | 10.122 | 11.538 | 10.566 | 9.484 | 10.982 | 11.009 | 10.210 | 11.325 | 9.798 | 9.733 | 10.647 | 11.641 | 9.963 | 11.704 | 10.999 | 11.504 | 10.732 | 11.534 | -0.064 | 0.1044 | 0.3027 | 0 | 0 | 0 |
| 'Abhd15' | 1.350 | 1.062 | 1.340 | 1.434 | 1.320 | 1.060 | 1.244 | 1.153 | 1.377 | 1.112 | 1.193 | 1.352 | 1.093 | 1.153 | 1.275 | 1.140 | 0.924 | 1.357 | 0.080 | 0.4334 | 0.6782 | 0 | 0 | 0 |
| 'Abhd16a' | 11.939 | 11.843 | 14.563 | 11.273 | 12.609 | 13.664 | 11.476 | 12.807 | 13.749 | 12.460 | 10.187 | 12.937 | 12.347 | 11.810 | 12.191 | 11.958 | 11.678 | 11.436 | 0.080 | 0.0523 | 0.1941 | 0 | 0 | 0 |
| 'Abhd17a' | 34.909 | 36.159 | 37.076 | 30.396 | 35.556 | 42.013 | 32.321 | 35.206 | 36.967 | 33.420 | 30.004 | 39.369 | 35.908 | 34.421 | 39.758 | 38.737 | 30.291 | 40.162 | -0.015 | 0.7516 | 0.8894 | 0 | 0 | 0 |
| 'Abhd17b' | 12.315 | 12.969 | 11.918 | 11.765 | 12.612 | 10.889 | 12.637 | 15.350 | 10.407 | 11.525 | 13.627 | 11.472 | 11.504 | 14.749 | 11.306 | 12.306 | 13.555 | 11.501 | -0.016 | 0.7285 | 0.8755 | 0 | 0 | 0 |
| 'Abhd17c' | 44.374 | 39.439 | 44.274 | 39.436 | 42.452 | 40.993 | 39.438 | 43.938 | 35.110 | 47.770 | 47.712 | 39.917 | 44.957 | 48.443 | 42.917 | 41.820 | 41.082 | 46.482 | -0.128 | 0.0114 | 0.0690 | 0 | 0 | 0 |
| 'Abhd18' | 1.625 | 1.648 | 1.519 | 1.785 | 1.502 | 1.462 | 1.757 | 1.733 | 1.419 | 1.504 | 1.883 | 1.618 | 1.703 | 1.889 | 1.698 | 1.746 | 1.889 | 1.516 | -0.106 | 0.0383 | 0.1590 | 0 | 0 | 0 |
| 'Abhd2' | 24.225 | 26.089 | 24.919 | 23.707 | 25.661 | 26.558 | 23.066 | 24.446 | 26.457 | 26.440 | 25.345 | 27.626 | 23.582 | 25.159 | 27.592 | 25.992 | 24.682 | 28.468 | -0.072 | 0.0155 | 0.0858 | 0 | 0 | 0 |
| 'Abhd4' | 9.958 | 12.322 | 10.996 | 10.051 | 11.148 | 11.417 | 10.479 | 11.458 | 10.807 | 11.105 | 10.294 | 10.985 | 10.607 | 10.954 | 10.180 | 10.259 | 10.912 | 9.719 | 0.041 | 0.2418 | 0.4924 | 0 | 0 | 0 |
| 'Abhd5' | 2.601 | 3.327 | 3.207 | 2.763 | 3.130 | 3.094 | 2.984 | 3.196 | 2.959 | 3.044 | 3.107 | 2.978 | 3.178 | 3.334 | 2.676 | 3.036 | 3.005 | 2.997 | -0.014 | 0.7934 | 0.9078 | 0 | 0 | 0 |
| 'Abhd6' | 7.670 | 7.821 | 7.465 | 8.578 | 8.053 | 7.008 | 8.631 | 6.792 | 7.445 | 8.009 | 7.725 | 9.090 | 8.396 | 7.902 | 7.530 | 7.637 | 9.136 | 8.222 | -0.108 | 0.0657 | 0.2235 | 0 | 0 | 0 |
| 'Abhd8' | 124.447 | 137.148 | 139.237 | 114.973 | 131.228 | 157.457 | 117.977 | 128.498 | 143.498 | 127.261 | 112.603 | 145.267 | 128.551 | 122.650 | 150.888 | 144.575 | 105.091 | 154.595 | -0.003 | 0.9473 | 0.9796 | 0 | 0 | 0 |
| 'Abi1' | 13.854 | 14.831 | 13.338 | 13.896 | 14.438 | 12.820 | 14.464 | 14.522 | 13.018 | 14.253 | 14.061 | 13.602 | 13.768 | 14.510 | 13.067 | 14.062 | 14.816 | 13.394 | -0.016 | 0.4013 | 0.6501 | 0 | 0 | 0 |
| 'Abi2' | 3.893 | 4.216 | 4.111 | 3.963 | 4.017 | 3.814 | 4.295 | 4.082 | 3.966 | 4.394 | 4.526 | 4.318 | 3.950 | 4.609 | 4.190 | 3.991 | 5.055 | 4.285 | -0.122 | 0.0012 | 0.0120 | 1 | 0 | 0 |
| 'Abi3' | 3.381 | 3.806 | 3.504 | 3.883 | 3.938 | 3.718 | 3.544 | 3.992 | 3.795 | 3.086 | 3.216 | 3.339 | 4.474 | 3.023 | 3.235 | 3.162 | 2.837 | 3.275 | 0.166 | 0.0138 | 0.0791 | 0 | 0 | 0 |
| 'Abl1' | 7.931 | 7.481 | 8.356 | 7.835 | 7.386 | 8.249 | 7.362 | 7.467 | 8.155 | 8.702 | 8.260 | 8.289 | 8.463 | 7.987 | 7.964 | 8.149 | 7.281 | 8.013 | -0.067 | 0.0259 | 0.1236 | 0 | 0 | 0 |
| 'Abl2' | 5.203 | 4.754 | 4.370 | 4.287 | 4.246 | 4.075 | 4.237 | 4.854 | 4.365 | 4.780 | 3.904 | 4.453 | 4.321 | 4.218 | 4.006 | 5.109 | 4.289 | 4.479 | 0.021 | 0.6727 | 0.8425 | 0 | 0 | 0 |
| 'Ablim1' | 21.722 | 21.689 | 20.641 | 21.405 | 21.140 | 20.682 | 20.121 | 21.516 | 20.696 | 21.573 | 22.360 | 21.310 | 20.674 | 21.682 | 21.731 | 20.725 | 21.145 | 22.230 | -0.040 | 0.1096 | 0.3122 | 0 | 0 | 0 |
| 'Ablim2' | 2.798 | 3.792 | 3.001 | 2.679 | 3.111 | 3.578 | 3.533 | 2.973 | 3.636 | 3.434 | 2.434 | 2.650 | 4.235 | 3.186 | 2.499 | 3.129 | 3.100 | 2.563 | 0.097 | 0.3583 | 0.6109 | 0 | 0 | 0 |
| 'Abr' | 9.970 | 9.609 | 10.368 | 9.671 | 9.781 | 10.499 | 9.258 | 9.980 | 10.369 | 10.489 | 9.872 | 10.818 | 9.969 | 10.362 | 10.836 | 9.753 | 10.077 | 11.092 | -0.069 | 0.0031 | 0.0258 | 1 | 0 | 0 |
| 'Abracl' | 80.866 | 89.356 | 90.900 | 91.402 | 89.720 | 86.118 | 94.980 | 88.230 | 80.968 | 85.728 | 85.568 | 83.360 | 87.612 | 93.180 | 90.798 | 85.259 | 96.246 | 86.660 | -0.014 | 0.7085 | 0.8650 | 0 | 0 | 0 |
| 'Abt1' | 27.179 | 20.320 | 18.144 | 24.244 | 20.882 | 20.959 | 24.045 | 21.768 | 18.282 | 21.131 | 19.036 | 18.617 | 19.813 | 22.483 | 18.788 | 20.980 | 20.015 | 17.005 | 0.125 | 0.0211 | 0.1066 | 0 | 0 | 0 |
| 'Abtb1' | 6.571 | 7.690 | 8.109 | 6.432 | 7.503 | 8.130 | 6.877 | 8.114 | 8.316 | 7.583 | 7.299 | 7.444 | 7.780 | 6.773 | 7.208 | 7.420 | 6.738 | 6.959 | 0.042 | 0.3553 | 0.6086 | 0 | 0 | 0 |
| 'Abtb2' | 45.033 | 38.890 | 40.243 | 42.212 | 38.090 | 39.930 | 35.746 | 41.558 | 39.952 | 45.112 | 39.182 | 40.634 | 42.299 | 38.738 | 38.063 | 42.906 | 37.246 | 41.829 | -0.026 | 0.5060 | 0.7322 | 0 | 0 | 0 |
| 'Acaa1b' | 7.679 | 6.942 | 8.850 | 9.659 | 6.901 | 8.586 | 8.737 | 7.908 | 9.362 | 7.838 | 6.872 | 8.929 | 7.875 | 8.073 | 9.312 | 8.461 | 7.450 | 7.290 | 0.039 | 0.5468 | 0.7610 | 0 | 0 | 0 |
| 'Acaa2' | 46.964 | 44.587 | 44.095 | 44.838 | 45.601 | 44.980 | 44.006 | 45.801 | 43.573 | 39.296 | 32.116 | 39.719 | 35.805 | 35.409 | 42.580 | 38.482 | 31.422 | 39.690 | 0.265 | 0.0000 | 0.0000 | 1 | 0 | 0 |
| 'Acaca' | 4.214 | 3.780 | 4.260 | 4.374 | 3.743 | 3.998 | 4.453 | 3.867 | 4.149 | 4.038 | 4.038 | 4.386 | 3.725 | 3.622 | 4.298 | 3.583 | 3.889 | 4.571 | 0.016 | 0.7214 | 0.8716 | 0 | 0 | 0 |
| 'Acacb' | 0.895 | 0.718 | 0.953 | 0.840 | 0.791 | 1.000 | 0.891 | 0.954 | 1.039 | 0.950 | 0.824 | 0.833 | 0.818 | 0.842 | 0.852 | 0.676 | 0.837 | 0.881 | 0.101 | 0.1300 | 0.3461 | 0 | 0 | 0 |
| 'Acad10' | 3.110 | 2.794 | 3.432 | 3.103 | 3.009 | 3.222 | 3.206 | 3.051 | 3.122 | 3.215 | 2.838 | 2.987 | 2.873 | 2.737 | 3.024 | 2.954 | 2.611 | 2.836 | 0.101 | 0.0182 | 0.0968 | 0 | 0 | 0 |
| 'Acad11' | 1.865 | 2.147 | 1.896 | 1.922 | 2.135 | 1.875 | 1.987 | 2.133 | 1.990 | 2.237 | 2.283 | 2.163 | 2.012 | 2.039 | 1.980 | 2.001 | 2.048 | 1.986 | -0.080 | 0.0967 | 0.2894 | 0 | 0 | 0 |
| 'Acad8' | 2.506 | 2.423 | 2.809 | 2.523 | 2.808 | 2.691 | 2.486 | 2.922 | 2.922 | 2.570 | 2.271 | 2.565 | 2.830 | 2.662 | 2.505 | 2.470 | 2.748 | 2.359 | 0.060 | 0.2371 | 0.4868 | 0 | 0 | 0 |
| 'Acad9' | 4.714 | 5.128 | 5.102 | 4.519 | 5.214 | 5.439 | 5.008 | 5.392 | 5.339 | 5.545 | 4.990 | 5.182 | 5.635 | 4.948 | 5.017 | 5.506 | 5.147 | 5.122 | -0.047 | 0.2546 | 0.5066 | 0 | 0 | 0 |
| 'Acadl' | 21.748 | 20.768 | 19.543 | 22.122 | 23.074 | 21.222 | 24.558 | 22.365 | 20.055 | 8.717 | 10.896 | 9.807 | 10.638 | 10.142 | 9.988 | 10.782 | 11.059 | 10.408 | 1.068 | 0.0000 | 0.0000 | 1 | 1 | 1 |
| 'Acadm' | 4.706 | 5.528 | 4.730 | 4.634 | 5.419 | 5.368 | 5.122 | 5.118 | 5.265 | 5.033 | 5.147 | 4.941 | 5.318 | 4.608 | 4.769 | 5.008 | 4.817 | 4.309 | 0.050 | 0.2632 | 0.5172 | 0 | 0 | 0 |
| 'Acads' | 5.919 | 5.947 | 7.013 | 5.515 | 5.793 | 7.144 | 6.782 | 6.327 | 6.992 | 12.269 | 10.359 | 12.577 | 11.163 | 10.412 | 12.048 | 11.465 | 10.371 | 11.294 | -0.837 | 0.0000 | 0.0000 | 1 | 1 | 0 |
| 'Acadsb' | 3.584 | 3.886 | 3.237 | 3.233 | 4.279 | 3.448 | 3.540 | 3.721 | 3.496 | 3.618 | 3.738 | 3.751 | 2.996 | 3.149 | 3.253 | 3.192 | 3.594 | 3.358 | 0.063 | 0.2335 | 0.4820 | 0 | 0 | 0 |
| 'Acadvl' | 18.585 | 17.779 | 17.681 | 18.528 | 16.447 | 16.953 | 17.460 | 17.507 | 17.259 | 17.346 | 16.195 | 17.641 | 18.162 | 16.253 | 17.037 | 17.806 | 16.395 | 16.505 | 0.034 | 0.1935 | 0.4310 | 0 | 0 | 0 |
| 'Acap1' | 88.389 | 78.313 | 98.863 | 84.552 | 79.307 | 96.193 | 87.407 | 79.653 | 96.719 | 88.742 | 82.756 | 98.389 | 89.577 | 83.111 | 94.366 | 83.171 | 87.596 | 96.902 | -0.040 | 0.0873 | 0.2695 | 0 | 0 | 0 |
| 'Acap2' | 24.942 | 26.486 | 24.261 | 24.359 | 25.433 | 23.794 | 23.616 | 26.505 | 24.549 | 25.460 | 24.553 | 25.308 | 21.835 | 23.760 | 24.564 | 23.283 | 22.679 | 26.106 | 0.030 | 0.4086 | 0.6576 | 0 | 0 | 0 |
| 'Acap3' | 3.965 | 3.701 | 4.857 | 3.519 | 3.537 | 4.082 | 3.658 | 3.993 | 4.339 | 4.025 | 3.876 | 4.245 | 4.286 | 3.801 | 4.087 | 4.306 | 3.813 | 4.204 | -0.048 | 0.2809 | 0.5379 | 0 | 0 | 0 |
| 'Acat2' | 10.343 | 10.593 | 10.774 | 9.965 | 10.622 | 11.003 | 11.052 | 11.395 | 12.167 | 9.713 | 10.630 | 11.807 | 10.404 | 9.894 | 11.811 | 9.991 | 10.373 | 10.962 | 0.021 | 0.6344 | 0.8189 | 0 | 0 | 0 |
| 'Acat3' | 2.746 | 2.603 | 2.805 | 3.246 | 2.584 | 2.867 | 2.729 | 2.597 | 2.822 | 2.126 | 2.995 | 2.665 | 2.473 | 3.034 | 2.673 | 2.418 | 2.652 | 2.960 | 0.052 | 0.5389 | 0.7568 | 0 | 0 | 0 |
| 'Acbd3' | 11.540 | 11.590 | 11.310 | 11.876 | 11.321 | 10.972 | 11.299 | 12.031 | 10.813 | 11.499 | 11.362 | 12.070 | 11.829 | 12.829 | 12.182 | 12.275 | 12.445 | 12.081 | -0.091 | 0.0025 | 0.0220 | 1 | 0 | 0 |
| 'Acbd4' | 4.351 | 4.743 | 4.695 | 4.229 | 4.416 | 4.937 | 4.702 | 4.669 | 4.882 | 4.132 | 4.014 | 4.377 | 4.568 | 4.399 | 4.453 | 4.097 | 3.818 | 3.904 | 0.129 | 0.0012 | 0.0123 | 1 | 0 | 0 |
| 'Acbd5' | 4.354 | 4.962 | 4.787 | 5.230 | 5.060 | 4.738 | 5.025 | 5.118 | 4.503 | 4.487 | 5.412 | 5.499 | 4.282 | 5.222 | 5.314 | 4.538 | 5.822 | 4.788 | -0.062 | 0.2096 | 0.4547 | 0 | 0 | 0 |
| 'Acbd6' | 6.293 | 6.003 | 6.717 | 5.816 | 6.000 | 6.762 | 6.459 | 6.069 | 6.155 | 6.075 | 5.748 | 6.098 | 6.149 | 5.502 | 6.196 | 5.912 | 5.348 | 5.904 | 0.081 | 0.0135 | 0.0776 | 0 | 0 | 0 |
| 'Accs' | 2.998 | 3.288 | 3.540 | 3.183 | 3.140 | 3.534 | 3.389 | 3.486 | 4.070 | 3.427 | 3.022 | 3.143 | 3.385 | 3.184 | 2.867 | 3.061 | 2.894 | 2.635 | 0.140 | 0.0124 | 0.0732 | 0 | 0 | 0 |
| 'Acd' | 39.106 | 39.034 | 38.250 | 37.908 | 36.096 | 38.968 | 38.557 | 38.780 | 38.215 | 38.422 | 34.310 | 40.935 | 40.708 | 37.182 | 40.513 | 40.160 | 37.955 | 38.934 | -0.030 | 0.3433 | 0.5993 | 0 | 0 | 0 |
| 'Acer2' | 0.503 | 0.626 | 0.569 | 0.622 | 0.463 | 0.588 | 0.674 | 0.734 | 0.648 | 0.577 | 0.425 | 0.704 | 0.551 | 0.517 | 0.714 | 0.602 | 0.679 | 0.588 | 0.000 | 0.9969 | 0.9994 | 0 | 0 | 0 |
| 'Acer3' | 0.688 | 0.828 | 0.842 | 0.817 | 0.804 | 0.633 | 0.792 | 0.901 | 0.739 | 0.785 | 1.090 | 0.790 | 0.870 | 0.985 | 0.833 | 0.808 | 0.827 | 0.746 | -0.139 | 0.0857 | 0.2662 | 0 | 0 | 0 |
| 'Acin1' | 37.444 | 31.306 | 37.502 | 36.474 | 31.553 | 38.002 | 34.620 | 32.723 | 37.928 | 36.424 | 34.439 | 39.106 | 36.029 | 33.514 | 37.133 | 34.015 | 34.039 | 38.447 | -0.037 | 0.1547 | 0.3831 | 0 | 0 | 0 |
| 'Acly' | 36.480 | 36.072 | 38.726 | 37.465 | 37.904 | 37.848 | 36.449 | 36.845 | 37.987 | 39.493 | 38.633 | 39.341 | 38.763 | 37.997 | 40.391 | 36.815 | 38.582 | 38.683 | -0.065 | 0.0007 | 0.0083 | 1 | 0 | 0 |
| 'Aco1' | 11.040 | 11.865 | 12.361 | 11.091 | 11.606 | 11.935 | 11.767 | 12.153 | 12.497 | 11.967 | 11.668 | 11.699 | 11.902 | 11.127 | 11.789 | 11.682 | 11.662 | 11.123 | 0.015 | 0.6157 | 0.8082 | 0 | 0 | 0 |
| 'Aco2' | 43.640 | 43.653 | 44.780 | 43.664 | 44.192 | 46.252 | 44.527 | 42.908 | 44.651 | 44.459 | 42.801 | 44.889 | 47.457 | 43.391 | 43.923 | 43.645 | 42.019 | 44.199 | -0.005 | 0.7915 | 0.9070 | 0 | 0 | 0 |
| 'Acod1' | 1.791 | 2.722 | 2.006 | 1.992 | 2.407 | 2.020 | 2.220 | 1.973 | 2.059 | 3.440 | 2.881 | 2.039 | 4.078 | 2.774 | 2.103 | 3.183 | 2.551 | 1.773 | -0.351 | 0.0019 | 0.0176 | 1 | 0 | 0 |
| 'Acot11' | 1.786 | 1.345 | 1.801 | 1.740 | 1.466 | 1.541 | 1.436 | 1.270 | 1.627 | 1.675 | 1.683 | 1.875 | 1.295 | 1.433 | 1.731 | 1.147 | 1.208 | 1.949 | -0.006 | 0.9466 | 0.9794 | 0 | 0 | 0 |
| 'Acot13' | 27.184 | 31.338 | 26.596 | 30.724 | 29.552 | 32.677 | 27.634 | 27.892 | 26.135 | 26.621 | 26.316 | 29.674 | 28.383 | 26.499 | 25.257 | 24.545 | 32.190 | 24.419 | 0.074 | 0.2266 | 0.4732 | 0 | 0 | 0 |
| 'Acot2' | 9.559 | 7.991 | 7.823 | 8.023 | 9.309 | 8.800 | 8.788 | 8.542 | 7.866 | 10.390 | 8.286 | 7.128 | 9.943 | 8.233 | 6.717 | 9.421 | 8.019 | 6.227 | 0.053 | 0.3890 | 0.6384 | 0 | 0 | 0 |
| 'Acot6' | 1.002 | 1.520 | 1.041 | 1.272 | 1.027 | 1.064 | 1.165 | 1.205 | 1.030 | 1.044 | 1.197 | 1.158 | 0.765 | 1.314 | 1.113 | 1.027 | 1.246 | 0.825 | 0.073 | 0.5267 | 0.7483 | 0 | 0 | 0 |
| 'Acot7' | 26.125 | 22.516 | 29.684 | 28.381 | 25.431 | 29.723 | 25.927 | 23.715 | 28.117 | 26.923 | 22.890 | 29.946 | 28.505 | 23.188 | 31.033 | 25.721 | 24.084 | 29.821 | -0.025 | 0.4310 | 0.6762 | 0 | 0 | 0 |
| 'Acot8' | 6.256 | 6.118 | 7.491 | 6.141 | 6.975 | 7.427 | 5.757 | 6.461 | 7.233 | 6.040 | 5.710 | 7.024 | 6.312 | 6.017 | 6.704 | 5.914 | 5.816 | 7.360 | 0.063 | 0.1636 | 0.3954 | 0 | 0 | 0 |
| 'Acot9' | 5.238 | 5.716 | 5.606 | 5.596 | 6.350 | 5.243 | 5.044 | 5.433 | 5.203 | 5.839 | 4.842 | 4.839 | 5.400 | 4.754 | 5.372 | 5.616 | 4.904 | 4.992 | 0.078 | 0.1440 | 0.3681 | 0 | 0 | 0 |
| 'Acox1' | 4.100 | 3.623 | 4.452 | 4.210 | 4.518 | 4.364 | 3.886 | 4.084 | 4.312 | 4.137 | 4.488 | 4.585 | 4.050 | 4.545 | 4.375 | 3.703 | 4.547 | 4.410 | -0.057 | 0.1824 | 0.4185 | 0 | 0 | 0 |
| 'Acox3' | 4.219 | 4.483 | 5.026 | 4.116 | 4.158 | 4.789 | 4.546 | 4.267 | 4.745 | 4.141 | 4.782 | 4.986 | 4.522 | 4.623 | 4.604 | 3.993 | 4.711 | 4.421 | -0.027 | 0.4828 | 0.7146 | 0 | 0 | 0 |
| 'Acp1' | 10.108 | 11.201 | 10.086 | 10.042 | 10.623 | 10.021 | 10.253 | 11.116 | 10.260 | 9.324 | 9.671 | 8.846 | 10.218 | 9.662 | 9.555 | 9.284 | 10.413 | 8.828 | 0.119 | 0.0033 | 0.0273 | 1 | 0 | 0 |
| 'Acp2' | 4.479 | 5.321 | 5.387 | 4.246 | 5.685 | 5.182 | 4.881 | 5.015 | 5.421 | 5.249 | 4.802 | 4.622 | 5.121 | 5.209 | 4.417 | 5.557 | 4.834 | 4.287 | 0.042 | 0.4609 | 0.6973 | 0 | 0 | 0 |
| 'Acp5' | 61.801 | 59.522 | 67.076 | 61.452 | 61.282 | 71.574 | 61.799 | 64.851 | 68.615 | 66.844 | 53.334 | 73.844 | 66.419 | 58.671 | 71.449 | 74.709 | 55.819 | 67.377 | -0.032 | 0.4210 | 0.6677 | 0 | 0 | 0 |
| 'Acp6' | 3.010 | 2.798 | 2.885 | 2.778 | 2.751 | 2.861 | 2.852 | 2.938 | 3.217 | 2.623 | 2.767 | 3.240 | 2.922 | 2.850 | 2.887 | 2.686 | 2.968 | 3.176 | -0.016 | 0.7203 | 0.8713 | 0 | 0 | 0 |
| 'Acpp' | 2.397 | 2.519 | 2.523 | 2.560 | 2.750 | 2.746 | 2.238 | 2.750 | 2.671 | 1.996 | 1.649 | 1.723 | 1.641 | 1.862 | 1.548 | 1.818 | 1.886 | 1.806 | 0.534 | 0.0000 | 0.0000 | 1 | 1 | 0 |
| 'Acrbp' | 2.487 | 2.086 | 2.360 | 2.386 | 1.686 | 2.510 | 2.984 | 1.785 | 2.791 | 1.872 | 2.323 | 3.922 | 1.835 | 2.231 | 3.311 | 1.866 | 2.642 | 3.890 | -0.162 | 0.1673 | 0.3998 | 0 | 0 | 0 |
| 'Acsbg1' | 0.985 | 0.666 | 1.232 | 0.811 | 0.906 | 1.216 | 0.967 | 1.163 | 1.196 | 0.757 | 0.741 | 1.257 | 1.090 | 0.877 | 1.239 | 0.914 | 0.960 | 1.007 | 0.041 | 0.7088 | 0.8650 | 0 | 0 | 0 |
| 'Acsf2' | 7.454 | 6.535 | 7.767 | 7.436 | 6.862 | 7.806 | 8.170 | 7.166 | 7.936 | 7.650 | 6.527 | 7.170 | 6.927 | 6.414 | 6.777 | 6.259 | 5.764 | 6.767 | 0.146 | 0.0004 | 0.0055 | 1 | 0 | 0 |
| 'Acsf3' | 15.378 | 16.182 | 17.173 | 15.580 | 16.878 | 16.398 | 16.743 | 17.059 | 16.628 | 18.225 | 17.381 | 16.538 | 19.889 | 16.523 | 15.806 | 18.892 | 17.345 | 14.943 | -0.075 | 0.0949 | 0.2862 | 0 | 0 | 0 |
| 'Acsl1' | 1.271 | 1.173 | 1.092 | 0.931 | 1.101 | 1.057 | 1.073 | 1.173 | 1.207 | 1.161 | 1.096 | 1.171 | 1.329 | 1.296 | 1.109 | 1.357 | 1.174 | 1.136 | -0.108 | 0.0859 | 0.2668 | 0 | 0 | 0 |
| 'Acsl3' | 2.922 | 2.983 | 3.335 | 2.931 | 2.915 | 2.870 | 3.339 | 3.304 | 3.079 | 2.889 | 3.627 | 2.872 | 3.196 | 3.336 | 3.038 | 2.556 | 3.583 | 2.650 | -0.006 | 0.9237 | 0.9708 | 0 | 0 | 0 |
| 'Acsl4' | 11.283 | 12.610 | 12.064 | 12.491 | 11.542 | 10.492 | 13.244 | 12.958 | 10.835 | 11.133 | 13.208 | 12.599 | 11.573 | 13.565 | 11.966 | 10.722 | 12.855 | 11.771 | -0.036 | 0.4310 | 0.6762 | 0 | 0 | 0 |
| 'Acsl5' | 50.587 | 48.393 | 49.115 | 50.326 | 49.148 | 50.672 | 51.671 | 52.015 | 51.384 | 45.539 | 51.887 | 54.401 | 51.661 | 50.321 | 53.158 | 49.152 | 53.657 | 53.121 | -0.041 | 0.1796 | 0.4148 | 0 | 0 | 0 |
| 'Acss1' | 9.129 | 9.678 | 10.007 | 9.838 | 9.590 | 9.542 | 9.480 | 9.873 | 10.229 | 8.308 | 9.135 | 9.505 | 9.392 | 9.040 | 9.305 | 8.982 | 7.575 | 8.526 | 0.119 | 0.0020 | 0.0181 | 1 | 0 | 0 |
| 'Acss2' | 0.566 | 0.636 | 0.594 | 0.637 | 0.591 | 0.657 | 0.648 | 0.563 | 0.548 | 0.634 | 0.665 | 0.612 | 0.581 | 0.646 | 0.537 | 0.526 | 0.704 | 0.542 | -0.013 | 0.8615 | 0.9401 | 0 | 0 | 0 |
| 'Actb' | 582.630 | 677.393 | 622.028 | 670.924 | 676.093 | 655.655 | 692.129 | 621.939 | 684.102 | 644.113 | 729.298 | 521.674 | 644.131 | 660.081 | 678.349 | 619.256 | 692.006 | 638.299 | 0.004 | 0.9333 | 0.9746 | 0 | 0 | 0 |
| 'Actl6a' | 12.082 | 12.658 | 12.749 | 13.049 | 12.300 | 12.625 | 11.612 | 12.606 | 12.580 | 11.418 | 11.700 | 12.619 | 12.488 | 11.738 | 11.632 | 12.753 | 11.337 | 11.741 | 0.052 | 0.0813 | 0.2571 | 0 | 0 | 0 |
| 'Actn1' | 24.688 | 15.660 | 28.093 | 24.297 | 15.408 | 25.778 | 22.087 | 17.358 | 25.312 | 28.793 | 23.431 | 36.113 | 23.495 | 21.664 | 31.290 | 27.341 | 19.490 | 33.295 | -0.318 | 0.0000 | 0.0000 | 1 | 0 | 0 |
| 'Actn4' | 50.170 | 43.024 | 50.617 | 51.151 | 42.243 | 50.298 | 51.090 | 43.915 | 48.904 | 48.876 | 44.555 | 48.864 | 50.737 | 44.492 | 48.630 | 47.372 | 42.973 | 49.502 | 0.006 | 0.7746 | 0.8994 | 0 | 0 | 0 |
| 'Actr10' | 20.244 | 22.831 | 20.410 | 20.385 | 21.667 | 20.542 | 20.612 | 22.373 | 19.804 | 19.699 | 18.416 | 19.158 | 20.473 | 20.483 | 19.119 | 19.033 | 20.929 | 18.330 | 0.092 | 0.0012 | 0.0120 | 1 | 0 | 0 |
| 'Actr1a' | 53.674 | 58.484 | 56.955 | 51.238 | 56.853 | 59.535 | 54.022 | 58.810 | 57.427 | 54.771 | 56.794 | 59.864 | 55.287 | 58.639 | 60.858 | 57.332 | 55.565 | 58.434 | -0.040 | 0.0742 | 0.2423 | 0 | 0 | 0 |
| 'Actr1b' | 18.407 | 18.423 | 19.304 | 18.339 | 17.190 | 18.275 | 17.955 | 17.215 | 18.659 | 17.830 | 16.751 | 20.026 | 19.114 | 17.482 | 18.057 | 18.357 | 18.178 | 19.639 | -0.027 | 0.3376 | 0.5955 | 0 | 0 | 0 |
| 'Actr2' | 113.733 | 130.805 | 120.292 | 124.743 | 130.604 | 116.386 | 126.433 | 130.637 | 114.779 | 121.125 | 128.659 | 110.287 | 116.182 | 132.615 | 115.782 | 122.870 | 129.583 | 112.525 | 0.015 | 0.5145 | 0.7384 | 0 | 0 | 0 |
| 'Actr3' | 80.494 | 88.428 | 79.872 | 88.054 | 88.680 | 81.684 | 88.572 | 86.524 | 80.641 | 86.970 | 88.983 | 77.293 | 85.576 | 91.811 | 82.698 | 86.406 | 91.845 | 79.018 | -0.024 | 0.2949 | 0.5505 | 0 | 0 | 0 |
| 'Actr5' | 6.492 | 5.486 | 7.388 | 6.678 | 5.305 | 7.408 | 6.857 | 6.313 | 6.684 | 6.746 | 5.931 | 6.724 | 6.367 | 5.454 | 6.121 | 6.133 | 6.334 | 6.494 | 0.048 | 0.2878 | 0.5444 | 0 | 0 | 0 |
| 'Actr6' | 8.751 | 9.858 | 9.792 | 9.940 | 11.795 | 9.234 | 11.834 | 10.954 | 9.073 | 8.463 | 10.402 | 9.941 | 8.940 | 10.658 | 9.647 | 7.917 | 10.968 | 9.203 | 0.065 | 0.2945 | 0.5504 | 0 | 0 | 0 |
| 'Actr8' | 22.255 | 22.823 | 22.603 | 20.854 | 23.271 | 23.872 | 20.305 | 24.631 | 22.329 | 23.242 | 21.901 | 21.626 | 19.528 | 21.343 | 23.373 | 22.755 | 23.265 | 20.764 | 0.030 | 0.4664 | 0.7021 | 0 | 0 | 0 |
| 'Acvr1b' | 10.923 | 11.916 | 12.264 | 10.809 | 11.370 | 12.007 | 9.402 | 13.328 | 12.204 | 11.396 | 10.629 | 10.224 | 9.782 | 10.709 | 10.415 | 9.923 | 9.855 | 9.674 | 0.162 | 0.0009 | 0.0096 | 1 | 0 | 0 |
| 'Acvr2a' | 0.811 | 0.814 | 0.999 | 0.700 | 1.034 | 0.880 | 0.729 | 1.080 | 0.887 | 1.242 | 0.851 | 0.713 | 1.093 | 0.996 | 0.662 | 1.038 | 0.864 | 0.760 | -0.033 | 0.7737 | 0.8993 | 0 | 0 | 0 |
| 'Acy1' | 1.771 | 1.514 | 2.039 | 1.420 | 1.999 | 2.161 | 1.807 | 1.512 | 2.210 | 1.865 | 1.322 | 1.825 | 2.008 | 1.352 | 1.642 | 1.697 | 1.407 | 1.755 | 0.135 | 0.0991 | 0.2943 | 0 | 0 | 0 |
| 'Acyp1' | 8.015 | 11.175 | 9.903 | 9.242 | 12.093 | 10.244 | 9.849 | 10.565 | 9.582 | 10.813 | 10.716 | 9.845 | 10.072 | 10.994 | 9.562 | 9.750 | 10.456 | 8.322 | -0.011 | 0.8329 | 0.9271 | 0 | 0 | 0 |
| 'Ada' | 1.503 | 1.577 | 1.607 | 1.611 | 1.987 | 1.802 | 1.781 | 1.710 | 1.487 | 1.414 | 1.724 | 1.714 | 1.756 | 1.773 | 1.823 | 2.020 | 1.869 | 1.564 | -0.065 | 0.5397 | 0.7573 | 0 | 0 | 0 |
| 'Adal' | 1.158 | 1.243 | 1.196 | 1.148 | 1.241 | 1.141 | 1.208 | 1.124 | 1.218 | 1.078 | 1.183 | 1.256 | 1.138 | 1.200 | 1.239 | 1.263 | 1.369 | 1.165 | -0.042 | 0.4534 | 0.6924 | 0 | 0 | 0 |
| 'Adam10' | 45.960 | 51.983 | 48.971 | 47.763 | 50.449 | 46.077 | 46.881 | 48.581 | 46.526 | 49.604 | 48.895 | 48.002 | 45.911 | 51.337 | 46.594 | 47.143 | 49.524 | 48.292 | -0.018 | 0.4268 | 0.6733 | 0 | 0 | 0 |
| 'Adam15' | 3.033 | 2.709 | 3.508 | 2.750 | 2.772 | 3.313 | 3.179 | 2.824 | 3.250 | 3.224 | 2.476 | 3.449 | 3.238 | 2.514 | 3.204 | 2.978 | 2.627 | 2.898 | 0.029 | 0.4923 | 0.7217 | 0 | 0 | 0 |
| 'Adam17' | 8.684 | 9.525 | 9.341 | 8.816 | 9.485 | 8.873 | 8.661 | 9.053 | 9.213 | 9.013 | 9.302 | 9.167 | 9.226 | 9.175 | 8.821 | 9.038 | 9.261 | 8.589 | -0.010 | 0.6711 | 0.8415 | 0 | 0 | 0 |
| 'Adam19' | 18.473 | 14.221 | 20.279 | 14.933 | 15.417 | 17.788 | 16.425 | 16.050 | 17.597 | 19.257 | 18.160 | 18.520 | 17.094 | 15.629 | 20.104 | 18.632 | 17.229 | 19.919 | -0.134 | 0.0046 | 0.0349 | 1 | 0 | 0 |
| 'Adam33' | 3.526 | 5.019 | 3.864 | 3.986 | 4.714 | 3.956 | 4.211 | 4.987 | 4.517 | 4.129 | 4.400 | 3.979 | 5.123 | 4.701 | 3.654 | 4.342 | 5.400 | 3.606 | -0.030 | 0.5985 | 0.7958 | 0 | 0 | 0 |
| 'Adam8' | 1.285 | 0.865 | 1.635 | 1.387 | 0.960 | 1.413 | 1.190 | 1.137 | 1.466 | 0.750 | 0.481 | 1.040 | 0.877 | 0.527 | 0.960 | 0.615 | 0.836 | 0.996 | 0.666 | 0.0000 | 0.0000 | 1 | 1 | 0 |
| 'Adam9' | 4.224 | 5.589 | 4.741 | 4.713 | 5.340 | 4.380 | 4.927 | 4.718 | 5.195 | 4.726 | 4.852 | 5.035 | 4.688 | 5.941 | 4.567 | 5.171 | 5.039 | 4.879 | -0.051 | 0.3691 | 0.6203 | 0 | 0 | 0 |
| 'Adamts10' | 14.823 | 12.967 | 17.030 | 14.675 | 13.595 | 16.307 | 15.307 | 13.442 | 17.169 | 16.187 | 14.553 | 18.524 | 17.295 | 13.900 | 16.737 | 14.945 | 16.196 | 17.896 | -0.125 | 0.0002 | 0.0025 | 1 | 0 | 0 |
| 'Adamts6' | 4.798 | 4.051 | 4.139 | 4.197 | 4.404 | 4.516 | 4.466 | 3.902 | 5.302 | 4.469 | 4.359 | 4.599 | 4.335 | 4.389 | 3.967 | 4.719 | 5.536 | 3.662 | -0.016 | 0.8238 | 0.9226 | 0 | 0 | 0 |
| 'Adamtsl2' | 1.592 | 1.914 | 1.562 | 1.842 | 1.544 | 1.686 | 2.070 | 1.422 | 2.126 | 1.510 | 1.240 | 1.656 | 1.710 | 1.317 | 1.200 | 1.338 | 1.563 | 1.468 | 0.258 | 0.0018 | 0.0165 | 1 | 0 | 0 |
| 'Adamtsl4' | 1.167 | 1.512 | 1.848 | 1.481 | 1.480 | 1.562 | 1.359 | 1.526 | 1.895 | 1.401 | 1.477 | 1.738 | 1.517 | 1.253 | 1.430 | 1.398 | 1.521 | 1.577 | 0.038 | 0.6204 | 0.8101 | 0 | 0 | 0 |
| 'Adap1' | 4.062 | 3.734 | 4.272 | 3.745 | 3.392 | 4.605 | 3.815 | 4.082 | 4.025 | 4.971 | 4.909 | 5.227 | 5.194 | 3.959 | 4.746 | 4.380 | 3.766 | 5.588 | -0.266 | 0.0000 | 0.0001 | 1 | 0 | 0 |
| 'Adar' | 22.919 | 21.529 | 24.057 | 20.643 | 20.885 | 24.779 | 22.192 | 23.806 | 25.858 | 23.619 | 24.344 | 27.209 | 24.151 | 22.304 | 26.389 | 22.981 | 23.659 | 27.033 | -0.111 | 0.0002 | 0.0030 | 1 | 0 | 0 |
| 'Adarb1' | 0.628 | 0.645 | 0.755 | 0.637 | 0.646 | 0.805 | 0.649 | 0.592 | 0.818 | 0.831 | 0.560 | 0.739 | 0.607 | 0.547 | 0.645 | 0.644 | 0.735 | 0.732 | 0.019 | 0.7855 | 0.9044 | 0 | 0 | 0 |
| 'Adat1' | 3.454 | 2.468 | 3.342 | 3.462 | 2.914 | 3.378 | 3.394 | 2.832 | 3.300 | 2.941 | 2.717 | 4.231 | 2.650 | 2.761 | 4.018 | 2.246 | 2.980 | 3.750 | 0.003 | 0.9678 | 0.9891 | 0 | 0 | 0 |
| 'Adat2' | 7.516 | 7.492 | 7.606 | 7.129 | 8.221 | 8.173 | 7.527 | 7.760 | 7.412 | 8.223 | 8.217 | 7.108 | 8.375 | 9.274 | 7.994 | 8.884 | 6.466 | 6.664 | -0.046 | 0.5137 | 0.7378 | 0 | 0 | 0 |
| 'Adat3' | 3.834 | 3.874 | 4.107 | 3.921 | 3.336 | 3.809 | 3.875 | 3.766 | 3.771 | 3.121 | 3.449 | 3.823 | 3.535 | 3.244 | 3.867 | 3.122 | 3.265 | 3.693 | 0.126 | 0.0120 | 0.0712 | 0 | 0 | 0 |
| 'Adck1' | 7.760 | 7.499 | 7.815 | 6.695 | 7.803 | 7.367 | 8.234 | 7.517 | 8.012 | 7.258 | 7.032 | 8.241 | 7.734 | 7.605 | 7.214 | 7.853 | 6.685 | 6.784 | 0.035 | 0.4921 | 0.7216 | 0 | 0 | 0 |
| 'Adck2' | 6.775 | 6.480 | 7.411 | 6.499 | 6.453 | 7.526 | 6.997 | 7.125 | 7.225 | 6.938 | 7.129 | 7.225 | 6.851 | 6.643 | 6.875 | 7.064 | 6.793 | 6.331 | 0.009 | 0.8082 | 0.9139 | 0 | 0 | 0 |
| 'Adck5' | 6.822 | 6.694 | 7.329 | 6.235 | 7.023 | 7.110 | 6.702 | 6.857 | 7.326 | 7.500 | 6.199 | 8.046 | 8.201 | 6.598 | 7.032 | 7.484 | 7.473 | 7.226 | -0.093 | 0.0259 | 0.1236 | 0 | 0 | 0 |
| 'Adcy3' | 1.815 | 1.609 | 2.380 | 1.727 | 2.184 | 2.225 | 2.036 | 2.029 | 2.174 | 2.326 | 1.825 | 2.141 | 2.188 | 2.253 | 1.927 | 1.938 | 1.739 | 2.007 | -0.024 | 0.7270 | 0.8750 | 0 | 0 | 0 |
| 'Adcy6' | 0.869 | 0.886 | 1.072 | 1.095 | 0.878 | 0.930 | 0.978 | 0.967 | 1.007 | 0.810 | 0.839 | 0.950 | 0.846 | 0.775 | 0.836 | 0.687 | 0.706 | 0.910 | 0.219 | 0.0038 | 0.0303 | 1 | 0 | 0 |
| 'Adcy7' | 89.220 | 90.769 | 99.462 | 84.822 | 89.797 | 96.441 | 84.992 | 90.027 | 102.442 | 91.911 | 91.958 | 100.642 | 87.384 | 90.706 | 98.379 | 93.485 | 91.434 | 103.222 | -0.047 | 0.0300 | 0.1349 | 0 | 0 | 0 |
| 'Adcy9' | 0.864 | 1.306 | 1.054 | 0.982 | 1.240 | 1.121 | 0.861 | 1.000 | 1.050 | 0.993 | 1.073 | 0.959 | 1.250 | 1.012 | 0.884 | 1.092 | 0.871 | 1.046 | 0.030 | 0.7162 | 0.8688 | 0 | 0 | 0 |
| 'Add1' | 12.329 | 12.958 | 13.097 | 11.974 | 12.737 | 13.951 | 12.107 | 13.674 | 13.671 | 13.628 | 13.087 | 13.402 | 12.884 | 13.134 | 13.887 | 13.821 | 12.667 | 12.846 | -0.045 | 0.1136 | 0.3197 | 0 | 0 | 0 |
| 'Add3' | 32.352 | 31.299 | 36.333 | 31.611 | 33.357 | 36.279 | 36.220 | 32.558 | 35.348 | 31.559 | 28.316 | 32.996 | 26.755 | 28.960 | 33.410 | 28.872 | 28.229 | 30.268 | 0.171 | 0.0000 | 0.0000 | 1 | 0 | 0 |
| 'Adgre5' | 10.372 | 8.809 | 9.929 | 9.467 | 8.380 | 10.849 | 10.282 | 10.092 | 10.427 | 9.931 | 9.472 | 11.276 | 10.159 | 9.978 | 11.457 | 10.414 | 9.881 | 10.847 | -0.085 | 0.0208 | 0.1056 | 0 | 0 | 0 |
| 'Adgrg5' | 0.759 | 0.695 | 0.931 | 0.448 | 0.493 | 0.579 | 0.527 | 0.499 | 0.607 | 0.579 | 0.760 | 0.711 | 0.621 | 0.632 | 0.813 | 0.568 | 0.811 | 0.767 | -0.177 | 0.1551 | 0.3835 | 0 | 0 | 0 |
| 'Adgrl1' | 6.024 | 5.153 | 7.243 | 6.027 | 5.067 | 6.466 | 5.408 | 5.507 | 7.259 | 5.706 | 5.229 | 6.165 | 5.698 | 5.233 | 5.798 | 4.842 | 5.022 | 6.076 | 0.107 | 0.0116 | 0.0695 | 0 | 0 | 0 |
| 'Adh5' | 9.352 | 10.409 | 9.780 | 10.002 | 10.267 | 9.854 | 10.592 | 10.175 | 9.741 | 9.725 | 9.320 | 9.620 | 9.968 | 8.756 | 9.583 | 9.533 | 9.095 | 9.226 | 0.074 | 0.0136 | 0.0781 | 0 | 0 | 0 |
| 'Adi1' | 21.372 | 21.212 | 22.510 | 22.809 | 23.310 | 22.311 | 21.868 | 22.408 | 22.700 | 22.318 | 21.340 | 22.822 | 19.276 | 19.994 | 22.033 | 21.105 | 20.532 | 21.285 | 0.061 | 0.0726 | 0.2384 | 0 | 0 | 0 |
| 'Adipor1' | 15.815 | 15.506 | 14.806 | 15.567 | 15.459 | 14.383 | 14.253 | 15.319 | 14.780 | 14.829 | 14.985 | 14.988 | 15.465 | 15.344 | 15.083 | 15.426 | 14.553 | 14.792 | -0.006 | 0.8075 | 0.9136 | 0 | 0 | 0 |
| 'Adipor2' | 25.880 | 27.913 | 24.999 | 25.967 | 27.363 | 25.758 | 25.329 | 27.504 | 25.860 | 26.566 | 27.846 | 26.651 | 26.513 | 27.742 | 26.897 | 27.782 | 26.883 | 27.050 | -0.056 | 0.0062 | 0.0439 | 1 | 0 | 0 |
| 'Adk' | 21.039 | 19.652 | 22.434 | 20.739 | 20.581 | 20.854 | 22.854 | 20.468 | 18.757 | 18.609 | 17.071 | 20.856 | 18.091 | 19.501 | 20.369 | 18.650 | 17.022 | 18.546 | 0.139 | 0.0012 | 0.0123 | 1 | 0 | 0 |
| 'Adnp' | 3.640 | 2.848 | 4.073 | 4.293 | 3.344 | 3.589 | 3.608 | 3.367 | 2.927 | 3.342 | 4.094 | 2.788 | 3.192 | 3.320 | 3.078 | 3.716 | 3.436 | 4.066 | 0.022 | 0.8019 | 0.9114 | 0 | 0 | 0 |
| 'Adnp2' | 6.849 | 7.889 | 7.714 | 6.549 | 7.744 | 7.965 | 6.055 | 8.283 | 7.832 | 8.371 | 7.132 | 7.088 | 6.947 | 7.272 | 6.522 | 8.590 | 6.633 | 6.588 | 0.030 | 0.6441 | 0.8256 | 0 | 0 | 0 |
| 'Ado' | 5.191 | 5.488 | 6.997 | 5.016 | 5.890 | 7.281 | 4.912 | 6.039 | 6.527 | 5.302 | 5.596 | 6.730 | 5.364 | 5.190 | 6.589 | 5.790 | 5.126 | 6.991 | 0.010 | 0.8232 | 0.9223 | 0 | 0 | 0 |
| 'Adora2a' | 36.637 | 36.007 | 34.444 | 35.462 | 34.200 | 38.735 | 37.257 | 33.702 | 36.610 | 32.982 | 32.592 | 36.193 | 30.387 | 31.871 | 36.180 | 34.556 | 32.208 | 35.337 | 0.085 | 0.0138 | 0.0792 | 0 | 0 | 0 |
| 'Adpgk' | 16.890 | 16.503 | 17.306 | 15.481 | 17.267 | 18.339 | 14.818 | 16.641 | 16.973 | 15.692 | 16.625 | 18.249 | 17.373 | 17.093 | 17.614 | 17.138 | 17.368 | 17.118 | -0.048 | 0.1921 | 0.4292 | 0 | 0 | 0 |
| 'Adprh' | 79.417 | 86.186 | 83.908 | 80.244 | 88.735 | 89.843 | 84.618 | 83.340 | 85.188 | 85.554 | 88.579 | 86.991 | 89.975 | 90.802 | 86.910 | 85.273 | 89.687 | 81.746 | -0.055 | 0.0334 | 0.1453 | 0 | 0 | 0 |
| 'Adprhl2' | 9.484 | 10.358 | 11.347 | 9.531 | 9.601 | 11.236 | 9.054 | 10.468 | 10.849 | 9.571 | 8.508 | 10.311 | 9.908 | 9.874 | 10.440 | 10.170 | 8.524 | 10.558 | 0.056 | 0.1536 | 0.3812 | 0 | 0 | 0 |
| 'Adprm' | 14.061 | 15.538 | 14.190 | 14.187 | 15.134 | 13.493 | 14.545 | 16.085 | 14.021 | 12.326 | 16.326 | 13.384 | 14.030 | 15.238 | 13.225 | 12.855 | 15.544 | 13.298 | 0.048 | 0.1451 | 0.3695 | 0 | 0 | 0 |
| 'Adra1b' | 1.543 | 1.376 | 1.607 | 1.305 | 1.403 | 1.440 | 1.560 | 0.880 | 1.545 | 0.855 | 0.905 | 1.032 | 0.858 | 0.548 | 0.822 | 1.035 | 0.815 | 0.958 | 0.677 | 0.0000 | 0.0000 | 1 | 1 | 0 |
| 'Adra2a' | 4.272 | 2.374 | 3.510 | 3.001 | 2.699 | 3.379 | 2.802 | 3.192 | 3.470 | 3.738 | 2.092 | 3.179 | 2.512 | 2.475 | 2.856 | 4.071 | 2.714 | 3.079 | 0.107 | 0.2688 | 0.5231 | 0 | 0 | 0 |
| 'Adrb2' | 6.542 | 3.182 | 1.919 | 4.495 | 3.077 | 2.444 | 5.037 | 3.494 | 1.786 | 5.023 | 3.890 | 2.270 | 5.750 | 4.420 | 2.089 | 5.322 | 4.484 | 1.851 | -0.141 | 0.1385 | 0.3592 | 0 | 0 | 0 |
| 'Adrbk1' | 76.978 | 73.607 | 79.600 | 75.943 | 75.083 | 81.781 | 76.529 | 77.041 | 77.930 | 77.133 | 76.033 | 82.578 | 83.641 | 79.848 | 81.790 | 76.894 | 77.194 | 85.685 | -0.064 | 0.0024 | 0.0208 | 1 | 0 | 0 |
| 'Adrm1' | 8.667 | 9.125 | 9.155 | 7.679 | 8.799 | 9.543 | 8.201 | 8.760 | 8.830 | 9.193 | 7.785 | 9.097 | 8.560 | 8.589 | 9.432 | 9.328 | 7.663 | 8.829 | -0.004 | 0.9178 | 0.9687 | 0 | 0 | 0 |
| 'Adsl' | 8.150 | 6.676 | 8.431 | 7.558 | 7.557 | 8.207 | 7.909 | 7.561 | 8.211 | 7.598 | 7.162 | 8.274 | 8.327 | 7.265 | 7.876 | 7.318 | 7.623 | 7.924 | 0.008 | 0.7975 | 0.9101 | 0 | 0 | 0 |
| 'Adss' | 13.904 | 14.466 | 14.230 | 13.447 | 15.193 | 14.398 | 14.783 | 15.215 | 13.655 | 13.688 | 14.266 | 14.148 | 13.491 | 14.679 | 12.602 | 14.464 | 14.304 | 12.852 | 0.045 | 0.1177 | 0.3264 | 0 | 0 | 0 |
| 'Adssl1' | 3.019 | 3.718 | 2.834 | 3.831 | 3.317 | 3.133 | 2.828 | 3.582 | 3.270 | 2.747 | 2.661 | 1.931 | 2.375 | 3.026 | 2.299 | 1.756 | 2.572 | 1.685 | 0.491 | 0.0000 | 0.0000 | 1 | 1 | 0 |
| 'Aebp2' | 21.957 | 24.212 | 22.792 | 22.255 | 23.744 | 21.959 | 22.668 | 23.420 | 21.857 | 22.001 | 22.541 | 24.247 | 22.703 | 23.639 | 24.202 | 23.308 | 22.451 | 23.530 | -0.038 | 0.1131 | 0.3192 | 0 | 0 | 0 |
| 'Aen' | 3.007 | 2.723 | 2.986 | 2.842 | 2.590 | 2.887 | 2.639 | 2.939 | 2.876 | 2.943 | 2.530 | 3.102 | 2.887 | 2.407 | 3.039 | 2.691 | 2.414 | 3.306 | 0.002 | 0.9716 | 0.9899 | 0 | 0 | 0 |
| 'Aes' | 381.051 | 359.746 | 414.840 | 380.759 | 359.462 | 460.659 | 389.997 | 379.120 | 395.201 | 397.631 | 334.955 | 363.485 | 383.892 | 346.026 | 423.602 | 402.053 | 334.116 | 422.233 | 0.038 | 0.3056 | 0.5634 | 0 | 0 | 0 |
| 'Afap1' | 0.897 | 1.043 | 1.039 | 0.995 | 0.822 | 1.003 | 0.834 | 1.158 | 1.033 | 0.969 | 0.925 | 0.892 | 0.997 | 1.048 | 0.876 | 0.909 | 0.823 | 0.844 | 0.087 | 0.1988 | 0.4387 | 0 | 0 | 0 |
| 'Aff1' | 16.628 | 16.060 | 15.208 | 15.654 | 15.357 | 16.015 | 14.524 | 16.063 | 16.601 | 16.179 | 17.340 | 17.679 | 16.809 | 15.404 | 16.577 | 14.986 | 14.958 | 18.275 | -0.071 | 0.0748 | 0.2434 | 0 | 0 | 0 |
| 'Aff3' | 12.471 | 10.771 | 12.517 | 13.209 | 11.409 | 12.238 | 14.074 | 11.187 | 11.978 | 12.767 | 13.021 | 14.413 | 11.931 | 12.020 | 13.384 | 10.571 | 13.510 | 14.713 | -0.094 | 0.0950 | 0.2862 | 0 | 0 | 0 |
| 'Aff4' | 9.997 | 8.480 | 8.417 | 8.769 | 8.337 | 7.755 | 7.990 | 8.584 | 7.942 | 9.273 | 8.871 | 7.899 | 8.535 | 8.237 | 7.669 | 8.723 | 7.845 | 8.195 | 0.010 | 0.7975 | 0.9101 | 0 | 0 | 0 |
| 'Afg3l1' | 11.020 | 11.271 | 11.170 | 10.714 | 11.574 | 11.207 | 11.406 | 11.667 | 11.859 | 11.517 | 11.071 | 11.107 | 11.766 | 9.953 | 10.096 | 10.937 | 10.766 | 10.058 | 0.057 | 0.0812 | 0.2571 | 0 | 0 | 0 |
| 'Afg3l2' | 38.552 | 36.854 | 36.835 | 35.643 | 37.576 | 36.694 | 38.477 | 36.838 | 37.247 | 37.279 | 35.991 | 35.964 | 37.958 | 36.493 | 35.013 | 38.615 | 35.311 | 34.969 | 0.023 | 0.3319 | 0.5898 | 0 | 0 | 0 |
| 'Afmid' | 1.998 | 2.160 | 2.543 | 2.101 | 2.169 | 2.500 | 2.032 | 2.413 | 2.185 | 2.673 | 2.274 | 2.224 | 2.415 | 2.284 | 2.231 | 2.497 | 1.883 | 1.985 | -0.025 | 0.7230 | 0.8727 | 0 | 0 | 0 |
| 'Aftph' | 12.214 | 14.412 | 11.881 | 11.923 | 12.797 | 11.814 | 11.490 | 13.168 | 11.295 | 13.174 | 13.790 | 11.935 | 12.035 | 13.257 | 11.571 | 13.479 | 13.334 | 11.919 | -0.055 | 0.0749 | 0.2436 | 0 | 0 | 0 |
| 'Aga' | 14.351 | 18.503 | 15.079 | 16.386 | 17.030 | 18.054 | 15.695 | 17.149 | 16.519 | 15.726 | 15.330 | 16.344 | 16.215 | 17.388 | 15.954 | 15.096 | 15.680 | 13.983 | 0.054 | 0.3009 | 0.5577 | 0 | 0 | 0 |
| 'Agap2' | 37.145 | 33.133 | 38.897 | 32.980 | 32.958 | 37.807 | 32.295 | 34.014 | 39.615 | 34.318 | 31.360 | 42.454 | 35.778 | 33.142 | 39.596 | 33.398 | 31.733 | 42.526 | -0.031 | 0.3434 | 0.5993 | 0 | 0 | 0 |
| 'Agap3' | 8.616 | 6.921 | 8.928 | 8.635 | 7.221 | 8.804 | 8.096 | 7.673 | 8.571 | 8.052 | 7.271 | 9.956 | 7.691 | 7.162 | 10.006 | 8.020 | 7.548 | 10.616 | -0.059 | 0.1816 | 0.4173 | 0 | 0 | 0 |
| 'Agbl5' | 2.352 | 2.180 | 2.544 | 2.273 | 2.242 | 2.595 | 2.185 | 2.278 | 2.742 | 2.440 | 2.058 | 2.552 | 2.694 | 2.212 | 2.591 | 2.408 | 2.187 | 2.296 | -0.012 | 0.7828 | 0.9025 | 0 | 0 | 0 |
| 'Ager' | 0.743 | 0.631 | 0.760 | 0.762 | 0.596 | 0.824 | 0.825 | 0.742 | 0.736 | 0.688 | 0.816 | 0.829 | 0.625 | 0.591 | 0.666 | 0.606 | 0.598 | 0.840 | 0.067 | 0.4914 | 0.7209 | 0 | 0 | 0 |
| 'Agfg1' | 31.182 | 28.488 | 30.558 | 28.139 | 28.127 | 27.747 | 26.314 | 30.846 | 26.466 | 28.320 | 31.127 | 29.735 | 25.823 | 30.261 | 29.490 | 27.599 | 29.140 | 29.859 | -0.028 | 0.4771 | 0.7107 | 0 | 0 | 0 |
| 'Agfg2' | 6.336 | 5.497 | 6.217 | 6.047 | 5.625 | 6.392 | 5.786 | 4.776 | 6.111 | 5.362 | 5.678 | 6.530 | 6.069 | 6.047 | 6.819 | 5.544 | 5.246 | 6.398 | -0.036 | 0.4332 | 0.6782 | 0 | 0 | 0 |
| 'Aggf1' | 15.975 | 16.507 | 15.943 | 13.901 | 17.923 | 16.801 | 14.144 | 17.461 | 16.871 | 16.175 | 15.277 | 16.777 | 16.668 | 17.043 | 16.517 | 17.543 | 15.648 | 16.578 | -0.039 | 0.3493 | 0.6043 | 0 | 0 | 0 |
| 'Agl' | 1.624 | 1.919 | 1.651 | 1.697 | 1.870 | 1.604 | 1.784 | 1.840 | 1.563 | 1.920 | 2.079 | 1.727 | 1.918 | 1.828 | 1.650 | 1.871 | 1.940 | 1.587 | -0.098 | 0.0070 | 0.0480 | 1 | 0 | 0 |
| 'Ago1' | 12.750 | 12.226 | 13.264 | 12.501 | 11.735 | 12.853 | 11.585 | 12.563 | 13.078 | 12.819 | 12.667 | 13.896 | 12.475 | 12.525 | 13.024 | 12.276 | 12.015 | 14.070 | -0.050 | 0.0677 | 0.2280 | 0 | 0 | 0 |
| 'Ago2' | 19.644 | 17.312 | 17.321 | 19.071 | 15.563 | 16.272 | 18.319 | 16.564 | 17.110 | 18.016 | 18.258 | 20.122 | 17.315 | 17.621 | 18.273 | 16.340 | 17.887 | 22.003 | -0.089 | 0.0967 | 0.2894 | 0 | 0 | 0 |
| 'Ago3' | 4.255 | 3.921 | 3.961 | 4.140 | 3.798 | 3.417 | 4.147 | 3.748 | 3.691 | 4.263 | 4.777 | 4.286 | 4.087 | 3.767 | 3.958 | 4.163 | 4.420 | 4.266 | -0.128 | 0.0061 | 0.0436 | 1 | 0 | 0 |
| 'Ago4' | 0.497 | 0.739 | 0.667 | 0.445 | 0.615 | 0.664 | 0.550 | 0.804 | 0.601 | 0.568 | 0.520 | 0.600 | 0.614 | 0.682 | 0.574 | 0.679 | 0.725 | 0.507 | 0.022 | 0.8343 | 0.9283 | 0 | 0 | 0 |
| 'Agpat1' | 11.563 | 10.713 | 12.233 | 11.744 | 10.943 | 13.390 | 10.599 | 10.028 | 12.431 | 11.458 | 10.719 | 14.656 | 11.510 | 11.040 | 14.178 | 12.007 | 11.094 | 14.522 | -0.110 | 0.0014 | 0.0140 | 1 | 0 | 0 |
| 'Agpat2' | 2.205 | 2.522 | 2.261 | 2.329 | 2.782 | 2.616 | 2.347 | 2.284 | 2.564 | 2.079 | 2.290 | 2.491 | 2.089 | 2.194 | 2.624 | 2.326 | 2.028 | 2.533 | 0.067 | 0.3482 | 0.6036 | 0 | 0 | 0 |
| 'Agpat3' | 30.565 | 32.404 | 31.185 | 30.717 | 32.226 | 31.228 | 29.672 | 32.132 | 31.591 | 29.126 | 32.080 | 33.043 | 32.041 | 31.827 | 33.408 | 30.579 | 30.834 | 35.039 | -0.042 | 0.1137 | 0.3198 | 0 | 0 | 0 |
| 'Agpat5' | 1.859 | 1.810 | 1.731 | 1.726 | 1.991 | 1.670 | 1.756 | 1.588 | 1.777 | 1.818 | 1.877 | 1.824 | 1.788 | 1.938 | 1.974 | 1.838 | 1.860 | 1.872 | -0.089 | 0.0692 | 0.2315 | 0 | 0 | 0 |
| 'Agps' | 8.051 | 8.220 | 8.380 | 8.448 | 8.664 | 8.064 | 8.385 | 8.536 | 7.563 | 8.489 | 8.706 | 8.700 | 7.928 | 8.829 | 7.795 | 8.239 | 8.947 | 7.832 | -0.033 | 0.2737 | 0.5290 | 0 | 0 | 0 |
| 'Agrn' | 5.154 | 4.814 | 5.903 | 4.710 | 5.192 | 5.219 | 4.783 | 5.287 | 5.912 | 5.046 | 4.937 | 5.550 | 5.297 | 4.753 | 4.970 | 4.409 | 4.766 | 5.503 | 0.043 | 0.2818 | 0.5381 | 0 | 0 | 0 |
| 'Agtpbp1' | 2.856 | 2.719 | 2.974 | 2.941 | 2.776 | 2.896 | 3.184 | 3.087 | 2.993 | 2.697 | 3.190 | 2.860 | 2.598 | 2.843 | 2.872 | 2.476 | 2.955 | 2.847 | 0.052 | 0.2289 | 0.4756 | 0 | 0 | 0 |
| 'Agtrap' | 8.189 | 9.190 | 9.837 | 8.238 | 10.057 | 9.666 | 8.785 | 9.428 | 9.862 | 8.700 | 9.010 | 9.296 | 8.840 | 10.042 | 9.461 | 9.944 | 8.729 | 9.146 | -0.007 | 0.8618 | 0.9401 | 0 | 0 | 0 |
| 'Ahctf1' | 6.937 | 6.723 | 6.676 | 7.513 | 6.481 | 5.980 | 6.886 | 6.716 | 6.433 | 6.830 | 6.871 | 6.747 | 6.330 | 6.600 | 6.492 | 7.002 | 6.818 | 6.543 | -0.009 | 0.7788 | 0.9003 | 0 | 0 | 0 |
| 'Ahcyl1' | 15.721 | 16.795 | 17.545 | 15.802 | 17.126 | 16.749 | 16.460 | 16.723 | 17.221 | 15.333 | 16.023 | 15.569 | 14.975 | 16.139 | 15.648 | 14.937 | 16.902 | 15.123 | 0.085 | 0.0002 | 0.0027 | 1 | 0 | 0 |
| 'Ahcyl2' | 2.740 | 2.339 | 2.699 | 2.690 | 2.805 | 2.604 | 2.613 | 2.771 | 2.547 | 2.735 | 2.659 | 2.871 | 2.452 | 2.523 | 2.588 | 2.464 | 2.917 | 2.684 | -0.016 | 0.7093 | 0.8653 | 0 | 0 | 0 |
| 'Ahdc1' | 3.882 | 3.060 | 3.272 | 4.038 | 2.835 | 3.194 | 3.496 | 2.923 | 3.437 | 3.298 | 2.687 | 4.089 | 3.272 | 2.989 | 3.766 | 3.317 | 3.119 | 4.429 | -0.048 | 0.4618 | 0.6980 | 0 | 0 | 0 |
| 'Ahi1' | 10.079 | 10.026 | 8.774 | 11.077 | 10.018 | 9.263 | 9.837 | 10.042 | 9.917 | 11.732 | 12.569 | 11.388 | 10.397 | 12.488 | 10.710 | 10.594 | 13.347 | 11.262 | -0.241 | 0.0000 | 0.0000 | 1 | 0 | 0 |
| 'Ahnak' | 8.829 | 6.049 | 8.432 | 7.808 | 6.539 | 8.215 | 7.338 | 7.151 | 7.901 | 5.865 | 6.846 | 8.858 | 5.073 | 7.242 | 8.044 | 6.710 | 6.742 | 8.863 | 0.087 | 0.2427 | 0.4934 | 0 | 0 | 0 |
| 'Ahr' | 1.729 | 1.026 | 1.923 | 1.333 | 1.524 | 1.428 | 1.534 | 1.703 | 1.437 | 0.963 | 0.965 | 1.288 | 0.817 | 0.971 | 1.142 | 1.012 | 0.816 | 1.010 | 0.597 | 0.0000 | 0.0000 | 1 | 1 | 0 |
| 'Ahsa1' | 96.474 | 90.395 | 94.005 | 103.483 | 91.978 | 96.432 | 91.934 | 89.697 | 93.259 | 85.595 | 88.492 | 122.348 | 96.866 | 95.745 | 110.154 | 92.137 | 101.882 | 126.320 | -0.121 | 0.0271 | 0.1268 | 0 | 0 | 0 |
| 'Ahsa2' | 6.494 | 5.497 | 5.808 | 6.739 | 5.116 | 5.233 | 6.028 | 5.390 | 5.222 | 4.767 | 5.474 | 8.413 | 5.202 | 6.044 | 7.215 | 5.079 | 6.581 | 8.705 | -0.149 | 0.1981 | 0.4380 | 0 | 0 | 0 |
| 'Aifm1' | 11.000 | 11.376 | 11.688 | 12.037 | 12.079 | 11.802 | 11.593 | 11.975 | 11.382 | 11.064 | 10.911 | 11.543 | 12.360 | 10.860 | 11.526 | 11.590 | 11.532 | 10.390 | 0.033 | 0.2904 | 0.5472 | 0 | 0 | 0 |
| 'Aifm2' | 0.718 | 0.890 | 0.982 | 0.886 | 0.826 | 0.915 | 0.896 | 0.771 | 0.866 | 0.858 | 0.783 | 0.630 | 0.743 | 0.628 | 0.581 | 0.481 | 0.803 | 0.556 | 0.352 | 0.0018 | 0.0168 | 1 | 0 | 0 |
| 'Aig1' | 1.069 | 0.915 | 0.983 | 1.075 | 0.958 | 0.995 | 1.105 | 1.013 | 0.833 | 0.953 | 0.939 | 0.995 | 0.947 | 0.805 | 0.849 | 0.825 | 0.813 | 0.925 | 0.131 | 0.0580 | 0.2076 | 0 | 0 | 0 |
| 'Aim1' | 2.310 | 2.177 | 2.410 | 2.313 | 2.153 | 2.392 | 2.365 | 2.310 | 2.508 | 2.301 | 2.088 | 2.543 | 2.601 | 2.011 | 2.261 | 2.343 | 2.059 | 2.288 | 0.017 | 0.7069 | 0.8642 | 0 | 0 | 0 |
| 'Aim1l' | 1.130 | 1.288 | 1.547 | 1.239 | 1.328 | 1.578 | 1.521 | 1.258 | 1.733 | 1.416 | 1.199 | 1.186 | 1.277 | 1.140 | 1.088 | 1.183 | 1.344 | 1.117 | 0.194 | 0.0107 | 0.0657 | 0 | 0 | 0 |
| 'Aim2' | 20.740 | 23.405 | 21.484 | 21.272 | 23.122 | 20.381 | 21.720 | 22.068 | 21.970 | 22.060 | 19.986 | 19.033 | 22.919 | 21.729 | 17.710 | 22.398 | 22.964 | 15.704 | 0.086 | 0.0606 | 0.2123 | 0 | 0 | 0 |
| 'Aimp1' | 7.567 | 7.178 | 7.347 | 7.422 | 8.029 | 7.378 | 7.533 | 7.991 | 7.323 | 7.422 | 7.549 | 7.408 | 7.516 | 7.224 | 7.032 | 7.552 | 7.355 | 6.743 | 0.033 | 0.3015 | 0.5584 | 0 | 0 | 0 |
| 'Aimp2' | 6.221 | 5.338 | 6.292 | 5.393 | 5.176 | 5.340 | 5.242 | 6.127 | 6.833 | 5.061 | 5.062 | 5.895 | 5.406 | 5.934 | 6.013 | 5.270 | 5.081 | 5.957 | 0.059 | 0.3416 | 0.5981 | 0 | 0 | 0 |
| 'Aip' | 43.083 | 44.821 | 47.594 | 44.448 | 44.231 | 46.857 | 45.116 | 45.600 | 46.991 | 39.889 | 41.262 | 42.483 | 44.779 | 41.602 | 45.038 | 42.776 | 42.004 | 42.192 | 0.088 | 0.0005 | 0.0055 | 1 | 0 | 0 |
| 'Ak2' | 54.781 | 52.696 | 59.888 | 53.503 | 55.479 | 53.680 | 55.170 | 55.759 | 55.280 | 49.261 | 50.163 | 52.243 | 51.832 | 51.285 | 54.671 | 52.437 | 49.845 | 52.042 | 0.088 | 0.0005 | 0.0060 | 1 | 0 | 0 |
| 'Ak3' | 15.875 | 18.211 | 16.200 | 15.883 | 16.847 | 15.492 | 15.821 | 16.436 | 15.793 | 15.185 | 15.827 | 16.830 | 14.446 | 15.993 | 16.247 | 12.857 | 16.031 | 16.384 | 0.053 | 0.1906 | 0.4271 | 0 | 0 | 0 |
| 'Ak4' | 2.631 | 2.414 | 1.364 | 2.533 | 1.787 | 1.608 | 2.784 | 1.770 | 1.654 | 1.261 | 3.534 | 3.666 | 2.277 | 3.311 | 4.246 | 2.345 | 3.693 | 3.862 | -0.598 | 0.0075 | 0.0505 | 0 | 0 | 0 |
| 'Akap1' | 1.437 | 1.092 | 1.497 | 1.307 | 1.242 | 1.264 | 1.361 | 1.049 | 1.279 | 1.029 | 1.042 | 1.179 | 1.210 | 0.815 | 1.196 | 1.075 | 1.269 | 1.158 | 0.197 | 0.0096 | 0.0608 | 0 | 0 | 0 |
| 'Akap10' | 8.060 | 7.493 | 7.655 | 8.266 | 7.708 | 6.863 | 8.112 | 7.993 | 7.322 | 7.703 | 7.769 | 7.902 | 7.778 | 7.743 | 7.117 | 6.635 | 8.151 | 6.911 | 0.023 | 0.5667 | 0.7748 | 0 | 0 | 0 |
| 'Akap11' | 10.533 | 12.673 | 11.535 | 11.286 | 12.198 | 11.568 | 10.591 | 11.454 | 11.136 | 11.751 | 12.269 | 11.978 | 10.863 | 11.832 | 11.519 | 12.852 | 12.155 | 11.391 | -0.061 | 0.0640 | 0.2205 | 0 | 0 | 0 |
| 'Akap12' | 16.036 | 16.068 | 15.904 | 16.923 | 15.221 | 16.677 | 16.516 | 16.420 | 17.188 | 16.208 | 16.553 | 18.718 | 16.481 | 15.933 | 18.636 | 16.242 | 16.105 | 18.571 | -0.074 | 0.0206 | 0.1053 | 0 | 0 | 0 |
| 'Akap13' | 41.677 | 38.900 | 38.103 | 39.526 | 37.821 | 37.355 | 36.887 | 38.774 | 39.393 | 42.003 | 42.104 | 37.818 | 40.306 | 40.330 | 37.170 | 40.997 | 38.878 | 39.033 | -0.051 | 0.0725 | 0.2384 | 0 | 0 | 0 |
| 'Akap5' | 0.416 | 0.513 | 0.490 | 0.565 | 0.483 | 0.538 | 0.472 | 0.479 | 0.542 | 0.607 | 0.580 | 0.745 | 0.554 | 0.494 | 0.653 | 0.467 | 0.867 | 0.723 | -0.355 | 0.0002 | 0.0031 | 1 | 0 | 0 |

| | | | | | | | | | | | | | | | | | | | | | | | | |
|---|---|---|---|---|---|---|---|---|---|---|---|---|---|---|---|---|---|---|---|---|---|---|---|---|
| 'Akap6' | 0.695 | 0.581 | 0.716 | 0.739 | 0.538 | 0.477 | 0.665 | 0.620 | 0.569 | 0.728 | 0.967 | 0.846 | 1.164 | 1.002 | 0.929 | 0.500 | 1.032 | 1.049 | -0.566 | 0.0000 | 0.0002 | 1 | 1 | 0 |
| 'Akap7' | 0.328 | 0.614 | 0.514 | 0.328 | 0.553 | 0.527 | 0.452 | 0.506 | 0.469 | 0.608 | 0.436 | 0.492 | 0.407 | 0.495 | 0.429 | 0.387 | 0.453 | 0.403 | 0.037 | 0.7558 | 0.8912 | 0 | 0 | 0 |
| 'Akap8' | 73.744 | 76.225 | 75.237 | 70.330 | 74.808 | 72.647 | 71.674 | 78.807 | 76.460 | 73.688 | 73.958 | 77.916 | 68.882 | 72.929 | 73.892 | 74.785 | 76.206 | 73.422 | -0.001 | 0.9473 | 0.9796 | 0 | 0 | 0 |
| 'Akap8l' | 103.065 | 90.554 | 105.749 | 94.664 | 89.027 | 100.548 | 94.616 | 91.408 | 103.553 | 100.710 | 87.040 | 109.444 | 96.834 | 89.742 | 98.637 | 96.362 | 97.006 | 100.834 | -0.016 | 0.5549 | 0.7675 | 0 | 0 | 0 |
| 'Akap9' | 6.589 | 5.675 | 5.687 | 5.878 | 5.518 | 5.312 | 5.662 | 5.503 | 5.522 | 6.165 | 6.547 | 5.837 | 5.560 | 5.476 | 5.138 | 5.924 | 5.966 | 6.072 | -0.048 | 0.2918 | 0.5486 | 0 | 0 | 0 |
| 'Akip1' | 5.963 | 6.673 | 7.011 | 6.613 | 6.654 | 7.363 | 6.702 | 6.840 | 6.127 | 5.922 | 5.751 | 5.492 | 7.500 | 6.497 | 6.542 | 7.252 | 5.191 | 5.598 | 0.104 | 0.1515 | 0.3786 | 0 | 0 | 0 |
| 'Akirin1' | 23.896 | 32.111 | 25.558 | 23.900 | 26.591 | 26.286 | 24.114 | 26.052 | 25.497 | 25.528 | 24.994 | 27.377 | 25.269 | 26.580 | 28.594 | 27.710 | 25.639 | 26.297 | -0.040 | 0.3775 | 0.6293 | 0 | 0 | 0 |
| 'Akirin2' | 34.054 | 32.325 | 32.714 | 32.508 | 32.208 | 34.453 | 35.791 | 35.487 | 27.653 | 30.332 | 36.943 | 34.164 | 30.980 | 36.398 | 33.235 | 32.064 | 37.448 | 36.316 | -0.059 | 0.2473 | 0.4979 | 0 | 0 | 0 |
| 'Akna' | 158.709 | 152.671 | 157.251 | 154.733 | 142.043 | 162.054 | 148.581 | 143.878 | 168.391 | 149.499 | 155.355 | 191.840 | 154.717 | 149.627 | 186.980 | 151.320 | 148.117 | 206.996 | -0.108 | 0.0077 | 0.0512 | 0 | 0 | 0 |
| 'Akr1a1' | 53.012 | 61.105 | 51.146 | 53.187 | 59.616 | 55.923 | 56.052 | 58.195 | 54.548 | 52.740 | 52.139 | 53.151 | 55.722 | 56.138 | 56.450 | 56.763 | 53.112 | 55.573 | 0.019 | 0.5468 | 0.7610 | 0 | 0 | 0 |
| 'Akr1b10' | 4.106 | 4.949 | 4.955 | 4.792 | 5.450 | 4.630 | 4.736 | 4.542 | 5.291 | 5.599 | 4.330 | 4.882 | 5.269 | 4.974 | 4.754 | 5.256 | 4.999 | 4.401 | -0.043 | 0.4433 | 0.6868 | 0 | 0 | 0 |
| 'Akr1b3' | 13.206 | 12.751 | 12.415 | 12.896 | 13.195 | 12.664 | 13.539 | 13.012 | 12.646 | 13.132 | 12.532 | 12.842 | 13.239 | 12.863 | 12.816 | 12.340 | 12.902 | 12.854 | -0.002 | 0.9142 | 0.9671 | 0 | 0 | 0 |
| 'Akr1c18' | 10.447 | 11.463 | 10.927 | 10.418 | 15.532 | 10.902 | 12.298 | 10.326 | 8.701 | 7.647 | 4.947 | 6.628 | 6.238 | 8.845 | 6.653 | 6.843 | 11.555 | 5.576 | 0.631 | 0.0000 | 0.0000 | 1 | 1 | 0 |
| 'Akr1e1' | 1.512 | 1.274 | 1.470 | 0.991 | 1.196 | 1.404 | 1.441 | 1.348 | 1.544 | 1.799 | 1.632 | 1.661 | 1.796 | 1.790 | 1.535 | 1.539 | 2.041 | 1.641 | -0.341 | 0.0001 | 0.0017 | 1 | 0 | 0 |
| 'Akr7a5' | 7.926 | 8.043 | 7.725 | 7.175 | 8.111 | 9.408 | 6.508 | 7.803 | 8.333 | 8.122 | 6.884 | 8.926 | 7.185 | 7.201 | 8.573 | 9.009 | 6.550 | 9.293 | -0.022 | 0.7395 | 0.8819 | 0 | 0 | 0 |
| 'Akt1' | 34.943 | 36.662 | 37.557 | 31.640 | 36.505 | 41.600 | 34.135 | 37.719 | 38.576 | 34.646 | 31.060 | 40.247 | 36.749 | 33.836 | 42.346 | 37.544 | 31.185 | 41.154 | -0.004 | 0.9173 | 0.9686 | 0 | 0 | 0 |
| 'Akt2' | 13.983 | 14.163 | 13.811 | 12.540 | 14.440 | 14.598 | 12.314 | 15.659 | 13.721 | 14.117 | 13.571 | 14.620 | 13.085 | 14.401 | 14.710 | 14.176 | 12.737 | 14.986 | -0.025 | 0.5218 | 0.7444 | 0 | 0 | 0 |
| 'Akt3' | 1.684 | 2.075 | 1.697 | 1.644 | 1.965 | 1.524 | 1.964 | 1.952 | 1.632 | 1.781 | 2.379 | 2.382 | 1.431 | 1.944 | 2.208 | 1.729 | 2.229 | 2.420 | -0.208 | 0.0117 | 0.0699 | 0 | 0 | 0 |
| 'Aktip' | 4.796 | 6.041 | 4.578 | 5.747 | 5.797 | 5.685 | 5.444 | 6.125 | 5.245 | 5.204 | 5.636 | 6.027 | 4.706 | 6.496 | 6.152 | 5.422 | 6.218 | 5.751 | -0.078 | 0.1847 | 0.4203 | 0 | 0 | 0 |
| 'Alad' | 45.221 | 52.150 | 44.026 | 44.947 | 51.746 | 48.180 | 44.807 | 46.971 | 47.102 | 45.291 | 45.984 | 40.664 | 55.103 | 48.513 | 41.976 | 48.657 | 47.032 | 38.294 | 0.041 | 0.3785 | 0.6301 | 0 | 0 | 0 |
| 'Alas1' | 6.130 | 6.078 | 7.101 | 6.371 | 6.713 | 6.795 | 6.553 | 6.315 | 6.500 | 6.058 | 6.256 | 6.243 | 6.418 | 5.976 | 6.581 | 6.158 | 5.757 | 6.120 | 0.068 | 0.0461 | 0.1789 | 0 | 0 | 0 |
| 'Alcam' | 2.647 | 2.511 | 2.606 | 2.322 | 2.432 | 2.275 | 2.470 | 2.413 | 2.340 | 2.811 | 3.030 | 3.043 | 2.821 | 2.944 | 2.918 | 2.606 | 3.353 | 2.788 | -0.268 | 0.0000 | 0.0000 | 1 | 0 | 0 |
| 'Aldh16a1' | 4.489 | 4.082 | 5.265 | 4.319 | 4.000 | 5.304 | 4.620 | 4.404 | 5.242 | 4.383 | 4.355 | 4.975 | 4.647 | 3.980 | 5.024 | 4.156 | 4.307 | 5.121 | 0.018 | 0.5941 | 0.7933 | 0 | 0 | 0 |
| 'Aldh18a1' | 10.143 | 9.197 | 10.663 | 9.317 | 9.584 | 10.015 | 9.418 | 9.644 | 10.448 | 10.517 | 9.874 | 10.492 | 10.115 | 9.032 | 10.442 | 9.588 | 9.256 | 10.455 | -0.030 | 0.3327 | 0.5908 | 0 | 0 | 0 |
| 'Aldh1b1' | 3.382 | 3.489 | 3.629 | 3.273 | 3.871 | 3.864 | 3.017 | 3.735 | 4.419 | 4.248 | 3.186 | 2.649 | 3.899 | 3.456 | 2.681 | 3.547 | 2.820 | 2.114 | 0.207 | 0.0532 | 0.1968 | 0 | 0 | 0 |
| 'Aldh2' | 2.438 | 2.028 | 2.497 | 2.385 | 2.065 | 2.656 | 2.890 | 2.222 | 2.589 | 1.731 | 1.870 | 2.083 | 1.845 | 1.673 | 2.319 | 1.959 | 1.753 | 1.905 | 0.335 | 0.0000 | 0.0000 | 1 | 0 | 0 |
| 'Aldh3a2' | 5.846 | 7.004 | 6.828 | 5.981 | 6.887 | 6.945 | 6.228 | 7.111 | 7.130 | 6.724 | 6.651 | 6.954 | 6.434 | 6.788 | 6.999 | 6.365 | 6.483 | 6.422 | -0.009 | 0.7891 | 0.9059 | 0 | 0 | 0 |
| 'Aldh4a1' | 4.449 | 5.244 | 4.307 | 4.164 | 4.866 | 4.491 | 4.071 | 4.649 | 4.679 | 5.217 | 5.107 | 4.819 | 5.906 | 5.538 | 4.336 | 5.016 | 5.175 | 4.615 | -0.170 | 0.0006 | 0.0067 | 1 | 0 | 0 |
| 'Aldh5a1' | 2.742 | 2.959 | 3.248 | 2.463 | 3.011 | 2.958 | 3.406 | 3.132 | 2.803 | 2.958 | 2.886 | 2.386 | 2.503 | 2.629 | 2.502 | 2.767 | 2.296 | 2.146 | 0.211 | 0.0022 | 0.0194 | 1 | 0 | 0 |
| 'Aldh6a1' | 4.254 | 5.050 | 4.525 | 4.099 | 5.434 | 4.622 | 4.657 | 5.634 | 5.115 | 5.605 | 5.304 | 4.268 | 4.705 | 4.972 | 4.369 | 5.574 | 4.424 | 4.036 | 0.002 | 0.9730 | 0.9906 | 0 | 0 | 0 |
| 'Aldh7a1' | 5.018 | 6.329 | 5.072 | 6.208 | 5.812 | 5.905 | 5.874 | 5.818 | 5.897 | 4.791 | 4.569 | 4.320 | 5.776 | 4.788 | 4.304 | 4.983 | 4.734 | 4.401 | 0.274 | 0.0000 | 0.0000 | 1 | 0 | 0 |
| 'Aldh9a1' | 15.209 | 15.116 | 15.071 | 14.830 | 15.909 | 14.922 | 14.880 | 15.547 | 14.890 | 16.416 | 15.034 | 15.339 | 15.960 | 14.998 | 15.450 | 14.381 | 14.375 | 15.425 | -0.022 | 0.4732 | 0.7088 | 0 | 0 | 0 |
| 'Aldoa' | 142.984 | 142.774 | 133.576 | 148.249 | 143.565 | 140.183 | 148.083 | 131.465 | 136.167 | 139.344 | 149.233 | 130.930 | 155.325 | 150.302 | 149.974 | 145.293 | 150.182 | 139.589 | -0.059 | 0.0674 | 0.2275 | 0 | 0 | 0 |
| 'Aldoc' | 3.896 | 3.967 | 3.398 | 3.903 | 4.549 | 3.273 | 4.935 | 3.498 | 3.372 | 4.004 | 3.864 | 3.507 | 4.907 | 4.369 | 4.002 | 4.441 | 3.914 | 3.407 | -0.082 | 0.1822 | 0.4183 | 0 | 0 | 0 |
| 'Alg1' | 6.063 | 6.611 | 6.344 | 5.562 | 6.699 | 6.846 | 5.537 | 6.731 | 6.606 | 7.251 | 5.868 | 6.749 | 6.530 | 5.736 | 6.565 | 7.296 | 6.046 | 6.046 | -0.036 | 0.5142 | 0.7381 | 0 | 0 | 0 |
| 'Alg10b' | 14.268 | 15.030 | 15.016 | 13.270 | 15.322 | 14.915 | 13.159 | 14.524 | 14.089 | 15.233 | 15.484 | 15.651 | 15.521 | 15.478 | 15.012 | 14.286 | 13.352 | 15.377 | -0.075 | 0.0451 | 0.1759 | 0 | 0 | 0 |
| 'Alg11' | 1.740 | 2.388 | 1.986 | 1.822 | 2.286 | 1.950 | 1.853 | 2.492 | 1.751 | 1.705 | 2.027 | 2.286 | 1.801 | 1.997 | 2.247 | 2.180 | 2.161 | 2.051 | -0.038 | 0.5709 | 0.7777 | 0 | 0 | 0 |
| 'Alg12' | 6.030 | 6.448 | 7.921 | 6.199 | 7.169 | 7.639 | 6.539 | 7.031 | 7.552 | 7.289 | 5.759 | 6.776 | 7.298 | 5.877 | 6.525 | 6.806 | 6.579 | 7.006 | 0.052 | 0.3200 | 0.5782 | 0 | 0 | 0 |
| 'Alg13' | 1.199 | 0.968 | 1.393 | 1.275 | 1.108 | 0.998 | 1.253 | 1.331 | 1.227 | 1.194 | 1.486 | 1.408 | 1.249 | 1.267 | 1.183 | 1.277 | 1.356 | 1.042 | -0.098 | 0.2060 | 0.4499 | 0 | 0 | 0 |
| 'Alg14' | 0.950 | 1.210 | 1.222 | 1.111 | 1.129 | 1.351 | 1.108 | 1.576 | 1.352 | 1.320 | 1.467 | 1.337 | 1.473 | 1.464 | 1.218 | 1.083 | 1.455 | 1.326 | -0.147 | 0.1689 | 0.4013 | 0 | 0 | 0 |
| 'Alg2' | 5.916 | 5.827 | 6.463 | 6.546 | 5.954 | 6.273 | 6.078 | 5.228 | 6.293 | 5.635 | 5.595 | 6.767 | 5.317 | 6.132 | 6.282 | 6.024 | 5.848 | 6.443 | 0.000 | 0.9968 | 0.9994 | 0 | 0 | 0 |
| 'Alg3' | 3.117 | 2.878 | 3.157 | 3.156 | 3.397 | 3.305 | 2.959 | 3.485 | 3.363 | 2.873 | 2.989 | 3.407 | 3.201 | 2.822 | 3.345 | 2.762 | 2.676 | 3.009 | 0.074 | 0.2125 | 0.4576 | 0 | 0 | 0 |
| 'Alg5' | 5.366 | 6.283 | 6.325 | 5.501 | 6.626 | 6.274 | 5.892 | 6.054 | 6.159 | 5.693 | 5.717 | 5.815 | 6.322 | 5.964 | 5.613 | 6.163 | 5.929 | 5.123 | 0.048 | 0.2836 | 0.5397 | 0 | 0 | 0 |
| 'Alg6' | 1.032 | 1.130 | 1.056 | 1.134 | 1.013 | 0.988 | 1.377 | 0.939 | 1.054 | 0.963 | 1.119 | 1.087 | 0.906 | 1.193 | 1.038 | 0.934 | 1.023 | 0.877 | 0.073 | 0.3555 | 0.6086 | 0 | 0 | 0 |
| 'Alg8' | 1.644 | 1.508 | 1.993 | 1.568 | 1.652 | 2.093 | 1.473 | 2.082 | 2.079 | 2.191 | 1.663 | 1.694 | 1.692 | 1.860 | 1.771 | 1.793 | 1.803 | 1.600 | 0.005 | 0.9536 | 0.9829 | 0 | 0 | 0 |
| 'Alg9' | 2.419 | 2.313 | 2.455 | 2.377 | 2.607 | 2.439 | 2.353 | 2.190 | 2.513 | 2.563 | 2.283 | 2.309 | 2.290 | 2.149 | 2.411 | 2.062 | 2.413 | 2.263 | 0.053 | 0.3137 | 0.5715 | 0 | 0 | 0 |
| 'Alkbh1' | 2.175 | 2.197 | 2.400 | 2.273 | 2.105 | 2.134 | 2.306 | 2.371 | 2.171 | 2.010 | 2.048 | 2.477 | 2.237 | 2.189 | 2.174 | 2.100 | 2.177 | 2.219 | 0.022 | 0.5973 | 0.7953 | 0 | 0 | 0 |
| 'Alkbh2' | 4.999 | 4.345 | 4.084 | 5.191 | 4.648 | 4.845 | 4.444 | 4.704 | 4.529 | 3.976 | 3.949 | 5.009 | 4.020 | 4.914 | 5.192 | 4.003 | 4.261 | 3.794 | 0.078 | 0.3438 | 0.5995 | 0 | 0 | 0 |
| 'Alkbh3' | 4.710 | 5.611 | 5.268 | 4.797 | 5.014 | 5.391 | 4.668 | 5.514 | 4.788 | 5.163 | 4.657 | 4.384 | 5.468 | 3.990 | 4.023 | 5.648 | 5.179 | 3.897 | 0.109 | 0.1010 | 0.2978 | 0 | 0 | 0 |
| 'Alkbh4' | 4.361 | 4.623 | 4.758 | 4.285 | 4.253 | 5.239 | 3.946 | 4.866 | 4.713 | 3.981 | 3.664 | 4.755 | 3.988 | 4.565 | 4.900 | 4.135 | 4.317 | 4.659 | 0.066 | 0.1662 | 0.3984 | 0 | 0 | 0 |
| 'Alkbh5' | 42.475 | 35.453 | 31.531 | 38.890 | 33.967 | 32.189 | 35.450 | 34.423 | 31.222 | 35.937 | 41.355 | 40.334 | 36.879 | 41.528 | 40.358 | 39.204 | 41.154 | 43.079 | -0.204 | 0.0002 | 0.0032 | 1 | 0 | 0 |
| 'Alkbh6' | 4.788 | 5.144 | 5.125 | 4.675 | 5.299 | 5.289 | 5.014 | 5.369 | 5.434 | 4.828 | 4.755 | 5.437 | 5.293 | 4.439 | 5.189 | 5.065 | 4.456 | 5.188 | 0.033 | 0.4190 | 0.6660 | 0 | 0 | 0 |
| 'Alkbh7' | 7.461 | 8.679 | 7.085 | 8.070 | 9.055 | 7.186 | 6.535 | 7.540 | 8.387 | 5.514 | 6.791 | 7.586 | 7.845 | 7.735 | 7.446 | 7.243 | 5.740 | 7.027 | 0.135 | 0.0766 | 0.2467 | 0 | 0 | 0 |
| 'Alkbh8' | 5.613 | 5.777 | 5.898 | 6.514 | 5.580 | 5.153 | 5.960 | 5.848 | 4.885 | 5.225 | 5.308 | 5.626 | 5.478 | 5.674 | 5.615 | 5.194 | 5.697 | 5.559 | 0.037 | 0.4505 | 0.6903 | 0 | 0 | 0 |
| 'Alox12' | 1.057 | 1.413 | 1.518 | 0.966 | 1.723 | 1.473 | 1.159 | 1.391 | 1.187 | 1.378 | 1.978 | 1.391 | 0.971 | 1.945 | 1.558 | 1.278 | 1.782 | 1.382 | -0.192 | 0.0395 | 0.1621 | 0 | 0 | 0 |
| 'Alpk2' | 0.877 | 0.857 | 0.711 | 0.821 | 0.743 | 0.734 | 0.854 | 0.859 | 0.650 | 0.682 | 0.717 | 0.692 | 0.515 | 0.652 | 0.601 | 0.453 | 0.864 | 0.654 | 0.267 | 0.0070 | 0.0480 | 1 | 0 | 0 |
| 'Alpl' | 2.969 | 1.823 | 3.160 | 3.132 | 2.006 | 2.830 | 3.182 | 2.361 | 2.789 | 2.271 | 2.363 | 3.060 | 2.895 | 2.161 | 2.894 | 2.867 | 1.920 | 2.932 | 0.041 | 0.5516 | 0.7649 | 0 | 0 | 0 |
| 'Als2' | 8.664 | 7.557 | 8.914 | 9.151 | 8.697 | 8.307 | 8.660 | 8.118 | 8.551 | 7.025 | 7.425 | 7.306 | 6.870 | 7.540 | 7.129 | 6.541 | 7.714 | 7.428 | 0.228 | 0.0000 | 0.0000 | 1 | 0 | 0 |
| 'Als2cl' | 12.244 | 11.493 | 13.674 | 13.646 | 11.032 | 12.911 | 13.434 | 11.239 | 13.542 | 11.276 | 10.299 | 12.320 | 12.801 | 10.495 | 11.364 | 12.742 | 11.564 | 11.672 | 0.103 | 0.0021 | 0.0189 | 1 | 0 | 0 |
| 'Alyref' | 22.497 | 19.682 | 21.283 | 22.725 | 19.920 | 22.136 | 21.984 | 22.528 | 18.353 | 21.346 | 21.717 | 17.393 | 23.070 | 23.488 | 20.077 | 20.834 | 20.979 | 20.643 | 0.005 | 0.9262 | 0.9719 | 0 | 0 | 0 |
| 'Alyref2' | 6.473 | 6.661 | 8.890 | 6.133 | 7.726 | 8.966 | 8.193 | 8.392 | 9.086 | 7.730 | 8.542 | 8.985 | 6.564 | 7.972 | 6.594 | 7.589 | 7.123 | 6.818 | 0.045 | 0.5915 | 0.7921 | 0 | 0 | 0 |
| 'Amacr' | 1.576 | 1.950 | 1.740 | 1.600 | 1.449 | 1.790 | 1.666 | 1.801 | 1.464 | 1.464 | 1.988 | 1.852 | 2.013 | 1.941 | 1.584 | 1.547 | 1.272 | 1.907 | -0.067 | 0.5322 | 0.7526 | 0 | 0 | 0 |
| 'Ambra1' | 10.819 | 10.527 | 11.302 | 10.606 | 9.911 | 10.752 | 10.499 | 10.232 | 11.426 | 11.331 | 10.402 | 11.401 | 9.965 | 9.838 | 10.998 | 10.198 | 10.459 | 11.291 | -0.008 | 0.7962 | 0.9092 | 0 | 0 | 0 |
| 'Amdhd2' | 4.169 | 4.693 | 5.055 | 4.639 | 5.131 | 5.664 | 4.671 | 4.782 | 5.381 | 4.982 | 4.933 | 4.842 | 5.221 | 4.604 | 4.651 | 5.422 | 4.972 | 4.279 | 0.003 | 0.9640 | 0.9869 | 0 | 0 | 0 |
| 'Amer1' | 1.080 | 1.322 | 1.117 | 0.963 | 1.530 | 1.446 | 1.156 | 1.410 | 1.443 | 1.278 | 1.317 | 1.605 | 1.215 | 1.420 | 1.476 | 1.480 | 1.183 | 1.600 | -0.152 | 0.0510 | 0.1912 | 0 | 0 | 0 |
| 'Amfr' | 41.842 | 44.782 | 44.046 | 39.564 | 44.816 | 46.960 | 41.547 | 46.762 | 42.610 | 42.968 | 40.148 | 44.128 | 42.882 | 43.086 | 42.192 | 43.165 | 38.231 | 43.431 | 0.036 | 0.2648 | 0.5184 | 0 | 0 | 0 |
| 'Amigo1' | 0.821 | 1.134 | 0.848 | 0.941 | 0.831 | 0.837 | 0.719 | 0.918 | 1.046 | 0.739 | 0.846 | 0.936 | 0.814 | 1.018 | 0.813 | 0.846 | 0.823 | 1.036 | 0.024 | 0.7617 | 0.8943 | 0 | 0 | 0 |
| 'Amigo3' | 6.020 | 4.936 | 5.369 | 4.196 | 4.570 | 4.123 | 3.707 | 5.364 | 4.495 | 6.064 | 5.077 | 4.504 | 5.279 | 4.219 | 4.088 | 4.609 | 4.193 | 4.232 | 0.018 | 0.8455 | 0.9343 | 0 | 0 | 0 |
| 'Ammecr1' | 5.976 | 5.371 | 5.840 | 6.311 | 5.545 | 5.894 | 5.570 | 5.983 | 5.363 | 6.060 | 5.535 | 5.632 | 6.146 | 6.113 | 5.782 | 5.400 | 6.670 | 6.114 | -0.052 | 0.2813 | 0.5379 | 0 | 0 | 0 |
| 'Ammecr1l' | 17.367 | 17.820 | 16.820 | 18.044 | 18.025 | 16.889 | 16.538 | 16.848 | 16.980 | 17.331 | 17.800 | 20.314 | 16.824 | 17.801 | 18.566 | 17.530 | 17.936 | 19.996 | -0.092 | 0.0137 | 0.0785 | 0 | 0 | 0 |
| 'Amn1' | 0.260 | 0.429 | 0.398 | 0.299 | 0.449 | 0.385 | 0.327 | 0.467 | 0.391 | 0.394 | 0.434 | 0.330 | 0.371 | 0.387 | 0.336 | 0.391 | 0.384 | 0.279 | 0.035 | 0.6937 | 0.8558 | 0 | 0 | 0 |
| 'Ampd2' | 5.941 | 5.298 | 6.693 | 5.770 | 5.582 | 5.897 | 6.019 | 5.251 | 6.426 | 5.867 | 5.071 | 6.411 | 5.440 | 5.242 | 5.924 | 5.196 | 5.822 | 5.784 | 0.047 | 0.2133 | 0.4581 | 0 | 0 | 0 |
| 'Ampd3' | 2.697 | 3.352 | 3.531 | 2.874 | 3.304 | 3.083 | 2.587 | 3.059 | 3.506 | 3.071 | 3.047 | 3.127 | 2.751 | 3.258 | 2.936 | 2.640 | 2.690 | 2.976 | 0.067 | 0.1928 | 0.4302 | 0 | 0 | 0 |
| 'Amz2' | 6.042 | 7.180 | 5.566 | 6.060 | 6.882 | 6.524 | 6.223 | 6.424 | 6.317 | 6.323 | 6.180 | 5.972 | 7.164 | 6.850 | 6.903 | 6.595 | 6.900 | 5.734 | -0.047 | 0.3031 | 0.5606 | 0 | 0 | 0 |
| 'Anapc1' | 9.957 | 10.336 | 10.893 | 10.235 | 9.957 | 10.559 | 9.722 | 10.290 | 10.416 | 10.495 | 11.312 | 10.746 | 9.893 | 10.503 | 10.411 | 10.165 | 10.992 | 9.988 | -0.042 | 0.1207 | 0.3311 | 0 | 0 | 0 |
| 'Anapc10' | 16.056 | 17.553 | 16.682 | 18.104 | 19.686 | 17.159 | 21.218 | 18.727 | 15.606 | 18.813 | 17.420 | 15.968 | 15.042 | 15.896 | 15.834 | 16.951 | 19.471 | 14.272 | 0.091 | 0.1563 | 0.3855 | 0 | 0 | 0 |
| 'Anapc11' | 6.234 | 8.191 | 6.647 | 5.379 | 7.771 | 7.189 | 6.238 | 7.449 | 7.172 | 6.042 | 5.857 | 6.392 | 6.356 | 6.965 | 6.400 | 6.449 | 6.663 | 6.034 | 0.107 | 0.0276 | 0.1282 | 0 | 0 | 0 |
| 'Anapc13' | 10.423 | 10.891 | 10.539 | 12.224 | 11.989 | 10.943 | 12.075 | 11.750 | 9.961 | 9.394 | 11.010 | 9.646 | 11.353 | 11.153 | 10.247 | 10.504 | 10.364 | 9.586 | 0.101 | 0.0262 | 0.1243 | 0 | 0 | 0 |
| 'Anapc15' | 1.426 | 1.605 | 1.599 | 1.610 | 1.757 | 1.580 | 1.509 | 1.408 | 1.654 | 1.539 | 1.268 | 1.532 | 1.473 | 1.607 | 1.457 | 1.492 | 1.669 | 1.393 | 0.061 | 0.3182 | 0.5767 | 0 | 0 | 0 |
| 'Anapc16' | 5.951 | 7.698 | 6.223 | 6.344 | 7.162 | 6.185 | 6.232 | 6.925 | 6.201 | 6.600 | 6.853 | 6.530 | 6.066 | 6.477 | 6.868 | 6.911 | 6.347 | 6.347 | -0.018 | 0.6628 | 0.8371 | 0 | 0 | 0 |
| 'Anapc2' | 25.250 | 25.572 | 25.920 | 24.367 | 24.619 | 26.224 | 25.281 | 25.292 | 27.350 | 25.525 | 24.446 | 24.790 | 25.902 | 24.534 | 24.092 | 24.621 | 24.953 | 24.383 | 0.033 | 0.1835 | 0.4192 | 0 | 0 | 0 |
| 'Anapc4' | 7.645 | 8.018 | 8.126 | 7.342 | 8.946 | 7.606 | 8.055 | 7.956 | 7.998 | 7.872 | 8.046 | 7.536 | 8.058 | 7.393 | 7.130 | 7.577 | 8.501 | 6.913 | 0.045 | 0.2124 | 0.4575 | 0 | 0 | 0 |
| 'Anapc5' | 15.880 | 16.126 | 17.260 | 16.134 | 15.777 | 16.873 | 16.661 | 17.042 | 17.620 | 15.170 | 15.966 | 16.300 | 17.620 | 16.294 | 16.398 | 16.258 | 16.147 | 15.792 | 0.023 | 0.3889 | 0.6384 | 0 | 0 | 0 |
| 'Anapc7' | 9.335 | 8.742 | 10.031 | 9.592 | 9.360 | 10.032 | 9.219 | 9.838 | 9.824 | 9.183 | 8.852 | 9.872 | 9.278 | 9.325 | 9.396 | 8.964 | 8.715 | 9.218 | 0.045 | 0.1262 | 0.3401 | 0 | 0 | 0 |
| 'Angel1' | 3.226 | 3.502 | 3.827 | 2.881 | 3.501 | 3.483 | 2.821 | 3.718 | 3.954 | 3.679 | 3.331 | 3.803 | 3.290 | 3.231 | 3.358 | 3.828 | 2.821 | 3.359 | 0.000 | 0.9943 | 0.9990 | 0 | 0 | 0 |
| 'Angel2' | 9.145 | 11.229 | 10.381 | 9.988 | 10.965 | 10.670 | 9.485 | 11.370 | 10.467 | 10.517 | 10.939 | 10.177 | 10.262 | 11.150 | 10.440 | 10.945 | 11.063 | 9.570 | -0.031 | 0.3142 | 0.5719 | 0 | 0 | 0 |
| 'Angptl2' | 30.157 | 27.480 | 31.847 | 24.923 | 31.097 | 30.987 | 23.673 | 32.485 | 29.196 | 37.787 | 39.187 | 37.417 | 31.466 | 38.557 | 38.061 | 35.532 | 35.946 | 37.581 | -0.353 | 0.0000 | 0.0000 | 1 | 0 | 0 |
| 'Ank' | 7.815 | 8.440 | 9.010 | 7.059 | 9.076 | 9.708 | 7.443 | 9.217 | 7.565 | 9.060 | 8.603 | 8.014 | 8.322 | 9.208 | 8.174 | 8.595 | 8.513 | 8.614 | -0.041 | 0.4435 | 0.6869 | 0 | 0 | 0 |
| 'Ankfy1' | 10.113 | 9.143 | 9.610 | 9.891 | 9.397 | 9.047 | 9.749 | 9.486 | 9.129 | 10.681 | 10.854 | 10.461 | 10.171 | 10.078 | 10.134 | 10.174 | 10.961 | 10.659 | -0.149 | 0.0000 | 0.0000 | 1 | 0 | 0 |
| 'Ankhd1' | 17.866 | 15.304 | 16.888 | 18.075 | 15.632 | 15.953 | 16.243 | 15.867 | 16.991 | 16.924 | 17.934 | 18.227 | 16.128 | 16.964 | 16.711 | 16.763 | 17.468 | 18.227 | -0.073 | 0.0546 | 0.2001 | 0 | 0 | 0 |
| 'Ankib1' | 10.494 | 10.375 | 9.963 | 10.346 | 9.392 | 10.080 | 9.945 | 10.493 | 10.033 | 9.621 | 10.314 | 11.176 | 9.027 | 9.927 | 10.177 | 10.697 | 10.347 | 11.271 | -0.033 | 0.4181 | 0.6655 | 0 | 0 | 0 |
| 'Ankle1' | 1.744 | 1.962 | 2.735 | 1.894 | 2.188 | 2.840 | 2.681 | 1.988 | 3.142 | 1.470 | 2.026 | 2.571 | 2.935 | 2.444 | 2.940 | 2.126 | 2.277 | 2.424 | -0.026 | 0.8005 | 0.9111 | 0 | 0 | 0 |
| 'Ankle2' | 8.609 | 8.910 | 9.244 | 8.994 | 8.906 | 8.876 | 9.018 | 9.112 | 8.719 | 8.677 | 8.922 | 9.364 | 7.864 | 8.679 | 9.386 | 8.309 | 9.178 | 9.285 | 0.002 | 0.9359 | 0.9755 | 0 | 0 | 0 |
| 'Ankmy2' | 25.506 | 26.772 | 24.157 | 25.385 | 24.716 | 26.822 | 24.331 | 27.619 | 27.164 | 25.902 | 26.029 | 26.686 | 24.070 | 24.957 | 26.513 | 27.052 | 25.666 | 25.565 | -0.009 | 0.7878 | 0.9048 | 0 | 0 | 0 |
| 'Ankra2' | 4.307 | 6.239 | 5.391 | 4.478 | 6.248 | 5.844 | 4.934 | 5.803 | 5.499 | 5.387 | 5.167 | 5.115 | 5.039 | 5.798 | 5.331 | 5.655 | 5.715 | 4.776 | 0.008 | 0.8792 | 0.9490 | 0 | 0 | 0 |
| 'Ankrd10' | 30.312 | 29.842 | 29.583 | 29.650 | 28.866 | 29.513 | 30.020 | 28.503 | 29.744 | 28.347 | 30.220 | 34.397 | 26.422 | 29.642 | 31.959 | 27.664 | 28.820 | 32.216 | -0.029 | 0.4367 | 0.6801 | 0 | 0 | 0 |
| 'Ankrd12' | 21.516 | 23.384 | 18.562 | 22.780 | 22.040 | 19.547 | 21.562 | 22.670 | 18.945 | 22.062 | 24.448 | 21.860 | 22.781 | 24.219 | 20.261 | 22.223 | 25.879 | 21.502 | -0.116 | 0.0000 | 0.0006 | 1 | 0 | 0 |
| 'Ankrd13a' | 25.188 | 23.380 | 26.667 | 23.531 | 24.908 | 28.903 | 23.269 | 25.910 | 25.186 | 27.273 | 24.206 | 26.511 | 24.283 | 25.352 | 27.428 | 27.161 | 23.023 | 28.031 | -0.050 | 0.1829 | 0.4190 | 0 | 0 | 0 |
| 'Ankrd13b' | 3.728 | 2.900 | 3.496 | 3.230 | 2.589 | 3.204 | 3.017 | 2.828 | 3.585 | 2.742 | 2.334 | 3.424 | 2.631 | 2.122 | 3.262 | 2.345 | 2.408 | 3.384 | 0.208 | 0.0005 | 0.0061 | 1 | 0 | 0 |
| 'Ankrd13c' | 5.419 | 5.241 | 4.796 | 5.251 | 5.417 | 4.777 | 4.998 | 5.244 | 4.725 | 4.964 | 5.210 | 5.218 | 4.909 | 5.301 | 5.071 | 5.092 | 5.283 | 5.552 | -0.036 | 0.3157 | 0.5735 | 0 | 0 | 0 |
| 'Ankrd13d' | 2.133 | 1.586 | 2.016 | 2.414 | 1.759 | 1.779 | 2.050 | 1.837 | 1.730 | 1.619 | 1.579 | 2.237 | 1.903 | 1.603 | 1.994 | 1.622 | 1.842 | 2.237 | 0.037 | 0.6498 | 0.8294 | 0 | 0 | 0 |
| 'Ankrd16' | 5.452 | 5.838 | 6.130 | 4.711 | 5.738 | 5.840 | 5.291 | 6.056 | 6.805 | 5.387 | 5.734 | 5.602 | 6.235 | 5.599 | 5.575 | 5.526 | 5.488 | 5.585 | 0.019 | 0.6801 | 0.8471 | 0 | 0 | 0 |
| 'Ankrd17' | 7.387 | 6.594 | 6.305 | 7.420 | 6.333 | 6.205 | 6.912 | 6.644 | 6.290 | 6.879 | 7.090 | 7.111 | 6.595 | 6.664 | 6.739 | 6.364 | 7.009 | 7.243 | -0.051 | 0.2105 | 0.4559 | 0 | 0 | 0 |
| 'Ankrd26' | 5.751 | 4.731 | 5.337 | 5.841 | 5.149 | 5.297 | 5.910 | 5.099 | 5.689 | 4.766 | 5.219 | 5.872 | 5.473 | 4.862 | 5.590 | 5.422 | 5.566 | 5.390 | 0.005 | 0.9120 | 0.9662 | 0 | 0 | 0 |
| 'Ankrd28' | 7.300 | 8.702 | 8.078 | 6.797 | 8.781 | 8.139 | 7.431 | 8.510 | 7.942 | 8.546 | 8.645 | 7.488 | 8.343 | 9.011 | 7.470 | 7.662 | 8.013 | 6.823 | -0.015 | 0.7299 | 0.8766 | 0 | 0 | 0 |
| 'Ankrd37' | 3.883 | 3.776 | 3.131 | 5.126 | 3.024 | 2.927 | 4.238 | 3.304 | 2.835 | 2.537 | 4.919 | 3.967 | 3.556 | 5.888 | 4.379 | 2.886 | 6.626 | 3.280 | -0.221 | 0.2013 | 0.4426 | 0 | 0 | 0 |
| 'Ankrd39' | 24.736 | 21.637 | 23.262 | 21.173 | 20.639 | 24.343 | 17.940 | 20.966 | 24.177 | 21.514 | 18.901 | 24.228 | 19.597 | 20.560 | 24.163 | 23.712 | 18.532 | 23.115 | 0.029 | 0.6712 | 0.8415 | 0 | 0 | 0 |
| 'Ankrd40' | 28.024 | 31.295 | 27.610 | 27.762 | 29.063 | 27.427 | 28.836 | 29.300 | 28.443 | 26.677 | 28.700 | 28.657 | 28.412 | 30.313 | 26.291 | 28.810 | 29.666 | 26.886 | 0.007 | 0.7918 | 0.9070 | 0 | 0 | 0 |
| 'Ankrd44' | 13.360 | 12.924 | 13.203 | 13.932 | 12.756 | 13.296 | 13.617 | 12.795 | 13.190 | 10.960 | 12.872 | 12.275 | 10.761 | 11.796 | 11.784 | 11.313 | 11.517 | 11.943 | 0.168 | 0.0000 | 0.0000 | 1 | 0 | 0 |
| 'Ankrd46' | 51.862 | 62.479 | 54.882 | 49.202 | 58.896 | 52.934 | 48.420 | 59.719 | 50.932 | 51.514 | 58.413 | 54.112 | 50.104 | 56.715 | 52.696 | 52.793 | 58.175 | 52.516 | -0.005 | 0.8309 | 0.9266 | 0 | 0 | 0 |
| 'Ankrd49' | 4.863 | 6.982 | 6.658 | 4.744 | 6.998 | 6.703 | 5.814 | 6.335 | 5.983 | 5.878 | 5.614 | 5.728 | 5.043 | 6.330 | 5.046 | 6.206 | 5.924 | 5.305 | 0.099 | 0.1753 | 0.4100 | 0 | 0 | 0 |
| 'Ankrd52' | 16.161 | 15.708 | 19.004 | 15.886 | 15.783 | 17.405 | 15.298 | 15.838 | 19.112 | 17.345 | 15.702 | 17.390 | 17.985 | 15.812 | 18.183 | 17.165 | 14.594 | 17.964 | -0.029 | 0.3839 | 0.6343 | 0 | 0 | 0 |
| 'Ankrd54' | 5.553 | 5.765 | 6.151 | 4.986 | 5.709 | 6.283 | 5.007 | 5.367 | 6.116 | 5.452 | 4.800 | 6.785 | 5.852 | 5.451 | 6.471 | 5.480 | 5.274 | 6.820 | -0.052 | 0.1985 | 0.4383 | 0 | 0 | 0 |
| 'Ankrd55' | 2.069 | 3.042 | 1.950 | 1.841 | 2.768 | 2.200 | 2.139 | 2.739 | 2.289 | 2.520 | 2.015 | 2.172 | 2.634 | 2.465 | 1.826 | 2.659 | 2.703 | 1.722 | 0.006 | 0.9390 | 0.9765 | 0 | 0 | 0 |
| 'Ankrd6' | 0.575 | 0.768 | 0.510 | 0.527 | 0.630 | 0.644 | 0.702 | 0.670 | 0.583 | 0.888 | 0.731 | 0.782 | 0.802 | 0.996 | 0.815 | 0.741 | 0.849 | 0.767 | -0.414 | 0.0000 | 0.0001 | 1 | 1 | 0 |
| 'Ankrd9' | 1.318 | 0.731 | 0.934 | 0.912 | 0.926 | 0.861 | 1.083 | 1.044 | 0.913 | 1.062 | 0.951 | 0.922 | 1.285 | 0.978 | 0.806 | 1.036 | 0.928 | 0.749 | 0.001 | 0.9937 | 0.9990 | 0 | 0 | 0 |
| 'Anks1' | 4.416 | 3.661 | 3.984 | 4.183 | 3.961 | 4.268 | 3.536 | 3.986 | 3.852 | 4.404 | 3.877 | 4.296 | 4.039 | 3.808 | 3.650 | 3.586 | 3.854 | 4.198 | -0.005 | 0.9165 | 0.9681 | 0 | 0 | 0 |
| 'Anks3' | 24.241 | 25.435 | 30.346 | 25.016 | 24.876 | 29.973 | 24.536 | 24.601 | 28.079 | 24.573 | 25.303 | 30.077 | 28.549 | 25.245 | 26.166 | 25.115 | 27.537 | 27.120 | -0.029 | 0.4551 | 0.6934 | 0 | 0 | 0 |
| 'Ankzf1' | 4.865 | 4.985 | 5.110 | 4.942 | 4.631 | 5.195 | 5.273 | 4.859 | 5.255 | 4.851 | 5.244 | 6.006 | 5.385 | 5.085 | 5.561 | 5.072 | 5.974 | 5.332 | -0.117 | 0.0022 | 0.0198 | 1 | 0 | 0 |
| 'Anln' | 1.508 | 2.287 | 2.759 | 2.077 | 2.170 | 2.561 | 2.545 | 2.138 | 3.070 | 1.627 | 1.641 | 2.313 | 2.305 | 2.494 | 2.681 | 2.049 | 2.301 | 2.400 | 0.073 | 0.4279 | 0.6739 | 0 | 0 | 0 |
| 'Ano10' | 12.762 | 13.825 | 13.333 | 13.026 | 12.808 | 15.315 | 14.182 | 14.874 | 15.104 | 13.474 | 13.511 | 13.542 | 12.304 | 13.515 | 12.863 | 12.822 | 13.945 | 13.508 | 0.060 | 0.1634 | 0.3954 | 0 | 0 | 0 |
| 'Ano6' | 13.727 | 13.313 | 14.882 | 14.380 | 13.034 | 14.682 | 13.625 | 14.437 | 14.510 | 14.104 | 14.421 | 14.146 | 13.672 | 13.762 | 13.920 | 13.490 | 13.308 | 13.775 | 0.015 | 0.5949 | 0.7937 | 0 | 0 | 0 |
| 'Ano8' | 14.084 | 12.028 | 16.916 | 13.540 | 13.063 | 15.121 | 13.583 | 13.326 | 14.731 | 13.839 | 14.188 | 15.038 | 14.061 | 13.519 | 13.967 | 13.290 | 12.804 | 14.388 | 0.007 | 0.8703 | 0.9446 | 0 | 0 | 0 |
| 'Anp32a' | 80.394 | 83.330 | 79.844 | 89.135 | 86.085 | 82.774 | 87.297 | 82.874 | 80.504 | 78.040 | 76.894 | 74.845 | 82.311 | 80.992 | 82.230 | 81.386 | 85.534 | 75.327 | 0.057 | 0.0389 | 0.1608 | 0 | 0 | 0 |
| 'Anp32b' | 241.616 | 249.583 | 242.203 | 258.101 | 253.765 | 255.990 | 259.410 | 247.359 | 250.252 | 229.470 | 235.505 | 234.879 | 256.700 | 249.536 | 254.580 | 245.277 | 248.329 | 238.196 | 0.032 | 0.2173 | 0.4639 | 0 | 0 | 0 |
| 'Anp32e' | 25.655 | 29.141 | 28.396 | 29.268 | 29.475 | 28.147 | 29.880 | 28.744 | 28.122 | 25.358 | 27.746 | 27.841 | 27.431 | 29.927 | 29.613 | 26.739 | 29.584 | 26.954 | 0.021 | 0.4882 | 0.7184 | 0 | 0 | 0 |
| 'Antxr2' | 10.430 | 11.703 | 11.332 | 10.238 | 11.694 | 11.632 | 11.010 | 11.669 | 11.026 | 12.034 | 11.034 | 10.358 | 11.169 | 11.432 | 9.761 | 10.887 | 10.060 | 10.113 | 0.048 | 0.2153 | 0.4611 | 0 | 0 | 0 |
| 'Anxa11' | 20.563 | 22.861 | 21.435 | 19.818 | 21.270 | 21.825 | 20.813 | 23.136 | 21.924 | 20.124 | 19.558 | 18.662 | 20.887 | 18.994 | 18.399 | 18.628 | 19.060 | 18.243 | 0.156 | 0.0000 | 0.0000 | 1 | 0 | 0 |
| 'Anxa2' | 10.528 | 12.080 | 10.215 | 11.550 | 14.405 | 11.898 | 11.692 | 13.372 | 10.855 | 7.349 | 8.888 | 8.535 | 9.615 | 9.463 | 9.107 | 8.588 | 10.372 | 7.712 | 0.406 | 0.0000 | 0.0000 | 1 | 1 | 0 |
| 'Anxa4' | 0.474 | 0.399 | 0.432 | 0.471 | 0.527 | 0.525 | 0.354 | 0.530 | 0.498 | 0.526 | 0.466 | 0.463 | 0.355 | 0.357 | 0.435 | 0.499 | 0.536 | 0.445 | 0.044 | 0.6974 | 0.8579 | 0 | 0 | 0 |
| 'Anxa5' | 29.006 | 28.891 | 30.381 | 28.357 | 29.651 | 30.825 | 30.428 | 29.026 | 30.755 | 30.952 | 28.673 | 33.015 | 28.449 | 30.343 | 32.209 | 30.887 | 31.552 | 33.959 | -0.078 | 0.0110 | 0.0668 | 0 | 0 | 0 |
| 'Anxa6' | 11.003 | 8.436 | 12.638 | 9.615 | 9.591 | 13.694 | 11.816 | 11.371 | 10.768 | 9.724 | 9.255 | 15.831 | 8.831 | 9.216 | 14.120 | 12.197 | 9.349 | 14.493 | -0.048 | 0.5160 | 0.7399 | 0 | 0 | 0 |
| 'Anxa7' | 10.334 | 9.573 | 11.074 | 11.269 | 10.646 | 10.587 | 11.701 | 11.516 | 9.954 | 10.586 | 11.515 | 12.093 | 10.486 | 10.921 | 11.641 | 10.810 | 11.076 | 10.901 | -0.061 | 0.1510 | 0.3777 | 0 | 0 | 0 |
| 'Aoc2' | 4.590 | 4.164 | 5.237 | 4.415 | 3.470 | 4.306 | 4.068 | 3.708 | 4.301 | 4.484 | 4.204 | 5.404 | 5.005 | 4.824 | 4.745 | 3.730 | 5.232 | 5.333 | -0.181 | 0.0135 | 0.0779 | 0 | 0 | 0 |
| 'Ap1ar' | 4.757 | 4.970 | 4.628 | 4.363 | 4.565 | 4.417 | 4.368 | 4.834 | 4.431 | 4.742 | 4.538 | 4.890 | 4.570 | 4.760 | 4.721 | 5.148 | 4.748 | 4.591 | -0.058 | 0.0845 | 0.2638 | 0 | 0 | 0 |
| 'Ap1b1' | 35.135 | 33.291 | 36.737 | 35.681 | 33.186 | 37.635 | 34.404 | 34.800 | 36.983 | 36.812 | 33.879 | 36.943 | 34.356 | 32.421 | 36.426 | 34.724 | 32.516 | 36.887 | 0.004 | 0.8644 | 0.9410 | 0 | 0 | 0 |
| 'Ap1g1' | 14.549 | 13.457 | 13.812 | 14.241 | 13.763 | 12.815 | 13.927 | 13.155 | 13.275 | 13.825 | 14.033 | 14.909 | 12.477 | 13.369 | 13.377 | 13.126 | 13.942 | 14.640 | -0.020 | 0.6022 | 0.7982 | 0 | 0 | 0 |
| 'Ap1g2' | 10.154 | 8.964 | 12.104 | 10.271 | 9.779 | 12.308 | 10.438 | 9.705 | 12.101 | 10.757 | 9.259 | 11.679 | 11.409 | 9.380 | 11.205 | 10.139 | 9.581 | 10.639 | 0.015 | 0.6278 | 0.8154 | 0 | 0 | 0 |
| 'Ap1m1' | 19.914 | 19.061 | 22.101 | 18.698 | 20.501 | 22.735 | 18.775 | 20.902 | 22.320 | 20.826 | 19.906 | 21.474 | 20.685 | 19.382 | 21.502 | 20.552 | 19.659 | 19.845 | 0.000 | 0.9891 | 0.9968 | 0 | 0 | 0 |
| 'Ap1m2' | 2.343 | 1.835 | 1.995 | 2.035 | 1.945 | 2.450 | 2.444 | 2.035 | 2.538 | 1.647 | 1.884 | 1.864 | 2.031 | 1.914 | 1.710 | 2.240 | 1.445 | 1.250 | 0.296 | 0.0103 | 0.0639 | 0 | 0 | 0 |
| 'Ap1s1' | 20.709 | 25.683 | 22.708 | 20.986 | 25.664 | 24.962 | 22.619 | 24.067 | 24.109 | 22.558 | 20.107 | 23.414 | 23.714 | 23.655 | 23.689 | 25.196 | 21.398 | 22.956 | 0.020 | 0.6583 | 0.8350 | 0 | 0 | 0 |
| 'Ap1s2' | 4.330 | 5.687 | 4.913 | 5.164 | 5.862 | 4.789 | 5.486 | 4.952 | 4.964 | 5.460 | 5.646 | 5.875 | 5.363 | 6.712 | 5.416 | 5.848 | 6.697 | 5.583 | -0.203 | 0.0000 | 0.0005 | 1 | 0 | 0 |
| 'Ap2a1' | 30.998 | 33.701 | 34.872 | 31.420 | 31.657 | 35.297 | 32.899 | 31.993 | 34.673 | 35.920 | 31.240 | 34.559 | 34.523 | 30.372 | 34.650 | 32.286 | 31.190 | 33.737 | -0.015 | 0.5886 | 0.7893 | 0 | 0 | 0 |
| 'Ap2a2' | 13.189 | 14.407 | 15.935 | 13.492 | 13.970 | 14.784 | 13.186 | 14.412 | 15.337 | 14.334 | 13.894 | 14.909 | 13.699 | 12.667 | 14.656 | 13.998 | 13.021 | 15.085 | 0.018 | 0.5613 | 0.7712 | 0 | 0 | 0 |
| 'Ap2b1' | 17.464 | 17.475 | 18.306 | 17.608 | 18.416 | 19.493 | 17.418 | 18.597 | 18.960 | 19.055 | 19.310 | 19.176 | 17.693 | 18.062 | 19.158 | 17.848 | 19.013 | 18.789 | -0.048 | 0.0450 | 0.1759 | 0 | 0 | 0 |
| 'Ap2m1' | 41.557 | 44.321 | 41.582 | 42.668 | 44.965 | 44.337 | 41.335 | 43.233 | 42.573 | 40.805 | 42.409 | 41.187 | 43.698 | 42.429 | 43.204 | 43.318 | 42.537 | 42.030 | 0.008 | 0.7161 | 0.8688 | 0 | 0 | 0 |
| 'Ap2s1' | 35.507 | 42.327 | 38.850 | 36.627 | 41.232 | 41.518 | 36.386 | 37.857 | 39.554 | 36.881 | 34.609 | 37.464 | 39.163 | 37.597 | 40.202 | 38.958 | 36.766 | 38.241 | 0.030 | 0.4003 | 0.6491 | 0 | 0 | 0 |
| 'Ap3b1' | 32.707 | 32.636 | 31.792 | 34.157 | 31.496 | 32.960 | 33.909 | 32.575 | 33.877 | 34.483 | 32.366 | 32.366 | 32.872 | 33.356 | 31.194 | 32.525 | 34.423 | 31.341 | -0.004 | 0.8784 | 0.9485 | 0 | 0 | 0 |
| 'Ap3d1' | 53.145 | 47.205 | 53.827 | 51.408 | 48.064 | 54.680 | 50.239 | 50.723 | 55.387 | 56.012 | 51.433 | 59.070 | 54.513 | 50.788 | 54.990 | 51.732 | 51.868 | 56.561 | -0.078 | 0.0006 | 0.0073 | 1 | 0 | 0 |
| 'Ap3m1' | 3.014 | 3.749 | 3.338 | 2.774 | 3.421 | 3.169 | 2.845 | 3.449 | 3.223 | 2.904 | 3.200 | 3.319 | 3.065 | 3.169 | 2.975 | 3.253 | 3.112 | 3.242 | 0.022 | 0.6016 | 0.7979 | 0 | 0 | 0 |
| 'Ap3m2' | 14.794 | 15.332 | 16.312 | 15.475 | 15.469 | 14.969 | 16.751 | 17.037 | 17.349 | 17.301 | 16.158 | 15.406 | 17.164 | 14.143 | 14.655 | 14.312 | 16.333 | 13.639 | 0.036 | 0.4622 | 0.6981 | 0 | 0 | 0 |
| 'Ap3s1' | 8.566 | 10.352 | 9.874 | 9.528 | 9.803 | 9.401 | 9.730 | 10.216 | 8.948 | 8.104 | 10.563 | 7.936 | 9.163 | 10.632 | 8.041 | 7.841 | 10.315 | 8.078 | 0.098 | 0.0435 | 0.1726 | 0 | 0 | 0 |
| 'Ap3s2' | 7.045 | 7.984 | 7.827 | 6.688 | 9.078 | 8.252 | 7.545 | 7.996 | 8.087 | 7.502 | 7.491 | 7.118 | 7.603 | 7.096 | 7.369 | 7.651 | 7.577 | 6.660 | 0.083 | 0.0599 | 0.2110 | 0 | 0 | 0 |
| 'Ap4e1' | 2.928 | 2.466 | 2.877 | 3.028 | 3.005 | 2.756 | 2.713 | 3.084 | 2.908 | 2.989 | 2.967 | 2.925 | 2.788 | 3.338 | 2.685 | 2.876 | 3.422 | 2.569 | -0.048 | 0.3459 | 0.6012 | 0 | 0 | 0 |
| 'Ap4m1' | 3.983 | 3.987 | 4.546 | 4.094 | 3.926 | 4.375 | 4.194 | 3.722 | 4.630 | 3.979 | 3.922 | 4.328 | 4.370 | 3.833 | 4.087 | 4.159 | 4.391 | 4.172 | -0.002 | 0.9599 | 0.9852 | 0 | 0 | 0 |
| 'Ap4s1' | 15.890 | 19.314 | 17.837 | 17.523 | 19.208 | 18.994 | 19.691 | 21.314 | 18.172 | 20.452 | 18.392 | 18.611 | 17.945 | 18.686 | 19.199 | 19.066 | 20.665 | 20.559 | -0.059 | 0.2657 | 0.5192 | 0 | 0 | 0 |
| 'Ap5b1' | 8.271 | 8.120 | 10.617 | 8.402 | 8.376 | 10.358 | 8.749 | 7.376 | 10.949 | 8.777 | 7.486 | 10.363 | 8.295 | 7.415 | 10.292 | 7.433 | 6.943 | 10.036 | 0.062 | 0.1518 | 0.3789 | 0 | 0 | 0 |
| 'Ap5m1' | 11.191 | 13.034 | 11.916 | 11.065 | 11.579 | 12.045 | 10.432 | 10.967 | 11.110 | 10.657 | 11.762 | 11.539 | 11.652 | 11.211 | 10.687 | 10.865 | 12.768 | 10.198 | 0.018 | 0.6929 | 0.8555 | 0 | 0 | 0 |
| 'Ap5s1' | 2.685 | 2.955 | 2.964 | 3.069 | 3.019 | 3.398 | 2.606 | 3.346 | 3.331 | 3.650 | 3.313 | 3.302 | 3.607 | 3.022 | 2.671 | 3.121 | 3.121 | 2.982 | -0.080 | 0.2361 | 0.4859 | 0 | 0 | 0 |
| 'Ap5z1' | 4.667 | 4.284 | 5.987 | 4.661 | 4.278 | 5.494 | 4.773 | 4.662 | 5.806 | 5.593 | 4.732 | 5.992 | 5.856 | 4.717 | 5.495 | 4.794 | 4.923 | 5.225 | -0.098 | 0.0279 | 0.1292 | 0 | 0 | 0 |
| 'Apaf1' | 5.099 | 5.299 | 5.343 | 5.227 | 5.146 | 5.337 | 4.854 | 4.901 | 5.069 | 5.001 | 5.332 | 4.897 | 5.282 | 5.148 | 4.759 | 4.768 | 5.153 | 4.689 | 0.031 | 0.3466 | 0.6017 | 0 | 0 | 0 |
| 'Apba3' | 31.555 | 28.089 | 31.696 | 29.284 | 27.728 | 31.291 | 29.110 | 27.546 | 30.093 | 28.812 | 30.244 | 34.215 | 31.457 | 28.508 | 30.986 | 28.447 | 27.667 | 33.861 | -0.053 | 0.1364 | 0.3555 | 0 | 0 | 0 |
| 'Apbb1ip' | 16.454 | 16.591 | 16.462 | 17.383 | 16.098 | 16.339 | 17.876 | 16.317 | 17.011 | 16.922 | 16.153 | 18.275 | 15.663 | 15.292 | 16.629 | 15.890 | 16.024 | 18.113 | 0.003 | 0.9271 | 0.9720 | 0 | 0 | 0 |
| 'Apc' | 6.180 | 5.446 | 5.383 | 5.660 | 5.712 | 5.363 | 5.559 | 5.850 | 5.349 | 6.582 | 6.193 | 5.660 | 6.230 | 6.190 | 5.391 | 6.192 | 5.770 | 5.790 | -0.105 | 0.0003 | 0.0042 | 1 | 0 | 0 |
| 'Apeh' | 13.359 | 13.135 | 14.432 | 14.092 | 13.571 | 14.744 | 13.270 | 13.444 | 14.277 | 13.921 | 13.189 | 14.628 | 14.204 | 12.507 | 13.729 | 12.818 | 13.085 | 13.117 | 0.025 | 0.3333 | 0.5915 | 0 | 0 | 0 |
| 'Apex1' | 16.548 | 14.087 | 14.181 | 14.424 | 16.160 | 15.445 | 16.186 | 15.839 | 14.777 | 14.603 | 14.011 | 14.309 | 15.726 | 11.870 | 13.446 | 13.817 | 14.498 | 13.589 | 0.117 | 0.0104 | 0.0643 | 0 | 0 | 0 |
| 'Apex2' | 1.434 | 1.147 | 1.292 | 1.329 | 0.899 | 1.103 | 1.539 | 1.203 | 1.456 | 1.162 | 1.459 | 1.738 | 1.225 | 1.154 | 1.611 | 1.105 | 1.367 | 1.581 | -0.131 | 0.1572 | 0.3868 | 0 | 0 | 0 |
| 'Aph1a' | 12.614 | 14.724 | 12.461 | 11.989 | 14.150 | 14.184 | 12.694 | 13.666 | 13.572 | 12.737 | 12.928 | 13.668 | 13.200 | 14.090 | 14.428 | 14.178 | 12.515 | 13.568 | -0.028 | 0.4658 | 0.7014 | 0 | 0 | 0 |
| 'Api5' | 24.596 | 26.241 | 24.767 | 25.525 | 25.431 | 23.635 | 25.059 | 25.866 | 23.283 | 24.247 | 24.304 | 25.316 | 23.310 | 25.597 | 25.002 | 24.040 | 25.316 | 25.576 | -0.002 | 0.9404 | 0.9769 | 0 | 0 | 0 |
| 'Apip' | 2.078 | 3.007 | 2.367 | 2.576 | 2.421 | 2.880 | 2.874 | 2.898 | 2.748 | 2.413 | 2.854 | 2.660 | 2.888 | 2.450 | 2.920 | 2.905 | 2.643 | 2.155 | -0.015 | 0.8650 | 0.9414 | 0 | 0 | 0 |
| 'Apitd1' | 4.041 | 4.236 | 4.213 | 3.552 | 3.376 | 4.135 | 4.184 | 4.001 | 3.676 | 3.169 | 3.895 | 3.608 | 4.576 | 4.034 | 4.055 | 3.374 | 3.637 | 3.751 | 0.046 | 0.5268 | 0.7483 | 0 | 0 | 0 |
| 'Aplf' | 1.254 | 1.383 | 1.335 | 1.297 | 1.332 | 1.517 | 1.369 | 1.521 | 1.432 | 0.982 | 1.009 | 1.349 | 1.206 | 1.218 | 1.350 | 1.126 | 1.163 | 1.130 | 0.221 | 0.0011 | 0.0113 | 1 | 0 | 0 |
| 'Aplp1' | 19.135 | 19.228 | 21.140 | 18.253 | 21.063 | 20.743 | 18.269 | 19.685 | 19.676 | 16.675 | 16.793 | 19.553 | 17.300 | 18.072 | 17.875 | 16.950 | 15.945 | 19.118 | 0.150 | 0.0000 | 0.0008 | 1 | 0 | 0 |
| 'Aplp2' | 28.539 | 32.134 | 33.079 | 32.554 | 32.296 | 35.226 | 30.142 | 31.350 | 33.207 | 35.824 | 31.309 | 33.426 | 29.871 | 30.775 | 32.019 | 33.162 | 27.962 | 32.754 | -0.003 | 0.9342 | 0.9748 | 0 | 0 | 0 |
| 'Apmap' | 12.913 | 14.946 | 14.392 | 11.864 | 15.290 | 14.571 | 13.256 | 16.385 | 13.927 | 13.903 | 13.694 | 13.342 | 13.334 | 14.047 | 13.048 | 14.790 | 11.924 | 12.782 | 0.067 | 0.1777 | 0.4131 | 0 | 0 | 0 |
| 'Apoa1bp' | 46.748 | 55.040 | 49.010 | 44.093 | 52.145 | 52.654 | 45.151 | 52.846 | 52.987 | 47.246 | 41.620 | 47.598 | 50.304 | 49.449 | 50.169 | 50.202 | 44.444 | 46.949 | 0.063 | 0.1734 | 0.4069 | 0 | 0 | 0 |
| 'Apobec1' | 1.809 | 2.504 | 1.752 | 1.646 | 2.484 | 1.764 | 1.954 | 2.320 | 2.049 | 1.823 | 1.918 | 1.542 | 2.091 | 2.031 | 1.490 | 1.834 | 1.698 | 1.392 | 0.195 | 0.0063 | 0.0445 | 1 | 0 | 0 |
| 'Apobec3' | 4.899 | 4.505 | 5.308 | 5.183 | 4.622 | 4.822 | 5.037 | 4.667 | 5.080 | 4.385 | 4.497 | 5.390 | 4.875 | 4.712 | 5.193 | 4.576 | 4.747 | 4.960 | 0.013 | 0.7115 | 0.8659 | 0 | 0 | 0 |
| 'Apobr' | 3.804 | 3.100 | 4.335 | 3.701 | 3.377 | 4.008 | 4.057 | 3.476 | 3.765 | 3.552 | 3.096 | 4.384 | 3.333 | 3.446 | 4.138 | 3.911 | 2.926 | 3.141 | 0.065 | 0.2998 | 0.5560 | 0 | 0 | 0 |
| 'Apoe' | 17.597 | 28.144 | 13.195 | 14.767 | 23.310 | 19.665 | 16.129 | 22.291 | 16.118 | 21.846 | 18.190 | 16.283 | 16.834 | 26.283 | 14.135 | 28.578 | 27.720 | 15.564 | -0.126 | 0.3405 | 0.5975 | 0 | 0 | 0 |
| 'Apol10b' | 3.460 | 4.641 | 3.688 | 3.311 | 3.663 | 3.961 | 3.990 | 3.543 | 3.620 | 4.528 | 5.692 | 4.120 | 4.249 | 6.724 | 3.253 | 3.408 | 5.145 | 3.433 | -0.247 | 0.0138 | 0.0790 | 0 | 0 | 0 |
| 'Apol9b' | 3.003 | 3.351 | 3.204 | 2.763 | 4.039 | 3.542 | 2.926 | 3.804 | 3.456 | 3.618 | 3.313 | 4.050 | 3.461 | 4.186 | 3.123 | 4.413 | 4.376 | 3.010 | -0.168 | 0.0844 | 0.2636 | 0 | 0 | 0 |
| 'Apoo' | 5.922 | 6.833 | 7.179 | 6.759 | 6.930 | 6.750 | 6.735 | 7.599 | 5.570 | 6.185 | 5.984 | 5.913 | 7.605 | 7.506 | 5.913 | 6.490 | 7.063 | 5.501 | 0.047 | 0.5452 | 0.7606 | 0 | 0 | 0 |
| 'Apool' | 4.507 | 4.110 | 4.190 | 4.609 | 4.755 | 4.632 | 4.668 | 4.808 | 4.392 | 4.480 | 4.527 | 4.777 | 4.587 | 4.835 | 4.339 | 4.032 | 5.214 | 4.213 | -0.026 | 0.6307 | 0.8173 | 0 | 0 | 0 |
| 'Apopt1' | 5.753 | 5.586 | 6.317 | 6.499 | 6.534 | 6.207 | 5.473 | 5.925 | 5.734 | 5.990 | 5.033 | 5.842 | 6.242 | 6.599 | 5.760 | 6.454 | 5.603 | 5.418 | 0.022 | 0.6698 | 0.8408 | 0 | 0 | 0 |
| 'App' | 0.703 | 1.201 | 0.781 | 0.680 | 1.166 | 0.881 | 0.880 | 1.495 | 0.941 | 1.610 | 2.485 | 1.327 | 1.705 | 2.249 | 1.373 | 1.588 | 2.155 | 1.416 | -0.863 | 0.0000 | 0.0000 | 1 | 1 | 0 |
| 'Appbp2' | 11.188 | 10.257 | 10.973 | 11.604 | 10.590 | 9.884 | 12.035 | 10.728 | 10.165 | 9.534 | 11.353 | 11.653 | 10.293 | 11.393 | 10.675 | 10.894 | 11.873 | 10.476 | -0.023 | 0.6276 | 0.8154 | 0 | 0 | 0 |
| 'Appl1' | 11.345 | 11.987 | 11.772 | 11.879 | 12.918 | 11.182 | 12.329 | 12.058 | 11.825 | 11.277 | 12.627 | 12.814 | 11.192 | 12.260 | 12.152 | 11.234 | 13.815 | 11.330 | -0.030 | 0.3858 | 0.6354 | 0 | 0 | 0 |
| 'Appl2' | 2.645 | 2.555 | 2.899 | 2.967 | 2.542 | 2.957 | 2.870 | 2.468 | 2.796 | 1.763 | 1.696 | 1.966 | 1.708 | 2.053 | 1.813 | 1.586 | 2.000 | 2.005 | 0.562 | 0.0000 | 0.0000 | 1 | 1 | 0 |
| 'Aprt' | 104.204 | 100.806 | 105.781 | 98.467 | 109.472 | 109.426 | 107.743 | 102.606 | 99.394 | 100.513 | 90.264 | 105.227 | 106.687 | 100.525 | 105.811 | 103.472 | 102.979 | 104.484 | 0.016 | 0.5988 | 0.7958 | 0 | 0 | 0 |
| 'Aptx' | 3.666 | 4.022 | 3.895 | 3.538 | 4.016 | 3.800 | 3.903 | 4.149 | 3.861 | 3.593 | 4.070 | 3.841 | 3.540 | 3.525 | 3.613 | 3.296 | 3.833 | 3.349 | 0.080 | 0.0432 | 0.1721 | 0 | 0 | 0 |
| 'Aqr' | 8.485 | 8.623 | 9.135 | 8.595 | 9.370 | 8.873 | 8.354 | 8.955 | 8.981 | 9.204 | 8.439 | 9.010 | 8.430 | 8.489 | 8.844 | 8.579 | 8.769 | 8.894 | 0.002 | 0.9230 | 0.9705 | 0 | 0 | 0 |
| 'Ar' | 1.507 | 1.093 | 1.469 | 1.703 | 1.338 | 1.137 | 1.109 | 1.372 | 1.536 | 1.130 | 1.062 | 1.287 | 1.029 | 1.104 | 0.970 | 0.891 | 1.032 | 1.283 | 0.312 | 0.0009 | 0.0093 | 1 | 0 | 0 |
| 'Araf' | 14.392 | 14.192 | 15.653 | 13.081 | 13.898 | 14.640 | 13.485 | 14.547 | 16.192 | 15.356 | 13.683 | 15.498 | 14.556 | 13.400 | 13.914 | 14.025 | 14.925 | 14.449 | -0.008 | 0.8132 | 0.9157 | 0 | 0 | 0 |
| 'Arap1' | 14.270 | 13.783 | 15.433 | 14.351 | 13.664 | 15.273 | 13.456 | 14.498 | 15.554 | 15.428 | 13.919 | 16.365 | 15.108 | 14.882 | 15.089 | 14.932 | 14.150 | 15.990 | -0.071 | 0.0037 | 0.0295 | 1 | 0 | 0 |
| 'Arap2' | 14.170 | 15.997 | 13.294 | 14.783 | 14.803 | 13.981 | 13.507 | 15.629 | 13.251 | 15.830 | 14.381 | 12.489 | 13.479 | 14.111 | 11.922 | 19.155 | 13.430 | 12.280 | 0.023 | 0.6857 | 0.8513 | 0 | 0 | 0 |
| 'Arc' | 23.650 | 16.588 | 15.007 | 24.395 | 15.402 | 14.916 | 25.563 | 15.849 | 15.370 | 18.255 | 19.630 | 20.600 | 21.774 | 19.863 | 17.936 | 18.128 | 22.379 | 20.332 | -0.148 | 0.1324 | 0.3501 | 0 | 0 | 0 |
| 'Arcn1' | 19.873 | 20.814 | 19.730 | 20.635 | 20.290 | 20.139 | 20.583 | 21.242 | 19.180 | 18.457 | 21.260 | 20.392 | 19.204 | 20.045 | 19.753 | 19.818 | 20.133 | 18.970 | 0.024 | 0.3438 | 0.5995 | 0 | 0 | 0 |
| 'Arel1' | 11.137 | 13.018 | 12.718 | 12.160 | 13.359 | 13.115 | 12.566 | 13.163 | 13.098 | 13.173 | 14.215 | 12.832 | 12.976 | 13.365 | 13.043 | 13.594 | 13.053 | 12.445 | -0.064 | 0.0315 | 0.1394 | 0 | 0 | 0 |
| 'Arf1' | 156.932 | 170.088 | 156.531 | 162.085 | 171.528 | 163.642 | 159.773 | 171.117 | 157.148 | 157.052 | 155.528 | 154.503 | 158.969 | 164.045 | 163.472 | 160.633 | 153.466 | 154.238 | 0.035 | 0.1078 | 0.3092 | 0 | 0 | 0 |
| 'Arf2' | 6.128 | 8.123 | 6.605 | 6.362 | 7.954 | 7.002 | 6.811 | 7.592 | 6.964 | 6.500 | 6.878 | 5.549 | 6.634 | 7.228 | 5.878 | 6.996 | 5.301 | 5.282 | 0.169 | 0.0066 | 0.0462 | 1 | 0 | 0 |
| 'Arf3' | 102.753 | 113.015 | 107.736 | 104.800 | 113.550 | 113.919 | 97.454 | 110.565 | 113.403 | 108.640 | 103.012 | 112.592 | 102.264 | 109.765 | 116.617 | 111.752 | 105.647 | 117.551 | -0.027 | 0.3528 | 0.6063 | 0 | 0 | 0 |
| 'Arf4' | 18.505 | 16.959 | 15.443 | 16.397 | 17.151 | 16.307 | 15.767 | 17.359 | 15.972 | 17.369 | 17.707 | 17.190 | 16.526 | 18.975 | 16.114 | 17.980 | 17.745 | 16.333 | -0.066 | 0.0446 | 0.1750 | 0 | 0 | 0 |
| 'Arf5' | 79.838 | 93.474 | 84.014 | 77.698 | 89.501 | 89.103 | 79.262 | 92.303 | 84.855 | 80.837 | 83.914 | 81.550 | 85.493 | 88.890 | 86.282 | 89.822 | 84.645 | 84.145 | -0.004 | 0.9035 | 0.9619 | 0 | 0 | 0 |
| 'Arf6' | 112.727 | 111.763 | 111.271 | 103.894 | 113.138 | 113.801 | 107.152 | 115.560 | 103.011 | 110.973 | 108.843 | 103.321 | 106.290 | 114.316 | 112.316 | 111.434 | 111.449 | 112.732 | -0.009 | 0.7395 | 0.8819 | 0 | 0 | 0 |
| 'Arfgap1' | 4.806 | 4.831 | 5.319 | 4.712 | 5.235 | 5.447 | 4.767 | 4.935 | 5.134 | 5.501 | 4.897 | 5.219 | 4.943 | 4.801 | 4.990 | 5.075 | 4.756 | 5.029 | -0.010 | 0.7574 | 0.8914 | 0 | 0 | 0 |
| 'Arfgap2' | 16.003 | 15.967 | 16.914 | 15.862 | 16.209 | 16.219 | 16.025 | 17.565 | 16.516 | 17.776 | 16.149 | 17.364 | 17.386 | 15.868 | 16.427 | 16.236 | 16.542 | 17.783 | -0.052 | 0.0954 | 0.2874 | 0 | 0 | 0 |
| 'Arfgap3' | 7.792 | 9.232 | 8.429 | 8.424 | 8.518 | 9.502 | 8.091 | 8.471 | 8.931 | 7.613 | 7.585 | 9.032 | 8.284 | 8.203 | 7.862 | 7.462 | 8.201 | 8.301 | 0.074 | 0.1147 | 0.3213 | 0 | 0 | 0 |
| 'Arfgef1' | 15.191 | 15.384 | 14.474 | 14.928 | 15.135 | 14.086 | 14.947 | 15.049 | 13.922 | 15.554 | 15.443 | 16.056 | 14.360 | 15.564 | 15.318 | 15.460 | 15.598 | 15.915 | -0.077 | 0.0048 | 0.0361 | 1 | 0 | 0 |
| 'Arfgef2' | 13.387 | 13.353 | 13.696 | 13.180 | 13.324 | 13.065 | 11.947 | 13.897 | 13.603 | 14.183 | 13.984 | 13.098 | 13.815 | 13.615 | 13.313 | 14.151 | 13.518 | 13.284 | -0.049 | 0.0997 | 0.2954 | 0 | 0 | 0 |
| 'Arfip2' | 3.931 | 4.155 | 4.227 | 3.802 | 3.948 | 4.347 | 3.800 | 4.207 | 4.343 | 3.774 | 3.583 | 4.012 | 4.208 | 4.097 | 4.075 | 4.117 | 3.901 | 3.804 | 0.039 | 0.2761 | 0.5318 | 0 | 0 | 0 |
| 'Arfrp1' | 9.272 | 10.142 | 9.901 | 9.284 | 9.579 | 9.540 | 8.872 | 9.517 | 10.037 | 9.153 | 9.752 | 10.796 | 9.587 | 9.690 | 10.122 | 9.593 | 10.197 | 10.339 | -0.064 | 0.0170 | 0.0920 | 0 | 0 | 0 |
| 'Arglu1' | 90.268 | 104.115 | 93.906 | 88.765 | 100.343 | 97.076 | 89.230 | 105.043 | 93.496 | 85.576 | 91.075 | 89.854 | 88.602 | 101.966 | 93.570 | 95.864 | 91.962 | 87.981 | 0.050 | 0.0807 | 0.2562 | 0 | 0 | 0 |
| 'Arhgap1' | 15.414 | 18.698 | 17.320 | 15.330 | 17.755 | 18.491 | 15.480 | 17.920 | 18.845 | 16.623 | 16.052 | 18.209 | 15.909 | 16.200 | 17.875 | 16.271 | 14.934 | 17.716 | 0.038 | 0.2633 | 0.5173 | 0 | 0 | 0 |
| 'Arhgap10' | 22.122 | 18.800 | 20.737 | 20.616 | 20.167 | 20.743 | 21.113 | 18.261 | 19.620 | 22.877 | 20.732 | 21.385 | 25.830 | 19.109 | 20.156 | 20.804 | 20.258 | 20.980 | -0.087 | 0.0269 | 0.1263 | 0 | 0 | 0 |

| | | | | | | | | | | | | | | | | | | | | | | | | |
|---|---|---|---|---|---|---|---|---|---|---|---|---|---|---|---|---|---|---|---|---|---|---|---|---|
| 'Arhgap11a' | 7.288 | 7.739 | 9.366 | 7.542 | 8.032 | 8.808 | 7.838 | 7.211 | 9.060 | 6.005 | 7.878 | 7.599 | 9.545 | 9.177 | 9.699 | 7.774 | 8.703 | 7.841 | -0.038 | 0.5941 | 0.7933 | 0 | 0 | 0 |
| 'Arhgap12' | 3.154 | 3.333 | 3.157 | 3.494 | 3.214 | 3.062 | 3.438 | 3.167 | 3.266 | 3.174 | 3.304 | 3.431 | 3.082 | 3.344 | 3.041 | 3.135 | 4.021 | 3.317 | -0.039 | 0.4008 | 0.6495 | 0 | 0 | 0 |
| 'Arhgap15' | 11.584 | 11.773 | 11.304 | 13.274 | 11.518 | 10.672 | 13.541 | 11.439 | 10.654 | 11.367 | 10.903 | 10.633 | 10.365 | 10.996 | 10.437 | 11.189 | 11.660 | 9.388 | 0.113 | 0.0032 | 0.0263 | 1 | 0 | 0 |
| 'Arhgap17' | 2.603 | 2.136 | 2.573 | 2.162 | 2.150 | 2.253 | 2.287 | 2.100 | 2.292 | 2.387 | 2.625 | 2.587 | 2.424 | 2.373 | 2.378 | 2.448 | 2.434 | 2.729 | -0.132 | 0.0064 | 0.0453 | 1 | 0 | 0 |
| 'Arhgap18' | 3.252 | 3.647 | 3.484 | 3.791 | 3.250 | 3.417 | 4.105 | 3.390 | 3.708 | 3.518 | 3.218 | 3.947 | 3.366 | 3.796 | 3.696 | 3.414 | 3.960 | 3.315 | -0.025 | 0.6519 | 0.8310 | 0 | 0 | 0 |
| 'Arhgap19' | 2.827 | 3.363 | 4.029 | 3.672 | 3.459 | 3.837 | 3.784 | 3.608 | 4.190 | 0.481 | 0.753 | 0.628 | 0.860 | 0.687 | 0.803 | 0.797 | 0.732 | 0.626 | 2.361 | 0.0000 | 0.0000 | 1 | 1 | 1 |
| 'Arhgap21' | 0.294 | 0.314 | 0.387 | 0.452 | 0.333 | 0.415 | 0.413 | 0.345 | 0.469 | 0.345 | 0.454 | 0.421 | 0.475 | 0.447 | 0.380 | 0.352 | 0.368 | 0.471 | -0.133 | 0.1731 | 0.4066 | 0 | 0 | 0 |
| 'Arhgap25' | 28.088 | 23.252 | 25.376 | 29.509 | 23.441 | 26.089 | 30.069 | 24.542 | 25.968 | 24.066 | 26.785 | 28.222 | 23.366 | 23.342 | 27.768 | 21.716 | 26.736 | 27.799 | 0.028 | 0.6325 | 0.8178 | 0 | 0 | 0 |
| 'Arhgap26' | 1.913 | 1.923 | 1.986 | 1.719 | 1.740 | 1.787 | 2.046 | 1.951 | 1.893 | 2.032 | 1.671 | 2.058 | 1.552 | 1.651 | 1.943 | 1.858 | 1.873 | 2.126 | 0.007 | 0.9020 | 0.9612 | 0 | 0 | 0 |
| 'Arhgap27' | 9.908 | 8.795 | 10.425 | 9.175 | 8.835 | 10.072 | 9.358 | 9.269 | 10.568 | 9.665 | 9.104 | 11.171 | 9.574 | 8.334 | 10.259 | 9.701 | 8.830 | 10.831 | -0.027 | 0.3286 | 0.5868 | 0 | 0 | 0 |
| 'Arhgap30' | 36.731 | 37.164 | 36.994 | 36.308 | 36.442 | 37.800 | 34.509 | 36.002 | 39.128 | 36.445 | 37.426 | 41.341 | 35.651 | 37.088 | 42.408 | 37.763 | 36.545 | 43.205 | -0.080 | 0.0038 | 0.0304 | 1 | 0 | 0 |
| 'Arhgap31' | 1.362 | 1.497 | 1.620 | 1.351 | 1.437 | 1.662 | 1.652 | 1.521 | 1.374 | 1.835 | 1.584 | 2.178 | 1.776 | 1.649 | 2.044 | 2.026 | 1.793 | 1.951 | -0.334 | 0.0000 | 0.0000 | 1 | 0 | 0 |
| 'Arhgap33' | 1.301 | 1.558 | 2.228 | 1.152 | 1.039 | 1.763 | 1.396 | 1.486 | 2.201 | 1.275 | 1.564 | 1.604 | 1.632 | 1.545 | 1.825 | 1.360 | 1.384 | 1.428 | 0.037 | 0.7263 | 0.8747 | 0 | 0 | 0 |
| 'Arhgap35' | 2.911 | 3.078 | 2.854 | 2.987 | 2.776 | 3.015 | 2.924 | 3.108 | 3.175 | 3.296 | 3.047 | 2.821 | 2.989 | 3.207 | 2.744 | 3.618 | 2.810 | 3.073 | -0.044 | 0.3754 | 0.6267 | 0 | 0 | 0 |
| 'Arhgap39' | 6.068 | 5.602 | 5.935 | 5.592 | 6.181 | 6.014 | 4.993 | 5.777 | 5.291 | 6.060 | 6.166 | 6.227 | 6.243 | 5.866 | 5.834 | 5.726 | 6.336 | 5.828 | -0.087 | 0.0262 | 0.1242 | 0 | 0 | 0 |
| 'Arhgap4' | 33.713 | 33.359 | 38.500 | 36.253 | 33.943 | 39.088 | 36.428 | 34.380 | 39.905 | 36.106 | 32.143 | 38.957 | 35.570 | 32.205 | 36.933 | 33.972 | 34.372 | 36.288 | 0.029 | 0.1595 | 0.3900 | 0 | 0 | 0 |
| 'Arhgap5' | 2.922 | 3.224 | 2.902 | 2.866 | 3.215 | 2.895 | 2.907 | 3.574 | 2.913 | 3.076 | 3.133 | 3.067 | 2.484 | 3.588 | 2.673 | 2.874 | 3.802 | 3.141 | -0.030 | 0.5895 | 0.7903 | 0 | 0 | 0 |
| 'Arhgap9' | 138.934 | 134.599 | 145.134 | 130.612 | 142.371 | 154.170 | 135.320 | 141.856 | 145.389 | 139.414 | 132.030 | 148.246 | 138.763 | 135.651 | 147.654 | 146.832 | 132.122 | 141.813 | -0.003 | 0.9003 | 0.9605 | 0 | 0 | 0 |
| 'Arhgdia' | 159.159 | 157.940 | 162.454 | 148.980 | 155.984 | 178.665 | 151.720 | 154.876 | 166.254 | 162.251 | 152.382 | 161.346 | 159.750 | 154.095 | 172.572 | 160.774 | 147.904 | 171.962 | -0.017 | 0.5094 | 0.7342 | 0 | 0 | 0 |
| 'Arhgdib' | 305.099 | 316.255 | 298.924 | 322.163 | 320.602 | 320.721 | 325.493 | 305.382 | 317.155 | 323.304 | 315.374 | 286.869 | 327.694 | 312.467 | 329.527 | 325.436 | 310.139 | 309.506 | -0.015 | 0.5896 | 0.7903 | 0 | 0 | 0 |
| 'Arhgef1' | 87.238 | 75.435 | 100.262 | 83.860 | 77.020 | 95.934 | 86.240 | 79.262 | 99.546 | 90.271 | 81.351 | 95.320 | 89.853 | 76.587 | 92.487 | 85.361 | 82.176 | 92.835 | -0.017 | 0.4845 | 0.7160 | 0 | 0 | 0 |
| 'Arhgef10' | 2.888 | 3.319 | 3.347 | 2.894 | 3.498 | 3.413 | 3.162 | 3.347 | 3.441 | 2.989 | 3.278 | 3.209 | 3.387 | 3.376 | 3.514 | 2.922 | 3.435 | 3.204 | -0.011 | 0.7732 | 0.8993 | 0 | 0 | 0 |
| 'Arhgef11' | 2.670 | 2.231 | 2.894 | 2.726 | 2.520 | 2.727 | 2.513 | 2.689 | 2.831 | 2.944 | 2.891 | 2.751 | 2.717 | 2.760 | 2.642 | 2.579 | 2.776 | 2.717 | -0.065 | 0.1399 | 0.3619 | 0 | 0 | 0 |
| 'Arhgef12' | 9.949 | 9.394 | 9.652 | 10.131 | 9.121 | 9.089 | 9.094 | 9.530 | 9.365 | 8.404 | 8.716 | 8.335 | 8.102 | 7.707 | 7.920 | 7.844 | 8.459 | 8.963 | 0.186 | 0.0000 | 0.0000 | 1 | 0 | 0 |
| 'Arhgef16' | 1.255 | 1.224 | 1.276 | 1.078 | 1.201 | 1.318 | 1.174 | 1.234 | 1.413 | 1.191 | 1.154 | 1.209 | 1.243 | 1.174 | 1.189 | 1.412 | 1.226 | 1.022 | 0.046 | 0.4763 | 0.7107 | 0 | 0 | 0 |
| 'Arhgef18' | 2.933 | 2.168 | 2.912 | 2.630 | 2.233 | 2.971 | 2.872 | 2.643 | 2.985 | 2.579 | 2.504 | 3.614 | 2.474 | 2.090 | 3.271 | 2.814 | 2.295 | 3.428 | -0.043 | 0.4458 | 0.6878 | 0 | 0 | 0 |
| 'Arhgef2' | 10.095 | 8.945 | 8.996 | 9.825 | 8.114 | 9.025 | 9.150 | 8.457 | 9.321 | 8.806 | 8.636 | 10.917 | 8.668 | 8.633 | 10.240 | 8.786 | 8.745 | 10.707 | -0.046 | 0.3110 | 0.5692 | 0 | 0 | 0 |
| 'Arhgef3' | 66.132 | 60.089 | 61.702 | 64.093 | 61.365 | 60.376 | 66.455 | 61.514 | 61.031 | 63.138 | 63.908 | 63.134 | 53.625 | 59.541 | 61.760 | 60.757 | 62.556 | 60.450 | 0.025 | 0.4836 | 0.7151 | 0 | 0 | 0 |
| 'Arhgef39' | 0.755 | 0.959 | 1.011 | 0.916 | 0.797 | 0.969 | 0.928 | 0.732 | 0.951 | 0.574 | 0.768 | 0.928 | 0.814 | 0.750 | 0.992 | 0.731 | 0.735 | 1.008 | 0.118 | 0.1584 | 0.3886 | 0 | 0 | 0 |
| 'Arhgef6' | 16.922 | 18.233 | 17.813 | 17.667 | 17.613 | 17.277 | 18.599 | 18.680 | 17.706 | 18.612 | 18.589 | 17.567 | 18.258 | 17.607 | 17.199 | 18.054 | 18.337 | 16.853 | -0.015 | 0.4956 | 0.7247 | 0 | 0 | 0 |
| 'Arhgef7' | 6.281 | 6.530 | 6.781 | 6.100 | 6.292 | 6.762 | 6.199 | 6.413 | 6.883 | 6.989 | 6.949 | 6.686 | 6.487 | 6.205 | 6.910 | 6.996 | 6.389 | 6.747 | -0.059 | 0.0562 | 0.2036 | 0 | 0 | 0 |
| 'Arid1a' | 43.074 | 37.609 | 38.805 | 39.701 | 34.171 | 38.173 | 34.368 | 37.761 | 39.496 | 39.870 | 37.824 | 43.285 | 39.110 | 38.212 | 41.957 | 39.805 | 37.610 | 45.610 | -0.091 | 0.0227 | 0.1125 | 0 | 0 | 0 |
| 'Arid1b' | 16.366 | 13.856 | 15.834 | 15.660 | 14.187 | 16.549 | 15.111 | 14.895 | 15.486 | 16.732 | 16.432 | 14.718 | 15.214 | 16.096 | 15.075 | 16.748 | 15.080 | 15.357 | -0.046 | 0.2296 | 0.4766 | 0 | 0 | 0 |
| 'Arid2' | 9.834 | 9.100 | 9.753 | 9.332 | 9.097 | 8.887 | 8.204 | 9.605 | 9.440 | 9.369 | 9.859 | 9.959 | 8.477 | 9.644 | 9.561 | 9.047 | 10.052 | 10.715 | -0.067 | 0.1066 | 0.3067 | 0 | 0 | 0 |
| 'Arid3a' | 1.583 | 1.358 | 1.739 | 1.519 | 1.470 | 1.811 | 1.480 | 1.340 | 1.678 | 1.400 | 1.491 | 1.727 | 1.252 | 1.115 | 1.747 | 1.253 | 1.171 | 1.573 | 0.126 | 0.0261 | 0.1241 | 0 | 0 | 0 |
| 'Arid3b' | 0.784 | 0.642 | 0.933 | 0.691 | 0.833 | 0.940 | 0.846 | 0.695 | 0.941 | 0.791 | 0.740 | 1.020 | 0.717 | 0.583 | 0.974 | 0.836 | 0.748 | 0.932 | -0.017 | 0.7944 | 0.9082 | 0 | 0 | 0 |
| 'Arid4a' | 21.648 | 21.327 | 19.493 | 21.681 | 20.174 | 21.368 | 21.741 | 20.957 | 21.847 | 18.930 | 22.416 | 24.505 | 18.620 | 21.152 | 22.853 | 20.155 | 21.700 | 24.375 | -0.040 | 0.3900 | 0.6393 | 0 | 0 | 0 |
| 'Arid4b' | 7.234 | 6.816 | 6.609 | 7.707 | 6.754 | 6.306 | 7.164 | 7.024 | 6.519 | 7.184 | 6.829 | 6.752 | 6.596 | 6.877 | 6.312 | 6.930 | 7.100 | 5.816 | 0.030 | 0.3333 | 0.5915 | 0 | 0 | 0 |
| 'Arid5a' | 6.761 | 5.371 | 7.661 | 5.737 | 6.315 | 6.744 | 6.110 | 7.324 | 6.321 | 5.775 | 6.983 | 6.678 | 5.531 | 5.807 | 6.295 | 4.538 | 5.902 | 6.705 | 0.100 | 0.1187 | 0.3278 | 0 | 0 | 0 |
| 'Arid5b' | 27.299 | 19.984 | 20.107 | 21.707 | 20.713 | 18.053 | 19.707 | 21.571 | 19.168 | 30.163 | 25.007 | 18.568 | 25.945 | 24.746 | 18.182 | 28.454 | 23.253 | 17.546 | -0.159 | 0.0093 | 0.0594 | 0 | 0 | 0 |
| 'Arih1' | 26.563 | 24.714 | 24.937 | 27.189 | 26.118 | 24.515 | 26.218 | 25.914 | 23.576 | 26.553 | 27.008 | 25.804 | 25.108 | 26.928 | 25.933 | 24.788 | 26.980 | 27.113 | -0.052 | 0.0985 | 0.2933 | 0 | 0 | 0 |
| 'Arih2' | 5.317 | 5.737 | 5.005 | 4.757 | 5.562 | 5.371 | 5.029 | 5.734 | 5.036 | 5.281 | 5.245 | 5.305 | 5.311 | 5.690 | 5.290 | 5.999 | 5.204 | 4.825 | -0.028 | 0.4409 | 0.6843 | 0 | 0 | 0 |
| 'Arl1' | 19.885 | 24.335 | 21.604 | 22.046 | 25.490 | 19.906 | 22.243 | 22.973 | 20.066 | 23.333 | 20.414 | 20.831 | 21.804 | 22.566 | 20.232 | 22.542 | 21.619 | 19.672 | 0.026 | 0.5211 | 0.7440 | 0 | 0 | 0 |
| 'Arl10' | 1.925 | 2.702 | 1.924 | 1.741 | 2.351 | 2.653 | 1.622 | 2.617 | 2.163 | 2.124 | 1.568 | 1.853 | 1.812 | 1.982 | 2.026 | 2.193 | 1.533 | 1.907 | 0.200 | 0.0709 | 0.2347 | 0 | 0 | 0 |
| 'Arl13b' | 3.468 | 3.686 | 3.376 | 3.186 | 3.336 | 3.353 | 3.220 | 3.493 | 3.344 | 3.974 | 3.598 | 3.621 | 3.448 | 4.136 | 3.325 | 3.391 | 3.867 | 2.865 | -0.087 | 0.1079 | 0.3092 | 0 | 0 | 0 |
| 'Arl14ep' | 4.098 | 5.759 | 4.797 | 5.552 | 5.640 | 4.995 | 5.194 | 5.056 | 4.929 | 3.941 | 5.017 | 4.911 | 4.014 | 5.150 | 4.599 | 4.794 | 5.056 | 4.477 | 0.119 | 0.0291 | 0.1322 | 0 | 0 | 0 |
| 'Arl15' | 2.810 | 2.589 | 3.036 | 3.238 | 2.815 | 3.271 | 3.690 | 3.285 | 2.988 | 3.523 | 3.487 | 3.142 | 2.742 | 3.168 | 3.313 | 4.076 | 3.439 | 3.199 | -0.119 | 0.1260 | 0.3400 | 0 | 0 | 0 |
| 'Arl16' | 2.139 | 2.381 | 2.318 | 2.099 | 2.252 | 2.140 | 2.336 | 2.328 | 2.103 | 1.921 | 2.286 | 2.215 | 2.299 | 1.952 | 2.067 | 1.754 | 2.334 | 1.948 | 0.080 | 0.1645 | 0.3968 | 0 | 0 | 0 |
| 'Arl2' | 5.998 | 6.197 | 5.956 | 6.050 | 5.968 | 6.157 | 6.242 | 6.069 | 6.884 | 5.819 | 5.053 | 6.203 | 6.116 | 6.276 | 5.800 | 6.638 | 5.609 | 5.648 | 0.051 | 0.2829 | 0.5394 | 0 | 0 | 0 |
| 'Arl2bp' | 9.862 | 12.726 | 10.477 | 8.422 | 10.469 | 11.339 | 8.809 | 13.528 | 10.682 | 13.610 | 10.787 | 9.880 | 12.375 | 11.225 | 10.795 | 13.593 | 7.756 | 8.830 | -0.041 | 0.6852 | 0.8511 | 0 | 0 | 0 |
| 'Arl3' | 23.491 | 25.865 | 24.977 | 27.934 | 26.706 | 25.999 | 23.586 | 26.563 | 26.515 | 22.612 | 22.509 | 23.634 | 22.506 | 23.333 | 22.502 | 27.494 | 22.300 | 22.753 | 0.135 | 0.0092 | 0.0590 | 0 | 0 | 0 |
| 'Arl4a' | 3.813 | 2.909 | 2.529 | 3.482 | 3.089 | 2.480 | 3.113 | 2.933 | 2.545 | 3.637 | 3.470 | 2.515 | 3.400 | 3.484 | 2.351 | 3.004 | 3.277 | 2.186 | -0.024 | 0.6662 | 0.8390 | 0 | 0 | 0 |
| 'Arl4c' | 4.227 | 4.757 | 4.301 | 4.344 | 5.433 | 5.710 | 5.054 | 5.744 | 4.928 | 12.201 | 9.207 | 8.555 | 15.160 | 13.004 | 9.660 | 17.140 | 10.186 | 9.348 | -1.214 | 0.0000 | 0.0000 | 1 | 1 | 1 |
| 'Arl5a' | 12.319 | 12.462 | 13.423 | 12.712 | 13.224 | 12.237 | 12.711 | 13.159 | 11.942 | 12.414 | 11.441 | 13.216 | 11.622 | 13.158 | 11.729 | 13.664 | 13.325 | 13.100 | -0.004 | 0.9147 | 0.9671 | 0 | 0 | 0 |
| 'Arl5b' | 1.444 | 1.171 | 1.058 | 1.236 | 1.075 | 0.949 | 1.279 | 1.209 | 0.993 | 1.131 | 1.391 | 1.313 | 1.108 | 1.234 | 1.152 | 1.338 | 1.361 | 1.115 | -0.115 | 0.0932 | 0.2826 | 0 | 0 | 0 |
| 'Arl5c' | 50.334 | 57.538 | 50.606 | 51.048 | 59.323 | 54.884 | 52.550 | 54.197 | 52.570 | 57.014 | 54.162 | 53.835 | 71.792 | 61.121 | 54.456 | 62.413 | 56.629 | 52.676 | -0.126 | 0.0074 | 0.0497 | 1 | 0 | 0 |
| 'Arl6' | 0.977 | 1.437 | 1.257 | 1.241 | 1.480 | 1.071 | 1.180 | 1.520 | 1.150 | 0.998 | 1.007 | 0.865 | 1.015 | 1.089 | 0.835 | 1.113 | 1.337 | 0.881 | 0.303 | 0.0043 | 0.0338 | 1 | 0 | 0 |
| 'Arl6ip1' | 60.551 | 70.600 | 68.868 | 66.928 | 71.460 | 70.586 | 68.696 | 72.597 | 70.280 | 62.149 | 68.464 | 72.431 | 61.487 | 73.522 | 72.664 | 66.564 | 70.004 | 73.050 | -0.010 | 0.7215 | 0.8716 | 0 | 0 | 0 |
| 'Arl6ip4' | 16.450 | 18.215 | 18.224 | 15.458 | 16.916 | 20.203 | 16.804 | 16.950 | 18.731 | 16.347 | 13.520 | 18.898 | 17.644 | 15.314 | 18.762 | 16.749 | 16.182 | 17.615 | 0.052 | 0.2067 | 0.4509 | 0 | 0 | 0 |
| 'Arl6ip5' | 167.931 | 189.227 | 162.506 | 174.956 | 187.956 | 167.456 | 173.084 | 173.184 | 166.825 | 161.966 | 158.827 | 161.284 | 167.270 | 169.650 | 165.001 | 166.343 | 163.273 | 160.603 | 0.071 | 0.0045 | 0.0349 | 1 | 0 | 0 |
| 'Arl6ip6' | 12.376 | 12.558 | 13.585 | 14.625 | 12.446 | 11.484 | 13.147 | 12.436 | 11.719 | 11.167 | 12.954 | 13.848 | 13.198 | 13.288 | 13.654 | 10.451 | 14.051 | 13.904 | -0.039 | 0.5175 | 0.7407 | 0 | 0 | 0 |
| 'Arl8a' | 14.577 | 15.216 | 14.873 | 14.249 | 14.012 | 15.134 | 14.177 | 13.960 | 14.428 | 13.872 | 15.080 | 19.001 | 13.962 | 16.097 | 17.482 | 14.252 | 14.381 | 17.911 | -0.131 | 0.0049 | 0.0367 | 1 | 0 | 0 |
| 'Arl8b' | 17.215 | 20.836 | 17.076 | 18.147 | 19.835 | 17.928 | 17.972 | 18.517 | 17.031 | 17.870 | 19.012 | 20.430 | 18.882 | 20.862 | 19.822 | 18.482 | 20.061 | 20.485 | -0.113 | 0.0017 | 0.0157 | 1 | 0 | 0 |
| 'Armc1' | 11.234 | 14.257 | 12.640 | 11.526 | 13.506 | 11.945 | 13.131 | 12.960 | 11.654 | 12.220 | 12.653 | 13.183 | 11.599 | 13.915 | 12.366 | 11.685 | 13.198 | 12.615 | -0.025 | 0.5188 | 0.7416 | 0 | 0 | 0 |
| 'Armc10' | 8.044 | 9.014 | 9.643 | 7.561 | 9.416 | 8.718 | 8.432 | 9.960 | 9.140 | 9.102 | 8.790 | 8.950 | 8.746 | 9.217 | 8.564 | 9.199 | 8.708 | 8.305 | -0.003 | 0.9352 | 0.9752 | 0 | 0 | 0 |
| 'Armc5' | 12.475 | 13.131 | 15.091 | 10.248 | 13.178 | 15.960 | 10.145 | 13.052 | 15.339 | 13.851 | 11.428 | 12.758 | 14.849 | 12.143 | 13.268 | 14.634 | 11.517 | 12.289 | 0.008 | 0.9133 | 0.9669 | 0 | 0 | 0 |
| 'Armc6' | 2.737 | 2.843 | 2.933 | 2.963 | 2.700 | 2.852 | 2.612 | 2.677 | 2.935 | 2.929 | 2.926 | 3.732 | 3.055 | 3.026 | 3.491 | 2.942 | 2.892 | 3.400 | -0.188 | 0.0002 | 0.0027 | 1 | 0 | 0 |
| 'Armc7' | 12.845 | 11.794 | 13.036 | 14.076 | 10.948 | 12.511 | 13.576 | 12.225 | 13.056 | 11.288 | 11.137 | 14.993 | 11.468 | 10.278 | 14.763 | 10.977 | 11.619 | 14.032 | 0.039 | 0.4780 | 0.7111 | 0 | 0 | 0 |
| 'Armc8' | 8.799 | 8.966 | 10.188 | 8.821 | 9.726 | 9.772 | 9.039 | 9.654 | 9.180 | 10.703 | 9.379 | 9.387 | 8.938 | 9.445 | 8.277 | 9.655 | 9.368 | 8.349 | 0.004 | 0.9246 | 0.9715 | 0 | 0 | 0 |
| 'Armc9' | 0.462 | 0.350 | 0.471 | 0.387 | 0.515 | 0.426 | 0.368 | 0.420 | 0.504 | 0.559 | 0.485 | 0.532 | 0.515 | 0.473 | 0.399 | 0.489 | 0.473 | 0.466 | -0.176 | 0.0406 | 0.1649 | 0 | 0 | 0 |
| 'Armcx2' | 4.537 | 4.116 | 3.546 | 4.022 | 4.126 | 3.818 | 3.641 | 4.331 | 3.647 | 4.486 | 4.553 | 4.425 | 4.432 | 4.540 | 3.991 | 4.406 | 4.658 | 3.973 | -0.153 | 0.0003 | 0.0036 | 1 | 0 | 0 |
| 'Armcx5' | 5.869 | 4.946 | 3.581 | 4.524 | 4.424 | 3.755 | 4.171 | 4.201 | 3.738 | 5.577 | 5.503 | 3.127 | 5.384 | 4.747 | 2.710 | 5.459 | 4.489 | 2.700 | 0.015 | 0.8579 | 0.9390 | 0 | 0 | 0 |
| 'Armcx6' | 2.512 | 1.978 | 2.006 | 1.957 | 2.006 | 1.637 | 2.125 | 2.064 | 1.900 | 2.099 | 1.859 | 1.527 | 2.214 | 1.971 | 1.777 | 2.090 | 1.917 | 1.437 | 0.114 | 0.1349 | 0.3536 | 0 | 0 | 0 |
| 'Armt1' | 4.456 | 5.357 | 5.231 | 5.304 | 5.723 | 5.024 | 4.470 | 5.201 | 5.081 | 4.750 | 5.288 | 5.115 | 4.935 | 5.193 | 5.179 | 4.372 | 4.925 | 4.747 | 0.028 | 0.4974 | 0.7258 | 0 | 0 | 0 |
| 'Arnt' | 6.823 | 7.492 | 7.194 | 6.971 | 7.591 | 7.029 | 6.959 | 7.408 | 7.290 | 7.225 | 7.286 | 6.796 | 6.876 | 7.079 | 6.778 | 7.257 | 7.111 | 6.452 | 0.035 | 0.1722 | 0.4060 | 0 | 0 | 0 |
| 'Arnt2' | 10.632 | 15.301 | 10.667 | 12.327 | 12.579 | 11.647 | 11.378 | 13.879 | 12.250 | 9.472 | 9.589 | 7.825 | 9.951 | 10.384 | 7.169 | 10.046 | 10.802 | 8.254 | 0.401 | 0.0000 | 0.0000 | 1 | 1 | 0 |
| 'Arntl' | 16.793 | 18.722 | 14.857 | 17.765 | 19.368 | 17.138 | 16.862 | 17.786 | 17.238 | 18.367 | 17.839 | 17.454 | 16.819 | 17.815 | 16.003 | 17.015 | 18.145 | 16.874 | -0.014 | 0.7103 | 0.8654 | 0 | 0 | 0 |
| 'Arntl2' | 1.036 | 1.020 | 1.343 | 1.481 | 1.314 | 1.146 | 1.256 | 1.457 | 1.321 | 1.553 | 1.851 | 1.771 | 1.993 | 1.819 | 1.370 | 1.290 | 1.718 | 1.866 | -0.434 | 0.0000 | 0.0000 | 1 | 1 | 0 |
| 'Arpc1a' | 16.997 | 18.444 | 17.514 | 16.094 | 18.126 | 17.510 | 16.740 | 18.295 | 17.031 | 16.938 | 18.454 | 19.386 | 18.935 | 18.142 | 19.799 | 19.321 | 19.446 | 18.815 | -0.123 | 0.0001 | 0.0010 | 1 | 0 | 0 |
| 'Arpc1b' | 74.613 | 75.277 | 75.544 | 77.187 | 74.320 | 80.732 | 77.961 | 73.176 | 75.110 | 73.871 | 72.996 | 74.720 | 78.532 | 75.070 | 83.461 | 80.556 | 73.478 | 79.413 | -0.028 | 0.2928 | 0.5493 | 0 | 0 | 0 |
| 'Arpc2' | 252.667 | 283.952 | 249.769 | 271.286 | 276.341 | 254.900 | 272.846 | 279.419 | 250.888 | 263.964 | 277.266 | 231.971 | 272.569 | 283.531 | 262.617 | 275.962 | 276.695 | 248.233 | -0.011 | 0.6749 | 0.8440 | 0 | 0 | 0 |
| 'Arpc3' | 27.979 | 29.079 | 27.623 | 30.178 | 29.276 | 29.477 | 29.590 | 28.387 | 27.758 | 24.814 | 24.579 | 23.606 | 26.299 | 26.278 | 27.537 | 25.360 | 26.889 | 23.999 | 0.167 | 0.0000 | 0.0000 | 1 | 0 | 0 |
| 'Arpc4' | 119.612 | 131.308 | 129.152 | 123.047 | 136.621 | 135.897 | 127.516 | 132.539 | 127.108 | 133.156 | 127.834 | 118.449 | 133.268 | 130.148 | 132.750 | 132.656 | 125.640 | 123.617 | -0.004 | 0.9040 | 0.9624 | 0 | 0 | 0 |
| 'Arpc5' | 104.260 | 120.310 | 105.717 | 106.793 | 120.430 | 110.235 | 111.304 | 112.065 | 107.423 | 101.905 | 103.971 | 100.514 | 107.994 | 111.015 | 109.266 | 113.101 | 108.617 | 100.215 | 0.050 | 0.0753 | 0.2440 | 0 | 0 | 0 |
| 'Arpc5l' | 17.337 | 19.838 | 17.370 | 17.944 | 20.951 | 18.376 | 17.783 | 20.116 | 18.447 | 16.126 | 17.927 | 18.280 | 16.990 | 21.140 | 20.213 | 17.917 | 19.042 | 18.894 | 0.002 | 0.9525 | 0.9823 | 0 | 0 | 0 |
| 'Arpp19' | 4.325 | 4.546 | 4.524 | 4.858 | 4.877 | 4.705 | 4.749 | 4.633 | 4.463 | 4.154 | 4.009 | 4.340 | 4.522 | 4.517 | 4.680 | 4.490 | 4.667 | 4.551 | 0.050 | 0.1718 | 0.4055 | 0 | 0 | 0 |
| 'Arpp21' | 0.250 | 0.345 | 0.209 | 0.270 | 0.333 | 0.255 | 0.263 | 0.273 | 0.244 | 0.225 | 0.315 | 0.201 | 0.412 | 0.283 | 0.178 | 0.223 | 0.324 | 0.143 | 0.096 | 0.4141 | 0.6620 | 0 | 0 | 0 |
| 'Arrb1' | 4.040 | 4.197 | 4.349 | 3.515 | 4.536 | 4.478 | 3.492 | 4.250 | 4.154 | 4.250 | 3.948 | 4.274 | 3.071 | 3.704 | 4.113 | 4.079 | 3.419 | 4.047 | 0.073 | 0.2011 | 0.4425 | 0 | 0 | 0 |
| 'Arrb2' | 61.533 | 64.627 | 64.246 | 59.711 | 61.700 | 68.003 | 64.333 | 60.635 | 68.780 | 63.368 | 60.493 | 66.399 | 71.652 | 64.372 | 67.477 | 63.791 | 65.737 | 67.596 | -0.054 | 0.0572 | 0.2055 | 0 | 0 | 0 |
| 'Arrdc1' | 11.225 | 10.880 | 13.299 | 10.978 | 10.710 | 12.911 | 11.292 | 10.960 | 12.825 | 10.979 | 10.765 | 13.293 | 12.359 | 10.457 | 12.845 | 10.932 | 11.985 | 11.939 | -0.019 | 0.5616 | 0.7712 | 0 | 0 | 0 |
| 'Arrdc2' | 6.427 | 6.841 | 8.551 | 6.101 | 7.889 | 7.423 | 6.979 | 7.133 | 8.472 | 6.914 | 7.170 | 7.445 | 7.665 | 6.913 | 6.761 | 7.174 | 7.575 | 6.373 | 0.032 | 0.6121 | 0.8056 | 0 | 0 | 0 |
| 'Arrdc3' | 1.716 | 1.548 | 1.860 | 1.346 | 2.039 | 1.694 | 1.381 | 2.031 | 1.510 | 2.331 | 1.391 | 1.181 | 1.825 | 1.456 | 1.020 | 1.827 | 1.344 | 1.040 | 0.204 | 0.1391 | 0.3605 | 0 | 0 | 0 |
| 'Arrdc4' | 3.026 | 3.202 | 3.170 | 2.706 | 3.591 | 2.972 | 3.024 | 2.969 | 3.072 | 3.799 | 4.335 | 5.916 | 2.942 | 3.466 | 5.087 | 4.174 | 4.307 | 6.930 | -0.549 | 0.0000 | 0.0000 | 1 | 1 | 0 |
| 'Arsa' | 4.590 | 4.759 | 5.124 | 4.084 | 4.528 | 4.850 | 4.221 | 4.393 | 5.194 | 4.864 | 4.251 | 4.612 | 4.973 | 4.358 | 4.336 | 4.783 | 4.143 | 4.356 | 0.030 | 0.5278 | 0.7486 | 0 | 0 | 0 |
| 'Arsb' | 5.829 | 6.923 | 6.884 | 6.587 | 7.068 | 7.394 | 6.839 | 6.677 | 7.212 | 7.040 | 6.000 | 6.338 | 7.305 | 7.074 | 6.466 | 7.221 | 7.094 | 6.640 | -0.002 | 0.9712 | 0.9898 | 0 | 0 | 0 |
| 'Arsg' | 0.871 | 1.150 | 0.905 | 0.812 | 1.026 | 1.043 | 1.096 | 1.220 | 1.035 | 0.780 | 0.670 | 0.644 | 0.632 | 0.678 | 0.561 | 0.666 | 0.770 | 0.615 | 0.596 | 0.0000 | 0.0000 | 1 | 1 | 0 |
| 'Arsk' | 0.918 | 0.664 | 0.890 | 0.842 | 0.922 | 0.939 | 0.769 | 1.065 | 0.906 | 1.045 | 1.012 | 0.890 | 0.924 | 1.129 | 0.954 | 0.997 | 0.994 | 0.794 | -0.129 | 0.1635 | 0.3954 | 0 | 0 | 0 |
| 'Arv1' | 9.583 | 12.965 | 12.390 | 9.494 | 12.521 | 12.871 | 11.307 | 10.910 | 12.445 | 10.902 | 10.223 | 11.701 | 10.261 | 11.215 | 11.673 | 12.402 | 12.518 | 11.056 | 0.024 | 0.7099 | 0.8653 | 0 | 0 | 0 |
| 'Arvcf' | 0.492 | 0.595 | 0.656 | 0.485 | 0.587 | 0.738 | 0.517 | 0.580 | 0.643 | 0.638 | 0.460 | 0.566 | 0.459 | 0.368 | 0.575 | 0.447 | 0.407 | 0.622 | 0.212 | 0.0190 | 0.0998 | 0 | 0 | 0 |
| 'As3mt' | 5.133 | 5.332 | 5.803 | 6.136 | 5.531 | 5.922 | 6.958 | 5.050 | 5.990 | 6.077 | 5.980 | 4.913 | 5.516 | 5.288 | 5.118 | 4.988 | 5.413 | 4.236 | 0.122 | 0.1166 | 0.3243 | 0 | 0 | 0 |
| 'Asah1' | 2.889 | 3.358 | 3.071 | 2.844 | 3.769 | 3.275 | 3.182 | 3.487 | 3.049 | 3.044 | 3.759 | 2.796 | 3.399 | 3.685 | 3.137 | 3.156 | 3.239 | 2.903 | -0.013 | 0.8088 | 0.9139 | 0 | 0 | 0 |
| 'Asah2' | 2.019 | 2.162 | 2.026 | 2.106 | 1.992 | 2.190 | 2.521 | 2.464 | 2.670 | 2.632 | 2.506 | 2.673 | 2.739 | 2.590 | 2.159 | 2.831 | 2.680 | 2.488 | -0.219 | 0.0023 | 0.0205 | 1 | 0 | 0 |
| 'Asap1' | 31.935 | 31.910 | 31.703 | 30.412 | 31.055 | 30.542 | 29.554 | 30.972 | 30.657 | 34.656 | 31.037 | 31.481 | 31.750 | 30.679 | 31.911 | 33.299 | 28.560 | 30.916 | -0.038 | 0.1883 | 0.4246 | 0 | 0 | 0 |
| 'Asap2' | 2.760 | 2.433 | 2.824 | 2.204 | 2.570 | 2.926 | 2.609 | 2.979 | 2.967 | 2.917 | 3.733 | 3.130 | 3.191 | 3.642 | 3.031 | 2.677 | 3.292 | 3.203 | -0.247 | 0.0001 | 0.0016 | 1 | 0 | 0 |
| 'Asb1' | 4.025 | 4.104 | 4.119 | 3.557 | 4.214 | 4.087 | 3.723 | 4.539 | 3.964 | 4.025 | 4.159 | 3.823 | 3.907 | 4.364 | 4.046 | 4.205 | 3.727 | 3.680 | 0.012 | 0.7805 | 0.9013 | 0 | 0 | 0 |
| 'Asb13' | 11.146 | 13.205 | 12.667 | 10.006 | 12.949 | 12.207 | 10.832 | 12.550 | 13.383 | 11.051 | 10.926 | 13.414 | 12.229 | 12.213 | 13.519 | 13.163 | 12.352 | 13.850 | -0.063 | 0.1677 | 0.4000 | 0 | 0 | 0 |
| 'Asb2' | 79.238 | 81.033 | 82.896 | 81.355 | 82.459 | 83.029 | 81.242 | 79.784 | 86.775 | 82.733 | 78.268 | 82.349 | 81.441 | 75.435 | 82.235 | 76.172 | 73.084 | 82.511 | 0.036 | 0.1037 | 0.3020 | 0 | 0 | 0 |
| 'Asb3' | 2.347 | 2.667 | 2.304 | 2.798 | 2.241 | 2.213 | 3.072 | 2.468 | 2.271 | 2.223 | 2.669 | 2.747 | 2.075 | 2.433 | 2.579 | 2.168 | 2.664 | 2.515 | 0.004 | 0.9568 | 0.9839 | 0 | 0 | 0 |
| 'Asb6' | 6.904 | 6.803 | 7.172 | 6.160 | 6.510 | 7.290 | 6.611 | 6.491 | 7.309 | 6.375 | 5.709 | 7.769 | 6.790 | 6.537 | 7.302 | 6.772 | 6.270 | 7.325 | -0.003 | 0.9304 | 0.9737 | 0 | 0 | 0 |
| 'Asb7' | 2.312 | 2.986 | 2.376 | 2.169 | 2.826 | 2.440 | 2.332 | 2.824 | 2.355 | 2.634 | 2.501 | 2.274 | 2.717 | 2.846 | 2.241 | 2.901 | 2.852 | 2.073 | -0.033 | 0.5362 | 0.7552 | 0 | 0 | 0 |
| 'Asb8' | 8.267 | 9.454 | 10.152 | 8.633 | 9.624 | 10.152 | 9.768 | 10.332 | 10.797 | 9.152 | 9.418 | 9.414 | 7.764 | 9.708 | 10.091 | 8.764 | 8.985 | 9.541 | 0.067 | 0.1308 | 0.3473 | 0 | 0 | 0 |
| 'Ascc1' | 22.747 | 25.406 | 22.521 | 25.978 | 24.777 | 25.370 | 24.184 | 24.839 | 22.449 | 24.412 | 22.901 | 21.489 | 27.286 | 25.892 | 23.189 | 26.216 | 26.413 | 20.990 | -0.009 | 0.8458 | 0.9343 | 0 | 0 | 0 |
| 'Ascc2' | 13.816 | 12.485 | 15.447 | 14.211 | 13.734 | 14.168 | 15.177 | 13.291 | 14.457 | 13.907 | 15.376 | 15.727 | 14.664 | 13.651 | 14.920 | 14.554 | 14.162 | 14.614 | -0.064 | 0.0625 | 0.2169 | 0 | 0 | 0 |
| 'Ascc3' | 13.355 | 12.340 | 12.426 | 13.180 | 13.058 | 11.333 | 12.883 | 11.970 | 12.323 | 13.285 | 12.836 | 12.634 | 11.692 | 12.777 | 11.732 | 11.978 | 13.666 | 12.751 | -0.018 | 0.6495 | 0.8292 | 0 | 0 | 0 |
| 'Ascl2' | 27.813 | 30.728 | 25.026 | 27.384 | 29.080 | 25.140 | 26.679 | 28.088 | 26.827 | 27.173 | 31.375 | 29.042 | 34.079 | 30.907 | 29.052 | 29.839 | 34.232 | 29.052 | -0.168 | 0.0000 | 0.0002 | 1 | 0 | 0 |
| 'Asf1a' | 11.731 | 13.532 | 12.645 | 13.538 | 13.854 | 12.894 | 14.307 | 13.033 | 14.204 | 12.388 | 12.821 | 13.333 | 12.316 | 11.528 | 13.070 | 11.667 | 12.756 | 13.133 | 0.065 | 0.1563 | 0.3855 | 0 | 0 | 0 |
| 'Asf1b' | 13.359 | 13.054 | 16.410 | 15.815 | 14.546 | 17.325 | 15.460 | 13.674 | 17.495 | 10.389 | 13.710 | 15.874 | 17.866 | 15.940 | 17.955 | 12.566 | 14.625 | 15.942 | 0.012 | 0.8742 | 0.9463 | 0 | 0 | 0 |
| 'Ash1l' | 10.881 | 9.036 | 9.224 | 10.767 | 8.676 | 8.658 | 9.562 | 9.617 | 9.707 | 10.275 | 10.615 | 10.486 | 9.334 | 9.887 | 9.620 | 10.191 | 9.870 | 11.390 | -0.103 | 0.0380 | 0.1585 | 0 | 0 | 0 |
| 'Ash2l' | 5.944 | 6.639 | 7.077 | 5.800 | 6.761 | 6.562 | 6.655 | 6.860 | 6.471 | 6.447 | 5.929 | 5.917 | 6.603 | 6.304 | 6.040 | 6.475 | 6.311 | 5.410 | 0.077 | 0.0655 | 0.2234 | 0 | 0 | 0 |
| 'Asic1' | 2.258 | 2.328 | 2.816 | 2.184 | 2.261 | 2.762 | 1.989 | 1.928 | 2.263 | 1.802 | 2.190 | 2.626 | 1.966 | 2.258 | 2.298 | 1.786 | 1.902 | 2.394 | 0.099 | 0.1991 | 0.4393 | 0 | 0 | 0 |
| 'Asl' | 8.834 | 8.752 | 9.148 | 9.307 | 8.527 | 9.180 | 9.615 | 8.919 | 9.021 | 8.895 | 8.661 | 9.558 | 9.488 | 8.518 | 9.462 | 9.451 | 9.951 | 8.510 | -0.032 | 0.3417 | 0.5981 | 0 | 0 | 0 |
| 'Asna1' | 49.684 | 52.345 | 47.898 | 50.163 | 52.054 | 54.101 | 52.231 | 52.491 | 52.810 | 51.614 | 45.349 | 52.257 | 55.444 | 47.207 | 52.434 | 52.034 | 46.168 | 48.355 | 0.029 | 0.4266 | 0.6730 | 0 | 0 | 0 |
| 'Asns' | 1.090 | 0.924 | 1.372 | 0.812 | 1.043 | 1.011 | 1.057 | 1.027 | 1.260 | 0.929 | 0.612 | 0.791 | 0.848 | 0.792 | 0.893 | 0.845 | 0.611 | 0.877 | 0.421 | 0.0002 | 0.0031 | 1 | 1 | 0 |
| 'Aspm' | 1.700 | 2.223 | 2.587 | 2.379 | 2.132 | 2.443 | 2.206 | 2.208 | 2.569 | 1.463 | 2.504 | 2.418 | 2.352 | 2.725 | 2.992 | 1.957 | 2.689 | 2.388 | -0.074 | 0.3615 | 0.6135 | 0 | 0 | 0 |
| 'Aspscr1' | 10.953 | 11.043 | 11.913 | 10.373 | 10.787 | 12.680 | 10.364 | 11.345 | 12.391 | 11.803 | 10.310 | 11.764 | 11.906 | 10.768 | 11.375 | 11.662 | 10.720 | 11.042 | -0.003 | 0.9355 | 0.9752 | 0 | 0 | 0 |
| 'Asrgl1' | 11.323 | 10.571 | 10.889 | 10.909 | 10.000 | 10.205 | 11.799 | 11.254 | 10.878 | 9.313 | 10.191 | 10.255 | 10.825 | 10.714 | 10.507 | 9.900 | 10.454 | 9.590 | 0.082 | 0.0377 | 0.1575 | 0 | 0 | 0 |
| 'Aste1' | 1.264 | 1.259 | 1.291 | 1.189 | 1.159 | 1.200 | 1.230 | 1.398 | 1.282 | 1.317 | 1.138 | 1.250 | 1.386 | 1.203 | 1.130 | 1.270 | 1.334 | 1.244 | -0.007 | 0.9080 | 0.9639 | 0 | 0 | 0 |
| 'Asun' | 6.677 | 6.479 | 6.471 | 7.185 | 6.573 | 6.227 | 7.187 | 6.588 | 6.730 | 6.310 | 6.911 | 7.288 | 6.437 | 7.121 | 7.258 | 6.357 | 7.406 | 7.119 | -0.061 | 0.1413 | 0.3636 | 0 | 0 | 0 |
| 'Asxl1' | 23.066 | 19.209 | 22.895 | 21.354 | 19.639 | 21.002 | 22.229 | 20.913 | 21.938 | 23.870 | 22.443 | 21.081 | 24.556 | 21.361 | 19.641 | 21.664 | 22.302 | 21.506 | -0.055 | 0.1470 | 0.3726 | 0 | 0 | 0 |
| 'Asxl2' | 6.454 | 6.357 | 6.384 | 6.532 | 6.497 | 6.218 | 6.602 | 6.025 | 6.595 | 6.346 | 6.032 | 6.254 | 5.915 | 5.740 | 5.998 | 6.129 | 6.113 | 6.195 | 0.064 | 0.0098 | 0.0616 | 0 | 0 | 0 |
| 'Atad1' | 33.393 | 39.781 | 37.446 | 35.798 | 39.115 | 34.617 | 38.170 | 39.782 | 35.046 | 36.349 | 36.454 | 35.176 | 33.490 | 40.143 | 34.205 | 35.472 | 39.393 | 33.373 | 0.030 | 0.3168 | 0.5748 | 0 | 0 | 0 |
| 'Atad2' | 23.254 | 22.558 | 26.087 | 26.224 | 22.647 | 24.134 | 23.557 | 22.237 | 24.879 | 21.331 | 24.895 | 26.376 | 25.696 | 25.616 | 25.897 | 20.234 | 24.430 | 28.507 | -0.058 | 0.2455 | 0.4959 | 0 | 0 | 0 |
| 'Atad2b' | 11.721 | 11.304 | 11.842 | 11.819 | 11.528 | 10.922 | 12.377 | 11.793 | 10.861 | 12.008 | 13.023 | 12.118 | 11.531 | 12.340 | 11.180 | 11.841 | 12.395 | 11.950 | -0.068 | 0.0305 | 0.1363 | 0 | 0 | 0 |
| 'Atad3a' | 7.619 | 6.880 | 8.250 | 7.603 | 7.812 | 7.106 | 7.225 | 7.725 | 7.470 | 7.874 | 6.284 | 8.109 | 7.554 | 6.223 | 7.605 | 7.475 | 7.021 | 7.186 | 0.041 | 0.3427 | 0.5990 | 0 | 0 | 0 |
| 'Atad5' | 3.061 | 2.782 | 3.578 | 3.522 | 2.897 | 2.999 | 3.467 | 2.909 | 3.243 | 3.052 | 3.191 | 2.995 | 3.854 | 3.000 | 2.845 | 2.800 | 3.058 | 2.883 | 0.029 | 0.6176 | 0.8087 | 0 | 0 | 0 |
| 'Atat1' | 1.408 | 1.411 | 1.589 | 1.523 | 1.599 | 1.624 | 1.478 | 1.525 | 1.934 | 1.252 | 1.283 | 1.876 | 1.434 | 1.528 | 1.642 | 1.302 | 1.407 | 1.687 | 0.052 | 0.3923 | 0.6418 | 0 | 0 | 0 |
| 'Atcay' | 12.476 | 13.637 | 11.831 | 13.235 | 13.051 | 14.096 | 14.152 | 12.396 | 13.431 | 13.309 | 13.139 | 14.618 | 13.989 | 13.119 | 13.854 | 12.855 | 14.439 | 15.449 | -0.089 | 0.0328 | 0.1434 | 0 | 0 | 0 |
| 'Ate1' | 6.815 | 8.215 | 7.574 | 7.637 | 7.720 | 7.735 | 8.184 | 7.541 | 7.688 | 8.024 | 8.537 | 8.428 | 7.786 | 8.234 | 8.376 | 7.450 | 8.718 | 8.013 | -0.104 | 0.0025 | 0.0216 | 1 | 0 | 0 |
| 'Atf2' | 6.981 | 6.974 | 6.675 | 6.696 | 6.906 | 6.320 | 7.190 | 7.173 | 6.560 | 6.751 | 7.584 | 7.206 | 6.950 | 7.546 | 6.594 | 6.696 | 7.609 | 6.463 | -0.055 | 0.0630 | 0.2181 | 0 | 0 | 0 |
| 'Atf4' | 194.077 | 144.987 | 114.192 | 158.334 | 136.988 | 122.657 | 151.795 | 143.980 | 110.901 | 173.560 | 146.319 | 100.496 | 178.812 | 165.046 | 106.891 | 159.708 | 152.980 | 97.242 | 0.005 | 0.9287 | 0.9728 | 0 | 0 | 0 |
| 'Atf5' | 3.445 | 2.968 | 3.650 | 2.992 | 2.961 | 3.927 | 3.091 | 3.186 | 3.516 | 3.496 | 3.028 | 4.110 | 3.737 | 3.263 | 4.203 | 2.803 | 3.465 | 4.574 | -0.146 | 0.0325 | 0.1423 | 0 | 0 | 0 |
| 'Atf6' | 8.978 | 8.285 | 8.741 | 8.875 | 8.290 | 7.896 | 8.896 | 8.413 | 8.014 | 4.915 | 5.798 | 4.719 | 4.760 | 4.913 | 4.679 | 4.725 | 4.707 | 4.420 | 0.799 | 0.0000 | 0.0000 | 1 | 1 | 0 |
| 'Atf6b' | 39.944 | 35.544 | 45.536 | 36.499 | 37.982 | 46.360 | 38.821 | 37.521 | 43.345 | 41.160 | 35.429 | 44.620 | 41.067 | 36.258 | 41.858 | 39.355 | 35.362 | 43.756 | 0.001 | 0.9756 | 0.9918 | 0 | 0 | 0 |
| 'Atf7ip' | 10.141 | 9.460 | 9.347 | 9.705 | 9.476 | 8.876 | 9.551 | 9.382 | 9.667 | 9.866 | 10.889 | 10.706 | 8.905 | 9.661 | 10.302 | 9.838 | 10.216 | 10.517 | -0.097 | 0.0115 | 0.0693 | 0 | 0 | 0 |
| 'Atg10' | 4.525 | 4.444 | 3.607 | 5.836 | 4.244 | 4.169 | 4.871 | 4.580 | 3.755 | 4.704 | 5.054 | 4.763 | 4.224 | 4.440 | 4.519 | 3.471 | 5.527 | 4.471 | -0.066 | 0.4788 | 0.7114 | 0 | 0 | 0 |
| 'Atg101' | 26.259 | 26.151 | 22.454 | 26.267 | 25.897 | 25.289 | 25.294 | 27.395 | 24.154 | 24.529 | 23.428 | 26.030 | 25.511 | 27.423 | 25.406 | 27.716 | 26.009 | 27.311 | -0.040 | 0.3711 | 0.6230 | 0 | 0 | 0 |
| 'Atg12' | 27.226 | 25.564 | 22.939 | 24.010 | 27.378 | 23.207 | 23.401 | 24.988 | 21.802 | 23.728 | 24.801 | 22.476 | 23.825 | 26.805 | 20.700 | 25.362 | 25.312 | 22.578 | 0.024 | 0.5165 | 0.7404 | 0 | 0 | 0 |
| 'Atg13' | 6.835 | 7.465 | 7.494 | 6.529 | 7.512 | 7.436 | 6.956 | 7.611 | 7.324 | 7.150 | 7.200 | 7.611 | 6.619 | 7.000 | 7.381 | 7.111 | 7.155 | 7.365 | 0.002 | 0.9567 | 0.9839 | 0 | 0 | 0 |
| 'Atg14' | 7.453 | 7.779 | 7.728 | 5.949 | 8.365 | 7.772 | 7.532 | 8.528 | 7.415 | 8.205 | 7.280 | 7.696 | 7.363 | 7.650 | 6.648 | 8.042 | 8.147 | 7.094 | 0.000 | 0.9926 | 0.9986 | 0 | 0 | 0 |
| 'Atg16l1' | 4.209 | 4.099 | 4.387 | 3.791 | 3.949 | 4.488 | 3.952 | 4.436 | 4.425 | 4.137 | 4.146 | 4.395 | 3.898 | 3.755 | 4.121 | 4.062 | 4.162 | 4.133 | 0.027 | 0.4036 | 0.6527 | 0 | 0 | 0 |
| 'Atg16l2' | 6.480 | 5.306 | 7.139 | 5.431 | 5.466 | 7.334 | 6.524 | 5.812 | 7.152 | 6.682 | 6.499 | 8.431 | 6.948 | 5.965 | 7.272 | 6.084 | 7.083 | 7.639 | -0.158 | 0.0005 | 0.0064 | 1 | 0 | 0 |
| 'Atg2a' | 12.044 | 10.956 | 13.008 | 11.309 | 11.533 | 12.251 | 10.878 | 10.903 | 13.021 | 12.669 | 12.083 | 13.120 | 12.345 | 11.863 | 12.422 | 12.757 | 12.396 | 13.097 | -0.100 | 0.0009 | 0.0095 | 1 | 0 | 0 |
| 'Atg2b' | 9.801 | 8.818 | 8.910 | 9.049 | 8.432 | 8.413 | 8.629 | 8.903 | 9.010 | 8.983 | 9.326 | 9.639 | 9.104 | 8.860 | 9.713 | 8.093 | 8.933 | 9.610 | -0.051 | 0.1804 | 0.4156 | 0 | 0 | 0 |
| 'Atg3' | 14.237 | 16.330 | 16.561 | 16.355 | 16.082 | 15.819 | 15.560 | 15.682 | 15.603 | 15.195 | 15.662 | 15.210 | 15.516 | 15.439 | 14.953 | 13.549 | 17.551 | 14.562 | 0.037 | 0.3294 | 0.5875 | 0 | 0 | 0 |
| 'Atg4a' | 7.794 | 10.446 | 8.759 | 7.969 | 8.919 | 8.504 | 8.322 | 9.571 | 8.066 | 8.564 | 8.946 | 8.736 | 8.726 | 9.669 | 7.703 | 8.319 | 9.214 | 7.761 | -0.001 | 0.9763 | 0.9918 | 0 | 0 | 0 |
| 'Atg4b' | 6.622 | 6.745 | 6.905 | 6.672 | 6.703 | 6.918 | 6.162 | 7.491 | 6.523 | 6.310 | 6.789 | 6.785 | 6.227 | 6.049 | 6.470 | 6.732 | 6.216 | 6.542 | 0.054 | 0.1186 | 0.3278 | 0 | 0 | 0 |
| 'Atg4d' | 18.051 | 17.630 | 18.211 | 18.189 | 21.316 | 20.380 | 17.431 | 18.332 | 20.146 | 17.519 | 18.064 | 20.564 | 17.482 | 18.856 | 20.211 | 19.145 | 17.116 | 20.231 | -0.007 | 0.8622 | 0.9403 | 0 | 0 | 0 |
| 'Atg5' | 14.966 | 15.944 | 14.312 | 15.051 | 16.196 | 14.597 | 13.977 | 17.169 | 13.886 | 15.605 | 16.737 | 14.299 | 15.290 | 16.028 | 13.433 | 15.635 | 15.824 | 13.638 | -0.010 | 0.7872 | 0.9048 | 0 | 0 | 0 |
| 'Atg7' | 4.549 | 4.365 | 4.193 | 4.267 | 4.165 | 4.078 | 4.115 | 4.204 | 3.991 | 4.420 | 4.298 | 4.205 | 4.176 | 4.527 | 4.238 | 4.324 | 4.108 | 4.301 | -0.034 | 0.3194 | 0.5780 | 0 | 0 | 0 |
| 'Atg9a' | 4.288 | 4.746 | 4.719 | 4.226 | 4.119 | 4.879 | 4.142 | 4.408 | 5.239 | 4.450 | 4.351 | 5.381 | 4.305 | 4.703 | 4.684 | 4.470 | 4.451 | 4.926 | -0.046 | 0.1916 | 0.4285 | 0 | 0 | 0 |
| 'Athl1' | 6.667 | 6.895 | 7.273 | 6.224 | 7.091 | 7.671 | 7.210 | 7.135 | 7.780 | 7.276 | 7.253 | 7.333 | 7.414 | 6.253 | 6.628 | 6.590 | 7.091 | 6.498 | 0.026 | 0.5451 | 0.7606 | 0 | 0 | 0 |
| 'Atic' | 14.729 | 13.889 | 15.289 | 14.736 | 14.967 | 14.164 | 14.769 | 15.032 | 15.038 | 14.596 | 13.731 | 15.945 | 14.276 | 13.955 | 15.126 | 13.407 | 13.637 | 15.740 | 0.013 | 0.6686 | 0.8400 | 0 | 0 | 0 |
| 'Atl2' | 7.030 | 8.269 | 7.073 | 7.150 | 7.170 | 6.746 | 7.735 | 8.119 | 7.076 | 5.909 | 7.908 | 6.650 | 6.705 | 7.309 | 6.348 | 6.444 | 7.426 | 6.962 | 0.095 | 0.0252 | 0.1213 | 0 | 0 | 0 |
| 'Atl3' | 25.314 | 24.914 | 26.039 | 25.689 | 26.830 | 23.710 | 26.051 | 26.112 | 24.513 | 26.363 | 28.199 | 25.882 | 25.615 | 27.977 | 25.763 | 27.199 | 29.781 | 24.639 | -0.083 | 0.0031 | 0.0261 | 1 | 0 | 0 |
| 'Atm' | 9.175 | 9.337 | 9.603 | 10.194 | 9.763 | 8.864 | 9.175 | 9.139 | 9.665 | 10.093 | 9.530 | 10.234 | 9.881 | 9.189 | 9.213 | 9.540 | 10.020 | 10.142 | -0.061 | 0.0572 | 0.2055 | 0 | 0 | 0 |
| 'Atmin' | 25.754 | 26.270 | 24.714 | 23.647 | 25.271 | 26.894 | 23.553 | 26.112 | 25.873 | 24.773 | 21.309 | 23.580 | 23.252 | 23.521 | 22.462 | 24.238 | 20.822 | 24.316 | 0.121 | 0.0009 | 0.0096 | 1 | 0 | 0 |
| 'Atox1' | 39.750 | 44.017 | 39.922 | 40.190 | 42.069 | 45.525 | 40.787 | 43.541 | 43.128 | 36.078 | 38.276 | 42.176 | 43.066 | 45.124 | 44.076 | 39.805 | 40.791 | 37.075 | 0.036 | 0.4189 | 0.6660 | 0 | 0 | 0 |
| 'Atp10a' | 9.150 | 8.894 | 10.881 | 9.743 | 8.605 | 10.132 | 9.430 | 8.291 | 9.670 | 9.777 | 8.681 | 11.469 | 10.727 | 9.675 | 11.218 | 9.876 | 9.851 | 12.208 | -0.153 | 0.0001 | 0.0010 | 1 | 0 | 0 |
| 'Atp10d' | 5.386 | 5.931 | 5.622 | 5.550 | 5.865 | 5.584 | 5.197 | 5.585 | 5.631 | 5.749 | 6.221 | 6.381 | 5.986 | 5.715 | 6.376 | 5.643 | 5.837 | 6.224 | -0.118 | 0.0000 | 0.0008 | 1 | 0 | 0 |
| 'Atp11a' | 5.773 | 4.841 | 6.374 | 5.446 | 4.907 | 5.470 | 5.461 | 5.635 | 6.184 | 5.508 | 5.783 | 6.250 | 5.364 | 5.011 | 5.851 | 4.892 | 5.724 | 6.472 | -0.032 | 0.4960 | 0.7249 | 0 | 0 | 0 |
| 'Atp11b' | 11.968 | 12.588 | 11.972 | 11.532 | 12.167 | 11.232 | 12.614 | 12.471 | 11.750 | 12.138 | 12.524 | 12.353 | 12.188 | 12.586 | 11.549 | 12.046 | 12.401 | 11.739 | -0.028 | 0.1907 | 0.4271 | 0 | 0 | 0 |
| 'Atp11c' | 2.359 | 2.384 | 2.300 | 2.202 | 2.033 | 2.014 | 2.542 | 2.244 | 2.255 | 2.220 | 2.252 | 2.471 | 2.065 | 2.407 | 2.168 | 2.236 | 2.404 | 2.216 | -0.020 | 0.6747 | 0.8439 | 0 | 0 | 0 |
| 'Atp13a1' | 33.030 | 29.518 | 39.611 | 31.638 | 28.879 | 36.386 | 32.355 | 29.966 | 37.140 | 30.516 | 27.933 | 40.819 | 34.499 | 28.475 | 36.799 | 32.738 | 32.419 | 38.191 | -0.029 | 0.3715 | 0.6235 | 0 | 0 | 0 |
| 'Atp13a2' | 11.019 | 10.800 | 12.475 | 10.035 | 10.591 | 13.327 | 10.757 | 10.551 | 12.802 | 11.078 | 9.937 | 11.734 | 12.273 | 9.114 | 11.483 | 10.807 | 9.838 | 12.050 | 0.047 | 0.2185 | 0.4655 | 0 | 0 | 0 |
| 'Atp13a3' | 18.017 | 19.191 | 18.550 | 18.515 | 18.021 | 16.883 | 19.293 | 18.523 | 17.156 | 17.418 | 20.304 | 19.249 | 18.404 | 20.106 | 18.416 | 17.593 | 19.759 | 20.152 | -0.073 | 0.0406 | 0.1649 | 0 | 0 | 0 |
| 'Atp1a1' | 88.970 | 88.979 | 89.687 | 82.328 | 88.262 | 90.066 | 83.739 | 89.025 | 90.744 | 88.671 | 83.625 | 90.615 | 84.037 | 85.732 | 89.568 | 89.622 | 84.380 | 90.953 | -0.002 | 0.9324 | 0.9746 | 0 | 0 | 0 |
| 'Atp1a3' | 21.307 | 24.336 | 18.461 | 20.710 | 24.620 | 21.395 | 20.093 | 22.943 | 20.485 | 24.252 | 25.296 | 18.402 | 28.806 | 25.124 | 17.781 | 21.624 | 24.024 | 19.314 | -0.076 | 0.1690 | 0.4013 | 0 | 0 | 0 |
| 'Atp1b1' | 21.378 | 31.435 | 19.792 | 23.527 | 31.442 | 24.044 | 21.932 | 27.761 | 21.663 | 22.165 | 25.855 | 18.138 | 27.385 | 26.824 | 18.140 | 25.456 | 25.340 | 17.524 | 0.101 | 0.0895 | 0.2744 | 0 | 0 | 0 |
| 'Atp1b3' | 87.977 | 98.076 | 91.893 | 93.307 | 99.532 | 95.433 | 93.830 | 102.744 | 91.786 | 93.690 | 101.332 | 96.038 | 95.844 | 98.933 | 100.435 | 104.844 | 97.645 | 101.292 | -0.070 | 0.0109 | 0.0667 | 0 | 0 | 0 |
| 'Atp2a2' | 30.730 | 30.386 | 32.151 | 30.262 | 32.719 | 31.890 | 30.660 | 32.155 | 30.642 | 30.950 | 31.744 | 31.272 | 31.837 | 31.121 | 31.102 | 30.659 | 30.262 | 32.677 | -0.011 | 0.6318 | 0.8176 | 0 | 0 | 0 |
| 'Atp2a3' | 60.755 | 63.988 | 69.624 | 59.930 | 64.713 | 70.155 | 60.257 | 63.930 | 69.860 | 67.521 | 62.720 | 63.849 | 66.983 | 60.744 | 65.464 | 66.239 | 58.343 | 65.328 | 0.003 | 0.9190 | 0.9693 | 0 | 0 | 0 |
| 'Atp2b1' | 12.859 | 13.413 | 13.769 | 13.933 | 13.736 | 13.186 | 13.180 | 13.528 | 13.284 | 14.676 | 14.810 | 14.678 | 13.235 | 13.769 | 13.444 | 14.169 | 15.254 | 14.174 | -0.095 | 0.0012 | 0.0123 | 1 | 0 | 0 |
| 'Atp2b4' | 3.582 | 3.676 | 3.834 | 3.696 | 3.434 | 3.616 | 2.740 | 3.659 | 3.447 | 7.567 | 8.327 | 7.435 | 7.086 | 8.362 | 6.768 | 7.234 | 7.785 | 7.646 | -1.114 | 0.0000 | 0.0000 | 1 | 1 | 1 |
| 'Atp2c1' | 5.403 | 5.686 | 5.704 | 5.395 | 6.004 | 5.470 | 5.626 | 5.586 | 5.618 | 5.509 | 5.792 | 5.704 | 5.039 | 5.464 | 5.385 | 5.484 | 5.570 | 5.604 | 0.016 | 0.5690 | 0.7768 | 0 | 0 | 0 |
| 'Atp5a1' | 121.143 | 127.428 | 117.902 | 119.836 | 127.359 | 123.008 | 117.862 | 126.423 | 121.361 | 121.255 | 119.389 | 111.813 | 124.040 | 123.027 | 120.202 | 122.003 | 119.620 | 116.530 | 0.022 | 0.3000 | 0.5564 | 0 | 0 | 0 |
| 'Atp5b' | 102.247 | 105.364 | 101.135 | 109.777 | 108.558 | 103.067 | 110.686 | 105.095 | 101.103 | 97.238 | 103.369 | 94.585 | 106.997 | 103.788 | 101.680 | 102.722 | 105.253 | 99.110 | 0.039 | 0.1044 | 0.3027 | 0 | 0 | 0 |
| 'Atp5c1' | 59.878 | 67.341 | 63.409 | 63.883 | 63.290 | 58.569 | 65.034 | 63.518 | 58.006 | 57.567 | 60.668 | 54.792 | 59.484 | 62.175 | 58.927 | 58.836 | 60.185 | 52.529 | 0.090 | 0.0011 | 0.0112 | 1 | 0 | 0 |
| 'Atp5d' | 34.997 | 36.466 | 36.283 | 33.132 | 35.804 | 36.410 | 33.885 | 34.522 | 33.788 | 32.488 | 31.467 | 33.711 | 33.845 | 32.192 | 35.167 | 33.841 | 31.616 | 33.201 | 0.073 | 0.0051 | 0.0380 | 1 | 0 | 0 |
| 'Atp5e' | 53.133 | 55.448 | 51.819 | 58.614 | 56.481 | 55.271 | 56.774 | 54.428 | 53.762 | 49.013 | 53.706 | 44.283 | 53.965 | 52.461 | 48.075 | 54.904 | 54.099 | 48.013 | 0.106 | 0.0021 | 0.0188 | 1 | 0 | 0 |
| 'Atp5f1' | 27.669 | 32.507 | 29.090 | 28.478 | 32.765 | 29.577 | 31.659 | 30.767 | 29.276 | 27.456 | 27.951 | 27.555 | 30.771 | 30.549 | 28.722 | 30.758 | 29.923 | 27.592 | 0.045 | 0.1507 | 0.3777 | 0 | 0 | 0 |
| 'Atp5g1' | 6.092 | 6.895 | 6.225 | 6.288 | 6.848 | 6.205 | 6.406 | 6.369 | 6.104 | 6.387 | 6.204 | 5.961 | 6.458 | 6.331 | 6.352 | 5.982 | 6.325 | 6.199 | 0.018 | 0.5603 | 0.7706 | 0 | 0 | 0 |
| 'Atp5g2' | 75.238 | 77.526 | 73.510 | 76.524 | 77.687 | 77.222 | 74.707 | 78.969 | 72.361 | 69.930 | 72.726 | 68.639 | 79.026 | 73.973 | 78.750 | 71.275 | 72.835 | 73.674 | 0.038 | 0.2239 | 0.4711 | 0 | 0 | 0 |
| 'Atp5g3' | 68.655 | 72.662 | 66.930 | 68.918 | 74.526 | 70.954 | 66.938 | 71.084 | 67.932 | 63.281 | 66.827 | 60.658 | 67.587 | 69.570 | 67.550 | 67.978 | 69.698 | 63.177 | 0.067 | 0.0133 | 0.0769 | 0 | 0 | 0 |
| 'Atp5h' | 51.061 | 56.941 | 52.280 | 55.854 | 55.594 | 53.638 | 57.515 | 55.073 | 53.752 | 47.039 | 50.547 | 48.908 | 52.345 | 52.024 | 53.615 | 52.366 | 51.119 | 48.475 | 0.096 | 0.0013 | 0.0129 | 1 | 0 | 0 |
| 'Atp5j' | 18.525 | 21.691 | 18.661 | 20.424 | 21.401 | 19.321 | 20.111 | 19.719 | 19.478 | 18.071 | 18.246 | 17.883 | 19.723 | 19.744 | 18.560 | 20.982 | 19.693 | 17.450 | 0.063 | 0.0567 | 0.2047 | 0 | 0 | 0 |
| 'Atp5j2' | 23.147 | 26.854 | 23.777 | 25.815 | 26.036 | 25.544 | 26.106 | 24.069 | 24.885 | 23.020 | 24.564 | 22.214 | 24.600 | 25.338 | 24.920 | 24.166 | 24.928 | 22.795 | 0.053 | 0.1235 | 0.3356 | 0 | 0 | 0 |
| 'Atp5k' | 45.074 | 47.627 | 44.912 | 50.666 | 47.161 | 47.180 | 50.704 | 46.294 | 45.527 | 40.812 | 43.411 | 41.816 | 45.515 | 45.084 | 47.726 | 48.475 | 47.535 | 44.355 | 0.061 | 0.1509 | 0.3777 | 0 | 0 | 0 |
| 'Atp5l' | 155.806 | 169.039 | 149.188 | 165.916 | 168.470 | 153.757 | 162.859 | 164.065 | 158.315 | 137.065 | 157.307 | 134.395 | 148.442 | 162.503 | 144.452 | 146.323 | 157.264 | 146.708 | 0.108 | 0.0009 | 0.0096 | 1 | 0 | 0 |
| 'Atp5o' | 27.302 | 29.639 | 27.687 | 29.743 | 31.250 | 29.444 | 28.373 | 28.733 | 28.644 | 25.968 | 27.055 | 27.299 | 28.775 | 26.992 | 27.865 | 28.592 | 28.786 | 27.154 | 0.058 | 0.0478 | 0.1828 | 0 | 0 | 0 |
| 'Atp5s' | 3.132 | 4.185 | 5.032 | 3.528 | 4.360 | 3.695 | 4.916 | 3.640 | 4.029 | 3.411 | 3.136 | 4.003 | 3.427 | 4.069 | 3.918 | 3.777 | 3.718 | 3.774 | 0.119 | 0.2162 | 0.4622 | 0 | 0 | 0 |
| 'Atp5sl' | 8.777 | 8.975 | 10.400 | 7.953 | 9.166 | 9.945 | 8.294 | 9.565 | 9.905 | 10.075 | 8.729 | 9.560 | 9.076 | 8.342 | 8.381 | 9.438 | 7.985 | 8.165 | 0.048 | 0.3516 | 0.6056 | 0 | 0 | 0 |
| 'Atp6ap1' | 14.893 | 16.280 | 14.994 | 14.647 | 15.933 | 15.760 | 14.058 | 15.479 | 15.236 | 17.304 | 17.126 | 16.764 | 16.778 | 17.570 | 17.117 | 16.573 | 17.172 | 16.859 | -0.169 | 0.0000 | 0.0000 | 1 | 0 | 0 |
| 'Atp6ap2' | 25.620 | 29.379 | 26.257 | 28.130 | 27.421 | 25.430 | 27.642 | 27.604 | 26.555 | 27.570 | 29.177 | 27.196 | 26.997 | 30.524 | 25.947 | 26.203 | 28.238 | 26.048 | -0.035 | 0.2447 | 0.4952 | 0 | 0 | 0 |
| 'Atp6v0a1' | 1.090 | 1.128 | 1.281 | 1.183 | 1.086 | 1.215 | 1.082 | 1.105 | 1.448 | 1.146 | 1.165 | 1.009 | 0.901 | 1.175 | 0.884 | 1.004 | 1.304 | 0.809 | 0.180 | 0.0380 | 0.1583 | 0 | 0 | 0 |
| 'Atp6v0a2' | 24.929 | 25.650 | 23.434 | 24.691 | 23.718 | 23.462 | 23.334 | 23.940 | 23.345 | 24.670 | 25.992 | 26.532 | 25.771 | 25.382 | 25.365 | 24.209 | 25.075 | 26.490 | -0.096 | 0.0005 | 0.0061 | 1 | 0 | 0 |
| 'Atp6v0b' | 18.218 | 19.833 | 19.089 | 18.729 | 20.363 | 20.780 | 18.219 | 20.770 | 19.261 | 18.462 | 18.718 | 21.680 | 19.567 | 20.930 | 20.388 | 20.351 | 21.068 | 20.671 | -0.066 | 0.0316 | 0.1395 | 0 | 0 | 0 |
| 'Atp6v0c' | 10.745 | 9.922 | 10.047 | 10.454 | 10.650 | 10.494 | 9.382 | 10.405 | 10.453 | 11.628 | 10.058 | 9.961 | 10.712 | 10.856 | 9.781 | 10.306 | 9.798 | 10.293 | -0.020 | 0.6069 | 0.8023 | 0 | 0 | 0 |
| 'Atp6v0d1' | 34.825 | 39.449 | 37.147 | 36.867 | 38.668 | 40.820 | 34.642 | 40.537 | 38.253 | 36.996 | 37.738 | 40.086 | 39.518 | 40.473 | 40.303 | 41.257 | 37.797 | 36.531 | -0.050 | 0.1358 | 0.3545 | 0 | 0 | 0 |
| 'Atp6v0e' | 25.106 | 29.225 | 26.135 | 25.863 | 29.245 | 27.595 | 25.572 | 28.844 | 26.028 | 25.181 | 27.225 | 26.216 | 25.022 | 27.643 | 26.178 | 26.104 | 25.728 | 24.498 | 0.047 | 0.0717 | 0.2363 | 0 | 0 | 0 |
| 'Atp6v0e2' | 5.617 | 6.950 | 6.143 | 6.242 | 6.948 | 6.627 | 5.847 | 7.184 | 6.633 | 5.171 | 4.055 | 5.525 | 5.466 | 5.850 | 5.387 | 4.868 | 5.071 | 4.850 | 0.313 | 0.0000 | 0.0000 | 1 | 0 | 0 |
| 'Atp6v1a' | 8.637 | 9.629 | 9.440 | 9.480 | 9.456 | 8.850 | 8.924 | 9.971 | 9.519 | 9.320 | 9.325 | 9.088 | 9.255 | 9.583 | 8.957 | 9.311 | 9.583 | 8.547 | 0.007 | 0.7927 | 0.9074 | 0 | 0 | 0 |
| 'Atp6v1b2' | 14.260 | 15.994 | 14.624 | 13.603 | 15.875 | 14.495 | 14.215 | 15.374 | 14.610 | 15.799 | 15.421 | 15.504 | 15.460 | 15.519 | 15.754 | 15.506 | 14.394 | 15.607 | -0.075 | 0.0107 | 0.0657 | 0 | 0 | 0 |
| 'Atp6v1c1' | 32.216 | 34.121 | 31.385 | 33.201 | 36.093 | 30.848 | 31.720 | 36.020 | 34.685 | 31.698 | 34.165 | 35.156 | 35.834 | 36.060 | 35.173 | 37.519 | 35.331 | 32.138 | -0.072 | 0.0422 | 0.1695 | 0 | 0 | 0 |
| 'Atp6v1d' | 51.353 | 55.984 | 47.294 | 52.150 | 54.047 | 52.321 | 51.468 | 55.026 | 50.757 | 51.311 | 49.913 | 49.348 | 52.166 | 55.729 | 48.993 | 52.652 | 54.589 | 50.498 | 0.005 | 0.8679 | 0.9429 | 0 | 0 | 0 |
| 'Atp6v1e1' | 8.600 | 8.955 | 8.352 | 8.303 | 8.836 | 8.968 | 8.080 | 9.412 | 9.408 | 8.716 | 8.107 | 8.726 | 8.989 | 8.388 | 8.830 | 9.247 | 8.470 | 7.739 | 0.023 | 0.5626 | 0.7718 | 0 | 0 | 0 |

| | | | | | | | | | | | | | | | | | | | | | | | | |
|---|---|---|---|---|---|---|---|---|---|---|---|---|---|---|---|---|---|---|---|---|---|---|---|---|
| 'Atp6v1f' | 96.131 | 107.328 | 95.256 | 99.604 | 105.543 | 93.264 | 97.452 | 106.994 | 96.781 | 93.809 | 95.181 | 87.405 | 92.719 | 102.236 | 96.217 | 96.701 | 99.483 | 88.359 | 0.067 | 0.0117 | 0.0699 | 0 | 0 | 0 |
| 'Atp6v1g1' | 66.760 | 81.629 | 68.562 | 71.460 | 76.914 | 73.409 | 71.778 | 76.391 | 68.279 | 60.238 | 67.791 | 65.372 | 71.658 | 73.421 | 65.528 | 67.245 | 74.785 | 62.764 | 0.093 | 0.0051 | 0.0379 | 1 | 0 | 0 |
| 'Atp6v1g2' | 1.493 | 1.959 | 2.082 | 1.608 | 1.993 | 2.239 | 1.806 | 2.028 | 2.137 | 1.818 | 1.819 | 2.040 | 2.206 | 2.179 | 2.017 | 1.877 | 2.633 | 2.063 | -0.117 | 0.1098 | 0.3127 | 0 | 0 | 0 |
| 'Atp6v1g3' | 5.562 | 7.068 | 5.385 | 6.435 | 7.109 | 5.067 | 5.563 | 6.400 | 4.560 | 7.209 | 5.800 | 5.790 | 8.387 | 7.711 | 5.023 | 7.145 | 7.868 | 5.703 | -0.205 | 0.0051 | 0.0380 | 1 | 0 | 0 |
| 'Atp6v1h' | 4.953 | 5.475 | 5.079 | 5.466 | 5.728 | 4.550 | 5.715 | 5.276 | 4.675 | 5.090 | 4.867 | 4.919 | 4.876 | 5.094 | 4.706 | 4.546 | 5.360 | 4.882 | 0.065 | 0.1028 | 0.3006 | 0 | 0 | 0 |
| 'Atp7a' | 1.662 | 1.337 | 1.444 | 1.661 | 1.411 | 1.144 | 1.385 | 1.555 | 1.329 | 1.539 | 1.511 | 1.692 | 1.278 | 1.397 | 1.517 | 1.585 | 1.442 | 1.546 | -0.079 | 0.2841 | 0.5403 | 0 | 0 | 0 |
| 'Atp8a1' | 4.169 | 3.960 | 4.311 | 3.886 | 4.004 | 4.015 | 4.135 | 4.441 | 4.274 | 4.577 | 5.071 | 4.454 | 4.621 | 4.768 | 3.900 | 4.397 | 5.217 | 3.987 | -0.145 | 0.0010 | 0.0101 | 1 | 0 | 0 |
| 'Atp8b2' | 8.485 | 8.080 | 8.601 | 7.971 | 8.317 | 8.512 | 7.279 | 8.885 | 8.562 | 8.353 | 7.877 | 8.442 | 8.067 | 8.031 | 8.628 | 8.210 | 7.642 | 8.262 | 0.014 | 0.6607 | 0.8362 | 0 | 0 | 0 |
| 'Atp9a' | 19.707 | 22.282 | 19.991 | 18.703 | 20.336 | 20.878 | 19.391 | 21.679 | 19.422 | 26.861 | 24.370 | 20.803 | 24.783 | 22.451 | 23.673 | 24.812 | 23.558 | 23.788 | -0.248 | 0.0000 | 0.0000 | 1 | 0 | 0 |
| 'Atp9b' | 40.802 | 41.588 | 42.588 | 39.999 | 41.698 | 40.949 | 41.166 | 41.070 | 42.297 | 43.694 | 42.359 | 41.299 | 43.817 | 41.075 | 37.898 | 42.800 | 44.219 | 38.839 | -0.023 | 0.3798 | 0.6314 | 0 | 0 | 0 |
| 'Atpaf1' | 3.741 | 4.727 | 4.239 | 4.054 | 4.515 | 4.492 | 4.024 | 4.428 | 4.193 | 3.551 | 4.152 | 4.246 | 3.729 | 4.353 | 4.144 | 3.859 | 3.910 | 3.880 | 0.085 | 0.0240 | 0.1173 | 0 | 0 | 0 |
| 'Atpaf2' | 9.537 | 9.631 | 10.854 | 8.976 | 10.227 | 10.690 | 10.560 | 10.148 | 10.363 | 9.209 | 9.187 | 10.331 | 10.358 | 9.894 | 10.084 | 9.667 | 9.439 | 9.516 | 0.043 | 0.2687 | 0.5230 | 0 | 0 | 0 |
| 'Atpif1' | 24.842 | 26.601 | 23.190 | 27.778 | 25.804 | 26.393 | 27.870 | 26.401 | 28.455 | 22.256 | 23.899 | 21.935 | 27.129 | 24.074 | 24.971 | 23.255 | 25.987 | 22.860 | 0.123 | 0.0116 | 0.0695 | 0 | 0 | 0 |
| 'Atr' | 1.829 | 1.953 | 2.035 | 2.029 | 2.205 | 1.987 | 2.200 | 2.073 | 1.951 | 2.066 | 1.880 | 2.069 | 1.792 | 1.909 | 1.884 | 2.063 | 2.231 | 1.759 | 0.037 | 0.4050 | 0.6541 | 0 | 0 | 0 |
| 'Atraid' | 6.397 | 7.043 | 6.774 | 6.814 | 7.526 | 6.666 | 6.067 | 7.732 | 6.820 | 7.138 | 6.540 | 6.483 | 6.776 | 7.408 | 5.952 | 6.430 | 6.604 | 5.537 | 0.063 | 0.1792 | 0.4145 | 0 | 0 | 0 |
| 'Atrn' | 3.132 | 3.886 | 3.477 | 3.167 | 3.408 | 3.253 | 2.967 | 3.591 | 3.280 | 3.633 | 3.824 | 3.499 | 3.449 | 3.454 | 3.045 | 3.468 | 3.483 | 3.167 | -0.052 | 0.2132 | 0.4581 | 0 | 0 | 0 |
| 'Atrnl1' | 1.507 | 1.466 | 1.623 | 1.331 | 1.352 | 1.540 | 1.805 | 1.344 | 1.479 | 1.719 | 1.304 | 1.399 | 1.750 | 1.697 | 1.126 | 1.560 | 1.599 | 1.373 | -0.010 | 0.9020 | 0.9612 | 0 | 0 | 0 |
| 'Atrx' | 10.774 | 9.793 | 10.619 | 11.011 | 9.836 | 9.853 | 10.187 | 10.331 | 10.161 | 10.369 | 10.347 | 10.244 | 9.509 | 10.260 | 9.543 | 10.127 | 10.214 | 10.163 | 0.018 | 0.5644 | 0.7735 | 0 | 0 | 0 |
| 'Atxn1' | 3.454 | 2.989 | 3.323 | 3.341 | 3.086 | 3.158 | 3.345 | 2.648 | 3.363 | 4.081 | 3.966 | 4.397 | 3.104 | 3.662 | 3.831 | 3.896 | 3.765 | 3.973 | -0.285 | 0.0000 | 0.0000 | 1 | 0 | 0 |
| 'Atxn10' | 44.334 | 44.452 | 44.652 | 46.403 | 45.647 | 46.652 | 45.789 | 45.733 | 43.850 | 48.656 | 43.115 | 47.454 | 43.477 | 44.152 | 46.737 | 42.459 | 48.515 | 44.358 | -0.017 | 0.5903 | 0.7909 | 0 | 0 | 0 |
| 'Atxn1l' | 8.617 | 10.429 | 10.048 | 9.070 | 10.051 | 9.937 | 8.945 | 10.346 | 9.751 | 9.883 | 10.079 | 10.511 | 9.479 | 9.906 | 10.215 | 9.645 | 9.099 | 10.955 | -0.055 | 0.1070 | 0.3075 | 0 | 0 | 0 |
| 'Atxn2' | 2.524 | 2.091 | 2.397 | 2.289 | 2.061 | 2.231 | 2.218 | 2.162 | 2.250 | 2.375 | 2.235 | 2.491 | 2.189 | 2.388 | 2.401 | 2.340 | 2.319 | 2.749 | -0.097 | 0.0298 | 0.1341 | 0 | 0 | 0 |
| 'Atxn2l' | 75.860 | 62.201 | 80.640 | 73.871 | 61.868 | 74.167 | 71.321 | 63.971 | 77.834 | 74.206 | 70.963 | 79.981 | 75.640 | 70.339 | 74.941 | 68.690 | 72.953 | 78.689 | -0.069 | 0.0459 | 0.1783 | 0 | 0 | 0 |
| 'Atxn3' | 6.184 | 7.268 | 7.106 | 6.540 | 6.747 | 6.088 | 7.027 | 6.987 | 7.113 | 6.101 | 6.674 | 7.226 | 6.137 | 6.530 | 6.514 | 6.699 | 7.312 | 6.467 | 0.019 | 0.6463 | 0.8273 | 0 | 0 | 0 |
| 'Atxn7' | 14.393 | 14.237 | 14.793 | 14.260 | 14.589 | 14.090 | 12.561 | 14.918 | 14.233 | 15.029 | 15.373 | 16.066 | 14.428 | 14.084 | 15.222 | 13.948 | 15.135 | 15.952 | -0.089 | 0.0069 | 0.0479 | 1 | 0 | 0 |
| 'Atxn7l1' | 4.035 | 3.520 | 3.847 | 3.684 | 3.538 | 3.822 | 3.479 | 3.535 | 3.932 | 3.909 | 3.833 | 4.379 | 3.884 | 3.850 | 3.952 | 3.576 | 3.987 | 4.031 | -0.095 | 0.0073 | 0.0495 | 1 | 0 | 0 |
| 'Atxn7l2' | 2.500 | 2.075 | 2.684 | 2.057 | 2.302 | 2.585 | 2.228 | 2.047 | 2.629 | 3.199 | 1.816 | 2.229 | 2.715 | 2.054 | 2.325 | 2.496 | 1.983 | 2.017 | 0.023 | 0.7657 | 0.8956 | 0 | 0 | 0 |
| 'Atxn7l3' | 15.241 | 14.808 | 14.856 | 14.174 | 15.000 | 15.921 | 13.867 | 14.356 | 15.072 | 14.374 | 15.211 | 16.472 | 14.376 | 15.246 | 16.603 | 15.072 | 14.263 | 15.639 | -0.052 | 0.0675 | 0.2276 | 0 | 0 | 0 |
| 'Atxn7l3b' | 24.543 | 27.974 | 25.102 | 26.021 | 26.882 | 26.243 | 25.950 | 27.165 | 26.537 | 24.566 | 26.191 | 28.369 | 23.137 | 28.981 | 26.785 | 26.134 | 27.713 | 27.415 | -0.029 | 0.3692 | 0.6203 | 0 | 0 | 0 |
| 'Auh' | 3.477 | 3.412 | 3.048 | 3.135 | 3.273 | 3.191 | 3.445 | 3.527 | 3.147 | 3.038 | 3.696 | 3.065 | 3.469 | 3.282 | 3.216 | 3.275 | 3.615 | 2.941 | -0.004 | 0.9140 | 0.9671 | 0 | 0 | 0 |
| 'Aup1' | 14.112 | 14.157 | 16.122 | 14.041 | 14.198 | 15.569 | 14.933 | 14.465 | 15.634 | 14.394 | 13.653 | 15.866 | 15.219 | 13.874 | 15.461 | 14.636 | 14.206 | 15.024 | -0.001 | 0.9618 | 0.9864 | 0 | 0 | 0 |
| 'Aurka' | 5.255 | 5.746 | 6.067 | 5.810 | 6.002 | 6.464 | 5.460 | 5.260 | 6.269 | 3.695 | 5.317 | 6.032 | 6.598 | 5.893 | 6.845 | 5.249 | 5.866 | 5.616 | 0.022 | 0.7645 | 0.8953 | 0 | 0 | 0 |
| 'Aurkaip1' | 21.760 | 22.849 | 23.508 | 21.805 | 23.485 | 22.422 | 23.476 | 24.323 | 23.041 | 19.953 | 20.548 | 21.832 | 21.532 | 21.834 | 23.052 | 22.011 | 21.645 | 20.730 | 0.087 | 0.0036 | 0.0288 | 1 | 0 | 0 |
| 'Aurkb' | 4.742 | 4.893 | 6.819 | 5.151 | 5.307 | 6.850 | 6.381 | 5.482 | 7.241 | 3.373 | 5.341 | 6.390 | 6.200 | 6.375 | 7.660 | 5.400 | 5.364 | 6.196 | 0.004 | 0.9637 | 0.9869 | 0 | 0 | 0 |
| 'Aven' | 4.772 | 4.926 | 5.111 | 5.339 | 5.268 | 5.487 | 4.988 | 4.992 | 4.523 | 5.650 | 4.748 | 4.851 | 5.469 | 5.526 | 5.267 | 5.422 | 5.316 | 4.828 | -0.058 | 0.2237 | 0.4711 | 0 | 0 | 0 |
| 'Avl9' | 6.830 | 7.229 | 6.549 | 6.568 | 7.308 | 6.653 | 6.274 | 7.379 | 6.815 | 7.369 | 7.434 | 7.371 | 6.506 | 7.793 | 7.434 | 7.533 | 7.974 | 7.330 | -0.125 | 0.0001 | 0.0016 | 1 | 0 | 0 |
| 'Avpr2' | 0.994 | 0.966 | 1.265 | 1.298 | 1.169 | 1.092 | 1.542 | 1.121 | 1.369 | 0.896 | 1.349 | 1.666 | 0.944 | 0.782 | 1.529 | 0.688 | 1.264 | 1.260 | 0.063 | 0.6345 | 0.8189 | 0 | 0 | 0 |
| 'Axin1' | 30.914 | 26.524 | 27.819 | 27.911 | 25.751 | 28.938 | 27.884 | 28.456 | 26.382 | 28.952 | 27.543 | 27.729 | 27.695 | 26.587 | 28.041 | 28.545 | 24.740 | 28.802 | 0.002 | 0.9504 | 0.9809 | 0 | 0 | 0 |
| 'Axin2' | 6.264 | 5.043 | 6.471 | 6.145 | 4.496 | 5.784 | 6.114 | 5.547 | 5.997 | 4.675 | 5.555 | 6.699 | 3.539 | 4.430 | 6.063 | 4.830 | 4.480 | 6.041 | 0.162 | 0.0427 | 0.1704 | 0 | 0 | 0 |
| 'Axl' | 2.195 | 2.203 | 2.877 | 2.458 | 2.586 | 3.000 | 2.886 | 2.757 | 3.170 | 2.562 | 1.920 | 3.418 | 2.552 | 2.314 | 2.817 | 3.023 | 2.420 | 2.830 | 0.008 | 0.9057 | 0.9629 | 0 | 0 | 0 |
| 'Azi2' | 8.304 | 8.632 | 8.375 | 8.280 | 8.654 | 8.244 | 8.379 | 8.891 | 8.613 | 8.682 | 9.390 | 9.750 | 7.829 | 8.814 | 8.859 | 8.357 | 9.182 | 9.522 | -0.085 | 0.0105 | 0.0649 | 0 | 0 | 0 |
| 'Azin1' | 21.055 | 21.671 | 21.998 | 22.065 | 22.877 | 20.015 | 23.966 | 22.031 | 20.047 | 20.423 | 24.887 | 21.261 | 23.194 | 23.878 | 21.155 | 18.179 | 24.792 | 20.121 | -0.023 | 0.6122 | 0.8056 | 0 | 0 | 0 |
| 'B230219D22Rik' | 22.252 | 26.584 | 23.698 | 23.294 | 24.164 | 23.442 | 25.880 | 23.387 | 24.046 | 19.977 | 25.657 | 27.456 | 22.331 | 25.393 | 25.726 | 20.568 | 26.839 | 26.979 | -0.034 | 0.4903 | 0.7202 | 0 | 0 | 0 |
| 'B2m' | 980.189 | ####### | 964.469 | ####### | ####### | 995.275 | ####### | ####### | ####### | ####### | ####### | 877.357 | ####### | ####### | ####### | ####### | ####### | ####### | -0.018 | 0.6460 | 0.8271 | 0 | 0 | 0 |
| 'B3galnt2' | 10.240 | 11.957 | 11.735 | 12.276 | 11.849 | 11.032 | 11.137 | 11.703 | 10.391 | 10.593 | 11.700 | 12.331 | 10.733 | 11.359 | 11.330 | 11.322 | 11.626 | 12.035 | -0.026 | 0.5381 | 0.7566 | 0 | 0 | 0 |
| 'B3galt4' | 12.151 | 13.910 | 15.790 | 12.764 | 11.934 | 14.670 | 16.304 | 14.495 | 15.469 | 12.896 | 11.320 | 16.121 | 11.576 | 11.513 | 15.037 | 10.965 | 11.954 | 14.551 | 0.124 | 0.0413 | 0.1670 | 0 | 0 | 0 |
| 'B3galt5' | 0.569 | 0.756 | 0.580 | 0.534 | 0.717 | 0.713 | 0.488 | 0.586 | 0.677 | 0.595 | 0.668 | 0.725 | 0.786 | 0.695 | 0.660 | 0.657 | 0.610 | 0.624 | -0.121 | 0.1682 | 0.4005 | 0 | 0 | 0 |
| 'B3galt6' | 3.622 | 3.980 | 3.701 | 3.570 | 3.457 | 3.908 | 3.863 | 3.778 | 4.401 | 3.865 | 3.170 | 3.920 | 3.755 | 3.308 | 4.025 | 3.473 | 3.270 | 3.296 | 0.080 | 0.2137 | 0.4588 | 0 | 0 | 0 |
| 'B3gat3' | 39.806 | 44.985 | 39.513 | 37.924 | 41.673 | 47.601 | 39.631 | 39.737 | 39.899 | 39.709 | 37.128 | 46.158 | 42.402 | 42.044 | 47.182 | 47.468 | 39.964 | 46.336 | -0.077 | 0.1047 | 0.3032 | 0 | 0 | 0 |
| 'B3glct' | 0.935 | 0.988 | 1.096 | 0.859 | 0.914 | 0.955 | 0.953 | 1.018 | 1.176 | 0.920 | 0.967 | 1.069 | 0.779 | 1.060 | 1.093 | 1.070 | 0.974 | 1.075 | -0.020 | 0.8079 | 0.9138 | 0 | 0 | 0 |
| 'B3gnt2' | 5.454 | 5.123 | 5.439 | 5.320 | 5.289 | 5.793 | 5.235 | 5.431 | 5.339 | 5.618 | 5.566 | 5.648 | 5.514 | 5.067 | 5.832 | 5.368 | 5.382 | 5.262 | -0.032 | 0.4004 | 0.6492 | 0 | 0 | 0 |
| 'B3gnt5' | 0.173 | 0.315 | 0.155 | 0.286 | 0.394 | 0.201 | 0.268 | 0.321 | 0.199 | 0.344 | 0.362 | 0.248 | 0.392 | 0.390 | 0.253 | 0.273 | 0.374 | 0.240 | -0.357 | 0.0015 | 0.0143 | 1 | 0 | 0 |
| 'B3gnt9' | 1.190 | 1.097 | 1.127 | 0.961 | 1.311 | 1.246 | 1.372 | 1.287 | 1.417 | 1.030 | 0.910 | 1.158 | 1.333 | 1.083 | 1.029 | 1.157 | 1.066 | 0.848 | 0.183 | 0.0563 | 0.2036 | 0 | 0 | 0 |
| 'B3gntl1' | 7.566 | 6.847 | 8.280 | 7.280 | 7.280 | 7.532 | 7.297 | 8.996 | 7.106 | 7.696 | 7.072 | 7.358 | 7.942 | 6.434 | 6.466 | 6.163 | 7.984 | 5.732 | 0.110 | 0.0988 | 0.2937 | 0 | 0 | 0 |
| 'B4galnt1' | 122.187 | 124.215 | 144.827 | 107.836 | 123.095 | 147.825 | 114.791 | 121.754 | 147.581 | 133.957 | 103.840 | 138.052 | 117.590 | 114.893 | 139.755 | 137.830 | 99.589 | 136.540 | 0.031 | 0.5121 | 0.7373 | 0 | 0 | 0 |
| 'B4galnt4' | 70.239 | 67.817 | 80.051 | 64.305 | 63.538 | 72.908 | 61.897 | 68.795 | 73.835 | 69.211 | 68.208 | 75.154 | 72.568 | 65.392 | 75.926 | 66.593 | 62.201 | 76.622 | -0.030 | 0.3691 | 0.6203 | 0 | 0 | 0 |
| 'B4galt1' | 28.493 | 30.491 | 29.119 | 27.731 | 29.817 | 30.731 | 28.367 | 30.958 | 29.881 | 30.557 | 30.617 | 29.781 | 30.805 | 32.090 | 30.333 | 33.926 | 27.945 | 29.515 | -0.062 | 0.0628 | 0.2176 | 0 | 0 | 0 |
| 'B4galt3' | 2.246 | 2.006 | 2.687 | 2.119 | 2.292 | 2.245 | 1.947 | 2.302 | 2.370 | 2.458 | 1.929 | 2.395 | 2.375 | 2.208 | 2.025 | 2.333 | 2.065 | 2.326 | 0.000 | 0.9938 | 0.9990 | 0 | 0 | 0 |
| 'B4galt5' | 6.659 | 5.536 | 6.437 | 5.610 | 5.903 | 6.475 | 6.275 | 6.715 | 6.819 | 6.141 | 5.651 | 6.856 | 6.124 | 6.419 | 6.597 | 5.994 | 6.412 | 7.352 | -0.036 | 0.4500 | 0.6898 | 0 | 0 | 0 |
| 'B4galt7' | 5.473 | 5.095 | 6.097 | 5.037 | 4.952 | 6.246 | 5.305 | 5.635 | 5.561 | 5.279 | 5.435 | 6.283 | 5.363 | 5.460 | 6.542 | 4.825 | 5.110 | 6.703 | -0.053 | 0.2500 | 0.5010 | 0 | 0 | 0 |
| 'B4gat1' | 9.147 | 10.988 | 10.473 | 10.597 | 10.752 | 9.653 | 8.925 | 10.120 | 10.556 | 10.392 | 9.595 | 10.563 | 9.788 | 10.508 | 9.287 | 11.407 | 9.487 | 9.755 | -0.005 | 0.9221 | 0.9705 | 0 | 0 | 0 |
| 'B630019K06Rik' | 1.908 | 1.853 | 1.977 | 1.966 | 1.805 | 1.962 | 2.077 | 2.150 | 2.140 | 1.930 | 1.646 | 1.807 | 1.648 | 2.532 | 1.557 | 1.750 | 1.754 | 1.517 | 0.144 | 0.1526 | 0.3797 | 0 | 0 | 0 |
| 'B9d2' | 16.122 | 16.513 | 17.982 | 15.490 | 18.506 | 18.412 | 15.979 | 16.151 | 17.340 | 17.438 | 16.494 | 15.721 | 17.336 | 17.949 | 16.140 | 17.812 | 15.318 | 13.681 | 0.040 | 0.4545 | 0.6931 | 0 | 0 | 0 |
| 'BC003331' | 3.644 | 4.032 | 3.929 | 3.792 | 4.358 | 3.813 | 3.938 | 4.344 | 3.982 | 4.430 | 4.013 | 4.073 | 4.146 | 4.095 | 3.923 | 4.033 | 4.244 | 3.692 | -0.044 | 0.2974 | 0.5532 | 0 | 0 | 0 |
| 'BC003965' | 7.466 | 8.237 | 8.499 | 7.775 | 9.426 | 9.014 | 7.893 | 8.526 | 8.336 | 6.937 | 8.136 | 8.621 | 7.332 | 8.710 | 7.984 | 7.638 | 8.053 | 7.813 | 0.064 | 0.1484 | 0.3748 | 0 | 0 | 0 |
| 'BC004004' | 7.096 | 7.952 | 7.121 | 6.570 | 8.336 | 7.903 | 6.936 | 8.493 | 7.686 | 9.009 | 7.321 | 6.466 | 7.990 | 7.963 | 6.738 | 8.102 | 7.170 | 5.683 | 0.033 | 0.6009 | 0.7972 | 0 | 0 | 0 |
| 'BC005537' | 91.705 | 102.175 | 99.156 | 93.523 | 100.829 | 96.285 | 98.760 | 95.962 | 97.200 | 96.254 | 104.735 | 105.868 | 94.281 | 107.773 | 104.719 | 92.952 | 104.128 | 108.727 | -0.081 | 0.0020 | 0.0183 | 1 | 0 | 0 |
| 'BC005561' | 1.384 | 1.713 | 1.514 | 1.771 | 1.978 | 1.780 | 1.617 | 1.656 | 1.546 | 1.521 | 1.195 | 1.634 | 1.515 | 1.510 | 1.371 | 1.716 | 1.793 | 1.441 | 0.106 | 0.1933 | 0.4308 | 0 | 0 | 0 |
| 'BC005624' | 18.199 | 19.951 | 19.491 | 19.146 | 21.502 | 21.249 | 19.329 | 21.428 | 21.765 | 19.977 | 19.642 | 19.075 | 21.934 | 19.952 | 18.542 | 20.062 | 21.058 | 17.959 | 0.022 | 0.5878 | 0.7885 | 0 | 0 | 0 |
| 'BC017158' | 3.537 | 3.409 | 3.805 | 3.054 | 3.552 | 3.919 | 3.316 | 3.677 | 3.708 | 3.772 | 3.674 | 3.714 | 3.567 | 3.187 | 3.795 | 3.507 | 3.538 | 3.280 | -0.010 | 0.8298 | 0.9262 | 0 | 0 | 0 |
| 'BC017643' | 16.724 | 17.070 | 18.065 | 17.670 | 17.838 | 18.107 | 17.405 | 16.394 | 19.224 | 14.749 | 19.397 | 26.089 | 15.801 | 17.749 | 22.894 | 14.715 | 20.041 | 25.704 | -0.145 | 0.0315 | 0.1392 | 0 | 0 | 0 |
| 'BC021614' | 97.156 | 113.545 | 92.010 | 94.197 | 109.502 | 107.144 | 97.693 | 105.696 | 105.427 | 105.322 | 78.571 | 87.973 | 90.989 | 91.224 | 84.826 | 101.162 | 91.965 | 85.040 | 0.165 | 0.0010 | 0.0103 | 1 | 0 | 0 |
| 'BC025920' | 1.046 | 1.249 | 1.322 | 1.199 | 1.071 | 1.476 | 1.268 | 1.073 | 1.687 | 1.072 | 1.067 | 1.563 | 1.235 | 1.014 | 1.450 | 0.838 | 1.482 | 1.228 | 0.036 | 0.6880 | 0.8525 | 0 | 0 | 0 |
| 'BC029214' | 4.526 | 5.198 | 4.983 | 4.351 | 4.973 | 5.204 | 4.540 | 5.295 | 5.003 | 5.380 | 4.093 | 4.692 | 5.829 | 4.561 | 4.667 | 4.736 | 4.438 | 4.270 | 0.039 | 0.5133 | 0.7375 | 0 | 0 | 0 |
| 'BC029722' | 32.207 | 32.535 | 34.004 | 31.940 | 34.619 | 33.930 | 31.397 | 33.101 | 34.456 | 31.157 | 29.414 | 35.222 | 32.314 | 32.991 | 33.300 | 31.818 | 32.766 | 35.262 | 0.006 | 0.8615 | 0.9401 | 0 | 0 | 0 |
| 'BC030336' | 3.270 | 3.597 | 3.594 | 3.162 | 3.852 | 3.474 | 3.292 | 3.563 | 3.762 | 2.935 | 4.114 | 4.195 | 2.840 | 4.083 | 3.725 | 3.750 | 3.642 | 4.003 | -0.086 | 0.1454 | 0.3699 | 0 | 0 | 0 |
| 'BC030867' | 0.814 | 0.897 | 1.055 | 0.870 | 0.920 | 1.005 | 1.060 | 0.843 | 1.058 | 0.540 | 0.789 | 0.789 | 1.028 | 0.869 | 0.933 | 0.768 | 0.908 | 0.807 | 0.188 | 0.0866 | 0.2680 | 0 | 0 | 0 |
| 'BC031181' | 55.939 | 65.791 | 56.937 | 62.701 | 63.997 | 54.632 | 63.237 | 63.235 | 50.752 | 52.552 | 58.503 | 54.774 | 59.991 | 59.309 | 52.670 | 53.415 | 64.839 | 55.286 | 0.055 | 0.1555 | 0.3842 | 0 | 0 | 0 |
| 'BC037034' | 11.662 | 11.728 | 14.033 | 11.584 | 11.414 | 14.509 | 11.008 | 11.396 | 13.744 | 11.000 | 11.458 | 15.698 | 11.869 | 11.740 | 14.918 | 10.868 | 12.054 | 15.447 | -0.058 | 0.0471 | 0.1808 | 0 | 0 | 0 |
| 'BC048403' | 1.412 | 1.689 | 1.702 | 1.312 | 1.492 | 1.668 | 1.411 | 1.785 | 1.735 | 1.430 | 1.298 | 1.450 | 1.564 | 1.539 | 1.433 | 1.661 | 1.325 | 1.187 | 0.137 | 0.0754 | 0.2441 | 0 | 0 | 0 |
| 'BC052040' | 3.198 | 3.341 | 3.446 | 3.076 | 3.477 | 3.301 | 3.173 | 3.470 | 3.414 | 3.410 | 2.889 | 3.377 | 3.652 | 3.648 | 3.168 | 3.172 | 3.365 | 2.963 | 0.001 | 0.9792 | 0.9926 | 0 | 0 | 0 |
| 'BC055324' | 1.734 | 2.057 | 2.324 | 1.844 | 2.061 | 2.117 | 2.010 | 1.747 | 2.297 | 1.789 | 2.104 | 2.035 | 2.033 | 2.025 | 2.197 | 1.727 | 1.863 | 1.885 | 0.032 | 0.6244 | 0.8127 | 0 | 0 | 0 |
| 'BC068281' | 1.577 | 1.481 | 1.227 | 1.643 | 1.587 | 1.473 | 1.337 | 1.302 | 1.555 | 0.984 | 1.422 | 1.756 | 1.119 | 1.521 | 1.807 | 1.244 | 1.641 | 1.834 | -0.021 | 0.8414 | 0.9326 | 0 | 0 | 0 |
| 'BC094916' | 8.492 | 6.028 | 6.180 | 6.351 | 7.350 | 6.800 | 5.675 | 7.145 | 6.365 | 7.162 | 8.592 | 10.045 | 7.474 | 7.079 | 9.148 | 8.446 | 8.227 | 9.919 | -0.341 | 0.0001 | 0.0010 | 1 | 0 | 0 |
| 'Babam1' | 55.001 | 55.244 | 59.539 | 51.683 | 62.277 | 60.171 | 57.184 | 54.727 | 59.891 | 53.132 | 52.866 | 56.688 | 57.313 | 53.361 | 57.552 | 56.832 | 50.697 | 57.362 | 0.046 | 0.1578 | 0.3878 | 0 | 0 | 0 |
| 'Bace1' | 1.404 | 1.617 | 1.576 | 1.286 | 1.670 | 1.481 | 1.369 | 1.331 | 1.733 | 1.509 | 1.857 | 1.645 | 1.629 | 1.555 | 1.508 | 1.549 | 1.458 | 1.593 | -0.099 | 0.1004 | 0.2965 | 0 | 0 | 0 |
| 'Bach1' | 2.812 | 3.110 | 3.039 | 2.997 | 3.014 | 2.980 | 2.739 | 3.465 | 3.005 | 2.640 | 2.993 | 2.891 | 2.729 | 3.119 | 2.866 | 3.012 | 2.891 | 3.007 | 0.048 | 0.2773 | 0.5334 | 0 | 0 | 0 |
| 'Bach2' | 1.006 | 0.894 | 0.881 | 0.834 | 0.855 | 0.890 | 0.862 | 1.067 | 0.813 | 0.728 | 0.683 | 0.852 | 0.618 | 0.676 | 0.862 | 0.811 | 0.866 | 0.699 | 0.243 | 0.0022 | 0.0199 | 1 | 0 | 0 |
| 'Bad' | 6.232 | 7.055 | 6.626 | 5.952 | 7.462 | 7.248 | 7.074 | 7.186 | 6.985 | 6.073 | 6.702 | 7.108 | 7.143 | 7.057 | 6.602 | 6.257 | 6.972 | 6.476 | 0.017 | 0.7087 | 0.8650 | 0 | 0 | 0 |
| 'Bag1' | 38.289 | 36.717 | 36.677 | 38.733 | 38.237 | 38.556 | 39.663 | 38.887 | 35.624 | 37.926 | 39.036 | 38.255 | 40.220 | 40.399 | 37.988 | 38.266 | 38.830 | 37.681 | -0.041 | 0.1019 | 0.2993 | 0 | 0 | 0 |
| 'Bag3' | 47.667 | 12.218 | 15.336 | 28.209 | 15.317 | 13.558 | 24.975 | 13.742 | 14.293 | 18.190 | 19.268 | 17.913 | 16.870 | 14.915 | 15.157 | 15.041 | 18.853 | 19.555 | 0.105 | 0.6067 | 0.8021 | 0 | 0 | 0 |
| 'Bag4' | 8.990 | 8.942 | 9.273 | 9.125 | 9.343 | 9.062 | 9.102 | 9.235 | 8.821 | 8.855 | 7.990 | 8.759 | 8.312 | 9.264 | 8.702 | 8.493 | 8.177 | 8.883 | 0.070 | 0.0497 | 0.1881 | 0 | 0 | 0 |
| 'Bag5' | 6.377 | 6.668 | 6.350 | 5.848 | 6.317 | 6.299 | 6.481 | 6.581 | 6.341 | 6.786 | 5.788 | 7.048 | 6.616 | 6.486 | 6.811 | 6.715 | 6.866 | 6.633 | -0.076 | 0.0615 | 0.2145 | 0 | 0 | 0 |
| 'Bag6' | 32.107 | 29.543 | 35.245 | 30.778 | 31.100 | 35.190 | 31.875 | 30.967 | 36.543 | 34.002 | 29.252 | 33.093 | 33.616 | 30.699 | 32.824 | 33.204 | 29.403 | 32.729 | 0.012 | 0.6529 | 0.8315 | 0 | 0 | 0 |
| 'Bahd1' | 12.471 | 13.089 | 13.878 | 11.972 | 11.977 | 13.722 | 11.518 | 12.238 | 13.480 | 12.538 | 12.754 | 14.440 | 12.417 | 12.946 | 14.692 | 12.609 | 12.419 | 14.812 | -0.075 | 0.0054 | 0.0396 | 1 | 0 | 0 |
| 'Baiap2' | 7.188 | 6.766 | 7.951 | 6.962 | 6.504 | 8.131 | 7.079 | 6.822 | 8.141 | 8.538 | 6.916 | 8.434 | 7.736 | 7.060 | 8.546 | 7.379 | 7.561 | 8.254 | -0.113 | 0.0006 | 0.0073 | 1 | 0 | 0 |
| 'Bak1' | 20.825 | 21.786 | 22.456 | 20.342 | 23.065 | 24.590 | 21.115 | 21.333 | 23.376 | 21.220 | 18.816 | 23.003 | 24.147 | 20.861 | 23.466 | 21.421 | 20.933 | 23.664 | -0.001 | 0.9694 | 0.9898 | 0 | 0 | 0 |
| 'Banf1' | 77.718 | 90.223 | 78.989 | 86.610 | 90.024 | 88.301 | 83.827 | 87.644 | 89.313 | 75.888 | 78.635 | 81.998 | 87.048 | 84.599 | 86.585 | 86.233 | 75.648 | 84.544 | 0.047 | 0.2336 | 0.4820 | 0 | 0 | 0 |
| 'Banp' | 4.337 | 2.978 | 3.434 | 4.612 | 3.128 | 3.265 | 3.688 | 3.411 | 3.791 | 2.942 | 2.921 | 5.860 | 2.575 | 2.946 | 4.807 | 2.868 | 3.712 | 6.534 | -0.057 | 0.7310 | 0.8773 | 0 | 0 | 0 |
| 'Bap1' | 11.354 | 10.742 | 11.545 | 11.210 | 11.608 | 12.076 | 11.212 | 11.225 | 12.006 | 12.397 | 10.384 | 12.040 | 11.937 | 11.620 | 11.286 | 12.203 | 10.880 | 11.357 | -0.025 | 0.3843 | 0.6347 | 0 | 0 | 0 |
| 'Bard1' | 3.184 | 2.803 | 3.824 | 3.834 | 3.081 | 3.619 | 4.167 | 3.169 | 3.526 | 2.904 | 3.838 | 3.854 | 3.131 | 3.738 | 3.832 | 2.535 | 3.876 | 3.246 | 0.004 | 0.9640 | 0.9869 | 0 | 0 | 0 |
| 'Batf' | 155.080 | 166.439 | 145.523 | 158.058 | 167.218 | 154.472 | 150.213 | 171.002 | 148.771 | 143.295 | 163.480 | 143.983 | 157.999 | 163.594 | 154.165 | 153.380 | 158.006 | 141.409 | 0.028 | 0.2708 | 0.5261 | 0 | 0 | 0 |
| 'Bax' | 132.058 | 145.353 | 139.777 | 136.945 | 138.774 | 136.284 | 140.475 | 145.799 | 140.220 | 130.197 | 124.846 | 139.535 | 137.087 | 138.093 | 133.992 | 136.910 | 133.220 | 137.910 | 0.038 | 0.1338 | 0.3522 | 0 | 0 | 0 |
| 'Baz1a' | 58.465 | 55.559 | 53.694 | 64.117 | 57.529 | 50.521 | 55.512 | 59.074 | 50.200 | 53.943 | 60.796 | 51.435 | 53.657 | 56.633 | 51.007 | 51.724 | 59.316 | 54.326 | 0.023 | 0.5532 | 0.7659 | 0 | 0 | 0 |
| 'Baz1b' | 24.967 | 20.867 | 23.771 | 26.443 | 21.605 | 23.079 | 23.969 | 22.298 | 23.317 | 21.984 | 24.411 | 26.100 | 23.609 | 23.844 | 24.934 | 21.826 | 24.178 | 26.625 | -0.060 | 0.1838 | 0.4193 | 0 | 0 | 0 |
| 'Baz2a' | 54.234 | 52.908 | 53.866 | 53.968 | 50.558 | 50.893 | 51.136 | 51.358 | 55.983 | 52.486 | 56.791 | 57.926 | 55.061 | 52.468 | 55.054 | 51.386 | 54.243 | 60.298 | -0.072 | 0.0211 | 0.1067 | 0 | 0 | 0 |
| 'Baz2b' | 1.557 | 1.481 | 1.317 | 1.395 | 1.411 | 1.433 | 1.428 | 1.577 | 1.475 | 1.755 | 1.855 | 1.772 | 1.648 | 1.787 | 1.616 | 1.729 | 1.797 | 1.494 | -0.250 | 0.0000 | 0.0000 | 1 | 0 | 0 |
| 'Bbc3' | 9.313 | 8.907 | 8.495 | 9.069 | 9.014 | 8.893 | 9.257 | 8.976 | 8.153 | 7.600 | 8.612 | 8.070 | 8.368 | 7.580 | 8.590 | 7.266 | 8.516 | 8.897 | 0.110 | 0.0289 | 0.1317 | 0 | 0 | 0 |
| 'Bbip1' | 10.162 | 11.857 | 11.080 | 11.418 | 12.082 | 10.604 | 10.164 | 10.683 | 10.527 | 10.502 | 10.474 | 10.364 | 10.068 | 11.873 | 9.628 | 10.315 | 11.234 | 9.386 | 0.060 | 0.1828 | 0.4190 | 0 | 0 | 0 |
| 'Bbof1' | 1.625 | 1.596 | 1.440 | 1.723 | 1.735 | 1.437 | 2.000 | 1.449 | 1.423 | 1.629 | 1.420 | 1.182 | 2.263 | 1.298 | 1.411 | 1.117 | 1.873 | 1.484 | 0.060 | 0.6178 | 0.8087 | 0 | 0 | 0 |
| 'Bbs1' | 0.573 | 0.747 | 0.683 | 0.727 | 0.889 | 0.693 | 0.660 | 0.716 | 0.846 | 0.762 | 0.778 | 0.516 | 0.598 | 0.567 | 0.598 | 0.385 | 0.751 | 0.544 | 0.245 | 0.0448 | 0.1754 | 0 | 0 | 0 |
| 'Bbs12' | 1.516 | 2.100 | 1.461 | 1.401 | 2.026 | 1.615 | 2.100 | 2.086 | 2.102 | 1.465 | 1.713 | 2.116 | 1.541 | 1.937 | 1.682 | 1.618 | 1.881 | 1.738 | 0.036 | 0.6915 | 0.8550 | 0 | 0 | 0 |
| 'Bbs2' | 3.705 | 4.180 | 3.619 | 3.977 | 4.801 | 3.651 | 4.180 | 4.248 | 3.981 | 4.830 | 3.897 | 4.033 | 3.748 | 4.083 | 3.844 | 3.811 | 4.611 | 3.272 | -0.008 | 0.9006 | 0.9605 | 0 | 0 | 0 |
| 'Bbs4' | 5.886 | 6.697 | 6.970 | 6.485 | 6.886 | 5.847 | 8.079 | 7.254 | 7.437 | 7.047 | 5.482 | 5.965 | 6.333 | 5.955 | 5.989 | 6.841 | 7.184 | 4.932 | 0.136 | 0.0566 | 0.2046 | 0 | 0 | 0 |
| 'Bbs9' | 1.408 | 1.472 | 1.321 | 1.525 | 1.321 | 1.317 | 1.799 | 1.367 | 1.391 | 1.234 | 1.190 | 1.331 | 1.296 | 1.431 | 1.328 | 1.254 | 1.463 | 1.138 | 0.128 | 0.0475 | 0.1822 | 0 | 0 | 0 |
| 'Bbx' | 2.952 | 2.684 | 2.606 | 3.153 | 2.749 | 2.570 | 3.031 | 2.763 | 2.568 | 2.863 | 3.054 | 3.150 | 2.591 | 2.651 | 2.837 | 2.566 | 2.943 | 2.956 | -0.046 | 0.3832 | 0.6337 | 0 | 0 | 0 |
| 'Bcap29' | 19.065 | 19.669 | 20.123 | 18.866 | 20.111 | 17.693 | 19.320 | 18.588 | 17.726 | 20.178 | 18.109 | 17.623 | 19.277 | 18.527 | 15.996 | 18.558 | 19.637 | 15.702 | 0.058 | 0.1619 | 0.3936 | 0 | 0 | 0 |
| 'Bcap31' | 93.951 | 95.124 | 98.490 | 93.882 | 94.103 | 92.004 | 95.930 | 94.693 | 95.941 | 92.041 | 98.430 | 101.562 | 89.847 | 93.923 | 97.593 | 99.486 | 95.942 | 96.514 | -0.029 | 0.2659 | 0.5194 | 0 | 0 | 0 |
| 'Bcas2' | 10.684 | 10.811 | 9.622 | 9.829 | 10.844 | 9.861 | 9.746 | 11.084 | 9.405 | 10.384 | 10.889 | 9.090 | 11.252 | 11.836 | 8.875 | 11.332 | 10.958 | 8.795 | -0.026 | 0.4525 | 0.6915 | 0 | 0 | 0 |
| 'Bcas3' | 4.673 | 4.401 | 4.480 | 4.313 | 4.605 | 4.431 | 4.483 | 4.552 | 4.117 | 4.959 | 5.092 | 4.414 | 4.397 | 4.051 | 4.169 | 4.682 | 4.716 | 4.064 | -0.024 | 0.5753 | 0.7806 | 0 | 0 | 0 |
| 'Bcat1' | 1.063 | 0.671 | 0.998 | 0.804 | 0.865 | 0.859 | 0.912 | 0.980 | 0.798 | 0.878 | 0.905 | 1.071 | 0.818 | 0.820 | 1.161 | 0.762 | 1.034 | 1.094 | -0.107 | 0.2373 | 0.4868 | 0 | 0 | 0 |
| 'Bcat2' | 9.323 | 9.344 | 11.847 | 9.802 | 10.346 | 11.573 | 10.133 | 9.622 | 11.699 | 9.559 | 8.164 | 9.754 | 9.282 | 8.375 | 10.066 | 10.043 | 8.580 | 9.741 | 0.157 | 0.0000 | 0.0005 | 1 | 0 | 0 |
| 'Bccip' | 27.292 | 27.694 | 27.226 | 28.078 | 28.826 | 27.143 | 27.912 | 29.397 | 27.047 | 25.934 | 25.524 | 25.597 | 27.061 | 25.577 | 24.247 | 26.013 | 27.298 | 24.115 | 0.105 | 0.0000 | 0.0004 | 1 | 0 | 0 |
| 'Bcdin3d' | 8.441 | 9.876 | 11.028 | 7.991 | 9.877 | 10.832 | 9.345 | 9.629 | 10.773 | 7.842 | 9.415 | 9.231 | 7.600 | 7.921 | 8.903 | 7.595 | 8.939 | 8.138 | 0.208 | 0.0009 | 0.0094 | 1 | 0 | 0 |
| 'Bckdha' | 7.514 | 7.695 | 7.313 | 6.691 | 7.466 | 8.222 | 6.875 | 7.524 | 8.189 | 7.223 | 7.252 | 6.801 | 7.376 | 6.886 | 6.688 | 7.423 | 6.003 | 5.728 | 0.134 | 0.0253 | 0.1216 | 0 | 0 | 0 |
| 'Bckdhb' | 6.655 | 7.022 | 6.450 | 7.224 | 6.576 | 6.319 | 6.914 | 7.160 | 5.757 | 6.975 | 6.478 | 6.792 | 6.401 | 5.994 | 5.742 | 6.718 | 6.364 | 6.316 | 0.038 | 0.4785 | 0.7112 | 0 | 0 | 0 |
| 'Bckdk' | 8.250 | 10.542 | 9.782 | 8.636 | 10.076 | 10.332 | 9.202 | 9.736 | 10.521 | 9.172 | 8.389 | 8.510 | 9.962 | 9.450 | 9.456 | 9.747 | 8.867 | 7.957 | 0.084 | 0.1027 | 0.3004 | 0 | 0 | 0 |
| 'Bcl10' | 20.110 | 21.438 | 17.914 | 18.943 | 20.756 | 19.363 | 18.569 | 20.669 | 19.511 | 20.094 | 19.979 | 18.777 | 19.649 | 23.344 | 18.628 | 22.150 | 21.552 | 19.492 | -0.059 | 0.0945 | 0.2852 | 0 | 0 | 0 |
| 'Bcl11b' | 50.453 | 46.531 | 49.080 | 47.143 | 44.728 | 46.633 | 47.488 | 47.601 | 47.139 | 48.789 | 43.871 | 45.980 | 44.715 | 45.990 | 44.710 | 47.114 | 41.515 | 45.883 | 0.053 | 0.0424 | 0.1700 | 0 | 0 | 0 |
| 'Bcl2' | 6.177 | 7.447 | 6.685 | 7.457 | 6.788 | 6.619 | 8.155 | 7.046 | 6.966 | 6.918 | 7.348 | 6.928 | 6.581 | 7.313 | 6.888 | 7.104 | 8.239 | 6.927 | -0.032 | 0.4985 | 0.7264 | 0 | 0 | 0 |
| 'Bcl2l1' | 1.699 | 1.737 | 2.224 | 1.957 | 2.095 | 2.303 | 1.877 | 1.874 | 2.167 | 2.114 | 1.763 | 1.735 | 2.072 | 1.843 | 1.992 | 1.986 | 1.584 | 1.645 | 0.095 | 0.1517 | 0.3788 | 0 | 0 | 0 |
| 'Bcl2l11' | 2.094 | 2.176 | 1.925 | 1.644 | 1.885 | 2.182 | 1.911 | 2.060 | 2.172 | 1.983 | 1.809 | 2.041 | 1.770 | 2.159 | 2.143 | 2.601 | 1.738 | 2.115 | -0.032 | 0.6455 | 0.8268 | 0 | 0 | 0 |
| 'Bcl2l12' | 16.877 | 18.443 | 21.874 | 15.379 | 19.340 | 20.096 | 16.656 | 20.721 | 21.444 | 19.035 | 17.332 | 19.394 | 18.886 | 14.121 | 17.630 | 17.723 | 18.465 | 16.326 | 0.092 | 0.1618 | 0.3935 | 0 | 0 | 0 |
| 'Bcl2l13' | 7.033 | 7.386 | 7.283 | 7.456 | 7.883 | 7.661 | 7.377 | 7.807 | 7.099 | 7.387 | 7.182 | 7.317 | 7.764 | 7.764 | 7.425 | 8.102 | 7.089 | 7.370 | -0.018 | 0.5610 | 0.7711 | 0 | 0 | 0 |
| 'Bcl2l15' | 6.220 | 6.821 | 5.318 | 6.545 | 7.233 | 5.522 | 6.606 | 6.317 | 5.772 | 6.206 | 5.948 | 5.950 | 7.178 | 7.209 | 6.176 | 6.092 | 7.441 | 6.030 | -0.067 | 0.1844 | 0.4199 | 0 | 0 | 0 |
| 'Bcl2l2' | 0.610 | 0.759 | 0.902 | 0.614 | 0.842 | 0.697 | 0.658 | 0.678 | 0.790 | 0.644 | 0.676 | 0.586 | 0.673 | 0.596 | 0.709 | 0.760 | 0.590 | 0.645 | 0.157 | 0.1057 | 0.3050 | 0 | 0 | 0 |
| 'Bcl3' | 15.942 | 13.593 | 17.895 | 15.077 | 15.426 | 15.772 | 14.666 | 17.455 | 16.337 | 13.901 | 18.669 | 15.333 | 16.172 | 17.085 | 14.650 | 13.968 | 16.189 | 15.147 | 0.005 | 0.9228 | 0.9705 | 0 | 0 | 0 |
| 'Bcl6' | 65.662 | 76.666 | 62.624 | 67.438 | 71.740 | 63.612 | 68.431 | 70.707 | 69.623 | 71.464 | 70.275 | 60.420 | 77.715 | 72.875 | 60.656 | 69.101 | 70.464 | 62.785 | -0.007 | 0.8458 | 0.9343 | 0 | 0 | 0 |
| 'Bcl7a' | 4.290 | 4.742 | 5.383 | 4.371 | 5.105 | 4.888 | 4.101 | 4.747 | 5.119 | 4.356 | 4.665 | 5.144 | 5.150 | 5.125 | 4.849 | 4.630 | 4.622 | 4.981 | -0.039 | 0.3255 | 0.5828 | 0 | 0 | 0 |
| 'Bcl7b' | 16.579 | 16.656 | 17.284 | 16.698 | 17.838 | 17.258 | 17.283 | 17.762 | 17.972 | 17.381 | 17.351 | 16.561 | 16.692 | 17.283 | 16.591 | 18.155 | 16.705 | 16.320 | 0.016 | 0.5958 | 0.7944 | 0 | 0 | 0 |
| 'Bcl7c' | 7.577 | 8.046 | 8.338 | 7.648 | 7.894 | 9.054 | 7.503 | 7.564 | 8.641 | 7.464 | 6.522 | 7.812 | 8.046 | 7.212 | 7.962 | 7.796 | 6.990 | 7.845 | 0.085 | 0.0132 | 0.0767 | 0 | 0 | 0 |
| 'Bcl9' | 5.955 | 6.843 | 7.068 | 5.924 | 7.235 | 7.184 | 5.981 | 6.847 | 7.345 | 7.381 | 5.629 | 5.947 | 6.110 | 6.724 | 6.004 | 6.394 | 5.440 | 5.328 | 0.125 | 0.0396 | 0.1621 | 0 | 0 | 0 |
| 'Bcl9l' | 16.178 | 15.307 | 17.268 | 15.593 | 14.673 | 17.004 | 15.326 | 15.712 | 16.836 | 20.248 | 15.553 | 18.158 | 17.426 | 17.205 | 18.462 | 18.542 | 14.623 | 17.795 | -0.142 | 0.0002 | 0.0032 | 1 | 0 | 0 |
| 'Bclaf1' | 18.437 | 17.969 | 19.124 | 18.385 | 18.316 | 17.756 | 19.409 | 18.710 | 18.196 | 17.834 | 19.224 | 18.440 | 17.894 | 18.511 | 17.496 | 17.471 | 18.940 | 17.294 | 0.018 | 0.4692 | 0.7053 | 0 | 0 | 0 |
| 'Bcor' | 26.420 | 25.582 | 25.447 | 24.803 | 25.629 | 26.035 | 23.454 | 25.715 | 26.611 | 28.257 | 24.019 | 24.661 | 24.737 | 25.197 | 24.172 | 27.919 | 24.545 | 22.827 | 0.014 | 0.7084 | 0.8650 | 0 | 0 | 0 |
| 'Bcorl1' | 0.613 | 0.458 | 0.695 | 0.867 | 0.545 | 0.596 | 0.658 | 0.493 | 0.512 | 0.490 | 0.624 | 0.901 | 0.511 | 0.696 | 0.810 | 0.481 | 0.653 | 0.898 | -0.161 | 0.2221 | 0.4694 | 0 | 0 | 0 |
| 'Bcr' | 5.952 | 4.933 | 6.457 | 6.800 | 5.773 | 7.146 | 5.650 | 5.584 | 6.197 | 6.202 | 5.576 | 6.303 | 5.910 | 6.217 | 6.257 | 6.399 | 5.256 | 6.585 | -0.014 | 0.7736 | 0.8993 | 0 | 0 | 0 |
| 'Bcs1l' | 4.044 | 4.156 | 4.274 | 4.295 | 4.015 | 3.810 | 3.968 | 4.050 | 4.493 | 3.991 | 4.124 | 3.897 | 3.659 | 4.223 | 3.370 | 4.150 | 3.880 | 3.503 | 0.089 | 0.1458 | 0.3706 | 0 | 0 | 0 |
| 'Bdh1' | 11.017 | 11.255 | 11.479 | 11.308 | 11.223 | 11.745 | 11.325 | 11.010 | 10.845 | 11.368 | 11.319 | 11.572 | 11.456 | 10.553 | 12.077 | 11.724 | 11.388 | 12.219 | -0.045 | 0.1352 | 0.3537 | 0 | 0 | 0 |
| 'Bdp1' | 6.425 | 5.598 | 5.914 | 6.917 | 6.043 | 5.455 | 5.833 | 5.676 | 5.929 | 5.934 | 6.349 | 6.390 | 5.812 | 5.922 | 5.781 | 6.304 | 6.051 | 6.243 | -0.039 | 0.3545 | 0.6080 | 0 | 0 | 0 |
| 'Becn1' | 14.638 | 16.267 | 16.124 | 15.113 | 15.716 | 16.190 | 15.715 | 16.942 | 15.210 | 15.947 | 15.831 | 15.522 | 16.831 | 16.326 | 15.754 | 15.829 | 16.770 | 14.693 | -0.026 | 0.3397 | 0.5969 | 0 | 0 | 0 |
| 'Begain' | 1.594 | 1.551 | 2.174 | 1.721 | 1.574 | 1.834 | 1.903 | 1.768 | 2.045 | 1.608 | 1.348 | 1.880 | 1.987 | 1.347 | 2.092 | 1.755 | 1.496 | 1.663 | 0.083 | 0.3133 | 0.5715 | 0 | 0 | 0 |
| 'Bend3' | 2.102 | 2.100 | 2.454 | 1.908 | 2.162 | 2.106 | 1.978 | 2.164 | 2.310 | 2.689 | 2.389 | 1.909 | 2.357 | 2.060 | 2.215 | 2.147 | 2.049 | 1.783 | -0.023 | 0.7435 | 0.8843 | 0 | 0 | 0 |
| 'Bet1' | 3.735 | 5.054 | 4.492 | 3.980 | 4.931 | 4.403 | 4.835 | 4.799 | 4.364 | 4.148 | 4.155 | 4.212 | 4.130 | 4.416 | 3.862 | 4.518 | 4.358 | 3.895 | 0.092 | 0.0662 | 0.2250 | 0 | 0 | 0 |
| 'Bet1l' | 21.577 | 21.134 | 19.894 | 18.399 | 21.225 | 21.806 | 18.068 | 19.260 | 20.612 | 17.620 | 19.683 | 20.422 | 19.565 | 19.606 | 19.742 | 20.786 | 19.676 | 21.262 | 0.019 | 0.6754 | 0.8443 | 0 | 0 | 0 |
| 'Bex2' | 7.642 | 6.168 | 7.674 | 6.683 | 7.627 | 7.336 | 7.043 | 7.225 | 5.635 | 7.756 | 6.713 | 5.672 | 6.406 | 6.348 | 5.598 | 4.751 | 6.252 | 5.670 | 0.192 | 0.0419 | 0.1686 | 0 | 0 | 0 |
| 'Bfar' | 7.129 | 7.757 | 7.421 | 7.605 | 6.924 | 7.310 | 7.814 | 7.431 | 7.426 | 6.976 | 8.067 | 7.822 | 6.810 | 7.913 | 7.725 | 7.375 | 8.041 | 7.485 | -0.040 | 0.2862 | 0.5428 | 0 | 0 | 0 |
| 'Bhlhb9' | 6.053 | 5.613 | 5.448 | 5.299 | 6.101 | 5.359 | 6.141 | 6.034 | 5.949 | 6.100 | 6.539 | 6.306 | 6.023 | 6.227 | 6.270 | 6.233 | 6.730 | 6.167 | -0.133 | 0.0007 | 0.0082 | 1 | 0 | 0 |
| 'Bhlhe40' | 26.994 | 19.932 | 13.401 | 16.412 | 18.293 | 14.230 | 16.288 | 20.159 | 12.370 | 28.628 | 22.435 | 11.481 | 23.649 | 28.440 | 12.711 | 32.387 | 21.307 | 12.441 | -0.239 | 0.0496 | 0.1878 | 0 | 0 | 0 |
| 'Bicd2' | 9.383 | 9.141 | 9.864 | 8.881 | 9.959 | 10.067 | 8.859 | 10.169 | 9.986 | 10.335 | 9.031 | 10.581 | 10.459 | 9.704 | 9.541 | 9.530 | 9.612 | 9.666 | -0.046 | 0.1809 | 0.4165 | 0 | 0 | 0 |
| 'Bid' | 4.206 | 4.569 | 3.961 | 4.367 | 4.507 | 3.735 | 4.098 | 3.979 | 4.038 | 3.947 | 3.691 | 3.905 | 3.838 | 3.863 | 3.745 | 4.392 | 3.970 | 3.789 | 0.076 | 0.1139 | 0.3199 | 0 | 0 | 0 |
| 'Bin1' | 51.387 | 50.543 | 52.619 | 48.523 | 50.402 | 54.712 | 51.275 | 49.559 | 56.301 | 59.680 | 51.620 | 62.082 | 53.458 | 55.017 | 61.475 | 55.199 | 53.887 | 59.873 | -0.149 | 0.0000 | 0.0000 | 1 | 0 | 0 |
| 'Bin2' | 94.791 | 82.220 | 95.030 | 94.370 | 85.942 | 100.121 | 92.715 | 86.054 | 97.503 | 95.366 | 82.510 | 97.495 | 89.708 | 85.548 | 99.359 | 92.995 | 85.818 | 98.482 | -0.007 | 0.7250 | 0.8737 | 0 | 0 | 0 |
| 'Bin3' | 32.008 | 36.570 | 36.586 | 33.703 | 35.870 | 36.574 | 34.197 | 36.971 | 35.074 | 36.859 | 35.916 | 35.119 | 35.860 | 36.357 | 34.862 | 35.441 | 36.518 | 31.736 | -0.013 | 0.6988 | 0.8590 | 0 | 0 | 0 |
| 'Birc2' | 12.680 | 13.437 | 12.165 | 11.400 | 13.198 | 11.744 | 11.505 | 12.764 | 11.939 | 12.546 | 12.542 | 11.844 | 11.973 | 12.855 | 11.136 | 13.074 | 12.542 | 11.239 | 0.006 | 0.8534 | 0.9372 | 0 | 0 | 0 |
| 'Birc3' | 15.871 | 16.019 | 15.948 | 15.240 | 16.792 | 15.540 | 16.847 | 16.680 | 15.267 | 16.653 | 16.806 | 14.583 | 16.868 | 16.188 | 14.146 | 17.477 | 16.952 | 12.782 | 0.014 | 0.6898 | 0.8538 | 0 | 0 | 0 |
| 'Birc5' | 6.784 | 8.295 | 10.394 | 9.281 | 8.153 | 10.206 | 10.086 | 8.712 | 9.894 | 5.344 | 8.041 | 8.614 | 9.407 | 10.219 | 10.191 | 7.898 | 8.355 | 9.532 | 0.066 | 0.4541 | 0.6929 | 0 | 0 | 0 |
| 'Birc6' | 11.266 | 9.842 | 11.015 | 11.175 | 9.685 | 9.991 | 10.034 | 9.782 | 10.594 | 10.076 | 10.723 | 11.251 | 9.257 | 10.299 | 10.378 | 9.915 | 10.655 | 12.246 | -0.031 | 0.5010 | 0.7282 | 0 | 0 | 0 |
| 'Bivm' | 2.772 | 2.863 | 2.797 | 3.118 | 2.834 | 2.564 | 2.709 | 2.716 | 2.877 | 2.381 | 2.862 | 2.463 | 2.944 | 3.166 | 2.434 | 2.357 | 2.823 | 2.647 | 0.061 | 0.3064 | 0.5643 | 0 | 0 | 0 |
| 'Blcap' | 6.072 | 7.033 | 6.621 | 5.960 | 6.150 | 6.080 | 5.650 | 6.369 | 6.104 | 6.789 | 7.321 | 7.793 | 5.898 | 6.444 | 7.170 | 6.054 | 7.182 | 7.513 | -0.164 | 0.0013 | 0.0130 | 1 | 0 | 0 |
| 'Blm' | 1.676 | 1.832 | 2.024 | 1.831 | 1.756 | 1.753 | 1.987 | 1.648 | 1.965 | 1.608 | 1.880 | 1.559 | 2.011 | 1.638 | 1.672 | 1.607 | 1.771 | 1.465 | 0.108 | 0.0810 | 0.2565 | 0 | 0 | 0 |
| 'Blmh' | 12.706 | 13.004 | 14.104 | 13.947 | 13.529 | 13.581 | 14.285 | 13.151 | 13.331 | 12.971 | 13.221 | 13.400 | 13.499 | 12.197 | 13.036 | 11.951 | 14.376 | 13.121 | 0.035 | 0.3127 | 0.5712 | 0 | 0 | 0 |
| 'Bloc1s1' | 20.948 | 23.146 | 22.675 | 20.423 | 22.343 | 24.687 | 21.475 | 24.877 | 21.693 | 21.785 | 19.823 | 22.206 | 21.805 | 23.048 | 23.424 | 23.062 | 23.233 | 20.559 | 0.015 | 0.7186 | 0.8703 | 0 | 0 | 0 |
| 'Bloc1s2' | 56.271 | 59.511 | 54.601 | 54.820 | 59.729 | 58.522 | 54.811 | 58.642 | 53.367 | 50.854 | 54.431 | 55.892 | 59.730 | 58.253 | 52.929 | 57.937 | 55.329 | 54.920 | 0.016 | 0.6475 | 0.8282 | 0 | 0 | 0 |
| 'Bloc1s3' | 8.075 | 9.118 | 10.627 | 9.885 | 10.383 | 10.264 | 7.940 | 11.084 | 9.573 | 9.452 | 6.950 | 8.576 | 9.245 | 8.662 | 8.724 | 9.996 | 7.795 | 8.965 | 0.143 | 0.0477 | 0.1827 | 0 | 0 | 0 |
| 'Bloc1s4' | 20.635 | 20.954 | 19.321 | 17.922 | 21.922 | 19.910 | 15.591 | 19.440 | 19.637 | 19.045 | 17.441 | 18.987 | 18.862 | 20.089 | 18.636 | 22.267 | 19.874 | 19.452 | -0.002 | 0.9649 | 0.9874 | 0 | 0 | 0 |
| 'Bloc1s5' | 7.651 | 10.529 | 8.882 | 8.502 | 9.709 | 9.328 | 9.503 | 10.532 | 10.172 | 9.305 | 10.209 | 8.872 | 9.672 | 10.570 | 8.685 | 8.980 | 9.788 | 8.254 | -0.001 | 0.9890 | 0.9968 | 0 | 0 | 0 |
| 'Bloc1s6' | 7.876 | 9.064 | 8.465 | 7.436 | 9.437 | 8.730 | 8.187 | 8.945 | 8.574 | 8.431 | 7.965 | 8.884 | 8.127 | 8.814 | 8.048 | 9.147 | 8.447 | 8.011 | 0.003 | 0.9382 | 0.9764 | 0 | 0 | 0 |
| 'Blvra' | 9.670 | 12.236 | 11.876 | 10.296 | 12.806 | 12.204 | 12.732 | 11.838 | 11.910 | 11.585 | 11.501 | 11.431 | 12.943 | 11.960 | 11.460 | 11.238 | 11.774 | 10.503 | 0.003 | 0.9450 | 0.9789 | 0 | 0 | 0 |
| 'Blvrb' | 1.607 | 2.073 | 1.355 | 1.851 | 1.498 | 1.473 | 1.829 | 2.068 | 1.687 | 0.832 | 1.477 | 1.613 | 1.489 | 1.159 | 1.592 | 1.763 | 1.426 | 1.292 | 0.259 | 0.0458 | 0.1782 | 0 | 0 | 0 |
| 'Blzf1' | 2.566 | 2.668 | 2.820 | 2.476 | 3.165 | 2.647 | 2.699 | 3.121 | 2.564 | 3.094 | 2.850 | 2.469 | 2.820 | 2.481 | 2.184 | 2.512 | 2.667 | 2.311 | 0.074 | 0.2103 | 0.4556 | 0 | 0 | 0 |
| 'Bmf' | 1.127 | 1.725 | 0.880 | 1.063 | 1.686 | 1.299 | 1.171 | 1.318 | 1.126 | 1.360 | 1.077 | 1.302 | 1.154 | 1.200 | 0.997 | 1.887 | 1.322 | 0.887 | 0.007 | 0.9500 | 0.9809 | 0 | 0 | 0 |
| 'Bmi1' | 19.400 | 19.240 | 18.440 | 18.913 | 20.478 | 18.203 | 18.572 | 20.403 | 16.650 | 19.576 | 19.431 | 20.677 | 18.143 | 20.401 | 19.158 | 19.467 | 21.989 | 19.415 | -0.079 | 0.0281 | 0.1296 | 0 | 0 | 0 |
| 'Bmp1' | 1.474 | 1.486 | 1.572 | 1.541 | 1.466 | 1.622 | 1.375 | 1.331 | 2.067 | 1.813 | 2.021 | 1.767 | 1.877 | 1.868 | 1.375 | 1.241 | 2.171 | 1.836 | -0.204 | 0.0522 | 0.1938 | 0 | 0 | 0 |
| 'Bmp2k' | 8.640 | 9.067 | 9.354 | 8.830 | 8.799 | 8.248 | 8.225 | 8.984 | 8.411 | 10.214 | 10.250 | 8.792 | 8.529 | 9.815 | 8.859 | 10.223 | 9.745 | 8.954 | -0.126 | 0.0021 | 0.0187 | 1 | 0 | 0 |
| 'Bmp7' | 1.794 | 1.821 | 1.983 | 1.570 | 1.905 | 2.216 | 1.282 | 2.288 | 2.167 | 2.070 | 1.448 | 2.511 | 1.940 | 1.921 | 2.210 | 1.922 | 2.186 | 2.217 | -0.128 | 0.1378 | 0.3581 | 0 | 0 | 0 |
| 'Bmpr2' | 1.258 | 1.127 | 1.377 | 1.385 | 1.189 | 1.154 | 1.502 | 1.353 | 1.379 | 0.630 | 0.602 | 0.893 | 0.613 | 0.704 | 0.730 | 0.751 | 0.806 | 0.805 | 0.829 | 0.0000 | 0.0000 | 1 | 1 | 0 |
| 'Bms1' | 20.399 | 19.050 | 21.419 | 21.113 | 20.082 | 20.830 | 19.411 | 18.839 | 20.045 | 19.770 | 20.614 | 20.916 | 19.819 | 20.108 | 19.755 | 19.416 | 20.561 | 21.200 | -0.018 | 0.5457 | 0.7606 | 0 | 0 | 0 |
| 'Bmt2' | 3.726 | 3.313 | 3.575 | 4.025 | 3.333 | 3.694 | 5.100 | 3.373 | 3.344 | 3.243 | 4.925 | 4.332 | 3.414 | 4.055 | 4.150 | 2.929 | 4.740 | 4.242 | -0.111 | 0.3357 | 0.5944 | 0 | 0 | 0 |
| 'Bmyc' | 10.028 | 10.056 | 11.609 | 11.530 | 9.561 | 13.419 | 10.447 | 11.628 | 11.502 | 9.591 | 9.051 | 11.217 | 9.771 | 8.747 | 10.051 | 10.599 | 7.445 | 12.011 | 0.165 | 0.0213 | 0.1071 | 0 | 0 | 0 |
| 'Bnip1' | 2.200 | 2.358 | 2.691 | 2.148 | 2.575 | 2.572 | 2.162 | 2.722 | 2.682 | 2.738 | 2.014 | 2.145 | 2.845 | 2.194 | 2.358 | 2.750 | 2.190 | 1.926 | 0.063 | 0.4116 | 0.6601 | 0 | 0 | 0 |
| 'Bnip2' | 13.694 | 15.150 | 13.810 | 13.684 | 15.420 | 15.418 | 15.037 | 15.558 | 15.166 | 13.183 | 15.191 | 15.148 | 13.092 | 15.383 | 15.653 | 13.228 | 15.535 | 14.701 | 0.009 | 0.7696 | 0.8975 | 0 | 0 | 0 |
| 'Bnip3' | 4.676 | 4.449 | 3.094 | 4.761 | 4.005 | 3.037 | 4.244 | 4.228 | 3.479 | 3.253 | 5.581 | 4.706 | 4.231 | 5.405 | 4.658 | 3.700 | 4.817 | 4.398 | -0.201 | 0.0418 | 0.1683 | 0 | 0 | 0 |
| 'Bnip3l' | 20.788 | 24.563 | 23.314 | 23.653 | 24.213 | 21.082 | 23.932 | 23.843 | 20.796 | 22.593 | 23.614 | 23.161 | 23.202 | 25.734 | 22.041 | 22.510 | 25.018 | 21.517 | -0.034 | 0.2607 | 0.5143 | 0 | 0 | 0 |
| 'Bod1' | 8.160 | 8.081 | 7.131 | 7.544 | 8.326 | 7.019 | 5.780 | 8.577 | 6.439 | 8.630 | 7.157 | 6.478 | 8.471 | 9.061 | 7.278 | 8.780 | 8.122 | 6.618 | -0.074 | 0.2748 | 0.5304 | 0 | 0 | 0 |
| 'Bod1l' | 13.216 | 12.268 | 12.719 | 13.344 | 12.772 | 12.908 | 13.942 | 12.672 | 12.155 | 12.368 | 14.157 | 13.525 | 12.382 | 12.766 | 12.667 | 12.374 | 13.001 | 14.101 | -0.028 | 0.4655 | 0.7014 | 0 | 0 | 0 |
| 'Bola1' | 6.720 | 6.538 | 6.785 | 6.085 | 7.753 | 6.713 | 6.137 | 7.477 | 6.583 | 5.788 | 6.202 | 5.437 | 6.822 | 6.746 | 6.059 | 5.936 | 5.854 | 5.408 | 0.163 | 0.0194 | 0.1011 | 0 | 0 | 0 |
| 'Bola2' | 5.630 | 5.493 | 5.679 | 6.216 | 5.236 | 5.494 | 5.354 | 5.715 | 5.537 | 4.715 | 4.887 | 5.521 | 5.723 | 5.051 | 5.316 | 5.201 | 5.581 | 5.560 | 0.069 | 0.1127 | 0.3189 | 0 | 0 | 0 |
| 'Bola3' | 5.820 | 6.859 | 5.651 | 5.605 | 6.224 | 5.627 | 5.914 | 6.544 | 6.009 | 5.774 | 5.903 | 5.230 | 6.175 | 6.017 | 5.650 | 5.976 | 6.046 | 5.033 | 0.059 | 0.1447 | 0.3691 | 0 | 0 | 0 |
| 'Bop1' | 43.327 | 38.847 | 45.816 | 41.048 | 37.435 | 42.271 | 43.484 | 38.345 | 44.774 | 41.289 | 33.419 | 47.189 | 40.796 | 32.885 | 42.271 | 38.614 | 35.722 | 45.406 | 0.061 | 0.0736 | 0.2407 | 0 | 0 | 0 |
| 'Bora' | 12.865 | 12.327 | 14.562 | 13.106 | 13.312 | 13.792 | 13.789 | 11.799 | 13.370 | 9.877 | 12.341 | 12.063 | 13.801 | 13.471 | 12.615 | 11.579 | 12.412 | 12.000 | 0.101 | 0.0694 | 0.2316 | 0 | 0 | 0 |
| 'Borcs5' | 3.573 | 3.368 | 3.803 | 3.327 | 3.641 | 4.015 | 3.522 | 3.921 | 3.727 | 3.792 | 3.320 | 3.288 | 4.065 | 3.285 | 3.568 | 3.211 | 3.555 | 3.259 | 0.067 | 0.2812 | 0.5379 | 0 | 0 | 0 |
| 'Borcs6' | 18.281 | 18.708 | 19.226 | 17.786 | 16.837 | 19.490 | 16.956 | 17.026 | 17.507 | 17.325 | 17.547 | 17.900 | 17.262 | 19.062 | 17.091 | 16.910 | 16.171 | 17.732 | 0.036 | 0.3997 | 0.6486 | 0 | 0 | 0 |
| 'Borcs7' | 4.320 | 4.466 | 4.967 | 3.882 | 5.478 | 4.665 | 5.284 | 5.289 | 4.476 | 5.205 | 4.590 | 3.969 | 5.025 | 4.625 | 3.642 | 4.857 | 3.870 | 3.940 | 0.109 | 0.1692 | 0.4016 | 0 | 0 | 0 |
| 'Bpgm' | 6.256 | 9.104 | 6.925 | 6.800 | 8.594 | 7.380 | 7.106 | 8.110 | 7.802 | 7.105 | 7.194 | 5.684 | 7.921 | 7.440 | 6.930 | 7.404 | 7.495 | 5.600 | 0.108 | 0.0782 | 0.2506 | 0 | 0 | 0 |
| 'Bphl' | 17.363 | 21.220 | 16.939 | 18.871 | 19.166 | 19.725 | 17.494 | 18.819 | 17.404 | 17.825 | 16.736 | 17.947 | 19.996 | 18.721 | 18.151 | 17.616 | 17.410 | 16.377 | 0.038 | 0.4575 | 0.6947 | 0 | 0 | 0 |
| 'Bpnt1' | 2.699 | 2.826 | 2.556 | 2.998 | 3.021 | 2.633 | 2.930 | 3.287 | 2.906 | 2.370 | 3.163 | 2.838 | 3.115 | 2.810 | 2.833 | 2.356 | 2.646 | 2.938 | 0.025 | 0.7343 | 0.8791 | 0 | 0 | 0 |
| 'Bptf' | 18.196 | 16.133 | 17.450 | 17.656 | 15.759 | 16.960 | 17.054 | 16.557 | 17.494 | 17.227 | 17.252 | 17.822 | 15.324 | 15.821 | 16.814 | 16.333 | 16.025 | 18.771 | 0.007 | 0.8463 | 0.9343 | 0 | 0 | 0 |
| 'Braf' | 3.373 | 3.589 | 3.584 | 3.438 | 3.851 | 3.373 | 3.898 | 3.767 | 3.227 | 3.478 | 3.955 | 3.411 | 3.632 | 3.875 | 2.944 | 3.755 | 3.831 | 3.196 | -0.005 | 0.8880 | 0.9540 | 0 | 0 | 0 |
| 'Brap' | 9.209 | 8.825 | 8.873 | 8.420 | 9.315 | 9.154 | 8.139 | 8.989 | 8.770 | 8.958 | 9.533 | 9.514 | 9.018 | 9.097 | 9.405 | 8.843 | 9.086 | 9.815 | -0.073 | 0.0092 | 0.0587 | 0 | 0 | 0 |
| 'Brat1' | 2.964 | 2.918 | 3.604 | 2.685 | 3.185 | 3.665 | 3.275 | 3.443 | 3.658 | 3.347 | 2.791 | 3.518 | 3.057 | 2.785 | 3.350 | 2.995 | 2.763 | 3.365 | 0.062 | 0.1732 | 0.4066 | 0 | 0 | 0 |
| 'Brca1' | 2.189 | 1.808 | 2.531 | 2.574 | 1.984 | 2.362 | 2.357 | 2.134 | 2.486 | 1.980 | 2.465 | 2.330 | 2.797 | 2.371 | 2.401 | 1.881 | 2.210 | 2.275 | -0.034 | 0.6326 | 0.8178 | 0 | 0 | 0 |
| 'Brca2' | 0.966 | 0.767 | 1.111 | 1.170 | 0.978 | 1.018 | 1.143 | 1.051 | 1.108 | 1.110 | 1.050 | 0.942 | 1.185 | 1.165 | 1.079 | 1.077 | 0.974 | 0.912 | -0.033 | 0.6235 | 0.8120 | 0 | 0 | 0 |
| 'Brcc3' | 3.221 | 3.364 | 3.107 | 3.155 | 3.595 | 3.145 | 3.032 | 3.219 | 2.772 | 3.099 | 3.264 | 3.090 | 3.099 | 3.266 | 3.019 | 3.140 | 3.109 | 2.663 | 0.032 | 0.4134 | 0.6616 | 0 | 0 | 0 |
| 'Brd1' | 18.333 | 17.752 | 16.291 | 17.542 | 16.979 | 17.007 | 16.319 | 18.577 | 16.367 | 17.585 | 18.064 | 18.088 | 16.825 | 18.365 | 17.487 | 18.720 | 16.570 | 18.720 | -0.059 | 0.0962 | 0.2885 | 0 | 0 | 0 |

| | | | | | | | | | | | | | | | | | | | | | | | | |
|---|---|---|---|---|---|---|---|---|---|---|---|---|---|---|---|---|---|---|---|---|---|---|---|---|
| 'Brd2' | 45.620 | 37.893 | 38.719 | 43.131 | 36.998 | 39.349 | 42.019 | 38.787 | 38.298 | 41.023 | 39.041 | 39.240 | 41.890 | 39.719 | 38.888 | 41.247 | 39.661 | 38.459 | -0.006 | 0.8222 | 0.9217 | 0 | 0 | 0 |
| 'Brd3' | 4.555 | 4.146 | 4.823 | 4.341 | 4.164 | 4.666 | 4.863 | 4.262 | 4.806 | 3.880 | 4.439 | 5.854 | 4.034 | 4.329 | 5.167 | 4.059 | 4.549 | 5.642 | -0.050 | 0.3357 | 0.5944 | 0 | 0 | 0 |
| 'Brd4' | 23.587 | 20.692 | 22.544 | 22.713 | 20.071 | 22.366 | 21.836 | 21.481 | 22.276 | 24.790 | 22.857 | 23.212 | 24.177 | 22.189 | 22.527 | 22.568 | 22.055 | 23.513 | -0.084 | 0.0011 | 0.0117 | 1 | 0 | 0 |
| 'Brd7' | 11.416 | 12.202 | 13.434 | 11.371 | 12.526 | 13.081 | 12.860 | 12.892 | 12.261 | 12.009 | 12.164 | 11.911 | 12.567 | 12.050 | 11.765 | 11.611 | 11.786 | 11.991 | 0.046 | 0.1504 | 0.3777 | 0 | 0 | 0 |
| 'Brd8' | 12.242 | 13.428 | 13.881 | 12.750 | 13.471 | 13.370 | 13.422 | 13.403 | 14.520 | 13.114 | 13.073 | 12.634 | 13.519 | 13.188 | 12.261 | 13.017 | 13.431 | 11.273 | 0.052 | 0.1281 | 0.3429 | 0 | 0 | 0 |
| 'Brd9' | 26.975 | 28.672 | 31.497 | 27.254 | 28.988 | 30.912 | 28.844 | 28.047 | 30.116 | 27.622 | 27.504 | 29.810 | 29.899 | 28.711 | 27.857 | 27.602 | 31.109 | 26.526 | 0.015 | 0.6544 | 0.8326 | 0 | 0 | 0 |
| 'Bre' | 6.714 | 7.229 | 6.459 | 6.430 | 6.791 | 6.582 | 6.965 | 6.791 | 6.340 | 6.739 | 5.988 | 6.356 | 6.243 | 6.208 | 6.051 | 6.192 | 6.536 | 6.298 | 0.077 | 0.0219 | 0.1095 | 0 | 0 | 0 |
| 'Brf1' | 48.516 | 44.083 | 51.768 | 47.737 | 45.648 | 48.633 | 45.172 | 47.143 | 51.502 | 44.196 | 46.038 | 47.053 | 46.331 | 42.891 | 49.354 | 45.855 | 45.310 | 47.672 | 0.045 | 0.1064 | 0.3065 | 0 | 0 | 0 |
| 'Brf2' | 7.505 | 7.091 | 7.159 | 7.876 | 8.403 | 8.580 | 6.261 | 7.563 | 8.175 | 7.937 | 5.750 | 8.301 | 7.773 | 5.579 | 7.518 | 7.090 | 6.148 | 8.084 | 0.082 | 0.2831 | 0.5394 | 0 | 0 | 0 |
| 'Bri3' | 4.470 | 4.646 | 4.735 | 3.907 | 4.870 | 5.352 | 4.225 | 5.449 | 4.457 | 4.744 | 4.542 | 4.637 | 4.318 | 4.126 | 4.779 | 4.497 | 5.337 | 4.507 | 0.017 | 0.8120 | 0.9155 | 0 | 0 | 0 |
| 'Bri3bp' | 9.741 | 9.512 | 10.121 | 9.336 | 10.017 | 10.581 | 10.614 | 8.829 | 9.951 | 9.885 | 10.894 | 10.516 | 10.905 | 10.787 | 11.407 | 10.052 | 10.358 | 10.711 | -0.117 | 0.0019 | 0.0175 | 1 | 0 | 0 |
| 'Brip1' | 1.769 | 1.558 | 2.491 | 2.433 | 1.774 | 2.074 | 2.347 | 1.720 | 2.157 | 1.588 | 2.351 | 2.257 | 2.147 | 2.185 | 2.503 | 1.806 | 2.394 | 2.089 | -0.089 | 0.3368 | 0.5954 | 0 | 0 | 0 |
| 'Brix1' | 13.652 | 13.992 | 13.188 | 15.118 | 14.166 | 13.629 | 13.418 | 14.454 | 13.235 | 13.477 | 12.859 | 13.692 | 13.201 | 14.100 | 13.355 | 14.768 | 14.050 | 13.391 | 0.011 | 0.7502 | 0.8886 | 0 | 0 | 0 |
| 'Brk1' | 132.551 | 153.399 | 135.007 | 137.849 | 148.262 | 144.770 | 137.306 | 150.517 | 140.477 | 137.275 | 139.702 | 131.381 | 140.290 | 139.724 | 136.606 | 144.405 | 142.961 | 127.767 | 0.036 | 0.1618 | 0.3935 | 0 | 0 | 0 |
| 'Brms1' | 40.479 | 40.694 | 44.164 | 41.584 | 39.999 | 48.217 | 40.377 | 41.826 | 44.784 | 42.884 | 39.542 | 42.201 | 45.883 | 41.412 | 43.105 | 41.225 | 40.110 | 39.552 | 0.016 | 0.6699 | 0.8408 | 0 | 0 | 0 |
| 'Brms1l' | 8.090 | 9.203 | 9.585 | 8.117 | 9.506 | 9.069 | 9.138 | 9.119 | 9.228 | 7.060 | 9.604 | 9.872 | 7.954 | 9.880 | 10.746 | 7.255 | 9.262 | 9.508 | -0.012 | 0.8185 | 0.9196 | 0 | 0 | 0 |
| 'Brox' | 7.501 | 8.655 | 7.968 | 7.606 | 8.456 | 8.028 | 8.454 | 8.498 | 7.638 | 7.661 | 8.423 | 8.568 | 7.884 | 8.417 | 8.169 | 7.917 | 8.947 | 7.917 | -0.036 | 0.2077 | 0.4523 | 0 | 0 | 0 |
| 'Brpf1' | 8.319 | 7.514 | 8.316 | 7.701 | 7.461 | 8.153 | 7.472 | 7.555 | 8.635 | 7.896 | 7.915 | 8.540 | 8.064 | 7.614 | 7.997 | 8.061 | 7.558 | 7.965 | -0.019 | 0.4793 | 0.7118 | 0 | 0 | 0 |
| 'Brpf3' | 4.272 | 3.829 | 4.456 | 3.597 | 3.711 | 4.622 | 3.655 | 3.993 | 4.455 | 3.830 | 4.278 | 4.221 | 4.439 | 4.165 | 4.295 | 3.933 | 3.592 | 4.175 | -0.023 | 0.6154 | 0.8081 | 0 | 0 | 0 |
| 'Brwd1' | 6.829 | 6.037 | 6.083 | 6.918 | 5.642 | 5.495 | 6.560 | 5.836 | 6.203 | 5.999 | 6.577 | 6.458 | 5.406 | 5.450 | 5.714 | 5.642 | 6.254 | 6.591 | 0.026 | 0.6292 | 0.8163 | 0 | 0 | 0 |
| 'Brwd3' | 2.371 | 1.882 | 1.989 | 2.658 | 1.960 | 1.719 | 2.110 | 1.775 | 2.087 | 1.913 | 2.119 | 2.512 | 2.161 | 1.884 | 2.042 | 2.034 | 2.102 | 2.415 | -0.067 | 0.3540 | 0.6074 | 0 | 0 | 0 |
| 'Bscl2' | 9.241 | 9.735 | 10.023 | 9.557 | 9.387 | 10.363 | 9.458 | 9.428 | 9.875 | 9.128 | 9.114 | 9.078 | 9.462 | 9.094 | 9.197 | 9.064 | 8.792 | 8.972 | 0.080 | 0.0016 | 0.0152 | 1 | 0 | 0 |
| 'Bsdc1' | 7.781 | 7.075 | 7.400 | 8.207 | 7.377 | 7.771 | 7.363 | 6.870 | 7.617 | 7.656 | 8.058 | 8.675 | 7.335 | 7.787 | 7.950 | 8.054 | 7.209 | 8.307 | -0.086 | 0.0259 | 0.1236 | 0 | 0 | 0 |
| 'Bsg' | 44.950 | 47.148 | 44.652 | 45.248 | 49.382 | 48.513 | 44.622 | 45.330 | 44.479 | 44.777 | 45.199 | 48.634 | 45.543 | 46.511 | 49.110 | 47.686 | 46.676 | 47.219 | -0.036 | 0.1686 | 0.4009 | 0 | 0 | 0 |
| 'Bsn' | 0.712 | 0.738 | 0.679 | 0.726 | 0.636 | 0.768 | 0.699 | 0.604 | 0.803 | 0.557 | 0.577 | 0.601 | 0.752 | 0.521 | 0.711 | 0.474 | 0.532 | 0.730 | 0.209 | 0.0079 | 0.0523 | 0 | 0 | 0 |
| 'Bspry' | 1.035 | 1.018 | 1.395 | 1.055 | 1.067 | 1.386 | 1.223 | 1.108 | 1.410 | 1.329 | 1.043 | 0.976 | 1.505 | 1.073 | 1.056 | 1.140 | 0.716 | 0.706 | 0.176 | 0.1718 | 0.4055 | 0 | 0 | 0 |
| 'Bst2' | 26.075 | 28.117 | 28.188 | 23.115 | 28.215 | 30.077 | 23.554 | 31.697 | 27.400 | 24.588 | 19.656 | 41.689 | 32.889 | 24.660 | 41.251 | 31.467 | 26.008 | 39.760 | -0.182 | 0.0570 | 0.2053 | 0 | 0 | 0 |
| 'Btaf1' | 20.027 | 18.982 | 20.694 | 20.963 | 20.305 | 18.575 | 19.583 | 19.960 | 19.844 | 20.651 | 20.065 | 19.476 | 19.604 | 19.279 | 18.300 | 18.153 | 19.950 | 19.159 | 0.025 | 0.4241 | 0.6709 | 0 | 0 | 0 |
| 'Btbd1' | 26.771 | 30.414 | 26.072 | 27.844 | 28.385 | 26.747 | 28.481 | 28.161 | 24.433 | 25.255 | 28.186 | 29.275 | 25.685 | 28.782 | 30.752 | 25.664 | 29.343 | 30.196 | -0.046 | 0.2935 | 0.5497 | 0 | 0 | 0 |
| 'Btbd10' | 3.689 | 3.861 | 3.489 | 3.647 | 3.770 | 3.558 | 3.780 | 3.933 | 3.265 | 3.632 | 3.865 | 3.832 | 3.931 | 3.879 | 3.707 | 3.717 | 4.145 | 3.709 | -0.077 | 0.0486 | 0.1851 | 0 | 0 | 0 |
| 'Btbd11' | 1.795 | 2.013 | 1.825 | 1.457 | 1.924 | 2.043 | 1.517 | 1.633 | 1.979 | 3.110 | 2.849 | 2.869 | 2.407 | 3.205 | 3.002 | 2.414 | 2.635 | 2.715 | -0.652 | 0.0000 | 0.0000 | 1 | 1 | 0 |
| 'Btbd19' | 5.229 | 6.216 | 6.551 | 4.837 | 6.059 | 6.281 | 5.148 | 6.450 | 6.100 | 7.327 | 6.190 | 6.193 | 6.747 | 6.877 | 6.159 | 7.244 | 6.438 | 5.747 | -0.166 | 0.0055 | 0.0401 | 1 | 0 | 0 |
| 'Btbd2' | 12.733 | 11.079 | 13.673 | 11.309 | 11.881 | 14.687 | 10.834 | 12.283 | 13.112 | 13.058 | 10.711 | 13.755 | 12.415 | 12.135 | 13.950 | 12.725 | 11.543 | 13.641 | -0.038 | 0.3264 | 0.5842 | 0 | 0 | 0 |
| 'Btbd6' | 18.450 | 17.099 | 20.126 | 15.272 | 17.613 | 21.959 | 17.342 | 17.424 | 18.830 | 17.884 | 17.192 | 21.111 | 17.271 | 16.508 | 20.627 | 17.593 | 16.820 | 19.610 | -0.013 | 0.7352 | 0.8795 | 0 | 0 | 0 |
| 'Btbd7' | 9.471 | 9.912 | 9.530 | 9.812 | 9.815 | 10.065 | 9.396 | 9.258 | 9.002 | 9.166 | 10.057 | 10.443 | 9.446 | 9.387 | 10.238 | 9.679 | 8.530 | 11.158 | -0.043 | 0.3303 | 0.5880 | 0 | 0 | 0 |
| 'Btbd9' | 7.078 | 6.666 | 6.641 | 6.969 | 6.927 | 7.066 | 7.125 | 6.751 | 7.008 | 5.475 | 5.460 | 5.649 | 5.097 | 5.134 | 5.606 | 5.117 | 5.268 | 5.557 | 0.351 | 0.0000 | 0.0000 | 1 | 0 | 0 |
| 'Btd' | 6.601 | 6.718 | 7.385 | 6.717 | 6.119 | 7.590 | 6.206 | 6.496 | 7.171 | 7.443 | 7.526 | 8.122 | 6.520 | 8.097 | 6.806 | 6.636 | 7.264 | 7.248 | -0.115 | 0.0329 | 0.1438 | 0 | 0 | 0 |
| 'Btf3' | 143.212 | 141.487 | 132.937 | 151.338 | 143.655 | 136.053 | 148.616 | 144.080 | 136.582 | 134.650 | 136.393 | 121.222 | 143.963 | 137.927 | 134.587 | 139.521 | 142.042 | 130.078 | 0.057 | 0.0297 | 0.1339 | 0 | 0 | 0 |
| 'Btf3l4' | 16.288 | 19.711 | 17.864 | 18.340 | 19.793 | 18.057 | 18.286 | 21.411 | 17.643 | 17.019 | 17.170 | 16.477 | 17.923 | 18.399 | 16.793 | 16.819 | 17.655 | 15.456 | 0.111 | 0.0013 | 0.0126 | 1 | 0 | 0 |
| 'Btg1' | 113.241 | 107.682 | 74.518 | 82.500 | 106.216 | 86.240 | 81.574 | 114.952 | 74.168 | 111.661 | 98.618 | 60.788 | 94.651 | 117.026 | 70.123 | 119.301 | 83.295 | 64.044 | 0.048 | 0.6131 | 0.8059 | 0 | 0 | 0 |
| 'Btg2' | 117.858 | 104.578 | 80.349 | 116.084 | 92.357 | 83.673 | 126.796 | 91.528 | 84.274 | 81.714 | 111.578 | 105.797 | 94.813 | 111.677 | 102.632 | 97.648 | 116.824 | 103.894 | -0.069 | 0.4602 | 0.6967 | 0 | 0 | 0 |
| 'Btla' | 48.052 | 58.142 | 55.976 | 53.765 | 58.644 | 53.091 | 58.484 | 54.732 | 56.927 | 54.929 | 55.322 | 54.209 | 46.841 | 52.933 | 51.694 | 51.305 | 57.664 | 55.025 | 0.042 | 0.2659 | 0.5194 | 0 | 0 | 0 |
| 'Btrc' | 3.513 | 4.296 | 3.840 | 3.811 | 3.977 | 4.042 | 4.383 | 4.097 | 4.238 | 4.234 | 3.673 | 3.572 | 3.689 | 3.557 | 3.425 | 4.174 | 3.477 | 3.143 | 0.129 | 0.0157 | 0.0866 | 0 | 0 | 0 |
| 'Bub1' | 3.398 | 4.456 | 5.246 | 5.229 | 4.310 | 4.705 | 5.118 | 3.627 | 5.038 | 2.745 | 4.627 | 4.564 | 5.315 | 5.052 | 5.473 | 3.963 | 4.621 | 4.490 | -0.001 | 0.9929 | 0.9986 | 0 | 0 | 0 |
| 'Bub1b' | 11.694 | 13.227 | 15.925 | 15.815 | 13.369 | 15.939 | 16.467 | 13.155 | 17.047 | 9.890 | 13.853 | 14.696 | 16.090 | 14.806 | 17.273 | 12.800 | 14.082 | 15.008 | 0.034 | 0.6559 | 0.8333 | 0 | 0 | 0 |
| 'Bub3' | 57.367 | 62.955 | 61.060 | 61.635 | 61.965 | 62.620 | 60.857 | 62.126 | 60.945 | 50.108 | 60.494 | 63.904 | 58.023 | 62.043 | 64.796 | 54.018 | 60.676 | 63.479 | 0.027 | 0.4278 | 0.6739 | 0 | 0 | 0 |
| 'Bud13' | 3.957 | 3.769 | 4.333 | 4.070 | 4.222 | 4.217 | 4.139 | 3.870 | 4.384 | 4.116 | 3.794 | 4.662 | 3.545 | 3.646 | 4.287 | 4.018 | 3.646 | 4.760 | 0.007 | 0.8860 | 0.9530 | 0 | 0 | 0 |
| 'Bud31' | 24.260 | 24.017 | 21.732 | 24.492 | 23.614 | 21.881 | 23.332 | 23.302 | 21.874 | 23.188 | 23.790 | 21.535 | 24.573 | 23.995 | 21.518 | 22.696 | 23.240 | 21.427 | 0.007 | 0.7905 | 0.9064 | 0 | 0 | 0 |
| 'Bysl' | 7.884 | 6.840 | 8.058 | 7.447 | 6.496 | 7.608 | 7.635 | 6.743 | 6.845 | 7.287 | 6.680 | 7.581 | 7.292 | 6.407 | 7.915 | 7.077 | 6.185 | 7.564 | 0.026 | 0.5584 | 0.7697 | 0 | 0 | 0 |
| 'Bzrap1' | 1.000 | 0.857 | 1.346 | 1.070 | 0.682 | 1.083 | 0.897 | 0.857 | 1.221 | 0.941 | 0.849 | 1.104 | 0.941 | 0.730 | 1.108 | 0.908 | 0.702 | 0.982 | 0.117 | 0.0744 | 0.2425 | 0 | 0 | 0 |
| 'Bzw1' | 51.948 | 56.020 | 51.892 | 55.494 | 53.461 | 48.474 | 55.101 | 53.510 | 48.712 | 50.097 | 53.318 | 51.125 | 48.160 | 52.301 | 49.681 | 47.899 | 53.844 | 50.580 | 0.043 | 0.1507 | 0.3777 | 0 | 0 | 0 |
| 'Bzw2' | 219.274 | 238.803 | 228.991 | 238.152 | 248.644 | 224.645 | 232.380 | 235.642 | 228.740 | 214.537 | 234.787 | 201.227 | 215.065 | 222.566 | 211.529 | 207.361 | 220.144 | 201.025 | 0.110 | 0.0000 | 0.0001 | 1 | 0 | 0 |
| 'C1d' | 7.317 | 9.850 | 8.123 | 7.838 | 9.968 | 8.437 | 8.242 | 9.059 | 8.083 | 7.835 | 8.237 | 8.357 | 7.719 | 9.149 | 7.501 | 8.961 | 8.823 | 7.816 | 0.032 | 0.4528 | 0.6916 | 0 | 0 | 0 |
| 'C1galt1' | 17.303 | 20.270 | 19.002 | 19.691 | 21.230 | 18.606 | 20.541 | 20.607 | 17.794 | 20.294 | 20.445 | 19.154 | 21.073 | 22.072 | 19.044 | 20.221 | 21.026 | 18.648 | -0.068 | 0.0207 | 0.1056 | 0 | 0 | 0 |
| 'C1galt1c1' | 7.324 | 7.851 | 6.761 | 7.415 | 7.525 | 6.639 | 8.515 | 7.553 | 6.589 | 5.856 | 9.019 | 9.263 | 6.666 | 8.520 | 8.325 | 6.873 | 9.235 | 8.833 | -0.149 | 0.0728 | 0.2387 | 0 | 0 | 0 |
| 'C1qbp' | 14.057 | 12.950 | 14.187 | 14.135 | 13.834 | 14.068 | 14.340 | 13.335 | 14.296 | 13.459 | 12.567 | 15.225 | 14.291 | 12.226 | 14.193 | 12.825 | 13.024 | 14.074 | 0.023 | 0.5006 | 0.7278 | 0 | 0 | 0 |
| 'C2cd2' | 5.622 | 5.159 | 5.299 | 5.774 | 6.144 | 5.039 | 5.344 | 5.541 | 5.043 | 5.325 | 5.767 | 5.541 | 6.259 | 5.720 | 5.113 | 5.542 | 6.630 | 5.153 | -0.071 | 0.1129 | 0.3192 | 0 | 0 | 0 |
| 'C2cd2l' | 6.048 | 5.358 | 6.805 | 5.926 | 5.429 | 6.685 | 4.986 | 6.170 | 6.449 | 6.060 | 6.921 | 7.416 | 6.573 | 5.901 | 7.104 | 6.699 | 5.500 | 6.823 | -0.137 | 0.0068 | 0.0473 | 1 | 0 | 0 |
| 'C2cd3' | 8.638 | 7.806 | 8.725 | 9.337 | 7.710 | 8.423 | 8.078 | 7.920 | 8.537 | 8.827 | 8.992 | 9.301 | 8.769 | 8.210 | 8.835 | 7.821 | 8.492 | 9.069 | -0.071 | 0.0556 | 0.2021 | 0 | 0 | 0 |
| 'C2cd5' | 8.893 | 9.831 | 8.839 | 9.176 | 9.532 | 8.648 | 8.931 | 9.399 | 9.320 | 8.975 | 9.172 | 8.752 | 8.591 | 8.722 | 8.259 | 8.620 | 8.884 | 8.357 | 0.064 | 0.0044 | 0.0340 | 1 | 0 | 0 |
| 'C3' | 8.000 | 11.318 | 8.334 | 8.596 | 10.489 | 10.422 | 9.594 | 10.565 | 10.634 | 19.964 | 18.086 | 15.627 | 16.424 | 20.016 | 15.229 | 20.520 | 18.680 | 17.960 | -0.898 | 0.0000 | 0.0000 | 1 | 1 | 0 |
| 'C330007P06Rik' | 4.733 | 6.176 | 5.809 | 5.015 | 6.335 | 5.289 | 5.376 | 6.105 | 5.538 | 5.751 | 6.208 | 5.164 | 5.533 | 6.072 | 5.405 | 5.848 | 5.452 | 5.106 | -0.014 | 0.7495 | 0.8879 | 0 | 0 | 0 |
| 'C330018D20Rik' | 3.321 | 3.855 | 3.921 | 3.926 | 4.431 | 3.525 | 3.593 | 4.177 | 4.267 | 3.517 | 3.824 | 3.662 | 3.386 | 3.775 | 3.854 | 3.377 | 4.031 | 3.725 | 0.067 | 0.2562 | 0.5084 | 0 | 0 | 0 |
| 'C330027C09Rik' | 1.998 | 2.454 | 3.149 | 2.471 | 2.447 | 2.438 | 3.135 | 2.386 | 2.597 | 1.803 | 2.414 | 2.436 | 2.444 | 2.415 | 2.705 | 2.146 | 2.691 | 2.428 | 0.093 | 0.2298 | 0.4766 | 0 | 0 | 0 |
| 'C530008M17Rik' | 0.936 | 0.975 | 0.980 | 1.009 | 0.870 | 1.145 | 1.047 | 0.906 | 0.905 | 0.731 | 0.789 | 0.948 | 0.696 | 0.834 | 0.695 | 0.793 | 0.805 | 0.757 | 0.302 | 0.0000 | 0.0001 | 1 | 0 | 0 |
| 'C87436' | 3.753 | 3.446 | 3.213 | 3.435 | 3.513 | 3.244 | 3.401 | 3.910 | 3.249 | 3.628 | 3.422 | 3.660 | 3.106 | 3.655 | 3.578 | 3.306 | 3.530 | 3.078 | -0.004 | 0.9404 | 0.9769 | 0 | 0 | 0 |
| 'C8g' | 1.353 | 0.883 | 1.846 | 1.314 | 1.458 | 1.313 | 1.094 | 1.288 | 1.587 | 1.516 | 1.146 | 1.268 | 1.334 | 1.191 | 1.530 | 1.473 | 1.416 | 1.090 | 0.028 | 0.8101 | 0.9142 | 0 | 0 | 0 |
| 'Caap1' | 4.383 | 4.973 | 5.210 | 4.858 | 5.216 | 4.423 | 5.829 | 4.836 | 4.882 | 4.671 | 4.425 | 4.260 | 4.960 | 5.167 | 3.586 | 4.306 | 5.535 | 3.786 | 0.127 | 0.0569 | 0.2050 | 0 | 0 | 0 |
| 'Cab39' | 39.815 | 43.447 | 42.252 | 39.989 | 42.997 | 41.177 | 42.532 | 41.473 | 39.218 | 39.806 | 37.996 | 38.161 | 39.708 | 41.570 | 37.638 | 38.353 | 39.869 | 38.612 | 0.073 | 0.0012 | 0.0119 | 1 | 0 | 0 |
| 'Cab39l' | 12.105 | 13.020 | 12.817 | 12.698 | 12.779 | 12.101 | 13.600 | 14.397 | 11.685 | 12.440 | 12.755 | 13.201 | 12.668 | 12.098 | 12.276 | 12.108 | 15.009 | 11.233 | 0.005 | 0.9194 | 0.9695 | 0 | 0 | 0 |
| 'Cabin1' | 29.476 | 24.472 | 30.455 | 27.002 | 24.426 | 28.438 | 26.430 | 26.162 | 29.729 | 31.181 | 26.975 | 31.933 | 27.900 | 25.691 | 29.654 | 28.611 | 25.702 | 30.183 | -0.074 | 0.0154 | 0.0854 | 0 | 0 | 0 |
| 'Cables2' | 8.432 | 8.788 | 9.542 | 8.189 | 9.065 | 9.209 | 8.290 | 8.979 | 9.125 | 9.542 | 8.625 | 8.912 | 9.729 | 8.904 | 8.225 | 9.264 | 8.928 | 8.345 | -0.023 | 0.5472 | 0.7612 | 0 | 0 | 0 |
| 'Cacfd1' | 6.149 | 6.681 | 6.140 | 5.409 | 6.771 | 6.337 | 5.480 | 6.730 | 6.338 | 5.939 | 6.319 | 6.093 | 6.474 | 6.447 | 6.192 | 6.578 | 5.879 | 5.790 | -0.002 | 0.9694 | 0.9898 | 0 | 0 | 0 |
| 'Cacna1a' | 0.188 | 0.227 | 0.221 | 0.213 | 0.162 | 0.174 | 0.160 | 0.203 | 0.204 | 0.324 | 0.370 | 0.266 | 0.266 | 0.230 | 0.283 | 0.293 | 0.275 | 0.285 | -0.563 | 0.0000 | 0.0000 | 1 | 1 | 0 |
| 'Cacna1d' | 0.820 | 0.721 | 0.633 | 0.606 | 0.735 | 0.657 | 0.777 | 0.655 | 0.670 | 0.519 | 0.574 | 0.517 | 0.444 | 0.334 | 0.607 | 0.526 | 0.405 | 0.397 | 0.524 | 0.0000 | 0.0000 | 1 | 1 | 0 |
| 'Cacna1g' | 0.449 | 0.546 | 0.473 | 0.577 | 0.619 | 0.543 | 0.469 | 0.561 | 0.545 | 0.611 | 0.610 | 0.559 | 0.883 | 0.689 | 0.507 | 0.590 | 0.646 | 0.500 | -0.232 | 0.0086 | 0.0559 | 0 | 0 | 0 |
| 'Cacna2d2' | 0.916 | 0.743 | 1.081 | 0.896 | 0.624 | 1.107 | 0.811 | 0.837 | 1.119 | 0.694 | 0.532 | 0.755 | 0.536 | 0.425 | 0.728 | 0.517 | 0.432 | 0.662 | 0.617 | 0.0000 | 0.0000 | 1 | 1 | 0 |
| 'Cacna2d4' | 1.262 | 1.243 | 1.286 | 1.204 | 1.283 | 1.438 | 1.388 | 1.323 | 1.431 | 1.411 | 1.259 | 1.293 | 1.299 | 1.277 | 1.170 | 1.410 | 1.349 | 1.064 | 0.037 | 0.5223 | 0.7449 | 0 | 0 | 0 |
| 'Cacnb1' | 4.485 | 3.663 | 4.866 | 4.554 | 3.689 | 4.661 | 4.544 | 3.930 | 4.282 | 4.562 | 3.721 | 5.352 | 3.931 | 3.368 | 4.920 | 4.203 | 3.883 | 5.223 | -0.027 | 0.5975 | 0.7954 | 0 | 0 | 0 |
| 'Cactin' | 40.021 | 33.352 | 40.861 | 34.907 | 32.815 | 43.477 | 36.290 | 34.934 | 41.666 | 38.395 | 31.388 | 42.171 | 41.309 | 33.856 | 41.459 | 37.967 | 31.011 | 39.182 | -0.001 | 0.9735 | 0.9908 | 0 | 0 | 0 |
| 'Cacul1' | 9.769 | 9.127 | 8.313 | 8.909 | 9.205 | 9.043 | 9.201 | 9.649 | 8.577 | 9.752 | 9.785 | 9.348 | 8.942 | 9.675 | 8.262 | 9.665 | 9.484 | 8.729 | -0.042 | 0.1919 | 0.4290 | 0 | 0 | 0 |
| 'Cacybp' | 25.359 | 21.289 | 20.129 | 26.530 | 22.466 | 19.738 | 25.999 | 20.943 | 19.476 | 19.372 | 22.816 | 28.665 | 20.041 | 23.193 | 24.927 | 17.234 | 25.039 | 28.497 | -0.061 | 0.5789 | 0.7820 | 0 | 0 | 0 |
| 'Cad' | 8.032 | 7.235 | 8.858 | 8.644 | 7.390 | 8.439 | 8.039 | 7.767 | 8.256 | 8.668 | 7.793 | 8.223 | 8.791 | 7.162 | 7.638 | 7.286 | 7.535 | 8.650 | 0.007 | 0.8515 | 0.9362 | 0 | 0 | 0 |
| 'Calcoco1' | 22.041 | 22.824 | 21.799 | 20.770 | 23.341 | 22.323 | 22.807 | 21.978 | 23.464 | 21.045 | 23.033 | 22.079 | 22.107 | 22.404 | 21.959 | 22.134 | 21.906 | 20.870 | 0.017 | 0.5094 | 0.7342 | 0 | 0 | 0 |
| 'Calcrl' | 1.387 | 1.280 | 1.739 | 1.824 | 1.443 | 1.668 | 1.833 | 1.356 | 1.572 | 1.099 | 1.333 | 1.798 | 1.292 | 1.370 | 1.170 | 1.222 | 1.590 | 1.544 | 0.165 | 0.0743 | 0.2424 | 0 | 0 | 0 |
| 'Calhm2' | 16.974 | 17.888 | 15.738 | 16.671 | 15.234 | 16.448 | 16.986 | 15.041 | 17.762 | 13.503 | 15.070 | 20.177 | 13.181 | 14.268 | 17.836 | 13.271 | 16.923 | 20.000 | 0.047 | 0.4890 | 0.7189 | 0 | 0 | 0 |
| 'Calm1' | 104.780 | 117.530 | 106.661 | 114.860 | 119.673 | 107.233 | 117.538 | 112.414 | 108.920 | 105.430 | 109.716 | 94.675 | 108.404 | 111.026 | 104.904 | 108.421 | 115.001 | 101.736 | 0.064 | 0.0198 | 0.1021 | 0 | 0 | 0 |
| 'Calm2' | 197.379 | 230.049 | 180.757 | 207.691 | 218.460 | 185.765 | 207.810 | 218.823 | 193.537 | 194.312 | 208.029 | 169.158 | 199.157 | 217.737 | 183.073 | 215.574 | 210.479 | 172.064 | 0.048 | 0.0834 | 0.2617 | 0 | 0 | 0 |
| 'Calm3' | 100.350 | 108.904 | 107.010 | 104.319 | 111.298 | 114.500 | 103.282 | 106.219 | 111.244 | 101.629 | 101.974 | 103.187 | 107.527 | 106.563 | 114.287 | 107.692 | 102.635 | 108.404 | 0.009 | 0.7329 | 0.8781 | 0 | 0 | 0 |
| 'Calr' | 163.118 | 160.148 | 164.010 | 178.494 | 174.450 | 169.279 | 166.729 | 164.920 | 165.177 | 190.316 | 159.355 | 150.307 | 168.183 | 165.844 | 155.734 | 179.104 | 164.309 | 156.849 | 0.008 | 0.8090 | 0.9139 | 0 | 0 | 0 |
| 'Calu' | 10.343 | 11.358 | 10.718 | 10.830 | 11.114 | 10.726 | 10.543 | 11.316 | 10.223 | 10.861 | 10.813 | 10.774 | 10.814 | 11.360 | 10.103 | 10.635 | 10.838 | 10.525 | -0.005 | 0.8268 | 0.9244 | 0 | 0 | 0 |
| 'Camk1d' | 4.481 | 3.610 | 4.458 | 4.303 | 3.997 | 4.631 | 4.086 | 3.895 | 4.199 | 4.454 | 4.509 | 4.864 | 4.335 | 4.209 | 4.157 | 3.736 | 3.958 | 4.451 | -0.050 | 0.3022 | 0.5592 | 0 | 0 | 0 |
| 'Camk2d' | 2.995 | 3.475 | 3.073 | 3.125 | 3.310 | 3.036 | 3.247 | 3.356 | 3.108 | 3.471 | 3.739 | 3.007 | 3.479 | 3.538 | 2.927 | 3.499 | 3.678 | 2.801 | -0.074 | 0.0212 | 0.1068 | 0 | 0 | 0 |
| 'Camk2g' | 8.357 | 7.079 | 9.119 | 6.992 | 7.488 | 7.968 | 8.587 | 8.157 | 8.581 | 8.086 | 6.808 | 8.292 | 6.906 | 7.545 | 7.877 | 7.148 | 7.144 | 6.701 | 0.112 | 0.0295 | 0.1336 | 0 | 0 | 0 |
| 'Camk4' | 30.336 | 29.409 | 32.271 | 30.588 | 30.091 | 29.849 | 30.935 | 30.793 | 30.915 | 31.518 | 29.340 | 29.467 | 27.011 | 27.571 | 28.787 | 27.884 | 26.093 | 29.549 | 0.087 | 0.0052 | 0.0388 | 1 | 0 | 0 |
| 'Camkk1' | 1.663 | 1.761 | 1.618 | 1.662 | 1.640 | 1.437 | 1.519 | 1.278 | 1.610 | 1.428 | 1.560 | 1.905 | 1.180 | 1.405 | 1.799 | 1.709 | 1.671 | 1.926 | -0.051 | 0.5521 | 0.7650 | 0 | 0 | 0 |
| 'Camkk2' | 1.767 | 1.683 | 2.087 | 1.920 | 1.774 | 2.072 | 1.846 | 1.832 | 1.974 | 1.773 | 1.708 | 1.839 | 1.972 | 1.489 | 2.011 | 1.552 | 1.462 | 2.109 | 0.083 | 0.1274 | 0.3420 | 0 | 0 | 0 |
| 'Camkmt' | 8.352 | 9.443 | 7.505 | 6.890 | 6.200 | 7.407 | 8.514 | 7.433 | 8.077 | 8.730 | 6.318 | 7.787 | 7.243 | 7.636 | 8.097 | 7.408 | 6.310 | 6.235 | 0.072 | 0.4096 | 0.6588 | 0 | 0 | 0 |
| 'Caml' | 8.498 | 10.404 | 9.426 | 9.534 | 9.294 | 9.010 | 8.338 | 9.331 | 9.077 | 8.816 | 9.396 | 9.763 | 8.573 | 9.620 | 9.825 | 8.544 | 9.693 | 9.379 | -0.027 | 0.5242 | 0.7465 | 0 | 0 | 0 |
| 'Camsap2' | 2.919 | 2.471 | 2.830 | 2.861 | 2.785 | 2.830 | 2.944 | 2.807 | 2.825 | 2.851 | 2.974 | 3.079 | 2.624 | 2.730 | 3.010 | 3.283 | 2.962 | 3.018 | -0.079 | 0.0619 | 0.2156 | 0 | 0 | 0 |
| 'Camta1' | 0.358 | 0.334 | 0.329 | 0.277 | 0.313 | 0.311 | 0.275 | 0.327 | 0.330 | 0.260 | 0.330 | 0.288 | 0.337 | 0.354 | 0.263 | 0.283 | 0.368 | 0.254 | 0.062 | 0.4361 | 0.6799 | 0 | 0 | 0 |
| 'Camta2' | 11.840 | 10.435 | 11.872 | 10.496 | 10.548 | 11.012 | 10.518 | 10.701 | 11.981 | 11.225 | 10.538 | 12.193 | 11.282 | 10.674 | 11.529 | 10.730 | 10.610 | 12.155 | -0.032 | 0.2786 | 0.5352 | 0 | 0 | 0 |
| 'Cand1' | 20.763 | 21.117 | 20.571 | 21.073 | 20.940 | 20.791 | 20.225 | 20.864 | 20.693 | 21.174 | 21.124 | 20.995 | 19.586 | 20.352 | 20.593 | 20.089 | 21.385 | 20.016 | 0.003 | 0.8910 | 0.9554 | 0 | 0 | 0 |
| 'Cand2' | 7.810 | 9.059 | 10.126 | 7.891 | 8.835 | 9.455 | 8.624 | 8.800 | 9.528 | 8.257 | 7.218 | 6.837 | 7.489 | 7.355 | 6.895 | 7.668 | 6.632 | 6.768 | 0.292 | 0.0000 | 0.0000 | 1 | 0 | 0 |
| 'Cant1' | 20.179 | 21.338 | 21.487 | 18.629 | 19.293 | 21.843 | 18.387 | 21.169 | 21.012 | 21.175 | 20.485 | 22.679 | 20.103 | 20.497 | 21.360 | 21.240 | 18.763 | 22.455 | -0.052 | 0.1116 | 0.3165 | 0 | 0 | 0 |
| 'Canx' | 42.929 | 46.286 | 47.900 | 49.428 | 50.769 | 47.041 | 47.380 | 48.999 | 45.518 | 48.113 | 46.479 | 47.750 | 45.888 | 48.879 | 45.532 | 44.682 | 48.244 | 48.003 | -0.002 | 0.9341 | 0.9748 | 0 | 0 | 0 |
| 'Cap1' | 157.545 | 164.367 | 162.684 | 167.433 | 167.340 | 160.585 | 167.566 | 161.306 | 162.486 | 165.730 | 171.472 | 160.631 | 163.735 | 167.303 | 171.120 | 162.598 | 170.389 | 167.855 | -0.039 | 0.0676 | 0.2278 | 0 | 0 | 0 |
| 'Cap2' | 2.525 | 2.409 | 3.142 | 2.795 | 2.804 | 2.637 | 2.953 | 2.709 | 3.507 | 2.783 | 2.725 | 3.085 | 2.102 | 3.405 | 2.566 | 2.567 | 2.374 | 2.654 | 0.062 | 0.5402 | 0.7577 | 0 | 0 | 0 |
| 'Capg' | 5.640 | 4.924 | 5.937 | 4.858 | 6.264 | 6.016 | 5.484 | 5.481 | 5.731 | 5.240 | 4.194 | 6.912 | 4.840 | 4.850 | 6.093 | 6.121 | 5.617 | 5.662 | 0.014 | 0.8326 | 0.9270 | 0 | 0 | 0 |
| 'Capn1' | 93.321 | 87.971 | 100.688 | 95.090 | 91.246 | 104.153 | 92.107 | 91.760 | 101.042 | 97.687 | 92.894 | 100.913 | 92.190 | 91.221 | 100.427 | 94.206 | 85.399 | 99.185 | -0.004 | 0.8381 | 0.9309 | 0 | 0 | 0 |
| 'Capn10' | 2.307 | 2.620 | 2.483 | 2.076 | 2.460 | 2.539 | 2.195 | 2.561 | 2.704 | 2.219 | 2.209 | 2.348 | 2.633 | 2.389 | 2.285 | 1.944 | 1.955 | 2.102 | 0.114 | 0.0726 | 0.2384 | 0 | 0 | 0 |
| 'Capn15' | 10.907 | 9.782 | 12.023 | 10.368 | 10.290 | 10.757 | 9.389 | 10.032 | 12.145 | 10.578 | 10.326 | 12.695 | 10.773 | 9.856 | 11.340 | 9.355 | 9.941 | 12.738 | -0.038 | 0.3505 | 0.6046 | 0 | 0 | 0 |
| 'Capn2' | 4.741 | 3.929 | 5.180 | 4.304 | 4.348 | 5.390 | 4.742 | 4.632 | 4.795 | 5.242 | 4.420 | 5.685 | 4.526 | 4.477 | 6.102 | 4.855 | 4.232 | 5.816 | -0.113 | 0.0113 | 0.0681 | 0 | 0 | 0 |
| 'Capn3' | 1.631 | 1.653 | 1.622 | 2.063 | 1.403 | 1.669 | 1.766 | 1.182 | 1.914 | 1.691 | 1.081 | 1.618 | 1.576 | 1.189 | 1.583 | 1.578 | 1.160 | 1.558 | 0.178 | 0.0061 | 0.0436 | 1 | 0 | 0 |
| 'Capn5' | 0.755 | 1.022 | 0.452 | 0.730 | 0.779 | 0.813 | 0.737 | 0.769 | 0.711 | 0.762 | 0.610 | 0.669 | 0.618 | 0.854 | 0.579 | 0.736 | 0.670 | 0.567 | 0.134 | 0.2373 | 0.4868 | 0 | 0 | 0 |
| 'Capn7' | 11.286 | 12.771 | 10.920 | 10.754 | 12.484 | 11.505 | 11.401 | 13.308 | 11.817 | 10.495 | 11.631 | 12.266 | 10.741 | 12.575 | 11.706 | 10.848 | 13.033 | 11.486 | 0.006 | 0.8294 | 0.9261 | 0 | 0 | 0 |
| 'Capns1' | 72.574 | 74.528 | 73.280 | 73.897 | 76.027 | 82.300 | 74.752 | 75.874 | 75.818 | 74.070 | 74.046 | 73.970 | 75.014 | 71.366 | 75.120 | 72.619 | 71.585 | 72.787 | 0.029 | 0.1924 | 0.4295 | 0 | 0 | 0 |
| 'Caprin1' | 24.355 | 23.307 | 23.432 | 25.141 | 23.279 | 22.212 | 24.011 | 23.018 | 22.887 | 22.706 | 23.672 | 25.469 | 22.687 | 23.671 | 24.592 | 21.599 | 24.325 | 26.138 | -0.032 | 0.3904 | 0.6397 | 0 | 0 | 0 |
| 'Caprin2' | 0.492 | 0.513 | 0.487 | 0.624 | 0.436 | 0.459 | 0.533 | 0.507 | 0.584 | 0.551 | 0.459 | 0.512 | 0.483 | 0.473 | 0.505 | 0.474 | 0.645 | 0.419 | 0.024 | 0.7732 | 0.8993 | 0 | 0 | 0 |
| 'Capza1' | 101.600 | 110.989 | 108.382 | 111.780 | 112.921 | 104.935 | 114.496 | 108.230 | 105.196 | 103.305 | 110.850 | 105.637 | 104.323 | 113.589 | 104.479 | 96.962 | 114.206 | 104.823 | 0.020 | 0.4719 | 0.7075 | 0 | 0 | 0 |
| 'Capza2' | 20.665 | 24.681 | 21.584 | 22.512 | 24.467 | 20.810 | 23.365 | 22.775 | 20.105 | 21.904 | 22.606 | 20.932 | 21.748 | 23.235 | 19.880 | 21.490 | 23.771 | 18.874 | 0.036 | 0.1796 | 0.4148 | 0 | 0 | 0 |
| 'Capzb' | 65.828 | 68.452 | 63.743 | 68.947 | 68.555 | 67.487 | 67.664 | 64.970 | 66.229 | 63.602 | 65.382 | 61.204 | 70.424 | 65.722 | 66.356 | 67.668 | 64.393 | 63.837 | 0.022 | 0.3833 | 0.6338 | 0 | 0 | 0 |
| 'Car12' | 0.899 | 0.652 | 0.790 | 0.761 | 0.564 | 0.615 | 0.887 | 0.794 | 0.854 | 0.794 | 0.849 | 1.185 | 0.963 | 0.644 | 0.942 | 0.749 | 0.807 | 1.109 | -0.253 | 0.0159 | 0.0875 | 0 | 0 | 0 |
| 'Card11' | 53.105 | 45.997 | 55.530 | 51.618 | 46.914 | 53.687 | 51.273 | 47.626 | 52.146 | 54.559 | 49.929 | 53.984 | 51.266 | 45.717 | 52.811 | 48.833 | 47.312 | 55.269 | -0.016 | 0.5451 | 0.7606 | 0 | 0 | 0 |
| 'Card19' | 20.771 | 26.444 | 23.657 | 20.138 | 20.701 | 21.709 | 22.303 | 26.823 | 23.827 | 20.522 | 20.270 | 21.653 | 22.060 | 20.291 | 20.863 | 24.679 | 19.332 | 19.494 | 0.115 | 0.0702 | 0.2334 | 0 | 0 | 0 |
| 'Card6' | 6.857 | 7.653 | 6.581 | 6.238 | 7.197 | 6.933 | 7.132 | 6.915 | 7.137 | 7.244 | 6.798 | 8.428 | 6.223 | 7.062 | 7.966 | 7.998 | 6.588 | 8.154 | -0.096 | 0.0701 | 0.2332 | 0 | 0 | 0 |
| 'Card9' | 1.062 | 1.462 | 2.110 | 1.561 | 1.084 | 1.962 | 1.182 | 1.109 | 1.453 | 1.299 | 0.835 | 1.613 | 1.333 | 1.015 | 1.550 | 1.108 | 0.896 | 1.544 | 0.199 | 0.0779 | 0.2503 | 0 | 0 | 0 |
| 'Carf' | 0.590 | 0.802 | 0.729 | 0.659 | 0.833 | 0.726 | 0.707 | 0.808 | 0.849 | 1.013 | 1.114 | 1.039 | 1.096 | 1.040 | 0.998 | 1.060 | 1.108 | 0.883 | -0.493 | 0.0000 | 0.0000 | 1 | 1 | 0 |
| 'Carhsp1' | 22.552 | 22.663 | 21.601 | 21.363 | 22.159 | 22.807 | 22.325 | 24.071 | 22.311 | 23.804 | 22.301 | 25.115 | 25.859 | 25.458 | 26.353 | 26.191 | 21.259 | 24.113 | -0.137 | 0.0005 | 0.0061 | 1 | 0 | 0 |
| 'Carkd' | 4.291 | 4.989 | 5.907 | 4.740 | 5.063 | 5.883 | 4.611 | 5.603 | 5.884 | 4.677 | 4.789 | 5.363 | 5.174 | 5.306 | 5.297 | 5.569 | 5.282 | 5.208 | 0.000 | 0.9984 | 0.9994 | 0 | 0 | 0 |
| 'Carm1' | 6.631 | 5.907 | 6.327 | 5.563 | 5.809 | 6.326 | 5.805 | 5.474 | 6.228 | 6.088 | 5.644 | 6.411 | 5.918 | 5.397 | 6.000 | 5.738 | 5.590 | 5.906 | 0.027 | 0.4310 | 0.6762 | 0 | 0 | 0 |
| 'Carnmt1' | 13.811 | 12.103 | 14.349 | 14.291 | 13.154 | 13.796 | 13.947 | 13.424 | 12.196 | 12.485 | 13.210 | 13.503 | 12.258 | 14.419 | 12.825 | 12.026 | 13.298 | 11.629 | 0.058 | 0.2849 | 0.5413 | 0 | 0 | 0 |
| 'Carns1' | 9.543 | 8.443 | 8.969 | 8.291 | 7.684 | 8.405 | 8.633 | 9.321 | 8.643 | 9.505 | 7.897 | 10.555 | 9.146 | 7.945 | 9.019 | 10.252 | 8.308 | 8.071 | -0.059 | 0.2721 | 0.5273 | 0 | 0 | 0 |
| 'Cars' | 5.080 | 5.185 | 5.987 | 5.434 | 5.296 | 5.630 | 6.283 | 5.413 | 5.706 | 5.209 | 5.381 | 5.553 | 5.579 | 5.343 | 5.154 | 5.247 | 6.126 | 5.326 | 0.022 | 0.6310 | 0.8173 | 0 | 0 | 0 |
| 'Cars2' | 18.115 | 15.968 | 18.924 | 18.177 | 14.725 | 19.606 | 17.575 | 16.100 | 19.297 | 18.865 | 17.718 | 18.890 | 20.523 | 18.245 | 17.945 | 18.889 | 18.864 | 18.973 | -0.100 | 0.0185 | 0.0977 | 0 | 0 | 0 |
| 'Casc3' | 19.694 | 19.351 | 20.140 | 18.864 | 18.698 | 20.249 | 17.875 | 18.832 | 20.986 | 19.807 | 20.063 | 22.119 | 19.078 | 19.459 | 20.971 | 20.355 | 19.265 | 21.046 | -0.070 | 0.0058 | 0.0421 | 1 | 0 | 0 |
| 'Casc4' | 1.428 | 1.733 | 1.430 | 1.620 | 1.657 | 1.339 | 1.787 | 1.609 | 1.419 | 1.883 | 1.973 | 1.475 | 1.668 | 1.815 | 1.406 | 1.820 | 1.635 | 1.286 | -0.099 | 0.0653 | 0.2228 | 0 | 0 | 0 |
| 'Casc5' | 1.640 | 1.618 | 2.189 | 2.243 | 1.677 | 2.105 | 2.061 | 1.852 | 2.110 | 1.282 | 1.987 | 2.218 | 1.923 | 2.184 | 2.373 | 1.442 | 2.269 | 2.086 | -0.024 | 0.7938 | 0.9080 | 0 | 0 | 0 |
| 'Cask' | 0.585 | 0.531 | 0.520 | 0.590 | 0.567 | 0.572 | 0.574 | 0.577 | 0.630 | 0.574 | 0.460 | 0.533 | 0.415 | 0.612 | 0.543 | 0.543 | 0.667 | 0.529 | 0.072 | 0.3579 | 0.6103 | 0 | 0 | 0 |
| 'Caskin1' | 1.682 | 1.946 | 1.672 | 1.885 | 1.942 | 1.839 | 1.975 | 1.931 | 1.894 | 2.085 | 2.641 | 1.889 | 3.122 | 2.585 | 1.676 | 1.639 | 2.496 | 1.905 | -0.258 | 0.0059 | 0.0427 | 1 | 0 | 0 |
| 'Caskin2' | 6.915 | 5.702 | 7.226 | 6.754 | 5.803 | 6.695 | 6.186 | 5.562 | 6.795 | 6.452 | 6.290 | 6.938 | 6.398 | 5.550 | 6.859 | 6.479 | 5.895 | 7.568 | -0.028 | 0.5012 | 0.7283 | 0 | 0 | 0 |
| 'Casp1' | 77.704 | 93.234 | 81.870 | 93.408 | 95.845 | 77.343 | 95.418 | 84.815 | 83.872 | 79.852 | 81.992 | 76.112 | 81.875 | 84.826 | 70.798 | 79.629 | 87.390 | 71.645 | 0.122 | 0.0009 | 0.0098 | 1 | 0 | 0 |
| 'Casp2' | 6.694 | 7.423 | 7.582 | 6.304 | 7.210 | 7.052 | 7.424 | 7.723 | 7.621 | 7.433 | 7.970 | 7.950 | 7.110 | 7.858 | 7.792 | 6.840 | 8.072 | 7.564 | -0.087 | 0.0053 | 0.0393 | 1 | 0 | 0 |
| 'Casp3' | 36.479 | 37.727 | 38.739 | 39.231 | 35.641 | 36.467 | 38.498 | 37.353 | 38.186 | 35.953 | 39.961 | 39.303 | 36.700 | 41.340 | 39.776 | 35.436 | 37.709 | 38.027 | -0.035 | 0.2887 | 0.5451 | 0 | 0 | 0 |
| 'Casp4' | 14.225 | 15.670 | 16.302 | 14.563 | 16.448 | 14.957 | 16.897 | 16.466 | 15.663 | 15.395 | 15.393 | 12.830 | 13.949 | 14.566 | 11.706 | 15.133 | 14.886 | 11.634 | 0.167 | 0.0004 | 0.0049 | 1 | 0 | 0 |
| 'Casp6' | 3.598 | 4.229 | 4.234 | 3.730 | 4.253 | 4.273 | 4.061 | 3.983 | 4.129 | 4.168 | 4.099 | 3.984 | 4.342 | 3.860 | 4.138 | 3.659 | 3.993 | 4.277 | -0.013 | 0.7789 | 0.9003 | 0 | 0 | 0 |
| 'Casp7' | 21.986 | 26.975 | 25.256 | 25.336 | 27.353 | 25.125 | 26.870 | 25.833 | 25.659 | 24.445 | 27.137 | 24.994 | 29.585 | 26.674 | 25.520 | 25.493 | 27.434 | 24.143 | -0.043 | 0.2910 | 0.5477 | 0 | 0 | 0 |
| 'Casp8' | 31.880 | 33.192 | 31.839 | 33.590 | 34.320 | 32.877 | 34.672 | 34.527 | 33.243 | 35.677 | 37.664 | 37.985 | 39.665 | 36.734 | 37.738 | 36.763 | 38.180 | 36.669 | -0.180 | 0.0000 | 0.0000 | 1 | 0 | 0 |
| 'Casp8ap2' | 11.802 | 10.223 | 10.034 | 12.248 | 11.019 | 9.146 | 10.797 | 10.435 | 9.609 | 10.585 | 10.987 | 10.503 | 11.012 | 10.998 | 9.694 | 11.050 | 11.539 | 10.356 | -0.035 | 0.3611 | 0.6135 | 0 | 0 | 0 |
| 'Casp9' | 3.724 | 4.422 | 4.437 | 3.922 | 4.407 | 4.444 | 3.798 | 4.145 | 4.536 | 4.325 | 3.735 | 4.004 | 4.537 | 4.328 | 4.156 | 3.940 | 3.789 | 3.494 | 0.049 | 0.3597 | 0.6121 | 0 | 0 | 0 |
| 'Cass4' | 1.070 | 1.167 | 1.117 | 1.231 | 1.200 | 1.135 | 0.997 | 1.182 | 1.074 | 1.452 | 1.561 | 1.354 | 1.204 | 1.165 | 1.402 | 1.262 | 1.364 | 1.152 | -0.238 | 0.0010 | 0.0109 | 1 | 0 | 0 |
| 'Cast' | 43.264 | 40.854 | 43.547 | 43.219 | 42.718 | 45.208 | 45.331 | 44.317 | 41.726 | 45.635 | 45.926 | 46.867 | 42.679 | 45.656 | 46.724 | 43.610 | 43.220 | 45.089 | -0.065 | 0.0215 | 0.1080 | 0 | 0 | 0 |
| 'Casz1' | 0.921 | 0.711 | 0.828 | 0.698 | 0.570 | 0.753 | 0.803 | 0.633 | 0.662 | 0.795 | 0.686 | 0.718 | 0.552 | 0.702 | 0.682 | 0.646 | 0.627 | 0.621 | 0.117 | 0.1786 | 0.4140 | 0 | 0 | 0 |
| 'Cat' | 23.545 | 25.769 | 26.063 | 25.952 | 25.867 | 27.107 | 25.560 | 24.720 | 25.304 | 26.168 | 24.258 | 26.249 | 25.021 | 27.525 | 25.973 | 25.455 | 24.954 | 25.914 | -0.021 | 0.4810 | 0.7131 | 0 | 0 | 0 |
| 'Catsper2' | 1.314 | 1.504 | 1.489 | 1.166 | 1.256 | 1.561 | 1.511 | 1.669 | 1.563 | 1.303 | 1.320 | 1.362 | 1.539 | 1.178 | 1.302 | 0.974 | 1.760 | 1.319 | 0.100 | 0.2812 | 0.5379 | 0 | 0 | 0 |
| 'Cbarp' | 4.542 | 3.399 | 4.121 | 3.501 | 3.425 | 3.949 | 3.086 | 3.440 | 3.709 | 4.217 | 2.990 | 4.227 | 3.329 | 3.034 | 3.997 | 3.209 | 2.901 | 3.866 | 0.059 | 0.3692 | 0.6203 | 0 | 0 | 0 |
| 'Cbfa2t2' | 4.877 | 4.644 | 4.599 | 5.004 | 4.586 | 4.921 | 4.565 | 5.366 | 4.509 | 4.752 | 4.414 | 4.423 | 4.362 | 4.468 | 4.249 | 5.188 | 4.832 | 4.423 | 0.061 | 0.1431 | 0.3663 | 0 | 0 | 0 |
| 'Cbfa2t3' | 6.062 | 6.254 | 5.362 | 5.834 | 5.511 | 4.999 | 5.324 | 5.587 | 5.639 | 6.888 | 6.348 | 6.857 | 8.231 | 6.957 | 6.094 | 6.288 | 6.876 | 7.425 | -0.306 | 0.0000 | 0.0000 | 1 | 0 | 0 |
| 'Cbfb' | 29.510 | 32.205 | 31.478 | 30.743 | 31.920 | 30.022 | 31.237 | 31.603 | 30.738 | 27.862 | 30.963 | 30.929 | 28.924 | 32.377 | 31.265 | 27.709 | 31.436 | 31.805 | 0.022 | 0.3975 | 0.6467 | 0 | 0 | 0 |
| 'Cbl' | 23.287 | 19.039 | 21.968 | 22.296 | 19.152 | 20.752 | 21.808 | 19.623 | 21.117 | 21.749 | 21.863 | 23.926 | 19.369 | 19.286 | 21.686 | 19.743 | 20.263 | 23.247 | -0.027 | 0.5268 | 0.7483 | 0 | 0 | 0 |
| 'Cblb' | 20.834 | 20.628 | 19.026 | 19.481 | 19.622 | 19.788 | 20.239 | 21.170 | 19.680 | 20.781 | 21.612 | 21.675 | 18.082 | 21.519 | 20.445 | 23.488 | 21.673 | 21.036 | -0.087 | 0.0236 | 0.1157 | 0 | 0 | 0 |
| 'Cbll1' | 3.467 | 3.497 | 3.592 | 3.585 | 3.767 | 3.394 | 3.966 | 3.517 | 3.524 | 3.507 | 3.766 | 4.079 | 3.133 | 3.651 | 4.091 | 3.332 | 3.645 | 3.862 | -0.047 | 0.3476 | 0.6031 | 0 | 0 | 0 |
| 'Cbr1' | 18.845 | 17.695 | 18.577 | 19.039 | 17.746 | 17.975 | 16.912 | 14.383 | 17.568 | 16.932 | 15.212 | 18.133 | 15.396 | 16.040 | 18.018 | 15.421 | 14.255 | 16.259 | 0.109 | 0.0463 | 0.1794 | 0 | 0 | 0 |
| 'Cbr4' | 2.398 | 2.768 | 2.842 | 2.125 | 2.659 | 2.679 | 2.322 | 2.902 | 2.922 | 2.485 | 2.484 | 2.230 | 2.616 | 3.052 | 2.109 | 2.976 | 2.745 | 2.002 | 0.061 | 0.4315 | 0.6763 | 0 | 0 | 0 |
| 'Cbwd1' | 6.705 | 7.418 | 7.497 | 7.450 | 6.857 | 5.979 | 6.143 | 6.095 | 6.649 | 5.848 | 5.826 | 6.042 | 6.702 | 6.873 | 5.697 | 6.063 | 6.877 | 5.262 | 0.130 | 0.0582 | 0.2077 | 0 | 0 | 0 |
| 'Cbx1' | 15.461 | 17.691 | 16.367 | 16.117 | 16.669 | 16.671 | 15.862 | 16.851 | 16.273 | 14.078 | 15.501 | 18.153 | 15.709 | 16.999 | 16.969 | 14.781 | 16.374 | 17.080 | 0.009 | 0.7830 | 0.9025 | 0 | 0 | 0 |
| 'Cbx2' | 2.767 | 2.713 | 3.182 | 3.093 | 1.994 | 3.442 | 3.036 | 2.490 | 3.382 | 2.270 | 2.929 | 3.622 | 2.711 | 2.525 | 3.747 | 2.113 | 2.243 | 3.785 | 0.003 | 0.9742 | 0.9912 | 0 | 0 | 0 |
| 'Cbx4' | 28.866 | 26.628 | 24.367 | 22.902 | 25.230 | 26.717 | 22.265 | 26.004 | 23.662 | 28.825 | 22.397 | 24.342 | 28.159 | 26.291 | 24.342 | 30.744 | 21.223 | 23.169 | -0.021 | 0.7274 | 0.8750 | 0 | 0 | 0 |
| 'Cbx5' | 9.398 | 8.595 | 10.171 | 10.156 | 9.449 | 9.000 | 10.011 | 8.787 | 9.674 | 8.328 | 9.947 | 10.952 | 9.668 | 9.893 | 10.072 | 8.616 | 10.526 | 10.622 | -0.066 | 0.1826 | 0.4189 | 0 | 0 | 0 |
| 'Cbx6' | 11.244 | 11.826 | 13.177 | 12.041 | 11.528 | 12.415 | 12.808 | 10.975 | 13.083 | 10.863 | 11.867 | 13.681 | 11.185 | 11.240 | 13.763 | 11.031 | 11.895 | 13.946 | -0.014 | 0.7047 | 0.8623 | 0 | 0 | 0 |
| 'Cbx7' | 10.030 | 11.684 | 11.893 | 8.552 | 10.827 | 11.711 | 9.968 | 12.217 | 12.304 | 10.479 | 9.803 | 10.566 | 10.710 | 9.302 | 10.494 | 10.437 | 9.412 | 9.656 | 0.111 | 0.0311 | 0.1383 | 0 | 0 | 0 |
| 'Cbx8' | 4.576 | 3.312 | 3.421 | 3.962 | 3.335 | 3.735 | 4.094 | 3.551 | 3.753 | 3.090 | 3.436 | 4.028 | 2.704 | 3.207 | 3.713 | 3.343 | 2.885 | 3.920 | 0.142 | 0.0673 | 0.2275 | 0 | 0 | 0 |
| 'Cby1' | 14.769 | 18.580 | 15.926 | 16.047 | 17.118 | 17.708 | 15.614 | 15.948 | 16.054 | 14.264 | 13.439 | 14.633 | 16.581 | 17.074 | 16.273 | 17.321 | 13.578 | 12.745 | 0.111 | 0.0895 | 0.2744 | 0 | 0 | 0 |
| 'Cc2d1a' | 17.345 | 15.385 | 19.027 | 15.880 | 15.803 | 17.646 | 14.959 | 15.806 | 19.025 | 17.702 | 14.360 | 18.248 | 18.400 | 15.445 | 17.472 | 17.967 | 14.737 | 18.200 | -0.023 | 0.5334 | 0.7533 | 0 | 0 | 0 |
| 'Cc2d1b' | 15.907 | 15.617 | 18.933 | 15.265 | 16.025 | 17.678 | 15.768 | 15.834 | 18.204 | 17.029 | 15.611 | 17.937 | 17.546 | 16.067 | 16.478 | 17.112 | 16.305 | 16.740 | -0.026 | 0.3875 | 0.6374 | 0 | 0 | 0 |
| 'Cc2d2a' | 0.769 | 0.877 | 0.669 | 0.661 | 0.959 | 0.824 | 0.664 | 0.817 | 0.715 | 0.760 | 0.666 | 0.841 | 0.940 | 0.829 | 0.726 | 1.040 | 0.895 | 0.622 | -0.087 | 0.3561 | 0.6090 | 0 | 0 | 0 |
| 'Ccar1' | 40.374 | 39.070 | 38.256 | 43.610 | 37.790 | 37.820 | 42.895 | 37.225 | 38.871 | 36.541 | 40.770 | 41.013 | 40.003 | 41.611 | 38.151 | 37.058 | 41.673 | 37.935 | -0.007 | 0.8459 | 0.9343 | 0 | 0 | 0 |
| 'Ccar2' | 56.037 | 57.976 | 62.219 | 55.851 | 60.195 | 61.742 | 57.597 | 58.107 | 63.621 | 59.245 | 56.252 | 56.754 | 59.094 | 57.488 | 56.470 | 56.633 | 55.631 | 52.003 | 0.056 | 0.0557 | 0.2021 | 0 | 0 | 0 |
| 'Ccdc102a' | 3.630 | 2.541 | 4.001 | 3.267 | 3.133 | 3.144 | 3.040 | 2.629 | 2.994 | 2.965 | 3.094 | 3.535 | 3.356 | 2.781 | 3.993 | 2.648 | 3.297 | 3.783 | -0.062 | 0.4640 | 0.6999 | 0 | 0 | 0 |
| 'Ccdc107' | 3.355 | 4.539 | 4.403 | 3.486 | 4.189 | 4.263 | 3.774 | 4.121 | 4.530 | 4.150 | 3.771 | 4.162 | 3.715 | 3.678 | 4.097 | 3.896 | 3.189 | 3.402 | 0.092 | 0.1592 | 0.3895 | 0 | 0 | 0 |
| 'Ccdc109b' | 6.111 | 7.520 | 5.964 | 6.844 | 7.287 | 5.933 | 6.249 | 7.276 | 6.649 | 6.712 | 6.833 | 6.762 | 8.588 | 7.569 | 7.122 | 6.722 | 8.376 | 6.552 | -0.143 | 0.0217 | 0.1086 | 0 | 0 | 0 |
| 'Ccdc115' | 14.408 | 16.248 | 13.671 | 14.384 | 15.734 | 14.553 | 13.658 | 15.986 | 13.322 | 13.387 | 14.513 | 13.652 | 14.605 | 13.608 | 12.870 | 14.378 | 13.545 | 13.140 | 0.079 | 0.0262 | 0.1242 | 0 | 0 | 0 |
| 'Ccdc117' | 23.006 | 16.272 | 15.038 | 20.040 | 16.261 | 15.139 | 18.255 | 17.316 | 15.260 | 15.006 | 21.321 | 30.231 | 16.040 | 21.708 | 24.935 | 16.343 | 24.265 | 32.646 | -0.351 | 0.0247 | 0.1197 | 0 | 0 | 0 |
| 'Ccdc12' | 13.246 | 14.699 | 14.147 | 14.199 | 14.320 | 14.673 | 14.467 | 15.346 | 15.026 | 12.872 | 13.474 | 13.772 | 14.884 | 13.185 | 13.391 | 14.546 | 13.698 | 12.696 | 0.076 | 0.0237 | 0.1161 | 0 | 0 | 0 |
| 'Ccdc120' | 1.012 | 2.031 | 1.279 | 1.496 | 1.706 | 1.370 | 1.132 | 1.416 | 1.746 | 1.459 | 1.721 | 1.640 | 1.999 | 1.775 | 1.476 | 1.880 | 1.630 | 1.573 | -0.225 | 0.0297 | 0.1339 | 0 | 0 | 0 |
| 'Ccdc124' | 81.857 | 87.497 | 90.032 | 86.845 | 84.969 | 96.742 | 86.389 | 83.623 | 94.112 | 80.652 | 68.533 | 84.692 | 90.632 | 73.889 | 90.106 | 89.056 | 75.971 | 82.578 | 0.097 | 0.0098 | 0.0614 | 0 | 0 | 0 |
| 'Ccdc125' | 15.548 | 18.896 | 18.375 | 14.494 | 21.479 | 19.201 | 17.182 | 19.335 | 18.429 | 19.895 | 22.001 | 18.767 | 21.131 | 19.313 | 17.734 | 19.471 | 20.960 | 16.927 | -0.123 | 0.0298 | 0.1341 | 0 | 0 | 0 |
| 'Ccdc126' | 15.830 | 18.657 | 13.878 | 15.115 | 18.083 | 14.401 | 15.879 | 17.595 | 12.857 | 15.473 | 19.013 | 15.760 | 16.781 | 19.774 | 15.920 | 16.351 | 18.386 | 15.932 | -0.125 | 0.0003 | 0.0035 | 1 | 0 | 0 |
| 'Ccdc127' | 2.237 | 2.198 | 2.373 | 2.677 | 2.415 | 2.268 | 2.518 | 2.368 | 2.220 | 1.976 | 2.500 | 2.698 | 2.101 | 2.490 | 2.338 | 2.034 | 2.632 | 2.552 | -0.014 | 0.8248 | 0.9227 | 0 | 0 | 0 |
| 'Ccdc130' | 6.040 | 6.930 | 6.532 | 6.238 | 6.538 | 7.000 | 6.671 | 6.373 | 6.687 | 6.757 | 6.090 | 6.299 | 6.534 | 6.379 | 6.379 | 6.122 | 5.765 | 6.068 | 0.055 | 0.2236 | 0.4711 | 0 | 0 | 0 |
| 'Ccdc134' | 24.880 | 27.165 | 26.042 | 25.233 | 27.341 | 25.301 | 26.199 | 25.012 | 26.645 | 25.927 | 30.921 | 30.447 | 33.721 | 28.469 | 27.456 | 26.340 | 31.086 | 29.650 | -0.189 | 0.0000 | 0.0006 | 1 | 0 | 0 |
| 'Ccdc137' | 4.953 | 4.454 | 5.644 | 4.985 | 4.531 | 4.562 | 4.702 | 4.675 | 5.005 | 4.154 | 5.230 | 5.973 | 4.679 | 5.098 | 5.949 | 4.111 | 4.983 | 5.983 | -0.091 | 0.1287 | 0.3438 | 0 | 0 | 0 |
| 'Ccdc14' | 1.807 | 2.082 | 2.436 | 1.649 | 2.131 | 2.062 | 1.959 | 2.547 | 2.279 | 2.033 | 2.396 | 1.738 | 2.392 | 1.754 | 1.846 | 2.145 | 1.969 | 1.323 | 0.111 | 0.2811 | 0.5379 | 0 | 0 | 0 |
| 'Ccdc141' | 0.176 | 0.145 | 0.125 | 0.173 | 0.157 | 0.126 | 0.141 | 0.183 | 0.166 | 0.222 | 0.173 | 0.279 | 0.169 | 0.209 | 0.298 | 0.260 | 0.254 | 0.258 | -0.620 | 0.0000 | 0.0000 | 1 | 1 | 0 |
| 'Ccdc142' | 0.958 | 1.107 | 1.208 | 0.905 | 0.882 | 1.138 | 0.850 | 1.141 | 0.810 | 1.045 | 0.857 | 0.921 | 1.155 | 0.829 | 1.065 | 0.834 | 0.730 | 0.990 | 0.090 | 0.4099 | 0.6590 | 0 | 0 | 0 |
| 'Ccdc15' | 1.546 | 1.402 | 1.582 | 1.921 | 1.649 | 1.405 | 1.870 | 1.331 | 1.426 | 1.184 | 1.943 | 1.503 | 1.409 | 1.918 | 1.287 | 1.413 | 1.562 | 1.645 | 0.017 | 0.8755 | 0.9470 | 0 | 0 | 0 |
| 'Ccdc157' | 1.580 | 1.691 | 1.969 | 1.477 | 1.466 | 1.967 | 1.487 | 1.605 | 1.887 | 1.854 | 1.822 | 1.818 | 2.032 | 1.714 | 1.871 | 1.873 | 1.678 | 1.497 | -0.101 | 0.1347 | 0.3536 | 0 | 0 | 0 |
| 'Ccdc163' | 2.390 | 2.256 | 2.389 | 2.199 | 2.396 | 2.853 | 1.849 | 2.334 | 2.509 | 2.550 | 2.340 | 2.846 | 2.668 | 2.513 | 2.551 | 2.277 | 2.828 | 2.337 | -0.124 | 0.0556 | 0.2019 | 0 | 0 | 0 |
| 'Ccdc166' | 2.349 | 2.213 | 2.739 | 1.686 | 2.699 | 2.764 | 2.162 | 2.548 | 2.333 | 2.550 | 2.240 | 2.525 | 2.605 | 2.091 | 2.203 | 2.328 | 2.050 | 2.633 | 0.009 | 0.9079 | 0.9639 | 0 | 0 | 0 |
| 'Ccdc167' | 3.519 | 3.441 | 3.873 | 3.753 | 3.695 | 3.554 | 3.165 | 3.258 | 3.515 | 3.280 | 3.352 | 3.741 | 3.513 | 3.541 | 3.398 | 3.594 | 3.981 | 3.603 | -0.021 | 0.6669 | 0.8396 | 0 | 0 | 0 |
| 'Ccdc17' | 1.501 | 1.563 | 1.535 | 1.391 | 1.600 | 1.471 | 1.423 | 1.442 | 1.829 | 1.583 | 1.247 | 1.684 | 1.431 | 1.439 | 1.360 | 1.644 | 1.317 | 1.424 | 0.051 | 0.4982 | 0.7261 | 0 | 0 | 0 |
| 'Ccdc171' | 1.103 | 0.917 | 0.892 | 1.103 | 0.884 | 0.741 | 1.173 | 0.802 | 0.868 | 0.959 | 1.029 | 1.056 | 0.997 | 1.015 | 0.977 | 0.904 | 1.104 | 0.994 | -0.119 | 0.1490 | 0.3757 | 0 | 0 | 0 |
| 'Ccdc173' | 1.351 | 1.603 | 1.644 | 1.952 | 2.006 | 1.652 | 2.280 | 2.377 | 2.161 | 1.666 | 1.862 | 1.772 | 1.956 | 1.765 | 1.627 | 2.032 | 2.024 | 1.573 | 0.053 | 0.6192 | 0.8095 | 0 | 0 | 0 |
| 'Ccdc174' | 3.427 | 3.298 | 3.267 | 3.484 | 3.654 | 3.221 | 3.509 | 3.807 | 3.204 | 3.612 | 3.213 | 3.153 | 3.715 | 3.539 | 3.090 | 3.602 | 3.305 | 2.832 | 0.032 | 0.4022 | 0.6511 | 0 | 0 | 0 |
| 'Ccdc18' | 0.451 | 0.658 | 0.656 | 0.817 | 0.620 | 0.704 | 0.643 | 0.692 | 0.713 | 0.540 | 0.690 | 0.510 | 0.606 | 0.789 | 0.517 | 0.567 | 0.688 | 0.612 | 0.113 | 0.2965 | 0.5524 | 0 | 0 | 0 |
| 'Ccdc181' | 6.531 | 7.233 | 7.370 | 6.939 | 7.929 | 7.030 | 6.102 | 8.105 | 7.666 | 7.197 | 6.434 | 6.873 | 7.118 | 7.507 | 6.379 | 7.464 | 7.594 | 6.020 | 0.047 | 0.4464 | 0.6880 | 0 | 0 | 0 |
| 'Ccdc186' | 5.005 | 5.080 | 4.034 | 4.668 | 4.843 | 4.048 | 4.868 | 5.231 | 4.689 | 4.629 | 4.608 | 4.646 | 4.108 | 4.661 | 4.447 | 4.992 | 4.898 | 4.498 | 0.018 | 0.6935 | 0.8558 | 0 | 0 | 0 |
| 'Ccdc191' | 2.588 | 2.284 | 2.035 | 2.258 | 2.251 | 2.069 | 2.779 | 2.397 | 2.518 | 2.642 | 2.745 | 2.380 | 2.540 | 2.460 | 2.203 | 2.444 | 2.722 | 2.140 | -0.083 | 0.1866 | 0.4223 | 0 | 0 | 0 |
| 'Ccdc22' | 4.181 | 4.441 | 5.047 | 4.942 | 4.015 | 5.480 | 4.354 | 4.572 | 4.718 | 4.450 | 4.474 | 4.676 | 4.628 | 4.476 | 4.550 | 4.221 | 4.231 | 4.442 | 0.049 | 0.3387 | 0.5961 | 0 | 0 | 0 |
| 'Ccdc25' | 19.321 | 20.732 | 20.448 | 22.049 | 20.733 | 22.323 | 22.654 | 19.584 | 19.484 | 20.058 | 22.927 | 20.993 | 21.148 | 21.497 | 19.952 | 18.687 | 21.775 | 19.068 | -0.002 | 0.9708 | 0.9898 | 0 | 0 | 0 |
| 'Ccdc28a' | 8.073 | 9.925 | 8.062 | 6.859 | 8.644 | 8.899 | 8.436 | 9.863 | 8.701 | 8.146 | 9.049 | 8.105 | 9.289 | 9.313 | 8.781 | 8.788 | 8.221 | 7.310 | -0.002 | 0.9667 | 0.9887 | 0 | 0 | 0 |
| 'Ccdc28b' | 3.674 | 3.368 | 3.530 | 3.881 | 3.753 | 3.494 | 3.785 | 3.427 | 3.325 | 3.102 | 3.101 | 3.266 | 3.560 | 3.676 | 3.436 | 3.529 | 3.544 | 2.983 | 0.084 | 0.1240 | 0.3366 | 0 | 0 | 0 |
| 'Ccdc32' | 8.157 | 9.911 | 8.646 | 8.267 | 10.116 | 9.322 | 9.000 | 10.105 | 8.385 | 8.094 | 7.473 | 8.286 | 8.179 | 8.555 | 8.492 | 8.527 | 8.566 | 7.545 | 0.136 | 0.0038 | 0.0302 | 1 | 0 | 0 |
| 'Ccdc34' | 1.390 | 1.572 | 1.804 | 1.510 | 1.630 | 1.591 | 1.778 | 1.602 | 1.692 | 1.369 | 1.472 | 1.647 | 1.742 | 1.592 | 1.641 | 1.381 | 1.550 | 1.588 | 0.043 | 0.4567 | 0.6943 | 0 | 0 | 0 |
| 'Ccdc43' | 4.690 | 5.237 | 5.237 | 5.393 | 5.770 | 4.835 | 4.637 | 5.050 | 4.832 | 4.649 | 4.669 | 5.041 | 5.012 | 5.039 | 4.594 | 5.195 | 5.362 | 4.639 | 0.034 | 0.4460 | 0.6878 | 0 | 0 | 0 |

| | | | | | | | | | | | | | | | | | | | | | | | | |
|---|---|---|---|---|---|---|---|---|---|---|---|---|---|---|---|---|---|---|---|---|---|---|---|---|
| 'Ccdc47' | 18.890 | 21.243 | 20.704 | 21.567 | 21.871 | 19.322 | 21.008 | 20.014 | 19.515 | 19.466 | 19.468 | 20.879 | 19.244 | 20.722 | 20.504 | 20.344 | 20.949 | 20.204 | 0.006 | 0.8605 | 0.9397 | 0 | 0 | 0 |
| 'Ccdc50' | 15.965 | 17.933 | 15.711 | 16.030 | 17.011 | 15.053 | 18.213 | 16.693 | 16.211 | 13.936 | 15.033 | 15.091 | 13.734 | 15.361 | 14.221 | 13.514 | 16.132 | 13.791 | 0.174 | 0.0000 | 0.0000 | 1 | 0 | 0 |
| 'Ccdc51' | 6.422 | 7.249 | 7.786 | 6.945 | 7.616 | 7.248 | 6.879 | 6.678 | 7.244 | 5.781 | 5.933 | 7.258 | 7.223 | 6.887 | 7.152 | 5.171 | 6.029 | 6.710 | 0.118 | 0.0876 | 0.2701 | 0 | 0 | 0 |
| 'Ccdc53' | 6.391 | 6.993 | 6.044 | 6.882 | 6.059 | 5.521 | 6.675 | 5.959 | 5.893 | 5.677 | 6.034 | 6.022 | 6.551 | 6.160 | 6.600 | 6.709 | 5.832 | 5.766 | 0.012 | 0.7906 | 0.9064 | 0 | 0 | 0 |
| 'Ccdc58' | 11.171 | 14.090 | 13.485 | 14.780 | 14.582 | 14.013 | 13.700 | 14.612 | 14.664 | 12.479 | 13.882 | 12.667 | 12.919 | 12.358 | 13.552 | 14.071 | 14.281 | 12.411 | 0.068 | 0.2383 | 0.4883 | 0 | 0 | 0 |
| 'Ccdc59' | 8.791 | 8.335 | 8.512 | 7.153 | 8.940 | 8.229 | 7.818 | 9.595 | 7.894 | 9.540 | 7.688 | 7.532 | 9.186 | 8.316 | 7.003 | 9.132 | 7.594 | 6.731 | 0.048 | 0.4069 | 0.6562 | 0 | 0 | 0 |
| 'Ccdc6' | 9.512 | 9.571 | 8.858 | 8.500 | 8.851 | 9.322 | 7.875 | 8.262 | 9.235 | 8.486 | 9.027 | 10.654 | 9.163 | 9.015 | 10.376 | 8.985 | 9.602 | 10.268 | -0.108 | 0.0093 | 0.0592 | 0 | 0 | 0 |
| 'Ccdc61' | 7.790 | 8.463 | 8.663 | 7.338 | 7.790 | 8.636 | 8.822 | 7.858 | 8.916 | 8.204 | 7.779 | 8.947 | 7.924 | 7.953 | 8.321 | 7.180 | 8.403 | 8.704 | 0.003 | 0.9452 | 0.9790 | 0 | 0 | 0 |
| 'Ccdc62' | 0.353 | 0.477 | 0.398 | 0.345 | 0.324 | 0.473 | 0.510 | 0.545 | 0.485 | 0.404 | 0.369 | 0.375 | 0.345 | 0.356 | 0.335 | 0.346 | 0.327 | 0.331 | 0.284 | 0.0095 | 0.0601 | 0 | 0 | 0 |
| 'Ccdc64' | 8.959 | 6.751 | 8.957 | 7.628 | 6.765 | 9.513 | 7.691 | 7.360 | 8.578 | 7.907 | 6.839 | 8.968 | 7.728 | 7.000 | 8.891 | 8.227 | 6.905 | 9.340 | 0.000 | 0.9998 | 0.9999 | 0 | 0 | 0 |
| 'Ccdc66' | 12.663 | 14.303 | 14.977 | 13.993 | 14.348 | 15.870 | 14.388 | 15.643 | 15.040 | 16.378 | 13.657 | 14.314 | 15.796 | 14.322 | 13.267 | 15.426 | 14.058 | 13.754 | -0.004 | 0.9336 | 0.9746 | 0 | 0 | 0 |
| 'Ccdc67' | 2.026 | 2.332 | 2.060 | 1.939 | 2.572 | 1.898 | 2.959 | 2.503 | 2.473 | 1.713 | 3.341 | 2.984 | 2.784 | 2.409 | 2.711 | 2.606 | 2.862 | 2.913 | -0.242 | 0.0143 | 0.0807 | 0 | 0 | 0 |
| 'Ccdc69' | 24.093 | 23.563 | 25.122 | 25.277 | 24.534 | 27.122 | 23.316 | 22.717 | 24.373 | 23.725 | 26.424 | 25.848 | 24.652 | 26.284 | 28.455 | 24.398 | 26.465 | 26.467 | -0.088 | 0.0226 | 0.1121 | 0 | 0 | 0 |
| 'Ccdc71' | 7.818 | 7.256 | 8.815 | 7.111 | 7.570 | 9.409 | 7.498 | 7.269 | 8.622 | 8.036 | 6.617 | 8.641 | 7.315 | 6.447 | 8.238 | 7.685 | 6.542 | 8.638 | 0.058 | 0.1114 | 0.3160 | 0 | 0 | 0 |
| 'Ccdc71l' | 7.871 | 8.205 | 8.488 | 8.287 | 7.807 | 7.805 | 7.924 | 7.853 | 7.295 | 8.553 | 8.810 | 8.261 | 7.627 | 9.281 | 9.018 | 8.735 | 8.194 | 8.559 | -0.114 | 0.0115 | 0.0692 | 0 | 0 | 0 |
| 'Ccdc77' | 1.963 | 1.796 | 2.129 | 1.942 | 1.810 | 1.981 | 1.926 | 1.776 | 2.210 | 1.626 | 1.809 | 2.102 | 1.881 | 1.834 | 1.913 | 1.905 | 1.996 | 1.904 | 0.036 | 0.4764 | 0.7107 | 0 | 0 | 0 |
| 'Ccdc80' | 0.633 | 0.567 | 0.593 | 0.611 | 0.551 | 0.735 | 0.613 | 0.580 | 0.589 | 0.580 | 0.520 | 0.659 | 0.503 | 0.678 | 0.539 | 0.707 | 0.654 | 0.482 | 0.035 | 0.7493 | 0.8877 | 0 | 0 | 0 |
| 'Ccdc82' | 21.923 | 19.397 | 21.631 | 23.010 | 20.739 | 21.501 | 21.858 | 22.043 | 21.843 | 19.987 | 23.404 | 22.038 | 19.030 | 21.597 | 21.718 | 23.363 | 25.311 | 21.711 | -0.038 | 0.4512 | 0.6904 | 0 | 0 | 0 |
| 'Ccdc84' | 30.647 | 31.233 | 34.874 | 30.951 | 29.427 | 31.438 | 34.279 | 32.172 | 35.202 | 25.472 | 25.088 | 30.307 | 28.416 | 25.637 | 25.150 | 26.463 | 27.024 | 25.309 | 0.266 | 0.0000 | 0.0000 | 1 | 0 | 0 |
| 'Ccdc85b' | 56.941 | 51.788 | 59.895 | 52.085 | 56.897 | 57.887 | 55.113 | 53.118 | 52.806 | 54.739 | 44.213 | 58.166 | 55.736 | 50.921 | 57.350 | 52.332 | 53.103 | 57.131 | 0.026 | 0.5440 | 0.7606 | 0 | 0 | 0 |
| 'Ccdc86' | 2.511 | 1.960 | 2.274 | 2.116 | 2.143 | 2.399 | 2.136 | 1.844 | 2.120 | 2.916 | 2.054 | 2.191 | 2.545 | 1.893 | 1.933 | 2.960 | 1.749 | 2.083 | -0.051 | 0.4522 | 0.6912 | 0 | 0 | 0 |
| 'Ccdc88a' | 0.807 | 0.716 | 0.844 | 0.789 | 0.884 | 0.902 | 0.903 | 0.890 | 0.879 | 1.019 | 0.874 | 0.894 | 0.780 | 1.347 | 0.791 | 0.833 | 1.229 | 0.981 | -0.197 | 0.0317 | 0.1400 | 0 | 0 | 0 |
| 'Ccdc88b' | 87.490 | 72.886 | 91.114 | 80.037 | 74.517 | 83.483 | 79.505 | 75.789 | 86.533 | 86.929 | 76.588 | 89.285 | 88.883 | 75.116 | 84.642 | 81.538 | 76.880 | 89.072 | -0.045 | 0.0932 | 0.2826 | 0 | 0 | 0 |
| 'Ccdc88c' | 34.503 | 25.767 | 30.424 | 31.927 | 24.373 | 28.873 | 30.037 | 27.367 | 31.016 | 30.990 | 27.736 | 31.002 | 30.762 | 26.720 | 29.051 | 28.117 | 27.460 | 31.635 | -0.010 | 0.8082 | 0.9139 | 0 | 0 | 0 |
| 'Ccdc9' | 12.233 | 11.686 | 13.132 | 10.722 | 11.321 | 13.417 | 11.101 | 11.864 | 13.474 | 12.979 | 10.791 | 11.797 | 12.409 | 11.107 | 11.484 | 12.589 | 11.178 | 10.495 | 0.046 | 0.3106 | 0.5689 | 0 | 0 | 0 |
| 'Ccdc90b' | 3.624 | 4.595 | 4.213 | 3.699 | 4.641 | 5.010 | 4.287 | 3.983 | 3.706 | 4.062 | 4.233 | 4.069 | 4.317 | 3.437 | 3.339 | 4.018 | 4.265 | 3.197 | 0.100 | 0.2271 | 0.4734 | 0 | 0 | 0 |
| 'Ccdc91' | 8.793 | 8.609 | 8.448 | 7.934 | 8.624 | 9.243 | 8.050 | 9.692 | 8.676 | 9.228 | 8.856 | 8.723 | 8.411 | 7.963 | 8.742 | 9.357 | 7.783 | 8.772 | -0.002 | 0.9710 | 0.9898 | 0 | 0 | 0 |
| 'Ccdc92' | 13.196 | 11.442 | 12.573 | 11.633 | 11.383 | 14.602 | 11.907 | 11.593 | 12.364 | 11.870 | 11.341 | 14.959 | 13.574 | 13.040 | 15.076 | 11.881 | 10.996 | 15.378 | -0.102 | 0.0381 | 0.1587 | 0 | 0 | 0 |
| 'Ccdc93' | 5.410 | 4.842 | 5.044 | 5.157 | 4.890 | 4.744 | 5.212 | 4.969 | 4.892 | 5.184 | 5.493 | 6.169 | 5.058 | 5.188 | 5.673 | 5.182 | 5.264 | 6.040 | -0.137 | 0.0013 | 0.0132 | 1 | 0 | 0 |
| 'Ccdc94' | 20.921 | 20.646 | 23.371 | 17.964 | 20.192 | 26.187 | 20.108 | 21.467 | 24.386 | 22.809 | 17.535 | 20.996 | 22.062 | 20.179 | 20.638 | 23.024 | 19.564 | 18.052 | 0.072 | 0.2451 | 0.4957 | 0 | 0 | 0 |
| 'Ccdc97' | 13.398 | 13.682 | 14.680 | 12.685 | 13.714 | 15.226 | 12.799 | 13.860 | 15.198 | 14.422 | 12.990 | 15.376 | 14.632 | 14.458 | 15.060 | 14.372 | 12.775 | 15.182 | -0.056 | 0.0620 | 0.2156 | 0 | 0 | 0 |
| 'Cchcr1' | 5.254 | 4.559 | 6.704 | 5.491 | 4.360 | 6.169 | 5.747 | 4.549 | 6.934 | 5.469 | 4.805 | 5.766 | 6.592 | 4.951 | 5.744 | 5.356 | 5.103 | 5.601 | -0.007 | 0.8967 | 0.9586 | 0 | 0 | 0 |
| 'Ccl25' | 0.743 | 0.874 | 1.029 | 0.634 | 0.995 | 0.811 | 0.837 | 0.850 | 0.699 | 0.948 | 0.744 | 0.837 | 0.940 | 0.664 | 0.603 | 0.789 | 0.864 | 0.703 | 0.063 | 0.6000 | 0.7967 | 0 | 0 | 0 |
| 'Ccm2' | 37.446 | 37.198 | 38.751 | 40.195 | 35.761 | 38.103 | 39.235 | 37.204 | 38.861 | 35.654 | 37.648 | 42.414 | 34.455 | 34.506 | 40.892 | 35.559 | 36.357 | 40.988 | 0.009 | 0.7927 | 0.9074 | 0 | 0 | 0 |
| 'Ccna2' | 6.257 | 7.204 | 9.095 | 7.498 | 7.568 | 8.254 | 8.105 | 6.880 | 8.588 | 4.712 | 7.409 | 8.273 | 7.849 | 9.201 | 9.548 | 6.624 | 8.297 | 8.404 | -0.022 | 0.7835 | 0.9028 | 0 | 0 | 0 |
| 'Ccnb1' | 3.408 | 4.528 | 5.261 | 4.423 | 4.750 | 5.419 | 4.860 | 4.542 | 5.642 | 3.020 | 4.795 | 4.867 | 4.735 | 4.904 | 5.930 | 3.954 | 4.661 | 4.741 | 0.032 | 0.6659 | 0.8389 | 0 | 0 | 0 |
| 'Ccnb2' | 22.842 | 27.187 | 31.143 | 28.894 | 29.096 | 33.979 | 28.693 | 27.430 | 35.665 | 19.317 | 26.720 | 30.142 | 28.790 | 29.661 | 36.047 | 25.802 | 27.590 | 30.749 | 0.046 | 0.4806 | 0.7127 | 0 | 0 | 0 |
| 'Ccnc' | 3.036 | 3.589 | 3.312 | 3.170 | 3.591 | 3.104 | 3.330 | 3.411 | 3.054 | 2.965 | 3.255 | 3.330 | 2.873 | 3.650 | 3.018 | 2.889 | 3.609 | 3.030 | 0.033 | 0.4195 | 0.6664 | 0 | 0 | 0 |
| 'Ccnd2' | 46.572 | 50.167 | 47.307 | 48.948 | 49.566 | 49.265 | 46.214 | 46.946 | 48.653 | 41.761 | 42.364 | 45.846 | 40.508 | 41.510 | 46.406 | 41.413 | 40.584 | 49.102 | 0.146 | 0.0000 | 0.0001 | 1 | 0 | 0 |
| 'Ccnd3' | 9.886 | 9.385 | 9.918 | 9.963 | 9.138 | 10.018 | 10.174 | 9.654 | 10.021 | 9.563 | 9.894 | 11.790 | 9.625 | 9.499 | 10.872 | 11.197 | 10.500 | 11.249 | -0.106 | 0.0033 | 0.0270 | 1 | 0 | 0 |
| 'Ccndbp1' | 21.643 | 25.411 | 22.345 | 22.398 | 25.072 | 22.816 | 21.724 | 24.617 | 22.961 | 23.351 | 22.982 | 22.648 | 23.151 | 24.755 | 22.186 | 23.077 | 23.937 | 21.304 | -0.001 | 0.9772 | 0.9920 | 0 | 0 | 0 |
| 'Ccne1' | 14.056 | 15.622 | 15.182 | 15.508 | 16.133 | 13.963 | 15.910 | 16.180 | 14.999 | 12.161 | 13.081 | 13.440 | 18.008 | 14.614 | 14.620 | 13.104 | 13.681 | 13.511 | 0.111 | 0.0522 | 0.1938 | 0 | 0 | 0 |
| 'Ccne2' | 1.342 | 1.140 | 1.520 | 1.513 | 1.404 | 1.255 | 1.905 | 1.407 | 1.353 | 1.161 | 1.337 | 1.615 | 1.840 | 1.827 | 1.817 | 1.471 | 1.441 | 1.672 | -0.160 | 0.0927 | 0.2819 | 0 | 0 | 0 |
| 'Ccnf' | 6.605 | 10.278 | 11.367 | 8.526 | 8.982 | 10.874 | 7.764 | 9.116 | 12.424 | 5.824 | 8.943 | 11.243 | 8.928 | 10.671 | 12.660 | 8.702 | 8.321 | 11.261 | -0.023 | 0.7473 | 0.8861 | 0 | 0 | 0 |
| 'Ccng1' | 51.512 | 54.169 | 50.032 | 49.450 | 54.437 | 48.274 | 52.395 | 51.616 | 47.791 | 49.464 | 51.131 | 53.255 | 46.567 | 52.493 | 49.165 | 47.642 | 54.922 | 52.533 | -0.004 | 0.8975 | 0.9589 | 0 | 0 | 0 |
| 'Ccng2' | 8.814 | 9.953 | 7.755 | 10.052 | 8.411 | 7.991 | 9.436 | 8.880 | 8.491 | 6.241 | 10.981 | 9.190 | 8.397 | 10.113 | 8.925 | 7.272 | 10.933 | 8.936 | -0.020 | 0.7898 | 0.9061 | 0 | 0 | 0 |
| 'Ccnh' | 7.682 | 9.022 | 7.472 | 8.115 | 8.560 | 7.430 | 8.781 | 8.208 | 7.520 | 7.731 | 7.774 | 7.640 | 7.084 | 8.647 | 6.901 | 7.722 | 7.628 | 6.943 | 0.080 | 0.0486 | 0.1851 | 0 | 0 | 0 |
| 'Ccni' | 59.197 | 64.697 | 57.913 | 58.266 | 64.294 | 60.838 | 60.524 | 61.886 | 58.077 | 58.632 | 56.586 | 53.450 | 55.554 | 58.893 | 56.246 | 56.041 | 55.758 | 53.059 | 0.103 | 0.0000 | 0.0002 | 1 | 0 | 0 |
| 'Ccnj' | 0.940 | 1.172 | 1.015 | 0.963 | 1.408 | 0.909 | 0.839 | 1.281 | 0.956 | 1.111 | 0.985 | 1.033 | 0.962 | 0.972 | 0.989 | 1.013 | 0.882 | 1.040 | 0.051 | 0.5680 | 0.7756 | 0 | 0 | 0 |
| 'Ccnk' | 23.107 | 21.924 | 22.486 | 20.770 | 20.843 | 23.361 | 21.047 | 22.749 | 22.906 | 23.354 | 20.214 | 23.168 | 23.865 | 22.240 | 22.514 | 22.490 | 20.531 | 22.860 | -0.023 | 0.4636 | 0.6997 | 0 | 0 | 0 |
| 'Ccnl1' | 20.439 | 18.391 | 15.551 | 17.456 | 18.213 | 15.531 | 17.073 | 19.110 | 14.734 | 22.815 | 17.528 | 14.474 | 18.305 | 20.654 | 13.715 | 22.807 | 17.868 | 14.106 | -0.047 | 0.3822 | 0.6330 | 0 | 0 | 0 |
| 'Ccnl2' | 42.156 | 44.205 | 44.680 | 40.548 | 43.577 | 42.026 | 40.510 | 44.172 | 44.444 | 41.233 | 42.580 | 44.548 | 43.553 | 43.394 | 43.074 | 41.190 | 46.941 | 43.910 | -0.026 | 0.2314 | 0.4788 | 0 | 0 | 0 |
| 'Ccnt1' | 6.466 | 6.484 | 6.887 | 7.485 | 6.762 | 6.504 | 6.812 | 6.843 | 6.830 | 5.607 | 7.146 | 7.911 | 5.986 | 6.996 | 7.422 | 6.229 | 7.757 | 7.823 | -0.046 | 0.3940 | 0.6434 | 0 | 0 | 0 |
| 'Ccnt2' | 6.065 | 6.585 | 5.759 | 6.047 | 6.380 | 5.728 | 6.488 | 6.141 | 6.034 | 5.359 | 6.199 | 6.846 | 5.801 | 6.498 | 6.551 | 5.890 | 6.447 | 6.787 | -0.041 | 0.3372 | 0.5955 | 0 | 0 | 0 |
| 'Ccny' | 28.676 | 31.296 | 29.207 | 28.093 | 30.957 | 29.848 | 28.221 | 32.864 | 29.215 | 31.057 | 31.092 | 30.124 | 29.133 | 29.717 | 28.865 | 29.969 | 29.486 | 31.129 | -0.023 | 0.4180 | 0.6655 | 0 | 0 | 0 |
| 'Ccnyl1' | 2.841 | 3.404 | 2.723 | 2.461 | 3.124 | 2.992 | 2.646 | 3.136 | 3.087 | 4.001 | 3.575 | 3.198 | 4.642 | 4.183 | 3.448 | 4.778 | 3.436 | 3.079 | -0.377 | 0.0000 | 0.0000 | 1 | 0 | 0 |
| 'Ccp110' | 2.385 | 2.540 | 2.697 | 2.574 | 2.816 | 2.686 | 2.485 | 2.533 | 2.546 | 2.485 | 2.630 | 2.610 | 3.131 | 3.138 | 2.737 | 2.920 | 2.783 | 2.508 | -0.108 | 0.0259 | 0.1236 | 0 | 0 | 0 |
| 'Ccpg1' | 0.777 | 0.672 | 0.641 | 0.679 | 0.769 | 0.731 | 0.752 | 0.828 | 0.756 | 0.815 | 0.788 | 0.820 | 0.778 | 0.759 | 0.781 | 0.761 | 0.679 | 0.769 | -0.087 | 0.1856 | 0.4213 | 0 | 0 | 0 |
| 'Ccr4' | 22.704 | 19.924 | 24.950 | 25.642 | 18.966 | 23.587 | 27.773 | 18.029 | 24.704 | 19.110 | 20.474 | 29.109 | 18.092 | 18.240 | 26.638 | 19.923 | 20.103 | 28.260 | 0.042 | 0.5581 | 0.7697 | 0 | 0 | 0 |
| 'Ccr7' | 2.581 | 3.968 | 3.585 | 3.983 | 3.737 | 5.798 | 4.651 | 4.827 | 4.734 | 2.623 | 2.708 | 5.500 | 4.941 | 3.382 | 6.421 | 6.704 | 2.962 | 5.576 | -0.082 | 0.6547 | 0.8326 | 0 | 0 | 0 |
| 'Ccr9' | 0.498 | 0.365 | 0.502 | 0.512 | 0.400 | 0.521 | 0.483 | 0.364 | 0.474 | 0.243 | 0.155 | 0.301 | 0.240 | 0.207 | 0.347 | 0.256 | 0.192 | 0.319 | 0.853 | 0.0000 | 0.0000 | 1 | 1 | 0 |
| 'Ccrl2' | 4.562 | 1.382 | 0.678 | 3.119 | 1.731 | 0.930 | 3.231 | 1.911 | 0.843 | 2.656 | 2.576 | 0.672 | 2.622 | 2.991 | 0.711 | 3.005 | 2.807 | 0.471 | 0.007 | 0.9704 | 0.9898 | 0 | 0 | 0 |
| 'Ccs' | 39.500 | 42.456 | 43.383 | 39.827 | 40.248 | 44.414 | 36.612 | 42.590 | 42.428 | 42.752 | 39.999 | 42.774 | 38.101 | 36.291 | 43.803 | 44.574 | 41.080 | 42.505 | -0.009 | 0.8230 | 0.9223 | 0 | 0 | 0 |
| 'Ccsap' | 5.364 | 5.300 | 6.044 | 5.951 | 5.294 | 6.501 | 6.101 | 5.525 | 6.018 | 4.451 | 5.679 | 5.963 | 5.569 | 5.988 | 6.078 | 5.685 | 5.925 | 5.712 | 0.020 | 0.6968 | 0.8575 | 0 | 0 | 0 |
| 'Ccser2' | 5.379 | 5.344 | 4.579 | 5.214 | 5.171 | 4.593 | 4.494 | 5.370 | 4.501 | 5.055 | 5.205 | 5.061 | 4.632 | 5.217 | 4.554 | 5.242 | 5.002 | 4.932 | -0.021 | 0.5880 | 0.7887 | 0 | 0 | 0 |
| 'Cct2' | 184.722 | 172.825 | 170.589 | 186.166 | 173.426 | 165.703 | 184.664 | 176.966 | 160.995 | 159.783 | 166.451 | 169.744 | 178.065 | 165.812 | 166.242 | 164.309 | 176.021 | 169.295 | 0.044 | 0.1338 | 0.3522 | 0 | 0 | 0 |
| 'Cct3' | 18.354 | 17.224 | 18.094 | 19.064 | 18.951 | 17.579 | 18.798 | 18.205 | 17.727 | 18.286 | 17.549 | 20.114 | 18.234 | 17.649 | 19.041 | 17.599 | 18.826 | 20.347 | -0.043 | 0.1960 | 0.4350 | 0 | 0 | 0 |
| 'Cct4' | 28.990 | 26.929 | 26.458 | 30.113 | 26.728 | 26.344 | 27.266 | 27.706 | 25.498 | 27.630 | 28.420 | 29.606 | 27.236 | 28.241 | 28.758 | 26.724 | 29.579 | 30.334 | -0.072 | 0.0517 | 0.1926 | 0 | 0 | 0 |
| 'Cct5' | 53.928 | 50.683 | 49.597 | 55.426 | 51.086 | 51.887 | 55.706 | 49.293 | 52.248 | 50.807 | 52.682 | 55.033 | 53.190 | 50.246 | 55.026 | 50.828 | 52.536 | 58.315 | -0.039 | 0.2490 | 0.5000 | 0 | 0 | 0 |
| 'Cct7' | 28.530 | 27.940 | 26.511 | 29.451 | 27.576 | 28.524 | 28.483 | 27.831 | 26.525 | 25.260 | 25.864 | 29.725 | 26.538 | 26.895 | 28.760 | 26.435 | 27.968 | 31.004 | 0.007 | 0.8563 | 0.9385 | 0 | 0 | 0 |
| 'Cct8' | 36.744 | 35.476 | 36.157 | 39.660 | 38.060 | 35.489 | 37.018 | 36.080 | 34.711 | 33.792 | 38.200 | 37.050 | 36.427 | 36.353 | 34.805 | 35.152 | 38.221 | 36.160 | 0.003 | 0.9197 | 0.9698 | 0 | 0 | 0 |
| 'Ccz1' | 9.092 | 10.229 | 10.204 | 9.465 | 10.353 | 9.237 | 10.153 | 10.846 | 9.909 | 9.784 | 10.087 | 9.342 | 9.970 | 9.070 | 9.062 | 9.244 | 9.821 | 8.495 | 0.066 | 0.0653 | 0.2228 | 0 | 0 | 0 |
| 'Cd101' | 3.423 | 5.460 | 4.289 | 4.604 | 4.138 | 4.352 | 4.856 | 4.599 | 4.558 | 3.994 | 4.690 | 4.721 | 3.207 | 3.980 | 4.542 | 4.232 | 4.209 | 4.201 | 0.077 | 0.3048 | 0.5622 | 0 | 0 | 0 |
| 'Cd151' | 4.853 | 6.714 | 6.258 | 4.489 | 6.547 | 6.509 | 6.159 | 6.500 | 7.305 | 4.585 | 5.398 | 7.122 | 4.877 | 5.246 | 6.030 | 5.796 | 6.096 | 7.410 | 0.056 | 0.4468 | 0.6880 | 0 | 0 | 0 |
| 'Cd160' | 18.813 | 17.347 | 20.495 | 18.239 | 17.964 | 18.743 | 19.608 | 18.550 | 18.497 | 15.736 | 14.723 | 16.444 | 14.227 | 14.640 | 17.157 | 13.833 | 15.276 | 15.856 | 0.277 | 0.0000 | 0.0000 | 1 | 0 | 0 |
| 'Cd164' | 57.027 | 59.720 | 56.531 | 57.060 | 60.764 | 53.906 | 54.279 | 59.352 | 52.231 | 56.776 | 54.433 | 52.516 | 53.411 | 57.016 | 49.834 | 51.540 | 55.992 | 49.942 | 0.075 | 0.0013 | 0.0131 | 1 | 0 | 0 |
| 'Cd2' | 145.971 | 168.537 | 142.025 | 153.281 | 163.712 | 149.664 | 148.300 | 161.491 | 150.179 | 153.442 | 145.199 | 133.293 | 146.655 | 144.408 | 140.737 | 153.082 | 146.742 | 140.077 | 0.074 | 0.0073 | 0.0495 | 1 | 0 | 0 |
| 'Cd200' | 19.334 | 19.932 | 18.031 | 20.671 | 19.034 | 18.241 | 20.095 | 18.728 | 19.258 | 16.785 | 16.445 | 18.729 | 14.303 | 17.970 | 18.421 | 17.810 | 19.352 | 17.734 | 0.127 | 0.0087 | 0.0564 | 0 | 0 | 0 |
| 'Cd200r1' | 18.139 | 21.145 | 15.537 | 19.566 | 19.991 | 17.948 | 19.298 | 20.909 | 18.230 | 15.194 | 14.306 | 14.769 | 15.107 | 17.520 | 13.765 | 16.677 | 16.065 | 13.474 | 0.303 | 0.0000 | 0.0000 | 1 | 0 | 0 |
| 'Cd22' | 4.170 | 5.354 | 5.314 | 4.503 | 6.068 | 4.829 | 4.293 | 5.964 | 4.395 | 4.328 | 5.438 | 4.359 | 4.882 | 5.516 | 3.864 | 3.143 | 5.603 | 4.274 | 0.108 | 0.0781 | 0.2506 | 0 | 0 | 0 |
| 'Cd226' | 4.005 | 3.700 | 4.741 | 4.099 | 4.316 | 5.326 | 5.492 | 4.685 | 4.548 | 3.901 | 4.093 | 5.397 | 3.080 | 3.302 | 4.266 | 3.442 | 3.571 | 4.210 | 0.202 | 0.0183 | 0.0968 | 0 | 0 | 0 |
| 'Cd247' | 31.783 | 30.854 | 30.011 | 31.853 | 30.711 | 31.282 | 32.106 | 30.211 | 30.961 | 29.656 | 28.748 | 29.709 | 29.207 | 29.680 | 30.555 | 30.418 | 29.026 | 29.882 | 0.057 | 0.0038 | 0.0301 | 1 | 0 | 0 |
| 'Cd27' | 98.337 | 107.225 | 103.654 | 95.360 | 104.692 | 110.399 | 96.986 | 102.848 | 111.741 | 95.003 | 97.164 | 107.231 | 96.686 | 100.124 | 112.781 | 96.186 | 93.111 | 108.500 | 0.028 | 0.2609 | 0.5143 | 0 | 0 | 0 |
| 'Cd274' | 31.041 | 29.166 | 22.922 | 24.417 | 28.866 | 25.434 | 25.255 | 29.321 | 25.048 | 31.723 | 27.033 | 25.237 | 29.513 | 31.190 | 24.334 | 33.657 | 28.333 | 24.551 | -0.087 | 0.0738 | 0.2411 | 0 | 0 | 0 |
| 'Cd28' | 62.883 | 66.190 | 63.303 | 64.346 | 66.973 | 63.909 | 65.046 | 63.884 | 64.911 | 65.947 | 68.685 | 77.571 | 56.758 | 65.929 | 72.696 | 70.515 | 64.769 | 76.928 | -0.100 | 0.0176 | 0.0943 | 0 | 0 | 0 |
| 'Cd2ap' | 26.838 | 26.576 | 22.398 | 25.316 | 26.066 | 23.946 | 27.162 | 26.336 | 25.815 | 25.775 | 27.453 | 22.786 | 26.828 | 26.701 | 23.248 | 27.154 | 28.025 | 23.605 | -0.013 | 0.7247 | 0.8736 | 0 | 0 | 0 |
| 'Cd2bp2' | 12.625 | 12.718 | 13.755 | 12.729 | 13.548 | 13.480 | 13.040 | 13.402 | 14.476 | 12.197 | 12.568 | 12.921 | 13.237 | 12.227 | 12.923 | 12.664 | 12.600 | 12.058 | 0.069 | 0.0074 | 0.0499 | 1 | 0 | 0 |
| 'Cd320' | 2.030 | 1.674 | 1.938 | 1.550 | 2.001 | 1.844 | 1.865 | 1.725 | 1.752 | 1.695 | 1.830 | 2.126 | 1.791 | 1.467 | 1.775 | 1.970 | 1.411 | 1.758 | 0.034 | 0.6953 | 0.8567 | 0 | 0 | 0 |
| 'Cd37' | 39.529 | 41.323 | 45.668 | 37.072 | 41.362 | 45.208 | 39.415 | 41.812 | 46.657 | 44.074 | 41.678 | 42.620 | 42.713 | 41.744 | 43.564 | 36.808 | 40.287 | 41.674 | -0.002 | 0.9496 | 0.9809 | 0 | 0 | 0 |
| 'Cd38' | 7.580 | 11.857 | 8.199 | 9.375 | 10.667 | 8.744 | 8.928 | 11.522 | 8.336 | 8.205 | 10.361 | 9.793 | 7.339 | 11.693 | 8.554 | 7.668 | 10.425 | 9.194 | 0.017 | 0.7526 | 0.8897 | 0 | 0 | 0 |
| 'Cd3d' | 283.530 | 330.567 | 274.745 | 301.682 | 316.937 | 290.805 | 298.286 | 301.692 | 292.083 | 286.666 | 294.302 | 282.230 | 282.770 | 304.737 | 303.961 | 310.383 | 304.012 | 285.098 | 0.007 | 0.7930 | 0.9075 | 0 | 0 | 0 |
| 'Cd3e' | 346.476 | 372.538 | 363.929 | 347.983 | 385.122 | 369.675 | 357.167 | 371.406 | 361.234 | 324.140 | 334.128 | 319.769 | 319.349 | 327.700 | 329.015 | 332.261 | 328.777 | 330.553 | 0.142 | 0.0000 | 0.0000 | 1 | 0 | 0 |
| 'Cd3eap' | 8.738 | 8.267 | 9.197 | 7.484 | 8.467 | 10.046 | 8.035 | 8.351 | 8.670 | 9.580 | 7.798 | 9.744 | 9.085 | 6.582 | 8.505 | 9.970 | 6.720 | 8.202 | 0.016 | 0.8129 | 0.9157 | 0 | 0 | 0 |
| 'Cd3g' | 406.786 | 481.221 | 416.832 | 459.491 | 485.201 | 426.503 | 453.567 | 458.657 | 420.653 | 406.605 | 410.927 | 364.349 | 384.950 | 419.495 | 397.273 | 431.859 | 429.813 | 387.088 | 0.131 | 0.0000 | 0.0002 | 1 | 0 | 0 |
| 'Cd4' | 161.583 | 160.343 | 160.110 | 161.426 | 161.486 | 167.506 | 157.948 | 153.593 | 168.376 | 171.286 | 166.896 | 160.830 | 157.156 | 165.195 | 176.142 | 168.633 | 159.955 | 175.834 | -0.059 | 0.0302 | 0.1352 | 0 | 0 | 0 |
| 'Cd40lg' | 127.712 | 149.252 | 121.518 | 122.758 | 138.027 | 130.044 | 122.303 | 144.146 | 132.002 | 147.838 | 124.646 | 133.034 | 128.482 | 134.979 | 129.593 | 149.420 | 117.093 | 134.033 | -0.026 | 0.5941 | 0.7933 | 0 | 0 | 0 |
| 'Cd44' | 29.447 | 32.078 | 28.752 | 29.505 | 31.405 | 28.986 | 29.902 | 30.532 | 29.791 | 30.959 | 29.295 | 28.524 | 27.899 | 30.081 | 27.986 | 31.222 | 29.265 | 27.357 | 0.032 | 0.1822 | 0.4183 | 0 | 0 | 0 |
| 'Cd46' | 0.492 | 0.638 | 0.585 | 0.506 | 0.510 | 0.594 | 0.624 | 0.636 | 0.648 | 0.517 | 0.632 | 0.393 | 0.507 | 0.556 | 0.500 | 0.764 | 0.519 | 0.467 | 0.127 | 0.2650 | 0.5184 | 0 | 0 | 0 |
| 'Cd47' | 14.888 | 15.978 | 15.401 | 15.262 | 16.476 | 15.262 | 16.116 | 16.304 | 14.536 | 15.381 | 15.757 | 14.791 | 15.414 | 17.402 | 14.614 | 16.486 | 16.723 | 14.710 | -0.019 | 0.4584 | 0.6955 | 0 | 0 | 0 |
| 'Cd48' | 27.917 | 30.469 | 29.966 | 29.806 | 31.915 | 30.126 | 29.464 | 29.660 | 29.335 | 26.939 | 28.526 | 31.562 | 25.940 | 28.417 | 32.461 | 25.806 | 29.230 | 30.304 | 0.038 | 0.3104 | 0.5687 | 0 | 0 | 0 |
| 'Cd5' | 411.128 | 420.599 | 424.502 | 426.245 | 431.514 | 450.522 | 443.674 | 402.051 | 445.621 | 373.836 | 328.090 | 362.612 | 370.920 | 339.372 | 365.208 | 386.456 | 320.665 | 375.594 | 0.250 | 0.0000 | 0.0000 | 1 | 0 | 0 |
| 'Cd52' | 203.744 | 248.646 | 219.304 | 223.521 | 225.263 | 225.021 | 226.882 | 203.950 | 232.666 | 222.097 | 195.534 | 176.323 | 204.914 | 205.184 | 216.317 | 220.965 | 200.531 | 198.596 | 0.117 | 0.0101 | 0.0630 | 0 | 0 | 0 |
| 'Cd53' | 256.018 | 288.013 | 266.317 | 267.049 | 287.289 | 248.003 | 281.977 | 284.459 | 253.980 | 243.751 | 281.931 | 253.764 | 207.132 | 282.681 | 262.388 | 238.494 | 274.711 | 252.935 | 0.074 | 0.0599 | 0.2110 | 0 | 0 | 0 |
| 'Cd6' | 157.942 | 164.082 | 160.429 | 162.071 | 161.759 | 174.370 | 165.803 | 155.080 | 167.345 | 154.793 | 132.747 | 142.358 | 154.263 | 135.836 | 142.909 | 158.202 | 126.622 | 143.793 | 0.178 | 0.0000 | 0.0000 | 1 | 0 | 0 |
| 'Cd68' | 6.313 | 4.935 | 4.917 | 5.223 | 5.727 | 4.738 | 5.531 | 5.804 | 4.761 | 5.513 | 5.556 | 4.741 | 7.833 | 6.015 | 5.119 | 5.591 | 6.717 | 4.108 | -0.094 | 0.2179 | 0.4647 | 0 | 0 | 0 |
| 'Cd69' | 24.842 | 18.598 | 11.852 | 20.151 | 20.902 | 11.734 | 16.801 | 24.601 | 10.281 | 29.786 | 20.672 | 8.951 | 27.706 | 21.977 | 8.162 | 24.672 | 18.573 | 8.709 | 0.004 | 0.9706 | 0.9898 | 0 | 0 | 0 |
| 'Cd7' | 13.447 | 12.062 | 15.792 | 12.858 | 11.668 | 12.846 | 12.269 | 13.256 | 13.288 | 8.471 | 8.353 | 11.461 | 9.084 | 7.610 | 10.584 | 9.394 | 8.015 | 9.360 | 0.501 | 0.0000 | 0.0000 | 1 | 1 | 0 |
| 'Cd72' | 0.643 | 1.051 | 0.525 | 0.880 | 0.899 | 0.606 | 0.740 | 0.885 | 0.805 | 0.514 | 0.782 | 0.563 | 0.723 | 0.699 | 0.596 | 0.452 | 0.863 | 0.474 | 0.284 | 0.0098 | 0.0615 | 0 | 0 | 0 |
| 'Cd81' | 9.292 | 9.970 | 9.145 | 9.037 | 10.345 | 10.525 | 9.184 | 11.337 | 9.712 | 8.468 | 8.530 | 9.929 | 8.438 | 9.712 | 9.220 | 7.829 | 9.043 | 9.043 | 0.127 | 0.0037 | 0.0298 | 1 | 0 | 0 |
| 'Cd82' | 109.940 | 125.132 | 111.169 | 112.695 | 121.512 | 115.653 | 110.988 | 120.454 | 115.961 | 120.948 | 119.595 | 109.602 | 107.235 | 121.873 | 118.278 | 120.717 | 119.623 | 115.856 | -0.025 | 0.3559 | 0.6089 | 0 | 0 | 0 |
| 'Cd83' | 21.350 | 20.611 | 20.012 | 17.961 | 20.923 | 21.181 | 22.028 | 19.372 | 22.158 | 22.846 | 15.540 | 20.273 | 17.520 | 18.929 | 17.986 | 21.199 | 20.913 | 18.649 | 0.085 | 0.1510 | 0.3777 | 0 | 0 | 0 |
| 'Cd84' | 22.214 | 23.479 | 23.029 | 22.160 | 22.862 | 22.455 | 22.510 | 22.637 | 23.072 | 22.949 | 21.823 | 22.848 | 19.689 | 22.518 | 22.193 | 22.723 | 21.172 | 22.245 | 0.035 | 0.2458 | 0.4962 | 0 | 0 | 0 |
| 'Cd8b1' | 2.620 | 3.333 | 2.460 | 2.332 | 3.483 | 2.913 | 2.480 | 3.127 | 2.541 | 2.089 | 2.548 | 2.059 | 2.392 | 2.726 | 2.363 | 2.118 | 2.115 | 1.726 | 0.317 | 0.0035 | 0.0285 | 1 | 0 | 0 |
| 'Cd9' | 9.082 | 9.007 | 9.054 | 8.626 | 9.195 | 10.680 | 8.430 | 8.684 | 9.471 | 11.678 | 9.363 | 12.235 | 9.800 | 10.502 | 11.949 | 10.265 | 10.963 | 10.850 | -0.259 | 0.0000 | 0.0000 | 1 | 0 | 0 |
| 'Cd96' | 55.625 | 55.105 | 61.820 | 56.478 | 51.998 | 59.897 | 58.982 | 52.951 | 59.963 | 57.047 | 49.351 | 58.801 | 47.799 | 47.578 | 55.892 | 49.600 | 42.827 | 55.041 | 0.135 | 0.0002 | 0.0025 | 1 | 0 | 0 |
| 'Cd99l2' | 6.446 | 7.210 | 6.548 | 6.491 | 6.479 | 6.109 | 6.555 | 7.070 | 6.161 | 6.618 | 6.443 | 6.288 | 6.144 | 6.442 | 6.386 | 5.795 | 6.122 | 6.390 | 0.047 | 0.2359 | 0.4858 | 0 | 0 | 0 |
| 'Cdadc1' | 14.797 | 15.501 | 14.482 | 15.136 | 16.856 | 15.426 | 15.721 | 16.346 | 14.600 | 15.279 | 15.118 | 13.208 | 14.284 | 14.965 | 12.888 | 16.491 | 14.581 | 13.965 | 0.083 | 0.0554 | 0.2019 | 0 | 0 | 0 |
| 'Cdan1' | 3.427 | 2.872 | 3.572 | 3.002 | 2.730 | 3.144 | 3.026 | 2.865 | 3.548 | 3.001 | 3.483 | 3.971 | 3.309 | 3.135 | 3.817 | 3.075 | 3.327 | 3.954 | -0.149 | 0.0012 | 0.0122 | 1 | 0 | 0 |
| 'Cdc123' | 12.619 | 12.868 | 13.461 | 13.980 | 14.728 | 12.666 | 13.933 | 13.162 | 12.883 | 12.117 | 13.077 | 13.131 | 13.705 | 13.111 | 13.436 | 12.580 | 13.874 | 12.813 | 0.017 | 0.6411 | 0.8239 | 0 | 0 | 0 |
| 'Cdc14a' | 3.484 | 3.283 | 3.127 | 3.755 | 3.274 | 3.325 | 3.475 | 3.411 | 3.159 | 3.623 | 3.665 | 3.901 | 3.473 | 3.767 | 3.680 | 3.168 | 3.972 | 3.943 | -0.145 | 0.0022 | 0.0194 | 1 | 0 | 0 |
| 'Cdc14b' | 0.512 | 0.663 | 0.539 | 0.538 | 0.669 | 0.697 | 0.580 | 0.734 | 0.641 | 0.560 | 0.516 | 0.754 | 0.468 | 0.574 | 0.616 | 0.608 | 0.571 | 0.586 | 0.057 | 0.5789 | 0.7820 | 0 | 0 | 0 |
| 'Cdc16' | 9.133 | 10.739 | 10.670 | 9.309 | 10.526 | 9.791 | 9.823 | 10.622 | 10.652 | 9.754 | 9.155 | 9.459 | 10.008 | 9.670 | 9.417 | 10.603 | 9.185 | 8.505 | 0.081 | 0.0538 | 0.1983 | 0 | 0 | 0 |
| 'Cdc20' | 18.163 | 20.841 | 24.248 | 19.413 | 20.105 | 25.402 | 25.123 | 19.109 | 25.910 | 14.685 | 18.613 | 22.007 | 21.100 | 19.934 | 25.892 | 18.973 | 19.585 | 20.944 | 0.114 | 0.0821 | 0.2590 | 0 | 0 | 0 |
| 'Cdc23' | 9.868 | 10.868 | 11.282 | 11.229 | 11.841 | 11.581 | 11.081 | 11.469 | 11.685 | 10.185 | 11.714 | 11.284 | 10.536 | 11.224 | 10.701 | 10.201 | 11.828 | 10.528 | 0.029 | 0.3230 | 0.5806 | 0 | 0 | 0 |
| 'Cdc25a' | 5.485 | 4.867 | 5.516 | 5.608 | 4.975 | 5.314 | 5.350 | 4.802 | 5.234 | 4.411 | 5.039 | 5.775 | 5.387 | 5.214 | 5.617 | 5.018 | 4.925 | 6.148 | -0.023 | 0.6223 | 0.8114 | 0 | 0 | 0 |
| 'Cdc25b' | 33.583 | 46.033 | 32.763 | 35.890 | 44.475 | 40.466 | 34.458 | 40.973 | 38.395 | 34.288 | 40.589 | 41.622 | 47.440 | 47.155 | 42.116 | 42.601 | 43.600 | 42.475 | -0.155 | 0.0049 | 0.0367 | 1 | 0 | 0 |
| 'Cdc25c' | 2.988 | 3.719 | 4.725 | 2.920 | 3.327 | 3.828 | 4.066 | 3.306 | 4.660 | 2.116 | 3.691 | 3.263 | 4.301 | 3.900 | 4.232 | 2.842 | 4.381 | 3.122 | 0.064 | 0.5934 | 0.7931 | 0 | 0 | 0 |
| 'Cdc26' | 4.794 | 5.825 | 5.295 | 4.958 | 5.830 | 5.093 | 5.464 | 4.979 | 5.324 | 4.726 | 5.072 | 4.596 | 5.055 | 4.837 | 4.724 | 4.989 | 5.498 | 4.729 | 0.094 | 0.0306 | 0.1366 | 0 | 0 | 0 |
| 'Cdc27' | 8.596 | 8.299 | 8.888 | 9.356 | 9.165 | 8.091 | 9.846 | 8.491 | 8.621 | 8.553 | 9.028 | 8.835 | 8.453 | 9.134 | 8.579 | 8.198 | 9.377 | 8.312 | 0.004 | 0.9147 | 0.9671 | 0 | 0 | 0 |
| 'Cdc34' | 50.950 | 46.343 | 57.189 | 47.417 | 46.036 | 60.540 | 49.620 | 51.025 | 51.578 | 52.129 | 46.354 | 50.721 | 56.086 | 49.861 | 56.924 | 57.964 | 42.161 | 52.917 | -0.021 | 0.6614 | 0.8364 | 0 | 0 | 0 |
| 'Cdc37' | 110.262 | 106.821 | 112.683 | 108.926 | 106.733 | 119.357 | 112.793 | 103.197 | 109.329 | 109.629 | 100.497 | 113.371 | 108.498 | 103.108 | 116.548 | 113.137 | 103.809 | 110.908 | 0.005 | 0.7937 | 0.9079 | 0 | 0 | 0 |
| 'Cdc37l1' | 7.211 | 8.142 | 8.770 | 7.578 | 8.197 | 7.720 | 8.076 | 7.769 | 7.928 | 7.125 | 8.573 | 8.811 | 7.211 | 8.792 | 8.154 | 7.229 | 8.554 | 7.999 | -0.031 | 0.4103 | 0.6592 | 0 | 0 | 0 |
| 'Cdc40' | 27.639 | 29.342 | 27.272 | 30.612 | 29.605 | 27.508 | 31.555 | 29.981 | 26.294 | 26.429 | 28.040 | 28.406 | 25.751 | 28.074 | 28.047 | 25.529 | 30.235 | 26.191 | 0.061 | 0.1019 | 0.2993 | 0 | 0 | 0 |
| 'Cdc42' | 158.934 | 174.206 | 160.001 | 172.796 | 182.615 | 161.940 | 175.584 | 172.522 | 158.013 | 160.819 | 169.580 | 150.621 | 159.613 | 172.173 | 156.036 | 156.593 | 175.249 | 154.565 | 0.049 | 0.0324 | 0.1421 | 0 | 0 | 0 |
| 'Cdc42bpg' | 4.672 | 4.691 | 5.603 | 4.389 | 4.597 | 5.052 | 4.655 | 4.908 | 5.609 | 5.650 | 4.745 | 4.410 | 5.167 | 4.079 | 4.136 | 4.216 | 4.145 | 3.903 | 0.121 | 0.0765 | 0.2465 | 0 | 0 | 0 |
| 'Cdc42ep3' | 8.166 | 8.336 | 7.894 | 7.969 | 8.571 | 8.551 | 7.558 | 8.826 | 7.217 | 7.293 | 8.285 | 9.069 | 7.047 | 9.863 | 9.053 | 7.244 | 8.469 | 9.083 | -0.057 | 0.3466 | 0.6017 | 0 | 0 | 0 |
| 'Cdc42se1' | 49.346 | 52.936 | 52.350 | 49.168 | 54.510 | 54.571 | 52.463 | 54.001 | 51.157 | 45.128 | 48.090 | 54.846 | 46.296 | 50.813 | 56.783 | 50.270 | 50.626 | 55.940 | 0.027 | 0.4010 | 0.6496 | 0 | 0 | 0 |
| 'Cdc42se2' | 59.419 | 70.312 | 58.769 | 58.059 | 68.546 | 62.896 | 58.461 | 66.615 | 59.516 | 62.879 | 59.456 | 58.939 | 63.566 | 66.037 | 60.722 | 64.219 | 58.532 | 58.125 | 0.013 | 0.7022 | 0.8611 | 0 | 0 | 0 |
| 'Cdc45' | 9.284 | 9.897 | 11.451 | 11.000 | 9.739 | 11.848 | 12.479 | 9.204 | 12.101 | 8.011 | 10.958 | 10.552 | 12.497 | 10.505 | 11.678 | 10.033 | 11.134 | 10.423 | 0.004 | 0.9584 | 0.9847 | 0 | 0 | 0 |
| 'Cdc5l' | 34.178 | 33.642 | 35.706 | 39.418 | 34.353 | 34.815 | 38.910 | 33.480 | 34.532 | 34.574 | 37.363 | 37.116 | 34.589 | 35.051 | 34.534 | 33.055 | 36.491 | 35.112 | -0.007 | 0.8471 | 0.9343 | 0 | 0 | 0 |
| 'Cdc6' | 3.168 | 2.320 | 3.289 | 3.323 | 2.966 | 2.946 | 3.467 | 2.240 | 3.024 | 2.529 | 2.831 | 2.958 | 3.619 | 2.510 | 3.328 | 2.693 | 2.973 | 3.199 | -0.013 | 0.8630 | 0.9405 | 0 | 0 | 0 |
| 'Cdc7' | 5.610 | 5.471 | 6.376 | 5.785 | 5.944 | 5.736 | 6.031 | 5.726 | 6.451 | 5.618 | 6.263 | 6.014 | 6.115 | 6.242 | 6.084 | 5.081 | 6.183 | 6.200 | -0.028 | 0.4935 | 0.7228 | 0 | 0 | 0 |
| 'Cdc73' | 4.759 | 4.580 | 4.816 | 5.240 | 4.594 | 4.205 | 4.884 | 4.630 | 4.291 | 4.575 | 4.512 | 4.491 | 4.378 | 4.473 | 4.037 | 4.412 | 4.888 | 4.168 | 0.061 | 0.1001 | 0.2959 | 0 | 0 | 0 |
| 'Cdca2' | 2.584 | 2.895 | 3.740 | 3.224 | 3.097 | 3.375 | 3.015 | 2.895 | 3.620 | 2.156 | 2.964 | 3.252 | 3.006 | 3.446 | 3.616 | 2.444 | 3.142 | 3.389 | 0.045 | 0.4805 | 0.7127 | 0 | 0 | 0 |
| 'Cdca3' | 3.612 | 4.887 | 5.223 | 3.794 | 4.056 | 5.188 | 4.420 | 4.133 | 5.140 | 3.214 | 4.104 | 4.569 | 5.135 | 4.746 | 5.890 | 4.199 | 4.515 | 4.355 | -0.025 | 0.7473 | 0.8861 | 0 | 0 | 0 |
| 'Cdca4' | 34.908 | 39.244 | 38.915 | 36.631 | 38.081 | 40.825 | 40.075 | 36.604 | 38.876 | 33.436 | 36.483 | 41.431 | 32.219 | 39.373 | 39.191 | 34.644 | 36.421 | 40.030 | 0.036 | 0.3186 | 0.5771 | 0 | 0 | 0 |
| 'Cdca5' | 4.763 | 4.772 | 5.987 | 5.605 | 5.054 | 5.540 | 5.498 | 5.480 | 5.633 | 4.132 | 4.666 | 5.568 | 6.639 | 6.399 | 5.943 | 5.297 | 5.345 | 5.133 | -0.034 | 0.6445 | 0.8259 | 0 | 0 | 0 |
| 'Cdca7' | 13.630 | 11.185 | 15.449 | 15.644 | 13.919 | 14.674 | 14.744 | 12.260 | 14.137 | 13.379 | 10.530 | 15.264 | 14.825 | 11.992 | 15.672 | 12.431 | 12.150 | 14.290 | 0.050 | 0.2958 | 0.5518 | 0 | 0 | 0 |
| 'Cdca7l' | 14.462 | 11.934 | 15.099 | 14.830 | 13.786 | 15.164 | 14.149 | 14.469 | 15.343 | 14.585 | 13.612 | 14.431 | 15.122 | 12.462 | 14.448 | 13.937 | 13.184 | 13.438 | 0.039 | 0.2987 | 0.5550 | 0 | 0 | 0 |
| 'Cdca8' | 6.148 | 7.488 | 9.308 | 7.720 | 7.677 | 9.437 | 8.567 | 7.464 | 9.670 | 5.440 | 7.748 | 8.263 | 9.071 | 8.516 | 9.548 | 6.567 | 8.628 | 8.249 | 0.015 | 0.8470 | 0.9343 | 0 | 0 | 0 |
| 'Cdh11' | 12.965 | 12.797 | 12.762 | 13.712 | 13.349 | 11.961 | 14.565 | 14.467 | 12.438 | 12.468 | 11.753 | 11.134 | 13.122 | 11.367 | 10.514 | 12.586 | 13.280 | 11.301 | 0.137 | 0.0002 | 0.0031 | 1 | 0 | 0 |
| 'Cdh23' | 3.103 | 3.029 | 3.003 | 2.642 | 2.999 | 2.551 | 2.873 | 2.947 | 2.872 | 3.297 | 3.922 | 3.423 | 3.728 | 3.274 | 3.157 | 3.354 | 3.552 | 3.143 | -0.255 | 0.0000 | 0.0000 | 1 | 0 | 0 |
| 'Cdh24' | 12.180 | 10.271 | 14.058 | 13.143 | 9.643 | 13.252 | 11.737 | 10.639 | 13.831 | 12.369 | 9.657 | 13.461 | 10.682 | 8.920 | 12.047 | 9.664 | 9.435 | 12.002 | 0.134 | 0.0035 | 0.0286 | 1 | 0 | 0 |
| 'Cdip1' | 3.678 | 4.523 | 3.898 | 3.590 | 3.770 | 3.948 | 3.764 | 4.426 | 4.447 | 4.018 | 3.861 | 3.686 | 4.032 | 3.930 | 3.620 | 3.955 | 3.937 | 3.161 | 0.068 | 0.1807 | 0.4162 | 0 | 0 | 0 |
| 'Cdipt' | 22.554 | 24.862 | 22.097 | 21.469 | 25.877 | 24.919 | 20.588 | 24.566 | 23.845 | 23.316 | 21.585 | 25.365 | 23.866 | 23.366 | 24.474 | 25.143 | 21.864 | 25.248 | -0.036 | 0.3937 | 0.6431 | 0 | 0 | 0 |
| 'Cdk1' | 4.167 | 4.933 | 6.404 | 5.395 | 5.323 | 6.334 | 5.590 | 5.298 | 6.015 | 3.448 | 5.021 | 5.571 | 5.863 | 6.337 | 6.223 | 5.519 | 5.510 | 5.425 | 0.003 | 0.9704 | 0.9898 | 0 | 0 | 0 |
| 'Cdk11b' | 29.793 | 26.317 | 31.013 | 28.891 | 27.448 | 30.772 | 29.688 | 27.494 | 31.238 | 28.417 | 27.455 | 30.321 | 29.605 | 26.639 | 28.579 | 28.789 | 30.138 | 29.166 | 0.009 | 0.7519 | 0.8895 | 0 | 0 | 0 |
| 'Cdk12' | 7.341 | 6.735 | 7.578 | 7.830 | 6.731 | 6.779 | 8.065 | 7.077 | 7.539 | 7.115 | 7.563 | 8.132 | 6.955 | 7.470 | 7.324 | 6.987 | 8.004 | 7.767 | -0.047 | 0.2801 | 0.5371 | 0 | 0 | 0 |
| 'Cdk13' | 28.783 | 25.378 | 29.581 | 27.111 | 25.772 | 29.149 | 26.488 | 27.441 | 28.076 | 30.213 | 27.368 | 30.215 | 27.425 | 28.126 | 29.881 | 28.563 | 27.502 | 32.157 | -0.087 | 0.0028 | 0.0235 | 1 | 0 | 0 |
| 'Cdk16' | 7.823 | 7.943 | 8.396 | 6.784 | 7.780 | 8.594 | 7.768 | 7.715 | 8.926 | 7.454 | 7.228 | 9.525 | 7.376 | 7.604 | 9.491 | 7.671 | 7.983 | 9.683 | -0.054 | 0.1428 | 0.3662 | 0 | 0 | 0 |
| 'Cdk17' | 61.517 | 59.651 | 55.989 | 59.737 | 58.523 | 53.736 | 59.535 | 62.045 | 55.095 | 58.049 | 66.386 | 64.179 | 56.010 | 67.915 | 62.642 | 63.078 | 66.213 | 63.585 | -0.122 | 0.0008 | 0.0085 | 1 | 0 | 0 |
| 'Cdk19' | 16.736 | 18.347 | 17.785 | 17.933 | 18.321 | 17.875 | 16.535 | 18.002 | 18.031 | 16.932 | 18.385 | 16.128 | 17.855 | 19.248 | 16.391 | 17.137 | 18.484 | 15.312 | 0.028 | 0.3920 | 0.6414 | 0 | 0 | 0 |
| 'Cdk2' | 28.851 | 29.969 | 32.825 | 30.715 | 29.262 | 32.212 | 29.934 | 28.287 | 34.582 | 26.916 | 29.718 | 32.534 | 32.065 | 30.609 | 33.280 | 26.933 | 30.607 | 31.921 | -0.001 | 0.9689 | 0.9897 | 0 | 0 | 0 |
| 'Cdk20' | 2.064 | 3.050 | 2.996 | 2.297 | 2.868 | 2.524 | 2.980 | 2.510 | 3.233 | 2.065 | 2.458 | 2.321 | 2.809 | 2.616 | 2.454 | 1.972 | 2.176 | 2.461 | 0.183 | 0.0801 | 0.2549 | 0 | 0 | 0 |
| 'Cdk2ap2' | 86.460 | 99.903 | 93.252 | 85.972 | 105.330 | 98.474 | 86.118 | 101.009 | 92.314 | 84.647 | 85.174 | 81.552 | 88.400 | 91.527 | 90.867 | 85.612 | 84.553 | 84.894 | 0.114 | 0.0005 | 0.0057 | 1 | 0 | 0 |
| 'Cdk4' | 15.180 | 17.313 | 16.935 | 15.403 | 17.897 | 17.205 | 16.440 | 17.565 | 17.643 | 14.694 | 14.819 | 17.423 | 17.678 | 16.082 | 18.564 | 16.987 | 15.883 | 17.382 | 0.008 | 0.8368 | 0.9300 | 0 | 0 | 0 |
| 'Cdk5' | 20.376 | 22.786 | 23.584 | 21.639 | 22.437 | 24.900 | 21.096 | 24.530 | 24.323 | 22.324 | 21.267 | 23.974 | 25.112 | 21.255 | 24.557 | 20.785 | 24.047 | 23.082 | -0.018 | 0.6612 | 0.8362 | 0 | 0 | 0 |
| 'Cdk5r1' | 18.327 | 20.550 | 16.760 | 19.627 | 18.005 | 17.706 | 19.518 | 18.488 | 16.400 | 21.020 | 26.737 | 25.673 | 23.785 | 26.003 | 23.839 | 23.706 | 25.901 | 25.965 | -0.442 | 0.0000 | 0.0000 | 1 | 1 | 0 |
| 'Cdk5rap1' | 2.319 | 2.721 | 2.975 | 2.797 | 2.869 | 2.476 | 2.848 | 2.827 | 2.908 | 2.962 | 2.508 | 2.407 | 2.681 | 2.683 | 2.240 | 2.555 | 2.521 | 2.211 | 0.115 | 0.0674 | 0.2275 | 0 | 0 | 0 |
| 'Cdk5rap2' | 4.406 | 3.898 | 4.486 | 4.820 | 4.104 | 4.583 | 4.767 | 3.869 | 4.697 | 4.603 | 4.465 | 4.546 | 4.332 | 4.567 | 4.535 | 4.408 | 4.244 | 4.523 | -0.034 | 0.3682 | 0.6201 | 0 | 0 | 0 |
| 'Cdk5rap3' | 9.689 | 10.150 | 11.373 | 10.630 | 10.898 | 11.990 | 10.548 | 11.036 | 11.716 | 12.136 | 10.330 | 10.880 | 12.356 | 10.335 | 10.840 | 11.065 | 10.652 | 10.297 | -0.022 | 0.6080 | 0.8028 | 0 | 0 | 0 |
| 'Cdk6' | 6.350 | 4.750 | 5.112 | 6.342 | 5.074 | 5.257 | 5.070 | 5.299 | 5.556 | 5.862 | 6.591 | 6.035 | 6.302 | 5.765 | 5.888 | 6.077 | 5.798 | 6.592 | -0.181 | 0.0011 | 0.0112 | 1 | 0 | 0 |
| 'Cdk7' | 5.482 | 6.250 | 6.038 | 6.463 | 6.574 | 5.992 | 6.377 | 6.396 | 5.797 | 5.968 | 6.728 | 6.259 | 5.698 | 6.241 | 5.846 | 6.092 | 5.983 | 5.524 | 0.014 | 0.7135 | 0.8672 | 0 | 0 | 0 |
| 'Cdkal1' | 1.467 | 1.193 | 1.560 | 1.515 | 1.576 | 1.426 | 1.664 | 1.432 | 1.468 | 1.413 | 1.542 | 1.467 | 1.340 | 1.444 | 1.331 | 1.399 | 1.929 | 1.373 | 0.001 | 0.9911 | 0.9976 | 0 | 0 | 0 |
| 'Cdkl3' | 0.607 | 0.645 | 0.653 | 0.576 | 0.681 | 0.547 | 0.667 | 0.654 | 0.652 | 0.567 | 0.586 | 0.536 | 0.566 | 0.539 | 0.450 | 0.607 | 0.505 | 0.486 | 0.223 | 0.0093 | 0.0590 | 0 | 0 | 0 |
| 'Cdkn1a' | 6.111 | 5.657 | 5.627 | 6.723 | 6.296 | 5.394 | 6.526 | 6.456 | 4.672 | 5.757 | 8.297 | 5.721 | 7.175 | 7.936 | 6.353 | 7.623 | 7.163 | 5.131 | -0.195 | 0.0031 | 0.0260 | 1 | 0 | 0 |
| 'Cdkn1b' | 35.038 | 38.133 | 29.908 | 32.493 | 36.406 | 31.326 | 32.233 | 35.285 | 29.558 | 31.078 | 39.485 | 32.367 | 31.372 | 41.984 | 32.287 | 33.682 | 38.373 | 32.001 | -0.065 | 0.0642 | 0.2209 | 0 | 0 | 0 |
| 'Cdkn2aip' | 14.837 | 15.336 | 14.347 | 14.555 | 14.373 | 13.601 | 15.923 | 15.785 | 13.069 | 14.051 | 16.044 | 13.987 | 15.353 | 16.048 | 13.777 | 16.467 | 15.141 | 13.666 | -0.037 | 0.3070 | 0.5653 | 0 | 0 | 0 |
| 'Cdkn2aipnl' | 40.561 | 48.013 | 42.042 | 41.314 | 45.909 | 43.885 | 40.814 | 45.398 | 42.000 | 38.596 | 41.358 | 40.793 | 41.486 | 41.788 | 41.855 | 42.571 | 38.279 | 40.710 | 0.073 | 0.0202 | 0.1038 | 0 | 0 | 0 |
| 'Cdkn2c' | 23.552 | 27.039 | 28.292 | 27.134 | 25.232 | 30.237 | 27.792 | 26.490 | 29.018 | 19.663 | 28.470 | 28.628 | 27.315 | 30.496 | 31.453 | 23.831 | 28.209 | 29.935 | -0.025 | 0.6576 | 0.8349 | 0 | 0 | 0 |
| 'Cdkn2d' | 33.762 | 34.258 | 34.644 | 37.175 | 36.688 | 39.338 | 35.814 | 39.449 | 34.939 | 26.493 | 32.769 | 32.476 | 32.545 | 36.136 | 35.705 | 31.223 | 33.012 | 27.786 | 0.170 | 0.0010 | 0.0106 | 1 | 0 | 0 |
| 'Cdkn3' | 5.841 | 6.450 | 6.748 | 6.505 | 6.562 | 7.341 | 6.143 | 5.098 | 6.520 | 4.568 | 6.069 | 5.731 | 5.832 | 5.891 | 6.848 | 5.433 | 6.590 | 4.188 | 0.155 | 0.1212 | 0.3319 | 0 | 0 | 0 |
| 'Cdon' | 4.209 | 4.341 | 4.086 | 4.131 | 4.061 | 4.259 | 4.136 | 4.206 | 4.128 | 4.248 | 4.507 | 4.639 | 4.270 | 4.072 | 4.275 | 4.319 | 4.363 | 4.480 | -0.074 | 0.0230 | 0.1135 | 0 | 0 | 0 |
| 'Cdpf1' | 4.295 | 5.596 | 4.500 | 5.225 | 5.044 | 5.013 | 4.342 | 4.711 | 4.652 | 4.482 | 5.078 | 5.132 | 4.544 | 4.941 | 5.352 | 5.125 | 4.425 | 5.082 | -0.043 | 0.4857 | 0.7168 | 0 | 0 | 0 |
| 'Cdr2' | 2.473 | 2.052 | 2.159 | 2.265 | 1.992 | 1.924 | 2.389 | 2.072 | 1.971 | 1.842 | 2.487 | 3.492 | 2.020 | 1.863 | 2.947 | 1.695 | 2.398 | 3.889 | -0.210 | 0.0883 | 0.2714 | 0 | 0 | 0 |
| 'Cds2' | 28.271 | 27.660 | 28.152 | 25.838 | 27.357 | 27.026 | 25.667 | 28.024 | 28.713 | 27.094 | 30.063 | 32.517 | 26.605 | 28.817 | 31.825 | 27.590 | 27.824 | 34.154 | -0.119 | 0.0010 | 0.0101 | 1 | 0 | 0 |
| 'Cdt1' | 32.133 | 28.529 | 33.969 | 30.629 | 27.726 | 32.393 | 28.951 | 29.531 | 34.289 | 32.027 | 29.636 | 31.154 | 37.619 | 30.046 | 31.543 | 31.253 | 26.332 | 32.434 | -0.028 | 0.5249 | 0.7470 | 0 | 0 | 0 |
| 'Cdv3' | 16.268 | 13.390 | 14.371 | 14.458 | 13.695 | 13.563 | 13.778 | 13.721 | 12.796 | 13.738 | 14.499 | 15.127 | 13.268 | 15.029 | 15.599 | 14.291 | 14.651 | 15.664 | -0.075 | 0.1054 | 0.3042 | 0 | 0 | 0 |
| 'Cdyl' | 3.349 | 3.749 | 3.317 | 3.630 | 3.418 | 3.416 | 3.737 | 3.720 | 3.460 | 3.729 | 3.319 | 3.709 | 3.612 | 4.296 | 3.842 | 3.542 | 3.732 | 3.840 | -0.095 | 0.0491 | 0.1868 | 0 | 0 | 0 |
| 'Cdyl2' | 7.580 | 8.738 | 7.729 | 7.831 | 8.043 | 8.130 | 8.706 | 7.422 | 8.686 | 6.540 | 7.401 | 7.569 | 6.497 | 6.653 | 7.602 | 7.069 | 7.106 | 7.438 | 0.177 | 0.0000 | 0.0005 | 1 | 0 | 0 |
| 'Cebpa' | 117.855 | 123.926 | 112.665 | 109.355 | 122.803 | 117.435 | 115.099 | 115.846 | 116.687 | 132.903 | 118.467 | 108.541 | 141.227 | 116.355 | 117.014 | 128.912 | 113.840 | 112.658 | -0.058 | 0.1296 | 0.3451 | 0 | 0 | 0 |
| 'Cebpb' | 109.228 | 37.819 | 59.766 | 89.059 | 46.078 | 55.267 | 74.440 | 58.001 | 31.781 | 84.182 | 67.319 | 39.506 | 82.510 | 68.160 | 47.781 | 71.364 | 70.346 | 57.765 | -0.117 | 0.4292 | 0.6749 | 0 | 0 | 0 |
| 'Cebpg' | 9.568 | 9.480 | 9.015 | 10.504 | 9.101 | 8.365 | 11.168 | 8.451 | 8.734 | 6.982 | 11.103 | 11.243 | 8.926 | 10.132 | 10.728 | 7.540 | 11.702 | 10.353 | -0.073 | 0.4772 | 0.7108 | 0 | 0 | 0 |
| 'Cebpz' | 20.681 | 19.623 | 19.672 | 19.040 | 22.733 | 19.922 | 19.769 | 19.957 | 19.133 | 20.895 | 19.654 | 20.307 | 17.092 | 20.401 | 18.034 | 20.196 | 18.425 | 19.825 | 0.035 | 0.4137 | 0.6620 | 0 | 0 | 0 |
| 'Cebpzos' | 8.115 | 11.171 | 8.656 | 9.456 | 10.391 | 8.768 | 9.218 | 9.883 | 9.149 | 9.920 | 7.989 | 9.146 | 9.244 | 9.682 | 8.321 | 8.714 | 9.468 | 7.525 | 0.061 | 0.3653 | 0.6175 | 0 | 0 | 0 |
| 'Cecr5' | 11.204 | 9.486 | 12.088 | 10.426 | 10.375 | 11.489 | 10.525 | 10.595 | 12.180 | 10.275 | 9.846 | 11.229 | 10.384 | 10.462 | 10.781 | 9.793 | 9.838 | 10.329 | 0.074 | 0.0289 | 0.1317 | 0 | 0 | 0 |
| 'Cela1' | 12.302 | 12.376 | 12.236 | 12.411 | 12.618 | 12.481 | 14.589 | 14.126 | 12.073 | 15.509 | 11.920 | 11.574 | 14.842 | 12.059 | 11.874 | 12.016 | 11.374 | 11.888 | 0.020 | 0.7504 | 0.8887 | 0 | 0 | 0 |
| 'Celf1' | 10.250 | 10.497 | 11.084 | 10.762 | 10.419 | 10.267 | 10.904 | 9.905 | 10.735 | 9.304 | 10.731 | 12.598 | 10.076 | 10.149 | 12.041 | 9.419 | 11.068 | 12.526 | -0.051 | 0.2536 | 0.5051 | 0 | 0 | 0 |
| 'Celf2' | 6.519 | 6.809 | 7.125 | 6.874 | 6.847 | 6.703 | 7.297 | 6.750 | 6.890 | 6.987 | 7.367 | 6.936 | 6.854 | 7.188 | 6.624 | 6.522 | 7.004 | 6.474 | -0.013 | 0.6321 | 0.8176 | 0 | 0 | 0 |
| 'Celf3' | 0.352 | 0.440 | 0.454 | 0.343 | 0.518 | 0.494 | 0.373 | 0.644 | 0.590 | 0.691 | 0.521 | 0.588 | 0.626 | 0.700 | 0.582 | 0.647 | 0.654 | 0.561 | -0.417 | 0.0001 | 0.0009 | 1 | 1 | 0 |
| 'Celf5' | 0.338 | 0.463 | 0.440 | 0.268 | 0.413 | 0.365 | 0.396 | 0.385 | 0.453 | 0.378 | 0.361 | 0.322 | 0.418 | 0.359 | 0.398 | 0.363 | 0.340 | 0.297 | 0.118 | 0.2630 | 0.5169 | 0 | 0 | 0 |
| 'Celsr1' | 20.686 | 19.669 | 22.671 | 17.968 | 19.674 | 22.003 | 16.816 | 20.355 | 21.610 | 20.353 | 18.551 | 18.134 | 17.531 | 17.485 | 18.168 | 17.915 | 15.633 | 20.098 | 0.134 | 0.0069 | 0.0479 | 1 | 0 | 0 |
| 'Celsr2' | 0.603 | 0.643 | 0.647 | 0.632 | 0.641 | 0.723 | 0.499 | 0.633 | 0.738 | 0.669 | 0.492 | 0.591 | 0.634 | 0.424 | 0.586 | 0.458 | 0.426 | 0.607 | 0.226 | 0.0084 | 0.0551 | 0 | 0 | 0 |
| 'Cenpa' | 6.680 | 7.788 | 8.716 | 6.690 | 7.828 | 8.551 | 7.317 | 8.487 | 8.786 | 5.907 | 7.125 | 8.219 | 7.693 | 8.582 | 8.830 | 7.155 | 7.619 | 7.849 | 0.026 | 0.5585 | 0.7697 | 0 | 0 | 0 |
| 'Cenpb' | 38.370 | 38.682 | 44.633 | 34.207 | 40.378 | 49.020 | 38.174 | 39.581 | 44.993 | 42.660 | 34.660 | 36.389 | 44.095 | 37.330 | 38.786 | 44.922 | 32.781 | 35.637 | 0.075 | 0.2196 | 0.4665 | 0 | 0 | 0 |
| 'Cenpc1' | 4.164 | 4.642 | 4.754 | 4.062 | 4.437 | 4.033 | 5.297 | 3.968 | 4.888 | 3.872 | 4.386 | 4.675 | 3.954 | 4.770 | 4.506 | 5.103 | 4.526 | [illegible] | 0.003 | 0.9634 | 0.9869 | 0 | 0 | 0 |
| 'Cenpe' | 5.963 | 5.989 | 7.614 | 6.936 | 6.335 | 6.690 | 7.371 | 5.930 | 7.378 | 6.596 | 6.823 | 6.836 | 6.812 | 7.587 | 7.935 | 5.864 | 6.426 | 7.184 | -0.004 | 0.9531 | 0.9826 | 0 | 0 | 0 |
| 'Cenpf' | 1.939 | 2.549 | 2.804 | 2.666 | 2.507 | 2.746 | 2.438 | 2.357 | 2.956 | 1.662 | 2.554 | 2.645 | 2.312 | 2.695 | 2.803 | 2.258 | 2.517 | 2.870 | -0.034 | 0.5954 | 0.7941 | 0 | 0 | 0 |
| 'Cenph' | 4.999 | 6.075 | 7.263 | 6.839 | 6.074 | 6.334 | 8.189 | 6.123 | 6.972 | 5.424 | 6.260 | 6.148 | 7.635 | 5.918 | 6.940 | 6.605 | 5.541 | 5.421 | 0.063 | 0.4485 | 0.6888 | 0 | 0 | 0 |
| 'Cenpi' | 0.698 | 0.793 | 0.846 | 1.013 | 0.717 | 0.873 | 0.791 | 0.690 | 0.756 | 0.507 | 0.626 | 0.806 | 0.974 | 0.814 | 0.828 | 0.791 | 0.937 | 0.831 | -0.008 | 0.9428 | 0.9785 | 0 | 0 | 0 |
| 'Cenpj' | 4.606 | 4.174 | 4.030 | 5.314 | 4.103 | 4.170 | 4.703 | 4.200 | 4.714 | 4.069 | 4.518 | 4.592 | 4.531 | 4.440 | 4.093 | 4.469 | 4.489 | 4.765 | -0.015 | 0.7916 | 0.9070 | 0 | 0 | 0 |
| 'Cenpk' | 1.376 | 1.901 | 2.630 | 2.303 | 1.847 | 2.210 | 2.553 | 1.742 | 2.188 | 1.938 | 2.504 | 2.012 | 2.231 | 2.040 | 2.117 | 1.586 | 2.217 | 1.635 | 0.028 | 0.8260 | 0.9237 | 0 | 0 | 0 |
| 'Cenpl' | 1.199 | 1.393 | 1.383 | 1.250 | 1.337 | 1.345 | 1.438 | 1.509 | 1.279 | 1.148 | 1.277 | 1.211 | 1.247 | 1.351 | 1.162 | 1.107 | 1.455 | 1.104 | 0.121 | 0.0182 | 0.0966 | 0 | 0 | 0 |
| 'Cenpm' | 4.412 | 4.670 | 5.125 | 4.724 | 4.605 | 6.321 | 4.482 | 4.856 | 6.047 | 4.078 | 5.153 | 4.751 | 5.222 | 5.195 | 4.690 | 4.732 | [illegible] | [illegible] | -0.083 | 0.3288 | 0.5870 | 0 | 0 | 0 |
| 'Cenpn' | 8.954 | 8.476 | 11.760 | 11.156 | 10.225 | 11.501 | 10.879 | 10.490 | 10.907 | 7.771 | 11.266 | 9.318 | 12.593 | 11.721 | 11.567 | 8.580 | 8.924 | [illegible] | 0.041 | 0.6436 | 0.8251 | 0 | 0 | 0 |

| | | | | | | | | | | | | | | | | | | | | | | | | |
|---|---|---|---|---|---|---|---|---|---|---|---|---|---|---|---|---|---|---|---|---|---|---|---|---|
| 'Cenpo' | 3.298 | 3.717 | 3.815 | 2.925 | 3.197 | 3.739 | 3.651 | 3.023 | 3.430 | 3.362 | 3.498 | 3.552 | 3.571 | 3.314 | 3.498 | 3.465 | 3.395 | 3.398 | -0.022 | 0.6610 | 0.8362 | 0 | 0 | 0 |
| 'Cenpq' | 11.675 | 12.773 | 10.224 | 12.021 | 10.980 | 10.976 | 11.454 | 11.990 | 9.484 | 10.550 | 11.403 | 10.017 | 11.015 | 11.512 | 9.913 | 10.034 | 11.778 | 9.337 | 0.075 | 0.1195 | 0.3292 | 0 | 0 | 0 |
| 'Cenpt' | 11.059 | 11.855 | 15.217 | 12.370 | 12.546 | 13.431 | 12.620 | 11.228 | 14.076 | 12.997 | 13.278 | 12.851 | 16.739 | 13.433 | 11.963 | 13.960 | 12.425 | 12.418 | -0.077 | 0.2552 | 0.5073 | 0 | 0 | 0 |
| 'Cenpu' | 0.875 | 0.842 | 1.239 | 1.110 | 0.884 | 0.894 | 1.111 | 0.860 | 1.011 | 0.977 | 0.982 | 0.889 | 1.211 | 1.023 | 0.808 | 1.089 | 0.849 | 0.970 | 0.005 | 0.9612 | 0.9860 | 0 | 0 | 0 |
| 'Cenpv' | 2.190 | 1.535 | 2.067 | 1.720 | 1.715 | 1.690 | 1.624 | 2.022 | 1.963 | 1.391 | 1.091 | 1.693 | 1.720 | 1.518 | 1.858 | 1.499 | 1.212 | 1.335 | 0.305 | 0.0021 | 0.0187 | 1 | 0 | 0 |
| 'Cenpw' | 3.994 | 4.313 | 4.175 | 3.899 | 3.896 | 4.273 | 4.009 | 4.177 | 3.785 | 3.301 | 3.702 | 4.026 | 4.807 | 3.991 | 4.907 | 4.008 | 4.523 | 3.564 | -0.023 | 0.7989 | 0.9107 | 0 | 0 | 0 |
| 'Cep104' | 5.629 | 5.357 | 5.852 | 5.284 | 5.163 | 6.228 | 5.271 | 5.360 | 6.108 | 5.636 | 5.640 | 6.359 | 5.736 | 5.867 | 6.268 | 5.488 | 5.494 | 5.674 | -0.063 | 0.1053 | 0.3042 | 0 | 0 | 0 |
| 'Cep120' | 24.033 | 26.295 | 24.744 | 23.778 | 26.944 | 26.379 | 25.145 | 25.518 | 24.873 | 27.157 | 27.754 | 25.679 | 25.602 | 27.584 | 24.085 | 26.030 | 27.279 | 25.771 | -0.067 | 0.0167 | 0.0911 | 0 | 0 | 0 |
| 'Cep128' | 1.462 | 1.264 | 1.618 | 1.546 | 1.409 | 1.406 | 1.505 | 1.343 | 1.261 | 1.306 | 1.498 | 1.538 | 1.437 | 1.343 | 1.498 | 1.106 | 1.313 | 1.395 | 0.030 | 0.6308 | 0.8173 | 0 | 0 | 0 |
| 'Cep131' | 4.099 | 4.213 | 4.761 | 3.928 | 4.209 | 5.180 | 4.729 | 4.139 | 5.326 | 4.901 | 3.937 | 4.072 | 5.301 | 4.205 | 4.189 | 4.777 | 3.530 | 3.961 | 0.058 | 0.3849 | 0.6348 | 0 | 0 | 0 |
| 'Cep135' | 4.337 | 4.758 | 4.538 | 4.430 | 4.427 | 4.268 | 4.624 | 4.577 | 4.712 | 4.356 | 4.649 | 4.266 | 4.450 | 4.485 | 4.008 | 4.565 | 4.534 | 3.851 | 0.047 | 0.1678 | 0.4000 | 0 | 0 | 0 |
| 'Cep152' | 7.072 | 5.822 | 7.808 | 7.619 | 6.637 | 7.034 | 7.012 | 6.568 | 6.951 | 6.740 | 6.248 | 6.302 | 7.038 | 6.840 | 6.784 | 6.671 | 7.544 | 6.939 | 0.025 | 0.6214 | 0.8108 | 0 | 0 | 0 |
| 'Cep162' | 2.539 | 2.690 | 3.057 | 3.253 | 2.590 | 2.736 | 2.826 | 2.692 | 2.891 | 2.742 | 2.793 | 3.291 | 2.801 | 2.694 | 2.856 | 2.921 | 2.843 | 3.258 | -0.065 | 0.1666 | 0.3991 | 0 | 0 | 0 |
| 'Cep164' | 5.358 | 5.567 | 6.418 | 5.842 | 5.551 | 6.801 | 5.791 | 5.825 | 6.370 | 6.378 | 6.297 | 6.504 | 6.623 | 5.811 | 6.592 | 5.935 | 5.483 | 6.346 | -0.074 | 0.0439 | 0.1733 | 0 | 0 | 0 |
| 'Cep170' | 0.866 | 0.877 | 0.739 | 0.953 | 0.847 | 0.719 | 0.841 | 0.808 | 0.862 | 0.700 | 0.838 | 0.932 | 0.824 | 0.770 | 0.930 | 0.737 | 0.861 | 0.976 | -0.024 | 0.7226 | 0.8724 | 0 | 0 | 0 |
| 'Cep170b' | 4.956 | 4.032 | 5.288 | 4.439 | 4.015 | 5.106 | 3.888 | 4.139 | 4.917 | 4.752 | 3.849 | 4.596 | 4.421 | 3.921 | 4.341 | 4.346 | 3.863 | 4.494 | 0.074 | 0.1044 | 0.3027 | 0 | 0 | 0 |
| 'Cep19' | 6.866 | 8.024 | 6.975 | 6.682 | 8.494 | 6.935 | 6.805 | 7.527 | 7.437 | 6.337 | 6.226 | 7.295 | 6.642 | 7.727 | 7.093 | 6.803 | 6.297 | 7.536 | 0.068 | 0.1771 | 0.4124 | 0 | 0 | 0 |
| 'Cep192' | 16.176 | 14.545 | 17.279 | 16.573 | 15.144 | 15.015 | 16.726 | 14.720 | 17.032 | 15.773 | 18.060 | 17.425 | 16.599 | 15.268 | 17.188 | 14.307 | 18.710 | 16.652 | -0.077 | 0.1229 | 0.3344 | 0 | 0 | 0 |
| 'Cep250' | 14.669 | 12.944 | 15.925 | 14.231 | 12.297 | 16.429 | 13.799 | 13.341 | 15.836 | 15.612 | 14.774 | 16.828 | 15.753 | 14.556 | 15.911 | 14.694 | 14.494 | 16.793 | -0.120 | 0.0000 | 0.0004 | 1 | 0 | 0 |
| 'Cep290' | 0.938 | 1.251 | 1.082 | 1.251 | 1.077 | 0.804 | 0.953 | 0.860 | 0.987 | 0.932 | 0.988 | 0.710 | 0.979 | 0.715 | 0.707 | 1.276 | 0.817 | 0.760 | 0.229 | 0.0269 | 0.1262 | 0 | 0 | 0 |
| 'Cep295' | 2.947 | 2.765 | 3.316 | 2.725 | 2.783 | 3.082 | 2.813 | 2.731 | 3.198 | 2.906 | 3.133 | 2.885 | 2.868 | 2.881 | 2.826 | 2.940 | 2.881 | 2.701 | 0.012 | 0.7809 | 0.9017 | 0 | 0 | 0 |
| 'Cep350' | 7.473 | 6.694 | 6.666 | 7.406 | 6.981 | 6.652 | 6.375 | 6.543 | 7.020 | 8.225 | 8.249 | 8.414 | 7.426 | 7.552 | 8.083 | 8.549 | 7.632 | 9.156 | -0.256 | 0.0000 | 0.0000 | 1 | 0 | 0 |
| 'Cep41' | 2.375 | 2.407 | 2.402 | 2.216 | 2.342 | 2.449 | 2.082 | 2.248 | 2.306 | 2.105 | 2.171 | 2.283 | 2.155 | 2.093 | 2.072 | 2.082 | 2.428 | 1.773 | 0.111 | 0.0286 | 0.1309 | 0 | 0 | 0 |
| 'Cep44' | 4.399 | 4.899 | 4.858 | 4.126 | 5.237 | 4.686 | 4.431 | 4.860 | 4.668 | 5.005 | 4.414 | 4.094 | 4.752 | 4.896 | 4.850 | 5.087 | 5.103 | 4.308 | -0.014 | 0.8025 | 0.9115 | 0 | 0 | 0 |
| 'Cep55' | 7.945 | 9.492 | 11.127 | 10.397 | 9.525 | 11.396 | 10.274 | 8.374 | 10.968 | 6.078 | 9.294 | 9.818 | 10.029 | 10.718 | 11.835 | 8.643 | 9.618 | 9.533 | 0.055 | 0.4994 | 0.7273 | 0 | 0 | 0 |
| 'Cep57' | 7.217 | 7.174 | 6.862 | 7.298 | 6.661 | 6.447 | 7.476 | 6.992 | 6.878 | 7.233 | 6.835 | 6.283 | 6.754 | 7.103 | 6.411 | 6.421 | 7.003 | 5.569 | 0.072 | 0.0470 | 0.1806 | 0 | 0 | 0 |
| 'Cep57l1' | 1.488 | 1.255 | 1.485 | 1.359 | 1.290 | 1.445 | 1.386 | 1.439 | 1.564 | 1.103 | 1.476 | 1.359 | 1.612 | 1.308 | 1.256 | 1.356 | 1.452 | 1.286 | 0.053 | 0.4319 | 0.6767 | 0 | 0 | 0 |
| 'Cep63' | 3.165 | 3.114 | 3.329 | 3.618 | 2.878 | 3.295 | 3.223 | 3.294 | 3.402 | 3.311 | 3.185 | 3.384 | 3.025 | 3.035 | 3.067 | 3.015 | 3.692 | 2.838 | 0.028 | 0.5772 | 0.7819 | 0 | 0 | 0 |
| 'Cep68' | 7.043 | 6.712 | 7.140 | 6.150 | 7.147 | 6.396 | 6.569 | 5.978 | 6.621 | 6.464 | 6.514 | 7.536 | 6.466 | 6.615 | 6.977 | 6.846 | 6.540 | 7.590 | -0.055 | 0.2176 | 0.4642 | 0 | 0 | 0 |
| 'Cep70' | 2.047 | 2.112 | 2.355 | 2.143 | 2.286 | 2.350 | 2.653 | 2.200 | 2.390 | 2.525 | 2.217 | 2.124 | 2.219 | 2.342 | 1.878 | 2.589 | 2.221 | 1.765 | 0.047 | 0.4826 | 0.7146 | 0 | 0 | 0 |
| 'Cep72' | 2.283 | 2.851 | 2.869 | 2.574 | 2.311 | 2.807 | 2.764 | 2.687 | 2.784 | 2.540 | 2.905 | 2.404 | 2.434 | 2.630 | 2.325 | 2.173 | 2.514 | 2.086 | 0.118 | 0.1130 | 0.3192 | 0 | 0 | 0 |
| 'Cep76' | 5.742 | 5.324 | 5.409 | 5.123 | 5.157 | 5.284 | 5.662 | 4.824 | 4.925 | 3.842 | 5.540 | 5.935 | 4.869 | 5.541 | 5.486 | 4.890 | 4.887 | 5.252 | 0.025 | 0.7018 | 0.8609 | 0 | 0 | 0 |
| 'Cep78' | 7.100 | 7.174 | 8.507 | 6.788 | 7.799 | 7.573 | 6.346 | 7.533 | 7.075 | 7.136 | 6.643 | 6.634 | 7.196 | 6.230 | 6.335 | 5.713 | 7.112 | 6.294 | 0.145 | 0.0150 | 0.0838 | 0 | 0 | 0 |
| 'Cep83' | 19.284 | 18.655 | 20.156 | 19.471 | 20.157 | 19.908 | 19.684 | 18.977 | 18.910 | 18.539 | 18.641 | 19.696 | 19.884 | 19.491 | 19.603 | 18.883 | 21.031 | 19.158 | -0.008 | 0.8002 | 0.9111 | 0 | 0 | 0 |
| 'Cep85' | 4.988 | 5.100 | 4.999 | 5.051 | 5.191 | 5.311 | 4.713 | 5.443 | 5.601 | 4.284 | 4.291 | 4.888 | 5.070 | 4.623 | 4.847 | 4.301 | 4.385 | 4.457 | 0.160 | 0.0001 | 0.0015 | 1 | 0 | 0 |
| 'Cep85l' | 3.685 | 2.653 | 3.156 | 3.637 | 3.066 | 3.147 | 3.132 | 2.725 | 3.221 | 4.129 | 3.247 | 3.066 | 2.941 | 3.277 | 3.095 | 2.961 | 3.496 | 3.514 | -0.070 | 0.3995 | 0.6485 | 0 | 0 | 0 |
| 'Cep95' | 9.048 | 8.462 | 8.656 | 9.116 | 7.795 | 8.370 | 8.020 | 8.136 | 9.036 | 7.846 | 8.693 | 10.098 | 8.995 | 8.514 | 9.523 | 7.993 | 9.547 | 9.631 | -0.088 | 0.0571 | 0.2053 | 0 | 0 | 0 |
| 'Cep97' | 8.420 | 8.452 | 8.149 | 7.881 | 8.553 | 7.555 | 8.214 | 8.580 | 8.411 | 7.907 | 9.286 | 9.959 | 7.744 | 8.621 | 9.344 | 8.645 | 9.705 | 9.788 | -0.136 | 0.0020 | 0.0179 | 1 | 0 | 0 |
| 'Cept1' | 16.475 | 18.795 | 16.855 | 18.097 | 17.841 | 16.398 | 18.815 | 17.308 | 16.382 | 14.162 | 17.731 | 17.803 | 15.098 | 16.481 | 17.274 | 14.625 | 18.945 | 16.372 | 0.070 | 0.1395 | 0.3611 | 0 | 0 | 0 |
| 'Cerk' | 2.928 | 4.151 | 2.689 | 3.099 | 3.463 | 3.199 | 2.855 | 3.833 | 3.076 | 2.870 | 3.283 | 2.932 | 3.213 | 4.148 | 3.011 | 3.504 | 3.822 | 3.007 | -0.037 | 0.4909 | 0.7208 | 0 | 0 | 0 |
| 'Cerkl' | 0.414 | 0.577 | 0.496 | 0.438 | 0.605 | 0.405 | 0.621 | 0.519 | 0.462 | 0.473 | 0.741 | 0.469 | 0.531 | 0.501 | 0.528 | 0.571 | 0.603 | 0.559 | -0.146 | 0.1601 | 0.3904 | 0 | 0 | 0 |
| 'Cers2' | 19.878 | 21.351 | 20.969 | 19.699 | 21.501 | 21.785 | 19.042 | 21.718 | 21.531 | 20.212 | 18.650 | 22.169 | 20.517 | 20.143 | 21.870 | 20.259 | 20.509 | 20.818 | 0.006 | 0.8245 | 0.9227 | 0 | 0 | 0 |
| 'Cers4' | 5.570 | 6.919 | 6.062 | 6.053 | 7.189 | 6.223 | 5.944 | 6.632 | 6.479 | 6.446 | 6.209 | 6.719 | 5.880 | 6.399 | 6.747 | 6.749 | 7.055 | 6.504 | -0.055 | 0.1290 | 0.3443 | 0 | 0 | 0 |
| 'Cers5' | 13.364 | 13.887 | 14.960 | 13.655 | 14.423 | 14.560 | 13.394 | 14.387 | 15.509 | 14.583 | 14.708 | 15.888 | 13.854 | 13.798 | 14.774 | 12.760 | 14.663 | 15.279 | -0.036 | 0.2077 | 0.4523 | 0 | 0 | 0 |
| 'Cers6' | 3.322 | 3.698 | 2.922 | 3.727 | 3.557 | 3.255 | 3.565 | 2.966 | 2.757 | 3.628 | 4.269 | 3.592 | 4.183 | 3.978 | 3.372 | 3.537 | 4.524 | 3.556 | -0.232 | 0.0000 | 0.0004 | 1 | 0 | 0 |
| 'Cetn2' | 31.166 | 36.065 | 30.776 | 31.812 | 37.449 | 30.366 | 34.701 | 35.666 | 32.213 | 31.959 | 30.610 | 31.390 | 33.229 | 34.999 | 32.154 | 35.042 | 32.796 | 31.214 | 0.018 | 0.6203 | 0.8101 | 0 | 0 | 0 |
| 'Cetn3' | 43.360 | 50.261 | 45.589 | 48.843 | 51.992 | 45.552 | 52.161 | 52.457 | 44.703 | 45.504 | 43.266 | 46.571 | 48.836 | 50.168 | 44.679 | 48.236 | 47.930 | 40.842 | 0.048 | 0.2140 | 0.4593 | 0 | 0 | 0 |
| 'Cfap157' | 8.036 | 9.419 | 9.879 | 8.652 | 9.582 | 10.575 | 8.089 | 10.276 | 9.929 | 10.545 | 9.531 | 9.857 | 11.602 | 9.117 | 9.828 | 11.114 | 9.902 | 10.323 | -0.128 | 0.0391 | 0.1613 | 0 | 0 | 0 |
| 'Cfap20' | 27.557 | 28.115 | 25.046 | 25.690 | 27.208 | 26.037 | 26.156 | 27.625 | 26.106 | 28.773 | 27.422 | 24.605 | 27.327 | 27.074 | 25.076 | 26.920 | 26.595 | 22.886 | 0.012 | 0.7461 | 0.8854 | 0 | 0 | 0 |
| 'Cfap36' | 8.307 | 8.999 | 8.731 | 8.792 | 9.091 | 8.598 | 10.248 | 9.735 | 8.431 | 9.580 | 8.459 | 8.113 | 9.047 | 8.889 | 7.466 | 9.228 | 7.994 | 7.732 | 0.073 | 0.0934 | 0.2829 | 0 | 0 | 0 |
| 'Cfap43' | 8.594 | 7.926 | 7.578 | 8.259 | 7.357 | 8.385 | 7.726 | 7.863 | 8.069 | 8.128 | 8.671 | 8.357 | 8.380 | 8.553 | 8.091 | 8.528 | 8.606 | 7.815 | -0.075 | 0.0245 | 0.1188 | 0 | 0 | 0 |
| 'Cfdp1' | 51.020 | 54.370 | 50.393 | 58.211 | 57.333 | 54.616 | 53.799 | 56.178 | 54.073 | 56.814 | 47.821 | 50.502 | 56.637 | 49.717 | 53.142 | 55.677 | 50.137 | 48.501 | 0.053 | 0.1449 | 0.3693 | 0 | 0 | 0 |
| 'Cfl1' | 873.340 | 968.766 | 854.820 | 934.836 | 945.509 | 903.894 | 951.034 | 880.694 | 903.549 | 874.911 | 931.718 | 755.043 | 918.329 | 906.968 | 927.677 | 942.595 | 905.832 | 863.858 | 0.023 | 0.5252 | 0.7470 | 0 | 0 | 0 |
| 'Cfl2' | 3.969 | 4.808 | 4.773 | 3.633 | 4.372 | 4.825 | 3.992 | 4.883 | 4.491 | 3.256 | 4.611 | 5.026 | 3.326 | 5.655 | 4.518 | 3.965 | 5.149 | 4.678 | -0.020 | 0.7565 | 0.8913 | 0 | 0 | 0 |
| 'Cflar' | 8.513 | 8.201 | 8.165 | 8.096 | 8.304 | 7.326 | 8.820 | 8.193 | 8.207 | 8.054 | 9.284 | 9.221 | 7.849 | 9.068 | 8.378 | 8.382 | 9.382 | 8.760 | -0.096 | 0.0148 | 0.0827 | 0 | 0 | 0 |
| 'Cfp' | 6.063 | 5.084 | 6.676 | 5.058 | 5.001 | 5.740 | 5.229 | 5.267 | 5.766 | 7.373 | 5.218 | 7.580 | 5.920 | 5.303 | 6.430 | 5.491 | 6.980 | 6.643 | -0.202 | 0.0065 | 0.0457 | 1 | 0 | 0 |
| 'Cggbp1' | 20.679 | 20.533 | 19.416 | 23.122 | 19.633 | 19.830 | 20.957 | 20.297 | 18.849 | 17.340 | 21.501 | 23.639 | 18.429 | 22.039 | 21.947 | 19.858 | 23.524 | 22.831 | -0.068 | 0.2310 | 0.4782 | 0 | 0 | 0 |
| 'Cgrrf1' | 2.055 | 2.509 | 1.974 | 2.204 | 2.338 | 2.124 | 2.201 | 2.614 | 1.804 | 2.689 | 2.213 | 2.184 | 2.290 | 2.470 | 2.024 | 2.544 | 2.215 | 1.724 | -0.050 | 0.4750 | 0.7099 | 0 | 0 | 0 |
| 'Chac2' | 2.461 | 3.161 | 2.323 | 2.745 | 3.044 | 1.852 | 2.759 | 2.473 | 2.181 | 1.876 | 2.632 | 2.357 | 2.550 | 2.835 | 2.315 | 2.167 | 2.868 | 1.881 | 0.068 | 0.4742 | 0.7094 | 0 | 0 | 0 |
| 'Chaf1a' | 15.163 | 13.518 | 19.706 | 15.885 | 13.793 | 17.030 | 15.914 | 14.131 | 19.386 | 14.292 | 15.089 | 16.459 | 21.528 | 16.461 | 18.320 | 14.544 | 14.685 | 16.582 | -0.052 | 0.4473 | 0.6880 | 0 | 0 | 0 |
| 'Chaf1b' | 4.099 | 3.360 | 4.877 | 4.043 | 3.904 | 4.641 | 4.010 | 3.713 | 4.571 | 3.884 | 4.028 | 3.829 | 4.859 | 3.750 | 4.479 | 4.368 | 4.110 | 3.706 | 0.000 | 0.9975 | 0.9994 | 0 | 0 | 0 |
| 'Champ1' | 6.563 | 8.001 | 7.218 | 6.681 | 7.176 | 7.493 | 7.056 | 7.293 | 7.609 | 6.606 | 7.467 | 6.965 | 7.027 | 6.993 | 6.912 | 6.917 | 6.406 | 6.906 | 0.054 | 0.1510 | 0.3777 | 0 | 0 | 0 |
| 'Chchd1' | 37.934 | 43.438 | 42.521 | 34.102 | 41.675 | 41.509 | 39.295 | 40.504 | 40.554 | 34.991 | 35.811 | 38.426 | 36.072 | 39.935 | 43.450 | 37.065 | 39.065 | 35.600 | 0.078 | 0.1079 | 0.3092 | 0 | 0 | 0 |
| 'Chchd10' | 19.391 | 20.052 | 18.258 | 15.597 | 15.509 | 19.324 | 17.137 | 18.374 | 16.255 | 21.014 | 18.865 | 18.540 | 22.200 | 17.295 | 16.578 | 21.049 | 18.037 | 17.748 | -0.102 | 0.1120 | 0.3174 | 0 | 0 | 0 |
| 'Chchd2' | 66.472 | 59.690 | 55.688 | 67.510 | 59.874 | 60.314 | 67.369 | 58.648 | 54.739 | 53.980 | 63.807 | 56.014 | 57.349 | 62.734 | 60.513 | 56.239 | 66.772 | 56.828 | 0.031 | 0.5333 | 0.7533 | 0 | 0 | 0 |
| 'Chchd3' | 10.402 | 10.977 | 11.265 | 11.847 | 11.257 | 11.303 | 11.547 | 10.865 | 11.051 | 10.536 | 10.893 | 10.897 | 11.874 | 11.348 | 10.958 | 11.149 | 10.708 | 10.691 | 0.010 | 0.7445 | 0.8847 | 0 | 0 | 0 |
| 'Chchd4' | 3.139 | 3.046 | 2.698 | 2.951 | 2.864 | 2.856 | 2.933 | 2.899 | 2.805 | 2.148 | 2.741 | 3.146 | 2.626 | 2.826 | 3.005 | 2.304 | 2.287 | 2.705 | 0.121 | 0.0809 | 0.2564 | 0 | 0 | 0 |
| 'Chchd5' | 6.502 | 7.017 | 7.026 | 6.244 | 6.889 | 7.179 | 5.997 | 7.677 | 7.291 | 6.821 | 5.851 | 6.231 | 6.783 | 6.404 | 6.441 | 6.349 | 6.372 | 5.892 | 0.108 | 0.0403 | 0.1641 | 0 | 0 | 0 |
| 'Chchd6' | 0.853 | 1.074 | 1.170 | 0.931 | 1.185 | 1.026 | 0.791 | 1.016 | 1.184 | 0.956 | 0.915 | 0.990 | 0.888 | 1.025 | 1.157 | 1.009 | 0.900 | 0.836 | 0.084 | 0.4333 | 0.6782 | 0 | 0 | 0 |
| 'Chchd7' | 1.992 | 2.364 | 1.669 | 1.896 | 2.288 | 2.183 | 1.755 | 2.528 | 2.303 | 2.056 | 1.835 | 1.871 | 1.964 | 1.993 | 1.981 | 2.172 | 1.929 | 1.702 | 0.104 | 0.1438 | 0.3679 | 0 | 0 | 0 |
| 'Chd1l' | 11.741 | 11.178 | 12.005 | 11.969 | 12.513 | 12.414 | 12.891 | 12.496 | 13.000 | 11.705 | 12.634 | 12.056 | 13.114 | 11.940 | 11.747 | 12.840 | 13.324 | 10.805 | -0.007 | 0.8544 | 0.9375 | 0 | 0 | 0 |
| 'Chd2' | 12.605 | 10.190 | 10.227 | 12.169 | 9.909 | 9.683 | 11.480 | 10.898 | 10.607 | 12.201 | 11.404 | 11.016 | 10.908 | 10.596 | 10.543 | 11.853 | 10.716 | 11.073 | -0.051 | 0.1657 | 0.3981 | 0 | 0 | 0 |
| 'Chd3' | 95.963 | 84.037 | 100.566 | 90.450 | 86.040 | 99.777 | 88.993 | 89.266 | 101.155 | 97.373 | 86.129 | 96.696 | 87.142 | 83.386 | 92.594 | 88.241 | 87.756 | 95.129 | 0.026 | 0.3244 | 0.5817 | 0 | 0 | 0 |
| 'Chd4' | 106.608 | 103.557 | 103.254 | 107.737 | 101.454 | 105.928 | 102.349 | 99.589 | 102.696 | 105.493 | 105.670 | 102.338 | 103.509 | 103.384 | 101.488 | 107.674 | 103.490 | 104.679 | -0.018 | 0.3940 | 0.6434 | 0 | 0 | 0 |
| 'Chd6' | 7.357 | 6.802 | 6.737 | 7.667 | 6.743 | 6.762 | 6.918 | 6.796 | 7.331 | 6.760 | 7.042 | 7.257 | 6.081 | 6.507 | 6.437 | 6.946 | 6.376 | 6.775 | 0.057 | 0.1333 | 0.3518 | 0 | 0 | 0 |
| 'Chd7' | 2.121 | 1.475 | 2.205 | 1.788 | 1.568 | 2.010 | 1.713 | 1.699 | 1.984 | 2.255 | 1.984 | 2.252 | 1.697 | 1.673 | 2.189 | 2.054 | 2.018 | 2.221 | -0.158 | 0.0085 | 0.0555 | 0 | 0 | 0 |
| 'Chd9' | 5.755 | 6.042 | 5.419 | 5.805 | 5.774 | 5.697 | 5.655 | 5.538 | 5.791 | 6.163 | 7.379 | 6.467 | 6.096 | 6.769 | 5.891 | 6.238 | 7.177 | 6.323 | -0.193 | 0.0000 | 0.0000 | 1 | 0 | 0 |
| 'Chdh' | 6.210 | 5.821 | 7.262 | 6.169 | 4.730 | 6.637 | 5.493 | 5.477 | 7.096 | 4.154 | 3.132 | 4.508 | 2.512 | 3.080 | 4.420 | 3.671 | 3.566 | 4.105 | 0.721 | 0.0000 | 0.0000 | 1 | 1 | 0 |
| 'Chek1' | 1.659 | 1.611 | 2.325 | 2.267 | 2.008 | 1.986 | 2.510 | 1.911 | 2.096 | 1.828 | 2.059 | 2.202 | 2.835 | 2.057 | 2.289 | 1.910 | 2.096 | 2.344 | -0.110 | 0.1898 | 0.4262 | 0 | 0 | 0 |
| 'Chek2' | 1.115 | 1.404 | 1.696 | 1.695 | 1.297 | 1.722 | 1.430 | 1.375 | 1.698 | 0.833 | 1.769 | 1.590 | 1.475 | 1.735 | 1.870 | 1.050 | 1.327 | 1.561 | 0.021 | 0.8493 | 0.9351 | 0 | 0 | 0 |
| 'Cherp' | 34.376 | 30.370 | 36.650 | 31.349 | 30.027 | 35.792 | 31.751 | 31.177 | 37.815 | 34.973 | 30.206 | 38.372 | 35.929 | 30.953 | 35.785 | 33.200 | 32.472 | 37.089 | -0.056 | 0.0398 | 0.1628 | 0 | 0 | 0 |
| 'Chfr' | 9.932 | 12.306 | 11.470 | 9.635 | 11.861 | 11.507 | 10.532 | 11.940 | 11.360 | 10.986 | 11.302 | 11.199 | 11.341 | 11.247 | 11.290 | 11.205 | 10.986 | 10.440 | -0.005 | 0.8657 | 0.9418 | 0 | 0 | 0 |
| 'Chic1' | 2.805 | 2.413 | 2.747 | 2.343 | 2.518 | 2.201 | 2.409 | 2.257 | 2.392 | 2.117 | 2.571 | 2.482 | 2.640 | 2.031 | 2.008 | 2.813 | 2.739 | 2.258 | 0.021 | 0.7739 | 0.8993 | 0 | 0 | 0 |
| 'Chic2' | 10.297 | 11.205 | 9.207 | 10.238 | 11.415 | 9.807 | 10.622 | 11.137 | 8.921 | 9.697 | 9.707 | 8.650 | 10.169 | 10.933 | 9.331 | 11.165 | 10.233 | 8.920 | 0.054 | 0.1047 | 0.3032 | 0 | 0 | 0 |
| 'Chid1' | 2.189 | 2.187 | 2.192 | 2.030 | 2.320 | 2.398 | 1.839 | 2.237 | 2.215 | 2.286 | 2.016 | 2.389 | 2.312 | 2.177 | 2.333 | 2.304 | 2.208 | 2.458 | -0.073 | 0.1040 | 0.3023 | 0 | 0 | 0 |
| 'Chil5' | 1.623 | 1.484 | 1.499 | 1.479 | 1.018 | 1.203 | 2.477 | 1.488 | 1.040 | 2.363 | 2.172 | 1.870 | 1.893 | 1.622 | 1.630 | 1.691 | 1.758 | 1.359 | -0.323 | 0.0118 | 0.0706 | 0 | 0 | 0 |
| 'Chka' | 1.932 | 1.383 | 1.573 | 1.722 | 1.271 | 1.401 | 1.723 | 1.303 | 1.334 | 1.894 | 1.589 | 1.432 | 1.684 | 1.423 | 1.214 | 1.328 | 1.631 | 1.331 | -0.002 | 0.9805 | 0.9926 | 0 | 0 | 0 |
| 'Chkb' | 7.416 | 6.820 | 7.680 | 7.355 | 6.725 | 7.307 | 7.136 | 6.933 | 7.671 | 7.837 | 7.125 | 7.990 | 7.713 | 7.325 | 7.046 | 6.974 | 8.262 | 7.690 | -0.074 | 0.0481 | 0.1838 | 0 | 0 | 0 |
| 'Chm' | 3.910 | 4.018 | 4.090 | 4.552 | 4.113 | 3.693 | 4.493 | 4.099 | 3.743 | 4.281 | 4.337 | 4.129 | 4.008 | 3.898 | 3.645 | 3.554 | 4.520 | 3.798 | 0.007 | 0.8870 | 0.9534 | 0 | 0 | 0 |
| 'Chml' | 1.899 | 1.644 | 1.177 | 1.723 | 1.210 | 1.339 | 1.882 | 1.377 | 1.409 | 1.257 | 1.753 | 1.718 | 1.644 | 1.516 | 1.712 | 1.580 | 1.917 | 1.736 | -0.146 | 0.1452 | 0.3696 | 0 | 0 | 0 |
| 'Chmp1a' | 26.888 | 34.550 | 30.734 | 25.044 | 33.230 | 34.963 | 26.640 | 32.460 | 31.451 | 30.203 | 28.355 | 30.622 | 31.048 | 31.316 | 31.907 | 34.903 | 27.513 | 29.620 | -0.014 | 0.8050 | 0.9125 | 0 | 0 | 0 |
| 'Chmp2a' | 131.301 | 147.516 | 134.685 | 142.667 | 142.861 | 141.987 | 136.233 | 134.466 | 137.749 | 131.859 | 128.246 | 134.697 | 132.219 | 139.600 | 133.478 | 141.649 | 138.118 | 135.198 | 0.029 | 0.2987 | 0.5550 | 0 | 0 | 0 |
| 'Chmp2b' | 17.005 | 19.763 | 17.804 | 18.579 | 20.530 | 18.710 | 19.471 | 20.122 | 16.984 | 18.357 | 18.076 | 20.083 | 17.962 | 19.751 | 18.846 | 17.339 | 19.188 | 18.930 | -0.017 | 0.6910 | 0.8544 | 0 | 0 | 0 |
| 'Chmp3' | 11.586 | 14.458 | 12.962 | 13.296 | 14.391 | 13.435 | 13.474 | 14.203 | 13.359 | 12.262 | 13.558 | 13.527 | 13.013 | 14.253 | 13.710 | 13.825 | 13.392 | 13.542 | -0.013 | 0.6985 | 0.8588 | 0 | 0 | 0 |
| 'Chmp4b' | 47.228 | 48.504 | 46.157 | 45.690 | 48.790 | 49.217 | 44.709 | 48.579 | 47.397 | 46.518 | 44.326 | 45.967 | 46.878 | 45.495 | 49.610 | 46.900 | 42.509 | 45.133 | 0.034 | 0.2262 | 0.4729 | 0 | 0 | 0 |
| 'Chmp5' | 41.305 | 44.470 | 38.867 | 44.720 | 43.592 | 43.117 | 40.762 | 46.425 | 42.838 | 38.084 | 41.724 | 43.235 | 42.141 | 45.766 | 43.111 | 40.499 | 46.710 | 42.269 | -0.003 | 0.9328 | 0.9746 | 0 | 0 | 0 |
| 'Chmp6' | 12.822 | 14.310 | 14.240 | 12.741 | 14.509 | 16.166 | 12.972 | 13.871 | 14.110 | 13.857 | 11.186 | 13.459 | 14.110 | 13.051 | 14.062 | 15.984 | 11.821 | 12.505 | 0.060 | 0.2799 | 0.5369 | 0 | 0 | 0 |
| 'Chmp7' | 23.487 | 22.759 | 23.955 | 24.178 | 25.319 | 24.927 | 22.825 | 22.853 | 24.627 | 21.826 | 25.313 | 27.309 | 23.555 | 24.523 | 26.981 | 23.866 | 26.143 | 27.028 | -0.087 | 0.0151 | 0.0842 | 0 | 0 | 0 |
| 'Chn1' | 2.063 | 2.838 | 1.594 | 1.876 | 2.467 | 2.131 | 1.926 | 2.666 | 1.929 | 3.226 | 4.360 | 2.989 | 3.507 | 4.523 | 2.732 | 3.198 | 4.529 | 2.658 | -0.707 | 0.0000 | 0.0000 | 1 | 1 | 0 |
| 'Chn2' | 2.797 | 3.665 | 3.276 | 2.762 | 3.402 | 3.319 | 2.791 | 3.563 | 3.505 | 3.425 | 3.511 | 3.633 | 2.582 | 3.678 | 3.472 | 3.533 | 3.594 | 3.395 | -0.096 | 0.0571 | 0.2053 | 0 | 0 | 0 |
| 'Chordc1' | 48.731 | 46.788 | 45.255 | 53.971 | 46.298 | 43.496 | 50.635 | 43.351 | 42.842 | 39.949 | 51.131 | 71.079 | 39.920 | 48.591 | 61.378 | 39.087 | 57.149 | 76.503 | -0.180 | 0.1599 | 0.3902 | 0 | 0 | 0 |
| 'Chp1' | 25.971 | 28.809 | 28.587 | 26.343 | 27.909 | 29.404 | 28.223 | 29.732 | 28.228 | 27.461 | 28.338 | 28.155 | 27.941 | 29.171 | 28.665 | 26.426 | 27.898 | 27.013 | 0.002 | 0.9262 | 0.9719 | 0 | 0 | 0 |
| 'Chpf' | 6.920 | 5.935 | 7.039 | 6.222 | 6.135 | 7.072 | 6.172 | 6.171 | 6.974 | 6.174 | 5.179 | 6.685 | 6.054 | 5.552 | 6.081 | 5.975 | 5.252 | 6.909 | 0.114 | 0.0011 | 0.0114 | 1 | 0 | 0 |
| 'Chpf2' | 5.042 | 5.187 | 6.147 | 4.384 | 5.908 | 6.276 | 4.085 | 5.574 | 5.726 | 5.428 | 5.067 | 5.325 | 5.028 | 4.902 | 5.551 | 5.740 | 4.574 | 5.197 | 0.031 | 0.6099 | 0.8041 | 0 | 0 | 0 |
| 'Chpt1' | 0.683 | 0.950 | 0.719 | 0.658 | 0.888 | 0.775 | 0.840 | 0.772 | 0.782 | 0.847 | 0.964 | 0.664 | 0.804 | 0.900 | 0.684 | 0.809 | 0.872 | 0.770 | -0.053 | 0.4557 | 0.6936 | 0 | 0 | 0 |
| 'Chrac1' | 13.176 | 16.112 | 14.214 | 13.208 | 15.875 | 15.587 | 13.896 | 15.111 | 15.173 | 14.608 | 13.638 | 12.702 | 16.402 | 15.092 | 13.232 | 16.331 | 13.596 | 12.653 | 0.039 | 0.4770 | 0.7107 | 0 | 0 | 0 |
| 'Chrnb2' | 2.814 | 2.860 | 2.800 | 2.538 | 2.581 | 2.643 | 2.596 | 2.609 | 2.988 | 2.496 | 2.545 | 2.896 | 2.522 | 2.440 | 2.709 | 2.539 | 2.503 | 2.790 | 0.046 | 0.2404 | 0.4911 | 0 | 0 | 0 |
| 'Chst10' | 0.770 | 0.751 | 0.982 | 0.984 | 0.922 | 1.004 | 0.829 | 0.832 | 0.926 | 0.778 | 0.686 | 0.665 | 0.795 | 0.542 | 0.679 | 0.661 | 0.598 | 0.665 | 0.393 | 0.0000 | 0.0000 | 1 | 1 | 0 |
| 'Chst11' | 7.470 | 7.041 | 7.052 | 5.925 | 7.489 | 7.275 | 5.477 | 8.379 | 6.568 | 7.079 | 7.521 | 6.103 | 4.789 | 6.834 | 6.527 | 5.697 | 5.536 | 5.823 | 0.156 | 0.0375 | 0.1572 | 0 | 0 | 0 |
| 'Chst12' | 9.407 | 9.341 | 11.228 | 10.033 | 10.193 | 12.101 | 8.986 | 9.658 | 10.935 | 10.441 | 8.524 | 9.847 | 9.697 | 8.961 | 10.187 | 9.810 | 9.051 | 10.317 | 0.076 | 0.1072 | 0.3080 | 0 | 0 | 0 |
| 'Chst14' | 1.691 | 1.961 | 1.762 | 1.551 | 1.740 | 1.877 | 1.502 | 2.167 | 1.662 | 1.188 | 1.659 | 1.960 | 1.591 | 1.458 | 1.562 | 1.295 | 1.469 | 1.549 | 0.186 | 0.0381 | 0.1585 | 0 | 0 | 0 |
| 'Chst15' | 5.739 | 7.695 | 5.139 | 5.830 | 6.443 | 5.277 | 4.680 | 5.888 | 5.246 | 7.089 | 6.590 | 7.474 | 5.548 | 7.339 | 6.975 | 7.239 | 6.573 | 7.029 | -0.272 | 0.0001 | 0.0017 | 1 | 0 | 0 |
| 'Chst2' | 7.615 | 6.490 | 9.544 | 6.729 | 7.073 | 9.102 | 6.910 | 7.565 | 7.463 | 9.288 | 7.265 | 8.813 | 7.810 | 7.927 | 9.165 | 9.114 | 6.520 | 8.981 | -0.136 | 0.0192 | 0.1004 | 0 | 0 | 0 |
| 'Chsy1' | 4.083 | 3.852 | 4.056 | 3.367 | 4.399 | 3.918 | 3.451 | 4.455 | 3.566 | 2.597 | 2.753 | 2.943 | 3.411 | 3.327 | 2.862 | 2.991 | 2.805 | 2.490 | 0.415 | 0.0000 | 0.0000 | 1 | 1 | 0 |
| 'Chtf18' | 2.374 | 2.134 | 3.402 | 2.459 | 1.951 | 3.145 | 2.599 | 2.054 | 3.064 | 2.044 | 2.173 | 2.750 | 3.202 | 2.103 | 3.353 | 2.458 | 2.236 | 2.554 | 0.000 | 0.9989 | 0.9997 | 0 | 0 | 0 |
| 'Chtop' | 48.754 | 51.938 | 56.539 | 51.078 | 54.928 | 53.528 | 51.980 | 53.820 | 54.033 | 50.034 | 53.404 | 53.127 | 49.154 | 50.253 | 54.015 | 48.620 | 53.039 | 49.657 | 0.037 | 0.1160 | 0.3235 | 0 | 0 | 0 |
| 'Chuk' | 10.900 | 11.776 | 11.817 | 11.193 | 11.359 | 10.497 | 10.727 | 11.979 | 11.125 | 11.303 | 11.719 | 10.824 | 11.134 | 11.835 | 10.522 | 10.572 | 12.403 | 10.087 | 0.007 | 0.8290 | 0.9260 | 0 | 0 | 0 |
| 'Churc1' | 3.586 | 4.428 | 4.000 | 4.167 | 4.733 | 4.090 | 4.436 | 4.662 | 4.110 | 3.777 | 3.439 | 3.968 | 4.071 | 4.399 | 3.945 | 4.269 | 4.059 | 3.080 | 0.112 | 0.0562 | 0.2034 | 0 | 0 | 0 |
| 'Ciao1' | 10.221 | 10.362 | 10.593 | 9.617 | 10.622 | 10.925 | 10.860 | 10.884 | 10.082 | 9.101 | 10.380 | 11.308 | 9.106 | 11.329 | 10.827 | 9.957 | 10.713 | 10.964 | -0.003 | 0.9416 | 0.9778 | 0 | 0 | 0 |
| 'Ciapin1' | 10.746 | 11.746 | 11.131 | 10.315 | 11.265 | 10.899 | 11.138 | 11.046 | 10.581 | 9.932 | 10.339 | 10.490 | 11.238 | 10.915 | 11.200 | 10.602 | 10.057 | 10.432 | 0.043 | 0.1724 | 0.4062 | 0 | 0 | 0 |
| 'Ciart' | 11.329 | 4.346 | 3.889 | 8.340 | 4.256 | 3.057 | 7.175 | 4.448 | 2.635 | 8.678 | 7.937 | 3.222 | 8.625 | 7.749 | 3.408 | 5.859 | 8.318 | 2.833 | -0.222 | 0.1778 | 0.4131 | 0 | 0 | 0 |
| 'Cib1' | 10.748 | 11.892 | 11.363 | 10.580 | 11.222 | 11.905 | 10.273 | 11.450 | 11.377 | 10.535 | 10.271 | 11.750 | 10.817 | 10.906 | 11.338 | 11.078 | 11.432 | 11.260 | 0.008 | 0.8019 | 0.9114 | 0 | 0 | 0 |
| 'Cic' | 21.579 | 20.278 | 24.052 | 19.481 | 19.580 | 23.156 | 18.244 | 21.401 | 23.491 | 23.338 | 20.504 | 21.975 | 21.852 | 20.875 | 21.574 | 21.770 | 19.611 | 21.778 | -0.028 | 0.4857 | 0.7168 | 0 | 0 | 0 |
| 'Cinp' | 10.289 | 12.064 | 11.146 | 12.405 | 10.855 | 10.987 | 9.933 | 10.230 | 11.763 | 9.360 | 8.847 | 12.087 | 10.839 | 10.576 | 12.231 | 9.779 | 10.649 | 10.994 | 0.046 | 0.4239 | 0.6707 | 0 | 0 | 0 |
| 'Cipc' | 5.915 | 6.741 | 6.462 | 6.363 | 6.452 | 6.602 | 6.235 | 7.013 | 6.632 | 6.437 | 6.384 | 7.076 | 5.704 | 6.281 | 6.784 | 6.199 | 6.312 | 6.642 | 0.002 | 0.9546 | 0.9832 | 0 | 0 | 0 |
| 'Cirbp' | 28.387 | 40.947 | 37.346 | 26.047 | 41.408 | 37.838 | 32.334 | 44.310 | 39.292 | 36.967 | 34.206 | 24.015 | 36.129 | 33.086 | 24.920 | 37.370 | 28.867 | 19.527 | 0.255 | 0.0404 | 0.1643 | 0 | 0 | 0 |
| 'Cirh1a' | 26.414 | 22.964 | 25.591 | 24.587 | 26.360 | 22.139 | 26.112 | 25.758 | 23.079 | 22.049 | 22.310 | 23.592 | 20.464 | 21.935 | 23.535 | 23.617 | 23.780 | 25.215 | 0.100 | 0.0395 | 0.1621 | 0 | 0 | 0 |
| 'Cisd1' | 26.426 | 28.299 | 24.852 | 27.157 | 29.318 | 28.111 | 25.803 | 26.012 | 25.669 | 21.683 | 22.588 | 28.256 | 30.407 | 29.625 | 30.470 | 25.372 | 28.291 | 29.302 | -0.040 | 0.5227 | 0.7450 | 0 | 0 | 0 |
| 'Cisd2' | 7.562 | 9.260 | 8.392 | 7.434 | 9.266 | 7.538 | 8.158 | 8.615 | 7.920 | 7.361 | 7.881 | 7.978 | 8.510 | 8.549 | 7.634 | 7.095 | 8.519 | 7.494 | 0.046 | 0.2148 | 0.4603 | 0 | 0 | 0 |
| 'Cisd3' | 6.134 | 8.299 | 6.460 | 7.664 | 6.997 | 7.426 | 6.334 | 7.575 | 6.332 | 5.678 | 4.684 | 6.089 | 7.036 | 6.438 | 5.792 | 5.572 | 6.912 | 6.476 | 0.184 | 0.0855 | 0.2658 | 0 | 0 | 0 |
| 'Cish' | 1.244 | 0.639 | 0.873 | 1.019 | 0.617 | 1.111 | 1.089 | 0.683 | 1.012 | 0.524 | 0.413 | 0.761 | 0.702 | 0.393 | 0.832 | 0.874 | 0.555 | 0.771 | 0.498 | 0.0000 | 0.0000 | 1 | 1 | 0 |
| 'Cit' | 1.272 | 0.929 | 1.773 | 1.476 | 1.113 | 1.471 | 1.414 | 1.147 | 1.584 | 1.031 | 1.425 | 1.548 | 1.615 | 1.330 | 1.780 | 1.151 | 1.407 | 1.623 | -0.104 | 0.2128 | 0.4579 | 0 | 0 | 0 |
| 'Cited2' | 76.138 | 55.134 | 45.410 | 64.060 | 57.888 | 49.027 | 65.339 | 58.708 | 42.767 | 69.502 | 62.609 | 48.524 | 62.224 | 61.425 | 46.294 | 68.786 | 63.005 | 47.517 | -0.055 | 0.1882 | 0.4246 | 0 | 0 | 0 |
| 'Ciz1' | 6.981 | 6.114 | 7.748 | 6.410 | 6.035 | 7.350 | 6.404 | 6.382 | 7.803 | 7.673 | 6.481 | 7.292 | 7.528 | 6.008 | 7.055 | 6.623 | 6.314 | 6.898 | -0.026 | 0.4973 | 0.7258 | 0 | 0 | 0 |
| 'Ckap2' | 5.462 | 6.093 | 8.635 | 8.319 | 6.757 | 7.685 | 7.483 | 6.695 | 8.935 | 4.583 | 7.174 | 7.324 | 7.532 | 7.148 | 8.481 | 5.773 | 6.672 | 7.110 | 0.084 | 0.3392 | 0.5966 | 0 | 0 | 0 |
| 'Ckap2l' | 7.975 | 7.885 | 11.755 | 10.004 | 9.019 | 10.330 | 10.049 | 8.911 | 11.544 | 6.404 | 8.365 | 9.878 | 10.256 | 10.130 | 12.171 | 7.529 | 9.963 | 10.352 | 0.030 | 0.7088 | 0.8650 | 0 | 0 | 0 |
| 'Ckap5' | 8.047 | 8.497 | 9.288 | 9.631 | 8.333 | 8.681 | 10.063 | 8.576 | 9.225 | 8.023 | 9.380 | 9.526 | 8.489 | 9.307 | 9.238 | 7.002 | 9.442 | 9.492 | 0.000 | 0.9982 | 0.9994 | 0 | 0 | 0 |
| 'Ckb' | 12.594 | 12.209 | 11.599 | 12.299 | 12.572 | 14.849 | 14.828 | 10.031 | 14.449 | 10.389 | 7.618 | 13.332 | 8.418 | 8.999 | 12.787 | 9.227 | 8.865 | 11.947 | 0.326 | 0.0000 | 0.0006 | 1 | 0 | 0 |
| 'Cklf' | 8.409 | 11.377 | 12.519 | 7.647 | 8.541 | 9.476 | 9.621 | 10.037 | 10.061 | 7.143 | 10.736 | 11.141 | 10.868 | 9.543 | 11.065 | 8.126 | 7.996 | 8.593 | 0.024 | 0.8272 | 0.9247 | 0 | 0 | 0 |
| 'Cks1b' | 12.514 | 15.182 | 15.073 | 14.801 | 12.647 | 17.012 | 14.907 | 13.137 | 14.764 | 9.723 | 15.515 | 13.791 | 15.758 | 15.300 | 16.034 | 12.885 | 14.017 | 13.603 | 0.029 | 0.7084 | 0.8650 | 0 | 0 | 0 |
| 'Cks2' | 61.591 | 58.501 | 56.477 | 60.941 | 56.293 | 54.605 | 61.681 | 57.108 | 54.731 | 58.490 | 56.432 | 48.893 | 61.905 | 70.194 | 58.523 | 56.461 | 58.316 | 50.581 | 0.001 | 0.9878 | 0.9964 | 0 | 0 | 0 |
| 'Clasp1' | 1.608 | 1.436 | 1.566 | 1.630 | 1.382 | 1.437 | 1.525 | 1.337 | 1.455 | 1.502 | 1.559 | 1.608 | 1.404 | 1.627 | 1.459 | 1.450 | 1.606 | 1.623 | -0.061 | 0.1963 | 0.4353 | 0 | 0 | 0 |
| 'Clasp2' | 7.984 | 8.092 | 9.062 | 8.254 | 9.446 | 9.296 | 8.543 | 9.376 | 8.690 | 7.811 | 8.317 | 8.666 | 7.987 | 8.757 | 8.609 | 8.157 | 8.584 | 8.422 | 0.055 | 0.0979 | 0.2920 | 0 | 0 | 0 |
| 'Clasrp' | 47.753 | 41.140 | 54.190 | 44.814 | 40.052 | 51.904 | 44.704 | 43.247 | 55.196 | 47.608 | 41.248 | 53.648 | 53.481 | 42.085 | 47.172 | 47.325 | 43.865 | 52.194 | -0.031 | 0.3965 | 0.6455 | 0 | 0 | 0 |
| 'Clcc1' | 5.223 | 6.074 | 5.909 | 5.326 | 6.209 | 6.192 | 6.256 | 5.803 | 6.178 | 5.687 | 5.841 | 6.112 | 5.804 | 6.009 | 5.561 | 5.382 | 5.396 | 5.682 | 0.032 | 0.4218 | 0.6682 | 0 | 0 | 0 |
| 'Clcf1' | 4.868 | 5.101 | 5.908 | 4.516 | 5.045 | 6.036 | 5.031 | 5.350 | 6.439 | 5.254 | 4.681 | 5.619 | 4.430 | 4.435 | 5.526 | 4.062 | 4.892 | 5.679 | 0.104 | 0.0172 | 0.0929 | 0 | 0 | 0 |
| 'Clcn2' | 0.697 | 0.660 | 0.780 | 0.646 | 0.513 | 0.616 | 0.729 | 0.672 | 0.746 | 0.661 | 0.668 | 0.744 | 0.680 | 0.591 | 0.555 | 0.486 | 0.679 | 0.636 | 0.073 | 0.4023 | 0.6511 | 0 | 0 | 0 |
| 'Clcn3' | 3.250 | 2.972 | 2.999 | 3.356 | 3.127 | 2.645 | 3.419 | 3.193 | 2.950 | 2.834 | 3.488 | 3.155 | 3.390 | 3.346 | 3.037 | 3.190 | 3.497 | 3.117 | -0.070 | 0.1028 | 0.3006 | 0 | 0 | 0 |
| 'Clcn4' | 26.126 | 28.377 | 27.775 | 27.233 | 28.388 | 27.519 | 25.112 | 28.240 | 29.566 | 28.141 | 27.337 | 28.351 | 26.478 | 28.950 | 27.670 | 27.928 | 27.182 | 27.032 | -0.014 | 0.5876 | 0.7884 | 0 | 0 | 0 |
| 'Clcn5' | 1.469 | 1.452 | 1.587 | 1.357 | 1.274 | 1.362 | 1.646 | 1.413 | 1.442 | 1.360 | 1.723 | 1.501 | 1.478 | 1.415 | 1.447 | 1.335 | 1.530 | 1.528 | -0.044 | 0.4512 | 0.6904 | 0 | 0 | 0 |
| 'Clcn6' | 3.531 | 3.883 | 4.009 | 3.578 | 3.074 | 3.890 | 3.754 | 3.695 | 3.896 | 4.008 | 3.966 | 3.689 | 3.658 | 3.094 | 3.750 | 3.792 | 3.450 | 3.306 | 0.020 | 0.7114 | 0.8659 | 0 | 0 | 0 |
| 'Clcn7' | 7.177 | 7.202 | 8.210 | 7.016 | 8.084 | 8.004 | 7.051 | 7.817 | 8.060 | 7.827 | 7.820 | 8.993 | 8.461 | 7.895 | 7.962 | 7.811 | 7.663 | 8.380 | -0.098 | 0.0032 | 0.0264 | 1 | 0 | 0 |
| 'Cldnd1' | 8.171 | 6.929 | 7.594 | 7.652 | 8.222 | 6.752 | 6.676 | 7.651 | 7.174 | 6.989 | 8.107 | 7.865 | 7.251 | 8.953 | 7.241 | 7.581 | 7.485 | 7.655 | -0.057 | 0.2627 | 0.5166 | 0 | 0 | 0 |
| 'Clec16a' | 4.799 | 4.557 | 5.113 | 4.621 | 4.496 | 5.363 | 4.313 | 4.592 | 5.093 | 5.056 | 4.978 | 5.074 | 4.824 | 5.088 | 4.693 | 4.744 | 4.502 | 4.948 | -0.041 | 0.2712 | 0.5262 | 0 | 0 | 0 |
| 'Clec2d' | 88.741 | 89.415 | 102.398 | 97.001 | 96.071 | 105.146 | 100.015 | 90.584 | 111.303 | 83.130 | 78.948 | 105.953 | 91.186 | 90.845 | 109.925 | 88.439 | 99.486 | 106.069 | 0.035 | 0.3880 | 0.6376 | 0 | 0 | 0 |
| 'Clic1' | 120.073 | 134.057 | 124.253 | 120.650 | 133.746 | 131.660 | 119.514 | 130.404 | 126.791 | 117.469 | 116.748 | 118.189 | 127.522 | 129.389 | 128.682 | 127.417 | 123.475 | 119.747 | 0.030 | 0.2874 | 0.5438 | 0 | 0 | 0 |
| 'Clic3' | 1.798 | 1.649 | 2.065 | 2.333 | 1.434 | 1.727 | 1.796 | 1.356 | 2.294 | 1.578 | 1.709 | 2.738 | 1.674 | 1.823 | 2.276 | 1.505 | 1.777 | 2.934 | -0.143 | 0.2010 | 0.4424 | 0 | 0 | 0 |
| 'Clic4' | 1.773 | 1.861 | 1.966 | 1.988 | 2.040 | 2.190 | 1.741 | 1.872 | 1.957 | 1.838 | 1.665 | 2.261 | 2.340 | 2.027 | 2.250 | 1.894 | 2.042 | 2.046 | -0.096 | 0.1282 | 0.3432 | 0 | 0 | 0 |
| 'Clint1' | 28.334 | 27.704 | 28.061 | 25.988 | 28.526 | 28.174 | 27.823 | 27.944 | 27.796 | 29.028 | 27.886 | 28.528 | 26.287 | 27.427 | 28.467 | 26.910 | 27.143 | 28.067 | -0.007 | 0.7714 | 0.8985 | 0 | 0 | 0 |
| 'Clip1' | 11.070 | 9.934 | 10.949 | 10.932 | 10.312 | 10.831 | 10.500 | 10.477 | 10.504 | 10.897 | 10.866 | 11.451 | 10.198 | 10.664 | 11.015 | 10.589 | 10.341 | 11.552 | -0.041 | 0.1343 | 0.3530 | 0 | 0 | 0 |
| 'Clip2' | 5.795 | 6.122 | 5.766 | 4.728 | 5.702 | 6.475 | 5.924 | 5.802 | 6.313 | 7.091 | 5.675 | 6.561 | 5.762 | 6.107 | 5.965 | 5.555 | 5.772 | 6.304 | -0.070 | 0.1814 | 0.4171 | 0 | 0 | 0 |
| 'Clip3' | 7.510 | 7.629 | 7.746 | 8.133 | 8.804 | 8.424 | 8.180 | 8.397 | 8.049 | 7.584 | 7.098 | 6.626 | 7.972 | 6.703 | 6.868 | 7.750 | 6.777 | 6.388 | 0.187 | 0.0000 | 0.0002 | 1 | 0 | 0 |
| 'Clk1' | 37.297 | 29.648 | 26.642 | 36.956 | 28.769 | 25.875 | 37.235 | 31.355 | 28.116 | 29.836 | 34.456 | 30.188 | 28.519 | 31.736 | 27.863 | 28.885 | 36.174 | 25.935 | 0.025 | 0.6929 | 0.8555 | 0 | 0 | 0 |
| 'Clk2' | 14.633 | 13.776 | 15.149 | 14.166 | 13.828 | 13.914 | 13.693 | 13.753 | 15.189 | 13.624 | 13.644 | 16.249 | 14.615 | 13.677 | 15.014 | 13.242 | 14.205 | 15.797 | -0.033 | 0.2830 | 0.5394 | 0 | 0 | 0 |
| 'Clk3' | 41.131 | 39.888 | 40.597 | 39.985 | 37.251 | 39.033 | 41.849 | 39.344 | 43.342 | 39.479 | 40.186 | 45.898 | 40.480 | 41.163 | 41.804 | 40.148 | 42.842 | 40.826 | -0.053 | 0.0989 | 0.2940 | 0 | 0 | 0 |
| 'Clk4' | 13.813 | 14.553 | 13.582 | 15.377 | 14.021 | 13.254 | 14.942 | 14.653 | 14.028 | 13.531 | 14.088 | 15.087 | 13.816 | 15.495 | 14.328 | 13.359 | 17.003 | 13.517 | -0.033 | 0.4215 | 0.6681 | 0 | 0 | 0 |
| 'Clmn' | 0.010 | 0.013 | 0.004 | 0.018 | 0.024 | 0.014 | 0.013 | 0.027 | 0.016 | 0.800 | 1.456 | 1.151 | 0.653 | 0.775 | 0.764 | 0.474 | 1.423 | 0.437 | -5.849 | 0.0000 | 0.0000 | 1 | 1 | 1 |
| 'Cln3' | 5.719 | 5.581 | 6.812 | 5.852 | 5.757 | 5.948 | 6.199 | 5.808 | 6.649 | 6.614 | 5.876 | 6.449 | 6.265 | 5.084 | 6.034 | 5.836 | 5.978 | 5.696 | 0.003 | 0.9485 | 0.9803 | 0 | 0 | 0 |
| 'Cln5' | 14.667 | 14.178 | 12.679 | 13.576 | 13.953 | 12.910 | 12.601 | 14.086 | 12.845 | 12.902 | 13.866 | 13.057 | 13.916 | 15.458 | 13.402 | 14.133 | 14.632 | 14.805 | -0.063 | 0.1572 | 0.3868 | 0 | 0 | 0 |
| 'Cln6' | 12.021 | 12.188 | 12.968 | 11.675 | 12.763 | 13.780 | 11.865 | 13.445 | 12.941 | 14.541 | 12.729 | 12.807 | 14.217 | 13.465 | 12.898 | 11.513 | 11.975 | 12.008 | -0.040 | 0.3909 | 0.6403 | 0 | 0 | 0 |
| 'Cln8' | 1.763 | 1.628 | 1.725 | 1.585 | 2.109 | 1.874 | 1.436 | 1.730 | 1.536 | 1.715 | 1.784 | 1.698 | 1.653 | 1.822 | 1.476 | 1.805 | 1.799 | 1.682 | -0.010 | 0.8747 | 0.9463 | 0 | 0 | 0 |
| 'Clns1a' | 13.500 | 15.400 | 14.226 | 14.281 | 15.999 | 14.327 | 14.322 | 14.730 | 13.982 | 14.332 | 13.558 | 13.730 | 14.108 | 13.120 | 13.327 | 14.287 | 13.624 | 13.953 | 0.063 | 0.0513 | 0.1917 | 0 | 0 | 0 |
| 'Clock' | 2.001 | 1.954 | 1.803 | 2.086 | 1.899 | 1.735 | 2.159 | 2.064 | 1.876 | 1.783 | 1.925 | 1.787 | 1.546 | 1.753 | 1.618 | 1.666 | 1.969 | 1.772 | 0.139 | 0.0045 | 0.0346 | 1 | 0 | 0 |
| 'Clp1' | 6.240 | 6.255 | 5.350 | 6.610 | 5.904 | 5.732 | 6.989 | 6.143 | 5.977 | 5.516 | 6.432 | 6.389 | 6.110 | 6.077 | 6.386 | 5.335 | 6.108 | 6.346 | -0.005 | 0.9249 | 0.9715 | 0 | 0 | 0 |
| 'Clpb' | 3.838 | 3.425 | 3.799 | 3.270 | 3.328 | 3.911 | 3.267 | 3.349 | 3.715 | 3.488 | 2.950 | 3.720 | 3.659 | 3.041 | 3.379 | 3.608 | 3.293 | 3.682 | 0.041 | 0.3078 | 0.5662 | 0 | 0 | 0 |
| 'Clpp' | 29.307 | 29.835 | 27.717 | 29.451 | 29.837 | 27.680 | 29.395 | 29.123 | 29.193 | 28.345 | 25.505 | 28.721 | 28.845 | 23.988 | 26.937 | 29.133 | 26.621 | 27.095 | 0.077 | 0.0511 | 0.1915 | 0 | 0 | 0 |
| 'Clptm1' | 33.346 | 32.237 | 34.356 | 32.004 | 33.195 | 36.445 | 32.842 | 33.967 | 33.504 | 34.162 | 31.950 | 35.243 | 34.236 | 31.690 | 33.242 | 36.246 | 31.238 | 33.938 | -0.009 | 0.7364 | 0.8800 | 0 | 0 | 0 |
| 'Clptm1l' | 73.692 | 78.224 | 76.958 | 75.020 | 81.363 | 80.494 | 71.088 | 82.232 | 71.632 | 81.817 | 73.897 | 77.261 | 79.677 | 82.262 | 78.315 | 83.504 | 77.611 | 78.451 | -0.056 | 0.0765 | 0.2465 | 0 | 0 | 0 |
| 'Clpx' | 9.010 | 10.417 | 9.419 | 9.057 | 9.043 | 9.014 | 9.731 | 9.454 | 9.348 | 9.570 | 9.351 | 9.465 | 9.314 | 9.126 | 8.356 | 8.999 | 10.204 | 8.653 | 0.012 | 0.7392 | 0.8819 | 0 | 0 | 0 |
| 'Clspn' | 4.786 | 4.197 | 6.446 | 5.856 | 4.400 | 5.921 | 6.259 | 4.594 | 5.778 | 4.304 | 5.024 | 4.989 | 6.834 | 5.118 | 5.844 | 4.801 | 4.854 | 4.756 | 0.031 | 0.6926 | 0.8555 | 0 | 0 | 0 |
| 'Clstn1' | 12.470 | 11.618 | 14.759 | 11.505 | 11.580 | 14.644 | 11.674 | 12.565 | 14.422 | 14.195 | 10.966 | 12.874 | 12.718 | 11.549 | 12.132 | 11.882 | 10.589 | 12.506 | 0.064 | 0.1669 | 0.3994 | 0 | 0 | 0 |
| 'Clta' | 44.042 | 47.851 | 45.663 | 44.401 | 46.838 | 47.383 | 44.416 | 47.127 | 45.282 | 44.471 | 42.706 | 43.955 | 46.468 | 44.861 | 48.171 | 45.396 | 42.548 | 45.133 | 0.022 | 0.4123 | 0.6608 | 0 | 0 | 0 |
| 'Cltb' | 10.236 | 11.413 | 10.768 | 11.975 | 11.483 | 11.376 | 11.350 | 10.907 | 10.766 | 9.912 | 9.889 | 9.934 | 11.480 | 9.595 | 9.115 | 10.146 | 9.598 | 9.482 | 0.157 | 0.0000 | 0.0004 | 1 | 0 | 0 |
| 'Cltc' | 24.526 | 26.317 | 24.677 | 24.769 | 25.545 | 24.751 | 25.652 | 25.336 | 24.905 | 23.858 | 26.640 | 27.604 | 22.967 | 26.250 | 26.320 | 23.434 | 27.263 | 27.624 | -0.044 | 0.1693 | 0.4017 | 0 | 0 | 0 |
| 'Cluap1' | 7.484 | 7.579 | 6.793 | 8.111 | 7.061 | 7.509 | 7.422 | 7.523 | 7.646 | 7.475 | 7.337 | 7.483 | 6.357 | 8.157 | 7.323 | 7.440 | 8.046 | 7.344 | -0.006 | 0.9051 | 0.9626 | 0 | 0 | 0 |
| 'Cluh' | 7.349 | 6.964 | 8.293 | 6.842 | 7.046 | 7.356 | 6.986 | 7.196 | 7.521 | 7.408 | 7.118 | 7.761 | 7.317 | 6.297 | 6.935 | 7.099 | 6.793 | 7.506 | 0.020 | 0.5675 | 0.7754 | 0 | 0 | 0 |
| 'Cmah' | 1.295 | 1.369 | 1.333 | 1.404 | 1.736 | 1.514 | 1.390 | 1.518 | 1.549 | 1.441 | 1.923 | 2.166 | 1.955 | 1.761 | 2.004 | 1.622 | 1.799 | 1.969 | -0.370 | 0.0000 | 0.0002 | 1 | 0 | 0 |
| 'Cmas' | 10.377 | 12.737 | 10.708 | 10.911 | 11.848 | 11.333 | 10.793 | 11.870 | 10.607 | 10.943 | 11.894 | 10.833 | 11.917 | 12.279 | 11.012 | 12.437 | 12.576 | 10.967 | -0.061 | 0.0553 | 0.2017 | 0 | 0 | 0 |
| 'Cmc1' | 21.986 | 24.222 | 22.452 | 22.410 | 23.975 | 22.914 | 23.269 | 24.191 | 23.601 | 21.306 | 23.105 | 20.864 | 22.726 | 20.880 | 19.489 | 23.198 | 23.824 | 19.807 | 0.092 | 0.0347 | 0.1491 | 0 | 0 | 0 |
| 'Cmc2' | 5.937 | 6.186 | 5.954 | 6.300 | 6.167 | 6.087 | 6.504 | 6.445 | 5.870 | 4.966 | 5.805 | 5.135 | 6.773 | 5.834 | 6.406 | 5.677 | 5.813 | 5.397 | 0.088 | 0.1000 | 0.2958 | 0 | 0 | 0 |
| 'Cmc4' | 1.370 | 2.378 | 1.793 | 2.117 | 1.658 | 2.012 | 2.519 | 2.022 | 2.129 | 1.902 | 2.265 | 2.177 | 1.878 | 2.454 | 2.183 | 1.850 | 2.148 | 2.174 | -0.102 | 0.3643 | 0.6162 | 0 | 0 | 0 |
| 'Cmip' | 61.518 | 57.881 | 60.350 | 58.448 | 58.088 | 60.349 | 58.264 | 60.088 | 58.211 | 65.368 | 59.610 | 60.165 | 63.887 | 62.396 | 59.669 | 57.205 | 58.378 | 64.752 | -0.059 | 0.0462 | 0.1789 | 0 | 0 | 0 |
| 'Cmpk1' | 21.127 | 23.222 | 23.042 | 23.216 | 23.718 | 21.234 | 24.174 | 25.084 | 21.491 | 21.610 | 23.874 | 20.332 | 21.586 | 22.699 | 20.097 | 20.700 | 24.274 | 19.660 | 0.075 | 0.0174 | 0.0936 | 0 | 0 | 0 |
| 'Cmpk2' | 3.757 | 3.487 | 3.833 | 2.511 | 4.760 | 4.381 | 3.358 | 4.691 | 3.519 | 3.625 | 3.721 | 5.149 | 4.509 | 4.048 | 5.304 | 3.932 | 3.992 | 5.085 | -0.211 | 0.0200 | 0.1029 | 0 | 0 | 0 |
| 'Cmss1' | 18.269 | 22.187 | 20.227 | 21.129 | 22.447 | 19.511 | 20.270 | 21.624 | 21.342 | 21.580 | 17.862 | 17.634 | 17.839 | 18.174 | 17.171 | 17.711 | 18.811 | 14.605 | 0.202 | 0.0004 | 0.0055 | 1 | 0 | 0 |
| 'Cmtm3' | 9.664 | 9.713 | 10.050 | 10.864 | 10.505 | 11.520 | 8.211 | 10.205 | 10.992 | 10.743 | 8.811 | 10.963 | 9.095 | 9.099 | 11.215 | 10.580 | 9.210 | 9.329 | 0.034 | 0.6555 | 0.8331 | 0 | 0 | 0 |
| 'Cmtm4' | 2.362 | 2.374 | 2.722 | 2.798 | 1.957 | 2.756 | 2.556 | 2.433 | 2.859 | 1.890 | 2.372 | 3.487 | 2.124 | 2.736 | 3.357 | 2.189 | 2.307 | 3.675 | -0.073 | 0.3717 | 0.6236 | 0 | 0 | 0 |
| 'Cmtm6' | 51.976 | 60.875 | 56.063 | 52.377 | 60.166 | 57.930 | 52.156 | 59.810 | 56.148 | 55.313 | 57.984 | 59.913 | 54.327 | 61.564 | 59.984 | 57.017 | 55.475 | 57.908 | -0.046 | 0.0570 | 0.2051 | 0 | 0 | 0 |
| 'Cmtm7' | 32.442 | 37.396 | 36.132 | 32.551 | 35.522 | 40.753 | 35.379 | 37.954 | 38.018 | 31.804 | 25.804 | 33.033 | 32.454 | 29.718 | 34.922 | 30.239 | 30.766 | 32.773 | 0.200 | 0.0000 | 0.0002 | 1 | 0 | 0 |
| 'Cmtr1' | 3.071 | 2.947 | 3.359 | 3.112 | 2.962 | 2.993 | 2.688 | 3.127 | 3.362 | 3.065 | 3.127 | 3.209 | 3.188 | 2.846 | 3.208 | 3.296 | 2.941 | 3.078 | -0.023 | 0.5734 | 0.7795 | 0 | 0 | 0 |
| 'Cmtr2' | 3.462 | 3.147 | 2.684 | 3.187 | 3.286 | 2.783 | 2.567 | 3.563 | 2.963 | 3.392 | 2.655 | 2.904 | 2.612 | 2.713 | 2.854 | 3.055 | 2.986 | 2.810 | 0.077 | 0.2748 | 0.5304 | 0 | 0 | 0 |
| 'Cmya5' | 0.636 | 0.775 | 0.814 | 0.734 | 0.733 | 0.789 | 0.757 | 0.690 | 0.715 | 0.800 | 0.615 | 0.870 | 0.693 | 0.732 | 0.701 | 0.694 | 0.706 | 0.664 | 0.014 | 0.8494 | 0.9351 | 0 | 0 | 0 |
| 'Cnbd2' | 1.912 | 1.801 | 2.030 | 1.973 | 1.591 | 1.655 | 2.099 | 1.778 | 1.660 | 1.805 | 1.868 | 1.708 | 1.958 | 1.998 | 1.595 | 1.699 | 1.358 | 1.753 | 0.058 | 0.4474 | 0.6880 | 0 | 0 | 0 |
| 'Cnbp' | 49.934 | 55.794 | 46.927 | 48.825 | 53.554 | 47.962 | 50.151 | 54.817 | 46.871 | 51.473 | 49.988 | 45.023 | 48.014 | 52.667 | 45.617 | 53.178 | 48.518 | 44.546 | 0.040 | 0.1181 | 0.3270 | 0 | 0 | 0 |
| 'Cndp2' | 39.717 | 37.398 | 40.880 | 37.181 | 38.127 | 41.931 | 41.767 | 40.540 | 40.205 | 40.944 | 39.134 | 39.884 | 41.009 | 36.327 | 40.333 | 40.750 | 38.840 | 38.928 | -0.001 | 0.9763 | 0.9918 | 0 | 0 | 0 |
| 'Cnep1r1' | 3.110 | 3.612 | 3.599 | 3.294 | 4.097 | 3.245 | 3.686 | 3.252 | 3.603 | 3.326 | 3.514 | 3.732 | 3.180 | 3.232 | 3.606 | 3.143 | 3.891 | 3.736 | -0.010 | 0.8458 | 0.9343 | 0 | 0 | 0 |
| 'Cnga1' | 1.725 | 1.686 | 1.464 | 2.033 | 1.567 | 1.320 | 1.873 | 1.426 | 1.431 | 1.511 | 1.329 | 1.838 | 1.585 | 1.753 | 1.384 | 1.275 | 1.687 | 1.511 | 0.036 | 0.6826 | 0.8492 | 0 | 0 | 0 |
| 'Cnih1' | 14.398 | 14.491 | 14.074 | 13.646 | 15.327 | 13.150 | 14.510 | 14.697 | 12.522 | 12.776 | 13.586 | 13.647 | 12.848 | 12.812 | 13.217 | 12.372 | 14.268 | 12.972 | 0.082 | 0.0391 | 0.1613 | 0 | 0 | 0 |
| 'Cnih4' | 5.349 | 5.409 | 5.092 | 5.579 | 5.290 | 5.366 | 4.671 | 5.699 | 5.550 | 4.813 | 4.997 | 5.541 | 5.171 | 5.134 | 5.220 | 5.335 | 5.490 | 5.069 | 0.025 | 0.5965 | 0.7949 | 0 | 0 | 0 |
| 'Cnn2' | 221.855 | 214.888 | 236.512 | 219.575 | 223.473 | 259.331 | 221.550 | 213.010 | 243.152 | 237.584 | 216.050 | 241.662 | 244.951 | 222.149 | 265.105 | 245.825 | 215.604 | 254.843 | -0.072 | 0.0056 | 0.0411 | 1 | 0 | 0 |
| 'Cnn3' | 0.326 | 0.315 | 0.446 | 0.325 | 0.422 | 0.362 | 0.420 | 0.368 | 0.521 | 0.504 | 0.505 | 0.519 | 0.460 | 0.609 | 0.553 | 0.512 | 0.447 | 0.510 | -0.401 | 0.0007 | 0.0076 | 1 | 1 | 0 |
| 'Cnnm2' | 4.074 | 3.748 | 4.101 | 3.641 | 4.053 | 4.242 | 3.640 | 3.470 | 3.849 | 3.748 | 3.740 | 3.822 | 4.356 | 3.718 | 3.584 | 3.488 | 3.879 | 3.480 | 0.034 | 0.5856 | 0.7869 | 0 | 0 | 0 |
| 'Cnnm3' | 5.830 | 6.223 | 6.716 | 5.701 | 6.189 | 7.179 | 5.247 | 6.609 | 6.996 | 6.453 | 6.361 | 7.367 | 6.665 | 6.258 | 6.930 | 7.227 | 5.522 | 7.755 | -0.105 | 0.0266 | 0.1254 | 0 | 0 | 0 |
| 'Cnnm4' | 4.134 | 4.284 | 4.540 | 4.353 | 4.413 | 4.621 | 4.636 | 4.279 | 4.393 | 4.426 | 4.060 | 4.396 | 4.800 | 3.988 | 4.424 | 4.402 | 3.993 | 4.446 | 0.015 | 0.7282 | 0.8753 | 0 | 0 | 0 |
| 'Cnot1' | 57.797 | 54.687 | 54.457 | 57.949 | 54.703 | 54.048 | 55.142 | 55.546 | 55.205 | 53.542 | 56.570 | 57.805 | 52.445 | 54.216 | 55.988 | 53.617 | 55.518 | 58.494 | -0.007 | 0.8087 | 0.9139 | 0 | 0 | 0 |

| | | | | | | | | | | | | | | | | | | | | | | | | |
|---|---|---|---|---|---|---|---|---|---|---|---|---|---|---|---|---|---|---|---|---|---|---|---|---|
| 'Cnot10' | 29.743 | 28.680 | 29.159 | 29.177 | 26.996 | 28.482 | 28.682 | 28.251 | 29.476 | 28.733 | 28.675 | 28.415 | 28.705 | 27.332 | 28.057 | 27.260 | 29.690 | 27.708 | 0.014 | 0.6273 | 0.8154 | 0 | 0 | 0 |
| 'Cnot11' | 8.977 | 9.675 | 10.138 | 8.737 | 9.255 | 10.286 | 8.847 | 9.731 | 10.127 | 10.141 | 8.956 | 9.945 | 10.286 | 9.320 | 10.062 | 9.976 | 8.679 | 9.879 | -0.034 | 0.3337 | 0.5919 | 0 | 0 | 0 |
| 'Cnot2' | 11.110 | 10.585 | 10.788 | 11.362 | 11.130 | 10.426 | 11.614 | 10.849 | 10.790 | 10.310 | 10.965 | 10.720 | 9.915 | 10.950 | 10.015 | 10.251 | 10.248 | 10.441 | 0.061 | 0.0402 | 0.1639 | 0 | 0 | 0 |
| 'Cnot3' | 14.103 | 13.238 | 14.902 | 13.668 | 12.574 | 14.516 | 13.658 | 13.251 | 14.781 | 13.168 | 12.763 | 16.335 | 13.841 | 13.390 | 15.541 | 13.744 | 13.601 | 15.644 | -0.048 | 0.0607 | 0.2124 | 0 | 0 | 0 |
| 'Cnot4' | 3.309 | 3.058 | 3.056 | 3.376 | 3.037 | 2.992 | 3.539 | 2.940 | 3.140 | 3.243 | 3.218 | 3.502 | 2.945 | 3.152 | 3.336 | 2.899 | 3.515 | 3.474 | -0.056 | 0.2571 | 0.5093 | 0 | 0 | 0 |
| 'Cnot6' | 16.972 | 18.321 | 17.939 | 17.912 | 17.964 | 16.412 | 18.593 | 17.953 | 16.687 | 15.484 | 17.774 | 18.032 | 17.054 | 17.809 | 17.396 | 16.104 | 17.102 | 17.431 | 0.030 | 0.3463 | 0.6015 | 0 | 0 | 0 |
| 'Cnot6l' | 14.770 | 13.928 | 13.929 | 14.892 | 12.829 | 13.054 | 15.936 | 13.385 | 12.751 | 13.502 | 14.792 | 15.938 | 12.336 | 15.042 | 14.712 | 13.693 | 14.045 | 15.511 | -0.058 | 0.2765 | 0.5323 | 0 | 0 | 0 |
| 'Cnot7' | 4.833 | 5.747 | 5.070 | 5.166 | 5.898 | 4.851 | 5.746 | 5.528 | 5.018 | 5.130 | 5.506 | 5.357 | 5.280 | 5.925 | 4.997 | 5.622 | 5.587 | 5.128 | -0.035 | 0.2844 | 0.5407 | 0 | 0 | 0 |
| 'Cnot8' | 37.191 | 38.302 | 36.396 | 36.358 | 39.046 | 36.669 | 36.311 | 37.639 | 37.364 | 33.892 | 36.418 | 41.162 | 36.269 | 39.771 | 39.690 | 36.485 | 41.128 | 40.222 | -0.053 | 0.0938 | 0.2835 | 0 | 0 | 0 |
| 'Cnot9' | 14.071 | 15.301 | 15.886 | 14.414 | 15.214 | 15.255 | 15.134 | 14.998 | 14.635 | 13.259 | 14.191 | 14.617 | 14.878 | 15.026 | 15.968 | 14.470 | 12.907 | 14.854 | 0.041 | 0.2504 | 0.5014 | 0 | 0 | 0 |
| 'Cnp' | 22.346 | 23.800 | 24.457 | 23.420 | 23.581 | 27.239 | 23.648 | 24.090 | 25.547 | 22.194 | 22.088 | 24.941 | 22.815 | 21.512 | 25.423 | 23.652 | 20.828 | 23.802 | 0.063 | 0.0185 | 0.0976 | 0 | 0 | 0 |
| 'Cnppd1' | 14.926 | 13.777 | 12.506 | 12.965 | 13.344 | 13.623 | 12.913 | 13.903 | 13.251 | 14.176 | 13.818 | 14.339 | 14.284 | 14.433 | 14.438 | 13.591 | 13.801 | 13.338 | -0.069 | 0.0306 | 0.1365 | 0 | 0 | 0 |
| 'Cnpy2' | 26.627 | 34.637 | 32.587 | 28.295 | 34.777 | 34.267 | 32.476 | 34.883 | 32.486 | 32.183 | 29.263 | 29.597 | 35.092 | 32.227 | 32.140 | 32.391 | 30.771 | 29.280 | 0.029 | 0.5786 | 0.7820 | 0 | 0 | 0 |
| 'Cnpy3' | 8.625 | 8.397 | 8.501 | 7.396 | 7.803 | 8.600 | 8.025 | 9.138 | 9.100 | 7.467 | 7.549 | 8.882 | 8.683 | 8.162 | 8.036 | 8.906 | 7.795 | 8.884 | 0.014 | 0.7622 | 0.8944 | 0 | 0 | 0 |
| 'Cnpy4' | 11.907 | 12.581 | 12.229 | 13.242 | 12.112 | 12.298 | 11.714 | 12.635 | 12.243 | 10.976 | 13.011 | 12.056 | 12.469 | 12.240 | 12.264 | 11.337 | 13.154 | 11.630 | 0.014 | 0.7211 | 0.8716 | 0 | 0 | 0 |
| 'Cnr2' | 3.976 | 4.118 | 4.266 | 3.512 | 4.108 | 4.224 | 3.522 | 3.764 | 4.417 | 3.230 | 2.966 | 3.041 | 2.576 | 2.510 | 3.390 | 3.001 | 2.626 | 2.690 | 0.459 | 0.0000 | 0.0000 | 1 | 1 | 0 |
| 'Cnst' | 13.949 | 13.241 | 14.084 | 14.481 | 13.495 | 14.538 | 12.601 | 12.715 | 14.232 | 12.675 | 12.302 | 12.454 | 12.434 | 12.034 | 13.177 | 12.236 | 11.575 | 12.602 | 0.136 | 0.0000 | 0.0001 | 1 | 0 | 0 |
| 'Cntln' | 2.486 | 2.608 | 2.310 | 2.541 | 2.839 | 2.622 | 2.730 | 2.765 | 2.502 | 3.195 | 2.538 | 3.002 | 2.714 | 2.962 | 2.720 | 2.680 | 2.994 | 2.660 | -0.139 | 0.0045 | 0.0349 | 1 | 0 | 0 |
| 'Cntrl' | 26.095 | 23.137 | 26.682 | 25.576 | 22.487 | 25.748 | 25.837 | 22.472 | 27.200 | 23.757 | 25.815 | 26.899 | 24.252 | 23.303 | 25.545 | 24.509 | 25.338 | 26.427 | -0.017 | 0.6262 | 0.8144 | 0 | 0 | 0 |
| 'Cntrob' | 4.192 | 3.781 | 5.395 | 4.625 | 4.097 | 4.652 | 4.126 | 4.635 | 5.167 | 4.672 | 4.565 | 4.547 | 4.759 | 4.391 | 4.543 | 4.368 | 4.091 | 4.284 | 0.006 | 0.8987 | 0.9598 | 0 | 0 | 0 |
| 'Coa3' | 29.906 | 37.794 | 33.868 | 35.391 | 37.607 | 38.516 | 33.660 | 34.859 | 32.499 | 31.585 | 35.586 | 34.083 | 34.495 | 34.839 | 32.699 | 33.846 | 33.694 | 31.203 | 0.044 | 0.3084 | 0.5668 | 0 | 0 | 0 |
| 'Coa4' | 5.406 | 6.761 | 6.214 | 4.667 | 6.494 | 5.651 | 6.147 | 6.283 | 5.617 | 5.074 | 5.078 | 5.607 | 5.590 | 5.576 | 5.853 | 4.941 | 5.780 | 5.153 | 0.111 | 0.1011 | 0.2979 | 0 | 0 | 0 |
| 'Coa5' | 26.166 | 29.333 | 27.203 | 27.241 | 30.186 | 26.767 | 26.732 | 29.959 | 26.312 | 28.526 | 27.053 | 27.295 | 25.206 | 28.093 | 27.713 | 27.169 | 27.322 | 26.785 | 0.015 | 0.5944 | 0.7935 | 0 | 0 | 0 |
| 'Coa6' | 18.949 | 20.386 | 18.806 | 22.504 | 21.371 | 18.445 | 20.113 | 20.236 | 19.782 | 19.091 | 20.045 | 20.264 | 18.835 | 20.542 | 19.148 | 18.399 | 19.573 | 18.236 | 0.033 | 0.5577 | 0.7693 | 0 | 0 | 0 |
| 'Coa7' | 1.746 | 2.407 | 2.168 | 2.355 | 2.241 | 2.223 | 2.279 | 1.725 | 2.423 | 2.307 | 1.615 | 1.914 | 1.982 | 2.296 | 1.849 | 2.163 | 1.797 | 1.858 | 0.127 | 0.1274 | 0.3421 | 0 | 0 | 0 |
| 'Coasy' | 6.080 | 6.767 | 6.911 | 6.932 | 5.978 | 6.641 | 6.274 | 6.542 | 7.136 | 6.287 | 5.689 | 6.350 | 6.884 | 6.262 | 6.683 | 6.544 | 6.492 | 6.012 | 0.042 | 0.3400 | 0.5970 | 0 | 0 | 0 |
| 'Cobll1' | 0.524 | 0.573 | 0.790 | 0.563 | 0.563 | 0.621 | 0.543 | 0.656 | 0.572 | 0.679 | 0.832 | 0.768 | 0.770 | 0.889 | 0.699 | 0.703 | 0.693 | 0.712 | -0.321 | 0.0001 | 0.0009 | 1 | 0 | 0 |
| 'Cog1' | 16.366 | 16.069 | 18.755 | 15.581 | 16.467 | 19.391 | 16.832 | 15.925 | 19.231 | 16.720 | 15.695 | 18.529 | 16.545 | 15.510 | 18.261 | 16.139 | 15.735 | 17.083 | 0.031 | 0.2015 | 0.4426 | 0 | 0 | 0 |
| 'Cog2' | 11.363 | 12.348 | 13.802 | 12.466 | 12.374 | 12.732 | 11.854 | 13.051 | 12.952 | 13.215 | 12.253 | 12.232 | 12.769 | 12.448 | 11.752 | 12.326 | 11.768 | 11.484 | 0.026 | 0.4611 | 0.6975 | 0 | 0 | 0 |
| 'Cog3' | 16.921 | 18.285 | 17.563 | 17.623 | 18.606 | 16.822 | 17.969 | 18.504 | 17.555 | 17.817 | 17.865 | 18.376 | 16.325 | 17.545 | 17.946 | 18.563 | 17.928 | 17.224 | -0.010 | 0.7357 | 0.8797 | 0 | 0 | 0 |
| 'Cog4' | 7.247 | 7.134 | 7.685 | 6.847 | 6.967 | 7.279 | 7.057 | 7.146 | 7.205 | 7.035 | 7.361 | 7.986 | 7.272 | 6.862 | 7.607 | 6.663 | 7.483 | 7.406 | -0.037 | 0.2434 | 0.4935 | 0 | 0 | 0 |
| 'Cog5' | 16.873 | 16.648 | 16.503 | 18.773 | 15.223 | 15.755 | 17.551 | 15.191 | 16.979 | 16.794 | 18.832 | 18.274 | 15.879 | 17.134 | 16.058 | 16.232 | 19.461 | 17.475 | -0.075 | 0.1482 | 0.3745 | 0 | 0 | 0 |
| 'Cog6' | 5.168 | 6.187 | 5.928 | 5.431 | 6.628 | 5.858 | 5.794 | 6.681 | 6.160 | 6.190 | 6.162 | 6.073 | 5.945 | 5.911 | 6.334 | 5.607 | 6.288 | 5.697 | -0.024 | 0.5278 | 0.7486 | 0 | 0 | 0 |
| 'Cog7' | 2.922 | 2.711 | 2.951 | 2.551 | 3.259 | 3.324 | 2.335 | 2.897 | 2.854 | 2.647 | 2.670 | 2.597 | 2.750 | 2.817 | 2.325 | 2.755 | 2.679 | 2.104 | 0.140 | 0.0304 | 0.1361 | 0 | 0 | 0 |
| 'Cog8' | 8.813 | 9.066 | 9.012 | 8.132 | 8.664 | 9.239 | 7.915 | 9.230 | 8.306 | 8.385 | 7.963 | 9.592 | 8.149 | 8.865 | 9.651 | 8.454 | 8.789 | 9.104 | -0.021 | 0.5555 | 0.7676 | 0 | 0 | 0 |
| 'Coil' | 5.205 | 5.528 | 6.476 | 5.115 | 5.713 | 5.644 | 5.215 | 6.336 | 6.502 | 5.802 | 5.282 | 5.486 | 5.465 | 5.639 | 5.822 | 4.825 | 5.399 | 5.333 | 0.070 | 0.1762 | 0.4114 | 0 | 0 | 0 |
| 'Col11a2' | 2.158 | 1.827 | 2.723 | 2.181 | 1.870 | 2.394 | 2.160 | 2.066 | 2.491 | 1.606 | 1.474 | 2.535 | 1.812 | 1.523 | 2.362 | 1.775 | 1.887 | 2.689 | 0.165 | 0.0087 | 0.0564 | 0 | 0 | 0 |
| 'Col16a1' | 0.487 | 0.439 | 0.675 | 0.438 | 0.588 | 0.657 | 0.660 | 0.479 | 0.721 | 0.490 | 0.187 | 0.615 | 0.340 | 0.417 | 0.543 | 0.552 | 0.473 | 0.439 | 0.333 | 0.0092 | 0.0587 | 0 | 0 | 0 |
| 'Col1a1' | 1.050 | 0.877 | 1.202 | 0.667 | 0.904 | 0.954 | 0.607 | 1.007 | 1.289 | 1.668 | 1.371 | 1.203 | 1.282 | 1.217 | 1.341 | 1.453 | 1.203 | 1.486 | -0.519 | 0.0000 | 0.0001 | 1 | 1 | 0 |
| 'Col20a1' | 1.818 | 1.793 | 2.340 | 2.069 | 1.669 | 1.806 | 1.658 | 1.658 | 2.109 | 2.397 | 1.784 | 2.572 | 1.876 | 1.754 | 1.952 | 1.782 | 1.931 | 1.934 | -0.104 | 0.2079 | 0.4525 | 0 | 0 | 0 |
| 'Col4a3bp' | 15.326 | 16.856 | 15.657 | 15.702 | 16.014 | 15.147 | 16.281 | 17.279 | 15.541 | 16.049 | 16.857 | 16.522 | 16.030 | 16.502 | 15.380 | 16.116 | 16.901 | 16.758 | -0.046 | 0.0841 | 0.2631 | 0 | 0 | 0 |
| 'Colgalt1' | 19.542 | 19.355 | 20.922 | 17.160 | 19.188 | 20.192 | 17.409 | 22.071 | 21.286 | 19.348 | 18.506 | 20.435 | 19.249 | 18.360 | 19.720 | 20.394 | 18.532 | 18.658 | 0.024 | 0.5676 | 0.7754 | 0 | 0 | 0 |
| 'Colq' | 1.765 | 2.273 | 2.374 | 1.596 | 2.105 | 2.446 | 1.885 | 1.945 | 2.198 | 2.810 | 2.152 | 2.464 | 2.645 | 2.343 | 2.219 | 2.373 | 2.997 | 2.037 | -0.257 | 0.0032 | 0.0266 | 1 | 0 | 0 |
| 'Commd10' | 17.682 | 20.807 | 19.261 | 19.660 | 19.578 | 18.783 | 20.814 | 21.379 | 18.067 | 19.328 | 18.964 | 19.008 | 18.529 | 18.783 | 17.885 | 21.152 | 20.851 | 18.307 | 0.015 | 0.7268 | 0.8750 | 0 | 0 | 0 |
| 'Commd2' | 6.303 | 6.681 | 6.711 | 6.403 | 6.788 | 6.421 | 6.102 | 6.844 | 6.610 | 6.264 | 6.432 | 6.784 | 6.083 | 6.869 | 6.112 | 6.648 | 6.868 | 5.920 | 0.012 | 0.7749 | 0.8994 | 0 | 0 | 0 |
| 'Commd3' | 9.150 | 11.629 | 10.061 | 10.964 | 12.171 | 10.482 | 10.962 | 11.778 | 10.656 | 10.402 | 10.628 | 10.298 | 11.498 | 11.697 | 9.955 | 10.493 | 11.284 | 10.110 | 0.008 | 0.8354 | 0.9289 | 0 | 0 | 0 |
| 'Commd4' | 89.937 | 104.448 | 101.436 | 89.584 | 100.819 | 108.285 | 96.254 | 99.650 | 99.627 | 91.047 | 88.763 | 101.288 | 104.042 | 97.120 | 100.527 | 94.740 | 90.506 | 92.058 | 0.036 | 0.3121 | 0.5708 | 0 | 0 | 0 |
| 'Commd5' | 8.948 | 9.822 | 9.266 | 8.612 | 9.421 | 10.361 | 9.202 | 9.477 | 8.248 | 8.293 | 8.046 | 9.411 | 9.314 | 8.261 | 8.483 | 8.456 | 8.405 | 9.477 | 0.076 | 0.1710 | 0.4043 | 0 | 0 | 0 |
| 'Commd6' | 11.592 | 13.674 | 12.599 | 12.167 | 14.403 | 12.968 | 13.126 | 13.081 | 12.664 | 11.680 | 11.267 | 12.868 | 12.611 | 14.000 | 12.563 | 13.025 | 13.725 | 12.698 | 0.008 | 0.8638 | 0.9407 | 0 | 0 | 0 |
| 'Commd7' | 8.432 | 10.882 | 9.249 | 9.241 | 10.564 | 9.694 | 9.481 | 9.796 | 9.199 | 8.961 | 9.378 | 10.296 | 9.765 | 9.855 | 10.196 | 10.526 | 9.283 | 9.849 | -0.042 | 0.3093 | 0.5675 | 0 | 0 | 0 |
| 'Commd8' | 9.542 | 10.724 | 9.604 | 9.068 | 10.945 | 9.637 | 10.597 | 10.620 | 10.314 | 9.713 | 9.800 | 9.758 | 9.547 | 9.541 | 9.768 | 9.606 | 9.830 | 9.205 | 0.056 | 0.0684 | 0.2294 | 0 | 0 | 0 |
| 'Commd9' | 6.896 | 7.636 | 7.225 | 6.015 | 7.576 | 6.612 | 5.770 | 8.119 | 7.556 | 7.699 | 5.820 | 6.522 | 7.191 | 6.343 | 7.728 | 7.070 | 5.760 | 6.189 | 0.063 | 0.4021 | 0.6510 | 0 | 0 | 0 |
| 'Comt' | 16.047 | 18.066 | 16.717 | 15.111 | 15.956 | 16.787 | 16.023 | 16.394 | 15.343 | 17.164 | 14.283 | 16.171 | 16.630 | 14.637 | 15.426 | 18.060 | 13.239 | 15.939 | 0.041 | 0.3897 | 0.6389 | 0 | 0 | 0 |
| 'Comtd1' | 5.644 | 4.446 | 6.677 | 5.622 | 4.891 | 6.127 | 5.602 | 5.668 | 6.707 | 5.476 | 5.173 | 6.486 | 5.558 | 5.492 | 5.402 | 5.065 | 5.095 | 5.407 | 0.054 | 0.3857 | 0.6354 | 0 | 0 | 0 |
| 'Copa' | 20.935 | 20.873 | 20.949 | 21.731 | 22.382 | 22.016 | 21.690 | 22.026 | 21.901 | 21.991 | 21.700 | 20.805 | 20.480 | 21.129 | 21.188 | 20.838 | 21.756 | 21.248 | 0.015 | 0.5017 | 0.7288 | 0 | 0 | 0 |
| 'Copb1' | 55.854 | 58.575 | 55.866 | 60.719 | 59.058 | 55.188 | 56.751 | 60.127 | 54.776 | 53.958 | 57.024 | 57.379 | 53.567 | 58.080 | 55.590 | 52.391 | 58.702 | 54.779 | 0.032 | 0.2074 | 0.4519 | 0 | 0 | 0 |
| 'Copb2' | 21.957 | 23.858 | 23.683 | 22.897 | 24.481 | 22.760 | 23.577 | 23.547 | 23.879 | 23.678 | 24.485 | 22.560 | 22.635 | 24.026 | 21.282 | 21.165 | 23.491 | 20.502 | 0.038 | 0.1830 | 0.4190 | 0 | 0 | 0 |
| 'Cope' | 13.630 | 14.245 | 14.937 | 13.764 | 14.642 | 15.671 | 13.623 | 14.514 | 14.682 | 13.674 | 13.810 | 15.087 | 14.905 | 13.822 | 15.015 | 15.037 | 13.432 | 14.281 | -0.003 | 0.9134 | 0.9669 | 0 | 0 | 0 |
| 'Copg1' | 12.037 | 12.169 | 13.205 | 11.996 | 12.318 | 12.833 | 11.877 | 12.216 | 13.281 | 12.362 | 12.428 | 13.437 | 12.422 | 12.102 | 12.754 | 12.232 | 12.094 | 13.162 | -0.024 | 0.1888 | 0.4250 | 0 | 0 | 0 |
| 'Copg2' | 7.513 | 8.457 | 8.259 | 8.517 | 8.609 | 8.746 | 9.032 | 8.389 | 8.841 | 8.386 | 8.170 | 8.518 | 8.443 | 7.986 | 7.941 | 7.661 | 8.522 | 7.629 | 0.046 | 0.1907 | 0.4271 | 0 | 0 | 0 |
| 'Cops2' | 15.802 | 17.928 | 16.200 | 16.642 | 16.712 | 15.629 | 16.962 | 16.945 | 14.878 | 15.987 | 16.004 | 16.186 | 15.271 | 17.732 | 14.606 | 15.821 | 16.331 | 14.406 | 0.041 | 0.1873 | 0.4232 | 0 | 0 | 0 |
| 'Cops3' | 9.970 | 11.026 | 10.967 | 10.230 | 10.658 | 10.778 | 9.946 | 11.642 | 10.756 | 10.476 | 10.568 | 10.879 | 10.760 | 10.579 | 10.924 | 10.127 | 10.057 | 9.436 | 0.022 | 0.5155 | 0.7392 | 0 | 0 | 0 |
| 'Cops4' | 9.740 | 9.796 | 9.937 | 9.047 | 10.111 | 9.428 | 10.407 | 10.086 | 9.366 | 9.576 | 8.816 | 9.391 | 10.217 | 8.728 | 9.497 | 9.799 | 9.669 | 9.180 | 0.040 | 0.2277 | 0.4743 | 0 | 0 | 0 |
| 'Cops5' | 8.986 | 10.236 | 9.414 | 9.401 | 10.364 | 9.942 | 9.509 | 9.748 | 9.685 | 8.459 | 8.429 | 9.461 | 9.419 | 9.923 | 9.600 | 9.425 | 9.168 | 9.208 | 0.057 | 0.0941 | 0.2841 | 0 | 0 | 0 |
| 'Cops6' | 26.390 | 27.433 | 26.289 | 26.662 | 28.070 | 26.768 | 27.823 | 27.264 | 26.473 | 25.691 | 24.837 | 23.923 | 26.712 | 26.196 | 25.797 | 25.715 | 26.170 | 24.235 | 0.076 | 0.0020 | 0.0179 | 1 | 0 | 0 |
| 'Cops7a' | 12.259 | 13.379 | 13.458 | 10.519 | 12.869 | 13.092 | 11.851 | 13.980 | 14.096 | 11.695 | 11.602 | 13.958 | 12.275 | 11.977 | 13.131 | 12.466 | 11.214 | 12.581 | 0.046 | 0.2730 | 0.5281 | 0 | 0 | 0 |
| 'Cops7b' | 2.360 | 2.565 | 2.920 | 2.480 | 2.514 | 2.757 | 2.641 | 2.471 | 2.836 | 2.377 | 2.726 | 2.780 | 2.464 | 2.538 | 2.676 | 2.285 | 2.666 | 2.579 | 0.017 | 0.6764 | 0.8451 | 0 | 0 | 0 |
| 'Cops8' | 9.566 | 9.593 | 8.634 | 8.580 | 9.179 | 9.074 | 8.163 | 9.585 | 8.778 | 9.638 | 7.951 | 8.210 | 9.249 | 9.681 | 7.886 | 9.981 | 8.298 | 7.636 | 0.043 | 0.3599 | 0.6123 | 0 | 0 | 0 |
| 'Copz1' | 63.825 | 72.579 | 67.626 | 61.911 | 76.296 | 71.304 | 65.887 | 73.620 | 69.885 | 69.848 | 66.749 | 65.154 | 67.911 | 67.278 | 68.768 | 72.708 | 65.313 | 63.542 | 0.026 | 0.4729 | 0.7086 | 0 | 0 | 0 |
| 'Coq10a' | 9.155 | 10.034 | 10.497 | 9.413 | 11.383 | 10.890 | 10.475 | 11.335 | 10.242 | 10.228 | 8.653 | 9.650 | 9.946 | 10.632 | 9.827 | 9.645 | 9.973 | 9.647 | 0.073 | 0.1287 | 0.3438 | 0 | 0 | 0 |
| 'Coq10b' | 4.434 | 3.377 | 2.597 | 3.849 | 3.251 | 2.675 | 3.356 | 3.532 | 2.585 | 3.916 | 3.329 | 3.005 | 3.503 | 3.760 | 2.744 | 3.587 | 3.816 | 2.495 | -0.038 | 0.5405 | 0.7579 | 0 | 0 | 0 |
| 'Coq2' | 10.214 | 11.440 | 11.732 | 10.367 | 11.716 | 11.675 | 10.526 | 10.865 | 11.540 | 12.000 | 10.608 | 12.067 | 11.315 | 12.558 | 10.755 | 10.568 | 11.057 | 11.061 | -0.040 | 0.3329 | 0.5910 | 0 | 0 | 0 |
| 'Coq3' | 4.297 | 5.383 | 4.863 | 5.319 | 5.470 | 5.059 | 5.609 | 5.229 | 5.612 | 4.580 | 5.475 | 5.224 | 5.009 | 4.729 | 4.870 | 4.603 | 5.314 | 5.210 | 0.038 | 0.5047 | 0.7312 | 0 | 0 | 0 |
| 'Coq4' | 0.714 | 0.686 | 0.780 | 0.556 | 0.691 | 0.683 | 0.511 | 0.631 | 0.736 | 0.479 | 0.479 | 0.647 | 0.744 | 0.643 | 0.639 | 0.693 | 0.558 | 0.573 | 0.128 | 0.2393 | 0.4895 | 0 | 0 | 0 |
| 'Coq5' | 9.296 | 10.090 | 9.576 | 8.536 | 10.132 | 9.502 | 8.315 | 9.490 | 9.470 | 9.442 | 9.294 | 9.370 | 9.367 | 9.015 | 9.490 | 10.050 | 9.772 | 8.954 | -0.014 | 0.7220 | 0.8719 | 0 | 0 | 0 |
| 'Coq6' | 1.791 | 1.910 | 2.197 | 1.906 | 1.953 | 2.308 | 2.061 | 2.166 | 2.266 | 1.960 | 1.795 | 2.133 | 2.029 | 1.594 | 1.844 | 2.120 | 1.967 | 1.977 | 0.082 | 0.1523 | 0.3794 | 0 | 0 | 0 |
| 'Coq7' | 1.852 | 1.454 | 1.720 | 1.411 | 1.543 | 1.754 | 1.599 | 1.722 | 1.645 | 1.778 | 1.559 | 1.649 | 1.627 | 1.940 | 1.847 | 1.648 | 1.469 | 1.502 | -0.028 | 0.7282 | 0.8753 | 0 | 0 | 0 |
| 'Coq8a' | 11.745 | 15.395 | 11.923 | 11.996 | 15.318 | 12.848 | 11.616 | 14.385 | 13.049 | 15.745 | 13.888 | 11.442 | 16.870 | 15.699 | 11.368 | 15.577 | 14.805 | 10.877 | -0.095 | 0.1305 | 0.3469 | 0 | 0 | 0 |
| 'Coq8b' | 4.078 | 3.490 | 3.952 | 3.906 | 3.898 | 4.356 | 3.880 | 4.277 | 4.344 | 4.280 | 3.903 | 3.872 | 4.370 | 3.907 | 4.101 | 4.390 | 3.699 | 3.846 | -0.010 | 0.8300 | 0.9262 | 0 | 0 | 0 |
| 'Coq9' | 9.776 | 10.246 | 10.101 | 10.000 | 10.766 | 10.099 | 9.565 | 9.375 | 9.804 | 10.107 | 8.805 | 10.382 | 9.725 | 9.897 | 10.665 | 9.655 | 10.460 | 10.190 | -0.017 | 0.6686 | 0.8400 | 0 | 0 | 0 |
| 'Coro1a' | 277.509 | 300.500 | 292.582 | 285.378 | 295.159 | 312.295 | 296.635 | 284.073 | 302.571 | 287.816 | 281.999 | 273.053 | 297.765 | 286.565 | 313.561 | 301.584 | 286.525 | 295.624 | 0.001 | 0.9686 | 0.9896 | 0 | 0 | 0 |
| 'Coro1b' | 77.416 | 78.736 | 81.895 | 75.793 | 81.378 | 86.655 | 78.209 | 77.204 | 82.690 | 82.121 | 75.473 | 79.371 | 81.991 | 78.583 | 83.832 | 82.502 | 77.851 | 78.482 | -0.011 | 0.6538 | 0.8322 | 0 | 0 | 0 |
| 'Coro1c' | 11.006 | 12.670 | 11.405 | 10.946 | 12.775 | 12.695 | 10.754 | 11.789 | 12.715 | 10.545 | 10.918 | 11.450 | 10.994 | 12.036 | 12.357 | 10.906 | 10.511 | 12.039 | 0.057 | 0.0984 | 0.2931 | 0 | 0 | 0 |
| 'Coro2b' | 32.325 | 36.988 | 30.882 | 30.885 | 37.907 | 34.328 | 29.506 | 36.017 | 34.061 | 37.744 | 36.746 | 31.035 | 44.147 | 37.958 | 31.274 | 35.899 | 37.836 | 29.048 | -0.092 | 0.0912 | 0.2786 | 0 | 0 | 0 |
| 'Coro7' | 22.221 | 18.561 | 24.914 | 21.542 | 18.566 | 22.302 | 22.097 | 19.775 | 22.812 | 23.637 | 22.227 | 23.920 | 21.995 | 19.838 | 23.342 | 19.172 | 21.737 | 23.658 | -0.064 | 0.1299 | 0.3459 | 0 | 0 | 0 |
| 'Cotl1' | 452.298 | 517.699 | 455.256 | 450.846 | 510.002 | 513.588 | 455.481 | 473.059 | 492.788 | 525.731 | 494.236 | 460.498 | 532.321 | 522.217 | 555.332 | 554.775 | 517.300 | 516.954 | -0.127 | 0.0014 | 0.0139 | 1 | 0 | 0 |
| 'Cox10' | 8.630 | 8.551 | 9.384 | 7.655 | 9.843 | 9.449 | 8.585 | 9.202 | 9.139 | 9.240 | 7.688 | 8.571 | 8.921 | 8.274 | 8.772 | 9.727 | 7.731 | 7.876 | 0.061 | 0.2366 | 0.4863 | 0 | 0 | 0 |
| 'Cox11' | 12.608 | 15.282 | 14.117 | 13.225 | 14.712 | 14.144 | 12.982 | 15.531 | 13.872 | 14.319 | 12.420 | 13.342 | 15.309 | 14.112 | 12.608 | 16.551 | 12.429 | 10.857 | 0.047 | 0.4595 | 0.6963 | 0 | 0 | 0 |
| 'Cox14' | 70.259 | 83.989 | 71.136 | 72.476 | 78.090 | 78.795 | 69.000 | 78.820 | 78.968 | 69.396 | 64.357 | 70.003 | 72.158 | 69.320 | 73.891 | 75.021 | 70.566 | 66.477 | 0.099 | 0.0138 | 0.0790 | 0 | 0 | 0 |
| 'Cox15' | 7.555 | 8.135 | 7.965 | 6.927 | 8.002 | 8.045 | 7.374 | 8.613 | 7.907 | 8.349 | 8.284 | 7.704 | 7.861 | 7.578 | 8.023 | 7.753 | 7.614 | 7.016 | 0.000 | 0.9997 | 0.9999 | 0 | 0 | 0 |
| 'Cox16' | 2.579 | 2.526 | 2.409 | 2.440 | 2.653 | 2.617 | 2.309 | 2.510 | 2.133 | 1.986 | 2.422 | 2.208 | 2.218 | 2.460 | 2.170 | 2.394 | 2.411 | 1.674 | 0.149 | 0.0193 | 0.1007 | 0 | 0 | 0 |
| 'Cox18' | 4.740 | 5.833 | 5.888 | 4.383 | 5.466 | 5.721 | 4.719 | 5.873 | 5.193 | 5.082 | 5.839 | 5.296 | 5.935 | 5.613 | 5.238 | 5.129 | 4.550 | 4.792 | -0.001 | 0.9801 | 0.9926 | 0 | 0 | 0 |
| 'Cox19' | 3.527 | 3.447 | 3.430 | 3.056 | 3.683 | 3.702 | 3.169 | 3.325 | 3.355 | 3.527 | 3.161 | 3.517 | 3.441 | 3.963 | 3.494 | 3.062 | 3.230 | 3.294 | -0.010 | 0.8501 | 0.9354 | 0 | 0 | 0 |
| 'Cox20' | 41.013 | 41.256 | 39.276 | 42.313 | 43.069 | 36.954 | 46.610 | 42.952 | 40.901 | 36.948 | 40.435 | 42.013 | 43.045 | 43.602 | 39.471 | 36.660 | 44.714 | 34.614 | 0.036 | 0.4394 | 0.6829 | 0 | 0 | 0 |
| 'Cox4i1' | 96.154 | 106.329 | 92.153 | 99.262 | 107.466 | 99.919 | 102.184 | 100.861 | 100.377 | 91.470 | 94.036 | 85.931 | 99.964 | 95.931 | 96.067 | 102.546 | 97.050 | 91.572 | 0.072 | 0.0220 | 0.1096 | 0 | 0 | 0 |
| 'Cox5a' | 97.739 | 104.948 | 104.822 | 104.314 | 105.426 | 105.020 | 99.399 | 96.109 | 100.223 | 89.891 | 98.242 | 99.019 | 97.912 | 103.413 | 105.146 | 89.822 | 97.678 | 97.594 | 0.052 | 0.1234 | 0.3356 | 0 | 0 | 0 |
| 'Cox5b' | 45.177 | 50.517 | 44.305 | 45.975 | 48.363 | 47.131 | 47.789 | 50.308 | 45.784 | 41.303 | 48.230 | 40.067 | 47.369 | 45.880 | 44.978 | 46.277 | 48.151 | 41.556 | 0.066 | 0.0396 | 0.1621 | 0 | 0 | 0 |
| 'Cox6b1' | 197.273 | 226.974 | 209.673 | 216.853 | 221.612 | 210.450 | 218.208 | 219.715 | 204.723 | 198.703 | 202.830 | 194.648 | 213.747 | 208.962 | 214.420 | 220.727 | 227.863 | 197.176 | 0.025 | 0.4367 | 0.6801 | 0 | 0 | 0 |
| 'Cox6c' | 20.324 | 24.727 | 20.724 | 22.201 | 23.522 | 20.828 | 22.822 | 23.073 | 20.962 | 21.375 | 21.446 | 18.892 | 21.450 | 24.421 | 20.597 | 22.754 | 22.544 | 19.080 | 0.040 | 0.2216 | 0.4691 | 0 | 0 | 0 |
| 'Cox7a2' | 75.292 | 86.987 | 72.704 | 80.332 | 84.966 | 77.008 | 73.175 | 77.082 | 72.306 | 68.597 | 76.538 | 68.162 | 75.126 | 77.969 | 75.195 | 73.460 | 78.017 | 67.623 | 0.072 | 0.0471 | 0.1808 | 0 | 0 | 0 |
| 'Cox7a2l' | 147.775 | 147.197 | 142.970 | 152.069 | 150.985 | 147.648 | 149.721 | 153.235 | 141.172 | 151.190 | 146.342 | 144.078 | 147.965 | 154.405 | 150.595 | 152.685 | 147.599 | 141.514 | -0.013 | 0.5367 | 0.7556 | 0 | 0 | 0 |
| 'Cox7b' | 46.438 | 50.143 | 44.660 | 48.547 | 51.861 | 48.728 | 50.369 | 48.545 | 44.486 | 41.780 | 43.816 | 42.922 | 46.803 | 45.848 | 44.757 | 42.411 | 47.731 | 42.847 | 0.107 | 0.0010 | 0.0102 | 1 | 0 | 0 |
| 'Cox7c' | 39.075 | 46.181 | 37.773 | 43.438 | 45.097 | 40.725 | 44.301 | 42.627 | 39.483 | 39.359 | 41.540 | 35.662 | 42.658 | 44.842 | 36.654 | 41.307 | 42.640 | 35.781 | 0.062 | 0.0586 | 0.2086 | 0 | 0 | 0 |
| 'Cox8a' | 281.321 | 299.712 | 286.839 | 294.789 | 291.040 | 281.673 | 277.699 | 304.492 | 275.097 | 269.754 | 285.453 | 241.099 | 291.380 | 291.923 | 286.481 | 275.514 | 294.521 | 257.952 | 0.048 | 0.1465 | 0.3719 | 0 | 0 | 0 |
| 'Cpd' | 1.822 | 2.188 | 2.106 | 2.061 | 2.181 | 2.098 | 1.998 | 2.381 | 2.009 | 2.154 | 2.779 | 2.065 | 1.901 | 2.660 | 2.034 | 2.224 | 1.904 | 2.076 | -0.074 | 0.2435 | 0.4936 | 0 | 0 | 0 |
| 'Cpeb1' | 1.810 | 3.132 | 2.305 | 2.320 | 2.686 | 2.715 | 1.864 | 2.657 | 2.771 | 2.218 | 2.372 | 1.865 | 2.543 | 2.349 | 1.960 | 2.102 | 1.890 | 1.945 | 0.192 | 0.0270 | 0.1265 | 0 | 0 | 0 |
| 'Cpeb2' | 1.730 | 1.619 | 1.387 | 1.760 | 1.462 | 1.430 | 1.661 | 1.525 | 1.272 | 1.598 | 2.086 | 1.898 | 1.930 | 2.197 | 1.636 | 1.764 | 2.026 | 2.150 | -0.336 | 0.0000 | 0.0000 | 1 | 0 | 0 |
| 'Cpeb4' | 7.615 | 7.826 | 6.726 | 7.765 | 7.758 | 6.410 | 8.162 | 8.195 | 6.352 | 8.220 | 7.794 | 6.839 | 8.097 | 8.122 | 6.299 | 8.947 | 7.913 | 6.671 | -0.053 | 0.0973 | 0.2907 | 0 | 0 | 0 |
| 'Cpm' | 1.294 | 2.013 | 1.256 | 1.405 | 2.037 | 1.402 | 1.417 | 1.859 | 1.162 | 1.366 | 1.184 | 1.084 | 1.309 | 1.394 | 1.145 | 1.316 | 1.647 | 1.122 | 0.232 | 0.0013 | 0.0131 | 1 | 0 | 0 |
| 'Cpne1' | 0.287 | 0.329 | 0.319 | 0.273 | 0.232 | 0.277 | 0.353 | 0.219 | 0.281 | 0.236 | 0.282 | 0.324 | 0.240 | 0.278 | 0.257 | 0.369 | 0.259 | 0.248 | 0.028 | 0.8129 | 0.9157 | 0 | 0 | 0 |
| 'Cpne2' | 1.005 | 1.299 | 1.238 | 1.404 | 1.146 | 1.245 | 1.298 | 0.976 | 1.284 | 1.207 | 1.058 | 1.273 | 1.222 | 1.216 | 1.331 | 1.149 | 1.181 | 1.251 | -0.020 | 0.8129 | 0.9157 | 0 | 0 | 0 |
| 'Cpne3' | 5.736 | 6.809 | 6.671 | 6.660 | 6.565 | 6.117 | 6.631 | 6.716 | 6.413 | 6.318 | 7.034 | 7.007 | 5.858 | 7.353 | 6.261 | 5.906 | 6.850 | 6.681 | -0.035 | 0.3778 | 0.6296 | 0 | 0 | 0 |
| 'Cpne5' | 0.580 | 0.779 | 0.550 | 0.619 | 0.633 | 0.675 | 0.640 | 0.632 | 0.724 | 0.479 | 0.407 | 0.305 | 0.625 | 0.390 | 0.424 | 0.560 | 0.495 | 0.342 | 0.549 | 0.0000 | 0.0000 | 1 | 1 | 0 |
| 'Cpne7' | 0.317 | 0.451 | 0.383 | 0.250 | 0.466 | 0.499 | 0.397 | 0.518 | 0.420 | 0.432 | 0.378 | 0.452 | 0.303 | 0.395 | 0.460 | 0.346 | 0.486 | 0.359 | 0.016 | 0.8829 | 0.9507 | 0 | 0 | 0 |
| 'Cpox' | 12.605 | 13.026 | 13.258 | 13.595 | 13.866 | 13.853 | 12.747 | 14.179 | 13.254 | 12.334 | 11.900 | 14.222 | 13.418 | 13.761 | 14.634 | 12.331 | 15.592 | 14.766 | -0.043 | 0.3375 | 0.5955 | 0 | 0 | 0 |
| 'Cpped1' | 1.322 | 1.272 | 1.377 | 1.494 | 1.326 | 1.150 | 1.390 | 1.320 | 1.360 | 1.596 | 1.348 | 1.382 | 1.522 | 1.150 | 1.440 | 1.468 | 1.257 | 1.475 | -0.084 | 0.1845 | 0.4199 | 0 | 0 | 0 |
| 'Cpq' | 4.332 | 5.140 | 5.301 | 5.160 | 5.071 | 5.131 | 4.096 | 4.710 | 4.224 | 3.213 | 2.986 | 3.335 | 3.199 | 3.379 | 3.235 | 2.862 | 2.524 | 3.152 | 0.614 | 0.0000 | 0.0000 | 1 | 1 | 0 |
| 'Cpsf1' | 44.862 | 43.337 | 48.173 | 42.968 | 42.774 | 47.881 | 42.533 | 44.111 | 47.076 | 46.598 | 42.426 | 49.153 | 48.093 | 41.292 | 45.039 | 47.069 | 44.103 | 47.868 | -0.038 | 0.1343 | 0.3530 | 0 | 0 | 0 |
| 'Cpsf2' | 19.704 | 18.717 | 19.294 | 20.300 | 19.113 | 19.150 | 21.569 | 20.582 | 18.672 | 18.031 | 19.198 | 20.151 | 19.169 | 19.937 | 18.557 | 17.383 | 18.542 | 19.393 | 0.043 | 0.2300 | 0.4768 | 0 | 0 | 0 |
| 'Cpsf3' | 47.591 | 46.521 | 48.180 | 48.700 | 49.686 | 48.985 | 47.834 | 49.807 | 49.782 | 49.746 | 46.070 | 47.770 | 47.451 | 46.273 | 45.506 | 46.923 | 49.796 | 42.569 | 0.041 | 0.1327 | 0.3506 | 0 | 0 | 0 |
| 'Cpsf3l' | 15.006 | 15.271 | 16.715 | 16.573 | 15.741 | 17.068 | 15.429 | 15.770 | 16.149 | 15.821 | 15.588 | 17.701 | 16.924 | 15.890 | 17.108 | 16.658 | 15.081 | 17.718 | -0.058 | 0.0295 | 0.1335 | 0 | 0 | 0 |
| 'Cpsf4' | 7.438 | 7.383 | 7.639 | 7.117 | 7.701 | 7.947 | 7.139 | 7.922 | 7.537 | 7.094 | 7.362 | 7.591 | 7.671 | 7.245 | 7.497 | 7.599 | 7.135 | 7.695 | 0.011 | 0.7147 | 0.8683 | 0 | 0 | 0 |
| 'Cpsf6' | 11.840 | 12.539 | 12.768 | 11.957 | 11.974 | 11.489 | 13.486 | 11.938 | 12.402 | 10.152 | 12.679 | 13.922 | 11.330 | 12.754 | 12.875 | 10.381 | 13.588 | 13.623 | -0.017 | 0.7427 | 0.8839 | 0 | 0 | 0 |
| 'Cpsf7' | 46.242 | 46.784 | 52.581 | 44.804 | 46.925 | 48.155 | 48.084 | 47.942 | 49.127 | 47.638 | 47.618 | 49.252 | 48.635 | 47.006 | 46.248 | 44.707 | 49.578 | 44.088 | 0.010 | 0.7397 | 0.8819 | 0 | 0 | 0 |
| 'Cpt1a' | 94.884 | 89.110 | 70.383 | 95.418 | 83.819 | 69.525 | 92.955 | 90.087 | 72.305 | 84.683 | 78.731 | 84.216 | 91.184 | 80.271 | 78.404 | 84.431 | 82.714 | 84.863 | -0.004 | 0.9340 | 0.9748 | 0 | 0 | 0 |
| 'Cpt2' | 2.718 | 3.050 | 3.790 | 2.711 | 3.390 | 3.750 | 2.851 | 4.132 | 3.528 | 2.668 | 3.061 | 3.248 | 3.340 | 3.171 | 3.515 | 2.765 | 3.539 | 3.032 | 0.067 | 0.3048 | 0.5622 | 0 | 0 | 0 |
| 'Cptp' | 3.331 | 3.997 | 3.375 | 3.351 | 3.402 | 3.372 | 4.091 | 3.565 | 3.421 | 3.084 | 3.222 | 3.583 | 3.106 | 2.987 | 3.363 | 3.633 | 3.154 | 4.021 | 0.063 | 0.3693 | 0.6203 | 0 | 0 | 0 |
| 'Cr1l' | 5.691 | 6.459 | 6.111 | 5.692 | 6.154 | 6.040 | 6.087 | 6.253 | 5.629 | 6.171 | 6.407 | 6.610 | 5.789 | 6.368 | 6.517 | 5.897 | 6.347 | 6.444 | -0.077 | 0.0089 | 0.0575 | 0 | 0 | 0 |
| 'Cracr2a' | 7.702 | 7.149 | 8.317 | 6.882 | 7.221 | 8.543 | 6.390 | 7.546 | 8.641 | 10.990 | 8.581 | 8.054 | 9.393 | 7.965 | 7.869 | 8.122 | 8.593 | 7.747 | -0.177 | 0.0205 | 0.1051 | 0 | 0 | 0 |
| 'Cradd' | 4.769 | 6.312 | 5.465 | 5.570 | 5.900 | 6.454 | 6.165 | 7.358 | 6.251 | 6.743 | 6.569 | 5.203 | 5.789 | 5.912 | 5.888 | 6.163 | 5.524 | 5.199 | 0.032 | 0.6879 | 0.8525 | 0 | 0 | 0 |
| 'Cramp1l' | 13.074 | 10.885 | 12.699 | 12.647 | 10.694 | 11.235 | 11.854 | 11.412 | 12.907 | 12.449 | 12.559 | 14.583 | 12.509 | 11.831 | 13.591 | 13.278 | 12.734 | 15.526 | -0.158 | 0.0003 | 0.0042 | 1 | 0 | 0 |
| 'Crat' | 1.746 | 1.928 | 1.806 | 1.594 | 1.844 | 1.881 | 1.758 | 2.048 | 1.726 | 1.855 | 1.688 | 1.727 | 1.798 | 1.811 | 1.636 | 1.768 | 1.603 | 1.551 | 0.072 | 0.1344 | 0.3530 | 0 | 0 | 0 |
| 'Crbn' | 18.277 | 21.650 | 19.093 | 17.076 | 21.537 | 18.612 | 19.421 | 22.144 | 18.506 | 18.860 | 20.012 | 17.184 | 17.855 | 19.952 | 17.188 | 19.055 | 19.768 | 16.712 | 0.072 | 0.0107 | 0.0656 | 0 | 0 | 0 |
| 'Crcp' | 9.613 | 9.424 | 9.954 | 10.117 | 10.447 | 10.533 | 9.303 | 10.258 | 9.579 | 9.550 | 8.835 | 9.485 | 9.784 | 9.315 | 9.772 | 9.359 | 9.705 | 9.113 | 0.063 | 0.0961 | 0.2883 | 0 | 0 | 0 |
| 'Creb1' | 6.226 | 6.083 | 5.845 | 6.404 | 6.142 | 5.620 | 6.646 | 6.269 | 5.576 | 5.567 | 6.130 | 6.608 | 5.339 | 6.078 | 5.677 | 5.375 | 6.523 | 6.157 | 0.024 | 0.5997 | 0.7967 | 0 | 0 | 0 |
| 'Creb3' | 8.874 | 8.580 | 8.466 | 8.222 | 9.271 | 8.668 | 8.652 | 8.961 | 8.092 | 8.478 | 8.192 | 8.928 | 8.437 | 7.993 | 8.757 | 10.021 | 7.924 | 8.169 | 0.006 | 0.8785 | 0.9485 | 0 | 0 | 0 |
| 'Creb3l2' | 1.266 | 1.650 | 1.351 | 1.616 | 1.528 | 1.776 | 1.430 | 1.648 | 1.961 | 1.622 | 1.604 | 1.652 | 1.574 | 1.478 | 1.414 | 1.612 | 1.927 | 1.493 | -0.027 | 0.7325 | 0.8781 | 0 | 0 | 0 |
| 'Crebbp' | 9.472 | 8.257 | 7.972 | 9.355 | 7.845 | 7.492 | 8.500 | 8.831 | 8.406 | 9.149 | 8.883 | 8.988 | 8.355 | 9.006 | 8.562 | 8.967 | 8.481 | 9.313 | -0.079 | 0.0592 | 0.2100 | 0 | 0 | 0 |
| 'Crebl2' | 5.846 | 7.513 | 6.073 | 5.384 | 6.870 | 6.559 | 5.595 | 7.419 | 6.279 | 5.428 | 6.157 | 6.837 | 5.004 | 6.769 | 7.025 | 6.196 | 6.107 | 6.668 | 0.020 | 0.6922 | 0.8554 | 0 | 0 | 0 |
| 'Crebrf' | 4.576 | 4.698 | 4.967 | 4.732 | 5.130 | 4.737 | 5.908 | 4.804 | 5.466 | 4.361 | 6.092 | 6.466 | 4.267 | 5.142 | 5.792 | 4.560 | 5.889 | 5.837 | -0.106 | 0.1206 | 0.3311 | 0 | 0 | 0 |
| 'Crebzf' | 9.032 | 10.261 | 12.023 | 8.647 | 10.110 | 11.676 | 10.078 | 9.963 | 11.408 | 8.144 | 10.646 | 12.670 | 9.324 | 10.336 | 12.158 | 9.091 | 10.717 | 11.950 | -0.036 | 0.2896 | 0.5461 | 0 | 0 | 0 |
| 'Creg1' | 4.758 | 4.460 | 4.697 | 4.677 | 5.223 | 4.859 | 4.392 | 4.635 | 4.587 | 5.105 | 4.149 | 5.627 | 4.212 | 5.044 | 4.965 | 5.099 | 4.269 | 5.011 | -0.054 | 0.3578 | 0.6103 | 0 | 0 | 0 |
| 'Creld1' | 2.165 | 2.783 | 2.529 | 2.679 | 2.372 | 2.596 | 2.248 | 2.473 | 2.387 | 2.398 | 2.163 | 2.322 | 2.212 | 2.166 | 1.946 | 2.237 | 2.286 | 2.034 | 0.157 | 0.0067 | 0.0463 | 1 | 0 | 0 |
| 'Creld2' | 40.174 | 38.364 | 40.037 | 40.521 | 43.023 | 46.665 | 35.787 | 44.151 | 45.726 | 41.326 | 35.499 | 40.713 | 42.968 | 41.699 | 40.430 | 44.540 | 39.141 | 38.224 | 0.031 | 0.5344 | 0.7542 | 0 | 0 | 0 |
| 'Crem' | 0.388 | 0.381 | 0.288 | 0.361 | 0.334 | 0.376 | 0.344 | 0.385 | 0.334 | 0.431 | 0.477 | 0.409 | 0.351 | 0.470 | 0.408 | 0.426 | 0.492 | 0.386 | -0.276 | 0.0002 | 0.0032 | 1 | 0 | 0 |
| 'Crim1' | 2.892 | 3.311 | 3.207 | 3.081 | 3.411 | 3.776 | 3.196 | 3.453 | 3.573 | 3.213 | 2.876 | 2.500 | 2.704 | 3.150 | 2.156 | 3.233 | 2.848 | 2.457 | 0.253 | 0.0005 | 0.0057 | 1 | 0 | 0 |
| 'Crip1' | 2.566 | 2.480 | 2.570 | 2.683 | 2.525 | 3.011 | 2.600 | 2.886 | 2.740 | 2.498 | 2.721 | 3.169 | 2.956 | 3.159 | 3.172 | 2.535 | 2.761 | 2.785 | -0.111 | 0.0743 | 0.2424 | 0 | 0 | 0 |
| 'Cript' | 11.307 | 12.347 | 11.693 | 10.850 | 10.732 | 10.858 | 11.629 | 10.489 | 10.998 | 10.657 | 11.301 | 13.046 | 10.003 | 12.041 | 12.095 | 9.970 | 13.168 | 11.406 | -0.053 | 0.3316 | 0.5897 | 0 | 0 | 0 |
| 'Crk' | 8.620 | 9.310 | 8.505 | 8.782 | 7.995 | 7.907 | 9.471 | 8.398 | 8.615 | 7.564 | 8.955 | 10.081 | 7.539 | 8.897 | 9.137 | 8.454 | 9.736 | 9.676 | -0.053 | 0.3318 | 0.5898 | 0 | 0 | 0 |
| 'Crkl' | 29.587 | 28.006 | 28.031 | 28.844 | 26.761 | 27.333 | 27.851 | 25.429 | 28.454 | 27.057 | 29.471 | 31.853 | 26.544 | 29.143 | 30.752 | 28.240 | 27.984 | 33.566 | -0.090 | 0.0313 | 0.1386 | 0 | 0 | 0 |
| 'Crlf2' | 34.850 | 35.767 | 40.927 | 29.253 | 33.802 | 38.754 | 31.717 | 36.602 | 39.139 | 38.983 | 33.248 | 38.679 | 34.964 | 35.410 | 38.405 | 40.309 | 32.600 | 39.216 | -0.059 | 0.2032 | 0.4456 | 0 | 0 | 0 |
| 'Crlf3' | 36.564 | 35.529 | 34.899 | 36.474 | 35.325 | 35.253 | 36.944 | 35.239 | 33.848 | 29.438 | 34.709 | 38.590 | 30.850 | 35.014 | 38.116 | 33.674 | 35.783 | 40.384 | 0.009 | 0.8454 | 0.9343 | 0 | 0 | 0 |
| 'Crls1' | 3.717 | 3.882 | 3.696 | 3.462 | 4.004 | 3.692 | 3.754 | 4.008 | 3.505 | 3.381 | 3.147 | 4.046 | 3.152 | 3.851 | 4.396 | 3.510 | 3.448 | 3.718 | 0.031 | 0.6231 | 0.8119 | 0 | 0 | 0 |
| 'Crmp1' | 0.559 | 0.426 | 0.581 | 0.421 | 0.472 | 0.571 | 0.475 | 0.566 | 0.653 | 0.443 | 0.390 | 0.669 | 0.395 | 0.434 | 0.619 | 0.365 | 0.372 | 0.508 | 0.160 | 0.1125 | 0.3183 | 0 | 0 | 0 |
| 'Crnkl1' | 18.203 | 18.937 | 18.580 | 18.314 | 19.287 | 18.944 | 21.040 | 18.892 | 19.248 | 18.594 | 21.738 | 19.739 | 18.690 | 18.836 | 18.588 | 17.700 | 18.892 | 18.145 | -0.007 | 0.8513 | 0.9360 | 0 | 0 | 0 |
| 'Crocc' | 2.539 | 2.191 | 2.907 | 2.476 | 2.277 | 2.617 | 2.594 | 2.130 | 2.883 | 2.729 | 2.434 | 2.681 | 2.683 | 2.577 | 2.347 | 2.452 | 2.630 | 2.483 | -0.037 | 0.4712 | 0.7069 | 0 | 0 | 0 |
| 'Crot' | 9.439 | 11.324 | 10.574 | 10.253 | 10.312 | 10.278 | 10.653 | 11.195 | 9.918 | 9.700 | 10.016 | 10.954 | 9.246 | 10.303 | 9.987 | 10.006 | 11.479 | 9.630 | 0.028 | 0.4459 | 0.6878 | 0 | 0 | 0 |
| 'Crtam' | 1.086 | 0.648 | 1.564 | 1.171 | 1.113 | 1.300 | 1.173 | 1.249 | 1.025 | 0.895 | 0.698 | 1.093 | 1.044 | 0.592 | 1.076 | 0.710 | 0.505 | 0.979 | 0.446 | 0.0003 | 0.0040 | 1 | 1 | 0 |
| 'Crtap' | 6.333 | 6.341 | 6.693 | 6.012 | 6.895 | 6.694 | 5.222 | 6.194 | 6.174 | 5.472 | 4.919 | 6.349 | 6.437 | 5.458 | 6.851 | 6.727 | 5.359 | 6.870 | 0.046 | 0.4870 | 0.7175 | 0 | 0 | 0 |
| 'Crtc1' | 3.799 | 3.577 | 3.693 | 3.106 | 3.473 | 3.804 | 3.366 | 3.551 | 3.981 | 3.846 | 3.313 | 3.935 | 3.657 | 3.300 | 3.492 | 3.639 | 3.110 | 3.603 | 0.011 | 0.7895 | 0.9061 | 0 | 0 | 0 |
| 'Crtc2' | 16.354 | 15.728 | 16.314 | 15.664 | 15.001 | 16.394 | 16.301 | 15.772 | 16.571 | 16.550 | 15.976 | 17.099 | 17.084 | 15.152 | 16.818 | 16.082 | 16.583 | 16.825 | -0.051 | 0.0289 | 0.1317 | 0 | 0 | 0 |
| 'Crtc3' | 12.176 | 11.027 | 12.426 | 10.406 | 11.456 | 12.477 | 10.379 | 11.710 | 12.611 | 13.161 | 12.552 | 12.993 | 12.029 | 12.335 | 13.142 | 12.579 | 11.877 | 12.387 | -0.121 | 0.0005 | 0.0064 | 1 | 0 | 0 |
| 'Cry1' | 5.849 | 5.232 | 5.947 | 5.274 | 5.760 | 5.122 | 5.543 | 5.528 | 5.481 | 5.184 | 5.982 | 6.530 | 5.214 | 5.339 | 6.347 | 5.567 | 5.596 | 6.032 | -0.072 | 0.2074 | 0.4519 | 0 | 0 | 0 |
| 'Cry2' | 3.539 | 4.149 | 3.772 | 3.056 | 3.563 | 3.507 | 3.085 | 3.820 | 3.613 | 4.422 | 3.722 | 3.671 | 3.904 | 3.815 | 3.691 | 4.127 | 3.422 | 3.276 | -0.091 | 0.1348 | 0.3536 | 0 | 0 | 0 |
| 'Cryl1' | 11.187 | 12.909 | 11.900 | 12.735 | 14.246 | 11.479 | 11.467 | 14.760 | 13.186 | 11.559 | 12.452 | 13.743 | 12.385 | 11.362 | 13.363 | 12.986 | 12.209 | 13.450 | -0.015 | 0.8066 | 0.9134 | 0 | 0 | 0 |
| 'Cryz' | 0.498 | 0.511 | 0.482 | 0.514 | 0.549 | 0.560 | 0.449 | 0.535 | 0.504 | 0.604 | 0.401 | 0.429 | 0.536 | 0.437 | 0.477 | 0.550 | 0.559 | 0.345 | 0.091 | 0.3644 | 0.6162 | 0 | 0 | 0 |
| 'Cryzl1' | 4.106 | 5.659 | 4.731 | 4.681 | 5.261 | 4.677 | 4.915 | 5.255 | 4.933 | 4.562 | 4.283 | 4.383 | 4.446 | 4.762 | 4.006 | 4.423 | 4.516 | 3.782 | 0.160 | 0.0004 | 0.0047 | 1 | 0 | 0 |
| 'Cs' | 87.740 | 91.133 | 93.991 | 90.434 | 92.998 | 93.676 | 90.559 | 91.681 | 92.039 | 87.064 | 87.306 | 91.544 | 94.400 | 88.610 | 97.134 | 89.315 | 89.907 | 90.648 | 0.004 | 0.8522 | 0.9367 | 0 | 0 | 0 |
| 'Csad' | 9.076 | 8.745 | 10.574 | 7.567 | 8.813 | 10.273 | 8.786 | 9.493 | 10.173 | 9.437 | 9.958 | 10.032 | 9.534 | 9.027 | 8.833 | 8.783 | 9.608 | 8.628 | -0.012 | 0.8325 | 0.9270 | 0 | 0 | 0 |
| 'Csde1' | 18.392 | 20.394 | 19.088 | 20.215 | 20.243 | 18.543 | 20.767 | 20.898 | 17.968 | 19.265 | 20.290 | 17.984 | 18.964 | 19.999 | 17.222 | 19.581 | 20.525 | 17.207 | 0.036 | 0.1155 | 0.3228 | 0 | 0 | 0 |
| 'Cse1l' | 7.793 | 8.222 | 8.715 | 8.441 | 8.453 | 8.075 | 8.902 | 8.187 | 7.967 | 7.031 | 7.517 | 8.543 | 7.992 | 8.573 | 8.188 | 7.172 | 8.465 | 8.122 | 0.050 | 0.2088 | 0.4535 | 0 | 0 | 0 |
| 'Csf1' | 26.167 | 21.878 | 27.856 | 22.457 | 22.626 | 24.122 | 23.030 | 23.341 | 25.781 | 24.368 | 24.233 | 29.100 | 17.884 | 22.043 | 26.690 | 23.085 | 22.280 | 29.203 | -0.014 | 0.8014 | 0.9114 | 0 | 0 | 0 |
| 'Csf2ra' | 9.073 | 7.382 | 9.379 | 6.522 | 8.124 | 9.800 | 7.546 | 8.288 | 9.208 | 10.278 | 6.546 | 10.282 | 7.792 | 7.156 | 9.710 | 8.424 | 8.388 | 9.717 | -0.061 | 0.3848 | 0.6347 | 0 | 0 | 0 |
| 'Csgalnact2' | 2.324 | 3.331 | 2.736 | 2.433 | 3.247 | 3.018 | 2.914 | 3.180 | 2.524 | 3.252 | 3.011 | 2.657 | 3.089 | 3.422 | 2.464 | 2.764 | 2.944 | 2.274 | -0.019 | 0.7660 | 0.8956 | 0 | 0 | 0 |
| 'Csk' | 253.740 | 274.084 | 265.670 | 261.936 | 261.829 | 277.204 | 259.447 | 260.570 | 273.270 | 250.644 | 238.027 | 268.291 | 246.276 | 246.518 | 272.722 | 254.419 | 232.697 | 276.176 | 0.053 | 0.0283 | 0.1303 | 0 | 0 | 0 |
| 'Csnk1a1' | 37.111 | 41.312 | 37.166 | 38.979 | 40.959 | 36.465 | 39.759 | 40.654 | 36.342 | 39.930 | 39.873 | 36.220 | 38.624 | 41.365 | 36.022 | 39.457 | 40.871 | 36.329 | -0.010 | 0.5387 | 0.7567 | 0 | 0 | 0 |
| 'Csnk1d' | 36.781 | 38.371 | 35.828 | 34.805 | 37.233 | 37.436 | 34.102 | 37.882 | 36.911 | 36.004 | 37.007 | 39.017 | 34.928 | 36.820 | 39.248 | 38.012 | 36.968 | 39.131 | -0.044 | 0.0778 | 0.2501 | 0 | 0 | 0 |
| 'Csnk1e' | 3.948 | 4.213 | 4.589 | 4.063 | 4.014 | 4.634 | 4.099 | 3.984 | 4.223 | 4.260 | 3.408 | 3.750 | 3.757 | 3.891 | 3.751 | 3.571 | 3.406 | 3.860 | 0.158 | 0.0003 | 0.0041 | 1 | 0 | 0 |
| 'Csnk1g1' | 2.620 | 2.650 | 2.876 | 2.653 | 2.709 | 2.574 | 2.771 | 2.869 | 2.718 | 2.629 | 2.967 | 2.987 | 2.750 | 2.742 | 2.839 | 2.616 | 3.047 | 2.938 | -0.073 | 0.0288 | 0.1315 | 0 | 0 | 0 |
| 'Csnk1g2' | 165.616 | 175.101 | 168.390 | 158.164 | 168.654 | 193.824 | 160.363 | 169.879 | 173.959 | 174.236 | 161.009 | 171.214 | 178.066 | 172.302 | 182.810 | 193.770 | 161.173 | 183.728 | -0.052 | 0.1629 | 0.3947 | 0 | 0 | 0 |
| 'Csnk1g3' | 20.389 | 21.501 | 19.021 | 21.324 | 21.406 | 20.332 | 21.907 | 22.010 | 20.070 | 19.345 | 21.505 | 22.005 | 19.845 | 22.395 | 22.405 | 19.026 | 22.334 | 23.109 | -0.042 | 0.2941 | 0.5500 | 0 | 0 | 0 |
| 'Csnk2a1' | 27.914 | 31.360 | 28.479 | 28.973 | 31.046 | 28.145 | 28.866 | 31.479 | 28.241 | 28.643 | 30.271 | 30.296 | 28.244 | 31.246 | 29.043 | 28.804 | 30.454 | 30.187 | -0.028 | 0.1786 | 0.4140 | 0 | 0 | 0 |
| 'Csnk2a2' | 15.358 | 15.965 | 15.450 | 16.550 | 15.814 | 14.521 | 14.424 | 15.522 | 14.717 | 14.714 | 14.994 | 15.753 | 14.878 | 15.546 | 15.756 | 14.441 | 16.235 | 16.041 | -0.014 | 0.7005 | 0.8602 | 0 | 0 | 0 |
| 'Csnk2b' | 32.018 | 31.801 | 33.022 | 32.378 | 33.051 | 34.268 | 34.168 | 32.141 | 33.765 | 30.177 | 32.166 | 32.333 | 32.829 | 31.894 | 33.975 | 30.182 | 33.161 | 32.539 | 0.026 | 0.3384 | 0.5957 | 0 | 0 | 0 |
| 'Cspp1' | 1.616 | 1.570 | 1.886 | 1.696 | 1.755 | 1.542 | 1.792 | 1.645 | 1.941 | 1.939 | 1.863 | 1.770 | 1.856 | 1.569 | 1.603 | 1.715 | 1.732 | 1.673 | -0.035 | 0.4946 | 0.7237 | 0 | 0 | 0 |
| 'Csrnp1' | 74.747 | 39.294 | 27.038 | 44.693 | 39.102 | 30.535 | 33.300 | 45.298 | 26.034 | 79.665 | 44.760 | 26.465 | 62.773 | 59.794 | 27.228 | 85.310 | 38.344 | 24.727 | -0.241 | 0.1091 | 0.3110 | 0 | 0 | 0 |
| 'Csrnp2' | 3.499 | 2.702 | 3.500 | 3.434 | 3.189 | 3.489 | 3.425 | 3.503 | 3.436 | 3.603 | 3.442 | 4.283 | 3.119 | 3.913 | 3.536 | 3.462 | 3.575 | 3.905 | -0.132 | 0.0396 | 0.1621 | 0 | 0 | 0 |
| 'Csrp1' | 57.310 | 58.791 | 57.222 | 56.834 | 55.473 | 63.401 | 56.217 | 55.066 | 60.671 | 78.541 | 74.288 | 71.855 | 81.870 | 74.682 | 80.077 | 76.493 | 73.028 | 73.383 | -0.402 | 0.0000 | 0.0000 | 1 | 1 | 0 |
| 'Csrp2bp' | 4.875 | 5.270 | 5.526 | 5.167 | 5.078 | 5.641 | 5.156 | 5.244 | 5.479 | 5.154 | 5.253 | 5.596 | 5.197 | 5.541 | 5.619 | 5.578 | 5.475 | 5.334 | -0.047 | 0.1531 | 0.3805 | 0 | 0 | 0 |
| 'Cst3' | 31.105 | 35.504 | 30.936 | 32.196 | 35.311 | 34.660 | 29.482 | 36.394 | 32.020 | 31.255 | 34.482 | 32.453 | 30.945 | 33.846 | 34.535 | 38.017 | 33.466 | 32.169 | -0.028 | 0.5084 | 0.7336 | 0 | 0 | 0 |
| 'Cst7' | 69.280 | 57.210 | 78.005 | 67.102 | 57.261 | 66.307 | 58.911 | 65.014 | 71.600 | 59.218 | 55.016 | 64.560 | 41.232 | 51.872 | 66.183 | 50.271 | 49.482 | 64.534 | 0.229 | 0.0002 | 0.0032 | 1 | 0 | 0 |
| 'Cstb' | 68.442 | 72.429 | 61.591 | 63.766 | 68.027 | 62.027 | 59.871 | 60.215 | 60.503 | 63.748 | 61.776 | 61.117 | 64.593 | 68.701 | 60.705 | 58.833 | 69.349 | 53.027 | 0.027 | 0.5466 | 0.7610 | 0 | 0 | 0 |
| 'Cstf1' | 12.914 | 14.761 | 15.392 | 12.535 | 13.636 | 14.914 | 13.178 | 14.506 | 15.057 | 12.579 | 13.669 | 16.040 | 13.022 | 13.041 | 15.409 | 14.057 | 13.382 | 15.518 | -0.010 | 0.7603 | 0.8934 | 0 | 0 | 0 |
| 'Cstf2' | 6.823 | 7.130 | 7.450 | 6.765 | 7.150 | 6.954 | 6.670 | 6.908 | 6.851 | 6.350 | 6.770 | 7.111 | 7.429 | 7.273 | 6.967 | 7.080 | 7.131 | 7.010 | -0.020 | 0.4993 | 0.7273 | 0 | 0 | 0 |
| 'Cstf2t' | 15.448 | 16.507 | 16.175 | 16.348 | 16.644 | 15.084 | 16.430 | 16.519 | 17.067 | 13.828 | 15.851 | 16.793 | 13.839 | 15.657 | 15.747 | 14.117 | 16.486 | 16.522 | 0.062 | 0.1066 | 0.3067 | 0 | 0 | 0 |
| 'Cstf3' | 6.317 | 7.542 | 7.226 | 7.578 | 7.021 | 6.942 | 8.291 | 7.556 | 7.546 | 6.993 | 7.608 | 7.433 | 6.814 | 7.531 | 7.037 | 6.888 | 8.035 | 6.698 | 0.010 | 0.8133 | 0.9157 | 0 | 0 | 0 |
| 'Ctage5' | 12.374 | 11.194 | 12.288 | 12.740 | 11.628 | 11.328 | 12.975 | 11.715 | 11.202 | 11.980 | 11.422 | 11.879 | 11.692 | 11.834 | 11.675 | 11.404 | 11.570 | 12.009 | 0.015 | 0.6204 | 0.8101 | 0 | 0 | 0 |
| 'Ctbp1' | 30.672 | 33.727 | 31.666 | 29.871 | 33.610 | 33.824 | 30.129 | 33.264 | 31.345 | 30.453 | 30.604 | 32.725 | 32.504 | 33.111 | 34.630 | 35.579 | 30.848 | 33.546 | -0.041 | 0.2286 | 0.4752 | 0 | 0 | 0 |
| 'Ctbs' | 0.724 | 1.042 | 0.805 | 0.859 | 0.944 | 0.745 | 0.763 | 0.846 | 0.863 | 0.971 | 0.766 | 0.722 | 1.012 | 0.976 | 0.829 | 0.868 | 0.678 | 0.628 | 0.019 | 0.8420 | 0.9331 | 0 | 0 | 0 |
| 'Ctc1' | 3.519 | 3.505 | 4.186 | 3.310 | 3.573 | 4.332 | 3.617 | 3.629 | 4.167 | 4.272 | 3.552 | 3.753 | 3.974 | 3.611 | 3.824 | 3.633 | 3.212 | 3.396 | 0.017 | 0.7397 | 0.8819 | 0 | 0 | 0 |
| 'Ctcf' | 42.884 | 45.022 | 44.697 | 45.830 | 45.550 | 44.525 | 43.889 | 45.307 | 44.757 | 44.784 | 44.616 | 41.739 | 45.300 | 46.318 | 43.248 | 44.587 | 46.201 | 40.682 | 0.010 | 0.6663 | 0.8391 | 0 | 0 | 0 |
| 'Ctdnep1' | 35.948 | 37.369 | 37.279 | 35.741 | 36.017 | 38.824 | 33.928 | 37.082 | 37.719 | 35.896 | 35.621 | 38.019 | 35.092 | 35.688 | 36.360 | 36.662 | 33.354 | 37.257 | 0.016 | 0.5355 | 0.7551 | 0 | 0 | 0 |
| 'Ctdp1' | 16.772 | 17.118 | 17.583 | 16.316 | 16.633 | 18.054 | 15.314 | 17.919 | 16.230 | 17.099 | 18.054 | 17.372 | 17.072 | 16.694 | 16.749 | 18.136 | 16.976 | 17.006 | -0.038 | 0.2436 | 0.4936 | 0 | 0 | 0 |
| 'Ctdsp1' | 36.678 | 40.047 | 42.653 | 32.330 | 39.429 | 45.683 | 32.813 | 44.976 | 43.705 | 37.058 | 35.871 | 39.014 | 36.725 | 32.762 | 39.858 | 38.305 | 33.029 | 38.022 | 0.103 | 0.0468 | 0.1803 | 0 | 0 | 0 |
| 'Ctdsp2' | 39.722 | 42.644 | 36.319 | 37.699 | 42.147 | 40.890 | 36.339 | 43.224 | 38.760 | 38.821 | 41.924 | 45.962 | 35.708 | 44.140 | 44.549 | 42.639 | 41.626 | 45.760 | -0.102 | 0.0128 | 0.0750 | 0 | 0 | 0 |
| 'Ctdspl' | 1.255 | 1.475 | 1.695 | 1.511 | 1.145 | 1.583 | 1.465 | 1.260 | 1.835 | 1.085 | 1.169 | 1.722 | 1.526 | 1.056 | 1.797 | 1.319 | 1.211 | 1.650 | 0.060 | 0.4343 | 0.6788 | 0 | 0 | 0 |
| 'Ctdspl2' | 4.327 | 4.258 | 4.592 | 4.908 | 4.595 | 4.188 | 4.817 | 4.224 | 4.348 | 3.735 | 4.468 | 4.754 | 4.253 | 4.433 | 4.554 | 3.808 | 4.768 | 4.558 | 0.023 | 0.6424 | 0.8247 | 0 | 0 | 0 |
| 'Ctif' | 0.580 | 0.688 | 0.667 | 0.658 | 0.629 | 0.930 | 0.739 | 0.719 | 0.838 | 0.792 | 1.008 | 0.531 | 0.708 | 0.503 | 0.574 | 0.712 | 0.538 | 0.486 | 0.153 | 0.2794 | 0.5363 | 0 | 0 | 0 |
| 'Ctla4' | 119.827 | 102.933 | 61.986 | 87.521 | 111.617 | 64.696 | 90.971 | 104.919 | 57.726 | 129.870 | 107.380 | 63.054 | 155.993 | 130.044 | 62.133 | 127.277 | 111.092 | 61.874 | -0.216 | 0.0062 | 0.0440 | 1 | 0 | 0 |
| 'Ctnna1' | 6.398 | 5.473 | 7.863 | 8.259 | 7.605 | 8.053 | 7.087 | 6.998 | 8.102 | 7.831 | 8.942 | 9.149 | 7.511 | 9.201 | 8.808 | 6.795 | 8.161 | 9.338 | -0.211 | 0.0013 | 0.0129 | 1 | 0 | 0 |
| 'Ctnnb1' | 14.700 | 16.416 | 16.529 | 15.634 | 15.473 | 17.310 | 15.892 | 15.926 | 16.919 | 12.643 | 14.921 | 18.739 | 13.636 | 15.300 | 18.021 | 14.223 | 15.705 | 18.833 | 0.026 | 0.5270 | 0.7483 | 0 | 0 | 0 |
| 'Ctnnbip1' | 5.505 | 6.789 | 6.343 | 6.036 | 5.861 | 7.496 | 6.062 | 6.528 | 6.226 | 5.871 | 5.542 | 5.735 | 5.286 | 6.051 | 5.768 | 6.135 | 4.658 | 4.993 | 0.177 | 0.0050 | 0.0375 | 1 | 0 | 0 |
| 'Ctnnbl1' | 23.449 | 22.294 | 25.238 | 27.017 | 25.153 | 25.456 | 25.254 | 26.025 | 24.109 | 25.705 | 24.266 | 24.126 | 25.561 | 23.642 | 25.867 | 25.702 | 23.345 | 23.567 | 0.006 | 0.8592 | 0.9390 | 0 | 0 | 0 |
| 'Ctns' | 2.906 | 3.386 | 3.355 | 3.701 | 3.461 | 3.601 | 3.255 | 3.252 | 3.521 | 3.433 | 2.969 | 3.412 | 3.310 | 3.483 | 3.357 | 3.280 | 3.185 | 3.231 | 0.024 | 0.6410 | 0.8239 | 0 | 0 | 0 |
| 'Ctps' | 9.284 | 7.817 | 8.998 | 8.917 | 8.553 | 8.205 | 9.081 | 9.140 | 8.355 | 8.648 | 8.282 | 8.701 | 8.974 | 8.157 | 8.280 | 7.411 | 8.192 | 8.490 | 0.048 | 0.2667 | 0.5201 | 0 | 0 | 0 |

| | | | | | | | | | | | | | | | | | | | | | | | | |
|---|---|---|---|---|---|---|---|---|---|---|---|---|---|---|---|---|---|---|---|---|---|---|---|---|
| 'Ctps2' | 7.737 | 8.670 | 7.873 | 8.450 | 9.068 | 7.595 | 7.869 | 8.648 | 7.741 | 8.755 | 8.603 | 8.094 | 8.336 | 8.831 | 7.705 | 8.136 | 8.443 | 7.411 | -0.025 | 0.3607 | 0.6132 | 0 | 0 | 0 |
| 'Ctr9' | 7.461 | 7.515 | 8.068 | 7.938 | 7.618 | 8.406 | 7.631 | 7.982 | 8.241 | 8.249 | 8.402 | 8.217 | 8.074 | 7.926 | 7.983 | 7.642 | 7.690 | 7.946 | -0.034 | 0.2306 | 0.4777 | 0 | 0 | 0 |
| 'Ctsa' | 27.475 | 26.998 | 30.195 | 26.224 | 27.654 | 28.737 | 25.145 | 28.464 | 28.032 | 25.725 | 26.223 | 27.334 | 25.470 | 26.953 | 27.762 | 24.753 | 25.274 | 26.680 | 0.066 | 0.0056 | 0.0407 | 1 | 0 | 0 |
| 'Ctsb' | 137.059 | 144.295 | 130.267 | 139.116 | 147.628 | 134.727 | 132.919 | 142.596 | 130.325 | 153.280 | 162.496 | 140.589 | 147.120 | 160.789 | 150.549 | 160.058 | 157.545 | 147.985 | -0.166 | 0.0000 | 0.0000 | 1 | 0 | 0 |
| 'Ctsc' | 6.295 | 6.245 | 6.612 | 6.064 | 5.711 | 6.125 | 6.379 | 6.366 | 5.492 | 5.295 | 5.672 | 6.696 | 5.153 | 5.140 | 5.842 | 5.777 | 5.646 | 6.125 | 0.095 | 0.0567 | 0.2046 | 0 | 0 | 0 |
| 'Ctsd' | 56.722 | 56.957 | 54.965 | 54.128 | 55.469 | 59.434 | 52.487 | 56.204 | 55.076 | 57.600 | 53.701 | 59.008 | 55.982 | 55.877 | 59.065 | 60.016 | 52.121 | 59.158 | -0.041 | 0.1411 | 0.3636 | 0 | 0 | 0 |
| 'Ctse' | 88.339 | 102.616 | 88.598 | 86.431 | 100.837 | 94.462 | 87.564 | 98.438 | 91.825 | 110.918 | 102.610 | 99.577 | 104.856 | 106.308 | 103.276 | 112.034 | 103.905 | 95.791 | -0.174 | 0.0000 | 0.0000 | 1 | 0 | 0 |
| 'Ctsl' | 35.499 | 46.794 | 27.404 | 38.292 | 43.262 | 33.714 | 36.954 | 37.412 | 35.072 | 44.544 | 43.122 | 34.759 | 53.214 | 50.247 | 36.478 | 50.321 | 49.390 | 35.376 | -0.255 | 0.0000 | 0.0002 | 1 | 0 | 0 |
| 'Ctso' | 8.564 | 10.970 | 8.785 | 8.592 | 9.971 | 9.008 | 8.933 | 10.361 | 9.801 | 10.239 | 10.284 | 9.486 | 9.739 | 9.869 | 9.233 | 10.097 | 10.375 | 8.816 | -0.065 | 0.0828 | 0.2605 | 0 | 0 | 0 |
| 'Ctss' | 49.442 | 58.199 | 48.753 | 48.408 | 56.977 | 54.490 | 50.572 | 57.946 | 52.010 | 59.185 | 55.696 | 61.519 | 58.741 | 62.977 | 60.674 | 62.225 | 62.051 | 59.885 | -0.202 | 0.0000 | 0.0000 | 1 | 0 | 0 |
| 'Ctsw' | 6.495 | 7.196 | 7.951 | 5.197 | 6.172 | 9.304 | 8.521 | 7.089 | 8.973 | 7.817 | 5.368 | 10.199 | 5.271 | 7.695 | 9.978 | 10.074 | 8.317 | 9.880 | -0.163 | 0.1288 | 0.3439 | 0 | 0 | 0 |
| 'Ctsz' | 152.636 | 169.160 | 141.645 | 157.253 | 172.355 | 141.908 | 146.189 | 173.715 | 141.486 | 140.768 | 155.276 | 118.972 | 175.794 | 154.061 | 127.564 | 142.640 | 154.486 | 120.727 | 0.107 | 0.0078 | 0.0522 | 0 | 0 | 0 |
| 'Cttn' | 2.715 | 2.834 | 3.087 | 2.595 | 3.286 | 3.038 | 3.255 | 2.781 | 2.514 | 3.182 | 2.559 | 2.723 | 2.862 | 3.015 | 2.664 | 2.557 | 3.021 | 2.378 | 0.055 | 0.3852 | 0.6349 | 0 | 0 | 0 |
| 'Ctu1' | 5.272 | 5.230 | 5.874 | 3.657 | 5.623 | 6.703 | 4.211 | 5.990 | 5.583 | 5.116 | 4.263 | 5.403 | 4.498 | 4.780 | 5.594 | 5.479 | 4.534 | 5.108 | 0.096 | 0.2195 | 0.4665 | 0 | 0 | 0 |
| 'Ctu2' | 3.737 | 3.805 | 4.301 | 3.931 | 4.009 | 4.607 | 3.355 | 3.904 | 3.930 | 4.035 | 3.183 | 4.076 | 3.742 | 3.680 | 4.171 | 3.874 | 3.565 | 3.726 | 0.054 | 0.2412 | 0.4920 | 0 | 0 | 0 |
| 'Cuedc1' | 0.446 | 0.537 | 0.499 | 0.519 | 0.627 | 0.564 | 0.532 | 0.516 | 0.509 | 0.510 | 0.671 | 0.367 | 0.551 | 0.410 | 0.405 | 0.693 | 0.602 | 0.302 | 0.101 | 0.4129 | 0.6611 | 0 | 0 | 0 |
| 'Cuedc2' | 11.412 | 11.439 | 11.759 | 13.005 | 12.419 | 13.592 | 14.331 | 12.611 | 12.006 | 10.305 | 10.384 | 10.386 | 12.286 | 11.699 | 9.914 | 11.485 | 10.449 | 10.981 | 0.192 | 0.0004 | 0.0050 | 1 | 0 | 0 |
| 'Cul1' | 20.764 | 22.112 | 21.340 | 22.726 | 21.856 | 19.692 | 21.739 | 22.475 | 20.530 | 21.136 | 21.129 | 20.885 | 19.909 | 23.333 | 20.130 | 20.126 | 21.938 | 19.979 | 0.024 | 0.3992 | 0.6482 | 0 | 0 | 0 |
| 'Cul2' | 3.851 | 4.106 | 4.006 | 4.332 | 4.659 | 3.799 | 4.545 | 4.278 | 4.164 | 4.389 | 4.165 | 4.291 | 4.226 | 3.978 | 4.028 | 3.880 | 4.683 | 4.186 | -0.020 | 0.6478 | 0.8283 | 0 | 0 | 0 |
| 'Cul3' | 23.938 | 25.382 | 25.438 | 24.494 | 25.948 | 24.907 | 26.162 | 27.352 | 24.062 | 25.259 | 24.584 | 25.124 | 23.865 | 27.283 | 24.547 | 26.479 | 27.221 | 24.598 | -0.018 | 0.5152 | 0.7390 | 0 | 0 | 0 |
| 'Cul4a' | 9.184 | 9.553 | 9.529 | 8.585 | 9.666 | 9.154 | 8.698 | 10.389 | 9.114 | 9.103 | 9.126 | 9.314 | 9.579 | 9.840 | 9.038 | 9.569 | 9.260 | 9.242 | -0.013 | 0.6635 | 0.8375 | 0 | 0 | 0 |
| 'Cul4b' | 10.321 | 10.349 | 10.488 | 10.661 | 10.823 | 10.049 | 10.404 | 11.391 | 9.941 | 10.825 | 11.417 | 11.769 | 10.428 | 11.302 | 10.225 | 10.283 | 11.106 | 11.110 | -0.074 | 0.0287 | 0.1312 | 0 | 0 | 0 |
| 'Cul5' | 5.128 | 5.401 | 5.162 | 5.174 | 5.213 | 4.994 | 5.249 | 5.169 | 4.944 | 5.263 | 4.895 | 5.235 | 4.724 | 5.263 | 4.924 | 5.136 | 5.394 | 4.732 | 0.015 | 0.6188 | 0.8093 | 0 | 0 | 0 |
| 'Cul7' | 2.560 | 2.634 | 2.824 | 2.594 | 2.622 | 2.745 | 2.436 | 2.515 | 2.900 | 2.413 | 2.410 | 2.502 | 2.770 | 2.345 | 2.342 | 2.326 | 2.386 | 2.468 | 0.107 | 0.0080 | 0.0530 | 0 | 0 | 0 |
| 'Cul9' | 1.095 | 0.906 | 1.127 | 0.955 | 1.034 | 1.076 | 1.051 | 1.104 | 1.183 | 0.880 | 1.219 | 0.845 | 0.865 | 0.819 | 0.927 | 0.678 | 0.848 | 0.824 | 0.266 | 0.0006 | 0.0073 | 1 | 0 | 0 |
| 'Cuta' | 38.063 | 42.989 | 42.234 | 44.611 | 44.305 | 46.161 | 40.240 | 42.282 | 40.582 | 39.362 | 39.815 | 45.766 | 46.285 | 39.763 | 45.378 | 39.179 | 42.169 | 43.595 | -0.016 | 0.7227 | 0.8724 | 0 | 0 | 0 |
| 'Cutc' | 3.080 | 3.199 | 3.554 | 2.875 | 4.524 | 3.454 | 3.020 | 4.157 | 3.638 | 3.485 | 3.831 | 2.862 | 2.955 | 3.803 | 3.209 | 3.964 | 3.440 | 2.381 | 0.079 | 0.3741 | 0.6253 | 0 | 0 | 0 |
| 'Cux1' | 3.549 | 3.004 | 3.725 | 3.501 | 3.381 | 3.630 | 3.522 | 3.277 | 3.607 | 3.803 | 3.868 | 3.743 | 3.926 | 3.511 | 3.377 | 3.393 | 3.659 | 3.693 | -0.091 | 0.0210 | 0.1063 | 0 | 0 | 0 |
| 'Cwc15' | 78.553 | 81.633 | 84.425 | 79.491 | 87.302 | 86.378 | 81.528 | 85.797 | 82.880 | 79.409 | 76.959 | 73.851 | 81.587 | 77.719 | 77.557 | 80.885 | 77.540 | 75.472 | 0.087 | 0.0033 | 0.0269 | 1 | 0 | 0 |
| 'Cwc22' | 4.269 | 4.583 | 4.676 | 4.784 | 4.609 | 4.908 | 4.847 | 4.426 | 4.924 | 4.652 | 4.599 | 4.230 | 4.455 | 4.560 | 4.255 | 4.150 | 4.509 | 3.854 | 0.090 | 0.0264 | 0.1247 | 0 | 0 | 0 |
| 'Cwc25' | 7.532 | 6.757 | 7.362 | 6.828 | 7.023 | 7.409 | 7.649 | 7.134 | 7.363 | 7.352 | 6.952 | 6.923 | 7.249 | 6.517 | 6.844 | 7.069 | 7.189 | 6.626 | 0.047 | 0.1569 | 0.3866 | 0 | 0 | 0 |
| 'Cwc27' | 8.493 | 8.367 | 7.539 | 8.024 | 6.906 | 7.598 | 8.720 | 7.320 | 7.184 | 7.243 | 6.760 | 7.308 | 7.214 | 6.790 | 8.089 | 7.192 | 8.303 | 7.434 | 0.066 | 0.2294 | 0.4764 | 0 | 0 | 0 |
| 'Cwf19l1' | 5.273 | 5.905 | 5.949 | 6.198 | 5.841 | 6.495 | 5.553 | 6.200 | 6.661 | 5.333 | 5.595 | 6.438 | 5.955 | 5.572 | 5.802 | 6.473 | 5.909 | 5.774 | 0.021 | 0.6836 | 0.8500 | 0 | 0 | 0 |
| 'Cwf19l2' | 7.863 | 10.221 | 9.491 | 9.309 | 9.891 | 9.326 | 9.382 | 9.015 | 8.993 | 9.319 | 10.635 | 9.376 | 8.780 | 8.867 | 8.109 | 7.965 | 9.609 | 8.009 | 0.037 | 0.4625 | 0.6985 | 0 | 0 | 0 |
| 'Cxcr4' | 96.281 | 99.196 | 69.440 | 87.759 | 100.042 | 72.084 | 84.785 | 108.868 | 78.158 | 99.583 | 101.209 | 82.782 | 86.925 | 114.275 | 77.912 | 107.629 | 106.731 | 82.402 | -0.124 | 0.0031 | 0.0262 | 1 | 0 | 0 |
| 'Cxcr5' | 286.946 | 293.602 | 275.005 | 320.241 | 286.469 | 280.297 | 326.977 | 254.389 | 296.298 | 249.658 | 299.667 | 294.130 | 284.530 | 257.754 | 296.917 | 239.072 | 285.866 | 310.258 | 0.047 | 0.3894 | 0.6388 | 0 | 0 | 0 |
| 'Cxcr6' | 24.506 | 23.416 | 24.940 | 27.487 | 19.442 | 21.666 | 28.823 | 22.044 | 22.762 | 14.515 | 17.759 | 21.432 | 14.702 | 15.722 | 18.572 | 15.338 | 16.016 | 19.456 | 0.475 | 0.0000 | 0.0000 | 1 | 1 | 0 |
| 'Cxx1b' | 37.437 | 40.132 | 36.000 | 35.132 | 40.557 | 43.879 | 38.217 | 38.030 | 40.658 | 38.878 | 35.163 | 37.167 | 36.909 | 36.962 | 41.305 | 37.880 | 34.760 | 35.468 | 0.056 | 0.2084 | 0.4530 | 0 | 0 | 0 |
| 'Cxx1c' | 43.960 | 53.217 | 47.172 | 48.451 | 46.693 | 46.973 | 43.571 | 46.252 | 52.943 | 52.903 | 48.888 | 50.900 | 49.127 | 50.212 | 50.424 | 49.673 | 47.623 | 47.943 | -0.073 | 0.1500 | 0.3772 | 0 | 0 | 0 |
| 'Cxxc1' | 76.523 | 70.718 | 79.663 | 72.833 | 68.244 | 79.861 | 71.720 | 70.114 | 79.633 | 79.363 | 70.876 | 77.588 | 79.490 | 70.356 | 75.254 | 76.169 | 71.410 | 76.379 | -0.025 | 0.2965 | 0.5524 | 0 | 0 | 0 |
| 'Cxxc5' | 49.357 | 69.093 | 43.917 | 51.300 | 59.325 | 52.631 | 49.468 | 57.191 | 50.502 | 49.286 | 48.986 | 51.449 | 55.340 | 62.840 | 51.019 | 57.145 | 57.084 | 52.587 | -0.027 | 0.6075 | 0.8026 | 0 | 0 | 0 |
| 'Cyb561a3' | 11.645 | 14.478 | 14.671 | 12.033 | 14.163 | 14.577 | 12.113 | 14.229 | 15.255 | 14.940 | 14.795 | 14.977 | 14.328 | 13.611 | 13.211 | 13.098 | 12.862 | 13.043 | -0.033 | 0.5562 | 0.7679 | 0 | 0 | 0 |
| 'Cyb561d1' | 3.244 | 4.011 | 3.596 | 3.186 | 4.081 | 3.941 | 3.148 | 4.129 | 3.889 | 3.678 | 3.218 | 4.078 | 3.686 | 3.998 | 3.976 | 3.732 | 3.582 | 3.917 | -0.044 | 0.3506 | 0.6047 | 0 | 0 | 0 |
| 'Cyb561d2' | 2.764 | 2.906 | 3.343 | 3.275 | 3.517 | 3.747 | 3.047 | 3.836 | 3.788 | 2.982 | 3.148 | 3.465 | 3.168 | 3.087 | 3.244 | 3.375 | 3.070 | 3.021 | 0.072 | 0.2511 | 0.5023 | 0 | 0 | 0 |
| 'Cyb5a' | 16.710 | 18.724 | 16.659 | 17.870 | 17.360 | 17.252 | 18.374 | 18.399 | 16.258 | 15.551 | 17.593 | 17.590 | 17.612 | 17.762 | 17.888 | 16.186 | 16.403 | 16.629 | 0.026 | 0.4794 | 0.7118 | 0 | 0 | 0 |
| 'Cyb5b' | 37.425 | 41.787 | 38.729 | 39.936 | 38.060 | 38.478 | 39.654 | 39.054 | 38.766 | 35.195 | 39.989 | 41.619 | 38.271 | 41.332 | 41.664 | 35.681 | 39.656 | 40.728 | -0.020 | 0.5320 | 0.7526 | 0 | 0 | 0 |
| 'Cyb5d2' | 1.408 | 1.197 | 1.528 | 1.455 | 1.389 | 1.551 | 1.525 | 1.273 | 1.441 | 1.608 | 1.452 | 1.543 | 1.361 | 1.462 | 1.387 | 1.392 | 1.572 | 1.358 | -0.048 | 0.4782 | 0.7112 | 0 | 0 | 0 |
| 'Cyb5r1' | 3.030 | 3.594 | 4.130 | 3.381 | 3.341 | 3.744 | 3.066 | 3.518 | 3.726 | 3.162 | 3.418 | 3.824 | 3.471 | 3.311 | 3.784 | 3.268 | 3.184 | 3.481 | 0.018 | 0.6919 | 0.8552 | 0 | 0 | 0 |
| 'Cyb5r3' | 18.698 | 21.689 | 19.816 | 17.121 | 20.351 | 20.480 | 18.348 | 20.491 | 21.368 | 21.554 | 20.213 | 20.433 | 21.231 | 18.561 | 19.605 | 22.337 | 17.986 | 19.638 | -0.036 | 0.4426 | 0.6862 | 0 | 0 | 0 |
| 'Cyb5rl' | 0.534 | 0.477 | 0.646 | 0.504 | 0.559 | 0.551 | 0.525 | 0.592 | 0.675 | 0.580 | 0.497 | 0.570 | 0.528 | 0.516 | 0.485 | 0.624 | 0.523 | 0.509 | 0.067 | 0.3799 | 0.6316 | 0 | 0 | 0 |
| 'Cyba' | 146.423 | 158.255 | 156.602 | 153.017 | 154.080 | 160.877 | 152.958 | 154.274 | 161.334 | 157.613 | 140.843 | 155.541 | 148.162 | 154.632 | 158.236 | 166.460 | 147.488 | 155.487 | 0.004 | 0.8777 | 0.9485 | 0 | 0 | 0 |
| 'Cybb' | 11.904 | 15.105 | 11.176 | 11.813 | 17.034 | 11.244 | 11.948 | 14.184 | 10.408 | 14.962 | 19.359 | 14.004 | 16.670 | 18.734 | 12.567 | 16.078 | 21.528 | 12.319 | -0.354 | 0.0000 | 0.0000 | 1 | 0 | 0 |
| 'Cyc1' | 76.103 | 82.134 | 90.820 | 78.527 | 88.503 | 90.215 | 73.936 | 87.915 | 88.323 | 78.740 | 71.807 | 85.685 | 92.323 | 80.318 | 88.379 | 88.480 | 76.644 | 78.714 | 0.019 | 0.6821 | 0.8487 | 0 | 0 | 0 |
| 'Cyfip1' | 9.901 | 13.008 | 9.394 | 10.180 | 12.013 | 9.877 | 10.145 | 11.179 | 10.278 | 11.596 | 11.767 | 10.205 | 12.171 | 12.601 | 9.720 | 11.908 | 12.217 | 9.763 | -0.098 | 0.0063 | 0.0446 | 1 | 0 | 0 |
| 'Cyfip2' | 61.955 | 56.587 | 60.235 | 58.622 | 57.083 | 58.396 | 56.543 | 58.613 | 57.053 | 61.451 | 59.923 | 57.684 | 58.929 | 58.782 | 59.140 | 58.387 | 57.045 | 58.793 | -0.024 | 0.3148 | 0.5724 | 0 | 0 | 0 |
| 'Cyhr1' | 19.251 | 18.172 | 20.209 | 18.258 | 18.958 | 21.074 | 16.964 | 18.706 | 18.725 | 19.496 | 18.607 | 21.183 | 19.258 | 18.637 | 20.884 | 20.483 | 17.591 | 22.432 | -0.077 | 0.0264 | 0.1245 | 0 | 0 | 0 |
| 'Cyld' | 32.279 | 34.747 | 32.412 | 35.391 | 34.988 | 30.914 | 32.992 | 34.644 | 31.039 | 32.681 | 35.624 | 34.312 | 33.597 | 37.036 | 31.332 | 32.444 | 35.143 | 33.177 | -0.040 | 0.1242 | 0.3369 | 0 | 0 | 0 |
| 'Cyp20a1' | 2.081 | 2.482 | 2.634 | 2.172 | 2.422 | 2.385 | 2.766 | 2.874 | 2.441 | 2.832 | 2.976 | 3.188 | 2.688 | 2.669 | 2.671 | 2.847 | 2.901 | 3.040 | -0.229 | 0.0005 | 0.0061 | 1 | 0 | 0 |
| 'Cyp2s1' | 0.822 | 0.785 | 0.844 | 0.680 | 0.681 | 0.941 | 0.786 | 0.669 | 0.914 | 0.887 | 0.738 | 1.290 | 0.594 | 0.670 | 1.091 | 0.795 | 0.547 | 1.104 | -0.116 | 0.2538 | 0.5055 | 0 | 0 | 0 |
| 'Cyp4f13' | 4.023 | 4.465 | 4.154 | 3.694 | 4.530 | 4.431 | 4.500 | 4.501 | 4.821 | 6.208 | 5.294 | 4.986 | 6.283 | 5.062 | 4.504 | 5.893 | 5.178 | 4.224 | -0.284 | 0.0000 | 0.0003 | 1 | 0 | 0 |
| 'Cyp4f17' | 4.555 | 4.310 | 4.402 | 3.565 | 4.025 | 4.593 | 3.757 | 4.426 | 4.415 | 4.535 | 4.464 | 4.069 | 4.255 | 3.994 | 3.665 | 4.118 | 4.076 | 4.320 | 0.023 | 0.7741 | 0.8993 | 0 | 0 | 0 |
| 'Cyp4v3' | 3.183 | 4.043 | 3.512 | 3.557 | 4.178 | 3.267 | 4.113 | 4.151 | 3.696 | 3.347 | 3.685 | 2.946 | 3.045 | 3.428 | 2.619 | 3.492 | 3.485 | 2.581 | 0.229 | 0.0000 | 0.0004 | 1 | 0 | 0 |
| 'Cyp51' | 8.915 | 9.919 | 10.164 | 10.670 | 9.578 | 10.720 | 10.748 | 8.661 | 10.197 | 10.150 | 11.693 | 10.157 | 10.791 | 11.174 | 11.204 | 9.156 | 10.527 | 9.943 | -0.092 | 0.0798 | 0.2544 | 0 | 0 | 0 |
| 'Cysltr2' | 3.508 | 5.311 | 5.060 | 4.452 | 5.415 | 3.644 | 4.380 | 4.997 | 5.194 | 5.158 | 4.631 | 3.542 | 4.573 | 4.200 | 3.028 | 4.634 | 4.719 | 3.131 | 0.162 | 0.1065 | 0.3065 | 0 | 0 | 0 |
| 'Cyth1' | 28.392 | 35.003 | 31.001 | 31.133 | 33.433 | 30.074 | 33.064 | 32.515 | 30.324 | 51.693 | 52.643 | 54.079 | 53.454 | 51.501 | 54.453 | 56.247 | 50.707 | 53.606 | -0.761 | 0.0000 | 0.0000 | 1 | 1 | 0 |
| 'Cyth2' | 22.628 | 22.686 | 21.686 | 20.858 | 22.029 | 23.020 | 22.898 | 22.821 | 21.523 | 21.562 | 19.981 | 22.147 | 22.372 | 21.300 | 21.479 | 22.540 | 21.627 | 20.038 | 0.041 | 0.1419 | 0.3648 | 0 | 0 | 0 |
| 'Cyth3' | 12.885 | 13.295 | 11.913 | 12.692 | 12.736 | 12.538 | 12.309 | 12.927 | 12.831 | 12.023 | 12.933 | 12.872 | 10.774 | 12.153 | 11.976 | 12.187 | 11.288 | 12.815 | 0.055 | 0.1377 | 0.3579 | 0 | 0 | 0 |
| 'Cyth4' | 50.446 | 48.456 | 47.128 | 52.229 | 47.648 | 48.453 | 53.674 | 48.511 | 49.038 | 53.800 | 53.215 | 59.561 | 59.128 | 51.393 | 54.116 | 58.724 | 56.546 | 57.812 | -0.192 | 0.0000 | 0.0000 | 1 | 0 | 0 |
| 'Cytip' | 23.142 | 24.465 | 23.189 | 23.083 | 24.547 | 22.324 | 23.307 | 26.134 | 21.920 | 20.483 | 22.718 | 22.978 | 16.405 | 21.644 | 23.201 | 21.173 | 21.630 | 22.796 | 0.127 | 0.0088 | 0.0568 | 0 | 0 | 0 |
| 'D030056L22Rik' | 6.816 | 6.502 | 6.722 | 6.011 | 6.032 | 6.250 | 6.721 | 6.857 | 6.531 | 5.501 | 7.047 | 7.414 | 6.283 | 7.863 | 7.046 | 6.603 | 7.058 | 7.432 | -0.099 | 0.0747 | 0.2432 | 0 | 0 | 0 |
| 'D10Jhu81e' | 53.819 | 61.820 | 55.244 | 53.456 | 54.423 | 61.575 | 53.186 | 57.118 | 56.589 | 54.746 | 52.911 | 55.939 | 55.572 | 52.837 | 55.758 | 54.296 | 49.704 | 54.696 | 0.049 | 0.1489 | 0.3756 | 0 | 0 | 0 |
| 'D10Wsu102e' | 22.177 | 24.281 | 21.102 | 21.812 | 23.675 | 21.931 | 23.127 | 22.893 | 22.034 | 21.899 | 22.482 | 21.908 | 21.944 | 23.288 | 23.099 | 22.291 | 22.921 | 23.255 | -0.013 | 0.6556 | 0.8332 | 0 | 0 | 0 |
| 'D11Wsu47e' | 5.194 | 5.063 | 5.189 | 4.517 | 5.273 | 5.266 | 5.047 | 5.411 | 5.444 | 5.038 | 4.397 | 5.776 | 4.471 | 5.111 | 5.478 | 5.752 | 4.957 | 4.941 | 0.004 | 0.9376 | 0.9764 | 0 | 0 | 0 |
| 'D130040H23Rik' | 4.469 | 4.571 | 4.506 | 4.482 | 4.720 | 4.179 | 5.013 | 4.297 | 4.486 | 5.519 | 5.564 | 4.866 | 5.580 | 5.056 | 4.233 | 4.810 | 5.476 | 4.026 | -0.157 | 0.0140 | 0.0800 | 0 | 0 | 0 |
| 'D16Ertd472e' | 8.787 | 8.060 | 8.730 | 8.777 | 9.759 | 8.951 | 8.803 | 8.990 | 8.188 | 8.136 | 7.743 | 8.043 | 7.395 | 8.555 | 7.340 | 8.543 | 7.592 | 7.039 | 0.159 | 0.0004 | 0.0048 | 1 | 0 | 0 |
| 'D17H6S53E' | 7.257 | 7.577 | 8.598 | 7.730 | 7.737 | 9.087 | 7.524 | 7.854 | 10.237 | 6.396 | 8.713 | 11.503 | 6.720 | 7.925 | 11.027 | 6.813 | 7.601 | 11.668 | -0.088 | 0.1680 | 0.4004 | 0 | 0 | 0 |
| 'D17Wsu92e' | 22.934 | 24.094 | 21.961 | 21.626 | 22.874 | 23.985 | 20.341 | 24.121 | 22.644 | 22.587 | 22.684 | 24.586 | 21.836 | 23.351 | 25.614 | 24.134 | 21.861 | 25.386 | -0.062 | 0.0870 | 0.2690 | 0 | 0 | 0 |
| 'D1Ertd622e' | 0.210 | 0.135 | 0.207 | 0.247 | 0.187 | 0.235 | 0.223 | 0.194 | 0.225 | 0.153 | 0.229 | 0.273 | 0.192 | 0.222 | 0.300 | 0.182 | 0.236 | 0.265 | -0.140 | 0.1707 | 0.4037 | 0 | 0 | 0 |
| 'D230025D16Rik' | 3.566 | 3.648 | 3.540 | 3.414 | 3.877 | 3.739 | 3.370 | 4.010 | 3.369 | 3.497 | 3.763 | 3.854 | 3.526 | 3.712 | 3.388 | 3.677 | 4.468 | 3.833 | -0.062 | 0.1529 | 0.3803 | 0 | 0 | 0 |
| 'D2Wsu81e' | 3.630 | 3.744 | 4.136 | 3.498 | 3.950 | 4.225 | 3.888 | 3.814 | 4.045 | 3.762 | 3.639 | 4.005 | 4.167 | 3.278 | 4.034 | 3.987 | 3.718 | 3.595 | 0.021 | 0.6133 | 0.8061 | 0 | 0 | 0 |
| 'D3Ertd254e' | 0.816 | 0.791 | 0.663 | 0.748 | 0.772 | 0.702 | 0.813 | 0.678 | 0.723 | 0.715 | 0.711 | 0.822 | 0.670 | 0.686 | 0.627 | 0.686 | 0.609 | 0.672 | 0.086 | 0.2578 | 0.5105 | 0 | 0 | 0 |
| 'D3Ertd751e' | 0.721 | 0.910 | 0.878 | 0.805 | 1.018 | 0.856 | 0.953 | 1.026 | 0.865 | 0.785 | 0.937 | 0.847 | 0.923 | 0.932 | 0.843 | 0.805 | 0.909 | 0.710 | 0.049 | 0.4278 | 0.6739 | 0 | 0 | 0 |
| 'D430042O09Rik' | 1.888 | 1.539 | 1.857 | 1.724 | 1.777 | 1.846 | 2.009 | 1.744 | 2.143 | 1.853 | 1.901 | 2.103 | 1.647 | 1.737 | 1.957 | 1.862 | 1.803 | 2.036 | -0.043 | 0.3746 | 0.6259 | 0 | 0 | 0 |
| 'D5Ertd579e' | 3.834 | 4.282 | 3.360 | 3.576 | 3.617 | 3.719 | 3.456 | 3.911 | 3.664 | 3.408 | 3.605 | 4.016 | 3.603 | 3.988 | 3.677 | 3.940 | 3.919 | 4.000 | -0.044 | 0.3020 | 0.5591 | 0 | 0 | 0 |
| 'D630039A03Rik' | 3.196 | 4.547 | 3.259 | 3.433 | 3.219 | 4.010 | 3.385 | 3.182 | 3.507 | 2.665 | 3.672 | 3.548 | 2.642 | 3.428 | 3.735 | 3.763 | 3.670 | 3.028 | 0.064 | 0.4565 | 0.6942 | 0 | 0 | 0 |
| 'D6Wsu163e' | 1.698 | 1.660 | 2.042 | 1.764 | 1.772 | 1.843 | 1.973 | 2.163 | 1.807 | 2.092 | 2.133 | 1.977 | 2.178 | 1.930 | 1.796 | 1.741 | 2.157 | 1.884 | -0.103 | 0.0914 | 0.2786 | 0 | 0 | 0 |
| 'D8Ertd738e' | 154.997 | 153.471 | 140.099 | 154.448 | 155.493 | 146.847 | 142.709 | 151.872 | 134.074 | 148.920 | 146.743 | 137.956 | 155.002 | 149.158 | 141.947 | 164.609 | 156.629 | 137.923 | -0.013 | 0.6545 | 0.8326 | 0 | 0 | 0 |
| 'D8Ertd82e' | 28.897 | 25.410 | 25.930 | 25.396 | 24.035 | 25.759 | 22.671 | 25.579 | 25.386 | 29.430 | 30.222 | 29.895 | 27.885 | 28.004 | 29.111 | 29.292 | 24.722 | 29.317 | -0.181 | 0.0000 | 0.0004 | 1 | 0 | 0 |
| 'D930015E06Rik' | 11.890 | 12.418 | 11.733 | 11.769 | 11.565 | 11.490 | 11.832 | 11.794 | 12.691 | 12.138 | 11.436 | 12.507 | 13.288 | 11.967 | 11.772 | 11.673 | 12.057 | 12.173 | -0.036 | 0.1965 | 0.4354 | 0 | 0 | 0 |
| 'Daam1' | 1.564 | 1.542 | 1.762 | 1.789 | 1.434 | 1.995 | 1.787 | 1.598 | 1.663 | 1.581 | 1.521 | 1.752 | 1.027 | 1.946 | 1.502 | 1.250 | 1.834 | 1.749 | 0.093 | 0.3354 | 0.5941 | 0 | 0 | 0 |
| 'Dab2ip' | 1.156 | 1.126 | 1.296 | 1.133 | 1.207 | 1.451 | 1.008 | 1.384 | 1.382 | 1.471 | 1.224 | 1.196 | 1.347 | 1.312 | 1.182 | 1.267 | 1.092 | 1.150 | -0.020 | 0.7742 | 0.8993 | 0 | 0 | 0 |
| 'Dad1' | 20.110 | 20.657 | 19.756 | 20.256 | 20.502 | 21.508 | 18.543 | 21.158 | 18.785 | 20.822 | 19.538 | 18.581 | 19.550 | 19.922 | 19.130 | 20.611 | 20.014 | 18.321 | 0.032 | 0.3303 | 0.5880 | 0 | 0 | 0 |
| 'Dag1' | 1.669 | 1.669 | 1.665 | 1.617 | 1.491 | 1.652 | 1.523 | 1.642 | 1.700 | 1.570 | 1.603 | 1.808 | 1.583 | 1.831 | 1.799 | 1.790 | 1.535 | 1.877 | -0.082 | 0.0697 | 0.2323 | 0 | 0 | 0 |
| 'Daglb' | 1.275 | 1.140 | 1.312 | 1.264 | 1.394 | 1.329 | 1.096 | 1.219 | 1.225 | 1.177 | 1.205 | 1.315 | 1.200 | 1.313 | 1.241 | 1.202 | 1.229 | 1.258 | 0.005 | 0.9249 | 0.9715 | 0 | 0 | 0 |
| 'Dalrd3' | 8.321 | 7.309 | 9.779 | 8.010 | 8.467 | 10.315 | 8.120 | 8.176 | 9.913 | 8.639 | 7.643 | 10.162 | 8.933 | 7.721 | 8.737 | 8.661 | 8.405 | 8.702 | 0.003 | 0.9499 | 0.9809 | 0 | 0 | 0 |
| 'Dap' | 60.806 | 68.716 | 64.216 | 59.802 | 65.764 | 64.753 | 59.797 | 66.573 | 63.887 | 66.878 | 62.978 | 64.667 | 68.609 | 69.675 | 65.630 | 66.693 | 62.868 | 60.306 | -0.045 | 0.1315 | 0.3484 | 0 | 0 | 0 |
| 'Dap3' | 19.674 | 18.847 | 20.354 | 19.922 | 19.612 | 21.011 | 19.334 | 19.335 | 20.443 | 19.365 | 19.381 | 21.044 | 19.173 | 20.089 | 21.316 | 18.653 | 20.055 | 20.917 | -0.022 | 0.3878 | 0.6374 | 0 | 0 | 0 |
| 'Dapk3' | 34.558 | 31.815 | 41.055 | 31.697 | 33.729 | 42.115 | 30.244 | 33.673 | 36.131 | 35.185 | 28.626 | 36.383 | 38.633 | 31.070 | 35.537 | 37.719 | 29.708 | 35.902 | 0.021 | 0.6769 | 0.8454 | 0 | 0 | 0 |
| 'Dapp1' | 23.024 | 23.129 | 20.553 | 24.117 | 23.398 | 20.455 | 23.294 | 23.147 | 20.882 | 19.798 | 21.597 | 20.167 | 20.640 | 21.506 | 20.200 | 20.193 | 23.255 | 19.323 | 0.101 | 0.0005 | 0.0057 | 1 | 0 | 0 |
| 'Dars' | 27.115 | 29.677 | 27.343 | 29.750 | 31.772 | 27.066 | 29.465 | 30.765 | 27.895 | 28.930 | 30.992 | 28.899 | 30.369 | 31.370 | 27.397 | 28.661 | 33.025 | 27.000 | -0.043 | 0.0786 | 0.2516 | 0 | 0 | 0 |
| 'Dars2' | 2.348 | 2.153 | 2.549 | 2.619 | 2.503 | 2.525 | 2.559 | 2.618 | 2.560 | 2.184 | 2.623 | 2.497 | 2.511 | 2.802 | 2.357 | 2.364 | 2.444 | 2.363 | 0.012 | 0.8085 | 0.9139 | 0 | 0 | 0 |
| 'Daxx' | 16.448 | 15.864 | 15.719 | 14.626 | 16.342 | 18.134 | 14.162 | 16.745 | 17.977 | 14.474 | 15.301 | 18.945 | 16.478 | 15.282 | 18.380 | 15.793 | 15.497 | 17.997 | -0.030 | 0.4473 | 0.6880 | 0 | 0 | 0 |
| 'Dazap1' | 42.722 | 43.019 | 47.604 | 41.945 | 42.762 | 46.410 | 44.789 | 44.466 | 45.830 | 42.432 | 41.593 | 43.793 | 45.083 | 41.450 | 44.183 | 42.113 | 42.832 | 41.958 | 0.042 | 0.0599 | 0.2110 | 0 | 0 | 0 |
| 'Dazap2' | 185.269 | 209.198 | 184.664 | 174.519 | 207.951 | 185.963 | 184.023 | 208.542 | 185.094 | 203.587 | 199.415 | 170.062 | 187.263 | 205.125 | 178.528 | 192.112 | 185.505 | 168.772 | 0.019 | 0.5383 | 0.7566 | 0 | 0 | 0 |
| 'Dbf4' | 3.706 | 3.969 | 4.403 | 4.411 | 3.518 | 4.339 | 4.467 | 3.753 | 4.079 | 3.273 | 3.650 | 4.397 | 4.002 | 4.309 | 4.720 | 3.803 | 4.020 | 4.712 | -0.019 | 0.7336 | 0.8786 | 0 | 0 | 0 |
| 'Dbh' | 1.304 | 1.906 | 1.493 | 2.109 | 1.275 | 1.254 | 1.362 | 1.506 | 1.868 | 1.637 | 1.199 | 1.473 | 1.504 | 1.306 | 1.000 | 1.417 | 1.542 | 1.459 | 0.147 | 0.2386 | 0.4887 | 0 | 0 | 0 |
| 'Dbi' | 13.670 | 16.256 | 14.611 | 15.084 | 15.592 | 15.503 | 15.735 | 14.308 | 15.042 | 12.615 | 14.511 | 14.396 | 18.006 | 16.220 | 14.943 | 15.357 | 14.678 | 13.551 | 0.005 | 0.9314 | 0.9746 | 0 | 0 | 0 |
| 'Dbnl' | 69.520 | 67.521 | 74.324 | 65.652 | 71.740 | 79.682 | 66.188 | 69.997 | 75.576 | 76.089 | 66.238 | 74.566 | 76.979 | 68.356 | 74.490 | 76.196 | 67.148 | 71.810 | -0.036 | 0.2593 | 0.5126 | 0 | 0 | 0 |
| 'Dbp' | 2.579 | 2.695 | 3.042 | 2.382 | 2.550 | 2.814 | 2.435 | 2.708 | 2.881 | 2.059 | 2.064 | 2.806 | 2.102 | 2.043 | 2.703 | 2.191 | 1.951 | 2.552 | 0.219 | 0.0003 | 0.0042 | 1 | 0 | 0 |
| 'Dbr1' | 8.133 | 7.945 | 7.960 | 7.808 | 8.094 | 7.774 | 7.263 | 8.417 | 7.842 | 8.343 | 8.447 | 8.030 | 8.113 | 8.963 | 7.600 | 8.035 | 8.802 | 7.427 | -0.054 | 0.1631 | 0.3949 | 0 | 0 | 0 |
| 'Dbt' | 6.251 | 7.286 | 6.720 | 6.458 | 7.573 | 7.133 | 6.673 | 7.517 | 5.987 | 7.724 | 7.961 | 6.501 | 7.244 | 7.447 | 6.755 | 7.013 | 6.582 | 5.748 | -0.038 | 0.4510 | 0.6904 | 0 | 0 | 0 |
| 'Dcaf10' | 3.876 | 3.788 | 3.467 | 3.630 | 3.603 | 3.400 | 3.285 | 3.756 | 3.251 | 3.958 | 3.649 | 3.638 | 3.481 | 3.874 | 3.399 | 3.952 | 3.815 | 3.560 | -0.065 | 0.1005 | 0.2967 | 0 | 0 | 0 |
| 'Dcaf11' | 6.942 | 7.249 | 6.990 | 6.453 | 6.897 | 7.238 | 6.726 | 7.334 | 7.452 | 7.572 | 6.833 | 7.273 | 7.217 | 7.081 | 7.024 | 7.127 | 7.414 | 7.008 | -0.039 | 0.1633 | 0.3953 | 0 | 0 | 0 |
| 'Dcaf12' | 21.194 | 20.902 | 21.422 | 22.066 | 20.342 | 20.584 | 23.466 | 20.347 | 22.135 | 16.807 | 23.720 | 25.846 | 20.214 | 23.079 | 25.534 | 20.410 | 23.793 | 24.588 | -0.088 | 0.1382 | 0.3586 | 0 | 0 | 0 |
| 'Dcaf13' | 41.911 | 47.194 | 43.172 | 41.614 | 43.248 | 38.848 | 41.848 | 41.261 | 41.914 | 41.206 | 45.431 | 41.367 | 39.547 | 42.453 | 38.999 | 38.231 | 46.375 | 37.927 | 0.026 | 0.4751 | 0.7100 | 0 | 0 | 0 |
| 'Dcaf15' | 72.463 | 74.957 | 82.731 | 69.192 | 70.154 | 80.850 | 72.279 | 73.803 | 81.505 | 76.226 | 70.300 | 79.614 | 80.110 | 67.997 | 77.759 | 72.581 | 69.592 | 71.813 | 0.015 | 0.6240 | 0.8125 | 0 | 0 | 0 |
| 'Dcaf17' | 3.751 | 4.806 | 4.771 | 4.017 | 5.165 | 4.209 | 4.513 | 4.727 | 4.509 | 4.270 | 4.010 | 4.191 | 4.023 | 3.601 | 3.716 | 3.721 | 4.389 | 3.761 | 0.163 | 0.0039 | 0.0309 | 1 | 0 | 0 |
| 'Dcaf4' | 10.728 | 11.310 | 14.073 | 10.645 | 11.514 | 11.238 | 12.373 | 11.529 | 12.194 | 11.521 | 10.631 | 10.827 | 11.145 | 10.382 | 10.407 | 12.112 | 10.751 | 8.837 | 0.124 | 0.0388 | 0.1606 | 0 | 0 | 0 |
| 'Dcaf5' | 33.927 | 30.629 | 30.825 | 32.007 | 30.094 | 31.976 | 33.569 | 29.125 | 30.134 | 33.360 | 28.432 | 31.490 | 30.838 | 29.410 | 30.186 | 32.365 | 28.749 | 31.463 | 0.020 | 0.4285 | 0.6742 | 0 | 0 | 0 |
| 'Dcaf6' | 3.825 | 4.945 | 4.353 | 3.892 | 4.478 | 4.227 | 4.176 | 4.815 | 4.392 | 4.737 | 4.978 | 4.489 | 4.026 | 4.617 | 4.046 | 4.354 | 4.803 | 4.146 | -0.050 | 0.2153 | 0.4610 | 0 | 0 | 0 |
| 'Dcaf7' | 32.586 | 34.162 | 33.907 | 33.554 | 33.479 | 34.520 | 32.077 | 34.079 | 36.774 | 31.506 | 34.271 | 38.734 | 28.922 | 34.123 | 38.329 | 33.693 | 33.052 | 39.138 | -0.038 | 0.2928 | 0.5493 | 0 | 0 | 0 |
| 'Dcaf8' | 13.841 | 14.557 | 14.520 | 13.712 | 14.086 | 14.513 | 13.641 | 14.520 | 14.484 | 14.404 | 14.215 | 13.647 | 13.686 | 13.188 | 12.813 | 14.103 | 13.810 | 12.731 | 0.052 | 0.0486 | 0.1851 | 0 | 0 | 0 |
| 'Dcakd' | 5.399 | 5.127 | 6.321 | 4.978 | 5.445 | 6.327 | 5.623 | 4.636 | 6.294 | 5.652 | 4.549 | 5.924 | 4.905 | 4.937 | 5.822 | 5.217 | 4.731 | 5.750 | 0.069 | 0.1414 | 0.3636 | 0 | 0 | 0 |
| 'Dck' | 38.212 | 40.184 | 40.395 | 40.075 | 43.464 | 39.791 | 43.969 | 40.937 | 36.796 | 35.790 | 47.578 | 47.430 | 43.424 | 48.878 | 45.239 | 39.101 | 48.147 | 47.926 | -0.157 | 0.0012 | 0.0122 | 1 | 0 | 0 |
| 'Dclk2' | 1.880 | 1.300 | 2.225 | 1.629 | 1.592 | 1.843 | 1.760 | 1.612 | 1.980 | 1.710 | 1.565 | 2.031 | 1.586 | 1.546 | 2.022 | 1.627 | 1.173 | 2.058 | 0.042 | 0.4696 | 0.7055 | 0 | 0 | 0 |
| 'Dclre1a' | 1.130 | 1.285 | 1.230 | 1.275 | 1.362 | 1.401 | 1.432 | 1.439 | 1.337 | 1.215 | 1.017 | 0.981 | 1.150 | 1.236 | 0.970 | 1.222 | 1.016 | 0.924 | 0.288 | 0.0000 | 0.0002 | 1 | 0 | 0 |
| 'Dclre1b' | 1.118 | 1.211 | 1.183 | 1.244 | 1.072 | 1.233 | 1.114 | 0.969 | 1.149 | 1.034 | 0.971 | 1.150 | 0.918 | 1.044 | 1.178 | 1.020 | 0.973 | 1.097 | 0.118 | 0.0534 | 0.1973 | 0 | 0 | 0 |
| 'Dclre1c' | 5.108 | 5.271 | 5.269 | 5.389 | 5.009 | 4.810 | 5.490 | 4.733 | 5.654 | 5.402 | 5.913 | 6.138 | 6.515 | 5.619 | 5.731 | 4.931 | 7.220 | 5.620 | -0.195 | 0.0008 | 0.0089 | 1 | 0 | 0 |
| 'Dcp1a' | 14.194 | 13.719 | 14.490 | 13.395 | 14.145 | 14.059 | 13.640 | 15.029 | 14.991 | 13.913 | 14.522 | 15.681 | 14.109 | 13.273 | 15.102 | 14.344 | 13.791 | 14.427 | -0.028 | 0.3601 | 0.6124 | 0 | 0 | 0 |
| 'Dcp1b' | 3.578 | 3.398 | 3.921 | 2.958 | 3.413 | 3.833 | 3.486 | 3.429 | 3.744 | 3.587 | 3.007 | 3.620 | 3.041 | 2.869 | 3.321 | 3.362 | 3.065 | 3.090 | 0.124 | 0.0079 | 0.0523 | 0 | 0 | 0 |
| 'Dcp2' | 9.885 | 9.243 | 10.001 | 10.356 | 8.444 | 9.284 | 10.249 | 9.322 | 9.373 | 9.267 | 11.679 | 11.683 | 9.073 | 9.950 | 10.300 | 9.271 | 11.796 | 11.194 | -0.137 | 0.0195 | 0.1011 | 0 | 0 | 0 |
| 'Dcps' | 20.995 | 22.912 | 23.799 | 23.107 | 23.435 | 23.759 | 21.996 | 22.155 | 24.167 | 21.922 | 22.235 | 23.143 | 23.937 | 23.496 | 22.566 | 25.055 | 22.232 | 21.910 | -0.010 | 0.7744 | 0.8994 | 0 | 0 | 0 |
| 'Dctd' | 0.590 | 0.309 | 0.517 | 0.534 | 0.520 | 0.447 | 0.488 | 0.556 | 0.495 | 0.386 | 0.501 | 0.652 | 0.493 | 0.449 | 0.577 | 0.429 | 0.399 | 0.532 | -0.002 | 0.9899 | 0.9972 | 0 | 0 | 0 |
| 'Dctn1' | 23.846 | 22.649 | 25.368 | 22.962 | 21.804 | 27.054 | 22.207 | 23.063 | 26.184 | 25.804 | 23.229 | 25.403 | 25.773 | 21.556 | 24.730 | 24.846 | 21.225 | 24.721 | -0.024 | 0.4472 | 0.6880 | 0 | 0 | 0 |
| 'Dctn2' | 64.367 | 63.238 | 68.138 | 59.667 | 63.361 | 68.715 | 64.565 | 67.154 | 66.085 | 65.696 | 57.632 | 64.470 | 65.013 | 60.919 | 64.632 | 64.547 | 60.286 | 60.032 | 0.047 | 0.0938 | 0.2835 | 0 | 0 | 0 |
| 'Dctn3' | 28.364 | 25.819 | 24.550 | 24.571 | 30.300 | 27.443 | 25.056 | 28.400 | 26.495 | 25.447 | 25.743 | 25.030 | 26.837 | 28.273 | 25.815 | 26.755 | 25.707 | 23.809 | 0.038 | 0.3518 | 0.6056 | 0 | 0 | 0 |
| 'Dctn4' | 7.827 | 7.203 | 7.530 | 7.217 | 8.078 | 7.594 | 6.903 | 8.525 | 7.574 | 7.552 | 7.573 | 8.018 | 7.442 | 7.884 | 8.238 | 8.735 | 7.968 | 7.811 | -0.065 | 0.1155 | 0.3228 | 0 | 0 | 0 |
| 'Dctn5' | 17.157 | 18.540 | 18.843 | 16.938 | 19.069 | 18.631 | 18.035 | 18.093 | 19.189 | 17.512 | 18.165 | 19.929 | 17.335 | 18.216 | 19.133 | 17.262 | 19.802 | 19.656 | -0.035 | 0.1872 | 0.4231 | 0 | 0 | 0 |
| 'Dctn6' | 8.196 | 9.085 | 8.382 | 8.535 | 9.361 | 8.324 | 8.746 | 9.671 | 8.081 | 7.248 | 8.500 | 7.603 | 8.050 | 8.007 | 7.860 | 8.105 | 7.874 | 7.366 | 0.139 | 0.0000 | 0.0007 | 1 | 0 | 0 |
| 'Dctpp1' | 46.407 | 45.611 | 45.788 | 46.706 | 47.543 | 43.883 | 40.695 | 50.006 | 43.019 | 40.049 | 37.618 | 41.503 | 49.001 | 42.396 | 45.462 | 42.725 | 39.386 | 44.978 | 0.085 | 0.1037 | 0.3020 | 0 | 0 | 0 |
| 'Dcun1d1' | 6.089 | 5.705 | 5.638 | 6.844 | 5.839 | 5.641 | 6.779 | 5.661 | 5.481 | 5.400 | 5.935 | 5.891 | 5.526 | 6.108 | 5.748 | 5.550 | 5.775 | 6.001 | 0.032 | 0.5096 | 0.7343 | 0 | 0 | 0 |
| 'Dcun1d2' | 3.961 | 4.069 | 3.903 | 4.092 | 3.482 | 3.839 | 4.145 | 3.846 | 3.747 | 3.653 | 3.658 | 4.479 | 3.598 | 4.057 | 4.170 | 3.493 | 3.897 | 4.096 | -0.014 | 0.7687 | 0.8967 | 0 | 0 | 0 |
| 'Dcun1d3' | 1.465 | 1.184 | 1.595 | 1.497 | 1.677 | 1.341 | 1.759 | 1.554 | 1.620 | 1.236 | 1.519 | 1.640 | 1.436 | 1.533 | 1.620 | 1.381 | 1.622 | 1.402 | 0.020 | 0.7769 | 0.9002 | 0 | 0 | 0 |
| 'Dcun1d4' | 2.733 | 3.397 | 3.060 | 2.729 | 3.686 | 2.983 | 2.850 | 3.088 | 3.507 | 2.937 | 3.963 | 3.993 | 2.945 | 3.448 | 3.665 | 3.436 | 3.545 | 3.620 | -0.186 | 0.0002 | 0.0033 | 1 | 0 | 0 |
| 'Dcun1d5' | 65.423 | 76.365 | 71.807 | 72.738 | 79.027 | 74.294 | 75.546 | 79.727 | 68.719 | 66.505 | 70.405 | 74.178 | 68.293 | 72.936 | 71.601 | 70.359 | 78.809 | 66.640 | 0.039 | 0.2205 | 0.4679 | 0 | 0 | 0 |
| 'Dcxr' | 4.841 | 5.324 | 5.026 | 4.293 | 5.579 | 5.226 | 4.178 | 5.344 | 4.909 | 4.698 | 5.163 | 3.857 | 5.243 | 4.063 | 4.051 | 4.761 | 3.746 | 3.366 | 0.200 | 0.0142 | 0.0806 | 0 | 0 | 0 |
| 'Dda1' | 9.641 | 11.330 | 9.746 | 9.383 | 11.360 | 11.846 | 9.696 | 11.094 | 10.383 | 10.736 | 9.045 | 10.531 | 10.908 | 9.801 | 10.165 | 12.011 | 9.221 | 10.154 | 0.019 | 0.7240 | 0.8731 | 0 | 0 | 0 |
| 'Ddb1' | 98.977 | 95.807 | 100.292 | 98.128 | 99.832 | 101.800 | 95.166 | 97.773 | 97.699 | 102.161 | 97.983 | 98.625 | 95.575 | 95.452 | 99.237 | 92.632 | 97.544 | 97.242 | 0.005 | 0.8321 | 0.9270 | 0 | 0 | 0 |
| 'Ddb2' | 6.026 | 5.620 | 6.933 | 5.875 | 5.709 | 6.378 | 5.731 | 5.950 | 6.630 | 6.540 | 6.166 | 6.810 | 6.249 | 6.024 | 6.549 | 5.604 | 5.898 | 6.457 | -0.046 | 0.1910 | 0.4276 | 0 | 0 | 0 |
| 'Ddc' | 2.223 | 2.622 | 2.013 | 2.449 | 2.036 | 2.447 | 2.402 | 2.155 | 2.016 | 0.921 | 0.924 | 0.952 | 0.859 | 0.895 | 0.836 | 0.899 | 0.797 | 0.784 | 1.351 | 0.0000 | 0.0000 | 1 | 1 | 1 |
| 'Ddhd1' | 3.252 | 2.973 | 3.766 | 3.182 | 3.357 | 3.602 | 2.976 | 3.654 | 3.697 | 3.695 | 3.728 | 3.681 | 3.486 | 3.385 | 3.391 | 3.346 | 3.727 | 3.753 | -0.089 | 0.0425 | 0.1701 | 0 | 0 | 0 |
| 'Ddhd2' | 10.081 | 12.544 | 12.559 | 10.723 | 12.743 | 12.207 | 10.820 | 12.318 | 12.122 | 11.660 | 12.327 | 10.461 | 11.565 | 11.072 | 10.883 | 11.162 | 11.886 | 9.070 | 0.077 | 0.1251 | 0.3385 | 0 | 0 | 0 |
| 'Ddias' | 1.974 | 1.874 | 2.194 | 2.000 | 1.949 | 2.305 | 1.899 | 1.658 | 2.138 | 1.870 | 1.787 | 2.034 | 2.021 | 1.799 | 2.041 | 1.761 | 1.706 | 1.856 | 0.082 | 0.3099 | 0.5684 | 0 | 0 | 0 |
| 'Ddit3' | 19.178 | 12.752 | 10.893 | 14.297 | 11.629 | 11.566 | 15.419 | 14.152 | 11.574 | 17.590 | 15.534 | 11.688 | 16.767 | 13.733 | 11.658 | 15.482 | 14.812 | 11.329 | -0.088 | 0.1291 | 0.3443 | 0 | 0 | 0 |
| 'Ddit4' | 184.597 | 157.561 | 134.308 | 233.367 | 143.778 | 133.495 | 234.440 | 137.138 | 133.485 | 114.588 | 208.999 | 174.410 | 148.783 | 174.274 | 189.854 | 127.532 | 246.942 | 187.359 | -0.098 | 0.5539 | 0.7667 | 0 | 0 | 0 |
| 'Ddost' | 73.053 | 80.197 | 77.880 | 71.024 | 81.635 | 79.778 | 72.866 | 77.548 | 79.217 | 79.657 | 72.297 | 77.323 | 78.364 | 76.333 | 78.229 | 80.555 | 71.213 | 76.969 | -0.006 | 0.8407 | 0.9323 | 0 | 0 | 0 |
| 'Ddr1' | 0.721 | 0.518 | 0.963 | 0.742 | 0.546 | 0.774 | 0.661 | 0.574 | 0.732 | 0.385 | 0.608 | 0.681 | 0.532 | 0.593 | 0.734 | 0.498 | 0.507 | 0.685 | 0.251 | 0.0130 | 0.0756 | 0 | 0 | 0 |
| 'Ddrgk1' | 29.511 | 31.051 | 32.160 | 31.729 | 30.347 | 32.340 | 30.628 | 31.615 | 33.614 | 32.483 | 28.838 | 30.219 | 32.106 | 28.513 | 30.183 | 30.197 | 29.980 | 27.857 | 0.057 | 0.0680 | 0.2287 | 0 | 0 | 0 |
| 'Ddt' | 22.617 | 27.007 | 22.880 | 22.506 | 24.402 | 25.764 | 23.371 | 26.097 | 22.751 | 22.577 | 22.531 | 23.580 | 24.051 | 22.352 | 24.190 | 20.922 | 23.822 | 21.700 | 0.063 | 0.1549 | 0.3832 | 0 | 0 | 0 |
| 'Ddx1' | 37.749 | 37.866 | 38.224 | 38.310 | 41.207 | 40.324 | 41.021 | 39.699 | 40.838 | 36.695 | 37.905 | 34.449 | 40.640 | 36.259 | 35.126 | 38.657 | 39.811 | 32.842 | 0.089 | 0.0125 | 0.0735 | 0 | 0 | 0 |
| 'Ddx10' | 19.748 | 21.501 | 19.015 | 20.278 | 21.343 | 18.403 | 22.662 | 20.954 | 19.003 | 20.516 | 20.548 | 19.934 | 19.443 | 18.546 | 18.539 | 21.154 | 20.632 | 19.606 | 0.016 | 0.6435 | 0.8251 | 0 | 0 | 0 |
| 'Ddx11' | 13.236 | 10.886 | 14.871 | 12.357 | 12.659 | 14.054 | 13.451 | 11.097 | 13.669 | 12.896 | 11.744 | 14.045 | 13.845 | 11.744 | 13.444 | 11.871 | 12.685 | 12.940 | 0.001 | 0.9796 | 0.9926 | 0 | 0 | 0 |
| 'Ddx17' | 107.885 | 123.065 | 113.503 | 109.659 | 126.310 | 117.094 | 107.129 | 123.610 | 118.084 | 115.517 | 115.342 | 118.357 | 110.426 | 118.344 | 115.865 | 115.889 | 119.551 | 115.374 | -0.010 | 0.6112 | 0.8049 | 0 | 0 | 0 |
| 'Ddx18' | 23.464 | 21.742 | 23.636 | 23.338 | 22.609 | 22.825 | 24.451 | 23.972 | 22.377 | 21.112 | 21.268 | 23.812 | 22.036 | 20.365 | 22.730 | 22.109 | 20.757 | 22.673 | 0.070 | 0.0242 | 0.1178 | 0 | 0 | 0 |
| 'Ddx19a' | 22.793 | 23.215 | 23.506 | 22.301 | 21.340 | 24.498 | 22.659 | 22.786 | 23.432 | 23.162 | 20.094 | 22.633 | 22.420 | 22.382 | 22.273 | 21.426 | 19.524 | 22.672 | 0.061 | 0.0399 | 0.1628 | 0 | 0 | 0 |
| 'Ddx19b' | 10.823 | 11.308 | 11.832 | 9.933 | 11.295 | 11.933 | 10.146 | 11.254 | 11.968 | 11.516 | 10.944 | 10.484 | 11.258 | 11.373 | 10.833 | 11.650 | 10.080 | 9.929 | 0.027 | 0.5221 | 0.7447 | 0 | 0 | 0 |
| 'Ddx20' | 6.027 | 6.864 | 6.658 | 5.906 | 6.323 | 6.275 | 5.717 | 6.177 | 5.961 | 5.925 | 5.836 | 6.253 | 5.645 | 5.980 | 5.776 | 5.580 | 5.503 | 5.379 | 0.095 | 0.0193 | 0.1007 | 0 | 0 | 0 |
| 'Ddx21' | 91.115 | 77.628 | 83.570 | 85.531 | 83.344 | 77.865 | 85.454 | 82.793 | 78.983 | 84.857 | 79.627 | 82.155 | 81.039 | 80.320 | 83.137 | 80.310 | 81.838 | 83.332 | 0.007 | 0.8005 | 0.9111 | 0 | 0 | 0 |
| 'Ddx23' | 22.618 | 22.220 | 23.987 | 22.679 | 21.644 | 23.503 | 21.920 | 21.263 | 24.480 | 22.535 | 21.487 | 24.374 | 23.654 | 20.798 | 22.313 | 21.891 | 22.280 | 24.038 | -0.005 | 0.8429 | 0.9337 | 0 | 0 | 0 |
| 'Ddx24' | 89.765 | 87.542 | 84.125 | 80.851 | 88.969 | 88.043 | 79.333 | 90.995 | 86.673 | 99.398 | 87.546 | 91.342 | 88.718 | 86.008 | 89.352 | 95.442 | 82.436 | 92.974 | -0.077 | 0.0331 | 0.1442 | 0 | 0 | 0 |
| 'Ddx26b' | 4.500 | 4.811 | 5.023 | 4.486 | 5.244 | 5.200 | 4.409 | 4.760 | 5.221 | 4.757 | 4.691 | 5.107 | 4.664 | 4.961 | 4.668 | 4.546 | 5.157 | 4.697 | 0.002 | 0.9476 | 0.9796 | 0 | 0 | 0 |
| 'Ddx27' | 18.184 | 16.414 | 19.269 | 17.566 | 17.731 | 18.681 | 18.793 | 17.969 | 19.425 | 18.489 | 17.027 | 19.503 | 19.157 | 16.634 | 19.090 | 18.027 | 17.840 | 18.657 | -0.014 | 0.5845 | 0.7863 | 0 | 0 | 0 |
| 'Ddx28' | 14.869 | 12.625 | 15.504 | 13.372 | 13.891 | 13.217 | 15.143 | 13.467 | 14.689 | 15.154 | 14.862 | 14.600 | 13.848 | 14.749 | 14.094 | 13.818 | 15.191 | 14.296 | -0.050 | 0.2821 | 0.5383 | 0 | 0 | 0 |
| 'Ddx31' | 3.599 | 3.603 | 3.525 | 3.677 | 3.557 | 3.763 | 4.123 | 3.761 | 3.790 | 3.287 | 3.419 | 4.630 | 3.565 | 2.895 | 4.239 | 3.474 | 3.763 | 3.905 | -0.007 | 0.9143 | 0.9671 | 0 | 0 | 0 |
| 'Ddx39' | 15.950 | 17.312 | 17.546 | 16.375 | 16.869 | 17.096 | 17.454 | 17.138 | 17.252 | 15.331 | 15.226 | 17.370 | 17.325 | 16.215 | 17.481 | 16.097 | 16.729 | 16.228 | 0.035 | 0.1965 | 0.4354 | 0 | 0 | 0 |
| 'Ddx39b' | 84.515 | 80.518 | 88.044 | 82.151 | 79.767 | 84.156 | 80.880 | 83.102 | 89.775 | 78.412 | 81.909 | 82.102 | 87.104 | 78.767 | 82.480 | 77.110 | 81.980 | 81.251 | 0.032 | 0.2256 | 0.4722 | 0 | 0 | 0 |
| 'Ddx3x' | 75.363 | 68.123 | 55.997 | 68.056 | 68.458 | 54.968 | 67.054 | 72.706 | 52.402 | 79.863 | 67.460 | 47.035 | 71.778 | 73.234 | 45.987 | 77.266 | 64.286 | 44.212 | 0.043 | 0.3542 | 0.6075 | 0 | 0 | 0 |
| 'Ddx41' | 59.697 | 61.874 | 61.562 | 57.642 | 61.263 | 63.193 | 54.771 | 59.611 | 59.307 | 62.887 | 57.272 | 59.740 | 66.401 | 59.489 | 58.434 | 63.338 | 56.707 | 58.118 | -0.018 | 0.5868 | 0.7880 | 0 | 0 | 0 |
| 'Ddx42' | 57.415 | 58.595 | 58.763 | 57.840 | 57.894 | 56.181 | 59.692 | 58.730 | 57.256 | 57.688 | 59.154 | 54.752 | 55.722 | 55.906 | 53.963 | 55.171 | 58.248 | 50.097 | 0.052 | 0.0248 | 0.1199 | 0 | 0 | 0 |
| 'Ddx43' | 5.885 | 6.719 | 7.414 | 5.502 | 4.843 | 4.698 | 7.222 | 6.229 | 6.034 | 5.767 | 5.886 | 7.179 | 5.761 | 5.461 | 5.622 | 6.353 | 7.200 | 8.309 | -0.089 | 0.3671 | 0.6192 | 0 | 0 | 0 |
| 'Ddx46' | 38.702 | 40.231 | 38.813 | 37.051 | 41.816 | 40.607 | 37.136 | 41.773 | 39.913 | 37.428 | 38.210 | 38.272 | 37.617 | 38.309 | 38.278 | 40.074 | 36.838 | 39.321 | 0.037 | 0.1358 | 0.3545 | 0 | 0 | 0 |
| 'Ddx47' | 10.278 | 10.228 | 10.609 | 10.325 | 10.481 | 10.151 | 9.996 | 10.336 | 10.255 | 10.021 | 10.502 | 10.352 | 9.998 | 9.827 | 10.206 | 9.807 | 10.558 | 10.187 | 0.009 | 0.7102 | 0.8654 | 0 | 0 | 0 |
| 'Ddx49' | 6.250 | 5.156 | 6.603 | 5.769 | 6.400 | 6.910 | 6.214 | 6.203 | 6.303 | 5.791 | 5.108 | 6.330 | 6.205 | 5.475 | 5.881 | 6.109 | 6.100 | 5.958 | 0.067 | 0.1302 | 0.3464 | 0 | 0 | 0 |
| 'Ddx5' | 258.474 | 265.244 | 230.394 | 241.587 | 262.998 | 222.941 | 236.618 | 261.661 | 225.003 | 269.094 | 252.637 | 192.745 | 244.036 | 275.265 | 202.203 | 258.891 | 252.705 | 192.516 | 0.042 | 0.2609 | 0.5143 | 0 | 0 | 0 |
| 'Ddx50' | 38.129 | 42.178 | 39.168 | 38.709 | 42.520 | 38.257 | 41.777 | 42.205 | 37.405 | 38.305 | 35.611 | 36.547 | 35.384 | 38.559 | 33.559 | 36.605 | 40.081 | 33.595 | 0.123 | 0.0000 | 0.0002 | 1 | 0 | 0 |
| 'Ddx51' | 4.728 | 4.395 | 5.518 | 4.566 | 4.513 | 5.274 | 4.337 | 4.728 | 5.112 | 4.941 | 3.892 | 5.361 | 4.559 | 4.222 | 4.831 | 4.929 | 4.025 | 4.387 | 0.061 | 0.1861 | 0.4218 | 0 | 0 | 0 |
| 'Ddx52' | 18.664 | 20.561 | 20.095 | 19.543 | 20.046 | 20.369 | 21.024 | 21.612 | 19.730 | 19.843 | 19.373 | 18.517 | 20.666 | 20.122 | 17.477 | 20.241 | 20.591 | 17.721 | 0.051 | 0.1221 | 0.3333 | 0 | 0 | 0 |
| 'Ddx54' | 20.621 | 18.405 | 22.556 | 17.535 | 19.670 | 23.594 | 18.986 | 19.470 | 21.868 | 21.894 | 18.325 | 22.501 | 21.283 | 18.417 | 21.588 | 22.295 | 17.944 | 21.563 | -0.033 | 0.3811 | 0.6324 | 0 | 0 | 0 |
| 'Ddx55' | 5.910 | 5.694 | 5.850 | 5.994 | 5.933 | 5.530 | 6.111 | 5.857 | 5.880 | 5.884 | 5.982 | 6.727 | 6.028 | 5.701 | 5.731 | 5.525 | 6.739 | 6.870 | -0.079 | 0.0855 | 0.2658 | 0 | 0 | 0 |
| 'Ddx56' | 13.857 | 13.543 | 15.252 | 14.052 | 14.021 | 13.921 | 13.021 | 13.855 | 13.708 | 13.950 | 12.101 | 13.861 | 14.089 | 12.200 | 14.003 | 13.409 | 12.345 | 13.754 | 0.056 | 0.1245 | 0.3375 | 0 | 0 | 0 |
| 'Ddx58' | 6.861 | 7.403 | 7.304 | 6.668 | 7.736 | 7.583 | 7.141 | 7.920 | 7.132 | 7.605 | 7.460 | 8.577 | 7.303 | 7.366 | 8.227 | 7.805 | 7.183 | 8.538 | -0.104 | 0.0040 | 0.0316 | 1 | 0 | 0 |
| 'Ddx59' | 3.827 | 4.168 | 3.961 | 3.174 | 4.137 | 4.337 | 4.072 | 4.406 | 4.594 | 6.210 | 5.215 | 4.599 | 5.890 | 6.093 | 4.284 | 5.202 | 4.916 | 4.257 | -0.347 | 0.0000 | 0.0002 | 1 | 0 | 0 |
| 'Ddx6' | 150.617 | 150.885 | 140.651 | 156.853 | 155.078 | 134.078 | 146.687 | 152.893 | 140.683 | 135.799 | 140.212 | 122.395 | 136.397 | 138.666 | 124.925 | 133.848 | 130.309 | 124.800 | 0.151 | 0.0000 | 0.0000 | 1 | 0 | 0 |
| 'Ddx60' | 3.693 | 4.515 | 3.922 | 3.915 | 4.456 | 3.951 | 4.042 | 4.588 | 4.203 | 4.979 | 5.022 | 5.284 | 4.260 | 4.098 | 4.798 | 4.323 | 5.247 | 4.655 | -0.211 | 0.0000 | 0.0005 | 1 | 0 | 0 |
| 'Deaf1' | 24.312 | 24.231 | 28.146 | 24.409 | 22.288 | 27.120 | 24.961 | 25.949 | 28.368 | 24.149 | 24.172 | 29.098 | 25.493 | 22.193 | 26.259 | 22.053 | 23.370 | 27.337 | 0.024 | 0.5189 | 0.7416 | 0 | 0 | 0 |
| 'Deb1' | 15.029 | 21.397 | 18.041 | 19.503 | 20.675 | 16.044 | 19.418 | 16.373 | 15.429 | 16.934 | 20.710 | 16.739 | 17.839 | 18.193 | 17.778 | 18.522 | 19.143 | 14.652 | -0.004 | 0.9623 | 0.9866 | 0 | 0 | 0 |
| 'Decr1' | 7.922 | 9.354 | 8.853 | 8.774 | 9.195 | 8.742 | 8.089 | 9.173 | 8.643 | 10.254 | 8.741 | 8.560 | 8.905 | 8.652 | 8.119 | 10.237 | 9.400 | 7.552 | -0.035 | 0.4816 | 0.7136 | 0 | 0 | 0 |
| 'Decr2' | 4.612 | 4.393 | 4.976 | 5.584 | 5.800 | 5.124 | 5.233 | 5.121 | 5.587 | 5.727 | 5.191 | 4.964 | 4.569 | 5.114 | 4.945 | 5.592 | 5.726 | 4.497 | -0.002 | 0.9791 | 0.9926 | 0 | 0 | 0 |
| 'Dedd' | 2.506 | 2.562 | 2.754 | 2.310 | 2.513 | 2.410 | 2.182 | 2.467 | 2.811 | 2.334 | 2.628 | 3.075 | 2.318 | 2.407 | 2.775 | 2.516 | 2.561 | 3.046 | -0.083 | 0.0848 | 0.2643 | 0 | 0 | 0 |
| 'Dedd2' | 13.906 | 12.483 | 13.618 | 13.710 | 12.540 | 13.451 | 12.906 | 12.885 | 13.900 | 12.417 | 12.506 | 18.082 | 11.430 | 11.920 | 15.651 | 12.441 | 11.992 | 18.397 | -0.056 | 0.3515 | 0.6056 | 0 | 0 | 0 |
| 'Def6' | 97.935 | 84.550 | 102.233 | 88.002 | 85.891 | 108.559 | 91.834 | 87.667 | 100.745 | 95.607 | 82.359 | 104.476 | 95.676 | 84.764 | 104.125 | 95.223 | 82.624 | 98.835 | -0.002 | 0.9180 | 0.9687 | 0 | 0 | 0 |
| 'Def8' | 5.145 | 5.397 | 5.217 | 5.151 | 5.785 | 6.052 | 4.870 | 5.996 | 5.921 | 5.673 | 5.440 | 4.995 | 5.963 | 5.166 | 5.243 | 5.293 | 5.150 | 5.074 | 0.038 | 0.4455 | 0.6878 | 0 | 0 | 0 |
| 'Degs1' | 17.799 | 20.967 | 18.186 | 18.364 | 19.817 | 16.837 | 18.633 | 17.746 | 17.365 | 16.534 | 19.348 | 19.549 | 18.594 | 19.830 | 17.659 | 16.908 | 20.866 | 17.937 | -0.027 | 0.4889 | 0.7189 | 0 | 0 | 0 |
| 'Degs2' | 15.008 | 14.049 | 17.467 | 14.085 | 14.252 | 15.150 | 16.188 | 14.361 | 17.117 | 12.813 | 14.759 | 18.255 | 14.276 | 13.612 | 16.957 | 12.347 | 14.880 | 18.886 | 0.004 | 0.9459 | 0.9791 | 0 | 0 | 0 |
| 'Dek' | 45.681 | 47.927 | 47.108 | 49.816 | 48.103 | 45.127 | 50.779 | 47.701 | 45.429 | 41.942 | 47.107 | 43.377 | 48.173 | 48.708 | 44.483 | 43.091 | 47.026 | 41.873 | 0.066 | 0.0316 | 0.1395 | 0 | 0 | 0 |
| 'Dennd1a' | 2.528 | 2.265 | 2.491 | 2.493 | 2.230 | 2.645 | 2.229 | 2.385 | 2.550 | 2.632 | 2.315 | 2.390 | 2.488 | 2.303 | 2.317 | 2.462 | 2.089 | 2.300 | 0.030 | 0.4397 | 0.6831 | 0 | 0 | 0 |
| 'Dennd1b' | 1.845 | 1.794 | 1.704 | 1.605 | 1.620 | 1.729 | 1.681 | 1.613 | 1.591 | 1.892 | 1.916 | 1.706 | 1.739 | 1.771 | 1.495 | 1.748 | 1.730 | 1.650 | -0.051 | 0.2362 | 0.4860 | 0 | 0 | 0 |
| 'Dennd1c' | 100.627 | 97.695 | 122.580 | 97.807 | 96.735 | 114.497 | 110.233 | 99.271 | 126.418 | 107.379 | 99.913 | 130.386 | 108.298 | 97.457 | 122.410 | 101.690 | 107.562 | 127.579 | -0.065 | 0.0223 | 0.1109 | 0 | 0 | 0 |
| 'Dennd2d' | 178.370 | 197.937 | 187.993 | 176.997 | 188.106 | 178.866 | 180.162 | 190.832 | 192.831 | 172.185 | 182.103 | 180.676 | 175.462 | 178.040 | 181.866 | 169.141 | 186.825 | 182.709 | 0.044 | 0.0199 | 0.1028 | 0 | 0 | 0 |
| 'Dennd3' | 1.857 | 1.829 | 1.976 | 1.716 | 1.557 | 1.813 | 1.820 | 1.842 | 1.977 | 1.411 | 1.582 | 1.797 | 1.665 | 1.437 | 1.716 | 1.868 | 1.311 | 1.314 | 0.209 | 0.0021 | 0.0187 | 1 | 0 | 0 |
| 'Dennd4a' | 2.535 | 2.850 | 2.736 | 2.707 | 2.755 | 2.719 | 2.925 | 3.025 | 2.608 | 2.524 | 2.568 | 2.583 | 2.411 | 2.559 | 2.718 | 2.322 | 2.435 | 2.526 | 0.120 | 0.0102 | 0.0634 | 0 | 0 | 0 |
| 'Dennd4b' | 27.032 | 26.031 | 29.962 | 27.053 | 26.440 | 27.773 | 25.874 | 25.458 | 30.302 | 28.391 | 29.092 | 31.692 | 30.343 | 27.512 | 30.071 | 26.724 | 28.762 | 31.488 | -0.114 | 0.0000 | 0.0008 | 1 | 0 | 0 |
| 'Dennd4c' | 13.857 | 13.737 | 13.233 | 14.915 | 13.763 | 12.651 | 14.063 | 13.187 | 13.402 | 13.207 | 14.490 | 15.644 | 12.757 | 13.494 | 14.385 | 13.804 | 13.581 | 15.642 | -0.060 | 0.1728 | 0.4066 | 0 | 0 | 0 |
| 'Dennd5a' | 0.560 | 0.560 | 0.524 | 0.520 | 0.874 | 0.622 | 0.666 | 0.801 | 0.556 | 0.333 | 0.415 | 0.489 | 0.585 | 0.453 | 0.480 | 0.616 | 0.430 | 0.496 | 0.379 | 0.0013 | 0.0130 | 1 | 1 | 0 |
| 'Dennd6a' | 4.734 | 5.568 | 5.348 | 5.003 | 4.808 | 5.076 | 5.218 | 5.611 | 5.128 | 4.691 | 5.777 | 5.926 | 4.861 | 5.603 | 5.516 | 4.142 | 5.223 | 5.844 | -0.044 | 0.3668 | 0.6189 | 0 | 0 | 0 |

| | | | | | | | | | | | | | | | | | | | | | | | | |
|---|---|---|---|---|---|---|---|---|---|---|---|---|---|---|---|---|---|---|---|---|---|---|---|---|
| 'Dennd6b' | 8.232 | 8.140 | 8.924 | 7.572 | 7.474 | 8.977 | 7.668 | 8.566 | 9.639 | 7.879 | 7.348 | 9.507 | 8.114 | 6.588 | 8.329 | 7.666 | 7.504 | 8.493 | 0.062 | 0.1541 | 0.3820 | 0 | 0 | 0 |
| 'Denr' | 8.579 | 9.347 | 8.813 | 8.482 | 10.047 | 9.464 | 10.044 | 10.094 | 9.107 | 8.917 | 8.557 | 7.729 | 9.800 | 9.169 | 7.914 | 9.671 | 9.144 | 7.575 | 0.093 | 0.0346 | 0.1489 | 0 | 0 | 0 |
| 'Depdc1a' | 2.405 | 2.618 | 3.468 | 2.602 | 3.414 | 3.253 | 3.375 | 2.726 | 3.622 | 1.982 | 3.793 | 3.303 | 3.750 | 3.918 | 3.448 | 3.328 | 3.496 | 2.837 | -0.128 | 0.2081 | 0.4526 | 0 | 0 | 0 |
| 'Depdc1b' | 1.440 | 1.634 | 2.494 | 2.048 | 1.997 | 2.341 | 2.025 | 1.954 | 2.591 | 1.308 | 2.160 | 2.622 | 2.273 | 2.546 | 2.680 | 2.010 | 2.470 | 2.458 | -0.160 | 0.0816 | 0.2578 | 0 | 0 | 0 |
| 'Depdc5' | 3.143 | 2.862 | 3.426 | 3.724 | 3.043 | 3.352 | 3.176 | 3.261 | 3.346 | 3.178 | 3.638 | 3.279 | 3.542 | 3.410 | 3.332 | 3.391 | 3.306 | 3.335 | -0.061 | 0.1602 | 0.3904 | 0 | 0 | 0 |
| 'Dera' | 5.760 | 5.320 | 5.832 | 5.674 | 5.687 | 5.682 | 5.988 | 5.805 | 5.606 | 5.432 | 5.606 | 5.468 | 6.074 | 5.634 | 5.606 | 5.701 | 5.568 | 5.339 | 0.019 | 0.5308 | 0.7514 | 0 | 0 | 0 |
| 'Derl1' | 101.361 | 111.350 | 97.439 | 88.554 | 103.152 | 99.417 | 86.866 | 110.208 | 96.732 | 103.873 | 100.220 | 98.137 | 104.642 | 111.808 | 101.744 | 107.396 | 102.726 | 94.898 | -0.058 | 0.1308 | 0.3473 | 0 | 0 | 0 |
| 'Derl2' | 5.636 | 7.436 | 6.734 | 6.298 | 7.176 | 6.594 | 6.334 | 7.158 | 6.549 | 5.893 | 6.332 | 7.278 | 6.198 | 6.660 | 7.145 | 6.402 | 6.859 | 6.889 | -0.011 | 0.7619 | 0.8944 | 0 | 0 | 0 |
| 'Desi1' | 10.804 | 10.946 | 10.897 | 10.764 | 9.939 | 11.364 | 10.600 | 10.467 | 10.530 | 9.660 | 11.019 | 11.177 | 11.234 | 10.780 | 10.796 | 9.948 | 10.723 | 10.880 | -0.010 | 0.7685 | 0.8967 | 0 | 0 | 0 |
| 'Desi2' | 8.943 | 9.929 | 8.565 | 9.204 | 9.604 | 8.745 | 9.507 | 9.547 | 9.181 | 9.083 | 9.885 | 9.931 | 9.595 | 9.630 | 9.852 | 9.848 | 9.551 | 10.167 | -0.087 | 0.0047 | 0.0361 | 1 | 0 | 0 |
| 'Det1' | 5.193 | 5.827 | 5.422 | 5.581 | 6.158 | 5.305 | 4.730 | 5.578 | 5.748 | 5.318 | 5.431 | 5.315 | 5.358 | 5.011 | 5.189 | 5.269 | 5.204 | 5.528 | 0.044 | 0.3361 | 0.5947 | 0 | 0 | 0 |
| 'Dexi' | 7.073 | 6.528 | 6.900 | 5.839 | 7.061 | 7.782 | 5.874 | 7.605 | 7.081 | 6.568 | 7.185 | 7.028 | 6.819 | 6.597 | 7.281 | 7.152 | 6.340 | 7.525 | -0.022 | 0.6788 | 0.8464 | 0 | 0 | 0 |
| 'Dffa' | 14.192 | 15.014 | 15.918 | 14.936 | 15.841 | 17.944 | 15.008 | 15.477 | 17.519 | 15.031 | 13.867 | 16.568 | 16.369 | 14.662 | 17.185 | 15.467 | 15.391 | 16.082 | 0.001 | 0.9762 | 0.9918 | 0 | 0 | 0 |
| 'Dffb' | 4.124 | 4.059 | 4.373 | 4.507 | 4.630 | 4.856 | 4.470 | 4.775 | 4.114 | 4.346 | 4.471 | 4.836 | 4.505 | 3.888 | 4.598 | 4.053 | 4.566 | 4.597 | -0.013 | 0.7969 | 0.9097 | 0 | 0 | 0 |
| 'Dfna5' | 0.554 | 0.586 | 0.663 | 0.615 | 0.490 | 0.543 | 0.582 | 0.552 | 0.620 | 0.643 | 0.603 | 0.515 | 0.497 | 0.562 | 0.461 | 0.441 | 0.630 | 0.482 | 0.105 | 0.2215 | 0.4691 | 0 | 0 | 0 |
| 'Dgat1' | 4.923 | 5.031 | 5.721 | 4.914 | 5.499 | 5.071 | 4.902 | 5.511 | 5.291 | 5.373 | 5.153 | 5.199 | 5.344 | 5.100 | 4.848 | 4.484 | 5.550 | 4.384 | 0.036 | 0.4562 | 0.6941 | 0 | 0 | 0 |
| 'Dgcr14' | 17.937 | 18.813 | 21.701 | 17.022 | 17.916 | 22.191 | 18.023 | 19.546 | 21.323 | 20.545 | 17.986 | 20.147 | 19.165 | 18.013 | 20.092 | 20.790 | 15.675 | 17.810 | 0.027 | 0.5829 | 0.7850 | 0 | 0 | 0 |
| 'Dgcr6' | 2.983 | 3.948 | 3.407 | 3.314 | 3.280 | 3.862 | 3.333 | 3.698 | 3.555 | 3.098 | 2.788 | 3.084 | 3.811 | 2.776 | 3.291 | 3.620 | 2.963 | 3.219 | 0.122 | 0.0588 | 0.2090 | 0 | 0 | 0 |
| 'Dgcr8' | 4.857 | 5.009 | 5.701 | 4.720 | 5.417 | 5.621 | 4.425 | 5.409 | 5.550 | 4.304 | 4.675 | 5.241 | 5.029 | 4.779 | 5.076 | 5.124 | 5.025 | 5.203 | 0.061 | 0.1031 | 0.3009 | 0 | 0 | 0 |
| 'Dgka' | 298.290 | 264.024 | 320.502 | 298.841 | 269.164 | 301.230 | 299.175 | 280.539 | 320.219 | 307.082 | 311.592 | 339.637 | 296.073 | 281.536 | 326.332 | 280.655 | 307.445 | 327.265 | -0.078 | 0.0132 | 0.0766 | 0 | 0 | 0 |
| 'Dgkd' | 31.706 | 34.043 | 31.923 | 31.084 | 32.387 | 32.090 | 29.222 | 32.410 | 31.939 | 31.662 | 30.702 | 30.538 | 31.639 | 30.298 | 30.263 | 31.579 | 29.388 | 30.796 | 0.040 | 0.1132 | 0.3194 | 0 | 0 | 0 |
| 'Dgke' | 2.231 | 2.478 | 2.885 | 2.335 | 2.516 | 2.413 | 2.399 | 2.476 | 2.542 | 2.658 | 2.465 | 2.309 | 2.472 | 2.494 | 2.153 | 2.415 | 2.335 | 2.359 | 0.035 | 0.4862 | 0.7170 | 0 | 0 | 0 |
| 'Dgkq' | 1.762 | 1.660 | 2.016 | 1.657 | 1.814 | 1.834 | 1.883 | 1.749 | 2.034 | 1.819 | 1.815 | 1.919 | 1.687 | 1.786 | 1.767 | 1.750 | 1.921 | 1.771 | 0.007 | 0.8629 | 0.9405 | 0 | 0 | 0 |
| 'Dgkz' | 112.595 | 103.566 | 119.560 | 111.426 | 102.141 | 120.258 | 108.795 | 99.291 | 123.315 | 124.078 | 112.554 | 128.606 | 129.457 | 115.499 | 126.521 | 123.629 | 122.231 | 131.217 | -0.167 | 0.0000 | 0.0000 | 1 | 0 | 0 |
| 'Dguok' | 8.639 | 10.085 | 8.956 | 9.092 | 8.903 | 9.974 | 8.283 | 9.212 | 9.951 | 9.822 | 8.825 | 9.282 | 8.910 | 9.246 | 8.755 | 8.577 | 9.098 | 8.344 | 0.028 | 0.4849 | 0.7162 | 0 | 0 | 0 |
| 'Dhcr24' | 15.764 | 17.607 | 19.496 | 17.723 | 16.011 | 20.078 | 16.570 | 15.915 | 19.788 | 17.321 | 17.982 | 18.854 | 17.138 | 18.280 | 21.235 | 16.585 | 15.677 | 18.975 | -0.039 | 0.3275 | 0.5852 | 0 | 0 | 0 |
| 'Dhcr7' | 9.680 | 10.077 | 10.535 | 10.083 | 9.977 | 10.304 | 9.606 | 9.430 | 10.680 | 10.229 | 11.088 | 11.441 | 10.365 | 10.479 | 11.419 | 9.050 | 10.576 | 11.117 | -0.096 | 0.0041 | 0.0321 | 1 | 0 | 0 |
| 'Dhdds' | 2.807 | 3.000 | 2.934 | 2.896 | 2.907 | 3.044 | 3.057 | 2.982 | 2.901 | 2.874 | 2.908 | 2.966 | 2.817 | 2.760 | 2.869 | 2.816 | 2.910 | 2.938 | 0.025 | 0.4480 | 0.6883 | 0 | 0 | 0 |
| 'Dhdh' | 1.944 | 1.628 | 1.816 | 1.625 | 2.176 | 1.870 | 2.304 | 1.821 | 1.888 | 1.648 | 1.738 | 1.678 | 1.742 | 1.877 | 1.834 | 1.467 | 1.669 | 1.723 | 0.138 | 0.1032 | 0.3013 | 0 | 0 | 0 |
| 'Dhfr' | 4.138 | 4.084 | 5.129 | 4.806 | 4.440 | 4.601 | 5.425 | 3.960 | 5.148 | 4.207 | 4.188 | 4.823 | 4.989 | 4.173 | 5.263 | 4.379 | 4.341 | 4.262 | 0.024 | 0.6683 | 0.8400 | 0 | 0 | 0 |
| 'Dhodh' | 1.797 | 1.534 | 2.006 | 1.767 | 1.779 | 1.836 | 1.800 | 1.704 | 1.980 | 1.870 | 1.450 | 1.837 | 1.834 | 1.495 | 1.730 | 1.632 | 1.698 | 1.605 | 0.087 | 0.0913 | 0.2786 | 0 | 0 | 0 |
| 'Dhps' | 7.809 | 8.295 | 9.286 | 8.597 | 9.190 | 9.614 | 8.354 | 8.899 | 9.787 | 8.627 | 8.195 | 9.266 | 8.447 | 8.056 | 8.656 | 8.005 | 8.728 | 7.958 | 0.060 | 0.1264 | 0.3405 | 0 | 0 | 0 |
| 'Dhrs1' | 10.609 | 14.354 | 10.220 | 10.952 | 14.166 | 11.130 | 11.319 | 12.258 | 10.608 | 11.720 | 11.611 | 10.572 | 12.663 | 11.575 | 11.432 | 11.418 | 12.223 | 9.336 | 0.024 | 0.6185 | 0.8092 | 0 | 0 | 0 |
| 'Dhrs11' | 7.262 | 7.931 | 7.807 | 5.496 | 7.514 | 8.367 | 6.421 | 7.709 | 7.580 | 7.299 | 5.521 | 6.744 | 6.014 | 6.659 | 7.089 | 7.117 | 6.506 | 6.138 | 0.154 | 0.0246 | 0.1192 | 0 | 0 | 0 |
| 'Dhrs13' | 1.019 | 1.299 | 1.470 | 1.158 | 1.232 | 1.648 | 1.175 | 1.113 | 1.399 | 1.322 | 1.030 | 1.362 | 1.160 | 1.096 | 1.444 | 1.112 | 1.040 | 1.463 | 0.051 | 0.5543 | 0.7669 | 0 | 0 | 0 |
| 'Dhrs3' | 7.412 | 7.587 | 7.613 | 7.712 | 7.012 | 8.056 | 7.617 | 6.399 | 8.563 | 8.175 | 6.715 | 9.382 | 7.338 | 7.655 | 8.130 | 8.998 | 8.273 | 8.348 | -0.117 | 0.0197 | 0.1020 | 0 | 0 | 0 |
| 'Dhrs4' | 16.687 | 18.852 | 17.770 | 16.890 | 19.354 | 20.182 | 17.201 | 18.346 | 18.120 | 16.958 | 18.125 | 18.678 | 19.905 | 18.398 | 18.432 | 18.629 | 17.299 | 17.853 | -0.020 | 0.6680 | 0.8400 | 0 | 0 | 0 |
| 'Dhrs7' | 25.540 | 30.634 | 28.838 | 28.397 | 29.115 | 27.716 | 26.565 | 29.713 | 28.098 | 25.572 | 28.018 | 26.433 | 27.264 | 26.039 | 25.786 | 28.442 | 28.394 | 25.539 | 0.067 | 0.0822 | 0.2593 | 0 | 0 | 0 |
| 'Dhrs7b' | 19.031 | 19.291 | 18.978 | 18.096 | 17.714 | 19.793 | 17.975 | 19.580 | 20.097 | 19.330 | 16.686 | 19.312 | 19.257 | 17.086 | 20.152 | 18.207 | 19.719 | 18.812 | 0.007 | 0.8839 | 0.9514 | 0 | 0 | 0 |
| 'Dhx15' | 31.811 | 33.208 | 32.024 | 34.555 | 34.136 | 30.810 | 34.649 | 34.356 | 31.316 | 31.103 | 33.548 | 33.131 | 31.942 | 32.582 | 31.318 | 30.158 | 34.782 | 32.465 | 0.017 | 0.5529 | 0.7658 | 0 | 0 | 0 |
| 'Dhx16' | 19.165 | 18.973 | 23.294 | 19.162 | 19.309 | 22.279 | 19.573 | 19.646 | 22.646 | 21.213 | 18.359 | 22.796 | 21.563 | 19.325 | 21.043 | 20.041 | 20.433 | 20.779 | -0.024 | 0.4244 | 0.6710 | 0 | 0 | 0 |
| 'Dhx29' | 14.046 | 14.444 | 16.727 | 14.853 | 14.657 | 15.195 | 15.013 | 15.079 | 15.371 | 15.701 | 16.247 | 15.379 | 14.024 | 14.130 | 13.927 | 14.174 | 14.936 | 15.560 | 0.005 | 0.9023 | 0.9612 | 0 | 0 | 0 |
| 'Dhx30' | 5.670 | 5.729 | 6.459 | 6.155 | 5.398 | 5.888 | 5.356 | 5.670 | 6.229 | 5.718 | 5.493 | 6.334 | 5.943 | 5.098 | 5.964 | 5.355 | 5.608 | 6.065 | 0.016 | 0.6174 | 0.8087 | 0 | 0 | 0 |
| 'Dhx32' | 5.336 | 5.834 | 4.967 | 5.191 | 5.351 | 5.370 | 5.245 | 5.234 | 5.215 | 5.119 | 4.335 | 4.644 | 5.561 | 4.764 | 4.606 | 5.571 | 4.686 | 4.590 | 0.114 | 0.0058 | 0.0418 | 1 | 0 | 0 |
| 'Dhx33' | 3.175 | 3.011 | 2.916 | 2.908 | 2.704 | 2.956 | 2.840 | 3.106 | 2.878 | 2.835 | 2.967 | 3.006 | 2.541 | 2.580 | 2.826 | 2.751 | 2.714 | 3.025 | 0.060 | 0.1718 | 0.4055 | 0 | 0 | 0 |
| 'Dhx34' | 30.423 | 35.875 | 33.826 | 30.181 | 33.025 | 34.191 | 30.647 | 31.492 | 35.159 | 35.946 | 30.798 | 34.530 | 39.746 | 32.308 | 32.528 | 36.607 | 35.100 | 33.288 | -0.086 | 0.0539 | 0.1984 | 0 | 0 | 0 |
| 'Dhx35' | 4.418 | 3.849 | 5.025 | 4.386 | 4.994 | 5.092 | 4.223 | 4.738 | 4.700 | 5.033 | 4.865 | 4.701 | 4.779 | 4.226 | 4.593 | 4.973 | 4.961 | 4.209 | -0.038 | 0.4747 | 0.7098 | 0 | 0 | 0 |
| 'Dhx36' | 8.523 | 8.797 | 9.402 | 9.444 | 8.884 | 8.127 | 9.941 | 9.381 | 8.972 | 8.654 | 9.860 | 9.189 | 8.549 | 9.217 | 8.332 | 8.631 | 9.993 | 7.987 | 0.010 | 0.8102 | 0.9142 | 0 | 0 | 0 |
| 'Dhx37' | 9.273 | 8.470 | 10.091 | 8.812 | 8.324 | 9.816 | 8.483 | 8.891 | 10.430 | 9.585 | 9.160 | 9.967 | 9.643 | 8.360 | 9.047 | 9.902 | 9.009 | 9.459 | -0.034 | 0.3603 | 0.6126 | 0 | 0 | 0 |
| 'Dhx38' | 35.356 | 32.675 | 36.130 | 34.457 | 33.571 | 36.549 | 32.299 | 33.849 | 35.274 | 36.762 | 33.428 | 35.101 | 37.655 | 33.075 | 35.025 | 36.525 | 33.595 | 33.372 | -0.028 | 0.3128 | 0.5712 | 0 | 0 | 0 |
| 'Dhx40' | 10.302 | 11.358 | 11.296 | 11.815 | 12.648 | 11.254 | 11.572 | 12.401 | 11.000 | 11.873 | 12.226 | 9.996 | 10.884 | 11.383 | 9.937 | 11.896 | 11.501 | 9.546 | 0.059 | 0.1404 | 0.3626 | 0 | 0 | 0 |
| 'Dhx57' | 13.224 | 14.865 | 14.332 | 12.228 | 14.159 | 15.122 | 12.023 | 14.348 | 15.799 | 14.204 | 13.095 | 13.623 | 13.633 | 13.722 | 13.770 | 13.399 | 13.803 | 13.374 | 0.029 | 0.4601 | 0.6967 | 0 | 0 | 0 |
| 'Dhx58' | 5.913 | 5.923 | 6.591 | 5.609 | 6.674 | 7.337 | 5.593 | 7.860 | 7.551 | 5.819 | 5.958 | 7.551 | 6.755 | 5.942 | 7.830 | 6.373 | 5.311 | 7.131 | 0.000 | 0.9983 | 0.9994 | 0 | 0 | 0 |
| 'Dhx8' | 17.293 | 17.175 | 18.219 | 16.696 | 17.426 | 18.785 | 16.943 | 17.540 | 18.653 | 18.258 | 17.971 | 19.750 | 16.002 | 17.416 | 17.856 | 17.798 | 16.746 | 19.096 | -0.030 | 0.3203 | 0.5784 | 0 | 0 | 0 |
| 'Dhx9' | 24.954 | 25.443 | 24.823 | 25.907 | 26.309 | 24.356 | 25.459 | 25.829 | 23.876 | 24.151 | 24.943 | 23.978 | 24.840 | 25.654 | 23.850 | 24.090 | 25.120 | 23.718 | 0.032 | 0.0612 | 0.2139 | 0 | 0 | 0 |
| 'Diablo' | 4.831 | 6.215 | 5.127 | 5.280 | 5.994 | 5.702 | 5.220 | 5.616 | 5.702 | 5.660 | 5.126 | 5.670 | 5.283 | 5.733 | 5.098 | 5.561 | 5.420 | 5.707 | -0.004 | 0.9330 | 0.9746 | 0 | 0 | 0 |
| 'Diaph1' | 19.817 | 19.298 | 21.141 | 20.252 | 19.830 | 21.624 | 19.775 | 20.133 | 21.074 | 20.498 | 19.909 | 21.220 | 19.860 | 20.385 | 20.830 | 19.955 | 18.589 | 21.161 | -0.006 | 0.7780 | 0.9002 | 0 | 0 | 0 |
| 'Diaph2' | 1.009 | 1.023 | 0.911 | 0.861 | 0.997 | 0.831 | 0.993 | 0.976 | 0.850 | 1.005 | 0.918 | 1.025 | 0.983 | 0.968 | 0.927 | 1.061 | 1.165 | 1.082 | -0.129 | 0.0215 | 0.1082 | 0 | 0 | 0 |
| 'Diaph3' | 4.732 | 4.098 | 5.283 | 5.172 | 4.987 | 4.734 | 5.106 | 4.917 | 4.619 | 3.853 | 5.624 | 4.966 | 5.302 | 5.023 | 5.559 | 4.192 | 5.179 | 5.036 | -0.044 | 0.5140 | 0.7381 | 0 | 0 | 0 |
| 'Dicer1' | 18.168 | 19.580 | 18.806 | 18.118 | 17.720 | 19.527 | 16.725 | 19.700 | 18.882 | 19.329 | 19.576 | 20.186 | 18.069 | 19.039 | 18.832 | 19.151 | 19.527 | 20.457 | -0.069 | 0.0181 | 0.0964 | 0 | 0 | 0 |
| 'Dido1' | 10.380 | 9.425 | 9.952 | 11.101 | 9.274 | 8.972 | 11.338 | 9.407 | 10.001 | 9.623 | 10.268 | 10.542 | 9.248 | 9.909 | 9.983 | 9.572 | 10.665 | 10.013 | -0.013 | 0.7749 | 0.8994 | 0 | 0 | 0 |
| 'Diexf' | 2.702 | 2.718 | 2.996 | 2.783 | 2.561 | 2.721 | 2.810 | 2.657 | 2.804 | 2.497 | 2.734 | 3.106 | 2.281 | 2.307 | 2.733 | 2.623 | 2.392 | 3.237 | 0.040 | 0.4499 | 0.6898 | 0 | 0 | 0 |
| 'Dimt1' | 8.961 | 10.023 | 9.520 | 8.745 | 8.622 | 7.398 | 8.684 | 10.516 | 9.534 | 8.973 | 8.435 | 9.444 | 9.407 | 8.227 | 9.508 | 8.787 | 9.430 | 9.867 | -0.017 | 0.8003 | 0.9111 | 0 | 0 | 0 |
| 'Dip2a' | 1.421 | 1.613 | 1.727 | 1.470 | 1.716 | 1.932 | 1.724 | 1.828 | 1.794 | 1.610 | 1.531 | 1.736 | 1.706 | 1.658 | 1.546 | 1.607 | 1.675 | 1.550 | 0.047 | 0.3518 | 0.6056 | 0 | 0 | 0 |
| 'Dip2b' | 19.344 | 20.333 | 19.510 | 19.853 | 20.796 | 18.711 | 18.046 | 21.105 | 19.671 | 22.258 | 24.702 | 20.226 | 23.595 | 23.036 | 19.478 | 21.574 | 21.982 | 20.279 | -0.160 | 0.0000 | 0.0000 | 1 | 0 | 0 |
| 'Dip2c' | 1.188 | 1.286 | 1.108 | 1.272 | 1.192 | 1.238 | 1.138 | 1.320 | 1.134 | 1.216 | 1.055 | 1.118 | 1.028 | 1.205 | 1.254 | 1.018 | 1.336 | 1.331 | 0.031 | 0.6654 | 0.8385 | 0 | 0 | 0 |
| 'Dirc2' | 1.727 | 1.922 | 1.857 | 1.731 | 2.007 | 1.748 | 1.589 | 2.229 | 1.600 | 1.997 | 2.016 | 1.823 | 1.717 | 2.015 | 1.743 | 1.877 | 2.037 | 1.709 | -0.051 | 0.3942 | 0.6434 | 0 | 0 | 0 |
| 'Dis3' | 12.183 | 10.940 | 12.292 | 12.200 | 11.989 | 11.344 | 11.674 | 12.023 | 11.813 | 11.544 | 10.492 | 11.680 | 10.776 | 10.247 | 12.285 | 10.786 | 10.711 | 12.225 | 0.069 | 0.0819 | 0.2586 | 0 | 0 | 0 |
| 'Dis3l' | 4.933 | 4.486 | 4.882 | 4.973 | 5.112 | 5.515 | 4.465 | 4.970 | 5.034 | 4.569 | 4.954 | 5.453 | 5.285 | 4.795 | 5.068 | 4.674 | 4.195 | 5.374 | -0.011 | 0.8084 | 0.9139 | 0 | 0 | 0 |
| 'Dis3l2' | 3.925 | 3.916 | 3.923 | 4.066 | 3.876 | 3.643 | 4.184 | 4.167 | 3.658 | 3.775 | 3.860 | 3.970 | 3.861 | 3.371 | 3.702 | 3.626 | 3.744 | 4.123 | 0.040 | 0.3387 | 0.5961 | 0 | 0 | 0 |
| 'Disp1' | 3.468 | 3.992 | 3.737 | 3.640 | 3.274 | 3.592 | 3.736 | 3.431 | 3.952 | 4.028 | 3.615 | 3.651 | 3.821 | 3.478 | 3.688 | 4.300 | 3.526 | 3.243 | -0.029 | 0.5710 | 0.7777 | 0 | 0 | 0 |
| 'Dkc1' | 8.845 | 8.049 | 8.431 | 8.863 | 8.044 | 7.478 | 8.415 | 7.637 | 8.085 | 7.762 | 8.238 | 9.005 | 8.428 | 7.500 | 8.723 | 8.371 | 8.431 | 8.722 | -0.040 | 0.3347 | 0.5932 | 0 | 0 | 0 |
| 'Dlat' | 8.193 | 8.987 | 8.723 | 7.494 | 9.293 | 8.640 | 8.568 | 8.538 | 8.035 | 8.203 | 8.118 | 8.341 | 9.047 | 8.134 | 8.042 | 8.985 | 8.354 | 8.198 | 0.009 | 0.8197 | 0.9202 | 0 | 0 | 0 |
| 'Dld' | 38.426 | 42.956 | 40.715 | 40.657 | 41.424 | 38.710 | 40.623 | 43.618 | 38.202 | 39.608 | 40.086 | 39.281 | 40.715 | 43.285 | 38.347 | 42.116 | 42.142 | 36.413 | 0.004 | 0.8866 | 0.9533 | 0 | 0 | 0 |
| 'Dlg1' | 6.017 | 6.376 | 6.272 | 6.183 | 6.866 | 6.202 | 6.091 | 7.060 | 6.064 | 6.371 | 6.254 | 6.277 | 6.036 | 6.360 | 5.682 | 6.190 | 6.303 | 5.985 | 0.031 | 0.2452 | 0.4958 | 0 | 0 | 0 |
| 'Dlg3' | 2.594 | 2.733 | 2.603 | 2.452 | 2.722 | 2.527 | 2.764 | 2.810 | 2.433 | 2.633 | 2.356 | 2.197 | 2.469 | 2.566 | 2.367 | 2.608 | 2.610 | 2.160 | 0.103 | 0.0110 | 0.0668 | 0 | 0 | 0 |
| 'Dlg4' | 12.315 | 11.918 | 12.265 | 11.727 | 11.510 | 12.162 | 11.651 | 11.910 | 12.742 | 12.614 | 11.320 | 12.509 | 12.589 | 11.527 | 11.898 | 12.070 | 10.780 | 11.864 | 0.004 | 0.8825 | 0.9506 | 0 | 0 | 0 |
| 'Dlgap4' | 5.423 | 4.867 | 5.511 | 4.901 | 5.150 | 5.702 | 5.007 | 5.150 | 5.551 | 5.875 | 4.872 | 6.048 | 5.367 | 4.933 | 5.655 | 5.251 | 5.091 | 5.808 | -0.059 | 0.0582 | 0.2077 | 0 | 0 | 0 |
| 'Dlgap5' | 2.840 | 3.106 | 3.789 | 3.174 | 3.208 | 3.831 | 3.325 | 2.989 | 3.521 | 2.469 | 2.942 | 3.483 | 3.755 | 3.280 | 3.647 | 2.863 | 3.179 | 3.155 | 0.035 | 0.5595 | 0.7702 | 0 | 0 | 0 |
| 'Dlst' | 58.575 | 58.182 | 62.391 | 56.866 | 59.840 | 60.267 | 59.266 | 60.734 | 63.786 | 59.801 | 60.037 | 62.469 | 63.052 | 56.834 | 60.744 | 57.728 | 60.932 | 57.434 | -0.008 | 0.7429 | 0.8839 | 0 | 0 | 0 |
| 'Dmap1' | 5.677 | 7.387 | 7.740 | 6.216 | 7.523 | 7.812 | 7.072 | 7.807 | 7.985 | 6.799 | 6.543 | 6.569 | 6.947 | 7.069 | 6.551 | 6.849 | 6.219 | 5.789 | 0.124 | 0.0259 | 0.1236 | 0 | 0 | 0 |
| 'Dmtf1' | 6.326 | 6.589 | 6.145 | 6.085 | 6.601 | 6.074 | 6.189 | 6.524 | 6.211 | 5.829 | 6.116 | 6.154 | 5.664 | 5.966 | 5.468 | 6.104 | 6.595 | 5.746 | 0.070 | 0.0082 | 0.0537 | 0 | 0 | 0 |
| 'Dmwd' | 2.603 | 2.257 | 2.664 | 2.616 | 2.458 | 2.153 | 2.106 | 2.312 | 2.394 | 2.505 | 2.164 | 1.891 | 2.930 | 2.262 | 2.148 | 1.867 | 2.337 | 2.058 | 0.095 | 0.2364 | 0.4861 | 0 | 0 | 0 |
| 'Dmxl1' | 2.859 | 3.082 | 2.853 | 3.067 | 2.917 | 2.764 | 2.954 | 3.107 | 2.767 | 3.227 | 3.358 | 2.846 | 2.775 | 3.255 | 2.633 | 2.878 | 3.157 | 2.811 | -0.038 | 0.2769 | 0.5326 | 0 | 0 | 0 |
| 'Dna2' | 1.372 | 1.165 | 1.764 | 1.736 | 1.351 | 1.584 | 1.837 | 1.284 | 1.712 | 1.021 | 1.320 | 1.656 | 1.839 | 1.531 | 1.636 | 1.455 | 1.414 | 1.764 | 0.000 | 0.9959 | 0.9994 | 0 | 0 | 0 |
| 'Dnaaf2' | 5.337 | 5.786 | 6.179 | 6.296 | 6.031 | 6.542 | 6.126 | 5.837 | 6.281 | 5.404 | 5.504 | 6.177 | 6.069 | 6.135 | 6.676 | 5.604 | 5.489 | 5.825 | 0.028 | 0.5725 | 0.7792 | 0 | 0 | 0 |
| 'Dnaaf5' | 6.694 | 6.236 | 8.353 | 6.457 | 6.862 | 9.366 | 5.914 | 6.935 | 7.478 | 8.818 | 5.972 | 7.306 | 7.649 | 6.423 | 7.048 | 7.302 | 5.770 | 6.789 | 0.021 | 0.7719 | 0.8988 | 0 | 0 | 0 |
| 'Dnah12' | 0.388 | 0.611 | 0.471 | 0.324 | 1.015 | 0.629 | 0.645 | 0.407 | 0.222 | 0.281 | 0.415 | 0.684 | 0.326 | 0.699 | 0.763 | 0.512 | 0.444 | 0.271 | 0.083 | 0.7620 | 0.8944 | 0 | 0 | 0 |
| 'Dnah17' | 39.531 | 31.779 | 40.671 | 38.930 | 30.861 | 34.459 | 35.259 | 32.766 | 41.555 | 0.298 | 0.332 | 0.410 | 0.360 | 0.277 | 0.294 | 0.308 | 0.266 | 0.272 | 6.828 | 0.0000 | 0.0000 | 1 | 1 | 1 |
| 'Dnah8' | 26.484 | 17.082 | 29.171 | 27.893 | 20.088 | 27.446 | 25.989 | 20.751 | 28.286 | 15.439 | 16.143 | 19.297 | 12.802 | 14.079 | 18.603 | 14.394 | 15.414 | 19.169 | 0.600 | 0.0000 | 0.0000 | 1 | 1 | 0 |
| 'Dnaja1' | 34.489 | 20.590 | 19.040 | 34.767 | 21.344 | 19.027 | 29.985 | 19.834 | 19.000 | 22.864 | 27.788 | 33.826 | 23.465 | 25.247 | 29.363 | 22.341 | 32.806 | 38.178 | -0.273 | 0.1005 | 0.2967 | 0 | 0 | 0 |
| 'Dnaja2' | 75.587 | 80.584 | 74.389 | 80.237 | 79.882 | 71.186 | 82.519 | 79.399 | 71.732 | 73.874 | 76.816 | 75.121 | 69.786 | 78.796 | 75.424 | 72.829 | 79.992 | 75.813 | 0.023 | 0.4445 | 0.6875 | 0 | 0 | 0 |
| 'Dnaja3' | 8.203 | 7.381 | 7.623 | 7.869 | 7.910 | 7.927 | 7.606 | 7.500 | 7.910 | 7.493 | 7.656 | 8.691 | 7.264 | 7.735 | 8.221 | 7.808 | 7.848 | 8.096 | -0.031 | 0.4075 | 0.6564 | 0 | 0 | 0 |
| 'Dnaja4' | 7.378 | 6.600 | 6.859 | 7.999 | 6.467 | 7.022 | 6.677 | 6.760 | 7.103 | 6.727 | 5.492 | 7.742 | 6.070 | 6.642 | 7.335 | 5.859 | 5.847 | 8.039 | 0.063 | 0.2934 | 0.5497 | 0 | 0 | 0 |
| 'Dnajb1' | 211.095 | 49.614 | 56.317 | 218.872 | 45.969 | 49.799 | 248.594 | 50.560 | 47.925 | 65.830 | 132.203 | 105.319 | 74.801 | 79.194 | 86.419 | 49.901 | 158.650 | 105.153 | -0.238 | 0.5250 | 0.7470 | 0 | 0 | 0 |
| 'Dnajb11' | 27.175 | 29.894 | 30.736 | 27.886 | 32.903 | 30.804 | 28.580 | 32.225 | 31.185 | 30.974 | 27.706 | 28.615 | 29.914 | 29.157 | 27.116 | 30.641 | 27.787 | 28.693 | 0.048 | 0.1961 | 0.4350 | 0 | 0 | 0 |
| 'Dnajb12' | 6.224 | 7.119 | 7.159 | 5.254 | 6.386 | 7.117 | 5.736 | 7.053 | 6.784 | 6.569 | 6.510 | 6.528 | 6.730 | 6.406 | 6.390 | 7.366 | 5.602 | 6.293 | 0.000 | 0.9956 | 0.9994 | 0 | 0 | 0 |
| 'Dnajb13' | 2.082 | 1.127 | 1.085 | 1.955 | 1.339 | 0.972 | 1.651 | 1.433 | 1.218 | 2.104 | 1.329 | 1.567 | 1.548 | 1.399 | 1.155 | 2.063 | 1.276 | 1.801 | -0.177 | 0.1424 | 0.3655 | 0 | 0 | 0 |
| 'Dnajb14' | 1.472 | 1.795 | 1.729 | 1.631 | 1.882 | 1.604 | 1.629 | 1.815 | 1.802 | 1.698 | 1.891 | 1.795 | 1.644 | 1.923 | 1.672 | 1.620 | 1.749 | 1.824 | -0.056 | 0.1870 | 0.4227 | 0 | 0 | 0 |
| 'Dnajb2' | 2.841 | 2.689 | 2.473 | 2.708 | 2.987 | 2.443 | 2.315 | 2.861 | 2.677 | 2.848 | 2.378 | 3.204 | 2.302 | 2.583 | 2.739 | 2.600 | 2.511 | 3.047 | -0.027 | 0.6838 | 0.8501 | 0 | 0 | 0 |
| 'Dnajb5' | 2.984 | 3.420 | 3.414 | 2.956 | 2.985 | 3.793 | 3.222 | 3.210 | 3.579 | 2.944 | 2.971 | 3.046 | 2.953 | 2.895 | 3.045 | 3.255 | 2.915 | 2.758 | 0.136 | 0.0047 | 0.0356 | 1 | 0 | 0 |
| 'Dnajb6' | 15.555 | 16.436 | 15.544 | 16.026 | 16.231 | 15.319 | 15.061 | 15.118 | 15.782 | 14.596 | 15.492 | 15.524 | 14.632 | 15.943 | 15.237 | 15.285 | 15.513 | 14.859 | 0.030 | 0.2298 | 0.4766 | 0 | 0 | 0 |
| 'Dnajb9' | 46.626 | 35.646 | 15.218 | 34.382 | 34.932 | 18.422 | 31.626 | 37.852 | 13.816 | 46.491 | 38.924 | 15.168 | 38.284 | 48.651 | 15.488 | 47.447 | 36.951 | 16.122 | -0.148 | 0.0555 | 0.2019 | 0 | 0 | 0 |
| 'Dnajc1' | 6.249 | 5.822 | 7.191 | 6.574 | 5.892 | 6.730 | 7.122 | 6.106 | 6.911 | 6.957 | 6.974 | 7.056 | 6.564 | 6.426 | 6.617 | 6.496 | 6.979 | 6.565 | -0.061 | 0.1117 | 0.3167 | 0 | 0 | 0 |
| 'Dnajc10' | 15.985 | 19.348 | 18.534 | 17.042 | 19.744 | 17.991 | 17.391 | 19.112 | 18.485 | 17.582 | 18.970 | 17.226 | 17.515 | 18.495 | 16.999 | 18.573 | 17.791 | 16.223 | 0.028 | 0.3613 | 0.6135 | 0 | 0 | 0 |
| 'Dnajc11' | 3.689 | 3.298 | 3.947 | 3.488 | 3.515 | 3.711 | 3.909 | 3.426 | 3.847 | 3.815 | 3.690 | 3.763 | 3.864 | 3.106 | 3.710 | 3.763 | 3.387 | 3.509 | 0.002 | 0.9615 | 0.9862 | 0 | 0 | 0 |
| 'Dnajc13' | 7.918 | 8.016 | 7.668 | 7.978 | 8.017 | 7.779 | 7.529 | 8.064 | 7.725 | 7.751 | 8.762 | 8.390 | 7.344 | 7.856 | 7.651 | 8.238 | 8.624 | 8.431 | -0.058 | 0.0816 | 0.2579 | 0 | 0 | 0 |
| 'Dnajc14' | 51.293 | 55.139 | 52.475 | 51.756 | 57.852 | 55.681 | 51.266 | 54.790 | 52.994 | 52.830 | 55.471 | 53.672 | 49.157 | 56.550 | 56.284 | 56.734 | 52.564 | 54.375 | -0.023 | 0.4429 | 0.6865 | 0 | 0 | 0 |
| 'Dnajc15' | 18.434 | 19.130 | 19.863 | 18.059 | 22.696 | 19.113 | 19.394 | 21.049 | 17.640 | 17.460 | 15.552 | 18.806 | 17.938 | 20.866 | 17.730 | 18.798 | 19.576 | 17.818 | 0.077 | 0.2173 | 0.4639 | 0 | 0 | 0 |
| 'Dnajc16' | 4.914 | 5.801 | 5.015 | 4.108 | 5.387 | 5.527 | 4.884 | 6.221 | 5.702 | 4.941 | 4.938 | 5.314 | 5.629 | 5.085 | 5.529 | 6.448 | 4.612 | 5.269 | -0.018 | 0.7860 | 0.9045 | 0 | 0 | 0 |
| 'Dnajc17' | 11.547 | 13.694 | 15.000 | 14.620 | 13.836 | 15.748 | 15.075 | 12.972 | 16.535 | 15.457 | 14.767 | 13.795 | 15.313 | 13.712 | 13.987 | 15.662 | 14.450 | 11.587 | -0.004 | 0.9518 | 0.9818 | 0 | 0 | 0 |
| 'Dnajc18' | 5.386 | 6.270 | 6.009 | 5.881 | 5.844 | 5.439 | 5.546 | 5.706 | 5.615 | 5.734 | 5.941 | 5.922 | 5.850 | 5.176 | 5.735 | 5.277 | 5.273 | 6.156 | 0.002 | 0.9698 | 0.9898 | 0 | 0 | 0 |
| 'Dnajc19' | 5.212 | 6.653 | 5.773 | 5.393 | 6.364 | 5.414 | 5.121 | 5.699 | 5.203 | 4.908 | 5.738 | 5.092 | 5.598 | 5.844 | 5.185 | 5.329 | 5.257 | 4.920 | 0.074 | 0.1075 | 0.3084 | 0 | 0 | 0 |
| 'Dnajc2' | 12.652 | 11.718 | 12.982 | 12.789 | 12.281 | 11.815 | 12.995 | 12.754 | 12.007 | 11.222 | 11.856 | 12.279 | 11.994 | 11.936 | 11.782 | 11.423 | 12.963 | 11.901 | 0.050 | 0.1232 | 0.3352 | 0 | 0 | 0 |
| 'Dnajc21' | 8.723 | 8.194 | 7.588 | 8.970 | 8.537 | 8.230 | 8.740 | 9.198 | 8.078 | 7.616 | 8.126 | 8.068 | 8.523 | 8.684 | 7.687 | 8.685 | 8.915 | 8.002 | 0.026 | 0.4636 | 0.6997 | 0 | 0 | 0 |
| 'Dnajc24' | 7.508 | 7.900 | 7.375 | 7.001 | 8.156 | 7.110 | 7.894 | 7.381 | 7.782 | 6.875 | 8.184 | 7.587 | 7.344 | 8.525 | 6.904 | 8.532 | 7.827 | 6.885 | -0.020 | 0.7233 | 0.8730 | 0 | 0 | 0 |
| 'Dnajc25' | 1.586 | 1.673 | 1.224 | 1.189 | 1.174 | 1.253 | 1.585 | 1.591 | 1.313 | 1.407 | 1.306 | 1.098 | 1.352 | 1.330 | 1.101 | 1.617 | 1.036 | 1.035 | 0.160 | 0.0851 | 0.2650 | 0 | 0 | 0 |
| 'Dnajc27' | 2.815 | 2.951 | 3.107 | 2.779 | 2.637 | 2.780 | 2.513 | 3.130 | 3.165 | 3.164 | 2.816 | 2.568 | 3.277 | 2.681 | 2.723 | 2.882 | 2.661 | 2.440 | 0.037 | 0.5521 | 0.7650 | 0 | 0 | 0 |
| 'Dnajc3' | 50.713 | 55.534 | 55.407 | 55.681 | 56.822 | 54.509 | 51.465 | 56.076 | 54.484 | 54.117 | 53.086 | 56.757 | 51.755 | 54.202 | 54.733 | 54.551 | 53.868 | 56.271 | -0.008 | 0.7350 | 0.8795 | 0 | 0 | 0 |
| 'Dnajc30' | 8.972 | 10.202 | 10.164 | 7.621 | 9.977 | 11.930 | 7.474 | 9.998 | 10.058 | 9.569 | 8.437 | 8.780 | 10.514 | 8.096 | 7.643 | 11.438 | 8.393 | 8.020 | 0.091 | 0.3131 | 0.5715 | 0 | 0 | 0 |
| 'Dnajc4' | 30.718 | 35.253 | 36.442 | 34.042 | 33.657 | 41.806 | 30.551 | 38.298 | 37.077 | 38.271 | 31.042 | 35.456 | 34.978 | 33.749 | 35.175 | 32.145 | 34.235 | 34.291 | 0.029 | 0.5771 | 0.7818 | 0 | 0 | 0 |
| 'Dnajc5' | 17.357 | 17.920 | 17.135 | 17.065 | 17.579 | 17.613 | 16.633 | 17.949 | 17.761 | 17.220 | 19.455 | 19.149 | 18.255 | 18.383 | 18.631 | 18.247 | 19.043 | 18.905 | -0.102 | 0.0000 | 0.0000 | 1 | 0 | 0 |
| 'Dnajc6' | 12.437 | 13.487 | 11.411 | 13.050 | 12.386 | 11.115 | 13.796 | 12.001 | 12.564 | 11.739 | 14.470 | 10.727 | 14.379 | 12.743 | 10.799 | 12.075 | 13.585 | 10.128 | 0.016 | 0.7407 | 0.8827 | 0 | 0 | 0 |
| 'Dnajc7' | 34.250 | 37.367 | 37.125 | 36.314 | 37.514 | 37.007 | 36.855 | 37.323 | 38.025 | 35.690 | 36.108 | 40.176 | 34.704 | 35.918 | 37.866 | 36.295 | 37.760 | 36.554 | -0.009 | 0.6737 | 0.8432 | 0 | 0 | 0 |
| 'Dnajc8' | 20.649 | 23.534 | 22.268 | 22.482 | 24.613 | 23.484 | 23.962 | 23.726 | 22.855 | 22.277 | 22.482 | 20.918 | 22.626 | 22.813 | 21.405 | 22.894 | 22.395 | 20.287 | 0.058 | 0.0404 | 0.1646 | 0 | 0 | 0 |
| 'Dnajc9' | 70.716 | 76.435 | 74.665 | 75.581 | 82.472 | 78.122 | 74.798 | 78.243 | 77.559 | 63.465 | 72.456 | 70.431 | 76.486 | 76.776 | 75.054 | 70.222 | 71.322 | 68.770 | 0.083 | 0.0111 | 0.0671 | 0 | 0 | 0 |
| 'Dnal4' | 3.386 | 3.310 | 3.825 | 2.893 | 3.753 | 4.379 | 2.946 | 4.098 | 3.540 | 3.523 | 3.163 | 3.055 | 3.612 | 3.568 | 3.403 | 3.720 | 3.183 | 2.975 | 0.090 | 0.2569 | 0.5093 | 0 | 0 | 0 |
| 'Dnase1' | 0.692 | 0.606 | 0.657 | 0.610 | 0.619 | 0.615 | 0.705 | 0.576 | 0.699 | 0.719 | 0.617 | 0.854 | 0.638 | 0.709 | 0.757 | 0.609 | 0.602 | 0.789 | -0.147 | 0.0960 | 0.2881 | 0 | 0 | 0 |
| 'Dnase1l1' | 2.772 | 2.592 | 2.465 | 2.579 | 3.017 | 3.057 | 2.551 | 2.861 | 3.097 | 2.647 | 2.700 | 2.785 | 2.923 | 2.814 | 2.703 | 2.857 | 3.071 | 2.733 | -0.021 | 0.7510 | 0.8892 | 0 | 0 | 0 |
| 'Dnase1l2' | 0.721 | 0.598 | 0.717 | 1.098 | 0.642 | 0.745 | 1.067 | 0.661 | 0.843 | 0.424 | 0.692 | 1.254 | 0.525 | 0.633 | 0.933 | 0.470 | 0.670 | 0.944 | 0.127 | 0.4436 | 0.6869 | 0 | 0 | 0 |
| 'Dnase1l3' | 7.717 | 10.366 | 8.286 | 8.489 | 10.847 | 9.489 | 9.497 | 8.460 | 9.089 | 13.097 | 14.230 | 10.693 | 17.136 | 13.352 | 11.127 | 13.175 | 14.763 | 10.429 | -0.522 | 0.0000 | 0.0000 | 1 | 1 | 0 |
| 'Dnase2a' | 3.775 | 3.782 | 3.938 | 3.374 | 3.969 | 4.018 | 3.634 | 3.851 | 4.083 | 3.999 | 2.769 | 3.562 | 3.461 | 3.136 | 3.416 | 3.625 | 3.264 | 3.303 | 0.165 | 0.0009 | 0.0099 | 1 | 0 | 0 |
| 'Dnd1' | 0.666 | 0.488 | 0.511 | 0.749 | 0.500 | 0.684 | 0.563 | 0.579 | 0.710 | 0.660 | 0.771 | 1.008 | 0.668 | 0.609 | 1.007 | 0.650 | 0.735 | 0.906 | -0.379 | 0.0015 | 0.0147 | 1 | 1 | 0 |
| 'Dnhd1' | 1.754 | 1.937 | 1.802 | 1.665 | 1.796 | 1.940 | 1.839 | 1.893 | 1.860 | 1.769 | 1.661 | 1.998 | 1.951 | 1.800 | 1.848 | 1.582 | 1.728 | 1.917 | 0.003 | 0.9378 | 0.9764 | 0 | 0 | 0 |
| 'Dnlz' | 5.848 | 7.001 | 6.383 | 5.388 | 6.780 | 6.894 | 5.961 | 6.661 | 6.522 | 6.353 | 5.930 | 6.692 | 6.262 | 6.441 | 6.480 | 6.768 | 5.888 | 6.510 | -0.010 | 0.8153 | 0.9170 | 0 | 0 | 0 |
| 'Dnm1l' | 32.530 | 34.443 | 32.123 | 34.991 | 35.996 | 31.489 | 34.254 | 34.528 | 33.981 | 34.551 | 35.267 | 33.514 | 33.179 | 33.729 | 32.347 | 31.814 | 36.692 | 30.731 | 0.000 | 0.9915 | 0.9979 | 0 | 0 | 0 |
| 'Dnm2' | 34.663 | 31.449 | 36.061 | 32.408 | 30.241 | 35.791 | 32.483 | 30.997 | 35.686 | 33.293 | 31.358 | 34.601 | 33.673 | 29.879 | 34.183 | 31.573 | 30.012 | 34.054 | 0.024 | 0.2616 | 0.5153 | 0 | 0 | 0 |
| 'Dnmbp' | 6.748 | 6.912 | 6.874 | 6.549 | 6.479 | 7.028 | 6.671 | 7.264 | 7.031 | 7.269 | 7.332 | 6.993 | 6.892 | 7.061 | 6.668 | 6.443 | 7.017 | 6.729 | -0.027 | 0.4396 | 0.6830 | 0 | 0 | 0 |
| 'Dnmt1' | 71.654 | 75.351 | 86.617 | 82.133 | 72.654 | 81.573 | 75.522 | 70.148 | 88.729 | 63.758 | 72.303 | 81.792 | 82.694 | 74.011 | 85.218 | 72.913 | 70.438 | 82.926 | 0.027 | 0.5026 | 0.7297 | 0 | 0 | 0 |
| 'Dnmt3a' | 5.537 | 5.070 | 5.880 | 5.329 | 5.155 | 5.495 | 4.957 | 4.998 | 5.640 | 5.281 | 5.391 | 5.711 | 5.306 | 5.142 | 5.403 | 5.016 | 4.851 | 5.898 | -0.008 | 0.7884 | 0.9053 | 0 | 0 | 0 |
| 'Dnmt3b' | 1.063 | 1.187 | 1.330 | 1.130 | 0.967 | 1.400 | 1.151 | 1.376 | 1.249 | 1.242 | 1.174 | 1.112 | 0.946 | 0.950 | 1.085 | 1.041 | 0.908 | 1.114 | 0.178 | 0.0216 | 0.1084 | 0 | 0 | 0 |
| 'Dnpep' | 5.154 | 5.517 | 5.238 | 4.945 | 5.462 | 5.777 | 4.927 | 5.479 | 5.028 | 4.955 | 4.718 | 4.649 | 5.095 | 4.971 | 4.766 | 5.422 | 4.504 | 4.503 | 0.119 | 0.0046 | 0.0352 | 1 | 0 | 0 |
| 'Dnph1' | 3.513 | 3.831 | 4.665 | 3.393 | 3.308 | 3.854 | 3.118 | 3.704 | 4.600 | 4.567 | 3.488 | 4.654 | 5.181 | 5.249 | 5.817 | 3.948 | 4.285 | 4.156 | -0.288 | 0.0072 | 0.0493 | 1 | 0 | 0 |
| 'Dnttip1' | 26.507 | 29.861 | 27.651 | 25.716 | 28.131 | 28.470 | 27.431 | 29.290 | 27.265 | 28.234 | 26.604 | 26.392 | 28.917 | 24.559 | 25.896 | 27.680 | 26.658 | 24.850 | 0.052 | 0.1327 | 0.3506 | 0 | 0 | 0 |
| 'Dnttip2' | 12.079 | 12.538 | 11.838 | 13.053 | 13.130 | 12.292 | 11.581 | 12.370 | 11.886 | 13.266 | 11.973 | 12.263 | 11.892 | 12.825 | 11.890 | 12.836 | 12.467 | 11.671 | -0.014 | 0.6582 | 0.8350 | 0 | 0 | 0 |
| 'Doc2a' | 4.934 | 4.658 | 4.497 | 4.255 | 4.358 | 4.722 | 4.788 | 4.867 | 4.911 | 4.126 | 4.036 | 4.829 | 4.785 | 4.173 | 4.947 | 4.244 | 3.922 | 4.851 | 0.060 | 0.2257 | 0.4722 | 0 | 0 | 0 |
| 'Dock10' | 12.554 | 10.743 | 11.937 | 12.123 | 10.869 | 11.068 | 12.238 | 10.754 | 11.655 | 10.388 | 10.594 | 10.456 | 9.386 | 9.445 | 10.052 | 9.752 | 9.623 | 9.996 | 0.200 | 0.0000 | 0.0000 | 1 | 0 | 0 |
| 'Dock11' | 22.897 | 21.330 | 21.935 | 24.722 | 23.091 | 21.657 | 24.176 | 23.044 | 21.775 | 23.569 | 24.321 | 24.300 | 22.578 | 23.284 | 22.734 | 21.960 | 24.564 | 23.612 | -0.056 | 0.0934 | 0.2829 | 0 | 0 | 0 |
| 'Dock2' | 43.545 | 40.431 | 42.649 | 45.188 | 40.460 | 42.485 | 43.567 | 40.145 | 42.814 | 44.551 | 46.497 | 44.092 | 42.355 | 42.036 | 44.285 | 41.113 | 44.492 | 44.857 | -0.060 | 0.0574 | 0.2060 | 0 | 0 | 0 |
| 'Dock4' | 1.302 | 1.917 | 1.596 | 1.533 | 2.393 | 1.757 | 1.720 | 1.805 | 1.550 | 1.896 | 2.129 | 1.812 | 1.814 | 2.707 | 1.952 | 1.862 | 2.896 | 1.840 | -0.287 | 0.0001 | 0.0010 | 1 | 0 | 0 |
| 'Dock5' | 0.432 | 0.300 | 0.474 | 0.569 | 0.330 | 0.378 | 0.398 | 0.306 | 0.295 | 0.567 | 0.609 | 0.618 | 0.476 | 0.554 | 0.546 | 0.660 | 0.348 | 0.645 | -0.544 | 0.0000 | 0.0003 | 1 | 1 | 0 |
| 'Dock6' | 4.089 | 3.913 | 4.098 | 3.903 | 3.589 | 4.010 | 3.729 | 3.607 | 4.268 | 4.099 | 3.120 | 4.030 | 4.021 | 3.167 | 3.644 | 3.278 | 3.171 | 3.558 | 0.120 | 0.0154 | 0.0853 | 0 | 0 | 0 |
| 'Dock8' | 65.254 | 60.031 | 66.932 | 67.385 | 62.281 | 66.387 | 64.414 | 62.295 | 66.540 | 64.979 | 67.523 | 68.157 | 63.320 | 63.402 | 64.178 | 59.779 | 63.852 | 68.464 | -0.016 | 0.5721 | 0.7788 | 0 | 0 | 0 |
| 'Dock9' | 2.775 | 2.390 | 3.124 | 2.814 | 2.643 | 2.875 | 2.823 | 2.755 | 2.714 | 3.521 | 2.722 | 3.649 | 3.024 | 2.866 | 3.506 | 3.327 | 3.079 | 3.448 | -0.235 | 0.0000 | 0.0000 | 1 | 0 | 0 |
| 'Dohh' | 4.810 | 4.912 | 4.421 | 4.127 | 5.127 | 5.037 | 3.963 | 5.177 | 4.763 | 4.539 | 3.946 | 4.922 | 4.400 | 4.953 | 5.057 | 5.231 | 3.675 | 5.054 | 0.009 | 0.8909 | 0.9554 | 0 | 0 | 0 |
| 'Dok1' | 12.501 | 13.314 | 12.752 | 12.811 | 12.366 | 13.803 | 12.539 | 12.199 | 13.099 | 11.968 | 12.099 | 12.330 | 12.414 | 11.579 | 12.998 | 11.550 | 11.963 | 11.922 | 0.074 | 0.0278 | 0.1286 | 0 | 0 | 0 |
| 'Dok2' | 18.257 | 19.040 | 21.871 | 17.349 | 19.242 | 24.538 | 17.922 | 20.152 | 22.399 | 19.016 | 15.271 | 22.409 | 18.701 | 18.278 | 21.112 | 19.712 | 14.651 | 22.218 | 0.069 | 0.1570 | 0.3866 | 0 | 0 | 0 |
| 'Dok3' | 15.641 | 19.131 | 14.763 | 17.328 | 17.309 | 15.798 | 17.398 | 16.042 | 16.129 | 14.854 | 16.754 | 17.838 | 20.049 | 17.635 | 15.011 | 13.875 | 20.693 | 16.321 | -0.051 | 0.4374 | 0.6805 | 0 | 0 | 0 |
| 'Dolk' | 4.079 | 4.475 | 5.597 | 4.058 | 4.671 | 5.339 | 3.861 | 4.202 | 5.732 | 2.945 | 3.518 | 5.276 | 3.968 | 3.667 | 5.187 | 4.624 | 4.373 | 5.537 | 0.092 | 0.2142 | 0.4596 | 0 | 0 | 0 |
| 'Dolpp1' | 3.768 | 4.354 | 3.983 | 3.602 | 3.630 | 4.031 | 3.367 | 3.873 | 4.042 | 3.667 | 3.410 | 4.172 | 4.006 | 3.895 | 4.091 | 3.622 | 3.542 | 4.178 | -0.011 | 0.8028 | 0.9115 | 0 | 0 | 0 |
| 'Donson' | 7.367 | 8.981 | 7.371 | 7.466 | 8.396 | 8.021 | 7.372 | 7.746 | 7.073 | 7.482 | 7.520 | 7.482 | 8.005 | 8.532 | 7.919 | 8.030 | 8.416 | 7.360 | -0.032 | 0.3905 | 0.6397 | 0 | 0 | 0 |
| 'Dopey2' | 15.740 | 13.388 | 15.712 | 16.089 | 14.266 | 15.557 | 14.162 | 14.152 | 15.909 | 16.732 | 17.016 | 16.989 | 16.393 | 14.878 | 15.664 | 15.491 | 17.429 | 16.219 | -0.132 | 0.0010 | 0.0104 | 1 | 0 | 0 |
| 'Dot1l' | 10.088 | 9.195 | 10.415 | 9.347 | 8.782 | 9.589 | 8.674 | 8.774 | 10.517 | 9.058 | 9.597 | 10.753 | 10.619 | 9.105 | 10.046 | 9.084 | 9.339 | 10.067 | -0.049 | 0.1793 | 0.4146 | 0 | 0 | 0 |
| 'Dpagt1' | 13.120 | 12.231 | 12.938 | 12.208 | 11.787 | 12.337 | 12.028 | 11.236 | 12.991 | 12.302 | 11.241 | 11.867 | 12.519 | 11.534 | 13.328 | 10.617 | 11.232 | 11.491 | 0.054 | 0.2665 | 0.5199 | 0 | 0 | 0 |
| 'Dpf2' | 14.699 | 14.504 | 15.531 | 14.312 | 14.440 | 15.404 | 14.096 | 14.834 | 15.351 | 14.862 | 13.565 | 16.825 | 14.114 | 14.935 | 15.989 | 15.396 | 14.252 | 16.369 | -0.044 | 0.1000 | 0.2958 | 0 | 0 | 0 |
| 'Dph1' | 2.303 | 2.237 | 2.876 | 2.038 | 2.250 | 2.487 | 2.512 | 2.723 | 2.699 | 2.592 | 2.116 | 2.291 | 2.256 | 2.065 | 2.152 | 2.527 | 2.171 | 2.255 | 0.116 | 0.1001 | 0.2960 | 0 | 0 | 0 |
| 'Dph2' | 2.161 | 2.248 | 2.655 | 2.202 | 2.346 | 2.497 | 2.234 | 2.112 | 2.826 | 2.092 | 2.265 | 2.654 | 2.407 | 2.180 | 2.548 | 2.175 | 2.067 | 2.722 | -0.003 | 0.9606 | 0.9857 | 0 | 0 | 0 |
| 'Dph3' | 2.736 | 2.920 | 2.328 | 2.495 | 2.802 | 2.685 | 2.256 | 2.791 | 2.316 | 2.453 | 2.561 | 2.381 | 3.081 | 2.878 | 2.513 | 2.896 | 2.459 | 2.190 | -0.013 | 0.8302 | 0.9262 | 0 | 0 | 0 |
| 'Dph5' | 1.557 | 1.880 | 1.848 | 1.877 | 1.844 | 1.931 | 1.925 | 2.089 | 2.093 | 1.796 | 1.803 | 1.743 | 1.935 | 1.945 | 1.626 | 1.835 | 1.643 | 1.615 | 0.089 | 0.1277 | 0.3423 | 0 | 0 | 0 |
| 'Dph6' | 2.098 | 2.553 | 2.334 | 1.994 | 2.191 | 2.321 | 2.267 | 2.260 | 2.327 | 2.172 | 2.186 | 2.483 | 2.142 | 2.363 | 2.364 | 2.508 | 2.382 | 2.270 | -0.049 | 0.2937 | 0.5497 | 0 | 0 | 0 |
| 'Dph7' | 3.310 | 3.239 | 4.280 | 3.949 | 3.668 | 4.164 | 3.792 | 4.114 | 4.026 | 3.835 | 3.732 | 4.159 | 3.929 | 3.330 | 3.990 | 3.723 | 3.750 | 4.003 | -0.005 | 0.9178 | 0.9687 | 0 | 0 | 0 |
| 'Dpm2' | 9.490 | 11.353 | 11.861 | 9.134 | 11.622 | 11.767 | 9.637 | 12.450 | 12.158 | 9.856 | 9.490 | 10.856 | 10.968 | 10.734 | 11.309 | 11.149 | 10.162 | 9.754 | 0.064 | 0.2190 | 0.4658 | 0 | 0 | 0 |
| 'Dpm3' | 14.639 | 16.308 | 14.101 | 14.463 | 15.292 | 15.674 | 13.728 | 15.934 | 13.435 | 13.235 | 12.616 | 14.799 | 13.571 | 12.553 | 15.674 | 15.461 | 13.539 | 13.031 | 0.087 | 0.1289 | 0.3439 | 0 | 0 | 0 |
| 'Dpp3' | 49.213 | 49.827 | 53.449 | 47.372 | 50.296 | 56.401 | 49.726 | 51.182 | 53.867 | 53.264 | 48.536 | 53.263 | 53.310 | 50.890 | 54.254 | 52.139 | 49.253 | 53.060 | -0.030 | 0.2379 | 0.4879 | 0 | 0 | 0 |
| 'Dpp7' | 1.924 | 2.067 | 2.124 | 2.112 | 1.891 | 2.329 | 1.962 | 2.117 | 2.124 | 2.206 | 1.966 | 2.350 | 2.189 | 1.741 | 2.029 | 2.011 | 2.231 | 2.430 | -0.052 | 0.3508 | 0.6049 | 0 | 0 | 0 |
| 'Dpp8' | 28.517 | 30.944 | 29.503 | 28.422 | 30.738 | 30.135 | 29.477 | 32.330 | 30.480 | 29.100 | 30.691 | 31.321 | 28.903 | 30.530 | 29.799 | 29.425 | 31.143 | 31.347 | -0.021 | 0.3310 | 0.5888 | 0 | 0 | 0 |
| 'Dpp9' | 26.778 | 25.095 | 29.099 | 25.866 | 26.496 | 30.173 | 28.338 | 25.410 | 28.480 | 28.237 | 24.854 | 30.610 | 28.357 | 24.871 | 30.757 | 26.017 | 25.872 | 30.362 | -0.035 | 0.1809 | 0.4165 | 0 | 0 | 0 |
| 'Dpy19l1' | 0.995 | 0.863 | 1.240 | 1.294 | 1.205 | 1.210 | 1.429 | 0.986 | 1.216 | 0.938 | 1.007 | 1.139 | 1.248 | 1.140 | 1.268 | 0.985 | 1.265 | 1.247 | 0.012 | 0.8820 | 0.9503 | 0 | 0 | 0 |
| 'Dpy19l3' | 5.457 | 5.499 | 5.323 | 5.016 | 5.522 | 4.773 | 5.563 | 5.235 | 5.362 | 5.590 | 5.732 | 5.447 | 5.749 | 5.569 | 4.951 | 5.333 | 5.011 | 5.388 | -0.042 | 0.2428 | 0.4935 | 0 | 0 | 0 |
| 'Dpy19l4' | 4.362 | 5.424 | 5.157 | 4.731 | 4.841 | 4.636 | 5.140 | 4.974 | 4.971 | 5.219 | 4.907 | 4.915 | 5.231 | 5.490 | 4.197 | 4.988 | 5.430 | 4.048 | -0.013 | 0.7921 | 0.9070 | 0 | 0 | 0 |
| 'Dpy30' | 46.081 | 49.600 | 47.490 | 51.854 | 56.586 | 49.860 | 50.597 | 51.446 | 50.378 | 46.654 | 47.331 | 46.399 | 51.024 | 48.086 | 46.328 | 47.753 | 48.592 | 43.368 | 0.080 | 0.0464 | 0.1795 | 0 | 0 | 0 |
| 'Dpysl2' | 5.638 | 5.673 | 6.225 | 6.336 | 6.017 | 5.164 | 6.222 | 5.613 | 5.575 | 5.064 | 5.745 | 6.186 | 4.929 | 5.755 | 6.645 | 5.301 | 5.241 | 6.788 | 0.013 | 0.8432 | 0.9337 | 0 | 0 | 0 |
| 'Dqx1' | 3.958 | 3.381 | 4.660 | 4.565 | 3.557 | 4.822 | 4.181 | 3.788 | 5.430 | 4.178 | 3.491 | 4.513 | 3.807 | 3.442 | 3.907 | 3.340 | 3.612 | 4.061 | 0.144 | 0.0161 | 0.0883 | 0 | 0 | 0 |
| 'Dr1' | 26.528 | 27.637 | 26.040 | 27.120 | 27.861 | 25.095 | 27.166 | 28.586 | 25.922 | 26.475 | 29.364 | 29.685 | 26.492 | 28.704 | 28.015 | 26.644 | 29.798 | 28.051 | -0.079 | 0.0065 | 0.0455 | 1 | 0 | 0 |
| 'Dram2' | 3.240 | 5.033 | 4.188 | 4.146 | 4.272 | 3.845 | 3.954 | 4.622 | 3.894 | 3.959 | 4.038 | 3.812 | 4.319 | 4.410 | 3.825 | 3.917 | 4.567 | 3.231 | 0.031 | 0.5835 | 0.7856 | 0 | 0 | 0 |
| 'Drap1' | 33.689 | 36.383 | 32.034 | 32.241 | 36.921 | 35.486 | 34.078 | 35.693 | 32.702 | 32.052 | 30.920 | 30.632 | 35.567 | 32.743 | 34.012 | 36.429 | 31.278 | 32.094 | 0.054 | 0.1545 | 0.3825 | 0 | 0 | 0 |
| 'Drg1' | 14.882 | 16.393 | 15.217 | 15.018 | 17.108 | 15.549 | 15.308 | 15.808 | 16.194 | 15.994 | 15.466 | 14.987 | 14.837 | 15.858 | 14.619 | 15.559 | 15.708 | 14.092 | 0.036 | 0.2250 | 0.4716 | 0 | 0 | 0 |
| 'Drg2' | 9.869 | 10.199 | 11.366 | 10.702 | 10.313 | 11.010 | 10.405 | 11.270 | 11.313 | 10.034 | 9.237 | 11.639 | 11.362 | 9.330 | 10.821 | 10.528 | 10.289 | 10.100 | 0.034 | 0.4107 | 0.6595 | 0 | 0 | 0 |
| 'Drosha' | 12.574 | 12.228 | 13.593 | 12.438 | 12.586 | 13.276 | 12.712 | 12.914 | 13.356 | 13.409 | 12.738 | 11.841 | 13.023 | 12.147 | 11.650 | 12.530 | 12.472 | 11.139 | 0.054 | 0.1149 | 0.3217 | 0 | 0 | 0 |
| 'Dscc1' | 1.190 | 0.737 | 1.289 | 1.369 | 0.936 | 1.227 | 1.132 | 1.158 | 1.084 | 0.982 | 1.012 | 1.171 | 1.476 | 0.804 | 1.102 | 0.867 | 0.963 | 1.002 | 0.098 | 0.3554 | 0.6086 | 0 | 0 | 0 |
| 'Dscr3' | 15.413 | 15.974 | 14.934 | 15.393 | 16.070 | 16.143 | 14.813 | 15.430 | 15.562 | 14.886 | 15.656 | 16.036 | 15.402 | 16.316 | 17.111 | 15.082 | 15.433 | 14.941 | -0.023 | 0.4549 | 0.6934 | 0 | 0 | 0 |
| 'Dse' | 0.769 | 0.780 | 0.567 | 0.436 | 0.849 | 1.036 | 0.961 | 0.779 | 0.684 | 1.722 | 1.566 | 1.457 | 1.072 | 1.809 | 1.387 | 1.886 | 1.567 | 1.410 | -1.020 | 0.0000 | 0.0000 | 1 | 1 | 1 |
| 'Dsel' | 1.375 | 1.501 | 1.058 | 1.092 | 1.162 | 1.096 | 1.428 | 1.510 | 1.269 | 1.551 | 1.106 | 1.335 | 1.401 | 1.564 | 1.048 | 1.593 | 1.265 | 1.157 | -0.076 | 0.3462 | 0.6014 | 0 | 0 | 0 |
| 'Dsn1' | 3.498 | 4.045 | 4.935 | 3.417 | 4.341 | 4.444 | 4.203 | 3.952 | 4.777 | 3.657 | 4.260 | 3.957 | 5.068 | 5.078 | 4.064 | 4.160 | 4.409 | 3.839 | -0.043 | 0.5655 | 0.7741 | 0 | 0 | 0 |
| 'Dsp' | 1.313 | 0.945 | 1.350 | 0.887 | 1.079 | 1.106 | 0.889 | 0.699 | 1.201 | 2.008 | 1.334 | 2.343 | 1.567 | 1.718 | 2.176 | 2.443 | 1.585 | 2.405 | -0.895 | 0.0000 | 0.0000 | 1 | 1 | 0 |
| 'Dst' | 1.252 | 1.187 | 1.280 | 1.452 | 1.187 | 1.069 | 1.211 | 1.303 | 1.265 | 1.091 | 0.991 | 1.238 | 0.860 | 0.990 | 1.176 | 0.974 | 1.173 | 1.264 | 0.193 | 0.0045 | 0.0349 | 1 | 0 | 0 |
| 'Dstn' | 6.776 | 8.544 | 6.608 | 6.990 | 8.883 | 8.650 | 7.164 | 8.767 | 7.152 | 6.966 | 7.321 | 6.511 | 8.874 | 8.514 | 6.599 | 6.972 | 7.555 | 6.546 | 0.067 | 0.3301 | 0.5880 | 0 | 0 | 0 |
| 'Dstyk' | 1.681 | 2.168 | 1.551 | 1.677 | 1.970 | 1.719 | 1.388 | 1.802 | 1.693 | 2.016 | 2.089 | 1.882 | 1.847 | 2.043 | 1.715 | 2.293 | 1.914 | 1.759 | -0.178 | 0.0036 | 0.0288 | 1 | 0 | 0 |
| 'Dtd1' | 9.559 | 10.505 | 9.396 | 8.881 | 10.594 | 9.058 | 9.441 | 10.009 | 8.816 | 9.756 | 9.678 | 9.844 | 10.070 | 9.819 | 9.582 | 9.848 | 9.450 | 9.096 | -0.031 | 0.4289 | 0.6747 | 0 | 0 | 0 |
| 'Dtd2' | 2.071 | 2.335 | 2.348 | 2.003 | 2.399 | 2.694 | 2.520 | 2.598 | 2.225 | 2.348 | 1.473 | 1.931 | 2.568 | 2.208 | 2.068 | 2.389 | 2.489 | 1.599 | 0.152 | 0.1182 | 0.3272 | 0 | 0 | 0 |
| 'Dtl' | 2.205 | 1.967 | 2.407 | 2.294 | 2.158 | 2.061 | 2.634 | 1.972 | 2.147 | 2.029 | 2.242 | 2.012 | 2.649 | 2.155 | 2.101 | 2.001 | 1.995 | 1.950 | 0.041 | 0.4755 | 0.7102 | 0 | 0 | 0 |
| 'Dtnb' | 4.056 | 4.029 | 4.107 | 3.807 | 4.154 | 4.509 | 3.782 | 4.095 | 4.144 | 4.493 | 3.682 | 3.950 | 4.125 | 4.007 | 3.921 | 4.145 | 3.745 | 3.703 | 0.029 | 0.4608 | 0.6973 | 0 | 0 | 0 |
| 'Dtnbp1' | 18.768 | 19.403 | 18.518 | 18.518 | 18.398 | 19.975 | 17.881 | 20.030 | 19.417 | 17.814 | 16.296 | 17.946 | 19.334 | 18.213 | 18.561 | 18.123 | 16.525 | 17.835 | 0.080 | 0.0470 | 0.1806 | 0 | 0 | 0 |
| 'Dtwd1' | 3.194 | 4.534 | 3.764 | 3.688 | 4.210 | 3.849 | 3.944 | 4.114 | 3.148 | 3.375 | 2.843 | 3.258 | 3.424 | 3.284 | 2.749 | 3.148 | 3.870 | 2.878 | 0.234 | 0.0035 | 0.0280 | 1 | 0 | 0 |
| 'Dtwd2' | 0.642 | 0.654 | 0.679 | 0.636 | 0.500 | 0.672 | 0.534 | 0.695 | 0.581 | 0.602 | 0.501 | 0.627 | 0.529 | 0.546 | 0.592 | 0.697 | 0.634 | 0.475 | 0.103 | 0.3431 | 0.5992 | 0 | 0 | 0 |
| 'Dtx1' | 26.136 | 21.014 | 30.195 | 19.957 | 20.050 | 28.208 | 20.863 | 20.935 | 26.868 | 29.459 | 25.672 | 39.732 | 23.258 | 22.892 | 37.495 | 27.465 | 23.516 | 40.352 | -0.325 | 0.0000 | 0.0000 | 1 | 0 | 0 |
| 'Dtx2' | 3.992 | 3.534 | 3.876 | 3.491 | 3.727 | 3.848 | 3.484 | 3.555 | 3.869 | 3.895 | 3.224 | 4.395 | 3.636 | 3.655 | 4.072 | 3.490 | 3.354 | 4.381 | -0.041 | 0.3499 | 0.6044 | 0 | 0 | 0 |
| 'Dtx3' | 33.985 | 32.164 | 39.076 | 31.690 | 33.896 | 37.698 | 33.875 | 33.598 | 38.670 | 36.789 | 31.128 | 35.211 | 36.407 | 30.387 | 35.175 | 32.525 | 33.251 | 33.514 | 0.037 | 0.2933 | 0.5497 | 0 | 0 | 0 |
| 'Dtx3l' | 17.769 | 19.267 | 18.657 | 18.958 | 19.301 | 19.567 | 18.809 | 18.797 | 18.727 | 17.571 | 21.214 | 23.817 | 18.303 | 19.375 | 24.933 | 18.170 | 21.771 | 25.336 | -0.167 | 0.0008 | 0.0087 | 1 | 0 | 0 |
| 'Dtx4' | 2.886 | 3.932 | 3.681 | 2.958 | 3.458 | 3.719 | 3.320 | 3.668 | 3.781 | 3.621 | 3.014 | 3.833 | 2.621 | 2.992 | 3.219 | 3.599 | 2.692 | 3.382 | 0.102 | 0.1486 | 0.3750 | 0 | 0 | 0 |
| 'Dtymk' | 20.562 | 21.339 | 23.177 | 20.286 | 21.931 | 22.643 | 22.328 | 21.860 | 22.713 | 18.677 | 19.821 | 21.785 | 21.758 | 21.205 | 23.646 | 20.471 | 21.483 | 22.092 | 0.033 | 0.3070 | 0.5653 | 0 | 0 | 0 |
| 'Dus1l' | 7.294 | 7.558 | 8.021 | 7.246 | 7.486 | 8.100 | 7.292 | 7.531 | 7.597 | 7.495 | 7.084 | 8.032 | 7.706 | 7.478 | 7.990 | 7.967 | 7.166 | 7.439 | -0.014 | 0.6312 | 0.8173 | 0 | 0 | 0 |
| 'Dus2' | 9.528 | 10.252 | 10.046 | 10.266 | 10.878 | 9.666 | 9.577 | 10.493 | 9.328 | 11.226 | 10.375 | 9.246 | 10.892 | 10.116 | 8.552 | 10.039 | 10.466 | 8.142 | 0.011 | 0.7858 | 0.9044 | 0 | 0 | 0 |
| 'Dus3l' | 35.578 | 35.906 | 41.294 | 31.126 | 33.487 | 39.294 | 33.441 | 34.718 | 40.501 | 36.231 | 33.708 | 40.798 | 39.714 | 33.759 | 37.389 | 34.862 | 34.622 | 37.906 | -0.028 | 0.4454 | 0.6878 | 0 | 0 | 0 |
| 'Dus4l' | 2.363 | 2.309 | 3.185 | 2.707 | 2.883 | 2.804 | 2.819 | 2.685 | 2.524 | 2.325 | 3.074 | 3.113 | 3.304 | 2.204 | 2.406 | 2.818 | 2.761 | 2.454 | -0.026 | 0.8063 | 0.9133 | 0 | 0 | 0 |

| | | | | | | | | | | | | | | | | | | | | | | | | |
|---|---|---|---|---|---|---|---|---|---|---|---|---|---|---|---|---|---|---|---|---|---|---|---|---|
| 'Dusp1' | 234.082 | 122.763 | 80.859 | 168.203 | 130.756 | 78.742 | 149.476 | 157.654 | 65.045 | 213.511 | 137.741 | 46.491 | 180.248 | 161.321 | 50.651 | 186.713 | 131.900 | 43.037 | 0.168 | 0.1886 | 0.4247 | 0 | 0 | 0 |
| 'Dusp10' | 32.797 | 32.230 | 28.701 | 30.108 | 30.512 | 31.120 | 29.715 | 32.026 | 30.020 | 33.855 | 25.400 | 24.850 | 31.702 | 28.788 | 25.135 | 35.156 | 22.843 | 25.148 | 0.135 | 0.0146 | 0.0819 | 0 | 0 | 0 |
| 'Dusp11' | 17.005 | 18.301 | 17.808 | 16.450 | 17.670 | 16.777 | 17.534 | 18.027 | 16.959 | 15.839 | 17.289 | 19.929 | 14.939 | 18.006 | 18.982 | 16.893 | 19.232 | 19.935 | -0.048 | 0.2370 | 0.4866 | 0 | 0 | 0 |
| 'Dusp12' | 1.590 | 1.970 | 1.857 | 1.731 | 2.422 | 2.103 | 1.935 | 2.124 | 2.112 | 2.012 | 1.636 | 1.682 | 2.008 | 1.866 | 1.724 | 1.791 | 1.991 | 1.583 | 0.121 | 0.0959 | 0.2881 | 0 | 0 | 0 |
| 'Dusp16' | 1.371 | 1.538 | 1.373 | 1.183 | 1.454 | 1.345 | 1.384 | 1.678 | 1.513 | 1.627 | 1.827 | 1.200 | 1.462 | 1.840 | 1.227 | 1.535 | 1.684 | 1.020 | -0.050 | 0.4960 | 0.7249 | 0 | 0 | 0 |
| 'Dusp19' | 2.061 | 2.317 | 2.243 | 2.296 | 2.322 | 2.441 | 2.199 | 2.240 | 2.410 | 2.584 | 2.548 | 2.412 | 2.130 | 2.893 | 2.377 | 2.198 | 2.302 | 2.502 | -0.103 | 0.0912 | 0.2786 | 0 | 0 | 0 |
| 'Dusp2' | 64.928 | 43.377 | 37.265 | 55.825 | 40.813 | 42.577 | 59.613 | 43.804 | 33.298 | 52.361 | 44.491 | 33.662 | 49.535 | 49.362 | 36.868 | 60.500 | 51.047 | 34.652 | 0.017 | 0.7721 | 0.8988 | 0 | 0 | 0 |
| 'Dusp23' | 2.289 | 2.816 | 3.105 | 2.747 | 2.781 | 2.812 | 4.071 | 2.709 | 2.750 | 2.756 | 3.561 | 2.844 | 2.312 | 3.165 | 2.804 | 3.245 | 2.302 | 2.813 | 0.008 | 0.9446 | 0.9789 | 0 | 0 | 0 |
| 'Dusp28' | 3.972 | 5.135 | 4.441 | 3.865 | 5.296 | 3.507 | 4.432 | 4.842 | 4.302 | 4.100 | 4.014 | 4.244 | 4.070 | 5.089 | 4.984 | 4.722 | 4.615 | 5.248 | -0.064 | 0.4887 | 0.7187 | 0 | 0 | 0 |
| 'Dusp4' | 28.729 | 30.098 | 22.866 | 25.921 | 29.514 | 26.160 | 23.126 | 26.632 | 25.564 | 29.699 | 24.220 | 23.460 | 32.220 | 26.393 | 22.625 | 35.996 | 25.643 | 22.695 | -0.023 | 0.7274 | 0.8750 | 0 | 0 | 0 |
| 'Dusp5' | 11.630 | 7.043 | 5.689 | 6.636 | 6.353 | 6.569 | 5.918 | 7.696 | 4.902 | 10.430 | 5.826 | 5.438 | 9.820 | 9.004 | 5.319 | 18.990 | 5.609 | 5.239 | -0.194 | 0.2933 | 0.5497 | 0 | 0 | 0 |
| 'Dusp6' | 39.219 | 43.492 | 34.348 | 34.403 | 44.671 | 34.693 | 32.943 | 48.380 | 34.640 | 46.463 | 33.368 | 31.357 | 39.640 | 42.446 | 29.852 | 42.607 | 33.378 | 32.025 | 0.058 | 0.4104 | 0.6592 | 0 | 0 | 0 |
| 'Dusp7' | 5.385 | 4.842 | 5.313 | 5.111 | 5.343 | 5.561 | 5.604 | 5.007 | 5.444 | 4.599 | 5.270 | 6.833 | 4.804 | 5.061 | 6.612 | 4.935 | 5.016 | 7.186 | -0.083 | 0.1809 | 0.4165 | 0 | 0 | 0 |
| 'Dut' | 21.496 | 23.978 | 25.282 | 23.044 | 23.538 | 25.396 | 26.109 | 24.468 | 24.033 | 19.366 | 22.801 | 24.749 | 24.561 | 24.587 | 26.872 | 21.325 | 23.439 | 22.641 | 0.036 | 0.4263 | 0.6727 | 0 | 0 | 0 |
| 'Dvl1' | 9.248 | 8.175 | 10.558 | 9.178 | 8.414 | 9.863 | 8.933 | 8.626 | 9.818 | 8.537 | 7.959 | 10.517 | 8.970 | 8.427 | 9.604 | 8.292 | 8.628 | 9.968 | 0.023 | 0.4243 | 0.6710 | 0 | 0 | 0 |
| 'Dvl2' | 7.486 | 6.990 | 7.788 | 7.024 | 6.810 | 7.754 | 7.219 | 6.987 | 7.356 | 7.275 | 6.837 | 7.807 | 7.061 | 6.780 | 7.662 | 7.296 | 7.422 | 7.701 | -0.018 | 0.5248 | 0.7470 | 0 | 0 | 0 |
| 'Dvl3' | 13.398 | 11.064 | 13.560 | 11.359 | 11.748 | 13.098 | 11.845 | 11.838 | 13.661 | 13.983 | 13.472 | 13.183 | 14.714 | 11.578 | 12.929 | 12.581 | 12.806 | 14.902 | -0.116 | 0.0149 | 0.0832 | 0 | 0 | 0 |
| 'Dxo' | 7.471 | 6.657 | 8.043 | 7.650 | 6.761 | 7.872 | 7.222 | 7.031 | 7.738 | 7.591 | 6.421 | 7.978 | 8.193 | 6.500 | 7.194 | 7.328 | 6.645 | 7.320 | 0.018 | 0.6164 | 0.8085 | 0 | 0 | 0 |
| 'Dym' | 10.631 | 9.176 | 11.809 | 10.097 | 11.032 | 12.308 | 10.327 | 11.411 | 11.362 | 10.221 | 10.642 | 10.578 | 11.175 | 9.933 | 9.741 | 9.797 | 9.365 | 10.191 | 0.093 | 0.0646 | 0.2217 | 0 | 0 | 0 |
| 'Dync1h1' | 34.442 | 30.668 | 34.503 | 33.807 | 30.006 | 33.634 | 29.776 | 31.106 | 34.362 | 33.980 | 33.639 | 34.674 | 30.061 | 30.880 | 33.270 | 31.439 | 31.192 | 37.266 | -0.031 | 0.4166 | 0.6643 | 0 | 0 | 0 |
| 'Dync1i2' | 6.756 | 7.179 | 6.847 | 7.093 | 7.284 | 7.020 | 8.058 | 7.182 | 6.972 | 6.481 | 6.848 | 7.504 | 6.768 | 7.113 | 6.792 | 6.498 | 7.688 | 6.985 | 0.026 | 0.4939 | 0.7233 | 0 | 0 | 0 |
| 'Dync1li1' | 35.955 | 33.968 | 34.632 | 35.794 | 35.085 | 33.681 | 34.503 | 36.143 | 34.433 | 33.789 | 35.660 | 37.375 | 34.780 | 35.402 | 35.795 | 34.405 | 36.924 | 35.564 | -0.037 | 0.1851 | 0.4207 | 0 | 0 | 0 |
| 'Dync1li2' | 13.855 | 17.120 | 15.600 | 15.011 | 16.801 | 14.957 | 16.027 | 16.338 | 17.828 | 14.303 | 16.178 | 16.207 | 14.971 | 17.051 | 15.486 | 14.336 | 16.759 | 16.824 | 0.001 | 0.9698 | 0.9898 | 0 | 0 | 0 |
| 'Dync2h1' | 0.976 | 0.963 | 0.915 | 1.059 | 0.805 | 0.875 | 1.019 | 0.901 | 1.010 | 0.907 | 0.794 | 0.886 | 0.767 | 0.791 | 0.807 | 0.640 | 0.971 | 0.869 | 0.182 | 0.0092 | 0.0588 | 0 | 0 | 0 |
| 'Dynll1' | 18.293 | 15.197 | 16.016 | 20.619 | 15.814 | 15.813 | 18.789 | 13.919 | 15.036 | 14.300 | 18.065 | 18.732 | 15.189 | 16.940 | 18.376 | 14.337 | 18.931 | 20.734 | -0.069 | 0.4702 | 0.7061 | 0 | 0 | 0 |
| 'Dynll2' | 52.524 | 54.235 | 49.856 | 56.040 | 51.997 | 51.450 | 50.977 | 51.382 | 52.773 | 47.366 | 50.954 | 52.873 | 52.008 | 53.684 | 54.861 | 51.468 | 51.016 | 55.938 | -0.008 | 0.8132 | 0.9157 | 0 | 0 | 0 |
| 'Dynlrb1' | 42.080 | 48.261 | 39.259 | 45.791 | 46.026 | 44.236 | 46.937 | 44.513 | 41.901 | 40.597 | 43.369 | 38.584 | 46.761 | 42.551 | 42.748 | 42.606 | 43.120 | 38.325 | 0.064 | 0.0756 | 0.2441 | 0 | 0 | 0 |
| 'Dynlt1c' | 73.899 | 95.697 | 84.364 | 83.358 | 92.823 | 93.932 | 79.222 | 94.928 | 95.155 | 73.563 | 71.462 | 80.152 | 87.356 | 89.159 | 76.268 | 82.787 | 75.498 | 76.087 | 0.141 | 0.0056 | 0.0407 | 1 | 0 | 0 |
| 'Dyrk1a' | 13.451 | 12.470 | 11.985 | 13.303 | 11.894 | 11.845 | 12.578 | 13.166 | 11.622 | 13.170 | 12.846 | 12.575 | 12.691 | 13.432 | 12.599 | 13.427 | 12.975 | 13.775 | -0.077 | 0.0212 | 0.1069 | 0 | 0 | 0 |
| 'Dyrk1b' | 6.728 | 6.676 | 7.577 | 6.000 | 6.912 | 8.376 | 6.326 | 6.352 | 7.769 | 6.884 | 6.700 | 7.905 | 7.168 | 5.822 | 7.604 | 6.539 | 7.214 | 7.357 | -0.023 | 0.6283 | 0.8156 | 0 | 0 | 0 |
| 'Dyrk2' | 30.463 | 27.537 | 30.427 | 25.593 | 27.915 | 30.395 | 27.788 | 28.148 | 30.076 | 28.022 | 30.233 | 31.834 | 24.863 | 29.689 | 33.542 | 30.487 | 28.320 | 33.371 | -0.072 | 0.0970 | 0.2901 | 0 | 0 | 0 |
| 'Dzip1' | 11.159 | 11.164 | 12.041 | 9.787 | 9.881 | 11.876 | 11.748 | 10.695 | 12.481 | 11.397 | 9.950 | 13.746 | 10.465 | 9.760 | 13.079 | 11.317 | 9.682 | 11.956 | -0.015 | 0.7109 | 0.8658 | 0 | 0 | 0 |
| 'Dzip3' | 1.445 | 1.367 | 1.532 | 1.716 | 1.690 | 1.382 | 1.481 | 1.482 | 1.600 | 1.314 | 1.362 | 1.985 | 1.306 | 1.520 | 1.741 | 1.316 | 1.458 | 1.904 | -0.027 | 0.7171 | 0.8693 | 0 | 0 | 0 |
| 'E130012A19Rik' | 11.963 | 7.991 | 10.143 | 9.180 | 7.678 | 11.063 | 11.113 | 9.447 | 10.193 | 9.330 | 7.645 | 9.430 | 9.798 | 9.515 | 10.016 | 8.894 | 7.430 | 9.154 | 0.119 | 0.0549 | 0.2008 | 0 | 0 | 0 |
| 'E130308A19Rik' | 1.726 | 1.967 | 1.739 | 1.590 | 1.735 | 1.626 | 1.803 | 2.144 | 1.731 | 2.144 | 1.869 | 1.858 | 2.435 | 2.134 | 1.845 | 2.101 | 2.121 | 1.571 | -0.176 | 0.0032 | 0.0264 | 1 | 0 | 0 |
| 'E130309D02Rik' | 6.771 | 7.331 | 7.102 | 6.194 | 7.395 | 7.597 | 6.403 | 7.252 | 6.976 | 6.622 | 6.682 | 7.280 | 7.126 | 6.770 | 7.706 | 7.642 | 5.998 | 7.406 | -0.015 | 0.7356 | 0.8796 | 0 | 0 | 0 |
| 'E130311K13Rik' | 2.629 | 3.641 | 4.167 | 2.759 | 4.354 | 4.051 | 3.392 | 4.446 | 3.902 | 4.039 | 2.951 | 2.711 | 4.033 | 3.463 | 2.666 | 4.048 | 2.737 | 2.499 | 0.197 | 0.0990 | 0.2941 | 0 | 0 | 0 |
| 'E2f1' | 6.279 | 5.295 | 7.461 | 5.958 | 5.114 | 6.407 | 6.944 | 5.215 | 6.260 | 5.749 | 6.026 | 6.610 | 7.975 | 5.701 | 7.706 | 5.638 | 5.745 | 6.788 | -0.090 | 0.1597 | 0.3901 | 0 | 0 | 0 |
| 'E2f2' | 15.201 | 12.324 | 16.603 | 15.576 | 13.934 | 16.671 | 15.552 | 13.862 | 16.040 | 12.560 | 17.181 | 15.854 | 16.966 | 16.449 | 16.759 | 14.273 | 14.819 | 15.140 | -0.057 | 0.3370 | 0.5955 | 0 | 0 | 0 |
| 'E2f3' | 5.092 | 4.963 | 6.375 | 5.712 | 5.135 | 6.077 | 5.788 | 5.317 | 6.312 | 5.675 | 5.897 | 6.105 | 6.154 | 5.122 | 6.156 | 5.499 | 5.366 | 6.207 | -0.048 | 0.2810 | 0.5379 | 0 | 0 | 0 |
| 'E2f4' | 105.438 | 111.381 | 106.337 | 104.409 | 111.967 | 109.538 | 103.713 | 106.259 | 109.823 | 106.462 | 102.880 | 114.321 | 123.333 | 110.746 | 116.531 | 113.069 | 109.935 | 116.628 | -0.077 | 0.0090 | 0.0578 | 0 | 0 | 0 |
| 'E2f5' | 3.544 | 3.269 | 3.311 | 2.888 | 3.736 | 3.400 | 3.060 | 4.287 | 3.032 | 3.882 | 3.253 | 3.495 | 3.619 | 3.700 | 3.231 | 3.432 | 3.509 | 2.703 | -0.021 | 0.7820 | 0.9025 | 0 | 0 | 0 |
| 'E2f6' | 12.688 | 14.088 | 13.815 | 13.164 | 14.607 | 12.665 | 14.221 | 13.969 | 14.498 | 11.117 | 12.329 | 15.259 | 13.001 | 13.274 | 13.828 | 12.971 | 13.649 | 14.349 | 0.030 | 0.5188 | 0.7416 | 0 | 0 | 0 |
| 'E2f7' | 0.752 | 0.573 | 0.877 | 0.838 | 0.642 | 0.722 | 0.807 | 0.749 | 0.841 | 0.617 | 0.818 | 0.732 | 1.129 | 0.795 | 0.827 | 0.883 | 0.709 | 0.823 | -0.115 | 0.2194 | 0.4664 | 0 | 0 | 0 |
| 'E2f8' | 4.827 | 4.493 | 6.906 | 5.700 | 4.984 | 6.309 | 6.187 | 4.884 | 6.113 | 4.057 | 5.371 | 6.044 | 7.506 | 6.060 | 6.948 | 5.382 | 5.355 | 5.853 | -0.079 | 0.3503 | 0.6045 | 0 | 0 | 0 |
| 'E330009J07Rik' | 4.975 | 5.246 | 4.948 | 5.237 | 4.797 | 5.336 | 5.000 | 5.326 | 5.367 | 4.933 | 6.014 | 5.978 | 5.144 | 5.144 | 5.768 | 6.163 | 5.583 | 5.859 | -0.140 | 0.0017 | 0.0160 | 1 | 0 | 0 |
| 'E430025E21Rik' | 24.852 | 25.568 | 26.507 | 25.061 | 26.226 | 25.258 | 26.969 | 25.286 | 26.623 | 29.297 | 27.688 | 26.037 | 28.192 | 25.735 | 26.656 | 27.057 | 25.117 | 25.072 | -0.060 | 0.0502 | 0.1893 | 0 | 0 | 0 |
| 'E4f1' | 27.025 | 26.332 | 32.145 | 24.923 | 25.954 | 32.350 | 27.816 | 27.816 | 32.688 | 28.796 | 24.201 | 30.727 | 27.614 | 23.009 | 28.774 | 28.982 | 27.414 | 27.926 | 0.044 | 0.2663 | 0.5199 | 0 | 0 | 0 |
| 'Eaf1' | 24.355 | 26.250 | 22.550 | 21.898 | 26.627 | 23.500 | 21.499 | 26.245 | 23.683 | 24.361 | 26.176 | 25.641 | 25.444 | 25.767 | 26.052 | 26.483 | 25.195 | 26.392 | -0.109 | 0.0018 | 0.0166 | 1 | 0 | 0 |
| 'Eapp' | 12.894 | 13.811 | 12.719 | 13.269 | 14.680 | 13.053 | 12.060 | 13.657 | 13.417 | 13.224 | 12.985 | 12.997 | 12.057 | 14.256 | 12.311 | 14.030 | 13.439 | 12.345 | 0.014 | 0.6944 | 0.8560 | 0 | 0 | 0 |
| 'Ears2' | 2.299 | 2.422 | 2.496 | 2.328 | 2.743 | 2.371 | 2.239 | 2.452 | 2.689 | 2.704 | 2.211 | 3.026 | 2.440 | 2.334 | 2.605 | 2.511 | 2.409 | 2.821 | -0.083 | 0.1358 | 0.3545 | 0 | 0 | 0 |
| 'Ebag9' | 26.796 | 32.747 | 27.381 | 31.632 | 35.024 | 31.627 | 31.985 | 32.215 | 31.678 | 29.950 | 26.896 | 27.205 | 29.896 | 33.046 | 28.258 | 27.884 | 28.399 | 25.699 | 0.114 | 0.0284 | 0.1304 | 0 | 0 | 0 |
| 'Ebi3' | 4.015 | 3.688 | 5.968 | 3.774 | 3.618 | 6.253 | 5.698 | 3.963 | 5.583 | 8.519 | 3.903 | 7.656 | 7.183 | 4.145 | 6.677 | 6.548 | 4.289 | 6.251 | -0.375 | 0.0002 | 0.0032 | 1 | 0 | 0 |
| 'Ebna1bp2' | 12.825 | 12.634 | 13.899 | 13.671 | 13.710 | 13.367 | 13.245 | 13.237 | 13.684 | 11.603 | 12.229 | 13.325 | 13.177 | 12.512 | 13.277 | 12.676 | 12.638 | 12.673 | 0.064 | 0.0302 | 0.1352 | 0 | 0 | 0 |
| 'Ebp' | 8.841 | 9.785 | 9.873 | 9.468 | 9.852 | 9.751 | 9.858 | 9.293 | 10.228 | 9.419 | 8.429 | 10.374 | 8.840 | 9.309 | 10.159 | 9.018 | 8.693 | 10.601 | 0.019 | 0.6352 | 0.8194 | 0 | 0 | 0 |
| 'Ebpl' | 7.311 | 9.665 | 10.332 | 8.004 | 9.390 | 9.342 | 7.929 | 9.885 | 9.158 | 9.075 | 9.244 | 8.645 | 7.878 | 9.265 | 10.007 | 10.102 | 8.620 | 7.668 | 0.002 | 0.9700 | 0.9898 | 0 | 0 | 0 |
| 'Ecd' | 23.505 | 26.177 | 25.410 | 24.390 | 26.081 | 24.459 | 25.741 | 25.250 | 24.147 | 25.782 | 23.941 | 22.935 | 25.435 | 23.480 | 22.712 | 24.970 | 23.501 | 21.511 | 0.063 | 0.0434 | 0.1726 | 0 | 0 | 0 |
| 'Ece1' | 79.663 | 82.555 | 81.620 | 74.673 | 76.047 | 80.202 | 73.379 | 78.384 | 84.403 | 93.855 | 81.678 | 82.166 | 83.849 | 83.918 | 82.333 | 89.232 | 79.734 | 85.324 | -0.110 | 0.0013 | 0.0127 | 1 | 0 | 0 |
| 'Ece2' | 0.447 | 0.526 | 0.440 | 0.482 | 0.464 | 0.334 | 0.451 | 0.397 | 0.416 | 0.413 | 0.509 | 0.502 | 0.524 | 0.457 | 0.485 | 0.413 | 0.404 | 0.359 | -0.069 | 0.4921 | 0.7216 | 0 | 0 | 0 |
| 'Ech1' | 17.186 | 17.466 | 17.798 | 19.031 | 18.551 | 19.157 | 19.198 | 16.998 | 18.138 | 17.539 | 15.913 | 17.537 | 17.306 | 17.173 | 17.396 | 18.239 | 17.059 | 15.809 | 0.076 | 0.0183 | 0.0968 | 0 | 0 | 0 |
| 'Echdc1' | 1.347 | 1.411 | 1.269 | 1.544 | 1.183 | 1.489 | 1.349 | 1.262 | 1.283 | 1.400 | 1.298 | 1.550 | 1.370 | 1.487 | 1.399 | 1.249 | 1.541 | 1.605 | -0.107 | 0.1814 | 0.4171 | 0 | 0 | 0 |
| 'Echdc2' | 0.813 | 0.782 | 0.986 | 0.850 | 0.663 | 1.032 | 0.731 | 0.903 | 0.989 | 0.768 | 0.728 | 0.922 | 0.962 | 0.648 | 0.797 | 0.708 | 0.624 | 0.800 | 0.145 | 0.1206 | 0.3311 | 0 | 0 | 0 |
| 'Echs1' | 5.088 | 5.796 | 5.792 | 4.940 | 6.011 | 5.670 | 5.066 | 6.151 | 5.482 | 5.962 | 5.373 | 5.058 | 5.661 | 5.256 | 4.962 | 5.168 | 5.251 | 4.625 | 0.071 | 0.1324 | 0.3501 | 0 | 0 | 0 |
| 'Eci1' | 18.758 | 22.840 | 20.564 | 17.889 | 21.217 | 21.135 | 19.193 | 23.220 | 21.827 | 20.104 | 19.065 | 17.909 | 18.913 | 20.986 | 16.405 | 21.524 | 20.677 | 14.533 | 0.132 | 0.0317 | 0.1400 | 0 | 0 | 0 |
| 'Eci2' | 7.608 | 7.325 | 7.823 | 7.527 | 8.291 | 7.842 | 8.048 | 8.175 | 8.370 | 7.017 | 8.117 | 7.606 | 8.439 | 7.516 | 7.616 | 7.415 | 7.587 | 6.976 | 0.047 | 0.2311 | 0.4784 | 0 | 0 | 0 |
| 'Ecm1' | 3.850 | 2.416 | 3.622 | 3.293 | 2.971 | 3.381 | 3.668 | 2.973 | 4.052 | 3.686 | 2.917 | 4.495 | 3.044 | 2.462 | 4.077 | 2.981 | 2.974 | 4.306 | -0.040 | 0.5854 | 0.7868 | 0 | 0 | 0 |
| 'Ecsit' | 14.459 | 14.197 | 15.136 | 12.298 | 13.665 | 15.261 | 12.871 | 15.287 | 14.520 | 15.102 | 13.735 | 14.363 | 13.607 | 12.439 | 12.807 | 15.189 | 14.302 | 13.705 | 0.022 | 0.6561 | 0.8334 | 0 | 0 | 0 |
| 'Ect2' | 1.987 | 1.984 | 2.496 | 2.472 | 2.267 | 2.294 | 2.865 | 2.410 | 2.339 | 1.492 | 2.428 | 2.381 | 2.435 | 2.770 | 2.782 | 1.838 | 2.590 | 2.359 | 0.000 | 0.9972 | 0.9994 | 0 | 0 | 0 |
| 'Edc3' | 9.027 | 8.542 | 8.102 | 8.430 | 8.474 | 8.315 | 8.572 | 8.291 | 8.524 | 8.276 | 8.432 | 9.555 | 8.175 | 8.950 | 8.931 | 8.402 | 8.454 | 9.072 | -0.050 | 0.1521 | 0.3793 | 0 | 0 | 0 |
| 'Edc4' | 18.144 | 17.874 | 19.015 | 16.937 | 17.515 | 18.717 | 18.122 | 17.224 | 19.364 | 18.212 | 17.446 | 18.891 | 17.547 | 16.638 | 17.641 | 17.421 | 17.390 | 17.273 | 0.029 | 0.2268 | 0.4734 | 0 | 0 | 0 |
| 'Edem1' | 22.385 | 22.456 | 24.273 | 23.012 | 22.414 | 23.765 | 20.193 | 22.542 | 23.429 | 24.018 | 26.662 | 27.565 | 26.539 | 26.687 | 27.076 | 24.279 | 27.728 | 28.822 | -0.237 | 0.0000 | 0.0000 | 1 | 0 | 0 |
| 'Edem2' | 20.139 | 22.228 | 20.967 | 19.277 | 20.821 | 21.495 | 19.371 | 22.582 | 21.128 | 22.278 | 19.727 | 21.726 | 23.308 | 21.640 | 20.450 | 22.935 | 21.177 | 20.450 | -0.053 | 0.1469 | 0.3726 | 0 | 0 | 0 |
| 'Edem3' | 7.893 | 8.825 | 8.054 | 8.131 | 8.651 | 7.305 | 7.746 | 8.339 | 7.523 | 7.884 | 8.449 | 8.662 | 7.742 | 8.646 | 8.089 | 7.879 | 8.299 | 8.670 | -0.052 | 0.1411 | 0.3636 | 0 | 0 | 0 |
| 'Edf1' | 43.455 | 42.968 | 44.492 | 41.785 | 45.524 | 44.467 | 44.012 | 42.347 | 46.142 | 42.693 | 38.205 | 42.166 | 41.767 | 40.339 | 44.632 | 45.202 | 41.473 | 41.099 | 0.057 | 0.0438 | 0.1733 | 0 | 0 | 0 |
| 'Edrf1' | 4.582 | 5.083 | 4.614 | 4.561 | 4.994 | 4.830 | 4.533 | 4.844 | 4.727 | 4.522 | 5.222 | 5.435 | 4.428 | 5.072 | 4.627 | 4.702 | 5.385 | 5.167 | -0.074 | 0.0579 | 0.2074 | 0 | 0 | 0 |
| 'Eea1' | 36.350 | 37.035 | 36.468 | 37.108 | 37.481 | 34.873 | 37.474 | 37.807 | 37.168 | 38.374 | 40.189 | 37.114 | 33.114 | 38.712 | 34.770 | 36.819 | 40.426 | 35.275 | -0.022 | 0.4785 | 0.7112 | 0 | 0 | 0 |
| 'Eed' | 36.424 | 36.656 | 35.070 | 41.783 | 39.248 | 37.349 | 41.527 | 37.635 | 36.808 | 35.077 | 39.725 | 39.304 | 38.137 | 37.312 | 38.547 | 37.681 | 41.785 | 39.249 | -0.031 | 0.4311 | 0.6762 | 0 | 0 | 0 |
| 'Eef1akmt1' | 19.839 | 24.963 | 21.878 | 24.121 | 25.572 | 21.016 | 21.633 | 22.507 | 21.713 | 26.342 | 21.660 | 23.510 | 25.262 | 20.218 | 23.087 | 21.993 | 23.093 | 21.496 | -0.043 | 0.4583 | 0.6954 | 0 | 0 | 0 |
| 'Eef1b2' | 68.906 | 66.514 | 63.258 | 69.526 | 65.648 | 62.948 | 71.017 | 64.728 | 62.474 | 62.295 | 66.910 | 58.625 | 66.118 | 63.934 | 65.897 | 63.659 | 66.371 | 61.922 | 0.036 | 0.2200 | 0.4672 | 0 | 0 | 0 |
| 'Eef1d' | 10.057 | 9.542 | 9.960 | 10.197 | 10.023 | 10.034 | 10.093 | 9.881 | 9.873 | 9.191 | 9.703 | 10.340 | 9.846 | 9.034 | 10.369 | 9.559 | 9.405 | 10.319 | 0.019 | 0.4886 | 0.7187 | 0 | 0 | 0 |
| 'Eef1e1' | 19.105 | 19.158 | 19.490 | 21.609 | 19.672 | 17.079 | 18.341 | 19.643 | 18.685 | 16.053 | 15.868 | 20.340 | 17.662 | 21.078 | 21.041 | 18.410 | 18.370 | 18.274 | 0.033 | 0.5920 | 0.7921 | 0 | 0 | 0 |
| 'Eef1g' | 95.329 | 81.971 | 90.444 | 102.148 | 88.221 | 85.834 | 101.088 | 86.216 | 86.352 | 85.119 | 90.548 | 92.263 | 91.236 | 86.891 | 95.320 | 83.813 | 96.490 | 94.356 | -0.011 | 0.8062 | 0.9133 | 0 | 0 | 0 |
| 'Eef2' | 795.872 | 743.661 | 748.286 | 785.577 | 756.802 | 770.153 | 792.939 | 738.629 | 751.606 | 786.919 | 794.604 | 682.471 | 802.334 | 760.196 | 785.449 | 776.992 | 756.050 | 771.628 | -0.018 | 0.5298 | 0.7503 | 0 | 0 | 0 |
| 'Eef2k' | 0.378 | 0.606 | 0.439 | 0.628 | 0.547 | 0.491 | 0.431 | 0.622 | 0.536 | 0.419 | 0.664 | 0.547 | 0.548 | 0.620 | 0.444 | 0.561 | 0.595 | 0.551 | -0.099 | 0.3230 | 0.5806 | 0 | 0 | 0 |
| 'Eef2kmt' | 4.172 | 3.778 | 4.151 | 3.953 | 4.080 | 4.418 | 3.660 | 4.156 | 4.357 | 4.297 | 4.026 | 4.635 | 4.348 | 4.063 | 4.201 | 4.388 | 4.365 | 3.986 | -0.069 | 0.1552 | 0.3836 | 0 | 0 | 0 |
| 'Eefsec' | 12.277 | 10.063 | 13.805 | 12.202 | 11.171 | 13.228 | 11.403 | 10.902 | 12.548 | 12.348 | 11.304 | 12.827 | 12.257 | 10.208 | 12.109 | 11.815 | 10.059 | 12.436 | 0.023 | 0.5144 | 0.7384 | 0 | 0 | 0 |
| 'Efcab14' | 3.872 | 4.246 | 4.544 | 4.151 | 4.494 | 4.384 | 4.051 | 4.412 | 4.374 | 4.075 | 4.330 | 4.554 | 4.213 | 4.490 | 4.401 | 4.219 | 4.218 | 4.690 | -0.036 | 0.2423 | 0.4928 | 0 | 0 | 0 |
| 'Efcab2' | 1.137 | 1.048 | 1.066 | 1.062 | 1.097 | 0.918 | 1.040 | 1.045 | 1.163 | 1.059 | 1.017 | 1.152 | 0.952 | 0.966 | 0.891 | 0.627 | 1.156 | 0.833 | 0.122 | 0.2496 | 0.5007 | 0 | 0 | 0 |
| 'Efcab7' | 1.835 | 1.663 | 1.922 | 1.966 | 2.083 | 1.639 | 2.136 | 1.675 | 2.006 | 1.682 | 1.600 | 1.480 | 1.933 | 2.048 | 1.601 | 1.668 | 1.821 | 1.062 | 0.185 | 0.0597 | 0.2110 | 0 | 0 | 0 |
| 'Efhd2' | 69.678 | 77.961 | 77.255 | 66.023 | 78.248 | 85.158 | 66.911 | 76.988 | 78.954 | 79.367 | 74.564 | 81.720 | 82.326 | 76.171 | 84.948 | 85.587 | 72.918 | 82.302 | -0.100 | 0.0109 | 0.0664 | 0 | 0 | 0 |
| 'Efl1' | 6.144 | 6.264 | 6.516 | 6.764 | 6.561 | 6.182 | 6.856 | 6.412 | 6.597 | 6.642 | 5.878 | 6.903 | 6.798 | 5.935 | 6.104 | 6.281 | 7.065 | 6.229 | -0.004 | 0.9156 | 0.9677 | 0 | 0 | 0 |
| 'Efnb2' | 2.397 | 2.711 | 2.519 | 1.968 | 3.379 | 2.879 | 2.033 | 3.393 | 2.571 | 3.629 | 2.364 | 2.320 | 2.391 | 3.159 | 2.079 | 3.293 | 2.451 | 1.902 | 0.013 | 0.9085 | 0.9641 | 0 | 0 | 0 |
| 'Efr3a' | 7.824 | 8.131 | 7.733 | 8.281 | 8.289 | 7.533 | 8.169 | 8.381 | 7.502 | 7.327 | 8.730 | 7.978 | 7.535 | 8.199 | 7.856 | 7.438 | 7.728 | 7.536 | 0.019 | 0.5361 | 0.7552 | 0 | 0 | 0 |
| 'Eftud2' | 17.135 | 15.786 | 16.971 | 16.759 | 16.512 | 16.832 | 16.106 | 16.435 | 17.030 | 17.394 | 16.559 | 16.316 | 17.660 | 15.737 | 16.187 | 16.113 | 16.058 | 15.722 | 0.009 | 0.7209 | 0.8716 | 0 | 0 | 0 |
| 'Egln1' | 52.042 | 51.874 | 51.473 | 49.253 | 47.988 | 52.604 | 51.605 | 50.639 | 47.426 | 46.001 | 52.892 | 55.861 | 53.168 | 58.463 | 58.847 | 52.231 | 49.848 | 60.790 | -0.110 | 0.0117 | 0.0699 | 0 | 0 | 0 |
| 'Egln2' | 37.041 | 43.455 | 41.834 | 36.776 | 42.457 | 46.193 | 37.185 | 40.980 | 41.369 | 38.437 | 37.327 | 43.632 | 42.349 | 40.614 | 42.555 | 44.901 | 36.968 | 43.797 | -0.025 | 0.5488 | 0.7626 | 0 | 0 | 0 |
| 'Egln3' | 92.952 | 128.294 | 86.617 | 85.704 | 117.013 | 95.093 | 88.248 | 115.961 | 91.017 | 100.481 | 105.982 | 75.087 | 107.877 | 116.456 | 91.775 | 111.131 | 101.025 | 81.559 | 0.002 | 0.9771 | 0.9920 | 0 | 0 | 0 |
| 'Egr1' | 33.910 | 27.581 | 31.652 | 34.777 | 27.623 | 31.184 | 44.518 | 28.616 | 33.870 | 30.156 | 22.311 | 27.104 | 27.015 | 22.975 | 27.540 | 26.821 | 24.318 | 26.256 | 0.308 | 0.0000 | 0.0000 | 1 | 0 | 0 |
| 'Egr2' | 12.173 | 11.298 | 12.004 | 11.631 | 11.383 | 11.471 | 12.697 | 11.115 | 12.949 | 12.205 | 10.903 | 13.410 | 9.767 | 11.494 | 13.772 | 12.597 | 11.530 | 13.972 | -0.047 | 0.3490 | 0.6042 | 0 | 0 | 0 |
| 'Egr3' | 15.684 | 18.506 | 21.565 | 12.630 | 15.554 | 22.130 | 13.908 | 15.877 | 22.489 | 20.451 | 13.956 | 18.388 | 15.531 | 13.026 | 15.903 | 19.694 | 13.093 | 18.078 | 0.080 | 0.3830 | 0.6336 | 0 | 0 | 0 |
| 'Egr4' | 9.422 | 2.739 | 4.270 | 8.737 | 2.472 | 3.728 | 11.916 | 3.012 | 3.947 | 4.139 | 3.739 | 2.937 | 4.116 | 2.771 | 1.926 | 3.174 | 4.153 | 1.581 | 0.644 | 0.0046 | 0.0350 | 1 | 1 | 0 |
| 'Ehbp1' | 0.812 | 0.718 | 0.589 | 0.666 | 0.579 | 0.682 | 0.891 | 0.772 | 0.561 | 0.544 | 0.577 | 0.647 | 0.546 | 0.625 | 0.507 | 0.660 | 0.525 | 0.664 | 0.220 | 0.0289 | 0.1317 | 0 | 0 | 0 |
| 'Ehbp1l1' | 20.660 | 18.990 | 23.078 | 20.609 | 19.845 | 22.755 | 22.559 | 19.334 | 22.902 | 19.358 | 20.187 | 23.925 | 19.771 | 18.990 | 22.834 | 18.981 | 20.378 | 22.879 | 0.015 | 0.6242 | 0.8126 | 0 | 0 | 0 |
| 'Ehd1' | 16.325 | 15.256 | 17.366 | 14.685 | 16.107 | 17.589 | 15.709 | 16.010 | 18.357 | 18.526 | 15.310 | 16.270 | 19.604 | 16.485 | 16.509 | 17.560 | 15.526 | 16.534 | -0.055 | 0.2229 | 0.4704 | 0 | 0 | 0 |
| 'Ehd3' | 77.185 | 84.846 | 72.907 | 74.401 | 78.486 | 78.090 | 73.656 | 79.014 | 74.744 | 78.742 | 77.504 | 78.367 | 75.308 | 78.147 | 79.492 | 77.636 | 71.089 | 80.477 | -0.019 | 0.5442 | 0.7606 | 0 | 0 | 0 |
| 'Ehd4' | 5.058 | 5.802 | 6.094 | 5.629 | 6.167 | 6.396 | 5.571 | 6.327 | 6.233 | 5.648 | 6.254 | 5.646 | 7.232 | 6.999 | 5.931 | 6.311 | 6.638 | 5.273 | -0.075 | 0.2041 | 0.4471 | 0 | 0 | 0 |
| 'Ehhadh' | 3.920 | 3.899 | 4.316 | 4.048 | 5.135 | 4.504 | 4.520 | 4.877 | 4.454 | 4.641 | 4.435 | 3.369 | 4.159 | 4.164 | 3.712 | 4.290 | 3.695 | 3.029 | 0.166 | 0.0268 | 0.1260 | 0 | 0 | 0 |
| 'Ehmt1' | 5.710 | 4.987 | 5.562 | 5.624 | 5.260 | 5.542 | 5.442 | 5.229 | 5.568 | 5.755 | 5.400 | 5.770 | 5.112 | 4.955 | 5.343 | 4.726 | 5.268 | 5.884 | 0.010 | 0.7998 | 0.9111 | 0 | 0 | 0 |
| 'Ehmt2' | 30.316 | 27.009 | 31.409 | 29.078 | 26.900 | 31.746 | 27.875 | 27.427 | 30.883 | 29.057 | 26.922 | 33.185 | 29.090 | 26.935 | 31.933 | 29.293 | 27.443 | 32.678 | -0.031 | 0.1364 | 0.3555 | 0 | 0 | 0 |
| 'Ei24' | 18.870 | 20.718 | 18.875 | 19.257 | 21.196 | 19.475 | 18.510 | 20.700 | 19.454 | 20.507 | 20.284 | 19.841 | 18.988 | 20.470 | 20.022 | 20.385 | 19.695 | 20.142 | -0.038 | 0.1661 | 0.3983 | 0 | 0 | 0 |
| 'Eid1' | 35.115 | 40.689 | 38.766 | 37.821 | 40.982 | 35.325 | 39.516 | 38.961 | 35.101 | 38.552 | 39.389 | 35.420 | 37.822 | 40.807 | 35.227 | 34.296 | 40.228 | 33.737 | 0.019 | 0.5821 | 0.7842 | 0 | 0 | 0 |
| 'Eid2' | 9.873 | 10.371 | 11.118 | 9.041 | 9.225 | 11.504 | 9.689 | 11.619 | 8.934 | 9.241 | 7.918 | 8.175 | 8.703 | 9.186 | 11.348 | 9.081 | 9.013 | 10.246 | 0.138 | 0.0718 | 0.2363 | 0 | 0 | 0 |
| 'Eid2b' | 2.871 | 3.399 | 3.913 | 3.340 | 3.433 | 3.707 | 3.256 | 3.492 | 3.835 | 2.862 | 3.183 | 2.842 | 3.129 | 3.001 | 3.033 | 3.224 | 2.645 | 2.974 | 0.218 | 0.0048 | 0.0363 | 1 | 0 | 0 |
| 'Eif1' | 122.404 | 115.423 | 94.445 | 114.633 | 108.499 | 96.937 | 114.818 | 112.018 | 89.545 | 121.249 | 105.519 | 81.749 | 126.120 | 122.854 | 88.359 | 122.253 | 111.923 | 82.842 | 0.010 | 0.7774 | 0.9002 | 0 | 0 | 0 |
| 'Eif1a' | 9.447 | 9.377 | 8.589 | 9.657 | 9.566 | 7.939 | 9.893 | 9.528 | 7.614 | 8.264 | 9.276 | 8.666 | 6.662 | 9.393 | 7.855 | 6.973 | 9.279 | 7.444 | 0.133 | 0.0264 | 0.1245 | 0 | 0 | 0 |
| 'Eif1ad' | 20.142 | 20.558 | 21.198 | 21.427 | 21.148 | 19.939 | 19.429 | 22.106 | 20.446 | 20.669 | 19.356 | 20.455 | 22.420 | 21.662 | 20.859 | 21.344 | 21.166 | 19.900 | -0.020 | 0.5631 | 0.7720 | 0 | 0 | 0 |
| 'Eif1ax' | 15.721 | 16.525 | 15.648 | 16.121 | 16.972 | 15.225 | 16.154 | 16.257 | 15.669 | 14.535 | 14.625 | 15.866 | 16.905 | 16.123 | 15.481 | 14.820 | 16.689 | 15.117 | 0.027 | 0.3838 | 0.6342 | 0 | 0 | 0 |
| 'Eif1b' | 65.196 | 70.327 | 62.906 | 73.757 | 64.621 | 69.475 | 61.087 | 60.558 | 66.167 | 57.725 | 59.156 | 71.206 | 57.190 | 66.515 | 75.547 | 61.906 | 66.734 | 73.606 | 0.003 | 0.9589 | 0.9847 | 0 | 0 | 0 |
| 'Eif2a' | 5.033 | 5.281 | 5.477 | 5.155 | 5.451 | 4.949 | 5.569 | 5.410 | 5.281 | 5.170 | 4.709 | 4.940 | 5.502 | 5.054 | 4.969 | 5.284 | 5.167 | 4.550 | 0.061 | 0.0820 | 0.2590 | 0 | 0 | 0 |
| 'Eif2ak1' | 12.701 | 13.985 | 13.260 | 13.197 | 13.785 | 14.033 | 13.308 | 13.810 | 13.431 | 12.920 | 13.541 | 12.748 | 13.461 | 13.817 | 12.757 | 12.181 | 13.211 | 12.400 | 0.045 | 0.1035 | 0.3017 | 0 | 0 | 0 |
| 'Eif2ak2' | 2.476 | 2.882 | 2.746 | 2.028 | 3.121 | 3.156 | 2.803 | 3.041 | 3.218 | 2.691 | 2.639 | 3.947 | 3.306 | 2.700 | 3.665 | 3.210 | 2.864 | 3.485 | -0.177 | 0.0147 | 0.0823 | 0 | 0 | 0 |
| 'Eif2ak3' | 5.249 | 5.574 | 4.522 | 4.614 | 5.465 | 5.072 | 4.698 | 5.394 | 4.819 | 5.599 | 5.535 | 4.479 | 5.395 | 5.583 | 4.653 | 5.345 | 5.094 | 4.651 | -0.034 | 0.4227 | 0.6693 | 0 | 0 | 0 |
| 'Eif2ak4' | 4.877 | 5.343 | 5.524 | 5.305 | 5.383 | 5.335 | 5.462 | 5.642 | 5.461 | 5.381 | 5.559 | 4.877 | 5.346 | 5.238 | 4.942 | 5.235 | 5.332 | 4.098 | 0.067 | 0.1284 | 0.3432 | 0 | 0 | 0 |
| 'Eif2b1' | 8.127 | 7.983 | 8.604 | 7.647 | 8.642 | 7.962 | 8.000 | 8.940 | 7.657 | 7.580 | 7.690 | 7.786 | 7.910 | 6.948 | 8.533 | 7.262 | 6.992 | 7.259 | 0.103 | 0.0250 | 0.1207 | 0 | 0 | 0 |
| 'Eif2b2' | 14.403 | 13.963 | 14.184 | 13.698 | 14.372 | 14.668 | 13.575 | 14.912 | 13.344 | 15.011 | 12.648 | 13.508 | 14.102 | 13.187 | 14.064 | 14.131 | 13.817 | 12.932 | 0.035 | 0.2965 | 0.5524 | 0 | 0 | 0 |
| 'Eif2b3' | 2.145 | 2.278 | 2.539 | 2.481 | 2.590 | 2.094 | 2.564 | 2.621 | 2.157 | 2.134 | 2.018 | 2.169 | 2.094 | 2.078 | 2.016 | 2.031 | 1.857 | 1.919 | 0.212 | 0.0003 | 0.0034 | 1 | 0 | 0 |
| 'Eif2b4' | 9.982 | 9.000 | 10.213 | 8.973 | 9.127 | 10.094 | 9.569 | 9.448 | 9.499 | 9.947 | 9.252 | 9.827 | 9.755 | 8.808 | 9.981 | 8.856 | 9.391 | 9.639 | -0.001 | 0.9794 | 0.9926 | 0 | 0 | 0 |
| 'Eif2b5' | 16.366 | 16.999 | 17.636 | 17.723 | 16.823 | 18.027 | 16.751 | 17.819 | 17.330 | 17.492 | 15.483 | 18.117 | 16.479 | 15.036 | 17.803 | 17.713 | 16.937 | 17.209 | 0.019 | 0.5622 | 0.7715 | 0 | 0 | 0 |
| 'Eif2d' | 6.367 | 6.213 | 6.743 | 6.279 | 6.504 | 6.760 | 5.790 | 6.260 | 6.662 | 6.312 | 6.302 | 6.949 | 6.686 | 5.573 | 6.543 | 6.257 | 6.299 | 6.850 | -0.016 | 0.6251 | 0.8133 | 0 | 0 | 0 |
| 'Eif2s1' | 27.641 | 28.715 | 27.454 | 28.600 | 28.342 | 26.426 | 30.499 | 27.123 | 25.828 | 25.203 | 24.773 | 27.148 | 26.333 | 26.401 | 26.033 | 26.286 | 27.576 | 25.822 | 0.074 | 0.0176 | 0.0942 | 0 | 0 | 0 |
| 'Eif2s2' | 4.821 | 4.751 | 4.499 | 5.260 | 4.730 | 4.620 | 5.033 | 4.777 | 4.515 | 4.553 | 4.556 | 4.479 | 4.794 | 4.758 | 4.395 | 4.851 | 5.052 | 4.576 | 0.022 | 0.5001 | 0.7275 | 0 | 0 | 0 |
| 'Eif2s3x' | 39.443 | 40.761 | 38.450 | 41.585 | 40.506 | 38.984 | 41.598 | 41.971 | 37.436 | 38.529 | 39.778 | 38.915 | 38.821 | 38.446 | 38.975 | 38.886 | 41.289 | 39.040 | 0.020 | 0.3897 | 0.6389 | 0 | 0 | 0 |
| 'Eif3a' | 115.606 | 107.817 | 103.575 | 117.613 | 107.239 | 106.300 | 116.059 | 106.080 | 106.350 | 108.563 | 103.732 | 106.473 | 107.774 | 105.785 | 109.779 | 107.858 | 108.596 | 106.948 | 0.019 | 0.4218 | 0.6682 | 0 | 0 | 0 |
| 'Eif3b' | 93.677 | 86.150 | 97.688 | 86.308 | 88.752 | 95.927 | 87.699 | 87.504 | 92.413 | 91.627 | 82.826 | 93.082 | 94.584 | 82.674 | 93.155 | 91.102 | 82.131 | 94.827 | 0.009 | 0.6998 | 0.8597 | 0 | 0 | 0 |
| 'Eif3c' | 94.912 | 88.987 | 91.998 | 93.338 | 91.619 | 93.275 | 90.182 | 86.632 | 91.803 | 93.420 | 89.526 | 91.110 | 91.474 | 86.341 | 90.956 | 90.707 | 88.161 | 92.484 | 0.005 | 0.8101 | 0.9142 | 0 | 0 | 0 |
| 'Eif3d' | 102.374 | 96.446 | 102.799 | 105.809 | 100.098 | 107.907 | 108.206 | 97.723 | 103.193 | 103.275 | 93.529 | 101.515 | 105.199 | 92.300 | 100.782 | 101.919 | 92.651 | 96.881 | 0.048 | 0.0134 | 0.0776 | 0 | 0 | 0 |
| 'Eif3e' | 165.910 | 164.495 | 155.109 | 181.339 | 174.112 | 150.493 | 177.280 | 162.478 | 151.136 | 157.897 | 154.652 | 152.216 | 157.622 | 158.656 | 151.605 | 151.629 | 164.453 | 145.065 | 0.075 | 0.0068 | 0.0472 | 1 | 0 | 0 |
| 'Eif3f' | 97.094 | 86.675 | 91.371 | 94.047 | 89.629 | 94.709 | 96.714 | 87.807 | 94.721 | 97.643 | 94.006 | 96.289 | 95.164 | 87.889 | 101.228 | 92.923 | 93.141 | 98.598 | -0.052 | 0.0366 | 0.1546 | 0 | 0 | 0 |
| 'Eif3g' | 105.295 | 94.365 | 98.886 | 94.477 | 99.568 | 107.275 | 98.943 | 101.409 | 100.204 | 97.729 | 89.570 | 98.832 | 99.040 | 89.414 | 98.697 | 102.551 | 94.271 | 99.382 | 0.042 | 0.1496 | 0.3765 | 0 | 0 | 0 |
| 'Eif3h' | 281.239 | 268.863 | 262.998 | 286.513 | 278.429 | 276.291 | 296.004 | 272.110 | 278.660 | 271.748 | 263.023 | 260.658 | 280.607 | 268.140 | 281.488 | 274.707 | 273.730 | 269.342 | 0.023 | 0.3246 | 0.5818 | 0 | 0 | 0 |
| 'Eif3i' | 29.583 | 28.055 | 28.271 | 30.569 | 28.902 | 28.862 | 30.208 | 29.878 | 28.388 | 25.821 | 28.021 | 27.990 | 28.198 | 26.823 | 28.613 | 28.174 | 27.932 | 26.906 | 0.069 | 0.0091 | 0.0583 | 0 | 0 | 0 |
| 'Eif3k' | 199.994 | 202.886 | 196.301 | 191.464 | 205.753 | 201.647 | 196.781 | 202.684 | 194.524 | 197.332 | 187.508 | 188.492 | 202.400 | 197.643 | 203.320 | 194.547 | 192.323 | 195.994 | 0.016 | 0.5037 | 0.7306 | 0 | 0 | 0 |
| 'Eif3l' | 92.388 | 84.464 | 92.684 | 93.424 | 87.505 | 89.523 | 92.175 | 86.554 | 91.212 | 90.719 | 89.836 | 91.977 | 91.182 | 85.404 | 90.854 | 85.959 | 89.396 | 92.480 | -0.007 | 0.7660 | 0.8956 | 0 | 0 | 0 |
| 'Eif3m' | 35.788 | 35.808 | 35.800 | 37.082 | 38.997 | 35.167 | 38.857 | 37.443 | 34.260 | 36.016 | 34.443 | 35.387 | 35.920 | 35.773 | 36.188 | 33.250 | 36.010 | 33.259 | 0.045 | 0.0993 | 0.2945 | 0 | 0 | 0 |
| 'Eif4a1' | 57.055 | 59.219 | 55.821 | 60.213 | 59.210 | 53.996 | 59.883 | 58.951 | 55.846 | 56.819 | 60.064 | 52.978 | 61.658 | 60.350 | 55.883 | 57.778 | 61.996 | 54.611 | -0.015 | 0.4951 | 0.7242 | 0 | 0 | 0 |
| 'Eif4a2' | 64.040 | 66.207 | 57.733 | 59.724 | 67.661 | 55.590 | 57.045 | 70.190 | 55.353 | 65.412 | 61.867 | 52.341 | 61.670 | 67.746 | 50.617 | 63.184 | 66.169 | 51.888 | 0.026 | 0.3947 | 0.6439 | 0 | 0 | 0 |
| 'Eif4a3' | 47.439 | 45.197 | 47.428 | 50.030 | 47.854 | 48.381 | 47.577 | 48.888 | 47.664 | 45.228 | 44.999 | 45.327 | 47.628 | 44.396 | 47.791 | 49.474 | 45.089 | 44.574 | 0.046 | 0.0598 | 0.2110 | 0 | 0 | 0 |
| 'Eif4b' | 168.267 | 153.809 | 162.052 | 167.588 | 159.187 | 160.212 | 173.316 | 158.533 | 156.643 | 157.183 | 159.848 | 160.100 | 159.224 | 158.329 | 160.454 | 158.756 | 157.976 | 160.559 | 0.016 | 0.4773 | 0.7108 | 0 | 0 | 0 |
| 'Eif4e' | 13.279 | 14.082 | 12.328 | 13.260 | 13.760 | 12.357 | 13.091 | 13.666 | 12.429 | 11.651 | 13.354 | 13.600 | 11.847 | 13.677 | 13.256 | 11.855 | 14.339 | 12.939 | 0.009 | 0.8013 | 0.9113 | 0 | 0 | 0 |
| 'Eif4e2' | 10.821 | 11.136 | 10.404 | 10.993 | 10.543 | 11.013 | 10.665 | 10.606 | 10.549 | 9.210 | 10.529 | 10.798 | 9.604 | 10.298 | 11.037 | 9.557 | 10.182 | 10.931 | 0.060 | 0.0789 | 0.2525 | 0 | 0 | 0 |
| 'Eif4e3' | 1.568 | 1.248 | 1.720 | 1.814 | 1.769 | 2.107 | 1.767 | 1.650 | 1.423 | 0.857 | 1.221 | 1.916 | 1.063 | 1.519 | 1.804 | 1.127 | 1.330 | 1.454 | 0.284 | 0.0262 | 0.1243 | 0 | 0 | 0 |
| 'Eif4ebp1' | 8.866 | 7.917 | 6.693 | 6.827 | 7.606 | 6.401 | 8.280 | 7.419 | 7.876 | 6.041 | 8.106 | 7.510 | 7.120 | 6.293 | 8.373 | 7.888 | 6.749 | 8.148 | 0.020 | 0.8222 | 0.9217 | 0 | 0 | 0 |
| 'Eif4ebp2' | 98.511 | 108.024 | 103.517 | 93.698 | 110.326 | 116.207 | 89.352 | 112.304 | 110.648 | 107.610 | 101.118 | 112.807 | 99.930 | 106.788 | 118.045 | 116.397 | 94.285 | 119.369 | -0.063 | 0.1688 | 0.4013 | 0 | 0 | 0 |
| 'Eif4enif1' | 12.868 | 13.367 | 13.519 | 12.794 | 13.298 | 13.845 | 12.672 | 14.148 | 13.509 | 14.629 | 14.127 | 14.379 | 14.257 | 14.245 | 13.844 | 14.195 | 13.820 | 14.307 | -0.100 | 0.0000 | 0.0003 | 1 | 0 | 0 |
| 'Eif4g1' | 31.550 | 30.018 | 34.522 | 32.940 | 30.460 | 32.976 | 31.043 | 30.911 | 34.736 | 33.167 | 31.399 | 34.449 | 33.111 | 30.230 | 33.176 | 30.825 | 31.587 | 34.991 | -0.030 | 0.1862 | 0.4219 | 0 | 0 | 0 |
| 'Eif4g2' | 110.876 | 112.341 | 105.239 | 113.339 | 109.074 | 104.703 | 110.304 | 108.464 | 104.782 | 106.866 | 111.412 | 106.169 | 102.542 | 113.240 | 110.365 | 105.629 | 114.257 | 111.861 | -0.016 | 0.5662 | 0.7744 | 0 | 0 | 0 |
| 'Eif4g3' | 24.696 | 20.687 | 25.231 | 25.193 | 22.063 | 24.772 | 23.840 | 22.315 | 24.385 | 23.674 | 24.568 | 24.120 | 25.263 | 22.617 | 22.918 | 22.949 | 23.150 | 24.205 | -0.014 | 0.6605 | 0.8362 | 0 | 0 | 0 |
| 'Eif4h' | 33.631 | 36.543 | 35.436 | 32.673 | 37.022 | 37.561 | 33.287 | 35.361 | 35.318 | 32.261 | 33.437 | 34.658 | 34.728 | 33.994 | 37.042 | 34.049 | 31.994 | 35.200 | 0.033 | 0.2272 | 0.4734 | 0 | 0 | 0 |
| 'Eif5' | 105.844 | 89.312 | 77.496 | 90.222 | 92.367 | 72.727 | 88.131 | 92.021 | 75.709 | 97.136 | 87.960 | 76.336 | 89.865 | 94.615 | 73.107 | 93.868 | 90.508 | 77.645 | -0.006 | 0.8708 | 0.9447 | 0 | 0 | 0 |
| 'Eif5a' | 359.427 | 360.716 | 344.874 | 352.421 | 376.080 | 348.634 | 352.904 | 376.089 | 344.310 | 337.359 | 338.860 | 317.206 | 359.127 | 352.033 | 335.528 | 365.635 | 338.775 | 326.746 | 0.057 | 0.0115 | 0.0692 | 0 | 0 | 0 |
| 'Eif5a2' | 3.495 | 4.417 | 4.540 | 3.602 | 4.681 | 4.274 | 3.994 | 4.730 | 4.198 | 4.341 | 3.900 | 4.085 | 3.485 | 4.166 | 3.931 | 4.176 | 4.423 | 3.658 | 0.059 | 0.2841 | 0.5403 | 0 | 0 | 0 |
| 'Eif5b' | 15.891 | 14.704 | 14.898 | 17.126 | 15.017 | 14.451 | 17.509 | 15.331 | 14.294 | 15.388 | 14.585 | 15.512 | 15.541 | 14.748 | 14.069 | 15.149 | 15.749 | 15.585 | 0.016 | 0.6325 | 0.8178 | 0 | 0 | 0 |
| 'Eif6' | 22.080 | 22.682 | 21.529 | 20.493 | 22.285 | 21.803 | 20.725 | 22.452 | 22.036 | 19.678 | 20.244 | 21.262 | 21.607 | 20.116 | 21.955 | 21.772 | 19.707 | 21.360 | 0.052 | 0.0803 | 0.2553 | 0 | 0 | 0 |
| 'Elac1' | 2.426 | 3.488 | 3.050 | 2.058 | 3.536 | 3.013 | 2.960 | 3.655 | 3.310 | 3.577 | 3.595 | 3.279 | 2.898 | 3.681 | 2.727 | 3.236 | 3.323 | 2.964 | -0.106 | 0.1451 | 0.3695 | 0 | 0 | 0 |
| 'Elac2' | 7.532 | 7.632 | 8.420 | 6.807 | 7.956 | 7.932 | 7.044 | 8.046 | 8.495 | 8.175 | 7.558 | 7.833 | 8.436 | 7.491 | 7.482 | 7.916 | 6.877 | 7.017 | 0.014 | 0.7638 | 0.8953 | 0 | 0 | 0 |
| 'Elavl1' | 24.272 | 22.873 | 25.657 | 26.229 | 24.948 | 23.996 | 25.696 | 25.133 | 23.520 | 22.125 | 26.479 | 26.477 | 21.376 | 25.470 | 25.845 | 21.141 | 26.423 | 25.792 | 0.001 | 0.9789 | 0.9926 | 0 | 0 | 0 |
| 'Elf1' | 57.180 | 54.366 | 54.807 | 58.687 | 56.002 | 52.135 | 58.891 | 55.278 | 53.833 | 55.610 | 60.601 | 54.013 | 52.093 | 57.131 | 54.148 | 52.566 | 56.845 | 53.595 | 0.003 | 0.9316 | 0.9746 | 0 | 0 | 0 |
| 'Elf2' | 6.038 | 6.517 | 5.966 | 6.120 | 6.390 | 6.036 | 5.640 | 6.604 | 6.237 | 6.225 | 6.383 | 6.248 | 6.147 | 6.282 | 5.791 | 6.824 | 5.717 | 5.833 | -0.008 | 0.8204 | 0.9206 | 0 | 0 | 0 |
| 'Elf4' | 24.515 | 22.931 | 22.903 | 22.872 | 21.080 | 22.701 | 21.149 | 22.028 | 23.822 | 22.897 | 23.653 | 28.035 | 22.456 | 22.966 | 26.905 | 21.522 | 24.915 | 28.625 | -0.127 | 0.0037 | 0.0295 | 1 | 0 | 0 |
| 'Elk1' | 5.690 | 5.230 | 5.595 | 5.299 | 5.612 | 5.520 | 4.346 | 5.617 | 6.041 | 6.441 | 5.641 | 6.122 | 5.782 | 5.618 | 5.688 | 5.366 | 4.993 | 5.721 | -0.079 | 0.1405 | 0.3628 | 0 | 0 | 0 |
| 'Elk3' | 17.488 | 17.281 | 16.428 | 17.442 | 17.399 | 18.335 | 16.763 | 18.049 | 17.882 | 18.876 | 17.037 | 18.413 | 17.755 | 18.808 | 17.959 | 18.750 | 17.415 | 17.308 | -0.057 | 0.0442 | 0.1742 | 0 | 0 | 0 |
| 'Elk4' | 25.395 | 29.413 | 27.462 | 26.811 | 28.513 | 26.181 | 26.733 | 28.371 | 27.910 | 26.592 | 28.229 | 28.706 | 24.618 | 26.506 | 26.853 | 27.633 | 27.097 | 30.230 | -0.010 | 0.7643 | 0.8953 | 0 | 0 | 0 |
| 'Ell' | 17.161 | 17.242 | 17.854 | 15.220 | 16.883 | 17.234 | 16.469 | 17.564 | 18.426 | 18.163 | 16.588 | 18.082 | 16.805 | 15.007 | 17.298 | 17.184 | 17.645 | 17.459 | -0.011 | 0.7605 | 0.8935 | 0 | 0 | 0 |
| 'Ell2' | 7.600 | 7.627 | 7.563 | 7.752 | 7.324 | 6.676 | 7.097 | 8.336 | 6.888 | 8.589 | 7.952 | 7.194 | 7.418 | 8.119 | 6.844 | 7.426 | 9.002 | 6.790 | -0.056 | 0.2406 | 0.4911 | 0 | 0 | 0 |
| 'Elmo1' | 21.462 | 19.964 | 20.242 | 22.269 | 21.255 | 20.675 | 22.449 | 20.608 | 20.683 | 21.422 | 22.686 | 22.132 | 21.850 | 21.705 | 21.108 | 20.854 | 23.090 | 21.499 | -0.062 | 0.0294 | 0.1332 | 0 | 0 | 0 |
| 'Elmo2' | 7.802 | 7.869 | 8.314 | 7.822 | 8.177 | 8.053 | 7.256 | 7.303 | 8.058 | 7.786 | 7.875 | 8.072 | 8.064 | 7.783 | 8.078 | 7.753 | 7.382 | 8.075 | -0.015 | 0.6212 | 0.8107 | 0 | 0 | 0 |
| 'Elmo3' | 8.698 | 10.102 | 8.804 | 9.548 | 9.237 | 9.168 | 8.364 | 9.620 | 9.093 | 9.618 | 7.976 | 7.214 | 10.023 | 7.997 | 7.077 | 8.888 | 9.247 | 7.171 | 0.135 | 0.0191 | 0.1002 | 0 | 0 | 0 |
| 'Elmod2' | 1.342 | 1.603 | 1.511 | 1.352 | 1.609 | 1.611 | 1.255 | 1.626 | 1.490 | 1.446 | 1.776 | 1.582 | 1.368 | 1.421 | 1.477 | 1.447 | 1.568 | 1.499 | -0.029 | 0.5362 | 0.7552 | 0 | 0 | 0 |
| 'Elmod3' | 4.474 | 4.737 | 4.788 | 4.691 | 4.813 | 4.992 | 4.598 | 5.064 | 4.834 | 4.731 | 4.138 | 4.650 | 4.587 | 4.547 | 4.978 | 4.911 | 4.312 | 3.933 | 0.068 | 0.1582 | 0.3883 | 0 | 0 | 0 |
| 'Elmsan1' | 15.069 | 14.458 | 13.935 | 14.566 | 14.310 | 13.679 | 12.636 | 14.945 | 14.393 | 14.184 | 14.995 | 14.968 | 15.089 | 15.156 | 14.400 | 15.308 | 14.052 | 15.149 | -0.069 | 0.0375 | 0.1572 | 0 | 0 | 0 |
| 'Elof1' | 15.903 | 18.080 | 16.681 | 16.998 | 16.345 | 18.891 | 15.782 | 18.468 | 18.339 | 15.625 | 16.666 | 16.706 | 16.787 | 17.241 | 18.479 | 17.655 | 17.152 | 16.315 | 0.020 | 0.6521 | 0.8310 | 0 | 0 | 0 |
| 'Elovl1' | 15.159 | 16.453 | 16.615 | 15.247 | 16.137 | 16.740 | 15.341 | 15.548 | 16.579 | 14.721 | 14.687 | 18.939 | 15.824 | 15.457 | 18.298 | 15.580 | 16.562 | 18.663 | -0.063 | 0.0808 | 0.2562 | 0 | 0 | 0 |
| 'Elovl5' | 33.723 | 35.053 | 35.057 | 35.025 | 35.879 | 35.161 | 36.430 | 35.619 | 35.362 | 33.560 | 33.695 | 37.666 | 31.469 | 34.632 | 34.739 | 33.439 | 35.253 | 38.297 | 0.009 | 0.7674 | 0.8961 | 0 | 0 | 0 |
| 'Elovl6' | 5.852 | 4.392 | 5.623 | 5.338 | 4.877 | 4.870 | 4.688 | 5.170 | 5.136 | 4.874 | 6.247 | 5.400 | 4.542 | 5.127 | 5.630 | 4.463 | 5.474 | 5.137 | -0.034 | 0.6061 | 0.8014 | 0 | 0 | 0 |
| 'Elp2' | 40.793 | 41.801 | 43.010 | 43.540 | 43.882 | 41.307 | 43.137 | 43.920 | 43.397 | 41.436 | 43.190 | 43.214 | 42.518 | 41.815 | 40.884 | 40.324 | 45.507 | 39.486 | 0.013 | 0.6153 | 0.8081 | 0 | 0 | 0 |
| 'Elp3' | 23.629 | 25.835 | 25.054 | 22.199 | 26.180 | 24.727 | 25.605 | 26.631 | 26.310 | 24.390 | 25.300 | 24.192 | 24.841 | 24.478 | 23.287 | 24.512 | 24.663 | 23.239 | 0.038 | 0.2238 | 0.4711 | 0 | 0 | 0 |
| 'Elp4' | 3.138 | 4.323 | 3.384 | 4.037 | 4.077 | 3.769 | 4.166 | 4.204 | 3.847 | 4.042 | 4.009 | 3.712 | 3.786 | 3.688 | 3.674 | 3.807 | 4.201 | 3.290 | 0.014 | 0.8139 | 0.9160 | 0 | 0 | 0 |
| 'Elp5' | 6.926 | 8.428 | 7.837 | 7.943 | 8.326 | 8.264 | 7.934 | 8.118 | 8.649 | 7.360 | 6.939 | 7.559 | 7.035 | 7.397 | 8.037 | 7.362 | 7.770 | 7.009 | 0.111 | 0.0042 | 0.0328 | 1 | 0 | 0 |
| 'Elp6' | 2.566 | 2.946 | 3.317 | 3.275 | 3.371 | 2.893 | 3.068 | 3.043 | 3.326 | 2.880 | 2.995 | 3.086 | 3.256 | 3.106 | 3.056 | 3.009 | 2.651 | 2.845 | 0.035 | 0.5874 | 0.7882 | 0 | 0 | 0 |
| 'Emb' | 44.656 | 47.184 | 46.720 | 44.654 | 48.750 | 47.555 | 46.393 | 51.474 | 46.446 | 47.318 | 50.076 | 53.115 | 41.298 | 48.664 | 51.503 | 52.502 | 47.888 | 48.787 | -0.068 | 0.0731 | 0.2395 | 0 | 0 | 0 |
| 'Emc1' | 6.122 | 6.068 | 6.885 | 5.500 | 6.516 | 6.521 | 5.767 | 6.712 | 6.841 | 6.863 | 7.151 | 6.717 | 7.119 | 6.014 | 6.275 | 6.462 | 6.783 | 6.171 | -0.074 | 0.1055 | 0.3045 | 0 | 0 | 0 |
| 'Emc10' | 37.188 | 38.819 | 36.926 | 34.092 | 39.253 | 38.381 | 33.625 | 39.878 | 37.003 | 39.036 | 34.359 | 35.587 | 36.570 | 36.120 | 36.671 | 38.927 | 31.053 | 35.362 | 0.040 | 0.3392 | 0.5966 | 0 | 0 | 0 |
| 'Emc2' | 36.516 | 40.961 | 37.167 | 40.610 | 42.237 | 37.615 | 43.701 | 39.277 | 35.214 | 38.622 | 44.014 | 36.680 | 39.581 | 41.435 | 38.850 | 40.797 | 41.931 | 34.073 | -0.021 | 0.5918 | 0.7921 | 0 | 0 | 0 |
| 'Emc3' | 14.017 | 18.127 | 17.116 | 16.161 | 17.192 | 17.285 | 16.148 | 17.766 | 16.656 | 18.051 | 15.607 | 16.312 | 16.432 | 17.266 | 16.684 | 16.523 | 14.869 | 14.915 | 0.025 | 0.5748 | 0.7803 | 0 | 0 | 0 |
| 'Emc4' | 13.566 | 17.960 | 14.955 | 14.300 | 16.824 | 15.374 | 13.136 | 15.955 | 15.782 | 14.891 | 14.067 | 15.482 | 15.905 | 16.864 | 15.877 | 17.195 | 15.449 | 14.293 | -0.037 | 0.4812 | 0.7131 | 0 | 0 | 0 |
| 'Emc6' | 18.177 | 21.197 | 18.424 | 19.224 | 19.761 | 19.028 | 17.793 | 18.436 | 17.666 | 18.942 | 17.865 | 19.572 | 19.265 | 19.816 | 19.491 | 21.265 | 17.691 | 19.132 | -0.042 | 0.3290 | 0.5872 | 0 | 0 | 0 |
| 'Emc7' | 9.271 | 10.418 | 8.929 | 8.663 | 10.793 | 9.229 | 8.299 | 10.596 | 8.406 | 8.343 | 8.985 | 9.671 | 9.762 | 10.061 | 9.272 | 9.307 | 9.960 | 9.292 | -0.018 | 0.6602 | 0.8362 | 0 | 0 | 0 |
| 'Emc8' | 2.911 | 3.328 | 3.518 | 3.019 | 3.526 | 3.536 | 3.183 | 3.036 | 3.400 | 2.797 | 2.912 | 3.570 | 3.274 | 3.110 | 3.371 | 3.089 | 2.816 | 3.399 | 0.040 | 0.3485 | 0.6039 | 0 | 0 | 0 |
| 'Emc9' | 3.692 | 4.950 | 4.513 | 3.862 | 4.320 | 4.493 | 4.373 | 5.040 | 4.882 | 3.843 | 4.425 | 4.334 | 3.834 | 3.583 | 4.069 | 3.581 | 4.118 | 3.746 | 0.159 | 0.0079 | 0.0523 | 0 | 0 | 0 |
| 'Emd' | 14.928 | 16.053 | 14.475 | 13.994 | 15.170 | 15.640 | 14.001 | 15.757 | 15.162 | 15.585 | 14.847 | 14.550 | 15.799 | 15.195 | 15.125 | 14.911 | 15.557 | 13.725 | -0.010 | 0.7633 | 0.8952 | 0 | 0 | 0 |
| 'Eme1' | 1.835 | 1.713 | 2.914 | 2.312 | 2.062 | 2.747 | 2.390 | 1.862 | 2.897 | 1.741 | 2.088 | 2.614 | 2.626 | 2.669 | 2.470 | 1.929 | 2.066 | 2.200 | 0.002 | 0.9863 | 0.9955 | 0 | 0 | 0 |
| 'Eme2' | 3.030 | 2.429 | 3.173 | 2.962 | 2.514 | 3.100 | 2.607 | 2.727 | 3.054 | 2.730 | 3.096 | 3.042 | 3.003 | 2.200 | 2.804 | 2.229 | 2.650 | 2.768 | 0.050 | 0.4260 | 0.6724 | 0 | 0 | 0 |
| 'Emg1' | 18.716 | 19.604 | 19.472 | 18.528 | 19.132 | 19.856 | 18.802 | 19.815 | 19.744 | 18.171 | 18.340 | 19.476 | 18.270 | 18.215 | 20.133 | 18.701 | 18.622 | 19.234 | 0.027 | 0.1945 | 0.4326 | 0 | 0 | 0 |
| 'Emid1' | 0.691 | 0.674 | 0.712 | 0.722 | 0.742 | 0.939 | 0.806 | 0.765 | 0.892 | 0.638 | 0.317 | 0.517 | 0.765 | 0.605 | 0.453 | 0.564 | 0.599 | 0.550 | 0.468 | 0.0002 | 0.0023 | 1 | 1 | 0 |
| 'Eml2' | 11.377 | 11.629 | 12.907 | 11.313 | 11.932 | 12.426 | 11.099 | 11.779 | 12.809 | 10.951 | 9.954 | 11.554 | 11.191 | 10.145 | 11.817 | 11.566 | 10.759 | 10.764 | 0.111 | 0.0003 | 0.0044 | 1 | 0 | 0 |
| 'Eml3' | 54.777 | 56.440 | 61.720 | 55.273 | 54.242 | 58.951 | 56.000 | 55.569 | 61.101 | 55.330 | 54.746 | 56.582 | 55.659 | 49.634 | 53.101 | 53.222 | 53.668 | 53.680 | 0.072 | 0.0059 | 0.0426 | 1 | 0 | 0 |
| 'Eml4' | 33.302 | 36.843 | 33.189 | 33.642 | 37.339 | 34.086 | 32.763 | 37.032 | 33.983 | 31.403 | 33.401 | 34.613 | 30.408 | 32.121 | 33.875 | 32.445 | 34.766 | 34.512 | 0.056 | 0.0398 | 0.1628 | 0 | 0 | 0 |
| 'Eml5' | 4.546 | 4.993 | 4.804 | 4.183 | 4.491 | 4.091 | 4.275 | 4.920 | 4.478 | 5.248 | 5.383 | 4.318 | 4.949 | 4.843 | 4.166 | 5.107 | 4.943 | 4.046 | -0.078 | 0.0988 | 0.2937 | 0 | 0 | 0 |
| 'Emp3' | 5.727 | 4.924 | 5.201 | 4.771 | 5.609 | 6.400 | 5.440 | 6.712 | 5.040 | 2.633 | 4.779 | 4.724 | 4.773 | 4.990 | 4.610 | 4.256 | 4.374 | 4.999 | 0.303 | 0.0043 | 0.0335 | 1 | 0 | 0 |
| 'Emsy' | 5.139 | 4.702 | 4.653 | 5.144 | 4.436 | 4.316 | 5.052 | 4.753 | 4.600 | 4.497 | 5.300 | 5.539 | 4.396 | 4.882 | 5.096 | 4.526 | 4.784 | 5.462 | -0.066 | 0.2115 | 0.4566 | 0 | 0 | 0 |
| 'Enc1' | 10.343 | 9.048 | 8.449 | 9.729 | 9.075 | 9.376 | 8.708 | 8.991 | 8.250 | 9.158 | 8.848 | 9.309 | 8.365 | 10.148 | 8.744 | 9.562 | 7.965 | 9.472 | -0.004 | 0.9345 | 0.9749 | 0 | 0 | 0 |
| 'Endod1' | 41.201 | 47.038 | 43.133 | 39.366 | 45.403 | 46.636 | 41.813 | 43.324 | 45.948 | 45.922 | 41.162 | 46.255 | 41.020 | 46.788 | 44.079 | 50.045 | 44.126 | 47.509 | -0.059 | 0.1412 | 0.3636 | 0 | 0 | 0 |
| 'Endog' | 7.473 | 6.126 | 8.137 | 5.608 | 7.897 | 8.128 | 6.705 | 7.760 | 8.509 | 6.779 | 5.999 | 7.334 | 7.701 | 6.084 | 6.376 | 6.919 | 5.180 | 6.143 | 0.175 | 0.0450 | 0.1759 | 0 | 0 | 0 |
| 'Endov' | 4.896 | 5.200 | 5.803 | 4.696 | 4.761 | 5.682 | 4.934 | 5.145 | 5.845 | 5.409 | 4.837 | 5.205 | 5.504 | 5.384 | 5.039 | 5.612 | 4.453 | 4.990 | 0.007 | 0.8733 | 0.9459 | 0 | 0 | 0 |
| 'Engase' | 1.324 | 1.466 | 1.682 | 1.472 | 1.489 | 1.522 | 1.385 | 1.525 | 1.681 | 1.477 | 1.216 | 1.343 | 1.536 | 1.389 | 1.288 | 1.474 | 1.245 | 1.344 | 0.133 | 0.0194 | 0.1009 | 0 | 0 | 0 |

| | | | | | | | | | | | | | | | | | | | | | | | | |
|---|---|---|---|---|---|---|---|---|---|---|---|---|---|---|---|---|---|---|---|---|---|---|---|---|
| 'Enkd1' | 10.166 | 9.930 | 11.553 | 8.621 | 8.911 | 11.955 | 10.483 | 7.884 | 12.690 | 9.636 | 8.913 | 11.366 | 12.528 | 9.532 | 10.432 | 9.576 | 8.986 | 11.048 | -0.013 | 0.8529 | 0.9370 | 0 | 0 | 0 |
| 'Eno1' | 71.288 | 66.562 | 68.844 | 68.975 | 71.173 | 71.921 | 71.861 | 65.083 | 68.071 | 64.392 | 70.249 | 71.632 | 74.684 | 73.826 | 80.697 | 73.198 | 72.214 | 73.447 | -0.079 | 0.0216 | 0.1086 | 0 | 0 | 0 |
| 'Eno2' | 0.876 | 0.908 | 0.851 | 0.810 | 0.898 | 0.737 | 0.838 | 0.903 | 0.949 | 0.886 | 0.793 | 0.753 | 1.111 | 0.808 | 0.888 | 0.925 | 0.742 | 0.672 | 0.033 | 0.6696 | 0.8408 | 0 | 0 | 0 |
| 'Eno3' | 3.882 | 3.400 | 4.096 | 4.267 | 3.345 | 3.960 | 4.512 | 3.867 | 3.923 | 3.522 | 3.200 | 4.403 | 4.159 | 3.185 | 4.292 | 3.778 | 3.562 | 3.763 | 0.039 | 0.5383 | 0.7566 | 0 | 0 | 0 |
| 'Enoph1' | 4.741 | 5.124 | 4.900 | 4.246 | 5.302 | 5.402 | 5.212 | 5.493 | 5.043 | 5.060 | 5.103 | 5.197 | 5.292 | 4.348 | 5.430 | 5.003 | 5.271 | 5.631 | -0.040 | 0.4637 | 0.6997 | 0 | 0 | 0 |
| 'Enox1' | 2.981 | 4.705 | 3.660 | 3.089 | 5.388 | 3.964 | 3.808 | 4.665 | 3.336 | 4.615 | 5.373 | 3.833 | 4.379 | 5.855 | 3.477 | 5.097 | 5.613 | 3.237 | -0.222 | 0.0032 | 0.0265 | 1 | 0 | 0 |
| 'Enox2' | 3.267 | 3.527 | 3.255 | 3.910 | 2.932 | 3.178 | 3.601 | 2.848 | 3.208 | 3.342 | 3.585 | 3.999 | 3.154 | 2.745 | 3.806 | 3.265 | 3.601 | 3.995 | -0.104 | 0.1727 | 0.4066 | 0 | 0 | 0 |
| 'Enpp1' | 0.874 | 1.029 | 0.979 | 0.980 | 1.004 | 0.935 | 1.088 | 1.243 | 0.900 | 1.179 | 1.208 | 0.940 | 1.037 | 1.525 | 0.804 | 1.228 | 1.244 | 0.837 | -0.136 | 0.0677 | 0.2279 | 0 | 0 | 0 |
| 'Enpp4' | 1.865 | 2.070 | 2.050 | 2.368 | 2.371 | 2.150 | 1.878 | 1.904 | 2.036 | 1.701 | 2.275 | 2.356 | 1.954 | 1.865 | 2.108 | 1.868 | 1.988 | 2.383 | -0.002 | 0.9813 | 0.9931 | 0 | 0 | 0 |
| 'Enpp5' | 0.653 | 1.220 | 0.862 | 0.838 | 1.186 | 0.868 | 0.967 | 0.944 | 0.977 | 0.915 | 1.041 | 0.825 | 0.822 | 0.855 | 0.802 | 0.901 | 0.709 | 0.710 | 0.145 | 0.1319 | 0.3493 | 0 | 0 | 0 |
| 'Ensa' | 23.662 | 25.478 | 23.398 | 22.654 | 24.004 | 26.571 | 21.959 | 24.152 | 23.665 | 23.791 | 23.200 | 25.330 | 23.456 | 24.830 | 24.562 | 24.654 | 25.031 | 23.539 | -0.030 | 0.3204 | 0.5785 | 0 | 0 | 0 |
| 'Enthd2' | 7.193 | 6.335 | 8.924 | 7.594 | 6.590 | 8.269 | 7.757 | 7.256 | 9.053 | 7.435 | 6.618 | 9.936 | 7.515 | 6.292 | 8.717 | 6.888 | 6.892 | 8.947 | -0.018 | 0.6729 | 0.8425 | 0 | 0 | 0 |
| 'Entpd5' | 9.070 | 10.962 | 9.427 | 9.586 | 10.130 | 9.855 | 9.414 | 10.174 | 9.330 | 9.091 | 9.640 | 9.443 | 10.877 | 10.582 | 9.249 | 9.401 | 9.860 | 9.182 | -0.002 | 0.9525 | 0.9823 | 0 | 0 | 0 |
| 'Entpd6' | 10.320 | 10.114 | 10.837 | 10.621 | 10.375 | 10.855 | 9.234 | 10.151 | 10.400 | 10.285 | 9.412 | 11.035 | 10.959 | 9.718 | 10.396 | 11.445 | 8.918 | 10.010 | 0.003 | 0.9391 | 0.9765 | 0 | 0 | 0 |
| 'Entpd7' | 1.152 | 0.903 | 1.253 | 1.244 | 1.286 | 1.161 | 0.954 | 1.041 | 1.283 | 1.101 | 1.199 | 1.197 | 1.100 | 1.256 | 1.286 | 1.217 | 1.100 | 1.378 | -0.080 | 0.2646 | 0.5184 | 0 | 0 | 0 |
| 'Eny2' | 14.634 | 16.408 | 15.231 | 14.967 | 17.514 | 14.490 | 17.825 | 18.068 | 13.938 | 15.138 | 16.952 | 14.542 | 16.872 | 17.033 | 14.473 | 15.297 | 15.696 | 13.614 | 0.023 | 0.5814 | 0.7838 | 0 | 0 | 0 |
| 'Eogt' | 4.649 | 5.214 | 5.375 | 4.810 | 4.907 | 5.081 | 4.966 | 5.241 | 4.966 | 4.849 | 5.571 | 4.999 | 5.388 | 5.047 | 4.510 | 4.360 | 4.864 | 4.883 | 0.014 | 0.7612 | 0.8940 | 0 | 0 | 0 |
| 'Eomes' | 0.748 | 0.770 | 0.716 | 0.692 | 0.795 | 1.277 | 0.710 | 0.840 | 0.705 | 0.520 | 0.298 | 1.717 | 1.447 | 0.753 | 1.507 | 1.032 | 0.518 | 1.305 | -0.233 | 0.3402 | 0.5970 | 0 | 0 | 0 |
| 'Ep300' | 13.286 | 11.309 | 12.232 | 12.314 | 11.251 | 12.028 | 10.978 | 11.550 | 12.112 | 13.015 | 13.675 | 13.529 | 11.617 | 12.358 | 12.863 | 12.524 | 11.828 | 13.750 | -0.115 | 0.0053 | 0.0393 | 1 | 0 | 0 |
| 'Ep400' | 17.079 | 14.678 | 17.680 | 16.430 | 15.714 | 16.920 | 15.975 | 15.836 | 17.581 | 18.416 | 17.560 | 17.344 | 16.543 | 15.942 | 16.862 | 16.949 | 16.030 | 17.200 | -0.058 | 0.0644 | 0.2213 | 0 | 0 | 0 |
| 'Epas1' | 47.082 | 38.990 | 48.818 | 44.307 | 39.932 | 43.828 | 44.551 | 39.971 | 48.096 | 41.568 | 33.542 | 41.934 | 32.215 | 33.141 | 41.591 | 35.386 | 30.387 | 42.722 | 0.244 | 0.0000 | 0.0000 | 1 | 0 | 0 |
| 'Epb41' | 12.565 | 13.339 | 12.508 | 12.406 | 12.722 | 13.288 | 11.615 | 13.017 | 12.446 | 14.470 | 13.508 | 12.721 | 13.176 | 12.892 | 12.713 | 13.356 | 12.027 | 12.585 | -0.054 | 0.1089 | 0.3109 | 0 | 0 | 0 |
| 'Epb41l2' | 9.946 | 9.668 | 11.224 | 9.470 | 9.696 | 10.898 | 9.827 | 9.573 | 10.829 | 9.598 | 11.232 | 11.419 | 10.734 | 10.462 | 11.951 | 10.636 | 11.350 | 11.107 | -0.120 | 0.0003 | 0.0044 | 1 | 0 | 0 |
| 'Epc1' | 22.609 | 20.489 | 19.276 | 19.146 | 20.650 | 18.679 | 20.245 | 21.211 | 18.754 | 21.420 | 22.821 | 20.386 | 21.607 | 22.744 | 20.228 | 22.785 | 22.414 | 20.501 | -0.115 | 0.0002 | 0.0023 | 1 | 0 | 0 |
| 'Epc2' | 16.403 | 15.753 | 14.439 | 15.951 | 15.348 | 14.860 | 16.105 | 17.699 | 14.977 | 18.165 | 16.875 | 15.540 | 15.958 | 18.211 | 15.345 | 17.428 | 17.425 | 14.549 | -0.085 | 0.0119 | 0.0708 | 0 | 0 | 0 |
| 'Epg5' | 6.259 | 6.092 | 7.214 | 6.118 | 6.275 | 6.521 | 6.180 | 6.494 | 7.053 | 6.308 | 7.920 | 6.594 | 6.545 | 6.139 | 6.135 | 5.806 | 6.745 | 6.158 | -0.012 | 0.8137 | 0.9160 | 0 | 0 | 0 |
| 'Ephb6' | 15.419 | 15.257 | 16.228 | 13.510 | 15.309 | 17.456 | 14.136 | 15.048 | 17.239 | 17.029 | 15.915 | 18.089 | 17.667 | 14.933 | 16.464 | 17.702 | 14.238 | 15.850 | -0.093 | 0.0332 | 0.1446 | 0 | 0 | 0 |
| 'Ephx1' | 135.134 | 144.480 | 131.120 | 133.853 | 141.480 | 141.902 | 134.711 | 141.494 | 134.085 | 149.373 | 136.540 | 141.990 | 122.952 | 140.453 | 140.030 | 148.559 | 136.275 | 139.445 | -0.031 | 0.3532 | 0.6066 | 0 | 0 | 0 |
| 'Ephx4' | 3.051 | 2.381 | 2.451 | 2.569 | 2.613 | 2.564 | 3.140 | 2.400 | 1.906 | 3.474 | 2.150 | 2.246 | 3.147 | 2.528 | 2.179 | 2.596 | 2.609 | 2.114 | -0.002 | 0.9841 | 0.9943 | 0 | 0 | 0 |
| 'Epm2aip1' | 5.727 | 4.844 | 5.634 | 5.768 | 4.939 | 4.688 | 5.737 | 4.536 | 5.400 | 4.743 | 6.532 | 6.233 | 5.143 | 5.292 | 5.236 | 4.391 | 6.394 | 4.845 | -0.054 | 0.4805 | 0.7127 | 0 | 0 | 0 |
| 'Epn1' | 27.889 | 26.654 | 28.402 | 25.092 | 26.032 | 29.150 | 26.277 | 25.694 | 28.357 | 26.240 | 25.391 | 30.071 | 26.394 | 24.388 | 30.433 | 27.568 | 24.891 | 30.821 | -0.024 | 0.4033 | 0.6522 | 0 | 0 | 0 |
| 'Eprs' | 7.215 | 6.751 | 7.325 | 7.386 | 7.401 | 6.902 | 7.395 | 6.853 | 7.036 | 7.168 | 7.251 | 6.964 | 6.998 | 6.721 | 6.707 | 7.033 | 7.299 | 6.827 | 0.019 | 0.4694 | 0.7053 | 0 | 0 | 0 |
| 'Eps15' | 10.952 | 11.930 | 12.765 | 12.100 | 12.776 | 12.005 | 12.290 | 12.857 | 11.808 | 12.230 | 13.179 | 13.159 | 11.526 | 12.830 | 12.274 | 12.066 | 12.596 | 12.582 | -0.049 | 0.0807 | 0.2562 | 0 | 0 | 0 |
| 'Eps15l1' | 34.671 | 31.272 | 35.353 | 34.935 | 32.684 | 33.946 | 33.400 | 34.003 | 34.115 | 35.592 | 34.039 | 36.235 | 35.703 | 34.509 | 35.187 | 34.073 | 35.548 | 36.375 | -0.069 | 0.0074 | 0.0497 | 1 | 0 | 0 |
| 'Epsti1' | 25.758 | 23.948 | 25.840 | 25.387 | 26.869 | 27.507 | 27.294 | 27.220 | 24.771 | 25.728 | 26.096 | 30.106 | 26.693 | 27.000 | 31.540 | 27.488 | 26.913 | 28.117 | -0.101 | 0.0079 | 0.0526 | 0 | 0 | 0 |
| 'Eral1' | 6.506 | 7.458 | 7.134 | 6.128 | 7.203 | 7.271 | 6.090 | 6.853 | 7.299 | 6.807 | 6.029 | 6.465 | 6.585 | 6.413 | 6.356 | 6.396 | 6.672 | 5.931 | 0.094 | 0.0360 | 0.1527 | 0 | 0 | 0 |
| 'Erap1' | 33.062 | 33.607 | 35.410 | 36.587 | 36.362 | 33.919 | 35.155 | 35.576 | 35.060 | 37.189 | 35.883 | 36.390 | 35.791 | 35.544 | 31.835 | 36.806 | 35.570 | 35.943 | -0.039 | 0.1885 | 0.4247 | 0 | 0 | 0 |
| 'Erbb3' | 4.877 | 5.125 | 4.885 | 3.927 | 4.639 | 5.372 | 4.713 | 5.610 | 4.792 | 4.176 | 4.580 | 3.867 | 4.182 | 4.410 | 3.762 | 4.393 | 3.858 | 3.595 | 0.252 | 0.0000 | 0.0003 | 1 | 0 | 0 |
| 'Erbin' | 36.664 | 35.705 | 35.131 | 36.957 | 36.695 | 32.656 | 35.875 | 36.381 | 33.129 | 38.180 | 39.625 | 36.423 | 34.293 | 38.754 | 33.857 | 35.133 | 40.301 | 36.335 | -0.071 | 0.0243 | 0.1181 | 0 | 0 | 0 |
| 'Erc1' | 1.413 | 1.371 | 1.438 | 1.518 | 1.352 | 1.326 | 1.436 | 1.218 | 1.627 | 1.435 | 1.473 | 1.481 | 1.289 | 1.408 | 1.337 | 1.322 | 1.499 | 1.441 | -0.010 | 0.8395 | 0.9315 | 0 | 0 | 0 |
| 'Ercc1' | 5.222 | 5.551 | 6.086 | 5.344 | 5.246 | 6.250 | 5.331 | 6.105 | 6.049 | 6.668 | 5.117 | 6.064 | 6.383 | 5.436 | 6.359 | 5.835 | 5.639 | 5.332 | -0.053 | 0.2986 | 0.5549 | 0 | 0 | 0 |
| 'Ercc2' | 8.245 | 7.088 | 9.021 | 7.639 | 7.032 | 8.095 | 7.439 | 7.220 | 9.680 | 7.245 | 6.573 | 10.264 | 7.069 | 6.760 | 8.832 | 7.174 | 7.704 | 9.927 | -0.007 | 0.8926 | 0.9562 | 0 | 0 | 0 |
| 'Ercc4' | 5.335 | 5.442 | 6.209 | 5.338 | 5.678 | 5.625 | 4.903 | 5.767 | 5.736 | 6.125 | 5.498 | 5.647 | 5.628 | 5.668 | 5.075 | 5.483 | 5.376 | 5.297 | -0.001 | 0.9795 | 0.9926 | 0 | 0 | 0 |
| 'Ercc5' | 5.310 | 5.097 | 5.393 | 5.664 | 5.128 | 4.884 | 5.425 | 4.796 | 5.364 | 5.130 | 5.217 | 5.391 | 5.254 | 4.908 | 4.721 | 5.063 | 5.485 | 4.720 | 0.024 | 0.5392 | 0.7570 | 0 | 0 | 0 |
| 'Ercc6' | 15.293 | 14.009 | 15.805 | 14.156 | 13.959 | 14.602 | 13.445 | 13.547 | 15.727 | 16.803 | 15.477 | 14.851 | 15.533 | 13.972 | 13.974 | 14.685 | 15.080 | 14.719 | -0.057 | 0.1765 | 0.4118 | 0 | 0 | 0 |
| 'Ercc6l' | 2.405 | 1.944 | 2.552 | 3.307 | 2.053 | 2.467 | 2.634 | 2.174 | 2.678 | 1.772 | 2.767 | 2.912 | 2.421 | 2.230 | 3.144 | 2.075 | 2.757 | 2.809 | -0.055 | 0.5840 | 0.7860 | 0 | 0 | 0 |
| 'Ercc6l2' | 4.773 | 5.214 | 4.835 | 4.983 | 5.197 | 4.763 | 5.336 | 5.067 | 4.448 | 5.052 | 4.727 | 4.439 | 4.681 | 4.987 | 4.097 | 4.622 | 4.462 | 3.884 | 0.114 | 0.0012 | 0.0121 | 1 | 0 | 0 |
| 'Ercc8' | 1.228 | 1.175 | 1.209 | 1.091 | 1.292 | 1.269 | 1.380 | 1.222 | 1.233 | 1.181 | 1.112 | 1.212 | 1.131 | 1.106 | 1.262 | 1.159 | 1.120 | 1.168 | 0.074 | 0.1765 | 0.4118 | 0 | 0 | 0 |
| 'Erf' | 16.125 | 14.892 | 15.184 | 14.834 | 13.955 | 15.852 | 13.905 | 12.965 | 16.842 | 13.012 | 14.287 | 19.249 | 11.669 | 13.527 | 19.993 | 14.182 | 14.595 | 19.315 | -0.049 | 0.4455 | 0.6878 | 0 | 0 | 0 |
| 'Ergic1' | 9.143 | 8.762 | 8.765 | 8.597 | 9.188 | 8.716 | 8.905 | 9.405 | 9.373 | 9.746 | 7.782 | 9.413 | 8.838 | 8.477 | 8.917 | 10.327 | 8.296 | 8.825 | -0.004 | 0.9179 | 0.9687 | 0 | 0 | 0 |
| 'Ergic2' | 6.522 | 7.173 | 7.008 | 6.929 | 7.113 | 6.858 | 7.571 | 7.359 | 6.596 | 6.778 | 6.322 | 6.899 | 7.238 | 7.164 | 6.547 | 6.609 | 7.619 | 6.164 | 0.029 | 0.4096 | 0.6588 | 0 | 0 | 0 |
| 'Ergic3' | 21.473 | 23.851 | 23.522 | 21.237 | 24.488 | 24.297 | 22.245 | 24.962 | 24.041 | 24.130 | 22.736 | 22.735 | 23.135 | 22.025 | 22.849 | 25.460 | 21.653 | 22.148 | 0.013 | 0.7274 | 0.8750 | 0 | 0 | 0 |
| 'Eri1' | 17.280 | 16.946 | 19.133 | 18.595 | 18.069 | 18.175 | 19.452 | 16.448 | 18.420 | 15.728 | 17.441 | 20.208 | 17.617 | 18.466 | 19.020 | 16.986 | 18.146 | 20.160 | -0.022 | 0.5795 | 0.7822 | 0 | 0 | 0 |
| 'Eri2' | 0.478 | 0.592 | 0.543 | 0.496 | 0.573 | 0.503 | 0.617 | 0.535 | 0.580 | 0.487 | 0.540 | 0.523 | 0.501 | 0.651 | 0.455 | 0.511 | 0.548 | 0.492 | 0.050 | 0.5039 | 0.7309 | 0 | 0 | 0 |
| 'Eri3' | 6.369 | 5.988 | 7.139 | 6.318 | 6.272 | 6.361 | 6.229 | 6.691 | 7.048 | 6.422 | 6.025 | 6.613 | 6.621 | 5.740 | 6.264 | 7.063 | 6.149 | 6.608 | 0.017 | 0.6811 | 0.8482 | 0 | 0 | 0 |
| 'Erich1' | 15.314 | 14.409 | 13.761 | 13.750 | 14.039 | 13.753 | 14.462 | 14.478 | 14.320 | 14.502 | 13.163 | 15.324 | 13.548 | 14.101 | 14.370 | 15.186 | 12.359 | 13.433 | 0.012 | 0.8005 | 0.9111 | 0 | 0 | 0 |
| 'Erlec1' | 8.808 | 9.500 | 10.195 | 8.827 | 9.999 | 9.296 | 10.876 | 10.985 | 9.380 | 10.120 | 9.953 | 8.254 | 9.552 | 9.256 | 8.738 | 8.461 | 8.827 | 8.153 | 0.105 | 0.0386 | 0.1602 | 0 | 0 | 0 |
| 'Erlin1' | 1.640 | 1.919 | 1.827 | 1.529 | 1.850 | 1.553 | 1.863 | 2.169 | 1.638 | 1.568 | 1.717 | 2.193 | 1.817 | 1.743 | 1.913 | 1.648 | 1.623 | 2.225 | -0.061 | 0.4460 | 0.6878 | 0 | 0 | 0 |
| 'Erlin2' | 5.956 | 7.163 | 8.498 | 6.869 | 7.678 | 7.947 | 6.666 | 7.192 | 8.283 | 6.900 | 7.009 | 8.401 | 7.308 | 6.728 | 7.386 | 7.713 | 7.513 | 7.963 | -0.029 | 0.5426 | 0.7595 | 0 | 0 | 0 |
| 'Ermp1' | 4.046 | 4.294 | 4.530 | 4.588 | 4.295 | 4.562 | 4.375 | 4.534 | 4.485 | 4.540 | 4.699 | 5.039 | 4.605 | 4.506 | 4.816 | 4.105 | 4.762 | 4.967 | -0.095 | 0.0032 | 0.0263 | 1 | 0 | 0 |
| 'Ern1' | 3.938 | 4.150 | 4.835 | 4.273 | 4.084 | 4.494 | 4.109 | 4.323 | 4.895 | 4.215 | 4.311 | 4.733 | 4.292 | 3.875 | 4.593 | 4.516 | 4.199 | 4.839 | -0.026 | 0.4437 | 0.6869 | 0 | 0 | 0 |
| 'Ero1l' | 19.264 | 18.187 | 16.909 | 18.962 | 17.622 | 16.499 | 20.855 | 17.043 | 17.108 | 15.658 | 21.409 | 20.370 | 18.017 | 20.549 | 20.905 | 16.961 | 20.785 | 20.362 | -0.117 | 0.0596 | 0.2110 | 0 | 0 | 0 |
| 'Ero1lb' | 3.278 | 4.030 | 3.814 | 3.556 | 3.652 | 3.067 | 3.380 | 3.567 | 3.079 | 3.365 | 3.675 | 3.409 | 2.829 | 4.103 | 3.072 | 3.666 | 4.137 | 3.419 | -0.018 | 0.7829 | 0.9025 | 0 | 0 | 0 |
| 'Erp29' | 6.732 | 6.277 | 6.816 | 5.862 | 6.989 | 7.072 | 6.463 | 6.697 | 6.544 | 6.555 | 6.122 | 6.905 | 6.552 | 6.635 | 7.145 | 6.911 | 6.142 | 6.776 | -0.016 | 0.6476 | 0.8282 | 0 | 0 | 0 |
| 'Erp44' | 17.559 | 18.470 | 17.448 | 17.289 | 18.881 | 16.512 | 18.176 | 18.298 | 17.101 | 17.911 | 18.337 | 18.081 | 18.122 | 18.932 | 17.133 | 17.785 | 20.331 | 16.663 | -0.044 | 0.1256 | 0.3395 | 0 | 0 | 0 |
| 'Errfi1' | 3.251 | 2.830 | 2.337 | 2.559 | 2.541 | 2.583 | 2.378 | 2.663 | 2.052 | 2.714 | 2.891 | 2.239 | 2.944 | 2.985 | 2.278 | 2.863 | 3.296 | 2.598 | -0.097 | 0.1588 | 0.3889 | 0 | 0 | 0 |
| 'Esco1' | 13.778 | 13.448 | 13.067 | 14.968 | 14.000 | 12.305 | 15.946 | 13.279 | 12.088 | 12.913 | 14.328 | 14.021 | 12.260 | 13.443 | 12.581 | 12.915 | 14.954 | 13.537 | 0.008 | 0.8626 | 0.9405 | 0 | 0 | 0 |
| 'Esco2' | 3.500 | 3.816 | 4.424 | 4.408 | 4.033 | 4.296 | 4.291 | 3.983 | 4.381 | 2.148 | 4.336 | 4.423 | 4.495 | 4.974 | 4.983 | 3.393 | 4.686 | 3.920 | -0.010 | 0.9221 | 0.9705 | 0 | 0 | 0 |
| 'Esd' | 11.887 | 14.568 | 13.148 | 12.496 | 12.995 | 12.288 | 12.852 | 13.859 | 12.270 | 12.429 | 12.533 | 12.851 | 12.871 | 13.034 | 12.156 | 12.094 | 12.520 | 11.728 | 0.037 | 0.2418 | 0.4924 | 0 | 0 | 0 |
| 'Esf1' | 6.636 | 6.431 | 5.927 | 6.911 | 7.064 | 6.087 | 7.385 | 6.672 | 5.634 | 7.140 | 6.136 | 6.151 | 6.935 | 6.166 | 6.141 | 7.490 | 7.451 | 6.034 | -0.034 | 0.4098 | 0.6589 | 0 | 0 | 0 |
| 'Esm1' | 13.828 | 10.680 | 15.055 | 12.015 | 16.322 | 11.323 | 12.218 | 16.936 | 11.360 | 14.649 | 17.875 | 13.990 | 12.364 | 13.711 | 16.624 | 11.267 | 11.047 | 14.695 | -0.086 | 0.3807 | 0.6321 | 0 | 0 | 0 |
| 'Espl1' | 5.362 | 5.479 | 8.331 | 6.827 | 5.032 | 7.967 | 6.889 | 5.367 | 8.071 | 4.703 | 6.531 | 6.621 | 7.721 | 6.540 | 7.781 | 5.720 | 6.308 | 6.375 | -0.002 | 0.9787 | 0.9926 | 0 | 0 | 0 |
| 'Espn' | 0.525 | 0.609 | 0.793 | 0.543 | 0.731 | 0.714 | 0.623 | 0.791 | 0.855 | 0.656 | 0.515 | 0.752 | 0.635 | 0.559 | 0.616 | 0.437 | 0.678 | 0.783 | 0.114 | 0.2043 | 0.4472 | 0 | 0 | 0 |
| 'Esrra' | 14.855 | 11.718 | 16.036 | 14.515 | 13.320 | 15.556 | 13.909 | 14.360 | 15.641 | 14.092 | 14.199 | 16.246 | 14.346 | 14.069 | 15.898 | 15.641 | 14.326 | 15.232 | -0.052 | 0.1510 | 0.3777 | 0 | 0 | 0 |
| 'Esyt1' | 17.741 | 13.406 | 20.776 | 15.949 | 14.671 | 20.248 | 18.560 | 18.147 | 17.478 | 13.078 | 14.713 | 21.993 | 16.177 | 13.553 | 20.971 | 15.522 | 14.896 | 20.857 | 0.052 | 0.4128 | 0.6610 | 0 | 0 | 0 |
| 'Esyt2' | 49.252 | 49.963 | 49.655 | 49.338 | 48.987 | 47.959 | 46.728 | 49.489 | 49.620 | 49.632 | 53.250 | 56.865 | 46.793 | 52.288 | 53.837 | 48.913 | 51.893 | 55.136 | -0.097 | 0.0007 | 0.0078 | 1 | 0 | 0 |
| 'Etaa1' | 7.113 | 7.236 | 7.191 | 7.881 | 7.891 | 7.079 | 8.280 | 7.466 | 6.955 | 7.097 | 8.067 | 8.412 | 6.641 | 7.565 | 7.549 | 7.010 | 7.921 | 8.137 | -0.044 | 0.4026 | 0.6514 | 0 | 0 | 0 |
| 'Etf1' | 55.542 | 58.188 | 53.911 | 55.626 | 60.444 | 50.450 | 58.743 | 57.544 | 52.295 | 53.284 | 55.464 | 52.662 | 55.829 | 57.199 | 51.334 | 53.930 | 56.971 | 51.832 | 0.029 | 0.1818 | 0.4175 | 0 | 0 | 0 |
| 'Etfa' | 30.018 | 35.018 | 30.305 | 34.137 | 33.590 | 29.857 | 33.225 | 34.184 | 30.676 | 30.875 | 31.726 | 29.889 | 31.149 | 32.158 | 28.925 | 30.793 | 33.478 | 26.903 | 0.064 | 0.0205 | 0.1050 | 0 | 0 | 0 |
| 'Etfb' | 6.055 | 7.379 | 6.667 | 6.952 | 7.377 | 6.314 | 6.903 | 7.113 | 6.634 | 6.603 | 6.809 | 6.758 | 7.243 | 6.824 | 5.792 | 6.931 | 6.417 | 6.142 | 0.027 | 0.6169 | 0.8087 | 0 | 0 | 0 |
| 'Etfdh' | 4.942 | 5.392 | 4.903 | 5.799 | 5.526 | 5.210 | 5.581 | 5.515 | 4.973 | 4.836 | 4.963 | 5.272 | 5.608 | 5.181 | 5.212 | 4.979 | 5.394 | 4.881 | 0.030 | 0.4351 | 0.6791 | 0 | 0 | 0 |
| 'Ethe1' | 8.362 | 8.987 | 9.270 | 6.943 | 8.073 | 8.966 | 8.004 | 8.755 | 10.454 | 8.214 | 6.260 | 8.688 | 8.053 | 8.963 | 8.825 | 9.292 | 7.749 | 7.745 | 0.070 | 0.3279 | 0.5857 | 0 | 0 | 0 |
| 'Etnk1' | 18.328 | 22.469 | 19.142 | 18.396 | 22.045 | 19.680 | 17.833 | 21.365 | 19.817 | 20.032 | 21.553 | 21.679 | 19.695 | 21.241 | 20.877 | 21.621 | 21.532 | 21.761 | -0.100 | 0.0005 | 0.0065 | 1 | 0 | 0 |
| 'Ets1' | 261.096 | 285.501 | 254.130 | 261.580 | 281.513 | 258.570 | 254.889 | 277.604 | 255.273 | 272.657 | 282.534 | 228.156 | 251.545 | 276.466 | 254.309 | 275.306 | 254.607 | 249.095 | 0.017 | 0.5792 | 0.7822 | 0 | 0 | 0 |
| 'Ets2' | 6.478 | 4.542 | 4.689 | 5.813 | 4.481 | 4.985 | 4.943 | 4.453 | 4.921 | 6.866 | 5.351 | 5.375 | 5.976 | 5.411 | 5.956 | 6.075 | 5.329 | 5.228 | -0.196 | 0.0002 | 0.0022 | 1 | 0 | 0 |
| 'Etv3' | 3.832 | 3.870 | 3.725 | 3.274 | 3.645 | 3.656 | 3.189 | 3.856 | 3.593 | 4.017 | 3.873 | 4.174 | 3.516 | 3.727 | 4.169 | 4.196 | 3.653 | 4.166 | -0.130 | 0.0030 | 0.0250 | 1 | 0 | 0 |
| 'Etv5' | 1.196 | 1.340 | 1.411 | 1.331 | 1.607 | 1.465 | 1.464 | 1.569 | 1.732 | 1.380 | 1.036 | 1.325 | 1.443 | 1.264 | 1.250 | 1.667 | 1.340 | 1.058 | 0.152 | 0.0750 | 0.2437 | 0 | 0 | 0 |
| 'Etv6' | 7.274 | 6.452 | 6.555 | 6.701 | 6.289 | 6.328 | 6.199 | 7.051 | 6.273 | 7.050 | 7.618 | 7.049 | 7.449 | 7.497 | 6.935 | 7.228 | 6.452 | 6.982 | -0.129 | 0.0009 | 0.0093 | 1 | 0 | 0 |
| 'Eva1b' | 6.314 | 6.281 | 7.387 | 6.453 | 6.923 | 8.688 | 5.916 | 7.129 | 7.992 | 6.900 | 6.265 | 7.923 | 6.813 | 6.058 | 7.719 | 7.447 | 5.530 | 7.257 | 0.019 | 0.7241 | 0.8732 | 0 | 0 | 0 |
| 'Evi2a' | 16.799 | 19.633 | 16.671 | 17.057 | 19.885 | 17.763 | 17.170 | 18.494 | 17.967 | 19.134 | 17.504 | 16.906 | 16.366 | 18.757 | 17.648 | 19.746 | 18.219 | 15.296 | 0.007 | 0.8695 | 0.9441 | 0 | 0 | 0 |
| 'Evl' | 153.838 | 161.512 | 152.688 | 145.676 | 157.958 | 160.198 | 153.743 | 155.606 | 159.394 | 145.932 | 138.329 | 136.749 | 148.032 | 146.408 | 141.383 | 149.341 | 137.060 | 141.386 | 0.115 | 0.0000 | 0.0001 | 1 | 0 | 0 |
| 'Ewsr1' | 64.417 | 66.741 | 67.350 | 63.349 | 65.521 | 65.463 | 67.983 | 66.126 | 69.534 | 65.039 | 60.825 | 63.277 | 68.751 | 60.167 | 62.862 | 65.801 | 63.293 | 60.562 | 0.054 | 0.0188 | 0.0990 | 0 | 0 | 0 |
| 'Exd2' | 12.803 | 13.221 | 13.806 | 13.044 | 14.094 | 12.683 | 12.752 | 13.199 | 14.169 | 13.412 | 12.676 | 13.369 | 13.163 | 11.965 | 12.079 | 11.944 | 12.771 | 11.958 | 0.067 | 0.0814 | 0.2574 | 0 | 0 | 0 |
| 'Exo1' | 0.831 | 1.009 | 1.038 | 0.892 | 0.659 | 1.000 | 1.006 | 0.961 | 1.125 | 0.730 | 0.947 | 1.024 | 1.123 | 1.019 | 1.213 | 0.830 | 0.877 | 0.892 | -0.035 | 0.7016 | 0.8608 | 0 | 0 | 0 |
| 'Exo5' | 3.682 | 4.843 | 4.692 | 4.355 | 4.471 | 4.711 | 5.408 | 4.462 | 5.099 | 3.773 | 4.550 | 5.461 | 3.915 | 4.940 | 5.273 | 3.840 | 4.987 | 5.114 | -0.015 | 0.8140 | 0.9160 | 0 | 0 | 0 |
| 'Exoc1' | 3.879 | 3.971 | 4.596 | 3.883 | 4.456 | 4.272 | 4.356 | 4.516 | 4.661 | 4.525 | 4.232 | 3.928 | 4.563 | 3.961 | 3.634 | 4.297 | 4.253 | 3.336 | 0.066 | 0.2225 | 0.4701 | 0 | 0 | 0 |
| 'Exoc2' | 23.077 | 25.381 | 22.707 | 23.946 | 25.621 | 23.244 | 25.076 | 24.544 | 22.927 | 23.927 | 24.631 | 23.241 | 24.425 | 24.149 | 22.088 | 24.483 | 25.506 | 21.396 | 0.006 | 0.8029 | 0.9115 | 0 | 0 | 0 |
| 'Exoc3' | 23.089 | 23.352 | 23.571 | 21.554 | 22.743 | 23.249 | 22.826 | 23.720 | 23.301 | 24.779 | 23.743 | 23.688 | 22.560 | 22.864 | 22.223 | 23.136 | 22.505 | 21.995 | -0.009 | 0.7354 | 0.8796 | 0 | 0 | 0 |
| 'Exoc3l' | 2.740 | 2.796 | 2.922 | 3.183 | 2.672 | 3.091 | 2.885 | 3.066 | 2.767 | 3.048 | 2.901 | 2.960 | 3.541 | 3.031 | 2.991 | 2.669 | 3.350 | 3.159 | -0.092 | 0.2423 | 0.4928 | 0 | 0 | 0 |
| 'Exoc4' | 5.494 | 5.572 | 5.326 | 5.734 | 5.427 | 5.263 | 5.552 | 5.146 | 5.298 | 5.457 | 5.986 | 5.322 | 5.895 | 5.495 | 5.429 | 5.868 | 5.443 | 5.137 | -0.045 | 0.1352 | 0.3537 | 0 | 0 | 0 |
| 'Exoc5' | 7.627 | 7.346 | 7.390 | 7.837 | 7.944 | 6.624 | 8.590 | 7.477 | 6.592 | 7.836 | 7.808 | 7.543 | 7.077 | 7.902 | 7.293 | 7.050 | 8.261 | 7.379 | -0.029 | 0.4924 | 0.7217 | 0 | 0 | 0 |
| 'Exoc6' | 25.869 | 26.919 | 27.220 | 26.412 | 28.059 | 25.613 | 28.315 | 27.471 | 25.741 | 27.699 | 29.495 | 24.268 | 25.645 | 30.386 | 23.879 | 24.600 | 30.710 | 21.027 | 0.022 | 0.6176 | 0.8087 | 0 | 0 | 0 |
| 'Exoc6b' | 1.464 | 1.293 | 1.399 | 1.628 | 1.287 | 1.390 | 1.568 | 1.320 | 1.500 | 1.581 | 1.714 | 1.667 | 1.534 | 1.544 | 1.502 | 1.509 | 1.724 | 1.387 | -0.153 | 0.0049 | 0.0367 | 1 | 0 | 0 |
| 'Exoc7' | 9.138 | 8.934 | 10.363 | 8.406 | 9.349 | 10.952 | 8.955 | 8.986 | 9.833 | 9.435 | 8.675 | 9.672 | 10.049 | 8.892 | 9.508 | 9.552 | 8.867 | 9.164 | 0.009 | 0.7997 | 0.9111 | 0 | 0 | 0 |
| 'Exoc8' | 4.362 | 4.785 | 5.516 | 5.121 | 4.799 | 5.239 | 4.375 | 4.932 | 5.172 | 3.827 | 4.833 | 6.114 | 3.753 | 4.847 | 5.595 | 4.281 | 4.962 | 6.162 | -0.004 | 0.9484 | 0.9802 | 0 | 0 | 0 |
| 'Exog' | 2.401 | 2.218 | 2.254 | 2.502 | 2.065 | 2.249 | 2.360 | 1.869 | 2.019 | 1.553 | 2.491 | 3.151 | 2.241 | 2.449 | 2.987 | 1.612 | 2.522 | 3.104 | -0.143 | 0.1810 | 0.4165 | 0 | 0 | 0 |
| 'Exosc1' | 13.030 | 15.401 | 14.217 | 12.720 | 16.828 | 14.578 | 12.251 | 16.341 | 13.299 | 12.929 | 12.927 | 15.372 | 13.778 | 13.835 | 13.723 | 13.620 | 14.671 | 12.641 | 0.039 | 0.4641 | 0.6999 | 0 | 0 | 0 |
| 'Exosc10' | 18.948 | 17.013 | 20.076 | 18.558 | 17.822 | 19.394 | 17.714 | 17.207 | 19.140 | 19.247 | 16.586 | 20.045 | 19.119 | 17.432 | 19.207 | 17.729 | 17.593 | 19.050 | -0.012 | 0.6310 | 0.8173 | 0 | 0 | 0 |
| 'Exosc2' | 6.698 | 7.261 | 7.707 | 7.067 | 6.640 | 7.098 | 8.068 | 6.841 | 7.231 | 6.181 | 5.678 | 7.142 | 6.703 | 6.532 | 6.447 | 7.283 | 6.505 | 6.639 | 0.115 | 0.0095 | 0.0603 | 0 | 0 | 0 |
| 'Exosc3' | 31.873 | 36.608 | 32.903 | 31.348 | 31.816 | 32.747 | 31.131 | 33.004 | 30.410 | 29.608 | 31.647 | 30.631 | 32.752 | 30.326 | 31.697 | 29.942 | 31.030 | 30.549 | 0.057 | 0.0849 | 0.2646 | 0 | 0 | 0 |
| 'Exosc4' | 11.041 | 12.681 | 11.847 | 8.930 | 12.111 | 11.631 | 11.136 | 12.851 | 12.200 | 11.789 | 9.969 | 11.835 | 11.045 | 10.256 | 11.103 | 11.280 | 9.527 | 10.323 | 0.091 | 0.1238 | 0.3364 | 0 | 0 | 0 |
| 'Exosc5' | 4.668 | 5.257 | 5.508 | 4.817 | 4.982 | 5.229 | 5.008 | 5.484 | 5.702 | 4.870 | 4.335 | 5.761 | 4.628 | 4.900 | 5.304 | 5.184 | 4.424 | 5.424 | 0.045 | 0.2727 | 0.5277 | 0 | 0 | 0 |
| 'Exosc7' | 11.294 | 12.006 | 12.302 | 11.985 | 13.532 | 11.950 | 12.228 | 13.037 | 12.182 | 11.518 | 11.114 | 11.655 | 12.259 | 12.628 | 12.434 | 12.099 | 12.263 | 11.671 | 0.028 | 0.5062 | 0.7324 | 0 | 0 | 0 |
| 'Exosc8' | 7.603 | 9.345 | 8.453 | 8.619 | 9.083 | 8.094 | 8.561 | 8.746 | 8.669 | 7.207 | 8.283 | 8.816 | 8.814 | 8.547 | 8.485 | 7.876 | 9.034 | 8.015 | 0.025 | 0.5066 | 0.7326 | 0 | 0 | 0 |
| 'Exosc9' | 3.346 | 3.897 | 3.802 | 3.428 | 4.427 | 4.106 | 3.678 | 4.464 | 4.160 | 4.173 | 3.565 | 3.525 | 4.278 | 3.974 | 3.641 | 3.528 | 3.841 | 3.251 | 0.054 | 0.3678 | 0.6198 | 0 | 0 | 0 |
| 'Exph5' | 0.769 | 0.648 | 0.688 | 0.707 | 0.665 | 0.686 | 0.643 | 0.701 | 0.680 | 0.643 | 0.650 | 0.652 | 0.465 | 0.661 | 0.587 | 0.855 | 0.566 | 0.604 | 0.121 | 0.1577 | 0.3877 | 0 | 0 | 0 |
| 'Ext1' | 16.434 | 16.905 | 15.371 | 17.619 | 17.466 | 15.987 | 16.683 | 17.018 | 15.909 | 19.394 | 18.729 | 15.619 | 16.955 | 17.399 | 14.696 | 18.784 | 17.294 | 15.855 | -0.057 | 0.0869 | 0.2686 | 0 | 0 | 0 |
| 'Ext2' | 8.643 | 10.184 | 10.291 | 9.444 | 9.796 | 10.192 | 9.701 | 9.947 | 10.276 | 10.758 | 9.326 | 9.744 | 10.396 | 9.986 | 9.828 | 10.228 | 9.523 | 9.321 | -0.020 | 0.5699 | 0.7774 | 0 | 0 | 0 |
| 'Extl3' | 16.654 | 17.370 | 17.906 | 16.368 | 17.081 | 17.707 | 15.073 | 17.455 | 17.829 | 16.384 | 17.208 | 16.805 | 16.137 | 16.270 | 17.610 | 16.041 | 14.815 | 17.298 | 0.037 | 0.2424 | 0.4929 | 0 | 0 | 0 |
| 'Eya2' | 4.661 | 3.712 | 4.354 | 4.182 | 4.076 | 4.286 | 3.923 | 3.964 | 4.342 | 4.419 | 3.531 | 3.883 | 4.416 | 4.007 | 3.577 | 4.501 | 3.335 | 3.740 | 0.083 | 0.1082 | 0.3097 | 0 | 0 | 0 |
| 'Eya3' | 4.353 | 4.486 | 4.516 | 4.046 | 4.348 | 4.220 | 3.910 | 4.228 | 4.660 | 4.031 | 4.753 | 5.043 | 4.137 | 4.121 | 4.749 | 4.007 | 4.503 | 4.761 | -0.060 | 0.1079 | 0.3092 | 0 | 0 | 0 |
| 'Ezh1' | 15.817 | 15.375 | 17.195 | 16.542 | 15.838 | 16.095 | 15.192 | 15.889 | 17.246 | 15.790 | 15.466 | 17.766 | 14.605 | 14.960 | 17.121 | 14.547 | 15.586 | 16.971 | 0.014 | 0.6320 | 0.8176 | 0 | 0 | 0 |
| 'Ezh2' | 16.710 | 16.781 | 16.291 | 18.138 | 17.417 | 16.187 | 18.548 | 16.693 | 16.683 | 16.276 | 17.411 | 16.181 | 20.744 | 18.346 | 16.527 | 17.328 | 18.231 | 15.203 | -0.035 | 0.3595 | 0.6118 | 0 | 0 | 0 |
| 'Ezr' | 133.024 | 126.437 | 128.707 | 126.269 | 129.068 | 136.068 | 123.920 | 125.036 | 128.832 | 127.327 | 115.629 | 118.951 | 131.984 | 122.304 | 126.940 | 127.944 | 113.842 | 118.871 | 0.059 | 0.0240 | 0.1173 | 0 | 0 | 0 |
| 'F2r' | 2.157 | 2.657 | 3.120 | 2.515 | 2.625 | 2.770 | 2.989 | 2.543 | 2.507 | 1.935 | 1.690 | 2.319 | 1.617 | 1.666 | 1.980 | 1.830 | 1.370 | 2.133 | 0.510 | 0.0000 | 0.0000 | 1 | 1 | 0 |
| 'F2rl1' | 1.053 | 0.940 | 1.078 | 0.695 | 0.990 | 1.464 | 1.002 | 1.057 | 1.049 | 1.394 | 1.143 | 1.731 | 1.447 | 1.281 | 2.023 | 1.558 | 1.544 | 1.859 | -0.585 | 0.0000 | 0.0000 | 1 | 1 | 0 |
| 'F8a' | 9.028 | 7.955 | 8.258 | 7.228 | 8.048 | 7.956 | 7.128 | 8.006 | 8.801 | 7.582 | 7.794 | 8.360 | 7.646 | 9.446 | 8.714 | 8.380 | 7.401 | 8.822 | -0.039 | 0.4951 | 0.7242 | 0 | 0 | 0 |
| 'Faah' | 15.343 | 14.665 | 16.934 | 14.470 | 13.363 | 16.799 | 15.240 | 13.736 | 18.146 | 16.970 | 12.864 | 18.738 | 14.957 | 13.693 | 17.211 | 16.561 | 14.596 | 18.108 | -0.059 | 0.0775 | 0.2494 | 0 | 0 | 0 |
| 'Faap100' | 1.882 | 2.066 | 2.293 | 2.003 | 2.055 | 2.296 | 2.140 | 2.188 | 2.279 | 2.379 | 1.754 | 2.224 | 2.076 | 1.994 | 2.253 | 1.969 | 1.969 | 2.066 | 0.029 | 0.5494 | 0.7630 | 0 | 0 | 0 |
| 'Faap20' | 6.243 | 6.674 | 6.779 | 6.477 | 6.991 | 7.565 | 5.986 | 6.747 | 7.092 | 6.417 | 6.406 | 7.165 | 6.474 | 6.740 | 7.330 | 6.186 | 6.619 | 6.828 | -0.002 | 0.9475 | 0.9796 | 0 | 0 | 0 |
| 'Faap24' | 3.620 | 3.726 | 4.171 | 4.000 | 3.846 | 4.160 | 3.714 | 4.382 | 3.921 | 3.676 | 3.238 | 4.465 | 4.101 | 4.210 | 4.289 | 4.238 | 3.788 | 4.080 | -0.034 | 0.5521 | 0.7650 | 0 | 0 | 0 |
| 'Fabp5' | 5.989 | 6.530 | 7.139 | 5.935 | 7.866 | 6.587 | 6.794 | 6.979 | 6.605 | 5.367 | 6.442 | 6.502 | 8.235 | 6.810 | 7.016 | 6.900 | 6.787 | 6.171 | -0.008 | 0.9043 | 0.9624 | 0 | 0 | 0 |
| 'Fadd' | 7.317 | 8.670 | 9.214 | 6.942 | 9.292 | 9.193 | 8.075 | 9.173 | 10.024 | 9.416 | 8.235 | 7.321 | 9.551 | 8.216 | 7.884 | 8.230 | 7.237 | 6.364 | 0.097 | 0.2107 | 0.4560 | 0 | 0 | 0 |
| 'Fads1' | 10.590 | 11.240 | 12.925 | 11.098 | 12.936 | 12.621 | 10.824 | 11.282 | 12.326 | 11.306 | 11.212 | 13.166 | 12.936 | 12.209 | 13.534 | 11.863 | 11.256 | 12.535 | -0.068 | 0.0907 | 0.2776 | 0 | 0 | 0 |
| 'Fads2' | 11.459 | 14.047 | 15.377 | 13.958 | 14.557 | 15.241 | 15.194 | 12.728 | 15.910 | 10.942 | 12.164 | 14.037 | 15.995 | 15.009 | 15.922 | 12.904 | 11.739 | 14.862 | 0.042 | 0.5467 | 0.7610 | 0 | 0 | 0 |
| 'Faf1' | 8.422 | 8.990 | 9.101 | 9.440 | 8.891 | 9.044 | 9.763 | 8.810 | 8.285 | 8.324 | 8.627 | 8.149 | 9.201 | 8.434 | 8.054 | 8.633 | 9.423 | 7.813 | 0.066 | 0.0852 | 0.2652 | 0 | 0 | 0 |
| 'Faf2' | 13.478 | 14.266 | 14.326 | 13.730 | 14.091 | 13.715 | 13.542 | 13.935 | 14.243 | 13.527 | 14.200 | 14.472 | 13.243 | 14.106 | 13.902 | 13.952 | 13.407 | 14.511 | -0.011 | 0.6639 | 0.8377 | 0 | 0 | 0 |
| 'Fah' | 1.370 | 1.651 | 1.694 | 1.305 | 1.431 | 1.723 | 1.726 | 1.812 | 1.740 | 1.616 | 1.105 | 1.640 | 1.431 | 1.221 | 1.209 | 1.471 | 1.287 | 1.428 | 0.202 | 0.0619 | 0.2156 | 0 | 0 | 0 |
| 'Fahd2a' | 2.669 | 3.201 | 2.852 | 3.002 | 3.587 | 3.124 | 2.669 | 3.259 | 3.219 | 3.015 | 2.331 | 2.787 | 3.122 | 2.961 | 3.115 | 3.278 | 2.913 | 2.625 | 0.066 | 0.3557 | 0.6089 | 0 | 0 | 0 |
| 'Faim' | 5.342 | 5.704 | 4.895 | 5.240 | 5.778 | 5.376 | 4.649 | 5.957 | 5.115 | 5.424 | 4.824 | 4.656 | 5.299 | 5.564 | 4.851 | 5.363 | 4.816 | 4.616 | 0.074 | 0.1053 | 0.3042 | 0 | 0 | 0 |
| 'Fam101b' | 64.245 | 71.284 | 72.673 | 60.415 | 68.819 | 74.431 | 56.229 | 75.560 | 69.588 | 68.420 | 65.325 | 79.749 | 50.236 | 63.359 | 79.721 | 67.317 | 54.536 | 86.009 | -0.006 | 0.9207 | 0.9702 | 0 | 0 | 0 |
| 'Fam102a' | 33.213 | 28.869 | 31.213 | 29.453 | 26.898 | 31.136 | 27.555 | 28.849 | 30.076 | 26.958 | 28.226 | 31.194 | 25.181 | 26.562 | 32.429 | 26.966 | 24.645 | 32.053 | 0.066 | 0.1334 | 0.3518 | 0 | 0 | 0 |
| 'Fam102b' | 1.659 | 1.289 | 1.780 | 1.888 | 1.563 | 1.563 | 1.542 | 1.652 | 1.589 | 1.853 | 1.570 | 1.882 | 1.521 | 1.546 | 1.516 | 1.641 | 1.660 | 1.832 | -0.060 | 0.4381 | 0.6813 | 0 | 0 | 0 |
| 'Fam105a' | 16.729 | 17.838 | 15.305 | 16.178 | 16.423 | 15.824 | 16.706 | 17.434 | 16.062 | 16.574 | 17.225 | 19.060 | 16.454 | 18.639 | 18.934 | 17.439 | 18.331 | 18.171 | -0.129 | 0.0005 | 0.0060 | 1 | 0 | 0 |
| 'Fam107b' | 34.593 | 35.155 | 34.476 | 33.082 | 35.718 | 34.378 | 35.744 | 36.231 | 33.142 | 36.935 | 38.108 | 35.383 | 33.003 | 37.842 | 36.355 | 35.783 | 36.007 | 33.887 | -0.059 | 0.0255 | 0.1221 | 0 | 0 | 0 |
| 'Fam109a' | 1.726 | 1.697 | 2.069 | 1.841 | 1.922 | 2.287 | 1.612 | 2.055 | 2.119 | 1.952 | 1.610 | 1.948 | 1.430 | 1.532 | 1.858 | 1.992 | 1.729 | 1.794 | 0.127 | 0.0958 | 0.2881 | 0 | 0 | 0 |
| 'Fam110a' | 8.729 | 7.095 | 6.369 | 7.987 | 6.718 | 6.251 | 7.600 | 7.030 | 6.450 | 8.192 | 6.477 | 6.705 | 8.246 | 7.181 | 6.854 | 7.643 | 7.482 | 7.041 | -0.049 | 0.2997 | 0.5560 | 0 | 0 | 0 |
| 'Fam111a' | 4.487 | 4.807 | 5.504 | 5.694 | 5.180 | 5.210 | 5.875 | 4.761 | 5.254 | 3.894 | 4.800 | 5.708 | 4.626 | 5.091 | 5.655 | 4.609 | 5.259 | 5.497 | 0.043 | 0.4766 | 0.7107 | 0 | 0 | 0 |
| 'Fam114a2' | 14.201 | 14.248 | 13.633 | 13.596 | 12.968 | 14.728 | 14.004 | 14.519 | 14.116 | 12.881 | 15.080 | 15.679 | 12.667 | 14.388 | 13.872 | 13.478 | 15.565 | 14.069 | -0.029 | 0.4879 | 0.7183 | 0 | 0 | 0 |
| 'Fam117a' | 15.519 | 15.743 | 14.515 | 14.820 | 14.498 | 16.128 | 13.357 | 14.775 | 15.498 | 15.017 | 15.630 | 17.351 | 15.370 | 14.397 | 15.892 | 15.518 | 13.589 | 15.806 | -0.051 | 0.2006 | 0.4418 | 0 | 0 | 0 |
| 'Fam117b' | 11.216 | 10.798 | 11.491 | 11.165 | 10.912 | 11.574 | 11.858 | 11.329 | 10.372 | 14.816 | 15.912 | 16.307 | 13.384 | 15.500 | 16.296 | 15.322 | 15.664 | 17.157 | -0.488 | 0.0000 | 0.0000 | 1 | 1 | 0 |
| 'Fam118a' | 13.613 | 17.215 | 15.676 | 14.847 | 16.080 | 14.678 | 12.777 | 15.816 | 15.388 | 16.976 | 18.112 | 15.874 | 15.708 | 17.858 | 15.654 | 16.720 | 17.075 | 14.381 | -0.132 | 0.0022 | 0.0195 | 1 | 0 | 0 |
| 'Fam118b' | 3.229 | 3.344 | 3.325 | 3.781 | 3.942 | 3.442 | 3.389 | 3.647 | 3.312 | 3.410 | 3.628 | 3.649 | 3.240 | 3.422 | 3.567 | 3.143 | 4.096 | 3.070 | -0.005 | 0.9209 | 0.9703 | 0 | 0 | 0 |
| 'Fam120a' | 78.157 | 79.560 | 80.304 | 77.743 | 82.102 | 79.352 | 76.722 | 81.321 | 78.849 | 78.490 | 76.075 | 83.740 | 79.258 | 80.995 | 82.678 | 80.273 | 76.336 | 86.189 | -0.031 | 0.1778 | 0.4131 | 0 | 0 | 0 |
| 'Fam120b' | 15.504 | 15.351 | 16.401 | 15.068 | 16.006 | 16.868 | 15.214 | 15.561 | 15.960 | 17.038 | 14.725 | 17.663 | 15.147 | 15.891 | 16.452 | 15.854 | 15.905 | 15.784 | -0.037 | 0.2395 | 0.4897 | 0 | 0 | 0 |
| 'Fam120c' | 1.834 | 1.710 | 1.879 | 2.035 | 2.193 | 1.858 | 2.017 | 1.705 | 1.914 | 2.136 | 2.072 | 2.441 | 1.891 | 2.279 | 2.133 | 2.070 | 1.982 | 2.402 | -0.194 | 0.0012 | 0.0119 | 1 | 0 | 0 |
| 'Fam122a' | 29.842 | 25.601 | 25.292 | 22.639 | 24.249 | 27.959 | 20.524 | 25.727 | 29.018 | 26.938 | 23.389 | 25.023 | 24.636 | 26.386 | 26.068 | 25.537 | 21.358 | 24.625 | 0.039 | 0.5441 | 0.7606 | 0 | 0 | 0 |
| 'Fam122b' | 1.460 | 1.740 | 1.699 | 1.485 | 1.807 | 1.917 | 1.898 | 1.825 | 1.755 | 1.595 | 2.200 | 1.894 | 1.566 | 1.826 | 1.442 | 1.342 | 1.808 | 1.812 | 0.000 | 0.9983 | 0.9994 | 0 | 0 | 0 |
| 'Fam126a' | 2.826 | 2.933 | 2.985 | 3.085 | 3.078 | 2.859 | 3.015 | 2.754 | 3.065 | 3.032 | 3.658 | 3.912 | 2.895 | 3.208 | 3.495 | 2.690 | 3.349 | 3.692 | -0.180 | 0.0009 | 0.0093 | 1 | 0 | 0 |
| 'Fam126b' | 1.231 | 1.251 | 1.163 | 1.355 | 1.321 | 0.987 | 1.247 | 1.231 | 1.100 | 1.282 | 1.364 | 1.155 | 1.169 | 1.374 | 0.970 | 1.327 | 1.376 | 1.076 | -0.035 | 0.4771 | 0.7107 | 0 | 0 | 0 |
| 'Fam129a' | 3.824 | 3.138 | 4.402 | 3.082 | 3.512 | 4.634 | 3.271 | 3.507 | 4.266 | 3.200 | 2.705 | 4.421 | 2.422 | 2.716 | 3.819 | 3.133 | 3.012 | 3.586 | 0.208 | 0.0004 | 0.0049 | 1 | 0 | 0 |
| 'Fam129c' | 0.867 | 0.679 | 0.766 | 0.900 | 0.683 | 0.890 | 0.729 | 0.693 | 0.900 | 0.716 | 0.856 | 1.300 | 0.829 | 0.708 | 0.880 | 0.653 | 0.810 | 0.930 | -0.132 | 0.2018 | 0.4432 | 0 | 0 | 0 |
| 'Fam131a' | 1.065 | 1.737 | 1.213 | 1.114 | 1.317 | 1.167 | 0.798 | 1.235 | 0.998 | 0.918 | 1.028 | 1.270 | 1.157 | 1.446 | 1.402 | 1.131 | 1.082 | 1.434 | -0.056 | 0.5971 | 0.7953 | 0 | 0 | 0 |
| 'Fam133b' | 4.556 | 3.802 | 4.466 | 4.572 | 4.319 | 4.080 | 4.867 | 4.222 | 4.104 | 3.574 | 4.384 | 4.474 | 3.946 | 4.361 | 4.168 | 3.453 | 4.451 | 4.132 | 0.067 | 0.2413 | 0.4920 | 0 | 0 | 0 |
| 'Fam134a' | 14.594 | 15.210 | 14.852 | 13.320 | 14.460 | 15.869 | 13.505 | 15.834 | 16.229 | 15.156 | 13.568 | 17.069 | 14.550 | 15.583 | 16.323 | 15.699 | 14.418 | 17.899 | -0.077 | 0.0464 | 0.1794 | 0 | 0 | 0 |
| 'Fam134b' | 38.875 | 36.489 | 36.860 | 38.669 | 39.613 | 35.884 | 36.554 | 40.000 | 35.309 | 38.727 | 41.156 | 41.923 | 33.106 | 39.648 | 38.566 | 37.848 | 38.062 | 40.538 | -0.058 | 0.1524 | 0.3796 | 0 | 0 | 0 |
| 'Fam134c' | 20.422 | 22.221 | 20.156 | 20.095 | 21.673 | 20.914 | 20.258 | 20.887 | 21.169 | 20.053 | 20.932 | 22.273 | 19.466 | 21.333 | 22.144 | 20.569 | 21.169 | 21.916 | -0.028 | 0.2669 | 0.5202 | 0 | 0 | 0 |
| 'Fam136a' | 7.272 | 9.307 | 7.363 | 8.211 | 8.173 | 8.360 | 8.076 | 7.586 | 7.784 | 6.721 | 6.490 | 7.620 | 7.615 | 7.204 | 7.339 | 6.969 | 7.120 | 7.582 | 0.137 | 0.0072 | 0.0490 | 1 | 0 | 0 |
| 'Fam13b' | 27.736 | 27.012 | 27.948 | 28.047 | 27.683 | 25.226 | 27.668 | 26.369 | 27.394 | 29.191 | 27.814 | 27.608 | 26.347 | 27.434 | 26.428 | 27.408 | 27.646 | 25.832 | -0.014 | 0.6428 | 0.8250 | 0 | 0 | 0 |
| 'Fam149b' | 12.230 | 12.528 | 14.632 | 12.351 | 13.596 | 13.954 | 13.276 | 15.075 | 14.837 | 15.523 | 14.090 | 13.691 | 13.663 | 13.639 | 13.638 | 13.849 | 14.049 | 13.238 | -0.040 | 0.3736 | 0.6246 | 0 | 0 | 0 |
| 'Fam160a2' | 2.149 | 1.954 | 2.134 | 1.882 | 1.945 | 2.124 | 1.899 | 2.271 | 2.067 | 2.286 | 1.769 | 2.198 | 2.315 | 2.248 | 1.816 | 2.175 | 2.217 | 2.016 | -0.054 | 0.3244 | 0.5817 | 0 | 0 | 0 |
| 'Fam160b1' | 11.035 | 10.371 | 12.120 | 10.788 | 10.962 | 11.586 | 10.471 | 11.064 | 11.573 | 10.662 | 12.227 | 14.498 | 9.706 | 11.712 | 13.646 | 11.135 | 12.050 | 14.063 | -0.139 | 0.0016 | 0.0151 | 1 | 0 | 0 |
| 'Fam160b2' | 21.916 | 24.451 | 26.654 | 21.924 | 23.184 | 26.583 | 21.717 | 24.678 | 26.362 | 25.032 | 23.710 | 25.697 | 25.320 | 23.050 | 24.785 | 25.007 | 23.256 | 24.096 | -0.027 | 0.4325 | 0.6775 | 0 | 0 | 0 |
| 'Fam162a' | 83.375 | 80.679 | 74.839 | 83.912 | 88.235 | 75.474 | 88.096 | 78.961 | 71.712 | 79.118 | 84.682 | 81.690 | 86.798 | 88.531 | 89.197 | 93.258 | 87.039 | 84.904 | -0.110 | 0.0071 | 0.0483 | 1 | 0 | 0 |
| 'Fam163b' | 2.289 | 2.387 | 3.529 | 2.106 | 1.897 | 3.109 | 2.498 | 2.460 | 3.981 | 2.317 | 1.597 | 2.212 | 1.324 | 1.384 | 2.330 | 1.804 | 1.691 | 1.892 | 0.542 | 0.0000 | 0.0000 | 1 | 1 | 0 |
| 'Fam166a' | 9.858 | 9.415 | 9.420 | 10.247 | 10.044 | 10.723 | 10.021 | 10.907 | 11.595 | 10.413 | 11.568 | 11.024 | 9.403 | 11.815 | 12.372 | 10.313 | 11.688 | 10.167 | -0.103 | 0.1023 | 0.3000 | 0 | 0 | 0 |
| 'Fam168a' | 32.044 | 29.099 | 30.520 | 29.549 | 30.102 | 29.727 | 26.221 | 27.757 | 29.532 | 31.097 | 28.513 | 31.012 | 28.555 | 28.656 | 30.560 | 28.152 | 26.439 | 30.076 | -0.002 | 0.9656 | 0.9878 | 0 | 0 | 0 |
| 'Fam168b' | 31.812 | 33.552 | 30.545 | 31.511 | 33.855 | 32.441 | 30.817 | 34.264 | 30.772 | 31.602 | 34.063 | 35.469 | 29.760 | 34.759 | 35.679 | 32.220 | 31.997 | 34.416 | -0.061 | 0.0512 | 0.1916 | 0 | 0 | 0 |
| 'Fam171b' | 0.729 | 0.815 | 0.786 | 0.728 | 1.094 | 0.898 | 0.938 | 0.660 | 0.926 | 1.355 | 1.201 | 1.083 | 1.399 | 1.441 | 1.194 | 1.400 | 1.390 | 1.410 | -0.657 | 0.0000 | 0.0000 | 1 | 1 | 0 |
| 'Fam172a' | 11.428 | 12.095 | 11.038 | 10.442 | 11.589 | 10.304 | 11.637 | 11.825 | 10.101 | 11.745 | 11.364 | 11.414 | 10.558 | 10.958 | 10.699 | 10.239 | 11.893 | 11.406 | -0.012 | 0.7571 | 0.8913 | 0 | 0 | 0 |
| 'Fam173a' | 96.188 | 118.172 | 119.720 | 104.599 | 111.825 | 119.381 | 106.037 | 114.299 | 114.000 | 102.029 | 95.265 | 103.129 | 106.823 | 106.188 | 103.974 | 106.966 | 108.555 | 99.456 | 0.096 | 0.0083 | 0.0544 | 0 | 0 | 0 |
| 'Fam173b' | 2.241 | 2.599 | 2.806 | 2.717 | 2.570 | 2.742 | 2.106 | 2.520 | 2.951 | 2.578 | 2.685 | 2.919 | 2.805 | 2.822 | 2.685 | 2.340 | 2.770 | 3.102 | -0.101 | 0.0604 | 0.2119 | 0 | 0 | 0 |
| 'Fam174a' | 4.527 | 5.759 | 4.124 | 4.279 | 5.717 | 5.035 | 4.387 | 5.622 | 4.408 | 4.810 | 4.889 | 5.153 | 5.485 | 5.816 | 4.979 | 6.036 | 5.354 | 4.894 | -0.131 | 0.0237 | 0.1162 | 0 | 0 | 0 |
| 'Fam175a' | 2.230 | 2.957 | 2.876 | 2.191 | 3.098 | 2.502 | 3.064 | 2.886 | 2.560 | 2.297 | 2.223 | 2.305 | 2.384 | 2.591 | 2.363 | 2.398 | 2.422 | 2.296 | 0.180 | 0.0028 | 0.0242 | 1 | 0 | 0 |
| 'Fam175b' | 3.475 | 3.873 | 3.686 | 4.100 | 3.683 | 3.668 | 3.803 | 3.558 | 3.601 | 3.306 | 4.070 | 4.030 | 3.489 | 3.595 | 3.560 | 3.611 | 3.921 | 3.892 | -0.015 | 0.7412 | 0.8828 | 0 | 0 | 0 |
| 'Fam178a' | 8.240 | 8.307 | 8.174 | 8.880 | 7.876 | 7.757 | 8.523 | 8.082 | 7.919 | 8.414 | 9.522 | 8.820 | 8.009 | 8.607 | 8.321 | 7.462 | 9.379 | 9.305 | -0.088 | 0.0552 | 0.2016 | 0 | 0 | 0 |
| 'Fam179b' | 9.193 | 8.316 | 8.860 | 8.979 | 8.485 | 7.735 | 8.349 | 8.648 | 8.005 | 8.238 | 8.994 | 8.403 | 7.311 | 7.561 | 7.910 | 7.418 | 9.652 | 7.884 | 0.052 | 0.3136 | 0.5715 | 0 | 0 | 0 |
| 'Fam185a' | 1.742 | 1.750 | 1.828 | 1.827 | 1.872 | 1.679 | 1.475 | 1.756 | 1.413 | 1.660 | 1.898 | 1.675 | 1.895 | 1.944 | 1.571 | 1.599 | 1.442 | 1.777 | -0.021 | 0.7829 | 0.9025 | 0 | 0 | 0 |
| 'Fam188a' | 1.808 | 2.627 | 2.332 | 2.240 | 2.548 | 2.288 | 2.369 | 2.686 | 2.461 | 2.096 | 2.498 | 2.480 | 2.120 | 2.486 | 2.383 | 2.020 | 2.707 | 2.366 | -0.003 | 0.9430 | 0.9785 | 0 | 0 | 0 |
| 'Fam189b' | 15.425 | 14.458 | 17.001 | 13.916 | 15.175 | 16.563 | 14.369 | 15.579 | 17.283 | 13.553 | 12.932 | 14.093 | 12.841 | 12.969 | 13.909 | 12.629 | 13.237 | 13.689 | 0.211 | 0.0000 | 0.0000 | 1 | 0 | 0 |
| 'Fam192a' | 34.980 | 35.538 | 38.074 | 38.129 | 35.570 | 39.332 | 40.256 | 36.213 | 38.308 | 35.078 | 35.444 | 37.599 | 36.411 | 36.273 | 36.250 | 36.791 | 36.691 | 35.293 | 0.035 | 0.2484 | 0.4993 | 0 | 0 | 0 |
| 'Fam193a' | 19.649 | 15.322 | 19.369 | 19.690 | 16.451 | 18.854 | 20.296 | 16.403 | 18.861 | 20.364 | 19.073 | 20.576 | 18.898 | 17.657 | 19.759 | 17.015 | 19.881 | 21.178 | -0.096 | 0.0390 | 0.1610 | 0 | 0 | 0 |
| 'Fam193b' | 60.725 | 48.882 | 69.244 | 57.929 | 52.720 | 66.539 | 57.175 | 55.483 | 66.739 | 64.614 | 56.089 | 69.614 | 62.824 | 57.876 | 63.393 | 54.543 | 62.223 | 68.703 | -0.079 | 0.0431 | 0.1717 | 0 | 0 | 0 |
| 'Fam195a' | 5.898 | 5.576 | 5.491 | 4.419 | 3.781 | 4.729 | 4.132 | 4.097 | 4.695 | 5.034 | 4.825 | 4.422 | 4.548 | 4.322 | 4.294 | 4.224 | 3.587 | 4.098 | 0.121 | 0.2868 | 0.5433 | 0 | 0 | 0 |
| 'Fam195b' | 22.102 | 28.436 | 24.900 | 22.638 | 29.121 | 28.345 | 21.921 | 28.304 | 27.207 | 24.300 | 22.138 | 25.476 | 25.944 | 25.889 | 27.403 | 27.083 | 23.082 | 24.688 | 0.027 | 0.6164 | 0.8085 | 0 | 0 | 0 |
| 'Fam199x' | 4.491 | 4.845 | 4.142 | 5.007 | 4.206 | 3.676 | 4.400 | 4.357 | 3.879 | 4.785 | 4.682 | 4.395 | 4.022 | 4.290 | 3.984 | 4.094 | 4.700 | 4.531 | -0.034 | 0.5256 | 0.7473 | 0 | 0 | 0 |
| 'Fam204a' | 33.682 | 35.235 | 35.311 | 31.950 | 39.306 | 34.206 | 37.704 | 39.172 | 32.846 | 36.428 | 32.765 | 25.004 | 39.649 | 33.737 | 25.913 | 34.405 | 35.614 | 20.907 | 0.180 | 0.0108 | 0.0661 | 0 | 0 | 0 |
| 'Fam206a' | 3.670 | 3.803 | 4.118 | 3.540 | 3.945 | 4.036 | 3.469 | 4.022 | 3.619 | 3.735 | 3.416 | 3.576 | 3.407 | 3.562 | 3.505 | 3.584 | 3.642 | 3.360 | 0.101 | 0.0106 | 0.0652 | 0 | 0 | 0 |
| 'Fam207a' | 27.668 | 25.117 | 27.611 | 26.897 | 25.628 | 28.813 | 25.133 | 27.057 | 26.708 | 26.084 | 25.270 | 30.386 | 26.397 | 25.131 | 28.284 | 26.957 | 24.869 | 29.705 | -0.025 | 0.4524 | 0.6913 | 0 | 0 | 0 |
| 'Fam208a' | 8.797 | 12.844 | 11.242 | 9.678 | 12.673 | 12.062 | 10.927 | 12.471 | 11.469 | 9.816 | 10.073 | 11.782 | 9.475 | 10.954 | 10.959 | 12.047 | 10.150 | 10.650 | 0.073 | 0.1785 | 0.4140 | 0 | 0 | 0 |
| 'Fam208b' | 11.687 | 12.668 | 12.415 | 12.140 | 11.842 | 11.174 | 12.042 | 12.082 | 12.026 | 12.409 | 12.580 | 14.703 | 10.664 | 11.716 | 12.307 | 11.920 | 12.784 | 14.013 | -0.076 | 0.0998 | 0.2955 | 0 | 0 | 0 |
| 'Fam20a' | 1.801 | 2.474 | 2.432 | 1.764 | 2.158 | 2.280 | 1.899 | 2.368 | 2.001 | 2.546 | 2.572 | 2.635 | 1.475 | 2.388 | 2.606 | 2.083 | 2.658 | 2.169 | -0.149 | 0.0603 | 0.2118 | 0 | 0 | 0 |
| 'Fam20b' | 6.914 | 7.256 | 7.407 | 6.545 | 7.579 | 7.286 | 6.439 | 7.295 | 7.346 | 6.871 | 7.074 | 7.148 | 7.478 | 7.711 | 6.890 | 7.134 | 6.977 | 6.850 | -0.011 | 0.7239 | 0.8731 | 0 | 0 | 0 |
| 'Fam21' | 7.965 | 7.670 | 8.273 | 8.212 | 7.808 | 8.183 | 8.083 | 7.843 | 8.092 | 8.853 | 8.442 | 8.546 | 8.616 | 8.590 | 7.898 | 7.865 | 9.176 | 8.024 | -0.085 | 0.0073 | 0.0495 | 1 | 0 | 0 |
| 'Fam210a' | 7.973 | 9.057 | 7.630 | 8.054 | 8.986 | 7.931 | 8.386 | 8.559 | 7.709 | 8.122 | 8.600 | 9.160 | 8.038 | 9.404 | 8.350 | 9.306 | 8.642 | 8.821 | -0.094 | 0.0108 | 0.0661 | 0 | 0 | 0 |
| 'Fam210b' | 6.108 | 7.288 | 7.088 | 5.534 | 6.902 | 7.103 | 5.614 | 7.304 | 6.269 | 7.764 | 5.812 | 6.375 | 6.623 | 7.639 | 6.098 | 6.439 | 6.320 | 5.763 | 0.000 | 0.9999 | 0.9999 | 0 | 0 | 0 |
| 'Fam212b' | 0.819 | 0.691 | 0.728 | 0.793 | 0.730 | 0.875 | 0.791 | 0.737 | 0.916 | 0.788 | 0.825 | 0.958 | 0.854 | 0.882 | 0.864 | 0.826 | 1.005 | 1.222 | -0.227 | 0.0190 | 0.0998 | 0 | 0 | 0 |
| 'Fam213a' | 1.868 | 1.750 | 1.717 | 1.642 | 2.045 | 1.999 | 1.785 | 1.871 | 1.525 | 2.454 | 1.951 | 1.682 | 2.056 | 1.894 | 1.409 | 2.126 | 1.572 | 1.450 | -0.027 | 0.7669 | 0.8961 | 0 | 0 | 0 |
| 'Fam213b' | 5.099 | 6.448 | 5.360 | 5.287 | 6.226 | 5.992 | 4.536 | 6.352 | 5.018 | 5.225 | 5.520 | 3.973 | 4.266 | 4.983 | 5.018 | 4.579 | 3.845 | 2.943 | 0.316 | 0.0008 | 0.0092 | 1 | 0 | 0 |
| 'Fam214a' | 22.481 | 20.988 | 21.618 | 22.631 | 19.869 | 21.620 | 22.996 | 22.440 | 22.404 | 22.131 | 22.831 | 21.167 | 20.482 | 20.022 | 19.741 | 20.925 | 20.790 | 19.972 | 0.058 | 0.0880 | 0.2707 | 0 | 0 | 0 |
| 'Fam214b' | 1.945 | 2.427 | 2.242 | 1.727 | 2.695 | 2.471 | 2.180 | 2.000 | 2.248 | 2.011 | 2.193 | 1.935 | 2.652 | 2.613 | 1.824 | 1.925 | 1.867 | 1.715 | 0.080 | 0.3616 | 0.6135 | 0 | 0 | 0 |
| 'Fam216a' | 14.246 | 14.187 | 12.314 | 13.839 | 13.594 | 14.027 | 12.864 | 12.663 | 12.987 | 11.063 | 13.784 | 14.669 | 14.036 | 12.971 | 14.485 | 12.253 | 13.006 | 14.039 | -0.013 | 0.8220 | 0.9217 | 0 | 0 | 0 |
| 'Fam219a' | 1.841 | 2.225 | 2.105 | 1.675 | 2.286 | 2.360 | 1.369 | 2.159 | 2.287 | 2.170 | 2.114 | 2.073 | 2.376 | 2.145 | 2.219 | 2.057 | 1.912 | 1.976 | -0.072 | 0.3399 | 0.5970 | 0 | 0 | 0 |
| 'Fam219b' | 16.694 | 18.094 | 15.264 | 13.877 | 19.566 | 16.776 | 17.545 | 15.340 | 19.879 | 15.192 | 13.387 | 15.715 | 14.791 | 16.577 | 17.988 | 18.185 | 14.785 | 17.062 | 0.079 | 0.3234 | 0.5808 | 0 | 0 | 0 |
| 'Fam221a' | 0.863 | 0.885 | 1.001 | 0.848 | 0.655 | 0.913 | 0.938 | 0.958 | 1.025 | 0.905 | 1.096 | 0.921 | 1.140 | 0.935 | 0.993 | 0.975 | 0.854 | 0.878 | -0.101 | 0.3558 | 0.6089 | 0 | 0 | 0 |

| | | | | | | | | | | | | | | | | | | | | | | | |
|---|---|---|---|---|---|---|---|---|---|---|---|---|---|---|---|---|---|---|---|---|---|---|---|
| 'Fam222a' | 12.759 | 10.451 | 10.801 | 10.263 | 10.471 | 11.814 | 8.711 | 10.765 | 10.940 | 11.017 | 9.889 | 9.925 | 10.970 | 12.141 | 10.239 | 12.657 | 9.870 | 11.098 | -0.017 | 0.7917 | 0.9070 | 0 | 0 | 0 |
| 'Fam222b' | 5.329 | 5.105 | 4.535 | 4.789 | 4.957 | 5.226 | 4.241 | 4.656 | 4.940 | 5.229 | 4.859 | 5.852 | 4.393 | 4.806 | 5.616 | 4.587 | 4.687 | 5.747 | -0.075 | 0.1882 | 0.4246 | 0 | 0 | 0 |
| 'Fam234a' | 9.639 | 8.831 | 9.350 | 9.396 | 9.608 | 9.435 | 8.852 | 9.196 | 9.132 | 9.651 | 9.532 | 9.589 | 11.707 | 9.408 | 9.768 | 9.728 | 9.516 | 9.565 | -0.093 | 0.0164 | 0.0898 | 0 | 0 | 0 |
| 'Fam234b' | 6.827 | 7.152 | 7.561 | 6.731 | 7.093 | 7.359 | 7.039 | 7.196 | 7.514 | 6.855 | 6.774 | 7.669 | 7.503 | 7.605 | 7.443 | 7.275 | 6.605 | 7.272 | -0.023 | 0.4503 | 0.6902 | 0 | 0 | 0 |
| 'Fam26f' | 9.665 | 13.875 | 9.698 | 9.268 | 10.362 | 11.574 | 10.076 | 12.081 | 10.818 | 13.781 | 15.308 | 15.021 | 14.603 | 15.990 | 15.212 | 15.322 | 15.746 | 16.785 | -0.516 | 0.0000 | 0.0000 | 1 | 1 | 0 |
| 'Fam32a' | 58.818 | 63.848 | 58.372 | 54.585 | 64.547 | 63.437 | 57.110 | 62.822 | 60.401 | 61.593 | 53.924 | 58.449 | 61.187 | 57.849 | 59.824 | 61.327 | 55.972 | 56.571 | 0.036 | 0.3092 | 0.5675 | 0 | 0 | 0 |
| 'Fam35a' | 5.923 | 6.398 | 6.578 | 6.172 | 6.370 | 5.734 | 6.671 | 5.429 | 6.215 | 6.351 | 7.014 | 5.396 | 5.812 | 6.803 | 5.868 | 5.847 | 6.513 | 5.261 | 0.015 | 0.8035 | 0.9117 | 0 | 0 | 0 |
| 'Fam3a' | 4.466 | 5.219 | 4.491 | 3.859 | 4.796 | 4.389 | 4.319 | 5.018 | 4.539 | 5.065 | 4.322 | 5.151 | 4.596 | 4.864 | 5.026 | 4.632 | 5.265 | 4.571 | -0.098 | 0.0425 | 0.1701 | 0 | 0 | 0 |
| 'Fam3c' | 12.816 | 12.273 | 16.066 | 13.082 | 13.461 | 14.711 | 13.244 | 15.100 | 15.706 | 16.082 | 17.183 | 19.709 | 11.738 | 14.122 | 18.981 | 16.902 | 15.446 | 18.594 | -0.240 | 0.0001 | 0.0009 | 1 | 0 | 0 |
| 'Fam43a' | 88.940 | 98.062 | 60.507 | 74.748 | 93.203 | 68.935 | 73.481 | 94.003 | 64.388 | 107.973 | 93.398 | 68.101 | 94.346 | 108.107 | 68.801 | 123.451 | 92.142 | 70.838 | -0.210 | 0.0008 | 0.0083 | 1 | 0 | 0 |
| 'Fam45a' | 4.412 | 5.506 | 5.148 | 4.722 | 5.400 | 4.656 | 4.712 | 5.326 | 5.012 | 4.999 | 4.732 | 5.102 | 4.689 | 5.259 | 4.457 | 4.800 | 4.988 | 4.568 | 0.027 | 0.4880 | 0.7184 | 0 | 0 | 0 |
| 'Fam46c' | 29.461 | 24.550 | 25.906 | 27.730 | 26.513 | 26.444 | 26.532 | 27.673 | 25.257 | 29.097 | 33.690 | 36.112 | 24.784 | 30.622 | 32.545 | 28.912 | 29.718 | 38.358 | -0.246 | 0.0001 | 0.0012 | 1 | 0 | 0 |
| 'Fam49a' | 8.185 | 8.224 | 8.409 | 8.702 | 8.201 | 8.273 | 9.504 | 8.600 | 8.114 | 9.200 | 10.138 | 11.282 | 9.088 | 9.917 | 10.194 | 9.406 | 10.931 | 10.101 | -0.258 | 0.0000 | 0.0000 | 1 | 0 | 0 |
| 'Fam49b' | 46.239 | 51.724 | 45.738 | 49.048 | 50.584 | 45.807 | 49.556 | 52.299 | 44.356 | 45.775 | 50.333 | 44.718 | 49.677 | 48.767 | 44.446 | 45.949 | 49.386 | 43.189 | 0.032 | 0.1212 | 0.3319 | 0 | 0 | 0 |
| 'Fam50a' | 18.900 | 15.390 | 17.341 | 19.626 | 16.847 | 17.968 | 18.648 | 17.513 | 18.507 | 18.181 | 18.770 | 19.298 | 18.100 | 18.367 | 18.549 | 18.538 | 17.936 | 18.446 | -0.059 | 0.1026 | 0.3002 | 0 | 0 | 0 |
| 'Fam53a' | 5.714 | 6.186 | 6.095 | 5.576 | 5.905 | 6.266 | 6.108 | 5.907 | 6.041 | 5.872 | 5.866 | 6.872 | 6.344 | 5.782 | 6.647 | 6.137 | 6.083 | 6.567 | -0.077 | 0.0091 | 0.0583 | 0 | 0 | 0 |
| 'Fam53b' | 13.303 | 12.023 | 12.691 | 11.483 | 11.980 | 12.453 | 11.622 | 11.199 | 12.050 | 12.662 | 12.627 | 14.788 | 11.054 | 12.049 | 13.527 | 13.118 | 10.838 | 14.644 | -0.091 | 0.0569 | 0.2050 | 0 | 0 | 0 |
| 'Fam53c' | 9.420 | 9.856 | 9.547 | 8.880 | 8.872 | 8.996 | 9.106 | 9.152 | 9.908 | 8.952 | 8.842 | 9.976 | 9.185 | 9.778 | 9.714 | 9.093 | 8.419 | 9.435 | -0.006 | 0.8546 | 0.9376 | 0 | 0 | 0 |
| 'Fam57a' | 0.463 | 0.571 | 0.479 | 0.404 | 0.556 | 0.467 | 0.512 | 0.630 | 0.435 | 0.410 | 0.535 | 0.494 | 0.462 | 0.579 | 0.548 | 0.384 | 0.557 | 0.517 | -0.008 | 0.9339 | 0.9747 | 0 | 0 | 0 |
| 'Fam58b' | 15.871 | 17.883 | 17.157 | 16.355 | 16.645 | 19.907 | 16.215 | 17.927 | 16.934 | 16.616 | 15.502 | 16.264 | 16.585 | 15.970 | 17.599 | 15.961 | 15.546 | 18.268 | 0.054 | 0.2899 | 0.5465 | 0 | 0 | 0 |
| 'Fam60a' | 7.414 | 6.067 | 5.653 | 6.359 | 5.252 | 5.643 | 6.321 | 5.782 | 5.531 | 4.972 | 6.202 | 7.196 | 5.222 | 6.122 | 7.238 | 5.735 | 6.195 | 7.089 | -0.059 | 0.4518 | 0.6910 | 0 | 0 | 0 |
| 'Fam63a' | 1.896 | 1.723 | 1.995 | 1.579 | 1.841 | 2.293 | 1.772 | 2.142 | 2.181 | 2.202 | 2.060 | 1.835 | 2.018 | 1.849 | 1.991 | 1.844 | 1.935 | 1.939 | -0.027 | 0.6718 | 0.8418 | 0 | 0 | 0 |
| 'Fam63b' | 10.383 | 10.661 | 10.046 | 11.130 | 10.107 | 9.660 | 11.532 | 10.253 | 9.977 | 9.731 | 11.927 | 12.584 | 10.405 | 11.343 | 12.064 | 9.120 | 12.063 | 12.067 | -0.120 | 0.0338 | 0.1465 | 0 | 0 | 0 |
| 'Fam64a' | 1.494 | 2.339 | 3.278 | 2.008 | 2.066 | 3.042 | 1.956 | 2.547 | 3.015 | 1.344 | 2.148 | 2.690 | 2.551 | 2.395 | 2.787 | 2.129 | 2.184 | 2.547 | 0.041 | 0.6658 | 0.8389 | 0 | 0 | 0 |
| 'Fam65a' | 8.284 | 7.160 | 8.928 | 8.185 | 7.141 | 8.047 | 8.933 | 8.188 | 8.667 | 8.326 | 7.415 | 10.131 | 7.876 | 7.279 | 9.297 | 7.625 | 8.159 | 9.739 | -0.056 | 0.2601 | 0.5137 | 0 | 0 | 0 |
| 'Fam65b' | 3.552 | 2.998 | 4.057 | 3.382 | 3.654 | 4.270 | 3.496 | 3.888 | 3.826 | 3.564 | 3.954 | 4.564 | 3.079 | 3.647 | 4.495 | 3.407 | 3.271 | 4.277 | -0.053 | 0.3022 | 0.5592 | 0 | 0 | 0 |
| 'Fam69a' | 4.249 | 5.174 | 4.626 | 4.543 | 5.035 | 4.935 | 5.400 | 4.407 | 5.169 | 4.707 | 4.771 | 5.120 | 4.230 | 4.577 | 5.068 | 4.945 | 4.341 | 4.947 | 0.011 | 0.8509 | 0.9358 | 0 | 0 | 0 |
| 'Fam69b' | 2.680 | 2.409 | 2.866 | 2.458 | 2.445 | 3.148 | 2.430 | 3.110 | 3.009 | 2.681 | 2.151 | 2.417 | 2.522 | 1.934 | 2.422 | 2.688 | 1.763 | 2.274 | 0.240 | 0.0048 | 0.0361 | 1 | 0 | 0 |
| 'Fam71b' | 17.712 | 23.812 | 15.132 | 19.924 | 19.782 | 19.993 | 22.431 | 17.403 | 19.766 | 15.116 | 14.422 | 16.269 | 19.888 | 17.118 | 14.998 | 16.368 | 17.364 | 14.970 | 0.250 | 0.0004 | 0.0046 | 1 | 0 | 0 |
| 'Fam73a' | 2.009 | 2.287 | 1.976 | 1.910 | 2.101 | 1.952 | 1.818 | 2.307 | 1.875 | 1.907 | 2.202 | 2.061 | 2.050 | 2.137 | 1.938 | 2.025 | 1.876 | 1.770 | 0.008 | 0.8588 | 0.9390 | 0 | 0 | 0 |
| 'Fam73b' | 2.012 | 1.936 | 2.076 | 1.873 | 2.145 | 2.463 | 1.674 | 2.139 | 2.309 | 2.026 | 1.693 | 2.385 | 2.170 | 2.128 | 1.969 | 1.786 | 2.043 | 2.225 | 0.001 | 0.9814 | 0.9931 | 0 | 0 | 0 |
| 'Fam76a' | 7.810 | 9.494 | 8.055 | 7.808 | 10.525 | 8.475 | 6.574 | 9.451 | 9.111 | 7.954 | 8.744 | 8.208 | 8.820 | 8.984 | 8.189 | 10.001 | 8.261 | 6.885 | 0.012 | 0.8627 | 0.9405 | 0 | 0 | 0 |
| 'Fam76b' | 8.364 | 11.435 | 9.159 | 8.194 | 10.095 | 9.369 | 9.034 | 10.197 | 9.210 | 8.271 | 8.785 | 8.876 | 8.711 | 10.637 | 8.923 | 9.070 | 9.983 | 8.405 | 0.045 | 0.2292 | 0.4761 | 0 | 0 | 0 |
| 'Fam78a' | 33.556 | 35.379 | 34.412 | 33.045 | 32.725 | 36.789 | 32.482 | 32.737 | 34.974 | 30.647 | 32.358 | 41.843 | 32.898 | 33.013 | 40.140 | 34.184 | 32.293 | 42.531 | -0.067 | 0.0924 | 0.2812 | 0 | 0 | 0 |
| 'Fam83d' | 1.171 | 0.982 | 0.829 | 0.802 | 1.103 | 0.844 | 1.055 | 0.848 | 1.213 | 0.915 | 1.084 | 1.010 | 1.239 | 1.169 | 1.099 | 1.123 | 0.984 | 0.953 | -0.125 | 0.2193 | 0.4661 | 0 | 0 | 0 |
| 'Fam83g' | 3.841 | 3.511 | 4.033 | 3.547 | 3.418 | 4.077 | 3.229 | 3.764 | 4.325 | 4.609 | 3.762 | 3.985 | 4.651 | 3.693 | 4.026 | 4.979 | 3.724 | 3.647 | -0.135 | 0.0272 | 0.1270 | 0 | 0 | 0 |
| 'Fam84b' | 2.155 | 1.681 | 1.492 | 1.922 | 1.921 | 1.842 | 2.065 | 1.814 | 1.546 | 2.041 | 1.950 | 1.626 | 2.183 | 1.771 | 1.904 | 2.244 | 1.902 | 1.583 | -0.069 | 0.2994 | 0.5557 | 0 | 0 | 0 |
| 'Fam8a1' | 13.619 | 19.530 | 18.190 | 16.372 | 18.987 | 17.841 | 16.024 | 18.151 | 17.477 | 13.376 | 19.486 | 24.851 | 14.079 | 19.642 | 22.106 | 17.883 | 20.939 | 26.570 | -0.188 | 0.0066 | 0.0462 | 1 | 0 | 0 |
| 'Fam91a1' | 14.402 | 13.971 | 15.037 | 13.748 | 14.588 | 13.998 | 14.642 | 16.603 | 15.320 | 15.503 | 14.217 | 13.228 | 14.903 | 14.041 | 13.224 | 16.532 | 14.073 | 13.430 | 0.030 | 0.4794 | 0.7118 | 0 | 0 | 0 |
| 'Fam92a' | 4.148 | 4.523 | 4.486 | 4.214 | 4.691 | 4.234 | 5.020 | 4.747 | 4.996 | 4.077 | 3.827 | 3.901 | 4.369 | 3.543 | 3.809 | 4.557 | 3.918 | 3.224 | 0.214 | 0.0001 | 0.0014 | 1 | 0 | 0 |
| 'Fam96a' | 32.987 | 37.502 | 35.976 | 34.881 | 37.780 | 33.568 | 36.542 | 37.409 | 33.477 | 32.815 | 31.575 | 33.771 | 33.949 | 34.914 | 31.614 | 35.955 | 35.857 | 33.800 | 0.060 | 0.0755 | 0.2441 | 0 | 0 | 0 |
| 'Fam96b' | 19.837 | 22.303 | 20.611 | 23.887 | 23.254 | 19.302 | 19.871 | 22.104 | 20.717 | 18.024 | 18.760 | 20.732 | 21.346 | 22.467 | 20.154 | 19.027 | 20.943 | 20.160 | 0.063 | 0.1691 | 0.4013 | 0 | 0 | 0 |
| 'Fam98a' | 10.534 | 10.771 | 11.029 | 9.314 | 10.741 | 11.040 | 10.688 | 10.399 | 10.844 | 10.034 | 10.636 | 11.892 | 10.433 | 10.841 | 11.136 | 11.376 | 11.064 | 11.771 | -0.068 | 0.0592 | 0.2100 | 0 | 0 | 0 |
| 'Fam98b' | 12.067 | 12.763 | 13.080 | 11.507 | 14.458 | 13.756 | 13.470 | 13.351 | 12.417 | 12.705 | 12.078 | 12.558 | 13.402 | 13.598 | 12.408 | 12.324 | 12.503 | 11.544 | 0.036 | 0.3603 | 0.6126 | 0 | 0 | 0 |
| 'Fam98c' | 5.489 | 4.886 | 5.946 | 4.891 | 4.833 | 5.483 | 5.003 | 4.625 | 5.773 | 4.845 | 4.770 | 5.824 | 5.422 | 4.404 | 5.220 | 4.961 | 4.769 | 5.061 | 0.040 | 0.3326 | 0.5908 | 0 | 0 | 0 |
| 'Fan1' | 1.051 | 1.146 | 1.302 | 0.907 | 1.089 | 1.161 | 1.201 | 1.333 | 1.215 | 1.083 | 1.009 | 0.951 | 1.265 | 1.032 | 0.886 | 1.194 | 0.939 | 0.802 | 0.187 | 0.0261 | 0.1241 | 0 | 0 | 0 |
| 'Fanca' | 1.358 | 1.315 | 1.787 | 1.611 | 1.098 | 1.473 | 1.590 | 1.217 | 1.731 | 1.278 | 1.374 | 1.410 | 1.871 | 1.367 | 1.517 | 1.407 | 1.467 | 1.464 | -0.013 | 0.8619 | 0.9402 | 0 | 0 | 0 |
| 'Fancb' | 1.140 | 1.167 | 1.589 | 1.335 | 1.320 | 1.452 | 1.255 | 1.413 | 1.504 | 1.205 | 1.290 | 1.469 | 1.389 | 1.704 | 1.607 | 1.236 | 1.563 | 1.488 | -0.091 | 0.3137 | 0.5715 | 0 | 0 | 0 |
| 'Fancc' | 0.531 | 0.554 | 0.519 | 0.773 | 0.502 | 0.492 | 0.624 | 0.562 | 0.651 | 0.435 | 0.641 | 0.625 | 0.441 | 0.663 | 0.653 | 0.578 | 0.530 | 0.591 | 0.003 | 0.9756 | 0.9918 | 0 | 0 | 0 |
| 'Fancd2' | 1.618 | 1.738 | 2.100 | 1.996 | 1.470 | 2.130 | 2.004 | 1.620 | 2.151 | 1.334 | 1.890 | 1.662 | 2.194 | 1.829 | 1.855 | 1.540 | 1.931 | 1.504 | 0.083 | 0.3657 | 0.6178 | 0 | 0 | 0 |
| 'Fance' | 3.120 | 3.391 | 3.993 | 2.466 | 3.210 | 3.678 | 3.254 | 3.542 | 3.839 | 3.646 | 2.827 | 3.550 | 3.687 | 3.473 | 3.593 | 3.244 | 3.180 | 3.530 | -0.024 | 0.6993 | 0.8592 | 0 | 0 | 0 |
| 'Fancf' | 4.252 | 3.944 | 5.516 | 3.873 | 3.946 | 4.633 | 5.665 | 4.878 | 4.086 | 4.859 | 4.004 | 3.820 | 3.711 | 3.911 | 4.892 | 4.304 | 4.987 | 4.603 | 0.062 | 0.5265 | 0.7481 | 0 | 0 | 0 |
| 'Fancg' | 3.654 | 3.681 | 4.433 | 3.652 | 3.520 | 3.902 | 3.794 | 3.294 | 4.431 | 3.372 | 3.366 | 3.915 | 4.225 | 3.729 | 4.247 | 3.552 | 3.435 | 3.674 | 0.024 | 0.6309 | 0.8173 | 0 | 0 | 0 |
| 'Fanci' | 1.830 | 1.841 | 2.133 | 1.903 | 1.784 | 1.904 | 1.714 | 1.647 | 2.001 | 1.607 | 1.895 | 1.998 | 2.240 | 1.944 | 2.088 | 1.874 | 1.990 | 1.727 | -0.061 | 0.3249 | 0.5821 | 0 | 0 | 0 |
| 'Fancl' | 3.441 | 4.003 | 4.123 | 3.987 | 3.564 | 4.143 | 4.323 | 3.914 | 3.614 | 3.693 | 3.663 | 3.947 | 3.312 | 3.733 | 3.538 | 3.082 | 4.245 | 3.394 | 0.093 | 0.1329 | 0.3509 | 0 | 0 | 0 |
| 'Fancm' | 5.582 | 5.169 | 6.279 | 5.387 | 5.656 | 5.280 | 5.737 | 5.494 | 5.539 | 5.111 | 6.066 | 5.563 | 5.575 | 5.530 | 5.375 | 4.900 | 5.941 | 5.471 | 0.009 | 0.8372 | 0.9302 | 0 | 0 | 0 |
| 'Far1' | 7.309 | 8.668 | 8.274 | 7.839 | 8.304 | 8.280 | 8.110 | 8.676 | 7.898 | 7.502 | 7.641 | 7.874 | 7.591 | 8.545 | 7.679 | 7.558 | 8.418 | 7.558 | 0.049 | 0.0640 | 0.2205 | 0 | 0 | 0 |
| 'Farp1' | 0.563 | 0.508 | 0.698 | 0.574 | 0.739 | 0.924 | 0.553 | 0.579 | 0.610 | 0.843 | 0.858 | 0.832 | 0.659 | 0.595 | 0.702 | 0.575 | 0.657 | 0.938 | -0.222 | 0.0540 | 0.1988 | 0 | 0 | 0 |
| 'Farp2' | 0.875 | 0.916 | 1.030 | 0.846 | 0.747 | 1.107 | 0.798 | 0.881 | 1.218 | 1.064 | 1.061 | 0.965 | 0.882 | 0.752 | 1.022 | 0.826 | 0.814 | 1.023 | 0.001 | 0.9887 | 0.9968 | 0 | 0 | 0 |
| 'Fars2' | 18.095 | 18.290 | 15.570 | 18.577 | 18.367 | 15.684 | 19.111 | 16.824 | 15.967 | 17.972 | 19.161 | 17.874 | 19.955 | 18.239 | 17.822 | 17.768 | 19.944 | 16.743 | -0.100 | 0.0198 | 0.1022 | 0 | 0 | 0 |
| 'Farsa' | 13.828 | 12.807 | 13.983 | 13.911 | 13.187 | 13.473 | 13.905 | 13.499 | 13.422 | 12.988 | 12.500 | 14.410 | 13.927 | 12.387 | 14.204 | 13.009 | 12.993 | 14.187 | 0.004 | 0.8726 | 0.9454 | 0 | 0 | 0 |
| 'Farsb' | 9.879 | 10.613 | 10.559 | 11.226 | 11.050 | 9.824 | 10.337 | 10.125 | 9.145 | 10.032 | 10.946 | 10.717 | 10.638 | 9.486 | 10.605 | 9.300 | 11.141 | 10.086 | -0.018 | 0.6882 | 0.8526 | 0 | 0 | 0 |
| 'Fas' | 36.802 | 45.348 | 36.369 | 40.455 | 41.904 | 36.110 | 43.315 | 41.790 | 39.723 | 49.397 | 52.196 | 45.445 | 50.966 | 53.167 | 44.717 | 49.046 | 54.566 | 43.221 | -0.303 | 0.0000 | 0.0000 | 1 | 0 | 0 |
| 'Fasn' | 22.306 | 19.233 | 24.138 | 21.627 | 19.114 | 22.975 | 20.471 | 20.202 | 23.308 | 22.855 | 22.221 | 25.652 | 21.428 | 20.812 | 24.363 | 21.675 | 21.471 | 26.112 | -0.106 | 0.0004 | 0.0049 | 1 | 0 | 0 |
| 'Fastk' | 6.596 | 6.662 | 7.501 | 5.699 | 6.401 | 7.621 | 6.314 | 6.326 | 7.533 | 7.133 | 6.091 | 7.619 | 7.200 | 6.399 | 7.397 | 6.924 | 6.319 | 7.344 | -0.052 | 0.1275 | 0.3421 | 0 | 0 | 0 |
| 'Fastkd1' | 1.696 | 1.650 | 1.751 | 1.783 | 1.763 | 1.718 | 1.585 | 1.707 | 1.565 | 1.608 | 1.547 | 1.863 | 1.484 | 1.579 | 1.372 | 1.374 | 1.649 | 1.678 | 0.085 | 0.1863 | 0.4219 | 0 | 0 | 0 |
| 'Fastkd2' | 1.789 | 1.938 | 2.477 | 2.225 | 2.438 | 2.065 | 2.292 | 2.388 | 2.311 | 2.176 | 2.281 | 2.175 | 2.099 | 2.122 | 2.048 | 2.108 | 1.918 | 2.020 | 0.063 | 0.2930 | 0.5494 | 0 | 0 | 0 |
| 'Fastkd3' | 6.838 | 6.645 | 6.811 | 6.744 | 7.753 | 6.351 | 7.414 | 6.988 | 7.017 | 5.459 | 6.266 | 7.341 | 6.116 | 6.439 | 7.361 | 5.230 | 6.234 | 6.981 | 0.106 | 0.0883 | 0.2714 | 0 | 0 | 0 |
| 'Fastkd5' | 0.871 | 0.846 | 0.645 | 0.819 | 0.875 | 0.824 | 0.785 | 0.733 | 0.777 | 0.488 | 1.029 | 1.007 | 0.646 | 0.870 | 0.781 | 0.582 | 0.788 | 0.932 | -0.003 | 0.9798 | 0.9926 | 0 | 0 | 0 |
| 'Fau' | 10.361 | 12.469 | 11.368 | 12.350 | 12.818 | 12.112 | 11.120 | 12.029 | 13.207 | 10.903 | 12.668 | 9.863 | 11.528 | 12.115 | 12.022 | 12.403 | 10.803 | 10.005 | 0.076 | 0.2733 | 0.5284 | 0 | 0 | 0 |
| 'Fbf1' | 2.049 | 1.652 | 2.440 | 2.175 | 2.008 | 2.445 | 2.039 | 1.947 | 2.320 | 2.293 | 2.226 | 2.231 | 2.182 | 1.939 | 2.211 | 1.787 | 1.676 | 2.203 | 0.013 | 0.8178 | 0.9193 | 0 | 0 | 0 |
| 'Fbl' | 138.390 | 122.793 | 130.430 | 132.524 | 132.833 | 123.754 | 128.798 | 133.088 | 124.482 | 126.964 | 120.304 | 124.061 | 135.698 | 120.977 | 128.518 | 131.803 | 120.814 | 125.039 | 0.032 | 0.2298 | 0.4766 | 0 | 0 | 0 |
| 'Fbln1' | 1.322 | 1.454 | 2.198 | 1.568 | 1.421 | 2.000 | 1.943 | 1.907 | 2.301 | 1.521 | 1.560 | 2.123 | 1.779 | 1.628 | 2.166 | 1.534 | 1.678 | 2.226 | -0.020 | 0.8042 | 0.9121 | 0 | 0 | 0 |
| 'Fbn2' | 1.062 | 1.452 | 1.596 | 1.412 | 1.088 | 1.182 | 1.609 | 1.487 | 1.295 | 4.569 | 4.583 | 3.599 | 3.171 | 4.207 | 3.289 | 5.256 | 5.261 | 4.113 | -1.646 | 0.0000 | 0.0000 | 1 | 1 | 1 |
| 'Fbrsl1' | 7.452 | 6.148 | 7.226 | 6.392 | 5.832 | 6.531 | 6.224 | 6.732 | 6.929 | 8.084 | 6.584 | 7.092 | 7.412 | 6.817 | 6.870 | 7.219 | 6.700 | 6.951 | -0.107 | 0.0099 | 0.0619 | 0 | 0 | 0 |
| 'Fbxl12' | 2.191 | 2.425 | 2.469 | 1.769 | 2.309 | 2.535 | 1.920 | 2.496 | 2.724 | 2.471 | 2.258 | 2.478 | 2.454 | 2.220 | 2.247 | 2.495 | 2.247 | 2.216 | -0.030 | 0.6174 | 0.8087 | 0 | 0 | 0 |
| 'Fbxl14' | 51.625 | 52.471 | 52.903 | 51.041 | 52.742 | 56.959 | 51.278 | 51.596 | 51.473 | 52.545 | 52.683 | 55.749 | 52.354 | 54.424 | 57.035 | 54.247 | 47.535 | 56.983 | -0.045 | 0.1252 | 0.3386 | 0 | 0 | 0 |
| 'Fbxl15' | 6.636 | 6.419 | 6.287 | 6.175 | 5.907 | 6.950 | 5.794 | 5.277 | 6.993 | 5.998 | 5.382 | 6.063 | 5.991 | 5.252 | 5.295 | 5.951 | 5.306 | 5.682 | 0.138 | 0.0312 | 0.1385 | 0 | 0 | 0 |
| 'Fbxl17' | 4.152 | 3.573 | 4.020 | 3.844 | 3.542 | 4.062 | 3.593 | 3.702 | 3.577 | 4.118 | 4.057 | 3.767 | 3.640 | 3.946 | 3.897 | 3.709 | 3.894 | 3.834 | -0.040 | 0.3502 | 0.6045 | 0 | 0 | 0 |
| 'Fbxl18' | 1.407 | 1.317 | 1.469 | 1.304 | 1.464 | 1.310 | 1.309 | 1.419 | 1.649 | 1.402 | 1.427 | 1.424 | 1.529 | 1.470 | 1.367 | 1.268 | 1.316 | 1.356 | 0.001 | 0.9820 | 0.9933 | 0 | 0 | 0 |
| 'Fbxl19' | 3.472 | 3.217 | 3.619 | 3.405 | 3.162 | 3.925 | 3.317 | 3.252 | 3.921 | 3.323 | 3.408 | 3.824 | 3.709 | 3.607 | 3.662 | 3.094 | 3.237 | 3.867 | -0.031 | 0.4417 | 0.6852 | 0 | 0 | 0 |
| 'Fbxl20' | 8.177 | 8.574 | 8.090 | 7.197 | 8.286 | 7.852 | 7.405 | 8.600 | 7.881 | 8.795 | 9.218 | 8.007 | 7.833 | 8.679 | 7.378 | 9.341 | 7.959 | 7.408 | -0.058 | 0.1854 | 0.4209 | 0 | 0 | 0 |
| 'Fbxl3' | 8.326 | 8.832 | 7.842 | 7.964 | 8.807 | 8.381 | 8.055 | 9.168 | 7.676 | 8.684 | 8.878 | 8.360 | 8.009 | 9.275 | 8.248 | 9.168 | 8.848 | 8.190 | -0.058 | 0.0484 | 0.1848 | 0 | 0 | 0 |
| 'Fbxl4' | 1.564 | 2.235 | 1.666 | 1.952 | 2.262 | 1.746 | 2.114 | 1.961 | 2.104 | 2.205 | 2.112 | 1.472 | 1.935 | 1.749 | 1.497 | 1.684 | 1.944 | 1.101 | 0.167 | 0.0607 | 0.2124 | 0 | 0 | 0 |
| 'Fbxl5' | 5.367 | 6.103 | 5.742 | 5.733 | 6.068 | 5.149 | 5.606 | 6.068 | 5.364 | 5.771 | 6.119 | 5.845 | 5.396 | 5.863 | 5.145 | 5.577 | 6.022 | 5.693 | -0.020 | 0.5269 | 0.7483 | 0 | 0 | 0 |
| 'Fbxl6' | 44.575 | 41.290 | 51.105 | 41.784 | 44.586 | 47.374 | 39.971 | 45.474 | 47.915 | 42.642 | 39.734 | 47.940 | 40.864 | 37.258 | 43.016 | 42.511 | 40.263 | 44.339 | 0.085 | 0.0154 | 0.0853 | 0 | 0 | 0 |
| 'Fbxl8' | 6.643 | 6.994 | 7.638 | 6.208 | 6.009 | 8.133 | 7.060 | 6.370 | 8.323 | 5.983 | 6.288 | 8.340 | 6.579 | 6.727 | 8.403 | 6.394 | 6.734 | 7.845 | -0.011 | 0.8078 | 0.9138 | 0 | 0 | 0 |
| 'Fbxo10' | 1.934 | 1.897 | 2.561 | 1.745 | 2.381 | 2.212 | 1.865 | 1.960 | 2.214 | 2.022 | 2.053 | 1.950 | 1.777 | 1.905 | 1.816 | 1.937 | 1.862 | 1.478 | 0.156 | 0.0431 | 0.1717 | 0 | 0 | 0 |
| 'Fbxo11' | 13.199 | 14.644 | 12.679 | 14.122 | 13.181 | 12.172 | 13.656 | 14.084 | 12.109 | 13.424 | 14.268 | 13.219 | 12.981 | 13.717 | 12.508 | 13.313 | 14.658 | 13.316 | -0.032 | 0.2966 | 0.5524 | 0 | 0 | 0 |
| 'Fbxo17' | 3.414 | 3.095 | 3.929 | 3.040 | 2.638 | 3.689 | 3.372 | 4.121 | 3.750 | 2.198 | 2.505 | 2.694 | 2.018 | 2.201 | 3.073 | 2.121 | 2.304 | 2.493 | 0.521 | 0.0000 | 0.0000 | 1 | 1 | 0 |
| 'Fbxo18' | 8.023 | 8.179 | 8.010 | 7.965 | 8.519 | 8.047 | 7.727 | 8.287 | 8.411 | 8.865 | 8.737 | 8.139 | 8.419 | 8.489 | 7.866 | 8.052 | 8.105 | 7.892 | -0.036 | 0.2216 | 0.4691 | 0 | 0 | 0 |
| 'Fbxo21' | 3.605 | 3.941 | 3.892 | 3.486 | 4.035 | 3.982 | 3.890 | 3.854 | 4.085 | 3.377 | 3.900 | 4.411 | 3.673 | 3.583 | 4.146 | 4.269 | 4.005 | 4.321 | -0.050 | 0.2399 | 0.4902 | 0 | 0 | 0 |
| 'Fbxo22' | 14.279 | 15.644 | 14.443 | 13.995 | 14.891 | 15.100 | 15.856 | 15.697 | 14.495 | 14.268 | 14.616 | 14.404 | 15.454 | 15.222 | 13.812 | 14.157 | 15.815 | 13.874 | 0.019 | 0.5379 | 0.7566 | 0 | 0 | 0 |
| 'Fbxo24' | 1.067 | 1.152 | 1.131 | 1.064 | 1.053 | 1.237 | 0.907 | 1.170 | 1.432 | 0.871 | 0.966 | 1.421 | 1.052 | 0.934 | 1.322 | 1.293 | 1.069 | 1.179 | -0.001 | 0.9946 | 0.9991 | 0 | 0 | 0 |
| 'Fbxo25' | 10.981 | 12.820 | 11.470 | 11.720 | 11.171 | 12.421 | 11.074 | 11.522 | 11.552 | 10.983 | 11.024 | 13.161 | 10.631 | 10.690 | 11.816 | 11.021 | 11.769 | 12.320 | 0.001 | 0.9770 | 0.9920 | 0 | 0 | 0 |
| 'Fbxo27' | 1.115 | 0.972 | 1.410 | 1.080 | 1.016 | 1.238 | 0.795 | 1.309 | 1.144 | 1.022 | 1.314 | 1.372 | 1.214 | 1.004 | 1.438 | 0.659 | 0.934 | 1.136 | -0.010 | 0.9348 | 0.9749 | 0 | 0 | 0 |
| 'Fbxo28' | 8.577 | 9.064 | 9.146 | 8.926 | 9.955 | 8.949 | 9.294 | 9.297 | 8.966 | 8.643 | 9.307 | 8.927 | 8.278 | 9.307 | 9.135 | 8.904 | 9.063 | 8.812 | 0.022 | 0.4350 | 0.6791 | 0 | 0 | 0 |
| 'Fbxo3' | 3.338 | 4.241 | 3.666 | 3.679 | 4.302 | 3.879 | 4.181 | 3.997 | 3.966 | 3.543 | 3.768 | 3.901 | 3.604 | 4.202 | 3.996 | 3.684 | 4.324 | 3.695 | 0.008 | 0.8320 | 0.9270 | 0 | 0 | 0 |
| 'Fbxo30' | 0.664 | 0.771 | 0.872 | 0.897 | 0.757 | 0.681 | 0.975 | 0.784 | 0.857 | 0.649 | 0.644 | 0.656 | 0.437 | 0.922 | 0.763 | 0.488 | 0.558 | 0.674 | 0.319 | 0.0114 | 0.0690 | 0 | 0 | 0 |
| 'Fbxo31' | 7.840 | 7.560 | 8.894 | 7.350 | 7.709 | 7.948 | 7.239 | 8.116 | 7.318 | 7.562 | 6.541 | 7.474 | 6.749 | 6.663 | 7.406 | 7.662 | 6.594 | 7.147 | 0.126 | 0.0028 | 0.0240 | 1 | 0 | 0 |
| 'Fbxo32' | 5.094 | 5.251 | 4.924 | 4.257 | 5.382 | 5.061 | 4.787 | 5.180 | 5.636 | 5.314 | 5.349 | 5.092 | 4.015 | 6.000 | 5.262 | 6.112 | 5.419 | 4.876 | -0.064 | 0.3059 | 0.5638 | 0 | 0 | 0 |
| 'Fbxo33' | 22.266 | 20.029 | 15.587 | 20.303 | 18.905 | 15.940 | 19.360 | 18.861 | 15.009 | 19.696 | 20.081 | 17.186 | 18.787 | 21.723 | 17.134 | 22.211 | 20.105 | 18.080 | -0.090 | 0.0350 | 0.1496 | 0 | 0 | 0 |
| 'Fbxo34' | 3.581 | 4.113 | 3.954 | 4.138 | 4.578 | 3.676 | 4.441 | 4.315 | 3.120 | 3.368 | 4.128 | 3.420 | 3.878 | 3.878 | 3.567 | 3.515 | 3.751 | 3.532 | 0.104 | 0.1182 | 0.3272 | 0 | 0 | 0 |
| 'Fbxo38' | 26.977 | 25.889 | 25.407 | 27.559 | 25.538 | 26.417 | 25.375 | 27.750 | 27.373 | 26.309 | 27.973 | 28.165 | 25.027 | 27.166 | 25.388 | 26.556 | 26.449 | 26.770 | -0.019 | 0.5392 | 0.7570 | 0 | 0 | 0 |
| 'Fbxo4' | 6.366 | 7.646 | 7.126 | 6.322 | 7.655 | 7.500 | 7.114 | 8.390 | 6.896 | 7.861 | 7.703 | 6.930 | 7.820 | 8.237 | 7.302 | 7.997 | 7.491 | 6.560 | -0.066 | 0.2210 | 0.4683 | 0 | 0 | 0 |
| 'Fbxo42' | 10.381 | 9.890 | 9.417 | 9.571 | 8.647 | 9.030 | 8.989 | 8.642 | 9.644 | 8.251 | 9.018 | 10.564 | 8.672 | 8.891 | 9.596 | 8.563 | 8.882 | 10.275 | 0.015 | 0.7521 | 0.8895 | 0 | 0 | 0 |
| 'Fbxo44' | 1.115 | 1.520 | 1.429 | 1.353 | 1.258 | 1.384 | 1.398 | 1.364 | 1.154 | 0.990 | 0.957 | 0.902 | 1.342 | 1.049 | 1.023 | 1.040 | 0.897 | 0.827 | 0.401 | 0.0000 | 0.0001 | 1 | 1 | 0 |
| 'Fbxo45' | 7.659 | 7.871 | 7.490 | 7.854 | 8.476 | 7.622 | 7.075 | 7.688 | 7.435 | 7.218 | 7.503 | 7.876 | 7.457 | 9.219 | 8.375 | 7.913 | 8.394 | 8.655 | -0.081 | 0.0823 | 0.2594 | 0 | 0 | 0 |
| 'Fbxo46' | 6.518 | 6.697 | 7.246 | 6.592 | 6.670 | 7.156 | 6.323 | 6.046 | 7.342 | 6.426 | 6.795 | 8.001 | 6.149 | 6.473 | 8.089 | 7.383 | 7.010 | 8.230 | -0.101 | 0.0125 | 0.0735 | 0 | 0 | 0 |
| 'Fbxo5' | 10.227 | 11.117 | 13.072 | 12.161 | 11.547 | 12.182 | 13.909 | 12.024 | 15.353 | 8.101 | 10.077 | 12.810 | 12.054 | 12.300 | 14.386 | 9.708 | 11.427 | 12.613 | 0.101 | 0.1799 | 0.4151 | 0 | 0 | 0 |
| 'Fbxo6' | 5.636 | 6.270 | 5.653 | 4.872 | 6.146 | 6.236 | 5.089 | 5.801 | 5.242 | 6.023 | 5.089 | 5.463 | 5.110 | 5.190 | 5.418 | 5.817 | 4.644 | 4.915 | 0.086 | 0.1413 | 0.3636 | 0 | 0 | 0 |
| 'Fbxo7' | 3.238 | 3.164 | 3.286 | 3.375 | 3.275 | 3.286 | 3.256 | 3.261 | 3.079 | 3.420 | 3.058 | 3.374 | 3.378 | 3.439 | 3.246 | 3.146 | 3.262 | 3.234 | -0.029 | 0.3824 | 0.6330 | 0 | 0 | 0 |
| 'Fbxo8' | 5.669 | 5.457 | 5.351 | 5.522 | 5.402 | 4.953 | 6.106 | 5.300 | 5.435 | 5.791 | 5.454 | 5.485 | 5.843 | 5.393 | 5.615 | 5.014 | 5.630 | 5.121 | -0.020 | 0.6833 | 0.8498 | 0 | 0 | 0 |
| 'Fbxo9' | 8.538 | 10.208 | 9.581 | 10.141 | 9.702 | 10.001 | 9.247 | 10.029 | 9.236 | 9.028 | 8.896 | 9.888 | 8.852 | 9.547 | 9.702 | 8.355 | 9.700 | 9.790 | 0.034 | 0.3637 | 0.6157 | 0 | 0 | 0 |
| 'Fbxw11' | 14.798 | 14.629 | 13.374 | 13.800 | 14.212 | 14.216 | 13.646 | 15.693 | 13.570 | 15.575 | 14.193 | 14.159 | 14.069 | 15.515 | 14.340 | 14.880 | 14.254 | 14.034 | -0.044 | 0.1559 | 0.3849 | 0 | 0 | 0 |
| 'Fbxw17' | 2.315 | 2.282 | 2.616 | 2.371 | 2.469 | 2.432 | 2.602 | 2.395 | 2.629 | 2.689 | 2.399 | 2.258 | 2.354 | 2.246 | 2.267 | 2.348 | 2.392 | 1.932 | 0.079 | 0.1314 | 0.3484 | 0 | 0 | 0 |
| 'Fbxw2' | 5.309 | 6.265 | 5.380 | 5.311 | 6.094 | 5.520 | 5.680 | 6.427 | 5.332 | 5.874 | 5.703 | 5.542 | 5.867 | 5.960 | 5.511 | 5.993 | 5.607 | 5.476 | -0.019 | 0.5275 | 0.7486 | 0 | 0 | 0 |
| 'Fbxw4' | 11.063 | 9.695 | 12.519 | 10.110 | 9.388 | 12.806 | 9.001 | 10.997 | 11.921 | 12.105 | 9.574 | 11.084 | 10.873 | 9.762 | 10.348 | 11.641 | 9.808 | 11.460 | 0.004 | 0.9444 | 0.9789 | 0 | 0 | 0 |
| 'Fbxw5' | 12.044 | 13.680 | 13.483 | 10.976 | 12.805 | 14.088 | 11.440 | 12.852 | 13.852 | 13.869 | 12.009 | 13.123 | 13.089 | 11.729 | 12.456 | 12.951 | 12.492 | 12.446 | 0.001 | 0.9728 | 0.9906 | 0 | 0 | 0 |
| 'Fbxw7' | 2.939 | 2.816 | 2.983 | 3.112 | 2.998 | 2.900 | 3.190 | 2.777 | 2.834 | 2.994 | 3.009 | 3.260 | 2.734 | 2.863 | 2.964 | 2.807 | 3.382 | 3.442 | -0.062 | 0.2233 | 0.4710 | 0 | 0 | 0 |
| 'Fbxw8' | 17.680 | 19.139 | 19.252 | 18.441 | 18.121 | 21.157 | 18.959 | 19.398 | 20.472 | 19.505 | 17.017 | 17.737 | 18.211 | 17.313 | 18.251 | 19.164 | 15.978 | 16.885 | 0.100 | 0.0061 | 0.0436 | 1 | 0 | 0 |
| 'Fbxw9' | 1.349 | 1.612 | 1.479 | 1.288 | 1.677 | 1.911 | 1.225 | 1.624 | 1.646 | 1.494 | 1.284 | 1.385 | 1.543 | 1.670 | 1.362 | 1.741 | 1.288 | 1.193 | 0.085 | 0.3268 | 0.5848 | 0 | 0 | 0 |
| 'Fcf1' | 24.663 | 28.331 | 25.540 | 23.896 | 28.696 | 24.535 | 26.537 | 27.922 | 27.334 | 22.324 | 25.806 | 24.327 | 27.286 | 26.174 | 24.183 | 30.171 | 27.879 | 24.831 | 0.018 | 0.7131 | 0.8669 | 0 | 0 | 0 |
| 'Fcho1' | 27.779 | 25.106 | 32.339 | 28.099 | 24.943 | 31.099 | 29.015 | 24.709 | 32.033 | 28.224 | 25.664 | 32.123 | 30.102 | 25.183 | 30.008 | 25.760 | 26.885 | 30.670 | -0.011 | 0.6981 | 0.8585 | 0 | 0 | 0 |
| 'Fcho2' | 9.227 | 9.625 | 7.570 | 8.939 | 8.975 | 7.164 | 8.647 | 8.716 | 7.576 | 8.417 | 9.305 | 9.662 | 8.690 | 9.247 | 8.945 | 7.742 | 10.760 | 10.026 | -0.131 | 0.0268 | 0.1260 | 0 | 0 | 0 |
| 'Fchsd1' | 3.475 | 3.181 | 3.703 | 2.962 | 3.384 | 3.596 | 2.900 | 3.279 | 3.732 | 4.218 | 3.248 | 3.783 | 3.270 | 3.017 | 3.511 | 3.729 | 3.176 | 3.692 | -0.073 | 0.1867 | 0.4225 | 0 | 0 | 0 |
| 'Fchsd2' | 4.758 | 3.995 | 4.637 | 4.907 | 4.051 | 4.288 | 4.356 | 4.511 | 4.773 | 4.560 | 4.213 | 4.890 | 4.001 | 3.867 | 4.237 | 3.970 | 3.982 | 4.587 | 0.061 | 0.1833 | 0.4191 | 0 | 0 | 0 |
| 'Fdft1' | 30.242 | 28.806 | 28.747 | 30.552 | 27.575 | 29.864 | 28.324 | 26.825 | 29.628 | 26.963 | 32.538 | 26.766 | 29.669 | 30.190 | 30.135 | 26.856 | 29.362 | 25.093 | 0.010 | 0.8314 | 0.9267 | 0 | 0 | 0 |
| 'Fdps' | 4.919 | 5.211 | 5.030 | 5.442 | 5.053 | 5.358 | 5.308 | 4.501 | 5.223 | 4.349 | 5.677 | 5.504 | 5.597 | 5.299 | 6.122 | 4.768 | 4.828 | 5.087 | -0.050 | 0.3819 | 0.6330 | 0 | 0 | 0 |
| 'Fdx1' | 16.905 | 20.776 | 18.338 | 17.438 | 21.997 | 16.993 | 17.178 | 19.850 | 19.186 | 14.705 | 16.002 | 16.527 | 15.782 | 19.118 | 18.214 | 15.215 | 18.619 | 14.865 | 0.163 | 0.0044 | 0.0339 | 1 | 0 | 0 |
| 'Fdx1l' | 29.142 | 24.981 | 31.119 | 24.752 | 28.990 | 29.317 | 25.871 | 29.233 | 28.593 | 26.611 | 24.162 | 28.747 | 28.193 | 23.870 | 30.632 | 29.488 | 27.095 | 27.324 | 0.029 | 0.5756 | 0.7807 | 0 | 0 | 0 |
| 'Fdxacb1' | 0.633 | 0.914 | 1.062 | 0.777 | 1.071 | 0.912 | 0.684 | 0.707 | 1.098 | 1.153 | 0.688 | 0.877 | 0.924 | 0.789 | 0.617 | 0.837 | 0.901 | 0.694 | 0.059 | 0.6611 | 0.8362 | 0 | 0 | 0 |
| 'Fdxr' | 10.827 | 10.013 | 11.723 | 10.755 | 10.046 | 12.717 | 10.554 | 10.645 | 11.186 | 11.626 | 10.429 | 12.122 | 11.807 | 10.553 | 11.607 | 10.309 | 10.463 | 11.061 | -0.032 | 0.3768 | 0.6285 | 0 | 0 | 0 |
| 'Fech' | 10.614 | 11.818 | 10.814 | 10.072 | 12.192 | 12.033 | 9.615 | 12.972 | 11.973 | 12.652 | 11.534 | 10.715 | 12.660 | 11.576 | 10.594 | 12.623 | 10.331 | 9.256 | -0.003 | 0.9583 | 0.9847 | 0 | 0 | 0 |
| 'Fem1b' | 9.903 | 9.843 | 10.505 | 10.890 | 9.808 | 10.272 | 11.291 | 9.806 | 10.688 | 8.937 | 11.103 | 13.167 | 9.866 | 10.590 | 11.882 | 9.612 | 11.215 | 13.656 | -0.109 | 0.0591 | 0.2099 | 0 | 0 | 0 |
| 'Fem1c' | 11.925 | 11.878 | 10.489 | 11.091 | 11.250 | 10.470 | 12.030 | 11.125 | 10.277 | 10.549 | 11.591 | 11.658 | 10.414 | 11.511 | 10.932 | 11.813 | 10.443 | 10.643 | 0.000 | 0.9974 | 0.9994 | 0 | 0 | 0 |
| 'Fen1' | 9.145 | 8.857 | 10.307 | 9.453 | 8.221 | 10.227 | 10.072 | 8.402 | 10.273 | 8.006 | 8.551 | 9.643 | 10.272 | 8.568 | 10.379 | 9.370 | 9.290 | 10.006 | 0.003 | 0.9472 | 0.9796 | 0 | 0 | 0 |
| 'Fer1l5' | 0.555 | 0.542 | 0.694 | 0.606 | 0.547 | 0.607 | 0.761 | 0.581 | 0.631 | 0.555 | 0.553 | 0.962 | 0.568 | 0.614 | 0.717 | 0.603 | 0.650 | 0.968 | -0.181 | 0.0911 | 0.2785 | 0 | 0 | 0 |
| 'Fermt3' | 204.089 | 195.485 | 214.478 | 192.624 | 195.974 | 224.889 | 205.247 | 190.816 | 211.155 | 200.007 | 188.595 | 216.880 | 208.053 | 187.082 | 211.256 | 198.795 | 185.439 | 217.935 | 0.006 | 0.7544 | 0.8906 | 0 | 0 | 0 |
| 'Fez2' | 11.540 | 13.555 | 11.660 | 11.222 | 11.791 | 12.376 | 11.419 | 13.692 | 13.122 | 10.902 | 10.562 | 9.305 | 10.914 | 10.732 | 9.498 | 11.833 | 11.421 | 9.776 | 0.216 | 0.0001 | 0.0011 | 1 | 0 | 0 |
| 'Fgd1' | 0.821 | 0.656 | 0.872 | 0.666 | 0.955 | 1.042 | 1.138 | 0.810 | 0.793 | 0.805 | 0.699 | 0.754 | 1.025 | 0.765 | 0.902 | 1.059 | 0.571 | 0.633 | 0.110 | 0.3588 | 0.6113 | 0 | 0 | 0 |
| 'Fgd3' | 21.502 | 18.793 | 20.364 | 21.550 | 19.080 | 19.335 | 19.768 | 18.667 | 20.913 | 19.191 | 20.332 | 23.846 | 18.115 | 19.838 | 21.850 | 18.471 | 19.712 | 23.401 | -0.046 | 0.3297 | 0.5877 | 0 | 0 | 0 |
| 'Fgd6' | 0.511 | 0.533 | 0.592 | 0.476 | 0.664 | 0.526 | 0.589 | 0.673 | 0.763 | 0.578 | 0.781 | 0.619 | 0.560 | 0.737 | 0.538 | 0.427 | 0.622 | 0.559 | -0.029 | 0.7762 | 0.9001 | 0 | 0 | 0 |
| 'Fgf13' | 6.179 | 6.192 | 6.634 | 5.872 | 4.973 | 6.138 | 5.930 | 6.078 | 6.714 | 7.777 | 7.376 | 8.134 | 6.667 | 7.591 | 8.082 | 7.309 | 6.865 | 8.148 | -0.321 | 0.0000 | 0.0000 | 1 | 0 | 0 |
| 'Fgf2' | 3.932 | 5.556 | 4.090 | 4.433 | 5.089 | 4.227 | 5.365 | 4.740 | 4.434 | 4.018 | 4.853 | 5.038 | 3.935 | 4.692 | 4.816 | 4.169 | 5.852 | 4.544 | -0.016 | 0.8033 | 0.9117 | 0 | 0 | 0 |
| 'Fgfr1op' | 10.692 | 12.196 | 10.946 | 11.768 | 12.147 | 10.345 | 11.348 | 11.963 | 11.076 | 12.072 | 13.533 | 11.192 | 12.547 | 12.959 | 11.152 | 11.267 | 13.257 | 11.019 | -0.098 | 0.0017 | 0.0157 | 1 | 0 | 0 |
| 'Fgfr1op2' | 21.181 | 24.717 | 22.568 | 22.049 | 24.098 | 22.055 | 23.503 | 22.475 | 22.011 | 20.291 | 22.554 | 21.874 | 22.758 | 22.573 | 21.062 | 22.550 | 23.649 | 20.568 | 0.037 | 0.1898 | 0.4262 | 0 | 0 | 0 |
| 'Fh1' | 15.074 | 17.285 | 17.887 | 17.192 | 17.567 | 18.080 | 17.485 | 16.424 | 16.328 | 15.271 | 14.352 | 16.214 | 17.425 | 16.043 | 15.869 | 15.553 | 15.735 | 16.419 | 0.089 | 0.0167 | 0.0909 | 0 | 0 | 0 |
| 'Fhdc1' | 1.584 | 1.097 | 1.423 | 1.377 | 1.275 | 1.424 | 1.164 | 1.258 | 1.268 | 1.578 | 1.684 | 1.967 | 1.627 | 1.579 | 1.707 | 1.575 | 1.768 | 2.056 | -0.397 | 0.0000 | 0.0000 | 1 | 1 | 0 |
| 'Fhl3' | 18.924 | 20.898 | 21.145 | 20.192 | 19.360 | 23.173 | 19.110 | 20.369 | 22.224 | 21.372 | 17.824 | 21.586 | 20.536 | 19.942 | 22.191 | 18.990 | 18.844 | 20.443 | 0.018 | 0.6523 | 0.8310 | 0 | 0 | 0 |
| 'Fhod1' | 6.753 | 5.589 | 7.273 | 6.878 | 5.473 | 6.540 | 6.620 | 6.351 | 7.435 | 5.712 | 5.424 | 7.861 | 6.368 | 5.673 | 7.341 | 5.948 | 6.053 | 8.235 | 0.002 | 0.9726 | 0.9905 | 0 | 0 | 0 |
| 'Fibp' | 33.547 | 39.818 | 38.754 | 33.192 | 40.466 | 42.044 | 34.224 | 41.249 | 41.465 | 37.945 | 31.255 | 34.755 | 41.065 | 36.835 | 34.567 | 40.374 | 33.579 | 32.750 | 0.084 | 0.1579 | 0.3880 | 0 | 0 | 0 |
| 'Ficd' | 4.908 | 6.189 | 5.792 | 4.889 | 5.866 | 5.832 | 4.650 | 5.528 | 5.717 | 5.894 | 4.904 | 5.520 | 5.351 | 5.797 | 4.972 | 5.775 | 5.276 | 5.773 | -0.008 | 0.8980 | 0.9593 | 0 | 0 | 0 |
| 'Fig4' | 8.264 | 10.291 | 10.412 | 9.518 | 11.147 | 10.645 | 10.333 | 10.806 | 10.722 | 10.485 | 10.158 | 9.835 | 10.402 | 10.607 | 10.035 | 9.838 | 10.317 | 9.206 | 0.010 | 0.8325 | 0.9270 | 0 | 0 | 0 |
| 'Fignl1' | 2.419 | 2.317 | 3.447 | 3.322 | 2.722 | 3.307 | 3.315 | 2.666 | 3.321 | 2.183 | 2.630 | 2.749 | 3.242 | 2.463 | 2.936 | 2.170 | 2.691 | 2.728 | 0.157 | 0.0441 | 0.1739 | 0 | 0 | 0 |
| 'Filip1l' | 9.578 | 12.035 | 10.500 | 9.821 | 11.981 | 12.298 | 10.818 | 11.237 | 11.338 | 10.494 | 11.060 | 13.747 | 8.730 | 10.925 | 13.145 | 12.000 | 10.515 | 14.570 | -0.083 | 0.1726 | 0.4064 | 0 | 0 | 0 |
| 'Fip1l1' | 10.514 | 9.954 | 9.879 | 10.433 | 9.675 | 10.011 | 10.211 | 10.008 | 9.820 | 10.504 | 9.783 | 10.479 | 10.133 | 10.057 | 10.259 | 9.622 | 10.105 | 10.038 | -0.019 | 0.4063 | 0.6556 | 0 | 0 | 0 |
| 'Fis1' | 37.696 | 45.525 | 35.318 | 37.546 | 43.128 | 42.248 | 36.453 | 42.830 | 39.777 | 37.511 | 37.313 | 37.968 | 38.353 | 41.732 | 39.999 | 43.687 | 39.581 | 38.054 | 0.012 | 0.7733 | 0.8993 | 0 | 0 | 0 |
| 'Fitm2' | 1.272 | 1.839 | 1.668 | 1.391 | 1.610 | 1.379 | 1.120 | 1.454 | 1.742 | 1.052 | 1.290 | 1.717 | 1.073 | 1.278 | 1.521 | 1.642 | 1.458 | 1.765 | 0.057 | 0.5691 | 0.7768 | 0 | 0 | 0 |
| 'Fiz1' | 16.267 | 16.915 | 16.732 | 13.556 | 15.897 | 18.395 | 14.547 | 16.132 | 16.999 | 14.923 | 14.029 | 16.648 | 15.328 | 15.065 | 16.662 | 16.112 | 12.715 | 16.369 | 0.068 | 0.1419 | 0.3648 | 0 | 0 | 0 |
| 'Fkbp15' | 10.056 | 9.035 | 10.963 | 9.710 | 9.893 | 11.280 | 9.478 | 10.099 | 10.459 | 10.739 | 10.147 | 10.760 | 10.426 | 9.802 | 9.776 | 10.134 | 9.647 | 9.340 | -0.006 | 0.8641 | 0.9410 | 0 | 0 | 0 |
| 'Fkbp1a' | 32.027 | 34.931 | 33.773 | 33.091 | 35.420 | 33.051 | 33.455 | 34.253 | 33.513 | 33.573 | 33.863 | 34.225 | 32.617 | 34.382 | 35.069 | 33.297 | 34.349 | 35.542 | -0.028 | 0.1860 | 0.4217 | 0 | 0 | 0 |
| 'Fkbp2' | 37.075 | 42.932 | 41.676 | 43.622 | 44.073 | 45.020 | 39.579 | 43.733 | 43.231 | 39.534 | 36.724 | 37.342 | 45.073 | 41.466 | 38.819 | 42.968 | 41.628 | 35.304 | 0.079 | 0.1039 | 0.3023 | 0 | 0 | 0 |
| 'Fkbp3' | 75.746 | 85.994 | 76.760 | 85.719 | 89.012 | 78.710 | 90.306 | 85.113 | 80.850 | 74.457 | 70.251 | 74.891 | 80.560 | 75.582 | 69.879 | 78.794 | 81.180 | 72.378 | 0.127 | 0.0003 | 0.0035 | 1 | 0 | 0 |
| 'Fkbp4' | 20.753 | 16.371 | 21.402 | 21.643 | 18.733 | 21.891 | 19.676 | 18.315 | 19.303 | 17.662 | 16.772 | 28.207 | 18.735 | 19.300 | 25.388 | 19.584 | 20.523 | 29.866 | -0.129 | 0.0599 | 0.2110 | 0 | 0 | 0 |
| 'Fkbp5' | 25.552 | 28.760 | 26.714 | 28.743 | 25.953 | 26.784 | 25.079 | 26.573 | 25.712 | 24.039 | 23.872 | 27.110 | 25.909 | 24.443 | 28.133 | 33.409 | 24.171 | 31.383 | -0.023 | 0.6919 | 0.8552 | 0 | 0 | 0 |
| 'Fkbp8' | 29.174 | 28.429 | 33.206 | 26.580 | 28.910 | 35.077 | 27.845 | 28.028 | 32.779 | 29.849 | 24.983 | 33.887 | 30.724 | 28.014 | 32.842 | 32.133 | 26.395 | 32.976 | -0.019 | 0.5737 | 0.7796 | 0 | 0 | 0 |
| 'Fkrp' | 7.700 | 7.741 | 8.096 | 7.193 | 8.555 | 8.250 | 7.090 | 7.142 | 7.944 | 7.017 | 7.393 | 9.255 | 7.372 | 8.233 | 8.952 | 7.994 | 7.144 | 9.393 | -0.071 | 0.1211 | 0.3319 | 0 | 0 | 0 |
| 'Fktn' | 1.112 | 1.216 | 1.367 | 1.268 | 1.368 | 1.461 | 1.395 | 1.490 | 1.436 | 1.467 | 1.382 | 1.326 | 1.325 | 1.131 | 1.258 | 1.270 | 1.579 | 1.274 | 0.005 | 0.9385 | 0.9764 | 0 | 0 | 0 |
| 'Flad1' | 7.755 | 8.087 | 9.086 | 8.660 | 8.912 | 8.826 | 7.661 | 8.031 | 9.593 | 8.697 | 7.600 | 9.117 | 8.067 | 8.057 | 9.480 | 7.709 | 8.418 | 8.741 | 0.002 | 0.9624 | 0.9866 | 0 | 0 | 0 |
| 'Flcn' | 10.393 | 9.831 | 10.389 | 9.071 | 9.446 | 10.608 | 9.744 | 9.846 | 10.264 | 10.978 | 11.378 | 13.323 | 10.526 | 10.914 | 12.691 | 10.930 | 11.891 | 13.769 | -0.256 | 0.0000 | 0.0000 | 1 | 0 | 0 |
| 'Fli1' | 15.953 | 14.276 | 15.394 | 16.700 | 13.540 | 13.976 | 16.476 | 15.884 | 15.137 | 15.227 | 16.906 | 16.431 | 12.176 | 13.153 | 15.368 | 12.799 | 14.627 | 16.254 | 0.037 | 0.5728 | 0.7794 | 0 | 0 | 0 |
| 'Flii' | 49.491 | 46.882 | 55.519 | 50.731 | 48.173 | 54.601 | 49.173 | 48.349 | 52.519 | 51.894 | 51.216 | 54.035 | 54.198 | 46.605 | 51.830 | 48.816 | 50.121 | 52.747 | -0.030 | 0.2434 | 0.4935 | 0 | 0 | 0 |
| 'Flna' | 43.355 | 37.393 | 46.036 | 45.367 | 41.636 | 44.990 | 42.493 | 40.696 | 44.715 | 48.288 | 48.755 | 47.522 | 45.178 | 46.125 | 51.029 | 43.736 | 46.604 | 50.604 | -0.158 | 0.0000 | 0.0002 | 1 | 0 | 0 |
| 'Flnb' | 13.456 | 11.202 | 14.073 | 13.386 | 11.747 | 14.071 | 12.726 | 12.470 | 13.816 | 14.303 | 15.120 | 13.814 | 13.766 | 13.432 | 13.921 | 13.333 | 14.143 | 14.489 | -0.124 | 0.0024 | 0.0207 | 1 | 0 | 0 |
| 'Flot1' | 7.339 | 7.169 | 8.200 | 6.587 | 7.077 | 7.690 | 6.509 | 8.165 | 7.365 | 6.746 | 6.781 | 7.262 | 6.334 | 6.876 | 7.105 | 7.039 | 6.368 | 6.833 | 0.102 | 0.0152 | 0.0848 | 0 | 0 | 0 |
| 'Flot2' | 9.631 | 10.124 | 9.632 | 9.234 | 10.448 | 11.199 | 9.340 | 9.981 | 11.052 | 10.078 | 9.033 | 9.915 | 9.790 | 8.968 | 10.363 | 10.006 | 7.678 | 10.105 | 0.068 | 0.1420 | 0.3649 | 0 | 0 | 0 |
| 'Flrt2' | 0.449 | 0.578 | 0.666 | 0.428 | 0.500 | 0.553 | 0.609 | 0.595 | 0.523 | 0.470 | 0.564 | 0.557 | 0.159 | 0.534 | 0.412 | 0.349 | 0.560 | 0.405 | 0.302 | 0.0362 | 0.1530 | 0 | 0 | 0 |
| 'Flywch1' | 9.924 | 9.907 | 10.424 | 9.494 | 10.493 | 11.251 | 10.164 | 9.286 | 11.206 | 10.890 | 9.226 | 10.470 | 10.579 | 9.680 | 9.675 | 9.799 | 8.978 | 9.833 | 0.037 | 0.3379 | 0.5955 | 0 | 0 | 0 |
| 'Fmc1' | 6.931 | 7.052 | 8.303 | 8.194 | 7.035 | 6.943 | 7.201 | 9.527 | 6.416 | 7.016 | 6.273 | 8.066 | 8.696 | 7.431 | 7.451 | 7.278 | 7.048 | 6.137 | 0.033 | 0.7162 | 0.8688 | 0 | 0 | 0 |
| 'Fmnl1' | 85.888 | 77.400 | 85.749 | 86.688 | 77.801 | 87.296 | 82.975 | 77.461 | 86.497 | 81.075 | 80.643 | 94.908 | 86.270 | 78.740 | 92.809 | 80.305 | 82.380 | 100.160 | -0.064 | 0.0373 | 0.1565 | 0 | 0 | 0 |
| 'Fmnl3' | 30.042 | 30.845 | 32.177 | 28.232 | 28.811 | 32.603 | 29.099 | 30.171 | 32.070 | 33.081 | 28.108 | 31.540 | 30.871 | 30.113 | 31.216 | 29.865 | 28.590 | 30.683 | -0.010 | 0.7153 | 0.8687 | 0 | 0 | 0 |
| 'Fmo5' | 1.058 | 1.776 | 1.303 | 1.741 | 1.236 | 1.518 | 1.527 | 1.220 | 1.316 | 1.889 | 1.729 | 1.551 | 1.364 | 1.465 | 1.378 | 1.605 | 1.766 | 1.430 | -0.175 | 0.1182 | 0.3272 | 0 | 0 | 0 |
| 'Fmr1' | 6.223 | 6.926 | 6.724 | 6.569 | 6.772 | 6.057 | 6.447 | 6.942 | 6.383 | 6.362 | 7.382 | 6.286 | 6.681 | 6.709 | 6.113 | 6.461 | 7.059 | 6.132 | -0.012 | 0.6905 | 0.8542 | 0 | 0 | 0 |
| 'Fn3krp' | 2.325 | 3.114 | 3.806 | 3.886 | 3.731 | 3.362 | 3.133 | 4.002 | 3.266 | 2.780 | 3.657 | 3.606 | 3.076 | 3.938 | 3.740 | 3.011 | 3.293 | 3.629 | -0.016 | 0.8468 | 0.9343 | 0 | 0 | 0 |
| 'Fnbp1' | 27.170 | 26.589 | 26.609 | 25.800 | 26.648 | 26.757 | 25.900 | 26.379 | 27.809 | 27.434 | 26.251 | 26.199 | 25.337 | 24.988 | 26.328 | 27.250 | 25.262 | 27.409 | 0.009 | 0.7071 | 0.8643 | 0 | 0 | 0 |
| 'Fnbp4' | 22.757 | 20.322 | 21.901 | 20.969 | 19.061 | 21.287 | 21.640 | 20.617 | 21.720 | 21.476 | 22.100 | 23.303 | 21.892 | 20.959 | 21.692 | 20.649 | 21.912 | 22.036 | -0.054 | 0.0690 | 0.2308 | 0 | 0 | 0 |
| 'Fndc3a' | 8.576 | 7.928 | 8.199 | 8.059 | 8.397 | 7.798 | 9.262 | 9.042 | 8.305 | 8.709 | 8.746 | 9.021 | 8.161 | 8.462 | 8.114 | 8.663 | 9.262 | 8.757 | -0.055 | 0.1309 | 0.3475 | 0 | 0 | 0 |
| 'Fndc3b' | 0.623 | 0.538 | 0.475 | 0.470 | 0.598 | 0.529 | 0.465 | 0.613 | 0.484 | 0.388 | 0.405 | 0.413 | 0.378 | 0.544 | 0.224 | 0.364 | 0.445 | 0.370 | 0.435 | 0.0000 | 0.0005 | 1 | 1 | 0 |
| 'Fndc9' | 6.663 | 3.702 | 4.735 | 6.913 | 3.479 | 4.127 | 6.160 | 4.405 | 4.445 | 3.678 | 5.159 | 6.095 | 5.078 | 4.527 | 5.847 | 3.514 | 6.486 | 6.345 | -0.100 | 0.5508 | 0.7646 | 0 | 0 | 0 |
| 'Fnip1' | 4.777 | 4.606 | 4.520 | 4.623 | 4.848 | 3.931 | 4.074 | 4.962 | 4.552 | 5.254 | 5.102 | 4.931 | 4.546 | 4.742 | 4.414 | 4.762 | 4.470 | 4.573 | -0.077 | 0.1091 | 0.3110 | 0 | 0 | 0 |
| 'Fnip2' | 0.202 | 0.263 | 0.322 | 0.261 | 0.263 | 0.238 | 0.311 | 0.260 | 0.227 | 0.295 | 0.249 | 0.332 | 0.380 | 0.331 | 0.340 | 0.266 | 0.292 | 0.326 | -0.282 | 0.0096 | 0.0607 | 0 | 0 | 0 |
| 'Fnta' | 29.613 | 35.497 | 30.035 | 32.788 | 35.027 | 29.324 | 32.380 | 33.524 | 33.620 | 28.744 | 30.954 | 36.026 | 32.183 | 33.838 | 37.590 | 34.008 | 34.663 | 36.307 | -0.075 | 0.1251 | 0.3385 | 0 | 0 | 0 |
| 'Fntb' | 5.628 | 5.808 | 5.783 | 5.392 | 5.400 | 6.179 | 5.342 | 5.516 | 6.126 | 5.080 | 4.953 | 5.834 | 5.111 | 4.407 | 5.487 | 4.642 | 4.858 | 5.030 | 0.157 | 0.0001 | 0.0019 | 1 | 0 | 0 |
| 'Focad' | 2.775 | 2.410 | 2.567 | 2.968 | 2.722 | 2.327 | 2.650 | 2.419 | 2.560 | 2.774 | 2.602 | 2.565 | 2.862 | 2.217 | 2.380 | 2.307 | 2.611 | 2.344 | 0.032 | 0.5042 | 0.7311 | 0 | 0 | 0 |
| 'Fopnl' | 28.733 | 32.101 | 32.118 | 27.902 | 35.170 | 30.877 | 30.607 | 34.162 | 29.330 | 32.055 | 32.982 | 29.473 | 31.505 | 32.219 | 29.369 | 33.186 | 32.233 | 26.278 | 0.003 | 0.9443 | 0.9789 | 0 | 0 | 0 |
| 'Fos' | 391.557 | 114.829 | 154.971 | 383.153 | 139.696 | 98.682 | 405.099 | 190.012 | 103.700 | 252.320 | 201.644 | 74.408 | 226.183 | 156.070 | 68.221 | 146.264 | 206.324 | 63.013 | 0.451 | 0.0350 | 0.1496 | 0 | 0 | 0 |
| 'Fosb' | 222.165 | 31.290 | 54.109 | 179.938 | 43.071 | 24.263 | 185.601 | 70.932 | 25.988 | 146.021 | 68.698 | 10.096 | 132.405 | 44.844 | 10.092 | 51.958 | 67.765 | 8.217 | 0.787 | 0.0135 | 0.0778 | 0 | 0 | 0 |
| 'Fosl2' | 1.375 | 0.602 | 0.582 | 0.810 | 0.606 | 0.521 | 0.588 | 0.777 | 0.509 | 1.172 | 1.021 | 0.734 | 0.852 | 1.122 | 0.554 | 1.274 | 1.064 | 0.584 | -0.389 | 0.0033 | 0.0269 | 1 | 1 | 0 |
| 'Foxj2' | 7.675 | 7.677 | 7.324 | 6.813 | 6.784 | 8.076 | 6.959 | 7.256 | 7.767 | 7.593 | 7.085 | 8.570 | 7.242 | 7.579 | 8.229 | 8.584 | 7.229 | 8.054 | -0.090 | 0.0218 | 0.1089 | 0 | 0 | 0 |
| 'Foxk1' | 14.244 | 12.966 | 15.384 | 13.966 | 12.589 | 14.929 | 12.467 | 13.243 | 14.678 | 13.940 | 14.281 | 17.207 | 13.801 | 13.447 | 15.679 | 14.858 | 13.089 | 17.635 | -0.114 | 0.0017 | 0.0162 | 1 | 0 | 0 |
| 'Foxk2' | 16.289 | 13.993 | 15.098 | 16.838 | 13.592 | 15.671 | 15.643 | 14.141 | 15.351 | 14.305 | 15.894 | 18.470 | 15.432 | 16.204 | 18.103 | 15.740 | 15.946 | 19.993 | -0.145 | 0.0046 | 0.0350 | 1 | 0 | 0 |
| 'Foxm1' | 3.601 | 3.527 | 5.099 | 3.886 | 3.316 | 4.722 | 4.128 | 3.531 | 4.978 | 2.832 | 3.539 | 4.703 | 4.917 | 4.500 | 5.521 | 3.080 | 3.924 | 4.859 | -0.055 | 0.5022 | 0.7294 | 0 | 0 | 0 |
| 'Foxn2' | 16.810 | 16.616 | 15.933 | 16.905 | 16.171 | 14.488 | 16.285 | 16.722 | 15.270 | 15.139 | 18.496 | 17.622 | 14.798 | 17.279 | 16.488 | 14.845 | 18.835 | 17.460 | -0.064 | 0.1592 | 0.3895 | 0 | 0 | 0 |
| 'Foxn3' | 4.212 | 3.229 | 3.608 | 3.910 | 3.396 | 3.270 | 4.206 | 3.326 | 2.910 | 3.934 | 3.776 | 3.913 | 3.558 | 3.608 | 3.731 | 3.886 | 3.764 | 3.979 | -0.108 | 0.0584 | 0.2082 | 0 | 0 | 0 |
| 'Foxo1' | 23.706 | 24.088 | 23.475 | 21.644 | 23.900 | 24.293 | 21.669 | 23.865 | 23.203 | 25.065 | 23.609 | 24.365 | 23.305 | 23.955 | 24.405 | 26.911 | 23.694 | 24.465 | -0.076 | 0.0149 | 0.0833 | 0 | 0 | 0 |
| 'Foxo3' | 9.753 | 9.381 | 8.802 | 8.833 | 8.520 | 9.344 | 8.465 | 8.905 | 8.894 | 9.014 | 8.950 | 10.061 | 8.385 | 8.505 | 9.462 | 10.083 | 7.886 | 10.363 | -0.040 | 0.3824 | 0.6330 | 0 | 0 | 0 |
| 'Foxo4' | 14.084 | 14.573 | 15.225 | 13.374 | 14.596 | 15.165 | 11.916 | 14.351 | 14.608 | 15.120 | 14.818 | 16.264 | 14.881 | 15.659 | 15.149 | 16.084 | 13.368 | 16.201 | -0.115 | 0.0046 | 0.0352 | 1 | 0 | 0 |
| 'Foxp1' | 6.742 | 6.200 | 5.688 | 6.287 | 5.821 | 5.971 | 6.138 | 6.022 | 5.901 | 6.724 | 6.482 | 5.961 | 5.841 | 5.983 | 5.445 | 6.533 | 6.309 | 5.353 | -0.004 | 0.9024 | 0.9612 | 0 | 0 | 0 |
| 'Foxp3' | 3.243 | 2.106 | 3.473 | 2.875 | 3.241 | 4.063 | 4.114 | 3.417 | 3.765 | 1.848 | 0.798 | 4.270 | 2.539 | 0.931 | 3.461 | 2.730 | 0.977 | 3.389 | 0.716 | 0.0006 | 0.0067 | 1 | 1 | 0 |
| 'Foxp4' | 16.472 | 13.941 | 17.869 | 15.095 | 13.872 | 17.598 | 14.762 | 14.220 | 19.234 | 17.006 | 12.943 | 18.076 | 15.535 | 13.823 | 17.111 | 16.243 | 13.664 | 18.480 | -0.007 | 0.8313 | 0.9267 | 0 | 0 | 0 |
| 'Foxred1' | 2.873 | 3.064 | 3.157 | 2.862 | 3.062 | 3.198 | 2.912 | 2.949 | 3.461 | 2.766 | 2.903 | 3.420 | 3.007 | 3.125 | 3.193 | 2.918 | 3.149 | 3.199 | -0.022 | 0.5486 | 0.7624 | 0 | 0 | 0 |
| 'Foxred2' | 1.386 | 1.199 | 1.564 | 1.214 | 1.235 | 1.149 | 1.251 | 1.297 | 1.309 | 1.235 | 1.007 | 1.039 | 1.225 | 0.910 | 1.049 | 0.937 | 0.880 | 0.920 | 0.330 | 0.0000 | 0.0001 | 1 | 0 | 0 |
| 'Fpgs' | 1.854 | 1.853 | 2.020 | 1.668 | 1.951 | 1.903 | 1.938 | 1.916 | 1.985 | 2.121 | 1.634 | 1.760 | 2.005 | 1.411 | 1.761 | 1.891 | 1.422 | 1.782 | 0.110 | 0.0861 | 0.2671 | 0 | 0 | 0 |
| 'Fpgt' | 2.412 | 3.264 | 2.983 | 2.762 | 2.884 | 2.565 | 2.690 | 3.114 | 2.653 | 2.622 | 2.654 | 3.123 | 2.305 | 2.786 | 2.506 | 2.280 | 2.757 | 2.858 | 0.063 | 0.2774 | 0.5335 | 0 | 0 | 0 |

| | | | | | | | | | | | | | | | | | | | | | | | | |
|---|---|---|---|---|---|---|---|---|---|---|---|---|---|---|---|---|---|---|---|---|---|---|---|---|
| 'Fra10ac1' | 14.559 | 17.459 | 15.461 | 14.121 | 16.221 | 14.750 | 15.305 | 17.805 | 15.367 | 15.948 | 14.095 | 12.993 | 12.456 | 14.939 | 12.947 | 15.519 | 14.616 | 11.626 | 0.169 | 0.0023 | 0.0201 | 1 | 0 | 0 |
| 'Frat2' | 45.233 | 25.011 | 24.543 | 29.302 | 27.531 | 26.970 | 25.923 | 29.997 | 21.831 | 40.001 | 32.479 | 20.430 | 31.831 | 36.794 | 21.298 | 37.399 | 31.275 | 22.448 | -0.080 | 0.4769 | 0.7107 | 0 | 0 | 0 |
| 'Frg1' | 60.165 | 62.617 | 63.167 | 70.832 | 67.900 | 61.984 | 74.976 | 65.862 | 62.574 | 58.493 | 60.028 | 62.187 | 59.438 | 65.689 | 62.891 | 59.777 | 64.855 | 62.054 | 0.074 | 0.0768 | 0.2473 | 0 | 0 | 0 |
| 'Frmd4b' | 2.970 | 2.481 | 2.423 | 2.594 | 2.651 | 2.310 | 2.718 | 2.962 | 2.168 | 3.019 | 3.134 | 2.524 | 2.821 | 2.909 | 2.495 | 2.821 | 3.021 | 2.484 | -0.123 | 0.0022 | 0.0197 | 1 | 0 | 0 |
| 'Frmd6' | 1.851 | 1.609 | 1.697 | 1.987 | 1.713 | 1.884 | 1.651 | 1.868 | 1.948 | 1.748 | 1.573 | 2.050 | 1.678 | 1.736 | 1.573 | 1.790 | 1.945 | 1.973 | -0.002 | 0.9804 | 0.9926 | 0 | 0 | 0 |
| 'Frmd8' | 36.689 | 41.255 | 41.497 | 39.767 | 39.697 | 46.792 | 39.703 | 40.866 | 43.669 | 41.534 | 39.936 | 53.488 | 39.774 | 41.631 | 51.476 | 48.484 | 38.888 | 54.453 | -0.151 | 0.0006 | 0.0072 | 1 | 0 | 0 |
| 'Frrs1' | 1.220 | 1.555 | 1.771 | 1.491 | 1.469 | 1.263 | 1.442 | 1.247 | 1.272 | 1.685 | 1.395 | 1.307 | 1.571 | 1.497 | 1.355 | 1.577 | 1.373 | 1.203 | -0.028 | 0.7211 | 0.8716 | 0 | 0 | 0 |
| 'Frs2' | 8.675 | 9.279 | 8.410 | 8.316 | 8.750 | 8.262 | 8.524 | 9.716 | 9.053 | 8.190 | 7.856 | 8.850 | 8.471 | 9.558 | 7.997 | 8.899 | 8.648 | 8.762 | 0.020 | 0.6171 | 0.8087 | 0 | 0 | 0 |
| 'Fry' | 0.157 | 0.129 | 0.144 | 0.160 | 0.191 | 0.209 | 0.196 | 0.198 | 0.182 | 0.183 | 0.186 | 0.167 | 0.161 | 0.176 | 0.153 | 0.140 | 0.227 | 0.201 | -0.030 | 0.7760 | 0.9000 | 0 | 0 | 0 |
| 'Fryl' | 13.934 | 12.698 | 13.875 | 13.809 | 12.425 | 13.330 | 12.880 | 12.341 | 14.312 | 15.268 | 14.887 | 14.392 | 13.573 | 13.808 | 14.316 | 14.029 | 13.729 | 15.347 | -0.124 | 0.0003 | 0.0037 | 1 | 0 | 0 |
| 'Frzb' | 8.347 | 9.595 | 10.293 | 9.426 | 10.579 | 8.277 | 9.738 | 11.437 | 8.519 | 7.524 | 7.231 | 6.050 | 7.264 | 6.694 | 6.375 | 6.824 | 7.401 | 5.942 | 0.484 | 0.0000 | 0.0000 | 1 | 1 | 0 |
| 'Fth1' | 645.164 | 622.927 | 593.836 | 646.133 | 639.777 | 633.764 | 646.053 | 636.450 | 625.271 | 649.351 | 619.574 | 526.890 | 621.644 | 633.712 | 579.811 | 646.319 | 610.461 | 557.231 | 0.056 | 0.0463 | 0.1793 | 0 | 0 | 0 |
| 'Ftl1-ps1' | 170.855 | 132.442 | 128.585 | 170.145 | 133.209 | 130.021 | 181.985 | 135.891 | 121.395 | 164.077 | 153.299 | 121.994 | 167.325 | 148.188 | 131.150 | 148.287 | 162.888 | 126.246 | -0.038 | 0.3993 | 0.6482 | 0 | 0 | 0 |
| 'Fto' | 5.658 | 5.929 | 5.507 | 5.867 | 6.073 | 5.539 | 6.281 | 6.000 | 5.566 | 6.571 | 6.439 | 5.694 | 6.878 | 6.498 | 5.328 | 6.191 | 6.444 | 5.279 | -0.085 | 0.0057 | 0.0413 | 1 | 0 | 0 |
| 'Ftsj1' | 2.210 | 2.108 | 2.403 | 2.101 | 2.447 | 2.150 | 2.099 | 2.340 | 2.175 | 2.184 | 2.166 | 2.474 | 2.143 | 2.212 | 2.478 | 2.375 | 2.359 | 2.288 | -0.057 | 0.2814 | 0.5379 | 0 | 0 | 0 |
| 'Ftsj3' | 17.297 | 14.335 | 16.104 | 16.195 | 15.424 | 14.563 | 16.538 | 16.680 | 15.920 | 15.763 | 15.079 | 16.832 | 15.940 | 14.026 | 16.601 | 15.210 | 15.201 | 16.588 | 0.007 | 0.8472 | 0.9343 | 0 | 0 | 0 |
| 'Fubp1' | 5.863 | 5.556 | 5.686 | 6.031 | 5.232 | 5.185 | 5.904 | 5.366 | 5.518 | 4.918 | 5.436 | 6.744 | 5.247 | 5.536 | 5.984 | 5.100 | 6.249 | 6.521 | -0.046 | 0.3991 | 0.6482 | 0 | 0 | 0 |
| 'Fubp3' | 3.747 | 4.863 | 4.660 | 3.791 | 4.583 | 4.852 | 4.106 | 4.697 | 4.810 | 4.202 | 4.426 | 4.186 | 4.521 | 4.555 | 4.413 | 4.723 | 4.424 | 3.953 | 0.014 | 0.7641 | 0.8953 | 0 | 0 | 0 |
| 'Fuca1' | 13.973 | 15.360 | 14.911 | 13.778 | 14.245 | 15.682 | 12.837 | 14.728 | 14.596 | 14.028 | 13.189 | 14.071 | 15.590 | 14.575 | 14.987 | 15.070 | 13.613 | 14.138 | 0.001 | 0.9879 | 0.9964 | 0 | 0 | 0 |
| 'Fuca2' | 6.857 | 8.331 | 7.307 | 7.301 | 7.602 | 7.091 | 7.420 | 7.883 | 7.187 | 7.857 | 7.650 | 7.281 | 7.499 | 7.737 | 7.492 | 8.066 | 8.069 | 6.496 | -0.035 | 0.3103 | 0.5686 | 0 | 0 | 0 |
| 'Fuk' | 8.249 | 7.931 | 8.825 | 8.207 | 8.112 | 7.954 | 7.887 | 7.515 | 9.181 | 7.422 | 7.528 | 9.447 | 7.266 | 7.467 | 8.307 | 7.885 | 7.554 | 9.147 | 0.022 | 0.6224 | 0.8114 | 0 | 0 | 0 |
| 'Fundc1' | 16.206 | 19.423 | 17.201 | 18.119 | 18.656 | 16.857 | 19.025 | 19.068 | 18.120 | 19.101 | 16.543 | 16.657 | 16.689 | 19.008 | 17.130 | 18.754 | 18.947 | 16.448 | 0.021 | 0.6541 | 0.8324 | 0 | 0 | 0 |
| 'Fundc2' | 16.447 | 17.791 | 15.464 | 15.498 | 17.036 | 15.809 | 17.190 | 16.807 | 14.531 | 15.066 | 16.266 | 14.435 | 16.878 | 16.305 | 15.256 | 15.242 | 16.913 | 14.019 | 0.052 | 0.1352 | 0.3537 | 0 | 0 | 0 |
| 'Fuom' | 1.138 | 1.241 | 1.344 | 1.013 | 1.442 | 1.449 | 1.019 | 1.792 | 1.083 | 1.375 | 1.146 | 1.266 | 1.192 | 1.349 | 1.427 | 1.200 | 1.158 | 1.232 | 0.010 | 0.9121 | 0.9662 | 0 | 0 | 0 |
| 'Furin' | 7.332 | 7.021 | 8.776 | 7.276 | 7.574 | 8.013 | 6.406 | 8.006 | 8.402 | 7.558 | 7.251 | 8.142 | 7.364 | 7.151 | 7.670 | 7.513 | 6.709 | 7.955 | 0.021 | 0.5755 | 0.7807 | 0 | 0 | 0 |
| 'Fus' | 29.577 | 35.155 | 32.818 | 27.663 | 34.309 | 30.194 | 31.854 | 35.854 | 33.260 | 34.203 | 30.701 | 25.780 | 32.788 | 30.514 | 27.516 | 33.452 | 29.118 | 24.242 | 0.110 | 0.0421 | 0.1692 | 0 | 0 | 0 |
| 'Fut10' | 3.224 | 3.348 | 3.101 | 3.307 | 3.106 | 3.025 | 3.050 | 3.075 | 2.964 | 2.363 | 2.681 | 2.303 | 2.508 | 2.638 | 1.897 | 2.483 | 2.474 | 1.865 | 0.407 | 0.0000 | 0.0000 | 1 | 1 | 0 |
| 'Fut11' | 7.212 | 7.130 | 7.493 | 6.997 | 7.463 | 7.474 | 7.170 | 7.028 | 7.210 | 6.302 | 7.202 | 8.010 | 6.558 | 7.887 | 8.003 | 6.688 | 7.314 | 8.512 | -0.040 | 0.4122 | 0.6607 | 0 | 0 | 0 |
| 'Fut8' | 25.553 | 26.831 | 23.168 | 25.218 | 24.740 | 23.277 | 26.174 | 25.087 | 23.901 | 25.034 | 26.960 | 23.524 | 28.373 | 26.831 | 22.694 | 25.255 | 26.568 | 22.779 | -0.035 | 0.2247 | 0.4716 | 0 | 0 | 0 |
| 'Fuz' | 20.639 | 22.699 | 25.548 | 20.470 | 25.238 | 27.155 | 24.114 | 25.511 | 27.130 | 24.843 | 21.666 | 23.461 | 22.349 | 21.002 | 21.969 | 24.649 | 20.823 | 21.118 | 0.104 | 0.0546 | 0.2000 | 0 | 0 | 0 |
| 'Fv1' | 6.197 | 4.911 | 5.689 | 4.214 | 5.585 | 4.793 | 5.265 | 5.227 | 5.610 | 3.393 | 6.551 | 6.798 | 4.082 | 5.105 | 5.481 | 4.094 | 5.746 | 5.902 | 0.010 | 0.9239 | 0.9709 | 0 | 0 | 0 |
| 'Fxn' | 3.409 | 3.557 | 4.176 | 3.466 | 3.819 | 3.993 | 3.609 | 3.706 | 3.582 | 3.703 | 3.454 | 3.535 | 3.671 | 3.460 | 4.268 | 3.742 | 3.506 | 3.209 | 0.031 | 0.6267 | 0.8148 | 0 | 0 | 0 |
| 'Fxr1' | 9.912 | 10.158 | 9.997 | 10.828 | 10.834 | 9.189 | 10.414 | 10.202 | 9.161 | 9.579 | 10.478 | 11.294 | 9.531 | 10.313 | 10.126 | 9.031 | 10.216 | 11.565 | -0.034 | 0.4741 | 0.7094 | 0 | 0 | 0 |
| 'Fxr2' | 35.074 | 32.721 | 33.719 | 32.229 | 32.196 | 33.808 | 32.628 | 33.304 | 32.731 | 32.422 | 32.808 | 34.646 | 34.679 | 30.516 | 34.600 | 32.813 | 31.040 | 34.895 | -0.010 | 0.7034 | 0.8612 | 0 | 0 | 0 |
| 'Fxyd5' | 35.354 | 32.298 | 36.263 | 33.690 | 36.835 | 37.704 | 35.472 | 35.887 | 36.588 | 35.698 | 34.146 | 37.629 | 32.098 | 33.401 | 40.410 | 35.307 | 33.375 | 36.864 | -0.004 | 0.9104 | 0.9652 | 0 | 0 | 0 |
| 'Fyb' | 183.003 | 191.463 | 183.245 | 187.280 | 189.418 | 179.491 | 195.321 | 185.849 | 192.597 | 201.216 | 199.269 | 170.670 | 199.298 | 191.952 | 173.468 | 198.242 | 194.113 | 174.289 | -0.021 | 0.4474 | 0.6880 | 0 | 0 | 0 |
| 'Fyco1' | 7.086 | 7.110 | 7.714 | 6.808 | 6.930 | 7.487 | 7.083 | 7.117 | 7.768 | 7.673 | 7.863 | 7.159 | 7.195 | 6.978 | 7.226 | 7.914 | 7.440 | 6.931 | -0.035 | 0.3402 | 0.5970 | 0 | 0 | 0 |
| 'Fyn' | 172.814 | 183.188 | 168.679 | 172.242 | 180.217 | 169.867 | 167.659 | 182.396 | 167.007 | 186.565 | 199.580 | 161.197 | 180.466 | 190.090 | 177.101 | 186.081 | 188.692 | 171.317 | -0.079 | 0.0014 | 0.0140 | 1 | 0 | 0 |
| 'Fyttd1' | 5.913 | 6.922 | 6.122 | 5.908 | 6.676 | 5.786 | 6.805 | 6.850 | 5.815 | 6.153 | 6.213 | 6.040 | 6.271 | 7.086 | 5.798 | 6.033 | 6.665 | 6.029 | -0.001 | 0.9853 | 0.9949 | 0 | 0 | 0 |
| 'Fzd5' | 1.395 | 0.776 | 0.648 | 1.095 | 0.731 | 0.658 | 1.039 | 0.747 | 0.634 | 0.934 | 1.132 | 0.692 | 0.979 | 1.146 | 0.796 | 0.836 | 0.803 | 0.692 | -0.084 | 0.4490 | 0.6890 | 0 | 0 | 0 |
| 'Fzr1' | 11.277 | 11.414 | 13.195 | 10.192 | 11.055 | 13.391 | 11.139 | 11.260 | 13.314 | 10.656 | 10.106 | 13.096 | 12.202 | 11.407 | 13.268 | 11.816 | 10.645 | 12.850 | -0.008 | 0.8050 | 0.9125 | 0 | 0 | 0 |
| 'G2e3' | 2.441 | 2.683 | 2.768 | 2.671 | 2.696 | 2.633 | 2.948 | 2.382 | 2.719 | 1.794 | 2.731 | 3.287 | 2.336 | 2.981 | 2.885 | 2.478 | 3.079 | 3.052 | -0.043 | 0.5616 | 0.7712 | 0 | 0 | 0 |
| 'G3bp1' | 22.619 | 22.874 | 23.462 | 22.899 | 22.913 | 22.948 | 22.426 | 22.778 | 22.991 | 22.311 | 21.394 | 22.811 | 22.495 | 21.123 | 22.541 | 22.551 | 21.659 | 21.706 | 0.042 | 0.0100 | 0.0622 | 0 | 0 | 0 |
| 'G3bp2' | 17.445 | 19.438 | 17.275 | 18.133 | 19.192 | 17.800 | 17.904 | 18.854 | 17.376 | 17.797 | 18.611 | 17.952 | 18.037 | 19.213 | 18.132 | 17.593 | 18.793 | 17.466 | -0.014 | 0.4194 | 0.6664 | 0 | 0 | 0 |
| 'G6pc3' | 6.229 | 6.507 | 6.689 | 5.770 | 6.474 | 6.893 | 6.416 | 7.016 | 6.648 | 5.785 | 5.239 | 6.293 | 5.291 | 5.166 | 5.944 | 6.339 | 5.684 | 5.653 | 0.180 | 0.0001 | 0.0011 | 1 | 0 | 0 |
| 'G6pdx' | 13.161 | 13.383 | 13.516 | 13.925 | 13.001 | 14.453 | 13.072 | 13.788 | 12.933 | 13.134 | 13.487 | 14.108 | 14.075 | 13.229 | 14.024 | 13.216 | 13.255 | 13.196 | -0.017 | 0.5952 | 0.7939 | 0 | 0 | 0 |
| 'Gaa' | 10.476 | 8.969 | 10.975 | 9.171 | 9.891 | 10.355 | 9.823 | 9.338 | 10.345 | 10.681 | 8.637 | 10.485 | 8.703 | 7.941 | 9.906 | 10.665 | 7.885 | 9.095 | 0.083 | 0.0779 | 0.2502 | 0 | 0 | 0 |
| 'Gab3' | 1.468 | 1.514 | 1.508 | 1.819 | 1.622 | 1.528 | 2.102 | 1.512 | 1.534 | 2.051 | 1.780 | 1.765 | 1.596 | 2.016 | 1.715 | 1.755 | 1.744 | 1.985 | -0.185 | 0.0469 | 0.1806 | 0 | 0 | 0 |
| 'Gabarap' | 52.832 | 58.243 | 51.375 | 57.249 | 57.783 | 55.034 | 56.461 | 58.439 | 53.116 | 55.647 | 55.698 | 52.037 | 53.594 | 55.395 | 53.247 | 54.648 | 56.589 | 51.859 | 0.023 | 0.2908 | 0.5477 | 0 | 0 | 0 |
| 'Gabarapl1' | 12.775 | 13.497 | 11.393 | 11.476 | 12.660 | 11.604 | 11.354 | 12.626 | 11.014 | 14.725 | 12.146 | 12.083 | 14.960 | 14.233 | 10.775 | 12.201 | 13.266 | 11.684 | -0.108 | 0.0426 | 0.1704 | 0 | 0 | 0 |
| 'Gabarapl2' | 32.722 | 34.384 | 35.139 | 33.172 | 34.975 | 31.805 | 32.260 | 33.131 | 33.113 | 28.179 | 31.054 | 31.671 | 28.228 | 31.877 | 32.875 | 29.825 | 29.476 | 30.182 | 0.127 | 0.0000 | 0.0007 | 1 | 0 | 0 |
| 'Gabbr1' | 5.527 | 4.828 | 6.431 | 5.109 | 4.518 | 5.520 | 4.956 | 4.788 | 6.481 | 5.538 | 5.127 | 6.014 | 5.364 | 4.260 | 5.217 | 5.139 | 5.101 | 5.503 | 0.013 | 0.7851 | 0.9042 | 0 | 0 | 0 |
| 'Gabpa' | 14.279 | 15.952 | 14.647 | 14.468 | 15.723 | 14.123 | 15.126 | 15.354 | 14.040 | 14.227 | 14.605 | 15.287 | 14.057 | 15.602 | 14.600 | 14.120 | 14.678 | 14.450 | 0.008 | 0.7706 | 0.8977 | 0 | 0 | 0 |
| 'Gabpb1' | 4.588 | 4.880 | 4.780 | 4.376 | 4.743 | 4.747 | 4.904 | 5.000 | 4.512 | 5.075 | 4.286 | 4.530 | 4.420 | 4.467 | 4.437 | 4.669 | 4.471 | 3.843 | 0.073 | 0.0646 | 0.2217 | 0 | 0 | 0 |
| 'Gabpb2' | 11.830 | 12.017 | 12.336 | 11.519 | 11.158 | 11.888 | 11.386 | 11.566 | 12.402 | 11.565 | 13.317 | 13.356 | 11.974 | 12.233 | 13.057 | 12.212 | 12.555 | 13.628 | -0.112 | 0.0001 | 0.0010 | 1 | 0 | 0 |
| 'Gabrr2' | 2.097 | 2.011 | 1.903 | 1.241 | 1.688 | 1.800 | 1.991 | 2.120 | 1.897 | 1.726 | 1.824 | 1.896 | 1.935 | 1.548 | 1.371 | 1.345 | 1.664 | 1.541 | 0.156 | 0.1453 | 0.3699 | 0 | 0 | 0 |
| 'Gadd45a' | 13.513 | 11.077 | 9.170 | 13.242 | 10.502 | 8.229 | 12.208 | 11.789 | 8.136 | 13.870 | 12.477 | 7.708 | 14.148 | 12.732 | 7.871 | 12.731 | 12.787 | 6.898 | -0.033 | 0.5176 | 0.7407 | 0 | 0 | 0 |
| 'Gadd45b' | 248.532 | 167.445 | 125.621 | 213.056 | 166.146 | 119.255 | 183.405 | 200.028 | 114.110 | 248.424 | 184.825 | 104.129 | 277.769 | 216.396 | 106.221 | 248.226 | 184.265 | 96.314 | -0.068 | 0.3223 | 0.5800 | 0 | 0 | 0 |
| 'Gadd45g' | 23.777 | 14.523 | 12.960 | 25.382 | 15.023 | 13.671 | 26.468 | 16.503 | 11.367 | 19.187 | 15.632 | 11.467 | 13.685 | 16.947 | 11.109 | 17.231 | 19.388 | 9.292 | 0.218 | 0.0209 | 0.1061 | 0 | 0 | 0 |
| 'Gadd45gip1' | 9.973 | 8.641 | 10.584 | 9.221 | 9.046 | 11.056 | 9.969 | 9.810 | 9.474 | 8.949 | 7.666 | 10.853 | 8.776 | 8.621 | 10.315 | 9.140 | 8.733 | 10.465 | 0.061 | 0.2098 | 0.4550 | 0 | 0 | 0 |
| 'Gak' | 16.454 | 17.254 | 17.138 | 16.553 | 16.853 | 17.233 | 16.729 | 16.931 | 16.888 | 16.576 | 16.888 | 17.446 | 16.979 | 17.057 | 17.218 | 17.037 | 17.582 | 17.440 | -0.031 | 0.0697 | 0.2323 | 0 | 0 | 0 |
| 'Gal' | 17.591 | 20.935 | 13.174 | 16.952 | 24.183 | 18.445 | 21.848 | 24.080 | 13.056 | 12.390 | 15.371 | 8.895 | 16.938 | 15.482 | 8.875 | 11.190 | 15.663 | 8.145 | 0.595 | 0.0000 | 0.0000 | 1 | 1 | 0 |
| 'Gale' | 3.449 | 3.277 | 4.385 | 3.943 | 3.556 | 4.580 | 3.895 | 3.605 | 3.988 | 3.267 | 3.371 | 4.066 | 3.706 | 3.038 | 3.938 | 3.329 | 3.572 | 3.846 | 0.098 | 0.0663 | 0.2252 | 0 | 0 | 0 |
| 'Galk1' | 30.231 | 28.625 | 29.686 | 26.119 | 29.033 | 31.751 | 27.142 | 29.258 | 27.742 | 27.350 | 26.598 | 27.743 | 31.206 | 26.865 | 30.029 | 29.968 | 24.786 | 26.381 | 0.043 | 0.3534 | 0.6069 | 0 | 0 | 0 |
| 'Galk2' | 4.872 | 5.084 | 5.272 | 5.156 | 4.980 | 5.315 | 5.122 | 5.247 | 5.156 | 5.535 | 4.950 | 5.395 | 5.797 | 5.164 | 5.442 | 5.162 | 4.898 | 5.308 | -0.055 | 0.1046 | 0.3032 | 0 | 0 | 0 |
| 'Galm' | 14.961 | 14.224 | 15.811 | 14.631 | 15.812 | 15.634 | 14.722 | 15.014 | 14.732 | 15.345 | 14.714 | 17.058 | 13.429 | 14.719 | 16.040 | 15.402 | 14.168 | 16.280 | -0.028 | 0.4782 | 0.7112 | 0 | 0 | 0 |
| 'Galns' | 10.233 | 9.682 | 11.880 | 9.903 | 10.178 | 11.494 | 9.811 | 10.302 | 11.012 | 10.409 | 11.079 | 11.679 | 12.756 | 10.793 | 11.749 | 10.869 | 10.451 | 11.810 | -0.112 | 0.0140 | 0.0800 | 0 | 0 | 0 |
| 'Galnt1' | 24.104 | 26.548 | 27.296 | 26.179 | 27.294 | 25.037 | 25.555 | 26.672 | 23.739 | 25.035 | 24.049 | 22.771 | 23.237 | 25.684 | 22.125 | 23.577 | 24.550 | 22.832 | 0.111 | 0.0001 | 0.0021 | 1 | 0 | 0 |
| 'Galnt10' | 5.710 | 5.825 | 6.910 | 5.997 | 6.184 | 7.756 | 6.076 | 6.607 | 6.778 | 7.246 | 6.725 | 6.775 | 6.619 | 5.792 | 6.506 | 7.351 | 5.679 | 6.336 | -0.037 | 0.4996 | 0.7273 | 0 | 0 | 0 |
| 'Galnt11' | 3.612 | 4.628 | 4.349 | 3.990 | 4.504 | 4.103 | 3.798 | 4.672 | 4.551 | 4.632 | 4.596 | 3.792 | 4.441 | 3.894 | 4.106 | 4.510 | 4.572 | 2.883 | 0.025 | 0.7199 | 0.8711 | 0 | 0 | 0 |
| 'Galnt12' | 25.292 | 25.971 | 27.424 | 23.256 | 25.240 | 27.620 | 24.304 | 24.450 | 26.488 | 24.355 | 22.878 | 28.364 | 24.811 | 19.783 | 25.818 | 23.191 | 22.307 | 26.830 | 0.062 | 0.1272 | 0.3416 | 0 | 0 | 0 |
| 'Galnt2' | 79.418 | 88.936 | 78.115 | 79.517 | 88.229 | 82.508 | 75.079 | 88.014 | 78.880 | 84.576 | 89.922 | 81.949 | 95.514 | 87.509 | 81.832 | 87.072 | 85.238 | 87.012 | -0.092 | 0.0038 | 0.0302 | 1 | 0 | 0 |
| 'Galnt4' | 4.193 | 4.209 | 4.598 | 4.816 | 5.568 | 4.426 | 4.954 | 4.206 | 4.369 | 3.798 | 3.466 | 3.500 | 4.182 | 3.798 | 3.680 | 4.110 | 3.862 | 3.665 | 0.269 | 0.0000 | 0.0001 | 1 | 0 | 0 |
| 'Galnt6' | 52.384 | 45.590 | 50.338 | 50.295 | 43.546 | 48.764 | 49.280 | 46.418 | 50.102 | 46.514 | 46.311 | 50.860 | 48.308 | 44.808 | 51.837 | 47.270 | 43.649 | 51.735 | 0.008 | 0.7826 | 0.9025 | 0 | 0 | 0 |
| 'Galnt7' | 5.597 | 5.967 | 5.725 | 5.952 | 6.500 | 6.102 | 5.476 | 6.449 | 6.599 | 6.272 | 5.790 | 6.914 | 5.327 | 6.554 | 6.098 | 6.145 | 6.651 | 6.649 | -0.065 | 0.1350 | 0.3537 | 0 | 0 | 0 |
| 'Galnt9' | 10.846 | 10.229 | 9.904 | 10.460 | 11.053 | 11.058 | 10.310 | 10.411 | 10.855 | 11.601 | 10.608 | 12.015 | 10.476 | 11.252 | 11.615 | 11.948 | 11.029 | 11.553 | -0.113 | 0.0015 | 0.0144 | 1 | 0 | 0 |
| 'Galt' | 5.548 | 5.119 | 5.871 | 5.385 | 5.447 | 5.884 | 5.396 | 5.961 | 5.532 | 5.844 | 4.907 | 5.232 | 5.513 | 4.675 | 4.585 | 5.184 | 5.123 | 4.423 | 0.136 | 0.0055 | 0.0406 | 1 | 0 | 0 |
| 'Gan' | 1.546 | 1.428 | 1.645 | 1.892 | 1.345 | 1.473 | 1.395 | 1.647 | 1.456 | 1.607 | 1.423 | 1.636 | 1.511 | 1.564 | 1.516 | 1.520 | 1.407 | 1.724 | -0.019 | 0.7568 | 0.8913 | 0 | 0 | 0 |
| 'Ganab' | 58.433 | 60.654 | 61.466 | 56.942 | 63.064 | 60.375 | 54.657 | 60.532 | 64.309 | 62.075 | 59.020 | 62.853 | 59.636 | 60.097 | 61.916 | 60.560 | 58.014 | 62.046 | -0.026 | 0.2793 | 0.5363 | 0 | 0 | 0 |
| 'Ganc' | 2.999 | 3.130 | 3.108 | 3.271 | 3.291 | 3.181 | 3.286 | 2.970 | 2.919 | 2.888 | 2.967 | 2.793 | 3.046 | 3.020 | 2.590 | 2.891 | 2.941 | 2.649 | 0.117 | 0.0005 | 0.0061 | 1 | 0 | 0 |
| 'Gapvd1' | 9.283 | 8.700 | 8.970 | 8.900 | 8.778 | 9.031 | 8.380 | 9.292 | 8.799 | 8.812 | 8.975 | 8.746 | 8.191 | 8.986 | 8.348 | 8.602 | 9.073 | 8.849 | 0.019 | 0.4862 | 0.7170 | 0 | 0 | 0 |
| 'Gar1' | 11.254 | 11.392 | 13.489 | 12.495 | 12.701 | 12.241 | 11.727 | 12.466 | 13.085 | 11.603 | 10.905 | 12.964 | 12.115 | 10.609 | 12.993 | 11.510 | 10.938 | 11.467 | 0.065 | 0.0987 | 0.2936 | 0 | 0 | 0 |
| 'Garem' | 0.883 | 1.004 | 0.762 | 0.658 | 0.831 | 0.761 | 0.685 | 0.821 | 0.893 | 1.190 | 0.991 | 0.779 | 1.042 | 1.224 | 0.735 | 1.027 | 1.133 | 0.700 | -0.257 | 0.0202 | 0.1038 | 0 | 0 | 0 |
| 'Gars' | 25.720 | 26.593 | 27.407 | 25.892 | 27.065 | 26.609 | 25.763 | 26.101 | 25.612 | 27.274 | 24.385 | 25.249 | 27.684 | 23.909 | 25.653 | 26.393 | 24.925 | 24.844 | 0.031 | 0.2731 | 0.5281 | 0 | 0 | 0 |
| 'Gas2' | 0.868 | 0.862 | 0.813 | 0.832 | 1.039 | 0.825 | 1.012 | 1.098 | 0.810 | 0.493 | 0.927 | 0.707 | 0.782 | 0.795 | 0.795 | 0.829 | 1.021 | 0.856 | 0.172 | 0.0916 | 0.2792 | 0 | 0 | 0 |
| 'Gas2l1' | 0.758 | 0.715 | 0.875 | 0.700 | 0.824 | 0.986 | 0.777 | 0.739 | 1.000 | 0.720 | 0.586 | 0.933 | 0.791 | 0.652 | 0.933 | 0.843 | 0.784 | 0.811 | 0.051 | 0.4676 | 0.7033 | 0 | 0 | 0 |
| 'Gas2l3' | 0.834 | 1.200 | 1.310 | 1.092 | 1.130 | 1.239 | 0.975 | 1.217 | 1.275 | 0.938 | 1.264 | 1.058 | 1.198 | 1.235 | 1.140 | 0.835 | 1.256 | 1.218 | 0.012 | 0.8902 | 0.9551 | 0 | 0 | 0 |
| 'Gas8' | 6.770 | 7.570 | 7.200 | 5.961 | 6.965 | 7.205 | 6.846 | 7.225 | 8.192 | 9.145 | 5.158 | 6.762 | 7.573 | 6.871 | 7.219 | 7.985 | 6.185 | 6.752 | 0.005 | 0.9507 | 0.9810 | 0 | 0 | 0 |
| 'Gata3' | 41.963 | 33.431 | 33.446 | 36.052 | 32.592 | 33.556 | 36.734 | 35.122 | 34.541 | 35.981 | 30.169 | 28.000 | 29.497 | 29.527 | 27.145 | 30.274 | 27.954 | 29.528 | 0.234 | 0.0000 | 0.0000 | 1 | 0 | 0 |
| 'Gatad1' | 14.014 | 15.602 | 16.512 | 14.196 | 16.727 | 17.201 | 14.227 | 15.659 | 16.839 | 13.372 | 14.988 | 16.537 | 15.952 | 16.117 | 17.877 | 16.802 | 15.462 | 16.422 | -0.037 | 0.3666 | 0.6187 | 0 | 0 | 0 |
| 'Gatad2a' | 26.438 | 25.430 | 26.661 | 26.139 | 25.063 | 27.877 | 25.813 | 26.643 | 26.451 | 26.307 | 26.037 | 27.364 | 26.574 | 25.491 | 28.567 | 25.806 | 24.867 | 28.897 | -0.030 | 0.2181 | 0.4649 | 0 | 0 | 0 |
| 'Gatad2b' | 10.359 | 9.543 | 9.075 | 9.360 | 9.305 | 9.266 | 9.534 | 9.478 | 9.335 | 9.788 | 10.500 | 9.789 | 9.638 | 9.888 | 9.678 | 9.265 | 9.910 | 9.800 | -0.060 | 0.0469 | 0.1806 | 0 | 0 | 0 |
| 'Gatb' | 1.859 | 1.728 | 1.780 | 1.683 | 1.510 | 1.590 | 1.570 | 1.721 | 1.699 | 1.643 | 1.877 | 1.801 | 1.826 | 1.885 | 1.674 | 1.858 | 1.739 | 1.690 | -0.083 | 0.1779 | 0.4131 | 0 | 0 | 0 |
| 'Gatc' | 8.954 | 10.985 | 8.997 | 7.901 | 10.163 | 8.639 | 8.609 | 10.413 | 10.011 | 8.386 | 8.540 | 8.978 | 9.211 | 8.744 | 8.837 | 8.479 | 8.023 | 8.124 | 0.112 | 0.0546 | 0.2001 | 0 | 0 | 0 |
| 'Gatm' | 0.977 | 1.121 | 1.563 | 1.219 | 1.578 | 1.530 | 1.904 | 1.555 | 1.597 | 1.212 | 0.697 | 0.980 | 0.904 | 1.553 | 1.263 | 1.094 | 1.234 | 1.059 | 0.379 | 0.0058 | 0.0418 | 1 | 1 | 0 |
| 'Gatsl2' | 2.941 | 3.440 | 3.327 | 3.355 | 3.279 | 3.524 | 3.244 | 3.223 | 3.616 | 3.384 | 3.773 | 3.433 | 2.977 | 3.026 | 2.966 | 3.217 | 2.975 | 2.971 | 0.050 | 0.3493 | 0.6043 | 0 | 0 | 0 |
| 'Gatsl3' | 19.194 | 18.243 | 22.644 | 17.534 | 19.644 | 20.823 | 16.991 | 21.990 | 19.154 | 20.666 | 16.407 | 16.558 | 16.537 | 15.593 | 17.189 | 16.451 | 14.480 | 15.036 | 0.237 | 0.0003 | 0.0038 | 1 | 0 | 0 |
| 'Gba' | 16.722 | 15.902 | 17.406 | 16.875 | 17.158 | 17.804 | 16.903 | 17.692 | 17.950 | 18.152 | 16.495 | 17.028 | 17.689 | 16.884 | 18.012 | 17.187 | 16.621 | 16.787 | -0.011 | 0.7181 | 0.8700 | 0 | 0 | 0 |
| 'Gba2' | 8.193 | 7.654 | 8.788 | 8.076 | 7.558 | 8.543 | 7.946 | 7.736 | 8.979 | 8.423 | 8.177 | 8.797 | 8.720 | 7.697 | 8.061 | 7.977 | 8.009 | 8.544 | -0.028 | 0.3293 | 0.5875 | 0 | 0 | 0 |
| 'Gbas' | 12.887 | 16.032 | 13.846 | 12.577 | 15.164 | 14.508 | 13.537 | 15.061 | 14.214 | 14.121 | 13.958 | 12.671 | 13.977 | 13.560 | 13.452 | 14.561 | 13.723 | 11.466 | 0.064 | 0.1340 | 0.3525 | 0 | 0 | 0 |
| 'Gbe1' | 1.150 | 1.256 | 1.160 | 1.202 | 1.381 | 1.156 | 1.258 | 1.305 | 1.097 | 1.198 | 1.228 | 1.267 | 1.100 | 1.176 | 1.270 | 1.071 | 1.246 | 1.257 | -0.001 | 0.9856 | 0.9951 | 0 | 0 | 0 |
| 'Gbf1' | 10.220 | 9.397 | 10.792 | 9.915 | 9.354 | 10.721 | 9.849 | 9.551 | 10.620 | 11.463 | 10.927 | 10.674 | 10.369 | 10.153 | 10.092 | 10.019 | 10.035 | 10.646 | -0.073 | 0.0297 | 0.1339 | 0 | 0 | 0 |
| 'Gbp2' | 14.061 | 14.955 | 15.700 | 15.706 | 13.604 | 15.852 | 15.645 | 13.073 | 16.413 | 17.309 | 15.788 | 20.829 | 15.955 | 15.974 | 20.685 | 17.843 | 17.499 | 21.591 | -0.287 | 0.0000 | 0.0000 | 1 | 0 | 0 |
| 'Gbp2b' | 8.508 | 8.727 | 9.495 | 9.398 | 9.397 | 9.969 | 8.748 | 8.640 | 10.434 | 10.694 | 9.535 | 13.801 | 10.403 | 11.024 | 14.061 | 10.764 | 11.094 | 14.700 | -0.359 | 0.0000 | 0.0000 | 1 | 0 | 0 |
| 'Gbp3' | 4.391 | 4.141 | 4.884 | 3.783 | 4.747 | 5.283 | 5.189 | 4.323 | 5.084 | 5.514 | 4.616 | 5.735 | 6.206 | 5.134 | 5.534 | 5.553 | 4.784 | 4.969 | -0.210 | 0.0002 | 0.0027 | 1 | 0 | 0 |
| 'Gbp5' | 4.136 | 4.445 | 4.673 | 3.950 | 4.085 | 3.969 | 3.905 | 3.538 | 4.497 | 5.292 | 4.606 | 5.078 | 5.857 | 4.226 | 4.980 | 4.766 | 5.417 | 4.968 | -0.292 | 0.0000 | 0.0000 | 1 | 0 | 0 |
| 'Gbp7' | 8.072 | 7.681 | 9.071 | 8.140 | 8.154 | 8.741 | 8.640 | 8.586 | 9.378 | 8.572 | 7.263 | 10.988 | 7.204 | 7.150 | 10.255 | 9.129 | 7.677 | 10.317 | -0.040 | 0.4458 | 0.6878 | 0 | 0 | 0 |
| 'Gcat' | 6.027 | 6.032 | 6.813 | 5.816 | 6.708 | 6.725 | 6.074 | 6.640 | 7.497 | 6.153 | 6.053 | 6.951 | 5.983 | 6.295 | 7.893 | 5.981 | 6.500 | 6.484 | -0.008 | 0.8653 | 0.9415 | 0 | 0 | 0 |
| 'Gcc1' | 8.263 | 6.546 | 7.346 | 7.244 | 6.515 | 7.148 | 6.779 | 6.981 | 7.117 | 6.992 | 7.113 | 8.080 | 7.181 | 7.018 | 7.451 | 6.446 | 6.689 | 7.291 | -0.018 | 0.6830 | 0.8495 | 0 | 0 | 0 |
| 'Gcc2' | 3.587 | 3.619 | 4.138 | 3.847 | 4.079 | 3.999 | 3.793 | 3.790 | 4.203 | 3.752 | 3.998 | 3.936 | 3.634 | 3.920 | 3.934 | 4.056 | 3.935 | 3.980 | -0.011 | 0.7571 | 0.8913 | 0 | 0 | 0 |
| 'Gcdh' | 7.750 | 9.015 | 9.633 | 8.952 | 9.295 | 9.042 | 9.892 | 9.681 | 8.999 | 8.677 | 8.611 | 8.733 | 9.353 | 9.521 | 8.465 | 7.716 | 9.617 | 7.736 | 0.058 | 0.2279 | 0.4743 | 0 | 0 | 0 |
| 'Gcfc2' | 1.297 | 1.166 | 1.253 | 1.381 | 0.989 | 1.107 | 1.315 | 1.152 | 1.150 | 1.073 | 1.242 | 1.334 | 1.287 | 1.008 | 1.065 | 1.127 | 1.287 | 1.076 | 0.024 | 0.7336 | 0.8786 | 0 | 0 | 0 |
| 'Gch1' | 7.374 | 7.744 | 6.720 | 7.370 | 6.952 | 7.076 | 6.543 | 6.972 | 6.742 | 7.472 | 7.090 | 8.080 | 7.234 | 7.836 | 8.505 | 8.334 | 7.683 | 7.019 | -0.138 | 0.0090 | 0.0580 | 0 | 0 | 0 |
| 'Gclc' | 15.687 | 16.067 | 14.949 | 12.556 | 15.237 | 14.102 | 13.768 | 16.664 | 14.111 | 14.854 | 14.237 | 14.793 | 15.863 | 15.951 | 15.053 | 16.472 | 14.172 | 15.003 | -0.045 | 0.3351 | 0.5939 | 0 | 0 | 0 |
| 'Gclm' | 10.027 | 9.523 | 11.410 | 9.996 | 10.753 | 10.815 | 9.811 | 9.849 | 11.473 | 11.855 | 10.233 | 10.064 | 12.205 | 11.731 | 10.942 | 10.616 | 10.719 | 9.400 | -0.066 | 0.2436 | 0.4936 | 0 | 0 | 0 |
| 'Gcn1l1' | 15.265 | 13.288 | 16.549 | 14.403 | 14.140 | 15.169 | 14.412 | 13.509 | 16.605 | 16.115 | 14.748 | 15.607 | 15.830 | 13.404 | 14.588 | 14.289 | 14.843 | 15.667 | -0.031 | 0.3770 | 0.6288 | 0 | 0 | 0 |
| 'Gcnt1' | 0.529 | 0.440 | 0.624 | 0.445 | 0.418 | 0.431 | 0.387 | 0.604 | 0.400 | 0.303 | 0.372 | 0.530 | 0.402 | 0.485 | 0.436 | 0.429 | 0.421 | 0.511 | 0.132 | 0.2816 | 0.5380 | 0 | 0 | 0 |
| 'Gcsh' | 8.475 | 8.331 | 8.642 | 8.016 | 8.246 | 8.418 | 8.347 | 8.440 | 8.079 | 8.662 | 8.101 | 8.051 | 9.319 | 8.366 | 8.615 | 8.009 | 7.897 | 7.606 | 0.001 | 0.9868 | 0.9959 | 0 | 0 | 0 |
| 'Gdap2' | 5.000 | 4.768 | 4.723 | 4.494 | 4.898 | 5.187 | 4.409 | 5.109 | 5.000 | 4.816 | 5.283 | 4.856 | 5.163 | 4.974 | 4.970 | 4.712 | 5.077 | 4.697 | -0.036 | 0.3791 | 0.6307 | 0 | 0 | 0 |
| 'Gde1' | 24.651 | 23.612 | 21.197 | 22.318 | 25.107 | 22.930 | 21.167 | 25.611 | 23.018 | 23.693 | 21.603 | 23.855 | 25.473 | 25.281 | 22.630 | 24.741 | 24.712 | 23.285 | -0.050 | 0.2132 | 0.4581 | 0 | 0 | 0 |
| 'Gdi1' | 24.929 | 26.451 | 26.694 | 23.592 | 26.843 | 26.732 | 23.891 | 25.806 | 27.428 | 26.495 | 27.302 | 25.887 | 27.361 | 26.493 | 25.790 | 24.980 | 28.117 | 24.852 | -0.041 | 0.1627 | 0.3947 | 0 | 0 | 0 |
| 'Gdi2' | 187.623 | 202.059 | 180.682 | 198.570 | 211.797 | 183.028 | 204.016 | 199.923 | 182.835 | 200.531 | 216.268 | 181.839 | 209.579 | 218.733 | 195.133 | 204.160 | 218.757 | 189.186 | -0.077 | 0.0007 | 0.0075 | 1 | 0 | 0 |
| 'Gdpd3' | 6.115 | 5.057 | 4.378 | 4.942 | 5.640 | 3.907 | 4.784 | 6.604 | 4.398 | 6.295 | 5.978 | 4.308 | 7.596 | 6.422 | 3.867 | 5.054 | 5.245 | 3.861 | -0.081 | 0.2980 | 0.5539 | 0 | 0 | 0 |
| 'Gdpd5' | 5.415 | 6.012 | 7.551 | 5.859 | 5.805 | 7.286 | 5.545 | 6.208 | 6.940 | 7.090 | 6.429 | 7.514 | 5.898 | 6.561 | 7.298 | 6.541 | 6.366 | 6.569 | -0.100 | 0.0305 | 0.1363 | 0 | 0 | 0 |
| 'Gdpgp1' | 2.933 | 3.329 | 3.698 | 3.600 | 3.715 | 3.480 | 3.583 | 3.115 | 3.976 | 3.132 | 2.756 | 3.926 | 3.179 | 3.101 | 3.471 | 3.178 | 3.526 | 3.758 | 0.044 | 0.4775 | 0.7109 | 0 | 0 | 0 |
| 'Gem' | 13.873 | 2.475 | 1.601 | 10.432 | 2.851 | 1.345 | 9.669 | 3.269 | 1.364 | 6.373 | 6.676 | 1.881 | 5.857 | 5.118 | 1.836 | 5.932 | 6.110 | 1.669 | -0.160 | 0.4945 | 0.7237 | 0 | 0 | 0 |
| 'Gemin2' | 3.943 | 4.347 | 4.867 | 3.526 | 5.051 | 4.195 | 4.320 | 5.127 | 4.346 | 4.008 | 4.305 | 4.096 | 4.859 | 4.018 | 3.836 | 4.097 | 3.997 | 4.083 | 0.079 | 0.2256 | 0.4722 | 0 | 0 | 0 |
| 'Gemin5' | 9.003 | 7.327 | 9.053 | 8.917 | 8.145 | 9.204 | 7.968 | 8.412 | 8.746 | 9.029 | 8.288 | 9.291 | 8.198 | 7.945 | 8.896 | 8.441 | 7.715 | 9.393 | -0.016 | 0.6626 | 0.8371 | 0 | 0 | 0 |
| 'Gemin6' | 3.763 | 3.643 | 4.243 | 4.775 | 4.108 | 4.361 | 5.508 | 4.802 | 4.475 | 3.342 | 3.706 | 4.751 | 4.339 | 4.768 | 4.495 | 4.240 | 4.788 | 4.420 | 0.012 | 0.8923 | 0.9561 | 0 | 0 | 0 |
| 'Gemin7' | 8.175 | 9.340 | 8.402 | 7.356 | 8.811 | 8.840 | 7.533 | 9.367 | 8.529 | 8.001 | 7.387 | 7.928 | 8.298 | 8.350 | 8.314 | 8.289 | 7.312 | 8.030 | 0.075 | 0.1030 | 0.3009 | 0 | 0 | 0 |
| 'Gemin8' | 10.324 | 9.312 | 10.174 | 9.303 | 9.239 | 11.179 | 10.142 | 10.175 | 10.065 | 11.231 | 9.766 | 9.688 | 10.947 | 11.819 | 9.614 | 10.503 | 10.217 | 8.433 | -0.035 | 0.5721 | 0.7788 | 0 | 0 | 0 |
| 'Gen1' | 2.717 | 2.936 | 3.329 | 2.799 | 2.837 | 3.382 | 3.077 | 2.690 | 2.828 | 1.926 | 3.210 | 3.147 | 3.237 | 2.571 | 3.635 | 3.310 | 3.138 | 3.226 | -0.052 | 0.5466 | 0.7610 | 0 | 0 | 0 |
| 'Get4' | 16.901 | 18.603 | 17.571 | 15.412 | 18.305 | 18.916 | 16.277 | 19.303 | 18.117 | 17.458 | 15.844 | 17.232 | 18.894 | 17.188 | 17.421 | 19.021 | 16.811 | 16.954 | 0.013 | 0.7572 | 0.8913 | 0 | 0 | 0 |
| 'Gfer' | 14.152 | 14.007 | 16.431 | 13.863 | 14.219 | 17.785 | 13.305 | 14.880 | 15.947 | 12.907 | 13.025 | 14.319 | 15.777 | 15.708 | 15.432 | 15.121 | 13.414 | 15.232 | 0.036 | 0.5350 | 0.7546 | 0 | 0 | 0 |
| 'Gfi1' | 16.300 | 14.321 | 17.488 | 14.887 | 13.248 | 16.472 | 15.047 | 13.747 | 16.454 | 16.097 | 12.877 | 15.923 | 15.729 | 13.146 | 14.856 | 14.659 | 12.373 | 15.201 | 0.068 | 0.0346 | 0.1487 | 0 | 0 | 0 |
| 'Gfm1' | 9.861 | 10.967 | 10.431 | 9.975 | 11.009 | 10.086 | 10.529 | 10.700 | 9.764 | 9.128 | 9.438 | 9.023 | 9.316 | 9.004 | 8.594 | 9.072 | 9.003 | 8.475 | 0.191 | 0.0000 | 0.0000 | 1 | 0 | 0 |
| 'Gfm2' | 4.245 | 4.092 | 4.703 | 4.497 | 4.460 | 4.247 | 4.367 | 4.374 | 4.624 | 4.532 | 4.186 | 4.367 | 4.587 | 4.613 | 4.256 | 4.165 | 4.670 | 4.204 | -0.008 | 0.8246 | 0.9227 | 0 | 0 | 0 |
| 'Gfod1' | 6.582 | 5.885 | 5.969 | 5.997 | 5.446 | 6.122 | 5.518 | 5.090 | 5.949 | 7.077 | 5.302 | 7.002 | 6.801 | 6.485 | 7.012 | 7.221 | 6.171 | 7.170 | -0.206 | 0.0000 | 0.0001 | 1 | 0 | 0 |
| 'Gfod2' | 0.864 | 0.969 | 0.871 | 0.802 | 0.977 | 0.858 | 0.919 | 0.881 | 0.924 | 0.844 | 0.901 | 1.114 | 0.535 | 0.774 | 1.043 | 0.904 | 0.633 | 1.148 | 0.020 | 0.8608 | 0.9399 | 0 | 0 | 0 |
| 'Gfpt1' | 11.018 | 11.996 | 10.826 | 11.971 | 12.248 | 10.788 | 10.338 | 11.572 | 10.915 | 10.696 | 11.962 | 11.369 | 11.743 | 11.877 | 11.107 | 10.599 | 12.022 | 11.422 | -0.029 | 0.3441 | 0.5996 | 0 | 0 | 0 |
| 'Gfra4' | 32.208 | 53.081 | 33.817 | 34.279 | 49.194 | 39.813 | 35.529 | 47.015 | 37.818 | 38.480 | 39.856 | 30.647 | 44.924 | 43.160 | 29.660 | 40.987 | 42.900 | 30.104 | 0.077 | 0.2441 | 0.4944 | 0 | 0 | 0 |
| 'Gga1' | 35.932 | 34.521 | 44.054 | 34.615 | 33.837 | 40.952 | 34.577 | 35.345 | 41.818 | 38.778 | 34.176 | 40.714 | 39.516 | 31.390 | 36.636 | 38.987 | 34.275 | 35.712 | 0.013 | 0.7528 | 0.8898 | 0 | 0 | 0 |
| 'Gga2' | 1.144 | 1.104 | 1.071 | 0.965 | 1.205 | 1.159 | 1.093 | 1.150 | 1.063 | 1.170 | 1.108 | 1.131 | 1.202 | 1.193 | 1.068 | 1.307 | 1.162 | 1.024 | -0.063 | 0.2390 | 0.4894 | 0 | 0 | 0 |
| 'Gga3' | 7.187 | 7.026 | 8.202 | 7.578 | 6.957 | 7.754 | 7.244 | 7.083 | 8.319 | 7.478 | 7.584 | 8.730 | 7.405 | 7.238 | 7.944 | 6.833 | 8.257 | 8.396 | -0.064 | 0.0626 | 0.2171 | 0 | 0 | 0 |
| 'Ggact' | 8.184 | 9.761 | 7.596 | 8.360 | 8.704 | 8.861 | 8.752 | 8.543 | 8.204 | 8.587 | 8.699 | 7.988 | 8.631 | 7.710 | 8.107 | 8.607 | 7.593 | 8.198 | 0.040 | 0.3929 | 0.6426 | 0 | 0 | 0 |
| 'Ggct' | 1.850 | 1.702 | 1.478 | 1.750 | 1.653 | 1.802 | 1.750 | 1.945 | 1.723 | 1.531 | 1.609 | 1.726 | 1.673 | 1.770 | 1.605 | 1.822 | 1.541 | 1.389 | 0.083 | 0.1732 | 0.4066 | 0 | 0 | 0 |
| 'Ggcx' | 3.310 | 3.129 | 3.617 | 3.405 | 3.254 | 3.209 | 3.044 | 3.233 | 3.346 | 4.131 | 3.487 | 3.015 | 3.688 | 3.020 | 2.789 | 3.055 | 3.278 | 2.879 | 0.010 | 0.8813 | 0.9499 | 0 | 0 | 0 |
| 'Ggh' | 2.094 | 2.634 | 2.014 | 2.209 | 2.554 | 1.934 | 2.494 | 2.605 | 2.204 | 3.152 | 2.003 | 1.925 | 1.860 | 2.392 | 2.162 | 2.961 | 1.631 | 1.461 | 0.086 | 0.4800 | 0.7122 | 0 | 0 | 0 |
| 'Ggnbp1' | 0.676 | 0.712 | 0.865 | 0.798 | 0.642 | 0.753 | 0.659 | 0.680 | 0.910 | 0.682 | 0.640 | 0.868 | 0.598 | 0.615 | 0.798 | 0.857 | 0.681 | 0.646 | 0.058 | 0.5212 | 0.7440 | 0 | 0 | 0 |
| 'Ggnbp2' | 30.106 | 29.742 | 27.665 | 31.708 | 29.570 | 27.750 | 32.071 | 30.402 | 28.925 | 29.481 | 31.038 | 29.410 | 29.888 | 30.557 | 28.698 | 30.022 | 32.948 | 28.404 | -0.025 | 0.3510 | 0.6052 | 0 | 0 | 0 |
| 'Ggps1' | 12.903 | 13.474 | 12.851 | 13.729 | 12.533 | 12.500 | 14.198 | 13.980 | 12.806 | 11.937 | 13.854 | 15.113 | 12.535 | 13.833 | 13.365 | 11.731 | 15.921 | 13.436 | -0.047 | 0.3753 | 0.6267 | 0 | 0 | 0 |
| 'Ggt5' | 2.195 | 1.827 | 2.234 | 2.274 | 2.077 | 2.059 | 1.835 | 2.084 | 2.154 | 2.179 | 2.496 | 2.306 | 2.083 | 2.282 | 2.137 | 2.057 | 2.277 | 2.400 | -0.114 | 0.0693 | 0.2315 | 0 | 0 | 0 |
| 'Ggt7' | 4.283 | 4.111 | 4.773 | 3.902 | 4.751 | 4.397 | 4.869 | 4.683 | 4.730 | 5.349 | 4.483 | 4.581 | 5.562 | 4.200 | 4.361 | 4.204 | 4.213 | 4.301 | -0.035 | 0.5752 | 0.7806 | 0 | 0 | 0 |
| 'Ggta1' | 7.557 | 7.278 | 7.961 | 8.214 | 7.805 | 7.258 | 8.122 | 7.411 | 7.414 | 6.389 | 7.543 | 8.002 | 6.084 | 6.723 | 7.211 | 5.593 | 8.146 | 7.267 | 0.125 | 0.0333 | 0.1447 | 0 | 0 | 0 |
| 'Ghdc' | 3.339 | 3.178 | 3.303 | 3.143 | 3.499 | 3.603 | 3.463 | 3.165 | 3.254 | 3.711 | 2.582 | 3.740 | 3.158 | 2.977 | 3.546 | 3.145 | 3.495 | 3.484 | -0.013 | 0.8467 | 0.9343 | 0 | 0 | 0 |
| 'Ghitm' | 65.836 | 74.100 | 67.567 | 70.324 | 70.203 | 67.960 | 71.532 | 72.813 | 66.520 | 68.133 | 75.230 | 67.829 | 73.381 | 75.084 | 68.910 | 70.145 | 75.059 | 64.214 | -0.035 | 0.1239 | 0.3364 | 0 | 0 | 0 |
| 'Gid4' | 6.860 | 8.125 | 7.261 | 6.762 | 8.719 | 7.239 | 6.710 | 7.444 | 6.900 | 7.156 | 6.865 | 7.711 | 7.182 | 7.835 | 7.013 | 7.493 | 6.767 | 7.646 | -0.010 | 0.8346 | 0.9283 | 0 | 0 | 0 |
| 'Gid8' | 17.094 | 19.598 | 18.942 | 17.380 | 18.449 | 19.588 | 18.724 | 18.746 | 19.629 | 17.392 | 16.747 | 18.948 | 17.040 | 18.858 | 18.046 | 17.361 | 18.779 | 17.894 | 0.050 | 0.0983 | 0.2930 | 0 | 0 | 0 |
| 'Gigyf2' | 6.193 | 5.681 | 5.984 | 6.022 | 5.842 | 5.908 | 5.664 | 5.807 | 6.165 | 5.909 | 5.731 | 6.054 | 5.543 | 5.318 | 5.663 | 6.341 | 5.649 | 6.155 | 0.015 | 0.6252 | 0.8133 | 0 | 0 | 0 |
| 'Gimap1' | 51.552 | 60.923 | 54.603 | 50.229 | 58.608 | 57.678 | 53.870 | 54.456 | 57.102 | 50.842 | 54.027 | 58.067 | 50.079 | 55.122 | 57.936 | 56.052 | 52.059 | 55.307 | 0.016 | 0.5979 | 0.7954 | 0 | 0 | 0 |
| 'Gimap3' | 204.528 | 242.743 | 233.281 | 197.216 | 239.609 | 231.881 | 207.155 | 234.210 | 241.639 | 212.675 | 231.773 | 240.346 | 221.242 | 234.084 | 246.862 | 221.005 | 228.064 | 238.931 | -0.043 | 0.0406 | 0.1649 | 0 | 0 | 0 |
| 'Gimap4' | 21.413 | 18.247 | 24.377 | 21.138 | 22.026 | 25.321 | 23.187 | 20.478 | 23.214 | 18.187 | 18.559 | 31.369 | 14.521 | 18.596 | 29.680 | 20.955 | 20.422 | 28.820 | 0.011 | 0.9005 | 0.9605 | 0 | 0 | 0 |
| 'Gimap5' | 24.312 | 27.764 | 23.759 | 23.330 | 28.033 | 24.977 | 23.093 | 26.329 | 24.789 | 25.175 | 26.293 | 25.817 | 23.591 | 27.580 | 26.085 | 27.111 | 26.223 | 25.435 | -0.056 | 0.0638 | 0.2204 | 0 | 0 | 0 |
| 'Gimap6' | 121.908 | 140.473 | 125.768 | 121.788 | 145.210 | 128.444 | 129.354 | 143.710 | 125.050 | 143.890 | 133.013 | 109.181 | 138.056 | 137.122 | 116.419 | 142.020 | 124.124 | 107.195 | 0.032 | 0.4543 | 0.6930 | 0 | 0 | 0 |
| 'Gimap7' | 36.318 | 28.003 | 32.170 | 35.866 | 31.059 | 30.154 | 33.861 | 28.645 | 31.990 | 22.181 | 28.248 | 38.419 | 21.959 | 24.778 | 34.008 | 24.148 | 23.943 | 37.196 | 0.189 | 0.0580 | 0.2074 | 0 | 0 | 0 |
| 'Gimap8' | 53.075 | 56.051 | 50.944 | 52.822 | 55.111 | 53.668 | 55.096 | 55.718 | 53.179 | 46.641 | 55.043 | 53.166 | 44.651 | 54.466 | 53.179 | 52.722 | 54.251 | 53.018 | 0.047 | 0.1674 | 0.3998 | 0 | 0 | 0 |
| 'Gimap9' | 56.458 | 64.696 | 61.069 | 67.268 | 65.887 | 61.936 | 67.422 | 60.788 | 59.312 | 50.435 | 55.616 | 62.293 | 51.532 | 60.024 | 63.253 | 53.481 | 62.957 | 61.252 | 0.108 | 0.0233 | 0.1147 | 0 | 0 | 0 |
| 'Gin1' | 1.692 | 1.855 | 1.837 | 1.772 | 1.719 | 1.618 | 1.953 | 1.661 | 1.928 | 1.855 | 1.605 | 1.267 | 2.024 | 1.395 | 1.196 | 1.678 | 1.701 | 1.297 | 0.202 | 0.0087 | 0.0564 | 0 | 0 | 0 |
| 'Ginm1' | 4.643 | 5.401 | 4.533 | 4.682 | 5.261 | 4.903 | 5.262 | 4.943 | 4.524 | 5.317 | 4.802 | 4.192 | 4.635 | 4.812 | 3.923 | 5.146 | 4.851 | 3.871 | 0.084 | 0.0868 | 0.2683 | 0 | 0 | 0 |
| 'Gins1' | 4.887 | 4.677 | 6.213 | 5.410 | 4.695 | 5.448 | 5.290 | 5.969 | 5.800 | 4.166 | 4.411 | 4.761 | 5.748 | 4.480 | 5.253 | 3.915 | 5.123 | 4.613 | 0.180 | 0.0196 | 0.1016 | 0 | 0 | 0 |
| 'Gins2' | 1.994 | 1.773 | 2.368 | 2.229 | 1.656 | 1.987 | 2.284 | 1.808 | 2.008 | 1.684 | 1.732 | 1.920 | 2.375 | 1.740 | 2.175 | 1.805 | 1.743 | 1.978 | 0.066 | 0.3206 | 0.5786 | 0 | 0 | 0 |
| 'Gins3' | 7.405 | 7.449 | 8.944 | 7.781 | 8.269 | 8.573 | 9.182 | 8.807 | 8.854 | 6.869 | 8.030 | 8.536 | 8.303 | 8.805 | 8.383 | 6.924 | 7.253 | 8.101 | 0.068 | 0.2240 | 0.4711 | 0 | 0 | 0 |
| 'Gins4' | 26.188 | 29.334 | 26.628 | 28.003 | 27.928 | 29.139 | 27.754 | 28.538 | 26.800 | 23.848 | 25.051 | 26.026 | 29.046 | 26.830 | 27.416 | 29.599 | 26.265 | 26.045 | 0.049 | 0.2491 | 0.5000 | 0 | 0 | 0 |
| 'Gipc1' | 32.634 | 35.564 | 37.833 | 31.723 | 39.885 | 40.967 | 32.795 | 38.210 | 41.330 | 34.818 | 29.165 | 34.404 | 38.169 | 32.812 | 34.388 | 38.328 | 29.913 | 30.434 | 0.120 | 0.0454 | 0.1770 | 0 | 0 | 0 |
| 'Git1' | 18.814 | 17.291 | 19.213 | 16.395 | 15.069 | 18.777 | 16.867 | 17.146 | 19.396 | 17.072 | 15.449 | 21.872 | 19.108 | 16.432 | 19.993 | 18.864 | 17.376 | 21.130 | -0.081 | 0.0352 | 0.1501 | 0 | 0 | 0 |
| 'Git2' | 10.758 | 10.163 | 10.135 | 10.616 | 9.837 | 10.391 | 9.812 | 9.960 | 10.359 | 10.173 | 10.403 | 11.515 | 9.817 | 10.621 | 10.952 | 10.238 | 9.840 | 11.291 | -0.053 | 0.0868 | 0.2684 | 0 | 0 | 0 |
| 'Gjb2' | 2.049 | 2.120 | 2.296 | 1.866 | 2.147 | 2.155 | 2.198 | 2.039 | 2.167 | 2.715 | 2.192 | 3.177 | 1.973 | 2.352 | 2.793 | 2.420 | 2.694 | 3.127 | -0.318 | 0.0005 | 0.0064 | 1 | 0 | 0 |
| 'Gk' | 1.371 | 1.430 | 1.352 | 1.219 | 1.465 | 1.152 | 1.461 | 1.331 | 1.178 | 1.340 | 1.468 | 1.354 | 1.546 | 1.615 | 1.215 | 1.540 | 1.968 | 1.339 | -0.169 | 0.0120 | 0.0716 | 0 | 0 | 0 |
| 'Gla' | 17.039 | 5.039 | 3.761 | 11.775 | 5.321 | 3.372 | 8.819 | 5.699 | 2.891 | 11.444 | 9.799 | 2.911 | 11.987 | 8.562 | 2.824 | 7.729 | 9.250 | 2.702 | -0.094 | 0.5949 | 0.7937 | 0 | 0 | 0 |
| 'Glb1' | 24.164 | 23.324 | 26.065 | 22.954 | 24.892 | 25.350 | 22.557 | 24.200 | 23.656 | 24.098 | 22.255 | 23.524 | 24.563 | 23.411 | 23.481 | 26.582 | 23.019 | 21.311 | 0.028 | 0.4723 | 0.7079 | 0 | 0 | 0 |
| 'Glb1l' | 0.864 | 0.918 | 1.129 | 0.957 | 0.889 | 0.958 | 0.943 | 0.897 | 1.110 | 1.080 | 0.880 | 1.021 | 1.064 | 1.012 | 1.010 | 0.830 | 0.918 | 0.788 | 0.000 | 0.9975 | 0.9994 | 0 | 0 | 0 |
| 'Glcci1' | 1.410 | 1.167 | 1.439 | 1.596 | 1.284 | 1.361 | 1.237 | 1.468 | 1.051 | 1.664 | 1.891 | 1.224 | 1.603 | 1.401 | 1.352 | 1.662 | 1.437 | 1.545 | -0.196 | 0.0164 | 0.0898 | 0 | 0 | 0 |
| 'Glce' | 1.413 | 1.863 | 1.834 | 1.405 | 1.940 | 1.974 | 1.527 | 1.937 | 1.777 | 1.766 | 1.877 | 1.430 | 1.763 | 1.745 | 1.536 | 2.151 | 1.648 | 1.197 | 0.052 | 0.5622 | 0.7715 | 0 | 0 | 0 |
| 'Gle1' | 7.852 | 7.927 | 8.700 | 7.858 | 8.783 | 9.229 | 8.455 | 8.480 | 9.176 | 8.330 | 7.745 | 8.563 | 8.097 | 8.426 | 8.955 | 8.677 | 7.992 | 8.211 | 0.019 | 0.5555 | 0.7676 | 0 | 0 | 0 |
| 'Glg1' | 33.209 | 32.099 | 33.171 | 33.496 | 32.523 | 32.236 | 30.853 | 33.305 | 32.897 | 35.662 | 36.355 | 34.483 | 33.287 | 34.072 | 33.914 | 34.890 | 32.784 | 35.832 | -0.094 | 0.0006 | 0.0071 | 1 | 0 | 0 |
| 'Glipr1' | 14.948 | 16.817 | 14.715 | 13.914 | 16.874 | 15.715 | 13.440 | 16.515 | 15.117 | 14.505 | 13.835 | 15.587 | 14.552 | 16.481 | 16.025 | 15.872 | 15.489 | 14.717 | -0.003 | 0.9495 | 0.9809 | 0 | 0 | 0 |
| 'Glipr2' | 6.784 | 6.859 | 7.441 | 6.925 | 6.421 | 8.897 | 6.381 | 7.227 | 7.662 | 8.019 | 5.933 | 8.708 | 7.088 | 8.169 | 10.116 | 7.623 | 6.471 | 7.699 | -0.119 | 0.0756 | 0.2441 | 0 | 0 | 0 |
| 'Glis3' | 0.152 | 0.186 | 0.194 | 0.245 | 0.216 | 0.184 | 0.182 | 0.186 | 0.145 | 0.263 | 0.302 | 0.260 | 0.283 | 0.343 | 0.283 | 0.275 | 0.286 | 0.244 | -0.599 | 0.0000 | 0.0000 | 1 | 1 | 0 |
| 'Glmn' | 1.505 | 1.771 | 1.778 | 1.870 | 2.002 | 1.710 | 2.138 | 1.926 | 1.574 | 1.783 | 1.833 | 1.668 | 1.755 | 1.659 | 1.725 | 1.457 | 1.660 | 1.393 | 0.105 | 0.1559 | 0.3849 | 0 | 0 | 0 |
| 'Glmp' | 7.674 | 7.635 | 8.369 | 7.764 | 8.281 | 8.415 | 7.895 | 7.903 | 8.561 | 7.988 | 7.640 | 8.682 | 8.072 | 8.157 | 8.497 | 7.764 | 8.114 | 8.586 | -0.031 | 0.1772 | 0.4124 | 0 | 0 | 0 |
| 'Glo1' | 52.276 | 58.727 | 60.426 | 59.951 | 60.039 | 59.472 | 56.940 | 58.631 | 57.024 | 34.863 | 34.553 | 36.552 | 36.186 | 37.275 | 37.591 | 34.976 | 35.596 | 36.503 | 0.680 | 0.0000 | 0.0000 | 1 | 1 | 0 |
| 'Glod4' | 5.013 | 5.327 | 5.279 | 5.259 | 5.297 | 4.685 | 5.358 | 5.840 | 5.213 | 5.085 | 5.093 | 5.016 | 4.919 | 5.260 | 4.440 | 4.628 | 4.823 | 4.143 | 0.111 | 0.0071 | 0.0485 | 1 | 0 | 0 |
| 'Glrx' | 46.132 | 43.398 | 45.367 | 45.635 | 46.031 | 44.283 | 46.605 | 43.933 | 40.909 | 48.118 | 50.165 | 44.401 | 47.999 | 47.206 | 45.058 | 49.555 | 43.425 | 40.085 | -0.054 | 0.1248 | 0.3381 | 0 | 0 | 0 |
| 'Glrx2' | 1.792 | 2.167 | 2.023 | 2.112 | 2.175 | 1.976 | 2.092 | 2.136 | 1.884 | 1.849 | 1.734 | 2.143 | 1.822 | 1.923 | 2.016 | 1.947 | 2.038 | 1.803 | 0.069 | 0.1357 | 0.3545 | 0 | 0 | 0 |
| 'Glrx5' | 61.480 | 64.198 | 64.212 | 63.151 | 66.031 | 66.831 | 59.245 | 70.958 | 62.810 | 63.581 | 60.022 | 60.010 | 64.605 | 63.688 | 62.880 | 62.359 | 61.604 | 61.455 | 0.040 | 0.2394 | 0.4896 | 0 | 0 | 0 |
| 'Gls' | 12.421 | 14.004 | 12.325 | 12.373 | 13.499 | 11.710 | 13.294 | 14.134 | 11.441 | 13.233 | 13.935 | 12.112 | 12.002 | 13.638 | 11.821 | 12.948 | 14.080 | 12.143 | -0.020 | 0.4170 | 0.6648 | 0 | 0 | 0 |
| 'Glt8d1' | 7.511 | 11.301 | 9.531 | 8.580 | 10.780 | 10.476 | 7.663 | 10.028 | 10.902 | 9.840 | 9.709 | 10.536 | 10.236 | 10.813 | 10.120 | 10.360 | 12.856 | 9.609 | -0.133 | 0.0342 | 0.1478 | 0 | 0 | 0 |
| 'Gltp' | 99.395 | 102.585 | 96.019 | 99.440 | 105.869 | 100.943 | 95.984 | 108.595 | 97.428 | 93.680 | 107.108 | 90.493 | 97.840 | 97.588 | 97.290 | 91.861 | 96.263 | 95.473 | 0.052 | 0.0563 | 0.2038 | 0 | 0 | 0 |
| 'Gltscr1' | 18.814 | 16.086 | 19.016 | 17.987 | 15.700 | 17.560 | 15.512 | 17.582 | 20.179 | 19.312 | 17.700 | 18.376 | 19.395 | 17.069 | 17.287 | 18.224 | 15.812 | 17.320 | -0.027 | 0.5428 | 0.7596 | 0 | 0 | 0 |
| 'Gltscr1l' | 16.606 | 15.097 | 14.550 | 15.437 | 15.266 | 14.736 | 14.443 | 15.579 | 14.406 | 17.351 | 16.634 | 14.523 | 15.204 | 15.571 | 13.726 | 16.500 | 14.874 | 14.150 | -0.031 | 0.3813 | 0.6326 | 0 | 0 | 0 |
| 'Gltscr2' | 88.384 | 76.747 | 86.339 | 80.556 | 80.895 | 89.190 | 84.681 | 80.075 | 83.364 | 88.679 | 76.997 | 84.557 | 88.073 | 76.941 | 84.777 | 89.927 | 75.559 | 82.322 | -0.003 | 0.8927 | 0.9562 | 0 | 0 | 0 |
| 'Glud1' | 27.013 | 27.286 | 28.217 | 26.028 | 27.395 | 27.970 | 28.551 | 27.867 | 28.512 | 28.078 | 26.188 | 27.567 | 26.952 | 27.091 | 27.209 | 28.268 | 27.025 | 26.117 | 0.016 | 0.4562 | 0.6941 | 0 | 0 | 0 |
| 'Glul' | 10.168 | 9.164 | 8.293 | 10.027 | 9.148 | 9.139 | 9.009 | 9.507 | 8.776 | 9.494 | 8.726 | 7.667 | 9.745 | 9.810 | 8.041 | 10.132 | 8.402 | 7.052 | 0.075 | 0.0978 | 0.2920 | 0 | 0 | 0 |
| 'Glyr1' | 51.416 | 55.358 | 55.137 | 53.684 | 54.936 | 52.129 | 47.947 | 51.078 | 51.310 | 52.197 | 51.686 | 54.852 | 52.145 | 49.397 | 53.170 | 53.834 | 52.319 | 53.578 | -0.012 | 0.6784 | 0.8461 | 0 | 0 | 0 |
| 'Gm10053' | 16.191 | 13.299 | 14.252 | 17.613 | 16.556 | 14.278 | 17.793 | 15.360 | 14.584 | 12.567 | 14.131 | 13.406 | 15.470 | 16.225 | 13.665 | 14.406 | 16.977 | 12.577 | 0.102 | 0.1503 | 0.3777 | 0 | 0 | 0 |
| 'Gm10076' | 17.223 | 21.591 | 15.760 | 24.466 | 20.134 | 19.150 | 20.670 | 22.477 | 19.341 | 15.414 | 16.948 | 14.918 | 17.101 | 17.621 | 18.000 | 21.708 | 21.460 | 16.798 | 0.167 | 0.0668 | 0.2260 | 0 | 0 | 0 |
| 'Gm10146' | 66.412 | 72.754 | 74.378 | 66.677 | 80.325 | 74.466 | 75.647 | 79.835 | 70.729 | 63.893 | 69.310 | 68.791 | 72.878 | 71.917 | 69.350 | 73.500 | 72.974 | 70.660 | 0.052 | 0.2063 | 0.4504 | 0 | 0 | 0 |
| 'Gm10184' | 9.458 | 13.007 | 13.341 | 10.998 | 12.572 | 13.777 | 10.249 | 12.003 | 14.090 | 8.110 | 10.650 | 13.366 | 11.477 | 13.981 | 14.717 | 10.699 | 10.921 | 13.270 | 0.020 | 0.7371 | 0.8802 | 0 | 0 | 0 |
| 'Gm10250' | 17.163 | 18.137 | 15.058 | 19.977 | 23.996 | 19.370 | 20.399 | 18.790 | 15.070 | 14.152 | 16.270 | 17.630 | 18.063 | 18.275 | 17.203 | 22.004 | 19.285 | 14.364 | 0.076 | 0.3803 | 0.6318 | 0 | 0 | 0 |
| 'Gm10263' | 53.350 | 62.503 | 55.583 | 69.180 | 60.023 | 62.020 | 74.077 | 68.290 | 49.830 | 50.283 | 61.673 | 55.337 | 59.483 | 61.730 | 60.507 | 77.120 | 60.173 | 52.940 | 0.030 | 0.6895 | 0.8534 | 0 | 0 | 0 |
| 'Gm10269' | 73.731 | 63.903 | 64.296 | 78.938 | 65.954 | 66.118 | 82.605 | 72.610 | 59.772 | 60.648 | 66.245 | 60.226 | 65.427 | 70.129 | 64.702 | 78.462 | 68.949 | 59.546 | 0.070 | 0.1955 | 0.4340 | 0 | 0 | 0 |
| 'Gm10382' | 57.142 | 26.066 | 32.009 | 51.469 | 28.138 | 27.541 | 56.573 | 28.049 | 28.860 | 39.676 | 41.662 | 26.621 | 39.668 | 33.885 | 27.511 | 34.580 | 36.882 | 28.619 | 0.053 | 0.6590 | 0.8353 | 0 | 0 | 0 |
| 'Gm10521' | 3.160 | 3.625 | 3.910 | 3.168 | 3.173 | 3.117 | 3.148 | 4.046 | 3.498 | 4.309 | 3.607 | 3.119 | 3.147 | 3.250 | 2.564 | 4.175 | 3.055 | 3.342 | 0.018 | 0.8495 | 0.9351 | 0 | 0 | 0 |
| 'Gm10553' | 14.338 | 19.397 | 14.724 | 18.684 | 22.962 | 16.404 | 18.777 | 17.852 | 11.508 | 14.570 | 14.077 | 12.031 | 16.744 | 15.574 | 13.442 | 15.391 | 18.002 | 12.916 | 0.200 | 0.0255 | 0.1221 | 0 | 0 | 0 |
| 'Gm10638' | 4.806 | 4.768 | 4.318 | 4.473 | 4.360 | 4.455 | 5.335 | 5.183 | 5.344 | 3.796 | 3.131 | 3.736 | 5.367 | 4.141 | 4.570 | 4.740 | 4.544 | 5.657 | 0.109 | 0.3482 | 0.6036 | 0 | 0 | 0 |

| | | | | | | | | | | | | | | | | | | | | | | | | |
|---|---|---|---|---|---|---|---|---|---|---|---|---|---|---|---|---|---|---|---|---|---|---|---|---|
| 'Gm10709' | 237.037 | 227.967 | 220.092 | 256.294 | 236.930 | 236.096 | 252.412 | 229.136 | 228.513 | 224.930 | 233.430 | 184.412 | 227.857 | 225.454 | 231.539 | 246.463 | 225.447 | 216.989 | 0.067 | 0.1128 | 0.3190 | 0 | 0 | 0 |
| 'Gm12258' | 2.782 | 3.075 | 3.956 | 2.620 | 3.378 | 3.361 | 3.418 | 3.146 | 3.934 | 2.792 | 3.228 | 4.166 | 2.619 | 2.533 | 3.792 | 2.382 | 3.847 | 3.578 | 0.025 | 0.7348 | 0.8795 | 0 | 0 | 0 |
| 'Gm12355' | 7.096 | 10.758 | 9.674 | 10.170 | 8.173 | 8.629 | 10.508 | 10.066 | 6.758 | 8.667 | 8.056 | 8.224 | 9.153 | 9.225 | 8.604 | 7.112 | 8.382 | 7.054 | 0.116 | 0.2019 | 0.4432 | 0 | 0 | 0 |
| 'Gm14085' | 15.834 | 15.691 | 17.173 | 16.516 | 16.911 | 17.345 | 17.275 | 16.071 | 16.872 | 17.465 | 18.098 | 18.524 | 16.454 | 18.983 | 17.610 | 16.868 | 17.892 | 16.405 | -0.090 | 0.0064 | 0.0448 | 1 | 0 | 0 |
| 'Gm15013' | 6.096 | 6.763 | 6.811 | 6.181 | 6.464 | 6.600 | 7.032 | 6.177 | 6.777 | 5.993 | 6.218 | 5.080 | 5.797 | 6.744 | 6.294 | 5.411 | 5.892 | 6.218 | 0.134 | 0.0541 | 0.1989 | 0 | 0 | 0 |
| 'Gm15440' | 3.062 | 2.613 | 2.286 | 2.630 | 3.555 | 2.667 | 2.677 | 2.803 | 2.245 | 3.173 | 3.517 | 3.259 | 3.058 | 2.507 | 2.894 | 3.423 | 2.831 | 3.268 | -0.209 | 0.0668 | 0.2261 | 0 | 0 | 0 |
| 'Gm15800' | 13.069 | 9.663 | 11.661 | 12.647 | 9.629 | 11.008 | 10.969 | 10.002 | 12.061 | 10.833 | 12.123 | 13.344 | 10.437 | 11.043 | 11.935 | 9.958 | 12.146 | 14.617 | -0.090 | 0.1756 | 0.4105 | 0 | 0 | 0 |
| 'Gm15821' | 4.564 | 5.915 | 6.994 | 4.390 | 4.933 | 5.775 | 3.428 | 5.807 | 7.650 | 5.468 | 4.370 | 6.396 | 6.149 | 5.175 | 6.425 | 4.570 | 5.948 | 5.756 | -0.044 | 0.6590 | 0.8353 | 0 | 0 | 0 |
| 'Gm16286' | 20.340 | 18.371 | 16.257 | 16.108 | 19.159 | 16.359 | 15.098 | 17.881 | 16.243 | 19.687 | 17.640 | 15.164 | 19.380 | 18.552 | 15.679 | 19.906 | 15.750 | 14.737 | -0.010 | 0.8596 | 0.9390 | 0 | 0 | 0 |
| 'Gm16372' | 9.514 | 7.133 | 7.741 | 9.302 | 9.386 | 8.814 | 9.175 | 9.324 | 8.494 | 7.616 | 8.729 | 9.678 | 8.106 | 9.288 | 10.317 | 11.721 | 10.200 | 8.109 | -0.094 | 0.3873 | 0.6372 | 0 | 0 | 0 |
| 'Gm17018' | 6.878 | 8.560 | 7.928 | 7.293 | 8.618 | 8.707 | 9.013 | 8.068 | 7.702 | 6.809 | 7.093 | 7.412 | 7.839 | 7.246 | 8.625 | 7.400 | 6.558 | 6.668 | 0.134 | 0.0379 | 0.1583 | 0 | 0 | 0 |
| 'Gm18025' | 5.863 | 4.517 | 5.092 | 6.053 | 5.380 | 5.323 | 5.450 | 6.312 | 4.680 | 4.594 | 6.133 | 4.846 | 5.439 | 5.582 | 5.975 | 6.375 | 5.530 | 5.105 | -0.028 | 0.7197 | 0.8711 | 0 | 0 | 0 |
| 'Gm20518' | 17.331 | 18.775 | 19.334 | 15.971 | 19.239 | 19.453 | 14.679 | 18.617 | 19.530 | 17.830 | 17.477 | 20.192 | 16.653 | 17.181 | 19.716 | 18.695 | 16.258 | 20.507 | -0.025 | 0.5522 | 0.7650 | 0 | 0 | 0 |
| 'Gm20604' | 2.759 | 1.937 | 2.181 | 2.528 | 2.336 | 2.391 | 2.577 | 2.257 | 2.180 | 2.192 | 2.225 | 2.302 | 2.421 | 2.524 | 2.117 | 2.111 | 2.394 | 2.150 | 0.037 | 0.6589 | 0.8353 | 0 | 0 | 0 |
| 'Gm20716' | 6.349 | 7.611 | 6.809 | 6.499 | 7.394 | 6.906 | 7.014 | 7.280 | 6.390 | 7.221 | 7.161 | 6.460 | 6.997 | 7.185 | 6.344 | 6.867 | 6.603 | 6.076 | 0.022 | 0.5757 | 0.7807 | 0 | 0 | 0 |
| 'Gm20721' | 3.936 | 3.638 | 3.292 | 4.155 | 3.808 | 3.600 | 3.811 | 3.950 | 4.517 | 2.625 | 3.651 | 3.866 | 3.081 | 3.951 | 2.972 | 2.935 | 3.419 | 3.406 | 0.195 | 0.0466 | 0.1799 | 0 | 0 | 0 |
| 'Gm21685' | 13.927 | 16.298 | 15.943 | 13.054 | 15.608 | 16.544 | 13.410 | 16.158 | 17.871 | 15.614 | 15.676 | 14.665 | 16.387 | 16.163 | 13.762 | 15.801 | 14.583 | 14.118 | 0.011 | 0.8405 | 0.9322 | 0 | 0 | 0 |
| 'Gm21975' | 26.856 | 26.985 | 28.927 | 26.297 | 26.129 | 27.358 | 28.850 | 25.654 | 29.383 | 25.234 | 27.834 | 31.755 | 27.109 | 27.682 | 30.123 | 24.473 | 28.988 | 29.909 | -0.049 | 0.1724 | 0.4062 | 0 | 0 | 0 |
| 'Gm21987' | 8.655 | 9.811 | 8.391 | 9.559 | 10.923 | 8.695 | 9.001 | 9.302 | 8.240 | 8.413 | 9.711 | 8.138 | 9.929 | 10.304 | 8.934 | 9.580 | 10.065 | 8.180 | -0.022 | 0.5720 | 0.7788 | 0 | 0 | 0 |
| 'Gm21992' | 6.319 | 6.260 | 6.297 | 7.041 | 5.836 | 6.195 | 6.847 | 5.868 | 6.236 | 5.287 | 6.425 | 7.926 | 5.855 | 5.551 | 7.224 | 5.638 | 7.208 | 8.209 | -0.064 | 0.3682 | 0.6201 | 0 | 0 | 0 |
| 'Gm28035' | 3.740 | 3.101 | 3.641 | 4.083 | 3.345 | 3.378 | 3.283 | 3.136 | 3.187 | 3.002 | 3.535 | 4.124 | 3.589 | 3.311 | 4.213 | 3.573 | 3.636 | 4.385 | -0.122 | 0.0679 | 0.2284 | 0 | 0 | 0 |
| 'Gm28042' | 0.930 | 0.758 | 1.173 | 0.924 | 0.936 | 1.039 | 1.115 | 0.908 | 1.251 | 1.315 | 0.996 | 1.315 | 1.212 | 1.003 | 1.093 | 0.901 | 1.139 | 1.213 | -0.189 | 0.0155 | 0.0856 | 0 | 0 | 0 |
| 'Gm28043' | 1.636 | 2.009 | 2.246 | 1.862 | 2.343 | 1.935 | 1.939 | 2.050 | 2.215 | 1.787 | 1.847 | 1.846 | 1.967 | 1.983 | 1.568 | 1.997 | 2.031 | 1.859 | 0.102 | 0.1336 | 0.3522 | 0 | 0 | 0 |
| 'Gm28048' | 6.948 | 9.628 | 8.273 | 8.282 | 9.484 | 7.971 | 8.902 | 7.795 | 8.715 | 8.780 | 7.397 | 8.214 | 8.863 | 9.795 | 7.928 | 9.625 | 8.605 | 6.697 | -0.009 | 0.8987 | 0.9598 | 0 | 0 | 0 |
| 'Gm28053' | 4.102 | 4.317 | 5.325 | 4.242 | 4.198 | 5.081 | 4.752 | 4.897 | 5.541 | 4.059 | 4.126 | 4.825 | 3.955 | 4.364 | 4.515 | 4.885 | 4.383 | 4.061 | 0.113 | 0.2148 | 0.4603 | 0 | 0 | 0 |
| 'Gm28285' | 2.279 | 2.208 | 1.582 | 1.972 | 1.794 | 1.698 | 2.054 | 1.902 | 1.739 | 1.774 | 1.758 | 1.904 | 2.096 | 1.740 | 2.001 | 1.677 | 2.522 | 1.937 | -0.042 | 0.6934 | 0.8558 | 0 | 0 | 0 |
| 'Gm28551' | 22.508 | 23.385 | 22.035 | 22.154 | 25.237 | 21.849 | 22.942 | 22.904 | 21.787 | 23.029 | 23.957 | 24.362 | 23.630 | 26.549 | 23.444 | 23.061 | 26.756 | 22.588 | -0.099 | 0.0012 | 0.0126 | 1 | 0 | 0 |
| 'Gm29094' | 93.640 | 110.644 | 98.483 | 103.879 | 104.950 | 95.638 | 102.755 | 103.636 | 99.098 | 95.864 | 96.812 | 89.754 | 104.116 | 98.011 | 88.914 | 108.500 | 101.795 | 89.925 | 0.054 | 0.1007 | 0.2971 | 0 | 0 | 0 |
| 'Gm29394' | 9.701 | 9.835 | 10.682 | 8.393 | 10.345 | 10.967 | 9.859 | 9.579 | 11.455 | 11.076 | 8.830 | 11.498 | 10.287 | 10.770 | 9.989 | 12.167 | 9.262 | 9.158 | -0.043 | 0.4890 | 0.7189 | 0 | 0 | 0 |
| 'Gm29427' | 4.192 | 4.028 | 3.985 | 4.472 | 3.758 | 4.104 | 2.930 | 3.672 | 4.343 | 4.445 | 3.547 | 4.190 | 3.145 | 3.927 | 3.671 | 3.170 | 4.089 | 4.296 | 0.031 | 0.7313 | 0.8773 | 0 | 0 | 0 |
| 'Gm2a' | 32.627 | 30.308 | 33.693 | 31.172 | 32.486 | 34.027 | 31.346 | 32.213 | 31.527 | 34.259 | 31.576 | 32.206 | 30.507 | 32.086 | 32.474 | 32.004 | 29.167 | 32.490 | 0.005 | 0.8768 | 0.9479 | 0 | 0 | 0 |
| 'Gm340' | 2.892 | 2.419 | 2.907 | 2.901 | 2.097 | 2.302 | 3.596 | 2.386 | 2.543 | 1.932 | 2.407 | 2.382 | 2.371 | 2.069 | 2.110 | 1.746 | 2.486 | 2.522 | 0.241 | 0.0125 | 0.0735 | 0 | 0 | 0 |
| 'Gm3550' | 28.675 | 26.593 | 26.915 | 27.322 | 28.106 | 26.033 | 28.448 | 28.559 | 28.387 | 25.051 | 24.838 | 24.934 | 27.506 | 24.315 | 27.816 | 28.328 | 26.642 | 25.527 | 0.076 | 0.0587 | 0.2089 | 0 | 0 | 0 |
| 'Gm42688' | 3.679 | 3.124 | 3.807 | 2.931 | 3.901 | 4.208 | 3.620 | 3.347 | 3.933 | 4.014 | 3.265 | 3.624 | 3.193 | 3.371 | 3.376 | 3.219 | 3.446 | 3.544 | 0.062 | 0.3564 | 0.6092 | 0 | 0 | 0 |
| 'Gm42715' | 0.335 | 0.350 | 0.366 | 0.247 | 0.351 | 0.299 | 0.392 | 0.430 | 0.326 | 0.364 | 0.386 | 0.332 | 0.395 | 0.397 | 0.311 | 0.403 | 0.377 | 0.256 | -0.053 | 0.6015 | 0.7979 | 0 | 0 | 0 |
| 'Gm4353' | 1.970 | 1.973 | 1.821 | 1.936 | 1.956 | 2.234 | 1.721 | 2.091 | 2.351 | 1.797 | 1.814 | 2.325 | 1.865 | 1.989 | 1.875 | 1.853 | 1.980 | 2.076 | 0.021 | 0.7636 | 0.8953 | 0 | 0 | 0 |
| 'Gm4924' | 2.011 | 2.936 | 1.989 | 1.781 | 3.383 | 2.170 | 2.160 | 3.054 | 2.084 | 2.386 | 2.004 | 1.832 | 1.988 | 2.878 | 1.648 | 2.504 | 2.087 | 1.681 | 0.162 | 0.0785 | 0.2513 | 0 | 0 | 0 |
| 'Gm5160' | 12.016 | 12.369 | 13.378 | 15.250 | 13.572 | 11.027 | 17.432 | 16.635 | 12.448 | 9.101 | 11.349 | 12.331 | 11.554 | 13.572 | 11.514 | 16.047 | 13.658 | 8.119 | 0.199 | 0.0753 | 0.2440 | 0 | 0 | 0 |
| 'Gm5218' | 12.856 | 9.807 | 9.340 | 10.415 | 10.321 | 10.868 | 7.410 | 11.189 | 9.333 | 9.689 | 10.585 | 10.592 | 10.167 | 8.493 | 15.038 | 8.854 | 8.547 | 8.132 | 0.018 | 0.8895 | 0.9547 | 0 | 0 | 0 |
| 'Gm5239' | 44.474 | 46.989 | 40.388 | 50.062 | 42.967 | 37.621 | 47.141 | 49.970 | 45.959 | 36.867 | 45.447 | 46.767 | 45.995 | 42.818 | 42.344 | 51.073 | 42.309 | 46.862 | 0.000 | 0.9997 | 0.9999 | 0 | 0 | 0 |
| 'Gm527' | 2.576 | 2.842 | 2.956 | 3.119 | 2.893 | 2.953 | 3.824 | 3.096 | 2.576 | 3.038 | 3.283 | 2.500 | 3.618 | 3.968 | 2.003 | 2.042 | 4.002 | 2.928 | -0.027 | 0.8231 | 0.9223 | 0 | 0 | 0 |
| 'Gm5422' | 1.377 | 1.161 | 1.327 | 1.621 | 1.333 | 1.308 | 1.978 | 1.651 | 0.908 | 1.066 | 1.409 | 1.354 | 1.258 | 1.163 | 1.641 | 1.371 | 1.247 | 1.330 | 0.074 | 0.5612 | 0.7711 | 0 | 0 | 0 |
| 'Gm561' | 12.805 | 11.826 | 12.611 | 13.907 | 14.318 | 14.554 | 12.202 | 14.079 | 13.603 | 10.295 | 12.202 | 14.839 | 13.788 | 11.389 | 13.781 | 11.548 | 13.318 | 13.643 | 0.041 | 0.6379 | 0.8219 | 0 | 0 | 0 |
| 'Gm5612' | 26.852 | 27.732 | 24.481 | 27.826 | 24.716 | 23.454 | 23.143 | 32.445 | 21.499 | 25.763 | 30.470 | 25.456 | 26.248 | 27.620 | 25.188 | 31.535 | 30.248 | 21.738 | -0.077 | 0.2641 | 0.5180 | 0 | 0 | 0 |
| 'Gm5617' | 11.597 | 13.540 | 12.860 | 13.652 | 14.926 | 13.750 | 12.235 | 14.435 | 13.981 | 13.252 | 11.556 | 10.458 | 11.795 | 11.497 | 12.231 | 13.468 | 11.916 | 10.027 | 0.190 | 0.0164 | 0.0898 | 0 | 0 | 0 |
| 'Gm5819' | 138.902 | 169.682 | 147.708 | 140.726 | 155.530 | 140.956 | 144.747 | 144.886 | 159.866 | 125.990 | 150.951 | 178.827 | 154.124 | 146.592 | 164.315 | 128.475 | 158.127 | 170.090 | -0.049 | 0.3259 | 0.5834 | 0 | 0 | 0 |
| 'Gm6563' | 14.439 | 16.789 | 15.709 | 20.999 | 18.635 | 15.350 | 18.977 | 18.794 | 16.062 | 13.958 | 16.424 | 16.198 | 16.599 | 16.023 | 13.143 | 17.631 | 18.479 | 13.429 | 0.117 | 0.1088 | 0.3107 | 0 | 0 | 0 |
| 'Gm6576' | 36.252 | 36.168 | 35.856 | 39.970 | 40.925 | 34.908 | 42.142 | 41.966 | 35.917 | 30.311 | 36.620 | 37.725 | 40.910 | 37.397 | 37.012 | 35.947 | 39.187 | 35.662 | 0.040 | 0.4537 | 0.6927 | 0 | 0 | 0 |
| 'Gm684' | 3.122 | 3.727 | 2.599 | 3.963 | 3.124 | 1.870 | 3.337 | 2.774 | 2.835 | 3.253 | 4.052 | 4.055 | 3.543 | 3.142 | 3.230 | 3.093 | 4.525 | 4.343 | -0.322 | 0.0115 | 0.0692 | 0 | 0 | 0 |
| 'Gm6904' | 9.463 | 9.337 | 9.974 | 11.378 | 15.317 | 11.520 | 11.941 | 11.180 | 10.713 | 13.552 | 12.945 | 12.074 | 11.450 | 10.730 | 11.560 | 12.032 | 11.545 | 9.487 | -0.078 | 0.3727 | 0.6242 | 0 | 0 | 0 |
| 'Gm7293' | 17.793 | 18.602 | 16.870 | 18.041 | 18.435 | 17.091 | 19.498 | 17.500 | 17.521 | 17.022 | 17.603 | 16.294 | 19.859 | 17.517 | 15.927 | 17.938 | 17.115 | 16.604 | 0.038 | 0.3035 | 0.5608 | 0 | 0 | 0 |
| 'Gm7334' | 8.237 | 5.604 | 6.937 | 7.049 | 7.104 | 6.125 | 8.521 | 6.476 | 6.321 | 6.231 | 6.774 | 6.696 | 6.113 | 7.157 | 6.248 | 7.400 | 7.128 | 7.718 | 0.009 | 0.9155 | 0.9677 | 0 | 0 | 0 |
| 'Gm8186' | 19.213 | 17.407 | 21.225 | 17.672 | 17.009 | 15.757 | 18.866 | 18.353 | 19.663 | 15.967 | 14.119 | 18.097 | 17.416 | 16.082 | 16.587 | 22.587 | 15.888 | 15.562 | 0.113 | 0.2143 | 0.4597 | 0 | 0 | 0 |
| 'Gm8225' | 28.285 | 24.197 | 25.488 | 29.799 | 29.632 | 23.792 | 28.529 | 30.986 | 23.554 | 25.072 | 30.380 | 23.447 | 30.770 | 30.452 | 27.785 | 30.186 | 28.220 | 24.346 | -0.042 | 0.4694 | 0.7053 | 0 | 0 | 0 |
| 'Gm8369' | 7.533 | 7.815 | 6.274 | 6.561 | 7.516 | 6.262 | 7.883 | 7.473 | 6.051 | 7.440 | 7.361 | 5.818 | 7.356 | 7.859 | 5.776 | 7.464 | 8.420 | 4.501 | 0.035 | 0.4995 | 0.7273 | 0 | 0 | 0 |
| 'Gm8394' | 3.884 | 4.425 | 3.858 | 4.670 | 4.383 | 4.061 | 4.953 | 6.146 | 5.536 | 4.431 | 4.904 | 5.336 | 5.259 | 5.613 | 4.995 | 5.671 | 5.816 | 3.929 | -0.146 | 0.1930 | 0.4305 | 0 | 0 | 0 |
| 'Gm8797' | 80.841 | 61.834 | 55.907 | 81.811 | 59.647 | 61.670 | 94.610 | 70.494 | 53.367 | 60.289 | 76.759 | 52.298 | 66.132 | 63.442 | 56.920 | 72.421 | 68.904 | 48.196 | 0.116 | 0.1025 | 0.3002 | 0 | 0 | 0 |
| 'Gm8909' | 15.847 | 18.132 | 16.438 | 17.979 | 16.390 | 16.451 | 17.778 | 16.960 | 16.282 | 18.286 | 19.095 | 19.134 | 20.141 | 19.437 | 18.929 | 18.446 | 20.211 | 18.409 | -0.193 | 0.0000 | 0.0008 | 1 | 0 | 0 |
| 'Gm9803' | 9.744 | 9.924 | 9.510 | 9.263 | 10.249 | 9.897 | 10.272 | 10.654 | 9.171 | 9.748 | 8.671 | 10.067 | 9.581 | 9.296 | 9.291 | 9.643 | 9.694 | 10.126 | 0.025 | 0.6433 | 0.8251 | 0 | 0 | 0 |
| 'Gm9821' | 99.476 | 84.217 | 98.731 | 99.035 | 81.430 | 93.372 | 91.312 | 79.085 | 109.167 | 114.803 | 106.309 | 119.076 | 118.644 | 102.700 | 108.869 | 104.704 | 112.798 | 118.840 | -0.283 | 0.0000 | 0.0000 | 1 | 0 | 0 |
| 'Gm9887' | 13.474 | 9.806 | 13.361 | 10.732 | 13.681 | 13.109 | 10.316 | 13.220 | 13.085 | 13.276 | 11.103 | 13.383 | 11.631 | 10.481 | 11.599 | 13.014 | 11.280 | 13.562 | 0.016 | 0.8446 | 0.9342 | 0 | 0 | 0 |
| 'Gm9938' | 2.705 | 3.570 | 3.468 | 3.809 | 4.081 | 3.539 | 3.444 | 3.864 | 3.537 | 3.639 | 3.896 | 3.531 | 2.639 | 3.135 | 3.464 | 3.007 | 3.885 | 3.923 | 0.026 | 0.7717 | 0.8987 | 0 | 0 | 0 |
| 'Gm996' | 0.455 | 0.290 | 0.538 | 0.550 | 0.304 | 0.537 | 0.454 | 0.345 | 0.643 | 0.384 | 0.249 | 0.373 | 0.294 | 0.209 | 0.380 | 0.271 | 0.343 | 0.330 | 0.522 | 0.0000 | 0.0000 | 1 | 1 | 0 |
| 'Gmcl1' | 8.320 | 9.524 | 8.158 | 7.723 | 9.264 | 9.059 | 8.203 | 9.381 | 8.544 | 8.763 | 9.059 | 9.110 | 8.459 | 9.417 | 9.179 | 8.654 | 9.059 | 8.958 | -0.058 | 0.0639 | 0.2205 | 0 | 0 | 0 |
| 'Gmds' | 2.658 | 2.932 | 3.396 | 3.118 | 2.634 | 3.091 | 3.113 | 2.952 | 2.756 | 2.999 | 3.267 | 3.239 | 3.343 | 3.496 | 3.083 | 3.096 | 2.726 | 3.053 | -0.095 | 0.1319 | 0.3493 | 0 | 0 | 0 |
| 'Gmeb1' | 3.324 | 4.010 | 3.921 | 3.462 | 3.805 | 4.068 | 3.414 | 3.904 | 3.821 | 3.981 | 3.540 | 3.621 | 3.952 | 3.675 | 3.419 | 4.016 | 3.528 | 3.055 | 0.033 | 0.5199 | 0.7425 | 0 | 0 | 0 |
| 'Gmeb2' | 12.336 | 12.349 | 12.690 | 10.677 | 12.061 | 11.968 | 10.881 | 12.682 | 13.100 | 13.018 | 11.622 | 10.974 | 11.865 | 12.561 | 10.596 | 11.638 | 11.083 | 10.281 | 0.063 | 0.1886 | 0.4247 | 0 | 0 | 0 |
| 'Gmfb' | 21.962 | 26.456 | 25.607 | 24.927 | 27.231 | 24.293 | 27.156 | 25.912 | 24.187 | 21.084 | 26.233 | 27.539 | 23.292 | 24.703 | 27.365 | 21.193 | 26.308 | 26.149 | 0.016 | 0.7307 | 0.8772 | 0 | 0 | 0 |
| 'Gmfg' | 25.422 | 25.638 | 27.428 | 24.981 | 26.107 | 26.911 | 27.825 | 25.696 | 25.864 | 26.663 | 27.168 | 27.185 | 26.713 | 28.323 | 29.850 | 27.116 | 28.325 | 26.627 | -0.081 | 0.0075 | 0.0501 | 0 | 0 | 0 |
| 'Gmip' | 29.678 | 25.351 | 31.291 | 27.187 | 25.280 | 29.975 | 25.714 | 27.244 | 30.670 | 28.741 | 26.386 | 34.669 | 29.104 | 25.688 | 30.528 | 27.868 | 27.094 | 33.664 | -0.072 | 0.0254 | 0.1220 | 0 | 0 | 0 |
| 'Gmnn' | 12.963 | 12.437 | 16.693 | 15.509 | 13.619 | 13.205 | 14.512 | 15.430 | 14.070 | 10.995 | 14.274 | 12.920 | 16.887 | 14.090 | 14.313 | 13.708 | 13.827 | 11.678 | 0.058 | 0.4203 | 0.6675 | 0 | 0 | 0 |
| 'Gmppa' | 9.040 | 9.381 | 11.390 | 9.415 | 10.285 | 11.045 | 9.151 | 10.277 | 11.528 | 10.369 | 9.402 | 9.773 | 10.709 | 9.212 | 10.230 | 9.727 | 9.488 | 8.724 | 0.052 | 0.2866 | 0.5433 | 0 | 0 | 0 |
| 'Gmppb' | 4.418 | 4.287 | 4.572 | 4.395 | 4.548 | 4.671 | 4.053 | 4.277 | 4.455 | 4.564 | 4.302 | 4.582 | 4.855 | 4.530 | 4.497 | 4.360 | 4.199 | 4.382 | -0.030 | 0.4852 | 0.7166 | 0 | 0 | 0 |
| 'Gmpr2' | 25.180 | 26.338 | 27.118 | 27.044 | 28.547 | 26.207 | 25.279 | 25.787 | 26.826 | 26.231 | 24.645 | 30.936 | 26.985 | 25.861 | 29.187 | 25.769 | 27.960 | 27.933 | -0.060 | 0.1225 | 0.3338 | 0 | 0 | 0 |
| 'Gmps' | 17.178 | 16.874 | 18.133 | 19.194 | 18.043 | 16.398 | 18.811 | 17.868 | 17.754 | 17.632 | 16.913 | 17.255 | 16.289 | 16.433 | 16.208 | 16.458 | 17.424 | 16.040 | 0.077 | 0.0221 | 0.1102 | 0 | 0 | 0 |
| 'Gna11' | 7.465 | 5.986 | 7.041 | 6.542 | 6.175 | 6.571 | 5.746 | 5.502 | 6.705 | 5.645 | 5.800 | 7.497 | 6.400 | 5.745 | 6.607 | 5.774 | 6.117 | 8.336 | -0.013 | 0.8370 | 0.9301 | 0 | 0 | 0 |
| 'Gna12' | 33.040 | 35.995 | 31.623 | 31.953 | 36.841 | 35.588 | 32.665 | 36.981 | 31.434 | 34.216 | 30.616 | 28.667 | 33.566 | 35.213 | 29.761 | 36.639 | 30.515 | 31.164 | 0.068 | 0.1201 | 0.3301 | 0 | 0 | 0 |
| 'Gna13' | 109.821 | 134.503 | 91.834 | 101.672 | 127.414 | 99.190 | 97.997 | 121.696 | 97.390 | 119.185 | 123.690 | 102.210 | 129.685 | 141.468 | 105.711 | 134.916 | 134.010 | 106.693 | -0.175 | 0.0000 | 0.0003 | 1 | 0 | 0 |
| 'Gna14' | 6.753 | 8.442 | 7.209 | 7.919 | 8.673 | 7.407 | 7.747 | 8.415 | 7.575 | 8.341 | 8.890 | 7.741 | 6.388 | 8.298 | 7.250 | 7.722 | 10.754 | 7.258 | -0.058 | 0.3218 | 0.5798 | 0 | 0 | 0 |
| 'Gnai2' | 117.484 | 124.875 | 119.691 | 119.786 | 122.987 | 131.843 | 120.147 | 118.108 | 123.700 | 120.080 | 118.888 | 117.269 | 126.113 | 120.295 | 127.327 | 121.190 | 114.737 | 124.953 | -0.001 | 0.9813 | 0.9931 | 0 | 0 | 0 |
| 'Gnai3' | 91.799 | 100.029 | 93.099 | 94.533 | 97.889 | 93.256 | 97.022 | 95.962 | 91.307 | 91.997 | 98.522 | 91.570 | 91.543 | 102.885 | 93.051 | 95.492 | 100.594 | 92.153 | -0.015 | 0.4832 | 0.7148 | 0 | 0 | 0 |
| 'Gnal' | 1.883 | 2.643 | 2.299 | 1.879 | 2.529 | 2.281 | 2.222 | 2.828 | 2.651 | 2.561 | 2.119 | 2.309 | 2.199 | 2.029 | 2.145 | 2.435 | 2.421 | 2.114 | 0.046 | 0.5389 | 0.7568 | 0 | 0 | 0 |
| 'Gnaq' | 4.868 | 5.338 | 4.919 | 4.639 | 5.652 | 5.042 | 4.490 | 5.066 | 5.506 | 5.750 | 5.131 | 4.749 | 5.151 | 5.737 | 4.592 | 5.696 | 6.000 | 4.580 | -0.061 | 0.2542 | 0.5061 | 0 | 0 | 0 |
| 'Gnas' | 20.808 | 18.959 | 21.289 | 21.035 | 20.728 | 22.097 | 21.691 | 20.808 | 19.576 | 19.511 | 17.664 | 16.363 | 19.859 | 17.916 | 18.601 | 19.214 | 18.674 | 18.696 | 0.158 | 0.0000 | 0.0001 | 1 | 0 | 0 |
| 'Gnb1' | 40.682 | 43.965 | 39.735 | 42.387 | 44.789 | 42.782 | 41.316 | 44.517 | 39.893 | 41.024 | 43.011 | 39.131 | 42.863 | 43.672 | 40.864 | 41.814 | 43.259 | 40.831 | 0.003 | 0.8841 | 0.9514 | 0 | 0 | 0 |
| 'Gnb1l' | 1.592 | 1.575 | 1.487 | 1.478 | 1.557 | 1.713 | 1.672 | 1.714 | 1.691 | 1.564 | 1.384 | 1.375 | 1.504 | 1.451 | 1.403 | 1.505 | 1.172 | 1.253 | 0.195 | 0.0014 | 0.0139 | 1 | 0 | 0 |
| 'Gnb2' | 36.019 | 37.652 | 36.600 | 34.223 | 36.785 | 41.252 | 35.731 | 36.417 | 38.800 | 33.945 | 32.761 | 38.132 | 38.977 | 36.474 | 38.772 | 36.079 | 35.342 | 37.644 | 0.012 | 0.6855 | 0.8512 | 0 | 0 | 0 |
| 'Gnb4' | 0.268 | 0.250 | 0.217 | 0.222 | 0.262 | 0.236 | 0.192 | 0.221 | 0.211 | 0.180 | 0.247 | 0.240 | 0.196 | 0.138 | 0.219 | 0.214 | 0.274 | 0.235 | 0.080 | 0.4964 | 0.7252 | 0 | 0 | 0 |
| 'Gnb5' | 5.028 | 6.802 | 5.220 | 4.274 | 5.556 | 5.091 | 4.064 | 5.446 | 4.705 | 5.839 | 5.434 | 4.162 | 4.364 | 4.639 | 4.545 | 4.772 | 4.576 | 3.782 | 0.131 | 0.1742 | 0.4083 | 0 | 0 | 0 |
| 'Gne' | 2.994 | 3.089 | 3.206 | 3.251 | 3.274 | 3.247 | 3.310 | 2.923 | 3.278 | 3.360 | 2.984 | 3.236 | 2.970 | 2.862 | 3.086 | 3.208 | 2.720 | 3.207 | 0.036 | 0.3402 | 0.5970 | 0 | 0 | 0 |
| 'Gng10' | 16.322 | 17.199 | 16.145 | 15.676 | 17.860 | 17.209 | 17.647 | 18.195 | 16.115 | 15.090 | 15.136 | 15.197 | 16.832 | 14.941 | 15.811 | 16.901 | 15.954 | 15.535 | 0.097 | 0.0344 | 0.1481 | 0 | 0 | 0 |
| 'Gng12' | 1.065 | 1.123 | 1.158 | 1.099 | 1.141 | 0.977 | 1.050 | 1.099 | 0.977 | 0.810 | 0.982 | 0.818 | 0.940 | 0.788 | 0.802 | 0.861 | 1.114 | 0.698 | 0.306 | 0.0000 | 0.0003 | 1 | 0 | 0 |
| 'Gng2' | 38.182 | 46.399 | 42.423 | 42.255 | 44.130 | 43.055 | 43.337 | 45.017 | 42.509 | 40.095 | 38.976 | 33.915 | 38.983 | 38.559 | 35.346 | 39.414 | 36.672 | 32.587 | 0.203 | 0.0000 | 0.0000 | 1 | 0 | 0 |
| 'Gng3' | 3.789 | 3.417 | 3.190 | 3.509 | 3.547 | 3.763 | 3.699 | 3.268 | 4.178 | 1.982 | 1.992 | 1.812 | 3.148 | 2.509 | 2.164 | 1.849 | 2.215 | 2.029 | 0.713 | 0.0000 | 0.0000 | 1 | 1 | 0 |
| 'Gng5' | 19.279 | 20.491 | 19.024 | 18.075 | 20.311 | 19.284 | 18.798 | 19.527 | 20.085 | 16.462 | 19.722 | 18.808 | 19.340 | 19.893 | 19.281 | 18.705 | 19.371 | 18.907 | 0.026 | 0.4196 | 0.6665 | 0 | 0 | 0 |
| 'Gngt2' | 6.723 | 6.950 | 6.155 | 6.466 | 7.114 | 6.342 | 6.397 | 7.626 | 6.984 | 5.819 | 6.142 | 5.859 | 7.600 | 6.644 | 6.324 | 5.223 | 6.251 | 5.809 | 0.112 | 0.0676 | 0.2279 | 0 | 0 | 0 |
| 'Gnl1' | 77.862 | 74.425 | 79.008 | 73.038 | 75.123 | 86.950 | 72.888 | 76.411 | 80.053 | 75.654 | 67.560 | 86.335 | 74.446 | 69.928 | 84.098 | 83.513 | 65.267 | 86.975 | -0.002 | 0.9560 | 0.9836 | 0 | 0 | 0 |
| 'Gnl2' | 12.737 | 12.355 | 13.162 | 12.066 | 13.328 | 12.621 | 12.961 | 13.040 | 12.227 | 12.699 | 11.932 | 12.340 | 12.502 | 10.512 | 11.980 | 13.255 | 12.356 | 11.514 | 0.061 | 0.0836 | 0.2621 | 0 | 0 | 0 |
| 'Gnl3' | 40.250 | 34.832 | 31.216 | 31.697 | 35.089 | 29.826 | 32.557 | 39.340 | 30.455 | 42.704 | 36.961 | 29.625 | 37.733 | 43.514 | 29.347 | 39.668 | 33.794 | 27.855 | -0.072 | 0.2058 | 0.4494 | 0 | 0 | 0 |
| 'Gnl3l' | 13.358 | 14.373 | 13.132 | 13.768 | 14.439 | 13.336 | 13.457 | 14.491 | 13.352 | 14.375 | 14.536 | 14.022 | 13.134 | 14.263 | 12.838 | 13.859 | 13.678 | 12.349 | -0.004 | 0.8921 | 0.9560 | 0 | 0 | 0 |
| 'Gnpat' | 12.275 | 12.837 | 13.326 | 12.350 | 12.892 | 13.834 | 12.372 | 13.625 | 13.454 | 12.835 | 12.460 | 14.143 | 13.480 | 13.501 | 13.180 | 13.074 | 13.715 | 14.034 | -0.053 | 0.0579 | 0.2074 | 0 | 0 | 0 |
| 'Gnpda2' | 1.940 | 2.127 | 2.041 | 2.102 | 2.238 | 1.859 | 2.168 | 2.552 | 2.170 | 1.991 | 2.420 | 2.386 | 2.053 | 2.292 | 2.229 | 1.876 | 2.269 | 2.264 | -0.060 | 0.2569 | 0.5093 | 0 | 0 | 0 |
| 'Gnpnat1' | 6.631 | 7.527 | 6.647 | 6.881 | 7.057 | 6.987 | 6.526 | 6.990 | 6.200 | 5.829 | 7.498 | 7.665 | 6.032 | 7.714 | 6.649 | 5.432 | 6.759 | 6.628 | 0.013 | 0.8213 | 0.9213 | 0 | 0 | 0 |
| 'Gnptab' | 15.988 | 15.721 | 15.570 | 16.054 | 16.722 | 14.534 | 14.851 | 16.682 | 14.927 | 15.661 | 16.179 | 14.546 | 14.813 | 15.244 | 14.293 | 14.117 | 15.382 | 14.302 | 0.058 | 0.0336 | 0.1458 | 0 | 0 | 0 |
| 'Gnptg' | 26.461 | 35.384 | 30.377 | 28.949 | 32.279 | 30.258 | 31.932 | 31.201 | 29.883 | 27.300 | 31.582 | 35.172 | 31.175 | 32.987 | 32.552 | 30.568 | 34.763 | 32.875 | -0.077 | 0.0508 | 0.1908 | 0 | 0 | 0 |
| 'Gns' | 19.745 | 21.587 | 21.068 | 18.437 | 20.465 | 21.706 | 18.442 | 21.462 | 21.127 | 22.460 | 21.520 | 21.573 | 20.393 | 21.020 | 19.858 | 21.775 | 18.888 | 19.442 | -0.032 | 0.4256 | 0.6723 | 0 | 0 | 0 |
| 'Golga1' | 1.303 | 1.701 | 1.807 | 1.591 | 1.710 | 1.772 | 1.781 | 1.556 | 1.874 | 1.473 | 1.486 | 1.944 | 1.429 | 1.598 | 1.883 | 1.301 | 2.089 | 1.907 | -0.012 | 0.8537 | 0.9373 | 0 | 0 | 0 |
| 'Golga2' | 7.935 | 7.902 | 8.731 | 7.470 | 8.002 | 8.730 | 7.351 | 7.576 | 9.064 | 8.728 | 7.939 | 8.480 | 8.627 | 7.742 | 8.111 | 8.272 | 7.591 | 8.046 | -0.026 | 0.4552 | 0.6934 | 0 | 0 | 0 |
| 'Golga3' | 6.929 | 6.739 | 7.185 | 7.212 | 7.119 | 7.390 | 6.840 | 6.893 | 7.020 | 7.158 | 7.563 | 7.516 | 6.855 | 7.137 | 6.800 | 7.020 | 7.990 | 7.233 | -0.053 | 0.1010 | 0.2976 | 0 | 0 | 0 |
| 'Golga4' | 21.757 | 19.169 | 21.550 | 22.356 | 19.208 | 20.516 | 20.148 | 19.842 | 20.992 | 21.177 | 20.665 | 23.140 | 20.539 | 20.956 | 21.301 | 21.533 | 21.325 | 23.769 | -0.078 | 0.0200 | 0.1030 | 0 | 0 | 0 |
| 'Golga5' | 28.113 | 30.558 | 27.080 | 24.854 | 29.454 | 28.503 | 26.064 | 29.824 | 28.304 | 28.533 | 27.866 | 24.803 | 27.436 | 26.897 | 25.469 | 29.284 | 26.546 | 24.335 | 0.061 | 0.1190 | 0.3284 | 0 | 0 | 0 |
| 'Golga7' | 34.123 | 41.720 | 34.708 | 35.124 | 39.884 | 36.704 | 34.194 | 39.122 | 36.076 | 34.961 | 36.495 | 36.142 | 35.008 | 37.698 | 36.810 | 33.935 | 37.113 | 34.816 | 0.023 | 0.3817 | 0.6330 | 0 | 0 | 0 |
| 'Golgb1' | 8.708 | 7.759 | 9.071 | 9.281 | 7.331 | 8.089 | 8.701 | 7.806 | 8.828 | 9.461 | 9.352 | 8.882 | 7.938 | 7.036 | 7.947 | 8.992 | 8.733 | 8.354 | -0.034 | 0.5151 | 0.7389 | 0 | 0 | 0 |
| 'Golim4' | 1.373 | 1.486 | 1.583 | 1.398 | 1.433 | 1.347 | 1.429 | 1.590 | 1.414 | 1.476 | 1.526 | 1.504 | 1.330 | 1.493 | 1.467 | 1.425 | 1.480 | 1.469 | -0.021 | 0.6137 | 0.8065 | 0 | 0 | 0 |
| 'Golm1' | 2.968 | 2.677 | 3.796 | 3.069 | 2.720 | 3.666 | 3.113 | 3.066 | 3.112 | 2.936 | 2.750 | 4.311 | 3.576 | 2.726 | 3.671 | 3.361 | 3.416 | 3.791 | -0.130 | 0.0548 | 0.2006 | 0 | 0 | 0 |
| 'Golph3' | 58.233 | 67.707 | 57.308 | 58.374 | 63.536 | 59.106 | 58.576 | 65.045 | 57.652 | 62.617 | 65.438 | 63.066 | 55.472 | 66.013 | 61.764 | 60.765 | 62.744 | 61.424 | -0.049 | 0.0794 | 0.2537 | 0 | 0 | 0 |
| 'Golph3l' | 2.637 | 2.757 | 2.959 | 3.049 | 2.929 | 2.852 | 3.037 | 3.094 | 2.885 | 2.462 | 3.238 | 3.062 | 2.445 | 3.094 | 2.918 | 2.801 | 3.107 | 3.076 | -0.008 | 0.8743 | 0.9463 | 0 | 0 | 0 |
| 'Golt1b' | 7.028 | 7.735 | 7.575 | 6.817 | 7.838 | 8.121 | 6.544 | 8.176 | 8.026 | 6.449 | 8.503 | 8.151 | 7.299 | 8.214 | 7.665 | 7.745 | 7.593 | 8.021 | -0.045 | 0.3201 | 0.5782 | 0 | 0 | 0 |
| 'Gon4l' | 16.915 | 13.971 | 16.752 | 16.380 | 13.900 | 15.976 | 15.582 | 14.332 | 16.607 | 16.619 | 16.511 | 16.949 | 15.638 | 14.838 | 15.585 | 14.586 | 14.895 | 16.828 | -0.034 | 0.3848 | 0.6347 | 0 | 0 | 0 |
| 'Gopc' | 8.523 | 8.039 | 8.297 | 8.927 | 7.700 | 8.170 | 8.306 | 8.123 | 7.941 | 7.716 | 9.021 | 9.105 | 7.999 | 9.243 | 8.965 | 7.759 | 10.024 | 8.917 | -0.097 | 0.0425 | 0.1700 | 0 | 0 | 0 |
| 'Gorab' | 2.310 | 1.999 | 2.137 | 1.888 | 2.155 | 2.081 | 2.069 | 2.375 | 2.322 | 2.103 | 1.781 | 1.874 | 2.174 | 1.922 | 1.697 | 2.121 | 1.635 | 1.614 | 0.192 | 0.0052 | 0.0384 | 1 | 0 | 0 |
| 'Gorasp1' | 3.313 | 3.604 | 3.905 | 3.698 | 4.243 | 4.118 | 3.630 | 3.854 | 4.202 | 3.867 | 3.323 | 4.042 | 4.387 | 3.059 | 3.546 | 3.761 | 3.402 | 3.525 | 0.056 | 0.4060 | 0.6552 | 0 | 0 | 0 |
| 'Gorasp2' | 10.971 | 11.536 | 11.086 | 11.299 | 11.429 | 11.670 | 10.879 | 12.344 | 11.591 | 11.327 | 10.589 | 11.550 | 11.325 | 11.158 | 11.638 | 12.831 | 11.173 | 11.895 | -0.019 | 0.5798 | 0.7822 | 0 | 0 | 0 |
| 'Gosr1' | 19.173 | 21.370 | 20.908 | 19.858 | 23.018 | 22.365 | 20.856 | 22.538 | 23.185 | 20.033 | 22.366 | 22.602 | 19.970 | 21.141 | 21.009 | 20.541 | 21.311 | 20.539 | 0.016 | 0.5776 | 0.7819 | 0 | 0 | 0 |
| 'Gosr2' | 28.317 | 31.770 | 30.256 | 28.842 | 29.412 | 31.585 | 29.452 | 31.736 | 30.801 | 31.729 | 29.484 | 31.841 | 28.898 | 29.472 | 32.235 | 28.356 | 30.650 | 29.376 | -0.011 | 0.6971 | 0.8577 | 0 | 0 | 0 |
| 'Got1' | 8.879 | 8.715 | 9.327 | 9.688 | 8.930 | 8.966 | 8.512 | 10.023 | 9.240 | 8.663 | 8.746 | 9.638 | 10.120 | 9.742 | 9.339 | 9.342 | 8.354 | 9.495 | -0.031 | 0.4723 | 0.7079 | 0 | 0 | 0 |
| 'Got2' | 49.850 | 53.078 | 51.759 | 51.005 | 53.649 | 54.189 | 50.751 | 49.664 | 51.203 | 50.552 | 48.379 | 49.402 | 54.251 | 47.950 | 51.984 | 49.967 | 46.472 | 51.278 | 0.036 | 0.2013 | 0.4426 | 0 | 0 | 0 |
| 'Gpaa1' | 6.833 | 6.472 | 7.473 | 6.223 | 6.324 | 7.624 | 6.302 | 6.180 | 7.555 | 6.586 | 5.726 | 8.002 | 7.805 | 6.572 | 7.168 | 7.039 | 6.663 | 7.291 | -0.056 | 0.1717 | 0.4055 | 0 | 0 | 0 |
| 'Gpalpp1' | 9.622 | 11.434 | 11.019 | 10.496 | 12.036 | 11.081 | 10.226 | 10.856 | 11.575 | 10.730 | 11.084 | 11.068 | 10.488 | 10.818 | 11.124 | 10.195 | 10.490 | 10.577 | 0.013 | 0.6938 | 0.8558 | 0 | 0 | 0 |
| 'Gpam' | 4.131 | 5.057 | 4.583 | 3.515 | 5.036 | 4.720 | 3.900 | 4.502 | 4.147 | 4.657 | 4.561 | 3.740 | 4.158 | 3.931 | 3.253 | 4.358 | 3.401 | 3.449 | 0.151 | 0.0508 | 0.1908 | 0 | 0 | 0 |
| 'Gpank1' | 17.285 | 18.460 | 19.364 | 18.399 | 17.245 | 19.320 | 19.114 | 17.321 | 19.610 | 16.303 | 16.654 | 19.251 | 17.995 | 16.294 | 19.300 | 17.939 | 18.016 | 18.645 | 0.037 | 0.3492 | 0.6043 | 0 | 0 | 0 |
| 'Gpat4' | 21.172 | 21.106 | 20.730 | 20.750 | 21.173 | 20.257 | 19.449 | 20.259 | 20.066 | 20.627 | 20.182 | 22.844 | 21.031 | 20.645 | 23.580 | 20.435 | 21.613 | 24.028 | -0.088 | 0.0143 | 0.0806 | 0 | 0 | 0 |
| 'Gpatch1' | 4.014 | 4.490 | 4.459 | 3.631 | 4.389 | 4.693 | 3.813 | 4.796 | 4.374 | 4.899 | 4.306 | 4.193 | 4.185 | 3.947 | 3.871 | 4.428 | 4.745 | 4.161 | -0.011 | 0.8504 | 0.9357 | 0 | 0 | 0 |
| 'Gpatch11' | 8.723 | 9.335 | 9.628 | 10.003 | 9.768 | 9.765 | 10.211 | 8.801 | 9.858 | 7.948 | 9.006 | 10.348 | 8.974 | 8.886 | 9.252 | 8.652 | 9.490 | 9.842 | 0.047 | 0.2914 | 0.5480 | 0 | 0 | 0 |
| 'Gpatch2' | 1.925 | 1.840 | 2.038 | 1.806 | 2.042 | 1.925 | 2.072 | 1.955 | 1.874 | 2.086 | 2.075 | 2.050 | 2.038 | 1.996 | 1.949 | 1.849 | 2.103 | 1.913 | -0.057 | 0.1510 | 0.3777 | 0 | 0 | 0 |
| 'Gpatch2l' | 15.207 | 16.571 | 17.576 | 13.983 | 16.240 | 17.130 | 15.508 | 16.400 | 16.853 | 16.573 | 16.033 | 15.037 | 15.842 | 16.264 | 15.370 | 15.543 | 15.584 | 14.176 | 0.044 | 0.2739 | 0.5293 | 0 | 0 | 0 |
| 'Gpatch3' | 6.969 | 5.487 | 6.535 | 5.561 | 6.661 | 6.760 | 5.421 | 5.981 | 6.589 | 6.957 | 4.968 | 5.516 | 7.436 | 6.439 | 5.195 | 7.944 | 4.706 | 5.018 | 0.058 | 0.5049 | 0.7314 | 0 | 0 | 0 |
| 'Gpatch4' | 5.558 | 4.242 | 5.173 | 5.110 | 4.735 | 4.281 | 4.033 | 4.670 | 5.043 | 4.567 | 3.858 | 5.005 | 4.697 | 3.893 | 4.986 | 4.807 | 4.134 | 4.894 | 0.062 | 0.2938 | 0.5498 | 0 | 0 | 0 |
| 'Gpatch8' | 7.940 | 7.358 | 8.001 | 8.162 | 7.216 | 7.560 | 7.724 | 7.462 | 8.129 | 7.535 | 7.831 | 8.394 | 7.112 | 7.258 | 7.938 | 7.443 | 7.715 | 8.667 | -0.018 | 0.6279 | 0.8154 | 0 | 0 | 0 |
| 'Gpbp1' | 28.555 | 27.243 | 25.535 | 29.013 | 26.947 | 24.180 | 29.020 | 27.552 | 24.131 | 26.531 | 27.792 | 27.168 | 23.872 | 27.652 | 24.913 | 25.475 | 29.080 | 26.558 | 0.006 | 0.8866 | 0.9533 | 0 | 0 | 0 |
| 'Gpbp1l1' | 39.194 | 38.104 | 37.787 | 39.823 | 38.187 | 36.615 | 38.954 | 40.075 | 36.138 | 36.100 | 42.749 | 38.996 | 34.044 | 39.557 | 37.116 | 35.745 | 41.446 | 38.323 | -0.005 | 0.8833 | 0.9509 | 0 | 0 | 0 |
| 'Gpc1' | 0.913 | 0.831 | 1.127 | 1.018 | 1.151 | 1.111 | 0.880 | 1.111 | 1.211 | 0.872 | 0.718 | 0.884 | 0.948 | 0.627 | 0.845 | 0.688 | 0.564 | 0.907 | 0.395 | 0.0000 | 0.0007 | 1 | 1 | 0 |
| 'Gpcpd1' | 6.167 | 7.979 | 7.517 | 5.752 | 8.400 | 8.037 | 6.728 | 9.132 | 7.634 | 8.336 | 7.226 | 5.544 | 7.660 | 7.910 | 5.091 | 8.310 | 6.791 | 4.882 | 0.129 | 0.2294 | 0.4764 | 0 | 0 | 0 |
| 'Gpd1l' | 9.065 | 9.875 | 9.913 | 9.009 | 11.163 | 10.635 | 9.673 | 10.198 | 9.887 | 10.320 | 9.495 | 9.609 | 9.155 | 10.315 | 9.293 | 10.837 | 8.532 | 9.787 | 0.024 | 0.6125 | 0.8056 | 0 | 0 | 0 |
| 'Gpd2' | 2.609 | 2.530 | 3.101 | 3.108 | 2.512 | 2.930 | 3.108 | 2.593 | 2.986 | 2.866 | 2.661 | 3.272 | 3.006 | 2.623 | 3.355 | 2.878 | 2.750 | 3.271 | -0.081 | 0.0712 | 0.2354 | 0 | 0 | 0 |
| 'Gphn' | 2.287 | 2.747 | 2.355 | 2.489 | 2.734 | 2.714 | 2.796 | 2.732 | 2.607 | 3.014 | 2.779 | 2.657 | 3.015 | 2.562 | 2.503 | 2.625 | 3.068 | 2.570 | -0.093 | 0.0966 | 0.2893 | 0 | 0 | 0 |
| 'Gpi1' | 60.332 | 54.410 | 55.650 | 61.944 | 57.508 | 57.217 | 59.197 | 55.601 | 54.570 | 57.242 | 61.384 | 58.728 | 64.528 | 59.656 | 61.547 | 57.679 | 60.509 | 61.159 | -0.083 | 0.0076 | 0.0511 | 0 | 0 | 0 |
| 'Gpkow' | 16.394 | 16.268 | 15.649 | 15.576 | 16.980 | 17.078 | 16.633 | 16.495 | 16.808 | 15.074 | 17.640 | 17.573 | 15.995 | 18.537 | 17.031 | 17.649 | 16.921 | 17.658 | -0.069 | 0.0500 | 0.1886 | 0 | 0 | 0 |
| 'Gpld1' | 3.638 | 4.402 | 3.868 | 3.755 | 3.930 | 3.856 | 3.538 | 4.156 | 3.996 | 3.392 | 3.704 | 3.840 | 3.930 | 3.977 | 3.820 | 4.451 | 3.556 | 3.693 | 0.022 | 0.6939 | 0.8558 | 0 | 0 | 0 |
| 'Gpm6b' | 20.158 | 27.009 | 20.296 | 21.493 | 25.101 | 18.622 | 21.391 | 22.252 | 20.049 | 25.050 | 27.175 | 23.008 | 21.958 | 27.407 | 21.429 | 24.231 | 28.551 | 20.476 | -0.170 | 0.0000 | 0.0001 | 1 | 0 | 0 |
| 'Gpn1' | 3.342 | 3.417 | 3.590 | 2.776 | 3.579 | 3.412 | 2.844 | 3.492 | 3.361 | 3.312 | 3.104 | 3.230 | 3.441 | 3.008 | 3.299 | 3.244 | 3.442 | 3.226 | 0.016 | 0.7399 | 0.8820 | 0 | 0 | 0 |
| 'Gpn2' | 7.136 | 6.883 | 7.718 | 6.513 | 7.411 | 8.291 | 7.072 | 7.945 | 7.681 | 6.829 | 5.651 | 7.110 | 7.337 | 7.061 | 7.265 | 6.634 | 6.459 | 6.825 | 0.116 | 0.0308 | 0.1370 | 0 | 0 | 0 |
| 'Gpn3' | 7.301 | 8.400 | 7.708 | 6.752 | 8.666 | 8.003 | 6.868 | 8.241 | 7.620 | 7.530 | 7.070 | 7.082 | 7.928 | 7.518 | 7.283 | 7.460 | 7.879 | 7.139 | 0.046 | 0.2759 | 0.5317 | 0 | 0 | 0 |
| 'Gpr107' | 5.213 | 5.074 | 5.341 | 5.249 | 5.147 | 5.114 | 4.962 | 5.554 | 4.956 | 5.021 | 5.508 | 5.450 | 5.138 | 4.884 | 5.497 | 5.013 | 5.511 | 5.845 | -0.049 | 0.2004 | 0.4416 | 0 | 0 | 0 |
| 'Gpr108' | 41.961 | 38.235 | 46.662 | 40.574 | 41.503 | 47.622 | 40.528 | 42.308 | 46.830 | 45.069 | 40.587 | 44.114 | 46.183 | 42.057 | 44.365 | 44.430 | 39.523 | 43.116 | -0.020 | 0.5551 | 0.7675 | 0 | 0 | 0 |
| 'Gpr132' | 40.563 | 29.637 | 43.523 | 36.316 | 31.206 | 40.930 | 39.537 | 33.856 | 44.193 | 40.997 | 35.020 | 47.067 | 30.900 | 31.756 | 42.883 | 36.457 | 32.227 | 43.977 | -0.016 | 0.7512 | 0.8893 | 0 | 0 | 0 |
| 'Gpr137' | 6.893 | 5.805 | 7.623 | 6.124 | 6.300 | 8.273 | 6.061 | 7.063 | 7.588 | 6.748 | 5.826 | 7.529 | 7.083 | 5.990 | 7.038 | 6.245 | 5.461 | 6.654 | 0.068 | 0.1845 | 0.4199 | 0 | 0 | 0 |
| 'Gpr137b' | 6.230 | 5.708 | 6.349 | 5.230 | 6.798 | 7.261 | 5.122 | 6.750 | 5.526 | 6.382 | 5.529 | 6.852 | 5.541 | 8.393 | 6.269 | 6.611 | 5.602 | 5.834 | -0.057 | 0.5287 | 0.7495 | 0 | 0 | 0 |
| 'Gpr146' | 1.272 | 1.238 | 1.608 | 1.304 | 1.519 | 1.383 | 1.219 | 1.772 | 1.491 | 0.909 | 1.400 | 1.519 | 0.785 | 1.159 | 1.721 | 1.060 | 0.815 | 1.818 | 0.220 | 0.0422 | 0.1695 | 0 | 0 | 0 |
| 'Gpr153' | 1.739 | 2.071 | 1.642 | 1.363 | 1.596 | 1.879 | 1.430 | 1.774 | 1.962 | 2.132 | 2.057 | 1.782 | 1.878 | 2.265 | 1.969 | 2.107 | 2.033 | 1.957 | -0.236 | 0.0013 | 0.0130 | 1 | 0 | 0 |
| 'Gpr155' | 5.280 | 6.503 | 5.253 | 5.400 | 6.158 | 5.193 | 4.986 | 5.956 | 5.202 | 5.750 | 7.157 | 6.059 | 5.154 | 6.563 | 5.691 | 5.916 | 7.131 | 5.633 | -0.151 | 0.0000 | 0.0003 | 1 | 0 | 0 |
| 'Gpr162' | 1.194 | 1.303 | 1.261 | 1.051 | 1.357 | 1.327 | 1.134 | 0.985 | 1.059 | 0.945 | 0.598 | 1.088 | 0.892 | 1.031 | 1.028 | 0.986 | 1.229 | 1.183 | 0.234 | 0.0183 | 0.0968 | 0 | 0 | 0 |
| 'Gpr171' | 7.893 | 6.274 | 7.491 | 7.736 | 7.219 | 6.624 | 8.572 | 6.079 | 6.969 | 6.535 | 7.392 | 8.274 | 6.227 | 7.164 | 7.690 | 5.335 | 7.796 | 7.275 | 0.012 | 0.8880 | 0.9540 | 0 | 0 | 0 |
| 'Gpr174' | 4.478 | 4.507 | 4.352 | 4.416 | 4.249 | 3.917 | 5.004 | 4.454 | 4.133 | 4.656 | 5.131 | 4.969 | 4.700 | 4.967 | 4.423 | 4.719 | 5.516 | 4.800 | -0.163 | 0.0001 | 0.0011 | 1 | 0 | 0 |
| 'Gpr179' | 0.517 | 0.714 | 0.644 | 0.670 | 0.626 | 0.552 | 0.521 | 0.372 | 0.530 | 0.503 | 0.377 | 0.488 | 0.371 | 0.465 | 0.495 | 0.433 | 0.321 | 0.571 | 0.339 | 0.0053 | 0.0392 | 1 | 0 | 0 |
| 'Gpr18' | 3.455 | 2.757 | 2.866 | 2.418 | 3.507 | 3.080 | 2.517 | 2.842 | 2.687 | 1.646 | 1.985 | 3.681 | 2.377 | 1.572 | 3.248 | 2.048 | 2.875 | 3.672 | 0.175 | 0.2248 | 0.4716 | 0 | 0 | 0 |
| 'Gpr180' | 6.110 | 5.983 | 6.340 | 5.740 | 6.366 | 6.389 | 6.162 | 6.896 | 6.161 | 6.629 | 5.322 | 5.457 | 5.240 | 5.578 | 5.436 | 6.957 | 6.040 | 5.143 | 0.117 | 0.0543 | 0.1995 | 0 | 0 | 0 |
| 'Gpr183' | 49.414 | 44.454 | 57.547 | 48.280 | 46.794 | 53.002 | 48.839 | 47.555 | 53.758 | 47.843 | 46.885 | 58.616 | 34.798 | 41.889 | 57.151 | 40.961 | 42.335 | 56.859 | 0.073 | 0.1797 | 0.4150 | 0 | 0 | 0 |
| 'Gpr19' | 1.157 | 0.974 | 1.127 | 0.996 | 0.986 | 1.099 | 0.960 | 1.034 | 1.142 | 0.956 | 0.978 | 0.769 | 0.847 | 0.856 | 0.720 | 0.858 | 0.981 | 0.550 | 0.347 | 0.0001 | 0.0021 | 1 | 0 | 0 |
| 'Gpr35' | 0.482 | 0.449 | 0.513 | 0.329 | 0.361 | 0.491 | 0.396 | 0.519 | 0.535 | 0.437 | 0.599 | 0.493 | 0.282 | 0.575 | 0.413 | 0.385 | 0.521 | 0.430 | -0.008 | 0.9444 | 0.9789 | 0 | 0 | 0 |
| 'Gpr65' | 3.028 | 2.527 | 3.764 | 3.305 | 3.921 | 3.486 | 3.366 | 3.841 | 3.617 | 2.728 | 2.767 | 3.312 | 2.743 | 2.475 | 3.140 | 2.804 | 2.415 | 2.728 | 0.284 | 0.0014 | 0.0135 | 1 | 0 | 0 |
| 'Gpr68' | 10.783 | 9.839 | 9.998 | 10.120 | 8.890 | 10.378 | 10.548 | 9.082 | 10.902 | 9.352 | 10.721 | 13.837 | 7.776 | 9.502 | 12.305 | 8.814 | 10.888 | 13.391 | -0.088 | 0.2248 | 0.4716 | 0 | 0 | 0 |
| 'Gpr83' | 2.554 | 2.387 | 3.313 | 2.343 | 2.291 | 3.122 | 2.246 | 2.339 | 2.490 | 2.024 | 1.908 | 3.306 | 1.640 | 1.822 | 2.881 | 1.770 | 1.637 | 3.386 | 0.205 | 0.0109 | 0.0664 | 0 | 0 | 0 |
| 'Gpr89' | 3.686 | 3.327 | 3.522 | 3.649 | 3.444 | 3.463 | 3.159 | 3.294 | 3.541 | 3.472 | 3.579 | 3.989 | 3.286 | 3.702 | 3.199 | 3.362 | 3.881 | 3.572 | -0.055 | 0.2614 | 0.5150 | 0 | 0 | 0 |
| 'Gprasp1' | 6.125 | 4.465 | 4.564 | 5.975 | 4.621 | 4.236 | 5.163 | 4.386 | 4.492 | 5.610 | 5.667 | 6.717 | 4.818 | 5.077 | 4.987 | 4.504 | 5.284 | 5.616 | -0.153 | 0.0522 | 0.1938 | 0 | 0 | 0 |
| 'Gprin3' | 4.228 | 4.384 | 4.824 | 4.828 | 4.293 | 3.758 | 4.834 | 4.804 | 4.346 | 4.547 | 5.023 | 5.032 | 4.869 | 4.644 | 4.360 | 4.401 | 5.157 | 5.112 | -0.112 | 0.0368 | 0.1550 | 0 | 0 | 0 |
| 'Gps1' | 23.143 | 22.755 | 24.992 | 22.294 | 22.856 | 24.584 | 22.588 | 22.999 | 24.150 | 22.598 | 21.432 | 23.984 | 21.592 | 21.061 | 22.889 | 22.746 | 21.921 | 22.577 | 0.058 | 0.0145 | 0.0817 | 0 | 0 | 0 |
| 'Gps2' | 31.932 | 27.467 | 35.637 | 29.520 | 29.572 | 38.195 | 28.823 | 31.241 | 34.217 | 31.313 | 26.580 | 34.729 | 34.386 | 28.533 | 32.449 | 32.005 | 29.650 | 32.230 | 0.014 | 0.7304 | 0.8771 | 0 | 0 | 0 |
| 'Gpsm1' | 2.928 | 2.522 | 3.209 | 2.516 | 2.453 | 3.019 | 2.403 | 3.150 | 3.001 | 2.783 | 2.121 | 2.625 | 2.521 | 2.396 | 2.504 | 2.576 | 2.541 | 2.354 | 0.164 | 0.0074 | 0.0497 | 1 | 0 | 0 |
| 'Gpsm2' | 3.056 | 3.351 | 4.352 | 3.753 | 3.354 | 4.394 | 4.038 | 3.941 | 4.084 | 2.619 | 3.564 | 4.155 | 4.418 | 3.763 | 4.419 | 3.451 | 3.747 | 3.874 | 0.001 | 0.9943 | 0.9990 | 0 | 0 | 0 |
| 'Gpsm3' | 73.934 | 86.983 | 81.732 | 71.196 | 81.383 | 89.889 | 74.993 | 82.415 | 86.568 | 82.474 | 76.905 | 83.826 | 78.950 | 78.332 | 89.308 | 84.299 | 77.198 | 85.367 | -0.027 | 0.4256 | 0.6723 | 0 | 0 | 0 |
| 'Gpt' | 7.062 | 6.684 | 7.746 | 7.196 | 6.686 | 7.114 | 7.010 | 7.213 | 7.864 | 4.967 | 6.617 | 7.270 | 6.084 | 6.176 | 7.620 | 4.731 | 7.332 | 6.512 | 0.161 | 0.0229 | 0.1130 | 0 | 0 | 0 |
| 'Gpx1' | 37.576 | 31.363 | 32.209 | 34.900 | 34.777 | 33.123 | 34.924 | 32.393 | 30.853 | 36.180 | 39.458 | 34.192 | 38.273 | 38.100 | 37.140 | 36.374 | 38.975 | 34.052 | -0.149 | 0.0001 | 0.0009 | 1 | 0 | 0 |
| 'Gpx4' | 32.110 | 34.302 | 32.185 | 33.738 | 33.658 | 33.721 | 35.043 | 35.480 | 32.719 | 31.273 | 33.866 | 30.396 | 32.317 | 33.646 | 30.428 | 29.357 | 30.986 | 30.140 | 0.091 | 0.0011 | 0.0110 | 1 | 0 | 0 |
| 'Gramd1a' | 81.545 | 68.936 | 83.480 | 77.145 | 67.632 | 83.835 | 77.369 | 66.014 | 86.843 | 75.419 | 64.664 | 92.662 | 66.967 | 66.536 | 88.591 | 73.523 | 72.839 | 95.182 | -0.012 | 0.7622 | 0.8944 | 0 | 0 | 0 |
| 'Gramd1b' | 20.668 | 18.005 | 19.785 | 18.503 | 18.760 | 18.517 | 19.310 | 18.738 | 21.443 | 20.195 | 18.873 | 20.455 | 17.978 | 16.860 | 18.835 | 17.365 | 18.052 | 20.927 | 0.025 | 0.5280 | 0.7488 | 0 | 0 | 0 |
| 'Gramd1c' | 0.509 | 0.594 | 0.475 | 0.582 | 0.617 | 0.714 | 0.527 | 0.659 | 0.519 | 1.008 | 0.596 | 0.686 | 0.685 | 0.797 | 0.593 | 0.817 | 0.887 | 0.593 | -0.358 | 0.0032 | 0.0264 | 1 | 0 | 0 |
| 'Gramd3' | 22.149 | 27.949 | 22.564 | 22.008 | 26.906 | 23.218 | 22.474 | 27.568 | 22.882 | 22.983 | 27.473 | 27.671 | 27.945 | 28.294 | 25.844 | 26.850 | 30.662 | 26.753 | -0.185 | 0.0000 | 0.0000 | 1 | 0 | 0 |
| 'Gramd4' | 13.639 | 10.587 | 12.162 | 11.874 | 10.437 | 11.786 | 11.917 | 11.540 | 12.421 | 11.812 | 10.874 | 12.822 | 10.754 | 10.389 | 12.152 | 11.770 | 10.887 | 12.144 | 0.028 | 0.4645 | 0.7004 | 0 | 0 | 0 |
| 'Grap' | 63.704 | 64.738 | 66.641 | 65.619 | 63.925 | 75.988 | 62.481 | 66.071 | 69.892 | 59.606 | 54.889 | 71.252 | 59.327 | 59.517 | 69.051 | 64.259 | 57.057 | 63.624 | 0.091 | 0.0046 | 0.0350 | 1 | 0 | 0 |
| 'Grap2' | 45.607 | 41.728 | 45.400 | 45.734 | 38.545 | 43.780 | 42.882 | 40.565 | 44.916 | 40.455 | 40.695 | 45.202 | 39.476 | 37.469 | 45.911 | 43.212 | 34.627 | 47.598 | 0.047 | 0.2243 | 0.4713 | 0 | 0 | 0 |
| 'Grasp' | 2.372 | 1.839 | 2.547 | 1.872 | 1.503 | 2.499 | 2.331 | 1.855 | 1.535 | 2.351 | 1.806 | 1.787 | 1.987 | 1.649 | 2.063 | 1.872 | 1.773 | 2.111 | 0.083 | 0.4735 | 0.7089 | 0 | 0 | 0 |
| 'Grb2' | 62.983 | 64.000 | 59.236 | 63.392 | 62.592 | 60.265 | 60.371 | 65.139 | 58.970 | 61.602 | 60.338 | 55.374 | 61.129 | 62.741 | 58.035 | 60.016 | 58.028 | 56.600 | 0.052 | 0.0175 | 0.0940 | 0 | 0 | 0 |
| 'Grcc10' | 36.490 | 46.732 | 38.787 | 38.782 | 41.900 | 41.758 | 38.671 | 43.850 | 40.781 | 38.338 | 38.909 | 36.170 | 39.165 | 41.131 | 41.176 | 40.503 | 41.940 | 37.448 | 0.040 | 0.2499 | 0.5009 | 0 | 0 | 0 |
| 'Grhl1' | 11.655 | 10.120 | 11.460 | 11.090 | 10.301 | 10.169 | 10.641 | 10.783 | 10.394 | 12.380 | 12.485 | 13.003 | 12.044 | 12.625 | 12.271 | 12.436 | 12.895 | 12.303 | -0.228 | 0.0000 | 0.0000 | 1 | 0 | 0 |
| 'Grhpr' | 14.454 | 14.455 | 13.125 | 13.007 | 14.414 | 13.568 | 12.717 | 13.369 | 12.968 | 12.783 | 13.555 | 13.369 | 14.886 | 13.756 | 12.911 | 13.765 | 12.910 | 12.152 | 0.013 | 0.7509 | 0.8892 | 0 | 0 | 0 |
| 'Grik5' | 10.777 | 10.179 | 12.840 | 10.716 | 10.652 | 11.613 | 10.391 | 11.329 | 12.240 | 10.787 | 8.545 | 10.571 | 10.629 | 8.129 | 9.491 | 9.439 | 8.988 | 9.336 | 0.221 | 0.0000 | 0.0000 | 1 | 0 | 0 |
| 'Grina' | 83.197 | 76.184 | 89.690 | 72.292 | 78.435 | 87.068 | 75.589 | 81.174 | 92.750 | 87.037 | 76.639 | 90.802 | 84.516 | 77.740 | 86.488 | 75.343 | 82.337 | 86.063 | -0.032 | 0.3511 | 0.6052 | 0 | 0 | 0 |
| 'Gripap1' | 12.327 | 11.229 | 14.368 | 12.837 | 12.506 | 13.675 | 13.736 | 12.575 | 13.898 | 13.945 | 12.714 | 13.872 | 13.946 | 12.477 | 13.304 | 12.904 | 13.732 | 13.840 | -0.055 | 0.0967 | 0.2894 | 0 | 0 | 0 |
| 'Grk4' | 0.939 | 1.216 | 1.109 | 1.022 | 1.046 | 1.160 | 0.922 | 0.749 | 1.192 | 1.060 | 1.093 | 1.144 | 1.023 | 0.986 | 1.130 | 0.922 | 1.048 | 0.798 | 0.006 | 0.9542 | 0.9832 | 0 | 0 | 0 |
| 'Grk5' | 2.677 | 2.382 | 2.818 | 2.491 | 2.632 | 2.951 | 2.593 | 2.882 | 2.960 | 2.769 | 3.088 | 3.024 | 3.124 | 2.905 | 2.889 | 2.883 | 2.711 | 2.974 | -0.115 | 0.1001 | 0.2959 | 0 | 0 | 0 |
| 'Grk6' | 95.169 | 95.439 | 103.408 | 90.045 | 99.386 | 110.905 | 91.396 | 102.006 | 103.209 | 99.947 | 97.827 | 101.749 | 102.975 | 106.225 | 110.426 | 104.271 | 97.046 | 104.930 | -0.066 | 0.0261 | 0.1241 | 0 | 0 | 0 |
| 'Grm4' | 1.173 | 1.066 | 2.031 | 1.098 | 1.091 | 1.931 | 1.256 | 1.146 | 1.855 | 0.845 | 0.593 | 0.876 | 0.637 | 0.467 | 1.043 | 0.939 | 0.587 | 0.825 | 0.893 | 0.0000 | 0.0000 | 1 | 1 | 0 |
| 'Grm6' | 0.564 | 0.722 | 0.632 | 0.807 | 0.571 | 0.688 | 0.576 | 0.725 | 0.622 | 0.655 | 0.620 | 0.614 | 0.917 | 0.826 | 0.513 | 0.644 | 0.808 | 0.689 | -0.095 | 0.3478 | 0.6032 | 0 | 0 | 0 |
| 'Grn' | 14.067 | 15.584 | 14.435 | 14.816 | 15.012 | 15.310 | 14.295 | 14.764 | 15.451 | 16.411 | 16.484 | 16.073 | 19.347 | 16.759 | 16.074 | 16.321 | 15.827 | 16.136 | -0.171 | 0.0000 | 0.0000 | 1 | 0 | 0 |
| 'Grpel1' | 12.623 | 13.654 | 12.230 | 13.976 | 14.742 | 13.552 | 12.728 | 13.891 | 12.498 | 12.942 | 11.738 | 13.009 | 13.132 | 13.581 | 14.377 | 15.483 | 12.288 | 12.631 | -0.003 | 0.9567 | 0.9839 | 0 | 0 | 0 |
| 'Grpel2' | 6.000 | 6.346 | 4.862 | 5.826 | 5.501 | 5.532 | 5.571 | 5.900 | 5.564 | 4.913 | 5.587 | 6.549 | 5.119 | 5.270 | 5.616 | 6.064 | 6.365 | 6.197 | -0.033 | 0.6054 | 0.8013 | 0 | 0 | 0 |
| 'Grsf1' | 9.810 | 10.307 | 9.862 | 10.805 | 10.576 | 10.729 | 9.975 | 10.681 | 9.391 | 9.807 | 10.107 | 10.205 | 10.713 | 11.164 | 10.751 | 9.927 | 9.836 | 10.671 | -0.028 | 0.4461 | 0.6878 | 0 | 0 | 0 |
| 'Grtp1' | 12.040 | 16.949 | 13.843 | 13.427 | 14.567 | 15.215 | 13.981 | 14.446 | 15.247 | 13.161 | 12.561 | 11.460 | 14.872 | 12.621 | 11.840 | 14.153 | 12.131 | 11.565 | 0.175 | 0.0073 | 0.0496 | 1 | 0 | 0 |
| 'Grwd1' | 4.373 | 4.006 | 4.646 | 4.119 | 4.255 | 4.763 | 4.214 | 4.680 | 4.428 | 4.021 | 3.881 | 5.148 | 4.265 | 3.940 | 4.930 | 4.438 | 4.052 | 5.099 | -0.022 | 0.6647 | 0.8381 | 0 | 0 | 0 |
| 'Gsap' | 3.810 | 4.372 | 4.189 | 3.966 | 4.002 | 3.932 | 4.284 | 4.193 | 4.295 | 4.129 | 4.626 | 4.475 | 3.988 | 4.702 | 4.191 | 4.238 | 5.027 | 4.165 | -0.103 | 0.0049 | 0.0367 | 1 | 0 | 0 |
| 'Gsdmd' | 41.434 | 38.918 | 46.739 | 38.290 | 39.021 | 44.726 | 38.273 | 40.163 | 44.985 | 40.688 | 36.130 | 45.204 | 41.727 | 38.523 | 43.956 | 37.917 | 40.029 | 42.859 | 0.011 | 0.7027 | 0.8611 | 0 | 0 | 0 |
| 'Gse1' | 11.657 | 12.140 | 10.531 | 11.526 | 10.443 | 11.216 | 10.479 | 10.843 | 11.682 | 12.489 | 12.211 | 12.435 | 13.793 | 12.045 | 12.100 | 12.850 | 12.646 | 12.568 | -0.181 | 0.0000 | 0.0000 | 1 | 0 | 0 |
| 'Gsg2' | 4.213 | 5.308 | 5.753 | 5.399 | 4.165 | 5.928 | 5.349 | 5.050 | 5.950 | 3.837 | 4.542 | 6.038 | 5.150 | 5.496 | 6.178 | 4.583 | 4.766 | 5.340 | 0.022 | 0.7472 | 0.8861 | 0 | 0 | 0 |
| 'Gsk3a' | 47.629 | 47.212 | 51.260 | 46.013 | 46.446 | 51.705 | 44.697 | 47.888 | 50.645 | 48.436 | 45.319 | 51.954 | 50.141 | 47.976 | 51.860 | 47.690 | 44.178 | 51.765 | -0.029 | 0.2120 | 0.4573 | 0 | 0 | 0 |

| | | | | | | | | | | | | | | | | | | | | | | | | |
|---|---|---|---|---|---|---|---|---|---|---|---|---|---|---|---|---|---|---|---|---|---|---|---|---|
| 'Gsk3b' | 11.650 | 12.267 | 11.555 | 10.965 | 11.635 | 11.543 | 10.445 | 13.008 | 10.966 | 12.846 | 11.976 | 11.697 | 11.807 | 12.665 | 11.416 | 11.706 | 11.049 | 12.655 | -0.063 | 0.1112 | 0.3156 | 0 | 0 | 0 |
| 'Gskip' | 13.375 | 15.234 | 14.669 | 15.321 | 14.274 | 12.565 | 15.501 | 13.727 | 13.192 | 14.591 | 13.431 | 15.804 | 12.808 | 14.730 | 13.716 | 12.138 | 15.845 | 14.764 | -0.015 | 0.7869 | 0.9047 | 0 | 0 | 0 |
| 'Gsn' | 24.573 | 21.463 | 26.119 | 24.033 | 22.175 | 24.900 | 24.705 | 20.607 | 24.725 | 20.173 | 18.628 | 21.472 | 19.135 | 18.700 | 20.799 | 17.617 | 17.736 | 21.032 | 0.271 | 0.0000 | 0.0000 | 1 | 0 | 0 |
| 'Gspt1' | 28.127 | 26.631 | 28.503 | 28.281 | 26.038 | 26.180 | 29.593 | 26.737 | 27.284 | 27.452 | 28.361 | 28.575 | 26.826 | 26.662 | 27.739 | 24.985 | 28.402 | 28.945 | -0.015 | 0.6673 | 0.8396 | 0 | 0 | 0 |
| 'Gspt2' | 1.668 | 2.079 | 1.699 | 2.183 | 1.673 | 1.409 | 1.718 | 1.807 | 1.678 | 1.723 | 1.426 | 1.486 | 1.582 | 1.814 | 1.171 | 2.082 | 2.036 | 1.443 | 0.102 | 0.2594 | 0.5126 | 0 | 0 | 0 |
| 'Gsr' | 14.606 | 16.169 | 15.133 | 15.476 | 16.147 | 15.169 | 15.985 | 15.763 | 14.576 | 13.766 | 14.884 | 13.880 | 15.864 | 14.878 | 13.860 | 13.414 | 13.897 | 14.224 | 0.098 | 0.0055 | 0.0403 | 1 | 0 | 0 |
| 'Gss' | 2.049 | 2.474 | 2.138 | 1.901 | 2.110 | 2.087 | 2.117 | 1.977 | 2.219 | 2.274 | 2.139 | 1.839 | 2.319 | 2.005 | 1.805 | 1.970 | 1.730 | 1.904 | 0.078 | 0.2217 | 0.4691 | 0 | 0 | 0 |
| 'Gstcd' | 2.053 | 2.242 | 2.498 | 1.882 | 1.978 | 2.166 | 2.113 | 1.952 | 1.959 | 1.844 | 2.100 | 1.996 | 2.129 | 2.159 | 1.555 | 1.921 | 1.943 | 1.745 | 0.109 | 0.1034 | 0.3016 | 0 | 0 | 0 |
| 'Gstk1' | 1.562 | 2.139 | 1.951 | 2.610 | 1.767 | 1.797 | 2.164 | 2.018 | 2.310 | 1.880 | 1.939 | 1.873 | 1.828 | 1.738 | 1.886 | 2.239 | 1.717 | 1.887 | 0.096 | 0.3154 | 0.5732 | 0 | 0 | 0 |
| 'Gstm4' | 6.196 | 6.380 | 6.724 | 6.331 | 6.605 | 7.587 | 6.747 | 6.957 | 6.079 | 5.476 | 5.450 | 5.392 | 6.528 | 6.222 | 5.714 | 7.722 | 6.979 | 5.266 | 0.127 | 0.1083 | 0.3098 | 0 | 0 | 0 |
| 'Gstp1' | 126.582 | 137.709 | 123.279 | 131.117 | 133.568 | 130.592 | 128.255 | 132.516 | 126.529 | 119.272 | 129.699 | 120.574 | 130.468 | 119.801 | 126.754 | 137.192 | 133.801 | 121.670 | 0.029 | 0.3525 | 0.6063 | 0 | 0 | 0 |
| 'Gstp2' | 47.510 | 60.227 | 48.499 | 49.798 | 52.342 | 46.098 | 46.958 | 55.452 | 51.545 | 48.221 | 48.030 | 44.770 | 51.754 | 47.427 | 49.326 | 50.796 | 50.190 | 46.137 | 0.058 | 0.1968 | 0.4357 | 0 | 0 | 0 |
| 'Gstt2' | 40.083 | 43.064 | 41.573 | 35.777 | 38.855 | 42.466 | 40.553 | 40.770 | 43.829 | 41.685 | 42.641 | 44.236 | 47.999 | 44.092 | 42.349 | 48.147 | 42.480 | 38.019 | -0.102 | 0.0226 | 0.1122 | 0 | 0 | 0 |
| 'Gstt3' | 1.920 | 2.334 | 2.350 | 2.295 | 2.582 | 2.612 | 2.151 | 2.561 | 3.014 | 2.472 | 2.399 | 2.772 | 2.920 | 2.856 | 2.632 | 2.959 | 2.592 | 2.231 | -0.138 | 0.0808 | 0.2562 | 0 | 0 | 0 |
| 'Gstz1' | 7.618 | 9.046 | 9.387 | 8.241 | 8.630 | 8.972 | 8.113 | 8.787 | 9.261 | 8.997 | 7.244 | 7.790 | 7.290 | 8.595 | 7.674 | 10.150 | 8.384 | 7.732 | 0.079 | 0.2413 | 0.4920 | 0 | 0 | 0 |
| 'Gtdc1' | 2.987 | 3.273 | 2.395 | 3.006 | 2.943 | 2.403 | 2.884 | 3.175 | 2.359 | 3.090 | 2.629 | 2.761 | 2.636 | 3.021 | 2.616 | 2.684 | 3.113 | 2.646 | -0.010 | 0.8495 | 0.9351 | 0 | 0 | 0 |
| 'Gtf2a1' | 4.004 | 4.976 | 4.666 | 4.525 | 5.309 | 4.733 | 4.853 | 5.750 | 4.801 | 5.135 | 4.772 | 4.051 | 4.828 | 5.353 | 3.799 | 4.836 | 4.888 | 3.628 | 0.077 | 0.1684 | 0.4008 | 0 | 0 | 0 |
| 'Gtf2a2' | 6.966 | 8.204 | 8.582 | 7.114 | 8.432 | 7.567 | 7.464 | 8.060 | 7.762 | 7.001 | 7.329 | 7.933 | 7.317 | 8.265 | 7.943 | 7.216 | 8.080 | 7.507 | 0.020 | 0.5376 | 0.7564 | 0 | 0 | 0 |
| 'Gtf2b' | 10.756 | 11.430 | 10.675 | 10.073 | 11.671 | 10.561 | 10.867 | 11.776 | 10.043 | 10.652 | 10.301 | 10.110 | 11.661 | 11.338 | 10.433 | 10.798 | 11.309 | 10.365 | 0.003 | 0.9300 | 0.9736 | 0 | 0 | 0 |
| 'Gtf2e1' | 4.727 | 4.948 | 5.217 | 4.887 | 5.494 | 5.382 | 5.701 | 5.688 | 5.093 | 4.609 | 5.171 | 5.298 | 4.441 | 4.421 | 5.142 | 5.098 | 5.195 | 4.722 | 0.084 | 0.0736 | 0.2407 | 0 | 0 | 0 |
| 'Gtf2e2' | 8.664 | 8.708 | 8.267 | 9.931 | 9.252 | 8.075 | 9.118 | 9.024 | 7.426 | 7.543 | 8.720 | 8.339 | 8.957 | 10.008 | 8.296 | 7.491 | 9.771 | 7.897 | 0.014 | 0.7955 | 0.9092 | 0 | 0 | 0 |
| 'Gtf2f1' | 76.123 | 71.007 | 74.871 | 78.097 | 71.991 | 78.772 | 77.836 | 72.095 | 74.662 | 74.759 | 67.084 | 74.471 | 77.868 | 69.805 | 76.313 | 74.602 | 66.491 | 73.693 | 0.034 | 0.1357 | 0.3545 | 0 | 0 | 0 |
| 'Gtf2f2' | 17.432 | 18.022 | 16.091 | 17.609 | 18.877 | 17.159 | 19.171 | 18.449 | 16.484 | 16.618 | 15.671 | 15.943 | 16.702 | 17.733 | 16.055 | 15.875 | 17.104 | 14.815 | 0.105 | 0.0150 | 0.0838 | 0 | 0 | 0 |
| 'Gtf2h1' | 14.208 | 15.389 | 14.387 | 13.088 | 15.168 | 13.938 | 13.834 | 14.508 | 13.246 | 13.247 | 13.322 | 13.601 | 13.714 | 13.253 | 13.373 | 13.830 | 13.985 | 13.129 | 0.061 | 0.0394 | 0.1621 | 0 | 0 | 0 |
| 'Gtf2h2' | 6.131 | 6.254 | 6.778 | 6.962 | 7.485 | 5.930 | 6.375 | 6.450 | 6.248 | 7.270 | 7.360 | 7.368 | 7.486 | 7.289 | 6.841 | 6.576 | 8.067 | 6.992 | -0.168 | 0.0001 | 0.0016 | 1 | 0 | 0 |
| 'Gtf2h3' | 4.418 | 4.071 | 4.572 | 4.114 | 4.747 | 4.823 | 4.492 | 4.527 | 4.993 | 4.797 | 4.355 | 4.829 | 4.344 | 4.385 | 4.851 | 3.948 | 4.061 | 4.455 | 0.015 | 0.7187 | 0.8703 | 0 | 0 | 0 |
| 'Gtf2h4' | 2.271 | 2.199 | 2.266 | 2.188 | 2.255 | 2.286 | 2.168 | 2.281 | 2.394 | 2.029 | 2.052 | 2.293 | 2.280 | 2.214 | 2.227 | 1.947 | 2.136 | 2.169 | 0.056 | 0.2782 | 0.5348 | 0 | 0 | 0 |
| 'Gtf2h5' | 8.005 | 9.956 | 7.618 | 8.346 | 9.519 | 8.940 | 8.296 | 9.708 | 8.267 | 8.720 | 7.612 | 7.675 | 8.926 | 8.491 | 8.373 | 9.246 | 8.621 | 6.910 | 0.066 | 0.1964 | 0.4354 | 0 | 0 | 0 |
| 'Gtf2i' | 23.785 | 23.930 | 23.489 | 24.075 | 23.948 | 24.444 | 23.377 | 22.800 | 24.540 | 26.032 | 24.526 | 26.342 | 25.058 | 24.596 | 24.913 | 25.192 | 24.174 | 25.931 | -0.092 | 0.0000 | 0.0000 | 1 | 0 | 0 |
| 'Gtf2ird1' | 0.428 | 0.474 | 0.468 | 0.444 | 0.351 | 0.443 | 0.497 | 0.428 | 0.433 | 0.356 | 0.385 | 0.456 | 0.447 | 0.477 | 0.447 | 0.478 | 0.334 | 0.506 | 0.017 | 0.8368 | 0.9300 | 0 | 0 | 0 |
| 'Gtf2ird2' | 3.190 | 2.441 | 2.641 | 2.475 | 2.404 | 2.719 | 2.660 | 2.383 | 2.806 | 2.564 | 2.582 | 3.097 | 2.458 | 2.701 | 2.918 | 2.613 | 2.562 | 2.817 | -0.047 | 0.4126 | 0.6609 | 0 | 0 | 0 |
| 'Gtf3a' | 5.005 | 4.588 | 5.706 | 5.313 | 5.512 | 5.192 | 5.115 | 5.240 | 5.193 | 5.174 | 4.318 | 5.210 | 5.425 | 5.083 | 5.398 | 5.608 | 4.638 | 5.594 | 0.006 | 0.9059 | 0.9631 | 0 | 0 | 0 |
| 'Gtf3c1' | 10.571 | 9.598 | 10.746 | 10.982 | 9.437 | 10.331 | 10.146 | 10.098 | 10.309 | 10.612 | 10.928 | 10.964 | 10.191 | 9.652 | 10.072 | 9.748 | 9.901 | 10.829 | -0.022 | 0.5173 | 0.7406 | 0 | 0 | 0 |
| 'Gtf3c2' | 5.322 | 5.908 | 6.284 | 5.171 | 5.711 | 5.523 | 5.365 | 5.688 | 6.302 | 6.053 | 5.841 | 5.881 | 6.813 | 5.884 | 5.595 | 5.398 | 5.895 | 5.733 | -0.062 | 0.1689 | 0.4013 | 0 | 0 | 0 |
| 'Gtf3c3' | 4.985 | 5.955 | 5.637 | 4.763 | 5.820 | 5.253 | 5.116 | 5.993 | 5.447 | 5.853 | 5.418 | 5.327 | 4.682 | 5.455 | 4.757 | 5.558 | 5.719 | 4.862 | 0.031 | 0.5166 | 0.7404 | 0 | 0 | 0 |
| 'Gtf3c4' | 1.767 | 2.047 | 1.833 | 1.837 | 2.081 | 2.195 | 2.014 | 2.101 | 2.143 | 1.810 | 1.934 | 2.301 | 1.630 | 2.083 | 2.326 | 2.049 | 1.888 | 2.525 | -0.051 | 0.3859 | 0.6354 | 0 | 0 | 0 |
| 'Gtf3c5' | 5.308 | 5.446 | 6.262 | 5.077 | 5.851 | 5.773 | 5.785 | 5.863 | 6.485 | 6.223 | 5.442 | 5.677 | 6.274 | 5.292 | 5.499 | 5.484 | 5.473 | 5.545 | 0.018 | 0.7052 | 0.8628 | 0 | 0 | 0 |
| 'Gtf3c6' | 25.413 | 28.767 | 25.339 | 24.965 | 27.360 | 25.187 | 27.497 | 28.331 | 26.171 | 24.231 | 26.012 | 25.354 | 27.339 | 25.936 | 24.611 | 25.462 | 24.941 | 22.484 | 0.064 | 0.1395 | 0.3611 | 0 | 0 | 0 |
| 'Gtpbp1' | 53.960 | 56.197 | 57.553 | 50.941 | 53.372 | 58.891 | 52.613 | 54.311 | 56.174 | 58.744 | 54.588 | 58.843 | 56.787 | 54.477 | 58.171 | 57.491 | 51.237 | 57.670 | -0.050 | 0.0502 | 0.1891 | 0 | 0 | 0 |
| 'Gtpbp10' | 1.064 | 1.560 | 1.485 | 1.464 | 1.419 | 1.258 | 1.367 | 1.702 | 1.520 | 1.443 | 1.509 | 0.985 | 1.281 | 1.228 | 1.013 | 1.437 | 1.129 | 0.702 | 0.272 | 0.0085 | 0.0558 | 0 | 0 | 0 |
| 'Gtpbp2' | 18.244 | 17.758 | 18.633 | 16.673 | 17.143 | 18.326 | 16.785 | 17.290 | 19.653 | 18.063 | 16.686 | 19.758 | 19.186 | 17.231 | 18.861 | 17.224 | 18.588 | 20.043 | -0.056 | 0.0572 | 0.2057 | 0 | 0 | 0 |
| 'Gtpbp3' | 4.520 | 3.811 | 4.779 | 4.015 | 4.141 | 4.939 | 4.354 | 4.184 | 5.087 | 4.625 | 4.046 | 4.665 | 4.401 | 3.968 | 4.568 | 4.093 | 3.645 | 4.733 | 0.033 | 0.4119 | 0.6604 | 0 | 0 | 0 |
| 'Gtpbp4' | 30.542 | 29.147 | 29.704 | 31.396 | 31.231 | 28.598 | 30.426 | 31.639 | 27.344 | 30.427 | 28.483 | 29.298 | 26.580 | 28.215 | 26.609 | 31.781 | 29.546 | 28.025 | 0.050 | 0.1555 | 0.3842 | 0 | 0 | 0 |
| 'Gtpbp6' | 4.372 | 4.128 | 4.914 | 4.059 | 4.014 | 4.980 | 4.155 | 3.770 | 4.719 | 4.288 | 3.545 | 5.687 | 3.925 | 3.984 | 4.691 | 4.286 | 4.122 | 5.115 | -0.032 | 0.4860 | 0.7170 | 0 | 0 | 0 |
| 'Gtpbp8' | 6.098 | 7.642 | 7.280 | 6.876 | 7.378 | 6.709 | 6.581 | 6.945 | 7.657 | 6.701 | 6.333 | 6.358 | 6.493 | 6.911 | 5.643 | 7.157 | 7.052 | 5.559 | 0.111 | 0.0338 | 0.1462 | 0 | 0 | 0 |
| 'Gtse1' | 5.882 | 5.997 | 9.671 | 6.681 | 6.512 | 8.785 | 7.238 | 7.232 | 8.633 | 5.247 | 7.089 | 8.735 | 8.283 | 9.750 | 9.014 | 6.025 | 6.890 | 8.121 | -0.069 | 0.4210 | 0.6677 | 0 | 0 | 0 |
| 'Gucd1' | 2.770 | 3.963 | 3.329 | 3.113 | 3.630 | 3.990 | 3.167 | 3.218 | 3.559 | 3.040 | 3.346 | 3.732 | 3.268 | 3.317 | 3.418 | 3.563 | 3.411 | 3.277 | 0.000 | 0.9927 | 0.9986 | 0 | 0 | 0 |
| 'Guf1' | 2.708 | 3.259 | 3.038 | 2.593 | 3.371 | 2.916 | 3.152 | 3.250 | 3.226 | 2.977 | 2.814 | 2.870 | 3.023 | 3.048 | 2.601 | 3.077 | 2.962 | 2.520 | 0.077 | 0.0954 | 0.2872 | 0 | 0 | 0 |
| 'Guk1' | 8.213 | 10.158 | 8.424 | 8.712 | 9.724 | 9.642 | 8.195 | 9.005 | 8.065 | 8.671 | 7.518 | 9.141 | 9.076 | 8.148 | 9.492 | 8.576 | 7.403 | 8.670 | 0.046 | 0.4207 | 0.6675 | 0 | 0 | 0 |
| 'Gusb' | 4.753 | 4.330 | 5.413 | 4.308 | 4.680 | 4.985 | 4.727 | 5.256 | 4.798 | 4.598 | 4.649 | 5.639 | 4.817 | 4.421 | 5.047 | 4.865 | 4.943 | 5.364 | -0.046 | 0.2813 | 0.5379 | 0 | 0 | 0 |
| 'Gxylt1' | 7.742 | 7.430 | 7.032 | 7.076 | 8.631 | 6.991 | 7.708 | 7.902 | 6.999 | 7.997 | 7.806 | 7.524 | 7.742 | 8.793 | 7.234 | 7.886 | 8.049 | 7.630 | -0.077 | 0.0267 | 0.1256 | 0 | 0 | 0 |
| 'Gyg' | 10.918 | 11.182 | 10.968 | 11.332 | 12.507 | 11.900 | 11.518 | 12.150 | 11.180 | 11.278 | 12.176 | 11.971 | 12.223 | 12.458 | 12.459 | 12.374 | 12.453 | 12.287 | -0.092 | 0.0065 | 0.0457 | 1 | 0 | 0 |
| 'Gypc' | 8.321 | 7.871 | 8.624 | 8.019 | 7.507 | 9.379 | 8.334 | 7.615 | 9.117 | 8.931 | 7.478 | 9.389 | 7.539 | 7.509 | 9.330 | 8.175 | 8.239 | 8.936 | -0.025 | 0.5295 | 0.7499 | 0 | 0 | 0 |
| 'Gys1' | 4.210 | 4.697 | 5.088 | 4.919 | 4.872 | 4.945 | 4.824 | 4.420 | 4.938 | 4.763 | 5.051 | 4.849 | 5.424 | 4.449 | 4.732 | 4.453 | 4.313 | 4.772 | -0.009 | 0.8513 | 0.9360 | 0 | 0 | 0 |
| 'Gzf1' | 10.171 | 10.591 | 9.728 | 9.987 | 10.697 | 10.480 | 9.912 | 10.617 | 9.942 | 9.044 | 11.086 | 9.434 | 9.803 | 9.842 | 9.731 | 10.545 | 10.589 | 9.486 | 0.034 | 0.3271 | 0.5849 | 0 | 0 | 0 |
| 'Gzma' | 64.460 | 38.780 | 92.585 | 55.155 | 58.013 | 56.749 | 57.255 | 69.634 | 54.583 | 29.761 | 37.303 | 34.339 | 21.132 | 26.616 | 40.106 | 22.337 | 30.007 | 42.901 | 0.952 | 0.0000 | 0.0000 | 1 | 1 | 0 |
| 'H13' | 16.593 | 15.700 | 16.151 | 16.451 | 16.334 | 16.494 | 15.639 | 16.614 | 15.891 | 17.134 | 16.179 | 17.158 | 16.357 | 15.434 | 15.992 | 16.720 | 16.225 | 16.143 | -0.025 | 0.2869 | 0.5436 | 0 | 0 | 0 |
| 'H1f0' | 104.008 | 142.056 | 130.512 | 106.870 | 142.351 | 146.966 | 121.564 | 141.587 | 142.840 | 143.585 | 115.910 | 98.008 | 139.804 | 123.291 | 111.195 | 137.443 | 114.345 | 91.829 | 0.127 | 0.1769 | 0.4122 | 0 | 0 | 0 |
| 'H1fx' | 2.430 | 3.094 | 3.168 | 3.205 | 3.007 | 3.740 | 3.337 | 3.987 | 3.509 | 4.095 | 3.364 | 2.695 | 4.126 | 5.253 | 3.918 | 3.399 | 3.608 | 3.477 | -0.196 | 0.1159 | 0.3233 | 0 | 0 | 0 |
| 'H2-Ab1' | 13.821 | 21.360 | 12.795 | 12.922 | 20.658 | 14.633 | 14.657 | 20.121 | 14.384 | 9.711 | 19.232 | 8.528 | 9.581 | 18.231 | 9.428 | 9.032 | 19.715 | 9.039 | 0.416 | 0.0000 | 0.0000 | 1 | 1 | 0 |
| 'H2-DMa' | 9.712 | 11.505 | 9.196 | 8.598 | 11.283 | 9.412 | 9.391 | 10.241 | 10.237 | 9.078 | 10.259 | 7.379 | 9.318 | 9.516 | 9.222 | 9.107 | 8.627 | 7.085 | 0.166 | 0.0060 | 0.0430 | 1 | 0 | 0 |
| 'H2-Ke6' | 20.172 | 22.052 | 22.737 | 19.644 | 21.289 | 23.699 | 20.371 | 22.957 | 23.481 | 19.754 | 19.763 | 21.990 | 21.339 | 19.725 | 21.771 | 20.497 | 20.878 | 21.572 | 0.057 | 0.0411 | 0.1663 | 0 | 0 | 0 |
| 'H2-M3' | 49.560 | 51.394 | 47.108 | 49.081 | 52.682 | 47.909 | 50.945 | 51.515 | 47.129 | 46.927 | 49.056 | 52.987 | 51.763 | 51.410 | 50.773 | 53.169 | 52.588 | 47.774 | -0.044 | 0.1514 | 0.3784 | 0 | 0 | 0 |
| 'H2-Oa' | 2.551 | 3.451 | 3.101 | 2.856 | 3.774 | 3.652 | 2.941 | 3.092 | 3.602 | 3.253 | 2.411 | 3.025 | 3.036 | 2.915 | 2.824 | 3.138 | 2.555 | 2.478 | 0.164 | 0.0664 | 0.2253 | 0 | 0 | 0 |
| 'H2-Ob' | 1.315 | 1.971 | 1.145 | 1.451 | 2.052 | 1.778 | 1.617 | 2.107 | 1.720 | 1.222 | 1.318 | 1.074 | 0.876 | 1.487 | 1.433 | 1.149 | 2.055 | 0.866 | 0.400 | 0.0003 | 0.0037 | 1 | 1 | 0 |
| 'H2-Q4' | 192.424 | 215.751 | 189.363 | 207.486 | 214.445 | 195.445 | 204.395 | 215.076 | 198.540 | 211.606 | 208.934 | 183.259 | 220.787 | 219.732 | 198.935 | 234.102 | 214.234 | 198.633 | -0.053 | 0.0806 | 0.2562 | 0 | 0 | 0 |
| 'H2-T23' | 57.225 | 64.836 | 60.640 | 54.804 | 61.011 | 64.206 | 58.575 | 62.032 | 65.184 | 61.480 | 58.984 | 68.351 | 65.837 | 62.964 | 67.474 | 63.422 | 62.568 | 67.299 | -0.088 | 0.0012 | 0.0119 | 1 | 0 | 0 |
| 'H2-T24' | 23.078 | 23.007 | 25.161 | 19.801 | 22.100 | 25.671 | 24.603 | 24.961 | 25.745 | 24.151 | 26.954 | 35.280 | 26.054 | 26.141 | 35.371 | 28.110 | 26.203 | 35.677 | -0.303 | 0.0000 | 0.0000 | 1 | 0 | 0 |
| 'H2afj' | 45.334 | 44.539 | 48.419 | 44.219 | 46.341 | 46.893 | 43.853 | 43.176 | 48.401 | 40.101 | 41.305 | 43.952 | 46.255 | 41.444 | 47.049 | 42.477 | 45.773 | 43.957 | 0.057 | 0.0710 | 0.2349 | 0 | 0 | 0 |
| 'H2afv' | 46.146 | 60.213 | 51.627 | 47.928 | 56.019 | 54.136 | 50.102 | 54.358 | 54.894 | 43.269 | 45.724 | 47.826 | 50.304 | 53.225 | 55.730 | 54.699 | 51.091 | 49.286 | 0.062 | 0.1898 | 0.4262 | 0 | 0 | 0 |
| 'H2afx' | 70.725 | 73.976 | 87.689 | 75.085 | 74.591 | 94.603 | 73.585 | 76.275 | 91.810 | 63.210 | 66.785 | 78.316 | 75.548 | 76.884 | 89.652 | 69.137 | 67.533 | 78.099 | 0.099 | 0.0125 | 0.0735 | 0 | 0 | 0 |
| 'H2afy' | 26.963 | 26.161 | 28.838 | 24.019 | 25.225 | 25.790 | 23.967 | 25.907 | 28.466 | 43.712 | 41.232 | 41.569 | 43.116 | 40.241 | 43.508 | 42.678 | 41.416 | 42.350 | -0.701 | 0.0000 | 0.0000 | 1 | 1 | 0 |
| 'H2afz' | 101.440 | 96.544 | 93.594 | 99.073 | 100.804 | 91.110 | 96.142 | 102.164 | 91.567 | 90.593 | 95.665 | 74.959 | 108.387 | 101.080 | 91.861 | 100.431 | 95.020 | 82.442 | 0.048 | 0.2432 | 0.4935 | 0 | 0 | 0 |
| 'H3f3a' | 53.493 | 59.231 | 52.607 | 55.446 | 57.761 | 53.245 | 55.393 | 58.093 | 52.589 | 54.706 | 56.660 | 46.408 | 54.527 | 55.617 | 53.220 | 55.724 | 54.693 | 48.983 | 0.043 | 0.1223 | 0.3336 | 0 | 0 | 0 |
| 'H3f3b' | 127.365 | 144.822 | 106.878 | 105.046 | 143.722 | 117.450 | 103.529 | 142.502 | 103.719 | 147.715 | 117.921 | 70.309 | 137.695 | 147.600 | 84.321 | 158.726 | 110.257 | 70.329 | 0.103 | 0.3756 | 0.6268 | 0 | 0 | 0 |
| 'H6pd' | 6.615 | 6.625 | 7.073 | 6.848 | 6.146 | 7.809 | 5.789 | 6.887 | 7.262 | 6.545 | 6.794 | 7.690 | 6.066 | 6.770 | 7.241 | 5.940 | 6.378 | 6.946 | 0.007 | 0.8711 | 0.9447 | 0 | 0 | 0 |
| 'Habp4' | 26.654 | 28.966 | 29.152 | 26.081 | 28.528 | 30.362 | 22.765 | 31.004 | 30.732 | 30.535 | 27.594 | 31.437 | 28.168 | 29.251 | 27.031 | 29.449 | 27.227 | 27.862 | -0.037 | 0.4277 | 0.6739 | 0 | 0 | 0 |
| 'Hacd2' | 4.635 | 4.703 | 4.411 | 4.147 | 5.575 | 4.317 | 4.105 | 4.444 | 4.428 | 3.682 | 4.064 | 4.223 | 3.696 | 4.181 | 4.002 | 3.700 | 4.140 | 3.871 | 0.179 | 0.0018 | 0.0169 | 1 | 0 | 0 |
| 'Hacd3' | 60.153 | 75.553 | 60.256 | 62.601 | 72.134 | 63.403 | 65.715 | 69.451 | 65.271 | 57.235 | 62.241 | 66.775 | 59.706 | 66.356 | 64.911 | 64.211 | 68.626 | 65.862 | 0.031 | 0.3642 | 0.6162 | 0 | 0 | 0 |
| 'Hace1' | 2.320 | 2.332 | 2.352 | 2.397 | 2.780 | 2.221 | 2.278 | 2.413 | 2.101 | 2.414 | 2.649 | 2.287 | 2.511 | 2.602 | 2.001 | 2.713 | 2.661 | 2.321 | -0.071 | 0.1141 | 0.3201 | 0 | 0 | 0 |
| 'Hacl1' | 1.880 | 2.441 | 2.093 | 1.807 | 1.934 | 2.194 | 1.982 | 2.130 | 1.926 | 2.065 | 2.363 | 2.065 | 1.966 | 2.294 | 1.893 | 2.115 | 2.191 | 1.642 | -0.021 | 0.7260 | 0.8745 | 0 | 0 | 0 |
| 'Hadh' | 10.321 | 10.196 | 10.591 | 11.544 | 11.149 | 10.346 | 10.264 | 10.929 | 10.759 | 10.442 | 9.053 | 10.520 | 10.047 | 9.949 | 10.016 | 10.178 | 9.775 | 9.223 | 0.095 | 0.0067 | 0.0463 | 1 | 0 | 0 |
| 'Hadha' | 22.676 | 23.035 | 22.930 | 23.714 | 23.575 | 24.266 | 23.871 | 23.514 | 22.955 | 24.451 | 22.719 | 22.842 | 23.268 | 22.857 | 22.463 | 22.922 | 22.486 | 22.822 | 0.015 | 0.4442 | 0.6872 | 0 | 0 | 0 |
| 'Hadhb' | 12.104 | 13.853 | 12.064 | 13.175 | 14.165 | 12.723 | 13.188 | 13.349 | 11.886 | 11.691 | 12.677 | 12.723 | 11.530 | 12.655 | 12.266 | 12.434 | 12.737 | 12.156 | 0.057 | 0.0576 | 0.2066 | 0 | 0 | 0 |
| 'Hagh' | 8.670 | 8.890 | 9.250 | 9.005 | 10.447 | 9.828 | 8.287 | 10.151 | 8.930 | 10.243 | 9.125 | 9.816 | 10.933 | 10.259 | 8.514 | 9.605 | 10.330 | 9.274 | -0.088 | 0.1008 | 0.2973 | 0 | 0 | 0 |
| 'Haghl' | 5.051 | 5.837 | 6.693 | 5.115 | 5.986 | 6.623 | 5.702 | 6.133 | 6.779 | 5.753 | 4.930 | 5.071 | 6.750 | 5.265 | 5.083 | 5.759 | 5.367 | 4.079 | 0.160 | 0.0361 | 0.1527 | 0 | 0 | 0 |
| 'Hap1' | 5.750 | 7.637 | 8.544 | 6.671 | 7.064 | 9.863 | 6.582 | 8.042 | 10.308 | 7.186 | 6.103 | 9.370 | 6.372 | 6.599 | 8.705 | 6.591 | 6.434 | 9.193 | 0.068 | 0.1839 | 0.4193 | 0 | 0 | 0 |
| 'Harbi1' | 1.228 | 1.404 | 1.335 | 1.067 | 1.342 | 1.442 | 1.344 | 1.413 | 1.256 | 1.142 | 1.225 | 1.384 | 1.085 | 1.384 | 1.532 | 1.156 | 1.174 | 1.332 | 0.040 | 0.5554 | 0.7676 | 0 | 0 | 0 |
| 'Hars' | 45.963 | 48.411 | 51.005 | 48.346 | 51.543 | 51.209 | 49.721 | 49.426 | 47.599 | 47.978 | 49.932 | 49.951 | 51.983 | 48.756 | 49.619 | 50.635 | 49.654 | 49.949 | -0.027 | 0.3123 | 0.5709 | 0 | 0 | 0 |
| 'Hars2' | 6.748 | 7.518 | 8.232 | 6.928 | 7.994 | 7.962 | 7.871 | 7.998 | 7.946 | 7.399 | 7.422 | 7.830 | 7.585 | 7.510 | 7.461 | 7.182 | 8.219 | 7.085 | 0.021 | 0.5605 | 0.7707 | 0 | 0 | 0 |
| 'Hat1' | 15.480 | 16.052 | 18.010 | 19.009 | 16.648 | 16.246 | 18.105 | 16.460 | 16.332 | 14.945 | 16.011 | 16.479 | 18.820 | 18.017 | 16.940 | 15.902 | 16.968 | 15.962 | 0.010 | 0.8281 | 0.9252 | 0 | 0 | 0 |
| 'Haus1' | 17.096 | 18.129 | 18.165 | 18.546 | 18.006 | 20.253 | 19.082 | 20.191 | 20.394 | 19.184 | 17.331 | 17.389 | 20.600 | 18.225 | 18.080 | 19.305 | 18.813 | 14.934 | 0.049 | 0.4007 | 0.6494 | 0 | 0 | 0 |
| 'Haus2' | 9.682 | 8.442 | 7.622 | 8.053 | 10.281 | 8.360 | 8.646 | 9.296 | 7.444 | 11.919 | 8.579 | 5.854 | 11.110 | 9.930 | 6.573 | 9.446 | 7.975 | 5.348 | 0.049 | 0.5573 | 0.7689 | 0 | 0 | 0 |
| 'Haus3' | 10.013 | 10.999 | 9.591 | 10.802 | 10.628 | 8.534 | 11.069 | 10.226 | 9.150 | 8.906 | 12.300 | 10.677 | 10.507 | 12.055 | 9.848 | 10.625 | 12.417 | 10.438 | -0.114 | 0.0195 | 0.1014 | 0 | 0 | 0 |
| 'Haus4' | 24.883 | 25.410 | 26.826 | 22.955 | 25.270 | 26.823 | 24.433 | 23.543 | 25.607 | 24.698 | 27.187 | 25.103 | 26.271 | 27.227 | 27.835 | 25.681 | 25.078 | 23.241 | -0.047 | 0.2724 | 0.5275 | 0 | 0 | 0 |
| 'Haus5' | 4.117 | 4.063 | 5.526 | 4.400 | 3.771 | 5.152 | 4.487 | 4.226 | 5.444 | 3.954 | 3.836 | 4.645 | 4.520 | 3.524 | 4.838 | 4.132 | 3.781 | 4.368 | 0.123 | 0.0068 | 0.0473 | 1 | 0 | 0 |
| 'Haus6' | 4.125 | 4.546 | 4.794 | 5.052 | 4.839 | 4.359 | 4.978 | 4.557 | 4.440 | 4.499 | 5.115 | 4.617 | 4.536 | 4.671 | 4.440 | 4.000 | 4.957 | 4.388 | 0.004 | 0.9235 | 0.9708 | 0 | 0 | 0 |
| 'Haus7' | 12.694 | 13.799 | 13.982 | 12.866 | 13.418 | 13.131 | 13.478 | 14.095 | 13.516 | 12.453 | 14.851 | 13.259 | 13.106 | 12.573 | 13.918 | 12.497 | 13.369 | 12.158 | 0.025 | 0.5745 | 0.7800 | 0 | 0 | 0 |
| 'Hax1' | 6.226 | 7.099 | 6.699 | 7.206 | 6.782 | 6.363 | 7.089 | 6.931 | 6.523 | 5.957 | 6.326 | 6.739 | 6.500 | 6.520 | 7.222 | 6.300 | 6.984 | 6.716 | 0.025 | 0.5421 | 0.7590 | 0 | 0 | 0 |
| 'Hbp1' | 11.088 | 12.233 | 11.745 | 11.718 | 12.106 | 10.733 | 12.164 | 12.830 | 12.025 | 11.011 | 13.900 | 13.191 | 10.241 | 12.912 | 11.678 | 11.777 | 13.097 | 12.081 | -0.052 | 0.2089 | 0.4535 | 0 | 0 | 0 |
| 'Hbs1l' | 23.022 | 24.296 | 24.216 | 23.092 | 24.344 | 22.445 | 23.245 | 23.423 | 23.523 | 23.872 | 22.529 | 23.713 | 23.574 | 22.351 | 22.042 | 23.395 | 23.779 | 23.038 | 0.011 | 0.6685 | 0.8400 | 0 | 0 | 0 |
| 'Hc' | 2.700 | 2.630 | 3.249 | 3.216 | 2.676 | 3.073 | 2.948 | 2.215 | 3.097 | 2.222 | 1.705 | 2.498 | 1.658 | 1.543 | 2.270 | 2.108 | 1.673 | 2.364 | 0.505 | 0.0000 | 0.0000 | 1 | 1 | 0 |
| 'Hccs' | 8.713 | 8.577 | 6.814 | 7.341 | 7.766 | 6.695 | 7.889 | 7.948 | 6.521 | 7.123 | 7.310 | 6.804 | 7.957 | 7.968 | 7.033 | 7.330 | 7.768 | 6.637 | 0.035 | 0.3941 | 0.6434 | 0 | 0 | 0 |
| 'Hcfc1' | 24.140 | 22.391 | 25.953 | 23.030 | 22.525 | 24.711 | 23.758 | 22.735 | 25.523 | 24.009 | 23.844 | 26.763 | 24.320 | 23.397 | 26.254 | 22.223 | 24.137 | 27.557 | -0.061 | 0.0171 | 0.0927 | 0 | 0 | 0 |
| 'Hcfc1r1' | 11.981 | 13.249 | 12.458 | 11.697 | 11.450 | 12.818 | 11.621 | 12.140 | 13.602 | 11.462 | 11.366 | 11.964 | 11.818 | 10.464 | 11.489 | 11.771 | 11.170 | 11.244 | 0.102 | 0.0452 | 0.1764 | 0 | 0 | 0 |
| 'Hcfc2' | 5.136 | 6.182 | 5.361 | 5.896 | 5.255 | 5.388 | 5.725 | 5.554 | 5.701 | 4.678 | 5.825 | 6.200 | 4.818 | 6.134 | 5.851 | 4.896 | 6.326 | 6.000 | -0.025 | 0.6464 | 0.8273 | 0 | 0 | 0 |
| 'Hcls1' | 170.425 | 169.675 | 169.896 | 169.917 | 172.065 | 184.240 | 171.233 | 165.764 | 178.268 | 170.969 | 167.579 | 162.616 | 180.429 | 169.347 | 174.704 | 176.356 | 164.204 | 167.932 | 0.006 | 0.8002 | 0.9111 | 0 | 0 | 0 |
| 'Hcn2' | 1.343 | 1.473 | 1.631 | 1.567 | 1.370 | 1.875 | 1.295 | 1.282 | 1.474 | 1.159 | 1.380 | 1.445 | 1.298 | 1.518 | 1.672 | 1.175 | 0.807 | 1.357 | 0.164 | 0.1269 | 0.3414 | 0 | 0 | 0 |
| 'Hcst' | 115.046 | 139.135 | 135.936 | 108.794 | 126.735 | 150.796 | 117.314 | 138.673 | 138.124 | 132.689 | 127.014 | 146.357 | 130.870 | 132.343 | 150.510 | 146.032 | 129.188 | 132.936 | -0.081 | 0.0664 | 0.2253 | 0 | 0 | 0 |
| 'Hdac10' | 6.673 | 5.586 | 7.191 | 5.879 | 5.789 | 7.295 | 5.873 | 6.423 | 6.944 | 6.222 | 5.979 | 8.105 | 6.134 | 5.428 | 6.923 | 5.774 | 6.382 | 7.215 | -0.022 | 0.6150 | 0.8079 | 0 | 0 | 0 |
| 'Hdac2' | 16.000 | 16.844 | 16.126 | 17.282 | 17.436 | 14.734 | 17.249 | 16.296 | 15.908 | 15.193 | 16.863 | 16.195 | 15.698 | 16.418 | 15.532 | 14.739 | 16.238 | 15.229 | 0.043 | 0.1792 | 0.4145 | 0 | 0 | 0 |
| 'Hdac3' | 12.493 | 13.310 | 13.301 | 12.794 | 13.715 | 13.829 | 12.689 | 13.472 | 13.908 | 13.181 | 12.538 | 12.974 | 14.700 | 13.135 | 12.891 | 12.851 | 13.733 | 12.017 | 0.008 | 0.8086 | 0.9139 | 0 | 0 | 0 |
| 'Hdac4' | 3.611 | 3.535 | 3.910 | 3.598 | 3.604 | 3.470 | 3.542 | 3.621 | 3.705 | 4.272 | 4.515 | 4.059 | 3.895 | 4.086 | 4.055 | 4.312 | 4.108 | 4.191 | -0.209 | 0.0000 | 0.0000 | 1 | 0 | 0 |
| 'Hdac5' | 5.024 | 4.999 | 5.514 | 4.811 | 4.907 | 5.705 | 5.257 | 5.149 | 5.736 | 5.388 | 4.927 | 5.919 | 4.902 | 4.840 | 5.701 | 5.538 | 4.597 | 5.432 | -0.014 | 0.6559 | 0.8333 | 0 | 0 | 0 |
| 'Hdac6' | 2.573 | 2.266 | 2.818 | 2.478 | 2.399 | 2.791 | 2.597 | 2.637 | 3.087 | 2.724 | 2.602 | 2.741 | 3.119 | 2.553 | 2.593 | 2.526 | 2.792 | 2.399 | -0.033 | 0.5438 | 0.7606 | 0 | 0 | 0 |
| 'Hdac7' | 42.028 | 32.986 | 42.890 | 36.120 | 32.444 | 40.771 | 35.204 | 33.530 | 42.700 | 39.289 | 32.156 | 42.202 | 37.206 | 32.914 | 39.938 | 36.162 | 34.806 | 41.257 | 0.000 | 0.9968 | 0.9994 | 0 | 0 | 0 |
| 'Hdac8' | 2.655 | 2.299 | 2.521 | 2.651 | 2.693 | 2.646 | 3.087 | 2.764 | 2.367 | 2.795 | 2.823 | 2.999 | 2.448 | 2.478 | 2.849 | 2.724 | 2.797 | 2.551 | -0.062 | 0.3059 | 0.5638 | 0 | 0 | 0 |
| 'Hddc2' | 3.335 | 3.629 | 3.869 | 2.826 | 3.661 | 3.805 | 3.623 | 3.892 | 3.506 | 3.483 | 3.292 | 3.350 | 3.819 | 3.575 | 3.902 | 3.949 | 3.795 | 3.254 | -0.017 | 0.7771 | 0.9002 | 0 | 0 | 0 |
| 'Hddc3' | 4.007 | 4.620 | 4.434 | 4.434 | 4.932 | 5.705 | 4.460 | 5.505 | 5.003 | 4.768 | 4.046 | 4.199 | 5.089 | 4.402 | 4.517 | 4.850 | 4.080 | 3.788 | 0.112 | 0.1352 | 0.3537 | 0 | 0 | 0 |
| 'Hdgf' | 55.066 | 50.530 | 56.167 | 57.281 | 51.167 | 59.862 | 58.183 | 54.047 | 53.368 | 51.592 | 49.057 | 52.515 | 56.800 | 52.833 | 57.193 | 53.055 | 49.331 | 57.449 | 0.036 | 0.2561 | 0.5084 | 0 | 0 | 0 |
| 'Hdgfrp2' | 52.919 | 47.806 | 64.761 | 51.014 | 49.676 | 63.508 | 54.744 | 50.055 | 63.443 | 55.112 | 47.975 | 58.755 | 60.126 | 49.590 | 56.493 | 55.418 | 47.811 | 54.309 | 0.023 | 0.5046 | 0.7312 | 0 | 0 | 0 |
| 'Hdhd2' | 3.050 | 3.416 | 3.203 | 3.189 | 3.527 | 3.036 | 3.009 | 3.257 | 2.925 | 2.918 | 3.223 | 3.167 | 2.922 | 3.299 | 3.106 | 3.104 | 3.057 | 2.949 | 0.030 | 0.4616 | 0.6979 | 0 | 0 | 0 |
| 'Hdhd3' | 2.140 | 2.121 | 2.782 | 2.426 | 2.648 | 2.088 | 2.677 | 2.523 | 2.567 | 2.086 | 2.128 | 1.896 | 2.338 | 0.872 | 1.666 | 2.071 | 1.470 | 1.653 | 0.436 | 0.0009 | 0.0098 | 1 | 1 | 0 |
| 'Hdlbp' | 17.895 | 16.887 | 18.580 | 18.128 | 17.549 | 17.842 | 17.020 | 16.612 | 18.441 | 18.238 | 17.331 | 18.790 | 18.329 | 17.578 | 17.575 | 16.979 | 17.551 | 18.135 | -0.025 | 0.2876 | 0.5442 | 0 | 0 | 0 |
| 'Heatr1' | 15.062 | 12.074 | 16.094 | 15.447 | 13.553 | 13.288 | 14.513 | 14.054 | 14.105 | 14.670 | 14.103 | 16.052 | 12.797 | 13.557 | 14.376 | 12.071 | 14.511 | 15.576 | -0.005 | 0.9255 | 0.9715 | 0 | 0 | 0 |
| 'Heatr3' | 6.717 | 6.373 | 6.900 | 6.656 | 6.717 | 6.418 | 6.451 | 6.596 | 6.490 | 6.810 | 6.096 | 6.703 | 6.703 | 5.925 | 6.239 | 6.101 | 6.107 | 6.257 | 0.047 | 0.1609 | 0.3919 | 0 | 0 | 0 |
| 'Heatr5a' | 4.155 | 4.334 | 4.257 | 4.390 | 4.051 | 3.994 | 4.205 | 3.890 | 4.185 | 3.778 | 3.800 | 4.184 | 3.997 | 3.618 | 3.800 | 3.228 | 3.628 | 3.673 | 0.134 | 0.0030 | 0.0255 | 1 | 0 | 0 |
| 'Heatr5b' | 12.324 | 11.591 | 12.135 | 13.148 | 12.237 | 12.063 | 12.298 | 10.754 | 11.885 | 12.168 | 14.017 | 12.644 | 11.377 | 12.444 | 12.601 | 10.557 | 12.972 | 13.050 | -0.054 | 0.2751 | 0.5307 | 0 | 0 | 0 |
| 'Heatr6' | 6.217 | 5.735 | 7.059 | 6.750 | 5.935 | 6.430 | 6.213 | 6.176 | 6.550 | 6.276 | 6.420 | 6.681 | 6.662 | 6.464 | 6.380 | 5.930 | 6.469 | 6.459 | -0.027 | 0.4506 | 0.6903 | 0 | 0 | 0 |
| 'Heca' | 29.341 | 29.458 | 31.947 | 30.381 | 29.087 | 31.281 | 28.731 | 31.453 | 27.931 | 30.645 | 31.471 | 31.919 | 26.795 | 32.498 | 31.492 | 30.498 | 30.918 | 34.790 | -0.069 | 0.0828 | 0.2604 | 0 | 0 | 0 |
| 'Hectd1' | 49.077 | 47.820 | 45.470 | 50.414 | 50.294 | 46.443 | 46.593 | 48.846 | 46.157 | 49.229 | 48.624 | 44.994 | 46.361 | 49.579 | 44.063 | 49.434 | 47.691 | 45.137 | 0.011 | 0.6321 | 0.8176 | 0 | 0 | 0 |
| 'Hectd2' | 1.716 | 2.077 | 1.661 | 1.639 | 1.818 | 1.571 | 1.793 | 1.994 | 1.927 | 1.756 | 2.044 | 2.037 | 1.696 | 1.945 | 1.863 | 2.041 | 2.098 | 1.861 | -0.113 | 0.0211 | 0.1066 | 0 | 0 | 0 |
| 'Hectd3' | 21.377 | 20.542 | 23.455 | 20.876 | 20.766 | 24.326 | 21.103 | 21.801 | 23.132 | 22.563 | 20.947 | 24.023 | 20.876 | 22.060 | 23.036 | 22.122 | 20.923 | 22.357 | -0.020 | 0.4294 | 0.6749 | 0 | 0 | 0 |
| 'Heg1' | 10.198 | 9.776 | 10.230 | 9.844 | 9.694 | 10.094 | 10.214 | 10.026 | 10.132 | 10.170 | 9.968 | 10.225 | 10.105 | 10.212 | 10.489 | 10.730 | 9.129 | 11.585 | -0.048 | 0.1321 | 0.3496 | 0 | 0 | 0 |
| 'Helb' | 12.457 | 13.226 | 14.105 | 11.980 | 13.402 | 12.879 | 12.691 | 13.145 | 14.044 | 11.458 | 13.472 | 13.195 | 13.467 | 13.427 | 13.413 | 13.467 | 13.396 | 12.752 | -0.011 | 0.7446 | 0.8847 | 0 | 0 | 0 |
| 'Hells' | 7.299 | 7.241 | 8.421 | 8.393 | 7.041 | 7.679 | 9.464 | 6.738 | 7.198 | 6.807 | 7.106 | 7.631 | 8.806 | 7.453 | 7.613 | 6.811 | 7.532 | 6.935 | 0.044 | 0.4456 | 0.6878 | 0 | 0 | 0 |
| 'Helq' | 5.232 | 4.733 | 5.251 | 5.528 | 4.936 | 4.902 | 5.056 | 5.057 | 5.148 | 5.183 | 4.844 | 5.296 | 5.356 | 4.721 | 5.070 | 4.672 | 5.144 | 4.844 | 0.010 | 0.7878 | 0.9048 | 0 | 0 | 0 |
| 'Helz' | 9.366 | 8.532 | 9.165 | 9.290 | 8.191 | 8.726 | 8.679 | 8.668 | 9.283 | 9.685 | 9.561 | 9.979 | 8.871 | 8.928 | 9.063 | 8.667 | 8.589 | 10.197 | -0.075 | 0.0410 | 0.1658 | 0 | 0 | 0 |
| 'Helz2' | 15.339 | 14.423 | 16.545 | 14.150 | 14.879 | 15.968 | 12.979 | 15.094 | 16.738 | 15.485 | 15.377 | 17.933 | 16.474 | 14.890 | 17.093 | 15.126 | 15.345 | 18.245 | -0.111 | 0.0004 | 0.0047 | 1 | 0 | 0 |
| 'Hemk1' | 3.001 | 3.126 | 3.295 | 2.879 | 2.964 | 3.380 | 3.054 | 3.187 | 3.389 | 3.080 | 2.732 | 2.653 | 3.161 | 2.950 | 2.686 | 2.771 | 2.593 | 2.481 | 0.170 | 0.0023 | 0.0201 | 1 | 0 | 0 |
| 'Herc1' | 12.400 | 10.988 | 11.827 | 12.808 | 11.142 | 11.064 | 11.675 | 10.662 | 11.676 | 11.923 | 12.717 | 12.177 | 10.867 | 11.381 | 11.316 | 11.243 | 11.486 | 12.757 | -0.034 | 0.4181 | 0.6655 | 0 | 0 | 0 |
| 'Herc2' | 15.231 | 12.645 | 14.570 | 16.011 | 13.039 | 13.935 | 13.327 | 13.673 | 14.709 | 13.960 | 14.998 | 15.181 | 12.860 | 13.681 | 14.036 | 13.505 | 13.622 | 16.268 | -0.022 | 0.6483 | 0.8285 | 0 | 0 | 0 |
| 'Herc3' | 7.509 | 7.926 | 7.597 | 7.001 | 7.826 | 7.536 | 7.367 | 7.865 | 7.349 | 6.882 | 6.750 | 6.893 | 6.999 | 7.173 | 6.918 | 7.466 | 7.279 | 6.721 | 0.099 | 0.0002 | 0.0030 | 1 | 0 | 0 |
| 'Herc4' | 20.839 | 20.873 | 20.795 | 20.108 | 19.404 | 19.035 | 21.536 | 18.647 | 19.496 | 18.600 | 21.578 | 22.108 | 20.038 | 20.746 | 19.965 | 18.295 | 23.973 | 21.113 | -0.055 | 0.2485 | 0.4993 | 0 | 0 | 0 |
| 'Herc6' | 5.618 | 5.251 | 5.365 | 5.282 | 5.718 | 5.372 | 5.790 | 5.570 | 5.935 | 5.652 | 6.448 | 7.066 | 5.906 | 5.682 | 6.359 | 5.647 | 6.273 | 6.731 | -0.172 | 0.0000 | 0.0008 | 1 | 0 | 0 |
| 'Herpud1' | 19.511 | 16.559 | 13.436 | 16.776 | 17.272 | 15.911 | 15.552 | 15.733 | 15.847 | 17.466 | 17.038 | 16.229 | 14.910 | 18.619 | 14.775 | 17.021 | 16.542 | 16.060 | -0.031 | 0.5529 | 0.7658 | 0 | 0 | 0 |
| 'Herpud2' | 29.924 | 29.106 | 28.770 | 27.652 | 29.726 | 27.248 | 29.323 | 30.808 | 27.788 | 31.317 | 30.040 | 29.064 | 29.911 | 30.825 | 29.685 | 28.477 | 32.639 | 29.224 | -0.068 | 0.0211 | 0.1066 | 0 | 0 | 0 |
| 'Hes5' | 14.720 | 12.858 | 11.911 | 10.656 | 10.697 | 12.493 | 14.495 | 11.428 | 13.274 | 12.241 | 11.982 | 14.395 | 10.054 | 11.320 | 13.025 | 11.140 | 11.853 | 12.811 | 0.036 | 0.6182 | 0.8090 | 0 | 0 | 0 |
| 'Hes6' | 10.927 | 13.679 | 11.622 | 10.688 | 12.817 | 12.194 | 12.163 | 11.571 | 11.185 | 11.229 | 10.588 | 11.614 | 12.160 | 12.224 | 11.397 | 11.680 | 10.735 | 11.253 | 0.040 | 0.3551 | 0.6084 | 0 | 0 | 0 |
| 'Hexa' | 29.787 | 31.845 | 33.109 | 30.859 | 34.427 | 33.326 | 29.297 | 34.199 | 31.759 | 37.266 | 33.561 | 32.102 | 32.362 | 34.123 | 30.678 | 34.176 | 31.168 | 33.362 | -0.058 | 0.1606 | 0.3913 | 0 | 0 | 0 |
| 'Hexb' | 14.923 | 19.037 | 21.675 | 18.670 | 19.329 | 20.693 | 19.762 | 20.076 | 16.923 | 19.454 | 18.815 | 20.327 | 17.598 | 20.379 | 19.188 | 18.106 | 17.937 | 19.015 | -0.010 | 0.8537 | 0.9373 | 0 | 0 | 0 |
| 'Hexdc' | 2.967 | 2.589 | 3.298 | 2.306 | 2.607 | 3.116 | 3.137 | 2.838 | 3.070 | 2.658 | 2.713 | 3.241 | 2.975 | 2.901 | 3.059 | 3.152 | 2.678 | 2.924 | -0.027 | 0.6632 | 0.8374 | 0 | 0 | 0 |
| 'Hexim1' | 55.397 | 38.466 | 31.779 | 45.133 | 38.136 | 33.177 | 41.019 | 40.872 | 30.312 | 45.430 | 38.747 | 32.220 | 47.474 | 45.609 | 28.731 | 47.855 | 38.769 | 29.446 | -0.004 | 0.9367 | 0.9759 | 0 | 0 | 0 |
| 'Hexim2' | 1.129 | 1.035 | 1.240 | 0.939 | 1.273 | 1.309 | 1.260 | 1.038 | 1.264 | 0.743 | 0.732 | 0.994 | 1.027 | 0.859 | 1.019 | 0.995 | 0.980 | 0.825 | 0.347 | 0.0005 | 0.0060 | 1 | 0 | 0 |
| 'Hey1' | 3.478 | 3.036 | 1.837 | 2.640 | 2.340 | 1.634 | 2.422 | 2.841 | 1.820 | 2.839 | 4.994 | 3.113 | 3.726 | 5.301 | 3.422 | 3.865 | 5.282 | 3.144 | -0.702 | 0.0000 | 0.0000 | 1 | 1 | 0 |
| 'Heyl' | 0.796 | 1.134 | 0.923 | 0.891 | 0.869 | 0.989 | 0.693 | 0.750 | 1.067 | 1.607 | 1.964 | 2.234 | 1.568 | 1.806 | 2.232 | 1.814 | 1.801 | 2.606 | -1.137 | 0.0000 | 0.0000 | 1 | 1 | 1 |
| 'Hgh1' | 11.899 | 11.974 | 12.662 | 10.932 | 11.865 | 12.938 | 10.992 | 11.697 | 12.408 | 12.204 | 9.003 | 12.892 | 11.505 | 9.828 | 13.196 | 11.380 | 10.104 | 12.374 | 0.056 | 0.2668 | 0.5202 | 0 | 0 | 0 |
| 'Hgs' | 14.543 | 13.656 | 15.438 | 13.412 | 14.004 | 16.892 | 13.952 | 13.589 | 15.329 | 15.240 | 13.774 | 17.773 | 15.712 | 14.462 | 16.386 | 15.254 | 13.395 | 17.835 | -0.106 | 0.0009 | 0.0099 | 1 | 0 | 0 |
| 'Hgsnat' | 3.831 | 4.627 | 4.804 | 3.461 | 4.660 | 4.257 | 4.262 | 4.005 | 4.713 | 4.047 | 4.291 | 4.502 | 4.176 | 4.608 | 3.881 | 4.003 | 4.189 | 3.822 | 0.029 | 0.6684 | 0.8400 | 0 | 0 | 0 |
| 'Hhat' | 0.418 | 0.418 | 0.386 | 0.341 | 0.467 | 0.421 | 0.484 | 0.362 | 0.236 | 0.465 | 0.334 | 0.545 | 0.496 | 0.406 | 0.386 | 0.439 | 0.482 | 0.387 | -0.190 | 0.1262 | 0.3401 | 0 | 0 | 0 |
| 'Hibadh' | 5.446 | 5.573 | 5.094 | 5.747 | 6.114 | 5.521 | 5.643 | 5.383 | 5.008 | 5.452 | 5.592 | 5.690 | 5.325 | 5.751 | 5.290 | 5.208 | 5.745 | 5.562 | -0.018 | 0.6185 | 0.8092 | 0 | 0 | 0 |
| 'Hibch' | 2.055 | 2.016 | 2.632 | 2.152 | 2.278 | 2.254 | 2.411 | 2.206 | 2.189 | 3.213 | 3.744 | 3.879 | 3.514 | 3.451 | 3.521 | 3.383 | 3.901 | 3.457 | -0.677 | 0.0000 | 0.0000 | 1 | 1 | 0 |
| 'Hic1' | 3.348 | 2.502 | 3.626 | 3.041 | 2.236 | 3.103 | 3.166 | 1.967 | 2.990 | 2.766 | 2.054 | 4.468 | 2.136 | 2.095 | 4.844 | 2.702 | 1.617 | 4.582 | -0.021 | 0.8495 | 0.9351 | 0 | 0 | 0 |
| 'Hic2' | 0.739 | 0.661 | 0.693 | 0.693 | 0.565 | 0.607 | 0.561 | 0.675 | 0.801 | 0.739 | 0.707 | 0.629 | 0.690 | 0.717 | 0.548 | 0.645 | 0.664 | 0.643 | 0.007 | 0.9395 | 0.9766 | 0 | 0 | 0 |
| 'Hid1' | 1.829 | 2.143 | 2.063 | 1.677 | 2.148 | 2.221 | 2.149 | 2.173 | 2.110 | 1.928 | 2.070 | 1.835 | 1.913 | 1.943 | 1.787 | 1.645 | 2.056 | 1.602 | 0.135 | 0.0297 | 0.1339 | 0 | 0 | 0 |
| 'Hif1a' | 220.594 | 274.126 | 204.222 | 234.885 | 263.894 | 204.849 | 242.228 | 247.550 | 213.004 | 269.353 | 274.927 | 190.888 | 288.797 | 291.753 | 201.258 | 259.144 | 276.279 | 196.924 | -0.095 | 0.0159 | 0.0873 | 0 | 0 | 0 |
| 'Hif1an' | 10.852 | 12.870 | 11.914 | 10.822 | 12.155 | 12.510 | 10.932 | 11.815 | 13.213 | 11.668 | 11.298 | 13.253 | 10.382 | 12.243 | 12.866 | 12.025 | 11.216 | 13.728 | -0.033 | 0.3844 | 0.6347 | 0 | 0 | 0 |
| 'Higd1a' | 23.710 | 23.292 | 21.141 | 24.960 | 22.511 | 21.422 | 24.370 | 23.639 | 22.032 | 19.994 | 23.707 | 20.832 | 23.754 | 25.155 | 22.334 | 22.380 | 25.221 | 21.305 | 0.007 | 0.8627 | 0.9405 | 0 | 0 | 0 |
| 'Higd2a' | 77.585 | 86.446 | 82.506 | 76.662 | 82.367 | 84.981 | 82.071 | 86.029 | 78.610 | 82.623 | 82.011 | 80.752 | 88.931 | 84.486 | 79.782 | 79.920 | 81.099 | 76.590 | -0.008 | 0.8175 | 0.9190 | 0 | 0 | 0 |
| 'Hilpda' | 8.415 | 6.362 | 6.450 | 7.046 | 7.174 | 5.725 | 7.616 | 7.101 | 5.668 | 8.829 | 6.673 | 5.000 | 7.787 | 7.438 | 5.885 | 7.988 | 6.502 | 4.523 | 0.038 | 0.5983 | 0.7958 | 0 | 0 | 0 |
| 'Hinfp' | 12.426 | 11.395 | 12.308 | 10.584 | 11.376 | 11.850 | 11.385 | 11.990 | 11.994 | 11.930 | 12.222 | 12.940 | 11.694 | 11.303 | 13.157 | 12.841 | 11.749 | 12.439 | -0.073 | 0.1015 | 0.2986 | 0 | 0 | 0 |
| 'Hint1' | 72.818 | 71.035 | 72.336 | 73.051 | 77.934 | 75.698 | 72.976 | 73.689 | 71.229 | 63.691 | 64.746 | 67.785 | 74.183 | 71.129 | 73.222 | 68.168 | 69.010 | 66.420 | 0.085 | 0.0095 | 0.0600 | 0 | 0 | 0 |
| 'Hint2' | 4.600 | 6.051 | 5.544 | 4.731 | 5.926 | 5.294 | 5.638 | 5.899 | 6.185 | 5.839 | 4.479 | 4.693 | 5.150 | 4.301 | 4.777 | 5.260 | 5.262 | 4.501 | 0.162 | 0.0168 | 0.0913 | 0 | 0 | 0 |
| 'Hint3' | 7.012 | 8.697 | 8.772 | 7.024 | 8.462 | 9.424 | 7.768 | 9.811 | 8.303 | 8.503 | 8.634 | 8.106 | 8.346 | 8.015 | 8.751 | 7.902 | 8.387 | 8.547 | -0.005 | 0.9392 | 0.9765 | 0 | 0 | 0 |
| 'Hip1' | 0.694 | 0.685 | 0.691 | 0.584 | 0.626 | 0.669 | 0.703 | 0.538 | 0.692 | 0.457 | 0.632 | 0.656 | 0.542 | 0.619 | 0.576 | 0.565 | 0.535 | 0.558 | 0.181 | 0.0118 | 0.0706 | 0 | 0 | 0 |
| 'Hip1r' | 30.586 | 30.199 | 30.615 | 28.484 | 28.008 | 31.508 | 30.452 | 28.836 | 32.778 | 31.774 | 27.903 | 30.640 | 34.199 | 30.110 | 28.437 | 30.945 | 29.544 | 29.976 | -0.021 | 0.5248 | 0.7470 | 0 | 0 | 0 |
| 'Hipk1' | 24.953 | 25.313 | 23.234 | 27.750 | 23.118 | 22.954 | 27.030 | 24.261 | 23.884 | 22.061 | 25.096 | 26.100 | 23.575 | 23.894 | 23.818 | 22.271 | 25.192 | 25.745 | 0.019 | 0.6730 | 0.8425 | 0 | 0 | 0 |
| 'Hipk2' | 4.939 | 3.450 | 4.863 | 5.568 | 3.550 | 4.200 | 5.241 | 3.485 | 4.949 | 5.340 | 4.586 | 5.044 | 4.710 | 3.747 | 4.893 | 4.501 | 3.676 | 5.942 | -0.097 | 0.1421 | 0.3651 | 0 | 0 | 0 |
| 'Hipk3' | 16.252 | 16.972 | 17.345 | 18.397 | 16.159 | 16.454 | 17.743 | 17.050 | 16.636 | 16.787 | 18.085 | 18.085 | 16.461 | 17.566 | 16.595 | 16.597 | 17.507 | 17.961 | -0.036 | 0.2733 | 0.5284 | 0 | 0 | 0 |
| 'Hira' | 3.381 | 3.560 | 3.541 | 3.552 | 3.326 | 3.387 | 3.408 | 3.718 | 3.631 | 3.364 | 3.541 | 3.753 | 3.524 | 3.466 | 3.807 | 3.041 | 3.301 | 3.795 | -0.018 | 0.6687 | 0.8401 | 0 | 0 | 0 |
| 'Hirip3' | 8.619 | 6.483 | 9.138 | 8.887 | 6.969 | 8.520 | 8.185 | 6.932 | 8.540 | 7.586 | 7.189 | 8.781 | 9.686 | 7.797 | 9.052 | 7.943 | 8.287 | 8.656 | -0.068 | 0.1628 | 0.3947 | 0 | 0 | 0 |
| 'Hist1h1c' | 6.452 | 7.188 | 9.071 | 4.961 | 7.418 | 8.558 | 3.360 | 7.868 | 6.659 | 6.435 | 5.101 | 4.458 | 6.250 | 6.637 | 3.496 | 7.420 | 3.969 | 3.863 | 0.369 | 0.0620 | 0.2157 | 0 | 0 | 0 |
| 'Hist1h2ab' | 6.742 | 8.958 | 8.704 | 7.624 | 10.163 | 14.613 | 8.586 | 10.600 | 10.490 | 4.698 | 7.822 | 7.254 | 9.345 | 10.564 | 9.108 | 8.260 | 9.431 | 7.510 | 0.217 | 0.1084 | 0.3099 | 0 | 0 | 0 |
| 'Hist1h2ac' | 22.660 | 24.889 | 28.134 | 20.872 | 26.074 | 26.019 | 19.158 | 25.485 | 25.996 | 22.212 | 26.132 | 24.595 | 25.681 | 28.561 | 28.842 | 28.649 | 22.548 | 26.670 | -0.095 | 0.1835 | 0.4192 | 0 | 0 | 0 |
| 'Hist1h2ad' | 161.142 | 173.460 | 186.661 | 132.860 | 156.140 | 195.359 | 116.112 | 159.118 | 197.477 | 159.280 | 166.564 | 150.203 | 184.302 | 178.722 | 193.304 | 170.158 | 144.909 | 169.442 | -0.059 | 0.4290 | 0.6747 | 0 | 0 | 0 |
| 'Hist1h2ae' | 67.315 | 73.339 | 77.030 | 60.957 | 71.585 | 82.898 | 58.139 | 71.730 | 87.661 | 62.515 | 76.163 | 62.363 | 79.365 | 84.350 | 77.050 | 72.946 | 65.759 | 68.937 | -0.010 | 0.8927 | 0.9562 | 0 | 0 | 0 |
| 'Hist1h2af' | 27.624 | 25.945 | 29.337 | 18.110 | 28.809 | 27.894 | 21.571 | 23.774 | 32.888 | 25.719 | 28.827 | 22.449 | 27.394 | 27.774 | 32.906 | 27.447 | 24.285 | 27.150 | -0.056 | 0.5366 | 0.7555 | 0 | 0 | 0 |
| 'Hist2h2ac' | 10.371 | 10.260 | 16.033 | 9.058 | 12.831 | 15.287 | 8.694 | 12.131 | 14.977 | 12.821 | 7.721 | 8.540 | 10.704 | 9.187 | 11.267 | 9.429 | 11.662 | 6.229 | 0.312 | 0.0262 | 0.1242 | 0 | 0 | 0 |
| 'Hist2h2be' | 2.743 | 3.113 | 3.161 | 2.808 | 3.586 | 3.648 | 3.391 | 3.394 | 2.971 | 1.850 | 3.034 | 2.435 | 1.353 | 3.084 | 2.696 | 1.023 | 2.367 | 2.252 | 0.541 | 0.0000 | 0.0001 | 1 | 1 | 0 |
| 'Hist3h2a' | 4.105 | 3.033 | 2.233 | 4.045 | 3.241 | 3.005 | 2.320 | 3.315 | 2.457 | 3.450 | 3.630 | 2.903 | 2.770 | 3.697 | 2.782 | 3.319 | 3.214 | 2.677 | -0.045 | 0.6609 | 0.8362 | 0 | 0 | 0 |
| 'Hist3h2ba' | 9.067 | 6.772 | 8.487 | 8.391 | 7.127 | 8.443 | 6.989 | 6.761 | 8.287 | 6.691 | 6.923 | 6.081 | 7.643 | 6.941 | 6.362 | 5.490 | 6.697 | 5.871 | 0.265 | 0.0104 | 0.0640 | 0 | 0 | 0 |
| 'Hivep1' | 5.990 | 5.445 | 6.128 | 5.617 | 5.516 | 6.408 | 5.336 | 5.221 | 6.553 | 7.011 | 6.245 | 6.516 | 5.832 | 5.871 | 6.199 | 6.606 | 5.874 | 5.957 | -0.112 | 0.0121 | 0.0717 | 0 | 0 | 0 |
| 'Hivep2' | 32.499 | 28.146 | 29.302 | 29.134 | 28.239 | 28.411 | 27.888 | 28.365 | 28.601 | 36.039 | 31.793 | 27.414 | 31.293 | 30.578 | 27.464 | 32.839 | 29.236 | 27.505 | -0.079 | 0.0450 | 0.1759 | 0 | 0 | 0 |
| 'Hivep3' | 5.233 | 4.579 | 5.978 | 4.891 | 5.013 | 5.498 | 4.846 | 4.984 | 6.361 | 5.757 | 4.829 | 5.606 | 5.078 | 5.084 | 5.233 | 5.425 | 4.592 | 5.193 | 0.006 | 0.8893 | 0.9547 | 0 | 0 | 0 |
| 'Hjurp' | 20.995 | 20.836 | 23.636 | 21.687 | 20.636 | 22.241 | 21.573 | 20.662 | 23.437 | 19.899 | 23.855 | 25.450 | 21.805 | 22.964 | 25.993 | 22.242 | 24.628 | 25.827 | -0.128 | 0.0002 | 0.0027 | 1 | 0 | 0 |
| 'Hk1' | 30.132 | 28.934 | 28.820 | 29.413 | 28.682 | 30.240 | 29.642 | 28.781 | 28.983 | 30.222 | 29.444 | 31.016 | 29.882 | 29.228 | 30.942 | 30.705 | 28.110 | 31.906 | -0.053 | 0.0142 | 0.0806 | 0 | 0 | 0 |
| 'Hk2' | 2.000 | 1.270 | 1.477 | 1.883 | 1.185 | 1.194 | 2.072 | 1.362 | 1.637 | 1.308 | 2.894 | 2.034 | 1.794 | 1.713 | 2.651 | 1.340 | 2.020 | 2.182 | -0.364 | 0.0425 | 0.1700 | 0 | 0 | 0 |
| 'Hlcs' | 7.910 | 8.086 | 8.525 | 7.078 | 7.834 | 8.056 | 6.841 | 7.292 | 8.014 | 8.567 | 7.541 | 7.144 | 7.820 | 6.926 | 6.691 | 7.984 | 6.997 | 7.146 | 0.058 | 0.3137 | 0.5715 | 0 | 0 | 0 |
| 'Hltf' | 7.454 | 7.546 | 7.884 | 7.758 | 8.373 | 7.404 | 7.801 | 8.145 | 7.707 | 7.547 | 8.381 | 8.074 | 7.127 | 8.702 | 7.744 | 7.284 | 8.531 | 7.965 | -0.035 | 0.2620 | 0.5159 | 0 | 0 | 0 |
| 'Hmbox1' | 3.847 | 3.234 | 3.671 | 3.400 | 3.665 | 3.653 | 3.437 | 3.593 | 3.561 | 3.856 | 4.309 | 4.084 | 3.598 | 3.505 | 3.774 | 3.575 | 3.880 | 3.969 | -0.116 | 0.0085 | 0.0556 | 0 | 0 | 0 |
| 'Hmbs' | 20.066 | 23.537 | 24.359 | 23.773 | 25.792 | 25.190 | 24.327 | 22.122 | 24.625 | 22.075 | 18.183 | 21.726 | 22.998 | 22.454 | 23.114 | 23.591 | 20.135 | 20.026 | 0.127 | 0.0161 | 0.0883 | 0 | 0 | 0 |
| 'Hmces' | 17.436 | 17.642 | 16.669 | 18.539 | 18.132 | 15.921 | 16.844 | 17.406 | 17.148 | 18.088 | 17.555 | 18.219 | 18.263 | 18.604 | 16.981 | 17.939 | 20.162 | 16.370 | -0.071 | 0.0260 | 0.1236 | 0 | 0 | 0 |
| 'Hmg20a' | 30.258 | 31.969 | 31.455 | 29.610 | 32.750 | 31.572 | 30.021 | 32.033 | 32.690 | 31.219 | 33.297 | 32.325 | 28.535 | 30.243 | 31.967 | 30.511 | 30.515 | 31.082 | 0.003 | 0.9003 | 0.9605 | 0 | 0 | 0 |
| 'Hmg20b' | 9.834 | 9.135 | 11.826 | 9.147 | 9.308 | 11.395 | 10.048 | 9.223 | 11.910 | 10.624 | 8.504 | 12.038 | 11.758 | 8.891 | 10.651 | 10.010 | 9.638 | 11.137 | -0.034 | 0.3833 | 0.6338 | 0 | 0 | 0 |
| 'Hmgb1' | 79.833 | 90.632 | 81.925 | 82.270 | 90.241 | 83.190 | 84.684 | 88.077 | 84.867 | 77.247 | 88.750 | 71.628 | 86.677 | 89.887 | 81.548 | 84.790 | 88.395 | 75.700 | 0.032 | 0.3086 | 0.5669 | 0 | 0 | 0 |
| 'Hmgb3' | 9.022 | 10.695 | 9.739 | 10.422 | 10.712 | 10.766 | 10.188 | 10.051 | 10.196 | 8.921 | 9.822 | 9.103 | 10.383 | 11.009 | 11.040 | 9.484 | 9.837 | 8.506 | 0.050 | 0.3755 | 0.6267 | 0 | 0 | 0 |
| 'Hmgcl' | 12.628 | 13.197 | 12.607 | 13.494 | 13.854 | 12.713 | 13.713 | 13.966 | 13.218 | 12.792 | 12.571 | 14.365 | 12.844 | 13.689 | 13.814 | 12.569 | 13.579 | 12.997 | -0.015 | 0.6741 | 0.8433 | 0 | 0 | 0 |
| 'Hmgcr' | 10.086 | 11.043 | 11.764 | 10.837 | 10.760 | 11.527 | 10.111 | 10.138 | 10.909 | 10.735 | 12.435 | 11.044 | 11.018 | 12.229 | 11.583 | 10.908 | 11.306 | 10.163 | -0.070 | 0.0707 | 0.2345 | 0 | 0 | 0 |
| 'Hmgcs1' | 17.362 | 20.967 | 22.597 | 19.850 | 20.237 | 21.274 | 20.005 | 19.695 | 22.225 | 18.602 | 23.305 | 18.893 | 18.939 | 22.048 | 22.263 | 19.455 | 20.804 | 18.680 | 0.002 | 0.9643 | 0.9869 | 0 | 0 | 0 |
| 'Hmgn1' | 23.149 | 22.472 | 19.519 | 21.741 | 23.555 | 21.558 | 20.573 | 22.366 | 21.104 | 22.001 | 22.105 | 19.682 | 22.620 | 23.664 | 20.600 | 24.364 | 20.543 | 19.430 | 0.001 | 0.9696 | 0.9898 | 0 | 0 | 0 |
| 'Hmgn5' | 14.212 | 10.020 | 9.484 | 14.194 | 10.869 | 10.097 | 12.719 | 10.844 | 9.873 | 11.874 | 12.131 | 10.932 | 14.983 | 12.843 | 11.762 | 11.012 | 13.983 | 11.156 | -0.134 | 0.0494 | 0.1875 | 0 | 0 | 0 |
| 'Hmgxb3' | 14.085 | 13.212 | 13.770 | 13.663 | 13.234 | 13.506 | 13.280 | 12.778 | 14.505 | 14.681 | 12.764 | 15.298 | 13.565 | 13.599 | 13.525 | 13.337 | 13.008 | 14.338 | -0.037 | 0.2525 | 0.5040 | 0 | 0 | 0 |
| 'Hmgxb4' | 4.666 | 5.486 | 5.521 | 4.942 | 4.703 | 5.372 | 5.407 | 5.344 | 5.570 | 4.616 | 5.270 | 5.120 | 4.810 | 5.506 | 5.223 | 4.989 | 5.119 | 4.700 | 0.044 | 0.2812 | 0.5379 | 0 | 0 | 0 |
| 'Hmha1' | 170.899 | 155.218 | 188.366 | 170.898 | 157.524 | 183.095 | 166.932 | 157.187 | 187.482 | 177.251 | 164.989 | 192.447 | 177.012 | 162.842 | 184.653 | 168.517 | 175.047 | 193.686 | -0.066 | 0.0021 | 0.0189 | 1 | 0 | 0 |
| 'Hmmr' | 3.637 | 4.629 | 5.417 | 4.956 | 4.942 | 4.956 | 5.315 | 4.306 | 5.520 | 2.934 | 4.463 | 5.249 | 4.796 | 5.993 | 6.051 | 4.331 | 5.523 | 5.221 | -0.032 | 0.7240 | 0.8731 | 0 | 0 | 0 |

| | | | | | | | | | | | | | | | | | | | | | | | | |
|---|---|---|---|---|---|---|---|---|---|---|---|---|---|---|---|---|---|---|---|---|---|---|---|---|
| 'Hmox1' | 21.409 | 15.284 | 12.874 | 19.481 | 13.330 | 13.892 | 20.126 | 14.924 | 11.519 | 16.615 | 21.331 | 14.466 | 20.526 | 19.910 | 15.166 | 16.833 | 22.381 | 13.304 | -0.187 | 0.0237 | 0.1162 | 0 | 0 | 0 |
| 'Hmox2' | 19.685 | 20.746 | 20.044 | 19.153 | 20.031 | 20.808 | 19.965 | 18.799 | 19.185 | 17.733 | 17.064 | 18.495 | 18.023 | 18.961 | 19.260 | 18.356 | 17.361 | 19.119 | 0.107 | 0.0009 | 0.0096 | 1 | 0 | 0 |
| 'Hn1' | 60.192 | 69.694 | 63.240 | 63.968 | 66.836 | 65.732 | 62.790 | 64.875 | 66.863 | 58.774 | 60.197 | 60.682 | 60.697 | 65.237 | 65.246 | 65.933 | 64.708 | 60.770 | 0.044 | 0.1197 | 0.3293 | 0 | 0 | 0 |
| 'Hn1l' | 10.706 | 10.698 | 10.476 | 10.139 | 10.421 | 9.880 | 10.306 | 10.914 | 12.185 | 9.404 | 10.884 | 11.456 | 10.561 | 10.951 | 12.194 | 9.190 | 11.104 | 11.335 | -0.033 | 0.5278 | 0.7486 | 0 | 0 | 0 |
| 'Hnrnpa0' | 296.780 | 256.462 | 302.407 | 290.553 | 262.643 | 299.448 | 291.972 | 279.284 | 257.498 | 290.798 | 283.254 | 239.365 | 296.299 | 281.752 | 281.496 | 293.299 | 267.567 | 286.198 | -0.001 | 0.9732 | 0.9906 | 0 | 0 | 0 |
| 'Hnrnpa1' | 250.195 | 267.883 | 215.592 | 233.559 | 248.717 | 220.200 | 240.416 | 239.687 | 237.361 | 225.242 | 243.762 | 191.762 | 235.248 | 225.504 | 217.943 | 242.167 | 230.238 | 193.838 | 0.095 | 0.0076 | 0.0509 | 0 | 0 | 0 |
| 'Hnrnpa2b1' | 104.473 | 106.773 | 98.307 | 109.930 | 106.215 | 97.426 | 115.189 | 103.761 | 99.080 | 94.761 | 104.419 | 93.311 | 106.215 | 109.786 | 100.371 | 98.729 | 115.902 | 97.008 | 0.022 | 0.5318 | 0.7525 | 0 | 0 | 0 |
| 'Hnrnpa3' | 130.709 | 144.777 | 131.907 | 143.263 | 144.064 | 137.106 | 142.379 | 142.706 | 138.558 | 128.174 | 134.667 | 119.123 | 131.791 | 135.204 | 133.438 | 134.380 | 137.280 | 124.081 | 0.081 | 0.0014 | 0.0136 | 1 | 0 | 0 |
| 'Hnrnpab' | 139.409 | 142.294 | 148.506 | 143.357 | 148.900 | 141.921 | 148.410 | 143.270 | 141.102 | 130.902 | 134.794 | 132.363 | 148.428 | 137.839 | 146.977 | 136.553 | 137.979 | 141.836 | 0.045 | 0.1013 | 0.2981 | 0 | 0 | 0 |
| 'Hnrnpc' | 79.774 | 80.365 | 81.708 | 89.999 | 85.110 | 80.444 | 87.555 | 82.089 | 80.496 | 84.669 | 81.797 | 76.045 | 79.975 | 82.749 | 75.825 | 80.029 | 86.684 | 73.114 | 0.043 | 0.1277 | 0.3423 | 0 | 0 | 0 |
| 'Hnrnpd' | 31.600 | 33.758 | 32.611 | 32.461 | 33.479 | 30.837 | 36.046 | 33.310 | 32.307 | 29.738 | 31.097 | 27.913 | 32.658 | 32.264 | 29.523 | 30.621 | 31.558 | 27.959 | 0.108 | 0.0001 | 0.0015 | 1 | 0 | 0 |
| 'Hnrnpdl' | 43.375 | 44.702 | 37.791 | 42.135 | 42.581 | 37.563 | 45.358 | 44.270 | 38.435 | 40.349 | 39.878 | 36.134 | 42.426 | 40.150 | 36.240 | 44.389 | 42.902 | 34.348 | 0.067 | 0.0029 | 0.0247 | 1 | 0 | 0 |
| 'Hnrnph1' | 54.260 | 54.191 | 51.273 | 53.213 | 53.627 | 48.786 | 54.472 | 53.493 | 49.997 | 53.123 | 55.823 | 47.478 | 52.859 | 53.926 | 46.400 | 51.587 | 55.110 | 44.854 | 0.030 | 0.1869 | 0.4227 | 0 | 0 | 0 |
| 'Hnrnph3' | 10.495 | 12.203 | 12.212 | 11.165 | 11.659 | 11.932 | 11.972 | 11.779 | 12.485 | 9.453 | 12.067 | 13.154 | 10.676 | 12.775 | 11.649 | 10.219 | 14.730 | 12.170 | -0.019 | 0.6901 | 0.8539 | 0 | 0 | 0 |
| 'Hnrnpk' | 58.546 | 66.059 | 60.144 | 61.491 | 65.668 | 59.340 | 64.828 | 64.243 | 61.404 | 58.991 | 61.484 | 56.631 | 60.573 | 62.643 | 58.990 | 60.381 | 63.341 | 56.249 | 0.048 | 0.0121 | 0.0716 | 0 | 0 | 0 |
| 'Hnrnpl' | 41.857 | 42.718 | 42.939 | 42.462 | 42.730 | 41.993 | 44.607 | 42.879 | 44.498 | 38.119 | 42.386 | 43.354 | 42.012 | 40.340 | 43.226 | 38.155 | 43.035 | 42.942 | 0.039 | 0.1473 | 0.3732 | 0 | 0 | 0 |
| 'Hnrnpll' | 16.152 | 14.821 | 15.667 | 16.178 | 15.228 | 14.338 | 16.859 | 15.762 | 15.377 | 14.757 | 14.104 | 15.410 | 15.469 | 15.336 | 15.331 | 14.899 | 15.448 | 14.726 | 0.039 | 0.1580 | 0.3880 | 0 | 0 | 0 |
| 'Hnrnpm' | 62.028 | 61.584 | 63.654 | 66.706 | 65.850 | 65.113 | 65.079 | 64.037 | 65.183 | 62.222 | 63.102 | 62.054 | 63.818 | 65.035 | 65.051 | 62.353 | 65.270 | 63.617 | 0.006 | 0.7703 | 0.8977 | 0 | 0 | 0 |
| 'Hnrnpu' | 106.539 | 99.550 | 105.138 | 111.582 | 100.555 | 99.951 | 112.413 | 98.452 | 104.242 | 100.225 | 102.245 | 99.705 | 102.446 | 101.298 | 102.198 | 97.733 | 104.106 | 100.958 | 0.031 | 0.2704 | 0.5253 | 0 | 0 | 0 |
| 'Hnrnpul1' | 61.954 | 63.326 | 65.630 | 61.185 | 62.623 | 70.958 | 62.472 | 62.714 | 68.842 | 66.862 | 60.208 | 62.748 | 66.228 | 61.232 | 66.287 | 66.357 | 58.368 | 63.621 | 0.009 | 0.7372 | 0.8802 | 0 | 0 | 0 |
| 'Hnrnpul2' | 101.932 | 104.268 | 103.726 | 103.019 | 103.078 | 106.691 | 98.952 | 100.245 | 106.820 | 101.201 | 103.149 | 101.767 | 102.572 | 104.676 | 106.005 | 102.539 | 102.665 | 104.512 | -0.011 | 0.5694 | 0.7769 | 0 | 0 | 0 |
| 'Homer1' | 1.805 | 1.704 | 1.842 | 1.732 | 1.661 | 1.574 | 1.639 | 1.586 | 1.568 | 1.711 | 1.563 | 1.566 | 1.752 | 1.636 | 1.377 | 1.571 | 1.836 | 1.591 | 0.042 | 0.4138 | 0.6620 | 0 | 0 | 0 |
| 'Homez' | 1.539 | 1.767 | 1.822 | 1.670 | 2.030 | 1.862 | 1.752 | 2.050 | 2.168 | 1.747 | 1.973 | 1.693 | 1.750 | 1.642 | 1.671 | 1.530 | 1.515 | 1.524 | 0.136 | 0.0293 | 0.1329 | 0 | 0 | 0 |
| 'Hook1' | 4.120 | 4.385 | 4.152 | 4.142 | 4.098 | 4.037 | 4.555 | 4.081 | 4.207 | 4.099 | 4.746 | 4.351 | 4.183 | 4.207 | 4.122 | 4.144 | 4.287 | 3.984 | -0.025 | 0.4745 | 0.7095 | 0 | 0 | 0 |
| 'Hook2' | 5.892 | 4.645 | 6.429 | 5.678 | 4.823 | 4.752 | 5.796 | 5.951 | 5.830 | 5.351 | 5.052 | 5.493 | 7.004 | 5.277 | 4.937 | 5.384 | 5.830 | 4.590 | 0.021 | 0.7869 | 0.9047 | 0 | 0 | 0 |
| 'Hook3' | 6.057 | 7.314 | 6.860 | 6.609 | 7.381 | 6.667 | 6.426 | 7.066 | 6.700 | 7.159 | 7.012 | 7.041 | 6.138 | 7.069 | 6.111 | 6.874 | 6.958 | 6.567 | -0.009 | 0.8003 | 0.9111 | 0 | 0 | 0 |
| 'Hp1bp3' | 18.961 | 20.483 | 19.943 | 19.438 | 20.501 | 20.364 | 19.528 | 20.217 | 19.780 | 19.285 | 20.827 | 21.194 | 19.970 | 21.148 | 21.192 | 20.689 | 21.617 | 20.818 | -0.070 | 0.0001 | 0.0009 | 1 | 0 | 0 |
| 'Hpcal1' | 108.390 | 99.524 | 94.189 | 99.008 | 101.428 | 94.918 | 96.873 | 98.809 | 93.741 | 99.494 | 100.172 | 100.223 | 104.681 | 97.889 | 94.981 | 97.184 | 95.517 | 104.353 | -0.024 | 0.4349 | 0.6790 | 0 | 0 | 0 |
| 'Hpf1' | 14.368 | 15.336 | 16.185 | 15.338 | 15.435 | 14.769 | 15.516 | 16.469 | 14.394 | 13.745 | 14.477 | 14.544 | 15.140 | 15.771 | 14.218 | 13.508 | 16.067 | 13.480 | 0.064 | 0.0764 | 0.2462 | 0 | 0 | 0 |
| 'Hpgds' | 7.176 | 9.174 | 7.265 | 7.292 | 8.140 | 7.661 | 7.819 | 7.628 | 7.853 | 7.408 | 7.163 | 6.572 | 7.517 | 7.196 | 6.304 | 7.580 | 7.201 | 5.867 | 0.148 | 0.0008 | 0.0085 | 1 | 0 | 0 |
| 'Hpn' | 2.096 | 2.251 | 2.984 | 2.259 | 2.497 | 2.479 | 2.769 | 1.984 | 3.138 | 2.988 | 2.709 | 2.705 | 3.437 | 2.929 | 2.366 | 2.667 | 2.926 | 2.658 | -0.189 | 0.0258 | 0.1234 | 0 | 0 | 0 |
| 'Hprt' | 73.148 | 83.689 | 76.728 | 79.991 | 84.763 | 69.445 | 80.792 | 80.265 | 71.581 | 68.756 | 80.808 | 73.479 | 75.824 | 80.555 | 72.551 | 73.460 | 80.188 | 69.751 | 0.039 | 0.1956 | 0.4341 | 0 | 0 | 0 |
| 'Hps1' | 5.072 | 4.938 | 5.977 | 4.458 | 5.025 | 5.804 | 4.959 | 4.661 | 5.494 | 5.879 | 4.582 | 5.848 | 5.434 | 4.880 | 5.345 | 5.496 | 4.617 | 5.330 | -0.041 | 0.3393 | 0.5966 | 0 | 0 | 0 |
| 'Hps3' | 2.372 | 2.527 | 2.791 | 2.397 | 2.697 | 2.843 | 2.584 | 2.837 | 2.585 | 2.708 | 2.686 | 2.540 | 2.575 | 2.563 | 2.229 | 2.622 | 2.700 | 2.295 | 0.038 | 0.3947 | 0.6439 | 0 | 0 | 0 |
| 'Hps4' | 3.551 | 3.200 | 3.910 | 3.581 | 3.322 | 4.051 | 3.368 | 3.516 | 3.871 | 4.103 | 2.760 | 3.688 | 3.631 | 3.215 | 3.776 | 3.610 | 3.270 | 3.471 | 0.033 | 0.4719 | 0.7075 | 0 | 0 | 0 |
| 'Hps5' | 4.613 | 4.518 | 5.320 | 4.925 | 4.519 | 4.902 | 4.408 | 4.863 | 5.149 | 4.704 | 4.828 | 5.180 | 4.780 | 4.433 | 4.681 | 4.187 | 5.154 | 5.041 | -0.003 | 0.9362 | 0.9756 | 0 | 0 | 0 |
| 'Hps6' | 6.173 | 4.802 | 7.132 | 5.678 | 5.615 | 6.004 | 6.310 | 5.639 | 6.803 | 6.120 | 4.176 | 6.626 | 5.749 | 5.261 | 6.435 | 5.172 | 4.964 | 6.414 | 0.078 | 0.1832 | 0.4191 | 0 | 0 | 0 |
| 'Hpse' | 0.904 | 1.223 | 1.088 | 0.944 | 1.154 | 1.043 | 1.118 | 1.247 | 1.042 | 0.915 | 0.891 | 1.142 | 1.002 | 1.293 | 0.924 | 0.982 | 1.122 | 1.025 | 0.045 | 0.6562 | 0.8334 | 0 | 0 | 0 |
| 'Hras' | 4.924 | 4.765 | 5.071 | 5.004 | 4.570 | 5.195 | 5.000 | 5.093 | 4.457 | 4.764 | 4.462 | 4.620 | 4.819 | 4.754 | 5.111 | 4.940 | 4.463 | 5.314 | 0.020 | 0.6420 | 0.8243 | 0 | 0 | 0 |
| 'Hrh2' | 3.411 | 3.489 | 2.785 | 3.036 | 3.474 | 3.016 | 3.930 | 3.036 | 3.289 | 3.817 | 3.050 | 2.226 | 3.602 | 3.400 | 2.349 | 3.380 | 3.400 | 1.743 | 0.148 | 0.1016 | 0.2987 | 0 | 0 | 0 |
| 'Hrh3' | 1.218 | 1.797 | 1.213 | 1.317 | 1.395 | 1.347 | 1.207 | 1.538 | 1.447 | 2.041 | 1.813 | 1.542 | 3.250 | 2.376 | 1.584 | 2.423 | 1.979 | 1.434 | -0.548 | 0.0000 | 0.0000 | 1 | 1 | 0 |
| 'Hs1bp3' | 6.342 | 6.742 | 6.319 | 5.950 | 7.042 | 7.200 | 5.102 | 6.739 | 6.849 | 7.984 | 7.060 | 7.773 | 6.821 | 6.191 | 7.629 | 8.431 | 6.463 | 6.708 | -0.167 | 0.0125 | 0.0735 | 0 | 0 | 0 |
| 'Hs2st1' | 2.321 | 2.570 | 2.400 | 2.128 | 2.973 | 2.343 | 2.468 | 2.412 | 2.734 | 2.532 | 2.424 | 2.510 | 2.352 | 2.448 | 2.438 | 2.485 | 2.483 | 2.452 | 0.000 | 0.9983 | 0.9994 | 0 | 0 | 0 |
| 'Hs3st3b1' | 5.832 | 6.677 | 5.702 | 5.450 | 6.327 | 5.730 | 6.030 | 6.269 | 5.664 | 4.466 | 5.639 | 7.094 | 3.884 | 5.864 | 6.298 | 5.182 | 6.286 | 6.767 | 0.058 | 0.4346 | 0.6788 | 0 | 0 | 0 |
| 'Hs6st1' | 13.186 | 11.248 | 14.131 | 12.347 | 11.851 | 13.994 | 11.948 | 12.090 | 13.476 | 12.393 | 9.165 | 11.291 | 10.502 | 10.756 | 11.800 | 11.453 | 8.670 | 12.036 | 0.214 | 0.0000 | 0.0000 | 1 | 0 | 0 |
| 'Hsbp1' | 98.824 | 117.243 | 97.228 | 104.287 | 112.884 | 103.269 | 107.468 | 106.901 | 98.671 | 97.110 | 100.462 | 91.939 | 104.591 | 100.025 | 97.946 | 104.408 | 103.318 | 91.053 | 0.076 | 0.0097 | 0.0610 | 0 | 0 | 0 |
| 'Hscb' | 12.278 | 13.678 | 14.062 | 12.746 | 13.663 | 13.257 | 13.757 | 12.843 | 13.227 | 12.438 | 13.545 | 13.878 | 15.040 | 12.727 | 13.684 | 13.312 | 13.074 | 12.354 | -0.020 | 0.6689 | 0.8402 | 0 | 0 | 0 |
| 'Hsd11b1' | 6.375 | 8.518 | 6.595 | 6.622 | 7.767 | 7.012 | 7.184 | 7.501 | 7.332 | 9.084 | 8.715 | 8.138 | 9.869 | 9.923 | 8.400 | 8.886 | 9.201 | 8.082 | -0.319 | 0.0000 | 0.0000 | 1 | 0 | 0 |
| 'Hsd17b10' | 15.293 | 15.714 | 14.864 | 16.707 | 16.471 | 14.978 | 16.202 | 16.338 | 14.537 | 14.888 | 14.426 | 15.263 | 14.027 | 13.411 | 16.151 | 15.089 | 13.547 | 14.301 | 0.091 | 0.0336 | 0.1456 | 0 | 0 | 0 |
| 'Hsd17b11' | 5.998 | 6.734 | 5.686 | 6.138 | 6.022 | 6.692 | 6.267 | 6.497 | 6.378 | 6.529 | 6.289 | 7.214 | 6.871 | 7.661 | 6.829 | 7.572 | 6.360 | 6.762 | -0.151 | 0.0025 | 0.0217 | 1 | 0 | 0 |
| 'Hsd17b12' | 10.166 | 11.517 | 11.682 | 10.791 | 10.913 | 11.159 | 12.281 | 11.342 | 11.189 | 10.829 | 11.419 | 10.812 | 12.080 | 11.582 | 11.413 | 11.599 | 11.170 | 10.917 | -0.020 | 0.5735 | 0.7795 | 0 | 0 | 0 |
| 'Hsd17b4' | 14.456 | 16.252 | 16.020 | 16.216 | 16.449 | 16.340 | 15.911 | 16.893 | 16.457 | 15.616 | 15.235 | 15.430 | 15.236 | 14.669 | 15.186 | 14.626 | 15.291 | 14.069 | 0.088 | 0.0045 | 0.0346 | 1 | 0 | 0 |
| 'Hsd17b7' | 1.749 | 1.970 | 2.031 | 1.752 | 1.954 | 1.972 | 1.949 | 1.925 | 1.905 | 4.865 | 5.315 | 4.032 | 5.472 | 5.703 | 4.009 | 5.196 | 4.864 | 3.518 | -1.312 | 0.0000 | 0.0000 | 1 | 1 | 1 |
| 'Hsdl1' | 48.305 | 61.763 | 51.665 | 48.460 | 55.300 | 58.676 | 51.281 | 54.799 | 57.616 | 54.718 | 53.481 | 57.429 | 53.245 | 57.574 | 59.910 | 62.560 | 51.063 | 58.542 | -0.073 | 0.1043 | 0.3027 | 0 | 0 | 0 |
| 'Hsdl2' | 6.625 | 6.317 | 7.405 | 7.207 | 7.338 | 6.205 | 7.538 | 7.298 | 6.534 | 7.138 | 6.751 | 6.940 | 6.960 | 6.909 | 6.326 | 7.286 | 7.417 | 5.764 | 0.014 | 0.7676 | 0.8963 | 0 | 0 | 0 |
| 'Hsf1' | 28.524 | 27.282 | 32.668 | 27.335 | 27.987 | 32.906 | 25.629 | 28.536 | 30.140 | 29.546 | 26.462 | 35.177 | 30.063 | 26.134 | 32.218 | 27.977 | 28.261 | 32.752 | -0.052 | 0.1243 | 0.3372 | 0 | 0 | 0 |
| 'Hsf2' | 5.205 | 6.922 | 5.397 | 6.183 | 5.751 | 4.949 | 6.408 | 6.183 | 4.987 | 5.193 | 6.674 | 5.594 | 5.463 | 6.145 | 5.266 | 4.908 | 5.534 | 5.290 | 0.032 | 0.6113 | 0.8049 | 0 | 0 | 0 |
| 'Hsf2bp' | 1.031 | 0.923 | 1.254 | 0.828 | 1.088 | 0.994 | 0.902 | 1.424 | 0.887 | 1.303 | 2.450 | 1.432 | 2.183 | 1.768 | 1.427 | 1.262 | 2.078 | 1.571 | -0.719 | 0.0000 | 0.0000 | 1 | 1 | 0 |
| 'Hsf4' | 1.870 | 1.686 | 2.166 | 1.828 | 2.159 | 2.215 | 1.702 | 1.790 | 2.295 | 1.930 | 1.579 | 2.736 | 2.098 | 1.702 | 2.099 | 2.141 | 2.064 | 2.346 | -0.098 | 0.2177 | 0.4642 | 0 | 0 | 0 |
| 'Hsh2d' | 4.943 | 6.012 | 5.740 | 4.159 | 5.463 | 5.876 | 4.441 | 7.075 | 6.498 | 5.354 | 4.525 | 6.996 | 5.306 | 4.540 | 6.300 | 5.907 | 4.874 | 7.006 | -0.030 | 0.7363 | 0.8800 | 0 | 0 | 0 |
| 'Hsp90aa1' | 148.523 | 68.202 | 72.988 | 148.664 | 72.603 | 69.017 | 152.589 | 72.353 | 65.597 | 87.663 | 116.906 | 104.189 | 94.178 | 90.950 | 87.912 | 78.191 | 127.591 | 109.274 | -0.153 | 0.4151 | 0.6630 | 0 | 0 | 0 |
| 'Hsp90ab1' | 452.913 | 269.229 | 308.886 | 445.473 | 278.045 | 305.153 | 432.537 | 270.137 | 296.175 | 330.723 | 380.012 | 342.303 | 347.331 | 324.460 | 361.886 | 316.010 | 410.464 | 395.440 | -0.112 | 0.3139 | 0.5715 | 0 | 0 | 0 |
| 'Hsp90b1' | 40.719 | 36.859 | 45.323 | 47.039 | 43.078 | 42.788 | 40.448 | 41.878 | 41.295 | 38.292 | 37.372 | 41.453 | 35.024 | 37.975 | 37.843 | 34.770 | 40.740 | 42.261 | 0.124 | 0.0052 | 0.0386 | 1 | 0 | 0 |
| 'Hspa12a' | 2.975 | 3.819 | 2.955 | 3.288 | 3.467 | 3.186 | 2.688 | 3.048 | 2.784 | 4.031 | 4.822 | 3.619 | 4.685 | 4.731 | 3.207 | 3.623 | 4.901 | 3.573 | -0.402 | 0.0000 | 0.0000 | 1 | 1 | 0 |
| 'Hspa13' | 6.021 | 5.836 | 4.885 | 5.517 | 5.073 | 4.879 | 4.997 | 5.178 | 4.384 | 5.774 | 6.298 | 5.127 | 5.579 | 6.199 | 4.385 | 5.628 | 5.680 | 5.030 | -0.094 | 0.0537 | 0.1980 | 0 | 0 | 0 |
| 'Hspa1l' | 161.554 | 2.793 | 23.820 | 211.499 | 3.367 | 4.264 | 242.757 | 2.944 | 12.933 | 11.960 | 103.350 | 56.227 | 26.667 | 26.963 | 43.083 | 2.913 | 145.063 | 53.637 | -1.538 | 0.0640 | 0.2206 | 0 | 0 | 0 |
| 'Hspa2' | 10.861 | 6.507 | 6.886 | 10.455 | 6.007 | 5.920 | 10.765 | 6.975 | 6.331 | 6.628 | 7.373 | 4.715 | 7.363 | 5.920 | 5.676 | 5.350 | 7.800 | 5.038 | 0.305 | 0.0025 | 0.0218 | 1 | 0 | 0 |
| 'Hspa4' | 36.793 | 35.021 | 34.258 | 37.224 | 35.489 | 33.815 | 38.610 | 34.833 | 35.163 | 34.252 | 35.125 | 37.284 | 34.426 | 33.861 | 36.953 | 33.515 | 36.477 | 37.520 | -0.004 | 0.9114 | 0.9659 | 0 | 0 | 0 |
| 'Hspa4l' | 3.468 | 3.209 | 3.422 | 3.672 | 3.184 | 3.023 | 3.535 | 3.149 | 3.213 | 3.516 | 3.462 | 5.122 | 3.134 | 3.492 | 4.229 | 3.347 | 3.882 | 5.389 | -0.248 | 0.0018 | 0.0170 | 1 | 0 | 0 |
| 'Hspa5' | 181.185 | 143.830 | 130.698 | 176.859 | 151.767 | 130.577 | 169.226 | 148.344 | 125.576 | 166.398 | 151.776 | 119.208 | 153.091 | 156.259 | 122.277 | 161.583 | 152.461 | 127.290 | 0.039 | 0.2378 | 0.4879 | 0 | 0 | 0 |
| 'Hspa9' | 36.314 | 35.777 | 35.688 | 34.276 | 36.457 | 33.682 | 37.352 | 37.839 | 32.888 | 37.584 | 34.658 | 36.553 | 37.593 | 35.344 | 34.202 | 37.328 | 36.481 | 35.899 | -0.036 | 0.1630 | 0.3947 | 0 | 0 | 0 |
| 'Hspb11' | 2.587 | 2.682 | 2.644 | 2.887 | 3.021 | 3.026 | 3.502 | 2.968 | 3.187 | 2.326 | 2.720 | 3.096 | 2.494 | 2.300 | 2.563 | 2.591 | 2.633 | 2.669 | 0.155 | 0.0564 | 0.2039 | 0 | 0 | 0 |
| 'Hspb6' | 0.917 | 0.861 | 1.176 | 0.636 | 0.787 | 0.826 | 0.792 | 1.001 | 0.768 | 0.710 | 0.947 | 0.841 | 0.909 | 0.837 | 0.836 | 0.921 | 0.529 | 0.793 | 0.087 | 0.4842 | 0.7158 | 0 | 0 | 0 |
| 'Hspbap1' | 2.249 | 2.376 | 2.407 | 2.886 | 2.454 | 2.472 | 2.705 | 2.408 | 2.751 | 2.612 | 2.642 | 3.056 | 2.711 | 2.658 | 2.728 | 2.303 | 3.113 | 3.119 | -0.156 | 0.0241 | 0.1176 | 0 | 0 | 0 |
| 'Hspbp1' | 10.650 | 10.274 | 11.642 | 10.056 | 11.975 | 12.760 | 9.914 | 11.144 | 11.634 | 10.774 | 8.844 | 12.141 | 10.349 | 10.482 | 11.692 | 11.942 | 9.149 | 11.540 | 0.037 | 0.4586 | 0.6955 | 0 | 0 | 0 |
| 'Hspd1' | 57.587 | 39.194 | 44.948 | 55.807 | 43.693 | 41.240 | 52.986 | 42.840 | 41.421 | 41.269 | 46.363 | 58.427 | 46.034 | 43.656 | 50.989 | 38.272 | 52.248 | 60.716 | -0.074 | 0.4780 | 0.7111 | 0 | 0 | 0 |
| 'Hspe1' | 82.044 | 47.940 | 49.530 | 86.088 | 43.135 | 46.831 | 89.946 | 44.453 | 44.560 | 44.794 | 66.002 | 59.591 | 52.813 | 52.472 | 55.923 | 46.900 | 88.626 | 59.757 | -0.046 | 0.8066 | 0.9134 | 0 | 0 | 0 |
| 'Hsph1' | 51.794 | 7.038 | 9.914 | 34.686 | 8.261 | 9.055 | 31.074 | 8.284 | 8.841 | 14.884 | 26.240 | 28.331 | 16.014 | 14.067 | 19.018 | 8.618 | 27.615 | 32.750 | -0.593 | 0.1504 | 0.3777 | 0 | 0 | 0 |
| 'Htatip2' | 13.896 | 16.124 | 16.318 | 13.242 | 18.037 | 17.337 | 15.034 | 17.628 | 16.685 | 16.221 | 16.533 | 15.874 | 18.464 | 16.566 | 16.309 | 17.301 | 14.749 | 14.578 | -0.034 | 0.5470 | 0.7610 | 0 | 0 | 0 |
| 'Htatsf1' | 15.203 | 15.683 | 15.921 | 14.897 | 16.094 | 15.961 | 15.601 | 15.841 | 15.058 | 16.089 | 14.025 | 14.632 | 15.002 | 14.468 | 14.570 | 15.892 | 14.586 | 13.574 | 0.071 | 0.0184 | 0.0974 | 0 | 0 | 0 |
| 'Htra2' | 4.263 | 4.605 | 4.936 | 4.479 | 4.349 | 4.707 | 4.599 | 4.659 | 5.140 | 4.489 | 3.985 | 4.434 | 4.643 | 4.198 | 4.244 | 3.924 | 4.234 | 4.250 | 0.109 | 0.0071 | 0.0485 | 1 | 0 | 0 |
| 'Htt' | 6.441 | 5.699 | 6.909 | 6.360 | 5.752 | 6.229 | 5.969 | 5.989 | 6.512 | 6.450 | 7.127 | 6.811 | 6.080 | 6.182 | 6.795 | 5.940 | 6.671 | 6.695 | -0.083 | 0.0354 | 0.1504 | 0 | 0 | 0 |
| 'Hus1' | 1.390 | 1.580 | 1.684 | 1.471 | 1.659 | 1.650 | 1.638 | 1.693 | 1.612 | 1.388 | 1.652 | 1.623 | 1.455 | 1.773 | 1.593 | 1.549 | 1.601 | 1.717 | -0.005 | 0.9123 | 0.9663 | 0 | 0 | 0 |
| 'Huwe1' | 19.138 | 16.502 | 18.753 | 19.425 | 16.061 | 16.945 | 16.895 | 16.781 | 17.860 | 20.067 | 19.245 | 19.483 | 16.734 | 17.012 | 17.661 | 17.856 | 18.302 | 20.042 | -0.084 | 0.0650 | 0.2224 | 0 | 0 | 0 |
| 'Hvcn1' | 9.614 | 9.956 | 10.110 | 8.239 | 9.444 | 10.111 | 9.252 | 10.090 | 10.548 | 10.591 | 7.731 | 9.569 | 7.417 | 8.837 | 9.361 | 9.376 | 7.902 | 7.918 | 0.142 | 0.0212 | 0.1067 | 0 | 0 | 0 |
| 'Hyal2' | 1.065 | 1.371 | 1.125 | 1.078 | 1.299 | 1.302 | 1.309 | 1.344 | 1.501 | 1.289 | 1.133 | 1.371 | 1.367 | 1.029 | 1.343 | 1.309 | 1.003 | 1.262 | 0.020 | 0.8065 | 0.9134 | 0 | 0 | 0 |
| 'Hyi' | 7.123 | 7.996 | 8.668 | 6.970 | 7.743 | 8.170 | 6.945 | 8.683 | 7.927 | 7.107 | 7.116 | 7.739 | 8.068 | 6.925 | 7.664 | 7.661 | 6.786 | 6.832 | 0.082 | 0.0984 | 0.2931 | 0 | 0 | 0 |
| 'Hyls1' | 2.063 | 2.263 | 2.096 | 2.146 | 2.207 | 1.994 | 2.186 | 2.125 | 2.053 | 1.705 | 2.245 | 2.418 | 1.749 | 2.437 | 2.320 | 2.001 | 2.312 | 2.327 | -0.043 | 0.5093 | 0.7342 | 0 | 0 | 0 |
| 'Hyou1' | 16.131 | 14.775 | 16.810 | 16.095 | 16.480 | 17.468 | 14.476 | 15.426 | 16.669 | 16.798 | 14.815 | 16.293 | 15.638 | 15.282 | 15.131 | 15.734 | 14.981 | 16.511 | 0.022 | 0.5042 | 0.7311 | 0 | 0 | 0 |
| 'Hypk' | 2.702 | 2.975 | 2.555 | 2.412 | 2.740 | 2.739 | 2.335 | 2.724 | 2.918 | 2.445 | 2.325 | 2.652 | 2.875 | 2.249 | 2.404 | 2.553 | 2.596 | 2.342 | 0.088 | 0.1269 | 0.3414 | 0 | 0 | 0 |
| 'Iah1' | 14.450 | 17.595 | 16.315 | 16.106 | 17.876 | 19.534 | 16.688 | 17.640 | 16.672 | 14.885 | 15.439 | 17.029 | 14.603 | 14.411 | 16.804 | 15.521 | 15.782 | 16.018 | 0.105 | 0.0628 | 0.2176 | 0 | 0 | 0 |
| 'Iars' | 9.229 | 8.270 | 9.329 | 8.121 | 9.797 | 9.183 | 9.441 | 8.973 | 9.565 | 9.049 | 8.667 | 10.376 | 8.544 | 8.519 | 9.575 | 8.672 | 9.654 | 10.309 | -0.036 | 0.3827 | 0.6332 | 0 | 0 | 0 |
| 'Iars2' | 6.433 | 7.300 | 6.939 | 6.362 | 6.621 | 6.564 | 6.682 | 6.752 | 6.642 | 6.846 | 6.411 | 6.638 | 7.299 | 6.481 | 6.483 | 6.449 | 6.584 | 5.887 | 0.018 | 0.5935 | 0.7931 | 0 | 0 | 0 |
| 'Iba57' | 2.034 | 2.440 | 2.007 | 1.264 | 1.725 | 2.229 | 1.926 | 2.077 | 2.056 | 2.667 | 2.007 | 2.149 | 2.102 | 2.060 | 2.275 | 2.161 | 1.664 | 1.946 | -0.108 | 0.2640 | 0.5179 | 0 | 0 | 0 |
| 'Ibtk' | 6.852 | 6.760 | 6.836 | 7.261 | 6.544 | 6.654 | 7.273 | 6.982 | 6.429 | 6.928 | 7.411 | 6.651 | 6.491 | 7.274 | 6.196 | 6.417 | 7.802 | 6.204 | -0.002 | 0.9562 | 0.9837 | 0 | 0 | 0 |
| 'Ica1l' | 3.212 | 3.999 | 2.816 | 3.751 | 3.714 | 3.314 | 3.865 | 4.046 | 3.126 | 9.514 | 10.722 | 9.434 | 11.089 | 10.686 | 9.422 | 11.345 | 12.420 | 9.058 | -1.572 | 0.0000 | 0.0000 | 1 | 1 | 1 |
| 'Icam1' | 19.862 | 17.359 | 19.223 | 17.356 | 20.505 | 19.889 | 21.300 | 19.549 | 19.041 | 17.420 | 19.652 | 18.317 | 20.239 | 18.256 | 16.691 | 18.972 | 17.781 | 16.187 | 0.083 | 0.0712 | 0.2354 | 0 | 0 | 0 |
| 'Icam2' | 11.505 | 13.239 | 13.746 | 11.691 | 13.112 | 13.578 | 11.048 | 12.933 | 13.664 | 13.168 | 11.804 | 14.229 | 12.011 | 12.166 | 12.716 | 13.220 | 11.939 | 13.181 | -0.011 | 0.7777 | 0.9002 | 0 | 0 | 0 |
| 'Icam5' | 4.084 | 3.973 | 4.561 | 4.245 | 3.825 | 4.906 | 4.626 | 3.742 | 5.140 | 3.681 | 4.726 | 4.696 | 4.648 | 4.805 | 5.066 | 4.290 | 4.277 | 4.538 | -0.068 | 0.3047 | 0.5622 | 0 | 0 | 0 |
| 'Ice1' | 19.372 | 20.761 | 19.750 | 18.315 | 18.423 | 19.416 | 18.122 | 18.608 | 19.706 | 18.980 | 19.306 | 21.131 | 18.366 | 19.149 | 20.276 | 20.223 | 19.052 | 20.822 | -0.051 | 0.0851 | 0.2650 | 0 | 0 | 0 |
| 'Ice2' | 3.128 | 3.159 | 3.188 | 3.236 | 3.425 | 3.190 | 3.051 | 3.397 | 3.070 | 3.088 | 2.896 | 3.131 | 2.803 | 3.098 | 2.638 | 2.922 | 3.591 | 2.778 | 0.088 | 0.0451 | 0.1759 | 0 | 0 | 0 |
| 'Ick' | 0.709 | 0.668 | 0.758 | 0.623 | 0.715 | 0.664 | 0.657 | 0.662 | 0.633 | 0.757 | 0.575 | 0.640 | 0.683 | 0.664 | 0.516 | 0.701 | 0.727 | 0.539 | 0.069 | 0.3435 | 0.5994 | 0 | 0 | 0 |
| 'Icmt' | 5.648 | 6.053 | 5.748 | 5.803 | 5.467 | 6.359 | 5.421 | 5.800 | 5.952 | 5.576 | 5.940 | 6.372 | 5.583 | 5.946 | 6.120 | 5.780 | 5.797 | 6.225 | -0.041 | 0.1984 | 0.4383 | 0 | 0 | 0 |
| 'Icos' | 98.298 | 130.266 | 98.799 | 102.950 | 119.934 | 103.964 | 110.514 | 111.300 | 108.137 | 94.399 | 94.814 | 96.320 | 87.454 | 104.765 | 93.432 | 102.998 | 100.993 | 100.540 | 0.154 | 0.0002 | 0.0022 | 1 | 0 | 0 |
| 'Icosl' | 6.123 | 5.608 | 7.979 | 6.409 | 5.675 | 7.545 | 6.312 | 6.984 | 7.612 | 6.769 | 5.628 | 7.607 | 4.198 | 3.840 | 6.107 | 6.115 | 4.931 | 6.559 | 0.214 | 0.0108 | 0.0661 | 0 | 0 | 0 |
| 'Ict1' | 5.281 | 6.630 | 6.437 | 5.952 | 6.197 | 6.481 | 6.098 | 6.256 | 6.199 | 6.046 | 5.463 | 5.471 | 6.319 | 5.641 | 5.696 | 5.935 | 5.441 | 5.039 | 0.113 | 0.0133 | 0.0769 | 0 | 0 | 0 |
| 'Id1' | 6.791 | 5.513 | 5.630 | 6.895 | 6.687 | 4.929 | 8.132 | 5.309 | 5.284 | 6.825 | 10.876 | 8.551 | 7.617 | 9.871 | 10.022 | 6.665 | 10.728 | 8.366 | -0.537 | 0.0000 | 0.0000 | 1 | 1 | 0 |
| 'Id2' | 71.983 | 100.535 | 72.101 | 67.405 | 91.197 | 74.856 | 74.703 | 94.392 | 69.230 | 70.893 | 76.936 | 62.837 | 62.765 | 92.263 | 60.695 | 81.669 | 69.752 | 58.403 | 0.158 | 0.0048 | 0.0365 | 1 | 0 | 0 |
| 'Id3' | 18.801 | 17.094 | 20.581 | 22.967 | 17.966 | 19.426 | 22.517 | 15.655 | 19.901 | 13.124 | 15.104 | 23.281 | 12.743 | 14.448 | 20.220 | 13.433 | 18.877 | 21.677 | 0.203 | 0.0645 | 0.2216 | 0 | 0 | 0 |
| 'Ide' | 16.885 | 16.793 | 17.373 | 16.930 | 18.632 | 16.875 | 17.364 | 18.082 | 16.858 | 18.871 | 17.519 | 17.040 | 17.909 | 17.688 | 15.286 | 16.502 | 17.417 | 16.528 | 0.000 | 0.9952 | 0.9994 | 0 | 0 | 0 |
| 'Idh1' | 0.821 | 1.371 | 1.141 | 1.075 | 1.238 | 1.007 | 0.848 | 1.050 | 1.014 | 0.745 | 1.132 | 1.217 | 0.978 | 0.927 | 1.360 | 1.169 | 1.295 | 1.121 | -0.080 | 0.4510 | 0.6904 | 0 | 0 | 0 |
| 'Idh2' | 12.482 | 14.012 | 13.208 | 13.003 | 13.224 | 13.930 | 12.674 | 12.795 | 13.182 | 12.543 | 11.804 | 12.659 | 13.709 | 12.026 | 12.844 | 13.523 | 12.133 | 12.271 | 0.052 | 0.1147 | 0.3213 | 0 | 0 | 0 |
| 'Idh3a' | 26.713 | 27.588 | 28.429 | 29.065 | 30.481 | 28.645 | 29.881 | 29.593 | 28.203 | 26.764 | 26.932 | 27.677 | 28.228 | 25.963 | 28.160 | 26.206 | 28.221 | 26.154 | 0.069 | 0.0270 | 0.1265 | 0 | 0 | 0 |
| 'Idh3b' | 34.182 | 34.135 | 35.962 | 33.522 | 34.171 | 34.953 | 35.257 | 32.384 | 36.143 | 32.610 | 33.098 | 35.355 | 33.820 | 32.746 | 35.386 | 32.190 | 34.329 | 33.992 | 0.022 | 0.3342 | 0.5925 | 0 | 0 | 0 |
| 'Idh3g' | 16.449 | 17.198 | 17.403 | 15.623 | 18.075 | 17.972 | 16.312 | 17.696 | 17.623 | 17.157 | 16.817 | 16.424 | 17.807 | 16.651 | 16.939 | 16.952 | 16.008 | 15.583 | 0.029 | 0.3574 | 0.6103 | 0 | 0 | 0 |
| 'Idnk' | 19.664 | 20.180 | 19.607 | 18.903 | 20.135 | 20.905 | 20.457 | 20.364 | 19.509 | 17.833 | 19.569 | 22.709 | 20.001 | 19.724 | 21.241 | 19.871 | 22.654 | 21.786 | -0.058 | 0.1340 | 0.3525 | 0 | 0 | 0 |
| 'Ids' | 7.614 | 8.453 | 8.220 | 7.546 | 8.311 | 8.553 | 6.984 | 8.734 | 8.089 | 9.532 | 9.440 | 9.008 | 8.324 | 9.961 | 8.219 | 8.846 | 8.963 | 8.241 | -0.158 | 0.0001 | 0.0021 | 1 | 0 | 0 |
| 'Idua' | 2.070 | 2.382 | 2.388 | 2.065 | 2.370 | 2.454 | 2.221 | 2.312 | 2.491 | 2.702 | 2.470 | 2.340 | 2.295 | 2.191 | 2.320 | 2.601 | 2.418 | 2.322 | -0.067 | 0.1898 | 0.4262 | 0 | 0 | 0 |
| 'Ier2' | 126.124 | 65.293 | 66.987 | 105.213 | 67.539 | 71.890 | 116.559 | 74.026 | 58.761 | 88.480 | 65.109 | 53.266 | 85.088 | 71.020 | 51.624 | 101.357 | 68.276 | 50.009 | 0.228 | 0.0001 | 0.0013 | 1 | 0 | 0 |
| 'Ier3ip1' | 10.999 | 13.434 | 11.490 | 12.466 | 12.946 | 11.108 | 11.789 | 12.655 | 10.656 | 11.035 | 11.179 | 11.319 | 10.581 | 11.983 | 10.850 | 10.454 | 12.026 | 10.721 | 0.084 | 0.0228 | 0.1128 | 0 | 0 | 0 |
| 'Ier5' | 90.398 | 69.737 | 64.611 | 73.753 | 69.200 | 70.597 | 72.070 | 73.268 | 62.868 | 92.775 | 66.474 | 61.558 | 92.851 | 83.558 | 62.739 | 102.778 | 70.911 | 61.986 | -0.096 | 0.0921 | 0.2805 | 0 | 0 | 0 |
| 'Ier5l' | 25.099 | 15.475 | 16.057 | 24.623 | 14.177 | 16.378 | 23.129 | 16.032 | 15.358 | 20.527 | 18.606 | 17.850 | 17.588 | 19.281 | 17.958 | 19.473 | 21.296 | 20.123 | -0.096 | 0.2446 | 0.4952 | 0 | 0 | 0 |
| 'Iffo1' | 5.408 | 5.197 | 6.481 | 5.206 | 5.416 | 6.044 | 5.693 | 5.762 | 6.153 | 5.685 | 5.068 | 5.528 | 5.668 | 5.218 | 5.353 | 5.116 | 5.196 | 5.279 | 0.085 | 0.0177 | 0.0944 | 0 | 0 | 0 |
| 'Iffo2' | 13.164 | 13.540 | 9.230 | 11.188 | 11.741 | 11.105 | 9.919 | 11.032 | 11.080 | 11.292 | 13.330 | 15.722 | 11.743 | 14.154 | 15.833 | 13.919 | 12.650 | 18.612 | -0.321 | 0.0018 | 0.0170 | 1 | 0 | 0 |
| 'Ifi202b' | 4.021 | 3.872 | 2.786 | 3.459 | 5.733 | 3.960 | 4.683 | 3.896 | 3.543 | 5.223 | 5.135 | 6.434 | 5.579 | 5.570 | 5.907 | 5.536 | 7.043 | 5.065 | -0.548 | 0.0000 | 0.0000 | 1 | 1 | 0 |
| 'Ifi27' | 37.992 | 44.291 | 41.026 | 39.864 | 46.739 | 44.912 | 42.279 | 44.833 | 43.835 | 42.150 | 40.867 | 45.156 | 41.515 | 40.925 | 44.630 | 43.834 | 44.835 | 43.012 | -0.018 | 0.5729 | 0.7795 | 0 | 0 | 0 |
| 'Ifi27l2a' | 312.983 | 339.745 | 292.957 | 324.585 | 346.305 | 350.346 | 326.881 | 365.056 | 323.683 | 348.220 | 318.665 | 399.868 | 329.665 | 365.050 | 478.408 | 405.553 | 397.112 | 395.823 | -0.213 | 0.0003 | 0.0043 | 1 | 0 | 0 |
| 'Ifi30' | 11.420 | 11.528 | 10.315 | 10.363 | 12.810 | 10.462 | 11.572 | 12.158 | 9.357 | 7.834 | 11.730 | 10.041 | 10.692 | 12.898 | 10.434 | 8.130 | 12.019 | 8.530 | 0.104 | 0.1621 | 0.3938 | 0 | 0 | 0 |
| 'Ifi35' | 13.764 | 13.406 | 12.521 | 11.364 | 13.757 | 13.740 | 11.675 | 14.287 | 13.504 | 12.360 | 11.902 | 14.685 | 13.627 | 13.163 | 14.783 | 13.767 | 11.951 | 14.889 | -0.048 | 0.3446 | 0.5997 | 0 | 0 | 0 |
| 'Ifi44' | 0.862 | 0.817 | 0.741 | 0.710 | 1.021 | 0.958 | 0.496 | 0.966 | 0.933 | 1.266 | 1.180 | 1.779 | 1.262 | 1.251 | 1.481 | 1.304 | 1.452 | 2.017 | -0.801 | 0.0000 | 0.0000 | 1 | 1 | 0 |
| 'Ifi44l' | 2.194 | 2.227 | 2.503 | 1.768 | 3.185 | 1.635 | 2.008 | 2.472 | 1.737 | 3.529 | 5.271 | 3.654 | 3.495 | 3.904 | 3.261 | 4.126 | 6.493 | 3.194 | -0.900 | 0.0000 | 0.0000 | 1 | 1 | 0 |
| 'Ifi47' | 28.081 | 30.013 | 29.619 | 29.729 | 29.082 | 29.695 | 29.960 | 28.403 | 30.144 | 26.313 | 31.670 | 38.346 | 27.476 | 29.808 | 36.823 | 28.773 | 33.785 | 38.739 | -0.141 | 0.0058 | 0.0421 | 1 | 0 | 0 |
| 'Ifih1' | 6.239 | 6.103 | 6.242 | 5.499 | 6.537 | 6.263 | 5.880 | 6.650 | 6.722 | 8.008 | 6.970 | 8.451 | 7.630 | 7.087 | 7.577 | 7.611 | 7.741 | 7.503 | -0.300 | 0.0000 | 0.0000 | 1 | 0 | 0 |
| 'Ifit1' | 4.426 | 4.461 | 3.704 | 2.175 | 4.616 | 5.819 | 3.354 | 5.184 | 4.747 | 3.804 | 3.564 | 6.919 | 4.308 | 3.849 | 6.920 | 4.844 | 3.285 | 6.782 | -0.183 | 0.1650 | 0.3974 | 0 | 0 | 0 |
| 'Ifit2' | 6.495 | 7.727 | 7.352 | 6.608 | 7.624 | 8.431 | 8.197 | 7.891 | 7.697 | 7.790 | 7.880 | 7.501 | 7.587 | 8.231 | 7.407 | 7.282 | 7.487 | 6.868 | -0.009 | 0.8444 | 0.9341 | 0 | 0 | 0 |
| 'Ifitm2' | 20.624 | 18.394 | 23.520 | 20.190 | 19.566 | 22.876 | 22.244 | 18.831 | 21.238 | 15.252 | 15.445 | 15.974 | 15.928 | 16.643 | 18.872 | 17.228 | 13.673 | 16.748 | 0.364 | 0.0000 | 0.0000 | 1 | 0 | 0 |
| 'Ifnar1' | 13.464 | 12.992 | 14.395 | 12.854 | 13.763 | 14.051 | 13.140 | 13.462 | 14.940 | 14.369 | 12.771 | 14.581 | 14.603 | 13.213 | 13.876 | 13.799 | 13.773 | 13.797 | -0.030 | 0.2628 | 0.5168 | 0 | 0 | 0 |
| 'Ifnar2' | 16.583 | 16.457 | 17.569 | 17.387 | 17.244 | 18.218 | 18.149 | 16.591 | 18.941 | 16.564 | 17.488 | 18.988 | 16.890 | 17.123 | 18.607 | 17.130 | 16.712 | 17.414 | -0.011 | 0.7384 | 0.8814 | 0 | 0 | 0 |
| 'Ifngr1' | 9.393 | 11.586 | 10.708 | 10.311 | 10.595 | 10.952 | 9.890 | 11.056 | 9.398 | 10.194 | 10.894 | 10.260 | 10.541 | 11.569 | 10.897 | 10.717 | 10.172 | 9.989 | -0.030 | 0.4408 | 0.6843 | 0 | 0 | 0 |
| 'Ifngr2' | 1.206 | 1.248 | 1.158 | 1.160 | 1.164 | 1.434 | 1.231 | 1.223 | 1.593 | 1.475 | 1.042 | 1.395 | 1.204 | 1.030 | 1.086 | 1.280 | 1.076 | 1.383 | 0.045 | 0.5948 | 0.7937 | 0 | 0 | 0 |
| 'Ifrd1' | 6.572 | 3.069 | 2.405 | 4.306 | 3.358 | 2.509 | 3.936 | 3.236 | 2.491 | 6.480 | 3.924 | 2.827 | 5.215 | 4.889 | 2.788 | 6.051 | 3.670 | 2.593 | -0.268 | 0.0013 | 0.0129 | 1 | 0 | 0 |
| 'Ifrd2' | 4.584 | 3.652 | 4.622 | 3.799 | 3.621 | 4.648 | 3.668 | 4.210 | 4.463 | 3.982 | 4.012 | 4.921 | 4.029 | 3.409 | 4.091 | 4.117 | 3.882 | 4.941 | -0.012 | 0.8302 | 0.9262 | 0 | 0 | 0 |
| 'Ift122' | 0.408 | 0.538 | 0.434 | 0.451 | 0.601 | 0.557 | 0.556 | 0.446 | 0.484 | 0.519 | 0.580 | 0.448 | 0.510 | 0.484 | 0.436 | 0.560 | 0.482 | 0.341 | 0.034 | 0.7098 | 0.8653 | 0 | 0 | 0 |
| 'Ift140' | 5.332 | 5.017 | 6.009 | 5.472 | 5.046 | 5.954 | 5.629 | 5.276 | 5.639 | 5.574 | 5.511 | 5.528 | 5.058 | 5.344 | 5.469 | 5.362 | 5.049 | 5.350 | 0.026 | 0.4361 | 0.6799 | 0 | 0 | 0 |
| 'Ift172' | 2.010 | 2.073 | 2.240 | 1.957 | 1.921 | 2.161 | 1.825 | 2.086 | 2.312 | 2.237 | 2.016 | 2.047 | 2.429 | 2.066 | 2.073 | 1.818 | 2.178 | 1.979 | -0.027 | 0.6279 | 0.8154 | 0 | 0 | 0 |
| 'Ift20' | 17.169 | 17.824 | 15.982 | 16.289 | 18.980 | 18.174 | 16.932 | 17.721 | 17.055 | 17.892 | 15.954 | 15.985 | 16.857 | 17.638 | 16.250 | 15.945 | 16.703 | 14.932 | 0.066 | 0.0844 | 0.2636 | 0 | 0 | 0 |
| 'Ift22' | 1.441 | 1.931 | 1.781 | 1.917 | 1.680 | 1.886 | 2.124 | 1.726 | 1.936 | 1.641 | 2.051 | 1.706 | 2.360 | 1.510 | 1.373 | 1.831 | 1.849 | 1.498 | 0.045 | 0.6414 | 0.8241 | 0 | 0 | 0 |
| 'Ift27' | 23.035 | 26.048 | 25.488 | 24.435 | 25.520 | 25.956 | 21.883 | 25.199 | 23.833 | 22.892 | 21.083 | 23.659 | 23.996 | 24.834 | 23.216 | 22.862 | 22.521 | 21.682 | 0.087 | 0.0567 | 0.2047 | 0 | 0 | 0 |
| 'Ift46' | 3.599 | 4.179 | 4.389 | 3.574 | 4.443 | 4.402 | 3.987 | 4.368 | 4.092 | 3.746 | 3.199 | 2.944 | 3.718 | 3.559 | 2.974 | 3.217 | 3.263 | 2.356 | 0.351 | 0.0000 | 0.0000 | 1 | 0 | 0 |
| 'Ift52' | 12.606 | 13.398 | 14.652 | 12.505 | 13.851 | 13.544 | 13.590 | 14.435 | 14.104 | 13.716 | 13.445 | 13.712 | 12.691 | 13.873 | 12.948 | 13.324 | 13.636 | 12.314 | 0.029 | 0.4217 | 0.6681 | 0 | 0 | 0 |
| 'Ift57' | 2.866 | 3.323 | 3.196 | 3.798 | 3.807 | 3.239 | 4.178 | 3.688 | 3.364 | 3.607 | 3.584 | 2.960 | 2.896 | 3.291 | 2.964 | 3.245 | 3.376 | 2.569 | 0.135 | 0.0508 | 0.1908 | 0 | 0 | 0 |
| 'Ift74' | 1.829 | 2.079 | 2.301 | 2.136 | 1.991 | 2.744 | 2.369 | 2.364 | 2.397 | 2.146 | 2.260 | 1.902 | 2.358 | 2.457 | 2.048 | 2.392 | 2.111 | 1.928 | 0.052 | 0.5657 | 0.7742 | 0 | 0 | 0 |
| 'Ift80' | 3.614 | 3.576 | 4.126 | 3.957 | 3.325 | 3.904 | 3.879 | 3.580 | 3.929 | 3.784 | 3.701 | 4.353 | 3.279 | 3.884 | 3.735 | 3.591 | 3.532 | 3.757 | 0.000 | 0.9964 | 0.9994 | 0 | 0 | 0 |
| 'Ift81' | 0.681 | 0.780 | 0.771 | 0.640 | 0.964 | 0.690 | 0.736 | 0.852 | 0.752 | 0.859 | 0.774 | 0.638 | 0.690 | 0.805 | 0.662 | 0.640 | 0.889 | 0.555 | 0.076 | 0.3845 | 0.6347 | 0 | 0 | 0 |
| 'Ift88' | 0.845 | 0.693 | 0.827 | 0.729 | 0.762 | 0.894 | 1.034 | 0.965 | 1.102 | 1.037 | 0.836 | 0.783 | 0.814 | 0.877 | 0.658 | 0.768 | 0.815 | 0.786 | 0.090 | 0.3621 | 0.6142 | 0 | 0 | 0 |
| 'Igbp1' | 8.664 | 9.488 | 8.992 | 9.126 | 9.034 | 8.875 | 9.502 | 9.087 | 8.738 | 9.791 | 8.724 | 9.211 | 9.970 | 9.986 | 9.839 | 9.549 | 9.673 | 8.864 | -0.082 | 0.0129 | 0.0751 | 0 | 0 | 0 |
| 'Igf1r' | 13.643 | 12.027 | 12.156 | 11.593 | 13.819 | 14.090 | 11.135 | 13.212 | 12.601 | 17.671 | 16.792 | 15.673 | 16.040 | 16.180 | 16.400 | 17.736 | 16.786 | 16.238 | -0.395 | 0.0000 | 0.0000 | 1 | 1 | 0 |
| 'Igf2bp2' | 2.640 | 2.870 | 3.251 | 2.238 | 3.341 | 2.483 | 2.803 | 3.261 | 2.642 | 2.224 | 3.380 | 2.186 | 2.656 | 3.206 | 2.266 | 2.064 | 3.131 | 2.306 | 0.130 | 0.0575 | 0.2064 | 0 | 0 | 0 |
| 'Igf2r' | 5.764 | 5.956 | 5.564 | 5.595 | 5.416 | 6.059 | 5.495 | 6.097 | 5.893 | 5.765 | 4.626 | 7.192 | 5.568 | 5.130 | 6.170 | 6.462 | 5.350 | 6.970 | -0.043 | 0.4424 | 0.6861 | 0 | 0 | 0 |
| 'Igfbp4' | 0.665 | 1.344 | 1.638 | 2.070 | 1.770 | 3.196 | 2.770 | 2.855 | 2.172 | 0.773 | 0.866 | 1.447 | 2.101 | 1.455 | 2.320 | 2.408 | 1.446 | 2.044 | 0.318 | 0.2624 | 0.5162 | 0 | 0 | 0 |
| 'Ighd' | 6.774 | 11.173 | 6.460 | 7.918 | 12.496 | 7.259 | 8.008 | 11.711 | 7.980 | 8.349 | 11.536 | 6.178 | 11.462 | 12.650 | 5.297 | 7.614 | 12.497 | 5.406 | -0.010 | 0.9142 | 0.9671 | 0 | 0 | 0 |
| 'Ighm' | 437.048 | 574.033 | 420.428 | 460.269 | 548.814 | 475.453 | 484.480 | 493.320 | 464.972 | 521.412 | 528.020 | 411.766 | 638.307 | 558.333 | 431.747 | 499.755 | 552.568 | 402.131 | -0.063 | 0.2465 | 0.4972 | 0 | 0 | 0 |
| 'Ighmbp2' | 5.392 | 5.046 | 5.589 | 5.113 | 4.941 | 5.549 | 5.498 | 4.997 | 5.469 | 5.537 | 5.315 | 5.435 | 5.075 | 4.615 | 5.403 | 4.939 | 4.631 | 5.318 | 0.032 | 0.4157 | 0.6634 | 0 | 0 | 0 |
| 'Igsf23' | 6.395 | 7.576 | 6.159 | 6.564 | 7.307 | 6.809 | 5.702 | 6.089 | 6.231 | 4.122 | 4.238 | 4.700 | 5.731 | 5.003 | 4.972 | 4.811 | 4.193 | 4.466 | 0.460 | 0.0000 | 0.0000 | 1 | 1 | 0 |
| 'Igtp' | 19.245 | 21.012 | 19.336 | 19.650 | 19.972 | 21.123 | 20.730 | 19.536 | 22.135 | 19.573 | 21.674 | 24.859 | 21.018 | 20.992 | 25.185 | 20.804 | 22.250 | 24.930 | -0.149 | 0.0001 | 0.0012 | 1 | 0 | 0 |
| 'Iigp1' | 4.456 | 5.254 | 4.757 | 4.669 | 6.119 | 4.650 | 5.510 | 4.618 | 4.848 | 6.647 | 6.744 | 10.165 | 8.098 | 8.263 | 9.071 | 7.156 | 11.104 | 9.539 | -0.788 | 0.0000 | 0.0000 | 1 | 1 | 0 |
| 'Ik' | 83.594 | 84.429 | 76.463 | 87.325 | 89.198 | 81.623 | 83.084 | 84.282 | 81.663 | 80.316 | 80.063 | 77.479 | 87.023 | 81.895 | 79.894 | 88.216 | 84.778 | 75.080 | 0.023 | 0.3686 | 0.6201 | 0 | 0 | 0 |
| 'Ikbip' | 1.468 | 1.759 | 1.470 | 1.718 | 1.707 | 1.362 | 1.613 | 1.743 | 1.562 | 1.130 | 1.441 | 1.351 | 1.232 | 1.480 | 1.560 | 1.472 | 1.680 | 1.275 | 0.177 | 0.0155 | 0.0856 | 0 | 0 | 0 |
| 'Ikbkap' | 5.354 | 5.670 | 6.415 | 5.237 | 5.596 | 5.813 | 5.730 | 6.353 | 5.834 | 5.745 | 5.610 | 5.438 | 5.729 | 5.569 | 4.741 | 5.671 | 5.675 | 4.529 | 0.089 | 0.0706 | 0.2342 | 0 | 0 | 0 |
| 'Ikbkb' | 6.509 | 6.880 | 7.495 | 6.064 | 6.533 | 7.232 | 6.546 | 6.948 | 7.084 | 7.271 | 6.776 | 7.605 | 7.376 | 6.915 | 6.996 | 7.073 | 6.712 | 7.548 | -0.080 | 0.0066 | 0.0458 | 1 | 0 | 0 |
| 'Ikbke' | 34.391 | 29.502 | 36.800 | 32.216 | 30.935 | 35.185 | 32.245 | 30.739 | 36.846 | 35.164 | 33.128 | 38.765 | 32.260 | 30.764 | 34.960 | 32.919 | 32.951 | 37.647 | -0.058 | 0.0485 | 0.1851 | 0 | 0 | 0 |
| 'Ikbkg' | 2.847 | 3.086 | 3.184 | 2.753 | 3.000 | 3.073 | 2.913 | 3.299 | 3.135 | 2.995 | 3.141 | 3.155 | 2.972 | 3.272 | 3.131 | 3.292 | 3.196 | 2.843 | -0.044 | 0.1840 | 0.4194 | 0 | 0 | 0 |
| 'Ikzf1' | 101.996 | 104.298 | 98.501 | 103.857 | 100.159 | 95.371 | 99.980 | 101.780 | 96.656 | 98.141 | 107.612 | 95.848 | 93.612 | 102.625 | 98.253 | 94.740 | 98.831 | 100.863 | 0.009 | 0.7549 | 0.8907 | 0 | 0 | 0 |
| 'Ikzf2' | 8.692 | 8.962 | 7.647 | 8.758 | 9.447 | 7.716 | 8.920 | 8.420 | 8.509 | 10.824 | 9.932 | 9.157 | 12.402 | 9.868 | 8.847 | 10.727 | 9.963 | 9.070 | -0.245 | 0.0000 | 0.0000 | 1 | 0 | 0 |
| 'Ikzf3' | 53.727 | 57.860 | 48.173 | 52.662 | 57.836 | 50.695 | 51.244 | 56.836 | 50.111 | 51.125 | 55.846 | 43.515 | 58.798 | 53.041 | 45.495 | 49.838 | 52.931 | 44.980 | 0.065 | 0.0479 | 0.1831 | 0 | 0 | 0 |
| 'Ikzf4' | 0.595 | 0.455 | 0.625 | 0.565 | 0.472 | 0.700 | 0.565 | 0.723 | 0.709 | 0.422 | 0.290 | 0.649 | 0.368 | 0.415 | 0.599 | 0.420 | 0.284 | 0.561 | 0.440 | 0.0001 | 0.0009 | 1 | 1 | 0 |
| 'Ikzf5' | 7.421 | 7.728 | 7.566 | 7.553 | 8.221 | 6.732 | 7.734 | 7.713 | 7.332 | 7.478 | 7.722 | 6.957 | 7.232 | 7.940 | 6.893 | 7.666 | 7.352 | 6.291 | 0.046 | 0.1858 | 0.4214 | 0 | 0 | 0 |
| 'Il10ra' | 5.975 | 6.101 | 5.534 | 5.309 | 5.345 | 5.742 | 6.269 | 5.016 | 5.484 | 6.819 | 8.237 | 8.120 | 6.874 | 8.349 | 8.392 | 8.170 | 8.921 | 7.403 | -0.498 | 0.0000 | 0.0000 | 1 | 1 | 0 |
| 'Il10rb' | 4.334 | 4.921 | 4.793 | 4.903 | 5.351 | 4.792 | 4.906 | 4.950 | 4.909 | 4.662 | 4.810 | 4.821 | 5.063 | 4.800 | 4.527 | 4.992 | 4.779 | 4.219 | 0.027 | 0.5326 | 0.7527 | 0 | 0 | 0 |
| 'Il12rb1' | 11.767 | 10.677 | 13.585 | 10.631 | 11.212 | 12.812 | 11.256 | 10.622 | 13.292 | 13.619 | 9.180 | 11.908 | 13.163 | 9.617 | 11.697 | 13.786 | 9.946 | 11.056 | 0.026 | 0.6495 | 0.8292 | 0 | 0 | 0 |
| 'Il13ra1' | 1.453 | 2.430 | 1.179 | 1.308 | 2.551 | 1.210 | 1.445 | 2.004 | 1.100 | 2.213 | 2.821 | 2.433 | 2.146 | 2.475 | 1.780 | 2.642 | 2.741 | 1.689 | -0.571 | 0.0000 | 0.0000 | 1 | 1 | 0 |
| 'Il15ra' | 1.178 | 1.294 | 1.213 | 1.094 | 1.320 | 1.125 | 1.323 | 1.130 | 1.125 | 1.297 | 1.092 | 1.261 | 1.128 | 1.186 | 1.187 | 1.193 | 1.288 | 1.124 | -0.012 | 0.8314 | 0.9267 | 0 | 0 | 0 |
| 'Il16' | 29.256 | 30.804 | 31.203 | 31.166 | 30.067 | 32.577 | 32.043 | 29.860 | 32.308 | 30.302 | 33.105 | 35.365 | 29.920 | 32.454 | 35.603 | 29.956 | 33.161 | 35.787 | -0.091 | 0.0025 | 0.0220 | 1 | 0 | 0 |
| 'Il17ra' | 10.712 | 10.138 | 11.072 | 9.556 | 10.233 | 11.516 | 9.190 | 10.511 | 11.199 | 10.530 | 11.077 | 12.521 | 10.504 | 10.846 | 12.593 | 10.333 | 10.384 | 12.631 | -0.116 | 0.0001 | 0.0022 | 1 | 0 | 0 |
| 'Il17rb' | 7.430 | 6.015 | 9.479 | 6.931 | 5.764 | 8.230 | 7.488 | 6.726 | 9.457 | 3.725 | 3.350 | 4.787 | 3.010 | 2.834 | 4.854 | 3.519 | 2.827 | 4.190 | 1.024 | 0.0000 | 0.0000 | 1 | 1 | 1 |
| 'Il18bp' | 5.964 | 4.035 | 4.355 | 4.521 | 3.792 | 3.497 | 4.598 | 4.506 | 3.903 | 5.003 | 5.005 | 4.704 | 5.224 | 5.249 | 5.334 | 5.034 | 4.231 | 4.030 | -0.169 | 0.0652 | 0.2228 | 0 | 0 | 0 |
| 'Il1r1' | 19.711 | 21.196 | 20.033 | 21.396 | 20.783 | 17.860 | 19.198 | 21.279 | 19.411 | 22.374 | 26.833 | 22.883 | 21.963 | 24.778 | 24.120 | 20.752 | 26.697 | 26.103 | -0.268 | 0.0000 | 0.0000 | 1 | 0 | 0 |
| 'Il1r2' | 23.733 | 27.493 | 22.611 | 25.427 | 29.432 | 20.621 | 26.123 | 27.755 | 22.857 | 33.208 | 32.429 | 25.297 | 40.614 | 32.250 | 26.192 | 31.984 | 34.734 | 24.833 | -0.321 | 0.0000 | 0.0000 | 1 | 0 | 0 |
| 'Il1rap' | 1.189 | 1.578 | 1.469 | 1.447 | 1.513 | 1.616 | 1.282 | 1.646 | 1.484 | 1.438 | 1.399 | 1.425 | 1.396 | 1.416 | 1.331 | 1.626 | 1.583 | 1.442 | 0.009 | 0.8747 | 0.9463 | 0 | 0 | 0 |
| 'Il1rn' | 0.451 | 0.474 | 0.553 | 0.509 | 0.514 | 0.562 | 0.549 | 0.592 | 0.628 | 0.798 | 0.797 | 0.632 | 0.775 | 0.871 | 0.709 | 0.492 | 0.895 | 0.626 | -0.444 | 0.0000 | 0.0006 | 1 | 1 | 0 |
| 'Il21' | 48.147 | 61.216 | 41.391 | 46.847 | 57.349 | 42.722 | 43.063 | 52.807 | 40.257 | 56.460 | 61.708 | 51.975 | 65.307 | 65.129 | 54.147 | 59.857 | 66.028 | 55.836 | -0.326 | 0.0000 | 0.0000 | 1 | 0 | 0 |
| 'Il21r' | 47.946 | 42.739 | 48.224 | 46.097 | 40.239 | 45.785 | 43.860 | 43.280 | 49.481 | 44.179 | 42.312 | 53.807 | 40.940 | 39.579 | 50.598 | 40.452 | 38.276 | 54.036 | 0.009 | 0.8434 | 0.9337 | 0 | 0 | 0 |
| 'Il27ra' | 138.717 | 121.713 | 149.090 | 131.459 | 129.278 | 144.842 | 131.371 | 128.709 | 141.931 | 136.466 | 128.838 | 152.900 | 118.631 | 125.704 | 148.658 | 128.221 | 122.004 | 145.952 | 0.003 | 0.9206 | 0.9702 | 0 | 0 | 0 |
| 'Il2ra' | 2.387 | 1.536 | 2.778 | 2.359 | 1.525 | 2.702 | 2.521 | 2.001 | 2.723 | 1.613 | 0.731 | 2.861 | 1.565 | 0.856 | 2.413 | 2.068 | 0.694 | 2.168 | 0.532 | 0.0000 | 0.0001 | 1 | 1 | 0 |
| 'Il2rb' | 85.342 | 76.754 | 71.128 | 73.212 | 72.882 | 73.252 | 72.278 | 73.710 | 72.342 | 78.213 | 80.157 | 88.209 | 81.901 | 81.129 | 85.976 | 80.667 | 79.204 | 92.012 | -0.166 | 0.0000 | 0.0007 | 1 | 0 | 0 |
| 'Il2rg' | 74.388 | 71.004 | 79.191 | 71.457 | 71.014 | 75.837 | 77.188 | 68.673 | 76.842 | 76.974 | 69.947 | 76.280 | 70.770 | 69.711 | 77.208 | 72.177 | 71.108 | 77.293 | -0.002 | 0.9476 | 0.9796 | 0 | 0 | 0 |
| 'Il3ra' | 12.521 | 12.382 | 15.626 | 10.612 | 13.535 | 15.578 | 8.740 | 13.227 | 15.139 | 13.962 | 10.448 | 15.185 | 10.413 | 11.070 | 16.648 | 12.585 | 11.300 | 15.665 | -0.005 | 0.9438 | 0.9788 | 0 | 0 | 0 |
| 'Il4' | 107.841 | 95.419 | 84.733 | 112.933 | 93.075 | 77.071 | 108.083 | 95.577 | 76.027 | 85.266 | 84.755 | 67.836 | 90.730 | 80.856 | 70.098 | 75.677 | 87.759 | 67.203 | 0.244 | 0.0000 | 0.0000 | 1 | 0 | 0 |
| 'Il4i1' | 9.279 | 9.120 | 10.361 | 9.184 | 9.089 | 10.731 | 9.530 | 9.105 | 10.382 | 8.795 | 8.720 | 10.454 | 9.302 | 9.194 | 10.683 | 9.479 | 9.315 | 10.189 | 0.001 | 0.9711 | 0.9898 | 0 | 0 | 0 |
| 'Il4ra' | 7.490 | 6.274 | 9.516 | 7.874 | 6.829 | 9.323 | 8.268 | 7.883 | 8.468 | 6.187 | 5.256 | 7.093 | 5.497 | 4.565 | 7.049 | 6.806 | 4.758 | 6.959 | 0.406 | 0.0000 | 0.0000 | 1 | 1 | 0 |
| 'Il6ra' | 30.498 | 38.071 | 25.605 | 30.141 | 33.820 | 29.347 | 29.623 | 33.722 | 29.699 | 32.737 | 34.231 | 29.778 | 35.495 | 35.577 | 29.606 | 37.522 | 34.426 | 30.148 | -0.107 | 0.0068 | 0.0473 | 1 | 0 | 0 |
| 'Il6st' | 36.252 | 42.454 | 36.311 | 36.630 | 41.260 | 37.838 | 38.862 | 41.656 | 38.536 | 43.624 | 47.464 | 41.907 | 39.947 | 45.990 | 41.513 | 44.230 | 47.645 | 41.809 | -0.183 | 0.0000 | 0.0000 | 1 | 0 | 0 |
| 'Il7r' | 0.831 | 1.369 | 1.511 | 1.399 | 2.370 | 2.424 | 2.054 | 2.639 | 2.095 | 0.987 | 1.423 | 1.927 | 1.548 | 1.924 | 1.916 | 3.487 | 2.129 | 2.036 | -0.086 | 0.7208 | 0.8716 | 0 | 0 | 0 |
| 'Il9r' | 19.563 | 22.915 | 18.033 | 22.089 | 19.523 | 18.914 | 22.992 | 20.104 | 20.837 | 14.578 | 15.097 | 14.923 | 15.667 | 14.184 | 14.225 | 14.251 | 15.538 | 13.475 | 0.474 | 0.0000 | 0.0000 | 1 | 1 | 0 |
| 'Ilf2' | 8.006 | 8.066 | 7.971 | 7.973 | 7.917 | 8.154 | 7.928 | 8.500 | 8.400 | 7.934 | 7.993 | 7.905 | 8.308 | 8.062 | 7.735 | 7.686 | 8.468 | 7.352 | 0.021 | 0.4519 | 0.6910 | 0 | 0 | 0 |
| 'Ilf3' | 33.204 | 30.134 | 34.787 | 32.714 | 28.704 | 32.994 | 32.989 | 32.755 | 32.765 | 31.038 | 31.501 | 34.044 | 32.310 | 30.031 | 34.773 | 31.001 | 31.970 | 33.431 | -0.005 | 0.8548 | 0.9376 | 0 | 0 | 0 |
| 'Ilkap' | 9.314 | 10.647 | 9.956 | 9.056 | 10.802 | 10.198 | 9.293 | 9.929 | 9.996 | 9.485 | 9.309 | 9.567 | 9.656 | 9.873 | 9.593 | 9.780 | 10.108 | 8.916 | 0.037 | 0.2156 | 0.4613 | 0 | 0 | 0 |
| 'Ilvbl' | 37.846 | 42.612 | 39.856 | 34.544 | 39.606 | 41.148 | 37.339 | 39.478 | 40.685 | 42.328 | 40.122 | 36.384 | 44.325 | 40.308 | 35.488 | 45.890 | 37.108 | 31.995 | -0.007 | 0.9018 | 0.9612 | 0 | 0 | 0 |
| 'Immp1l' | 23.620 | 29.082 | 25.591 | 29.515 | 29.372 | 23.705 | 29.208 | 25.574 | 21.244 | 23.554 | 25.309 | 27.397 | 23.428 | 29.395 | 22.617 | 23.264 | 28.609 | 23.414 | 0.042 | 0.5017 | 0.7288 | 0 | 0 | 0 |
| 'Immt' | 8.365 | 7.668 | 8.210 | 7.957 | 7.756 | 7.741 | 8.178 | 8.440 | 8.361 | 7.427 | 8.021 | 7.950 | 8.067 | 7.940 | 7.844 | 7.761 | 8.298 | 7.742 | 0.024 | 0.4124 | 0.6608 | 0 | 0 | 0 |
| 'Imp3' | 29.935 | 31.076 | 27.182 | 32.179 | 31.211 | 31.295 | 27.617 | 33.083 | 30.317 | 27.114 | 28.032 | 29.763 | 28.998 | 31.733 | 30.898 | 31.755 | 31.907 | 26.912 | 0.026 | 0.5873 | 0.7882 | 0 | 0 | 0 |
| 'Imp4' | 6.385 | 6.522 | 6.719 | 6.371 | 7.198 | 6.750 | 6.030 | 7.558 | 6.689 | 7.019 | 6.167 | 6.523 | 7.096 | 6.661 | 6.726 | 7.041 | 6.564 | 6.178 | -0.003 | 0.9390 | 0.9765 | 0 | 0 | 0 |
| 'Impa1' | 9.349 | 10.607 | 11.525 | 10.941 | 10.843 | 11.007 | 11.262 | 10.900 | 10.979 | 9.356 | 11.811 | 11.552 | 9.989 | 10.966 | 10.912 | 9.508 | 11.047 | 10.958 | 0.010 | 0.7936 | 0.9079 | 0 | 0 | 0 |
| 'Impa2' | 18.607 | 17.665 | 19.921 | 17.635 | 19.306 | 20.184 | 19.654 | 16.197 | 20.004 | 17.232 | 15.358 | 17.514 | 19.778 | 16.856 | 21.615 | 18.219 | 15.694 | 17.676 | 0.072 | 0.1595 | 0.3899 | 0 | 0 | 0 |
| 'Impact' | 13.264 | 14.898 | 12.590 | 10.406 | 13.686 | 12.200 | 12.940 | 14.774 | 12.257 | 15.130 | 15.848 | 12.744 | 12.881 | 14.491 | 11.361 | 14.346 | 13.459 | 11.431 | -0.063 | 0.2383 | 0.4883 | 0 | 0 | 0 |

| | | | | | | | | | | | | | | | | | | | | | | | | |
|---|---|---|---|---|---|---|---|---|---|---|---|---|---|---|---|---|---|---|---|---|---|---|---|---|
| 'Impad1' | 8.260 | 8.597 | 8.338 | 7.806 | 8.552 | 7.871 | 8.819 | 9.006 | 7.142 | 8.996 | 9.181 | 8.668 | 9.049 | 9.749 | 7.708 | 9.636 | 8.618 | 8.559 | -0.117 | 0.0025 | 0.0214 | 1 | 0 | 0 |
| 'Impdh1' | 9.287 | 8.746 | 9.122 | 8.158 | 8.007 | 9.350 | 8.377 | 8.387 | 9.115 | 8.717 | 7.573 | 7.474 | 9.011 | 7.069 | 7.772 | 8.433 | 6.598 | 7.162 | 0.167 | 0.0003 | 0.0036 | 1 | 0 | 0 |
| 'Impdh2' | 20.063 | 16.498 | 18.769 | 20.662 | 17.164 | 16.786 | 20.626 | 18.291 | 17.317 | 18.504 | 18.837 | 18.571 | 20.859 | 18.007 | 18.930 | 17.298 | 19.502 | 19.051 | -0.044 | 0.3184 | 0.5768 | 0 | 0 | 0 |
| 'Inadl' | 4.631 | 4.391 | 4.359 | 4.470 | 4.730 | 4.044 | 4.622 | 4.888 | 4.107 | 5.025 | 4.359 | 4.506 | 4.315 | 4.406 | 3.995 | 4.071 | 4.814 | 4.405 | -0.001 | 0.9853 | 0.9949 | 0 | 0 | 0 |
| 'Incenp' | 53.157 | 47.678 | 64.565 | 56.141 | 47.856 | 65.624 | 59.004 | 47.429 | 64.509 | 44.716 | 52.104 | 59.879 | 62.629 | 55.828 | 64.193 | 51.700 | 56.482 | 59.631 | -0.022 | 0.6888 | 0.8532 | 0 | 0 | 0 |
| 'Inf2' | 4.366 | 3.841 | 5.692 | 4.402 | 3.811 | 5.707 | 4.212 | 4.557 | 5.933 | 4.448 | 3.509 | 4.999 | 3.756 | 3.818 | 5.121 | 3.839 | 3.927 | 4.935 | 0.144 | 0.0056 | 0.0411 | 1 | 0 | 0 |
| 'Ing1' | 36.038 | 34.483 | 31.232 | 35.472 | 34.450 | 34.093 | 34.556 | 33.833 | 32.828 | 30.081 | 38.762 | 38.525 | 31.870 | 37.805 | 37.667 | 33.645 | 37.149 | 40.221 | -0.093 | 0.0632 | 0.2190 | 0 | 0 | 0 |
| 'Ing2' | 6.292 | 5.339 | 4.939 | 6.159 | 4.994 | 4.830 | 5.907 | 5.064 | 4.678 | 5.028 | 6.055 | 5.243 | 5.722 | 6.239 | 5.435 | 6.358 | 6.220 | 5.152 | -0.106 | 0.0555 | 0.2019 | 0 | 0 | 0 |
| 'Ing3' | 4.545 | 4.302 | 3.772 | 3.946 | 4.209 | 3.971 | 4.016 | 4.185 | 3.937 | 4.435 | 4.670 | 4.308 | 4.160 | 4.503 | 4.073 | 4.645 | 4.043 | 3.938 | -0.081 | 0.0418 | 0.1683 | 0 | 0 | 0 |
| 'Ing4' | 6.192 | 6.599 | 6.942 | 6.135 | 6.962 | 7.350 | 6.163 | 6.508 | 7.029 | 6.066 | 5.981 | 7.098 | 6.427 | 6.101 | 6.756 | 6.579 | 6.518 | 6.565 | 0.031 | 0.2927 | 0.5493 | 0 | 0 | 0 |
| 'Ing5' | 4.859 | 6.093 | 6.198 | 4.730 | 5.635 | 5.483 | 5.455 | 5.661 | 5.796 | 4.960 | 5.429 | 5.937 | 5.572 | 5.469 | 5.641 | 5.505 | 5.833 | 5.404 | -0.009 | 0.8168 | 0.9185 | 0 | 0 | 0 |
| 'Inip' | 5.149 | 5.323 | 4.843 | 4.465 | 4.768 | 5.084 | 5.114 | 4.654 | 4.712 | 4.160 | 4.386 | 5.058 | 4.696 | 4.783 | 4.515 | 4.733 | 4.825 | 4.855 | 0.053 | 0.3239 | 0.5814 | 0 | 0 | 0 |
| 'Ino80' | 13.246 | 12.020 | 12.755 | 12.213 | 12.720 | 11.402 | 12.568 | 12.517 | 12.677 | 12.678 | 12.171 | 12.736 | 11.997 | 11.930 | 11.522 | 11.514 | 12.513 | 11.745 | 0.032 | 0.3316 | 0.5897 | 0 | 0 | 0 |
| 'Ino80b' | 13.367 | 12.620 | 16.155 | 13.744 | 13.331 | 16.083 | 13.224 | 13.325 | 15.775 | 13.345 | 11.694 | 13.987 | 15.035 | 11.889 | 14.852 | 13.528 | 11.319 | 13.974 | 0.086 | 0.0312 | 0.1384 | 0 | 0 | 0 |
| 'Ino80c' | 9.700 | 11.290 | 10.800 | 9.994 | 12.804 | 11.127 | 11.190 | 11.372 | 11.526 | 10.472 | 10.509 | 9.879 | 9.877 | 10.476 | 10.441 | 10.033 | 9.828 | 9.102 | 0.128 | 0.0017 | 0.0162 | 1 | 0 | 0 |
| 'Ino80d' | 3.825 | 2.826 | 3.167 | 3.371 | 2.598 | 2.616 | 2.910 | 2.791 | 3.024 | 4.086 | 3.144 | 3.398 | 3.327 | 2.891 | 3.086 | 3.756 | 2.994 | 3.569 | -0.166 | 0.0047 | 0.0355 | 1 | 0 | 0 |
| 'Inpp1' | 11.831 | 12.328 | 13.030 | 12.234 | 12.757 | 12.482 | 13.004 | 12.457 | 13.490 | 16.567 | 15.808 | 16.706 | 15.584 | 14.861 | 16.673 | 15.474 | 16.720 | 16.244 | -0.360 | 0.0000 | 0.0000 | 1 | 0 | 0 |
| 'Inpp4a' | 6.615 | 6.424 | 6.013 | 6.482 | 5.825 | 5.969 | 5.861 | 6.218 | 6.450 | 6.398 | 6.978 | 7.279 | 6.243 | 6.618 | 6.770 | 6.064 | 7.062 | 7.406 | -0.133 | 0.0018 | 0.0163 | 1 | 0 | 0 |
| 'Inpp4b' | 26.803 | 22.536 | 25.055 | 28.704 | 22.817 | 22.281 | 27.715 | 22.981 | 23.227 | 22.961 | 28.966 | 27.278 | 19.996 | 23.515 | 24.686 | 19.861 | 26.956 | 26.070 | 0.003 | 0.9748 | 0.9916 | 0 | 0 | 0 |
| 'Inpp5a' | 1.666 | 1.598 | 1.762 | 1.760 | 1.573 | 1.646 | 1.637 | 1.838 | 1.555 | 1.820 | 2.150 | 1.653 | 1.737 | 1.840 | 1.560 | 1.875 | 1.913 | 2.018 | -0.139 | 0.0429 | 0.1713 | 0 | 0 | 0 |
| 'Inpp5b' | 28.059 | 26.449 | 28.879 | 27.893 | 27.571 | 27.916 | 27.114 | 27.479 | 28.255 | 28.191 | 30.048 | 30.069 | 23.851 | 27.359 | 30.186 | 25.362 | 28.576 | 28.444 | -0.023 | 0.5425 | 0.7594 | 0 | 0 | 0 |
| 'Inpp5d' | 26.498 | 24.175 | 25.808 | 26.816 | 24.708 | 24.919 | 25.446 | 24.906 | 26.609 | 26.480 | 26.776 | 27.159 | 26.746 | 24.564 | 26.055 | 25.187 | 24.706 | 27.221 | -0.042 | 0.1084 | 0.3099 | 0 | 0 | 0 |
| 'Inpp5e' | 6.510 | 6.665 | 6.945 | 5.905 | 6.006 | 7.153 | 5.813 | 6.761 | 6.973 | 6.471 | 6.264 | 6.870 | 6.647 | 6.317 | 6.642 | 6.388 | 6.375 | 6.381 | 0.001 | 0.9794 | 0.9926 | 0 | 0 | 0 |
| 'Inpp5f' | 4.273 | 4.548 | 4.193 | 4.281 | 4.525 | 4.069 | 4.260 | 4.697 | 4.246 | 4.636 | 4.051 | 4.157 | 4.546 | 4.545 | 4.025 | 4.872 | 4.451 | 3.859 | -0.009 | 0.7778 | 0.9002 | 0 | 0 | 0 |
| 'Inpp5k' | 9.732 | 10.653 | 10.566 | 9.145 | 10.601 | 10.358 | 9.641 | 11.358 | 10.285 | 10.826 | 11.120 | 9.948 | 10.249 | 9.739 | 10.287 | 10.806 | 10.464 | 10.233 | -0.030 | 0.3890 | 0.6384 | 0 | 0 | 0 |
| 'Inppl1' | 19.646 | 18.007 | 20.140 | 19.111 | 16.905 | 20.135 | 17.757 | 17.625 | 19.640 | 18.717 | 18.234 | 21.386 | 18.300 | 17.656 | 19.873 | 17.699 | 17.559 | 20.174 | -0.016 | 0.5720 | 0.7788 | 0 | 0 | 0 |
| 'Insig1' | 35.605 | 27.446 | 21.169 | 27.385 | 26.633 | 22.885 | 23.992 | 26.973 | 21.282 | 36.927 | 37.452 | 16.703 | 39.507 | 35.510 | 21.059 | 34.404 | 29.727 | 15.570 | -0.144 | 0.1954 | 0.4339 | 0 | 0 | 0 |
| 'Insig2' | 4.808 | 5.586 | 5.093 | 4.656 | 5.664 | 4.746 | 5.376 | 5.353 | 5.316 | 4.284 | 4.981 | 6.237 | 5.409 | 5.941 | 5.971 | 4.701 | 6.392 | 6.043 | -0.115 | 0.0498 | 0.1882 | 0 | 0 | 0 |
| 'Insl3' | 21.035 | 20.526 | 26.515 | 21.854 | 19.345 | 23.116 | 24.600 | 21.636 | 29.843 | 21.512 | 23.252 | 28.673 | 21.372 | 20.642 | 26.371 | 13.772 | 22.822 | 26.666 | 0.012 | 0.8784 | 0.9485 | 0 | 0 | 0 |
| 'Insr' | 5.320 | 6.992 | 5.234 | 4.808 | 6.582 | 5.718 | 5.041 | 6.664 | 5.819 | 6.301 | 6.255 | 5.609 | 6.765 | 6.097 | 5.354 | 7.313 | 5.745 | 5.431 | -0.086 | 0.1730 | 0.4066 | 0 | 0 | 0 |
| 'Ints1' | 14.850 | 14.007 | 17.397 | 14.168 | 13.340 | 16.362 | 14.468 | 14.196 | 17.078 | 16.192 | 14.404 | 15.998 | 16.062 | 13.994 | 15.600 | 15.157 | 14.455 | 15.767 | -0.031 | 0.3162 | 0.5741 | 0 | 0 | 0 |
| 'Ints10' | 4.790 | 5.086 | 5.654 | 5.326 | 5.396 | 5.647 | 5.051 | 5.145 | 5.257 | 5.303 | 5.156 | 5.829 | 5.340 | 5.509 | 5.226 | 5.184 | 5.535 | 5.099 | -0.037 | 0.2746 | 0.5303 | 0 | 0 | 0 |
| 'Ints12' | 2.888 | 2.520 | 2.663 | 2.580 | 2.460 | 2.719 | 2.844 | 3.007 | 2.645 | 2.587 | 2.426 | 2.604 | 2.908 | 2.600 | 2.419 | 2.976 | 2.027 | 2.256 | 0.089 | 0.1139 | 0.3199 | 0 | 0 | 0 |
| 'Ints2' | 2.748 | 2.258 | 2.897 | 2.848 | 3.084 | 2.624 | 2.994 | 2.453 | 3.004 | 2.680 | 2.873 | 2.846 | 2.954 | 2.520 | 2.490 | 2.266 | 2.864 | 2.706 | 0.029 | 0.6381 | 0.8220 | 0 | 0 | 0 |
| 'Ints3' | 12.568 | 11.131 | 13.038 | 12.667 | 11.689 | 12.899 | 12.251 | 11.472 | 13.415 | 12.124 | 12.367 | 14.419 | 12.024 | 11.795 | 12.936 | 11.448 | 12.378 | 14.368 | -0.046 | 0.1762 | 0.4114 | 0 | 0 | 0 |
| 'Ints4' | 5.289 | 4.970 | 5.448 | 5.311 | 4.989 | 5.070 | 4.991 | 5.115 | 5.205 | 4.847 | 5.246 | 5.320 | 4.671 | 4.683 | 5.046 | 4.601 | 4.945 | 4.932 | 0.056 | 0.1123 | 0.3181 | 0 | 0 | 0 |
| 'Ints5' | 2.812 | 3.111 | 3.291 | 2.776 | 2.798 | 3.240 | 2.725 | 2.975 | 3.113 | 3.106 | 2.919 | 3.464 | 3.023 | 2.613 | 3.442 | 3.071 | 3.079 | 3.603 | -0.089 | 0.1179 | 0.3268 | 0 | 0 | 0 |
| 'Ints6' | 22.146 | 18.487 | 18.627 | 23.562 | 17.386 | 18.141 | 20.673 | 18.810 | 18.058 | 17.352 | 21.299 | 21.667 | 18.265 | 19.372 | 20.212 | 18.509 | 21.807 | 20.897 | -0.042 | 0.4996 | 0.7273 | 0 | 0 | 0 |
| 'Ints7' | 7.340 | 7.363 | 8.067 | 7.642 | 7.699 | 7.808 | 7.092 | 7.691 | 7.905 | 7.642 | 7.736 | 7.600 | 7.761 | 7.019 | 7.300 | 7.567 | 7.510 | 7.150 | 0.020 | 0.4961 | 0.7249 | 0 | 0 | 0 |
| 'Ints8' | 6.475 | 6.887 | 6.621 | 7.021 | 6.591 | 6.218 | 6.698 | 6.564 | 6.529 | 5.637 | 7.249 | 8.202 | 5.824 | 6.961 | 7.163 | 5.807 | 7.418 | 8.522 | -0.077 | 0.2110 | 0.4562 | 0 | 0 | 0 |
| 'Ints9' | 20.153 | 17.092 | 20.290 | 19.004 | 19.168 | 20.363 | 20.933 | 19.180 | 19.036 | 20.284 | 19.271 | 19.969 | 20.229 | 19.333 | 20.586 | 19.553 | 19.430 | 19.093 | -0.028 | 0.4103 | 0.6592 | 0 | 0 | 0 |
| 'Invs' | 2.262 | 2.246 | 2.385 | 2.202 | 2.353 | 2.460 | 2.204 | 2.273 | 2.446 | 2.558 | 2.749 | 2.589 | 2.304 | 2.380 | 2.504 | 2.198 | 2.195 | 2.698 | -0.100 | 0.0246 | 0.1191 | 0 | 0 | 0 |
| 'Ip6k1' | 17.681 | 17.642 | 17.002 | 16.615 | 16.969 | 17.635 | 16.072 | 17.168 | 17.899 | 16.643 | 18.279 | 20.999 | 17.232 | 18.167 | 19.504 | 18.223 | 18.006 | 19.747 | -0.118 | 0.0001 | 0.0019 | 1 | 0 | 0 |
| 'Ip6k2' | 7.714 | 7.759 | 8.012 | 7.760 | 8.113 | 7.634 | 7.335 | 7.616 | 7.720 | 7.683 | 7.591 | 7.994 | 7.373 | 7.924 | 7.487 | 7.645 | 8.255 | 7.639 | -0.009 | 0.7578 | 0.8916 | 0 | 0 | 0 |
| 'Ipcef1' | 13.423 | 11.260 | 10.973 | 12.189 | 10.864 | 10.495 | 11.925 | 10.709 | 11.106 | 12.964 | 12.698 | 12.796 | 11.426 | 12.599 | 12.808 | 13.157 | 12.990 | 13.474 | -0.172 | 0.0002 | 0.0028 | 1 | 0 | 0 |
| 'Ipmk' | 10.742 | 10.533 | 8.879 | 9.147 | 10.140 | 8.391 | 9.231 | 10.566 | 8.722 | 9.765 | 11.513 | 9.582 | 9.351 | 10.747 | 9.815 | 10.198 | 10.794 | 9.697 | -0.094 | 0.0139 | 0.0793 | 0 | 0 | 0 |
| 'Ipo11' | 2.043 | 2.403 | 2.229 | 2.162 | 2.223 | 2.271 | 2.524 | 2.152 | 2.117 | 2.082 | 2.060 | 2.178 | 2.033 | 2.323 | 2.040 | 2.229 | 2.398 | 1.881 | 0.055 | 0.2553 | 0.5074 | 0 | 0 | 0 |
| 'Ipo13' | 10.001 | 8.947 | 9.794 | 9.520 | 9.900 | 9.973 | 9.287 | 9.397 | 9.588 | 10.157 | 9.502 | 10.698 | 9.630 | 8.759 | 10.663 | 10.203 | 9.352 | 10.402 | -0.059 | 0.0824 | 0.2594 | 0 | 0 | 0 |
| 'Ipo4' | 4.459 | 3.925 | 4.867 | 4.156 | 4.230 | 4.251 | 3.886 | 4.795 | 4.844 | 4.756 | 3.879 | 4.789 | 4.655 | 3.888 | 4.226 | 4.297 | 4.278 | 4.508 | -0.004 | 0.9400 | 0.9769 | 0 | 0 | 0 |
| 'Ipo5' | 34.488 | 33.601 | 38.646 | 37.070 | 35.137 | 36.049 | 37.516 | 37.079 | 36.810 | 34.844 | 33.703 | 37.187 | 37.875 | 35.705 | 37.634 | 33.111 | 34.379 | 36.270 | 0.014 | 0.6174 | 0.8087 | 0 | 0 | 0 |
| 'Ipo7' | 30.585 | 28.744 | 28.069 | 31.970 | 29.230 | 27.979 | 31.419 | 28.920 | 26.357 | 26.446 | 28.088 | 34.523 | 28.958 | 29.965 | 31.698 | 27.601 | 32.848 | 34.711 | -0.073 | 0.1892 | 0.4258 | 0 | 0 | 0 |
| 'Ipo8' | 4.466 | 4.553 | 4.466 | 4.670 | 4.372 | 4.445 | 4.419 | 4.462 | 4.431 | 4.386 | 4.732 | 4.541 | 4.395 | 4.503 | 4.650 | 4.285 | 4.691 | 4.497 | -0.024 | 0.3537 | 0.6072 | 0 | 0 | 0 |
| 'Ipo9' | 6.359 | 5.884 | 6.944 | 6.061 | 6.662 | 6.560 | 6.316 | 6.418 | 6.291 | 7.470 | 7.633 | 7.489 | 7.760 | 7.120 | 7.431 | 6.777 | 7.179 | 7.488 | -0.216 | 0.0000 | 0.0000 | 1 | 0 | 0 |
| 'Ipp' | 10.494 | 11.631 | 11.535 | 12.090 | 10.947 | 10.213 | 11.872 | 10.999 | 11.178 | 10.996 | 10.883 | 10.160 | 10.732 | 12.391 | 10.837 | 10.871 | 12.055 | 11.007 | 0.007 | 0.8824 | 0.9506 | 0 | 0 | 0 |
| 'Ippk' | 5.139 | 6.020 | 4.993 | 5.409 | 5.401 | 4.980 | 4.638 | 5.593 | 5.589 | 5.361 | 5.332 | 5.081 | 4.639 | 5.665 | 4.801 | 5.492 | 6.177 | 5.401 | -0.013 | 0.8243 | 0.9227 | 0 | 0 | 0 |
| 'Iqcb1' | 1.463 | 1.600 | 1.260 | 1.597 | 1.360 | 1.288 | 1.723 | 1.241 | 1.224 | 1.206 | 1.400 | 1.187 | 1.227 | 1.480 | 1.112 | 1.272 | 1.522 | 0.970 | 0.153 | 0.0470 | 0.1807 | 0 | 0 | 0 |
| 'Iqce' | 2.518 | 2.575 | 3.089 | 2.499 | 2.628 | 3.060 | 2.606 | 2.581 | 2.876 | 2.824 | 2.378 | 2.908 | 2.954 | 2.412 | 2.744 | 2.371 | 2.378 | 2.658 | 0.036 | 0.3887 | 0.6382 | 0 | 0 | 0 |
| 'Iqgap1' | 31.548 | 28.188 | 33.935 | 31.712 | 30.245 | 32.841 | 31.103 | 30.614 | 32.487 | 34.034 | 34.159 | 32.345 | 30.850 | 32.786 | 32.046 | 30.358 | 31.171 | 33.483 | -0.055 | 0.1038 | 0.3022 | 0 | 0 | 0 |
| 'Iqgap2' | 5.332 | 4.641 | 5.804 | 5.299 | 5.140 | 5.636 | 5.810 | 5.257 | 5.721 | 5.012 | 5.163 | 5.651 | 5.198 | 4.603 | 5.631 | 5.569 | 4.982 | 5.665 | 0.027 | 0.5080 | 0.7336 | 0 | 0 | 0 |
| 'Iqgap3' | 0.569 | 0.564 | 0.781 | 0.684 | 0.568 | 0.735 | 0.642 | 0.684 | 0.740 | 0.449 | 0.515 | 0.708 | 0.685 | 0.755 | 0.864 | 0.616 | 0.652 | 0.724 | -0.008 | 0.9266 | 0.9719 | 0 | 0 | 0 |
| 'Iqsec1' | 6.205 | 5.836 | 6.601 | 6.224 | 5.681 | 6.451 | 5.425 | 6.247 | 6.881 | 6.023 | 5.387 | 5.540 | 5.369 | 5.088 | 5.532 | 5.327 | 5.418 | 5.760 | 0.159 | 0.0000 | 0.0006 | 1 | 0 | 0 |
| 'Iqsec2' | 0.248 | 0.307 | 0.297 | 0.238 | 0.275 | 0.325 | 0.196 | 0.324 | 0.373 | 0.445 | 0.302 | 0.379 | 0.443 | 0.375 | 0.371 | 0.414 | 0.393 | 0.290 | -0.404 | 0.0007 | 0.0078 | 1 | 1 | 0 |
| 'Irak1' | 8.722 | 8.182 | 9.926 | 7.680 | 8.122 | 9.351 | 8.632 | 8.661 | 8.820 | 9.110 | 8.884 | 9.882 | 9.469 | 9.127 | 9.607 | 8.928 | 9.296 | 9.397 | -0.110 | 0.0003 | 0.0043 | 1 | 0 | 0 |
| 'Irak2' | 3.075 | 3.118 | 3.072 | 3.128 | 3.068 | 3.054 | 3.126 | 3.277 | 3.215 | 3.420 | 2.952 | 2.862 | 3.016 | 2.998 | 2.381 | 2.893 | 2.770 | 2.502 | 0.120 | 0.0204 | 0.1047 | 0 | 0 | 0 |
| 'Irak4' | 13.475 | 13.851 | 13.182 | 12.496 | 13.882 | 13.699 | 12.153 | 14.242 | 13.551 | 14.030 | 16.026 | 14.614 | 14.299 | 16.496 | 13.964 | 14.055 | 13.832 | 14.377 | -0.136 | 0.0001 | 0.0010 | 1 | 0 | 0 |
| 'Ireb2' | 11.791 | 14.643 | 15.010 | 14.040 | 15.452 | 13.577 | 14.738 | 14.927 | 14.124 | 12.604 | 14.342 | 13.801 | 13.774 | 15.272 | 13.204 | 13.336 | 14.988 | 12.575 | 0.040 | 0.2936 | 0.5497 | 0 | 0 | 0 |
| 'Irf1' | 36.848 | 30.394 | 29.642 | 33.290 | 29.801 | 30.275 | 34.720 | 30.311 | 31.300 | 35.915 | 32.967 | 32.458 | 35.255 | 32.595 | 30.791 | 36.821 | 33.678 | 31.094 | -0.084 | 0.0014 | 0.0139 | 1 | 0 | 0 |
| 'Irf2' | 53.550 | 50.177 | 49.979 | 52.637 | 52.277 | 51.016 | 57.040 | 50.609 | 51.477 | 49.865 | 54.016 | 56.252 | 51.771 | 53.070 | 55.421 | 49.363 | 52.001 | 59.011 | -0.048 | 0.2042 | 0.4471 | 0 | 0 | 0 |
| 'Irf2bp1' | 29.884 | 28.102 | 31.920 | 28.432 | 27.768 | 33.159 | 29.073 | 26.011 | 31.129 | 26.707 | 27.884 | 36.121 | 28.742 | 28.229 | 33.236 | 27.777 | 26.060 | 35.344 | -0.034 | 0.3418 | 0.5981 | 0 | 0 | 0 |
| 'Irf2bp2' | 30.555 | 30.954 | 28.149 | 22.945 | 30.989 | 32.678 | 23.905 | 33.567 | 27.618 | 39.856 | 26.723 | 25.593 | 31.893 | 36.039 | 27.599 | 44.957 | 24.505 | 26.843 | -0.115 | 0.3515 | 0.6056 | 0 | 0 | 0 |
| 'Irf3' | 63.099 | 61.745 | 69.010 | 60.000 | 61.059 | 71.426 | 66.256 | 60.213 | 71.367 | 60.728 | 56.516 | 74.722 | 63.663 | 59.353 | 69.630 | 62.652 | 64.318 | 69.318 | -0.004 | 0.8870 | 0.9534 | 0 | 0 | 0 |
| 'Irf4' | 12.476 | 12.018 | 13.303 | 11.016 | 12.895 | 12.773 | 11.502 | 12.888 | 12.880 | 15.959 | 11.493 | 13.937 | 12.494 | 12.255 | 13.435 | 16.546 | 12.025 | 13.789 | -0.131 | 0.0255 | 0.1221 | 0 | 0 | 0 |
| 'Irf5' | 2.508 | 3.218 | 3.039 | 2.669 | 3.163 | 3.166 | 2.844 | 2.948 | 3.382 | 2.574 | 2.790 | 2.591 | 2.558 | 2.563 | 2.901 | 2.950 | 2.820 | 2.551 | 0.145 | 0.0350 | 0.1496 | 0 | 0 | 0 |
| 'Irf7' | 7.847 | 8.431 | 10.243 | 7.138 | 8.709 | 11.742 | 7.905 | 11.110 | 10.791 | 9.487 | 8.209 | 13.458 | 9.718 | 8.595 | 13.416 | 9.130 | 8.624 | 12.613 | -0.154 | 0.0139 | 0.0796 | 0 | 0 | 0 |
| 'Irf8' | 4.150 | 4.264 | 3.741 | 3.779 | 3.931 | 4.084 | 3.543 | 4.009 | 3.916 | 3.780 | 4.165 | 3.491 | 4.427 | 4.092 | 3.684 | 4.104 | 4.446 | 3.437 | -0.010 | 0.8303 | 0.9262 | 0 | 0 | 0 |
| 'Irf9' | 11.779 | 12.900 | 12.295 | 11.701 | 12.047 | 12.991 | 11.923 | 12.775 | 13.554 | 11.988 | 13.007 | 16.694 | 12.758 | 12.765 | 15.750 | 12.484 | 14.086 | 15.465 | -0.166 | 0.0000 | 0.0002 | 1 | 0 | 0 |
| 'Irgm1' | 7.955 | 8.206 | 7.955 | 7.406 | 7.926 | 8.770 | 8.928 | 8.134 | 9.133 | 8.105 | 8.580 | 12.594 | 8.271 | 8.824 | 12.607 | 9.020 | 10.048 | 12.524 | -0.278 | 0.0000 | 0.0002 | 1 | 0 | 0 |
| 'Irgm2' | 4.414 | 6.052 | 5.402 | 5.012 | 5.147 | 5.863 | 5.721 | 5.126 | 6.548 | 4.625 | 5.832 | 7.604 | 4.814 | 4.865 | 7.606 | 4.848 | 5.519 | 7.443 | -0.105 | 0.1220 | 0.3333 | 0 | 0 | 0 |
| 'Irgq' | 5.386 | 5.894 | 5.140 | 5.408 | 5.331 | 5.776 | 5.782 | 5.451 | 6.206 | 5.004 | 5.112 | 6.346 | 4.988 | 5.738 | 6.648 | 5.881 | 5.019 | 6.819 | -0.042 | 0.4542 | 0.6929 | 0 | 0 | 0 |
| 'Irs2' | 3.069 | 1.384 | 1.204 | 1.811 | 1.373 | 1.417 | 1.390 | 1.327 | 1.670 | 3.012 | 1.761 | 1.317 | 1.926 | 1.461 | 1.570 | 2.875 | 1.628 | 1.441 | -0.195 | 0.1346 | 0.3535 | 0 | 0 | 0 |
| 'Isca2' | 27.324 | 34.709 | 35.359 | 29.846 | 37.357 | 34.044 | 30.724 | 35.653 | 35.603 | 31.500 | 31.747 | 30.752 | 32.756 | 37.335 | 31.303 | 33.098 | 30.272 | 27.566 | 0.061 | 0.2641 | 0.5180 | 0 | 0 | 0 |
| 'Iscu' | 10.982 | 13.385 | 11.028 | 11.754 | 13.403 | 11.363 | 10.871 | 11.898 | 11.822 | 11.490 | 11.096 | 11.095 | 11.511 | 11.941 | 11.278 | 11.348 | 10.275 | 9.712 | 0.080 | 0.0649 | 0.2222 | 0 | 0 | 0 |
| 'Isg15' | 58.140 | 58.096 | 57.913 | 54.498 | 64.970 | 64.322 | 49.815 | 66.210 | 61.081 | 58.519 | 68.619 | 81.775 | 66.300 | 69.152 | 88.763 | 62.954 | 72.529 | 84.628 | -0.294 | 0.0000 | 0.0000 | 1 | 0 | 0 |
| 'Isg20' | 9.708 | 9.603 | 9.175 | 8.644 | 10.005 | 10.213 | 8.868 | 10.559 | 9.707 | 9.809 | 8.547 | 10.868 | 11.524 | 8.650 | 11.236 | 10.628 | 9.669 | 9.960 | -0.081 | 0.1352 | 0.3537 | 0 | 0 | 0 |
| 'Isg20l2' | 21.665 | 24.409 | 23.643 | 21.830 | 23.930 | 23.411 | 22.227 | 24.612 | 23.033 | 22.348 | 22.320 | 23.740 | 23.140 | 23.015 | 23.426 | 23.200 | 21.468 | 22.549 | 0.012 | 0.6782 | 0.8461 | 0 | 0 | 0 |
| 'Isoc1' | 12.263 | 15.366 | 14.720 | 13.869 | 15.367 | 15.036 | 12.886 | 15.293 | 15.660 | 15.188 | 13.516 | 16.075 | 14.755 | 15.139 | 16.518 | 18.622 | 14.059 | 14.941 | -0.101 | 0.0716 | 0.2363 | 0 | 0 | 0 |
| 'Isoc2a' | 21.752 | 22.218 | 20.853 | 21.731 | 24.246 | 22.998 | 22.270 | 22.978 | 23.328 | 17.759 | 16.375 | 19.913 | 19.356 | 17.604 | 21.068 | 20.984 | 18.961 | 19.500 | 0.225 | 0.0000 | 0.0002 | 1 | 0 | 0 |
| 'Ispd' | 4.053 | 3.383 | 3.681 | 4.596 | 4.000 | 4.047 | 4.900 | 3.367 | 3.199 | 4.707 | 4.654 | 4.211 | 3.996 | 3.907 | 3.694 | 4.243 | 5.493 | 4.389 | -0.174 | 0.0426 | 0.1704 | 0 | 0 | 0 |
| 'Ist1' | 53.826 | 55.096 | 51.241 | 50.678 | 53.506 | 51.353 | 50.280 | 54.145 | 51.300 | 51.926 | 55.795 | 58.565 | 50.981 | 56.033 | 55.039 | 50.397 | 57.342 | 55.501 | -0.072 | 0.0103 | 0.0638 | 0 | 0 | 0 |
| 'Isy1' | 10.058 | 10.764 | 10.526 | 9.504 | 10.016 | 10.661 | 9.579 | 10.394 | 10.490 | 10.457 | 9.506 | 10.057 | 9.811 | 10.175 | 9.686 | 9.453 | 9.258 | 9.687 | 0.053 | 0.0851 | 0.2650 | 0 | 0 | 0 |
| 'Isyna1' | 90.867 | 75.635 | 97.208 | 86.583 | 80.102 | 103.346 | 80.888 | 76.614 | 95.310 | 90.271 | 70.840 | 100.495 | 87.088 | 71.445 | 91.485 | 88.043 | 77.063 | 94.668 | 0.019 | 0.5253 | 0.7470 | 0 | 0 | 0 |
| 'Itch' | 8.633 | 8.865 | 8.208 | 9.110 | 9.312 | 8.703 | 9.013 | 9.070 | 8.229 | 8.602 | 9.065 | 9.053 | 8.274 | 9.204 | 8.375 | 8.447 | 9.411 | 8.936 | -0.017 | 0.5707 | 0.7776 | 0 | 0 | 0 |
| 'Itfg1' | 18.286 | 20.204 | 19.585 | 19.940 | 19.245 | 18.663 | 19.711 | 20.810 | 18.422 | 19.551 | 21.449 | 20.101 | 18.917 | 20.699 | 19.279 | 19.227 | 19.715 | 18.888 | -0.035 | 0.2530 | 0.5047 | 0 | 0 | 0 |
| 'Itfg2' | 2.091 | 2.170 | 2.221 | 1.952 | 2.122 | 2.455 | 1.965 | 2.444 | 2.292 | 2.358 | 1.936 | 2.020 | 2.302 | 2.084 | 1.870 | 2.360 | 1.732 | 1.683 | 0.102 | 0.1338 | 0.3522 | 0 | 0 | 0 |
| 'Itga3' | 0.905 | 0.861 | 0.961 | 0.833 | 0.812 | 0.977 | 0.850 | 0.976 | 0.910 | 0.876 | 0.873 | 0.911 | 0.722 | 0.914 | 1.027 | 0.953 | 0.852 | 0.957 | 0.003 | 0.9716 | 0.9899 | 0 | 0 | 0 |
| 'Itga4' | 5.517 | 4.551 | 5.847 | 5.013 | 4.446 | 5.177 | 5.426 | 5.170 | 5.156 | 6.817 | 5.676 | 7.840 | 6.152 | 5.479 | 6.714 | 7.028 | 5.041 | 7.240 | -0.328 | 0.0000 | 0.0000 | 1 | 0 | 0 |
| 'Itga6' | 5.966 | 6.045 | 6.266 | 6.051 | 5.655 | 6.499 | 6.467 | 5.922 | 6.052 | 4.977 | 5.106 | 5.385 | 5.284 | 5.033 | 5.344 | 5.482 | 5.173 | 5.291 | 0.213 | 0.0000 | 0.0000 | 1 | 0 | 0 |
| 'Itga7' | 3.771 | 3.656 | 3.610 | 2.806 | 4.058 | 3.359 | 4.201 | 4.138 | 3.991 | 7.686 | 8.122 | 7.796 | 7.303 | 7.087 | 8.388 | 6.643 | 7.242 | 7.134 | -1.017 | 0.0000 | 0.0000 | 1 | 1 | 1 |
| 'Itgae' | 2.722 | 1.674 | 2.757 | 2.041 | 1.607 | 2.863 | 2.992 | 1.771 | 2.895 | 1.480 | 0.722 | 2.783 | 1.572 | 1.144 | 2.439 | 1.880 | 0.937 | 2.475 | 0.500 | 0.0000 | 0.0000 | 1 | 1 | 0 |
| 'Itgal' | 247.724 | 245.902 | 261.523 | 249.997 | 249.849 | 256.169 | 237.469 | 245.772 | 264.794 | 280.229 | 269.787 | 249.992 | 263.673 | 260.156 | 264.975 | 259.788 | 255.788 | 267.430 | -0.081 | 0.0024 | 0.0213 | 1 | 0 | 0 |
| 'Itgav' | 4.292 | 5.101 | 4.297 | 3.952 | 4.879 | 4.291 | 4.643 | 4.795 | 4.593 | 4.236 | 4.201 | 4.690 | 4.195 | 4.682 | 4.498 | 4.662 | 4.370 | 4.778 | 0.003 | 0.9335 | 0.9746 | 0 | 0 | 0 |
| 'Itgb1' | 15.335 | 20.324 | 15.257 | 16.856 | 20.248 | 16.317 | 18.074 | 21.078 | 15.146 | 16.119 | 18.194 | 17.800 | 18.558 | 20.400 | 15.628 | 17.162 | 25.696 | 16.501 | -0.079 | 0.1206 | 0.3311 | 0 | 0 | 0 |
| 'Itgb1bp1' | 9.499 | 11.156 | 9.839 | 9.779 | 10.505 | 9.924 | 9.971 | 9.825 | 9.770 | 9.076 | 10.094 | 10.164 | 9.610 | 9.686 | 10.177 | 9.984 | 9.999 | 9.809 | 0.011 | 0.7428 | 0.8839 | 0 | 0 | 0 |
| 'Itgb2' | 117.503 | 125.209 | 119.898 | 122.274 | 121.354 | 122.868 | 116.265 | 119.499 | 122.847 | 119.034 | 116.776 | 118.425 | 117.341 | 118.274 | 122.410 | 122.265 | 112.946 | 122.234 | 0.013 | 0.5191 | 0.7417 | 0 | 0 | 0 |
| 'Itgb3' | 0.279 | 0.417 | 0.423 | 0.340 | 0.468 | 0.659 | 0.546 | 0.595 | 0.542 | 0.345 | 0.482 | 0.575 | 0.482 | 0.516 | 0.530 | 0.361 | 0.482 | 0.540 | -0.034 | 0.7736 | 0.8993 | 0 | 0 | 0 |
| 'Itgb3bp' | 1.725 | 2.126 | 1.841 | 1.737 | 2.074 | 1.529 | 2.216 | 2.056 | 1.737 | 1.862 | 2.155 | 1.910 | 1.966 | 1.878 | 1.532 | 2.033 | 1.772 | 1.525 | 0.016 | 0.8113 | 0.9151 | 0 | 0 | 0 |
| 'Itgb7' | 64.667 | 50.766 | 69.699 | 60.155 | 60.194 | 68.931 | 63.735 | 59.981 | 63.879 | 60.083 | 55.992 | 78.883 | 59.782 | 55.158 | 73.075 | 65.611 | 56.503 | 73.837 | -0.047 | 0.2346 | 0.4837 | 0 | 0 | 0 |
| 'Itgb8' | 2.567 | 2.618 | 1.695 | 2.419 | 2.584 | 1.978 | 2.717 | 2.730 | 2.126 | 2.544 | 2.605 | 2.232 | 2.138 | 2.596 | 1.748 | 2.942 | 3.188 | 2.229 | -0.066 | 0.3243 | 0.5817 | 0 | 0 | 0 |
| 'Itih5' | 1.782 | 1.836 | 1.232 | 1.396 | 1.443 | 1.545 | 1.522 | 1.671 | 1.645 | 1.286 | 1.404 | 1.803 | 1.022 | 1.348 | 1.424 | 1.667 | 0.996 | 1.586 | 0.157 | 0.1421 | 0.3651 | 0 | 0 | 0 |
| 'Itk' | 151.653 | 160.754 | 139.857 | 162.973 | 152.499 | 143.258 | 165.070 | 145.298 | 153.438 | 150.438 | 146.608 | 140.660 | 151.753 | 141.438 | 134.066 | 144.565 | 155.516 | 140.174 | 0.064 | 0.0277 | 0.1284 | 0 | 0 | 0 |
| 'Itm2a' | 3.824 | 5.399 | 5.468 | 4.346 | 5.306 | 5.450 | 4.833 | 6.677 | 5.330 | 4.634 | 3.611 | 6.640 | 3.192 | 3.160 | 5.204 | 4.876 | 4.356 | 5.392 | 0.174 | 0.1208 | 0.3314 | 0 | 0 | 0 |
| 'Itm2b' | 392.472 | 512.235 | 390.861 | 430.950 | 474.867 | 432.460 | 433.247 | 462.362 | 420.045 | 445.288 | 447.252 | 384.913 | 477.227 | 488.955 | 419.408 | 494.572 | 458.796 | 410.100 | -0.039 | 0.3171 | 0.5753 | 0 | 0 | 0 |
| 'Itm2c' | 76.948 | 81.627 | 75.180 | 72.432 | 82.658 | 81.819 | 69.818 | 79.206 | 79.325 | 80.538 | 73.175 | 79.562 | 73.744 | 76.790 | 80.814 | 88.057 | 68.770 | 82.221 | -0.020 | 0.6414 | 0.8241 | 0 | 0 | 0 |
| 'Itpa' | 13.814 | 14.665 | 13.805 | 12.983 | 15.131 | 14.656 | 14.324 | 15.545 | 15.717 | 12.750 | 13.086 | 13.406 | 15.211 | 13.868 | 13.374 | 14.136 | 13.007 | 11.803 | 0.105 | 0.0189 | 0.0994 | 0 | 0 | 0 |
| 'Itpk1' | 21.951 | 17.117 | 17.778 | 17.674 | 17.464 | 16.873 | 16.565 | 18.366 | 16.401 | 18.095 | 18.172 | 20.464 | 18.618 | 18.334 | 19.423 | 19.094 | 18.181 | 20.673 | -0.105 | 0.0405 | 0.1647 | 0 | 0 | 0 |
| 'Itpkb' | 51.379 | 45.777 | 45.073 | 47.666 | 44.178 | 45.495 | 43.333 | 44.787 | 45.611 | 45.838 | 43.583 | 50.386 | 43.181 | 44.867 | 49.274 | 47.121 | 43.140 | 54.452 | -0.038 | 0.3718 | 0.6237 | 0 | 0 | 0 |
| 'Itpkc' | 2.898 | 3.071 | 2.844 | 2.811 | 2.643 | 2.809 | 2.501 | 2.746 | 2.816 | 2.873 | 2.837 | 2.959 | 2.698 | 2.956 | 2.560 | 3.072 | 2.539 | 2.753 | -0.015 | 0.7697 | 0.8975 | 0 | 0 | 0 |
| 'Itpr1' | 12.247 | 10.259 | 10.906 | 11.698 | 9.773 | 9.892 | 11.818 | 9.629 | 10.669 | 12.363 | 12.371 | 12.440 | 11.437 | 10.959 | 11.767 | 11.533 | 11.327 | 12.622 | -0.156 | 0.0003 | 0.0038 | 1 | 0 | 0 |
| 'Itpr2' | 31.695 | 27.854 | 29.933 | 32.986 | 28.757 | 29.351 | 30.991 | 28.056 | 30.034 | 33.955 | 36.450 | 30.522 | 33.431 | 31.259 | 29.995 | 32.204 | 33.014 | 32.289 | -0.132 | 0.0005 | 0.0062 | 1 | 0 | 0 |
| 'Itpr3' | 29.613 | 25.082 | 30.468 | 27.180 | 25.667 | 28.173 | 25.867 | 26.741 | 28.754 | 28.129 | 26.033 | 27.548 | 27.720 | 24.075 | 24.765 | 25.981 | 25.173 | 25.304 | 0.067 | 0.0561 | 0.2032 | 0 | 0 | 0 |
| 'Itpripl1' | 4.202 | 4.519 | 4.602 | 4.114 | 4.345 | 4.965 | 4.891 | 4.502 | 4.702 | 4.060 | 4.557 | 4.770 | 3.638 | 4.014 | 4.902 | 4.109 | 4.439 | 4.680 | 0.050 | 0.2786 | 0.5352 | 0 | 0 | 0 |
| 'Itpripl2' | 0.730 | 0.746 | 0.762 | 0.678 | 0.777 | 0.758 | 0.706 | 0.655 | 0.974 | 0.493 | 0.415 | 0.580 | 0.503 | 0.581 | 0.529 | 0.404 | 0.581 | 0.550 | 0.531 | 0.0000 | 0.0000 | 1 | 1 | 0 |
| 'Itsn1' | 0.117 | 0.133 | 0.104 | 0.158 | 0.143 | 0.121 | 0.142 | 0.125 | 0.108 | 0.082 | 0.083 | 0.090 | 0.125 | 0.081 | 0.114 | 0.082 | 0.103 | 0.062 | 0.456 | 0.0002 | 0.0025 | 1 | 1 | 0 |
| 'Itsn2' | 46.021 | 38.414 | 42.529 | 45.948 | 40.701 | 39.744 | 41.237 | 40.667 | 41.593 | 43.266 | 44.424 | 43.779 | 41.109 | 39.473 | 42.016 | 40.438 | 42.399 | 45.342 | -0.032 | 0.4072 | 0.6562 | 0 | 0 | 0 |
| 'Ivd' | 5.678 | 6.859 | 7.107 | 6.444 | 6.541 | 7.274 | 6.253 | 6.382 | 7.443 | 6.943 | 6.863 | 6.581 | 6.333 | 6.501 | 6.852 | 6.688 | 6.373 | 6.157 | 0.007 | 0.8666 | 0.9424 | 0 | 0 | 0 |
| 'Ivns1abp' | 47.476 | 54.688 | 47.986 | 43.783 | 51.998 | 48.637 | 42.098 | 50.765 | 49.062 | 42.448 | 39.943 | 39.745 | 40.048 | 45.890 | 40.793 | 42.256 | 42.417 | 40.891 | 0.208 | 0.0000 | 0.0000 | 1 | 0 | 0 |
| 'Iws1' | 10.889 | 12.066 | 12.146 | 11.334 | 11.637 | 11.565 | 11.467 | 11.463 | 12.297 | 11.246 | 11.168 | 12.024 | 11.563 | 11.180 | 11.575 | 10.875 | 10.828 | 11.129 | 0.033 | 0.1896 | 0.4262 | 0 | 0 | 0 |
| 'Izumo1r' | 188.120 | 186.547 | 200.602 | 178.754 | 193.280 | 201.305 | 170.395 | 202.322 | 192.687 | 195.439 | 194.260 | 201.274 | 162.854 | 184.388 | 206.691 | 187.157 | 182.809 | 203.364 | -0.014 | 0.6882 | 0.8526 | 0 | 0 | 0 |
| 'Izumo4' | 23.120 | 37.889 | 36.554 | 20.007 | 31.434 | 35.311 | 25.162 | 32.647 | 37.097 | 30.012 | 28.034 | 33.972 | 28.158 | 27.577 | 34.785 | 32.098 | 23.109 | 30.577 | 0.029 | 0.7328 | 0.8781 | 0 | 0 | 0 |
| 'Jade1' | 2.792 | 2.766 | 2.425 | 2.547 | 2.761 | 2.513 | 2.494 | 2.539 | 2.449 | 2.554 | 2.796 | 3.076 | 2.458 | 2.824 | 2.706 | 2.802 | 2.686 | 3.013 | -0.111 | 0.0195 | 0.1015 | 0 | 0 | 0 |
| 'Jade2' | 25.201 | 26.443 | 24.380 | 23.199 | 24.946 | 24.704 | 21.518 | 26.171 | 24.548 | 26.468 | 24.434 | 26.159 | 24.091 | 24.741 | 24.576 | 25.697 | 24.103 | 25.190 | -0.039 | 0.2450 | 0.4957 | 0 | 0 | 0 |
| 'Jade3' | 1.403 | 1.224 | 1.390 | 1.813 | 1.255 | 1.444 | 1.452 | 1.305 | 1.355 | 1.512 | 1.354 | 1.570 | 1.289 | 1.136 | 1.273 | 1.362 | 1.595 | 1.432 | -0.005 | 0.9514 | 0.9816 | 0 | 0 | 0 |
| 'Jag2' | 1.044 | 1.181 | 1.170 | 0.853 | 0.832 | 1.066 | 0.945 | 1.144 | 1.193 | 1.296 | 0.854 | 1.207 | 1.279 | 0.888 | 0.879 | 0.907 | 1.117 | 1.070 | -0.025 | 0.8248 | 0.9227 | 0 | 0 | 0 |
| 'Jagn1' | 4.554 | 5.453 | 4.574 | 4.420 | 4.987 | 4.863 | 4.506 | 5.163 | 4.854 | 4.130 | 4.431 | 5.160 | 4.320 | 4.676 | 5.440 | 4.472 | 4.640 | 5.003 | 0.019 | 0.7021 | 0.8611 | 0 | 0 | 0 |
| 'Jak1' | 87.018 | 90.286 | 86.705 | 88.341 | 90.911 | 87.570 | 88.488 | 92.653 | 86.805 | 90.320 | 93.725 | 87.845 | 80.584 | 88.460 | 86.931 | 89.594 | 89.511 | 87.428 | -0.003 | 0.9042 | 0.9624 | 0 | 0 | 0 |
| 'Jak2' | 163.680 | 171.353 | 162.211 | 167.340 | 173.873 | 158.242 | 163.378 | 174.455 | 155.452 | 172.354 | 172.754 | 146.586 | 154.622 | 170.289 | 148.143 | 166.516 | 170.573 | 153.227 | 0.025 | 0.3013 | 0.5581 | 0 | 0 | 0 |
| 'Jak3' | 77.533 | 64.924 | 87.677 | 70.627 | 68.685 | 78.850 | 71.721 | 73.324 | 83.920 | 81.298 | 77.920 | 81.067 | 75.257 | 69.828 | 77.024 | 67.002 | 74.780 | 78.363 | -0.026 | 0.5405 | 0.7579 | 0 | 0 | 0 |
| 'Jakmip1' | 12.527 | 10.810 | 11.124 | 11.856 | 10.791 | 11.781 | 11.544 | 10.938 | 11.414 | 11.131 | 11.189 | 11.568 | 11.625 | 11.421 | 11.044 | 11.644 | 10.307 | 11.176 | 0.013 | 0.6337 | 0.8184 | 0 | 0 | 0 |
| 'Jarid2' | 12.286 | 12.906 | 12.534 | 12.053 | 13.362 | 13.009 | 12.208 | 12.530 | 13.265 | 13.464 | 12.543 | 12.523 | 13.174 | 13.043 | 12.824 | 13.192 | 12.811 | 12.502 | -0.034 | 0.1699 | 0.4027 | 0 | 0 | 0 |
| 'Jazf1' | 2.531 | 2.374 | 2.674 | 2.308 | 2.407 | 2.451 | 2.353 | 2.483 | 2.658 | 2.983 | 2.901 | 2.755 | 2.764 | 2.772 | 2.545 | 3.075 | 2.507 | 2.635 | -0.164 | 0.0090 | 0.0578 | 0 | 0 | 0 |
| 'Jdp2' | 14.742 | 16.131 | 12.983 | 15.380 | 15.186 | 14.022 | 14.695 | 15.162 | 12.808 | 14.051 | 13.848 | 12.402 | 21.077 | 16.226 | 13.478 | 13.579 | 14.754 | 14.427 | -0.040 | 0.5515 | 0.7649 | 0 | 0 | 0 |
| 'Jkamp' | 4.036 | 5.622 | 4.706 | 4.219 | 4.879 | 5.176 | 4.811 | 5.715 | 4.444 | 4.596 | 4.871 | 4.913 | 4.928 | 4.864 | 5.214 | 4.949 | 4.677 | 4.730 | -0.020 | 0.7155 | 0.8688 | 0 | 0 | 0 |
| 'Jmjd1c' | 8.478 | 7.750 | 7.843 | 8.844 | 7.795 | 7.242 | 8.406 | 8.019 | 7.653 | 8.316 | 7.835 | 7.104 | 7.216 | 7.263 | 6.737 | 7.439 | 7.919 | 6.814 | 0.102 | 0.0027 | 0.0229 | 1 | 0 | 0 |
| 'Jmjd4' | 1.666 | 1.773 | 1.808 | 1.634 | 1.633 | 1.746 | 1.616 | 1.750 | 1.744 | 1.618 | 1.660 | 2.116 | 1.507 | 1.590 | 2.020 | 1.756 | 1.719 | 2.203 | -0.086 | 0.1162 | 0.3236 | 0 | 0 | 0 |
| 'Jmjd6' | 9.315 | 7.203 | 6.736 | 8.547 | 6.780 | 6.706 | 8.685 | 6.402 | 6.942 | 6.852 | 9.097 | 9.686 | 7.633 | 8.477 | 9.660 | 6.804 | 9.904 | 9.138 | -0.211 | 0.0473 | 0.1814 | 0 | 0 | 0 |
| 'Jmjd8' | 4.990 | 5.251 | 5.936 | 5.074 | 5.140 | 5.856 | 5.758 | 5.307 | 5.692 | 4.942 | 4.506 | 6.333 | 5.071 | 4.948 | 5.755 | 4.715 | 4.965 | 5.522 | 0.051 | 0.2032 | 0.4456 | 0 | 0 | 0 |
| 'Jmy' | 5.722 | 5.251 | 4.856 | 5.295 | 5.973 | 5.092 | 5.881 | 5.431 | 5.158 | 5.592 | 5.548 | 5.357 | 5.485 | 4.893 | 4.960 | 5.512 | 5.142 | 5.191 | 0.014 | 0.7079 | 0.8650 | 0 | 0 | 0 |
| 'Josd1' | 13.446 | 11.627 | 10.456 | 10.929 | 12.112 | 10.370 | 10.409 | 11.183 | 9.198 | 13.969 | 12.457 | 11.916 | 12.895 | 12.601 | 11.661 | 14.288 | 11.253 | 11.637 | -0.185 | 0.0003 | 0.0035 | 1 | 0 | 0 |
| 'Josd2' | 5.962 | 6.494 | 7.770 | 5.956 | 7.027 | 7.855 | 5.844 | 6.487 | 7.527 | 7.251 | 6.165 | 7.698 | 6.863 | 6.733 | 7.213 | 7.019 | 6.617 | 6.941 | -0.051 | 0.2241 | 0.4711 | 0 | 0 | 0 |
| 'Jrk' | 1.714 | 1.555 | 2.154 | 1.320 | 2.059 | 2.116 | 1.571 | 2.165 | 2.125 | 1.760 | 1.520 | 1.384 | 1.706 | 1.394 | 1.757 | 1.584 | 1.631 | 1.107 | 0.272 | 0.0093 | 0.0593 | 0 | 0 | 0 |
| 'Jrkl' | 6.482 | 6.378 | 9.003 | 6.706 | 8.384 | 8.059 | 8.177 | 8.036 | 8.341 | 6.913 | 7.361 | 8.436 | 6.618 | 7.145 | 6.869 | 5.424 | 9.137 | 6.955 | 0.091 | 0.2130 | 0.4581 | 0 | 0 | 0 |
| 'Jtb' | 16.377 | 18.113 | 15.257 | 14.008 | 17.260 | 16.773 | 14.461 | 17.412 | 15.479 | 14.631 | 15.904 | 16.002 | 15.377 | 16.465 | 15.386 | 16.122 | 14.712 | 16.059 | 0.032 | 0.4391 | 0.6828 | 0 | 0 | 0 |
| 'Jun' | 416.992 | 95.854 | 150.704 | 409.880 | 106.800 | 112.170 | 469.616 | 145.525 | 107.002 | 257.490 | 223.326 | 115.973 | 265.289 | 170.508 | 95.301 | 171.075 | 255.530 | 95.799 | 0.086 | 0.7362 | 0.8799 | 0 | 0 | 0 |
| 'Junb' | 960.090 | 619.542 | 458.875 | 667.761 | 616.004 | 497.399 | 599.394 | 717.228 | 382.933 | 977.594 | 563.390 | 296.430 | 833.506 | 744.762 | 336.581 | 971.794 | 475.771 | 304.799 | 0.097 | 0.4752 | 0.7100 | 0 | 0 | 0 |
| 'Jund' | 424.787 | 303.344 | 263.153 | 360.303 | 297.810 | 292.641 | 379.930 | 323.842 | 256.198 | 413.395 | 288.315 | 219.430 | 400.289 | 313.696 | 251.886 | 403.507 | 278.041 | 244.082 | 0.053 | 0.2370 | 0.4866 | 0 | 0 | 0 |
| 'Kansl1' | 22.226 | 19.945 | 19.941 | 21.394 | 20.214 | 20.227 | 20.842 | 21.066 | 20.353 | 21.986 | 20.677 | 19.826 | 21.436 | 20.222 | 19.972 | 21.440 | 19.910 | 20.010 | -0.003 | 0.8932 | 0.9564 | 0 | 0 | 0 |
| 'Kansl1l' | 4.965 | 5.169 | 5.174 | 5.666 | 5.380 | 4.544 | 6.009 | 5.256 | 5.265 | 5.690 | 6.069 | 6.055 | 6.030 | 6.167 | 5.459 | 5.433 | 6.822 | 5.450 | -0.178 | 0.0001 | 0.0021 | 1 | 0 | 0 |
| 'Kansl2' | 29.985 | 30.243 | 29.504 | 29.234 | 27.463 | 27.187 | 27.903 | 28.641 | 28.806 | 25.619 | 27.596 | 31.945 | 26.513 | 28.994 | 31.497 | 26.423 | 29.676 | 31.198 | -0.012 | 0.7725 | 0.8992 | 0 | 0 | 0 |
| 'Kansl3' | 10.206 | 9.482 | 9.786 | 9.461 | 9.147 | 9.816 | 9.904 | 10.003 | 9.781 | 9.740 | 9.330 | 10.400 | 9.543 | 9.235 | 10.176 | 9.520 | 9.922 | 10.564 | -0.024 | 0.4071 | 0.6562 | 0 | 0 | 0 |
| 'Kars' | 53.179 | 53.792 | 54.485 | 54.704 | 55.461 | 53.347 | 52.945 | 57.511 | 54.192 | 52.090 | 49.025 | 52.296 | 55.085 | 49.028 | 52.247 | 53.046 | 48.129 | 51.809 | 0.071 | 0.0067 | 0.0469 | 1 | 0 | 0 |
| 'Kat2a' | 11.275 | 9.204 | 12.863 | 11.591 | 10.861 | 12.240 | 10.894 | 11.163 | 12.417 | 10.854 | 9.877 | 13.100 | 10.766 | 9.848 | 12.292 | 10.547 | 9.662 | 13.037 | 0.028 | 0.4257 | 0.6724 | 0 | 0 | 0 |
| 'Kat2b' | 20.037 | 20.292 | 18.989 | 19.163 | 20.967 | 18.973 | 19.468 | 20.964 | 19.014 | 19.213 | 20.838 | 21.669 | 17.429 | 21.058 | 19.858 | 20.009 | 21.707 | 20.256 | -0.045 | 0.1821 | 0.4183 | 0 | 0 | 0 |
| 'Kat5' | 22.121 | 26.178 | 24.575 | 22.064 | 25.080 | 25.387 | 21.982 | 23.708 | 26.619 | 22.941 | 21.722 | 22.767 | 24.867 | 23.898 | 21.570 | 24.189 | 24.367 | 20.372 | 0.065 | 0.1581 | 0.3881 | 0 | 0 | 0 |
| 'Kat6a' | 38.705 | 32.486 | 35.081 | 38.067 | 32.848 | 34.239 | 36.465 | 33.286 | 36.126 | 35.539 | 37.374 | 38.971 | 35.928 | 34.485 | 37.363 | 36.127 | 36.204 | 42.188 | -0.087 | 0.0277 | 0.1284 | 0 | 0 | 0 |
| 'Kat6b' | 2.287 | 2.052 | 2.175 | 2.125 | 2.102 | 2.179 | 2.239 | 2.266 | 2.210 | 2.309 | 2.480 | 2.243 | 2.238 | 2.225 | 2.059 | 2.190 | 2.210 | 2.130 | -0.040 | 0.2406 | 0.4912 | 0 | 0 | 0 |
| 'Kat7' | 15.082 | 14.943 | 14.020 | 14.138 | 15.175 | 15.040 | 14.209 | 15.396 | 14.546 | 14.305 | 14.181 | 14.200 | 14.035 | 15.081 | 14.144 | 14.736 | 14.488 | 13.693 | 0.032 | 0.1967 | 0.4356 | 0 | 0 | 0 |
| 'Kat8' | 6.445 | 7.779 | 6.998 | 7.077 | 8.110 | 7.312 | 7.202 | 7.505 | 7.648 | 7.485 | 6.563 | 8.529 | 8.280 | 8.728 | 8.873 | 8.974 | 8.354 | 7.457 | -0.163 | 0.0034 | 0.0276 | 1 | 0 | 0 |
| 'Katna1' | 15.193 | 16.124 | 14.936 | 16.298 | 15.958 | 14.705 | 16.379 | 15.534 | 15.410 | 15.048 | 15.456 | 15.776 | 15.085 | 15.505 | 14.879 | 15.040 | 17.949 | 14.975 | -0.005 | 0.8843 | 0.9514 | 0 | 0 | 0 |
| 'Katnal1' | 0.550 | 0.485 | 0.505 | 0.498 | 0.536 | 0.506 | 0.556 | 0.464 | 0.517 | 0.452 | 0.473 | 0.513 | 0.476 | 0.506 | 0.439 | 0.498 | 0.517 | 0.470 | 0.074 | 0.2922 | 0.5491 | 0 | 0 | 0 |
| 'Katnb1' | 10.260 | 10.514 | 13.601 | 10.379 | 11.060 | 14.060 | 11.657 | 11.192 | 13.332 | 12.447 | 10.666 | 11.764 | 13.193 | 10.192 | 11.874 | 11.609 | 10.521 | 10.982 | 0.028 | 0.6048 | 0.8011 | 0 | 0 | 0 |
| 'Katnbl1' | 2.535 | 3.515 | 3.497 | 2.724 | 3.168 | 3.030 | 3.019 | 2.894 | 3.154 | 2.755 | 3.153 | 3.091 | 2.858 | 3.006 | 2.735 | 2.968 | 3.244 | 2.963 | 0.028 | 0.6465 | 0.8274 | 0 | 0 | 0 |
| 'Kbtbd11' | 10.453 | 9.455 | 12.394 | 10.823 | 10.041 | 11.597 | 10.510 | 11.074 | 10.946 | 10.576 | 12.050 | 14.541 | 11.257 | 10.524 | 14.878 | 11.272 | 10.951 | 14.952 | -0.191 | 0.0000 | 0.0005 | 1 | 0 | 0 |
| 'Kbtbd2' | 10.942 | 12.245 | 9.475 | 10.639 | 11.410 | 10.235 | 10.334 | 11.324 | 9.592 | 12.045 | 10.943 | 10.501 | 10.827 | 13.172 | 10.229 | 12.036 | 11.482 | 10.392 | -0.091 | 0.0140 | 0.0799 | 0 | 0 | 0 |
| 'Kbtbd3' | 1.556 | 2.581 | 2.346 | 2.326 | 2.423 | 2.095 | 2.256 | 2.427 | 2.094 | 2.595 | 1.874 | 2.108 | 1.654 | 2.577 | 2.089 | 2.388 | 2.082 | 1.696 | 0.065 | 0.5454 | 0.7606 | 0 | 0 | 0 |
| 'Kbtbd4' | 9.125 | 9.135 | 8.919 | 8.843 | 9.076 | 9.506 | 9.392 | 9.039 | 9.967 | 8.166 | 9.495 | 11.188 | 7.895 | 9.200 | 10.825 | 9.272 | 9.976 | 11.343 | -0.081 | 0.1089 | 0.3109 | 0 | 0 | 0 |
| 'Kbtbd7' | 5.130 | 5.394 | 4.768 | 4.651 | 5.041 | 4.645 | 5.676 | 5.342 | 4.573 | 4.767 | 4.677 | 5.134 | 4.552 | 5.220 | 4.342 | 4.706 | 4.704 | 4.437 | 0.068 | 0.2030 | 0.4453 | 0 | 0 | 0 |
| 'Kbtbd8' | 1.256 | 1.860 | 2.173 | 2.076 | 1.760 | 1.955 | 2.068 | 2.028 | 2.080 | 1.294 | 1.738 | 2.414 | 1.339 | 1.622 | 1.879 | 1.316 | 1.453 | 2.242 | 0.154 | 0.1750 | 0.4097 | 0 | 0 | 0 |
| 'Kcmf1' | 17.761 | 16.590 | 14.836 | 18.067 | 17.656 | 15.539 | 16.045 | 17.744 | 14.950 | 16.658 | 17.138 | 16.864 | 16.669 | 18.711 | 18.059 | 17.455 | 17.529 | 16.792 | -0.077 | 0.0496 | 0.1879 | 0 | 0 | 0 |
| 'Kcna2' | 3.098 | 2.547 | 3.360 | 3.063 | 2.213 | 3.035 | 2.667 | 2.517 | 3.332 | 2.295 | 2.412 | 3.065 | 1.755 | 1.702 | 2.669 | 1.959 | 1.634 | 3.223 | 0.324 | 0.0001 | 0.0012 | 1 | 0 | 0 |
| 'Kcna3' | 88.201 | 82.579 | 92.903 | 83.772 | 80.227 | 86.556 | 80.378 | 80.503 | 91.309 | 90.432 | 86.182 | 95.615 | 85.099 | 76.616 | 90.152 | 93.003 | 77.760 | 100.063 | -0.061 | 0.0744 | 0.2425 | 0 | 0 | 0 |
| 'Kcnab2' | 16.132 | 15.241 | 17.346 | 15.668 | 16.853 | 18.942 | 15.816 | 16.375 | 17.927 | 16.915 | 14.901 | 18.321 | 17.863 | 16.244 | 18.717 | 17.056 | 15.247 | 18.509 | -0.042 | 0.1532 | 0.3806 | 0 | 0 | 0 |
| 'Kcnab3' | 0.935 | 1.264 | 1.084 | 1.201 | 0.907 | 1.293 | 1.094 | 1.221 | 0.999 | 1.044 | 0.931 | 0.741 | 0.779 | 0.929 | 0.609 | 0.688 | 0.981 | 0.778 | 0.422 | 0.0001 | 0.0020 | 1 | 1 | 0 |
| 'Kcnf1' | 9.997 | 10.811 | 9.853 | 9.788 | 10.753 | 11.059 | 9.633 | 10.014 | 9.637 | 8.648 | 6.828 | 8.526 | 7.451 | 6.802 | 7.317 | 8.464 | 6.887 | 7.841 | 0.401 | 0.0000 | 0.0000 | 1 | 1 | 0 |
| 'Kcnh2' | 4.083 | 4.470 | 3.746 | 3.831 | 4.345 | 3.927 | 3.635 | 4.005 | 4.085 | 3.827 | 3.840 | 3.635 | 4.200 | 3.946 | 3.445 | 4.057 | 3.982 | 3.327 | 0.068 | 0.1430 | 0.3663 | 0 | 0 | 0 |
| 'Kcnh3' | 1.554 | 1.642 | 2.181 | 1.500 | 1.722 | 2.363 | 1.540 | 1.881 | 2.104 | 1.673 | 1.362 | 1.366 | 1.980 | 1.313 | 1.459 | 1.390 | 1.206 | 1.356 | 0.329 | 0.0024 | 0.0211 | 1 | 0 | 0 |
| 'Kcnk5' | 12.532 | 9.235 | 11.726 | 11.333 | 9.255 | 11.182 | 12.529 | 9.456 | 10.531 | 10.250 | 11.772 | 12.600 | 11.177 | 10.070 | 11.782 | 10.157 | 11.748 | 12.331 | -0.075 | 0.2184 | 0.4654 | 0 | 0 | 0 |
| 'Kcnk6' | 1.863 | 1.929 | 1.649 | 1.308 | 2.096 | 2.075 | 2.276 | 1.759 | 2.217 | 1.328 | 1.399 | 2.500 | 1.714 | 1.950 | 2.293 | 2.070 | 1.849 | 2.187 | -0.021 | 0.8432 | 0.9337 | 0 | 0 | 0 |
| 'Kcnn4' | 46.828 | 48.068 | 50.435 | 47.910 | 45.564 | 52.447 | 49.989 | 42.901 | 51.575 | 45.380 | 43.455 | 56.113 | 48.433 | 42.772 | 53.777 | 50.140 | 44.885 | 55.353 | -0.023 | 0.4437 | 0.6869 | 0 | 0 | 0 |
| 'Kcnq5' | 1.272 | 1.105 | 0.954 | 0.975 | 1.232 | 0.952 | 1.271 | 1.032 | 0.963 | 1.181 | 1.123 | 1.142 | 1.125 | 1.205 | 0.925 | 1.120 | 1.234 | 1.180 | -0.089 | 0.1732 | 0.4066 | 0 | 0 | 0 |
| 'Kctd1' | 3.024 | 2.538 | 3.095 | 3.222 | 2.581 | 3.534 | 3.077 | 3.473 | 3.783 | 3.193 | 3.140 | 3.426 | 2.998 | 2.787 | 3.108 | 2.606 | 2.876 | 3.642 | 0.020 | 0.7845 | 0.9037 | 0 | 0 | 0 |
| 'Kctd10' | 8.445 | 10.823 | 9.270 | 8.829 | 10.555 | 10.054 | 7.919 | 10.086 | 9.974 | 9.700 | 9.318 | 9.637 | 9.870 | 10.019 | 9.271 | 9.646 | 9.174 | 9.715 | -0.022 | 0.6060 | 0.8014 | 0 | 0 | 0 |
| 'Kctd11' | 1.307 | 1.594 | 1.357 | 1.612 | 1.389 | 1.481 | 1.289 | 1.369 | 1.635 | 1.432 | 1.600 | 2.066 | 1.386 | 1.703 | 2.152 | 1.630 | 2.168 | 2.073 | -0.332 | 0.0008 | 0.0089 | 1 | 0 | 0 |
| 'Kctd13' | 11.387 | 12.531 | 11.704 | 9.948 | 12.495 | 12.512 | 10.333 | 11.711 | 12.228 | 11.707 | 10.536 | 11.122 | 12.349 | 10.749 | 11.126 | 11.128 | 11.085 | 10.241 | 0.057 | 0.2379 | 0.4879 | 0 | 0 | 0 |
| 'Kctd17' | 7.872 | 9.439 | 9.776 | 6.749 | 8.549 | 10.763 | 6.823 | 8.840 | 9.556 | 8.603 | 8.046 | 9.337 | 9.385 | 8.504 | 10.207 | 8.954 | 7.349 | 9.452 | -0.043 | 0.4780 | 0.7111 | 0 | 0 | 0 |
| 'Kctd18' | 1.863 | 2.276 | 2.410 | 1.689 | 2.449 | 2.264 | 1.834 | 2.326 | 2.393 | 2.533 | 2.133 | 2.012 | 2.366 | 2.155 | 1.940 | 2.373 | 2.168 | 1.849 | -0.013 | 0.8526 | 0.9370 | 0 | 0 | 0 |
| 'Kctd20' | 11.764 | 12.763 | 11.991 | 11.347 | 12.397 | 12.473 | 11.474 | 12.635 | 11.980 | 12.027 | 12.697 | 12.612 | 12.226 | 12.439 | 12.520 | 12.564 | 12.496 | 12.624 | -0.055 | 0.0176 | 0.0942 | 0 | 0 | 0 |
| 'Kctd3' | 2.645 | 2.544 | 2.688 | 2.437 | 2.510 | 2.320 | 2.261 | 2.849 | 2.508 | 2.661 | 2.520 | 2.533 | 2.546 | 2.633 | 2.321 | 2.346 | 2.773 | 2.213 | 0.007 | 0.8776 | 0.9485 | 0 | 0 | 0 |
| 'Kctd5' | 10.841 | 10.893 | 10.768 | 10.765 | 10.679 | 10.602 | 10.714 | 12.134 | 10.688 | 10.590 | 10.830 | 11.560 | 10.890 | 11.817 | 11.129 | 10.340 | 10.510 | 11.302 | -0.026 | 0.5169 | 0.7404 | 0 | 0 | 0 |
| 'Kctd6' | 1.967 | 1.526 | 2.027 | 1.972 | 2.093 | 1.879 | 1.937 | 2.155 | 2.131 | 2.619 | 2.705 | 2.044 | 2.688 | 2.461 | 1.974 | 2.450 | 2.726 | 2.224 | -0.298 | 0.0001 | 0.0018 | 1 | 0 | 0 |
| 'Kctd7' | 2.190 | 2.977 | 2.438 | 2.481 | 3.039 | 2.780 | 2.188 | 2.383 | 2.129 | 1.997 | 2.654 | 2.291 | 2.282 | 2.815 | 2.603 | 2.272 | 2.568 | 2.127 | 0.053 | 0.4373 | 0.6804 | 0 | 0 | 0 |
| 'Kctd9' | 4.009 | 4.570 | 4.453 | 3.767 | 4.384 | 4.203 | 3.827 | 4.258 | 4.222 | 3.801 | 4.744 | 3.939 | 3.861 | 4.470 | 4.120 | 3.833 | 4.439 | 3.876 | 0.020 | 0.5988 | 0.7958 | 0 | 0 | 0 |
| 'Kdelc1' | 2.785 | 3.403 | 2.759 | 2.789 | 3.043 | 2.703 | 3.282 | 3.150 | 2.707 | 2.589 | 2.941 | 2.831 | 2.904 | 3.006 | 2.664 | 2.893 | 3.633 | 2.802 | 0.003 | 0.9523 | 0.9822 | 0 | 0 | 0 |
| 'Kdelc2' | 5.842 | 7.473 | 6.894 | 6.465 | 7.188 | 7.273 | 6.416 | 6.674 | 6.914 | 7.801 | 7.175 | 8.519 | 6.351 | 8.109 | 7.947 | 7.339 | 7.710 | 8.260 | -0.195 | 0.0000 | 0.0008 | 1 | 0 | 0 |
| 'Kdelr1' | 17.189 | 18.565 | 18.157 | 17.470 | 19.640 | 18.980 | 17.393 | 18.746 | 17.966 | 17.937 | 18.191 | 19.260 | 18.488 | 18.754 | 20.485 | 17.388 | 16.855 | 19.601 | -0.038 | 0.2945 | 0.5504 | 0 | 0 | 0 |
| 'Kdelr2' | 15.800 | 17.893 | 17.655 | 16.583 | 17.306 | 18.538 | 17.477 | 18.770 | 18.010 | 14.658 | 16.488 | 18.992 | 14.899 | 17.405 | 20.187 | 14.406 | 16.737 | 18.388 | 0.042 | 0.3472 | 0.6025 | 0 | 0 | 0 |
| 'Kdm1a' | 22.816 | 20.878 | 22.105 | 23.341 | 21.152 | 21.749 | 23.767 | 21.943 | 20.338 | 21.886 | 22.306 | 20.735 | 23.312 | 22.216 | 20.966 | 22.442 | 23.131 | 20.798 | -0.007 | 0.7941 | 0.9080 | 0 | 0 | 0 |
| 'Kdm1b' | 4.685 | 4.938 | 5.156 | 4.727 | 5.295 | 5.177 | 5.129 | 4.689 | 4.930 | 4.964 | 5.505 | 5.700 | 5.035 | 5.035 | 5.168 | 5.326 | 5.652 | 5.659 | -0.116 | 0.0019 | 0.0174 | 1 | 0 | 0 |
| 'Kdm2a' | 13.592 | 12.868 | 12.556 | 13.645 | 12.096 | 12.554 | 13.230 | 12.315 | 13.061 | 12.232 | 13.846 | 15.443 | 11.631 | 12.728 | 14.542 | 12.536 | 13.377 | 15.417 | -0.077 | 0.1082 | 0.3097 | 0 | 0 | 0 |
| 'Kdm2b' | 6.965 | 5.683 | 6.959 | 6.460 | 5.901 | 7.080 | 6.991 | 6.032 | 7.264 | 6.877 | 6.413 | 7.275 | 6.645 | 6.413 | 7.262 | 6.383 | 6.519 | 7.045 | -0.048 | 0.1191 | 0.3284 | 0 | 0 | 0 |
| 'Kdm3a' | 8.094 | 8.398 | 7.611 | 8.566 | 7.711 | 7.415 | 8.303 | 7.649 | 7.686 | 7.423 | 9.762 | 8.764 | 8.055 | 9.501 | 8.962 | 8.024 | 10.284 | 8.561 | -0.158 | 0.0016 | 0.0150 | 1 | 0 | 0 |
| 'Kdm3b' | 24.205 | 24.441 | 25.237 | 23.776 | 24.815 | 25.519 | 22.512 | 26.217 | 26.370 | 25.113 | 24.763 | 24.584 | 23.874 | 24.199 | 23.607 | 25.360 | 23.321 | 23.241 | 0.023 | 0.4207 | 0.6675 | 0 | 0 | 0 |
| 'Kdm4a' | 20.662 | 19.829 | 21.142 | 19.080 | 19.919 | 21.269 | 18.465 | 20.860 | 21.322 | 21.734 | 20.851 | 20.528 | 19.901 | 19.955 | 20.403 | 20.479 | 19.157 | 20.381 | -0.016 | 0.5999 | 0.7967 | 0 | 0 | 0 |
| 'Kdm4b' | 13.286 | 12.080 | 11.921 | 11.517 | 11.161 | 12.386 | 11.158 | 10.926 | 12.959 | 11.665 | 12.239 | 14.198 | 12.522 | 11.264 | 12.609 | 12.322 | 11.841 | 13.514 | -0.073 | 0.0645 | 0.2217 | 0 | 0 | 0 |
| 'Kdm4c' | 11.148 | 11.402 | 10.811 | 11.266 | 10.863 | 10.125 | 11.084 | 11.171 | 9.945 | 10.449 | 12.118 | 10.804 | 10.801 | 11.636 | 10.201 | 10.752 | 11.875 | 11.114 | -0.039 | 0.2473 | 0.4979 | 0 | 0 | 0 |
| 'Kdm5a' | 10.127 | 9.550 | 9.303 | 10.198 | 9.523 | 8.752 | 10.445 | 9.517 | 8.956 | 9.693 | 10.421 | 9.391 | 8.712 | 9.349 | 8.828 | 9.560 | 10.192 | 8.661 | 0.015 | 0.6728 | 0.8425 | 0 | 0 | 0 |

| | | | | | | | | | | | | | | | | | | | | | | | | |
|---|---|---|---|---|---|---|---|---|---|---|---|---|---|---|---|---|---|---|---|---|---|---|---|---|
| 'Kdm5b' | 5.232 | 4.758 | 4.063 | 5.266 | 4.701 | 4.282 | 5.225 | 4.394 | 4.172 | 4.660 | 5.956 | 6.519 | 4.595 | 5.200 | 6.096 | 4.814 | 5.431 | 6.693 | -0.254 | 0.0069 | 0.0476 | 1 | 0 | 0 |
| 'Kdm5c' | 15.170 | 13.118 | 15.018 | 15.276 | 13.323 | 14.774 | 14.284 | 13.777 | 14.512 | 14.052 | 15.343 | 16.411 | 14.081 | 14.281 | 15.528 | 13.744 | 14.607 | 16.222 | -0.065 | 0.0747 | 0.2432 | 0 | 0 | 0 |
| 'Kdm6a' | 7.797 | 6.917 | 7.339 | 8.086 | 7.127 | 6.990 | 7.818 | 7.066 | 6.999 | 6.661 | 7.122 | 7.606 | 6.796 | 6.675 | 6.784 | 6.845 | 6.909 | 7.400 | 0.061 | 0.1149 | 0.3218 | 0 | 0 | 0 |
| 'Kdm6b' | 41.320 | 32.714 | 35.617 | 36.034 | 31.511 | 34.014 | 33.826 | 33.549 | 38.274 | 43.178 | 37.147 | 38.467 | 38.285 | 36.968 | 37.890 | 41.400 | 36.746 | 37.822 | -0.145 | 0.0002 | 0.0023 | 1 | 0 | 0 |
| 'Kdm7a' | 13.042 | 12.729 | 11.406 | 13.175 | 12.763 | 10.914 | 12.888 | 12.595 | 11.950 | 13.207 | 13.640 | 13.504 | 12.640 | 13.737 | 11.938 | 12.957 | 14.721 | 12.543 | -0.106 | 0.0017 | 0.0156 | 1 | 0 | 0 |
| 'Kdm8' | 3.076 | 3.687 | 4.360 | 3.956 | 3.550 | 3.876 | 4.037 | 3.743 | 4.137 | 3.644 | 3.503 | 3.808 | 4.546 | 3.610 | 4.458 | 3.218 | 3.350 | 3.218 | 0.033 | 0.6812 | 0.8482 | 0 | 0 | 0 |
| 'Kdsr' | 3.803 | 4.053 | 4.132 | 4.058 | 4.073 | 3.751 | 4.083 | 4.369 | 3.879 | 3.893 | 4.131 | 4.168 | 3.294 | 4.189 | 3.988 | 4.278 | 3.900 | 3.833 | 0.011 | 0.8061 | 0.9133 | 0 | 0 | 0 |
| 'Keap1' | 15.980 | 15.865 | 16.853 | 15.568 | 16.649 | 17.506 | 15.047 | 16.041 | 16.909 | 16.042 | 15.459 | 17.540 | 16.717 | 15.695 | 18.176 | 16.694 | 15.001 | 17.436 | -0.033 | 0.2208 | 0.4681 | 0 | 0 | 0 |
| 'Khdc3' | 1.526 | 0.758 | 0.919 | 1.564 | 0.728 | 0.793 | 1.147 | 0.690 | 0.865 | 1.025 | 1.521 | 1.682 | 1.067 | 1.384 | 1.373 | 0.851 | 1.751 | 1.690 | -0.498 | 0.0054 | 0.0396 | 1 | 1 | 0 |
| 'Khdrbs1' | 36.920 | 37.355 | 38.263 | 37.610 | 38.631 | 36.143 | 39.758 | 38.083 | 37.046 | 37.694 | 35.231 | 35.525 | 35.484 | 35.968 | 36.023 | 35.310 | 36.161 | 34.340 | 0.068 | 0.0010 | 0.0101 | 1 | 0 | 0 |
| 'Khk' | 4.320 | 4.413 | 4.189 | 4.399 | 3.929 | 4.936 | 4.558 | 4.652 | 4.587 | 4.422 | 3.593 | 3.908 | 4.065 | 3.981 | 4.026 | 4.828 | 3.448 | 3.723 | 0.151 | 0.0142 | 0.0806 | 0 | 0 | 0 |
| 'Khnyn' | 29.099 | 30.726 | 31.190 | 26.849 | 29.392 | 32.190 | 27.837 | 30.059 | 33.739 | 30.007 | 30.591 | 34.747 | 29.834 | 30.335 | 33.805 | 32.850 | 29.544 | 33.858 | -0.086 | 0.0028 | 0.0239 | 1 | 0 | 0 |
| 'Khsrp' | 67.022 | 65.153 | 74.313 | 67.155 | 63.201 | 72.569 | 66.072 | 66.399 | 67.381 | 66.824 | 64.989 | 73.142 | 72.870 | 68.194 | 74.021 | 69.694 | 65.219 | 78.932 | -0.067 | 0.0133 | 0.0769 | 0 | 0 | 0 |
| 'Kidins220' | 13.859 | 13.401 | 12.439 | 12.503 | 12.574 | 12.230 | 13.176 | 12.300 | 12.681 | 13.682 | 14.096 | 13.172 | 13.128 | 12.265 | 12.612 | 13.020 | 14.126 | 12.983 | -0.059 | 0.0745 | 0.2429 | 0 | 0 | 0 |
| 'Kif11' | 13.032 | 13.317 | 17.690 | 15.730 | 15.229 | 15.659 | 15.743 | 13.739 | 16.905 | 10.383 | 14.377 | 16.801 | 16.363 | 18.147 | 18.584 | 15.277 | 16.676 | 16.912 | -0.075 | 0.3051 | 0.5627 | 0 | 0 | 0 |
| 'Kif13a' | 2.983 | 3.607 | 3.451 | 2.964 | 3.816 | 3.808 | 2.765 | 3.834 | 3.539 | 4.460 | 3.748 | 3.809 | 3.373 | 4.320 | 3.758 | 4.031 | 4.324 | 3.746 | -0.220 | 0.0002 | 0.0027 | 1 | 0 | 0 |
| 'Kif13b' | 5.109 | 4.800 | 5.300 | 5.629 | 4.938 | 5.023 | 5.351 | 4.589 | 5.459 | 4.428 | 5.646 | 5.841 | 4.678 | 5.065 | 5.375 | 4.893 | 5.420 | 5.547 | -0.032 | 0.5345 | 0.7542 | 0 | 0 | 0 |
| 'Kif14' | 1.116 | 1.293 | 1.727 | 1.374 | 1.294 | 1.449 | 1.545 | 1.276 | 1.879 | 0.955 | 1.542 | 1.343 | 1.476 | 1.604 | 1.600 | 1.280 | 1.585 | 1.338 | 0.014 | 0.8732 | 0.9459 | 0 | 0 | 0 |
| 'Kif15' | 9.269 | 10.097 | 11.650 | 11.292 | 10.229 | 11.084 | 12.235 | 9.405 | 11.515 | 8.355 | 11.968 | 11.281 | 11.151 | 11.171 | 11.268 | 9.777 | 12.431 | 10.610 | -0.028 | 0.6741 | 0.8433 | 0 | 0 | 0 |
| 'Kif16b' | 5.102 | 5.216 | 5.019 | 5.399 | 5.159 | 5.128 | 5.502 | 5.250 | 5.067 | 5.503 | 5.324 | 5.142 | 4.921 | 5.146 | 4.748 | 5.157 | 5.259 | 5.163 | 0.004 | 0.9118 | 0.9661 | 0 | 0 | 0 |
| 'Kif18a' | 0.742 | 0.807 | 0.964 | 0.947 | 0.812 | 0.908 | 1.100 | 0.834 | 0.929 | 0.708 | 0.920 | 0.805 | 0.937 | 0.913 | 0.901 | 0.981 | 0.830 | 0.738 | 0.051 | 0.5239 | 0.7463 | 0 | 0 | 0 |
| 'Kif18b' | 7.909 | 7.698 | 11.882 | 9.681 | 8.005 | 11.272 | 9.605 | 8.472 | 11.970 | 6.317 | 7.873 | 10.507 | 9.664 | 9.151 | 11.891 | 9.272 | 8.548 | 10.265 | 0.035 | 0.6004 | 0.7968 | 0 | 0 | 0 |
| 'Kif1b' | 2.210 | 2.261 | 2.319 | 2.408 | 2.090 | 2.089 | 2.235 | 2.390 | 2.325 | 2.208 | 2.506 | 2.560 | 1.988 | 2.274 | 2.401 | 2.310 | 2.090 | 2.737 | -0.061 | 0.2471 | 0.4979 | 0 | 0 | 0 |
| 'Kif1bp' | 4.851 | 5.227 | 4.702 | 4.721 | 5.041 | 4.968 | 5.033 | 4.973 | 4.819 | 4.700 | 5.004 | 5.422 | 4.623 | 4.723 | 5.427 | 4.564 | 5.331 | 5.389 | -0.043 | 0.2934 | 0.5497 | 0 | 0 | 0 |
| 'Kif1c' | 12.085 | 11.600 | 12.229 | 11.283 | 11.912 | 13.315 | 10.582 | 11.906 | 12.155 | 12.849 | 12.007 | 12.895 | 12.553 | 11.754 | 13.044 | 12.235 | 10.869 | 12.825 | -0.062 | 0.0645 | 0.2217 | 0 | 0 | 0 |
| 'Kif20a' | 20.786 | 20.259 | 26.988 | 24.873 | 23.852 | 26.589 | 26.151 | 23.407 | 27.362 | 17.243 | 21.540 | 24.547 | 22.914 | 24.325 | 28.071 | 19.951 | 24.142 | 24.440 | 0.080 | 0.1886 | 0.4247 | 0 | 0 | 0 |
| 'Kif20b' | 5.518 | 6.040 | 6.459 | 7.149 | 5.885 | 6.041 | 7.361 | 5.906 | 6.406 | 5.439 | 6.756 | 6.999 | 6.067 | 7.006 | 6.851 | 5.898 | 7.084 | 6.372 | -0.054 | 0.3803 | 0.6318 | 0 | 0 | 0 |
| 'Kif21b' | 30.377 | 25.199 | 30.684 | 27.196 | 26.275 | 29.261 | 25.821 | 27.421 | 30.449 | 35.237 | 34.165 | 37.137 | 33.528 | 32.302 | 35.756 | 34.062 | 29.908 | 39.089 | -0.310 | 0.0000 | 0.0000 | 1 | 0 | 0 |
| 'Kif22' | 5.605 | 5.837 | 7.994 | 6.225 | 5.805 | 7.484 | 6.658 | 6.214 | 7.222 | 4.325 | 6.146 | 6.778 | 6.934 | 6.724 | 7.619 | 5.637 | 6.103 | 6.653 | 0.041 | 0.5414 | 0.7584 | 0 | 0 | 0 |
| 'Kif23' | 10.454 | 11.934 | 14.297 | 13.600 | 12.388 | 14.084 | 13.362 | 11.694 | 15.110 | 9.618 | 12.043 | 12.977 | 14.296 | 14.014 | 14.159 | 12.775 | 13.056 | 12.141 | 0.009 | 0.8910 | 0.9554 | 0 | 0 | 0 |
| 'Kif24' | 0.526 | 0.562 | 0.792 | 0.809 | 0.535 | 0.732 | 0.674 | 0.572 | 0.792 | 0.592 | 0.572 | 0.677 | 0.761 | 0.730 | 0.757 | 0.498 | 0.732 | 0.539 | 0.017 | 0.8765 | 0.9477 | 0 | 0 | 0 |
| 'Kif2a' | 8.480 | 8.688 | 8.453 | 8.560 | 8.471 | 8.001 | 8.468 | 8.095 | 8.204 | 8.478 | 7.918 | 9.266 | 8.284 | 8.721 | 8.067 | 7.424 | 8.452 | 8.178 | -0.003 | 0.9332 | 0.9746 | 0 | 0 | 0 |
| 'Kif2c' | 2.857 | 3.298 | 4.205 | 4.071 | 3.069 | 3.943 | 4.103 | 3.046 | 4.040 | 2.027 | 3.635 | 3.729 | 4.057 | 3.566 | 4.288 | 3.176 | 3.557 | 3.645 | 0.030 | 0.7645 | 0.8953 | 0 | 0 | 0 |
| 'Kif3a' | 7.589 | 7.931 | 7.829 | 8.918 | 8.407 | 7.777 | 8.728 | 8.670 | 7.839 | 6.400 | 5.887 | 6.175 | 6.083 | 6.246 | 5.926 | 6.259 | 6.298 | 5.977 | 0.401 | 0.0000 | 0.0000 | 1 | 1 | 0 |
| 'Kif3b' | 19.460 | 20.160 | 19.549 | 20.427 | 18.244 | 20.322 | 19.039 | 19.432 | 19.272 | 19.564 | 18.904 | 19.723 | 19.185 | 19.483 | 18.930 | 20.379 | 19.135 | 19.237 | 0.002 | 0.9455 | 0.9790 | 0 | 0 | 0 |
| 'Kif3c' | 5.283 | 4.868 | 5.904 | 6.222 | 5.436 | 6.030 | 6.098 | 5.652 | 5.548 | 5.241 | 5.621 | 6.959 | 5.713 | 5.483 | 6.592 | 5.537 | 5.689 | 6.804 | -0.083 | 0.0925 | 0.2812 | 0 | 0 | 0 |
| 'Kif4' | 4.457 | 4.544 | 5.474 | 5.288 | 4.470 | 5.852 | 5.854 | 4.321 | 5.860 | 3.500 | 4.523 | 5.237 | 5.698 | 5.694 | 6.340 | 4.473 | 5.564 | 5.471 | -0.025 | 0.7660 | 0.8956 | 0 | 0 | 0 |
| 'Kif5b' | 23.134 | 23.886 | 20.978 | 23.308 | 22.741 | 20.482 | 22.487 | 22.454 | 20.379 | 22.593 | 24.166 | 23.327 | 22.367 | 23.794 | 21.185 | 22.671 | 23.642 | 22.282 | -0.057 | 0.0364 | 0.1537 | 0 | 0 | 0 |
| 'Kif5c' | 2.409 | 2.523 | 2.827 | 2.497 | 2.608 | 2.482 | 2.625 | 2.335 | 2.381 | 2.619 | 3.141 | 2.891 | 2.514 | 2.761 | 2.621 | 2.861 | 2.865 | 3.129 | -0.172 | 0.0016 | 0.0149 | 1 | 0 | 0 |
| 'Kif7' | 1.565 | 1.818 | 2.159 | 1.998 | 1.948 | 1.993 | 1.569 | 1.977 | 2.313 | 1.860 | 1.700 | 1.438 | 2.131 | 1.750 | 1.778 | 1.727 | 1.615 | 1.518 | 0.159 | 0.0713 | 0.2355 | 0 | 0 | 0 |
| 'Kifap3' | 1.429 | 1.759 | 1.680 | 1.745 | 2.074 | 1.740 | 2.527 | 1.938 | 1.862 | 2.184 | 2.042 | 2.155 | 1.951 | 2.083 | 1.926 | 2.158 | 2.131 | 1.955 | -0.169 | 0.0345 | 0.1485 | 0 | 0 | 0 |
| 'Kifc2' | 6.679 | 7.439 | 8.837 | 6.369 | 6.482 | 8.742 | 6.997 | 7.239 | 9.312 | 8.378 | 6.763 | 8.758 | 7.345 | 6.360 | 7.475 | 6.289 | 7.850 | 7.430 | 0.014 | 0.8206 | 0.9207 | 0 | 0 | 0 |
| 'Kifc3' | 5.875 | 5.643 | 5.390 | 6.384 | 4.756 | 4.983 | 5.458 | 5.019 | 5.566 | 4.559 | 3.831 | 4.768 | 4.002 | 3.990 | 4.456 | 5.060 | 4.473 | 4.102 | 0.308 | 0.0000 | 0.0000 | 1 | 0 | 0 |
| 'Kin' | 16.102 | 16.534 | 16.788 | 17.377 | 16.227 | 15.847 | 17.220 | 15.982 | 17.325 | 14.440 | 15.839 | 17.306 | 14.812 | 14.946 | 16.437 | 15.525 | 18.328 | 17.581 | 0.029 | 0.5323 | 0.7526 | 0 | 0 | 0 |
| 'Kirrel3' | 1.181 | 1.381 | 1.484 | 1.169 | 1.535 | 1.422 | 0.714 | 1.363 | 1.274 | 1.343 | 1.379 | 1.692 | 0.881 | 1.268 | 1.329 | 1.415 | 1.504 | 1.574 | -0.119 | 0.2109 | 0.4561 | 0 | 0 | 0 |
| 'Kit' | 0.351 | 0.333 | 0.387 | 0.255 | 0.331 | 0.295 | 0.270 | 0.303 | 0.267 | 0.558 | 0.561 | 0.529 | 0.476 | 0.439 | 0.401 | 0.618 | 0.517 | 0.561 | -0.740 | 0.0000 | 0.0000 | 1 | 1 | 0 |
| 'Kiz' | 8.930 | 10.051 | 9.066 | 10.369 | 9.426 | 8.833 | 10.219 | 9.839 | 9.096 | 9.599 | 9.596 | 9.077 | 10.156 | 10.161 | 8.903 | 9.262 | 9.751 | 7.977 | 0.010 | 0.8138 | 0.9160 | 0 | 0 | 0 |
| 'Klc1' | 16.050 | 15.215 | 15.638 | 15.368 | 15.938 | 16.558 | 14.173 | 15.453 | 16.600 | 15.803 | 15.644 | 17.715 | 17.404 | 16.161 | 17.892 | 16.880 | 16.766 | 17.857 | -0.120 | 0.0001 | 0.0009 | 1 | 0 | 0 |
| 'Klc2' | 6.151 | 6.037 | 7.425 | 6.674 | 5.697 | 7.313 | 6.570 | 6.912 | 7.413 | 7.116 | 6.318 | 7.429 | 6.508 | 6.078 | 6.796 | 6.263 | 6.129 | 6.805 | 0.008 | 0.8347 | 0.9283 | 0 | 0 | 0 |
| 'Klc3' | 6.262 | 4.918 | 7.266 | 5.390 | 5.554 | 7.576 | 5.578 | 5.988 | 7.487 | 6.117 | 4.242 | 6.203 | 6.037 | 4.929 | 6.228 | 5.897 | 4.598 | 5.894 | 0.155 | 0.0049 | 0.0368 | 1 | 0 | 0 |
| 'Klc4' | 15.897 | 16.789 | 18.656 | 14.786 | 16.995 | 18.902 | 16.869 | 17.626 | 17.810 | 18.490 | 15.624 | 18.076 | 17.349 | 15.275 | 16.735 | 18.149 | 16.398 | 15.070 | 0.021 | 0.6554 | 0.8331 | 0 | 0 | 0 |
| 'Klf10' | 14.588 | 13.810 | 11.432 | 12.641 | 12.392 | 11.747 | 13.464 | 13.580 | 11.902 | 15.017 | 13.212 | 12.967 | 13.298 | 16.604 | 12.809 | 15.850 | 15.794 | 12.799 | -0.159 | 0.0009 | 0.0099 | 1 | 0 | 0 |
| 'Klf11' | 4.891 | 1.797 | 1.442 | 2.893 | 1.547 | 1.154 | 2.033 | 2.171 | 0.983 | 4.627 | 2.284 | 1.343 | 2.843 | 2.230 | 1.392 | 3.728 | 2.261 | 1.156 | -0.207 | 0.1206 | 0.3311 | 0 | 0 | 0 |
| 'Klf13' | 27.943 | 29.342 | 28.779 | 26.042 | 28.717 | 28.855 | 25.911 | 31.531 | 29.148 | 30.946 | 31.748 | 29.660 | 30.232 | 32.251 | 31.167 | 34.441 | 28.020 | 29.822 | -0.130 | 0.0009 | 0.0100 | 1 | 0 | 0 |
| 'Klf16' | 10.713 | 9.037 | 8.536 | 9.550 | 8.156 | 9.651 | 7.938 | 8.347 | 9.372 | 8.211 | 7.390 | 9.982 | 9.102 | 8.629 | 9.667 | 7.879 | 7.324 | 10.823 | 0.032 | 0.6294 | 0.8165 | 0 | 0 | 0 |
| 'Klf2' | 51.587 | 12.015 | 13.986 | 51.055 | 13.781 | 13.840 | 68.296 | 18.059 | 10.706 | 26.202 | 19.597 | 12.151 | 27.108 | 17.974 | 11.408 | 20.562 | 26.815 | 10.714 | 0.264 | 0.2269 | 0.4734 | 0 | 0 | 0 |
| 'Klf3' | 17.071 | 21.298 | 15.030 | 16.280 | 19.513 | 16.196 | 16.627 | 18.756 | 16.449 | 18.954 | 20.026 | 18.953 | 20.361 | 21.565 | 18.197 | 22.017 | 19.271 | 19.058 | -0.199 | 0.0000 | 0.0000 | 1 | 0 | 0 |
| 'Klf4' | 25.063 | 5.841 | 4.706 | 16.402 | 6.613 | 2.958 | 14.343 | 11.764 | 2.697 | 23.072 | 10.583 | 1.537 | 16.238 | 8.781 | 1.796 | 10.799 | 8.995 | 1.378 | 0.334 | 0.1548 | 0.3832 | 0 | 0 | 0 |
| 'Klf6' | 55.801 | 15.795 | 15.014 | 38.325 | 18.917 | 13.455 | 38.633 | 21.862 | 11.886 | 40.005 | 31.421 | 12.504 | 31.485 | 26.505 | 12.301 | 25.597 | 30.576 | 10.914 | -0.014 | 0.9336 | 0.9746 | 0 | 0 | 0 |
| 'Klf7' | 4.955 | 5.397 | 4.949 | 5.204 | 5.577 | 5.418 | 5.636 | 6.061 | 5.452 | 6.144 | 5.933 | 4.789 | 6.271 | 6.604 | 4.661 | 6.694 | 5.932 | 4.428 | -0.076 | 0.1733 | 0.4066 | 0 | 0 | 0 |
| 'Klf9' | 1.332 | 1.465 | 1.101 | 1.194 | 1.164 | 1.076 | 1.150 | 1.007 | 1.308 | 1.361 | 1.133 | 0.850 | 0.953 | 0.923 | 0.827 | 1.220 | 0.960 | 0.928 | 0.245 | 0.0359 | 0.1520 | 0 | 0 | 0 |
| 'Klhdc1' | 3.096 | 3.462 | 2.988 | 3.656 | 3.356 | 3.471 | 3.200 | 2.978 | 2.872 | 2.545 | 3.849 | 3.841 | 2.682 | 2.932 | 3.567 | 3.138 | 3.168 | 3.452 | -0.019 | 0.8121 | 0.9155 | 0 | 0 | 0 |
| 'Klhdc10' | 3.634 | 4.190 | 3.913 | 3.249 | 4.172 | 3.979 | 3.450 | 4.603 | 4.020 | 4.220 | 4.037 | 3.623 | 3.916 | 4.215 | 3.731 | 4.352 | 3.822 | 3.559 | -0.020 | 0.7027 | 0.8611 | 0 | 0 | 0 |
| 'Klhdc2' | 7.273 | 8.335 | 7.678 | 7.362 | 8.627 | 7.608 | 7.481 | 8.687 | 7.495 | 7.634 | 7.740 | 6.847 | 7.778 | 8.828 | 7.013 | 8.851 | 8.069 | 6.492 | 0.022 | 0.5968 | 0.7950 | 0 | 0 | 0 |
| 'Klhdc3' | 39.502 | 41.961 | 40.448 | 37.878 | 40.212 | 41.628 | 38.681 | 40.485 | 42.234 | 37.626 | 41.416 | 43.454 | 41.058 | 36.635 | 43.557 | 40.413 | 40.781 | 45.324 | -0.040 | 0.2094 | 0.4546 | 0 | 0 | 0 |
| 'Klhdc4' | 6.308 | 5.710 | 7.829 | 6.870 | 6.312 | 7.279 | 7.092 | 6.782 | 7.492 | 7.318 | 7.098 | 6.973 | 7.495 | 6.843 | 6.596 | 6.714 | 6.586 | 6.590 | -0.022 | 0.6360 | 0.8202 | 0 | 0 | 0 |
| 'Klhdc8b' | 0.356 | 0.303 | 0.411 | 0.345 | 0.370 | 0.401 | 0.352 | 0.314 | 0.424 | 0.382 | 0.268 | 0.333 | 0.374 | 0.269 | 0.329 | 0.296 | 0.226 | 0.302 | 0.232 | 0.0072 | 0.0488 | 1 | 0 | 0 |
| 'Klhl11' | 8.433 | 5.672 | 6.141 | 5.074 | 6.056 | 6.211 | 5.732 | 6.962 | 5.302 | 7.233 | 6.120 | 6.147 | 6.636 | 6.879 | 5.449 | 9.124 | 5.825 | 5.807 | -0.086 | 0.3326 | 0.5908 | 0 | 0 | 0 |
| 'Klhl12' | 1.767 | 2.368 | 2.181 | 1.651 | 2.391 | 2.033 | 1.853 | 2.298 | 2.168 | 1.906 | 1.293 | 1.573 | 1.850 | 1.982 | 1.574 | 2.174 | 1.836 | 1.646 | 0.229 | 0.0061 | 0.0437 | 1 | 0 | 0 |
| 'Klhl15' | 0.789 | 0.913 | 0.957 | 0.968 | 0.985 | 0.864 | 0.984 | 1.059 | 1.010 | 0.865 | 0.976 | 0.848 | 1.002 | 1.014 | 0.807 | 0.784 | 1.037 | 0.681 | 0.085 | 0.2187 | 0.4655 | 0 | 0 | 0 |
| 'Klhl17' | 22.593 | 22.128 | 27.573 | 22.130 | 23.841 | 26.893 | 22.992 | 24.417 | 27.211 | 25.342 | 23.071 | 25.911 | 24.714 | 21.877 | 22.457 | 24.491 | 22.629 | 24.616 | 0.020 | 0.6102 | 0.8042 | 0 | 0 | 0 |
| 'Klhl18' | 3.190 | 3.145 | 3.345 | 2.995 | 3.244 | 3.246 | 2.772 | 3.359 | 3.298 | 3.695 | 3.512 | 3.929 | 3.632 | 3.616 | 3.916 | 3.790 | 3.669 | 3.762 | -0.238 | 0.0000 | 0.0000 | 1 | 0 | 0 |
| 'Klhl2' | 12.537 | 12.242 | 10.611 | 12.408 | 11.526 | 11.521 | 13.654 | 11.603 | 10.735 | 12.735 | 12.857 | 13.120 | 12.682 | 11.903 | 12.286 | 12.283 | 13.179 | 12.780 | -0.110 | 0.0115 | 0.0692 | 0 | 0 | 0 |
| 'Klhl20' | 1.955 | 2.353 | 2.517 | 2.125 | 2.515 | 2.376 | 2.206 | 2.627 | 2.390 | 2.820 | 3.335 | 3.114 | 3.065 | 3.333 | 3.018 | 3.112 | 3.063 | 3.103 | -0.419 | 0.0000 | 0.0000 | 1 | 1 | 0 |
| 'Klhl21' | 8.677 | 7.619 | 6.900 | 6.968 | 7.410 | 7.844 | 7.222 | 7.603 | 7.624 | 7.238 | 8.914 | 9.427 | 7.214 | 7.347 | 8.887 | 6.865 | 8.245 | 9.178 | -0.119 | 0.0552 | 0.2017 | 0 | 0 | 0 |
| 'Klhl22' | 2.307 | 2.379 | 2.712 | 2.224 | 2.878 | 2.472 | 2.126 | 2.586 | 2.303 | 2.245 | 2.184 | 2.005 | 2.347 | 2.063 | 1.971 | 2.076 | 2.099 | 1.949 | 0.209 | 0.0003 | 0.0039 | 1 | 0 | 0 |
| 'Klhl24' | 14.620 | 16.756 | 14.503 | 15.324 | 16.176 | 15.457 | 15.408 | 16.835 | 14.630 | 14.363 | 18.549 | 17.532 | 13.742 | 17.193 | 15.543 | 15.530 | 18.350 | 18.054 | -0.100 | 0.0207 | 0.1055 | 0 | 0 | 0 |
| 'Klhl25' | 4.164 | 3.979 | 4.450 | 3.749 | 3.861 | 4.604 | 3.764 | 4.477 | 4.483 | 4.677 | 5.358 | 5.502 | 4.491 | 4.954 | 5.643 | 5.181 | 5.512 | 5.540 | -0.325 | 0.0000 | 0.0000 | 1 | 0 | 0 |
| 'Klhl26' | 11.915 | 11.522 | 13.795 | 12.462 | 11.680 | 12.923 | 10.505 | 12.149 | 12.497 | 13.311 | 11.639 | 14.295 | 11.817 | 11.773 | 12.551 | 11.800 | 11.666 | 12.878 | -0.040 | 0.3428 | 0.5990 | 0 | 0 | 0 |
| 'Klhl28' | 6.999 | 6.111 | 6.399 | 7.205 | 4.758 | 5.982 | 7.932 | 5.789 | 6.435 | 5.719 | 7.348 | 7.985 | 5.648 | 5.801 | 7.155 | 4.865 | 7.165 | 7.571 | -0.053 | 0.5799 | 0.7822 | 0 | 0 | 0 |
| 'Klhl3' | 2.139 | 2.017 | 1.864 | 1.645 | 1.804 | 2.028 | 1.884 | 1.898 | 2.211 | 2.503 | 1.914 | 2.418 | 2.104 | 1.478 | 2.408 | 1.948 | 1.460 | 2.554 | -0.111 | 0.2282 | 0.4748 | 0 | 0 | 0 |
| 'Klhl36' | 19.663 | 20.374 | 22.246 | 17.840 | 20.708 | 23.568 | 17.968 | 21.606 | 22.296 | 20.320 | 17.173 | 19.751 | 21.844 | 17.470 | 20.096 | 19.430 | 18.393 | 18.154 | 0.100 | 0.0528 | 0.1953 | 0 | 0 | 0 |
| 'Klhl4' | 2.600 | 3.306 | 3.088 | 3.498 | 2.982 | 2.837 | 2.920 | 3.257 | 2.889 | 4.445 | 5.090 | 4.432 | 4.610 | 4.878 | 3.692 | 3.862 | 4.359 | 4.684 | -0.562 | 0.0000 | 0.0000 | 1 | 1 | 0 |
| 'Klhl42' | 3.698 | 3.390 | 3.432 | 3.959 | 3.294 | 3.610 | 4.085 | 3.531 | 3.665 | 2.802 | 3.412 | 5.196 | 3.299 | 3.474 | 4.483 | 2.943 | 4.179 | 4.740 | -0.075 | 0.3934 | 0.6428 | 0 | 0 | 0 |
| 'Klhl5' | 3.684 | 3.333 | 3.335 | 3.787 | 3.645 | 3.756 | 3.657 | 3.819 | 3.338 | 3.989 | 3.676 | 3.746 | 4.125 | 4.146 | 3.843 | 4.021 | 4.278 | 3.563 | -0.136 | 0.0006 | 0.0068 | 1 | 0 | 0 |
| 'Klhl6' | 30.044 | 30.857 | 31.470 | 31.357 | 32.039 | 31.838 | 31.837 | 32.946 | 29.210 | 28.414 | 33.872 | 31.376 | 30.728 | 31.836 | 31.246 | 31.333 | 34.696 | 31.152 | -0.025 | 0.4092 | 0.6583 | 0 | 0 | 0 |
| 'Klhl7' | 2.487 | 2.504 | 2.697 | 2.382 | 2.807 | 2.209 | 2.684 | 2.438 | 2.409 | 2.769 | 2.897 | 2.685 | 2.601 | 2.501 | 2.400 | 2.547 | 2.719 | 2.397 | -0.068 | 0.1852 | 0.4208 | 0 | 0 | 0 |
| 'Klhl9' | 9.477 | 9.781 | 9.377 | 11.236 | 9.167 | 10.241 | 12.836 | 9.973 | 9.604 | 7.026 | 9.985 | 12.509 | 8.083 | 9.531 | 10.419 | 7.498 | 11.491 | 11.201 | 0.066 | 0.5445 | 0.7606 | 0 | 0 | 0 |
| 'Klk8' | 15.344 | 15.001 | 17.537 | 15.743 | 15.934 | 16.689 | 15.763 | 16.114 | 16.377 | 14.503 | 12.355 | 15.094 | 13.228 | 11.213 | 16.476 | 13.909 | 11.577 | 15.312 | 0.217 | 0.0000 | 0.0001 | 1 | 0 | 0 |
| 'Klrb1f' | 1.151 | 1.592 | 1.333 | 1.136 | 1.297 | 1.308 | 1.436 | 1.320 | 1.163 | 1.262 | 1.493 | 1.068 | 1.420 | 1.005 | 1.520 | 0.946 | 1.245 | 1.118 | 0.071 | 0.5005 | 0.7278 | 0 | 0 | 0 |
| 'Kmt2a' | 18.971 | 15.356 | 16.871 | 17.596 | 14.592 | 15.333 | 15.349 | 15.110 | 16.733 | 16.461 | 16.718 | 17.930 | 15.912 | 15.229 | 16.537 | 16.307 | 14.793 | 20.090 | -0.050 | 0.3271 | 0.5849 | 0 | 0 | 0 |
| 'Kmt2b' | 23.854 | 20.363 | 23.489 | 22.402 | 20.060 | 22.291 | 20.141 | 21.354 | 23.908 | 23.750 | 22.255 | 25.031 | 22.298 | 21.118 | 23.678 | 22.405 | 21.153 | 26.355 | -0.082 | 0.0203 | 0.1041 | 0 | 0 | 0 |
| 'Kmt2c' | 7.746 | 6.339 | 6.922 | 7.653 | 6.122 | 6.536 | 6.602 | 6.384 | 7.144 | 6.791 | 7.705 | 7.569 | 5.922 | 6.515 | 6.854 | 6.408 | 6.648 | 8.099 | -0.036 | 0.5492 | 0.7629 | 0 | 0 | 0 |
| 'Kmt2d' | 23.146 | 17.829 | 21.811 | 22.182 | 17.437 | 19.103 | 18.570 | 19.342 | 22.198 | 20.606 | 21.710 | 24.224 | 19.374 | 19.535 | 22.535 | 20.724 | 19.140 | 28.467 | -0.119 | 0.0494 | 0.1875 | 0 | 0 | 0 |
| 'Kmt2e' | 25.306 | 21.533 | 22.099 | 24.890 | 20.247 | 20.687 | 24.967 | 21.505 | 21.631 | 24.023 | 25.113 | 23.699 | 22.268 | 22.734 | 22.303 | 23.187 | 24.068 | 24.123 | -0.075 | 0.0796 | 0.2541 | 0 | 0 | 0 |
| 'Kmt5a' | 7.311 | 7.162 | 7.632 | 7.767 | 6.995 | 7.744 | 7.121 | 7.376 | 7.158 | 6.876 | 6.968 | 8.108 | 7.685 | 7.208 | 8.127 | 7.621 | 7.567 | 8.469 | -0.061 | 0.0752 | 0.2439 | 0 | 0 | 0 |
| 'Kndc1' | 0.258 | 0.258 | 0.279 | 0.255 | 0.260 | 0.256 | 0.265 | 0.262 | 0.279 | 0.233 | 0.201 | 0.170 | 0.166 | 0.163 | 0.170 | 0.158 | 0.184 | 0.168 | 0.558 | 0.0000 | 0.0002 | 1 | 1 | 0 |
| 'Knop1' | 2.221 | 2.566 | 2.437 | 2.421 | 2.346 | 2.717 | 2.676 | 2.487 | 2.635 | 2.294 | 2.334 | 2.894 | 2.232 | 2.379 | 2.728 | 2.545 | 2.484 | 2.985 | -0.036 | 0.4472 | 0.6880 | 0 | 0 | 0 |
| 'Knstrn' | 3.215 | 3.579 | 4.608 | 4.204 | 3.786 | 3.905 | 4.356 | 3.715 | 4.434 | 2.996 | 3.978 | 4.111 | 4.105 | 4.156 | 4.870 | 3.645 | 4.108 | 3.761 | -0.008 | 0.9140 | 0.9671 | 0 | 0 | 0 |
| 'Kntc1' | 4.547 | 3.732 | 5.642 | 5.363 | 3.776 | 4.562 | 4.749 | 4.049 | 5.176 | 4.213 | 4.634 | 5.064 | 5.351 | 4.018 | 5.320 | 4.047 | 4.496 | 4.778 | -0.028 | 0.6523 | 0.8310 | 0 | 0 | 0 |
| 'Kpna1' | 6.137 | 5.934 | 5.957 | 6.270 | 5.976 | 5.156 | 6.656 | 5.681 | 5.627 | 5.395 | 6.307 | 6.553 | 5.278 | 5.848 | 5.944 | 4.981 | 6.583 | 6.508 | -0.010 | 0.8636 | 0.9407 | 0 | 0 | 0 |
| 'Kpna3' | 19.802 | 21.219 | 20.971 | 22.483 | 20.866 | 20.514 | 21.277 | 20.171 | 19.475 | 19.006 | 21.445 | 22.324 | 21.163 | 22.979 | 20.788 | 21.014 | 22.932 | 22.178 | -0.065 | 0.0805 | 0.2559 | 0 | 0 | 0 |
| 'Kpna4' | 16.504 | 15.746 | 16.168 | 18.346 | 16.985 | 15.527 | 18.244 | 16.372 | 15.817 | 15.333 | 17.550 | 19.297 | 15.845 | 16.051 | 17.656 | 14.228 | 19.487 | 18.668 | -0.050 | 0.3818 | 0.6330 | 0 | 0 | 0 |
| 'Kpna6' | 9.838 | 9.970 | 10.193 | 10.231 | 9.964 | 10.124 | 9.698 | 10.679 | 9.841 | 9.869 | 9.668 | 10.807 | 9.726 | 10.082 | 10.610 | 9.536 | 10.449 | 10.863 | -0.029 | 0.3380 | 0.5956 | 0 | 0 | 0 |
| 'Kptn' | 5.193 | 5.530 | 6.224 | 5.400 | 5.308 | 6.310 | 5.483 | 5.650 | 6.516 | 5.425 | 5.021 | 5.685 | 6.219 | 5.852 | 5.503 | 5.201 | 5.518 | 4.989 | 0.052 | 0.2909 | 0.5477 | 0 | 0 | 0 |
| 'Kras' | 25.585 | 34.058 | 27.567 | 29.666 | 31.902 | 27.130 | 29.591 | 30.234 | 27.438 | 27.180 | 29.470 | 27.289 | 28.761 | 31.049 | 27.234 | 27.417 | 29.987 | 27.195 | 0.028 | 0.3483 | 0.6037 | 0 | 0 | 0 |
| 'Krba1' | 3.218 | 3.065 | 3.940 | 3.060 | 3.007 | 3.376 | 2.976 | 3.003 | 3.741 | 3.442 | 3.122 | 3.327 | 3.472 | 2.887 | 3.164 | 3.103 | 3.099 | 3.365 | 0.008 | 0.8597 | 0.9390 | 0 | 0 | 0 |
| 'Krcc1' | 19.734 | 24.065 | 20.127 | 21.330 | 22.622 | 20.640 | 23.465 | 20.636 | 19.688 | 19.102 | 23.770 | 23.034 | 21.298 | 23.291 | 23.777 | 19.425 | 24.019 | 21.554 | -0.063 | 0.1728 | 0.4066 | 0 | 0 | 0 |
| 'Kremen1' | 9.971 | 9.345 | 10.306 | 8.977 | 9.650 | 10.393 | 9.028 | 9.833 | 9.973 | 10.921 | 10.528 | 10.089 | 10.067 | 9.741 | 10.080 | 9.785 | 9.666 | 10.033 | -0.062 | 0.0735 | 0.2405 | 0 | 0 | 0 |
| 'Kri1' | 12.486 | 10.669 | 13.874 | 10.747 | 10.578 | 13.048 | 12.278 | 11.981 | 13.966 | 12.781 | 12.183 | 13.215 | 12.390 | 10.855 | 12.122 | 12.464 | 11.421 | 12.242 | -0.012 | 0.7775 | 0.9002 | 0 | 0 | 0 |
| 'Krit1' | 3.869 | 4.013 | 3.903 | 4.101 | 3.967 | 3.699 | 4.079 | 3.904 | 4.040 | 3.932 | 3.795 | 4.230 | 3.737 | 4.033 | 3.738 | 3.991 | 4.246 | 4.056 | -0.020 | 0.5358 | 0.7552 | 0 | 0 | 0 |
| 'Krr1' | 6.040 | 6.510 | 5.884 | 5.657 | 6.552 | 5.343 | 5.633 | 6.296 | 6.018 | 5.753 | 5.707 | 5.872 | 5.889 | 6.000 | 5.468 | 5.686 | 6.330 | 5.718 | 0.027 | 0.4332 | 0.6782 | 0 | 0 | 0 |
| 'Krt17' | 5.122 | 5.868 | 3.158 | 5.979 | 4.255 | 4.234 | 4.941 | 4.519 | 4.373 | 6.513 | 6.244 | 4.815 | 6.157 | 6.491 | 5.296 | 6.619 | 6.785 | 4.244 | -0.336 | 0.0001 | 0.0009 | 1 | 0 | 0 |
| 'Krt83' | 2.967 | 4.739 | 2.526 | 2.577 | 4.140 | 3.169 | 3.067 | 4.066 | 3.021 | 4.716 | 4.847 | 3.932 | 5.921 | 6.588 | 3.647 | 5.365 | 5.417 | 3.457 | -0.547 | 0.0000 | 0.0000 | 1 | 1 | 0 |
| 'Krtcap2' | 16.802 | 17.132 | 17.659 | 15.788 | 18.824 | 17.773 | 14.016 | 18.202 | 17.018 | 15.829 | 16.991 | 16.939 | 16.605 | 16.896 | 18.092 | 15.977 | 15.906 | 15.896 | 0.030 | 0.4737 | 0.7090 | 0 | 0 | 0 |
| 'Krtcap3' | 2.047 | 2.108 | 2.634 | 1.942 | 1.962 | 2.015 | 2.522 | 1.980 | 2.127 | 1.574 | 1.540 | 2.278 | 1.622 | 1.460 | 1.936 | 2.083 | 1.972 | 2.226 | 0.201 | 0.0143 | 0.0806 | 0 | 0 | 0 |
| 'Ksr1' | 34.997 | 30.360 | 33.671 | 33.640 | 30.580 | 29.478 | 31.825 | 32.827 | 31.177 | 35.348 | 38.550 | 33.168 | 37.793 | 33.241 | 33.196 | 33.939 | 34.512 | 35.819 | -0.140 | 0.0003 | 0.0043 | 1 | 0 | 0 |
| 'Kti12' | 11.284 | 11.222 | 10.556 | 10.165 | 10.349 | 11.139 | 9.314 | 11.054 | 10.796 | 10.046 | 9.425 | 12.251 | 10.955 | 10.511 | 11.602 | 11.005 | 10.012 | 12.443 | -0.049 | 0.3779 | 0.6297 | 0 | 0 | 0 |
| 'Ktn1' | 5.367 | 5.735 | 5.756 | 5.600 | 5.426 | 5.123 | 5.747 | 5.581 | 5.410 | 5.642 | 5.765 | 4.886 | 5.496 | 5.722 | 4.671 | 5.864 | 5.987 | 4.452 | 0.035 | 0.3857 | 0.6354 | 0 | 0 | 0 |
| 'Kxd1' | 132.684 | 133.122 | 124.946 | 142.434 | 133.752 | 129.139 | 142.919 | 128.158 | 130.444 | 126.828 | 131.350 | 107.018 | 129.022 | 124.461 | 128.566 | 136.356 | 129.493 | 122.762 | 0.066 | 0.0591 | 0.2099 | 0 | 0 | 0 |
| 'Kyat1' | 1.241 | 1.468 | 1.544 | 1.223 | 1.321 | 1.629 | 1.181 | 1.307 | 1.495 | 1.411 | 1.001 | 1.280 | 1.323 | 1.320 | 1.418 | 1.284 | 1.325 | 1.251 | 0.091 | 0.2336 | 0.4820 | 0 | 0 | 0 |
| 'L2hgdh' | 2.345 | 3.572 | 3.715 | 2.844 | 3.482 | 2.675 | 2.939 | 2.939 | 3.575 | 2.824 | 3.231 | 3.091 | 2.747 | 2.664 | 2.925 | 2.593 | 2.986 | 2.801 | 0.101 | 0.1953 | 0.4337 | 0 | 0 | 0 |
| 'L3mbtl2' | 3.473 | 3.596 | 3.978 | 3.314 | 3.345 | 3.757 | 3.808 | 3.901 | 3.996 | 3.706 | 3.278 | 3.883 | 3.541 | 3.549 | 3.555 | 3.659 | 3.385 | 3.606 | 0.035 | 0.3518 | 0.6056 | 0 | 0 | 0 |
| 'L3mbtl3' | 7.323 | 6.536 | 7.292 | 7.421 | 7.345 | 7.140 | 7.062 | 6.859 | 7.298 | 8.693 | 8.672 | 9.288 | 7.831 | 7.809 | 7.905 | 8.367 | 9.666 | 8.267 | -0.262 | 0.0000 | 0.0000 | 1 | 0 | 0 |
| 'Lace1' | 1.900 | 1.661 | 1.765 | 2.034 | 1.855 | 1.913 | 1.640 | 1.887 | 2.128 | 2.026 | 1.965 | 1.895 | 2.141 | 1.867 | 1.741 | 1.725 | 1.879 | 1.642 | -0.015 | 0.8476 | 0.9344 | 0 | 0 | 0 |
| 'Lactb' | 13.904 | 16.649 | 16.446 | 15.871 | 15.189 | 15.429 | 13.583 | 15.158 | 14.504 | 13.595 | 14.149 | 15.218 | 13.811 | 14.799 | 14.312 | 15.465 | 14.276 | 15.176 | 0.052 | 0.2392 | 0.4895 | 0 | 0 | 0 |
| 'Lactb2' | 2.916 | 3.707 | 3.243 | 2.891 | 3.458 | 3.071 | 3.737 | 3.090 | 3.313 | 2.926 | 3.451 | 2.959 | 2.961 | 3.402 | 2.932 | 2.558 | 3.346 | 2.458 | 0.114 | 0.0726 | 0.2384 | 0 | 0 | 0 |
| 'Lag3' | 46.914 | 43.475 | 42.949 | 39.871 | 49.986 | 42.322 | 37.198 | 48.853 | 41.430 | 44.664 | 52.466 | 48.354 | 40.683 | 47.180 | 52.670 | 45.605 | 50.178 | 48.151 | -0.139 | 0.0034 | 0.0275 | 1 | 0 | 0 |
| 'Lage3' | 6.620 | 7.120 | 6.484 | 6.427 | 6.736 | 7.019 | 6.780 | 7.309 | 6.543 | 6.016 | 6.606 | 6.783 | 6.977 | 6.229 | 7.312 | 5.750 | 6.902 | 5.814 | 0.049 | 0.3375 | 0.5955 | 0 | 0 | 0 |
| 'Lamc1' | 2.198 | 1.784 | 2.500 | 2.422 | 2.036 | 2.428 | 2.316 | 1.879 | 2.438 | 2.222 | 2.045 | 2.250 | 2.280 | 1.732 | 1.901 | 2.429 | 1.733 | 2.226 | 0.081 | 0.1201 | 0.3301 | 0 | 0 | 0 |
| 'Lamp1' | 103.375 | 113.907 | 107.222 | 106.266 | 117.838 | 115.206 | 105.754 | 115.145 | 107.209 | 120.685 | 107.849 | 107.488 | 119.766 | 118.621 | 111.321 | 114.830 | 111.826 | 109.813 | -0.054 | 0.0728 | 0.2387 | 0 | 0 | 0 |
| 'Lamp2' | 8.296 | 9.351 | 8.753 | 8.304 | 9.434 | 8.448 | 9.409 | 9.782 | 8.456 | 9.513 | 10.667 | 10.728 | 9.198 | 11.160 | 10.368 | 9.385 | 11.113 | 11.136 | -0.229 | 0.0000 | 0.0000 | 1 | 0 | 0 |
| 'Lamtor1' | 47.789 | 61.798 | 46.087 | 42.438 | 58.040 | 56.227 | 40.915 | 54.660 | 55.156 | 47.717 | 44.310 | 51.696 | 50.457 | 49.021 | 55.784 | 57.073 | 44.043 | 50.667 | 0.022 | 0.7544 | 0.8906 | 0 | 0 | 0 |
| 'Lamtor2' | 21.194 | 24.517 | 21.045 | 21.788 | 23.572 | 21.769 | 22.491 | 21.282 | 21.717 | 18.978 | 18.762 | 20.223 | 20.304 | 20.816 | 20.030 | 19.292 | 20.914 | 18.923 | 0.145 | 0.0000 | 0.0004 | 1 | 0 | 0 |
| 'Lamtor4' | 27.198 | 36.249 | 34.992 | 28.138 | 36.468 | 36.913 | 30.743 | 34.898 | 34.838 | 32.277 | 32.857 | 31.412 | 33.383 | 32.766 | 33.254 | 31.519 | 32.803 | 31.480 | 0.028 | 0.5207 | 0.7435 | 0 | 0 | 0 |
| 'Lamtor5' | 27.589 | 31.024 | 28.148 | 27.522 | 31.697 | 27.525 | 31.098 | 30.952 | 28.839 | 26.258 | 27.970 | 27.571 | 29.402 | 29.873 | 28.643 | 28.172 | 28.429 | 26.990 | 0.047 | 0.1934 | 0.4308 | 0 | 0 | 0 |
| 'Lancl1' | 6.716 | 7.973 | 6.851 | 6.537 | 7.585 | 7.180 | 6.949 | 7.967 | 6.946 | 6.551 | 5.866 | 6.424 | 6.395 | 5.896 | 6.002 | 6.128 | 6.242 | 5.725 | 0.212 | 0.0000 | 0.0000 | 1 | 0 | 0 |
| 'Lancl2' | 2.234 | 3.228 | 2.881 | 2.771 | 2.686 | 3.197 | 2.953 | 2.770 | 2.652 | 2.389 | 2.913 | 2.817 | 3.077 | 2.847 | 2.638 | 2.952 | 3.565 | 2.252 | -0.013 | 0.8681 | 0.9429 | 0 | 0 | 0 |
| 'Lancl3' | 1.133 | 1.219 | 1.280 | 0.922 | 0.899 | 0.953 | 0.960 | 1.000 | 1.070 | 1.822 | 1.973 | 1.593 | 2.133 | 1.897 | 1.439 | 2.188 | 1.956 | 1.793 | -0.822 | 0.0000 | 0.0000 | 1 | 1 | 0 |
| 'Lap3' | 4.316 | 4.506 | 4.894 | 4.593 | 4.581 | 4.312 | 4.735 | 5.187 | 4.167 | 4.706 | 3.885 | 4.373 | 4.797 | 4.312 | 4.597 | 4.606 | 4.147 | 4.203 | 0.050 | 0.3396 | 0.5969 | 0 | 0 | 0 |
| 'Laptm4a' | 51.283 | 65.312 | 57.223 | 53.920 | 64.457 | 56.238 | 57.700 | 64.253 | 58.098 | 56.862 | 54.397 | 57.561 | 51.924 | 62.213 | 54.852 | 60.191 | 58.023 | 55.071 | 0.034 | 0.3412 | 0.5981 | 0 | 0 | 0 |
| 'Laptm5' | 334.125 | 339.726 | 340.026 | 328.741 | 342.055 | 357.416 | 329.530 | 328.215 | 353.059 | 372.055 | 341.025 | 314.393 | 322.661 | 332.960 | 347.526 | 352.297 | 327.160 | 354.055 | -0.017 | 0.6166 | 0.8086 | 0 | 0 | 0 |
| 'Larp1' | 29.804 | 25.138 | 28.197 | 30.007 | 23.651 | 27.185 | 28.074 | 25.156 | 26.995 | 27.608 | 26.480 | 30.634 | 27.556 | 25.301 | 29.344 | 26.397 | 25.570 | 29.530 | -0.037 | 0.2886 | 0.5451 | 0 | 0 | 0 |
| 'Larp1b' | 3.871 | 3.745 | 3.627 | 3.748 | 3.692 | 4.177 | 4.095 | 3.503 | 3.711 | 3.850 | 3.566 | 4.038 | 4.035 | 3.683 | 3.873 | 3.730 | 4.463 | 3.700 | -0.045 | 0.3102 | 0.5686 | 0 | 0 | 0 |
| 'Larp4' | 11.813 | 11.744 | 11.469 | 12.255 | 10.972 | 10.017 | 12.557 | 11.342 | 10.784 | 11.006 | 11.634 | 12.150 | 10.618 | 11.488 | 11.677 | 11.128 | 12.594 | 11.861 | -0.030 | 0.4980 | 0.7259 | 0 | 0 | 0 |
| 'Larp4b' | 9.368 | 9.905 | 10.222 | 10.442 | 10.175 | 9.677 | 10.353 | 9.943 | 10.201 | 9.122 | 10.605 | 10.574 | 9.550 | 9.924 | 10.178 | 9.610 | 10.386 | 10.398 | -0.012 | 0.7015 | 0.8608 | 0 | 0 | 0 |
| 'Lars' | 28.227 | 30.255 | 30.717 | 29.068 | 30.735 | 29.474 | 30.823 | 32.913 | 29.206 | 31.053 | 28.217 | 28.031 | 30.682 | 28.339 | 26.249 | 31.570 | 29.535 | 26.284 | 0.055 | 0.0960 | 0.2881 | 0 | 0 | 0 |
| 'Lars2' | 7.827 | 7.869 | 9.458 | 9.143 | 8.777 | 9.464 | 9.623 | 8.495 | 9.441 | 8.819 | 9.294 | 8.921 | 9.362 | 8.655 | 8.881 | 8.116 | 8.733 | 8.597 | 0.003 | 0.9447 | 0.9789 | 0 | 0 | 0 |
| 'Las1l' | 4.852 | 5.022 | 4.681 | 4.674 | 4.532 | 4.623 | 4.758 | 5.100 | 4.753 | 5.119 | 4.619 | 4.788 | 4.671 | 4.446 | 4.337 | 4.942 | 4.560 | 4.402 | 0.028 | 0.4186 | 0.6658 | 0 | 0 | 0 |
| 'Lasp1' | 55.953 | 53.262 | 54.811 | 51.761 | 50.223 | 58.015 | 49.832 | 51.238 | 55.031 | 51.627 | 49.397 | 57.100 | 50.713 | 49.363 | 58.765 | 52.437 | 46.755 | 59.751 | 0.005 | 0.8671 | 0.9426 | 0 | 0 | 0 |
| 'Lat' | 149.425 | 155.557 | 155.672 | 152.116 | 157.462 | 156.551 | 149.126 | 153.943 | 155.483 | 148.461 | 140.922 | 155.168 | 159.121 | 143.158 | 161.857 | 157.556 | 147.338 | 159.968 | 0.002 | 0.9202 | 0.9700 | 0 | 0 | 0 |
| 'Lats1' | 6.406 | 7.387 | 6.707 | 7.255 | 7.315 | 6.996 | 7.510 | 7.269 | 6.981 | 6.211 | 7.217 | 7.518 | 6.603 | 6.578 | 6.899 | 7.447 | 6.547 | 7.109 | 0.025 | 0.5556 | 0.7677 | 0 | 0 | 0 |
| 'Lats2' | 5.226 | 4.763 | 4.704 | 4.762 | 5.182 | 5.119 | 4.814 | 5.011 | 4.606 | 5.129 | 5.712 | 5.424 | 5.063 | 5.752 | 5.136 | 5.444 | 5.210 | 5.552 | -0.141 | 0.0002 | 0.0023 | 1 | 0 | 0 |
| 'Lax1' | 119.526 | 126.888 | 114.112 | 118.065 | 119.584 | 114.135 | 112.265 | 116.867 | 118.570 | 106.001 | 129.229 | 147.767 | 101.286 | 125.655 | 142.981 | 112.235 | 138.812 | 153.662 | -0.126 | 0.0239 | 0.1170 | 0 | 0 | 0 |
| 'Lbh' | 143.332 | 173.463 | 138.672 | 133.831 | 167.828 | 156.773 | 131.347 | 160.303 | 151.953 | 163.274 | 166.760 | 157.011 | 152.315 | 185.300 | 170.657 | 173.872 | 163.983 | 166.909 | -0.159 | 0.0001 | 0.0013 | 1 | 0 | 0 |
| 'Lbhd1' | 1.274 | 1.107 | 1.229 | 1.481 | 1.543 | 1.319 | 1.684 | 1.297 | 1.293 | 1.199 | 0.892 | 1.558 | 0.985 | 1.377 | 1.421 | 0.957 | 1.483 | 1.607 | 0.073 | 0.5347 | 0.7544 | 0 | 0 | 0 |
| 'Lbr' | 34.425 | 36.402 | 37.597 | 37.726 | 35.997 | 36.857 | 37.181 | 36.048 | 36.737 | 32.659 | 38.209 | 39.049 | 35.004 | 38.933 | 39.570 | 33.828 | 37.877 | 39.158 | -0.032 | 0.3180 | 0.5764 | 0 | 0 | 0 |
| 'Lca5' | 0.345 | 0.467 | 0.496 | 0.660 | 0.504 | 0.517 | 0.456 | 0.402 | 0.482 | 0.561 | 0.502 | 0.472 | 0.567 | 0.485 | 0.441 | 0.447 | 0.550 | 0.429 | -0.053 | 0.6397 | 0.8228 | 0 | 0 | 0 |
| 'Lck' | 235.704 | 258.952 | 237.220 | 241.498 | 254.338 | 258.193 | 235.383 | 242.474 | 253.768 | 262.186 | 249.237 | 238.833 | 262.233 | 260.032 | 260.121 | 272.175 | 256.724 | 256.671 | -0.075 | 0.0089 | 0.0574 | 0 | 0 | 0 |
| 'Lclat1' | 4.771 | 5.574 | 5.342 | 5.332 | 5.500 | 5.636 | 5.624 | 4.948 | 5.539 | 5.541 | 5.250 | 5.944 | 5.365 | 5.977 | 5.424 | 5.121 | 5.810 | 5.348 | -0.060 | 0.1592 | 0.3895 | 0 | 0 | 0 |
| 'Lcmt1' | 4.362 | 4.278 | 4.726 | 4.896 | 4.800 | 4.812 | 4.979 | 4.661 | 4.749 | 4.102 | 4.644 | 4.391 | 4.316 | 4.619 | 4.255 | 4.593 | 4.771 | 3.910 | 0.088 | 0.0679 | 0.2284 | 0 | 0 | 0 |
| 'Lcmt2' | 1.966 | 1.891 | 2.267 | 1.471 | 1.968 | 2.540 | 1.847 | 2.113 | 2.634 | 2.072 | 2.222 | 1.651 | 1.807 | 1.800 | 1.812 | 1.772 | 1.448 | 1.464 | 0.215 | 0.0333 | 0.1447 | 0 | 0 | 0 |
| 'Lcor' | 2.051 | 1.965 | 1.770 | 1.938 | 1.889 | 1.449 | 2.158 | 1.814 | 1.647 | 1.933 | 1.887 | 1.749 | 1.795 | 1.611 | 1.593 | 1.853 | 1.806 | 1.825 | 0.035 | 0.5535 | 0.7663 | 0 | 0 | 0 |
| 'Lcorl' | 0.390 | 0.321 | 0.374 | 0.358 | 0.349 | 0.359 | 0.494 | 0.408 | 0.356 | 0.431 | 0.440 | 0.409 | 0.411 | 0.441 | 0.400 | 0.348 | 0.364 | 0.404 | -0.110 | 0.1540 | 0.3819 | 0 | 0 | 0 |
| 'Lcp1' | 121.069 | 117.441 | 125.337 | 132.138 | 121.801 | 121.530 | 136.149 | 115.887 | 124.713 | 134.896 | 133.707 | 118.695 | 129.663 | 132.103 | 131.155 | 123.440 | 129.085 | 126.372 | -0.066 | 0.0500 | 0.1886 | 0 | 0 | 0 |
| 'Lcp2' | 83.435 | 82.762 | 83.300 | 84.887 | 83.209 | 82.207 | 84.303 | 84.554 | 84.254 | 82.816 | 80.924 | 80.053 | 80.913 | 78.846 | 82.768 | 79.270 | 77.276 | 82.262 | 0.044 | 0.0191 | 0.1002 | 0 | 0 | 0 |
| 'Ldah' | 30.661 | 30.613 | 29.222 | 28.856 | 30.156 | 30.132 | 32.285 | 30.286 | 27.054 | 31.800 | 35.025 | 31.864 | 29.729 | 31.999 | 28.586 | 31.121 | 33.305 | 29.937 | -0.084 | 0.0459 | 0.1783 | 0 | 0 | 0 |
| 'Ldb1' | 59.216 | 54.809 | 61.590 | 56.514 | 53.835 | 61.814 | 58.698 | 55.517 | 60.502 | 56.587 | 53.224 | 65.562 | 58.262 | 54.414 | 63.491 | 56.825 | 54.739 | 67.408 | -0.031 | 0.1945 | 0.4326 | 0 | 0 | 0 |
| 'Ldha' | 127.110 | 129.444 | 118.279 | 130.188 | 132.393 | 118.101 | 131.283 | 126.382 | 117.852 | 111.233 | 131.965 | 119.699 | 129.354 | 133.121 | 136.074 | 121.856 | 130.438 | 127.577 | -0.025 | 0.5078 | 0.7336 | 0 | 0 | 0 |
| 'Ldhb' | 15.366 | 18.357 | 15.342 | 18.019 | 19.566 | 16.781 | 18.524 | 20.357 | 18.038 | 17.567 | 18.300 | 16.225 | 18.886 | 15.904 | 14.955 | 21.012 | 16.345 | 15.435 | 0.043 | 0.4469 | 0.6880 | 0 | 0 | 0 |
| 'Ldhd' | 1.260 | 1.648 | 1.427 | 1.646 | 1.679 | 1.415 | 1.441 | 1.544 | 1.536 | 1.034 | 0.954 | 0.812 | 0.895 | 0.945 | 0.696 | 0.889 | 1.020 | 0.808 | 0.750 | 0.0000 | 0.0000 | 1 | 1 | 0 |
| 'Ldlr' | 4.134 | 4.883 | 6.279 | 4.321 | 4.908 | 6.128 | 4.228 | 3.974 | 6.917 | 5.440 | 7.524 | 4.739 | 6.227 | 6.244 | 6.200 | 6.002 | 4.646 | 4.335 | -0.193 | 0.1474 | 0.3732 | 0 | 0 | 0 |
| 'Ldlrad4' | 27.633 | 24.987 | 22.867 | 25.004 | 25.036 | 23.776 | 27.388 | 23.962 | 22.625 | 27.689 | 29.718 | 27.629 | 28.019 | 29.005 | 26.890 | 28.732 | 31.788 | 28.081 | -0.218 | 0.0000 | 0.0000 | 1 | 0 | 0 |
| 'Ldlrap1' | 14.861 | 17.265 | 13.438 | 14.186 | 16.009 | 15.943 | 13.645 | 16.528 | 13.853 | 15.278 | 16.256 | 16.919 | 16.487 | 16.590 | 15.324 | 16.973 | 16.759 | 16.900 | -0.135 | 0.0008 | 0.0089 | 1 | 0 | 0 |
| 'Ldoc1l' | 5.496 | 5.963 | 6.683 | 4.461 | 5.874 | 5.850 | 5.183 | 5.512 | 6.294 | 5.804 | 5.663 | 4.982 | 5.493 | 4.671 | 5.522 | 4.817 | 4.982 | 4.845 | 0.125 | 0.0526 | 0.1947 | 0 | 0 | 0 |
| 'Lef1' | 12.477 | 11.168 | 13.147 | 12.461 | 11.271 | 12.798 | 13.298 | 11.463 | 13.288 | 11.829 | 12.355 | 13.859 | 9.805 | 10.887 | 13.913 | 10.529 | 11.985 | 14.346 | 0.019 | 0.7196 | 0.8711 | 0 | 0 | 0 |
| 'Lemd2' | 46.837 | 44.382 | 54.880 | 41.577 | 42.049 | 53.585 | 42.051 | 44.918 | 44.947 | 46.818 | 41.019 | 48.069 | 46.475 | 43.894 | 51.179 | 49.125 | 40.395 | 52.705 | -0.024 | 0.5781 | 0.7820 | 0 | 0 | 0 |
| 'Lemd3' | 10.504 | 10.623 | 11.151 | 10.101 | 10.430 | 10.037 | 10.047 | 11.645 | 10.057 | 10.690 | 10.346 | 10.245 | 9.835 | 10.029 | 10.194 | 11.090 | 10.293 | 10.370 | 0.015 | 0.6628 | 0.8371 | 0 | 0 | 0 |
| 'Leng1' | 9.598 | 8.826 | 9.057 | 10.349 | 7.607 | 8.578 | 9.362 | 8.078 | 9.209 | 8.321 | 7.986 | 10.283 | 8.856 | 8.504 | 9.852 | 9.254 | 8.483 | 8.647 | -0.010 | 0.8610 | 0.9399 | 0 | 0 | 0 |
| 'Leng8' | 15.138 | 16.538 | 19.787 | 12.432 | 15.588 | 17.886 | 14.292 | 16.037 | 20.264 | 18.606 | 17.219 | 17.194 | 18.182 | 15.948 | 16.908 | 15.750 | 16.375 | 16.522 | -0.067 | 0.2637 | 0.5178 | 0 | 0 | 0 |
| 'Leng9' | 8.180 | 7.973 | 8.219 | 8.826 | 6.970 | 8.711 | 8.130 | 7.649 | 7.813 | 7.967 | 7.393 | 6.738 | 7.254 | 6.900 | 6.889 | 8.471 | 7.026 | 6.456 | 0.157 | 0.0039 | 0.0309 | 1 | 0 | 0 |
| 'Leo1' | 17.116 | 15.641 | 17.330 | 17.425 | 17.051 | 17.686 | 18.354 | 16.996 | 17.021 | 15.516 | 15.696 | 17.161 | 15.709 | 15.086 | 15.488 | 16.112 | 15.453 | 15.587 | 0.111 | 0.0029 | 0.0248 | 1 | 0 | 0 |
| 'Leprot' | 43.633 | 55.830 | 48.724 | 45.925 | 55.005 | 49.070 | 50.195 | 55.082 | 48.406 | 50.476 | 51.260 | 47.307 | 53.309 | 57.249 | 48.200 | 54.404 | 52.719 | 44.349 | -0.034 | 0.3576 | 0.6103 | 0 | 0 | 0 |
| 'Leprotl1' | 50.070 | 56.434 | 51.551 | 50.217 | 54.853 | 52.366 | 50.619 | 55.126 | 50.760 | 48.187 | 52.741 | 53.739 | 46.837 | 51.540 | 51.328 | 49.917 | 52.303 | 51.267 | 0.031 | 0.1853 | 0.4208 | 0 | 0 | 0 |
| 'Letm1' | 10.814 | 10.690 | 11.336 | 10.742 | 9.950 | 11.760 | 10.834 | 10.743 | 11.332 | 11.115 | 10.684 | 11.121 | 11.072 | 10.082 | 11.066 | 10.970 | 10.217 | 11.155 | 0.002 | 0.9280 | 0.9723 | 0 | 0 | 0 |
| 'Letm2' | 1.962 | 2.094 | 2.583 | 2.245 | 2.051 | 1.730 | 2.520 | 2.467 | 2.215 | 2.086 | 2.514 | 2.381 | 1.925 | 2.557 | 2.355 | 2.191 | 2.392 | 2.042 | -0.048 | 0.6383 | 0.8220 | 0 | 0 | 0 |
| 'Letmd1' | 19.142 | 18.360 | 20.553 | 18.164 | 19.429 | 20.153 | 19.299 | 21.056 | 20.466 | 19.491 | 18.303 | 20.176 | 19.537 | 17.026 | 19.697 | 19.145 | 19.308 | 18.905 | 0.033 | 0.3521 | 0.6059 | 0 | 0 | 0 |
| 'Lfng' | 10.242 | 11.869 | 12.507 | 9.714 | 11.644 | 12.921 | 10.115 | 12.985 | 12.341 | 9.056 | 8.721 | 15.427 | 8.361 | 8.910 | 14.088 | 11.759 | 8.047 | 13.978 | 0.099 | 0.2206 | 0.4680 | 0 | 0 | 0 |
| 'Lgals1' | 256.089 | 283.026 | 245.824 | 293.740 | 290.995 | 263.678 | 271.369 | 267.954 | 256.892 | 249.264 | 291.402 | 219.124 | 343.389 | 299.640 | 270.416 | 267.081 | 305.451 | 250.110 | -0.046 | 0.3972 | 0.6462 | 0 | 0 | 0 |
| 'Lgals3' | 3.970 | 4.358 | 3.138 | 3.667 | 4.346 | 3.778 | 3.488 | 5.314 | 3.084 | 4.200 | 6.659 | 3.780 | 6.181 | 5.873 | 4.221 | 4.037 | 6.520 | 4.344 | -0.382 | 0.0000 | 0.0001 | 1 | 1 | 0 |
| 'Lgals3bp' | 26.344 | 25.706 | 28.543 | 21.420 | 27.844 | 30.936 | 23.854 | 31.272 | 29.577 | 29.707 | 29.108 | 36.732 | 30.797 | 27.704 | 37.342 | 32.504 | 26.053 | 36.955 | -0.231 | 0.0000 | 0.0007 | 1 | 0 | 0 |
| 'Lgals4' | 1.865 | 1.623 | 2.285 | 2.449 | 1.936 | 2.319 | 2.275 | 1.855 | 1.977 | 1.742 | 1.593 | 1.963 | 1.767 | 1.818 | 2.015 | 1.585 | 1.802 | 1.962 | 0.180 | 0.0192 | 0.1004 | 0 | 0 | 0 |
| 'Lgals7' | 13.216 | 11.694 | 12.493 | 11.608 | 15.841 | 10.584 | 12.268 | 13.776 | 8.767 | 8.870 | 10.183 | 9.233 | 10.674 | 10.953 | 10.555 | 10.090 | 12.643 | 9.106 | 0.243 | 0.0014 | 0.0135 | 1 | 0 | 0 |
| 'Lgals8' | 10.797 | 11.467 | 10.669 | 9.918 | 12.097 | 11.147 | 10.202 | 11.583 | 12.278 | 10.917 | 11.597 | 12.746 | 10.818 | 11.147 | 11.173 | 11.260 | 12.161 | 12.100 | -0.067 | 0.0646 | 0.2217 | 0 | 0 | 0 |
| 'Lgals9' | 34.333 | 35.721 | 35.743 | 32.346 | 36.020 | 35.307 | 35.187 | 37.785 | 35.672 | 33.974 | 32.327 | 36.928 | 36.782 | 33.437 | 37.605 | 35.383 | 33.998 | 35.773 | -0.003 | 0.9217 | 0.9705 | 0 | 0 | 0 |
| 'Lgalsl' | 11.924 | 13.475 | 12.377 | 12.320 | 14.041 | 11.137 | 13.328 | 14.237 | 12.281 | 12.074 | 13.912 | 10.521 | 10.475 | 13.815 | 11.004 | 10.502 | 13.890 | 10.515 | 0.106 | 0.0096 | 0.0608 | 0 | 0 | 0 |
| 'Lgmn' | 6.987 | 7.763 | 6.825 | 7.711 | 8.519 | 6.957 | 8.029 | 7.673 | 6.106 | 6.272 | 7.905 | 6.006 | 7.290 | 6.031 | 5.433 | 6.354 | 6.775 | 4.972 | 0.211 | 0.0002 | 0.0026 | 1 | 0 | 0 |
| 'Lias' | 7.054 | 7.983 | 8.447 | 7.403 | 9.532 | 8.004 | 8.765 | 8.172 | 7.596 | 8.050 | 8.140 | 6.769 | 8.412 | 8.028 | 7.175 | 7.760 | 7.646 | 6.611 | 0.083 | 0.1176 | 0.3263 | 0 | 0 | 0 |
| 'Lif' | 5.695 | 2.512 | 2.322 | 4.091 | 3.461 | 2.118 | 4.475 | 4.490 | 1.698 | 4.152 | 3.781 | 2.128 | 4.934 | 3.990 | 2.348 | 4.259 | 3.332 | 2.000 | -0.023 | 0.8104 | 0.9142 | 0 | 0 | 0 |
| 'Lig1' | 21.585 | 20.352 | 23.538 | 23.675 | 20.786 | 23.399 | 24.288 | 20.074 | 25.677 | 21.418 | 21.186 | 22.267 | 28.946 | 22.754 | 23.711 | 22.625 | 21.484 | 21.812 | -0.032 | 0.5093 | 0.7342 | 0 | 0 | 0 |
| 'Lig3' | 4.100 | 3.868 | 4.104 | 3.945 | 3.775 | 3.863 | 4.016 | 3.706 | 3.801 | 3.518 | 3.710 | 3.836 | 3.448 | 3.611 | 3.727 | 3.505 | 3.645 | 3.574 | 0.100 | 0.0022 | 0.0193 | 1 | 0 | 0 |
| 'Lig4' | 4.745 | 4.695 | 4.220 | 5.530 | 4.782 | 4.633 | 5.065 | 4.884 | 4.475 | 3.852 | 4.645 | 4.326 | 4.131 | 4.837 | 3.965 | 4.030 | 4.510 | 4.195 | 0.147 | 0.0033 | 0.0271 | 1 | 0 | 0 |
| 'Lima1' | 0.793 | 0.943 | 1.015 | 0.864 | 0.922 | 0.788 | 0.857 | 1.043 | 0.807 | 0.763 | 0.696 | 0.704 | 0.852 | 1.025 | 0.635 | 0.667 | 0.799 | 0.642 | 0.239 | 0.0102 | 0.0633 | 0 | 0 | 0 |
| 'Limd1' | 22.619 | 21.459 | 23.088 | 21.359 | 20.332 | 23.764 | 20.232 | 21.630 | 22.789 | 21.032 | 21.079 | 25.121 | 19.282 | 22.222 | 23.828 | 20.101 | 19.998 | 25.399 | -0.013 | 0.7215 | 0.8716 | 0 | 0 | 0 |
| 'Limd2' | 139.495 | 166.427 | 152.514 | 145.594 | 163.004 | 174.296 | 155.511 | 156.595 | 164.540 | 158.502 | 142.612 | 149.702 | 149.737 | 154.507 | 167.195 | 173.111 | 143.691 | 154.999 | 0.013 | 0.7568 | 0.8913 | 0 | 0 | 0 |
| 'Lime1' | 1.672 | 1.693 | 1.899 | 1.451 | 1.668 | 2.057 | 1.639 | 1.675 | 1.741 | 1.866 | 1.845 | 2.046 | 1.847 | 1.859 | 1.955 | 1.866 | 1.542 | 2.070 | -0.131 | 0.0546 | 0.2000 | 0 | 0 | 0 |
| 'Limk1' | 5.794 | 5.550 | 6.459 | 5.152 | 5.616 | 6.781 | 5.415 | 5.498 | 6.729 | 5.990 | 4.718 | 5.723 | 4.979 | 4.705 | 5.632 | 5.376 | 4.291 | 5.816 | 0.158 | 0.0003 | 0.0041 | 1 | 0 | 0 |
| 'Limk2' | 10.647 | 9.770 | 10.997 | 10.696 | 9.883 | 11.054 | 10.401 | 10.365 | 11.561 | 10.596 | 9.832 | 11.998 | 10.819 | 9.773 | 10.958 | 9.574 | 10.517 | 11.468 | -0.014 | 0.6301 | 0.8169 | 0 | 0 | 0 |
| 'Lims1' | 36.116 | 38.839 | 35.501 | 37.914 | 37.601 | 33.378 | 40.923 | 36.477 | 35.062 | 32.916 | 40.918 | 41.317 | 33.748 | 40.179 | 39.767 | 34.145 | 44.012 | 40.625 | -0.074 | 0.1477 | 0.3736 | 0 | 0 | 0 |
| 'Lin37' | 10.135 | 10.527 | 12.189 | 10.520 | 9.929 | 11.921 | 11.776 | 10.457 | 11.407 | 10.284 | 9.822 | 12.295 | 10.318 | 10.147 | 12.280 | 9.632 | 11.481 | 11.711 | 0.000 | 0.9956 | 0.9994 | 0 | 0 | 0 |
| 'Lin52' | 2.048 | 1.784 | 1.765 | 1.673 | 2.326 | 2.062 | 1.965 | 2.139 | 2.005 | 1.955 | 2.095 | 2.356 | 1.505 | 2.737 | 1.890 | 1.816 | 1.888 | 1.718 | -0.028 | 0.8006 | 0.9111 | 0 | 0 | 0 |
| 'Lin54' | 1.350 | 1.248 | 1.396 | 1.429 | 1.363 | 1.249 | 1.325 | 1.147 | 1.249 | 1.203 | 1.382 | 1.415 | 1.237 | 1.214 | 1.367 | 1.376 | 1.411 | 1.292 | -0.030 | 0.5745 | 0.7800 | 0 | 0 | 0 |
| 'Lin7c' | 14.065 | 17.846 | 15.480 | 15.009 | 17.560 | 14.795 | 16.661 | 18.553 | 15.033 | 16.389 | 15.391 | 14.772 | 15.157 | 17.958 | 13.529 | 15.532 | 15.518 | 13.691 | 0.059 | 0.1287 | 0.3438 | 0 | 0 | 0 |
| 'Lin9' | 2.436 | 2.395 | 2.153 | 2.354 | 2.262 | 2.149 | 2.410 | 2.277 | 2.203 | 2.248 | 2.264 | 2.124 | 2.431 | 2.538 | 2.183 | 2.132 | 2.192 | 2.134 | 0.016 | 0.6989 | 0.8590 | 0 | 0 | 0 |
| 'Lins1' | 0.639 | 0.465 | 0.554 | 0.552 | 0.518 | 0.590 | 0.467 | 0.570 | 0.492 | 0.542 | 0.581 | 0.655 | 0.537 | 0.588 | 0.607 | 0.647 | 0.550 | 0.593 | -0.134 | 0.0705 | 0.2340 | 0 | 0 | 0 |

| | | | | | | | | | | | | | | | | | | | | | | | | |
|---|---|---|---|---|---|---|---|---|---|---|---|---|---|---|---|---|---|---|---|---|---|---|---|---|
| 'Lipa' | 28.493 | 29.628 | 31.598 | 30.911 | 32.749 | 32.377 | 33.484 | 31.042 | 30.187 | 32.609 | 34.863 | 35.071 | 30.566 | 33.331 | 33.616 | 34.199 | 34.633 | 33.943 | -0.121 | 0.0003 | 0.0040 | 1 | 0 | 0 |
| 'Lipe' | 2.179 | 2.454 | 2.976 | 2.272 | 2.411 | 2.967 | 2.460 | 2.652 | 2.926 | 2.456 | 2.194 | 2.495 | 2.266 | 2.118 | 2.229 | 2.805 | 2.247 | 2.091 | 0.150 | 0.0231 | 0.1141 | 0 | 0 | 0 |
| 'Liph' | 1.009 | 1.427 | 1.360 | 1.048 | 1.437 | 1.078 | 0.950 | 1.223 | 1.341 | 1.438 | 2.034 | 1.417 | 1.899 | 2.073 | 0.878 | 1.434 | 1.782 | 1.639 | -0.425 | 0.0005 | 0.0056 | 1 | 1 | 0 |
| 'Litaf' | 6.522 | 7.644 | 6.971 | 6.248 | 6.630 | 6.533 | 6.242 | 7.516 | 6.946 | 6.456 | 8.065 | 7.311 | 6.924 | 8.127 | 8.314 | 6.312 | 7.621 | 7.659 | -0.133 | 0.0070 | 0.0480 | 1 | 0 | 0 |
| 'Lix1l' | 1.281 | 1.733 | 1.852 | 1.495 | 1.986 | 1.712 | 1.401 | 1.684 | 1.563 | 1.488 | 1.300 | 1.384 | 1.303 | 1.530 | 1.459 | 1.167 | 1.203 | 1.340 | 0.258 | 0.0040 | 0.0312 | 1 | 0 | 0 |
| 'Llgl1' | 15.517 | 14.624 | 16.802 | 13.593 | 14.962 | 16.645 | 13.405 | 14.945 | 16.126 | 15.256 | 14.199 | 15.836 | 16.065 | 15.451 | 15.843 | 14.505 | 13.456 | 15.242 | -0.002 | 0.9581 | 0.9847 | 0 | 0 | 0 |
| 'Llgl2' | 7.319 | 6.516 | 7.725 | 6.625 | 5.940 | 7.686 | 6.209 | 6.696 | 7.300 | 7.085 | 7.356 | 7.972 | 6.722 | 6.429 | 7.323 | 6.750 | 6.953 | 7.897 | -0.065 | 0.1141 | 0.3201 | 0 | 0 | 0 |
| 'Llph' | 49.276 | 48.424 | 46.873 | 50.003 | 52.449 | 46.965 | 49.278 | 49.167 | 47.122 | 42.792 | 40.157 | 44.256 | 44.388 | 41.912 | 42.987 | 47.359 | 43.419 | 40.944 | 0.165 | 0.0000 | 0.0000 | 1 | 0 | 0 |
| 'Lman1' | 7.042 | 7.790 | 7.544 | 7.299 | 7.858 | 7.414 | 7.463 | 7.754 | 7.927 | 7.176 | 7.443 | 7.873 | 7.927 | 7.481 | 7.236 | 7.388 | 7.585 | 7.321 | 0.000 | 0.9890 | 0.9968 | 0 | 0 | 0 |
| 'Lman2' | 49.777 | 53.000 | 53.534 | 47.501 | 55.164 | 54.426 | 48.873 | 52.602 | 52.679 | 51.395 | 50.121 | 54.489 | 51.283 | 52.539 | 54.941 | 53.068 | 50.202 | 55.414 | -0.029 | 0.2253 | 0.4720 | 0 | 0 | 0 |
| 'Lman2l' | 2.526 | 2.294 | 2.769 | 2.660 | 2.345 | 2.673 | 2.300 | 2.293 | 2.925 | 2.823 | 2.567 | 3.767 | 2.561 | 2.516 | 3.057 | 2.451 | 2.639 | 4.344 | -0.231 | 0.0006 | 0.0074 | 1 | 0 | 0 |
| 'Lmbr1' | 0.539 | 0.701 | 0.485 | 0.598 | 0.604 | 0.489 | 0.724 | 0.593 | 0.566 | 0.338 | 0.616 | 0.483 | 0.403 | 0.478 | 0.496 | 0.481 | 0.582 | 0.490 | 0.261 | 0.0070 | 0.0479 | 1 | 0 | 0 |
| 'Lmbr1l' | 17.333 | 16.609 | 17.731 | 15.848 | 16.402 | 16.490 | 15.451 | 16.716 | 17.119 | 17.143 | 17.646 | 18.164 | 17.680 | 17.510 | 17.983 | 16.826 | 18.608 | 16.399 | -0.086 | 0.0066 | 0.0460 | 1 | 0 | 0 |
| 'Lmbrd1' | 8.249 | 8.240 | 7.654 | 9.116 | 8.033 | 6.893 | 9.858 | 8.162 | 7.589 | 7.483 | 9.501 | 8.395 | 7.075 | 8.644 | 7.032 | 7.347 | 9.299 | 8.022 | 0.008 | 0.9022 | 0.9612 | 0 | 0 | 0 |
| 'Lmbrd2' | 6.320 | 5.727 | 5.692 | 6.039 | 5.460 | 5.637 | 6.705 | 5.652 | 5.886 | 5.823 | 6.135 | 6.322 | 6.399 | 5.288 | 5.799 | 6.346 | 5.665 | 5.639 | -0.024 | 0.6084 | 0.8029 | 0 | 0 | 0 |
| 'Lmf1' | 1.393 | 1.607 | 1.561 | 1.447 | 1.550 | 1.519 | 1.520 | 1.476 | 1.719 | 1.522 | 1.372 | 1.497 | 1.484 | 1.345 | 1.410 | 1.682 | 1.526 | 1.356 | 0.055 | 0.2716 | 0.5264 | 0 | 0 | 0 |
| 'Lmf2' | 28.236 | 25.459 | 34.141 | 26.237 | 26.687 | 31.576 | 26.607 | 26.456 | 32.615 | 25.400 | 25.033 | 31.256 | 29.464 | 27.033 | 29.572 | 26.415 | 25.615 | 29.195 | 0.040 | 0.2452 | 0.4957 | 0 | 0 | 0 |
| 'Lmln' | 0.859 | 1.055 | 0.804 | 0.742 | 0.893 | 0.796 | 0.668 | 0.631 | 0.970 | 0.889 | 0.931 | 0.823 | 0.944 | 0.741 | 0.872 | 0.896 | 0.651 | 0.832 | -0.043 | 0.6706 | 0.8413 | 0 | 0 | 0 |
| 'Lmnb1' | 64.081 | 63.198 | 72.926 | 68.678 | 63.177 | 75.203 | 66.631 | 61.874 | 75.221 | 55.442 | 62.769 | 68.522 | 73.925 | 67.695 | 73.125 | 64.075 | 66.275 | 68.160 | 0.013 | 0.7514 | 0.8894 | 0 | 0 | 0 |
| 'Lmnb2' | 10.542 | 12.079 | 11.197 | 11.138 | 11.478 | 12.057 | 11.964 | 11.060 | 11.904 | 10.640 | 11.258 | 11.607 | 13.295 | 11.439 | 11.002 | 11.168 | 12.821 | 10.886 | -0.021 | 0.6189 | 0.8093 | 0 | 0 | 0 |
| 'Lmntd2' | 6.204 | 5.618 | 7.112 | 5.642 | 6.044 | 6.211 | 5.681 | 5.564 | 6.615 | 4.965 | 4.809 | 6.738 | 5.563 | 4.303 | 6.388 | 5.131 | 5.431 | 6.074 | 0.131 | 0.0315 | 0.1394 | 0 | 0 | 0 |
| 'Lmo4' | 18.286 | 23.306 | 23.936 | 21.908 | 22.396 | 23.475 | 20.159 | 25.621 | 22.816 | 24.250 | 27.624 | 24.151 | 21.481 | 24.984 | 25.510 | 25.930 | 25.318 | 26.784 | -0.176 | 0.0001 | 0.0012 | 1 | 0 | 0 |
| 'Lmtk2' | 15.036 | 13.585 | 14.026 | 13.941 | 14.023 | 13.716 | 13.222 | 14.452 | 13.322 | 13.952 | 14.788 | 13.806 | 14.177 | 14.563 | 13.975 | 15.154 | 12.833 | 14.081 | -0.031 | 0.3554 | 0.6086 | 0 | 0 | 0 |
| 'Lnpep' | 28.854 | 30.324 | 30.267 | 31.545 | 28.990 | 27.105 | 28.894 | 28.118 | 28.026 | 27.917 | 30.309 | 29.748 | 26.532 | 30.647 | 27.573 | 25.616 | 30.884 | 30.354 | 0.004 | 0.9161 | 0.9680 | 0 | 0 | 0 |
| 'Lnpk1' | 1.410 | 1.587 | 1.709 | 1.476 | 1.634 | 1.562 | 1.750 | 1.618 | 1.472 | 1.454 | 1.723 | 1.624 | 1.525 | 1.517 | 1.407 | 1.350 | 1.633 | 1.432 | 0.044 | 0.3847 | 0.6347 | 0 | 0 | 0 |
| 'Lnx2' | 59.457 | 59.278 | 55.097 | 60.150 | 58.688 | 57.570 | 59.360 | 58.506 | 57.246 | 57.402 | 61.857 | 59.410 | 61.516 | 60.858 | 57.520 | 60.789 | 63.072 | 61.498 | -0.061 | 0.0096 | 0.0606 | 0 | 0 | 0 |
| 'Lonp1' | 44.778 | 42.736 | 45.338 | 40.884 | 43.509 | 45.802 | 41.018 | 43.706 | 44.834 | 41.662 | 41.239 | 44.065 | 47.093 | 40.004 | 43.466 | 44.549 | 44.082 | 43.532 | 0.001 | 0.9636 | 0.9869 | 0 | 0 | 0 |
| 'Lonp2' | 19.393 | 20.839 | 18.816 | 20.221 | 20.458 | 20.164 | 20.218 | 19.906 | 20.344 | 19.342 | 18.323 | 18.099 | 18.431 | 18.745 | 17.256 | 19.104 | 18.292 | 16.629 | 0.126 | 0.0000 | 0.0000 | 1 | 0 | 0 |
| 'Lonrf1' | 1.522 | 1.672 | 1.585 | 1.741 | 1.511 | 1.797 | 1.574 | 1.738 | 1.976 | 2.031 | 1.903 | 1.849 | 1.759 | 1.719 | 1.911 | 1.991 | 1.576 | 1.643 | -0.119 | 0.1523 | 0.3794 | 0 | 0 | 0 |
| 'Lpar2' | 0.698 | 0.549 | 0.699 | 0.658 | 0.609 | 0.667 | 0.580 | 0.587 | 0.655 | 0.579 | 0.569 | 0.790 | 0.526 | 0.548 | 0.777 | 0.826 | 0.655 | 0.791 | -0.093 | 0.3272 | 0.5850 | 0 | 0 | 0 |
| 'Lpar3' | 3.654 | 3.467 | 3.731 | 2.288 | 2.906 | 3.323 | 2.414 | 3.309 | 3.238 | 3.885 | 5.090 | 4.532 | 3.059 | 4.991 | 4.959 | 3.417 | 4.392 | 4.040 | -0.431 | 0.0000 | 0.0000 | 1 | 1 | 0 |
| 'Lpar5' | 7.610 | 6.565 | 7.434 | 7.874 | 6.134 | 8.665 | 7.043 | 6.631 | 9.047 | 6.005 | 7.416 | 9.425 | 6.088 | 6.888 | 9.403 | 6.130 | 6.310 | 8.576 | 0.010 | 0.8811 | 0.9499 | 0 | 0 | 0 |
| 'Lpar6' | 2.695 | 3.313 | 3.860 | 3.285 | 3.069 | 3.926 | 3.661 | 2.925 | 4.199 | 2.981 | 3.208 | 4.266 | 2.910 | 3.089 | 3.351 | 2.766 | 3.588 | 3.173 | 0.058 | 0.4057 | 0.6550 | 0 | 0 | 0 |
| 'Lpcat1' | 10.145 | 9.000 | 10.513 | 9.817 | 9.234 | 9.996 | 9.365 | 9.889 | 10.068 | 10.123 | 9.399 | 10.635 | 9.843 | 9.220 | 10.006 | 9.936 | 9.433 | 10.852 | -0.032 | 0.2775 | 0.5335 | 0 | 0 | 0 |
| 'Lpcat3' | 4.303 | 5.194 | 4.937 | 4.191 | 4.841 | 4.610 | 4.646 | 4.917 | 5.433 | 4.519 | 4.284 | 4.951 | 4.845 | 4.483 | 5.268 | 4.460 | 4.385 | 4.685 | 0.026 | 0.5603 | 0.7706 | 0 | 0 | 0 |
| 'Lpcat4' | 25.647 | 23.375 | 25.462 | 23.097 | 24.256 | 25.719 | 22.426 | 22.394 | 24.730 | 25.045 | 23.284 | 29.184 | 22.028 | 22.463 | 29.273 | 23.587 | 25.719 | 30.370 | -0.094 | 0.0339 | 0.1467 | 0 | 0 | 0 |
| 'Lpgat1' | 4.994 | 4.862 | 5.112 | 5.016 | 4.842 | 4.643 | 5.380 | 4.946 | 4.804 | 4.155 | 5.653 | 5.442 | 4.618 | 4.981 | 4.684 | 4.317 | 5.624 | 5.277 | -0.013 | 0.8098 | 0.9142 | 0 | 0 | 0 |
| 'Lpin1' | 8.202 | 7.980 | 8.322 | 8.007 | 7.974 | 7.482 | 8.518 | 8.306 | 8.062 | 8.533 | 8.825 | 8.449 | 8.516 | 8.255 | 8.538 | 7.910 | 8.429 | 8.276 | -0.067 | 0.0327 | 0.1432 | 0 | 0 | 0 |
| 'Lpin2' | 7.075 | 7.666 | 6.119 | 7.689 | 6.882 | 6.535 | 7.213 | 6.559 | 6.944 | 7.546 | 7.816 | 6.702 | 8.753 | 7.283 | 6.324 | 7.349 | 7.850 | 6.274 | -0.080 | 0.0579 | 0.2074 | 0 | 0 | 0 |
| 'Lpp' | 14.561 | 15.289 | 12.182 | 12.928 | 14.317 | 11.506 | 13.461 | 13.687 | 12.189 | 16.026 | 15.515 | 11.888 | 16.701 | 14.711 | 10.616 | 16.276 | 15.675 | 11.213 | -0.095 | 0.0284 | 0.1306 | 0 | 0 | 0 |
| 'Lpxn' | 127.746 | 137.378 | 132.994 | 136.471 | 134.614 | 133.054 | 138.942 | 137.827 | 139.082 | 126.007 | 139.413 | 132.040 | 122.500 | 136.619 | 133.750 | 122.439 | 136.567 | 128.781 | 0.038 | 0.1374 | 0.3575 | 0 | 0 | 0 |
| 'Lrba' | 10.082 | 9.610 | 10.276 | 10.601 | 10.229 | 9.418 | 10.237 | 9.706 | 9.572 | 10.691 | 12.215 | 10.398 | 10.739 | 10.713 | 10.043 | 10.273 | 11.052 | 10.994 | -0.124 | 0.0006 | 0.0073 | 1 | 0 | 0 |
| 'Lrch1' | 13.193 | 13.266 | 12.242 | 13.086 | 12.602 | 12.620 | 12.334 | 12.967 | 12.478 | 13.206 | 12.863 | 12.024 | 12.680 | 12.720 | 10.911 | 11.893 | 12.632 | 11.938 | 0.041 | 0.1776 | 0.4131 | 0 | 0 | 0 |
| 'Lrch3' | 9.105 | 9.256 | 9.078 | 9.050 | 9.039 | 8.978 | 8.686 | 9.388 | 9.330 | 8.624 | 9.583 | 10.846 | 8.374 | 9.312 | 9.931 | 9.025 | 9.409 | 10.935 | -0.080 | 0.0386 | 0.1599 | 0 | 0 | 0 |
| 'Lrfn4' | 1.869 | 1.887 | 2.293 | 2.408 | 1.948 | 2.007 | 1.906 | 1.888 | 1.870 | 1.950 | 1.932 | 2.031 | 1.902 | 2.008 | 1.833 | 1.667 | 1.722 | 2.137 | 0.060 | 0.3851 | 0.6349 | 0 | 0 | 0 |
| 'Lrif1' | 2.820 | 3.710 | 3.137 | 2.955 | 3.559 | 2.680 | 3.507 | 3.788 | 3.011 | 3.148 | 3.144 | 3.081 | 2.934 | 3.622 | 2.691 | 3.329 | 3.657 | 2.798 | 0.024 | 0.6003 | 0.7968 | 0 | 0 | 0 |
| 'Lrig1' | 46.709 | 48.610 | 42.842 | 40.932 | 46.112 | 43.807 | 41.843 | 47.824 | 41.451 | 54.185 | 49.213 | 44.170 | 50.241 | 50.458 | 44.839 | 57.026 | 49.140 | 44.654 | -0.157 | 0.0001 | 0.0009 | 1 | 0 | 0 |
| 'Lrig2' | 1.523 | 1.838 | 1.823 | 1.712 | 1.843 | 1.850 | 1.657 | 1.919 | 1.860 | 1.901 | 1.938 | 1.864 | 1.625 | 1.839 | 1.584 | 1.764 | 1.804 | 1.701 | -0.010 | 0.8218 | 0.9217 | 0 | 0 | 0 |
| 'Lrmp' | 31.238 | 29.472 | 31.285 | 29.411 | 29.796 | 30.997 | 31.141 | 30.160 | 31.474 | 33.473 | 31.857 | 32.838 | 33.474 | 31.591 | 31.653 | 31.766 | 31.211 | 31.602 | -0.084 | 0.0000 | 0.0003 | 1 | 0 | 0 |
| 'Lrp1' | 1.859 | 2.431 | 2.093 | 2.386 | 2.323 | 2.209 | 1.999 | 1.884 | 2.094 | 2.605 | 2.436 | 2.352 | 2.852 | 2.404 | 2.461 | 2.894 | 2.842 | 2.376 | -0.278 | 0.0000 | 0.0000 | 1 | 0 | 0 |
| 'Lrp10' | 160.410 | 157.003 | 172.470 | 146.929 | 158.912 | 177.424 | 149.771 | 156.574 | 172.657 | 168.951 | 150.665 | 160.894 | 155.215 | 150.565 | 164.303 | 160.839 | 144.383 | 166.944 | 0.018 | 0.5348 | 0.7544 | 0 | 0 | 0 |
| 'Lrp12' | 1.933 | 1.786 | 2.129 | 1.900 | 2.304 | 2.002 | 2.306 | 2.061 | 2.196 | 2.574 | 2.577 | 1.857 | 2.141 | 2.047 | 2.266 | 2.196 | 1.941 | 1.593 | -0.041 | 0.6329 | 0.8179 | 0 | 0 | 0 |
| 'Lrp3' | 2.860 | 2.362 | 3.188 | 2.398 | 2.547 | 2.368 | 2.770 | 2.641 | 2.542 | 2.732 | 3.390 | 2.962 | 2.844 | 2.807 | 2.895 | 2.381 | 2.730 | 3.022 | -0.129 | 0.0515 | 0.1919 | 0 | 0 | 0 |
| 'Lrp5' | 0.521 | 0.564 | 0.519 | 0.501 | 0.479 | 0.525 | 0.467 | 0.540 | 0.641 | 0.467 | 0.399 | 0.457 | 0.519 | 0.472 | 0.553 | 0.457 | 0.385 | 0.453 | 0.186 | 0.0232 | 0.1143 | 0 | 0 | 0 |
| 'Lrp6' | 6.491 | 6.413 | 6.158 | 5.689 | 6.198 | 5.840 | 5.790 | 6.481 | 5.822 | 6.549 | 7.091 | 6.063 | 5.965 | 6.405 | 6.035 | 6.092 | 6.184 | 6.481 | -0.060 | 0.1024 | 0.3002 | 0 | 0 | 0 |
| 'Lrp8' | 1.483 | 1.171 | 1.587 | 1.239 | 1.357 | 1.703 | 1.313 | 1.369 | 1.673 | 1.550 | 1.405 | 1.582 | 1.665 | 1.534 | 1.585 | 1.622 | 1.438 | 1.637 | -0.125 | 0.0193 | 0.1008 | 0 | 0 | 0 |
| 'Lrpap1' | 8.723 | 8.380 | 9.099 | 7.582 | 8.312 | 9.518 | 8.160 | 8.891 | 8.973 | 9.288 | 7.982 | 8.292 | 8.784 | 8.140 | 8.093 | 8.523 | 7.169 | 7.635 | 0.065 | 0.1575 | 0.3874 | 0 | 0 | 0 |
| 'Lrpprc' | 5.912 | 5.809 | 6.033 | 5.736 | 5.673 | 5.066 | 5.705 | 5.659 | 5.754 | 5.819 | 6.265 | 6.259 | 5.967 | 5.788 | 5.707 | 5.697 | 5.540 | 6.057 | -0.061 | 0.1031 | 0.3009 | 0 | 0 | 0 |
| 'Lrrc1' | 1.787 | 1.612 | 1.573 | 1.893 | 1.604 | 1.453 | 1.655 | 1.598 | 1.765 | 1.633 | 1.601 | 1.684 | 1.495 | 1.484 | 1.734 | 1.654 | 1.505 | 1.848 | 0.016 | 0.7798 | 0.9008 | 0 | 0 | 0 |
| 'Lrrc14' | 1.493 | 1.573 | 1.837 | 1.412 | 1.724 | 1.683 | 1.594 | 1.705 | 1.730 | 1.645 | 1.696 | 1.583 | 1.644 | 1.431 | 1.625 | 1.779 | 1.518 | 1.376 | 0.040 | 0.4731 | 0.7088 | 0 | 0 | 0 |
| 'Lrrc20' | 11.514 | 13.121 | 12.256 | 12.842 | 12.271 | 13.900 | 12.172 | 12.393 | 12.807 | 12.574 | 11.544 | 12.312 | 11.585 | 11.504 | 11.756 | 11.028 | 12.002 | 11.250 | 0.089 | 0.0331 | 0.1442 | 0 | 0 | 0 |
| 'Lrrc24' | 5.184 | 5.890 | 5.890 | 5.397 | 5.541 | 6.624 | 5.221 | 6.188 | 6.146 | 5.888 | 5.125 | 5.222 | 5.707 | 5.022 | 5.372 | 6.167 | 4.582 | 4.625 | 0.123 | 0.0452 | 0.1763 | 0 | 0 | 0 |
| 'Lrrc28' | 1.421 | 1.465 | 1.598 | 1.751 | 1.737 | 1.684 | 1.611 | 1.521 | 1.578 | 1.775 | 1.680 | 1.630 | 1.718 | 1.929 | 1.579 | 1.674 | 1.587 | 1.516 | -0.078 | 0.1852 | 0.4208 | 0 | 0 | 0 |
| 'Lrrc32' | 0.896 | 0.805 | 1.130 | 0.764 | 0.695 | 1.182 | 0.875 | 0.808 | 1.079 | 0.870 | 0.309 | 1.106 | 0.812 | 0.364 | 0.941 | 0.981 | 0.249 | 0.976 | 0.399 | 0.0036 | 0.0288 | 1 | 1 | 0 |
| 'Lrrc40' | 2.354 | 2.776 | 2.591 | 2.429 | 2.531 | 2.226 | 2.446 | 2.675 | 2.434 | 2.325 | 2.564 | 2.106 | 2.381 | 2.816 | 2.120 | 2.179 | 2.134 | 2.064 | 0.113 | 0.0328 | 0.1433 | 0 | 0 | 0 |
| 'Lrrc41' | 6.761 | 6.761 | 6.895 | 6.027 | 6.519 | 6.411 | 5.993 | 6.232 | 6.721 | 6.739 | 5.659 | 7.136 | 6.218 | 5.657 | 7.038 | 6.367 | 5.508 | 7.126 | 0.010 | 0.8222 | 0.9217 | 0 | 0 | 0 |
| 'Lrrc42' | 13.278 | 13.594 | 13.650 | 13.749 | 13.769 | 13.308 | 13.434 | 13.117 | 14.334 | 12.994 | 12.112 | 13.270 | 12.440 | 12.245 | 12.551 | 12.197 | 12.255 | 12.510 | 0.105 | 0.0009 | 0.0100 | 1 | 0 | 0 |
| 'Lrrc45' | 12.395 | 10.543 | 14.567 | 12.763 | 11.234 | 14.166 | 11.900 | 11.603 | 14.451 | 11.932 | 10.696 | 14.959 | 13.456 | 11.736 | 13.432 | 12.109 | 12.041 | 13.189 | -0.012 | 0.7338 | 0.8786 | 0 | 0 | 0 |
| 'Lrrc47' | 15.774 | 14.937 | 16.050 | 13.876 | 15.662 | 16.950 | 13.678 | 15.173 | 14.793 | 16.351 | 13.907 | 15.706 | 16.202 | 14.915 | 16.151 | 16.360 | 13.086 | 16.327 | -0.030 | 0.4906 | 0.7205 | 0 | 0 | 0 |
| 'Lrrc49' | 0.239 | 0.359 | 0.247 | 0.246 | 0.290 | 0.327 | 0.256 | 0.247 | 0.317 | 0.256 | 0.251 | 0.302 | 0.275 | 0.240 | 0.326 | 0.297 | 0.336 | 0.256 | -0.028 | 0.8051 | 0.9125 | 0 | 0 | 0 |
| 'Lrrc56' | 0.813 | 0.706 | 0.961 | 0.923 | 0.889 | 0.884 | 0.904 | 0.859 | 1.214 | 0.775 | 0.720 | 0.890 | 0.853 | 0.846 | 1.017 | 0.644 | 0.781 | 0.863 | 0.130 | 0.1592 | 0.3895 | 0 | 0 | 0 |
| 'Lrrc57' | 8.333 | 7.618 | 6.949 | 8.442 | 7.746 | 6.041 | 7.520 | 7.594 | 6.278 | 7.830 | 7.217 | 6.161 | 8.902 | 7.611 | 6.381 | 6.744 | 8.735 | 5.992 | 0.012 | 0.8233 | 0.9223 | 0 | 0 | 0 |
| 'Lrrc58' | 6.197 | 6.535 | 5.098 | 5.798 | 5.858 | 4.986 | 5.912 | 5.945 | 5.602 | 6.454 | 5.768 | 6.023 | 6.140 | 6.582 | 5.750 | 6.794 | 6.137 | 6.389 | -0.125 | 0.0041 | 0.0325 | 1 | 0 | 0 |
| 'Lrrc59' | 21.323 | 19.842 | 22.802 | 21.264 | 22.029 | 24.286 | 21.788 | 22.848 | 22.229 | 19.859 | 20.210 | 23.898 | 20.968 | 19.978 | 24.437 | 22.695 | 19.533 | 24.133 | 0.012 | 0.7461 | 0.8854 | 0 | 0 | 0 |
| 'Lrrc61' | 5.498 | 6.353 | 5.872 | 5.196 | 6.523 | 6.076 | 5.046 | 6.199 | 6.154 | 6.550 | 5.990 | 5.849 | 5.583 | 5.866 | 5.422 | 5.851 | 5.795 | 5.729 | -0.003 | 0.9434 | 0.9788 | 0 | 0 | 0 |
| 'Lrrc8b' | 3.507 | 4.267 | 4.483 | 3.652 | 4.946 | 4.571 | 3.792 | 4.454 | 4.134 | 4.393 | 3.720 | 3.411 | 4.654 | 5.023 | 3.764 | 4.326 | 4.453 | 3.113 | 0.034 | 0.6585 | 0.8350 | 0 | 0 | 0 |
| 'Lrrc8c' | 4.912 | 5.366 | 5.810 | 5.265 | 5.291 | 5.872 | 4.696 | 4.981 | 5.401 | 5.422 | 4.889 | 5.541 | 6.225 | 5.077 | 5.809 | 5.588 | 4.612 | 5.271 | -0.034 | 0.4665 | 0.7022 | 0 | 0 | 0 |
| 'Lrrc8d' | 4.138 | 3.913 | 4.193 | 3.820 | 3.999 | 3.895 | 3.714 | 4.188 | 3.844 | 4.122 | 3.947 | 3.426 | 3.894 | 3.751 | 3.275 | 3.907 | 3.836 | 3.277 | 0.092 | 0.0219 | 0.1095 | 0 | 0 | 0 |
| 'Lrrcc1' | 4.754 | 4.878 | 4.892 | 5.066 | 5.372 | 4.917 | 5.105 | 5.436 | 4.787 | 5.246 | 4.979 | 4.556 | 4.746 | 5.131 | 4.475 | 4.893 | 4.662 | 3.740 | 0.086 | 0.0394 | 0.1621 | 0 | 0 | 0 |
| 'Lrrfip1' | 33.389 | 31.567 | 32.035 | 33.148 | 32.049 | 32.343 | 34.276 | 31.271 | 32.601 | 32.004 | 31.559 | 31.287 | 30.807 | 32.072 | 30.467 | 31.519 | 30.920 | 30.648 | 0.047 | 0.0217 | 0.1086 | 0 | 0 | 0 |
| 'Lrrfip2' | 3.740 | 3.931 | 3.609 | 3.732 | 4.098 | 3.620 | 4.125 | 3.888 | 4.056 | 4.063 | 3.760 | 3.764 | 4.011 | 4.319 | 3.774 | 3.965 | 4.574 | 3.569 | -0.050 | 0.2150 | 0.4606 | 0 | 0 | 0 |
| 'Lrsam1' | 1.732 | 2.011 | 2.242 | 1.735 | 2.096 | 2.190 | 1.840 | 2.255 | 2.416 | 2.202 | 2.130 | 1.952 | 2.205 | 2.100 | 1.910 | 1.923 | 1.878 | 1.924 | 0.016 | 0.8075 | 0.9136 | 0 | 0 | 0 |
| 'Lrwd1' | 7.133 | 6.138 | 7.782 | 7.138 | 6.736 | 7.675 | 6.812 | 6.783 | 7.269 | 6.982 | 6.547 | 7.352 | 7.371 | 6.419 | 7.079 | 6.405 | 6.242 | 7.406 | 0.029 | 0.3590 | 0.6115 | 0 | 0 | 0 |
| 'Lsg1' | 5.778 | 5.476 | 5.642 | 5.541 | 5.866 | 5.841 | 5.687 | 6.250 | 6.511 | 5.725 | 5.688 | 5.664 | 5.619 | 5.223 | 5.447 | 5.828 | 5.330 | 5.626 | 0.060 | 0.1003 | 0.2964 | 0 | 0 | 0 |
| 'Lsm1' | 9.656 | 10.771 | 9.848 | 10.478 | 11.612 | 10.056 | 9.903 | 10.600 | 10.247 | 9.959 | 8.907 | 11.140 | 10.058 | 9.995 | 11.464 | 11.134 | 10.661 | 9.563 | -0.013 | 0.8049 | 0.9125 | 0 | 0 | 0 |
| 'Lsm10' | 7.590 | 7.954 | 8.383 | 7.847 | 8.946 | 7.951 | 7.477 | 8.135 | 8.371 | 7.244 | 6.595 | 7.950 | 7.688 | 8.255 | 7.847 | 7.840 | 6.967 | 8.106 | 0.072 | 0.1869 | 0.4227 | 0 | 0 | 0 |
| 'Lsm11' | 1.737 | 1.484 | 1.700 | 1.821 | 1.420 | 1.769 | 1.644 | 1.471 | 1.786 | 1.342 | 1.699 | 2.155 | 1.427 | 1.716 | 2.304 | 1.420 | 1.637 | 2.288 | -0.109 | 0.1764 | 0.4117 | 0 | 0 | 0 |
| 'Lsm12' | 16.351 | 17.116 | 16.920 | 17.773 | 18.656 | 16.210 | 18.010 | 16.491 | 16.740 | 15.601 | 16.578 | 17.073 | 17.106 | 16.982 | 17.490 | 16.261 | 16.577 | 16.729 | 0.021 | 0.5361 | 0.7552 | 0 | 0 | 0 |
| 'Lsm14a' | 27.681 | 28.880 | 26.835 | 27.441 | 29.106 | 26.185 | 27.889 | 28.517 | 26.251 | 25.300 | 27.867 | 26.979 | 26.738 | 28.198 | 27.616 | 27.829 | 28.751 | 27.425 | 0.001 | 0.9827 | 0.9934 | 0 | 0 | 0 |
| 'Lsm14b' | 18.773 | 20.400 | 18.458 | 18.948 | 20.075 | 19.275 | 16.822 | 20.419 | 18.233 | 17.267 | 20.085 | 19.449 | 22.203 | 21.504 | 20.935 | 19.006 | 18.940 | 21.267 | -0.088 | 0.0504 | 0.1897 | 0 | 0 | 0 |
| 'Lsm2' | 3.963 | 3.963 | 4.552 | 4.100 | 4.093 | 4.450 | 4.212 | 3.805 | 4.330 | 3.742 | 3.532 | 4.341 | 3.819 | 4.138 | 4.702 | 3.847 | 3.893 | 4.719 | 0.018 | 0.6870 | 0.8521 | 0 | 0 | 0 |
| 'Lsm3' | 9.196 | 10.278 | 9.651 | 9.554 | 10.558 | 9.121 | 10.148 | 9.819 | 9.483 | 8.054 | 9.490 | 8.962 | 9.965 | 9.435 | 8.631 | 8.798 | 9.429 | 8.806 | 0.092 | 0.0197 | 0.1020 | 0 | 0 | 0 |
| 'Lsm4' | 81.534 | 86.019 | 83.609 | 81.313 | 84.159 | 87.316 | 81.376 | 86.278 | 80.490 | 73.886 | 69.257 | 77.217 | 84.795 | 77.158 | 80.850 | 83.241 | 75.338 | 78.749 | 0.092 | 0.0057 | 0.0415 | 1 | 0 | 0 |
| 'Lsm5' | 14.983 | 17.403 | 15.687 | 15.966 | 17.531 | 15.235 | 16.625 | 17.117 | 15.463 | 12.927 | 14.393 | 13.407 | 16.761 | 16.838 | 14.002 | 15.920 | 17.754 | 13.307 | 0.102 | 0.0464 | 0.1795 | 0 | 0 | 0 |
| 'Lsm6' | 7.309 | 8.795 | 8.068 | 7.994 | 7.934 | 8.421 | 8.416 | 8.323 | 8.392 | 6.907 | 7.860 | 7.648 | 8.044 | 7.715 | 7.884 | 8.043 | 8.046 | 7.503 | 0.070 | 0.0599 | 0.2110 | 0 | 0 | 0 |
| 'Lsm8' | 7.484 | 7.864 | 7.869 | 7.802 | 8.912 | 7.863 | 6.940 | 8.493 | 7.546 | 6.363 | 7.863 | 7.259 | 7.378 | 8.241 | 7.510 | 7.295 | 7.816 | 7.151 | 0.074 | 0.0569 | 0.2050 | 0 | 0 | 0 |
| 'Lsp1' | 32.236 | 29.706 | 32.273 | 30.657 | 30.137 | 36.332 | 33.173 | 29.173 | 32.694 | 35.252 | 29.652 | 34.582 | 35.438 | 31.419 | 36.782 | 34.894 | 30.727 | 34.215 | -0.091 | 0.0007 | 0.0080 | 1 | 0 | 0 |
| 'Lsr' | 0.924 | 1.157 | 1.401 | 0.998 | 0.836 | 1.096 | 0.912 | 1.039 | 1.046 | 0.737 | 0.543 | 0.654 | 0.662 | 0.699 | 0.670 | 0.802 | 0.597 | 0.659 | 0.651 | 0.0000 | 0.0000 | 1 | 1 | 0 |
| 'Lss' | 1.409 | 1.448 | 1.818 | 1.480 | 1.628 | 1.640 | 1.690 | 1.456 | 1.877 | 1.528 | 1.691 | 1.544 | 1.824 | 1.614 | 1.644 | 1.630 | 1.381 | 1.477 | 0.002 | 0.9799 | 0.9926 | 0 | 0 | 0 |
| 'Lta' | 9.644 | 9.984 | 12.095 | 9.260 | 10.322 | 11.204 | 10.362 | 9.044 | 12.490 | 13.148 | 10.304 | 11.821 | 14.126 | 11.265 | 10.301 | 11.743 | 12.247 | 10.161 | -0.162 | 0.0304 | 0.1360 | 0 | 0 | 0 |
| 'Lta4h' | 33.243 | 30.919 | 34.271 | 35.064 | 31.802 | 33.708 | 31.033 | 30.238 | 33.080 | 31.772 | 29.723 | 35.095 | 32.471 | 32.043 | 33.586 | 31.779 | 31.294 | 33.494 | -0.002 | 0.9502 | 0.9809 | 0 | 0 | 0 |
| 'Ltb' | 106.990 | 119.697 | 113.772 | 105.294 | 116.801 | 127.057 | 105.216 | 110.443 | 120.105 | 120.198 | 108.890 | 123.009 | 117.144 | 120.403 | 125.503 | 133.149 | 110.017 | 122.533 | -0.087 | 0.0198 | 0.1024 | 0 | 0 | 0 |
| 'Ltbp3' | 1.598 | 1.799 | 1.586 | 1.886 | 1.520 | 1.465 | 1.166 | 1.267 | 1.585 | 0.753 | 0.592 | 0.642 | 0.641 | 0.666 | 0.553 | 0.576 | 0.658 | 0.765 | 1.233 | 0.0000 | 0.0000 | 1 | 1 | 1 |
| 'Ltk' | 1.677 | 1.574 | 2.002 | 1.579 | 1.692 | 1.918 | 1.619 | 1.600 | 2.031 | 1.517 | 1.420 | 1.691 | 1.858 | 1.427 | 1.521 | 1.496 | 1.451 | 1.539 | 0.163 | 0.0069 | 0.0479 | 1 | 0 | 0 |
| 'Ltn1' | 14.180 | 14.653 | 15.100 | 15.077 | 15.642 | 15.263 | 15.193 | 16.160 | 15.205 | 14.371 | 15.142 | 15.571 | 14.671 | 14.741 | 15.547 | 14.952 | 15.927 | 14.963 | -0.005 | 0.8497 | 0.9352 | 0 | 0 | 0 |
| 'Ltv1' | 19.997 | 20.613 | 24.461 | 20.170 | 22.871 | 22.082 | 22.087 | 22.408 | 22.976 | 24.902 | 22.781 | 20.001 | 23.237 | 22.845 | 20.264 | 23.946 | 21.509 | 19.140 | -0.009 | 0.8711 | 0.9447 | 0 | 0 | 0 |
| 'Luc7l' | 8.132 | 8.455 | 7.831 | 7.678 | 7.968 | 7.859 | 7.828 | 8.246 | 8.088 | 6.799 | 7.893 | 8.997 | 7.511 | 7.796 | 8.487 | 7.223 | 8.358 | 8.794 | -0.005 | 0.8992 | 0.9600 | 0 | 0 | 0 |
| 'Luc7l2' | 17.739 | 18.291 | 17.056 | 16.577 | 17.816 | 16.015 | 17.863 | 18.561 | 17.059 | 17.982 | 18.034 | 16.108 | 17.972 | 17.616 | 15.854 | 17.754 | 18.328 | 16.272 | 0.001 | 0.9775 | 0.9920 | 0 | 0 | 0 |
| 'Luc7l3' | 11.582 | 11.450 | 11.765 | 13.490 | 11.166 | 10.783 | 13.985 | 10.374 | 11.781 | 8.639 | 12.635 | 14.509 | 10.295 | 11.060 | 12.637 | 9.256 | 14.163 | 13.206 | 0.001 | 0.9957 | 0.9994 | 0 | 0 | 0 |
| 'Luzp1' | 0.843 | 0.859 | 0.742 | 0.735 | 0.862 | 0.773 | 0.579 | 0.773 | 0.810 | 0.973 | 0.927 | 1.148 | 1.145 | 1.015 | 0.952 | 1.121 | 1.067 | 1.142 | -0.458 | 0.0000 | 0.0000 | 1 | 1 | 0 |
| 'Lxn' | 2.104 | 1.809 | 1.989 | 2.197 | 1.871 | 2.177 | 2.520 | 1.773 | 1.875 | 1.375 | 1.607 | 1.669 | 1.382 | 1.111 | 1.563 | 1.390 | 1.265 | 1.421 | 0.494 | 0.0000 | 0.0000 | 1 | 1 | 0 |
| 'Ly6e' | 15.268 | 14.666 | 17.398 | 15.600 | 15.566 | 18.245 | 15.794 | 14.647 | 18.538 | 15.577 | 15.906 | 20.753 | 17.349 | 16.819 | 21.831 | 18.259 | 16.601 | 20.836 | -0.177 | 0.0000 | 0.0000 | 1 | 0 | 0 |
| 'Ly75' | 12.255 | 14.277 | 12.648 | 12.298 | 14.838 | 12.087 | 13.485 | 13.403 | 12.768 | 13.920 | 14.241 | 12.896 | 14.535 | 13.503 | 12.649 | 14.538 | 13.854 | 13.743 | -0.082 | 0.0182 | 0.0966 | 0 | 0 | 0 |
| 'Ly9' | 27.245 | 26.931 | 29.330 | 28.601 | 27.415 | 29.245 | 27.408 | 26.565 | 28.525 | 29.731 | 25.677 | 31.319 | 28.413 | 27.977 | 29.388 | 29.784 | 27.618 | 30.052 | -0.060 | 0.0070 | 0.0480 | 1 | 0 | 0 |
| 'Ly96' | 5.780 | 7.211 | 5.430 | 5.764 | 6.222 | 5.928 | 5.605 | 6.909 | 5.976 | 5.449 | 6.013 | 5.373 | 5.757 | 7.182 | 4.796 | 5.206 | 7.263 | 5.377 | 0.057 | 0.4166 | 0.6643 | 0 | 0 | 0 |
| 'Lyar' | 10.053 | 9.698 | 9.872 | 10.632 | 10.240 | 9.523 | 10.788 | 10.717 | 10.275 | 9.759 | 9.117 | 10.173 | 10.279 | 8.988 | 9.547 | 10.852 | 10.160 | 9.886 | 0.036 | 0.3370 | 0.5955 | 0 | 0 | 0 |
| 'Lypla1' | 7.140 | 8.277 | 8.336 | 7.710 | 8.680 | 7.547 | 8.170 | 7.914 | 7.693 | 7.930 | 7.547 | 7.637 | 7.570 | 8.333 | 7.334 | 6.959 | 8.220 | 6.680 | 0.056 | 0.1336 | 0.3522 | 0 | 0 | 0 |
| 'Lypla2' | 44.527 | 49.297 | 50.013 | 43.467 | 49.922 | 54.256 | 46.077 | 50.461 | 52.362 | 49.423 | 44.408 | 50.285 | 46.849 | 46.120 | 51.540 | 48.861 | 44.435 | 49.622 | 0.018 | 0.5733 | 0.7795 | 0 | 0 | 0 |
| 'Lyrm1' | 1.315 | 1.611 | 1.341 | 1.060 | 1.363 | 1.183 | 1.292 | 1.223 | 1.501 | 1.341 | 1.448 | 1.230 | 1.305 | 1.036 | 1.180 | 1.362 | 1.346 | 1.025 | 0.070 | 0.5124 | 0.7374 | 0 | 0 | 0 |
| 'Lyrm2' | 11.794 | 15.072 | 14.121 | 13.226 | 15.075 | 14.085 | 13.513 | 14.701 | 14.119 | 14.152 | 12.020 | 12.736 | 15.341 | 13.662 | 13.463 | 15.538 | 13.586 | 11.655 | 0.034 | 0.5583 | 0.7697 | 0 | 0 | 0 |
| 'Lyrm4' | 12.637 | 13.658 | 12.431 | 14.378 | 14.431 | 12.368 | 13.686 | 13.035 | 11.582 | 13.070 | 15.178 | 11.940 | 15.305 | 15.588 | 12.111 | 14.868 | 14.335 | 11.688 | -0.074 | 0.1429 | 0.3662 | 0 | 0 | 0 |
| 'Lyrm5' | 1.203 | 1.651 | 1.739 | 1.378 | 1.666 | 1.547 | 1.479 | 1.675 | 1.696 | 1.691 | 1.691 | 1.439 | 1.315 | 1.515 | 1.442 | 1.301 | 1.619 | 1.274 | 0.074 | 0.3137 | 0.5715 | 0 | 0 | 0 |
| 'Lyrm9' | 2.222 | 2.284 | 2.603 | 2.160 | 2.105 | 2.113 | 2.677 | 2.160 | 2.355 | 2.243 | 2.338 | 2.224 | 2.339 | 1.968 | 2.114 | 2.213 | 1.948 | 1.691 | 0.108 | 0.0937 | 0.2834 | 0 | 0 | 0 |
| 'Lysmd2' | 9.520 | 8.826 | 9.727 | 9.812 | 9.446 | 10.224 | 10.217 | 10.603 | 10.416 | 7.015 | 4.987 | 10.050 | 6.321 | 6.308 | 7.468 | 7.598 | 6.331 | 8.663 | 0.441 | 0.0000 | 0.0000 | 1 | 1 | 0 |
| 'Lysmd3' | 6.905 | 6.329 | 6.706 | 6.973 | 6.958 | 5.993 | 6.721 | 6.317 | 6.414 | 6.661 | 7.762 | 7.083 | 6.603 | 6.986 | 7.208 | 7.136 | 7.117 | 7.325 | -0.117 | 0.0205 | 0.1048 | 0 | 0 | 0 |
| 'Lysmd4' | 3.790 | 4.665 | 4.525 | 3.661 | 4.266 | 3.747 | 3.609 | 4.031 | 4.234 | 3.370 | 3.847 | 4.174 | 3.775 | 3.721 | 4.278 | 3.789 | 3.529 | 3.906 | 0.071 | 0.2468 | 0.4977 | 0 | 0 | 0 |
| 'Lyst' | 11.872 | 11.742 | 11.123 | 13.602 | 11.205 | 10.758 | 11.476 | 10.928 | 12.125 | 11.694 | 13.871 | 12.354 | 11.334 | 12.312 | 12.111 | 11.174 | 13.637 | 12.797 | -0.096 | 0.0635 | 0.2198 | 0 | 0 | 0 |
| 'Lzic' | 8.904 | 9.540 | 9.512 | 9.777 | 9.254 | 10.072 | 10.270 | 10.512 | 9.217 | 8.070 | 8.782 | 9.607 | 9.332 | 8.827 | 9.007 | 8.681 | 9.527 | 9.567 | 0.082 | 0.0642 | 0.2209 | 0 | 0 | 0 |
| 'Lztfl1' | 1.466 | 2.199 | 1.798 | 2.063 | 2.120 | 1.768 | 1.943 | 1.692 | 1.966 | 1.416 | 1.391 | 1.514 | 1.488 | 1.276 | 1.466 | 1.657 | 1.332 | 1.357 | 0.383 | 0.0000 | 0.0000 | 1 | 1 | 0 |
| 'Lztr1' | 5.928 | 6.064 | 6.585 | 6.311 | 6.189 | 6.877 | 5.676 | 6.268 | 6.634 | 5.857 | 6.113 | 7.218 | 6.216 | 5.866 | 6.683 | 6.202 | 6.578 | 6.770 | -0.037 | 0.2285 | 0.4751 | 0 | 0 | 0 |
| 'Lzts1' | 1.672 | 1.991 | 1.947 | 1.927 | 1.886 | 1.927 | 1.927 | 1.990 | 1.862 | 1.592 | 2.070 | 1.973 | 1.756 | 1.831 | 1.868 | 1.944 | 1.921 | 1.964 | 0.005 | 0.9253 | 0.9715 | 0 | 0 | 0 |
| 'Lzts2' | 2.492 | 1.867 | 2.119 | 2.165 | 1.352 | 2.399 | 1.495 | 1.248 | 2.139 | 2.204 | 1.874 | 2.452 | 1.804 | 1.992 | 2.108 | 1.976 | 1.636 | 2.072 | -0.084 | 0.4302 | 0.6758 | 0 | 0 | 0 |
| 'Lzts3' | 2.792 | 3.331 | 3.337 | 3.271 | 2.893 | 3.288 | 3.080 | 3.415 | 3.327 | 3.730 | 2.918 | 2.809 | 3.844 | 3.068 | 2.708 | 3.242 | 3.058 | 2.975 | 0.018 | 0.7828 | 0.9025 | 0 | 0 | 0 |
| 'M6pr' | 40.007 | 39.932 | 37.403 | 38.110 | 40.212 | 36.682 | 37.427 | 37.857 | 37.954 | 41.133 | 38.638 | 38.400 | 40.818 | 40.560 | 38.779 | 41.008 | 39.866 | 38.655 | -0.060 | 0.0099 | 0.0619 | 0 | 0 | 0 |
| 'Macf1' | 22.368 | 16.948 | 22.771 | 23.148 | 17.132 | 20.374 | 19.629 | 17.823 | 22.096 | 19.952 | 20.746 | 22.791 | 16.469 | 17.426 | 20.122 | 18.914 | 17.403 | 24.712 | 0.017 | 0.7795 | 0.9008 | 0 | 0 | 0 |
| 'Macrod1' | 15.294 | 12.932 | 14.561 | 12.016 | 13.965 | 14.559 | 12.992 | 13.754 | 12.572 | 10.943 | 10.568 | 11.387 | 14.392 | 12.286 | 11.453 | 12.748 | 10.012 | 11.605 | 0.214 | 0.0012 | 0.0121 | 1 | 0 | 0 |
| 'Mad1l1' | 7.398 | 6.345 | 8.143 | 8.305 | 6.735 | 8.920 | 8.071 | 7.141 | 8.358 | 6.629 | 7.268 | 7.727 | 8.280 | 7.164 | 7.275 | 7.097 | 7.243 | 7.411 | 0.058 | 0.2396 | 0.4899 | 0 | 0 | 0 |
| 'Mad2l1' | 4.234 | 4.564 | 5.284 | 4.750 | 4.702 | 4.912 | 5.236 | 4.640 | 5.118 | 3.354 | 4.282 | 4.813 | 4.838 | 5.168 | 4.990 | 4.304 | 5.242 | 4.720 | 0.049 | 0.4238 | 0.6706 | 0 | 0 | 0 |
| 'Mad2l1bp' | 23.168 | 22.071 | 20.901 | 23.024 | 20.296 | 19.882 | 20.712 | 19.671 | 19.403 | 19.848 | 21.404 | 19.703 | 18.191 | 20.210 | 21.728 | 20.768 | 22.024 | 19.225 | 0.038 | 0.4732 | 0.7088 | 0 | 0 | 0 |
| 'Mad2l2' | 6.260 | 5.454 | 6.601 | 6.691 | 5.778 | 7.181 | 5.968 | 5.965 | 6.550 | 5.605 | 5.605 | 6.518 | 6.365 | 6.631 | 6.412 | 6.376 | 5.847 | 6.076 | 0.020 | 0.7075 | 0.8645 | 0 | 0 | 0 |
| 'Madd' | 13.639 | 12.912 | 13.099 | 12.729 | 13.119 | 13.218 | 11.901 | 12.838 | 13.537 | 14.059 | 14.842 | 15.645 | 14.107 | 14.363 | 15.292 | 13.419 | 14.338 | 16.010 | -0.184 | 0.0000 | 0.0000 | 1 | 0 | 0 |
| 'Maea' | 27.448 | 26.372 | 27.386 | 27.914 | 26.861 | 28.183 | 26.929 | 27.261 | 26.158 | 24.664 | 24.043 | 29.066 | 26.700 | 26.539 | 29.593 | 27.006 | 26.220 | 28.127 | 0.004 | 0.8891 | 0.9546 | 0 | 0 | 0 |
| 'Maf' | 158.312 | 164.874 | 155.848 | 161.881 | 164.197 | 153.481 | 168.028 | 154.476 | 150.928 | 155.946 | 177.348 | 157.996 | 166.583 | 173.868 | 165.008 | 155.666 | 176.677 | 175.787 | -0.082 | 0.0125 | 0.0734 | 0 | 0 | 0 |
| 'Maf1' | 15.252 | 17.106 | 14.730 | 14.216 | 15.977 | 16.520 | 14.070 | 16.870 | 15.125 | 15.282 | 14.429 | 16.806 | 14.002 | 15.876 | 16.238 | 16.389 | 14.926 | 16.312 | -0.016 | 0.6879 | 0.8525 | 0 | 0 | 0 |
| 'Maff' | 29.761 | 4.630 | 2.903 | 8.618 | 4.776 | 2.829 | 4.776 | 7.891 | 2.373 | 24.229 | 9.888 | 1.811 | 17.907 | 11.727 | 1.654 | 20.509 | 5.759 | 1.751 | -0.211 | 0.5007 | 0.7278 | 0 | 0 | 0 |
| 'Mafg' | 19.060 | 21.940 | 20.096 | 19.520 | 21.180 | 20.640 | 19.464 | 20.826 | 19.723 | 19.702 | 24.139 | 23.362 | 22.850 | 24.605 | 23.547 | 21.826 | 24.057 | 24.297 | -0.203 | 0.0000 | 0.0000 | 1 | 0 | 0 |
| 'Mafk' | 17.323 | 10.102 | 7.824 | 10.683 | 10.570 | 7.957 | 10.083 | 11.478 | 7.205 | 17.649 | 11.723 | 8.356 | 15.016 | 14.016 | 8.383 | 18.020 | 11.455 | 8.195 | -0.256 | 0.0077 | 0.0512 | 0 | 0 | 0 |
| 'Maged1' | 0.853 | 0.731 | 0.896 | 1.029 | 0.921 | 0.966 | 1.262 | 0.994 | 0.859 | 0.858 | 0.611 | 0.896 | 0.972 | 0.890 | 0.702 | 0.912 | 0.881 | 0.839 | 0.152 | 0.0837 | 0.2622 | 0 | 0 | 0 |
| 'Maged2' | 1.599 | 1.326 | 1.644 | 1.737 | 1.702 | 1.632 | 1.685 | 1.715 | 1.670 | 1.707 | 1.369 | 1.962 | 1.648 | 1.680 | 1.641 | 1.647 | 1.585 | 1.402 | -0.010 | 0.8889 | 0.9545 | 0 | 0 | 0 |
| 'Magee1' | 6.975 | 5.348 | 6.415 | 6.758 | 5.699 | 6.085 | 6.476 | 6.303 | 5.591 | 5.919 | 6.786 | 6.993 | 5.380 | 6.252 | 5.858 | 6.043 | 6.793 | 6.372 | -0.030 | 0.6436 | 0.8251 | 0 | 0 | 0 |
| 'Magoh' | 49.253 | 52.949 | 46.876 | 55.369 | 54.339 | 45.690 | 50.870 | 52.308 | 46.453 | 47.113 | 49.919 | 43.460 | 50.345 | 52.160 | 49.231 | 48.965 | 49.115 | 40.905 | 0.064 | 0.0780 | 0.2504 | 0 | 0 | 0 |
| 'Magohb' | 3.519 | 4.087 | 3.513 | 3.428 | 4.221 | 3.760 | 3.992 | 4.511 | 3.426 | 3.158 | 3.692 | 3.379 | 3.529 | 3.040 | 3.063 | 3.569 | 3.567 | 2.962 | 0.185 | 0.0042 | 0.0331 | 1 | 0 | 0 |
| 'Magt1' | 6.856 | 7.589 | 7.775 | 7.792 | 7.924 | 7.594 | 7.842 | 7.502 | 7.340 | 7.415 | 7.805 | 8.672 | 7.101 | 7.780 | 7.665 | 6.745 | 8.296 | 8.118 | -0.042 | 0.2804 | 0.5374 | 0 | 0 | 0 |
| 'Mak16' | 31.263 | 31.126 | 31.605 | 32.238 | 33.882 | 29.146 | 31.818 | 31.851 | 31.712 | 27.817 | 29.439 | 32.511 | 28.858 | 29.634 | 31.345 | 29.228 | 31.289 | 32.112 | 0.050 | 0.1777 | 0.4131 | 0 | 0 | 0 |
| 'Malsu1' | 2.263 | 2.588 | 2.136 | 2.138 | 2.316 | 2.288 | 2.097 | 2.421 | 2.336 | 2.193 | 2.420 | 1.997 | 2.160 | 2.011 | 1.975 | 2.258 | 2.181 | 1.774 | 0.114 | 0.0456 | 0.1774 | 0 | 0 | 0 |
| 'Malt1' | 110.180 | 113.586 | 101.968 | 105.855 | 120.167 | 97.042 | 106.503 | 122.214 | 99.915 | 117.151 | 120.758 | 104.818 | 106.899 | 116.999 | 102.416 | 109.571 | 123.941 | 109.284 | -0.062 | 0.0169 | 0.0918 | 0 | 0 | 0 |
| 'Mamdc4' | 0.962 | 0.669 | 1.172 | 1.016 | 1.001 | 1.018 | 1.046 | 0.886 | 1.276 | 0.836 | 0.928 | 1.229 | 1.041 | 0.980 | 1.016 | 1.057 | 1.146 | 1.207 | -0.074 | 0.4166 | 0.6643 | 0 | 0 | 0 |
| 'Maml1' | 16.861 | 17.180 | 16.425 | 15.698 | 15.927 | 16.106 | 15.169 | 15.575 | 15.349 | 17.898 | 17.420 | 18.157 | 16.674 | 17.540 | 17.617 | 16.936 | 15.909 | 17.962 | -0.125 | 0.0002 | 0.0032 | 1 | 0 | 0 |
| 'Man1a' | 5.169 | 4.979 | 5.636 | 5.368 | 5.471 | 5.754 | 4.824 | 5.726 | 5.331 | 5.251 | 5.111 | 5.572 | 4.916 | 4.896 | 5.272 | 5.444 | 4.661 | 5.259 | 0.049 | 0.1922 | 0.4294 | 0 | 0 | 0 |
| 'Man1a2' | 8.031 | 8.521 | 8.577 | 8.776 | 8.277 | 8.663 | 8.123 | 8.635 | 8.267 | 8.148 | 8.566 | 9.951 | 8.360 | 8.955 | 9.594 | 9.209 | 8.608 | 10.388 | -0.118 | 0.0016 | 0.0155 | 1 | 0 | 0 |
| 'Man1b1' | 13.599 | 14.033 | 14.448 | 13.426 | 13.992 | 14.527 | 13.034 | 14.048 | 14.050 | 13.043 | 14.654 | 15.596 | 13.575 | 13.245 | 14.896 | 13.650 | 13.900 | 15.560 | -0.045 | 0.1112 | 0.3157 | 0 | 0 | 0 |
| 'Man1c1' | 3.437 | 3.763 | 2.797 | 3.311 | 3.663 | 3.688 | 3.074 | 4.107 | 2.766 | 3.206 | 3.102 | 3.096 | 3.371 | 4.102 | 2.557 | 4.158 | 3.476 | 3.040 | 0.015 | 0.8289 | 0.9259 | 0 | 0 | 0 |
| 'Man2a1' | 28.960 | 34.477 | 34.906 | 29.701 | 33.650 | 34.268 | 30.258 | 32.999 | 32.963 | 34.273 | 31.208 | 32.787 | 35.486 | 32.962 | 32.613 | 35.237 | 31.397 | 33.817 | -0.049 | 0.1885 | 0.4247 | 0 | 0 | 0 |
| 'Man2a2' | 18.672 | 17.065 | 19.665 | 17.616 | 17.018 | 17.755 | 17.192 | 17.063 | 19.434 | 20.035 | 19.844 | 20.532 | 19.215 | 18.047 | 19.040 | 18.290 | 18.272 | 20.390 | -0.116 | 0.0002 | 0.0027 | 1 | 0 | 0 |
| 'Man2b1' | 54.850 | 49.848 | 62.784 | 55.048 | 52.250 | 58.215 | 53.012 | 51.845 | 61.706 | 56.181 | 52.445 | 63.032 | 56.804 | 50.633 | 58.533 | 50.802 | 54.276 | 59.369 | -0.019 | 0.4813 | 0.7131 | 0 | 0 | 0 |
| 'Man2b2' | 32.223 | 30.012 | 36.363 | 28.329 | 28.932 | 36.515 | 31.252 | 29.587 | 36.431 | 34.500 | 31.582 | 36.789 | 32.836 | 30.530 | 36.469 | 33.308 | 30.984 | 35.497 | -0.073 | 0.0090 | 0.0578 | 0 | 0 | 0 |
| 'Man2c1' | 6.522 | 6.249 | 7.264 | 5.949 | 5.973 | 6.784 | 6.542 | 5.773 | 6.823 | 6.331 | 6.031 | 6.932 | 6.586 | 5.638 | 6.393 | 6.102 | 6.227 | 6.623 | 0.014 | 0.6583 | 0.8350 | 0 | 0 | 0 |
| 'Manba' | 5.498 | 5.054 | 5.345 | 5.234 | 5.639 | 4.924 | 5.200 | 5.316 | 5.048 | 5.721 | 5.488 | 5.732 | 5.001 | 5.058 | 5.518 | 5.279 | 5.165 | 5.345 | -0.044 | 0.2513 | 0.5025 | 0 | 0 | 0 |
| 'Manbal' | 18.956 | 22.756 | 19.049 | 18.297 | 23.847 | 23.709 | 17.782 | 24.624 | 21.400 | 20.650 | 17.399 | 19.074 | 21.959 | 22.127 | 19.941 | 22.926 | 14.954 | 18.809 | 0.087 | 0.2825 | 0.5390 | 0 | 0 | 0 |
| 'Manea' | 2.136 | 2.890 | 2.618 | 1.996 | 2.874 | 2.787 | 2.728 | 2.916 | 2.459 | 2.595 | 2.366 | 2.220 | 2.254 | 2.548 | 1.992 | 2.373 | 2.406 | 1.868 | 0.175 | 0.0116 | 0.0695 | 0 | 0 | 0 |
| 'Maneal' | 1.106 | 1.381 | 2.083 | 1.215 | 1.499 | 1.204 | 1.264 | 1.779 | 1.543 | 1.111 | 0.895 | 1.092 | 0.801 | 0.997 | 1.348 | 0.843 | 0.767 | 1.188 | 0.527 | 0.0001 | 0.0010 | 1 | 1 | 0 |
| 'Manf' | 19.412 | 20.382 | 19.760 | 20.546 | 23.202 | 20.829 | 21.153 | 21.279 | 19.551 | 18.603 | 18.296 | 17.189 | 19.326 | 18.645 | 17.442 | 18.778 | 19.389 | 17.612 | 0.161 | 0.0000 | 0.0000 | 1 | 0 | 0 |
| 'Mansc1' | 3.229 | 3.975 | 3.862 | 4.654 | 3.155 | 3.787 | 4.043 | 3.432 | 3.577 | 4.117 | 3.495 | 3.951 | 3.154 | 3.293 | 3.945 | 2.695 | 2.919 | 3.304 | 0.107 | 0.2314 | 0.4788 | 0 | 0 | 0 |
| 'Map1a' | 0.402 | 0.261 | 0.403 | 0.350 | 0.331 | 0.412 | 0.418 | 0.452 | 0.407 | 0.353 | 0.398 | 0.454 | 0.443 | 0.431 | 0.462 | 0.330 | 0.365 | 0.455 | -0.109 | 0.2124 | 0.4575 | 0 | 0 | 0 |
| 'Map1b' | 6.522 | 5.179 | 6.726 | 6.968 | 6.163 | 6.153 | 6.092 | 6.641 | 6.180 | 5.377 | 6.724 | 6.254 | 5.204 | 5.979 | 6.635 | 6.207 | 5.600 | 7.554 | 0.021 | 0.7432 | 0.8842 | 0 | 0 | 0 |
| 'Map1lc3a' | 5.087 | 2.570 | 2.688 | 3.408 | 2.485 | 2.256 | 3.174 | 3.106 | 2.412 | 3.550 | 2.846 | 2.099 | 4.006 | 3.486 | 2.401 | 3.750 | 2.887 | 1.790 | 0.034 | 0.7396 | 0.8819 | 0 | 0 | 0 |
| 'Map1lc3b' | 21.628 | 16.814 | 13.103 | 16.120 | 16.905 | 13.645 | 13.947 | 17.604 | 13.003 | 20.547 | 18.693 | 13.003 | 20.415 | 20.680 | 13.310 | 19.986 | 17.041 | 12.691 | -0.125 | 0.0460 | 0.1783 | 0 | 0 | 0 |
| 'Map1s' | 30.049 | 31.180 | 34.952 | 29.047 | 28.643 | 32.773 | 28.534 | 29.092 | 33.988 | 31.521 | 29.894 | 37.116 | 33.967 | 28.689 | 33.560 | 32.672 | 30.321 | 37.453 | -0.096 | 0.0031 | 0.0256 | 1 | 0 | 0 |
| 'Map2k1' | 23.779 | 26.553 | 22.590 | 24.150 | 24.594 | 24.555 | 24.005 | 26.349 | 23.308 | 24.622 | 25.128 | 27.701 | 24.315 | 27.237 | 26.123 | 26.146 | 24.749 | 25.932 | -0.091 | 0.0061 | 0.0437 | 1 | 0 | 0 |
| 'Map2k2' | 19.264 | 20.118 | 20.805 | 18.089 | 18.025 | 21.969 | 18.846 | 18.898 | 21.209 | 18.562 | 17.502 | 22.014 | 20.514 | 17.226 | 21.264 | 21.003 | 16.915 | 21.792 | -0.005 | 0.8946 | 0.9571 | 0 | 0 | 0 |
| 'Map2k3' | 17.930 | 17.898 | 15.907 | 17.429 | 16.989 | 17.160 | 16.930 | 16.715 | 16.819 | 18.827 | 19.302 | 19.937 | 19.313 | 20.267 | 19.784 | 20.953 | 19.610 | 20.594 | -0.227 | 0.0000 | 0.0000 | 1 | 0 | 0 |
| 'Map2k4' | 6.979 | 8.299 | 7.305 | 6.879 | 7.794 | 7.738 | 7.519 | 8.960 | 7.380 | 7.043 | 7.560 | 6.900 | 6.798 | 7.719 | 6.948 | 7.839 | 6.808 | 6.280 | 0.099 | 0.0210 | 0.1065 | 0 | 0 | 0 |
| 'Map2k5' | 18.609 | 17.023 | 16.474 | 14.981 | 16.715 | 17.563 | 16.760 | 19.333 | 16.648 | 17.733 | 16.162 | 16.674 | 17.064 | 14.481 | 15.822 | 16.888 | 15.515 | 16.585 | 0.060 | 0.2056 | 0.4492 | 0 | 0 | 0 |
| 'Map2k6' | 3.916 | 3.588 | 3.510 | 4.642 | 3.679 | 3.386 | 4.066 | 3.776 | 3.748 | 3.730 | 3.334 | 3.809 | 3.510 | 3.571 | 3.531 | 3.600 | 3.449 | 3.603 | 0.074 | 0.1640 | 0.3960 | 0 | 0 | 0 |
| 'Map3k1' | 18.229 | 17.273 | 17.020 | 17.322 | 17.933 | 16.439 | 16.335 | 18.020 | 16.714 | 17.810 | 20.601 | 18.856 | 16.339 | 19.168 | 18.521 | 19.132 | 18.766 | 19.671 | -0.131 | 0.0006 | 0.0070 | 1 | 0 | 0 |
| 'Map3k10' | 3.810 | 3.782 | 4.290 | 3.376 | 3.800 | 4.658 | 3.366 | 4.459 | 3.750 | 3.683 | 3.323 | 4.037 | 4.239 | 3.533 | 3.978 | 4.049 | 3.180 | 3.694 | 0.058 | 0.3528 | 0.6063 | 0 | 0 | 0 |
| 'Map3k11' | 42.163 | 39.176 | 42.100 | 39.434 | 38.232 | 46.418 | 38.325 | 36.811 | 46.856 | 40.662 | 38.198 | 48.189 | 38.920 | 39.317 | 46.260 | 41.310 | 40.107 | 47.842 | -0.053 | 0.0795 | 0.2537 | 0 | 0 | 0 |
| 'Map3k12' | 2.887 | 2.584 | 3.013 | 2.993 | 2.714 | 3.141 | 2.982 | 2.527 | 3.007 | 3.028 | 2.683 | 3.007 | 2.963 | 2.876 | 2.992 | 2.635 | 2.931 | 2.749 | -0.012 | 0.7749 | 0.8994 | 0 | 0 | 0 |
| 'Map3k14' | 8.454 | 7.779 | 8.914 | 7.705 | 7.403 | 8.731 | 8.589 | 8.315 | 8.744 | 7.945 | 7.698 | 9.087 | 8.510 | 7.855 | 8.627 | 8.702 | 7.780 | 8.726 | -0.015 | 0.6278 | 0.8154 | 0 | 0 | 0 |
| 'Map3k2' | 10.532 | 10.818 | 9.793 | 10.421 | 10.225 | 9.673 | 9.992 | 10.142 | 9.103 | 10.310 | 11.382 | 11.267 | 9.796 | 11.168 | 10.221 | 10.280 | 11.529 | 11.682 | -0.118 | 0.0033 | 0.0270 | 1 | 0 | 0 |
| 'Map3k3' | 22.224 | 20.723 | 22.103 | 21.647 | 19.610 | 21.558 | 21.390 | 22.233 | 20.546 | 21.035 | 21.083 | 22.533 | 20.002 | 21.866 | 23.283 | 21.959 | 20.605 | 24.845 | -0.047 | 0.2241 | 0.4711 | 0 | 0 | 0 |
| 'Map3k4' | 17.282 | 17.126 | 16.571 | 16.495 | 15.700 | 17.992 | 15.586 | 16.240 | 17.073 | 16.719 | 15.850 | 17.621 | 16.315 | 15.944 | 16.762 | 16.217 | 15.411 | 16.071 | 0.020 | 0.5036 | 0.7306 | 0 | 0 | 0 |
| 'Map3k5' | 22.488 | 22.501 | 23.426 | 22.506 | 23.343 | 22.920 | 20.807 | 24.400 | 23.078 | 25.743 | 27.113 | 24.219 | 22.763 | 24.822 | 22.311 | 24.688 | 27.046 | 24.539 | -0.128 | 0.0003 | 0.0035 | 1 | 0 | 0 |
| 'Map3k7' | 10.986 | 11.886 | 11.812 | 10.623 | 11.708 | 11.359 | 10.493 | 12.661 | 11.202 | 12.671 | 11.377 | 10.755 | 11.395 | 11.551 | 10.827 | 12.086 | 11.323 | 10.474 | -0.005 | 0.9014 | 0.9611 | 0 | 0 | 0 |
| 'Map3k8' | 4.586 | 3.661 | 3.494 | 4.374 | 3.568 | 3.212 | 3.745 | 3.867 | 3.479 | 4.086 | 3.635 | 3.566 | 3.679 | 3.826 | 3.351 | 4.092 | 3.937 | 3.497 | 0.003 | 0.9589 | 0.9847 | 0 | 0 | 0 |
| 'Map4' | 6.184 | 5.731 | 6.870 | 6.170 | 6.184 | 7.286 | 6.409 | 6.139 | 6.886 | 6.823 | 6.857 | 7.055 | 6.928 | 6.432 | 7.208 | 6.155 | 6.651 | 7.198 | -0.095 | 0.0014 | 0.0140 | 1 | 0 | 0 |
| 'Map4k1' | 136.740 | 143.331 | 146.357 | 138.090 | 138.937 | 144.370 | 140.710 | 144.153 | 142.531 | 152.042 | 143.183 | 138.373 | 164.111 | 135.886 | 135.724 | 137.776 | 133.757 | 138.816 | -0.014 | 0.6605 | 0.8362 | 0 | 0 | 0 |
| 'Map4k2' | 26.140 | 22.997 | 28.077 | 24.261 | 23.166 | 27.093 | 25.239 | 23.255 | 28.401 | 25.278 | 22.025 | 27.086 | 26.518 | 22.693 | 25.693 | 23.745 | 23.825 | 26.284 | 0.023 | 0.3476 | 0.6031 | 0 | 0 | 0 |
| 'Map4k3' | 2.529 | 2.869 | 3.669 | 3.092 | 2.965 | 3.258 | 3.603 | 3.367 | 3.638 | 2.753 | 2.950 | 3.043 | 2.677 | 2.854 | 2.609 | 2.732 | 3.198 | 2.551 | 0.184 | 0.0097 | 0.0613 | 0 | 0 | 0 |
| 'Map4k4' | 31.719 | 29.572 | 31.545 | 30.800 | 29.879 | 30.677 | 28.913 | 30.561 | 29.812 | 29.286 | 29.548 | 29.718 | 27.399 | 28.060 | 30.550 | 27.652 | 27.780 | 30.779 | 0.059 | 0.0393 | 0.1619 | 0 | 0 | 0 |
| 'Map4k5' | 0.453 | 0.417 | 0.480 | 0.423 | 0.431 | 0.398 | 0.522 | 0.531 | 0.481 | 0.348 | 0.444 | 0.344 | 0.360 | 0.344 | 0.278 | 0.289 | 0.326 | 0.272 | 0.456 | 0.0000 | 0.0000 | 1 | 1 | 0 |
| 'Map6' | 0.638 | 0.718 | 0.810 | 0.483 | 0.738 | 0.881 | 0.608 | 0.650 | 0.707 | 0.889 | 0.688 | 0.823 | 0.840 | 0.922 | 1.135 | 0.868 | 0.672 | 0.901 | -0.320 | 0.0006 | 0.0065 | 1 | 0 | 0 |
| 'Map7' | 10.587 | 12.284 | 9.611 | 11.401 | 11.337 | 10.649 | 10.166 | 11.895 | 10.828 | 12.226 | 13.834 | 10.823 | 13.266 | 14.395 | 10.882 | 12.028 | 12.189 | 10.635 | -0.165 | 0.0001 | 0.0014 | 1 | 0 | 0 |
| 'Map7d1' | 40.037 | 33.181 | 40.605 | 36.264 | 32.879 | 42.280 | 39.157 | 31.955 | 42.150 | 37.346 | 35.965 | 51.428 | 39.577 | 36.469 | 47.788 | 38.590 | 38.209 | 52.651 | -0.164 | 0.0000 | 0.0007 | 1 | 0 | 0 |
| 'Map9' | 0.510 | 0.566 | 0.486 | 0.715 | 0.678 | 0.696 | 0.614 | 0.538 | 0.701 | 0.663 | 0.692 | 0.530 | 0.652 | 0.685 | 0.535 | 0.654 | 0.604 | 0.412 | 0.024 | 0.8292 | 0.9261 | 0 | 0 | 0 |
| 'Mapk1' | 38.576 | 42.997 | 37.221 | 40.896 | 42.788 | 38.594 | 42.275 | 40.733 | 38.840 | 36.999 | 41.901 | 39.457 | 39.318 | 41.803 | 38.597 | 39.347 | 43.665 | 39.285 | -0.001 | 0.9594 | 0.9850 | 0 | 0 | 0 |
| 'Mapk11' | 16.232 | 13.086 | 18.857 | 14.810 | 13.084 | 17.942 | 14.478 | 15.268 | 18.045 | 13.800 | 12.423 | 17.258 | 11.920 | 12.408 | 16.668 | 13.030 | 12.311 | 15.609 | 0.171 | 0.0001 | 0.0017 | 1 | 0 | 0 |
| 'Mapk14' | 26.329 | 28.310 | 25.022 | 25.633 | 28.655 | 26.066 | 24.116 | 28.522 | 26.111 | 27.401 | 29.278 | 29.399 | 29.842 | 29.786 | 29.653 | 28.155 | 27.197 | 30.504 | -0.143 | 0.0000 | 0.0003 | 1 | 0 | 0 |
| 'Mapk1ip1' | 6.356 | 5.401 | 6.262 | 5.580 | 5.500 | 5.778 | 6.814 | 5.169 | 6.181 | 6.130 | 6.152 | 7.151 | 6.143 | 5.804 | 6.466 | 4.805 | 5.947 | 6.112 | -0.060 | 0.3089 | 0.5671 | 0 | 0 | 0 |
| 'Mapk3' | 22.729 | 23.978 | 23.579 | 20.248 | 24.062 | 23.664 | 23.364 | 24.704 | 24.619 | 22.839 | 20.703 | 21.900 | 22.148 | 23.007 | 23.798 | 21.419 | 20.922 | 21.112 | 0.082 | 0.0208 | 0.1056 | 0 | 0 | 0 |
| 'Mapk6' | 8.734 | 7.504 | 7.246 | 7.628 | 7.107 | 7.039 | 7.163 | 7.702 | 6.759 | 7.529 | 7.641 | 7.230 | 7.101 | 8.050 | 6.917 | 8.071 | 7.181 | 7.106 | -0.008 | 0.8457 | 0.9343 | 0 | 0 | 0 |
| 'Mapk7' | 2.526 | 2.311 | 2.850 | 2.449 | 2.404 | 2.849 | 2.417 | 2.442 | 2.774 | 2.629 | 2.104 | 2.532 | 2.595 | 2.252 | 2.499 | 2.403 | 2.203 | 2.475 | 0.080 | 0.0340 | 0.1469 | 0 | 0 | 0 |
| 'Mapk8' | 3.200 | 3.276 | 3.370 | 2.979 | 3.412 | 3.196 | 3.194 | 3.540 | 3.011 | 3.198 | 3.223 | 3.148 | 3.246 | 3.285 | 3.126 | 3.147 | 3.268 | 2.977 | 0.020 | 0.5643 | 0.7734 | 0 | 0 | 0 |
| 'Mapk8ip1' | 1.242 | 1.836 | 1.572 | 1.674 | 1.431 | 1.529 | 1.389 | 1.564 | 1.698 | 1.365 | 1.363 | 1.476 | 1.457 | 1.308 | 1.420 | 1.856 | 1.436 | 1.429 | 0.077 | 0.3376 | 0.5955 | 0 | 0 | 0 |
| 'Mapk8ip3' | 9.292 | 7.441 | 10.078 | 8.797 | 8.032 | 9.055 | 8.311 | 8.195 | 9.869 | 9.387 | 8.881 | 10.832 | 8.455 | 8.135 | 9.396 | 8.455 | 9.372 | 10.308 | -0.085 | 0.0438 | 0.1733 | 0 | 0 | 0 |

| | | | | | | | | | | | | | | | | | | | | | | | | |
|---|---|---|---|---|---|---|---|---|---|---|---|---|---|---|---|---|---|---|---|---|---|---|---|---|
| 'Mapk9' | 5.647 | 6.462 | 6.056 | 5.337 | 6.423 | 5.978 | 5.680 | 5.897 | 6.020 | 5.925 | 5.277 | 5.423 | 5.571 | 5.575 | 5.313 | 5.817 | 5.512 | 4.701 | 0.114 | 0.0041 | 0.0326 | 1 | 0 | 0 |
| 'Mapkap1' | 15.084 | 16.850 | 16.333 | 15.050 | 17.285 | 17.340 | 16.681 | 17.682 | 16.570 | 17.884 | 16.039 | 16.339 | 16.501 | 16.580 | 16.524 | 17.701 | 15.253 | 16.658 | -0.016 | 0.6685 | 0.8400 | 0 | 0 | 0 |
| 'Mapkapk2' | 57.797 | 56.662 | 61.085 | 53.032 | 57.661 | 66.730 | 54.463 | 61.252 | 56.409 | 60.084 | 58.259 | 62.337 | 61.657 | 62.253 | 65.469 | 61.298 | 58.655 | 65.380 | -0.091 | 0.0058 | 0.0422 | 1 | 0 | 0 |
| 'Mapkapk3' | 4.900 | 5.068 | 5.211 | 4.342 | 4.882 | 5.424 | 4.804 | 5.417 | 5.210 | 4.447 | 4.232 | 5.647 | 4.690 | 4.870 | 5.635 | 4.940 | 5.101 | 5.660 | -0.009 | 0.8294 | 0.9261 | 0 | 0 | 0 |
| 'Mapkapk5' | 3.095 | 2.943 | 3.334 | 3.082 | 2.858 | 3.367 | 2.986 | 2.822 | 2.685 | 3.241 | 2.587 | 3.239 | 3.055 | 3.013 | 3.299 | 3.041 | 2.666 | 3.662 | -0.042 | 0.4466 | 0.6880 | 0 | 0 | 0 |
| 'Mapkbp1' | 3.732 | 3.890 | 4.439 | 3.830 | 3.740 | 4.573 | 3.800 | 4.143 | 4.743 | 4.310 | 4.063 | 3.757 | 4.227 | 4.154 | 3.555 | 4.438 | 3.830 | 3.224 | 0.048 | 0.4504 | 0.6902 | 0 | 0 | 0 |
| 'Mapre1' | 25.492 | 27.520 | 27.244 | 27.264 | 29.074 | 26.635 | 28.324 | 27.970 | 27.395 | 27.775 | 27.289 | 25.717 | 26.898 | 27.951 | 26.040 | 26.115 | 28.155 | 25.744 | 0.021 | 0.3647 | 0.6165 | 0 | 0 | 0 |
| 'Mapre2' | 20.057 | 22.573 | 19.812 | 19.834 | 22.216 | 20.981 | 20.848 | 21.811 | 20.806 | 21.612 | 19.448 | 18.803 | 22.932 | 21.066 | 19.459 | 22.438 | 20.022 | 19.271 | 0.021 | 0.5374 | 0.7562 | 0 | 0 | 0 |
| 'Mapre3' | 4.751 | 4.925 | 5.876 | 4.560 | 5.030 | 5.690 | 5.039 | 5.289 | 5.573 | 5.481 | 5.188 | 5.173 | 5.702 | 5.212 | 4.870 | 4.705 | 5.275 | 4.855 | 0.000 | 0.9982 | 0.9994 | 0 | 0 | 0 |
| 'Marc2' | 5.981 | 6.511 | 6.333 | 6.330 | 6.990 | 6.540 | 6.487 | 6.368 | 6.857 | 6.037 | 6.027 | 6.261 | 6.179 | 6.214 | 6.277 | 6.446 | 5.792 | 6.376 | 0.059 | 0.0749 | 0.2436 | 0 | 0 | 0 |
| 'March2' | 6.066 | 7.247 | 6.559 | 6.119 | 7.163 | 6.767 | 6.056 | 6.761 | 6.587 | 6.630 | 6.766 | 6.990 | 6.747 | 7.138 | 7.102 | 7.077 | 6.900 | 6.893 | -0.082 | 0.0062 | 0.0442 | 1 | 0 | 0 |
| 'March5' | 22.166 | 22.545 | 20.960 | 21.403 | 22.379 | 21.421 | 21.373 | 23.405 | 20.920 | 20.853 | 22.140 | 21.401 | 22.097 | 21.428 | 20.722 | 23.398 | 22.974 | 20.351 | 0.000 | 0.9980 | 0.9994 | 0 | 0 | 0 |
| 'March6' | 33.858 | 33.113 | 32.182 | 32.964 | 32.927 | 30.434 | 32.896 | 33.479 | 30.513 | 31.840 | 37.008 | 35.069 | 31.162 | 35.487 | 32.784 | 32.026 | 34.765 | 34.345 | -0.069 | 0.0371 | 0.1561 | 0 | 0 | 0 |
| 'March7' | 9.584 | 11.549 | 9.075 | 9.056 | 11.052 | 9.144 | 9.585 | 11.019 | 9.047 | 10.353 | 9.922 | 8.360 | 9.704 | 10.231 | 7.968 | 10.476 | 9.933 | 7.207 | 0.077 | 0.0568 | 0.2049 | 0 | 0 | 0 |
| 'March9' | 7.485 | 6.613 | 7.206 | 7.294 | 6.402 | 7.492 | 6.737 | 6.320 | 6.577 | 5.699 | 6.196 | 7.710 | 6.558 | 6.727 | 7.641 | 6.937 | 6.242 | 8.068 | -0.003 | 0.9568 | 0.9839 | 0 | 0 | 0 |
| 'Marcks' | 2.491 | 1.782 | 3.075 | 2.308 | 2.214 | 2.944 | 2.990 | 2.543 | 2.885 | 3.342 | 2.780 | 3.370 | 2.951 | 3.012 | 3.489 | 3.291 | 3.476 | 3.109 | -0.315 | 0.0000 | 0.0005 | 1 | 0 | 0 |
| 'Marcksl1' | 85.680 | 84.911 | 83.101 | 80.570 | 83.055 | 83.855 | 73.956 | 82.519 | 90.769 | 92.327 | 85.750 | 93.701 | 88.627 | 92.820 | 90.929 | 98.055 | 90.338 | 88.590 | -0.143 | 0.0000 | 0.0003 | 1 | 0 | 0 |
| 'Marf1' | 19.147 | 16.255 | 18.422 | 20.506 | 16.168 | 16.532 | 18.268 | 16.405 | 17.929 | 16.515 | 19.855 | 22.530 | 16.971 | 17.282 | 19.894 | 16.071 | 21.855 | 22.295 | -0.125 | 0.0681 | 0.2290 | 0 | 0 | 0 |
| 'Mark2' | 17.192 | 16.473 | 17.307 | 16.456 | 16.439 | 17.809 | 16.108 | 16.265 | 17.442 | 16.712 | 16.802 | 17.835 | 16.813 | 16.583 | 17.270 | 16.358 | 16.262 | 17.611 | -0.017 | 0.3691 | 0.6203 | 0 | 0 | 0 |
| 'Mark3' | 36.045 | 36.750 | 34.052 | 37.710 | 36.653 | 37.166 | 35.239 | 36.473 | 36.151 | 36.436 | 36.368 | 36.073 | 36.080 | 37.329 | 35.877 | 35.501 | 35.384 | 37.254 | -0.011 | 0.6624 | 0.8371 | 0 | 0 | 0 |
| 'Mark4' | 20.716 | 17.587 | 20.037 | 17.501 | 17.803 | 20.374 | 16.196 | 18.143 | 20.708 | 19.082 | 17.477 | 20.414 | 19.302 | 18.677 | 20.051 | 18.782 | 16.289 | 21.284 | -0.027 | 0.5083 | 0.7336 | 0 | 0 | 0 |
| 'Mars' | 8.847 | 8.012 | 9.601 | 9.037 | 9.141 | 10.253 | 8.451 | 10.015 | 9.685 | 8.984 | 8.822 | 9.939 | 9.929 | 9.413 | 10.092 | 9.459 | 8.662 | 9.477 | -0.038 | 0.3199 | 0.5782 | 0 | 0 | 0 |
| 'Mars2' | 1.119 | 1.257 | 1.545 | 1.244 | 1.359 | 1.249 | 1.364 | 1.293 | 1.312 | 1.514 | 1.485 | 1.713 | 1.391 | 1.191 | 1.515 | 1.350 | 1.345 | 1.572 | -0.171 | 0.0062 | 0.0440 | 1 | 0 | 0 |
| 'Marveld1' | 5.486 | 4.967 | 5.707 | 5.187 | 5.130 | 6.553 | 5.020 | 5.108 | 6.034 | 4.579 | 4.110 | 5.582 | 4.189 | 4.911 | 5.418 | 4.738 | 4.403 | 5.783 | 0.162 | 0.0003 | 0.0037 | 1 | 0 | 0 |
| 'Marveld2' | 1.282 | 1.190 | 0.857 | 1.268 | 1.365 | 1.303 | 1.399 | 1.247 | 1.356 | 1.489 | 0.708 | 0.947 | 1.249 | 1.058 | 1.093 | 1.162 | 1.093 | 0.671 | 0.249 | 0.0369 | 0.1553 | 0 | 0 | 0 |
| 'Mast2' | 5.895 | 4.971 | 6.356 | 5.733 | 5.077 | 6.348 | 5.441 | 5.402 | 6.346 | 6.275 | 6.009 | 6.582 | 6.539 | 5.849 | 6.516 | 5.916 | 6.293 | 6.733 | -0.148 | 0.0000 | 0.0002 | 1 | 0 | 0 |
| 'Mast3' | 50.780 | 47.425 | 54.399 | 47.292 | 46.018 | 52.312 | 46.351 | 48.108 | 53.854 | 52.637 | 47.533 | 54.900 | 52.152 | 44.200 | 52.222 | 47.524 | 44.616 | 51.247 | -0.011 | 0.6929 | 0.8555 | 0 | 0 | 0 |
| 'Mast4' | 0.592 | 0.630 | 0.530 | 0.674 | 0.548 | 0.597 | 0.620 | 0.608 | 0.568 | 0.679 | 0.680 | 0.623 | 0.700 | 0.670 | 0.611 | 0.749 | 0.667 | 0.625 | -0.170 | 0.0001 | 0.0010 | 1 | 0 | 0 |
| 'Mastl' | 1.084 | 1.001 | 1.361 | 1.212 | 1.427 | 1.257 | 1.444 | 1.296 | 1.426 | 0.850 | 1.215 | 1.303 | 1.434 | 1.254 | 1.313 | 1.228 | 1.076 | 0.915 | 0.108 | 0.2759 | 0.5317 | 0 | 0 | 0 |
| 'Mat2a' | 22.387 | 16.844 | 15.174 | 20.882 | 17.123 | 14.769 | 21.081 | 17.371 | 15.209 | 15.834 | 19.974 | 19.917 | 17.465 | 17.865 | 17.441 | 16.191 | 21.576 | 19.836 | -0.069 | 0.4585 | 0.6955 | 0 | 0 | 0 |
| 'Mat2b' | 15.772 | 17.585 | 15.041 | 15.172 | 18.002 | 16.266 | 15.859 | 17.337 | 14.974 | 15.625 | 16.441 | 16.441 | 15.410 | 16.197 | 15.723 | 16.539 | 16.338 | 14.815 | 0.011 | 0.7122 | 0.8661 | 0 | 0 | 0 |
| 'Matk' | 21.924 | 21.801 | 23.661 | 19.586 | 22.251 | 25.205 | 21.402 | 21.103 | 24.050 | 24.529 | 20.095 | 25.615 | 24.319 | 22.613 | 23.367 | 24.448 | 22.560 | 23.437 | -0.081 | 0.0232 | 0.1143 | 0 | 0 | 0 |
| 'Matn2' | 0.942 | 1.100 | 1.287 | 1.114 | 0.942 | 1.090 | 1.347 | 0.972 | 1.440 | 0.929 | 0.859 | 0.938 | 0.777 | 1.325 | 0.895 | 1.040 | 1.039 | 0.907 | 0.232 | 0.0446 | 0.1751 | 0 | 0 | 0 |
| 'Matr3' | 13.325 | 14.286 | 13.653 | 15.312 | 14.271 | 12.538 | 15.198 | 14.051 | 12.575 | 13.594 | 14.509 | 14.283 | 12.631 | 14.689 | 12.666 | 12.678 | 15.339 | 12.945 | 0.011 | 0.7926 | 0.9074 | 0 | 0 | 0 |
| 'Mau2' | 31.776 | 28.561 | 32.904 | 30.675 | 28.992 | 31.695 | 30.091 | 29.488 | 33.438 | 29.869 | 28.908 | 36.473 | 30.908 | 29.137 | 33.162 | 29.086 | 32.303 | 34.275 | -0.044 | 0.1528 | 0.3800 | 0 | 0 | 0 |
| 'Mavs' | 6.310 | 6.208 | 6.691 | 5.823 | 6.077 | 6.303 | 6.087 | 6.533 | 6.409 | 5.938 | 6.087 | 6.772 | 5.930 | 6.191 | 6.909 | 6.131 | 6.009 | 6.724 | -0.016 | 0.6559 | 0.8333 | 0 | 0 | 0 |
| 'Max' | 115.628 | 122.828 | 101.021 | 117.533 | 112.887 | 106.874 | 111.946 | 113.072 | 103.535 | 101.237 | 118.626 | 111.604 | 106.529 | 118.274 | 114.830 | 103.930 | 115.584 | 113.448 | -0.010 | 0.7925 | 0.9074 | 0 | 0 | 0 |
| 'Maz' | 88.568 | 81.346 | 100.616 | 81.672 | 80.728 | 104.281 | 86.131 | 82.170 | 98.535 | 87.673 | 81.426 | 99.445 | 91.938 | 85.606 | 104.031 | 88.918 | 83.567 | 107.103 | -0.056 | 0.0178 | 0.0950 | 0 | 0 | 0 |
| 'Mb21d1' | 1.687 | 1.398 | 1.885 | 1.794 | 1.128 | 1.767 | 1.908 | 1.611 | 2.025 | 1.536 | 1.692 | 1.806 | 1.588 | 1.560 | 1.701 | 1.667 | 1.206 | 1.780 | 0.059 | 0.5045 | 0.7312 | 0 | 0 | 0 |
| 'Mbd1' | 25.113 | 27.206 | 26.161 | 25.683 | 23.454 | 25.527 | 24.860 | 25.217 | 26.905 | 24.520 | 25.392 | 27.686 | 23.152 | 23.539 | 26.469 | 25.480 | 23.159 | 26.080 | 0.018 | 0.5957 | 0.7944 | 0 | 0 | 0 |
| 'Mbd2' | 36.400 | 36.840 | 34.959 | 35.070 | 37.101 | 36.279 | 40.583 | 36.704 | 32.404 | 33.396 | 33.240 | 32.073 | 33.853 | 36.055 | 34.268 | 35.425 | 34.974 | 34.531 | 0.073 | 0.0290 | 0.1319 | 0 | 0 | 0 |
| 'Mbd3' | 41.963 | 38.848 | 43.835 | 38.417 | 37.904 | 46.704 | 38.920 | 36.718 | 44.097 | 41.767 | 37.024 | 48.164 | 46.872 | 39.030 | 48.021 | 43.910 | 37.481 | 49.887 | -0.102 | 0.0010 | 0.0104 | 1 | 0 | 0 |
| 'Mbd4' | 1.662 | 1.919 | 1.937 | 1.976 | 1.933 | 1.971 | 2.073 | 1.762 | 1.843 | 1.487 | 2.047 | 2.081 | 1.836 | 1.945 | 2.157 | 1.757 | 2.331 | 2.212 | -0.075 | 0.2482 | 0.4991 | 0 | 0 | 0 |
| 'Mbd5' | 1.123 | 1.125 | 1.170 | 1.096 | 1.150 | 1.084 | 1.350 | 1.030 | 1.192 | 1.171 | 1.402 | 1.297 | 1.091 | 1.237 | 1.053 | 1.114 | 1.174 | 1.263 | -0.078 | 0.1904 | 0.4268 | 0 | 0 | 0 |
| 'Mbd6' | 21.289 | 19.041 | 23.535 | 20.319 | 18.401 | 21.283 | 21.695 | 20.352 | 23.593 | 21.995 | 22.600 | 23.626 | 21.608 | 20.868 | 22.179 | 19.820 | 21.475 | 21.214 | -0.056 | 0.1408 | 0.3634 | 0 | 0 | 0 |
| 'Mbip' | 13.412 | 15.909 | 15.781 | 13.704 | 17.685 | 14.200 | 15.844 | 16.185 | 14.027 | 16.825 | 14.974 | 12.740 | 16.129 | 15.535 | 13.172 | 14.520 | 15.658 | 11.252 | 0.061 | 0.3040 | 0.5614 | 0 | 0 | 0 |
| 'Mblac2' | 3.024 | 3.345 | 3.293 | 3.403 | 2.893 | 3.122 | 3.346 | 3.462 | 3.049 | 2.854 | 3.438 | 3.026 | 2.888 | 3.285 | 2.833 | 2.558 | 3.384 | 3.449 | 0.053 | 0.4272 | 0.6735 | 0 | 0 | 0 |
| 'Mbnl1' | 27.949 | 29.572 | 29.626 | 29.676 | 29.321 | 27.806 | 30.146 | 29.732 | 28.427 | 29.871 | 32.020 | 27.926 | 28.214 | 30.210 | 27.599 | 27.678 | 29.073 | 27.877 | 0.000 | 0.9978 | 0.9994 | 0 | 0 | 0 |
| 'Mbnl2' | 14.481 | 16.680 | 15.290 | 14.241 | 15.536 | 14.208 | 15.585 | 16.815 | 15.465 | 13.016 | 17.015 | 15.937 | 12.501 | 16.786 | 15.027 | 14.302 | 17.941 | 14.710 | 0.005 | 0.9044 | 0.9624 | 0 | 0 | 0 |
| 'Mbnl3' | 1.976 | 1.676 | 2.140 | 2.156 | 1.774 | 1.836 | 1.907 | 1.770 | 1.988 | 1.925 | 1.855 | 1.947 | 2.053 | 1.997 | 1.935 | 1.769 | 1.905 | 1.953 | -0.019 | 0.7095 | 0.8653 | 0 | 0 | 0 |
| 'Mboat1' | 4.779 | 3.761 | 5.289 | 4.698 | 4.421 | 5.156 | 4.086 | 5.010 | 5.253 | 5.715 | 4.409 | 6.392 | 4.097 | 5.985 | 6.181 | 5.602 | 5.318 | 5.545 | -0.220 | 0.0021 | 0.0191 | 1 | 0 | 0 |
| 'Mboat7' | 5.943 | 5.775 | 6.221 | 5.321 | 5.769 | 6.081 | 5.594 | 5.937 | 6.136 | 6.267 | 5.541 | 6.546 | 5.925 | 5.850 | 6.256 | 6.336 | 5.999 | 6.406 | -0.072 | 0.0093 | 0.0594 | 0 | 0 | 0 |
| 'Mbp' | 15.490 | 14.951 | 15.116 | 14.025 | 14.760 | 15.335 | 14.334 | 14.603 | 14.595 | 15.114 | 14.846 | 17.203 | 13.267 | 14.622 | 16.260 | 14.995 | 14.344 | 16.866 | -0.054 | 0.1311 | 0.3479 | 0 | 0 | 0 |
| 'Mbtd1' | 3.379 | 3.230 | 3.231 | 3.347 | 3.389 | 2.900 | 3.609 | 3.194 | 3.052 | 3.297 | 3.552 | 3.633 | 3.043 | 3.553 | 3.195 | 3.218 | 3.577 | 3.395 | -0.068 | 0.1227 | 0.3341 | 0 | 0 | 0 |
| 'Mbtps1' | 60.628 | 62.010 | 62.376 | 60.578 | 59.822 | 59.290 | 60.095 | 58.763 | 62.537 | 61.520 | 62.300 | 63.439 | 56.544 | 62.875 | 58.666 | 59.219 | 61.022 | 62.571 | -0.016 | 0.5444 | 0.7606 | 0 | 0 | 0 |
| 'Mbtps2' | 1.361 | 1.687 | 1.659 | 1.618 | 1.658 | 1.492 | 1.498 | 1.605 | 1.504 | 1.532 | 1.712 | 1.536 | 1.543 | 1.401 | 1.498 | 1.461 | 1.660 | 1.406 | 0.022 | 0.6460 | 0.8271 | 0 | 0 | 0 |
| 'Mcat' | 15.623 | 15.417 | 14.617 | 11.975 | 15.538 | 17.314 | 13.319 | 16.462 | 15.661 | 14.942 | 15.014 | 15.287 | 16.259 | 15.567 | 14.687 | 15.824 | 14.654 | 14.434 | -0.016 | 0.7759 | 0.9000 | 0 | 0 | 0 |
| 'Mccc1' | 2.650 | 2.875 | 2.684 | 2.824 | 2.520 | 2.556 | 2.581 | 2.652 | 2.920 | 3.020 | 2.818 | 2.612 | 2.627 | 2.587 | 2.678 | 2.373 | 2.697 | 2.757 | -0.005 | 0.9187 | 0.9692 | 0 | 0 | 0 |
| 'Mccc2' | 9.014 | 10.200 | 9.695 | 9.221 | 11.219 | 10.039 | 9.890 | 9.625 | 9.556 | 9.771 | 9.531 | 8.949 | 8.817 | 7.684 | 8.415 | 10.104 | 8.503 | 7.678 | 0.146 | 0.0113 | 0.0681 | 0 | 0 | 0 |
| 'Mcee' | 3.183 | 3.286 | 3.261 | 3.083 | 3.972 | 3.328 | 2.841 | 3.738 | 3.387 | 3.664 | 3.487 | 3.328 | 2.732 | 3.573 | 2.600 | 3.135 | 2.953 | 3.018 | 0.069 | 0.4165 | 0.6643 | 0 | 0 | 0 |
| 'Mcf2l' | 0.649 | 0.860 | 0.492 | 0.654 | 0.752 | 0.673 | 0.577 | 0.849 | 0.673 | 0.456 | 0.568 | 0.392 | 0.594 | 0.539 | 0.337 | 0.409 | 0.569 | 0.294 | 0.573 | 0.0000 | 0.0000 | 1 | 1 | 0 |
| 'Mcfd2' | 5.484 | 6.294 | 5.728 | 5.324 | 5.465 | 5.831 | 6.054 | 6.112 | 5.952 | 5.715 | 5.727 | 6.561 | 5.888 | 5.420 | 5.908 | 5.681 | 5.901 | 6.397 | -0.044 | 0.3240 | 0.5814 | 0 | 0 | 0 |
| 'Mcl1' | 112.607 | 104.204 | 85.149 | 99.797 | 94.145 | 91.967 | 100.202 | 101.471 | 80.680 | 91.204 | 103.535 | 98.710 | 90.419 | 114.444 | 100.912 | 105.405 | 108.527 | 108.350 | -0.096 | 0.0796 | 0.2541 | 0 | 0 | 0 |
| 'Mcm10' | 1.720 | 1.420 | 2.402 | 2.039 | 1.705 | 2.182 | 1.989 | 1.591 | 2.023 | 1.581 | 1.689 | 1.919 | 2.321 | 1.758 | 2.243 | 1.832 | 1.900 | 2.100 | -0.039 | 0.5799 | 0.7822 | 0 | 0 | 0 |
| 'Mcm2' | 24.886 | 21.695 | 25.912 | 26.650 | 23.350 | 25.109 | 25.508 | 22.907 | 25.273 | 23.680 | 22.843 | 25.984 | 29.445 | 23.312 | 26.227 | 23.179 | 23.230 | 25.783 | -0.027 | 0.4616 | 0.6979 | 0 | 0 | 0 |
| 'Mcm3' | 22.727 | 18.867 | 24.973 | 23.615 | 20.587 | 25.014 | 24.934 | 20.160 | 24.110 | 20.841 | 21.158 | 22.736 | 28.343 | 20.385 | 24.945 | 22.241 | 20.339 | 22.879 | -0.006 | 0.9047 | 0.9624 | 0 | 0 | 0 |
| 'Mcm3ap' | 23.142 | 21.820 | 25.148 | 23.825 | 23.437 | 24.984 | 22.826 | 22.767 | 25.441 | 24.102 | 23.661 | 25.009 | 24.259 | 23.743 | 22.871 | 23.132 | 24.747 | 23.256 | -0.020 | 0.4973 | 0.7258 | 0 | 0 | 0 |
| 'Mcm4' | 49.941 | 41.993 | 52.312 | 53.521 | 45.375 | 49.722 | 50.432 | 47.154 | 48.249 | 45.579 | 49.621 | 50.446 | 57.886 | 48.261 | 51.266 | 46.726 | 47.734 | 49.928 | -0.041 | 0.3327 | 0.5908 | 0 | 0 | 0 |
| 'Mcm5' | 63.571 | 54.806 | 71.553 | 65.864 | 57.274 | 70.195 | 66.831 | 55.918 | 70.953 | 58.398 | 60.505 | 66.786 | 77.457 | 60.948 | 72.591 | 62.485 | 60.580 | 66.476 | -0.038 | 0.3782 | 0.6299 | 0 | 0 | 0 |
| 'Mcm6' | 43.831 | 41.032 | 45.706 | 45.924 | 43.865 | 45.375 | 47.238 | 43.041 | 45.102 | 44.677 | 46.773 | 43.185 | 57.037 | 44.945 | 47.014 | 43.036 | 45.901 | 42.110 | -0.058 | 0.1887 | 0.4247 | 0 | 0 | 0 |
| 'Mcm7' | 28.580 | 25.564 | 31.445 | 31.194 | 27.247 | 30.919 | 31.280 | 26.347 | 32.019 | 26.237 | 26.655 | 28.763 | 31.644 | 28.013 | 30.904 | 26.198 | 26.818 | 28.333 | 0.048 | 0.2210 | 0.4683 | 0 | 0 | 0 |
| 'Mcm8' | 1.763 | 1.675 | 2.382 | 2.095 | 1.640 | 2.162 | 1.801 | 1.674 | 2.208 | 1.627 | 2.139 | 1.932 | 2.366 | 2.472 | 2.118 | 1.817 | 1.533 | 2.039 | -0.062 | 0.5167 | 0.7404 | 0 | 0 | 0 |
| 'Mcm9' | 5.484 | 4.575 | 5.450 | 5.764 | 4.935 | 5.196 | 6.085 | 5.217 | 5.901 | 5.896 | 5.194 | 5.743 | 5.494 | 5.009 | 5.038 | 5.186 | 5.644 | 4.769 | 0.007 | 0.8895 | 0.9547 | 0 | 0 | 0 |
| 'Mcmbp' | 19.648 | 19.159 | 20.683 | 19.460 | 19.388 | 20.216 | 19.860 | 19.438 | 19.617 | 17.283 | 19.544 | 21.940 | 19.585 | 19.387 | 21.037 | 16.770 | 20.245 | 22.410 | -0.015 | 0.7022 | 0.8611 | 0 | 0 | 0 |
| 'Mcoln1' | 8.072 | 9.127 | 8.733 | 7.172 | 8.599 | 9.265 | 8.667 | 8.613 | 8.994 | 9.436 | 9.207 | 9.126 | 9.648 | 8.447 | 9.124 | 8.877 | 9.060 | 8.972 | -0.095 | 0.0171 | 0.0927 | 0 | 0 | 0 |
| 'Mcph1' | 1.740 | 1.520 | 1.796 | 1.870 | 1.720 | 1.679 | 1.880 | 1.811 | 1.586 | 1.802 | 1.882 | 1.611 | 1.989 | 2.013 | 1.688 | 1.460 | 1.599 | 1.580 | -0.009 | 0.8800 | 0.9494 | 0 | 0 | 0 |
| 'Mcrs1' | 31.585 | 32.153 | 36.174 | 26.507 | 34.351 | 39.385 | 32.006 | 33.453 | 37.627 | 33.902 | 28.521 | 32.075 | 36.485 | 32.614 | 31.950 | 36.883 | 30.823 | 30.477 | 0.036 | 0.5493 | 0.7629 | 0 | 0 | 0 |
| 'Mctp2' | 1.148 | 1.041 | 0.934 | 0.870 | 0.822 | 1.015 | 1.006 | 0.837 | 1.075 | 0.997 | 0.971 | 1.159 | 0.930 | 1.066 | 1.008 | 0.784 | 1.052 | 1.289 | -0.093 | 0.2570 | 0.5093 | 0 | 0 | 0 |
| 'Mcts1' | 13.819 | 15.829 | 14.779 | 14.956 | 15.601 | 14.190 | 14.172 | 14.897 | 14.233 | 13.341 | 14.446 | 14.588 | 14.042 | 14.316 | 13.605 | 13.860 | 16.049 | 14.206 | 0.030 | 0.4157 | 0.6634 | 0 | 0 | 0 |
| 'Mcts2' | 5.811 | 3.030 | 5.221 | 5.915 | 3.819 | 4.269 | 4.959 | 4.446 | 3.737 | 5.132 | 6.164 | 5.557 | 5.013 | 5.983 | 5.936 | 4.638 | 5.747 | 5.889 | -0.289 | 0.0125 | 0.0734 | 0 | 0 | 0 |
| 'Mcu' | 4.681 | 5.669 | 4.673 | 4.211 | 5.050 | 5.237 | 4.594 | 5.103 | 4.883 | 5.445 | 5.552 | 4.977 | 5.097 | 5.795 | 5.123 | 5.622 | 5.427 | 4.938 | -0.128 | 0.0029 | 0.0245 | 1 | 0 | 0 |
| 'Mcur1' | 34.619 | 38.740 | 34.595 | 34.446 | 40.075 | 35.899 | 36.483 | 38.756 | 34.755 | 34.973 | 35.818 | 32.260 | 38.024 | 37.368 | 34.308 | 34.835 | 34.464 | 31.129 | 0.058 | 0.0510 | 0.1912 | 0 | 0 | 0 |
| 'Mdc1' | 9.881 | 10.005 | 10.569 | 9.304 | 10.287 | 10.585 | 8.749 | 10.345 | 10.715 | 10.766 | 10.854 | 10.929 | 10.269 | 9.733 | 10.555 | 10.100 | 10.400 | 10.591 | -0.069 | 0.0295 | 0.1335 | 0 | 0 | 0 |
| 'Mdfic' | 6.063 | 6.981 | 6.300 | 5.983 | 6.014 | 6.002 | 6.630 | 6.229 | 6.549 | 6.344 | 7.387 | 6.548 | 5.633 | 6.867 | 6.508 | 6.531 | 7.803 | 6.413 | -0.088 | 0.0510 | 0.1912 | 0 | 0 | 0 |
| 'Mdh1' | 42.988 | 46.320 | 42.981 | 46.234 | 46.727 | 44.571 | 48.032 | 44.470 | 45.110 | 40.920 | 43.174 | 42.987 | 43.611 | 42.620 | 44.318 | 41.341 | 45.653 | 41.647 | 0.064 | 0.0196 | 0.1017 | 0 | 0 | 0 |
| 'Mdh2' | 37.842 | 38.274 | 38.600 | 38.818 | 40.191 | 39.804 | 37.767 | 39.400 | 37.885 | 37.203 | 36.366 | 36.833 | 37.791 | 37.501 | 38.700 | 37.132 | 35.939 | 36.883 | 0.050 | 0.0159 | 0.0877 | 0 | 0 | 0 |
| 'Mdk' | 4.219 | 4.368 | 4.510 | 4.381 | 3.164 | 3.938 | 3.379 | 4.666 | 4.120 | 3.787 | 4.948 | 5.119 | 3.339 | 4.173 | 5.352 | 4.899 | 4.496 | 5.155 | -0.169 | 0.0687 | 0.2303 | 0 | 0 | 0 |
| 'Mdm1' | 5.688 | 6.750 | 6.418 | 6.075 | 5.352 | 5.888 | 6.398 | 6.110 | 6.640 | 5.067 | 5.349 | 5.855 | 6.618 | 5.996 | 6.182 | 5.472 | 5.797 | 5.290 | 0.086 | 0.1448 | 0.3693 | 0 | 0 | 0 |
| 'Mdm2' | 12.814 | 12.481 | 12.044 | 12.401 | 12.850 | 11.880 | 12.198 | 12.868 | 11.473 | 12.258 | 13.494 | 12.408 | 11.697 | 13.218 | 12.094 | 12.209 | 13.108 | 12.751 | -0.039 | 0.1795 | 0.4147 | 0 | 0 | 0 |
| 'Mdm4' | 10.742 | 12.077 | 11.838 | 10.078 | 11.753 | 11.276 | 9.552 | 11.845 | 12.305 | 10.154 | 10.638 | 11.181 | 10.078 | 11.243 | 10.843 | 10.431 | 10.277 | 11.473 | 0.062 | 0.0438 | 0.1733 | 0 | 0 | 0 |
| 'Mdn1' | 8.805 | 6.856 | 8.532 | 8.677 | 6.994 | 7.468 | 6.974 | 7.246 | 8.209 | 8.023 | 7.804 | 8.839 | 6.867 | 6.870 | 7.820 | 7.344 | 7.651 | 9.505 | -0.030 | 0.6077 | 0.8027 | 0 | 0 | 0 |
| 'Mdp1' | 47.749 | 49.436 | 52.891 | 48.115 | 50.876 | 53.115 | 50.233 | 51.394 | 47.914 | 45.972 | 48.883 | 54.158 | 47.045 | 49.528 | 51.643 | 46.019 | 51.479 | 52.363 | 0.003 | 0.9211 | 0.9703 | 0 | 0 | 0 |
| 'Me2' | 24.305 | 24.255 | 25.916 | 26.321 | 26.913 | 26.631 | 26.969 | 26.419 | 24.049 | 24.113 | 23.860 | 26.545 | 23.785 | 22.570 | 26.277 | 26.650 | 26.318 | 25.685 | 0.026 | 0.5012 | 0.7283 | 0 | 0 | 0 |
| 'Mea1' | 55.364 | 67.340 | 59.739 | 57.300 | 63.968 | 65.420 | 58.202 | 65.468 | 61.613 | 55.961 | 52.956 | 57.429 | 59.899 | 61.676 | 59.495 | 63.568 | 60.156 | 54.543 | 0.066 | 0.1059 | 0.3053 | 0 | 0 | 0 |
| 'Meaf6' | 10.082 | 11.549 | 11.485 | 10.913 | 11.745 | 11.535 | 11.146 | 11.883 | 11.744 | 11.044 | 11.760 | 11.456 | 11.592 | 10.977 | 11.068 | 10.676 | 11.957 | 11.246 | -0.007 | 0.7961 | 0.9092 | 0 | 0 | 0 |
| 'Mecp2' | 4.217 | 4.335 | 4.495 | 4.098 | 4.013 | 4.428 | 4.453 | 4.210 | 4.534 | 4.722 | 4.169 | 4.641 | 4.139 | 4.066 | 4.247 | 4.612 | 4.055 | 4.250 | -0.014 | 0.6488 | 0.8287 | 0 | 0 | 0 |
| 'Mecr' | 3.361 | 3.167 | 3.317 | 2.934 | 2.956 | 3.097 | 3.350 | 3.089 | 3.461 | 2.922 | 2.888 | 3.457 | 3.528 | 2.486 | 2.973 | 3.388 | 2.849 | 3.352 | 0.032 | 0.5710 | 0.7777 | 0 | 0 | 0 |
| 'Med1' | 9.553 | 10.564 | 10.153 | 9.687 | 10.038 | 9.684 | 10.153 | 10.230 | 10.109 | 9.581 | 10.649 | 10.637 | 10.005 | 10.569 | 10.328 | 10.548 | 9.909 | 10.220 | -0.048 | 0.0652 | 0.2228 | 0 | 0 | 0 |
| 'Med10' | 66.800 | 66.441 | 64.804 | 65.144 | 66.913 | 64.019 | 64.007 | 68.863 | 62.300 | 63.595 | 61.012 | 62.139 | 71.094 | 69.053 | 64.108 | 66.034 | 59.914 | 57.455 | 0.028 | 0.4293 | 0.6749 | 0 | 0 | 0 |
| 'Med11' | 10.230 | 10.794 | 10.616 | 10.619 | 9.870 | 11.239 | 9.919 | 9.718 | 11.472 | 8.523 | 9.134 | 11.067 | 12.688 | 9.371 | 10.877 | 10.503 | 9.957 | 10.419 | 0.016 | 0.7865 | 0.9047 | 0 | 0 | 0 |
| 'Med12' | 26.900 | 23.231 | 27.795 | 25.585 | 23.322 | 24.845 | 25.905 | 24.159 | 27.467 | 27.054 | 26.922 | 27.693 | 25.982 | 23.824 | 26.532 | 23.941 | 26.021 | 26.818 | -0.047 | 0.1877 | 0.4239 | 0 | 0 | 0 |
| 'Med13' | 18.710 | 17.528 | 16.591 | 18.169 | 17.788 | 15.249 | 18.483 | 17.031 | 16.166 | 18.334 | 19.561 | 17.528 | 17.730 | 19.062 | 16.070 | 18.644 | 19.027 | 17.950 | -0.086 | 0.0088 | 0.0568 | 0 | 0 | 0 |
| 'Med14' | 6.470 | 6.466 | 6.902 | 6.590 | 6.768 | 6.693 | 6.530 | 6.477 | 6.836 | 6.096 | 6.785 | 7.043 | 6.207 | 6.190 | 6.786 | 6.121 | 6.687 | 7.323 | 0.001 | 0.9669 | 0.9887 | 0 | 0 | 0 |
| 'Med15' | 47.429 | 46.660 | 48.458 | 42.335 | 45.678 | 46.161 | 43.559 | 45.085 | 47.827 | 50.624 | 49.431 | 48.223 | 50.264 | 44.031 | 46.542 | 47.850 | 44.625 | 48.088 | -0.066 | 0.0439 | 0.1733 | 0 | 0 | 0 |
| 'Med16' | 8.517 | 8.623 | 9.406 | 7.698 | 8.339 | 9.980 | 7.818 | 8.442 | 9.940 | 9.674 | 7.930 | 8.831 | 9.441 | 7.668 | 9.411 | 8.885 | 7.921 | 8.072 | 0.008 | 0.8697 | 0.9441 | 0 | 0 | 0 |
| 'Med17' | 19.096 | 20.806 | 18.871 | 19.083 | 20.994 | 17.773 | 17.590 | 19.193 | 18.993 | 18.153 | 17.447 | 19.523 | 17.971 | 17.434 | 18.122 | 18.352 | 17.304 | 18.893 | 0.064 | 0.0860 | 0.2669 | 0 | 0 | 0 |
| 'Med18' | 2.490 | 3.776 | 3.435 | 2.936 | 3.920 | 3.072 | 3.091 | 3.554 | 4.113 | 2.657 | 2.697 | 3.600 | 2.982 | 3.091 | 3.861 | 2.658 | 3.325 | 4.033 | 0.047 | 0.5794 | 0.7822 | 0 | 0 | 0 |
| 'Med19' | 8.380 | 9.762 | 9.514 | 8.146 | 9.129 | 9.137 | 7.779 | 9.464 | 9.321 | 8.427 | 8.325 | 8.626 | 9.079 | 8.729 | 8.727 | 8.614 | 8.181 | 8.158 | 0.060 | 0.1928 | 0.4302 | 0 | 0 | 0 |
| 'Med20' | 2.435 | 2.488 | 2.403 | 2.461 | 2.530 | 2.507 | 2.592 | 2.310 | 2.305 | 2.006 | 2.708 | 2.877 | 2.071 | 2.711 | 2.710 | 2.054 | 2.444 | 2.781 | -0.030 | 0.6373 | 0.8213 | 0 | 0 | 0 |
| 'Med21' | 4.319 | 4.323 | 3.955 | 4.457 | 4.402 | 4.232 | 5.074 | 3.842 | 4.222 | 3.725 | 4.466 | 4.889 | 4.142 | 3.990 | 4.476 | 3.387 | 4.460 | 4.651 | 0.006 | 0.9271 | 0.9720 | 0 | 0 | 0 |
| 'Med22' | 13.552 | 13.913 | 12.847 | 13.789 | 12.722 | 13.681 | 13.543 | 12.362 | 13.318 | 11.645 | 14.009 | 14.561 | 11.604 | 12.751 | 15.326 | 11.910 | 12.919 | 14.567 | -0.003 | 0.9552 | 0.9834 | 0 | 0 | 0 |
| 'Med23' | 5.419 | 6.204 | 5.927 | 5.673 | 6.069 | 5.936 | 5.901 | 6.490 | 6.191 | 6.342 | 6.081 | 5.889 | 5.811 | 5.788 | 5.502 | 6.040 | 6.508 | 5.529 | -0.001 | 0.9877 | 0.9964 | 0 | 0 | 0 |
| 'Med24' | 9.686 | 8.846 | 10.994 | 8.687 | 8.676 | 10.490 | 8.984 | 9.084 | 10.683 | 10.121 | 9.527 | 10.068 | 10.240 | 9.199 | 10.028 | 8.902 | 9.580 | 9.171 | -0.024 | 0.5476 | 0.7614 | 0 | 0 | 0 |
| 'Med25' | 35.242 | 35.112 | 42.127 | 34.443 | 34.199 | 43.033 | 34.696 | 37.226 | 42.247 | 40.490 | 33.110 | 41.739 | 39.129 | 32.920 | 40.288 | 37.922 | 34.470 | 39.551 | -0.015 | 0.6623 | 0.8370 | 0 | 0 | 0 |
| 'Med26' | 11.331 | 10.142 | 11.090 | 9.902 | 10.195 | 10.952 | 9.377 | 9.098 | 10.764 | 12.184 | 9.893 | 10.241 | 11.309 | 10.539 | 9.750 | 11.895 | 8.896 | 9.847 | -0.028 | 0.5978 | 0.7954 | 0 | 0 | 0 |
| 'Med27' | 5.196 | 5.266 | 5.539 | 5.077 | 5.043 | 5.140 | 4.942 | 5.347 | 4.828 | 5.505 | 5.136 | 4.929 | 5.310 | 4.739 | 4.760 | 4.797 | 4.978 | 4.711 | 0.041 | 0.3501 | 0.6044 | 0 | 0 | 0 |
| 'Med28' | 24.700 | 29.880 | 26.587 | 26.808 | 29.917 | 27.404 | 26.902 | 29.114 | 27.930 | 24.245 | 26.467 | 28.140 | 26.751 | 27.393 | 28.429 | 29.633 | 29.303 | 27.055 | -0.003 | 0.9258 | 0.9717 | 0 | 0 | 0 |
| 'Med29' | 11.272 | 11.147 | 9.437 | 9.192 | 10.137 | 9.713 | 8.315 | 12.247 | 10.002 | 11.198 | 8.912 | 10.025 | 11.119 | 9.006 | 9.686 | 9.329 | 8.162 | 9.461 | 0.062 | 0.4216 | 0.6681 | 0 | 0 | 0 |
| 'Med30' | 33.555 | 37.381 | 35.716 | 35.995 | 36.158 | 37.697 | 39.503 | 37.202 | 34.109 | 31.623 | 34.529 | 33.514 | 36.652 | 36.448 | 35.416 | 33.071 | 35.529 | 31.429 | 0.075 | 0.0806 | 0.2562 | 0 | 0 | 0 |
| 'Med31' | 9.374 | 7.029 | 6.337 | 7.404 | 7.145 | 5.334 | 6.942 | 8.095 | 5.847 | 7.146 | 9.260 | 6.794 | 8.057 | 8.275 | 5.894 | 6.595 | 8.860 | 6.275 | -0.089 | 0.2391 | 0.4895 | 0 | 0 | 0 |
| 'Med4' | 30.284 | 31.707 | 30.770 | 31.380 | 30.699 | 29.288 | 29.735 | 31.257 | 28.178 | 28.663 | 31.275 | 28.926 | 31.010 | 31.446 | 27.400 | 32.137 | 29.504 | 26.682 | 0.026 | 0.4744 | 0.7095 | 0 | 0 | 0 |
| 'Med6' | 6.491 | 7.202 | 7.127 | 7.446 | 7.359 | 6.367 | 6.533 | 7.348 | 7.237 | 6.782 | 6.822 | 7.020 | 6.645 | 7.882 | 6.565 | 7.105 | 7.639 | 6.411 | -0.004 | 0.9386 | 0.9764 | 0 | 0 | 0 |
| 'Med7' | 3.499 | 4.460 | 4.243 | 3.773 | 4.219 | 4.522 | 4.025 | 4.534 | 4.527 | 4.083 | 3.937 | 4.286 | 4.078 | 4.055 | 3.571 | 3.542 | 4.352 | 3.948 | 0.061 | 0.2081 | 0.4526 | 0 | 0 | 0 |
| 'Med8' | 10.724 | 12.125 | 11.119 | 11.025 | 12.032 | 11.684 | 10.753 | 11.287 | 11.119 | 10.586 | 11.075 | 11.401 | 11.290 | 11.938 | 11.975 | 11.867 | 11.347 | 10.872 | -0.018 | 0.5700 | 0.7774 | 0 | 0 | 0 |
| 'Med9' | 11.537 | 13.117 | 12.810 | 10.311 | 11.620 | 13.560 | 10.417 | 11.766 | 12.398 | 10.804 | 11.851 | 10.710 | 12.847 | 11.444 | 11.698 | 12.010 | 9.474 | 10.204 | 0.083 | 0.1906 | 0.4271 | 0 | 0 | 0 |
| 'Mef2a' | 3.967 | 3.959 | 3.836 | 3.868 | 4.553 | 3.952 | 4.022 | 4.444 | 3.999 | 4.297 | 4.282 | 3.734 | 4.058 | 4.176 | 3.428 | 4.514 | 4.022 | 3.620 | 0.014 | 0.7207 | 0.8716 | 0 | 0 | 0 |
| 'Mef2d' | 16.758 | 17.017 | 15.643 | 14.623 | 16.242 | 16.348 | 15.021 | 17.022 | 16.827 | 16.428 | 16.194 | 16.140 | 16.680 | 16.609 | 16.011 | 17.330 | 15.212 | 15.864 | -0.019 | 0.5588 | 0.7698 | 0 | 0 | 0 |
| 'Megf9' | 1.181 | 1.466 | 1.300 | 1.037 | 1.276 | 1.257 | 1.209 | 1.333 | 1.230 | 1.273 | 1.445 | 1.163 | 1.441 | 1.373 | 1.131 | 1.243 | 1.443 | 1.169 | -0.051 | 0.4430 | 0.6866 | 0 | 0 | 0 |
| 'Meis3' | 8.640 | 9.112 | 11.123 | 9.018 | 9.338 | 10.771 | 8.888 | 9.107 | 10.708 | 9.038 | 6.999 | 9.384 | 9.660 | 7.917 | 9.319 | 8.075 | 7.997 | 8.614 | 0.159 | 0.0003 | 0.0036 | 1 | 0 | 0 |
| 'Melk' | 3.252 | 3.845 | 5.230 | 3.896 | 4.286 | 4.529 | 5.221 | 3.970 | 4.757 | 3.477 | 4.312 | 4.384 | 4.505 | 4.583 | 5.765 | 4.276 | 4.712 | 4.189 | -0.055 | 0.5062 | 0.7324 | 0 | 0 | 0 |
| 'Memo1' | 23.475 | 24.914 | 25.589 | 22.984 | 27.760 | 24.531 | 25.946 | 23.471 | 23.248 | 24.856 | 25.084 | 26.000 | 24.790 | 23.877 | 24.706 | 24.731 | 25.358 | 23.073 | -0.019 | 0.6893 | 0.8534 | 0 | 0 | 0 |
| 'Men1' | 5.602 | 5.427 | 5.829 | 6.003 | 5.292 | 5.977 | 5.597 | 5.166 | 5.672 | 4.823 | 4.804 | 6.244 | 5.420 | 4.999 | 6.146 | 4.910 | 5.066 | 6.079 | 0.050 | 0.2014 | 0.4426 | 0 | 0 | 0 |
| 'Mepce' | 29.953 | 24.879 | 28.550 | 28.274 | 24.636 | 28.126 | 28.567 | 26.337 | 29.223 | 25.725 | 28.970 | 33.552 | 24.959 | 28.103 | 32.555 | 25.839 | 29.129 | 32.206 | -0.077 | 0.1139 | 0.3199 | 0 | 0 | 0 |
| 'Mesdc1' | 10.863 | 9.869 | 11.330 | 10.365 | 9.411 | 11.684 | 10.035 | 8.949 | 11.653 | 9.500 | 10.257 | 14.310 | 10.689 | 11.043 | 13.419 | 10.652 | 10.661 | 13.999 | -0.159 | 0.0005 | 0.0060 | 1 | 0 | 0 |
| 'Mesdc2' | 8.533 | 9.679 | 9.350 | 8.973 | 9.289 | 9.772 | 9.196 | 9.726 | 9.838 | 9.888 | 9.600 | 9.845 | 8.863 | 9.243 | 9.818 | 10.436 | 9.058 | 9.064 | -0.034 | 0.3531 | 0.6066 | 0 | 0 | 0 |
| 'Metap1' | 7.905 | 8.230 | 7.702 | 7.945 | 7.647 | 7.579 | 8.144 | 7.782 | 7.523 | 7.690 | 7.628 | 8.999 | 7.635 | 8.259 | 8.539 | 7.390 | 9.097 | 8.526 | -0.079 | 0.0555 | 0.2019 | 0 | 0 | 0 |
| 'Metap1d' | 6.023 | 6.571 | 6.808 | 6.025 | 6.045 | 6.454 | 5.955 | 6.733 | 6.445 | 6.487 | 6.198 | 6.453 | 6.410 | 6.293 | 5.945 | 6.002 | 6.714 | 6.176 | 0.001 | 0.9784 | 0.9925 | 0 | 0 | 0 |
| 'Metap2' | 10.112 | 10.772 | 9.716 | 10.717 | 10.724 | 10.377 | 10.917 | 11.058 | 10.348 | 10.307 | 10.056 | 10.100 | 10.733 | 10.404 | 9.857 | 10.587 | 10.286 | 9.744 | 0.029 | 0.1748 | 0.4095 | 0 | 0 | 0 |
| 'Metrn' | 17.865 | 19.732 | 20.738 | 16.652 | 18.116 | 22.895 | 17.803 | 18.271 | 21.762 | 15.986 | 14.752 | 16.922 | 18.631 | 16.553 | 18.539 | 18.069 | 14.397 | 17.810 | 0.187 | 0.0003 | 0.0039 | 1 | 0 | 0 |
| 'Metrnl' | 5.104 | 3.836 | 2.760 | 4.139 | 2.923 | 3.099 | 3.997 | 3.603 | 2.560 | 2.854 | 3.721 | 3.393 | 3.870 | 4.491 | 3.511 | 3.559 | 3.124 | 3.108 | -0.011 | 0.9278 | 0.9722 | 0 | 0 | 0 |
| 'Mettl1' | 1.828 | 2.031 | 2.312 | 2.034 | 2.080 | 2.253 | 2.207 | 2.476 | 2.188 | 2.504 | 1.946 | 1.984 | 2.453 | 1.931 | 2.080 | 2.058 | 1.845 | 1.617 | 0.074 | 0.3528 | 0.6063 | 0 | 0 | 0 |
| 'Mettl10' | 6.088 | 5.553 | 6.645 | 5.814 | 6.752 | 6.063 | 6.623 | 6.475 | 5.661 | 5.731 | 6.721 | 5.946 | 5.729 | 6.721 | 5.483 | 5.606 | 5.839 | 5.568 | 0.056 | 0.2839 | 0.5401 | 0 | 0 | 0 |
| 'Mettl13' | 1.141 | 1.106 | 1.157 | 0.977 | 1.121 | 1.015 | 0.814 | 1.076 | 1.025 | 0.795 | 0.772 | 0.788 | 0.708 | 0.729 | 0.843 | 0.742 | 0.667 | 0.702 | 0.475 | 0.0000 | 0.0000 | 1 | 1 | 0 |
| 'Mettl14' | 4.205 | 4.842 | 4.822 | 4.072 | 5.713 | 4.563 | 4.312 | 4.949 | 5.213 | 4.636 | 5.008 | 4.109 | 4.861 | 4.828 | 4.196 | 4.752 | 4.638 | 3.909 | 0.053 | 0.3303 | 0.5880 | 0 | 0 | 0 |
| 'Mettl15' | 1.588 | 1.256 | 1.361 | 1.969 | 1.889 | 1.626 | 1.986 | 1.863 | 1.323 | 1.419 | 1.543 | 1.467 | 1.564 | 1.533 | 1.561 | 1.082 | 1.704 | 1.310 | 0.151 | 0.1245 | 0.3375 | 0 | 0 | 0 |
| 'Mettl16' | 2.670 | 2.595 | 2.582 | 2.813 | 2.573 | 2.265 | 2.804 | 2.437 | 2.481 | 2.603 | 2.418 | 2.952 | 2.506 | 2.413 | 2.701 | 2.530 | 2.597 | 2.849 | -0.038 | 0.4311 | 0.6762 | 0 | 0 | 0 |
| 'Mettl17' | 6.049 | 5.744 | 7.026 | 6.102 | 6.569 | 6.857 | 6.215 | 6.403 | 7.471 | 6.083 | 6.228 | 6.775 | 6.636 | 6.128 | 6.549 | 5.922 | 6.117 | 6.368 | 0.030 | 0.3760 | 0.6273 | 0 | 0 | 0 |
| 'Mettl18' | 4.091 | 3.428 | 4.196 | 4.135 | 3.762 | 4.747 | 4.401 | 4.106 | 2.862 | 4.120 | 4.865 | 4.142 | 4.088 | 3.945 | 3.191 | 3.443 | 3.655 | 4.370 | -0.013 | 0.8995 | 0.9600 | 0 | 0 | 0 |
| 'Mettl2' | 4.793 | 4.592 | 5.009 | 4.438 | 4.899 | 4.863 | 4.499 | 4.590 | 4.523 | 4.850 | 4.510 | 4.909 | 4.752 | 4.445 | 4.850 | 4.457 | 4.228 | 4.791 | 0.003 | 0.9378 | 0.9764 | 0 | 0 | 0 |
| 'Mettl20' | 0.883 | 0.962 | 0.871 | 0.820 | 0.880 | 0.867 | 0.868 | 0.811 | 0.957 | 0.613 | 0.965 | 0.745 | 0.943 | 0.669 | 0.811 | 0.968 | 0.803 | 0.728 | 0.126 | 0.2269 | 0.4734 | 0 | 0 | 0 |
| 'Mettl21a' | 4.754 | 5.563 | 4.926 | 4.652 | 5.854 | 4.873 | 4.902 | 6.017 | 5.267 | 5.319 | 5.701 | 5.211 | 5.241 | 6.120 | 5.334 | 5.231 | 5.622 | 4.910 | -0.067 | 0.1068 | 0.3071 | 0 | 0 | 0 |
| 'Mettl22' | 3.130 | 3.004 | 3.838 | 2.877 | 2.995 | 3.078 | 2.873 | 3.082 | 3.440 | 3.205 | 2.533 | 3.059 | 3.511 | 3.068 | 3.207 | 3.218 | 2.854 | 3.153 | 0.024 | 0.7186 | 0.8703 | 0 | 0 | 0 |
| 'Mettl25' | 1.681 | 1.490 | 1.551 | 1.355 | 1.588 | 1.354 | 1.621 | 1.785 | 1.464 | 1.319 | 1.758 | 1.527 | 1.496 | 1.367 | 1.395 | 1.515 | 1.597 | 1.207 | 0.066 | 0.4079 | 0.6567 | 0 | 0 | 0 |
| 'Mettl26' | 13.850 | 13.487 | 12.789 | 13.029 | 12.569 | 13.969 | 14.519 | 12.088 | 13.279 | 12.316 | 12.671 | 13.815 | 13.673 | 12.337 | 12.417 | 12.469 | 13.792 | 12.822 | 0.024 | 0.5690 | 0.7768 | 0 | 0 | 0 |
| 'Mettl3' | 3.805 | 3.302 | 3.865 | 3.794 | 3.860 | 3.636 | 3.962 | 3.521 | 3.927 | 3.573 | 3.639 | 4.101 | 3.699 | 3.435 | 4.058 | 3.771 | 3.929 | 3.812 | -0.027 | 0.4849 | 0.7162 | 0 | 0 | 0 |
| 'Mettl4' | 7.552 | 8.685 | 9.331 | 8.729 | 8.408 | 8.814 | 8.483 | 8.200 | 9.086 | 7.424 | 9.081 | 9.822 | 7.335 | 9.775 | 9.004 | 8.124 | 9.855 | 9.136 | -0.052 | 0.3382 | 0.5957 | 0 | 0 | 0 |
| 'Mettl5' | 3.963 | 4.815 | 4.071 | 3.644 | 4.662 | 4.560 | 4.802 | 4.588 | 4.098 | 3.821 | 4.365 | 3.832 | 4.573 | 4.681 | 4.073 | 3.832 | 4.062 | 3.597 | 0.079 | 0.1897 | 0.4262 | 0 | 0 | 0 |
| 'Mettl6' | 15.163 | 15.459 | 14.952 | 16.193 | 13.724 | 14.070 | 17.773 | 13.877 | 15.187 | 14.265 | 17.000 | 17.286 | 13.329 | 15.325 | 15.516 | 15.001 | 16.766 | 16.020 | -0.056 | 0.3444 | 0.5996 | 0 | 0 | 0 |
| 'Mettl8' | 2.234 | 2.066 | 2.089 | 2.283 | 2.036 | 2.247 | 2.377 | 2.061 | 2.150 | 2.115 | 1.920 | 2.119 | 1.911 | 1.699 | 2.079 | 1.719 | 1.842 | 1.712 | 0.175 | 0.0011 | 0.0112 | 1 | 0 | 0 |
| 'Mettl9' | 31.381 | 32.338 | 31.746 | 29.487 | 32.760 | 30.288 | 32.469 | 33.408 | 31.198 | 27.426 | 29.988 | 32.113 | 30.311 | 32.710 | 31.433 | 29.811 | 28.920 | 28.788 | 0.057 | 0.1035 | 0.3017 | 0 | 0 | 0 |
| 'Mex3b' | 9.103 | 8.950 | 8.018 | 9.540 | 7.292 | 7.896 | 9.957 | 7.743 | 8.630 | 8.144 | 9.667 | 11.709 | 8.583 | 10.256 | 11.089 | 9.740 | 9.728 | 11.579 | -0.242 | 0.0010 | 0.0102 | 1 | 0 | 0 |
| 'Mex3c' | 12.586 | 11.289 | 9.858 | 10.570 | 10.496 | 10.171 | 10.411 | 11.051 | 9.745 | 11.784 | 11.919 | 12.399 | 10.704 | 13.252 | 12.806 | 13.754 | 12.139 | 13.383 | -0.233 | 0.0000 | 0.0007 | 1 | 0 | 0 |
| 'Mex3d' | 2.809 | 2.442 | 2.270 | 2.488 | 2.276 | 2.608 | 2.216 | 2.434 | 2.461 | 2.466 | 2.469 | 2.694 | 2.278 | 2.493 | 2.643 | 2.384 | 2.144 | 2.735 | -0.030 | 0.6082 | 0.8029 | 0 | 0 | 0 |
| 'Mfap3' | 4.676 | 4.971 | 4.784 | 4.587 | 4.822 | 4.737 | 5.532 | 4.869 | 4.941 | 3.799 | 5.232 | 5.352 | 4.242 | 4.725 | 5.608 | 4.351 | 5.057 | 5.370 | 0.000 | 0.9985 | 0.9994 | 0 | 0 | 0 |
| 'Mff' | 4.875 | 5.738 | 4.905 | 5.039 | 5.550 | 5.222 | 4.980 | 5.377 | 5.136 | 4.841 | 5.176 | 5.040 | 4.896 | 5.908 | 5.133 | 5.503 | 5.556 | 4.954 | -0.016 | 0.6212 | 0.8107 | 0 | 0 | 0 |
| 'Mfge8' | 2.493 | 2.487 | 2.861 | 2.267 | 2.547 | 2.690 | 2.291 | 2.663 | 2.600 | 2.417 | 2.496 | 2.586 | 2.172 | 2.329 | 2.602 | 2.459 | 2.188 | 2.466 | 0.072 | 0.1185 | 0.3277 | 0 | 0 | 0 |
| 'Mfhas1' | 6.827 | 6.235 | 6.789 | 5.859 | 6.643 | 7.229 | 5.755 | 6.720 | 6.668 | 7.843 | 7.291 | 8.167 | 6.278 | 6.950 | 7.793 | 7.667 | 6.464 | 8.230 | -0.191 | 0.0001 | 0.0017 | 1 | 0 | 0 |
| 'Mfn1' | 5.327 | 5.817 | 5.772 | 5.304 | 6.535 | 5.579 | 5.581 | 6.424 | 5.374 | 5.571 | 5.426 | 5.230 | 5.742 | 5.919 | 4.964 | 5.588 | 5.607 | 4.912 | 0.070 | 0.0584 | 0.2082 | 0 | 0 | 0 |
| 'Mfn2' | 12.968 | 14.038 | 14.103 | 14.353 | 13.862 | 14.751 | 12.892 | 14.758 | 14.248 | 14.852 | 14.513 | 13.527 | 14.896 | 13.963 | 14.281 | 13.717 | 13.315 | 14.186 | -0.024 | 0.4836 | 0.7151 | 0 | 0 | 0 |
| 'Mfng' | 29.398 | 32.325 | 30.488 | 28.528 | 29.725 | 33.150 | 26.723 | 33.203 | 31.496 | 29.449 | 29.330 | 36.739 | 29.916 | 30.043 | 34.593 | 31.787 | 29.243 | 34.783 | -0.066 | 0.0875 | 0.2697 | 0 | 0 | 0 |
| 'Mfsd1' | 21.745 | 25.221 | 22.735 | 22.142 | 24.675 | 23.775 | 21.475 | 25.327 | 22.663 | 23.650 | 24.003 | 23.731 | 23.882 | 25.118 | 22.925 | 23.910 | 24.141 | 22.401 | -0.038 | 0.1513 | 0.3781 | 0 | 0 | 0 |
| 'Mfsd10' | 4.263 | 4.380 | 4.633 | 4.215 | 4.030 | 4.517 | 4.629 | 4.160 | 4.750 | 3.903 | 3.870 | 4.235 | 4.159 | 3.662 | 4.296 | 4.277 | 3.661 | 3.951 | 0.126 | 0.0002 | 0.0027 | 1 | 0 | 0 |
| 'Mfsd11' | 1.921 | 1.771 | 1.828 | 1.786 | 1.755 | 1.834 | 1.948 | 1.720 | 1.787 | 1.716 | 1.873 | 2.086 | 2.062 | 2.027 | 2.027 | 2.082 | 2.167 | 2.101 | -0.162 | 0.0006 | 0.0067 | 1 | 0 | 0 |
| 'Mfsd12' | 2.540 | 2.824 | 2.916 | 2.570 | 2.832 | 2.923 | 2.443 | 2.666 | 2.983 | 2.644 | 2.664 | 2.998 | 2.237 | 2.279 | 2.741 | 2.509 | 2.373 | 2.628 | 0.083 | 0.1762 | 0.4114 | 0 | 0 | 0 |
| 'Mfsd13a' | 4.666 | 4.575 | 4.650 | 4.691 | 3.748 | 4.659 | 4.372 | 4.439 | 5.237 | 4.391 | 3.307 | 4.424 | 4.379 | 4.331 | 4.118 | 4.323 | 4.035 | 3.767 | 0.137 | 0.0142 | 0.0806 | 0 | 0 | 0 |
| 'Mfsd14a' | 17.954 | 17.330 | 15.708 | 16.382 | 17.231 | 14.848 | 16.554 | 17.445 | 15.820 | 18.399 | 17.749 | 17.441 | 17.769 | 18.951 | 16.352 | 17.418 | 18.670 | 16.489 | -0.105 | 0.0001 | 0.0011 | 1 | 0 | 0 |
| 'Mfsd14b' | 10.135 | 11.326 | 10.441 | 10.536 | 10.918 | 10.327 | 10.163 | 11.159 | 10.571 | 10.057 | 10.524 | 11.310 | 10.145 | 10.498 | 10.429 | 9.630 | 11.415 | 10.444 | 0.001 | 0.9730 | 0.9906 | 0 | 0 | 0 |
| 'Mfsd2a' | 2.190 | 0.821 | 1.162 | 2.060 | 0.930 | 1.153 | 1.482 | 0.896 | 1.172 | 1.777 | 1.612 | 1.045 | 1.366 | 1.154 | 1.111 | 1.290 | 1.269 | 1.106 | -0.025 | 0.8456 | 0.9343 | 0 | 0 | 0 |
| 'Mfsd3' | 3.422 | 3.798 | 3.940 | 3.393 | 3.539 | 3.904 | 3.462 | 4.126 | 4.691 | 3.198 | 3.339 | 3.490 | 3.698 | 3.117 | 3.549 | 3.426 | 3.824 | 3.252 | 0.144 | 0.0640 | 0.2205 | 0 | 0 | 0 |
| 'Mfsd4a' | 6.306 | 7.458 | 7.090 | 6.343 | 7.426 | 6.960 | 6.889 | 7.539 | 7.041 | 6.633 | 6.681 | 7.454 | 5.794 | 6.755 | 7.078 | 6.298 | 6.747 | 7.148 | 0.044 | 0.1803 | 0.4156 | 0 | 0 | 0 |
| 'Mfsd4b4' | 4.435 | 4.438 | 4.957 | 4.117 | 4.988 | 4.624 | 4.505 | 5.333 | 4.705 | 4.448 | 4.628 | 4.011 | 4.415 | 4.111 | 4.176 | 4.431 | 4.339 | 3.715 | 0.139 | 0.0207 | 0.1056 | 0 | 0 | 0 |
| 'Mfsd5' | 27.762 | 29.229 | 26.892 | 23.405 | 28.457 | 28.933 | 22.347 | 29.031 | 24.590 | 28.183 | 24.180 | 28.750 | 24.936 | 27.842 | 26.875 | 28.674 | 23.126 | 27.501 | -0.008 | 0.8862 | 0.9530 | 0 | 0 | 0 |
| 'Mfsd6' | 7.456 | 7.403 | 7.602 | 7.274 | 7.928 | 7.672 | 6.601 | 7.437 | 7.257 | 7.699 | 8.215 | 8.081 | 7.735 | 7.762 | 8.119 | 7.302 | 7.826 | 8.458 | -0.106 | 0.0006 | 0.0073 | 1 | 0 | 0 |
| 'Mfsd7b' | 2.566 | 2.579 | 2.459 | 2.379 | 2.869 | 2.312 | 2.225 | 2.814 | 2.456 | 2.283 | 2.551 | 2.851 | 2.463 | 2.647 | 2.671 | 2.372 | 2.630 | 2.769 | -0.052 | 0.2588 | 0.5119 | 0 | 0 | 0 |
| 'Mfsd8' | 2.906 | 2.827 | 3.049 | 2.756 | 2.788 | 2.767 | 3.079 | 3.196 | 3.305 | 3.154 | 2.931 | 2.785 | 2.985 | 3.017 | 2.515 | 3.036 | 2.907 | 2.527 | 0.043 | 0.4058 | 0.6551 | 0 | 0 | 0 |
| 'Mga' | 3.810 | 3.452 | 3.711 | 3.551 | 3.521 | 3.180 | 3.797 | 3.595 | 3.543 | 3.820 | 4.135 | 3.831 | 3.313 | 3.560 | 3.329 | 3.582 | 3.843 | 3.809 | -0.058 | 0.2101 | 0.4553 | 0 | 0 | 0 |
| 'Mgat1' | 7.806 | 7.942 | 8.489 | 7.865 | 8.528 | 8.517 | 7.820 | 8.075 | 8.640 | 7.960 | 8.348 | 8.674 | 7.728 | 7.812 | 8.525 | 7.913 | 7.879 | 8.482 | -0.003 | 0.8817 | 0.9501 | 0 | 0 | 0 |
| 'Mgat2' | 32.388 | 32.835 | 31.855 | 32.012 | 31.287 | 32.327 | 34.358 | 30.546 | 30.828 | 29.319 | 29.275 | 35.075 | 28.914 | 32.273 | 33.359 | 30.245 | 30.396 | 34.635 | 0.013 | 0.7409 | 0.8827 | 0 | 0 | 0 |
| 'Mgat4b' | 2.651 | 2.566 | 2.568 | 2.608 | 2.888 | 3.346 | 2.796 | 2.532 | 2.700 | 2.104 | 2.130 | 2.222 | 2.037 | 1.941 | 2.108 | 2.169 | 1.710 | 1.862 | 0.421 | 0.0000 | 0.0000 | 1 | 1 | 0 |
| 'Mgat5' | 9.598 | 9.320 | 9.326 | 8.650 | 9.635 | 9.421 | 8.682 | 9.706 | 9.458 | 9.344 | 10.019 | 10.114 | 8.748 | 9.419 | 9.994 | 9.803 | 8.368 | 9.825 | -0.041 | 0.2581 | 0.5107 | 0 | 0 | 0 |
| 'Mgea5' | 57.134 | 61.059 | 64.763 | 54.948 | 61.087 | 64.413 | 55.612 | 61.373 | 63.810 | 70.617 | 60.349 | 58.084 | 64.010 | 61.770 | 57.364 | 64.685 | 64.246 | 52.294 | -0.033 | 0.4911 | 0.7208 | 0 | 0 | 0 |
| 'Mgme1' | 7.942 | 8.266 | 7.915 | 7.172 | 7.495 | 7.838 | 7.670 | 6.817 | 8.327 | 7.273 | 6.221 | 7.531 | 6.659 | 8.129 | 6.153 | 5.852 | 7.160 | 6.284 | 0.165 | 0.0349 | 0.1496 | 0 | 0 | 0 |
| 'Mgrn1' | 33.784 | 37.974 | 37.952 | 32.779 | 37.125 | 37.222 | 31.776 | 35.855 | 34.549 | 39.051 | 34.946 | 36.997 | 35.213 | 36.263 | 34.009 | 37.897 | 35.899 | 33.154 | -0.030 | 0.4293 | 0.6749 | 0 | 0 | 0 |
| 'Mgst2' | 3.268 | 3.645 | 3.989 | 3.122 | 4.274 | 3.793 | 3.544 | 3.452 | 4.440 | 2.952 | 2.401 | 2.976 | 3.020 | 2.479 | 3.167 | 2.244 | 2.844 | 2.881 | 0.408 | 0.0000 | 0.0000 | 1 | 1 | 0 |
| 'Mib1' | 4.836 | 4.786 | 4.701 | 4.593 | 4.792 | 4.610 | 4.110 | 4.942 | 4.619 | 4.344 | 4.818 | 5.440 | 4.381 | 4.680 | 4.838 | 4.650 | 5.157 | 5.404 | -0.068 | 0.0997 | 0.2954 | 0 | 0 | 0 |
| 'Mib2' | 4.420 | 4.363 | 4.818 | 4.504 | 4.265 | 4.843 | 4.248 | 4.533 | 4.990 | 4.904 | 3.946 | 5.350 | 4.479 | 4.247 | 4.564 | 4.789 | 4.393 | 5.067 | -0.037 | 0.3233 | 0.5808 | 0 | 0 | 0 |
| 'Mical1' | 6.055 | 4.808 | 7.143 | 5.318 | 5.290 | 6.538 | 6.441 | 5.310 | 6.294 | 5.755 | 5.208 | 6.380 | 4.564 | 5.097 | 6.145 | 5.510 | 4.892 | 5.455 | 0.106 | 0.0371 | 0.1560 | 0 | 0 | 0 |
| 'Mical2' | 0.441 | 0.811 | 0.504 | 0.564 | 0.474 | 0.557 | 0.521 | 0.525 | 0.616 | 0.759 | 0.667 | 0.718 | 0.917 | 0.760 | 0.644 | 0.736 | 0.595 | 0.627 | -0.373 | 0.0002 | 0.0032 | 1 | 0 | 0 |
| 'Mical3' | 0.364 | 0.285 | 0.422 | 0.284 | 0.250 | 0.337 | 0.334 | 0.349 | 0.375 | 0.345 | 0.246 | 0.345 | 0.290 | 0.275 | 0.351 | 0.269 | 0.312 | 0.373 | 0.092 | 0.2392 | 0.4895 | 0 | 0 | 0 |
| 'Micall1' | 13.027 | 11.318 | 12.374 | 11.505 | 11.887 | 12.443 | 10.644 | 11.502 | 12.745 | 12.242 | 11.440 | 14.491 | 11.821 | 12.001 | 13.162 | 11.599 | 10.781 | 14.840 | -0.072 | 0.1000 | 0.2958 | 0 | 0 | 0 |
| 'Micu1' | 4.643 | 4.746 | 4.359 | 4.437 | 4.366 | 4.344 | 4.565 | 4.641 | 4.669 | 4.909 | 4.695 | 5.067 | 4.494 | 4.377 | 5.065 | 4.915 | 4.921 | 4.887 | -0.100 | 0.0048 | 0.0363 | 1 | 0 | 0 |
| 'Micu2' | 46.767 | 53.797 | 51.929 | 47.598 | 57.516 | 53.563 | 52.580 | 56.295 | 52.695 | 52.506 | 52.624 | 45.993 | 48.665 | 53.031 | 47.348 | 53.295 | 53.224 | 41.523 | 0.070 | 0.0814 | 0.2576 | 0 | 0 | 0 |
| 'Micu3' | 1.977 | 2.106 | 1.877 | 2.249 | 2.268 | 2.289 | 2.355 | 2.190 | 1.941 | 2.307 | 2.211 | 2.117 | 2.072 | 2.090 | 1.907 | 1.921 | 2.671 | 1.698 | 0.007 | 0.9164 | 0.9681 | 0 | 0 | 0 |
| 'Mid1ip1' | 8.741 | 8.406 | 8.604 | 9.148 | 9.117 | 9.311 | 9.206 | 9.040 | 9.279 | 8.918 | 9.732 | 8.946 | 10.390 | 9.595 | 8.952 | 8.599 | 9.634 | 8.105 | -0.044 | 0.3135 | 0.5715 | 0 | 0 | 0 |
| 'Mid2' | 0.786 | 1.099 | 0.951 | 0.831 | 0.953 | 0.708 | 1.032 | 1.087 | 0.789 | 0.809 | 0.953 | 0.908 | 0.715 | 0.910 | 0.659 | 1.025 | 0.945 | 0.788 | 0.077 | 0.3683 | 0.6201 | 0 | 0 | 0 |
| 'Midn' | 26.040 | 21.930 | 22.901 | 23.607 | 21.885 | 23.250 | 22.624 | 21.973 | 23.170 | 23.264 | 22.877 | 24.753 | 22.489 | 23.392 | 24.449 | 23.718 | 21.296 | 25.005 | -0.037 | 0.2766 | 0.5323 | 0 | 0 | 0 |
| 'Mief1' | 12.075 | 11.470 | 12.025 | 11.596 | 12.120 | 11.602 | 13.247 | 11.249 | 12.611 | 10.667 | 12.682 | 13.111 | 10.855 | 12.369 | 12.413 | 11.802 | 12.712 | 13.028 | -0.032 | 0.4698 | 0.7056 | 0 | 0 | 0 |
| 'Mief2' | 6.549 | 6.372 | 7.363 | 5.220 | 6.266 | 7.691 | 5.585 | 6.584 | 7.740 | 7.268 | 5.344 | 5.818 | 6.694 | 5.540 | 5.912 | 6.436 | 5.370 | 5.231 | 0.142 | 0.0586 | 0.2088 | 0 | 0 | 0 |
| 'Mien1' | 63.810 | 75.724 | 65.712 | 70.050 | 72.509 | 70.327 | 70.783 | 73.257 | 67.435 | 65.363 | 67.032 | 65.017 | 70.351 | 67.683 | 67.979 | 69.226 | 65.406 | 63.712 | 0.052 | 0.1260 | 0.3400 | 0 | 0 | 0 |
| 'Mier1' | 7.517 | 8.144 | 7.620 | 7.649 | 7.947 | 7.048 | 7.861 | 8.371 | 7.415 | 7.357 | 8.000 | 7.251 | 7.146 | 7.944 | 7.114 | 7.277 | 7.773 | 7.702 | 0.031 | 0.2456 | 0.4960 | 0 | 0 | 0 |
| 'Mier2' | 5.950 | 5.011 | 6.498 | 5.305 | 5.261 | 6.436 | 5.999 | 5.672 | 6.884 | 5.598 | 5.170 | 6.491 | 5.889 | 4.735 | 6.250 | 5.307 | 4.954 | 5.932 | 0.065 | 0.0755 | 0.2441 | 0 | 0 | 0 |
| 'Mier3' | 2.862 | 3.103 | 2.708 | 3.307 | 2.768 | 2.549 | 3.073 | 2.895 | 2.822 | 2.808 | 3.522 | 3.555 | 2.494 | 3.348 | 3.176 | 2.676 | 3.387 | 3.606 | -0.139 | 0.0468 | 0.1803 | 0 | 0 | 0 |
| 'Mif' | 312.652 | 282.948 | 289.045 | 291.859 | 306.643 | 279.072 | 301.819 | 280.666 | 280.491 | 280.810 | 314.213 | 280.356 | 332.155 | 321.536 | 310.341 | 300.356 | 297.789 | 291.953 | -0.066 | 0.0647 | 0.2218 | 0 | 0 | 0 |
| 'Mif4gd' | 14.894 | 15.814 | 15.516 | 14.269 | 15.850 | 14.795 | 14.627 | 16.450 | 15.230 | 13.721 | 14.805 | 14.806 | 16.085 | 15.119 | 14.773 | 14.163 | 15.865 | 14.151 | 0.031 | 0.3052 | 0.5628 | 0 | 0 | 0 |

| | | | | | | | | | | | | | | | | | | | | | | | | |
|---|---|---|---|---|---|---|---|---|---|---|---|---|---|---|---|---|---|---|---|---|---|---|---|---|
| 'Miip' | 8.405 | 9.214 | 12.175 | 9.137 | 8.631 | 11.614 | 10.583 | 9.320 | 11.864 | 8.987 | 8.298 | 11.831 | 9.650 | 8.027 | 10.866 | 9.668 | 9.034 | 10.955 | 0.046 | 0.2351 | 0.4844 | 0 | 0 | 0 |
| 'Milr1' | 1.869 | 2.340 | 2.090 | 2.157 | 2.496 | 2.300 | 2.456 | 2.172 | 2.239 | 2.078 | 2.038 | 2.249 | 2.750 | 2.570 | 2.107 | 2.245 | 2.497 | 1.872 | -0.035 | 0.6229 | 0.8118 | 0 | 0 | 0 |
| 'Mina' | 3.430 | 3.713 | 3.699 | 3.493 | 3.754 | 3.552 | 3.466 | 3.834 | 3.742 | 3.459 | 3.309 | 3.389 | 3.304 | 3.304 | 3.382 | 3.785 | 2.988 | 3.250 | 0.108 | 0.0179 | 0.0956 | 0 | 0 | 0 |
| 'Mink1' | 50.216 | 41.909 | 51.301 | 46.363 | 43.172 | 47.935 | 44.386 | 45.973 | 50.158 | 48.902 | 46.601 | 52.288 | 46.824 | 44.707 | 48.437 | 43.988 | 45.871 | 51.660 | -0.038 | 0.2446 | 0.4952 | 0 | 0 | 0 |
| 'Minos1' | 29.474 | 34.089 | 28.210 | 33.258 | 32.358 | 30.393 | 32.952 | 31.502 | 29.458 | 27.939 | 29.792 | 27.098 | 30.829 | 30.693 | 28.504 | 30.171 | 31.731 | 26.744 | 0.085 | 0.0066 | 0.0462 | 1 | 0 | 0 |
| 'Minpp1' | 18.953 | 20.700 | 21.130 | 17.984 | 20.708 | 20.586 | 18.346 | 19.310 | 19.594 | 18.926 | 17.405 | 20.616 | 19.212 | 18.154 | 19.784 | 19.481 | 18.868 | 19.449 | 0.032 | 0.3576 | 0.6103 | 0 | 0 | 0 |
| 'Mios' | 3.860 | 4.887 | 4.272 | 4.167 | 5.009 | 4.095 | 4.951 | 4.457 | 4.327 | 4.608 | 4.922 | 3.771 | 4.176 | 4.356 | 3.705 | 5.020 | 4.140 | 3.315 | 0.071 | 0.2219 | 0.4694 | 0 | 0 | 0 |
| 'Mipep' | 9.218 | 9.044 | 8.869 | 11.730 | 10.449 | 9.541 | 10.364 | 9.199 | 8.809 | 10.117 | 8.945 | 9.061 | 8.335 | 8.716 | 9.132 | 9.437 | 8.876 | 8.479 | 0.090 | 0.1025 | 0.3002 | 0 | 0 | 0 |
| 'Mipol1' | 0.443 | 0.409 | 0.412 | 0.390 | 0.482 | 0.426 | 0.386 | 0.444 | 0.433 | 0.436 | 0.410 | 0.384 | 0.321 | 0.340 | 0.391 | 0.393 | 0.480 | 0.342 | 0.127 | 0.1899 | 0.4262 | 0 | 0 | 0 |
| 'Mis12' | 9.394 | 9.626 | 10.066 | 11.379 | 8.987 | 9.852 | 10.166 | 9.400 | 10.024 | 8.014 | 10.212 | 11.153 | 9.172 | 9.409 | 11.351 | 8.608 | 11.316 | 11.441 | -0.035 | 0.5745 | 0.7800 | 0 | 0 | 0 |
| 'Mis18a' | 6.031 | 7.148 | 6.774 | 5.299 | 6.329 | 6.349 | 5.671 | 6.158 | 6.448 | 5.627 | 6.718 | 6.549 | 6.136 | 6.369 | 6.664 | 6.112 | 6.288 | 6.223 | -0.023 | 0.6083 | 0.8029 | 0 | 0 | 0 |
| 'Mis18bp1' | 1.095 | 1.359 | 1.457 | 1.389 | 1.449 | 1.302 | 1.530 | 1.370 | 1.508 | 0.837 | 1.297 | 1.315 | 1.386 | 1.633 | 1.531 | 1.250 | 1.402 | 1.100 | 0.077 | 0.3851 | 0.6349 | 0 | 0 | 0 |
| 'Mitd1' | 8.081 | 8.717 | 8.089 | 8.761 | 8.926 | 7.691 | 8.461 | 9.263 | 8.753 | 6.382 | 8.399 | 9.771 | 7.235 | 9.074 | 10.164 | 7.958 | 9.713 | 9.951 | -0.040 | 0.5359 | 0.7552 | 0 | 0 | 0 |
| 'Mki67' | 44.334 | 42.537 | 55.447 | 50.986 | 42.637 | 56.405 | 51.012 | 41.432 | 55.053 | 36.514 | 47.971 | 49.209 | 52.839 | 52.250 | 57.008 | 46.105 | 47.166 | 53.627 | -0.026 | 0.6760 | 0.8448 | 0 | 0 | 0 |
| 'Mkks' | 5.178 | 5.644 | 4.841 | 5.219 | 4.604 | 4.592 | 5.335 | 4.660 | 4.600 | 4.340 | 4.910 | 5.766 | 4.533 | 5.230 | 4.757 | 4.501 | 5.844 | 4.881 | -0.021 | 0.7552 | 0.8907 | 0 | 0 | 0 |
| 'Mkl1' | 19.152 | 17.175 | 20.495 | 17.913 | 17.727 | 20.269 | 18.068 | 17.896 | 20.215 | 18.940 | 18.143 | 21.933 | 18.231 | 17.493 | 21.016 | 18.079 | 17.501 | 21.563 | -0.043 | 0.0769 | 0.2475 | 0 | 0 | 0 |
| 'Mkl2' | 4.246 | 3.798 | 4.509 | 4.272 | 3.748 | 4.571 | 3.810 | 4.238 | 4.353 | 4.651 | 4.319 | 4.686 | 4.072 | 4.073 | 4.488 | 4.553 | 4.449 | 4.695 | -0.097 | 0.0132 | 0.0767 | 0 | 0 | 0 |
| 'Mkln1' | 2.637 | 2.643 | 2.534 | 2.667 | 2.597 | 2.466 | 2.740 | 2.633 | 2.441 | 2.572 | 2.885 | 2.716 | 2.522 | 2.571 | 2.386 | 2.508 | 2.707 | 2.492 | -0.013 | 0.7035 | 0.8613 | 0 | 0 | 0 |
| 'Mknk1' | 7.485 | 7.532 | 8.176 | 6.942 | 7.448 | 7.854 | 6.466 | 8.200 | 7.974 | 7.940 | 7.620 | 8.101 | 8.489 | 7.933 | 7.877 | 8.262 | 7.618 | 7.450 | -0.075 | 0.0648 | 0.2221 | 0 | 0 | 0 |
| 'Mknk2' | 92.717 | 85.567 | 95.543 | 83.154 | 88.229 | 106.498 | 84.494 | 89.827 | 97.918 | 94.713 | 89.446 | 105.537 | 97.996 | 92.627 | 107.922 | 99.716 | 81.317 | 110.457 | -0.103 | 0.0028 | 0.0238 | 1 | 0 | 0 |
| 'Mkrn1' | 25.390 | 29.587 | 24.606 | 24.983 | 27.747 | 27.356 | 25.193 | 28.220 | 25.798 | 26.403 | 26.204 | 27.640 | 24.979 | 27.880 | 27.116 | 28.395 | 26.789 | 26.889 | -0.033 | 0.2865 | 0.5432 | 0 | 0 | 0 |
| 'Mkrn2' | 6.784 | 6.032 | 5.688 | 5.687 | 5.682 | 5.399 | 5.468 | 5.781 | 5.297 | 6.494 | 6.610 | 5.751 | 6.202 | 6.075 | 5.502 | 6.117 | 6.479 | 5.817 | -0.094 | 0.0288 | 0.1316 | 0 | 0 | 0 |
| 'Mks1' | 2.183 | 3.172 | 2.935 | 2.359 | 2.890 | 3.134 | 2.411 | 2.798 | 3.057 | 2.941 | 2.709 | 2.816 | 3.081 | 2.680 | 2.534 | 3.008 | 2.531 | 2.305 | 0.007 | 0.9267 | 0.9719 | 0 | 0 | 0 |
| 'Mlec' | 63.731 | 61.943 | 65.946 | 64.325 | 58.995 | 70.394 | 62.519 | 61.432 | 68.409 | 59.457 | 63.453 | 78.802 | 57.316 | 61.841 | 77.978 | 62.621 | 63.022 | 82.082 | -0.072 | 0.0708 | 0.2347 | 0 | 0 | 0 |
| 'Mlf2' | 20.783 | 21.087 | 20.649 | 19.556 | 21.893 | 23.804 | 20.447 | 21.121 | 21.668 | 19.647 | 18.642 | 21.499 | 21.105 | 20.469 | 23.029 | 21.966 | 19.771 | 22.092 | 0.011 | 0.7436 | 0.8843 | 0 | 0 | 0 |
| 'Mlh1' | 3.918 | 4.108 | 3.927 | 4.085 | 3.837 | 4.201 | 4.463 | 4.085 | 4.320 | 4.161 | 4.306 | 4.355 | 4.356 | 4.071 | 4.160 | 3.585 | 4.568 | 3.932 | -0.035 | 0.4316 | 0.6763 | 0 | 0 | 0 |
| 'Mlh3' | 6.005 | 7.275 | 7.147 | 6.095 | 7.219 | 7.023 | 6.177 | 7.494 | 7.577 | 7.271 | 6.751 | 6.977 | 6.423 | 6.194 | 6.392 | 6.795 | 6.745 | 6.091 | 0.045 | 0.3522 | 0.6059 | 0 | 0 | 0 |
| 'Mllt1' | 22.260 | 19.396 | 25.332 | 21.336 | 20.142 | 24.746 | 22.004 | 21.413 | 24.720 | 21.764 | 19.115 | 27.181 | 22.778 | 20.327 | 24.901 | 23.017 | 20.670 | 26.208 | -0.042 | 0.1449 | 0.3693 | 0 | 0 | 0 |
| 'Mllt10' | 13.521 | 11.836 | 12.206 | 12.719 | 12.118 | 12.034 | 12.417 | 12.484 | 11.980 | 12.809 | 12.153 | 12.931 | 12.202 | 12.066 | 12.218 | 11.630 | 11.754 | 12.195 | 0.006 | 0.8552 | 0.9380 | 0 | 0 | 0 |
| 'Mllt11' | 3.225 | 3.190 | 3.417 | 3.456 | 4.103 | 3.566 | 4.141 | 3.847 | 3.444 | 2.562 | 4.069 | 4.315 | 3.006 | 3.759 | 3.797 | 2.954 | 4.352 | 3.851 | -0.020 | 0.8084 | 0.9139 | 0 | 0 | 0 |
| 'Mllt3' | 8.581 | 7.675 | 8.412 | 9.103 | 8.508 | 7.622 | 9.123 | 7.908 | 7.297 | 8.455 | 8.351 | 8.536 | 8.916 | 8.354 | 8.304 | 8.169 | 8.744 | 8.763 | -0.060 | 0.1255 | 0.3391 | 0 | 0 | 0 |
| 'Mllt4' | 0.535 | 0.584 | 0.545 | 0.593 | 0.531 | 0.682 | 0.591 | 0.538 | 0.643 | 0.519 | 0.459 | 0.444 | 0.466 | 0.429 | 0.483 | 0.347 | 0.431 | 0.501 | 0.351 | 0.0000 | 0.0000 | 1 | 0 | 0 |
| 'Mllt6' | 26.757 | 24.401 | 26.942 | 24.517 | 24.285 | 26.051 | 22.334 | 25.308 | 27.446 | 23.323 | 25.082 | 28.064 | 23.964 | 24.799 | 27.708 | 23.291 | 23.930 | 31.811 | -0.031 | 0.4538 | 0.6927 | 0 | 0 | 0 |
| 'Mlst8' | 7.489 | 7.995 | 8.904 | 7.813 | 8.593 | 8.453 | 7.570 | 8.432 | 9.093 | 8.660 | 8.716 | 8.307 | 8.953 | 8.823 | 8.202 | 8.198 | 8.032 | 8.304 | -0.041 | 0.3565 | 0.6092 | 0 | 0 | 0 |
| 'Mlx' | 15.524 | 14.751 | 15.049 | 13.684 | 14.778 | 15.211 | 15.406 | 14.447 | 14.754 | 13.171 | 14.024 | 15.401 | 14.197 | 14.506 | 15.707 | 12.988 | 15.373 | 15.605 | 0.017 | 0.6534 | 0.8320 | 0 | 0 | 0 |
| 'Mlxip' | 12.078 | 10.984 | 11.945 | 10.359 | 10.753 | 10.745 | 10.003 | 10.634 | 11.945 | 12.613 | 11.741 | 12.578 | 11.760 | 11.035 | 11.244 | 11.777 | 11.078 | 12.968 | -0.113 | 0.0054 | 0.0396 | 1 | 0 | 0 |
| 'Mlycd' | 7.734 | 7.952 | 9.565 | 7.373 | 7.853 | 9.606 | 7.631 | 8.118 | 9.010 | 8.556 | 7.451 | 8.510 | 7.774 | 7.710 | 8.842 | 8.780 | 6.968 | 8.423 | 0.030 | 0.4920 | 0.7216 | 0 | 0 | 0 |
| 'Mmaa' | 7.194 | 6.791 | 7.006 | 8.069 | 6.748 | 6.216 | 5.616 | 6.066 | 7.212 | 7.228 | 6.360 | 7.600 | 6.770 | 5.682 | 5.602 | 5.305 | 6.998 | 6.029 | 0.063 | 0.4460 | 0.6878 | 0 | 0 | 0 |
| 'Mmab' | 4.371 | 6.349 | 5.321 | 4.928 | 5.304 | 5.744 | 5.131 | 4.701 | 6.006 | 5.158 | 5.423 | 5.378 | 6.151 | 4.974 | 5.852 | 5.277 | 5.254 | 4.970 | -0.033 | 0.6129 | 0.8058 | 0 | 0 | 0 |
| 'Mmachc' | 2.379 | 2.611 | 2.723 | 2.079 | 2.693 | 2.697 | 2.106 | 2.963 | 2.406 | 1.877 | 2.529 | 2.568 | 1.711 | 1.991 | 2.201 | 2.194 | 2.212 | 2.183 | 0.207 | 0.0178 | 0.0950 | 0 | 0 | 0 |
| 'Mmadhc' | 10.410 | 12.657 | 11.262 | 10.809 | 11.904 | 11.478 | 11.658 | 12.149 | 10.849 | 11.456 | 11.735 | 10.525 | 11.400 | 12.452 | 11.408 | 11.235 | 11.718 | 10.272 | 0.006 | 0.8726 | 0.9454 | 0 | 0 | 0 |
| 'Mmd' | 12.344 | 12.705 | 10.496 | 10.987 | 12.245 | 12.105 | 12.352 | 13.109 | 11.024 | 11.343 | 10.485 | 12.136 | 10.531 | 11.868 | 12.297 | 12.776 | 12.012 | 11.750 | 0.016 | 0.7273 | 0.8750 | 0 | 0 | 0 |
| 'Mmgt1' | 6.218 | 7.171 | 6.683 | 6.049 | 7.455 | 6.503 | 7.239 | 6.125 | 6.513 | 6.156 | 7.289 | 7.049 | 6.316 | 6.248 | 7.068 | 7.473 | 7.556 | 6.392 | -0.051 | 0.3457 | 0.6009 | 0 | 0 | 0 |
| 'Mmgt2' | 3.565 | 3.212 | 3.883 | 3.656 | 3.729 | 3.577 | 3.513 | 3.406 | 4.006 | 3.263 | 3.440 | 3.814 | 3.422 | 3.572 | 3.629 | 3.406 | 3.281 | 3.917 | 0.025 | 0.5900 | 0.7907 | 0 | 0 | 0 |
| 'Mms19' | 6.130 | 5.798 | 6.354 | 5.638 | 5.988 | 5.566 | 5.561 | 6.015 | 6.295 | 6.544 | 6.010 | 5.970 | 6.753 | 5.672 | 5.520 | 5.407 | 6.269 | 5.391 | -0.014 | 0.7564 | 0.8913 | 0 | 0 | 0 |
| 'Mms22l' | 2.974 | 2.878 | 3.474 | 2.894 | 3.280 | 3.364 | 3.177 | 3.245 | 3.239 | 2.936 | 3.387 | 3.145 | 3.712 | 3.475 | 3.314 | 3.159 | 3.286 | 3.256 | -0.065 | 0.1660 | 0.3983 | 0 | 0 | 0 |
| 'Mnat1' | 3.147 | 3.704 | 3.114 | 3.599 | 3.458 | 3.499 | 3.953 | 3.632 | 3.355 | 3.256 | 3.163 | 2.994 | 3.738 | 3.542 | 2.988 | 3.952 | 3.438 | 2.746 | 0.072 | 0.1788 | 0.4143 | 0 | 0 | 0 |
| 'Mndal' | 21.700 | 21.308 | 23.281 | 21.534 | 21.573 | 23.426 | 22.898 | 23.907 | 23.385 | 21.891 | 22.586 | 28.698 | 22.662 | 24.975 | 27.803 | 24.757 | 26.282 | 27.256 | -0.171 | 0.0000 | 0.0005 | 1 | 0 | 0 |
| 'Mns1' | 0.739 | 0.763 | 0.874 | 1.185 | 0.915 | 1.073 | 0.977 | 0.868 | 0.938 | 0.881 | 0.515 | 0.800 | 0.971 | 0.777 | 0.858 | 0.938 | 0.795 | 0.754 | 0.184 | 0.0731 | 0.2396 | 0 | 0 | 0 |
| 'Mnt' | 11.929 | 11.027 | 13.114 | 10.319 | 13.550 | 13.291 | 10.705 | 13.146 | 13.605 | 13.510 | 13.062 | 12.192 | 12.925 | 12.404 | 12.404 | 13.361 | 11.658 | 12.080 | -0.048 | 0.3681 | 0.6200 | 0 | 0 | 0 |
| 'Mob1a' | 40.627 | 44.005 | 42.160 | 44.197 | 46.186 | 42.476 | 41.994 | 44.505 | 40.686 | 41.459 | 46.656 | 44.262 | 40.024 | 46.040 | 43.358 | 40.793 | 46.476 | 44.839 | -0.036 | 0.1701 | 0.4029 | 0 | 0 | 0 |
| 'Mob1b' | 2.115 | 2.492 | 2.086 | 2.076 | 2.448 | 1.940 | 1.994 | 2.460 | 2.074 | 2.309 | 2.242 | 2.245 | 2.325 | 2.644 | 1.938 | 2.358 | 2.298 | 2.215 | -0.079 | 0.0649 | 0.2221 | 0 | 0 | 0 |
| 'Mob2' | 30.468 | 30.444 | 29.311 | 27.851 | 32.963 | 30.265 | 27.000 | 32.512 | 28.989 | 28.770 | 29.902 | 30.394 | 31.708 | 30.030 | 31.247 | 31.465 | 29.918 | 30.284 | -0.031 | 0.4117 | 0.6601 | 0 | 0 | 0 |
| 'Mob3a' | 78.340 | 79.256 | 79.346 | 71.787 | 73.693 | 83.071 | 71.298 | 76.385 | 80.698 | 69.002 | 66.671 | 85.462 | 62.290 | 67.288 | 85.231 | 69.337 | 65.217 | 88.559 | 0.072 | 0.0882 | 0.2713 | 0 | 0 | 0 |
| 'Mob3b' | 0.754 | 0.713 | 0.678 | 0.738 | 0.810 | 0.650 | 0.769 | 0.617 | 0.709 | 0.697 | 0.790 | 0.771 | 0.496 | 0.449 | 0.675 | 0.576 | 0.743 | 0.770 | 0.088 | 0.4469 | 0.6880 | 0 | 0 | 0 |
| 'Mob3c' | 6.937 | 7.581 | 7.353 | 6.095 | 6.803 | 7.127 | 6.876 | 6.285 | 7.081 | 7.282 | 6.440 | 7.817 | 6.780 | 5.936 | 6.652 | 6.580 | 7.251 | 7.639 | -0.021 | 0.6852 | 0.8511 | 0 | 0 | 0 |
| 'Mob4' | 4.795 | 5.558 | 4.597 | 4.903 | 5.332 | 4.454 | 5.076 | 5.101 | 4.485 | 4.796 | 5.141 | 5.301 | 4.553 | 5.579 | 4.572 | 4.729 | 5.638 | 4.732 | -0.041 | 0.2988 | 0.5551 | 0 | 0 | 0 |
| 'Mocos' | 1.971 | 2.068 | 2.132 | 1.557 | 1.965 | 2.197 | 2.498 | 1.944 | 2.113 | 2.896 | 2.457 | 2.301 | 2.157 | 1.962 | 2.370 | 2.577 | 1.474 | 2.790 | -0.189 | 0.0692 | 0.2315 | 0 | 0 | 0 |
| 'Mocs1' | 4.871 | 4.965 | 5.343 | 4.630 | 4.711 | 5.642 | 5.439 | 5.053 | 5.380 | 5.200 | 4.765 | 5.026 | 5.265 | 4.488 | 4.636 | 5.115 | 4.231 | 4.753 | 0.075 | 0.1025 | 0.3002 | 0 | 0 | 0 |
| 'Mocs2' | 2.400 | 2.972 | 2.370 | 2.590 | 2.797 | 2.580 | 2.750 | 2.506 | 2.612 | 2.296 | 2.361 | 2.358 | 2.635 | 2.667 | 2.239 | 2.588 | 2.717 | 2.125 | 0.087 | 0.0782 | 0.2506 | 0 | 0 | 0 |
| 'Mocs3' | 10.741 | 9.251 | 10.706 | 8.380 | 9.476 | 11.719 | 9.346 | 8.813 | 10.489 | 10.361 | 9.283 | 11.070 | 8.727 | 9.074 | 12.210 | 10.003 | 8.606 | 12.294 | -0.047 | 0.4188 | 0.6660 | 0 | 0 | 0 |
| 'Mogs' | 42.821 | 42.489 | 48.423 | 41.642 | 43.056 | 47.675 | 40.859 | 43.144 | 45.874 | 40.244 | 41.388 | 50.089 | 40.408 | 41.622 | 49.699 | 45.648 | 39.879 | 49.769 | -0.018 | 0.5386 | 0.7567 | 0 | 0 | 0 |
| 'Mon1a' | 7.611 | 6.145 | 6.629 | 6.236 | 6.011 | 7.766 | 6.930 | 6.091 | 6.366 | 6.202 | 6.298 | 8.222 | 6.878 | 6.035 | 7.826 | 6.128 | 5.972 | 8.432 | -0.060 | 0.3119 | 0.5704 | 0 | 0 | 0 |
| 'Mon1b' | 10.416 | 11.739 | 12.880 | 10.415 | 12.020 | 13.064 | 11.495 | 11.607 | 12.669 | 11.309 | 11.557 | 13.582 | 11.439 | 12.091 | 12.876 | 11.766 | 11.309 | 14.086 | -0.062 | 0.0514 | 0.1917 | 0 | 0 | 0 |
| 'Mon2' | 17.943 | 17.187 | 17.917 | 16.923 | 17.834 | 17.953 | 16.731 | 18.785 | 16.678 | 19.210 | 18.900 | 17.248 | 17.781 | 18.839 | 16.655 | 17.404 | 18.032 | 17.295 | -0.039 | 0.2044 | 0.4472 | 0 | 0 | 0 |
| 'Morc2a' | 11.517 | 11.782 | 12.440 | 11.915 | 11.481 | 11.510 | 11.743 | 12.316 | 12.200 | 11.966 | 11.952 | 11.626 | 10.480 | 10.842 | 10.855 | 10.589 | 12.682 | 10.622 | 0.064 | 0.0834 | 0.2617 | 0 | 0 | 0 |
| 'Morc3' | 9.770 | 9.470 | 9.762 | 10.119 | 9.745 | 9.450 | 9.273 | 9.883 | 9.658 | 9.162 | 9.843 | 10.154 | 9.076 | 9.801 | 9.436 | 8.987 | 10.013 | 9.749 | 0.004 | 0.8936 | 0.9565 | 0 | 0 | 0 |
| 'Morf4l1' | 29.764 | 32.706 | 28.557 | 31.199 | 33.763 | 29.179 | 32.537 | 32.848 | 29.748 | 28.153 | 30.600 | 26.932 | 29.554 | 30.848 | 27.087 | 29.846 | 31.293 | 26.173 | 0.096 | 0.0000 | 0.0000 | 1 | 0 | 0 |
| 'Morn1' | 1.833 | 2.052 | 2.089 | 1.956 | 1.845 | 2.028 | 2.213 | 1.883 | 2.238 | 1.800 | 2.127 | 1.650 | 1.796 | 1.563 | 1.643 | 1.816 | 1.713 | 1.426 | 0.222 | 0.0016 | 0.0149 | 1 | 0 | 0 |
| 'Mospd1' | 3.082 | 3.157 | 3.576 | 3.531 | 3.702 | 3.002 | 3.243 | 3.366 | 2.236 | 3.892 | 3.453 | 3.684 | 3.463 | 3.526 | 3.438 | 3.439 | 4.028 | 3.343 | -0.176 | 0.0444 | 0.1747 | 0 | 0 | 0 |
| 'Mospd2' | 1.152 | 1.517 | 1.170 | 1.025 | 1.356 | 1.344 | 1.183 | 1.306 | 1.203 | 1.135 | 1.453 | 1.251 | 1.346 | 1.343 | 1.269 | 1.162 | 1.348 | 0.992 | -0.018 | 0.7759 | 0.9000 | 0 | 0 | 0 |
| 'Mospd3' | 23.582 | 24.670 | 26.489 | 23.884 | 24.500 | 27.161 | 24.314 | 25.133 | 25.563 | 24.272 | 22.551 | 26.199 | 26.796 | 24.380 | 27.104 | 23.800 | 21.895 | 25.044 | 0.010 | 0.7565 | 0.8913 | 0 | 0 | 0 |
| 'Mov10' | 5.287 | 5.061 | 5.964 | 4.990 | 4.779 | 5.783 | 5.430 | 4.824 | 5.785 | 5.336 | 5.365 | 7.336 | 5.100 | 4.392 | 6.385 | 4.942 | 5.357 | 7.094 | -0.103 | 0.0364 | 0.1538 | 0 | 0 | 0 |
| 'Mpc1' | 11.531 | 14.391 | 11.968 | 11.741 | 13.574 | 12.941 | 12.196 | 13.525 | 12.828 | 12.639 | 12.045 | 11.803 | 12.954 | 12.290 | 11.622 | 12.606 | 11.630 | 10.811 | 0.069 | 0.1150 | 0.3219 | 0 | 0 | 0 |
| 'Mpc2' | 3.262 | 3.423 | 3.058 | 2.877 | 3.670 | 3.266 | 2.775 | 3.517 | 3.297 | 2.961 | 3.426 | 3.071 | 3.584 | 3.174 | 2.939 | 3.495 | 3.195 | 2.800 | 0.017 | 0.7475 | 0.8862 | 0 | 0 | 0 |
| 'Mpdu1' | 8.374 | 10.756 | 11.118 | 7.983 | 10.512 | 10.885 | 9.347 | 10.662 | 11.594 | 10.569 | 8.860 | 8.604 | 10.872 | 9.620 | 9.940 | 11.001 | 9.045 | 8.980 | 0.048 | 0.4915 | 0.7210 | 0 | 0 | 0 |
| 'Mpg' | 3.706 | 4.236 | 4.527 | 3.795 | 4.182 | 4.488 | 3.937 | 4.412 | 4.636 | 3.976 | 3.942 | 4.233 | 3.895 | 3.680 | 4.283 | 3.736 | 3.803 | 3.680 | 0.096 | 0.0442 | 0.1742 | 0 | 0 | 0 |
| 'Mphosph10' | 7.365 | 6.267 | 6.559 | 6.697 | 7.243 | 6.466 | 7.634 | 6.734 | 6.449 | 7.031 | 6.606 | 7.148 | 5.635 | 6.997 | 6.878 | 7.456 | 7.357 | 6.856 | -0.024 | 0.6513 | 0.8307 | 0 | 0 | 0 |
| 'Mphosph6' | 21.731 | 30.816 | 25.214 | 24.788 | 32.739 | 27.380 | 24.719 | 29.924 | 26.102 | 27.602 | 25.323 | 21.879 | 29.875 | 29.004 | 24.526 | 32.199 | 28.368 | 20.867 | 0.017 | 0.8130 | 0.9157 | 0 | 0 | 0 |
| 'Mphosph8' | 14.989 | 14.780 | 13.431 | 15.885 | 14.473 | 13.009 | 16.085 | 13.946 | 13.300 | 13.012 | 15.022 | 15.490 | 13.509 | 15.002 | 14.422 | 13.282 | 16.465 | 14.637 | -0.024 | 0.6364 | 0.8206 | 0 | 0 | 0 |
| 'Mphosph9' | 3.604 | 3.190 | 3.525 | 3.643 | 3.228 | 3.042 | 4.012 | 3.157 | 3.400 | 3.078 | 3.507 | 3.478 | 3.414 | 3.483 | 3.233 | 2.915 | 3.632 | 3.426 | 0.017 | 0.7580 | 0.8917 | 0 | 0 | 0 |
| 'Mpi' | 2.636 | 2.761 | 2.973 | 2.670 | 2.211 | 2.934 | 2.647 | 2.557 | 2.773 | 2.738 | 2.598 | 2.879 | 2.732 | 2.939 | 2.737 | 2.529 | 2.940 | 2.901 | -0.056 | 0.3671 | 0.6192 | 0 | 0 | 0 |
| 'Mplkip' | 7.683 | 9.425 | 9.283 | 8.676 | 9.129 | 8.623 | 8.833 | 8.091 | 9.802 | 7.788 | 8.427 | 9.235 | 7.904 | 8.031 | 8.535 | 7.684 | 8.425 | 7.449 | 0.097 | 0.0594 | 0.2105 | 0 | 0 | 0 |
| 'Mpnd' | 10.048 | 10.034 | 11.147 | 10.214 | 10.295 | 12.263 | 10.934 | 10.559 | 10.831 | 9.332 | 9.262 | 10.691 | 9.836 | 9.278 | 10.342 | 9.625 | 9.272 | 9.395 | 0.135 | 0.0001 | 0.0016 | 1 | 0 | 0 |
| 'Mpp1' | 8.023 | 8.136 | 8.626 | 9.035 | 8.410 | 8.882 | 8.647 | 8.494 | 8.253 | 8.597 | 9.138 | 9.334 | 8.871 | 9.334 | 9.176 | 7.905 | 8.179 | 9.348 | -0.074 | 0.0441 | 0.1739 | 0 | 0 | 0 |
| 'Mpp2' | 1.420 | 1.552 | 1.269 | 1.338 | 1.411 | 1.297 | 1.270 | 1.447 | 1.383 | 1.058 | 1.402 | 1.757 | 1.316 | 1.509 | 1.414 | 1.383 | 1.440 | 1.609 | -0.078 | 0.2969 | 0.5529 | 0 | 0 | 0 |
| 'Mpp4' | 0.282 | 0.298 | 0.337 | 0.224 | 0.327 | 0.254 | 0.361 | 0.246 | 0.325 | 0.307 | 0.293 | 0.284 | 0.394 | 0.301 | 0.284 | 0.279 | 0.334 | 0.270 | -0.060 | 0.6052 | 0.8013 | 0 | 0 | 0 |
| 'Mpp5' | 5.799 | 6.424 | 5.850 | 6.846 | 6.470 | 5.793 | 6.810 | 6.348 | 5.665 | 7.128 | 6.818 | 6.198 | 5.951 | 6.420 | 5.300 | 5.724 | 6.772 | 5.382 | -0.005 | 0.9222 | 0.9705 | 0 | 0 | 0 |
| 'Mpp6' | 3.442 | 3.044 | 3.708 | 3.642 | 3.391 | 3.385 | 3.746 | 3.331 | 3.089 | 3.401 | 3.420 | 3.382 | 3.432 | 3.044 | 3.416 | 2.803 | 3.393 | 3.249 | 0.047 | 0.3486 | 0.6039 | 0 | 0 | 0 |
| 'Mppe1' | 23.143 | 25.096 | 26.379 | 20.132 | 26.487 | 25.056 | 22.375 | 27.293 | 25.819 | 26.979 | 21.693 | 23.282 | 25.776 | 23.580 | 23.470 | 27.827 | 23.997 | 20.772 | 0.022 | 0.7196 | 0.8711 | 0 | 0 | 0 |
| 'Mpst' | 29.590 | 31.871 | 32.253 | 30.092 | 30.474 | 34.356 | 30.520 | 29.905 | 31.842 | 30.705 | 27.443 | 30.146 | 28.714 | 27.710 | 29.693 | 28.887 | 26.235 | 27.558 | 0.118 | 0.0009 | 0.0097 | 1 | 0 | 0 |
| 'Mpv17' | 2.184 | 2.438 | 2.405 | 2.228 | 2.664 | 2.480 | 2.557 | 2.553 | 2.603 | 2.473 | 2.690 | 2.692 | 2.643 | 2.501 | 2.519 | 2.443 | 2.654 | 2.193 | -0.059 | 0.2624 | 0.5162 | 0 | 0 | 0 |
| 'Mpv17l2' | 26.110 | 35.736 | 30.174 | 26.510 | 36.358 | 32.813 | 27.121 | 35.115 | 33.346 | 29.336 | 27.452 | 30.065 | 32.514 | 27.855 | 30.316 | 36.260 | 26.784 | 30.962 | 0.045 | 0.5245 | 0.7468 | 0 | 0 | 0 |
| 'Mpzl3' | 2.572 | 3.347 | 2.896 | 2.511 | 3.387 | 2.837 | 2.805 | 3.314 | 2.898 | 4.907 | 4.264 | 3.712 | 4.355 | 4.654 | 3.858 | 4.927 | 4.557 | 3.127 | -0.534 | 0.0000 | 0.0000 | 1 | 1 | 0 |
| 'Mr1' | 2.640 | 3.455 | 2.842 | 2.629 | 3.725 | 2.915 | 2.804 | 3.351 | 2.788 | 3.284 | 3.322 | 2.587 | 3.803 | 3.153 | 2.655 | 2.904 | 2.923 | 2.543 | -0.012 | 0.8475 | 0.9343 | 0 | 0 | 0 |
| 'Mre11a' | 8.612 | 8.698 | 9.943 | 8.925 | 8.356 | 8.720 | 9.167 | 8.892 | 9.018 | 8.379 | 8.610 | 9.642 | 8.381 | 7.755 | 8.499 | 7.969 | 8.815 | 8.416 | 0.059 | 0.1191 | 0.3284 | 0 | 0 | 0 |
| 'Mrfap1' | 69.601 | 69.126 | 62.871 | 67.181 | 67.025 | 63.371 | 65.429 | 69.508 | 62.417 | 58.796 | 68.595 | 64.175 | 63.646 | 68.099 | 68.177 | 62.363 | 66.489 | 70.318 | 0.003 | 0.9305 | 0.9737 | 0 | 0 | 0 |
| 'Mrgbp' | 6.949 | 7.431 | 8.490 | 7.435 | 7.625 | 8.228 | 7.625 | 7.752 | 8.605 | 6.883 | 6.492 | 8.301 | 7.349 | 6.583 | 7.897 | 7.588 | 7.098 | 8.073 | 0.070 | 0.0299 | 0.1345 | 0 | 0 | 0 |
| 'Mrgpre' | 8.647 | 10.275 | 8.469 | 9.022 | 8.915 | 9.796 | 9.715 | 8.536 | 9.349 | 9.949 | 8.904 | 9.441 | 8.549 | 8.147 | 8.967 | 7.894 | 7.844 | 9.045 | 0.056 | 0.2873 | 0.5438 | 0 | 0 | 0 |
| 'Mri1' | 4.016 | 3.553 | 4.372 | 4.033 | 4.054 | 4.176 | 3.849 | 3.836 | 4.457 | 3.690 | 3.295 | 4.162 | 4.215 | 3.286 | 3.694 | 3.804 | 3.448 | 3.936 | 0.104 | 0.0357 | 0.1516 | 0 | 0 | 0 |
| 'Mrm1' | 4.767 | 5.592 | 4.756 | 4.739 | 4.777 | 5.294 | 4.767 | 5.730 | 5.360 | 4.884 | 4.897 | 5.118 | 4.776 | 4.571 | 5.136 | 5.508 | 5.425 | 4.491 | 0.022 | 0.7086 | 0.8650 | 0 | 0 | 0 |
| 'Mrm2' | 7.193 | 8.319 | 7.836 | 6.834 | 7.109 | 7.744 | 6.426 | 7.099 | 6.965 | 7.421 | 6.220 | 7.658 | 5.938 | 6.416 | 7.723 | 7.167 | 5.990 | 8.254 | 0.050 | 0.4856 | 0.7168 | 0 | 0 | 0 |
| 'Mroh1' | 4.777 | 4.546 | 5.358 | 4.551 | 4.850 | 5.393 | 4.697 | 4.665 | 5.414 | 5.387 | 5.360 | 5.168 | 5.341 | 4.940 | 4.757 | 4.969 | 5.046 | 4.751 | -0.058 | 0.1519 | 0.3790 | 0 | 0 | 0 |
| 'Mrpl1' | 3.130 | 2.875 | 3.035 | 3.544 | 3.271 | 2.856 | 3.701 | 2.941 | 2.882 | 2.985 | 2.874 | 2.720 | 3.294 | 2.975 | 2.946 | 2.906 | 3.475 | 2.612 | 0.064 | 0.2498 | 0.5008 | 0 | 0 | 0 |
| 'Mrpl10' | 26.996 | 32.186 | 31.676 | 29.158 | 31.155 | 31.204 | 27.755 | 30.550 | 31.035 | 29.155 | 27.088 | 28.993 | 31.837 | 27.282 | 28.461 | 27.921 | 28.530 | 28.324 | 0.065 | 0.0848 | 0.2643 | 0 | 0 | 0 |
| 'Mrpl11' | 8.068 | 9.392 | 9.113 | 7.712 | 9.808 | 9.294 | 8.429 | 9.634 | 8.739 | 8.459 | 8.032 | 8.921 | 10.074 | 8.618 | 8.531 | 9.357 | 8.997 | 8.266 | 0.004 | 0.9325 | 0.9746 | 0 | 0 | 0 |
| 'Mrpl12' | 18.782 | 18.673 | 19.429 | 17.551 | 18.761 | 20.647 | 18.018 | 18.058 | 17.933 | 16.449 | 15.716 | 19.685 | 20.477 | 18.407 | 19.681 | 20.145 | 15.117 | 18.550 | 0.022 | 0.6603 | 0.8362 | 0 | 0 | 0 |
| 'Mrpl13' | 5.041 | 5.541 | 5.086 | 5.578 | 6.228 | 5.404 | 5.729 | 5.182 | 5.547 | 4.895 | 4.864 | 5.091 | 5.371 | 5.359 | 5.079 | 5.375 | 4.930 | 4.650 | 0.099 | 0.0168 | 0.0915 | 0 | 0 | 0 |
| 'Mrpl14' | 14.093 | 14.928 | 16.124 | 15.356 | 16.202 | 15.092 | 14.172 | 15.053 | 15.563 | 12.096 | 14.701 | 14.910 | 14.459 | 15.943 | 15.765 | 15.485 | 13.717 | 14.090 | 0.050 | 0.3102 | 0.5686 | 0 | 0 | 0 |
| 'Mrpl15' | 2.669 | 3.097 | 2.936 | 2.888 | 3.172 | 3.224 | 3.065 | 2.770 | 2.971 | 2.719 | 2.782 | 2.995 | 3.050 | 2.867 | 3.103 | 2.764 | 2.872 | 2.774 | 0.032 | 0.4343 | 0.6788 | 0 | 0 | 0 |
| 'Mrpl16' | 20.820 | 15.719 | 20.650 | 18.399 | 19.525 | 19.048 | 20.176 | 20.314 | 20.881 | 17.851 | 16.891 | 17.670 | 20.561 | 19.348 | 18.180 | 18.338 | 19.216 | 16.395 | 0.091 | 0.1075 | 0.3084 | 0 | 0 | 0 |
| 'Mrpl17' | 4.272 | 5.612 | 4.765 | 4.773 | 5.263 | 5.136 | 4.760 | 5.282 | 4.808 | 4.531 | 4.472 | 4.395 | 4.882 | 4.710 | 4.266 | 4.701 | 4.340 | 4.023 | 0.135 | 0.0031 | 0.0259 | 1 | 0 | 0 |
| 'Mrpl18' | 26.667 | 26.833 | 25.279 | 28.245 | 26.524 | 28.015 | 27.477 | 26.221 | 26.730 | 24.342 | 26.732 | 26.816 | 28.346 | 26.927 | 27.890 | 26.582 | 27.614 | 26.445 | -0.010 | 0.7643 | 0.8953 | 0 | 0 | 0 |
| 'Mrpl19' | 4.859 | 5.082 | 5.563 | 4.683 | 5.623 | 5.554 | 5.115 | 5.650 | 5.387 | 4.948 | 4.700 | 4.543 | 5.199 | 5.290 | 4.578 | 4.988 | 5.189 | 4.352 | 0.115 | 0.0160 | 0.0881 | 0 | 0 | 0 |
| 'Mrpl2' | 15.756 | 16.729 | 16.221 | 14.964 | 18.040 | 17.102 | 14.961 | 19.475 | 14.890 | 16.208 | 13.883 | 15.106 | 18.534 | 15.768 | 17.287 | 19.471 | 15.426 | 14.500 | 0.011 | 0.8592 | 0.9390 | 0 | 0 | 0 |
| 'Mrpl20' | 9.328 | 9.090 | 8.931 | 8.926 | 10.029 | 8.733 | 9.123 | 10.033 | 8.540 | 7.158 | 7.413 | 8.891 | 8.918 | 8.843 | 10.194 | 9.096 | 8.520 | 8.888 | 0.073 | 0.2054 | 0.4490 | 0 | 0 | 0 |
| 'Mrpl21' | 8.077 | 8.550 | 9.706 | 9.269 | 9.334 | 9.224 | 8.266 | 8.858 | 9.552 | 8.051 | 8.055 | 8.480 | 9.163 | 8.397 | 8.656 | 7.497 | 8.991 | 8.119 | 0.090 | 0.0488 | 0.1856 | 0 | 0 | 0 |
| 'Mrpl22' | 7.521 | 9.107 | 8.482 | 7.780 | 8.514 | 8.629 | 9.542 | 8.671 | 8.228 | 8.298 | 9.191 | 8.427 | 7.821 | 8.816 | 7.488 | 8.247 | 8.678 | 7.250 | 0.033 | 0.5918 | 0.7921 | 0 | 0 | 0 |
| 'Mrpl23' | 12.400 | 12.336 | 12.433 | 11.456 | 12.436 | 12.680 | 11.386 | 12.758 | 11.555 | 11.304 | 12.101 | 12.962 | 12.871 | 12.652 | 11.922 | 11.504 | 11.178 | 11.377 | 0.008 | 0.8539 | 0.9373 | 0 | 0 | 0 |
| 'Mrpl24' | 27.223 | 25.710 | 25.719 | 25.794 | 27.374 | 25.884 | 27.043 | 26.697 | 25.363 | 25.119 | 26.771 | 26.771 | 26.189 | 26.445 | 26.090 | 24.859 | 25.922 | 25.659 | 0.006 | 0.8221 | 0.9217 | 0 | 0 | 0 |
| 'Mrpl27' | 11.925 | 14.844 | 14.280 | 12.714 | 14.938 | 13.327 | 13.002 | 13.416 | 16.082 | 12.811 | 11.294 | 12.595 | 13.776 | 12.468 | 13.911 | 13.707 | 11.646 | 11.953 | 0.114 | 0.0459 | 0.1783 | 0 | 0 | 0 |
| 'Mrpl28' | 18.558 | 21.470 | 20.535 | 19.288 | 20.187 | 22.106 | 18.060 | 19.650 | 20.141 | 18.127 | 15.665 | 21.704 | 21.330 | 22.382 | 23.192 | 22.692 | 19.612 | 21.395 | -0.059 | 0.3019 | 0.5590 | 0 | 0 | 0 |
| 'Mrpl3' | 6.551 | 6.732 | 7.084 | 6.967 | 7.744 | 7.149 | 7.016 | 6.891 | 7.306 | 7.302 | 6.643 | 6.821 | 7.504 | 7.572 | 6.359 | 7.362 | 6.606 | 6.683 | 0.005 | 0.8989 | 0.9598 | 0 | 0 | 0 |
| 'Mrpl30' | 9.136 | 9.021 | 8.828 | 8.560 | 8.847 | 8.418 | 9.013 | 9.098 | 8.904 | 8.498 | 8.276 | 9.174 | 8.407 | 9.289 | 9.505 | 9.095 | 8.244 | 8.655 | 0.001 | 0.9849 | 0.9949 | 0 | 0 | 0 |
| 'Mrpl32' | 30.353 | 28.398 | 28.971 | 32.389 | 30.452 | 29.702 | 33.826 | 31.941 | 27.561 | 29.902 | 30.857 | 32.523 | 33.184 | 35.212 | 30.293 | 29.527 | 34.940 | 32.379 | -0.093 | 0.0605 | 0.2120 | 0 | 0 | 0 |
| 'Mrpl33' | 4.561 | 4.862 | 5.382 | 5.065 | 5.250 | 5.155 | 5.182 | 4.780 | 5.301 | 4.280 | 4.500 | 4.911 | 4.747 | 4.404 | 5.220 | 4.208 | 4.746 | 4.651 | 0.115 | 0.0021 | 0.0187 | 1 | 0 | 0 |
| 'Mrpl34' | 39.710 | 41.298 | 37.626 | 34.856 | 37.986 | 36.762 | 34.745 | 41.775 | 35.404 | 30.849 | 32.275 | 37.415 | 39.233 | 39.368 | 39.337 | 36.997 | 34.849 | 37.154 | 0.041 | 0.4540 | 0.6929 | 0 | 0 | 0 |
| 'Mrpl35' | 3.426 | 3.522 | 3.873 | 4.041 | 3.789 | 4.137 | 3.600 | 3.962 | 4.078 | 3.304 | 3.172 | 3.814 | 3.566 | 3.682 | 3.634 | 3.455 | 3.856 | 3.316 | 0.103 | 0.0554 | 0.2018 | 0 | 0 | 0 |
| 'Mrpl36' | 30.768 | 33.162 | 36.227 | 32.745 | 36.868 | 32.229 | 33.073 | 34.843 | 34.863 | 34.235 | 31.177 | 33.228 | 34.478 | 33.596 | 34.505 | 32.467 | 34.423 | 31.438 | 0.015 | 0.7266 | 0.8749 | 0 | 0 | 0 |
| 'Mrpl37' | 8.330 | 8.908 | 9.778 | 7.959 | 8.859 | 8.670 | 7.447 | 8.028 | 9.371 | 8.828 | 7.709 | 8.819 | 8.374 | 7.786 | 8.747 | 9.079 | 8.759 | 8.047 | 0.013 | 0.7802 | 0.9011 | 0 | 0 | 0 |
| 'Mrpl38' | 6.630 | 6.662 | 7.182 | 5.871 | 7.310 | 7.394 | 6.991 | 7.027 | 7.653 | 6.902 | 5.986 | 7.747 | 6.735 | 5.892 | 6.591 | 6.808 | 6.262 | 7.479 | 0.041 | 0.3802 | 0.6318 | 0 | 0 | 0 |
| 'Mrpl39' | 10.420 | 11.562 | 11.399 | 11.160 | 12.228 | 12.723 | 10.831 | 12.215 | 12.491 | 10.690 | 9.869 | 10.491 | 12.364 | 12.187 | 9.608 | 10.882 | 10.395 | 9.814 | 0.117 | 0.0313 | 0.1386 | 0 | 0 | 0 |
| 'Mrpl4' | 49.349 | 54.098 | 54.182 | 47.131 | 54.672 | 59.086 | 47.820 | 51.475 | 55.902 | 45.966 | 48.639 | 53.825 | 53.720 | 50.156 | 52.381 | 51.153 | 49.583 | 48.738 | 0.049 | 0.1830 | 0.4190 | 0 | 0 | 0 |
| 'Mrpl40' | 13.340 | 13.351 | 14.876 | 13.177 | 15.151 | 14.592 | 14.060 | 13.359 | 13.266 | 13.066 | 11.391 | 12.755 | 13.401 | 13.565 | 13.915 | 12.655 | 14.063 | 12.626 | 0.083 | 0.0868 | 0.2683 | 0 | 0 | 0 |
| 'Mrpl41' | 26.555 | 31.929 | 25.501 | 27.700 | 26.819 | 28.141 | 26.824 | 26.751 | 25.882 | 24.489 | 26.222 | 26.709 | 28.746 | 28.533 | 27.136 | 29.616 | 29.076 | 29.428 | -0.036 | 0.4607 | 0.6973 | 0 | 0 | 0 |
| 'Mrpl42' | 61.408 | 62.431 | 62.593 | 72.038 | 67.398 | 62.221 | 65.617 | 62.805 | 64.711 | 60.723 | 61.340 | 60.422 | 62.496 | 58.700 | 60.660 | 63.629 | 60.419 | 62.517 | 0.065 | 0.0809 | 0.2564 | 0 | 0 | 0 |
| 'Mrpl43' | 32.556 | 36.091 | 31.446 | 31.732 | 34.421 | 37.278 | 31.335 | 33.246 | 36.195 | 32.369 | 30.051 | 34.569 | 31.627 | 31.321 | 34.002 | 31.263 | 28.500 | 32.904 | 0.072 | 0.0815 | 0.2576 | 0 | 0 | 0 |
| 'Mrpl44' | 8.438 | 10.533 | 10.314 | 9.243 | 10.688 | 10.054 | 10.155 | 10.017 | 10.718 | 8.575 | 8.763 | 10.785 | 9.052 | 8.625 | 10.126 | 9.356 | 9.055 | 9.819 | 0.079 | 0.1089 | 0.3109 | 0 | 0 | 0 |
| 'Mrpl45' | 10.910 | 11.306 | 11.245 | 11.945 | 11.304 | 10.816 | 11.740 | 11.222 | 11.451 | 11.666 | 9.827 | 11.679 | 12.134 | 10.295 | 11.664 | 11.653 | 10.627 | 10.779 | 0.007 | 0.8580 | 0.9390 | 0 | 0 | 0 |
| 'Mrpl46' | 13.750 | 14.850 | 13.671 | 11.802 | 13.578 | 14.144 | 12.812 | 14.542 | 12.813 | 13.313 | 13.784 | 13.769 | 12.797 | 14.025 | 11.960 | 13.131 | 12.056 | 13.074 | 0.038 | 0.4272 | 0.6735 | 0 | 0 | 0 |
| 'Mrpl47' | 2.388 | 2.533 | 2.929 | 2.335 | 2.489 | 2.672 | 2.360 | 2.691 | 2.622 | 2.174 | 2.366 | 2.560 | 2.237 | 2.304 | 2.270 | 2.374 | 2.483 | 2.200 | 0.127 | 0.0304 | 0.1361 | 0 | 0 | 0 |
| 'Mrpl49' | 19.936 | 20.046 | 19.972 | 21.369 | 19.497 | 22.044 | 20.203 | 20.695 | 20.499 | 17.794 | 18.837 | 21.274 | 20.916 | 18.606 | 20.749 | 18.107 | 20.278 | 19.538 | 0.050 | 0.2264 | 0.4730 | 0 | 0 | 0 |
| 'Mrpl50' | 9.930 | 11.542 | 10.500 | 10.897 | 12.149 | 10.463 | 11.419 | 10.814 | 10.845 | 10.353 | 12.104 | 11.863 | 9.964 | 11.610 | 10.625 | 9.676 | 11.539 | 11.714 | -0.032 | 0.5163 | 0.7403 | 0 | 0 | 0 |
| 'Mrpl51' | 7.244 | 9.350 | 8.423 | 9.405 | 8.432 | 8.779 | 8.331 | 8.438 | 8.619 | 7.349 | 8.059 | 8.643 | 8.451 | 8.848 | 8.684 | 8.345 | 8.321 | 8.520 | 0.020 | 0.6598 | 0.8361 | 0 | 0 | 0 |
| 'Mrpl52' | 25.877 | 28.481 | 29.746 | 25.823 | 27.334 | 28.115 | 25.992 | 27.552 | 27.831 | 23.751 | 24.579 | 24.765 | 26.120 | 26.016 | 26.099 | 25.571 | 25.929 | 24.370 | 0.112 | 0.0008 | 0.0084 | 1 | 0 | 0 |
| 'Mrpl54' | 34.108 | 34.869 | 36.118 | 29.724 | 32.011 | 38.402 | 30.679 | 35.277 | 35.061 | 31.951 | 32.145 | 32.840 | 36.718 | 30.911 | 33.332 | 38.372 | 30.362 | 32.110 | 0.030 | 0.5586 | 0.7697 | 0 | 0 | 0 |
| 'Mrpl55' | 4.437 | 5.311 | 5.886 | 5.036 | 5.859 | 5.574 | 4.962 | 5.514 | 4.945 | 5.724 | 4.490 | 5.003 | 5.220 | 4.948 | 4.882 | 5.440 | 4.487 | 4.267 | 0.089 | 0.1899 | 0.4262 | 0 | 0 | 0 |
| 'Mrpl57' | 17.489 | 20.315 | 18.432 | 18.201 | 19.658 | 20.376 | 20.734 | 18.049 | 18.700 | 16.887 | 16.011 | 17.762 | 20.314 | 18.617 | 16.731 | 19.705 | 18.198 | 16.251 | 0.089 | 0.0641 | 0.2209 | 0 | 0 | 0 |
| 'Mrpl9' | 20.963 | 21.413 | 19.996 | 21.212 | 21.340 | 22.187 | 19.456 | 22.229 | 20.822 | 19.897 | 21.171 | 21.350 | 20.279 | 20.714 | 21.280 | 21.683 | 20.109 | 21.373 | 0.003 | 0.9172 | 0.9686 | 0 | 0 | 0 |
| 'Mrps11' | 6.040 | 5.639 | 6.380 | 5.353 | 5.527 | 6.136 | 5.376 | 5.499 | 5.839 | 6.522 | 5.066 | 5.923 | 5.531 | 5.377 | 5.995 | 6.314 | 5.430 | 5.902 | -0.011 | 0.8181 | 0.9194 | 0 | 0 | 0 |
| 'Mrps12' | 9.257 | 8.995 | 9.335 | 8.731 | 9.025 | 9.356 | 9.609 | 8.692 | 9.676 | 8.579 | 8.400 | 10.496 | 8.689 | 8.864 | 9.467 | 9.302 | 10.016 | 10.440 | -0.043 | 0.3777 | 0.6296 | 0 | 0 | 0 |
| 'Mrps14' | 10.566 | 11.838 | 10.953 | 11.332 | 12.493 | 10.851 | 12.199 | 11.998 | 11.429 | 7.062 | 6.594 | 6.955 | 6.512 | 7.096 | 6.487 | 7.616 | 7.424 | 5.800 | 0.740 | 0.0000 | 0.0000 | 1 | 1 | 0 |
| 'Mrps15' | 16.329 | 15.740 | 16.455 | 17.249 | 17.007 | 18.890 | 17.133 | 16.880 | 17.320 | 16.420 | 14.910 | 17.864 | 17.036 | 15.967 | 16.874 | 17.645 | 16.178 | 16.897 | 0.019 | 0.6242 | 0.8126 | 0 | 0 | 0 |
| 'Mrps16' | 57.885 | 66.396 | 70.246 | 58.772 | 60.205 | 68.799 | 62.415 | 67.631 | 64.141 | 57.708 | 55.640 | 60.133 | 65.080 | 63.489 | 61.097 | 58.608 | 59.353 | 56.304 | 0.093 | 0.0358 | 0.1520 | 0 | 0 | 0 |
| 'Mrps17' | 2.942 | 3.294 | 2.911 | 2.893 | 3.111 | 3.081 | 3.207 | 3.070 | 3.173 | 2.984 | 3.027 | 3.297 | 3.068 | 2.767 | 3.371 | 2.879 | 3.037 | 3.107 | -0.009 | 0.8145 | 0.9164 | 0 | 0 | 0 |
| 'Mrps18a' | 4.805 | 5.599 | 4.936 | 4.701 | 4.910 | 6.003 | 5.077 | 5.934 | 5.250 | 5.285 | 4.728 | 5.116 | 6.026 | 4.989 | 5.001 | 5.453 | 5.152 | 5.263 | -0.004 | 0.9504 | 0.9809 | 0 | 0 | 0 |
| 'Mrps18b' | 5.619 | 5.109 | 5.810 | 5.502 | 4.896 | 5.072 | 4.950 | 5.323 | 5.399 | 4.456 | 4.204 | 5.468 | 4.734 | 4.881 | 4.713 | 4.869 | 4.502 | 4.736 | 0.150 | 0.0050 | 0.0372 | 1 | 0 | 0 |
| 'Mrps18c' | 10.600 | 13.416 | 10.855 | 11.402 | 12.335 | 11.809 | 10.316 | 12.071 | 10.889 | 9.098 | 10.265 | 9.915 | 10.981 | 12.601 | 10.962 | 11.471 | 11.260 | 9.748 | 0.097 | 0.0689 | 0.2306 | 0 | 0 | 0 |
| 'Mrps2' | 4.973 | 6.014 | 6.585 | 5.418 | 6.248 | 5.804 | 4.711 | 6.241 | 5.828 | 5.559 | 4.990 | 5.424 | 5.333 | 5.062 | 5.766 | 5.088 | 5.489 | 4.907 | 0.110 | 0.0345 | 0.1486 | 0 | 0 | 0 |
| 'Mrps21' | 33.984 | 38.242 | 35.252 | 38.323 | 35.825 | 37.774 | 37.112 | 35.119 | 35.824 | 31.685 | 33.085 | 33.994 | 37.981 | 36.374 | 37.382 | 35.610 | 36.513 | 32.881 | 0.042 | 0.2948 | 0.5505 | 0 | 0 | 0 |
| 'Mrps22' | 3.196 | 2.801 | 3.034 | 3.268 | 2.993 | 2.982 | 3.043 | 2.923 | 2.901 | 2.840 | 3.092 | 3.102 | 2.983 | 2.574 | 2.973 | 2.940 | 3.359 | 2.937 | 0.006 | 0.9222 | 0.9705 | 0 | 0 | 0 |
| 'Mrps23' | 18.773 | 22.033 | 20.301 | 17.663 | 23.315 | 20.833 | 20.243 | 21.858 | 18.895 | 19.109 | 21.356 | 19.963 | 20.237 | 18.862 | 20.727 | 20.206 | 19.051 | 18.851 | 0.030 | 0.4737 | 0.7090 | 0 | 0 | 0 |
| 'Mrps24' | 11.378 | 11.472 | 12.716 | 11.630 | 12.851 | 12.465 | 10.872 | 12.202 | 11.369 | 11.269 | 9.679 | 12.274 | 11.402 | 11.202 | 11.824 | 12.273 | 11.490 | 12.002 | 0.038 | 0.3787 | 0.6303 | 0 | 0 | 0 |
| 'Mrps25' | 2.770 | 2.879 | 3.481 | 3.593 | 3.170 | 2.968 | 3.168 | 2.802 | 3.518 | 2.844 | 2.582 | 3.416 | 2.934 | 2.905 | 3.355 | 2.811 | 3.017 | 3.231 | 0.049 | 0.3878 | 0.6374 | 0 | 0 | 0 |
| 'Mrps26' | 15.114 | 16.573 | 16.451 | 14.168 | 15.393 | 17.746 | 14.722 | 15.979 | 15.500 | 14.121 | 12.804 | 15.920 | 14.607 | 14.892 | 16.172 | 16.303 | 14.479 | 16.778 | 0.048 | 0.2471 | 0.4979 | 0 | 0 | 0 |
| 'Mrps27' | 69.629 | 85.290 | 68.435 | 75.826 | 87.889 | 70.570 | 75.221 | 77.137 | 70.899 | 67.637 | 77.666 | 62.748 | 93.403 | 77.940 | 62.728 | 67.697 | 82.220 | 61.543 | 0.052 | 0.3249 | 0.5821 | 0 | 0 | 0 |
| 'Mrps28' | 16.885 | 19.138 | 15.572 | 18.140 | 17.683 | 14.861 | 18.497 | 17.146 | 17.246 | 15.310 | 14.379 | 17.561 | 18.330 | 14.734 | 15.259 | 15.263 | 15.133 | 15.413 | 0.104 | 0.1124 | 0.3181 | 0 | 0 | 0 |
| 'Mrps30' | 12.608 | 13.961 | 12.276 | 12.266 | 12.766 | 13.534 | 11.716 | 13.242 | 11.689 | 12.237 | 11.691 | 11.809 | 12.070 | 14.412 | 10.474 | 12.816 | 11.442 | 10.508 | 0.080 | 0.1472 | 0.3731 | 0 | 0 | 0 |
| 'Mrps31' | 6.914 | 6.742 | 7.328 | 7.686 | 7.463 | 6.537 | 8.287 | 5.986 | 6.766 | 6.452 | 6.248 | 6.453 | 7.517 | 7.010 | 6.803 | 6.962 | 6.821 | 6.509 | 0.054 | 0.3551 | 0.6084 | 0 | 0 | 0 |
| 'Mrps33' | 11.793 | 14.228 | 12.683 | 12.968 | 13.985 | 14.295 | 13.982 | 13.637 | 12.820 | 12.559 | 12.833 | 12.393 | 12.641 | 13.030 | 13.180 | 14.259 | 12.921 | 12.581 | 0.039 | 0.3633 | 0.6153 | 0 | 0 | 0 |
| 'Mrps34' | 6.167 | 6.183 | 7.048 | 5.359 | 6.417 | 6.912 | 6.144 | 6.513 | 6.869 | 6.575 | 5.818 | 5.917 | 6.905 | 5.331 | 6.000 | 7.187 | 5.512 | 5.257 | 0.079 | 0.2076 | 0.4522 | 0 | 0 | 0 |
| 'Mrps35' | 2.516 | 2.810 | 2.860 | 2.836 | 3.011 | 2.912 | 2.729 | 2.774 | 3.156 | 2.860 | 2.772 | 2.805 | 2.808 | 2.676 | 2.880 | 2.898 | 2.560 | 2.704 | 0.027 | 0.5406 | 0.7579 | 0 | 0 | 0 |
| 'Mrps36' | 6.158 | 6.287 | 6.501 | 7.144 | 6.800 | 7.130 | 7.588 | 6.708 | 6.997 | 5.989 | 5.821 | 6.936 | 6.737 | 6.654 | 6.493 | 6.488 | 6.516 | 6.688 | 0.055 | 0.2801 | 0.5371 | 0 | 0 | 0 |
| 'Mrps5' | 3.742 | 4.116 | 3.773 | 3.560 | 4.005 | 3.772 | 3.345 | 3.915 | 4.020 | 4.021 | 3.420 | 3.553 | 3.900 | 3.439 | 3.665 | 3.714 | 3.540 | 3.240 | 0.067 | 0.1492 | 0.3758 | 0 | 0 | 0 |
| 'Mrps6' | 41.505 | 38.366 | 42.313 | 40.791 | 40.782 | 41.640 | 40.642 | 40.366 | 38.749 | 39.546 | 39.874 | 46.011 | 38.108 | 41.490 | 44.864 | 40.780 | 42.143 | 42.354 | -0.050 | 0.1491 | 0.3758 | 0 | 0 | 0 |
| 'Mrps7' | 14.755 | 14.024 | 15.222 | 13.535 | 14.388 | 15.046 | 14.306 | 15.300 | 13.717 | 12.893 | 13.168 | 14.880 | 15.919 | 14.534 | 15.482 | 14.970 | 13.177 | 15.840 | -0.016 | 0.7312 | 0.8773 | 0 | 0 | 0 |
| 'Mrps9' | 6.836 | 6.859 | 6.825 | 6.911 | 6.508 | 6.568 | 7.196 | 7.128 | 6.305 | 6.898 | 6.767 | 6.551 | 6.775 | 6.299 | 5.904 | 6.915 | 6.466 | 5.880 | 0.056 | 0.0864 | 0.2677 | 0 | 0 | 0 |
| 'Mrrf' | 2.974 | 2.336 | 2.681 | 3.043 | 2.637 | 2.901 | 3.028 | 2.923 | 2.759 | 2.711 | 2.662 | 2.824 | 3.039 | 2.715 | 2.882 | 2.464 | 2.703 | 3.082 | -0.001 | 0.9877 | 0.9964 | 0 | 0 | 0 |
| 'Mrs2' | 19.434 | 17.850 | 18.795 | 17.974 | 16.957 | 17.182 | 16.456 | 17.300 | 18.025 | 17.937 | 17.797 | 17.630 | 19.281 | 18.995 | 17.332 | 18.108 | 18.323 | 17.611 | -0.033 | 0.4248 | 0.6715 | 0 | 0 | 0 |
| 'Mrto4' | 7.466 | 7.285 | 7.362 | 7.095 | 7.905 | 7.321 | 6.940 | 7.201 | 6.921 | 8.685 | 7.065 | 6.818 | 8.188 | 7.146 | 6.305 | 7.898 | 6.996 | 6.890 | -0.014 | 0.7647 | 0.8953 | 0 | 0 | 0 |
| 'Ms4a4b' | 28.023 | 35.600 | 28.441 | 28.662 | 36.104 | 32.242 | 31.275 | 30.233 | 30.381 | 26.178 | 26.450 | 29.481 | 29.461 | 30.743 | 29.028 | 29.370 | 30.201 | 26.028 | 0.115 | 0.0088 | 0.0568 | 0 | 0 | 0 |
| 'Ms4a4c' | 1.648 | 2.013 | 1.505 | 1.619 | 2.433 | 2.138 | 2.213 | 2.174 | 1.900 | 1.574 | 1.335 | 1.642 | 1.251 | 1.482 | 1.580 | 1.755 | 2.012 | 0.958 | 0.358 | 0.0046 | 0.0349 | 1 | 0 | 0 |
| 'Ms4a6b' | 132.988 | 148.138 | 147.759 | 146.305 | 150.688 | 145.343 | 153.374 | 144.152 | 148.615 | 134.076 | 137.528 | 143.103 | 134.902 | 135.562 | 142.004 | 135.446 | 144.183 | 136.132 | 0.072 | 0.0020 | 0.0184 | 1 | 0 | 0 |
| 'Ms4a6c' | 14.522 | 18.196 | 17.355 | 17.006 | 16.700 | 17.673 | 19.224 | 15.404 | 17.297 | 16.605 | 17.176 | 20.204 | 17.588 | 19.636 | 18.118 | 17.340 | 20.333 | 16.882 | -0.109 | 0.0382 | 0.1588 | 0 | 0 | 0 |
| 'Msantd2' | 10.741 | 10.591 | 10.996 | 11.736 | 11.126 | 9.286 | 9.809 | 11.141 | 11.580 | 9.297 | 12.107 | 12.147 | 9.608 | 11.493 | 11.318 | 10.107 | 12.606 | 11.160 | -0.051 | 0.3824 | 0.6330 | 0 | 0 | 0 |
| 'Msantd4' | 10.978 | 11.003 | 12.097 | 12.154 | 11.295 | 10.862 | 13.918 | 10.858 | 11.431 | 10.308 | 11.589 | 12.121 | 10.810 | 11.059 | 11.674 | 10.099 | 11.127 | 12.262 | 0.034 | 0.5173 | 0.7406 | 0 | 0 | 0 |
| 'Msh2' | 8.857 | 9.175 | 9.295 | 9.538 | 9.060 | 9.295 | 9.080 | 8.713 | 9.305 | 8.648 | 8.724 | 8.879 | 8.748 | 8.395 | 9.365 | 8.218 | 8.579 | 8.850 | 0.059 | 0.0427 | 0.1704 | 0 | 0 | 0 |
| 'Msh3' | 2.096 | 2.178 | 1.978 | 2.173 | 1.938 | 1.776 | 2.098 | 1.981 | 1.989 | 2.095 | 1.973 | 2.156 | 2.079 | 1.882 | 2.022 | 2.041 | 1.951 | 1.881 | -0.007 | 0.8527 | 0.9370 | 0 | 0 | 0 |
| 'Msh5' | 1.662 | 1.774 | 2.827 | 2.096 | 1.610 | 2.263 | 2.196 | 1.841 | 2.688 | 1.772 | 1.738 | 2.401 | 2.046 | 2.026 | 2.386 | 1.753 | 2.022 | 2.222 | 0.027 | 0.6846 | 0.8509 | 0 | 0 | 0 |
| 'Msh6' | 40.376 | 40.518 | 39.847 | 44.825 | 39.057 | 40.203 | 42.621 | 38.706 | 41.947 | 36.412 | 44.288 | 43.652 | 41.422 | 41.286 | 44.345 | 36.882 | 42.320 | 43.609 | -0.034 | 0.4303 | 0.6760 | 0 | 0 | 0 |
| 'Msi1' | 1.116 | 0.965 | 0.962 | 0.954 | 0.977 | 0.905 | 1.023 | 1.207 | 0.929 | 0.829 | 0.804 | 0.664 | 0.857 | 0.568 | 0.714 | 0.852 | 0.648 | 0.417 | 0.513 | 0.0000 | 0.0000 | 1 | 1 | 0 |
| 'Msi2' | 9.357 | 11.826 | 10.137 | 9.499 | 11.490 | 9.866 | 10.181 | 11.695 | 9.842 | 10.602 | 11.284 | 10.318 | 10.023 | 11.385 | 9.487 | 11.196 | 10.613 | 10.508 | -0.037 | 0.2649 | 0.5184 | 0 | 0 | 0 |
| 'Msl1' | 26.665 | 26.501 | 25.607 | 26.513 | 24.485 | 26.041 | 24.393 | 24.462 | 25.865 | 23.425 | 25.478 | 32.107 | 23.896 | 26.207 | 30.952 | 24.767 | 27.345 | 33.157 | -0.103 | 0.0382 | 0.1588 | 0 | 0 | 0 |
| 'Msl2' | 37.409 | 34.019 | 35.844 | 35.274 | 34.490 | 36.019 | 35.895 | 35.969 | 33.112 | 34.435 | 36.324 | 35.688 | 32.832 | 35.696 | 35.548 | 36.144 | 34.951 | 36.197 | -0.009 | 0.7640 | 0.8953 | 0 | 0 | 0 |
| 'Msl3' | 5.789 | 6.881 | 6.631 | 6.408 | 6.437 | 5.991 | 6.886 | 6.323 | 6.294 | 5.260 | 6.965 | 7.263 | 6.641 | 6.176 | 6.737 | 5.993 | 6.580 | 6.530 | -0.027 | 0.5889 | 0.7896 | 0 | 0 | 0 |
| 'Msl3l2' | 13.223 | 12.392 | 11.177 | 10.558 | 11.700 | 12.543 | 11.065 | 11.866 | 11.417 | 12.493 | 11.000 | 10.858 | 10.557 | 11.263 | 10.254 | 11.456 | 10.455 | 9.741 | 0.106 | 0.0436 | 0.1729 | 0 | 0 | 0 |
| 'Msmo1' | 2.350 | 3.466 | 3.238 | 2.912 | 3.507 | 3.342 | 2.701 | 3.029 | 3.300 | 3.105 | 4.023 | 3.095 | 3.418 | 3.706 | 3.195 | 3.319 | 3.548 | 2.665 | -0.119 | 0.0605 | 0.2120 | 0 | 0 | 0 |
| 'Msn' | 117.793 | 126.407 | 121.913 | 124.293 | 126.331 | 124.782 | 122.257 | 124.507 | 123.573 | 124.821 | 127.343 | 122.463 | 124.707 | 128.132 | 128.802 | 121.512 | 127.528 | 126.397 | -0.036 | 0.0381 | 0.1585 | 0 | 0 | 0 |
| 'Msra' | 50.339 | 56.706 | 49.398 | 52.642 | 57.084 | 54.174 | 51.443 | 57.759 | 52.572 | 51.638 | 54.077 | 51.705 | 48.280 | 54.297 | 54.980 | 52.251 | 52.283 | 51.150 | 0.023 | 0.4371 | 0.6803 | 0 | 0 | 0 |
| 'Msrb1' | 16.482 | 17.378 | 18.933 | 18.791 | 19.726 | 18.195 | 17.559 | 18.765 | 17.031 | 17.562 | 18.641 | 17.274 | 19.262 | 17.360 | 17.362 | 18.348 | 17.339 | 15.533 | 0.030 | 0.4969 | 0.7258 | 0 | 0 | 0 |
| 'Msrb2' | 3.183 | 3.634 | 4.303 | 3.669 | 3.432 | 4.119 | 3.666 | 3.231 | 3.985 | 4.544 | 4.202 | 4.675 | 3.764 | 3.626 | 4.686 | 4.475 | 4.632 | 3.998 | -0.224 | 0.0042 | 0.0331 | 1 | 0 | 0 |
| 'Mss51' | 2.473 | 2.659 | 2.450 | 2.173 | 2.157 | 1.827 | 2.723 | 2.594 | 2.438 | 1.915 | 3.530 | 2.841 | 2.656 | 3.054 | 2.481 | 2.304 | 3.201 | 2.742 | -0.213 | 0.0449 | 0.1758 | 0 | 0 | 0 |
| 'Mst1' | 1.616 | 1.244 | 1.903 | 1.379 | 1.258 | 1.370 | 1.786 | 1.529 | 1.932 | 1.683 | 1.967 | 1.552 | 1.718 | 0.918 | 1.297 | 1.049 | 1.490 | 1.368 | 0.092 | 0.4692 | 0.7053 | 0 | 0 | 0 |
| 'Msto1' | 6.600 | 6.405 | 7.140 | 6.956 | 6.370 | 7.345 | 6.483 | 6.903 | 7.015 | 6.744 | 6.299 | 7.071 | 7.104 | 6.119 | 6.885 | 6.560 | 6.267 | 6.406 | 0.033 | 0.2952 | 0.5508 | 0 | 0 | 0 |
| 'Mta1' | 22.696 | 22.061 | 23.557 | 21.759 | 20.067 | 23.932 | 21.955 | 22.150 | 23.057 | 22.476 | 19.790 | 22.863 | 23.073 | 20.614 | 22.353 | 21.378 | 19.646 | 22.673 | 0.037 | 0.1429 | 0.3662 | 0 | 0 | 0 |
| 'Mta2' | 80.846 | 80.996 | 84.035 | 80.166 | 79.529 | 83.509 | 80.849 | 79.863 | 81.646 | 77.924 | 76.036 | 80.161 | 80.350 | 76.557 | 82.893 | 78.256 | 78.283 | 79.788 | 0.032 | 0.0381 | 0.1585 | 0 | 0 | 0 |
| 'Mta3' | 10.476 | 9.931 | 9.905 | 9.881 | 10.009 | 10.070 | 10.046 | 9.425 | 9.749 | 9.851 | 9.408 | 10.013 | 10.269 | 9.879 | 10.277 | 9.666 | 9.090 | 9.713 | 0.010 | 0.6993 | 0.8592 | 0 | 0 | 0 |
| 'Mtap' | 17.158 | 17.697 | 16.925 | 17.302 | 17.058 | 16.872 | 17.810 | 17.639 | 17.836 | 16.034 | 15.014 | 16.870 | 15.074 | 16.423 | 15.679 | 16.048 | 15.784 | 16.018 | 0.115 | 0.0003 | 0.0042 | 1 | 0 | 0 |
| 'Mtbp' | 6.240 | 5.965 | 6.880 | 6.566 | 6.577 | 6.127 | 6.552 | 6.149 | 6.483 | 6.239 | 6.149 | 6.361 | 6.576 | 6.394 | 6.465 | 6.919 | 6.706 | 5.736 | -0.007 | 0.8493 | 0.9351 | 0 | 0 | 0 |
| 'Mtch1' | 20.240 | 18.815 | 22.153 | 19.769 | 20.023 | 21.574 | 21.624 | 21.002 | 20.734 | 18.312 | 19.966 | 22.127 | 22.268 | 19.210 | 23.346 | 20.358 | 21.718 | 22.608 | -0.040 | 0.2877 | 0.5442 | 0 | 0 | 0 |
| 'Mtch2' | 4.821 | 5.551 | 5.456 | 4.920 | 5.895 | 5.183 | 5.416 | 5.621 | 5.199 | 5.352 | 5.174 | 5.317 | 5.632 | 5.702 | 5.173 | 5.898 | 5.497 | 5.242 | -0.039 | 0.2697 | 0.5244 | 0 | 0 | 0 |
| 'Mtcp1' | 1.064 | 1.956 | 1.397 | 0.912 | 1.710 | 1.319 | 1.093 | 1.573 | 1.389 | 1.162 | 1.309 | 1.523 | 1.361 | 1.613 | 1.288 | 1.562 | 1.344 | 1.385 | -0.052 | 0.5731 | 0.7795 | 0 | 0 | 0 |
| 'Mtdh' | 8.138 | 8.152 | 8.553 | 7.769 | 8.269 | 8.041 | 8.173 | 8.723 | 8.036 | 8.469 | 8.009 | 7.686 | 7.868 | 8.044 | 7.145 | 8.720 | 8.057 | 7.103 | 0.048 | 0.1366 | 0.3560 | 0 | 0 | 0 |
| 'Mterf2' | 6.929 | 8.121 | 7.858 | 7.524 | 7.334 | 7.766 | 6.659 | 8.165 | 7.585 | 6.627 | 7.697 | 6.578 | 6.695 | 7.513 | 7.194 | 6.364 | 6.413 | 6.309 | 0.144 | 0.0242 | 0.1177 | 0 | 0 | 0 |
| 'Mterf3' | 5.812 | 6.675 | 6.129 | 6.289 | 6.412 | 6.236 | 6.376 | 6.381 | 6.248 | 5.919 | 6.404 | 6.671 | 6.000 | 6.403 | 5.994 | 5.959 | 6.985 | 5.695 | -0.002 | 0.9533 | 0.9827 | 0 | 0 | 0 |

| | | | | | | | | | | | | | | | | | | | | | | | | |
|---|---|---|---|---|---|---|---|---|---|---|---|---|---|---|---|---|---|---|---|---|---|---|---|---|
| 'Mterf4' | 1.620 | 1.807 | 1.568 | 1.505 | 1.715 | 1.617 | 1.898 | 1.630 | 1.610 | 1.435 | 1.564 | 1.693 | 1.438 | 1.534 | 1.563 | 1.158 | 1.686 | 1.723 | 0.095 | 0.1817 | 0.4174 | 0 | 0 | 0 |
| 'Mtf1' | 4.180 | 4.062 | 3.516 | 3.958 | 3.772 | 3.768 | 3.963 | 3.725 | 3.735 | 3.703 | 3.682 | 4.133 | 3.668 | 4.453 | 4.038 | 3.952 | 4.140 | 4.349 | -0.071 | 0.1350 | 0.3537 | 0 | 0 | 0 |
| 'Mtf2' | 6.365 | 6.542 | 6.372 | 6.752 | 6.091 | 5.981 | 7.101 | 6.505 | 6.162 | 5.852 | 6.344 | 6.552 | 5.784 | 6.433 | 6.240 | 5.793 | 6.955 | 6.427 | 0.026 | 0.5254 | 0.7470 | 0 | 0 | 0 |
| 'Mtfmt' | 6.726 | 6.832 | 7.137 | 6.577 | 6.462 | 6.813 | 6.686 | 6.871 | 7.496 | 7.087 | 6.509 | 8.125 | 6.839 | 6.529 | 7.291 | 5.847 | 6.819 | 7.450 | -0.036 | 0.3826 | 0.6330 | 0 | 0 | 0 |
| 'Mtfr1' | 4.274 | 4.289 | 3.904 | 4.341 | 4.048 | 3.701 | 4.531 | 4.345 | 3.926 | 4.018 | 4.336 | 4.870 | 3.948 | 4.146 | 4.118 | 4.643 | 4.403 | 4.372 | -0.075 | 0.1477 | 0.3736 | 0 | 0 | 0 |
| 'Mtfr1l' | 6.399 | 7.329 | 7.027 | 5.768 | 7.280 | 6.850 | 6.785 | 7.351 | 7.313 | 6.922 | 7.049 | 6.716 | 7.169 | 6.455 | 6.322 | 6.958 | 6.206 | 6.602 | 0.029 | 0.5066 | 0.7326 | 0 | 0 | 0 |
| 'Mtfr2' | 2.547 | 2.550 | 3.135 | 2.618 | 2.394 | 2.732 | 3.062 | 2.486 | 2.924 | 2.011 | 2.835 | 3.003 | 3.044 | 2.679 | 3.759 | 2.321 | 3.826 | 2.845 | -0.115 | 0.2652 | 0.5185 | 0 | 0 | 0 |
| 'Mtg1' | 2.238 | 2.639 | 2.777 | 2.241 | 2.782 | 2.825 | 2.293 | 2.549 | 2.836 | 2.564 | 2.230 | 2.345 | 2.773 | 2.329 | 2.482 | 2.469 | 2.209 | 2.155 | 0.098 | 0.1440 | 0.3682 | 0 | 0 | 0 |
| 'Mthfd1' | 24.912 | 20.322 | 24.220 | 25.108 | 22.014 | 22.303 | 24.114 | 22.756 | 23.280 | 23.664 | 22.170 | 25.443 | 25.775 | 23.412 | 25.162 | 23.278 | 22.834 | 24.581 | -0.061 | 0.0643 | 0.2213 | 0 | 0 | 0 |
| 'Mthfd1l' | 17.005 | 14.742 | 17.014 | 17.555 | 17.923 | 15.892 | 18.204 | 15.704 | 15.661 | 16.174 | 16.703 | 18.472 | 18.605 | 16.602 | 17.052 | 16.692 | 16.545 | 17.520 | -0.058 | 0.1462 | 0.3715 | 0 | 0 | 0 |
| 'Mthfd2' | 24.579 | 27.078 | 25.900 | 25.811 | 27.223 | 25.245 | 24.740 | 28.203 | 25.585 | 22.917 | 27.286 | 24.786 | 24.544 | 28.137 | 25.763 | 22.323 | 25.421 | 25.196 | 0.040 | 0.1586 | 0.3887 | 0 | 0 | 0 |
| 'Mthfr' | 3.178 | 2.772 | 3.137 | 2.986 | 2.728 | 2.976 | 3.023 | 2.793 | 2.930 | 2.768 | 2.670 | 3.005 | 2.684 | 2.751 | 2.628 | 2.707 | 2.807 | 2.664 | 0.092 | 0.0208 | 0.1057 | 0 | 0 | 0 |
| 'Mthfsd' | 1.245 | 1.426 | 1.748 | 1.453 | 1.571 | 1.637 | 1.471 | 1.381 | 1.667 | 1.489 | 1.312 | 1.595 | 1.457 | 1.415 | 1.493 | 1.450 | 1.406 | 1.547 | 0.034 | 0.4587 | 0.6955 | 0 | 0 | 0 |
| 'Mtif2' | 7.962 | 8.198 | 9.087 | 8.705 | 9.297 | 8.892 | 9.378 | 9.189 | 8.882 | 9.600 | 9.002 | 9.217 | 9.444 | 9.246 | 7.821 | 9.250 | 9.684 | 8.424 | -0.046 | 0.2609 | 0.5143 | 0 | 0 | 0 |
| 'Mtif3' | 4.111 | 3.634 | 4.184 | 4.242 | 4.135 | 4.247 | 4.664 | 3.906 | 4.178 | 3.535 | 3.870 | 4.944 | 4.058 | 3.848 | 4.619 | 3.519 | 4.152 | 4.631 | -0.008 | 0.8802 | 0.9494 | 0 | 0 | 0 |
| 'Mtm1' | 0.349 | 0.466 | 0.415 | 0.414 | 0.423 | 0.422 | 0.385 | 0.481 | 0.403 | 0.451 | 0.427 | 0.416 | 0.359 | 0.401 | 0.362 | 0.388 | 0.458 | 0.331 | 0.054 | 0.4500 | 0.6898 | 0 | 0 | 0 |
| 'Mtmr1' | 8.170 | 8.863 | 8.250 | 8.291 | 8.718 | 8.092 | 8.388 | 8.897 | 8.364 | 8.482 | 7.417 | 7.760 | 7.981 | 8.294 | 7.099 | 8.173 | 7.600 | 7.544 | 0.101 | 0.0010 | 0.0102 | 1 | 0 | 0 |
| 'Mtmr10' | 1.159 | 1.308 | 1.347 | 1.199 | 1.379 | 1.462 | 1.263 | 1.460 | 1.487 | 1.441 | 1.163 | 1.058 | 1.271 | 1.253 | 0.995 | 1.139 | 1.313 | 0.891 | 0.199 | 0.0121 | 0.0716 | 0 | 0 | 0 |
| 'Mtmr12' | 9.385 | 9.607 | 8.656 | 8.867 | 9.318 | 9.042 | 8.673 | 9.598 | 9.281 | 9.398 | 9.076 | 9.131 | 9.065 | 9.685 | 8.613 | 8.988 | 8.750 | 8.627 | 0.009 | 0.7444 | 0.8847 | 0 | 0 | 0 |
| 'Mtmr14' | 3.650 | 3.828 | 4.113 | 3.843 | 4.035 | 4.645 | 4.169 | 4.578 | 4.153 | 3.903 | 4.255 | 4.281 | 3.981 | 4.447 | 4.114 | 4.495 | 4.387 | 3.945 | -0.036 | 0.4285 | 0.6742 | 0 | 0 | 0 |
| 'Mtmr2' | 5.934 | 6.607 | 6.390 | 5.869 | 6.057 | 6.092 | 5.847 | 6.853 | 6.166 | 6.352 | 6.311 | 5.600 | 6.272 | 6.105 | 5.472 | 6.468 | 6.160 | 5.629 | 0.034 | 0.3760 | 0.6273 | 0 | 0 | 0 |
| 'Mtmr3' | 13.440 | 13.691 | 13.669 | 13.542 | 13.513 | 13.774 | 12.557 | 13.914 | 13.530 | 13.909 | 12.839 | 13.371 | 12.330 | 13.127 | 12.891 | 13.676 | 12.502 | 12.853 | 0.040 | 0.1348 | 0.3536 | 0 | 0 | 0 |
| 'Mtmr4' | 12.925 | 12.148 | 12.459 | 13.078 | 12.424 | 12.502 | 11.560 | 12.311 | 12.747 | 12.764 | 11.521 | 12.323 | 11.061 | 11.476 | 11.780 | 10.905 | 11.144 | 12.158 | 0.083 | 0.0173 | 0.0931 | 0 | 0 | 0 |
| 'Mtmr6' | 14.717 | 15.616 | 14.625 | 15.146 | 16.066 | 14.638 | 15.354 | 16.422 | 13.434 | 14.193 | 16.372 | 16.430 | 15.380 | 17.026 | 15.611 | 15.729 | 15.827 | 15.860 | -0.080 | 0.0144 | 0.0814 | 0 | 0 | 0 |
| 'Mtmr7' | 0.850 | 0.832 | 0.805 | 0.815 | 0.604 | 0.782 | 0.813 | 0.858 | 1.029 | 0.746 | 0.590 | 0.705 | 0.800 | 0.734 | 0.538 | 0.717 | 0.565 | 0.538 | 0.312 | 0.0034 | 0.0276 | 1 | 0 | 0 |
| 'Mtmr9' | 10.333 | 13.522 | 13.147 | 12.501 | 14.515 | 14.149 | 13.112 | 13.916 | 14.225 | 12.084 | 11.603 | 11.953 | 12.018 | 12.758 | 11.455 | 11.386 | 12.390 | 12.637 | 0.128 | 0.0084 | 0.0549 | 0 | 0 | 0 |
| 'Mto1' | 3.571 | 3.276 | 3.940 | 4.181 | 3.562 | 3.676 | 3.851 | 3.730 | 3.578 | 3.274 | 3.217 | 4.108 | 3.613 | 3.014 | 3.426 | 2.845 | 3.253 | 3.654 | 0.116 | 0.0448 | 0.1754 | 0 | 0 | 0 |
| 'Mtor' | 9.486 | 9.088 | 9.868 | 10.042 | 9.266 | 10.073 | 8.773 | 9.167 | 10.158 | 9.343 | 9.721 | 10.180 | 9.260 | 8.899 | 8.730 | 9.349 | 9.942 | 9.543 | 0.005 | 0.8783 | 0.9485 | 0 | 0 | 0 |
| 'Mtpap' | 15.530 | 16.307 | 16.334 | 15.796 | 15.317 | 15.840 | 17.864 | 16.799 | 15.708 | 16.894 | 16.720 | 15.494 | 14.636 | 15.634 | 13.131 | 16.212 | 14.224 | 14.209 | 0.077 | 0.0886 | 0.2723 | 0 | 0 | 0 |
| 'Mtpn' | 36.346 | 40.656 | 41.038 | 37.637 | 42.707 | 38.095 | 40.216 | 41.003 | 37.391 | 36.815 | 39.985 | 38.773 | 37.013 | 39.446 | 37.139 | 39.052 | 41.671 | 38.639 | 0.016 | 0.5452 | 0.7606 | 0 | 0 | 0 |
| 'Mtr' | 11.774 | 10.398 | 12.475 | 12.052 | 12.149 | 11.721 | 12.105 | 11.464 | 12.641 | 11.870 | 13.526 | 12.401 | 11.713 | 11.592 | 11.471 | 12.358 | 11.352 | 11.633 | -0.023 | 0.5706 | 0.7776 | 0 | 0 | 0 |
| 'Mtrf1' | 2.870 | 3.598 | 3.299 | 2.624 | 3.028 | 3.480 | 3.737 | 3.728 | 2.560 | 3.186 | 2.579 | 2.699 | 3.336 | 3.049 | 2.216 | 3.011 | 3.500 | 2.282 | 0.158 | 0.1506 | 0.3777 | 0 | 0 | 0 |
| 'Mtrf1l' | 4.692 | 4.273 | 4.434 | 3.912 | 4.433 | 4.284 | 4.776 | 4.024 | 4.128 | 3.562 | 4.082 | 4.611 | 4.158 | 4.483 | 4.828 | 4.373 | 4.873 | 4.884 | -0.045 | 0.4812 | 0.7131 | 0 | 0 | 0 |
| 'Mtrr' | 6.959 | 7.365 | 6.919 | 6.448 | 6.832 | 6.379 | 7.548 | 7.321 | 6.905 | 7.202 | 6.406 | 7.703 | 7.261 | 6.279 | 7.690 | 6.462 | 7.017 | 7.634 | -0.040 | 0.4342 | 0.6788 | 0 | 0 | 0 |
| 'Mtss1' | 3.420 | 3.495 | 3.164 | 3.837 | 3.883 | 3.577 | 3.662 | 3.469 | 3.752 | 3.956 | 3.792 | 3.523 | 3.762 | 3.960 | 3.624 | 3.632 | 3.773 | 3.368 | -0.060 | 0.1838 | 0.4193 | 0 | 0 | 0 |
| 'Mttp' | 0.863 | 0.723 | 0.920 | 0.858 | 0.765 | 0.895 | 1.004 | 1.042 | 0.879 | 0.905 | 0.830 | 0.801 | 0.703 | 0.857 | 0.689 | 0.749 | 0.813 | 0.777 | 0.157 | 0.0618 | 0.2155 | 0 | 0 | 0 |
| 'Mturn' | 8.238 | 11.939 | 8.441 | 10.002 | 10.827 | 9.294 | 9.664 | 9.679 | 9.965 | 7.556 | 9.667 | 8.293 | 10.685 | 9.104 | 7.498 | 8.402 | 10.045 | 7.967 | 0.140 | 0.0211 | 0.1066 | 0 | 0 | 0 |
| 'Mtx1' | 5.821 | 4.732 | 4.905 | 5.144 | 4.520 | 4.516 | 4.754 | 4.976 | 4.850 | 5.512 | 4.062 | 4.557 | 5.548 | 4.543 | 4.238 | 5.132 | 4.579 | 4.507 | 0.047 | 0.2881 | 0.5447 | 0 | 0 | 0 |
| 'Mtx2' | 6.773 | 8.215 | 7.897 | 6.451 | 8.348 | 7.101 | 7.378 | 8.043 | 6.658 | 7.008 | 7.258 | 6.400 | 7.399 | 7.266 | 6.226 | 7.310 | 7.087 | 6.282 | 0.095 | 0.0399 | 0.1628 | 0 | 0 | 0 |
| 'Mtx3' | 2.920 | 4.201 | 4.393 | 2.885 | 4.870 | 3.901 | 3.554 | 4.201 | 4.414 | 3.949 | 4.304 | 3.458 | 3.713 | 3.947 | 3.131 | 4.529 | 3.688 | 3.205 | 0.045 | 0.6021 | 0.7982 | 0 | 0 | 0 |
| 'Mul1' | 3.713 | 4.132 | 4.340 | 3.266 | 4.035 | 3.695 | 3.812 | 4.029 | 3.757 | 4.097 | 3.364 | 3.762 | 3.584 | 3.992 | 3.744 | 4.240 | 3.458 | 3.505 | 0.036 | 0.5146 | 0.7385 | 0 | 0 | 0 |
| 'Mum1' | 8.338 | 8.120 | 9.972 | 8.498 | 8.172 | 9.599 | 8.762 | 8.179 | 9.602 | 8.233 | 7.986 | 9.180 | 8.489 | 7.669 | 8.863 | 7.626 | 7.628 | 8.432 | 0.085 | 0.0005 | 0.0059 | 1 | 0 | 0 |
| 'Mus81' | 6.252 | 6.684 | 8.207 | 6.405 | 6.505 | 7.914 | 6.748 | 6.816 | 7.948 | 6.699 | 6.213 | 6.881 | 7.622 | 6.731 | 7.125 | 6.676 | 6.957 | 6.611 | 0.033 | 0.4690 | 0.7052 | 0 | 0 | 0 |
| 'Mut' | 6.927 | 9.116 | 8.238 | 7.890 | 8.784 | 8.245 | 9.029 | 8.186 | 8.648 | 7.477 | 8.955 | 8.788 | 8.583 | 8.179 | 8.816 | 8.109 | 9.672 | 8.814 | -0.060 | 0.2135 | 0.4586 | 0 | 0 | 0 |
| 'Mutyh' | 1.384 | 1.637 | 2.215 | 1.431 | 1.508 | 2.094 | 1.514 | 1.772 | 2.134 | 1.831 | 1.642 | 1.631 | 2.079 | 1.092 | 1.562 | 1.816 | 1.454 | 1.148 | 0.131 | 0.2558 | 0.5081 | 0 | 0 | 0 |
| 'Mvb12a' | 40.592 | 43.373 | 40.901 | 37.908 | 44.194 | 44.343 | 38.605 | 42.826 | 40.649 | 36.659 | 37.137 | 42.225 | 38.818 | 36.605 | 41.241 | 39.678 | 36.492 | 41.055 | 0.080 | 0.0391 | 0.1612 | 0 | 0 | 0 |
| 'Mvb12b' | 24.196 | 24.457 | 28.918 | 24.455 | 24.746 | 27.845 | 23.021 | 26.108 | 27.553 | 25.903 | 23.060 | 21.619 | 23.030 | 21.223 | 21.826 | 21.906 | 19.724 | 21.585 | 0.199 | 0.0000 | 0.0004 | 1 | 0 | 0 |
| 'Mvd' | 6.948 | 7.128 | 8.256 | 7.313 | 7.211 | 8.635 | 7.963 | 7.501 | 8.311 | 6.693 | 7.779 | 7.794 | 7.902 | 6.802 | 9.296 | 6.917 | 6.491 | 9.434 | -0.004 | 0.9500 | 0.9809 | 0 | 0 | 0 |
| 'Mvk' | 3.795 | 4.105 | 3.935 | 3.546 | 3.702 | 4.360 | 3.665 | 2.918 | 4.054 | 3.548 | 3.850 | 3.956 | 3.787 | 3.766 | 4.090 | 3.829 | 3.258 | 4.105 | -0.016 | 0.7810 | 0.9017 | 0 | 0 | 0 |
| 'Mvp' | 9.340 | 8.795 | 10.021 | 8.751 | 9.776 | 10.174 | 9.565 | 9.301 | 9.528 | 9.566 | 9.807 | 10.453 | 9.043 | 9.078 | 9.548 | 9.228 | 9.650 | 9.839 | -0.027 | 0.3991 | 0.6482 | 0 | 0 | 0 |
| 'Mxd1' | 4.338 | 3.661 | 3.276 | 3.461 | 3.295 | 3.060 | 3.417 | 3.396 | 3.137 | 3.659 | 4.038 | 3.655 | 3.411 | 4.145 | 3.415 | 4.004 | 3.770 | 3.883 | -0.141 | 0.0183 | 0.0971 | 0 | 0 | 0 |
| 'Mxd3' | 3.581 | 5.132 | 5.852 | 4.463 | 5.035 | 7.293 | 5.099 | 5.192 | 5.978 | 3.573 | 4.200 | 5.412 | 5.178 | 5.199 | 7.144 | 4.796 | 4.970 | 5.881 | 0.024 | 0.7623 | 0.8944 | 0 | 0 | 0 |
| 'Mxd4' | 32.865 | 33.030 | 29.139 | 31.534 | 30.424 | 32.956 | 28.939 | 29.585 | 30.418 | 30.082 | 36.162 | 41.768 | 33.236 | 37.664 | 42.671 | 35.195 | 34.058 | 46.853 | -0.277 | 0.0000 | 0.0000 | 1 | 0 | 0 |
| 'Mxi1' | 4.455 | 4.142 | 4.370 | 4.362 | 4.840 | 4.359 | 4.322 | 4.646 | 3.750 | 3.672 | 4.213 | 4.029 | 3.203 | 4.440 | 3.785 | 3.946 | 3.891 | 3.675 | 0.163 | 0.0020 | 0.0181 | 1 | 0 | 0 |
| 'Mxra8' | 1.288 | 1.030 | 1.166 | 1.289 | 1.256 | 1.252 | 0.820 | 1.373 | 1.344 | 1.398 | 1.315 | 1.281 | 1.389 | 1.227 | 1.202 | 1.354 | 1.277 | 1.178 | -0.102 | 0.2347 | 0.4838 | 0 | 0 | 0 |
| 'Myb' | 11.720 | 9.496 | 12.540 | 10.310 | 10.407 | 12.371 | 10.855 | 11.410 | 12.176 | 11.839 | 11.534 | 12.735 | 10.680 | 10.284 | 12.990 | 10.940 | 10.597 | 12.988 | -0.054 | 0.1654 | 0.3978 | 0 | 0 | 0 |
| 'Mybbp1a' | 15.510 | 12.073 | 16.219 | 14.173 | 13.271 | 14.930 | 14.579 | 14.333 | 14.574 | 15.496 | 13.163 | 15.858 | 14.043 | 11.729 | 14.524 | 14.421 | 12.106 | 14.458 | 0.036 | 0.3383 | 0.5957 | 0 | 0 | 0 |
| 'Mybl2' | 5.932 | 5.216 | 7.482 | 5.967 | 5.852 | 7.192 | 6.836 | 5.811 | 7.103 | 5.070 | 6.188 | 7.074 | 7.935 | 6.324 | 7.010 | 5.907 | 6.056 | 6.808 | -0.039 | 0.5289 | 0.7495 | 0 | 0 | 0 |
| 'Myc' | 3.760 | 2.168 | 3.474 | 3.143 | 2.882 | 3.092 | 3.186 | 2.918 | 2.775 | 3.710 | 2.457 | 2.450 | 3.172 | 2.677 | 2.348 | 3.398 | 2.898 | 2.263 | 0.117 | 0.1178 | 0.3266 | 0 | 0 | 0 |
| 'Mycbp' | 1.575 | 1.676 | 1.480 | 1.572 | 1.487 | 1.469 | 1.462 | 1.543 | 1.602 | 1.438 | 1.714 | 1.595 | 1.618 | 1.447 | 1.596 | 1.239 | 1.648 | 1.636 | -0.025 | 0.7527 | 0.8897 | 0 | 0 | 0 |
| 'Mycbp2' | 24.838 | 21.763 | 22.781 | 24.726 | 21.476 | 22.095 | 21.922 | 21.559 | 23.120 | 24.187 | 23.151 | 23.719 | 20.045 | 21.516 | 21.206 | 21.807 | 21.934 | 24.884 | 0.001 | 0.9750 | 0.9916 | 0 | 0 | 0 |
| 'Myd88' | 17.714 | 18.359 | 17.254 | 17.069 | 19.269 | 18.497 | 17.709 | 19.368 | 17.289 | 16.700 | 17.767 | 18.658 | 16.830 | 18.865 | 19.152 | 17.704 | 17.617 | 18.027 | -0.001 | 0.9645 | 0.9870 | 0 | 0 | 0 |
| 'Mydgf' | 14.869 | 15.966 | 14.842 | 15.664 | 19.261 | 15.891 | 14.491 | 16.825 | 16.057 | 15.120 | 14.613 | 15.128 | 15.777 | 15.555 | 14.953 | 17.081 | 13.750 | 15.998 | 0.047 | 0.3490 | 0.6042 | 0 | 0 | 0 |
| 'Myef2' | 3.758 | 4.800 | 4.082 | 4.024 | 4.874 | 4.103 | 4.876 | 4.843 | 4.076 | 3.864 | 4.803 | 3.650 | 4.242 | 4.744 | 3.806 | 4.661 | 4.547 | 3.483 | 0.056 | 0.2280 | 0.4744 | 0 | 0 | 0 |
| 'Myg1' | 6.702 | 6.537 | 7.735 | 6.847 | 7.051 | 7.199 | 6.208 | 6.688 | 7.114 | 7.025 | 6.524 | 7.237 | 6.963 | 6.985 | 7.915 | 6.490 | 6.281 | 6.763 | -0.011 | 0.7849 | 0.9041 | 0 | 0 | 0 |
| 'Myh10' | 0.482 | 0.585 | 0.449 | 0.549 | 0.517 | 0.532 | 0.496 | 0.458 | 0.549 | 0.667 | 0.573 | 0.547 | 0.504 | 0.579 | 0.446 | 0.473 | 0.447 | 0.524 | -0.060 | 0.4864 | 0.7172 | 0 | 0 | 0 |
| 'Myh3' | 2.514 | 2.046 | 2.396 | 2.339 | 2.170 | 2.071 | 2.377 | 1.888 | 2.384 | 2.743 | 3.195 | 1.869 | 3.023 | 1.792 | 2.659 | 2.151 | 2.785 | 2.628 | -0.187 | 0.0653 | 0.2228 | 0 | 0 | 0 |
| 'Myh9' | 206.221 | 187.273 | 210.310 | 213.119 | 192.174 | 212.396 | 207.928 | 191.941 | 215.571 | 232.164 | 221.524 | 213.174 | 217.662 | 213.206 | 229.578 | 220.064 | 209.509 | 232.126 | -0.127 | 0.0000 | 0.0001 | 1 | 0 | 0 |
| 'Myl12a' | 55.513 | 62.256 | 53.561 | 57.451 | 61.816 | 56.374 | 57.130 | 58.720 | 56.002 | 59.188 | 59.644 | 54.180 | 64.496 | 63.573 | 60.893 | 60.437 | 62.119 | 56.404 | -0.071 | 0.0108 | 0.0662 | 0 | 0 | 0 |
| 'Myl12b' | 121.514 | 123.576 | 119.346 | 120.796 | 130.478 | 122.638 | 119.110 | 126.136 | 120.555 | 118.941 | 128.990 | 118.451 | 118.106 | 127.509 | 127.712 | 119.370 | 125.551 | 122.845 | -0.014 | 0.5290 | 0.7495 | 0 | 0 | 0 |
| 'Myl6' | 732.293 | 768.227 | 691.694 | 761.829 | 757.746 | 730.821 | 746.848 | 728.811 | 713.349 | 733.171 | 739.416 | 629.377 | 723.987 | 761.010 | 769.187 | 755.329 | 762.578 | 692.042 | 0.004 | 0.8989 | 0.9598 | 0 | 0 | 0 |
| 'Mylip' | 16.709 | 19.395 | 15.373 | 16.122 | 19.521 | 17.239 | 16.184 | 20.160 | 16.824 | 17.553 | 16.783 | 15.796 | 14.491 | 16.747 | 15.160 | 18.463 | 16.706 | 15.088 | 0.089 | 0.0514 | 0.1917 | 0 | 0 | 0 |
| 'Mylpf' | 4.685 | 6.119 | 6.967 | 6.150 | 4.546 | 5.897 | 5.336 | 5.143 | 5.642 | 5.065 | 5.044 | 5.781 | 5.244 | 5.885 | 6.481 | 4.695 | 5.048 | 5.930 | 0.028 | 0.7329 | 0.8781 | 0 | 0 | 0 |
| 'Mynn' | 3.045 | 3.019 | 2.646 | 2.932 | 3.018 | 2.469 | 3.011 | 2.714 | 2.603 | 2.791 | 3.074 | 2.734 | 2.400 | 2.801 | 2.441 | 2.451 | 2.777 | 2.443 | 0.076 | 0.1152 | 0.3222 | 0 | 0 | 0 |
| 'Myo10' | 0.240 | 0.182 | 0.310 | 0.242 | 0.185 | 0.242 | 0.238 | 0.265 | 0.276 | 0.322 | 0.376 | 0.280 | 0.342 | 0.242 | 0.308 | 0.213 | 0.315 | 0.277 | -0.292 | 0.0165 | 0.0899 | 0 | 0 | 0 |
| 'Myo18a' | 13.724 | 13.212 | 13.596 | 13.958 | 13.073 | 13.678 | 13.716 | 13.811 | 14.259 | 14.979 | 14.028 | 13.962 | 15.060 | 13.534 | 13.296 | 13.483 | 13.040 | 14.294 | -0.041 | 0.1284 | 0.3432 | 0 | 0 | 0 |
| 'Myo19' | 0.966 | 0.971 | 1.063 | 1.001 | 0.932 | 0.876 | 0.902 | 0.861 | 1.027 | 0.803 | 0.927 | 1.128 | 0.943 | 0.730 | 0.869 | 0.820 | 1.012 | 0.958 | 0.050 | 0.4968 | 0.7257 | 0 | 0 | 0 |
| 'Myo1b' | 0.290 | 0.368 | 0.255 | 0.368 | 0.286 | 0.307 | 0.343 | 0.296 | 0.251 | 0.247 | 0.277 | 0.231 | 0.331 | 0.309 | 0.199 | 0.224 | 0.375 | 0.189 | 0.206 | 0.0525 | 0.1947 | 0 | 0 | 0 |
| 'Myo1c' | 7.754 | 7.195 | 8.205 | 7.286 | 7.397 | 8.384 | 6.939 | 7.995 | 8.021 | 7.591 | 7.786 | 8.076 | 7.499 | 7.523 | 7.312 | 6.590 | 7.275 | 7.763 | 0.028 | 0.4462 | 0.6878 | 0 | 0 | 0 |
| 'Myo1e' | 2.135 | 2.253 | 2.076 | 2.322 | 3.062 | 2.260 | 2.031 | 2.759 | 2.439 | 1.594 | 2.150 | 3.144 | 2.053 | 2.616 | 2.870 | 1.720 | 2.763 | 2.747 | -0.029 | 0.7588 | 0.8921 | 0 | 0 | 0 |
| 'Myo1f' | 9.838 | 9.334 | 10.381 | 10.362 | 10.555 | 11.057 | 11.374 | 9.362 | 12.285 | 14.342 | 11.196 | 14.043 | 14.009 | 12.571 | 13.770 | 13.708 | 10.050 | 13.350 | -0.316 | 0.0000 | 0.0000 | 1 | 0 | 0 |
| 'Myo1g' | 17.763 | 14.547 | 19.312 | 16.583 | 16.001 | 18.745 | 16.893 | 16.459 | 18.240 | 15.793 | 15.916 | 18.725 | 16.331 | 14.814 | 18.311 | 15.556 | 15.642 | 18.117 | 0.041 | 0.1508 | 0.3777 | 0 | 0 | 0 |
| 'Myo1h' | 2.087 | 2.256 | 2.441 | 1.922 | 1.845 | 2.146 | 2.443 | 2.025 | 2.506 | 3.003 | 3.049 | 2.714 | 2.829 | 2.595 | 2.281 | 2.952 | 3.224 | 2.187 | -0.339 | 0.0000 | 0.0002 | 1 | 0 | 0 |
| 'Myo5a' | 1.942 | 1.508 | 2.042 | 2.051 | 1.674 | 1.907 | 1.893 | 1.903 | 1.851 | 1.845 | 1.978 | 1.992 | 1.919 | 1.792 | 1.919 | 1.829 | 1.996 | 1.982 | -0.051 | 0.2860 | 0.5428 | 0 | 0 | 0 |
| 'Myo6' | 0.547 | 0.412 | 0.540 | 0.488 | 0.542 | 0.546 | 0.549 | 0.585 | 0.478 | 0.284 | 0.456 | 0.363 | 0.370 | 0.447 | 0.336 | 0.402 | 0.404 | 0.339 | 0.468 | 0.0000 | 0.0000 | 1 | 1 | 0 |
| 'Myo9a' | 2.097 | 2.286 | 2.215 | 2.459 | 2.258 | 2.021 | 2.612 | 2.173 | 2.191 | 2.615 | 2.589 | 2.464 | 2.236 | 2.323 | 2.100 | 2.227 | 2.497 | 2.538 | -0.103 | 0.0581 | 0.2077 | 0 | 0 | 0 |
| 'Myo9b' | 46.455 | 39.254 | 50.060 | 43.293 | 41.176 | 45.943 | 40.494 | 40.828 | 48.010 | 49.678 | 45.871 | 48.531 | 46.536 | 44.916 | 46.014 | 45.126 | 44.242 | 48.695 | -0.098 | 0.0044 | 0.0340 | 1 | 0 | 0 |
| 'Mypop' | 4.490 | 5.389 | 5.740 | 4.929 | 5.064 | 5.414 | 4.593 | 5.037 | 5.676 | 4.754 | 4.164 | 4.987 | 4.797 | 3.929 | 4.634 | 5.014 | 4.164 | 4.960 | 0.151 | 0.0025 | 0.0218 | 1 | 0 | 0 |
| 'Mysm1' | 4.358 | 4.662 | 3.913 | 3.951 | 4.289 | 3.931 | 4.228 | 4.084 | 3.933 | 3.910 | 4.497 | 4.207 | 4.069 | 4.376 | 4.086 | 4.421 | 4.451 | 4.115 | -0.042 | 0.2233 | 0.4710 | 0 | 0 | 0 |
| 'Mzt1' | 23.519 | 31.785 | 25.292 | 26.196 | 29.478 | 25.023 | 28.703 | 30.160 | 25.892 | 26.747 | 24.848 | 23.653 | 26.197 | 27.492 | 23.400 | 26.284 | 27.372 | 21.328 | 0.101 | 0.0109 | 0.0665 | 0 | 0 | 0 |
| 'Mzt2' | 5.493 | 4.862 | 4.591 | 4.919 | 5.684 | 5.176 | 4.546 | 5.322 | 5.255 | 4.800 | 4.322 | 4.445 | 4.662 | 5.010 | 4.475 | 5.019 | 4.289 | 4.463 | 0.139 | 0.0344 | 0.1483 | 0 | 0 | 0 |
| 'N4bp1' | 12.040 | 11.841 | 10.119 | 10.142 | 11.027 | 11.462 | 10.749 | 11.711 | 10.712 | 12.329 | 10.849 | 12.707 | 11.036 | 11.285 | 12.468 | 14.120 | 10.918 | 11.961 | -0.119 | 0.0191 | 0.1001 | 0 | 0 | 0 |
| 'N4bp2' | 5.544 | 5.697 | 6.026 | 5.462 | 5.901 | 5.868 | 6.055 | 5.599 | 5.985 | 5.581 | 5.436 | 5.618 | 5.667 | 5.955 | 5.120 | 5.383 | 5.399 | 5.167 | 0.070 | 0.0245 | 0.1191 | 0 | 0 | 0 |
| 'N4bp2l1' | 8.740 | 9.980 | 8.538 | 8.542 | 9.323 | 8.567 | 8.971 | 9.886 | 8.516 | 8.061 | 8.873 | 9.650 | 9.167 | 8.796 | 8.945 | 8.777 | 9.223 | 9.645 | -0.018 | 0.6416 | 0.8242 | 0 | 0 | 0 |
| 'N4bp2l2' | 4.458 | 4.475 | 4.606 | 4.948 | 4.601 | 4.115 | 5.576 | 4.454 | 4.287 | 4.265 | 4.848 | 4.747 | 4.084 | 3.957 | 3.996 | 3.998 | 4.867 | 4.132 | 0.081 | 0.1519 | 0.3789 | 0 | 0 | 0 |
| 'N4bp3' | 2.657 | 2.162 | 3.356 | 3.327 | 1.932 | 3.299 | 2.467 | 3.005 | 2.781 | 2.377 | 2.730 | 3.351 | 2.585 | 3.072 | 3.806 | 2.372 | 2.460 | 3.979 | -0.096 | 0.3214 | 0.5793 | 0 | 0 | 0 |
| 'N6amt1' | 1.815 | 2.667 | 2.400 | 1.827 | 2.529 | 2.445 | 1.888 | 2.415 | 2.565 | 2.350 | 1.912 | 1.779 | 2.160 | 1.864 | 1.736 | 2.660 | 1.705 | 1.532 | 0.214 | 0.0376 | 0.1573 | 0 | 0 | 0 |
| 'Naa10' | 8.707 | 8.146 | 9.151 | 9.168 | 8.709 | 8.771 | 9.113 | 8.781 | 9.172 | 8.096 | 7.729 | 9.640 | 8.400 | 8.063 | 9.952 | 7.900 | 8.354 | 9.205 | 0.033 | 0.3746 | 0.6259 | 0 | 0 | 0 |
| 'Naa15' | 9.678 | 8.912 | 9.504 | 10.618 | 8.910 | 8.450 | 11.502 | 9.339 | 9.116 | 8.481 | 9.271 | 9.599 | 8.424 | 9.031 | 9.067 | 8.508 | 10.090 | 9.120 | 0.062 | 0.2432 | 0.4935 | 0 | 0 | 0 |
| 'Naa16' | 6.365 | 6.081 | 6.166 | 6.478 | 6.145 | 5.070 | 6.572 | 6.246 | 5.244 | 5.664 | 6.321 | 6.137 | 5.962 | 6.120 | 5.643 | 6.254 | 6.839 | 5.934 | -0.027 | 0.5445 | 0.7606 | 0 | 0 | 0 |
| 'Naa20' | 10.090 | 10.739 | 9.827 | 9.821 | 10.420 | 9.195 | 9.174 | 10.850 | 9.298 | 10.273 | 10.084 | 10.225 | 10.467 | 10.721 | 9.120 | 10.914 | 9.974 | 9.415 | -0.040 | 0.2751 | 0.5307 | 0 | 0 | 0 |
| 'Naa25' | 3.564 | 3.523 | 3.624 | 3.558 | 3.052 | 3.228 | 3.056 | 3.371 | 3.386 | 3.250 | 2.893 | 3.659 | 2.996 | 3.115 | 3.517 | 3.429 | 3.353 | 3.493 | 0.021 | 0.6500 | 0.8295 | 0 | 0 | 0 |
| 'Naa30' | 4.757 | 5.374 | 5.618 | 5.377 | 5.345 | 4.846 | 5.576 | 5.399 | 5.203 | 4.442 | 5.381 | 5.879 | 4.335 | 5.023 | 5.445 | 4.843 | 5.284 | 5.961 | 0.020 | 0.6960 | 0.8571 | 0 | 0 | 0 |
| 'Naa38' | 11.339 | 14.201 | 12.285 | 10.361 | 11.542 | 11.585 | 10.788 | 11.475 | 11.369 | 10.045 | 10.864 | 11.769 | 11.616 | 9.360 | 13.126 | 11.042 | 11.526 | 11.207 | 0.047 | 0.4400 | 0.6834 | 0 | 0 | 0 |
| 'Naa40' | 24.523 | 28.028 | 28.847 | 24.360 | 25.607 | 28.582 | 25.926 | 28.151 | 27.521 | 26.034 | 26.780 | 28.142 | 26.977 | 27.442 | 29.520 | 26.246 | 26.155 | 27.118 | -0.027 | 0.3504 | 0.6045 | 0 | 0 | 0 |
| 'Naa50' | 10.904 | 10.960 | 11.328 | 11.940 | 11.211 | 10.265 | 12.056 | 11.263 | 10.689 | 9.923 | 10.779 | 10.893 | 10.591 | 10.672 | 10.367 | 9.422 | 10.707 | 11.202 | 0.077 | 0.0434 | 0.1725 | 0 | 0 | 0 |
| 'Naa60' | 15.724 | 15.851 | 16.493 | 15.270 | 16.684 | 16.949 | 15.635 | 15.748 | 16.834 | 16.217 | 15.284 | 16.450 | 16.219 | 16.278 | 16.429 | 14.455 | 15.531 | 15.803 | 0.014 | 0.6060 | 0.8014 | 0 | 0 | 0 |
| 'Naaa' | 3.384 | 3.570 | 3.900 | 3.796 | 4.049 | 4.052 | 3.340 | 3.895 | 4.505 | 3.825 | 2.789 | 3.685 | 3.241 | 3.841 | 3.362 | 3.885 | 3.389 | 3.484 | 0.122 | 0.0536 | 0.1977 | 0 | 0 | 0 |
| 'Nab1' | 40.293 | 44.234 | 38.975 | 41.186 | 46.051 | 37.051 | 41.450 | 46.659 | 36.133 | 45.067 | 50.793 | 38.287 | 46.373 | 49.117 | 40.193 | 44.801 | 47.777 | 39.041 | -0.118 | 0.0000 | 0.0000 | 1 | 0 | 0 |
| 'Nab2' | 22.574 | 18.558 | 21.745 | 20.256 | 15.949 | 23.448 | 20.444 | 17.692 | 22.301 | 15.227 | 14.896 | 26.296 | 13.734 | 15.684 | 23.499 | 16.964 | 17.417 | 24.309 | 0.136 | 0.0568 | 0.2049 | 0 | 0 | 0 |
| 'Nabp1' | 2.310 | 2.710 | 2.673 | 2.303 | 2.610 | 2.740 | 3.006 | 2.492 | 2.668 | 2.281 | 2.351 | 2.440 | 1.833 | 2.294 | 2.425 | 2.184 | 2.488 | 2.280 | 0.182 | 0.0007 | 0.0076 | 1 | 0 | 0 |
| 'Nabp2' | 14.725 | 16.070 | 14.832 | 15.361 | 16.089 | 16.313 | 15.712 | 15.500 | 15.115 | 15.186 | 14.145 | 15.101 | 16.861 | 15.338 | 15.151 | 15.864 | 15.374 | 15.223 | 0.003 | 0.9144 | 0.9671 | 0 | 0 | 0 |
| 'Naca' | 57.015 | 57.765 | 54.013 | 61.857 | 59.236 | 53.926 | 61.499 | 54.305 | 54.702 | 55.014 | 56.110 | 50.120 | 57.348 | 55.099 | 55.427 | 56.330 | 57.362 | 52.473 | 0.043 | 0.1270 | 0.3414 | 0 | 0 | 0 |
| 'Nacc1' | 46.101 | 43.275 | 46.647 | 42.070 | 41.426 | 49.015 | 41.430 | 44.018 | 46.672 | 44.866 | 42.071 | 46.558 | 47.325 | 40.809 | 46.110 | 45.863 | 39.257 | 49.443 | -0.015 | 0.6453 | 0.8266 | 0 | 0 | 0 |
| 'Nadk' | 20.750 | 22.396 | 21.305 | 21.336 | 22.402 | 22.453 | 21.766 | 22.862 | 22.575 | 22.535 | 20.162 | 20.363 | 22.507 | 21.705 | 22.070 | 23.484 | 22.154 | 21.839 | -0.002 | 0.9554 | 0.9835 | 0 | 0 | 0 |
| 'Nadk2' | 3.394 | 3.444 | 3.447 | 3.170 | 3.175 | 3.769 | 3.519 | 3.919 | 3.183 | 3.277 | 3.728 | 3.728 | 3.136 | 3.637 | 3.779 | 3.500 | 3.989 | 3.611 | -0.068 | 0.1636 | 0.3955 | 0 | 0 | 0 |
| 'Nadsyn1' | 4.246 | 3.873 | 4.430 | 3.926 | 3.797 | 3.886 | 4.760 | 3.911 | 4.444 | 4.045 | 4.491 | 4.881 | 4.228 | 3.859 | 4.206 | 4.086 | 4.683 | 4.377 | -0.074 | 0.1663 | 0.3986 | 0 | 0 | 0 |
| 'Nae1' | 3.724 | 3.902 | 4.015 | 4.101 | 4.227 | 3.690 | 4.247 | 4.209 | 4.009 | 3.617 | 4.185 | 4.316 | 3.695 | 4.181 | 4.006 | 3.667 | 4.280 | 3.814 | 0.001 | 0.9775 | 0.9920 | 0 | 0 | 0 |
| 'Naf1' | 6.759 | 4.938 | 6.681 | 6.176 | 6.689 | 6.781 | 5.046 | 6.755 | 6.295 | 7.260 | 7.157 | 6.440 | 6.932 | 5.439 | 5.892 | 4.420 | 7.270 | 6.119 | -0.027 | 0.7697 | 0.8975 | 0 | 0 | 0 |
| 'Naga' | 32.768 | 31.456 | 32.660 | 30.893 | 31.388 | 34.086 | 30.604 | 32.129 | 33.227 | 29.325 | 28.192 | 30.822 | 28.583 | 31.192 | 29.428 | 29.074 | 27.689 | 27.200 | 0.137 | 0.0000 | 0.0006 | 1 | 0 | 0 |
| 'Nagk' | 1.873 | 2.019 | 2.123 | 1.813 | 2.153 | 2.088 | 2.098 | 1.948 | 2.338 | 1.910 | 2.224 | 2.089 | 1.930 | 2.157 | 2.082 | 1.988 | 1.969 | 2.054 | -0.004 | 0.9429 | 0.9785 | 0 | 0 | 0 |
| 'Naglu' | 7.262 | 7.615 | 9.291 | 7.230 | 8.634 | 8.662 | 7.224 | 8.182 | 8.670 | 8.387 | 7.459 | 7.482 | 7.469 | 7.441 | 8.023 | 7.151 | 6.993 | 6.360 | 0.116 | 0.0384 | 0.1594 | 0 | 0 | 0 |
| 'Nagpa' | 10.866 | 10.762 | 12.354 | 9.968 | 11.980 | 13.270 | 10.341 | 10.960 | 12.784 | 12.016 | 10.585 | 14.120 | 11.554 | 10.592 | 12.320 | 12.242 | 11.068 | 13.762 | -0.079 | 0.0544 | 0.1996 | 0 | 0 | 0 |
| 'Naif1' | 2.893 | 3.543 | 3.149 | 2.944 | 3.379 | 3.617 | 2.835 | 3.328 | 3.593 | 3.273 | 3.149 | 3.684 | 3.133 | 3.682 | 3.755 | 3.147 | 3.252 | 3.146 | -0.061 | 0.2882 | 0.5447 | 0 | 0 | 0 |
| 'Naip2' | 1.609 | 2.665 | 2.059 | 2.173 | 2.612 | 2.212 | 1.735 | 2.095 | 1.799 | 2.392 | 3.374 | 2.073 | 2.847 | 2.888 | 2.258 | 2.194 | 3.281 | 2.563 | -0.339 | 0.0001 | 0.0011 | 1 | 0 | 0 |
| 'Nampt' | 12.880 | 13.626 | 11.551 | 13.387 | 12.738 | 11.689 | 13.764 | 13.269 | 11.626 | 13.020 | 14.879 | 14.645 | 13.156 | 14.895 | 14.243 | 13.883 | 16.469 | 13.538 | -0.183 | 0.0000 | 0.0003 | 1 | 0 | 0 |
| 'Nanos1' | 18.321 | 21.155 | 17.551 | 19.078 | 18.487 | 19.343 | 19.241 | 17.734 | 17.816 | 18.297 | 20.288 | 23.041 | 17.214 | 21.797 | 23.600 | 17.774 | 23.032 | 24.732 | -0.174 | 0.0033 | 0.0271 | 1 | 0 | 0 |
| 'Nans' | 18.938 | 18.164 | 18.388 | 18.347 | 19.095 | 19.273 | 17.714 | 18.383 | 18.995 | 18.524 | 17.766 | 19.227 | 20.071 | 18.476 | 19.342 | 19.234 | 17.808 | 17.795 | -0.018 | 0.5047 | 0.7312 | 0 | 0 | 0 |
| 'Nap1l1' | 52.668 | 52.376 | 52.652 | 53.725 | 53.282 | 48.930 | 56.090 | 52.588 | 50.938 | 51.566 | 51.251 | 51.973 | 57.043 | 54.327 | 53.681 | 52.093 | 57.023 | 52.361 | -0.036 | 0.1835 | 0.4192 | 0 | 0 | 0 |
| 'Nap1l4' | 122.630 | 125.500 | 121.405 | 123.692 | 125.537 | 124.995 | 120.636 | 128.799 | 123.336 | 121.535 | 121.424 | 125.443 | 123.221 | 119.870 | 124.298 | 125.334 | 121.048 | 121.899 | 0.005 | 0.7634 | 0.8953 | 0 | 0 | 0 |
| 'Napa' | 21.688 | 19.593 | 21.169 | 19.694 | 20.018 | 21.384 | 20.346 | 20.772 | 20.922 | 21.814 | 19.580 | 21.828 | 21.337 | 19.906 | 21.136 | 22.266 | 18.506 | 20.462 | -0.018 | 0.5217 | 0.7444 | 0 | 0 | 0 |
| 'Napg' | 9.622 | 11.283 | 10.745 | 9.517 | 11.022 | 9.584 | 8.799 | 11.141 | 10.367 | 10.149 | 10.470 | 10.553 | 9.423 | 10.332 | 9.641 | 10.367 | 10.749 | 9.992 | -0.006 | 0.8673 | 0.9426 | 0 | 0 | 0 |
| 'Naprt' | 6.137 | 7.430 | 8.244 | 7.204 | 7.592 | 7.711 | 7.136 | 7.987 | 7.079 | 7.732 | 8.081 | 6.221 | 6.757 | 6.804 | 6.474 | 5.675 | 6.712 | 5.252 | 0.154 | 0.0375 | 0.1572 | 0 | 0 | 0 |
| 'Narf' | 14.911 | 14.182 | 12.777 | 14.334 | 13.598 | 13.188 | 14.697 | 13.552 | 13.477 | 12.212 | 14.497 | 17.159 | 12.738 | 16.042 | 16.917 | 12.675 | 16.087 | 16.887 | -0.120 | 0.0589 | 0.2095 | 0 | 0 | 0 |
| 'Narfl' | 3.934 | 4.615 | 4.805 | 3.774 | 4.114 | 4.621 | 4.147 | 4.267 | 4.921 | 3.771 | 3.805 | 4.204 | 4.488 | 4.323 | 3.843 | 3.847 | 3.894 | 4.409 | 0.087 | 0.0812 | 0.2570 | 0 | 0 | 0 |
| 'Nars' | 50.475 | 50.584 | 50.802 | 51.171 | 54.132 | 49.921 | 51.446 | 54.058 | 49.303 | 49.281 | 48.160 | 48.649 | 50.756 | 48.470 | 50.329 | 50.792 | 49.652 | 47.975 | 0.046 | 0.0450 | 0.1759 | 0 | 0 | 0 |
| 'Nars2' | 3.062 | 3.512 | 2.801 | 3.411 | 3.551 | 2.833 | 3.612 | 3.331 | 3.030 | 3.057 | 3.227 | 3.269 | 3.157 | 2.861 | 2.807 | 3.405 | 3.450 | 3.055 | 0.019 | 0.7351 | 0.8795 | 0 | 0 | 0 |
| 'Nasp' | 10.180 | 10.042 | 11.225 | 11.472 | 9.895 | 10.416 | 11.370 | 10.072 | 10.250 | 9.419 | 10.077 | 10.156 | 10.665 | 9.860 | 10.731 | 10.023 | 10.879 | 9.712 | 0.041 | 0.2763 | 0.5319 | 0 | 0 | 0 |
| 'Nat10' | 3.603 | 3.221 | 4.082 | 3.702 | 3.397 | 3.294 | 3.792 | 3.462 | 3.276 | 3.522 | 2.970 | 3.964 | 3.516 | 2.876 | 3.361 | 3.327 | 3.315 | 3.733 | 0.046 | 0.3612 | 0.6135 | 0 | 0 | 0 |
| 'Nat2' | 3.701 | 4.236 | 4.860 | 4.077 | 4.502 | 5.211 | 5.078 | 4.858 | 4.933 | 3.542 | 4.362 | 4.512 | 3.540 | 3.917 | 4.693 | 3.626 | 4.228 | 4.286 | 0.166 | 0.0094 | 0.0598 | 0 | 0 | 0 |
| 'Nat6' | 2.020 | 1.696 | 2.315 | 1.742 | 1.994 | 2.243 | 1.854 | 1.868 | 2.187 | 2.010 | 1.729 | 2.368 | 1.741 | 2.143 | 2.050 | 1.985 | 1.889 | 1.954 | -0.004 | 0.9518 | 0.9818 | 0 | 0 | 0 |
| 'Nat9' | 3.696 | 3.897 | 4.066 | 3.514 | 4.481 | 4.065 | 3.774 | 4.333 | 3.843 | 3.546 | 3.676 | 3.582 | 3.839 | 3.807 | 2.951 | 3.511 | 3.774 | 3.297 | 0.149 | 0.0067 | 0.0469 | 1 | 0 | 0 |
| 'Natd1' | 8.684 | 10.088 | 9.252 | 8.587 | 9.114 | 10.431 | 8.383 | 9.216 | 9.296 | 8.674 | 8.127 | 9.710 | 8.931 | 9.028 | 10.034 | 9.783 | 9.024 | 9.875 | -0.014 | 0.7421 | 0.8835 | 0 | 0 | 0 |
| 'Nav2' | 13.281 | 13.829 | 13.208 | 14.532 | 13.677 | 13.002 | 13.177 | 14.082 | 14.549 | 13.717 | 14.088 | 10.307 | 14.825 | 12.707 | 9.868 | 13.174 | 12.207 | 10.750 | 0.144 | 0.0061 | 0.0435 | 1 | 0 | 0 |
| 'Nbas' | 7.450 | 7.554 | 6.831 | 6.435 | 6.888 | 6.740 | 7.061 | 7.910 | 6.915 | 7.573 | 7.722 | 7.476 | 6.619 | 6.933 | 7.396 | 6.752 | 7.481 | 7.159 | -0.041 | 0.3240 | 0.5814 | 0 | 0 | 0 |
| 'Nbeal1' | 0.945 | 0.846 | 0.935 | 1.013 | 0.840 | 0.829 | 0.965 | 0.927 | 0.846 | 1.093 | 0.926 | 1.010 | 0.813 | 1.003 | 0.873 | 0.985 | 1.120 | 0.965 | -0.121 | 0.0422 | 0.1695 | 0 | 0 | 0 |
| 'Nbeal2' | 20.457 | 18.630 | 23.522 | 19.959 | 19.146 | 22.343 | 19.300 | 18.977 | 23.414 | 21.375 | 20.321 | 23.827 | 19.495 | 18.270 | 21.711 | 19.542 | 18.912 | 23.485 | -0.020 | 0.4873 | 0.7176 | 0 | 0 | 0 |
| 'Nbn' | 18.616 | 17.032 | 17.013 | 17.714 | 17.757 | 18.512 | 18.182 | 17.234 | 18.145 | 19.294 | 19.272 | 18.502 | 18.769 | 20.049 | 19.226 | 18.779 | 19.849 | 18.468 | -0.111 | 0.0011 | 0.0118 | 1 | 0 | 0 |
| 'Nbr1' | 11.178 | 11.887 | 11.697 | 11.257 | 11.245 | 11.558 | 10.948 | 12.331 | 11.603 | 11.343 | 12.789 | 12.744 | 11.488 | 12.438 | 12.066 | 12.148 | 12.315 | 12.039 | -0.087 | 0.0002 | 0.0032 | 1 | 0 | 0 |
| 'Ncapd2' | 15.773 | 14.909 | 20.262 | 19.043 | 14.993 | 19.101 | 18.329 | 15.479 | 19.544 | 13.649 | 17.857 | 17.510 | 19.706 | 18.009 | 19.916 | 14.645 | 17.276 | 18.429 | -0.012 | 0.8578 | 0.9390 | 0 | 0 | 0 |
| 'Ncapd3' | 17.779 | 17.433 | 20.331 | 19.528 | 17.353 | 19.096 | 19.595 | 18.799 | 19.862 | 17.334 | 18.500 | 18.763 | 18.651 | 18.329 | 20.090 | 17.488 | 20.055 | 18.731 | 0.006 | 0.8697 | 0.9441 | 0 | 0 | 0 |
| 'Ncapg' | 2.712 | 2.852 | 3.808 | 3.649 | 2.926 | 2.979 | 3.669 | 2.689 | 3.262 | 2.219 | 3.112 | 3.458 | 3.133 | 3.627 | 3.654 | 2.558 | 3.301 | 3.281 | 0.003 | 0.9699 | 0.9898 | 0 | 0 | 0 |
| 'Ncapg2' | 8.595 | 8.932 | 11.408 | 9.972 | 8.982 | 10.308 | 11.266 | 8.963 | 10.232 | 7.800 | 10.192 | 10.325 | 11.154 | 10.536 | 10.706 | 8.386 | 10.563 | 9.816 | -0.025 | 0.7029 | 0.8611 | 0 | 0 | 0 |
| 'Ncaph' | 5.082 | 5.543 | 7.173 | 6.113 | 5.631 | 6.507 | 6.205 | 5.464 | 6.603 | 4.257 | 5.357 | 6.126 | 6.386 | 6.315 | 6.459 | 5.303 | 5.462 | 6.045 | 0.058 | 0.3483 | 0.6036 | 0 | 0 | 0 |
| 'Ncaph2' | 14.149 | 14.279 | 15.409 | 13.811 | 14.586 | 16.297 | 14.522 | 14.719 | 15.640 | 14.129 | 13.249 | 16.028 | 15.227 | 14.784 | 16.055 | 15.065 | 14.035 | 16.073 | -0.024 | 0.3441 | 0.5995 | 0 | 0 | 0 |
| 'Ncbp1' | 32.821 | 33.755 | 34.259 | 34.805 | 34.079 | 32.846 | 36.096 | 33.575 | 32.208 | 30.333 | 32.748 | 33.790 | 33.293 | 32.530 | 33.173 | 32.225 | 33.893 | 32.534 | 0.035 | 0.1973 | 0.4366 | 0 | 0 | 0 |
| 'Ncbp2' | 12.016 | 12.574 | 11.435 | 13.677 | 12.156 | 13.281 | 13.410 | 12.355 | 12.457 | 11.168 | 11.503 | 14.003 | 11.611 | 12.431 | 14.233 | 12.087 | 13.223 | 13.481 | -0.018 | 0.7259 | 0.8744 | 0 | 0 | 0 |
| 'Ncbp3' | 4.931 | 5.246 | 5.215 | 5.161 | 5.058 | 5.115 | 5.564 | 4.951 | 4.949 | 5.173 | 5.225 | 5.002 | 5.087 | 5.194 | 4.745 | 4.915 | 4.837 | 4.581 | 0.035 | 0.2484 | 0.4993 | 0 | 0 | 0 |
| 'Ncdn' | 12.822 | 12.214 | 13.555 | 11.632 | 12.917 | 13.035 | 11.727 | 12.710 | 14.158 | 13.161 | 11.467 | 13.014 | 12.945 | 11.571 | 13.084 | 12.651 | 11.716 | 12.270 | 0.028 | 0.4049 | 0.6541 | 0 | 0 | 0 |
| 'Nceh1' | 6.416 | 7.482 | 7.357 | 6.050 | 6.469 | 6.289 | 6.611 | 7.577 | 6.245 | 7.211 | 7.509 | 6.321 | 6.889 | 6.626 | 5.926 | 6.435 | 6.160 | 5.475 | 0.040 | 0.4546 | 0.6931 | 0 | 0 | 0 |
| 'Ncf4' | 22.103 | 20.327 | 22.767 | 19.241 | 20.882 | 22.108 | 20.754 | 18.557 | 22.374 | 21.737 | 19.752 | 24.838 | 21.265 | 20.377 | 22.064 | 21.468 | 22.619 | 22.411 | -0.068 | 0.0662 | 0.2249 | 0 | 0 | 0 |
| 'Nck1' | 7.533 | 7.717 | 6.573 | 8.731 | 7.256 | 6.701 | 7.788 | 6.810 | 6.969 | 6.757 | 7.770 | 8.670 | 6.135 | 7.433 | 7.575 | 6.685 | 7.391 | 7.922 | -0.019 | 0.7761 | 0.9000 | 0 | 0 | 0 |
| 'Nck2' | 8.631 | 10.255 | 8.709 | 9.309 | 8.726 | 9.087 | 8.507 | 8.455 | 8.475 | 7.876 | 8.383 | 9.116 | 8.762 | 8.100 | 9.628 | 9.295 | 8.249 | 8.748 | 0.024 | 0.6009 | 0.7972 | 0 | 0 | 0 |
| 'Nckap1' | 0.703 | 0.783 | 0.842 | 0.818 | 0.743 | 0.939 | 0.832 | 0.824 | 0.762 | 0.425 | 0.636 | 0.666 | 0.510 | 0.678 | 0.682 | 0.581 | 0.616 | 0.680 | 0.397 | 0.0000 | 0.0003 | 1 | 1 | 0 |
| 'Nckap1l' | 97.314 | 95.985 | 98.893 | 98.731 | 99.967 | 101.675 | 98.920 | 98.251 | 100.973 | 99.056 | 101.531 | 98.984 | 99.112 | 95.827 | 98.898 | 91.795 | 98.792 | 99.434 | 0.001 | 0.9550 | 0.9834 | 0 | 0 | 0 |
| 'Nckap5l' | 7.770 | 6.150 | 8.382 | 7.611 | 6.648 | 7.248 | 6.802 | 6.934 | 7.973 | 7.276 | 6.741 | 7.191 | 6.644 | 6.414 | 7.154 | 6.346 | 5.632 | 7.058 | 0.109 | 0.0193 | 0.1007 | 0 | 0 | 0 |
| 'Nckipsd' | 3.676 | 2.625 | 3.999 | 2.889 | 2.989 | 3.280 | 2.983 | 3.365 | 3.789 | 2.416 | 3.002 | 3.487 | 2.941 | 2.908 | 3.188 | 2.946 | 2.613 | 3.345 | 0.135 | 0.0399 | 0.1628 | 0 | 0 | 0 |
| 'Ncl' | 55.171 | 48.842 | 50.891 | 53.335 | 52.370 | 49.461 | 55.979 | 51.315 | 49.932 | 49.886 | 49.688 | 46.830 | 50.435 | 48.272 | 49.303 | 51.131 | 47.823 | 47.577 | 0.073 | 0.0009 | 0.0098 | 1 | 0 | 0 |
| 'Ncln' | 11.182 | 10.622 | 12.771 | 10.207 | 10.170 | 11.787 | 10.059 | 10.714 | 12.541 | 11.557 | 10.260 | 13.587 | 11.263 | 10.097 | 11.822 | 11.177 | 10.908 | 12.594 | -0.056 | 0.0908 | 0.2777 | 0 | 0 | 0 |
| 'Ncoa1' | 13.008 | 13.304 | 11.334 | 12.531 | 13.537 | 11.923 | 12.252 | 12.853 | 11.363 | 13.030 | 12.577 | 11.681 | 13.285 | 13.224 | 10.730 | 12.353 | 12.790 | 11.991 | -0.006 | 0.8388 | 0.9311 | 0 | 0 | 0 |
| 'Ncoa2' | 6.098 | 5.270 | 5.859 | 5.586 | 5.561 | 5.608 | 5.883 | 5.757 | 5.998 | 5.860 | 5.357 | 6.014 | 5.184 | 5.499 | 5.690 | 5.612 | 5.051 | 5.803 | 0.034 | 0.2915 | 0.5482 | 0 | 0 | 0 |
| 'Ncoa3' | 32.047 | 28.225 | 28.037 | 29.481 | 28.547 | 28.733 | 27.344 | 27.576 | 29.322 | 26.882 | 27.327 | 29.722 | 23.721 | 26.294 | 29.198 | 26.631 | 25.926 | 30.810 | 0.065 | 0.1334 | 0.3518 | 0 | 0 | 0 |
| 'Ncoa5' | 8.981 | 9.002 | 9.976 | 8.324 | 8.552 | 9.720 | 8.191 | 8.817 | 10.128 | 9.828 | 8.150 | 9.815 | 8.601 | 8.854 | 9.336 | 8.951 | 8.194 | 8.863 | 0.009 | 0.7961 | 0.9092 | 0 | 0 | 0 |
| 'Ncoa6' | 7.501 | 6.565 | 7.251 | 7.036 | 6.690 | 7.189 | 5.996 | 7.068 | 6.855 | 7.959 | 7.110 | 7.256 | 7.213 | 7.129 | 7.366 | 7.426 | 6.775 | 7.210 | -0.082 | 0.0321 | 0.1412 | 0 | 0 | 0 |
| 'Ncoa7' | 9.672 | 10.307 | 12.031 | 10.251 | 9.984 | 11.402 | 11.840 | 10.488 | 11.744 | 10.660 | 11.449 | 12.371 | 9.437 | 10.825 | 12.937 | 9.432 | 11.444 | 12.203 | -0.052 | 0.2422 | 0.4928 | 0 | 0 | 0 |
| 'Ncor1' | 24.529 | 20.638 | 22.501 | 24.626 | 21.325 | 22.354 | 22.850 | 21.987 | 22.190 | 23.835 | 23.332 | 23.505 | 22.397 | 22.260 | 22.539 | 21.766 | 22.859 | 24.523 | -0.041 | 0.2414 | 0.4920 | 0 | 0 | 0 |
| 'Ncor2' | 17.758 | 16.120 | 19.391 | 16.607 | 15.706 | 19.005 | 15.213 | 17.259 | 18.718 | 19.641 | 17.389 | 18.305 | 19.332 | 17.239 | 17.817 | 17.671 | 16.276 | 18.830 | -0.073 | 0.0638 | 0.2204 | 0 | 0 | 0 |
| 'Ncstn' | 14.948 | 16.316 | 17.237 | 14.936 | 16.105 | 16.394 | 15.225 | 16.110 | 18.176 | 15.726 | 15.480 | 16.327 | 16.172 | 15.264 | 15.741 | 16.714 | 16.015 | 15.287 | 0.017 | 0.5970 | 0.7952 | 0 | 0 | 0 |
| 'Ndc1' | 1.697 | 1.720 | 2.441 | 1.905 | 1.975 | 1.909 | 1.962 | 1.969 | 2.136 | 1.861 | 2.031 | 2.020 | 1.782 | 2.037 | 1.992 | 1.636 | 2.013 | 1.963 | 0.025 | 0.6666 | 0.8393 | 0 | 0 | 0 |
| 'Ndc80' | 8.522 | 10.343 | 13.663 | 13.117 | 11.104 | 12.966 | 12.198 | 10.635 | 12.421 | 7.629 | 10.072 | 11.691 | 12.230 | 12.846 | 12.454 | 10.287 | 12.115 | 9.923 | 0.068 | 0.4408 | 0.6843 | 0 | 0 | 0 |
| 'Nde1' | 8.727 | 9.376 | 9.740 | 9.631 | 9.167 | 10.169 | 10.416 | 9.039 | 9.414 | 7.982 | 9.030 | 10.485 | 9.061 | 9.199 | 10.574 | 8.503 | 9.689 | 9.391 | 0.019 | 0.6583 | 0.8350 | 0 | 0 | 0 |
| 'Ndel1' | 6.736 | 6.806 | 6.122 | 6.275 | 7.456 | 5.482 | 6.192 | 6.949 | 6.424 | 5.953 | 6.544 | 6.945 | 6.114 | 6.800 | 7.033 | 5.671 | 7.129 | 6.776 | -0.029 | 0.5664 | 0.7746 | 0 | 0 | 0 |
| 'Ndfip1' | 305.652 | 366.544 | 318.648 | 304.197 | 360.088 | 312.170 | 319.789 | 351.892 | 312.024 | 317.973 | 318.818 | 281.302 | 296.796 | 334.560 | 301.055 | 327.199 | 326.456 | 288.961 | 0.068 | 0.0134 | 0.0775 | 0 | 0 | 0 |
| 'Ndfip2' | 2.557 | 3.183 | 2.738 | 2.804 | 3.340 | 3.021 | 2.843 | 3.272 | 2.763 | 2.785 | 3.062 | 2.692 | 3.132 | 2.985 | 2.515 | 2.742 | 2.853 | 2.542 | 0.055 | 0.1769 | 0.4122 | 0 | 0 | 0 |
| 'Ndnl2' | 5.289 | 5.295 | 4.955 | 4.397 | 5.662 | 5.196 | 4.514 | 5.181 | 4.630 | 4.386 | 4.372 | 4.925 | 5.189 | 5.281 | 5.008 | 5.596 | 4.704 | 4.528 | 0.026 | 0.6391 | 0.8223 | 0 | 0 | 0 |
| 'Ndor1' | 5.044 | 4.632 | 6.060 | 4.715 | 4.754 | 5.161 | 4.799 | 5.116 | 5.528 | 4.764 | 4.952 | 5.630 | 5.216 | 4.511 | 5.127 | 4.792 | 4.825 | 5.213 | 0.014 | 0.6963 | 0.8571 | 0 | 0 | 0 |
| 'Ndrg1' | 0.814 | 0.909 | 0.620 | 0.947 | 0.869 | 0.637 | 0.847 | 0.791 | 0.714 | 0.566 | 0.745 | 1.199 | 0.760 | 0.773 | 1.099 | 0.700 | 0.911 | 0.927 | -0.118 | 0.3495 | 0.6044 | 0 | 0 | 0 |
| 'Ndrg2' | 6.545 | 9.584 | 7.092 | 6.905 | 7.431 | 7.706 | 7.225 | 8.189 | 7.818 | 6.882 | 7.131 | 6.445 | 6.565 | 7.099 | 6.964 | 7.472 | 5.471 | 6.867 | 0.157 | 0.0223 | 0.1110 | 0 | 0 | 0 |
| 'Ndrg3' | 18.801 | 20.619 | 19.258 | 19.891 | 21.452 | 19.305 | 18.766 | 22.778 | 18.354 | 18.045 | 20.015 | 16.539 | 18.793 | 18.030 | 16.764 | 16.309 | 18.330 | 16.470 | 0.159 | 0.0000 | 0.0000 | 1 | 0 | 0 |
| 'Ndst1' | 22.252 | 24.549 | 22.639 | 19.114 | 21.881 | 23.071 | 19.385 | 22.713 | 23.295 | 23.791 | 23.145 | 21.328 | 22.091 | 21.954 | 21.494 | 23.127 | 20.857 | 21.957 | -0.017 | 0.6962 | 0.8571 | 0 | 0 | 0 |
| 'Ndst2' | 16.092 | 18.358 | 19.395 | 14.945 | 17.631 | 18.775 | 15.813 | 17.278 | 18.410 | 19.315 | 14.652 | 16.793 | 17.309 | 16.149 | 16.192 | 17.339 | 14.878 | 14.683 | 0.080 | 0.1485 | 0.3749 | 0 | 0 | 0 |
| 'Ndufa1' | 99.229 | 126.014 | 97.501 | 100.294 | 115.742 | 107.506 | 109.368 | 115.807 | 104.783 | 96.599 | 103.158 | 95.019 | 113.831 | 108.737 | 103.098 | 112.811 | 115.609 | 100.621 | 0.028 | 0.5278 | 0.7486 | 0 | 0 | 0 |
| 'Ndufa10' | 12.861 | 11.972 | 13.313 | 13.133 | 12.496 | 13.878 | 12.960 | 11.667 | 13.345 | 13.207 | 12.272 | 13.383 | 14.540 | 12.945 | 13.144 | 13.006 | 12.802 | 12.740 | -0.040 | 0.1951 | 0.4335 | 0 | 0 | 0 |
| 'Ndufa11' | 13.598 | 15.331 | 13.884 | 12.639 | 14.612 | 14.634 | 12.729 | 15.480 | 13.832 | 13.337 | 14.024 | 13.435 | 14.130 | 13.642 | 15.025 | 14.078 | 12.965 | 12.663 | 0.030 | 0.4868 | 0.7174 | 0 | 0 | 0 |
| 'Ndufa12' | 31.969 | 36.991 | 30.986 | 39.022 | 35.257 | 35.501 | 35.475 | 34.888 | 34.628 | 32.406 | 32.004 | 31.775 | 34.939 | 32.847 | 33.340 | 33.544 | 34.809 | 32.436 | 0.065 | 0.0990 | 0.2941 | 0 | 0 | 0 |
| 'Ndufa13' | 61.580 | 70.123 | 65.273 | 64.833 | 68.393 | 65.666 | 63.993 | 67.664 | 64.124 | 58.948 | 61.249 | 57.504 | 62.737 | 66.037 | 62.019 | 67.057 | 63.698 | 59.345 | 0.075 | 0.0172 | 0.0929 | 0 | 0 | 0 |
| 'Ndufa2' | 99.873 | 98.448 | 96.158 | 93.903 | 99.982 | 91.821 | 101.815 | 101.575 | 94.822 | 84.745 | 91.875 | 87.707 | 97.058 | 88.467 | 94.394 | 89.938 | 98.851 | 87.223 | 0.088 | 0.0123 | 0.0727 | 0 | 0 | 0 |
| 'Ndufa3' | 28.944 | 34.391 | 31.207 | 33.868 | 35.348 | 30.922 | 33.592 | 33.997 | 31.002 | 28.138 | 29.524 | 28.362 | 29.949 | 32.905 | 30.738 | 30.762 | 32.551 | 28.783 | 0.098 | 0.0048 | 0.0362 | 1 | 0 | 0 |
| 'Ndufa4' | 46.931 | 58.651 | 50.564 | 52.568 | 56.019 | 53.976 | 48.892 | 57.015 | 50.339 | 46.718 | 48.468 | 46.780 | 51.895 | 53.913 | 53.723 | 52.588 | 50.501 | 48.194 | 0.057 | 0.1512 | 0.3781 | 0 | 0 | 0 |
| 'Ndufa5' | 20.163 | 22.229 | 19.586 | 19.160 | 20.409 | 21.982 | 19.334 | 21.597 | 19.529 | 19.198 | 19.683 | 17.197 | 20.887 | 19.850 | 18.389 | 19.857 | 19.988 | 18.405 | 0.081 | 0.0637 | 0.2202 | 0 | 0 | 0 |
| 'Ndufa6' | 101.686 | 105.914 | 101.705 | 109.343 | 111.966 | 103.273 | 113.168 | 105.518 | 97.243 | 95.607 | 101.929 | 97.407 | 111.091 | 100.912 | 103.809 | 103.834 | 110.895 | 97.207 | 0.030 | 0.4070 | 0.6562 | 0 | 0 | 0 |
| 'Ndufa7' | 11.889 | 14.591 | 13.142 | 12.571 | 14.095 | 14.369 | 12.645 | 15.413 | 13.397 | 12.878 | 11.840 | 12.998 | 13.071 | 14.032 | 13.109 | 13.741 | 12.212 | 12.743 | 0.054 | 0.2171 | 0.4638 | 0 | 0 | 0 |
| 'Ndufa8' | 65.329 | 78.091 | 69.479 | 67.441 | 75.656 | 73.525 | 68.087 | 76.332 | 66.676 | 62.027 | 67.758 | 69.302 | 72.889 | 70.670 | 72.521 | 81.592 | 67.188 | 70.384 | 0.002 | 0.9658 | 0.9879 | 0 | 0 | 0 |
| 'Ndufa9' | 22.108 | 26.453 | 24.423 | 23.142 | 26.548 | 25.386 | 26.241 | 26.722 | 24.803 | 22.024 | 25.032 | 23.112 | 24.718 | 24.778 | 22.845 | 25.242 | 24.647 | 21.361 | 0.069 | 0.0513 | 0.1917 | 0 | 0 | 0 |
| 'Ndufab1' | 8.227 | 8.335 | 9.227 | 8.769 | 8.963 | 8.506 | 8.926 | 8.910 | 8.995 | 7.784 | 8.084 | 9.524 | 8.082 | 9.198 | 9.058 | 8.173 | 8.631 | 8.866 | 0.013 | 0.7402 | 0.8822 | 0 | 0 | 0 |
| 'Ndufaf1' | 7.012 | 8.030 | 8.307 | 7.533 | 7.830 | 6.913 | 6.976 | 7.057 | 7.880 | 7.046 | 7.987 | 6.799 | 6.995 | 7.855 | 6.790 | 6.384 | 7.986 | 6.933 | 0.056 | 0.3359 | 0.5946 | 0 | 0 | 0 |
| 'Ndufaf2' | 12.816 | 13.096 | 14.980 | 13.651 | 16.964 | 12.821 | 13.147 | 13.332 | 12.620 | 14.572 | 11.926 | 14.077 | 13.243 | 12.380 | 14.294 | 11.485 | 12.281 | 13.000 | 0.054 | 0.4792 | 0.7118 | 0 | 0 | 0 |
| 'Ndufaf3' | 10.369 | 10.906 | 11.499 | 11.334 | 11.864 | 11.640 | 10.534 | 11.870 | 11.231 | 11.207 | 9.765 | 10.209 | 10.519 | 10.558 | 11.429 | 10.412 | 10.770 | 10.637 | 0.078 | 0.1087 | 0.3105 | 0 | 0 | 0 |
| 'Ndufaf4' | 3.421 | 3.397 | 3.488 | 3.568 | 3.719 | 3.390 | 3.330 | 3.650 | 3.371 | 3.255 | 3.397 | 3.747 | 3.012 | 3.496 | 3.468 | 3.084 | 3.911 | 3.656 | 0.000 | 0.9982 | 0.9994 | 0 | 0 | 0 |
| 'Ndufaf5' | 4.397 | 4.830 | 4.495 | 4.582 | 5.165 | 4.267 | 4.851 | 4.913 | 4.784 | 4.849 | 4.902 | 4.862 | 4.575 | 4.422 | 4.815 | 3.864 | 4.461 | 3.943 | 0.035 | 0.5765 | 0.7816 | 0 | 0 | 0 |
| 'Ndufaf6' | 2.839 | 2.477 | 2.717 | 2.899 | 2.681 | 3.087 | 2.886 | 2.858 | 2.384 | 2.712 | 2.577 | 2.938 | 3.193 | 2.831 | 2.912 | 2.574 | 2.746 | 2.951 | -0.049 | 0.4856 | 0.7168 | 0 | 0 | 0 |
| 'Ndufaf7' | 9.850 | 13.190 | 11.946 | 10.606 | 12.374 | 10.590 | 11.846 | 11.842 | 11.931 | 10.410 | 10.624 | 9.696 | 10.724 | 9.659 | 9.482 | 10.605 | 10.652 | 9.034 | 0.186 | 0.0003 | 0.0039 | 1 | 0 | 0 |
| 'Ndufb10' | 19.273 | 19.122 | 19.902 | 19.569 | 20.936 | 20.572 | 20.152 | 20.387 | 20.784 | 18.437 | 19.648 | 17.705 | 19.331 | 19.432 | 18.891 | 18.873 | 19.650 | 18.034 | 0.083 | 0.0099 | 0.0618 | 0 | 0 | 0 |
| 'Ndufb11' | 117.062 | 126.581 | 112.883 | 119.958 | 123.569 | 117.172 | 121.023 | 127.601 | 113.445 | 111.847 | 119.396 | 103.894 | 119.960 | 116.704 | 116.532 | 121.074 | 126.193 | 106.383 | 0.043 | 0.1469 | 0.3725 | 0 | 0 | 0 |
| 'Ndufb2' | 2.949 | 3.521 | 3.472 | 3.182 | 3.326 | 3.086 | 3.189 | 3.189 | 3.066 | 2.866 | 3.041 | 3.283 | 3.525 | 3.497 | 3.195 | 3.302 | 2.976 | 2.664 | 0.019 | 0.7326 | 0.8781 | 0 | 0 | 0 |

| | | | | | | | | | | | | | | | | | | | | | | | | |
|---|---|---|---|---|---|---|---|---|---|---|---|---|---|---|---|---|---|---|---|---|---|---|---|---|
| 'Ndufb3' | 45.144 | 59.638 | 50.879 | 55.076 | 54.157 | 52.771 | 56.544 | 54.621 | 52.849 | 47.134 | 49.529 | 48.443 | 48.777 | 53.934 | 47.731 | 51.609 | 57.366 | 47.586 | 0.079 | 0.0674 | 0.2275 | 0 | 0 | 0 |
| 'Ndufb4' | 19.221 | 21.908 | 19.341 | 21.525 | 21.432 | 21.581 | 19.798 | 21.438 | 20.820 | 18.290 | 18.038 | 18.659 | 19.419 | 19.907 | 20.121 | 20.486 | 21.114 | 17.354 | 0.100 | 0.0260 | 0.1239 | 0 | 0 | 0 |
| 'Ndufb5' | 10.329 | 11.375 | 10.748 | 10.566 | 10.632 | 10.527 | 10.783 | 10.631 | 10.394 | 10.245 | 10.462 | 10.561 | 10.799 | 11.139 | 10.959 | 10.194 | 10.888 | 10.069 | -0.002 | 0.9571 | 0.9840 | 0 | 0 | 0 |
| 'Ndufb6' | 31.856 | 38.439 | 33.057 | 35.923 | 39.149 | 34.172 | 36.705 | 34.448 | 32.898 | 33.549 | 31.593 | 32.264 | 36.165 | 34.258 | 31.032 | 36.204 | 33.384 | 31.661 | 0.062 | 0.1302 | 0.3464 | 0 | 0 | 0 |
| 'Ndufb7' | 14.580 | 15.448 | 14.611 | 13.791 | 16.715 | 15.683 | 14.025 | 14.764 | 15.905 | 13.234 | 12.583 | 13.768 | 15.323 | 13.712 | 14.751 | 15.632 | 14.000 | 13.560 | 0.088 | 0.0455 | 0.1772 | 0 | 0 | 0 |
| 'Ndufb8' | 11.348 | 13.078 | 11.317 | 11.485 | 12.380 | 11.855 | 11.030 | 12.085 | 11.877 | 10.534 | 9.713 | 11.432 | 12.002 | 11.508 | 11.558 | 11.660 | 10.953 | 11.087 | 0.070 | 0.0751 | 0.2438 | 0 | 0 | 0 |
| 'Ndufb9' | 85.155 | 105.329 | 96.214 | 93.267 | 99.510 | 100.960 | 95.700 | 101.369 | 94.939 | 89.484 | 80.667 | 88.479 | 103.190 | 98.207 | 96.092 | 100.853 | 89.611 | 89.398 | 0.051 | 0.2709 | 0.5261 | 0 | 0 | 0 |
| 'Ndufc1' | 9.031 | 10.647 | 10.037 | 10.456 | 11.140 | 11.463 | 10.189 | 11.024 | 9.296 | 8.464 | 9.393 | 9.639 | 10.735 | 10.541 | 9.862 | 11.094 | 11.108 | 9.778 | 0.031 | 0.5784 | 0.7820 | 0 | 0 | 0 |
| 'Ndufc2' | 39.681 | 51.714 | 40.901 | 41.426 | 47.474 | 41.666 | 46.125 | 48.584 | 42.001 | 39.193 | 44.862 | 39.082 | 42.029 | 41.810 | 41.252 | 43.654 | 43.160 | 36.112 | 0.093 | 0.0161 | 0.0882 | 0 | 0 | 0 |
| 'Ndufs1' | 9.606 | 10.160 | 10.251 | 10.266 | 10.621 | 9.871 | 10.544 | 10.379 | 9.932 | 10.019 | 10.546 | 9.451 | 10.610 | 10.367 | 9.384 | 10.055 | 10.623 | 9.018 | 0.017 | 0.5236 | 0.7461 | 0 | 0 | 0 |
| 'Ndufs2' | 35.028 | 38.138 | 36.643 | 35.531 | 40.415 | 38.979 | 35.220 | 38.284 | 38.452 | 36.899 | 37.188 | 37.095 | 40.265 | 38.331 | 37.900 | 38.473 | 39.395 | 36.398 | -0.033 | 0.2260 | 0.4726 | 0 | 0 | 0 |
| 'Ndufs3' | 20.202 | 21.537 | 22.088 | 21.079 | 21.931 | 20.848 | 21.131 | 21.835 | 21.985 | 21.058 | 21.775 | 21.617 | 22.773 | 21.440 | 23.265 | 20.905 | 21.850 | 20.580 | -0.030 | 0.3681 | 0.6200 | 0 | 0 | 0 |
| 'Ndufs4' | 28.011 | 32.630 | 27.991 | 31.761 | 32.035 | 29.440 | 31.222 | 30.823 | 26.819 | 28.097 | 28.485 | 27.632 | 30.203 | 28.599 | 28.460 | 29.216 | 30.214 | 25.679 | 0.062 | 0.0877 | 0.2701 | 0 | 0 | 0 |
| 'Ndufs5' | 20.864 | 26.946 | 23.782 | 22.640 | 24.863 | 24.495 | 22.858 | 24.942 | 22.404 | 20.036 | 19.009 | 23.187 | 23.758 | 21.977 | 22.638 | 25.775 | 22.473 | 22.948 | 0.068 | 0.2047 | 0.4477 | 0 | 0 | 0 |
| 'Ndufs6' | 41.886 | 48.981 | 43.834 | 46.615 | 48.109 | 46.939 | 45.200 | 46.086 | 46.224 | 43.878 | 44.295 | 40.456 | 46.400 | 46.079 | 44.725 | 44.770 | 45.980 | 43.140 | 0.041 | 0.2620 | 0.5159 | 0 | 0 | 0 |
| 'Ndufs7' | 16.323 | 18.836 | 18.034 | 16.987 | 18.304 | 18.567 | 16.378 | 18.939 | 17.299 | 17.183 | 16.029 | 17.777 | 18.601 | 17.472 | 18.200 | 18.615 | 17.016 | 16.854 | 0.006 | 0.8654 | 0.9415 | 0 | 0 | 0 |
| 'Ndufs8' | 65.575 | 76.255 | 69.046 | 69.087 | 76.731 | 75.070 | 65.570 | 72.166 | 73.833 | 73.290 | 61.235 | 69.167 | 73.385 | 70.434 | 71.045 | 73.984 | 66.803 | 67.866 | 0.025 | 0.5293 | 0.7498 | 0 | 0 | 0 |
| 'Ndufv1' | 15.388 | 17.415 | 16.195 | 16.080 | 17.685 | 16.517 | 16.331 | 17.721 | 17.075 | 16.389 | 15.843 | 15.751 | 16.615 | 16.184 | 16.362 | 16.702 | 15.648 | 15.352 | 0.043 | 0.1241 | 0.3368 | 0 | 0 | 0 |
| 'Ndufv2' | 46.355 | 54.290 | 47.780 | 52.606 | 51.715 | 45.931 | 52.795 | 52.603 | 47.092 | 45.170 | 47.063 | 44.798 | 50.673 | 47.353 | 46.822 | 46.998 | 48.478 | 44.435 | 0.083 | 0.0076 | 0.0508 | 0 | 0 | 0 |
| 'Ndufv3' | 20.680 | 23.114 | 20.323 | 22.095 | 21.699 | 21.448 | 22.533 | 21.390 | 21.078 | 19.406 | 20.213 | 19.645 | 21.528 | 20.578 | 21.001 | 20.808 | 20.104 | 18.250 | 0.087 | 0.0070 | 0.0483 | 1 | 0 | 0 |
| 'Necap1' | 4.495 | 4.872 | 4.110 | 4.170 | 4.959 | 4.073 | 4.620 | 4.998 | 3.830 | 4.152 | 4.907 | 4.535 | 4.428 | 5.313 | 4.765 | 5.365 | 5.039 | 4.511 | -0.113 | 0.0313 | 0.1387 | 0 | 0 | 0 |
| 'Necap2' | 14.924 | 14.691 | 15.628 | 14.002 | 14.755 | 15.508 | 14.852 | 15.707 | 15.523 | 12.997 | 14.228 | 14.666 | 14.402 | 14.142 | 15.066 | 13.646 | 14.278 | 14.528 | 0.074 | 0.0041 | 0.0323 | 1 | 0 | 0 |
| 'Nedd1' | 7.792 | 8.988 | 9.360 | 9.556 | 9.214 | 8.522 | 9.278 | 9.563 | 9.097 | 8.287 | 8.753 | 9.193 | 8.367 | 10.207 | 9.035 | 7.954 | 10.055 | 8.335 | 0.010 | 0.8326 | 0.9270 | 0 | 0 | 0 |
| 'Nedd8' | 95.719 | 109.551 | 97.167 | 98.175 | 108.092 | 97.958 | 101.488 | 102.706 | 99.399 | 93.845 | 101.625 | 94.892 | 104.010 | 104.023 | 101.216 | 101.975 | 105.283 | 95.360 | 0.001 | 0.9573 | 0.9840 | 0 | 0 | 0 |
| 'Nedd9' | 53.927 | 53.829 | 48.139 | 52.838 | 52.876 | 49.265 | 51.000 | 51.935 | 50.613 | 57.212 | 53.133 | 49.159 | 52.521 | 53.571 | 48.130 | 59.799 | 54.063 | 47.486 | -0.040 | 0.1446 | 0.3690 | 0 | 0 | 0 |
| 'Nefh' | 9.866 | 5.569 | 11.328 | 8.758 | 7.152 | 10.387 | 9.559 | 6.824 | 9.998 | 7.294 | 6.287 | 8.296 | 7.249 | 6.060 | 7.330 | 6.397 | 5.384 | 8.521 | 0.320 | 0.0000 | 0.0000 | 1 | 0 | 0 |
| 'Neil1' | 12.715 | 17.057 | 16.604 | 14.571 | 16.535 | 17.322 | 14.542 | 17.018 | 16.846 | 17.585 | 16.529 | 14.684 | 19.488 | 18.694 | 14.635 | 17.085 | 16.168 | 13.934 | -0.062 | 0.3454 | 0.6007 | 0 | 0 | 0 |
| 'Neil2' | 3.433 | 4.391 | 3.982 | 2.892 | 4.286 | 3.813 | 3.588 | 3.569 | 3.786 | 4.386 | 4.195 | 4.115 | 3.025 | 4.153 | 3.988 | 3.629 | 3.151 | 3.431 | -0.028 | 0.7572 | 0.8913 | 0 | 0 | 0 |
| 'Neil3' | 2.135 | 2.257 | 2.734 | 2.554 | 2.084 | 2.316 | 2.745 | 2.296 | 2.985 | 1.719 | 2.587 | 2.516 | 3.017 | 3.078 | 3.022 | 2.096 | 2.979 | 2.660 | -0.107 | 0.2654 | 0.5188 | 0 | 0 | 0 |
| 'Nek1' | 1.401 | 1.205 | 1.335 | 1.637 | 1.596 | 1.335 | 1.644 | 1.374 | 1.318 | 1.525 | 1.470 | 1.620 | 1.380 | 1.489 | 1.426 | 1.359 | 1.601 | 1.453 | -0.072 | 0.2457 | 0.4961 | 0 | 0 | 0 |
| 'Nek2' | 4.100 | 4.804 | 6.457 | 4.766 | 5.072 | 5.938 | 5.430 | 5.238 | 6.585 | 3.444 | 5.276 | 5.060 | 5.581 | 5.643 | 5.838 | 4.581 | 4.961 | 5.117 | 0.074 | 0.3290 | 0.5872 | 0 | 0 | 0 |
| 'Nek3' | 1.361 | 1.516 | 1.920 | 2.253 | 1.727 | 1.763 | 1.640 | 2.115 | 1.530 | 1.741 | 1.661 | 1.882 | 1.589 | 1.944 | 1.997 | 1.480 | 1.678 | 1.630 | 0.008 | 0.9337 | 0.9746 | 0 | 0 | 0 |
| 'Nek4' | 5.748 | 7.150 | 6.603 | 5.879 | 7.163 | 6.547 | 6.010 | 6.950 | 7.047 | 7.092 | 7.805 | 6.779 | 7.153 | 7.605 | 6.279 | 7.205 | 6.605 | 5.862 | -0.084 | 0.1343 | 0.3530 | 0 | 0 | 0 |
| 'Nek6' | 0.406 | 0.329 | 0.320 | 0.447 | 0.326 | 0.304 | 0.432 | 0.292 | 0.299 | 0.418 | 0.364 | 0.366 | 0.398 | 0.318 | 0.315 | 0.360 | 0.342 | 0.264 | -0.024 | 0.8020 | 0.9114 | 0 | 0 | 0 |
| 'Nek7' | 44.381 | 51.518 | 46.569 | 49.542 | 53.232 | 44.087 | 49.693 | 49.409 | 44.448 | 45.224 | 50.517 | 43.914 | 47.836 | 52.549 | 41.940 | 44.109 | 52.204 | 41.320 | 0.036 | 0.1940 | 0.4317 | 0 | 0 | 0 |
| 'Nek8' | 1.831 | 1.719 | 1.909 | 1.946 | 1.793 | 2.026 | 1.954 | 1.689 | 2.123 | 1.936 | 1.617 | 1.989 | 2.055 | 1.761 | 2.080 | 1.737 | 1.992 | 1.824 | -0.013 | 0.7911 | 0.9067 | 0 | 0 | 0 |
| 'Nek9' | 45.133 | 43.670 | 45.898 | 44.054 | 44.115 | 43.999 | 41.531 | 43.494 | 45.632 | 44.989 | 46.440 | 45.766 | 42.078 | 41.873 | 43.206 | 44.336 | 46.172 | 42.649 | -0.010 | 0.7312 | 0.8773 | 0 | 0 | 0 |
| 'Nelfa' | 9.669 | 11.671 | 11.569 | 8.971 | 10.337 | 11.216 | 10.004 | 9.576 | 10.933 | 9.286 | 8.915 | 11.825 | 10.371 | 10.214 | 10.458 | 9.933 | 9.207 | 11.645 | 0.018 | 0.6933 | 0.8558 | 0 | 0 | 0 |
| 'Nelfb' | 17.591 | 17.605 | 17.862 | 17.154 | 17.636 | 17.920 | 17.068 | 17.306 | 17.992 | 16.933 | 18.623 | 20.506 | 16.984 | 17.601 | 19.280 | 16.862 | 18.055 | 20.758 | -0.076 | 0.0152 | 0.0847 | 0 | 0 | 0 |
| 'Nelfcd' | 15.205 | 17.057 | 17.769 | 16.896 | 15.930 | 17.718 | 15.817 | 16.396 | 17.767 | 15.877 | 15.248 | 15.730 | 17.597 | 15.438 | 15.528 | 16.434 | 14.656 | 15.282 | 0.077 | 0.0326 | 0.1428 | 0 | 0 | 0 |
| 'Nelfe' | 13.689 | 13.413 | 14.510 | 12.858 | 14.365 | 14.974 | 14.124 | 14.205 | 14.768 | 14.337 | 12.888 | 13.306 | 15.911 | 14.405 | 14.300 | 14.407 | 12.519 | 13.519 | 0.008 | 0.8499 | 0.9353 | 0 | 0 | 0 |
| 'Nemf' | 27.877 | 26.980 | 26.645 | 27.862 | 26.728 | 25.002 | 28.572 | 29.026 | 26.524 | 27.943 | 26.473 | 25.299 | 25.758 | 27.038 | 23.269 | 28.334 | 28.443 | 23.788 | 0.046 | 0.1283 | 0.3432 | 0 | 0 | 0 |
| 'Nemp2' | 10.419 | 12.456 | 12.701 | 10.476 | 12.578 | 11.404 | 11.381 | 13.337 | 12.402 | 12.869 | 13.594 | 10.760 | 13.283 | 12.769 | 10.843 | 11.577 | 13.011 | 10.007 | -0.028 | 0.5770 | 0.7818 | 0 | 0 | 0 |
| 'Nenf' | 13.208 | 14.485 | 12.922 | 13.985 | 16.919 | 13.430 | 11.877 | 13.007 | 12.867 | 13.070 | 11.353 | 11.799 | 11.326 | 12.280 | 12.000 | 11.977 | 10.520 | 11.406 | 0.199 | 0.0064 | 0.0451 | 1 | 0 | 0 |
| 'Nepro' | 3.564 | 3.427 | 3.626 | 3.328 | 3.793 | 3.283 | 3.167 | 3.879 | 3.350 | 3.243 | 3.245 | 3.191 | 3.573 | 3.409 | 3.023 | 3.626 | 3.315 | 3.168 | 0.071 | 0.1186 | 0.3278 | 0 | 0 | 0 |
| 'Net1' | 12.234 | 12.005 | 11.653 | 11.759 | 12.168 | 11.974 | 11.640 | 11.920 | 12.792 | 11.309 | 11.361 | 11.767 | 13.155 | 13.271 | 11.941 | 12.240 | 11.822 | 11.756 | -0.015 | 0.6963 | 0.8571 | 0 | 0 | 0 |
| 'Neu1' | 15.338 | 13.318 | 14.597 | 14.667 | 14.034 | 13.975 | 15.542 | 13.988 | 14.258 | 13.576 | 14.932 | 15.216 | 13.139 | 13.509 | 15.467 | 14.623 | 13.698 | 15.857 | -0.014 | 0.7358 | 0.8797 | 0 | 0 | 0 |
| 'Neu3' | 13.000 | 13.221 | 14.271 | 15.860 | 13.506 | 13.259 | 16.262 | 12.910 | 15.040 | 10.986 | 14.422 | 15.360 | 11.042 | 12.009 | 14.700 | 10.961 | 14.374 | 14.636 | 0.098 | 0.1679 | 0.4003 | 0 | 0 | 0 |
| 'Neurl3' | 28.628 | 32.545 | 27.397 | 25.441 | 32.419 | 31.542 | 26.168 | 32.485 | 28.053 | 26.680 | 26.057 | 25.721 | 20.184 | 25.133 | 24.871 | 24.832 | 22.921 | 23.457 | 0.253 | 0.0000 | 0.0000 | 1 | 0 | 0 |
| 'Neurl4' | 10.734 | 9.709 | 11.930 | 10.761 | 10.632 | 11.468 | 9.835 | 10.416 | 11.593 | 11.211 | 10.802 | 11.667 | 11.143 | 10.032 | 10.326 | 10.406 | 10.963 | 10.991 | -0.018 | 0.6027 | 0.7987 | 0 | 0 | 0 |
| 'Nf1' | 1.801 | 1.889 | 1.782 | 1.694 | 1.736 | 1.625 | 1.851 | 2.027 | 1.689 | 1.722 | 2.066 | 1.665 | 1.488 | 1.622 | 1.481 | 1.668 | 1.880 | 1.804 | 0.055 | 0.3061 | 0.5640 | 0 | 0 | 0 |
| 'Nf2' | 13.279 | 13.415 | 13.545 | 12.646 | 13.853 | 14.329 | 13.348 | 13.307 | 13.856 | 13.356 | 12.754 | 14.165 | 12.983 | 14.141 | 13.696 | 13.345 | 13.154 | 13.848 | -0.009 | 0.6948 | 0.8564 | 0 | 0 | 0 |
| 'Nfat5' | 6.690 | 5.877 | 6.299 | 6.118 | 5.414 | 5.575 | 5.891 | 5.704 | 6.343 | 6.978 | 6.255 | 6.698 | 6.484 | 5.836 | 5.642 | 6.470 | 6.303 | 6.739 | -0.101 | 0.0135 | 0.0776 | 0 | 0 | 0 |
| 'Nfatc1' | 80.911 | 78.342 | 84.504 | 70.855 | 75.234 | 85.149 | 69.228 | 78.847 | 86.714 | 89.461 | 80.270 | 87.947 | 77.199 | 84.714 | 89.716 | 81.980 | 78.521 | 93.252 | -0.116 | 0.0012 | 0.0119 | 1 | 0 | 0 |
| 'Nfatc2' | 12.147 | 10.838 | 10.889 | 12.155 | 10.487 | 10.740 | 11.060 | 10.998 | 11.825 | 12.401 | 11.506 | 13.187 | 11.091 | 10.990 | 12.446 | 11.244 | 10.913 | 14.293 | -0.104 | 0.0231 | 0.1139 | 0 | 0 | 0 |
| 'Nfatc2ip' | 11.846 | 13.319 | 13.849 | 12.455 | 12.704 | 12.973 | 13.851 | 13.341 | 13.801 | 11.908 | 12.501 | 12.948 | 12.776 | 12.764 | 13.268 | 12.424 | 12.543 | 13.160 | 0.037 | 0.2832 | 0.5394 | 0 | 0 | 0 |
| 'Nfatc3' | 39.375 | 38.086 | 37.777 | 38.134 | 39.149 | 36.833 | 36.656 | 38.643 | 36.447 | 38.232 | 40.347 | 39.225 | 35.074 | 39.090 | 38.290 | 36.680 | 38.157 | 37.953 | -0.019 | 0.4893 | 0.7192 | 0 | 0 | 0 |
| 'Nfe2l1' | 14.418 | 15.050 | 14.484 | 12.981 | 13.844 | 15.136 | 13.099 | 14.104 | 14.382 | 14.509 | 14.147 | 16.111 | 13.438 | 14.248 | 15.758 | 14.316 | 14.089 | 16.039 | -0.067 | 0.0282 | 0.1300 | 0 | 0 | 0 |
| 'Nfe2l2' | 11.045 | 10.502 | 11.453 | 12.066 | 9.676 | 11.250 | 12.446 | 10.145 | 10.458 | 9.530 | 11.583 | 13.213 | 8.724 | 10.886 | 12.318 | 9.435 | 11.960 | 12.881 | -0.026 | 0.7097 | 0.8653 | 0 | 0 | 0 |
| 'Nfia' | 1.556 | 1.694 | 1.097 | 1.472 | 1.562 | 1.048 | 1.557 | 1.571 | 1.174 | 1.495 | 1.962 | 1.404 | 1.973 | 1.948 | 1.346 | 1.637 | 1.943 | 1.475 | -0.273 | 0.0000 | 0.0000 | 1 | 0 | 0 |
| 'Nfic' | 4.596 | 4.409 | 5.037 | 4.083 | 3.957 | 4.553 | 4.850 | 4.492 | 5.122 | 4.663 | 3.727 | 4.065 | 3.921 | 3.962 | 4.313 | 4.442 | 3.554 | 3.688 | 0.172 | 0.0007 | 0.0083 | 1 | 0 | 0 |
| 'Nfil3' | 65.258 | 27.533 | 22.556 | 40.758 | 29.885 | 22.689 | 33.587 | 31.774 | 20.399 | 49.861 | 44.651 | 22.465 | 42.800 | 48.024 | 22.938 | 53.355 | 39.184 | 23.001 | -0.238 | 0.0513 | 0.1917 | 0 | 0 | 0 |
| 'Nfix' | 1.665 | 1.850 | 2.336 | 1.736 | 2.197 | 2.194 | 1.538 | 2.109 | 2.199 | 2.269 | 1.557 | 2.059 | 2.007 | 1.852 | 1.952 | 1.975 | 1.634 | 2.510 | -0.010 | 0.9049 | 0.9625 | 0 | 0 | 0 |
| 'Nfkb1' | 20.972 | 20.578 | 21.386 | 21.147 | 20.715 | 21.710 | 21.214 | 20.627 | 21.681 | 23.261 | 21.116 | 22.087 | 21.449 | 21.865 | 22.058 | 23.392 | 22.342 | 22.370 | -0.083 | 0.0001 | 0.0013 | 1 | 0 | 0 |
| 'Nfkb2' | 49.343 | 44.387 | 51.872 | 44.732 | 46.771 | 53.963 | 49.427 | 48.208 | 54.510 | 57.388 | 48.424 | 56.213 | 51.980 | 51.176 | 52.979 | 53.050 | 51.004 | 55.149 | -0.117 | 0.0001 | 0.0020 | 1 | 0 | 0 |
| 'Nfkbia' | 276.170 | 201.540 | 155.430 | 194.285 | 219.286 | 185.039 | 162.607 | 249.615 | 151.664 | 316.631 | 169.153 | 103.636 | 284.991 | 250.296 | 123.253 | 359.968 | 147.396 | 111.156 | 0.052 | 0.7639 | 0.8953 | 0 | 0 | 0 |
| 'Nfkbib' | 20.513 | 16.768 | 17.767 | 16.894 | 18.231 | 19.834 | 17.539 | 18.528 | 17.885 | 21.353 | 17.278 | 17.171 | 20.701 | 19.615 | 16.628 | 20.887 | 15.483 | 15.931 | -0.010 | 0.8576 | 0.9390 | 0 | 0 | 0 |
| 'Nfkbid' | 39.854 | 34.421 | 32.803 | 36.392 | 34.266 | 33.351 | 35.045 | 35.583 | 34.943 | 40.497 | 32.622 | 36.094 | 39.756 | 34.204 | 34.076 | 39.106 | 34.525 | 36.133 | -0.056 | 0.0704 | 0.2338 | 0 | 0 | 0 |
| 'Nfkbie' | 35.794 | 33.591 | 33.628 | 32.362 | 32.227 | 35.461 | 34.424 | 32.033 | 36.047 | 37.566 | 34.164 | 38.012 | 41.751 | 37.562 | 37.948 | 38.178 | 37.324 | 37.877 | -0.166 | 0.0000 | 0.0000 | 1 | 0 | 0 |
| 'Nfkbil1' | 16.580 | 14.657 | 17.493 | 15.354 | 16.736 | 19.037 | 15.588 | 16.949 | 17.541 | 17.116 | 14.671 | 16.007 | 15.913 | 14.844 | 16.833 | 16.601 | 14.244 | 14.541 | 0.087 | 0.0597 | 0.2110 | 0 | 0 | 0 |
| 'Nfrkb' | 6.935 | 6.278 | 7.118 | 6.765 | 6.291 | 6.646 | 6.781 | 6.920 | 7.185 | 6.398 | 6.935 | 7.092 | 6.517 | 6.465 | 6.565 | 6.196 | 6.371 | 7.043 | 0.022 | 0.5032 | 0.7302 | 0 | 0 | 0 |
| 'Nfs1' | 11.576 | 13.209 | 13.073 | 12.284 | 12.542 | 13.239 | 13.317 | 12.774 | 12.237 | 12.742 | 11.074 | 12.339 | 11.985 | 12.801 | 12.126 | 13.170 | 12.098 | 12.497 | 0.034 | 0.3501 | 0.6044 | 0 | 0 | 0 |
| 'Nfu1' | 4.127 | 4.733 | 4.337 | 4.420 | 5.273 | 4.763 | 4.602 | 5.131 | 4.504 | 4.451 | 3.703 | 4.068 | 4.362 | 4.162 | 3.907 | 4.296 | 4.702 | 3.689 | 0.154 | 0.0015 | 0.0143 | 1 | 0 | 0 |
| 'Nfx1' | 20.144 | 19.575 | 20.038 | 19.447 | 18.982 | 18.699 | 19.839 | 19.952 | 20.394 | 19.901 | 19.897 | 20.683 | 19.220 | 19.622 | 18.626 | 18.739 | 21.069 | 19.730 | -0.014 | 0.6205 | 0.8101 | 0 | 0 | 0 |
| 'Nfxl1' | 1.890 | 2.016 | 2.010 | 1.817 | 2.246 | 1.882 | 1.891 | 2.180 | 1.745 | 1.842 | 1.773 | 1.587 | 2.062 | 1.872 | 1.695 | 1.841 | 1.735 | 1.541 | 0.143 | 0.0054 | 0.0396 | 1 | 0 | 0 |
| 'Nfya' | 4.391 | 4.540 | 4.763 | 4.899 | 4.280 | 4.989 | 4.086 | 4.403 | 4.840 | 4.095 | 5.080 | 5.846 | 4.380 | 5.108 | 5.795 | 4.482 | 5.144 | 6.280 | -0.169 | 0.0008 | 0.0083 | 1 | 0 | 0 |
| 'Nfyb' | 6.162 | 6.867 | 7.134 | 6.093 | 7.275 | 6.634 | 6.309 | 7.007 | 6.377 | 6.612 | 6.600 | 6.935 | 6.922 | 7.053 | 6.563 | 6.183 | 7.012 | 6.059 | -0.014 | 0.6879 | 0.8525 | 0 | 0 | 0 |
| 'Nfyc' | 15.283 | 15.115 | 15.405 | 14.940 | 15.017 | 15.571 | 15.249 | 15.438 | 14.897 | 15.049 | 14.503 | 14.923 | 15.672 | 14.747 | 14.364 | 13.612 | 14.090 | 13.614 | 0.058 | 0.0543 | 0.1995 | 0 | 0 | 0 |
| 'Ngdn' | 53.437 | 56.255 | 55.415 | 54.367 | 55.391 | 53.805 | 57.441 | 59.270 | 50.909 | 49.638 | 53.758 | 55.641 | 48.968 | 57.354 | 53.512 | 51.606 | 54.164 | 51.326 | 0.048 | 0.1470 | 0.3726 | 0 | 0 | 0 |
| 'Ngfrap1' | 65.794 | 58.609 | 57.351 | 63.470 | 57.837 | 61.692 | 60.646 | 60.214 | 60.766 | 56.188 | 54.274 | 55.200 | 64.985 | 56.854 | 59.822 | 54.398 | 56.089 | 54.716 | 0.082 | 0.0409 | 0.1657 | 0 | 0 | 0 |
| 'Ngly1' | 12.945 | 16.013 | 12.218 | 13.348 | 14.274 | 11.781 | 14.249 | 13.920 | 11.566 | 14.875 | 16.299 | 13.815 | 17.244 | 17.037 | 13.079 | 15.042 | 19.193 | 12.713 | -0.220 | 0.0000 | 0.0000 | 1 | 0 | 0 |
| 'Ngrn' | 10.827 | 12.108 | 11.503 | 11.018 | 11.475 | 11.631 | 11.168 | 11.724 | 11.105 | 10.282 | 10.017 | 12.098 | 9.720 | 12.701 | 11.428 | 11.722 | 11.355 | 11.066 | 0.019 | 0.6792 | 0.8466 | 0 | 0 | 0 |
| 'Nhlrc2' | 11.733 | 11.148 | 11.484 | 12.029 | 11.014 | 10.850 | 11.103 | 10.813 | 10.729 | 10.477 | 12.288 | 12.519 | 11.625 | 11.675 | 11.594 | 11.431 | 10.541 | 12.098 | -0.060 | 0.1505 | 0.3777 | 0 | 0 | 0 |
| 'Nhlrc3' | 2.269 | 2.628 | 2.578 | 2.318 | 2.361 | 2.548 | 2.532 | 2.716 | 2.713 | 2.395 | 2.551 | 2.356 | 2.403 | 2.048 | 2.125 | 2.249 | 2.348 | 2.236 | 0.120 | 0.0103 | 0.0638 | 0 | 0 | 0 |
| 'Nhp2' | 21.183 | 21.221 | 23.166 | 21.208 | 20.019 | 22.554 | 22.903 | 22.265 | 21.901 | 21.246 | 19.946 | 20.838 | 23.062 | 18.893 | 22.953 | 23.143 | 18.292 | 20.847 | 0.048 | 0.2368 | 0.4865 | 0 | 0 | 0 |
| 'Nhsl1' | 1.005 | 1.782 | 1.253 | 1.194 | 1.442 | 1.342 | 1.003 | 1.637 | 1.303 | 1.300 | 1.599 | 1.357 | 1.634 | 1.860 | 1.347 | 1.609 | 1.731 | 1.223 | -0.209 | 0.0029 | 0.0248 | 1 | 0 | 0 |
| 'Nid2' | 1.142 | 1.175 | 1.186 | 1.132 | 0.883 | 1.111 | 1.383 | 1.073 | 0.992 | 0.677 | 0.517 | 0.735 | 0.806 | 0.859 | 0.707 | 0.723 | 0.743 | 0.611 | 0.643 | 0.0000 | 0.0000 | 1 | 1 | 0 |
| 'Nif3l1' | 2.126 | 2.102 | 2.384 | 2.207 | 2.458 | 2.194 | 2.114 | 2.150 | 2.202 | 2.457 | 2.102 | 2.225 | 2.287 | 2.243 | 2.086 | 2.195 | 1.743 | 2.366 | 0.007 | 0.8934 | 0.9565 | 0 | 0 | 0 |
| 'Nifk' | 19.230 | 18.848 | 17.727 | 19.323 | 19.420 | 18.361 | 20.086 | 19.498 | 17.245 | 18.808 | 16.685 | 19.342 | 17.933 | 18.096 | 18.976 | 18.438 | 18.329 | 17.862 | 0.028 | 0.4514 | 0.6906 | 0 | 0 | 0 |
| 'Nin' | 13.407 | 12.996 | 14.161 | 13.722 | 13.020 | 14.131 | 12.379 | 13.078 | 14.231 | 13.790 | 14.841 | 14.974 | 13.392 | 13.443 | 13.979 | 14.572 | 12.834 | 16.092 | -0.089 | 0.0125 | 0.0734 | 0 | 0 | 0 |
| 'Ninj1' | 6.191 | 6.949 | 6.408 | 5.791 | 6.439 | 7.038 | 5.757 | 7.270 | 6.234 | 7.456 | 5.728 | 6.435 | 7.898 | 5.783 | 5.792 | 7.089 | 6.366 | 6.197 | -0.022 | 0.7454 | 0.8851 | 0 | 0 | 0 |
| 'Ninl' | 0.522 | 0.660 | 0.615 | 0.783 | 0.738 | 0.700 | 0.663 | 0.813 | 0.638 | 0.559 | 0.635 | 0.660 | 0.580 | 0.567 | 0.584 | 0.618 | 0.538 | 0.537 | 0.193 | 0.0435 | 0.1728 | 0 | 0 | 0 |
| 'Nip7' | 14.205 | 14.453 | 13.419 | 14.530 | 13.097 | 13.350 | 13.054 | 13.367 | 11.737 | 13.296 | 12.217 | 13.877 | 13.512 | 12.834 | 13.503 | 13.557 | 13.415 | 13.867 | -0.001 | 0.9761 | 0.9918 | 0 | 0 | 0 |
| 'Nipa2' | 1.995 | 2.483 | 2.319 | 2.235 | 2.435 | 2.172 | 2.285 | 2.518 | 1.902 | 2.107 | 2.232 | 2.468 | 2.013 | 2.292 | 1.919 | 2.038 | 2.411 | 2.277 | 0.022 | 0.6843 | 0.8505 | 0 | 0 | 0 |
| 'Nipal1' | 5.008 | 6.823 | 6.803 | 6.011 | 8.533 | 6.405 | 6.430 | 7.247 | 7.138 | 4.993 | 6.061 | 4.892 | 4.412 | 5.164 | 5.195 | 4.431 | 5.011 | 4.277 | 0.429 | 0.0000 | 0.0000 | 1 | 1 | 0 |
| 'Nipal3' | 11.973 | 12.418 | 11.773 | 10.680 | 12.445 | 12.559 | 11.586 | 12.208 | 11.627 | 12.501 | 11.879 | 12.864 | 10.953 | 11.899 | 12.140 | 12.452 | 11.279 | 13.242 | -0.037 | 0.3215 | 0.5795 | 0 | 0 | 0 |
| 'Nipbl' | 40.588 | 33.696 | 36.615 | 41.319 | 34.295 | 33.268 | 37.842 | 35.138 | 35.491 | 39.162 | 42.121 | 38.651 | 36.234 | 37.798 | 35.496 | 34.720 | 40.048 | 39.891 | -0.081 | 0.0978 | 0.2920 | 0 | 0 | 0 |
| 'Nipsnap1' | 11.671 | 12.391 | 12.980 | 11.591 | 12.189 | 13.205 | 12.120 | 12.882 | 12.641 | 12.658 | 10.907 | 11.704 | 12.362 | 11.723 | 11.315 | 11.530 | 10.757 | 10.196 | 0.107 | 0.0082 | 0.0537 | 0 | 0 | 0 |
| 'Nipsnap3b' | 12.876 | 13.213 | 12.530 | 12.980 | 14.727 | 13.948 | 12.649 | 14.478 | 12.630 | 13.543 | 14.413 | 13.307 | 12.688 | 12.684 | 13.693 | 12.742 | 13.928 | 12.804 | -0.008 | 0.8539 | 0.9373 | 0 | 0 | 0 |
| 'Nisch' | 29.364 | 25.394 | 31.052 | 27.604 | 25.413 | 27.526 | 26.881 | 25.075 | 30.514 | 29.625 | 28.065 | 31.245 | 29.958 | 25.201 | 27.380 | 26.680 | 28.187 | 29.346 | -0.052 | 0.1601 | 0.3904 | 0 | 0 | 0 |
| 'Nit1' | 5.256 | 5.744 | 5.621 | 5.639 | 6.107 | 5.556 | 5.280 | 5.756 | 5.498 | 5.625 | 5.236 | 5.430 | 5.886 | 5.409 | 5.137 | 5.693 | 5.431 | 5.441 | 0.022 | 0.5132 | 0.7375 | 0 | 0 | 0 |
| 'Nit2' | 14.905 | 15.507 | 15.877 | 14.604 | 14.068 | 15.661 | 15.505 | 16.216 | 15.113 | 15.889 | 12.732 | 15.138 | 13.813 | 14.066 | 13.643 | 13.945 | 14.157 | 12.530 | 0.116 | 0.0289 | 0.1317 | 0 | 0 | 0 |
| 'Nkap' | 7.235 | 6.961 | 7.698 | 6.980 | 7.166 | 7.599 | 7.542 | 7.295 | 7.479 | 6.855 | 7.284 | 7.992 | 6.187 | 6.752 | 7.324 | 6.394 | 7.609 | 7.244 | 0.040 | 0.3494 | 0.6043 | 0 | 0 | 0 |
| 'Nkiras1' | 4.442 | 5.331 | 4.968 | 4.297 | 5.144 | 4.539 | 4.880 | 4.973 | 5.144 | 4.898 | 5.167 | 4.713 | 4.850 | 4.661 | 4.489 | 4.414 | 5.420 | 4.501 | 0.009 | 0.8346 | 0.9283 | 0 | 0 | 0 |
| 'Nkiras2' | 3.677 | 4.001 | 3.649 | 3.326 | 4.680 | 4.635 | 3.920 | 3.990 | 4.087 | 4.070 | 3.389 | 3.947 | 4.021 | 4.015 | 4.251 | 3.983 | 3.208 | 3.676 | 0.045 | 0.4857 | 0.7168 | 0 | 0 | 0 |
| 'Nkrf' | 3.975 | 3.577 | 3.173 | 4.031 | 3.973 | 3.384 | 3.845 | 3.233 | 3.344 | 3.442 | 3.565 | 3.389 | 3.306 | 3.508 | 2.777 | 3.727 | 3.690 | 3.197 | 0.072 | 0.2758 | 0.5317 | 0 | 0 | 0 |
| 'Nktr' | 15.841 | 16.269 | 15.769 | 15.225 | 16.012 | 15.052 | 15.410 | 16.293 | 15.805 | 16.597 | 16.713 | 15.007 | 16.849 | 16.312 | 14.666 | 15.562 | 16.152 | 14.619 | -0.017 | 0.4977 | 0.7258 | 0 | 0 | 0 |
| 'Nle1' | 5.137 | 4.941 | 6.003 | 5.170 | 4.992 | 5.218 | 5.003 | 5.489 | 5.617 | 4.909 | 4.451 | 5.630 | 4.997 | 4.348 | 4.776 | 4.749 | 4.251 | 5.017 | 0.129 | 0.0045 | 0.0349 | 1 | 0 | 0 |
| 'Nlgn2' | 1.170 | 1.117 | 1.397 | 1.271 | 1.249 | 1.329 | 1.303 | 1.326 | 1.345 | 1.340 | 1.099 | 0.948 | 1.071 | 1.077 | 0.984 | 1.006 | 0.954 | 1.133 | 0.264 | 0.0009 | 0.0101 | 1 | 0 | 0 |
| 'Nlk' | 14.336 | 13.362 | 13.119 | 14.169 | 14.416 | 13.872 | 13.257 | 13.111 | 13.995 | 12.762 | 14.747 | 14.598 | 13.353 | 12.553 | 13.534 | 13.331 | 12.926 | 14.190 | 0.007 | 0.8455 | 0.9343 | 0 | 0 | 0 |
| 'Nln' | 5.630 | 5.583 | 5.528 | 5.432 | 5.807 | 5.958 | 5.899 | 5.491 | 5.783 | 6.503 | 5.374 | 5.533 | 6.020 | 6.104 | 5.486 | 5.614 | 5.937 | 5.252 | -0.026 | 0.5923 | 0.7922 | 0 | 0 | 0 |
| 'Nlrc3' | 6.663 | 5.750 | 7.029 | 5.928 | 5.710 | 6.175 | 6.451 | 5.947 | 6.664 | 5.998 | 5.689 | 6.771 | 5.636 | 5.479 | 6.694 | 5.332 | 5.098 | 6.718 | 0.068 | 0.0910 | 0.2783 | 0 | 0 | 0 |
| 'Nlrc5' | 29.359 | 28.185 | 30.957 | 28.639 | 27.297 | 27.552 | 26.558 | 27.846 | 30.770 | 32.903 | 32.449 | 32.789 | 35.926 | 29.641 | 30.624 | 29.773 | 34.144 | 32.958 | -0.190 | 0.0000 | 0.0000 | 1 | 0 | 0 |
| 'Nlrx1' | 7.803 | 8.454 | 9.981 | 7.951 | 8.297 | 9.457 | 7.674 | 7.604 | 9.160 | 9.081 | 7.929 | 7.687 | 8.328 | 7.380 | 7.421 | 8.860 | 7.746 | 7.059 | 0.090 | 0.1378 | 0.3581 | 0 | 0 | 0 |
| 'Nmb' | 5.199 | 5.711 | 5.019 | 4.541 | 5.349 | 5.344 | 3.912 | 5.289 | 4.488 | 5.171 | 4.019 | 4.950 | 5.087 | 4.860 | 5.456 | 6.053 | 4.578 | 5.287 | -0.027 | 0.7291 | 0.8759 | 0 | 0 | 0 |
| 'Nmd3' | 3.846 | 4.104 | 4.046 | 3.717 | 3.958 | 3.981 | 4.536 | 4.155 | 3.672 | 3.893 | 3.821 | 3.492 | 3.754 | 4.302 | 3.283 | 4.039 | 4.337 | 3.007 | 0.084 | 0.0879 | 0.2706 | 0 | 0 | 0 |
| 'Nme1' | 14.627 | 14.162 | 15.654 | 15.197 | 16.484 | 14.560 | 15.430 | 15.890 | 14.568 | 13.888 | 14.255 | 14.489 | 14.636 | 15.314 | 15.127 | 15.363 | 15.449 | 13.474 | 0.041 | 0.2547 | 0.5066 | 0 | 0 | 0 |
| 'Nme2' | 21.021 | 19.882 | 23.280 | 20.863 | 21.367 | 21.070 | 21.423 | 22.327 | 21.538 | 19.971 | 19.592 | 21.259 | 20.068 | 19.440 | 23.263 | 20.066 | 20.759 | 21.156 | 0.047 | 0.2496 | 0.5007 | 0 | 0 | 0 |
| 'Nme6' | 0.985 | 1.131 | 0.903 | 1.048 | 1.084 | 1.078 | 1.158 | 1.073 | 1.164 | 1.098 | 0.797 | 1.035 | 1.030 | 1.072 | 0.976 | 1.118 | 1.029 | 0.771 | 0.092 | 0.3043 | 0.5618 | 0 | 0 | 0 |
| 'Nme7' | 1.176 | 1.574 | 1.213 | 1.233 | 1.348 | 1.310 | 1.464 | 1.289 | 1.421 | 1.251 | 1.504 | 1.508 | 1.287 | 1.294 | 1.416 | 1.456 | 1.540 | 1.165 | -0.065 | 0.3457 | 0.6009 | 0 | 0 | 0 |
| 'Nmi' | 17.080 | 17.254 | 16.901 | 17.668 | 19.991 | 18.406 | 18.657 | 20.042 | 17.681 | 18.537 | 17.047 | 17.691 | 17.979 | 16.432 | 16.822 | 17.544 | 18.967 | 16.604 | 0.042 | 0.2608 | 0.5143 | 0 | 0 | 0 |
| 'Nmnat1' | 2.346 | 2.049 | 2.568 | 1.990 | 2.414 | 2.902 | 2.015 | 2.191 | 2.680 | 2.006 | 1.824 | 2.324 | 2.338 | 2.095 | 2.197 | 2.113 | 1.961 | 2.488 | 0.121 | 0.0657 | 0.2235 | 0 | 0 | 0 |
| 'Nmnat3' | 0.981 | 1.214 | 1.312 | 0.805 | 0.874 | 1.153 | 0.996 | 1.068 | 0.972 | 0.834 | 0.829 | 0.994 | 0.867 | 0.849 | 0.936 | 1.028 | 0.805 | 0.809 | 0.232 | 0.0239 | 0.1170 | 0 | 0 | 0 |
| 'Nmral1' | 4.719 | 4.408 | 5.855 | 5.080 | 4.687 | 5.797 | 4.859 | 5.746 | 5.101 | 4.767 | 4.512 | 4.532 | 5.729 | 4.636 | 5.210 | 4.476 | 4.643 | 4.402 | 0.101 | 0.1044 | 0.3027 | 0 | 0 | 0 |
| 'Nmt1' | 22.151 | 20.783 | 21.956 | 22.478 | 20.994 | 24.071 | 22.502 | 22.296 | 22.303 | 21.990 | 21.066 | 21.962 | 22.949 | 20.700 | 21.724 | 22.241 | 20.860 | 20.781 | 0.030 | 0.2257 | 0.4723 | 0 | 0 | 0 |
| 'Nmt2' | 6.005 | 5.180 | 5.868 | 5.728 | 5.932 | 5.595 | 6.065 | 5.548 | 5.392 | 5.754 | 5.456 | 6.510 | 6.002 | 5.517 | 5.677 | 5.752 | 5.900 | 6.363 | -0.060 | 0.1475 | 0.3734 | 0 | 0 | 0 |
| 'Nnt' | 6.342 | 6.297 | 6.644 | 7.151 | 6.920 | 6.778 | 6.606 | 6.726 | 6.863 | 7.517 | 7.110 | 7.720 | 6.774 | 6.501 | 7.454 | 6.910 | 7.394 | 6.956 | -0.105 | 0.0034 | 0.0275 | 1 | 0 | 0 |
| 'Noa1' | 5.194 | 4.431 | 4.831 | 4.605 | 4.312 | 4.695 | 4.492 | 4.252 | 4.675 | 4.458 | 4.434 | 5.613 | 5.014 | 4.429 | 5.316 | 4.642 | 4.797 | 5.626 | -0.109 | 0.0239 | 0.1169 | 0 | 0 | 0 |
| 'Nob1' | 7.089 | 6.659 | 6.912 | 6.618 | 6.182 | 6.228 | 6.559 | 6.467 | 6.800 | 6.581 | 5.977 | 7.943 | 6.688 | 5.961 | 7.446 | 6.524 | 6.942 | 7.616 | -0.063 | 0.1659 | 0.3983 | 0 | 0 | 0 |
| 'Noc2l' | 35.803 | 30.825 | 35.203 | 31.898 | 31.563 | 33.448 | 32.018 | 34.142 | 34.037 | 32.054 | 27.882 | 35.683 | 33.254 | 27.634 | 33.385 | 31.199 | 29.788 | 35.043 | 0.056 | 0.1136 | 0.3198 | 0 | 0 | 0 |
| 'Noc3l' | 9.220 | 8.195 | 8.866 | 8.473 | 8.850 | 7.930 | 9.491 | 9.513 | 8.312 | 9.222 | 9.140 | 8.567 | 8.610 | 8.086 | 8.256 | 8.077 | 9.156 | 8.255 | 0.017 | 0.6906 | 0.8542 | 0 | 0 | 0 |
| 'Noc4l' | 10.601 | 9.072 | 11.677 | 9.548 | 9.866 | 11.198 | 10.055 | 9.947 | 10.556 | 9.745 | 8.755 | 11.249 | 10.493 | 8.760 | 10.865 | 9.568 | 8.250 | 11.245 | 0.048 | 0.2037 | 0.4464 | 0 | 0 | 0 |
| 'Noct' | 3.392 | 3.348 | 4.115 | 3.647 | 3.789 | 4.242 | 4.021 | 3.544 | 4.344 | 3.523 | 3.752 | 4.528 | 3.512 | 4.337 | 4.882 | 3.862 | 3.801 | 4.530 | -0.102 | 0.0457 | 0.1775 | 0 | 0 | 0 |
| 'Nod1' | 3.169 | 3.025 | 2.814 | 2.826 | 2.846 | 2.661 | 2.503 | 2.654 | 2.942 | 2.747 | 2.634 | 3.336 | 3.008 | 2.318 | 2.883 | 3.015 | 2.798 | 2.672 | -0.015 | 0.8131 | 0.9157 | 0 | 0 | 0 |
| 'Nod2' | 1.569 | 1.731 | 1.887 | 1.409 | 1.732 | 1.749 | 1.792 | 1.637 | 1.978 | 1.512 | 1.552 | 1.727 | 1.817 | 1.629 | 1.354 | 1.669 | 1.650 | 1.394 | 0.103 | 0.1251 | 0.3385 | 0 | 0 | 0 |
| 'Nol10' | 17.504 | 21.935 | 22.832 | 21.262 | 22.657 | 23.880 | 22.903 | 23.390 | 23.136 | 15.538 | 19.028 | 18.310 | 17.942 | 17.101 | 16.920 | 14.746 | 18.143 | 15.961 | 0.364 | 0.0000 | 0.0000 | 1 | 0 | 0 |
| 'Nol11' | 7.098 | 6.634 | 7.640 | 7.342 | 7.470 | 7.311 | 7.548 | 7.692 | 7.353 | 7.552 | 7.278 | 7.437 | 7.195 | 6.928 | 7.113 | 6.737 | 7.483 | 6.807 | 0.025 | 0.4796 | 0.7118 | 0 | 0 | 0 |
| 'Nol12' | 7.271 | 6.620 | 7.839 | 7.120 | 7.120 | 8.368 | 7.402 | 7.402 | 8.200 | 7.369 | 6.606 | 7.962 | 7.253 | 7.086 | 7.868 | 7.591 | 6.710 | 7.688 | 0.019 | 0.5703 | 0.7775 | 0 | 0 | 0 |
| 'Nol6' | 14.727 | 14.060 | 15.640 | 14.449 | 14.524 | 15.244 | 14.804 | 15.030 | 15.971 | 18.330 | 14.208 | 14.896 | 15.308 | 13.769 | 14.453 | 16.099 | 13.060 | 14.285 | -0.005 | 0.9069 | 0.9636 | 0 | 0 | 0 |
| 'Nol7' | 71.436 | 74.469 | 67.273 | 76.319 | 76.097 | 75.160 | 76.123 | 73.801 | 70.405 | 68.870 | 67.776 | 63.695 | 73.826 | 68.454 | 69.624 | 74.029 | 69.313 | 61.819 | 0.090 | 0.0027 | 0.0234 | 1 | 0 | 0 |
| 'Nol8' | 11.852 | 11.337 | 11.301 | 12.127 | 11.331 | 11.420 | 12.374 | 10.956 | 11.446 | 9.986 | 11.062 | 13.031 | 10.510 | 11.347 | 11.957 | 10.637 | 11.748 | 13.091 | -0.002 | 0.9748 | 0.9916 | 0 | 0 | 0 |
| 'Nol9' | 6.826 | 6.606 | 7.221 | 6.607 | 6.144 | 6.734 | 6.570 | 6.914 | 6.971 | 6.209 | 6.198 | 6.879 | 6.547 | 6.214 | 6.845 | 6.272 | 6.213 | 6.615 | 0.054 | 0.0664 | 0.2253 | 0 | 0 | 0 |
| 'Nolc1' | 19.434 | 16.165 | 19.618 | 18.566 | 17.096 | 19.265 | 18.235 | 16.338 | 18.267 | 18.152 | 15.452 | 19.147 | 18.651 | 15.051 | 17.949 | 18.417 | 15.126 | 18.857 | 0.047 | 0.0847 | 0.2642 | 0 | 0 | 0 |
| 'Nom1' | 5.793 | 6.006 | 5.861 | 5.657 | 6.115 | 6.159 | 5.144 | 5.917 | 6.275 | 6.058 | 5.400 | 5.852 | 5.698 | 5.025 | 5.739 | 5.696 | 5.591 | 5.920 | 0.044 | 0.2532 | 0.5048 | 0 | 0 | 0 |
| 'Nomo1' | 14.589 | 12.869 | 14.889 | 14.956 | 13.920 | 14.690 | 12.999 | 14.995 | 13.620 | 14.092 | 13.852 | 14.683 | 14.011 | 13.162 | 14.006 | 15.085 | 13.093 | 14.838 | 0.000 | 0.9943 | 0.9990 | 0 | 0 | 0 |
| 'Nono' | 56.248 | 59.147 | 61.162 | 62.629 | 61.757 | 61.018 | 61.255 | 59.617 | 60.805 | 59.094 | 56.471 | 57.111 | 59.644 | 59.952 | 61.145 | 59.210 | 58.861 | 57.645 | 0.029 | 0.2133 | 0.4581 | 0 | 0 | 0 |
| 'Nop10' | 49.203 | 54.685 | 46.564 | 51.032 | 56.602 | 48.876 | 56.040 | 49.684 | 45.479 | 49.431 | 51.541 | 46.729 | 48.568 | 50.757 | 47.965 | 47.320 | 52.666 | 48.427 | 0.032 | 0.3877 | 0.6374 | 0 | 0 | 0 |
| 'Nop14' | 14.796 | 13.758 | 13.755 | 14.134 | 15.475 | 13.597 | 13.578 | 15.262 | 14.635 | 14.180 | 13.042 | 15.126 | 13.593 | 13.489 | 13.472 | 13.853 | 13.301 | 12.981 | 0.054 | 0.1735 | 0.4069 | 0 | 0 | 0 |
| 'Nop16' | 9.201 | 7.907 | 8.640 | 8.344 | 8.793 | 8.218 | 8.688 | 9.435 | 8.020 | 8.393 | 8.471 | 9.046 | 8.751 | 7.206 | 8.860 | 8.477 | 8.377 | 9.001 | 0.000 | 0.9970 | 0.9994 | 0 | 0 | 0 |
| 'Nop2' | 20.403 | 16.279 | 21.659 | 22.023 | 18.425 | 18.980 | 21.772 | 18.031 | 21.067 | 18.492 | 17.501 | 22.044 | 18.699 | 16.215 | 21.251 | 18.264 | 18.084 | 20.461 | 0.050 | 0.2544 | 0.5063 | 0 | 0 | 0 |
| 'Nop58' | 6.778 | 6.685 | 6.460 | 7.254 | 6.742 | 6.406 | 7.350 | 6.681 | 6.413 | 6.881 | 6.536 | 6.685 | 6.945 | 6.902 | 6.395 | 6.923 | 7.072 | 6.116 | -0.005 | 0.8417 | 0.9329 | 0 | 0 | 0 |
| 'Nop9' | 13.830 | 14.145 | 15.460 | 13.587 | 15.459 | 15.874 | 14.314 | 16.025 | 14.946 | 14.950 | 13.875 | 14.336 | 14.271 | 14.323 | 14.431 | 14.165 | 14.094 | 14.256 | 0.047 | 0.1707 | 0.4037 | 0 | 0 | 0 |
| 'Nosip' | 29.704 | 32.960 | 33.533 | 29.641 | 32.215 | 33.864 | 29.773 | 31.120 | 35.668 | 30.721 | 27.750 | 33.225 | 32.471 | 28.845 | 33.976 | 31.026 | 29.716 | 33.015 | 0.027 | 0.4020 | 0.6509 | 0 | 0 | 0 |
| 'Notch1' | 25.433 | 20.420 | 24.156 | 22.103 | 20.659 | 21.871 | 18.709 | 21.574 | 22.369 | 23.968 | 25.777 | 27.664 | 21.459 | 24.409 | 26.274 | 23.799 | 23.556 | 30.195 | -0.210 | 0.0001 | 0.0018 | 1 | 0 | 0 |
| 'Notch2' | 8.936 | 7.359 | 8.981 | 8.377 | 8.015 | 8.616 | 7.980 | 8.308 | 8.792 | 8.383 | 8.148 | 8.379 | 7.586 | 7.528 | 8.319 | 7.767 | 7.809 | 8.780 | 0.043 | 0.2220 | 0.4694 | 0 | 0 | 0 |
| 'Notch3' | 2.869 | 4.057 | 2.734 | 3.266 | 3.674 | 2.943 | 2.658 | 3.117 | 2.803 | 5.367 | 6.871 | 5.429 | 10.434 | 6.998 | 5.591 | 8.171 | 5.987 | 5.858 | -1.114 | 0.0000 | 0.0000 | 1 | 1 | 1 |
| 'Npas4' | 2.247 | 2.226 | 2.465 | 1.909 | 1.686 | 2.303 | 2.759 | 2.188 | 2.722 | 2.918 | 2.234 | 3.734 | 2.964 | 2.041 | 3.453 | 3.250 | 3.105 | 3.188 | -0.404 | 0.0000 | 0.0001 | 1 | 1 | 0 |
| 'Npat' | 5.194 | 4.905 | 4.860 | 5.820 | 5.371 | 5.414 | 5.083 | 5.583 | 5.310 | 4.938 | 5.360 | 5.617 | 5.028 | 5.162 | 5.512 | 5.279 | 5.612 | 5.351 | -0.021 | 0.6022 | 0.7982 | 0 | 0 | 0 |
| 'Npc1' | 4.140 | 3.858 | 4.169 | 4.122 | 3.943 | 3.992 | 3.622 | 4.237 | 4.098 | 4.211 | 4.453 | 4.252 | 4.091 | 3.939 | 4.045 | 3.800 | 4.002 | 4.321 | -0.046 | 0.2228 | 0.4704 | 0 | 0 | 0 |
| 'Npc2' | 36.240 | 38.325 | 36.705 | 35.546 | 36.849 | 38.031 | 37.810 | 38.688 | 36.384 | 37.091 | 38.871 | 39.543 | 37.653 | 39.707 | 40.128 | 39.397 | 38.279 | 37.231 | -0.067 | 0.0051 | 0.0378 | 1 | 0 | 0 |
| 'Npepl1' | 37.860 | 39.318 | 42.480 | 34.112 | 39.646 | 46.350 | 39.331 | 42.419 | 45.811 | 38.576 | 30.685 | 41.524 | 35.483 | 33.299 | 38.410 | 42.396 | 34.139 | 39.016 | 0.130 | 0.0071 | 0.0483 | 1 | 0 | 0 |
| 'Npepps' | 11.006 | 10.213 | 10.994 | 11.664 | 10.549 | 9.992 | 11.594 | 10.349 | 10.866 | 10.167 | 10.388 | 11.664 | 10.779 | 10.724 | 10.693 | 9.567 | 11.319 | 11.627 | -0.007 | 0.8572 | 0.9388 | 0 | 0 | 0 |
| 'Nphp4' | 0.432 | 0.539 | 0.691 | 0.787 | 0.548 | 0.632 | 0.658 | 0.592 | 0.845 | 0.668 | 0.594 | 0.689 | 0.943 | 0.722 | 0.718 | 0.643 | 0.825 | 0.678 | -0.194 | 0.0853 | 0.2652 | 0 | 0 | 0 |
| 'Nploc4' | 16.189 | 17.274 | 16.806 | 15.298 | 17.275 | 17.251 | 14.410 | 17.405 | 17.294 | 17.100 | 16.731 | 16.811 | 17.129 | 17.308 | 17.902 | 17.088 | 15.881 | 17.667 | -0.052 | 0.1187 | 0.3278 | 0 | 0 | 0 |
| 'Npm1' | 110.519 | 102.185 | 101.668 | 118.897 | 113.202 | 96.504 | 124.786 | 111.379 | 97.511 | 101.427 | 106.001 | 97.153 | 108.715 | 105.372 | 102.860 | 105.948 | 109.470 | 96.590 | 0.051 | 0.1241 | 0.3368 | 0 | 0 | 0 |
| 'Npm3' | 38.961 | 37.301 | 39.741 | 41.946 | 39.820 | 39.259 | 39.513 | 39.112 | 37.995 | 35.827 | 36.905 | 37.076 | 38.249 | 34.337 | 39.236 | 35.569 | 37.442 | 39.959 | 0.068 | 0.0599 | 0.2110 | 0 | 0 | 0 |
| 'Nprl2' | 6.451 | 6.978 | 6.995 | 8.187 | 7.064 | 7.812 | 7.688 | 6.811 | 7.614 | 5.943 | 5.997 | 8.549 | 7.110 | 6.997 | 7.657 | 6.903 | 7.899 | 7.922 | 0.000 | 0.9940 | 0.9990 | 0 | 0 | 0 |
| 'Nprl3' | 3.014 | 3.471 | 3.653 | 2.992 | 3.091 | 3.506 | 3.385 | 3.542 | 3.678 | 3.432 | 3.238 | 2.948 | 3.425 | 3.122 | 3.173 | 3.442 | 3.008 | 2.623 | 0.089 | 0.1105 | 0.3142 | 0 | 0 | 0 |
| 'Nptn' | 8.328 | 9.645 | 8.566 | 8.952 | 9.601 | 8.734 | 9.300 | 9.229 | 8.962 | 8.292 | 8.484 | 8.927 | 7.974 | 8.487 | 8.624 | 8.341 | 8.471 | 8.997 | 0.071 | 0.0241 | 0.1177 | 0 | 0 | 0 |
| 'Nqo2' | 1.035 | 1.105 | 1.101 | 1.018 | 1.389 | 1.318 | 1.233 | 1.180 | 1.280 | 1.203 | 1.445 | 1.123 | 1.002 | 1.072 | 1.141 | 0.944 | 1.315 | 1.288 | 0.012 | 0.9087 | 0.9643 | 0 | 0 | 0 |
| 'Nr1d2' | 6.048 | 6.806 | 5.947 | 5.923 | 6.073 | 5.647 | 5.975 | 6.875 | 5.882 | 5.096 | 6.929 | 6.039 | 5.179 | 6.236 | 5.596 | 6.213 | 6.090 | 5.635 | 0.048 | 0.3122 | 0.5709 | 0 | 0 | 0 |
| 'Nr1h2' | 18.400 | 16.219 | 18.083 | 16.156 | 15.393 | 18.290 | 15.976 | 15.595 | 17.416 | 17.687 | 15.845 | 18.242 | 18.357 | 16.283 | 17.682 | 17.449 | 15.530 | 17.525 | -0.038 | 0.1879 | 0.4241 | 0 | 0 | 0 |
| 'Nr2c1' | 1.742 | 1.801 | 2.237 | 1.978 | 1.995 | 1.774 | 1.745 | 1.982 | 1.817 | 2.539 | 1.719 | 1.813 | 1.874 | 1.701 | 1.404 | 1.845 | 1.951 | 1.648 | 0.049 | 0.5785 | 0.7820 | 0 | 0 | 0 |
| 'Nr2c2' | 5.763 | 5.641 | 6.021 | 5.709 | 5.484 | 5.588 | 5.367 | 5.725 | 5.694 | 5.445 | 6.219 | 6.212 | 5.331 | 5.680 | 6.084 | 5.396 | 6.066 | 6.649 | -0.066 | 0.0754 | 0.2441 | 0 | 0 | 0 |
| 'Nr2c2ap' | 3.504 | 4.407 | 4.124 | 3.507 | 3.969 | 3.138 | 2.870 | 3.717 | 3.260 | 3.783 | 3.149 | 3.650 | 3.669 | 4.128 | 3.471 | 3.260 | 3.440 | 4.313 | -0.034 | 0.7464 | 0.8855 | 0 | 0 | 0 |
| 'Nr2f6' | 3.899 | 4.115 | 5.059 | 3.722 | 4.033 | 5.593 | 3.864 | 4.180 | 4.769 | 4.772 | 3.400 | 4.742 | 4.555 | 4.323 | 4.853 | 4.829 | 4.222 | 4.906 | -0.060 | 0.2963 | 0.5524 | 0 | 0 | 0 |
| 'Nr3c1' | 14.576 | 14.628 | 12.794 | 15.415 | 13.644 | 12.180 | 14.352 | 13.488 | 12.781 | 12.739 | 13.883 | 14.626 | 11.756 | 14.188 | 13.187 | 12.563 | 13.826 | 15.070 | 0.012 | 0.8255 | 0.9233 | 0 | 0 | 0 |
| 'Nr4a1' | 62.582 | 32.473 | 34.922 | 53.516 | 28.736 | 36.042 | 59.006 | 34.236 | 35.797 | 42.846 | 30.964 | 35.887 | 38.954 | 33.179 | 34.294 | 51.125 | 35.300 | 33.259 | 0.120 | 0.0702 | 0.2334 | 0 | 0 | 0 |
| 'Nr4a2' | 3.501 | 0.720 | 0.471 | 2.045 | 0.913 | 0.402 | 1.332 | 1.159 | 0.292 | 2.065 | 1.466 | 0.525 | 1.898 | 1.268 | 0.613 | 2.087 | 1.479 | 0.565 | -0.307 | 0.1039 | 0.3023 | 0 | 0 | 0 |
| 'Nr4a3' | 4.162 | 3.681 | 4.421 | 3.648 | 3.319 | 3.959 | 3.387 | 3.385 | 4.275 | 4.556 | 3.265 | 3.829 | 3.205 | 3.569 | 3.521 | 4.145 | 3.230 | 3.910 | 0.035 | 0.5782 | 0.7820 | 0 | 0 | 0 |
| 'Nrarp' | 2.268 | 1.854 | 1.615 | 2.708 | 1.473 | 1.673 | 2.137 | 1.483 | 1.703 | 1.487 | 1.616 | 2.169 | 2.135 | 1.992 | 2.158 | 1.483 | 2.375 | 1.801 | -0.063 | 0.6204 | 0.8101 | 0 | 0 | 0 |
| 'Nras' | 10.023 | 11.004 | 11.468 | 10.266 | 11.950 | 11.815 | 11.170 | 11.201 | 10.753 | 11.181 | 11.580 | 11.650 | 10.592 | 11.820 | 11.890 | 10.880 | 11.748 | 11.411 | -0.055 | 0.0520 | 0.1933 | 0 | 0 | 0 |
| 'Nrbf2' | 9.363 | 11.683 | 10.444 | 10.450 | 10.217 | 9.700 | 10.317 | 10.923 | 10.849 | 9.854 | 9.441 | 10.179 | 9.875 | 10.403 | 10.793 | 10.115 | 9.776 | 9.566 | 0.047 | 0.3139 | 0.5715 | 0 | 0 | 0 |
| 'Nrbp1' | 16.974 | 17.980 | 16.776 | 16.632 | 17.929 | 17.896 | 16.903 | 17.494 | 17.332 | 19.022 | 15.743 | 16.799 | 17.575 | 16.785 | 16.529 | 18.031 | 16.804 | 16.323 | 0.012 | 0.6836 | 0.8500 | 0 | 0 | 0 |
| 'Nrd1' | 20.779 | 21.415 | 22.493 | 23.041 | 21.892 | 22.063 | 21.596 | 21.723 | 22.301 | 22.214 | 22.320 | 21.832 | 21.247 | 21.756 | 21.170 | 19.543 | 22.561 | 21.767 | 0.011 | 0.7032 | 0.8611 | 0 | 0 | 0 |
| 'Nrde2' | 13.504 | 12.349 | 15.606 | 11.367 | 14.163 | 15.488 | 13.358 | 14.800 | 15.538 | 15.195 | 13.534 | 11.866 | 15.142 | 12.672 | 11.881 | 14.157 | 12.431 | 11.198 | 0.091 | 0.1856 | 0.4213 | 0 | 0 | 0 |
| 'Nrf1' | 4.315 | 4.249 | 4.154 | 3.793 | 3.810 | 4.259 | 4.001 | 4.151 | 4.298 | 4.326 | 4.127 | 4.517 | 4.296 | 4.269 | 4.479 | 4.358 | 4.309 | 4.677 | -0.097 | 0.0018 | 0.0165 | 1 | 0 | 0 |
| 'Nrg4' | 0.490 | 0.536 | 0.576 | 0.503 | 0.476 | 0.446 | 0.493 | 0.558 | 0.701 | 0.765 | 0.649 | 0.558 | 0.593 | 0.562 | 0.601 | 0.578 | 0.617 | 0.636 | -0.217 | 0.0294 | 0.1331 | 0 | 0 | 0 |
| 'Nrgn' | 14.759 | 17.050 | 16.059 | 13.999 | 15.514 | 18.579 | 16.857 | 14.238 | 20.158 | 14.350 | 10.124 | 16.208 | 11.279 | 11.129 | 17.856 | 15.428 | 11.666 | 14.213 | 0.265 | 0.0003 | 0.0035 | 1 | 0 | 0 |
| 'Nrip1' | 12.850 | 11.207 | 10.184 | 10.885 | 11.125 | 10.847 | 10.231 | 11.552 | 10.300 | 12.710 | 11.126 | 10.986 | 11.275 | 11.408 | 10.475 | 14.588 | 10.220 | 11.681 | -0.082 | 0.1139 | 0.3199 | 0 | 0 | 0 |
| 'Nrm' | 8.891 | 7.911 | 10.617 | 8.530 | 7.308 | 10.149 | 9.320 | 8.098 | 10.251 | 8.191 | 8.143 | 10.294 | 11.204 | 8.403 | 11.315 | 6.849 | 8.694 | 10.101 | -0.051 | 0.4568 | 0.6943 | 0 | 0 | 0 |
| 'Nrn1' | 3.482 | 3.552 | 5.488 | 3.331 | 4.871 | 4.842 | 5.805 | 4.657 | 4.845 | 1.711 | 1.673 | 2.657 | 2.294 | 2.104 | 2.759 | 2.300 | 2.028 | 2.244 | 1.034 | 0.0000 | 0.0000 | 1 | 1 | 1 |
| 'Nrp1' | 2.217 | 3.943 | 1.966 | 2.777 | 4.164 | 2.554 | 3.138 | 3.804 | 2.115 | 2.416 | 3.291 | 3.091 | 4.866 | 4.862 | 2.051 | 3.591 | 6.474 | 2.696 | -0.331 | 0.0226 | 0.1122 | 0 | 0 | 0 |
| 'Nrros' | 3.731 | 3.278 | 4.463 | 3.643 | 3.729 | 4.575 | 3.774 | 3.987 | 4.494 | 3.568 | 3.355 | 4.121 | 3.348 | 3.491 | 3.862 | 3.550 | 3.507 | 4.193 | 0.105 | 0.0048 | 0.0361 | 1 | 0 | 0 |
| 'Nsa2' | 64.331 | 62.815 | 61.803 | 66.393 | 63.608 | 59.530 | 71.254 | 61.714 | 60.482 | 62.838 | 64.877 | 57.233 | 63.074 | 62.644 | 60.131 | 66.327 | 63.378 | 57.638 | 0.026 | 0.3411 | 0.5981 | 0 | 0 | 0 |
| 'Nsd1' | 23.554 | 21.904 | 22.496 | 24.557 | 21.653 | 20.524 | 23.273 | 22.825 | 22.149 | 21.436 | 23.788 | 23.901 | 21.109 | 21.509 | 21.808 | 21.138 | 21.423 | 23.593 | 0.012 | 0.7698 | 0.8975 | 0 | 0 | 0 |
| 'Nsdhl' | 10.255 | 10.871 | 10.715 | 10.850 | 9.373 | 10.405 | 11.224 | 10.200 | 11.266 | 9.758 | 10.013 | 10.536 | 9.996 | 11.089 | 10.667 | 10.640 | 10.782 | 10.032 | 0.016 | 0.7140 | 0.8677 | 0 | 0 | 0 |
| 'Nsf' | 12.273 | 11.652 | 12.558 | 12.777 | 11.769 | 12.246 | 12.212 | 12.170 | 12.362 | 12.123 | 12.224 | 11.786 | 10.568 | 11.134 | 11.609 | 11.313 | 11.398 | 11.735 | 0.073 | 0.0191 | 0.1001 | 0 | 0 | 0 |
| 'Nsfl1c' | 12.263 | 13.387 | 12.837 | 13.242 | 14.012 | 13.516 | 12.461 | 13.130 | 13.291 | 12.734 | 12.402 | 13.092 | 12.651 | 11.389 | 13.025 | 13.043 | 11.795 | 12.172 | 0.061 | 0.0513 | 0.1917 | 0 | 0 | 0 |
| 'Nsg2' | 5.210 | 5.534 | 5.999 | 5.141 | 5.440 | 6.737 | 5.300 | 5.736 | 6.336 | 7.753 | 5.717 | 8.256 | 5.818 | 6.484 | 8.170 | 8.837 | 6.006 | 7.263 | -0.328 | 0.0000 | 0.0000 | 1 | 0 | 0 |
| 'Nsl1' | 2.126 | 1.696 | 2.731 | 2.482 | 1.961 | 2.660 | 2.439 | 1.930 | 2.640 | 1.782 | 2.070 | 2.396 | 2.804 | 2.339 | 2.534 | 1.882 | 2.038 | 2.386 | 0.014 | 0.8658 | 0.9418 | 0 | 0 | 0 |
| 'Nsmaf' | 18.103 | 18.385 | 18.265 | 18.394 | 19.130 | 18.013 | 18.899 | 19.158 | 18.089 | 19.222 | 19.485 | 17.901 | 19.362 | 19.192 | 17.580 | 18.874 | 20.534 | 16.807 | -0.030 | 0.1898 | 0.4262 | 0 | 0 | 0 |
| 'Nsmce1' | 32.998 | 34.729 | 32.126 | 33.304 | 36.843 | 35.655 | 34.394 | 36.929 | 34.271 | 30.434 | 29.415 | 28.839 | 30.381 | 31.351 | 27.589 | 27.725 | 27.286 | 28.504 | 0.239 | 0.0000 | 0.0000 | 1 | 0 | 0 |
| 'Nsmce2' | 7.833 | 9.798 | 7.615 | 8.656 | 9.482 | 7.563 | 8.812 | 8.256 | 7.936 | 7.548 | 8.251 | 8.366 | 9.199 | 8.600 | 7.776 | 8.163 | 8.705 | 7.985 | 0.003 | 0.9547 | 0.9832 | 0 | 0 | 0 |
| 'Nsmce4a' | 17.461 | 16.690 | 17.968 | 17.549 | 18.002 | 18.036 | 18.713 | 18.604 | 16.953 | 18.161 | 18.518 | 17.311 | 19.171 | 19.651 | 17.613 | 18.383 | 18.099 | 17.988 | -0.050 | 0.1132 | 0.3194 | 0 | 0 | 0 |
| 'Nsmf' | 6.516 | 5.993 | 7.467 | 6.432 | 6.141 | 7.587 | 6.085 | 6.520 | 7.396 | 7.474 | 6.284 | 7.548 | 6.796 | 6.189 | 7.585 | 7.346 | 6.524 | 6.902 | -0.067 | 0.0539 | 0.1985 | 0 | 0 | 0 |
| 'Nsrp1' | 14.107 | 12.686 | 13.872 | 14.585 | 12.979 | 14.085 | 14.809 | 13.090 | 14.073 | 14.126 | 13.429 | 13.912 | 13.249 | 12.679 | 13.395 | 13.162 | 14.534 | 12.542 | 0.028 | 0.4498 | 0.6898 | 0 | 0 | 0 |
| 'Nsun2' | 23.856 | 23.452 | 23.935 | 24.347 | 24.241 | 22.517 | 23.816 | 25.553 | 22.869 | 23.524 | 23.146 | 24.006 | 23.022 | 22.114 | 22.606 | 23.916 | 22.843 | 23.247 | 0.030 | 0.2346 | 0.4837 | 0 | 0 | 0 |
| 'Nsun3' | 1.989 | 2.511 | 2.510 | 2.074 | 2.648 | 2.240 | 2.452 | 2.662 | 2.758 | 2.604 | 2.502 | 1.797 | 2.354 | 2.424 | 1.568 | 2.544 | 2.131 | 1.466 | 0.185 | 0.0438 | 0.1733 | 0 | 0 | 0 |
| 'Nsun5' | 2.515 | 2.617 | 3.009 | 2.447 | 2.630 | 2.830 | 2.760 | 2.585 | 2.764 | 2.698 | 2.676 | 3.277 | 2.477 | 2.130 | 3.005 | 2.694 | 2.740 | 3.023 | -0.046 | 0.3559 | 0.6089 | 0 | 0 | 0 |
| 'Nt5c' | 34.574 | 36.131 | 38.228 | 36.266 | 37.762 | 40.343 | 35.651 | 37.217 | 39.306 | 32.915 | 32.397 | 38.036 | 36.368 | 35.669 | 38.418 | 35.873 | 36.055 | 38.151 | 0.039 | 0.1465 | 0.3719 | 0 | 0 | 0 |
| 'Nt5c2' | 7.688 | 8.224 | 7.969 | 8.645 | 8.261 | 7.823 | 8.056 | 8.245 | 7.998 | 8.523 | 8.641 | 8.008 | 7.352 | 8.360 | 7.704 | 7.735 | 8.391 | 7.304 | 0.008 | 0.8172 | 0.9188 | 0 | 0 | 0 |
| 'Nt5c3' | 6.824 | 6.889 | 6.423 | 7.240 | 7.707 | 6.362 | 7.423 | 7.215 | 6.141 | 6.747 | 6.664 | 7.111 | 6.883 | 7.040 | 6.873 | 6.924 | 6.520 | 6.195 | 0.011 | 0.7823 | 0.9025 | 0 | 0 | 0 |
| 'Nt5c3b' | 19.423 | 19.371 | 20.209 | 19.528 | 19.088 | 21.041 | 19.330 | 18.843 | 21.556 | 19.127 | 17.950 | 21.087 | 20.277 | 18.323 | 20.453 | 19.455 | 18.578 | 19.779 | 0.015 | 0.5654 | 0.7741 | 0 | 0 | 0 |
| 'Nt5dc1' | 16.420 | 17.700 | 15.456 | 15.913 | 16.973 | 15.485 | 17.343 | 17.223 | 15.817 | 16.780 | 16.591 | 14.471 | 15.075 | 16.631 | 14.821 | 15.778 | 16.218 | 12.989 | 0.083 | 0.0077 | 0.0512 | 0 | 0 | 0 |

| | | | | | | | | | | | | | | | | | | | | | | | | |
|---|---|---|---|---|---|---|---|---|---|---|---|---|---|---|---|---|---|---|---|---|---|---|---|---|
| 'Nt5dc2' | 4.342 | 6.136 | 5.747 | 6.051 | 4.987 | 5.545 | 5.942 | 4.217 | 5.626 | 5.537 | 4.894 | 5.195 | 7.770 | 5.698 | 7.295 | 7.374 | 5.265 | 7.334 | -0.222 | 0.0285 | 0.1307 | 0 | 0 | 0 |
| 'Nt5dc3' | 5.105 | 5.022 | 4.824 | 4.847 | 4.508 | 5.013 | 4.635 | 5.109 | 4.717 | 4.741 | 4.206 | 5.545 | 4.493 | 5.000 | 5.405 | 5.277 | 4.942 | 5.547 | -0.056 | 0.2727 | 0.5277 | 0 | 0 | 0 |
| 'Nt5e' | 11.231 | 14.331 | 10.221 | 10.649 | 13.507 | 11.015 | 11.571 | 13.312 | 11.202 | 12.580 | 14.645 | 15.789 | 12.593 | 14.913 | 13.985 | 14.861 | 16.633 | 15.715 | -0.317 | 0.0000 | 0.0000 | 1 | 0 | 0 |
| 'Nt5m' | 3.652 | 4.789 | 3.676 | 3.762 | 4.225 | 3.964 | 3.322 | 4.059 | 3.997 | 4.343 | 3.346 | 4.414 | 4.379 | 4.755 | 3.891 | 4.693 | 3.954 | 4.417 | -0.124 | 0.0674 | 0.2275 | 0 | 0 | 0 |
| 'Ntan1' | 12.113 | 13.685 | 12.136 | 12.015 | 14.440 | 12.306 | 14.040 | 12.838 | 13.046 | 12.995 | 12.498 | 12.207 | 13.163 | 12.975 | 11.754 | 12.362 | 14.377 | 11.645 | 0.020 | 0.5935 | 0.7931 | 0 | 0 | 0 |
| 'Ntf5' | 2.106 | 3.170 | 2.159 | 2.221 | 2.752 | 2.328 | 2.940 | 2.206 | 2.249 | 2.446 | 2.336 | 3.046 | 2.847 | 2.857 | 3.261 | 2.782 | 3.316 | 2.646 | -0.241 | 0.0185 | 0.0978 | 0 | 0 | 0 |
| 'Ntmt1' | 2.732 | 2.124 | 2.568 | 2.152 | 2.616 | 2.381 | 2.268 | 2.209 | 2.358 | 2.176 | 2.288 | 2.664 | 2.630 | 2.175 | 2.426 | 2.434 | 2.114 | 2.405 | -0.006 | 0.9284 | 0.9727 | 0 | 0 | 0 |
| 'Ntn1' | 3.889 | 4.893 | 3.586 | 4.491 | 3.982 | 3.987 | 4.398 | 4.341 | 3.765 | 4.888 | 5.735 | 4.422 | 6.748 | 5.492 | 4.586 | 4.428 | 5.312 | 4.059 | -0.299 | 0.0000 | 0.0001 | 1 | 0 | 0 |
| 'Ntng2' | 12.231 | 12.033 | 13.238 | 11.039 | 13.255 | 12.406 | 11.092 | 12.127 | 13.671 | 14.876 | 13.720 | 13.169 | 14.636 | 12.984 | 12.387 | 12.875 | 12.596 | 12.532 | -0.118 | 0.0129 | 0.0753 | 0 | 0 | 0 |
| 'Ntpcr' | 2.717 | 3.364 | 3.248 | 2.679 | 3.331 | 2.988 | 2.741 | 2.909 | 3.297 | 3.043 | 2.355 | 2.480 | 2.751 | 2.721 | 2.631 | 2.948 | 2.493 | 2.338 | 0.194 | 0.0048 | 0.0363 | 1 | 0 | 0 |
| 'Ntrk3' | 3.888 | 4.261 | 4.891 | 4.485 | 4.504 | 5.216 | 4.324 | 4.340 | 5.643 | 5.020 | 4.302 | 4.550 | 3.437 | 4.144 | 4.344 | 4.681 | 4.039 | 4.316 | 0.082 | 0.1722 | 0.4060 | 0 | 0 | 0 |
| 'Nub1' | 14.840 | 15.033 | 14.934 | 14.956 | 15.341 | 15.846 | 15.867 | 15.717 | 15.764 | 15.745 | 14.465 | 16.203 | 15.000 | 14.990 | 14.839 | 15.804 | 15.729 | 15.047 | -0.006 | 0.7987 | 0.9107 | 0 | 0 | 0 |
| 'Nubp1' | 6.082 | 6.720 | 6.673 | 5.097 | 6.185 | 6.929 | 5.784 | 6.310 | 6.792 | 6.054 | 6.065 | 6.362 | 6.502 | 5.578 | 6.284 | 6.973 | 5.655 | 5.627 | 0.029 | 0.5870 | 0.7881 | 0 | 0 | 0 |
| 'Nubp2' | 36.658 | 39.008 | 38.080 | 35.203 | 40.636 | 39.678 | 37.297 | 44.036 | 38.389 | 39.890 | 34.631 | 38.551 | 39.230 | 40.135 | 39.247 | 40.501 | 36.701 | 37.666 | 0.000 | 0.9958 | 0.9994 | 0 | 0 | 0 |
| 'Nubpl' | 0.914 | 1.019 | 0.657 | 0.901 | 1.091 | 0.641 | 0.810 | 1.080 | 0.780 | 0.837 | 0.891 | 1.095 | 0.881 | 0.859 | 1.151 | 0.597 | 0.892 | 0.765 | -0.055 | 0.6609 | 0.8362 | 0 | 0 | 0 |
| 'Nucb1' | 21.980 | 22.024 | 23.857 | 20.549 | 21.324 | 24.252 | 22.063 | 21.082 | 23.103 | 23.965 | 21.427 | 25.326 | 23.175 | 21.426 | 25.329 | 23.254 | 19.513 | 24.906 | -0.066 | 0.0196 | 0.1017 | 0 | 0 | 0 |
| 'Nucb2' | 8.262 | 9.662 | 8.176 | 8.148 | 7.896 | 7.415 | 9.137 | 8.300 | 7.468 | 9.022 | 7.943 | 7.446 | 10.761 | 8.304 | 6.416 | 9.862 | 9.040 | 6.845 | -0.023 | 0.7001 | 0.8599 | 0 | 0 | 0 |
| 'Nucks1' | 13.521 | 15.240 | 16.364 | 15.502 | 15.626 | 14.569 | 17.533 | 15.940 | 15.146 | 12.267 | 14.959 | 14.386 | 13.851 | 15.264 | 13.904 | 13.739 | 15.189 | 13.635 | 0.123 | 0.0042 | 0.0331 | 1 | 0 | 0 |
| 'Nudc' | 41.447 | 37.107 | 42.284 | 41.448 | 40.403 | 43.699 | 39.514 | 38.864 | 43.668 | 39.570 | 34.231 | 42.894 | 43.351 | 36.758 | 43.532 | 40.655 | 36.704 | 41.204 | 0.028 | 0.3219 | 0.5798 | 0 | 0 | 0 |
| 'Nudcd1' | 4.084 | 4.477 | 4.112 | 4.116 | 4.161 | 3.873 | 4.531 | 4.599 | 3.773 | 4.414 | 3.692 | 4.385 | 4.049 | 4.166 | 4.165 | 4.314 | 3.977 | 4.073 | 0.001 | 0.9873 | 0.9963 | 0 | 0 | 0 |
| 'Nudcd2' | 8.406 | 9.240 | 9.072 | 8.506 | 9.609 | 8.858 | 9.438 | 9.175 | 8.843 | 7.965 | 8.521 | 8.896 | 9.113 | 8.421 | 8.847 | 8.347 | 9.199 | 7.803 | 0.058 | 0.1413 | 0.3636 | 0 | 0 | 0 |
| 'Nudcd3' | 14.234 | 14.326 | 14.768 | 13.777 | 14.564 | 15.685 | 14.445 | 14.713 | 15.025 | 15.087 | 14.465 | 14.759 | 14.352 | 14.274 | 14.549 | 14.820 | 13.786 | 14.946 | -0.004 | 0.8800 | 0.9494 | 0 | 0 | 0 |
| 'Nudt1' | 6.392 | 8.238 | 8.145 | 7.994 | 8.281 | 7.826 | 6.849 | 8.125 | 7.779 | 7.153 | 7.636 | 7.059 | 7.130 | 7.658 | 8.265 | 7.001 | 7.542 | 6.428 | 0.072 | 0.1954 | 0.4339 | 0 | 0 | 0 |
| 'Nudt13' | 5.938 | 6.308 | 7.295 | 7.919 | 6.916 | 6.935 | 6.652 | 7.440 | 7.412 | 7.624 | 6.298 | 6.479 | 7.378 | 7.202 | 6.480 | 6.282 | 6.802 | 5.642 | 0.056 | 0.3864 | 0.6357 | 0 | 0 | 0 |
| 'Nudt14' | 27.202 | 33.736 | 31.081 | 24.409 | 28.317 | 30.218 | 29.860 | 32.362 | 32.791 | 27.941 | 27.700 | 28.348 | 26.907 | 31.230 | 31.294 | 30.528 | 24.874 | 25.759 | 0.076 | 0.1985 | 0.4383 | 0 | 0 | 0 |
| 'Nudt16' | 4.041 | 4.992 | 4.657 | 4.128 | 5.106 | 5.403 | 3.971 | 4.748 | 5.294 | 4.848 | 4.403 | 4.013 | 4.394 | 4.158 | 4.550 | 5.050 | 4.206 | 3.787 | 0.095 | 0.1630 | 0.3947 | 0 | 0 | 0 |
| 'Nudt16l1' | 46.244 | 52.612 | 49.891 | 45.526 | 53.028 | 56.241 | 43.961 | 53.150 | 54.297 | 49.143 | 49.199 | 52.926 | 47.174 | 47.254 | 54.592 | 53.790 | 48.992 | 52.036 | -0.012 | 0.7613 | 0.8940 | 0 | 0 | 0 |
| 'Nudt18' | 7.102 | 8.004 | 8.135 | 6.320 | 7.859 | 8.249 | 8.065 | 8.693 | 7.710 | 7.451 | 7.947 | 8.537 | 7.991 | 7.279 | 8.180 | 7.878 | 6.998 | 7.313 | -0.001 | 0.9820 | 0.9933 | 0 | 0 | 0 |
| 'Nudt19' | 6.662 | 7.265 | 7.106 | 6.756 | 6.871 | 7.107 | 7.035 | 7.925 | 7.099 | 6.325 | 6.998 | 7.430 | 6.936 | 7.058 | 7.468 | 7.073 | 5.992 | 7.194 | 0.018 | 0.6951 | 0.8566 | 0 | 0 | 0 |
| 'Nudt2' | 10.431 | 9.470 | 10.792 | 11.757 | 14.093 | 9.680 | 11.155 | 11.612 | 10.333 | 10.201 | 8.851 | 10.391 | 11.819 | 12.349 | 10.129 | 11.424 | 11.795 | 8.531 | 0.044 | 0.5827 | 0.7850 | 0 | 0 | 0 |
| 'Nudt21' | 34.490 | 37.046 | 35.981 | 38.705 | 37.702 | 34.675 | 37.808 | 36.616 | 35.362 | 34.195 | 35.597 | 36.036 | 35.802 | 38.008 | 36.927 | 34.377 | 37.315 | 34.769 | 0.011 | 0.7140 | 0.8677 | 0 | 0 | 0 |
| 'Nudt22' | 14.351 | 17.563 | 16.682 | 16.258 | 16.272 | 16.060 | 17.019 | 15.764 | 16.695 | 12.689 | 11.161 | 12.503 | 13.635 | 12.284 | 12.147 | 13.982 | 11.939 | 11.418 | 0.381 | 0.0000 | 0.0000 | 1 | 1 | 0 |
| 'Nudt3' | 23.732 | 25.762 | 24.465 | 22.879 | 26.024 | 25.807 | 22.101 | 25.869 | 24.258 | 22.676 | 22.895 | 23.951 | 23.787 | 21.866 | 25.051 | 24.136 | 21.626 | 25.441 | 0.052 | 0.1523 | 0.3794 | 0 | 0 | 0 |
| 'Nudt4' | 19.383 | 17.094 | 18.483 | 18.109 | 19.295 | 18.113 | 18.428 | 19.173 | 18.157 | 16.842 | 17.106 | 17.500 | 17.059 | 18.468 | 16.463 | 17.760 | 17.904 | 16.864 | 0.084 | 0.0101 | 0.0626 | 0 | 0 | 0 |
| 'Nudt5' | 6.722 | 7.640 | 7.014 | 6.878 | 7.462 | 7.010 | 7.387 | 7.508 | 6.757 | 7.203 | 6.816 | 7.312 | 7.378 | 7.283 | 6.793 | 6.915 | 7.173 | 6.685 | 0.002 | 0.9427 | 0.9785 | 0 | 0 | 0 |
| 'Nudt6' | 1.858 | 2.069 | 1.914 | 1.506 | 1.556 | 1.789 | 1.700 | 2.020 | 1.780 | 1.710 | 1.514 | 1.586 | 1.507 | 1.377 | 1.742 | 1.529 | 1.563 | 1.507 | 0.200 | 0.0165 | 0.0899 | 0 | 0 | 0 |
| 'Nudt7' | 0.472 | 0.470 | 0.606 | 0.503 | 0.602 | 0.525 | 0.615 | 0.736 | 0.452 | 0.589 | 0.670 | 0.532 | 0.562 | 0.546 | 0.592 | 0.597 | 0.568 | 0.551 | -0.064 | 0.5327 | 0.7528 | 0 | 0 | 0 |
| 'Nudt8' | 2.183 | 1.606 | 2.057 | 1.506 | 1.704 | 2.326 | 1.554 | 1.941 | 2.123 | 1.784 | 1.813 | 2.033 | 1.885 | 1.853 | 1.900 | 1.484 | 1.821 | 2.062 | 0.027 | 0.7248 | 0.8736 | 0 | 0 | 0 |
| 'Nudt9' | 13.532 | 14.601 | 13.316 | 13.294 | 14.850 | 13.998 | 12.302 | 13.792 | 14.002 | 13.800 | 12.536 | 13.040 | 14.416 | 13.293 | 13.810 | 14.358 | 12.462 | 13.446 | 0.020 | 0.6203 | 0.8101 | 0 | 0 | 0 |
| 'Nuf2' | 2.896 | 3.702 | 4.212 | 4.091 | 3.338 | 3.732 | 4.200 | 3.401 | 4.053 | 2.708 | 3.735 | 3.780 | 3.894 | 4.056 | 4.537 | 3.121 | 3.636 | 3.646 | 0.012 | 0.8783 | 0.9485 | 0 | 0 | 0 |
| 'Nufip1' | 6.858 | 6.574 | 6.399 | 6.556 | 6.571 | 6.030 | 6.961 | 6.688 | 6.243 | 6.694 | 6.201 | 6.266 | 6.427 | 5.880 | 6.404 | 6.015 | 6.138 | 5.692 | 0.066 | 0.1315 | 0.3484 | 0 | 0 | 0 |
| 'Nufip2' | 10.426 | 10.584 | 9.069 | 10.517 | 10.818 | 9.030 | 10.902 | 10.464 | 9.994 | 8.942 | 11.587 | 13.211 | 9.207 | 11.152 | 11.652 | 11.038 | 12.309 | 12.363 | -0.152 | 0.0171 | 0.0927 | 0 | 0 | 0 |
| 'Numa1' | 80.252 | 72.624 | 83.120 | 79.661 | 74.563 | 83.390 | 73.688 | 73.775 | 83.005 | 87.582 | 82.316 | 86.434 | 78.697 | 80.634 | 83.324 | 81.338 | 76.958 | 88.278 | -0.094 | 0.0008 | 0.0091 | 1 | 0 | 0 |
| 'Numb' | 3.844 | 3.538 | 3.605 | 3.692 | 3.937 | 4.173 | 4.164 | 4.091 | 4.175 | 3.677 | 4.092 | 4.229 | 3.909 | 3.846 | 3.849 | 3.739 | 3.907 | 4.040 | -0.015 | 0.7197 | 0.8711 | 0 | 0 | 0 |
| 'Numbl' | 0.336 | 0.230 | 0.246 | 0.232 | 0.216 | 0.243 | 0.252 | 0.251 | 0.323 | 0.328 | 0.366 | 0.236 | 0.297 | 0.357 | 0.242 | 0.241 | 0.209 | 0.288 | -0.131 | 0.3195 | 0.5781 | 0 | 0 | 0 |
| 'Nup107' | 18.672 | 20.553 | 20.941 | 19.780 | 20.602 | 21.093 | 21.594 | 21.178 | 21.168 | 17.852 | 20.223 | 18.256 | 19.241 | 19.813 | 19.083 | 18.789 | 20.134 | 18.169 | 0.106 | 0.0012 | 0.0120 | 1 | 0 | 0 |
| 'Nup133' | 8.304 | 8.145 | 9.541 | 9.382 | 8.291 | 8.786 | 8.737 | 8.848 | 9.125 | 8.306 | 8.854 | 8.551 | 9.417 | 9.038 | 8.256 | 7.861 | 8.115 | 8.257 | 0.037 | 0.3747 | 0.6259 | 0 | 0 | 0 |
| 'Nup153' | 44.441 | 44.159 | 42.147 | 43.420 | 44.787 | 41.688 | 42.615 | 44.625 | 42.237 | 44.517 | 44.936 | 46.333 | 41.167 | 44.541 | 43.912 | 43.726 | 45.139 | 45.273 | -0.046 | 0.0608 | 0.2127 | 0 | 0 | 0 |
| 'Nup155' | 7.735 | 8.382 | 8.913 | 8.403 | 8.393 | 8.440 | 8.738 | 8.573 | 9.232 | 7.353 | 8.584 | 8.781 | 8.473 | 8.586 | 9.296 | 7.934 | 9.091 | 8.715 | -0.011 | 0.7706 | 0.8977 | 0 | 0 | 0 |
| 'Nup160' | 4.915 | 5.638 | 5.696 | 5.799 | 5.589 | 5.753 | 5.514 | 5.816 | 5.592 | 5.271 | 5.658 | 5.623 | 5.407 | 5.549 | 5.405 | 5.553 | 5.839 | 5.232 | 0.012 | 0.7024 | 0.8611 | 0 | 0 | 0 |
| 'Nup188' | 11.765 | 10.319 | 12.052 | 11.530 | 10.597 | 11.435 | 11.039 | 10.791 | 12.258 | 11.072 | 10.780 | 11.584 | 10.778 | 10.436 | 11.146 | 10.724 | 10.757 | 11.860 | 0.029 | 0.3162 | 0.5741 | 0 | 0 | 0 |
| 'Nup205' | 7.626 | 6.813 | 8.282 | 8.007 | 7.084 | 7.696 | 7.688 | 7.300 | 7.837 | 6.799 | 7.281 | 7.665 | 7.675 | 7.381 | 7.661 | 6.426 | 7.846 | 7.784 | 0.028 | 0.4902 | 0.7201 | 0 | 0 | 0 |
| 'Nup210' | 32.955 | 30.653 | 36.424 | 32.259 | 31.432 | 33.553 | 32.214 | 32.215 | 34.743 | 33.003 | 33.240 | 35.318 | 32.909 | 31.418 | 35.295 | 31.373 | 30.283 | 36.154 | -0.022 | 0.3591 | 0.6115 | 0 | 0 | 0 |
| 'Nup214' | 8.073 | 7.484 | 8.911 | 7.846 | 8.239 | 8.618 | 7.590 | 7.356 | 8.345 | 8.707 | 8.010 | 8.454 | 7.984 | 7.506 | 8.039 | 7.763 | 7.411 | 7.939 | 0.003 | 0.9324 | 0.9746 | 0 | 0 | 0 |
| 'Nup37' | 9.310 | 12.251 | 12.416 | 11.924 | 12.360 | 11.867 | 11.911 | 13.202 | 12.848 | 9.924 | 11.025 | 10.266 | 9.842 | 11.637 | 11.919 | 10.279 | 11.855 | 9.345 | 0.164 | 0.0071 | 0.0483 | 1 | 0 | 0 |
| 'Nup43' | 1.870 | 2.166 | 2.381 | 2.355 | 2.384 | 2.407 | 2.565 | 2.094 | 2.164 | 1.991 | 2.212 | 2.429 | 2.013 | 1.975 | 2.147 | 1.759 | 1.771 | 2.336 | 0.106 | 0.1656 | 0.3980 | 0 | 0 | 0 |
| 'Nup50' | 30.052 | 30.862 | 29.214 | 30.069 | 28.569 | 30.448 | 30.371 | 28.726 | 29.318 | 30.740 | 29.892 | 32.053 | 30.814 | 31.667 | 32.945 | 31.543 | 30.931 | 32.209 | -0.091 | 0.0003 | 0.0037 | 1 | 0 | 0 |
| 'Nup85' | 12.314 | 13.559 | 15.321 | 12.186 | 12.980 | 14.073 | 12.820 | 13.255 | 14.586 | 12.825 | 13.185 | 13.351 | 14.698 | 12.853 | 13.329 | 13.068 | 13.983 | 12.718 | 0.002 | 0.9549 | 0.9833 | 0 | 0 | 0 |
| 'Nup88' | 5.977 | 5.926 | 6.094 | 6.197 | 5.973 | 5.960 | 5.940 | 6.221 | 6.384 | 5.938 | 5.860 | 5.773 | 6.315 | 6.131 | 5.503 | 5.799 | 5.834 | 5.685 | 0.041 | 0.1433 | 0.3668 | 0 | 0 | 0 |
| 'Nup93' | 22.620 | 22.918 | 25.043 | 24.994 | 24.042 | 23.588 | 25.177 | 24.300 | 25.345 | 23.273 | 24.480 | 24.183 | 24.319 | 22.696 | 24.690 | 23.386 | 24.065 | 24.030 | 0.009 | 0.7649 | 0.8953 | 0 | 0 | 0 |
| 'Nup98' | 17.122 | 17.329 | 17.221 | 17.715 | 17.487 | 17.439 | 16.716 | 16.865 | 16.756 | 16.652 | 18.303 | 17.742 | 15.273 | 17.610 | 16.752 | 16.478 | 18.104 | 17.345 | -0.006 | 0.8530 | 0.9370 | 0 | 0 | 0 |
| 'Nupl1' | 24.593 | 23.111 | 23.353 | 23.713 | 23.656 | 21.846 | 25.901 | 24.761 | 21.906 | 22.473 | 24.718 | 23.990 | 22.655 | 24.047 | 22.775 | 20.616 | 23.958 | 23.956 | 0.012 | 0.7651 | 0.8954 | 0 | 0 | 0 |
| 'Nupl2' | 2.297 | 2.948 | 3.144 | 2.207 | 3.319 | 3.071 | 2.581 | 3.243 | 3.220 | 3.004 | 2.565 | 2.195 | 3.202 | 2.190 | 2.243 | 2.850 | 2.735 | 2.001 | 0.176 | 0.0717 | 0.2363 | 0 | 0 | 0 |
| 'Nus1' | 12.554 | 12.727 | 12.121 | 10.930 | 12.693 | 12.368 | 11.456 | 13.237 | 10.539 | 12.331 | 11.426 | 13.067 | 11.977 | 13.265 | 11.923 | 12.885 | 11.903 | 12.630 | -0.049 | 0.2571 | 0.5093 | 0 | 0 | 0 |
| 'Nusap1' | 6.065 | 7.595 | 8.610 | 7.916 | 7.012 | 8.207 | 7.550 | 6.589 | 8.704 | 5.123 | 6.738 | 8.271 | 7.563 | 8.557 | 9.734 | 6.950 | 8.323 | 8.404 | -0.037 | 0.6162 | 0.8085 | 0 | 0 | 0 |
| 'Nutf2-ps1' | 19.481 | 21.988 | 20.408 | 21.829 | 21.279 | 20.398 | 21.013 | 20.854 | 20.607 | 18.304 | 19.301 | 22.328 | 19.985 | 19.395 | 20.655 | 21.146 | 18.985 | 22.137 | 0.026 | 0.5511 | 0.7647 | 0 | 0 | 0 |
| 'Nvl' | 6.057 | 6.484 | 6.683 | 6.244 | 6.591 | 6.223 | 6.641 | 6.660 | 6.473 | 6.327 | 6.713 | 6.964 | 6.611 | 6.663 | 6.150 | 6.839 | 7.098 | 6.520 | -0.056 | 0.0494 | 0.1875 | 0 | 0 | 0 |
| 'Nxf1' | 37.065 | 36.891 | 37.195 | 36.801 | 35.299 | 37.325 | 35.093 | 36.829 | 38.279 | 36.739 | 37.253 | 39.933 | 38.326 | 38.753 | 36.787 | 38.068 | 39.471 | 38.267 | -0.065 | 0.0026 | 0.0220 | 1 | 0 | 0 |
| 'Nxn' | 4.566 | 3.806 | 4.580 | 4.346 | 4.535 | 5.270 | 4.291 | 4.604 | 4.823 | 4.406 | 4.125 | 4.686 | 5.689 | 5.175 | 4.889 | 4.906 | 4.096 | 5.258 | -0.089 | 0.1766 | 0.4118 | 0 | 0 | 0 |
| 'Nxpe3' | 73.500 | 77.175 | 72.690 | 69.293 | 73.300 | 71.426 | 68.921 | 69.638 | 72.503 | 64.805 | 69.338 | 76.471 | 57.576 | 65.893 | 75.427 | 66.819 | 68.940 | 77.191 | 0.051 | 0.2036 | 0.4462 | 0 | 0 | 0 |
| 'Nxt1' | 23.483 | 24.673 | 21.063 | 23.200 | 23.102 | 23.588 | 22.480 | 19.801 | 22.689 | 18.842 | 20.841 | 22.072 | 21.812 | 22.278 | 21.790 | 21.146 | 23.548 | 23.048 | 0.049 | 0.3421 | 0.5981 | 0 | 0 | 0 |
| 'Nxt2' | 0.967 | 1.122 | 1.004 | 1.140 | 1.040 | 0.801 | 0.943 | 0.856 | 0.910 | 0.762 | 1.251 | 0.766 | 0.954 | 1.153 | 0.833 | 0.943 | 1.010 | 0.664 | 0.083 | 0.4345 | 0.6788 | 0 | 0 | 0 |
| 'Nyap1' | 0.542 | 0.407 | 0.639 | 0.400 | 0.417 | 0.772 | 0.610 | 0.511 | 0.619 | 0.570 | 0.418 | 0.689 | 0.379 | 0.399 | 0.559 | 0.513 | 0.379 | 0.594 | 0.121 | 0.2763 | 0.5319 | 0 | 0 | 0 |
| 'Nynrin' | 0.300 | 0.308 | 0.327 | 0.406 | 0.552 | 0.462 | 0.301 | 0.383 | 0.320 | 0.293 | 0.217 | 0.329 | 0.255 | 0.246 | 0.350 | 0.448 | 0.396 | 0.288 | 0.238 | 0.1063 | 0.3065 | 0 | 0 | 0 |
| 'Oaf' | 4.003 | 2.757 | 4.448 | 3.510 | 4.172 | 3.880 | 3.335 | 4.241 | 3.548 | 1.972 | 1.779 | 1.858 | 1.964 | 2.060 | 2.115 | 1.891 | 1.808 | 1.683 | 0.989 | 0.0000 | 0.0000 | 1 | 1 | 0 |
| 'Oard1' | 12.143 | 20.346 | 14.473 | 13.057 | 16.344 | 14.771 | 14.338 | 16.098 | 14.614 | 12.764 | 14.082 | 13.893 | 14.284 | 14.284 | 12.781 | 14.594 | 15.541 | 12.611 | 0.105 | 0.0632 | 0.2190 | 0 | 0 | 0 |
| 'Oas1b' | 5.602 | 7.363 | 7.808 | 6.007 | 6.733 | 7.973 | 4.986 | 6.973 | 9.065 | 5.391 | 6.954 | 10.526 | 5.853 | 7.099 | 10.239 | 6.374 | 6.843 | 10.094 | -0.157 | 0.0086 | 0.0559 | 0 | 0 | 0 |
| 'Oas1c' | 7.285 | 8.345 | 7.921 | 7.959 | 8.120 | 8.227 | 8.015 | 8.099 | 8.040 | 7.802 | 7.185 | 8.103 | 7.567 | 6.728 | 6.400 | 7.355 | 7.476 | 6.238 | 0.135 | 0.0086 | 0.0562 | 0 | 0 | 0 |
| 'Oas3' | 4.367 | 4.406 | 4.792 | 3.795 | 3.785 | 4.659 | 3.455 | 5.130 | 4.899 | 4.178 | 3.966 | 7.265 | 4.670 | 4.288 | 6.600 | 4.355 | 3.914 | 7.602 | -0.227 | 0.0064 | 0.0450 | 1 | 0 | 0 |
| 'Oat' | 26.532 | 28.287 | 28.630 | 26.774 | 27.065 | 28.334 | 27.381 | 27.845 | 26.465 | 26.313 | 27.620 | 27.045 | 28.134 | 26.632 | 26.763 | 26.982 | 27.847 | 23.927 | 0.027 | 0.4038 | 0.6528 | 0 | 0 | 0 |
| 'Oaz1' | 118.617 | 131.867 | 113.290 | 120.000 | 125.813 | 121.520 | 124.389 | 122.264 | 120.357 | 119.103 | 124.849 | 104.194 | 130.811 | 123.501 | 122.487 | 132.632 | 121.614 | 116.072 | -0.006 | 0.8609 | 0.9399 | 0 | 0 | 0 |
| 'Oaz2' | 13.123 | 15.377 | 12.448 | 12.277 | 16.284 | 13.903 | 12.370 | 15.579 | 12.720 | 13.442 | 14.176 | 13.284 | 13.490 | 14.601 | 12.974 | 14.723 | 13.270 | 12.550 | 0.003 | 0.9323 | 0.9746 | 0 | 0 | 0 |
| 'Obfc1' | 2.756 | 2.525 | 3.074 | 2.818 | 3.025 | 2.823 | 3.208 | 2.908 | 2.686 | 2.779 | 2.698 | 2.906 | 2.992 | 2.858 | 2.533 | 2.568 | 2.918 | 2.530 | 0.047 | 0.3932 | 0.6428 | 0 | 0 | 0 |
| 'Obscn' | 0.270 | 0.334 | 0.316 | 0.260 | 0.232 | 0.262 | 0.244 | 0.377 | 0.264 | 0.510 | 0.736 | 0.500 | 0.405 | 0.465 | 0.553 | 0.436 | 0.579 | 0.557 | -0.892 | 0.0000 | 0.0000 | 1 | 1 | 0 |
| 'Ocel1' | 2.915 | 3.346 | 3.022 | 2.734 | 3.189 | 3.542 | 2.839 | 2.984 | 3.556 | 2.908 | 2.883 | 3.709 | 3.159 | 3.181 | 3.879 | 3.470 | 3.024 | 3.610 | -0.097 | 0.0500 | 0.1886 | 0 | 0 | 0 |
| 'Ociad1' | 21.929 | 23.013 | 21.038 | 22.067 | 23.285 | 20.759 | 22.422 | 22.525 | 21.732 | 21.232 | 21.693 | 22.027 | 21.422 | 22.772 | 21.587 | 22.091 | 22.197 | 21.329 | 0.005 | 0.8195 | 0.9202 | 0 | 0 | 0 |
| 'Ociad2' | 0.865 | 0.710 | 0.708 | 0.827 | 0.878 | 0.773 | 0.694 | 0.823 | 0.600 | 0.858 | 0.607 | 0.998 | 0.860 | 0.924 | 0.722 | 0.963 | 0.596 | 0.894 | -0.131 | 0.2477 | 0.4985 | 0 | 0 | 0 |
| 'Ocrl' | 2.502 | 2.795 | 2.435 | 2.046 | 2.344 | 2.589 | 2.278 | 2.582 | 2.472 | 2.378 | 2.719 | 2.477 | 2.827 | 2.651 | 2.360 | 2.469 | 2.667 | 2.348 | -0.063 | 0.1969 | 0.4358 | 0 | 0 | 0 |
| 'Odc1' | 119.671 | 124.994 | 80.944 | 111.742 | 123.326 | 90.641 | 106.502 | 127.184 | 80.789 | 128.111 | 115.692 | 76.213 | 118.851 | 138.230 | 79.531 | 139.834 | 108.193 | 72.495 | -0.009 | 0.8589 | 0.9390 | 0 | 0 | 0 |
| 'Odf2' | 6.175 | 6.271 | 7.224 | 5.971 | 6.190 | 7.021 | 6.175 | 6.449 | 7.118 | 6.110 | 5.973 | 6.809 | 7.074 | 6.623 | 6.886 | 6.535 | 6.803 | 6.588 | -0.031 | 0.3549 | 0.6084 | 0 | 0 | 0 |
| 'Odf2l' | 0.801 | 0.859 | 0.905 | 0.865 | 0.764 | 0.871 | 0.889 | 0.880 | 0.952 | 0.923 | 0.842 | 0.804 | 0.852 | 0.871 | 0.741 | 0.846 | 0.980 | 0.800 | 0.022 | 0.7169 | 0.8692 | 0 | 0 | 0 |
| 'Ofd1' | 10.848 | 11.963 | 11.351 | 11.255 | 12.083 | 11.539 | 13.010 | 12.567 | 10.819 | 12.598 | 12.429 | 10.097 | 12.586 | 10.776 | 9.559 | 12.009 | 11.989 | 8.693 | 0.065 | 0.1874 | 0.4234 | 0 | 0 | 0 |
| 'Ogdh' | 42.120 | 41.535 | 43.925 | 43.562 | 42.726 | 43.442 | 41.177 | 41.286 | 43.244 | 44.485 | 43.871 | 44.222 | 44.095 | 42.157 | 42.588 | 41.175 | 42.335 | 42.634 | -0.028 | 0.1875 | 0.4235 | 0 | 0 | 0 |
| 'Ogfod1' | 5.426 | 5.896 | 5.764 | 5.815 | 6.186 | 5.512 | 5.549 | 5.581 | 5.619 | 5.422 | 5.398 | 6.530 | 5.439 | 5.111 | 6.433 | 5.386 | 5.750 | 6.468 | -0.030 | 0.4993 | 0.7273 | 0 | 0 | 0 |
| 'Ogfod2' | 3.781 | 4.123 | 4.000 | 3.643 | 4.142 | 4.247 | 3.619 | 4.624 | 3.951 | 4.103 | 3.646 | 4.120 | 4.198 | 3.582 | 4.287 | 4.096 | 3.960 | 4.027 | -0.007 | 0.8939 | 0.9566 | 0 | 0 | 0 |
| 'Ogfod3' | 3.635 | 4.378 | 4.303 | 3.489 | 3.291 | 3.215 | 3.984 | 4.040 | 3.457 | 2.369 | 1.986 | 3.742 | 3.338 | 4.101 | 3.601 | 4.393 | 3.836 | 3.954 | 0.096 | 0.4416 | 0.6851 | 0 | 0 | 0 |
| 'Ogfr' | 51.137 | 45.751 | 49.534 | 42.498 | 45.380 | 53.954 | 45.100 | 46.281 | 50.554 | 52.179 | 43.443 | 54.904 | 51.719 | 46.438 | 53.823 | 57.536 | 39.783 | 57.058 | -0.091 | 0.0472 | 0.1812 | 0 | 0 | 0 |
| 'Ogg1' | 2.354 | 2.846 | 3.156 | 2.305 | 2.809 | 2.624 | 2.156 | 2.542 | 2.610 | 2.368 | 2.328 | 2.528 | 2.852 | 2.913 | 2.635 | 2.424 | 2.651 | 2.163 | 0.024 | 0.7318 | 0.8778 | 0 | 0 | 0 |
| 'Ogt' | 15.713 | 19.167 | 19.535 | 15.487 | 17.723 | 17.553 | 16.228 | 18.042 | 19.832 | 14.262 | 17.936 | 21.198 | 15.410 | 17.326 | 19.538 | 16.272 | 18.660 | 20.407 | -0.022 | 0.5261 | 0.7477 | 0 | 0 | 0 |
| 'Oip5' | 1.305 | 1.296 | 1.942 | 1.553 | 1.211 | 1.904 | 1.482 | 1.611 | 1.745 | 1.137 | 1.550 | 1.535 | 1.957 | 1.550 | 1.757 | 1.374 | 1.691 | 1.745 | -0.029 | 0.7969 | 0.9097 | 0 | 0 | 0 |
| 'Ola1' | 10.738 | 10.033 | 10.853 | 11.337 | 11.074 | 10.010 | 12.370 | 11.143 | 9.958 | 9.780 | 10.700 | 10.465 | 10.517 | 10.577 | 10.324 | 9.982 | 11.150 | 9.992 | 0.048 | 0.2204 | 0.4678 | 0 | 0 | 0 |
| 'Olr1' | 1.226 | 1.414 | 1.452 | 1.043 | 1.396 | 0.983 | 1.399 | 1.257 | 1.539 | 1.731 | 2.119 | 1.800 | 2.085 | 1.925 | 1.684 | 1.638 | 2.529 | 1.920 | -0.582 | 0.0000 | 0.0000 | 1 | 1 | 0 |
| 'Oma1' | 8.398 | 8.152 | 9.282 | 10.579 | 9.766 | 9.781 | 9.267 | 8.491 | 8.983 | 10.704 | 9.806 | 9.738 | 10.136 | 9.232 | 8.382 | 8.515 | 9.034 | 7.963 | -0.025 | 0.6790 | 0.8465 | 0 | 0 | 0 |
| 'Opa1' | 6.377 | 7.152 | 7.176 | 6.480 | 7.778 | 6.975 | 7.331 | 7.274 | 7.225 | 6.972 | 7.281 | 6.596 | 7.032 | 7.248 | 6.531 | 6.907 | 6.787 | 6.236 | 0.041 | 0.2231 | 0.4707 | 0 | 0 | 0 |
| 'Opa3' | 8.837 | 8.623 | 8.368 | 9.028 | 8.762 | 9.838 | 8.563 | 8.953 | 9.068 | 8.704 | 7.735 | 10.509 | 8.724 | 7.903 | 10.073 | 9.691 | 8.385 | 10.313 | -0.045 | 0.3284 | 0.5865 | 0 | 0 | 0 |
| 'Optn' | 1.906 | 2.579 | 2.244 | 2.187 | 2.966 | 2.927 | 2.585 | 2.492 | 2.389 | 2.136 | 2.667 | 2.122 | 2.624 | 2.415 | 2.564 | 2.144 | 2.401 | 2.265 | 0.049 | 0.5091 | 0.7342 | 0 | 0 | 0 |
| 'Orai1' | 20.055 | 19.253 | 21.209 | 17.493 | 19.068 | 21.898 | 17.471 | 18.619 | 22.644 | 19.340 | 17.402 | 23.725 | 17.923 | 18.929 | 22.823 | 20.457 | 19.651 | 23.783 | -0.059 | 0.1173 | 0.3259 | 0 | 0 | 0 |
| 'Orai2' | 20.692 | 20.045 | 20.112 | 18.341 | 18.998 | 21.174 | 19.528 | 19.567 | 21.714 | 21.353 | 21.322 | 23.794 | 21.332 | 21.272 | 23.171 | 23.342 | 19.618 | 23.534 | -0.151 | 0.0000 | 0.0000 | 1 | 0 | 0 |
| 'Orai3' | 7.226 | 8.417 | 7.179 | 7.645 | 7.487 | 7.234 | 6.951 | 7.231 | 7.367 | 7.406 | 7.065 | 8.724 | 7.039 | 7.795 | 8.026 | 7.528 | 7.886 | 8.433 | -0.085 | 0.0728 | 0.2389 | 0 | 0 | 0 |
| 'Oraov1' | 3.577 | 3.840 | 4.068 | 4.008 | 3.590 | 3.744 | 4.800 | 4.124 | 4.181 | 3.265 | 4.151 | 4.376 | 3.891 | 3.539 | 4.213 | 3.665 | 4.102 | 4.308 | 0.004 | 0.9546 | 0.9832 | 0 | 0 | 0 |
| 'Orc1' | 2.199 | 1.831 | 2.345 | 2.039 | 1.749 | 1.656 | 2.650 | 1.651 | 1.963 | 1.515 | 1.788 | 2.308 | 2.467 | 2.205 | 2.429 | 2.282 | 1.931 | 2.090 | -0.092 | 0.3361 | 0.5947 | 0 | 0 | 0 |
| 'Orc2' | 8.419 | 8.719 | 8.795 | 8.702 | 8.932 | 8.279 | 8.964 | 9.371 | 8.096 | 8.737 | 8.733 | 8.728 | 7.615 | 8.305 | 8.028 | 8.242 | 9.114 | 7.912 | 0.042 | 0.2787 | 0.5352 | 0 | 0 | 0 |
| 'Orc3' | 8.279 | 8.983 | 8.890 | 8.506 | 9.103 | 8.693 | 8.865 | 9.174 | 8.660 | 9.212 | 8.415 | 8.123 | 8.639 | 8.629 | 7.751 | 7.980 | 8.874 | 7.316 | 0.071 | 0.0344 | 0.1481 | 0 | 0 | 0 |
| 'Orc4' | 4.466 | 4.935 | 4.655 | 5.214 | 4.487 | 3.988 | 5.511 | 4.716 | 4.523 | 3.899 | 5.114 | 5.311 | 4.042 | 5.154 | 4.783 | 3.785 | 6.202 | 4.952 | -0.028 | 0.7202 | 0.8713 | 0 | 0 | 0 |
| 'Orc5' | 3.961 | 3.918 | 3.711 | 3.675 | 3.735 | 3.386 | 4.136 | 3.978 | 3.833 | 3.604 | 3.896 | 3.754 | 3.623 | 4.081 | 4.039 | 3.251 | 4.456 | 3.469 | -0.003 | 0.9465 | 0.9794 | 0 | 0 | 0 |
| 'Orc6' | 12.168 | 11.320 | 13.282 | 12.629 | 11.559 | 12.035 | 13.987 | 12.049 | 12.665 | 11.372 | 12.543 | 11.001 | 13.562 | 11.774 | 11.321 | 11.368 | 11.294 | 10.819 | 0.082 | 0.1026 | 0.3002 | 0 | 0 | 0 |
| 'Ormdl1' | 6.842 | 8.248 | 7.395 | 7.914 | 8.061 | 6.438 | 7.239 | 7.543 | 6.679 | 7.621 | 8.118 | 7.805 | 7.196 | 8.951 | 7.725 | 7.299 | 8.028 | 7.592 | -0.100 | 0.0220 | 0.1099 | 0 | 0 | 0 |
| 'Ormdl2' | 17.100 | 19.967 | 17.902 | 19.196 | 21.330 | 20.098 | 19.316 | 19.939 | 18.217 | 18.319 | 20.344 | 20.614 | 18.482 | 20.955 | 20.685 | 19.535 | 20.960 | 20.312 | -0.074 | 0.0561 | 0.2033 | 0 | 0 | 0 |
| 'Ormdl3' | 26.191 | 28.391 | 24.351 | 24.803 | 26.357 | 26.812 | 23.885 | 28.223 | 24.494 | 23.348 | 22.329 | 24.022 | 23.969 | 26.458 | 24.570 | 25.854 | 25.479 | 26.722 | 0.057 | 0.1851 | 0.4207 | 0 | 0 | 0 |
| 'Os9' | 86.846 | 83.477 | 86.465 | 82.441 | 82.333 | 91.413 | 85.183 | 84.131 | 91.496 | 90.918 | 81.106 | 87.405 | 84.206 | 83.868 | 86.455 | 90.327 | 79.323 | 85.687 | -0.001 | 0.9637 | 0.9869 | 0 | 0 | 0 |
| 'Osbp' | 29.641 | 30.754 | 30.544 | 27.901 | 31.554 | 31.167 | 27.130 | 30.845 | 31.340 | 26.864 | 30.536 | 30.947 | 28.384 | 30.395 | 30.948 | 29.731 | 30.416 | 32.713 | -0.010 | 0.7125 | 0.8664 | 0 | 0 | 0 |
| 'Osbpl11' | 13.933 | 13.955 | 13.431 | 13.026 | 13.904 | 13.509 | 12.393 | 13.252 | 13.840 | 12.858 | 12.506 | 13.179 | 12.894 | 13.575 | 13.128 | 13.017 | 13.246 | 13.350 | 0.032 | 0.3039 | 0.5614 | 0 | 0 | 0 |
| 'Osbpl2' | 9.101 | 10.704 | 9.570 | 9.778 | 10.325 | 10.316 | 9.108 | 10.367 | 10.196 | 9.112 | 10.639 | 9.920 | 9.214 | 9.943 | 10.194 | 10.540 | 9.419 | 9.782 | 0.001 | 0.9767 | 0.9920 | 0 | 0 | 0 |
| 'Osbpl3' | 4.797 | 5.255 | 4.664 | 5.151 | 5.296 | 4.894 | 5.390 | 5.423 | 4.970 | 4.205 | 5.074 | 4.432 | 4.942 | 4.723 | 4.373 | 4.445 | 4.915 | 4.374 | 0.132 | 0.0001 | 0.0009 | 1 | 0 | 0 |
| 'Osbpl5' | 4.499 | 4.940 | 5.328 | 4.579 | 5.122 | 5.333 | 5.138 | 5.413 | 4.990 | 5.910 | 5.592 | 5.201 | 5.655 | 5.811 | 4.713 | 5.583 | 4.890 | 5.096 | -0.102 | 0.0390 | 0.1612 | 0 | 0 | 0 |
| 'Osbpl6' | 0.280 | 0.271 | 0.475 | 0.228 | 0.239 | 0.347 | 0.265 | 0.318 | 0.445 | 0.297 | 0.197 | 0.449 | 0.140 | 0.206 | 0.322 | 0.201 | 0.241 | 0.358 | 0.249 | 0.0445 | 0.1748 | 0 | 0 | 0 |
| 'Osbpl7' | 6.327 | 6.640 | 7.471 | 6.291 | 6.369 | 7.532 | 6.385 | 6.055 | 7.421 | 7.329 | 6.071 | 7.389 | 7.270 | 5.590 | 6.809 | 6.744 | 6.430 | 6.959 | -0.013 | 0.7327 | 0.8781 | 0 | 0 | 0 |
| 'Osbpl8' | 13.320 | 15.726 | 14.124 | 14.895 | 15.097 | 14.375 | 16.744 | 14.726 | 14.503 | 12.906 | 15.253 | 15.673 | 14.889 | 14.686 | 14.344 | 14.396 | 15.698 | 16.086 | -0.018 | 0.6864 | 0.8517 | 0 | 0 | 0 |
| 'Osbpl9' | 5.278 | 5.295 | 5.100 | 5.613 | 5.447 | 5.183 | 5.235 | 5.502 | 5.519 | 5.280 | 6.053 | 5.655 | 5.700 | 6.013 | 5.512 | 5.727 | 5.679 | 5.468 | -0.095 | 0.0016 | 0.0149 | 1 | 0 | 0 |
| 'Oser1' | 11.629 | 9.616 | 7.666 | 9.153 | 9.595 | 8.204 | 7.981 | 9.313 | 7.834 | 11.220 | 10.232 | 8.034 | 9.486 | 11.328 | 8.181 | 12.418 | 9.838 | 8.431 | -0.140 | 0.0244 | 0.1187 | 0 | 0 | 0 |
| 'Osgep' | 5.118 | 5.554 | 4.690 | 4.747 | 5.513 | 5.305 | 5.094 | 5.612 | 5.309 | 5.591 | 4.702 | 5.323 | 4.899 | 4.774 | 5.237 | 5.432 | 5.404 | 4.811 | 0.011 | 0.8070 | 0.9135 | 0 | 0 | 0 |
| 'Osgepl1' | 1.180 | 1.487 | 1.353 | 1.454 | 1.518 | 1.155 | 1.345 | 1.427 | 1.348 | 1.364 | 1.366 | 1.283 | 1.384 | 1.503 | 1.250 | 1.275 | 1.419 | 1.142 | 0.019 | 0.7563 | 0.8913 | 0 | 0 | 0 |
| 'Osgin1' | 3.949 | 3.650 | 3.783 | 3.534 | 3.109 | 4.009 | 3.939 | 3.590 | 3.876 | 4.017 | 3.382 | 3.857 | 3.450 | 3.419 | 3.373 | 3.837 | 3.569 | 3.156 | 0.054 | 0.3306 | 0.5884 | 0 | 0 | 0 |
| 'Osgin2' | 2.188 | 2.304 | 2.252 | 2.088 | 2.407 | 1.964 | 2.276 | 2.319 | 1.990 | 2.496 | 2.130 | 2.276 | 2.365 | 2.675 | 1.990 | 2.595 | 2.054 | 1.977 | -0.064 | 0.2557 | 0.5081 | 0 | 0 | 0 |
| 'Osm' | 4.799 | 4.200 | 5.082 | 4.810 | 4.241 | 4.805 | 5.335 | 4.391 | 4.290 | 3.561 | 3.361 | 4.549 | 3.925 | 4.103 | 4.528 | 3.547 | 4.166 | 3.946 | 0.216 | 0.0012 | 0.0119 | 1 | 0 | 0 |
| 'Ost4' | 33.916 | 40.824 | 35.991 | 38.766 | 39.540 | 34.495 | 35.152 | 36.298 | 34.399 | 31.794 | 36.070 | 34.750 | 35.059 | 37.982 | 37.673 | 35.766 | 38.091 | 35.628 | 0.016 | 0.6816 | 0.8485 | 0 | 0 | 0 |
| 'Ostc' | 18.986 | 24.721 | 21.465 | 20.597 | 24.598 | 21.814 | 20.787 | 24.231 | 19.773 | 21.857 | 19.801 | 20.785 | 21.634 | 22.587 | 20.898 | 21.660 | 22.441 | 20.128 | 0.023 | 0.5509 | 0.7646 | 0 | 0 | 0 |
| 'Ostf1' | 12.197 | 13.646 | 13.040 | 12.924 | 13.164 | 14.125 | 13.496 | 13.438 | 13.358 | 11.894 | 12.598 | 12.412 | 12.851 | 12.412 | 12.256 | 13.364 | 12.532 | 11.935 | 0.080 | 0.0095 | 0.0600 | 0 | 0 | 0 |
| 'Ostm1' | 6.886 | 8.499 | 7.906 | 7.079 | 8.872 | 8.113 | 7.160 | 8.637 | 7.602 | 7.619 | 8.781 | 7.656 | 7.486 | 8.491 | 7.071 | 7.658 | 8.109 | 7.412 | 0.000 | 0.9980 | 0.9994 | 0 | 0 | 0 |
| 'Otub1' | 21.712 | 23.065 | 23.478 | 21.302 | 22.929 | 24.230 | 21.263 | 23.504 | 22.706 | 20.621 | 21.902 | 21.370 | 22.341 | 20.408 | 22.086 | 24.320 | 20.140 | 22.038 | 0.056 | 0.1214 | 0.3322 | 0 | 0 | 0 |
| 'Otub2' | 1.197 | 1.188 | 1.113 | 1.054 | 1.299 | 1.194 | 1.077 | 1.069 | 1.007 | 1.019 | 1.078 | 0.978 | 0.993 | 1.086 | 1.134 | 1.146 | 1.156 | 1.097 | 0.070 | 0.2893 | 0.5459 | 0 | 0 | 0 |
| 'Otud1' | 13.432 | 9.447 | 7.213 | 9.823 | 7.650 | 8.807 | 8.659 | 9.481 | 7.186 | 11.391 | 9.702 | 7.840 | 9.365 | 10.924 | 7.728 | 11.254 | 10.386 | 6.826 | -0.065 | 0.4225 | 0.6692 | 0 | 0 | 0 |
| 'Otud3' | 1.907 | 1.595 | 2.213 | 1.629 | 1.502 | 1.619 | 1.421 | 1.451 | 1.164 | 8.539 | 8.947 | 8.797 | 7.898 | 8.087 | 8.246 | 7.743 | 6.878 | 8.311 | -2.340 | 0.0000 | 0.0000 | 1 | 1 | 1 |
| 'Otud4' | 14.386 | 14.571 | 15.467 | 14.131 | 15.072 | 13.618 | 13.253 | 15.336 | 15.271 | 14.356 | 14.573 | 14.447 | 13.473 | 14.192 | 13.842 | 14.796 | 14.453 | 14.014 | 0.023 | 0.4426 | 0.6862 | 0 | 0 | 0 |
| 'Otud5' | 18.705 | 16.828 | 19.463 | 17.316 | 18.058 | 20.467 | 17.508 | 17.051 | 19.434 | 19.174 | 18.033 | 19.861 | 19.933 | 17.979 | 19.817 | 18.908 | 18.321 | 19.671 | -0.069 | 0.0048 | 0.0363 | 1 | 0 | 0 |
| 'Otud6b' | 5.292 | 5.647 | 5.224 | 5.936 | 6.432 | 5.368 | 5.885 | 6.531 | 5.328 | 5.345 | 5.895 | 5.934 | 5.296 | 6.045 | 5.630 | 6.499 | 5.734 | 5.137 | -0.012 | 0.8325 | 0.9270 | 0 | 0 | 0 |
| 'Otud7b' | 12.333 | 12.545 | 12.116 | 12.388 | 12.749 | 12.340 | 11.981 | 12.216 | 11.788 | 13.562 | 13.569 | 13.326 | 11.303 | 12.750 | 13.467 | 11.714 | 13.461 | 13.594 | -0.090 | 0.0149 | 0.0833 | 0 | 0 | 0 |
| 'Otulin' | 29.830 | 26.678 | 27.679 | 26.408 | 27.233 | 30.404 | 30.066 | 29.072 | 25.739 | 27.733 | 29.359 | 24.977 | 30.195 | 29.227 | 27.085 | 29.288 | 29.170 | 27.435 | -0.011 | 0.8092 | 0.9140 | 0 | 0 | 0 |
| 'Ovca2' | 4.958 | 5.284 | 5.447 | 5.064 | 5.210 | 4.993 | 4.959 | 5.036 | 5.278 | 5.423 | 5.487 | 5.733 | 5.177 | 5.057 | 5.635 | 5.137 | 5.490 | 5.568 | -0.089 | 0.0734 | 0.2402 | 0 | 0 | 0 |
| 'Oxa1l' | 21.058 | 22.733 | 21.269 | 22.077 | 22.702 | 22.431 | 20.911 | 22.209 | 21.788 | 21.045 | 20.548 | 21.775 | 22.274 | 19.808 | 22.082 | 21.320 | 20.065 | 20.932 | 0.042 | 0.1349 | 0.3537 | 0 | 0 | 0 |
| 'Oxct1' | 21.726 | 23.091 | 24.470 | 24.590 | 23.001 | 24.684 | 25.017 | 22.857 | 23.739 | 23.129 | 23.575 | 24.885 | 22.309 | 23.373 | 23.315 | 22.072 | 22.970 | 23.989 | 0.012 | 0.6715 | 0.8417 | 0 | 0 | 0 |
| 'Oxld1' | 3.915 | 4.237 | 4.221 | 3.354 | 4.238 | 3.646 | 2.457 | 3.685 | 4.294 | 4.035 | 3.654 | 4.024 | 3.132 | 2.976 | 4.266 | 3.879 | 2.646 | 4.321 | 0.041 | 0.7122 | 0.8661 | 0 | 0 | 0 |
| 'Oxnad1' | 1.243 | 1.429 | 1.159 | 1.314 | 1.332 | 1.214 | 0.990 | 1.383 | 1.200 | 1.178 | 1.201 | 1.197 | 1.180 | 1.345 | 1.273 | 1.223 | 1.301 | 1.073 | 0.027 | 0.6585 | 0.8350 | 0 | 0 | 0 |
| 'Oxr1' | 10.700 | 11.854 | 11.080 | 10.713 | 12.158 | 11.522 | 11.663 | 11.721 | 11.082 | 11.808 | 13.081 | 12.062 | 10.589 | 12.389 | 11.121 | 11.872 | 12.398 | 11.693 | -0.072 | 0.0174 | 0.0937 | 0 | 0 | 0 |
| 'Oxsm' | 2.082 | 2.683 | 2.742 | 2.050 | 3.253 | 2.879 | 2.668 | 2.737 | 2.585 | 2.324 | 2.595 | 2.530 | 2.694 | 2.183 | 2.622 | 2.596 | 2.653 | 2.571 | 0.040 | 0.5587 | 0.7697 | 0 | 0 | 0 |
| 'Oxsr1' | 15.030 | 14.980 | 15.133 | 14.243 | 15.605 | 15.272 | 14.949 | 15.664 | 14.611 | 14.354 | 15.257 | 15.003 | 14.271 | 15.264 | 15.030 | 14.737 | 13.965 | 15.475 | 0.013 | 0.6316 | 0.8176 | 0 | 0 | 0 |
| 'P2rx4' | 1.842 | 2.329 | 2.169 | 1.546 | 2.314 | 2.148 | 1.685 | 2.579 | 2.150 | 2.179 | 2.112 | 1.879 | 2.126 | 2.015 | 1.643 | 1.745 | 2.076 | 1.721 | 0.086 | 0.2255 | 0.4722 | 0 | 0 | 0 |
| 'P2rx7' | 32.906 | 33.034 | 34.324 | 32.921 | 33.624 | 34.235 | 33.269 | 33.341 | 35.939 | 34.091 | 37.866 | 38.854 | 35.462 | 36.212 | 39.231 | 36.024 | 37.396 | 40.326 | -0.154 | 0.0000 | 0.0000 | 1 | 0 | 0 |
| 'P2ry10' | 10.453 | 9.966 | 9.582 | 9.960 | 9.746 | 9.298 | 11.205 | 10.790 | 8.912 | 10.394 | 9.539 | 10.792 | 8.998 | 10.110 | 10.216 | 10.619 | 9.287 | 10.889 | -0.028 | 0.5628 | 0.7719 | 0 | 0 | 0 |
| 'P3h1' | 0.725 | 0.837 | 0.779 | 0.707 | 0.897 | 0.786 | 0.830 | 0.715 | 0.790 | 0.888 | 0.762 | 0.871 | 0.936 | 0.700 | 0.705 | 0.993 | 0.723 | 0.665 | -0.047 | 0.5468 | 0.7610 | 0 | 0 | 0 |
| 'P4ha1' | 14.759 | 13.320 | 11.402 | 16.397 | 11.598 | 11.280 | 15.277 | 11.665 | 11.501 | 10.608 | 14.767 | 20.892 | 13.095 | 15.100 | 19.550 | 13.805 | 16.415 | 23.039 | -0.320 | 0.0287 | 0.1313 | 0 | 0 | 0 |
| 'P4hb' | 79.257 | 69.874 | 72.878 | 73.791 | 71.512 | 70.600 | 72.958 | 72.005 | 69.730 | 78.564 | 70.662 | 72.196 | 75.454 | 76.931 | 73.569 | 75.291 | 72.301 | 70.823 | -0.038 | 0.1118 | 0.3168 | 0 | 0 | 0 |
| 'P4htm' | 2.142 | 2.164 | 2.479 | 1.692 | 1.740 | 2.320 | 2.223 | 2.086 | 2.201 | 2.328 | 1.648 | 2.260 | 2.033 | 1.946 | 1.812 | 2.053 | 1.901 | 1.947 | 0.079 | 0.2715 | 0.5264 | 0 | 0 | 0 |
| 'Pa2g4' | 48.933 | 45.706 | 50.967 | 51.893 | 46.146 | 46.619 | 51.456 | 46.936 | 48.135 | 42.430 | 42.723 | 49.286 | 49.769 | 43.937 | 47.628 | 42.745 | 43.849 | 48.053 | 0.078 | 0.0228 | 0.1127 | 0 | 0 | 0 |
| 'Pabpc1' | 244.961 | 220.503 | 228.850 | 250.774 | 219.374 | 223.829 | 259.446 | 213.270 | 229.812 | 241.919 | 231.115 | 208.048 | 246.468 | 223.651 | 228.868 | 232.589 | 222.947 | 227.423 | 0.007 | 0.8192 | 0.9200 | 0 | 0 | 0 |
| 'Pabpc4' | 31.543 | 30.512 | 32.963 | 29.379 | 32.922 | 32.916 | 32.677 | 33.925 | 33.131 | 34.783 | 28.551 | 31.028 | 32.995 | 30.388 | 31.508 | 34.883 | 28.239 | 30.726 | 0.028 | 0.4601 | 0.6967 | 0 | 0 | 0 |
| 'Pabpn1' | 24.823 | 27.143 | 27.865 | 22.848 | 25.303 | 26.768 | 24.522 | 26.212 | 28.245 | 23.403 | 24.697 | 26.094 | 25.827 | 25.609 | 25.961 | 24.859 | 25.708 | 25.172 | 0.029 | 0.2886 | 0.5451 | 0 | 0 | 0 |
| 'Pacs1' | 36.538 | 34.825 | 38.514 | 34.873 | 36.259 | 40.451 | 35.481 | 36.543 | 39.446 | 40.417 | 35.473 | 38.691 | 38.687 | 35.058 | 38.697 | 36.442 | 31.644 | 37.758 | -0.009 | 0.7801 | 0.9010 | 0 | 0 | 0 |
| 'Pacs2' | 15.991 | 17.420 | 18.631 | 15.977 | 17.534 | 18.504 | 14.893 | 17.527 | 19.186 | 17.806 | 16.355 | 16.931 | 16.049 | 16.573 | 16.146 | 16.701 | 15.782 | 16.466 | 0.054 | 0.1522 | 0.3794 | 0 | 0 | 0 |
| 'Pacsin1' | 18.118 | 19.036 | 19.784 | 17.737 | 17.769 | 20.604 | 17.558 | 19.232 | 20.719 | 20.615 | 18.924 | 19.149 | 18.543 | 17.842 | 18.621 | 19.449 | 18.374 | 17.743 | 0.002 | 0.9643 | 0.9869 | 0 | 0 | 0 |
| 'Pacsin2' | 15.312 | 16.403 | 15.956 | 15.939 | 17.010 | 16.815 | 14.694 | 16.893 | 16.103 | 15.607 | 15.476 | 15.171 | 15.988 | 15.718 | 15.394 | 16.961 | 15.505 | 15.036 | 0.035 | 0.2570 | 0.5093 | 0 | 0 | 0 |
| 'Padi4' | 12.200 | 13.557 | 11.891 | 14.517 | 14.872 | 12.298 | 12.566 | 14.671 | 12.214 | 11.502 | 12.411 | 10.394 | 17.214 | 12.142 | 8.593 | 12.314 | 13.931 | 8.947 | 0.150 | 0.0373 | 0.1564 | 0 | 0 | 0 |
| 'Paf1' | 15.242 | 14.252 | 15.075 | 14.987 | 14.599 | 15.229 | 15.155 | 14.926 | 15.190 | 15.910 | 15.482 | 15.304 | 16.757 | 14.759 | 15.324 | 15.193 | 14.603 | 14.719 | -0.044 | 0.0798 | 0.2542 | 0 | 0 | 0 |
| 'Pafah1b1' | 12.736 | 13.512 | 12.708 | 12.911 | 13.177 | 11.686 | 12.772 | 13.500 | 12.341 | 12.632 | 13.010 | 12.120 | 12.603 | 13.658 | 12.152 | 12.890 | 13.065 | 12.144 | 0.003 | 0.8650 | 0.9414 | 0 | 0 | 0 |
| 'Pafah1b2' | 23.203 | 26.028 | 23.198 | 24.278 | 24.849 | 22.479 | 23.363 | 25.832 | 22.585 | 22.073 | 23.762 | 23.237 | 23.071 | 24.830 | 22.471 | 21.839 | 22.942 | 23.380 | 0.041 | 0.1375 | 0.3575 | 0 | 0 | 0 |
| 'Pafah1b3' | 3.082 | 4.190 | 3.444 | 3.374 | 3.866 | 3.385 | 3.924 | 3.589 | 3.124 | 2.989 | 3.091 | 3.168 | 3.247 | 3.494 | 3.530 | 3.232 | 2.906 | 3.210 | 0.127 | 0.0487 | 0.1853 | 0 | 0 | 0 |
| 'Pafah2' | 0.835 | 0.641 | 0.984 | 1.103 | 1.047 | 0.931 | 0.985 | 0.923 | 0.990 | 0.944 | 0.996 | 0.845 | 0.733 | 0.772 | 0.716 | 0.671 | 0.855 | 0.620 | 0.234 | 0.0387 | 0.1604 | 0 | 0 | 0 |
| 'Pag1' | 2.595 | 2.699 | 2.764 | 2.724 | 2.439 | 2.645 | 2.332 | 2.653 | 2.846 | 2.419 | 2.736 | 2.769 | 2.431 | 2.534 | 2.453 | 2.478 | 2.828 | 2.961 | -0.004 | 0.9324 | 0.9746 | 0 | 0 | 0 |
| 'Pagr1a' | 14.044 | 13.969 | 15.192 | 14.617 | 13.282 | 15.301 | 14.166 | 13.580 | 15.111 | 12.274 | 13.859 | 15.993 | 14.038 | 13.861 | 15.093 | 13.264 | 14.751 | 15.927 | -0.009 | 0.8035 | 0.9117 | 0 | 0 | 0 |
| 'Paics' | 19.255 | 18.656 | 19.542 | 20.754 | 19.335 | 19.003 | 20.229 | 19.430 | 19.482 | 16.803 | 18.067 | 19.646 | 18.927 | 17.814 | 18.826 | 16.365 | 19.386 | 17.629 | 0.092 | 0.0096 | 0.0607 | 0 | 0 | 0 |
| 'Paip1' | 11.096 | 10.259 | 10.582 | 11.393 | 10.664 | 9.903 | 11.495 | 11.405 | 9.675 | 10.063 | 10.568 | 12.693 | 10.662 | 11.518 | 11.157 | 10.637 | 11.715 | 12.066 | -0.081 | 0.1016 | 0.2987 | 0 | 0 | 0 |
| 'Paip2' | 38.571 | 42.975 | 38.333 | 41.298 | 41.433 | 37.403 | 43.319 | 40.291 | 39.021 | 36.306 | 39.210 | 38.099 | 37.183 | 40.572 | 38.231 | 37.302 | 41.094 | 36.915 | 0.060 | 0.0322 | 0.1417 | 0 | 0 | 0 |
| 'Paip2b' | 6.650 | 7.017 | 6.904 | 6.885 | 7.331 | 7.280 | 6.253 | 7.150 | 6.911 | 6.945 | 6.131 | 6.776 | 5.736 | 6.367 | 6.992 | 6.872 | 6.817 | 6.693 | 0.063 | 0.1217 | 0.3325 | 0 | 0 | 0 |
| 'Pak1ip1' | 8.927 | 9.214 | 9.570 | 8.683 | 9.226 | 9.692 | 8.457 | 9.777 | 9.362 | 9.331 | 8.742 | 8.779 | 9.167 | 9.051 | 8.546 | 9.195 | 8.812 | 7.516 | 0.061 | 0.1174 | 0.3260 | 0 | 0 | 0 |
| 'Pak2' | 71.018 | 71.772 | 70.171 | 71.891 | 73.364 | 68.289 | 74.108 | 71.119 | 67.788 | 68.578 | 74.973 | 71.819 | 70.711 | 73.870 | 70.892 | 67.509 | 75.100 | 71.780 | -0.024 | 0.3209 | 0.5789 | 0 | 0 | 0 |
| 'Pakap' | 0.339 | 0.362 | 0.322 | 0.336 | 0.328 | 0.345 | 0.365 | 0.469 | 0.358 | 0.367 | 0.398 | 0.307 | 0.359 | 0.393 | 0.299 | 0.308 | 0.390 | 0.326 | 0.035 | 0.6658 | 0.8389 | 0 | 0 | 0 |
| 'Palb2' | 0.710 | 0.599 | 0.928 | 0.850 | 0.789 | 0.697 | 0.859 | 0.890 | 0.967 | 0.620 | 0.844 | 0.860 | 0.788 | 0.864 | 0.810 | 0.822 | 0.762 | 0.941 | -0.012 | 0.8890 | 0.9545 | 0 | 0 | 0 |
| 'Palld' | 1.510 | 1.402 | 1.397 | 1.390 | 1.373 | 1.432 | 1.402 | 1.259 | 1.408 | 2.237 | 2.097 | 1.810 | 2.562 | 2.019 | 2.063 | 2.002 | 2.232 | 1.836 | -0.588 | 0.0000 | 0.0000 | 1 | 1 | 0 |
| 'Palm3' | 2.675 | 1.802 | 2.674 | 1.925 | 2.464 | 2.358 | 2.785 | 2.164 | 2.534 | 2.205 | 1.619 | 1.990 | 1.798 | 1.688 | 2.134 | 1.484 | 1.809 | 2.055 | 0.340 | 0.0000 | 0.0008 | 1 | 0 | 0 |
| 'Pan2' | 20.069 | 19.962 | 22.382 | 20.635 | 18.775 | 20.602 | 21.443 | 20.863 | 22.512 | 21.837 | 21.222 | 22.710 | 22.421 | 21.194 | 20.395 | 19.306 | 22.769 | 21.435 | -0.057 | 0.1226 | 0.3341 | 0 | 0 | 0 |
| 'Pan3' | 7.990 | 7.507 | 7.520 | 7.609 | 7.361 | 6.921 | 7.292 | 7.679 | 6.989 | 7.678 | 8.055 | 7.625 | 7.127 | 7.327 | 7.331 | 7.477 | 7.468 | 8.220 | -0.042 | 0.2523 | 0.5038 | 0 | 0 | 0 |
| 'Pank2' | 8.384 | 9.604 | 9.188 | 9.144 | 9.075 | 9.580 | 9.167 | 9.783 | 9.016 | 8.467 | 9.337 | 9.105 | 9.351 | 9.916 | 9.339 | 8.779 | 9.192 | 8.285 | 0.010 | 0.7366 | 0.8802 | 0 | 0 | 0 |
| 'Pank3' | 9.340 | 10.679 | 9.919 | 10.377 | 10.718 | 9.778 | 10.337 | 10.811 | 9.120 | 9.833 | 12.424 | 12.020 | 10.121 | 11.467 | 10.880 | 9.888 | 11.557 | 11.099 | -0.137 | 0.0005 | 0.0060 | 1 | 0 | 0 |
| 'Pank4' | 20.546 | 17.909 | 21.154 | 20.955 | 19.419 | 20.297 | 20.992 | 18.542 | 20.784 | 18.588 | 18.270 | 23.812 | 19.215 | 18.505 | 21.332 | 18.485 | 20.099 | 22.384 | -0.010 | 0.7898 | 0.9061 | 0 | 0 | 0 |
| 'Panx1' | 20.373 | 19.769 | 19.503 | 19.838 | 19.100 | 19.944 | 18.977 | 18.983 | 18.643 | 20.006 | 19.958 | 21.778 | 19.465 | 19.958 | 22.743 | 18.874 | 20.350 | 21.469 | -0.087 | 0.0119 | 0.0707 | 0 | 0 | 0 |
| 'Paox' | 4.415 | 4.129 | 5.631 | 3.667 | 4.900 | 5.386 | 5.278 | 5.313 | 4.747 | 3.997 | 4.364 | 4.436 | 4.522 | 5.102 | 4.696 | 4.333 | 5.658 | 4.417 | 0.067 | 0.4153 | 0.6631 | 0 | 0 | 0 |
| 'Papd4' | 34.458 | 34.892 | 36.752 | 34.932 | 35.927 | 33.461 | 38.295 | 34.974 | 33.736 | 38.042 | 34.973 | 32.275 | 36.332 | 34.722 | 29.013 | 32.763 | 35.426 | 29.427 | 0.061 | 0.1261 | 0.3401 | 0 | 0 | 0 |
| 'Papd5' | 11.259 | 11.484 | 10.293 | 11.120 | 11.098 | 10.510 | 10.381 | 11.413 | 9.462 | 10.993 | 11.161 | 12.100 | 11.485 | 11.740 | 11.385 | 10.232 | 11.793 | 12.768 | -0.109 | 0.0121 | 0.0716 | 0 | 0 | 0 |
| 'Papd7' | 5.669 | 5.924 | 5.244 | 5.366 | 5.436 | 5.466 | 4.698 | 5.860 | 5.060 | 5.315 | 5.235 | 6.239 | 5.365 | 5.458 | 6.118 | 6.360 | 5.596 | 6.370 | -0.107 | 0.0417 | 0.1681 | 0 | 0 | 0 |
| 'Papola' | 13.016 | 13.989 | 13.835 | 14.147 | 13.540 | 13.070 | 13.794 | 13.828 | 12.914 | 12.635 | 12.786 | 13.429 | 12.472 | 13.003 | 12.557 | 12.505 | 13.124 | 13.520 | 0.062 | 0.0175 | 0.0938 | 0 | 0 | 0 |
| 'Papolg' | 5.591 | 7.260 | 6.688 | 6.211 | 6.683 | 6.131 | 6.093 | 6.083 | 6.119 | 5.448 | 5.974 | 7.093 | 5.754 | 6.589 | 6.459 | 6.686 | 7.349 | 6.810 | -0.046 | 0.3500 | 0.6044 | 0 | 0 | 0 |
| 'Papss1' | 13.492 | 16.678 | 12.059 | 13.949 | 15.441 | 14.237 | 14.369 | 14.988 | 12.912 | 13.939 | 13.524 | 12.358 | 14.932 | 15.004 | 11.666 | 14.461 | 14.372 | 11.236 | 0.067 | 0.0832 | 0.2611 | 0 | 0 | 0 |
| 'Paqr4' | 2.569 | 2.411 | 2.787 | 2.350 | 2.719 | 3.195 | 2.660 | 2.409 | 3.183 | 2.580 | 2.513 | 2.879 | 2.787 | 2.844 | 2.705 | 2.415 | 2.670 | 2.606 | 0.008 | 0.9230 | 0.9705 | 0 | 0 | 0 |
| 'Paqr7' | 3.947 | 4.868 | 4.645 | 3.622 | 4.273 | 4.803 | 4.033 | 4.842 | 5.016 | 3.923 | 4.277 | 4.650 | 3.372 | 3.838 | 4.850 | 4.062 | 3.531 | 4.958 | 0.087 | 0.1064 | 0.3065 | 0 | 0 | 0 |
| 'Pard6a' | 4.734 | 4.202 | 3.872 | 4.673 | 4.184 | 3.334 | 4.469 | 4.192 | 4.206 | 4.393 | 3.707 | 3.651 | 4.691 | 3.968 | 3.697 | 3.839 | 4.035 | 3.904 | 0.062 | 0.3703 | 0.6219 | 0 | 0 | 0 |
| 'Pard6b' | 4.350 | 4.362 | 3.710 | 3.681 | 4.221 | 4.864 | 4.622 | 4.838 | 4.472 | 3.943 | 3.637 | 3.719 | 3.805 | 4.843 | 3.414 | 4.057 | 3.937 | 3.197 | 0.176 | 0.0134 | 0.0775 | 0 | 0 | 0 |
| 'Pard6g' | 2.306 | 3.195 | 2.073 | 2.225 | 3.150 | 2.932 | 2.901 | 3.225 | 2.838 | 3.386 | 2.731 | 3.363 | 2.552 | 3.367 | 3.271 | 3.802 | 3.304 | 3.690 | -0.267 | 0.0049 | 0.0372 | 1 | 0 | 0 |
| 'Parg' | 6.739 | 6.740 | 7.177 | 7.117 | 7.204 | 6.945 | 7.136 | 7.294 | 6.644 | 7.191 | 7.039 | 6.787 | 6.486 | 7.253 | 6.487 | 6.729 | 6.974 | 6.822 | 0.020 | 0.5064 | 0.7326 | 0 | 0 | 0 |
| 'Park7' | 28.341 | 33.780 | 29.270 | 29.923 | 32.654 | 31.312 | 31.157 | 33.781 | 30.488 | 28.548 | 28.301 | 27.166 | 29.743 | 29.971 | 28.003 | 30.350 | 29.254 | 27.853 | 0.104 | 0.0003 | 0.0038 | 1 | 0 | 0 |
| 'Parl' | 8.150 | 9.077 | 8.611 | 7.768 | 9.226 | 8.538 | 9.224 | 9.294 | 9.310 | 8.716 | 8.299 | 7.680 | 9.280 | 8.260 | 8.002 | 8.548 | 8.093 | 7.414 | 0.084 | 0.0469 | 0.1806 | 0 | 0 | 0 |
| 'Parn' | 16.925 | 20.141 | 16.053 | 16.966 | 18.046 | 16.733 | 18.828 | 17.055 | 17.726 | 17.591 | 19.615 | 17.553 | 16.812 | 17.498 | 16.924 | 16.975 | 18.205 | 15.284 | 0.005 | 0.9001 | 0.9605 | 0 | 0 | 0 |
| 'Parp1' | 31.932 | 29.624 | 34.801 | 32.923 | 30.842 | 34.901 | 33.815 | 30.857 | 35.102 | 31.346 | 31.767 | 34.560 | 36.157 | 31.049 | 33.894 | 32.551 | 31.161 | 33.531 | -0.017 | 0.4924 | 0.7217 | 0 | 0 | 0 |
| 'Parp10' | 13.361 | 12.879 | 14.458 | 12.776 | 12.727 | 15.031 | 12.674 | 13.445 | 15.001 | 14.118 | 12.297 | 16.587 | 13.971 | 13.540 | 15.364 | 13.505 | 13.422 | 15.914 | -0.084 | 0.0018 | 0.0168 | 1 | 0 | 0 |
| 'Parp11' | 4.686 | 5.678 | 5.442 | 5.521 | 5.333 | 5.474 | 5.381 | 5.155 | 5.895 | 4.049 | 5.466 | 6.841 | 4.651 | 5.650 | 6.394 | 4.868 | 6.145 | 6.461 | -0.058 | 0.3240 | 0.5814 | 0 | 0 | 0 |
| 'Parp12' | 2.448 | 2.683 | 3.195 | 2.118 | 2.944 | 3.369 | 2.368 | 2.861 | 3.244 | 2.255 | 2.433 | 3.892 | 2.561 | 2.582 | 3.265 | 2.927 | 2.515 | 3.672 | -0.064 | 0.3502 | 0.6045 | 0 | 0 | 0 |

| | | | | | | | | | | | | | | | | | | | | | | | | |
|---|---|---|---|---|---|---|---|---|---|---|---|---|---|---|---|---|---|---|---|---|---|---|---|---|
| 'Parp14' | 16.296 | 14.986 | 16.154 | 15.318 | 15.540 | 15.987 | 15.073 | 15.978 | 16.485 | 15.649 | 18.615 | 20.272 | 16.746 | 16.746 | 19.151 | 15.335 | 18.240 | 21.025 | -0.196 | 0.0000 | 0.0001 | 1 | 0 | 0 |
| 'Parp16' | 6.609 | 7.014 | 7.492 | 6.758 | 6.934 | 7.215 | 6.265 | 7.122 | 7.361 | 7.188 | 7.515 | 7.174 | 6.176 | 7.340 | 6.801 | 7.497 | 5.754 | 6.017 | 0.026 | 0.6626 | 0.8371 | 0 | 0 | 0 |
| 'Parp2' | 30.745 | 33.611 | 34.914 | 31.324 | 32.981 | 33.247 | 34.372 | 32.777 | 36.042 | 31.827 | 28.621 | 32.130 | 36.655 | 33.126 | 31.838 | 32.521 | 36.252 | 29.131 | 0.029 | 0.5326 | 0.7527 | 0 | 0 | 0 |
| 'Parp3' | 9.871 | 10.749 | 12.450 | 10.571 | 10.253 | 11.985 | 11.052 | 10.083 | 12.630 | 11.003 | 8.944 | 12.044 | 11.038 | 9.070 | 11.612 | 10.433 | 10.299 | 11.179 | 0.047 | 0.1618 | 0.3935 | 0 | 0 | 0 |
| 'Parp4' | 25.054 | 24.211 | 24.783 | 23.762 | 23.377 | 23.079 | 23.691 | 23.679 | 24.152 | 27.329 | 25.532 | 24.719 | 23.804 | 23.139 | 23.196 | 24.232 | 25.766 | 23.459 | -0.045 | 0.1704 | 0.4031 | 0 | 0 | 0 |
| 'Parp6' | 19.322 | 18.314 | 19.129 | 19.023 | 19.814 | 20.015 | 17.564 | 20.062 | 19.138 | 19.612 | 18.227 | 19.797 | 18.877 | 19.589 | 18.012 | 17.884 | 19.089 | 19.198 | 0.007 | 0.8326 | 0.9270 | 0 | 0 | 0 |
| 'Parp8' | 27.745 | 29.710 | 27.008 | 29.229 | 28.980 | 28.811 | 29.317 | 30.634 | 27.855 | 30.259 | 31.344 | 31.948 | 27.206 | 27.804 | 27.745 | 28.291 | 31.677 | 30.343 | -0.053 | 0.1510 | 0.3777 | 0 | 0 | 0 |
| 'Parp9' | 20.302 | 20.632 | 19.184 | 19.999 | 21.730 | 21.206 | 19.357 | 21.735 | 20.490 | 19.343 | 23.187 | 24.692 | 21.791 | 21.817 | 25.084 | 19.561 | 23.753 | 24.307 | -0.150 | 0.0002 | 0.0026 | 1 | 0 | 0 |
| 'Parpbp' | 1.724 | 2.184 | 2.740 | 2.346 | 1.758 | 2.837 | 2.644 | 2.039 | 2.301 | 1.586 | 2.416 | 2.605 | 2.714 | 2.780 | 3.007 | 2.206 | 2.852 | 2.175 | -0.131 | 0.2252 | 0.4719 | 0 | 0 | 0 |
| 'Pars2' | 1.254 | 1.377 | 1.763 | 1.464 | 1.787 | 1.570 | 1.633 | 1.613 | 1.820 | 1.350 | 1.480 | 1.771 | 1.402 | 1.520 | 1.872 | 1.428 | 1.463 | 1.645 | 0.021 | 0.7649 | 0.8953 | 0 | 0 | 0 |
| 'Parvg' | 36.180 | 38.338 | 37.070 | 35.059 | 38.951 | 39.248 | 35.603 | 38.298 | 37.754 | 37.618 | 35.362 | 38.100 | 39.340 | 36.478 | 37.395 | 38.379 | 33.912 | 36.780 | 0.003 | 0.9198 | 0.9698 | 0 | 0 | 0 |
| 'Pask' | 0.872 | 0.737 | 1.032 | 0.845 | 0.688 | 1.009 | 0.978 | 0.667 | 0.885 | 0.795 | 1.199 | 0.966 | 1.174 | 0.704 | 1.050 | 1.206 | 0.810 | 0.942 | -0.211 | 0.0273 | 0.1271 | 0 | 0 | 0 |
| 'Patl1' | 24.428 | 23.371 | 21.692 | 23.928 | 23.783 | 22.834 | 21.975 | 23.900 | 22.905 | 21.768 | 23.714 | 23.784 | 23.054 | 24.327 | 22.336 | 22.096 | 23.601 | 23.740 | -0.009 | 0.7719 | 0.8988 | 0 | 0 | 0 |
| 'Patz1' | 5.091 | 4.352 | 5.133 | 4.491 | 4.805 | 5.477 | 4.282 | 4.542 | 5.224 | 4.475 | 4.320 | 5.604 | 4.349 | 4.475 | 5.604 | 4.834 | 4.137 | 5.607 | -0.006 | 0.8826 | 0.9507 | 0 | 0 | 0 |
| 'Pawr' | 8.411 | 8.299 | 6.677 | 8.106 | 7.745 | 7.695 | 9.025 | 7.878 | 6.952 | 7.528 | 7.469 | 6.232 | 7.736 | 7.092 | 6.828 | 6.725 | 7.107 | 8.318 | 0.108 | 0.1128 | 0.3190 | 0 | 0 | 0 |
| 'Pax5' | 4.229 | 5.657 | 3.504 | 4.555 | 5.859 | 3.722 | 3.897 | 5.634 | 4.043 | 5.650 | 7.267 | 3.526 | 7.556 | 6.008 | 3.555 | 5.278 | 6.755 | 3.479 | -0.236 | 0.0029 | 0.0244 | 1 | 0 | 0 |
| 'Paxbp1' | 25.946 | 22.683 | 26.684 | 24.984 | 24.390 | 24.306 | 24.780 | 25.372 | 24.688 | 26.172 | 25.165 | 23.525 | 23.812 | 24.118 | 22.428 | 25.845 | 24.882 | 23.964 | 0.017 | 0.6123 | 0.8056 | 0 | 0 | 0 |
| 'Paxip1' | 6.456 | 5.760 | 7.228 | 7.125 | 6.573 | 6.910 | 6.642 | 6.786 | 6.682 | 6.649 | 6.560 | 6.248 | 7.744 | 6.445 | 6.707 | 6.346 | 6.430 | 6.442 | 0.006 | 0.8949 | 0.9571 | 0 | 0 | 0 |
| 'Pbdc1' | 4.486 | 4.413 | 4.600 | 5.009 | 4.469 | 4.345 | 4.559 | 4.171 | 4.803 | 4.354 | 4.579 | 4.779 | 4.624 | 4.404 | 4.401 | 4.298 | 4.455 | 4.049 | 0.018 | 0.6852 | 0.8511 | 0 | 0 | 0 |
| 'Pbk' | 4.588 | 5.483 | 6.184 | 6.211 | 5.865 | 5.039 | 6.198 | 5.261 | 6.157 | 2.516 | 5.018 | 6.321 | 6.141 | 6.493 | 7.171 | 4.445 | 5.821 | 5.603 | 0.040 | 0.7389 | 0.8819 | 0 | 0 | 0 |
| 'Pbrm1' | 12.852 | 13.489 | 13.437 | 13.742 | 13.062 | 12.727 | 13.993 | 13.157 | 13.191 | 14.282 | 15.183 | 13.348 | 13.646 | 13.555 | 12.780 | 12.909 | 14.759 | 12.953 | -0.054 | 0.0928 | 0.2820 | 0 | 0 | 0 |
| 'Pbx2' | 36.981 | 37.190 | 37.600 | 38.272 | 37.076 | 37.448 | 35.132 | 34.943 | 39.513 | 33.490 | 32.999 | 40.559 | 33.992 | 36.634 | 38.750 | 36.373 | 34.349 | 40.357 | 0.019 | 0.5654 | 0.7741 | 0 | 0 | 0 |
| 'Pbx3' | 2.192 | 2.575 | 2.477 | 2.264 | 2.233 | 2.227 | 2.189 | 2.563 | 2.413 | 2.497 | 2.776 | 2.693 | 2.096 | 2.537 | 2.717 | 2.263 | 2.466 | 2.502 | -0.104 | 0.0472 | 0.1812 | 0 | 0 | 0 |
| 'Pbx4' | 0.586 | 0.688 | 0.884 | 0.743 | 0.820 | 0.812 | 0.774 | 0.832 | 1.005 | 0.732 | 0.690 | 0.755 | 0.686 | 0.546 | 0.668 | 0.744 | 0.770 | 0.737 | 0.167 | 0.1246 | 0.3375 | 0 | 0 | 0 |
| 'Pbxip1' | 59.027 | 62.113 | 62.377 | 57.438 | 59.016 | 65.268 | 59.456 | 57.336 | 64.875 | 69.441 | 66.428 | 69.433 | 64.240 | 65.310 | 68.278 | 64.773 | 62.016 | 68.506 | -0.141 | 0.0000 | 0.0000 | 1 | 0 | 0 |
| 'Pcbd2' | 34.868 | 39.023 | 38.168 | 36.416 | 37.640 | 38.900 | 38.514 | 37.740 | 36.136 | 37.171 | 35.378 | 36.862 | 40.184 | 38.365 | 36.929 | 40.892 | 41.366 | 34.195 | -0.024 | 0.5753 | 0.7806 | 0 | 0 | 0 |
| 'Pcbp1' | 188.746 | 161.115 | 177.478 | 178.233 | 164.554 | 176.716 | 184.199 | 164.751 | 164.619 | 165.380 | 164.525 | 162.340 | 174.609 | 168.480 | 176.511 | 163.278 | 166.619 | 178.533 | 0.026 | 0.4346 | 0.6788 | 0 | 0 | 0 |
| 'Pcbp2' | 133.541 | 132.641 | 135.769 | 136.885 | 131.003 | 135.857 | 136.166 | 133.671 | 138.043 | 134.596 | 130.049 | 129.639 | 137.108 | 134.114 | 134.786 | 134.600 | 132.217 | 130.289 | 0.010 | 0.5649 | 0.7737 | 0 | 0 | 0 |
| 'Pcbp3' | 0.796 | 0.984 | 0.789 | 0.705 | 0.577 | 0.736 | 0.880 | 0.957 | 0.877 | 0.837 | 0.820 | 0.782 | 0.859 | 0.939 | 0.904 | 0.870 | 0.668 | 0.958 | -0.073 | 0.4609 | 0.6973 | 0 | 0 | 0 |
| 'Pcbp4' | 3.983 | 4.092 | 5.171 | 3.989 | 4.227 | 5.098 | 3.969 | 4.361 | 4.800 | 3.927 | 3.745 | 4.517 | 4.006 | 3.425 | 4.591 | 4.393 | 3.130 | 4.112 | 0.140 | 0.0038 | 0.0302 | 1 | 0 | 0 |
| 'Pcca' | 3.320 | 3.432 | 3.361 | 3.310 | 3.411 | 3.157 | 3.832 | 3.340 | 3.281 | 3.337 | 3.269 | 3.562 | 3.258 | 3.297 | 3.174 | 3.138 | 3.345 | 2.944 | 0.036 | 0.4204 | 0.6675 | 0 | 0 | 0 |
| 'Pccb' | 11.000 | 11.740 | 11.526 | 11.212 | 12.543 | 11.847 | 11.312 | 11.465 | 11.703 | 11.881 | 10.453 | 10.344 | 10.364 | 10.120 | 9.914 | 11.050 | 9.269 | 8.958 | 0.168 | 0.0001 | 0.0018 | 1 | 0 | 0 |
| 'Pcdhga8' | 0.325 | 0.344 | 0.425 | 0.351 | 0.303 | 0.337 | 0.354 | 0.348 | 0.372 | 0.316 | 0.369 | 0.353 | 0.436 | 0.328 | 0.420 | 0.371 | 0.284 | 0.349 | -0.033 | 0.7421 | 0.8835 | 0 | 0 | 0 |
| 'Pcdhgc3' | 0.582 | 0.693 | 0.760 | 0.600 | 0.702 | 0.712 | 0.611 | 0.564 | 0.684 | 0.722 | 0.628 | 0.684 | 0.599 | 0.738 | 0.764 | 0.585 | 0.789 | 0.780 | -0.097 | 0.2237 | 0.4711 | 0 | 0 | 0 |
| 'Pced1a' | 2.016 | 2.247 | 2.380 | 2.034 | 2.262 | 2.468 | 2.190 | 2.113 | 2.606 | 2.004 | 2.264 | 2.540 | 2.444 | 2.109 | 2.431 | 2.036 | 2.210 | 2.288 | -0.016 | 0.7032 | 0.8611 | 0 | 0 | 0 |
| 'Pced1b' | 29.047 | 28.649 | 29.793 | 27.111 | 26.526 | 32.724 | 33.342 | 27.253 | 31.663 | 25.824 | 33.248 | 35.159 | 27.660 | 29.824 | 35.338 | 25.840 | 32.141 | 32.544 | -0.068 | 0.2158 | 0.4615 | 0 | 0 | 0 |
| 'Pcf11' | 22.604 | 19.780 | 17.194 | 21.237 | 18.241 | 16.188 | 21.484 | 19.798 | 17.237 | 19.035 | 20.749 | 21.312 | 19.166 | 21.146 | 20.561 | 20.335 | 22.295 | 20.519 | -0.109 | 0.0407 | 0.1650 | 0 | 0 | 0 |
| 'Pcgf1' | 5.752 | 6.845 | 7.738 | 6.487 | 8.102 | 7.383 | 6.617 | 7.313 | 7.488 | 6.680 | 5.975 | 6.558 | 6.962 | 7.509 | 5.835 | 6.844 | 6.455 | 5.869 | 0.109 | 0.0751 | 0.2438 | 0 | 0 | 0 |
| 'Pcgf2' | 5.274 | 5.009 | 5.353 | 5.250 | 4.632 | 4.921 | 5.016 | 4.798 | 5.276 | 4.408 | 4.788 | 4.846 | 4.604 | 3.866 | 4.610 | 3.553 | 4.612 | 5.190 | 0.157 | 0.0059 | 0.0426 | 1 | 0 | 0 |
| 'Pcgf3' | 2.198 | 2.271 | 2.406 | 2.442 | 2.396 | 2.116 | 2.479 | 2.280 | 2.324 | 2.104 | 2.328 | 2.296 | 2.057 | 2.393 | 2.273 | 1.984 | 2.240 | 2.173 | 0.062 | 0.1627 | 0.3947 | 0 | 0 | 0 |
| 'Pcgf5' | 6.744 | 7.740 | 7.255 | 6.943 | 7.077 | 6.363 | 6.751 | 7.424 | 6.633 | 7.224 | 7.789 | 7.390 | 7.575 | 8.669 | 7.064 | 7.302 | 8.009 | 6.954 | -0.121 | 0.0005 | 0.0061 | 1 | 0 | 0 |
| 'Pcgf6' | 11.056 | 14.547 | 13.263 | 12.418 | 13.370 | 12.410 | 11.250 | 13.312 | 12.065 | 11.235 | 11.446 | 13.132 | 9.988 | 10.788 | 11.828 | 10.735 | 12.192 | 11.504 | 0.125 | 0.0120 | 0.0712 | 0 | 0 | 0 |
| 'Pcid2' | 9.467 | 9.783 | 9.556 | 9.027 | 9.387 | 9.954 | 8.706 | 9.381 | 9.418 | 9.524 | 8.690 | 9.796 | 9.523 | 8.487 | 9.456 | 9.998 | 9.119 | 9.874 | -0.007 | 0.8489 | 0.9351 | 0 | 0 | 0 |
| 'Pcif1' | 20.381 | 21.577 | 22.480 | 20.231 | 21.571 | 23.581 | 21.796 | 22.419 | 23.038 | 22.150 | 20.204 | 21.457 | 22.545 | 20.457 | 20.668 | 23.602 | 20.976 | 21.036 | 0.020 | 0.5331 | 0.7530 | 0 | 0 | 0 |
| 'Pck2' | 13.508 | 15.091 | 14.560 | 13.810 | 13.947 | 15.993 | 14.334 | 13.748 | 15.647 | 14.024 | 14.913 | 16.348 | 18.586 | 16.578 | 15.881 | 16.696 | 15.744 | 15.903 | -0.159 | 0.0006 | 0.0071 | 1 | 0 | 0 |
| 'Pcm1' | 15.854 | 15.807 | 14.679 | 17.044 | 15.383 | 13.419 | 17.164 | 15.231 | 14.311 | 15.321 | 16.444 | 15.273 | 16.043 | 15.399 | 14.152 | 15.305 | 17.226 | 14.907 | -0.026 | 0.4641 | 0.6999 | 0 | 0 | 0 |
| 'Pcmt1' | 5.428 | 6.057 | 5.303 | 6.185 | 5.727 | 5.948 | 5.721 | 5.671 | 5.659 | 4.868 | 5.152 | 6.124 | 4.950 | 5.415 | 5.619 | 5.484 | 5.668 | 5.908 | 0.059 | 0.1831 | 0.4191 | 0 | 0 | 0 |
| 'Pcmtd1' | 4.492 | 5.780 | 4.958 | 4.964 | 5.423 | 4.872 | 5.125 | 5.478 | 4.752 | 5.072 | 5.600 | 4.807 | 4.674 | 5.871 | 4.359 | 5.215 | 5.154 | 4.605 | 0.006 | 0.8579 | 0.9390 | 0 | 0 | 0 |
| 'Pcmtd2' | 6.159 | 8.065 | 7.179 | 7.307 | 7.997 | 7.224 | 7.651 | 7.050 | 7.059 | 6.183 | 6.967 | 7.834 | 5.715 | 7.093 | 7.013 | 6.596 | 7.994 | 7.655 | 0.049 | 0.3417 | 0.5981 | 0 | 0 | 0 |
| 'Pcna' | 108.481 | 97.944 | 94.318 | 108.742 | 98.152 | 91.296 | 105.285 | 97.726 | 91.250 | 94.602 | 98.402 | 82.568 | 116.556 | 108.156 | 93.359 | 99.434 | 99.531 | 87.221 | 0.013 | 0.7457 | 0.8853 | 0 | 0 | 0 |
| 'Pcnp' | 18.494 | 21.187 | 18.826 | 18.765 | 21.236 | 18.204 | 19.538 | 21.198 | 18.656 | 17.425 | 18.772 | 20.016 | 17.262 | 20.730 | 19.395 | 18.144 | 20.854 | 19.791 | 0.017 | 0.5850 | 0.7865 | 0 | 0 | 0 |
| 'Pcnt' | 14.623 | 14.304 | 16.825 | 15.641 | 13.464 | 16.629 | 14.989 | 15.090 | 15.707 | 15.252 | 15.618 | 15.996 | 14.716 | 15.357 | 15.452 | 15.132 | 15.398 | 15.745 | -0.025 | 0.4266 | 0.6730 | 0 | 0 | 0 |
| 'Pcnx' | 4.322 | 3.950 | 4.447 | 3.786 | 3.913 | 4.168 | 3.889 | 4.130 | 4.314 | 4.490 | 4.172 | 4.189 | 3.792 | 3.966 | 4.047 | 4.029 | 3.873 | 4.317 | -0.006 | 0.8783 | 0.9485 | 0 | 0 | 0 |
| 'Pcnx3' | 13.962 | 13.183 | 15.663 | 13.132 | 12.929 | 14.409 | 12.385 | 13.586 | 15.421 | 14.134 | 13.798 | 14.592 | 13.868 | 12.667 | 14.197 | 13.418 | 13.419 | 14.707 | -0.013 | 0.6797 | 0.8470 | 0 | 0 | 0 |
| 'Pcnx4' | 2.362 | 2.450 | 2.028 | 2.031 | 2.759 | 2.189 | 1.976 | 2.293 | 1.933 | 2.331 | 2.298 | 2.329 | 2.346 | 2.627 | 2.079 | 2.808 | 2.097 | 2.317 | -0.099 | 0.1840 | 0.4193 | 0 | 0 | 0 |
| 'Pcolce' | 0.447 | 0.482 | 0.546 | 0.505 | 0.441 | 0.555 | 0.536 | 0.481 | 0.522 | 0.454 | 0.503 | 0.699 | 0.487 | 0.505 | 0.727 | 0.415 | 0.647 | 0.636 | -0.187 | 0.0674 | 0.2275 | 0 | 0 | 0 |
| 'Pcsk7' | 23.860 | 20.878 | 26.082 | 23.981 | 22.618 | 23.717 | 24.666 | 23.823 | 25.463 | 23.884 | 25.092 | 25.377 | 24.589 | 21.149 | 24.821 | 22.756 | 23.447 | 23.802 | -0.009 | 0.8006 | 0.9111 | 0 | 0 | 0 |
| 'Pctp' | 4.392 | 4.442 | 4.617 | 3.409 | 5.486 | 4.744 | 3.437 | 5.748 | 4.872 | 4.605 | 3.850 | 3.590 | 3.832 | 4.512 | 4.097 | 3.859 | 3.280 | 3.227 | 0.230 | 0.0127 | 0.0742 | 0 | 0 | 0 |
| 'Pcx' | 2.602 | 3.624 | 3.576 | 3.772 | 3.503 | 3.358 | 2.984 | 2.906 | 3.606 | 3.600 | 2.702 | 3.591 | 3.025 | 2.910 | 3.197 | 3.269 | 3.005 | 2.953 | 0.067 | 0.3595 | 0.6118 | 0 | 0 | 0 |
| 'Pcyox1' | 3.247 | 3.736 | 3.232 | 3.375 | 3.796 | 3.361 | 3.289 | 2.789 | 3.009 | 3.217 | 3.425 | 3.575 | 3.104 | 3.002 | 3.276 | 3.209 | 3.053 | 3.471 | 0.006 | 0.9216 | 0.9705 | 0 | 0 | 0 |
| 'Pcyox1l' | 6.884 | 7.107 | 7.420 | 6.906 | 7.730 | 7.077 | 5.921 | 7.230 | 7.438 | 6.499 | 6.806 | 9.374 | 8.045 | 7.570 | 8.724 | 6.078 | 7.139 | 9.851 | -0.147 | 0.0225 | 0.1119 | 0 | 0 | 0 |
| 'Pcyt1a' | 2.396 | 2.621 | 2.523 | 2.411 | 2.998 | 2.621 | 2.461 | 2.923 | 2.574 | 2.760 | 2.681 | 2.268 | 2.739 | 2.656 | 2.503 | 2.547 | 2.621 | 1.964 | 0.047 | 0.4304 | 0.6760 | 0 | 0 | 0 |
| 'Pcyt1b' | 1.771 | 1.695 | 2.084 | 2.287 | 2.165 | 1.996 | 2.135 | 1.704 | 2.142 | 2.117 | 2.011 | 2.005 | 2.534 | 2.034 | 2.052 | 2.189 | 1.742 | 1.968 | -0.062 | 0.3820 | 0.6330 | 0 | 0 | 0 |
| 'Pcyt2' | 10.239 | 10.331 | 12.164 | 10.389 | 9.963 | 11.276 | 11.421 | 9.692 | 11.528 | 11.314 | 9.864 | 11.794 | 11.131 | 9.341 | 10.910 | 10.578 | 10.794 | 10.901 | -0.007 | 0.8439 | 0.9338 | 0 | 0 | 0 |
| 'Pdap1' | 14.852 | 15.656 | 15.629 | 16.089 | 15.420 | 17.251 | 16.045 | 15.799 | 15.921 | 14.713 | 13.831 | 15.138 | 15.792 | 13.567 | 15.908 | 15.097 | 13.960 | 14.766 | 0.093 | 0.0016 | 0.0152 | 1 | 0 | 0 |
| 'Pdcd1' | 439.603 | 526.617 | 447.117 | 487.721 | 507.857 | 456.472 | 493.482 | 470.893 | 472.651 | 418.742 | 463.792 | 433.037 | 469.628 | 441.564 | 449.179 | 423.039 | 466.215 | 477.532 | 0.078 | 0.0281 | 0.1296 | 0 | 0 | 0 |
| 'Pdcd10' | 30.951 | 30.107 | 30.353 | 32.428 | 32.552 | 27.578 | 30.988 | 32.652 | 28.461 | 29.590 | 33.461 | 29.724 | 28.786 | 29.532 | 29.404 | 29.495 | 34.544 | 27.199 | 0.013 | 0.7394 | 0.8819 | 0 | 0 | 0 |
| 'Pdcd11' | 18.391 | 18.174 | 20.403 | 18.597 | 18.711 | 18.746 | 17.637 | 19.323 | 20.035 | 19.473 | 19.190 | 19.043 | 18.636 | 16.512 | 18.209 | 19.968 | 18.363 | 19.166 | 0.004 | 0.9140 | 0.9671 | 0 | 0 | 0 |
| 'Pdcd1lg2' | 12.021 | 14.356 | 13.811 | 12.840 | 13.432 | 13.412 | 14.488 | 12.389 | 14.128 | 15.066 | 13.497 | 12.791 | 12.340 | 12.061 | 11.593 | 13.785 | 14.364 | 12.100 | 0.032 | 0.5780 | 0.7820 | 0 | 0 | 0 |
| 'Pdcd2l' | 6.282 | 7.570 | 6.834 | 7.295 | 7.520 | 7.200 | 7.327 | 7.378 | 6.887 | 6.882 | 4.627 | 6.938 | 7.492 | 5.898 | 6.979 | 6.821 | 6.358 | 7.034 | 0.106 | 0.1133 | 0.3194 | 0 | 0 | 0 |
| 'Pdcd4' | 163.703 | 183.415 | 162.671 | 172.318 | 184.522 | 176.582 | 176.259 | 180.514 | 173.650 | 163.318 | 169.542 | 166.228 | 141.935 | 169.340 | 169.284 | 175.502 | 171.329 | 175.046 | 0.056 | 0.1109 | 0.3152 | 0 | 0 | 0 |
| 'Pdcd5' | 10.743 | 11.466 | 10.980 | 11.568 | 10.571 | 10.464 | 10.648 | 10.584 | 10.868 | 9.261 | 9.428 | 10.136 | 11.132 | 10.017 | 10.766 | 10.093 | 10.855 | 9.553 | 0.089 | 0.0296 | 0.1338 | 0 | 0 | 0 |
| 'Pdcd6' | 41.703 | 45.920 | 41.680 | 41.869 | 46.461 | 44.525 | 39.572 | 45.318 | 42.203 | 41.406 | 43.240 | 43.110 | 42.432 | 46.724 | 45.457 | 44.940 | 43.081 | 42.816 | -0.025 | 0.4284 | 0.6742 | 0 | 0 | 0 |
| 'Pdcd6ip' | 31.162 | 31.012 | 33.144 | 31.603 | 32.668 | 32.343 | 31.939 | 31.688 | 32.396 | 32.374 | 31.927 | 33.904 | 31.538 | 32.415 | 32.509 | 31.099 | 31.003 | 33.315 | -0.022 | 0.2381 | 0.4880 | 0 | 0 | 0 |
| 'Pdcd7' | 15.596 | 15.985 | 17.370 | 14.725 | 15.085 | 16.830 | 15.676 | 17.247 | 15.664 | 14.847 | 15.400 | 17.283 | 15.477 | 15.721 | 17.041 | 14.048 | 15.621 | 17.435 | 0.002 | 0.9571 | 0.9840 | 0 | 0 | 0 |
| 'Pdcl' | 4.790 | 5.197 | 5.576 | 5.614 | 6.237 | 6.013 | 5.372 | 5.778 | 5.393 | 5.418 | 5.240 | 4.796 | 4.997 | 5.504 | 4.506 | 5.335 | 5.586 | 4.679 | 0.113 | 0.0224 | 0.1111 | 0 | 0 | 0 |
| 'Pdcl3' | 8.307 | 8.588 | 8.776 | 8.691 | 9.078 | 8.891 | 8.901 | 8.929 | 8.257 | 7.633 | 7.468 | 8.086 | 8.749 | 8.445 | 8.886 | 8.211 | 8.292 | 8.201 | 0.073 | 0.0518 | 0.1929 | 0 | 0 | 0 |
| 'Pddc1' | 7.109 | 9.462 | 7.886 | 6.886 | 9.093 | 8.752 | 7.594 | 8.937 | 8.627 | 8.339 | 7.560 | 8.159 | 8.057 | 8.016 | 8.481 | 8.721 | 7.635 | 7.657 | 0.018 | 0.7248 | 0.8736 | 0 | 0 | 0 |
| 'Pde11a' | 0.950 | 0.566 | 0.720 | 0.785 | 0.625 | 0.727 | 0.875 | 0.762 | 0.717 | 0.921 | 0.970 | 0.974 | 0.836 | 0.850 | 0.915 | 0.829 | 0.806 | 0.926 | -0.267 | 0.0052 | 0.0386 | 1 | 0 | 0 |
| 'Pde12' | 8.489 | 7.514 | 7.291 | 7.906 | 7.531 | 7.180 | 7.846 | 7.404 | 6.894 | 7.139 | 7.531 | 7.269 | 7.925 | 7.847 | 7.443 | 7.675 | 7.497 | 7.256 | -0.001 | 0.9709 | 0.9898 | 0 | 0 | 0 |
| 'Pde1b' | 11.778 | 10.624 | 11.625 | 10.663 | 8.306 | 10.003 | 10.454 | 10.531 | 11.312 | 10.259 | 11.807 | 11.599 | 11.675 | 9.055 | 10.345 | 8.792 | 11.019 | 11.106 | -0.017 | 0.7952 | 0.9089 | 0 | 0 | 0 |
| 'Pde3b' | 27.310 | 31.424 | 27.420 | 28.022 | 30.846 | 29.394 | 27.700 | 30.490 | 26.901 | 31.432 | 28.399 | 26.644 | 31.935 | 32.824 | 25.946 | 31.689 | 29.735 | 28.726 | -0.052 | 0.1647 | 0.3970 | 0 | 0 | 0 |
| 'Pde4a' | 0.398 | 0.357 | 0.414 | 0.397 | 0.359 | 0.275 | 0.331 | 0.283 | 0.378 | 0.270 | 0.176 | 0.288 | 0.269 | 0.261 | 0.310 | 0.289 | 0.177 | 0.234 | 0.474 | 0.0000 | 0.0008 | 1 | 1 | 0 |
| 'Pde4b' | 8.918 | 7.341 | 7.113 | 7.329 | 7.869 | 7.154 | 7.905 | 7.562 | 7.137 | 8.419 | 7.180 | 7.388 | 7.257 | 7.700 | 7.344 | 8.337 | 7.170 | 7.065 | 0.000 | 0.9966 | 0.9994 | 0 | 0 | 0 |
| 'Pde4d' | 1.652 | 1.826 | 1.744 | 1.846 | 1.744 | 1.594 | 1.772 | 1.902 | 1.533 | 1.742 | 1.918 | 2.136 | 1.581 | 2.007 | 2.097 | 1.930 | 1.856 | 2.251 | -0.177 | 0.0034 | 0.0274 | 1 | 0 | 0 |
| 'Pde4dip' | 3.756 | 3.765 | 3.438 | 4.062 | 3.473 | 3.663 | 3.615 | 3.711 | 3.796 | 3.410 | 4.194 | 4.200 | 3.442 | 3.915 | 3.846 | 3.650 | 3.945 | 4.239 | -0.076 | 0.1110 | 0.3154 | 0 | 0 | 0 |
| 'Pde6d' | 8.630 | 8.949 | 8.338 | 9.373 | 9.037 | 8.371 | 8.044 | 8.690 | 8.249 | 7.870 | 8.172 | 8.157 | 8.045 | 7.937 | 7.933 | 7.559 | 8.763 | 8.238 | 0.082 | 0.0395 | 0.1621 | 0 | 0 | 0 |
| 'Pde7a' | 11.242 | 13.432 | 11.554 | 11.606 | 11.614 | 11.417 | 11.941 | 11.946 | 11.329 | 11.194 | 11.689 | 13.086 | 10.138 | 11.716 | 12.857 | 11.744 | 11.761 | 13.586 | -0.032 | 0.4749 | 0.7099 | 0 | 0 | 0 |
| 'Pdha1' | 34.564 | 35.729 | 33.674 | 35.315 | 36.224 | 34.631 | 32.170 | 35.511 | 34.125 | 34.785 | 34.993 | 33.031 | 33.194 | 34.663 | 33.525 | 32.737 | 34.251 | 32.595 | 0.029 | 0.2220 | 0.4694 | 0 | 0 | 0 |
| 'Pdhb' | 72.006 | 78.138 | 73.339 | 74.587 | 80.967 | 75.291 | 74.567 | 77.985 | 73.046 | 72.380 | 75.126 | 69.438 | 80.331 | 79.226 | 67.656 | 74.288 | 77.583 | 65.329 | 0.032 | 0.2629 | 0.5169 | 0 | 0 | 0 |
| 'Pdhx' | 3.906 | 4.478 | 4.425 | 4.269 | 4.044 | 4.460 | 3.854 | 4.729 | 4.555 | 3.796 | 4.181 | 4.025 | 4.315 | 4.194 | 4.085 | 4.355 | 4.092 | 3.642 | 0.074 | 0.1709 | 0.4040 | 0 | 0 | 0 |
| 'Pdia3' | 132.222 | 141.134 | 145.165 | 155.140 | 151.649 | 147.300 | 151.755 | 143.057 | 141.514 | 146.514 | 136.329 | 135.477 | 136.687 | 145.312 | 137.639 | 146.971 | 142.766 | 138.172 | 0.038 | 0.2002 | 0.4413 | 0 | 0 | 0 |
| 'Pdia4' | 53.352 | 51.037 | 55.415 | 55.703 | 57.481 | 58.891 | 53.532 | 55.397 | 56.384 | 7.335 | 31.902 | 22.751 | 6.464 | 37.538 | 17.084 | 6.770 | 40.485 | 37.913 | 1.524 | 0.0000 | 0.0000 | 1 | 1 | 1 |
| 'Pdia6' | 47.035 | 50.161 | 54.539 | 53.958 | 54.909 | 56.690 | 52.554 | 54.186 | 55.437 | 53.365 | 49.869 | 47.321 | 49.324 | 49.366 | 46.180 | 52.154 | 50.887 | 46.928 | 0.097 | 0.0078 | 0.0519 | 0 | 0 | 0 |
| 'Pdik1l' | 2.176 | 2.578 | 2.693 | 2.700 | 2.395 | 2.601 | 2.974 | 2.455 | 2.970 | 1.913 | 2.440 | 3.018 | 2.118 | 2.501 | 2.962 | 2.096 | 2.949 | 2.984 | 0.034 | 0.6335 | 0.8184 | 0 | 0 | 0 |
| 'Pdk1' | 5.244 | 5.342 | 4.805 | 4.909 | 4.984 | 4.568 | 5.218 | 4.810 | 4.797 | 4.591 | 6.013 | 5.422 | 5.498 | 5.923 | 5.433 | 5.116 | 5.414 | 4.826 | -0.121 | 0.0068 | 0.0472 | 1 | 0 | 0 |
| 'Pdk2' | 1.374 | 1.894 | 1.410 | 1.333 | 1.251 | 1.867 | 1.456 | 1.941 | 1.587 | 1.686 | 1.672 | 1.630 | 1.830 | 1.528 | 1.721 | 2.141 | 1.683 | 1.238 | -0.103 | 0.3232 | 0.5807 | 0 | 0 | 0 |
| 'Pdk3' | 14.008 | 14.189 | 14.094 | 15.509 | 15.025 | 14.155 | 13.982 | 14.029 | 14.023 | 14.084 | 14.719 | 14.394 | 15.654 | 14.872 | 14.448 | 13.978 | 15.516 | 14.164 | -0.043 | 0.2277 | 0.4743 | 0 | 0 | 0 |
| 'Pdlim2' | 12.772 | 13.362 | 13.313 | 11.728 | 13.701 | 13.977 | 11.636 | 12.592 | 13.510 | 12.946 | 11.709 | 12.844 | 13.404 | 12.044 | 13.929 | 12.700 | 11.784 | 12.611 | 0.023 | 0.5477 | 0.7614 | 0 | 0 | 0 |
| 'Pdlim5' | 3.194 | 3.396 | 3.510 | 3.668 | 3.465 | 3.392 | 3.417 | 3.244 | 3.622 | 3.688 | 3.606 | 3.748 | 3.460 | 4.268 | 3.712 | 3.589 | 4.273 | 3.726 | -0.149 | 0.0008 | 0.0090 | 1 | 0 | 0 |
| 'Pdlim7' | 6.058 | 6.665 | 6.655 | 6.337 | 5.985 | 6.497 | 6.202 | 6.004 | 6.796 | 7.042 | 6.123 | 6.920 | 6.729 | 6.791 | 6.704 | 6.164 | 6.448 | 6.462 | -0.066 | 0.0487 | 0.1854 | 0 | 0 | 0 |
| 'Pdp1' | 0.767 | 0.883 | 0.990 | 0.829 | 0.867 | 0.881 | 1.037 | 0.813 | 0.934 | 0.960 | 1.130 | 0.873 | 0.761 | 0.911 | 0.949 | 0.942 | 0.993 | 0.745 | -0.047 | 0.5851 | 0.7865 | 0 | 0 | 0 |
| 'Pdp2' | 3.333 | 3.156 | 3.234 | 2.982 | 2.618 | 3.261 | 3.347 | 2.742 | 3.375 | 3.116 | 3.801 | 3.306 | 3.152 | 3.231 | 3.004 | 3.324 | 3.325 | 3.633 | -0.097 | 0.1672 | 0.3997 | 0 | 0 | 0 |
| 'Pdpk1' | 14.270 | 14.828 | 13.098 | 13.968 | 14.408 | 13.289 | 13.727 | 14.273 | 12.976 | 15.045 | 15.392 | 15.014 | 15.226 | 16.210 | 14.411 | 15.635 | 15.494 | 14.746 | -0.148 | 0.0000 | 0.0000 | 1 | 0 | 0 |
| 'Pdpr' | 8.341 | 9.328 | 9.409 | 7.694 | 8.762 | 9.240 | 8.129 | 9.569 | 9.259 | 9.445 | 9.051 | 9.236 | 8.688 | 10.105 | 9.077 | 8.258 | 9.620 | 9.228 | -0.061 | 0.0930 | 0.2822 | 0 | 0 | 0 |
| 'Pdrg1' | 11.748 | 14.385 | 13.980 | 13.396 | 13.728 | 13.710 | 13.073 | 13.817 | 13.418 | 12.849 | 12.472 | 14.388 | 12.718 | 13.208 | 13.617 | 12.311 | 14.054 | 14.513 | -0.002 | 0.9604 | 0.9857 | 0 | 0 | 0 |
| 'Pds5a' | 10.029 | 9.196 | 10.205 | 10.860 | 9.881 | 9.601 | 9.873 | 10.103 | 9.647 | 10.017 | 9.715 | 9.810 | 10.034 | 9.936 | 9.289 | 9.600 | 10.234 | 9.644 | 0.008 | 0.7822 | 0.9025 | 0 | 0 | 0 |
| 'Pds5b' | 5.836 | 5.544 | 5.922 | 5.981 | 5.864 | 5.660 | 5.962 | 5.737 | 5.613 | 5.767 | 5.799 | 6.237 | 5.591 | 5.999 | 5.558 | 5.243 | 5.887 | 5.854 | -0.007 | 0.8240 | 0.9226 | 0 | 0 | 0 |
| 'Pdss1' | 1.234 | 1.536 | 1.498 | 1.361 | 1.282 | 1.253 | 1.631 | 1.625 | 1.318 | 1.308 | 1.407 | 1.388 | 1.543 | 1.176 | 1.312 | 1.429 | 1.376 | 0.982 | 0.083 | 0.3242 | 0.5816 | 0 | 0 | 0 |
| 'Pdss2' | 1.187 | 1.095 | 1.037 | 1.128 | 1.091 | 1.186 | 1.411 | 0.962 | 1.192 | 0.963 | 1.126 | 1.062 | 1.017 | 1.126 | 0.920 | 0.885 | 1.244 | 1.231 | 0.090 | 0.2973 | 0.5531 | 0 | 0 | 0 |
| 'Pdxdc1' | 3.493 | 4.116 | 4.146 | 3.641 | 4.018 | 4.231 | 4.057 | 4.042 | 4.150 | 3.893 | 3.671 | 3.936 | 3.989 | 3.702 | 3.764 | 3.774 | 3.710 | 3.739 | 0.059 | 0.0922 | 0.2806 | 0 | 0 | 0 |
| 'Pdxk' | 7.942 | 8.805 | 8.563 | 8.511 | 8.340 | 8.515 | 8.061 | 8.729 | 8.161 | 8.479 | 8.092 | 8.439 | 7.760 | 8.218 | 8.989 | 8.721 | 6.967 | 8.406 | 0.020 | 0.6336 | 0.8184 | 0 | 0 | 0 |
| 'Pdzd11' | 5.575 | 6.515 | 5.924 | 5.509 | 6.914 | 7.182 | 5.503 | 5.852 | 6.406 | 6.317 | 7.219 | 6.079 | 6.421 | 6.790 | 7.189 | 5.770 | 7.451 | 5.964 | -0.104 | 0.0642 | 0.2209 | 0 | 0 | 0 |
| 'Pdzd8' | 11.768 | 13.693 | 12.534 | 11.919 | 13.555 | 13.562 | 12.530 | 13.471 | 12.915 | 12.516 | 12.740 | 13.770 | 12.386 | 13.345 | 13.444 | 13.479 | 13.238 | 13.843 | -0.048 | 0.0972 | 0.2906 | 0 | 0 | 0 |
| 'Pea15a' | 9.671 | 11.099 | 10.398 | 9.526 | 11.027 | 11.674 | 9.569 | 10.255 | 11.647 | 10.728 | 10.158 | 11.629 | 10.606 | 10.838 | 11.927 | 11.648 | 10.641 | 11.541 | -0.084 | 0.0280 | 0.1292 | 0 | 0 | 0 |
| 'Peak1' | 6.422 | 5.684 | 5.928 | 6.249 | 5.517 | 5.556 | 5.215 | 5.546 | 5.852 | 6.840 | 6.649 | 6.471 | 6.437 | 5.791 | 5.985 | 6.585 | 5.639 | 6.871 | -0.150 | 0.0007 | 0.0082 | 1 | 0 | 0 |
| 'Pear1' | 18.189 | 17.592 | 18.116 | 17.711 | 17.029 | 17.141 | 16.511 | 16.923 | 18.194 | 18.358 | 18.782 | 20.516 | 19.619 | 18.310 | 18.771 | 16.898 | 19.063 | 19.904 | -0.123 | 0.0001 | 0.0014 | 1 | 0 | 0 |
| 'Pebp1' | 49.792 | 53.548 | 49.870 | 49.787 | 54.314 | 51.405 | 48.139 | 55.657 | 48.936 | 51.036 | 47.557 | 52.306 | 50.657 | 50.183 | 53.446 | 52.805 | 45.913 | 49.793 | 0.013 | 0.7102 | 0.8654 | 0 | 0 | 0 |
| 'Pecam1' | 32.368 | 34.476 | 34.147 | 33.556 | 34.524 | 33.507 | 34.321 | 34.063 | 34.411 | 33.633 | 34.504 | 30.497 | 35.456 | 32.773 | 29.493 | 31.492 | 33.790 | 27.932 | 0.069 | 0.0322 | 0.1414 | 0 | 0 | 0 |
| 'Pef1' | 14.818 | 15.680 | 15.388 | 14.938 | 15.752 | 14.739 | 12.692 | 14.709 | 14.877 | 13.949 | 13.938 | 15.747 | 13.322 | 15.017 | 15.016 | 14.543 | 14.219 | 14.867 | 0.020 | 0.6094 | 0.8038 | 0 | 0 | 0 |
| 'Peli1' | 25.277 | 25.374 | 23.927 | 24.685 | 24.931 | 24.218 | 26.401 | 25.965 | 22.807 | 23.493 | 26.587 | 23.872 | 21.687 | 24.833 | 21.816 | 23.114 | 24.782 | 23.675 | 0.054 | 0.1052 | 0.3040 | 0 | 0 | 0 |
| 'Pelo' | 11.197 | 8.903 | 8.200 | 10.013 | 8.141 | 8.307 | 9.705 | 8.721 | 10.396 | 7.600 | 8.293 | 9.760 | 9.545 | 7.648 | 8.644 | 7.637 | 8.652 | 9.008 | 0.101 | 0.1662 | 0.3984 | 0 | 0 | 0 |
| 'Pelp1' | 16.399 | 13.800 | 16.779 | 14.869 | 15.357 | 17.103 | 15.262 | 14.979 | 16.926 | 15.895 | 14.405 | 16.519 | 16.659 | 14.405 | 16.996 | 15.598 | 14.337 | 15.129 | 0.008 | 0.8034 | 0.9117 | 0 | 0 | 0 |
| 'Penk' | 9.591 | 10.523 | 11.194 | 9.631 | 10.301 | 11.307 | 10.223 | 10.328 | 14.453 | 4.260 | 5.552 | 7.135 | 4.860 | 5.262 | 6.536 | 4.734 | 5.118 | 5.564 | 0.979 | 0.0000 | 0.0000 | 1 | 1 | 0 |
| 'Peo1' | 9.726 | 8.705 | 7.978 | 7.280 | 9.358 | 7.878 | 7.514 | 8.393 | 9.148 | 8.341 | 8.397 | 8.955 | 8.417 | 8.469 | 8.284 | 7.951 | 8.013 | 9.046 | -0.011 | 0.8508 | 0.9358 | 0 | 0 | 0 |
| 'Pepd' | 7.265 | 7.200 | 6.996 | 7.529 | 7.311 | 6.802 | 7.421 | 7.890 | 7.429 | 7.926 | 6.934 | 7.890 | 8.060 | 6.973 | 7.264 | 7.121 | 7.334 | 7.828 | -0.048 | 0.2407 | 0.4912 | 0 | 0 | 0 |
| 'Per1' | 18.888 | 12.045 | 10.175 | 14.594 | 11.416 | 10.509 | 14.535 | 11.653 | 10.771 | 18.866 | 13.547 | 9.789 | 16.852 | 14.461 | 9.181 | 18.335 | 13.187 | 9.878 | -0.102 | 0.0761 | 0.2453 | 0 | 0 | 0 |
| 'Per2' | 4.571 | 4.515 | 4.761 | 4.636 | 5.040 | 4.512 | 4.979 | 4.108 | 4.609 | 4.320 | 4.233 | 4.697 | 4.689 | 4.496 | 4.092 | 4.359 | 4.345 | 4.006 | 0.073 | 0.1222 | 0.3336 | 0 | 0 | 0 |
| 'Per3' | 0.431 | 0.435 | 0.346 | 0.350 | 0.406 | 0.396 | 0.331 | 0.447 | 0.578 | 0.332 | 0.374 | 0.389 | 0.362 | 0.380 | 0.332 | 0.224 | 0.369 | 0.372 | 0.231 | 0.0274 | 0.1274 | 0 | 0 | 0 |
| 'Pes1' | 23.913 | 24.493 | 24.618 | 23.414 | 26.761 | 25.334 | 23.467 | 24.243 | 23.235 | 24.965 | 22.989 | 23.996 | 24.114 | 23.551 | 24.878 | 23.342 | 22.589 | 23.586 | 0.025 | 0.4111 | 0.6597 | 0 | 0 | 0 |
| 'Pet100' | 15.265 | 17.881 | 15.295 | 17.146 | 18.523 | 18.119 | 15.082 | 16.278 | 18.119 | 14.557 | 15.759 | 15.045 | 14.228 | 15.513 | 16.330 | 17.010 | 15.183 | 14.016 | 0.132 | 0.0199 | 0.1029 | 0 | 0 | 0 |
| 'Pex1' | 2.376 | 2.295 | 2.545 | 2.574 | 2.351 | 2.521 | 2.496 | 2.403 | 2.456 | 2.683 | 2.848 | 2.879 | 2.719 | 2.644 | 2.495 | 2.623 | 2.749 | 2.837 | -0.164 | 0.0000 | 0.0005 | 1 | 0 | 0 |
| 'Pex10' | 2.410 | 2.446 | 3.045 | 2.382 | 2.090 | 2.964 | 2.650 | 2.259 | 2.868 | 2.237 | 2.186 | 3.198 | 2.203 | 2.303 | 2.985 | 2.597 | 2.755 | 3.106 | -0.040 | 0.5054 | 0.7316 | 0 | 0 | 0 |
| 'Pex11a' | 1.755 | 2.344 | 1.997 | 2.054 | 2.355 | 2.389 | 1.958 | 1.971 | 2.000 | 2.078 | 2.042 | 1.874 | 1.716 | 1.704 | 2.021 | 1.506 | 1.753 | 1.540 | 0.197 | 0.0123 | 0.0729 | 0 | 0 | 0 |
| 'Pex11b' | 1.955 | 2.551 | 2.210 | 2.227 | 2.521 | 2.100 | 1.972 | 2.389 | 2.080 | 1.997 | 2.007 | 2.065 | 1.976 | 1.900 | 2.039 | 2.099 | 2.225 | 1.967 | 0.112 | 0.0358 | 0.1518 | 0 | 0 | 0 |
| 'Pex11g' | 1.435 | 1.922 | 2.167 | 1.127 | 1.849 | 2.146 | 1.662 | 1.775 | 2.260 | 1.873 | 1.627 | 1.513 | 1.713 | 1.594 | 1.627 | 1.662 | 1.636 | 1.360 | 0.154 | 0.1437 | 0.3678 | 0 | 0 | 0 |
| 'Pex12' | 1.987 | 2.114 | 1.906 | 1.785 | 1.943 | 2.048 | 1.937 | 2.193 | 2.160 | 1.997 | 2.193 | 2.004 | 1.975 | 2.103 | 1.806 | 1.750 | 2.111 | 1.716 | 0.026 | 0.6434 | 0.8251 | 0 | 0 | 0 |
| 'Pex13' | 3.032 | 3.225 | 3.442 | 3.552 | 3.962 | 3.192 | 3.107 | 3.828 | 3.354 | 3.312 | 3.265 | 3.440 | 3.424 | 3.479 | 3.647 | 3.556 | 3.172 | 3.326 | -0.009 | 0.8709 | 0.9447 | 0 | 0 | 0 |
| 'Pex14' | 4.554 | 4.965 | 4.922 | 4.062 | 4.708 | 5.014 | 4.576 | 4.545 | 4.665 | 4.872 | 4.140 | 4.863 | 4.720 | 4.086 | 4.440 | 4.218 | 4.000 | 4.404 | 0.067 | 0.1446 | 0.3690 | 0 | 0 | 0 |
| 'Pex16' | 8.299 | 11.015 | 9.718 | 7.295 | 10.483 | 10.240 | 7.804 | 10.380 | 9.595 | 9.963 | 7.970 | 8.552 | 9.947 | 8.259 | 9.052 | 10.802 | 7.728 | 7.996 | 0.068 | 0.3980 | 0.6470 | 0 | 0 | 0 |
| 'Pex19' | 8.821 | 10.073 | 9.219 | 8.958 | 10.490 | 10.013 | 9.031 | 10.249 | 9.964 | 9.785 | 9.315 | 8.877 | 9.765 | 9.891 | 9.077 | 10.347 | 9.273 | 8.279 | 0.030 | 0.4690 | 0.7052 | 0 | 0 | 0 |
| 'Pex2' | 3.857 | 5.270 | 4.810 | 4.009 | 5.204 | 4.661 | 4.686 | 5.238 | 4.663 | 4.506 | 4.474 | 3.898 | 4.087 | 4.782 | 4.114 | 4.996 | 4.417 | 3.646 | 0.116 | 0.0394 | 0.1621 | 0 | 0 | 0 |
| 'Pex26' | 2.373 | 1.861 | 1.785 | 2.095 | 1.968 | 2.007 | 1.875 | 1.663 | 1.984 | 1.418 | 2.020 | 2.514 | 1.586 | 1.972 | 2.453 | 2.033 | 1.943 | 2.781 | -0.087 | 0.3811 | 0.6324 | 0 | 0 | 0 |
| 'Pex3' | 4.273 | 5.584 | 4.682 | 4.630 | 5.396 | 4.732 | 5.189 | 5.321 | 4.604 | 5.195 | 4.468 | 4.286 | 5.470 | 4.962 | 3.923 | 4.746 | 5.377 | 3.980 | 0.057 | 0.2953 | 0.5510 | 0 | 0 | 0 |
| 'Pex5' | 6.531 | 7.056 | 7.135 | 7.000 | 7.105 | 7.336 | 6.698 | 7.270 | 7.334 | 6.925 | 7.081 | 7.040 | 6.831 | 7.430 | 6.972 | 7.093 | 7.400 | 6.545 | -0.003 | 0.9130 | 0.9668 | 0 | 0 | 0 |
| 'Pex6' | 11.887 | 11.177 | 13.978 | 11.476 | 12.056 | 13.773 | 11.145 | 12.058 | 12.876 | 13.464 | 11.703 | 14.836 | 12.542 | 11.214 | 13.195 | 12.136 | 11.506 | 14.344 | -0.067 | 0.0493 | 0.1872 | 0 | 0 | 0 |
| 'Pex7' | 3.369 | 3.896 | 3.414 | 3.353 | 3.617 | 3.597 | 3.687 | 3.974 | 3.360 | 3.551 | 4.796 | 3.481 | 3.743 | 3.273 | 3.514 | 4.118 | 3.693 | 3.129 | -0.050 | 0.4509 | 0.6904 | 0 | 0 | 0 |
| 'Pfas' | 3.041 | 2.544 | 3.538 | 3.074 | 2.951 | 3.054 | 3.112 | 2.955 | 3.275 | 2.949 | 2.998 | 3.288 | 3.217 | 2.950 | 3.158 | 2.965 | 2.718 | 3.357 | -0.012 | 0.7534 | 0.8901 | 0 | 0 | 0 |
| 'Pfdn1' | 31.713 | 33.582 | 36.045 | 32.322 | 35.437 | 34.872 | 33.798 | 37.545 | 34.292 | 31.703 | 28.217 | 32.984 | 34.556 | 30.568 | 33.762 | 35.069 | 32.876 | 28.135 | 0.096 | 0.0405 | 0.1646 | 0 | 0 | 0 |
| 'Pfdn2' | 20.923 | 21.173 | 20.778 | 21.244 | 22.954 | 21.506 | 20.794 | 21.816 | 19.541 | 18.671 | 21.393 | 19.480 | 20.414 | 19.589 | 19.839 | 19.604 | 21.505 | 18.543 | 0.082 | 0.0138 | 0.0791 | 0 | 0 | 0 |
| 'Pfdn4' | 11.427 | 12.845 | 12.976 | 11.183 | 14.137 | 11.464 | 15.552 | 12.527 | 13.239 | 11.158 | 10.806 | 13.035 | 12.412 | 12.536 | 12.897 | 12.693 | 15.033 | 11.677 | 0.022 | 0.7579 | 0.8916 | 0 | 0 | 0 |
| 'Pfdn5' | 64.371 | 68.604 | 62.518 | 66.065 | 68.991 | 63.238 | 67.353 | 66.860 | 64.731 | 63.969 | 60.854 | 62.124 | 61.089 | 62.402 | 62.806 | 67.272 | 63.276 | 59.760 | 0.061 | 0.0098 | 0.0615 | 0 | 0 | 0 |
| 'Pfdn6' | 18.780 | 19.049 | 19.161 | 21.717 | 18.954 | 20.373 | 19.494 | 20.468 | 18.711 | 16.092 | 17.873 | 18.485 | 17.865 | 18.385 | 19.591 | 18.951 | 20.592 | 17.964 | 0.082 | 0.0598 | 0.2110 | 0 | 0 | 0 |
| 'Pfkfb1' | 1.032 | 0.848 | 1.035 | 1.018 | 0.695 | 0.952 | 1.035 | 0.951 | 1.198 | 1.027 | 1.189 | 0.830 | 1.197 | 0.969 | 1.032 | 0.938 | 1.075 | 0.722 | -0.028 | 0.7958 | 0.9092 | 0 | 0 | 0 |
| 'Pfkfb3' | 8.322 | 8.029 | 6.832 | 8.173 | 7.342 | 6.796 | 7.822 | 7.062 | 7.102 | 7.286 | 8.811 | 8.606 | 8.091 | 8.780 | 8.949 | 7.561 | 9.034 | 8.570 | -0.177 | 0.0007 | 0.0081 | 1 | 0 | 0 |
| 'Pfkfb4' | 1.608 | 2.214 | 1.783 | 1.787 | 1.876 | 1.994 | 1.718 | 2.009 | 1.865 | 2.033 | 2.121 | 1.950 | 1.765 | 1.980 | 1.593 | 1.770 | 1.953 | 1.741 | -0.018 | 0.7569 | 0.8913 | 0 | 0 | 0 |
| 'Pfkl' | 49.877 | 44.038 | 42.813 | 49.232 | 43.844 | 43.368 | 48.111 | 45.115 | 45.100 | 41.545 | 51.529 | 54.218 | 54.147 | 51.092 | 56.209 | 48.864 | 52.590 | 53.946 | -0.185 | 0.0002 | 0.0033 | 1 | 0 | 0 |
| 'Pfkm' | 15.296 | 16.715 | 16.142 | 15.232 | 13.295 | 15.604 | 14.227 | 15.042 | 14.996 | 14.234 | 14.191 | 15.981 | 16.259 | 13.743 | 15.755 | 15.648 | 14.166 | 14.717 | 0.009 | 0.8335 | 0.9276 | 0 | 0 | 0 |
| 'Pfkp' | 39.148 | 33.585 | 33.700 | 39.278 | 34.582 | 31.549 | 36.625 | 33.979 | 33.276 | 34.388 | 39.133 | 36.632 | 37.385 | 36.741 | 37.633 | 34.021 | 39.219 | 38.332 | -0.092 | 0.0399 | 0.1628 | 0 | 0 | 0 |
| 'Pfn1' | 344.019 | 428.755 | 362.571 | 369.422 | 419.365 | 401.224 | 373.999 | 382.528 | 403.305 | 387.516 | 388.575 | 299.253 | 391.699 | 399.547 | 398.043 | 410.270 | 383.114 | 367.319 | 0.014 | 0.7866 | 0.9047 | 0 | 0 | 0 |
| 'Pfn2' | 2.595 | 3.620 | 2.713 | 2.794 | 2.848 | 3.159 | 2.815 | 3.333 | 2.734 | 2.380 | 3.194 | 2.592 | 2.509 | 2.802 | 2.491 | 2.735 | 2.743 | 2.429 | 0.147 | 0.0122 | 0.0723 | 0 | 0 | 0 |
| 'Pgam1' | 162.429 | 148.639 | 132.125 | 161.208 | 152.791 | 133.824 | 154.748 | 139.196 | 133.335 | 138.007 | 163.413 | 155.935 | 168.706 | 178.435 | 164.248 | 151.201 | 171.143 | 162.312 | -0.154 | 0.0017 | 0.0162 | 1 | 0 | 0 |
| 'Pgam2' | 11.943 | 13.449 | 11.725 | 13.843 | 11.358 | 11.110 | 12.133 | 11.361 | 11.067 | 13.950 | 14.621 | 11.294 | 21.374 | 15.573 | 12.583 | 16.325 | 16.926 | 14.490 | -0.345 | 0.0001 | 0.0010 | 1 | 0 | 0 |
| 'Pgam5' | 11.846 | 15.403 | 14.178 | 12.548 | 15.036 | 13.739 | 12.574 | 15.038 | 13.330 | 13.535 | 12.310 | 13.607 | 12.486 | 13.762 | 13.904 | 14.331 | 13.561 | 13.257 | 0.020 | 0.6252 | 0.8133 | 0 | 0 | 0 |
| 'Pgap1' | 1.261 | 1.181 | 1.201 | 0.995 | 1.226 | 0.934 | 1.217 | 1.205 | 1.018 | 1.423 | 1.643 | 1.511 | 1.428 | 1.352 | 1.299 | 1.502 | 1.466 | 1.448 | -0.366 | 0.0000 | 0.0000 | 1 | 0 | 0 |
| 'Pgap2' | 1.732 | 1.646 | 1.785 | 1.846 | 1.954 | 1.797 | 1.714 | 1.682 | 1.907 | 1.848 | 1.767 | 1.956 | 1.900 | 1.664 | 1.792 | 1.703 | 1.792 | 1.579 | -0.009 | 0.8544 | 0.9375 | 0 | 0 | 0 |
| 'Pgap3' | 2.233 | 2.715 | 2.569 | 2.294 | 2.504 | 2.779 | 2.124 | 2.682 | 2.503 | 2.563 | 2.064 | 2.348 | 2.191 | 2.279 | 2.530 | 2.594 | 2.057 | 2.325 | 0.090 | 0.1496 | 0.3765 | 0 | 0 | 0 |
| 'Pgd' | 14.867 | 14.444 | 15.971 | 14.934 | 14.566 | 16.208 | 15.046 | 14.725 | 15.885 | 14.255 | 15.428 | 16.896 | 16.121 | 16.095 | 16.385 | 14.519 | 15.561 | 16.510 | -0.064 | 0.0172 | 0.0929 | 0 | 0 | 0 |
| 'Pggt1b' | 11.195 | 10.269 | 10.250 | 11.113 | 9.705 | 10.194 | 10.787 | 10.152 | 9.760 | 10.268 | 10.622 | 9.778 | 10.236 | 9.485 | 9.094 | 9.580 | 9.913 | 9.379 | 0.071 | 0.0971 | 0.2904 | 0 | 0 | 0 |
| 'Pgls' | 9.193 | 8.343 | 9.383 | 7.914 | 7.615 | 10.191 | 7.725 | 8.302 | 9.498 | 8.253 | 8.295 | 9.690 | 8.759 | 7.829 | 8.857 | 8.268 | 8.225 | 9.377 | 0.002 | 0.9606 | 0.9857 | 0 | 0 | 0 |
| 'Pglyrp1' | 10.017 | 9.795 | 9.727 | 9.821 | 10.748 | 11.483 | 10.822 | 10.218 | 10.721 | 10.834 | 8.250 | 9.532 | 9.739 | 9.656 | 10.840 | 8.843 | 8.616 | 9.195 | 0.116 | 0.0301 | 0.1350 | 0 | 0 | 0 |
| 'Pglyrp2' | 34.520 | 33.373 | 44.121 | 32.321 | 36.365 | 40.200 | 30.391 | 35.528 | 40.039 | 32.362 | 28.670 | 39.544 | 26.113 | 26.969 | 36.758 | 30.041 | 25.833 | 37.582 | 0.199 | 0.0000 | 0.0005 | 1 | 0 | 0 |
| 'Pgm1' | 6.909 | 7.319 | 7.694 | 7.790 | 7.698 | 7.698 | 7.759 | 6.931 | 7.650 | 7.242 | 7.382 | 7.542 | 7.768 | 7.974 | 7.010 | 6.406 | 7.647 | 6.628 | 0.028 | 0.5564 | 0.7680 | 0 | 0 | 0 |
| 'Pgm2' | 27.912 | 26.617 | 24.137 | 26.581 | 28.219 | 24.256 | 28.596 | 26.173 | 25.567 | 26.883 | 28.274 | 28.253 | 29.462 | 27.900 | 29.420 | 27.932 | 24.624 | 27.579 | -0.087 | 0.0272 | 0.1270 | 0 | 0 | 0 |
| 'Pgm2l1' | 2.445 | 2.384 | 2.367 | 2.383 | 2.240 | 2.677 | 2.594 | 2.386 | 2.589 | 2.422 | 3.253 | 3.836 | 2.605 | 2.751 | 3.346 | 2.763 | 2.873 | 3.543 | -0.320 | 0.0000 | 0.0000 | 1 | 0 | 0 |
| 'Pgm3' | 2.165 | 2.583 | 2.692 | 2.372 | 2.688 | 2.531 | 2.301 | 2.800 | 2.630 | 2.579 | 2.830 | 2.290 | 2.789 | 2.804 | 2.334 | 2.385 | 2.650 | 2.087 | 0.000 | 0.9994 | 0.9999 | 0 | 0 | 0 |
| 'Pgp' | 46.639 | 41.175 | 48.354 | 44.286 | 38.070 | 49.021 | 42.084 | 38.638 | 48.438 | 43.018 | 39.097 | 49.133 | 47.345 | 38.067 | 53.752 | 42.893 | 38.140 | 54.278 | -0.040 | 0.3200 | 0.5782 | 0 | 0 | 0 |
| 'Pgpep1' | 1.447 | 1.925 | 1.225 | 1.518 | 1.923 | 1.544 | 1.549 | 1.612 | 1.373 | 1.236 | 1.250 | 1.571 | 1.103 | 1.637 | 1.528 | 1.248 | 1.423 | 1.530 | 0.140 | 0.1262 | 0.3401 | 0 | 0 | 0 |
| 'Pgrmc1' | 23.108 | 25.814 | 21.887 | 20.370 | 25.845 | 22.675 | 22.036 | 25.840 | 22.357 | 25.386 | 21.241 | 22.214 | 21.481 | 23.941 | 21.159 | 22.814 | 21.844 | 20.396 | 0.054 | 0.2322 | 0.4801 | 0 | 0 | 0 |
| 'Pgrmc2' | 15.837 | 19.859 | 17.803 | 17.526 | 18.358 | 17.858 | 17.568 | 17.867 | 17.241 | 18.449 | 18.027 | 18.914 | 18.327 | 18.886 | 19.658 | 17.180 | 18.919 | 18.531 | -0.077 | 0.0185 | 0.0978 | 0 | 0 | 0 |
| 'Pgs1' | 15.142 | 13.278 | 14.685 | 15.372 | 13.818 | 13.487 | 14.167 | 14.283 | 14.587 | 12.514 | 14.456 | 15.049 | 14.095 | 13.771 | 15.082 | 12.930 | 14.668 | 15.407 | -0.001 | 0.9875 | 0.9964 | 0 | 0 | 0 |
| 'Phactr2' | 7.798 | 6.834 | 7.318 | 7.454 | 6.855 | 7.759 | 6.822 | 7.017 | 7.603 | 9.853 | 8.691 | 9.690 | 8.758 | 9.613 | 8.805 | 9.988 | 9.383 | 9.382 | -0.372 | 0.0000 | 0.0000 | 1 | 0 | 0 |
| 'Phactr4' | 4.026 | 3.658 | 3.888 | 3.796 | 4.039 | 3.986 | 3.988 | 3.836 | 3.804 | 3.629 | 3.381 | 3.930 | 3.889 | 4.188 | 3.822 | 3.801 | 4.036 | 3.958 | 0.005 | 0.9085 | 0.9641 | 0 | 0 | 0 |
| 'Phax' | 20.011 | 19.256 | 19.841 | 20.632 | 21.781 | 19.610 | 20.829 | 20.675 | 18.680 | 17.992 | 18.469 | 18.157 | 18.938 | 20.003 | 19.088 | 20.912 | 18.431 | 19.467 | 0.071 | 0.0592 | 0.2100 | 0 | 0 | 0 |
| 'Phb' | 24.838 | 24.532 | 24.205 | 25.292 | 24.339 | 24.626 | 24.529 | 24.734 | 22.824 | 23.402 | 22.935 | 25.085 | 25.498 | 22.757 | 22.984 | 24.396 | 22.806 | 24.468 | 0.024 | 0.4138 | 0.6620 | 0 | 0 | 0 |
| 'Phb2' | 35.485 | 36.604 | 34.615 | 33.607 | 36.944 | 34.855 | 35.373 | 34.320 | 35.707 | 31.985 | 32.815 | 35.159 | 35.783 | 31.596 | 36.468 | 34.944 | 32.232 | 35.052 | 0.042 | 0.1642 | 0.3963 | 0 | 0 | 0 |
| 'Phc1' | 2.636 | 2.891 | 3.013 | 2.204 | 2.776 | 3.255 | 2.489 | 2.980 | 3.091 | 3.113 | 2.533 | 2.863 | 3.163 | 2.844 | 2.840 | 2.947 | 2.676 | 2.735 | -0.032 | 0.5872 | 0.7882 | 0 | 0 | 0 |
| 'Phc2' | 14.414 | 14.263 | 14.182 | 13.727 | 13.167 | 15.142 | 13.759 | 13.300 | 13.716 | 14.223 | 12.902 | 15.465 | 15.062 | 13.760 | 15.675 | 15.368 | 13.769 | 16.618 | -0.089 | 0.0064 | 0.0449 | 1 | 0 | 0 |
| 'Phc3' | 6.201 | 6.219 | 6.607 | 6.210 | 5.929 | 5.528 | 6.285 | 6.127 | 6.113 | 6.670 | 6.849 | 6.977 | 6.224 | 6.392 | 6.101 | 6.225 | 6.439 | 6.752 | -0.098 | 0.0074 | 0.0500 | 0 | 0 | 0 |
| 'Phf1' | 60.400 | 61.297 | 73.742 | 59.319 | 59.596 | 74.661 | 59.777 | 64.284 | 73.002 | 69.454 | 60.637 | 74.890 | 61.561 | 56.961 | 69.226 | 63.610 | 64.761 | 67.627 | -0.019 | 0.5707 | 0.7776 | 0 | 0 | 0 |
| 'Phf10' | 8.630 | 9.504 | 8.455 | 9.067 | 9.644 | 8.992 | 9.138 | 9.558 | 8.652 | 8.276 | 8.293 | 9.612 | 9.580 | 9.983 | 9.167 | 9.232 | 8.653 | 9.179 | -0.022 | 0.5807 | 0.7829 | 0 | 0 | 0 |
| 'Phf11a' | 4.006 | 3.852 | 4.264 | 3.278 | 3.065 | 3.770 | 3.205 | 4.215 | 3.997 | 3.795 | 3.865 | 5.229 | 4.117 | 3.206 | 5.170 | 4.735 | 4.182 | 5.176 | -0.237 | 0.0215 | 0.1080 | 0 | 0 | 0 |
| 'Phf11c' | 2.347 | 2.133 | 2.377 | 2.179 | 1.842 | 2.175 | 2.408 | 2.231 | 2.201 | 2.625 | 2.593 | 3.369 | 2.325 | 2.376 | 3.211 | 2.847 | 1.958 | 2.841 | -0.291 | 0.0005 | 0.0059 | 1 | 0 | 0 |
| 'Phf11d' | 0.560 | 0.517 | 0.623 | 0.666 | 0.682 | 0.441 | 0.693 | 0.587 | 0.684 | 0.733 | 0.664 | 0.646 | 0.770 | 0.710 | 0.555 | 0.634 | 0.844 | 0.641 | -0.196 | 0.0697 | 0.2322 | 0 | 0 | 0 |
| 'Phf12' | 28.313 | 27.502 | 29.524 | 27.418 | 26.121 | 28.650 | 26.976 | 27.369 | 30.209 | 29.159 | 26.980 | 28.598 | 28.816 | 27.620 | 27.360 | 28.707 | 25.465 | 27.098 | 0.004 | 0.8828 | 0.9507 | 0 | 0 | 0 |
| 'Phf13' | 9.463 | 7.613 | 6.829 | 9.231 | 6.178 | 6.901 | 8.512 | 6.157 | 7.271 | 7.249 | 9.138 | 8.218 | 7.611 | 9.171 | 7.563 | 7.816 | 8.481 | 8.150 | -0.127 | 0.1161 | 0.3235 | 0 | 0 | 0 |
| 'Phf14' | 5.199 | 5.043 | 4.903 | 5.722 | 5.302 | 5.096 | 6.302 | 5.120 | 5.271 | 5.433 | 5.257 | 4.755 | 5.037 | 4.875 | 4.370 | 4.825 | 5.355 | 4.232 | 0.110 | 0.0094 | 0.0598 | 0 | 0 | 0 |
| 'Phf19' | 1.997 | 1.874 | 2.533 | 2.154 | 1.994 | 2.575 | 1.951 | 1.732 | 2.562 | 1.729 | 1.897 | 2.247 | 2.549 | 1.801 | 2.337 | 2.048 | 1.981 | 2.092 | 0.037 | 0.5661 | 0.7744 | 0 | 0 | 0 |
| 'Phf2' | 43.166 | 40.698 | 43.159 | 41.362 | 40.167 | 44.326 | 37.264 | 40.352 | 44.122 | 40.946 | 38.940 | 44.275 | 42.879 | 40.378 | 42.539 | 42.257 | 37.933 | 44.965 | -0.012 | 0.6818 | 0.8486 | 0 | 0 | 0 |
| 'Phf20' | 6.387 | 6.020 | 5.931 | 6.323 | 5.611 | 6.000 | 6.447 | 5.823 | 6.170 | 5.846 | 6.433 | 6.559 | 5.700 | 5.865 | 6.467 | 6.350 | 6.021 | 6.519 | -0.039 | 0.2947 | 0.5504 | 0 | 0 | 0 |
| 'Phf20l1' | 54.728 | 53.460 | 57.632 | 54.960 | 55.230 | 50.789 | 54.848 | 57.285 | 52.329 | 51.090 | 54.297 | 49.526 | 49.040 | 52.355 | 44.655 | 49.515 | 55.978 | 46.760 | 0.108 | 0.0007 | 0.0079 | 1 | 0 | 0 |
| 'Phf21a' | 5.781 | 5.971 | 5.554 | 5.588 | 5.817 | 5.878 | 5.314 | 5.934 | 5.815 | 6.347 | 5.704 | 5.656 | 5.918 | 6.139 | 5.751 | 6.129 | 5.709 | 5.546 | -0.042 | 0.1742 | 0.4083 | 0 | 0 | 0 |
| 'Phf21b' | 1.010 | 0.742 | 1.128 | 0.964 | 0.915 | 1.174 | 0.881 | 0.964 | 1.195 | 0.621 | 0.610 | 0.836 | 0.771 | 0.736 | 0.844 | 0.576 | 0.706 | 0.815 | 0.453 | 0.0000 | 0.0000 | 1 | 1 | 0 |
| 'Phf23' | 21.566 | 19.826 | 19.642 | 20.549 | 18.726 | 20.063 | 21.931 | 19.154 | 20.503 | 17.814 | 19.957 | 22.652 | 18.437 | 19.665 | 23.909 | 18.041 | 21.066 | 22.148 | -0.021 | 0.6780 | 0.8461 | 0 | 0 | 0 |
| 'Phf3' | 8.001 | 7.189 | 6.854 | 8.245 | 7.143 | 7.206 | 7.466 | 7.073 | 6.831 | 7.434 | 7.708 | 7.746 | 7.004 | 7.267 | 7.218 | 7.379 | 8.034 | 8.111 | -0.054 | 0.2106 | 0.4559 | 0 | 0 | 0 |
| 'Phf5a' | 37.935 | 41.823 | 37.667 | 41.438 | 41.110 | 43.795 | 42.477 | 41.529 | 38.894 | 38.507 | 40.988 | 38.869 | 41.150 | 39.449 | 40.806 | 41.700 | 42.027 | 36.915 | 0.015 | 0.7180 | 0.8700 | 0 | 0 | 0 |
| 'Phf6' | 13.278 | 15.337 | 14.218 | 15.772 | 14.992 | 13.563 | 15.139 | 13.863 | 13.746 | 13.690 | 14.767 | 14.845 | 12.871 | 15.167 | 13.914 | 14.061 | 15.271 | 14.776 | -0.006 | 0.8809 | 0.9499 | 0 | 0 | 0 |
| 'Phf7' | 10.025 | 13.524 | 11.215 | 11.796 | 12.150 | 10.800 | 11.858 | 12.799 | 12.030 | 12.424 | 11.151 | 9.754 | 13.010 | 12.408 | 10.047 | 12.395 | 12.225 | 8.400 | 0.059 | 0.3309 | 0.5887 | 0 | 0 | 0 |
| 'Phf8' | 6.847 | 7.484 | 7.347 | 7.856 | 7.293 | 7.332 | 6.808 | 7.332 | 7.135 | 7.218 | 8.288 | 8.330 | 7.103 | 7.584 | 7.858 | 6.575 | 7.660 | 7.978 | -0.079 | 0.0379 | 0.1582 | 0 | 0 | 0 |
| 'Phip' | 4.120 | 3.563 | 3.577 | 4.524 | 3.644 | 3.168 | 3.941 | 3.823 | 3.392 | 3.688 | 4.102 | 4.087 | 3.301 | 3.844 | 3.546 | 3.941 | 3.730 | 4.049 | -0.039 | 0.5028 | 0.7298 | 0 | 0 | 0 |
| 'Phka2' | 2.823 | 3.082 | 2.889 | 2.734 | 2.641 | 2.834 | 2.749 | 2.892 | 3.079 | 2.704 | 2.912 | 2.674 | 2.717 | 2.379 | 2.610 | 2.715 | 2.715 | 2.860 | 0.074 | 0.0961 | 0.2882 | 0 | 0 | 0 |
| 'Phkb' | 4.583 | 4.958 | 4.487 | 4.819 | 4.641 | 4.579 | 4.969 | 4.673 | 4.454 | 4.845 | 5.173 | 4.952 | 4.562 | 4.997 | 4.635 | 5.107 | 5.036 | 4.071 | -0.052 | 0.1672 | 0.3997 | 0 | 0 | 0 |
| 'Phkg2' | 5.374 | 5.321 | 6.648 | 5.571 | 5.720 | 5.858 | 5.729 | 5.225 | 5.945 | 6.526 | 5.702 | 6.050 | 6.281 | 5.701 | 5.626 | 5.900 | 5.433 | 5.896 | -0.057 | 0.1600 | 0.3904 | 0 | 0 | 0 |
| 'Phlda1' | 54.249 | 22.576 | 22.526 | 47.260 | 22.599 | 21.934 | 47.451 | 26.082 | 19.297 | 36.835 | 30.423 | 22.708 | 35.656 | 32.276 | 21.923 | 38.984 | 34.924 | 19.373 | -0.021 | 0.8481 | 0.9348 | 0 | 0 | 0 |
| 'Phlda3' | 1.736 | 1.958 | 2.132 | 1.726 | 1.969 | 2.012 | 1.759 | 1.802 | 2.194 | 3.226 | 4.675 | 4.520 | 2.696 | 3.300 | 4.701 | 3.479 | 2.887 | 4.562 | -0.985 | 0.0000 | 0.0000 | 1 | 1 | 0 |

| | | | | | | | | | | | | | | | | | | | | | | | | |
|---|---|---|---|---|---|---|---|---|---|---|---|---|---|---|---|---|---|---|---|---|---|---|---|---|
| 'Phldb1' | 0.622 | 0.596 | 0.705 | 0.598 | 0.574 | 0.756 | 0.659 | 0.561 | 0.820 | 0.758 | 0.525 | 0.883 | 0.771 | 0.622 | 0.750 | 0.873 | 0.729 | 0.780 | -0.196 | 0.0060 | 0.0430 | 1 | 0 | 0 |
| 'Phldb3' | 5.922 | 6.094 | 6.478 | 5.700 | 5.524 | 6.992 | 5.657 | 5.775 | 6.756 | 5.673 | 4.315 | 6.199 | 5.586 | 5.408 | 5.411 | 6.488 | 5.193 | 5.560 | 0.131 | 0.0131 | 0.0760 | 0 | 0 | 0 |
| 'Phlpp1' | 4.015 | 3.994 | 4.249 | 4.776 | 4.254 | 4.777 | 3.998 | 4.849 | 4.392 | 5.934 | 6.310 | 5.840 | 5.297 | 6.434 | 6.146 | 5.732 | 5.995 | 5.977 | -0.455 | 0.0000 | 0.0000 | 1 | 1 | 0 |
| 'Phlpp2' | 2.356 | 1.994 | 2.716 | 2.621 | 2.236 | 2.426 | 2.146 | 1.840 | 2.633 | 3.011 | 2.848 | 2.452 | 2.743 | 2.876 | 2.227 | 2.572 | 2.457 | 2.597 | -0.186 | 0.0187 | 0.0985 | 0 | 0 | 0 |
| 'Phospho2' | 4.331 | 4.448 | 3.926 | 4.078 | 4.393 | 3.915 | 3.988 | 4.397 | 3.830 | 4.359 | 4.209 | 4.360 | 4.027 | 4.376 | 3.941 | 4.264 | 4.818 | 4.071 | -0.057 | 0.1618 | 0.3935 | 0 | 0 | 0 |
| 'Phpt1' | 8.045 | 10.736 | 8.930 | 7.900 | 8.737 | 8.823 | 7.937 | 8.562 | 9.305 | 8.370 | 7.166 | 7.871 | 8.050 | 7.704 | 8.006 | 10.148 | 7.394 | 7.739 | 0.117 | 0.0959 | 0.2881 | 0 | 0 | 0 |
| 'Phrf1' | 10.710 | 9.307 | 10.998 | 10.212 | 9.230 | 10.698 | 9.802 | 9.581 | 11.060 | 10.258 | 10.480 | 11.964 | 9.978 | 9.633 | 11.352 | 9.362 | 9.326 | 12.265 | -0.055 | 0.1255 | 0.3391 | 0 | 0 | 0 |
| 'Phtf1' | 6.735 | 8.096 | 7.462 | 7.392 | 8.065 | 7.372 | 7.201 | 7.527 | 7.388 | 7.248 | 7.420 | 7.537 | 7.157 | 7.759 | 7.137 | 7.069 | 7.506 | 7.304 | 0.010 | 0.7116 | 0.8659 | 0 | 0 | 0 |
| 'Phtf2' | 3.017 | 3.479 | 2.985 | 2.757 | 3.136 | 2.956 | 3.669 | 3.250 | 3.115 | 3.504 | 3.976 | 3.560 | 3.388 | 3.664 | 3.197 | 3.924 | 3.747 | 3.035 | -0.183 | 0.0002 | 0.0031 | 1 | 0 | 0 |
| 'Phyh' | 32.574 | 35.952 | 30.702 | 33.091 | 35.603 | 32.222 | 34.846 | 34.407 | 31.442 | 33.581 | 33.023 | 33.973 | 32.409 | 34.078 | 31.886 | 32.044 | 35.628 | 29.592 | 0.007 | 0.8249 | 0.9227 | 0 | 0 | 0 |
| 'Phykpl' | 1.786 | 1.525 | 1.496 | 1.733 | 1.732 | 1.404 | 1.512 | 1.652 | 1.446 | 1.807 | 1.848 | 1.343 | 1.758 | 1.555 | 1.258 | 1.638 | 1.448 | 1.222 | 0.044 | 0.4369 | 0.6802 | 0 | 0 | 0 |
| 'Pi4k2a' | 22.620 | 21.013 | 24.196 | 22.600 | 21.935 | 23.615 | 21.093 | 21.631 | 22.186 | 24.145 | 22.423 | 22.119 | 22.980 | 22.332 | 23.291 | 22.225 | 20.109 | 23.343 | -0.022 | 0.5126 | 0.7375 | 0 | 0 | 0 |
| 'Pi4ka' | 16.258 | 15.776 | 17.070 | 17.397 | 16.349 | 16.736 | 15.991 | 16.450 | 16.913 | 17.672 | 18.108 | 17.422 | 16.552 | 16.481 | 16.922 | 16.716 | 16.602 | 17.904 | -0.062 | 0.0248 | 0.1201 | 0 | 0 | 0 |
| 'Pi4kb' | 12.091 | 11.945 | 12.392 | 12.352 | 11.284 | 12.693 | 11.416 | 12.354 | 12.390 | 11.974 | 11.516 | 13.431 | 11.673 | 11.559 | 12.839 | 11.412 | 11.780 | 12.644 | -0.010 | 0.7114 | 0.8659 | 0 | 0 | 0 |
| 'Pianp' | 16.407 | 18.451 | 18.825 | 14.544 | 18.050 | 19.179 | 15.086 | 17.361 | 19.621 | 19.169 | 15.693 | 18.863 | 17.614 | 17.532 | 19.712 | 18.307 | 16.142 | 18.292 | -0.047 | 0.3047 | 0.5622 | 0 | 0 | 0 |
| 'Pias1' | 13.478 | 14.734 | 13.936 | 13.746 | 13.845 | 13.881 | 15.227 | 14.195 | 14.354 | 14.081 | 15.358 | 14.876 | 13.264 | 15.555 | 13.262 | 12.639 | 15.693 | 13.471 | -0.020 | 0.6227 | 0.8116 | 0 | 0 | 0 |
| 'Pias2' | 10.567 | 10.628 | 9.428 | 10.295 | 10.101 | 9.680 | 10.776 | 11.065 | 10.243 | 8.665 | 10.289 | 10.291 | 9.703 | 11.024 | 10.045 | 9.955 | 10.494 | 9.520 | 0.032 | 0.4151 | 0.6630 | 0 | 0 | 0 |
| 'Pias3' | 3.638 | 4.351 | 4.193 | 3.854 | 3.926 | 4.352 | 3.602 | 3.892 | 4.391 | 3.711 | 4.309 | 4.129 | 4.237 | 3.608 | 3.802 | 4.040 | 4.360 | 3.681 | 0.002 | 0.9672 | 0.9887 | 0 | 0 | 0 |
| 'Pias4' | 9.385 | 9.413 | 9.901 | 9.569 | 9.219 | 10.559 | 9.824 | 10.047 | 10.911 | 9.915 | 8.595 | 10.664 | 9.364 | 8.940 | 10.723 | 9.798 | 8.691 | 9.370 | 0.035 | 0.3063 | 0.5643 | 0 | 0 | 0 |
| 'Picalm' | 60.736 | 63.440 | 56.578 | 61.156 | 64.584 | 56.732 | 66.652 | 66.605 | 59.380 | 60.339 | 64.108 | 55.881 | 62.823 | 68.760 | 55.058 | 60.963 | 68.377 | 54.435 | 0.006 | 0.8272 | 0.9247 | 0 | 0 | 0 |
| 'Pick1' | 2.603 | 2.921 | 2.955 | 2.236 | 2.787 | 3.141 | 2.471 | 2.856 | 2.975 | 2.943 | 2.877 | 3.099 | 3.119 | 2.868 | 3.004 | 2.804 | 2.851 | 2.839 | -0.094 | 0.0319 | 0.1405 | 0 | 0 | 0 |
| 'Pidd1' | 1.541 | 1.310 | 1.966 | 1.458 | 1.417 | 1.636 | 1.752 | 1.351 | 1.830 | 1.568 | 1.334 | 1.427 | 1.806 | 1.272 | 1.674 | 1.275 | 1.325 | 1.331 | 0.120 | 0.0799 | 0.2546 | 0 | 0 | 0 |
| 'Piezo1' | 6.267 | 6.078 | 7.105 | 5.914 | 6.103 | 6.714 | 6.217 | 6.299 | 7.036 | 6.885 | 6.232 | 6.793 | 6.611 | 6.331 | 6.570 | 6.261 | 6.188 | 6.769 | -0.032 | 0.2593 | 0.5126 | 0 | 0 | 0 |
| 'Pif1' | 0.832 | 0.938 | 1.446 | 1.061 | 1.052 | 1.204 | 1.133 | 1.133 | 1.384 | 0.627 | 1.080 | 1.235 | 1.178 | 1.115 | 1.421 | 0.911 | 1.163 | 1.200 | 0.027 | 0.7664 | 0.8958 | 0 | 0 | 0 |
| 'Piga' | 1.050 | 1.263 | 0.983 | 1.088 | 1.263 | 1.061 | 1.224 | 1.038 | 1.165 | 0.923 | 1.089 | 1.147 | 1.105 | 1.105 | 1.077 | 1.185 | 1.175 | 1.091 | 0.013 | 0.8247 | 0.9227 | 0 | 0 | 0 |
| 'Pigb' | 1.840 | 1.996 | 2.002 | 1.902 | 1.912 | 1.868 | 2.103 | 1.905 | 1.932 | 1.906 | 1.985 | 1.952 | 1.947 | 1.920 | 1.892 | 1.770 | 1.849 | 1.861 | 0.018 | 0.6501 | 0.8295 | 0 | 0 | 0 |
| 'Pigc' | 2.155 | 2.584 | 2.526 | 2.363 | 2.270 | 2.554 | 2.340 | 2.257 | 2.537 | 1.629 | 2.151 | 2.240 | 1.747 | 2.089 | 2.315 | 1.811 | 2.186 | 2.384 | 0.209 | 0.0001 | 0.0009 | 1 | 0 | 0 |
| 'Pigf' | 9.919 | 13.658 | 10.337 | 10.354 | 12.344 | 9.911 | 12.255 | 12.715 | 10.172 | 9.543 | 9.164 | 12.334 | 9.832 | 12.867 | 12.326 | 11.243 | 11.993 | 11.580 | -0.016 | 0.8383 | 0.9309 | 0 | 0 | 0 |
| 'Pigg' | 1.602 | 1.972 | 2.090 | 1.960 | 1.934 | 1.927 | 2.474 | 1.884 | 2.185 | 1.737 | 2.214 | 1.877 | 2.062 | 2.214 | 1.855 | 1.963 | 2.111 | 1.721 | 0.015 | 0.8384 | 0.9309 | 0 | 0 | 0 |
| 'Pigh' | 4.989 | 5.112 | 5.948 | 5.799 | 6.080 | 5.961 | 5.614 | 6.154 | 6.447 | 5.593 | 6.304 | 6.200 | 6.182 | 5.571 | 6.043 | 4.601 | 5.877 | 5.362 | -0.003 | 0.9621 | 0.9866 | 0 | 0 | 0 |
| 'Pigk' | 5.206 | 5.315 | 5.478 | 5.290 | 5.581 | 5.589 | 5.508 | 5.978 | 5.444 | 5.627 | 5.413 | 5.690 | 5.384 | 5.785 | 5.730 | 5.369 | 5.883 | 5.211 | -0.030 | 0.3194 | 0.5780 | 0 | 0 | 0 |
| 'Pigl' | 5.341 | 6.283 | 6.146 | 4.608 | 5.275 | 6.270 | 5.468 | 5.829 | 6.197 | 5.982 | 6.019 | 6.310 | 5.841 | 5.163 | 5.583 | 4.853 | 6.419 | 5.192 | -0.011 | 0.8584 | 0.9390 | 0 | 0 | 0 |
| 'Pigm' | 8.429 | 9.014 | 9.433 | 8.476 | 8.530 | 8.673 | 8.170 | 8.925 | 8.585 | 7.774 | 8.878 | 9.539 | 8.362 | 8.600 | 8.995 | 8.567 | 8.753 | 9.039 | -0.017 | 0.5967 | 0.7950 | 0 | 0 | 0 |
| 'Pign' | 3.278 | 3.188 | 3.323 | 3.345 | 3.218 | 2.735 | 3.237 | 3.262 | 3.036 | 3.267 | 3.874 | 3.261 | 3.225 | 3.238 | 2.937 | 2.932 | 3.571 | 3.054 | -0.046 | 0.3278 | 0.5857 | 0 | 0 | 0 |
| 'Pigo' | 10.472 | 10.896 | 13.282 | 10.283 | 11.737 | 12.498 | 11.116 | 11.395 | 12.798 | 12.191 | 10.846 | 10.679 | 12.476 | 10.825 | 10.906 | 12.351 | 10.434 | 10.550 | 0.036 | 0.4719 | 0.7075 | 0 | 0 | 0 |
| 'Pigp' | 7.972 | 10.169 | 8.698 | 7.776 | 8.650 | 9.231 | 8.428 | 10.376 | 8.018 | 8.703 | 7.391 | 6.927 | 8.938 | 6.870 | 6.723 | 8.125 | 7.802 | 6.607 | 0.214 | 0.0013 | 0.0127 | 1 | 0 | 0 |
| 'Pigq' | 5.423 | 6.076 | 6.779 | 5.211 | 6.446 | 6.397 | 5.573 | 6.660 | 6.714 | 6.035 | 5.757 | 6.631 | 6.431 | 6.050 | 6.494 | 6.043 | 6.050 | 6.006 | -0.019 | 0.6201 | 0.8101 | 0 | 0 | 0 |
| 'Pigs' | 44.733 | 48.393 | 50.506 | 42.845 | 45.731 | 51.479 | 42.433 | 48.099 | 49.676 | 52.030 | 45.037 | 49.724 | 48.770 | 47.159 | 48.944 | 48.955 | 44.647 | 47.482 | -0.040 | 0.2504 | 0.5015 | 0 | 0 | 0 |
| 'Pigt' | 13.553 | 13.549 | 15.885 | 13.181 | 14.115 | 16.433 | 13.117 | 14.325 | 15.162 | 15.008 | 12.711 | 15.822 | 13.732 | 13.387 | 14.896 | 15.112 | 12.126 | 15.278 | 0.005 | 0.8938 | 0.9566 | 0 | 0 | 0 |
| 'Pigu' | 3.280 | 3.015 | 2.985 | 2.914 | 3.190 | 2.899 | 2.813 | 2.918 | 2.771 | 3.037 | 3.189 | 3.282 | 3.106 | 3.069 | 3.410 | 3.110 | 3.288 | 3.030 | -0.102 | 0.0173 | 0.0933 | 0 | 0 | 0 |
| 'Pigv' | 4.842 | 5.704 | 5.704 | 5.213 | 5.588 | 6.102 | 5.205 | 5.645 | 5.897 | 4.826 | 5.348 | 6.483 | 4.738 | 5.831 | 6.289 | 5.634 | 5.444 | 5.654 | -0.023 | 0.6052 | 0.8013 | 0 | 0 | 0 |
| 'Pigw' | 1.370 | 1.283 | 1.420 | 1.387 | 1.173 | 1.230 | 1.845 | 1.102 | 1.398 | 0.964 | 1.413 | 1.708 | 1.103 | 1.001 | 1.592 | 1.027 | 1.567 | 1.625 | 0.012 | 0.9228 | 0.9705 | 0 | 0 | 0 |
| 'Pigx' | 9.645 | 9.949 | 9.316 | 8.726 | 11.245 | 9.746 | 8.609 | 10.001 | 9.613 | 7.860 | 9.150 | 9.734 | 8.388 | 9.899 | 9.535 | 9.431 | 9.143 | 9.348 | 0.062 | 0.1292 | 0.3445 | 0 | 0 | 0 |
| 'Pigyl' | 11.759 | 12.704 | 11.745 | 12.722 | 12.663 | 13.109 | 11.123 | 12.820 | 12.874 | 10.954 | 11.083 | 10.527 | 12.180 | 11.417 | 11.061 | 13.187 | 12.099 | 9.336 | 0.130 | 0.0285 | 0.1307 | 0 | 0 | 0 |
| 'Pih1d1' | 32.411 | 34.348 | 33.261 | 32.823 | 32.348 | 35.307 | 34.114 | 35.384 | 35.003 | 31.248 | 33.116 | 34.501 | 35.637 | 36.021 | 33.893 | 33.101 | 32.169 | 30.354 | 0.013 | 0.7331 | 0.8782 | 0 | 0 | 0 |
| 'Pik3ap1' | 1.949 | 1.948 | 1.453 | 1.209 | 2.603 | 1.496 | 1.317 | 2.451 | 1.416 | 0.849 | 1.864 | 1.176 | 1.105 | 1.909 | 1.207 | 0.849 | 2.103 | 1.197 | 0.375 | 0.0012 | 0.0119 | 1 | 0 | 0 |
| 'Pik3c2a' | 5.506 | 5.228 | 5.566 | 5.523 | 5.727 | 5.166 | 5.555 | 5.767 | 5.395 | 6.233 | 5.745 | 5.063 | 5.585 | 5.374 | 4.882 | 5.565 | 5.586 | 4.985 | 0.007 | 0.8470 | 0.9343 | 0 | 0 | 0 |
| 'Pik3c2b' | 1.206 | 1.073 | 0.948 | 0.931 | 0.885 | 0.724 | 0.709 | 0.895 | 0.934 | 1.443 | 1.382 | 1.398 | 1.411 | 1.580 | 1.417 | 1.441 | 1.444 | 1.660 | -0.676 | 0.0000 | 0.0000 | 1 | 1 | 0 |
| 'Pik3c3' | 8.993 | 9.464 | 9.112 | 8.915 | 10.457 | 9.939 | 8.186 | 9.823 | 9.709 | 9.794 | 9.877 | 9.706 | 8.708 | 9.146 | 9.329 | 9.471 | 10.194 | 8.866 | -0.018 | 0.6346 | 0.8190 | 0 | 0 | 0 |
| 'Pik3ca' | 17.806 | 19.784 | 16.279 | 18.158 | 20.009 | 16.963 | 17.295 | 19.732 | 16.263 | 17.743 | 19.466 | 20.112 | 17.211 | 20.649 | 18.847 | 20.741 | 19.828 | 21.283 | -0.131 | 0.0033 | 0.0269 | 1 | 0 | 0 |
| 'Pik3cb' | 1.483 | 1.587 | 1.545 | 1.438 | 1.437 | 1.652 | 1.527 | 1.703 | 1.722 | 1.408 | 1.347 | 1.366 | 1.565 | 1.586 | 1.318 | 1.333 | 1.775 | 1.260 | 0.118 | 0.0682 | 0.2291 | 0 | 0 | 0 |
| 'Pik3cd' | 49.171 | 45.179 | 50.536 | 46.737 | 47.564 | 51.389 | 44.220 | 48.411 | 50.655 | 48.530 | 49.072 | 51.488 | 48.853 | 47.642 | 48.988 | 45.273 | 45.917 | 51.865 | -0.023 | 0.3794 | 0.6308 | 0 | 0 | 0 |
| 'Pik3cg' | 11.582 | 14.014 | 12.268 | 12.759 | 14.170 | 13.313 | 12.899 | 13.973 | 13.139 | 13.147 | 12.236 | 12.007 | 13.598 | 12.725 | 12.299 | 14.086 | 11.864 | 11.449 | 0.049 | 0.2249 | 0.4716 | 0 | 0 | 0 |
| 'Pik3ip1' | 4.095 | 4.204 | 4.573 | 3.992 | 4.111 | 4.709 | 4.717 | 4.060 | 4.703 | 4.050 | 4.219 | 5.662 | 3.461 | 3.738 | 4.884 | 4.457 | 4.024 | 5.286 | -0.028 | 0.6128 | 0.8058 | 0 | 0 | 0 |
| 'Pik3r1' | 7.706 | 7.507 | 7.382 | 7.858 | 7.975 | 7.762 | 7.441 | 8.003 | 7.856 | 7.662 | 7.317 | 8.703 | 7.548 | 7.868 | 8.300 | 8.271 | 7.706 | 8.878 | -0.067 | 0.0555 | 0.2019 | 0 | 0 | 0 |
| 'Pik3r2' | 11.662 | 11.824 | 12.733 | 11.010 | 12.398 | 14.140 | 11.452 | 11.546 | 13.091 | 12.024 | 10.439 | 12.106 | 12.432 | 11.438 | 11.434 | 11.534 | 10.527 | 11.835 | 0.071 | 0.0622 | 0.2160 | 0 | 0 | 0 |
| 'Pik3r3' | 0.599 | 0.590 | 0.776 | 0.586 | 0.719 | 0.620 | 0.520 | 0.557 | 0.759 | 0.606 | 0.627 | 0.562 | 0.499 | 0.509 | 0.364 | 0.520 | 0.432 | 0.485 | 0.311 | 0.0092 | 0.0587 | 0 | 0 | 0 |
| 'Pik3r4' | 3.767 | 3.646 | 4.285 | 3.965 | 4.365 | 3.916 | 4.028 | 3.928 | 4.201 | 4.352 | 4.082 | 4.110 | 3.677 | 4.020 | 4.141 | 3.955 | 4.011 | 4.004 | -0.019 | 0.6089 | 0.8033 | 0 | 0 | 0 |
| 'Pik3r5' | 1.861 | 1.410 | 2.012 | 1.897 | 1.189 | 2.292 | 1.795 | 1.892 | 1.733 | 1.671 | 1.331 | 2.321 | 1.757 | 1.682 | 2.071 | 1.745 | 1.469 | 2.313 | -0.028 | 0.7174 | 0.8695 | 0 | 0 | 0 |
| 'Pikfyve' | 2.886 | 2.972 | 3.024 | 2.957 | 3.172 | 2.907 | 2.745 | 2.956 | 2.901 | 2.988 | 3.050 | 3.103 | 2.801 | 3.303 | 2.809 | 3.025 | 3.414 | 2.924 | -0.057 | 0.0928 | 0.2820 | 0 | 0 | 0 |
| 'Pim1' | 107.429 | 89.722 | 70.768 | 72.209 | 98.105 | 82.746 | 58.462 | 107.402 | 70.078 | 144.541 | 84.666 | 53.655 | 133.496 | 119.855 | 60.475 | 157.378 | 76.686 | 56.279 | -0.144 | 0.4372 | 0.6804 | 0 | 0 | 0 |
| 'Pim2' | 17.959 | 18.979 | 16.267 | 16.818 | 18.789 | 18.979 | 18.759 | 17.886 | 18.443 | 21.440 | 17.489 | 18.794 | 23.007 | 21.275 | 17.258 | 19.433 | 22.965 | 17.030 | -0.140 | 0.0074 | 0.0498 | 1 | 0 | 0 |
| 'Pim3' | 47.344 | 23.027 | 18.246 | 28.933 | 21.561 | 19.217 | 22.429 | 25.319 | 16.800 | 36.499 | 28.055 | 18.117 | 33.907 | 31.837 | 17.549 | 36.801 | 27.977 | 16.241 | -0.137 | 0.2372 | 0.4868 | 0 | 0 | 0 |
| 'Pin1' | 10.812 | 11.208 | 11.371 | 9.876 | 11.718 | 11.481 | 10.746 | 11.666 | 10.919 | 10.275 | 9.345 | 10.733 | 10.404 | 10.923 | 12.400 | 10.703 | 10.213 | 10.701 | 0.051 | 0.2363 | 0.4861 | 0 | 0 | 0 |
| 'Pin4' | 17.982 | 19.248 | 18.307 | 20.801 | 19.149 | 19.333 | 18.801 | 17.089 | 19.293 | 15.201 | 18.466 | 17.802 | 15.959 | 16.154 | 18.152 | 17.743 | 21.341 | 15.688 | 0.109 | 0.0791 | 0.2530 | 0 | 0 | 0 |
| 'Pink1' | 26.599 | 27.889 | 27.543 | 23.375 | 29.032 | 29.852 | 24.867 | 29.889 | 26.153 | 28.181 | 27.959 | 27.704 | 24.751 | 26.780 | 27.655 | 28.366 | 25.999 | 26.481 | -0.003 | 0.9491 | 0.9806 | 0 | 0 | 0 |
| 'Pinx1' | 8.113 | 8.072 | 9.064 | 9.315 | 7.169 | 8.868 | 9.756 | 8.221 | 8.973 | 8.075 | 7.925 | 8.385 | 7.796 | 7.180 | 7.993 | 7.258 | 7.016 | 9.861 | 0.104 | 0.1284 | 0.3432 | 0 | 0 | 0 |
| 'Pip4k2a' | 46.841 | 48.354 | 47.414 | 46.433 | 47.980 | 46.987 | 44.602 | 46.959 | 47.071 | 52.349 | 48.947 | 51.758 | 46.638 | 51.724 | 50.357 | 50.001 | 49.076 | 51.413 | -0.108 | 0.0000 | 0.0003 | 1 | 0 | 0 |
| 'Pip4k2b' | 38.422 | 40.417 | 38.640 | 38.358 | 38.842 | 41.907 | 35.128 | 41.643 | 39.723 | 40.071 | 38.032 | 42.744 | 36.778 | 39.605 | 41.618 | 40.622 | 35.857 | 45.319 | -0.040 | 0.2820 | 0.5382 | 0 | 0 | 0 |
| 'Pip4k2c' | 28.513 | 31.159 | 30.889 | 30.669 | 31.936 | 32.045 | 28.958 | 31.352 | 32.689 | 30.639 | 30.592 | 31.945 | 30.601 | 32.496 | 31.293 | 33.741 | 30.573 | 32.423 | -0.041 | 0.1555 | 0.3842 | 0 | 0 | 0 |
| 'Pip5k1a' | 8.707 | 8.877 | 8.458 | 9.292 | 9.442 | 8.431 | 8.968 | 9.369 | 8.846 | 8.788 | 9.105 | 8.264 | 8.787 | 9.209 | 8.016 | 8.490 | 9.077 | 8.403 | 0.031 | 0.2219 | 0.4694 | 0 | 0 | 0 |
| 'Pip5k1c' | 11.617 | 11.336 | 11.466 | 10.529 | 10.895 | 12.402 | 10.853 | 10.967 | 12.493 | 12.067 | 11.163 | 11.964 | 11.951 | 11.647 | 12.286 | 12.055 | 10.328 | 11.335 | -0.041 | 0.2106 | 0.4559 | 0 | 0 | 0 |
| 'Pipox' | 6.571 | 11.125 | 7.463 | 6.978 | 10.356 | 8.496 | 8.401 | 11.237 | 8.245 | 7.747 | 7.644 | 5.489 | 8.935 | 7.533 | 5.152 | 7.591 | 7.660 | 4.864 | 0.329 | 0.0002 | 0.0025 | 1 | 0 | 0 |
| 'Pithd1' | 13.720 | 16.506 | 13.968 | 13.818 | 17.144 | 14.823 | 13.858 | 16.468 | 15.325 | 13.479 | 14.050 | 14.386 | 15.343 | 14.988 | 14.754 | 15.479 | 14.315 | 13.891 | 0.038 | 0.3391 | 0.5966 | 0 | 0 | 0 |
| 'Pitpna' | 28.001 | 30.351 | 29.248 | 28.479 | 31.259 | 29.994 | 28.944 | 29.940 | 30.493 | 29.103 | 29.315 | 29.079 | 30.082 | 30.162 | 30.494 | 29.447 | 29.280 | 29.474 | -0.009 | 0.6401 | 0.8230 | 0 | 0 | 0 |
| 'Pitpnb' | 10.473 | 11.226 | 9.766 | 10.094 | 11.561 | 9.879 | 10.743 | 10.690 | 9.900 | 10.104 | 10.520 | 10.885 | 9.993 | 11.100 | 10.730 | 10.159 | 10.669 | 10.411 | -0.020 | 0.5328 | 0.7528 | 0 | 0 | 0 |
| 'Pitpnc1' | 9.745 | 9.801 | 8.866 | 9.669 | 10.025 | 9.666 | 8.829 | 9.385 | 9.088 | 9.259 | 10.290 | 8.769 | 9.079 | 10.079 | 8.499 | 9.019 | 8.812 | 8.744 | 0.035 | 0.3172 | 0.5754 | 0 | 0 | 0 |
| 'Pitpnm1' | 38.230 | 31.990 | 39.922 | 35.091 | 32.180 | 38.002 | 35.317 | 33.473 | 38.936 | 34.567 | 34.775 | 41.440 | 34.346 | 31.517 | 38.171 | 32.147 | 32.960 | 40.875 | 0.001 | 0.9675 | 0.9890 | 0 | 0 | 0 |
| 'Pitpnm2' | 28.500 | 28.506 | 28.834 | 26.942 | 28.633 | 28.618 | 24.600 | 29.801 | 29.185 | 34.225 | 32.436 | 30.882 | 36.807 | 32.905 | 29.763 | 32.430 | 30.493 | 32.254 | -0.214 | 0.0000 | 0.0000 | 1 | 0 | 0 |
| 'Pitrm1' | 16.763 | 15.695 | 17.383 | 17.102 | 16.443 | 17.965 | 17.889 | 16.814 | 17.958 | 17.720 | 16.480 | 17.936 | 18.132 | 15.558 | 17.032 | 16.808 | 15.402 | 17.137 | 0.006 | 0.8355 | 0.9289 | 0 | 0 | 0 |
| 'Pja1' | 6.116 | 4.841 | 4.371 | 5.770 | 4.663 | 4.696 | 5.569 | 4.750 | 4.660 | 4.871 | 5.111 | 5.460 | 4.849 | 4.213 | 4.746 | 4.741 | 5.030 | 5.333 | 0.015 | 0.8007 | 0.9111 | 0 | 0 | 0 |
| 'Pja2' | 6.700 | 7.819 | 6.990 | 6.762 | 7.817 | 7.331 | 7.064 | 7.613 | 7.460 | 7.175 | 7.807 | 7.212 | 6.652 | 8.067 | 6.828 | 7.535 | 7.482 | 6.670 | -0.006 | 0.8565 | 0.9386 | 0 | 0 | 0 |
| 'Pkd1' | 15.033 | 13.589 | 16.058 | 13.388 | 13.060 | 14.444 | 12.393 | 14.329 | 15.472 | 16.891 | 15.435 | 16.262 | 13.480 | 15.036 | 14.872 | 14.405 | 14.114 | 16.540 | -0.112 | 0.0161 | 0.0885 | 0 | 0 | 0 |
| 'Pkd1l3' | 0.759 | 0.735 | 0.606 | 0.476 | 0.580 | 0.624 | 0.520 | 0.619 | 0.538 | 0.904 | 0.751 | 0.616 | 0.900 | 0.772 | 0.604 | 0.929 | 0.620 | 0.507 | -0.254 | 0.0296 | 0.1337 | 0 | 0 | 0 |
| 'Pkd2l2' | 5.764 | 6.474 | 6.782 | 6.294 | 6.808 | 6.331 | 6.094 | 6.633 | 6.903 | 6.341 | 6.644 | 6.587 | 6.018 | 7.249 | 7.153 | 6.351 | 6.248 | 5.467 | -0.007 | 0.9073 | 0.9636 | 0 | 0 | 0 |
| 'Pkig' | 0.712 | 0.905 | 0.662 | 0.591 | 0.849 | 0.564 | 0.683 | 0.677 | 0.404 | 0.580 | 0.449 | 0.564 | 0.749 | 0.870 | 0.613 | 0.761 | 0.682 | 0.519 | 0.040 | 0.7446 | 0.8847 | 0 | 0 | 0 |
| 'Pkm' | 720.865 | 705.580 | 667.711 | 731.854 | 716.432 | 681.371 | 736.204 | 667.052 | 668.677 | 671.929 | 772.132 | 629.443 | 814.576 | 751.234 | 748.515 | 714.957 | 721.785 | 703.190 | -0.063 | 0.1041 | 0.3023 | 0 | 0 | 0 |
| 'Pkmyt1' | 15.747 | 13.726 | 18.414 | 13.965 | 14.567 | 17.233 | 14.881 | 12.568 | 18.822 | 13.907 | 13.193 | 18.814 | 19.230 | 14.886 | 18.629 | 12.951 | 15.423 | 16.839 | -0.056 | 0.3712 | 0.6231 | 0 | 0 | 0 |
| 'Pkn1' | 22.377 | 19.071 | 22.866 | 21.527 | 19.134 | 22.816 | 19.451 | 20.037 | 22.472 | 21.984 | 19.321 | 21.582 | 22.676 | 19.107 | 21.483 | 21.908 | 18.253 | 21.778 | 0.004 | 0.8861 | 0.9530 | 0 | 0 | 0 |
| 'Pkn2' | 4.719 | 4.473 | 4.398 | 4.897 | 4.550 | 4.103 | 4.952 | 4.632 | 4.310 | 4.565 | 4.960 | 4.770 | 4.191 | 4.262 | 4.322 | 4.124 | 5.012 | 4.553 | -0.003 | 0.9391 | 0.9765 | 0 | 0 | 0 |
| 'Pknox1' | 4.197 | 4.087 | 4.866 | 4.002 | 4.312 | 5.067 | 4.263 | 4.849 | 4.585 | 4.783 | 4.192 | 5.221 | 4.500 | 4.845 | 4.994 | 4.509 | 4.537 | 5.286 | -0.100 | 0.0120 | 0.0714 | 0 | 0 | 0 |
| 'Pkp3' | 17.977 | 19.744 | 22.998 | 18.432 | 19.571 | 20.530 | 17.748 | 19.776 | 20.003 | 25.036 | 22.058 | 23.639 | 23.623 | 22.595 | 25.963 | 24.667 | 22.803 | 22.785 | -0.281 | 0.0000 | 0.0000 | 1 | 0 | 0 |
| 'Pkp4' | 3.851 | 4.218 | 3.804 | 3.917 | 3.635 | 3.856 | 3.573 | 3.827 | 4.063 | 5.012 | 4.920 | 4.115 | 4.431 | 4.784 | 4.132 | 4.146 | 4.177 | 3.863 | -0.194 | 0.0001 | 0.0010 | 1 | 0 | 0 |
| 'Pla2g12a' | 4.342 | 5.146 | 4.285 | 3.603 | 4.220 | 4.213 | 3.544 | 4.267 | 3.999 | 4.743 | 4.281 | 3.940 | 4.847 | 4.749 | 4.279 | 4.833 | 4.221 | 3.659 | -0.076 | 0.2368 | 0.4865 | 0 | 0 | 0 |
| 'Pla2g15' | 6.112 | 5.948 | 7.130 | 6.750 | 7.190 | 6.582 | 6.603 | 6.659 | 7.480 | 6.625 | 7.104 | 6.756 | 6.135 | 5.870 | 6.713 | 5.695 | 5.335 | 6.028 | 0.093 | 0.1130 | 0.3192 | 0 | 0 | 0 |
| 'Pla2g16' | 11.552 | 14.387 | 11.465 | 11.938 | 13.690 | 12.797 | 12.440 | 12.564 | 12.082 | 12.676 | 13.162 | 12.326 | 13.949 | 13.215 | 13.087 | 13.233 | 13.193 | 12.044 | -0.063 | 0.0736 | 0.2407 | 0 | 0 | 0 |
| 'Pla2g5' | 1.861 | 1.691 | 1.947 | 1.845 | 1.798 | 1.517 | 1.909 | 1.791 | 1.805 | 0.000 | 0.013 | 0.000 | 0.000 | 0.000 | 0.000 | 0.000 | 0.000 | 0.000 | 9.688 | 0.0000 | 0.0000 | 1 | 1 | 1 |
| 'Pla2g6' | 7.080 | 6.510 | 7.591 | 6.541 | 6.571 | 7.190 | 6.854 | 6.931 | 7.790 | 7.757 | 6.792 | 7.451 | 7.956 | 6.265 | 6.577 | 6.713 | 6.874 | 7.066 | -0.018 | 0.6641 | 0.8377 | 0 | 0 | 0 |
| 'Pla2g7' | 1.495 | 2.685 | 1.596 | 1.850 | 2.182 | 1.898 | 1.457 | 2.733 | 1.844 | 1.608 | 2.145 | 1.263 | 1.614 | 2.173 | 1.056 | 1.424 | 1.973 | 1.181 | 0.301 | 0.0006 | 0.0065 | 1 | 0 | 0 |
| 'Plaa' | 12.181 | 11.817 | 11.673 | 12.272 | 11.567 | 11.476 | 12.786 | 12.330 | 11.487 | 11.637 | 11.834 | 12.383 | 11.900 | 12.101 | 11.175 | 11.970 | 11.339 | 11.208 | 0.015 | 0.5843 | 0.7863 | 0 | 0 | 0 |
| 'Plagl1' | 0.421 | 0.707 | 0.522 | 0.345 | 0.539 | 0.497 | 0.441 | 0.516 | 0.652 | 0.788 | 0.704 | 0.788 | 0.707 | 0.694 | 0.748 | 0.772 | 0.981 | 0.771 | -0.607 | 0.0000 | 0.0000 | 1 | 1 | 0 |
| 'Plagl2' | 9.977 | 9.973 | 10.590 | 9.913 | 9.851 | 10.126 | 10.774 | 9.427 | 10.983 | 10.074 | 10.531 | 11.792 | 10.017 | 10.126 | 11.351 | 10.384 | 10.034 | 12.353 | -0.088 | 0.0101 | 0.0627 | 0 | 0 | 0 |
| 'Plbd2' | 6.787 | 7.540 | 7.333 | 6.209 | 6.590 | 7.587 | 6.768 | 7.096 | 7.510 | 7.889 | 6.897 | 7.917 | 7.668 | 6.661 | 7.462 | 7.983 | 6.445 | 7.478 | -0.075 | 0.0690 | 0.2309 | 0 | 0 | 0 |
| 'Plcb2' | 13.983 | 11.596 | 15.764 | 12.386 | 12.487 | 14.314 | 12.967 | 12.661 | 15.132 | 16.122 | 13.997 | 15.767 | 14.588 | 13.332 | 15.229 | 14.021 | 13.750 | 14.837 | -0.130 | 0.0004 | 0.0047 | 1 | 0 | 0 |
| 'Plcb3' | 13.279 | 12.905 | 12.764 | 13.279 | 13.269 | 12.519 | 12.696 | 13.334 | 13.994 | 12.264 | 11.788 | 12.822 | 13.367 | 12.910 | 11.254 | 11.452 | 11.697 | 12.510 | 0.088 | 0.0222 | 0.1105 | 0 | 0 | 0 |
| 'Plcb4' | 9.192 | 8.293 | 8.060 | 9.737 | 8.030 | 8.210 | 9.408 | 8.115 | 8.310 | 8.263 | 7.463 | 7.733 | 8.336 | 7.372 | 7.189 | 8.257 | 7.726 | 7.121 | 0.142 | 0.0000 | 0.0000 | 1 | 0 | 0 |
| 'Plcg1' | 69.292 | 69.052 | 67.497 | 68.278 | 68.256 | 66.326 | 67.839 | 66.935 | 68.959 | 71.898 | 76.045 | 72.412 | 75.248 | 71.645 | 71.220 | 72.171 | 77.072 | 73.156 | -0.120 | 0.0000 | 0.0000 | 1 | 0 | 0 |
| 'Plcl1' | 3.346 | 4.043 | 3.486 | 3.476 | 3.543 | 3.870 | 3.212 | 3.135 | 3.282 | 4.534 | 4.996 | 4.136 | 4.481 | 4.621 | 3.843 | 5.026 | 4.903 | 3.787 | -0.364 | 0.0000 | 0.0000 | 1 | 0 | 0 |
| 'Plcl2' | 25.571 | 22.987 | 26.167 | 25.630 | 25.119 | 26.473 | 25.695 | 24.588 | 25.124 | 27.374 | 23.110 | 25.115 | 23.608 | 22.957 | 24.371 | 23.189 | 23.295 | 24.206 | 0.056 | 0.0743 | 0.2424 | 0 | 0 | 0 |
| 'Plcxd1' | 4.087 | 3.455 | 3.028 | 4.085 | 3.141 | 3.317 | 4.114 | 2.887 | 3.740 | 3.097 | 4.585 | 4.477 | 3.882 | 3.906 | 4.753 | 3.653 | 4.223 | 4.227 | -0.224 | 0.0091 | 0.0583 | 0 | 0 | 0 |
| 'Plcxd2' | 2.558 | 1.978 | 2.525 | 2.618 | 1.949 | 2.687 | 2.286 | 2.235 | 2.613 | 2.349 | 1.974 | 2.696 | 1.911 | 1.966 | 2.510 | 2.275 | 1.796 | 2.224 | 0.113 | 0.0608 | 0.2127 | 0 | 0 | 0 |
| 'Pld2' | 1.541 | 1.698 | 1.783 | 1.852 | 1.738 | 1.755 | 1.893 | 1.642 | 2.120 | 1.616 | 1.880 | 1.751 | 1.547 | 1.521 | 1.554 | 1.712 | 1.992 | 1.776 | 0.052 | 0.4463 | 0.6878 | 0 | 0 | 0 |
| 'Pld3' | 27.609 | 30.431 | 27.257 | 26.273 | 29.416 | 28.538 | 26.608 | 27.434 | 27.632 | 27.026 | 25.081 | 26.877 | 28.411 | 24.949 | 26.100 | 27.785 | 21.877 | 26.052 | 0.091 | 0.0251 | 0.1210 | 0 | 0 | 0 |
| 'Plec' | 4.254 | 3.157 | 4.146 | 3.947 | 3.063 | 3.731 | 3.660 | 3.270 | 3.996 | 3.275 | 2.748 | 4.317 | 2.905 | 3.232 | 4.266 | 3.284 | 3.065 | 4.699 | 0.066 | 0.2866 | 0.5432 | 0 | 0 | 0 |
| 'Plek' | 1.555 | 1.970 | 1.588 | 1.597 | 2.152 | 2.205 | 1.626 | 2.301 | 1.685 | 1.445 | 1.824 | 1.476 | 1.371 | 2.222 | 1.569 | 1.364 | 2.566 | 1.172 | 0.164 | 0.0832 | 0.2611 | 0 | 0 | 0 |
| 'Plekha1' | 2.778 | 2.729 | 2.371 | 2.490 | 2.355 | 2.555 | 2.496 | 2.774 | 2.639 | 2.718 | 2.614 | 3.017 | 2.535 | 3.180 | 2.781 | 2.418 | 2.911 | 2.936 | -0.126 | 0.0145 | 0.0817 | 0 | 0 | 0 |
| 'Plekha2' | 27.906 | 26.465 | 24.221 | 26.909 | 25.279 | 25.035 | 26.325 | 24.379 | 24.719 | 22.999 | 25.862 | 28.223 | 24.214 | 25.996 | 28.660 | 24.333 | 25.607 | 29.867 | -0.036 | 0.4490 | 0.6890 | 0 | 0 | 0 |
| 'Plekha3' | 18.159 | 17.358 | 17.761 | 17.671 | 17.990 | 17.372 | 18.866 | 18.885 | 16.840 | 17.995 | 19.623 | 17.813 | 16.524 | 19.769 | 18.268 | 17.089 | 18.453 | 18.411 | -0.035 | 0.3415 | 0.5981 | 0 | 0 | 0 |
| 'Plekha5' | 3.840 | 3.729 | 4.021 | 4.000 | 3.558 | 3.980 | 3.906 | 4.256 | 4.037 | 4.147 | 4.076 | 3.984 | 3.952 | 3.830 | 3.834 | 3.865 | 3.864 | 3.969 | -0.015 | 0.6486 | 0.8286 | 0 | 0 | 0 |
| 'Plekha7' | 1.267 | 1.259 | 1.214 | 1.196 | 1.155 | 1.117 | 1.093 | 1.136 | 1.248 | 1.267 | 1.195 | 1.244 | 1.116 | 1.055 | 1.033 | 1.122 | 1.004 | 1.071 | 0.068 | 0.2064 | 0.4505 | 0 | 0 | 0 |
| 'Plekha8' | 7.843 | 9.153 | 9.086 | 8.963 | 9.244 | 9.896 | 8.154 | 8.917 | 8.294 | 6.951 | 8.479 | 7.925 | 7.661 | 7.443 | 8.322 | 7.415 | 9.025 | 8.338 | 0.144 | 0.0180 | 0.0961 | 0 | 0 | 0 |
| 'Plekhb2' | 16.879 | 19.336 | 16.053 | 16.253 | 17.876 | 16.862 | 16.055 | 17.628 | 16.318 | 16.534 | 17.062 | 18.157 | 17.031 | 18.528 | 16.734 | 17.434 | 16.898 | 17.297 | -0.037 | 0.2263 | 0.4730 | 0 | 0 | 0 |
| 'Plekhf2' | 16.106 | 18.967 | 16.320 | 15.002 | 17.738 | 16.363 | 16.105 | 18.129 | 16.048 | 15.745 | 16.049 | 17.252 | 15.862 | 18.089 | 16.444 | 16.143 | 17.233 | 16.634 | -0.003 | 0.9255 | 0.9715 | 0 | 0 | 0 |
| 'Plekhg2' | 26.436 | 22.234 | 27.704 | 24.371 | 22.018 | 26.281 | 23.809 | 22.828 | 27.762 | 26.607 | 27.004 | 30.509 | 25.807 | 23.703 | 29.539 | 23.964 | 26.497 | 29.673 | -0.134 | 0.0002 | 0.0031 | 1 | 0 | 0 |
| 'Plekhg3' | 41.653 | 42.704 | 40.612 | 38.383 | 39.182 | 42.377 | 37.781 | 40.682 | 41.818 | 40.706 | 40.829 | 36.262 | 39.673 | 39.316 | 35.718 | 36.121 | 40.195 | 33.463 | 0.086 | 0.0235 | 0.1155 | 0 | 0 | 0 |
| 'Plekhg5' | 2.141 | 2.302 | 2.497 | 1.860 | 2.263 | 2.564 | 1.996 | 2.121 | 2.554 | 2.198 | 1.731 | 2.074 | 1.991 | 1.664 | 2.039 | 1.979 | 1.800 | 1.935 | 0.213 | 0.0002 | 0.0030 | 1 | 0 | 0 |
| 'Plekhj1' | 93.584 | 106.626 | 102.807 | 98.089 | 111.722 | 106.263 | 96.826 | 107.443 | 103.567 | 96.676 | 92.834 | 96.983 | 107.318 | 104.495 | 99.291 | 96.006 | 99.885 | 97.205 | 0.045 | 0.1541 | 0.3820 | 0 | 0 | 0 |
| 'Plekhm1' | 7.903 | 7.969 | 8.525 | 7.279 | 8.297 | 9.011 | 7.207 | 8.113 | 8.987 | 8.165 | 8.399 | 9.161 | 7.954 | 8.383 | 8.742 | 8.646 | 8.442 | 9.307 | -0.084 | 0.0080 | 0.0526 | 0 | 0 | 0 |
| 'Plekhm2' | 9.508 | 9.372 | 11.162 | 9.341 | 9.126 | 10.131 | 9.099 | 10.473 | 10.767 | 11.255 | 9.134 | 10.128 | 10.718 | 8.947 | 9.287 | 11.109 | 9.306 | 9.631 | -0.014 | 0.7640 | 0.8953 | 0 | 0 | 0 |
| 'Plekhm3' | 0.687 | 0.909 | 0.647 | 0.801 | 0.684 | 0.793 | 0.815 | 1.022 | 0.824 | 0.893 | 0.958 | 0.890 | 0.936 | 0.988 | 0.831 | 0.947 | 0.910 | 0.987 | -0.230 | 0.0018 | 0.0170 | 1 | 0 | 0 |
| 'Plekhn1' | 5.987 | 4.632 | 7.484 | 5.776 | 5.739 | 6.956 | 6.160 | 5.730 | 6.747 | 6.183 | 5.219 | 7.632 | 6.050 | 5.375 | 5.930 | 4.524 | 6.001 | 6.289 | 0.037 | 0.5845 | 0.7863 | 0 | 0 | 0 |
| 'Plekho1' | 25.795 | 22.529 | 25.864 | 25.745 | 23.797 | 27.368 | 22.458 | 22.763 | 25.470 | 31.247 | 27.952 | 35.514 | 28.283 | 30.694 | 35.970 | 29.199 | 30.616 | 38.495 | -0.384 | 0.0000 | 0.0000 | 1 | 1 | 0 |
| 'Plekho2' | 38.469 | 41.950 | 42.519 | 38.542 | 40.289 | 44.730 | 40.506 | 42.327 | 44.818 | 46.860 | 43.469 | 43.225 | 44.895 | 43.269 | 43.377 | 43.633 | 39.803 | 41.685 | -0.071 | 0.0369 | 0.1553 | 0 | 0 | 0 |
| 'Plgrkt' | 24.956 | 29.898 | 27.968 | 25.665 | 29.335 | 26.435 | 27.308 | 28.594 | 28.379 | 26.718 | 26.640 | 26.937 | 28.357 | 27.150 | 24.931 | 25.769 | 30.705 | 23.718 | 0.034 | 0.3371 | 0.5955 | 0 | 0 | 0 |
| 'Plin2' | 2.696 | 2.232 | 2.287 | 2.573 | 2.148 | 2.249 | 2.251 | 2.015 | 2.207 | 2.680 | 2.079 | 4.025 | 2.628 | 2.487 | 3.162 | 2.806 | 2.644 | 4.197 | -0.371 | 0.0001 | 0.0011 | 1 | 0 | 0 |
| 'Plin3' | 17.694 | 16.995 | 17.448 | 15.589 | 17.757 | 18.492 | 16.968 | 18.376 | 18.826 | 15.275 | 15.779 | 15.639 | 14.523 | 14.816 | 16.097 | 15.883 | 14.743 | 15.141 | 0.191 | 0.0000 | 0.0000 | 1 | 0 | 0 |
| 'Plk1' | 11.531 | 13.013 | 16.192 | 14.282 | 12.373 | 17.502 | 14.434 | 12.969 | 16.089 | 8.868 | 13.602 | 15.037 | 14.011 | 15.010 | 16.937 | 12.845 | 13.860 | 14.604 | 0.030 | 0.6775 | 0.8457 | 0 | 0 | 0 |
| 'Plk2' | 8.964 | 2.577 | 1.526 | 6.411 | 3.278 | 1.357 | 4.760 | 3.908 | 1.233 | 6.804 | 4.750 | 1.369 | 4.481 | 4.906 | 0.882 | 5.224 | 5.723 | 1.212 | -0.055 | 0.7592 | 0.8924 | 0 | 0 | 0 |
| 'Plk3' | 32.252 | 26.888 | 27.396 | 28.843 | 27.159 | 27.571 | 27.972 | 28.536 | 26.109 | 33.540 | 26.506 | 30.064 | 30.021 | 30.271 | 28.955 | 35.726 | 28.580 | 30.080 | -0.123 | 0.0013 | 0.0129 | 1 | 0 | 0 |
| 'Plk4' | 2.106 | 2.539 | 2.770 | 2.331 | 2.526 | 2.495 | 2.432 | 2.167 | 2.586 | 1.914 | 2.466 | 2.595 | 2.663 | 2.282 | 2.516 | 2.256 | 2.599 | 2.415 | 0.002 | 0.9676 | 0.9890 | 0 | 0 | 0 |
| 'Plod1' | 2.226 | 2.310 | 2.644 | 2.450 | 2.023 | 2.629 | 2.335 | 2.339 | 2.795 | 2.738 | 2.586 | 3.083 | 2.472 | 2.097 | 2.726 | 2.794 | 2.060 | 3.473 | -0.151 | 0.0137 | 0.0785 | 0 | 0 | 0 |
| 'Plod3' | 4.944 | 4.414 | 5.045 | 4.849 | 4.842 | 5.168 | 4.733 | 4.368 | 4.952 | 4.406 | 3.933 | 5.496 | 4.865 | 4.741 | 4.746 | 4.565 | 4.888 | 5.130 | 0.004 | 0.9301 | 0.9737 | 0 | 0 | 0 |
| 'Plpp5' | 2.371 | 2.693 | 2.566 | 2.087 | 2.559 | 2.914 | 2.367 | 2.510 | 2.718 | 3.025 | 2.785 | 2.598 | 2.826 | 2.599 | 2.709 | 3.213 | 2.777 | 2.308 | -0.126 | 0.0580 | 0.2074 | 0 | 0 | 0 |
| 'Plpp6' | 8.864 | 9.604 | 9.879 | 8.546 | 9.887 | 9.570 | 8.336 | 9.580 | 9.974 | 8.998 | 8.482 | 8.639 | 9.277 | 9.742 | 9.082 | 9.137 | 9.483 | 8.150 | 0.053 | 0.2596 | 0.5127 | 0 | 0 | 0 |
| 'Plppr2' | 1.670 | 1.923 | 1.948 | 1.488 | 1.824 | 1.712 | 2.442 | 2.246 | 2.463 | 1.444 | 1.375 | 1.345 | 1.655 | 1.171 | 1.126 | 1.666 | 1.303 | 1.569 | 0.479 | 0.0000 | 0.0004 | 1 | 1 | 0 |
| 'Plppr3' | 2.927 | 1.433 | 1.277 | 1.580 | 1.277 | 1.147 | 1.238 | 1.484 | 1.395 | 2.936 | 1.568 | 1.329 | 2.569 | 1.834 | 1.149 | 2.264 | 1.414 | 1.266 | -0.212 | 0.1375 | 0.3575 | 0 | 0 | 0 |
| 'Plrg1' | 14.479 | 15.577 | 14.123 | 15.165 | 15.374 | 15.026 | 13.803 | 15.070 | 14.739 | 14.286 | 14.620 | 14.545 | 15.439 | 15.695 | 14.817 | 13.948 | 15.595 | 14.906 | -0.017 | 0.5922 | 0.7922 | 0 | 0 | 0 |
| 'Pls3' | 2.952 | 1.154 | 2.668 | 2.880 | 1.393 | 2.051 | 1.308 | 2.372 | 2.301 | 2.328 | 1.297 | 0.872 | 1.582 | 0.984 | 0.761 | 0.789 | 1.251 | 2.529 | 0.601 | 0.0277 | 0.1282 | 0 | 0 | 0 |
| 'Plscr1' | 5.231 | 4.832 | 5.964 | 5.247 | 6.541 | 5.224 | 4.608 | 6.354 | 4.965 | 5.846 | 5.269 | 6.081 | 5.531 | 7.021 | 5.850 | 5.573 | 6.016 | 5.371 | -0.111 | 0.1165 | 0.3241 | 0 | 0 | 0 |
| 'Plscr3' | 3.461 | 3.836 | 3.915 | 3.536 | 3.841 | 3.962 | 3.130 | 3.925 | 3.842 | 3.463 | 2.999 | 3.279 | 3.777 | 3.017 | 3.319 | 3.551 | 3.334 | 2.687 | 0.179 | 0.0052 | 0.0386 | 1 | 0 | 0 |
| 'Pltp' | 1.071 | 1.333 | 1.118 | 1.110 | 1.508 | 1.174 | 1.253 | 1.603 | 1.187 | 0.515 | 0.970 | 0.595 | 0.684 | 0.531 | 0.732 | 0.537 | 0.592 | 0.865 | 0.898 | 0.0000 | 0.0000 | 1 | 1 | 0 |
| 'Plxdc1' | 1.096 | 1.167 | 1.145 | 1.026 | 1.115 | 0.903 | 1.222 | 1.026 | 1.220 | 1.263 | 1.087 | 1.011 | 1.828 | 1.418 | 1.085 | 1.139 | 1.121 | 1.078 | -0.156 | 0.1137 | 0.3198 | 0 | 0 | 0 |
| 'Plxna1' | 1.547 | 1.817 | 2.079 | 1.591 | 1.752 | 2.079 | 1.414 | 1.673 | 1.920 | 1.852 | 1.332 | 1.849 | 2.142 | 1.389 | 1.642 | 1.739 | 1.400 | 1.942 | 0.046 | 0.5544 | 0.7670 | 0 | 0 | 0 |
| 'Plxna3' | 4.457 | 4.204 | 5.348 | 4.578 | 4.158 | 4.916 | 4.599 | 4.410 | 5.423 | 4.529 | 4.028 | 4.881 | 4.300 | 3.691 | 4.161 | 3.508 | 4.400 | 4.380 | 0.138 | 0.0039 | 0.0308 | 1 | 0 | 0 |
| 'Plxnb1' | 3.771 | 4.629 | 4.623 | 4.424 | 4.418 | 4.579 | 4.026 | 4.277 | 4.685 | 5.400 | 4.841 | 4.944 | 5.729 | 5.500 | 5.130 | 4.658 | 5.312 | 5.233 | -0.257 | 0.0000 | 0.0000 | 1 | 0 | 0 |
| 'Plxnb2' | 1.089 | 1.386 | 1.135 | 1.063 | 1.228 | 0.986 | 1.048 | 1.073 | 1.075 | 0.906 | 1.146 | 1.018 | 1.049 | 1.307 | 0.951 | 1.141 | 1.046 | 0.848 | 0.090 | 0.2259 | 0.4725 | 0 | 0 | 0 |
| 'Plxnc1' | 10.241 | 11.601 | 11.901 | 10.447 | 11.663 | 11.898 | 10.558 | 11.450 | 12.065 | 12.493 | 11.651 | 12.009 | 12.314 | 11.834 | 11.721 | 12.245 | 12.218 | 11.846 | -0.100 | 0.0015 | 0.0141 | 1 | 0 | 0 |
| 'Plxnd1' | 14.011 | 17.543 | 13.112 | 13.425 | 15.695 | 15.091 | 12.924 | 16.421 | 14.207 | 17.730 | 15.010 | 15.969 | 21.605 | 18.949 | 13.717 | 17.870 | 17.644 | 15.389 | -0.226 | 0.0004 | 0.0050 | 1 | 0 | 0 |
| 'Pmaip1' | 15.384 | 7.027 | 5.573 | 10.597 | 8.215 | 4.811 | 10.397 | 8.639 | 4.178 | 11.377 | 12.646 | 5.778 | 12.849 | 11.526 | 4.667 | 12.045 | 11.563 | 5.805 | -0.252 | 0.0084 | 0.0550 | 0 | 0 | 0 |
| 'Pmel' | 4.792 | 4.493 | 5.162 | 3.801 | 4.412 | 3.867 | 5.109 | 5.012 | 4.423 | 5.944 | 4.472 | 4.300 | 4.996 | 4.865 | 4.261 | 6.007 | 3.894 | 4.709 | -0.076 | 0.3644 | 0.6162 | 0 | 0 | 0 |
| 'Pmepa1' | 9.152 | 10.050 | 9.256 | 8.815 | 8.885 | 8.347 | 8.962 | 10.216 | 9.001 | 9.020 | 8.887 | 8.857 | 8.043 | 9.101 | 9.452 | 9.393 | 8.583 | 9.521 | 0.022 | 0.6223 | 0.8114 | 0 | 0 | 0 |
| 'Pmf1' | 4.407 | 4.491 | 5.495 | 4.576 | 4.343 | 5.799 | 5.030 | 4.550 | 5.652 | 4.033 | 4.494 | 4.765 | 5.790 | 4.570 | 5.830 | 4.398 | 4.300 | 4.692 | 0.036 | 0.5702 | 0.7775 | 0 | 0 | 0 |
| 'Pml' | 10.086 | 8.780 | 10.885 | 8.836 | 8.886 | 10.651 | 8.951 | 9.925 | 10.516 | 10.226 | 10.522 | 12.394 | 10.406 | 9.575 | 11.911 | 10.419 | 9.586 | 12.062 | -0.158 | 0.0000 | 0.0001 | 1 | 0 | 0 |
| 'Pmm1' | 1.975 | 1.492 | 1.829 | 1.646 | 1.649 | 1.806 | 1.874 | 2.028 | 1.680 | 1.731 | 1.843 | 1.903 | 1.802 | 1.812 | 1.945 | 1.793 | 1.719 | 1.430 | -0.004 | 0.9579 | 0.9846 | 0 | 0 | 0 |
| 'Pmm2' | 17.850 | 17.060 | 20.277 | 17.288 | 18.375 | 19.125 | 17.609 | 18.646 | 19.148 | 18.790 | 17.690 | 19.254 | 18.550 | 17.794 | 17.777 | 18.147 | 18.047 | 18.549 | -0.002 | 0.9403 | 0.9769 | 0 | 0 | 0 |
| 'Pmpca' | 17.114 | 19.058 | 18.757 | 16.847 | 20.676 | 19.091 | 17.449 | 19.565 | 18.797 | 18.245 | 16.253 | 17.937 | 18.464 | 18.054 | 16.983 | 18.000 | 17.268 | 17.062 | 0.068 | 0.0502 | 0.1891 | 0 | 0 | 0 |
| 'Pmpcb' | 7.674 | 8.871 | 8.014 | 7.992 | 8.608 | 8.296 | 7.833 | 8.555 | 8.048 | 7.807 | 8.026 | 7.692 | 8.452 | 8.479 | 7.275 | 7.591 | 8.203 | 7.354 | 0.049 | 0.1215 | 0.3323 | 0 | 0 | 0 |
| 'Pms1' | 2.757 | 2.254 | 3.135 | 2.822 | 2.758 | 2.560 | 3.295 | 2.591 | 2.662 | 2.666 | 3.060 | 2.832 | 2.747 | 2.772 | 2.506 | 2.502 | 3.112 | 2.392 | 0.005 | 0.9353 | 0.9752 | 0 | 0 | 0 |
| 'Pms2' | 2.392 | 2.443 | 2.773 | 2.364 | 2.585 | 2.539 | 2.246 | 2.571 | 2.461 | 2.581 | 2.474 | 2.437 | 2.559 | 2.631 | 2.485 | 2.412 | 2.683 | 2.383 | -0.022 | 0.5701 | 0.7775 | 0 | 0 | 0 |
| 'Pmvk' | 14.384 | 14.766 | 15.259 | 12.978 | 14.977 | 13.868 | 14.283 | 14.113 | 14.461 | 12.306 | 15.106 | 15.099 | 14.983 | 13.630 | 17.241 | 14.139 | 13.755 | 15.418 | -0.039 | 0.4229 | 0.6696 | 0 | 0 | 0 |
| 'Pnck' | 1.517 | 1.883 | 1.202 | 1.676 | 1.898 | 1.365 | 1.556 | 1.493 | 1.720 | 1.264 | 1.042 | 1.141 | 1.864 | 1.334 | 1.007 | 1.485 | 1.716 | 1.061 | 0.250 | 0.0274 | 0.1274 | 0 | 0 | 0 |
| 'Pnisr' | 13.245 | 13.982 | 14.178 | 12.754 | 14.124 | 14.140 | 13.858 | 13.292 | 14.750 | 12.550 | 14.137 | 15.466 | 13.150 | 15.481 | 14.463 | 13.527 | 15.309 | 14.419 | -0.057 | 0.0565 | 0.2041 | 0 | 0 | 0 |
| 'Pnkd' | 1.444 | 1.595 | 1.672 | 1.513 | 1.754 | 1.848 | 1.733 | 1.738 | 1.689 | 1.523 | 1.341 | 1.687 | 1.793 | 1.589 | 1.589 | 1.551 | 1.662 | 1.668 | 0.042 | 0.4365 | 0.6801 | 0 | 0 | 0 |
| 'Pnkp' | 9.780 | 10.453 | 11.837 | 10.106 | 11.008 | 12.185 | 10.476 | 10.860 | 11.408 | 10.183 | 9.782 | 11.225 | 11.276 | 10.323 | 10.539 | 10.590 | 9.948 | 10.795 | 0.040 | 0.2070 | 0.4515 | 0 | 0 | 0 |
| 'Pnn' | 78.782 | 74.474 | 80.139 | 79.706 | 73.167 | 77.861 | 84.915 | 75.968 | 80.192 | 72.025 | 77.869 | 84.047 | 75.714 | 76.982 | 82.067 | 70.559 | 84.405 | 81.654 | -0.010 | 0.7773 | 0.9002 | 0 | 0 | 0 |
| 'Pno1' | 13.286 | 12.801 | 13.517 | 12.979 | 13.914 | 11.988 | 14.974 | 15.302 | 12.979 | 13.390 | 12.846 | 14.458 | 13.223 | 12.421 | 12.993 | 14.858 | 13.585 | 12.735 | 0.001 | 0.9912 | 0.9976 | 0 | 0 | 0 |
| 'Pnp' | 17.935 | 17.131 | 16.736 | 17.690 | 18.511 | 17.850 | 19.308 | 18.096 | 17.462 | 18.236 | 18.765 | 17.178 | 19.992 | 18.600 | 17.850 | 18.504 | 18.025 | 17.444 | -0.043 | 0.1750 | 0.4097 | 0 | 0 | 0 |
| 'Pnpla2' | 9.117 | 8.807 | 9.433 | 8.381 | 8.880 | 10.284 | 8.708 | 8.700 | 9.584 | 9.306 | 8.542 | 10.086 | 10.481 | 9.391 | 10.464 | 10.337 | 9.492 | 9.078 | -0.099 | 0.0090 | 0.0578 | 0 | 0 | 0 |
| 'Pnpla6' | 18.367 | 16.850 | 19.954 | 18.075 | 16.727 | 18.529 | 16.595 | 18.493 | 19.633 | 18.017 | 16.894 | 19.098 | 18.999 | 16.713 | 17.945 | 17.090 | 17.476 | 18.302 | 0.015 | 0.6523 | 0.8310 | 0 | 0 | 0 |
| 'Pnpla7' | 3.888 | 3.385 | 4.288 | 3.398 | 3.467 | 3.832 | 3.744 | 3.962 | 4.277 | 3.817 | 3.810 | 4.404 | 3.619 | 3.557 | 4.153 | 3.788 | 3.652 | 3.908 | -0.028 | 0.4800 | 0.7122 | 0 | 0 | 0 |
| 'Pnpla8' | 7.279 | 8.847 | 6.809 | 7.503 | 8.077 | 6.945 | 8.152 | 8.059 | 7.361 | 7.379 | 8.557 | 7.636 | 7.141 | 7.570 | 6.702 | 8.897 | 8.613 | 6.688 | -0.016 | 0.7371 | 0.8802 | 0 | 0 | 0 |
| 'Pnpo' | 6.950 | 7.921 | 6.913 | 6.504 | 7.292 | 7.720 | 6.221 | 8.681 | 7.076 | 7.875 | 7.135 | 7.495 | 7.633 | 7.568 | 7.234 | 7.839 | 6.880 | 7.025 | -0.042 | 0.4250 | 0.6716 | 0 | 0 | 0 |
| 'Pnpt1' | 7.030 | 7.689 | 7.333 | 7.554 | 7.164 | 6.492 | 7.569 | 7.896 | 6.817 | 5.954 | 7.948 | 7.453 | 6.970 | 7.247 | 7.435 | 6.464 | 7.759 | 7.724 | 0.001 | 0.9803 | 0.9926 | 0 | 0 | 0 |
| 'Pnrc1' | 138.889 | 102.739 | 73.070 | 102.963 | 100.529 | 78.336 | 99.855 | 107.498 | 70.299 | 136.585 | 107.603 | 65.990 | 130.054 | 124.464 | 67.986 | 140.200 | 104.018 | 66.641 | -0.084 | 0.2834 | 0.5396 | 0 | 0 | 0 |
| 'Pnrc2' | 50.356 | 62.233 | 52.876 | 59.872 | 61.642 | 53.366 | 63.407 | 55.050 | 57.390 | 45.442 | 59.260 | 57.814 | 48.583 | 57.036 | 60.139 | 43.163 | 60.747 | 56.771 | 0.074 | 0.2098 | 0.4550 | 0 | 0 | 0 |
| 'Poc1a' | 1.960 | 2.011 | 2.707 | 2.337 | 1.969 | 2.313 | 2.329 | 2.128 | 2.807 | 1.604 | 2.536 | 2.157 | 2.389 | 2.333 | 2.358 | 2.085 | 2.830 | 2.405 | -0.014 | 0.8747 | 0.9463 | 0 | 0 | 0 |
| 'Poc1b' | 2.733 | 2.898 | 3.073 | 3.197 | 3.125 | 3.258 | 3.231 | 3.152 | 3.417 | 2.342 | 2.297 | 2.652 | 2.739 | 2.490 | 2.825 | 2.440 | 2.717 | 2.662 | 0.265 | 0.0000 | 0.0000 | 1 | 0 | 0 |
| 'Poc5' | 7.954 | 9.597 | 8.865 | 7.910 | 9.414 | 8.622 | 8.161 | 9.261 | 8.511 | 8.460 | 8.696 | 7.819 | 7.773 | 9.304 | 7.235 | 9.194 | 9.284 | 6.993 | 0.065 | 0.1848 | 0.4203 | 0 | 0 | 0 |
| 'Podnl1' | 81.501 | 78.655 | 103.267 | 77.944 | 79.498 | 103.900 | 83.354 | 77.681 | 101.236 | 83.829 | 72.287 | 94.409 | 74.257 | 71.799 | 88.347 | 81.286 | 76.004 | 89.104 | 0.091 | 0.0012 | 0.0125 | 1 | 0 | 0 |
| 'Pofut1' | 3.769 | 3.665 | 3.985 | 3.528 | 4.077 | 4.040 | 3.907 | 3.940 | 4.086 | 4.501 | 4.438 | 4.595 | 4.035 | 4.019 | 4.387 | 4.536 | 4.463 | 4.571 | -0.186 | 0.0000 | 0.0000 | 1 | 0 | 0 |
| 'Pofut2' | 15.732 | 14.885 | 16.415 | 14.427 | 15.615 | 15.809 | 15.678 | 16.522 | 17.153 | 14.932 | 14.440 | 17.442 | 15.075 | 16.426 | 16.335 | 15.812 | 15.925 | 18.670 | -0.039 | 0.3664 | 0.6184 | 0 | 0 | 0 |
| 'Pogk' | 0.605 | 0.579 | 0.620 | 0.523 | 0.597 | 0.545 | 0.520 | 0.535 | 0.582 | 0.605 | 0.488 | 0.571 | 0.514 | 0.461 | 0.577 | 0.589 | 0.492 | 0.593 | 0.054 | 0.3825 | 0.6330 | 0 | 0 | 0 |
| 'Poglut1' | 4.999 | 5.094 | 5.507 | 4.716 | 5.745 | 5.534 | 5.482 | 5.899 | 5.354 | 4.980 | 5.642 | 5.312 | 5.538 | 5.425 | 5.208 | 5.269 | 5.204 | 5.163 | 0.009 | 0.8088 | 0.9139 | 0 | 0 | 0 |
| 'Pogz' | 5.633 | 5.693 | 5.214 | 5.329 | 5.411 | 5.434 | 5.006 | 5.180 | 5.737 | 5.773 | 5.882 | 5.476 | 5.758 | 5.524 | 5.250 | 5.270 | 5.474 | 5.227 | -0.039 | 0.2605 | 0.5141 | 0 | 0 | 0 |
| 'Pola1' | 4.511 | 3.717 | 5.111 | 5.044 | 3.763 | 4.437 | 5.163 | 3.848 | 4.599 | 3.953 | 4.813 | 4.666 | 5.597 | 4.943 | 4.592 | 4.511 | 4.430 | 4.380 | -0.076 | 0.2501 | 0.5011 | 0 | 0 | 0 |
| 'Pola2' | 26.458 | 23.948 | 29.561 | 27.326 | 24.152 | 29.182 | 26.072 | 24.041 | 28.322 | 24.524 | 25.775 | 27.671 | 30.023 | 26.551 | 28.137 | 25.163 | 26.326 | 25.683 | -0.017 | 0.6861 | 0.8515 | 0 | 0 | 0 |
| 'Polb' | 22.637 | 26.983 | 25.694 | 27.531 | 22.978 | 22.769 | 26.293 | 27.902 | 25.339 | 25.962 | 25.859 | 28.520 | 26.107 | 26.951 | 24.157 | 23.414 | 27.550 | 26.440 | -0.059 | 0.2517 | 0.5032 | 0 | 0 | 0 |
| 'Pold1' | 11.954 | 10.117 | 13.920 | 11.789 | 10.134 | 12.761 | 11.875 | 10.230 | 12.820 | 10.719 | 10.637 | 12.080 | 13.077 | 10.545 | 12.646 | 10.822 | 9.800 | 12.314 | 0.028 | 0.4598 | 0.6965 | 0 | 0 | 0 |
| 'Pold2' | 12.599 | 13.080 | 14.207 | 12.092 | 12.889 | 15.844 | 12.384 | 13.677 | 15.121 | 13.842 | 11.420 | 13.401 | 14.156 | 11.787 | 14.780 | 13.741 | 10.613 | 13.033 | 0.055 | 0.2716 | 0.5264 | 0 | 0 | 0 |
| 'Pold3' | 9.380 | 8.878 | 10.575 | 9.792 | 8.942 | 9.367 | 9.831 | 8.676 | 10.117 | 8.819 | 8.965 | 10.534 | 9.791 | 9.417 | 10.185 | 9.356 | 9.674 | 9.922 | -0.031 | 0.3625 | 0.6146 | 0 | 0 | 0 |
| 'Pold4' | 57.223 | 55.867 | 57.491 | 52.568 | 60.388 | 60.704 | 54.652 | 54.625 | 60.049 | 50.324 | 55.458 | 54.786 | 54.211 | 55.963 | 54.540 | 55.900 | 59.052 | 49.314 | 0.062 | 0.1065 | 0.3065 | 0 | 0 | 0 |
| 'Poldip2' | 7.396 | 9.287 | 8.540 | 7.834 | 7.913 | 8.573 | 7.200 | 8.569 | 8.737 | 7.280 | 7.037 | 7.695 | 8.322 | 7.674 | 7.569 | 7.719 | 7.905 | 7.776 | 0.090 | 0.0395 | 0.1621 | 0 | 0 | 0 |
| 'Poldip3' | 22.927 | 24.203 | 22.520 | 21.378 | 24.319 | 24.365 | 21.978 | 23.598 | 23.843 | 23.033 | 24.039 | 25.114 | 21.203 | 23.877 | 25.792 | 23.875 | 23.846 | 25.249 | -0.057 | 0.0502 | 0.1893 | 0 | 0 | 0 |
| 'Pole' | 3.481 | 3.367 | 4.227 | 4.125 | 3.025 | 3.756 | 3.894 | 3.311 | 4.147 | 3.643 | 3.669 | 3.620 | 4.719 | 3.899 | 4.454 | 3.473 | 3.710 | 3.642 | -0.078 | 0.2116 | 0.4567 | 0 | 0 | 0 |
| 'Pole2' | 5.782 | 4.860 | 8.055 | 5.601 | 5.714 | 7.035 | 8.997 | 6.266 | 7.078 | 5.364 | 7.060 | 7.070 | 6.986 | 6.287 | 6.710 | 5.243 | 6.661 | 6.195 | 0.031 | 0.7482 | 0.8868 | 0 | 0 | 0 |
| 'Pole3' | 12.668 | 12.146 | 13.209 | 12.920 | 13.671 | 13.242 | 13.479 | 13.223 | 12.890 | 12.183 | 12.470 | 11.564 | 14.282 | 13.216 | 11.850 | 12.545 | 13.082 | 10.646 | 0.066 | 0.1351 | 0.3537 | 0 | 0 | 0 |
| 'Pole4' | 20.329 | 21.102 | 22.047 | 21.649 | 21.151 | 20.039 | 20.020 | 20.928 | 21.134 | 20.896 | 21.758 | 21.680 | 20.872 | 20.975 | 21.218 | 19.242 | 22.047 | 20.771 | -0.019 | 0.5383 | 0.7566 | 0 | 0 | 0 |
| 'Polg' | 10.899 | 10.491 | 12.438 | 10.346 | 10.767 | 12.464 | 10.131 | 10.523 | 12.043 | 11.400 | 10.584 | 12.029 | 11.374 | 10.733 | 11.506 | 10.465 | 10.987 | 11.820 | -0.023 | 0.3780 | 0.6297 | 0 | 0 | 0 |
| 'Polg2' | 3.233 | 2.230 | 2.919 | 2.991 | 2.699 | 2.420 | 3.272 | 2.887 | 2.423 | 2.968 | 2.789 | 2.217 | 2.967 | 2.654 | 2.003 | 2.482 | 2.565 | 1.879 | 0.157 | 0.0204 | 0.1047 | 0 | 0 | 0 |
| 'Polh' | 12.144 | 13.223 | 15.720 | 14.066 | 12.371 | 14.485 | 14.736 | 12.781 | 14.034 | 10.538 | 15.293 | 15.851 | 12.151 | 12.157 | 15.540 | 10.288 | 15.356 | 15.801 | 0.004 | 0.9528 | 0.9824 | 0 | 0 | 0 |
| 'Poli' | 2.973 | 3.146 | 3.238 | 3.215 | 3.778 | 3.321 | 3.577 | 3.818 | 3.521 | 3.278 | 2.796 | 3.023 | 2.994 | 3.086 | 3.020 | 3.171 | 2.850 | 2.595 | 0.180 | 0.0006 | 0.0065 | 1 | 0 | 0 |
| 'Polk' | 6.435 | 6.407 | 5.528 | 7.562 | 6.257 | 5.535 | 6.445 | 6.425 | 5.436 | 5.111 | 5.941 | 6.775 | 5.813 | 6.279 | 5.737 | 5.559 | 6.125 | 6.435 | 0.039 | 0.5554 | 0.7676 | 0 | 0 | 0 |
| 'Poll' | 13.140 | 13.053 | 15.578 | 12.787 | 14.143 | 15.249 | 12.982 | 12.497 | 14.831 | 14.590 | 10.643 | 13.490 | 14.210 | 12.927 | 12.221 | 12.514 | 12.231 | 13.038 | 0.091 | 0.0786 | 0.2516 | 0 | 0 | 0 |
| 'Polm' | 5.280 | 4.826 | 6.837 | 4.717 | 5.123 | 5.526 | 4.901 | 5.265 | 6.097 | 5.755 | 4.843 | 5.217 | 5.837 | 5.132 | 5.477 | 5.218 | 5.297 | 5.446 | 0.002 | 0.9778 | 0.9922 | 0 | 0 | 0 |
| 'Polq' | 1.092 | 0.906 | 1.204 | 1.161 | 0.950 | 1.149 | 1.273 | 0.953 | 1.200 | 1.008 | 1.333 | 1.154 | 1.326 | 1.094 | 1.154 | 1.069 | 1.208 | 1.105 | -0.093 | 0.1768 | 0.4121 | 0 | 0 | 0 |

| | | | | | | | | | | | | | | | | | | | | | | | | |
|---|---|---|---|---|---|---|---|---|---|---|---|---|---|---|---|---|---|---|---|---|---|---|---|---|
| 'Polr1a' | 5.621 | 4.872 | 5.689 | 5.583 | 4.758 | 5.437 | 4.958 | 4.969 | 5.193 | 5.791 | 5.068 | 5.627 | 5.041 | 4.822 | 5.319 | 4.715 | 4.762 | 5.398 | 0.006 | 0.8781 | 0.9485 | 0 | 0 | 0 |
| 'Polr1b' | 4.792 | 4.252 | 5.543 | 5.042 | 4.214 | 4.567 | 4.579 | 5.236 | 5.339 | 5.146 | 4.638 | 4.676 | 4.758 | 4.527 | 4.322 | 5.094 | 4.071 | 4.784 | 0.049 | 0.3816 | 0.6330 | 0 | 0 | 0 |
| 'Polr1c' | 14.490 | 13.575 | 12.669 | 14.808 | 14.017 | 12.503 | 14.562 | 14.565 | 12.887 | 12.374 | 13.440 | 14.102 | 13.763 | 13.143 | 13.578 | 12.988 | 14.592 | 14.113 | 0.007 | 0.8678 | 0.9428 | 0 | 0 | 0 |
| 'Polr1d' | 26.985 | 27.538 | 25.271 | 28.658 | 27.423 | 27.322 | 27.318 | 26.747 | 25.241 | 25.340 | 24.835 | 24.719 | 26.927 | 26.710 | 26.178 | 28.291 | 25.936 | 25.040 | 0.041 | 0.1346 | 0.3535 | 0 | 0 | 0 |
| 'Polr1e' | 4.731 | 5.017 | 5.264 | 4.885 | 5.203 | 4.765 | 4.987 | 4.766 | 5.267 | 4.972 | 4.576 | 5.012 | 4.797 | 4.509 | 4.761 | 4.328 | 4.401 | 4.973 | 0.069 | 0.0999 | 0.2958 | 0 | 0 | 0 |
| 'Polr2a' | 51.310 | 49.105 | 56.687 | 50.782 | 50.965 | 56.210 | 51.934 | 54.665 | 58.071 | 54.371 | 55.099 | 49.050 | 51.655 | 50.873 | 49.613 | 50.340 | 51.470 | 48.780 | 0.046 | 0.2073 | 0.4519 | 0 | 0 | 0 |
| 'Polr2b' | 23.800 | 24.667 | 23.350 | 25.027 | 25.512 | 22.858 | 24.663 | 25.732 | 23.202 | 23.806 | 24.656 | 23.133 | 23.102 | 25.197 | 22.277 | 22.841 | 24.404 | 21.895 | 0.039 | 0.0759 | 0.2449 | 0 | 0 | 0 |
| 'Polr2c' | 31.930 | 31.730 | 32.024 | 31.861 | 33.963 | 33.650 | 30.458 | 32.212 | 32.558 | 31.243 | 29.997 | 30.580 | 35.299 | 32.232 | 31.376 | 35.832 | 33.128 | 29.328 | 0.001 | 0.9783 | 0.9925 | 0 | 0 | 0 |
| 'Polr2d' | 43.127 | 42.182 | 38.155 | 46.190 | 42.323 | 41.756 | 40.480 | 43.199 | 43.165 | 35.857 | 40.354 | 38.874 | 40.700 | 43.431 | 39.237 | 43.594 | 43.379 | 40.165 | 0.049 | 0.2571 | 0.5093 | 0 | 0 | 0 |
| 'Polr2e' | 16.827 | 17.586 | 16.519 | 17.077 | 16.929 | 17.958 | 17.451 | 18.042 | 16.200 | 15.627 | 15.551 | 16.659 | 17.341 | 15.659 | 16.551 | 17.360 | 16.068 | 16.454 | 0.058 | 0.0546 | 0.2000 | 0 | 0 | 0 |
| 'Polr2f' | 66.317 | 74.409 | 68.134 | 64.921 | 71.740 | 73.516 | 66.188 | 78.042 | 64.199 | 62.508 | 65.392 | 66.630 | 69.646 | 67.667 | 68.552 | 69.696 | 67.131 | 68.503 | 0.040 | 0.3294 | 0.5875 | 0 | 0 | 0 |
| 'Polr2g' | 94.865 | 115.242 | 95.813 | 105.646 | 113.726 | 97.354 | 109.472 | 110.872 | 93.419 | 89.613 | 103.765 | 90.660 | 101.158 | 98.800 | 97.416 | 92.144 | 109.445 | 91.424 | 0.084 | 0.0153 | 0.0850 | 0 | 0 | 0 |
| 'Polr2h' | 13.407 | 13.235 | 13.171 | 13.962 | 12.221 | 12.075 | 13.648 | 13.806 | 11.569 | 11.808 | 12.367 | 11.791 | 12.888 | 12.722 | 12.582 | 12.543 | 13.878 | 11.124 | 0.060 | 0.2156 | 0.4613 | 0 | 0 | 0 |
| 'Polr2i' | 4.960 | 6.290 | 5.677 | 5.307 | 5.999 | 5.877 | 5.497 | 5.563 | 5.662 | 5.270 | 5.164 | 4.578 | 5.652 | 5.102 | 4.715 | 5.495 | 4.973 | 5.250 | 0.132 | 0.0328 | 0.1434 | 0 | 0 | 0 |
| 'Polr2j' | 19.855 | 21.997 | 21.272 | 21.706 | 21.295 | 21.473 | 20.467 | 21.566 | 22.858 | 19.309 | 17.939 | 19.306 | 19.618 | 20.558 | 21.123 | 20.704 | 21.130 | 20.864 | 0.084 | 0.0470 | 0.1806 | 0 | 0 | 0 |
| 'Polr2k' | 21.520 | 26.676 | 25.924 | 27.164 | 24.969 | 24.613 | 25.133 | 25.470 | 22.995 | 20.805 | 23.612 | 22.092 | 23.537 | 28.304 | 23.623 | 24.435 | 25.866 | 22.322 | 0.058 | 0.2795 | 0.5363 | 0 | 0 | 0 |
| 'Polr2m' | 59.293 | 64.359 | 61.453 | 61.601 | 63.381 | 63.457 | 65.727 | 61.390 | 62.628 | 65.048 | 61.870 | 60.259 | 59.899 | 61.216 | 60.384 | 61.983 | 60.338 | 57.533 | 0.028 | 0.2552 | 0.5074 | 0 | 0 | 0 |
| 'Polr3a' | 12.085 | 11.375 | 12.051 | 10.969 | 11.689 | 11.956 | 11.209 | 11.697 | 12.058 | 12.583 | 11.937 | 11.605 | 11.467 | 11.346 | 11.003 | 11.893 | 12.368 | 10.464 | 0.001 | 0.9814 | 0.9931 | 0 | 0 | 0 |
| 'Polr3b' | 12.901 | 12.688 | 13.062 | 13.718 | 12.271 | 12.338 | 12.504 | 13.180 | 13.479 | 13.656 | 13.614 | 13.716 | 12.854 | 12.022 | 12.351 | 13.359 | 13.440 | 12.803 | -0.031 | 0.3787 | 0.6303 | 0 | 0 | 0 |
| 'Polr3c' | 13.788 | 14.039 | 13.982 | 13.808 | 14.048 | 13.947 | 14.414 | 13.964 | 14.642 | 13.859 | 13.025 | 12.895 | 14.169 | 12.807 | 13.170 | 13.869 | 12.472 | 12.699 | 0.082 | 0.0026 | 0.0225 | 1 | 0 | 0 |
| 'Polr3d' | 10.471 | 10.052 | 11.493 | 9.524 | 10.826 | 10.475 | 8.796 | 11.149 | 11.611 | 11.437 | 9.207 | 9.513 | 10.511 | 8.214 | 9.088 | 11.132 | 8.660 | 8.018 | 0.138 | 0.0468 | 0.1803 | 0 | 0 | 0 |
| 'Polr3e' | 11.544 | 11.778 | 12.553 | 10.062 | 11.692 | 11.420 | 12.363 | 12.856 | 11.713 | 9.742 | 9.710 | 11.390 | 9.192 | 9.525 | 10.257 | 10.558 | 9.644 | 10.443 | 0.216 | 0.0000 | 0.0000 | 1 | 0 | 0 |
| 'Polr3f' | 4.220 | 4.770 | 4.325 | 4.141 | 4.805 | 4.342 | 4.179 | 4.886 | 4.292 | 4.277 | 4.157 | 4.354 | 4.524 | 4.873 | 4.014 | 4.584 | 4.635 | 3.963 | 0.010 | 0.7837 | 0.9030 | 0 | 0 | 0 |
| 'Polr3g' | 3.084 | 3.174 | 3.015 | 2.853 | 3.278 | 2.901 | 2.862 | 3.180 | 3.057 | 2.553 | 2.951 | 2.895 | 2.771 | 2.869 | 2.698 | 3.154 | 3.125 | 2.705 | 0.082 | 0.0883 | 0.2714 | 0 | 0 | 0 |
| 'Polr3gl' | 12.775 | 15.457 | 13.729 | 14.416 | 15.339 | 14.582 | 14.571 | 14.842 | 13.358 | 13.423 | 13.558 | 12.302 | 12.919 | 12.711 | 12.568 | 11.467 | 13.599 | 11.091 | 0.171 | 0.0000 | 0.0007 | 1 | 0 | 0 |
| 'Polr3h' | 5.213 | 6.554 | 6.280 | 4.655 | 6.193 | 5.591 | 5.517 | 6.176 | 5.496 | 5.599 | 4.792 | 4.958 | 4.797 | 5.145 | 5.301 | 6.003 | 4.935 | 4.946 | 0.146 | 0.0284 | 0.1304 | 0 | 0 | 0 |
| 'Polr3k' | 8.252 | 10.340 | 8.649 | 9.054 | 9.141 | 8.585 | 8.084 | 9.169 | 9.288 | 8.679 | 8.582 | 9.400 | 8.787 | 9.277 | 9.014 | 8.216 | 9.654 | 8.824 | -0.016 | 0.7028 | 0.8611 | 0 | 0 | 0 |
| 'Polrmt' | 4.652 | 4.398 | 4.800 | 4.097 | 4.482 | 4.671 | 4.039 | 4.275 | 4.818 | 3.986 | 3.595 | 4.857 | 4.331 | 3.683 | 4.698 | 4.181 | 3.864 | 4.992 | 0.066 | 0.1096 | 0.3122 | 0 | 0 | 0 |
| 'Pom121' | 21.874 | 17.949 | 23.370 | 19.307 | 19.132 | 22.559 | 18.394 | 20.678 | 22.391 | 22.839 | 18.696 | 22.008 | 21.436 | 21.889 | 22.973 | 22.608 | 17.706 | 23.079 | -0.066 | 0.1431 | 0.3663 | 0 | 0 | 0 |
| 'Pomgnt1' | 3.511 | 3.908 | 4.294 | 3.632 | 3.831 | 4.133 | 3.327 | 3.838 | 4.099 | 3.799 | 3.336 | 3.647 | 3.931 | 3.349 | 3.462 | 3.499 | 3.380 | 3.031 | 0.128 | 0.0135 | 0.0779 | 0 | 0 | 0 |
| 'Pomk' | 4.906 | 6.160 | 6.036 | 5.901 | 6.148 | 5.924 | 5.996 | 5.992 | 6.790 | 6.405 | 5.616 | 5.594 | 5.944 | 5.547 | 5.400 | 6.100 | 5.530 | 5.185 | 0.061 | 0.2756 | 0.5315 | 0 | 0 | 0 |
| 'Pomp' | 22.614 | 23.284 | 22.372 | 23.442 | 22.780 | 22.001 | 23.058 | 23.210 | 21.199 | 21.715 | 21.317 | 20.869 | 22.969 | 22.472 | 22.784 | 21.837 | 23.947 | 21.815 | 0.020 | 0.4941 | 0.7233 | 0 | 0 | 0 |
| 'Pomt1' | 1.611 | 1.527 | 2.093 | 1.914 | 1.872 | 2.059 | 1.742 | 1.960 | 2.311 | 2.181 | 1.783 | 2.326 | 2.211 | 1.911 | 1.753 | 1.911 | 2.207 | 1.904 | -0.103 | 0.1464 | 0.3719 | 0 | 0 | 0 |
| 'Pomt2' | 1.868 | 2.187 | 1.998 | 1.906 | 1.829 | 2.220 | 1.409 | 2.014 | 2.197 | 1.597 | 1.650 | 1.835 | 2.050 | 1.816 | 2.067 | 1.908 | 1.750 | 2.396 | 0.038 | 0.6449 | 0.8264 | 0 | 0 | 0 |
| 'Pon2' | 9.249 | 11.510 | 10.060 | 10.227 | 11.160 | 10.236 | 9.829 | 12.013 | 10.095 | 11.130 | 11.182 | 11.414 | 11.576 | 11.928 | 11.281 | 11.305 | 11.896 | 11.011 | -0.137 | 0.0000 | 0.0001 | 1 | 0 | 0 |
| 'Pop1' | 1.800 | 1.670 | 1.726 | 2.285 | 1.653 | 1.678 | 1.778 | 1.700 | 2.101 | 1.498 | 1.449 | 1.740 | 1.572 | 1.446 | 1.976 | 1.483 | 1.628 | 1.898 | 0.143 | 0.0654 | 0.2228 | 0 | 0 | 0 |
| 'Pop4' | 4.619 | 4.902 | 4.894 | 4.301 | 4.898 | 4.633 | 4.345 | 5.043 | 4.179 | 5.117 | 4.434 | 3.460 | 5.473 | 4.394 | 3.756 | 5.142 | 4.530 | 3.254 | 0.092 | 0.1946 | 0.4326 | 0 | 0 | 0 |
| 'Pop5' | 11.803 | 10.871 | 10.555 | 12.281 | 11.601 | 11.101 | 10.977 | 10.881 | 10.899 | 10.028 | 10.600 | 11.519 | 9.926 | 9.911 | 11.424 | 10.201 | 10.704 | 11.238 | 0.066 | 0.1358 | 0.3545 | 0 | 0 | 0 |
| 'Pop7' | 9.339 | 9.571 | 10.734 | 8.938 | 10.901 | 11.558 | 10.414 | 9.606 | 10.116 | 9.441 | 8.762 | 9.949 | 10.716 | 9.318 | 10.020 | 10.042 | 9.882 | 9.252 | 0.052 | 0.3417 | 0.5981 | 0 | 0 | 0 |
| 'Popdc2' | 0.359 | 0.485 | 0.429 | 0.543 | 0.591 | 0.425 | 0.513 | 0.509 | 0.431 | 0.647 | 1.109 | 0.784 | 0.815 | 0.946 | 0.750 | 0.737 | 0.945 | 0.655 | -0.802 | 0.0000 | 0.0000 | 1 | 1 | 0 |
| 'Por' | 24.899 | 24.553 | 25.817 | 24.913 | 24.703 | 24.742 | 25.070 | 24.826 | 23.966 | 24.529 | 24.875 | 25.370 | 26.694 | 23.997 | 24.760 | 23.575 | 23.281 | 25.641 | -0.006 | 0.8072 | 0.9136 | 0 | 0 | 0 |
| 'Pot1a' | 2.919 | 3.713 | 3.455 | 3.376 | 3.519 | 3.439 | 3.482 | 3.265 | 3.275 | 3.355 | 3.828 | 3.535 | 3.516 | 3.529 | 3.444 | 3.257 | 3.532 | 3.159 | -0.047 | 0.2568 | 0.5093 | 0 | 0 | 0 |
| 'Pot1b' | 5.579 | 5.583 | 5.963 | 5.677 | 5.493 | 6.067 | 5.890 | 5.955 | 6.240 | 5.770 | 6.566 | 6.500 | 6.136 | 6.213 | 6.181 | 5.212 | 6.389 | 5.785 | -0.073 | 0.1372 | 0.3572 | 0 | 0 | 0 |
| 'Pou2af1' | 124.699 | 176.111 | 120.748 | 132.593 | 167.384 | 137.632 | 136.926 | 159.222 | 134.775 | 138.208 | 150.558 | 113.231 | 157.907 | 166.217 | 120.820 | 145.189 | 150.913 | 112.954 | 0.030 | 0.5173 | 0.7406 | 0 | 0 | 0 |
| 'Pou2f1' | 2.272 | 2.280 | 2.350 | 2.451 | 2.196 | 2.078 | 2.613 | 2.361 | 2.397 | 2.480 | 2.493 | 2.447 | 2.498 | 2.370 | 2.248 | 2.448 | 2.347 | 2.391 | -0.061 | 0.0938 | 0.2835 | 0 | 0 | 0 |
| 'Pou2f2' | 16.142 | 15.399 | 16.785 | 15.663 | 14.853 | 16.108 | 16.412 | 14.867 | 17.513 | 16.488 | 14.471 | 18.248 | 15.275 | 15.101 | 15.767 | 16.304 | 14.758 | 16.422 | -0.002 | 0.9408 | 0.9772 | 0 | 0 | 0 |
| 'Pou6f1' | 5.498 | 5.910 | 4.807 | 4.730 | 5.514 | 5.187 | 4.551 | 5.672 | 5.679 | 5.859 | 5.065 | 5.434 | 5.021 | 5.109 | 4.706 | 5.097 | 4.842 | 5.005 | 0.031 | 0.5456 | 0.7606 | 0 | 0 | 0 |
| 'Ppa1' | 8.618 | 7.023 | 9.566 | 8.240 | 8.664 | 8.119 | 7.861 | 8.687 | 8.103 | 7.827 | 7.942 | 9.636 | 8.824 | 8.082 | 9.500 | 8.447 | 7.467 | 9.605 | -0.055 | 0.2952 | 0.5508 | 0 | 0 | 0 |
| 'Ppa2' | 1.411 | 1.328 | 1.410 | 1.152 | 1.617 | 1.225 | 1.467 | 1.399 | 1.448 | 1.644 | 1.413 | 1.500 | 1.295 | 1.294 | 1.466 | 1.499 | 1.417 | 1.267 | -0.050 | 0.4517 | 0.6909 | 0 | 0 | 0 |
| 'Ppan' | 10.334 | 8.589 | 11.027 | 9.074 | 8.494 | 10.054 | 9.582 | 9.419 | 10.249 | 10.105 | 8.771 | 11.393 | 9.515 | 7.698 | 9.685 | 9.419 | 8.593 | 10.169 | 0.016 | 0.7088 | 0.8650 | 0 | 0 | 0 |
| 'Ppard' | 5.076 | 4.557 | 5.246 | 4.517 | 4.535 | 5.398 | 4.764 | 4.755 | 5.275 | 4.850 | 4.542 | 5.445 | 4.741 | 4.510 | 5.337 | 4.259 | 4.809 | 5.774 | -0.015 | 0.7167 | 0.8692 | 0 | 0 | 0 |
| 'Ppargc1b' | 6.465 | 4.391 | 4.981 | 6.376 | 4.063 | 4.320 | 5.267 | 4.212 | 5.132 | 6.121 | 5.640 | 7.162 | 5.314 | 4.590 | 6.546 | 5.231 | 6.241 | 7.169 | -0.276 | 0.0018 | 0.0168 | 1 | 0 | 0 |
| 'Ppat' | 5.832 | 5.693 | 6.002 | 6.598 | 6.067 | 5.588 | 6.928 | 6.104 | 5.835 | 5.062 | 6.096 | 6.412 | 5.223 | 5.531 | 6.249 | 5.219 | 5.759 | 6.299 | 0.064 | 0.2421 | 0.4928 | 0 | 0 | 0 |
| 'Ppcdc' | 33.585 | 35.529 | 37.465 | 31.775 | 34.682 | 41.079 | 33.085 | 36.643 | 39.145 | 33.440 | 32.988 | 38.465 | 36.930 | 35.098 | 34.441 | 32.825 | 33.720 | 33.814 | 0.038 | 0.2840 | 0.5402 | 0 | 0 | 0 |
| 'Ppcs' | 9.660 | 10.108 | 10.047 | 9.874 | 11.442 | 10.364 | 11.404 | 10.203 | 11.440 | 9.335 | 8.849 | 11.242 | 9.358 | 9.794 | 11.050 | 10.448 | 10.961 | 10.954 | 0.023 | 0.6606 | 0.8362 | 0 | 0 | 0 |
| 'Ppdpf' | 14.649 | 16.714 | 15.342 | 14.232 | 16.766 | 16.601 | 13.598 | 16.276 | 16.179 | 14.683 | 13.207 | 15.359 | 14.872 | 13.955 | 15.812 | 15.501 | 13.305 | 14.315 | 0.087 | 0.0435 | 0.1726 | 0 | 0 | 0 |
| 'Ppfia1' | 15.804 | 16.330 | 15.214 | 16.054 | 16.100 | 14.935 | 14.896 | 16.370 | 14.767 | 17.277 | 18.939 | 15.219 | 19.950 | 18.057 | 14.934 | 17.969 | 18.631 | 15.442 | -0.160 | 0.0000 | 0.0001 | 1 | 0 | 0 |
| 'Ppfibp1' | 5.712 | 5.738 | 5.709 | 5.735 | 5.762 | 5.286 | 5.936 | 6.223 | 5.646 | 6.572 | 7.410 | 5.984 | 6.324 | 6.753 | 6.104 | 6.433 | 7.124 | 5.917 | -0.185 | 0.0000 | 0.0000 | 1 | 0 | 0 |
| 'Ppfibp2' | 1.017 | 0.895 | 0.967 | 0.900 | 0.994 | 1.174 | 1.296 | 0.857 | 1.306 | 0.906 | 0.786 | 0.720 | 0.534 | 1.078 | 0.597 | 0.874 | 0.767 | 0.745 | 0.428 | 0.0010 | 0.0108 | 1 | 1 | 0 |
| 'Pphln1' | 7.841 | 8.806 | 8.819 | 8.044 | 8.454 | 8.018 | 8.183 | 8.005 | 8.586 | 8.511 | 8.535 | 7.908 | 7.990 | 8.183 | 7.674 | 7.222 | 8.490 | 7.965 | 0.035 | 0.3499 | 0.6044 | 0 | 0 | 0 |
| 'Ppia' | 137.610 | 155.875 | 136.376 | 153.130 | 153.251 | 137.214 | 154.716 | 147.099 | 141.589 | 128.639 | 142.555 | 115.287 | 140.564 | 148.646 | 141.791 | 140.796 | 146.328 | 129.046 | 0.084 | 0.0242 | 0.1178 | 0 | 0 | 0 |
| 'Ppib' | 117.425 | 129.777 | 118.222 | 126.033 | 128.758 | 121.806 | 121.093 | 125.070 | 117.703 | 127.536 | 115.226 | 118.581 | 119.545 | 121.635 | 117.221 | 122.097 | 120.083 | 113.735 | 0.028 | 0.2645 | 0.5184 | 0 | 0 | 0 |
| 'Ppic' | 4.960 | 11.408 | 7.699 | 6.124 | 9.428 | 8.869 | 6.376 | 7.253 | 8.987 | 8.065 | 6.612 | 7.475 | 7.638 | 8.114 | 5.762 | 7.744 | 9.864 | 6.135 | 0.050 | 0.6650 | 0.8382 | 0 | 0 | 0 |
| 'Ppid' | 12.529 | 13.102 | 12.050 | 12.646 | 13.154 | 11.764 | 12.731 | 13.152 | 12.149 | 11.808 | 12.643 | 13.827 | 12.045 | 12.648 | 12.126 | 11.687 | 13.533 | 13.202 | -0.018 | 0.6118 | 0.8055 | 0 | 0 | 0 |
| 'Ppie' | 16.823 | 19.450 | 18.976 | 18.319 | 19.155 | 17.773 | 17.534 | 19.019 | 17.612 | 16.786 | 15.799 | 17.885 | 17.294 | 17.528 | 17.897 | 16.323 | 17.816 | 16.924 | 0.079 | 0.0193 | 0.1008 | 0 | 0 | 0 |
| 'Ppif' | 9.057 | 9.016 | 8.832 | 8.415 | 8.577 | 8.575 | 8.142 | 9.471 | 9.156 | 8.766 | 8.866 | 9.093 | 9.386 | 8.559 | 8.518 | 9.454 | 8.198 | 8.684 | -0.014 | 0.7151 | 0.8686 | 0 | 0 | 0 |
| 'Ppig' | 13.786 | 14.086 | 14.004 | 15.393 | 15.139 | 14.007 | 16.135 | 14.258 | 13.530 | 13.501 | 13.878 | 14.466 | 14.075 | 14.237 | 13.705 | 14.227 | 14.936 | 13.723 | 0.027 | 0.3932 | 0.6428 | 0 | 0 | 0 |
| 'Ppih' | 11.538 | 14.433 | 14.189 | 13.507 | 15.628 | 14.598 | 13.440 | 15.698 | 13.817 | 13.833 | 13.302 | 13.481 | 13.526 | 14.307 | 12.169 | 12.681 | 14.496 | 12.478 | 0.064 | 0.1491 | 0.3758 | 0 | 0 | 0 |
| 'Ppil1' | 17.960 | 18.066 | 16.831 | 17.986 | 18.764 | 17.888 | 17.795 | 17.981 | 17.071 | 16.832 | 15.733 | 17.716 | 18.714 | 17.350 | 18.713 | 18.236 | 16.687 | 17.525 | 0.013 | 0.7180 | 0.8700 | 0 | 0 | 0 |
| 'Ppil2' | 15.844 | 16.386 | 17.407 | 16.895 | 17.046 | 16.826 | 16.581 | 16.957 | 17.343 | 16.060 | 16.418 | 18.180 | 17.820 | 16.909 | 17.051 | 16.272 | 17.152 | 17.631 | -0.033 | 0.1629 | 0.3947 | 0 | 0 | 0 |
| 'Ppil3' | 4.735 | 5.345 | 4.919 | 5.120 | 4.716 | 4.484 | 4.843 | 4.901 | 4.939 | 4.331 | 4.349 | 4.651 | 4.161 | 4.855 | 4.042 | 4.317 | 4.848 | 4.516 | 0.121 | 0.0078 | 0.0520 | 0 | 0 | 0 |
| 'Ppil4' | 13.026 | 14.275 | 12.469 | 12.943 | 12.977 | 11.584 | 13.649 | 13.023 | 12.474 | 13.081 | 13.732 | 13.225 | 12.064 | 13.264 | 12.904 | 11.945 | 13.335 | 12.529 | -0.009 | 0.7869 | 0.9047 | 0 | 0 | 0 |
| 'Ppip5k1' | 2.531 | 2.461 | 2.826 | 2.610 | 2.482 | 2.650 | 2.491 | 2.581 | 2.773 | 2.887 | 2.742 | 2.614 | 2.573 | 2.526 | 2.554 | 2.574 | 2.709 | 2.482 | -0.022 | 0.5707 | 0.7776 | 0 | 0 | 0 |
| 'Ppip5k2' | 7.001 | 7.016 | 6.907 | 7.069 | 7.127 | 6.566 | 6.897 | 7.171 | 6.627 | 7.580 | 7.522 | 7.320 | 6.494 | 7.288 | 6.793 | 7.059 | 7.559 | 7.006 | -0.061 | 0.0437 | 0.1732 | 0 | 0 | 0 |
| 'Ppm1a' | 37.612 | 38.707 | 35.555 | 34.172 | 38.995 | 39.551 | 35.005 | 40.905 | 34.614 | 42.088 | 38.871 | 35.163 | 38.507 | 41.446 | 35.941 | 42.631 | 39.135 | 37.707 | -0.076 | 0.0559 | 0.2028 | 0 | 0 | 0 |
| 'Ppm1b' | 23.000 | 22.538 | 23.234 | 23.213 | 22.123 | 23.423 | 24.848 | 24.199 | 22.932 | 21.791 | 23.206 | 24.480 | 21.794 | 23.045 | 22.602 | 20.148 | 23.375 | 24.776 | 0.019 | 0.6078 | 0.8027 | 0 | 0 | 0 |
| 'Ppm1d' | 21.273 | 19.495 | 17.590 | 19.444 | 19.126 | 16.679 | 19.435 | 19.616 | 18.396 | 18.585 | 20.943 | 19.441 | 19.327 | 22.365 | 18.888 | 19.800 | 20.244 | 20.269 | -0.084 | 0.0421 | 0.1694 | 0 | 0 | 0 |
| 'Ppm1f' | 12.708 | 13.503 | 12.373 | 12.790 | 13.914 | 13.581 | 13.332 | 12.460 | 13.484 | 12.609 | 12.316 | 15.212 | 12.550 | 13.087 | 15.581 | 14.329 | 13.531 | 14.901 | -0.082 | 0.0520 | 0.1933 | 0 | 0 | 0 |
| 'Ppm1g' | 31.810 | 30.871 | 31.291 | 31.535 | 32.006 | 32.218 | 31.055 | 30.991 | 32.085 | 29.833 | 30.302 | 31.792 | 31.345 | 30.318 | 31.997 | 32.420 | 28.225 | 32.667 | 0.015 | 0.5338 | 0.7536 | 0 | 0 | 0 |
| 'Ppm1h' | 6.389 | 6.709 | 6.108 | 6.223 | 6.440 | 6.685 | 5.889 | 6.617 | 6.740 | 6.480 | 6.761 | 6.704 | 6.701 | 6.796 | 6.354 | 6.759 | 5.924 | 6.860 | -0.049 | 0.1647 | 0.3970 | 0 | 0 | 0 |
| 'Ppm1k' | 4.936 | 5.696 | 4.808 | 5.623 | 5.413 | 4.484 | 5.175 | 5.359 | 4.909 | 4.847 | 6.003 | 5.556 | 5.104 | 6.000 | 5.100 | 4.882 | 6.199 | 5.458 | -0.099 | 0.0346 | 0.1489 | 0 | 0 | 0 |
| 'Ppm1m' | 24.615 | 24.113 | 27.466 | 25.658 | 23.638 | 29.060 | 23.730 | 24.564 | 26.590 | 23.044 | 22.528 | 28.017 | 26.147 | 22.231 | 26.238 | 25.843 | 22.499 | 27.753 | 0.023 | 0.4565 | 0.6942 | 0 | 0 | 0 |
| 'Ppme1' | 40.631 | 38.696 | 34.857 | 38.481 | 34.655 | 35.215 | 35.428 | 37.480 | 35.873 | 33.844 | 40.166 | 39.928 | 36.412 | 38.268 | 40.030 | 35.645 | 36.944 | 41.360 | -0.059 | 0.1983 | 0.4382 | 0 | 0 | 0 |
| 'Ppox' | 6.854 | 7.129 | 7.861 | 7.218 | 7.327 | 8.023 | 7.147 | 7.324 | 8.106 | 7.623 | 7.245 | 8.073 | 8.403 | 6.724 | 7.785 | 6.994 | 7.577 | 7.162 | -0.025 | 0.5123 | 0.7374 | 0 | 0 | 0 |
| 'Ppp1ca' | 386.025 | 430.563 | 397.033 | 405.392 | 428.514 | 414.289 | 393.212 | 407.036 | 404.963 | 384.785 | 399.364 | 373.531 | 402.061 | 410.542 | 419.019 | 402.550 | 400.548 | 402.482 | 0.017 | 0.4958 | 0.7249 | 0 | 0 | 0 |
| 'Ppp1cb' | 16.277 | 19.366 | 16.743 | 18.108 | 19.248 | 17.066 | 19.062 | 19.011 | 16.238 | 16.345 | 17.995 | 17.815 | 16.064 | 19.389 | 17.448 | 16.847 | 18.750 | 17.973 | 0.010 | 0.7733 | 0.8993 | 0 | 0 | 0 |
| 'Ppp1cc' | 43.039 | 48.724 | 42.260 | 44.878 | 46.852 | 43.596 | 44.515 | 47.336 | 42.539 | 41.140 | 45.544 | 43.608 | 42.377 | 46.909 | 45.517 | 42.761 | 45.291 | 44.820 | 0.009 | 0.7223 | 0.8722 | 0 | 0 | 0 |
| 'Ppp1r10' | 11.659 | 10.089 | 11.983 | 12.252 | 10.062 | 11.691 | 11.301 | 10.478 | 12.693 | 10.350 | 11.770 | 12.428 | 10.689 | 11.414 | 11.630 | 10.288 | 11.647 | 11.695 | -0.007 | 0.8719 | 0.9452 | 0 | 0 | 0 |
| 'Ppp1r11' | 22.988 | 26.820 | 24.229 | 24.247 | 27.385 | 25.467 | 23.378 | 25.506 | 24.876 | 24.300 | 22.556 | 24.283 | 24.402 | 23.699 | 25.530 | 25.566 | 24.366 | 25.039 | 0.020 | 0.5647 | 0.7736 | 0 | 0 | 0 |
| 'Ppp1r12a' | 39.194 | 39.546 | 38.840 | 39.675 | 40.453 | 38.528 | 38.204 | 40.879 | 40.839 | 40.892 | 40.552 | 41.681 | 39.782 | 42.045 | 41.033 | 39.753 | 41.574 | 42.625 | -0.066 | 0.0021 | 0.0189 | 1 | 0 | 0 |
| 'Ppp1r12b' | 2.329 | 1.632 | 2.167 | 2.062 | 1.749 | 1.791 | 2.074 | 2.007 | 1.928 | 1.867 | 1.665 | 1.680 | 1.656 | 1.493 | 1.552 | 1.550 | 1.850 | 1.721 | 0.227 | 0.0002 | 0.0022 | 1 | 0 | 0 |
| 'Ppp1r12c' | 22.789 | 19.216 | 24.693 | 21.471 | 18.861 | 24.137 | 20.603 | 20.912 | 23.339 | 22.892 | 20.715 | 25.217 | 22.039 | 19.704 | 24.022 | 21.478 | 20.217 | 24.915 | -0.047 | 0.0897 | 0.2750 | 0 | 0 | 0 |
| 'Ppp1r13b' | 12.168 | 9.797 | 10.956 | 10.955 | 10.219 | 11.492 | 9.331 | 10.489 | 10.960 | 12.030 | 9.652 | 12.070 | 9.849 | 10.620 | 11.710 | 10.976 | 10.490 | 12.680 | -0.062 | 0.2182 | 0.4650 | 0 | 0 | 0 |
| 'Ppp1r14b' | 223.749 | 285.220 | 219.719 | 238.486 | 287.428 | 240.189 | 245.986 | 283.417 | 206.473 | 242.489 | 258.639 | 191.683 | 326.358 | 297.144 | 221.399 | 281.264 | 285.542 | 226.081 | -0.070 | 0.2226 | 0.4701 | 0 | 0 | 0 |
| 'Ppp1r15a' | 178.225 | 62.068 | 64.502 | 155.299 | 66.043 | 57.546 | 155.708 | 79.432 | 46.894 | 119.497 | 99.953 | 43.908 | 115.293 | 93.237 | 41.824 | 100.672 | 97.605 | 40.795 | 0.151 | 0.2913 | 0.5480 | 0 | 0 | 0 |
| 'Ppp1r15b' | 19.300 | 21.202 | 19.308 | 18.581 | 19.040 | 19.097 | 18.200 | 20.534 | 19.451 | 19.419 | 19.242 | 20.697 | 18.616 | 21.174 | 20.759 | 21.067 | 19.654 | 20.747 | -0.065 | 0.0368 | 0.1551 | 0 | 0 | 0 |
| 'Ppp1r16b' | 46.530 | 44.642 | 36.603 | 37.896 | 42.603 | 39.617 | 35.808 | 44.406 | 38.609 | 45.508 | 45.423 | 42.153 | 43.836 | 47.062 | 42.986 | 46.694 | 41.711 | 45.527 | -0.141 | 0.0019 | 0.0176 | 1 | 0 | 0 |
| 'Ppp1r18' | 29.477 | 28.149 | 28.829 | 28.034 | 26.887 | 31.751 | 27.260 | 27.352 | 30.449 | 26.197 | 26.152 | 32.119 | 26.441 | 28.004 | 32.993 | 28.204 | 26.276 | 32.787 | -0.012 | 0.7169 | 0.8692 | 0 | 0 | 0 |
| 'Ppp1r2' | 21.265 | 23.451 | 19.230 | 21.263 | 21.058 | 22.100 | 20.663 | 21.640 | 21.146 | 19.197 | 18.196 | 21.090 | 23.082 | 21.401 | 22.305 | 22.192 | 22.284 | 18.555 | 0.013 | 0.7961 | 0.9092 | 0 | 0 | 0 |
| 'Ppp1r21' | 7.280 | 7.739 | 8.133 | 7.237 | 7.664 | 8.048 | 7.620 | 7.880 | 7.777 | 7.957 | 7.135 | 7.677 | 8.440 | 7.874 | 7.158 | 8.003 | 8.180 | 7.315 | -0.015 | 0.6956 | 0.8570 | 0 | 0 | 0 |
| 'Ppp1r35' | 23.535 | 26.351 | 27.143 | 22.482 | 23.433 | 26.223 | 24.781 | 22.788 | 25.833 | 19.915 | 20.676 | 27.147 | 21.674 | 22.130 | 27.553 | 21.698 | 21.065 | 25.880 | 0.088 | 0.0361 | 0.1530 | 0 | 0 | 0 |
| 'Ppp1r37' | 22.309 | 21.654 | 27.361 | 21.031 | 21.829 | 26.407 | 22.414 | 21.540 | 25.644 | 23.348 | 19.875 | 26.045 | 22.941 | 20.282 | 22.873 | 22.331 | 20.081 | 25.901 | 0.035 | 0.3085 | 0.5669 | 0 | 0 | 0 |
| 'Ppp1r3f' | 1.502 | 1.433 | 1.646 | 1.226 | 1.202 | 1.523 | 1.655 | 1.385 | 1.228 | 1.266 | 1.590 | 1.271 | 1.167 | 1.437 | 1.129 | 0.922 | 1.108 | 1.123 | 0.212 | 0.0216 | 0.1085 | 0 | 0 | 0 |
| 'Ppp1r7' | 5.350 | 6.280 | 6.541 | 6.585 | 6.612 | 6.704 | 6.142 | 6.574 | 6.265 | 6.804 | 6.303 | 6.015 | 6.396 | 6.644 | 6.292 | 6.359 | 6.269 | 5.360 | 0.007 | 0.8678 | 0.9428 | 0 | 0 | 0 |
| 'Ppp1r8' | 15.094 | 16.471 | 14.473 | 14.726 | 14.916 | 15.471 | 14.544 | 14.709 | 14.887 | 13.762 | 15.172 | 16.003 | 14.217 | 16.727 | 16.868 | 14.180 | 15.693 | 15.797 | -0.044 | 0.2403 | 0.4909 | 0 | 0 | 0 |
| 'Ppp1r9b' | 71.839 | 65.418 | 71.029 | 65.482 | 62.213 | 75.287 | 57.898 | 66.001 | 71.261 | 70.323 | 63.433 | 76.083 | 67.005 | 68.710 | 73.985 | 69.729 | 60.150 | 78.907 | -0.060 | 0.1157 | 0.3230 | 0 | 0 | 0 |
| 'Ppp2ca' | 40.132 | 41.453 | 37.260 | 39.196 | 42.419 | 39.995 | 39.842 | 42.271 | 35.192 | 37.778 | 39.251 | 36.783 | 38.970 | 42.928 | 39.665 | 40.771 | 40.495 | 39.251 | -0.004 | 0.8918 | 0.9560 | 0 | 0 | 0 |
| 'Ppp2cb' | 32.893 | 36.845 | 29.813 | 34.403 | 35.753 | 33.675 | 34.118 | 37.015 | 32.086 | 35.097 | 34.797 | 33.221 | 36.312 | 37.620 | 34.516 | 38.804 | 35.076 | 34.762 | -0.075 | 0.0320 | 0.1407 | 0 | 0 | 0 |
| 'Ppp2r1a' | 49.730 | 48.983 | 49.404 | 48.234 | 51.068 | 52.057 | 49.295 | 50.019 | 49.900 | 51.177 | 48.037 | 49.106 | 49.652 | 47.582 | 50.029 | 49.363 | 47.451 | 47.606 | 0.019 | 0.3491 | 0.6042 | 0 | 0 | 0 |
| 'Ppp2r1b' | 10.432 | 11.206 | 10.948 | 10.664 | 10.861 | 10.612 | 11.726 | 10.647 | 11.021 | 10.281 | 10.358 | 11.766 | 10.747 | 11.265 | 11.406 | 10.348 | 11.526 | 11.767 | -0.032 | 0.3165 | 0.5745 | 0 | 0 | 0 |
| 'Ppp2r2a' | 30.874 | 27.479 | 27.847 | 31.734 | 26.032 | 24.838 | 31.866 | 26.624 | 26.990 | 26.138 | 29.505 | 28.921 | 25.408 | 28.121 | 25.224 | 24.936 | 30.933 | 27.091 | 0.033 | 0.5516 | 0.7649 | 0 | 0 | 0 |
| 'Ppp2r2d' | 8.173 | 7.829 | 7.943 | 7.687 | 7.944 | 7.812 | 8.129 | 7.577 | 6.620 | 8.093 | 7.017 | 6.891 | 8.117 | 7.675 | 7.178 | 7.991 | 8.280 | 7.042 | 0.024 | 0.5530 | 0.7658 | 0 | 0 | 0 |
| 'Ppp2r3a' | 0.933 | 0.863 | 1.059 | 0.841 | 0.689 | 0.763 | 1.023 | 0.868 | 0.869 | 0.625 | 0.591 | 0.740 | 0.465 | 0.532 | 0.795 | 0.485 | 0.464 | 0.715 | 0.546 | 0.0000 | 0.0000 | 1 | 1 | 0 |
| 'Ppp2r5a' | 37.999 | 34.892 | 38.904 | 37.102 | 36.384 | 38.371 | 36.642 | 37.508 | 34.276 | 38.233 | 38.847 | 37.147 | 35.620 | 39.306 | 37.361 | 35.579 | 38.700 | 41.157 | -0.052 | 0.1382 | 0.3586 | 0 | 0 | 0 |
| 'Ppp2r5b' | 7.462 | 7.436 | 8.501 | 6.607 | 7.457 | 8.837 | 6.113 | 7.567 | 8.065 | 7.655 | 6.586 | 8.071 | 7.947 | 7.720 | 8.350 | 7.601 | 7.090 | 8.044 | -0.029 | 0.5622 | 0.7715 | 0 | 0 | 0 |
| 'Ppp2r5c' | 62.028 | 69.046 | 61.113 | 65.092 | 66.239 | 61.818 | 64.007 | 65.401 | 60.776 | 62.087 | 67.056 | 63.336 | 61.029 | 66.488 | 61.828 | 62.246 | 65.647 | 60.913 | 0.001 | 0.9775 | 0.9920 | 0 | 0 | 0 |
| 'Ppp2r5d' | 37.179 | 35.769 | 40.266 | 36.898 | 34.668 | 42.484 | 36.665 | 36.281 | 41.709 | 39.036 | 35.329 | 44.338 | 42.560 | 37.698 | 41.777 | 38.802 | 38.089 | 40.976 | -0.080 | 0.0024 | 0.0207 | 1 | 0 | 0 |
| 'Ppp2r5e' | 22.180 | 21.784 | 22.012 | 22.731 | 22.172 | 20.822 | 21.692 | 21.900 | 21.019 | 23.392 | 22.614 | 21.609 | 23.483 | 22.483 | 20.872 | 23.407 | 23.281 | 22.209 | -0.059 | 0.0191 | 0.1000 | 0 | 0 | 0 |
| 'Ppp3ca' | 0.468 | 0.609 | 0.558 | 0.446 | 0.679 | 0.617 | 0.672 | 0.527 | 0.527 | 0.554 | 0.666 | 0.646 | 0.559 | 0.656 | 0.584 | 0.595 | 0.644 | 0.657 | -0.143 | 0.0822 | 0.2593 | 0 | 0 | 0 |
| 'Ppp3cb' | 14.337 | 15.180 | 15.224 | 14.727 | 14.673 | 14.401 | 15.589 | 15.354 | 14.141 | 15.045 | 14.958 | 14.141 | 14.200 | 15.138 | 13.251 | 14.442 | 15.359 | 12.878 | 0.038 | 0.1651 | 0.3974 | 0 | 0 | 0 |
| 'Ppp3cc' | 51.856 | 57.995 | 53.874 | 53.799 | 58.040 | 56.450 | 56.254 | 60.086 | 53.847 | 56.974 | 51.289 | 47.274 | 53.002 | 55.454 | 48.686 | 57.379 | 54.063 | 43.446 | 0.098 | 0.0091 | 0.0583 | 0 | 0 | 0 |
| 'Ppp3r1' | 34.246 | 37.239 | 33.230 | 33.924 | 36.037 | 31.082 | 36.478 | 35.741 | 32.171 | 32.778 | 33.831 | 32.483 | 31.570 | 33.209 | 29.186 | 31.240 | 33.350 | 30.695 | 0.091 | 0.0008 | 0.0091 | 1 | 0 | 0 |
| 'Ppp4c' | 20.728 | 22.014 | 22.221 | 20.651 | 22.188 | 23.081 | 19.808 | 20.689 | 22.651 | 20.947 | 20.515 | 21.302 | 21.646 | 21.344 | 22.383 | 21.941 | 20.519 | 21.500 | 0.004 | 0.8744 | 0.9463 | 0 | 0 | 0 |
| 'Ppp4r1' | 6.508 | 6.315 | 6.019 | 6.548 | 6.184 | 6.121 | 6.447 | 6.162 | 6.147 | 5.974 | 6.494 | 7.068 | 5.681 | 6.163 | 6.377 | 5.683 | 6.490 | 6.960 | -0.023 | 0.5993 | 0.7963 | 0 | 0 | 0 |
| 'Ppp5c' | 21.126 | 20.458 | 23.218 | 20.987 | 19.836 | 22.623 | 19.945 | 21.884 | 21.885 | 21.551 | 18.697 | 22.508 | 23.658 | 20.153 | 22.118 | 22.105 | 19.853 | 20.686 | -0.004 | 0.9093 | 0.9646 | 0 | 0 | 0 |
| 'Ppp6c' | 10.919 | 12.604 | 12.903 | 13.375 | 13.026 | 11.485 | 13.029 | 13.325 | 11.905 | 11.345 | 13.091 | 13.224 | 11.903 | 14.181 | 12.520 | 11.973 | 14.027 | 11.909 | -0.031 | 0.4542 | 0.6929 | 0 | 0 | 0 |
| 'Ppp6r1' | 77.011 | 69.446 | 80.595 | 74.685 | 70.215 | 82.519 | 73.131 | 72.949 | 76.225 | 78.859 | 72.778 | 78.090 | 75.613 | 69.560 | 80.490 | 73.570 | 69.477 | 81.415 | -0.017 | 0.4870 | 0.7175 | 0 | 0 | 0 |
| 'Ppp6r2' | 25.050 | 26.926 | 28.986 | 26.122 | 24.814 | 27.590 | 26.318 | 26.970 | 28.286 | 27.693 | 28.234 | 26.255 | 26.311 | 25.670 | 25.813 | 25.168 | 25.404 | 24.549 | 0.027 | 0.4339 | 0.6787 | 0 | 0 | 0 |
| 'Ppp6r3' | 27.222 | 31.753 | 27.575 | 27.822 | 31.009 | 27.407 | 27.062 | 30.364 | 27.714 | 25.748 | 28.479 | 27.543 | 29.006 | 29.112 | 27.380 | 27.775 | 29.095 | 28.982 | 0.013 | 0.6161 | 0.8085 | 0 | 0 | 0 |
| 'Pprc1' | 4.157 | 3.388 | 4.379 | 3.824 | 3.176 | 3.785 | 4.422 | 3.572 | 4.532 | 3.682 | 3.791 | 5.144 | 3.374 | 3.110 | 4.475 | 3.374 | 3.800 | 5.064 | -0.027 | 0.6902 | 0.8540 | 0 | 0 | 0 |
| 'Ppt1' | 22.482 | 29.425 | 22.293 | 23.927 | 26.044 | 23.250 | 24.328 | 27.163 | 23.832 | 25.805 | 25.498 | 22.186 | 26.012 | 26.415 | 21.186 | 24.213 | 25.325 | 21.323 | 0.020 | 0.5442 | 0.7606 | 0 | 0 | 0 |
| 'Ppt2' | 5.205 | 6.518 | 5.025 | 4.979 | 6.193 | 5.930 | 4.791 | 5.717 | 6.089 | 5.491 | 4.639 | 4.845 | 5.178 | 5.752 | 4.940 | 5.197 | 5.229 | 4.492 | 0.129 | 0.0242 | 0.1177 | 0 | 0 | 0 |
| 'Pptc7' | 80.424 | 91.236 | 74.896 | 82.352 | 85.811 | 82.422 | 75.235 | 85.293 | 79.775 | 84.309 | 94.595 | 87.643 | 99.433 | 101.605 | 93.153 | 103.214 | 88.150 | 92.838 | -0.208 | 0.0000 | 0.0000 | 1 | 0 | 0 |
| 'Pqbp1' | 33.844 | 33.407 | 31.695 | 35.433 | 33.806 | 31.875 | 33.327 | 31.936 | 32.308 | 32.143 | 30.992 | 31.667 | 33.338 | 33.100 | 31.461 | 34.159 | 30.470 | 31.579 | 0.031 | 0.2514 | 0.5026 | 0 | 0 | 0 |
| 'Pqlc1' | 0.376 | 0.290 | 0.223 | 0.293 | 0.312 | 0.238 | 0.322 | 0.399 | 0.237 | 0.321 | 0.252 | 0.235 | 0.307 | 0.246 | 0.229 | 0.398 | 0.327 | 0.253 | 0.060 | 0.5850 | 0.7865 | 0 | 0 | 0 |
| 'Pqlc2' | 1.791 | 1.583 | 1.798 | 1.511 | 1.770 | 1.769 | 1.216 | 1.646 | 1.625 | 1.754 | 1.915 | 1.784 | 1.553 | 1.483 | 1.684 | 1.850 | 1.695 | 1.874 | -0.086 | 0.2604 | 0.5140 | 0 | 0 | 0 |
| 'Pqlc3' | 4.527 | 5.163 | 5.872 | 4.936 | 5.658 | 5.593 | 4.603 | 4.975 | 5.098 | 5.165 | 5.181 | 4.917 | 4.886 | 5.156 | 4.906 | 5.041 | 4.626 | 5.103 | 0.036 | 0.4241 | 0.6709 | 0 | 0 | 0 |
| 'Pradc1' | 3.206 | 3.244 | 3.077 | 2.929 | 2.745 | 3.070 | 2.525 | 3.267 | 3.181 | 2.919 | 2.890 | 3.465 | 3.295 | 3.331 | 3.303 | 2.846 | 3.067 | 3.304 | -0.074 | 0.2217 | 0.4691 | 0 | 0 | 0 |
| 'Praf2' | 21.307 | 22.910 | 22.649 | 22.645 | 22.609 | 21.983 | 21.290 | 22.709 | 23.407 | 22.382 | 20.580 | 21.782 | 24.086 | 22.062 | 20.249 | 23.000 | 22.538 | 19.664 | 0.029 | 0.4971 | 0.7258 | 0 | 0 | 0 |
| 'Pramef8' | 5.896 | 6.103 | 6.438 | 5.384 | 6.710 | 6.700 | 6.231 | 6.601 | 6.212 | 7.297 | 5.935 | 5.771 | 6.550 | 6.397 | 5.671 | 6.456 | 6.003 | 5.680 | 0.009 | 0.8590 | 0.9390 | 0 | 0 | 0 |
| 'Prc1' | 1.905 | 2.260 | 2.613 | 2.335 | 2.302 | 2.731 | 2.500 | 2.273 | 2.970 | 1.711 | 2.455 | 2.484 | 2.652 | 2.954 | 3.083 | 2.394 | 2.498 | 2.664 | -0.077 | 0.3040 | 0.5614 | 0 | 0 | 0 |
| 'Prcc' | 46.994 | 48.444 | 44.892 | 50.083 | 45.805 | 47.672 | 45.708 | 44.851 | 45.143 | 44.608 | 44.441 | 48.291 | 45.606 | 44.429 | 48.171 | 45.951 | 40.973 | 47.527 | 0.021 | 0.5269 | 0.7483 | 0 | 0 | 0 |
| 'Prcp' | 5.515 | 5.935 | 4.805 | 5.435 | 5.737 | 5.893 | 6.404 | 5.943 | 4.616 | 5.313 | 5.445 | 4.644 | 5.839 | 5.374 | 5.028 | 4.609 | 4.771 | 4.235 | 0.139 | 0.0295 | 0.1334 | 0 | 0 | 0 |
| 'Prdm1' | 1.402 | 0.969 | 1.411 | 1.162 | 1.333 | 1.256 | 0.947 | 1.715 | 0.972 | 1.027 | 1.150 | 1.054 | 0.950 | 0.932 | 0.987 | 1.095 | 0.861 | 0.982 | 0.300 | 0.0039 | 0.0312 | 1 | 0 | 0 |
| 'Prdm10' | 3.300 | 3.250 | 3.846 | 3.794 | 3.494 | 3.577 | 3.527 | 3.230 | 3.685 | 3.156 | 3.307 | 4.066 | 3.507 | 3.610 | 3.552 | 3.455 | 3.604 | 3.948 | -0.037 | 0.3910 | 0.6403 | 0 | 0 | 0 |
| 'Prdm11' | 0.799 | 0.705 | 0.824 | 0.833 | 0.737 | 0.814 | 0.845 | 0.711 | 0.829 | 1.010 | 0.809 | 0.829 | 0.704 | 0.841 | 0.845 | 0.845 | 0.679 | 0.958 | -0.090 | 0.1919 | 0.4290 | 0 | 0 | 0 |
| 'Prdm15' | 2.900 | 2.156 | 2.498 | 2.678 | 1.990 | 2.119 | 2.072 | 2.190 | 2.316 | 2.296 | 2.316 | 3.059 | 2.440 | 2.288 | 2.640 | 2.260 | 2.773 | 3.385 | -0.172 | 0.0308 | 0.1370 | 0 | 0 | 0 |
| 'Prdm2' | 35.635 | 32.470 | 31.138 | 33.474 | 29.970 | 32.496 | 34.902 | 30.923 | 32.743 | 30.318 | 33.574 | 36.658 | 32.239 | 33.839 | 34.774 | 32.554 | 34.897 | 36.778 | -0.068 | 0.1035 | 0.3017 | 0 | 0 | 0 |
| 'Prdm4' | 11.690 | 12.689 | 11.363 | 10.573 | 11.336 | 11.194 | 10.480 | 12.483 | 11.282 | 12.228 | 11.757 | 12.135 | 12.239 | 12.081 | 12.354 | 12.757 | 11.310 | 11.069 | -0.076 | 0.0434 | 0.1725 | 0 | 0 | 0 |
| 'Prdm5' | 0.480 | 0.480 | 0.566 | 0.476 | 0.377 | 0.442 | 0.397 | 0.386 | 0.482 | 0.465 | 0.258 | 0.472 | 0.433 | 0.330 | 0.431 | 0.462 | 0.511 | 0.405 | 0.107 | 0.3154 | 0.5732 | 0 | 0 | 0 |
| 'Prdm9' | 2.976 | 2.769 | 2.964 | 2.935 | 3.482 | 2.617 | 3.143 | 3.375 | 2.915 | 2.829 | 3.461 | 2.577 | 3.193 | 2.786 | 2.863 | 2.216 | 2.631 | 2.775 | 0.090 | 0.1977 | 0.4373 | 0 | 0 | 0 |
| 'Prdx1' | 38.356 | 39.986 | 41.177 | 43.599 | 43.483 | 39.976 | 46.391 | 41.376 | 37.932 | 36.187 | 40.801 | 39.658 | 41.079 | 40.260 | 40.612 | 40.209 | 44.515 | 38.610 | 0.029 | 0.4550 | 0.6934 | 0 | 0 | 0 |
| 'Prdx2' | 17.761 | 18.345 | 17.121 | 18.132 | 18.487 | 18.152 | 17.160 | 18.076 | 18.306 | 15.914 | 16.211 | 15.638 | 16.319 | 15.118 | 16.640 | 15.300 | 16.236 | 15.235 | 0.169 | 0.0000 | 0.0000 | 1 | 0 | 0 |
| 'Prdx3' | 39.902 | 38.798 | 43.127 | 37.704 | 43.623 | 39.321 | 38.001 | 40.417 | 38.966 | 35.930 | 42.650 | 40.037 | 41.320 | 42.071 | 41.492 | 35.475 | 41.618 | 38.592 | -0.006 | 0.8709 | 0.9447 | 0 | 0 | 0 |
| 'Prdx4' | 5.019 | 5.961 | 6.183 | 4.994 | 6.675 | 6.249 | 5.636 | 6.901 | 5.896 | 5.446 | 5.438 | 6.254 | 6.541 | 6.510 | 6.069 | 6.102 | 6.457 | 5.880 | -0.045 | 0.4564 | 0.6942 | 0 | 0 | 0 |
| 'Prdx5' | 27.266 | 32.658 | 27.567 | 26.893 | 31.603 | 29.968 | 28.536 | 30.206 | 28.252 | 28.225 | 27.530 | 26.051 | 27.377 | 29.262 | 27.721 | 30.031 | 27.519 | 25.265 | 0.068 | 0.0488 | 0.1854 | 0 | 0 | 0 |
| 'Prdx6' | 8.670 | 8.222 | 8.106 | 8.113 | 8.717 | 8.819 | 8.243 | 8.790 | 8.457 | 9.074 | 8.291 | 8.856 | 8.500 | 8.749 | 8.105 | 8.390 | 7.619 | 7.746 | 0.005 | 0.8801 | 0.9494 | 0 | 0 | 0 |
| 'Preb' | 17.647 | 17.544 | 18.466 | 18.037 | 18.668 | 18.521 | 17.413 | 17.265 | 17.782 | 17.899 | 16.282 | 18.802 | 18.943 | 17.421 | 17.273 | 18.459 | 18.704 | 17.081 | -0.007 | 0.8153 | 0.9170 | 0 | 0 | 0 |
| 'Prelid1' | 145.539 | 165.008 | 140.409 | 138.509 | 161.314 | 157.639 | 137.871 | 157.389 | 147.013 | 136.394 | 144.082 | 147.235 | 156.853 | 157.448 | 170.718 | 164.572 | 149.038 | 162.120 | -0.052 | 0.2336 | 0.4820 | 0 | 0 | 0 |
| 'Prelid2' | 32.758 | 25.032 | 30.732 | 29.580 | 33.392 | 26.336 | 29.319 | 29.117 | 26.981 | 27.999 | 31.862 | 30.702 | 31.356 | 30.653 | 32.293 | 34.601 | 29.459 | 28.503 | -0.083 | 0.1508 | 0.3777 | 0 | 0 | 0 |
| 'Prep' | 19.676 | 17.882 | 20.107 | 21.089 | 19.768 | 18.575 | 21.846 | 19.993 | 18.689 | 17.151 | 17.569 | 19.933 | 18.257 | 16.022 | 19.639 | 16.984 | 18.440 | 18.320 | 0.116 | 0.0098 | 0.0615 | 0 | 0 | 0 |
| 'Prepl' | 6.440 | 7.204 | 8.068 | 7.168 | 7.314 | 7.475 | 7.541 | 7.392 | 6.960 | 7.411 | 7.688 | 8.316 | 7.289 | 7.473 | 7.600 | 7.500 | 8.163 | 7.982 | -0.095 | 0.0256 | 0.1227 | 0 | 0 | 0 |
| 'Prex1' | 51.117 | 51.737 | 51.786 | 51.561 | 50.615 | 51.052 | 47.505 | 51.163 | 51.367 | 51.828 | 51.417 | 50.787 | 52.336 | 51.979 | 49.956 | 50.472 | 50.865 | 51.449 | -0.020 | 0.3166 | 0.5745 | 0 | 0 | 0 |
| 'Prickle2' | 0.567 | 0.743 | 0.516 | 0.748 | 0.653 | 0.524 | 0.755 | 0.668 | 0.696 | 1.055 | 1.458 | 0.984 | 0.973 | 1.374 | 0.911 | 0.745 | 1.187 | 0.932 | -0.716 | 0.0000 | 0.0000 | 1 | 1 | 0 |
| 'Prickle3' | 5.257 | 5.355 | 5.605 | 5.155 | 4.828 | 5.078 | 5.139 | 5.202 | 5.439 | 5.295 | 5.439 | 5.455 | 5.161 | 4.657 | 5.262 | 4.937 | 5.878 | 5.203 | -0.017 | 0.6856 | 0.8513 | 0 | 0 | 0 |
| 'Prim1' | 6.806 | 7.273 | 8.594 | 8.110 | 7.637 | 7.648 | 8.238 | 6.699 | 8.236 | 6.253 | 6.910 | 7.141 | 8.829 | 7.384 | 7.945 | 6.678 | 7.608 | 7.210 | 0.057 | 0.3382 | 0.5957 | 0 | 0 | 0 |
| 'Prim2' | 4.492 | 4.451 | 5.345 | 5.690 | 4.689 | 5.316 | 5.194 | 5.016 | 5.416 | 4.177 | 4.931 | 5.031 | 5.540 | 5.247 | 5.258 | 4.706 | 4.481 | 4.766 | 0.036 | 0.5150 | 0.7389 | 0 | 0 | 0 |
| 'Primpol' | 4.600 | 5.363 | 5.190 | 4.704 | 4.956 | 4.677 | 4.922 | 5.135 | 4.788 | 4.771 | 5.407 | 4.593 | 4.667 | 4.857 | 4.182 | 4.493 | 5.118 | 3.966 | 0.070 | 0.0901 | 0.2760 | 0 | 0 | 0 |
| 'Prkaa1' | 12.113 | 11.095 | 11.949 | 11.799 | 11.764 | 10.523 | 14.625 | 11.865 | 11.380 | 12.838 | 13.672 | 12.873 | 11.787 | 12.966 | 11.704 | 10.982 | 13.316 | 11.924 | -0.078 | 0.1445 | 0.3689 | 0 | 0 | 0 |
| 'Prkab1' | 12.752 | 13.134 | 13.214 | 11.936 | 12.300 | 12.622 | 12.158 | 13.074 | 13.148 | 13.165 | 11.494 | 12.098 | 12.428 | 11.458 | 12.204 | 12.705 | 11.440 | 11.754 | 0.065 | 0.0388 | 0.1606 | 0 | 0 | 0 |
| 'Prkab2' | 0.519 | 0.391 | 0.463 | 0.518 | 0.505 | 0.464 | 0.471 | 0.424 | 0.458 | 0.487 | 0.509 | 0.521 | 0.461 | 0.494 | 0.540 | 0.505 | 0.638 | 0.434 | -0.131 | 0.1215 | 0.3323 | 0 | 0 | 0 |
| 'Prkaca' | 44.713 | 46.477 | 48.011 | 43.888 | 49.864 | 52.092 | 47.237 | 47.508 | 46.950 | 49.340 | 47.810 | 48.024 | 50.900 | 52.298 | 50.266 | 48.128 | 45.187 | 50.028 | -0.061 | 0.0707 | 0.2345 | 0 | 0 | 0 |
| 'Prkacb' | 29.136 | 33.265 | 30.684 | 29.764 | 31.909 | 28.723 | 29.009 | 31.363 | 29.271 | 28.268 | 28.432 | 26.616 | 25.952 | 28.159 | 27.223 | 25.694 | 25.427 | 26.794 | 0.159 | 0.0000 | 0.0000 | 1 | 0 | 0 |
| 'Prkag1' | 48.996 | 55.129 | 53.286 | 52.119 | 55.798 | 57.040 | 52.397 | 56.647 | 56.407 | 50.160 | 51.863 | 49.367 | 55.304 | 52.670 | 52.113 | 54.815 | 53.060 | 48.319 | 0.052 | 0.1211 | 0.3319 | 0 | 0 | 0 |
| 'Prkag2' | 2.794 | 3.064 | 2.798 | 2.726 | 3.076 | 3.167 | 2.976 | 3.290 | 3.020 | 3.202 | 2.807 | 3.256 | 3.353 | 3.151 | 3.261 | 3.260 | 3.173 | 3.177 | -0.103 | 0.0209 | 0.1061 | 0 | 0 | 0 |
| 'Prkar1a' | 116.073 | 136.390 | 120.595 | 120.585 | 137.987 | 127.368 | 121.985 | 134.359 | 125.461 | 124.578 | 127.404 | 118.471 | 121.619 | 137.037 | 122.872 | 127.026 | 129.046 | 122.104 | 0.002 | 0.9386 | 0.9764 | 0 | 0 | 0 |
| 'Prkar2a' | 6.674 | 6.895 | 7.061 | 6.796 | 7.065 | 7.112 | 6.013 | 7.206 | 6.703 | 6.421 | 6.569 | 6.794 | 7.066 | 7.256 | 6.814 | 6.302 | 6.837 | 6.990 | 0.002 | 0.9657 | 0.9878 | 0 | 0 | 0 |
| 'Prkca' | 35.515 | 34.298 | 33.850 | 35.108 | 33.843 | 32.466 | 34.654 | 34.809 | 33.710 | 36.879 | 37.051 | 35.809 | 33.355 | 34.823 | 33.625 | 34.375 | 35.530 | 34.106 | -0.045 | 0.0808 | 0.2562 | 0 | 0 | 0 |
| 'Prkcb' | 20.111 | 19.354 | 21.275 | 21.809 | 19.563 | 21.797 | 20.629 | 19.856 | 20.709 | 23.739 | 23.263 | 21.628 | 22.642 | 20.533 | 21.514 | 20.877 | 22.084 | 22.816 | -0.116 | 0.0006 | 0.0065 | 1 | 0 | 0 |
| 'Prkcd' | 8.247 | 8.042 | 8.609 | 8.187 | 8.377 | 8.700 | 7.503 | 8.062 | 9.145 | 8.857 | 8.418 | 9.568 | 8.831 | 8.195 | 8.789 | 7.760 | 8.715 | 9.323 | -0.080 | 0.0139 | 0.0794 | 0 | 0 | 0 |
| 'Prkce' | 2.625 | 2.457 | 2.364 | 2.221 | 2.111 | 2.192 | 2.145 | 2.548 | 2.176 | 2.659 | 2.473 | 2.445 | 2.558 | 2.239 | 2.268 | 2.661 | 2.189 | 2.534 | -0.086 | 0.1101 | 0.3132 | 0 | 0 | 0 |
| 'Prkch' | 90.794 | 90.701 | 86.808 | 86.767 | 90.620 | 90.479 | 82.376 | 92.925 | 86.536 | 91.905 | 91.020 | 90.575 | 85.497 | 91.724 | 91.654 | 90.174 | 90.546 | 93.176 | -0.043 | 0.0764 | 0.2464 | 0 | 0 | 0 |
| 'Prkci' | 4.204 | 4.924 | 4.069 | 4.760 | 4.674 | 4.011 | 4.099 | 4.503 | 4.433 | 4.420 | 4.589 | 4.150 | 4.096 | 4.295 | 3.834 | 3.683 | 4.252 | 4.196 | 0.064 | 0.1777 | 0.4131 | 0 | 0 | 0 |
| 'Prkcq' | 53.960 | 53.442 | 52.667 | 57.926 | 53.128 | 55.600 | 53.878 | 55.663 | 53.576 | 54.093 | 56.309 | 52.979 | 52.699 | 55.318 | 53.278 | 51.935 | 54.331 | 54.302 | 0.003 | 0.8994 | 0.9600 | 0 | 0 | 0 |
| 'Prkcsh' | 89.260 | 78.747 | 92.469 | 87.487 | 79.311 | 96.674 | 90.276 | 77.576 | 94.200 | 88.793 | 79.024 | 96.710 | 88.161 | 80.524 | 94.244 | 87.972 | 83.858 | 92.001 | -0.021 | 0.3144 | 0.5721 | 0 | 0 | 0 |
| 'Prkcz' | 2.471 | 2.666 | 2.506 | 2.268 | 2.924 | 3.091 | 2.303 | 2.763 | 2.766 | 2.966 | 2.960 | 3.014 | 3.083 | 3.172 | 2.925 | 3.106 | 2.917 | 3.066 | -0.205 | 0.0001 | 0.0010 | 1 | 0 | 0 |
| 'Prkd2' | 30.380 | 27.412 | 29.688 | 28.990 | 26.488 | 30.240 | 28.548 | 26.477 | 30.489 | 27.562 | 28.819 | 35.794 | 27.650 | 27.437 | 33.731 | 27.916 | 28.821 | 36.019 | -0.087 | 0.0265 | 0.1251 | 0 | 0 | 0 |
| 'Prkd3' | 0.983 | 1.043 | 1.254 | 0.997 | 1.105 | 1.244 | 1.057 | 1.181 | 1.092 | 0.662 | 0.745 | 0.759 | 0.655 | 0.705 | 0.824 | 0.597 | 0.763 | 0.911 | 0.583 | 0.0000 | 0.0000 | 1 | 1 | 0 |
| 'Prkdc' | 6.725 | 4.869 | 6.684 | 7.486 | 6.315 | 5.896 | 6.717 | 5.841 | 6.337 | 6.507 | 5.994 | 6.919 | 6.016 | 6.310 | 5.938 | 6.135 | 6.526 | 6.548 | -0.016 | 0.7744 | 0.8994 | 0 | 0 | 0 |
| 'Prkra' | 3.324 | 3.077 | 3.931 | 4.029 | 4.267 | 3.972 | 3.766 | 3.904 | 3.466 | 3.803 | 3.631 | 4.111 | 3.763 | 4.054 | 4.184 | 3.731 | 4.326 | 3.581 | -0.071 | 0.3354 | 0.5941 | 0 | 0 | 0 |
| 'Prkrip1' | 4.123 | 4.252 | 4.929 | 5.061 | 4.780 | 4.811 | 4.719 | 5.069 | 5.466 | 4.265 | 4.741 | 5.195 | 5.056 | 4.482 | 5.245 | 4.732 | 5.158 | 4.709 | -0.024 | 0.6419 | 0.8243 | 0 | 0 | 0 |
| 'Prkrir' | 16.260 | 17.945 | 16.731 | 16.435 | 16.409 | 16.475 | 15.298 | 17.975 | 16.136 | 17.201 | 16.863 | 16.131 | 16.962 | 18.235 | 16.322 | 17.266 | 15.760 | 15.634 | -0.015 | 0.6444 | 0.8259 | 0 | 0 | 0 |
| 'Prkx' | 7.455 | 7.866 | 7.682 | 6.998 | 7.509 | 7.749 | 7.659 | 8.405 | 7.611 | 7.652 | 8.327 | 8.370 | 7.410 | 8.234 | 7.728 | 7.248 | 7.525 | 8.301 | -0.050 | 0.1645 | 0.3968 | 0 | 0 | 0 |
| 'Prmt2' | 3.607 | 3.098 | 3.442 | 3.594 | 3.322 | 3.865 | 3.060 | 3.777 | 3.571 | 3.639 | 4.032 | 4.546 | 3.499 | 3.872 | 4.170 | 3.400 | 3.642 | 4.513 | -0.180 | 0.0011 | 0.0112 | 1 | 0 | 0 |
| 'Prmt3' | 12.679 | 13.893 | 12.923 | 12.994 | 15.473 | 13.702 | 13.771 | 15.046 | 12.461 | 13.337 | 14.932 | 14.192 | 15.115 | 14.746 | 14.246 | 14.329 | 16.189 | 14.123 | -0.107 | 0.0011 | 0.0114 | 1 | 0 | 0 |
| 'Prmt5' | 9.026 | 8.085 | 9.267 | 8.654 | 7.821 | 8.957 | 9.280 | 8.557 | 8.769 | 8.660 | 7.608 | 9.795 | 8.870 | 7.798 | 9.161 | 8.059 | 8.259 | 9.354 | 0.002 | 0.9449 | 0.9789 | 0 | 0 | 0 |
| 'Prmt6' | 4.864 | 5.233 | 5.667 | 4.540 | 5.339 | 5.599 | 4.933 | 5.051 | 5.268 | 5.145 | 4.810 | 5.073 | 5.221 | 4.936 | 4.624 | 4.752 | 4.731 | 4.901 | 0.063 | 0.1184 | 0.3276 | 0 | 0 | 0 |
| 'Prmt7' | 3.463 | 3.151 | 3.565 | 3.419 | 3.193 | 3.388 | 3.455 | 3.603 | 3.450 | 3.583 | 3.216 | 3.712 | 3.216 | 2.943 | 3.437 | 3.313 | 3.405 | 3.467 | 0.007 | 0.8662 | 0.9421 | 0 | 0 | 0 |
| 'Prmt9' | 6.097 | 5.482 | 5.465 | 6.026 | 5.838 | 5.350 | 5.570 | 5.510 | 5.737 | 5.265 | 6.236 | 6.730 | 5.793 | 5.492 | 6.208 | 5.324 | 6.304 | 6.006 | -0.076 | 0.1213 | 0.3321 | 0 | 0 | 0 |
| 'Prnp' | 14.370 | 19.018 | 16.943 | 14.260 | 18.793 | 16.359 | 15.931 | 19.257 | 16.871 | 18.398 | 16.499 | 16.806 | 15.021 | 17.111 | 14.976 | 17.106 | 18.200 | 14.380 | 0.018 | 0.7240 | 0.8731 | 0 | 0 | 0 |
| 'Prodh' | 8.940 | 9.084 | 9.013 | 9.799 | 9.215 | 9.281 | 8.638 | 9.127 | 10.594 | 7.399 | 7.428 | 7.373 | 9.773 | 7.893 | 7.707 | 7.806 | 7.671 | 7.254 | 0.243 | 0.0000 | 0.0001 | 1 | 0 | 0 |
| 'Prorsd1' | 5.943 | 5.413 | 4.913 | 5.827 | 5.892 | 5.658 | 6.042 | 6.181 | 5.943 | 5.044 | 5.173 | 5.238 | 5.396 | 5.866 | 5.246 | 5.487 | 5.839 | 5.561 | 0.074 | 0.1805 | 0.4159 | 0 | 0 | 0 |
| 'Pros1' | 1.146 | 0.791 | 1.282 | 1.499 | 1.230 | 0.988 | 1.221 | 1.196 | 1.115 | 0.978 | 0.842 | 1.115 | 0.694 | 0.712 | 1.217 | 0.974 | 0.745 | 1.086 | 0.317 | 0.0070 | 0.0482 | 1 | 0 | 0 |
| 'Prosc' | 12.233 | 11.493 | 12.109 | 11.412 | 12.538 | 12.362 | 11.128 | 12.575 | 11.919 | 12.635 | 12.478 | 13.099 | 13.254 | 12.623 | 11.713 | 12.519 | 13.256 | 12.129 | -0.086 | 0.0104 | 0.0643 | 0 | 0 | 0 |
| 'Proser1' | 5.393 | 5.380 | 6.322 | 4.827 | 5.232 | 5.403 | 5.202 | 5.564 | 5.863 | 5.915 | 5.092 | 5.341 | 5.688 | 5.696 | 5.204 | 5.455 | 5.179 | 5.343 | 0.001 | 0.9759 | 0.9918 | 0 | 0 | 0 |
| 'Prox2' | 0.918 | 1.087 | 0.966 | 0.821 | 1.214 | 1.161 | 1.214 | 0.999 | 1.209 | 1.066 | 1.204 | 1.180 | 0.972 | 0.741 | 1.137 | 0.620 | 0.865 | 0.837 | 0.131 | 0.2588 | 0.5119 | 0 | 0 | 0 |
| 'Proz' | 2.878 | 2.847 | 3.194 | 2.701 | 2.722 | 3.099 | 2.365 | 2.878 | 3.457 | 3.539 | 3.283 | 3.409 | 3.072 | 2.237 | 3.450 | 3.259 | 2.790 | 3.116 | -0.110 | 0.1839 | 0.4193 | 0 | 0 | 0 |
| 'Prpf18' | 14.924 | 17.593 | 15.136 | 15.171 | 17.107 | 15.208 | 16.271 | 16.384 | 15.575 | 16.537 | 17.448 | 15.733 | 17.978 | 17.565 | 15.330 | 16.109 | 18.225 | 15.415 | -0.080 | 0.0035 | 0.0282 | 1 | 0 | 0 |
| 'Prpf19' | 11.464 | 11.168 | 11.781 | 11.070 | 11.421 | 12.067 | 11.099 | 11.734 | 12.049 | 10.877 | 10.069 | 11.515 | 10.595 | 10.011 | 11.555 | 11.164 | 10.348 | 11.310 | 0.082 | 0.0007 | 0.0078 | 1 | 0 | 0 |
| 'Prpf3' | 5.406 | 5.051 | 5.139 | 6.100 | 5.272 | 5.102 | 5.753 | 4.900 | 5.148 | 4.774 | 5.603 | 6.260 | 5.397 | 5.004 | 6.098 | 5.046 | 5.460 | 6.406 | -0.076 | 0.1835 | 0.4192 | 0 | 0 | 0 |
| 'Prpf31' | 7.595 | 7.892 | 8.468 | 7.647 | 7.821 | 8.563 | 8.378 | 7.883 | 8.581 | 7.808 | 7.576 | 7.706 | 8.601 | 7.894 | 7.652 | 8.425 | 7.546 | 7.293 | 0.039 | 0.2645 | 0.5184 | 0 | 0 | 0 |
| 'Prpf38a' | 6.427 | 7.485 | 7.327 | 7.889 | 7.549 | 6.915 | 7.149 | 7.622 | 7.303 | 6.557 | 7.511 | 7.004 | 7.145 | 7.500 | 6.818 | 7.413 | 7.545 | 6.730 | 0.022 | 0.5545 | 0.7670 | 0 | 0 | 0 |
| 'Prpf38b' | 28.896 | 27.901 | 30.256 | 29.386 | 28.069 | 29.031 | 31.274 | 27.651 | 29.951 | 27.250 | 28.316 | 29.898 | 28.017 | 27.639 | 29.389 | 27.719 | 28.674 | 29.308 | 0.023 | 0.3337 | 0.5919 | 0 | 0 | 0 |
| 'Prpf39' | 16.946 | 16.876 | 16.149 | 17.158 | 17.145 | 14.806 | 17.990 | 17.963 | 15.753 | 15.391 | 17.278 | 17.700 | 14.746 | 17.078 | 15.822 | 16.050 | 18.469 | 16.534 | 0.005 | 0.9081 | 0.9639 | 0 | 0 | 0 |
| 'Prpf4' | 9.745 | 9.565 | 9.691 | 9.801 | 9.413 | 9.352 | 9.368 | 8.964 | 9.112 | 9.023 | 8.829 | 9.303 | 9.669 | 9.487 | 9.214 | 9.778 | 8.930 | 9.682 | 0.008 | 0.7982 | 0.9103 | 0 | 0 | 0 |

| | | | | | | | | | | | | | | | | | | | | | | | | |
|---|---|---|---|---|---|---|---|---|---|---|---|---|---|---|---|---|---|---|---|---|---|---|---|---|
| 'Prpf40a' | 13.414 | 12.944 | 13.715 | 14.653 | 14.130 | 12.867 | 16.180 | 13.983 | 12.983 | 13.345 | 13.393 | 13.877 | 13.365 | 13.217 | 13.180 | 13.190 | 14.854 | 13.885 | 0.017 | 0.6761 | 0.8449 | 0 | 0 | 0 |
| 'Prpf40b' | 1.876 | 1.688 | 1.800 | 1.664 | 1.424 | 1.663 | 1.688 | 1.713 | 1.982 | 1.719 | 1.351 | 1.770 | 1.662 | 1.277 | 1.595 | 1.554 | 1.510 | 1.455 | 0.147 | 0.0089 | 0.0574 | 0 | 0 | 0 |
| 'Prpf4b' | 43.468 | 42.434 | 47.464 | 43.350 | 43.156 | 44.083 | 44.588 | 40.801 | 45.325 | 40.604 | 44.435 | 47.544 | 42.166 | 44.503 | 45.608 | 43.787 | 47.126 | 47.387 | -0.041 | 0.1750 | 0.4097 | 0 | 0 | 0 |
| 'Prpf6' | 26.433 | 24.152 | 24.629 | 24.513 | 25.396 | 24.932 | 26.192 | 24.721 | 24.805 | 25.539 | 25.111 | 25.062 | 23.672 | 24.744 | 24.244 | 24.739 | 25.281 | 23.996 | 0.012 | 0.6400 | 0.8230 | 0 | 0 | 0 |
| 'Prpf8' | 82.302 | 75.978 | 82.763 | 82.137 | 80.241 | 82.582 | 79.536 | 78.906 | 82.052 | 83.838 | 80.621 | 80.643 | 80.771 | 78.257 | 79.985 | 80.740 | 77.624 | 82.156 | -0.007 | 0.7337 | 0.8786 | 0 | 0 | 0 |
| 'Prps1' | 16.772 | 16.761 | 16.271 | 15.874 | 15.621 | 14.484 | 15.175 | 18.053 | 16.154 | 16.463 | 15.623 | 15.435 | 16.187 | 14.431 | 15.694 | 15.076 | 15.492 | 14.531 | 0.053 | 0.2052 | 0.4489 | 0 | 0 | 0 |
| 'Prps1l3' | 34.995 | 33.434 | 33.144 | 34.659 | 33.326 | 29.203 | 32.613 | 33.759 | 33.048 | 29.248 | 32.408 | 34.559 | 30.077 | 33.026 | 35.034 | 29.441 | 35.457 | 35.294 | 0.007 | 0.8851 | 0.9521 | 0 | 0 | 0 |
| 'Prps2' | 36.505 | 38.735 | 35.882 | 36.764 | 38.573 | 35.312 | 35.660 | 37.301 | 35.276 | 36.837 | 39.787 | 35.962 | 37.870 | 38.811 | 35.642 | 36.042 | 40.214 | 36.096 | -0.041 | 0.0582 | 0.2077 | 0 | 0 | 0 |
| 'Prpsap1' | 9.705 | 10.042 | 10.155 | 9.531 | 10.337 | 10.330 | 9.017 | 9.587 | 10.641 | 10.066 | 9.611 | 11.153 | 9.373 | 9.992 | 9.637 | 10.185 | 9.698 | 10.375 | -0.024 | 0.4656 | 0.7014 | 0 | 0 | 0 |
| 'Prpsap2' | 7.743 | 8.416 | 8.298 | 8.709 | 8.707 | 9.078 | 8.580 | 9.122 | 9.447 | 8.993 | 8.097 | 9.103 | 8.856 | 8.878 | 8.513 | 8.959 | 7.342 | 8.623 | 0.002 | 0.9654 | 0.9878 | 0 | 0 | 0 |
| 'Prr11' | 4.655 | 5.683 | 6.350 | 6.082 | 5.347 | 6.025 | 6.172 | 5.738 | 6.406 | 3.417 | 5.435 | 5.984 | 5.994 | 6.053 | 6.504 | 4.588 | 6.142 | 5.424 | 0.075 | 0.3679 | 0.6199 | 0 | 0 | 0 |
| 'Prr12' | 24.831 | 22.412 | 25.156 | 22.073 | 21.632 | 24.600 | 20.721 | 23.401 | 26.367 | 23.608 | 22.995 | 25.575 | 23.368 | 24.392 | 25.869 | 24.359 | 20.377 | 26.285 | -0.047 | 0.2147 | 0.4602 | 0 | 0 | 0 |
| 'Prr13' | 51.583 | 56.721 | 55.684 | 54.183 | 54.737 | 60.058 | 53.709 | 55.778 | 57.260 | 55.322 | 53.196 | 51.759 | 53.252 | 54.045 | 54.936 | 54.670 | 52.002 | 51.210 | 0.047 | 0.0832 | 0.2611 | 0 | 0 | 0 |
| 'Prr14' | 13.245 | 12.105 | 13.886 | 13.263 | 12.209 | 14.176 | 13.761 | 11.889 | 13.355 | 13.522 | 12.635 | 14.703 | 13.529 | 11.597 | 13.819 | 12.793 | 13.155 | 14.484 | -0.041 | 0.1531 | 0.3804 | 0 | 0 | 0 |
| 'Prr14l' | 8.139 | 7.609 | 7.614 | 8.256 | 7.691 | 7.382 | 8.368 | 7.568 | 7.348 | 7.882 | 7.788 | 8.418 | 7.666 | 7.574 | 7.946 | 7.976 | 8.019 | 9.004 | -0.059 | 0.1165 | 0.3241 | 0 | 0 | 0 |
| 'Prr3' | 7.815 | 6.091 | 6.292 | 6.423 | 6.085 | 6.410 | 6.070 | 6.425 | 6.362 | 6.857 | 6.085 | 6.597 | 6.589 | 6.955 | 6.513 | 7.035 | 6.193 | 6.574 | -0.042 | 0.3660 | 0.6181 | 0 | 0 | 0 |
| 'Prr7' | 10.745 | 8.018 | 10.268 | 10.065 | 8.229 | 11.648 | 9.523 | 9.673 | 9.929 | 11.588 | 8.279 | 10.351 | 9.068 | 8.591 | 9.335 | 10.140 | 8.046 | 8.221 | 0.073 | 0.2862 | 0.5428 | 0 | 0 | 0 |
| 'Prrc1' | 5.049 | 5.494 | 6.129 | 5.246 | 5.874 | 5.830 | 5.988 | 5.961 | 6.030 | 5.654 | 4.630 | 5.605 | 5.205 | 5.269 | 4.942 | 5.222 | 5.953 | 5.255 | 0.101 | 0.0433 | 0.1723 | 0 | 0 | 0 |
| 'Prrc2a' | 90.694 | 81.301 | 97.435 | 89.003 | 81.794 | 93.924 | 80.794 | 85.110 | 98.050 | 91.277 | 85.608 | 93.909 | 90.185 | 85.372 | 91.459 | 86.714 | 83.593 | 95.348 | -0.022 | 0.4109 | 0.6596 | 0 | 0 | 0 |
| 'Prrc2b' | 27.370 | 25.053 | 28.141 | 27.774 | 24.199 | 26.734 | 25.574 | 25.206 | 28.523 | 27.778 | 27.658 | 29.655 | 27.763 | 27.335 | 28.062 | 26.571 | 26.164 | 31.345 | -0.092 | 0.0027 | 0.0228 | 1 | 0 | 0 |
| 'Prrc2c' | 25.816 | 22.590 | 25.733 | 26.079 | 23.330 | 25.414 | 24.629 | 23.420 | 26.566 | 23.692 | 26.199 | 26.819 | 22.900 | 23.795 | 26.308 | 24.490 | 24.174 | 28.321 | -0.030 | 0.4039 | 0.6528 | 0 | 0 | 0 |
| 'Prrg1' | 0.262 | 0.350 | 0.323 | 0.318 | 0.191 | 0.368 | 0.377 | 0.283 | 0.334 | 0.278 | 0.335 | 0.321 | 0.243 | 0.298 | 0.283 | 0.245 | 0.241 | 0.316 | 0.119 | 0.2947 | 0.5504 | 0 | 0 | 0 |
| 'Prrt1' | 4.278 | 5.312 | 4.216 | 4.537 | 5.113 | 4.988 | 4.370 | 6.283 | 4.992 | 5.064 | 3.703 | 4.658 | 4.979 | 4.897 | 3.767 | 4.560 | 4.819 | 3.556 | 0.125 | 0.1169 | 0.3249 | 0 | 0 | 0 |
| 'Prrt2' | 0.769 | 0.770 | 0.823 | 0.610 | 0.677 | 0.712 | 0.479 | 0.522 | 0.977 | 0.584 | 0.763 | 0.749 | 0.786 | 0.553 | 0.732 | 0.566 | 0.684 | 0.799 | 0.014 | 0.9025 | 0.9613 | 0 | 0 | 0 |
| 'Prss41' | 2.616 | 2.898 | 2.688 | 2.096 | 2.852 | 2.339 | 2.225 | 2.569 | 2.390 | 3.342 | 2.930 | 2.795 | 3.000 | 2.880 | 2.688 | 2.743 | 2.889 | 2.725 | -0.206 | 0.0042 | 0.0331 | 1 | 0 | 0 |
| 'Prune' | 5.289 | 5.332 | 4.833 | 4.720 | 5.609 | 5.239 | 4.797 | 5.179 | 5.344 | 4.956 | 4.928 | 4.718 | 5.535 | 5.219 | 4.667 | 5.304 | 4.968 | 4.098 | 0.057 | 0.2856 | 0.5425 | 0 | 0 | 0 |
| 'Prx' | 0.810 | 0.668 | 0.856 | 0.728 | 0.699 | 0.788 | 0.702 | 0.635 | 0.850 | 0.642 | 0.458 | 0.719 | 0.552 | 0.620 | 0.556 | 0.552 | 0.634 | 0.627 | 0.315 | 0.0000 | 0.0007 | 1 | 0 | 0 |
| 'Psap' | 138.998 | 150.828 | 144.274 | 142.529 | 147.954 | 150.054 | 139.211 | 149.554 | 143.893 | 145.517 | 145.569 | 138.086 | 140.041 | 146.571 | 144.830 | 148.595 | 140.452 | 141.411 | 0.008 | 0.7251 | 0.8737 | 0 | 0 | 0 |
| 'Psat1' | 19.170 | 19.330 | 19.676 | 20.317 | 19.173 | 18.282 | 18.978 | 19.339 | 19.185 | 17.358 | 18.157 | 18.000 | 19.296 | 17.928 | 18.068 | 16.395 | 18.736 | 16.662 | 0.100 | 0.0012 | 0.0119 | 1 | 0 | 0 |
| 'Psd' | 4.187 | 2.275 | 2.505 | 2.492 | 2.695 | 2.399 | 2.028 | 2.728 | 2.592 | 3.185 | 2.207 | 2.593 | 2.868 | 2.460 | 2.538 | 3.206 | 2.238 | 2.562 | 0.003 | 0.9755 | 0.9918 | 0 | 0 | 0 |
| 'Psd2' | 0.580 | 0.533 | 0.521 | 0.471 | 0.459 | 0.454 | 0.580 | 0.564 | 0.574 | 0.421 | 0.289 | 0.443 | 0.366 | 0.339 | 0.310 | 0.347 | 0.403 | 0.416 | 0.482 | 0.0000 | 0.0000 | 1 | 1 | 0 |
| 'Psd4' | 18.263 | 15.925 | 18.762 | 17.725 | 16.847 | 18.718 | 16.863 | 16.780 | 18.353 | 17.604 | 17.387 | 20.429 | 16.886 | 17.001 | 19.269 | 17.599 | 17.272 | 20.213 | -0.058 | 0.0327 | 0.1432 | 0 | 0 | 0 |
| 'Psen1' | 45.854 | 42.212 | 44.258 | 46.042 | 45.444 | 44.698 | 45.418 | 43.252 | 45.534 | 43.819 | 47.590 | 50.018 | 45.031 | 44.686 | 49.604 | 45.036 | 46.002 | 50.775 | -0.080 | 0.0103 | 0.0639 | 0 | 0 | 0 |
| 'Psen2' | 18.020 | 19.308 | 18.158 | 17.419 | 19.096 | 18.438 | 18.349 | 18.146 | 18.199 | 18.306 | 17.190 | 17.167 | 17.696 | 17.031 | 15.921 | 18.234 | 16.803 | 16.433 | 0.084 | 0.0013 | 0.0127 | 1 | 0 | 0 |
| 'Psenen' | 68.710 | 82.936 | 72.791 | 73.881 | 81.323 | 80.856 | 70.400 | 81.471 | 75.092 | 74.321 | 73.319 | 74.494 | 72.532 | 75.336 | 72.514 | 75.275 | 75.771 | 70.145 | 0.038 | 0.2156 | 0.4613 | 0 | 0 | 0 |
| 'Psip1' | 33.000 | 34.031 | 32.897 | 35.526 | 35.709 | 33.234 | 35.100 | 33.182 | 31.835 | 33.267 | 34.620 | 33.064 | 34.252 | 36.140 | 33.578 | 33.978 | 34.807 | 32.691 | -0.020 | 0.3917 | 0.6410 | 0 | 0 | 0 |
| 'Pskh1' | 15.000 | 17.908 | 16.809 | 13.679 | 17.433 | 18.935 | 13.749 | 18.949 | 17.984 | 17.897 | 16.351 | 16.161 | 16.162 | 16.458 | 16.695 | 18.782 | 13.083 | 15.617 | 0.017 | 0.7986 | 0.9106 | 0 | 0 | 0 |
| 'Psma1' | 34.967 | 37.493 | 36.014 | 35.639 | 39.383 | 36.333 | 38.658 | 37.838 | 35.334 | 33.216 | 33.872 | 34.226 | 36.218 | 35.756 | 35.037 | 32.694 | 36.853 | 33.163 | 0.080 | 0.0030 | 0.0249 | 1 | 0 | 0 |
| 'Psma2' | 149.085 | 169.709 | 151.193 | 156.703 | 164.531 | 154.258 | 161.461 | 157.231 | 147.643 | 143.538 | 144.535 | 142.668 | 153.539 | 146.117 | 154.867 | 156.142 | 158.002 | 137.523 | 0.067 | 0.0251 | 0.1210 | 0 | 0 | 0 |
| 'Psma3' | 18.820 | 19.559 | 17.485 | 20.125 | 19.305 | 17.106 | 20.191 | 19.221 | 16.862 | 17.185 | 18.149 | 18.151 | 18.548 | 20.029 | 17.624 | 18.335 | 20.172 | 17.062 | 0.016 | 0.6124 | 0.8056 | 0 | 0 | 0 |
| 'Psma4' | 33.170 | 35.625 | 34.726 | 36.186 | 37.641 | 35.440 | 39.403 | 35.984 | 37.262 | 33.130 | 35.730 | 35.809 | 36.060 | 34.741 | 34.490 | 35.140 | 37.229 | 33.251 | 0.032 | 0.2896 | 0.5461 | 0 | 0 | 0 |
| 'Psma5' | 39.827 | 46.135 | 43.068 | 46.625 | 48.433 | 43.417 | 47.950 | 45.226 | 43.424 | 39.515 | 41.386 | 42.641 | 45.103 | 42.593 | 41.058 | 48.634 | 45.378 | 43.519 | 0.039 | 0.3397 | 0.5969 | 0 | 0 | 0 |
| 'Psma6' | 40.172 | 43.776 | 41.456 | 44.556 | 44.135 | 41.727 | 44.432 | 43.014 | 40.817 | 37.071 | 38.565 | 37.541 | 40.209 | 40.174 | 38.995 | 41.478 | 39.835 | 38.336 | 0.114 | 0.0000 | 0.0007 | 1 | 0 | 0 |
| 'Psma7' | 33.293 | 35.474 | 33.451 | 32.375 | 35.076 | 33.152 | 32.785 | 32.862 | 32.510 | 28.733 | 28.968 | 31.191 | 32.432 | 31.556 | 31.924 | 32.307 | 33.871 | 30.759 | 0.084 | 0.0058 | 0.0418 | 1 | 0 | 0 |
| 'Psmb1' | 165.624 | 180.688 | 166.889 | 174.897 | 180.319 | 167.582 | 172.493 | 175.482 | 168.847 | 155.688 | 155.638 | 159.320 | 160.658 | 168.040 | 163.071 | 158.026 | 170.410 | 153.449 | 0.092 | 0.0000 | 0.0005 | 1 | 0 | 0 |
| 'Psmb10' | 77.393 | 76.354 | 78.820 | 73.712 | 80.439 | 83.592 | 70.356 | 77.687 | 78.636 | 82.512 | 72.283 | 88.624 | 84.693 | 77.537 | 89.739 | 87.528 | 78.918 | 90.909 | -0.120 | 0.0005 | 0.0061 | 1 | 0 | 0 |
| 'Psmb2' | 33.244 | 36.588 | 36.780 | 34.692 | 39.579 | 36.233 | 36.508 | 38.189 | 37.333 | 33.132 | 35.927 | 35.262 | 37.668 | 38.182 | 36.537 | 35.536 | 37.788 | 33.464 | 0.015 | 0.6401 | 0.8230 | 0 | 0 | 0 |
| 'Psmb3' | 36.820 | 40.783 | 37.824 | 39.472 | 39.307 | 36.616 | 36.392 | 38.946 | 36.817 | 34.512 | 35.593 | 36.069 | 36.719 | 37.556 | 37.906 | 35.817 | 37.821 | 36.601 | 0.049 | 0.0730 | 0.2395 | 0 | 0 | 0 |
| 'Psmb4' | 57.536 | 63.458 | 56.587 | 61.223 | 63.945 | 60.795 | 60.455 | 63.281 | 55.008 | 55.793 | 56.401 | 53.280 | 59.430 | 58.226 | 57.855 | 62.061 | 57.460 | 54.389 | 0.064 | 0.0277 | 0.1285 | 0 | 0 | 0 |
| 'Psmb5' | 97.068 | 106.378 | 97.637 | 97.001 | 111.276 | 102.971 | 97.042 | 105.106 | 98.073 | 86.676 | 94.821 | 101.239 | 95.922 | 100.063 | 107.171 | 93.937 | 95.625 | 101.785 | 0.043 | 0.1947 | 0.4328 | 0 | 0 | 0 |
| 'Psmb6' | 31.000 | 34.771 | 33.090 | 31.954 | 35.420 | 33.390 | 31.471 | 34.184 | 33.295 | 30.714 | 31.016 | 29.274 | 33.459 | 31.504 | 30.857 | 31.734 | 31.024 | 28.114 | 0.096 | 0.0020 | 0.0183 | 1 | 0 | 0 |
| 'Psmb7' | 22.132 | 25.169 | 23.046 | 23.776 | 24.596 | 22.476 | 24.181 | 23.777 | 21.881 | 20.458 | 22.576 | 19.935 | 23.247 | 21.238 | 20.126 | 21.987 | 22.395 | 20.201 | 0.125 | 0.0000 | 0.0006 | 1 | 0 | 0 |
| 'Psmb8' | 82.101 | 95.119 | 83.756 | 86.076 | 91.640 | 90.836 | 83.882 | 88.054 | 85.551 | 78.263 | 80.672 | 79.479 | 88.678 | 84.301 | 86.166 | 84.412 | 82.176 | 82.643 | 0.064 | 0.0349 | 0.1496 | 0 | 0 | 0 |
| 'Psmb9' | 38.403 | 41.828 | 40.720 | 39.913 | 43.940 | 41.904 | 39.786 | 39.834 | 41.816 | 36.759 | 38.028 | 39.590 | 41.342 | 40.394 | 39.572 | 39.944 | 41.767 | 38.637 | 0.037 | 0.1646 | 0.3969 | 0 | 0 | 0 |
| 'Psmc1' | 96.226 | 98.715 | 99.755 | 97.643 | 105.058 | 104.580 | 101.872 | 100.122 | 103.939 | 92.989 | 90.868 | 95.303 | 104.433 | 98.127 | 100.697 | 99.522 | 88.890 | 93.953 | 0.060 | 0.0436 | 0.1729 | 0 | 0 | 0 |
| 'Psmc2' | 28.165 | 28.689 | 28.691 | 29.774 | 29.308 | 27.498 | 29.755 | 30.883 | 27.542 | 26.471 | 27.556 | 27.778 | 28.693 | 27.747 | 27.240 | 27.954 | 28.999 | 26.826 | 0.051 | 0.0336 | 0.1458 | 0 | 0 | 0 |
| 'Psmc3' | 33.923 | 34.887 | 36.591 | 36.514 | 37.045 | 38.306 | 35.686 | 35.510 | 36.429 | 34.958 | 32.309 | 35.842 | 37.015 | 35.270 | 37.635 | 35.928 | 33.989 | 35.899 | 0.017 | 0.5102 | 0.7351 | 0 | 0 | 0 |
| 'Psmc3ip' | 2.479 | 2.348 | 2.450 | 2.454 | 2.310 | 2.249 | 2.795 | 2.581 | 2.273 | 2.284 | 2.087 | 2.488 | 2.799 | 2.366 | 2.718 | 2.292 | 2.324 | 2.308 | 0.000 | 0.9966 | 0.9994 | 0 | 0 | 0 |
| 'Psmc4' | 22.944 | 22.321 | 23.427 | 22.383 | 22.649 | 24.564 | 23.290 | 22.914 | 22.542 | 22.694 | 20.626 | 22.942 | 22.787 | 21.211 | 22.538 | 22.714 | 21.874 | 21.437 | 0.049 | 0.0438 | 0.1733 | 0 | 0 | 0 |
| 'Psmc5' | 75.709 | 76.149 | 76.357 | 79.188 | 80.732 | 76.580 | 75.951 | 77.149 | 77.093 | 73.023 | 76.591 | 72.906 | 82.053 | 73.486 | 77.930 | 79.074 | 74.994 | 72.554 | 0.016 | 0.5474 | 0.7613 | 0 | 0 | 0 |
| 'Psmc6' | 33.633 | 36.945 | 32.734 | 35.995 | 37.240 | 32.755 | 36.295 | 35.878 | 31.890 | 32.050 | 32.924 | 35.919 | 34.702 | 35.374 | 32.012 | 33.012 | 37.859 | 30.989 | 0.023 | 0.5092 | 0.7342 | 0 | 0 | 0 |
| 'Psmd1' | 12.422 | 12.371 | 12.459 | 13.149 | 12.798 | 11.636 | 13.151 | 13.037 | 11.961 | 12.368 | 12.486 | 12.382 | 13.265 | 12.711 | 12.451 | 12.253 | 12.932 | 11.825 | -0.008 | 0.7401 | 0.8822 | 0 | 0 | 0 |
| 'Psmd10' | 25.262 | 27.213 | 27.076 | 28.314 | 29.871 | 26.046 | 27.530 | 30.167 | 28.098 | 25.121 | 26.551 | 27.175 | 25.690 | 28.272 | 27.358 | 26.890 | 28.445 | 25.743 | 0.037 | 0.3229 | 0.5806 | 0 | 0 | 0 |
| 'Psmd11' | 8.280 | 9.002 | 8.893 | 9.206 | 8.950 | 8.970 | 9.062 | 9.471 | 8.738 | 8.009 | 8.883 | 9.197 | 8.862 | 8.911 | 9.022 | 8.599 | 8.887 | 8.994 | 0.009 | 0.7460 | 0.8854 | 0 | 0 | 0 |
| 'Psmd12' | 28.594 | 27.870 | 29.120 | 30.038 | 29.717 | 28.367 | 31.203 | 29.697 | 29.200 | 28.869 | 28.776 | 27.447 | 28.152 | 28.226 | 28.307 | 28.271 | 28.559 | 27.754 | 0.044 | 0.1028 | 0.3006 | 0 | 0 | 0 |
| 'Psmd13' | 13.056 | 15.119 | 14.156 | 13.434 | 14.404 | 14.035 | 14.248 | 14.780 | 14.453 | 13.022 | 13.541 | 14.037 | 13.552 | 13.836 | 13.974 | 14.024 | 14.003 | 13.974 | 0.030 | 0.2263 | 0.4730 | 0 | 0 | 0 |
| 'Psmd14' | 20.206 | 20.925 | 20.533 | 22.451 | 22.294 | 21.309 | 22.873 | 21.847 | 20.124 | 19.143 | 20.101 | 20.204 | 19.075 | 20.389 | 19.803 | 19.325 | 22.118 | 18.742 | 0.094 | 0.0038 | 0.0302 | 1 | 0 | 0 |
| 'Psmd2' | 37.777 | 40.622 | 40.234 | 40.502 | 41.069 | 39.560 | 39.311 | 39.744 | 40.009 | 38.505 | 39.700 | 38.807 | 39.052 | 38.762 | 40.132 | 37.127 | 39.658 | 39.249 | 0.021 | 0.2925 | 0.5493 | 0 | 0 | 0 |
| 'Psmd3' | 19.963 | 19.632 | 22.096 | 18.792 | 19.510 | 22.223 | 19.685 | 20.135 | 21.362 | 19.691 | 17.888 | 20.579 | 21.896 | 18.537 | 20.580 | 19.489 | 17.466 | 19.938 | 0.049 | 0.1070 | 0.3076 | 0 | 0 | 0 |
| 'Psmd4' | 19.429 | 20.710 | 20.173 | 20.107 | 20.470 | 21.533 | 20.037 | 21.028 | 20.986 | 19.705 | 19.122 | 19.511 | 19.968 | 19.244 | 20.519 | 19.647 | 19.961 | 18.358 | 0.058 | 0.0183 | 0.0970 | 0 | 0 | 0 |
| 'Psmd5' | 5.707 | 5.988 | 5.827 | 5.821 | 5.798 | 5.628 | 5.590 | 6.123 | 6.008 | 5.802 | 5.510 | 6.216 | 5.142 | 5.744 | 5.795 | 5.510 | 5.726 | 6.099 | 0.013 | 0.7068 | 0.8641 | 0 | 0 | 0 |
| 'Psmd6' | 21.574 | 22.879 | 23.182 | 21.669 | 25.319 | 22.241 | 22.886 | 23.046 | 23.050 | 20.852 | 22.008 | 22.953 | 22.022 | 21.967 | 22.422 | 23.440 | 21.403 | 22.081 | 0.035 | 0.2473 | 0.4979 | 0 | 0 | 0 |
| 'Psmd7' | 55.340 | 60.104 | 55.074 | 55.692 | 58.864 | 56.126 | 57.030 | 59.781 | 56.682 | 52.857 | 51.896 | 58.160 | 58.939 | 54.782 | 57.767 | 54.811 | 58.844 | 58.548 | 0.008 | 0.7899 | 0.9061 | 0 | 0 | 0 |
| 'Psmd8' | 75.122 | 80.565 | 72.710 | 73.865 | 80.245 | 76.699 | 73.695 | 79.532 | 73.493 | 72.689 | 70.940 | 69.834 | 77.315 | 72.749 | 75.492 | 75.212 | 71.223 | 71.348 | 0.052 | 0.0583 | 0.2079 | 0 | 0 | 0 |
| 'Psmd9' | 4.059 | 4.857 | 4.387 | 4.211 | 4.981 | 4.245 | 4.026 | 4.555 | 4.522 | 4.112 | 4.181 | 4.141 | 4.432 | 4.330 | 4.005 | 4.547 | 4.008 | 3.709 | 0.077 | 0.0882 | 0.2713 | 0 | 0 | 0 |
| 'Psme1' | 47.880 | 52.280 | 47.364 | 49.848 | 51.486 | 49.368 | 50.455 | 50.065 | 49.788 | 48.524 | 47.019 | 47.001 | 52.156 | 48.869 | 49.266 | 53.383 | 51.160 | 48.763 | -0.003 | 0.9100 | 0.9651 | 0 | 0 | 0 |
| 'Psme2' | 47.008 | 48.115 | 46.884 | 48.736 | 51.138 | 46.556 | 49.048 | 48.364 | 47.799 | 47.088 | 43.744 | 47.163 | 50.268 | 46.410 | 47.770 | 52.894 | 50.395 | 48.657 | -0.013 | 0.6540 | 0.8324 | 0 | 0 | 0 |
| 'Psme2b' | 32.423 | 36.556 | 32.374 | 36.403 | 37.112 | 36.406 | 42.847 | 40.195 | 33.919 | 36.851 | 36.308 | 42.503 | 38.011 | 39.596 | 37.266 | 49.975 | 36.306 | 38.636 | -0.128 | 0.0406 | 0.1649 | 0 | 0 | 0 |
| 'Psme3' | 36.622 | 38.761 | 39.686 | 36.772 | 40.041 | 39.869 | 35.931 | 39.151 | 38.921 | 38.279 | 35.742 | 40.200 | 38.191 | 36.700 | 40.656 | 37.207 | 37.570 | 40.396 | -0.008 | 0.7291 | 0.8759 | 0 | 0 | 0 |
| 'Psme4' | 11.280 | 12.700 | 12.065 | 12.702 | 12.472 | 11.364 | 12.738 | 12.677 | 11.626 | 12.530 | 12.887 | 12.170 | 12.264 | 13.173 | 11.564 | 11.459 | 12.709 | 12.306 | -0.031 | 0.2909 | 0.5477 | 0 | 0 | 0 |
| 'Psmf1' | 4.236 | 4.429 | 4.503 | 4.247 | 4.558 | 4.482 | 4.284 | 4.485 | 4.380 | 4.636 | 4.338 | 4.472 | 4.393 | 4.491 | 4.263 | 4.146 | 4.041 | 4.093 | 0.016 | 0.6337 | 0.8184 | 0 | 0 | 0 |
| 'Psmg1' | 16.688 | 20.726 | 18.837 | 18.894 | 19.458 | 17.817 | 18.626 | 19.199 | 19.298 | 18.377 | 17.018 | 17.384 | 18.236 | 17.862 | 17.863 | 18.145 | 17.028 | 16.830 | 0.082 | 0.0301 | 0.1350 | 0 | 0 | 0 |
| 'Psmg2' | 26.502 | 27.634 | 31.492 | 26.269 | 30.416 | 25.946 | 26.267 | 29.579 | 29.299 | 27.854 | 27.805 | 31.353 | 26.023 | 25.120 | 30.593 | 26.550 | 31.722 | 31.277 | -0.040 | 0.4164 | 0.6643 | 0 | 0 | 0 |
| 'Psmg3' | 2.700 | 3.404 | 2.881 | 3.407 | 2.828 | 3.322 | 2.635 | 2.995 | 3.057 | 2.708 | 2.437 | 3.658 | 2.815 | 2.877 | 3.556 | 2.047 | 2.747 | 3.414 | 0.036 | 0.6643 | 0.8378 | 0 | 0 | 0 |
| 'Psmg4' | 21.570 | 19.561 | 19.593 | 19.937 | 22.431 | 21.038 | 21.481 | 22.016 | 19.977 | 21.239 | 21.221 | 21.592 | 21.623 | 21.590 | 21.987 | 20.133 | 19.819 | 18.939 | -0.017 | 0.7578 | 0.8916 | 0 | 0 | 0 |
| 'Pspc1' | 13.975 | 14.296 | 16.384 | 16.420 | 15.670 | 15.101 | 17.284 | 16.107 | 15.699 | 16.299 | 15.211 | 12.669 | 16.170 | 13.600 | 12.594 | 13.275 | 14.713 | 10.731 | 0.168 | 0.0043 | 0.0336 | 1 | 0 | 0 |
| 'Psph' | 4.556 | 5.156 | 5.024 | 4.800 | 5.361 | 5.150 | 4.231 | 5.270 | 5.092 | 4.707 | 5.069 | 5.593 | 4.915 | 5.136 | 5.423 | 4.907 | 5.162 | 5.342 | -0.066 | 0.0956 | 0.2878 | 0 | 0 | 0 |
| 'Psrc1' | 0.494 | 0.488 | 0.677 | 0.619 | 0.580 | 0.629 | 0.519 | 0.371 | 0.699 | 0.441 | 0.430 | 0.694 | 0.590 | 0.449 | 0.671 | 0.560 | 0.609 | 0.650 | -0.027 | 0.7857 | 0.9044 | 0 | 0 | 0 |
| 'Pstk' | 3.197 | 4.067 | 3.835 | 3.347 | 4.050 | 3.556 | 3.215 | 4.086 | 4.360 | 4.172 | 3.355 | 3.614 | 3.884 | 3.778 | 3.433 | 3.989 | 3.537 | 2.823 | 0.041 | 0.5785 | 0.7820 | 0 | 0 | 0 |
| 'Pstpip1' | 40.169 | 37.554 | 41.655 | 38.830 | 37.600 | 43.472 | 39.861 | 39.286 | 41.381 | 41.830 | 38.123 | 44.591 | 45.545 | 38.225 | 44.056 | 41.294 | 39.971 | 44.494 | -0.082 | 0.0003 | 0.0041 | 1 | 0 | 0 |
| 'Ptar1' | 5.203 | 3.234 | 4.675 | 3.962 | 3.305 | 4.693 | 4.497 | 3.922 | 3.960 | 4.409 | 5.193 | 4.478 | 4.076 | 4.775 | 3.677 | 4.124 | 4.022 | 4.839 | -0.084 | 0.3687 | 0.6201 | 0 | 0 | 0 |
| 'Ptbp1' | 45.241 | 39.778 | 40.907 | 41.015 | 39.409 | 40.263 | 39.748 | 40.903 | 40.521 | 43.759 | 38.191 | 38.313 | 43.388 | 39.311 | 39.162 | 41.849 | 38.580 | 38.407 | 0.019 | 0.4659 | 0.7015 | 0 | 0 | 0 |
| 'Ptbp2' | 3.289 | 3.111 | 3.393 | 2.818 | 3.138 | 2.803 | 3.347 | 3.466 | 2.902 | 3.654 | 2.944 | 2.750 | 3.270 | 2.608 | 2.364 | 3.231 | 2.935 | 2.510 | 0.106 | 0.0718 | 0.2363 | 0 | 0 | 0 |
| 'Ptbp3' | 37.497 | 40.498 | 38.620 | 39.176 | 39.588 | 35.518 | 39.743 | 39.224 | 36.211 | 37.730 | 40.022 | 36.540 | 36.746 | 40.516 | 35.755 | 35.030 | 39.268 | 36.777 | 0.022 | 0.3884 | 0.6379 | 0 | 0 | 0 |
| 'Ptcd1' | 12.616 | 12.866 | 12.872 | 12.310 | 12.213 | 13.475 | 12.384 | 12.051 | 12.730 | 13.770 | 11.880 | 12.930 | 13.285 | 11.657 | 12.002 | 13.407 | 12.073 | 12.314 | -0.006 | 0.8722 | 0.9452 | 0 | 0 | 0 |
| 'Ptcd2' | 22.226 | 19.953 | 24.453 | 23.112 | 22.186 | 21.631 | 21.639 | 21.630 | 21.299 | 21.591 | 21.387 | 22.669 | 19.352 | 18.646 | 20.748 | 20.124 | 18.600 | 20.585 | 0.098 | 0.0259 | 0.1236 | 0 | 0 | 0 |
| 'Ptcd3' | 5.187 | 5.804 | 5.704 | 6.085 | 5.318 | 5.006 | 6.053 | 5.599 | 5.570 | 5.278 | 5.643 | 5.798 | 5.071 | 5.210 | 4.984 | 5.367 | 5.800 | 5.180 | 0.045 | 0.2946 | 0.5504 | 0 | 0 | 0 |
| 'Ptch1' | 4.058 | 3.985 | 3.812 | 3.728 | 3.743 | 3.860 | 3.705 | 3.887 | 3.982 | 3.345 | 3.376 | 3.671 | 3.344 | 3.311 | 3.581 | 3.179 | 3.117 | 3.580 | 0.175 | 0.0000 | 0.0000 | 1 | 0 | 0 |
| 'Ptdss1' | 18.378 | 20.499 | 19.674 | 18.798 | 20.383 | 19.134 | 18.047 | 22.393 | 18.545 | 19.273 | 20.893 | 19.784 | 19.757 | 20.061 | 20.073 | 19.635 | 19.190 | 20.566 | -0.039 | 0.2463 | 0.4969 | 0 | 0 | 0 |
| 'Ptdss2' | 2.436 | 2.792 | 3.019 | 2.462 | 2.524 | 2.728 | 2.563 | 2.979 | 2.891 | 2.703 | 2.549 | 3.018 | 2.716 | 2.596 | 2.605 | 2.552 | 2.453 | 2.817 | 0.009 | 0.8594 | 0.9390 | 0 | 0 | 0 |
| 'Pten' | 11.600 | 11.279 | 11.270 | 11.485 | 11.736 | 11.143 | 11.446 | 11.286 | 10.354 | 11.343 | 10.621 | 11.481 | 10.670 | 11.708 | 10.968 | 11.168 | 11.711 | 11.412 | -0.004 | 0.8802 | 0.9494 | 0 | 0 | 0 |
| 'Ptger1' | 19.075 | 17.303 | 21.960 | 18.593 | 16.712 | 20.450 | 17.612 | 16.296 | 20.992 | 20.600 | 19.466 | 22.018 | 19.613 | 16.392 | 20.480 | 19.009 | 18.088 | 23.110 | -0.093 | 0.0110 | 0.0669 | 0 | 0 | 0 |
| 'Ptger2' | 12.482 | 12.416 | 12.532 | 13.405 | 12.346 | 13.259 | 12.980 | 11.869 | 13.384 | 12.998 | 14.084 | 14.512 | 12.398 | 13.485 | 14.080 | 11.687 | 13.773 | 13.741 | -0.087 | 0.0311 | 0.1383 | 0 | 0 | 0 |
| 'Ptger4' | 4.259 | 2.835 | 3.020 | 3.126 | 2.525 | 3.021 | 3.291 | 2.771 | 3.162 | 3.856 | 2.744 | 3.490 | 3.022 | 3.009 | 3.100 | 4.027 | 3.402 | 2.967 | -0.092 | 0.1850 | 0.4207 | 0 | 0 | 0 |
| 'Ptges2' | 2.371 | 2.405 | 2.713 | 2.268 | 2.772 | 2.907 | 2.582 | 2.531 | 2.718 | 2.786 | 2.301 | 2.857 | 2.824 | 2.464 | 2.798 | 2.765 | 2.271 | 2.511 | -0.029 | 0.6074 | 0.8026 | 0 | 0 | 0 |
| 'Ptges3' | 38.489 | 36.851 | 35.868 | 40.126 | 36.447 | 35.266 | 36.757 | 36.180 | 34.006 | 33.092 | 36.336 | 46.717 | 31.126 | 41.141 | 42.549 | 33.742 | 41.937 | 48.924 | -0.107 | 0.1181 | 0.3270 | 0 | 0 | 0 |
| 'Ptges3l' | 1.239 | 1.525 | 1.579 | 1.144 | 1.561 | 1.455 | 1.519 | 1.299 | 1.684 | 1.230 | 1.622 | 1.599 | 1.760 | 1.542 | 1.697 | 1.667 | 1.653 | 1.340 | -0.128 | 0.2422 | 0.4928 | 0 | 0 | 0 |
| 'Ptgfrn' | 3.858 | 5.109 | 4.816 | 4.123 | 5.141 | 4.423 | 4.360 | 4.907 | 4.972 | 4.861 | 4.215 | 5.254 | 3.669 | 4.389 | 4.908 | 5.781 | 4.816 | 4.944 | -0.053 | 0.4367 | 0.6801 | 0 | 0 | 0 |
| 'Ptgir' | 3.625 | 3.682 | 3.681 | 4.113 | 3.105 | 3.487 | 3.952 | 3.192 | 3.573 | 2.111 | 1.515 | 2.038 | 1.656 | 1.743 | 2.277 | 1.969 | 1.560 | 1.611 | 0.962 | 0.0000 | 0.0000 | 1 | 1 | 0 |
| 'Ptgr1' | 1.465 | 1.470 | 1.839 | 1.700 | 2.157 | 1.832 | 2.137 | 1.832 | 1.341 | 1.467 | 1.544 | 1.726 | 2.235 | 1.601 | 1.677 | 1.335 | 1.488 | 1.366 | 0.103 | 0.3998 | 0.6487 | 0 | 0 | 0 |
| 'Ptgr2' | 5.392 | 6.621 | 5.958 | 5.852 | 6.618 | 6.103 | 6.134 | 6.862 | 6.354 | 5.848 | 6.160 | 5.539 | 6.245 | 6.342 | 5.543 | 5.815 | 6.570 | 5.080 | 0.065 | 0.0843 | 0.2636 | 0 | 0 | 0 |
| 'Ptk2' | 2.486 | 2.221 | 2.630 | 2.627 | 2.502 | 2.718 | 2.484 | 2.898 | 2.963 | 2.114 | 2.132 | 2.339 | 2.283 | 2.547 | 2.097 | 2.561 | 2.759 | 2.025 | 0.173 | 0.0155 | 0.0860 | 0 | 0 | 0 |
| 'Ptk2b' | 48.549 | 49.079 | 46.325 | 46.926 | 46.832 | 44.737 | 45.543 | 46.367 | 48.436 | 50.633 | 49.074 | 49.262 | 47.183 | 47.716 | 47.889 | 51.032 | 49.035 | 50.871 | -0.077 | 0.0032 | 0.0268 | 1 | 0 | 0 |
| 'Ptms' | 88.603 | 74.140 | 91.798 | 81.989 | 82.106 | 99.686 | 84.108 | 82.245 | 88.223 | 82.121 | 79.171 | 87.529 | 79.822 | 83.054 | 102.553 | 83.228 | 81.740 | 98.202 | -0.017 | 0.6468 | 0.8276 | 0 | 0 | 0 |
| 'Ptov1' | 10.865 | 7.459 | 10.668 | 8.376 | 7.717 | 9.713 | 10.529 | 8.755 | 9.677 | 7.665 | 8.414 | 11.363 | 7.006 | 6.681 | 9.687 | 6.752 | 8.416 | 11.167 | 0.115 | 0.1906 | 0.4271 | 0 | 0 | 0 |
| 'Ptp4a2' | 120.962 | 131.500 | 118.203 | 127.104 | 132.147 | 120.212 | 125.802 | 130.322 | 122.095 | 126.394 | 129.057 | 107.361 | 116.532 | 124.898 | 115.991 | 119.132 | 121.435 | 114.076 | 0.060 | 0.0174 | 0.0938 | 0 | 0 | 0 |
| 'Ptp4a3' | 20.906 | 25.187 | 22.640 | 20.840 | 25.547 | 25.061 | 20.455 | 25.444 | 23.160 | 23.740 | 21.647 | 20.685 | 24.770 | 22.856 | 21.274 | 23.183 | 21.817 | 19.274 | 0.060 | 0.2249 | 0.4716 | 0 | 0 | 0 |
| 'Ptpa' | 13.013 | 14.837 | 13.826 | 12.539 | 14.455 | 15.701 | 12.061 | 14.882 | 14.185 | 13.196 | 13.392 | 14.348 | 13.295 | 14.508 | 15.121 | 14.415 | 13.171 | 14.738 | -0.021 | 0.5669 | 0.7748 | 0 | 0 | 0 |
| 'Ptpdc1' | 1.049 | 1.074 | 1.016 | 0.939 | 1.415 | 1.256 | 1.084 | 1.276 | 1.224 | 1.438 | 1.187 | 1.130 | 1.317 | 1.654 | 1.053 | 1.315 | 1.488 | 0.958 | -0.153 | 0.1306 | 0.3469 | 0 | 0 | 0 |
| 'Ptpn1' | 53.126 | 50.514 | 50.476 | 47.949 | 52.555 | 50.304 | 46.863 | 55.619 | 48.718 | 53.401 | 60.597 | 53.559 | 51.505 | 56.582 | 55.884 | 53.021 | 54.393 | 56.492 | -0.130 | 0.0001 | 0.0009 | 1 | 0 | 0 |
| 'Ptpn11' | 66.934 | 83.143 | 65.524 | 73.286 | 78.421 | 69.562 | 68.375 | 77.609 | 70.962 | 75.496 | 88.559 | 69.916 | 79.232 | 83.711 | 75.204 | 76.141 | 83.726 | 76.650 | -0.128 | 0.0000 | 0.0002 | 1 | 0 | 0 |
| 'Ptpn12' | 11.589 | 10.754 | 9.456 | 10.955 | 9.689 | 9.112 | 10.184 | 10.186 | 8.645 | 9.850 | 10.803 | 10.973 | 9.526 | 10.347 | 10.193 | 9.026 | 11.516 | 11.447 | -0.062 | 0.2929 | 0.5493 | 0 | 0 | 0 |
| 'Ptpn13' | 28.161 | 27.128 | 26.562 | 29.596 | 28.212 | 25.007 | 27.334 | 27.767 | 26.997 | 28.386 | 30.254 | 26.071 | 29.653 | 28.328 | 26.291 | 27.048 | 29.662 | 27.598 | -0.048 | 0.1020 | 0.2994 | 0 | 0 | 0 |
| 'Ptpn18' | 9.769 | 10.383 | 10.486 | 9.549 | 9.867 | 10.281 | 8.938 | 10.323 | 11.039 | 9.454 | 9.319 | 10.348 | 9.267 | 9.480 | 10.388 | 10.461 | 9.405 | 10.003 | 0.031 | 0.3300 | 0.5880 | 0 | 0 | 0 |
| 'Ptpn2' | 15.093 | 16.856 | 15.088 | 16.811 | 16.526 | 16.158 | 16.514 | 16.447 | 16.118 | 16.441 | 16.068 | 15.437 | 15.273 | 16.074 | 14.209 | 14.629 | 15.815 | 14.183 | 0.064 | 0.0439 | 0.1733 | 0 | 0 | 0 |
| 'Ptpn21' | 1.677 | 2.050 | 2.170 | 1.487 | 1.737 | 1.806 | 1.856 | 2.048 | 2.221 | 2.417 | 2.236 | 1.890 | 2.053 | 2.074 | 2.042 | 1.829 | 1.579 | 1.811 | -0.078 | 0.3638 | 0.6158 | 0 | 0 | 0 |
| 'Ptpn22' | 50.544 | 51.665 | 48.720 | 52.882 | 51.656 | 47.121 | 49.456 | 52.980 | 49.661 | 55.300 | 54.003 | 52.420 | 50.561 | 53.912 | 51.170 | 52.401 | 58.380 | 55.422 | -0.100 | 0.0005 | 0.0061 | 1 | 0 | 0 |
| 'Ptpn23' | 11.355 | 10.404 | 12.130 | 10.255 | 10.115 | 11.897 | 10.625 | 10.847 | 12.783 | 12.522 | 11.501 | 12.274 | 12.306 | 10.220 | 11.817 | 11.044 | 11.695 | 12.614 | -0.088 | 0.0219 | 0.1092 | 0 | 0 | 0 |
| 'Ptpn3' | 3.027 | 3.238 | 3.451 | 3.621 | 3.245 | 3.380 | 3.200 | 3.086 | 3.392 | 3.135 | 3.287 | 3.702 | 2.802 | 3.016 | 3.467 | 2.831 | 3.379 | 3.698 | 0.002 | 0.9671 | 0.9887 | 0 | 0 | 0 |
| 'Ptpn4' | 3.387 | 3.849 | 3.769 | 3.977 | 3.868 | 3.645 | 4.090 | 3.703 | 3.520 | 3.623 | 4.346 | 3.689 | 3.224 | 3.905 | 3.450 | 3.573 | 4.055 | 3.534 | 0.010 | 0.8360 | 0.9293 | 0 | 0 | 0 |
| 'Ptpn5' | 2.107 | 3.086 | 1.837 | 1.392 | 2.637 | 2.254 | 1.861 | 3.048 | 2.066 | 2.157 | 2.166 | 2.213 | 1.666 | 2.717 | 1.852 | 2.948 | 2.083 | 2.167 | -0.009 | 0.9354 | 0.9752 | 0 | 0 | 0 |
| 'Ptpn6' | 53.531 | 51.296 | 54.266 | 53.550 | 54.303 | 54.776 | 54.965 | 51.010 | 54.334 | 50.513 | 46.028 | 61.845 | 46.747 | 48.456 | 57.554 | 51.323 | 52.477 | 60.334 | 0.014 | 0.7160 | 0.8688 | 0 | 0 | 0 |
| 'Ptpn7' | 42.047 | 38.061 | 43.862 | 42.674 | 38.131 | 44.563 | 43.516 | 38.338 | 43.408 | 44.553 | 45.396 | 56.079 | 43.503 | 45.505 | 56.458 | 46.342 | 45.211 | 57.938 | -0.241 | 0.0000 | 0.0000 | 1 | 0 | 0 |
| 'Ptpn9' | 9.054 | 7.818 | 9.872 | 9.041 | 9.149 | 9.776 | 8.544 | 8.179 | 9.002 | 8.572 | 8.467 | 9.142 | 8.305 | 8.810 | 9.224 | 8.390 | 8.080 | 8.918 | 0.039 | 0.3336 | 0.5919 | 0 | 0 | 0 |
| 'Ptpra' | 26.082 | 26.906 | 27.732 | 25.996 | 27.649 | 27.267 | 27.344 | 29.067 | 26.769 | 28.478 | 26.477 | 28.415 | 28.633 | 28.281 | 26.506 | 27.592 | 27.365 | 27.179 | -0.035 | 0.1830 | 0.4190 | 0 | 0 | 0 |
| 'Ptprc' | 121.316 | 134.891 | 123.564 | 133.171 | 133.387 | 119.518 | 131.308 | 128.689 | 123.868 | 135.453 | 143.964 | 112.853 | 130.398 | 139.605 | 121.087 | 131.996 | 138.850 | 120.680 | -0.039 | 0.1690 | 0.4013 | 0 | 0 | 0 |
| 'Ptprcap' | 466.467 | 534.542 | 493.882 | 464.838 | 524.313 | 544.524 | 478.219 | 500.700 | 527.618 | 512.685 | 457.693 | 450.250 | 506.033 | 478.730 | 518.420 | 538.335 | 459.465 | 478.070 | 0.033 | 0.4182 | 0.6655 | 0 | 0 | 0 |
| 'Ptpre' | 9.418 | 9.117 | 9.564 | 8.762 | 8.964 | 10.112 | 8.541 | 9.997 | 10.070 | 8.479 | 9.957 | 10.177 | 9.279 | 9.378 | 9.494 | 9.602 | 9.027 | 10.502 | -0.031 | 0.4206 | 0.6675 | 0 | 0 | 0 |
| 'Ptprf' | 0.744 | 0.843 | 0.842 | 0.752 | 0.707 | 1.026 | 0.810 | 0.723 | 0.802 | 1.148 | 0.720 | 0.980 | 1.063 | 0.764 | 1.095 | 0.932 | 0.840 | 0.921 | -0.230 | 0.0015 | 0.0146 | 1 | 0 | 0 |
| 'Ptprj' | 3.797 | 4.524 | 3.622 | 4.672 | 5.099 | 4.236 | 3.528 | 4.892 | 4.394 | 4.275 | 3.185 | 3.603 | 4.879 | 3.926 | 3.711 | 4.435 | 3.910 | 3.830 | 0.106 | 0.1737 | 0.4074 | 0 | 0 | 0 |
| 'Ptprs' | 4.151 | 3.419 | 4.474 | 3.504 | 3.660 | 4.013 | 4.346 | 3.822 | 4.408 | 4.709 | 3.390 | 4.778 | 3.873 | 3.261 | 4.323 | 4.277 | 3.556 | 4.412 | -0.037 | 0.4527 | 0.6916 | 0 | 0 | 0 |
| 'Ptrh1' | 38.439 | 42.324 | 41.328 | 37.700 | 41.110 | 39.125 | 39.706 | 42.050 | 44.315 | 43.176 | 45.474 | 42.333 | 49.495 | 44.012 | 45.194 | 47.029 | 46.613 | 42.358 | -0.157 | 0.0000 | 0.0004 | 1 | 0 | 0 |
| 'Ptrh2' | 5.536 | 5.386 | 5.790 | 5.828 | 5.818 | 5.278 | 5.848 | 4.906 | 6.197 | 4.623 | 4.676 | 5.574 | 5.070 | 5.021 | 4.805 | 4.696 | 6.164 | 5.822 | 0.107 | 0.1013 | 0.2981 | 0 | 0 | 0 |
| 'Ptrhd1' | 1.995 | 2.386 | 1.818 | 2.055 | 1.842 | 2.685 | 2.177 | 1.975 | 2.051 | 1.813 | 1.629 | 2.000 | 2.353 | 2.216 | 1.976 | 2.043 | 2.171 | 1.667 | 0.075 | 0.4070 | 0.6562 | 0 | 0 | 0 |
| 'Pts' | 23.643 | 33.883 | 27.187 | 26.523 | 35.004 | 32.168 | 29.245 | 32.088 | 28.704 | 29.151 | 30.427 | 28.326 | 30.475 | 35.430 | 29.193 | 30.792 | 29.764 | 26.045 | -0.020 | 0.7094 | 0.8653 | 0 | 0 | 0 |
| 'Pttg1' | 4.068 | 4.208 | 4.505 | 4.297 | 4.129 | 4.445 | 4.332 | 4.538 | 4.406 | 3.765 | 4.126 | 4.164 | 4.864 | 4.579 | 4.895 | 3.793 | 4.113 | 3.406 | 0.037 | 0.5617 | 0.7712 | 0 | 0 | 0 |
| 'Pttg1ip' | 14.757 | 16.233 | 14.179 | 14.467 | 15.597 | 14.464 | 13.105 | 15.537 | 13.879 | 15.211 | 13.144 | 14.977 | 14.035 | 15.193 | 13.792 | 14.995 | 14.905 | 14.109 | 0.007 | 0.8423 | 0.9332 | 0 | 0 | 0 |
| 'Puf60' | 125.074 | 124.042 | 132.534 | 118.824 | 127.758 | 140.844 | 121.920 | 121.589 | 134.490 | 130.923 | 120.197 | 128.539 | 129.064 | 123.028 | 127.476 | 139.276 | 125.325 | 128.119 | -0.016 | 0.5836 | 0.7856 | 0 | 0 | 0 |
| 'Pum1' | 19.238 | 17.636 | 16.043 | 17.939 | 16.407 | 15.943 | 15.964 | 17.681 | 16.924 | 17.862 | 16.402 | 17.946 | 15.837 | 17.632 | 16.781 | 17.244 | 17.129 | 18.312 | -0.024 | 0.5627 | 0.7718 | 0 | 0 | 0 |
| 'Pum2' | 16.743 | 17.158 | 15.376 | 16.655 | 17.287 | 15.282 | 16.267 | 17.165 | 15.311 | 16.717 | 17.298 | 16.171 | 15.820 | 17.946 | 15.713 | 16.478 | 16.873 | 15.963 | -0.028 | 0.1779 | 0.4131 | 0 | 0 | 0 |
| 'Pum3' | 19.128 | 17.631 | 19.793 | 19.196 | 20.163 | 19.301 | 20.453 | 19.825 | 18.669 | 18.752 | 19.671 | 19.057 | 19.404 | 16.111 | 17.804 | 19.429 | 19.790 | 17.761 | 0.044 | 0.2487 | 0.4994 | 0 | 0 | 0 |
| 'Pura' | 6.127 | 5.770 | 6.409 | 6.875 | 5.017 | 5.547 | 7.010 | 5.195 | 5.691 | 4.619 | 6.385 | 7.030 | 4.926 | 5.817 | 5.839 | 4.645 | 6.907 | 6.583 | 0.020 | 0.8451 | 0.9343 | 0 | 0 | 0 |
| 'Purb' | 35.257 | 36.544 | 36.825 | 36.917 | 34.770 | 35.334 | 36.128 | 35.353 | 34.097 | 35.328 | 38.869 | 37.756 | 33.567 | 37.013 | 36.024 | 35.252 | 35.900 | 38.627 | -0.042 | 0.1757 | 0.4107 | 0 | 0 | 0 |
| 'Purg' | 1.719 | 1.908 | 1.736 | 2.059 | 1.410 | 1.996 | 2.247 | 1.527 | 1.611 | 1.683 | 1.416 | 2.160 | 1.563 | 1.909 | 1.799 | 1.735 | 2.136 | 1.932 | -0.037 | 0.7255 | 0.8740 | 0 | 0 | 0 |
| 'Pus1' | 10.032 | 10.105 | 11.197 | 10.148 | 9.451 | 11.028 | 9.871 | 11.521 | 10.922 | 9.522 | 8.804 | 11.204 | 10.416 | 10.477 | 11.536 | 10.855 | 9.875 | 10.140 | 0.014 | 0.7583 | 0.8918 | 0 | 0 | 0 |
| 'Pus10' | 4.918 | 5.118 | 5.273 | 4.426 | 5.285 | 4.784 | 4.853 | 5.284 | 5.057 | 5.261 | 5.011 | 4.432 | 5.349 | 4.782 | 4.579 | 5.034 | 5.157 | 4.023 | 0.043 | 0.3861 | 0.6355 | 0 | 0 | 0 |
| 'Pus7' | 2.486 | 1.950 | 2.528 | 2.097 | 2.070 | 2.035 | 2.156 | 2.326 | 2.127 | 2.336 | 2.121 | 2.584 | 2.404 | 1.706 | 2.458 | 2.211 | 2.151 | 2.549 | -0.060 | 0.3300 | 0.5880 | 0 | 0 | 0 |
| 'Pus7l' | 4.107 | 4.230 | 4.504 | 3.669 | 4.896 | 4.424 | 3.968 | 4.729 | 4.502 | 4.769 | 4.144 | 4.070 | 4.156 | 3.868 | 4.387 | 3.885 | 4.280 | 3.543 | 0.065 | 0.2724 | 0.5275 | 0 | 0 | 0 |
| 'Pusl1' | 3.300 | 2.976 | 3.799 | 2.842 | 3.213 | 3.594 | 3.082 | 2.994 | 3.775 | 3.247 | 2.339 | 3.696 | 3.143 | 2.956 | 3.557 | 3.239 | 2.819 | 3.450 | 0.046 | 0.4271 | 0.6735 | 0 | 0 | 0 |
| 'Pvr' | 12.985 | 17.108 | 11.862 | 12.405 | 14.832 | 12.363 | 12.346 | 14.009 | 12.896 | 12.980 | 13.383 | 11.794 | 14.117 | 14.195 | 12.288 | 14.272 | 13.912 | 11.653 | 0.013 | 0.7623 | 0.8944 | 0 | 0 | 0 |
| 'Pwp1' | 8.558 | 9.198 | 9.031 | 8.032 | 9.591 | 8.568 | 8.913 | 9.831 | 8.470 | 9.019 | 8.646 | 8.774 | 9.040 | 8.752 | 8.438 | 7.863 | 8.333 | 8.570 | 0.036 | 0.3629 | 0.6149 | 0 | 0 | 0 |
| 'Pwp2' | 9.992 | 9.424 | 11.442 | 9.981 | 10.823 | 11.369 | 10.195 | 11.212 | 11.131 | 11.169 | 9.803 | 11.113 | 9.417 | 9.321 | 10.521 | 10.410 | 10.119 | 10.729 | 0.038 | 0.3612 | 0.6135 | 0 | 0 | 0 |
| 'Pwwp2a' | 4.979 | 4.813 | 4.847 | 4.648 | 4.476 | 5.072 | 5.066 | 4.927 | 4.620 | 4.330 | 5.175 | 5.950 | 4.576 | 5.405 | 6.275 | 5.080 | 5.532 | 6.016 | -0.159 | 0.0029 | 0.0243 | 1 | 0 | 0 |
| 'Pxk' | 21.827 | 19.706 | 19.614 | 21.252 | 23.127 | 21.166 | 20.317 | 22.508 | 20.139 | 21.476 | 19.599 | 21.154 | 21.899 | 21.111 | 20.020 | 20.673 | 23.406 | 20.991 | -0.017 | 0.6508 | 0.8303 | 0 | 0 | 0 |
| 'Pxmp2' | 2.984 | 4.504 | 3.115 | 3.875 | 3.692 | 3.648 | 3.398 | 4.098 | 3.556 | 3.706 | 5.298 | 3.935 | 4.213 | 4.422 | 4.776 | 3.913 | 4.884 | 3.743 | -0.255 | 0.0006 | 0.0073 | 1 | 0 | 0 |
| 'Pxmp4' | 13.370 | 16.230 | 14.974 | 12.877 | 15.167 | 14.331 | 13.059 | 16.164 | 14.252 | 12.520 | 16.741 | 13.443 | 13.444 | 13.751 | 14.274 | 12.683 | 14.037 | 14.441 | 0.048 | 0.2425 | 0.4930 | 0 | 0 | 0 |
| 'Pxn' | 8.906 | 8.411 | 8.513 | 8.553 | 8.263 | 9.274 | 7.955 | 8.474 | 8.998 | 8.600 | 8.030 | 8.581 | 8.972 | 8.178 | 8.355 | 8.242 | 8.019 | 8.735 | 0.021 | 0.4592 | 0.6960 | 0 | 0 | 0 |
| 'Pxylp1' | 0.486 | 0.651 | 0.531 | 0.396 | 0.544 | 0.565 | 0.545 | 0.469 | 0.521 | 0.479 | 0.548 | 0.527 | 0.513 | 0.616 | 0.554 | 0.551 | 0.518 | 0.537 | -0.051 | 0.5448 | 0.7606 | 0 | 0 | 0 |
| 'Pycard' | 11.444 | 11.501 | 11.994 | 11.884 | 11.957 | 11.803 | 11.578 | 12.295 | 11.892 | 12.003 | 10.966 | 11.287 | 11.710 | 12.369 | 12.178 | 11.538 | 10.415 | 10.643 | 0.037 | 0.3147 | 0.5723 | 0 | 0 | 0 |
| 'Pycr2' | 13.167 | 13.476 | 14.969 | 13.664 | 12.850 | 14.907 | 13.288 | 13.670 | 13.707 | 11.601 | 10.200 | 12.551 | 11.882 | 10.134 | 11.681 | 11.354 | 11.060 | 11.563 | 0.266 | 0.0000 | 0.0000 | 1 | 0 | 0 |
| 'Pycrl' | 17.658 | 16.758 | 18.910 | 18.548 | 16.702 | 19.670 | 18.747 | 16.537 | 20.101 | 17.172 | 15.362 | 17.696 | 18.109 | 15.860 | 18.299 | 16.137 | 16.999 | 16.217 | 0.098 | 0.0319 | 0.1405 | 0 | 0 | 0 |
| 'Pygb' | 26.292 | 24.664 | 26.677 | 26.126 | 25.488 | 27.563 | 24.983 | 24.115 | 25.814 | 28.855 | 26.761 | 29.744 | 25.745 | 26.043 | 30.257 | 26.349 | 25.231 | 29.944 | -0.113 | 0.0002 | 0.0027 | 1 | 0 | 0 |
| 'Pygo2' | 14.766 | 14.977 | 16.231 | 15.389 | 15.406 | 16.242 | 13.554 | 15.015 | 16.200 | 13.678 | 14.570 | 18.966 | 14.214 | 14.148 | 18.865 | 14.987 | 14.854 | 18.794 | -0.059 | 0.1497 | 0.3767 | 0 | 0 | 0 |
| 'Pym1' | 42.237 | 43.933 | 45.433 | 42.012 | 43.096 | 48.241 | 44.751 | 43.932 | 46.120 | 42.377 | 41.379 | 43.333 | 40.712 | 38.756 | 43.323 | 43.351 | 36.852 | 38.242 | 0.109 | 0.0008 | 0.0084 | 1 | 0 | 0 |
| 'Pyroxd1' | 2.031 | 2.434 | 2.604 | 2.063 | 2.862 | 2.381 | 1.919 | 2.539 | 2.554 | 2.545 | 2.085 | 2.293 | 2.455 | 2.068 | 2.069 | 2.318 | 2.363 | 1.893 | 0.077 | 0.2820 | 0.5382 | 0 | 0 | 0 |
| 'Pyurf' | 2.856 | 3.052 | 2.727 | 3.090 | 3.253 | 2.789 | 2.810 | 2.641 | 2.672 | 2.771 | 3.415 | 3.050 | 2.612 | 3.023 | 2.846 | 2.616 | 2.809 | 3.012 | -0.033 | 0.6327 | 0.8178 | 0 | 0 | 0 |
| 'Qars' | 12.398 | 12.312 | 13.188 | 12.826 | 12.714 | 12.670 | 12.849 | 12.526 | 12.973 | 12.416 | 13.170 | 12.676 | 13.450 | 12.014 | 12.658 | 12.158 | 12.831 | 11.953 | 0.004 | 0.8713 | 0.9448 | 0 | 0 | 0 |
| 'Qk' | 7.561 | 8.735 | 7.667 | 7.879 | 8.386 | 7.107 | 8.629 | 8.254 | 7.316 | 7.104 | 7.761 | 8.113 | 7.452 | 8.513 | 7.568 | 7.103 | 8.162 | 8.184 | 0.016 | 0.6884 | 0.8527 | 0 | 0 | 0 |
| 'Qpctl' | 4.129 | 4.405 | 4.743 | 3.585 | 4.635 | 4.851 | 4.128 | 4.574 | 5.082 | 4.462 | 4.063 | 3.947 | 4.336 | 4.368 | 3.841 | 4.333 | 4.172 | 3.386 | 0.117 | 0.0672 | 0.2272 | 0 | 0 | 0 |
| 'Qprt' | 5.266 | 4.568 | 5.669 | 4.333 | 5.083 | 5.404 | 5.201 | 5.022 | 5.579 | 5.130 | 4.164 | 4.877 | 4.629 | 4.083 | 4.357 | 5.250 | 3.988 | 3.615 | 0.200 | 0.0035 | 0.0280 | 1 | 0 | 0 |
| 'Qrich1' | 12.831 | 12.584 | 12.908 | 12.782 | 12.232 | 12.280 | 12.701 | 12.899 | 12.637 | 12.840 | 12.139 | 12.341 | 12.213 | 12.371 | 12.221 | 11.646 | 12.810 | 11.986 | 0.032 | 0.1725 | 0.4063 | 0 | 0 | 0 |
| 'Qrsl1' | 4.124 | 4.445 | 4.921 | 4.156 | 4.507 | 4.773 | 3.973 | 4.894 | 4.515 | 3.751 | 4.355 | 4.245 | 4.165 | 4.603 | 3.864 | 3.721 | 4.970 | 4.118 | 0.089 | 0.0752 | 0.2440 | 0 | 0 | 0 |
| 'Qsox1' | 8.466 | 8.011 | 8.578 | 7.430 | 7.726 | 7.809 | 7.936 | 8.375 | 7.883 | 8.618 | 8.399 | 8.878 | 8.272 | 7.589 | 8.820 | 7.945 | 8.061 | 9.020 | -0.076 | 0.0323 | 0.1417 | 0 | 0 | 0 |
| 'Qsox2' | 3.631 | 4.099 | 4.124 | 3.237 | 4.727 | 4.083 | 3.017 | 4.432 | 3.933 | 4.489 | 3.841 | 3.644 | 4.476 | 3.769 | 3.950 | 4.326 | 4.083 | 2.940 | -0.017 | 0.8286 | 0.9258 | 0 | 0 | 0 |
| 'Qtrt1' | 16.706 | 13.978 | 18.133 | 14.228 | 15.991 | 15.241 | 14.408 | 14.388 | 15.476 | 14.194 | 13.192 | 16.431 | 15.390 | 12.201 | 16.331 | 15.508 | 15.747 | 14.734 | 0.042 | 0.4656 | 0.7014 | 0 | 0 | 0 |
| 'Qtrtd1' | 2.121 | 2.145 | 2.174 | 1.884 | 2.278 | 2.106 | 2.006 | 2.290 | 2.146 | 2.194 | 1.989 | 2.250 | 2.200 | 2.107 | 2.120 | 2.330 | 2.265 | 2.149 | -0.044 | 0.3274 | 0.5852 | 0 | 0 | 0 |
| 'R3hcc1' | 6.849 | 8.149 | 8.546 | 7.662 | 8.422 | 8.654 | 7.367 | 8.413 | 8.663 | 9.028 | 6.846 | 7.337 | 8.634 | 7.650 | 7.396 | 8.364 | 7.206 | 7.079 | 0.059 | 0.3201 | 0.5782 | 0 | 0 | 0 |
| 'R3hcc1l' | 5.864 | 4.793 | 5.316 | 6.290 | 5.063 | 5.419 | 5.983 | 5.060 | 5.216 | 5.273 | 5.800 | 5.971 | 5.025 | 4.830 | 5.315 | 4.680 | 5.742 | 5.225 | 0.019 | 0.7443 | 0.8847 | 0 | 0 | 0 |
| 'R3hdm1' | 2.976 | 2.539 | 2.881 | 2.721 | 2.546 | 2.612 | 2.749 | 2.748 | 2.645 | 2.732 | 2.734 | 3.047 | 2.840 | 2.548 | 2.620 | 2.622 | 2.679 | 2.842 | -0.026 | 0.4770 | 0.7107 | 0 | 0 | 0 |
| 'R3hdm2' | 7.665 | 7.462 | 8.113 | 7.385 | 7.394 | 8.005 | 7.719 | 7.687 | 7.974 | 8.632 | 7.621 | 8.273 | 8.362 | 7.654 | 7.964 | 8.113 | 7.525 | 7.439 | -0.053 | 0.0498 | 0.1882 | 0 | 0 | 0 |
| 'R3hdm4' | 47.368 | 52.987 | 50.100 | 47.096 | 49.712 | 55.481 | 46.242 | 52.875 | 53.815 | 51.141 | 46.886 | 50.061 | 49.292 | 47.102 | 52.797 | 52.902 | 44.170 | 49.437 | 0.028 | 0.4554 | 0.6934 | 0 | 0 | 0 |
| 'Rab10' | 64.660 | 68.466 | 61.875 | 62.528 | 68.900 | 63.033 | 60.851 | 68.825 | 62.398 | 67.998 | 62.922 | 63.467 | 65.933 | 69.195 | 66.377 | 68.187 | 65.073 | 65.083 | -0.043 | 0.1110 | 0.3154 | 0 | 0 | 0 |
| 'Rab11a' | 23.939 | 27.400 | 21.484 | 24.578 | 26.941 | 21.817 | 24.810 | 25.019 | 22.172 | 21.985 | 25.195 | 21.799 | 23.465 | 26.099 | 22.928 | 23.447 | 27.146 | 20.642 | 0.024 | 0.4368 | 0.6801 | 0 | 0 | 0 |
| 'Rab11b' | 17.133 | 20.072 | 19.179 | 17.404 | 19.281 | 20.035 | 17.961 | 19.298 | 19.599 | 20.968 | 19.071 | 19.193 | 19.343 | 19.930 | 18.155 | 20.269 | 19.012 | 18.162 | -0.045 | 0.2113 | 0.4566 | 0 | 0 | 0 |
| 'Rab11fip1' | 7.870 | 7.397 | 7.347 | 6.727 | 6.764 | 7.152 | 6.542 | 7.168 | 7.231 | 7.288 | 6.237 | 8.426 | 6.858 | 6.910 | 8.395 | 7.695 | 7.132 | 8.376 | -0.076 | 0.1109 | 0.3152 | 0 | 0 | 0 |
| 'Rab11fip2' | 1.919 | 2.747 | 2.450 | 2.243 | 2.671 | 2.406 | 2.242 | 2.823 | 2.383 | 3.024 | 2.391 | 2.105 | 2.439 | 2.143 | 2.168 | 3.208 | 2.356 | 1.834 | 0.014 | 0.8667 | 0.9424 | 0 | 0 | 0 |
| 'Rab11fip3' | 3.744 | 3.232 | 4.197 | 3.869 | 3.237 | 3.967 | 3.640 | 3.220 | 3.596 | 3.955 | 4.024 | 4.782 | 3.628 | 3.654 | 4.433 | 3.957 | 3.461 | 4.802 | -0.175 | 0.0001 | 0.0020 | 1 | 0 | 0 |
| 'Rab11fip4' | 20.184 | 20.028 | 18.979 | 18.517 | 19.595 | 18.319 | 17.933 | 19.516 | 18.351 | 19.526 | 20.233 | 20.319 | 18.257 | 19.859 | 20.153 | 20.372 | 19.595 | 20.557 | -0.072 | 0.0218 | 0.1089 | 0 | 0 | 0 |
| 'Rab12' | 7.206 | 8.173 | 8.080 | 7.214 | 7.762 | 7.682 | 7.819 | 8.952 | 7.939 | 8.567 | 7.696 | 7.737 | 7.790 | 7.675 | 6.861 | 8.494 | 7.858 | 7.639 | 0.003 | 0.9446 | 0.9789 | 0 | 0 | 0 |
| 'Rab14' | 7.621 | 7.969 | 7.510 | 7.386 | 7.858 | 7.064 | 7.415 | 7.741 | 7.546 | 7.105 | 7.893 | 8.193 | 7.805 | 8.273 | 7.994 | 7.574 | 7.946 | 7.830 | -0.065 | 0.0167 | 0.0910 | 0 | 0 | 0 |
| 'Rab18' | 29.829 | 37.298 | 32.253 | 33.064 | 37.539 | 31.213 | 30.455 | 35.210 | 31.864 | 30.167 | 33.209 | 31.825 | 31.384 | 36.987 | 30.236 | 32.450 | 34.710 | 29.535 | 0.027 | 0.3750 | 0.6262 | 0 | 0 | 0 |
| 'Rab19' | 38.879 | 38.455 | 41.985 | 40.513 | 35.437 | 40.413 | 42.037 | 38.545 | 41.236 | 38.098 | 38.341 | 41.490 | 39.118 | 35.833 | 43.767 | 37.794 | 39.031 | 40.036 | 0.005 | 0.8737 | 0.9462 | 0 | 0 | 0 |
| 'Rab1a' | 24.060 | 28.408 | 25.046 | 26.086 | 28.668 | 25.043 | 27.931 | 27.163 | 25.565 | 22.905 | 27.285 | 26.165 | 23.908 | 27.733 | 25.399 | 25.103 | 27.986 | 26.508 | 0.018 | 0.5927 | 0.7924 | 0 | 0 | 0 |
| 'Rab1b' | 109.750 | 116.335 | 121.540 | 108.329 | 112.203 | 134.888 | 106.156 | 118.935 | 121.342 | 114.433 | 104.045 | 119.496 | 116.547 | 104.643 | 123.716 | 121.282 | 105.516 | 117.182 | 0.021 | 0.5275 | 0.7486 | 0 | 0 | 0 |
| 'Rab20' | 3.970 | 3.689 | 4.180 | 3.308 | 3.770 | 3.864 | 4.187 | 4.288 | 2.997 | 4.145 | 3.593 | 3.786 | 3.321 | 3.456 | 4.092 | 3.868 | 4.194 | 4.733 | -0.044 | 0.6520 | 0.8310 | 0 | 0 | 0 |
| 'Rab21' | 53.089 | 57.324 | 50.251 | 53.209 | 53.677 | 49.861 | 54.226 | 50.188 | 48.339 | 44.072 | 53.985 | 52.688 | 47.237 | 54.016 | 52.308 | 49.042 | 54.519 | 55.996 | 0.009 | 0.8484 | 0.9350 | 0 | 0 | 0 |
| 'Rab22a' | 6.481 | 6.725 | 6.043 | 6.370 | 6.234 | 6.241 | 6.659 | 6.738 | 6.419 | 6.149 | 6.391 | 7.128 | 5.882 | 6.591 | 6.705 | 6.031 | 6.880 | 7.157 | -0.036 | 0.3578 | 0.6103 | 0 | 0 | 0 |
| 'Rab23' | 0.741 | 0.704 | 0.730 | 0.795 | 0.728 | 0.734 | 0.713 | 0.715 | 0.883 | 0.633 | 0.829 | 0.894 | 0.726 | 0.718 | 0.885 | 0.745 | 0.822 | 0.798 | -0.077 | 0.2117 | 0.4569 | 0 | 0 | 0 |
| 'Rab24' | 44.649 | 47.536 | 50.237 | 45.578 | 50.968 | 52.105 | 45.527 | 50.966 | 49.035 | 46.077 | 44.284 | 50.542 | 52.230 | 51.736 | 48.542 | 48.143 | 47.947 | 49.469 | -0.019 | 0.5639 | 0.7730 | 0 | 0 | 0 |
| 'Rab27a' | 45.897 | 51.810 | 44.071 | 49.719 | 53.317 | 46.168 | 48.096 | 48.809 | 45.839 | 44.313 | 50.997 | 42.874 | 52.336 | 49.672 | 43.949 | 43.482 | 49.561 | 46.515 | 0.022 | 0.5171 | 0.7406 | 0 | 0 | 0 |
| 'Rab28' | 4.666 | 5.020 | 4.440 | 4.851 | 5.432 | 4.284 | 5.267 | 5.267 | 4.513 | 4.599 | 4.909 | 4.974 | 4.731 | 4.964 | 4.878 | 4.424 | 5.018 | 4.933 | -0.011 | 0.8070 | 0.9135 | 0 | 0 | 0 |
| 'Rab29' | 5.296 | 6.554 | 6.335 | 5.735 | 5.635 | 6.033 | 5.685 | 5.731 | 6.050 | 5.269 | 5.399 | 5.498 | 5.784 | 6.460 | 5.658 | 5.564 | 5.323 | 6.595 | 0.032 | 0.5820 | 0.7842 | 0 | 0 | 0 |
| 'Rab2a' | 43.831 | 49.099 | 44.158 | 44.952 | 46.143 | 43.811 | 45.547 | 47.637 | 44.262 | 39.617 | 45.354 | 46.670 | 40.403 | 46.334 | 47.708 | 44.168 | 47.414 | 44.418 | 0.014 | 0.6618 | 0.8369 | 0 | 0 | 0 |
| 'Rab2b' | 2.453 | 2.646 | 2.671 | 2.391 | 2.747 | 2.393 | 2.555 | 2.784 | 2.717 | 2.502 | 2.595 | 2.466 | 2.537 | 2.737 | 2.490 | 2.635 | 2.609 | 2.250 | 0.028 | 0.5225 | 0.7449 | 0 | 0 | 0 |
| 'Rab31' | 1.753 | 1.604 | 1.921 | 1.504 | 1.880 | 1.476 | 1.434 | 1.692 | 1.501 | 1.590 | 1.587 | 1.620 | 1.320 | 1.587 | 1.538 | 1.456 | 1.465 | 1.324 | 0.123 | 0.1825 | 0.4187 | 0 | 0 | 0 |

| | | | | | | | | | | | | | | | | | | | | | | | | |
|---|---|---|---|---|---|---|---|---|---|---|---|---|---|---|---|---|---|---|---|---|---|---|---|---|
| 'Rab32' | 1.101 | 1.303 | 2.176 | 1.397 | 1.804 | 1.503 | 1.418 | 1.918 | 2.205 | 2.492 | 2.951 | 2.705 | 2.640 | 2.370 | 3.188 | 2.798 | 2.215 | 2.156 | -0.672 | 0.0000 | 0.0000 | 1 | 1 | 0 |
| 'Rab33b' | 6.133 | 7.837 | 6.004 | 6.132 | 7.904 | 6.745 | 5.838 | 7.558 | 6.314 | 7.152 | 7.138 | 5.750 | 6.379 | 7.125 | 5.702 | 7.492 | 6.787 | 5.559 | 0.027 | 0.6124 | 0.8056 | 0 | 0 | 0 |
| 'Rab35' | 40.821 | 40.968 | 40.956 | 39.420 | 40.760 | 44.282 | 39.088 | 39.997 | 41.209 | 39.585 | 39.376 | 43.696 | 40.558 | 41.614 | 44.685 | 42.147 | 39.131 | 44.797 | -0.041 | 0.1131 | 0.3192 | 0 | 0 | 0 |
| 'Rab37' | 33.577 | 41.315 | 38.715 | 34.025 | 40.629 | 40.621 | 36.213 | 41.057 | 41.850 | 39.916 | 35.194 | 34.475 | 39.638 | 37.551 | 37.236 | 40.244 | 35.297 | 32.929 | 0.054 | 0.2649 | 0.5184 | 0 | 0 | 0 |
| 'Rab39b' | 3.444 | 3.899 | 3.846 | 3.895 | 4.493 | 3.540 | 3.853 | 4.344 | 3.620 | 3.228 | 3.982 | 3.509 | 3.690 | 4.084 | 3.446 | 3.514 | 4.058 | 3.241 | 0.082 | 0.1650 | 0.3974 | 0 | 0 | 0 |
| 'Rab3a' | 4.391 | 4.842 | 5.324 | 4.619 | 4.986 | 5.079 | 4.849 | 5.705 | 6.223 | 5.344 | 4.696 | 5.091 | 5.208 | 4.726 | 4.827 | 5.364 | 4.555 | 4.661 | 0.044 | 0.5427 | 0.7595 | 0 | 0 | 0 |
| 'Rab3d' | 1.587 | 1.872 | 1.853 | 1.498 | 1.762 | 2.224 | 1.739 | 2.107 | 1.962 | 1.599 | 1.271 | 1.629 | 1.290 | 1.496 | 1.801 | 1.497 | 1.566 | 1.479 | 0.276 | 0.0000 | 0.0008 | 1 | 0 | 0 |
| 'Rab3gap1' | 9.791 | 10.255 | 10.423 | 10.407 | 10.486 | 10.354 | 10.668 | 10.698 | 10.330 | 10.040 | 10.580 | 10.947 | 9.758 | 10.079 | 10.162 | 10.743 | 10.396 | 10.110 | -0.002 | 0.9336 | 0.9746 | 0 | 0 | 0 |
| 'Rab3gap2' | 7.193 | 7.402 | 7.111 | 7.084 | 7.172 | 6.743 | 7.220 | 7.260 | 6.774 | 7.220 | 7.322 | 6.603 | 7.129 | 6.312 | 6.239 | 6.154 | 6.852 | 6.373 | 0.076 | 0.0217 | 0.1088 | 0 | 0 | 0 |
| 'Rab3ip' | 5.314 | 5.346 | 5.068 | 5.356 | 5.102 | 4.802 | 5.374 | 6.010 | 4.675 | 5.013 | 5.547 | 6.009 | 5.036 | 5.251 | 6.100 | 5.474 | 5.718 | 6.170 | -0.113 | 0.0687 | 0.2303 | 0 | 0 | 0 |
| 'Rab40c' | 4.180 | 3.989 | 4.199 | 4.119 | 3.789 | 3.964 | 4.030 | 4.215 | 4.343 | 4.187 | 4.724 | 4.448 | 3.904 | 3.873 | 4.385 | 4.054 | 4.265 | 4.147 | -0.053 | 0.2126 | 0.4576 | 0 | 0 | 0 |
| 'Rab43' | 2.671 | 3.032 | 2.818 | 2.299 | 3.113 | 3.360 | 2.375 | 3.112 | 2.886 | 3.037 | 2.899 | 2.622 | 3.256 | 3.197 | 2.820 | 3.082 | 2.601 | 2.965 | -0.054 | 0.4259 | 0.6724 | 0 | 0 | 0 |
| 'Rab4a' | 9.723 | 10.082 | 9.182 | 10.086 | 10.228 | 9.716 | 9.771 | 9.229 | 9.112 | 8.993 | 7.588 | 8.473 | 8.657 | 8.374 | 9.199 | 8.499 | 7.242 | 7.684 | 0.206 | 0.0001 | 0.0017 | 1 | 0 | 0 |
| 'Rab4b' | 7.570 | 9.174 | 8.857 | 8.077 | 9.476 | 8.985 | 7.994 | 8.933 | 8.748 | 8.699 | 8.259 | 8.479 | 9.487 | 8.551 | 8.659 | 9.167 | 8.381 | 8.037 | -0.009 | 0.8331 | 0.9272 | 0 | 0 | 0 |
| 'Rab5a' | 32.408 | 34.082 | 31.602 | 33.632 | 34.272 | 30.400 | 35.954 | 33.750 | 28.090 | 30.283 | 32.088 | 32.081 | 29.966 | 35.510 | 30.562 | 31.925 | 34.232 | 32.321 | 0.011 | 0.7785 | 0.9002 | 0 | 0 | 0 |
| 'Rab5b' | 27.887 | 36.395 | 30.763 | 30.474 | 33.958 | 34.387 | 29.410 | 33.851 | 31.237 | 29.988 | 32.699 | 34.428 | 29.858 | 35.689 | 34.890 | 32.180 | 31.827 | 34.004 | -0.049 | 0.1538 | 0.3816 | 0 | 0 | 0 |
| 'Rab5c' | 61.667 | 68.787 | 61.741 | 59.741 | 67.827 | 66.136 | 60.557 | 63.654 | 63.815 | 63.408 | 59.203 | 61.521 | 62.369 | 61.617 | 62.986 | 64.116 | 61.091 | 60.831 | 0.031 | 0.2492 | 0.5001 | 0 | 0 | 0 |
| 'Rab6a' | 18.516 | 19.812 | 18.250 | 18.047 | 21.043 | 18.690 | 18.857 | 18.801 | 17.736 | 19.971 | 20.439 | 20.652 | 18.925 | 23.323 | 21.475 | 20.349 | 22.105 | 21.079 | -0.161 | 0.0000 | 0.0000 | 1 | 0 | 0 |
| 'Rab7' | 75.425 | 75.895 | 71.179 | 75.338 | 77.202 | 77.836 | 75.797 | 77.888 | 73.701 | 89.148 | 87.949 | 84.068 | 85.854 | 89.566 | 87.911 | 90.824 | 90.389 | 89.143 | -0.234 | 0.0000 | 0.0000 | 1 | 0 | 0 |
| 'Rab8a' | 34.701 | 37.106 | 35.515 | 37.527 | 38.292 | 36.422 | 36.701 | 37.865 | 35.303 | 36.751 | 35.752 | 35.270 | 37.214 | 36.057 | 37.073 | 36.750 | 36.264 | 35.706 | 0.000 | 0.9916 | 0.9979 | 0 | 0 | 0 |
| 'Rab8b' | 91.657 | 103.077 | 93.220 | 98.861 | 98.937 | 90.329 | 100.226 | 100.071 | 93.330 | 94.061 | 96.566 | 91.511 | 88.928 | 98.028 | 88.338 | 96.708 | 99.311 | 91.122 | 0.031 | 0.1675 | 0.3999 | 0 | 0 | 0 |
| 'Rab9' | 7.939 | 7.995 | 8.208 | 8.273 | 7.869 | 7.416 | 7.869 | 8.388 | 7.378 | 6.077 | 8.023 | 8.243 | 6.750 | 7.803 | 7.261 | 8.135 | 8.838 | 7.396 | 0.048 | 0.4174 | 0.6650 | 0 | 0 | 0 |
| 'Rabac1' | 26.151 | 30.876 | 23.880 | 25.130 | 28.254 | 25.913 | 21.916 | 29.238 | 24.878 | 24.939 | 24.700 | 25.025 | 22.516 | 25.175 | 24.571 | 27.339 | 24.482 | 24.233 | 0.068 | 0.1426 | 0.3660 | 0 | 0 | 0 |
| 'Rabep1' | 5.245 | 5.617 | 5.244 | 5.605 | 5.915 | 5.245 | 5.544 | 5.663 | 5.437 | 5.160 | 5.337 | 5.718 | 5.128 | 5.372 | 5.364 | 4.883 | 5.710 | 5.480 | 0.024 | 0.4598 | 0.6965 | 0 | 0 | 0 |
| 'Rabep2' | 8.566 | 8.950 | 11.664 | 7.877 | 8.923 | 12.391 | 8.444 | 9.206 | 11.042 | 9.184 | 8.319 | 9.767 | 9.565 | 8.685 | 9.120 | 9.489 | 8.713 | 8.596 | 0.075 | 0.1798 | 0.4151 | 0 | 0 | 0 |
| 'Rabepk' | 4.958 | 5.111 | 4.791 | 5.243 | 5.160 | 5.110 | 4.992 | 5.094 | 5.040 | 4.421 | 4.354 | 4.922 | 4.906 | 5.049 | 5.057 | 4.601 | 4.398 | 4.892 | 0.079 | 0.0656 | 0.2235 | 0 | 0 | 0 |
| 'Rabgap1' | 3.390 | 3.638 | 3.572 | 3.600 | 3.712 | 3.543 | 3.595 | 3.737 | 3.444 | 3.298 | 3.629 | 3.662 | 3.338 | 3.690 | 3.408 | 3.432 | 3.640 | 3.561 | 0.013 | 0.6623 | 0.8370 | 0 | 0 | 0 |
| 'Rabgap1l' | 11.162 | 13.173 | 10.072 | 10.951 | 12.114 | 10.285 | 11.210 | 12.178 | 10.651 | 13.516 | 13.136 | 11.000 | 12.917 | 13.372 | 10.658 | 13.073 | 12.810 | 10.955 | -0.140 | 0.0000 | 0.0000 | 1 | 0 | 0 |
| 'Rabgef1' | 5.048 | 5.611 | 4.894 | 5.289 | 4.918 | 5.124 | 5.207 | 5.000 | 5.110 | 5.399 | 5.957 | 6.221 | 5.255 | 6.031 | 5.525 | 5.406 | 6.489 | 5.570 | -0.180 | 0.0000 | 0.0004 | 1 | 0 | 0 |
| 'Rabggta' | 23.387 | 23.577 | 23.023 | 21.438 | 23.894 | 23.544 | 22.364 | 24.055 | 22.708 | 22.723 | 23.596 | 24.170 | 24.274 | 22.343 | 23.090 | 23.189 | 25.463 | 21.523 | -0.027 | 0.3728 | 0.6242 | 0 | 0 | 0 |
| 'Rabggtb' | 9.728 | 8.987 | 8.727 | 10.255 | 9.142 | 8.704 | 10.516 | 9.430 | 8.497 | 8.318 | 9.438 | 8.854 | 10.668 | 9.900 | 8.406 | 8.862 | 10.853 | 8.252 | -0.004 | 0.9383 | 0.9764 | 0 | 0 | 0 |
| 'Rabif' | 14.020 | 16.274 | 15.118 | 14.337 | 17.143 | 15.945 | 14.580 | 15.623 | 15.340 | 13.450 | 16.690 | 16.770 | 14.723 | 15.861 | 16.162 | 15.220 | 15.385 | 15.559 | -0.030 | 0.4002 | 0.6491 | 0 | 0 | 0 |
| 'Rabl3' | 1.765 | 2.143 | 2.177 | 2.078 | 2.532 | 1.927 | 1.871 | 2.203 | 2.333 | 2.237 | 2.269 | 1.974 | 2.172 | 2.194 | 1.788 | 2.190 | 2.065 | 1.826 | 0.017 | 0.7875 | 0.9048 | 0 | 0 | 0 |
| 'Rabl6' | 7.957 | 7.427 | 7.949 | 7.322 | 7.119 | 8.299 | 7.871 | 7.553 | 8.199 | 8.096 | 7.094 | 8.086 | 8.583 | 7.162 | 8.024 | 8.860 | 7.306 | 8.411 | -0.046 | 0.1386 | 0.3594 | 0 | 0 | 0 |
| 'Rac1' | 36.423 | 35.502 | 39.142 | 36.605 | 38.591 | 37.922 | 36.850 | 39.160 | 32.887 | 35.469 | 37.496 | 32.629 | 37.138 | 38.465 | 37.455 | 35.919 | 37.095 | 38.598 | 0.002 | 0.9552 | 0.9834 | 0 | 0 | 0 |
| 'Rac2' | 379.808 | 419.650 | 382.894 | 379.340 | 409.405 | 405.993 | 385.454 | 398.027 | 405.655 | 392.170 | 381.268 | 351.424 | 392.095 | 389.705 | 392.955 | 402.126 | 381.431 | 373.157 | 0.034 | 0.2124 | 0.4575 | 0 | 0 | 0 |
| 'Racgap1' | 6.819 | 6.961 | 8.414 | 7.568 | 7.578 | 8.224 | 7.409 | 7.415 | 8.535 | 6.031 | 7.109 | 7.302 | 8.114 | 8.063 | 8.206 | 6.861 | 7.265 | 6.996 | 0.052 | 0.3034 | 0.5608 | 0 | 0 | 0 |
| 'Rack1' | 111.100 | 110.631 | 105.806 | 114.833 | 110.271 | 107.176 | 115.462 | 107.469 | 105.964 | 106.135 | 110.839 | 97.482 | 108.991 | 105.526 | 110.118 | 108.778 | 107.044 | 105.398 | 0.031 | 0.2350 | 0.4842 | 0 | 0 | 0 |
| 'Rad1' | 1.413 | 1.538 | 1.551 | 1.502 | 1.561 | 1.636 | 1.371 | 1.619 | 1.553 | 1.489 | 1.628 | 1.878 | 1.452 | 1.592 | 1.751 | 1.722 | 1.692 | 1.724 | -0.133 | 0.0106 | 0.0654 | 0 | 0 | 0 |
| 'Rad17' | 11.456 | 11.757 | 10.454 | 12.000 | 11.746 | 10.484 | 11.251 | 12.001 | 9.668 | 11.642 | 12.252 | 11.898 | 11.132 | 13.101 | 11.563 | 11.533 | 12.567 | 11.134 | -0.098 | 0.0151 | 0.0843 | 0 | 0 | 0 |
| 'Rad18' | 2.963 | 3.127 | 3.115 | 3.121 | 3.003 | 3.080 | 3.222 | 3.203 | 3.003 | 3.010 | 3.591 | 3.081 | 3.252 | 3.157 | 2.872 | 3.193 | 3.421 | 2.832 | -0.035 | 0.3929 | 0.6426 | 0 | 0 | 0 |
| 'Rad21' | 81.656 | 88.860 | 88.892 | 88.646 | 90.431 | 89.553 | 91.302 | 86.945 | 87.279 | 78.819 | 89.282 | 87.465 | 84.915 | 93.244 | 88.780 | 84.262 | 89.482 | 86.292 | 0.010 | 0.7015 | 0.8608 | 0 | 0 | 0 |
| 'Rad23a' | 11.552 | 11.706 | 13.058 | 11.886 | 11.558 | 13.372 | 11.114 | 11.489 | 12.325 | 12.133 | 10.169 | 11.989 | 13.188 | 11.227 | 12.173 | 11.765 | 10.966 | 11.983 | 0.025 | 0.5073 | 0.7332 | 0 | 0 | 0 |
| 'Rad23b' | 20.994 | 20.223 | 20.169 | 20.199 | 20.664 | 19.909 | 21.320 | 20.215 | 20.365 | 20.250 | 19.850 | 20.349 | 20.623 | 19.862 | 19.869 | 20.150 | 20.291 | 19.588 | 0.014 | 0.4868 | 0.7174 | 0 | 0 | 0 |
| 'Rad50' | 10.894 | 10.163 | 10.872 | 11.956 | 10.568 | 10.864 | 11.672 | 10.338 | 10.817 | 9.971 | 11.085 | 11.327 | 10.614 | 10.338 | 10.928 | 9.703 | 11.000 | 11.619 | 0.011 | 0.7941 | 0.9080 | 0 | 0 | 0 |
| 'Rad51' | 2.441 | 2.550 | 3.331 | 2.732 | 2.676 | 3.113 | 3.150 | 2.650 | 3.099 | 2.317 | 2.587 | 2.926 | 3.557 | 3.220 | 3.391 | 2.677 | 3.003 | 2.790 | -0.054 | 0.4556 | 0.6936 | 0 | 0 | 0 |
| 'Rad51ap1' | 1.084 | 1.177 | 1.415 | 1.454 | 1.213 | 1.461 | 1.457 | 1.208 | 1.500 | 0.825 | 1.206 | 1.229 | 1.418 | 1.258 | 1.469 | 1.222 | 1.399 | 1.501 | 0.043 | 0.6312 | 0.8173 | 0 | 0 | 0 |
| 'Rad51c' | 1.688 | 1.808 | 1.963 | 1.837 | 1.908 | 2.170 | 1.426 | 1.888 | 1.800 | 1.522 | 1.652 | 1.819 | 1.846 | 1.943 | 1.808 | 1.867 | 1.771 | 1.914 | 0.025 | 0.7105 | 0.8655 | 0 | 0 | 0 |
| 'Rad51d' | 5.094 | 5.269 | 5.564 | 5.456 | 5.401 | 5.733 | 5.176 | 5.941 | 5.741 | 5.403 | 5.467 | 5.544 | 6.006 | 5.663 | 5.652 | 6.351 | 5.504 | 5.777 | -0.063 | 0.1190 | 0.3284 | 0 | 0 | 0 |
| 'Rad52' | 1.767 | 1.882 | 1.977 | 1.754 | 2.174 | 2.023 | 2.443 | 1.759 | 2.425 | 1.812 | 1.588 | 2.201 | 1.889 | 1.934 | 1.965 | 1.730 | 1.829 | 2.060 | 0.082 | 0.2186 | 0.4655 | 0 | 0 | 0 |
| 'Rad54b' | 1.270 | 1.125 | 1.529 | 1.563 | 1.349 | 1.496 | 1.855 | 1.203 | 1.504 | 1.011 | 1.352 | 1.501 | 1.720 | 1.401 | 1.647 | 1.223 | 1.329 | 1.406 | 0.015 | 0.8687 | 0.9434 | 0 | 0 | 0 |
| 'Rad54l' | 1.759 | 1.698 | 2.472 | 2.261 | 1.519 | 2.232 | 2.166 | 1.834 | 2.179 | 1.521 | 1.979 | 2.239 | 2.357 | 1.889 | 2.411 | 1.709 | 2.003 | 1.901 | -0.008 | 0.9224 | 0.9705 | 0 | 0 | 0 |
| 'Rad54l2' | 4.163 | 4.212 | 4.718 | 4.280 | 4.768 | 4.641 | 4.054 | 4.454 | 4.490 | 4.598 | 4.851 | 4.181 | 4.440 | 4.337 | 3.863 | 4.828 | 4.589 | 3.739 | 0.009 | 0.8487 | 0.9351 | 0 | 0 | 0 |
| 'Rad9a' | 21.653 | 19.173 | 21.241 | 22.230 | 18.654 | 22.917 | 21.124 | 20.844 | 22.687 | 18.334 | 19.702 | 26.821 | 20.751 | 20.365 | 25.381 | 19.209 | 23.198 | 25.253 | -0.071 | 0.1848 | 0.4204 | 0 | 0 | 0 |
| 'Rad9b' | 9.700 | 12.823 | 10.115 | 10.734 | 12.650 | 9.937 | 11.150 | 11.818 | 10.332 | 10.397 | 12.072 | 11.158 | 11.876 | 13.061 | 11.201 | 11.694 | 13.080 | 10.550 | -0.099 | 0.0043 | 0.0334 | 1 | 0 | 0 |
| 'Rae1' | 21.203 | 22.197 | 22.046 | 20.435 | 20.978 | 21.806 | 20.752 | 20.836 | 21.626 | 21.219 | 19.745 | 22.053 | 19.178 | 20.172 | 21.746 | 19.617 | 21.464 | 22.405 | 0.021 | 0.4911 | 0.7208 | 0 | 0 | 0 |
| 'Raf1' | 24.077 | 25.273 | 22.651 | 24.173 | 24.035 | 21.669 | 24.464 | 23.606 | 22.432 | 22.616 | 25.465 | 22.937 | 25.842 | 24.852 | 23.529 | 23.805 | 26.066 | 22.874 | -0.049 | 0.0723 | 0.2380 | 0 | 0 | 0 |
| 'Rai1' | 9.910 | 8.810 | 11.073 | 8.615 | 9.461 | 10.295 | 8.761 | 9.788 | 10.405 | 10.886 | 9.317 | 9.673 | 11.073 | 8.799 | 9.599 | 9.681 | 8.453 | 8.998 | 0.002 | 0.9696 | 0.9898 | 0 | 0 | 0 |
| 'Rala' | 11.649 | 12.603 | 12.139 | 13.578 | 14.225 | 13.182 | 13.852 | 13.464 | 13.274 | 12.199 | 11.884 | 13.109 | 12.000 | 13.394 | 12.868 | 11.479 | 12.028 | 12.189 | 0.069 | 0.1135 | 0.3197 | 0 | 0 | 0 |
| 'Ralb' | 3.870 | 4.355 | 4.646 | 3.918 | 5.114 | 4.534 | 4.137 | 5.002 | 4.299 | 4.955 | 4.505 | 4.346 | 5.229 | 4.610 | 4.292 | 4.282 | 4.267 | 4.271 | -0.042 | 0.4735 | 0.7089 | 0 | 0 | 0 |
| 'Ralbp1' | 33.308 | 33.934 | 32.095 | 32.800 | 33.561 | 35.200 | 34.135 | 33.779 | 31.833 | 32.453 | 32.873 | 35.724 | 33.271 | 32.191 | 34.413 | 32.340 | 33.330 | 35.889 | -0.021 | 0.4561 | 0.6941 | 0 | 0 | 0 |
| 'Ralgapa1' | 14.686 | 16.017 | 15.576 | 15.161 | 16.853 | 14.674 | 15.711 | 16.235 | 15.482 | 16.153 | 16.200 | 15.170 | 14.801 | 15.292 | 13.834 | 16.112 | 15.825 | 14.639 | 0.015 | 0.6152 | 0.8081 | 0 | 0 | 0 |
| 'Ralgapa2' | 3.762 | 4.249 | 3.748 | 3.828 | 4.015 | 3.865 | 3.934 | 3.996 | 3.895 | 4.288 | 4.371 | 3.913 | 4.005 | 3.968 | 3.688 | 3.982 | 4.159 | 3.781 | -0.045 | 0.1194 | 0.3289 | 0 | 0 | 0 |
| 'Ralgapb' | 8.823 | 9.180 | 8.978 | 8.757 | 9.471 | 8.905 | 8.044 | 8.858 | 8.686 | 9.583 | 9.703 | 8.815 | 8.556 | 9.053 | 8.600 | 8.805 | 9.317 | 8.985 | -0.040 | 0.1863 | 0.4219 | 0 | 0 | 0 |
| 'Ralgds' | 3.733 | 3.437 | 3.612 | 3.267 | 3.518 | 4.193 | 3.210 | 3.578 | 3.826 | 3.670 | 3.042 | 3.515 | 3.244 | 3.197 | 3.342 | 3.650 | 2.997 | 3.464 | 0.097 | 0.0333 | 0.1447 | 0 | 0 | 0 |
| 'Ralgps1' | 3.207 | 3.635 | 3.216 | 2.889 | 3.365 | 3.438 | 3.008 | 3.162 | 3.487 | 3.641 | 3.613 | 3.479 | 3.408 | 3.565 | 3.491 | 3.851 | 3.503 | 3.205 | -0.119 | 0.0060 | 0.0429 | 1 | 0 | 0 |
| 'Ralgps2' | 2.402 | 2.549 | 2.381 | 2.330 | 2.392 | 2.260 | 2.270 | 2.354 | 2.279 | 2.280 | 2.333 | 2.479 | 1.888 | 2.243 | 2.301 | 2.252 | 2.267 | 2.543 | 0.032 | 0.4738 | 0.7091 | 0 | 0 | 0 |
| 'Raly' | 57.564 | 53.768 | 59.780 | 56.039 | 54.235 | 63.017 | 56.680 | 54.669 | 58.382 | 55.985 | 53.063 | 58.839 | 59.276 | 54.834 | 61.315 | 58.954 | 54.904 | 58.795 | -0.015 | 0.4651 | 0.7012 | 0 | 0 | 0 |
| 'Ramp1' | 11.710 | 19.238 | 10.667 | 11.514 | 16.517 | 12.399 | 12.371 | 18.455 | 11.726 | 8.052 | 10.439 | 7.978 | 9.446 | 11.619 | 8.031 | 10.179 | 11.156 | 7.014 | 0.547 | 0.0000 | 0.0000 | 1 | 1 | 0 |
| 'Ramp3' | 6.057 | 5.259 | 6.478 | 5.708 | 5.368 | 6.375 | 5.226 | 5.799 | 6.585 | 6.535 | 5.523 | 6.145 | 3.658 | 4.496 | 6.581 | 5.703 | 4.509 | 5.986 | 0.103 | 0.2122 | 0.4574 | 0 | 0 | 0 |
| 'Ranbp1' | 46.260 | 51.862 | 52.841 | 49.822 | 49.607 | 49.728 | 50.513 | 47.554 | 49.519 | 44.992 | 46.663 | 45.842 | 52.221 | 47.202 | 50.828 | 44.581 | 48.162 | 47.718 | 0.053 | 0.1348 | 0.3536 | 0 | 0 | 0 |
| 'Ranbp10' | 14.344 | 12.017 | 13.665 | 13.629 | 12.191 | 14.390 | 13.181 | 13.265 | 14.283 | 14.223 | 13.334 | 15.001 | 13.324 | 13.692 | 13.971 | 14.722 | 13.984 | 13.740 | -0.067 | 0.0496 | 0.1878 | 0 | 0 | 0 |
| 'Ranbp2' | 28.734 | 26.880 | 28.042 | 30.209 | 28.834 | 25.018 | 29.425 | 26.721 | 25.437 | 27.821 | 30.489 | 27.570 | 24.674 | 28.667 | 25.658 | 26.976 | 29.190 | 27.072 | -0.004 | 0.9222 | 0.9705 | 0 | 0 | 0 |
| 'Ranbp3' | 58.634 | 57.084 | 67.569 | 58.795 | 56.826 | 64.508 | 59.205 | 56.856 | 64.086 | 60.024 | 56.689 | 66.718 | 64.112 | 58.180 | 62.294 | 56.931 | 58.484 | 64.393 | -0.023 | 0.3449 | 0.6000 | 0 | 0 | 0 |
| 'Ranbp6' | 5.666 | 8.180 | 6.538 | 6.772 | 7.433 | 7.529 | 5.779 | 6.611 | 6.768 | 5.815 | 6.918 | 6.882 | 5.356 | 7.565 | 6.542 | 6.129 | 6.322 | 6.440 | 0.066 | 0.2098 | 0.4550 | 0 | 0 | 0 |
| 'Ranbp9' | 66.999 | 70.687 | 62.373 | 69.447 | 70.577 | 64.504 | 70.933 | 69.967 | 64.403 | 66.254 | 65.387 | 63.626 | 69.512 | 71.751 | 63.212 | 70.752 | 70.017 | 64.557 | 0.000 | 0.9964 | 0.9994 | 0 | 0 | 0 |
| 'Rangap1' | 57.119 | 48.001 | 61.539 | 58.035 | 53.670 | 59.645 | 53.466 | 52.474 | 57.965 | 52.535 | 53.869 | 66.054 | 58.535 | 54.298 | 62.176 | 52.610 | 53.952 | 68.570 | -0.067 | 0.0688 | 0.2306 | 0 | 0 | 0 |
| 'Rap1a' | 74.300 | 90.168 | 72.718 | 84.506 | 86.884 | 72.630 | 87.951 | 85.094 | 72.006 | 73.731 | 84.031 | 72.869 | 74.095 | 87.460 | 72.444 | 73.359 | 85.215 | 72.329 | 0.050 | 0.1158 | 0.3232 | 0 | 0 | 0 |
| 'Rap1b' | 173.820 | 196.834 | 166.110 | 179.299 | 189.452 | 168.910 | 186.125 | 188.887 | 164.318 | 160.479 | 176.755 | 162.288 | 166.086 | 185.495 | 165.520 | 168.625 | 177.482 | 162.715 | 0.069 | 0.0021 | 0.0189 | 1 | 0 | 0 |
| 'Rap1gap' | 0.676 | 0.802 | 0.855 | 0.699 | 0.605 | 0.785 | 0.714 | 0.894 | 0.952 | 0.423 | 0.600 | 0.441 | 0.592 | 0.481 | 0.484 | 0.378 | 0.500 | 0.445 | 0.683 | 0.0000 | 0.0000 | 1 | 1 | 0 |
| 'Rap1gds1' | 15.008 | 15.038 | 14.585 | 15.537 | 14.845 | 14.558 | 14.648 | 15.178 | 14.674 | 14.473 | 16.636 | 15.930 | 14.153 | 15.509 | 16.311 | 13.100 | 15.654 | 16.256 | -0.051 | 0.1841 | 0.4195 | 0 | 0 | 0 |
| 'Rap2a' | 2.246 | 2.958 | 2.375 | 2.357 | 2.622 | 2.611 | 2.236 | 2.715 | 2.320 | 2.583 | 2.959 | 2.994 | 2.953 | 3.198 | 3.115 | 3.176 | 3.456 | 2.968 | -0.302 | 0.0000 | 0.0000 | 1 | 0 | 0 |
| 'Rap2b' | 3.907 | 4.084 | 3.911 | 4.059 | 3.988 | 4.383 | 4.056 | 3.816 | 3.825 | 3.992 | 4.110 | 4.576 | 3.862 | 4.800 | 4.243 | 3.863 | 4.460 | 4.445 | -0.103 | 0.0289 | 0.1317 | 0 | 0 | 0 |
| 'Rap2c' | 26.712 | 31.979 | 29.678 | 28.816 | 31.246 | 30.174 | 30.090 | 30.387 | 28.331 | 29.503 | 29.560 | 30.525 | 28.740 | 32.379 | 28.892 | 29.066 | 31.064 | 27.882 | -0.014 | 0.6157 | 0.8082 | 0 | 0 | 0 |
| 'Rapgef1' | 49.766 | 50.991 | 51.477 | 47.063 | 48.261 | 53.391 | 47.071 | 49.050 | 55.220 | 55.835 | 48.416 | 51.697 | 54.371 | 47.967 | 50.948 | 55.587 | 45.531 | 50.857 | -0.037 | 0.2884 | 0.5448 | 0 | 0 | 0 |
| 'Rapgef2' | 9.257 | 8.254 | 7.503 | 7.844 | 8.178 | 7.863 | 7.657 | 8.577 | 7.681 | 9.507 | 9.180 | 7.650 | 8.659 | 9.492 | 7.794 | 9.285 | 8.251 | 8.028 | -0.102 | 0.0119 | 0.0707 | 0 | 0 | 0 |
| 'Rapgef5' | 2.950 | 2.497 | 2.624 | 3.080 | 2.678 | 2.887 | 2.554 | 2.709 | 2.849 | 3.172 | 3.077 | 3.006 | 3.111 | 3.103 | 3.023 | 3.292 | 2.724 | 3.194 | -0.163 | 0.0009 | 0.0094 | 1 | 0 | 0 |
| 'Rapgef6' | 15.903 | 15.327 | 15.560 | 15.710 | 15.420 | 15.093 | 15.590 | 14.961 | 15.116 | 15.801 | 16.868 | 15.255 | 14.627 | 15.176 | 14.910 | 15.459 | 16.452 | 14.241 | -0.010 | 0.7307 | 0.8772 | 0 | 0 | 0 |
| 'Raph1' | 1.918 | 1.712 | 2.185 | 2.114 | 1.858 | 1.913 | 1.998 | 1.689 | 2.057 | 1.681 | 1.697 | 2.183 | 1.504 | 1.742 | 2.033 | 1.754 | 1.830 | 2.224 | 0.059 | 0.2872 | 0.5438 | 0 | 0 | 0 |
| 'Rapsn' | 1.685 | 1.401 | 1.293 | 1.047 | 0.956 | 1.709 | 1.392 | 0.979 | 1.768 | 1.631 | 1.215 | 1.443 | 1.503 | 1.446 | 1.738 | 1.252 | 1.097 | 1.281 | -0.053 | 0.6552 | 0.8329 | 0 | 0 | 0 |
| 'Rara' | 1.487 | 1.351 | 1.538 | 1.086 | 1.487 | 1.457 | 1.629 | 1.459 | 1.578 | 1.148 | 1.024 | 1.356 | 1.093 | 1.291 | 1.312 | 1.723 | 1.065 | 1.184 | 0.220 | 0.0175 | 0.0940 | 0 | 0 | 0 |
| 'Rarg' | 1.955 | 2.686 | 2.080 | 2.115 | 1.985 | 2.381 | 1.985 | 2.458 | 2.240 | 2.246 | 4.148 | 2.806 | 2.285 | 2.151 | 2.436 | 2.786 | 2.683 | 2.298 | -0.268 | 0.0059 | 0.0425 | 1 | 0 | 0 |
| 'Rars' | 14.636 | 14.917 | 15.352 | 15.408 | 14.330 | 15.045 | 14.378 | 15.816 | 15.630 | 14.986 | 14.525 | 15.193 | 14.264 | 13.988 | 14.259 | 14.150 | 15.131 | 15.151 | 0.032 | 0.2998 | 0.5560 | 0 | 0 | 0 |
| 'Rars2' | 6.219 | 6.193 | 6.381 | 6.022 | 6.989 | 5.634 | 6.418 | 6.737 | 6.204 | 6.146 | 6.018 | 6.179 | 6.230 | 6.791 | 6.094 | 6.770 | 6.575 | 5.602 | 0.001 | 0.9828 | 0.9934 | 0 | 0 | 0 |
| 'Rasa1' | 20.161 | 22.395 | 19.482 | 20.284 | 22.241 | 19.074 | 20.062 | 21.815 | 19.587 | 19.696 | 20.100 | 22.792 | 17.461 | 21.596 | 19.727 | 19.987 | 21.926 | 22.899 | -0.020 | 0.6393 | 0.8225 | 0 | 0 | 0 |
| 'Rasa2' | 5.510 | 6.044 | 5.214 | 5.490 | 5.333 | 5.099 | 5.892 | 5.509 | 5.123 | 5.077 | 5.513 | 5.812 | 5.080 | 5.383 | 5.455 | 4.871 | 6.058 | 5.575 | -0.001 | 0.9772 | 0.9920 | 0 | 0 | 0 |
| 'Rasa3' | 6.191 | 7.028 | 6.295 | 6.293 | 6.037 | 7.836 | 6.922 | 7.521 | 7.521 | 6.906 | 7.516 | 8.751 | 7.777 | 7.425 | 9.443 | 8.006 | 7.620 | 8.209 | -0.227 | 0.0000 | 0.0002 | 1 | 0 | 0 |
| 'Rasal1' | 3.441 | 3.120 | 5.004 | 3.579 | 3.163 | 3.929 | 3.648 | 3.411 | 3.909 | 4.802 | 4.494 | 5.818 | 3.835 | 3.362 | 4.853 | 4.199 | 4.050 | 5.791 | -0.319 | 0.0000 | 0.0001 | 1 | 0 | 0 |
| 'Rasal3' | 70.137 | 62.465 | 78.866 | 67.971 | 65.241 | 77.191 | 63.056 | 64.613 | 76.715 | 70.244 | 62.759 | 77.333 | 70.793 | 64.340 | 72.047 | 67.130 | 64.288 | 76.832 | -0.011 | 0.6642 | 0.8377 | 0 | 0 | 0 |
| 'Rasgef1a' | 4.309 | 4.823 | 4.281 | 3.989 | 4.327 | 4.854 | 4.358 | 4.408 | 4.174 | 4.785 | 4.794 | 6.107 | 5.309 | 5.249 | 6.182 | 4.401 | 4.578 | 6.493 | -0.289 | 0.0000 | 0.0001 | 1 | 0 | 0 |
| 'Rasgrf1' | 0.931 | 0.829 | 0.639 | 0.671 | 0.706 | 0.614 | 0.742 | 0.722 | 0.811 | 0.402 | 0.496 | 0.360 | 0.436 | 0.350 | 0.297 | 0.427 | 0.382 | 0.282 | 0.957 | 0.0000 | 0.0000 | 1 | 1 | 0 |
| 'Rasgrf2' | 1.173 | 0.940 | 1.156 | 1.101 | 0.872 | 0.994 | 1.146 | 1.040 | 0.929 | 1.208 | 1.104 | 1.132 | 1.062 | 1.287 | 1.084 | 0.957 | 1.060 | 1.075 | -0.101 | 0.1585 | 0.3886 | 0 | 0 | 0 |
| 'Rasgrp1' | 25.899 | 25.812 | 25.882 | 25.162 | 24.684 | 26.006 | 24.798 | 24.760 | 27.226 | 25.208 | 23.567 | 25.407 | 21.058 | 23.362 | 24.589 | 25.830 | 20.625 | 25.779 | 0.086 | 0.0249 | 0.1204 | 0 | 0 | 0 |
| 'Rasgrp2' | 1.000 | 0.695 | 1.049 | 0.809 | 0.782 | 1.190 | 1.013 | 0.874 | 1.104 | 0.804 | 0.854 | 1.317 | 1.047 | 0.811 | 1.263 | 1.071 | 0.902 | 1.223 | -0.133 | 0.0337 | 0.1460 | 0 | 0 | 0 |
| 'Rasl11a' | 10.524 | 10.869 | 9.142 | 11.710 | 11.067 | 8.782 | 12.285 | 10.339 | 8.892 | 7.619 | 11.669 | 10.226 | 6.503 | 9.913 | 10.113 | 6.396 | 12.734 | 9.987 | 0.139 | 0.1901 | 0.4265 | 0 | 0 | 0 |
| 'Rasl11b' | 3.049 | 1.683 | 1.496 | 3.139 | 1.693 | 1.298 | 2.857 | 2.097 | 1.398 | 2.240 | 1.880 | 1.279 | 2.093 | 1.616 | 1.310 | 1.870 | 2.412 | 0.911 | 0.233 | 0.0289 | 0.1317 | 0 | 0 | 0 |
| 'Rasl2-9' | 27.362 | 31.599 | 30.967 | 33.274 | 32.931 | 27.181 | 30.486 | 35.074 | 31.684 | 26.393 | 26.735 | 31.357 | 32.269 | 31.339 | 32.267 | 27.679 | 35.159 | 31.253 | 0.016 | 0.7787 | 0.9003 | 0 | 0 | 0 |
| 'Rassf1' | 4.234 | 3.461 | 3.582 | 3.975 | 3.798 | 3.943 | 4.079 | 3.832 | 3.563 | 3.774 | 3.855 | 3.941 | 4.196 | 3.869 | 3.665 | 4.011 | 3.723 | 3.525 | -0.015 | 0.6984 | 0.8587 | 0 | 0 | 0 |
| 'Rassf2' | 23.997 | 26.212 | 24.533 | 25.427 | 25.167 | 25.185 | 25.658 | 24.707 | 25.890 | 24.600 | 27.576 | 27.452 | 25.377 | 24.924 | 27.218 | 24.034 | 26.430 | 26.931 | -0.059 | 0.0347 | 0.1491 | 0 | 0 | 0 |
| 'Rassf3' | 23.222 | 25.396 | 23.712 | 25.112 | 25.128 | 26.128 | 26.572 | 24.581 | 24.365 | 23.746 | 28.013 | 27.491 | 24.880 | 27.777 | 28.571 | 28.399 | 26.956 | 27.951 | -0.131 | 0.0007 | 0.0075 | 1 | 0 | 0 |
| 'Rassf5' | 59.713 | 61.462 | 58.757 | 57.191 | 58.630 | 57.760 | 54.181 | 60.177 | 56.246 | 53.678 | 56.896 | 62.756 | 49.138 | 58.918 | 61.119 | 54.936 | 56.054 | 63.480 | 0.012 | 0.7492 | 0.8877 | 0 | 0 | 0 |
| 'Rassf7' | 2.217 | 1.932 | 2.493 | 1.679 | 2.269 | 2.733 | 2.270 | 2.212 | 2.502 | 1.995 | 2.001 | 2.470 | 2.359 | 2.000 | 2.438 | 2.392 | 1.841 | 2.416 | 0.022 | 0.7448 | 0.8848 | 0 | 0 | 0 |
| 'Raver1' | 68.945 | 65.797 | 76.951 | 62.920 | 64.295 | 75.436 | 67.453 | 66.233 | 76.121 | 69.891 | 65.662 | 74.928 | 74.210 | 66.458 | 74.677 | 70.018 | 62.343 | 73.291 | -0.027 | 0.2830 | 0.5394 | 0 | 0 | 0 |
| 'Raver2' | 0.713 | 0.426 | 0.768 | 0.747 | 0.534 | 0.709 | 0.691 | 0.486 | 0.800 | 0.665 | 0.574 | 0.502 | 0.687 | 0.599 | 0.707 | 0.482 | 0.498 | 0.538 | 0.160 | 0.1658 | 0.3982 | 0 | 0 | 0 |
| 'Rb1' | 18.246 | 20.984 | 18.091 | 17.508 | 19.976 | 17.256 | 18.758 | 19.333 | 16.968 | 19.872 | 19.705 | 16.997 | 19.100 | 19.593 | 16.311 | 18.052 | 19.280 | 14.164 | 0.029 | 0.4308 | 0.6762 | 0 | 0 | 0 |
| 'Rb1cc1' | 4.191 | 4.004 | 3.756 | 4.224 | 3.826 | 3.624 | 4.198 | 3.806 | 3.564 | 4.067 | 4.390 | 4.236 | 3.814 | 4.290 | 3.679 | 4.096 | 4.516 | 3.937 | -0.085 | 0.0357 | 0.1516 | 0 | 0 | 0 |
| 'Rbak' | 3.073 | 3.609 | 3.835 | 3.280 | 3.290 | 3.329 | 2.959 | 3.632 | 3.312 | 3.447 | 3.264 | 3.070 | 3.026 | 3.380 | 2.694 | 3.345 | 3.097 | 3.127 | 0.087 | 0.1147 | 0.3213 | 0 | 0 | 0 |
| 'Rbbp4' | 26.071 | 27.231 | 27.442 | 27.584 | 28.707 | 27.969 | 28.609 | 28.743 | 27.257 | 25.473 | 28.286 | 26.569 | 26.840 | 27.723 | 26.786 | 27.020 | 28.000 | 26.029 | 0.031 | 0.1538 | 0.3816 | 0 | 0 | 0 |
| 'Rbbp5' | 10.642 | 10.441 | 11.124 | 10.715 | 11.304 | 10.541 | 11.014 | 10.505 | 10.674 | 10.504 | 10.195 | 11.527 | 10.720 | 10.973 | 10.611 | 9.716 | 11.548 | 11.660 | -0.020 | 0.5604 | 0.7706 | 0 | 0 | 0 |
| 'Rbbp7' | 42.498 | 45.518 | 43.547 | 47.328 | 43.237 | 44.275 | 46.985 | 42.829 | 44.444 | 39.698 | 41.658 | 44.570 | 44.110 | 42.294 | 45.206 | 42.225 | 43.931 | 42.810 | 0.040 | 0.1790 | 0.4144 | 0 | 0 | 0 |
| 'Rbbp8' | 6.628 | 7.101 | 7.910 | 6.947 | 7.551 | 6.703 | 7.768 | 7.208 | 6.790 | 7.228 | 7.676 | 6.746 | 8.185 | 7.563 | 7.309 | 6.914 | 8.705 | 5.580 | -0.033 | 0.5860 | 0.7873 | 0 | 0 | 0 |
| 'Rbbp9' | 2.482 | 3.277 | 3.220 | 2.362 | 3.640 | 2.466 | 3.492 | 3.350 | 2.992 | 2.395 | 2.962 | 3.170 | 2.807 | 3.298 | 3.252 | 3.781 | 3.007 | 2.525 | -0.012 | 0.8961 | 0.9583 | 0 | 0 | 0 |
| 'Rbck1' | 45.073 | 44.981 | 46.762 | 39.610 | 43.344 | 48.117 | 41.755 | 42.138 | 47.319 | 43.495 | 41.640 | 47.366 | 46.739 | 41.873 | 46.907 | 44.589 | 42.268 | 47.461 | -0.022 | 0.4151 | 0.6630 | 0 | 0 | 0 |
| 'Rbfa' | 30.110 | 25.126 | 29.176 | 27.044 | 26.414 | 27.651 | 26.768 | 27.643 | 27.049 | 31.301 | 24.071 | 27.154 | 27.582 | 22.627 | 26.907 | 28.930 | 24.195 | 24.433 | 0.055 | 0.2116 | 0.4567 | 0 | 0 | 0 |
| 'Rbks' | 5.467 | 5.351 | 4.625 | 5.167 | 5.433 | 4.824 | 6.572 | 5.732 | 4.567 | 4.695 | 5.962 | 4.631 | 5.528 | 5.159 | 4.062 | 4.610 | 5.558 | 4.899 | 0.066 | 0.3505 | 0.6046 | 0 | 0 | 0 |
| 'Rbl1' | 12.314 | 15.298 | 15.152 | 13.865 | 15.194 | 14.054 | 14.227 | 14.819 | 14.994 | 12.969 | 15.317 | 14.728 | 14.316 | 16.097 | 14.602 | 14.374 | 14.195 | 14.422 | -0.024 | 0.4620 | 0.6981 | 0 | 0 | 0 |
| 'Rbl2' | 21.941 | 25.415 | 23.381 | 23.131 | 25.200 | 23.495 | 23.551 | 24.784 | 23.857 | 21.425 | 22.896 | 23.037 | 21.187 | 22.516 | 21.932 | 21.134 | 21.066 | 20.814 | 0.119 | 0.0000 | 0.0000 | 1 | 0 | 0 |
| 'Rbm10' | 24.033 | 19.305 | 24.495 | 24.130 | 20.170 | 24.818 | 23.434 | 20.689 | 24.241 | 21.707 | 21.191 | 27.301 | 22.371 | 21.610 | 25.527 | 22.331 | 22.447 | 25.683 | -0.045 | 0.1865 | 0.4221 | 0 | 0 | 0 |
| 'Rbm12' | 3.502 | 3.866 | 4.012 | 3.635 | 3.962 | 3.663 | 3.916 | 3.663 | 3.699 | 3.056 | 3.717 | 4.166 | 3.314 | 3.579 | 3.765 | 3.187 | 3.754 | 4.101 | 0.041 | 0.3959 | 0.6447 | 0 | 0 | 0 |
| 'Rbm14' | 10.547 | 8.243 | 10.277 | 10.220 | 8.424 | 9.414 | 9.063 | 8.275 | 9.453 | 8.424 | 9.681 | 11.789 | 8.810 | 9.078 | 11.973 | 7.962 | 9.174 | 13.018 | -0.099 | 0.1466 | 0.3719 | 0 | 0 | 0 |
| 'Rbm15' | 7.693 | 5.866 | 6.792 | 7.744 | 6.053 | 6.167 | 7.803 | 6.086 | 6.146 | 6.583 | 7.650 | 7.497 | 6.626 | 6.863 | 6.869 | 6.255 | 7.894 | 7.599 | -0.098 | 0.1414 | 0.3636 | 0 | 0 | 0 |
| 'Rbm15b' | 11.664 | 10.346 | 12.228 | 11.302 | 9.869 | 11.771 | 11.310 | 10.252 | 10.982 | 10.567 | 10.864 | 12.792 | 10.992 | 10.682 | 13.024 | 11.381 | 10.287 | 14.432 | -0.082 | 0.0573 | 0.2058 | 0 | 0 | 0 |
| 'Rbm17' | 29.393 | 32.059 | 31.988 | 31.886 | 32.127 | 32.615 | 32.065 | 34.582 | 31.305 | 29.589 | 31.317 | 31.170 | 30.942 | 30.691 | 31.126 | 30.920 | 31.471 | 31.199 | 0.038 | 0.1258 | 0.3398 | 0 | 0 | 0 |
| 'Rbm18' | 7.393 | 8.244 | 7.291 | 7.593 | 8.872 | 7.297 | 7.393 | 8.332 | 7.184 | 7.675 | 7.499 | 7.685 | 7.099 | 7.441 | 7.638 | 7.244 | 8.248 | 7.317 | 0.020 | 0.5691 | 0.7768 | 0 | 0 | 0 |
| 'Rbm19' | 4.197 | 3.550 | 4.192 | 3.737 | 3.791 | 4.472 | 4.122 | 3.715 | 4.092 | 4.198 | 3.732 | 4.152 | 3.709 | 3.663 | 4.039 | 4.056 | 3.455 | 4.237 | 0.019 | 0.6256 | 0.8137 | 0 | 0 | 0 |
| 'Rbm22' | 15.789 | 15.938 | 15.605 | 14.677 | 16.168 | 15.732 | 15.102 | 16.481 | 15.984 | 17.204 | 15.472 | 15.429 | 16.132 | 15.436 | 15.475 | 16.536 | 15.438 | 14.730 | -0.012 | 0.6925 | 0.8555 | 0 | 0 | 0 |
| 'Rbm25' | 15.907 | 16.624 | 16.312 | 17.019 | 16.389 | 15.673 | 16.724 | 16.870 | 16.810 | 16.759 | 17.376 | 17.220 | 16.172 | 16.797 | 15.270 | 16.681 | 17.051 | 15.876 | -0.019 | 0.4315 | 0.6763 | 0 | 0 | 0 |
| 'Rbm26' | 10.634 | 11.614 | 11.040 | 11.166 | 11.510 | 10.982 | 11.077 | 11.616 | 11.299 | 10.963 | 11.687 | 11.634 | 10.634 | 11.647 | 10.599 | 11.647 | 11.360 | 10.346 | -0.005 | 0.8556 | 0.9382 | 0 | 0 | 0 |
| 'Rbm27' | 14.671 | 13.273 | 13.645 | 15.381 | 14.134 | 13.489 | 14.668 | 13.875 | 13.439 | 14.368 | 14.826 | 14.135 | 13.475 | 14.181 | 13.559 | 13.461 | 13.569 | 14.533 | -0.007 | 0.8449 | 0.9343 | 0 | 0 | 0 |
| 'Rbm28' | 11.643 | 10.940 | 12.342 | 11.562 | 11.284 | 11.752 | 11.729 | 11.247 | 13.028 | 11.305 | 11.797 | 11.940 | 11.650 | 10.653 | 11.207 | 11.916 | 11.995 | 11.187 | 0.016 | 0.6109 | 0.8049 | 0 | 0 | 0 |
| 'Rbm3' | 54.947 | 57.252 | 49.249 | 52.150 | 55.517 | 47.436 | 59.274 | 57.111 | 47.148 | 47.688 | 56.029 | 40.481 | 55.865 | 52.916 | 47.613 | 49.107 | 50.486 | 43.928 | 0.102 | 0.0072 | 0.0491 | 1 | 0 | 0 |
| 'Rbm33' | 15.474 | 13.458 | 14.038 | 14.200 | 12.344 | 12.424 | 14.057 | 13.364 | 13.383 | 14.830 | 14.894 | 15.327 | 14.623 | 13.872 | 14.419 | 14.817 | 14.244 | 15.387 | -0.121 | 0.0008 | 0.0085 | 1 | 0 | 0 |
| 'Rbm34' | 10.299 | 11.188 | 10.136 | 11.010 | 11.612 | 9.978 | 11.174 | 11.525 | 9.891 | 10.743 | 10.700 | 10.897 | 10.710 | 10.287 | 9.750 | 10.635 | 11.728 | 10.055 | 0.001 | 0.9725 | 0.9905 | 0 | 0 | 0 |
| 'Rbm38' | 42.048 | 43.422 | 40.061 | 39.126 | 39.637 | 44.053 | 38.635 | 38.021 | 43.800 | 37.793 | 35.649 | 48.244 | 42.786 | 40.094 | 46.938 | 43.926 | 37.535 | 48.509 | -0.054 | 0.2087 | 0.4535 | 0 | 0 | 0 |
| 'Rbm39' | 36.885 | 38.545 | 36.441 | 38.997 | 37.925 | 34.893 | 39.269 | 38.958 | 35.207 | 35.725 | 36.849 | 35.906 | 33.985 | 36.552 | 35.171 | 36.975 | 39.031 | 35.279 | 0.039 | 0.1158 | 0.3231 | 0 | 0 | 0 |
| 'Rbm41' | 1.550 | 1.741 | 1.606 | 1.696 | 1.597 | 1.635 | 1.448 | 1.547 | 1.487 | 1.436 | 1.747 | 1.602 | 1.399 | 1.584 | 1.455 | 1.454 | 1.557 | 1.581 | 0.039 | 0.4569 | 0.6943 | 0 | 0 | 0 |
| 'Rbm42' | 39.684 | 37.158 | 41.053 | 36.363 | 38.362 | 44.286 | 37.664 | 37.300 | 41.499 | 40.536 | 37.062 | 41.551 | 41.168 | 34.423 | 41.779 | 38.493 | 32.488 | 42.890 | 0.004 | 0.9095 | 0.9647 | 0 | 0 | 0 |
| 'Rbm43' | 5.836 | 6.211 | 6.291 | 5.597 | 6.922 | 6.406 | 6.504 | 6.575 | 5.995 | 6.218 | 6.710 | 6.848 | 6.286 | 6.148 | 6.468 | 6.157 | 6.906 | 6.181 | -0.054 | 0.1660 | 0.3983 | 0 | 0 | 0 |
| 'Rbm45' | 30.522 | 32.985 | 33.611 | 29.376 | 38.180 | 34.918 | 36.501 | 36.871 | 33.728 | 33.404 | 30.252 | 32.145 | 31.859 | 31.600 | 33.659 | 36.246 | 33.384 | 31.974 | 0.047 | 0.3023 | 0.5594 | 0 | 0 | 0 |
| 'Rbm48' | 4.992 | 4.934 | 4.301 | 4.136 | 5.051 | 4.574 | 3.603 | 5.272 | 4.423 | 5.052 | 4.738 | 4.460 | 4.709 | 5.374 | 4.261 | 5.859 | 5.374 | 4.716 | -0.113 | 0.0714 | 0.2359 | 0 | 0 | 0 |
| 'Rbm4b' | 3.302 | 2.656 | 3.259 | 2.994 | 2.799 | 2.719 | 3.113 | 3.143 | 3.046 | 3.296 | 3.234 | 3.276 | 3.082 | 3.018 | 2.967 | 2.900 | 3.209 | 3.099 | -0.064 | 0.1651 | 0.3974 | 0 | 0 | 0 |
| 'Rbm5' | 13.043 | 13.596 | 13.363 | 13.935 | 12.874 | 13.431 | 13.510 | 13.085 | 14.178 | 12.302 | 13.877 | 15.016 | 11.699 | 14.057 | 13.794 | 13.458 | 14.900 | 14.247 | -0.036 | 0.3529 | 0.6063 | 0 | 0 | 0 |
| 'Rbm6' | 5.613 | 5.374 | 5.742 | 6.013 | 5.640 | 5.489 | 5.799 | 5.561 | 5.803 | 5.304 | 5.889 | 6.245 | 5.102 | 5.727 | 5.672 | 5.050 | 5.842 | 5.993 | -0.004 | 0.9201 | 0.9700 | 0 | 0 | 0 |
| 'Rbm7' | 41.152 | 45.555 | 40.516 | 40.564 | 45.246 | 40.047 | 42.066 | 46.028 | 39.615 | 40.565 | 43.131 | 41.283 | 40.943 | 43.201 | 38.434 | 40.521 | 43.920 | 36.669 | 0.033 | 0.2167 | 0.4630 | 0 | 0 | 0 |
| 'Rbm8a' | 14.083 | 14.968 | 13.477 | 14.262 | 15.317 | 13.880 | 15.321 | 15.561 | 13.629 | 13.210 | 13.766 | 12.629 | 14.200 | 13.950 | 13.253 | 14.616 | 15.244 | 12.597 | 0.071 | 0.0147 | 0.0824 | 0 | 0 | 0 |
| 'Rbms1' | 5.642 | 6.010 | 6.262 | 5.985 | 6.355 | 6.685 | 5.639 | 6.897 | 6.682 | 6.675 | 6.660 | 6.143 | 7.020 | 7.017 | 6.451 | 6.536 | 6.326 | 5.924 | -0.072 | 0.1066 | 0.3067 | 0 | 0 | 0 |
| 'Rbms2' | 34.237 | 37.480 | 37.798 | 33.620 | 38.049 | 40.384 | 34.076 | 38.089 | 35.940 | 41.943 | 39.790 | 38.594 | 40.578 | 37.586 | 38.684 | 41.252 | 36.871 | 37.578 | -0.106 | 0.0073 | 0.0495 | 1 | 0 | 0 |
| 'Rbmx' | 14.699 | 17.304 | 17.387 | 16.489 | 15.883 | 15.838 | 17.781 | 16.034 | 16.873 | 13.756 | 15.833 | 17.408 | 15.640 | 14.338 | 16.415 | 14.247 | 17.019 | 15.737 | 0.067 | 0.1212 | 0.3319 | 0 | 0 | 0 |
| 'Rbmx2' | 6.607 | 8.051 | 7.413 | 8.229 | 7.870 | 6.849 | 9.274 | 8.585 | 7.602 | 7.340 | 6.686 | 7.249 | 7.647 | 6.345 | 7.562 | 6.885 | 6.906 | 6.729 | 0.131 | 0.0647 | 0.2218 | 0 | 0 | 0 |
| 'Rbpj' | 5.987 | 7.090 | 6.416 | 6.646 | 6.577 | 6.189 | 6.527 | 6.732 | 6.321 | 6.043 | 6.142 | 6.036 | 5.887 | 6.503 | 5.561 | 6.420 | 6.506 | 5.148 | 0.099 | 0.0029 | 0.0245 | 1 | 0 | 0 |
| 'Rbsn' | 4.205 | 3.718 | 4.194 | 4.082 | 4.097 | 3.971 | 3.871 | 4.112 | 3.696 | 4.895 | 3.710 | 3.825 | 4.377 | 4.345 | 3.802 | 3.996 | 3.508 | 3.826 | -0.019 | 0.7040 | 0.8618 | 0 | 0 | 0 |
| 'Rbx1' | 40.259 | 50.676 | 45.623 | 46.781 | 45.341 | 45.769 | 47.738 | 48.800 | 44.214 | 37.477 | 43.480 | 41.923 | 42.206 | 46.223 | 43.525 | 44.068 | 45.988 | 41.415 | 0.093 | 0.0127 | 0.0745 | 0 | 0 | 0 |
| 'Rc3h1' | 9.565 | 7.993 | 8.071 | 9.158 | 7.815 | 7.145 | 9.076 | 8.031 | 7.622 | 7.588 | 8.558 | 8.993 | 7.274 | 7.767 | 7.605 | 7.910 | 8.883 | 8.795 | 0.006 | 0.9130 | 0.9668 | 0 | 0 | 0 |
| 'Rc3h2' | 2.746 | 2.648 | 2.722 | 2.915 | 2.918 | 2.566 | 3.020 | 2.851 | 2.628 | 2.628 | 2.932 | 2.985 | 2.899 | 2.644 | 2.467 | 2.801 | 2.818 | 2.904 | -0.019 | 0.6551 | 0.8329 | 0 | 0 | 0 |
| 'Rcan1' | 8.585 | 8.202 | 8.629 | 8.546 | 8.760 | 9.539 | 8.837 | 8.283 | 8.797 | 8.358 | 7.823 | 8.497 | 7.970 | 7.830 | 9.123 | 8.499 | 8.483 | 8.109 | 0.058 | 0.2187 | 0.4655 | 0 | 0 | 0 |
| 'Rcan3' | 8.151 | 8.008 | 7.289 | 7.562 | 8.733 | 8.917 | 7.185 | 8.593 | 8.509 | 7.856 | 8.009 | 8.521 | 7.495 | 7.600 | 8.824 | 7.793 | 7.457 | 8.131 | 0.014 | 0.7515 | 0.8894 | 0 | 0 | 0 |
| 'Rcbtb1' | 9.089 | 9.807 | 9.918 | 9.289 | 9.435 | 9.415 | 9.229 | 10.132 | 9.985 | 10.079 | 10.853 | 11.166 | 9.183 | 10.225 | 11.324 | 10.026 | 10.957 | 11.237 | -0.149 | 0.0000 | 0.0000 | 1 | 0 | 0 |
| 'Rcbtb2' | 1.844 | 2.072 | 2.006 | 2.002 | 2.101 | 2.019 | 2.257 | 2.152 | 2.020 | 2.043 | 2.117 | 2.023 | 2.027 | 1.968 | 1.854 | 1.903 | 2.093 | 1.782 | 0.040 | 0.3130 | 0.5715 | 0 | 0 | 0 |
| 'Rcc1' | 9.410 | 9.107 | 10.432 | 9.043 | 9.254 | 9.510 | 10.570 | 9.999 | 10.307 | 7.873 | 8.956 | 9.818 | 10.112 | 8.494 | 10.901 | 8.893 | 9.429 | 9.966 | 0.044 | 0.3587 | 0.6113 | 0 | 0 | 0 |
| 'Rcc2' | 54.908 | 48.443 | 56.504 | 54.124 | 50.166 | 53.649 | 51.508 | 51.496 | 51.379 | 49.805 | 51.335 | 51.084 | 50.747 | 48.067 | 51.816 | 48.106 | 47.578 | 54.056 | 0.051 | 0.0874 | 0.2697 | 0 | 0 | 0 |
| 'Rccd1' | 2.257 | 2.377 | 2.774 | 2.443 | 2.074 | 2.453 | 2.539 | 2.625 | 2.702 | 2.605 | 2.519 | 2.738 | 2.640 | 2.342 | 2.446 | 2.683 | 2.805 | 2.261 | -0.060 | 0.2649 | 0.5184 | 0 | 0 | 0 |
| 'Rce1' | 24.863 | 25.934 | 26.055 | 23.362 | 27.634 | 25.734 | 22.417 | 26.547 | 25.715 | 24.172 | 23.031 | 28.423 | 26.757 | 26.172 | 26.179 | 26.331 | 25.160 | 27.820 | -0.051 | 0.2282 | 0.4748 | 0 | 0 | 0 |
| 'Rchy1' | 9.634 | 11.640 | 10.319 | 10.126 | 11.341 | 10.138 | 10.264 | 11.323 | 10.101 | 10.023 | 10.414 | 10.333 | 9.439 | 11.059 | 10.198 | 10.703 | 11.188 | 9.544 | 0.018 | 0.5416 | 0.7586 | 0 | 0 | 0 |
| 'Rcl1' | 30.200 | 32.635 | 30.895 | 29.063 | 35.930 | 30.542 | 28.221 | 33.611 | 29.297 | 30.068 | 29.868 | 28.700 | 27.936 | 28.775 | 27.017 | 32.080 | 28.529 | 26.947 | 0.099 | 0.0123 | 0.0729 | 0 | 0 | 0 |
| 'Rcn1' | 6.061 | 8.758 | 5.954 | 6.094 | 7.965 | 6.068 | 6.812 | 8.415 | 6.081 | 6.739 | 6.647 | 6.473 | 7.731 | 7.651 | 5.900 | 6.733 | 7.031 | 5.811 | 0.011 | 0.8436 | 0.9337 | 0 | 0 | 0 |
| 'Rcn2' | 3.084 | 3.981 | 3.814 | 3.725 | 3.862 | 3.785 | 3.611 | 4.434 | 3.714 | 4.338 | 3.771 | 4.085 | 3.600 | 3.833 | 3.545 | 4.055 | 3.797 | 3.588 | -0.038 | 0.4811 | 0.7131 | 0 | 0 | 0 |
| 'Rcor1' | 27.787 | 25.805 | 26.285 | 27.514 | 24.666 | 24.738 | 26.969 | 25.746 | 24.823 | 26.380 | 28.245 | 27.670 | 24.833 | 27.838 | 26.278 | 25.819 | 26.660 | 27.612 | -0.054 | 0.1357 | 0.3545 | 0 | 0 | 0 |
| 'Rcor2' | 1.371 | 1.088 | 1.232 | 1.439 | 1.402 | 1.235 | 1.353 | 1.326 | 1.327 | 1.508 | 1.365 | 1.113 | 1.497 | 1.506 | 1.231 | 1.256 | 1.428 | 1.120 | -0.024 | 0.8130 | 0.9157 | 0 | 0 | 0 |
| 'Rcor3' | 0.995 | 1.353 | 1.080 | 1.035 | 1.221 | 1.050 | 1.044 | 1.136 | 1.142 | 1.149 | 1.032 | 1.070 | 0.990 | 1.074 | 1.037 | 1.047 | 0.983 | 1.000 | 0.082 | 0.1086 | 0.3103 | 0 | 0 | 0 |
| 'Rcsd1' | 9.363 | 8.759 | 9.559 | 8.984 | 8.388 | 10.352 | 9.868 | 8.488 | 10.137 | 9.695 | 9.358 | 12.400 | 10.088 | 10.114 | 11.846 | 11.543 | 9.712 | 11.081 | -0.202 | 0.0000 | 0.0000 | 1 | 0 | 0 |
| 'Rdh10' | 3.440 | 3.307 | 3.304 | 3.167 | 2.947 | 2.968 | 3.053 | 3.217 | 3.159 | 3.327 | 3.318 | 3.458 | 2.938 | 4.021 | 3.113 | 3.866 | 3.024 | 3.256 | -0.092 | 0.1415 | 0.3639 | 0 | 0 | 0 |
| 'Rdh11' | 6.347 | 5.857 | 6.312 | 6.471 | 6.496 | 6.583 | 6.537 | 6.769 | 6.344 | 5.908 | 6.875 | 7.234 | 5.907 | 6.683 | 6.866 | 5.996 | 6.072 | 7.484 | -0.043 | 0.3687 | 0.6201 | 0 | 0 | 0 |
| 'Rdh12' | 2.816 | 3.223 | 3.212 | 3.211 | 3.325 | 3.217 | 3.513 | 3.204 | 3.562 | 3.224 | 3.502 | 3.718 | 3.484 | 3.107 | 3.608 | 3.226 | 3.053 | 3.060 | -0.052 | 0.3414 | 0.5981 | 0 | 0 | 0 |
| 'Rdh13' | 1.929 | 1.719 | 1.811 | 1.621 | 1.876 | 1.771 | 1.652 | 1.830 | 1.729 | 1.730 | 1.914 | 1.722 | 1.621 | 1.669 | 1.752 | 1.828 | 1.623 | 1.576 | 0.044 | 0.4426 | 0.6862 | 0 | 0 | 0 |
| 'Rdh14' | 9.671 | 14.420 | 12.202 | 12.024 | 13.235 | 13.646 | 14.293 | 12.258 | 12.100 | 11.645 | 13.345 | 13.139 | 11.368 | 12.856 | 11.379 | 10.743 | 12.526 | 11.736 | 0.047 | 0.4457 | 0.6878 | 0 | 0 | 0 |
| 'Rdm1' | 8.955 | 9.555 | 11.149 | 10.142 | 10.623 | 11.977 | 9.981 | 9.581 | 11.085 | 9.382 | 10.350 | 11.513 | 10.522 | 11.637 | 11.058 | 9.099 | 9.647 | 10.278 | -0.020 | 0.6787 | 0.8463 | 0 | 0 | 0 |
| 'Rdx' | 23.806 | 22.256 | 23.600 | 24.710 | 23.256 | 23.981 | 24.763 | 23.749 | 24.683 | 23.682 | 23.375 | 24.949 | 23.790 | 23.792 | 23.488 | 22.476 | 22.767 | 23.174 | 0.010 | 0.6874 | 0.8525 | 0 | 0 | 0 |
| 'Reck' | 0.714 | 0.763 | 0.760 | 0.652 | 0.713 | 0.728 | 0.811 | 0.780 | 0.895 | 0.655 | 0.910 | 0.559 | 0.725 | 0.754 | 0.695 | 0.877 | 0.655 | 0.547 | 0.111 | 0.2666 | 0.5201 | 0 | 0 | 0 |
| 'Recql' | 4.385 | 4.895 | 4.847 | 4.480 | 4.585 | 5.369 | 4.978 | 4.861 | 4.952 | 4.631 | 4.703 | 4.977 | 5.170 | 4.718 | 5.141 | 4.590 | 4.819 | 4.502 | -0.008 | 0.8356 | 0.9289 | 0 | 0 | 0 |
| 'Recql4' | 4.421 | 3.841 | 5.766 | 4.879 | 4.006 | 5.419 | 4.689 | 3.890 | 5.605 | 4.533 | 4.978 | 5.301 | 5.997 | 4.946 | 5.229 | 4.219 | 3.945 | 4.715 | -0.062 | 0.3883 | 0.6378 | 0 | 0 | 0 |
| 'Recql5' | 2.974 | 3.306 | 3.688 | 3.024 | 3.157 | 3.658 | 3.310 | 3.452 | 3.979 | 3.757 | 3.194 | 3.314 | 3.789 | 3.433 | 2.993 | 3.412 | 3.365 | 3.067 | 0.002 | 0.9753 | 0.9918 | 0 | 0 | 0 |
| 'Reep2' | 4.456 | 7.047 | 5.790 | 6.152 | 5.848 | 6.191 | 6.247 | 5.963 | 6.294 | 6.119 | 6.172 | 5.253 | 7.084 | 6.258 | 6.085 | 5.131 | 5.971 | 5.344 | 0.003 | 0.9685 | 0.9896 | 0 | 0 | 0 |
| 'Reep3' | 10.730 | 12.122 | 12.212 | 10.905 | 13.261 | 11.741 | 11.711 | 12.525 | 11.423 | 11.521 | 11.197 | 11.248 | 11.052 | 10.560 | 11.153 | 11.252 | 11.833 | 11.232 | 0.065 | 0.0574 | 0.2060 | 0 | 0 | 0 |
| 'Reep4' | 24.916 | 25.762 | 28.415 | 22.632 | 25.225 | 27.154 | 24.250 | 23.840 | 28.485 | 21.994 | 25.806 | 29.485 | 26.408 | 26.145 | 31.324 | 24.243 | 26.874 | 29.452 | -0.078 | 0.0467 | 0.1802 | 0 | 0 | 0 |
| 'Reep5' | 25.238 | 26.341 | 24.371 | 24.507 | 29.514 | 28.374 | 26.894 | 26.277 | 24.733 | 25.264 | 24.530 | 28.176 | 27.012 | 27.973 | 29.605 | 27.468 | 27.494 | 26.978 | -0.062 | 0.1286 | 0.3437 | 0 | 0 | 0 |
| 'Rel' | 18.697 | 13.468 | 12.947 | 14.035 | 12.575 | 11.386 | 14.624 | 14.328 | 13.400 | 19.630 | 12.703 | 11.161 | 15.577 | 14.605 | 10.862 | 17.864 | 13.805 | 11.230 | -0.013 | 0.8439 | 0.9338 | 0 | 0 | 0 |
| 'Rela' | 65.119 | 65.283 | 66.882 | 61.592 | 66.087 | 68.228 | 59.928 | 68.445 | 67.736 | 71.388 | 60.874 | 66.652 | 66.687 | 62.988 | 66.496 | 67.697 | 60.665 | 65.534 | -0.008 | 0.7904 | 0.9064 | 0 | 0 | 0 |
| 'Relb' | 17.376 | 18.557 | 19.432 | 15.904 | 17.365 | 20.752 | 16.510 | 18.052 | 19.930 | 22.478 | 18.352 | 18.973 | 19.573 | 19.502 | 19.090 | 21.371 | 19.663 | 17.843 | -0.121 | 0.0172 | 0.0930 | 0 | 0 | 0 |
| 'Rell1' | 0.761 | 0.615 | 0.832 | 0.727 | 0.737 | 0.849 | 0.539 | 0.634 | 0.713 | 0.492 | 0.671 | 0.652 | 0.731 | 0.517 | 0.753 | 0.591 | 0.547 | 0.742 | 0.167 | 0.1020 | 0.2994 | 0 | 0 | 0 |
| 'Relt' | 5.298 | 4.384 | 5.932 | 5.941 | 5.282 | 5.859 | 4.645 | 5.165 | 5.590 | 4.719 | 4.749 | 6.871 | 5.051 | 4.758 | 5.588 | 4.213 | 4.369 | 5.839 | 0.047 | 0.4768 | 0.7107 | 0 | 0 | 0 |
| 'Renbp' | 0.766 | 0.935 | 0.763 | 0.803 | 0.762 | 0.868 | 0.704 | 0.767 | 0.631 | 0.716 | 0.701 | 0.732 | 0.987 | 0.844 | 0.685 | 0.825 | 0.601 | 0.720 | 0.028 | 0.7818 | 0.9024 | 0 | 0 | 0 |
| 'Repin1' | 1.287 | 1.533 | 1.821 | 1.402 | 1.396 | 1.655 | 1.409 | 1.465 | 1.727 | 1.417 | 1.395 | 1.647 | 1.383 | 1.321 | 1.437 | 1.199 | 1.114 | 1.545 | 0.121 | 0.0307 | 0.1368 | 0 | 0 | 0 |
| 'Reps1' | 18.259 | 16.854 | 16.729 | 17.551 | 17.535 | 17.334 | 17.660 | 18.640 | 16.681 | 18.679 | 16.782 | 15.474 | 20.574 | 18.506 | 15.280 | 17.751 | 17.479 | 15.105 | 0.011 | 0.7672 | 0.8961 | 0 | 0 | 0 |
| 'Rer1' | 25.490 | 29.008 | 27.729 | 24.545 | 27.227 | 26.250 | 26.469 | 28.419 | 25.466 | 26.076 | 24.821 | 27.939 | 24.623 | 27.443 | 28.775 | 24.517 | 26.821 | 26.142 | 0.008 | 0.8028 | 0.9115 | 0 | 0 | 0 |
| 'Rere' | 18.812 | 18.755 | 18.806 | 18.192 | 17.215 | 18.044 | 17.407 | 18.426 | 19.373 | 19.918 | 21.053 | 20.813 | 19.227 | 18.950 | 19.831 | 19.561 | 18.744 | 20.643 | -0.125 | 0.0000 | 0.0004 | 1 | 0 | 0 |
| 'Rest' | 14.003 | 14.937 | 13.002 | 15.520 | 14.682 | 12.538 | 15.725 | 13.826 | 13.933 | 12.281 | 17.028 | 16.182 | 12.861 | 16.250 | 15.391 | 13.567 | 16.049 | 15.770 | -0.086 | 0.1466 | 0.3719 | 0 | 0 | 0 |
| 'Retsat' | 1.910 | 1.818 | 1.274 | 1.455 | 1.570 | 1.494 | 1.205 | 1.639 | 1.455 | 1.660 | 1.606 | 1.282 | 1.693 | 1.250 | 1.099 | 2.003 | 1.111 | 1.010 | 0.131 | 0.2174 | 0.4639 | 0 | 0 | 0 |
| 'Rev1' | 2.775 | 2.948 | 3.070 | 2.821 | 2.744 | 2.678 | 2.729 | 2.727 | 2.798 | 2.376 | 2.456 | 2.565 | 2.421 | 2.193 | 2.207 | 2.240 | 2.791 | 2.509 | 0.204 | 0.0000 | 0.0005 | 1 | 0 | 0 |
| 'Rev3l' | 4.999 | 5.146 | 5.372 | 5.492 | 4.978 | 4.679 | 5.186 | 4.870 | 5.061 | 5.332 | 5.139 | 5.133 | 4.839 | 4.781 | 4.643 | 4.575 | 5.259 | 5.136 | 0.018 | 0.6377 | 0.8217 | 0 | 0 | 0 |
| 'Rexo1' | 14.613 | 12.963 | 16.828 | 13.118 | 14.309 | 16.044 | 13.758 | 14.558 | 16.305 | 15.660 | 13.416 | 14.899 | 15.826 | 12.774 | 14.231 | 15.035 | 12.788 | 13.939 | 0.034 | 0.4341 | 0.6788 | 0 | 0 | 0 |
| 'Rexo2' | 69.207 | 67.873 | 73.922 | 73.117 | 73.942 | 70.141 | 75.276 | 71.267 | 66.433 | 63.149 | 66.866 | 68.392 | 65.891 | 66.240 | 66.421 | 67.274 | 63.993 | 61.184 | 0.110 | 0.0005 | 0.0060 | 1 | 0 | 0 |
| 'Rexo4' | 22.709 | 24.053 | 23.981 | 22.525 | 22.320 | 23.552 | 24.060 | 22.715 | 24.012 | 9.490 | 11.421 | 14.192 | 11.375 | 11.832 | 13.419 | 11.431 | 12.595 | 14.114 | 0.928 | 0.0000 | 0.0000 | 1 | 1 | 0 |
| 'Rfc1' | 11.762 | 11.756 | 12.320 | 13.121 | 12.194 | 12.571 | 12.805 | 12.137 | 12.050 | 12.118 | 12.145 | 11.678 | 12.218 | 11.909 | 11.817 | 12.471 | 12.004 | 11.047 | 0.035 | 0.1800 | 0.4151 | 0 | 0 | 0 |
| 'Rfc2' | 11.041 | 11.267 | 11.593 | 12.198 | 11.487 | 12.098 | 11.720 | 11.369 | 13.019 | 11.767 | 11.928 | 11.724 | 12.781 | 11.433 | 11.982 | 11.836 | 11.686 | 10.910 | -0.013 | 0.6940 | 0.8558 | 0 | 0 | 0 |
| 'Rfc3' | 3.780 | 3.993 | 4.113 | 4.354 | 4.103 | 4.183 | 4.555 | 4.376 | 4.288 | 3.928 | 3.631 | 3.866 | 4.514 | 4.234 | 3.914 | 3.974 | 4.182 | 3.640 | 0.063 | 0.1759 | 0.4110 | 0 | 0 | 0 |

| | | | | | | | | | | | | | | | | | | | | | | | | |
|---|---|---|---|---|---|---|---|---|---|---|---|---|---|---|---|---|---|---|---|---|---|---|---|---|
| 'Rfc4' | 6.692 | 7.350 | 7.712 | 8.344 | 7.963 | 8.427 | 7.570 | 7.284 | 8.230 | 6.264 | 7.376 | 7.391 | 8.947 | 7.758 | 7.936 | 7.437 | 7.642 | 6.739 | 0.033 | 0.5977 | 0.7954 | 0 | 0 | 0 |
| 'Rfc5' | 5.226 | 5.914 | 6.053 | 4.874 | 5.595 | 5.753 | 5.084 | 5.186 | 5.929 | 4.652 | 4.456 | 5.210 | 6.404 | 5.330 | 5.621 | 5.265 | 4.964 | 5.534 | 0.053 | 0.3516 | 0.6056 | 0 | 0 | 0 |
| 'Rfesd' | 2.379 | 2.647 | 3.668 | 2.927 | 3.317 | 3.431 | 2.617 | 3.166 | 3.178 | 3.282 | 3.115 | 3.105 | 2.773 | 3.183 | 2.847 | 3.287 | 3.614 | 2.470 | -0.023 | 0.7751 | 0.8994 | 0 | 0 | 0 |
| 'Rffl' | 2.861 | 2.649 | 2.961 | 2.544 | 2.727 | 2.842 | 2.711 | 3.118 | 2.997 | 2.560 | 2.611 | 3.061 | 2.626 | 2.303 | 2.741 | 2.742 | 2.706 | 2.598 | 0.073 | 0.2215 | 0.4691 | 0 | 0 | 0 |
| 'Rfk' | 16.235 | 16.275 | 14.910 | 14.459 | 16.783 | 15.958 | 15.055 | 17.144 | 14.812 | 15.881 | 16.311 | 14.681 | 15.366 | 17.546 | 14.576 | 17.102 | 15.745 | 13.103 | 0.009 | 0.8103 | 0.9142 | 0 | 0 | 0 |
| 'Rfng' | 9.016 | 8.243 | 9.716 | 8.181 | 7.840 | 10.226 | 8.788 | 8.358 | 9.693 | 9.318 | 8.180 | 9.988 | 9.714 | 8.221 | 9.417 | 8.875 | 8.254 | 9.373 | -0.032 | 0.3958 | 0.6446 | 0 | 0 | 0 |
| 'Rft1' | 2.598 | 3.143 | 3.234 | 2.725 | 3.138 | 3.203 | 2.615 | 3.353 | 3.335 | 3.381 | 2.604 | 2.530 | 3.279 | 2.924 | 2.498 | 2.948 | 3.104 | 2.203 | 0.101 | 0.1898 | 0.4262 | 0 | 0 | 0 |
| 'Rftn1' | 24.747 | 23.000 | 24.555 | 24.299 | 24.496 | 26.091 | 24.316 | 25.029 | 27.004 | 25.564 | 21.571 | 21.916 | 26.614 | 21.825 | 22.195 | 23.825 | 19.851 | 21.659 | 0.119 | 0.0033 | 0.0271 | 1 | 0 | 0 |
| 'Rfwd2' | 2.596 | 2.822 | 2.465 | 2.766 | 2.597 | 2.385 | 3.137 | 2.977 | 2.334 | 2.491 | 2.661 | 2.526 | 2.425 | 2.464 | 2.399 | 2.516 | 2.719 | 2.548 | 0.065 | 0.1506 | 0.3777 | 0 | 0 | 0 |
| 'Rfwd3' | 22.709 | 19.861 | 24.779 | 23.345 | 20.729 | 22.655 | 23.922 | 20.435 | 23.385 | 20.439 | 22.881 | 25.865 | 21.554 | 22.159 | 25.998 | 19.586 | 23.200 | 27.166 | -0.058 | 0.2339 | 0.4825 | 0 | 0 | 0 |
| 'Rfx1' | 27.785 | 23.506 | 30.788 | 25.878 | 22.526 | 29.754 | 25.476 | 24.114 | 29.704 | 27.442 | 22.721 | 32.711 | 26.318 | 23.198 | 31.145 | 23.137 | 26.179 | 31.318 | -0.037 | 0.3282 | 0.5861 | 0 | 0 | 0 |
| 'Rfx2' | 1.729 | 1.661 | 1.787 | 1.844 | 1.797 | 1.838 | 1.714 | 1.946 | 1.892 | 2.109 | 1.691 | 1.603 | 2.034 | 2.082 | 1.885 | 2.079 | 1.268 | 1.854 | -0.029 | 0.7522 | 0.8895 | 0 | 0 | 0 |
| 'Rfx3' | 0.924 | 0.840 | 0.960 | 1.100 | 0.661 | 0.954 | 1.269 | 0.893 | 0.899 | 1.048 | 1.117 | 0.942 | 0.965 | 1.006 | 0.830 | 0.887 | 0.981 | 0.976 | -0.054 | 0.5520 | 0.7650 | 0 | 0 | 0 |
| 'Rfx5' | 3.344 | 3.531 | 3.728 | 2.934 | 3.720 | 3.852 | 3.412 | 3.618 | 3.870 | 3.654 | 3.153 | 3.539 | 3.610 | 3.743 | 3.573 | 3.356 | 3.396 | 3.463 | 0.012 | 0.7668 | 0.8961 | 0 | 0 | 0 |
| 'Rfx7' | 8.713 | 7.263 | 7.298 | 8.084 | 6.795 | 7.110 | 7.017 | 7.061 | 7.052 | 7.446 | 6.938 | 7.808 | 7.031 | 7.008 | 6.999 | 7.342 | 6.643 | 7.999 | 0.014 | 0.7543 | 0.8906 | 0 | 0 | 0 |
| 'Rfxank' | 11.378 | 13.206 | 11.741 | 11.317 | 13.276 | 12.532 | 11.449 | 12.933 | 12.096 | 12.169 | 11.266 | 11.682 | 12.468 | 12.245 | 10.839 | 12.691 | 11.461 | 10.631 | 0.049 | 0.2122 | 0.4574 | 0 | 0 | 0 |
| 'Rfxap' | 16.803 | 15.022 | 13.925 | 15.096 | 13.858 | 15.059 | 14.777 | 15.181 | 14.320 | 13.860 | 13.489 | 16.258 | 15.030 | 15.625 | 16.802 | 15.479 | 15.010 | 16.368 | -0.053 | 0.2685 | 0.5229 | 0 | 0 | 0 |
| 'Rgcc' | 32.089 | 45.124 | 36.897 | 37.520 | 36.026 | 38.025 | 33.891 | 35.297 | 38.247 | 28.758 | 29.074 | 40.429 | 27.939 | 30.019 | 39.172 | 29.626 | 34.014 | 42.112 | 0.139 | 0.0342 | 0.1477 | 0 | 0 | 0 |
| 'Rgl2' | 19.791 | 17.203 | 23.174 | 20.044 | 17.437 | 21.899 | 18.862 | 18.353 | 22.751 | 20.070 | 18.608 | 23.266 | 20.795 | 18.274 | 20.738 | 18.267 | 19.052 | 22.002 | -0.026 | 0.4092 | 0.6583 | 0 | 0 | 0 |
| 'Rgp1' | 14.469 | 14.616 | 15.604 | 13.288 | 14.379 | 15.380 | 13.671 | 14.899 | 15.785 | 14.635 | 13.339 | 14.213 | 14.283 | 13.999 | 15.338 | 14.731 | 13.386 | 13.605 | 0.042 | 0.1903 | 0.4268 | 0 | 0 | 0 |
| 'Rgs1' | 7.551 | 6.705 | 3.627 | 6.456 | 6.239 | 3.814 | 7.309 | 6.209 | 3.359 | 6.369 | 4.822 | 3.190 | 6.388 | 6.799 | 2.933 | 8.249 | 5.227 | 3.062 | 0.135 | 0.0311 | 0.1383 | 0 | 0 | 0 |
| 'Rgs10' | 140.706 | 167.209 | 132.481 | 152.713 | 160.513 | 145.736 | 151.623 | 151.834 | 143.337 | 143.811 | 145.443 | 124.173 | 157.061 | 156.427 | 141.024 | 155.461 | 155.003 | 132.291 | 0.029 | 0.3979 | 0.6470 | 0 | 0 | 0 |
| 'Rgs11' | 1.317 | 1.081 | 1.199 | 1.238 | 1.227 | 1.079 | 1.158 | 1.040 | 1.170 | 1.184 | 1.250 | 1.357 | 1.769 | 1.152 | 1.114 | 1.063 | 1.060 | 1.241 | -0.107 | 0.1963 | 0.4353 | 0 | 0 | 0 |
| 'Rgs12' | 0.669 | 0.692 | 0.688 | 0.713 | 0.695 | 0.714 | 0.663 | 0.544 | 0.742 | 0.672 | 0.740 | 0.736 | 0.654 | 0.669 | 0.785 | 0.623 | 0.639 | 0.693 | -0.036 | 0.5869 | 0.7880 | 0 | 0 | 0 |
| 'Rgs14' | 24.560 | 21.944 | 25.681 | 22.553 | 22.453 | 26.836 | 22.418 | 22.245 | 24.764 | 23.078 | 22.386 | 31.442 | 23.079 | 22.659 | 29.266 | 23.315 | 22.768 | 31.017 | -0.105 | 0.0064 | 0.0450 | 1 | 0 | 0 |
| 'Rgs16' | 97.422 | 97.667 | 54.932 | 77.395 | 83.370 | 65.491 | 79.377 | 84.874 | 60.720 | 108.575 | 88.740 | 57.123 | 112.401 | 114.744 | 56.635 | 130.211 | 104.725 | 54.315 | -0.201 | 0.0205 | 0.1048 | 0 | 0 | 0 |
| 'Rgs19' | 10.240 | 11.329 | 11.087 | 9.219 | 10.796 | 11.084 | 10.465 | 10.318 | 11.157 | 10.123 | 9.314 | 11.579 | 10.064 | 9.819 | 11.793 | 9.996 | 9.991 | 10.625 | 0.023 | 0.5178 | 0.7407 | 0 | 0 | 0 |
| 'Rgs2' | 32.599 | 13.436 | 7.562 | 29.148 | 13.497 | 7.826 | 28.441 | 15.526 | 6.568 | 15.485 | 14.829 | 5.692 | 12.716 | 13.947 | 5.120 | 14.839 | 15.710 | 4.345 | 0.503 | 0.0003 | 0.0041 | 1 | 1 | 0 |
| 'Rgs8' | 0.863 | 1.049 | 0.835 | 0.827 | 1.030 | 0.923 | 1.064 | 0.917 | 0.931 | 1.033 | 1.347 | 0.878 | 1.317 | 1.261 | 0.875 | 1.020 | 1.497 | 0.842 | -0.245 | 0.0016 | 0.0150 | 1 | 0 | 0 |
| 'Rgs9' | 1.588 | 1.965 | 1.285 | 1.603 | 1.965 | 1.473 | 1.309 | 1.660 | 1.688 | 1.511 | 1.191 | 1.214 | 1.722 | 1.438 | 0.817 | 1.729 | 1.373 | 1.015 | 0.279 | 0.0051 | 0.0379 | 1 | 0 | 0 |
| 'Rhbdd1' | 6.461 | 6.919 | 6.456 | 6.184 | 6.357 | 6.302 | 7.087 | 6.308 | 6.998 | 6.422 | 6.920 | 6.678 | 6.565 | 7.134 | 6.771 | 6.855 | 7.545 | 6.412 | -0.062 | 0.1161 | 0.3235 | 0 | 0 | 0 |
| 'Rhbdd2' | 2.860 | 3.194 | 3.312 | 2.739 | 3.080 | 3.150 | 2.579 | 3.141 | 3.406 | 3.032 | 2.634 | 2.800 | 2.772 | 2.869 | 2.702 | 2.954 | 2.618 | 2.573 | 0.130 | 0.0079 | 0.0526 | 0 | 0 | 0 |
| 'Rhbdd3' | 2.391 | 2.734 | 2.386 | 2.279 | 2.331 | 2.686 | 2.115 | 2.596 | 2.717 | 2.165 | 1.656 | 2.501 | 2.351 | 2.251 | 2.379 | 2.295 | 2.311 | 2.313 | 0.121 | 0.0625 | 0.2171 | 0 | 0 | 0 |
| 'Rhbdf1' | 0.336 | 0.532 | 0.281 | 0.264 | 0.375 | 0.334 | 0.313 | 0.440 | 0.451 | 0.363 | 0.270 | 0.290 | 0.352 | 0.300 | 0.277 | 0.250 | 0.302 | 0.281 | 0.279 | 0.0256 | 0.1227 | 0 | 0 | 0 |
| 'Rhbdf2' | 2.682 | 2.596 | 3.059 | 2.453 | 2.643 | 2.727 | 2.665 | 2.806 | 3.019 | 3.026 | 2.120 | 2.794 | 2.841 | 2.379 | 2.706 | 2.258 | 2.391 | 2.984 | 0.059 | 0.3109 | 0.5692 | 0 | 0 | 0 |
| 'Rhbdl3' | 3.337 | 3.172 | 2.852 | 3.350 | 3.184 | 3.348 | 2.378 | 3.103 | 3.024 | 3.052 | 2.932 | 3.270 | 2.997 | 3.165 | 3.338 | 3.084 | 2.674 | 3.547 | -0.026 | 0.6895 | 0.8534 | 0 | 0 | 0 |
| 'Rheb' | 8.084 | 9.223 | 7.469 | 8.240 | 8.619 | 7.062 | 8.365 | 8.471 | 7.335 | 7.427 | 9.450 | 8.350 | 7.584 | 9.405 | 7.629 | 8.062 | 9.123 | 7.863 | -0.052 | 0.1703 | 0.4030 | 0 | 0 | 0 |
| 'Rhebl1' | 2.030 | 1.982 | 1.974 | 1.833 | 1.942 | 2.079 | 2.047 | 2.082 | 2.176 | 2.090 | 1.790 | 1.639 | 1.969 | 1.353 | 1.285 | 1.797 | 1.735 | 1.339 | 0.276 | 0.0048 | 0.0363 | 1 | 0 | 0 |
| 'Rhoa' | 26.548 | 29.150 | 28.914 | 27.877 | 29.400 | 29.052 | 28.461 | 28.753 | 28.496 | 27.489 | 28.589 | 26.471 | 27.451 | 28.596 | 28.845 | 27.978 | 28.995 | 27.522 | 0.018 | 0.4591 | 0.6960 | 0 | 0 | 0 |
| 'Rhob' | 135.522 | 37.144 | 36.279 | 129.969 | 37.387 | 31.135 | 159.780 | 49.062 | 26.939 | 77.408 | 66.755 | 28.466 | 73.457 | 53.081 | 22.672 | 53.992 | 77.446 | 21.701 | 0.245 | 0.2523 | 0.5038 | 0 | 0 | 0 |
| 'Rhobtb2' | 10.984 | 9.295 | 10.550 | 10.145 | 10.315 | 10.293 | 9.682 | 9.675 | 10.717 | 9.529 | 11.185 | 11.382 | 9.050 | 10.144 | 11.542 | 10.076 | 10.075 | 11.718 | -0.054 | 0.2366 | 0.4864 | 0 | 0 | 0 |
| 'Rhof' | 19.361 | 18.362 | 18.333 | 19.425 | 17.670 | 19.878 | 18.851 | 16.696 | 19.253 | 17.191 | 17.038 | 23.364 | 17.013 | 17.155 | 22.004 | 18.113 | 17.031 | 23.767 | -0.042 | 0.3990 | 0.6482 | 0 | 0 | 0 |
| 'Rhog' | 133.070 | 135.014 | 145.429 | 133.376 | 139.588 | 148.139 | 124.764 | 133.112 | 148.754 | 131.375 | 123.238 | 140.828 | 136.291 | 130.398 | 151.174 | 136.760 | 121.188 | 148.971 | 0.015 | 0.6009 | 0.7972 | 0 | 0 | 0 |
| 'Rhoh' | 19.948 | 18.129 | 16.989 | 19.462 | 16.778 | 17.482 | 20.098 | 18.339 | 17.707 | 16.799 | 16.652 | 15.074 | 15.871 | 15.955 | 14.416 | 16.154 | 15.166 | 15.118 | 0.213 | 0.0000 | 0.0000 | 1 | 0 | 0 |
| 'Rhoq' | 1.333 | 1.370 | 1.382 | 1.302 | 1.491 | 1.044 | 1.423 | 1.502 | 1.055 | 1.102 | 1.532 | 1.562 | 1.182 | 1.348 | 1.346 | 0.932 | 1.433 | 1.328 | -0.009 | 0.9249 | 0.9715 | 0 | 0 | 0 |
| 'Rhot1' | 3.223 | 3.007 | 3.063 | 3.104 | 3.379 | 2.997 | 3.161 | 3.086 | 2.886 | 3.249 | 3.167 | 3.010 | 3.008 | 3.427 | 3.097 | 3.143 | 3.498 | 3.110 | -0.049 | 0.1768 | 0.4121 | 0 | 0 | 0 |
| 'Rhot2' | 6.734 | 7.065 | 7.507 | 6.520 | 6.797 | 7.538 | 6.497 | 7.191 | 7.838 | 6.877 | 6.375 | 7.363 | 7.415 | 6.409 | 6.917 | 6.638 | 6.406 | 6.908 | 0.044 | 0.1845 | 0.4199 | 0 | 0 | 0 |
| 'Ric1' | 5.969 | 6.177 | 5.478 | 6.410 | 6.405 | 5.289 | 5.607 | 5.825 | 5.407 | 5.703 | 6.698 | 6.109 | 6.224 | 6.619 | 5.855 | 5.816 | 6.794 | 6.231 | -0.105 | 0.0050 | 0.0373 | 1 | 0 | 0 |
| 'Ric8a' | 8.934 | 9.500 | 9.514 | 8.193 | 10.186 | 9.422 | 8.663 | 9.340 | 9.424 | 9.401 | 9.001 | 9.343 | 8.506 | 8.647 | 9.317 | 9.526 | 8.589 | 8.747 | 0.026 | 0.4568 | 0.6943 | 0 | 0 | 0 |
| 'Ric8b' | 9.026 | 9.280 | 8.964 | 9.432 | 9.726 | 8.984 | 8.401 | 9.524 | 8.865 | 9.235 | 10.358 | 9.522 | 8.561 | 9.509 | 9.544 | 9.014 | 8.782 | 9.390 | -0.040 | 0.3443 | 0.5996 | 0 | 0 | 0 |
| 'Rictor' | 15.017 | 15.455 | 14.776 | 14.150 | 15.273 | 13.680 | 15.162 | 15.497 | 15.139 | 15.147 | 15.975 | 14.474 | 14.179 | 14.997 | 13.395 | 15.133 | 14.913 | 14.461 | 0.007 | 0.8234 | 0.9223 | 0 | 0 | 0 |
| 'Rida' | 1.884 | 2.223 | 1.982 | 2.129 | 2.189 | 2.226 | 2.111 | 1.987 | 1.842 | 1.988 | 1.747 | 1.955 | 2.476 | 2.155 | 1.924 | 2.671 | 2.085 | 1.842 | -0.027 | 0.7127 | 0.8665 | 0 | 0 | 0 |
| 'Rif1' | 6.031 | 5.599 | 5.551 | 6.537 | 5.613 | 5.065 | 6.267 | 5.619 | 5.431 | 5.151 | 5.800 | 5.950 | 5.238 | 5.549 | 5.604 | 5.017 | 6.051 | 6.104 | 0.022 | 0.6699 | 0.8408 | 0 | 0 | 0 |
| 'Rilpl1' | 2.054 | 2.029 | 2.637 | 2.380 | 1.916 | 2.170 | 2.822 | 2.107 | 2.366 | 2.348 | 2.014 | 2.334 | 2.117 | 1.983 | 1.970 | 2.283 | 1.685 | 1.792 | 0.133 | 0.1075 | 0.3084 | 0 | 0 | 0 |
| 'Rilpl2' | 34.920 | 29.729 | 32.375 | 33.938 | 31.890 | 34.339 | 32.549 | 33.024 | 32.237 | 32.916 | 31.018 | 35.051 | 31.907 | 33.095 | 33.909 | 35.001 | 34.598 | 35.790 | -0.050 | 0.1310 | 0.3477 | 0 | 0 | 0 |
| 'Rims3' | 0.438 | 0.369 | 0.416 | 0.435 | 0.212 | 0.384 | 0.419 | 0.383 | 0.337 | 0.442 | 0.413 | 0.510 | 0.555 | 0.346 | 0.477 | 0.402 | 0.419 | 0.543 | -0.293 | 0.0111 | 0.0673 | 0 | 0 | 0 |
| 'Rin3' | 32.491 | 29.154 | 30.150 | 30.477 | 27.075 | 31.741 | 29.736 | 27.603 | 32.290 | 32.674 | 29.004 | 34.808 | 30.914 | 29.115 | 32.158 | 30.390 | 28.553 | 33.591 | -0.065 | 0.0217 | 0.1088 | 0 | 0 | 0 |
| 'Ring1' | 11.649 | 10.827 | 13.326 | 10.319 | 11.990 | 14.152 | 10.545 | 11.632 | 12.567 | 12.663 | 9.774 | 12.417 | 12.706 | 9.743 | 11.424 | 12.343 | 10.851 | 10.600 | 0.053 | 0.3364 | 0.5950 | 0 | 0 | 0 |
| 'Rinl' | 135.608 | 127.506 | 147.760 | 134.785 | 131.286 | 147.308 | 139.195 | 130.382 | 146.250 | 137.436 | 130.533 | 152.286 | 138.110 | 128.083 | 146.761 | 131.312 | 136.677 | 144.863 | -0.018 | 0.3132 | 0.5715 | 0 | 0 | 0 |
| 'Rint1' | 2.407 | 2.277 | 2.517 | 2.292 | 2.544 | 2.241 | 2.324 | 2.467 | 2.417 | 2.231 | 2.371 | 2.447 | 2.385 | 2.359 | 2.065 | 2.197 | 2.625 | 2.269 | 0.026 | 0.5559 | 0.7678 | 0 | 0 | 0 |
| 'Riok1' | 21.346 | 20.494 | 23.174 | 22.930 | 22.954 | 21.309 | 22.254 | 23.525 | 22.938 | 23.538 | 21.227 | 19.449 | 22.756 | 21.394 | 19.910 | 21.661 | 22.148 | 17.840 | 0.078 | 0.0646 | 0.2217 | 0 | 0 | 0 |
| 'Riok3' | 36.029 | 40.661 | 36.155 | 36.698 | 40.408 | 38.302 | 38.879 | 39.142 | 37.056 | 36.941 | 37.026 | 36.676 | 36.420 | 38.526 | 36.114 | 37.449 | 39.656 | 34.852 | 0.029 | 0.1971 | 0.4361 | 0 | 0 | 0 |
| 'Ripk1' | 7.303 | 6.220 | 6.660 | 7.117 | 6.510 | 6.702 | 6.864 | 6.941 | 6.390 | 6.733 | 7.135 | 7.721 | 6.438 | 7.433 | 7.039 | 6.853 | 6.887 | 7.422 | -0.080 | 0.0619 | 0.2156 | 0 | 0 | 0 |
| 'Ripk2' | 4.217 | 4.528 | 4.308 | 4.574 | 4.383 | 4.323 | 4.698 | 4.343 | 3.811 | 4.143 | 3.978 | 4.134 | 4.049 | 4.350 | 3.656 | 3.968 | 4.627 | 3.358 | 0.098 | 0.0621 | 0.2159 | 0 | 0 | 0 |
| 'Ripk3' | 39.695 | 33.191 | 41.779 | 37.948 | 35.054 | 39.565 | 40.931 | 35.825 | 38.878 | 36.958 | 36.314 | 38.327 | 39.776 | 34.518 | 38.901 | 35.336 | 38.249 | 38.462 | 0.015 | 0.6708 | 0.8413 | 0 | 0 | 0 |
| 'Rit1' | 3.557 | 3.882 | 3.623 | 3.784 | 3.743 | 4.049 | 3.288 | 4.179 | 4.097 | 3.913 | 4.278 | 4.074 | 3.465 | 3.969 | 4.124 | 4.625 | 3.802 | 4.230 | -0.101 | 0.0541 | 0.1989 | 0 | 0 | 0 |
| 'Rita1' | 1.224 | 1.260 | 1.267 | 1.249 | 1.218 | 1.255 | 1.283 | 0.940 | 0.936 | 1.081 | 1.102 | 1.302 | 0.976 | 1.062 | 1.278 | 0.896 | 1.199 | 1.246 | 0.045 | 0.6608 | 0.8362 | 0 | 0 | 0 |
| 'Rlf' | 9.234 | 8.766 | 7.707 | 9.064 | 8.379 | 7.494 | 8.625 | 8.880 | 7.449 | 8.971 | 10.217 | 8.799 | 9.309 | 9.483 | 8.739 | 9.823 | 10.234 | 8.630 | -0.167 | 0.0000 | 0.0000 | 1 | 0 | 0 |
| 'Rlim' | 6.549 | 6.487 | 7.042 | 7.343 | 6.479 | 6.424 | 8.354 | 6.687 | 6.458 | 6.067 | 7.307 | 7.708 | 5.700 | 6.444 | 6.644 | 5.235 | 7.614 | 7.116 | 0.039 | 0.5798 | 0.7822 | 0 | 0 | 0 |
| 'Rltpr' | 129.628 | 121.960 | 153.353 | 133.928 | 121.384 | 144.686 | 130.704 | 127.075 | 151.307 | 141.756 | 129.611 | 150.670 | 153.727 | 126.971 | 139.070 | 127.430 | 134.162 | 146.201 | -0.055 | 0.0901 | 0.2760 | 0 | 0 | 0 |
| 'Rmdn1' | 5.666 | 7.666 | 6.468 | 5.283 | 8.476 | 5.858 | 5.959 | 7.633 | 6.826 | 7.184 | 6.437 | 5.811 | 6.501 | 6.228 | 5.956 | 6.339 | 5.768 | 4.766 | 0.106 | 0.1765 | 0.4118 | 0 | 0 | 0 |
| 'Rmdn3' | 5.886 | 6.484 | 6.324 | 6.939 | 6.188 | 5.962 | 5.749 | 5.689 | 5.947 | 5.471 | 5.907 | 6.387 | 5.955 | 6.280 | 5.981 | 6.117 | 5.774 | 5.848 | 0.023 | 0.6110 | 0.8049 | 0 | 0 | 0 |
| 'Rmi1' | 4.215 | 4.255 | 4.350 | 4.952 | 4.420 | 4.236 | 4.286 | 4.700 | 4.601 | 3.729 | 4.024 | 4.928 | 3.734 | 4.194 | 4.322 | 3.763 | 4.670 | 4.880 | 0.048 | 0.4486 | 0.6888 | 0 | 0 | 0 |
| 'Rmi2' | 2.515 | 2.445 | 3.193 | 2.449 | 2.342 | 2.753 | 2.776 | 2.588 | 2.883 | 2.936 | 2.570 | 3.342 | 3.470 | 3.141 | 3.605 | 2.710 | 2.730 | 2.926 | -0.207 | 0.0016 | 0.0150 | 1 | 0 | 0 |
| 'Rmnd1' | 3.969 | 4.012 | 4.341 | 4.287 | 4.086 | 3.608 | 4.661 | 3.864 | 4.275 | 3.870 | 3.986 | 3.995 | 3.933 | 4.423 | 3.939 | 3.631 | 4.285 | 3.799 | 0.037 | 0.4571 | 0.6944 | 0 | 0 | 0 |
| 'Rmnd5a' | 17.031 | 17.729 | 17.321 | 17.155 | 16.834 | 16.868 | 16.749 | 17.049 | 17.272 | 16.459 | 18.382 | 18.914 | 16.283 | 18.884 | 17.560 | 17.372 | 17.822 | 19.055 | -0.072 | 0.0165 | 0.0900 | 0 | 0 | 0 |
| 'Rmnd5b' | 17.068 | 15.121 | 15.868 | 13.531 | 14.820 | 16.458 | 15.630 | 16.728 | 15.616 | 16.744 | 15.216 | 16.130 | 16.632 | 15.680 | 15.707 | 16.346 | 13.998 | 15.963 | -0.023 | 0.5786 | 0.7820 | 0 | 0 | 0 |
| 'Rnase4' | 6.556 | 6.111 | 5.944 | 5.954 | 6.958 | 5.789 | 6.816 | 6.258 | 6.281 | 7.107 | 6.409 | 7.277 | 7.377 | 7.533 | 7.765 | 7.738 | 6.760 | 7.351 | -0.220 | 0.0002 | 0.0033 | 1 | 0 | 0 |
| 'Rnaseh1' | 17.273 | 20.085 | 20.171 | 18.588 | 21.468 | 19.965 | 20.144 | 20.463 | 20.477 | 19.229 | 18.804 | 20.087 | 18.577 | 20.830 | 20.778 | 18.298 | 18.561 | 18.789 | 0.025 | 0.5592 | 0.7700 | 0 | 0 | 0 |
| 'Rnaseh2a' | 7.694 | 7.004 | 8.343 | 7.828 | 7.324 | 8.303 | 8.202 | 7.747 | 7.782 | 7.012 | 6.993 | 7.285 | 8.111 | 7.387 | 6.859 | 7.563 | 7.272 | 7.025 | 0.095 | 0.0234 | 0.1152 | 0 | 0 | 0 |
| 'Rnaseh2c' | 9.968 | 10.192 | 10.375 | 9.116 | 10.924 | 11.294 | 9.353 | 10.278 | 10.911 | 8.505 | 8.539 | 9.467 | 10.008 | 9.547 | 9.892 | 9.790 | 8.844 | 8.996 | 0.136 | 0.0027 | 0.0230 | 1 | 0 | 0 |
| 'Rnasek' | 5.158 | 5.244 | 5.141 | 5.356 | 5.221 | 5.341 | 4.213 | 4.708 | 5.079 | 5.228 | 5.433 | 5.141 | 5.031 | 4.001 | 5.419 | 4.708 | 5.072 | 4.534 | 0.019 | 0.7979 | 0.9103 | 0 | 0 | 0 |
| 'Rnasel' | 1.924 | 2.456 | 2.883 | 2.121 | 2.424 | 2.674 | 2.413 | 2.661 | 2.598 | 2.971 | 2.752 | 2.706 | 2.553 | 2.438 | 2.580 | 2.535 | 2.548 | 2.318 | -0.091 | 0.1445 | 0.3689 | 0 | 0 | 0 |
| 'Rnf10' | 25.040 | 24.551 | 26.032 | 24.670 | 25.121 | 27.267 | 25.303 | 25.421 | 26.609 | 24.148 | 24.334 | 27.560 | 24.330 | 24.979 | 27.187 | 24.257 | 24.589 | 27.533 | -0.003 | 0.8649 | 0.9414 | 0 | 0 | 0 |
| 'Rnf103' | 3.273 | 4.070 | 3.571 | 3.376 | 3.808 | 3.712 | 3.736 | 3.885 | 3.424 | 3.666 | 3.914 | 3.700 | 3.499 | 3.601 | 3.472 | 3.839 | 3.649 | 3.384 | -0.007 | 0.8532 | 0.9372 | 0 | 0 | 0 |
| 'Rnf11' | 5.607 | 6.659 | 6.260 | 6.045 | 6.472 | 5.411 | 5.959 | 6.991 | 5.748 | 6.577 | 6.034 | 6.519 | 5.964 | 7.034 | 5.990 | 6.131 | 6.141 | 6.107 | -0.050 | 0.2481 | 0.4990 | 0 | 0 | 0 |
| 'Rnf111' | 11.672 | 11.703 | 10.823 | 11.081 | 11.462 | 11.151 | 11.103 | 12.098 | 11.231 | 10.990 | 11.593 | 11.383 | 10.885 | 12.189 | 11.249 | 11.698 | 11.807 | 11.075 | -0.017 | 0.5119 | 0.7371 | 0 | 0 | 0 |
| 'Rnf113a1' | 5.168 | 5.260 | 5.710 | 5.777 | 6.570 | 5.699 | 5.639 | 4.629 | 4.982 | 3.910 | 5.343 | 5.941 | 4.998 | 5.047 | 6.858 | 4.197 | 5.870 | 5.383 | 0.039 | 0.6822 | 0.8487 | 0 | 0 | 0 |
| 'Rnf114' | 26.391 | 31.619 | 26.586 | 25.607 | 30.019 | 29.595 | 24.618 | 30.608 | 29.025 | 27.623 | 29.236 | 30.646 | 29.587 | 29.732 | 32.817 | 30.537 | 29.691 | 30.934 | -0.106 | 0.0035 | 0.0285 | 1 | 0 | 0 |
| 'Rnf115' | 16.440 | 17.659 | 16.514 | 16.650 | 17.613 | 15.517 | 16.810 | 17.117 | 16.128 | 16.033 | 15.632 | 15.780 | 15.450 | 17.410 | 15.836 | 15.255 | 16.593 | 15.818 | 0.052 | 0.0602 | 0.2116 | 0 | 0 | 0 |
| 'Rnf121' | 6.759 | 6.621 | 6.919 | 7.000 | 7.183 | 6.371 | 7.416 | 7.236 | 7.067 | 6.743 | 6.419 | 7.205 | 7.220 | 7.165 | 7.066 | 7.067 | 7.144 | 6.675 | -0.018 | 0.6740 | 0.8433 | 0 | 0 | 0 |
| 'Rnf122' | 1.490 | 2.137 | 1.426 | 1.617 | 2.148 | 1.532 | 1.566 | 1.898 | 1.501 | 1.515 | 1.730 | 1.618 | 1.115 | 1.813 | 1.255 | 1.605 | 1.442 | 1.336 | 0.164 | 0.0327 | 0.1432 | 0 | 0 | 0 |
| 'Rnf123' | 18.279 | 16.952 | 18.702 | 16.626 | 15.293 | 17.292 | 17.322 | 17.077 | 18.925 | 17.214 | 17.208 | 18.287 | 17.421 | 15.299 | 17.153 | 16.527 | 16.662 | 17.629 | 0.018 | 0.5792 | 0.7822 | 0 | 0 | 0 |
| 'Rnf125' | 56.501 | 58.114 | 55.895 | 57.661 | 59.477 | 57.766 | 55.890 | 56.295 | 58.873 | 57.177 | 63.702 | 53.155 | 61.256 | 53.701 | 56.708 | 51.292 | 55.004 | 61.172 | -0.001 | 0.9860 | 0.9953 | 0 | 0 | 0 |
| 'Rnf126' | 49.497 | 42.458 | 52.427 | 46.968 | 47.010 | 53.843 | 46.848 | 48.158 | 50.854 | 45.979 | 41.167 | 52.107 | 51.041 | 43.222 | 52.151 | 47.350 | 41.206 | 51.190 | 0.034 | 0.2927 | 0.5493 | 0 | 0 | 0 |
| 'Rnf128' | 3.975 | 4.940 | 4.061 | 4.116 | 4.556 | 3.876 | 3.913 | 4.805 | 4.102 | 7.436 | 6.775 | 6.131 | 5.887 | 7.729 | 5.430 | 6.275 | 7.167 | 6.023 | -0.627 | 0.0000 | 0.0000 | 1 | 1 | 0 |
| 'Rnf13' | 1.890 | 2.156 | 1.920 | 2.100 | 2.303 | 2.045 | 2.319 | 2.243 | 2.005 | 2.029 | 2.091 | 1.962 | 2.058 | 2.029 | 1.926 | 2.266 | 2.259 | 1.751 | 0.036 | 0.4027 | 0.6514 | 0 | 0 | 0 |
| 'Rnf130' | 2.516 | 4.050 | 2.776 | 2.425 | 3.900 | 3.511 | 3.114 | 3.375 | 2.863 | 2.996 | 2.983 | 2.924 | 2.937 | 3.309 | 2.082 | 2.650 | 2.870 | 2.266 | 0.164 | 0.1005 | 0.2967 | 0 | 0 | 0 |
| 'Rnf135' | 8.030 | 7.785 | 8.070 | 7.747 | 8.047 | 9.592 | 8.695 | 7.582 | 8.617 | 7.292 | 7.880 | 9.600 | 7.298 | 8.686 | 9.194 | 7.866 | 9.531 | 10.097 | -0.073 | 0.2162 | 0.4622 | 0 | 0 | 0 |
| 'Rnf138' | 13.742 | 15.287 | 13.991 | 13.583 | 16.216 | 13.206 | 14.661 | 15.914 | 13.532 | 15.202 | 12.708 | 13.645 | 13.722 | 15.595 | 12.893 | 15.100 | 15.240 | 13.136 | 0.020 | 0.6171 | 0.8087 | 0 | 0 | 0 |
| 'Rnf139' | 21.032 | 23.277 | 20.350 | 21.293 | 23.099 | 20.998 | 22.277 | 21.972 | 19.921 | 23.373 | 25.604 | 23.623 | 22.730 | 26.602 | 22.275 | 23.240 | 26.305 | 24.079 | -0.177 | 0.0000 | 0.0000 | 1 | 0 | 0 |
| 'Rnf14' | 18.840 | 22.751 | 20.416 | 17.746 | 23.433 | 21.530 | 18.946 | 25.044 | 21.028 | 19.236 | 20.931 | 21.282 | 20.730 | 22.187 | 22.122 | 22.907 | 21.185 | 20.560 | -0.026 | 0.5417 | 0.7586 | 0 | 0 | 0 |
| 'Rnf141' | 1.928 | 2.761 | 2.545 | 2.179 | 2.599 | 2.588 | 2.249 | 2.780 | 2.406 | 2.703 | 2.526 | 2.421 | 2.459 | 2.911 | 2.604 | 2.448 | 2.715 | 2.198 | -0.070 | 0.2239 | 0.4711 | 0 | 0 | 0 |
| 'Rnf144a' | 32.228 | 36.529 | 26.692 | 29.347 | 33.807 | 27.447 | 27.966 | 32.049 | 28.354 | 28.870 | 32.527 | 26.721 | 29.438 | 33.473 | 26.791 | 27.601 | 31.666 | 25.451 | 0.053 | 0.1025 | 0.3002 | 0 | 0 | 0 |
| 'Rnf145' | 45.150 | 43.833 | 44.374 | 44.181 | 45.443 | 44.120 | 46.509 | 48.811 | 45.093 | 47.954 | 45.918 | 41.484 | 45.848 | 42.898 | 40.464 | 47.749 | 46.831 | 39.258 | 0.026 | 0.3877 | 0.6374 | 0 | 0 | 0 |
| 'Rnf146' | 7.437 | 7.833 | 6.813 | 7.789 | 7.146 | 6.631 | 7.859 | 7.226 | 7.060 | 6.884 | 7.843 | 7.963 | 6.869 | 7.928 | 7.699 | 7.128 | 8.055 | 8.214 | -0.073 | 0.1215 | 0.3323 | 0 | 0 | 0 |
| 'Rnf149' | 31.839 | 34.393 | 34.325 | 29.717 | 36.743 | 31.731 | 31.964 | 38.264 | 31.687 | 37.132 | 41.756 | 33.879 | 31.739 | 40.518 | 34.558 | 36.882 | 40.287 | 36.528 | -0.157 | 0.0000 | 0.0004 | 1 | 0 | 0 |
| 'Rnf157' | 12.457 | 12.475 | 12.306 | 11.148 | 11.807 | 11.782 | 11.963 | 11.984 | 11.900 | 14.488 | 15.813 | 14.032 | 14.665 | 14.281 | 14.946 | 14.323 | 14.004 | 15.015 | -0.297 | 0.0000 | 0.0000 | 1 | 0 | 0 |
| 'Rnf166' | 24.342 | 25.577 | 25.795 | 22.223 | 22.106 | 27.446 | 21.191 | 24.373 | 26.173 | 22.533 | 21.420 | 25.742 | 21.708 | 21.836 | 27.686 | 23.949 | 21.501 | 25.735 | 0.040 | 0.3094 | 0.5676 | 0 | 0 | 0 |
| 'Rnf167' | 19.937 | 23.066 | 23.635 | 20.962 | 22.016 | 23.185 | 20.411 | 22.377 | 23.703 | 22.686 | 20.322 | 22.860 | 21.383 | 21.520 | 22.605 | 21.166 | 22.323 | 20.762 | 0.015 | 0.6146 | 0.8076 | 0 | 0 | 0 |
| 'Rnf168' | 4.638 | 5.311 | 5.690 | 4.816 | 5.612 | 5.900 | 5.264 | 5.448 | 5.606 | 5.040 | 6.070 | 5.643 | 5.295 | 6.006 | 5.833 | 5.455 | 5.522 | 6.123 | -0.086 | 0.0248 | 0.1200 | 0 | 0 | 0 |
| 'Rnf169' | 6.518 | 5.390 | 6.525 | 6.496 | 6.177 | 6.357 | 6.774 | 6.156 | 6.541 | 6.218 | 6.339 | 7.286 | 5.739 | 6.542 | 6.821 | 5.225 | 6.842 | 7.922 | -0.058 | 0.3206 | 0.5786 | 0 | 0 | 0 |
| 'Rnf181' | 7.607 | 9.304 | 8.671 | 7.607 | 9.177 | 9.190 | 7.901 | 8.801 | 8.621 | 7.821 | 8.949 | 8.792 | 8.466 | 9.356 | 9.149 | 8.458 | 8.806 | 8.505 | -0.038 | 0.1791 | 0.4145 | 0 | 0 | 0 |
| 'Rnf185' | 7.398 | 8.532 | 7.492 | 6.872 | 7.402 | 8.006 | 7.460 | 8.051 | 7.564 | 8.003 | 7.873 | 7.716 | 7.542 | 7.251 | 7.634 | 8.067 | 7.894 | 6.776 | -0.008 | 0.8570 | 0.9387 | 0 | 0 | 0 |
| 'Rnf19a' | 106.704 | 100.493 | 104.058 | 98.498 | 100.101 | 91.958 | 100.683 | 106.496 | 96.929 | 101.154 | 109.801 | 91.600 | 85.332 | 97.261 | 90.960 | 88.822 | 97.105 | 92.602 | 0.075 | 0.0659 | 0.2242 | 0 | 0 | 0 |
| 'Rnf19b' | 33.320 | 28.301 | 30.322 | 27.042 | 28.735 | 32.486 | 28.670 | 29.896 | 28.452 | 36.666 | 27.951 | 28.721 | 31.906 | 29.710 | 28.954 | 37.316 | 26.681 | 28.660 | -0.052 | 0.3194 | 0.5780 | 0 | 0 | 0 |
| 'Rnf2' | 9.546 | 10.059 | 9.673 | 9.511 | 10.148 | 9.021 | 10.635 | 9.871 | 8.957 | 7.280 | 8.841 | 8.898 | 7.094 | 8.234 | 7.804 | 7.013 | 8.422 | 8.275 | 0.271 | 0.0000 | 0.0000 | 1 | 0 | 0 |
| 'Rnf20' | 15.493 | 15.036 | 15.768 | 15.822 | 15.130 | 15.582 | 15.293 | 15.088 | 15.828 | 15.506 | 15.515 | 16.048 | 14.694 | 14.635 | 15.098 | 14.133 | 15.292 | 16.122 | 0.010 | 0.7113 | 0.8659 | 0 | 0 | 0 |
| 'Rnf213' | 14.488 | 12.078 | 15.095 | 13.566 | 12.444 | 15.102 | 11.591 | 14.243 | 16.154 | 14.260 | 14.067 | 17.425 | 12.926 | 12.535 | 16.462 | 13.040 | 12.851 | 19.274 | -0.093 | 0.0704 | 0.2338 | 0 | 0 | 0 |
| 'Rnf214' | 2.579 | 2.443 | 2.857 | 2.788 | 2.251 | 2.639 | 2.681 | 2.538 | 2.981 | 2.721 | 2.989 | 2.816 | 2.795 | 2.755 | 2.807 | 2.879 | 2.788 | 2.697 | -0.094 | 0.0388 | 0.1607 | 0 | 0 | 0 |
| 'Rnf215' | 13.045 | 13.028 | 15.978 | 11.238 | 13.329 | 15.966 | 11.818 | 13.127 | 15.560 | 15.641 | 12.500 | 15.388 | 15.722 | 11.222 | 14.317 | 12.861 | 11.849 | 14.099 | -0.017 | 0.7684 | 0.8967 | 0 | 0 | 0 |
| 'Rnf216' | 5.511 | 5.083 | 5.272 | 5.351 | 5.431 | 5.639 | 5.019 | 5.805 | 5.337 | 5.766 | 5.584 | 5.227 | 6.062 | 5.998 | 5.193 | 5.827 | 5.843 | 4.957 | -0.062 | 0.0958 | 0.2881 | 0 | 0 | 0 |
| 'Rnf219' | 5.493 | 6.822 | 6.257 | 6.145 | 5.185 | 5.722 | 6.928 | 6.489 | 6.185 | 5.158 | 6.461 | 6.131 | 5.468 | 5.904 | 5.508 | 5.426 | 5.599 | 5.611 | 0.093 | 0.1138 | 0.3199 | 0 | 0 | 0 |
| 'Rnf220' | 11.328 | 11.443 | 11.466 | 10.772 | 11.463 | 12.106 | 11.057 | 11.671 | 11.406 | 11.262 | 10.617 | 11.749 | 11.412 | 10.801 | 11.889 | 12.128 | 10.431 | 11.884 | -0.002 | 0.9466 | 0.9794 | 0 | 0 | 0 |
| 'Rnf24' | 0.624 | 0.723 | 0.569 | 0.633 | 0.628 | 0.581 | 0.631 | 0.666 | 0.637 | 0.596 | 0.754 | 0.605 | 0.582 | 0.727 | 0.529 | 0.572 | 0.743 | 0.570 | -0.004 | 0.9450 | 0.9789 | 0 | 0 | 0 |
| 'Rnf25' | 6.676 | 6.818 | 7.586 | 6.233 | 7.148 | 7.153 | 7.621 | 6.923 | 7.139 | 6.697 | 6.653 | 8.197 | 6.575 | 6.915 | 6.984 | 6.785 | 7.265 | 7.885 | -0.029 | 0.4978 | 0.7258 | 0 | 0 | 0 |
| 'Rnf26' | 18.554 | 22.334 | 26.243 | 19.599 | 21.775 | 25.475 | 19.112 | 22.202 | 22.950 | 20.116 | 19.483 | 19.926 | 22.762 | 20.139 | 21.767 | 19.624 | 19.033 | 19.367 | 0.108 | 0.0460 | 0.1785 | 0 | 0 | 0 |
| 'Rnf31' | 17.416 | 15.947 | 17.594 | 15.629 | 16.636 | 17.470 | 16.015 | 17.309 | 18.514 | 17.289 | 16.419 | 18.016 | 18.081 | 16.189 | 18.147 | 17.541 | 16.972 | 17.900 | -0.046 | 0.1040 | 0.3023 | 0 | 0 | 0 |
| 'Rnf34' | 9.799 | 10.771 | 10.921 | 10.418 | 11.011 | 10.903 | 10.957 | 11.405 | 11.173 | 10.361 | 12.028 | 11.802 | 10.465 | 10.991 | 10.996 | 10.463 | 11.405 | 10.902 | -0.042 | 0.1725 | 0.4063 | 0 | 0 | 0 |
| 'Rnf38' | 13.167 | 14.188 | 13.784 | 12.799 | 13.848 | 13.566 | 13.175 | 14.418 | 13.975 | 13.712 | 14.322 | 14.063 | 12.412 | 14.056 | 13.796 | 13.924 | 14.218 | 13.716 | -0.025 | 0.2970 | 0.5529 | 0 | 0 | 0 |
| 'Rnf4' | 8.772 | 8.673 | 7.799 | 8.304 | 8.608 | 8.014 | 8.055 | 8.642 | 7.998 | 8.621 | 8.771 | 8.755 | 8.552 | 8.854 | 8.381 | 9.282 | 8.758 | 8.235 | -0.074 | 0.0030 | 0.0250 | 1 | 0 | 0 |
| 'Rnf40' | 26.219 | 26.060 | 28.773 | 25.536 | 25.113 | 29.616 | 25.596 | 25.868 | 29.381 | 27.237 | 25.860 | 27.422 | 27.272 | 24.893 | 27.591 | 25.825 | 24.950 | 26.903 | 0.015 | 0.5368 | 0.7556 | 0 | 0 | 0 |
| 'Rnf41' | 5.521 | 5.062 | 5.437 | 4.936 | 4.629 | 5.476 | 5.495 | 4.471 | 5.548 | 5.071 | 5.740 | 7.308 | 5.016 | 6.238 | 6.458 | 5.987 | 5.582 | 7.273 | -0.239 | 0.0000 | 0.0008 | 1 | 0 | 0 |
| 'Rnf43' | 7.344 | 6.219 | 6.869 | 6.653 | 6.444 | 6.079 | 6.672 | 6.488 | 6.384 | 7.123 | 7.054 | 8.454 | 6.849 | 6.761 | 7.947 | 7.628 | 6.612 | 8.414 | -0.186 | 0.0001 | 0.0016 | 1 | 0 | 0 |
| 'Rnf44' | 9.576 | 9.601 | 10.525 | 8.909 | 9.786 | 10.097 | 9.162 | 10.014 | 10.425 | 9.894 | 9.620 | 11.146 | 10.048 | 9.585 | 10.697 | 9.508 | 9.324 | 10.505 | -0.047 | 0.0526 | 0.1949 | 0 | 0 | 0 |
| 'Rnf5' | 16.934 | 18.816 | 17.096 | 17.396 | 19.821 | 19.650 | 17.232 | 18.868 | 17.522 | 15.867 | 17.583 | 18.434 | 15.789 | 18.076 | 17.943 | 16.433 | 17.705 | 18.383 | 0.050 | 0.1693 | 0.4017 | 0 | 0 | 0 |
| 'Rnf6' | 7.617 | 8.312 | 8.068 | 7.456 | 7.575 | 7.660 | 7.833 | 7.484 | 8.050 | 6.988 | 8.026 | 8.718 | 7.277 | 8.047 | 8.915 | 7.427 | 8.178 | 8.407 | -0.050 | 0.1607 | 0.3916 | 0 | 0 | 0 |
| 'Rnf7' | 23.067 | 29.373 | 24.097 | 25.695 | 27.521 | 25.893 | 23.970 | 28.550 | 24.366 | 25.462 | 25.064 | 25.059 | 27.050 | 27.241 | 24.760 | 26.653 | 26.180 | 24.577 | -0.011 | 0.7657 | 0.8956 | 0 | 0 | 0 |
| 'Rnf8' | 1.477 | 1.407 | 1.591 | 1.549 | 1.417 | 1.631 | 1.464 | 1.387 | 1.501 | 1.251 | 1.565 | 1.698 | 1.560 | 1.585 | 1.478 | 1.244 | 1.440 | 1.459 | -0.001 | 0.9927 | 0.9986 | 0 | 0 | 0 |
| 'Rnft1' | 6.262 | 8.152 | 7.444 | 7.048 | 7.391 | 7.872 | 7.461 | 7.902 | 7.116 | 6.044 | 8.139 | 8.097 | 6.385 | 7.545 | 7.952 | 6.699 | 8.686 | 8.527 | -0.040 | 0.4103 | 0.6592 | 0 | 0 | 0 |
| 'Rngtt' | 5.694 | 5.670 | 5.609 | 6.345 | 5.909 | 5.441 | 6.775 | 5.317 | 5.526 | 5.703 | 5.724 | 5.858 | 5.496 | 5.900 | 5.590 | 5.486 | 5.718 | 5.566 | 0.019 | 0.6568 | 0.8340 | 0 | 0 | 0 |
| 'Rnh1' | 58.473 | 59.939 | 61.544 | 55.919 | 58.413 | 60.466 | 57.784 | 60.177 | 60.378 | 57.126 | 56.030 | 61.682 | 57.741 | 58.989 | 61.412 | 59.011 | 59.230 | 58.256 | 0.000 | 0.9910 | 0.9976 | 0 | 0 | 0 |
| 'Rnmt' | 3.329 | 3.838 | 3.427 | 3.369 | 4.003 | 3.463 | 3.760 | 3.771 | 3.551 | 3.324 | 3.615 | 3.742 | 3.365 | 3.783 | 3.513 | 3.658 | 3.604 | 3.351 | 0.008 | 0.8115 | 0.9153 | 0 | 0 | 0 |
| 'Rnmtl1' | 2.250 | 2.071 | 2.005 | 2.036 | 2.395 | 2.711 | 2.363 | 2.674 | 2.476 | 1.948 | 2.528 | 2.543 | 2.395 | 1.893 | 1.991 | 2.290 | 2.216 | 2.459 | 0.036 | 0.6870 | 0.8521 | 0 | 0 | 0 |
| 'Rnpc3' | 2.181 | 2.631 | 2.192 | 2.147 | 2.363 | 2.143 | 2.289 | 2.268 | 2.362 | 2.432 | 2.226 | 2.521 | 2.653 | 2.384 | 2.160 | 2.706 | 2.490 | 1.980 | -0.081 | 0.1404 | 0.3626 | 0 | 0 | 0 |
| 'Rnpep' | 11.963 | 10.739 | 12.849 | 12.481 | 11.778 | 13.219 | 12.523 | 10.784 | 12.373 | 4.281 | 4.280 | 4.902 | 5.017 | 4.098 | 4.087 | 4.115 | 4.121 | 4.153 | 1.463 | 0.0000 | 0.0000 | 1 | 1 | 1 |
| 'Rnpepl1' | 8.534 | 7.310 | 9.541 | 7.897 | 7.515 | 9.672 | 7.654 | 7.918 | 9.033 | 8.714 | 8.247 | 9.795 | 8.894 | 7.724 | 9.403 | 7.722 | 8.307 | 9.778 | -0.077 | 0.0188 | 0.0990 | 0 | 0 | 0 |
| 'Rnps1' | 74.025 | 75.841 | 76.965 | 73.285 | 72.689 | 78.104 | 73.348 | 79.261 | 75.996 | 70.907 | 66.907 | 73.742 | 77.080 | 72.010 | 76.490 | 74.179 | 71.209 | 72.410 | 0.043 | 0.0935 | 0.2831 | 0 | 0 | 0 |
| 'Rock1' | 45.841 | 44.926 | 43.579 | 49.417 | 46.883 | 42.620 | 50.352 | 45.001 | 42.520 | 45.721 | 46.739 | 45.979 | 42.495 | 45.409 | 41.359 | 41.787 | 46.619 | 45.414 | 0.021 | 0.5598 | 0.7704 | 0 | 0 | 0 |
| 'Rock2' | 12.318 | 12.387 | 12.409 | 12.888 | 12.840 | 12.201 | 12.157 | 12.920 | 12.637 | 12.071 | 12.352 | 13.273 | 12.696 | 12.410 | 12.188 | 12.230 | 12.048 | 13.387 | -0.012 | 0.6775 | 0.8457 | 0 | 0 | 0 |
| 'Rogdi' | 2.754 | 2.751 | 3.184 | 2.816 | 2.645 | 3.428 | 2.702 | 2.647 | 3.146 | 3.022 | 2.540 | 3.387 | 2.887 | 3.092 | 2.912 | 2.855 | 3.365 | 3.245 | -0.078 | 0.1934 | 0.4308 | 0 | 0 | 0 |
| 'Rom1' | 4.218 | 2.879 | 4.029 | 3.039 | 3.291 | 3.850 | 3.063 | 3.957 | 3.787 | 3.221 | 3.302 | 3.439 | 3.683 | 3.611 | 3.877 | 3.815 | 3.106 | 3.560 | 0.033 | 0.7284 | 0.8754 | 0 | 0 | 0 |
| 'Romo1' | 19.834 | 24.581 | 19.932 | 20.561 | 22.290 | 20.003 | 21.959 | 23.403 | 20.172 | 17.101 | 19.169 | 18.906 | 19.484 | 21.441 | 21.110 | 21.753 | 20.336 | 18.142 | 0.105 | 0.0264 | 0.1245 | 0 | 0 | 0 |
| 'Rora' | 1.392 | 1.884 | 1.309 | 1.409 | 1.634 | 1.327 | 1.403 | 1.795 | 1.272 | 1.858 | 2.003 | 1.549 | 2.129 | 1.955 | 1.331 | 2.034 | 1.755 | 1.343 | -0.256 | 0.0000 | 0.0004 | 1 | 0 | 0 |
| 'Rp2' | 2.907 | 3.217 | 3.278 | 2.925 | 3.073 | 2.936 | 3.282 | 3.104 | 2.884 | 2.913 | 3.317 | 2.838 | 2.920 | 3.354 | 3.235 | 2.969 | 3.230 | 2.942 | -0.010 | 0.8447 | 0.9343 | 0 | 0 | 0 |
| 'Rp9' | 52.707 | 58.051 | 52.884 | 51.234 | 52.328 | 58.763 | 50.327 | 54.198 | 53.515 | 48.711 | 54.602 | 59.032 | 52.947 | 54.954 | 60.109 | 56.954 | 51.816 | 57.229 | -0.047 | 0.1947 | 0.4328 | 0 | 0 | 0 |
| 'Rpa1' | 51.113 | 54.025 | 52.395 | 54.469 | 56.515 | 53.765 | 54.420 | 53.896 | 53.426 | 54.844 | 56.238 | 51.304 | 57.773 | 55.170 | 53.739 | 55.737 | 54.500 | 51.346 | -0.029 | 0.1780 | 0.4132 | 0 | 0 | 0 |
| 'Rpa2' | 13.170 | 14.137 | 15.031 | 14.009 | 15.354 | 14.992 | 13.653 | 13.764 | 15.299 | 14.201 | 14.152 | 15.221 | 15.741 | 14.452 | 15.611 | 14.755 | 13.943 | 14.315 | -0.044 | 0.1871 | 0.4230 | 0 | 0 | 0 |
| 'Rpa3' | 17.305 | 18.719 | 19.448 | 18.880 | 21.892 | 20.206 | 19.545 | 20.485 | 17.711 | 16.169 | 18.137 | 17.758 | 18.000 | 19.842 | 19.119 | 18.909 | 19.079 | 15.747 | 0.089 | 0.0718 | 0.2363 | 0 | 0 | 0 |
| 'Rpain' | 3.851 | 3.911 | 3.945 | 4.285 | 3.883 | 3.515 | 3.549 | 4.032 | 4.124 | 3.670 | 3.676 | 4.092 | 3.888 | 4.087 | 4.616 | 3.748 | 3.954 | 3.829 | -0.031 | 0.5623 | 0.7715 | 0 | 0 | 0 |
| 'Rpap1' | 5.531 | 5.246 | 6.597 | 5.284 | 5.418 | 6.073 | 5.250 | 5.667 | 6.907 | 5.938 | 5.563 | 5.809 | 5.953 | 5.188 | 5.442 | 5.651 | 5.308 | 5.148 | 0.044 | 0.3468 | 0.6020 | 0 | 0 | 0 |
| 'Rpap2' | 4.251 | 4.480 | 4.882 | 4.135 | 4.097 | 4.392 | 4.577 | 4.460 | 4.207 | 4.678 | 4.569 | 3.836 | 4.658 | 4.605 | 3.937 | 4.541 | 4.809 | 3.766 | 0.005 | 0.9224 | 0.9705 | 0 | 0 | 0 |
| 'Rpap3' | 12.983 | 14.164 | 14.687 | 12.798 | 14.856 | 13.993 | 14.478 | 13.908 | 13.423 | 14.601 | 12.611 | 12.124 | 13.281 | 11.933 | 13.065 | 13.424 | 13.292 | 10.888 | 0.115 | 0.0194 | 0.1008 | 0 | 0 | 0 |
| 'Rpe' | 7.443 | 8.327 | 8.216 | 8.999 | 9.091 | 7.837 | 8.780 | 8.165 | 7.699 | 7.029 | 8.252 | 8.561 | 7.642 | 8.660 | 8.215 | 7.032 | 7.935 | 8.128 | 0.047 | 0.3136 | 0.5715 | 0 | 0 | 0 |
| 'Rpf1' | 8.499 | 7.789 | 7.472 | 7.998 | 7.916 | 7.757 | 8.854 | 8.626 | 7.995 | 7.872 | 7.777 | 7.701 | 7.587 | 7.653 | 7.312 | 8.595 | 8.427 | 6.948 | 0.053 | 0.1612 | 0.3925 | 0 | 0 | 0 |
| 'Rpf2' | 6.318 | 6.214 | 6.780 | 6.335 | 6.236 | 5.601 | 7.450 | 6.352 | 6.933 | 5.662 | 5.426 | 6.327 | 5.883 | 5.716 | 5.092 | 5.411 | 6.776 | 5.468 | 0.154 | 0.0113 | 0.0681 | 0 | 0 | 0 |
| 'Rpgr' | 0.764 | 0.563 | 0.538 | 0.637 | 0.734 | 0.623 | 0.731 | 0.663 | 0.630 | 0.616 | 0.712 | 0.705 | 0.703 | 0.672 | 0.689 | 0.580 | 0.699 | 0.652 | -0.057 | 0.5087 | 0.7340 | 0 | 0 | 0 |
| 'Rpgrip1l' | 0.621 | 0.570 | 0.643 | 0.655 | 0.694 | 0.617 | 0.602 | 0.553 | 0.697 | 0.809 | 0.721 | 0.609 | 0.758 | 0.644 | 0.526 | 0.512 | 0.747 | 0.536 | -0.055 | 0.5399 | 0.7574 | 0 | 0 | 0 |
| 'Rph3a' | 0.773 | 0.692 | 0.939 | 0.815 | 0.821 | 0.724 | 0.676 | 1.022 | 0.908 | 0.670 | 0.724 | 0.601 | 0.399 | 0.731 | 0.690 | 0.359 | 0.654 | 0.716 | 0.419 | 0.0007 | 0.0076 | 1 | 1 | 0 |
| 'Rph3al' | 3.373 | 2.705 | 3.789 | 2.883 | 3.164 | 3.108 | 2.606 | 3.529 | 3.065 | 3.128 | 2.955 | 3.557 | 3.367 | 3.758 | 3.328 | 3.001 | 2.894 | 3.247 | -0.056 | 0.4308 | 0.6762 | 0 | 0 | 0 |
| 'Rpia' | 11.632 | 13.099 | 10.692 | 11.238 | 13.496 | 11.309 | 10.847 | 12.972 | 11.301 | 11.101 | 12.745 | 11.583 | 13.103 | 13.511 | 11.306 | 12.050 | 12.609 | 11.088 | -0.047 | 0.1575 | 0.3874 | 0 | 0 | 0 |
| 'Rpl10a' | 97.927 | 93.114 | 91.803 | 97.210 | 93.169 | 93.334 | 100.036 | 90.774 | 93.274 | 89.872 | 92.970 | 84.290 | 94.447 | 90.940 | 95.642 | 94.444 | 94.023 | 89.591 | 0.031 | 0.2508 | 0.5019 | 0 | 0 | 0 |
| 'Rpl11' | 101.949 | 109.461 | 98.530 | 111.750 | 106.430 | 100.234 | 111.440 | 103.642 | 98.221 | 95.709 | 103.763 | 82.058 | 100.710 | 101.233 | 101.312 | 105.809 | 103.746 | 92.883 | 0.076 | 0.0396 | 0.1624 | 0 | 0 | 0 |
| 'Rpl12' | 91.648 | 90.942 | 86.824 | 97.438 | 91.290 | 87.208 | 98.436 | 89.791 | 89.048 | 87.112 | 89.731 | 73.111 | 87.687 | 87.420 | 89.260 | 91.126 | 88.514 | 83.604 | 0.070 | 0.0394 | 0.1621 | 0 | 0 | 0 |
| 'Rpl13' | 189.693 | 181.778 | 174.966 | 196.092 | 187.217 | 181.204 | 197.025 | 180.694 | 178.767 | 177.675 | 177.787 | 159.342 | 185.872 | 176.689 | 185.341 | 190.293 | 179.183 | 175.402 | 0.042 | 0.1410 | 0.3636 | 0 | 0 | 0 |
| 'Rpl13a' | ####### | ####### | 985.946 | ####### | ####### | ####### | ####### | ####### | ####### | ####### | ####### | 836.586 | ####### | ####### | ####### | ####### | ####### | 976.022 | 0.061 | 0.0960 | 0.2881 | 0 | 0 | 0 |
| 'Rpl14' | 532.185 | 532.318 | 489.600 | 572.396 | 541.233 | 509.677 | 585.755 | 525.766 | 504.495 | 493.461 | 524.171 | 451.309 | 537.831 | 517.390 | 519.697 | 529.714 | 533.656 | 487.055 | 0.050 | 0.1150 | 0.3219 | 0 | 0 | 0 |
| 'Rpl15' | 114.386 | 120.056 | 108.223 | 121.499 | 120.528 | 109.876 | 125.364 | 116.923 | 109.973 | 109.431 | 117.172 | 98.425 | 117.357 | 120.115 | 112.210 | 115.324 | 117.941 | 104.801 | 0.038 | 0.1915 | 0.4284 | 0 | 0 | 0 |
| 'Rpl17' | 658.898 | 679.061 | 594.460 | 712.319 | 684.316 | 609.157 | 714.592 | 652.908 | 615.408 | 616.842 | 636.477 | 525.159 | 625.517 | 628.440 | 611.271 | 666.764 | 643.262 | 572.905 | 0.088 | 0.0044 | 0.0341 | 1 | 0 | 0 |
| 'Rpl18' | 97.875 | 100.169 | 89.812 | 104.618 | 102.050 | 93.864 | 105.909 | 97.204 | 92.265 | 91.123 | 95.170 | 80.074 | 94.821 | 93.322 | 94.410 | 99.325 | 98.679 | 90.460 | 0.066 | 0.0457 | 0.1775 | 0 | 0 | 0 |
| 'Rpl18a' | 411.451 | 410.557 | 377.233 | 439.450 | 413.270 | 397.901 | 433.696 | 402.563 | 396.429 | 392.061 | 416.008 | 324.505 | 412.711 | 393.973 | 408.998 | 423.954 | 402.777 | 378.133 | 0.041 | 0.2766 | 0.5323 | 0 | 0 | 0 |
| 'Rpl19' | 714.457 | 719.093 | 668.932 | 749.928 | 708.691 | 703.054 | 777.677 | 689.927 | 703.436 | 688.316 | 696.139 | 592.058 | 701.906 | 692.739 | 705.872 | 745.655 | 706.917 | 657.458 | 0.046 | 0.1624 | 0.3943 | 0 | 0 | 0 |
| 'Rpl21' | 144.779 | 152.983 | 135.715 | 156.152 | 154.465 | 142.474 | 160.185 | 141.824 | 143.670 | 137.577 | 143.653 | 115.208 | 139.883 | 142.250 | 142.641 | 149.759 | 144.245 | 133.182 | 0.083 | 0.0300 | 0.1349 | 0 | 0 | 0 |
| 'Rpl21-ps4' | 6.465 | 6.228 | 5.458 | 8.329 | 8.120 | 6.519 | 6.343 | 6.649 | 6.367 | 7.052 | 7.713 | 4.645 | 7.191 | 5.461 | 6.307 | 8.541 | 6.005 | 6.796 | 0.025 | 0.8351 | 0.9287 | 0 | 0 | 0 |
| 'Rpl22' | 49.874 | 51.862 | 47.098 | 53.671 | 52.404 | 49.236 | 54.592 | 50.466 | 47.981 | 49.266 | 50.119 | 43.883 | 48.610 | 50.666 | 48.969 | 51.411 | 53.412 | 46.186 | 0.037 | 0.2045 | 0.4475 | 0 | 0 | 0 |
| 'Rpl22l1' | 22.135 | 19.421 | 17.284 | 21.626 | 21.545 | 16.576 | 22.427 | 21.521 | 16.288 | 20.238 | 19.366 | 14.995 | 23.143 | 20.599 | 16.271 | 18.679 | 20.207 | 14.254 | 0.086 | 0.0262 | 0.1243 | 0 | 0 | 0 |
| 'Rpl23' | 187.219 | 199.929 | 171.756 | 209.226 | 200.757 | 179.790 | 215.855 | 189.569 | 182.012 | 182.427 | 199.497 | 146.364 | 184.778 | 194.821 | 183.348 | 199.717 | 195.364 | 169.118 | 0.059 | 0.1596 | 0.3901 | 0 | 0 | 0 |
| 'Rpl23a' | 253.482 | 262.213 | 227.625 | 271.784 | 261.757 | 240.285 | 278.303 | 239.071 | 245.001 | 243.032 | 259.798 | 198.372 | 252.560 | 239.242 | 246.058 | 266.766 | 249.663 | 231.752 | 0.049 | 0.2301 | 0.4769 | 0 | 0 | 0 |
| 'Rpl24' | 189.896 | 188.326 | 176.115 | 202.810 | 191.489 | 180.863 | 204.043 | 181.631 | 173.052 | 180.529 | 181.321 | 159.878 | 189.841 | 189.433 | 182.926 | 186.190 | 190.437 | 171.052 | 0.038 | 0.2095 | 0.4547 | 0 | 0 | 0 |
| 'Rpl26' | 149.984 | 147.517 | 134.283 | 159.444 | 148.088 | 141.741 | 159.640 | 141.804 | 138.106 | 140.323 | 143.398 | 121.634 | 144.507 | 142.504 | 149.515 | 154.588 | 147.714 | 136.295 | 0.033 | 0.3724 | 0.6241 | 0 | 0 | 0 |
| 'Rpl27a' | 102.777 | 105.647 | 95.270 | 109.296 | 103.137 | 98.497 | 107.669 | 98.388 | 95.206 | 99.300 | 103.002 | 80.605 | 99.517 | 101.897 | 100.888 | 104.315 | 102.090 | 94.430 | 0.037 | 0.3208 | 0.5788 | 0 | 0 | 0 |
| 'Rpl28' | 345.423 | 345.470 | 312.624 | 369.039 | 354.094 | 331.208 | 373.952 | 329.633 | 330.387 | 322.268 | 349.300 | 262.578 | 340.412 | 329.082 | 335.284 | 356.467 | 345.365 | 313.714 | 0.055 | 0.1899 | 0.4262 | 0 | 0 | 0 |
| 'Rpl3' | 136.684 | 131.718 | 124.003 | 143.290 | 133.854 | 128.463 | 142.959 | 132.829 | 123.801 | 131.048 | 135.200 | 112.198 | 136.020 | 129.720 | 127.140 | 136.557 | 130.944 | 120.237 | 0.037 | 0.1638 | 0.3958 | 0 | 0 | 0 |
| 'Rpl30' | 62.590 | 69.963 | 56.507 | 69.048 | 69.045 | 60.488 | 69.342 | 65.139 | 59.893 | 62.015 | 67.061 | 47.949 | 62.494 | 67.803 | 59.880 | 68.828 | 66.644 | 56.884 | 0.049 | 0.2413 | 0.4920 | 0 | 0 | 0 |
| 'Rpl31' | 289.763 | 318.827 | 276.621 | 336.663 | 319.831 | 287.970 | 337.327 | 295.914 | 295.245 | 289.802 | 321.325 | 229.986 | 306.361 | 307.403 | 302.366 | 320.761 | 312.740 | 277.131 | 0.038 | 0.4156 | 0.6634 | 0 | 0 | 0 |
| 'Rpl32' | 374.327 | 372.550 | 342.088 | 401.724 | 371.764 | 358.381 | 402.914 | 351.377 | 346.239 | 344.923 | 365.550 | 299.346 | 368.322 | 358.016 | 364.150 | 376.753 | 368.259 | 347.300 | 0.045 | 0.2272 | 0.4734 | 0 | 0 | 0 |
| 'Rpl34' | 78.337 | 80.222 | 71.627 | 80.845 | 77.729 | 74.395 | 82.554 | 76.930 | 71.646 | 73.131 | 75.236 | 61.237 | 72.661 | 77.105 | 75.369 | 79.638 | 77.825 | 69.433 | 0.059 | 0.1014 | 0.2981 | 0 | 0 | 0 |
| 'Rpl35' | 168.207 | 167.998 | 157.467 | 184.712 | 164.965 | 160.726 | 184.593 | 159.566 | 159.685 | 158.337 | 161.039 | 138.405 | 168.763 | 164.860 | 165.566 | 167.098 | 165.391 | 155.936 | 0.049 | 0.1667 | 0.3992 | 0 | 0 | 0 |
| 'Rpl35a' | 58.083 | 61.980 | 52.227 | 65.914 | 61.451 | 56.100 | 65.184 | 56.927 | 55.144 | 55.916 | 59.271 | 44.882 | 56.500 | 59.641 | 56.169 | 60.948 | 58.456 | 53.069 | 0.068 | 0.1160 | 0.3233 | 0 | 0 | 0 |
| 'Rpl36' | 592.079 | 582.780 | 575.649 | 592.230 | 549.592 | 575.314 | 588.010 | 557.158 | 554.252 | 549.101 | 548.973 | 452.205 | 552.869 | 554.604 | 562.210 | 570.782 | 546.198 | 519.837 | 0.081 | 0.0305 | 0.1362 | 0 | 0 | 0 |
| 'Rpl36a' | 67.679 | 68.349 | 62.578 | 73.663 | 70.247 | 64.970 | 75.452 | 68.601 | 65.065 | 63.364 | 70.870 | 56.077 | 65.502 | 68.255 | 70.267 | 66.996 | 68.037 | 62.848 | 0.047 | 0.2491 | 0.5000 | 0 | 0 | 0 |
| 'Rpl36al' | 69.998 | 72.568 | 68.847 | 75.839 | 73.702 | 70.569 | 76.631 | 71.759 | 70.492 | 66.793 | 67.809 | 59.541 | 69.648 | 68.461 | 72.125 | 71.730 | 73.674 | 66.860 | 0.067 | 0.0614 | 0.2145 | 0 | 0 | 0 |
| 'Rpl37' | 297.564 | 301.060 | 277.313 | 317.983 | 311.783 | 291.602 | 313.594 | 289.346 | 292.389 | 278.160 | 300.394 | 234.638 | 281.604 | 296.318 | 293.446 | 309.091 | 303.794 | 272.058 | 0.058 | 0.1530 | 0.3803 | 0 | 0 | 0 |
| 'Rpl37a' | 499.516 | 550.318 | 470.712 | 558.139 | 535.354 | 505.273 | 570.967 | 509.616 | 490.499 | 478.136 | 520.283 | 385.671 | 484.357 | 521.624 | 501.637 | 538.263 | 520.922 | 473.426 | 0.074 | 0.1091 | 0.3110 | 0 | 0 | 0 |
| 'Rpl38' | 249.525 | 259.142 | 233.325 | 270.434 | 257.501 | 242.721 | 271.633 | 261.816 | 241.679 | 226.398 | 253.676 | 204.661 | 237.719 | 252.891 | 240.205 | 251.658 | 256.500 | 228.596 | 0.079 | 0.0268 | 0.1260 | 0 | 0 | 0 |
| 'Rpl39' | 135.437 | 139.389 | 119.831 | 149.520 | 140.320 | 129.213 | 150.457 | 134.117 | 130.646 | 127.840 | 144.617 | 103.957 | 127.728 | 139.450 | 136.767 | 140.531 | 142.855 | 122.957 | 0.041 | 0.3990 | 0.6482 | 0 | 0 | 0 |
| 'Rpl4' | 930.251 | 906.037 | 864.523 | 968.238 | 911.581 | 877.653 | 973.890 | 878.023 | 879.473 | 916.557 | 888.408 | 781.567 | 919.073 | 882.093 | 906.012 | 947.927 | 903.132 | 864.276 | 0.021 | 0.4392 | 0.6829 | 0 | 0 | 0 |
| 'Rpl5' | 124.703 | 130.209 | 116.958 | 134.779 | 132.001 | 119.555 | 139.558 | 125.674 | 117.779 | 122.828 | 130.916 | 108.263 | 126.014 | 126.703 | 123.973 | 130.237 | 127.257 | 116.962 | 0.024 | 0.4171 | 0.6649 | 0 | 0 | 0 |
| 'Rpl7' | 146.412 | 149.425 | 138.035 | 156.037 | 150.604 | 141.887 | 157.631 | 144.334 | 142.484 | 141.430 | 147.531 | 125.168 | 145.147 | 145.590 | 144.086 | 145.754 | 148.538 | 135.672 | 0.042 | 0.1427 | 0.3660 | 0 | 0 | 0 |
| 'Rpl7a' | 180.535 | 164.319 | 163.843 | 181.989 | 167.523 | 167.962 | 184.054 | 167.282 | 166.606 | 166.553 | 165.416 | 148.781 | 181.987 | 163.924 | 164.719 | 172.603 | 163.533 | 157.071 | 0.046 | 0.0602 | 0.2116 | 0 | 0 | 0 |
| 'Rpl7l1' | 39.288 | 40.865 | 38.538 | 39.969 | 39.785 | 38.431 | 44.229 | 38.448 | 39.404 | 34.898 | 37.601 | 41.242 | 36.680 | 36.082 | 39.667 | 34.183 | 40.842 | 40.817 | 0.057 | 0.1671 | 0.3995 | 0 | 0 | 0 |
| 'Rpl8' | 912.063 | 893.675 | 887.966 | 933.355 | 927.458 | 909.015 | 920.637 | 879.079 | 889.469 | 891.589 | 902.027 | 774.261 | 897.510 | 893.514 | 934.827 | 898.345 | 868.798 | 885.877 | 0.026 | 0.4078 | 0.6567 | 0 | 0 | 0 |
| 'Rpl9' | 52.732 | 54.459 | 48.610 | 57.207 | 54.228 | 50.069 | 57.104 | 52.604 | 48.654 | 50.949 | 53.170 | 44.674 | 50.799 | 52.895 | 50.784 | 54.406 | 54.403 | 47.756 | 0.038 | 0.2073 | 0.4519 | 0 | 0 | 0 |
| 'Rplp0' | 474.713 | 454.351 | 440.860 | 479.411 | 460.643 | 448.358 | 497.943 | 444.944 | 450.274 | 458.133 | 474.974 | 385.652 | 476.736 | 454.510 | 471.643 | 468.769 | 468.957 | 452.005 | 0.003 | 0.9373 | 0.9763 | 0 | 0 | 0 |
| 'Rplp1' | ####### | ####### | ####### | ####### | ####### | ####### | ####### | ####### | ####### | ####### | ####### | ####### | ####### | ####### | ####### | ####### | ####### | ####### | 0.057 | 0.2335 | 0.4820 | 0 | 0 | 0 |

| | | | | | | | | | | | | | | | | | | | | | | | |
|---|---|---|---|---|---|---|---|---|---|---|---|---|---|---|---|---|---|---|---|---|---|---|---|
| 'Rplp2' | 125.890 | 131.387 | 120.149 | 131.748 | 126.934 | 124.251 | 134.428 | 124.408 | 120.866 | 120.719 | 127.806 | 98.007 | 122.457 | 124.178 | 125.800 | 130.207 | 125.576 | 115.417 | 0.055 | 0.1664 | 0.3987 | 0 | 0 | 0 |
| 'Rpn1' | 21.890 | 25.364 | 25.820 | 23.445 | 24.962 | 27.615 | 22.239 | 23.091 | 27.402 | 25.860 | 22.314 | 26.750 | 24.557 | 23.540 | 25.765 | 24.496 | 21.802 | 27.691 | -0.018 | 0.6234 | 0.8120 | 0 | 0 | 0 |
| 'Rpn2' | 26.261 | 25.637 | 27.139 | 26.936 | 25.668 | 26.789 | 25.661 | 25.366 | 26.384 | 27.046 | 25.395 | 26.978 | 25.702 | 25.565 | 27.145 | 27.022 | 24.803 | 27.625 | -0.018 | 0.3847 | 0.6347 | 0 | 0 | 0 |
| 'Rpp21' | 7.030 | 7.193 | 7.962 | 6.073 | 7.673 | 7.608 | 7.234 | 7.652 | 8.105 | 6.677 | 7.192 | 7.496 | 7.634 | 7.062 | 6.944 | 7.440 | 7.337 | 7.451 | 0.020 | 0.6797 | 0.8470 | 0 | 0 | 0 |
| 'Rpp25l' | 22.965 | 22.547 | 26.541 | 24.933 | 21.874 | 26.042 | 26.319 | 24.186 | 26.195 | 21.242 | 23.730 | 27.117 | 22.249 | 21.204 | 29.796 | 19.689 | 23.532 | 27.952 | 0.023 | 0.6756 | 0.8445 | 0 | 0 | 0 |
| 'Rpp30' | 9.891 | 12.294 | 12.407 | 12.298 | 13.168 | 12.019 | 11.412 | 12.137 | 11.039 | 10.204 | 10.157 | 11.184 | 12.757 | 9.989 | 11.290 | 9.883 | 10.922 | 9.816 | 0.130 | 0.0509 | 0.1910 | 0 | 0 | 0 |
| 'Rpp40' | 0.575 | 0.569 | 0.574 | 0.454 | 0.625 | 0.543 | 0.519 | 0.646 | 0.501 | 0.472 | 0.524 | 0.510 | 0.475 | 0.578 | 0.431 | 0.482 | 0.464 | 0.417 | 0.193 | 0.0543 | 0.1995 | 0 | 0 | 0 |
| 'Rprd1a' | 5.068 | 5.368 | 5.261 | 5.158 | 5.267 | 5.419 | 5.099 | 5.185 | 5.175 | 4.369 | 6.060 | 6.621 | 4.792 | 5.442 | 6.099 | 4.427 | 7.284 | 6.767 | -0.144 | 0.0352 | 0.1501 | 0 | 0 | 0 |
| 'Rprd1b' | 5.391 | 5.331 | 5.555 | 6.079 | 5.445 | 5.728 | 6.121 | 5.466 | 5.438 | 4.660 | 5.467 | 5.824 | 5.142 | 4.902 | 5.806 | 4.645 | 5.627 | 5.362 | 0.080 | 0.0895 | 0.2744 | 0 | 0 | 0 |
| 'Rprd2' | 6.558 | 5.500 | 6.119 | 6.744 | 5.632 | 5.844 | 6.873 | 5.868 | 5.919 | 5.968 | 6.746 | 6.920 | 5.854 | 6.278 | 6.406 | 5.516 | 6.377 | 6.620 | -0.055 | 0.2913 | 0.5480 | 0 | 0 | 0 |
| 'Rps10' | 134.689 | 137.555 | 131.318 | 144.152 | 136.092 | 133.812 | 141.435 | 132.481 | 130.485 | 127.099 | 129.163 | 115.664 | 131.950 | 132.785 | 137.727 | 132.386 | 131.238 | 123.982 | 0.062 | 0.0427 | 0.1704 | 0 | 0 | 0 |
| 'Rps11' | 781.583 | 829.118 | 764.234 | 860.781 | 814.759 | 795.819 | 875.399 | 792.774 | 796.706 | 790.138 | 829.551 | 657.199 | 808.322 | 797.118 | 812.610 | 853.909 | 839.965 | 760.899 | 0.022 | 0.5615 | 0.7712 | 0 | 0 | 0 |
| 'Rps12' | ####### | ####### | ####### | ####### | ####### | ####### | ####### | ####### | ####### | ####### | ####### | 908.763 | ####### | ####### | ####### | ####### | ####### | ####### | 0.016 | 0.6879 | 0.8525 | 0 | 0 | 0 |
| 'Rps13' | 55.721 | 56.901 | 50.656 | 58.854 | 56.032 | 51.559 | 58.045 | 53.479 | 49.803 | 53.583 | 55.962 | 44.522 | 54.984 | 52.991 | 52.877 | 55.573 | 55.310 | 50.104 | 0.034 | 0.2887 | 0.5451 | 0 | 0 | 0 |
| 'Rps14' | 186.014 | 203.586 | 172.930 | 204.174 | 192.890 | 183.217 | 202.305 | 191.464 | 183.607 | 174.727 | 187.832 | 148.980 | 175.380 | 186.706 | 182.473 | 195.663 | 186.534 | 168.256 | 0.088 | 0.0210 | 0.1064 | 0 | 0 | 0 |
| 'Rps15' | ####### | ####### | ####### | ####### | ####### | ####### | ####### | ####### | ####### | ####### | ####### | 895.989 | ####### | ####### | ####### | ####### | ####### | ####### | 0.052 | 0.1939 | 0.4316 | 0 | 0 | 0 |
| 'Rps15a' | 30.350 | 30.807 | 28.730 | 33.432 | 30.944 | 29.604 | 32.136 | 28.985 | 29.379 | 28.930 | 30.526 | 24.438 | 28.069 | 30.584 | 30.537 | 30.986 | 30.442 | 27.544 | 0.056 | 0.1703 | 0.4030 | 0 | 0 | 0 |
| 'Rps16' | 376.902 | 406.359 | 351.841 | 408.035 | 407.967 | 363.842 | 414.759 | 386.234 | 369.053 | 356.895 | 394.445 | 288.112 | 369.254 | 384.777 | 370.698 | 391.851 | 382.879 | 339.952 | 0.078 | 0.0555 | 0.2019 | 0 | 0 | 0 |
| 'Rps17' | 142.007 | 145.904 | 132.738 | 154.373 | 151.020 | 131.657 | 158.428 | 143.591 | 136.325 | 133.676 | 145.065 | 114.856 | 143.375 | 140.603 | 138.317 | 142.569 | 136.740 | 126.843 | 0.074 | 0.0353 | 0.1504 | 0 | 0 | 0 |
| 'Rps18' | 130.176 | 127.998 | 122.848 | 135.732 | 129.450 | 123.332 | 139.579 | 121.960 | 123.496 | 119.360 | 125.600 | 102.496 | 123.951 | 125.447 | 125.899 | 132.578 | 129.218 | 119.911 | 0.053 | 0.1649 | 0.3972 | 0 | 0 | 0 |
| 'Rps19' | 55.767 | 52.146 | 53.530 | 57.804 | 52.667 | 51.853 | 58.994 | 51.412 | 51.942 | 49.561 | 53.262 | 48.310 | 54.173 | 52.684 | 57.043 | 57.896 | 55.263 | 53.633 | 0.001 | 0.9827 | 0.9934 | 0 | 0 | 0 |
| 'Rps19bp1' | 19.190 | 25.488 | 23.097 | 20.176 | 22.007 | 22.192 | 19.343 | 24.157 | 19.040 | 19.764 | 17.686 | 22.548 | 19.970 | 18.017 | 22.132 | 21.558 | 19.192 | 21.199 | 0.079 | 0.2139 | 0.4592 | 0 | 0 | 0 |
| 'Rps2' | 392.256 | 364.664 | 352.525 | 396.137 | 373.397 | 342.887 | 407.826 | 357.713 | 343.595 | 355.830 | 383.944 | 294.041 | 394.699 | 365.383 | 360.715 | 379.580 | 369.715 | 336.571 | 0.029 | 0.4332 | 0.6782 | 0 | 0 | 0 |
| 'Rps20' | 77.645 | 77.434 | 72.607 | 83.735 | 77.721 | 74.779 | 86.885 | 75.038 | 72.688 | 73.252 | 80.143 | 61.623 | 76.296 | 75.326 | 77.154 | 79.494 | 79.698 | 72.134 | 0.038 | 0.3586 | 0.6113 | 0 | 0 | 0 |
| 'Rps21' | 143.440 | 164.247 | 143.768 | 165.326 | 158.748 | 142.384 | 163.920 | 147.586 | 132.436 | 141.515 | 156.896 | 112.779 | 140.026 | 150.409 | 142.783 | 160.339 | 153.937 | 131.639 | 0.069 | 0.1380 | 0.3585 | 0 | 0 | 0 |
| 'Rps23' | 695.777 | 739.983 | 644.580 | 744.711 | 730.477 | 660.645 | 760.538 | 677.099 | 662.036 | 650.239 | 723.762 | 532.867 | 683.976 | 717.000 | 671.568 | 741.609 | 723.956 | 625.596 | 0.048 | 0.2527 | 0.5042 | 0 | 0 | 0 |
| 'Rps24' | 440.345 | 444.486 | 403.603 | 482.973 | 442.479 | 414.821 | 478.854 | 421.580 | 421.860 | 414.363 | 438.943 | 342.738 | 411.051 | 424.502 | 426.573 | 444.319 | 437.226 | 401.250 | 0.068 | 0.0965 | 0.2890 | 0 | 0 | 0 |
| 'Rps25' | 515.787 | 512.709 | 476.446 | 554.661 | 518.330 | 485.750 | 550.868 | 502.649 | 485.585 | 361.344 | 376.159 | 306.523 | 378.009 | 392.572 | 372.461 | 382.266 | 388.535 | 348.740 | 0.467 | 0.0000 | 0.0000 | 1 | 1 | 0 |
| 'Rps26' | 788.868 | 774.895 | 727.192 | 819.271 | 772.609 | 750.512 | 847.054 | 739.604 | 752.315 | 720.013 | 780.264 | 628.271 | 749.588 | 766.591 | 782.752 | 767.128 | 768.237 | 746.235 | 0.044 | 0.2723 | 0.5275 | 0 | 0 | 0 |
| 'Rps27a' | 123.646 | 124.530 | 116.065 | 129.033 | 126.759 | 119.345 | 131.844 | 124.937 | 113.637 | 119.488 | 122.840 | 101.450 | 119.085 | 120.242 | 115.253 | 130.663 | 123.595 | 110.348 | 0.053 | 0.0704 | 0.2339 | 0 | 0 | 0 |
| 'Rps27l' | 11.373 | 17.570 | 13.389 | 14.678 | 16.979 | 14.964 | 14.343 | 15.430 | 14.659 | 9.938 | 12.881 | 14.477 | 15.360 | 16.630 | 15.627 | 14.520 | 14.262 | 15.518 | 0.028 | 0.7085 | 0.8650 | 0 | 0 | 0 |
| 'Rps27rt' | 121.584 | 127.904 | 110.132 | 131.549 | 124.111 | 115.186 | 132.933 | 117.514 | 113.682 | 117.437 | 126.371 | 86.213 | 111.194 | 119.680 | 116.971 | 127.829 | 123.458 | 106.941 | 0.070 | 0.1576 | 0.3874 | 0 | 0 | 0 |
| 'Rps28' | 100.391 | 109.543 | 93.688 | 116.282 | 108.477 | 98.789 | 111.147 | 103.515 | 98.585 | 99.419 | 111.798 | 74.450 | 100.229 | 104.014 | 99.569 | 111.148 | 106.522 | 94.711 | 0.052 | 0.3004 | 0.5570 | 0 | 0 | 0 |
| 'Rps29' | 277.386 | 302.651 | 253.127 | 304.514 | 295.821 | 268.768 | 305.892 | 267.161 | 266.832 | 275.564 | 298.807 | 195.905 | 264.573 | 283.311 | 271.946 | 306.079 | 297.652 | 247.752 | 0.051 | 0.3445 | 0.5997 | 0 | 0 | 0 |
| 'Rps3' | 122.261 | 125.164 | 114.847 | 131.815 | 128.782 | 118.058 | 131.021 | 124.523 | 117.498 | 113.483 | 119.844 | 101.600 | 117.948 | 120.910 | 119.894 | 125.496 | 122.378 | 109.455 | 0.073 | 0.0195 | 0.1013 | 0 | 0 | 0 |
| 'Rps3a1' | 703.758 | 743.853 | 676.333 | 768.934 | 755.923 | 683.668 | 791.915 | 711.373 | 689.344 | 692.380 | 726.903 | 589.962 | 703.889 | 729.792 | 687.272 | 741.582 | 741.997 | 642.932 | 0.051 | 0.1187 | 0.3278 | 0 | 0 | 0 |
| 'Rps4x' | 205.709 | 210.166 | 192.560 | 225.430 | 219.349 | 198.259 | 231.160 | 205.852 | 195.565 | 201.976 | 220.162 | 178.929 | 207.373 | 206.351 | 205.503 | 211.205 | 217.395 | 192.222 | 0.022 | 0.5072 | 0.7332 | 0 | 0 | 0 |
| 'Rps5' | 164.646 | 162.035 | 152.678 | 169.209 | 162.430 | 156.206 | 170.581 | 158.726 | 152.107 | 153.954 | 162.729 | 142.674 | 166.153 | 155.725 | 165.454 | 165.529 | 160.138 | 158.638 | 0.006 | 0.8394 | 0.9315 | 0 | 0 | 0 |
| 'Rps6' | 222.245 | 228.599 | 201.582 | 239.374 | 227.800 | 210.062 | 244.025 | 217.416 | 205.543 | 213.579 | 223.613 | 179.299 | 224.157 | 221.417 | 217.371 | 229.802 | 222.453 | 200.622 | 0.036 | 0.2939 | 0.5499 | 0 | 0 | 0 |
| 'Rps6ka1' | 32.866 | 30.262 | 37.181 | 34.818 | 32.583 | 37.310 | 33.187 | 31.909 | 36.308 | 29.511 | 28.152 | 35.158 | 30.138 | 29.552 | 35.076 | 28.955 | 29.510 | 34.887 | 0.116 | 0.0000 | 0.0000 | 1 | 0 | 0 |
| 'Rps6ka2' | 0.863 | 0.795 | 0.895 | 0.939 | 0.704 | 0.951 | 0.852 | 1.023 | 0.914 | 1.178 | 0.875 | 0.961 | 0.865 | 1.178 | 0.911 | 1.290 | 0.946 | 0.863 | -0.183 | 0.0676 | 0.2278 | 0 | 0 | 0 |
| 'Rps6ka3' | 8.748 | 9.961 | 10.348 | 9.663 | 9.699 | 9.430 | 10.007 | 10.158 | 9.276 | 9.468 | 10.288 | 10.932 | 8.967 | 9.720 | 9.668 | 9.530 | 10.305 | 10.277 | -0.042 | 0.2243 | 0.4713 | 0 | 0 | 0 |
| 'Rps6ka4' | 29.115 | 26.733 | 32.205 | 26.736 | 28.673 | 32.632 | 28.308 | 30.369 | 30.444 | 28.522 | 25.965 | 32.356 | 29.421 | 25.844 | 30.951 | 29.188 | 24.198 | 31.497 | 0.032 | 0.3539 | 0.6074 | 0 | 0 | 0 |
| 'Rps6ka5' | 1.578 | 1.437 | 1.861 | 1.727 | 1.628 | 1.753 | 1.601 | 1.668 | 1.620 | 1.650 | 1.787 | 1.841 | 1.492 | 1.687 | 1.911 | 1.482 | 1.753 | 1.698 | -0.047 | 0.4440 | 0.6872 | 0 | 0 | 0 |
| 'Rps6kb1' | 6.604 | 6.616 | 6.911 | 7.355 | 6.781 | 6.189 | 7.598 | 6.712 | 6.628 | 5.759 | 7.210 | 7.422 | 6.420 | 6.501 | 6.341 | 6.509 | 7.710 | 7.053 | 0.000 | 0.9966 | 0.9994 | 0 | 0 | 0 |
| 'Rps6kb2' | 9.309 | 8.716 | 10.101 | 9.285 | 9.008 | 10.116 | 9.241 | 9.517 | 10.244 | 9.370 | 8.687 | 9.702 | 10.052 | 8.529 | 9.612 | 8.892 | 9.393 | 8.946 | 0.031 | 0.3477 | 0.6032 | 0 | 0 | 0 |
| 'Rps6kc1' | 2.053 | 1.876 | 2.048 | 2.157 | 2.394 | 2.125 | 2.132 | 2.119 | 2.110 | 2.425 | 2.238 | 2.222 | 2.301 | 1.907 | 2.157 | 2.295 | 2.295 | 2.246 | -0.089 | 0.0467 | 0.1803 | 0 | 0 | 0 |
| 'Rps7' | 660.976 | 677.928 | 620.805 | 704.715 | 705.455 | 630.753 | 712.722 | 670.760 | 627.709 | 632.042 | 674.333 | 530.909 | 655.762 | 653.805 | 646.466 | 700.252 | 668.113 | 598.313 | 0.052 | 0.1342 | 0.3530 | 0 | 0 | 0 |
| 'Rps8' | 157.604 | 158.885 | 151.587 | 172.342 | 158.263 | 154.145 | 172.006 | 152.185 | 152.781 | 150.126 | 155.318 | 130.410 | 157.193 | 157.131 | 154.709 | 165.621 | 156.265 | 145.988 | 0.048 | 0.1602 | 0.3904 | 0 | 0 | 0 |
| 'Rps9' | 144.685 | 146.296 | 133.859 | 149.148 | 145.637 | 136.533 | 150.582 | 141.312 | 137.884 | 136.556 | 147.334 | 112.154 | 137.706 | 139.531 | 136.129 | 146.807 | 142.627 | 126.580 | 0.060 | 0.0832 | 0.2611 | 0 | 0 | 0 |
| 'Rpsa' | 984.484 | 994.070 | 926.340 | ####### | ####### | 947.831 | ####### | 945.138 | 946.133 | 996.761 | ####### | 791.963 | ####### | 976.368 | ####### | ####### | ####### | 948.783 | -0.007 | 0.8606 | 0.9398 | 0 | 0 | 0 |
| 'Rptor' | 3.462 | 2.962 | 3.339 | 3.224 | 2.776 | 3.098 | 3.185 | 2.888 | 3.126 | 3.220 | 3.169 | 3.542 | 2.979 | 3.051 | 3.451 | 2.958 | 3.102 | 3.277 | -0.046 | 0.2502 | 0.5012 | 0 | 0 | 0 |
| 'Rpusd1' | 9.228 | 9.628 | 11.107 | 8.427 | 9.478 | 12.650 | 10.349 | 10.289 | 10.991 | 10.361 | 7.397 | 11.260 | 10.863 | 8.932 | 11.469 | 9.986 | 8.308 | 10.584 | 0.038 | 0.5319 | 0.7526 | 0 | 0 | 0 |
| 'Rpusd2' | 2.061 | 2.585 | 2.585 | 2.261 | 2.497 | 2.197 | 1.901 | 2.116 | 2.581 | 2.494 | 2.099 | 2.597 | 2.142 | 2.149 | 2.467 | 2.893 | 2.116 | 2.379 | -0.049 | 0.5788 | 0.7820 | 0 | 0 | 0 |
| 'Rpusd3' | 1.846 | 2.668 | 2.886 | 2.294 | 2.565 | 2.763 | 2.005 | 2.374 | 2.588 | 2.320 | 2.420 | 2.448 | 2.808 | 2.547 | 2.404 | 2.672 | 2.188 | 1.982 | 0.003 | 0.9754 | 0.9918 | 0 | 0 | 0 |
| 'Rpusd4' | 9.086 | 9.739 | 9.991 | 9.136 | 8.826 | 9.471 | 8.688 | 9.415 | 9.790 | 9.103 | 9.296 | 10.600 | 9.047 | 7.967 | 10.697 | 9.172 | 10.229 | 9.584 | -0.038 | 0.3824 | 0.6330 | 0 | 0 | 0 |
| 'Rrad' | 11.634 | 6.376 | 7.841 | 12.803 | 8.062 | 7.424 | 13.623 | 6.606 | 7.817 | 7.483 | 4.949 | 8.042 | 7.484 | 6.112 | 7.039 | 8.950 | 7.789 | 6.766 | 0.295 | 0.0021 | 0.0189 | 1 | 0 | 0 |
| 'Rraga' | 33.668 | 31.983 | 31.928 | 29.684 | 35.139 | 32.768 | 31.571 | 36.165 | 29.407 | 30.532 | 33.943 | 33.801 | 35.999 | 33.952 | 33.069 | 29.811 | 35.645 | 31.720 | -0.042 | 0.3033 | 0.5608 | 0 | 0 | 0 |
| 'Rragb' | 3.698 | 4.013 | 3.706 | 3.404 | 3.645 | 3.492 | 3.652 | 4.232 | 4.204 | 4.413 | 4.828 | 3.969 | 4.226 | 3.939 | 3.704 | 4.268 | 3.273 | 4.137 | -0.116 | 0.1108 | 0.3150 | 0 | 0 | 0 |
| 'Rragc' | 11.381 | 12.102 | 10.637 | 11.953 | 11.916 | 11.386 | 10.346 | 12.416 | 11.206 | 11.985 | 10.811 | 12.297 | 11.391 | 11.561 | 11.673 | 11.782 | 12.156 | 13.077 | -0.060 | 0.1451 | 0.3696 | 0 | 0 | 0 |
| 'Rras' | 17.634 | 22.248 | 19.152 | 17.438 | 18.537 | 21.225 | 18.086 | 19.312 | 18.925 | 21.862 | 18.875 | 21.465 | 21.584 | 21.114 | 20.738 | 20.081 | 23.666 | 19.688 | -0.145 | 0.0063 | 0.0443 | 1 | 0 | 0 |
| 'Rras2' | 3.678 | 2.765 | 4.247 | 4.215 | 2.813 | 3.933 | 3.680 | 2.969 | 4.316 | 2.670 | 2.567 | 4.713 | 2.668 | 2.752 | 3.872 | 3.206 | 3.408 | 4.672 | 0.085 | 0.3576 | 0.6103 | 0 | 0 | 0 |
| 'Rrbp1' | 13.459 | 11.241 | 14.022 | 12.483 | 10.964 | 14.494 | 11.744 | 11.544 | 13.930 | 13.381 | 11.925 | 14.138 | 13.057 | 12.095 | 13.494 | 12.152 | 11.614 | 14.076 | -0.037 | 0.2334 | 0.4820 | 0 | 0 | 0 |
| 'Rreb1' | 0.679 | 0.512 | 0.587 | 0.607 | 0.540 | 0.609 | 0.483 | 0.494 | 0.648 | 0.531 | 0.451 | 0.581 | 0.371 | 0.380 | 0.510 | 0.379 | 0.366 | 0.509 | 0.331 | 0.0000 | 0.0006 | 1 | 0 | 0 |
| 'Rrm1' | 42.695 | 43.142 | 49.574 | 49.505 | 46.198 | 49.396 | 50.361 | 43.732 | 48.314 | 39.134 | 46.221 | 46.173 | 52.229 | 49.137 | 50.564 | 43.600 | 48.587 | 45.582 | -0.006 | 0.9093 | 0.9646 | 0 | 0 | 0 |
| 'Rrm2' | 30.729 | 30.557 | 40.465 | 38.821 | 34.149 | 36.301 | 38.191 | 32.378 | 37.788 | 25.377 | 35.376 | 37.539 | 40.509 | 37.535 | 42.363 | 30.933 | 35.995 | 37.358 | -0.026 | 0.7122 | 0.8661 | 0 | 0 | 0 |
| 'Rrm2b' | 3.457 | 4.480 | 3.947 | 3.998 | 4.277 | 3.769 | 3.951 | 4.533 | 3.769 | 4.184 | 4.288 | 3.996 | 3.780 | 4.187 | 3.751 | 3.656 | 4.432 | 3.418 | 0.005 | 0.9016 | 0.9612 | 0 | 0 | 0 |
| 'Rrn3' | 16.254 | 18.297 | 16.378 | 16.159 | 17.070 | 16.210 | 15.224 | 17.396 | 15.138 | 16.062 | 18.493 | 17.870 | 15.722 | 15.930 | 17.222 | 15.433 | 18.041 | 16.871 | -0.048 | 0.1988 | 0.4387 | 0 | 0 | 0 |
| 'Rrnad1' | 2.636 | 2.673 | 2.838 | 2.763 | 2.724 | 2.672 | 2.679 | 2.548 | 2.989 | 2.469 | 2.635 | 3.525 | 2.555 | 2.746 | 3.006 | 2.556 | 3.007 | 3.193 | -0.078 | 0.0991 | 0.2942 | 0 | 0 | 0 |
| 'Rrp1' | 153.720 | 145.618 | 155.341 | 151.187 | 148.589 | 159.211 | 148.988 | 153.205 | 153.178 | 156.089 | 140.524 | 148.001 | 159.908 | 144.447 | 153.450 | 157.569 | 141.930 | 149.468 | 0.010 | 0.6281 | 0.8154 | 0 | 0 | 0 |
| 'Rrp12' | 10.862 | 8.683 | 11.341 | 9.423 | 9.102 | 9.483 | 8.712 | 10.411 | 10.253 | 10.354 | 8.294 | 10.299 | 9.641 | 7.282 | 10.268 | 9.738 | 9.699 | 10.112 | 0.037 | 0.5181 | 0.7409 | 0 | 0 | 0 |
| 'Rrp15' | 5.926 | 5.582 | 6.176 | 6.637 | 5.820 | 5.495 | 6.286 | 6.035 | 5.565 | 6.562 | 5.317 | 5.523 | 5.812 | 4.829 | 5.454 | 5.139 | 5.480 | 5.564 | 0.095 | 0.0553 | 0.2017 | 0 | 0 | 0 |
| 'Rrp1b' | 4.533 | 3.655 | 4.608 | 3.774 | 3.815 | 3.904 | 4.002 | 4.394 | 4.189 | 4.121 | 4.246 | 4.541 | 4.385 | 3.633 | 4.366 | 3.663 | 4.123 | 4.116 | -0.022 | 0.6670 | 0.8396 | 0 | 0 | 0 |
| 'Rrp36' | 37.317 | 31.169 | 36.937 | 37.249 | 38.652 | 39.107 | 36.386 | 38.143 | 38.712 | 36.010 | 35.992 | 41.845 | 33.486 | 35.494 | 38.140 | 34.501 | 34.706 | 38.169 | 0.011 | 0.8119 | 0.9155 | 0 | 0 | 0 |
| 'Rrp8' | 5.584 | 5.971 | 5.893 | 5.325 | 5.973 | 5.650 | 4.765 | 6.049 | 5.791 | 5.788 | 4.909 | 5.883 | 5.850 | 5.764 | 5.922 | 5.585 | 5.492 | 5.526 | -0.004 | 0.9352 | 0.9752 | 0 | 0 | 0 |
| 'Rrp9' | 9.103 | 8.439 | 10.052 | 8.325 | 8.329 | 9.579 | 8.927 | 9.790 | 9.511 | 9.248 | 8.406 | 9.889 | 8.772 | 7.452 | 9.405 | 9.289 | 7.677 | 9.953 | 0.028 | 0.5137 | 0.7378 | 0 | 0 | 0 |
| 'Rrs1' | 10.419 | 9.615 | 11.704 | 8.948 | 10.019 | 9.900 | 11.188 | 10.030 | 9.937 | 9.799 | 9.918 | 10.422 | 9.566 | 9.688 | 9.873 | 9.875 | 9.938 | 10.561 | 0.026 | 0.6162 | 0.8085 | 0 | 0 | 0 |
| 'Rsad1' | 0.428 | 0.529 | 0.429 | 0.512 | 0.622 | 0.413 | 0.361 | 0.404 | 0.491 | 0.339 | 0.371 | 0.332 | 0.315 | 0.244 | 0.320 | 0.402 | 0.440 | 0.263 | 0.455 | 0.0002 | 0.0032 | 1 | 1 | 0 |
| 'Rsad2' | 0.896 | 0.976 | 0.680 | 0.684 | 1.074 | 0.995 | 0.546 | 1.334 | 0.902 | 1.000 | 0.979 | 1.502 | 1.344 | 1.056 | 1.519 | 1.027 | 0.985 | 1.426 | -0.442 | 0.0007 | 0.0080 | 1 | 1 | 0 |
| 'Rsbn1' | 9.356 | 10.856 | 8.706 | 9.526 | 9.828 | 8.373 | 10.386 | 9.129 | 8.335 | 8.776 | 10.127 | 9.903 | 8.582 | 9.992 | 9.051 | 9.643 | 10.556 | 9.674 | -0.045 | 0.3297 | 0.5877 | 0 | 0 | 0 |
| 'Rsbn1l' | 10.172 | 9.859 | 9.542 | 10.003 | 9.958 | 9.292 | 10.247 | 9.803 | 9.241 | 9.228 | 10.175 | 10.230 | 9.130 | 10.044 | 9.345 | 9.553 | 10.197 | 9.768 | -0.005 | 0.8824 | 0.9506 | 0 | 0 | 0 |
| 'Rsf1' | 5.864 | 4.886 | 5.048 | 6.003 | 4.957 | 4.650 | 5.693 | 5.000 | 4.854 | 5.005 | 5.231 | 5.430 | 4.796 | 5.117 | 5.034 | 4.901 | 5.452 | 5.140 | 0.011 | 0.8236 | 0.9224 | 0 | 0 | 0 |
| 'Rsl1d1' | 27.643 | 24.091 | 25.500 | 27.128 | 25.247 | 22.210 | 27.078 | 24.554 | 23.296 | 24.378 | 24.199 | 25.680 | 23.787 | 25.735 | 23.800 | 24.670 | 25.250 | 24.349 | 0.018 | 0.6483 | 0.8285 | 0 | 0 | 0 |
| 'Rsl24d1' | 18.014 | 18.109 | 17.013 | 17.445 | 16.982 | 15.873 | 17.902 | 18.105 | 16.429 | 16.151 | 16.876 | 17.549 | 15.190 | 16.094 | 17.328 | 16.469 | 17.322 | 17.560 | 0.037 | 0.3561 | 0.6090 | 0 | 0 | 0 |
| 'Rspry1' | 8.589 | 9.185 | 8.655 | 9.547 | 9.363 | 8.689 | 9.092 | 9.377 | 8.427 | 8.192 | 9.351 | 9.316 | 8.773 | 9.121 | 8.905 | 8.420 | 8.923 | 9.188 | -0.002 | 0.9539 | 0.9831 | 0 | 0 | 0 |
| 'Rsrc1' | 1.979 | 2.061 | 2.092 | 2.139 | 1.992 | 1.705 | 2.256 | 2.005 | 2.116 | 2.081 | 2.017 | 2.307 | 2.000 | 1.879 | 2.059 | 1.880 | 2.011 | 2.176 | -0.022 | 0.6718 | 0.8418 | 0 | 0 | 0 |
| 'Rsrc2' | 11.402 | 11.203 | 10.981 | 12.494 | 11.237 | 10.439 | 12.516 | 10.945 | 10.744 | 10.898 | 10.672 | 11.528 | 10.370 | 10.974 | 10.876 | 11.109 | 11.949 | 10.668 | 0.029 | 0.4300 | 0.6758 | 0 | 0 | 0 |
| 'Rsrp1' | 60.384 | 50.313 | 52.604 | 63.555 | 47.262 | 49.960 | 62.433 | 47.616 | 51.239 | 41.228 | 62.304 | 58.773 | 49.528 | 58.603 | 60.937 | 41.260 | 66.130 | 53.592 | -0.025 | 0.8165 | 0.9183 | 0 | 0 | 0 |
| 'Rsu1' | 4.240 | 4.869 | 4.536 | 4.579 | 4.734 | 4.760 | 5.222 | 4.529 | 4.627 | 4.837 | 4.867 | 4.695 | 5.563 | 5.414 | 4.949 | 5.159 | 5.280 | 4.685 | -0.121 | 0.0029 | 0.0248 | 1 | 0 | 0 |
| 'Rtca' | 10.982 | 11.976 | 11.102 | 12.096 | 11.342 | 11.646 | 10.405 | 12.969 | 10.973 | 10.995 | 10.540 | 10.682 | 11.820 | 11.638 | 10.801 | 11.403 | 10.962 | 10.903 | 0.044 | 0.2763 | 0.5319 | 0 | 0 | 0 |
| 'Rtcb' | 39.013 | 39.188 | 39.208 | 37.473 | 39.444 | 38.792 | 42.107 | 40.476 | 38.866 | 40.686 | 38.119 | 38.959 | 38.005 | 35.311 | 38.666 | 39.594 | 39.297 | 37.097 | 0.026 | 0.3437 | 0.5995 | 0 | 0 | 0 |
| 'Rtel1' | 4.099 | 4.312 | 5.373 | 4.187 | 4.293 | 4.794 | 4.558 | 4.431 | 5.117 | 4.602 | 4.664 | 4.760 | 5.192 | 4.627 | 4.735 | 4.349 | 4.994 | 4.518 | -0.056 | 0.2121 | 0.4574 | 0 | 0 | 0 |
| 'Rtf1' | 27.589 | 25.757 | 26.595 | 28.576 | 25.671 | 27.964 | 26.915 | 26.497 | 27.546 | 26.740 | 25.364 | 26.926 | 24.479 | 23.674 | 26.235 | 26.860 | 24.662 | 25.972 | 0.064 | 0.0131 | 0.0760 | 0 | 0 | 0 |
| 'Rtfdc1' | 29.333 | 29.193 | 29.325 | 27.854 | 30.217 | 30.748 | 27.617 | 30.441 | 29.282 | 29.210 | 27.662 | 29.590 | 28.098 | 29.156 | 30.309 | 28.821 | 29.328 | 27.919 | 0.012 | 0.6201 | 0.8101 | 0 | 0 | 0 |
| 'Rtn3' | 28.940 | 31.179 | 30.951 | 29.633 | 31.710 | 30.742 | 29.547 | 28.715 | 29.811 | 30.949 | 27.668 | 31.074 | 29.560 | 31.522 | 30.012 | 29.633 | 30.218 | 30.372 | -0.010 | 0.6905 | 0.8542 | 0 | 0 | 0 |
| 'Rtn4' | 8.917 | 8.476 | 9.366 | 8.812 | 9.606 | 9.187 | 8.876 | 9.245 | 8.560 | 9.895 | 9.022 | 9.065 | 9.181 | 9.735 | 9.484 | 9.726 | 9.048 | 9.445 | -0.070 | 0.0283 | 0.1303 | 0 | 0 | 0 |
| 'Rtn4ip1' | 3.963 | 4.642 | 4.255 | 3.728 | 4.260 | 3.786 | 3.698 | 4.450 | 4.265 | 4.235 | 4.074 | 4.104 | 4.349 | 4.445 | 4.064 | 4.215 | 4.641 | 3.639 | -0.036 | 0.4881 | 0.7184 | 0 | 0 | 0 |
| 'Rtn4rl1' | 1.677 | 0.979 | 1.788 | 1.774 | 1.261 | 1.699 | 1.521 | 1.085 | 1.941 | 1.604 | 1.161 | 2.582 | 1.377 | 1.125 | 1.840 | 1.412 | 1.087 | 2.222 | -0.083 | 0.4214 | 0.6681 | 0 | 0 | 0 |
| 'Rtp4' | 12.866 | 14.764 | 14.245 | 11.369 | 15.004 | 18.346 | 11.254 | 18.989 | 15.093 | 14.343 | 16.517 | 24.942 | 17.554 | 15.370 | 25.563 | 18.264 | 17.280 | 23.547 | -0.396 | 0.0000 | 0.0001 | 1 | 1 | 0 |
| 'Rttn' | 5.792 | 5.505 | 6.144 | 7.220 | 6.140 | 5.706 | 7.022 | 6.350 | 6.603 | 2.974 | 3.423 | 3.565 | 3.146 | 2.912 | 3.319 | 2.625 | 3.539 | 3.120 | 0.966 | 0.0000 | 0.0000 | 1 | 1 | 0 |
| 'Rubcn' | 2.801 | 2.907 | 2.909 | 3.020 | 2.815 | 2.889 | 2.736 | 2.852 | 2.834 | 2.844 | 3.121 | 3.149 | 3.049 | 2.631 | 3.196 | 2.781 | 2.944 | 3.042 | -0.067 | 0.0633 | 0.2191 | 0 | 0 | 0 |
| 'Rufy1' | 12.270 | 12.773 | 12.077 | 11.359 | 12.109 | 12.519 | 11.389 | 11.113 | 11.790 | 12.322 | 12.465 | 11.846 | 12.349 | 12.440 | 10.968 | 11.691 | 11.764 | 10.950 | 0.000 | 0.9992 | 0.9998 | 0 | 0 | 0 |
| 'Rufy2' | 4.867 | 5.388 | 4.871 | 5.049 | 4.970 | 4.978 | 4.797 | 5.437 | 5.187 | 4.706 | 5.212 | 5.081 | 4.601 | 5.196 | 4.847 | 4.600 | 5.431 | 4.754 | 0.025 | 0.4144 | 0.6624 | 0 | 0 | 0 |
| 'Rufy3' | 0.473 | 0.625 | 0.568 | 0.545 | 0.527 | 0.581 | 0.606 | 0.702 | 0.602 | 0.499 | 0.608 | 0.542 | 0.527 | 0.549 | 0.497 | 0.453 | 0.542 | 0.605 | 0.103 | 0.1720 | 0.4058 | 0 | 0 | 0 |
| 'Rundc1' | 18.891 | 18.695 | 20.501 | 19.090 | 18.119 | 18.963 | 19.354 | 19.120 | 20.190 | 18.702 | 18.150 | 21.465 | 16.390 | 16.576 | 19.998 | 18.398 | 16.998 | 19.856 | 0.044 | 0.2511 | 0.5023 | 0 | 0 | 0 |
| 'Rundc3a' | 3.309 | 2.896 | 3.594 | 2.414 | 3.092 | 3.423 | 2.795 | 3.156 | 3.400 | 3.566 | 3.242 | 4.093 | 3.186 | 3.210 | 3.650 | 3.133 | 3.034 | 3.750 | -0.145 | 0.0077 | 0.0516 | 0 | 0 | 0 |
| 'Rundc3b' | 8.222 | 7.643 | 8.507 | 7.046 | 7.508 | 8.056 | 8.038 | 8.450 | 7.482 | 9.273 | 8.296 | 9.455 | 6.803 | 9.427 | 9.198 | 8.313 | 8.050 | 9.272 | -0.146 | 0.0110 | 0.0668 | 0 | 0 | 0 |
| 'Runx1' | 10.688 | 9.670 | 10.148 | 10.352 | 10.149 | 9.481 | 9.934 | 10.599 | 9.741 | 10.823 | 9.317 | 9.233 | 9.973 | 9.836 | 8.921 | 10.258 | 9.510 | 8.579 | 0.064 | 0.0260 | 0.1239 | 0 | 0 | 0 |
| 'Runx2' | 0.998 | 0.871 | 1.087 | 0.995 | 0.961 | 0.979 | 0.920 | 1.022 | 1.028 | 1.072 | 1.214 | 1.294 | 1.004 | 1.187 | 0.950 | 0.941 | 1.083 | 1.008 | -0.148 | 0.0233 | 0.1146 | 0 | 0 | 0 |
| 'Runx3' | 3.684 | 2.321 | 4.200 | 2.722 | 2.204 | 3.856 | 3.546 | 3.129 | 3.538 | 3.410 | 2.867 | 5.426 | 2.960 | 2.885 | 5.131 | 3.661 | 2.485 | 5.192 | -0.203 | 0.0164 | 0.0897 | 0 | 0 | 0 |
| 'Rusc1' | 2.872 | 2.645 | 3.102 | 2.656 | 2.655 | 3.149 | 2.897 | 2.862 | 3.045 | 3.264 | 2.603 | 3.061 | 2.602 | 2.797 | 2.686 | 2.644 | 2.552 | 2.539 | 0.057 | 0.2993 | 0.5557 | 0 | 0 | 0 |
| 'Ruvbl1' | 28.007 | 27.780 | 29.679 | 33.039 | 30.287 | 29.194 | 29.556 | 30.366 | 29.633 | 27.155 | 28.299 | 29.795 | 29.170 | 28.415 | 28.499 | 29.623 | 28.137 | 28.713 | 0.041 | 0.2225 | 0.4701 | 0 | 0 | 0 |
| 'Ruvbl2' | 24.401 | 26.236 | 26.858 | 25.666 | 23.245 | 27.769 | 26.670 | 23.266 | 26.723 | 22.569 | 23.704 | 28.974 | 25.427 | 23.784 | 28.022 | 25.921 | 25.031 | 26.982 | -0.010 | 0.7652 | 0.8955 | 0 | 0 | 0 |
| 'Rwdd1' | 36.462 | 40.625 | 37.531 | 43.207 | 42.166 | 38.582 | 44.165 | 39.629 | 39.241 | 36.277 | 34.785 | 39.042 | 42.020 | 37.410 | 38.862 | 40.649 | 39.231 | 36.960 | 0.051 | 0.1999 | 0.4409 | 0 | 0 | 0 |
| 'Rwdd2b' | 7.349 | 8.512 | 8.643 | 7.396 | 7.847 | 8.529 | 7.973 | 8.223 | 8.071 | 8.804 | 9.137 | 8.773 | 8.226 | 8.986 | 8.965 | 7.017 | 8.027 | 8.316 | -0.082 | 0.1153 | 0.3225 | 0 | 0 | 0 |
| 'Rwdd3' | 0.639 | 0.783 | 0.660 | 0.659 | 0.768 | 0.629 | 0.647 | 0.672 | 0.731 | 0.686 | 0.681 | 0.548 | 0.742 | 0.613 | 0.491 | 0.710 | 0.567 | 0.436 | 0.178 | 0.0516 | 0.1922 | 0 | 0 | 0 |
| 'Rwdd4a' | 5.135 | 6.753 | 6.311 | 6.142 | 6.264 | 6.698 | 5.295 | 6.247 | 6.194 | 5.694 | 5.929 | 6.647 | 5.450 | 6.477 | 6.315 | 5.727 | 5.643 | 6.557 | 0.001 | 0.9740 | 0.9912 | 0 | 0 | 0 |
| 'Rxra' | 1.645 | 1.988 | 2.047 | 1.865 | 1.966 | 2.256 | 1.496 | 1.954 | 2.064 | 1.788 | 1.680 | 1.805 | 1.780 | 1.621 | 1.971 | 1.718 | 1.707 | 2.027 | 0.094 | 0.1202 | 0.3303 | 0 | 0 | 0 |
| 'Rxrb' | 14.859 | 13.910 | 14.545 | 14.395 | 14.041 | 15.858 | 13.842 | 13.994 | 15.660 | 12.759 | 12.701 | 16.322 | 13.655 | 14.477 | 15.469 | 14.103 | 14.177 | 16.207 | 0.004 | 0.9045 | 0.9624 | 0 | 0 | 0 |
| 'Rybp' | 6.280 | 6.110 | 5.386 | 6.460 | 5.587 | 5.637 | 6.249 | 6.175 | 5.502 | 5.097 | 6.723 | 6.320 | 5.230 | 6.195 | 6.032 | 6.289 | 6.207 | 5.945 | -0.030 | 0.5921 | 0.7921 | 0 | 0 | 0 |
| 'Ryk' | 2.406 | 2.821 | 2.522 | 2.332 | 2.439 | 2.277 | 2.782 | 2.777 | 2.577 | 2.282 | 2.268 | 2.044 | 2.245 | 2.337 | 2.125 | 2.101 | 2.393 | 2.089 | 0.197 | 0.0001 | 0.0015 | 1 | 0 | 0 |
| 'Ryr1' | 2.866 | 2.362 | 3.564 | 2.797 | 2.448 | 2.943 | 2.544 | 2.850 | 2.840 | 3.038 | 3.104 | 2.787 | 2.804 | 2.584 | 2.980 | 2.258 | 2.235 | 3.274 | -0.003 | 0.9685 | 0.9896 | 0 | 0 | 0 |
| 'Ryr3' | 0.393 | 0.374 | 0.352 | 0.402 | 0.358 | 0.274 | 0.408 | 0.369 | 0.343 | 0.438 | 0.426 | 0.397 | 0.339 | 0.375 | 0.357 | 0.380 | 0.373 | 0.345 | -0.086 | 0.2109 | 0.4561 | 0 | 0 | 0 |
| 'S100a10' | 64.856 | 60.644 | 68.439 | 66.967 | 69.217 | 72.919 | 65.333 | 68.416 | 61.183 | 57.998 | 57.215 | 68.297 | 48.242 | 60.793 | 71.194 | 56.666 | 62.044 | 64.102 | 0.124 | 0.0136 | 0.0782 | 0 | 0 | 0 |
| 'S100a11' | 366.536 | 394.270 | 367.515 | 387.135 | 407.697 | 389.243 | 382.855 | 386.896 | 382.759 | 364.274 | 365.767 | 311.699 | 344.299 | 363.548 | 384.119 | 347.667 | 366.358 | 348.213 | 0.107 | 0.0025 | 0.0214 | 1 | 0 | 0 |
| 'S100a13' | 48.824 | 53.656 | 51.088 | 52.392 | 49.666 | 53.172 | 54.231 | 49.842 | 53.147 | 48.508 | 52.303 | 49.860 | 51.290 | 51.389 | 55.034 | 54.537 | 55.856 | 48.077 | -0.010 | 0.8027 | 0.9115 | 0 | 0 | 0 |
| 'S100a4' | 5.632 | 7.902 | 8.498 | 6.265 | 6.710 | 9.048 | 7.811 | 9.115 | 10.770 | 8.156 | 6.703 | 7.896 | 7.132 | 7.137 | 9.849 | 6.246 | 7.187 | 9.145 | 0.037 | 0.7031 | 0.8611 | 0 | 0 | 0 |
| 'S100a6' | 6.369 | 8.903 | 7.776 | 8.847 | 8.549 | 9.133 | 9.569 | 8.436 | 8.754 | 8.165 | 8.852 | 8.366 | 8.794 | 9.785 | 10.120 | 9.010 | 7.584 | 9.393 | -0.082 | 0.2549 | 0.5070 | 0 | 0 | 0 |
| 'S100pbp' | 2.918 | 3.934 | 3.352 | 3.384 | 3.708 | 3.473 | 3.497 | 3.695 | 3.511 | 2.729 | 3.602 | 3.907 | 3.144 | 3.576 | 3.674 | 3.114 | 3.337 | 3.903 | 0.010 | 0.8395 | 0.9315 | 0 | 0 | 0 |
| 'S1pr1' | 4.788 | 4.833 | 6.266 | 5.667 | 5.876 | 7.606 | 6.754 | 6.695 | 7.183 | 5.230 | 5.882 | 8.399 | 5.775 | 6.670 | 7.905 | 6.878 | 7.171 | 7.189 | -0.144 | 0.0509 | 0.1910 | 0 | 0 | 0 |
| 'S1pr2' | 65.053 | 95.316 | 60.883 | 67.191 | 83.437 | 75.058 | 64.898 | 75.135 | 73.230 | 64.858 | 74.815 | 80.001 | 78.009 | 88.210 | 77.932 | 83.033 | 80.171 | 83.291 | -0.124 | 0.0338 | 0.1465 | 0 | 0 | 0 |
| 'S1pr4' | 22.427 | 20.912 | 26.043 | 22.876 | 21.453 | 27.259 | 22.139 | 21.943 | 25.378 | 17.951 | 23.045 | 28.944 | 18.666 | 22.161 | 29.800 | 24.395 | 22.606 | 28.754 | -0.041 | 0.4553 | 0.6934 | 0 | 0 | 0 |
| 'Saal1' | 1.847 | 2.007 | 2.266 | 1.883 | 2.220 | 2.091 | 1.954 | 2.051 | 2.187 | 2.039 | 1.929 | 2.046 | 2.220 | 2.007 | 1.962 | 2.250 | 2.157 | 2.069 | -0.021 | 0.6759 | 0.8448 | 0 | 0 | 0 |
| 'Sac3d1' | 8.399 | 8.336 | 9.026 | 8.356 | 8.406 | 8.301 | 8.980 | 8.122 | 9.325 | 7.254 | 7.449 | 10.577 | 8.499 | 7.060 | 11.121 | 7.973 | 8.218 | 11.103 | -0.051 | 0.4788 | 0.7114 | 0 | 0 | 0 |
| 'Sacm1l' | 25.388 | 29.069 | 26.350 | 28.405 | 28.838 | 25.118 | 30.603 | 29.038 | 25.651 | 24.188 | 29.217 | 26.862 | 24.401 | 28.320 | 24.564 | 24.480 | 30.149 | 25.124 | 0.055 | 0.1621 | 0.3938 | 0 | 0 | 0 |
| 'Sae1' | 68.022 | 72.209 | 75.287 | 70.354 | 74.596 | 69.955 | 69.838 | 74.719 | 70.878 | 63.092 | 68.367 | 69.477 | 72.269 | 70.327 | 70.951 | 65.794 | 71.955 | 67.438 | 0.049 | 0.0659 | 0.2240 | 0 | 0 | 0 |
| 'Safb2' | 22.520 | 18.032 | 21.496 | 20.869 | 17.690 | 19.998 | 20.571 | 18.389 | 21.000 | 19.421 | 21.353 | 25.028 | 21.180 | 20.643 | 22.313 | 20.090 | 22.295 | 25.021 | -0.139 | 0.0031 | 0.0261 | 1 | 0 | 0 |
| 'Sag' | 0.321 | 0.312 | 0.356 | 0.369 | 0.465 | 0.328 | 0.397 | 0.280 | 0.369 | 0.145 | 0.116 | 0.144 | 0.184 | 0.108 | 0.099 | 0.185 | 0.132 | 0.153 | 1.317 | 0.0000 | 0.0000 | 1 | 1 | 1 |
| 'Sall2' | 2.669 | 2.481 | 2.567 | 2.639 | 2.486 | 2.953 | 2.779 | 2.351 | 2.689 | 2.557 | 2.427 | 3.121 | 2.402 | 2.161 | 2.910 | 2.662 | 2.225 | 3.205 | -0.015 | 0.7784 | 0.9002 | 0 | 0 | 0 |
| 'Samd10' | 5.596 | 6.712 | 7.029 | 5.022 | 6.480 | 7.942 | 5.809 | 6.629 | 7.436 | 6.110 | 6.845 | 6.762 | 6.819 | 6.346 | 7.726 | 7.255 | 5.921 | 6.496 | -0.053 | 0.3722 | 0.6240 | 0 | 0 | 0 |
| 'Samd11' | 6.803 | 5.900 | 6.496 | 5.977 | 5.731 | 5.776 | 6.147 | 6.865 | 5.332 | 5.357 | 3.947 | 3.727 | 5.125 | 3.856 | 3.629 | 4.865 | 4.123 | 3.475 | 0.538 | 0.0000 | 0.0000 | 1 | 1 | 0 |
| 'Samd8' | 1.767 | 1.846 | 1.758 | 1.821 | 1.654 | 1.755 | 1.832 | 2.133 | 1.684 | 1.812 | 2.044 | 1.840 | 1.836 | 2.197 | 1.831 | 2.198 | 1.725 | 1.743 | -0.088 | 0.1262 | 0.3401 | 0 | 0 | 0 |
| 'Samd9l' | 10.452 | 15.489 | 12.957 | 10.275 | 16.219 | 17.354 | 10.765 | 16.758 | 14.800 | 15.366 | 12.700 | 18.321 | 13.171 | 16.371 | 18.742 | 18.877 | 14.015 | 18.714 | -0.253 | 0.0158 | 0.0870 | 0 | 0 | 0 |
| 'Samhd1' | 46.897 | 54.171 | 46.816 | 47.056 | 56.066 | 50.385 | 48.353 | 52.925 | 51.318 | 51.370 | 49.050 | 51.003 | 48.670 | 50.665 | 49.651 | 48.552 | 51.124 | 49.271 | 0.001 | 0.9558 | 0.9836 | 0 | 0 | 0 |
| 'Samm50' | 62.424 | 69.076 | 69.520 | 68.889 | 72.512 | 71.065 | 69.577 | 70.124 | 66.714 | 63.401 | 62.730 | 63.219 | 67.227 | 67.117 | 64.689 | 72.947 | 65.032 | 63.609 | 0.062 | 0.0536 | 0.1977 | 0 | 0 | 0 |
| 'Samsn1' | 22.323 | 24.186 | 20.484 | 22.745 | 23.546 | 20.400 | 24.165 | 23.203 | 20.336 | 19.574 | 21.349 | 19.755 | 18.740 | 22.395 | 19.815 | 19.317 | 21.380 | 19.310 | 0.136 | 0.0000 | 0.0004 | 1 | 0 | 0 |
| 'Sap130' | 20.976 | 19.927 | 22.504 | 21.528 | 20.594 | 23.181 | 19.575 | 20.032 | 24.341 | 22.808 | 21.642 | 22.856 | 22.179 | 21.296 | 21.708 | 21.505 | 21.515 | 22.059 | -0.047 | 0.1398 | 0.3618 | 0 | 0 | 0 |
| 'Sap18' | 31.057 | 29.128 | 26.692 | 32.233 | 28.517 | 27.739 | 33.681 | 29.990 | 26.601 | 27.130 | 29.406 | 26.593 | 31.083 | 29.375 | 26.551 | 26.876 | 32.695 | 25.227 | 0.047 | 0.2415 | 0.4920 | 0 | 0 | 0 |
| 'Sap25' | 23.985 | 21.482 | 25.695 | 23.593 | 20.762 | 23.454 | 23.269 | 20.476 | 26.581 | 22.084 | 22.820 | 29.499 | 23.191 | 20.515 | 25.333 | 19.784 | 24.345 | 25.923 | -0.039 | 0.4180 | 0.6655 | 0 | 0 | 0 |
| 'Sap30' | 19.641 | 13.320 | 12.519 | 17.743 | 12.058 | 11.784 | 19.971 | 11.716 | 11.997 | 11.377 | 18.246 | 14.586 | 14.129 | 15.408 | 15.458 | 12.291 | 20.471 | 14.280 | -0.090 | 0.5274 | 0.7486 | 0 | 0 | 0 |
| 'Sap30bp' | 10.035 | 10.303 | 10.105 | 9.913 | 10.933 | 10.078 | 10.419 | 11.194 | 11.611 | 11.243 | 9.576 | 9.206 | 9.973 | 9.511 | 9.336 | 9.528 | 9.594 | 8.722 | 0.119 | 0.0071 | 0.0483 | 1 | 0 | 0 |
| 'Sap30l' | 50.330 | 43.624 | 50.705 | 46.805 | 43.131 | 51.247 | 46.005 | 45.670 | 43.929 | 45.346 | 46.377 | 45.243 | 47.222 | 44.884 | 49.359 | 51.650 | 46.566 | 52.877 | -0.034 | 0.4448 | 0.6876 | 0 | 0 | 0 |
| 'Sapcd2' | 1.391 | 1.524 | 2.405 | 1.681 | 1.680 | 2.277 | 1.751 | 1.466 | 2.590 | 1.123 | 1.772 | 1.943 | 2.300 | 2.166 | 2.284 | 1.908 | 1.592 | 1.880 | -0.049 | 0.6474 | 0.8282 | 0 | 0 | 0 |
| 'Sar1a' | 33.314 | 34.495 | 34.087 | 36.615 | 36.323 | 35.056 | 34.162 | 36.652 | 32.524 | 32.783 | 37.541 | 38.082 | 32.772 | 36.279 | 36.059 | 36.823 | 37.185 | 37.316 | -0.064 | 0.0652 | 0.2228 | 0 | 0 | 0 |
| 'Sar1b' | 37.556 | 42.409 | 38.141 | 39.363 | 41.780 | 34.715 | 40.015 | 43.049 | 33.475 | 36.575 | 42.060 | 36.886 | 38.594 | 42.099 | 36.039 | 37.577 | 38.824 | 34.894 | 0.015 | 0.6387 | 0.8221 | 0 | 0 | 0 |
| 'Saraf' | 94.388 | 81.898 | 83.739 | 87.786 | 80.866 | 83.024 | 85.737 | 81.696 | 80.619 | 84.747 | 81.021 | 86.738 | 77.901 | 76.530 | 84.919 | 83.334 | 79.046 | 86.504 | 0.025 | 0.4477 | 0.6880 | 0 | 0 | 0 |
| 'Sarm1' | 0.368 | 0.311 | 0.331 | 0.434 | 0.310 | 0.304 | 0.317 | 0.404 | 0.415 | 0.261 | 0.284 | 0.396 | 0.298 | 0.259 | 0.263 | 0.250 | 0.209 | 0.359 | 0.283 | 0.0268 | 0.1260 | 0 | 0 | 0 |
| 'Sarnp' | 72.916 | 74.593 | 72.364 | 84.315 | 82.630 | 73.023 | 83.862 | 73.822 | 73.955 | 63.892 | 71.141 | 70.282 | 74.608 | 76.576 | 71.466 | 71.346 | 76.203 | 65.424 | 0.097 | 0.0175 | 0.0939 | 0 | 0 | 0 |
| 'Sars' | 26.572 | 24.864 | 25.864 | 25.714 | 25.284 | 26.439 | 26.961 | 26.201 | 26.254 | 25.953 | 25.684 | 25.692 | 26.487 | 25.877 | 26.655 | 24.793 | 25.680 | 25.691 | 0.001 | 0.9804 | 0.9926 | 0 | 0 | 0 |
| 'Sars2' | 13.692 | 9.488 | 16.315 | 11.791 | 11.193 | 14.838 | 12.518 | 10.981 | 15.127 | 11.601 | 10.594 | 12.654 | 13.343 | 9.926 | 12.210 | 11.712 | 10.874 | 11.980 | 0.135 | 0.0183 | 0.0971 | 0 | 0 | 0 |
| 'Sart1' | 46.062 | 41.177 | 49.918 | 44.221 | 42.176 | 49.049 | 44.284 | 43.991 | 47.413 | 46.551 | 41.693 | 47.581 | 47.068 | 42.192 | 46.646 | 45.987 | 43.344 | 46.434 | -0.006 | 0.7797 | 0.9008 | 0 | 0 | 0 |
| 'Sart3' | 14.463 | 14.363 | 14.912 | 13.801 | 14.170 | 15.548 | 13.936 | 14.106 | 15.482 | 14.193 | 13.340 | 15.136 | 14.697 | 13.884 | 15.488 | 14.853 | 13.928 | 14.645 | -0.003 | 0.9066 | 0.9635 | 0 | 0 | 0 |
| 'Sash3' | 185.400 | 194.422 | 189.807 | 184.829 | 190.806 | 199.490 | 189.731 | 187.200 | 194.390 | 177.509 | 195.548 | 206.733 | 178.411 | 188.368 | 211.793 | 186.960 | 182.537 | 202.568 | -0.021 | 0.4056 | 0.6548 | 0 | 0 | 0 |
| 'Sass6' | 1.708 | 1.999 | 2.095 | 1.883 | 1.511 | 1.994 | 2.026 | 1.785 | 2.147 | 1.390 | 1.853 | 1.934 | 1.730 | 2.125 | 1.964 | 1.771 | 1.746 | 1.893 | 0.055 | 0.4140 | 0.6620 | 0 | 0 | 0 |
| 'Sat1' | 27.022 | 20.808 | 17.271 | 22.467 | 20.176 | 17.410 | 22.888 | 21.823 | 15.954 | 20.846 | 24.637 | 17.780 | 22.332 | 23.009 | 17.577 | 20.564 | 26.438 | 17.082 | -0.045 | 0.4285 | 0.6742 | 0 | 0 | 0 |
| 'Satb1' | 6.651 | 5.659 | 7.129 | 6.440 | 5.395 | 6.547 | 6.622 | 5.930 | 6.824 | 6.231 | 5.780 | 6.846 | 4.941 | 5.320 | 6.794 | 6.081 | 5.034 | 7.157 | 0.071 | 0.1337 | 0.3522 | 0 | 0 | 0 |
| 'Sav1' | 22.952 | 20.453 | 20.831 | 24.085 | 20.834 | 23.210 | 22.558 | 23.689 | 21.053 | 20.385 | 21.523 | 21.202 | 23.415 | 23.700 | 20.802 | 19.820 | 19.413 | 21.899 | 0.044 | 0.3420 | 0.5981 | 0 | 0 | 0 |
| 'Saysd1' | 5.915 | 5.506 | 5.384 | 4.652 | 5.191 | 6.378 | 4.810 | 5.194 | 6.407 | 5.372 | 5.388 | 6.687 | 4.734 | 5.478 | 6.561 | 6.312 | 5.439 | 6.969 | -0.105 | 0.1223 | 0.3336 | 0 | 0 | 0 |
| 'Sbds' | 15.122 | 15.933 | 14.611 | 15.791 | 15.915 | 15.716 | 17.215 | 15.525 | 15.764 | 14.073 | 16.275 | 17.449 | 14.766 | 15.088 | 18.022 | 15.708 | 16.907 | 15.878 | -0.039 | 0.3889 | 0.6384 | 0 | 0 | 0 |
| 'Sbf1' | 36.155 | 34.625 | 37.624 | 35.350 | 34.009 | 35.731 | 33.543 | 34.662 | 37.947 | 37.275 | 35.421 | 38.592 | 38.254 | 33.486 | 35.553 | 34.730 | 34.994 | 37.594 | -0.039 | 0.1303 | 0.3465 | 0 | 0 | 0 |
| 'Sbf2' | 0.766 | 0.646 | 0.781 | 0.879 | 0.693 | 0.641 | 0.608 | 0.809 | 0.699 | 0.575 | 0.440 | 0.535 | 0.347 | 0.527 | 0.479 | 0.421 | 0.507 | 0.603 | 0.549 | 0.0000 | 0.0000 | 1 | 1 | 0 |
| 'Sbk1' | 8.777 | 9.203 | 9.084 | 8.316 | 8.269 | 10.006 | 8.377 | 8.039 | 9.365 | 8.867 | 8.412 | 9.464 | 9.166 | 8.711 | 9.790 | 8.563 | 7.790 | 9.739 | -0.030 | 0.4357 | 0.6798 | 0 | 0 | 0 |
| 'Sbno1' | 7.650 | 8.063 | 7.943 | 8.140 | 7.849 | 7.554 | 7.751 | 8.332 | 8.091 | 7.522 | 9.111 | 8.575 | 7.319 | 8.351 | 7.952 | 7.575 | 8.459 | 8.463 | -0.048 | 0.1522 | 0.3794 | 0 | 0 | 0 |
| 'Sbno2' | 39.959 | 34.663 | 43.849 | 36.114 | 38.187 | 38.026 | 36.766 | 41.885 | 39.164 | 39.385 | 40.113 | 38.181 | 38.797 | 36.243 | 34.761 | 33.320 | 38.570 | 36.510 | 0.044 | 0.3344 | 0.5927 | 0 | 0 | 0 |
| 'Sc5d' | 9.384 | 10.645 | 9.641 | 9.964 | 9.439 | 10.100 | 9.722 | 9.326 | 9.924 | 6.823 | 7.630 | 7.462 | 6.331 | 7.640 | 7.271 | 6.412 | 8.147 | 7.058 | 0.434 | 0.0000 | 0.0000 | 1 | 1 | 0 |
| 'Scaf1' | 16.931 | 15.401 | 19.132 | 16.728 | 15.339 | 18.122 | 16.362 | 15.031 | 19.492 | 17.955 | 16.210 | 19.572 | 17.625 | 15.321 | 18.636 | 16.993 | 15.874 | 19.117 | -0.056 | 0.0145 | 0.0817 | 0 | 0 | 0 |
| 'Scaf11' | 26.101 | 23.682 | 23.816 | 25.870 | 23.898 | 23.339 | 26.140 | 23.435 | 23.262 | 24.098 | 24.954 | 23.780 | 24.210 | 24.689 | 23.157 | 23.050 | 24.791 | 24.281 | 0.005 | 0.8769 | 0.9479 | 0 | 0 | 0 |
| 'Scaf4' | 20.426 | 18.682 | 21.468 | 20.786 | 20.009 | 20.116 | 21.206 | 18.240 | 21.334 | 18.908 | 20.082 | 23.503 | 20.513 | 19.706 | 21.574 | 19.284 | 20.270 | 22.137 | -0.041 | 0.2320 | 0.4798 | 0 | 0 | 0 |
| 'Scai' | 3.851 | 3.969 | 4.186 | 3.849 | 4.032 | 3.544 | 3.909 | 3.834 | 3.925 | 4.036 | 3.910 | 4.585 | 3.974 | 4.049 | 3.860 | 3.955 | 4.054 | 4.261 | -0.078 | 0.0395 | 0.1621 | 0 | 0 | 0 |
| 'Scamp2' | 31.310 | 36.764 | 36.839 | 31.406 | 34.353 | 37.426 | 31.687 | 36.642 | 38.126 | 37.312 | 35.313 | 39.133 | 34.174 | 34.091 | 37.543 | 37.405 | 33.316 | 37.134 | -0.061 | 0.0845 | 0.2638 | 0 | 0 | 0 |
| 'Scamp3' | 50.135 | 47.592 | 50.179 | 47.225 | 49.974 | 50.318 | 47.555 | 51.744 | 49.417 | 48.973 | 47.633 | 54.784 | 49.563 | 49.403 | 53.529 | 48.858 | 44.649 | 54.831 | -0.035 | 0.2437 | 0.4938 | 0 | 0 | 0 |
| 'Scand1' | 136.069 | 130.758 | 142.801 | 123.532 | 133.933 | 154.746 | 121.054 | 135.816 | 147.798 | 134.337 | 121.364 | 123.391 | 137.974 | 129.158 | 136.912 | 143.733 | 119.194 | 125.134 | 0.058 | 0.1935 | 0.4310 | 0 | 0 | 0 |
| 'Scap' | 16.370 | 14.181 | 16.578 | 15.362 | 14.919 | 16.287 | 15.250 | 15.518 | 16.640 | 16.211 | 14.239 | 17.354 | 16.100 | 14.616 | 16.121 | 16.449 | 14.140 | 16.534 | -0.015 | 0.5396 | 0.7573 | 0 | 0 | 0 |
| 'Scaper' | 7.589 | 6.874 | 7.260 | 7.875 | 7.148 | 6.071 | 7.883 | 7.579 | 6.360 | 8.149 | 7.845 | 8.194 | 7.319 | 7.520 | 7.611 | 6.657 | 8.884 | 7.494 | -0.123 | 0.0282 | 0.1300 | 0 | 0 | 0 |
| 'Scarb1' | 1.625 | 1.935 | 1.760 | 1.681 | 1.701 | 1.635 | 1.744 | 1.656 | 1.819 | 1.676 | 1.367 | 1.780 | 1.613 | 1.582 | 1.576 | 1.330 | 1.658 | 1.700 | 0.103 | 0.0597 | 0.2110 | 0 | 0 | 0 |
| 'Scarb2' | 15.759 | 19.188 | 15.153 | 16.633 | 17.493 | 16.100 | 15.645 | 17.192 | 15.689 | 14.990 | 14.986 | 14.358 | 16.898 | 16.040 | 14.280 | 16.233 | 15.421 | 14.384 | 0.101 | 0.0034 | 0.0275 | 1 | 0 | 0 |
| 'Sccpdh' | 6.063 | 7.537 | 5.314 | 5.856 | 7.460 | 5.897 | 6.536 | 6.486 | 5.956 | 6.046 | 7.466 | 7.380 | 6.997 | 7.866 | 7.026 | 7.221 | 8.611 | 6.599 | -0.208 | 0.0000 | 0.0002 | 1 | 0 | 0 |
| 'Scd1' | 0.983 | 1.670 | 0.937 | 0.989 | 1.527 | 0.881 | 0.824 | 1.861 | 0.870 | 0.905 | 1.871 | 0.824 | 0.756 | 1.823 | 0.784 | 0.887 | 2.031 | 0.745 | 0.019 | 0.8401 | 0.9319 | 0 | 0 | 0 |
| 'Scd2' | 83.130 | 95.918 | 87.997 | 81.910 | 88.782 | 84.777 | 90.573 | 85.221 | 89.700 | 76.672 | 95.148 | 84.436 | 76.945 | 88.732 | 86.514 | 78.133 | 96.618 | 83.010 | 0.033 | 0.3433 | 0.5993 | 0 | 0 | 0 |
| 'Scfd1' | 25.486 | 28.778 | 25.295 | 28.157 | 28.099 | 25.923 | 28.036 | 28.363 | 25.412 | 27.376 | 26.394 | 24.035 | 28.473 | 26.147 | 23.543 | 26.086 | 29.532 | 20.362 | 0.063 | 0.1091 | 0.3110 | 0 | 0 | 0 |
| 'Scfd2' | 1.513 | 1.298 | 1.179 | 1.327 | 1.192 | 1.275 | 1.262 | 1.254 | 1.083 | 1.276 | 1.397 | 1.454 | 1.200 | 1.169 | 1.296 | 1.468 | 1.306 | 1.566 | -0.105 | 0.1306 | 0.3470 | 0 | 0 | 0 |
| 'Scg5' | 18.353 | 17.512 | 17.765 | 17.367 | 17.885 | 18.519 | 16.765 | 18.890 | 18.728 | 19.557 | 16.830 | 19.459 | 13.635 | 15.709 | 17.415 | 17.908 | 16.409 | 17.172 | 0.061 | 0.2787 | 0.5352 | 0 | 0 | 0 |
| 'Sclt1' | 1.105 | 1.227 | 0.944 | 1.283 | 1.141 | 1.206 | 1.323 | 1.184 | 1.127 | 1.232 | 1.097 | 1.108 | 1.180 | 1.313 | 1.164 | 1.139 | 1.233 | 1.107 | -0.020 | 0.7451 | 0.8850 | 0 | 0 | 0 |
| 'Scly' | 9.494 | 10.212 | 10.387 | 9.332 | 9.997 | 10.583 | 9.661 | 10.770 | 10.595 | 8.766 | 8.260 | 8.199 | 9.110 | 8.461 | 7.427 | 8.518 | 7.852 | 7.501 | 0.290 | 0.0000 | 0.0000 | 1 | 0 | 0 |
| 'Scmh1' | 2.791 | 3.101 | 2.988 | 2.628 | 2.724 | 2.979 | 2.934 | 2.826 | 2.939 | 2.778 | 2.540 | 2.978 | 2.933 | 2.612 | 2.782 | 3.005 | 2.548 | 2.739 | 0.044 | 0.2711 | 0.5262 | 0 | 0 | 0 |
| 'Scml4' | 0.202 | 0.392 | 0.380 | 0.363 | 0.415 | 0.515 | 0.489 | 0.482 | 0.475 | 0.173 | 0.264 | 0.451 | 0.402 | 0.314 | 0.446 | 0.443 | 0.449 | 0.512 | 0.093 | 0.5569 | 0.7687 | 0 | 0 | 0 |
| 'Scn11a' | 2.613 | 2.836 | 2.634 | 2.950 | 2.483 | 2.290 | 3.006 | 2.522 | 2.852 | 3.184 | 2.630 | 3.488 | 2.384 | 2.304 | 3.405 | 2.625 | 2.567 | 3.168 | -0.104 | 0.1829 | 0.4190 | 0 | 0 | 0 |
| 'Scn1b' | 10.801 | 12.333 | 10.911 | 9.090 | 11.135 | 11.566 | 10.629 | 10.114 | 13.193 | 9.742 | 8.268 | 9.448 | 9.624 | 7.572 | 9.485 | 10.626 | 6.537 | 8.868 | 0.309 | 0.0000 | 0.0007 | 1 | 0 | 0 |
| 'Scn8a' | 0.320 | 0.339 | 0.407 | 0.236 | 0.328 | 0.289 | 0.304 | 0.326 | 0.336 | 0.329 | 0.290 | 0.327 | 0.256 | 0.248 | 0.326 | 0.205 | 0.292 | 0.349 | 0.127 | 0.1526 | 0.3797 | 0 | 0 | 0 |
| 'Scnm1' | 5.191 | 5.393 | 5.552 | 5.352 | 5.788 | 5.602 | 5.573 | 6.543 | 6.414 | 5.210 | 5.099 | 5.694 | 5.645 | 5.360 | 5.244 | 5.233 | 5.639 | 5.497 | 0.068 | 0.1625 | 0.3945 | 0 | 0 | 0 |
| 'Sco1' | 3.060 | 2.460 | 2.320 | 2.370 | 2.631 | 2.355 | 2.585 | 2.747 | 2.304 | 2.847 | 2.678 | 2.420 | 2.850 | 2.695 | 2.163 | 2.831 | 2.517 | 2.223 | -0.028 | 0.6038 | 0.8000 | 0 | 0 | 0 |
| 'Scp2' | 28.142 | 33.500 | 28.390 | 30.338 | 33.778 | 29.940 | 29.932 | 31.432 | 28.852 | 30.022 | 30.447 | 29.249 | 30.390 | 32.310 | 30.234 | 29.559 | 30.322 | 28.431 | 0.004 | 0.8720 | 0.9452 | 0 | 0 | 0 |
| 'Scpep1' | 16.785 | 15.938 | 16.401 | 17.092 | 16.729 | 15.712 | 17.431 | 17.637 | 15.336 | 17.182 | 17.008 | 16.892 | 17.436 | 16.190 | 17.312 | 14.358 | 14.624 | 15.919 | 0.006 | 0.8874 | 0.9536 | 0 | 0 | 0 |
| 'Scrib' | 32.146 | 28.722 | 35.942 | 29.520 | 29.769 | 35.330 | 28.876 | 30.089 | 35.615 | 32.068 | 33.139 | 36.211 | 33.690 | 32.004 | 34.891 | 33.912 | 31.434 | 36.392 | -0.098 | 0.0006 | 0.0066 | 1 | 0 | 0 |
| 'Scrn2' | 3.041 | 3.133 | 3.334 | 2.638 | 3.158 | 3.436 | 2.618 | 2.724 | 3.490 | 3.062 | 2.178 | 2.877 | 2.901 | 2.086 | 2.612 | 2.783 | 2.129 | 2.526 | 0.244 | 0.0009 | 0.0095 | 1 | 0 | 0 |
| 'Scrn3' | 3.433 | 4.350 | 3.724 | 3.599 | 4.318 | 3.828 | 4.014 | 4.146 | 3.596 | 3.772 | 3.927 | 3.677 | 3.429 | 3.870 | 3.913 | 3.906 | 4.272 | 3.355 | 0.025 | 0.5840 | 0.7860 | 0 | 0 | 0 |
| 'Scyl1' | 29.665 | 29.335 | 34.344 | 30.756 | 28.674 | 36.458 | 29.205 | 30.934 | 35.061 | 33.941 | 27.677 | 35.745 | 32.573 | 28.843 | 33.394 | 32.451 | 29.603 | 33.283 | -0.025 | 0.4282 | 0.6742 | 0 | 0 | 0 |
| 'Scyl2' | 4.101 | 4.093 | 4.120 | 3.864 | 4.413 | 4.062 | 4.070 | 4.122 | 3.893 | 4.188 | 4.234 | 4.314 | 3.998 | 4.427 | 3.653 | 3.844 | 4.390 | 4.158 | -0.030 | 0.4016 | 0.6504 | 0 | 0 | 0 |
| 'Scyl3' | 5.390 | 5.341 | 5.868 | 5.746 | 5.917 | 5.520 | 5.696 | 5.717 | 5.648 | 5.356 | 5.836 | 5.554 | 5.086 | 5.684 | 5.377 | 4.838 | 5.240 | 5.072 | 0.072 | 0.0352 | 0.1500 | 0 | 0 | 0 |
| 'Sdad1' | 6.700 | 6.362 | 7.360 | 6.172 | 6.358 | 6.649 | 6.477 | 7.269 | 6.868 | 6.767 | 6.027 | 7.035 | 6.208 | 6.000 | 6.338 | 6.314 | 6.547 | 6.655 | 0.047 | 0.2132 | 0.4581 | 0 | 0 | 0 |
| 'Sdc3' | 0.571 | 0.742 | 0.648 | 0.687 | 0.753 | 0.851 | 0.626 | 0.699 | 0.668 | 0.550 | 0.541 | 0.841 | 0.658 | 0.733 | 0.678 | 0.735 | 0.506 | 0.779 | 0.029 | 0.7784 | 0.9002 | 0 | 0 | 0 |
| 'Sdcbp' | 17.406 | 19.747 | 18.422 | 19.219 | 19.205 | 18.219 | 19.731 | 19.738 | 18.274 | 17.827 | 19.974 | 18.630 | 17.375 | 19.460 | 17.990 | 17.221 | 20.231 | 17.430 | 0.022 | 0.4182 | 0.6655 | 0 | 0 | 0 |
| 'Sdcbp2' | 11.274 | 10.823 | 9.424 | 9.995 | 11.384 | 10.005 | 11.010 | 11.946 | 11.269 | 13.720 | 14.914 | 13.310 | 10.610 | 12.607 | 12.688 | 15.059 | 12.605 | 12.211 | -0.286 | 0.0000 | 0.0003 | 1 | 0 | 0 |
| 'Sdccag3' | 9.903 | 10.051 | 10.915 | 9.908 | 10.476 | 11.075 | 9.427 | 9.564 | 11.302 | 8.909 | 10.391 | 12.592 | 9.947 | 10.641 | 12.407 | 9.206 | 10.675 | 12.236 | -0.074 | 0.0366 | 0.1546 | 0 | 0 | 0 |
| 'Sdccag8' | 3.306 | 3.312 | 3.656 | 3.901 | 3.346 | 3.444 | 3.966 | 3.350 | 3.347 | 3.355 | 3.378 | 3.764 | 3.738 | 3.692 | 3.617 | 3.467 | 3.897 | 3.420 | -0.047 | 0.3412 | 0.5981 | 0 | 0 | 0 |

| | | | | | | | | | | | | | | | | | | | | | | | | |
|---|---|---|---|---|---|---|---|---|---|---|---|---|---|---|---|---|---|---|---|---|---|---|---|---|
| 'Sde2' | 13.274 | 13.104 | 11.827 | 11.919 | 12.780 | 11.654 | 11.299 | 12.003 | 12.298 | 12.551 | 12.275 | 12.325 | 11.753 | 13.553 | 12.677 | 12.981 | 11.995 | 12.527 | -0.042 | 0.2926 | 0.5493 | 0 | 0 | 0 |
| 'Sdf2' | 20.413 | 20.825 | 20.119 | 20.867 | 21.690 | 22.015 | 20.667 | 21.007 | 22.645 | 20.385 | 22.546 | 23.815 | 20.082 | 21.697 | 24.115 | 19.846 | 22.878 | 23.285 | -0.074 | 0.0348 | 0.1492 | 0 | 0 | 0 |
| 'Sdf2l1' | 40.560 | 42.838 | 49.851 | 45.632 | 48.107 | 52.508 | 41.843 | 48.465 | 48.638 | 40.967 | 37.874 | 43.743 | 48.021 | 41.385 | 46.608 | 44.874 | 40.007 | 43.236 | 0.104 | 0.0260 | 0.1239 | 0 | 0 | 0 |
| 'Sdf4' | 38.594 | 42.623 | 38.865 | 38.506 | 44.143 | 42.424 | 38.216 | 42.436 | 40.566 | 40.087 | 39.467 | 40.472 | 39.243 | 41.444 | 40.934 | 42.828 | 39.960 | 39.949 | -0.004 | 0.8806 | 0.9497 | 0 | 0 | 0 |
| 'Sdha' | 52.169 | 56.064 | 51.923 | 53.554 | 54.298 | 52.772 | 50.975 | 54.244 | 52.658 | 55.386 | 55.178 | 51.171 | 54.617 | 54.223 | 52.714 | 52.834 | 53.806 | 50.557 | -0.015 | 0.4491 | 0.6890 | 0 | 0 | 0 |
| 'Sdhaf1' | 5.768 | 7.425 | 5.981 | 6.254 | 7.165 | 7.168 | 6.018 | 7.665 | 6.147 | 7.974 | 6.373 | 5.009 | 7.187 | 7.192 | 5.066 | 7.313 | 5.979 | 5.070 | 0.067 | 0.3992 | 0.6482 | 0 | 0 | 0 |
| 'Sdhaf2' | 20.790 | 21.438 | 21.443 | 22.416 | 22.268 | 21.750 | 22.598 | 21.594 | 21.119 | 20.643 | 19.937 | 22.419 | 21.398 | 22.478 | 22.005 | 20.241 | 21.409 | 21.380 | 0.011 | 0.7192 | 0.8709 | 0 | 0 | 0 |
| 'Sdhaf3' | 0.691 | 0.890 | 0.719 | 0.640 | 0.868 | 0.653 | 0.637 | 0.846 | 0.807 | 0.649 | 0.476 | 0.615 | 0.743 | 0.709 | 0.684 | 0.666 | 0.817 | 0.593 | 0.165 | 0.1395 | 0.3611 | 0 | 0 | 0 |
| 'Sdhaf4' | 8.620 | 11.521 | 9.952 | 8.486 | 10.987 | 10.327 | 9.941 | 10.005 | 10.638 | 9.283 | 9.079 | 10.069 | 10.029 | 8.408 | 10.247 | 10.629 | 9.859 | 7.154 | 0.079 | 0.3085 | 0.5669 | 0 | 0 | 0 |
| 'Sdhb' | 33.605 | 37.798 | 34.532 | 34.628 | 37.272 | 37.276 | 33.745 | 37.336 | 36.712 | 33.561 | 31.670 | 34.267 | 37.448 | 35.477 | 35.688 | 37.086 | 34.295 | 33.166 | 0.036 | 0.3010 | 0.5579 | 0 | 0 | 0 |
| 'Sdhc' | 18.769 | 20.275 | 18.911 | 17.015 | 19.957 | 18.875 | 18.789 | 19.458 | 18.713 | 20.283 | 19.422 | 18.628 | 19.832 | 19.928 | 20.000 | 20.392 | 20.085 | 19.299 | -0.067 | 0.0300 | 0.1349 | 0 | 0 | 0 |
| 'Sdhd' | 46.498 | 52.730 | 50.975 | 50.265 | 55.184 | 52.277 | 49.479 | 58.355 | 50.365 | 45.315 | 50.591 | 48.970 | 52.197 | 51.732 | 53.337 | 50.597 | 48.577 | 47.371 | 0.044 | 0.2359 | 0.4858 | 0 | 0 | 0 |
| 'Sdr39u1' | 50.373 | 56.761 | 53.767 | 48.984 | 57.989 | 57.979 | 55.004 | 55.102 | 55.484 | 53.228 | 51.888 | 59.166 | 52.568 | 54.186 | 59.506 | 55.364 | 56.622 | 55.311 | -0.032 | 0.3109 | 0.5692 | 0 | 0 | 0 |
| 'Sdr42e1' | 0.664 | 0.586 | 0.917 | 0.916 | 0.750 | 0.956 | 0.738 | 0.752 | 0.970 | 0.749 | 1.085 | 0.871 | 0.915 | 0.830 | 0.820 | 0.839 | 0.764 | 0.746 | -0.073 | 0.4971 | 0.7258 | 0 | 0 | 0 |
| 'Sec11a' | 13.173 | 12.962 | 12.665 | 14.624 | 13.210 | 13.070 | 13.583 | 13.825 | 12.855 | 12.891 | 13.851 | 13.831 | 12.722 | 13.036 | 14.083 | 12.525 | 14.107 | 13.472 | -0.019 | 0.6000 | 0.7967 | 0 | 0 | 0 |
| 'Sec11c' | 51.567 | 67.249 | 59.769 | 51.804 | 64.150 | 60.283 | 53.940 | 60.857 | 60.567 | 53.391 | 54.940 | 52.315 | 54.519 | 57.854 | 55.360 | 56.734 | 54.673 | 50.933 | 0.097 | 0.0063 | 0.0444 | 1 | 0 | 0 |
| 'Sec13' | 20.929 | 24.303 | 23.107 | 23.119 | 24.528 | 22.997 | 22.950 | 23.021 | 23.025 | 20.654 | 20.826 | 22.642 | 22.965 | 21.460 | 22.948 | 22.727 | 21.769 | 22.361 | 0.054 | 0.0782 | 0.2506 | 0 | 0 | 0 |
| 'Sec14l1' | 21.445 | 20.042 | 19.797 | 21.116 | 20.583 | 21.032 | 19.456 | 20.414 | 21.788 | 19.653 | 20.267 | 23.104 | 19.946 | 20.321 | 21.947 | 20.051 | 19.513 | 24.066 | -0.033 | 0.3527 | 0.6063 | 0 | 0 | 0 |
| 'Sec14l2' | 0.849 | 1.094 | 1.112 | 1.014 | 1.050 | 1.031 | 0.976 | 0.965 | 0.919 | 0.904 | 1.138 | 0.934 | 1.193 | 0.987 | 0.930 | 1.068 | 1.008 | 1.034 | -0.035 | 0.6948 | 0.8564 | 0 | 0 | 0 |
| 'Sec16a' | 15.163 | 14.889 | 16.771 | 15.861 | 14.661 | 15.598 | 15.162 | 15.328 | 16.600 | 17.033 | 16.866 | 16.708 | 15.601 | 15.223 | 15.362 | 14.714 | 15.380 | 16.437 | -0.045 | 0.1816 | 0.4173 | 0 | 0 | 0 |
| 'Sec16b' | 0.288 | 0.311 | 0.236 | 0.313 | 0.234 | 0.327 | 0.304 | 0.275 | 0.240 | 0.716 | 0.781 | 0.601 | 0.611 | 0.625 | 0.562 | 0.576 | 0.761 | 0.644 | -1.225 | 0.0000 | 0.0000 | 1 | 1 | 1 |
| 'Sec22a' | 2.517 | 2.575 | 2.678 | 2.642 | 2.696 | 2.624 | 3.068 | 2.779 | 2.630 | 2.676 | 2.802 | 2.547 | 2.662 | 2.705 | 2.221 | 2.750 | 2.619 | 2.372 | 0.045 | 0.3955 | 0.6444 | 0 | 0 | 0 |
| 'Sec22b' | 9.392 | 11.864 | 10.209 | 10.350 | 12.111 | 11.214 | 9.691 | 11.856 | 11.269 | 10.321 | 11.268 | 11.544 | 11.031 | 11.674 | 11.057 | 11.215 | 10.779 | 11.163 | -0.046 | 0.1945 | 0.4326 | 0 | 0 | 0 |
| 'Sec22c' | 4.536 | 5.909 | 4.803 | 4.594 | 5.896 | 5.591 | 4.934 | 4.958 | 5.299 | 5.507 | 5.635 | 5.176 | 5.410 | 5.680 | 5.199 | 6.017 | 5.467 | 5.022 | -0.090 | 0.0666 | 0.2256 | 0 | 0 | 0 |
| 'Sec23a' | 4.763 | 5.292 | 5.062 | 5.863 | 5.538 | 4.893 | 5.763 | 5.300 | 5.335 | 4.543 | 5.506 | 5.460 | 5.211 | 5.781 | 5.164 | 5.395 | 5.725 | 5.230 | -0.021 | 0.6700 | 0.8409 | 0 | 0 | 0 |
| 'Sec23b' | 8.852 | 9.544 | 9.581 | 9.269 | 10.217 | 9.948 | 9.416 | 9.377 | 9.742 | 9.557 | 9.183 | 9.951 | 10.100 | 10.406 | 9.639 | 9.915 | 9.342 | 9.872 | -0.045 | 0.1277 | 0.3423 | 0 | 0 | 0 |
| 'Sec23ip' | 13.850 | 14.470 | 13.720 | 14.808 | 13.958 | 13.558 | 14.674 | 13.738 | 13.993 | 14.563 | 14.834 | 14.344 | 14.508 | 14.628 | 13.797 | 13.902 | 15.271 | 14.039 | -0.047 | 0.0652 | 0.2228 | 0 | 0 | 0 |
| 'Sec24a' | 2.754 | 2.575 | 2.818 | 2.882 | 2.582 | 2.259 | 2.872 | 2.519 | 2.488 | 2.742 | 2.966 | 2.931 | 2.842 | 2.859 | 2.710 | 2.678 | 2.887 | 2.846 | -0.114 | 0.0129 | 0.0754 | 0 | 0 | 0 |
| 'Sec24b' | 15.307 | 13.527 | 14.925 | 14.912 | 14.085 | 14.179 | 14.402 | 14.368 | 14.782 | 15.030 | 14.656 | 15.940 | 14.251 | 14.497 | 15.176 | 13.394 | 15.073 | 16.426 | -0.054 | 0.1258 | 0.3398 | 0 | 0 | 0 |
| 'Sec24c' | 55.482 | 57.325 | 59.973 | 55.226 | 56.127 | 56.697 | 55.066 | 57.484 | 61.996 | 59.104 | 57.976 | 62.044 | 57.684 | 57.608 | 60.264 | 55.488 | 57.144 | 62.004 | -0.049 | 0.0173 | 0.0931 | 0 | 0 | 0 |
| 'Sec24d' | 4.364 | 4.033 | 4.354 | 4.702 | 4.387 | 4.616 | 4.286 | 4.814 | 4.539 | 4.838 | 4.215 | 4.279 | 4.350 | 4.676 | 4.196 | 4.488 | 4.155 | 4.312 | 0.015 | 0.7015 | 0.8608 | 0 | 0 | 0 |
| 'Sec31a' | 13.403 | 12.949 | 13.831 | 13.723 | 13.457 | 13.801 | 13.320 | 13.646 | 13.504 | 15.273 | 13.592 | 13.619 | 13.795 | 13.499 | 12.773 | 13.486 | 13.421 | 13.285 | -0.022 | 0.4258 | 0.6724 | 0 | 0 | 0 |
| 'Sec61a1' | 22.518 | 21.877 | 23.186 | 21.810 | 22.928 | 23.161 | 20.559 | 22.721 | 22.734 | 23.897 | 22.500 | 23.759 | 23.061 | 23.597 | 24.535 | 23.315 | 22.486 | 24.016 | -0.077 | 0.0010 | 0.0101 | 1 | 0 | 0 |
| 'Sec61a2' | 1.166 | 1.431 | 1.846 | 1.145 | 1.753 | 1.549 | 1.452 | 1.860 | 1.587 | 1.638 | 1.409 | 1.455 | 1.538 | 1.521 | 1.250 | 1.774 | 1.671 | 1.252 | 0.022 | 0.8094 | 0.9141 | 0 | 0 | 0 |
| 'Sec61g' | 8.752 | 9.284 | 8.174 | 9.212 | 8.129 | 8.353 | 9.627 | 8.715 | 8.048 | 7.478 | 8.383 | 7.967 | 8.100 | 9.165 | 8.854 | 8.332 | 9.646 | 7.456 | 0.044 | 0.3716 | 0.6236 | 0 | 0 | 0 |
| 'Sec62' | 34.698 | 34.715 | 32.321 | 37.468 | 33.300 | 31.211 | 38.553 | 32.407 | 31.247 | 32.812 | 35.447 | 34.264 | 30.879 | 33.570 | 31.692 | 33.068 | 34.687 | 31.786 | 0.023 | 0.5608 | 0.7710 | 0 | 0 | 0 |
| 'Sec63' | 5.901 | 5.623 | 6.241 | 5.846 | 6.017 | 5.478 | 5.912 | 5.564 | 5.547 | 5.679 | 5.740 | 5.859 | 5.631 | 6.000 | 5.298 | 5.355 | 5.937 | 5.572 | 0.019 | 0.5802 | 0.7824 | 0 | 0 | 0 |
| 'Secisbp2' | 10.843 | 10.665 | 11.168 | 9.440 | 11.236 | 11.112 | 10.306 | 10.845 | 10.927 | 10.888 | 10.213 | 11.384 | 11.301 | 10.223 | 11.239 | 11.334 | 10.543 | 11.181 | -0.036 | 0.2624 | 0.5162 | 0 | 0 | 0 |
| 'Secisbp2l' | 16.085 | 18.973 | 16.229 | 16.869 | 18.263 | 16.939 | 16.544 | 18.540 | 17.568 | 16.708 | 16.541 | 17.977 | 17.040 | 18.529 | 16.906 | 18.021 | 17.469 | 17.402 | -0.019 | 0.5050 | 0.7314 | 0 | 0 | 0 |
| 'Sectm1a' | 2.043 | 2.944 | 1.955 | 2.377 | 2.832 | 2.593 | 3.700 | 2.040 | 2.500 | 3.735 | 2.415 | 3.013 | 3.098 | 3.189 | 2.455 | 3.774 | 4.184 | 2.182 | -0.295 | 0.0143 | 0.0808 | 0 | 0 | 0 |
| 'Seh1l' | 9.499 | 10.432 | 9.381 | 9.687 | 9.810 | 8.925 | 10.802 | 8.940 | 9.495 | 8.222 | 9.664 | 10.764 | 7.922 | 10.538 | 10.418 | 8.996 | 11.418 | 10.402 | -0.031 | 0.6177 | 0.8087 | 0 | 0 | 0 |
| 'Sel1l' | 5.315 | 5.898 | 5.954 | 5.186 | 5.822 | 5.627 | 5.110 | 5.988 | 6.030 | 5.761 | 6.073 | 5.948 | 5.471 | 5.726 | 5.370 | 6.406 | 5.717 | 6.094 | -0.057 | 0.1359 | 0.3545 | 0 | 0 | 0 |
| 'Selk' | 61.641 | 60.983 | 49.671 | 60.925 | 63.236 | 50.991 | 56.409 | 67.166 | 46.941 | 67.346 | 58.545 | 45.928 | 61.311 | 67.435 | 45.546 | 62.750 | 58.437 | 44.160 | 0.017 | 0.6642 | 0.8377 | 0 | 0 | 0 |
| 'Sell' | 1.863 | 2.655 | 2.552 | 3.067 | 2.923 | 3.517 | 3.845 | 3.886 | 2.918 | 1.180 | 1.975 | 3.134 | 2.615 | 2.400 | 3.515 | 3.234 | 2.822 | 3.426 | 0.171 | 0.3601 | 0.6124 | 0 | 0 | 0 |
| 'Selplg' | 14.714 | 10.826 | 16.422 | 13.156 | 12.950 | 17.056 | 14.455 | 13.062 | 15.373 | 13.583 | 11.392 | 17.261 | 12.827 | 10.944 | 17.577 | 14.735 | 11.241 | 16.124 | 0.026 | 0.5572 | 0.7689 | 0 | 0 | 0 |
| 'Sema4a' | 1.409 | 1.355 | 1.737 | 1.072 | 1.275 | 1.608 | 1.370 | 1.549 | 1.602 | 0.927 | 1.075 | 1.102 | 0.893 | 0.982 | 1.217 | 1.008 | 0.945 | 1.217 | 0.472 | 0.0000 | 0.0000 | 1 | 1 | 0 |
| 'Sema4b' | 7.351 | 6.281 | 7.126 | 6.269 | 6.574 | 6.636 | 6.381 | 6.509 | 6.603 | 6.903 | 6.215 | 7.167 | 6.385 | 6.482 | 7.109 | 6.819 | 6.218 | 7.081 | -0.024 | 0.5080 | 0.7336 | 0 | 0 | 0 |
| 'Sema4d' | 29.357 | 29.738 | 28.679 | 28.980 | 28.231 | 29.163 | 29.221 | 30.193 | 29.778 | 29.844 | 29.691 | 30.157 | 29.747 | 29.098 | 30.331 | 29.737 | 27.115 | 31.020 | -0.029 | 0.2266 | 0.4732 | 0 | 0 | 0 |
| 'Sema4f' | 20.420 | 17.181 | 25.118 | 17.392 | 18.000 | 21.680 | 16.313 | 20.910 | 23.550 | 20.109 | 17.765 | 19.334 | 13.721 | 15.499 | 19.098 | 15.879 | 13.665 | 18.723 | 0.217 | 0.0014 | 0.0137 | 1 | 0 | 0 |
| 'Sema7a' | 7.020 | 7.379 | 8.853 | 6.208 | 6.530 | 8.231 | 6.649 | 7.573 | 8.823 | 7.070 | 7.381 | 8.887 | 5.729 | 6.377 | 10.073 | 6.415 | 6.386 | 8.559 | 0.006 | 0.9209 | 0.9703 | 0 | 0 | 0 |
| 'Senp1' | 6.716 | 6.721 | 6.830 | 6.991 | 6.553 | 6.653 | 6.495 | 6.198 | 7.026 | 6.671 | 7.023 | 6.768 | 7.214 | 6.764 | 7.074 | 6.210 | 7.035 | 6.495 | -0.036 | 0.2861 | 0.5428 | 0 | 0 | 0 |
| 'Senp2' | 23.556 | 23.685 | 25.782 | 24.035 | 25.568 | 24.758 | 23.441 | 25.812 | 25.585 | 23.402 | 25.327 | 26.229 | 25.094 | 25.782 | 25.789 | 24.406 | 26.825 | 24.860 | -0.045 | 0.0709 | 0.2347 | 0 | 0 | 0 |
| 'Senp3' | 19.151 | 20.073 | 19.697 | 17.773 | 18.746 | 19.716 | 17.657 | 19.152 | 19.852 | 19.594 | 18.018 | 19.893 | 19.615 | 19.168 | 18.819 | 19.780 | 18.202 | 18.444 | -0.007 | 0.7962 | 0.9092 | 0 | 0 | 0 |
| 'Senp5' | 4.955 | 4.918 | 4.912 | 4.332 | 4.897 | 4.407 | 4.537 | 4.901 | 4.295 | 4.894 | 4.981 | 5.102 | 4.492 | 4.999 | 4.748 | 4.945 | 4.960 | 4.919 | -0.074 | 0.0637 | 0.2202 | 0 | 0 | 0 |
| 'Senp6' | 13.359 | 13.227 | 12.052 | 13.494 | 12.542 | 12.036 | 13.666 | 13.445 | 11.958 | 13.180 | 12.325 | 12.304 | 12.034 | 13.127 | 11.774 | 12.272 | 13.159 | 11.745 | 0.037 | 0.1685 | 0.4008 | 0 | 0 | 0 |
| 'Senp7' | 4.741 | 5.165 | 4.482 | 5.240 | 4.751 | 4.431 | 4.962 | 4.553 | 4.359 | 4.512 | 4.748 | 4.981 | 4.326 | 4.522 | 4.419 | 4.044 | 4.878 | 4.749 | 0.038 | 0.4129 | 0.6611 | 0 | 0 | 0 |
| 'Senp8' | 1.575 | 1.717 | 1.883 | 2.057 | 2.157 | 2.051 | 2.041 | 1.521 | 2.054 | 1.403 | 1.640 | 2.249 | 1.468 | 1.822 | 1.606 | 1.758 | 2.426 | 2.039 | 0.039 | 0.7081 | 0.8650 | 0 | 0 | 0 |
| 'Sephs1' | 3.537 | 3.165 | 4.046 | 3.910 | 3.702 | 3.959 | 3.883 | 3.680 | 3.875 | 3.665 | 3.749 | 3.912 | 3.859 | 3.930 | 3.975 | 3.711 | 4.365 | 3.719 | -0.055 | 0.2420 | 0.4927 | 0 | 0 | 0 |
| 'Sepsecs' | 1.305 | 1.518 | 1.583 | 1.282 | 1.467 | 1.319 | 1.270 | 1.318 | 1.450 | 1.455 | 1.432 | 1.307 | 1.249 | 1.194 | 1.274 | 1.383 | 1.632 | 1.142 | 0.046 | 0.4762 | 0.7107 | 0 | 0 | 0 |
| 'Sept1' | 162.730 | 174.324 | 166.199 | 167.654 | 165.742 | 169.762 | 175.219 | 166.816 | 172.127 | 162.085 | 159.268 | 150.009 | 171.456 | 154.387 | 158.946 | 164.676 | 158.286 | 154.186 | 0.075 | 0.0010 | 0.0109 | 1 | 0 | 0 |
| 'Sept11' | 5.642 | 5.627 | 6.051 | 6.271 | 5.892 | 6.089 | 6.045 | 5.695 | 6.174 | 4.965 | 5.627 | 5.506 | 6.141 | 5.111 | 5.862 | 4.997 | 5.939 | 5.553 | 0.094 | 0.0276 | 0.1282 | 0 | 0 | 0 |
| 'Sept2' | 7.773 | 10.533 | 9.084 | 8.531 | 9.899 | 9.468 | 10.103 | 10.306 | 9.116 | 8.377 | 9.396 | 10.772 | 9.124 | 9.473 | 9.991 | 9.051 | 10.972 | 9.886 | -0.052 | 0.2534 | 0.5050 | 0 | 0 | 0 |
| 'Sept5' | 1.101 | 1.420 | 0.759 | 0.975 | 0.823 | 1.195 | 0.947 | 1.000 | 1.054 | 2.438 | 2.016 | 3.063 | 2.650 | 2.563 | 1.851 | 2.943 | 2.462 | 2.306 | -1.289 | 0.0000 | 0.0000 | 1 | 1 | 1 |
| 'Sept6' | 31.867 | 32.114 | 32.992 | 32.803 | 33.413 | 34.325 | 33.340 | 32.844 | 33.025 | 35.806 | 31.958 | 32.264 | 33.244 | 32.363 | 32.823 | 32.931 | 33.018 | 31.091 | -0.004 | 0.8765 | 0.9477 | 0 | 0 | 0 |
| 'Sept7' | 124.566 | 135.364 | 127.480 | 139.711 | 139.551 | 123.422 | 138.678 | 130.304 | 123.690 | 124.662 | 134.693 | 126.409 | 121.140 | 138.807 | 127.336 | 127.080 | 145.740 | 125.460 | 0.004 | 0.9044 | 0.9624 | 0 | 0 | 0 |
| 'Sept8' | 2.853 | 2.905 | 2.879 | 2.669 | 2.864 | 2.834 | 3.164 | 2.932 | 2.930 | 2.528 | 2.272 | 2.292 | 2.513 | 2.246 | 2.137 | 2.471 | 2.189 | 2.299 | 0.305 | 0.0000 | 0.0000 | 1 | 0 | 0 |
| 'Sept9' | 82.769 | 84.786 | 83.369 | 77.100 | 83.313 | 90.355 | 79.327 | 82.945 | 86.643 | 88.748 | 76.613 | 80.118 | 85.618 | 80.139 | 86.462 | 88.527 | 72.924 | 83.271 | 0.006 | 0.8530 | 0.9370 | 0 | 0 | 0 |
| 'Serac1' | 2.305 | 2.000 | 2.385 | 2.308 | 2.261 | 2.117 | 2.172 | 2.160 | 2.437 | 2.645 | 2.506 | 2.262 | 2.351 | 2.153 | 1.929 | 2.172 | 2.462 | 2.151 | -0.038 | 0.5558 | 0.7678 | 0 | 0 | 0 |
| 'Serbp1' | 23.633 | 24.203 | 22.652 | 24.906 | 24.190 | 22.807 | 24.917 | 23.584 | 23.028 | 22.735 | 23.287 | 21.273 | 22.908 | 23.131 | 22.466 | 22.916 | 23.375 | 22.522 | 0.053 | 0.0132 | 0.0767 | 0 | 0 | 0 |
| 'Serf2' | 27.601 | 31.233 | 28.931 | 29.698 | 30.437 | 29.165 | 27.527 | 28.657 | 28.534 | 28.158 | 26.714 | 25.625 | 27.709 | 28.045 | 28.857 | 30.195 | 29.120 | 26.957 | 0.048 | 0.1359 | 0.3545 | 0 | 0 | 0 |
| 'Sergef' | 1.978 | 1.731 | 1.767 | 1.940 | 2.052 | 2.130 | 2.505 | 1.817 | 1.976 | 2.036 | 1.737 | 2.180 | 1.728 | 1.922 | 1.913 | 2.410 | 1.979 | 1.985 | -0.015 | 0.8463 | 0.9343 | 0 | 0 | 0 |
| 'Serhl' | 3.212 | 3.577 | 4.155 | 3.220 | 3.888 | 4.002 | 3.613 | 3.608 | 4.146 | 3.606 | 3.366 | 3.862 | 3.877 | 3.707 | 3.781 | 3.710 | 3.701 | 3.429 | 0.005 | 0.9090 | 0.9645 | 0 | 0 | 0 |
| 'Serinc1' | 15.638 | 15.813 | 14.911 | 14.996 | 16.455 | 14.721 | 15.525 | 17.593 | 14.674 | 16.467 | 16.634 | 16.062 | 14.668 | 16.758 | 14.249 | 17.267 | 16.653 | 14.342 | -0.038 | 0.2531 | 0.5047 | 0 | 0 | 0 |
| 'Serinc3' | 126.056 | 149.927 | 131.017 | 142.916 | 145.314 | 128.619 | 141.935 | 145.630 | 126.496 | 119.479 | 142.001 | 123.893 | 131.742 | 145.692 | 128.037 | 135.950 | 142.328 | 132.008 | 0.032 | 0.2893 | 0.5459 | 0 | 0 | 0 |
| 'Serinc5' | 58.130 | 65.918 | 56.797 | 59.162 | 61.977 | 55.901 | 58.328 | 62.519 | 58.625 | 60.957 | 70.388 | 66.312 | 62.624 | 68.797 | 63.088 | 63.085 | 69.724 | 66.726 | -0.151 | 0.0000 | 0.0000 | 1 | 0 | 0 |
| 'Serp1' | 52.724 | 66.659 | 52.075 | 54.671 | 61.840 | 54.171 | 56.776 | 60.455 | 52.210 | 56.754 | 56.643 | 49.455 | 55.895 | 59.254 | 47.899 | 56.445 | 56.562 | 47.738 | 0.061 | 0.0271 | 0.1269 | 0 | 0 | 0 |
| 'Serpina9' | 2.351 | 4.204 | 2.760 | 2.954 | 3.354 | 3.015 | 3.361 | 3.156 | 3.261 | 3.051 | 3.290 | 2.677 | 3.624 | 4.287 | 2.805 | 3.819 | 3.446 | 3.069 | -0.090 | 0.2795 | 0.5363 | 0 | 0 | 0 |
| 'Serpinb6a' | 2.062 | 2.615 | 2.164 | 1.951 | 2.383 | 2.159 | 2.285 | 2.623 | 2.260 | 2.424 | 2.339 | 2.168 | 2.237 | 2.535 | 2.349 | 2.276 | 2.300 | 1.975 | -0.017 | 0.7351 | 0.8795 | 0 | 0 | 0 |
| 'Serpinb6b' | 3.075 | 3.216 | 3.089 | 3.784 | 3.057 | 3.108 | 3.141 | 3.022 | 2.938 | 3.494 | 4.321 | 4.154 | 3.577 | 3.982 | 4.151 | 3.460 | 4.342 | 4.489 | -0.353 | 0.0000 | 0.0000 | 1 | 0 | 0 |
| 'Serpinb9' | 2.703 | 2.685 | 2.786 | 2.939 | 2.993 | 3.023 | 2.723 | 2.987 | 3.143 | 2.678 | 3.362 | 3.758 | 3.086 | 3.365 | 3.099 | 2.759 | 3.612 | 3.976 | -0.206 | 0.0008 | 0.0089 | 1 | 0 | 0 |
| 'Serpinf1' | 2.667 | 3.375 | 2.430 | 2.649 | 3.529 | 2.551 | 2.970 | 3.180 | 3.130 | 2.854 | 3.118 | 2.445 | 3.486 | 3.255 | 2.508 | 3.181 | 3.117 | 2.236 | 0.009 | 0.8970 | 0.9586 | 0 | 0 | 0 |
| 'Sertad1' | 35.177 | 22.570 | 17.760 | 28.951 | 20.683 | 20.786 | 28.563 | 23.333 | 16.655 | 24.470 | 22.797 | 18.934 | 29.316 | 25.736 | 17.855 | 24.770 | 25.368 | 15.715 | 0.046 | 0.4998 | 0.7273 | 0 | 0 | 0 |
| 'Sertad2' | 10.612 | 9.410 | 9.642 | 9.771 | 8.129 | 8.947 | 9.377 | 9.212 | 9.832 | 9.915 | 10.602 | 11.024 | 9.051 | 10.292 | 10.264 | 9.519 | 9.319 | 10.669 | -0.104 | 0.0312 | 0.1385 | 0 | 0 | 0 |
| 'Sertad3' | 14.966 | 14.699 | 13.253 | 16.059 | 12.956 | 12.700 | 15.453 | 11.805 | 14.131 | 11.148 | 12.974 | 13.099 | 12.031 | 13.935 | 13.400 | 12.105 | 15.051 | 12.815 | 0.098 | 0.1590 | 0.3895 | 0 | 0 | 0 |
| 'Sesn1' | 8.523 | 9.291 | 8.708 | 7.941 | 9.509 | 9.072 | 8.983 | 9.221 | 9.364 | 7.915 | 7.076 | 8.888 | 6.840 | 8.128 | 8.149 | 8.243 | 7.709 | 8.385 | 0.161 | 0.0007 | 0.0078 | 1 | 0 | 0 |
| 'Sesn2' | 4.839 | 3.658 | 3.674 | 4.426 | 3.654 | 3.597 | 4.124 | 3.568 | 3.850 | 3.430 | 3.693 | 5.031 | 3.989 | 3.976 | 4.265 | 3.483 | 3.470 | 4.991 | -0.054 | 0.5181 | 0.7409 | 0 | 0 | 0 |
| 'Sesn3' | 59.517 | 48.045 | 55.387 | 57.495 | 51.532 | 51.235 | 49.941 | 51.495 | 52.322 | 53.079 | 60.778 | 61.383 | 47.605 | 54.724 | 59.649 | 48.510 | 56.803 | 63.333 | -0.092 | 0.1035 | 0.3017 | 0 | 0 | 0 |
| 'Set' | 86.727 | 84.392 | 92.775 | 91.647 | 89.190 | 88.486 | 92.112 | 90.033 | 86.937 | 85.138 | 83.412 | 85.859 | 87.664 | 86.297 | 89.465 | 80.415 | 88.535 | 89.006 | 0.038 | 0.1543 | 0.3822 | 0 | 0 | 0 |
| 'Setbp1' | 3.539 | 4.299 | 3.246 | 3.616 | 3.952 | 3.830 | 3.130 | 4.336 | 3.435 | 4.983 | 5.197 | 4.147 | 4.937 | 5.452 | 3.976 | 5.034 | 4.091 | 4.309 | -0.345 | 0.0000 | 0.0000 | 1 | 0 | 0 |
| 'Setd1a' | 23.062 | 21.302 | 25.054 | 22.198 | 21.561 | 23.554 | 20.862 | 22.118 | 24.451 | 23.857 | 22.572 | 21.843 | 24.444 | 20.652 | 20.847 | 23.809 | 21.283 | 20.494 | 0.023 | 0.5668 | 0.7748 | 0 | 0 | 0 |
| 'Setd2' | 10.903 | 8.980 | 8.757 | 10.362 | 8.971 | 8.813 | 9.563 | 9.150 | 8.849 | 10.856 | 10.282 | 9.212 | 9.263 | 9.734 | 8.576 | 11.194 | 9.588 | 9.162 | -0.069 | 0.0752 | 0.2439 | 0 | 0 | 0 |
| 'Setd4' | 9.452 | 10.535 | 10.833 | 8.446 | 11.372 | 10.614 | 8.384 | 11.871 | 10.097 | 9.889 | 10.513 | 9.345 | 10.544 | 10.507 | 10.370 | 9.249 | 10.728 | 9.185 | 0.007 | 0.8811 | 0.9499 | 0 | 0 | 0 |
| 'Setd5' | 10.157 | 9.821 | 9.728 | 9.930 | 9.490 | 9.484 | 9.623 | 9.497 | 9.904 | 9.991 | 10.060 | 9.874 | 9.587 | 9.725 | 9.412 | 9.671 | 9.431 | 9.525 | -0.004 | 0.8421 | 0.9331 | 0 | 0 | 0 |
| 'Setd6' | 7.568 | 6.956 | 8.323 | 6.488 | 7.302 | 9.115 | 7.401 | 7.704 | 8.446 | 7.705 | 6.473 | 7.573 | 7.325 | 6.169 | 7.269 | 7.285 | 7.146 | 7.336 | 0.102 | 0.0348 | 0.1492 | 0 | 0 | 0 |
| 'Setd7' | 11.257 | 11.282 | 10.383 | 11.835 | 10.528 | 10.703 | 10.827 | 10.298 | 11.147 | 10.672 | 11.554 | 14.044 | 11.181 | 11.282 | 12.797 | 12.000 | 11.916 | 14.735 | -0.172 | 0.0007 | 0.0075 | 1 | 0 | 0 |
| 'Setdb2' | 4.250 | 4.482 | 4.420 | 3.887 | 4.579 | 4.182 | 4.311 | 4.082 | 3.955 | 4.565 | 4.254 | 4.189 | 4.394 | 4.438 | 4.013 | 4.376 | 4.041 | 3.748 | -0.004 | 0.9380 | 0.9764 | 0 | 0 | 0 |
| 'Setx' | 24.314 | 23.322 | 23.332 | 24.852 | 22.595 | 22.676 | 22.846 | 22.894 | 23.042 | 24.192 | 25.559 | 23.911 | 21.526 | 24.122 | 23.192 | 23.677 | 23.574 | 24.468 | -0.040 | 0.2361 | 0.4859 | 0 | 0 | 0 |
| 'Sf1' | 30.790 | 30.587 | 32.068 | 29.369 | 30.063 | 31.589 | 29.506 | 30.073 | 33.079 | 33.086 | 30.391 | 31.983 | 32.945 | 30.464 | 31.014 | 31.484 | 30.180 | 30.645 | -0.036 | 0.1191 | 0.3284 | 0 | 0 | 0 |
| 'Sf3a1' | 25.070 | 24.546 | 25.527 | 25.623 | 24.356 | 26.684 | 24.853 | 24.200 | 26.210 | 25.250 | 24.716 | 28.083 | 26.079 | 24.626 | 25.990 | 25.265 | 24.535 | 28.160 | -0.047 | 0.0376 | 0.1573 | 0 | 0 | 0 |
| 'Sf3a2' | 30.888 | 30.055 | 33.833 | 31.946 | 29.782 | 32.336 | 33.005 | 31.828 | 34.856 | 31.658 | 29.344 | 33.925 | 32.533 | 30.126 | 34.000 | 30.681 | 30.381 | 34.589 | -0.004 | 0.8644 | 0.9410 | 0 | 0 | 0 |
| 'Sf3a3' | 27.738 | 28.709 | 28.109 | 28.239 | 28.824 | 28.050 | 28.349 | 26.578 | 27.866 | 27.560 | 26.417 | 28.059 | 27.872 | 27.586 | 26.473 | 26.079 | 27.963 | 27.762 | 0.025 | 0.3393 | 0.5966 | 0 | 0 | 0 |
| 'Sf3b1' | 50.951 | 52.861 | 51.105 | 53.340 | 52.685 | 50.218 | 54.005 | 51.564 | 50.571 | 50.759 | 52.751 | 50.403 | 49.291 | 51.757 | 48.154 | 50.968 | 55.274 | 48.120 | 0.021 | 0.3408 | 0.5979 | 0 | 0 | 0 |
| 'Sf3b2' | 150.470 | 150.985 | 157.359 | 153.365 | 152.295 | 164.964 | 151.151 | 150.780 | 161.117 | 159.517 | 152.223 | 152.896 | 158.359 | 154.842 | 157.077 | 160.641 | 149.750 | 153.251 | -0.016 | 0.4315 | 0.6763 | 0 | 0 | 0 |
| 'Sf3b3' | 77.074 | 73.783 | 73.854 | 77.636 | 71.406 | 68.871 | 75.151 | 70.669 | 70.860 | 72.594 | 74.720 | 77.903 | 69.728 | 71.076 | 74.481 | 68.983 | 75.032 | 81.529 | -0.026 | 0.4886 | 0.7187 | 0 | 0 | 0 |
| 'Sf3b4' | 34.999 | 32.430 | 38.786 | 34.716 | 35.439 | 39.411 | 33.793 | 34.222 | 38.134 | 36.928 | 30.616 | 37.615 | 36.535 | 32.832 | 38.381 | 33.306 | 30.580 | 38.676 | 0.020 | 0.5241 | 0.7464 | 0 | 0 | 0 |
| 'Sf3b5' | 17.803 | 19.793 | 18.836 | 18.011 | 19.052 | 19.531 | 18.753 | 18.253 | 18.792 | 18.587 | 18.898 | 18.701 | 20.779 | 18.060 | 18.968 | 19.227 | 18.401 | 17.986 | -0.017 | 0.5867 | 0.7880 | 0 | 0 | 0 |
| 'Sf3b6' | 55.231 | 57.557 | 56.082 | 58.943 | 63.054 | 57.964 | 57.868 | 58.218 | 58.191 | 56.897 | 54.653 | 58.290 | 57.263 | 56.548 | 57.804 | 56.410 | 56.696 | 52.711 | 0.030 | 0.3132 | 0.5715 | 0 | 0 | 0 |
| 'Sfmbt1' | 5.397 | 5.613 | 5.572 | 6.131 | 5.503 | 5.752 | 5.101 | 5.399 | 5.695 | 5.430 | 6.077 | 5.471 | 5.475 | 5.897 | 5.445 | 4.990 | 5.400 | 5.656 | 0.000 | 0.9981 | 0.9994 | 0 | 0 | 0 |
| 'Sfmbt2' | 1.027 | 1.091 | 1.006 | 1.151 | 1.189 | 0.978 | 1.260 | 1.105 | 0.932 | 1.112 | 1.116 | 1.199 | 1.039 | 0.890 | 1.132 | 1.015 | 1.345 | 0.977 | -0.035 | 0.6319 | 0.8176 | 0 | 0 | 0 |
| 'Sfn' | 6.964 | 5.496 | 5.513 | 4.840 | 5.949 | 4.488 | 5.587 | 4.702 | 5.163 | 5.487 | 5.965 | 7.376 | 5.383 | 5.988 | 6.282 | 5.585 | 5.530 | 7.095 | -0.185 | 0.0511 | 0.1914 | 0 | 0 | 0 |
| 'Sfpq' | 95.602 | 80.925 | 94.468 | 94.173 | 80.341 | 88.680 | 93.857 | 80.227 | 90.003 | 88.428 | 91.829 | 92.980 | 92.497 | 90.206 | 97.209 | 86.247 | 90.546 | 100.269 | -0.070 | 0.0591 | 0.2099 | 0 | 0 | 0 |
| 'Sfr1' | 49.486 | 43.633 | 43.243 | 49.097 | 44.048 | 40.364 | 45.917 | 46.119 | 39.029 | 47.011 | 43.416 | 38.346 | 45.800 | 43.884 | 38.592 | 46.677 | 45.934 | 38.225 | 0.043 | 0.1732 | 0.4066 | 0 | 0 | 0 |
| 'Sfswap' | 8.537 | 7.729 | 9.149 | 8.522 | 7.723 | 8.507 | 8.709 | 8.053 | 8.909 | 8.269 | 8.126 | 9.706 | 8.253 | 7.799 | 8.718 | 8.079 | 8.696 | 9.257 | -0.031 | 0.3152 | 0.5731 | 0 | 0 | 0 |
| 'Sft2d1' | 8.646 | 10.778 | 8.967 | 8.838 | 9.855 | 9.769 | 8.664 | 10.241 | 9.778 | 9.745 | 9.801 | 10.136 | 10.054 | 9.772 | 9.624 | 9.728 | 9.432 | 9.694 | -0.056 | 0.1157 | 0.3230 | 0 | 0 | 0 |
| 'Sft2d2' | 24.096 | 25.628 | 22.973 | 23.360 | 26.920 | 22.722 | 23.496 | 27.189 | 22.082 | 26.271 | 28.906 | 23.438 | 25.305 | 27.598 | 23.279 | 26.266 | 27.109 | 22.500 | -0.086 | 0.0001 | 0.0018 | 1 | 0 | 0 |
| 'Sft2d3' | 3.593 | 3.939 | 4.955 | 3.774 | 3.398 | 3.775 | 3.384 | 3.360 | 3.445 | 3.454 | 3.589 | 3.953 | 3.790 | 3.664 | 3.519 | 3.574 | 3.872 | 3.851 | 0.007 | 0.9268 | 0.9719 | 0 | 0 | 0 |
| 'Sfxn1' | 27.446 | 27.333 | 30.208 | 29.150 | 31.172 | 27.284 | 28.716 | 28.857 | 27.985 | 27.897 | 27.178 | 25.428 | 28.742 | 27.328 | 25.717 | 27.127 | 27.656 | 24.708 | 0.087 | 0.0080 | 0.0530 | 0 | 0 | 0 |
| 'Sfxn2' | 2.995 | 3.761 | 3.659 | 3.324 | 3.373 | 3.439 | 3.342 | 3.819 | 3.827 | 2.833 | 3.020 | 2.895 | 3.460 | 2.887 | 3.180 | 3.104 | 2.134 | 3.103 | 0.235 | 0.0022 | 0.0195 | 1 | 0 | 0 |
| 'Sfxn3' | 13.014 | 13.692 | 14.295 | 11.042 | 14.699 | 14.624 | 14.443 | 14.540 | 15.148 | 15.177 | 13.984 | 14.813 | 13.144 | 12.734 | 14.530 | 13.940 | 13.482 | 12.599 | 0.002 | 0.9742 | 0.9912 | 0 | 0 | 0 |
| 'Sfxn5' | 0.599 | 0.531 | 0.543 | 0.530 | 0.492 | 0.560 | 0.619 | 0.576 | 0.654 | 0.514 | 0.711 | 0.572 | 0.550 | 0.574 | 0.449 | 0.525 | 0.508 | 0.476 | 0.061 | 0.5052 | 0.7315 | 0 | 0 | 0 |
| 'Sgcb' | 2.581 | 3.196 | 3.571 | 2.530 | 3.142 | 3.092 | 2.923 | 3.643 | 3.697 | 3.076 | 2.595 | 3.000 | 3.199 | 3.063 | 2.813 | 3.482 | 2.544 | 2.534 | 0.102 | 0.2156 | 0.4613 | 0 | 0 | 0 |
| 'Sgf29' | 4.191 | 4.655 | 4.429 | 4.104 | 4.737 | 5.307 | 4.427 | 4.354 | 4.960 | 3.470 | 4.022 | 4.367 | 4.273 | 4.164 | 4.536 | 4.289 | 4.244 | 4.475 | 0.110 | 0.0379 | 0.1583 | 0 | 0 | 0 |
| 'Sgip1' | 0.283 | 0.289 | 0.278 | 0.225 | 0.337 | 0.232 | 0.191 | 0.357 | 0.235 | 0.290 | 0.263 | 0.147 | 0.300 | 0.354 | 0.159 | 0.273 | 0.213 | 0.166 | 0.199 | 0.1359 | 0.3546 | 0 | 0 | 0 |
| 'Sgk1' | 7.156 | 3.820 | 4.241 | 5.970 | 4.273 | 3.725 | 5.583 | 5.360 | 3.006 | 6.946 | 7.023 | 4.035 | 5.456 | 7.033 | 3.912 | 6.620 | 6.476 | 3.670 | -0.247 | 0.0274 | 0.1274 | 0 | 0 | 0 |
| 'Sgol1' | 2.371 | 2.987 | 4.041 | 3.269 | 2.950 | 3.723 | 3.476 | 3.028 | 3.616 | 2.184 | 2.461 | 3.374 | 3.412 | 3.420 | 3.780 | 3.007 | 3.505 | 3.413 | 0.031 | 0.7207 | 0.8716 | 0 | 0 | 0 |
| 'Sgol2a' | 0.746 | 0.792 | 1.091 | 0.984 | 1.005 | 1.083 | 1.247 | 0.891 | 1.236 | 0.518 | 0.904 | 1.108 | 1.060 | 0.936 | 1.065 | 0.918 | 1.090 | 1.180 | 0.034 | 0.7646 | 0.8953 | 0 | 0 | 0 |
| 'Sgpl1' | 8.035 | 8.801 | 8.464 | 7.872 | 8.915 | 9.429 | 8.071 | 9.148 | 8.600 | 8.941 | 8.893 | 9.659 | 7.883 | 9.409 | 9.031 | 8.528 | 9.281 | 9.177 | -0.074 | 0.0146 | 0.0818 | 0 | 0 | 0 |
| 'Sgpp1' | 21.056 | 22.210 | 20.677 | 19.174 | 23.115 | 22.233 | 21.025 | 22.513 | 19.935 | 20.564 | 21.957 | 22.270 | 20.292 | 21.931 | 21.913 | 21.503 | 21.136 | 22.726 | -0.030 | 0.3686 | 0.6201 | 0 | 0 | 0 |
| 'Sgpp2' | 1.144 | 1.411 | 1.172 | 1.140 | 1.039 | 1.238 | 1.531 | 1.414 | 1.412 | 1.312 | 0.787 | 0.845 | 0.968 | 0.881 | 0.990 | 0.900 | 0.609 | 0.822 | 0.500 | 0.0000 | 0.0004 | 1 | 1 | 0 |
| 'Sgsh' | 18.814 | 19.423 | 21.364 | 19.116 | 20.128 | 21.678 | 20.003 | 20.248 | 22.183 | 21.179 | 21.550 | 20.900 | 20.234 | 18.355 | 20.526 | 20.246 | 20.494 | 19.695 | -0.012 | 0.7030 | 0.8611 | 0 | 0 | 0 |
| 'Sgsm2' | 3.509 | 3.154 | 3.578 | 2.915 | 3.006 | 3.835 | 3.325 | 3.246 | 3.663 | 3.405 | 2.844 | 3.651 | 3.041 | 2.428 | 3.344 | 3.187 | 3.128 | 3.421 | 0.079 | 0.1051 | 0.3040 | 0 | 0 | 0 |
| 'Sgsm3' | 1.636 | 1.543 | 1.605 | 1.436 | 1.303 | 1.389 | 1.386 | 1.345 | 1.520 | 1.500 | 1.391 | 1.614 | 1.654 | 1.317 | 1.414 | 1.565 | 1.243 | 1.566 | -0.022 | 0.6966 | 0.8574 | 0 | 0 | 0 |
| 'Sgta' | 29.070 | 26.974 | 30.003 | 26.107 | 27.181 | 31.165 | 26.752 | 27.760 | 29.347 | 29.327 | 24.802 | 30.106 | 30.987 | 26.578 | 28.684 | 31.080 | 25.324 | 28.801 | -0.015 | 0.6749 | 0.8440 | 0 | 0 | 0 |
| 'Sh2b1' | 20.393 | 19.028 | 21.245 | 19.355 | 18.193 | 19.970 | 19.618 | 17.970 | 21.551 | 18.831 | 18.559 | 22.653 | 19.527 | 19.222 | 21.877 | 19.428 | 19.233 | 21.968 | -0.042 | 0.1142 | 0.3202 | 0 | 0 | 0 |
| 'Sh2b2' | 0.511 | 0.649 | 0.396 | 0.489 | 0.603 | 0.468 | 0.582 | 0.585 | 0.443 | 0.411 | 0.576 | 0.612 | 0.436 | 0.817 | 0.592 | 0.530 | 0.765 | 0.538 | -0.184 | 0.1048 | 0.3034 | 0 | 0 | 0 |
| 'Sh2b3' | 7.772 | 8.444 | 9.159 | 7.833 | 7.491 | 9.101 | 7.317 | 8.132 | 9.802 | 8.576 | 7.765 | 9.890 | 7.814 | 8.217 | 9.584 | 8.868 | 8.012 | 9.338 | -0.067 | 0.0421 | 0.1692 | 0 | 0 | 0 |
| 'Sh2d2a' | 43.613 | 38.885 | 42.055 | 41.434 | 39.110 | 44.465 | 39.914 | 37.267 | 42.305 | 41.028 | 38.398 | 49.281 | 43.537 | 39.480 | 49.281 | 43.849 | 39.092 | 49.165 | -0.098 | 0.0013 | 0.0132 | 1 | 0 | 0 |
| 'Sh2d3c' | 25.932 | 23.570 | 26.710 | 23.201 | 22.578 | 27.963 | 23.326 | 21.827 | 27.274 | 25.972 | 23.705 | 30.089 | 24.021 | 23.575 | 29.064 | 25.824 | 23.964 | 29.495 | -0.093 | 0.0013 | 0.0127 | 1 | 0 | 0 |
| 'Sh2d5' | 1.090 | 1.597 | 1.081 | 1.270 | 1.224 | 0.893 | 1.456 | 1.045 | 1.157 | 1.658 | 1.694 | 1.506 | 2.317 | 1.695 | 1.459 | 1.440 | 1.946 | 1.559 | -0.524 | 0.0000 | 0.0000 | 1 | 1 | 0 |
| 'Sh3bgrl' | 31.213 | 37.805 | 34.725 | 35.068 | 36.785 | 32.197 | 38.182 | 36.189 | 32.476 | 34.822 | 37.134 | 36.012 | 33.420 | 37.878 | 33.929 | 32.842 | 39.145 | 33.622 | -0.031 | 0.3577 | 0.6103 | 0 | 0 | 0 |
| 'Sh3bgrl3' | 109.247 | 107.015 | 115.222 | 111.501 | 109.481 | 124.046 | 113.511 | 108.623 | 118.063 | 107.737 | 98.401 | 103.538 | 111.085 | 100.192 | 116.721 | 107.367 | 97.123 | 107.756 | 0.089 | 0.0013 | 0.0130 | 1 | 0 | 0 |
| 'Sh3bp1' | 22.351 | 19.207 | 21.960 | 22.117 | 18.254 | 22.242 | 20.877 | 19.403 | 22.482 | 20.864 | 19.202 | 24.366 | 21.015 | 19.869 | 24.097 | 20.283 | 18.389 | 25.124 | -0.040 | 0.2309 | 0.4782 | 0 | 0 | 0 |
| 'Sh3bp2' | 2.515 | 2.413 | 2.349 | 2.164 | 2.391 | 2.604 | 2.338 | 2.556 | 2.432 | 2.657 | 2.122 | 2.937 | 2.126 | 2.152 | 2.724 | 3.276 | 2.374 | 2.528 | -0.079 | 0.2664 | 0.5199 | 0 | 0 | 0 |
| 'Sh3bp5' | 10.579 | 12.531 | 10.794 | 10.910 | 11.405 | 12.003 | 10.557 | 11.827 | 11.568 | 10.067 | 9.706 | 10.956 | 10.732 | 10.013 | 10.422 | 10.841 | 11.550 | 9.948 | 0.103 | 0.0102 | 0.0634 | 0 | 0 | 0 |
| 'Sh3bp5l' | 3.161 | 3.227 | 3.555 | 2.832 | 2.877 | 3.436 | 3.258 | 3.142 | 3.504 | 2.996 | 2.753 | 3.486 | 3.059 | 2.910 | 3.486 | 3.086 | 3.208 | 3.277 | 0.026 | 0.4979 | 0.7259 | 0 | 0 | 0 |
| 'Sh3d19' | 0.285 | 0.352 | 0.272 | 0.402 | 0.296 | 0.290 | 0.367 | 0.304 | 0.294 | 0.184 | 0.358 | 0.332 | 0.311 | 0.330 | 0.363 | 0.288 | 0.300 | 0.311 | 0.020 | 0.8629 | 0.9405 | 0 | 0 | 0 |
| 'Sh3d21' | 0.744 | 0.617 | 0.802 | 0.868 | 0.662 | 0.955 | 0.865 | 0.553 | 0.816 | 0.622 | 0.525 | 0.738 | 0.886 | 0.528 | 0.560 | 0.747 | 0.524 | 0.661 | 0.232 | 0.0304 | 0.1360 | 0 | 0 | 0 |
| 'Sh3gl1' | 63.389 | 59.523 | 67.425 | 62.727 | 57.960 | 68.278 | 60.506 | 56.777 | 66.326 | 60.000 | 55.401 | 71.721 | 59.944 | 56.852 | 65.290 | 60.009 | 59.035 | 71.153 | -0.002 | 0.9420 | 0.9781 | 0 | 0 | 0 |
| 'Sh3gl3' | 0.331 | 0.460 | 0.353 | 0.261 | 0.553 | 0.471 | 0.331 | 0.499 | 0.445 | 0.789 | 0.667 | 0.906 | 0.496 | 0.838 | 1.045 | 1.004 | 0.795 | 1.079 | -1.059 | 0.0000 | 0.0000 | 1 | 1 | 1 |
| 'Sh3glb1' | 13.631 | 13.322 | 12.838 | 14.244 | 14.219 | 12.461 | 14.759 | 14.535 | 12.081 | 12.901 | 15.054 | 13.713 | 14.086 | 14.223 | 13.196 | 13.591 | 14.861 | 13.160 | -0.044 | 0.1662 | 0.3984 | 0 | 0 | 0 |
| 'Sh3glb2' | 14.870 | 13.597 | 16.519 | 14.648 | 13.389 | 16.820 | 14.687 | 13.532 | 16.570 | 15.268 | 13.355 | 16.588 | 15.381 | 12.879 | 15.996 | 14.777 | 13.701 | 16.256 | -0.005 | 0.8006 | 0.9111 | 0 | 0 | 0 |
| 'Sh3kbp1' | 25.909 | 26.206 | 26.146 | 27.304 | 27.606 | 26.990 | 27.381 | 26.377 | 26.431 | 25.040 | 24.334 | 23.299 | 25.406 | 24.504 | 23.293 | 23.380 | 22.440 | 22.511 | 0.156 | 0.0000 | 0.0000 | 1 | 0 | 0 |
| 'Sh3pxd2a' | 0.499 | 0.788 | 0.742 | 0.619 | 0.549 | 0.766 | 0.696 | 0.810 | 0.751 | 0.464 | 0.376 | 0.376 | 0.355 | 0.583 | 0.412 | 0.234 | 0.533 | 0.365 | 0.750 | 0.0000 | 0.0000 | 1 | 1 | 0 |
| 'Sh3rf1' | 16.320 | 15.911 | 19.255 | 16.379 | 16.924 | 16.940 | 17.423 | 17.476 | 18.091 | 17.073 | 17.480 | 17.990 | 16.592 | 18.146 | 18.180 | 16.232 | 19.492 | 17.956 | -0.049 | 0.1598 | 0.3902 | 0 | 0 | 0 |
| 'Sh3tc1' | 0.323 | 0.576 | 0.294 | 0.327 | 0.451 | 0.488 | 0.383 | 0.513 | 0.431 | 0.413 | 0.445 | 0.275 | 0.418 | 0.456 | 0.320 | 0.457 | 0.399 | 0.291 | 0.128 | 0.2833 | 0.5395 | 0 | 0 | 0 |
| 'Sharpin' | 8.765 | 8.989 | 9.283 | 7.953 | 8.600 | 9.195 | 8.060 | 8.726 | 9.289 | 8.387 | 7.885 | 9.215 | 8.945 | 7.934 | 9.014 | 8.043 | 7.999 | 9.010 | 0.034 | 0.2664 | 0.5199 | 0 | 0 | 0 |
| 'Shb' | 1.149 | 1.151 | 0.814 | 0.897 | 1.038 | 1.031 | 1.036 | 1.150 | 0.909 | 1.514 | 1.170 | 0.894 | 2.069 | 1.316 | 0.974 | 1.051 | 1.553 | 0.926 | -0.308 | 0.0100 | 0.0622 | 0 | 0 | 0 |
| 'Shc1' | 16.078 | 18.366 | 17.848 | 15.703 | 17.801 | 17.681 | 16.924 | 18.265 | 18.432 | 17.836 | 17.325 | 17.823 | 17.312 | 17.890 | 18.324 | 17.473 | 16.931 | 16.671 | -0.015 | 0.5768 | 0.7816 | 0 | 0 | 0 |
| 'Shcbp1' | 6.745 | 7.691 | 9.532 | 8.319 | 8.985 | 7.997 | 11.322 | 7.713 | 8.691 | 5.560 | 8.309 | 8.189 | 9.138 | 8.996 | 10.158 | 6.791 | 9.186 | 7.253 | 0.056 | 0.5782 | 0.7820 | 0 | 0 | 0 |
| 'Shfm1' | 22.372 | 26.966 | 22.219 | 24.233 | 24.354 | 23.945 | 23.724 | 24.265 | 22.731 | 21.588 | 24.139 | 20.479 | 23.799 | 23.666 | 22.820 | 24.691 | 23.209 | 21.027 | 0.055 | 0.1231 | 0.3351 | 0 | 0 | 0 |
| 'Shisa5' | 132.911 | 161.188 | 141.467 | 129.424 | 158.359 | 152.081 | 134.622 | 152.914 | 153.087 | 156.939 | 146.954 | 131.555 | 136.982 | 149.086 | 148.060 | 158.148 | 140.446 | 142.237 | -0.008 | 0.8558 | 0.9383 | 0 | 0 | 0 |
| 'Shkbp1' | 11.072 | 11.584 | 12.647 | 10.290 | 11.314 | 12.846 | 10.333 | 11.664 | 12.325 | 11.334 | 10.258 | 11.299 | 11.563 | 10.531 | 11.151 | 10.801 | 10.608 | 10.611 | 0.074 | 0.0375 | 0.1572 | 0 | 0 | 0 |
| 'Shmt1' | 2.413 | 2.173 | 2.795 | 2.581 | 2.526 | 2.547 | 2.499 | 2.707 | 2.537 | 2.454 | 2.385 | 2.864 | 2.584 | 2.186 | 2.900 | 2.269 | 2.216 | 2.623 | 0.006 | 0.9071 | 0.9636 | 0 | 0 | 0 |
| 'Shmt2' | 37.938 | 35.194 | 45.598 | 39.368 | 37.908 | 41.959 | 40.792 | 36.115 | 38.504 | 34.067 | 34.840 | 40.532 | 40.911 | 32.583 | 41.843 | 36.793 | 31.991 | 38.757 | 0.079 | 0.0399 | 0.1629 | 0 | 0 | 0 |
| 'Shoc2' | 23.261 | 24.975 | 22.255 | 24.483 | 24.806 | 21.370 | 25.361 | 24.132 | 22.246 | 25.334 | 22.572 | 23.933 | 22.659 | 23.502 | 22.745 | 21.699 | 25.581 | 24.262 | -0.011 | 0.7673 | 0.8961 | 0 | 0 | 0 |
| 'Shpk' | 2.571 | 2.620 | 2.615 | 2.374 | 2.450 | 2.532 | 2.313 | 2.441 | 2.927 | 2.782 | 2.852 | 2.661 | 2.584 | 2.509 | 2.536 | 2.327 | 2.551 | 2.010 | -0.005 | 0.9437 | 0.9788 | 0 | 0 | 0 |
| 'Shprh' | 2.810 | 3.013 | 3.190 | 3.139 | 3.087 | 2.930 | 3.307 | 2.944 | 3.041 | 2.589 | 2.930 | 3.342 | 2.521 | 2.663 | 3.073 | 2.413 | 3.023 | 3.379 | 0.075 | 0.1568 | 0.3865 | 0 | 0 | 0 |
| 'Shq1' | 1.134 | 0.893 | 1.273 | 0.983 | 1.062 | 1.127 | 1.261 | 1.038 | 0.943 | 0.955 | 1.043 | 1.081 | 0.926 | 1.095 | 1.088 | 1.063 | 0.887 | 1.049 | 0.075 | 0.3179 | 0.5764 | 0 | 0 | 0 |
| 'Shtn1' | 8.717 | 11.090 | 8.602 | 8.307 | 8.741 | 8.476 | 8.285 | 9.412 | 10.203 | 7.246 | 7.599 | 7.053 | 5.209 | 6.938 | 6.362 | 6.505 | 7.756 | 4.519 | 0.461 | 0.0000 | 0.0000 | 1 | 1 | 0 |
| 'Siae' | 6.238 | 6.920 | 6.630 | 7.108 | 6.307 | 6.497 | 7.202 | 6.268 | 7.610 | 6.115 | 6.011 | 7.624 | 6.249 | 7.523 | 6.678 | 6.580 | 6.591 | 7.237 | -0.013 | 0.8206 | 0.9207 | 0 | 0 | 0 |
| 'Siah2' | 29.065 | 13.300 | 8.276 | 16.147 | 14.576 | 9.765 | 10.786 | 16.811 | 6.881 | 26.069 | 17.226 | 10.253 | 22.476 | 23.903 | 10.740 | 24.872 | 16.188 | 12.133 | -0.396 | 0.0123 | 0.0726 | 0 | 0 | 0 |
| 'Siah3' | 4.677 | 4.854 | 3.685 | 3.578 | 5.470 | 3.819 | 2.964 | 4.230 | 3.956 | 4.465 | 3.740 | 3.959 | 4.643 | 4.547 | 4.760 | 5.164 | 3.543 | 3.983 | -0.071 | 0.5432 | 0.7600 | 0 | 0 | 0 |
| 'Sidt1' | 6.505 | 6.312 | 6.392 | 6.038 | 5.920 | 6.682 | 6.445 | 6.048 | 6.030 | 5.642 | 5.709 | 7.118 | 5.603 | 4.874 | 5.879 | 5.738 | 5.828 | 6.972 | 0.067 | 0.1937 | 0.4313 | 0 | 0 | 0 |
| 'Sidt2' | 23.689 | 23.056 | 25.659 | 21.279 | 22.606 | 25.366 | 22.178 | 23.998 | 25.530 | 26.947 | 24.281 | 26.128 | 25.159 | 23.656 | 24.424 | 25.817 | 24.500 | 25.306 | -0.095 | 0.0028 | 0.0240 | 1 | 0 | 0 |
| 'Sigirr' | 66.438 | 66.391 | 74.392 | 67.420 | 71.271 | 72.799 | 66.657 | 73.665 | 75.151 | 72.084 | 66.585 | 70.721 | 71.079 | 66.913 | 69.135 | 70.412 | 66.406 | 64.161 | 0.029 | 0.3298 | 0.5877 | 0 | 0 | 0 |
| 'Sigmar1' | 3.550 | 2.977 | 3.627 | 3.284 | 3.311 | 3.570 | 3.317 | 3.043 | 3.521 | 2.851 | 2.943 | 3.507 | 2.924 | 3.195 | 3.505 | 3.360 | 2.924 | 3.686 | 0.056 | 0.3011 | 0.5579 | 0 | 0 | 0 |
| 'Sik1' | 29.285 | 7.337 | 3.530 | 12.960 | 6.938 | 3.968 | 9.561 | 9.814 | 3.396 | 28.250 | 11.907 | 3.505 | 22.643 | 14.600 | 2.935 | 25.548 | 10.795 | 3.420 | -0.364 | 0.0615 | 0.2147 | 0 | 0 | 0 |
| 'Sik2' | 2.421 | 2.192 | 2.302 | 2.216 | 2.291 | 2.186 | 2.276 | 2.229 | 2.469 | 2.342 | 2.498 | 2.661 | 2.330 | 2.202 | 2.402 | 2.146 | 2.249 | 2.608 | -0.073 | 0.1043 | 0.3027 | 0 | 0 | 0 |
| 'Sik3' | 14.636 | 12.673 | 13.620 | 13.634 | 12.789 | 13.909 | 13.856 | 13.874 | 13.166 | 15.115 | 15.036 | 14.175 | 15.107 | 14.081 | 14.385 | 14.317 | 14.423 | 15.141 | -0.119 | 0.0001 | 0.0010 | 1 | 0 | 0 |
| 'Sike1' | 2.399 | 3.359 | 3.241 | 2.745 | 3.561 | 3.327 | 2.825 | 3.572 | 3.450 | 2.874 | 3.045 | 3.382 | 2.752 | 3.146 | 3.435 | 3.041 | 3.348 | 3.378 | -0.013 | 0.7587 | 0.8921 | 0 | 0 | 0 |
| 'Sil1' | 14.744 | 13.646 | 14.799 | 16.246 | 15.201 | 15.375 | 16.479 | 14.323 | 14.739 | 17.166 | 13.894 | 18.911 | 14.118 | 13.665 | 16.205 | 16.039 | 14.504 | 16.522 | -0.074 | 0.1799 | 0.4151 | 0 | 0 | 0 |
| 'Simc1' | 2.285 | 1.982 | 2.303 | 2.221 | 2.039 | 2.037 | 2.197 | 2.288 | 2.189 | 2.288 | 2.038 | 2.215 | 2.137 | 2.079 | 1.923 | 1.974 | 1.972 | 2.105 | 0.051 | 0.3462 | 0.6014 | 0 | 0 | 0 |
| 'Sin3a' | 17.638 | 18.636 | 17.680 | 17.552 | 17.772 | 18.991 | 16.219 | 16.916 | 18.172 | 17.337 | 18.720 | 19.228 | 17.102 | 18.645 | 18.856 | 17.748 | 19.028 | 18.289 | -0.058 | 0.0373 | 0.1565 | 0 | 0 | 0 |
| 'Sin3b' | 33.994 | 33.558 | 32.718 | 31.822 | 32.055 | 32.186 | 33.028 | 31.405 | 32.041 | 30.719 | 31.753 | 37.549 | 31.610 | 32.989 | 35.239 | 30.059 | 33.981 | 37.422 | -0.051 | 0.1836 | 0.4193 | 0 | 0 | 0 |
| 'Sipa1' | 181.084 | 166.747 | 188.551 | 180.793 | 164.215 | 189.056 | 166.892 | 155.596 | 189.794 | 183.738 | 159.335 | 195.814 | 203.019 | 167.833 | 190.464 | 180.097 | 174.353 | 196.732 | -0.072 | 0.0094 | 0.0597 | 0 | 0 | 0 |
| 'Sipa1l1' | 23.759 | 21.462 | 24.582 | 23.789 | 23.108 | 24.465 | 22.065 | 23.031 | 25.127 | 24.077 | 24.429 | 24.255 | 21.842 | 21.789 | 23.150 | 22.286 | 21.363 | 24.182 | 0.017 | 0.5921 | 0.7921 | 0 | 0 | 0 |
| 'Sipa1l2' | 2.478 | 2.601 | 2.554 | 2.521 | 2.773 | 2.751 | 2.384 | 2.397 | 2.743 | 2.688 | 2.672 | 3.048 | 3.018 | 3.406 | 3.096 | 3.105 | 3.055 | 3.242 | -0.247 | 0.0000 | 0.0002 | 1 | 0 | 0 |
| 'Sipa1l3' | 1.438 | 1.294 | 1.725 | 1.386 | 1.365 | 1.664 | 1.385 | 1.634 | 1.572 | 1.670 | 1.431 | 1.470 | 1.724 | 1.447 | 1.328 | 1.336 | 1.255 | 1.432 | 0.032 | 0.6143 | 0.8072 | 0 | 0 | 0 |
| 'Sirt1' | 3.459 | 3.226 | 3.252 | 3.622 | 2.893 | 2.877 | 3.738 | 3.306 | 2.876 | 3.334 | 3.065 | 3.013 | 2.964 | 3.322 | 2.907 | 3.072 | 3.390 | 2.941 | 0.050 | 0.2943 | 0.5502 | 0 | 0 | 0 |
| 'Sirt2' | 15.786 | 14.420 | 16.011 | 15.492 | 16.186 | 16.908 | 15.320 | 15.146 | 16.094 | 17.562 | 16.028 | 16.505 | 17.572 | 15.275 | 15.957 | 16.166 | 15.532 | 16.012 | -0.061 | 0.0389 | 0.1607 | 0 | 0 | 0 |
| 'Sirt3' | 3.338 | 4.111 | 4.745 | 3.446 | 4.684 | 3.291 | 4.411 | 4.238 | 3.993 | 4.021 | 3.652 | 3.425 | 4.923 | 3.756 | 3.726 | 3.258 | 4.102 | 3.882 | 0.049 | 0.5965 | 0.7949 | 0 | 0 | 0 |
| 'Sirt4' | 2.726 | 2.475 | 2.895 | 2.741 | 2.716 | 2.743 | 3.088 | 2.666 | 2.633 | 2.919 | 2.499 | 2.809 | 2.843 | 2.677 | 2.677 | 2.510 | 3.078 | 2.339 | 0.008 | 0.8968 | 0.9586 | 0 | 0 | 0 |
| 'Sirt5' | 6.911 | 7.308 | 7.245 | 6.654 | 7.468 | 8.877 | 7.925 | 8.198 | 7.744 | 9.041 | 7.118 | 7.725 | 6.877 | 6.745 | 7.121 | 8.501 | 6.827 | 6.012 | 0.047 | 0.5456 | 0.7606 | 0 | 0 | 0 |
| 'Sirt6' | 15.786 | 14.831 | 16.943 | 13.799 | 16.230 | 16.764 | 15.250 | 15.319 | 17.485 | 15.442 | 14.038 | 15.962 | 15.944 | 12.877 | 16.982 | 14.100 | 11.706 | 15.304 | 0.097 | 0.0405 | 0.1647 | 0 | 0 | 0 |
| 'Sirt7' | 19.577 | 20.003 | 23.074 | 19.667 | 20.856 | 23.984 | 20.153 | 20.799 | 22.370 | 21.823 | 19.280 | 22.086 | 21.705 | 20.924 | 21.965 | 21.802 | 20.373 | 20.759 | -0.013 | 0.6504 | 0.8298 | 0 | 0 | 0 |
| 'Sit1' | 23.007 | 25.766 | 23.159 | 22.615 | 24.196 | 24.638 | 22.303 | 24.677 | 26.093 | 23.707 | 21.289 | 23.270 | 25.783 | 23.231 | 24.486 | 28.098 | 20.526 | 23.989 | 0.006 | 0.9029 | 0.9615 | 0 | 0 | 0 |
| 'Siva1' | 15.942 | 15.249 | 15.459 | 13.835 | 15.137 | 14.364 | 17.639 | 16.202 | 14.216 | 14.124 | 15.303 | 15.454 | 17.097 | 17.290 | 17.115 | 15.675 | 16.845 | 14.630 | -0.066 | 0.2994 | 0.5557 | 0 | 0 | 0 |
| 'Six5' | 5.804 | 5.714 | 6.481 | 6.266 | 4.843 | 6.341 | 5.020 | 4.590 | 6.538 | 5.599 | 5.263 | 7.123 | 5.680 | 5.547 | 6.113 | 5.146 | 5.470 | 8.027 | -0.078 | 0.2353 | 0.4846 | 0 | 0 | 0 |
| 'Ska1' | 2.745 | 2.847 | 3.601 | 3.103 | 3.594 | 3.145 | 3.681 | 2.999 | 3.407 | 2.467 | 2.965 | 3.156 | 3.847 | 3.250 | 3.632 | 3.468 | 3.396 | 2.997 | -0.014 | 0.8727 | 0.9454 | 0 | 0 | 0 |
| 'Ska2' | 5.609 | 7.035 | 7.815 | 7.403 | 7.086 | 8.023 | 7.412 | 7.914 | 7.690 | 6.054 | 7.472 | 8.096 | 6.657 | 8.897 | 8.406 | 6.294 | 7.741 | 8.013 | -0.043 | 0.4327 | 0.6777 | 0 | 0 | 0 |
| 'Ska3' | 4.054 | 3.715 | 5.234 | 4.754 | 3.968 | 3.925 | 4.340 | 4.103 | 5.725 | 3.635 | 4.354 | 4.443 | 5.336 | 4.877 | 5.084 | 3.525 | 4.472 | 4.485 | -0.026 | 0.7771 | 0.9002 | 0 | 0 | 0 |
| 'Skap1' | 39.268 | 39.731 | 38.618 | 40.794 | 39.988 | 39.164 | 41.424 | 39.646 | 38.225 | 37.041 | 37.856 | 36.586 | 34.883 | 36.000 | 37.133 | 36.397 | 37.555 | 34.660 | 0.110 | 0.0000 | 0.0000 | 1 | 0 | 0 |
| 'Skap2' | 2.430 | 2.377 | 2.312 | 2.470 | 2.309 | 2.247 | 2.711 | 2.591 | 2.208 | 2.348 | 2.745 | 2.358 | 2.433 | 2.653 | 2.449 | 2.288 | 2.673 | 2.178 | -0.040 | 0.3729 | 0.6242 | 0 | 0 | 0 |
| 'Ski' | 18.558 | 18.306 | 20.065 | 15.331 | 17.104 | 20.451 | 15.664 | 18.672 | 19.589 | 21.547 | 19.392 | 23.773 | 18.509 | 20.119 | 22.994 | 21.711 | 18.892 | 25.214 | -0.240 | 0.0000 | 0.0000 | 1 | 0 | 0 |
| 'Skil' | 3.282 | 3.966 | 3.125 | 3.131 | 3.893 | 3.530 | 3.331 | 3.801 | 3.324 | 3.758 | 3.751 | 3.461 | 3.680 | 4.292 | 3.232 | 3.946 | 4.306 | 3.199 | -0.108 | 0.0042 | 0.0331 | 1 | 0 | 0 |
| 'Skiv2l' | 16.889 | 16.036 | 20.144 | 16.988 | 16.087 | 19.175 | 17.792 | 16.240 | 20.156 | 17.658 | 17.136 | 20.215 | 19.396 | 16.545 | 18.821 | 17.713 | 16.516 | 18.768 | -0.042 | 0.1048 | 0.3033 | 0 | 0 | 0 |
| 'Skiv2l2' | 32.981 | 31.131 | 32.850 | 33.584 | 31.718 | 29.768 | 34.436 | 32.147 | 32.231 | 30.610 | 33.905 | 33.998 | 29.058 | 30.203 | 31.803 | 27.041 | 32.142 | 32.081 | 0.039 | 0.3591 | 0.6115 | 0 | 0 | 0 |
| 'Skp1a' | 41.178 | 47.582 | 41.653 | 44.392 | 48.423 | 41.978 | 43.759 | 46.594 | 42.074 | 40.238 | 43.776 | 42.396 | 40.096 | 46.268 | 39.446 | 42.413 | 46.835 | 42.780 | 0.036 | 0.1646 | 0.3969 | 0 | 0 | 0 |
| 'Skp2' | 0.573 | 0.769 | 0.716 | 0.797 | 0.684 | 0.723 | 0.803 | 0.676 | 0.815 | 0.602 | 0.785 | 0.887 | 0.839 | 0.821 | 0.814 | 0.757 | 0.892 | 0.952 | -0.185 | 0.0243 | 0.1182 | 0 | 0 | 0 |
| 'Sla' | 22.026 | 22.408 | 20.189 | 21.214 | 21.925 | 20.616 | 21.186 | 21.612 | 19.674 | 21.726 | 20.881 | 23.277 | 21.731 | 22.208 | 22.118 | 22.992 | 22.053 | 23.417 | -0.083 | 0.0062 | 0.0441 | 1 | 0 | 0 |
| 'Sla2' | 8.272 | 8.761 | 7.983 | 8.145 | 8.866 | 8.859 | 8.283 | 8.409 | 8.811 | 7.172 | 6.930 | 9.620 | 7.579 | 7.739 | 9.028 | 7.032 | 8.004 | 9.383 | 0.063 | 0.2038 | 0.4466 | 0 | 0 | 0 |
| 'Slain1' | 21.857 | 19.387 | 21.061 | 21.779 | 20.063 | 21.897 | 20.242 | 20.444 | 18.956 | 21.586 | 21.094 | 19.941 | 20.879 | 20.537 | 21.564 | 20.279 | 19.637 | 20.986 | -0.015 | 0.6569 | 0.8341 | 0 | 0 | 0 |
| 'Slain2' | 11.902 | 11.647 | 11.042 | 11.967 | 11.645 | 10.762 | 11.410 | 11.875 | 10.483 | 11.851 | 12.054 | 11.412 | 11.868 | 12.165 | 10.995 | 11.237 | 12.185 | 11.029 | -0.040 | 0.0822 | 0.2593 | 0 | 0 | 0 |
| 'Slamf1' | 5.355 | 4.830 | 5.460 | 5.529 | 4.569 | 5.319 | 5.140 | 5.283 | 5.197 | 5.154 | 5.325 | 5.547 | 4.639 | 5.105 | 4.972 | 4.814 | 4.901 | 4.906 | 0.032 | 0.4711 | 0.7068 | 0 | 0 | 0 |
| 'Slamf6' | 33.513 | 36.529 | 36.810 | 32.963 | 36.761 | 36.346 | 34.907 | 36.711 | 36.712 | 32.908 | 35.147 | 37.807 | 29.234 | 34.577 | 37.508 | 34.426 | 36.109 | 35.474 | 0.027 | 0.3398 | 0.5970 | 0 | 0 | 0 |

| | | | | | | | | | | | | | | | | | | | | | | | | |
|---|---|---|---|---|---|---|---|---|---|---|---|---|---|---|---|---|---|---|---|---|---|---|---|---|
| 'Slbp' | 13.199 | 11.016 | 11.328 | 13.391 | 12.002 | 11.422 | 13.050 | 11.451 | 11.561 | 10.997 | 12.585 | 12.321 | 13.561 | 13.510 | 12.405 | 11.097 | 13.309 | 11.880 | -0.055 | 0.2649 | 0.5184 | 0 | 0 | 0 |
| 'Slc10a3' | 5.897 | 6.726 | 6.842 | 6.007 | 7.074 | 6.051 | 5.959 | 6.552 | 6.194 | 6.953 | 6.669 | 6.593 | 5.761 | 6.168 | 6.448 | 6.903 | 6.281 | 5.555 | -0.010 | 0.8437 | 0.9337 | 0 | 0 | 0 |
| 'Slc10a7' | 7.358 | 7.986 | 7.647 | 8.770 | 7.639 | 7.834 | 8.266 | 8.254 | 8.404 | 7.171 | 7.623 | 8.289 | 8.220 | 7.420 | 8.684 | 7.629 | 7.845 | 7.797 | 0.014 | 0.7602 | 0.8934 | 0 | 0 | 0 |
| 'Slc11a2' | 18.655 | 18.479 | 17.060 | 18.376 | 18.320 | 17.236 | 18.930 | 17.481 | 17.905 | 17.564 | 18.056 | 19.600 | 18.604 | 19.964 | 19.794 | 20.029 | 19.948 | 19.870 | -0.107 | 0.0018 | 0.0165 | 1 | 0 | 0 |
| 'Slc12a2' | 4.492 | 4.537 | 5.459 | 5.080 | 4.215 | 4.224 | 4.349 | 4.516 | 4.663 | 5.765 | 6.111 | 5.705 | 5.190 | 6.155 | 5.675 | 6.134 | 5.763 | 6.107 | -0.347 | 0.0000 | 0.0000 | 1 | 0 | 0 |
| 'Slc12a4' | 10.160 | 10.509 | 11.050 | 10.169 | 10.009 | 10.358 | 9.500 | 10.491 | 10.366 | 10.610 | 10.652 | 10.323 | 10.881 | 10.154 | 10.252 | 10.512 | 10.031 | 10.386 | -0.025 | 0.4570 | 0.6944 | 0 | 0 | 0 |
| 'Slc12a6' | 6.916 | 7.646 | 7.838 | 6.498 | 7.191 | 7.744 | 7.376 | 7.446 | 8.106 | 6.598 | 7.812 | 8.246 | 7.015 | 7.849 | 8.372 | 6.871 | 8.242 | 8.093 | -0.059 | 0.0478 | 0.1828 | 0 | 0 | 0 |
| 'Slc12a7' | 25.519 | 23.440 | 27.623 | 23.236 | 24.272 | 27.729 | 22.567 | 25.470 | 28.363 | 27.786 | 27.251 | 32.704 | 22.494 | 25.336 | 30.653 | 26.428 | 24.695 | 30.840 | -0.128 | 0.0014 | 0.0137 | 1 | 0 | 0 |
| 'Slc12a8' | 1.575 | 1.765 | 1.846 | 1.446 | 1.855 | 1.892 | 1.704 | 1.715 | 1.884 | 2.438 | 1.792 | 1.592 | 1.990 | 1.634 | 1.396 | 1.763 | 1.766 | 1.351 | 0.003 | 0.9764 | 0.9918 | 0 | 0 | 0 |
| 'Slc12a9' | 6.334 | 5.269 | 7.640 | 6.383 | 5.663 | 6.531 | 5.492 | 5.674 | 7.078 | 6.633 | 5.942 | 6.630 | 7.066 | 5.164 | 6.165 | 5.755 | 5.936 | 6.647 | -0.007 | 0.8920 | 0.9560 | 0 | 0 | 0 |
| 'Slc14a1' | 5.403 | 7.311 | 6.339 | 5.232 | 7.102 | 6.896 | 6.695 | 7.353 | 6.899 | 9.863 | 8.541 | 6.139 | 8.489 | 8.556 | 7.768 | 8.623 | 6.643 | 4.635 | -0.224 | 0.0522 | 0.1938 | 0 | 0 | 0 |
| 'Slc15a3' | 1.151 | 1.127 | 1.106 | 1.048 | 1.326 | 1.158 | 1.110 | 1.340 | 1.278 | 1.236 | 1.042 | 1.194 | 1.424 | 1.294 | 1.040 | 1.598 | 1.149 | 1.089 | -0.058 | 0.4715 | 0.7072 | 0 | 0 | 0 |
| 'Slc15a4' | 2.538 | 2.345 | 2.418 | 2.552 | 2.425 | 2.454 | 2.680 | 2.349 | 2.399 | 2.184 | 2.623 | 3.451 | 2.340 | 2.484 | 3.283 | 2.335 | 2.976 | 3.481 | -0.184 | 0.0128 | 0.0750 | 0 | 0 | 0 |
| 'Slc16a1' | 9.603 | 6.966 | 9.089 | 8.946 | 6.435 | 7.444 | 11.219 | 7.260 | 7.720 | 7.532 | 8.254 | 8.511 | 7.570 | 7.272 | 8.280 | 6.162 | 8.189 | 7.873 | 0.077 | 0.3618 | 0.6137 | 0 | 0 | 0 |
| 'Slc16a10' | 7.386 | 8.785 | 6.888 | 7.586 | 9.669 | 8.244 | 8.126 | 8.966 | 7.487 | 9.064 | 9.721 | 8.129 | 9.156 | 10.231 | 8.200 | 8.721 | 9.368 | 7.787 | -0.148 | 0.0002 | 0.0025 | 1 | 0 | 0 |
| 'Slc16a13' | 1.523 | 1.826 | 1.665 | 1.694 | 1.845 | 1.637 | 1.411 | 1.518 | 1.955 | 1.586 | 1.470 | 1.626 | 1.743 | 1.293 | 2.006 | 2.095 | 1.415 | 1.485 | 0.028 | 0.7760 | 0.9000 | 0 | 0 | 0 |
| 'Slc16a3' | 4.233 | 3.765 | 3.485 | 4.407 | 3.363 | 3.808 | 3.831 | 3.438 | 3.224 | 2.566 | 5.825 | 5.219 | 5.081 | 5.266 | 6.727 | 4.205 | 5.157 | 5.839 | -0.448 | 0.0014 | 0.0135 | 1 | 1 | 0 |
| 'Slc16a5' | 4.859 | 5.146 | 6.274 | 5.453 | 4.597 | 5.670 | 5.356 | 4.640 | 5.946 | 3.851 | 3.663 | 4.492 | 3.938 | 4.131 | 4.250 | 3.853 | 3.250 | 4.068 | 0.421 | 0.0000 | 0.0000 | 1 | 1 | 0 |
| 'Slc16a6' | 3.496 | 4.390 | 3.392 | 3.862 | 4.348 | 4.027 | 3.283 | 4.271 | 3.890 | 2.561 | 2.262 | 2.472 | 3.643 | 2.807 | 2.814 | 2.570 | 2.477 | 2.585 | 0.515 | 0.0000 | 0.0000 | 1 | 1 | 0 |
| 'Slc17a5' | 2.354 | 3.673 | 3.196 | 2.670 | 3.358 | 3.284 | 2.740 | 3.111 | 3.526 | 3.118 | 3.305 | 3.103 | 2.951 | 3.275 | 3.394 | 3.229 | 3.407 | 2.635 | -0.039 | 0.5449 | 0.7606 | 0 | 0 | 0 |
| 'Slc17a9' | 3.737 | 3.445 | 3.983 | 2.798 | 3.369 | 3.782 | 3.443 | 3.485 | 3.894 | 3.204 | 2.673 | 4.273 | 3.672 | 3.376 | 3.994 | 3.862 | 3.216 | 3.813 | -0.017 | 0.7852 | 0.9042 | 0 | 0 | 0 |
| 'Slc19a1' | 3.603 | 2.941 | 3.846 | 3.360 | 3.480 | 3.213 | 3.608 | 3.714 | 3.642 | 3.031 | 3.238 | 3.672 | 2.796 | 2.869 | 3.659 | 2.922 | 3.191 | 3.673 | 0.105 | 0.0712 | 0.2354 | 0 | 0 | 0 |
| 'Slc19a2' | 1.889 | 1.473 | 1.237 | 1.523 | 1.178 | 1.248 | 1.185 | 1.629 | 1.578 | 1.450 | 1.456 | 1.678 | 1.612 | 1.754 | 1.538 | 1.512 | 1.549 | 1.745 | -0.155 | 0.0860 | 0.2670 | 0 | 0 | 0 |
| 'Slc1a4' | 3.989 | 6.771 | 4.236 | 4.465 | 6.864 | 4.539 | 4.936 | 6.204 | 5.381 | 7.960 | 8.863 | 6.627 | 5.836 | 7.813 | 6.430 | 7.570 | 8.325 | 6.894 | -0.507 | 0.0000 | 0.0000 | 1 | 1 | 0 |
| 'Slc1a5' | 23.443 | 20.840 | 23.124 | 22.193 | 21.155 | 21.671 | 21.540 | 20.194 | 22.876 | 21.421 | 21.391 | 26.421 | 20.329 | 20.447 | 24.753 | 20.933 | 21.771 | 27.239 | -0.061 | 0.1559 | 0.3849 | 0 | 0 | 0 |
| 'Slc20a1' | 12.275 | 9.724 | 11.056 | 10.684 | 9.428 | 9.892 | 9.994 | 9.693 | 10.700 | 11.008 | 9.690 | 10.498 | 10.738 | 9.214 | 9.906 | 10.394 | 10.143 | 10.163 | 0.016 | 0.6682 | 0.8400 | 0 | 0 | 0 |
| 'Slc20a2' | 5.244 | 6.300 | 6.298 | 5.869 | 6.979 | 7.088 | 5.103 | 6.338 | 6.592 | 6.995 | 6.408 | 5.324 | 6.491 | 5.860 | 5.746 | 7.117 | 5.596 | 5.526 | 0.016 | 0.8315 | 0.9267 | 0 | 0 | 0 |
| 'Slc22a15' | 8.108 | 9.068 | 7.813 | 7.652 | 9.646 | 7.793 | 8.013 | 9.530 | 8.049 | 8.627 | 8.584 | 7.651 | 8.491 | 8.228 | 7.110 | 8.501 | 9.008 | 7.058 | 0.036 | 0.2665 | 0.5199 | 0 | 0 | 0 |
| 'Slc22a17' | 1.653 | 2.085 | 2.643 | 1.778 | 2.669 | 2.199 | 1.851 | 1.797 | 2.579 | 2.180 | 0.913 | 1.743 | 1.793 | 1.669 | 1.360 | 2.036 | 1.549 | 1.489 | 0.379 | 0.0056 | 0.0411 | 1 | 1 | 0 |
| 'Slc22a5' | 9.920 | 10.563 | 10.936 | 9.239 | 9.831 | 10.378 | 7.637 | 9.765 | 9.962 | 10.617 | 8.224 | 8.885 | 10.224 | 8.592 | 7.569 | 9.618 | 9.652 | 8.111 | 0.111 | 0.1163 | 0.3237 | 0 | 0 | 0 |
| 'Slc23a2' | 7.122 | 7.167 | 7.072 | 7.480 | 7.041 | 6.862 | 7.084 | 7.328 | 6.943 | 6.939 | 6.916 | 6.879 | 6.910 | 7.403 | 6.522 | 6.898 | 7.056 | 6.166 | 0.046 | 0.1137 | 0.3198 | 0 | 0 | 0 |
| 'Slc24a3' | 0.405 | 0.477 | 0.429 | 0.390 | 0.534 | 0.448 | 0.311 | 0.425 | 0.410 | 0.675 | 0.745 | 0.370 | 0.529 | 0.737 | 0.375 | 0.466 | 0.565 | 0.412 | -0.318 | 0.0110 | 0.0670 | 0 | 0 | 0 |
| 'Slc25a1' | 10.855 | 12.479 | 11.648 | 9.897 | 12.539 | 12.085 | 11.391 | 12.875 | 12.356 | 11.548 | 10.727 | 11.346 | 10.735 | 11.689 | 11.720 | 12.457 | 9.635 | 10.705 | 0.066 | 0.1792 | 0.4145 | 0 | 0 | 0 |
| 'Slc25a10' | 3.767 | 4.055 | 4.354 | 3.817 | 3.968 | 4.535 | 3.935 | 3.641 | 4.221 | 3.783 | 3.730 | 4.264 | 4.269 | 4.090 | 4.444 | 4.250 | 3.687 | 4.203 | -0.026 | 0.5695 | 0.7769 | 0 | 0 | 0 |
| 'Slc25a11' | 31.484 | 38.228 | 35.424 | 34.816 | 39.629 | 38.538 | 34.823 | 37.742 | 36.808 | 35.539 | 33.585 | 33.998 | 37.212 | 37.370 | 35.670 | 34.854 | 35.133 | 32.495 | 0.040 | 0.2605 | 0.5141 | 0 | 0 | 0 |
| 'Slc25a12' | 9.932 | 10.410 | 10.042 | 10.442 | 10.777 | 10.052 | 10.415 | 10.722 | 10.085 | 10.437 | 9.991 | 10.191 | 10.201 | 9.478 | 9.796 | 10.092 | 10.054 | 9.781 | 0.032 | 0.1885 | 0.4247 | 0 | 0 | 0 |
| 'Slc25a13' | 3.642 | 4.177 | 3.772 | 3.881 | 4.205 | 3.521 | 3.977 | 3.921 | 3.534 | 3.292 | 3.226 | 3.365 | 3.584 | 3.209 | 3.396 | 2.908 | 3.641 | 3.527 | 0.178 | 0.0005 | 0.0062 | 1 | 0 | 0 |
| 'Slc25a14' | 1.410 | 1.922 | 1.738 | 1.554 | 2.123 | 1.922 | 1.589 | 2.032 | 1.764 | 1.701 | 1.588 | 1.740 | 1.903 | 2.008 | 1.981 | 1.834 | 1.918 | 1.654 | -0.038 | 0.6054 | 0.8013 | 0 | 0 | 0 |
| 'Slc25a15' | 3.058 | 3.023 | 3.216 | 2.858 | 3.929 | 3.121 | 3.042 | 3.468 | 2.930 | 3.298 | 2.783 | 3.129 | 3.194 | 3.215 | 3.196 | 3.752 | 2.888 | 2.846 | 0.009 | 0.8969 | 0.9586 | 0 | 0 | 0 |
| 'Slc25a16' | 3.759 | 4.975 | 4.305 | 4.123 | 4.890 | 4.414 | 4.441 | 4.976 | 4.503 | 4.603 | 4.313 | 3.738 | 4.806 | 4.647 | 4.007 | 4.993 | 4.309 | 3.193 | 0.063 | 0.3236 | 0.5811 | 0 | 0 | 0 |
| 'Slc25a17' | 17.816 | 19.136 | 19.172 | 18.199 | 21.100 | 20.179 | 19.351 | 21.877 | 19.062 | 20.640 | 19.026 | 19.312 | 21.305 | 21.724 | 19.844 | 19.404 | 21.152 | 18.830 | -0.053 | 0.1668 | 0.3992 | 0 | 0 | 0 |
| 'Slc25a19' | 7.582 | 7.316 | 7.342 | 6.991 | 7.334 | 6.756 | 7.046 | 7.729 | 7.158 | 6.514 | 7.285 | 7.829 | 7.049 | 8.193 | 7.852 | 6.967 | 8.276 | 8.026 | -0.069 | 0.1012 | 0.2979 | 0 | 0 | 0 |
| 'Slc25a20' | 10.414 | 10.455 | 9.082 | 10.183 | 10.688 | 9.263 | 10.400 | 11.102 | 9.319 | 10.157 | 9.785 | 10.452 | 10.602 | 11.023 | 10.574 | 11.601 | 10.227 | 10.406 | -0.076 | 0.0514 | 0.1917 | 0 | 0 | 0 |
| 'Slc25a22' | 3.335 | 3.039 | 3.695 | 2.892 | 2.948 | 3.512 | 3.257 | 2.779 | 3.514 | 2.940 | 2.864 | 3.774 | 3.344 | 2.902 | 3.499 | 3.110 | 3.022 | 3.523 | -0.013 | 0.7169 | 0.8692 | 0 | 0 | 0 |
| 'Slc25a23' | 5.167 | 5.467 | 5.737 | 4.795 | 5.101 | 5.924 | 4.972 | 5.181 | 5.605 | 5.364 | 4.140 | 5.532 | 4.863 | 4.235 | 5.639 | 4.902 | 3.873 | 5.433 | 0.117 | 0.0133 | 0.0770 | 0 | 0 | 0 |
| 'Slc25a24' | 7.418 | 8.476 | 8.346 | 7.599 | 8.708 | 8.907 | 8.124 | 8.648 | 7.861 | 7.479 | 7.911 | 7.516 | 7.778 | 8.124 | 8.028 | 7.408 | 7.740 | 7.025 | 0.095 | 0.0221 | 0.1103 | 0 | 0 | 0 |
| 'Slc25a25' | 2.823 | 2.425 | 1.924 | 2.568 | 2.273 | 2.281 | 2.443 | 2.399 | 2.019 | 2.614 | 2.474 | 2.171 | 2.835 | 2.588 | 2.244 | 2.893 | 2.213 | 2.103 | -0.073 | 0.1050 | 0.3038 | 0 | 0 | 0 |
| 'Slc25a26' | 0.900 | 0.918 | 0.886 | 1.079 | 1.025 | 0.844 | 1.151 | 1.003 | 2.700 | 1.053 | 1.105 | 1.075 | 1.015 | 0.905 | 0.955 | 0.887 | 1.176 | 1.010 | 0.163 | 0.4195 | 0.6664 | 0 | 0 | 0 |
| 'Slc25a28' | 22.636 | 19.225 | 23.336 | 18.569 | 21.086 | 22.904 | 19.386 | 20.441 | 23.647 | 19.928 | 19.137 | 21.097 | 21.699 | 18.104 | 20.640 | 19.431 | 20.946 | 19.186 | 0.079 | 0.1031 | 0.3009 | 0 | 0 | 0 |
| 'Slc25a29' | 1.932 | 1.905 | 2.385 | 1.581 | 1.876 | 2.529 | 1.670 | 2.081 | 1.897 | 1.866 | 2.402 | 2.254 | 2.311 | 1.899 | 2.343 | 2.314 | 2.495 | 1.724 | -0.135 | 0.1656 | 0.3980 | 0 | 0 | 0 |
| 'Slc25a3' | 60.346 | 54.185 | 47.628 | 52.960 | 52.584 | 47.378 | 52.290 | 54.617 | 47.809 | 58.466 | 53.009 | 43.430 | 58.473 | 55.147 | 45.191 | 56.958 | 51.526 | 43.783 | 0.008 | 0.8023 | 0.9114 | 0 | 0 | 0 |
| 'Slc25a30' | 10.639 | 10.209 | 11.454 | 9.879 | 9.840 | 9.893 | 10.556 | 10.808 | 10.198 | 10.303 | 12.460 | 10.471 | 11.580 | 11.458 | 11.516 | 9.475 | 13.441 | 11.055 | -0.125 | 0.0367 | 0.1547 | 0 | 0 | 0 |
| 'Slc25a32' | 10.200 | 11.429 | 11.713 | 10.946 | 11.400 | 10.366 | 11.683 | 11.515 | 10.600 | 10.199 | 10.426 | 11.788 | 10.507 | 11.453 | 10.397 | 9.937 | 12.117 | 11.140 | 0.011 | 0.8039 | 0.9119 | 0 | 0 | 0 |
| 'Slc25a33' | 3.411 | 2.952 | 2.872 | 2.925 | 3.384 | 3.297 | 2.315 | 3.571 | 3.029 | 3.640 | 2.921 | 3.102 | 2.632 | 2.843 | 3.075 | 3.735 | 2.548 | 2.713 | 0.029 | 0.7686 | 0.8967 | 0 | 0 | 0 |
| 'Slc25a35' | 1.034 | 1.093 | 1.337 | 1.025 | 1.182 | 1.321 | 1.289 | 1.118 | 1.665 | 1.053 | 1.471 | 1.106 | 1.094 | 0.887 | 1.405 | 1.002 | 0.805 | 1.074 | 0.153 | 0.1410 | 0.3636 | 0 | 0 | 0 |
| 'Slc25a36' | 7.393 | 9.728 | 8.255 | 7.694 | 9.442 | 9.041 | 8.145 | 8.902 | 8.665 | 7.801 | 7.935 | 8.230 | 6.737 | 8.647 | 7.271 | 8.825 | 8.323 | 7.322 | 0.107 | 0.0202 | 0.1038 | 0 | 0 | 0 |
| 'Slc25a37' | 6.207 | 6.276 | 6.128 | 5.678 | 6.170 | 5.871 | 5.527 | 6.200 | 6.213 | 5.601 | 6.196 | 6.725 | 6.063 | 6.643 | 6.294 | 6.129 | 6.176 | 6.572 | -0.067 | 0.0407 | 0.1651 | 0 | 0 | 0 |
| 'Slc25a38' | 6.824 | 6.625 | 7.603 | 6.415 | 6.328 | 7.453 | 6.760 | 6.908 | 7.037 | 6.715 | 6.155 | 7.536 | 6.570 | 6.467 | 7.156 | 7.142 | 7.066 | 7.546 | -0.017 | 0.6621 | 0.8370 | 0 | 0 | 0 |
| 'Slc25a39' | 10.855 | 13.546 | 11.451 | 11.063 | 11.913 | 12.852 | 11.550 | 12.363 | 12.797 | 11.084 | 10.383 | 11.754 | 12.293 | 11.501 | 12.283 | 12.361 | 10.796 | 12.120 | 0.039 | 0.3611 | 0.6135 | 0 | 0 | 0 |
| 'Slc25a4' | 40.936 | 45.332 | 42.029 | 42.962 | 47.797 | 41.618 | 44.613 | 46.451 | 41.592 | 39.903 | 40.768 | 38.058 | 43.077 | 42.691 | 40.498 | 44.530 | 41.448 | 36.635 | 0.088 | 0.0025 | 0.0215 | 1 | 0 | 0 |
| 'Slc25a40' | 1.443 | 1.889 | 1.911 | 1.665 | 1.759 | 1.661 | 2.008 | 1.776 | 1.954 | 1.661 | 1.827 | 1.716 | 1.642 | 1.845 | 1.783 | 1.514 | 1.842 | 1.545 | 0.053 | 0.3526 | 0.6063 | 0 | 0 | 0 |
| 'Slc25a42' | 4.407 | 5.555 | 4.940 | 4.448 | 5.173 | 5.162 | 4.465 | 4.651 | 4.821 | 4.580 | 4.903 | 6.751 | 4.425 | 5.490 | 5.856 | 5.267 | 5.501 | 6.269 | -0.179 | 0.0007 | 0.0075 | 1 | 0 | 0 |
| 'Slc25a44' | 4.548 | 4.324 | 4.623 | 4.159 | 4.385 | 4.546 | 4.055 | 4.681 | 4.304 | 4.487 | 4.426 | 4.303 | 4.689 | 4.418 | 4.643 | 4.759 | 4.106 | 4.567 | -0.032 | 0.4174 | 0.6650 | 0 | 0 | 0 |
| 'Slc25a45' | 15.785 | 16.217 | 17.718 | 15.362 | 16.341 | 18.723 | 14.672 | 16.771 | 17.273 | 16.092 | 16.274 | 18.958 | 14.966 | 16.832 | 18.278 | 15.446 | 17.239 | 17.989 | -0.040 | 0.1782 | 0.4135 | 0 | 0 | 0 |
| 'Slc25a46' | 10.202 | 12.995 | 11.361 | 10.877 | 12.868 | 11.544 | 10.940 | 11.767 | 11.079 | 10.770 | 10.501 | 12.006 | 9.975 | 11.970 | 11.762 | 11.217 | 11.433 | 11.390 | 0.020 | 0.6086 | 0.8031 | 0 | 0 | 0 |
| 'Slc25a5' | 155.850 | 148.713 | 148.077 | 161.197 | 154.837 | 144.744 | 161.635 | 149.529 | 141.289 | 138.389 | 150.244 | 135.565 | 155.423 | 154.337 | 151.510 | 148.442 | 154.071 | 142.079 | 0.027 | 0.3824 | 0.6330 | 0 | 0 | 0 |
| 'Slc25a51' | 15.306 | 16.416 | 14.605 | 13.909 | 14.552 | 14.447 | 13.693 | 15.494 | 14.793 | 13.086 | 14.068 | 15.909 | 15.625 | 15.402 | 14.550 | 14.805 | 14.930 | 15.311 | -0.020 | 0.6359 | 0.8201 | 0 | 0 | 0 |
| 'Slc25a53' | 0.577 | 0.640 | 0.603 | 0.603 | 0.684 | 0.710 | 0.683 | 0.611 | 0.551 | 0.520 | 0.466 | 0.519 | 0.530 | 0.522 | 0.449 | 0.563 | 0.512 | 0.510 | 0.288 | 0.0000 | 0.0002 | 1 | 0 | 0 |
| 'Slc26a10' | 2.178 | 1.762 | 1.873 | 1.953 | 2.051 | 1.811 | 2.190 | 1.849 | 1.924 | 1.925 | 1.511 | 1.813 | 1.604 | 1.771 | 1.712 | 2.158 | 1.366 | 1.706 | 0.165 | 0.0930 | 0.2822 | 0 | 0 | 0 |
| 'Slc26a11' | 5.543 | 5.112 | 6.027 | 5.812 | 5.193 | 5.791 | 5.585 | 5.473 | 5.580 | 5.684 | 6.347 | 6.945 | 6.366 | 6.094 | 6.124 | 5.861 | 6.456 | 6.360 | -0.178 | 0.0000 | 0.0003 | 1 | 0 | 0 |
| 'Slc26a2' | 3.771 | 4.411 | 4.639 | 3.964 | 4.399 | 4.113 | 4.156 | 4.507 | 4.509 | 4.508 | 5.039 | 4.279 | 4.133 | 4.408 | 4.135 | 4.395 | 4.643 | 3.904 | -0.043 | 0.2978 | 0.5536 | 0 | 0 | 0 |
| 'Slc26a6' | 1.061 | 0.860 | 1.163 | 1.109 | 0.948 | 1.119 | 1.217 | 0.995 | 1.256 | 0.977 | 1.104 | 1.320 | 0.910 | 1.001 | 1.232 | 0.895 | 1.017 | 1.057 | 0.019 | 0.7900 | 0.9061 | 0 | 0 | 0 |
| 'Slc27a1' | 6.862 | 9.060 | 9.930 | 8.025 | 7.772 | 8.897 | 7.069 | 8.636 | 8.713 | 8.427 | 8.529 | 8.461 | 8.409 | 7.488 | 8.277 | 7.998 | 7.226 | 7.677 | 0.038 | 0.5226 | 0.7449 | 0 | 0 | 0 |
| 'Slc27a4' | 8.390 | 9.725 | 9.219 | 7.997 | 9.178 | 10.167 | 7.910 | 8.816 | 9.679 | 8.968 | 7.583 | 9.255 | 8.352 | 9.124 | 9.153 | 9.272 | 7.793 | 9.298 | 0.030 | 0.5185 | 0.7413 | 0 | 0 | 0 |
| 'Slc29a1' | 17.500 | 16.011 | 19.207 | 16.634 | 16.498 | 18.916 | 15.454 | 15.976 | 18.584 | 18.587 | 16.321 | 21.559 | 17.260 | 17.741 | 20.695 | 19.456 | 19.061 | 20.786 | -0.157 | 0.0000 | 0.0000 | 1 | 0 | 0 |
| 'Slc29a3' | 3.533 | 3.404 | 3.848 | 2.747 | 3.257 | 3.789 | 2.706 | 3.655 | 3.663 | 3.873 | 3.161 | 3.620 | 3.469 | 3.321 | 3.187 | 3.354 | 2.574 | 3.595 | 0.014 | 0.8436 | 0.9337 | 0 | 0 | 0 |
| 'Slc2a1' | 18.248 | 18.311 | 15.042 | 19.702 | 15.531 | 15.255 | 17.147 | 15.344 | 15.254 | 14.148 | 25.667 | 17.227 | 19.257 | 23.125 | 21.242 | 16.599 | 25.490 | 17.791 | -0.264 | 0.0126 | 0.0736 | 0 | 0 | 0 |
| 'Slc2a3' | 6.853 | 6.554 | 6.183 | 7.060 | 5.923 | 5.995 | 6.814 | 5.616 | 5.979 | 6.197 | 6.994 | 6.352 | 6.800 | 6.199 | 6.854 | 5.969 | 5.430 | 5.944 | -0.008 | 0.8716 | 0.9450 | 0 | 0 | 0 |
| 'Slc2a6' | 6.282 | 8.292 | 7.918 | 6.658 | 7.278 | 7.884 | 7.426 | 6.385 | 8.560 | 7.631 | 6.374 | 6.268 | 7.813 | 6.799 | 6.122 | 7.537 | 6.836 | 5.713 | 0.120 | 0.0831 | 0.2611 | 0 | 0 | 0 |
| 'Slc2a8' | 1.975 | 2.158 | 2.544 | 1.727 | 2.136 | 2.769 | 1.972 | 2.374 | 2.655 | 1.759 | 1.937 | 2.371 | 2.079 | 1.984 | 1.971 | 2.001 | 1.565 | 2.096 | 0.179 | 0.0063 | 0.0443 | 1 | 0 | 0 |
| 'Slc2a9' | 3.322 | 3.661 | 3.299 | 2.928 | 3.146 | 3.177 | 2.982 | 3.482 | 3.103 | 3.219 | 3.211 | 3.190 | 3.094 | 3.537 | 3.211 | 3.339 | 2.866 | 3.416 | -0.009 | 0.8393 | 0.9315 | 0 | 0 | 0 |
| 'Slc30a1' | 8.221 | 6.244 | 5.838 | 6.566 | 6.316 | 5.597 | 7.782 | 6.614 | 5.264 | 4.734 | 5.760 | 5.760 | 4.832 | 5.141 | 5.407 | 5.126 | 5.500 | 5.570 | 0.268 | 0.0003 | 0.0037 | 1 | 0 | 0 |
| 'Slc30a4' | 0.744 | 1.006 | 0.769 | 0.821 | 1.030 | 0.746 | 0.796 | 1.023 | 0.693 | 0.941 | 0.818 | 0.861 | 0.686 | 0.952 | 0.803 | 1.061 | 1.368 | 0.858 | -0.146 | 0.1295 | 0.3450 | 0 | 0 | 0 |
| 'Slc30a5' | 18.265 | 21.823 | 21.152 | 20.356 | 21.129 | 19.267 | 20.930 | 22.207 | 20.535 | 19.199 | 19.125 | 21.237 | 19.369 | 19.401 | 19.503 | 18.297 | 19.760 | 19.135 | 0.068 | 0.0475 | 0.1822 | 0 | 0 | 0 |
| 'Slc30a6' | 7.517 | 8.592 | 8.363 | 7.271 | 9.172 | 7.931 | 7.913 | 8.484 | 8.086 | 7.286 | 9.151 | 7.985 | 7.367 | 7.753 | 7.986 | 8.322 | 7.665 | 7.356 | 0.040 | 0.4446 | 0.6875 | 0 | 0 | 0 |
| 'Slc30a7' | 3.521 | 3.676 | 3.672 | 3.465 | 3.315 | 3.566 | 3.657 | 3.385 | 3.625 | 2.872 | 3.888 | 4.098 | 3.188 | 3.414 | 3.652 | 3.030 | 3.647 | 3.773 | 0.005 | 0.9180 | 0.9687 | 0 | 0 | 0 |
| 'Slc30a9' | 4.112 | 4.145 | 3.756 | 4.160 | 4.383 | 3.768 | 4.224 | 4.224 | 3.886 | 3.754 | 4.320 | 4.774 | 3.955 | 4.718 | 4.559 | 3.906 | 4.848 | 4.700 | -0.119 | 0.0126 | 0.0738 | 0 | 0 | 0 |
| 'Slc31a1' | 1.863 | 2.437 | 2.272 | 1.934 | 2.134 | 1.920 | 2.023 | 2.196 | 2.183 | 1.842 | 2.248 | 2.116 | 2.171 | 2.133 | 1.990 | 2.142 | 1.965 | 1.879 | 0.024 | 0.6729 | 0.8425 | 0 | 0 | 0 |
| 'Slc31a2' | 4.188 | 5.344 | 5.171 | 3.803 | 3.581 | 4.127 | 3.735 | 4.243 | 4.794 | 3.638 | 4.115 | 5.334 | 4.134 | 4.746 | 5.768 | 3.734 | 4.584 | 5.491 | -0.104 | 0.2166 | 0.4628 | 0 | 0 | 0 |
| 'Slc33a1' | 4.645 | 5.910 | 5.420 | 4.731 | 5.498 | 6.058 | 5.886 | 5.554 | 5.870 | 4.573 | 4.840 | 5.681 | 5.109 | 5.099 | 5.122 | 5.164 | 5.296 | 5.141 | 0.091 | 0.0548 | 0.2005 | 0 | 0 | 0 |
| 'Slc35a1' | 7.605 | 8.749 | 7.819 | 8.082 | 8.939 | 7.555 | 7.412 | 8.809 | 8.588 | 7.853 | 6.967 | 7.872 | 6.945 | 8.480 | 7.673 | 7.002 | 7.333 | 7.992 | 0.095 | 0.0421 | 0.1692 | 0 | 0 | 0 |
| 'Slc35a2' | 3.617 | 4.096 | 4.026 | 3.361 | 4.155 | 4.387 | 3.595 | 4.113 | 4.066 | 3.816 | 3.847 | 4.468 | 3.814 | 4.072 | 4.526 | 3.968 | 3.813 | 4.298 | -0.061 | 0.1437 | 0.3678 | 0 | 0 | 0 |
| 'Slc35a3' | 2.771 | 2.984 | 3.247 | 2.762 | 3.249 | 3.077 | 3.097 | 3.413 | 2.672 | 2.780 | 3.162 | 3.052 | 3.072 | 3.098 | 2.918 | 3.220 | 3.226 | 3.020 | -0.023 | 0.6365 | 0.8206 | 0 | 0 | 0 |
| 'Slc35a4' | 10.118 | 11.394 | 11.220 | 9.891 | 11.367 | 11.659 | 10.594 | 11.233 | 11.836 | 10.821 | 10.780 | 10.809 | 10.640 | 11.087 | 10.714 | 11.081 | 10.557 | 10.729 | 0.022 | 0.4753 | 0.7100 | 0 | 0 | 0 |
| 'Slc35a5' | 1.708 | 2.137 | 1.764 | 1.936 | 2.034 | 1.687 | 2.322 | 1.890 | 1.956 | 1.646 | 2.151 | 2.167 | 1.718 | 1.996 | 2.052 | 1.788 | 2.118 | 2.097 | -0.041 | 0.5374 | 0.7562 | 0 | 0 | 0 |
| 'Slc35b1' | 10.106 | 11.138 | 11.536 | 9.854 | 12.222 | 12.353 | 10.649 | 11.997 | 11.701 | 11.924 | 10.123 | 10.210 | 12.005 | 10.687 | 10.887 | 10.770 | 11.085 | 10.347 | 0.042 | 0.3953 | 0.6444 | 0 | 0 | 0 |
| 'Slc35b2' | 26.520 | 27.320 | 25.313 | 25.244 | 25.190 | 26.044 | 27.708 | 26.253 | 23.330 | 23.217 | 24.446 | 27.676 | 26.953 | 26.528 | 26.338 | 28.768 | 24.514 | 25.227 | -0.018 | 0.6642 | 0.8377 | 0 | 0 | 0 |
| 'Slc35b3' | 19.184 | 23.035 | 21.764 | 21.858 | 23.518 | 22.435 | 22.989 | 24.140 | 22.952 | 21.090 | 22.169 | 22.830 | 22.074 | 23.731 | 20.386 | 24.164 | 22.901 | 19.876 | 0.008 | 0.8408 | 0.9323 | 0 | 0 | 0 |
| 'Slc35b4' | 4.004 | 3.971 | 4.399 | 3.861 | 4.265 | 4.460 | 3.861 | 4.709 | 4.154 | 4.278 | 4.306 | 4.562 | 4.267 | 4.504 | 4.342 | 4.566 | 4.128 | 4.502 | -0.074 | 0.0445 | 0.1750 | 0 | 0 | 0 |
| 'Slc35c1' | 10.661 | 9.322 | 9.783 | 9.950 | 9.755 | 10.401 | 10.229 | 9.382 | 10.382 | 8.989 | 10.055 | 12.357 | 8.909 | 8.949 | 12.200 | 9.652 | 9.586 | 12.782 | -0.059 | 0.2835 | 0.5397 | 0 | 0 | 0 |
| 'Slc35c2' | 6.921 | 6.872 | 7.531 | 6.536 | 7.370 | 7.937 | 6.085 | 7.660 | 7.258 | 7.306 | 6.127 | 7.933 | 7.066 | 6.858 | 7.347 | 7.768 | 6.198 | 7.990 | -0.018 | 0.7065 | 0.8639 | 0 | 0 | 0 |
| 'Slc35d1' | 2.116 | 1.975 | 2.068 | 1.876 | 1.949 | 1.937 | 1.868 | 1.937 | 1.757 | 1.862 | 1.851 | 2.107 | 1.922 | 2.014 | 1.954 | 2.002 | 2.121 | 2.185 | -0.055 | 0.2699 | 0.5246 | 0 | 0 | 0 |
| 'Slc35d2' | 7.869 | 10.336 | 9.909 | 7.498 | 10.145 | 9.866 | 9.255 | 10.110 | 9.296 | 8.991 | 9.612 | 8.709 | 10.401 | 10.263 | 8.375 | 9.522 | 10.132 | 7.365 | 0.009 | 0.8813 | 0.9499 | 0 | 0 | 0 |
| 'Slc35e1' | 21.812 | 19.958 | 21.966 | 21.401 | 20.952 | 21.776 | 19.815 | 19.975 | 19.001 | 20.965 | 22.615 | 23.546 | 21.395 | 23.968 | 24.684 | 21.733 | 20.195 | 25.378 | -0.140 | 0.0014 | 0.0137 | 1 | 0 | 0 |
| 'Slc35e2' | 5.633 | 5.186 | 5.813 | 5.577 | 5.342 | 5.224 | 6.355 | 4.906 | 5.696 | 5.137 | 6.355 | 7.052 | 5.317 | 5.435 | 6.286 | 4.995 | 6.335 | 7.112 | -0.127 | 0.0494 | 0.1875 | 0 | 0 | 0 |
| 'Slc35e3' | 1.639 | 2.155 | 1.807 | 1.668 | 2.066 | 1.882 | 1.645 | 2.207 | 1.818 | 1.778 | 1.669 | 1.790 | 1.645 | 1.734 | 1.586 | 1.532 | 1.924 | 1.804 | 0.107 | 0.0751 | 0.2439 | 0 | 0 | 0 |
| 'Slc35e4' | 1.975 | 1.775 | 1.425 | 1.207 | 1.605 | 1.890 | 1.276 | 1.635 | 1.202 | 2.092 | 1.487 | 1.568 | 2.238 | 1.571 | 1.285 | 1.657 | 1.906 | 1.519 | -0.130 | 0.2535 | 0.5050 | 0 | 0 | 0 |
| 'Slc35f2' | 9.529 | 10.693 | 9.192 | 10.432 | 8.325 | 9.399 | 9.430 | 9.741 | 8.365 | 7.212 | 10.056 | 8.400 | 8.904 | 7.927 | 8.836 | 8.114 | 7.599 | 7.954 | 0.169 | 0.0074 | 0.0496 | 1 | 0 | 0 |
| 'Slc35f6' | 1.406 | 1.814 | 1.723 | 1.359 | 1.837 | 1.868 | 1.630 | 1.766 | 2.052 | 1.951 | 1.656 | 1.869 | 1.682 | 1.657 | 1.936 | 1.666 | 1.673 | 1.774 | -0.053 | 0.3663 | 0.6184 | 0 | 0 | 0 |
| 'Slc35g1' | 5.680 | 5.289 | 5.044 | 5.683 | 5.208 | 4.780 | 5.829 | 5.235 | 4.415 | 3.783 | 5.262 | 4.935 | 4.952 | 4.636 | 4.568 | 4.569 | 4.839 | 5.539 | 0.112 | 0.1101 | 0.3133 | 0 | 0 | 0 |
| 'Slc36a1' | 6.747 | 6.360 | 7.397 | 6.237 | 6.294 | 6.702 | 5.826 | 6.816 | 7.265 | 6.670 | 6.833 | 8.672 | 6.610 | 6.878 | 8.288 | 6.502 | 6.229 | 8.690 | -0.139 | 0.0017 | 0.0158 | 1 | 0 | 0 |
| 'Slc36a4' | 16.536 | 20.479 | 17.376 | 17.313 | 19.755 | 16.740 | 19.154 | 18.321 | 17.547 | 18.704 | 17.150 | 15.109 | 19.455 | 18.262 | 14.713 | 17.369 | 17.864 | 13.931 | 0.092 | 0.0311 | 0.1384 | 0 | 0 | 0 |
| 'Slc37a1' | 19.143 | 16.552 | 18.438 | 15.588 | 15.848 | 18.301 | 16.632 | 16.593 | 16.785 | 17.421 | 16.089 | 19.998 | 15.938 | 15.970 | 17.616 | 17.219 | 16.377 | 19.566 | -0.030 | 0.4823 | 0.7144 | 0 | 0 | 0 |
| 'Slc37a2' | 3.744 | 4.688 | 3.788 | 3.800 | 4.349 | 4.031 | 4.211 | 4.006 | 4.342 | 4.780 | 5.200 | 5.077 | 4.566 | 5.080 | 4.544 | 5.494 | 5.694 | 4.532 | -0.295 | 0.0000 | 0.0000 | 1 | 0 | 0 |
| 'Slc37a3' | 5.200 | 4.840 | 5.453 | 5.213 | 4.850 | 5.124 | 5.356 | 5.021 | 5.573 | 4.473 | 5.174 | 5.674 | 4.674 | 4.742 | 5.404 | 4.171 | 5.382 | 5.582 | 0.034 | 0.4667 | 0.7023 | 0 | 0 | 0 |
| 'Slc37a4' | 11.764 | 13.660 | 12.347 | 12.239 | 13.994 | 13.991 | 13.055 | 13.769 | 13.646 | 12.863 | 13.561 | 13.368 | 14.398 | 14.656 | 13.629 | 12.669 | 14.097 | 11.611 | -0.040 | 0.3438 | 0.5995 | 0 | 0 | 0 |
| 'Slc38a1' | 38.880 | 40.924 | 40.390 | 39.118 | 40.081 | 37.695 | 39.672 | 40.428 | 39.418 | 40.555 | 41.706 | 42.116 | 39.106 | 41.118 | 40.747 | 39.223 | 40.003 | 43.221 | -0.056 | 0.0185 | 0.0977 | 0 | 0 | 0 |
| 'Slc38a10' | 16.904 | 15.673 | 18.550 | 15.589 | 15.837 | 18.655 | 16.069 | 16.268 | 17.649 | 17.913 | 16.808 | 18.936 | 17.140 | 16.685 | 18.208 | 18.177 | 16.226 | 17.869 | -0.073 | 0.0033 | 0.0269 | 1 | 0 | 0 |
| 'Slc38a2' | 60.043 | 42.501 | 32.025 | 52.007 | 38.646 | 31.030 | 43.517 | 42.971 | 32.437 | 45.887 | 43.944 | 35.914 | 44.166 | 47.037 | 32.646 | 44.449 | 44.220 | 36.274 | -0.026 | 0.6684 | 0.8400 | 0 | 0 | 0 |
| 'Slc38a6' | 2.480 | 3.184 | 3.147 | 2.492 | 2.812 | 3.068 | 2.746 | 3.256 | 3.308 | 3.040 | 3.180 | 2.822 | 2.669 | 3.332 | 2.616 | 3.049 | 3.174 | 2.213 | 0.022 | 0.7605 | 0.8935 | 0 | 0 | 0 |
| 'Slc38a7' | 5.728 | 6.516 | 7.121 | 5.330 | 6.038 | 6.468 | 5.550 | 6.472 | 6.970 | 6.906 | 6.574 | 7.002 | 6.430 | 5.648 | 6.489 | 6.125 | 6.098 | 6.850 | -0.059 | 0.2207 | 0.4680 | 0 | 0 | 0 |
| 'Slc38a9' | 4.859 | 5.818 | 5.565 | 4.425 | 5.279 | 4.995 | 5.081 | 5.377 | 5.517 | 5.485 | 5.591 | 5.678 | 5.822 | 5.243 | 5.039 | 5.165 | 5.591 | 5.691 | -0.085 | 0.0500 | 0.1886 | 0 | 0 | 0 |
| 'Slc39a1' | 15.285 | 16.989 | 15.330 | 13.986 | 17.861 | 18.184 | 14.248 | 17.660 | 15.552 | 13.998 | 15.825 | 18.222 | 16.379 | 16.685 | 18.265 | 17.044 | 15.188 | 17.685 | -0.055 | 0.2826 | 0.5390 | 0 | 0 | 0 |
| 'Slc39a10' | 2.822 | 2.773 | 3.428 | 3.236 | 3.579 | 3.112 | 2.803 | 2.955 | 3.246 | 2.870 | 3.321 | 3.686 | 2.753 | 2.738 | 2.898 | 3.102 | 3.236 | 3.810 | -0.035 | 0.5860 | 0.7873 | 0 | 0 | 0 |
| 'Slc39a11' | 9.507 | 9.727 | 9.466 | 9.583 | 10.131 | 9.182 | 8.987 | 10.233 | 9.351 | 9.575 | 9.629 | 9.632 | 9.587 | 10.114 | 9.844 | 10.525 | 8.458 | 9.666 | -0.024 | 0.5541 | 0.7667 | 0 | 0 | 0 |
| 'Slc39a13' | 4.632 | 4.688 | 4.771 | 4.334 | 4.759 | 5.596 | 4.246 | 4.675 | 5.525 | 4.711 | 4.186 | 4.987 | 4.488 | 4.547 | 4.753 | 5.197 | 4.589 | 4.855 | 0.022 | 0.6300 | 0.8169 | 0 | 0 | 0 |
| 'Slc39a14' | 1.923 | 1.747 | 1.948 | 1.724 | 1.774 | 2.011 | 1.840 | 2.138 | 2.071 | 2.046 | 1.932 | 2.014 | 1.821 | 1.793 | 1.723 | 1.780 | 1.823 | 1.777 | 0.033 | 0.5177 | 0.7407 | 0 | 0 | 0 |
| 'Slc39a3' | 2.509 | 3.045 | 3.417 | 2.653 | 3.079 | 3.649 | 2.543 | 3.160 | 3.497 | 3.004 | 2.936 | 3.180 | 3.126 | 2.734 | 2.911 | 3.035 | 2.823 | 3.360 | 0.008 | 0.8738 | 0.9462 | 0 | 0 | 0 |
| 'Slc39a4' | 17.913 | 16.545 | 19.939 | 16.910 | 17.483 | 19.866 | 18.089 | 16.557 | 19.798 | 18.290 | 16.603 | 18.514 | 16.219 | 16.918 | 16.815 | 18.705 | 17.143 | 17.054 | 0.055 | 0.1803 | 0.4156 | 0 | 0 | 0 |
| 'Slc39a6' | 39.264 | 44.854 | 38.287 | 42.680 | 41.979 | 37.628 | 38.837 | 40.730 | 38.804 | 34.207 | 37.757 | 38.106 | 38.294 | 37.888 | 37.985 | 35.125 | 33.802 | 39.496 | 0.113 | 0.0042 | 0.0331 | 1 | 0 | 0 |
| 'Slc39a7' | 21.256 | 20.440 | 22.623 | 20.680 | 20.803 | 22.938 | 19.806 | 19.487 | 22.094 | 20.748 | 18.597 | 22.950 | 20.456 | 18.780 | 20.611 | 20.108 | 19.893 | 22.038 | 0.034 | 0.1911 | 0.4278 | 0 | 0 | 0 |
| 'Slc39a8' | 0.605 | 0.488 | 0.686 | 0.592 | 0.604 | 0.597 | 0.579 | 0.602 | 0.594 | 0.534 | 0.797 | 0.639 | 0.572 | 0.538 | 0.574 | 0.537 | 0.561 | 0.570 | 0.003 | 0.9700 | 0.9898 | 0 | 0 | 0 |
| 'Slc39a9' | 12.414 | 14.499 | 13.857 | 12.559 | 14.109 | 13.290 | 12.080 | 13.481 | 14.300 | 13.297 | 13.752 | 13.906 | 12.382 | 14.425 | 13.210 | 14.525 | 13.294 | 13.323 | -0.028 | 0.4207 | 0.6675 | 0 | 0 | 0 |
| 'Slc3a2' | 52.194 | 42.992 | 30.500 | 36.815 | 39.296 | 32.193 | 36.699 | 42.079 | 31.332 | 51.860 | 38.088 | 29.745 | 51.889 | 48.088 | 30.719 | 53.631 | 36.400 | 28.943 | -0.087 | 0.2927 | 0.5493 | 0 | 0 | 0 |
| 'Slc41a1' | 40.817 | 38.551 | 38.106 | 37.184 | 39.691 | 37.637 | 34.206 | 40.462 | 37.040 | 42.829 | 50.294 | 48.944 | 41.304 | 51.034 | 48.127 | 45.696 | 50.585 | 48.953 | -0.323 | 0.0000 | 0.0000 | 1 | 0 | 0 |
| 'Slc41a3' | 4.283 | 4.177 | 5.245 | 3.539 | 4.190 | 4.670 | 3.791 | 4.445 | 4.402 | 5.182 | 4.841 | 4.316 | 5.136 | 3.989 | 4.736 | 5.500 | 3.711 | 4.110 | -0.096 | 0.2334 | 0.4820 | 0 | 0 | 0 |
| 'Slc43a2' | 12.460 | 11.820 | 11.341 | 12.560 | 12.128 | 11.474 | 12.127 | 11.551 | 11.855 | 9.968 | 12.291 | 12.466 | 11.780 | 12.324 | 12.773 | 10.850 | 12.532 | 13.427 | -0.023 | 0.6189 | 0.8093 | 0 | 0 | 0 |
| 'Slc43a3' | 2.992 | 2.994 | 3.608 | 3.586 | 3.052 | 3.311 | 3.537 | 2.806 | 3.544 | 2.702 | 3.159 | 3.174 | 4.380 | 2.871 | 3.302 | 2.912 | 2.755 | 2.971 | 0.047 | 0.5289 | 0.7495 | 0 | 0 | 0 |
| 'Slc44a1' | 2.129 | 2.214 | 2.431 | 2.358 | 2.095 | 2.099 | 2.771 | 2.660 | 2.022 | 2.454 | 2.433 | 2.450 | 2.499 | 2.688 | 2.370 | 2.726 | 2.797 | 2.400 | -0.144 | 0.0196 | 0.1017 | 0 | 0 | 0 |
| 'Slc44a2' | 74.377 | 70.453 | 79.921 | 73.919 | 72.112 | 76.516 | 70.436 | 71.305 | 76.688 | 74.430 | 71.314 | 79.029 | 70.866 | 72.944 | 78.128 | 70.784 | 72.786 | 79.031 | -0.018 | 0.3960 | 0.6448 | 0 | 0 | 0 |
| 'Slc45a4' | 8.808 | 8.872 | 10.121 | 7.929 | 9.262 | 9.925 | 6.921 | 9.502 | 9.257 | 9.638 | 9.674 | 10.361 | 9.001 | 9.542 | 10.250 | 9.956 | 8.560 | 10.984 | -0.139 | 0.0033 | 0.0269 | 1 | 0 | 0 |
| 'Slc46a1' | 1.691 | 1.428 | 1.816 | 1.936 | 2.234 | 1.894 | 1.260 | 1.939 | 1.900 | 1.668 | 1.433 | 1.466 | 1.771 | 1.627 | 1.587 | 2.032 | 1.314 | 1.535 | 0.159 | 0.1099 | 0.3127 | 0 | 0 | 0 |
| 'Slc46a3' | 3.467 | 5.044 | 4.361 | 3.885 | 5.616 | 5.159 | 4.114 | 4.147 | 4.633 | 4.178 | 4.813 | 3.916 | 4.382 | 4.443 | 4.211 | 4.631 | 4.151 | 3.901 | 0.053 | 0.4604 | 0.6970 | 0 | 0 | 0 |
| 'Slc48a1' | 16.884 | 19.267 | 17.777 | 16.196 | 17.935 | 19.985 | 16.265 | 19.970 | 18.929 | 16.630 | 16.710 | 17.541 | 16.409 | 17.697 | 18.405 | 17.850 | 15.244 | 16.828 | 0.080 | 0.0576 | 0.2065 | 0 | 0 | 0 |
| 'Slc4a1ap' | 3.678 | 3.662 | 3.857 | 4.092 | 3.690 | 3.812 | 3.718 | 3.775 | 3.455 | 3.587 | 3.793 | 4.217 | 3.539 | 3.724 | 3.673 | 3.793 | 4.253 | 3.754 | -0.038 | 0.3863 | 0.6356 | 0 | 0 | 0 |
| 'Slc4a2' | 10.194 | 9.657 | 10.673 | 9.655 | 9.703 | 10.607 | 10.053 | 9.491 | 10.705 | 9.878 | 10.125 | 11.223 | 10.584 | 9.992 | 10.783 | 9.405 | 10.003 | 10.704 | -0.043 | 0.0936 | 0.2833 | 0 | 0 | 0 |
| 'Slc4a7' | 10.550 | 10.822 | 11.555 | 10.780 | 11.275 | 10.714 | 12.044 | 11.247 | 10.807 | 10.038 | 10.146 | 11.329 | 9.087 | 9.908 | 10.024 | 9.956 | 10.728 | 9.682 | 0.122 | 0.0013 | 0.0131 | 1 | 0 | 0 |
| 'Slc4a8' | 8.620 | 8.329 | 9.614 | 8.366 | 7.187 | 8.622 | 8.018 | 8.355 | 9.445 | 7.885 | 8.936 | 8.376 | 8.960 | 8.316 | 8.422 | 8.290 | 8.089 | 7.954 | 0.018 | 0.7027 | 0.8611 | 0 | 0 | 0 |
| 'Slc50a1' | 17.606 | 18.887 | 19.748 | 17.924 | 19.758 | 20.540 | 18.283 | 19.597 | 19.787 | 19.534 | 17.140 | 19.091 | 19.847 | 17.976 | 18.678 | 19.070 | 18.535 | 17.605 | 0.030 | 0.3733 | 0.6243 | 0 | 0 | 0 |
| 'Slc52a2' | 14.669 | 16.417 | 16.173 | 14.746 | 15.677 | 17.750 | 14.077 | 16.254 | 17.284 | 14.442 | 14.463 | 16.119 | 13.865 | 14.915 | 15.829 | 14.896 | 13.785 | 15.363 | 0.088 | 0.0153 | 0.0849 | 0 | 0 | 0 |
| 'Slc5a3' | 7.630 | 5.646 | 6.231 | 7.931 | 4.754 | 5.381 | 6.272 | 5.621 | 5.839 | 6.078 | 8.133 | 11.141 | 4.925 | 6.986 | 9.479 | 5.761 | 8.670 | 12.554 | -0.387 | 0.0193 | 0.1007 | 0 | 0 | 0 |
| 'Slc5a6' | 1.942 | 1.671 | 1.711 | 1.579 | 1.808 | 1.723 | 1.757 | 1.981 | 1.705 | 1.839 | 1.382 | 1.587 | 1.792 | 1.266 | 1.616 | 1.656 | 1.326 | 1.376 | 0.191 | 0.0094 | 0.0598 | 0 | 0 | 0 |
| 'Slc6a6' | 9.439 | 9.982 | 9.060 | 8.778 | 8.869 | 8.795 | 8.731 | 9.077 | 9.054 | 9.382 | 9.610 | 8.902 | 9.784 | 9.356 | 9.115 | 9.377 | 8.937 | 8.869 | -0.036 | 0.2238 | 0.4711 | 0 | 0 | 0 |
| 'Slc6a8' | 0.580 | 0.457 | 0.571 | 0.522 | 0.464 | 0.622 | 0.417 | 0.506 | 0.564 | 0.472 | 0.515 | 0.503 | 0.499 | 0.533 | 0.394 | 0.489 | 0.445 | 0.362 | 0.162 | 0.0958 | 0.2881 | 0 | 0 | 0 |
| 'Slc7a1' | 9.571 | 9.509 | 10.305 | 8.693 | 9.200 | 9.973 | 9.215 | 9.667 | 9.562 | 10.583 | 10.496 | 9.579 | 9.958 | 10.285 | 9.773 | 10.037 | 9.805 | 9.574 | -0.079 | 0.0182 | 0.0968 | 0 | 0 | 0 |
| 'Slc7a10' | 10.748 | 11.419 | 11.351 | 10.586 | 10.494 | 8.946 | 10.835 | 11.573 | 10.246 | 12.417 | 12.901 | 10.332 | 14.166 | 13.513 | 11.026 | 9.485 | 13.940 | 9.849 | -0.165 | 0.0088 | 0.0571 | 0 | 0 | 0 |
| 'Slc7a11' | 0.638 | 0.620 | 0.639 | 0.678 | 0.665 | 0.658 | 0.546 | 0.690 | 0.601 | 0.564 | 0.692 | 0.581 | 0.722 | 0.668 | 0.595 | 0.642 | 0.638 | 0.672 | -0.012 | 0.8637 | 0.9407 | 0 | 0 | 0 |
| 'Slc7a5' | 19.568 | 21.709 | 21.416 | 18.874 | 20.769 | 20.054 | 18.137 | 20.753 | 21.429 | 19.262 | 19.238 | 20.656 | 23.444 | 19.958 | 20.280 | 20.984 | 19.935 | 20.680 | -0.025 | 0.5356 | 0.7551 | 0 | 0 | 0 |
| 'Slc7a6' | 18.469 | 17.815 | 19.712 | 17.205 | 17.271 | 17.760 | 18.634 | 17.403 | 18.978 | 16.629 | 16.949 | 20.387 | 17.410 | 16.313 | 18.450 | 17.869 | 18.555 | 18.530 | 0.007 | 0.8483 | 0.9349 | 0 | 0 | 0 |
| 'Slc7a6os' | 41.904 | 39.356 | 41.687 | 37.690 | 42.432 | 43.060 | 38.944 | 42.955 | 41.351 | 41.997 | 37.171 | 42.370 | 40.079 | 39.080 | 41.925 | 47.121 | 41.636 | 39.738 | -0.014 | 0.7184 | 0.8703 | 0 | 0 | 0 |
| 'Slc7a7' | 2.216 | 2.848 | 2.082 | 2.883 | 2.893 | 2.799 | 2.679 | 2.640 | 2.618 | 2.056 | 2.026 | 1.982 | 2.473 | 2.639 | 1.611 | 2.310 | 2.490 | 1.826 | 0.278 | 0.0004 | 0.0049 | 1 | 0 | 0 |
| 'Slc8b1' | 1.844 | 2.789 | 2.857 | 2.034 | 2.777 | 2.313 | 2.118 | 2.537 | 2.468 | 2.607 | 2.278 | 2.259 | 2.160 | 2.623 | 2.207 | 2.238 | 2.177 | 1.842 | 0.078 | 0.3031 | 0.5606 | 0 | 0 | 0 |
| 'Slc9a1' | 14.283 | 13.341 | 15.636 | 13.766 | 13.156 | 15.002 | 13.281 | 14.069 | 15.651 | 14.923 | 13.985 | 15.465 | 14.291 | 13.621 | 14.622 | 13.719 | 13.192 | 15.054 | -0.017 | 0.5470 | 0.7610 | 0 | 0 | 0 |
| 'Slc9a3r1' | 179.774 | 169.619 | 186.629 | 168.858 | 165.940 | 202.148 | 157.082 | 167.062 | 188.789 | 170.463 | 156.644 | 190.746 | 164.648 | 164.066 | 193.597 | 171.318 | 148.985 | 197.360 | 0.018 | 0.5649 | 0.7737 | 0 | 0 | 0 |
| 'Slc9a5' | 1.110 | 0.901 | 1.207 | 1.015 | 0.618 | 1.037 | 0.925 | 1.156 | 1.119 | 0.787 | 0.844 | 0.981 | 1.379 | 0.718 | 0.957 | 0.983 | 0.894 | 0.930 | 0.099 | 0.3978 | 0.6470 | 0 | 0 | 0 |
| 'Slc9a6' | 3.205 | 3.142 | 3.219 | 3.183 | 3.397 | 3.004 | 3.260 | 3.647 | 2.739 | 2.991 | 3.335 | 3.649 | 2.691 | 3.270 | 3.384 | 2.871 | 3.734 | 3.087 | -0.022 | 0.6908 | 0.8543 | 0 | 0 | 0 |
| 'Slc9a7' | 2.394 | 2.210 | 2.337 | 2.237 | 2.063 | 2.386 | 2.044 | 1.968 | 2.265 | 2.750 | 2.839 | 2.759 | 2.163 | 2.453 | 2.516 | 2.728 | 2.503 | 3.013 | -0.260 | 0.0001 | 0.0009 | 1 | 0 | 0 |
| 'Slc9a8' | 5.555 | 5.360 | 6.762 | 5.148 | 6.103 | 5.883 | 6.153 | 5.854 | 6.677 | 6.382 | 5.785 | 5.516 | 6.582 | 5.805 | 5.350 | 5.774 | 6.197 | 4.888 | 0.025 | 0.6623 | 0.8370 | 0 | 0 | 0 |
| 'Slc9a9' | 43.422 | 46.676 | 44.566 | 45.513 | 45.997 | 42.608 | 50.540 | 46.635 | 43.350 | 46.284 | 48.516 | 40.781 | 44.515 | 46.267 | 40.331 | 41.348 | 47.587 | 38.758 | 0.046 | 0.1581 | 0.3881 | 0 | 0 | 0 |
| 'Slc9b2' | 0.625 | 0.538 | 0.683 | 0.658 | 0.615 | 0.607 | 0.713 | 0.693 | 0.684 | 0.660 | 0.540 | 0.914 | 0.481 | 0.657 | 0.724 | 0.628 | 0.691 | 0.614 | -0.042 | 0.6943 | 0.8560 | 0 | 0 | 0 |
| 'Slco3a1' | 11.430 | 11.950 | 12.132 | 11.947 | 12.880 | 12.660 | 12.476 | 12.298 | 12.903 | 12.765 | 13.183 | 11.319 | 13.283 | 12.693 | 10.975 | 13.124 | 12.823 | 11.512 | -0.019 | 0.6086 | 0.8031 | 0 | 0 | 0 |
| 'Slco4a1' | 2.543 | 2.293 | 2.897 | 2.225 | 2.034 | 2.819 | 2.198 | 1.916 | 2.761 | 2.789 | 2.903 | 3.981 | 2.459 | 2.840 | 3.371 | 2.570 | 2.902 | 3.506 | -0.344 | 0.0000 | 0.0000 | 1 | 0 | 0 |
| 'Slf1' | 2.797 | 3.316 | 3.291 | 2.727 | 3.211 | 2.900 | 3.401 | 3.277 | 3.100 | 3.031 | 2.946 | 2.578 | 3.026 | 2.741 | 2.414 | 2.791 | 2.877 | 2.358 | 0.173 | 0.0015 | 0.0146 | 1 | 0 | 0 |
| 'Slfn1' | 42.557 | 43.100 | 47.636 | 41.220 | 44.530 | 53.462 | 42.344 | 47.378 | 46.939 | 39.815 | 46.166 | 61.370 | 37.591 | 39.424 | 62.775 | 46.607 | 46.403 | 63.257 | -0.108 | 0.0672 | 0.2272 | 0 | 0 | 0 |
| 'Slfn2' | 167.063 | 182.275 | 192.276 | 172.623 | 186.679 | 196.867 | 189.228 | 183.386 | 194.583 | 170.671 | 182.728 | 192.365 | 153.678 | 169.053 | 204.989 | 172.971 | 184.549 | 205.704 | 0.016 | 0.6339 | 0.8186 | 0 | 0 | 0 |
| 'Slfn3' | 3.981 | 3.423 | 5.154 | 2.912 | 3.881 | 4.658 | 4.280 | 3.889 | 5.006 | 3.918 | 3.572 | 4.491 | 4.127 | 3.548 | 4.211 | 3.432 | 3.162 | 3.890 | 0.106 | 0.2114 | 0.4566 | 0 | 0 | 0 |
| 'Slfn5' | 11.939 | 12.090 | 12.032 | 10.641 | 12.445 | 14.020 | 10.392 | 12.828 | 12.356 | 14.518 | 11.384 | 14.539 | 13.129 | 10.341 | 14.629 | 13.560 | 11.084 | 13.599 | -0.108 | 0.0637 | 0.2202 | 0 | 0 | 0 |
| 'Slfn8' | 7.299 | 6.410 | 7.840 | 6.673 | 7.982 | 8.245 | 6.591 | 8.380 | 7.948 | 7.918 | 7.138 | 9.770 | 6.923 | 6.858 | 9.539 | 8.575 | 6.736 | 10.653 | -0.138 | 0.0435 | 0.1726 | 0 | 0 | 0 |
| 'Slfn9' | 4.956 | 4.432 | 4.558 | 5.129 | 4.773 | 4.741 | 5.384 | 3.989 | 4.773 | 5.060 | 4.335 | 5.094 | 4.570 | 4.552 | 4.942 | 4.438 | 4.897 | 5.199 | -0.031 | 0.5902 | 0.7909 | 0 | 0 | 0 |
| 'Slirp' | 9.072 | 11.117 | 9.652 | 10.483 | 13.342 | 11.095 | 10.971 | 11.539 | 11.728 | 9.046 | 11.388 | 10.427 | 10.230 | 9.816 | 10.937 | 11.760 | 10.596 | 11.066 | 0.040 | 0.5445 | 0.7606 | 0 | 0 | 0 |
| 'Slk' | 18.010 | 18.022 | 18.347 | 17.717 | 17.961 | 19.193 | 17.083 | 18.580 | 17.728 | 18.286 | 17.675 | 16.910 | 17.378 | 19.064 | 16.829 | 17.583 | 18.102 | 16.636 | 0.030 | 0.2897 | 0.5462 | 0 | 0 | 0 |
| 'Slmap' | 4.991 | 5.165 | 4.987 | 5.331 | 4.976 | 4.876 | 4.731 | 5.310 | 5.127 | 5.076 | 5.030 | 5.220 | 4.801 | 5.266 | 4.939 | 5.051 | 5.331 | 5.071 | -0.020 | 0.4898 | 0.7197 | 0 | 0 | 0 |
| 'Slmo2' | 22.895 | 27.881 | 25.930 | 24.630 | 27.696 | 24.785 | 24.943 | 27.258 | 25.800 | 22.914 | 22.239 | 23.059 | 25.981 | 23.744 | 24.465 | 24.478 | 25.519 | 23.082 | 0.092 | 0.0143 | 0.0809 | 0 | 0 | 0 |
| 'Sltm' | 66.815 | 61.477 | 60.970 | 69.075 | 59.698 | 58.298 | 67.068 | 61.131 | 59.158 | 59.145 | 61.844 | 64.861 | 61.471 | 60.184 | 59.618 | 60.788 | 66.926 | 62.545 | 0.003 | 0.9368 | 0.9759 | 0 | 0 | 0 |
| 'Slu7' | 18.790 | 18.539 | 17.804 | 20.256 | 19.272 | 19.301 | 17.974 | 19.298 | 18.029 | 18.379 | 18.024 | 17.841 | 17.968 | 18.902 | 17.334 | 20.551 | 18.184 | 16.706 | 0.039 | 0.2295 | 0.4765 | 0 | 0 | 0 |
| 'Slx1b' | 1.266 | 1.131 | 1.373 | 1.364 | 1.211 | 1.589 | 1.325 | 1.377 | 1.586 | 1.165 | 1.340 | 1.337 | 1.221 | 0.948 | 1.388 | 1.241 | 1.196 | 1.291 | 0.131 | 0.0689 | 0.2306 | 0 | 0 | 0 |
| 'Slx4' | 5.252 | 4.263 | 5.227 | 5.329 | 4.771 | 4.929 | 5.011 | 4.159 | 4.957 | 4.730 | 5.156 | 5.858 | 4.912 | 4.728 | 5.368 | 3.989 | 4.836 | 6.031 | -0.064 | 0.2716 | 0.5264 | 0 | 0 | 0 |
| 'Slx4ip' | 1.059 | 1.076 | 1.192 | 1.361 | 1.091 | 1.172 | 1.519 | 1.077 | 1.320 | 0.959 | 1.168 | 1.322 | 1.127 | 1.145 | 1.139 | 0.857 | 1.216 | 1.125 | 0.094 | 0.2508 | 0.5019 | 0 | 0 | 0 |
| 'Smad1' | 2.178 | 2.830 | 2.403 | 1.879 | 2.387 | 2.040 | 2.045 | 2.566 | 2.183 | 2.396 | 2.205 | 2.415 | 2.373 | 2.711 | 2.242 | 2.208 | 2.583 | 2.495 | -0.094 | 0.1322 | 0.3498 | 0 | 0 | 0 |
| 'Smad2' | 22.500 | 20.677 | 20.120 | 21.831 | 20.211 | 21.492 | 21.755 | 20.949 | 18.547 | 19.302 | 21.711 | 20.879 | 18.541 | 21.081 | 20.700 | 21.666 | 21.204 | 20.384 | 0.010 | 0.8124 | 0.9157 | 0 | 0 | 0 |
| 'Smad3' | 15.363 | 16.029 | 15.547 | 14.428 | 15.621 | 16.730 | 15.630 | 16.783 | 16.760 | 15.465 | 14.851 | 17.437 | 13.518 | 13.621 | 15.724 | 15.467 | 13.303 | 16.100 | 0.067 | 0.1040 | 0.3023 | 0 | 0 | 0 |
| 'Smad4' | 12.657 | 13.887 | 13.712 | 12.587 | 13.998 | 13.489 | 12.973 | 14.171 | 12.888 | 13.671 | 13.973 | 14.705 | 12.822 | 14.304 | 14.166 | 14.128 | 13.347 | 14.899 | -0.078 | 0.0100 | 0.0622 | 0 | 0 | 0 |
| 'Smad5' | 2.620 | 3.078 | 2.795 | 2.907 | 2.815 | 2.717 | 3.077 | 2.649 | 3.174 | 2.584 | 3.115 | 3.715 | 2.602 | 2.927 | 3.343 | 2.824 | 3.342 | 3.461 | -0.120 | 0.0383 | 0.1590 | 0 | 0 | 0 |
| 'Smad7' | 0.643 | 0.758 | 0.547 | 0.623 | 0.642 | 0.914 | 0.905 | 0.971 | 0.667 | 0.562 | 0.554 | 0.722 | 1.022 | 0.614 | 0.857 | 1.517 | 0.715 | 0.785 | -0.126 | 0.4362 | 0.6799 | 0 | 0 | 0 |
| 'Smap1' | 29.772 | 32.059 | 27.336 | 29.560 | 30.127 | 29.027 | 30.554 | 31.567 | 28.044 | 32.591 | 30.687 | 27.807 | 33.606 | 32.667 | 30.656 | 32.258 | 31.928 | 29.639 | -0.082 | 0.0021 | 0.0190 | 1 | 0 | 0 |
| 'Smap2' | 42.874 | 45.343 | 42.229 | 40.479 | 45.866 | 45.312 | 42.148 | 44.889 | 44.782 | 48.368 | 45.668 | 48.964 | 48.386 | 50.533 | 49.642 | 50.160 | 46.380 | 47.644 | -0.157 | 0.0000 | 0.0000 | 1 | 0 | 0 |
| 'Smarca2' | 9.394 | 8.211 | 9.811 | 8.674 | 8.321 | 9.547 | 9.139 | 9.279 | 9.424 | 8.983 | 9.444 | 8.994 | 8.825 | 8.459 | 8.800 | 8.917 | 8.524 | 8.790 | 0.028 | 0.4005 | 0.6493 | 0 | 0 | 0 |
| 'Smarca4' | 40.576 | 35.586 | 42.068 | 39.604 | 36.220 | 42.078 | 40.516 | 36.612 | 43.294 | 40.964 | 39.884 | 43.846 | 41.263 | 37.410 | 40.703 | 39.113 | 38.725 | 43.681 | -0.048 | 0.0527 | 0.1953 | 0 | 0 | 0 |
| 'Smarca5' | 28.024 | 29.070 | 29.561 | 30.610 | 30.005 | 28.245 | 31.985 | 30.591 | 26.999 | 29.070 | 29.920 | 27.393 | 30.255 | 30.913 | 27.330 | 28.362 | 30.902 | 26.571 | 0.015 | 0.5945 | 0.7935 | 0 | 0 | 0 |
| 'Smarcal1' | 5.763 | 5.350 | 6.576 | 5.739 | 5.382 | 6.494 | 6.187 | 5.925 | 6.319 | 6.082 | 6.107 | 5.702 | 6.158 | 5.229 | 6.020 | 5.537 | 5.805 | 5.437 | 0.037 | 0.3846 | 0.6347 | 0 | 0 | 0 |
| 'Smarcb1' | 7.357 | 7.650 | 8.254 | 7.890 | 7.521 | 8.096 | 7.889 | 7.890 | 8.276 | 8.249 | 7.759 | 8.456 | 8.233 | 7.632 | 8.098 | 7.032 | 8.041 | 8.089 | -0.028 | 0.3497 | 0.6044 | 0 | 0 | 0 |
| 'Smarcc1' | 11.128 | 11.811 | 12.570 | 11.397 | 11.950 | 12.106 | 11.721 | 11.586 | 13.002 | 11.039 | 10.846 | 12.773 | 11.297 | 11.868 | 12.697 | 11.412 | 11.355 | 13.283 | -0.002 | 0.9459 | 0.9791 | 0 | 0 | 0 |
| 'Smarcc2' | 47.822 | 44.700 | 53.179 | 47.115 | 45.183 | 54.267 | 45.640 | 44.695 | 54.097 | 49.900 | 47.907 | 54.814 | 51.165 | 49.791 | 55.297 | 47.096 | 46.365 | 55.421 | -0.079 | 0.0006 | 0.0073 | 1 | 0 | 0 |
| 'Smarcd1' | 16.405 | 22.369 | 21.667 | 17.693 | 22.025 | 24.134 | 17.718 | 22.366 | 23.526 | 22.854 | 18.029 | 20.285 | 20.267 | 20.277 | 21.097 | 21.868 | 16.115 | 19.088 | 0.046 | 0.5072 | 0.7332 | 0 | 0 | 0 |
| 'Smarcd2' | 29.375 | 29.080 | 33.328 | 28.817 | 30.486 | 35.290 | 27.729 | 31.429 | 32.949 | 32.097 | 28.882 | 32.241 | 31.668 | 29.615 | 32.165 | 30.944 | 28.949 | 31.409 | -0.009 | 0.7782 | 0.9002 | 0 | 0 | 0 |
| 'Smarce1' | 14.426 | 15.268 | 14.830 | 15.002 | 14.527 | 14.946 | 15.827 | 15.164 | 15.111 | 14.413 | 13.899 | 14.339 | 15.105 | 14.453 | 15.024 | 14.773 | 13.831 | 14.114 | 0.045 | 0.0559 | 0.2027 | 0 | 0 | 0 |
| 'Smc1a' | 36.302 | 35.068 | 38.890 | 39.438 | 35.866 | 37.803 | 38.746 | 37.001 | 37.960 | 37.603 | 35.702 | 35.891 | 37.945 | 36.211 | 36.414 | 35.602 | 35.782 | 35.448 | 0.035 | 0.1152 | 0.3222 | 0 | 0 | 0 |
| 'Smc2' | 13.164 | 13.621 | 16.575 | 16.803 | 13.977 | 15.147 | 17.463 | 14.239 | 14.930 | 12.509 | 15.487 | 15.985 | 16.082 | 17.033 | 16.523 | 13.544 | 15.340 | 16.014 | -0.038 | 0.5330 | 0.7530 | 0 | 0 | 0 |
| 'Smc3' | 27.688 | 27.290 | 27.799 | 32.523 | 28.293 | 27.753 | 32.234 | 28.028 | 26.911 | 26.784 | 28.904 | 28.186 | 26.728 | 27.211 | 27.867 | 27.209 | 28.542 | 27.584 | 0.041 | 0.2836 | 0.5397 | 0 | 0 | 0 |
| 'Smc4' | 40.076 | 36.178 | 40.524 | 44.553 | 37.051 | 39.416 | 43.689 | 36.722 | 38.646 | 37.447 | 40.720 | 42.526 | 35.423 | 40.812 | 40.174 | 37.300 | 40.079 | 41.655 | -0.009 | 0.8430 | 0.9337 | 0 | 0 | 0 |
| 'Smc5' | 10.087 | 9.700 | 9.683 | 9.866 | 10.073 | 9.765 | 10.622 | 10.451 | 9.895 | 10.626 | 10.658 | 10.428 | 10.631 | 9.094 | 9.669 | 10.344 | 10.195 | 9.695 | -0.030 | 0.3228 | 0.5806 | 0 | 0 | 0 |
| 'Smc6' | 33.517 | 33.584 | 32.333 | 35.169 | 32.553 | 32.267 | 38.028 | 33.492 | 30.295 | 32.627 | 32.999 | 33.400 | 33.072 | 34.067 | 31.224 | 34.531 | 34.015 | 33.526 | -0.005 | 0.8832 | 0.9509 | 0 | 0 | 0 |
| 'Smchd1' | 8.811 | 8.208 | 8.928 | 8.953 | 8.737 | 8.420 | 9.277 | 8.955 | 8.581 | 8.776 | 9.085 | 9.413 | 8.988 | 9.510 | 8.987 | 8.743 | 8.903 | 9.398 | -0.064 | 0.0234 | 0.1149 | 0 | 0 | 0 |
| 'Smco4' | 134.388 | 165.918 | 127.449 | 144.088 | 165.250 | 139.468 | 135.933 | 155.584 | 136.217 | 127.551 | 136.292 | 116.425 | 146.610 | 150.115 | 130.132 | 134.909 | 142.581 | 124.041 | 0.097 | 0.0065 | 0.0458 | 1 | 0 | 0 |
| 'Smcr8' | 9.948 | 8.875 | 9.264 | 9.638 | 9.369 | 9.529 | 8.935 | 9.013 | 9.903 | 9.423 | 10.404 | 11.202 | 9.397 | 9.227 | 10.885 | 9.087 | 10.428 | 11.457 | -0.124 | 0.0031 | 0.0262 | 1 | 0 | 0 |

| | | | | | | | | | | | | | | | | | | | | | | | | |
|---|---|---|---|---|---|---|---|---|---|---|---|---|---|---|---|---|---|---|---|---|---|---|---|---|
| 'Smdt1' | 40.015 | 49.515 | 42.709 | 41.317 | 47.794 | 44.206 | 43.296 | 46.129 | 43.738 | 38.778 | 42.464 | 40.688 | 47.403 | 44.627 | 44.291 | 47.386 | 44.051 | 42.395 | 0.012 | 0.7749 | 0.8994 | 0 | 0 | 0 |
| 'Smek1' | 51.299 | 46.880 | 47.137 | 50.375 | 47.233 | 45.450 | 48.227 | 49.273 | 45.975 | 47.496 | 48.414 | 50.557 | 44.353 | 49.910 | 47.614 | 50.180 | 51.934 | 48.155 | -0.033 | 0.3124 | 0.5709 | 0 | 0 | 0 |
| 'Smek2' | 29.898 | 33.032 | 31.089 | 31.205 | 31.970 | 29.864 | 31.797 | 32.930 | 29.340 | 30.639 | 33.539 | 31.642 | 29.412 | 32.885 | 29.282 | 31.903 | 33.805 | 30.351 | -0.022 | 0.3431 | 0.5992 | 0 | 0 | 0 |
| 'Smg1' | 19.011 | 15.567 | 18.436 | 18.400 | 16.255 | 16.228 | 16.415 | 16.450 | 17.764 | 17.284 | 18.994 | 18.896 | 15.938 | 16.421 | 17.497 | 17.046 | 17.770 | 20.099 | -0.060 | 0.2104 | 0.4557 | 0 | 0 | 0 |
| 'Smg5' | 15.605 | 13.993 | 16.858 | 14.945 | 14.487 | 15.894 | 16.124 | 14.923 | 16.727 | 15.870 | 15.651 | 16.433 | 16.501 | 13.787 | 15.825 | 15.517 | 15.451 | 15.610 | -0.022 | 0.4522 | 0.6912 | 0 | 0 | 0 |
| 'Smg6' | 12.518 | 11.112 | 11.702 | 10.759 | 11.106 | 10.955 | 11.097 | 11.297 | 10.934 | 11.669 | 11.855 | 11.709 | 10.928 | 11.364 | 12.130 | 11.075 | 10.682 | 11.494 | -0.030 | 0.3933 | 0.6428 | 0 | 0 | 0 |
| 'Smg7' | 16.884 | 15.485 | 16.152 | 16.905 | 15.582 | 15.820 | 16.969 | 16.232 | 16.218 | 16.590 | 16.889 | 17.337 | 16.903 | 15.415 | 16.351 | 15.873 | 16.543 | 16.867 | -0.037 | 0.1722 | 0.4060 | 0 | 0 | 0 |
| 'Smg8' | 7.462 | 8.003 | 6.731 | 6.899 | 7.555 | 7.875 | 6.170 | 8.275 | 7.030 | 6.870 | 6.916 | 8.740 | 5.889 | 7.947 | 8.871 | 7.905 | 6.945 | 8.623 | -0.066 | 0.3228 | 0.5806 | 0 | 0 | 0 |
| 'Smg9' | 8.002 | 7.953 | 9.084 | 7.106 | 7.083 | 8.459 | 8.244 | 7.708 | 8.590 | 7.517 | 6.856 | 8.260 | 8.005 | 6.605 | 7.945 | 8.039 | 6.502 | 7.688 | 0.091 | 0.0178 | 0.0950 | 0 | 0 | 0 |
| 'Smim10l1' | 10.888 | 13.766 | 11.733 | 11.490 | 12.251 | 12.322 | 12.204 | 12.548 | 11.892 | 10.954 | 11.633 | 12.265 | 10.874 | 11.761 | 11.777 | 10.778 | 11.446 | 11.612 | 0.064 | 0.0754 | 0.2441 | 0 | 0 | 0 |
| 'Smim11' | 22.093 | 24.414 | 23.052 | 23.397 | 28.177 | 24.611 | 24.555 | 23.080 | 22.504 | 23.279 | 23.639 | 23.702 | 22.352 | 24.475 | 26.282 | 23.389 | 23.213 | 25.264 | -0.012 | 0.8010 | 0.9112 | 0 | 0 | 0 |
| 'Smim12' | 19.916 | 24.105 | 25.137 | 22.324 | 28.008 | 27.505 | 18.521 | 25.955 | 25.383 | 25.832 | 23.882 | 22.119 | 24.348 | 22.218 | 22.820 | 24.437 | 21.414 | 21.827 | 0.042 | 0.5308 | 0.7514 | 0 | 0 | 0 |
| 'Smim13' | 5.593 | 6.402 | 5.503 | 5.215 | 6.133 | 5.411 | 5.833 | 5.410 | 5.355 | 5.803 | 6.910 | 6.413 | 5.900 | 6.517 | 6.250 | 5.236 | 7.121 | 6.317 | -0.167 | 0.0006 | 0.0071 | 1 | 0 | 0 |
| 'Smim14' | 2.577 | 3.288 | 2.909 | 2.642 | 3.228 | 2.991 | 2.449 | 3.323 | 3.090 | 3.010 | 3.157 | 2.817 | 2.988 | 3.285 | 2.849 | 2.704 | 3.031 | 2.597 | -0.008 | 0.8404 | 0.9322 | 0 | 0 | 0 |
| 'Smim15' | 29.371 | 31.365 | 32.124 | 36.240 | 30.663 | 28.458 | 32.711 | 32.127 | 29.057 | 25.939 | 30.324 | 32.173 | 27.138 | 30.016 | 31.429 | 24.817 | 35.189 | 30.083 | 0.070 | 0.2395 | 0.4897 | 0 | 0 | 0 |
| 'Smim19' | 20.225 | 24.317 | 22.840 | 21.153 | 21.517 | 23.589 | 24.232 | 25.590 | 20.884 | 26.230 | 23.269 | 21.202 | 23.339 | 24.756 | 20.683 | 25.185 | 23.829 | 21.525 | -0.044 | 0.3811 | 0.6324 | 0 | 0 | 0 |
| 'Smim20' | 4.852 | 5.882 | 5.442 | 4.628 | 6.422 | 6.331 | 4.472 | 6.160 | 5.706 | 5.151 | 5.234 | 5.318 | 5.431 | 5.524 | 4.480 | 5.527 | 5.436 | 4.775 | 0.076 | 0.1974 | 0.4366 | 0 | 0 | 0 |
| 'Smim24' | 5.354 | 4.574 | 6.077 | 6.080 | 4.672 | 6.885 | 5.362 | 4.873 | 4.822 | 4.451 | 5.203 | 5.664 | 5.622 | 4.428 | 5.917 | 4.169 | 5.419 | 6.295 | 0.035 | 0.7158 | 0.8688 | 0 | 0 | 0 |
| 'Smim3' | 4.326 | 6.090 | 6.382 | 4.377 | 6.256 | 5.487 | 5.485 | 6.250 | 5.002 | 8.554 | 8.095 | 5.430 | 7.619 | 8.334 | 6.399 | 6.449 | 6.991 | 4.924 | -0.335 | 0.0004 | 0.0054 | 1 | 0 | 0 |
| 'Smim4' | 4.392 | 4.365 | 3.987 | 3.839 | 4.823 | 4.496 | 4.293 | 4.896 | 4.005 | 4.085 | 3.346 | 3.617 | 5.039 | 4.231 | 4.350 | 5.180 | 3.878 | 3.916 | 0.052 | 0.5090 | 0.7342 | 0 | 0 | 0 |
| 'Smim7' | 9.248 | 10.354 | 9.551 | 8.915 | 9.487 | 9.476 | 9.323 | 10.324 | 8.439 | 9.476 | 9.495 | 8.761 | 10.117 | 10.087 | 8.933 | 9.219 | 9.383 | 8.898 | 0.005 | 0.9008 | 0.9607 | 0 | 0 | 0 |
| 'Smim8' | 4.488 | 5.029 | 5.277 | 4.165 | 5.699 | 4.753 | 5.539 | 5.735 | 4.840 | 5.098 | 5.181 | 4.224 | 5.321 | 4.660 | 4.230 | 5.146 | 5.466 | 4.257 | 0.062 | 0.3496 | 0.6044 | 0 | 0 | 0 |
| 'Smn1' | 6.648 | 7.403 | 6.491 | 5.744 | 6.579 | 6.603 | 6.437 | 7.524 | 6.647 | 6.904 | 6.856 | 6.780 | 6.641 | 6.318 | 6.674 | 6.604 | 6.269 | 6.358 | 0.004 | 0.9215 | 0.9705 | 0 | 0 | 0 |
| 'Smndc1' | 34.306 | 32.961 | 33.109 | 31.850 | 35.967 | 31.245 | 33.301 | 33.940 | 30.753 | 36.894 | 32.615 | 31.981 | 32.358 | 37.804 | 31.703 | 37.856 | 32.144 | 32.769 | -0.049 | 0.2130 | 0.4581 | 0 | 0 | 0 |
| 'Smoc2' | 2.151 | 2.937 | 3.534 | 3.172 | 2.509 | 3.134 | 2.544 | 2.818 | 3.727 | 2.712 | 1.808 | 2.825 | 2.120 | 1.921 | 2.354 | 3.437 | 1.846 | 3.137 | 0.258 | 0.0164 | 0.0896 | 0 | 0 | 0 |
| 'Smox' | 18.915 | 24.131 | 15.727 | 20.074 | 22.834 | 18.436 | 18.983 | 21.316 | 18.559 | 23.918 | 22.545 | 20.075 | 30.745 | 25.777 | 19.492 | 27.051 | 25.940 | 19.308 | -0.269 | 0.0000 | 0.0000 | 1 | 0 | 0 |
| 'Smpd1' | 12.531 | 13.569 | 14.005 | 10.928 | 12.735 | 13.594 | 9.853 | 12.813 | 13.052 | 14.078 | 11.494 | 11.752 | 14.282 | 13.137 | 12.509 | 12.996 | 10.234 | 11.961 | 0.001 | 0.9900 | 0.9972 | 0 | 0 | 0 |
| 'Smpd2' | 2.644 | 2.397 | 2.863 | 2.482 | 2.391 | 2.554 | 2.718 | 2.527 | 2.849 | 2.479 | 2.231 | 2.636 | 2.648 | 2.397 | 2.674 | 2.553 | 2.566 | 2.440 | 0.042 | 0.3097 | 0.5681 | 0 | 0 | 0 |
| 'Smpd4' | 3.136 | 3.108 | 3.589 | 3.208 | 3.303 | 3.186 | 3.217 | 3.257 | 3.385 | 3.432 | 3.260 | 3.419 | 3.495 | 3.248 | 3.139 | 3.176 | 3.255 | 3.218 | -0.022 | 0.4956 | 0.7247 | 0 | 0 | 0 |
| 'Smpd5' | 0.839 | 0.979 | 1.027 | 0.808 | 1.038 | 1.035 | 0.784 | 0.955 | 0.988 | 0.908 | 0.550 | 0.693 | 0.994 | 0.642 | 0.688 | 0.860 | 0.730 | 0.522 | 0.365 | 0.0043 | 0.0336 | 1 | 0 | 0 |
| 'Smpdl3a' | 89.089 | 109.525 | 90.121 | 97.270 | 106.837 | 95.843 | 98.573 | 99.230 | 93.179 | 99.682 | 90.360 | 85.343 | 91.515 | 96.976 | 85.779 | 96.922 | 98.379 | 86.414 | 0.070 | 0.0253 | 0.1216 | 0 | 0 | 0 |
| 'Smpdl3b' | 11.303 | 10.755 | 13.456 | 9.539 | 10.729 | 11.400 | 11.155 | 10.240 | 12.618 | 12.321 | 8.792 | 13.659 | 9.195 | 9.818 | 12.800 | 12.870 | 10.111 | 14.506 | -0.044 | 0.5080 | 0.7336 | 0 | 0 | 0 |
| 'Sms' | 5.871 | 5.778 | 5.985 | 6.026 | 5.752 | 5.854 | 6.074 | 6.185 | 5.866 | 5.136 | 5.077 | 6.224 | 4.825 | 5.373 | 5.801 | 5.832 | 6.008 | 5.583 | 0.087 | 0.0508 | 0.1908 | 0 | 0 | 0 |
| 'Smtn' | 63.641 | 66.433 | 74.265 | 60.823 | 60.350 | 68.120 | 65.554 | 62.663 | 78.155 | 53.887 | 46.055 | 60.012 | 46.042 | 43.311 | 55.966 | 50.079 | 46.048 | 63.060 | 0.364 | 0.0000 | 0.0000 | 1 | 0 | 0 |
| 'Smu1' | 19.598 | 23.346 | 20.188 | 21.039 | 21.918 | 21.630 | 21.348 | 22.494 | 20.992 | 19.252 | 20.474 | 20.808 | 18.937 | 20.781 | 20.441 | 19.787 | 21.386 | 20.205 | 0.067 | 0.0143 | 0.0809 | 0 | 0 | 0 |
| 'Smug1' | 1.891 | 2.893 | 2.906 | 1.884 | 2.624 | 3.178 | 2.546 | 3.106 | 3.427 | 2.319 | 3.095 | 2.884 | 2.129 | 2.419 | 2.994 | 2.755 | 2.762 | 2.946 | -0.001 | 0.9881 | 0.9965 | 0 | 0 | 0 |
| 'Smurf1' | 4.610 | 5.530 | 5.337 | 4.148 | 5.264 | 5.302 | 4.606 | 5.005 | 5.052 | 4.917 | 4.863 | 5.237 | 4.740 | 5.095 | 4.902 | 5.229 | 4.782 | 5.219 | -0.017 | 0.6602 | 0.8362 | 0 | 0 | 0 |
| 'Smurf2' | 12.763 | 12.604 | 12.278 | 12.286 | 12.507 | 11.134 | 11.468 | 12.933 | 11.113 | 11.579 | 11.725 | 12.559 | 10.250 | 11.908 | 12.070 | 12.520 | 11.475 | 13.290 | 0.012 | 0.8002 | 0.9111 | 0 | 0 | 0 |
| 'Smyd2' | 2.326 | 2.153 | 2.502 | 2.402 | 2.162 | 2.455 | 2.517 | 2.353 | 2.402 | 2.114 | 1.620 | 2.370 | 2.008 | 2.001 | 2.639 | 1.752 | 1.434 | 2.291 | 0.210 | 0.0046 | 0.0351 | 1 | 0 | 0 |
| 'Smyd3' | 2.117 | 2.078 | 1.673 | 1.944 | 1.973 | 1.864 | 2.352 | 2.002 | 1.732 | 2.110 | 1.959 | 1.827 | 2.027 | 2.101 | 1.738 | 1.936 | 2.086 | 1.657 | 0.010 | 0.8245 | 0.9227 | 0 | 0 | 0 |
| 'Smyd4' | 5.964 | 6.011 | 5.888 | 6.025 | 6.412 | 5.804 | 6.354 | 5.555 | 6.010 | 6.067 | 6.486 | 6.214 | 6.481 | 6.607 | 6.149 | 6.499 | 5.974 | 6.561 | -0.091 | 0.0250 | 0.1208 | 0 | 0 | 0 |
| 'Smyd5' | 5.847 | 5.008 | 5.973 | 5.047 | 5.126 | 5.344 | 5.450 | 5.144 | 5.431 | 5.764 | 4.460 | 5.918 | 5.388 | 4.480 | 5.418 | 5.161 | 5.126 | 5.239 | 0.033 | 0.4362 | 0.6799 | 0 | 0 | 0 |
| 'Snap23' | 7.170 | 7.859 | 6.972 | 8.533 | 8.028 | 7.838 | 7.650 | 7.941 | 7.392 | 7.407 | 7.666 | 7.392 | 7.273 | 7.748 | 6.962 | 7.487 | 7.800 | 7.411 | 0.034 | 0.3674 | 0.6195 | 0 | 0 | 0 |
| 'Snap29' | 11.703 | 12.630 | 12.233 | 12.513 | 11.360 | 12.465 | 12.560 | 12.736 | 12.932 | 12.668 | 12.376 | 13.950 | 10.774 | 11.744 | 13.201 | 11.864 | 12.663 | 13.133 | -0.029 | 0.4926 | 0.7218 | 0 | 0 | 0 |
| 'Snap47' | 9.658 | 9.420 | 10.394 | 9.389 | 10.177 | 10.319 | 10.147 | 10.102 | 10.400 | 10.091 | 9.434 | 10.261 | 10.370 | 10.140 | 9.650 | 10.478 | 9.330 | 9.371 | 0.006 | 0.8596 | 0.9390 | 0 | 0 | 0 |
| 'Snapc1' | 20.002 | 19.937 | 18.808 | 21.403 | 19.547 | 18.135 | 20.303 | 18.952 | 19.360 | 16.321 | 19.636 | 19.531 | 18.875 | 20.161 | 19.896 | 17.665 | 22.139 | 19.200 | 0.013 | 0.7944 | 0.9082 | 0 | 0 | 0 |
| 'Snapc2' | 17.630 | 15.453 | 16.221 | 14.593 | 14.557 | 17.818 | 14.840 | 16.428 | 16.315 | 16.247 | 15.091 | 16.292 | 15.582 | 14.960 | 17.730 | 17.824 | 13.636 | 16.505 | -0.002 | 0.9631 | 0.9869 | 0 | 0 | 0 |
| 'Snapc3' | 3.803 | 3.525 | 3.616 | 3.752 | 3.670 | 3.475 | 4.459 | 3.622 | 3.761 | 3.254 | 4.147 | 4.155 | 3.224 | 3.747 | 3.445 | 3.163 | 3.808 | 3.984 | 0.022 | 0.7274 | 0.8750 | 0 | 0 | 0 |
| 'Snapc4' | 4.003 | 3.336 | 4.568 | 3.883 | 3.395 | 3.737 | 3.896 | 3.532 | 4.340 | 4.058 | 3.590 | 4.480 | 3.687 | 2.979 | 3.870 | 3.415 | 3.620 | 4.329 | 0.017 | 0.7325 | 0.8781 | 0 | 0 | 0 |
| 'Snapc5' | 38.684 | 41.871 | 37.446 | 37.625 | 37.054 | 40.329 | 42.719 | 39.670 | 36.176 | 39.945 | 38.481 | 38.062 | 37.198 | 37.762 | 37.334 | 40.481 | 41.616 | 34.645 | 0.015 | 0.7216 | 0.8716 | 0 | 0 | 0 |
| 'Snapin' | 5.377 | 7.051 | 5.937 | 5.379 | 7.371 | 5.783 | 5.133 | 6.914 | 5.800 | 6.490 | 5.918 | 5.263 | 6.146 | 6.375 | 5.394 | 6.078 | 6.055 | 4.613 | 0.053 | 0.3377 | 0.5955 | 0 | 0 | 0 |
| 'Snd1' | 14.512 | 13.488 | 15.269 | 14.804 | 13.831 | 14.965 | 14.462 | 13.400 | 14.530 | 14.732 | 13.874 | 14.417 | 14.813 | 14.357 | 14.147 | 14.650 | 13.906 | 14.409 | -0.009 | 0.6781 | 0.8461 | 0 | 0 | 0 |
| 'Snf8' | 11.672 | 13.815 | 13.622 | 12.523 | 14.167 | 13.662 | 13.148 | 13.966 | 13.643 | 12.275 | 12.704 | 12.825 | 12.981 | 12.993 | 13.323 | 13.545 | 13.598 | 12.080 | 0.038 | 0.2750 | 0.5307 | 0 | 0 | 0 |
| 'Snip1' | 18.076 | 17.578 | 18.467 | 16.761 | 17.544 | 17.840 | 16.346 | 18.110 | 17.195 | 17.758 | 17.111 | 17.925 | 18.038 | 17.970 | 16.845 | 18.220 | 17.864 | 16.215 | -0.007 | 0.8400 | 0.9319 | 0 | 0 | 0 |
| 'Snn' | 8.592 | 12.841 | 9.517 | 8.927 | 11.619 | 10.242 | 10.325 | 12.113 | 10.086 | 10.280 | 10.852 | 9.581 | 9.816 | 11.497 | 10.214 | 12.777 | 12.681 | 9.559 | -0.058 | 0.3188 | 0.5773 | 0 | 0 | 0 |
| 'Snrk' | 30.058 | 31.576 | 28.214 | 29.612 | 27.499 | 27.983 | 27.465 | 29.559 | 27.364 | 27.714 | 27.933 | 28.146 | 25.738 | 25.619 | 26.964 | 27.344 | 25.678 | 29.076 | 0.075 | 0.0394 | 0.1621 | 0 | 0 | 0 |
| 'Snrnp200' | 25.470 | 26.377 | 26.781 | 25.867 | 27.013 | 28.794 | 25.535 | 27.134 | 28.627 | 28.035 | 25.812 | 26.846 | 25.961 | 26.392 | 26.128 | 26.848 | 25.906 | 26.925 | 0.006 | 0.8120 | 0.9155 | 0 | 0 | 0 |
| 'Snrnp25' | 10.758 | 12.162 | 11.368 | 11.879 | 11.961 | 12.035 | 11.798 | 11.972 | 11.787 | 10.607 | 10.976 | 10.758 | 11.673 | 11.470 | 12.675 | 11.142 | 11.126 | 10.433 | 0.058 | 0.1840 | 0.4193 | 0 | 0 | 0 |
| 'Snrnp27' | 10.310 | 12.785 | 11.006 | 11.200 | 11.717 | 11.418 | 11.065 | 11.367 | 10.818 | 10.224 | 11.076 | 11.501 | 12.032 | 11.811 | 10.811 | 11.204 | 11.280 | 9.929 | 0.010 | 0.8275 | 0.9249 | 0 | 0 | 0 |
| 'Snrnp35' | 1.500 | 1.801 | 2.335 | 1.984 | 1.751 | 2.224 | 2.261 | 2.161 | 2.403 | 1.472 | 2.036 | 2.142 | 1.749 | 1.633 | 1.674 | 1.535 | 1.752 | 1.408 | 0.242 | 0.0214 | 0.1077 | 0 | 0 | 0 |
| 'Snrnp40' | 17.475 | 19.024 | 18.750 | 19.225 | 19.493 | 18.829 | 18.046 | 19.177 | 18.088 | 17.370 | 17.357 | 18.245 | 18.153 | 19.105 | 18.517 | 17.631 | 17.963 | 18.497 | 0.034 | 0.2203 | 0.4676 | 0 | 0 | 0 |
| 'Snrnp48' | 21.065 | 24.097 | 21.223 | 24.389 | 22.614 | 20.408 | 24.810 | 22.805 | 20.025 | 22.015 | 22.343 | 23.689 | 23.598 | 21.449 | 22.725 | 21.913 | 24.975 | 21.793 | -0.041 | 0.3446 | 0.5997 | 0 | 0 | 0 |
| 'Snrnp70' | 33.196 | 31.221 | 35.645 | 30.650 | 31.514 | 34.581 | 33.149 | 31.955 | 36.324 | 33.540 | 32.076 | 34.471 | 34.906 | 31.861 | 35.210 | 32.466 | 31.865 | 34.776 | -0.025 | 0.2530 | 0.5047 | 0 | 0 | 0 |
| 'Snrpa' | 18.318 | 18.429 | 20.066 | 18.093 | 19.600 | 19.634 | 19.158 | 17.952 | 19.951 | 19.461 | 16.925 | 18.575 | 20.249 | 15.830 | 19.089 | 19.270 | 18.187 | 19.419 | 0.027 | 0.4477 | 0.6880 | 0 | 0 | 0 |
| 'Snrpa1' | 10.673 | 10.263 | 11.304 | 11.459 | 10.662 | 10.727 | 11.417 | 10.638 | 10.693 | 11.073 | 10.156 | 10.727 | 10.936 | 10.508 | 10.531 | 10.119 | 10.852 | 8.903 | 0.051 | 0.1718 | 0.4055 | 0 | 0 | 0 |
| 'Snrpb' | 61.253 | 64.486 | 59.635 | 60.511 | 64.053 | 63.174 | 61.049 | 62.727 | 60.239 | 58.339 | 55.929 | 57.254 | 61.542 | 62.241 | 63.658 | 63.788 | 56.888 | 60.209 | 0.035 | 0.2518 | 0.5032 | 0 | 0 | 0 |
| 'Snrpb2' | 11.436 | 13.340 | 12.273 | 12.000 | 13.728 | 11.610 | 13.319 | 13.196 | 12.671 | 12.074 | 12.614 | 11.779 | 12.173 | 13.383 | 11.719 | 13.156 | 11.943 | 11.446 | 0.031 | 0.3628 | 0.6149 | 0 | 0 | 0 |
| 'Snrpc' | 7.617 | 9.212 | 7.603 | 7.571 | 9.028 | 8.395 | 7.719 | 8.435 | 7.923 | 7.681 | 7.099 | 7.470 | 8.254 | 8.888 | 7.484 | 8.925 | 7.640 | 7.278 | 0.044 | 0.3443 | 0.5996 | 0 | 0 | 0 |
| 'Snrpd1' | 54.939 | 54.553 | 59.455 | 58.912 | 60.769 | 54.161 | 62.989 | 59.131 | 58.158 | 50.047 | 53.710 | 55.464 | 53.268 | 57.244 | 56.895 | 52.538 | 59.205 | 49.746 | 0.089 | 0.0283 | 0.1304 | 0 | 0 | 0 |
| 'Snrpd2' | 34.797 | 34.865 | 34.216 | 37.311 | 36.316 | 33.763 | 38.561 | 34.283 | 33.074 | 32.137 | 30.626 | 31.212 | 35.153 | 34.375 | 35.055 | 35.230 | 33.680 | 31.822 | 0.072 | 0.0335 | 0.1456 | 0 | 0 | 0 |
| 'Snrpd3' | 19.486 | 20.139 | 21.278 | 22.081 | 21.796 | 20.963 | 20.501 | 21.694 | 19.950 | 18.236 | 18.953 | 18.781 | 20.968 | 20.098 | 19.646 | 19.500 | 19.628 | 18.744 | 0.097 | 0.0037 | 0.0295 | 1 | 0 | 0 |
| 'Snrpe' | 30.149 | 35.795 | 29.647 | 34.050 | 33.027 | 31.190 | 35.279 | 32.823 | 29.981 | 30.511 | 30.800 | 32.696 | 31.873 | 34.706 | 31.919 | 30.672 | 34.961 | 31.901 | -0.007 | 0.8588 | 0.9390 | 0 | 0 | 0 |
| 'Snrpf' | 62.304 | 61.690 | 65.120 | 68.383 | 69.759 | 69.497 | 67.707 | 65.629 | 64.469 | 53.441 | 60.773 | 58.880 | 66.101 | 62.900 | 62.340 | 66.401 | 65.682 | 53.497 | 0.104 | 0.0207 | 0.1056 | 0 | 0 | 0 |
| 'Snrpg' | 37.047 | 42.914 | 37.019 | 40.444 | 41.064 | 39.386 | 41.981 | 40.150 | 36.883 | 34.186 | 37.432 | 33.263 | 39.022 | 39.917 | 38.644 | 37.689 | 40.456 | 34.703 | 0.080 | 0.0344 | 0.1481 | 0 | 0 | 0 |
| 'Sntb1' | 4.785 | 4.412 | 3.724 | 4.012 | 4.052 | 4.371 | 4.723 | 3.840 | 4.489 | 5.458 | 4.814 | 5.292 | 5.709 | 5.143 | 5.050 | 5.550 | 5.494 | 5.235 | -0.328 | 0.0000 | 0.0000 | 1 | 0 | 0 |
| 'Sntb2' | 10.478 | 9.173 | 9.109 | 9.303 | 8.793 | 8.949 | 8.491 | 7.779 | 8.409 | 7.204 | 6.392 | 7.303 | 7.158 | 7.362 | 6.534 | 7.363 | 6.919 | 7.904 | 0.316 | 0.0000 | 0.0000 | 1 | 0 | 0 |
| 'Snupn' | 4.836 | 5.528 | 5.656 | 6.561 | 6.086 | 5.881 | 5.344 | 6.744 | 5.825 | 5.664 | 6.792 | 5.260 | 5.296 | 5.389 | 4.460 | 4.406 | 5.662 | 4.770 | 0.131 | 0.1133 | 0.3194 | 0 | 0 | 0 |
| 'Snw1' | 61.471 | 65.866 | 62.222 | 67.130 | 68.524 | 64.290 | 66.155 | 69.139 | 63.317 | 63.150 | 63.627 | 63.556 | 63.792 | 61.167 | 62.542 | 65.519 | 65.824 | 61.258 | 0.032 | 0.1814 | 0.4171 | 0 | 0 | 0 |
| 'Snx1' | 16.940 | 16.233 | 15.670 | 17.344 | 17.435 | 16.858 | 17.535 | 16.865 | 16.201 | 15.989 | 16.635 | 17.656 | 16.698 | 16.262 | 16.499 | 16.415 | 16.597 | 16.388 | 0.004 | 0.8884 | 0.9544 | 0 | 0 | 0 |
| 'Snx10' | 1.103 | 1.348 | 1.036 | 1.206 | 1.393 | 0.995 | 1.201 | 1.205 | 1.009 | 1.339 | 1.397 | 1.186 | 1.237 | 1.415 | 1.184 | 1.317 | 1.397 | 1.139 | -0.164 | 0.0025 | 0.0216 | 1 | 0 | 0 |
| 'Snx11' | 6.673 | 8.832 | 7.965 | 6.386 | 9.002 | 8.258 | 7.281 | 7.713 | 8.038 | 7.027 | 7.982 | 8.027 | 8.384 | 7.656 | 7.762 | 8.148 | 9.050 | 7.197 | -0.038 | 0.4629 | 0.6990 | 0 | 0 | 0 |
| 'Snx12' | 13.054 | 14.976 | 13.525 | 12.888 | 14.476 | 14.030 | 13.828 | 13.838 | 14.042 | 14.726 | 14.469 | 13.972 | 13.752 | 14.175 | 14.106 | 14.607 | 14.513 | 13.873 | -0.050 | 0.1019 | 0.2993 | 0 | 0 | 0 |
| 'Snx13' | 4.902 | 5.549 | 4.583 | 5.157 | 5.197 | 4.688 | 5.481 | 5.784 | 4.674 | 4.932 | 5.776 | 5.188 | 5.065 | 5.623 | 4.750 | 4.986 | 5.159 | 4.533 | -0.016 | 0.6868 | 0.8521 | 0 | 0 | 0 |
| 'Snx14' | 3.955 | 4.016 | 3.897 | 4.014 | 4.399 | 4.062 | 4.258 | 4.022 | 4.091 | 3.779 | 4.711 | 4.234 | 3.671 | 4.506 | 3.843 | 4.029 | 4.151 | 3.631 | -0.002 | 0.9590 | 0.9847 | 0 | 0 | 0 |
| 'Snx15' | 7.746 | 9.055 | 8.562 | 7.236 | 8.759 | 9.465 | 7.329 | 9.334 | 9.263 | 8.798 | 6.941 | 8.830 | 8.150 | 8.285 | 8.272 | 9.169 | 7.150 | 8.318 | 0.043 | 0.4657 | 0.7014 | 0 | 0 | 0 |
| 'Snx16' | 1.118 | 1.423 | 1.388 | 1.318 | 1.515 | 1.071 | 1.443 | 1.225 | 1.413 | 1.376 | 1.211 | 1.193 | 1.528 | 1.417 | 1.240 | 1.038 | 1.728 | 1.307 | -0.031 | 0.7579 | 0.8916 | 0 | 0 | 0 |
| 'Snx17' | 13.564 | 15.727 | 13.516 | 14.215 | 15.549 | 14.571 | 13.104 | 15.412 | 14.101 | 13.903 | 13.533 | 13.810 | 13.883 | 13.958 | 14.168 | 14.933 | 14.119 | 13.863 | 0.028 | 0.3807 | 0.6321 | 0 | 0 | 0 |
| 'Snx18' | 8.608 | 7.941 | 8.685 | 9.104 | 7.616 | 9.004 | 8.515 | 8.573 | 9.126 | 9.058 | 9.439 | 10.572 | 8.457 | 8.727 | 10.259 | 10.253 | 9.218 | 9.776 | -0.161 | 0.0002 | 0.0022 | 1 | 0 | 0 |
| 'Snx19' | 15.352 | 15.065 | 17.454 | 15.235 | 15.329 | 17.652 | 14.867 | 16.648 | 17.332 | 16.578 | 15.999 | 16.151 | 15.137 | 14.642 | 14.929 | 16.177 | 15.119 | 15.368 | 0.040 | 0.2701 | 0.5250 | 0 | 0 | 0 |
| 'Snx2' | 143.515 | 160.573 | 147.603 | 157.137 | 159.909 | 143.077 | 158.058 | 158.636 | 150.988 | 143.892 | 161.256 | 143.873 | 157.559 | 161.685 | 143.510 | 146.750 | 166.576 | 142.531 | 0.003 | 0.9151 | 0.9674 | 0 | 0 | 0 |
| 'Snx20' | 70.449 | 61.486 | 74.687 | 74.216 | 68.062 | 83.006 | 70.496 | 63.569 | 68.544 | 66.305 | 63.748 | 83.061 | 74.230 | 66.933 | 85.560 | 74.136 | 67.104 | 88.049 | -0.084 | 0.0447 | 0.1751 | 0 | 0 | 0 |
| 'Snx21' | 0.523 | 0.589 | 0.778 | 0.478 | 0.732 | 0.627 | 0.459 | 0.636 | 0.725 | 0.704 | 0.657 | 0.621 | 0.649 | 0.595 | 0.571 | 0.588 | 0.552 | 0.564 | 0.003 | 0.9719 | 0.9901 | 0 | 0 | 0 |
| 'Snx22' | 2.315 | 2.764 | 2.447 | 2.765 | 2.274 | 2.458 | 2.939 | 2.616 | 2.434 | 2.557 | 2.465 | 2.656 | 2.563 | 2.243 | 2.833 | 2.489 | 2.728 | 2.454 | -0.020 | 0.8135 | 0.9158 | 0 | 0 | 0 |
| 'Snx25' | 8.860 | 9.179 | 9.317 | 9.600 | 9.305 | 9.420 | 8.816 | 9.554 | 9.136 | 9.089 | 8.673 | 9.333 | 8.518 | 9.424 | 9.127 | 9.038 | 8.563 | 9.749 | 0.019 | 0.5591 | 0.7700 | 0 | 0 | 0 |
| 'Snx27' | 6.037 | 6.035 | 6.095 | 6.060 | 6.024 | 5.649 | 6.043 | 5.964 | 5.882 | 5.645 | 5.800 | 7.247 | 5.622 | 6.171 | 6.724 | 5.525 | 6.587 | 7.374 | -0.084 | 0.0831 | 0.2611 | 0 | 0 | 0 |
| 'Snx29' | 1.107 | 1.358 | 1.198 | 1.143 | 1.165 | 1.133 | 1.435 | 1.189 | 1.318 | 1.086 | 1.418 | 1.304 | 1.087 | 1.237 | 1.096 | 0.872 | 1.355 | 1.040 | 0.058 | 0.4672 | 0.7029 | 0 | 0 | 0 |
| 'Snx3' | 48.961 | 50.879 | 45.961 | 48.113 | 46.324 | 48.797 | 45.390 | 48.156 | 48.911 | 41.959 | 45.013 | 52.627 | 49.086 | 47.708 | 53.815 | 47.624 | 50.893 | 54.868 | -0.049 | 0.2283 | 0.4749 | 0 | 0 | 0 |
| 'Snx30' | 5.009 | 4.845 | 3.732 | 4.517 | 4.518 | 3.698 | 3.983 | 4.581 | 3.914 | 4.640 | 4.425 | 4.537 | 4.324 | 5.118 | 4.222 | 4.573 | 4.442 | 4.778 | -0.099 | 0.0651 | 0.2227 | 0 | 0 | 0 |
| 'Snx32' | 11.387 | 13.361 | 15.254 | 10.081 | 12.087 | 17.106 | 11.007 | 12.808 | 16.864 | 13.673 | 12.358 | 14.878 | 12.468 | 12.292 | 15.110 | 13.325 | 11.405 | 14.501 | -0.014 | 0.8071 | 0.9135 | 0 | 0 | 0 |
| 'Snx33' | 9.161 | 9.111 | 10.321 | 8.160 | 9.180 | 10.375 | 8.160 | 9.994 | 10.473 | 10.388 | 9.626 | 10.150 | 8.669 | 9.790 | 9.641 | 8.779 | 9.621 | 9.959 | -0.036 | 0.4018 | 0.6507 | 0 | 0 | 0 |
| 'Snx4' | 39.917 | 46.456 | 41.482 | 39.450 | 43.361 | 43.308 | 41.830 | 44.244 | 42.028 | 40.733 | 41.268 | 37.132 | 41.074 | 42.280 | 36.437 | 41.191 | 41.100 | 35.641 | 0.092 | 0.0032 | 0.0264 | 1 | 0 | 0 |
| 'Snx5' | 34.866 | 34.207 | 33.907 | 35.249 | 35.088 | 32.142 | 37.174 | 35.378 | 32.455 | 35.517 | 32.533 | 31.579 | 35.527 | 34.364 | 32.034 | 34.627 | 35.980 | 31.029 | 0.025 | 0.2602 | 0.5138 | 0 | 0 | 0 |
| 'Snx6' | 46.378 | 52.005 | 48.190 | 53.059 | 50.605 | 46.369 | 48.530 | 49.943 | 48.574 | 45.515 | 45.573 | 49.871 | 45.938 | 50.808 | 48.416 | 44.027 | 49.418 | 44.574 | 0.051 | 0.1270 | 0.3415 | 0 | 0 | 0 |
| 'Snx8' | 1.981 | 2.327 | 2.014 | 1.806 | 2.197 | 2.023 | 1.425 | 2.464 | 2.145 | 1.728 | 2.184 | 1.853 | 2.074 | 2.333 | 1.594 | 1.880 | 2.112 | 1.902 | 0.052 | 0.4546 | 0.6931 | 0 | 0 | 0 |
| 'Snx9' | 1.893 | 2.010 | 1.723 | 2.007 | 1.571 | 2.309 | 1.267 | 2.117 | 1.869 | 1.336 | 1.562 | 1.741 | 1.308 | 1.457 | 2.089 | 1.266 | 0.893 | 1.878 | 0.307 | 0.0149 | 0.0832 | 0 | 0 | 0 |
| 'Soat1' | 6.210 | 6.433 | 6.212 | 6.285 | 6.611 | 5.596 | 6.667 | 6.478 | 6.303 | 4.530 | 5.393 | 5.752 | 4.744 | 5.289 | 5.627 | 4.875 | 5.376 | 5.728 | 0.253 | 0.0000 | 0.0000 | 1 | 0 | 0 |
| 'Socs1' | 45.725 | 36.912 | 44.207 | 36.014 | 38.457 | 41.502 | 35.049 | 52.677 | 36.811 | 42.892 | 36.913 | 38.900 | 36.052 | 35.539 | 36.867 | 36.924 | 31.079 | 38.197 | 0.132 | 0.0709 | 0.2347 | 0 | 0 | 0 |
| 'Socs3' | 87.846 | 67.025 | 68.987 | 63.703 | 77.085 | 53.223 | 55.212 | 99.991 | 44.433 | 71.053 | 83.209 | 48.811 | 63.334 | 81.108 | 48.272 | 67.989 | 61.219 | 52.703 | 0.089 | 0.4274 | 0.6738 | 0 | 0 | 0 |
| 'Socs4' | 11.970 | 14.259 | 12.970 | 13.355 | 13.196 | 12.185 | 13.245 | 13.459 | 13.470 | 10.884 | 14.103 | 16.560 | 11.392 | 13.356 | 14.726 | 11.944 | 15.334 | 15.131 | -0.067 | 0.2254 | 0.4722 | 0 | 0 | 0 |
| 'Socs5' | 10.048 | 9.682 | 9.290 | 10.255 | 9.131 | 9.614 | 9.858 | 9.010 | 9.053 | 7.790 | 9.483 | 12.627 | 7.766 | 9.604 | 11.457 | 9.018 | 10.220 | 11.847 | -0.063 | 0.3801 | 0.6317 | 0 | 0 | 0 |
| 'Socs7' | 12.550 | 12.255 | 10.620 | 11.447 | 11.225 | 10.660 | 9.688 | 10.873 | 10.855 | 10.869 | 11.754 | 12.729 | 11.216 | 12.280 | 12.265 | 11.539 | 11.378 | 13.516 | -0.113 | 0.0276 | 0.1281 | 0 | 0 | 0 |
| 'Sod1' | 293.889 | 290.988 | 268.449 | 288.802 | 299.842 | 274.906 | 289.085 | 302.121 | 273.797 | 281.059 | 284.943 | 252.302 | 291.862 | 303.387 | 274.147 | 300.985 | 287.099 | 261.050 | 0.018 | 0.4704 | 0.7061 | 0 | 0 | 0 |
| 'Sod2' | 21.366 | 21.897 | 18.735 | 20.193 | 20.080 | 18.742 | 20.542 | 19.058 | 18.026 | 19.115 | 17.629 | 18.288 | 20.279 | 19.622 | 17.905 | 18.886 | 18.446 | 16.462 | 0.087 | 0.0090 | 0.0579 | 0 | 0 | 0 |
| 'Son' | 42.981 | 41.143 | 41.316 | 42.413 | 39.706 | 39.733 | 41.090 | 40.909 | 41.866 | 42.754 | 42.744 | 41.832 | 38.955 | 41.198 | 39.484 | 39.502 | 42.866 | 40.981 | -0.007 | 0.7932 | 0.9077 | 0 | 0 | 0 |
| 'Sorbs1' | 3.229 | 3.701 | 2.863 | 2.954 | 3.172 | 2.813 | 2.948 | 2.912 | 3.160 | 3.202 | 3.452 | 3.183 | 3.059 | 3.086 | 3.092 | 2.763 | 3.404 | 3.170 | -0.051 | 0.3750 | 0.6262 | 0 | 0 | 0 |
| 'Sord' | 4.428 | 4.083 | 4.957 | 4.552 | 4.346 | 5.308 | 4.185 | 3.910 | 4.636 | 4.540 | 4.467 | 4.155 | 4.330 | 4.734 | 4.779 | 3.465 | 4.133 | 3.945 | 0.065 | 0.3810 | 0.6324 | 0 | 0 | 0 |
| 'Sorl1' | 10.856 | 10.464 | 11.950 | 11.232 | 10.514 | 11.887 | 10.983 | 11.314 | 11.759 | 14.219 | 12.206 | 12.906 | 13.714 | 11.886 | 13.257 | 14.801 | 10.569 | 13.311 | -0.217 | 0.0000 | 0.0000 | 1 | 0 | 0 |
| 'Sos1' | 7.054 | 6.281 | 6.468 | 7.732 | 5.960 | 6.265 | 6.332 | 6.510 | 5.930 | 6.811 | 6.506 | 7.406 | 6.296 | 7.399 | 6.621 | 6.571 | 6.900 | 7.580 | -0.098 | 0.0682 | 0.2291 | 0 | 0 | 0 |
| 'Sos2' | 6.435 | 6.689 | 5.853 | 6.078 | 6.657 | 5.925 | 6.385 | 6.514 | 5.927 | 6.555 | 6.660 | 6.318 | 6.020 | 7.169 | 5.810 | 6.184 | 6.431 | 6.837 | -0.051 | 0.1682 | 0.4005 | 0 | 0 | 0 |
| 'Sostdc1' | 67.945 | 100.698 | 72.857 | 71.791 | 103.579 | 75.906 | 67.844 | 95.034 | 71.409 | 98.195 | 109.945 | 91.565 | 74.245 | 104.717 | 81.693 | 92.824 | 98.404 | 94.382 | -0.244 | 0.0000 | 0.0001 | 1 | 0 | 0 |
| 'Sox4' | 0.859 | 0.834 | 1.000 | 0.936 | 0.643 | 1.297 | 1.215 | 1.001 | 1.022 | 1.180 | 0.739 | 1.294 | 1.115 | 0.904 | 1.474 | 1.246 | 0.625 | 0.948 | -0.112 | 0.3378 | 0.5955 | 0 | 0 | 0 |
| 'Sox8' | 2.431 | 2.962 | 1.777 | 3.123 | 2.297 | 1.815 | 3.072 | 2.372 | 2.191 | 1.736 | 2.644 | 3.987 | 1.691 | 2.564 | 3.335 | 1.895 | 3.520 | 3.495 | -0.155 | 0.4182 | 0.6655 | 0 | 0 | 0 |
| 'Sp1' | 15.570 | 16.757 | 15.835 | 15.006 | 15.761 | 15.452 | 15.056 | 16.713 | 15.702 | 16.107 | 17.118 | 16.515 | 14.848 | 16.678 | 15.821 | 16.751 | 15.726 | 15.802 | -0.045 | 0.0913 | 0.2786 | 0 | 0 | 0 |
| 'Sp2' | 10.441 | 10.589 | 12.623 | 9.642 | 11.760 | 12.523 | 10.658 | 11.542 | 12.632 | 12.883 | 10.863 | 10.859 | 11.864 | 10.849 | 10.971 | 11.807 | 10.036 | 10.614 | 0.014 | 0.7907 | 0.9064 | 0 | 0 | 0 |
| 'Sp3' | 18.642 | 19.806 | 17.010 | 19.040 | 19.192 | 17.147 | 19.088 | 18.716 | 16.637 | 17.173 | 18.966 | 18.120 | 17.332 | 19.808 | 17.733 | 18.604 | 19.580 | 17.279 | -0.006 | 0.8183 | 0.9194 | 0 | 0 | 0 |
| 'Sp4' | 14.010 | 15.813 | 15.479 | 15.178 | 16.922 | 14.915 | 16.083 | 16.015 | 14.467 | 15.232 | 16.306 | 15.042 | 14.792 | 17.142 | 13.430 | 15.070 | 17.237 | 13.817 | 0.000 | 0.9935 | 0.9990 | 0 | 0 | 0 |
| 'Sp6' | 4.596 | 5.287 | 4.166 | 4.713 | 3.694 | 4.951 | 4.384 | 4.190 | 5.171 | 3.909 | 2.735 | 4.914 | 2.924 | 2.881 | 3.959 | 3.271 | 3.433 | 4.403 | 0.343 | 0.0000 | 0.0006 | 1 | 0 | 0 |
| 'Sp9' | 1.952 | 2.215 | 1.915 | 2.093 | 2.359 | 1.961 | 1.956 | 2.159 | 1.762 | 2.071 | 1.919 | 2.001 | 1.773 | 1.848 | 1.867 | 1.912 | 2.090 | 1.718 | 0.074 | 0.2712 | 0.5262 | 0 | 0 | 0 |
| 'Spag1' | 2.553 | 1.931 | 1.992 | 2.600 | 2.057 | 2.235 | 2.131 | 2.561 | 2.147 | 2.243 | 2.394 | 2.148 | 2.402 | 2.149 | 2.379 | 2.174 | 2.584 | 2.330 | -0.047 | 0.5595 | 0.7702 | 0 | 0 | 0 |
| 'Spag5' | 3.439 | 3.594 | 5.041 | 4.145 | 3.640 | 4.650 | 4.509 | 3.599 | 4.865 | 2.840 | 4.369 | 4.334 | 5.248 | 4.557 | 4.680 | 3.516 | 3.997 | 4.086 | -0.024 | 0.7857 | 0.9044 | 0 | 0 | 0 |
| 'Spag7' | 13.784 | 15.680 | 14.950 | 14.025 | 15.251 | 15.594 | 14.103 | 13.947 | 14.973 | 13.622 | 14.273 | 13.644 | 14.811 | 14.510 | 14.702 | 14.679 | 15.368 | 12.777 | 0.036 | 0.3790 | 0.6307 | 0 | 0 | 0 |
| 'Spag9' | 3.042 | 2.978 | 2.999 | 3.282 | 2.800 | 2.922 | 3.226 | 2.982 | 3.032 | 2.861 | 3.443 | 3.632 | 3.042 | 3.068 | 3.217 | 2.788 | 3.530 | 3.627 | -0.110 | 0.0382 | 0.1588 | 0 | 0 | 0 |
| 'Spast' | 8.004 | 7.445 | 7.803 | 7.537 | 7.763 | 8.058 | 7.688 | 8.166 | 7.191 | 8.142 | 7.772 | 7.839 | 7.842 | 8.504 | 7.625 | 7.563 | 7.860 | 8.132 | -0.041 | 0.2240 | 0.4711 | 0 | 0 | 0 |
| 'Spata1' | 1.020 | 1.532 | 1.374 | 1.316 | 1.678 | 1.322 | 1.496 | 1.534 | 1.395 | 1.967 | 1.408 | 1.421 | 1.724 | 1.796 | 1.310 | 1.427 | 1.714 | 1.035 | -0.130 | 0.1675 | 0.3999 | 0 | 0 | 0 |
| 'Spata13' | 47.342 | 50.230 | 43.612 | 46.731 | 50.436 | 45.190 | 44.168 | 49.340 | 45.151 | 51.950 | 52.998 | 45.773 | 51.860 | 52.834 | 46.776 | 51.291 | 49.884 | 47.051 | -0.104 | 0.0000 | 0.0002 | 1 | 0 | 0 |
| 'Spata2' | 8.767 | 8.190 | 8.148 | 8.208 | 8.045 | 8.569 | 7.115 | 8.471 | 7.712 | 8.842 | 8.566 | 9.101 | 8.665 | 10.028 | 8.947 | 8.812 | 7.085 | 9.145 | -0.122 | 0.0201 | 0.1036 | 0 | 0 | 0 |
| 'Spata24' | 4.525 | 6.197 | 6.777 | 5.601 | 5.183 | 5.605 | 5.537 | 5.869 | 5.446 | 4.736 | 4.551 | 4.400 | 6.098 | 4.602 | 5.755 | 6.347 | 5.372 | 3.976 | 0.150 | 0.1564 | 0.3855 | 0 | 0 | 0 |
| 'Spata2l' | 2.139 | 2.324 | 2.883 | 2.209 | 2.069 | 2.644 | 2.315 | 2.084 | 2.526 | 2.154 | 2.101 | 2.455 | 1.756 | 1.773 | 2.119 | 2.119 | 1.750 | 2.614 | 0.161 | 0.0187 | 0.0985 | 0 | 0 | 0 |
| 'Spata5' | 3.856 | 3.392 | 3.781 | 4.236 | 3.220 | 3.371 | 3.693 | 3.689 | 3.477 | 4.117 | 3.848 | 3.578 | 3.253 | 3.096 | 3.037 | 3.333 | 3.791 | 3.366 | 0.048 | 0.4343 | 0.6788 | 0 | 0 | 0 |
| 'Spata6' | 0.654 | 0.879 | 0.914 | 0.766 | 0.778 | 0.787 | 0.892 | 0.780 | 0.825 | 0.659 | 0.680 | 0.714 | 0.548 | 0.810 | 0.815 | 0.750 | 0.688 | 0.617 | 0.205 | 0.0130 | 0.0755 | 0 | 0 | 0 |
| 'Spc24' | 15.543 | 16.024 | 20.021 | 18.888 | 17.050 | 19.857 | 18.212 | 16.170 | 20.317 | 13.039 | 15.005 | 19.771 | 20.469 | 17.521 | 21.459 | 15.357 | 16.035 | 17.811 | 0.037 | 0.5799 | 0.7822 | 0 | 0 | 0 |
| 'Spc25' | 2.687 | 2.955 | 3.858 | 3.999 | 3.231 | 3.504 | 3.750 | 3.000 | 3.576 | 2.304 | 3.136 | 3.611 | 3.792 | 3.500 | 4.054 | 2.952 | 3.335 | 3.529 | 0.003 | 0.9711 | 0.9898 | 0 | 0 | 0 |
| 'Spcs1' | 16.962 | 24.526 | 17.280 | 19.220 | 23.909 | 19.144 | 19.306 | 22.413 | 19.160 | 19.935 | 20.372 | 17.250 | 21.015 | 22.797 | 17.862 | 21.166 | 22.033 | 17.209 | 0.004 | 0.9163 | 0.9681 | 0 | 0 | 0 |
| 'Spcs2' | 68.750 | 77.814 | 69.842 | 72.630 | 77.697 | 70.992 | 72.606 | 73.525 | 69.898 | 68.278 | 67.723 | 66.442 | 67.062 | 72.052 | 66.474 | 66.578 | 72.839 | 63.310 | 0.086 | 0.0000 | 0.0008 | 1 | 0 | 0 |
| 'Spcs3' | 54.809 | 63.379 | 61.971 | 58.138 | 66.855 | 58.999 | 60.425 | 63.325 | 57.990 | 60.195 | 59.832 | 57.914 | 59.121 | 62.525 | 57.593 | 56.850 | 61.473 | 57.081 | 0.024 | 0.3201 | 0.5782 | 0 | 0 | 0 |
| 'Spdl1' | 3.052 | 2.521 | 4.017 | 3.959 | 2.823 | 3.582 | 4.301 | 3.177 | 3.335 | 2.353 | 2.824 | 3.484 | 3.431 | 3.543 | 3.880 | 2.765 | 3.327 | 3.127 | 0.083 | 0.3821 | 0.6330 | 0 | 0 | 0 |
| 'Specc1l' | 25.703 | 25.271 | 25.485 | 25.256 | 25.203 | 25.827 | 25.283 | 26.832 | 25.415 | 26.648 | 26.931 | 24.098 | 24.102 | 25.606 | 23.601 | 27.477 | 23.718 | 22.982 | 0.025 | 0.4351 | 0.6791 | 0 | 0 | 0 |
| 'Spef1' | 4.533 | 3.529 | 3.996 | 3.745 | 3.650 | 4.546 | 4.352 | 3.897 | 3.798 | 3.029 | 3.367 | 3.330 | 3.864 | 4.064 | 3.473 | 3.051 | 2.878 | 3.968 | 0.211 | 0.0318 | 0.1400 | 0 | 0 | 0 |
| 'Spen' | 21.236 | 16.929 | 19.399 | 20.268 | 17.287 | 19.164 | 18.455 | 17.705 | 20.170 | 21.346 | 19.661 | 21.527 | 21.132 | 19.559 | 20.554 | 22.315 | 19.239 | 22.711 | -0.151 | 0.0000 | 0.0000 | 1 | 0 | 0 |
| 'Spg11' | 5.578 | 5.554 | 6.248 | 5.601 | 5.999 | 5.337 | 5.621 | 5.916 | 6.019 | 6.084 | 5.950 | 5.887 | 5.553 | 5.163 | 5.332 | 5.945 | 5.884 | 5.270 | 0.014 | 0.7323 | 0.8781 | 0 | 0 | 0 |
| 'Spg20' | 1.935 | 2.097 | 2.099 | 1.801 | 2.223 | 2.250 | 1.914 | 2.332 | 2.311 | 2.291 | 1.906 | 2.117 | 2.064 | 2.320 | 1.995 | 2.344 | 2.263 | 1.986 | -0.032 | 0.5459 | 0.7608 | 0 | 0 | 0 |
| 'Spg21' | 27.873 | 31.615 | 29.444 | 25.772 | 33.763 | 29.615 | 29.655 | 32.188 | 27.762 | 30.869 | 29.326 | 28.597 | 28.117 | 29.941 | 28.871 | 28.736 | 29.097 | 26.917 | 0.027 | 0.4433 | 0.6868 | 0 | 0 | 0 |
| 'Spg7' | 11.355 | 11.491 | 12.603 | 11.634 | 11.003 | 12.058 | 11.630 | 11.275 | 12.048 | 11.542 | 11.302 | 11.876 | 12.003 | 10.550 | 11.812 | 11.159 | 11.827 | 11.506 | 0.011 | 0.6778 | 0.8460 | 0 | 0 | 0 |
| 'Sphk2' | 8.558 | 8.613 | 8.679 | 7.773 | 8.252 | 9.486 | 7.258 | 8.719 | 8.771 | 8.000 | 7.552 | 9.538 | 8.604 | 7.682 | 8.687 | 9.101 | 7.258 | 8.666 | 0.010 | 0.8182 | 0.9194 | 0 | 0 | 0 |
| 'Spib' | 1.820 | 3.395 | 2.080 | 2.534 | 2.842 | 2.141 | 2.381 | 2.563 | 2.947 | 1.211 | 2.183 | 1.837 | 1.197 | 3.023 | 1.768 | 1.844 | 2.230 | 1.273 | 0.466 | 0.0001 | 0.0014 | 1 | 1 | 0 |
| 'Spice1' | 1.457 | 1.102 | 1.753 | 1.689 | 1.498 | 1.881 | 1.721 | 1.424 | 1.697 | 1.673 | 1.418 | 1.778 | 1.825 | 1.549 | 1.847 | 1.584 | 1.821 | 1.628 | -0.100 | 0.1329 | 0.3509 | 0 | 0 | 0 |
| 'Spidr' | 6.579 | 6.915 | 6.102 | 6.801 | 6.476 | 6.763 | 6.871 | 6.725 | 6.475 | 6.079 | 7.443 | 6.894 | 5.991 | 7.396 | 6.131 | 6.542 | 6.717 | 6.325 | -0.007 | 0.8913 | 0.9556 | 0 | 0 | 0 |
| 'Spin1' | 25.180 | 26.263 | 24.852 | 23.997 | 26.066 | 24.290 | 25.341 | 26.129 | 23.440 | 25.419 | 25.427 | 25.347 | 24.002 | 26.544 | 24.633 | 26.370 | 25.405 | 25.137 | -0.028 | 0.2613 | 0.5150 | 0 | 0 | 0 |
| 'Spint2' | 81.835 | 98.297 | 84.896 | 77.077 | 91.457 | 89.424 | 78.732 | 91.976 | 88.812 | 85.209 | 81.213 | 86.481 | 80.491 | 84.416 | 85.516 | 90.525 | 78.058 | 86.653 | 0.031 | 0.4192 | 0.6663 | 0 | 0 | 0 |
| 'Spn' | 79.954 | 75.314 | 81.333 | 78.129 | 74.082 | 87.486 | 81.339 | 71.415 | 83.382 | 75.808 | 81.180 | 97.024 | 69.697 | 80.480 | 100.381 | 82.131 | 80.484 | 101.291 | -0.113 | 0.0101 | 0.0630 | 0 | 0 | 0 |
| 'Spns1' | 8.344 | 7.465 | 9.091 | 8.511 | 8.028 | 8.475 | 8.328 | 7.763 | 9.418 | 7.734 | 8.207 | 10.451 | 8.365 | 8.209 | 9.709 | 6.896 | 8.468 | 9.563 | -0.049 | 0.2696 | 0.5244 | 0 | 0 | 0 |
| 'Spo11' | 0.820 | 1.730 | 1.272 | 0.551 | 1.532 | 1.518 | 0.871 | 1.301 | 1.324 | 1.121 | 0.775 | 1.169 | 1.302 | 1.335 | 1.075 | 1.700 | 1.140 | 1.290 | -0.055 | 0.7703 | 0.8977 | 0 | 0 | 0 |
| 'Spock2' | 14.161 | 19.566 | 13.321 | 14.220 | 18.415 | 16.654 | 14.812 | 16.800 | 16.359 | 16.077 | 14.821 | 12.666 | 17.349 | 15.326 | 12.395 | 17.067 | 14.604 | 12.428 | 0.114 | 0.0777 | 0.2498 | 0 | 0 | 0 |
| 'Spop' | 43.793 | 47.834 | 43.479 | 45.556 | 49.023 | 42.923 | 44.430 | 48.955 | 43.820 | 43.142 | 44.401 | 43.618 | 43.023 | 46.781 | 42.098 | 44.610 | 48.871 | 40.124 | 0.036 | 0.0934 | 0.2829 | 0 | 0 | 0 |
| 'Spopl' | 4.830 | 5.559 | 5.228 | 5.452 | 5.243 | 5.055 | 6.154 | 5.439 | 4.861 | 4.576 | 5.373 | 5.489 | 4.470 | 5.500 | 4.926 | 4.534 | 5.741 | 5.218 | 0.051 | 0.3220 | 0.5799 | 0 | 0 | 0 |
| 'Sppl2a' | 9.832 | 11.908 | 11.485 | 10.891 | 11.845 | 10.635 | 10.767 | 10.971 | 10.643 | 11.207 | 11.455 | 11.346 | 10.720 | 12.412 | 10.693 | 10.457 | 11.742 | 10.501 | -0.034 | 0.2647 | 0.5184 | 0 | 0 | 0 |
| 'Sppl2b' | 15.316 | 15.521 | 19.091 | 15.870 | 15.094 | 18.651 | 15.214 | 15.294 | 19.264 | 16.496 | 16.015 | 18.117 | 17.175 | 14.564 | 17.257 | 15.370 | 13.521 | 17.700 | 0.020 | 0.6096 | 0.8039 | 0 | 0 | 0 |
| 'Sppl3' | 7.019 | 6.577 | 6.896 | 6.879 | 7.071 | 7.030 | 7.154 | 7.338 | 6.747 | 6.578 | 6.722 | 6.996 | 6.206 | 6.991 | 6.851 | 6.609 | 6.190 | 7.475 | 0.039 | 0.3143 | 0.5719 | 0 | 0 | 0 |
| 'Spr' | 4.956 | 5.521 | 5.116 | 4.568 | 5.219 | 4.669 | 4.418 | 5.294 | 5.379 | 5.413 | 5.474 | 5.416 | 5.081 | 5.633 | 5.621 | 5.716 | 5.354 | 5.599 | -0.136 | 0.0020 | 0.0178 | 1 | 0 | 0 |
| 'Spred1' | 3.465 | 2.988 | 3.350 | 3.397 | 3.698 | 2.749 | 3.201 | 3.237 | 2.953 | 3.433 | 3.222 | 3.115 | 3.184 | 3.788 | 3.042 | 3.168 | 3.237 | 3.395 | -0.037 | 0.5111 | 0.7362 | 0 | 0 | 0 |
| 'Spred2' | 9.911 | 10.365 | 9.053 | 9.679 | 11.000 | 10.008 | 9.459 | 10.718 | 9.834 | 9.673 | 9.702 | 8.094 | 9.371 | 9.501 | 8.994 | 10.029 | 8.491 | 7.810 | 0.136 | 0.0017 | 0.0156 | 1 | 0 | 0 |
| 'Sprtn' | 13.702 | 15.683 | 15.859 | 12.765 | 15.629 | 15.653 | 14.522 | 15.058 | 16.059 | 15.213 | 14.176 | 16.109 | 13.849 | 14.169 | 15.365 | 13.722 | 14.822 | 15.776 | 0.005 | 0.9108 | 0.9654 | 0 | 0 | 0 |
| 'Spry1' | 19.550 | 14.946 | 13.011 | 13.246 | 16.685 | 13.933 | 12.825 | 20.250 | 13.117 | 21.551 | 14.762 | 11.417 | 19.743 | 17.709 | 11.582 | 21.339 | 14.890 | 10.721 | -0.035 | 0.7558 | 0.8912 | 0 | 0 | 0 |
| 'Spryd3' | 11.480 | 13.073 | 12.421 | 10.817 | 14.241 | 13.281 | 10.408 | 12.517 | 12.858 | 12.355 | 11.000 | 10.933 | 11.741 | 11.439 | 11.515 | 12.522 | 10.664 | 10.454 | 0.103 | 0.0508 | 0.1908 | 0 | 0 | 0 |
| 'Spryd4' | 6.279 | 7.496 | 6.904 | 6.234 | 7.311 | 6.641 | 5.808 | 6.971 | 6.921 | 5.791 | 5.366 | 6.894 | 6.257 | 6.272 | 6.476 | 6.783 | 6.215 | 6.955 | 0.069 | 0.3078 | 0.5662 | 0 | 0 | 0 |
| 'Spryd7' | 0.840 | 0.955 | 0.856 | 0.782 | 0.818 | 0.946 | 0.784 | 0.802 | 0.740 | 0.781 | 0.517 | 0.755 | 0.664 | 0.738 | 0.688 | 0.814 | 0.633 | 0.737 | 0.237 | 0.0253 | 0.1216 | 0 | 0 | 0 |
| 'Spsb1' | 14.989 | 15.454 | 13.214 | 13.497 | 15.450 | 13.725 | 12.758 | 15.204 | 13.019 | 15.106 | 17.795 | 16.564 | 16.653 | 17.618 | 15.876 | 14.566 | 16.846 | 16.390 | -0.224 | 0.0000 | 0.0000 | 1 | 0 | 0 |
| 'Spsb2' | 1.698 | 1.604 | 1.152 | 1.728 | 1.764 | 1.273 | 2.180 | 1.772 | 1.423 | 1.507 | 1.501 | 1.373 | 1.538 | 1.949 | 1.571 | 1.418 | 1.656 | 1.054 | 0.079 | 0.4207 | 0.6675 | 0 | 0 | 0 |
| 'Spsb3' | 1.730 | 1.773 | 2.096 | 1.903 | 1.855 | 1.990 | 1.963 | 2.117 | 2.072 | 1.587 | 1.960 | 2.119 | 1.694 | 1.758 | 2.152 | 2.129 | 1.954 | 2.013 | 0.004 | 0.9455 | 0.9790 | 0 | 0 | 0 |
| 'Sptan1' | 21.210 | 19.744 | 22.182 | 20.755 | 20.178 | 22.253 | 20.049 | 20.760 | 21.387 | 22.719 | 20.673 | 20.781 | 21.011 | 20.695 | 21.000 | 21.640 | 20.393 | 21.213 | -0.022 | 0.3863 | 0.6356 | 0 | 0 | 0 |
| 'Sptb' | 6.430 | 6.994 | 6.297 | 6.393 | 6.325 | 6.969 | 6.290 | 6.261 | 6.497 | 6.731 | 5.798 | 6.146 | 6.026 | 6.250 | 5.544 | 6.859 | 6.113 | 5.797 | 0.073 | 0.0610 | 0.2134 | 0 | 0 | 0 |
| 'Sptbn1' | 14.507 | 13.449 | 14.701 | 15.381 | 14.350 | 14.166 | 14.984 | 14.121 | 14.730 | 14.487 | 15.565 | 14.834 | 14.325 | 13.984 | 14.307 | 13.311 | 14.630 | 15.431 | -0.016 | 0.6278 | 0.8154 | 0 | 0 | 0 |
| 'Sptbn2' | 3.029 | 2.748 | 3.467 | 3.202 | 2.965 | 3.163 | 2.944 | 2.836 | 3.258 | 3.298 | 3.387 | 2.807 | 3.003 | 2.975 | 2.722 | 2.804 | 2.467 | 2.609 | 0.079 | 0.2005 | 0.4416 | 0 | 0 | 0 |
| 'Sptlc1' | 61.259 | 72.343 | 70.625 | 67.553 | 73.777 | 69.185 | 67.603 | 72.803 | 69.920 | 72.644 | 66.767 | 65.554 | 73.837 | 71.625 | 64.079 | 70.965 | 71.971 | 62.506 | 0.002 | 0.9423 | 0.9783 | 0 | 0 | 0 |
| 'Sptlc2' | 6.551 | 7.371 | 7.333 | 6.890 | 7.685 | 7.494 | 7.527 | 7.769 | 6.961 | 7.525 | 7.513 | 7.279 | 7.158 | 7.400 | 7.043 | 6.914 | 7.592 | 6.921 | -0.006 | 0.8455 | 0.9343 | 0 | 0 | 0 |
| 'Sptssa' | 110.436 | 126.992 | 125.888 | 122.588 | 123.871 | 109.746 | 124.659 | 127.437 | 118.790 | 109.802 | 122.175 | 109.262 | 103.248 | 115.979 | 122.021 | 107.119 | 119.321 | 111.471 | 0.086 | 0.0164 | 0.0897 | 0 | 0 | 0 |
| 'Spty2d1' | 32.441 | 18.851 | 11.553 | 20.361 | 18.708 | 13.135 | 17.973 | 20.893 | 12.134 | 28.780 | 21.556 | 12.395 | 25.253 | 24.706 | 12.098 | 32.607 | 17.841 | 11.563 | -0.142 | 0.1859 | 0.4216 | 0 | 0 | 0 |
| 'Sqle' | 3.602 | 4.238 | 4.502 | 3.813 | 4.177 | 4.244 | 3.962 | 3.660 | 4.449 | 3.504 | 4.650 | 3.553 | 4.129 | 4.093 | 3.870 | 3.704 | 3.435 | 3.152 | 0.097 | 0.1373 | 0.3573 | 0 | 0 | 0 |
| 'Sqrdl' | 3.493 | 4.303 | 3.564 | 3.980 | 4.045 | 4.153 | 3.860 | 3.754 | 3.784 | 4.109 | 3.712 | 3.868 | 3.565 | 4.506 | 3.372 | 4.414 | 4.080 | 3.726 | -0.027 | 0.6320 | 0.8176 | 0 | 0 | 0 |
| 'Sqstm1' | 35.551 | 31.657 | 30.686 | 32.077 | 31.020 | 30.744 | 30.180 | 32.219 | 29.729 | 34.055 | 30.408 | 29.572 | 31.314 | 30.546 | 29.810 | 34.831 | 28.615 | 29.270 | 0.020 | 0.5383 | 0.7566 | 0 | 0 | 0 |
| 'Sra1' | 13.812 | 14.268 | 15.067 | 14.869 | 14.676 | 15.348 | 14.898 | 15.114 | 15.552 | 14.180 | 14.177 | 14.496 | 14.654 | 14.528 | 14.550 | 14.390 | 15.154 | 13.662 | 0.033 | 0.2187 | 0.4655 | 0 | 0 | 0 |
| 'Srbd1' | 10.391 | 11.711 | 13.607 | 11.584 | 12.458 | 13.000 | 11.987 | 12.220 | 11.908 | 11.192 | 12.187 | 11.675 | 11.873 | 11.232 | 12.087 | 12.258 | 12.126 | 11.060 | 0.036 | 0.4114 | 0.6599 | 0 | 0 | 0 |
| 'Srcap' | 1.479 | 1.291 | 1.734 | 1.398 | 1.283 | 1.533 | 1.529 | 1.298 | 1.762 | 1.750 | 1.655 | 1.633 | 1.639 | 1.390 | 1.444 | 1.568 | 1.539 | 1.573 | -0.105 | 0.0557 | 0.2022 | 0 | 0 | 0 |
| 'Srd5a3' | 4.105 | 5.424 | 4.784 | 4.218 | 4.783 | 5.203 | 4.141 | 4.897 | 4.786 | 4.723 | 4.345 | 4.972 | 4.687 | 4.502 | 4.981 | 5.347 | 4.498 | 5.071 | -0.039 | 0.4476 | 0.6880 | 0 | 0 | 0 |
| 'Srebf1' | 15.694 | 16.606 | 19.178 | 16.071 | 15.621 | 17.880 | 16.700 | 16.388 | 19.199 | 17.468 | 16.616 | 17.714 | 16.949 | 16.093 | 17.959 | 16.380 | 15.538 | 16.641 | 0.007 | 0.8093 | 0.9140 | 0 | 0 | 0 |
| 'Srebf2' | 88.847 | 90.768 | 91.663 | 89.734 | 88.722 | 94.608 | 86.845 | 82.790 | 95.317 | 93.439 | 93.234 | 95.090 | 92.757 | 90.116 | 99.355 | 92.211 | 82.998 | 94.746 | -0.054 | 0.0387 | 0.1603 | 0 | 0 | 0 |
| 'Srek1' | 17.001 | 19.280 | 20.127 | 19.294 | 19.928 | 19.119 | 19.142 | 19.742 | 19.791 | 19.759 | 20.002 | 18.969 | 18.622 | 19.027 | 18.449 | 18.904 | 19.968 | 17.880 | 0.006 | 0.8384 | 0.9309 | 0 | 0 | 0 |
| 'Srek1ip1' | 2.094 | 2.554 | 2.483 | 2.567 | 2.802 | 2.568 | 2.843 | 2.576 | 2.700 | 2.540 | 2.418 | 2.557 | 2.534 | 2.699 | 2.385 | 2.550 | 2.611 | 2.060 | 0.040 | 0.4826 | 0.7146 | 0 | 0 | 0 |
| 'Srf' | 38.054 | 37.224 | 41.656 | 34.973 | 37.333 | 44.641 | 32.820 | 39.267 | 40.221 | 44.360 | 38.548 | 43.988 | 44.078 | 40.145 | 45.556 | 42.956 | 36.918 | 43.422 | -0.145 | 0.0003 | 0.0044 | 1 | 0 | 0 |
| 'Srfbp1' | 23.425 | 21.129 | 19.163 | 21.229 | 19.614 | 19.135 | 22.418 | 21.448 | 19.191 | 22.234 | 22.172 | 19.988 | 21.709 | 20.494 | 19.436 | 24.297 | 21.820 | 19.997 | -0.048 | 0.2432 | 0.4935 | 0 | 0 | 0 |
| 'Srgap2' | 6.790 | 5.725 | 7.254 | 7.227 | 6.096 | 7.099 | 6.673 | 6.676 | 7.428 | 7.027 | 7.362 | 7.183 | 7.195 | 6.941 | 6.656 | 6.487 | 7.358 | 7.040 | -0.064 | 0.1412 | 0.3636 | 0 | 0 | 0 |
| 'Srgn' | 150.589 | 173.710 | 143.708 | 155.687 | 173.190 | 144.422 | 159.231 | 161.951 | 146.465 | 166.824 | 165.537 | 124.523 | 157.092 | 169.850 | 143.535 | 171.417 | 169.335 | 135.749 | -0.001 | 0.9713 | 0.9898 | 0 | 0 | 0 |
| 'Sri' | 26.462 | 30.458 | 27.694 | 28.189 | 32.263 | 29.366 | 28.586 | 31.662 | 28.597 | 28.548 | 29.788 | 27.505 | 27.707 | 29.225 | 27.429 | 29.075 | 29.825 | 26.491 | 0.032 | 0.1753 | 0.4100 | 0 | 0 | 0 |
| 'Srm' | 12.234 | 9.815 | 11.709 | 11.516 | 10.184 | 10.775 | 10.638 | 11.354 | 10.887 | 10.194 | 9.417 | 13.070 | 10.936 | 9.971 | 11.714 | 10.725 | 9.539 | 12.333 | 0.007 | 0.8974 | 0.9589 | 0 | 0 | 0 |
| 'Srp14' | 40.228 | 45.507 | 36.190 | 42.148 | 43.721 | 40.072 | 41.427 | 40.259 | 38.036 | 37.579 | 39.244 | 37.406 | 43.661 | 41.403 | 37.763 | 42.956 | 43.393 | 37.212 | 0.015 | 0.6535 | 0.8320 | 0 | 0 | 0 |
| 'Srp19' | 7.669 | 8.678 | 7.931 | 8.209 | 9.036 | 8.272 | 8.594 | 8.354 | 7.979 | 7.563 | 8.045 | 8.504 | 8.222 | 8.535 | 8.646 | 7.969 | 8.886 | 7.847 | -0.006 | 0.8581 | 0.9390 | 0 | 0 | 0 |
| 'Srp68' | 31.178 | 31.258 | 31.479 | 31.397 | 32.656 | 32.406 | 31.019 | 32.892 | 31.320 | 29.290 | 28.494 | 29.957 | 32.692 | 30.186 | 29.743 | 29.132 | 30.259 | 29.017 | 0.077 | 0.0018 | 0.0170 | 1 | 0 | 0 |
| 'Srp72' | 19.893 | 19.793 | 18.775 | 20.145 | 20.746 | 19.922 | 20.482 | 19.993 | 19.212 | 19.852 | 18.696 | 19.025 | 19.158 | 19.779 | 19.230 | 19.463 | 19.435 | 18.745 | 0.034 | 0.0936 | 0.2832 | 0 | 0 | 0 |
| 'Srp9' | 14.188 | 16.174 | 13.237 | 13.680 | 16.202 | 14.078 | 14.255 | 15.540 | 13.382 | 14.648 | 14.125 | 12.115 | 15.322 | 15.984 | 13.081 | 15.826 | 15.060 | 12.595 | 0.015 | 0.6766 | 0.8452 | 0 | 0 | 0 |
| 'Srpk1' | 18.009 | 16.476 | 19.003 | 18.241 | 16.336 | 17.635 | 17.869 | 17.403 | 18.900 | 16.267 | 17.108 | 19.462 | 16.122 | 15.360 | 18.639 | 15.727 | 16.899 | 19.147 | 0.038 | 0.2689 | 0.5231 | 0 | 0 | 0 |
| 'Srpk2' | 5.627 | 6.021 | 5.783 | 5.878 | 5.851 | 5.878 | 6.027 | 5.940 | 5.882 | 5.603 | 6.346 | 6.020 | 5.534 | 5.760 | 5.926 | 5.422 | 5.813 | 6.176 | -0.004 | 0.9027 | 0.9613 | 0 | 0 | 0 |
| 'Srpr' | 32.921 | 35.079 | 34.520 | 34.264 | 35.741 | 33.597 | 34.022 | 36.238 | 33.680 | 31.566 | 35.610 | 37.226 | 33.831 | 36.197 | 35.418 | 34.909 | 35.774 | 37.806 | -0.050 | 0.0718 | 0.2363 | 0 | 0 | 0 |
| 'Srprb' | 4.008 | 4.714 | 4.023 | 3.932 | 4.253 | 4.210 | 3.924 | 4.376 | 4.272 | 3.406 | 3.769 | 4.736 | 3.789 | 3.996 | 4.681 | 3.656 | 4.287 | 4.804 | 0.007 | 0.8964 | 0.9584 | 0 | 0 | 0 |
| 'Srr' | 1.068 | 1.299 | 1.113 | 1.044 | 1.312 | 1.134 | 1.161 | 1.062 | 1.124 | 1.201 | 0.946 | 1.036 | 1.078 | 1.048 | 0.995 | 1.278 | 0.882 | 0.908 | 0.130 | 0.0533 | 0.1971 | 0 | 0 | 0 |
| 'Srrd' | 2.933 | 3.231 | 3.126 | 2.357 | 3.064 | 2.694 | 2.318 | 2.714 | 3.257 | 3.133 | 2.562 | 2.540 | 2.788 | 2.727 | 2.614 | 2.804 | 2.392 | 2.151 | 0.113 | 0.1683 | 0.4007 | 0 | 0 | 0 |

| | | | | | | | | | | | | | | | | | | | | | | | | |
|---|---|---|---|---|---|---|---|---|---|---|---|---|---|---|---|---|---|---|---|---|---|---|---|---|
| 'Srrm1' | 19.097 | 19.810 | 20.818 | 17.836 | 19.206 | 20.343 | 18.475 | 20.068 | 21.344 | 20.940 | 18.777 | 20.193 | 21.718 | 19.580 | 19.452 | 20.382 | 18.224 | 19.411 | -0.024 | 0.4840 | 0.7156 | 0 | 0 | 0 |
| 'Srrm2' | 69.431 | 60.803 | 65.026 | 61.354 | 58.495 | 63.972 | 59.441 | 58.498 | 66.955 | 66.925 | 66.619 | 65.321 | 63.242 | 61.405 | 64.611 | 63.628 | 60.721 | 67.981 | -0.053 | 0.1138 | 0.3199 | 0 | 0 | 0 |
| 'Srrt' | 33.813 | 29.755 | 37.165 | 33.880 | 31.645 | 35.210 | 34.070 | 31.358 | 37.033 | 32.816 | 30.853 | 35.027 | 35.336 | 30.148 | 33.394 | 31.283 | 31.450 | 34.130 | 0.034 | 0.1758 | 0.4109 | 0 | 0 | 0 |
| 'Srsf1' | 32.474 | 32.726 | 31.888 | 32.387 | 32.050 | 30.770 | 32.235 | 31.377 | 32.426 | 28.507 | 32.295 | 31.265 | 33.860 | 34.722 | 31.291 | 28.409 | 33.165 | 30.652 | 0.011 | 0.7410 | 0.8827 | 0 | 0 | 0 |
| 'Srsf11' | 17.503 | 16.421 | 17.350 | 17.378 | 16.974 | 16.943 | 16.748 | 17.001 | 17.413 | 18.083 | 16.475 | 17.053 | 17.649 | 16.288 | 15.928 | 16.458 | 17.182 | 16.095 | 0.014 | 0.5464 | 0.7610 | 0 | 0 | 0 |
| 'Srsf2' | 63.122 | 56.594 | 59.489 | 60.261 | 57.881 | 57.692 | 63.153 | 58.383 | 54.871 | 57.392 | 59.112 | 55.313 | 66.141 | 60.372 | 58.369 | 59.999 | 59.528 | 54.014 | -0.007 | 0.8188 | 0.9198 | 0 | 0 | 0 |
| 'Srsf3' | 32.290 | 32.007 | 28.581 | 30.897 | 32.105 | 28.446 | 33.003 | 31.657 | 29.222 | 29.719 | 30.537 | 27.938 | 32.499 | 32.841 | 29.306 | 30.422 | 30.791 | 28.459 | 0.019 | 0.4459 | 0.6878 | 0 | 0 | 0 |
| 'Srsf4' | 33.657 | 31.516 | 33.064 | 32.943 | 32.331 | 35.126 | 32.885 | 31.695 | 33.919 | 33.294 | 32.629 | 32.753 | 37.365 | 31.103 | 32.469 | 31.938 | 31.778 | 32.560 | -0.004 | 0.8843 | 0.9514 | 0 | 0 | 0 |
| 'Srsf5' | 51.572 | 55.889 | 42.728 | 45.326 | 55.159 | 44.829 | 46.752 | 54.529 | 42.855 | 54.380 | 46.520 | 33.254 | 52.674 | 55.278 | 35.912 | 54.707 | 47.228 | 29.317 | 0.118 | 0.0601 | 0.2114 | 0 | 0 | 0 |
| 'Srsf6' | 44.675 | 50.548 | 44.769 | 38.162 | 50.921 | 46.269 | 40.079 | 48.708 | 46.690 | 47.087 | 43.250 | 41.583 | 50.357 | 48.694 | 43.368 | 51.080 | 44.349 | 42.367 | -0.016 | 0.7460 | 0.8854 | 0 | 0 | 0 |
| 'Srsf7' | 113.787 | 109.024 | 101.131 | 116.126 | 109.672 | 95.996 | 109.280 | 110.644 | 99.465 | 107.032 | 103.708 | 103.666 | 108.783 | 113.276 | 101.856 | 105.957 | 116.979 | 103.682 | -0.012 | 0.6633 | 0.8375 | 0 | 0 | 0 |
| 'Srsf9' | 25.355 | 26.675 | 25.449 | 23.629 | 28.445 | 25.373 | 25.535 | 28.625 | 23.332 | 22.898 | 24.573 | 23.354 | 24.271 | 25.117 | 26.699 | 24.816 | 24.670 | 26.859 | 0.046 | 0.2698 | 0.5246 | 0 | 0 | 0 |
| 'Srxn1' | 3.958 | 4.298 | 4.088 | 4.457 | 4.107 | 4.202 | 3.290 | 4.363 | 4.219 | 2.818 | 3.987 | 4.632 | 3.049 | 3.378 | 4.538 | 3.269 | 4.585 | 4.745 | 0.076 | 0.3358 | 0.5944 | 0 | 0 | 0 |
| 'Ss18' | 17.836 | 19.197 | 17.394 | 18.313 | 20.047 | 18.878 | 16.869 | 19.469 | 19.060 | 16.330 | 18.266 | 21.193 | 18.878 | 20.110 | 20.886 | 20.648 | 19.381 | 21.113 | -0.094 | 0.0284 | 0.1306 | 0 | 0 | 0 |
| 'Ss18l1' | 3.072 | 3.596 | 3.036 | 3.271 | 3.106 | 2.746 | 3.795 | 2.608 | 3.139 | 2.714 | 3.663 | 3.957 | 2.878 | 3.842 | 3.651 | 2.699 | 3.676 | 4.392 | -0.158 | 0.0913 | 0.2786 | 0 | 0 | 0 |
| 'Ssb' | 42.864 | 46.008 | 42.582 | 46.243 | 48.366 | 41.400 | 48.942 | 44.999 | 41.685 | 41.924 | 42.857 | 41.586 | 43.097 | 45.456 | 41.318 | 44.668 | 46.426 | 38.745 | 0.051 | 0.0597 | 0.2110 | 0 | 0 | 0 |
| 'Ssbp1' | 3.207 | 3.700 | 3.606 | 3.139 | 3.617 | 3.412 | 3.239 | 3.845 | 3.504 | 2.879 | 3.351 | 3.313 | 3.463 | 3.614 | 3.486 | 3.598 | 3.347 | 2.699 | 0.063 | 0.2125 | 0.4575 | 0 | 0 | 0 |
| 'Ssbp2' | 1.777 | 2.029 | 1.730 | 1.439 | 2.075 | 1.771 | 1.943 | 2.086 | 1.765 | 1.893 | 2.084 | 2.171 | 2.245 | 2.282 | 1.983 | 1.833 | 2.115 | 2.101 | -0.192 | 0.0007 | 0.0081 | 1 | 0 | 0 |
| 'Ssbp3' | 9.777 | 9.190 | 10.715 | 10.486 | 10.321 | 11.159 | 9.204 | 10.420 | 10.465 | 11.442 | 9.589 | 10.773 | 10.409 | 9.898 | 10.606 | 10.120 | 9.186 | 10.671 | -0.024 | 0.5106 | 0.7356 | 0 | 0 | 0 |
| 'Ssbp4' | 91.309 | 70.897 | 103.935 | 77.740 | 78.408 | 98.011 | 81.518 | 81.676 | 94.514 | 79.956 | 69.223 | 92.670 | 80.265 | 66.756 | 92.206 | 77.753 | 63.409 | 92.399 | 0.117 | 0.0014 | 0.0140 | 1 | 0 | 0 |
| 'Ssfa2' | 3.347 | 3.374 | 3.216 | 3.061 | 3.028 | 3.228 | 3.581 | 3.265 | 3.186 | 2.872 | 3.469 | 3.324 | 3.242 | 3.473 | 3.413 | 3.213 | 3.399 | 3.280 | -0.030 | 0.4883 | 0.7185 | 0 | 0 | 0 |
| 'Ssh1' | 5.642 | 4.897 | 5.700 | 5.497 | 4.812 | 5.562 | 5.724 | 5.490 | 5.734 | 5.789 | 5.089 | 6.171 | 5.288 | 5.271 | 5.822 | 5.831 | 4.421 | 5.972 | -0.026 | 0.4998 | 0.7273 | 0 | 0 | 0 |
| 'Ssh2' | 18.194 | 18.288 | 15.917 | 16.524 | 17.414 | 14.916 | 16.076 | 18.540 | 15.422 | 18.483 | 20.370 | 16.841 | 16.570 | 19.281 | 16.605 | 18.865 | 19.488 | 18.102 | -0.133 | 0.0002 | 0.0025 | 1 | 0 | 0 |
| 'Ssh3' | 7.127 | 6.924 | 8.083 | 7.810 | 6.700 | 7.953 | 7.572 | 7.589 | 7.600 | 7.001 | 7.289 | 6.809 | 6.293 | 6.843 | 6.749 | 6.305 | 6.491 | 6.873 | 0.147 | 0.0017 | 0.0156 | 1 | 0 | 0 |
| 'Ssna1' | 14.662 | 15.628 | 16.743 | 15.364 | 17.255 | 16.750 | 14.603 | 16.691 | 15.998 | 14.183 | 13.959 | 16.291 | 15.464 | 15.431 | 16.314 | 14.977 | 14.851 | 16.657 | 0.044 | 0.2113 | 0.4566 | 0 | 0 | 0 |
| 'Ssr1' | 14.917 | 16.044 | 15.816 | 14.472 | 16.215 | 15.425 | 14.995 | 15.761 | 15.136 | 15.447 | 13.957 | 15.541 | 15.160 | 15.937 | 14.307 | 15.279 | 14.456 | 14.777 | 0.030 | 0.2858 | 0.5425 | 0 | 0 | 0 |
| 'Ssr2' | 32.994 | 36.039 | 36.904 | 32.672 | 37.440 | 35.753 | 34.434 | 35.149 | 38.487 | 32.283 | 32.523 | 35.968 | 34.485 | 34.063 | 37.450 | 35.223 | 33.876 | 35.169 | 0.029 | 0.3255 | 0.5828 | 0 | 0 | 0 |
| 'Ssr3' | 32.876 | 37.799 | 34.623 | 34.803 | 36.523 | 33.264 | 35.184 | 35.768 | 32.371 | 33.823 | 34.588 | 33.268 | 34.946 | 36.609 | 33.172 | 32.417 | 37.133 | 30.961 | 0.018 | 0.4653 | 0.7014 | 0 | 0 | 0 |
| 'Ssr4' | 11.861 | 13.524 | 12.134 | 11.105 | 13.293 | 11.954 | 11.259 | 13.867 | 12.036 | 11.977 | 12.046 | 11.518 | 12.877 | 12.932 | 12.014 | 12.740 | 13.754 | 11.205 | -0.011 | 0.7523 | 0.8895 | 0 | 0 | 0 |
| 'Ssrp1' | 44.253 | 42.194 | 47.053 | 46.404 | 43.658 | 46.817 | 47.519 | 43.977 | 47.618 | 44.101 | 43.688 | 45.228 | 48.267 | 44.384 | 46.528 | 44.439 | 43.346 | 44.704 | 0.006 | 0.7764 | 0.9001 | 0 | 0 | 0 |
| 'Sssca1' | 4.246 | 4.454 | 4.488 | 4.230 | 4.301 | 4.340 | 4.161 | 4.091 | 4.344 | 4.062 | 3.815 | 4.444 | 4.565 | 3.808 | 4.460 | 4.553 | 3.784 | 3.992 | 0.032 | 0.4910 | 0.7208 | 0 | 0 | 0 |
| 'Ssu72' | 21.037 | 23.863 | 21.032 | 21.302 | 23.316 | 21.633 | 21.584 | 22.255 | 21.164 | 21.476 | 20.864 | 21.985 | 22.905 | 23.066 | 22.339 | 24.153 | 23.387 | 20.708 | -0.039 | 0.2119 | 0.4573 | 0 | 0 | 0 |
| 'Ssx2ip' | 2.564 | 3.188 | 2.790 | 2.955 | 3.001 | 2.706 | 3.004 | 2.642 | 2.597 | 2.721 | 3.115 | 2.605 | 2.798 | 3.215 | 2.678 | 2.316 | 2.799 | 2.267 | 0.043 | 0.4479 | 0.6883 | 0 | 0 | 0 |
| 'St13' | 49.563 | 49.189 | 50.088 | 51.285 | 52.300 | 50.203 | 50.053 | 50.135 | 50.085 | 52.654 | 50.100 | 54.545 | 48.502 | 51.162 | 51.974 | 46.906 | 51.108 | 56.260 | -0.043 | 0.1495 | 0.3764 | 0 | 0 | 0 |
| 'St3gal1' | 124.756 | 117.634 | 117.885 | 125.135 | 116.748 | 115.065 | 115.462 | 120.603 | 118.503 | 140.756 | 134.169 | 119.718 | 133.618 | 123.689 | 122.371 | 128.639 | 124.199 | 121.393 | -0.109 | 0.0001 | 0.0010 | 1 | 0 | 0 |
| 'St3gal2' | 19.379 | 20.302 | 18.516 | 19.068 | 18.633 | 19.591 | 18.529 | 19.917 | 18.446 | 16.791 | 16.591 | 18.011 | 17.206 | 17.616 | 17.808 | 18.462 | 15.524 | 18.546 | 0.128 | 0.0002 | 0.0031 | 1 | 0 | 0 |
| 'St3gal3' | 3.275 | 3.290 | 3.342 | 3.311 | 3.338 | 3.695 | 3.242 | 3.530 | 3.584 | 3.507 | 3.557 | 3.439 | 3.960 | 3.081 | 3.123 | 3.359 | 3.189 | 3.411 | -0.009 | 0.8676 | 0.9428 | 0 | 0 | 0 |
| 'St3gal4' | 8.410 | 7.803 | 6.536 | 7.681 | 6.608 | 7.818 | 7.584 | 7.898 | 8.186 | 6.698 | 7.869 | 7.924 | 7.326 | 7.669 | 6.952 | 6.906 | 8.093 | 8.344 | 0.003 | 0.9612 | 0.9860 | 0 | 0 | 0 |
| 'St3gal5' | 2.388 | 3.048 | 2.942 | 3.006 | 3.097 | 2.851 | 2.825 | 3.957 | 3.150 | 2.997 | 2.353 | 2.723 | 2.229 | 2.245 | 2.919 | 3.613 | 2.477 | 2.204 | 0.191 | 0.0620 | 0.2156 | 0 | 0 | 0 |
| 'St6gal1' | 24.301 | 23.880 | 25.152 | 23.011 | 24.358 | 25.553 | 23.121 | 25.291 | 24.708 | 23.681 | 22.960 | 25.075 | 20.122 | 22.492 | 24.555 | 23.180 | 20.985 | 23.369 | 0.078 | 0.0158 | 0.0869 | 0 | 0 | 0 |
| 'St6galnac2' | 0.500 | 0.521 | 0.687 | 0.605 | 0.533 | 0.553 | 0.539 | 0.620 | 0.616 | 0.420 | 0.421 | 0.425 | 0.378 | 0.262 | 0.353 | 0.266 | 0.400 | 0.414 | 0.615 | 0.0000 | 0.0000 | 1 | 1 | 0 |
| 'St6galnac4' | 3.913 | 3.837 | 4.224 | 3.200 | 3.821 | 4.033 | 3.519 | 3.926 | 4.170 | 4.221 | 3.858 | 4.267 | 3.688 | 3.614 | 4.199 | 4.087 | 3.485 | 3.961 | -0.038 | 0.3955 | 0.6444 | 0 | 0 | 0 |
| 'St6galnac6' | 0.659 | 0.919 | 0.774 | 0.740 | 0.822 | 0.816 | 0.817 | 0.778 | 0.857 | 0.924 | 0.619 | 0.869 | 0.902 | 0.694 | 0.763 | 0.985 | 0.658 | 0.840 | -0.027 | 0.7466 | 0.8857 | 0 | 0 | 0 |
| 'St7' | 0.660 | 0.694 | 0.690 | 0.697 | 0.691 | 0.750 | 0.682 | 0.717 | 0.602 | 0.664 | 0.732 | 0.652 | 0.833 | 0.604 | 0.634 | 0.639 | 0.664 | 0.645 | 0.017 | 0.7980 | 0.9103 | 0 | 0 | 0 |
| 'St7l' | 2.051 | 1.890 | 1.827 | 2.090 | 1.858 | 1.795 | 2.181 | 1.876 | 1.807 | 1.773 | 1.817 | 2.036 | 1.811 | 1.939 | 1.824 | 1.622 | 2.020 | 1.778 | 0.046 | 0.3671 | 0.6192 | 0 | 0 | 0 |
| 'St8sia1' | 16.068 | 16.425 | 15.900 | 16.179 | 15.404 | 15.589 | 16.526 | 15.569 | 16.388 | 16.389 | 15.611 | 17.598 | 14.647 | 15.059 | 16.397 | 16.378 | 15.007 | 17.548 | -0.016 | 0.6203 | 0.8101 | 0 | 0 | 0 |
| 'St8sia4' | 6.051 | 6.581 | 6.240 | 6.672 | 6.464 | 5.719 | 6.732 | 6.081 | 5.611 | 5.134 | 6.292 | 6.153 | 5.028 | 5.995 | 5.237 | 4.760 | 5.898 | 5.789 | 0.150 | 0.0036 | 0.0288 | 1 | 0 | 0 |
| 'St8sia6' | 0.275 | 0.400 | 0.377 | 0.306 | 0.341 | 0.499 | 0.446 | 0.467 | 0.455 | 0.567 | 0.504 | 0.736 | 0.401 | 0.497 | 0.627 | 0.564 | 0.447 | 0.707 | -0.517 | 0.0000 | 0.0003 | 1 | 1 | 0 |
| 'Stag1' | 4.661 | 4.909 | 5.057 | 5.300 | 5.431 | 5.141 | 5.226 | 5.520 | 5.057 | 5.190 | 5.433 | 4.894 | 5.292 | 5.476 | 4.978 | 5.202 | 5.723 | 4.849 | -0.030 | 0.3217 | 0.5797 | 0 | 0 | 0 |
| 'Stag2' | 24.057 | 24.647 | 24.499 | 26.723 | 24.536 | 23.662 | 26.722 | 24.435 | 23.716 | 23.686 | 24.798 | 24.308 | 23.577 | 24.894 | 22.838 | 23.138 | 26.903 | 23.257 | 0.026 | 0.4210 | 0.6677 | 0 | 0 | 0 |
| 'Stag3' | 0.426 | 0.693 | 0.749 | 0.699 | 0.670 | 0.871 | 1.000 | 0.481 | 0.716 | 0.812 | 0.644 | 0.683 | 0.856 | 0.455 | 0.550 | 0.924 | 1.301 | 1.105 | -0.236 | 0.2634 | 0.5173 | 0 | 0 | 0 |
| 'Stam' | 2.135 | 2.440 | 2.070 | 2.445 | 2.259 | 2.175 | 2.449 | 2.340 | 2.372 | 2.274 | 2.264 | 2.330 | 2.014 | 2.314 | 2.034 | 2.275 | 2.676 | 2.018 | 0.021 | 0.6774 | 0.8457 | 0 | 0 | 0 |
| 'Stam2' | 5.562 | 5.859 | 6.077 | 5.313 | 5.997 | 6.212 | 5.352 | 6.096 | 5.958 | 6.336 | 5.818 | 5.636 | 5.785 | 6.172 | 5.314 | 5.832 | 5.963 | 5.284 | 0.003 | 0.9364 | 0.9758 | 0 | 0 | 0 |
| 'Stambp' | 3.522 | 3.700 | 3.440 | 3.427 | 3.695 | 3.541 | 3.690 | 3.730 | 3.600 | 4.140 | 4.388 | 3.866 | 4.341 | 4.289 | 3.546 | 3.855 | 4.068 | 3.552 | -0.164 | 0.0000 | 0.0001 | 1 | 0 | 0 |
| 'Stap1' | 5.073 | 6.689 | 5.286 | 5.170 | 6.117 | 5.216 | 5.683 | 5.498 | 5.987 | 5.215 | 4.631 | 5.076 | 4.959 | 5.325 | 4.521 | 5.122 | 4.800 | 4.410 | 0.188 | 0.0001 | 0.0018 | 1 | 0 | 0 |
| 'Star' | 1.373 | 1.774 | 1.739 | 1.664 | 1.905 | 1.612 | 1.430 | 1.683 | 1.758 | 1.957 | 1.469 | 1.611 | 2.095 | 1.687 | 1.896 | 2.034 | 1.875 | 1.631 | -0.127 | 0.2071 | 0.4517 | 0 | 0 | 0 |
| 'Stard10' | 7.487 | 6.461 | 8.715 | 6.162 | 7.003 | 8.755 | 7.045 | 6.697 | 7.532 | 9.143 | 7.269 | 8.914 | 7.846 | 7.389 | 9.338 | 8.767 | 7.362 | 9.024 | -0.194 | 0.0000 | 0.0008 | 1 | 0 | 0 |
| 'Stard3' | 8.804 | 9.297 | 9.812 | 8.555 | 9.020 | 10.022 | 8.422 | 9.104 | 9.602 | 8.742 | 8.427 | 10.339 | 9.149 | 8.542 | 10.003 | 8.686 | 8.301 | 9.971 | -0.005 | 0.8759 | 0.9473 | 0 | 0 | 0 |
| 'Stard3nl' | 3.177 | 3.843 | 3.667 | 3.286 | 4.233 | 3.448 | 3.185 | 3.681 | 3.675 | 3.673 | 3.847 | 4.063 | 4.239 | 4.190 | 3.744 | 3.866 | 3.724 | 3.431 | -0.127 | 0.0112 | 0.0680 | 0 | 0 | 0 |
| 'Stard4' | 8.751 | 8.517 | 9.354 | 8.971 | 8.691 | 9.461 | 9.658 | 8.696 | 9.609 | 7.560 | 9.970 | 10.141 | 8.333 | 9.893 | 10.260 | 7.886 | 10.106 | 9.494 | -0.040 | 0.4234 | 0.6703 | 0 | 0 | 0 |
| 'Stard5' | 9.953 | 13.193 | 9.259 | 10.137 | 11.955 | 10.236 | 9.720 | 11.628 | 9.592 | 10.937 | 12.333 | 10.387 | 13.149 | 13.670 | 9.616 | 11.885 | 13.682 | 9.931 | -0.152 | 0.0004 | 0.0047 | 1 | 0 | 0 |
| 'Stard7' | 10.675 | 11.226 | 10.507 | 10.089 | 10.489 | 10.843 | 10.098 | 10.672 | 10.953 | 9.901 | 10.320 | 11.294 | 10.733 | 10.761 | 11.623 | 10.820 | 9.973 | 11.186 | -0.027 | 0.3685 | 0.6201 | 0 | 0 | 0 |
| 'Stat1' | 37.376 | 37.705 | 38.358 | 39.093 | 39.428 | 39.549 | 38.947 | 36.041 | 40.418 | 43.149 | 43.065 | 49.143 | 45.915 | 41.800 | 48.267 | 43.747 | 46.925 | 49.613 | -0.257 | 0.0000 | 0.0000 | 1 | 0 | 0 |
| 'Stat2' | 23.598 | 21.613 | 25.648 | 22.286 | 21.102 | 23.841 | 22.989 | 21.344 | 26.568 | 23.111 | 23.807 | 30.528 | 24.149 | 23.114 | 28.747 | 21.654 | 24.266 | 29.272 | -0.138 | 0.0002 | 0.0025 | 1 | 0 | 0 |
| 'Stat3' | 35.318 | 33.637 | 34.497 | 34.511 | 34.355 | 33.885 | 33.987 | 34.021 | 33.698 | 32.807 | 38.287 | 36.138 | 33.484 | 37.007 | 35.176 | 32.881 | 35.446 | 37.376 | -0.059 | 0.0611 | 0.2134 | 0 | 0 | 0 |
| 'Stat4' | 52.674 | 58.814 | 51.820 | 58.070 | 57.840 | 52.450 | 61.102 | 54.885 | 53.276 | 50.193 | 51.495 | 50.096 | 50.856 | 51.919 | 47.985 | 50.093 | 56.608 | 46.628 | 0.124 | 0.0000 | 0.0004 | 1 | 0 | 0 |
| 'Stat5b' | 71.133 | 70.525 | 70.423 | 67.613 | 70.823 | 71.034 | 67.801 | 71.439 | 70.272 | 70.698 | 70.827 | 68.642 | 69.853 | 69.215 | 69.966 | 69.990 | 65.231 | 69.214 | 0.007 | 0.7275 | 0.8750 | 0 | 0 | 0 |
| 'Stat6' | 44.678 | 39.227 | 44.374 | 44.534 | 39.513 | 43.105 | 40.793 | 41.814 | 44.524 | 44.229 | 43.643 | 50.319 | 43.256 | 42.639 | 47.876 | 43.440 | 44.334 | 49.236 | -0.106 | 0.0003 | 0.0042 | 1 | 0 | 0 |
| 'Stau1' | 6.847 | 6.392 | 6.663 | 6.722 | 6.069 | 6.474 | 6.621 | 6.456 | 6.627 | 6.203 | 6.519 | 6.945 | 6.046 | 6.717 | 6.895 | 5.827 | 6.763 | 7.032 | -0.012 | 0.7566 | 0.8913 | 0 | 0 | 0 |
| 'Stau2' | 1.215 | 1.267 | 1.299 | 1.081 | 1.095 | 1.159 | 1.314 | 1.200 | 1.251 | 1.073 | 1.008 | 1.106 | 1.007 | 0.808 | 0.913 | 0.984 | 0.991 | 1.093 | 0.263 | 0.0000 | 0.0001 | 1 | 0 | 0 |
| 'Stc2' | 6.149 | 7.686 | 4.453 | 6.804 | 6.198 | 4.268 | 5.995 | 6.015 | 4.667 | 3.903 | 6.769 | 5.803 | 6.337 | 7.285 | 6.835 | 4.464 | 7.406 | 6.333 | -0.094 | 0.4411 | 0.6844 | 0 | 0 | 0 |
| 'Stil' | 1.893 | 1.767 | 2.243 | 1.937 | 1.797 | 1.776 | 2.044 | 1.731 | 2.181 | 1.504 | 1.922 | 1.805 | 2.250 | 2.240 | 2.244 | 1.836 | 1.740 | 1.753 | -0.004 | 0.9633 | 0.9869 | 0 | 0 | 0 |
| 'Stim1' | 66.516 | 65.564 | 66.407 | 63.325 | 68.329 | 66.939 | 61.036 | 67.629 | 67.403 | 65.286 | 62.972 | 68.396 | 62.399 | 64.803 | 68.735 | 60.253 | 59.603 | 68.582 | 0.019 | 0.4862 | 0.7170 | 0 | 0 | 0 |
| 'Stim2' | 16.684 | 17.273 | 17.072 | 16.911 | 16.560 | 16.685 | 16.193 | 16.984 | 17.147 | 15.707 | 16.153 | 18.982 | 16.896 | 17.602 | 17.441 | 17.028 | 16.894 | 18.967 | -0.051 | 0.0981 | 0.2925 | 0 | 0 | 0 |
| 'Stip1' | 99.297 | 87.806 | 99.230 | 106.516 | 90.789 | 99.088 | 96.344 | 86.144 | 94.952 | 88.203 | 91.231 | 136.508 | 100.980 | 93.371 | 122.131 | 90.007 | 102.767 | 149.803 | -0.172 | 0.0372 | 0.1561 | 0 | 0 | 0 |
| 'Stk10' | 54.651 | 49.441 | 54.225 | 50.822 | 47.312 | 53.708 | 50.307 | 49.860 | 52.044 | 51.237 | 47.086 | 55.876 | 46.306 | 48.723 | 54.503 | 51.646 | 44.772 | 57.010 | 0.008 | 0.8012 | 0.9113 | 0 | 0 | 0 |
| 'Stk11' | 21.736 | 19.467 | 21.856 | 20.956 | 20.234 | 22.327 | 20.436 | 20.500 | 21.739 | 20.755 | 19.787 | 23.000 | 21.125 | 19.451 | 21.343 | 19.911 | 20.785 | 22.506 | -0.006 | 0.7976 | 0.9101 | 0 | 0 | 0 |
| 'Stk11ip' | 5.757 | 6.481 | 6.749 | 5.525 | 6.269 | 7.085 | 5.720 | 6.315 | 6.962 | 6.077 | 5.270 | 5.361 | 5.756 | 5.338 | 5.178 | 5.475 | 5.629 | 4.994 | 0.202 | 0.0000 | 0.0006 | 1 | 0 | 0 |
| 'Stk16' | 4.483 | 5.275 | 5.135 | 4.436 | 4.854 | 5.063 | 4.612 | 4.689 | 5.114 | 4.359 | 4.137 | 5.418 | 4.727 | 4.435 | 5.000 | 4.058 | 4.411 | 5.131 | 0.051 | 0.1833 | 0.4191 | 0 | 0 | 0 |
| 'Stk17b' | 115.156 | 128.687 | 107.040 | 116.004 | 131.798 | 105.591 | 116.793 | 130.915 | 101.763 | 136.550 | 125.197 | 106.860 | 115.137 | 139.663 | 108.948 | 133.268 | 125.482 | 108.142 | -0.072 | 0.0292 | 0.1326 | 0 | 0 | 0 |
| 'Stk19' | 4.766 | 4.565 | 5.413 | 4.040 | 4.610 | 5.179 | 4.723 | 5.060 | 5.399 | 5.856 | 4.969 | 4.761 | 5.050 | 4.916 | 4.730 | 4.672 | 4.628 | 4.500 | -0.016 | 0.7862 | 0.9046 | 0 | 0 | 0 |
| 'Stk24' | 49.667 | 47.260 | 49.986 | 48.926 | 54.556 | 50.370 | 45.990 | 52.685 | 47.671 | 47.691 | 50.133 | 47.604 | 50.717 | 49.623 | 49.373 | 49.642 | 44.851 | 49.568 | 0.016 | 0.6420 | 0.8243 | 0 | 0 | 0 |
| 'Stk25' | 14.384 | 14.206 | 15.135 | 15.083 | 14.007 | 15.323 | 13.930 | 14.380 | 15.617 | 14.223 | 12.657 | 16.364 | 14.632 | 14.192 | 15.023 | 13.592 | 15.066 | 15.791 | -0.006 | 0.8466 | 0.9343 | 0 | 0 | 0 |
| 'Stk26' | 33.716 | 37.593 | 33.119 | 36.244 | 37.509 | 32.764 | 37.883 | 35.100 | 33.110 | 35.067 | 36.266 | 33.190 | 33.394 | 36.990 | 32.118 | 33.076 | 35.935 | 32.221 | 0.029 | 0.2673 | 0.5207 | 0 | 0 | 0 |
| 'Stk3' | 2.129 | 2.628 | 2.175 | 2.452 | 2.339 | 1.971 | 2.869 | 2.298 | 2.194 | 2.340 | 2.172 | 2.264 | 2.131 | 2.185 | 2.202 | 2.292 | 2.260 | 2.087 | 0.055 | 0.3988 | 0.6481 | 0 | 0 | 0 |
| 'Stk35' | 4.604 | 4.379 | 4.341 | 4.752 | 3.888 | 3.875 | 4.514 | 3.983 | 4.242 | 4.212 | 4.638 | 4.816 | 4.399 | 4.052 | 4.817 | 4.523 | 3.805 | 5.359 | -0.087 | 0.1283 | 0.3432 | 0 | 0 | 0 |
| 'Stk38' | 8.394 | 8.928 | 8.946 | 8.753 | 8.309 | 8.901 | 8.377 | 8.628 | 8.879 | 8.404 | 8.925 | 9.356 | 8.480 | 8.883 | 8.723 | 8.435 | 8.728 | 9.040 | -0.027 | 0.2561 | 0.5084 | 0 | 0 | 0 |
| 'Stk38l' | 2.538 | 2.528 | 2.703 | 2.346 | 2.287 | 2.511 | 2.839 | 2.536 | 2.533 | 2.356 | 3.270 | 2.730 | 2.313 | 2.305 | 2.336 | 2.535 | 2.702 | 2.269 | -0.006 | 0.9293 | 0.9732 | 0 | 0 | 0 |
| 'Stk39' | 25.788 | 23.870 | 25.492 | 28.740 | 26.732 | 23.902 | 28.120 | 27.480 | 22.049 | 26.461 | 28.797 | 23.329 | 29.892 | 27.215 | 22.652 | 24.762 | 27.993 | 24.178 | -0.028 | 0.5057 | 0.7319 | 0 | 0 | 0 |
| 'Stk4' | 89.924 | 89.975 | 86.004 | 89.408 | 87.227 | 84.975 | 86.725 | 86.723 | 82.577 | 85.148 | 92.893 | 91.519 | 73.945 | 89.634 | 88.851 | 89.880 | 84.827 | 93.284 | -0.022 | 0.5775 | 0.7819 | 0 | 0 | 0 |
| 'Stk40' | 22.616 | 22.395 | 20.589 | 20.883 | 21.660 | 23.226 | 21.073 | 21.056 | 20.827 | 23.264 | 22.433 | 23.970 | 22.560 | 22.408 | 23.988 | 23.541 | 20.852 | 24.397 | -0.104 | 0.0014 | 0.0137 | 1 | 0 | 0 |
| 'Stmn1' | 45.297 | 51.217 | 54.947 | 57.790 | 52.376 | 57.701 | 56.547 | 50.152 | 57.316 | 42.521 | 50.379 | 53.492 | 61.276 | 57.929 | 64.385 | 52.438 | 54.278 | 54.696 | -0.035 | 0.5590 | 0.7700 | 0 | 0 | 0 |
| 'Stom' | 11.055 | 10.071 | 10.657 | 11.209 | 10.926 | 10.794 | 8.941 | 11.716 | 9.293 | 9.231 | 14.795 | 11.390 | 9.513 | 12.431 | 11.998 | 7.584 | 12.454 | 10.386 | -0.069 | 0.3655 | 0.6177 | 0 | 0 | 0 |
| 'Stoml1' | 11.790 | 14.372 | 14.398 | 11.074 | 14.117 | 15.336 | 12.534 | 14.815 | 14.162 | 14.743 | 12.798 | 13.003 | 15.599 | 12.648 | 12.258 | 13.687 | 12.172 | 10.491 | 0.054 | 0.4466 | 0.6880 | 0 | 0 | 0 |
| 'Stoml2' | 8.412 | 9.701 | 9.297 | 8.682 | 10.281 | 10.084 | 8.828 | 9.431 | 9.889 | 8.994 | 8.726 | 8.746 | 9.180 | 9.119 | 9.312 | 8.993 | 8.557 | 8.666 | 0.065 | 0.0749 | 0.2436 | 0 | 0 | 0 |
| 'Ston1' | 0.629 | 0.655 | 0.480 | 0.514 | 0.662 | 0.632 | 0.458 | 0.685 | 0.571 | 0.512 | 0.510 | 0.704 | 0.517 | 0.531 | 0.494 | 0.802 | 0.443 | 0.651 | 0.016 | 0.8939 | 0.9566 | 0 | 0 | 0 |
| 'Ston2' | 0.875 | 0.638 | 1.142 | 0.902 | 0.753 | 0.876 | 0.822 | 0.686 | 0.844 | 0.825 | 0.826 | 1.020 | 0.875 | 0.788 | 1.022 | 0.731 | 0.649 | 1.133 | -0.070 | 0.4175 | 0.6650 | 0 | 0 | 0 |
| 'Strada' | 6.602 | 6.917 | 7.188 | 6.694 | 6.004 | 6.641 | 7.007 | 6.725 | 7.462 | 7.332 | 6.827 | 6.735 | 7.429 | 6.497 | 6.519 | 7.042 | 6.094 | 5.960 | 0.012 | 0.7741 | 0.8993 | 0 | 0 | 0 |
| 'Stradb' | 1.621 | 1.967 | 1.839 | 1.872 | 2.012 | 1.931 | 1.964 | 1.887 | 1.925 | 1.607 | 1.616 | 1.799 | 1.505 | 1.897 | 1.867 | 1.750 | 1.786 | 1.538 | 0.134 | 0.0132 | 0.0766 | 0 | 0 | 0 |
| 'Strap' | 39.124 | 39.250 | 35.053 | 36.524 | 40.480 | 34.684 | 35.744 | 39.392 | 32.522 | 36.803 | 36.859 | 33.821 | 36.866 | 39.568 | 33.105 | 39.272 | 36.848 | 34.304 | 0.013 | 0.6349 | 0.8193 | 0 | 0 | 0 |
| 'Strip1' | 8.227 | 8.864 | 8.591 | 8.241 | 8.980 | 9.347 | 8.589 | 8.474 | 9.062 | 9.070 | 8.194 | 8.511 | 8.517 | 8.111 | 8.150 | 8.831 | 8.669 | 7.746 | 0.039 | 0.2453 | 0.4958 | 0 | 0 | 0 |
| 'Strn' | 10.872 | 12.127 | 11.967 | 11.422 | 11.951 | 11.088 | 10.770 | 11.073 | 11.540 | 11.767 | 11.883 | 12.015 | 11.385 | 11.469 | 12.035 | 11.611 | 11.515 | 12.789 | -0.062 | 0.0446 | 0.1751 | 0 | 0 | 0 |
| 'Strn3' | 16.878 | 19.972 | 17.445 | 19.088 | 18.538 | 16.863 | 19.378 | 19.965 | 16.336 | 17.774 | 17.073 | 17.765 | 17.325 | 19.201 | 17.851 | 18.858 | 17.810 | 17.966 | 0.010 | 0.7899 | 0.9061 | 0 | 0 | 0 |
| 'Strn4' | 9.860 | 9.078 | 9.619 | 8.777 | 8.469 | 9.625 | 8.214 | 9.236 | 9.829 | 8.872 | 8.740 | 10.747 | 9.005 | 8.020 | 9.787 | 9.039 | 8.939 | 10.699 | -0.027 | 0.4801 | 0.7123 | 0 | 0 | 0 |
| 'Stt3a' | 20.181 | 22.751 | 21.250 | 21.214 | 22.298 | 20.542 | 21.821 | 23.311 | 21.235 | 22.171 | 22.546 | 21.896 | 20.494 | 21.755 | 20.346 | 22.113 | 22.534 | 21.157 | -0.015 | 0.5497 | 0.7633 | 0 | 0 | 0 |
| 'Stt3b' | 33.820 | 34.883 | 36.522 | 33.003 | 32.718 | 35.420 | 36.481 | 35.361 | 34.703 | 32.590 | 33.112 | 37.692 | 32.296 | 33.882 | 36.773 | 33.516 | 34.027 | 38.793 | -0.009 | 0.7701 | 0.8977 | 0 | 0 | 0 |
| 'Stub1' | 30.485 | 36.788 | 37.430 | 30.944 | 33.939 | 38.000 | 28.553 | 33.181 | 34.147 | 31.256 | 26.652 | 34.691 | 35.608 | 31.996 | 34.322 | 35.602 | 32.181 | 34.682 | 0.019 | 0.7162 | 0.8688 | 0 | 0 | 0 |
| 'Stx11' | 85.775 | 98.379 | 84.172 | 90.662 | 100.173 | 86.836 | 96.889 | 92.992 | 87.024 | 95.187 | 86.893 | 73.348 | 110.187 | 91.965 | 74.400 | 91.313 | 91.577 | 76.254 | 0.054 | 0.2248 | 0.4716 | 0 | 0 | 0 |
| 'Stx12' | 8.305 | 9.687 | 7.883 | 8.289 | 8.738 | 7.961 | 8.353 | 8.846 | 8.279 | 7.117 | 9.743 | 10.180 | 7.254 | 8.951 | 10.767 | 7.989 | 9.342 | 10.053 | -0.100 | 0.1094 | 0.3118 | 0 | 0 | 0 |
| 'Stx16' | 15.349 | 19.846 | 17.112 | 15.310 | 19.241 | 18.511 | 14.104 | 18.927 | 18.425 | 19.545 | 17.609 | 17.606 | 18.922 | 19.154 | 17.015 | 19.389 | 17.440 | 17.115 | -0.079 | 0.1370 | 0.3569 | 0 | 0 | 0 |
| 'Stx17' | 3.638 | 3.970 | 4.428 | 3.461 | 3.969 | 4.129 | 3.662 | 4.464 | 4.181 | 4.449 | 3.481 | 3.853 | 3.814 | 4.235 | 3.501 | 4.180 | 4.186 | 3.091 | 0.040 | 0.5345 | 0.7542 | 0 | 0 | 0 |
| 'Stx18' | 5.982 | 6.616 | 6.792 | 6.207 | 6.649 | 7.040 | 7.119 | 6.947 | 6.493 | 7.020 | 6.690 | 6.799 | 6.576 | 6.952 | 6.443 | 6.674 | 6.773 | 6.061 | -0.012 | 0.7751 | 0.8994 | 0 | 0 | 0 |
| 'Stx1a' | 0.579 | 0.542 | 0.521 | 0.498 | 0.518 | 0.558 | 0.848 | 0.655 | 0.666 | 0.548 | 0.424 | 0.647 | 0.545 | 0.282 | 0.750 | 0.706 | 0.343 | 0.666 | 0.139 | 0.3221 | 0.5799 | 0 | 0 | 0 |
| 'Stx2' | 1.816 | 2.175 | 2.113 | 2.058 | 2.287 | 2.449 | 2.213 | 2.224 | 2.213 | 1.712 | 1.628 | 1.769 | 1.759 | 1.605 | 1.792 | 1.894 | 1.401 | 1.787 | 0.338 | 0.0000 | 0.0000 | 1 | 0 | 0 |
| 'Stx4a' | 25.517 | 27.672 | 26.814 | 26.857 | 26.795 | 28.016 | 27.248 | 26.510 | 28.549 | 26.617 | 25.908 | 28.227 | 28.924 | 26.634 | 27.402 | 27.591 | 27.733 | 26.684 | -0.022 | 0.3631 | 0.6152 | 0 | 0 | 0 |
| 'Stx5a' | 7.371 | 7.505 | 7.390 | 7.467 | 7.612 | 7.645 | 7.415 | 8.071 | 7.370 | 7.429 | 7.469 | 7.883 | 7.766 | 7.272 | 7.540 | 7.666 | 7.978 | 7.144 | -0.018 | 0.4779 | 0.7111 | 0 | 0 | 0 |
| 'Stx6' | 14.114 | 15.455 | 13.362 | 13.639 | 15.211 | 15.012 | 13.203 | 15.572 | 14.211 | 15.303 | 14.820 | 14.603 | 15.006 | 15.587 | 14.489 | 15.304 | 14.111 | 13.517 | -0.044 | 0.1737 | 0.4074 | 0 | 0 | 0 |
| 'Stx7' | 10.311 | 10.508 | 8.811 | 10.263 | 10.817 | 9.439 | 9.707 | 10.965 | 9.745 | 9.450 | 10.247 | 9.866 | 9.370 | 11.230 | 9.012 | 10.305 | 9.801 | 9.304 | 0.019 | 0.6510 | 0.8304 | 0 | 0 | 0 |
| 'Stx8' | 3.057 | 3.355 | 2.907 | 3.391 | 3.391 | 3.111 | 3.232 | 3.078 | 3.028 | 3.055 | 3.395 | 3.162 | 3.109 | 2.768 | 3.191 | 3.113 | 3.410 | 3.323 | -0.015 | 0.7535 | 0.8901 | 0 | 0 | 0 |
| 'Stxbp1' | 2.215 | 2.945 | 2.441 | 2.615 | 2.665 | 2.786 | 2.541 | 2.674 | 2.680 | 2.982 | 3.045 | 2.937 | 3.049 | 3.296 | 3.084 | 3.108 | 3.040 | 3.088 | -0.242 | 0.0000 | 0.0000 | 1 | 0 | 0 |
| 'Stxbp2' | 17.319 | 16.731 | 19.275 | 16.424 | 17.761 | 20.037 | 17.368 | 17.674 | 19.509 | 22.388 | 19.466 | 21.511 | 21.973 | 19.715 | 21.178 | 21.571 | 19.774 | 20.195 | -0.222 | 0.0000 | 0.0000 | 1 | 0 | 0 |
| 'Stxbp3' | 6.973 | 7.702 | 6.810 | 7.595 | 7.650 | 6.257 | 6.845 | 7.274 | 7.428 | 7.075 | 7.750 | 8.084 | 6.677 | 7.292 | 7.499 | 7.420 | 8.390 | 7.745 | -0.088 | 0.0514 | 0.1917 | 0 | 0 | 0 |
| 'Stxbp4' | 0.446 | 0.402 | 0.397 | 0.457 | 0.373 | 0.402 | 0.524 | 0.452 | 0.458 | 0.455 | 0.281 | 0.394 | 0.369 | 0.371 | 0.370 | 0.404 | 0.417 | 0.373 | 0.172 | 0.0649 | 0.2221 | 0 | 0 | 0 |
| 'Stxbp5' | 4.239 | 4.100 | 4.424 | 4.253 | 4.620 | 4.445 | 4.260 | 4.378 | 4.355 | 4.640 | 4.296 | 4.125 | 4.563 | 4.333 | 4.190 | 4.365 | 4.387 | 4.054 | -0.002 | 0.9559 | 0.9836 | 0 | 0 | 0 |
| 'Sub1' | 30.902 | 32.929 | 26.116 | 29.580 | 31.495 | 26.437 | 29.059 | 31.632 | 25.084 | 30.793 | 30.682 | 23.608 | 27.315 | 31.726 | 25.822 | 33.255 | 28.357 | 23.842 | 0.037 | 0.3341 | 0.5924 | 0 | 0 | 0 |
| 'Sucla2' | 10.645 | 13.107 | 12.441 | 12.279 | 13.263 | 12.477 | 12.220 | 13.048 | 11.858 | 10.944 | 11.272 | 12.223 | 11.642 | 12.469 | 11.157 | 11.684 | 12.916 | 11.350 | 0.061 | 0.0825 | 0.2597 | 0 | 0 | 0 |
| 'Suclg1' | 17.269 | 19.337 | 17.629 | 18.731 | 19.517 | 18.326 | 17.367 | 19.345 | 17.644 | 17.206 | 16.926 | 17.092 | 18.775 | 17.799 | 16.316 | 17.852 | 16.927 | 16.646 | 0.075 | 0.0099 | 0.0618 | 0 | 0 | 0 |
| 'Suclg2' | 7.463 | 7.821 | 6.575 | 7.754 | 8.441 | 7.904 | 7.445 | 7.857 | 7.054 | 7.011 | 8.370 | 7.927 | 7.144 | 7.256 | 7.049 | 8.048 | 6.508 | 7.202 | 0.023 | 0.6804 | 0.8474 | 0 | 0 | 0 |
| 'Suco' | 9.286 | 10.424 | 9.524 | 8.988 | 10.102 | 8.951 | 9.049 | 10.182 | 8.754 | 10.945 | 10.477 | 9.817 | 9.579 | 10.751 | 9.573 | 11.952 | 10.427 | 8.873 | -0.124 | 0.0019 | 0.0177 | 1 | 0 | 0 |
| 'Suds3' | 9.166 | 8.544 | 9.249 | 9.091 | 9.280 | 10.252 | 8.547 | 9.212 | 9.681 | 9.446 | 8.419 | 9.914 | 9.420 | 9.486 | 10.165 | 10.034 | 8.912 | 9.825 | -0.053 | 0.0830 | 0.2609 | 0 | 0 | 0 |
| 'Sufu' | 13.461 | 13.265 | 12.809 | 13.236 | 13.764 | 13.733 | 13.189 | 13.398 | 13.650 | 14.417 | 13.276 | 13.488 | 13.133 | 13.827 | 13.322 | 12.867 | 12.965 | 14.437 | -0.025 | 0.4069 | 0.6562 | 0 | 0 | 0 |
| 'Sugp1' | 43.079 | 40.651 | 45.478 | 41.834 | 41.908 | 44.783 | 41.516 | 42.669 | 44.739 | 46.173 | 42.446 | 39.770 | 45.718 | 40.326 | 38.606 | 42.878 | 39.314 | 36.960 | 0.050 | 0.2001 | 0.4413 | 0 | 0 | 0 |
| 'Sugp2' | 4.054 | 4.057 | 4.725 | 3.886 | 4.119 | 4.681 | 4.099 | 4.065 | 4.836 | 4.224 | 4.013 | 4.558 | 4.461 | 4.216 | 4.516 | 4.095 | 4.390 | 4.418 | -0.025 | 0.4055 | 0.6548 | 0 | 0 | 0 |
| 'Sugt1' | 27.909 | 27.787 | 28.739 | 31.460 | 29.858 | 28.550 | 29.733 | 28.601 | 29.618 | 27.649 | 27.360 | 29.084 | 29.201 | 29.417 | 29.590 | 26.820 | 32.259 | 29.348 | -0.004 | 0.9173 | 0.9686 | 0 | 0 | 0 |
| 'Sumf1' | 6.601 | 8.699 | 7.582 | 7.195 | 8.572 | 8.864 | 7.474 | 8.589 | 7.832 | 8.169 | 8.124 | 7.639 | 7.619 | 8.082 | 7.254 | 8.807 | 7.650 | 6.951 | 0.013 | 0.8075 | 0.9136 | 0 | 0 | 0 |
| 'Sumf2' | 2.282 | 2.178 | 2.316 | 1.914 | 2.218 | 2.267 | 1.925 | 2.046 | 1.821 | 1.763 | 1.715 | 2.332 | 1.534 | 1.924 | 2.028 | 1.507 | 2.094 | 1.918 | 0.159 | 0.0378 | 0.1581 | 0 | 0 | 0 |
| 'Sumo1' | 38.889 | 44.395 | 38.963 | 39.970 | 42.782 | 37.944 | 42.343 | 40.936 | 37.897 | 37.047 | 39.834 | 38.667 | 39.083 | 41.481 | 38.464 | 38.171 | 40.295 | 35.951 | 0.047 | 0.0717 | 0.2363 | 0 | 0 | 0 |
| 'Sumo2' | 80.973 | 93.914 | 85.717 | 89.515 | 93.717 | 86.763 | 93.945 | 91.831 | 84.328 | 78.977 | 88.138 | 80.134 | 84.916 | 89.215 | 81.840 | 84.071 | 87.695 | 81.174 | 0.071 | 0.0065 | 0.0456 | 1 | 0 | 0 |
| 'Sumo3' | 12.986 | 13.926 | 12.623 | 12.841 | 13.324 | 13.257 | 12.597 | 13.496 | 12.788 | 12.110 | 12.829 | 12.861 | 13.220 | 13.426 | 13.510 | 12.825 | 12.995 | 12.249 | 0.011 | 0.6852 | 0.8511 | 0 | 0 | 0 |
| 'Sun1' | 4.484 | 4.234 | 4.535 | 3.992 | 4.237 | 4.169 | 4.151 | 4.120 | 4.328 | 4.059 | 4.143 | 4.140 | 3.818 | 4.017 | 4.463 | 3.895 | 3.781 | 3.925 | 0.070 | 0.0555 | 0.2019 | 0 | 0 | 0 |
| 'Sun2' | 53.788 | 56.014 | 59.000 | 54.280 | 54.441 | 61.783 | 54.700 | 54.181 | 61.286 | 59.663 | 55.408 | 59.311 | 52.753 | 55.164 | 60.183 | 58.563 | 54.621 | 59.143 | -0.026 | 0.2817 | 0.5380 | 0 | 0 | 0 |
| 'Suox' | 9.455 | 9.953 | 9.006 | 8.240 | 8.948 | 9.711 | 9.232 | 9.135 | 10.375 | 7.827 | 6.737 | 10.053 | 7.388 | 7.571 | 9.097 | 8.121 | 7.151 | 8.562 | 0.197 | 0.0007 | 0.0075 | 1 | 0 | 0 |
| 'Supt16' | 39.854 | 41.384 | 43.503 | 45.013 | 44.803 | 43.334 | 44.788 | 41.495 | 44.623 | 37.967 | 43.843 | 43.576 | 41.162 | 40.644 | 42.038 | 38.399 | 45.452 | 42.952 | 0.038 | 0.2828 | 0.5392 | 0 | 0 | 0 |
| 'Supt20' | 21.649 | 20.391 | 20.324 | 20.065 | 20.297 | 18.989 | 20.848 | 20.260 | 20.748 | 20.786 | 21.326 | 20.713 | 20.120 | 20.761 | 19.852 | 19.892 | 21.407 | 19.821 | -0.019 | 0.4704 | 0.7061 | 0 | 0 | 0 |
| 'Supt3' | 3.648 | 3.400 | 3.497 | 3.425 | 3.410 | 3.624 | 3.245 | 3.952 | 2.923 | 3.564 | 3.475 | 3.502 | 3.422 | 3.757 | 2.914 | 3.121 | 3.703 | 3.381 | 0.005 | 0.9316 | 0.9746 | 0 | 0 | 0 |
| 'Supt4a' | 65.612 | 66.722 | 58.049 | 66.453 | 64.574 | 62.967 | 65.128 | 67.271 | 59.975 | 60.087 | 64.247 | 61.097 | 62.055 | 66.229 | 63.862 | 68.698 | 67.338 | 59.701 | -0.002 | 0.9469 | 0.9795 | 0 | 0 | 0 |
| 'Supt5' | 97.251 | 92.438 | 100.047 | 92.901 | 93.021 | 103.034 | 95.184 | 95.252 | 101.549 | 101.809 | 93.221 | 100.679 | 100.959 | 90.221 | 97.064 | 96.241 | 95.793 | 99.974 | -0.019 | 0.3659 | 0.6181 | 0 | 0 | 0 |
| 'Supt6' | 29.426 | 28.215 | 29.571 | 29.704 | 27.450 | 29.264 | 28.692 | 27.645 | 29.092 | 29.324 | 28.436 | 29.038 | 28.900 | 27.223 | 28.861 | 28.788 | 28.232 | 28.561 | 0.000 | 0.9845 | 0.9946 | 0 | 0 | 0 |
| 'Supt7l' | 2.926 | 2.906 | 3.095 | 2.698 | 2.960 | 3.363 | 3.319 | 3.203 | 3.242 | 2.822 | 3.289 | 2.943 | 2.795 | 2.760 | 3.163 | 3.239 | 2.671 | 2.903 | 0.056 | 0.2932 | 0.5496 | 0 | 0 | 0 |
| 'Supv3l1' | 4.208 | 4.611 | 4.751 | 4.131 | 4.572 | 5.109 | 4.385 | 5.242 | 4.821 | 4.856 | 4.139 | 4.155 | 4.866 | 4.068 | 4.026 | 4.823 | 4.118 | 4.204 | 0.087 | 0.1223 | 0.3336 | 0 | 0 | 0 |
| 'Surf1' | 8.806 | 10.233 | 9.464 | 8.859 | 9.404 | 9.746 | 9.355 | 10.231 | 9.148 | 9.073 | 8.937 | 9.738 | 8.941 | 9.554 | 9.855 | 8.828 | 9.449 | 9.075 | 0.017 | 0.6306 | 0.8173 | 0 | 0 | 0 |
| 'Surf2' | 5.261 | 5.206 | 6.011 | 4.946 | 5.073 | 5.785 | 5.103 | 5.278 | 6.080 | 3.897 | 3.390 | 4.897 | 3.899 | 3.753 | 4.446 | 4.033 | 4.904 | 4.626 | 0.353 | 0.0000 | 0.0000 | 1 | 0 | 0 |
| 'Surf4' | 104.078 | 121.049 | 116.974 | 108.619 | 120.781 | 121.334 | 102.745 | 118.521 | 116.848 | 70.219 | 65.116 | 66.823 | 67.791 | 66.619 | 68.763 | 71.202 | 65.025 | 66.158 | 0.750 | 0.0000 | 0.0000 | 1 | 1 | 0 |
| 'Surf6' | 8.765 | 7.931 | 8.986 | 7.684 | 8.264 | 8.858 | 8.213 | 8.291 | 8.443 | 8.447 | 7.578 | 8.622 | 7.622 | 7.947 | 8.588 | 8.179 | 7.913 | 7.918 | 0.044 | 0.1965 | 0.4354 | 0 | 0 | 0 |
| 'Susd1' | 0.608 | 0.672 | 0.585 | 0.684 | 0.889 | 0.689 | 0.633 | 0.653 | 0.751 | 0.596 | 0.767 | 0.551 | 0.668 | 0.651 | 0.517 | 0.475 | 0.607 | 0.478 | 0.205 | 0.0553 | 0.2017 | 0 | 0 | 0 |
| 'Susd2' | 4.864 | 6.387 | 7.316 | 6.692 | 7.053 | 7.751 | 6.615 | 5.963 | 7.699 | 7.986 | 6.773 | 8.751 | 7.595 | 6.841 | 8.206 | 8.237 | 7.237 | 9.437 | -0.250 | 0.0000 | 0.0006 | 1 | 0 | 0 |
| 'Susd3' | 14.504 | 14.497 | 14.819 | 13.936 | 15.518 | 14.315 | 13.128 | 15.036 | 15.495 | 13.891 | 14.746 | 14.459 | 16.491 | 15.191 | 15.876 | 16.204 | 13.882 | 14.284 | -0.049 | 0.2400 | 0.4903 | 0 | 0 | 0 |
| 'Susd4' | 5.923 | 5.292 | 5.534 | 5.531 | 4.685 | 4.958 | 5.152 | 6.049 | 5.447 | 6.778 | 4.822 | 5.316 | 5.270 | 5.382 | 4.822 | 5.266 | 4.686 | 4.810 | 0.038 | 0.5387 | 0.7567 | 0 | 0 | 0 |
| 'Susd6' | 44.758 | 41.479 | 38.240 | 41.353 | 41.802 | 38.195 | 41.679 | 43.285 | 38.256 | 44.601 | 46.946 | 47.771 | 39.709 | 45.640 | 44.763 | 43.491 | 46.201 | 47.719 | -0.152 | 0.0002 | 0.0025 | 1 | 0 | 0 |
| 'Suv39h1' | 4.140 | 4.097 | 4.660 | 4.500 | 4.775 | 5.059 | 4.416 | 4.361 | 4.836 | 3.990 | 4.373 | 4.412 | 4.766 | 4.465 | 5.012 | 3.986 | 4.275 | 4.385 | 0.032 | 0.4735 | 0.7089 | 0 | 0 | 0 |
| 'Suv39h2' | 1.091 | 0.937 | 1.272 | 1.100 | 0.895 | 0.948 | 1.129 | 1.067 | 1.101 | 1.210 | 1.143 | 1.164 | 1.103 | 1.004 | 1.056 | 1.079 | 1.327 | 1.202 | -0.115 | 0.1318 | 0.3492 | 0 | 0 | 0 |
| 'Suv420h1' | 6.728 | 7.131 | 6.537 | 6.540 | 6.893 | 6.665 | 6.502 | 7.235 | 6.526 | 6.756 | 6.890 | 7.189 | 6.497 | 7.048 | 6.959 | 7.399 | 7.299 | 6.812 | -0.060 | 0.0203 | 0.1041 | 0 | 0 | 0 |
| 'Suv420h2' | 11.195 | 9.906 | 12.556 | 9.908 | 10.546 | 11.860 | 10.474 | 10.186 | 12.961 | 11.281 | 10.572 | 13.182 | 11.936 | 10.365 | 12.525 | 9.936 | 10.980 | 12.205 | -0.060 | 0.0862 | 0.2672 | 0 | 0 | 0 |
| 'Suz12' | 25.816 | 26.149 | 26.302 | 27.850 | 26.491 | 23.843 | 25.971 | 27.460 | 23.229 | 24.706 | 25.285 | 23.846 | 25.676 | 27.018 | 22.896 | 25.227 | 25.050 | 23.545 | 0.052 | 0.0558 | 0.2024 | 0 | 0 | 0 |
| 'Sv2a' | 5.172 | 5.401 | 6.143 | 4.933 | 4.835 | 5.556 | 4.997 | 5.132 | 5.967 | 5.371 | 5.034 | 4.927 | 5.085 | 3.991 | 5.603 | 4.523 | 5.355 | 4.508 | 0.108 | 0.0695 | 0.2318 | 0 | 0 | 0 |
| 'Svbp' | 12.473 | 15.071 | 12.958 | 11.926 | 13.697 | 13.645 | 12.960 | 14.392 | 13.213 | 12.263 | 13.222 | 12.533 | 13.956 | 13.338 | 12.965 | 13.137 | 12.650 | 11.544 | 0.047 | 0.3247 | 0.5819 | 0 | 0 | 0 |
| 'Svil' | 2.906 | 2.232 | 3.260 | 2.869 | 2.147 | 2.838 | 3.020 | 2.416 | 2.961 | 2.993 | 2.623 | 3.302 | 2.243 | 2.557 | 3.238 | 2.646 | 2.403 | 3.263 | -0.047 | 0.4141 | 0.6620 | 0 | 0 | 0 |
| 'Swap70' | 0.978 | 2.024 | 1.525 | 1.223 | 2.295 | 1.486 | 1.254 | 2.367 | 1.573 | 0.935 | 2.695 | 1.715 | 1.001 | 2.041 | 1.368 | 1.577 | 2.870 | 1.607 | -0.100 | 0.3079 | 0.5662 | 0 | 0 | 0 |
| 'Swi5' | 10.959 | 13.040 | 12.071 | 11.702 | 12.503 | 11.999 | 12.195 | 12.839 | 12.187 | 11.213 | 11.054 | 11.831 | 11.887 | 12.225 | 12.521 | 12.060 | 11.936 | 11.665 | 0.029 | 0.3491 | 0.6042 | 0 | 0 | 0 |
| 'Swsap1' | 4.460 | 6.333 | 5.815 | 5.462 | 6.840 | 5.835 | 5.479 | 6.951 | 5.172 | 6.385 | 5.160 | 4.688 | 4.937 | 4.777 | 5.726 | 5.520 | 5.248 | 4.500 | 0.146 | 0.0794 | 0.2537 | 0 | 0 | 0 |
| 'Swt1' | 2.230 | 2.139 | 2.131 | 2.635 | 2.160 | 1.971 | 2.561 | 2.080 | 1.979 | 2.113 | 2.631 | 2.905 | 2.257 | 2.519 | 2.342 | 2.160 | 2.758 | 2.735 | -0.188 | 0.0104 | 0.0641 | 0 | 0 | 0 |
| 'Syap1' | 15.594 | 18.654 | 15.482 | 17.280 | 18.148 | 14.187 | 16.173 | 19.078 | 15.912 | 16.387 | 17.843 | 16.134 | 16.821 | 17.390 | 15.107 | 17.163 | 17.502 | 14.107 | 0.005 | 0.8995 | 0.9600 | 0 | 0 | 0 |
| 'Syce2' | 3.036 | 3.461 | 3.589 | 3.153 | 2.937 | 3.126 | 3.672 | 3.165 | 2.970 | 2.116 | 2.648 | 2.898 | 3.602 | 2.869 | 3.245 | 3.234 | 2.322 | 2.951 | 0.157 | 0.0693 | 0.2315 | 0 | 0 | 0 |
| 'Sycp1' | 1.115 | 0.966 | 1.542 | 0.843 | 1.162 | 0.871 | 1.109 | 1.083 | 1.208 | 1.135 | 0.833 | 1.036 | 0.990 | 1.095 | 0.819 | 0.960 | 1.009 | 0.937 | 0.164 | 0.1525 | 0.3797 | 0 | 0 | 0 |
| 'Syde1' | 4.688 | 5.600 | 5.739 | 4.176 | 5.340 | 5.570 | 4.699 | 5.345 | 5.978 | 6.102 | 4.347 | 5.280 | 6.243 | 5.044 | 5.222 | 5.258 | 4.997 | 4.701 | -0.011 | 0.8797 | 0.9494 | 0 | 0 | 0 |
| 'Syf2' | 43.358 | 49.882 | 42.620 | 45.296 | 46.172 | 44.555 | 44.981 | 49.892 | 44.514 | 45.080 | 42.235 | 42.512 | 47.965 | 47.296 | 43.107 | 49.722 | 43.653 | 38.495 | 0.031 | 0.3860 | 0.6354 | 0 | 0 | 0 |
| 'Sympk' | 18.762 | 17.302 | 20.217 | 17.913 | 16.670 | 20.493 | 18.197 | 16.986 | 19.494 | 18.667 | 16.844 | 20.304 | 19.511 | 17.431 | 19.851 | 17.271 | 17.306 | 19.869 | -0.020 | 0.3836 | 0.6340 | 0 | 0 | 0 |
| 'Syncrip' | 10.345 | 10.662 | 10.650 | 11.192 | 11.014 | 9.705 | 11.425 | 10.991 | 10.419 | 9.548 | 10.670 | 11.003 | 10.074 | 10.882 | 10.306 | 9.763 | 11.092 | 10.648 | 0.025 | 0.4621 | 0.6981 | 0 | 0 | 0 |
| 'Syne2' | 2.327 | 2.280 | 2.390 | 2.472 | 2.294 | 2.284 | 2.182 | 2.489 | 2.458 | 2.487 | 2.579 | 2.137 | 2.448 | 2.416 | 2.238 | 2.090 | 2.278 | 2.059 | 0.024 | 0.5807 | 0.7829 | 0 | 0 | 0 |
| 'Syne3' | 4.171 | 3.512 | 4.614 | 3.623 | 4.082 | 4.549 | 3.944 | 4.051 | 4.297 | 3.165 | 2.869 | 4.307 | 3.332 | 3.313 | 5.068 | 4.188 | 3.298 | 4.877 | 0.099 | 0.1811 | 0.4165 | 0 | 0 | 0 |
| 'Syngr2' | 21.618 | 22.828 | 23.085 | 20.760 | 24.085 | 23.098 | 20.820 | 23.445 | 22.096 | 24.125 | 20.670 | 23.721 | 22.355 | 22.364 | 23.560 | 23.485 | 21.814 | 23.402 | -0.037 | 0.2688 | 0.5231 | 0 | 0 | 0 |
| 'Synj1' | 5.119 | 4.579 | 5.172 | 4.816 | 4.403 | 4.594 | 4.588 | 4.701 | 5.090 | 5.284 | 5.498 | 5.371 | 4.507 | 5.041 | 5.157 | 4.562 | 4.920 | 5.508 | -0.100 | 0.0272 | 0.1271 | 0 | 0 | 0 |
| 'Synj2' | 6.893 | 6.148 | 6.688 | 6.713 | 6.179 | 6.008 | 6.151 | 6.026 | 6.251 | 6.357 | 6.277 | 6.893 | 5.751 | 5.755 | 6.519 | 6.355 | 6.476 | 6.863 | -0.015 | 0.7063 | 0.8639 | 0 | 0 | 0 |
| 'Synj2bp' | 5.613 | 6.351 | 5.636 | 6.642 | 6.326 | 6.687 | 5.596 | 6.389 | 6.123 | 6.124 | 7.024 | 6.892 | 5.790 | 6.404 | 6.174 | 5.621 | 6.674 | 6.587 | -0.065 | 0.2326 | 0.4808 | 0 | 0 | 0 |
| 'Synpo' | 1.944 | 1.866 | 2.092 | 1.541 | 1.758 | 2.080 | 1.486 | 1.910 | 1.977 | 2.452 | 2.035 | 2.361 | 1.706 | 2.235 | 2.371 | 2.230 | 1.993 | 2.314 | -0.253 | 0.0000 | 0.0007 | 1 | 0 | 0 |
| 'Synrg' | 11.515 | 12.708 | 12.448 | 11.927 | 12.394 | 12.183 | 10.314 | 12.311 | 12.191 | 13.395 | 12.406 | 11.673 | 12.671 | 12.522 | 11.462 | 12.042 | 11.070 | 11.468 | -0.019 | 0.6355 | 0.8198 | 0 | 0 | 0 |
| 'Syp' | 1.869 | 2.318 | 1.812 | 1.713 | 2.334 | 1.754 | 1.761 | 2.470 | 1.650 | 2.040 | 2.996 | 2.259 | 1.612 | 2.566 | 2.280 | 2.469 | 2.528 | 2.214 | -0.257 | 0.0003 | 0.0035 | 1 | 0 | 0 |
| 'Sypl' | 8.359 | 9.975 | 8.974 | 8.760 | 9.219 | 8.586 | 9.443 | 9.401 | 8.391 | 8.813 | 10.156 | 8.937 | 8.854 | 10.169 | 9.028 | 8.189 | 9.563 | 8.226 | -0.025 | 0.4440 | 0.6872 | 0 | 0 | 0 |
| 'Sys1' | 3.927 | 4.379 | 3.880 | 3.780 | 4.091 | 4.435 | 3.973 | 4.430 | 3.778 | 3.751 | 4.072 | 3.946 | 4.044 | 3.943 | 4.112 | 4.580 | 3.832 | 4.135 | 0.001 | 0.9799 | 0.9926 | 0 | 0 | 0 |
| 'Syt3' | 0.900 | 0.961 | 0.821 | 0.805 | 0.800 | 0.693 | 0.670 | 0.743 | 0.687 | 0.825 | 0.603 | 0.502 | 0.776 | 0.563 | 0.415 | 0.620 | 0.457 | 0.594 | 0.406 | 0.0014 | 0.0140 | 1 | 1 | 0 |
| 'Sytl1' | 2.453 | 2.060 | 3.075 | 2.156 | 2.210 | 3.012 | 2.211 | 2.176 | 2.807 | 2.484 | 1.742 | 2.874 | 2.380 | 1.993 | 2.468 | 2.543 | 1.860 | 2.346 | 0.092 | 0.1783 | 0.4137 | 0 | 0 | 0 |
| 'Sytl2' | 1.180 | 1.273 | 1.141 | 1.076 | 1.071 | 1.119 | 1.300 | 1.423 | 1.197 | 0.984 | 0.840 | 1.043 | 1.202 | 0.780 | 0.925 | 1.016 | 0.787 | 0.896 | 0.336 | 0.0000 | 0.0001 | 1 | 0 | 0 |
| 'Syvn1' | 9.590 | 8.847 | 9.624 | 9.331 | 8.518 | 9.063 | 8.422 | 8.842 | 10.018 | 9.203 | 9.369 | 10.998 | 9.233 | 8.668 | 10.291 | 8.540 | 9.060 | 10.788 | -0.076 | 0.0447 | 0.1753 | 0 | 0 | 0 |
| 'Szrd1' | 23.082 | 24.599 | 23.223 | 23.950 | 24.608 | 24.586 | 22.807 | 24.261 | 25.580 | 22.245 | 22.762 | 25.882 | 24.130 | 23.780 | 26.989 | 24.171 | 23.109 | 26.836 | -0.032 | 0.3193 | 0.5780 | 0 | 0 | 0 |
| 'Szt2' | 12.118 | 9.899 | 13.609 | 11.581 | 9.838 | 11.384 | 11.120 | 10.501 | 13.419 | 12.231 | 12.629 | 12.714 | 12.522 | 9.901 | 11.550 | 10.595 | 12.301 | 11.700 | -0.053 | 0.3439 | 0.5995 | 0 | 0 | 0 |
| 'Tab1' | 12.642 | 11.456 | 15.698 | 11.902 | 11.651 | 14.759 | 13.256 | 12.692 | 14.444 | 14.591 | 9.966 | 11.552 | 12.940 | 9.452 | 10.858 | 13.749 | 9.464 | 10.588 | 0.202 | 0.0018 | 0.0165 | 1 | 0 | 0 |
| 'Tab2' | 25.312 | 26.498 | 23.158 | 25.045 | 24.728 | 24.044 | 24.018 | 26.242 | 23.216 | 24.900 | 25.270 | 24.426 | 23.270 | 25.562 | 24.069 | 24.459 | 25.329 | 23.370 | -0.001 | 0.9722 | 0.9903 | 0 | 0 | 0 |
| 'Tacc1' | 23.283 | 21.973 | 23.565 | 21.425 | 22.181 | 22.941 | 22.279 | 23.820 | 22.597 | 23.415 | 24.015 | 24.738 | 22.124 | 23.256 | 23.818 | 22.223 | 22.846 | 25.968 | -0.067 | 0.0252 | 0.1213 | 0 | 0 | 0 |
| 'Tacc3' | 8.810 | 8.878 | 11.296 | 10.065 | 9.284 | 11.128 | 10.911 | 9.320 | 12.301 | 7.305 | 10.189 | 10.052 | 11.404 | 10.498 | 11.519 | 9.543 | 10.883 | 9.749 | -0.001 | 0.9892 | 0.9968 | 0 | 0 | 0 |
| 'Taco1' | 15.765 | 15.090 | 15.938 | 16.017 | 16.599 | 17.035 | 16.845 | 16.490 | 15.169 | 14.706 | 16.680 | 16.711 | 13.456 | 13.231 | 16.656 | 15.565 | 14.716 | 18.056 | 0.042 | 0.4619 | 0.6981 | 0 | 0 | 0 |
| 'Tada1' | 9.698 | 9.807 | 9.446 | 9.959 | 9.191 | 9.342 | 8.341 | 9.658 | 9.514 | 8.707 | 8.217 | 9.638 | 8.309 | 9.243 | 8.623 | 8.666 | 9.024 | 9.043 | 0.084 | 0.0351 | 0.1497 | 0 | 0 | 0 |
| 'Tada2a' | 4.754 | 4.843 | 5.154 | 5.227 | 4.514 | 4.626 | 5.041 | 4.561 | 5.359 | 4.590 | 4.991 | 4.846 | 4.864 | 4.851 | 4.805 | 4.911 | 4.734 | 4.569 | 0.022 | 0.6578 | 0.8350 | 0 | 0 | 0 |
| 'Tada2b' | 8.909 | 10.840 | 9.141 | 8.463 | 9.291 | 9.497 | 8.219 | 10.328 | 9.135 | 8.958 | 8.758 | 10.381 | 8.767 | 9.396 | 9.256 | 9.060 | 9.173 | 9.494 | -0.007 | 0.8684 | 0.9431 | 0 | 0 | 0 |
| 'Tada3' | 6.970 | 7.081 | 8.582 | 7.556 | 7.110 | 9.043 | 8.501 | 7.611 | 8.786 | 8.220 | 7.106 | 9.124 | 8.225 | 7.346 | 8.764 | 7.832 | 7.831 | 8.529 | -0.047 | 0.1841 | 0.4195 | 0 | 0 | 0 |
| 'Taf1' | 10.126 | 10.992 | 10.348 | 10.486 | 11.159 | 10.621 | 10.061 | 10.538 | 10.819 | 10.126 | 11.365 | 10.660 | 10.445 | 11.859 | 10.427 | 10.513 | 10.973 | 10.626 | -0.037 | 0.1271 | 0.3416 | 0 | 0 | 0 |
| 'Taf10' | 20.370 | 21.019 | 21.366 | 19.167 | 20.742 | 22.855 | 21.350 | 21.300 | 20.485 | 20.764 | 20.171 | 20.317 | 21.208 | 20.099 | 21.524 | 20.863 | 19.975 | 20.338 | 0.017 | 0.5447 | 0.7606 | 0 | 0 | 0 |
| 'Taf11' | 8.460 | 10.321 | 8.722 | 7.741 | 10.795 | 10.005 | 8.939 | 10.152 | 8.875 | 8.782 | 8.281 | 9.003 | 9.807 | 10.489 | 8.479 | 10.298 | 9.009 | 8.071 | 0.018 | 0.7522 | 0.8895 | 0 | 0 | 0 |
| 'Taf12' | 10.444 | 11.411 | 11.635 | 11.298 | 11.341 | 11.757 | 10.845 | 11.627 | 11.162 | 10.764 | 10.232 | 12.059 | 11.055 | 11.778 | 12.113 | 11.476 | 11.079 | 11.449 | -0.019 | 0.5600 | 0.7704 | 0 | 0 | 0 |
| 'Taf13' | 3.971 | 4.448 | 4.005 | 4.143 | 4.540 | 3.956 | 3.702 | 4.661 | 3.787 | 4.168 | 4.417 | 3.737 | 4.183 | 4.577 | 3.624 | 3.961 | 4.272 | 3.598 | 0.021 | 0.6523 | 0.8310 | 0 | 0 | 0 |
| 'Taf15' | 44.363 | 45.043 | 45.525 | 47.921 | 48.053 | 44.321 | 47.530 | 46.269 | 48.197 | 46.026 | 46.364 | 43.863 | 48.821 | 45.325 | 43.073 | 46.881 | 46.653 | 42.027 | 0.020 | 0.4475 | 0.6880 | 0 | 0 | 0 |
| 'Taf1a' | 4.483 | 4.193 | 4.702 | 5.093 | 3.724 | 4.251 | 4.917 | 4.085 | 3.989 | 4.145 | 4.518 | 5.084 | 3.801 | 4.426 | 4.139 | 4.127 | 4.472 | 4.496 | -0.008 | 0.9103 | 0.9651 | 0 | 0 | 0 |
| 'Taf1b' | 20.392 | 22.422 | 20.593 | 23.478 | 20.790 | 20.719 | 21.857 | 24.641 | 21.116 | 23.760 | 25.658 | 20.678 | 22.379 | 23.359 | 21.155 | 22.641 | 21.636 | 18.192 | -0.030 | 0.5133 | 0.7375 | 0 | 0 | 0 |
| 'Taf1c' | 10.184 | 8.828 | 11.491 | 9.858 | 8.783 | 10.805 | 9.706 | 9.367 | 11.480 | 9.894 | 9.903 | 13.110 | 9.911 | 9.479 | 11.973 | 9.279 | 10.341 | 12.427 | -0.099 | 0.0073 | 0.0494 | 1 | 0 | 0 |
| 'Taf1d' | 37.628 | 30.318 | 27.443 | 34.879 | 30.384 | 24.863 | 41.157 | 33.330 | 25.110 | 33.712 | 29.649 | 26.604 | 35.633 | 30.868 | 25.760 | 30.893 | 32.224 | 26.091 | 0.051 | 0.2447 | 0.4952 | 0 | 0 | 0 |
| 'Taf2' | 13.906 | 14.996 | 15.313 | 14.359 | 16.620 | 14.524 | 15.393 | 16.518 | 15.550 | 16.298 | 14.407 | 14.688 | 15.277 | 14.411 | 14.849 | 15.039 | 15.912 | 15.560 | -0.004 | 0.9231 | 0.9705 | 0 | 0 | 0 |

| | | | | | | | | | | | | | | | | | | | | | | | | |
|---|---|---|---|---|---|---|---|---|---|---|---|---|---|---|---|---|---|---|---|---|---|---|---|---|
| 'Taf3' | 7.057 | 5.806 | 6.007 | 7.038 | 5.628 | 5.871 | 6.353 | 6.255 | 6.278 | 6.066 | 6.109 | 6.490 | 5.906 | 5.674 | 6.054 | 6.294 | 6.157 | 6.293 | 0.019 | 0.6525 | 0.8312 | 0 | 0 | 0 |
| 'Taf4' | 13.376 | 12.844 | 13.593 | 12.705 | 12.588 | 13.921 | 11.901 | 12.716 | 13.602 | 13.993 | 13.620 | 14.328 | 14.695 | 14.641 | 14.117 | 13.841 | 14.291 | 15.309 | -0.145 | 0.0000 | 0.0001 | 1 | 0 | 0 |
| 'Taf4b' | 1.211 | 1.115 | 1.371 | 1.188 | 0.982 | 1.273 | 1.398 | 1.173 | 1.245 | 1.441 | 1.395 | 1.151 | 1.331 | 1.302 | 0.957 | 1.344 | 1.296 | 1.219 | -0.055 | 0.5321 | 0.7526 | 0 | 0 | 0 |
| 'Taf5' | 6.546 | 6.811 | 6.777 | 6.208 | 6.932 | 6.697 | 6.770 | 7.409 | 6.339 | 6.611 | 6.400 | 6.315 | 6.880 | 7.078 | 5.903 | 7.026 | 6.307 | 5.641 | 0.052 | 0.2861 | 0.5428 | 0 | 0 | 0 |
| 'Taf5l' | 22.950 | 23.084 | 23.330 | 23.134 | 22.671 | 23.203 | 22.650 | 23.199 | 22.585 | 22.411 | 21.187 | 22.034 | 22.256 | 22.090 | 22.543 | 23.774 | 19.854 | 21.324 | 0.059 | 0.0481 | 0.1837 | 0 | 0 | 0 |
| 'Taf6' | 5.143 | 4.911 | 5.546 | 5.297 | 5.272 | 5.693 | 5.554 | 5.581 | 5.643 | 5.385 | 5.208 | 6.029 | 5.732 | 5.615 | 5.765 | 5.310 | 5.193 | 5.852 | -0.054 | 0.1197 | 0.3294 | 0 | 0 | 0 |
| 'Taf6l' | 4.425 | 5.070 | 4.905 | 4.470 | 4.988 | 5.324 | 4.201 | 4.958 | 4.891 | 4.567 | 4.383 | 4.856 | 4.891 | 4.955 | 4.974 | 4.689 | 4.852 | 4.523 | 0.007 | 0.8487 | 0.9351 | 0 | 0 | 0 |
| 'Taf7' | 12.640 | 9.074 | 6.311 | 9.715 | 9.665 | 7.220 | 10.115 | 9.467 | 5.971 | 11.153 | 9.582 | 5.194 | 11.681 | 11.436 | 5.530 | 12.641 | 8.525 | 4.887 | 0.029 | 0.6929 | 0.8555 | 0 | 0 | 0 |
| 'Taf8' | 18.112 | 20.326 | 21.451 | 19.968 | 18.200 | 22.220 | 18.282 | 20.720 | 23.732 | 20.284 | 20.483 | 23.660 | 20.167 | 21.816 | 21.456 | 18.884 | 20.691 | 22.521 | -0.066 | 0.0711 | 0.2352 | 0 | 0 | 0 |
| 'Taf9' | 27.634 | 31.367 | 28.453 | 31.883 | 32.827 | 28.125 | 32.921 | 31.044 | 27.551 | 27.777 | 29.844 | 29.124 | 29.403 | 32.764 | 29.500 | 27.311 | 31.578 | 29.752 | 0.008 | 0.8509 | 0.9358 | 0 | 0 | 0 |
| 'Taf9b' | 7.228 | 8.570 | 8.516 | 7.872 | 7.980 | 8.905 | 7.366 | 8.627 | 8.008 | 8.164 | 9.095 | 7.880 | 7.020 | 8.908 | 8.176 | 8.260 | 8.250 | 7.018 | 0.004 | 0.9434 | 0.9788 | 0 | 0 | 0 |
| 'Tagap1' | 58.553 | 48.910 | 54.031 | 54.276 | 48.617 | 54.745 | 55.377 | 51.125 | 53.887 | 56.693 | 49.983 | 51.969 | 51.146 | 48.399 | 50.403 | 54.680 | 53.737 | 50.160 | 0.029 | 0.3577 | 0.6103 | 0 | 0 | 0 |
| 'Tagln' | 3.144 | 3.216 | 3.745 | 2.692 | 3.438 | 3.124 | 3.721 | 3.637 | 3.089 | 4.076 | 3.624 | 2.704 | 4.026 | 2.317 | 2.427 | 3.236 | 3.208 | 2.470 | 0.096 | 0.3859 | 0.6354 | 0 | 0 | 0 |
| 'Tagln2' | 36.731 | 35.501 | 41.526 | 34.933 | 35.882 | 41.766 | 38.251 | 36.830 | 38.837 | 33.458 | 33.432 | 41.720 | 37.351 | 36.163 | 45.160 | 34.126 | 34.089 | 42.930 | 0.000 | 0.9936 | 0.9990 | 0 | 0 | 0 |
| 'Taldo1' | 63.019 | 60.439 | 62.549 | 65.522 | 66.097 | 64.978 | 63.683 | 62.947 | 64.228 | 57.939 | 62.596 | 60.419 | 64.985 | 63.299 | 65.711 | 59.586 | 59.766 | 61.401 | 0.035 | 0.2297 | 0.4766 | 0 | 0 | 0 |
| 'Tamm41' | 2.223 | 1.488 | 2.336 | 1.779 | 1.743 | 2.448 | 2.111 | 1.913 | 2.441 | 2.188 | 2.016 | 2.239 | 2.113 | 1.920 | 2.568 | 2.230 | 2.056 | 1.914 | -0.063 | 0.3574 | 0.6103 | 0 | 0 | 0 |
| 'Tanc1' | 0.544 | 0.618 | 0.650 | 0.803 | 0.610 | 0.725 | 0.676 | 0.690 | 0.692 | 1.016 | 0.999 | 0.739 | 1.141 | 0.965 | 0.739 | 0.940 | 0.753 | 0.817 | -0.432 | 0.0000 | 0.0000 | 1 | 1 | 0 |
| 'Tanc2' | 0.611 | 0.540 | 0.565 | 0.576 | 0.485 | 0.498 | 0.505 | 0.517 | 0.519 | 0.551 | 0.380 | 0.383 | 0.388 | 0.462 | 0.307 | 0.481 | 0.369 | 0.369 | 0.390 | 0.0000 | 0.0000 | 1 | 1 | 0 |
| 'Tango2' | 3.449 | 3.908 | 3.546 | 3.630 | 3.788 | 3.920 | 4.039 | 3.696 | 4.136 | 3.851 | 3.804 | 3.522 | 4.322 | 3.719 | 3.726 | 4.198 | 3.958 | 3.342 | -0.020 | 0.6767 | 0.8453 | 0 | 0 | 0 |
| 'Tango6' | 2.267 | 1.819 | 2.357 | 1.790 | 1.957 | 2.658 | 2.264 | 1.973 | 2.769 | 2.403 | 2.058 | 2.385 | 2.220 | 2.338 | 2.285 | 2.194 | 2.249 | 2.113 | -0.032 | 0.6708 | 0.8413 | 0 | 0 | 0 |
| 'Tank' | 17.167 | 20.111 | 16.644 | 17.287 | 19.732 | 15.933 | 18.623 | 17.899 | 16.433 | 16.710 | 17.200 | 15.058 | 18.142 | 17.905 | 14.390 | 17.191 | 18.216 | 13.740 | 0.097 | 0.0014 | 0.0135 | 1 | 0 | 0 |
| 'Taok1' | 19.426 | 14.781 | 15.785 | 17.982 | 14.910 | 13.481 | 16.738 | 15.173 | 15.032 | 16.673 | 17.568 | 16.637 | 15.904 | 16.116 | 15.642 | 17.591 | 17.830 | 16.625 | -0.086 | 0.0995 | 0.2952 | 0 | 0 | 0 |
| 'Taok2' | 24.248 | 21.256 | 25.466 | 23.132 | 21.063 | 23.518 | 23.511 | 21.659 | 24.895 | 22.527 | 23.458 | 26.288 | 23.680 | 21.914 | 25.044 | 21.566 | 23.338 | 25.794 | -0.044 | 0.1628 | 0.3947 | 0 | 0 | 0 |
| 'Taok3' | 12.132 | 10.662 | 11.081 | 11.292 | 11.446 | 11.186 | 11.267 | 11.151 | 10.891 | 12.168 | 11.653 | 11.807 | 12.402 | 11.416 | 11.489 | 11.759 | 11.262 | 11.958 | -0.077 | 0.0013 | 0.0127 | 1 | 0 | 0 |
| 'Tap1' | 33.674 | 36.136 | 38.099 | 31.889 | 35.386 | 38.062 | 34.036 | 34.181 | 38.268 | 36.411 | 35.512 | 37.916 | 38.450 | 34.802 | 37.924 | 36.708 | 36.220 | 36.046 | -0.058 | 0.0289 | 0.1317 | 0 | 0 | 0 |
| 'Tap2' | 74.184 | 76.820 | 75.685 | 70.582 | 72.503 | 73.899 | 71.665 | 71.855 | 77.159 | 68.536 | 65.824 | 70.978 | 75.239 | 68.281 | 71.367 | 71.525 | 67.415 | 70.700 | 0.067 | 0.0047 | 0.0356 | 1 | 0 | 0 |
| 'Tapbp' | 76.681 | 78.962 | 87.368 | 72.069 | 81.808 | 90.514 | 75.251 | 79.188 | 89.881 | 90.625 | 77.313 | 93.715 | 86.252 | 81.397 | 94.627 | 93.374 | 73.204 | 95.292 | -0.112 | 0.0024 | 0.0212 | 1 | 0 | 0 |
| 'Tapbpl' | 43.787 | 45.932 | 45.174 | 43.150 | 47.430 | 47.807 | 47.373 | 43.393 | 47.401 | 47.732 | 46.027 | 48.079 | 53.086 | 45.741 | 47.528 | 49.738 | 46.702 | 46.134 | -0.077 | 0.0124 | 0.0732 | 0 | 0 | 0 |
| 'Tapt1' | 12.626 | 13.689 | 14.024 | 12.757 | 13.899 | 12.501 | 12.944 | 14.482 | 12.809 | 12.603 | 14.653 | 12.827 | 14.558 | 14.177 | 12.342 | 12.230 | 15.212 | 12.274 | -0.022 | 0.5407 | 0.7580 | 0 | 0 | 0 |
| 'Tarbp1' | 7.405 | 6.305 | 7.569 | 6.497 | 6.382 | 6.399 | 6.237 | 6.447 | 6.408 | 6.014 | 6.174 | 7.587 | 5.580 | 6.293 | 7.099 | 5.735 | 7.047 | 7.414 | 0.011 | 0.8387 | 0.9311 | 0 | 0 | 0 |
| 'Tarbp2' | 4.619 | 4.730 | 5.456 | 4.373 | 4.832 | 5.574 | 4.975 | 5.231 | 5.541 | 4.865 | 4.994 | 5.470 | 5.223 | 4.867 | 5.614 | 5.454 | 4.707 | 5.119 | -0.039 | 0.2912 | 0.5480 | 0 | 0 | 0 |
| 'Tardbp' | 9.332 | 9.876 | 10.190 | 10.345 | 9.751 | 9.376 | 10.652 | 9.645 | 9.960 | 8.251 | 9.573 | 10.966 | 9.076 | 9.653 | 10.490 | 8.701 | 10.622 | 11.054 | 0.006 | 0.9076 | 0.9637 | 0 | 0 | 0 |
| 'Tars' | 30.226 | 29.555 | 32.440 | 30.153 | 31.597 | 30.987 | 29.216 | 30.720 | 31.333 | 31.016 | 30.196 | 30.877 | 31.426 | 28.463 | 30.640 | 29.324 | 31.940 | 29.847 | 0.004 | 0.8887 | 0.9544 | 0 | 0 | 0 |
| 'Tars2' | 5.915 | 5.093 | 6.474 | 5.970 | 5.217 | 6.111 | 5.885 | 5.499 | 6.069 | 6.140 | 5.606 | 6.724 | 6.165 | 5.316 | 6.176 | 5.171 | 5.632 | 6.376 | -0.041 | 0.2450 | 0.4957 | 0 | 0 | 0 |
| 'Tarsl2' | 0.900 | 1.115 | 1.139 | 0.973 | 1.043 | 1.039 | 0.905 | 0.890 | 1.071 | 1.045 | 0.921 | 0.927 | 0.907 | 0.779 | 0.948 | 1.130 | 1.028 | 0.879 | 0.080 | 0.3115 | 0.5700 | 0 | 0 | 0 |
| 'Tasp1' | 20.442 | 22.518 | 18.582 | 21.602 | 23.286 | 17.712 | 23.434 | 23.445 | 18.194 | 18.973 | 22.503 | 17.217 | 21.313 | 20.650 | 17.456 | 19.941 | 22.707 | 15.913 | 0.088 | 0.0068 | 0.0472 | 1 | 0 | 0 |
| 'Tatdn1' | 8.787 | 11.430 | 10.185 | 10.183 | 11.344 | 7.843 | 10.883 | 10.775 | 10.622 | 8.825 | 10.490 | 9.623 | 10.986 | 10.652 | 8.158 | 10.207 | 12.858 | 8.949 | 0.007 | 0.9157 | 0.9677 | 0 | 0 | 0 |
| 'Tatdn2' | 20.674 | 18.689 | 19.702 | 19.699 | 18.133 | 20.325 | 19.947 | 18.551 | 17.770 | 19.879 | 19.817 | 21.573 | 18.433 | 20.004 | 21.860 | 19.753 | 18.863 | 21.510 | -0.077 | 0.0425 | 0.1700 | 0 | 0 | 0 |
| 'Tatdn3' | 1.170 | 1.583 | 1.797 | 1.408 | 1.620 | 1.549 | 1.480 | 1.636 | 1.722 | 1.486 | 1.588 | 1.327 | 1.957 | 1.533 | 1.268 | 1.598 | 1.622 | 1.099 | 0.053 | 0.5873 | 0.7882 | 0 | 0 | 0 |
| 'Tax1bp1' | 17.625 | 19.269 | 17.567 | 18.589 | 18.819 | 17.851 | 18.874 | 18.371 | 17.429 | 18.177 | 18.672 | 17.633 | 17.414 | 18.961 | 17.183 | 18.743 | 18.662 | 16.608 | 0.011 | 0.6267 | 0.8148 | 0 | 0 | 0 |
| 'Tax1bp3' | 1.820 | 1.726 | 1.705 | 1.338 | 1.949 | 1.653 | 1.789 | 1.847 | 1.767 | 1.405 | 1.793 | 1.888 | 1.932 | 1.758 | 2.110 | 1.569 | 1.947 | 1.789 | -0.069 | 0.3728 | 0.6242 | 0 | 0 | 0 |
| 'Taz' | 3.795 | 4.658 | 4.496 | 3.756 | 4.708 | 4.531 | 3.952 | 4.694 | 4.746 | 4.143 | 4.487 | 4.585 | 4.329 | 4.472 | 4.478 | 4.329 | 4.516 | 4.414 | -0.028 | 0.3551 | 0.6084 | 0 | 0 | 0 |
| 'Tbc1d1' | 11.807 | 11.297 | 11.862 | 11.898 | 10.862 | 11.907 | 12.470 | 11.426 | 12.359 | 13.263 | 12.825 | 13.382 | 12.187 | 12.126 | 12.660 | 12.130 | 13.022 | 12.454 | -0.118 | 0.0001 | 0.0009 | 1 | 0 | 0 |
| 'Tbc1d10a' | 14.251 | 13.212 | 13.907 | 11.515 | 12.677 | 14.007 | 12.560 | 13.101 | 13.273 | 12.506 | 11.078 | 14.563 | 12.656 | 11.739 | 13.412 | 13.206 | 11.411 | 13.653 | 0.044 | 0.2672 | 0.5207 | 0 | 0 | 0 |
| 'Tbc1d10b' | 18.356 | 16.688 | 19.031 | 17.037 | 16.170 | 19.390 | 16.102 | 16.501 | 18.532 | 17.037 | 17.236 | 21.307 | 18.256 | 18.533 | 21.315 | 19.262 | 16.712 | 21.508 | -0.125 | 0.0001 | 0.0021 | 1 | 0 | 0 |
| 'Tbc1d10c' | 166.113 | 161.238 | 185.849 | 165.654 | 163.924 | 191.059 | 176.452 | 161.528 | 188.216 | 163.078 | 159.701 | 203.018 | 168.835 | 157.047 | 197.753 | 163.286 | 170.558 | 191.793 | -0.024 | 0.3077 | 0.5662 | 0 | 0 | 0 |
| 'Tbc1d12' | 1.519 | 1.420 | 1.564 | 1.490 | 1.640 | 1.482 | 1.643 | 1.682 | 1.406 | 1.780 | 1.650 | 1.580 | 1.833 | 1.803 | 1.696 | 1.277 | 1.378 | 1.232 | -0.050 | 0.5738 | 0.7796 | 0 | 0 | 0 |
| 'Tbc1d13' | 8.747 | 9.395 | 9.552 | 8.405 | 9.794 | 9.673 | 8.115 | 9.692 | 9.696 | 9.765 | 9.391 | 9.899 | 9.853 | 10.101 | 9.920 | 9.622 | 9.043 | 9.521 | -0.078 | 0.0228 | 0.1127 | 0 | 0 | 0 |
| 'Tbc1d14' | 9.900 | 10.283 | 9.082 | 9.698 | 10.092 | 9.442 | 8.731 | 10.174 | 9.325 | 9.875 | 9.757 | 9.314 | 9.369 | 9.679 | 9.139 | 9.321 | 9.696 | 9.548 | 0.006 | 0.8318 | 0.9270 | 0 | 0 | 0 |
| 'Tbc1d15' | 15.204 | 16.660 | 16.246 | 15.531 | 16.432 | 14.859 | 16.435 | 17.181 | 14.837 | 14.051 | 17.032 | 16.200 | 13.455 | 15.956 | 14.483 | 14.623 | 16.501 | 15.194 | 0.049 | 0.1934 | 0.4308 | 0 | 0 | 0 |
| 'Tbc1d17' | 8.355 | 7.472 | 8.797 | 7.498 | 7.290 | 8.622 | 7.719 | 8.029 | 8.673 | 8.132 | 7.378 | 8.219 | 7.786 | 6.727 | 7.888 | 8.234 | 6.736 | 7.780 | 0.067 | 0.0565 | 0.2043 | 0 | 0 | 0 |
| 'Tbc1d19' | 0.663 | 1.016 | 0.847 | 0.821 | 0.731 | 0.770 | 0.844 | 0.862 | 0.848 | 0.835 | 1.095 | 0.918 | 0.857 | 1.103 | 0.847 | 0.939 | 0.989 | 0.842 | -0.195 | 0.0232 | 0.1143 | 0 | 0 | 0 |
| 'Tbc1d2' | 0.596 | 0.598 | 1.108 | 0.887 | 0.559 | 0.929 | 0.720 | 0.869 | 0.890 | 0.955 | 0.781 | 0.671 | 0.851 | 0.677 | 0.692 | 0.869 | 0.667 | 0.602 | 0.092 | 0.4849 | 0.7162 | 0 | 0 | 0 |
| 'Tbc1d20' | 12.651 | 14.927 | 11.917 | 11.565 | 13.488 | 13.377 | 11.626 | 12.833 | 12.492 | 12.781 | 12.754 | 14.275 | 12.312 | 13.573 | 14.223 | 13.615 | 13.302 | 14.529 | -0.092 | 0.0258 | 0.1234 | 0 | 0 | 0 |
| 'Tbc1d22a' | 16.324 | 17.785 | 15.799 | 14.473 | 15.241 | 15.645 | 15.497 | 15.845 | 15.075 | 16.602 | 15.483 | 16.198 | 14.382 | 15.476 | 15.726 | 17.913 | 14.311 | 14.755 | -0.001 | 0.9830 | 0.9934 | 0 | 0 | 0 |
| 'Tbc1d22b' | 15.173 | 15.140 | 15.851 | 14.234 | 15.802 | 15.883 | 13.786 | 16.706 | 16.380 | 16.917 | 14.816 | 16.639 | 15.333 | 15.205 | 15.906 | 16.641 | 13.581 | 17.303 | -0.044 | 0.2961 | 0.5522 | 0 | 0 | 0 |
| 'Tbc1d23' | 8.786 | 10.680 | 9.167 | 8.581 | 10.345 | 8.760 | 8.791 | 10.123 | 9.682 | 9.218 | 10.422 | 9.626 | 8.894 | 9.479 | 8.564 | 9.522 | 10.037 | 8.661 | -0.005 | 0.8900 | 0.9550 | 0 | 0 | 0 |
| 'Tbc1d24' | 0.231 | 0.247 | 0.206 | 0.193 | 0.211 | 0.244 | 0.226 | 0.211 | 0.208 | 0.201 | 0.253 | 0.203 | 0.262 | 0.180 | 0.232 | 0.231 | 0.186 | 0.216 | 0.002 | 0.9823 | 0.9934 | 0 | 0 | 0 |
| 'Tbc1d25' | 2.885 | 2.726 | 2.854 | 2.610 | 2.657 | 3.171 | 2.378 | 2.875 | 3.125 | 2.667 | 2.859 | 2.958 | 2.661 | 3.078 | 3.055 | 2.651 | 3.229 | 2.824 | -0.042 | 0.4276 | 0.6739 | 0 | 0 | 0 |
| 'Tbc1d2b' | 29.055 | 30.499 | 27.367 | 28.280 | 29.270 | 27.908 | 26.917 | 29.654 | 29.485 | 26.953 | 27.584 | 28.333 | 29.144 | 29.279 | 27.705 | 29.431 | 26.303 | 28.307 | 0.019 | 0.5033 | 0.7303 | 0 | 0 | 0 |
| 'Tbc1d30' | 6.247 | 6.215 | 6.595 | 5.791 | 7.185 | 6.682 | 6.070 | 5.685 | 7.275 | 7.148 | 6.319 | 8.145 | 4.947 | 6.054 | 7.624 | 5.933 | 5.607 | 8.295 | -0.061 | 0.3663 | 0.6184 | 0 | 0 | 0 |
| 'Tbc1d31' | 4.178 | 4.466 | 4.301 | 4.680 | 4.262 | 4.340 | 4.471 | 3.755 | 4.441 | 4.163 | 4.636 | 4.165 | 5.160 | 4.719 | 4.123 | 4.088 | 5.152 | 4.447 | -0.073 | 0.1789 | 0.4144 | 0 | 0 | 0 |
| 'Tbc1d32' | 0.769 | 0.716 | 0.807 | 0.824 | 0.664 | 0.804 | 0.847 | 0.879 | 0.894 | 0.791 | 0.784 | 0.830 | 0.732 | 0.721 | 0.707 | 0.540 | 0.848 | 0.729 | 0.095 | 0.3088 | 0.5671 | 0 | 0 | 0 |
| 'Tbc1d4' | 40.388 | 46.240 | 39.328 | 40.364 | 44.753 | 39.396 | 41.139 | 43.025 | 41.615 | 48.371 | 49.756 | 41.951 | 45.571 | 49.743 | 43.457 | 48.537 | 52.014 | 44.285 | -0.181 | 0.0000 | 0.0000 | 1 | 0 | 0 |
| 'Tbc1d5' | 5.541 | 5.204 | 5.173 | 5.405 | 4.751 | 5.420 | 4.686 | 4.994 | 5.329 | 5.919 | 5.596 | 5.852 | 5.159 | 5.961 | 5.243 | 5.246 | 5.732 | 5.628 | -0.123 | 0.0056 | 0.0406 | 1 | 0 | 0 |
| 'Tbc1d7' | 4.654 | 5.913 | 5.491 | 4.725 | 7.009 | 6.391 | 5.294 | 5.832 | 5.391 | 5.405 | 5.604 | 4.842 | 7.045 | 6.154 | 4.971 | 6.682 | 5.156 | 4.193 | 0.018 | 0.8495 | 0.9351 | 0 | 0 | 0 |
| 'Tbc1d8' | 0.396 | 0.398 | 0.415 | 0.396 | 0.445 | 0.471 | 0.412 | 0.536 | 0.474 | 0.467 | 0.578 | 0.441 | 0.587 | 0.441 | 0.446 | 0.428 | 0.519 | 0.432 | -0.138 | 0.1862 | 0.4219 | 0 | 0 | 0 |
| 'Tbc1d8b' | 5.975 | 6.046 | 6.028 | 6.401 | 6.399 | 5.837 | 6.809 | 7.011 | 6.300 | 6.376 | 7.965 | 6.708 | 6.248 | 7.359 | 6.253 | 6.559 | 7.575 | 7.097 | -0.137 | 0.0032 | 0.0264 | 1 | 0 | 0 |
| 'Tbc1d9b' | 15.419 | 14.355 | 16.781 | 14.142 | 14.062 | 16.964 | 13.974 | 15.096 | 16.204 | 16.299 | 14.817 | 16.576 | 15.490 | 13.699 | 15.425 | 14.505 | 13.604 | 16.066 | -0.005 | 0.8897 | 0.9548 | 0 | 0 | 0 |
| 'Tbca' | 137.533 | 156.286 | 128.666 | 148.656 | 147.330 | 137.537 | 139.398 | 144.380 | 131.988 | 130.809 | 134.453 | 113.610 | 134.312 | 140.982 | 131.789 | 138.903 | 141.891 | 121.517 | 0.090 | 0.0116 | 0.0694 | 0 | 0 | 0 |
| 'Tbcb' | 19.414 | 21.676 | 22.329 | 21.078 | 23.734 | 25.545 | 20.801 | 23.126 | 22.355 | 21.747 | 19.016 | 20.801 | 23.792 | 22.372 | 21.557 | 22.110 | 20.729 | 19.123 | 0.055 | 0.2702 | 0.5251 | 0 | 0 | 0 |
| 'Tbcc' | 12.932 | 12.473 | 16.851 | 14.495 | 14.519 | 16.690 | 13.222 | 14.919 | 14.621 | 13.192 | 12.816 | 16.291 | 14.864 | 12.640 | 16.114 | 14.258 | 11.421 | 15.109 | 0.036 | 0.4784 | 0.7112 | 0 | 0 | 0 |
| 'Tbccd1' | 5.784 | 5.548 | 6.083 | 4.887 | 6.824 | 6.013 | 5.775 | 6.893 | 6.615 | 5.824 | 6.297 | 5.977 | 6.741 | 6.491 | 5.411 | 6.388 | 5.575 | 5.373 | 0.003 | 0.9635 | 0.9869 | 0 | 0 | 0 |
| 'Tbcd' | 7.464 | 7.107 | 7.605 | 7.363 | 7.567 | 7.403 | 7.717 | 7.409 | 7.714 | 7.356 | 7.825 | 7.792 | 7.228 | 7.256 | 7.916 | 6.918 | 7.064 | 7.522 | -0.001 | 0.9764 | 0.9918 | 0 | 0 | 0 |
| 'Tbce' | 6.186 | 6.641 | 7.713 | 6.791 | 7.011 | 6.717 | 6.879 | 7.521 | 7.920 | 7.037 | 6.389 | 6.894 | 6.667 | 6.813 | 6.090 | 6.627 | 6.675 | 5.852 | 0.093 | 0.0467 | 0.1803 | 0 | 0 | 0 |
| 'Tbcel' | 7.636 | 7.952 | 7.719 | 7.805 | 7.742 | 7.316 | 7.056 | 7.936 | 7.624 | 7.107 | 7.739 | 7.941 | 6.628 | 7.975 | 8.093 | 7.570 | 8.009 | 8.446 | -0.024 | 0.5197 | 0.7423 | 0 | 0 | 0 |
| 'Tbck' | 2.182 | 2.010 | 2.090 | 1.965 | 2.111 | 1.798 | 2.143 | 2.303 | 2.044 | 2.068 | 2.337 | 2.183 | 1.935 | 2.279 | 2.078 | 2.240 | 2.472 | 2.255 | -0.098 | 0.0438 | 0.1733 | 0 | 0 | 0 |
| 'Tbk1' | 65.880 | 64.925 | 66.741 | 68.636 | 66.076 | 61.466 | 64.479 | 66.632 | 67.167 | 62.562 | 67.313 | 66.155 | 67.245 | 68.218 | 65.243 | 64.310 | 75.615 | 62.287 | -0.026 | 0.4076 | 0.6565 | 0 | 0 | 0 |
| 'Tbl1x' | 21.289 | 21.521 | 20.387 | 20.550 | 21.328 | 19.949 | 20.522 | 21.285 | 19.922 | 20.999 | 21.165 | 20.057 | 19.872 | 20.041 | 19.815 | 19.781 | 20.370 | 20.541 | 0.021 | 0.3733 | 0.6243 | 0 | 0 | 0 |
| 'Tbl1xr1' | 6.626 | 6.717 | 6.344 | 7.247 | 6.462 | 5.598 | 7.476 | 6.757 | 6.595 | 6.242 | 6.933 | 6.650 | 5.765 | 6.646 | 5.860 | 6.130 | 6.860 | 6.087 | 0.053 | 0.2478 | 0.4986 | 0 | 0 | 0 |
| 'Tbl2' | 4.101 | 4.185 | 4.695 | 4.161 | 3.898 | 3.999 | 4.019 | 4.400 | 3.984 | 3.767 | 5.379 | 5.169 | 4.121 | 4.463 | 5.107 | 4.186 | 5.389 | 5.025 | -0.191 | 0.0007 | 0.0075 | 1 | 0 | 0 |
| 'Tbl3' | 4.582 | 4.260 | 4.863 | 4.143 | 4.703 | 4.885 | 4.975 | 4.555 | 4.772 | 4.656 | 4.680 | 4.747 | 5.059 | 4.183 | 4.829 | 4.869 | 4.569 | 4.342 | -0.014 | 0.7199 | 0.8711 | 0 | 0 | 0 |
| 'Tbp' | 2.994 | 3.251 | 3.119 | 2.924 | 3.268 | 3.222 | 3.148 | 3.145 | 2.985 | 2.630 | 3.113 | 3.202 | 2.804 | 2.987 | 3.111 | 2.982 | 3.085 | 3.070 | 0.043 | 0.2166 | 0.4628 | 0 | 0 | 0 |
| 'Tbpl1' | 9.964 | 11.271 | 10.501 | 10.975 | 12.108 | 9.584 | 11.394 | 10.912 | 10.464 | 10.175 | 11.896 | 12.158 | 10.533 | 11.809 | 10.987 | 11.124 | 11.566 | 11.509 | -0.083 | 0.0464 | 0.1794 | 0 | 0 | 0 |
| 'Tbrg1' | 13.426 | 12.715 | 12.689 | 12.867 | 13.152 | 13.972 | 13.832 | 12.789 | 14.088 | 12.676 | 12.822 | 14.771 | 13.838 | 13.623 | 13.689 | 12.899 | 13.931 | 13.881 | -0.044 | 0.1685 | 0.4008 | 0 | 0 | 0 |
| 'Tbrg4' | 8.210 | 6.923 | 8.607 | 8.339 | 7.423 | 8.212 | 8.937 | 7.898 | 8.175 | 8.243 | 6.531 | 8.300 | 7.699 | 7.198 | 7.730 | 7.918 | 7.401 | 7.627 | 0.072 | 0.0306 | 0.1365 | 0 | 0 | 0 |
| 'Tbx6' | 4.040 | 3.954 | 3.699 | 4.837 | 2.861 | 4.342 | 4.433 | 3.544 | 4.184 | 3.571 | 4.796 | 7.209 | 3.372 | 4.605 | 6.117 | 3.424 | 7.452 | 6.960 | -0.383 | 0.0061 | 0.0436 | 1 | 1 | 0 |
| 'Tbxa2r' | 5.855 | 6.497 | 5.675 | 6.763 | 6.673 | 7.090 | 6.302 | 4.985 | 6.235 | 5.653 | 5.761 | 8.858 | 7.212 | 6.919 | 6.777 | 8.039 | 5.930 | 8.001 | -0.190 | 0.0252 | 0.1212 | 0 | 0 | 0 |
| 'Tcaim' | 2.923 | 3.140 | 2.914 | 2.724 | 3.616 | 3.077 | 3.567 | 3.259 | 3.018 | 3.496 | 3.204 | 3.059 | 3.006 | 3.076 | 2.857 | 2.989 | 3.209 | 2.467 | 0.033 | 0.5726 | 0.7792 | 0 | 0 | 0 |
| 'Tcea1' | 33.381 | 37.741 | 35.154 | 37.805 | 38.564 | 33.525 | 37.468 | 37.104 | 32.925 | 35.818 | 37.047 | 32.848 | 34.795 | 36.172 | 32.426 | 33.533 | 38.672 | 31.755 | 0.038 | 0.1701 | 0.4029 | 0 | 0 | 0 |
| 'Tcea2' | 4.126 | 4.524 | 4.466 | 3.237 | 4.174 | 4.139 | 3.868 | 4.138 | 4.590 | 3.659 | 3.574 | 3.661 | 3.586 | 3.580 | 3.433 | 3.343 | 3.640 | 3.424 | 0.215 | 0.0000 | 0.0007 | 1 | 0 | 0 |
| 'Tceal3' | 4.396 | 5.098 | 3.668 | 4.251 | 6.410 | 4.419 | 3.665 | 5.742 | 5.216 | 6.382 | 5.542 | 5.124 | 5.615 | 5.239 | 5.547 | 7.122 | 5.662 | 4.998 | -0.269 | 0.0087 | 0.0565 | 0 | 0 | 0 |
| 'Tceanc' | 1.515 | 1.847 | 2.018 | 1.541 | 1.981 | 1.847 | 1.931 | 2.068 | 1.944 | 1.922 | 1.386 | 1.515 | 1.689 | 1.648 | 1.490 | 1.647 | 1.317 | 1.239 | 0.264 | 0.0022 | 0.0194 | 1 | 0 | 0 |
| 'Tceanc2' | 5.157 | 5.151 | 4.975 | 5.521 | 5.232 | 5.099 | 5.398 | 5.176 | 4.790 | 4.947 | 5.251 | 5.138 | 5.012 | 4.749 | 4.989 | 4.847 | 5.047 | 4.912 | 0.036 | 0.2968 | 0.5527 | 0 | 0 | 0 |
| 'Tceb1' | 10.797 | 12.155 | 11.138 | 12.441 | 13.321 | 10.831 | 12.207 | 11.723 | 10.797 | 10.502 | 11.603 | 11.066 | 11.279 | 11.978 | 11.236 | 10.840 | 11.653 | 9.867 | 0.061 | 0.0948 | 0.2857 | 0 | 0 | 0 |
| 'Tceb2' | 168.659 | 169.470 | 156.869 | 166.929 | 178.977 | 179.755 | 166.326 | 167.909 | 165.971 | 162.975 | 170.064 | 165.177 | 165.642 | 165.154 | 171.495 | 170.740 | 159.102 | 166.719 | 0.012 | 0.6935 | 0.8558 | 0 | 0 | 0 |
| 'Tceb3' | 20.623 | 17.984 | 18.432 | 19.998 | 19.204 | 18.283 | 19.666 | 19.907 | 18.281 | 19.372 | 19.470 | 20.465 | 17.437 | 18.688 | 18.635 | 18.115 | 19.605 | 20.488 | -0.011 | 0.7823 | 0.9025 | 0 | 0 | 0 |
| 'Tcerg1' | 35.873 | 32.560 | 36.811 | 34.480 | 33.115 | 33.225 | 37.621 | 34.598 | 35.415 | 35.881 | 33.286 | 34.939 | 35.608 | 32.511 | 32.476 | 33.281 | 32.023 | 31.575 | 0.047 | 0.0940 | 0.2840 | 0 | 0 | 0 |
| 'Tcf12' | 7.658 | 7.735 | 7.226 | 7.285 | 7.597 | 7.384 | 7.208 | 7.584 | 7.202 | 7.838 | 7.387 | 7.004 | 7.079 | 7.080 | 7.041 | 6.993 | 7.068 | 6.966 | 0.043 | 0.0918 | 0.2797 | 0 | 0 | 0 |
| 'Tcf19' | 10.112 | 10.223 | 13.228 | 11.539 | 10.695 | 13.374 | 12.052 | 10.193 | 12.601 | 8.986 | 11.131 | 11.795 | 13.879 | 11.522 | 14.444 | 9.908 | 10.542 | 10.846 | -0.001 | 0.9891 | 0.9968 | 0 | 0 | 0 |
| 'Tcf20' | 22.230 | 23.619 | 21.906 | 20.905 | 21.556 | 22.886 | 21.175 | 23.272 | 22.289 | 23.228 | 23.084 | 21.542 | 22.155 | 22.563 | 22.089 | 23.288 | 20.934 | 21.509 | -0.013 | 0.6586 | 0.8350 | 0 | 0 | 0 |
| 'Tcf25' | 113.290 | 114.864 | 115.318 | 101.376 | 115.358 | 113.230 | 102.200 | 114.873 | 112.414 | 119.975 | 118.895 | 116.829 | 114.770 | 114.819 | 117.433 | 120.620 | 115.896 | 116.071 | -0.084 | 0.0015 | 0.0146 | 1 | 0 | 0 |
| 'Tcf3' | 15.978 | 15.672 | 18.180 | 15.149 | 15.269 | 19.144 | 15.782 | 16.135 | 18.180 | 16.459 | 14.305 | 17.562 | 16.714 | 15.067 | 17.675 | 16.897 | 14.630 | 17.697 | 0.015 | 0.5847 | 0.7863 | 0 | 0 | 0 |
| 'Tcf4' | 0.076 | 0.090 | 0.060 | 0.076 | 0.082 | 0.052 | 0.043 | 0.081 | 0.056 | 0.074 | 0.119 | 0.075 | 0.091 | 0.130 | 0.077 | 0.116 | 0.095 | 0.091 | -0.500 | 0.0000 | 0.0004 | 1 | 1 | 0 |
| 'Tcf7' | 693.192 | 666.948 | 728.684 | 694.959 | 677.231 | 744.457 | 686.482 | 698.714 | 727.333 | 747.128 | 710.316 | 662.852 | 686.135 | 713.388 | 728.594 | 696.630 | 669.725 | 743.675 | -0.021 | 0.4766 | 0.7107 | 0 | 0 | 0 |
| 'Tchp' | 4.009 | 3.771 | 3.911 | 3.893 | 3.562 | 4.367 | 3.703 | 3.948 | 4.233 | 3.703 | 3.529 | 4.185 | 4.001 | 3.422 | 3.613 | 3.606 | 3.600 | 3.969 | 0.062 | 0.1131 | 0.3192 | 0 | 0 | 0 |
| 'Tcirg1' | 11.781 | 9.982 | 13.436 | 10.948 | 10.206 | 12.636 | 11.645 | 10.368 | 12.858 | 11.024 | 9.643 | 13.104 | 12.267 | 10.340 | 11.748 | 9.985 | 11.206 | 12.402 | 0.017 | 0.6650 | 0.8382 | 0 | 0 | 0 |
| 'Tcn2' | 7.041 | 7.099 | 6.879 | 7.264 | 8.037 | 7.694 | 6.454 | 7.645 | 7.012 | 7.544 | 7.524 | 7.461 | 6.712 | 6.736 | 6.908 | 5.882 | 6.689 | 6.917 | 0.047 | 0.3942 | 0.6434 | 0 | 0 | 0 |
| 'Tcp1' | 26.309 | 23.618 | 23.257 | 25.321 | 23.409 | 23.375 | 24.776 | 24.403 | 22.479 | 21.831 | 24.423 | 26.373 | 24.167 | 23.488 | 24.159 | 22.807 | 25.716 | 25.906 | -0.024 | 0.5562 | 0.7679 | 0 | 0 | 0 |
| 'Tcp11l2' | 15.392 | 16.991 | 15.174 | 14.734 | 16.079 | 15.797 | 13.916 | 16.858 | 14.887 | 15.824 | 16.567 | 16.516 | 13.369 | 16.230 | 15.236 | 15.682 | 16.291 | 15.715 | -0.027 | 0.4466 | 0.6880 | 0 | 0 | 0 |
| 'Tcta' | 3.352 | 4.345 | 3.344 | 2.495 | 4.321 | 4.254 | 3.224 | 4.259 | 3.669 | 3.628 | 3.455 | 3.199 | 3.502 | 3.514 | 3.599 | 4.406 | 3.354 | 3.131 | 0.053 | 0.5475 | 0.7613 | 0 | 0 | 0 |
| 'Tctex1d2' | 1.223 | 1.411 | 1.334 | 1.375 | 1.539 | 1.514 | 1.087 | 1.828 | 1.630 | 1.374 | 1.525 | 1.353 | 1.567 | 1.144 | 1.172 | 1.749 | 1.133 | 1.012 | 0.105 | 0.3845 | 0.6347 | 0 | 0 | 0 |
| 'Tctn1' | 1.305 | 1.236 | 1.336 | 1.274 | 1.338 | 1.267 | 1.497 | 1.130 | 1.409 | 1.156 | 1.211 | 1.220 | 1.261 | 1.179 | 1.114 | 1.343 | 1.201 | 1.091 | 0.120 | 0.0335 | 0.1455 | 0 | 0 | 0 |
| 'Tctn3' | 1.820 | 2.037 | 2.537 | 2.070 | 2.298 | 2.331 | 2.073 | 2.298 | 2.148 | 2.303 | 2.147 | 2.435 | 2.338 | 1.797 | 2.051 | 1.807 | 1.599 | 2.374 | 0.042 | 0.6001 | 0.7967 | 0 | 0 | 0 |
| 'Tdp1' | 4.050 | 3.901 | 4.213 | 3.942 | 3.954 | 4.397 | 4.248 | 4.193 | 4.885 | 4.030 | 3.802 | 4.169 | 4.022 | 4.074 | 4.663 | 3.581 | 3.772 | 3.718 | 0.067 | 0.1458 | 0.3707 | 0 | 0 | 0 |
| 'Tdp2' | 20.156 | 21.026 | 19.885 | 19.900 | 20.403 | 17.212 | 20.169 | 21.738 | 19.546 | 18.911 | 20.250 | 19.922 | 20.584 | 21.634 | 18.175 | 18.877 | 19.989 | 18.786 | 0.010 | 0.7905 | 0.9064 | 0 | 0 | 0 |
| 'Tdrd3' | 7.332 | 7.815 | 7.726 | 7.353 | 7.657 | 7.065 | 8.504 | 7.430 | 7.342 | 7.852 | 8.475 | 7.348 | 7.760 | 7.754 | 6.584 | 6.760 | 8.781 | 6.788 | -0.005 | 0.9103 | 0.9651 | 0 | 0 | 0 |
| 'Tdrd7' | 1.567 | 1.519 | 1.304 | 1.479 | 1.690 | 1.462 | 1.596 | 1.605 | 1.744 | 1.599 | 1.769 | 1.746 | 1.445 | 1.490 | 1.658 | 1.819 | 1.884 | 1.574 | -0.115 | 0.1316 | 0.3486 | 0 | 0 | 0 |
| 'Tdrp' | 0.662 | 0.688 | 0.681 | 0.585 | 0.570 | 0.763 | 0.901 | 0.509 | 0.748 | 0.535 | 0.521 | 0.810 | 0.614 | 0.353 | 0.884 | 0.824 | 0.761 | 0.891 | -0.031 | 0.8102 | 0.9142 | 0 | 0 | 0 |
| 'Tec' | 1.113 | 1.059 | 1.180 | 1.273 | 1.161 | 1.221 | 1.274 | 1.185 | 1.150 | 0.954 | 1.265 | 1.005 | 0.932 | 1.082 | 0.968 | 0.983 | 1.245 | 1.014 | 0.168 | 0.0208 | 0.1056 | 0 | 0 | 0 |
| 'Tecpr1' | 12.494 | 12.041 | 12.976 | 12.481 | 11.527 | 13.791 | 12.348 | 11.873 | 13.322 | 12.891 | 11.963 | 13.997 | 11.856 | 12.147 | 13.570 | 12.698 | 11.906 | 13.267 | -0.029 | 0.1929 | 0.4302 | 0 | 0 | 0 |
| 'Tecpr2' | 2.038 | 1.672 | 1.964 | 1.951 | 1.649 | 1.933 | 2.032 | 1.947 | 2.191 | 1.934 | 1.921 | 2.018 | 2.019 | 2.019 | 2.079 | 1.862 | 2.063 | 2.117 | -0.060 | 0.2432 | 0.4935 | 0 | 0 | 0 |
| 'Tecr' | 103.780 | 107.356 | 114.965 | 106.649 | 111.747 | 117.237 | 110.527 | 107.499 | 116.757 | 108.695 | 91.604 | 107.237 | 111.919 | 97.716 | 107.483 | 105.404 | 101.237 | 100.834 | 0.086 | 0.0025 | 0.0214 | 1 | 0 | 0 |
| 'Tef' | 5.906 | 7.358 | 6.139 | 5.446 | 6.087 | 6.238 | 5.890 | 6.566 | 6.471 | 6.064 | 6.011 | 6.097 | 5.858 | 5.616 | 5.947 | 6.326 | 5.408 | 5.573 | 0.072 | 0.1030 | 0.3009 | 0 | 0 | 0 |
| 'Tefm' | 2.079 | 2.369 | 2.577 | 1.710 | 3.000 | 2.524 | 1.933 | 2.756 | 2.397 | 2.548 | 1.934 | 1.951 | 1.992 | 2.146 | 1.830 | 2.568 | 1.966 | 2.254 | 0.148 | 0.1454 | 0.3699 | 0 | 0 | 0 |
| 'Telo2' | 2.543 | 2.135 | 2.832 | 2.464 | 2.315 | 2.662 | 2.557 | 2.345 | 2.656 | 2.279 | 2.337 | 2.635 | 2.370 | 1.897 | 2.603 | 2.380 | 2.308 | 2.474 | 0.072 | 0.0694 | 0.2318 | 0 | 0 | 0 |
| 'Ten1' | 5.687 | 7.096 | 5.906 | 5.970 | 6.306 | 6.268 | 5.263 | 6.431 | 6.320 | 5.748 | 5.074 | 6.154 | 6.571 | 5.961 | 6.251 | 6.583 | 5.733 | 6.172 | 0.012 | 0.8345 | 0.9283 | 0 | 0 | 0 |
| 'Tep1' | 8.789 | 6.887 | 9.908 | 8.974 | 7.905 | 9.027 | 8.096 | 8.350 | 9.482 | 9.432 | 8.347 | 9.375 | 8.739 | 9.006 | 8.707 | 8.648 | 8.593 | 9.287 | -0.060 | 0.1643 | 0.3965 | 0 | 0 | 0 |
| 'Tepp' | 1.752 | 1.333 | 1.732 | 1.251 | 1.490 | 1.196 | 1.620 | 1.283 | 1.494 | 1.373 | 1.397 | 1.572 | 1.133 | 1.526 | 1.801 | 1.475 | 1.764 | 1.934 | -0.091 | 0.4472 | 0.6880 | 0 | 0 | 0 |
| 'Terf1' | 4.926 | 5.872 | 5.504 | 5.858 | 5.348 | 4.955 | 5.970 | 5.471 | 5.269 | 4.227 | 5.291 | 6.195 | 4.920 | 5.129 | 5.865 | 4.444 | 6.252 | 5.910 | 0.019 | 0.7791 | 0.9004 | 0 | 0 | 0 |
| 'Terf2' | 6.310 | 6.303 | 6.081 | 5.939 | 5.835 | 6.409 | 6.344 | 6.424 | 6.635 | 5.954 | 6.218 | 6.879 | 6.106 | 6.224 | 6.459 | 6.426 | 6.055 | 6.267 | -0.020 | 0.5363 | 0.7553 | 0 | 0 | 0 |
| 'Terf2ip' | 13.653 | 13.994 | 12.893 | 13.883 | 13.779 | 13.184 | 14.172 | 13.509 | 13.551 | 13.127 | 13.639 | 15.330 | 12.288 | 13.128 | 15.230 | 13.621 | 13.148 | 15.267 | -0.038 | 0.3859 | 0.6354 | 0 | 0 | 0 |
| 'Tert' | 5.467 | 4.694 | 5.856 | 5.005 | 5.026 | 5.801 | 5.100 | 4.878 | 5.610 | 5.170 | 5.699 | 6.098 | 5.559 | 4.826 | 5.516 | 4.897 | 5.778 | 4.956 | -0.042 | 0.4580 | 0.6952 | 0 | 0 | 0 |
| 'Tes' | 10.521 | 11.718 | 10.923 | 10.301 | 11.588 | 11.438 | 11.056 | 11.033 | 10.847 | 11.228 | 11.159 | 11.417 | 11.512 | 10.984 | 11.584 | 11.765 | 11.905 | 11.118 | -0.058 | 0.0324 | 0.1423 | 0 | 0 | 0 |
| 'Tesc' | 14.362 | 13.462 | 17.023 | 16.368 | 14.562 | 16.933 | 14.739 | 13.654 | 16.525 | 14.995 | 12.148 | 14.880 | 15.200 | 11.886 | 14.288 | 14.715 | 11.959 | 14.301 | 0.138 | 0.0001 | 0.0020 | 1 | 0 | 0 |
| 'Tesk1' | 2.945 | 3.198 | 3.733 | 3.070 | 2.934 | 3.752 | 2.812 | 3.045 | 3.428 | 3.676 | 2.847 | 3.444 | 3.518 | 2.936 | 3.314 | 3.652 | 2.936 | 3.729 | -0.061 | 0.2644 | 0.5184 | 0 | 0 | 0 |
| 'Tesk2' | 1.707 | 2.209 | 1.766 | 1.892 | 1.927 | 2.135 | 1.827 | 2.130 | 1.976 | 1.915 | 1.990 | 1.739 | 1.858 | 1.963 | 1.642 | 1.849 | 1.762 | 1.403 | 0.118 | 0.0736 | 0.2407 | 0 | 0 | 0 |
| 'Tespa1' | 93.486 | 93.044 | 96.528 | 95.015 | 88.214 | 88.989 | 102.551 | 93.443 | 99.146 | 98.952 | 88.351 | 93.920 | 88.557 | 79.603 | 89.014 | 84.253 | 86.526 | 93.413 | 0.072 | 0.0418 | 0.1683 | 0 | 0 | 0 |
| 'Tet1' | 0.114 | 0.144 | 0.187 | 0.153 | 0.169 | 0.179 | 0.154 | 0.172 | 0.159 | 0.196 | 0.155 | 0.174 | 0.143 | 0.102 | 0.145 | 0.170 | 0.149 | 0.128 | 0.062 | 0.5908 | 0.7914 | 0 | 0 | 0 |
| 'Tet2' | 2.492 | 1.941 | 2.673 | 2.392 | 2.197 | 2.135 | 2.163 | 2.291 | 2.533 | 2.692 | 2.357 | 2.304 | 2.081 | 1.957 | 2.055 | 2.410 | 2.140 | 2.166 | 0.038 | 0.5225 | 0.7449 | 0 | 0 | 0 |
| 'Tet3' | 10.211 | 9.585 | 9.594 | 11.494 | 8.525 | 8.770 | 10.632 | 8.748 | 9.847 | 9.282 | 9.970 | 10.876 | 9.331 | 8.974 | 10.106 | 8.584 | 10.372 | 11.404 | -0.037 | 0.5397 | 0.7573 | 0 | 0 | 0 |
| 'Tex10' | 8.588 | 7.799 | 8.758 | 8.833 | 8.048 | 7.759 | 8.877 | 8.053 | 7.642 | 7.972 | 8.225 | 9.001 | 7.399 | 8.532 | 8.318 | 7.712 | 8.753 | 8.153 | -0.006 | 0.8924 | 0.9561 | 0 | 0 | 0 |
| 'Tex2' | 6.488 | 6.035 | 6.718 | 6.042 | 6.445 | 6.445 | 6.745 | 6.565 | 6.369 | 5.395 | 6.080 | 6.271 | 5.533 | 5.876 | 5.624 | 5.183 | 5.224 | 6.234 | 0.158 | 0.0002 | 0.0027 | 1 | 0 | 0 |
| 'Tex261' | 12.076 | 13.919 | 12.364 | 11.576 | 13.248 | 13.373 | 12.041 | 13.805 | 13.277 | 12.181 | 12.968 | 12.987 | 12.705 | 12.807 | 13.355 | 13.099 | 12.400 | 12.981 | -0.009 | 0.7647 | 0.8953 | 0 | 0 | 0 |
| 'Tex264' | 9.034 | 9.191 | 9.818 | 8.317 | 9.320 | 9.919 | 8.382 | 9.507 | 10.035 | 9.679 | 8.001 | 9.185 | 8.766 | 8.419 | 9.217 | 9.570 | 7.408 | 8.599 | 0.076 | 0.0908 | 0.2777 | 0 | 0 | 0 |
| 'Tex30' | 5.568 | 5.626 | 4.887 | 6.470 | 5.231 | 4.667 | 6.262 | 5.288 | 4.775 | 4.539 | 5.266 | 4.889 | 5.616 | 5.777 | 4.991 | 4.767 | 5.755 | 4.575 | 0.061 | 0.2847 | 0.5410 | 0 | 0 | 0 |
| 'Tex35' | 0.000 | 0.000 | 0.000 | 0.000 | 0.000 | 0.000 | 0.027 | 0.000 | 0.000 | 2.975 | 1.932 | 2.725 | 1.607 | 1.675 | 2.759 | 2.552 | 2.077 | 2.656 | -8.806 | 0.0000 | 0.0000 | 1 | 1 | 1 |
| 'Tex9' | 0.979 | 1.157 | 1.144 | 0.932 | 1.132 | 1.107 | 1.120 | 1.050 | 1.146 | 0.962 | 0.885 | 0.952 | 0.982 | 1.021 | 0.877 | 0.846 | 0.910 | 0.794 | 0.235 | 0.0001 | 0.0013 | 1 | 0 | 0 |
| 'Tfam' | 12.902 | 15.443 | 15.316 | 13.928 | 14.916 | 14.980 | 14.282 | 16.345 | 13.898 | 13.436 | 13.752 | 14.148 | 14.113 | 14.028 | 13.748 | 14.152 | 14.477 | 13.565 | 0.062 | 0.0553 | 0.2017 | 0 | 0 | 0 |
| 'Tfap4' | 6.790 | 6.423 | 7.490 | 6.270 | 6.897 | 5.950 | 7.112 | 7.416 | 7.085 | 6.571 | 6.970 | 6.881 | 7.013 | 5.456 | 7.000 | 6.527 | 7.198 | 6.515 | 0.021 | 0.7378 | 0.8809 | 0 | 0 | 0 |
| 'Tfb1m' | 7.008 | 7.940 | 8.265 | 7.560 | 7.694 | 7.048 | 7.756 | 7.656 | 7.970 | 8.037 | 8.266 | 7.546 | 10.057 | 7.770 | 7.375 | 8.390 | 8.054 | 6.708 | -0.072 | 0.3118 | 0.5704 | 0 | 0 | 0 |
| 'Tfb2m' | 6.317 | 6.933 | 6.771 | 7.014 | 7.800 | 6.650 | 6.949 | 7.843 | 6.619 | 6.842 | 6.346 | 7.165 | 6.690 | 7.296 | 6.499 | 7.976 | 7.052 | 6.295 | 0.004 | 0.9319 | 0.9746 | 0 | 0 | 0 |
| 'Tfcp2' | 5.151 | 4.985 | 6.060 | 5.834 | 6.816 | 5.863 | 5.413 | 6.369 | 6.045 | 5.276 | 6.023 | 5.720 | 5.252 | 5.871 | 5.564 | 6.002 | 6.234 | 5.414 | 0.028 | 0.6125 | 0.8056 | 0 | 0 | 0 |
| 'Tfdp1' | 57.376 | 55.447 | 61.459 | 61.640 | 58.603 | 62.123 | 60.981 | 58.467 | 58.976 | 54.550 | 57.630 | 60.835 | 64.839 | 58.175 | 64.588 | 55.093 | 59.302 | 62.743 | -0.018 | 0.5867 | 0.7880 | 0 | 0 | 0 |
| 'Tfdp2' | 1.528 | 1.695 | 1.696 | 1.725 | 1.769 | 1.535 | 1.544 | 1.772 | 1.592 | 1.749 | 1.429 | 1.600 | 1.538 | 1.666 | 1.672 | 1.402 | 1.580 | 1.404 | 0.067 | 0.2080 | 0.4525 | 0 | 0 | 0 |
| 'Tfe3' | 3.983 | 4.563 | 4.312 | 4.139 | 4.467 | 4.710 | 4.346 | 4.272 | 4.767 | 4.547 | 4.116 | 5.135 | 4.954 | 4.518 | 4.889 | 4.927 | 4.580 | 5.225 | -0.129 | 0.0003 | 0.0043 | 1 | 0 | 0 |
| 'Tfeb' | 4.804 | 4.525 | 5.215 | 4.525 | 4.263 | 5.836 | 5.028 | 4.183 | 5.605 | 4.391 | 3.785 | 5.156 | 4.650 | 4.086 | 4.789 | 5.003 | 4.192 | 5.330 | 0.077 | 0.0525 | 0.1947 | 0 | 0 | 0 |
| 'Tfg' | 13.322 | 14.737 | 13.484 | 13.442 | 13.672 | 13.470 | 14.856 | 12.947 | 13.054 | 13.185 | 13.517 | 14.363 | 13.598 | 14.448 | 13.811 | 13.132 | 14.333 | 13.475 | -0.026 | 0.4670 | 0.7026 | 0 | 0 | 0 |
| 'Tfip11' | 21.998 | 17.089 | 20.079 | 19.340 | 18.281 | 18.918 | 19.558 | 19.101 | 19.716 | 20.457 | 17.306 | 18.721 | 18.715 | 18.156 | 17.175 | 19.964 | 16.554 | 18.561 | 0.065 | 0.0695 | 0.2318 | 0 | 0 | 0 |
| 'Tfpi' | 0.344 | 0.446 | 0.330 | 0.359 | 0.513 | 0.267 | 0.405 | 0.491 | 0.403 | 0.421 | 0.275 | 0.414 | 0.381 | 0.293 | 0.369 | 0.417 | 0.385 | 0.301 | 0.090 | 0.4589 | 0.6958 | 0 | 0 | 0 |
| 'Tfpt' | 5.629 | 5.576 | 6.835 | 5.757 | 6.040 | 6.072 | 5.574 | 5.658 | 6.756 | 5.718 | 5.182 | 6.059 | 6.312 | 5.924 | 5.726 | 5.646 | 5.352 | 5.819 | 0.050 | 0.2995 | 0.5557 | 0 | 0 | 0 |
| 'Tfrc' | 19.030 | 18.643 | 19.694 | 20.161 | 18.486 | 16.349 | 19.954 | 19.703 | 16.720 | 15.012 | 19.951 | 16.235 | 16.413 | 19.690 | 17.700 | 14.926 | 18.128 | 17.008 | 0.115 | 0.0328 | 0.1433 | 0 | 0 | 0 |
| 'Tg' | 0.355 | 0.453 | 0.351 | 0.501 | 0.355 | 0.357 | 0.453 | 0.349 | 0.393 | 0.296 | 0.325 | 0.441 | 0.447 | 0.379 | 0.296 | 0.364 | 0.369 | 0.370 | 0.081 | 0.4494 | 0.6895 | 0 | 0 | 0 |
| 'Tgds' | 12.647 | 12.574 | 12.792 | 13.744 | 13.841 | 12.411 | 12.653 | 12.973 | 12.701 | 13.380 | 13.452 | 14.147 | 13.396 | 12.102 | 12.808 | 11.128 | 15.047 | 12.201 | -0.032 | 0.5179 | 0.7408 | 0 | 0 | 0 |
| 'Tgfb1' | 28.745 | 23.953 | 31.303 | 26.170 | 25.877 | 31.631 | 25.986 | 25.899 | 30.949 | 28.266 | 24.037 | 33.278 | 26.107 | 25.489 | 32.917 | 26.711 | 25.209 | 33.364 | -0.034 | 0.2459 | 0.4963 | 0 | 0 | 0 |
| 'Tgfb3' | 10.316 | 11.615 | 11.996 | 11.679 | 13.080 | 11.098 | 11.277 | 12.969 | 12.621 | 10.050 | 10.051 | 8.887 | 13.659 | 9.707 | 9.489 | 8.738 | 8.094 | 8.614 | 0.281 | 0.0002 | 0.0032 | 1 | 0 | 0 |
| 'Tgfbr1' | 4.736 | 5.825 | 4.789 | 4.876 | 5.437 | 5.009 | 5.410 | 5.545 | 5.033 | 4.279 | 5.170 | 5.801 | 4.733 | 5.041 | 5.239 | 5.319 | 5.462 | 5.441 | -0.011 | 0.8152 | 0.9170 | 0 | 0 | 0 |
| 'Tgfbr2' | 73.887 | 79.504 | 69.228 | 70.259 | 73.254 | 73.385 | 67.054 | 77.249 | 71.408 | 82.251 | 78.221 | 78.490 | 71.051 | 80.935 | 74.335 | 83.206 | 75.903 | 82.474 | -0.121 | 0.0010 | 0.0101 | 1 | 0 | 0 |
| 'Tgfbr3' | 3.628 | 3.197 | 3.572 | 3.354 | 2.978 | 2.942 | 3.751 | 3.618 | 3.736 | 2.861 | 3.246 | 3.790 | 2.640 | 2.760 | 3.478 | 3.477 | 2.941 | 3.692 | 0.084 | 0.2189 | 0.4657 | 0 | 0 | 0 |
| 'Tgfbr3l' | 4.660 | 4.088 | 6.550 | 4.743 | 4.851 | 5.499 | 4.298 | 4.954 | 6.021 | 5.652 | 5.512 | 4.697 | 4.689 | 3.654 | 4.798 | 4.662 | 5.101 | 4.031 | 0.087 | 0.3425 | 0.5988 | 0 | 0 | 0 |
| 'Tgfbrap1' | 4.971 | 5.488 | 5.234 | 4.897 | 5.136 | 5.254 | 4.865 | 5.319 | 5.394 | 5.418 | 5.251 | 5.261 | 5.307 | 5.049 | 4.977 | 5.484 | 5.368 | 5.174 | -0.032 | 0.2760 | 0.5317 | 0 | 0 | 0 |
| 'Tgif1' | 2.059 | 1.323 | 0.984 | 1.295 | 1.343 | 1.134 | 1.256 | 1.780 | 1.161 | 2.299 | 1.703 | 1.157 | 2.242 | 1.898 | 1.025 | 2.343 | 1.620 | 1.008 | -0.268 | 0.0065 | 0.0453 | 1 | 0 | 0 |
| 'Tgm2' | 4.163 | 3.480 | 4.279 | 3.938 | 4.239 | 4.418 | 4.139 | 4.144 | 3.850 | 5.359 | 5.066 | 5.250 | 6.412 | 5.176 | 5.116 | 5.811 | 4.284 | 5.829 | -0.403 | 0.0000 | 0.0000 | 1 | 1 | 0 |
| 'Tgoln1' | 51.228 | 50.577 | 48.629 | 51.139 | 50.796 | 46.050 | 48.283 | 51.985 | 46.684 | 52.883 | 56.880 | 52.715 | 47.500 | 54.222 | 49.955 | 53.065 | 51.092 | 52.691 | -0.092 | 0.0044 | 0.0341 | 1 | 0 | 0 |
| 'Tgs1' | 6.449 | 7.066 | 6.912 | 7.096 | 6.811 | 6.483 | 7.162 | 6.650 | 6.604 | 6.603 | 7.370 | 7.201 | 5.980 | 7.314 | 6.762 | 6.480 | 7.339 | 6.513 | -0.019 | 0.6452 | 0.8265 | 0 | 0 | 0 |
| 'Tgtp1' | 19.349 | 21.572 | 20.066 | 20.016 | 21.082 | 22.392 | 23.510 | 17.703 | 23.582 | 23.967 | 20.421 | 38.393 | 24.562 | 23.691 | 34.923 | 26.960 | 25.799 | 37.965 | -0.425 | 0.0000 | 0.0000 | 1 | 1 | 0 |
| 'Tha1' | 5.793 | 6.202 | 9.377 | 5.891 | 6.241 | 6.601 | 6.392 | 7.153 | 6.674 | 5.176 | 7.052 | 6.484 | 5.853 | 5.546 | 5.693 | 4.215 | 5.274 | 5.395 | 0.239 | 0.0025 | 0.0217 | 1 | 0 | 0 |
| 'Thada' | 13.239 | 11.494 | 11.011 | 13.059 | 11.129 | 11.880 | 12.461 | 11.728 | 11.321 | 13.915 | 15.479 | 14.087 | 11.885 | 13.245 | 12.685 | 13.847 | 12.969 | 12.955 | -0.186 | 0.0001 | 0.0019 | 1 | 0 | 0 |
| 'Thap1' | 6.259 | 5.852 | 5.259 | 5.748 | 6.548 | 5.763 | 5.683 | 6.222 | 5.457 | 5.556 | 5.527 | 6.033 | 6.417 | 5.804 | 5.444 | 5.715 | 5.749 | 5.112 | 0.020 | 0.7412 | 0.8828 | 0 | 0 | 0 |
| 'Thap11' | 20.992 | 17.067 | 19.287 | 17.483 | 18.284 | 19.731 | 18.702 | 18.963 | 18.830 | 15.670 | 17.339 | 20.930 | 18.338 | 18.372 | 21.168 | 18.945 | 14.785 | 20.976 | 0.017 | 0.7550 | 0.8907 | 0 | 0 | 0 |
| 'Thap2' | 7.725 | 7.136 | 6.254 | 8.314 | 6.884 | 6.795 | 7.308 | 6.518 | 6.229 | 5.810 | 7.670 | 8.803 | 6.453 | 8.604 | 8.344 | 6.249 | 8.528 | 8.477 | -0.133 | 0.1136 | 0.3197 | 0 | 0 | 0 |
| 'Thap3' | 18.467 | 20.965 | 20.646 | 17.879 | 20.913 | 22.161 | 18.853 | 20.170 | 19.920 | 18.086 | 17.666 | 21.608 | 18.078 | 17.690 | 22.506 | 20.472 | 19.025 | 21.391 | 0.016 | 0.6962 | 0.8571 | 0 | 0 | 0 |
| 'Thap4' | 3.156 | 2.707 | 2.774 | 2.851 | 2.356 | 2.774 | 2.573 | 2.566 | 2.880 | 2.659 | 2.389 | 3.144 | 2.547 | 2.529 | 3.248 | 2.707 | 2.533 | 3.504 | -0.043 | 0.4657 | 0.7014 | 0 | 0 | 0 |
| 'Thap7' | 5.266 | 4.964 | 5.595 | 4.081 | 5.460 | 5.885 | 5.019 | 4.755 | 5.902 | 4.950 | 5.113 | 5.934 | 5.118 | 4.458 | 5.302 | 5.175 | 4.812 | 5.667 | 0.001 | 0.9909 | 0.9976 | 0 | 0 | 0 |
| 'Them4' | 0.640 | 0.725 | 0.858 | 0.695 | 0.893 | 0.856 | 0.800 | 0.953 | 1.034 | 0.987 | 0.933 | 0.684 | 1.008 | 0.841 | 0.692 | 0.723 | 0.798 | 0.693 | 0.026 | 0.8304 | 0.9262 | 0 | 0 | 0 |
| 'Them6' | 14.530 | 16.271 | 14.751 | 13.490 | 14.517 | 17.081 | 13.702 | 15.292 | 13.520 | 12.193 | 11.518 | 14.439 | 16.580 | 13.099 | 14.232 | 13.656 | 11.135 | 13.734 | 0.131 | 0.0466 | 0.1801 | 0 | 0 | 0 |
| 'Themis' | 18.584 | 19.353 | 18.382 | 21.221 | 18.501 | 17.469 | 21.961 | 17.975 | 18.602 | 17.254 | 18.851 | 18.169 | 17.059 | 18.257 | 16.859 | 16.435 | 18.197 | 16.883 | 0.111 | 0.0082 | 0.0539 | 0 | 0 | 0 |
| 'Thg1l' | 3.361 | 4.035 | 4.038 | 3.686 | 3.610 | 3.739 | 3.460 | 3.924 | 3.485 | 3.646 | 3.678 | 3.355 | 3.849 | 3.219 | 3.522 | 3.373 | 3.225 | 3.497 | 0.080 | 0.1660 | 0.3983 | 0 | 0 | 0 |
| 'Thnsl1' | 0.427 | 0.517 | 0.585 | 0.589 | 0.556 | 0.521 | 0.455 | 0.606 | 0.462 | 0.478 | 0.569 | 0.554 | 0.479 | 0.674 | 0.401 | 0.505 | 0.486 | 0.574 | -0.010 | 0.9253 | 0.9715 | 0 | 0 | 0 |
| 'Thoc1' | 12.756 | 15.528 | 14.977 | 14.204 | 15.762 | 12.836 | 14.710 | 15.692 | 13.735 | 14.552 | 12.907 | 14.739 | 14.397 | 13.573 | 13.649 | 14.050 | 17.140 | 13.173 | 0.008 | 0.8744 | 0.9463 | 0 | 0 | 0 |
| 'Thoc2' | 7.040 | 7.886 | 7.969 | 7.961 | 7.741 | 7.297 | 8.134 | 7.605 | 7.664 | 7.471 | 7.682 | 7.434 | 7.273 | 7.632 | 6.683 | 7.694 | 7.499 | 6.758 | 0.058 | 0.0656 | 0.2235 | 0 | 0 | 0 |
| 'Thoc3' | 9.001 | 9.861 | 9.499 | 9.389 | 10.185 | 8.832 | 9.559 | 9.715 | 9.416 | 9.166 | 8.211 | 8.921 | 9.545 | 8.958 | 8.938 | 9.571 | 8.470 | 8.682 | 0.074 | 0.0371 | 0.1561 | 0 | 0 | 0 |
| 'Thoc5' | 17.287 | 16.601 | 18.543 | 16.491 | 17.243 | 18.623 | 17.189 | 17.166 | 16.679 | 17.433 | 16.203 | 15.329 | 17.606 | 16.041 | 16.036 | 16.176 | 16.348 | 15.655 | 0.080 | 0.0250 | 0.1206 | 0 | 0 | 0 |
| 'Thoc6' | 5.109 | 5.127 | 5.746 | 5.023 | 5.629 | 5.853 | 5.599 | 5.436 | 6.067 | 5.297 | 4.599 | 5.209 | 5.250 | 4.789 | 5.249 | 4.889 | 5.106 | 4.623 | 0.131 | 0.0038 | 0.0303 | 1 | 0 | 0 |
| 'Thoc7' | 44.274 | 48.445 | 47.275 | 46.844 | 45.226 | 41.614 | 49.475 | 48.503 | 43.417 | 39.434 | 41.071 | 44.897 | 44.030 | 43.980 | 44.338 | 41.771 | 47.543 | 40.880 | 0.083 | 0.0356 | 0.1512 | 0 | 0 | 0 |
| 'Thop1' | 3.620 | 3.454 | 4.086 | 3.263 | 2.877 | 4.152 | 3.721 | 3.081 | 3.911 | 3.666 | 2.908 | 4.052 | 3.686 | 2.981 | 4.478 | 3.777 | 3.237 | 3.789 | -0.026 | 0.5925 | 0.7923 | 0 | 0 | 0 |
| 'Thra' | 1.263 | 1.219 | 1.177 | 1.049 | 1.069 | 1.403 | 1.166 | 1.373 | 1.529 | 1.262 | 0.896 | 1.342 | 1.119 | 1.012 | 1.058 | 1.448 | 0.916 | 1.381 | 0.104 | 0.2649 | 0.5184 | 0 | 0 | 0 |
| 'Thrap3' | 37.572 | 36.677 | 40.363 | 38.725 | 38.674 | 39.119 | 39.843 | 36.728 | 40.503 | 40.234 | 38.107 | 35.042 | 39.581 | 37.311 | 36.306 | 37.740 | 35.951 | 35.891 | 0.041 | 0.1563 | 0.3855 | 0 | 0 | 0 |
| 'Thtpa' | 3.800 | 3.994 | 3.863 | 3.694 | 4.012 | 4.064 | 3.890 | 4.051 | 4.134 | 3.660 | 3.511 | 3.658 | 3.646 | 3.298 | 3.163 | 3.713 | 3.298 | 3.476 | 0.165 | 0.0002 | 0.0029 | 1 | 0 | 0 |
| 'Thumpd1' | 24.755 | 26.536 | 23.891 | 21.805 | 26.933 | 24.641 | 22.008 | 27.913 | 24.830 | 25.895 | 25.097 | 23.735 | 23.603 | 24.570 | 23.976 | 25.747 | 23.725 | 22.551 | 0.019 | 0.6342 | 0.8188 | 0 | 0 | 0 |
| 'Thumpd2' | 8.123 | 8.708 | 9.798 | 8.050 | 7.808 | 7.759 | 7.391 | 9.353 | 8.832 | 8.128 | 7.720 | 7.517 | 7.380 | 7.523 | 6.964 | 7.925 | 7.586 | 6.896 | 0.163 | 0.0090 | 0.0579 | 0 | 0 | 0 |
| 'Thumpd3' | 8.845 | 10.216 | 9.440 | 9.460 | 9.687 | 8.787 | 9.142 | 9.767 | 9.056 | 8.835 | 9.323 | 9.305 | 8.915 | 9.157 | 8.231 | 9.621 | 9.151 | 8.180 | 0.052 | 0.0871 | 0.2690 | 0 | 0 | 0 |
| 'Thy1' | 277.826 | 230.611 | 305.166 | 283.512 | 237.574 | 321.419 | 278.268 | 238.710 | 288.317 | 258.192 | 218.429 | 275.333 | 230.534 | 219.950 | 289.075 | 263.869 | 211.386 | 288.312 | 0.117 | 0.0001 | 0.0018 | 1 | 0 | 0 |
| 'Thyn1' | 11.670 | 11.699 | 9.455 | 13.046 | 11.669 | 10.091 | 10.547 | 11.750 | 9.366 | 8.708 | 11.247 | 10.997 | 10.243 | 10.498 | 10.662 | 12.462 | 11.919 | 10.460 | 0.011 | 0.8834 | 0.9510 | 0 | 0 | 0 |
| 'Tia1' | 2.165 | 3.128 | 2.677 | 2.028 | 2.817 | 2.740 | 2.122 | 2.890 | 2.862 | 2.393 | 2.400 | 2.533 | 2.595 | 2.722 | 2.479 | 2.580 | 2.481 | 2.357 | 0.033 | 0.5469 | 0.7610 | 0 | 0 | 0 |
| 'Tial1' | 11.811 | 13.215 | 12.481 | 12.566 | 13.125 | 11.658 | 12.562 | 12.538 | 12.340 | 11.449 | 12.635 | 12.847 | 12.787 | 13.386 | 12.239 | 12.029 | 14.372 | 12.461 | -0.035 | 0.2364 | 0.4861 | 0 | 0 | 0 |
| 'Tiam1' | 17.610 | 18.483 | 17.321 | 16.848 | 18.688 | 16.703 | 17.691 | 17.707 | 17.261 | 18.746 | 21.329 | 18.826 | 18.082 | 20.321 | 18.816 | 19.923 | 21.524 | 18.151 | -0.159 | 0.0000 | 0.0000 | 1 | 0 | 0 |
| 'Ticam1' | 9.652 | 8.837 | 10.136 | 10.208 | 9.495 | 10.743 | 9.528 | 9.226 | 9.929 | 10.061 | 8.473 | 10.212 | 10.141 | 8.449 | 10.022 | 8.322 | 8.013 | 9.411 | 0.065 | 0.1403 | 0.3625 | 0 | 0 | 0 |
| 'Ticrr' | 1.466 | 1.436 | 1.950 | 1.522 | 1.356 | 1.687 | 1.627 | 1.390 | 1.724 | 1.441 | 1.556 | 1.581 | 1.826 | 1.548 | 1.723 | 1.630 | 1.630 | 1.621 | -0.050 | 0.3730 | 0.6242 | 0 | 0 | 0 |
| 'Tigar' | 3.687 | 4.305 | 3.836 | 3.923 | 4.273 | 4.407 | 3.727 | 3.981 | 3.909 | 3.620 | 4.022 | 5.183 | 4.151 | 4.164 | 4.626 | 4.375 | 3.871 | 5.058 | -0.134 | 0.0235 | 0.1154 | 0 | 0 | 0 |
| 'Tigd2' | 15.794 | 15.550 | 13.881 | 15.419 | 14.219 | 15.498 | 15.392 | 16.047 | 13.807 | 15.364 | 16.600 | 15.724 | 14.700 | 16.911 | 14.347 | 15.312 | 16.758 | 15.150 | -0.064 | 0.1053 | 0.3042 | 0 | 0 | 0 |
| 'Tigit' | 271.119 | 266.283 | 273.503 | 259.486 | 296.485 | 268.105 | 246.608 | 304.511 | 258.521 | 300.182 | 318.326 | 264.212 | 272.453 | 286.511 | 293.091 | 282.464 | 260.871 | 293.526 | -0.085 | 0.0456 | 0.1775 | 0 | 0 | 0 |
| 'Timeless' | 5.178 | 4.331 | 6.543 | 5.539 | 4.652 | 6.186 | 5.813 | 4.514 | 6.148 | 4.945 | 5.224 | 5.287 | 7.267 | 5.064 | 5.740 | 5.651 | 4.739 | 4.984 | -0.018 | 0.7875 | 0.9048 | 0 | 0 | 0 |
| 'Timm10' | 7.484 | 6.130 | 7.091 | 7.366 | 6.991 | 6.929 | 6.728 | 7.337 | 7.831 | 7.391 | 6.498 | 6.645 | 6.037 | 5.291 | 6.405 | 5.784 | 6.670 | 6.192 | 0.159 | 0.0410 | 0.1658 | 0 | 0 | 0 |
| 'Timm13' | 44.733 | 45.407 | 41.772 | 42.596 | 50.306 | 46.176 | 41.537 | 45.596 | 42.506 | 37.878 | 40.104 | 44.220 | 44.771 | 45.112 | 46.940 | 45.340 | 42.445 | 41.709 | 0.032 | 0.4677 | 0.7034 | 0 | 0 | 0 |

| | | | | | | | | | | | | | | | | | | | | | | | | |
|---|---|---|---|---|---|---|---|---|---|---|---|---|---|---|---|---|---|---|---|---|---|---|---|---|
| 'Timm17a' | 21.573 | 21.903 | 22.275 | 21.875 | 24.607 | 20.315 | 21.344 | 23.533 | 19.317 | 18.324 | 19.442 | 17.617 | 21.322 | 20.989 | 19.629 | 18.868 | 17.885 | 18.155 | 0.182 | 0.0001 | 0.0017 | 1 | 0 | 0 |
| 'Timm17b' | 16.566 | 20.264 | 16.709 | 15.404 | 19.032 | 18.786 | 16.669 | 18.518 | 18.516 | 17.661 | 16.575 | 17.055 | 17.897 | 18.467 | 18.070 | 18.855 | 17.133 | 17.133 | 0.002 | 0.9590 | 0.9847 | 0 | 0 | 0 |
| 'Timm21' | 14.199 | 17.541 | 16.167 | 16.958 | 18.767 | 15.287 | 17.541 | 17.014 | 16.765 | 14.084 | 14.968 | 14.831 | 16.102 | 15.871 | 15.508 | 18.526 | 15.424 | 13.905 | 0.098 | 0.0806 | 0.2562 | 0 | 0 | 0 |
| 'Timm22' | 6.174 | 7.117 | 6.041 | 5.248 | 7.151 | 6.496 | 6.282 | 6.591 | 5.637 | 5.750 | 5.862 | 6.419 | 6.380 | 6.880 | 6.089 | 6.689 | 6.322 | 6.361 | -0.017 | 0.7452 | 0.8850 | 0 | 0 | 0 |
| 'Timm23' | 17.718 | 19.357 | 18.374 | 19.815 | 20.328 | 19.769 | 18.366 | 19.681 | 18.748 | 18.517 | 18.181 | 17.937 | 19.682 | 17.662 | 19.039 | 18.181 | 18.344 | 18.333 | 0.042 | 0.2239 | 0.4711 | 0 | 0 | 0 |
| 'Timm44' | 10.387 | 10.052 | 11.284 | 10.420 | 9.914 | 10.826 | 9.792 | 10.365 | 9.788 | 10.510 | 10.114 | 10.089 | 10.695 | 10.205 | 9.445 | 9.945 | 10.530 | 9.996 | 0.013 | 0.7104 | 0.8654 | 0 | 0 | 0 |
| 'Timm50' | 12.095 | 12.248 | 13.139 | 12.432 | 12.613 | 13.603 | 11.794 | 12.719 | 12.683 | 12.338 | 11.496 | 12.798 | 13.438 | 11.581 | 12.816 | 12.621 | 11.328 | 12.260 | 0.025 | 0.4698 | 0.7056 | 0 | 0 | 0 |
| 'Timm8b' | 42.142 | 47.787 | 43.901 | 44.689 | 47.645 | 43.393 | 38.389 | 42.763 | 41.237 | 36.932 | 46.313 | 40.598 | 43.905 | 43.507 | 45.780 | 42.011 | 46.157 | 39.653 | 0.017 | 0.7320 | 0.8779 | 0 | 0 | 0 |
| 'Timm9' | 8.829 | 7.784 | 9.162 | 10.555 | 9.021 | 9.280 | 7.979 | 9.798 | 12.155 | 9.923 | 10.750 | 9.691 | 9.702 | 9.601 | 9.931 | 8.202 | 11.365 | 10.623 | -0.091 | 0.2652 | 0.5185 | 0 | 0 | 0 |
| 'Timmdc1' | 11.849 | 12.832 | 12.195 | 11.092 | 13.133 | 11.490 | 11.141 | 12.475 | 13.030 | 11.493 | 10.539 | 13.096 | 13.425 | 13.453 | 13.317 | 12.157 | 12.729 | 13.477 | -0.073 | 0.1450 | 0.3694 | 0 | 0 | 0 |
| 'Timp2' | 94.956 | 112.021 | 94.048 | 92.791 | 112.592 | 104.121 | 93.905 | 109.395 | 90.880 | 118.390 | 116.822 | 102.257 | 122.335 | 122.936 | 109.360 | 122.622 | 116.684 | 106.650 | -0.210 | 0.0000 | 0.0000 | 1 | 0 | 0 |
| 'Tinf2' | 33.069 | 33.623 | 37.821 | 33.449 | 35.002 | 34.509 | 33.769 | 34.504 | 35.132 | 29.018 | 34.830 | 32.497 | 39.199 | 34.373 | 33.421 | 33.804 | 31.637 | 29.431 | 0.051 | 0.2880 | 0.5446 | 0 | 0 | 0 |
| 'Tiparp' | 6.595 | 4.151 | 3.791 | 5.557 | 4.297 | 3.503 | 4.984 | 5.081 | 3.693 | 5.555 | 5.511 | 3.643 | 5.434 | 5.292 | 3.641 | 4.870 | 5.058 | 3.484 | -0.034 | 0.6020 | 0.7982 | 0 | 0 | 0 |
| 'Tipin' | 28.395 | 32.654 | 32.174 | 28.887 | 33.153 | 31.467 | 35.823 | 32.794 | 33.880 | 28.340 | 27.529 | 25.829 | 34.321 | 32.161 | 26.327 | 29.132 | 28.689 | 22.344 | 0.179 | 0.0029 | 0.0248 | 1 | 0 | 0 |
| 'Tiprl' | 8.080 | 8.664 | 7.770 | 8.520 | 9.240 | 7.711 | 8.233 | 8.657 | 7.492 | 7.972 | 8.571 | 8.060 | 8.364 | 9.512 | 8.027 | 8.380 | 8.605 | 7.579 | -0.026 | 0.3896 | 0.6389 | 0 | 0 | 0 |
| 'Tirap' | 0.486 | 0.687 | 0.572 | 0.517 | 0.460 | 0.611 | 0.605 | 0.558 | 0.558 | 0.573 | 0.582 | 0.554 | 0.585 | 0.544 | 0.517 | 0.378 | 0.576 | 0.612 | 0.020 | 0.8473 | 0.9343 | 0 | 0 | 0 |
| 'Tjap1' | 18.124 | 18.005 | 20.420 | 16.715 | 18.506 | 21.166 | 19.093 | 17.618 | 20.823 | 18.880 | 18.148 | 21.304 | 19.127 | 16.989 | 19.219 | 18.049 | 17.247 | 19.268 | 0.006 | 0.8519 | 0.9365 | 0 | 0 | 0 |
| 'Tjp2' | 4.744 | 5.070 | 5.941 | 4.996 | 5.557 | 6.129 | 4.919 | 5.150 | 6.327 | 5.304 | 5.841 | 5.330 | 6.055 | 5.884 | 5.362 | 5.551 | 5.763 | 5.377 | -0.054 | 0.3377 | 0.5955 | 0 | 0 | 0 |
| 'Tjp3' | 14.113 | 12.298 | 15.061 | 14.262 | 12.360 | 15.239 | 13.177 | 13.015 | 14.810 | 12.427 | 11.377 | 12.749 | 15.014 | 11.422 | 12.518 | 13.731 | 11.547 | 11.437 | 0.143 | 0.0017 | 0.0160 | 1 | 0 | 0 |
| 'Tk1' | 5.172 | 4.572 | 6.251 | 5.426 | 4.848 | 5.552 | 5.841 | 4.886 | 5.511 | 4.225 | 4.887 | 5.406 | 6.160 | 5.206 | 5.905 | 5.267 | 5.358 | 5.699 | -0.014 | 0.8097 | 0.9142 | 0 | 0 | 0 |
| 'Tk2' | 24.452 | 27.034 | 25.837 | 23.612 | 27.530 | 26.329 | 25.033 | 28.148 | 26.904 | 28.370 | 28.704 | 24.652 | 27.422 | 27.159 | 23.360 | 28.825 | 27.095 | 20.527 | -0.011 | 0.8233 | 0.9223 | 0 | 0 | 0 |
| 'Tkfc' | 8.725 | 8.810 | 9.321 | 9.048 | 8.725 | 9.997 | 8.945 | 8.735 | 9.088 | 8.639 | 8.317 | 8.040 | 8.744 | 8.611 | 8.235 | 7.141 | 8.105 | 8.255 | 0.129 | 0.0115 | 0.0692 | 0 | 0 | 0 |
| 'Tkt' | 26.040 | 20.545 | 26.736 | 24.948 | 23.122 | 27.161 | 25.681 | 23.471 | 25.909 | 24.445 | 22.062 | 26.852 | 26.067 | 23.260 | 27.410 | 24.967 | 22.553 | 26.436 | -0.013 | 0.6309 | 0.8173 | 0 | 0 | 0 |
| 'Tlcd1' | 1.399 | 1.315 | 1.392 | 1.464 | 1.299 | 1.403 | 1.438 | 1.428 | 1.235 | 1.302 | 1.240 | 1.302 | 1.488 | 1.305 | 1.252 | 1.318 | 1.417 | 1.121 | 0.066 | 0.2870 | 0.5436 | 0 | 0 | 0 |
| 'Tlcd2' | 3.789 | 3.609 | 3.578 | 3.753 | 4.352 | 3.936 | 4.240 | 3.810 | 3.475 | 3.188 | 3.097 | 2.868 | 3.602 | 4.189 | 2.891 | 3.320 | 4.182 | 2.473 | 0.215 | 0.0050 | 0.0372 | 1 | 0 | 0 |
| 'Tldc1' | 5.745 | 5.852 | 6.761 | 5.775 | 6.996 | 6.535 | 5.835 | 5.697 | 6.955 | 6.567 | 5.995 | 7.596 | 6.551 | 6.823 | 6.941 | 6.875 | 6.120 | 7.307 | -0.129 | 0.0176 | 0.0942 | 0 | 0 | 0 |
| 'Tle3' | 20.550 | 18.193 | 19.823 | 19.138 | 17.977 | 18.975 | 17.718 | 19.164 | 19.682 | 21.433 | 20.567 | 21.558 | 18.566 | 20.086 | 20.903 | 19.626 | 20.175 | 22.047 | -0.121 | 0.0005 | 0.0060 | 1 | 0 | 0 |
| 'Tle4' | 10.699 | 9.745 | 9.210 | 11.154 | 9.324 | 9.791 | 10.742 | 9.676 | 9.691 | 10.042 | 9.666 | 10.360 | 9.810 | 9.741 | 9.648 | 10.519 | 9.650 | 11.166 | -0.023 | 0.5599 | 0.7704 | 0 | 0 | 0 |
| 'Tlk1' | 16.643 | 15.965 | 16.735 | 16.584 | 15.430 | 16.040 | 17.718 | 16.450 | 16.068 | 16.544 | 14.997 | 15.668 | 14.505 | 14.771 | 15.084 | 16.068 | 14.612 | 15.153 | 0.093 | 0.0006 | 0.0066 | 1 | 0 | 0 |
| 'Tlk2' | 6.290 | 6.381 | 6.273 | 6.098 | 6.348 | 6.416 | 6.659 | 6.919 | 6.227 | 6.280 | 6.544 | 6.206 | 6.002 | 6.585 | 6.109 | 6.510 | 6.379 | 6.210 | 0.012 | 0.6648 | 0.8381 | 0 | 0 | 0 |
| 'Tln1' | 129.546 | 119.111 | 131.963 | 126.850 | 118.579 | 137.059 | 118.752 | 117.482 | 133.419 | 147.715 | 135.029 | 133.699 | 136.708 | 130.535 | 138.098 | 133.332 | 124.654 | 141.085 | -0.120 | 0.0001 | 0.0010 | 1 | 0 | 0 |
| 'Tlr1' | 14.030 | 17.158 | 15.762 | 13.641 | 17.397 | 15.556 | 14.088 | 17.197 | 16.632 | 14.931 | 16.363 | 15.154 | 15.749 | 16.375 | 15.168 | 15.578 | 17.150 | 14.588 | -0.007 | 0.8304 | 0.9262 | 0 | 0 | 0 |
| 'Tlr4' | 1.030 | 1.451 | 0.941 | 1.371 | 1.223 | 0.840 | 1.373 | 1.192 | 1.216 | 0.944 | 1.115 | 1.224 | 1.040 | 1.327 | 0.962 | 0.930 | 1.329 | 0.973 | 0.080 | 0.4113 | 0.6599 | 0 | 0 | 0 |
| 'Tlr6' | 7.983 | 10.511 | 9.108 | 9.163 | 10.386 | 9.598 | 8.835 | 10.814 | 9.755 | 9.466 | 8.870 | 8.800 | 9.193 | 10.155 | 9.934 | 8.853 | 9.669 | 8.966 | 0.025 | 0.5738 | 0.7796 | 0 | 0 | 0 |
| 'Tlr9' | 4.568 | 7.106 | 4.704 | 4.698 | 6.662 | 5.167 | 4.649 | 5.472 | 5.309 | 5.577 | 5.591 | 4.207 | 5.617 | 5.557 | 3.733 | 5.000 | 4.669 | 3.935 | 0.129 | 0.0960 | 0.2881 | 0 | 0 | 0 |
| 'Tm2d1' | 10.543 | 14.265 | 11.508 | 12.096 | 15.254 | 11.420 | 12.280 | 14.289 | 11.969 | 11.138 | 10.392 | 11.080 | 11.800 | 13.248 | 9.972 | 12.355 | 11.384 | 9.740 | 0.149 | 0.0073 | 0.0494 | 1 | 0 | 0 |
| 'Tm2d2' | 12.096 | 14.220 | 10.055 | 12.132 | 12.723 | 12.645 | 11.614 | 12.173 | 12.415 | 11.207 | 10.395 | 11.905 | 12.236 | 13.951 | 12.341 | 12.988 | 13.994 | 12.489 | -0.034 | 0.5987 | 0.7958 | 0 | 0 | 0 |
| 'Tm2d3' | 5.466 | 6.910 | 5.639 | 5.195 | 6.112 | 5.430 | 5.223 | 6.049 | 6.004 | 5.603 | 5.140 | 6.314 | 5.688 | 5.708 | 5.949 | 6.391 | 5.984 | 5.769 | -0.032 | 0.5356 | 0.7551 | 0 | 0 | 0 |
| 'Tm6sf1' | 0.493 | 0.653 | 0.558 | 0.652 | 0.730 | 0.699 | 0.617 | 0.687 | 0.640 | 0.549 | 0.704 | 0.605 | 0.734 | 0.826 | 0.570 | 0.393 | 0.697 | 0.624 | -0.007 | 0.9380 | 0.9764 | 0 | 0 | 0 |
| 'Tm7sf2' | 4.322 | 5.401 | 4.741 | 4.777 | 5.363 | 5.354 | 4.370 | 4.762 | 5.205 | 4.629 | 4.900 | 4.375 | 5.610 | 5.178 | 4.534 | 4.484 | 5.041 | 4.075 | 0.040 | 0.4828 | 0.7146 | 0 | 0 | 0 |
| 'Tm7sf3' | 5.553 | 6.992 | 6.999 | 6.070 | 6.916 | 7.207 | 6.484 | 6.907 | 6.862 | 7.661 | 6.746 | 6.135 | 6.617 | 6.580 | 5.858 | 6.581 | 5.450 | 5.820 | 0.055 | 0.3722 | 0.6240 | 0 | 0 | 0 |
| 'Tm9sf1' | 6.155 | 6.963 | 7.196 | 5.915 | 6.828 | 6.881 | 6.051 | 6.829 | 6.982 | 6.573 | 5.748 | 6.654 | 6.032 | 6.240 | 6.777 | 6.397 | 6.169 | 6.474 | 0.056 | 0.0864 | 0.2678 | 0 | 0 | 0 |
| 'Tm9sf2' | 26.740 | 29.095 | 30.316 | 28.278 | 30.563 | 28.296 | 28.611 | 29.714 | 29.016 | 27.133 | 29.724 | 31.093 | 26.396 | 28.789 | 27.705 | 25.295 | 29.110 | 28.195 | 0.027 | 0.3497 | 0.6044 | 0 | 0 | 0 |
| 'Tm9sf3' | 51.126 | 57.998 | 54.831 | 52.634 | 57.121 | 53.103 | 53.482 | 56.643 | 52.781 | 50.433 | 54.845 | 53.902 | 48.478 | 57.040 | 52.834 | 51.111 | 57.276 | 55.694 | 0.013 | 0.5585 | 0.7697 | 0 | 0 | 0 |
| 'Tm9sf4' | 4.343 | 4.535 | 4.853 | 4.288 | 4.655 | 4.527 | 4.447 | 4.756 | 4.805 | 5.018 | 4.733 | 4.940 | 4.797 | 4.616 | 4.819 | 4.788 | 4.549 | 4.689 | -0.069 | 0.0273 | 0.1271 | 0 | 0 | 0 |
| 'Tma16' | 3.824 | 4.000 | 4.215 | 4.077 | 4.511 | 3.838 | 3.594 | 4.252 | 3.758 | 3.754 | 3.532 | 3.387 | 4.378 | 3.672 | 3.612 | 4.271 | 3.483 | 3.090 | 0.119 | 0.0839 | 0.2628 | 0 | 0 | 0 |
| 'Tma7' | 109.231 | 122.690 | 100.670 | 120.624 | 118.804 | 112.223 | 119.566 | 113.067 | 110.118 | 108.088 | 111.172 | 91.562 | 109.645 | 118.255 | 110.813 | 115.187 | 113.296 | 104.262 | 0.054 | 0.1635 | 0.3954 | 0 | 0 | 0 |
| 'Tmbim4' | 6.380 | 7.195 | 6.645 | 7.075 | 7.416 | 7.334 | 7.043 | 7.200 | 6.973 | 6.231 | 6.587 | 6.140 | 6.234 | 6.784 | 6.343 | 6.672 | 6.476 | 5.632 | 0.140 | 0.0005 | 0.0059 | 1 | 0 | 0 |
| 'Tmbim6' | 112.422 | 127.388 | 112.899 | 112.748 | 126.115 | 120.070 | 112.540 | 120.941 | 118.644 | 122.363 | 116.800 | 115.401 | 121.275 | 119.545 | 120.509 | 127.297 | 114.971 | 119.491 | -0.030 | 0.2882 | 0.5447 | 0 | 0 | 0 |
| 'Tmc3' | 0.740 | 1.050 | 0.804 | 0.789 | 0.972 | 0.821 | 0.636 | 0.983 | 1.062 | 0.949 | 0.887 | 1.104 | 1.030 | 1.145 | 0.949 | 0.858 | 0.933 | 0.781 | -0.160 | 0.0603 | 0.2118 | 0 | 0 | 0 |
| 'Tmc4' | 1.740 | 1.423 | 1.706 | 1.787 | 1.253 | 1.367 | 1.725 | 1.390 | 1.697 | 1.847 | 1.246 | 2.365 | 1.831 | 1.664 | 1.936 | 1.474 | 2.139 | 2.224 | -0.264 | 0.0051 | 0.0383 | 1 | 0 | 0 |
| 'Tmc6' | 48.189 | 44.238 | 52.142 | 46.334 | 43.685 | 48.434 | 45.453 | 44.873 | 51.525 | 48.573 | 46.479 | 55.668 | 48.435 | 44.000 | 51.248 | 45.792 | 48.832 | 55.016 | -0.074 | 0.0079 | 0.0523 | 0 | 0 | 0 |
| 'Tmc8' | 40.643 | 36.947 | 44.057 | 39.221 | 36.305 | 42.185 | 39.203 | 37.074 | 43.884 | 40.177 | 38.772 | 48.339 | 38.107 | 36.946 | 44.509 | 36.938 | 40.606 | 46.140 | -0.053 | 0.0844 | 0.2637 | 0 | 0 | 0 |
| 'Tmcc1' | 3.719 | 3.927 | 3.277 | 4.022 | 3.838 | 3.637 | 4.084 | 3.727 | 3.611 | 3.320 | 3.474 | 3.144 | 3.336 | 3.649 | 3.114 | 3.320 | 3.608 | 3.027 | 0.163 | 0.0000 | 0.0000 | 1 | 0 | 0 |
| 'Tmcc3' | 2.378 | 2.841 | 1.900 | 2.224 | 2.494 | 2.321 | 1.948 | 2.172 | 2.214 | 2.503 | 2.568 | 2.916 | 2.643 | 2.520 | 2.535 | 3.233 | 2.290 | 3.195 | -0.271 | 0.0017 | 0.0156 | 1 | 0 | 0 |
| 'Tmco1' | 7.317 | 8.585 | 7.918 | 7.226 | 8.309 | 7.836 | 7.842 | 8.295 | 7.188 | 7.579 | 7.662 | 7.580 | 7.641 | 8.066 | 7.417 | 7.495 | 8.048 | 6.746 | 0.036 | 0.3076 | 0.5661 | 0 | 0 | 0 |
| 'Tmco3' | 5.936 | 6.251 | 5.911 | 6.091 | 6.302 | 6.074 | 6.078 | 6.282 | 6.030 | 6.258 | 5.801 | 6.603 | 6.626 | 6.468 | 6.224 | 6.229 | 5.844 | 6.334 | -0.054 | 0.1599 | 0.3902 | 0 | 0 | 0 |
| 'Tmco4' | 10.599 | 9.312 | 10.321 | 9.614 | 9.676 | 10.788 | 9.519 | 9.675 | 10.133 | 10.407 | 10.022 | 10.714 | 11.489 | 10.563 | 10.611 | 10.451 | 9.902 | 10.657 | -0.089 | 0.0050 | 0.0375 | 1 | 0 | 0 |
| 'Tmco6' | 23.783 | 25.561 | 28.985 | 23.329 | 28.712 | 29.119 | 23.023 | 25.501 | 27.770 | 25.654 | 25.043 | 29.264 | 26.323 | 25.278 | 25.650 | 24.706 | 27.164 | 26.063 | -0.010 | 0.8051 | 0.9125 | 0 | 0 | 0 |
| 'Tmed10' | 40.895 | 46.437 | 43.557 | 42.026 | 46.387 | 45.407 | 41.799 | 45.321 | 45.186 | 45.666 | 45.609 | 41.603 | 42.265 | 41.501 | 40.421 | 43.415 | 40.211 | 40.667 | 0.048 | 0.1482 | 0.3745 | 0 | 0 | 0 |
| 'Tmed2' | 62.598 | 68.771 | 69.769 | 61.317 | 73.203 | 67.003 | 65.208 | 73.456 | 64.255 | 64.281 | 67.119 | 63.643 | 67.757 | 69.335 | 65.495 | 66.152 | 66.198 | 60.904 | 0.026 | 0.3690 | 0.6203 | 0 | 0 | 0 |
| 'Tmed3' | 8.564 | 9.602 | 9.139 | 8.580 | 9.467 | 9.945 | 8.881 | 8.939 | 9.684 | 9.739 | 8.479 | 9.928 | 9.664 | 9.346 | 9.761 | 10.398 | 8.959 | 9.127 | -0.056 | 0.2095 | 0.4547 | 0 | 0 | 0 |
| 'Tmed4' | 18.252 | 20.968 | 18.761 | 18.411 | 21.059 | 19.116 | 18.612 | 19.701 | 18.537 | 18.489 | 16.878 | 18.687 | 19.250 | 19.477 | 19.718 | 19.480 | 17.678 | 17.535 | 0.039 | 0.2948 | 0.5505 | 0 | 0 | 0 |
| 'Tmed5' | 8.425 | 8.150 | 8.087 | 8.087 | 7.909 | 7.083 | 8.253 | 7.706 | 6.919 | 8.067 | 8.579 | 8.145 | 7.726 | 8.735 | 7.209 | 8.390 | 8.677 | 7.790 | -0.065 | 0.0945 | 0.2851 | 0 | 0 | 0 |
| 'Tmed7' | 11.012 | 11.076 | 10.709 | 11.106 | 11.538 | 10.556 | 10.996 | 11.480 | 9.406 | 11.097 | 11.540 | 10.474 | 10.411 | 12.377 | 10.400 | 10.690 | 10.961 | 10.233 | -0.014 | 0.6552 | 0.8329 | 0 | 0 | 0 |
| 'Tmed8' | 12.948 | 13.169 | 12.485 | 14.629 | 13.051 | 11.915 | 14.037 | 12.527 | 12.511 | 11.435 | 15.299 | 15.207 | 12.261 | 14.679 | 13.595 | 10.885 | 14.573 | 14.598 | -0.069 | 0.2648 | 0.5184 | 0 | 0 | 0 |
| 'Tmed9' | 136.598 | 147.738 | 132.914 | 137.967 | 153.635 | 142.912 | 137.774 | 148.264 | 138.492 | 140.573 | 131.958 | 127.661 | 143.345 | 138.278 | 138.351 | 154.839 | 133.457 | 131.342 | 0.032 | 0.3314 | 0.5895 | 0 | 0 | 0 |
| 'Tmem101' | 6.315 | 8.142 | 8.446 | 6.099 | 8.340 | 7.541 | 5.985 | 7.612 | 7.785 | 7.335 | 6.698 | 8.077 | 7.765 | 6.979 | 7.036 | 6.490 | 6.185 | 7.874 | 0.021 | 0.7462 | 0.8854 | 0 | 0 | 0 |
| 'Tmem104' | 4.150 | 4.033 | 4.427 | 3.784 | 3.935 | 4.857 | 4.174 | 3.858 | 4.142 | 4.570 | 4.610 | 4.400 | 5.423 | 4.678 | 4.499 | 4.691 | 4.748 | 4.279 | -0.171 | 0.0011 | 0.0112 | 1 | 0 | 0 |
| 'Tmem106a' | 2.690 | 2.333 | 3.141 | 2.510 | 3.328 | 3.620 | 2.856 | 2.897 | 3.474 | 2.804 | 3.200 | 2.991 | 2.954 | 3.453 | 3.246 | 2.916 | 2.501 | 2.672 | -0.001 | 0.9900 | 0.9972 | 0 | 0 | 0 |
| 'Tmem106b' | 11.323 | 14.884 | 12.809 | 12.638 | 14.512 | 13.712 | 12.196 | 14.063 | 13.031 | 13.366 | 13.772 | 13.875 | 12.445 | 13.806 | 12.378 | 13.363 | 13.000 | 13.241 | -0.016 | 0.6432 | 0.8251 | 0 | 0 | 0 |
| 'Tmem106c' | 13.144 | 15.777 | 13.972 | 13.681 | 16.212 | 15.167 | 15.994 | 13.041 | 14.324 | 12.706 | 12.786 | 13.548 | 13.426 | 14.220 | 15.125 | 14.083 | 14.836 | 13.575 | 0.065 | 0.2478 | 0.4986 | 0 | 0 | 0 |
| 'Tmem109' | 6.534 | 5.501 | 7.260 | 5.976 | 5.864 | 6.841 | 6.702 | 5.950 | 6.875 | 5.637 | 6.083 | 6.746 | 6.071 | 6.309 | 6.678 | 5.769 | 5.809 | 6.700 | 0.035 | 0.3954 | 0.6444 | 0 | 0 | 0 |
| 'Tmem11' | 15.029 | 14.484 | 13.360 | 14.995 | 16.047 | 12.977 | 14.636 | 14.360 | 13.874 | 12.970 | 14.943 | 14.195 | 12.294 | 14.299 | 15.254 | 14.173 | 13.423 | 14.331 | 0.030 | 0.5449 | 0.7606 | 0 | 0 | 0 |
| 'Tmem110' | 4.019 | 5.047 | 4.624 | 4.269 | 4.432 | 4.478 | 4.237 | 4.777 | 4.467 | 4.668 | 4.199 | 4.493 | 4.317 | 4.750 | 4.561 | 4.028 | 4.337 | 4.314 | 0.011 | 0.7920 | 0.9070 | 0 | 0 | 0 |
| 'Tmem115' | 13.057 | 16.306 | 13.965 | 12.780 | 14.158 | 16.041 | 13.412 | 15.137 | 16.633 | 14.150 | 13.134 | 16.364 | 12.909 | 13.799 | 15.631 | 16.412 | 13.565 | 15.572 | -0.013 | 0.8060 | 0.9133 | 0 | 0 | 0 |
| 'Tmem120a' | 6.773 | 7.694 | 7.134 | 7.218 | 8.759 | 8.128 | 6.552 | 8.617 | 7.424 | 7.395 | 6.445 | 7.621 | 7.166 | 7.864 | 7.833 | 8.519 | 6.797 | 7.941 | 0.002 | 0.9704 | 0.9898 | 0 | 0 | 0 |
| 'Tmem120b' | 4.206 | 4.220 | 4.131 | 4.110 | 4.275 | 4.588 | 3.964 | 4.537 | 4.049 | 3.546 | 3.718 | 3.612 | 4.479 | 3.732 | 3.714 | 2.929 | 3.435 | 3.593 | 0.205 | 0.0007 | 0.0078 | 1 | 0 | 0 |
| 'Tmem121' | 17.029 | 21.649 | 15.765 | 16.489 | 18.071 | 17.099 | 14.696 | 16.891 | 15.742 | 17.518 | 16.195 | 17.869 | 19.515 | 17.348 | 18.687 | 19.395 | 17.734 | 16.367 | -0.082 | 0.1600 | 0.3904 | 0 | 0 | 0 |
| 'Tmem123' | 23.515 | 26.859 | 26.999 | 25.149 | 28.213 | 25.177 | 25.642 | 28.907 | 24.974 | 25.671 | 25.597 | 24.324 | 23.659 | 26.153 | 24.377 | 24.154 | 26.444 | 23.356 | 0.063 | 0.0173 | 0.0933 | 0 | 0 | 0 |
| 'Tmem126a' | 14.325 | 15.567 | 13.698 | 14.977 | 14.964 | 15.695 | 14.295 | 14.690 | 13.883 | 13.515 | 14.109 | 13.749 | 14.715 | 15.080 | 13.606 | 13.036 | 14.302 | 12.758 | 0.069 | 0.1220 | 0.3333 | 0 | 0 | 0 |
| 'Tmem126b' | 3.305 | 3.076 | 3.789 | 3.925 | 2.731 | 3.178 | 3.930 | 2.778 | 3.594 | 2.917 | 3.033 | 4.204 | 2.849 | 2.948 | 3.863 | 2.829 | 3.463 | 3.525 | 0.016 | 0.8412 | 0.9326 | 0 | 0 | 0 |
| 'Tmem127' | 21.829 | 25.757 | 24.617 | 22.038 | 24.635 | 27.724 | 23.178 | 25.017 | 26.732 | 23.556 | 25.002 | 28.654 | 22.658 | 24.508 | 28.226 | 25.074 | 23.993 | 29.175 | -0.070 | 0.0270 | 0.1265 | 0 | 0 | 0 |
| 'Tmem128' | 7.357 | 7.851 | 7.667 | 8.483 | 8.133 | 8.159 | 7.749 | 8.388 | 7.187 | 7.293 | 7.828 | 7.787 | 7.557 | 8.517 | 7.652 | 7.391 | 8.416 | 6.970 | 0.022 | 0.5940 | 0.7933 | 0 | 0 | 0 |
| 'Tmem129' | 4.394 | 4.459 | 4.457 | 4.494 | 4.038 | 4.386 | 4.200 | 3.867 | 4.471 | 3.481 | 4.952 | 5.592 | 4.045 | 4.501 | 5.124 | 4.086 | 4.730 | 5.054 | -0.110 | 0.0782 | 0.2506 | 0 | 0 | 0 |
| 'Tmem131' | 22.146 | 20.863 | 22.487 | 22.391 | 22.490 | 20.676 | 22.022 | 22.546 | 22.343 | 21.574 | 22.064 | 21.351 | 21.066 | 20.922 | 20.478 | 21.627 | 22.705 | 22.022 | 0.020 | 0.4499 | 0.6898 | 0 | 0 | 0 |
| 'Tmem135' | 5.014 | 5.712 | 5.230 | 5.455 | 5.428 | 5.193 | 5.525 | 5.808 | 4.857 | 5.647 | 6.004 | 5.280 | 5.918 | 5.420 | 4.667 | 4.978 | 5.555 | 5.181 | -0.025 | 0.5348 | 0.7544 | 0 | 0 | 0 |
| 'Tmem136' | 1.330 | 1.710 | 1.703 | 1.646 | 1.625 | 1.619 | 1.692 | 1.562 | 1.675 | 0.966 | 1.151 | 1.230 | 1.174 | 1.033 | 1.104 | 1.098 | 1.059 | 1.588 | 0.465 | 0.0000 | 0.0001 | 1 | 1 | 0 |
| 'Tmem138' | 4.494 | 5.636 | 4.728 | 4.081 | 4.882 | 5.227 | 5.263 | 4.756 | 5.422 | 4.117 | 3.721 | 4.738 | 4.539 | 4.677 | 4.380 | 4.007 | 4.129 | 4.246 | 0.184 | 0.0208 | 0.1058 | 0 | 0 | 0 |
| 'Tmem140' | 1.519 | 1.452 | 1.358 | 1.342 | 1.604 | 1.525 | 1.261 | 1.383 | 1.662 | 1.496 | 1.533 | 1.823 | 1.459 | 1.824 | 1.820 | 1.565 | 1.615 | 1.963 | -0.217 | 0.0015 | 0.0147 | 1 | 0 | 0 |
| 'Tmem143' | 1.057 | 1.423 | 1.458 | 1.255 | 1.509 | 1.476 | 1.224 | 1.333 | 1.434 | 1.398 | 1.201 | 1.344 | 1.349 | 1.035 | 1.364 | 1.231 | 1.227 | 1.235 | 0.081 | 0.2234 | 0.4710 | 0 | 0 | 0 |
| 'Tmem147' | 28.393 | 33.353 | 30.197 | 29.082 | 31.737 | 31.589 | 25.073 | 32.237 | 30.175 | 25.448 | 25.850 | 28.050 | 27.600 | 25.821 | 30.177 | 26.567 | 23.286 | 28.047 | 0.161 | 0.0017 | 0.0161 | 1 | 0 | 0 |
| 'Tmem14c' | 62.222 | 77.624 | 70.692 | 66.977 | 73.031 | 74.488 | 65.602 | 73.997 | 66.798 | 62.215 | 66.733 | 72.834 | 72.473 | 72.709 | 76.957 | 75.313 | 68.666 | 73.029 | -0.035 | 0.3951 | 0.6443 | 0 | 0 | 0 |
| 'Tmem154' | 5.320 | 5.377 | 6.519 | 5.338 | 6.179 | 7.019 | 5.766 | 5.611 | 6.845 | 6.228 | 5.268 | 7.379 | 5.611 | 5.810 | 6.833 | 6.279 | 5.826 | 6.868 | -0.068 | 0.1041 | 0.3023 | 0 | 0 | 0 |
| 'Tmem156' | 4.664 | 5.625 | 4.603 | 5.498 | 5.613 | 5.324 | 5.327 | 5.733 | 5.411 | 5.894 | 7.019 | 6.335 | 5.326 | 6.211 | 6.233 | 5.347 | 6.822 | 5.455 | -0.208 | 0.0001 | 0.0019 | 1 | 0 | 0 |
| 'Tmem158' | 4.352 | 3.488 | 4.814 | 4.801 | 3.148 | 3.891 | 5.309 | 3.144 | 4.170 | 5.094 | 4.182 | 4.720 | 4.852 | 4.115 | 5.339 | 4.865 | 3.943 | 5.309 | -0.205 | 0.0107 | 0.0659 | 0 | 0 | 0 |
| 'Tmem160' | 46.020 | 59.275 | 49.676 | 47.195 | 56.493 | 53.640 | 44.246 | 61.399 | 54.014 | 45.256 | 48.457 | 48.167 | 50.615 | 47.034 | 51.940 | 55.904 | 49.233 | 48.392 | 0.070 | 0.1922 | 0.4294 | 0 | 0 | 0 |
| 'Tmem161a' | 5.204 | 5.723 | 6.066 | 5.383 | 6.039 | 5.935 | 5.411 | 5.686 | 5.803 | 5.511 | 5.236 | 5.610 | 6.070 | 5.217 | 5.296 | 5.102 | 5.132 | 5.428 | 0.064 | 0.0911 | 0.2783 | 0 | 0 | 0 |
| 'Tmem161b' | 9.425 | 9.617 | 9.419 | 10.277 | 9.934 | 8.625 | 10.463 | 9.599 | 9.916 | 9.822 | 10.261 | 9.454 | 9.526 | 9.236 | 8.556 | 9.123 | 11.625 | 8.167 | 0.015 | 0.7778 | 0.9002 | 0 | 0 | 0 |
| 'Tmem164' | 15.223 | 16.908 | 16.175 | 15.915 | 16.473 | 16.324 | 17.067 | 15.294 | 17.024 | 16.226 | 16.389 | 18.166 | 15.785 | 17.435 | 17.705 | 16.004 | 16.434 | 19.264 | -0.079 | 0.0240 | 0.1173 | 0 | 0 | 0 |
| 'Tmem165' | 18.436 | 19.747 | 19.142 | 17.602 | 20.702 | 19.820 | 19.206 | 20.712 | 19.453 | 17.337 | 18.041 | 20.605 | 18.288 | 19.049 | 21.184 | 18.433 | 18.217 | 20.650 | 0.011 | 0.7549 | 0.8907 | 0 | 0 | 0 |
| 'Tmem167' | 5.698 | 6.801 | 6.395 | 6.878 | 6.771 | 6.290 | 6.317 | 6.307 | 6.260 | 6.057 | 6.213 | 6.363 | 6.060 | 6.486 | 6.023 | 6.109 | 6.552 | 6.102 | 0.030 | 0.3541 | 0.6075 | 0 | 0 | 0 |
| 'Tmem168' | 3.081 | 2.944 | 3.998 | 3.650 | 3.468 | 2.934 | 4.668 | 3.156 | 3.702 | 2.842 | 3.749 | 4.315 | 2.820 | 3.155 | 3.589 | 2.254 | 4.239 | 4.014 | 0.032 | 0.8004 | 0.9111 | 0 | 0 | 0 |
| 'Tmem170' | 2.865 | 2.059 | 2.119 | 3.371 | 2.732 | 2.164 | 2.800 | 2.525 | 2.715 | 2.080 | 2.674 | 3.064 | 1.713 | 2.723 | 3.104 | 2.642 | 2.511 | 2.635 | -0.004 | 0.9731 | 0.9906 | 0 | 0 | 0 |
| 'Tmem170b' | 3.923 | 3.629 | 3.802 | 3.610 | 3.870 | 3.591 | 3.813 | 3.820 | 3.889 | 3.680 | 4.820 | 4.432 | 3.775 | 4.365 | 4.143 | 3.747 | 4.296 | 4.563 | -0.165 | 0.0009 | 0.0096 | 1 | 0 | 0 |
| 'Tmem173' | 96.025 | 108.003 | 99.758 | 101.346 | 108.327 | 98.791 | 102.063 | 99.046 | 99.645 | 98.649 | 99.320 | 109.020 | 105.207 | 106.143 | 107.901 | 101.803 | 106.394 | 109.954 | -0.061 | 0.0271 | 0.1270 | 0 | 0 | 0 |
| 'Tmem175' | 3.351 | 4.053 | 4.050 | 3.246 | 4.256 | 3.695 | 2.852 | 4.084 | 4.008 | 4.224 | 3.292 | 3.615 | 4.092 | 3.274 | 3.603 | 3.891 | 3.616 | 3.465 | 0.010 | 0.8904 | 0.9552 | 0 | 0 | 0 |
| 'Tmem177' | 2.238 | 2.769 | 2.962 | 2.686 | 2.511 | 2.860 | 2.430 | 2.532 | 3.063 | 2.055 | 2.308 | 2.604 | 2.204 | 1.759 | 2.526 | 2.131 | 2.127 | 2.641 | 0.224 | 0.0023 | 0.0206 | 1 | 0 | 0 |
| 'Tmem18' | 9.200 | 12.099 | 9.600 | 10.345 | 10.818 | 10.161 | 11.582 | 11.110 | 10.222 | 9.214 | 11.121 | 11.087 | 9.775 | 9.678 | 11.250 | 9.645 | 9.512 | 10.427 | 0.034 | 0.5275 | 0.7486 | 0 | 0 | 0 |
| 'Tmem183a' | 10.455 | 13.505 | 11.852 | 10.685 | 13.620 | 11.968 | 11.167 | 12.664 | 11.346 | 11.807 | 12.840 | 11.047 | 12.344 | 12.037 | 10.988 | 12.223 | 11.157 | 10.354 | 0.021 | 0.6174 | 0.8087 | 0 | 0 | 0 |
| 'Tmem184b' | 10.510 | 9.824 | 11.153 | 9.429 | 9.510 | 11.296 | 9.988 | 9.910 | 10.998 | 10.572 | 9.361 | 12.387 | 10.366 | 8.699 | 10.990 | 9.108 | 9.501 | 11.905 | -0.015 | 0.7005 | 0.8602 | 0 | 0 | 0 |
| 'Tmem184c' | 18.202 | 18.185 | 17.563 | 18.967 | 18.245 | 16.552 | 17.952 | 18.188 | 17.331 | 18.000 | 19.634 | 19.304 | 15.984 | 17.714 | 18.013 | 17.923 | 20.587 | 18.833 | -0.054 | 0.2056 | 0.4492 | 0 | 0 | 0 |
| 'Tmem185a' | 6.160 | 7.795 | 6.534 | 6.320 | 7.773 | 6.664 | 5.982 | 7.066 | 6.731 | 7.138 | 6.687 | 7.383 | 6.081 | 7.406 | 6.999 | 7.177 | 6.398 | 7.638 | -0.061 | 0.2431 | 0.4935 | 0 | 0 | 0 |
| 'Tmem185b' | 2.382 | 2.641 | 2.409 | 2.016 | 2.415 | 2.284 | 1.880 | 2.292 | 2.294 | 2.229 | 1.932 | 2.469 | 1.837 | 2.055 | 2.508 | 2.251 | 2.108 | 2.489 | 0.038 | 0.6284 | 0.8157 | 0 | 0 | 0 |
| 'Tmem186' | 7.617 | 10.334 | 9.743 | 8.321 | 9.371 | 9.168 | 9.870 | 9.252 | 10.158 | 8.477 | 8.153 | 9.721 | 7.748 | 8.120 | 10.152 | 7.709 | 8.645 | 10.169 | 0.072 | 0.1838 | 0.4193 | 0 | 0 | 0 |
| 'Tmem19' | 12.426 | 17.255 | 14.369 | 12.641 | 16.350 | 14.563 | 13.595 | 15.100 | 14.816 | 14.610 | 15.066 | 14.116 | 13.561 | 14.697 | 14.599 | 15.856 | 14.704 | 13.816 | -0.013 | 0.7733 | 0.8993 | 0 | 0 | 0 |
| 'Tmem191c' | 9.373 | 9.354 | 11.235 | 8.679 | 9.508 | 10.288 | 8.712 | 9.947 | 10.804 | 9.481 | 8.979 | 10.374 | 11.103 | 8.415 | 9.901 | 8.953 | 9.615 | 10.026 | 0.005 | 0.9070 | 0.9636 | 0 | 0 | 0 |
| 'Tmem192' | 26.386 | 32.207 | 27.655 | 25.493 | 28.172 | 29.801 | 25.793 | 31.446 | 30.314 | 30.764 | 28.113 | 29.252 | 33.239 | 29.643 | 30.333 | 32.199 | 28.736 | 26.656 | -0.074 | 0.1567 | 0.3863 | 0 | 0 | 0 |
| 'Tmem198' | 0.923 | 0.805 | 1.004 | 0.725 | 0.872 | 1.014 | 0.963 | 0.757 | 0.855 | 0.748 | 0.808 | 0.904 | 1.018 | 0.862 | 0.686 | 1.043 | 0.730 | 0.898 | 0.036 | 0.7369 | 0.8802 | 0 | 0 | 0 |
| 'Tmem198b' | 1.619 | 1.686 | 1.647 | 1.535 | 1.749 | 1.543 | 1.500 | 1.790 | 1.904 | 1.685 | 1.642 | 1.392 | 1.681 | 1.580 | 1.336 | 1.598 | 1.599 | 1.207 | 0.130 | 0.0758 | 0.2448 | 0 | 0 | 0 |
| 'Tmem199' | 4.704 | 4.721 | 5.057 | 4.819 | 6.179 | 5.284 | 4.883 | 5.185 | 4.965 | 4.450 | 5.063 | 5.284 | 5.393 | 4.726 | 5.304 | 5.017 | 4.873 | 4.346 | 0.028 | 0.6385 | 0.8220 | 0 | 0 | 0 |
| 'Tmem2' | 5.636 | 6.425 | 6.057 | 6.060 | 7.059 | 6.886 | 6.218 | 6.930 | 6.557 | 7.505 | 7.688 | 7.402 | 7.203 | 8.609 | 7.082 | 8.296 | 7.297 | 7.687 | -0.259 | 0.0000 | 0.0000 | 1 | 0 | 0 |
| 'Tmem201' | 2.735 | 2.734 | 3.032 | 2.251 | 2.786 | 2.826 | 2.313 | 2.426 | 3.060 | 2.515 | 2.680 | 3.100 | 2.961 | 2.680 | 2.744 | 2.535 | 2.231 | 3.093 | -0.035 | 0.5234 | 0.7458 | 0 | 0 | 0 |
| 'Tmem203' | 15.345 | 17.731 | 16.841 | 12.151 | 17.977 | 17.875 | 13.909 | 19.255 | 17.283 | 16.324 | 13.552 | 14.329 | 17.317 | 14.457 | 13.924 | 15.878 | 14.527 | 12.968 | 0.146 | 0.0619 | 0.2156 | 0 | 0 | 0 |
| 'Tmem206' | 7.590 | 6.532 | 5.963 | 6.717 | 5.749 | 6.540 | 6.073 | 6.145 | 6.488 | 5.667 | 5.468 | 7.391 | 5.370 | 6.609 | 6.164 | 6.310 | 5.586 | 7.885 | 0.022 | 0.7765 | 0.9001 | 0 | 0 | 0 |
| 'Tmem208' | 9.541 | 9.306 | 9.654 | 10.255 | 10.706 | 9.570 | 10.605 | 9.215 | 9.674 | 7.774 | 8.335 | 11.432 | 8.596 | 9.186 | 10.587 | 9.112 | 9.379 | 10.345 | 0.050 | 0.3998 | 0.6487 | 0 | 0 | 0 |
| 'Tmem209' | 4.256 | 4.624 | 4.589 | 4.272 | 4.768 | 4.494 | 4.522 | 4.881 | 4.526 | 4.621 | 4.957 | 4.609 | 5.111 | 4.846 | 4.644 | 4.801 | 4.601 | 4.356 | -0.065 | 0.0480 | 0.1836 | 0 | 0 | 0 |
| 'Tmem214' | 5.689 | 4.959 | 6.027 | 5.263 | 5.684 | 6.003 | 5.414 | 5.730 | 5.916 | 6.056 | 5.060 | 6.715 | 5.644 | 5.563 | 6.006 | 6.004 | 5.365 | 6.639 | -0.074 | 0.0431 | 0.1717 | 0 | 0 | 0 |
| 'Tmem216' | 1.323 | 1.854 | 1.862 | 1.395 | 1.833 | 1.614 | 1.432 | 2.104 | 1.709 | 1.703 | 1.514 | 1.643 | 1.830 | 1.777 | 1.500 | 2.105 | 1.781 | 1.464 | -0.024 | 0.8031 | 0.9116 | 0 | 0 | 0 |
| 'Tmem218' | 5.520 | 6.005 | 6.085 | 6.572 | 8.659 | 7.292 | 6.543 | 7.936 | 7.212 | 5.948 | 6.413 | 5.769 | 5.569 | 8.149 | 6.370 | 6.766 | 5.335 | 5.332 | 0.148 | 0.1168 | 0.3247 | 0 | 0 | 0 |
| 'Tmem219' | 4.591 | 5.453 | 4.946 | 4.806 | 5.152 | 5.310 | 5.375 | 5.446 | 5.448 | 5.261 | 4.960 | 4.959 | 5.122 | 5.201 | 5.115 | 5.164 | 4.376 | 4.739 | 0.041 | 0.3964 | 0.6453 | 0 | 0 | 0 |
| 'Tmem220' | 1.756 | 2.509 | 2.178 | 2.165 | 2.264 | 2.284 | 2.255 | 2.713 | 2.196 | 2.183 | 2.652 | 1.603 | 2.560 | 2.194 | 1.592 | 2.058 | 1.843 | 1.559 | 0.165 | 0.1212 | 0.3319 | 0 | 0 | 0 |
| 'Tmem222' | 10.904 | 12.058 | 12.165 | 10.231 | 14.026 | 12.554 | 10.441 | 13.214 | 12.489 | 11.973 | 11.490 | 11.913 | 11.727 | 12.068 | 11.390 | 12.538 | 11.079 | 10.663 | 0.032 | 0.5118 | 0.7371 | 0 | 0 | 0 |
| 'Tmem229b' | 13.484 | 14.490 | 14.473 | 13.802 | 14.457 | 15.217 | 14.518 | 14.106 | 14.848 | 13.640 | 14.222 | 17.120 | 13.531 | 14.660 | 16.380 | 13.862 | 13.791 | 17.005 | -0.062 | 0.0594 | 0.2104 | 0 | 0 | 0 |
| 'Tmem230' | 8.567 | 8.169 | 7.465 | 7.695 | 8.682 | 8.166 | 7.947 | 8.773 | 7.440 | 8.726 | 8.484 | 7.239 | 8.191 | 8.045 | 6.657 | 8.518 | 8.762 | 6.590 | 0.033 | 0.4409 | 0.6843 | 0 | 0 | 0 |
| 'Tmem231' | 2.611 | 2.582 | 2.725 | 1.963 | 2.534 | 2.793 | 2.814 | 2.241 | 2.110 | 2.426 | 1.773 | 1.914 | 1.512 | 1.655 | 1.572 | 2.415 | 1.787 | 1.430 | 0.443 | 0.0000 | 0.0007 | 1 | 1 | 0 |
| 'Tmem234' | 18.621 | 20.189 | 20.697 | 18.966 | 19.341 | 22.032 | 19.533 | 19.444 | 20.599 | 20.522 | 16.801 | 20.461 | 20.310 | 19.247 | 20.744 | 20.590 | 18.973 | 19.855 | 0.005 | 0.8598 | 0.9390 | 0 | 0 | 0 |
| 'Tmem237' | 3.425 | 3.452 | 3.898 | 3.750 | 3.755 | 4.015 | 3.567 | 3.708 | 3.753 | 3.026 | 3.774 | 3.342 | 3.488 | 3.693 | 3.499 | 3.211 | 3.196 | 3.109 | 0.132 | 0.0211 | 0.1066 | 0 | 0 | 0 |
| 'Tmem238' | 5.670 | 5.049 | 5.739 | 5.192 | 5.283 | 5.347 | 4.363 | 5.939 | 4.509 | 4.918 | 5.506 | 4.403 | 4.390 | 4.593 | 5.581 | 4.835 | 4.011 | 5.561 | 0.106 | 0.2286 | 0.4752 | 0 | 0 | 0 |
| 'Tmem241' | 3.053 | 3.041 | 2.593 | 2.988 | 2.499 | 3.093 | 3.171 | 3.219 | 2.846 | 2.999 | 3.031 | 2.835 | 2.873 | 2.772 | 2.994 | 2.468 | 2.209 | 2.244 | 0.106 | 0.1677 | 0.4000 | 0 | 0 | 0 |
| 'Tmem242' | 12.680 | 16.023 | 13.979 | 14.799 | 16.075 | 14.621 | 14.546 | 14.852 | 13.564 | 14.210 | 14.092 | 13.946 | 16.115 | 14.843 | 14.659 | 13.851 | 13.732 | 11.565 | 0.032 | 0.5343 | 0.7542 | 0 | 0 | 0 |
| 'Tmem243' | 4.259 | 4.531 | 4.458 | 4.197 | 4.618 | 3.974 | 4.624 | 5.179 | 3.428 | 4.549 | 5.284 | 3.630 | 4.904 | 4.466 | 3.615 | 4.214 | 4.798 | 3.859 | -0.004 | 0.9386 | 0.9764 | 0 | 0 | 0 |
| 'Tmem245' | 3.228 | 3.020 | 3.410 | 2.873 | 2.797 | 2.595 | 2.897 | 3.210 | 3.119 | 3.481 | 3.258 | 3.370 | 3.138 | 2.909 | 3.081 | 2.949 | 3.171 | 3.530 | -0.099 | 0.0597 | 0.2110 | 0 | 0 | 0 |
| 'Tmem248' | 10.588 | 13.043 | 10.961 | 10.234 | 12.048 | 12.214 | 10.192 | 12.100 | 11.158 | 11.594 | 12.312 | 11.705 | 11.671 | 11.533 | 11.443 | 12.488 | 11.253 | 11.519 | -0.054 | 0.1634 | 0.3953 | 0 | 0 | 0 |
| 'Tmem251' | 7.005 | 8.197 | 6.976 | 6.772 | 7.687 | 7.267 | 6.710 | 7.169 | 7.196 | 6.776 | 7.081 | 7.159 | 6.123 | 7.482 | 7.144 | 7.641 | 7.096 | 6.420 | 0.036 | 0.4577 | 0.6949 | 0 | 0 | 0 |
| 'Tmem254b' | 8.370 | 8.800 | 8.878 | 8.324 | 10.355 | 9.787 | 8.196 | 9.798 | 9.809 | 8.765 | 8.868 | 8.920 | 9.917 | 10.225 | 8.673 | 9.246 | 8.180 | 10.038 | -0.019 | 0.7502 | 0.8886 | 0 | 0 | 0 |
| 'Tmem256' | 8.435 | 10.068 | 8.568 | 9.042 | 8.918 | 8.736 | 8.241 | 7.783 | 7.763 | 6.678 | 7.204 | 7.754 | 8.913 | 9.028 | 8.607 | 7.960 | 7.242 | 7.428 | 0.117 | 0.0682 | 0.2291 | 0 | 0 | 0 |
| 'Tmem258' | 20.421 | 25.370 | 20.640 | 22.413 | 25.131 | 23.411 | 22.460 | 23.106 | 23.070 | 19.341 | 22.000 | 19.800 | 22.239 | 22.732 | 20.802 | 21.747 | 22.203 | 19.844 | 0.100 | 0.0091 | 0.0583 | 0 | 0 | 0 |
| 'Tmem259' | 28.130 | 24.841 | 30.455 | 24.310 | 23.892 | 29.642 | 25.227 | 24.347 | 30.185 | 26.814 | 22.361 | 29.247 | 26.533 | 24.022 | 28.219 | 25.128 | 23.801 | 28.056 | 0.031 | 0.2721 | 0.5273 | 0 | 0 | 0 |
| 'Tmem260' | 2.976 | 3.490 | 3.406 | 3.219 | 3.457 | 3.147 | 3.371 | 3.332 | 3.312 | 3.674 | 3.359 | 3.360 | 3.373 | 3.347 | 3.432 | 3.174 | 2.899 | 3.363 | -0.026 | 0.5451 | 0.7606 | 0 | 0 | 0 |
| 'Tmem261' | 42.649 | 52.516 | 48.945 | 51.945 | 54.603 | 50.918 | 48.605 | 50.495 | 48.146 | 45.352 | 46.856 | 48.417 | 50.668 | 47.503 | 48.142 | 45.698 | 48.158 | 43.711 | 0.064 | 0.1272 | 0.3416 | 0 | 0 | 0 |
| 'Tmem263' | 7.220 | 7.004 | 6.819 | 8.208 | 6.993 | 6.744 | 8.526 | 6.798 | 6.639 | 6.030 | 8.155 | 8.795 | 6.259 | 7.534 | 8.400 | 5.155 | 7.921 | 8.760 | -0.048 | 0.5920 | 0.7921 | 0 | 0 | 0 |
| 'Tmem268' | 8.383 | 9.907 | 9.401 | 8.403 | 9.245 | 9.397 | 8.794 | 9.843 | 9.424 | 8.941 | 9.545 | 8.650 | 9.374 | 9.364 | 7.974 | 8.648 | 8.701 | 7.996 | 0.056 | 0.1409 | 0.3635 | 0 | 0 | 0 |
| 'Tmem29' | 1.671 | 1.872 | 1.613 | 1.775 | 1.755 | 1.547 | 1.606 | 1.766 | 1.692 | 1.900 | 1.865 | 1.651 | 2.128 | 2.030 | 1.572 | 1.965 | 1.937 | 1.604 | -0.127 | 0.0112 | 0.0680 | 0 | 0 | 0 |
| 'Tmem30a' | 25.561 | 30.357 | 26.658 | 27.925 | 29.295 | 25.604 | 28.010 | 28.420 | 25.366 | 26.334 | 29.723 | 27.856 | 28.238 | 31.170 | 26.379 | 26.289 | 30.077 | 26.513 | -0.043 | 0.0929 | 0.2822 | 0 | 0 | 0 |
| 'Tmem33' | 3.317 | 3.418 | 3.243 | 3.267 | 3.219 | 3.196 | 3.234 | 3.068 | 3.228 | 2.810 | 3.551 | 5.068 | 2.995 | 3.550 | 4.035 | 3.074 | 4.044 | 5.054 | -0.217 | 0.0052 | 0.0387 | 1 | 0 | 0 |
| 'Tmem35b' | 2.004 | 2.834 | 2.237 | 2.428 | 2.554 | 2.594 | 2.126 | 2.964 | 2.565 | 2.504 | 2.308 | 2.655 | 2.200 | 2.347 | 1.835 | 2.527 | 2.481 | 2.516 | 0.039 | 0.7021 | 0.8611 | 0 | 0 | 0 |
| 'Tmem38b' | 13.382 | 17.894 | 14.053 | 13.742 | 15.833 | 14.014 | 14.531 | 14.479 | 14.635 | 16.129 | 16.655 | 14.155 | 15.681 | 17.501 | 12.820 | 16.377 | 16.817 | 12.710 | -0.072 | 0.1162 | 0.3236 | 0 | 0 | 0 |
| 'Tmem39a' | 3.706 | 4.117 | 3.546 | 3.671 | 3.849 | 4.008 | 3.497 | 4.012 | 3.400 | 3.741 | 3.630 | 3.743 | 3.591 | 4.073 | 3.624 | 3.849 | 3.707 | 3.684 | -0.004 | 0.9230 | 0.9705 | 0 | 0 | 0 |
| 'Tmem39b' | 6.934 | 7.078 | 6.807 | 6.185 | 7.143 | 7.930 | 6.506 | 6.068 | 7.297 | 5.947 | 6.198 | 7.193 | 6.967 | 6.520 | 7.089 | 6.927 | 6.362 | 6.569 | 0.039 | 0.4083 | 0.6574 | 0 | 0 | 0 |
| 'Tmem41a' | 4.352 | 4.849 | 5.444 | 4.603 | 6.546 | 5.298 | 5.188 | 5.157 | 5.269 | 5.253 | 4.634 | 4.419 | 4.859 | 3.947 | 4.607 | 3.720 | 4.000 | 5.074 | 0.190 | 0.0363 | 0.1536 | 0 | 0 | 0 |
| 'Tmem43' | 15.813 | 16.265 | 16.658 | 15.523 | 15.906 | 16.019 | 14.447 | 15.489 | 16.286 | 15.429 | 16.039 | 17.034 | 16.583 | 15.911 | 15.515 | 14.366 | 15.272 | 16.888 | -0.019 | 0.5633 | 0.7722 | 0 | 0 | 0 |
| 'Tmem5' | 1.627 | 1.814 | 1.889 | 1.731 | 1.808 | 1.706 | 1.613 | 2.319 | 1.836 | 1.916 | 1.575 | 1.613 | 1.765 | 1.642 | 1.500 | 1.762 | 1.648 | 1.364 | 0.140 | 0.0498 | 0.1882 | 0 | 0 | 0 |
| 'Tmem50a' | 126.821 | 146.485 | 125.645 | 128.793 | 145.874 | 132.980 | 132.415 | 142.764 | 130.129 | 124.980 | 129.896 | 127.056 | 125.741 | 139.522 | 137.351 | 140.387 | 134.129 | 133.632 | 0.010 | 0.7345 | 0.8792 | 0 | 0 | 0 |
| 'Tmem50b' | 8.193 | 10.313 | 8.558 | 8.358 | 10.948 | 9.683 | 9.068 | 10.976 | 8.949 | 9.033 | 8.941 | 9.385 | 10.280 | 10.317 | 8.818 | 9.088 | 9.660 | 9.359 | -0.016 | 0.7396 | 0.8819 | 0 | 0 | 0 |
| 'Tmem55a' | 8.358 | 8.893 | 8.715 | 8.625 | 9.266 | 8.796 | 8.981 | 9.408 | 8.194 | 9.218 | 9.698 | 8.957 | 8.450 | 9.304 | 9.509 | 7.897 | 9.510 | 8.625 | -0.045 | 0.3440 | 0.5995 | 0 | 0 | 0 |
| 'Tmem55b' | 4.275 | 4.411 | 4.514 | 4.286 | 4.154 | 4.544 | 3.663 | 4.483 | 4.074 | 4.089 | 4.329 | 4.338 | 4.507 | 4.923 | 4.280 | 5.107 | 4.505 | 4.368 | -0.080 | 0.0979 | 0.2920 | 0 | 0 | 0 |
| 'Tmem57' | 4.381 | 4.350 | 4.619 | 4.259 | 4.435 | 4.517 | 4.125 | 4.441 | 4.362 | 4.411 | 4.126 | 4.202 | 4.118 | 3.798 | 4.092 | 4.303 | 3.608 | 3.858 | 0.106 | 0.0045 | 0.0345 | 1 | 0 | 0 |
| 'Tmem59' | 20.787 | 23.794 | 20.277 | 20.701 | 23.758 | 21.204 | 19.914 | 23.987 | 21.356 | 19.829 | 20.429 | 23.228 | 19.305 | 23.456 | 22.747 | 22.663 | 22.767 | 23.725 | -0.031 | 0.4486 | 0.6888 | 0 | 0 | 0 |
| 'Tmem60' | 3.383 | 3.870 | 4.059 | 3.165 | 3.495 | 3.707 | 3.296 | 4.070 | 3.419 | 3.241 | 3.741 | 3.794 | 3.102 | 3.890 | 3.645 | 3.779 | 3.878 | 3.567 | -0.011 | 0.8596 | 0.9390 | 0 | 0 | 0 |
| 'Tmem62' | 6.992 | 6.434 | 7.066 | 7.154 | 7.442 | 7.086 | 7.124 | 6.853 | 7.296 | 7.163 | 8.155 | 7.605 | 6.874 | 6.855 | 7.029 | 6.508 | 8.005 | 7.075 | -0.051 | 0.2497 | 0.5007 | 0 | 0 | 0 |
| 'Tmem63a' | 15.935 | 15.410 | 16.975 | 15.548 | 14.774 | 16.266 | 14.780 | 15.311 | 16.733 | 15.531 | 14.880 | 15.819 | 14.345 | 14.174 | 15.461 | 13.689 | 14.623 | 15.308 | 0.073 | 0.0115 | 0.0692 | 0 | 0 | 0 |
| 'Tmem63b' | 4.577 | 4.703 | 4.741 | 4.545 | 4.106 | 4.833 | 4.559 | 4.077 | 5.355 | 4.413 | 3.795 | 4.967 | 4.726 | 3.918 | 4.292 | 4.408 | 3.832 | 4.863 | 0.066 | 0.1422 | 0.3651 | 0 | 0 | 0 |
| 'Tmem64' | 7.326 | 8.719 | 8.413 | 7.533 | 8.219 | 8.055 | 7.515 | 7.978 | 7.263 | 7.715 | 7.317 | 7.591 | 6.532 | 7.931 | 7.897 | 6.381 | 7.701 | 7.976 | 0.071 | 0.1116 | 0.3165 | 0 | 0 | 0 |
| 'Tmem65' | 7.041 | 7.831 | 7.315 | 5.539 | 7.390 | 6.887 | 6.013 | 7.741 | 6.856 | 8.124 | 7.438 | 8.922 | 6.918 | 8.237 | 8.325 | 7.361 | 8.184 | 9.355 | -0.232 | 0.0000 | 0.0005 | 1 | 0 | 0 |
| 'Tmem67' | 1.279 | 1.055 | 1.483 | 1.531 | 1.122 | 1.013 | 1.696 | 1.066 | 1.301 | 1.085 | 1.391 | 1.558 | 1.118 | 1.212 | 1.320 | 0.892 | 1.455 | 1.548 | -0.018 | 0.8739 | 0.9462 | 0 | 0 | 0 |
| 'Tmem68' | 2.118 | 2.590 | 2.515 | 2.689 | 2.926 | 2.316 | 2.799 | 2.634 | 2.221 | 2.312 | 2.899 | 2.691 | 2.282 | 2.232 | 2.107 | 2.439 | 2.822 | 2.391 | 0.022 | 0.7487 | 0.8873 | 0 | 0 | 0 |
| 'Tmem69' | 3.450 | 4.177 | 4.105 | 3.299 | 3.820 | 4.303 | 3.435 | 4.116 | 4.744 | 4.062 | 3.893 | 4.428 | 3.528 | 4.392 | 4.260 | 3.996 | 3.863 | 4.017 | -0.049 | 0.4047 | 0.6539 | 0 | 0 | 0 |
| 'Tmem70' | 5.910 | 5.608 | 5.888 | 5.590 | 6.820 | 6.033 | 5.604 | 6.168 | 6.090 | 5.322 | 5.758 | 6.517 | 5.368 | 6.449 | 6.098 | 5.639 | 6.248 | 6.704 | -0.023 | 0.6299 | 0.8169 | 0 | 0 | 0 |
| 'Tmem71' | 20.571 | 24.159 | 22.884 | 21.203 | 24.812 | 20.277 | 21.868 | 27.436 | 20.737 | 19.587 | 22.172 | 22.014 | 18.688 | 21.900 | 20.398 | 21.496 | 22.558 | 19.836 | 0.098 | 0.0127 | 0.0744 | 0 | 0 | 0 |
| 'Tmem8' | 16.037 | 16.053 | 18.820 | 15.203 | 16.070 | 18.084 | 16.713 | 17.532 | 19.048 | 18.549 | 17.680 | 18.529 | 19.049 | 17.019 | 18.357 | 16.257 | 17.669 | 16.445 | -0.065 | 0.1143 | 0.3206 | 0 | 0 | 0 |
| 'Tmem80' | 2.433 | 3.351 | 3.003 | 2.628 | 2.772 | 3.166 | 2.717 | 2.874 | 3.068 | 2.662 | 2.500 | 2.770 | 2.777 | 2.568 | 2.791 | 2.687 | 2.844 | 2.865 | 0.075 | 0.1443 | 0.3687 | 0 | 0 | 0 |
| 'Tmem81' | 4.449 | 4.155 | 4.535 | 4.193 | 4.231 | 4.782 | 4.783 | 4.507 | 4.334 | 4.010 | 4.260 | 4.731 | 4.156 | 4.776 | 4.522 | 4.518 | 4.729 | 4.801 | -0.028 | 0.6176 | 0.8087 | 0 | 0 | 0 |
| 'Tmem86a' | 3.076 | 4.819 | 3.229 | 2.466 | 4.486 | 4.246 | 2.696 | 4.666 | 4.279 | 3.497 | 3.297 | 2.606 | 3.443 | 4.595 | 2.854 | 4.320 | 2.997 | 2.625 | 0.159 | 0.2073 | 0.4519 | 0 | 0 | 0 |
| 'Tmem86b' | 5.882 | 4.609 | 4.684 | 5.784 | 4.052 | 4.657 | 5.502 | 4.846 | 4.589 | 5.478 | 5.060 | 6.157 | 5.645 | 5.277 | 6.028 | 4.501 | 5.845 | 5.929 | -0.182 | 0.0105 | 0.0644 | 0 | 0 | 0 |
| 'Tmem87a' | 9.500 | 10.162 | 10.234 | 10.448 | 11.147 | 9.156 | 10.548 | 10.701 | 10.046 | 9.929 | 10.478 | 10.145 | 9.574 | 9.738 | 9.554 | 9.174 | 10.385 | 9.200 | 0.048 | 0.1422 | 0.3651 | 0 | 0 | 0 |
| 'Tmem87b' | 4.165 | 4.470 | 4.535 | 4.082 | 4.446 | 4.125 | 4.328 | 4.524 | 4.557 | 4.247 | 4.617 | 4.309 | 4.224 | 4.570 | 3.768 | 4.471 | 4.385 | 4.119 | 0.012 | 0.7396 | 0.8819 | 0 | 0 | 0 |
| 'Tmem9' | 7.756 | 8.880 | 7.210 | 7.208 | 8.932 | 8.716 | 7.718 | 8.917 | 7.936 | 7.436 | 6.766 | 6.749 | 8.471 | 7.758 | 6.900 | 8.176 | 6.923 | 6.167 | 0.157 | 0.0055 | 0.0405 | 1 | 0 | 0 |
| 'Tmem94' | 20.478 | 18.339 | 24.217 | 19.974 | 18.219 | 22.396 | 19.421 | 20.022 | 22.645 | 22.360 | 19.537 | 22.321 | 21.109 | 19.740 | 20.885 | 20.286 | 19.615 | 21.223 | -0.022 | 0.5080 | 0.7336 | 0 | 0 | 0 |
| 'Tmem9b' | 24.391 | 33.801 | 29.765 | 27.350 | 35.265 | 30.522 | 24.761 | 29.971 | 28.407 | 28.637 | 26.243 | 30.356 | 27.745 | 31.323 | 31.317 | 30.581 | 28.213 | 30.775 | -0.023 | 0.6545 | 0.8326 | 0 | 0 | 0 |
| 'Tmf1' | 15.418 | 14.426 | 14.007 | 15.128 | 14.498 | 13.610 | 14.804 | 14.335 | 13.583 | 14.415 | 14.772 | 15.506 | 14.092 | 15.238 | 14.603 | 14.726 | 15.770 | 15.357 | -0.064 | 0.0582 | 0.2077 | 0 | 0 | 0 |
| 'Tmie' | 5.555 | 5.441 | 5.559 | 6.401 | 5.565 | 5.840 | 6.035 | 4.230 | 6.903 | 3.938 | 4.187 | 4.930 | 3.720 | 4.254 | 4.794 | 3.962 | 3.973 | 4.728 | 0.405 | 0.0000 | 0.0000 | 1 | 1 | 0 |
| 'Tmod3' | 30.023 | 29.500 | 32.253 | 31.953 | 31.929 | 30.581 | 31.406 | 29.150 | 29.008 | 27.319 | 28.921 | 31.649 | 29.353 | 29.871 | 31.347 | 28.485 | 30.085 | 32.233 | 0.023 | 0.5005 | 0.7278 | 0 | 0 | 0 |
| 'Tmod4' | 3.417 | 2.688 | 4.155 | 3.983 | 2.596 | 3.523 | 3.509 | 3.315 | 3.982 | 3.386 | 3.000 | 5.405 | 3.495 | 3.224 | 4.617 | 2.576 | 3.816 | 4.249 | -0.125 | 0.1670 | 0.3995 | 0 | 0 | 0 |
| 'Tmpo' | 44.364 | 48.008 | 52.849 | 50.431 | 47.343 | 49.911 | 50.914 | 46.707 | 51.223 | 41.252 | 45.989 | 50.192 | 49.146 | 49.653 | 51.715 | 44.135 | 50.267 | 50.137 | 0.020 | 0.5798 | 0.7822 | 0 | 0 | 0 |
| 'Tmppe' | 4.591 | 5.029 | 5.711 | 5.240 | 4.461 | 5.275 | 5.433 | 4.651 | 5.761 | 3.949 | 6.291 | 7.375 | 4.208 | 5.015 | 6.796 | 4.232 | 5.611 | 7.755 | -0.134 | 0.1043 | 0.3027 | 0 | 0 | 0 |
| 'Tmprss3' | 1.094 | 1.376 | 1.363 | 1.194 | 1.330 | 1.106 | 1.674 | 1.339 | 1.414 | 1.048 | 0.626 | 0.885 | 0.849 | 0.657 | 0.837 | 1.040 | 0.854 | 0.847 | 0.621 | 0.0000 | 0.0000 | 1 | 1 | 0 |
| 'Tmsb15b1' | 20.450 | 20.933 | 19.102 | 24.248 | 23.970 | 20.448 | 23.469 | 21.891 | 17.780 | 17.188 | 19.181 | 19.494 | 19.752 | 19.267 | 18.260 | 18.485 | 18.111 | 14.789 | 0.199 | 0.0086 | 0.0563 | 0 | 0 | 0 |
| 'Tmsb4x' | 609.094 | 745.622 | 581.455 | 713.067 | 716.233 | 613.607 | 739.116 | 630.542 | 631.929 | 623.747 | 661.517 | 435.701 | 632.679 | 685.993 | 581.777 | 681.949 | 685.620 | 532.588 | 0.111 | 0.0391 | 0.1613 | 0 | 0 | 0 |
| 'Tmtc4' | 1.986 | 2.200 | 2.088 | 2.109 | 2.204 | 1.739 | 2.032 | 2.190 | 2.204 | 1.951 | 1.864 | 2.069 | 2.057 | 1.943 | 1.792 | 1.832 | 2.062 | 1.866 | 0.087 | 0.0785 | 0.2514 | 0 | 0 | 0 |
| 'Tmub1' | 7.146 | 7.330 | 7.264 | 7.099 | 6.359 | 7.520 | 7.083 | 6.998 | 7.153 | 7.704 | 6.541 | 7.652 | 7.648 | 7.163 | 8.007 | 6.694 | 6.776 | 7.810 | -0.057 | 0.1836 | 0.4192 | 0 | 0 | 0 |
| 'Tmub2' | 15.522 | 17.486 | 17.208 | 15.304 | 15.949 | 17.610 | 14.294 | 15.936 | 17.433 | 16.395 | 14.686 | 16.515 | 16.345 | 15.467 | 15.927 | 16.902 | 14.317 | 16.001 | 0.032 | 0.4125 | 0.6608 | 0 | 0 | 0 |
| 'Tmx1' | 20.306 | 23.134 | 20.422 | 20.462 | 24.418 | 19.925 | 21.440 | 22.065 | 19.784 | 20.857 | 21.157 | 22.509 | 20.989 | 24.495 | 21.646 | 20.337 | 21.919 | 22.825 | -0.052 | 0.1572 | 0.3868 | 0 | 0 | 0 |
| 'Tmx2' | 9.665 | 11.090 | 10.993 | 10.266 | 11.389 | 10.934 | 11.273 | 11.800 | 10.830 | 10.372 | 10.790 | 10.598 | 10.971 | 11.402 | 10.226 | 11.182 | 10.203 | 9.856 | 0.029 | 0.3891 | 0.6384 | 0 | 0 | 0 |
| 'Tmx3' | 19.996 | 23.906 | 21.946 | 22.265 | 23.929 | 20.051 | 23.234 | 23.642 | 19.942 | 20.389 | 22.914 | 22.877 | 20.190 | 25.563 | 20.880 | 20.885 | 23.429 | 21.425 | -0.010 | 0.7767 | 0.9001 | 0 | 0 | 0 |
| 'Tmx4' | 10.969 | 13.873 | 10.623 | 11.961 | 13.482 | 10.901 | 11.165 | 13.311 | 10.932 | 12.029 | 13.870 | 11.698 | 12.069 | 13.174 | 10.678 | 11.863 | 13.485 | 10.812 | -0.047 | 0.0548 | 0.2005 | 0 | 0 | 0 |
| 'Tnc' | 1.806 | 2.147 | 2.083 | 1.745 | 2.000 | 1.780 | 2.165 | 2.264 | 2.067 | 2.905 | 2.731 | 2.317 | 2.120 | 2.438 | 2.130 | 2.425 | 2.294 | 2.438 | -0.279 | 0.0000 | 0.0007 | 1 | 0 | 0 |
| 'Tnf' | 18.923 | 21.275 | 13.053 | 16.671 | 19.007 | 15.469 | 16.956 | 18.656 | 12.764 | 24.144 | 15.999 | 12.185 | 24.186 | 23.498 | 12.654 | 28.798 | 18.603 | 12.880 | -0.145 | 0.1582 | 0.3883 | 0 | 0 | 0 |
| 'Tnfaip1' | 5.349 | 5.016 | 5.847 | 6.175 | 6.146 | 6.434 | 5.771 | 6.485 | 5.270 | 5.480 | 5.591 | 5.745 | 5.185 | 5.609 | 5.574 | 5.034 | 5.002 | 5.624 | 0.094 | 0.0877 | 0.2701 | 0 | 0 | 0 |
| 'Tnfaip3' | 86.960 | 43.650 | 23.887 | 45.613 | 49.776 | 27.017 | 32.537 | 61.149 | 18.387 | 97.434 | 51.469 | 12.942 | 76.060 | 73.592 | 15.471 | 96.309 | 41.470 | 14.356 | -0.052 | 0.8172 | 0.9188 | 0 | 0 | 0 |
| 'Tnfaip8' | 25.325 | 29.059 | 26.029 | 26.080 | 28.697 | 26.901 | 28.627 | 27.230 | 26.967 | 27.477 | 28.365 | 25.866 | 27.259 | 27.669 | 26.814 | 26.887 | 29.481 | 25.753 | -0.014 | 0.5582 | 0.7697 | 0 | 0 | 0 |
| 'Tnfaip8l1' | 8.288 | 8.277 | 7.710 | 7.889 | 8.986 | 9.153 | 7.731 | 8.090 | 8.739 | 8.493 | 6.818 | 8.071 | 8.419 | 8.587 | 9.248 | 7.761 | 6.914 | 7.993 | 0.040 | 0.5002 | 0.7275 | 0 | 0 | 0 |
| 'Tnfaip8l2' | 55.249 | 52.094 | 62.837 | 55.492 | 55.755 | 67.306 | 62.015 | 53.434 | 63.034 | 50.660 | 49.119 | 61.771 | 46.741 | 50.104 | 59.134 | 53.873 | 48.515 | 59.599 | 0.127 | 0.0004 | 0.0045 | 1 | 0 | 0 |
| 'Tnfrsf12a' | 8.852 | 4.725 | 4.112 | 7.413 | 5.140 | 4.007 | 6.684 | 5.151 | 3.588 | 5.734 | 6.540 | 4.402 | 6.177 | 5.647 | 3.400 | 5.489 | 5.863 | 4.069 | 0.041 | 0.7046 | 0.8623 | 0 | 0 | 0 |
| 'Tnfrsf13b' | 12.605 | 14.254 | 11.487 | 12.051 | 13.824 | 13.144 | 12.798 | 13.468 | 12.770 | 12.154 | 12.972 | 12.811 | 13.498 | 13.487 | 11.702 | 13.513 | 13.630 | 11.271 | 0.005 | 0.8936 | 0.9565 | 0 | 0 | 0 |

| | | | | | | | | | | | | | | | | | | | | | | | | |
|---|---|---|---|---|---|---|---|---|---|---|---|---|---|---|---|---|---|---|---|---|---|---|---|---|
| 'Tnfrsf14' | 52.136 | 53.821 | 62.761 | 47.587 | 56.329 | 60.521 | 53.034 | 57.580 | 63.572 | 51.800 | 57.727 | 68.730 | 52.066 | 54.582 | 65.330 | 52.754 | 55.783 | 66.383 | -0.059 | 0.0286 | 0.1311 | 0 | 0 | 0 |
| 'Tnfrsf18' | 17.941 | 19.662 | 18.788 | 16.826 | 20.088 | 19.283 | 18.620 | 19.850 | 17.940 | 18.737 | 17.205 | 18.545 | 18.096 | 19.884 | 18.392 | 20.265 | 17.374 | 16.879 | 0.021 | 0.5830 | 0.7851 | 0 | 0 | 0 |
| 'Tnfrsf1a' | 5.978 | 7.446 | 6.334 | 6.262 | 5.903 | 7.338 | 5.714 | 6.238 | 6.519 | 6.202 | 6.036 | 7.266 | 6.391 | 6.613 | 7.090 | 6.698 | 5.670 | 6.826 | -0.040 | 0.4346 | 0.6788 | 0 | 0 | 0 |
| 'Tnfrsf1b' | 12.426 | 9.463 | 13.253 | 9.477 | 10.124 | 15.234 | 12.159 | 11.100 | 14.453 | 10.536 | 6.849 | 16.282 | 7.809 | 7.846 | 13.147 | 12.762 | 7.028 | 13.131 | 0.199 | 0.0211 | 0.1066 | 0 | 0 | 0 |
| 'Tnfrsf21' | 1.014 | 1.329 | 0.911 | 0.865 | 0.980 | 1.322 | 1.131 | 1.031 | 1.253 | 1.086 | 1.161 | 1.131 | 1.625 | 1.265 | 1.114 | 1.319 | 1.066 | 1.028 | -0.142 | 0.2267 | 0.4732 | 0 | 0 | 0 |
| 'Tnfrsf22' | 4.387 | 4.498 | 3.880 | 4.197 | 4.770 | 3.813 | 4.257 | 4.525 | 3.954 | 4.279 | 5.816 | 4.943 | 3.967 | 4.061 | 4.465 | 3.464 | 4.366 | 4.493 | -0.076 | 0.3034 | 0.5608 | 0 | 0 | 0 |
| 'Tnfrsf25' | 0.951 | 1.223 | 1.065 | 1.108 | 0.868 | 1.270 | 1.236 | 1.319 | 1.081 | 0.851 | 0.720 | 1.404 | 1.247 | 0.830 | 1.050 | 1.565 | 1.030 | 1.348 | 0.000 | 0.9974 | 0.9994 | 0 | 0 | 0 |
| 'Tnfrsf26' | 20.686 | 27.012 | 21.565 | 20.346 | 25.544 | 22.523 | 20.464 | 27.510 | 22.976 | 24.296 | 24.780 | 25.959 | 23.280 | 27.573 | 24.718 | 28.616 | 24.793 | 24.129 | -0.147 | 0.0022 | 0.0194 | 1 | 0 | 0 |
| 'Tnfrsf4' | 29.545 | 27.652 | 27.580 | 27.252 | 27.316 | 28.768 | 29.624 | 27.720 | 29.152 | 30.709 | 26.223 | 31.883 | 30.808 | 29.890 | 30.582 | 31.012 | 31.160 | 30.021 | -0.109 | 0.0004 | 0.0055 | 1 | 0 | 0 |
| 'Tnfrsf8' | 11.229 | 12.556 | 10.144 | 10.523 | 10.686 | 10.487 | 11.401 | 9.420 | 11.402 | 13.846 | 10.814 | 11.537 | 16.853 | 11.197 | 10.810 | 14.052 | 12.570 | 11.053 | -0.204 | 0.0010 | 0.0105 | 1 | 0 | 0 |
| 'Tnfrsf9' | 12.341 | 11.302 | 13.295 | 11.876 | 11.274 | 13.013 | 12.473 | 12.482 | 11.795 | 10.552 | 7.917 | 12.485 | 7.262 | 8.524 | 11.191 | 10.839 | 7.123 | 11.155 | 0.339 | 0.0000 | 0.0001 | 1 | 0 | 0 |
| 'Tnfsf10' | 4.703 | 5.197 | 5.188 | 4.638 | 4.997 | 5.219 | 4.844 | 5.290 | 5.241 | 3.931 | 4.130 | 4.793 | 4.416 | 4.000 | 4.402 | 4.101 | 4.181 | 4.406 | 0.223 | 0.0000 | 0.0000 | 1 | 0 | 0 |
| 'Tnfsf11' | 69.191 | 74.736 | 64.437 | 69.317 | 72.381 | 65.538 | 74.712 | 69.394 | 65.928 | 68.852 | 63.942 | 66.088 | 61.307 | 70.561 | 64.791 | 72.156 | 65.895 | 66.658 | 0.047 | 0.1675 | 0.3999 | 0 | 0 | 0 |
| 'Tnfsf13b' | 11.598 | 10.629 | 9.022 | 8.942 | 9.257 | 7.618 | 9.229 | 9.333 | 9.455 | 9.994 | 10.243 | 9.468 | 7.981 | 10.668 | 8.956 | 9.226 | 9.518 | 7.752 | 0.012 | 0.8795 | 0.9492 | 0 | 0 | 0 |
| 'Tnfsf14' | 40.332 | 41.570 | 44.513 | 45.567 | 39.246 | 44.186 | 42.706 | 39.529 | 46.761 | 36.190 | 41.843 | 54.125 | 40.745 | 36.130 | 52.005 | 38.119 | 41.616 | 54.181 | -0.040 | 0.4506 | 0.6903 | 0 | 0 | 0 |
| 'Tnfsf8' | 109.167 | 112.169 | 113.881 | 105.494 | 117.908 | 106.611 | 108.757 | 115.674 | 111.382 | 112.402 | 128.799 | 127.310 | 87.583 | 126.183 | 124.846 | 105.266 | 133.483 | 130.475 | -0.108 | 0.0267 | 0.1256 | 0 | 0 | 0 |
| 'Tnfsf9' | 7.449 | 4.146 | 4.999 | 6.761 | 5.483 | 5.480 | 7.102 | 5.085 | 5.362 | 4.052 | 5.441 | 8.164 | 3.815 | 4.959 | 7.612 | 3.301 | 7.593 | 10.279 | -0.053 | 0.7973 | 0.9100 | 0 | 0 | 0 |
| 'Tnik' | 4.968 | 4.856 | 5.238 | 5.678 | 4.678 | 5.010 | 5.407 | 4.831 | 5.461 | 5.101 | 5.566 | 5.194 | 5.007 | 4.897 | 5.120 | 5.087 | 5.177 | 4.979 | -0.012 | 0.7593 | 0.8924 | 0 | 0 | 0 |
| 'Tnip1' | 29.823 | 27.794 | 31.192 | 29.445 | 26.909 | 32.347 | 29.777 | 27.817 | 31.030 | 33.219 | 26.641 | 32.672 | 31.021 | 27.629 | 29.367 | 31.986 | 25.998 | 30.241 | -0.023 | 0.3905 | 0.6397 | 0 | 0 | 0 |
| 'Tnip2' | 10.249 | 9.046 | 10.021 | 8.895 | 8.644 | 9.712 | 8.935 | 9.300 | 10.079 | 9.548 | 10.259 | 11.585 | 9.353 | 10.078 | 10.228 | 9.713 | 9.249 | 10.723 | -0.106 | 0.0087 | 0.0564 | 0 | 0 | 0 |
| 'Tnk2' | 29.206 | 27.952 | 32.348 | 28.435 | 27.224 | 31.300 | 27.596 | 29.676 | 31.192 | 28.166 | 25.344 | 29.007 | 27.100 | 24.653 | 29.065 | 25.858 | 23.401 | 29.171 | 0.123 | 0.0000 | 0.0001 | 1 | 0 | 0 |
| 'Tnks' | 11.174 | 9.449 | 10.252 | 9.561 | 9.298 | 10.178 | 9.062 | 9.970 | 10.034 | 10.392 | 10.290 | 10.261 | 9.032 | 10.506 | 10.132 | 10.905 | 9.575 | 10.181 | -0.043 | 0.3269 | 0.5849 | 0 | 0 | 0 |
| 'Tnks1bp1' | 6.118 | 5.524 | 6.156 | 5.500 | 5.343 | 6.034 | 5.634 | 5.683 | 6.087 | 5.747 | 5.393 | 6.080 | 5.670 | 5.617 | 6.035 | 5.683 | 5.429 | 5.761 | 0.010 | 0.6940 | 0.8558 | 0 | 0 | 0 |
| 'Tnks2' | 29.708 | 32.090 | 28.186 | 29.782 | 31.872 | 29.563 | 28.942 | 31.199 | 26.790 | 28.962 | 31.727 | 30.670 | 27.945 | 32.232 | 30.110 | 29.080 | 32.131 | 31.713 | -0.046 | 0.1196 | 0.3292 | 0 | 0 | 0 |
| 'Tnnt1' | 1.090 | 0.957 | 1.298 | 1.141 | 1.021 | 0.995 | 1.266 | 1.124 | 0.862 | 0.976 | 1.341 | 1.314 | 0.998 | 0.654 | 1.372 | 1.188 | 0.921 | 1.394 | -0.066 | 0.5956 | 0.7942 | 0 | 0 | 0 |
| 'Tnpo1' | 32.128 | 28.962 | 32.808 | 33.003 | 28.727 | 31.611 | 34.388 | 29.120 | 31.531 | 29.783 | 33.453 | 35.349 | 30.657 | 30.177 | 32.111 | 30.559 | 36.068 | 36.031 | -0.071 | 0.1334 | 0.3518 | 0 | 0 | 0 |
| 'Tnpo2' | 18.590 | 17.003 | 18.898 | 17.791 | 17.901 | 17.852 | 17.341 | 17.741 | 18.093 | 16.096 | 18.677 | 19.421 | 17.427 | 16.890 | 19.192 | 15.755 | 18.191 | 18.925 | -0.004 | 0.9205 | 0.9702 | 0 | 0 | 0 |
| 'Tnpo3' | 6.117 | 6.195 | 6.212 | 6.190 | 6.129 | 6.257 | 6.321 | 6.519 | 6.339 | 6.024 | 6.144 | 6.307 | 6.152 | 5.965 | 6.020 | 5.955 | 6.416 | 5.974 | 0.023 | 0.3047 | 0.5622 | 0 | 0 | 0 |
| 'Tnrc6a' | 14.300 | 14.454 | 14.561 | 14.924 | 13.667 | 13.950 | 14.385 | 14.314 | 14.562 | 14.582 | 15.354 | 15.889 | 14.317 | 14.245 | 14.471 | 13.847 | 14.827 | 14.818 | -0.047 | 0.0837 | 0.2622 | 0 | 0 | 0 |
| 'Tnrc6b' | 12.594 | 9.547 | 10.955 | 12.143 | 9.321 | 9.878 | 12.043 | 10.200 | 10.594 | 11.395 | 11.609 | 11.723 | 10.691 | 10.591 | 10.545 | 10.603 | 11.149 | 11.624 | -0.056 | 0.2620 | 0.5159 | 0 | 0 | 0 |
| 'Tnrc6c' | 6.312 | 5.867 | 6.697 | 5.987 | 5.985 | 6.384 | 5.865 | 6.058 | 6.668 | 6.485 | 6.810 | 6.528 | 6.060 | 5.952 | 6.267 | 6.232 | 5.677 | 6.732 | -0.034 | 0.3212 | 0.5792 | 0 | 0 | 0 |
| 'Tns1' | 0.293 | 0.277 | 0.287 | 0.279 | 0.215 | 0.328 | 0.276 | 0.299 | 0.344 | 0.362 | 0.238 | 0.316 | 0.330 | 0.302 | 0.319 | 0.359 | 0.236 | 0.284 | -0.078 | 0.3420 | 0.5981 | 0 | 0 | 0 |
| 'Tns2' | 1.050 | 1.117 | 1.137 | 0.803 | 0.981 | 1.118 | 0.892 | 1.046 | 1.265 | 1.232 | 1.193 | 1.115 | 1.150 | 1.223 | 1.009 | 1.402 | 1.327 | 0.937 | -0.161 | 0.0979 | 0.2920 | 0 | 0 | 0 |
| 'Tob1' | 29.665 | 15.547 | 10.303 | 25.282 | 16.815 | 11.340 | 26.491 | 17.203 | 9.541 | 24.869 | 22.784 | 10.793 | 22.914 | 22.517 | 11.383 | 19.963 | 23.517 | 10.672 | -0.101 | 0.2296 | 0.4766 | 0 | 0 | 0 |
| 'Tob2' | 55.012 | 30.761 | 26.224 | 44.077 | 28.039 | 28.556 | 39.447 | 30.224 | 26.734 | 37.720 | 40.824 | 32.937 | 36.536 | 39.981 | 33.828 | 36.770 | 40.032 | 33.250 | -0.149 | 0.1551 | 0.3835 | 0 | 0 | 0 |
| 'Toe1' | 4.070 | 4.842 | 4.507 | 4.803 | 4.914 | 4.522 | 4.447 | 4.904 | 4.814 | 4.343 | 4.435 | 4.773 | 4.821 | 4.592 | 4.491 | 4.610 | 4.205 | 4.109 | 0.034 | 0.4461 | 0.6878 | 0 | 0 | 0 |
| 'Tollip' | 17.734 | 17.987 | 17.943 | 16.290 | 18.752 | 17.579 | 16.523 | 19.834 | 17.841 | 19.040 | 17.756 | 16.593 | 18.791 | 19.241 | 17.164 | 18.575 | 16.194 | 16.414 | -0.001 | 0.9827 | 0.9934 | 0 | 0 | 0 |
| 'Tom1' | 9.665 | 10.015 | 9.569 | 9.863 | 9.158 | 10.309 | 8.937 | 8.917 | 9.936 | 10.853 | 10.284 | 12.080 | 10.970 | 10.471 | 11.572 | 11.274 | 10.198 | 10.590 | -0.201 | 0.0000 | 0.0002 | 1 | 0 | 0 |
| 'Tom1l2' | 3.285 | 3.128 | 3.383 | 2.899 | 2.946 | 3.304 | 3.495 | 3.401 | 3.925 | 3.352 | 3.629 | 3.965 | 3.747 | 3.181 | 3.537 | 2.968 | 3.518 | 3.821 | -0.103 | 0.0436 | 0.1729 | 0 | 0 | 0 |
| 'Tomm20' | 24.744 | 23.364 | 20.522 | 22.799 | 22.501 | 20.370 | 23.272 | 22.809 | 19.738 | 21.601 | 22.056 | 20.810 | 21.968 | 22.735 | 21.795 | 23.437 | 22.879 | 19.334 | 0.014 | 0.6582 | 0.8350 | 0 | 0 | 0 |
| 'Tomm22' | 34.495 | 41.266 | 36.747 | 36.476 | 41.042 | 42.336 | 38.159 | 41.545 | 37.217 | 36.130 | 36.907 | 38.206 | 37.496 | 40.539 | 40.107 | 37.612 | 38.124 | 38.884 | 0.009 | 0.7959 | 0.9092 | 0 | 0 | 0 |
| 'Tomm34' | 20.186 | 19.901 | 22.366 | 19.792 | 21.377 | 20.920 | 20.625 | 20.902 | 21.500 | 21.415 | 20.496 | 23.417 | 19.560 | 19.004 | 21.813 | 19.729 | 20.184 | 22.525 | -0.016 | 0.6058 | 0.8014 | 0 | 0 | 0 |
| 'Tomm40' | 11.822 | 11.697 | 12.138 | 10.938 | 12.154 | 12.631 | 11.060 | 11.659 | 11.888 | 10.928 | 10.588 | 12.908 | 12.451 | 10.606 | 12.660 | 12.322 | 10.358 | 12.675 | -0.003 | 0.9329 | 0.9746 | 0 | 0 | 0 |
| 'Tomm40l' | 1.595 | 1.696 | 1.755 | 1.385 | 1.675 | 1.868 | 1.597 | 1.555 | 1.764 | 1.579 | 1.367 | 1.814 | 1.869 | 1.496 | 1.602 | 1.885 | 1.695 | 1.517 | -0.005 | 0.9409 | 0.9772 | 0 | 0 | 0 |
| 'Tomm6' | 2.592 | 2.888 | 1.872 | 2.773 | 2.326 | 2.045 | 2.586 | 2.103 | 2.394 | 2.243 | 3.002 | 2.576 | 2.846 | 2.770 | 2.014 | 3.055 | 2.688 | 2.106 | -0.132 | 0.1669 | 0.3994 | 0 | 0 | 0 |
| 'Tomm7' | 42.937 | 50.978 | 46.133 | 52.596 | 53.119 | 42.959 | 50.951 | 51.087 | 44.051 | 41.665 | 45.489 | 41.781 | 48.701 | 44.941 | 47.640 | 45.353 | 46.652 | 41.933 | 0.090 | 0.0425 | 0.1700 | 0 | 0 | 0 |
| 'Tomm70a' | 33.533 | 33.065 | 33.598 | 34.117 | 33.770 | 32.837 | 35.707 | 33.406 | 33.310 | 31.987 | 34.371 | 35.897 | 32.135 | 34.943 | 32.595 | 32.917 | 35.430 | 34.971 | -0.021 | 0.4867 | 0.7174 | 0 | 0 | 0 |
| 'Tonsl' | 16.773 | 17.264 | 18.693 | 16.289 | 17.148 | 19.076 | 17.173 | 17.870 | 18.509 | 17.476 | 16.616 | 17.947 | 18.967 | 17.827 | 17.865 | 17.403 | 17.342 | 17.084 | -0.008 | 0.7783 | 0.9002 | 0 | 0 | 0 |
| 'Top1' | 25.068 | 23.366 | 23.199 | 27.933 | 23.825 | 21.893 | 26.472 | 22.754 | 23.260 | 22.700 | 23.852 | 24.395 | 22.588 | 23.890 | 22.869 | 23.026 | 26.013 | 24.920 | 0.010 | 0.8273 | 0.9247 | 0 | 0 | 0 |
| 'Top1mt' | 9.446 | 11.791 | 11.446 | 11.099 | 10.668 | 10.957 | 11.637 | 11.417 | 12.113 | 11.962 | 10.219 | 10.474 | 11.184 | 10.117 | 10.233 | 10.185 | 9.559 | 9.252 | 0.100 | 0.0742 | 0.2422 | 0 | 0 | 0 |
| 'Top2a' | 18.223 | 19.077 | 24.182 | 22.661 | 19.809 | 23.321 | 20.852 | 19.365 | 23.711 | 14.754 | 21.996 | 22.896 | 23.102 | 25.264 | 25.668 | 20.884 | 21.997 | 23.021 | -0.072 | 0.2939 | 0.5499 | 0 | 0 | 0 |
| 'Top2b' | 52.370 | 55.062 | 51.432 | 56.431 | 53.306 | 49.641 | 59.103 | 52.735 | 49.725 | 53.031 | 56.821 | 53.396 | 54.824 | 56.834 | 49.854 | 54.935 | 59.002 | 52.062 | -0.044 | 0.1233 | 0.3353 | 0 | 0 | 0 |
| 'Top3a' | 4.504 | 3.959 | 4.643 | 4.701 | 4.412 | 4.391 | 4.925 | 4.351 | 4.936 | 4.762 | 4.646 | 4.722 | 4.669 | 4.307 | 4.394 | 4.410 | 4.662 | 4.758 | -0.029 | 0.4928 | 0.7220 | 0 | 0 | 0 |
| 'Top3b' | 4.827 | 5.136 | 5.770 | 4.565 | 5.171 | 5.444 | 4.809 | 5.392 | 5.713 | 5.191 | 5.006 | 4.916 | 5.573 | 4.958 | 4.836 | 4.914 | 5.093 | 4.430 | 0.050 | 0.2754 | 0.5313 | 0 | 0 | 0 |
| 'Topbp1' | 12.238 | 11.718 | 13.713 | 12.972 | 11.829 | 13.451 | 12.365 | 12.212 | 13.382 | 12.561 | 12.725 | 12.559 | 14.093 | 12.468 | 12.780 | 12.308 | 12.096 | 13.050 | -0.020 | 0.5177 | 0.7407 | 0 | 0 | 0 |
| 'Topors' | 28.407 | 22.503 | 18.379 | 23.591 | 21.061 | 19.136 | 24.921 | 21.043 | 18.911 | 22.705 | 23.440 | 20.663 | 23.266 | 22.983 | 19.545 | 24.716 | 22.720 | 19.645 | -0.030 | 0.4980 | 0.7259 | 0 | 0 | 0 |
| 'Tor1a' | 9.794 | 10.339 | 9.820 | 10.119 | 10.084 | 10.058 | 8.600 | 10.184 | 9.339 | 9.348 | 8.871 | 10.457 | 9.936 | 9.652 | 10.333 | 9.692 | 9.338 | 9.891 | 0.000 | 0.9902 | 0.9973 | 0 | 0 | 0 |
| 'Tor1aip1' | 9.277 | 9.138 | 10.407 | 9.246 | 9.013 | 9.976 | 9.161 | 9.853 | 9.500 | 9.914 | 10.012 | 10.673 | 10.232 | 10.094 | 9.928 | 10.277 | 9.978 | 10.467 | -0.107 | 0.0000 | 0.0006 | 1 | 0 | 0 |
| 'Tor1aip2' | 6.083 | 6.118 | 5.717 | 5.894 | 5.972 | 5.613 | 6.517 | 6.000 | 5.549 | 6.619 | 7.578 | 7.615 | 6.631 | 6.990 | 7.248 | 7.304 | 7.463 | 7.401 | -0.292 | 0.0000 | 0.0000 | 1 | 0 | 0 |
| 'Tor1b' | 6.951 | 7.089 | 7.705 | 6.594 | 7.374 | 7.617 | 6.726 | 7.322 | 7.802 | 7.518 | 7.576 | 8.111 | 7.091 | 7.391 | 7.201 | 7.267 | 6.883 | 7.649 | -0.044 | 0.1457 | 0.3705 | 0 | 0 | 0 |
| 'Tor2a' | 11.253 | 12.644 | 12.537 | 10.510 | 13.108 | 13.169 | 10.904 | 12.710 | 12.621 | 12.099 | 11.564 | 12.456 | 12.482 | 11.646 | 12.678 | 12.601 | 11.490 | 11.603 | -0.001 | 0.9750 | 0.9916 | 0 | 0 | 0 |
| 'Tor3a' | 4.963 | 5.593 | 5.062 | 4.639 | 5.114 | 5.342 | 4.923 | 5.006 | 5.707 | 5.720 | 5.994 | 6.908 | 6.075 | 5.796 | 6.768 | 6.161 | 6.111 | 6.753 | -0.295 | 0.0000 | 0.0000 | 1 | 0 | 0 |
| 'Tor4a' | 24.786 | 28.422 | 32.946 | 26.544 | 27.967 | 32.076 | 29.909 | 26.030 | 31.154 | 24.520 | 26.244 | 32.807 | 24.096 | 23.926 | 30.731 | 23.285 | 26.822 | 31.114 | 0.085 | 0.0237 | 0.1161 | 0 | 0 | 0 |
| 'Tox' | 36.763 | 40.600 | 36.604 | 39.290 | 40.546 | 38.011 | 41.068 | 38.930 | 39.609 | 40.981 | 43.132 | 36.387 | 38.210 | 40.306 | 35.416 | 37.819 | 42.171 | 35.730 | -0.003 | 0.9236 | 0.9708 | 0 | 0 | 0 |
| 'Tox2' | 112.444 | 125.568 | 116.624 | 110.927 | 124.139 | 118.530 | 108.843 | 120.618 | 124.684 | 133.200 | 118.666 | 106.900 | 139.007 | 115.941 | 112.478 | 122.974 | 109.831 | 113.362 | -0.022 | 0.6369 | 0.8209 | 0 | 0 | 0 |
| 'Tox4' | 13.978 | 13.444 | 14.112 | 14.447 | 14.046 | 13.839 | 14.457 | 13.603 | 13.940 | 14.960 | 14.307 | 14.975 | 14.471 | 14.277 | 14.656 | 13.598 | 14.628 | 14.453 | -0.062 | 0.0097 | 0.0610 | 0 | 0 | 0 |
| 'Tpcn1' | 10.497 | 10.830 | 10.604 | 10.785 | 10.738 | 10.923 | 9.565 | 10.722 | 11.574 | 12.005 | 11.377 | 12.295 | 12.154 | 10.579 | 11.695 | 11.843 | 10.214 | 12.389 | -0.129 | 0.0003 | 0.0037 | 1 | 0 | 0 |
| 'Tpcn2' | 8.711 | 8.307 | 9.762 | 7.564 | 7.744 | 10.356 | 9.905 | 10.472 | 10.386 | 8.959 | 9.175 | 9.558 | 9.530 | 8.437 | 9.202 | 8.901 | 9.320 | 8.997 | 0.012 | 0.8412 | 0.9326 | 0 | 0 | 0 |
| 'Tpd52' | 2.244 | 2.157 | 1.979 | 1.970 | 2.465 | 2.104 | 1.968 | 2.359 | 2.044 | 1.656 | 1.993 | 1.633 | 1.914 | 2.445 | 1.808 | 1.718 | 1.890 | 1.795 | 0.195 | 0.0291 | 0.1322 | 0 | 0 | 0 |
| 'Tpd52l2' | 3.483 | 3.606 | 3.148 | 3.450 | 3.333 | 3.516 | 3.438 | 3.478 | 3.227 | 3.171 | 3.351 | 3.709 | 3.278 | 3.419 | 3.662 | 3.415 | 3.480 | 3.649 | -0.036 | 0.3677 | 0.6197 | 0 | 0 | 0 |
| 'Tpgs1' | 38.448 | 36.056 | 43.968 | 35.963 | 37.923 | 44.523 | 35.845 | 34.620 | 40.219 | 37.006 | 33.752 | 40.607 | 39.751 | 32.037 | 38.557 | 37.087 | 30.893 | 37.960 | 0.076 | 0.0593 | 0.2104 | 0 | 0 | 0 |
| 'Tpgs2' | 2.582 | 2.922 | 2.788 | 2.814 | 3.266 | 2.770 | 2.482 | 2.839 | 2.626 | 2.979 | 2.910 | 2.725 | 2.842 | 2.842 | 2.951 | 2.755 | 2.844 | 2.684 | -0.036 | 0.4764 | 0.7107 | 0 | 0 | 0 |
| 'Tpi1' | 63.271 | 56.883 | 52.361 | 67.767 | 60.191 | 51.594 | 65.060 | 53.862 | 49.443 | 50.976 | 66.133 | 53.986 | 67.871 | 68.328 | 63.103 | 59.443 | 69.833 | 56.498 | -0.110 | 0.0656 | 0.2234 | 0 | 0 | 0 |
| 'Tpk1' | 0.797 | 0.875 | 0.610 | 0.729 | 0.687 | 0.610 | 0.768 | 0.838 | 0.750 | 0.781 | 0.594 | 0.725 | 0.609 | 0.654 | 0.537 | 0.552 | 0.542 | 0.600 | 0.221 | 0.0246 | 0.1191 | 0 | 0 | 0 |
| 'Tpm1' | 2.510 | 3.100 | 3.042 | 2.534 | 2.992 | 3.195 | 2.701 | 3.123 | 3.107 | 3.018 | 2.767 | 2.857 | 3.048 | 3.093 | 3.017 | 3.031 | 2.819 | 2.932 | -0.026 | 0.5460 | 0.7608 | 0 | 0 | 0 |
| 'Tpm3' | 93.170 | 98.630 | 99.191 | 97.879 | 100.382 | 100.529 | 101.613 | 98.308 | 99.955 | 95.873 | 96.715 | 88.342 | 98.694 | 96.847 | 96.393 | 94.895 | 96.049 | 95.248 | 0.040 | 0.0931 | 0.2826 | 0 | 0 | 0 |
| 'Tpm4' | 58.663 | 72.817 | 68.279 | 65.117 | 72.230 | 68.910 | 67.371 | 64.823 | 69.794 | 63.399 | 74.179 | 80.116 | 70.651 | 80.566 | 85.716 | 69.813 | 80.638 | 82.464 | -0.186 | 0.0000 | 0.0000 | 1 | 0 | 0 |
| 'Tpp1' | 26.438 | 27.865 | 28.963 | 26.561 | 28.478 | 29.049 | 26.472 | 30.907 | 28.888 | 31.032 | 30.307 | 29.837 | 29.550 | 29.508 | 28.903 | 29.600 | 27.069 | 27.347 | -0.062 | 0.0635 | 0.2196 | 0 | 0 | 0 |
| 'Tpp2' | 9.970 | 10.154 | 9.868 | 10.598 | 10.370 | 9.514 | 10.544 | 10.802 | 9.366 | 9.694 | 10.559 | 10.339 | 10.271 | 10.367 | 9.621 | 9.616 | 11.150 | 9.619 | -0.013 | 0.6431 | 0.8251 | 0 | 0 | 0 |
| 'Tpr' | 27.024 | 23.918 | 24.686 | 27.233 | 24.461 | 25.030 | 25.824 | 24.017 | 24.690 | 27.153 | 28.245 | 26.633 | 24.733 | 26.709 | 26.171 | 26.087 | 25.834 | 27.162 | -0.085 | 0.0124 | 0.0730 | 0 | 0 | 0 |
| 'Tpra1' | 5.238 | 5.137 | 5.720 | 4.945 | 4.851 | 5.252 | 5.208 | 4.616 | 5.431 | 5.258 | 5.060 | 6.556 | 5.186 | 4.969 | 5.760 | 5.049 | 5.244 | 6.210 | -0.100 | 0.0085 | 0.0559 | 0 | 0 | 0 |
| 'Tprgl' | 31.087 | 35.073 | 30.560 | 30.683 | 35.781 | 35.401 | 29.966 | 33.680 | 33.262 | 30.848 | 31.053 | 32.867 | 31.439 | 32.587 | 33.288 | 33.473 | 32.130 | 33.117 | 0.010 | 0.7469 | 0.8858 | 0 | 0 | 0 |
| 'Tprkb' | 2.656 | 3.196 | 2.948 | 3.087 | 3.141 | 3.101 | 3.378 | 2.996 | 3.337 | 2.695 | 2.967 | 2.783 | 2.949 | 3.045 | 2.643 | 2.498 | 2.980 | 2.547 | 0.137 | 0.0060 | 0.0433 | 1 | 0 | 0 |
| 'Tprn' | 6.750 | 5.880 | 6.743 | 6.306 | 6.075 | 6.619 | 5.169 | 6.047 | 6.433 | 7.133 | 5.723 | 6.681 | 6.884 | 6.040 | 7.146 | 6.488 | 4.858 | 7.126 | -0.056 | 0.3234 | 0.5808 | 0 | 0 | 0 |
| 'Tpst1' | 2.414 | 2.460 | 2.087 | 2.075 | 2.571 | 1.655 | 2.302 | 2.515 | 2.284 | 1.775 | 2.282 | 1.895 | 2.301 | 1.840 | 1.938 | 2.059 | 2.149 | 1.528 | 0.181 | 0.0276 | 0.1281 | 0 | 0 | 0 |
| 'Tpst2' | 9.883 | 8.767 | 9.836 | 7.908 | 8.632 | 10.177 | 8.712 | 9.820 | 9.603 | 8.891 | 7.692 | 10.081 | 8.652 | 8.254 | 10.160 | 8.964 | 7.516 | 9.729 | 0.054 | 0.2432 | 0.4935 | 0 | 0 | 0 |
| 'Tpt1' | 278.028 | 298.194 | 254.286 | 305.082 | 299.435 | 259.196 | 311.968 | 275.501 | 259.063 | 280.614 | 306.217 | 216.001 | 279.150 | 294.972 | 267.377 | 304.231 | 305.105 | 257.500 | 0.008 | 0.8469 | 0.9343 | 0 | 0 | 0 |
| 'Tpx2' | 6.757 | 7.307 | 9.209 | 8.623 | 7.535 | 8.736 | 8.573 | 7.139 | 9.290 | 5.568 | 7.262 | 8.787 | 9.143 | 8.999 | 10.203 | 6.819 | 7.941 | 8.719 | -0.015 | 0.8425 | 0.9333 | 0 | 0 | 0 |
| 'Tra2a' | 15.618 | 14.415 | 12.523 | 13.994 | 13.794 | 12.029 | 12.648 | 13.494 | 12.563 | 13.776 | 14.704 | 14.093 | 13.722 | 16.150 | 13.478 | 15.487 | 15.144 | 14.308 | -0.124 | 0.0036 | 0.0292 | 1 | 0 | 0 |
| 'Tra2b' | 33.897 | 28.242 | 26.268 | 31.451 | 27.568 | 24.675 | 29.692 | 28.326 | 25.095 | 28.047 | 30.294 | 26.879 | 27.034 | 31.503 | 26.801 | 28.271 | 29.131 | 28.393 | -0.021 | 0.6473 | 0.8281 | 0 | 0 | 0 |
| 'Trabd' | 10.444 | 11.346 | 11.049 | 10.193 | 10.771 | 11.539 | 9.904 | 10.663 | 11.307 | 10.197 | 9.487 | 11.704 | 10.107 | 10.359 | 11.556 | 10.624 | 10.115 | 11.579 | 0.011 | 0.6992 | 0.8592 | 0 | 0 | 0 |
| 'Tradd' | 19.524 | 19.097 | 21.694 | 18.848 | 21.287 | 21.959 | 20.372 | 20.711 | 22.291 | 20.486 | 18.295 | 21.516 | 20.786 | 20.324 | 21.058 | 19.794 | 19.095 | 19.135 | 0.031 | 0.3551 | 0.6084 | 0 | 0 | 0 |
| 'Traf1' | 11.348 | 10.640 | 12.572 | 10.827 | 10.638 | 13.150 | 10.847 | 10.420 | 12.724 | 12.434 | 11.138 | 14.773 | 12.534 | 11.823 | 13.375 | 12.758 | 12.582 | 13.385 | -0.166 | 0.0000 | 0.0000 | 1 | 0 | 0 |
| 'Traf2' | 14.414 | 13.461 | 13.714 | 13.777 | 12.946 | 13.886 | 13.963 | 12.930 | 14.184 | 13.575 | 12.102 | 14.466 | 12.412 | 12.205 | 14.142 | 13.442 | 12.821 | 13.586 | 0.043 | 0.1462 | 0.3714 | 0 | 0 | 0 |
| 'Traf3' | 8.725 | 9.441 | 8.848 | 8.852 | 8.618 | 8.841 | 8.779 | 8.701 | 9.344 | 9.075 | 9.400 | 10.363 | 8.837 | 8.894 | 9.711 | 9.611 | 8.664 | 10.428 | -0.095 | 0.0037 | 0.0295 | 1 | 0 | 0 |
| 'Traf3ip1' | 1.798 | 1.970 | 2.318 | 2.026 | 2.008 | 2.388 | 2.121 | 2.300 | 2.604 | 2.068 | 1.894 | 2.126 | 1.825 | 1.745 | 2.207 | 1.704 | 1.908 | 2.197 | 0.135 | 0.0257 | 0.1230 | 0 | 0 | 0 |
| 'Traf3ip2' | 20.369 | 20.712 | 19.283 | 19.717 | 19.079 | 19.603 | 19.420 | 20.321 | 19.765 | 20.230 | 19.505 | 19.843 | 22.192 | 19.328 | 19.009 | 18.890 | 19.641 | 18.321 | 0.000 | 0.9904 | 0.9974 | 0 | 0 | 0 |
| 'Traf3ip3' | 22.064 | 22.071 | 25.191 | 22.264 | 21.596 | 25.812 | 22.807 | 21.657 | 24.787 | 21.736 | 20.589 | 27.224 | 22.816 | 22.307 | 25.730 | 22.360 | 22.823 | 25.588 | -0.031 | 0.1490 | 0.3758 | 0 | 0 | 0 |
| 'Traf4' | 5.710 | 4.150 | 4.334 | 4.701 | 3.747 | 4.676 | 4.601 | 4.430 | 4.210 | 4.223 | 3.863 | 5.418 | 4.330 | 4.058 | 4.844 | 4.486 | 3.884 | 5.379 | -0.007 | 0.9165 | 0.9681 | 0 | 0 | 0 |
| 'Traf5' | 17.508 | 17.077 | 16.626 | 16.526 | 15.444 | 16.413 | 16.701 | 16.005 | 16.609 | 18.130 | 17.469 | 18.835 | 16.315 | 18.059 | 17.997 | 17.149 | 17.481 | 18.561 | -0.115 | 0.0004 | 0.0049 | 1 | 0 | 0 |
| 'Traf6' | 6.906 | 6.807 | 6.520 | 5.753 | 6.591 | 6.277 | 5.653 | 6.608 | 5.944 | 7.014 | 6.742 | 6.391 | 6.527 | 7.263 | 6.682 | 7.289 | 6.225 | 6.467 | -0.094 | 0.0330 | 0.1438 | 0 | 0 | 0 |
| 'Traf7' | 8.876 | 8.423 | 9.470 | 8.856 | 8.308 | 9.632 | 8.395 | 8.855 | 9.703 | 8.989 | 8.401 | 10.105 | 9.710 | 9.074 | 9.616 | 8.589 | 8.967 | 9.684 | -0.057 | 0.0281 | 0.1296 | 0 | 0 | 0 |
| 'Trafd1' | 9.780 | 9.784 | 10.573 | 9.391 | 9.683 | 11.251 | 10.052 | 10.212 | 11.000 | 10.397 | 9.836 | 11.646 | 10.452 | 9.631 | 11.636 | 10.913 | 10.231 | 11.055 | -0.072 | 0.0068 | 0.0472 | 1 | 0 | 0 |
| 'Traip' | 0.996 | 0.979 | 1.332 | 1.102 | 1.125 | 1.254 | 1.071 | 0.878 | 1.380 | 0.989 | 0.983 | 1.126 | 1.368 | 1.086 | 1.239 | 1.132 | 1.239 | 1.017 | -0.021 | 0.7980 | 0.9103 | 0 | 0 | 0 |
| 'Traj48' | 129.500 | 130.750 | 124.200 | 163.133 | 137.500 | 139.200 | 144.317 | 116.817 | 123.533 | 131.317 | 133.850 | 109.750 | 138.250 | 100.433 | 113.933 | 107.250 | 103.383 | 125.383 | 0.170 | 0.0405 | 0.1647 | 0 | 0 | 0 |
| 'Traj49' | 175.932 | 199.458 | 177.525 | 177.136 | 207.932 | 162.068 | 205.220 | 189.153 | 186.305 | 190.627 | 174.814 | 189.017 | 203.492 | 181.847 | 171.356 | 180.492 | 180.746 | 192.153 | -0.010 | 0.8774 | 0.9483 | 0 | 0 | 0 |
| 'Traj57' | 180.215 | 147.246 | 152.323 | 143.800 | 174.385 | 177.246 | 119.077 | 170.031 | 145.908 | 188.569 | 170.785 | 148.662 | 157.831 | 148.108 | 174.985 | 149.677 | 152.262 | 160.738 | -0.043 | 0.5995 | 0.7965 | 0 | 0 | 0 |
| 'Trak1' | 18.355 | 18.116 | 18.263 | 17.729 | 18.652 | 19.528 | 17.064 | 19.260 | 19.153 | 19.418 | 19.047 | 18.133 | 21.755 | 18.706 | 18.065 | 18.792 | 20.117 | 18.616 | -0.064 | 0.0801 | 0.2549 | 0 | 0 | 0 |
| 'Trak2' | 11.668 | 11.394 | 11.274 | 10.268 | 11.655 | 10.812 | 10.212 | 11.718 | 10.832 | 10.727 | 10.803 | 10.549 | 10.177 | 10.527 | 10.291 | 9.679 | 9.823 | 10.689 | 0.087 | 0.0098 | 0.0615 | 0 | 0 | 0 |
| 'Tram1' | 42.577 | 48.129 | 46.788 | 42.346 | 49.346 | 45.722 | 46.662 | 50.298 | 45.278 | 45.390 | 45.519 | 46.631 | 41.541 | 48.157 | 45.190 | 44.370 | 46.272 | 46.222 | 0.015 | 0.5440 | 0.7606 | 0 | 0 | 0 |
| 'Tram2' | 4.099 | 4.152 | 3.785 | 4.170 | 3.876 | 4.096 | 4.183 | 4.051 | 4.414 | 4.340 | 4.997 | 4.895 | 4.714 | 4.032 | 4.947 | 4.405 | 4.734 | 4.927 | -0.203 | 0.0000 | 0.0003 | 1 | 0 | 0 |
| 'Trap1' | 26.892 | 24.495 | 29.484 | 27.254 | 28.426 | 28.824 | 27.817 | 27.289 | 29.638 | 27.425 | 27.219 | 30.004 | 28.178 | 26.021 | 26.550 | 25.122 | 27.489 | 27.591 | 0.015 | 0.6515 | 0.8307 | 0 | 0 | 0 |
| 'Trappc1' | 18.329 | 21.908 | 19.920 | 22.530 | 22.318 | 22.250 | 20.293 | 22.033 | 21.478 | 19.342 | 18.772 | 19.491 | 21.133 | 21.325 | 19.701 | 20.897 | 21.857 | 17.657 | 0.074 | 0.0959 | 0.2881 | 0 | 0 | 0 |
| 'Trappc10' | 33.819 | 34.223 | 33.497 | 33.707 | 33.283 | 34.130 | 31.287 | 33.344 | 34.105 | 36.234 | 33.938 | 32.657 | 36.161 | 35.637 | 34.161 | 34.906 | 32.101 | 34.017 | -0.048 | 0.0932 | 0.2826 | 0 | 0 | 0 |
| 'Trappc11' | 7.233 | 7.701 | 7.730 | 7.576 | 7.441 | 7.592 | 7.282 | 7.520 | 7.500 | 7.096 | 7.913 | 7.774 | 7.457 | 7.519 | 7.566 | 7.538 | 7.814 | 7.327 | -0.019 | 0.4481 | 0.6883 | 0 | 0 | 0 |
| 'Trappc12' | 16.612 | 16.497 | 17.192 | 16.108 | 18.085 | 17.225 | 17.217 | 16.360 | 17.337 | 18.119 | 16.756 | 16.953 | 17.421 | 16.476 | 15.776 | 17.461 | 15.546 | 14.966 | 0.022 | 0.5322 | 0.7526 | 0 | 0 | 0 |
| 'Trappc13' | 4.398 | 4.819 | 4.719 | 4.725 | 4.331 | 3.979 | 4.839 | 3.992 | 4.302 | 3.628 | 4.370 | 5.030 | 4.082 | 4.760 | 4.664 | 4.406 | 4.894 | 4.791 | -0.030 | 0.6178 | 0.8087 | 0 | 0 | 0 |
| 'Trappc2' | 1.338 | 1.751 | 1.646 | 1.573 | 1.873 | 1.253 | 1.377 | 1.714 | 1.331 | 1.299 | 1.355 | 1.289 | 1.301 | 1.738 | 1.315 | 1.319 | 1.437 | 1.097 | 0.176 | 0.0278 | 0.1287 | 0 | 0 | 0 |
| 'Trappc2l' | 50.918 | 59.068 | 57.002 | 54.440 | 58.502 | 59.501 | 52.140 | 56.578 | 58.878 | 61.133 | 56.680 | 50.377 | 59.606 | 53.631 | 51.766 | 53.766 | 57.756 | 50.865 | 0.027 | 0.5656 | 0.7741 | 0 | 0 | 0 |
| 'Trappc3' | 34.127 | 38.262 | 37.253 | 34.118 | 37.860 | 38.235 | 33.381 | 38.388 | 38.086 | 35.021 | 35.017 | 35.685 | 37.144 | 35.804 | 35.414 | 37.252 | 36.412 | 35.433 | 0.020 | 0.5219 | 0.7444 | 0 | 0 | 0 |
| 'Trappc4' | 37.150 | 42.592 | 41.026 | 40.648 | 42.022 | 39.176 | 35.621 | 44.204 | 38.246 | 37.138 | 36.260 | 39.987 | 36.739 | 42.667 | 39.955 | 39.488 | 39.511 | 35.575 | 0.043 | 0.2744 | 0.5301 | 0 | 0 | 0 |
| 'Trappc5' | 16.718 | 18.489 | 16.496 | 15.812 | 16.174 | 18.055 | 14.387 | 16.390 | 16.677 | 14.919 | 15.555 | 17.768 | 15.690 | 17.578 | 17.847 | 16.098 | 14.509 | 17.708 | 0.003 | 0.9395 | 0.9766 | 0 | 0 | 0 |
| 'Trappc6a' | 9.126 | 9.916 | 8.881 | 7.802 | 8.892 | 9.488 | 8.436 | 8.954 | 8.246 | 8.073 | 7.875 | 9.018 | 8.811 | 7.910 | 8.728 | 7.871 | 7.635 | 7.484 | 0.105 | 0.0348 | 0.1492 | 0 | 0 | 0 |
| 'Trappc6b' | 42.404 | 47.736 | 44.172 | 42.256 | 46.339 | 42.674 | 45.099 | 48.325 | 42.925 | 41.997 | 48.469 | 42.815 | 44.127 | 46.530 | 44.343 | 43.132 | 47.355 | 40.911 | -0.001 | 0.9801 | 0.9926 | 0 | 0 | 0 |
| 'Trappc8' | 17.640 | 15.843 | 16.887 | 18.036 | 16.793 | 15.941 | 18.458 | 16.732 | 15.880 | 16.537 | 17.209 | 17.298 | 15.410 | 16.870 | 15.953 | 16.263 | 18.241 | 16.474 | 0.007 | 0.8595 | 0.9390 | 0 | 0 | 0 |
| 'Trappc9' | 14.524 | 14.107 | 14.533 | 13.938 | 14.274 | 14.889 | 14.597 | 13.292 | 13.769 | 15.567 | 13.390 | 14.691 | 14.517 | 13.559 | 13.874 | 14.797 | 12.576 | 13.651 | 0.005 | 0.8670 | 0.9425 | 0 | 0 | 0 |
| 'Trat1' | 11.784 | 12.617 | 16.379 | 13.557 | 13.215 | 18.358 | 14.341 | 12.297 | 19.560 | 14.276 | 11.050 | 21.836 | 9.813 | 10.421 | 21.415 | 16.352 | 10.291 | 19.477 | -0.022 | 0.8043 | 0.9122 | 0 | 0 | 0 |
| 'Trbj1-3' | 657.204 | 780.185 | 755.574 | 692.352 | 799.759 | 676.296 | 625.815 | 777.759 | 743.259 | 763.259 | 677.963 | 672.796 | 666.981 | 685.870 | 616.981 | 617.130 | 672.463 | 635.611 | 0.103 | 0.0291 | 0.1322 | 0 | 0 | 0 |
| 'Trbj1-7' | 239.689 | 256.289 | 264.022 | 194.622 | 304.489 | 261.133 | 194.133 | 287.533 | 312.644 | 257.400 | 234.444 | 257.356 | 271.644 | 279.244 | 254.022 | 204.289 | 202.889 | 226.378 | 0.064 | 0.4510 | 0.6904 | 0 | 0 | 0 |
| 'Trdmt1' | 1.678 | 2.027 | 1.971 | 1.750 | 2.076 | 2.013 | 1.933 | 2.140 | 1.909 | 1.963 | 1.563 | 1.341 | 1.816 | 1.717 | 1.508 | 1.726 | 1.671 | 1.210 | 0.277 | 0.0008 | 0.0085 | 1 | 0 | 0 |
| 'Treml2' | 2.082 | 1.704 | 2.483 | 2.283 | 2.433 | 2.936 | 2.344 | 2.928 | 2.520 | 1.955 | 2.303 | 3.310 | 2.322 | 2.166 | 3.412 | 2.530 | 2.023 | 3.288 | -0.098 | 0.2808 | 0.5378 | 0 | 0 | 0 |
| 'Triap1' | 25.156 | 27.690 | 23.803 | 28.410 | 25.320 | 25.572 | 24.300 | 27.750 | 22.573 | 23.150 | 23.518 | 24.325 | 23.273 | 25.396 | 25.607 | 22.673 | 23.894 | 24.008 | 0.079 | 0.0988 | 0.2937 | 0 | 0 | 0 |
| 'Trib1' | 37.326 | 29.178 | 16.175 | 22.373 | 30.402 | 18.812 | 18.616 | 32.308 | 15.286 | 48.394 | 27.376 | 15.259 | 36.479 | 40.191 | 15.194 | 55.599 | 23.788 | 15.356 | -0.242 | 0.1654 | 0.3978 | 0 | 0 | 0 |
| 'Trib2' | 51.001 | 58.110 | 57.857 | 51.927 | 57.644 | 59.441 | 49.513 | 59.980 | 56.270 | 48.323 | 48.058 | 53.998 | 44.044 | 48.375 | 54.854 | 50.111 | 41.238 | 53.378 | 0.171 | 0.0000 | 0.0006 | 1 | 0 | 0 |
| 'Trim11' | 16.070 | 15.102 | 17.620 | 15.623 | 15.071 | 16.874 | 15.709 | 14.971 | 17.700 | 14.121 | 16.854 | 20.041 | 16.616 | 16.387 | 18.769 | 13.579 | 17.420 | 19.087 | -0.086 | 0.0514 | 0.1917 | 0 | 0 | 0 |
| 'Trim14' | 5.770 | 6.423 | 6.593 | 4.738 | 5.987 | 7.472 | 5.908 | 6.942 | 6.729 | 5.717 | 5.869 | 7.893 | 5.916 | 5.992 | 7.533 | 6.540 | 5.916 | 7.358 | -0.064 | 0.2087 | 0.4535 | 0 | 0 | 0 |
| 'Trim2' | 0.276 | 0.379 | 0.313 | 0.253 | 0.388 | 0.321 | 0.263 | 0.343 | 0.323 | 0.386 | 0.389 | 0.321 | 0.366 | 0.459 | 0.312 | 0.375 | 0.404 | 0.289 | -0.211 | 0.0020 | 0.0182 | 1 | 0 | 0 |
| 'Trim21' | 20.476 | 22.163 | 22.915 | 21.219 | 22.590 | 23.192 | 24.019 | 23.591 | 24.740 | 21.525 | 22.556 | 27.398 | 22.925 | 22.097 | 25.867 | 23.008 | 23.952 | 27.005 | -0.091 | 0.0161 | 0.0883 | 0 | 0 | 0 |
| 'Trim23' | 4.531 | 4.856 | 4.971 | 4.631 | 5.516 | 4.671 | 5.509 | 5.333 | 4.630 | 4.689 | 5.070 | 4.553 | 4.518 | 5.074 | 4.215 | 4.577 | 5.129 | 4.124 | 0.081 | 0.0451 | 0.1760 | 0 | 0 | 0 |
| 'Trim24' | 6.087 | 6.021 | 6.483 | 5.572 | 6.203 | 6.324 | 5.874 | 5.942 | 6.479 | 6.884 | 6.909 | 6.291 | 6.625 | 6.873 | 6.178 | 6.651 | 6.962 | 5.924 | -0.114 | 0.0032 | 0.0268 | 1 | 0 | 0 |
| 'Trim25' | 13.057 | 14.645 | 13.802 | 10.575 | 13.391 | 14.020 | 11.775 | 15.346 | 14.164 | 14.203 | 15.156 | 16.839 | 13.197 | 14.589 | 17.318 | 16.183 | 13.321 | 18.794 | -0.219 | 0.0001 | 0.0022 | 1 | 0 | 0 |
| 'Trim26' | 15.188 | 15.209 | 15.366 | 14.514 | 15.128 | 16.550 | 13.343 | 15.991 | 16.582 | 15.550 | 14.181 | 15.616 | 15.214 | 15.575 | 15.469 | 15.289 | 13.740 | 14.879 | 0.014 | 0.6794 | 0.8468 | 0 | 0 | 0 |
| 'Trim27' | 17.168 | 16.932 | 17.854 | 15.250 | 16.006 | 17.506 | 16.453 | 16.738 | 16.857 | 14.988 | 14.297 | 18.596 | 15.564 | 14.912 | 18.650 | 16.663 | 15.681 | 18.505 | 0.020 | 0.5931 | 0.7929 | 0 | 0 | 0 |
| 'Trim28' | 39.096 | 34.801 | 39.426 | 36.671 | 34.839 | 37.559 | 37.347 | 34.442 | 36.342 | 36.227 | 33.982 | 38.950 | 37.420 | 34.683 | 38.690 | 35.312 | 35.410 | 40.752 | -0.014 | 0.6030 | 0.7991 | 0 | 0 | 0 |
| 'Trim3' | 1.756 | 1.536 | 1.923 | 1.752 | 1.727 | 2.065 | 1.744 | 1.911 | 2.210 | 1.612 | 1.666 | 1.852 | 1.501 | 1.594 | 1.748 | 1.834 | 1.623 | 1.658 | 0.137 | 0.0255 | 0.1221 | 0 | 0 | 0 |
| 'Trim30a' | 14.181 | 15.109 | 13.560 | 13.541 | 15.502 | 13.515 | 14.963 | 16.492 | 13.932 | 15.530 | 18.066 | 18.805 | 15.316 | 17.115 | 17.505 | 15.954 | 17.871 | 18.356 | -0.254 | 0.0000 | 0.0000 | 1 | 0 | 0 |
| 'Trim32' | 4.467 | 4.984 | 4.265 | 4.764 | 4.462 | 4.283 | 4.202 | 3.894 | 4.335 | 3.729 | 3.917 | 4.657 | 3.693 | 4.049 | 4.832 | 3.514 | 3.990 | 4.402 | 0.096 | 0.1056 | 0.3048 | 0 | 0 | 0 |
| 'Trim33' | 5.798 | 5.356 | 5.120 | 5.751 | 5.523 | 5.295 | 5.673 | 5.246 | 5.129 | 5.668 | 5.662 | 5.423 | 5.352 | 5.874 | 5.276 | 5.475 | 5.705 | 5.545 | -0.043 | 0.1776 | 0.4131 | 0 | 0 | 0 |
| 'Trim35' | 26.085 | 28.233 | 26.393 | 24.511 | 28.392 | 27.618 | 23.069 | 28.496 | 27.167 | 27.672 | 27.345 | 30.250 | 24.724 | 28.686 | 31.927 | 28.795 | 26.981 | 30.245 | -0.108 | 0.0051 | 0.0383 | 1 | 0 | 0 |
| 'Trim37' | 6.596 | 7.177 | 7.614 | 8.338 | 7.277 | 7.384 | 7.432 | 7.056 | 7.270 | 7.132 | 7.950 | 7.524 | 7.552 | 7.541 | 7.700 | 6.765 | 7.912 | 7.468 | -0.041 | 0.3213 | 0.5793 | 0 | 0 | 0 |
| 'Trim39' | 6.991 | 7.659 | 7.558 | 6.579 | 7.389 | 7.142 | 6.701 | 7.579 | 7.954 | 7.420 | 6.939 | 7.637 | 7.713 | 7.697 | 7.458 | 7.340 | 7.470 | 7.191 | -0.040 | 0.2079 | 0.4525 | 0 | 0 | 0 |
| 'Trim41' | 15.160 | 15.074 | 15.086 | 13.982 | 14.822 | 15.831 | 13.913 | 14.782 | 16.272 | 14.641 | 13.632 | 15.028 | 14.126 | 13.977 | 14.559 | 14.273 | 14.716 | 15.440 | 0.040 | 0.2023 | 0.4440 | 0 | 0 | 0 |
| 'Trim44' | 19.092 | 20.168 | 20.000 | 17.968 | 19.426 | 19.142 | 17.571 | 20.528 | 19.260 | 19.933 | 18.751 | 20.292 | 19.236 | 19.864 | 19.061 | 19.038 | 19.729 | 20.566 | -0.038 | 0.1749 | 0.4096 | 0 | 0 | 0 |
| 'Trim45' | 0.777 | 0.898 | 0.966 | 0.969 | 0.994 | 0.943 | 0.895 | 0.969 | 0.927 | 0.910 | 0.784 | 0.875 | 1.068 | 0.833 | 0.723 | 0.851 | 0.822 | 0.843 | 0.099 | 0.2174 | 0.4639 | 0 | 0 | 0 |
| 'Trim46' | 0.497 | 0.441 | 0.705 | 0.680 | 0.605 | 0.764 | 0.595 | 0.476 | 0.807 | 0.685 | 0.453 | 0.625 | 0.658 | 0.423 | 0.738 | 0.369 | 0.485 | 0.588 | 0.133 | 0.2454 | 0.4959 | 0 | 0 | 0 |
| 'Trim56' | 19.959 | 19.192 | 19.696 | 20.658 | 18.509 | 19.387 | 19.156 | 19.193 | 21.240 | 18.216 | 21.093 | 23.943 | 17.464 | 19.615 | 22.293 | 19.014 | 19.204 | 23.805 | -0.066 | 0.1510 | 0.3777 | 0 | 0 | 0 |
| 'Trim59' | 19.054 | 17.892 | 19.184 | 21.008 | 17.868 | 17.728 | 23.714 | 17.837 | 16.739 | 15.813 | 22.031 | 20.072 | 16.314 | 20.025 | 19.920 | 14.354 | 23.660 | 19.292 | -0.006 | 0.9477 | 0.9796 | 0 | 0 | 0 |
| 'Trim62' | 3.269 | 2.824 | 3.751 | 2.774 | 3.047 | 3.835 | 2.971 | 3.048 | 3.687 | 3.339 | 3.223 | 3.297 | 3.363 | 3.428 | 3.390 | 3.396 | 2.971 | 3.066 | -0.016 | 0.7823 | 0.9025 | 0 | 0 | 0 |
| 'Trim65' | 2.405 | 2.077 | 2.373 | 2.707 | 1.832 | 2.167 | 2.116 | 2.326 | 2.513 | 2.091 | 2.223 | 4.375 | 1.928 | 2.228 | 3.915 | 1.948 | 2.927 | 4.873 | -0.305 | 0.0403 | 0.1643 | 0 | 0 | 0 |
| 'Trim68' | 9.397 | 12.156 | 11.805 | 8.907 | 12.467 | 12.447 | 11.071 | 13.162 | 12.494 | 13.001 | 11.539 | 10.438 | 12.317 | 12.432 | 9.766 | 9.881 | 11.091 | 9.691 | 0.042 | 0.5599 | 0.7704 | 0 | 0 | 0 |
| 'Trim8' | 164.754 | 161.785 | 152.777 | 154.907 | 158.769 | 168.578 | 146.046 | 156.886 | 156.118 | 167.900 | 160.625 | 172.652 | 171.575 | 173.083 | 182.302 | 184.023 | 159.391 | 185.455 | -0.142 | 0.0000 | 0.0005 | 1 | 0 | 0 |
| 'Trio' | 2.961 | 2.410 | 2.743 | 2.408 | 2.540 | 2.786 | 3.167 | 2.580 | 2.738 | 2.707 | 2.432 | 2.382 | 2.588 | 2.551 | 2.249 | 2.539 | 2.541 | 2.173 | 0.131 | 0.0109 | 0.0667 | 0 | 0 | 0 |
| 'Triobp' | 7.771 | 6.864 | 8.344 | 6.851 | 7.001 | 9.178 | 7.675 | 7.057 | 8.616 | 7.573 | 6.394 | 8.685 | 8.436 | 7.400 | 8.813 | 8.097 | 6.696 | 9.062 | -0.046 | 0.1967 | 0.4356 | 0 | 0 | 0 |
| 'Trip10' | 1.595 | 1.473 | 1.524 | 1.896 | 1.379 | 1.534 | 1.807 | 1.907 | 1.344 | 1.265 | 1.066 | 1.675 | 1.359 | 1.155 | 1.957 | 1.505 | 1.826 | 1.620 | 0.088 | 0.4899 | 0.7197 | 0 | 0 | 0 |
| 'Trip11' | 11.646 | 9.752 | 10.160 | 10.827 | 10.150 | 9.886 | 11.177 | 10.247 | 10.039 | 11.374 | 11.348 | 10.230 | 10.797 | 10.265 | 9.648 | 10.881 | 10.912 | 9.876 | -0.031 | 0.3127 | 0.5712 | 0 | 0 | 0 |
| 'Trip12' | 27.181 | 26.934 | 27.485 | 29.636 | 28.155 | 26.877 | 29.148 | 26.676 | 27.188 | 28.122 | 29.090 | 27.334 | 26.013 | 27.354 | 26.619 | 25.854 | 28.974 | 27.569 | 0.003 | 0.9296 | 0.9734 | 0 | 0 | 0 |
| 'Trip13' | 2.915 | 2.923 | 4.176 | 3.721 | 3.479 | 3.968 | 3.716 | 3.140 | 3.937 | 3.106 | 3.445 | 3.260 | 3.711 | 4.063 | 3.601 | 2.899 | 3.833 | 3.352 | 0.025 | 0.7534 | 0.8901 | 0 | 0 | 0 |
| 'Trip4' | 5.229 | 5.743 | 5.928 | 5.983 | 5.535 | 5.706 | 5.768 | 6.330 | 5.682 | 6.199 | 5.732 | 5.397 | 5.842 | 6.039 | 5.225 | 5.706 | 6.010 | 5.381 | 0.004 | 0.9065 | 0.9635 | 0 | 0 | 0 |
| 'Trip6' | 2.500 | 2.185 | 2.512 | 2.543 | 2.113 | 2.437 | 2.422 | 2.267 | 2.271 | 2.922 | 2.024 | 3.378 | 2.411 | 2.558 | 3.031 | 1.518 | 2.706 | 3.302 | -0.177 | 0.0792 | 0.2532 | 0 | 0 | 0 |
| 'Trit1' | 5.727 | 5.099 | 5.541 | 5.595 | 5.075 | 5.186 | 5.826 | 5.275 | 5.294 | 5.831 | 5.045 | 6.079 | 5.577 | 4.844 | 5.106 | 5.261 | 5.635 | 4.836 | -0.005 | 0.9278 | 0.9722 | 0 | 0 | 0 |
| 'Trmo' | 1.340 | 1.312 | 1.347 | 1.522 | 1.440 | 1.349 | 1.259 | 0.918 | 1.223 | 1.085 | 1.149 | 1.427 | 1.046 | 1.542 | 1.310 | 1.305 | 1.484 | 1.271 | -0.005 | 0.9629 | 0.9869 | 0 | 0 | 0 |
| 'Trmt1' | 3.776 | 3.342 | 4.527 | 3.647 | 3.643 | 4.185 | 3.633 | 3.798 | 4.053 | 3.986 | 3.399 | 4.283 | 4.047 | 3.250 | 3.727 | 3.850 | 3.923 | 3.923 | -0.001 | 0.9780 | 0.9923 | 0 | 0 | 0 |
| 'Trmt10a' | 0.870 | 0.905 | 0.982 | 0.970 | 1.005 | 0.891 | 0.919 | 1.036 | 0.776 | 0.746 | 0.843 | 0.961 | 0.889 | 0.787 | 0.820 | 0.883 | 0.989 | 0.951 | 0.067 | 0.3686 | 0.6201 | 0 | 0 | 0 |
| 'Trmt10b' | 3.008 | 3.167 | 3.150 | 3.248 | 2.911 | 2.964 | 3.286 | 3.372 | 3.035 | 3.367 | 2.635 | 2.966 | 3.367 | 2.844 | 3.280 | 2.852 | 3.601 | 3.060 | -0.003 | 0.9622 | 0.9866 | 0 | 0 | 0 |
| 'Trmt10c' | 8.267 | 8.546 | 7.394 | 9.610 | 8.173 | 7.848 | 9.649 | 7.471 | 7.825 | 5.900 | 9.336 | 8.858 | 7.471 | 7.809 | 8.458 | 6.303 | 9.867 | 7.483 | 0.056 | 0.5134 | 0.7376 | 0 | 0 | 0 |
| 'Trmt11' | 6.170 | 6.685 | 6.980 | 6.333 | 6.873 | 5.787 | 6.334 | 7.176 | 5.751 | 6.052 | 6.441 | 6.043 | 6.084 | 6.511 | 5.454 | 5.846 | 7.209 | 5.005 | 0.080 | 0.1700 | 0.4027 | 0 | 0 | 0 |
| 'Trmt112' | 33.046 | 33.832 | 32.389 | 31.819 | 34.430 | 33.020 | 33.131 | 32.850 | 31.452 | 29.517 | 31.961 | 31.336 | 33.148 | 31.880 | 35.427 | 30.501 | 32.310 | 31.617 | 0.029 | 0.3744 | 0.6257 | 0 | 0 | 0 |
| 'Trmt12' | 2.652 | 2.828 | 3.253 | 2.742 | 3.055 | 3.275 | 3.189 | 2.909 | 3.115 | 2.200 | 2.514 | 3.113 | 2.139 | 2.472 | 2.877 | 2.132 | 2.816 | 2.895 | 0.205 | 0.0016 | 0.0155 | 1 | 0 | 0 |
| 'Trmt13' | 1.327 | 1.419 | 1.428 | 1.267 | 1.229 | 1.222 | 1.273 | 1.379 | 1.470 | 1.259 | 1.404 | 1.323 | 1.395 | 1.157 | 1.304 | 1.246 | 1.431 | 1.172 | 0.031 | 0.6090 | 0.8034 | 0 | 0 | 0 |
| 'Trmt1l' | 6.444 | 5.788 | 6.013 | 6.451 | 5.793 | 5.595 | 6.862 | 5.382 | 6.411 | 4.827 | 5.462 | 6.875 | 4.986 | 5.600 | 6.337 | 4.838 | 6.798 | 6.619 | 0.060 | 0.3976 | 0.6467 | 0 | 0 | 0 |
| 'Trmt2a' | 7.275 | 7.043 | 7.832 | 7.301 | 6.851 | 7.592 | 6.744 | 6.887 | 7.852 | 7.458 | 6.686 | 8.237 | 7.582 | 6.487 | 8.102 | 6.931 | 7.780 | 8.206 | -0.057 | 0.0955 | 0.2876 | 0 | 0 | 0 |
| 'Trmt2b' | 4.033 | 4.662 | 4.611 | 4.521 | 4.326 | 4.413 | 4.400 | 4.262 | 4.872 | 4.178 | 4.397 | 4.703 | 4.505 | 4.925 | 4.531 | 4.125 | 4.525 | 4.034 | -0.006 | 0.8886 | 0.9544 | 0 | 0 | 0 |
| 'Trmt44' | 2.065 | 1.823 | 1.876 | 1.486 | 1.694 | 1.931 | 1.503 | 1.540 | 1.627 | 1.436 | 1.739 | 2.388 | 1.507 | 2.043 | 2.374 | 1.809 | 2.004 | 2.079 | -0.165 | 0.0617 | 0.2151 | 0 | 0 | 0 |
| 'Trmt5' | 7.365 | 7.192 | 7.524 | 7.188 | 8.382 | 7.662 | 8.081 | 7.559 | 7.978 | 8.151 | 7.500 | 8.238 | 7.226 | 7.661 | 7.400 | 7.252 | 7.851 | 6.980 | -0.001 | 0.9829 | 0.9934 | 0 | 0 | 0 |
| 'Trmt6' | 5.183 | 5.851 | 5.769 | 5.103 | 5.456 | 5.731 | 5.250 | 6.073 | 5.602 | 5.895 | 5.343 | 5.137 | 5.530 | 5.142 | 4.923 | 5.727 | 5.493 | 4.554 | 0.061 | 0.1700 | 0.4027 | 0 | 0 | 0 |
| 'Trmt61a' | 3.519 | 3.767 | 4.031 | 3.068 | 4.067 | 3.765 | 3.577 | 3.847 | 3.519 | 3.975 | 3.570 | 3.929 | 3.366 | 3.725 | 3.735 | 3.755 | 2.882 | 3.954 | 0.001 | 0.9883 | 0.9965 | 0 | 0 | 0 |
| 'Trmu' | 2.915 | 2.986 | 3.430 | 2.564 | 3.265 | 3.429 | 3.310 | 3.189 | 3.408 | 2.909 | 2.328 | 3.195 | 3.263 | 2.872 | 3.176 | 2.433 | 2.984 | 2.969 | 0.110 | 0.0603 | 0.2118 | 0 | 0 | 0 |
| 'Trnau1ap' | 6.543 | 7.388 | 7.051 | 6.857 | 7.727 | 7.309 | 7.178 | 7.901 | 6.869 | 7.098 | 6.373 | 7.261 | 7.609 | 7.029 | 6.804 | 7.822 | 7.047 | 7.241 | -0.001 | 0.9817 | 0.9933 | 0 | 0 | 0 |
| 'Trnt1' | 10.808 | 11.752 | 10.505 | 11.263 | 11.765 | 10.274 | 11.675 | 12.391 | 10.257 | 10.698 | 10.716 | 10.458 | 11.077 | 10.970 | 10.088 | 10.650 | 11.191 | 10.765 | 0.044 | 0.1541 | 0.3820 | 0 | 0 | 0 |
| 'Troap' | 3.663 | 3.812 | 6.284 | 3.868 | 3.581 | 5.776 | 4.197 | 3.405 | 6.076 | 3.034 | 2.886 | 5.048 | 5.689 | 4.414 | 6.090 | 3.451 | 4.095 | 5.030 | 0.005 | 0.9584 | 0.9847 | 0 | 0 | 0 |
| 'Trove2' | 0.582 | 0.871 | 0.690 | 0.829 | 0.714 | 0.777 | 0.716 | 0.756 | 0.726 | 0.939 | 0.874 | 0.854 | 0.844 | 0.929 | 0.832 | 0.898 | 1.062 | 0.764 | -0.275 | 0.0001 | 0.0021 | 1 | 0 | 0 |
| 'Trp53' | 33.116 | 33.548 | 34.343 | 32.763 | 32.727 | 36.953 | 33.627 | 31.454 | 38.064 | 30.144 | 31.387 | 38.145 | 30.838 | 30.902 | 37.366 | 30.398 | 31.718 | 37.262 | 0.031 | 0.2899 | 0.5465 | 0 | 0 | 0 |
| 'Trp53bp1' | 6.768 | 5.938 | 7.380 | 6.917 | 6.329 | 6.634 | 6.289 | 6.169 | 7.117 | 7.395 | 6.252 | 6.929 | 6.452 | 6.324 | 6.667 | 6.498 | 6.288 | 7.019 | -0.017 | 0.6118 | 0.8055 | 0 | 0 | 0 |
| 'Trp53bp2' | 7.144 | 6.667 | 6.502 | 7.423 | 6.492 | 6.885 | 6.834 | 7.024 | 6.863 | 7.041 | 6.090 | 6.230 | 6.651 | 6.302 | 5.870 | 6.349 | 5.831 | 5.647 | 0.134 | 0.0001 | 0.0018 | 1 | 0 | 0 |
| 'Trp53i11' | 5.069 | 5.570 | 5.104 | 4.442 | 5.133 | 6.013 | 4.818 | 5.015 | 5.467 | 4.919 | 4.686 | 4.863 | 4.999 | 5.011 | 4.748 | 5.132 | 3.901 | 4.425 | 0.118 | 0.0291 | 0.1322 | 0 | 0 | 0 |
| 'Trp53i13' | 7.025 | 6.558 | 7.591 | 6.508 | 6.344 | 7.606 | 6.823 | 7.347 | 7.304 | 6.384 | 6.032 | 7.274 | 6.644 | 6.588 | 7.654 | 7.216 | 5.657 | 7.072 | 0.055 | 0.2527 | 0.5042 | 0 | 0 | 0 |
| 'Trp53inp1' | 43.909 | 49.003 | 40.664 | 45.319 | 48.206 | 39.547 | 46.801 | 46.782 | 39.870 | 46.138 | 51.995 | 46.263 | 43.084 | 53.030 | 43.861 | 46.310 | 54.831 | 45.381 | -0.118 | 0.0001 | 0.0019 | 1 | 0 | 0 |
| 'Trp53inp2' | 4.828 | 2.596 | 1.708 | 3.137 | 2.550 | 1.711 | 2.618 | 2.862 | 1.721 | 4.591 | 3.417 | 1.406 | 3.998 | 4.433 | 1.627 | 4.209 | 2.971 | 1.745 | -0.212 | 0.0494 | 0.1875 | 0 | 0 | 0 |
| 'Trpc4ap' | 15.648 | 15.335 | 16.098 | 15.498 | 15.332 | 16.320 | 15.722 | 15.909 | 15.686 | 16.055 | 15.108 | 16.689 | 16.107 | 14.579 | 15.849 | 15.778 | 15.686 | 16.306 | -0.017 | 0.3934 | 0.6428 | 0 | 0 | 0 |
| 'Trpm1' | 0.164 | 0.232 | 0.172 | 0.177 | 0.181 | 0.235 | 0.191 | 0.186 | 0.207 | 0.099 | 0.150 | 0.131 | 0.168 | 0.148 | 0.105 | 0.101 | 0.103 | 0.106 | 0.633 | 0.0000 | 0.0000 | 1 | 1 | 0 |
| 'Trpm4' | 5.191 | 5.064 | 5.315 | 5.635 | 3.971 | 4.776 | 4.314 | 4.750 | 4.824 | 4.997 | 4.822 | 5.308 | 5.168 | 5.072 | 5.007 | 4.458 | 4.942 | 4.714 | -0.032 | 0.5819 | 0.7842 | 0 | 0 | 0 |
| 'Trpm6' | 0.767 | 0.495 | 0.940 | 0.575 | 0.625 | 0.918 | 0.576 | 0.486 | 1.359 | 0.739 | 0.473 | 0.899 | 0.508 | 0.342 | 0.794 | 0.737 | 0.649 | 0.777 | 0.165 | 0.1981 | 0.4380 | 0 | 0 | 0 |
| 'Trpm7' | 7.819 | 8.436 | 8.270 | 7.972 | 8.618 | 7.498 | 8.132 | 8.518 | 7.769 | 8.942 | 8.900 | 8.075 | 7.931 | 8.791 | 7.677 | 8.145 | 8.236 | 7.798 | -0.038 | 0.1777 | 0.4131 | 0 | 0 | 0 |
| 'Trps1' | 0.695 | 0.586 | 0.608 | 0.720 | 0.640 | 0.561 | 0.834 | 0.711 | 0.652 | 0.553 | 0.661 | 0.699 | 0.545 | 0.611 | 0.572 | 0.419 | 0.761 | 0.529 | 0.154 | 0.0964 | 0.2888 | 0 | 0 | 0 |
| 'Trpt1' | 6.195 | 6.595 | 6.616 | 4.575 | 6.722 | 6.658 | 5.301 | 5.806 | 6.710 | 5.307 | 5.998 | 6.584 | 6.785 | 7.335 | 5.684 | 5.165 | 5.920 | 4.666 | 0.034 | 0.7160 | 0.8688 | 0 | 0 | 0 |
| 'Trpv2' | 19.928 | 18.852 | 19.807 | 20.011 | 19.600 | 19.501 | 19.550 | 19.249 | 19.962 | 20.520 | 17.721 | 21.904 | 19.770 | 18.796 | 21.647 | 18.449 | 18.960 | 20.920 | -0.029 | 0.3271 | 0.5849 | 0 | 0 | 0 |
| 'Trrap' | 11.625 | 9.610 | 11.807 | 11.376 | 10.048 | 10.951 | 10.059 | 10.010 | 11.497 | 11.853 | 11.665 | 11.631 | 10.972 | 10.079 | 11.117 | 10.304 | 10.513 | 12.537 | -0.066 | 0.1187 | 0.3278 | 0 | 0 | 0 |
| 'Trub1' | 1.760 | 1.852 | 2.143 | 1.767 | 2.100 | 1.942 | 1.808 | 2.189 | 2.016 | 1.862 | 1.784 | 1.854 | 1.766 | 1.645 | 1.526 | 1.705 | 1.786 | 1.663 | 0.163 | 0.0080 | 0.0526 | 0 | 0 | 0 |
| 'Trub2' | 2.394 | 3.354 | 3.202 | 2.379 | 3.046 | 3.158 | 2.492 | 3.329 | 3.144 | 3.050 | 2.300 | 2.747 | 2.753 | 2.462 | 2.903 | 2.999 | 2.545 | 2.355 | 0.121 | 0.1009 | 0.2976 | 0 | 0 | 0 |
| 'Tsc1' | 4.275 | 4.226 | 4.491 | 4.211 | 4.044 | 4.261 | 4.029 | 4.447 | 4.738 | 4.689 | 4.513 | 4.816 | 4.414 | 4.348 | 4.695 | 4.323 | 4.163 | 4.589 | -0.076 | 0.0138 | 0.0790 | 0 | 0 | 0 |
| 'Tsc2' | 15.997 | 15.428 | 17.556 | 16.176 | 15.478 | 17.546 | 17.188 | 15.920 | 18.517 | 17.710 | 15.739 | 17.247 | 17.707 | 16.391 | 16.569 | 15.660 | 15.860 | 17.154 | -0.013 | 0.6699 | 0.8408 | 0 | 0 | 0 |
| 'Tsc22d1' | 0.401 | 0.477 | 0.501 | 0.478 | 0.510 | 0.564 | 0.473 | 0.488 | 0.553 | 0.457 | 0.441 | 0.703 | 0.434 | 0.550 | 0.625 | 0.527 | 0.450 | 0.583 | -0.120 | 0.0992 | 0.2944 | 0 | 0 | 0 |
| 'Tsc22d2' | 7.496 | 5.970 | 6.050 | 6.610 | 5.832 | 5.836 | 6.505 | 6.247 | 5.683 | 7.148 | 6.971 | 6.946 | 6.870 | 7.015 | 6.288 | 6.948 | 6.530 | 6.591 | -0.137 | 0.0003 | 0.0041 | 1 | 0 | 0 |
| 'Tsc22d3' | 4.651 | 2.125 | 1.664 | 4.199 | 2.454 | 1.613 | 4.293 | 3.299 | 1.454 | 3.975 | 3.507 | 2.047 | 4.146 | 3.699 | 2.161 | 3.650 | 4.574 | 1.955 | -0.271 | 0.0057 | 0.0411 | 1 | 0 | 0 |
| 'Tsc22d4' | 29.603 | 30.227 | 29.588 | 27.410 | 29.371 | 33.261 | 26.480 | 28.394 | 31.785 | 29.057 | 29.062 | 35.199 | 31.105 | 30.745 | 36.615 | 31.953 | 29.295 | 37.349 | -0.133 | 0.0002 | 0.0025 | 1 | 0 | 0 |

| | | | | | | | | | | | | | | | | | | | | | | | | |
|---|---|---|---|---|---|---|---|---|---|---|---|---|---|---|---|---|---|---|---|---|---|---|---|---|
| 'Tsen15' | 2.514 | 2.602 | 2.689 | 2.083 | 2.530 | 2.777 | 2.206 | 2.974 | 2.621 | 2.451 | 2.318 | 2.669 | 2.163 | 2.770 | 2.565 | 2.578 | 2.345 | 2.172 | 0.056 | 0.3783 | 0.6301 | 0 | 0 | 0 |
| 'Tsen2' | 1.550 | 1.724 | 1.701 | 1.853 | 1.327 | 1.828 | 1.457 | 1.669 | 1.800 | 1.895 | 1.517 | 1.592 | 1.613 | 1.586 | 1.572 | 1.565 | 1.594 | 1.457 | 0.047 | 0.4944 | 0.7237 | 0 | 0 | 0 |
| 'Tsen34' | 1.743 | 1.875 | 1.481 | 1.666 | 1.687 | 1.818 | 1.841 | 1.797 | 1.730 | 1.525 | 1.614 | 1.577 | 1.620 | 1.654 | 1.806 | 1.879 | 1.613 | 1.649 | 0.057 | 0.3656 | 0.6178 | 0 | 0 | 0 |
| 'Tsen54' | 8.503 | 8.178 | 9.019 | 8.144 | 8.064 | 9.690 | 7.481 | 7.591 | 9.100 | 7.898 | 7.120 | 9.480 | 8.925 | 7.460 | 9.228 | 8.105 | 7.455 | 8.727 | 0.014 | 0.7120 | 0.8661 | 0 | 0 | 0 |
| 'Tsfm' | 4.164 | 4.639 | 4.531 | 4.321 | 4.535 | 4.619 | 4.405 | 4.666 | 3.829 | 3.844 | 3.656 | 4.031 | 4.577 | 3.924 | 4.113 | 4.153 | 3.736 | 3.709 | 0.139 | 0.0089 | 0.0576 | 0 | 0 | 0 |
| 'Tsg101' | 22.666 | 25.693 | 21.961 | 23.548 | 25.777 | 21.621 | 23.502 | 24.492 | 21.829 | 21.734 | 22.114 | 20.577 | 22.042 | 23.016 | 19.820 | 20.923 | 23.244 | 18.588 | 0.123 | 0.0000 | 0.0000 | 1 | 0 | 0 |
| 'Tshz1' | 5.736 | 5.160 | 5.536 | 5.511 | 4.854 | 6.307 | 5.283 | 5.530 | 5.753 | 4.670 | 4.891 | 5.934 | 4.847 | 5.168 | 5.699 | 5.183 | 5.019 | 6.222 | 0.052 | 0.2271 | 0.4734 | 0 | 0 | 0 |
| 'Tsn' | 9.216 | 10.569 | 10.196 | 9.414 | 10.553 | 10.171 | 10.044 | 11.043 | 9.951 | 9.798 | 10.013 | 9.655 | 9.971 | 10.108 | 9.627 | 9.793 | 10.027 | 9.466 | 0.033 | 0.1771 | 0.4124 | 0 | 0 | 0 |
| 'Tsnax' | 43.258 | 48.766 | 46.259 | 46.065 | 50.874 | 46.167 | 44.710 | 46.717 | 44.078 | 45.539 | 48.256 | 48.448 | 46.789 | 46.283 | 49.376 | 44.565 | 49.215 | 48.234 | -0.048 | 0.0859 | 0.2669 | 0 | 0 | 0 |
| 'Tspan13' | 72.179 | 79.269 | 73.500 | 77.902 | 79.751 | 75.270 | 76.510 | 80.456 | 71.760 | 70.841 | 71.153 | 65.651 | 65.866 | 70.920 | 65.846 | 68.826 | 72.855 | 65.931 | 0.142 | 0.0000 | 0.0000 | 1 | 0 | 0 |
| 'Tspan14' | 55.012 | 55.061 | 50.025 | 57.390 | 52.145 | 50.062 | 49.477 | 51.540 | 51.492 | 46.013 | 59.836 | 63.672 | 51.506 | 57.453 | 63.200 | 52.652 | 59.222 | 68.326 | -0.149 | 0.0096 | 0.0607 | 0 | 0 | 0 |
| 'Tspan3' | 20.906 | 27.196 | 20.325 | 21.678 | 26.612 | 21.768 | 22.972 | 23.400 | 21.227 | 25.138 | 24.512 | 26.325 | 28.170 | 26.536 | 26.794 | 28.241 | 23.435 | 24.307 | -0.199 | 0.0002 | 0.0024 | 1 | 0 | 0 |
| 'Tspan31' | 29.071 | 29.343 | 29.148 | 28.060 | 30.901 | 28.687 | 28.559 | 30.478 | 29.209 | 27.409 | 26.953 | 27.810 | 26.710 | 27.145 | 26.221 | 27.794 | 25.976 | 27.361 | 0.103 | 0.0023 | 0.0206 | 1 | 0 | 0 |
| 'Tspan32' | 24.958 | 31.274 | 27.410 | 25.022 | 29.044 | 28.245 | 25.189 | 30.290 | 29.625 | 30.698 | 28.463 | 26.523 | 31.735 | 30.085 | 26.252 | 30.211 | 29.908 | 25.210 | -0.056 | 0.2264 | 0.4730 | 0 | 0 | 0 |
| 'Tspan5' | 16.461 | 19.143 | 16.693 | 16.406 | 19.498 | 15.447 | 14.819 | 20.282 | 15.747 | 17.203 | 21.535 | 16.984 | 18.107 | 20.565 | 17.591 | 16.666 | 19.109 | 18.135 | -0.116 | 0.0004 | 0.0055 | 1 | 0 | 0 |
| 'Tspo' | 115.093 | 133.391 | 123.073 | 110.739 | 132.391 | 130.854 | 115.306 | 135.543 | 125.693 | 122.783 | 114.961 | 126.269 | 117.960 | 127.439 | 134.192 | 132.588 | 115.891 | 126.813 | -0.008 | 0.8280 | 0.9252 | 0 | 0 | 0 |
| 'Tspyl1' | 49.938 | 46.343 | 39.505 | 47.445 | 42.937 | 38.970 | 47.458 | 44.572 | 37.715 | 43.547 | 47.351 | 42.611 | 39.669 | 46.849 | 41.832 | 46.698 | 45.589 | 43.818 | -0.027 | 0.5508 | 0.7646 | 0 | 0 | 0 |
| 'Tspyl2' | 9.798 | 7.484 | 8.173 | 8.028 | 7.959 | 7.869 | 7.988 | 8.349 | 7.926 | 10.275 | 8.611 | 8.753 | 9.283 | 8.477 | 8.011 | 9.220 | 9.352 | 7.323 | -0.113 | 0.0171 | 0.0925 | 0 | 0 | 0 |
| 'Tspyl3' | 13.109 | 14.173 | 14.674 | 14.907 | 13.218 | 14.471 | 13.769 | 14.425 | 14.795 | 13.443 | 13.118 | 14.218 | 12.515 | 14.284 | 14.356 | 12.964 | 12.500 | 13.364 | 0.069 | 0.0684 | 0.2295 | 0 | 0 | 0 |
| 'Tspyl4' | 8.611 | 8.535 | 9.014 | 9.335 | 8.933 | 8.762 | 9.542 | 9.493 | 9.052 | 7.327 | 9.113 | 11.363 | 8.314 | 8.961 | 10.811 | 7.209 | 9.667 | 11.300 | -0.051 | 0.4441 | 0.6872 | 0 | 0 | 0 |
| 'Tsr1' | 7.653 | 6.545 | 7.339 | 6.554 | 6.965 | 6.607 | 6.953 | 7.063 | 7.280 | 7.491 | 7.342 | 6.977 | 6.783 | 6.853 | 6.562 | 6.905 | 6.525 | 6.763 | 0.011 | 0.7891 | 0.9059 | 0 | 0 | 0 |
| 'Tsr2' | 2.815 | 2.617 | 2.530 | 2.666 | 2.759 | 2.221 | 2.068 | 3.196 | 2.695 | 2.733 | 2.852 | 2.302 | 2.777 | 2.607 | 2.294 | 2.791 | 2.562 | 2.203 | 0.025 | 0.6851 | 0.8511 | 0 | 0 | 0 |
| 'Tsr3' | 23.302 | 23.628 | 23.519 | 23.215 | 23.412 | 26.143 | 21.665 | 25.690 | 26.189 | 22.204 | 22.457 | 26.881 | 23.715 | 22.379 | 23.228 | 20.952 | 22.044 | 24.850 | 0.039 | 0.3941 | 0.6434 | 0 | 0 | 0 |
| 'Tssc1' | 32.094 | 33.504 | 31.364 | 29.343 | 32.840 | 34.398 | 33.223 | 30.030 | 30.348 | 32.539 | 33.250 | 33.039 | 33.587 | 32.989 | 32.845 | 30.183 | 32.441 | 34.376 | -0.053 | 0.1624 | 0.3943 | 0 | 0 | 0 |
| 'Tssc4' | 4.661 | 5.572 | 5.419 | 4.550 | 5.186 | 6.000 | 5.248 | 5.496 | 5.586 | 3.961 | 4.718 | 5.708 | 4.896 | 4.823 | 5.546 | 4.785 | 4.486 | 5.532 | 0.088 | 0.0599 | 0.2110 | 0 | 0 | 0 |
| 'Tssk6' | 5.386 | 3.350 | 6.987 | 5.127 | 4.514 | 5.135 | 5.798 | 5.007 | 6.258 | 3.834 | 4.600 | 4.706 | 5.600 | 4.514 | 3.744 | 4.132 | 3.185 | 3.952 | 0.304 | 0.0058 | 0.0420 | 1 | 0 | 0 |
| 'Tsta3' | 30.785 | 28.964 | 33.984 | 31.942 | 30.644 | 35.282 | 31.622 | 32.406 | 35.214 | 30.719 | 29.307 | 32.678 | 31.302 | 30.549 | 32.687 | 29.978 | 26.529 | 29.608 | 0.081 | 0.0288 | 0.1315 | 0 | 0 | 0 |
| 'Tstd2' | 4.781 | 4.416 | 4.911 | 4.484 | 4.426 | 4.774 | 4.804 | 4.867 | 4.977 | 4.682 | 5.022 | 5.694 | 4.493 | 4.826 | 5.351 | 4.735 | 5.292 | 5.610 | -0.117 | 0.0040 | 0.0312 | 1 | 0 | 0 |
| 'Tstd3' | 13.839 | 14.673 | 15.816 | 15.247 | 16.670 | 14.328 | 15.538 | 14.225 | 15.544 | 14.843 | 14.031 | 14.204 | 15.258 | 14.531 | 12.914 | 13.508 | 14.528 | 13.785 | 0.078 | 0.1075 | 0.3084 | 0 | 0 | 0 |
| 'Ttbk1' | 3.153 | 3.265 | 3.045 | 3.073 | 3.462 | 3.366 | 2.982 | 3.297 | 3.040 | 3.826 | 4.312 | 3.520 | 4.365 | 4.055 | 3.699 | 3.947 | 3.468 | 3.697 | -0.288 | 0.0000 | 0.0000 | 1 | 0 | 0 |
| 'Ttbk2' | 1.805 | 2.008 | 1.655 | 1.858 | 1.841 | 1.719 | 1.722 | 1.579 | 1.866 | 1.953 | 2.222 | 2.088 | 2.004 | 2.282 | 1.971 | 2.146 | 1.957 | 1.956 | -0.224 | 0.0000 | 0.0002 | 1 | 0 | 0 |
| 'Ttc1' | 14.128 | 14.823 | 14.235 | 13.373 | 16.554 | 14.632 | 14.840 | 15.356 | 15.540 | 13.465 | 13.786 | 15.141 | 14.633 | 14.115 | 14.212 | 14.667 | 14.351 | 14.551 | 0.035 | 0.3429 | 0.5991 | 0 | 0 | 0 |
| 'Ttc13' | 7.706 | 7.508 | 8.451 | 7.640 | 8.280 | 8.198 | 7.766 | 8.019 | 8.581 | 8.347 | 7.579 | 8.516 | 8.501 | 7.276 | 8.107 | 8.379 | 8.005 | 8.190 | -0.025 | 0.3947 | 0.6439 | 0 | 0 | 0 |
| 'Ttc14' | 3.322 | 4.643 | 4.197 | 3.640 | 4.457 | 3.961 | 3.836 | 4.209 | 4.161 | 3.432 | 4.207 | 4.273 | 3.629 | 3.996 | 4.135 | 3.914 | 4.357 | 3.886 | 0.009 | 0.8010 | 0.9112 | 0 | 0 | 0 |
| 'Ttc16' | 1.776 | 1.551 | 1.618 | 1.531 | 1.507 | 1.538 | 1.633 | 1.456 | 1.540 | 2.070 | 2.200 | 1.668 | 2.778 | 1.757 | 1.444 | 1.673 | 2.236 | 1.714 | -0.308 | 0.0005 | 0.0056 | 1 | 0 | 0 |
| 'Ttc17' | 8.077 | 8.422 | 8.587 | 8.711 | 8.061 | 8.513 | 7.952 | 8.253 | 8.552 | 8.152 | 8.649 | 8.568 | 8.421 | 7.514 | 8.351 | 8.361 | 8.461 | 8.263 | -0.003 | 0.9234 | 0.9708 | 0 | 0 | 0 |
| 'Ttc19' | 3.119 | 3.323 | 3.450 | 3.377 | 3.342 | 3.505 | 3.300 | 3.958 | 3.059 | 3.271 | 3.123 | 3.231 | 3.459 | 3.424 | 3.049 | 3.861 | 3.169 | 3.083 | 0.032 | 0.5512 | 0.7647 | 0 | 0 | 0 |
| 'Ttc21b' | 1.819 | 1.956 | 2.130 | 2.059 | 1.807 | 1.988 | 2.057 | 2.138 | 2.091 | 1.561 | 1.866 | 2.021 | 1.907 | 1.845 | 1.777 | 1.923 | 1.944 | 1.785 | 0.106 | 0.0415 | 0.1678 | 0 | 0 | 0 |
| 'Ttc27' | 10.179 | 9.710 | 10.551 | 9.682 | 11.015 | 10.540 | 10.652 | 10.362 | 9.511 | 9.720 | 8.292 | 10.718 | 10.566 | 8.886 | 8.617 | 10.975 | 10.183 | 10.189 | 0.052 | 0.3256 | 0.5829 | 0 | 0 | 0 |
| 'Ttc28' | 2.021 | 2.098 | 1.983 | 1.664 | 1.887 | 2.065 | 1.867 | 1.906 | 1.972 | 2.326 | 1.443 | 2.202 | 1.902 | 1.726 | 1.857 | 2.549 | 1.514 | 2.119 | -0.012 | 0.8704 | 0.9446 | 0 | 0 | 0 |
| 'Ttc3' | 5.753 | 5.918 | 5.186 | 6.004 | 5.878 | 5.271 | 5.986 | 5.652 | 5.328 | 5.541 | 4.892 | 5.341 | 4.657 | 5.196 | 4.606 | 5.303 | 5.134 | 4.855 | 0.150 | 0.0000 | 0.0005 | 1 | 0 | 0 |
| 'Ttc32' | 7.538 | 7.532 | 7.399 | 6.256 | 7.246 | 7.373 | 5.747 | 6.955 | 6.783 | 6.735 | 7.558 | 8.402 | 6.767 | 7.275 | 8.157 | 6.665 | 7.458 | 7.993 | -0.106 | 0.1730 | 0.4066 | 0 | 0 | 0 |
| 'Ttc33' | 6.026 | 7.075 | 7.108 | 6.418 | 7.570 | 7.041 | 6.788 | 7.453 | 6.843 | 7.122 | 6.812 | 6.601 | 6.283 | 7.552 | 6.574 | 7.154 | 6.860 | 5.811 | 0.030 | 0.5130 | 0.7375 | 0 | 0 | 0 |
| 'Ttc37' | 6.796 | 8.147 | 8.273 | 7.499 | 7.999 | 7.598 | 8.130 | 7.791 | 7.050 | 7.357 | 7.797 | 7.580 | 7.419 | 7.399 | 6.620 | 7.162 | 7.781 | 6.820 | 0.059 | 0.1410 | 0.3636 | 0 | 0 | 0 |
| 'Ttc38' | 5.079 | 5.660 | 5.427 | 4.888 | 5.381 | 5.523 | 5.187 | 5.161 | 5.762 | 5.674 | 4.765 | 4.975 | 5.371 | 5.021 | 5.169 | 6.003 | 5.428 | 4.909 | 0.018 | 0.7074 | 0.8645 | 0 | 0 | 0 |
| 'Ttc39b' | 17.285 | 17.329 | 16.423 | 15.955 | 16.409 | 15.379 | 16.066 | 17.532 | 16.306 | 17.704 | 17.958 | 17.596 | 17.387 | 18.046 | 16.193 | 17.038 | 18.027 | 17.269 | -0.091 | 0.0003 | 0.0044 | 1 | 0 | 0 |
| 'Ttc39c' | 13.201 | 14.173 | 12.320 | 13.672 | 13.957 | 12.228 | 12.723 | 13.111 | 12.541 | 15.072 | 13.950 | 13.074 | 16.304 | 14.619 | 14.012 | 15.759 | 15.115 | 13.440 | -0.164 | 0.0000 | 0.0000 | 1 | 0 | 0 |
| 'Ttc4' | 5.734 | 6.450 | 6.302 | 5.597 | 6.707 | 6.693 | 5.740 | 6.431 | 6.743 | 6.625 | 5.937 | 6.498 | 6.131 | 6.230 | 5.813 | 5.807 | 6.651 | 6.050 | 0.004 | 0.9102 | 0.9651 | 0 | 0 | 0 |
| 'Ttc5' | 41.108 | 49.275 | 46.872 | 41.641 | 47.605 | 45.837 | 41.098 | 47.584 | 46.859 | 46.896 | 46.009 | 44.398 | 44.825 | 44.670 | 43.044 | 47.782 | 43.242 | 41.371 | 0.010 | 0.7895 | 0.9061 | 0 | 0 | 0 |
| 'Ttc7' | 7.851 | 7.739 | 8.700 | 7.204 | 8.176 | 8.887 | 7.181 | 7.754 | 8.569 | 7.652 | 7.292 | 8.583 | 7.878 | 7.611 | 8.753 | 8.333 | 6.794 | 8.872 | -0.003 | 0.9288 | 0.9728 | 0 | 0 | 0 |
| 'Ttc7b' | 6.963 | 6.559 | 6.552 | 6.219 | 6.778 | 6.760 | 6.476 | 7.280 | 6.418 | 6.683 | 6.564 | 6.882 | 6.120 | 6.027 | 6.780 | 6.942 | 6.032 | 6.202 | 0.034 | 0.4533 | 0.6923 | 0 | 0 | 0 |
| 'Ttc8' | 0.739 | 1.114 | 0.797 | 1.005 | 1.045 | 0.839 | 0.921 | 1.099 | 0.926 | 1.214 | 0.774 | 0.864 | 0.964 | 1.084 | 0.899 | 0.642 | 0.990 | 0.647 | 0.048 | 0.6661 | 0.8390 | 0 | 0 | 0 |
| 'Ttc9c' | 10.335 | 9.615 | 9.187 | 10.678 | 9.510 | 9.149 | 11.634 | 9.070 | 9.535 | 7.278 | 11.318 | 11.837 | 7.845 | 10.827 | 11.413 | 8.611 | 12.152 | 11.971 | -0.067 | 0.5212 | 0.7440 | 0 | 0 | 0 |
| 'Ttf1' | 5.845 | 5.505 | 6.225 | 5.896 | 5.457 | 5.823 | 6.406 | 5.878 | 6.204 | 5.695 | 6.171 | 6.708 | 5.823 | 5.802 | 6.657 | 5.361 | 6.291 | 6.150 | -0.049 | 0.2168 | 0.4632 | 0 | 0 | 0 |
| 'Ttf2' | 3.423 | 4.008 | 4.396 | 3.779 | 3.715 | 4.210 | 3.935 | 4.124 | 4.427 | 3.629 | 3.926 | 4.059 | 4.299 | 4.133 | 4.189 | 3.506 | 3.816 | 3.641 | 0.022 | 0.6381 | 0.8220 | 0 | 0 | 0 |
| 'Tti1' | 8.062 | 7.668 | 8.544 | 7.781 | 8.184 | 8.414 | 7.965 | 7.798 | 8.642 | 8.141 | 8.653 | 8.063 | 7.945 | 8.068 | 8.048 | 7.469 | 7.951 | 7.868 | 0.010 | 0.7680 | 0.8967 | 0 | 0 | 0 |
| 'Tti2' | 3.677 | 5.036 | 4.904 | 3.806 | 5.202 | 4.531 | 3.750 | 5.074 | 4.244 | 4.377 | 4.415 | 4.264 | 4.239 | 3.989 | 4.143 | 4.372 | 4.557 | 3.985 | 0.054 | 0.3379 | 0.5955 | 0 | 0 | 0 |
| 'Ttk' | 1.897 | 1.999 | 2.606 | 2.268 | 2.219 | 2.239 | 2.371 | 1.866 | 2.740 | 1.357 | 1.969 | 2.162 | 2.419 | 2.395 | 2.226 | 2.125 | 2.115 | 1.957 | 0.096 | 0.2805 | 0.5375 | 0 | 0 | 0 |
| 'Ttl' | 1.518 | 1.313 | 1.397 | 1.243 | 1.189 | 1.383 | 1.401 | 1.545 | 1.362 | 1.481 | 1.147 | 1.595 | 1.344 | 1.366 | 1.459 | 1.282 | 1.306 | 1.780 | -0.059 | 0.4614 | 0.6978 | 0 | 0 | 0 |
| 'Ttll1' | 3.736 | 3.640 | 4.477 | 3.857 | 4.715 | 4.280 | 4.543 | 4.470 | 5.180 | 4.773 | 3.517 | 4.086 | 4.759 | 3.441 | 4.071 | 4.212 | 3.443 | 3.837 | 0.101 | 0.2083 | 0.4529 | 0 | 0 | 0 |
| 'Ttll12' | 14.593 | 15.269 | 16.372 | 15.090 | 15.023 | 16.662 | 13.943 | 15.481 | 16.461 | 15.089 | 14.702 | 17.133 | 15.209 | 15.192 | 16.139 | 15.255 | 14.723 | 16.590 | -0.023 | 0.3388 | 0.5961 | 0 | 0 | 0 |
| 'Ttll3' | 1.746 | 1.272 | 1.820 | 1.885 | 1.390 | 1.490 | 1.874 | 1.498 | 1.698 | 1.899 | 1.860 | 3.011 | 1.843 | 1.669 | 2.274 | 1.790 | 2.180 | 2.959 | -0.411 | 0.0000 | 0.0002 | 1 | 1 | 0 |
| 'Ttll4' | 4.464 | 4.081 | 4.554 | 4.459 | 4.014 | 4.216 | 4.190 | 4.208 | 4.542 | 4.204 | 4.301 | 4.345 | 4.329 | 3.762 | 4.204 | 3.725 | 4.162 | 4.262 | 0.044 | 0.2210 | 0.4683 | 0 | 0 | 0 |
| 'Ttll5' | 1.601 | 1.606 | 1.734 | 1.955 | 1.624 | 1.744 | 1.764 | 1.711 | 1.692 | 1.603 | 1.661 | 1.501 | 1.832 | 1.627 | 1.487 | 1.409 | 1.607 | 1.364 | 0.123 | 0.0157 | 0.0866 | 0 | 0 | 0 |
| 'Ttn' | 0.230 | 0.157 | 0.190 | 0.225 | 0.172 | 0.134 | 0.274 | 0.152 | 0.195 | 0.265 | 0.226 | 0.222 | 0.202 | 0.163 | 0.155 | 0.244 | 0.213 | 0.256 | -0.201 | 0.0865 | 0.2679 | 0 | 0 | 0 |
| 'Ttyh1' | 0.264 | 0.267 | 0.198 | 0.263 | 0.284 | 0.215 | 0.219 | 0.267 | 0.242 | 0.170 | 0.208 | 0.198 | 0.189 | 0.238 | 0.155 | 0.227 | 0.222 | 0.146 | 0.323 | 0.0024 | 0.0209 | 1 | 0 | 0 |
| 'Ttyh2' | 1.234 | 1.219 | 1.221 | 1.020 | 1.234 | 1.239 | 1.117 | 1.108 | 1.171 | 1.175 | 0.978 | 1.640 | 1.017 | 1.107 | 1.236 | 1.062 | 0.933 | 1.286 | -0.009 | 0.9266 | 0.9719 | 0 | 0 | 0 |
| 'Ttyh3' | 38.272 | 38.177 | 42.854 | 34.843 | 37.912 | 43.318 | 34.614 | 38.939 | 41.848 | 43.488 | 39.232 | 42.376 | 38.071 | 37.547 | 42.581 | 42.974 | 35.318 | 43.425 | -0.069 | 0.0753 | 0.2440 | 0 | 0 | 0 |
| 'Tuba4a' | 13.693 | 8.574 | 8.474 | 10.802 | 9.621 | 8.497 | 9.825 | 10.147 | 7.883 | 11.515 | 10.210 | 8.275 | 11.323 | 11.048 | 8.170 | 10.642 | 10.770 | 7.867 | -0.044 | 0.4337 | 0.6784 | 0 | 0 | 0 |
| 'Tubb3' | 4.874 | 4.704 | 4.128 | 4.452 | 4.186 | 3.854 | 3.774 | 4.647 | 3.161 | 3.497 | 6.870 | 5.875 | 2.980 | 5.114 | 6.427 | 3.814 | 5.149 | 6.404 | -0.270 | 0.0350 | 0.1496 | 0 | 0 | 0 |
| 'Tubb4a' | 4.138 | 3.908 | 3.766 | 3.898 | 4.122 | 4.654 | 4.162 | 4.138 | 4.285 | 2.902 | 3.463 | 4.043 | 3.429 | 3.101 | 3.687 | 2.751 | 3.159 | 3.658 | 0.273 | 0.0004 | 0.0046 | 1 | 0 | 0 |
| 'Tubb5' | 148.577 | 144.679 | 166.109 | 161.457 | 145.299 | 166.388 | 149.646 | 145.867 | 166.662 | 133.950 | 151.147 | 149.402 | 161.546 | 160.739 | 176.363 | 143.115 | 152.426 | 163.792 | -0.009 | 0.8193 | 0.9201 | 0 | 0 | 0 |
| 'Tubb6' | 19.175 | 17.147 | 16.964 | 17.677 | 15.720 | 18.032 | 17.664 | 17.307 | 17.266 | 19.127 | 21.105 | 19.414 | 24.941 | 20.871 | 20.107 | 16.612 | 23.127 | 19.357 | -0.244 | 0.0001 | 0.0013 | 1 | 0 | 0 |
| 'Tubd1' | 3.488 | 3.775 | 4.084 | 3.971 | 3.557 | 4.032 | 4.191 | 4.560 | 4.832 | 3.781 | 3.616 | 4.581 | 3.502 | 4.126 | 4.838 | 3.519 | 3.938 | 4.381 | -0.004 | 0.9587 | 0.9847 | 0 | 0 | 0 |
| 'Tube1' | 2.653 | 2.973 | 3.558 | 2.714 | 3.209 | 2.989 | 3.637 | 3.295 | 3.452 | 2.598 | 3.360 | 2.929 | 4.229 | 3.161 | 3.138 | 2.354 | 3.236 | 2.872 | 0.019 | 0.8434 | 0.9337 | 0 | 0 | 0 |
| 'Tubg1' | 15.171 | 13.661 | 16.627 | 15.642 | 14.460 | 17.449 | 15.047 | 15.211 | 15.852 | 13.749 | 14.891 | 16.451 | 17.807 | 15.364 | 16.289 | 15.552 | 13.988 | 15.252 | -0.012 | 0.7767 | 0.9001 | 0 | 0 | 0 |
| 'Tubg2' | 7.207 | 8.633 | 8.764 | 6.755 | 8.221 | 8.079 | 7.296 | 8.637 | 9.096 | 8.639 | 5.880 | 6.300 | 7.934 | 6.620 | 5.698 | 6.333 | 6.596 | 4.706 | 0.306 | 0.0008 | 0.0091 | 1 | 0 | 0 |
| 'Tubgcp2' | 19.079 | 21.057 | 22.448 | 20.373 | 21.092 | 21.809 | 19.673 | 20.814 | 22.314 | 21.241 | 21.004 | 19.849 | 24.606 | 20.867 | 21.078 | 21.332 | 20.808 | 17.950 | -0.008 | 0.8634 | 0.9407 | 0 | 0 | 0 |
| 'Tubgcp3' | 6.065 | 6.457 | 6.924 | 6.382 | 6.229 | 6.448 | 6.481 | 6.405 | 6.777 | 6.104 | 6.084 | 6.903 | 6.386 | 6.215 | 6.335 | 6.460 | 6.479 | 5.970 | 0.019 | 0.4997 | 0.7273 | 0 | 0 | 0 |
| 'Tubgcp4' | 4.580 | 4.751 | 4.721 | 4.437 | 4.772 | 4.294 | 4.417 | 4.427 | 4.824 | 4.654 | 5.018 | 5.179 | 4.872 | 4.728 | 4.865 | 4.537 | 5.431 | 5.246 | -0.124 | 0.0007 | 0.0074 | 1 | 0 | 0 |
| 'Tubgcp5' | 2.382 | 2.555 | 2.553 | 2.511 | 2.668 | 2.650 | 2.803 | 2.732 | 2.580 | 2.492 | 2.644 | 2.681 | 2.347 | 2.887 | 2.649 | 2.573 | 2.675 | 2.115 | 0.014 | 0.7591 | 0.8924 | 0 | 0 | 0 |
| 'Tubgcp6' | 8.791 | 8.311 | 9.892 | 8.264 | 8.549 | 8.974 | 7.952 | 8.188 | 9.450 | 9.209 | 9.037 | 9.271 | 8.679 | 8.147 | 8.864 | 8.628 | 8.852 | 9.120 | -0.036 | 0.2744 | 0.5301 | 0 | 0 | 0 |
| 'Tufm' | 26.555 | 25.331 | 29.109 | 27.957 | 27.504 | 28.274 | 26.704 | 26.463 | 28.464 | 25.193 | 24.788 | 29.468 | 26.484 | 24.926 | 28.642 | 25.466 | 26.493 | 30.404 | 0.015 | 0.5973 | 0.7953 | 0 | 0 | 0 |
| 'Tuft1' | 0.797 | 0.453 | 0.806 | 0.839 | 0.650 | 0.837 | 0.598 | 0.733 | 0.716 | 0.799 | 0.566 | 0.642 | 0.767 | 0.689 | 0.772 | 0.670 | 0.599 | 0.579 | 0.090 | 0.4086 | 0.6576 | 0 | 0 | 0 |
| 'Tulp2' | 2.097 | 1.655 | 1.903 | 2.570 | 1.348 | 1.777 | 3.410 | 1.663 | 1.537 | 1.182 | 1.619 | 1.556 | 1.393 | 1.472 | 1.344 | 0.801 | 1.804 | 1.083 | 0.511 | 0.0055 | 0.0406 | 1 | 1 | 0 |
| 'Tulp3' | 0.957 | 1.097 | 1.112 | 0.943 | 1.042 | 1.115 | 0.872 | 1.005 | 1.137 | 1.105 | 0.960 | 0.984 | 0.983 | 0.909 | 0.994 | 0.927 | 0.818 | 1.049 | 0.081 | 0.2241 | 0.4711 | 0 | 0 | 0 |
| 'Tusc1' | 5.306 | 4.558 | 4.314 | 4.757 | 4.741 | 6.944 | 5.009 | 5.442 | 4.789 | 6.016 | 5.031 | 5.314 | 5.945 | 6.351 | 5.465 | 6.009 | 5.314 | 6.033 | -0.167 | 0.0664 | 0.2253 | 0 | 0 | 0 |
| 'Tusc2' | 12.129 | 15.487 | 12.278 | 10.540 | 15.065 | 14.443 | 11.361 | 13.813 | 14.271 | 12.756 | 11.937 | 14.061 | 12.743 | 13.333 | 14.114 | 14.093 | 12.207 | 14.152 | -0.018 | 0.7625 | 0.8944 | 0 | 0 | 0 |
| 'Tusc3' | 17.499 | 19.111 | 17.807 | 18.695 | 19.937 | 17.784 | 17.635 | 18.986 | 17.973 | 17.838 | 17.366 | 18.274 | 18.222 | 20.230 | 18.109 | 20.192 | 19.308 | 18.964 | -0.038 | 0.3123 | 0.5709 | 0 | 0 | 0 |
| 'Tut1' | 17.141 | 15.851 | 18.647 | 16.745 | 16.610 | 18.495 | 18.830 | 16.840 | 17.926 | 16.426 | 16.222 | 20.096 | 16.230 | 14.820 | 18.695 | 14.669 | 17.634 | 19.684 | 0.013 | 0.7763 | 0.9001 | 0 | 0 | 0 |
| 'Tvp23b' | 9.553 | 10.844 | 9.007 | 9.346 | 10.028 | 8.950 | 9.564 | 9.656 | 9.015 | 9.288 | 9.654 | 10.454 | 8.598 | 10.387 | 9.894 | 8.246 | 10.569 | 8.753 | -0.016 | 0.7409 | 0.8827 | 0 | 0 | 0 |
| 'Twf1' | 17.685 | 21.740 | 15.486 | 14.691 | 22.456 | 15.908 | 16.148 | 19.500 | 15.076 | 21.955 | 22.487 | 17.404 | 21.181 | 26.388 | 18.239 | 24.270 | 23.315 | 18.241 | -0.298 | 0.0000 | 0.0000 | 1 | 0 | 0 |
| 'Twf2' | 42.143 | 39.941 | 43.599 | 43.926 | 40.815 | 46.157 | 40.407 | 40.056 | 42.656 | 39.810 | 37.805 | 48.397 | 40.966 | 37.870 | 47.934 | 41.428 | 36.248 | 48.113 | -0.002 | 0.9543 | 0.9832 | 0 | 0 | 0 |
| 'Twistnb' | 12.313 | 14.695 | 14.169 | 13.849 | 13.902 | 13.002 | 14.321 | 13.868 | 12.632 | 13.225 | 14.747 | 14.167 | 13.025 | 13.860 | 14.355 | 13.821 | 12.965 | 13.574 | -0.026 | 0.5512 | 0.7647 | 0 | 0 | 0 |
| 'Twsg1' | 34.211 | 38.936 | 35.546 | 33.997 | 38.530 | 36.205 | 33.202 | 37.484 | 34.820 | 37.418 | 37.556 | 40.202 | 36.066 | 41.211 | 39.895 | 38.987 | 38.914 | 40.105 | -0.130 | 0.0000 | 0.0000 | 1 | 0 | 0 |
| 'Txk' | 1.224 | 1.152 | 1.446 | 1.205 | 1.369 | 1.391 | 1.369 | 1.240 | 1.300 | 1.075 | 1.172 | 1.152 | 1.012 | 0.905 | 1.162 | 1.026 | 1.052 | 0.894 | 0.299 | 0.0000 | 0.0000 | 1 | 0 | 0 |
| 'Txlna' | 11.629 | 11.165 | 12.663 | 11.321 | 11.842 | 12.410 | 11.057 | 11.174 | 12.156 | 12.040 | 10.848 | 12.752 | 11.607 | 10.609 | 12.607 | 11.325 | 10.559 | 12.536 | -0.002 | 0.9212 | 0.9704 | 0 | 0 | 0 |
| 'Txlnb' | 1.291 | 1.469 | 1.573 | 1.627 | 1.274 | 1.646 | 1.472 | 1.410 | 1.267 | 2.033 | 1.690 | 2.109 | 1.717 | 1.779 | 1.975 | 1.410 | 1.980 | 1.931 | -0.368 | 0.0000 | 0.0001 | 1 | 0 | 0 |
| 'Txlng' | 4.441 | 4.696 | 4.425 | 4.770 | 5.060 | 3.840 | 4.690 | 4.700 | 3.913 | 4.863 | 4.547 | 4.736 | 4.363 | 5.566 | 4.298 | 4.592 | 4.705 | 4.278 | -0.065 | 0.1517 | 0.3788 | 0 | 0 | 0 |
| 'Txn1' | 142.845 | 159.672 | 137.040 | 155.105 | 163.343 | 145.186 | 157.782 | 154.817 | 146.119 | 132.449 | 143.603 | 124.638 | 160.806 | 155.433 | 144.402 | 144.911 | 157.633 | 133.563 | 0.060 | 0.1195 | 0.3292 | 0 | 0 | 0 |
| 'Txn2' | 13.410 | 13.713 | 14.211 | 13.645 | 14.196 | 14.554 | 13.123 | 14.258 | 13.953 | 13.689 | 13.787 | 15.024 | 13.677 | 13.451 | 15.249 | 14.165 | 13.310 | 14.362 | -0.030 | 0.2711 | 0.5262 | 0 | 0 | 0 |
| 'Txndc11' | 4.677 | 4.681 | 5.065 | 4.609 | 4.739 | 4.646 | 4.840 | 4.852 | 5.066 | 5.318 | 5.163 | 5.188 | 5.047 | 4.844 | 5.054 | 4.988 | 5.087 | 5.113 | -0.094 | 0.0013 | 0.0126 | 1 | 0 | 0 |
| 'Txndc12' | 24.970 | 27.780 | 27.206 | 27.814 | 29.705 | 26.642 | 27.164 | 27.175 | 25.694 | 27.342 | 26.728 | 26.365 | 27.086 | 27.151 | 25.243 | 26.656 | 26.314 | 24.424 | 0.029 | 0.3418 | 0.5981 | 0 | 0 | 0 |
| 'Txndc15' | 37.210 | 44.617 | 39.364 | 37.548 | 42.732 | 38.554 | 38.089 | 39.204 | 38.983 | 38.078 | 33.698 | 42.963 | 40.650 | 42.468 | 41.695 | 39.287 | 43.120 | 44.686 | -0.056 | 0.2123 | 0.4575 | 0 | 0 | 0 |
| 'Txndc16' | 1.574 | 1.436 | 1.558 | 1.754 | 1.683 | 1.553 | 1.708 | 1.586 | 1.844 | 1.580 | 1.538 | 1.757 | 1.726 | 1.616 | 1.542 | 1.435 | 1.802 | 1.675 | -0.013 | 0.8090 | 0.9139 | 0 | 0 | 0 |
| 'Txndc17' | 35.053 | 36.512 | 34.038 | 37.836 | 39.181 | 34.119 | 39.396 | 39.364 | 34.103 | 38.465 | 36.237 | 34.254 | 41.033 | 36.460 | 35.166 | 35.064 | 37.083 | 33.656 | -0.004 | 0.9146 | 0.9671 | 0 | 0 | 0 |
| 'Txndc5' | 8.286 | 9.886 | 9.933 | 8.621 | 10.504 | 10.121 | 8.545 | 10.330 | 9.471 | 9.187 | 8.823 | 9.573 | 8.605 | 9.063 | 9.786 | 10.225 | 9.234 | 9.628 | 0.014 | 0.7292 | 0.8759 | 0 | 0 | 0 |
| 'Txndc9' | 4.295 | 4.953 | 4.480 | 4.328 | 5.063 | 4.856 | 4.542 | 4.885 | 4.990 | 4.216 | 4.296 | 4.899 | 4.443 | 4.500 | 4.599 | 4.372 | 4.570 | 4.507 | 0.052 | 0.1073 | 0.3081 | 0 | 0 | 0 |
| 'Txnip' | 31.392 | 37.517 | 37.434 | 35.759 | 37.891 | 37.459 | 37.300 | 36.755 | 38.480 | 31.727 | 35.813 | 39.788 | 31.179 | 34.076 | 36.765 | 31.341 | 40.195 | 37.258 | 0.044 | 0.2325 | 0.4806 | 0 | 0 | 0 |
| 'Txnl1' | 29.504 | 30.321 | 27.836 | 32.240 | 31.987 | 28.025 | 29.856 | 30.781 | 27.760 | 26.949 | 28.428 | 28.908 | 29.109 | 29.578 | 27.723 | 27.842 | 28.898 | 27.765 | 0.056 | 0.0701 | 0.2331 | 0 | 0 | 0 |
| 'Txnl4a' | 8.844 | 8.618 | 8.262 | 8.712 | 9.002 | 8.908 | 8.445 | 9.353 | 8.239 | 8.355 | 8.033 | 10.422 | 8.517 | 7.994 | 10.068 | 8.361 | 9.514 | 10.772 | -0.079 | 0.1719 | 0.4055 | 0 | 0 | 0 |
| 'Txnl4b' | 4.360 | 4.554 | 4.121 | 3.985 | 4.807 | 4.219 | 4.209 | 4.831 | 5.138 | 4.932 | 4.693 | 5.150 | 4.762 | 4.895 | 3.938 | 3.760 | 3.910 | 4.416 | -0.028 | 0.7090 | 0.8651 | 0 | 0 | 0 |
| 'Txnrd2' | 3.410 | 3.126 | 3.390 | 3.045 | 3.342 | 3.476 | 3.170 | 3.363 | 3.046 | 3.489 | 3.006 | 3.420 | 3.188 | 2.834 | 3.327 | 3.433 | 3.043 | 2.946 | 0.026 | 0.5915 | 0.7921 | 0 | 0 | 0 |
| 'Tyk2' | 28.181 | 26.451 | 31.781 | 29.056 | 26.985 | 31.640 | 27.180 | 27.890 | 32.069 | 31.547 | 28.912 | 31.796 | 30.094 | 27.780 | 30.571 | 28.409 | 29.569 | 30.070 | -0.052 | 0.0665 | 0.2254 | 0 | 0 | 0 |
| 'Tyro3' | 2.525 | 2.157 | 3.311 | 2.306 | 2.152 | 2.741 | 2.210 | 2.420 | 2.729 | 2.590 | 2.218 | 2.690 | 2.728 | 2.225 | 2.778 | 2.239 | 2.057 | 2.501 | 0.027 | 0.6429 | 0.8250 | 0 | 0 | 0 |
| 'Tysnd1' | 19.583 | 20.592 | 24.846 | 21.345 | 21.340 | 23.837 | 19.175 | 19.858 | 24.756 | 22.626 | 19.941 | 24.811 | 23.633 | 19.513 | 24.576 | 23.057 | 16.921 | 25.664 | -0.048 | 0.2927 | 0.5493 | 0 | 0 | 0 |
| 'Tyw1' | 6.913 | 6.303 | 6.801 | 7.260 | 6.817 | 7.055 | 6.659 | 6.811 | 6.863 | 7.237 | 6.556 | 7.213 | 6.919 | 6.668 | 7.120 | 6.813 | 7.261 | 6.682 | -0.033 | 0.2992 | 0.5556 | 0 | 0 | 0 |
| 'Tyw5' | 1.598 | 1.703 | 1.588 | 1.762 | 1.567 | 1.419 | 1.568 | 1.703 | 1.573 | 1.406 | 1.534 | 1.541 | 1.602 | 1.544 | 1.639 | 1.716 | 1.620 | 1.374 | 0.039 | 0.5394 | 0.7572 | 0 | 0 | 0 |
| 'U2af1l4' | 4.306 | 4.876 | 4.922 | 4.122 | 4.652 | 4.899 | 4.446 | 4.328 | 5.381 | 4.259 | 4.075 | 4.740 | 4.437 | 4.470 | 4.305 | 4.299 | 4.331 | 4.593 | 0.072 | 0.0409 | 0.1656 | 0 | 0 | 0 |
| 'U2af2' | 45.488 | 44.429 | 45.166 | 45.022 | 44.156 | 45.210 | 42.772 | 43.899 | 45.251 | 44.563 | 41.463 | 44.153 | 46.001 | 43.214 | 45.603 | 44.148 | 43.542 | 44.587 | 0.005 | 0.8006 | 0.9111 | 0 | 0 | 0 |
| 'U2surp' | 7.293 | 7.273 | 7.766 | 7.945 | 7.547 | 7.456 | 8.754 | 7.603 | 7.281 | 7.468 | 7.656 | 7.792 | 7.353 | 7.783 | 7.476 | 7.409 | 8.053 | 6.839 | 0.012 | 0.7423 | 0.8836 | 0 | 0 | 0 |
| 'Uap1' | 3.652 | 4.031 | 3.990 | 3.929 | 3.874 | 4.059 | 4.205 | 3.766 | 3.509 | 3.570 | 3.682 | 3.310 | 3.546 | 3.694 | 3.389 | 3.593 | 3.642 | 3.336 | 0.135 | 0.0020 | 0.0179 | 1 | 0 | 0 |
| 'Uap1l1' | 6.766 | 6.667 | 7.291 | 5.777 | 7.114 | 7.112 | 5.945 | 6.958 | 7.357 | 6.679 | 6.001 | 6.684 | 6.293 | 6.312 | 6.893 | 6.626 | 5.807 | 6.119 | 0.079 | 0.0711 | 0.2352 | 0 | 0 | 0 |
| 'Uba1' | 101.109 | 99.008 | 102.858 | 104.128 | 103.924 | 103.801 | 101.271 | 101.152 | 104.406 | 108.152 | 104.893 | 104.183 | 106.809 | 103.659 | 105.402 | 101.266 | 103.093 | 107.034 | -0.046 | 0.0141 | 0.0804 | 0 | 0 | 0 |
| 'Uba2' | 21.382 | 22.017 | 22.155 | 22.061 | 23.700 | 22.181 | 22.784 | 23.354 | 21.471 | 20.999 | 22.642 | 21.437 | 22.165 | 22.726 | 21.447 | 21.188 | 23.486 | 20.928 | 0.020 | 0.3589 | 0.6115 | 0 | 0 | 0 |
| 'Uba3' | 5.991 | 6.269 | 6.099 | 6.309 | 6.710 | 5.837 | 6.479 | 6.416 | 5.837 | 5.261 | 6.244 | 6.175 | 5.745 | 6.242 | 5.662 | 5.713 | 6.574 | 6.187 | 0.043 | 0.2473 | 0.4979 | 0 | 0 | 0 |
| 'Uba5' | 9.213 | 9.900 | 9.361 | 9.073 | 9.918 | 9.425 | 8.874 | 9.785 | 9.471 | 9.444 | 8.804 | 10.491 | 8.653 | 9.711 | 9.872 | 8.742 | 9.678 | 10.399 | -0.027 | 0.4394 | 0.6829 | 0 | 0 | 0 |
| 'Uba6' | 2.427 | 2.653 | 3.020 | 2.920 | 2.680 | 2.482 | 2.800 | 2.595 | 2.963 | 2.397 | 2.802 | 3.268 | 2.478 | 2.765 | 2.796 | 2.298 | 2.879 | 3.640 | -0.056 | 0.4130 | 0.6612 | 0 | 0 | 0 |
| 'Uba7' | 14.372 | 13.222 | 16.343 | 13.209 | 13.366 | 15.020 | 14.287 | 13.377 | 16.529 | 14.451 | 14.248 | 16.987 | 15.818 | 13.423 | 15.438 | 13.929 | 14.256 | 15.986 | -0.065 | 0.0324 | 0.1421 | 0 | 0 | 0 |
| 'Ubac1' | 6.878 | 7.068 | 7.647 | 6.900 | 7.198 | 7.863 | 7.100 | 7.104 | 7.381 | 7.027 | 7.453 | 8.272 | 7.370 | 7.547 | 7.968 | 7.266 | 7.974 | 7.380 | -0.079 | 0.0251 | 0.1208 | 0 | 0 | 0 |
| 'Ubac2' | 64.102 | 73.900 | 63.944 | 64.606 | 74.872 | 68.599 | 62.002 | 73.222 | 66.982 | 69.473 | 73.465 | 73.534 | 69.399 | 74.886 | 71.665 | 70.170 | 71.932 | 65.176 | -0.077 | 0.0033 | 0.0271 | 1 | 0 | 0 |
| 'Ubald1' | 13.155 | 10.988 | 12.723 | 10.699 | 11.049 | 12.867 | 11.780 | 11.797 | 12.553 | 13.183 | 13.154 | 12.592 | 13.407 | 12.233 | 13.380 | 13.357 | 12.089 | 12.169 | -0.110 | 0.0059 | 0.0426 | 1 | 0 | 0 |
| 'Ubald2' | 61.834 | 60.175 | 51.673 | 47.505 | 60.343 | 58.309 | 44.851 | 64.433 | 47.475 | 69.260 | 49.547 | 45.876 | 66.465 | 63.205 | 52.192 | 77.592 | 51.361 | 45.050 | -0.062 | 0.5421 | 0.7590 | 0 | 0 | 0 |
| 'Ubap1' | 11.057 | 10.790 | 9.752 | 10.421 | 10.411 | 11.197 | 10.783 | 10.718 | 9.992 | 10.356 | 10.912 | 10.704 | 9.644 | 10.483 | 10.750 | 9.712 | 10.447 | 10.478 | 0.014 | 0.7158 | 0.8688 | 0 | 0 | 0 |
| 'Ubap2' | 17.893 | 16.160 | 18.842 | 17.262 | 16.617 | 17.223 | 16.711 | 16.386 | 18.422 | 18.499 | 17.263 | 19.312 | 19.355 | 15.332 | 18.100 | 17.389 | 16.061 | 19.002 | -0.053 | 0.0871 | 0.2691 | 0 | 0 | 0 |
| 'Ubap2l' | 12.636 | 12.143 | 12.620 | 12.740 | 12.049 | 12.815 | 12.202 | 12.644 | 12.824 | 13.015 | 12.424 | 12.887 | 12.790 | 11.782 | 12.622 | 12.095 | 12.210 | 12.383 | -0.004 | 0.8303 | 0.9262 | 0 | 0 | 0 |
| 'Ubash3a' | 37.315 | 36.024 | 35.063 | 36.460 | 34.637 | 36.162 | 35.115 | 34.182 | 35.630 | 34.129 | 33.212 | 37.918 | 33.664 | 33.777 | 36.848 | 34.534 | 32.133 | 38.834 | 0.015 | 0.6219 | 0.8110 | 0 | 0 | 0 |
| 'Ubash3b' | 5.576 | 5.610 | 5.600 | 5.503 | 5.394 | 6.120 | 5.250 | 6.095 | 5.441 | 5.806 | 5.823 | 6.254 | 4.846 | 5.235 | 6.087 | 6.229 | 5.136 | 5.920 | -0.030 | 0.5484 | 0.7623 | 0 | 0 | 0 |
| 'Ubb' | ####### | ####### | ####### | ####### | ####### | ####### | ####### | ####### | ####### | ####### | ####### | 989.772 | ####### | ####### | ####### | ####### | ####### | ####### | 0.112 | 0.0487 | 0.1854 | 0 | 0 | 0 |
| 'Ube2a' | 10.578 | 10.423 | 10.114 | 10.101 | 10.479 | 10.064 | 9.539 | 10.381 | 9.216 | 9.466 | 9.401 | 10.078 | 10.941 | 10.977 | 10.074 | 10.194 | 10.718 | 9.277 | -0.014 | 0.7150 | 0.8686 | 0 | 0 | 0 |
| 'Ube2b' | 49.336 | 55.366 | 48.746 | 51.484 | 58.489 | 48.851 | 53.738 | 52.736 | 46.620 | 49.595 | 51.125 | 51.150 | 49.801 | 57.881 | 47.932 | 52.207 | 56.896 | 48.586 | -0.012 | 0.6738 | 0.8433 | 0 | 0 | 0 |
| 'Ube2c' | 9.957 | 11.607 | 14.931 | 12.425 | 12.731 | 15.676 | 13.461 | 12.537 | 16.189 | 7.927 | 9.466 | 13.442 | 12.415 | 14.285 | 16.191 | 11.748 | 13.323 | 13.164 | 0.083 | 0.3230 | 0.5806 | 0 | 0 | 0 |
| 'Ube2d1' | 7.330 | 8.434 | 7.961 | 6.868 | 7.599 | 7.122 | 7.061 | 7.186 | 7.519 | 7.028 | 6.852 | 8.107 | 7.059 | 7.668 | 7.917 | 7.074 | 7.726 | 8.332 | -0.029 | 0.4940 | 0.7233 | 0 | 0 | 0 |
| 'Ube2d2a' | 51.218 | 60.154 | 50.043 | 52.247 | 58.182 | 54.126 | 56.241 | 55.541 | 52.406 | 52.453 | 52.961 | 54.278 | 53.141 | 57.170 | 54.002 | 54.521 | 55.620 | 54.118 | -0.008 | 0.7683 | 0.8967 | 0 | 0 | 0 |
| 'Ube2d3' | 17.150 | 18.236 | 17.077 | 17.449 | 18.460 | 15.645 | 17.728 | 18.214 | 16.050 | 17.439 | 17.212 | 16.824 | 16.962 | 18.350 | 15.904 | 17.137 | 18.457 | 16.903 | -0.004 | 0.8474 | 0.9343 | 0 | 0 | 0 |
| 'Ube2e1' | 7.884 | 8.207 | 7.371 | 7.650 | 7.630 | 7.491 | 7.310 | 8.761 | 7.176 | 7.768 | 8.616 | 6.955 | 7.420 | 7.445 | 7.168 | 7.056 | 8.131 | 6.931 | 0.037 | 0.3721 | 0.6240 | 0 | 0 | 0 |
| 'Ube2e3' | 4.891 | 5.579 | 4.988 | 5.109 | 5.187 | 5.279 | 5.228 | 5.655 | 4.946 | 4.515 | 4.877 | 4.188 | 5.048 | 4.602 | 4.748 | 4.265 | 4.980 | 4.008 | 0.177 | 0.0001 | 0.0018 | 1 | 0 | 0 |
| 'Ube2f' | 4.691 | 4.991 | 4.355 | 4.935 | 4.948 | 4.242 | 4.565 | 4.815 | 4.068 | 4.763 | 5.069 | 4.752 | 4.840 | 5.113 | 4.515 | 4.710 | 5.127 | 4.482 | -0.075 | 0.0144 | 0.0810 | 0 | 0 | 0 |
| 'Ube2g1' | 13.290 | 15.125 | 12.558 | 12.490 | 14.529 | 12.257 | 13.713 | 14.559 | 11.766 | 12.558 | 14.075 | 12.897 | 13.197 | 14.845 | 12.576 | 12.714 | 15.247 | 12.670 | -0.020 | 0.4930 | 0.7221 | 0 | 0 | 0 |
| 'Ube2g2' | 24.878 | 27.741 | 24.628 | 24.899 | 26.874 | 27.463 | 22.828 | 27.489 | 25.708 | 25.700 | 26.060 | 25.993 | 26.122 | 24.067 | 24.700 | 26.455 | 24.689 | 26.420 | 0.003 | 0.9446 | 0.9789 | 0 | 0 | 0 |
| 'Ube2h' | 44.326 | 41.663 | 40.996 | 44.366 | 40.971 | 41.467 | 40.946 | 40.683 | 39.660 | 40.628 | 44.600 | 46.861 | 38.358 | 44.037 | 44.338 | 40.090 | 41.798 | 50.959 | -0.071 | 0.1267 | 0.3409 | 0 | 0 | 0 |
| 'Ube2i' | 28.011 | 31.344 | 28.938 | 27.584 | 29.947 | 27.661 | 27.659 | 30.486 | 29.774 | 27.751 | 30.027 | 28.089 | 29.940 | 29.560 | 28.696 | 29.024 | 28.876 | 28.631 | -0.007 | 0.7728 | 0.8993 | 0 | 0 | 0 |
| 'Ube2j1' | 45.564 | 49.392 | 41.822 | 43.815 | 50.093 | 43.097 | 45.192 | 45.538 | 39.243 | 48.026 | 44.267 | 41.333 | 51.449 | 51.704 | 40.803 | 47.242 | 47.925 | 41.094 | -0.045 | 0.1754 | 0.4103 | 0 | 0 | 0 |
| 'Ube2k' | 12.468 | 12.810 | 11.719 | 13.693 | 13.564 | 12.203 | 12.859 | 12.665 | 11.764 | 11.394 | 11.732 | 11.743 | 12.205 | 12.735 | 11.405 | 12.414 | 12.595 | 11.096 | 0.071 | 0.0153 | 0.0849 | 0 | 0 | 0 |
| 'Ube2l3' | 16.482 | 18.186 | 17.362 | 17.867 | 18.484 | 16.859 | 17.270 | 16.900 | 16.819 | 17.686 | 15.857 | 16.283 | 17.693 | 17.174 | 17.085 | 16.247 | 17.185 | 16.078 | 0.035 | 0.2227 | 0.4701 | 0 | 0 | 0 |
| 'Ube2m' | 42.077 | 36.272 | 45.704 | 43.150 | 39.729 | 46.411 | 42.863 | 42.242 | 39.914 | 41.853 | 39.813 | 37.702 | 45.469 | 43.571 | 42.426 | 41.839 | 40.496 | 46.056 | -0.014 | 0.7560 | 0.8913 | 0 | 0 | 0 |
| 'Ube2n' | 40.100 | 42.553 | 41.989 | 44.734 | 47.108 | 41.951 | 46.517 | 41.723 | 40.739 | 39.255 | 42.111 | 38.901 | 40.840 | 42.657 | 39.277 | 38.120 | 45.094 | 38.999 | 0.074 | 0.0293 | 0.1330 | 0 | 0 | 0 |
| 'Ube2o' | 9.424 | 8.409 | 9.169 | 9.333 | 8.545 | 9.342 | 8.787 | 8.493 | 9.178 | 9.147 | 9.011 | 10.141 | 9.721 | 8.554 | 9.653 | 8.294 | 9.217 | 10.434 | -0.073 | 0.0416 | 0.1680 | 0 | 0 | 0 |
| 'Ube2q1' | 30.796 | 34.561 | 32.050 | 31.522 | 33.816 | 32.729 | 31.371 | 33.490 | 32.235 | 30.408 | 32.687 | 33.756 | 32.339 | 33.349 | 32.972 | 33.503 | 32.470 | 32.770 | -0.021 | 0.3222 | 0.5799 | 0 | 0 | 0 |
| 'Ube2q2' | 11.077 | 11.776 | 11.293 | 12.359 | 11.800 | 11.083 | 12.509 | 11.737 | 10.965 | 11.729 | 12.884 | 11.968 | 11.744 | 12.503 | 11.727 | 12.241 | 13.140 | 11.690 | -0.079 | 0.0097 | 0.0610 | 0 | 0 | 0 |
| 'Ube2r2' | 29.005 | 26.061 | 26.555 | 29.676 | 27.536 | 28.636 | 29.767 | 28.458 | 26.508 | 27.854 | 26.546 | 27.875 | 27.076 | 26.824 | 27.983 | 28.567 | 27.582 | 27.766 | 0.012 | 0.6514 | 0.8307 | 0 | 0 | 0 |
| 'Ube2t' | 1.995 | 2.005 | 2.543 | 2.182 | 2.010 | 2.453 | 2.096 | 2.102 | 2.284 | 0.827 | 1.122 | 1.285 | 1.207 | 1.083 | 1.054 | 0.778 | 1.290 | 1.326 | 0.963 | 0.0000 | 0.0000 | 1 | 1 | 0 |
| 'Ube2v1' | 86.022 | 94.339 | 87.851 | 88.579 | 95.519 | 92.882 | 84.023 | 92.196 | 90.921 | 94.881 | 88.799 | 83.572 | 97.145 | 87.293 | 88.018 | 92.593 | 86.546 | 85.644 | 0.005 | 0.8841 | 0.9514 | 0 | 0 | 0 |
| 'Ube2v2' | 37.037 | 41.168 | 42.583 | 42.101 | 44.277 | 36.791 | 46.308 | 46.146 | 39.576 | 34.903 | 37.757 | 39.555 | 42.249 | 45.198 | 39.531 | 36.740 | 40.619 | 36.514 | 0.075 | 0.1857 | 0.4214 | 0 | 0 | 0 |
| 'Ube2w' | 4.053 | 4.840 | 4.771 | 4.878 | 4.390 | 4.026 | 4.702 | 4.591 | 4.338 | 4.497 | 4.431 | 4.620 | 3.716 | 4.365 | 4.015 | 4.299 | 5.009 | 4.200 | 0.039 | 0.4759 | 0.7107 | 0 | 0 | 0 |
| 'Ube2z' | 41.014 | 45.695 | 41.340 | 40.343 | 46.145 | 43.725 | 43.770 | 42.247 | 42.507 | 41.482 | 43.689 | 45.328 | 42.652 | 45.007 | 46.967 | 44.084 | 44.274 | 45.955 | -0.059 | 0.0350 | 0.1496 | 0 | 0 | 0 |
| 'Ube3a' | 4.978 | 4.920 | 4.880 | 5.737 | 4.667 | 4.246 | 5.507 | 5.100 | 4.375 | 4.515 | 5.292 | 5.701 | 4.239 | 4.794 | 4.841 | 4.403 | 5.270 | 5.590 | -0.019 | 0.7772 | 0.9002 | 0 | 0 | 0 |
| 'Ube3b' | 16.485 | 17.726 | 17.743 | 15.391 | 16.604 | 18.183 | 15.933 | 17.356 | 18.396 | 18.341 | 16.318 | 18.604 | 17.542 | 17.289 | 17.666 | 17.187 | 16.509 | 18.256 | -0.047 | 0.0963 | 0.2885 | 0 | 0 | 0 |
| 'Ube3c' | 6.602 | 6.349 | 6.744 | 7.180 | 5.982 | 6.577 | 6.673 | 6.814 | 6.025 | 6.543 | 6.817 | 6.388 | 5.963 | 6.351 | 5.995 | 5.943 | 6.783 | 6.423 | 0.034 | 0.4187 | 0.6660 | 0 | 0 | 0 |
| 'Ube4a' | 6.625 | 6.613 | 7.182 | 7.090 | 6.396 | 6.782 | 6.487 | 6.950 | 6.739 | 5.897 | 6.694 | 7.197 | 5.906 | 6.653 | 7.055 | 5.819 | 7.205 | 6.825 | 0.031 | 0.4309 | 0.6762 | 0 | 0 | 0 |
| 'Ube4b' | 12.598 | 11.836 | 13.006 | 12.661 | 12.086 | 12.337 | 12.064 | 12.349 | 12.947 | 12.823 | 11.867 | 13.547 | 12.141 | 12.735 | 13.527 | 12.120 | 12.210 | 13.949 | -0.049 | 0.0746 | 0.2430 | 0 | 0 | 0 |
| 'Ubfd1' | 4.201 | 3.998 | 3.870 | 4.010 | 3.932 | 3.814 | 3.798 | 4.012 | 4.185 | 3.620 | 4.154 | 5.714 | 3.640 | 4.046 | 5.086 | 3.841 | 4.392 | 5.993 | -0.170 | 0.0154 | 0.0856 | 0 | 0 | 0 |
| 'Ubiad1' | 8.286 | 8.309 | 8.468 | 7.960 | 9.045 | 9.135 | 8.653 | 8.626 | 9.627 | 9.322 | 7.699 | 9.706 | 8.395 | 7.886 | 9.609 | 8.801 | 8.319 | 8.549 | -0.016 | 0.7163 | 0.8688 | 0 | 0 | 0 |
| 'Ubl3' | 14.268 | 15.404 | 12.593 | 13.493 | 14.229 | 13.815 | 12.986 | 14.345 | 12.328 | 13.621 | 13.775 | 11.427 | 13.427 | 14.464 | 11.994 | 13.673 | 12.662 | 12.187 | 0.066 | 0.0598 | 0.2110 | 0 | 0 | 0 |
| 'Ubl4a' | 4.144 | 4.712 | 4.600 | 4.802 | 4.941 | 4.839 | 4.533 | 4.669 | 4.545 | 4.185 | 4.464 | 4.964 | 4.698 | 4.728 | 5.138 | 4.721 | 5.043 | 4.977 | -0.053 | 0.1924 | 0.4295 | 0 | 0 | 0 |
| 'Ubl5' | 13.816 | 15.759 | 13.100 | 14.823 | 15.392 | 14.715 | 14.552 | 15.423 | 14.258 | 14.367 | 14.791 | 11.926 | 14.384 | 15.109 | 13.951 | 14.683 | 15.482 | 13.437 | 0.034 | 0.3527 | 0.6063 | 0 | 0 | 0 |
| 'Ubl7' | 57.265 | 62.522 | 65.633 | 59.837 | 61.537 | 65.500 | 56.069 | 58.697 | 63.363 | 57.965 | 57.548 | 59.320 | 59.932 | 56.139 | 58.539 | 56.559 | 54.113 | 56.685 | 0.081 | 0.0074 | 0.0499 | 1 | 0 | 0 |
| 'Ublcp1' | 11.016 | 13.208 | 11.985 | 12.197 | 12.599 | 11.502 | 11.817 | 12.596 | 11.323 | 11.828 | 12.872 | 11.493 | 12.551 | 13.129 | 11.540 | 10.776 | 13.171 | 11.086 | -0.014 | 0.6773 | 0.8457 | 0 | 0 | 0 |
| 'Ubn1' | 43.930 | 43.225 | 45.099 | 43.532 | 41.410 | 44.888 | 41.679 | 43.059 | 43.841 | 44.561 | 47.250 | 48.552 | 41.823 | 44.646 | 46.415 | 40.223 | 46.179 | 48.945 | -0.074 | 0.0172 | 0.0928 | 0 | 0 | 0 |
| 'Ubox5' | 1.504 | 2.161 | 1.823 | 1.845 | 1.861 | 1.645 | 1.765 | 1.886 | 1.547 | 2.081 | 1.721 | 1.379 | 1.773 | 1.531 | 1.030 | 1.585 | 1.229 | 1.363 | 0.226 | 0.0148 | 0.0827 | 0 | 0 | 0 |
| 'Ubp1' | 30.221 | 34.490 | 34.804 | 28.349 | 34.384 | 34.691 | 30.159 | 34.074 | 35.945 | 34.044 | 32.929 | 34.543 | 36.220 | 32.581 | 34.266 | 33.966 | 34.265 | 32.510 | -0.052 | 0.1475 | 0.3734 | 0 | 0 | 0 |
| 'Ubqln1' | 59.089 | 58.426 | 60.167 | 61.437 | 58.349 | 60.135 | 58.315 | 57.293 | 56.062 | 57.991 | 58.504 | 64.088 | 57.591 | 59.232 | 62.976 | 55.618 | 58.901 | 67.053 | -0.044 | 0.1639 | 0.3960 | 0 | 0 | 0 |
| 'Ubqln2' | 24.149 | 18.388 | 21.342 | 20.264 | 19.195 | 21.130 | 20.549 | 20.196 | 18.984 | 23.173 | 21.889 | 20.275 | 20.947 | 22.892 | 18.976 | 21.918 | 20.753 | 19.835 | -0.056 | 0.2343 | 0.4833 | 0 | 0 | 0 |
| 'Ubqln4' | 11.979 | 10.866 | 11.711 | 11.611 | 11.148 | 12.718 | 10.686 | 11.444 | 12.499 | 11.987 | 10.457 | 12.563 | 12.083 | 10.549 | 11.558 | 12.522 | 10.149 | 11.988 | 0.003 | 0.9274 | 0.9720 | 0 | 0 | 0 |
| 'Ubr1' | 9.134 | 9.319 | 9.334 | 9.294 | 9.331 | 8.457 | 9.387 | 9.273 | 8.759 | 9.294 | 10.131 | 9.681 | 8.536 | 9.975 | 8.504 | 9.156 | 9.796 | 8.848 | -0.038 | 0.2455 | 0.4959 | 0 | 0 | 0 |
| 'Ubr2' | 34.391 | 30.645 | 33.324 | 33.791 | 30.949 | 31.581 | 32.049 | 33.004 | 33.230 | 33.584 | 35.655 | 36.526 | 31.707 | 31.804 | 35.107 | 32.550 | 34.180 | 37.993 | -0.087 | 0.0178 | 0.0951 | 0 | 0 | 0 |
| 'Ubr3' | 6.787 | 6.903 | 6.831 | 7.087 | 6.761 | 6.554 | 6.792 | 6.916 | 6.678 | 6.274 | 7.199 | 6.774 | 6.816 | 6.851 | 6.481 | 6.364 | 7.210 | 6.622 | 0.006 | 0.8218 | 0.9217 | 0 | 0 | 0 |
| 'Ubr4' | 26.945 | 22.868 | 26.960 | 26.208 | 23.943 | 26.120 | 23.322 | 24.027 | 26.813 | 27.748 | 26.410 | 26.787 | 24.796 | 24.225 | 25.057 | 24.951 | 24.898 | 28.507 | -0.050 | 0.1795 | 0.4147 | 0 | 0 | 0 |
| 'Ubr5' | 35.260 | 35.572 | 33.458 | 36.274 | 34.394 | 32.291 | 33.530 | 35.544 | 33.285 | 36.882 | 36.904 | 33.496 | 34.115 | 34.917 | 32.503 | 36.033 | 36.730 | 33.851 | -0.037 | 0.1336 | 0.3522 | 0 | 0 | 0 |
| 'Ubr7' | 24.074 | 25.491 | 25.196 | 26.496 | 25.695 | 23.648 | 24.686 | 24.774 | 25.244 | 24.454 | 25.411 | 27.895 | 23.260 | 25.755 | 26.024 | 25.293 | 26.853 | 26.451 | -0.051 | 0.1296 | 0.3451 | 0 | 0 | 0 |
| 'Ubtd1' | 8.924 | 8.837 | 9.145 | 8.097 | 7.423 | 9.016 | 7.377 | 6.744 | 8.473 | 9.217 | 9.023 | 12.276 | 9.578 | 8.518 | 12.189 | 9.382 | 8.764 | 13.548 | -0.330 | 0.0000 | 0.0000 | 1 | 0 | 0 |
| 'Ubtd2' | 3.954 | 3.798 | 3.450 | 3.345 | 4.183 | 4.212 | 4.467 | 4.208 | 3.786 | 3.594 | 2.957 | 5.435 | 3.559 | 4.647 | 4.598 | 4.466 | 3.994 | 3.769 | -0.090 | 0.4363 | 0.6800 | 0 | 0 | 0 |
| 'Ubtf' | 31.371 | 30.248 | 34.089 | 31.138 | 30.307 | 34.680 | 29.322 | 30.328 | 34.651 | 32.258 | 31.308 | 35.394 | 34.683 | 31.149 | 36.102 | 30.432 | 30.911 | 37.315 | -0.076 | 0.0026 | 0.0222 | 1 | 0 | 0 |
| 'Ubxn1' | 98.687 | 104.769 | 96.086 | 95.309 | 108.668 | 107.200 | 93.231 | 106.489 | 101.384 | 99.921 | 93.401 | 95.455 | 101.474 | 100.188 | 101.418 | 106.766 | 95.943 | 95.034 | 0.025 | 0.4575 | 0.6947 | 0 | 0 | 0 |
| 'Ubxn11' | 5.297 | 5.216 | 6.415 | 5.629 | 4.605 | 6.360 | 5.861 | 4.787 | 6.561 | 5.379 | 3.649 | 5.455 | 5.509 | 4.340 | 5.178 | 5.166 | 4.794 | 5.118 | 0.175 | 0.0002 | 0.0022 | 1 | 0 | 0 |
| 'Ubxn2a' | 8.294 | 9.374 | 8.846 | 8.779 | 8.744 | 8.216 | 8.576 | 8.819 | 8.721 | 8.087 | 8.418 | 9.175 | 8.643 | 9.001 | 8.663 | 9.001 | 8.384 | 9.254 | -0.020 | 0.5915 | 0.7921 | 0 | 0 | 0 |
| 'Ubxn2b' | 4.138 | 5.021 | 4.258 | 4.046 | 5.054 | 4.439 | 4.651 | 4.722 | 4.519 | 4.795 | 4.612 | 5.200 | 4.517 | 4.769 | 4.642 | 4.730 | 4.638 | 4.096 | -0.059 | 0.2131 | 0.4581 | 0 | 0 | 0 |
| 'Ubxn4' | 27.638 | 27.388 | 27.685 | 27.349 | 28.632 | 26.514 | 27.317 | 27.430 | 27.087 | 26.635 | 27.227 | 30.503 | 27.605 | 27.472 | 27.784 | 27.165 | 28.500 | 29.991 | -0.046 | 0.0974 | 0.2909 | 0 | 0 | 0 |
| 'Ubxn6' | 7.387 | 7.420 | 7.534 | 6.369 | 7.248 | 8.554 | 6.947 | 7.281 | 7.878 | 7.509 | 6.806 | 7.772 | 7.380 | 6.325 | 7.634 | 7.439 | 6.625 | 7.216 | 0.032 | 0.4072 | 0.6562 | 0 | 0 | 0 |
| 'Ubxn7' | 10.383 | 10.496 | 9.578 | 10.189 | 10.140 | 9.399 | 9.970 | 9.732 | 9.371 | 10.290 | 11.205 | 11.132 | 9.137 | 10.465 | 9.991 | 10.836 | 11.622 | 10.661 | -0.106 | 0.0089 | 0.0576 | 0 | 0 | 0 |
| 'Ubxn8' | 10.128 | 11.293 | 11.062 | 11.051 | 11.614 | 11.433 | 12.512 | 12.283 | 10.754 | 11.693 | 11.169 | 11.592 | 10.681 | 12.023 | 11.183 | 12.032 | 12.294 | 10.297 | -0.022 | 0.6059 | 0.8014 | 0 | 0 | 0 |
| 'Uchl3' | 20.327 | 23.278 | 21.130 | 20.946 | 22.232 | 17.534 | 23.832 | 21.388 | 22.225 | 18.427 | 20.158 | 19.546 | 19.610 | 19.462 | 19.505 | 21.215 | 16.926 | 18.543 | 0.138 | 0.0168 | 0.0914 | 0 | 0 | 0 |
| 'Uchl5' | 7.301 | 7.216 | 7.743 | 8.426 | 8.328 | 7.479 | 8.486 | 8.127 | 6.821 | 6.074 | 7.284 | 7.895 | 7.056 | 7.389 | 7.438 | 7.279 | 7.681 | 7.306 | 0.082 | 0.1080 | 0.3093 | 0 | 0 | 0 |
| 'Uck1' | 15.502 | 15.050 | 15.513 | 14.058 | 15.087 | 15.328 | 13.364 | 15.774 | 15.683 | 14.964 | 14.231 | 15.503 | 16.000 | 15.153 | 16.418 | 15.184 | 15.301 | 16.764 | -0.053 | 0.1371 | 0.3570 | 0 | 0 | 0 |
| 'Uck2' | 15.807 | 14.930 | 13.569 | 14.551 | 15.040 | 14.290 | 14.370 | 15.168 | 13.434 | 12.977 | 14.586 | 13.722 | 13.791 | 14.423 | 14.410 | 14.193 | 12.257 | 13.362 | 0.073 | 0.0596 | 0.2110 | 0 | 0 | 0 |
| 'Uckl1' | 7.734 | 7.955 | 9.223 | 8.198 | 8.251 | 8.829 | 8.505 | 8.631 | 9.511 | 8.357 | 8.007 | 10.222 | 8.573 | 7.898 | 9.640 | 8.281 | 8.533 | 9.171 | -0.047 | 0.1221 | 0.3333 | 0 | 0 | 0 |
| 'Ucp2' | 136.107 | 131.542 | 136.763 | 133.690 | 134.249 | 142.914 | 134.831 | 129.733 | 145.204 | 136.900 | 137.813 | 127.680 | 134.050 | 130.012 | 145.295 | 133.836 | 131.217 | 139.559 | -0.001 | 0.9704 | 0.9898 | 0 | 0 | 0 |
| 'Uevld' | 2.870 | 4.070 | 3.947 | 3.419 | 3.753 | 3.701 | 3.110 | 3.546 | 3.926 | 3.187 | 3.281 | 3.658 | 3.577 | 3.541 | 3.092 | 3.326 | 3.268 | 3.164 | 0.087 | 0.1550 | 0.3835 | 0 | 0 | 0 |
| 'Ufc1' | 13.187 | 14.240 | 13.452 | 13.250 | 13.339 | 12.966 | 12.538 | 14.157 | 12.909 | 12.947 | 13.113 | 12.384 | 14.490 | 13.509 | 13.185 | 14.260 | 13.385 | 12.354 | -0.002 | 0.9560 | 0.9836 | 0 | 0 | 0 |
| 'Ufd1l' | 9.888 | 10.767 | 10.040 | 9.763 | 10.520 | 9.794 | 9.910 | 10.748 | 10.050 | 10.251 | 9.850 | 9.582 | 10.298 | 10.421 | 9.447 | 9.964 | 10.557 | 9.135 | 0.023 | 0.3897 | 0.6389 | 0 | 0 | 0 |
| 'Ufl1' | 7.162 | 8.018 | 7.208 | 7.355 | 8.270 | 7.120 | 7.611 | 7.941 | 7.531 | 7.583 | 7.447 | 7.746 | 7.363 | 8.214 | 7.485 | 7.601 | 7.948 | 7.345 | -0.026 | 0.3615 | 0.6135 | 0 | 0 | 0 |
| 'Ufm1' | 8.337 | 9.744 | 8.511 | 7.922 | 10.017 | 7.956 | 8.624 | 9.415 | 8.227 | 8.618 | 8.356 | 8.461 | 8.873 | 9.874 | 8.280 | 8.535 | 8.210 | 7.865 | 0.015 | 0.7544 | 0.8906 | 0 | 0 | 0 |
| 'Ufsp2' | 6.191 | 6.522 | 6.498 | 6.718 | 6.607 | 6.338 | 7.260 | 6.338 | 6.360 | 6.477 | 6.080 | 7.048 | 6.805 | 6.902 | 6.601 | 6.275 | 7.328 | 5.950 | -0.032 | 0.4753 | 0.7100 | 0 | 0 | 0 |
| 'Ugcg' | 30.867 | 25.834 | 26.499 | 29.677 | 26.776 | 26.005 | 29.582 | 27.364 | 25.122 | 24.087 | 26.571 | 26.000 | 23.903 | 26.163 | 24.257 | 24.699 | 27.443 | 26.104 | 0.099 | 0.0219 | 0.1092 | 0 | 0 | 0 |
| 'Ugdh' | 8.129 | 9.345 | 8.227 | 8.445 | 8.553 | 8.729 | 8.166 | 8.653 | 8.462 | 7.647 | 9.052 | 8.509 | 8.598 | 8.705 | 8.248 | 8.557 | 8.957 | 8.322 | -0.009 | 0.7670 | 0.8961 | 0 | 0 | 0 |

| | | | | | | | | | | | | | | | | | | | | | | | | |
|---|---|---|---|---|---|---|---|---|---|---|---|---|---|---|---|---|---|---|---|---|---|---|---|---|
| 'Uggt1' | 16.733 | 15.863 | 17.672 | 18.081 | 16.339 | 15.712 | 17.334 | 16.242 | 16.092 | 16.120 | 17.100 | 16.170 | 15.778 | 14.636 | 15.210 | 14.611 | 16.312 | 16.330 | 0.065 | 0.0851 | 0.2650 | 0 | 0 | 0 |
| 'Ugp2' | 22.665 | 27.318 | 24.387 | 24.656 | 28.345 | 23.608 | 27.871 | 26.675 | 23.746 | 26.453 | 24.300 | 22.833 | 25.540 | 27.354 | 22.162 | 22.546 | 25.998 | 20.824 | 0.061 | 0.1266 | 0.3409 | 0 | 0 | 0 |
| 'Uhmk1' | 24.439 | 25.686 | 26.425 | 25.105 | 26.121 | 25.067 | 24.862 | 25.448 | 24.756 | 30.547 | 31.780 | 31.068 | 29.515 | 29.872 | 29.741 | 29.542 | 30.271 | 30.941 | -0.273 | 0.0000 | 0.0000 | 1 | 0 | 0 |
| 'Uhrf1' | 20.840 | 19.371 | 26.478 | 23.482 | 19.565 | 24.359 | 24.423 | 19.119 | 25.317 | 18.629 | 22.140 | 23.914 | 28.338 | 22.680 | 27.137 | 21.449 | 22.263 | 24.107 | -0.071 | 0.2473 | 0.4979 | 0 | 0 | 0 |
| 'Uhrf1bp1' | 1.794 | 1.613 | 1.973 | 1.527 | 1.865 | 1.839 | 1.734 | 1.662 | 1.813 | 1.771 | 1.642 | 1.751 | 1.872 | 1.431 | 1.839 | 1.416 | 1.956 | 1.802 | 0.021 | 0.7417 | 0.8832 | 0 | 0 | 0 |
| 'Uhrf1bp1l' | 13.289 | 14.011 | 13.769 | 14.491 | 13.780 | 13.094 | 13.935 | 14.731 | 13.137 | 14.201 | 14.329 | 12.445 | 13.861 | 13.454 | 12.959 | 13.587 | 14.534 | 12.118 | 0.026 | 0.3835 | 0.6340 | 0 | 0 | 0 |
| 'Uhrf2' | 9.022 | 8.112 | 8.143 | 9.381 | 8.774 | 7.679 | 8.588 | 8.399 | 7.920 | 7.830 | 8.563 | 9.821 | 7.549 | 8.912 | 8.874 | 8.327 | 9.584 | 9.069 | -0.057 | 0.2710 | 0.5262 | 0 | 0 | 0 |
| 'Uimc1' | 15.341 | 16.196 | 15.975 | 16.445 | 15.293 | 15.910 | 16.091 | 15.593 | 16.316 | 14.956 | 15.094 | 15.669 | 14.774 | 15.783 | 14.830 | 14.599 | 16.553 | 14.352 | 0.057 | 0.0442 | 0.1742 | 0 | 0 | 0 |
| 'Ulk1' | 7.817 | 8.030 | 8.083 | 7.037 | 7.329 | 7.949 | 7.492 | 7.203 | 8.415 | 8.184 | 8.817 | 9.267 | 8.230 | 9.261 | 8.803 | 8.608 | 8.679 | 9.071 | -0.196 | 0.0000 | 0.0000 | 1 | 0 | 0 |
| 'Ulk2' | 6.653 | 7.402 | 6.658 | 6.618 | 7.086 | 6.988 | 6.445 | 7.406 | 6.258 | 7.429 | 7.691 | 7.642 | 6.720 | 8.011 | 7.216 | 7.112 | 7.673 | 7.622 | -0.138 | 0.0000 | 0.0005 | 1 | 0 | 0 |
| 'Ulk3' | 19.953 | 19.433 | 22.491 | 18.595 | 18.354 | 21.249 | 19.633 | 20.232 | 23.370 | 19.903 | 20.115 | 20.956 | 21.310 | 19.296 | 20.335 | 20.035 | 18.672 | 20.049 | 0.013 | 0.7239 | 0.8731 | 0 | 0 | 0 |
| 'Umad1' | 1.181 | 1.365 | 1.168 | 1.465 | 1.486 | 1.307 | 1.402 | 1.277 | 1.155 | 1.075 | 1.539 | 1.353 | 1.263 | 1.516 | 1.381 | 1.189 | 1.432 | 1.425 | -0.060 | 0.3733 | 0.6243 | 0 | 0 | 0 |
| 'Umps' | 6.941 | 5.681 | 6.368 | 6.547 | 5.269 | 5.678 | 6.081 | 5.784 | 5.913 | 5.712 | 5.550 | 6.678 | 5.976 | 6.070 | 6.002 | 6.023 | 5.531 | 6.369 | -0.004 | 0.9377 | 0.9764 | 0 | 0 | 0 |
| 'Unc119' | 8.218 | 8.501 | 8.338 | 8.501 | 8.057 | 9.235 | 8.420 | 8.315 | 8.509 | 7.954 | 6.539 | 8.493 | 8.225 | 6.439 | 7.642 | 8.266 | 7.201 | 7.388 | 0.148 | 0.0002 | 0.0028 | 1 | 0 | 0 |
| 'Unc119b' | 27.899 | 27.528 | 24.782 | 26.470 | 25.790 | 27.137 | 25.407 | 27.111 | 25.906 | 22.968 | 26.321 | 30.218 | 24.861 | 25.860 | 30.778 | 24.713 | 25.321 | 31.497 | -0.032 | 0.5140 | 0.7381 | 0 | 0 | 0 |
| 'Unc13a' | 0.590 | 0.582 | 0.719 | 0.632 | 0.638 | 0.646 | 0.611 | 0.754 | 0.635 | 0.595 | 0.642 | 0.617 | 0.512 | 0.482 | 0.634 | 0.460 | 0.453 | 0.554 | 0.221 | 0.0046 | 0.0352 | 1 | 0 | 0 |
| 'Unc13d' | 38.930 | 34.837 | 42.151 | 36.392 | 35.227 | 40.968 | 37.620 | 34.559 | 41.799 | 39.485 | 37.058 | 42.383 | 39.288 | 35.772 | 39.158 | 36.456 | 36.499 | 39.623 | -0.026 | 0.3015 | 0.5584 | 0 | 0 | 0 |
| 'Unc45a' | 27.009 | 23.126 | 28.441 | 24.174 | 22.801 | 27.257 | 25.059 | 24.780 | 27.899 | 25.170 | 25.934 | 27.015 | 26.272 | 23.248 | 26.270 | 24.260 | 23.795 | 27.985 | -0.008 | 0.8130 | 0.9157 | 0 | 0 | 0 |
| 'Unc50' | 6.049 | 8.056 | 6.728 | 6.261 | 8.331 | 6.823 | 6.413 | 7.799 | 7.018 | 6.853 | 6.898 | 7.533 | 6.793 | 8.040 | 7.230 | 7.672 | 6.850 | 6.818 | -0.046 | 0.3426 | 0.5989 | 0 | 0 | 0 |
| 'Unc93b1' | 7.867 | 9.270 | 8.191 | 7.864 | 9.340 | 8.025 | 7.713 | 8.652 | 8.246 | 9.247 | 8.025 | 8.272 | 9.334 | 9.300 | 7.636 | 8.549 | 8.515 | 7.659 | -0.038 | 0.3806 | 0.6321 | 0 | 0 | 0 |
| 'Ung' | 6.196 | 5.335 | 6.675 | 6.567 | 5.409 | 6.288 | 6.258 | 6.107 | 6.391 | 6.098 | 5.340 | 6.409 | 7.412 | 5.553 | 6.280 | 6.054 | 5.232 | 5.763 | 0.020 | 0.6585 | 0.8350 | 0 | 0 | 0 |
| 'Unk' | 8.292 | 7.912 | 9.520 | 7.828 | 8.613 | 9.221 | 8.078 | 9.192 | 9.708 | 9.187 | 8.884 | 8.399 | 8.620 | 8.121 | 7.970 | 8.710 | 8.376 | 7.742 | 0.035 | 0.4398 | 0.6831 | 0 | 0 | 0 |
| 'Upf1' | 51.460 | 43.078 | 54.301 | 51.113 | 43.530 | 50.126 | 51.333 | 44.799 | 52.203 | 47.919 | 52.156 | 55.792 | 49.171 | 44.280 | 52.093 | 42.464 | 51.561 | 53.310 | -0.035 | 0.4592 | 0.6960 | 0 | 0 | 0 |
| 'Upf2' | 15.978 | 13.909 | 14.217 | 15.641 | 13.525 | 13.155 | 16.182 | 14.242 | 13.962 | 15.417 | 15.889 | 14.243 | 14.416 | 14.343 | 13.921 | 14.454 | 15.908 | 14.324 | -0.035 | 0.3721 | 0.6240 | 0 | 0 | 0 |
| 'Upf3a' | 38.424 | 37.325 | 39.212 | 38.478 | 38.377 | 42.730 | 39.901 | 38.917 | 43.188 | 39.725 | 37.850 | 42.650 | 41.882 | 36.743 | 41.382 | 42.185 | 39.671 | 39.983 | -0.032 | 0.3104 | 0.5687 | 0 | 0 | 0 |
| 'Upf3b' | 12.419 | 12.924 | 13.304 | 13.304 | 13.245 | 12.170 | 14.518 | 12.942 | 12.615 | 12.460 | 13.350 | 12.605 | 13.142 | 13.851 | 12.048 | 12.768 | 13.635 | 11.340 | 0.018 | 0.6046 | 0.8010 | 0 | 0 | 0 |
| 'Uprt' | 6.482 | 6.535 | 7.467 | 7.274 | 6.285 | 5.792 | 6.813 | 6.981 | 6.723 | 6.830 | 8.053 | 7.949 | 7.160 | 7.358 | 6.892 | 6.910 | 7.701 | 8.883 | -0.178 | 0.0062 | 0.0439 | 1 | 0 | 0 |
| 'Uqcc1' | 1.367 | 1.578 | 1.666 | 1.636 | 1.714 | 1.539 | 1.556 | 1.648 | 1.517 | 1.592 | 1.372 | 1.497 | 1.615 | 1.572 | 1.538 | 1.703 | 1.710 | 1.282 | 0.027 | 0.6173 | 0.8087 | 0 | 0 | 0 |
| 'Uqcc2' | 16.265 | 17.365 | 16.586 | 17.950 | 17.716 | 16.719 | 15.513 | 17.522 | 17.235 | 15.399 | 14.642 | 15.646 | 17.237 | 14.742 | 15.867 | 15.649 | 17.212 | 14.845 | 0.102 | 0.0246 | 0.1192 | 0 | 0 | 0 |
| 'Uqcc3' | 47.496 | 52.429 | 48.526 | 48.472 | 51.230 | 51.975 | 48.160 | 53.394 | 51.142 | 44.112 | 44.826 | 46.717 | 47.725 | 45.474 | 48.021 | 47.963 | 47.671 | 43.483 | 0.112 | 0.0020 | 0.0183 | 1 | 0 | 0 |
| 'Uqcr10' | 50.514 | 53.176 | 49.817 | 51.835 | 53.465 | 51.227 | 50.312 | 52.892 | 52.001 | 46.565 | 48.826 | 45.410 | 51.799 | 51.647 | 52.697 | 53.551 | 52.947 | 48.005 | 0.036 | 0.3219 | 0.5798 | 0 | 0 | 0 |
| 'Uqcr11' | 25.312 | 29.887 | 26.799 | 27.452 | 32.397 | 28.298 | 27.490 | 32.006 | 26.221 | 25.548 | 24.119 | 24.429 | 28.455 | 29.394 | 25.621 | 26.996 | 25.888 | 24.463 | 0.110 | 0.0158 | 0.0871 | 0 | 0 | 0 |
| 'Uqcrb' | 33.906 | 39.299 | 35.795 | 37.001 | 41.617 | 36.333 | 36.975 | 36.787 | 35.222 | 32.184 | 33.058 | 32.014 | 35.275 | 39.111 | 35.915 | 34.778 | 34.015 | 33.529 | 0.092 | 0.0314 | 0.1391 | 0 | 0 | 0 |
| 'Uqcrc1' | 28.182 | 28.985 | 30.365 | 27.344 | 29.551 | 31.179 | 27.409 | 28.685 | 29.926 | 28.845 | 27.456 | 28.821 | 29.685 | 26.689 | 29.749 | 27.698 | 25.594 | 28.488 | 0.038 | 0.1587 | 0.3887 | 0 | 0 | 0 |
| 'Uqcrc2' | 34.021 | 33.766 | 35.180 | 36.382 | 37.005 | 34.288 | 34.856 | 36.586 | 33.762 | 34.614 | 34.575 | 33.712 | 34.436 | 34.529 | 34.299 | 33.821 | 35.115 | 31.073 | 0.035 | 0.1477 | 0.3736 | 0 | 0 | 0 |
| 'Uqcrfs1' | 54.073 | 60.842 | 53.600 | 50.486 | 56.266 | 53.926 | 52.680 | 56.129 | 52.672 | 52.918 | 52.449 | 50.831 | 55.372 | 57.528 | 52.470 | 53.565 | 51.442 | 48.448 | 0.037 | 0.2520 | 0.5034 | 0 | 0 | 0 |
| 'Uqcrh' | 50.426 | 52.241 | 45.107 | 53.614 | 52.660 | 50.480 | 50.055 | 51.841 | 47.699 | 45.138 | 50.615 | 45.500 | 50.245 | 52.690 | 49.680 | 49.920 | 51.963 | 47.356 | 0.025 | 0.4443 | 0.6872 | 0 | 0 | 0 |
| 'Uqcrq' | 32.052 | 33.254 | 29.722 | 33.707 | 33.847 | 32.524 | 31.863 | 31.455 | 30.357 | 30.612 | 30.290 | 27.088 | 32.634 | 29.485 | 29.174 | 31.855 | 32.082 | 28.436 | 0.080 | 0.0142 | 0.0806 | 0 | 0 | 0 |
| 'Urb1' | 4.082 | 3.973 | 4.455 | 4.018 | 3.418 | 3.852 | 3.782 | 4.521 | 4.583 | 4.151 | 3.947 | 4.375 | 3.832 | 3.614 | 4.043 | 3.768 | 4.044 | 4.296 | 0.015 | 0.7607 | 0.8935 | 0 | 0 | 0 |
| 'Urb2' | 5.208 | 4.511 | 5.308 | 4.905 | 4.732 | 5.324 | 4.869 | 4.852 | 5.537 | 4.882 | 5.109 | 5.648 | 4.625 | 4.426 | 5.449 | 4.659 | 4.326 | 5.383 | 0.014 | 0.7011 | 0.8608 | 0 | 0 | 0 |
| 'Urgcp' | 15.006 | 14.712 | 14.108 | 14.088 | 13.602 | 14.727 | 12.596 | 13.251 | 14.637 | 13.031 | 14.231 | 15.807 | 13.072 | 13.493 | 16.135 | 12.991 | 14.103 | 14.878 | -0.022 | 0.6078 | 0.8027 | 0 | 0 | 0 |
| 'Uri1' | 13.505 | 14.226 | 13.962 | 13.667 | 14.701 | 13.603 | 15.040 | 14.846 | 13.901 | 13.180 | 12.843 | 12.954 | 12.979 | 12.970 | 12.385 | 12.239 | 12.693 | 11.897 | 0.147 | 0.0000 | 0.0000 | 1 | 0 | 0 |
| 'Urm1' | 5.389 | 5.272 | 5.879 | 4.707 | 5.451 | 5.984 | 5.575 | 5.592 | 5.412 | 5.398 | 4.575 | 5.763 | 5.442 | 5.346 | 5.525 | 5.573 | 5.855 | 5.637 | -0.005 | 0.9069 | 0.9636 | 0 | 0 | 0 |
| 'Urod' | 8.546 | 9.023 | 9.995 | 8.604 | 9.462 | 9.938 | 9.526 | 9.793 | 9.333 | 8.940 | 9.069 | 9.179 | 8.870 | 9.135 | 8.585 | 8.758 | 8.580 | 8.527 | 0.072 | 0.0353 | 0.1504 | 0 | 0 | 0 |
| 'Uros' | 3.566 | 3.478 | 3.816 | 3.916 | 3.725 | 3.913 | 3.584 | 3.956 | 3.908 | 3.265 | 3.536 | 3.785 | 3.340 | 3.395 | 3.755 | 4.098 | 3.778 | 3.429 | 0.058 | 0.2306 | 0.4777 | 0 | 0 | 0 |
| 'Usb1' | 13.694 | 15.655 | 14.855 | 17.356 | 17.034 | 15.693 | 15.416 | 17.709 | 16.013 | 14.442 | 14.665 | 17.213 | 14.991 | 16.735 | 17.064 | 16.454 | 15.827 | 16.450 | -0.018 | 0.7057 | 0.8633 | 0 | 0 | 0 |
| 'Use1' | 17.115 | 16.910 | 17.029 | 17.750 | 18.142 | 17.065 | 18.069 | 17.900 | 16.610 | 17.063 | 17.667 | 15.867 | 17.764 | 15.874 | 15.727 | 16.440 | 16.457 | 15.626 | 0.067 | 0.0117 | 0.0699 | 0 | 0 | 0 |
| 'Usf1' | 13.774 | 13.220 | 12.952 | 13.456 | 12.424 | 13.357 | 12.406 | 12.881 | 13.223 | 11.979 | 12.930 | 14.332 | 12.962 | 12.646 | 13.779 | 13.384 | 13.569 | 14.209 | -0.036 | 0.2831 | 0.5394 | 0 | 0 | 0 |
| 'Usf2' | 38.226 | 39.474 | 42.134 | 34.735 | 38.782 | 44.745 | 35.734 | 39.917 | 41.602 | 39.520 | 34.292 | 38.880 | 39.216 | 35.621 | 40.211 | 39.507 | 34.805 | 39.674 | 0.046 | 0.2013 | 0.4426 | 0 | 0 | 0 |
| 'Usf3' | 4.099 | 4.152 | 3.942 | 3.860 | 3.756 | 3.820 | 3.863 | 3.951 | 3.791 | 4.548 | 4.502 | 4.139 | 4.285 | 4.110 | 3.837 | 4.282 | 4.025 | 4.060 | -0.110 | 0.0006 | 0.0065 | 1 | 0 | 0 |
| 'Usmg5' | 143.015 | 161.742 | 137.683 | 161.868 | 170.830 | 135.138 | 163.598 | 153.680 | 136.487 | 129.754 | 143.839 | 124.598 | 139.387 | 154.085 | 134.933 | 141.985 | 165.150 | 129.762 | 0.098 | 0.0162 | 0.0888 | 0 | 0 | 0 |
| 'Uso1' | 10.124 | 10.963 | 10.546 | 10.572 | 11.110 | 10.301 | 11.060 | 10.503 | 10.341 | 10.681 | 10.560 | 10.422 | 10.627 | 10.175 | 9.604 | 9.533 | 11.404 | 9.645 | 0.032 | 0.2894 | 0.5460 | 0 | 0 | 0 |
| 'Usp1' | 27.792 | 28.932 | 28.345 | 27.987 | 28.017 | 28.404 | 27.682 | 27.792 | 26.212 | 26.414 | 26.612 | 27.342 | 28.759 | 30.283 | 27.664 | 27.658 | 26.943 | 25.924 | 0.010 | 0.7120 | 0.8661 | 0 | 0 | 0 |
| 'Usp10' | 7.904 | 7.786 | 8.135 | 7.279 | 7.489 | 7.846 | 7.441 | 7.242 | 8.152 | 6.863 | 7.024 | 8.689 | 7.270 | 7.381 | 8.367 | 7.110 | 7.401 | 8.505 | 0.003 | 0.9272 | 0.9720 | 0 | 0 | 0 |
| 'Usp11' | 2.681 | 2.459 | 2.897 | 2.820 | 2.704 | 2.687 | 2.369 | 2.728 | 2.969 | 1.620 | 1.664 | 1.815 | 1.602 | 1.461 | 1.800 | 1.380 | 1.582 | 1.539 | 0.737 | 0.0000 | 0.0000 | 1 | 1 | 0 |
| 'Usp12' | 26.517 | 29.052 | 24.226 | 25.570 | 27.452 | 25.645 | 23.825 | 29.472 | 23.886 | 26.390 | 29.216 | 27.107 | 24.266 | 28.236 | 28.397 | 27.986 | 27.384 | 28.599 | -0.084 | 0.0342 | 0.1476 | 0 | 0 | 0 |
| 'Usp14' | 6.343 | 6.995 | 5.872 | 7.220 | 7.087 | 5.469 | 7.233 | 7.270 | 6.075 | 7.431 | 6.857 | 6.922 | 6.945 | 7.330 | 6.576 | 6.430 | 7.710 | 6.419 | -0.092 | 0.0306 | 0.1366 | 0 | 0 | 0 |
| 'Usp15' | 34.681 | 30.609 | 33.280 | 35.943 | 33.413 | 33.523 | 34.124 | 36.037 | 31.746 | 35.165 | 34.303 | 32.986 | 32.925 | 33.635 | 30.244 | 30.649 | 34.687 | 31.860 | 0.024 | 0.5080 | 0.7336 | 0 | 0 | 0 |
| 'Usp16' | 6.168 | 6.403 | 6.457 | 6.655 | 6.435 | 6.459 | 6.762 | 6.587 | 6.402 | 6.644 | 6.517 | 6.288 | 6.285 | 6.704 | 5.696 | 6.537 | 6.754 | 6.177 | 0.010 | 0.7437 | 0.8844 | 0 | 0 | 0 |
| 'Usp18' | 2.906 | 2.888 | 2.700 | 1.959 | 3.412 | 2.903 | 2.213 | 3.129 | 3.076 | 2.898 | 3.121 | 4.750 | 3.430 | 2.289 | 4.554 | 3.145 | 3.374 | 4.777 | -0.360 | 0.0007 | 0.0078 | 1 | 0 | 0 |
| 'Usp19' | 22.463 | 21.251 | 22.715 | 20.841 | 21.194 | 22.203 | 20.283 | 21.522 | 22.829 | 21.980 | 21.264 | 23.936 | 21.912 | 20.057 | 23.217 | 21.531 | 21.280 | 23.959 | -0.038 | 0.1277 | 0.3423 | 0 | 0 | 0 |
| 'Usp20' | 2.721 | 3.041 | 3.161 | 2.656 | 2.710 | 3.099 | 2.918 | 2.810 | 3.173 | 2.862 | 2.883 | 2.955 | 2.706 | 2.411 | 2.551 | 2.847 | 2.850 | 2.663 | 0.078 | 0.0895 | 0.2744 | 0 | 0 | 0 |
| 'Usp21' | 3.315 | 3.546 | 3.900 | 3.305 | 3.635 | 3.882 | 3.407 | 3.645 | 3.898 | 3.585 | 3.236 | 4.104 | 3.769 | 3.563 | 3.880 | 3.636 | 3.333 | 3.845 | -0.031 | 0.3562 | 0.6090 | 0 | 0 | 0 |
| 'Usp22' | 5.947 | 5.507 | 5.301 | 4.754 | 4.771 | 5.569 | 5.222 | 6.236 | 5.438 | 6.208 | 5.279 | 4.696 | 5.418 | 5.308 | 4.699 | 5.377 | 5.273 | 4.573 | 0.057 | 0.3231 | 0.5806 | 0 | 0 | 0 |
| 'Usp24' | 11.955 | 10.856 | 11.767 | 11.864 | 11.386 | 11.458 | 11.074 | 11.906 | 11.890 | 12.249 | 11.537 | 12.352 | 10.976 | 11.548 | 11.816 | 11.493 | 12.293 | 12.215 | -0.042 | 0.1584 | 0.3886 | 0 | 0 | 0 |
| 'Usp25' | 44.900 | 45.174 | 44.670 | 47.847 | 46.266 | 44.367 | 48.517 | 47.965 | 41.461 | 45.630 | 47.510 | 46.132 | 48.054 | 48.691 | 45.247 | 47.514 | 47.397 | 45.210 | -0.047 | 0.0445 | 0.1749 | 0 | 0 | 0 |
| 'Usp27x' | 0.618 | 0.638 | 0.535 | 0.500 | 0.603 | 0.637 | 0.644 | 0.467 | 0.728 | 0.566 | 0.483 | 0.713 | 0.517 | 0.715 | 0.681 | 0.429 | 0.576 | 0.676 | -0.018 | 0.8673 | 0.9426 | 0 | 0 | 0 |
| 'Usp28' | 3.860 | 4.042 | 3.621 | 4.439 | 4.314 | 3.644 | 4.609 | 3.954 | 3.858 | 4.223 | 3.883 | 4.233 | 3.732 | 3.975 | 3.855 | 3.933 | 4.383 | 3.908 | -0.013 | 0.8023 | 0.9114 | 0 | 0 | 0 |
| 'Usp3' | 4.412 | 4.060 | 4.518 | 4.168 | 4.118 | 4.292 | 4.569 | 4.209 | 4.073 | 4.230 | 4.389 | 4.459 | 3.982 | 4.316 | 4.258 | 4.354 | 4.386 | 4.254 | -0.017 | 0.6231 | 0.8119 | 0 | 0 | 0 |
| 'Usp30' | 5.391 | 5.951 | 6.064 | 4.937 | 5.991 | 5.954 | 5.382 | 5.927 | 5.555 | 6.353 | 5.487 | 5.420 | 6.119 | 4.980 | 4.963 | 5.653 | 4.895 | 4.299 | 0.081 | 0.1827 | 0.4190 | 0 | 0 | 0 |
| 'Usp31' | 3.769 | 3.289 | 3.670 | 4.038 | 3.425 | 2.842 | 3.385 | 3.363 | 3.305 | 3.278 | 3.707 | 4.056 | 3.608 | 3.556 | 3.879 | 2.960 | 3.739 | 4.601 | -0.113 | 0.1268 | 0.3411 | 0 | 0 | 0 |
| 'Usp33' | 5.062 | 5.190 | 4.745 | 5.092 | 5.016 | 4.696 | 4.553 | 5.070 | 4.701 | 4.787 | 4.941 | 5.456 | 4.651 | 5.165 | 4.865 | 4.687 | 5.187 | 5.427 | -0.047 | 0.2271 | 0.4734 | 0 | 0 | 0 |
| 'Usp34' | 16.298 | 15.465 | 15.828 | 16.533 | 15.003 | 14.909 | 15.349 | 14.722 | 14.912 | 15.801 | 16.246 | 15.701 | 13.965 | 15.428 | 13.787 | 15.373 | 15.789 | 15.755 | 0.001 | 0.9689 | 0.9897 | 0 | 0 | 0 |
| 'Usp35' | 1.382 | 1.400 | 1.887 | 1.367 | 1.423 | 1.621 | 1.553 | 1.590 | 1.925 | 1.605 | 1.281 | 1.523 | 1.621 | 1.165 | 1.495 | 1.592 | 1.595 | 1.327 | 0.093 | 0.2112 | 0.4565 | 0 | 0 | 0 |
| 'Usp36' | 7.578 | 6.612 | 7.364 | 6.369 | 6.701 | 7.036 | 6.638 | 7.398 | 7.351 | 7.018 | 6.398 | 7.296 | 6.441 | 6.064 | 6.617 | 6.630 | 6.061 | 6.543 | 0.085 | 0.0267 | 0.1257 | 0 | 0 | 0 |
| 'Usp37' | 3.039 | 3.336 | 3.173 | 3.311 | 3.350 | 2.982 | 3.231 | 3.277 | 3.118 | 2.725 | 3.413 | 3.743 | 3.423 | 3.315 | 3.366 | 3.027 | 3.312 | 3.957 | -0.084 | 0.1081 | 0.3094 | 0 | 0 | 0 |
| 'Usp38' | 27.105 | 26.772 | 28.130 | 24.453 | 27.804 | 27.553 | 25.740 | 26.721 | 27.173 | 27.180 | 26.193 | 29.998 | 22.980 | 27.571 | 28.225 | 28.858 | 26.879 | 29.105 | -0.042 | 0.2563 | 0.5086 | 0 | 0 | 0 |
| 'Usp39' | 19.710 | 19.608 | 20.563 | 19.702 | 21.596 | 20.235 | 19.450 | 20.871 | 20.146 | 19.675 | 19.940 | 19.828 | 19.761 | 19.611 | 20.483 | 18.445 | 19.781 | 18.686 | 0.036 | 0.2101 | 0.4553 | 0 | 0 | 0 |
| 'Usp4' | 18.586 | 19.977 | 19.534 | 18.474 | 20.165 | 19.800 | 18.411 | 20.078 | 19.828 | 21.016 | 20.259 | 19.609 | 20.772 | 19.735 | 19.816 | 20.815 | 20.298 | 19.773 | -0.068 | 0.0042 | 0.0331 | 1 | 0 | 0 |
| 'Usp40' | 1.229 | 1.295 | 1.426 | 1.218 | 1.366 | 1.323 | 1.388 | 1.275 | 1.584 | 1.154 | 1.403 | 1.466 | 1.122 | 1.112 | 1.252 | 1.152 | 1.317 | 1.228 | 0.098 | 0.0716 | 0.2363 | 0 | 0 | 0 |
| 'Usp42' | 5.734 | 5.430 | 5.390 | 5.326 | 5.248 | 5.529 | 4.835 | 5.897 | 5.754 | 5.769 | 5.759 | 5.941 | 5.195 | 5.871 | 5.655 | 5.745 | 5.391 | 5.902 | -0.068 | 0.0781 | 0.2506 | 0 | 0 | 0 |
| 'Usp45' | 5.356 | 5.123 | 4.856 | 5.712 | 5.213 | 4.585 | 5.680 | 5.209 | 4.889 | 4.848 | 4.722 | 5.537 | 4.555 | 4.765 | 4.773 | 4.752 | 5.531 | 5.310 | 0.042 | 0.4124 | 0.6608 | 0 | 0 | 0 |
| 'Usp46' | 0.804 | 0.668 | 0.711 | 0.571 | 0.816 | 0.708 | 0.664 | 0.857 | 0.615 | 0.761 | 0.676 | 0.551 | 0.806 | 0.737 | 0.628 | 0.808 | 0.783 | 0.606 | 0.024 | 0.7735 | 0.8993 | 0 | 0 | 0 |
| 'Usp47' | 24.820 | 24.192 | 23.407 | 24.304 | 24.632 | 23.900 | 23.997 | 23.381 | 22.435 | 24.796 | 24.177 | 23.539 | 22.657 | 24.381 | 23.234 | 23.342 | 25.180 | 23.003 | -0.005 | 0.8471 | 0.9343 | 0 | 0 | 0 |
| 'Usp48' | 11.184 | 10.286 | 10.875 | 11.026 | 10.063 | 10.254 | 10.697 | 10.451 | 10.982 | 11.031 | 11.465 | 11.489 | 10.650 | 10.190 | 10.721 | 10.086 | 11.273 | 11.684 | -0.052 | 0.1358 | 0.3545 | 0 | 0 | 0 |
| 'Usp49' | 0.691 | 0.724 | 0.886 | 0.629 | 0.730 | 0.830 | 0.766 | 0.803 | 0.941 | 0.828 | 0.870 | 0.754 | 0.813 | 0.861 | 0.817 | 0.861 | 0.743 | 0.703 | -0.045 | 0.5846 | 0.7863 | 0 | 0 | 0 |
| 'Usp5' | 17.280 | 17.529 | 20.706 | 17.297 | 17.791 | 20.372 | 17.670 | 18.222 | 19.732 | 19.071 | 17.027 | 19.266 | 19.380 | 17.598 | 18.756 | 19.443 | 16.958 | 18.876 | -0.008 | 0.7962 | 0.9092 | 0 | 0 | 0 |
| 'Usp50' | 2.444 | 2.591 | 2.570 | 2.031 | 2.541 | 2.378 | 2.347 | 2.179 | 2.729 | 2.078 | 2.241 | 2.631 | 2.312 | 2.091 | 2.434 | 2.546 | 2.059 | 2.508 | 0.046 | 0.4976 | 0.7258 | 0 | 0 | 0 |
| 'Usp53' | 2.116 | 2.288 | 2.181 | 2.106 | 2.187 | 2.027 | 2.365 | 2.080 | 1.969 | 2.248 | 2.512 | 2.133 | 2.329 | 2.461 | 2.009 | 2.121 | 2.563 | 2.024 | -0.086 | 0.0446 | 0.1750 | 0 | 0 | 0 |
| 'Usp54' | 1.923 | 1.519 | 2.158 | 1.789 | 1.619 | 1.765 | 1.710 | 1.689 | 1.974 | 1.660 | 1.797 | 2.330 | 1.947 | 1.748 | 2.209 | 1.816 | 2.021 | 2.340 | -0.155 | 0.0064 | 0.0450 | 1 | 0 | 0 |
| 'Usp6nl' | 9.198 | 11.656 | 9.433 | 9.412 | 10.948 | 9.331 | 9.579 | 10.418 | 9.394 | 9.267 | 10.159 | 8.482 | 9.924 | 10.766 | 8.066 | 9.002 | 10.963 | 8.157 | 0.069 | 0.0350 | 0.1496 | 0 | 0 | 0 |
| 'Usp7' | 23.618 | 24.158 | 23.144 | 24.468 | 23.061 | 22.353 | 22.618 | 22.986 | 23.295 | 23.084 | 24.100 | 23.829 | 22.859 | 23.404 | 23.359 | 22.592 | 23.534 | 24.165 | -0.019 | 0.4366 | 0.6801 | 0 | 0 | 0 |
| 'Usp8' | 13.005 | 14.657 | 12.998 | 12.869 | 13.753 | 13.135 | 13.792 | 14.068 | 13.150 | 13.869 | 13.839 | 12.997 | 13.550 | 13.636 | 12.334 | 13.335 | 13.790 | 12.632 | 0.006 | 0.7856 | 0.9044 | 0 | 0 | 0 |
| 'Uspl1' | 3.768 | 3.381 | 3.544 | 3.538 | 3.257 | 3.312 | 3.824 | 3.614 | 3.203 | 3.350 | 3.409 | 4.518 | 3.355 | 3.244 | 3.778 | 3.097 | 3.599 | 4.634 | -0.074 | 0.2669 | 0.5202 | 0 | 0 | 0 |
| 'Utp11l' | 13.233 | 13.248 | 12.483 | 14.207 | 13.796 | 12.945 | 13.824 | 13.528 | 13.666 | 11.409 | 12.026 | 13.742 | 12.044 | 12.153 | 12.657 | 11.197 | 13.327 | 12.305 | 0.111 | 0.0054 | 0.0396 | 1 | 0 | 0 |
| 'Utp14a' | 10.597 | 9.547 | 10.913 | 10.830 | 10.064 | 10.165 | 10.564 | 9.678 | 9.764 | 10.755 | 9.485 | 10.756 | 9.895 | 8.378 | 10.195 | 9.416 | 9.933 | 10.043 | 0.039 | 0.3183 | 0.5768 | 0 | 0 | 0 |
| 'Utp14b' | 1.416 | 0.600 | 0.696 | 0.827 | 0.833 | 0.605 | 0.868 | 0.869 | 0.770 | 1.154 | 1.059 | 0.634 | 0.975 | 1.076 | 0.725 | 1.164 | 0.925 | 0.558 | -0.128 | 0.2568 | 0.5093 | 0 | 0 | 0 |
| 'Utp15' | 11.424 | 11.621 | 11.187 | 11.642 | 11.948 | 11.843 | 11.068 | 11.946 | 11.772 | 11.696 | 12.300 | 11.986 | 11.180 | 11.390 | 10.907 | 13.248 | 11.317 | 11.434 | -0.022 | 0.5561 | 0.7679 | 0 | 0 | 0 |
| 'Utp18' | 11.575 | 11.551 | 10.558 | 11.317 | 11.451 | 10.112 | 12.205 | 11.659 | 10.748 | 11.214 | 10.435 | 11.894 | 11.134 | 11.262 | 10.745 | 11.090 | 11.372 | 10.632 | 0.002 | 0.9469 | 0.9795 | 0 | 0 | 0 |
| 'Utp20' | 8.362 | 6.959 | 8.211 | 7.586 | 7.306 | 7.889 | 8.489 | 7.695 | 7.919 | 7.491 | 7.447 | 8.622 | 7.393 | 6.643 | 7.294 | 8.218 | 7.721 | 8.196 | 0.018 | 0.6702 | 0.8409 | 0 | 0 | 0 |
| 'Utp23' | 3.955 | 4.643 | 4.798 | 4.562 | 4.864 | 4.865 | 5.396 | 4.845 | 4.523 | 4.269 | 4.833 | 5.041 | 4.127 | 4.511 | 4.909 | 4.795 | 4.824 | 4.611 | 0.004 | 0.9335 | 0.9746 | 0 | 0 | 0 |
| 'Utp3' | 43.215 | 41.033 | 39.505 | 44.365 | 42.674 | 42.292 | 44.360 | 42.066 | 43.532 | 37.993 | 42.338 | 43.940 | 42.611 | 42.946 | 45.384 | 40.586 | 42.829 | 43.198 | -0.008 | 0.8202 | 0.9206 | 0 | 0 | 0 |
| 'Utp6' | 13.393 | 13.678 | 12.847 | 14.744 | 12.986 | 13.560 | 13.950 | 13.153 | 12.702 | 13.262 | 14.682 | 14.573 | 13.183 | 13.729 | 14.250 | 12.281 | 14.357 | 13.398 | -0.044 | 0.2622 | 0.5160 | 0 | 0 | 0 |
| 'Utrn' | 41.492 | 35.830 | 39.337 | 45.981 | 37.122 | 35.837 | 42.402 | 37.238 | 38.210 | 36.890 | 44.752 | 40.145 | 30.816 | 36.887 | 36.652 | 36.796 | 42.886 | 40.315 | 0.019 | 0.7685 | 0.8967 | 0 | 0 | 0 |
| 'Uvrag' | 36.390 | 38.085 | 35.903 | 38.448 | 40.215 | 37.423 | 35.838 | 38.214 | 34.551 | 42.890 | 37.623 | 34.811 | 39.935 | 39.825 | 34.125 | 40.224 | 35.499 | 34.488 | -0.026 | 0.4634 | 0.6995 | 0 | 0 | 0 |
| 'Uvssa' | 4.563 | 5.224 | 4.543 | 4.473 | 4.884 | 4.271 | 4.361 | 4.700 | 4.574 | 4.640 | 4.781 | 4.276 | 4.475 | 4.292 | 4.133 | 4.365 | 4.741 | 3.971 | 0.058 | 0.0830 | 0.2611 | 0 | 0 | 0 |
| 'Uxs1' | 1.919 | 2.576 | 2.372 | 2.069 | 2.659 | 2.329 | 2.297 | 2.408 | 2.180 | 2.170 | 2.145 | 2.451 | 2.233 | 2.176 | 2.194 | 2.163 | 2.410 | 2.171 | 0.026 | 0.6278 | 0.8154 | 0 | 0 | 0 |
| 'Uxt' | 4.118 | 5.227 | 4.318 | 3.310 | 4.537 | 4.656 | 4.224 | 4.813 | 4.206 | 4.304 | 4.127 | 3.703 | 4.189 | 4.647 | 4.215 | 4.560 | 3.913 | 3.516 | 0.080 | 0.2559 | 0.5082 | 0 | 0 | 0 |
| 'Vac14' | 27.060 | 25.962 | 28.619 | 25.593 | 24.403 | 29.981 | 25.940 | 25.782 | 28.543 | 28.224 | 27.041 | 27.705 | 28.817 | 25.482 | 28.191 | 28.615 | 25.043 | 27.243 | -0.033 | 0.2963 | 0.5524 | 0 | 0 | 0 |
| 'Vamp1' | 9.902 | 15.182 | 11.786 | 10.368 | 12.472 | 12.770 | 10.824 | 13.397 | 13.800 | 11.049 | 10.831 | 11.167 | 12.070 | 12.311 | 11.983 | 12.778 | 11.025 | 11.655 | 0.056 | 0.3444 | 0.5996 | 0 | 0 | 0 |
| 'Vamp2' | 25.566 | 28.794 | 22.261 | 20.222 | 25.825 | 23.933 | 21.295 | 25.087 | 23.015 | 29.889 | 29.280 | 18.815 | 27.963 | 30.489 | 21.711 | 28.895 | 24.763 | 19.184 | -0.091 | 0.2011 | 0.4424 | 0 | 0 | 0 |
| 'Vamp3' | 16.371 | 18.396 | 15.857 | 13.385 | 19.465 | 15.920 | 13.504 | 19.323 | 15.325 | 17.412 | 14.998 | 14.133 | 16.752 | 18.781 | 15.056 | 18.100 | 15.348 | 14.796 | 0.008 | 0.9018 | 0.9612 | 0 | 0 | 0 |
| 'Vamp4' | 4.972 | 5.229 | 4.850 | 5.020 | 5.212 | 4.787 | 4.905 | 5.251 | 4.993 | 5.014 | 5.011 | 4.963 | 4.738 | 5.321 | 4.567 | 4.977 | 5.106 | 4.818 | 0.012 | 0.7371 | 0.8802 | 0 | 0 | 0 |
| 'Vamp5' | 12.752 | 15.811 | 13.316 | 13.500 | 16.445 | 14.563 | 13.406 | 15.715 | 14.622 | 15.548 | 15.673 | 15.365 | 15.068 | 17.470 | 15.436 | 16.154 | 17.544 | 15.342 | -0.155 | 0.0000 | 0.0002 | 1 | 0 | 0 |
| 'Vamp8' | 32.398 | 37.060 | 31.201 | 34.079 | 35.895 | 33.197 | 34.929 | 35.893 | 33.249 | 32.836 | 32.569 | 32.958 | 34.408 | 37.022 | 34.474 | 36.321 | 37.256 | 33.427 | -0.028 | 0.3630 | 0.6150 | 0 | 0 | 0 |
| 'Vapa' | 51.884 | 58.673 | 48.601 | 53.103 | 56.944 | 51.010 | 53.135 | 54.706 | 48.780 | 48.634 | 55.651 | 49.379 | 50.797 | 50.819 | 47.495 | 53.308 | 54.043 | 46.775 | 0.049 | 0.0839 | 0.2628 | 0 | 0 | 0 |
| 'Vapb' | 12.838 | 15.645 | 13.253 | 12.953 | 14.278 | 13.995 | 13.304 | 14.369 | 13.348 | 13.020 | 12.898 | 13.845 | 13.518 | 14.373 | 13.470 | 13.990 | 12.700 | 13.487 | 0.018 | 0.5962 | 0.7947 | 0 | 0 | 0 |
| 'Vars' | 25.874 | 25.136 | 29.658 | 25.258 | 24.173 | 28.704 | 25.160 | 25.536 | 28.674 | 25.885 | 24.587 | 27.354 | 28.859 | 23.804 | 26.437 | 27.297 | 24.565 | 27.543 | 0.000 | 0.9928 | 0.9986 | 0 | 0 | 0 |
| 'Vars2' | 4.846 | 4.531 | 5.261 | 4.580 | 4.293 | 5.242 | 4.455 | 4.485 | 5.332 | 5.102 | 4.388 | 4.929 | 4.978 | 4.392 | 4.885 | 4.087 | 4.278 | 4.871 | 0.028 | 0.4796 | 0.7118 | 0 | 0 | 0 |
| 'Vasn' | 11.714 | 7.161 | 13.158 | 10.019 | 8.348 | 9.287 | 8.990 | 9.149 | 12.570 | 13.541 | 14.250 | 11.111 | 13.660 | 9.705 | 10.559 | 10.742 | 10.232 | 13.274 | -0.265 | 0.0191 | 0.1002 | 0 | 0 | 0 |
| 'Vasp' | 126.915 | 128.700 | 131.584 | 122.699 | 131.257 | 135.353 | 122.967 | 124.186 | 135.689 | 137.725 | 123.896 | 134.467 | 135.314 | 128.396 | 139.926 | 134.098 | 130.716 | 131.741 | -0.056 | 0.0130 | 0.0757 | 0 | 0 | 0 |
| 'Vat1' | 3.753 | 3.971 | 3.617 | 3.273 | 4.253 | 4.852 | 4.433 | 4.230 | 4.791 | 3.336 | 2.989 | 3.977 | 3.492 | 3.215 | 4.341 | 3.490 | 3.276 | 3.948 | 0.199 | 0.0082 | 0.0539 | 0 | 0 | 0 |
| 'Vat1l' | 1.445 | 1.599 | 1.452 | 1.417 | 1.747 | 1.233 | 1.310 | 2.039 | 1.337 | 1.380 | 2.131 | 1.617 | 1.551 | 2.114 | 1.581 | 1.431 | 1.792 | 1.443 | -0.156 | 0.0144 | 0.0814 | 0 | 0 | 0 |
| 'Vav1' | 52.571 | 49.991 | 50.597 | 53.403 | 49.641 | 50.877 | 51.434 | 51.026 | 50.548 | 52.174 | 49.572 | 56.534 | 51.254 | 50.408 | 53.745 | 51.459 | 49.413 | 57.131 | -0.047 | 0.0843 | 0.2635 | 0 | 0 | 0 |
| 'Vav2' | 2.820 | 2.492 | 2.698 | 2.752 | 2.462 | 2.819 | 2.669 | 2.721 | 3.023 | 2.833 | 2.352 | 3.265 | 2.495 | 2.404 | 2.794 | 2.523 | 2.397 | 3.266 | -0.002 | 0.9627 | 0.9868 | 0 | 0 | 0 |
| 'Vav3' | 2.344 | 2.687 | 2.386 | 2.516 | 2.568 | 2.477 | 2.495 | 2.421 | 2.493 | 2.886 | 2.998 | 2.882 | 2.804 | 2.895 | 2.597 | 3.018 | 2.972 | 2.504 | -0.202 | 0.0000 | 0.0000 | 1 | 0 | 0 |
| 'Vbp1' | 20.900 | 23.800 | 22.824 | 24.925 | 25.739 | 19.697 | 25.135 | 23.681 | 20.756 | 19.325 | 22.139 | 23.449 | 21.618 | 25.250 | 21.459 | 20.002 | 23.204 | 21.617 | 0.051 | 0.3138 | 0.5715 | 0 | 0 | 0 |
| 'Vcp' | 72.633 | 77.383 | 71.562 | 75.932 | 80.145 | 76.054 | 76.578 | 75.961 | 72.796 | 75.915 | 75.352 | 72.766 | 75.907 | 75.839 | 73.022 | 75.732 | 75.412 | 73.929 | 0.000 | 0.9898 | 0.9972 | 0 | 0 | 0 |
| 'Vcpip1' | 13.130 | 12.908 | 12.365 | 12.902 | 13.482 | 12.061 | 12.348 | 13.236 | 12.565 | 13.016 | 14.214 | 14.403 | 11.443 | 14.331 | 13.932 | 12.392 | 14.608 | 14.654 | -0.106 | 0.0090 | 0.0581 | 0 | 0 | 0 |
| 'Vcpkmt' | 12.614 | 15.022 | 12.181 | 13.718 | 15.605 | 12.160 | 13.117 | 14.135 | 12.685 | 11.756 | 11.932 | 12.726 | 11.667 | 13.126 | 12.456 | 12.259 | 12.308 | 12.972 | 0.104 | 0.0199 | 0.1028 | 0 | 0 | 0 |
| 'Vdac1' | 36.409 | 36.823 | 36.559 | 37.225 | 37.275 | 37.225 | 37.560 | 39.034 | 36.095 | 34.518 | 37.292 | 35.839 | 36.179 | 37.547 | 36.393 | 35.088 | 35.875 | 36.004 | 0.031 | 0.1703 | 0.4030 | 0 | 0 | 0 |
| 'Vdac2' | 27.872 | 31.530 | 27.433 | 29.055 | 30.711 | 27.619 | 29.166 | 30.734 | 27.217 | 28.978 | 28.142 | 28.223 | 30.878 | 30.490 | 28.484 | 30.180 | 30.355 | 26.646 | -0.018 | 0.4279 | 0.6739 | 0 | 0 | 0 |
| 'Vdac3-ps1' | 19.041 | 24.057 | 21.166 | 21.136 | 24.625 | 20.233 | 27.181 | 23.441 | 20.050 | 20.189 | 23.961 | 21.304 | 23.851 | 21.692 | 21.032 | 22.001 | 24.162 | 21.040 | -0.006 | 0.9108 | 0.9654 | 0 | 0 | 0 |
| 'Vdr' | 50.564 | 52.488 | 51.983 | 51.688 | 58.126 | 51.683 | 52.020 | 52.111 | 51.890 | 44.731 | 45.431 | 42.321 | 42.521 | 44.047 | 42.238 | 41.176 | 46.899 | 42.446 | 0.260 | 0.0000 | 0.0000 | 1 | 0 | 0 |
| 'Vegfb' | 7.325 | 6.433 | 7.768 | 7.305 | 6.727 | 7.231 | 8.027 | 8.968 | 7.121 | 7.794 | 6.919 | 7.691 | 9.377 | 7.899 | 7.054 | 6.852 | 6.699 | 7.665 | -0.032 | 0.6417 | 0.8243 | 0 | 0 | 0 |
| 'Vezf1' | 19.064 | 19.467 | 18.342 | 18.425 | 18.975 | 19.170 | 18.720 | 19.435 | 19.151 | 18.167 | 19.965 | 19.520 | 18.602 | 19.942 | 19.279 | 20.697 | 20.350 | 19.497 | -0.053 | 0.0584 | 0.2082 | 0 | 0 | 0 |
| 'Vezt' | 0.837 | 0.890 | 0.984 | 0.873 | 0.899 | 0.857 | 1.009 | 0.956 | 1.031 | 0.959 | 0.880 | 0.837 | 0.853 | 0.863 | 0.877 | 0.769 | 0.890 | 0.766 | 0.108 | 0.0511 | 0.1914 | 0 | 0 | 0 |
| 'Vgll4' | 13.355 | 14.140 | 11.095 | 12.958 | 12.427 | 11.211 | 12.051 | 12.781 | 10.847 | 11.954 | 14.343 | 14.725 | 11.690 | 14.493 | 13.655 | 12.873 | 13.882 | 15.279 | -0.161 | 0.0050 | 0.0376 | 1 | 0 | 0 |
| 'Vhl' | 9.187 | 11.084 | 8.860 | 9.264 | 9.628 | 9.838 | 8.931 | 9.344 | 9.048 | 8.325 | 10.070 | 9.662 | 9.568 | 10.381 | 8.811 | 9.354 | 10.093 | 9.489 | -0.023 | 0.5828 | 0.7850 | 0 | 0 | 0 |
| 'Vim' | 7.787 | 5.907 | 9.539 | 7.287 | 8.002 | 9.915 | 8.296 | 8.388 | 7.640 | 6.218 | 6.905 | 12.113 | 6.950 | 8.000 | 11.795 | 7.707 | 7.088 | 10.574 | -0.068 | 0.3883 | 0.6378 | 0 | 0 | 0 |
| 'Vimp' | 10.494 | 12.733 | 11.879 | 12.031 | 13.219 | 13.145 | 12.419 | 13.144 | 12.538 | 13.221 | 12.183 | 11.688 | 12.170 | 12.117 | 11.562 | 14.026 | 12.283 | 10.206 | 0.021 | 0.6713 | 0.8415 | 0 | 0 | 0 |
| 'Vipas39' | 20.160 | 20.884 | 22.051 | 22.098 | 22.470 | 21.007 | 21.250 | 23.893 | 21.185 | 22.665 | 20.812 | 20.924 | 22.366 | 21.711 | 20.715 | 21.358 | 20.773 | 20.019 | 0.019 | 0.5847 | 0.7863 | 0 | 0 | 0 |
| 'Vipr1' | 0.726 | 0.982 | 0.768 | 0.980 | 0.773 | 1.041 | 1.263 | 1.034 | 1.115 | 0.641 | 0.724 | 0.900 | 0.926 | 0.978 | 1.229 | 0.996 | 0.806 | 1.002 | 0.071 | 0.5612 | 0.7711 | 0 | 0 | 0 |
| 'Vipr2' | 1.593 | 2.512 | 1.923 | 1.824 | 2.080 | 1.922 | 1.950 | 2.181 | 2.202 | 1.884 | 1.513 | 1.425 | 1.913 | 1.548 | 1.278 | 1.692 | 1.363 | 1.074 | 0.407 | 0.0000 | 0.0004 | 1 | 1 | 0 |
| 'Vkorc1' | 5.262 | 5.868 | 5.367 | 5.189 | 7.170 | 5.933 | 5.356 | 5.983 | 6.200 | 5.079 | 4.682 | 4.780 | 5.487 | 4.821 | 5.248 | 6.482 | 4.615 | 3.908 | 0.209 | 0.0141 | 0.0801 | 0 | 0 | 0 |
| 'Vkorc1l1' | 2.571 | 3.133 | 3.027 | 2.978 | 3.497 | 3.116 | 3.070 | 3.211 | 3.162 | 2.878 | 3.039 | 3.369 | 2.922 | 3.190 | 3.161 | 2.762 | 2.997 | 3.509 | -0.023 | 0.6398 | 0.8229 | 0 | 0 | 0 |
| 'Vma21' | 2.781 | 3.084 | 2.721 | 2.797 | 3.440 | 2.517 | 2.595 | 2.911 | 2.663 | 2.927 | 3.058 | 2.528 | 2.662 | 2.802 | 2.394 | 2.567 | 2.616 | 2.536 | 0.070 | 0.1621 | 0.3938 | 0 | 0 | 0 |
| 'Vmac' | 3.144 | 3.652 | 3.500 | 2.649 | 3.614 | 3.787 | 3.181 | 3.816 | 3.933 | 3.506 | 2.966 | 3.384 | 3.326 | 3.279 | 3.377 | 3.400 | 3.060 | 3.146 | 0.078 | 0.2248 | 0.4716 | 0 | 0 | 0 |
| 'Vmp1' | 17.564 | 18.501 | 20.325 | 17.014 | 20.421 | 17.225 | 17.692 | 20.354 | 18.491 | 17.651 | 20.160 | 17.116 | 17.954 | 19.998 | 16.147 | 16.385 | 20.206 | 14.709 | 0.059 | 0.1670 | 0.3995 | 0 | 0 | 0 |
| 'Vopp1' | 4.317 | 4.813 | 3.860 | 4.081 | 4.858 | 3.948 | 3.887 | 4.739 | 3.911 | 4.218 | 4.752 | 4.284 | 4.107 | 4.749 | 4.287 | 4.658 | 4.663 | 4.497 | -0.080 | 0.0350 | 0.1496 | 0 | 0 | 0 |
| 'Vprbp' | 6.711 | 5.765 | 6.411 | 6.659 | 5.744 | 6.273 | 7.117 | 6.123 | 6.104 | 5.964 | 5.729 | 6.561 | 5.538 | 5.101 | 6.215 | 6.271 | 5.630 | 6.077 | 0.087 | 0.0330 | 0.1438 | 0 | 0 | 0 |
| 'Vps11' | 26.832 | 30.158 | 30.800 | 28.221 | 30.352 | 31.528 | 28.324 | 30.069 | 29.933 | 32.815 | 29.744 | 31.540 | 30.504 | 28.826 | 30.339 | 28.774 | 31.067 | 31.766 | -0.061 | 0.0514 | 0.1917 | 0 | 0 | 0 |
| 'Vps13a' | 11.419 | 11.308 | 11.176 | 12.345 | 11.211 | 9.931 | 11.854 | 11.208 | 10.306 | 11.175 | 12.028 | 11.035 | 10.459 | 11.488 | 10.093 | 9.997 | 12.042 | 11.318 | 0.005 | 0.9123 | 0.9663 | 0 | 0 | 0 |
| 'Vps13b' | 10.431 | 9.764 | 9.829 | 11.081 | 9.669 | 9.521 | 10.873 | 9.456 | 10.044 | 11.157 | 10.918 | 10.215 | 10.012 | 10.344 | 9.100 | 10.131 | 10.817 | 10.839 | -0.057 | 0.1597 | 0.3901 | 0 | 0 | 0 |
| 'Vps13c' | 28.155 | 27.328 | 27.086 | 29.528 | 25.981 | 25.035 | 26.101 | 25.362 | 26.457 | 28.397 | 30.209 | 25.196 | 28.600 | 28.538 | 24.863 | 28.587 | 28.245 | 27.814 | -0.065 | 0.0825 | 0.2597 | 0 | 0 | 0 |
| 'Vps13d' | 5.337 | 4.477 | 4.926 | 5.338 | 4.454 | 4.592 | 4.719 | 4.941 | 4.938 | 5.091 | 5.064 | 5.400 | 4.578 | 5.042 | 4.837 | 4.884 | 5.143 | 5.388 | -0.066 | 0.1275 | 0.3421 | 0 | 0 | 0 |
| 'Vps16' | 21.846 | 22.508 | 24.467 | 22.504 | 21.328 | 23.559 | 22.507 | 21.878 | 24.179 | 22.376 | 22.567 | 23.894 | 24.290 | 21.696 | 23.317 | 20.590 | 22.871 | 22.388 | -0.006 | 0.8322 | 0.9270 | 0 | 0 | 0 |
| 'Vps18' | 3.335 | 2.707 | 2.646 | 2.917 | 2.791 | 2.718 | 2.865 | 2.703 | 2.816 | 3.353 | 2.844 | 2.845 | 3.202 | 3.005 | 2.679 | 2.843 | 2.336 | 2.775 | -0.031 | 0.5298 | 0.7503 | 0 | 0 | 0 |
| 'Vps26a' | 14.678 | 16.449 | 14.995 | 14.961 | 16.524 | 15.735 | 15.880 | 17.136 | 14.906 | 14.018 | 15.670 | 16.575 | 15.173 | 17.261 | 14.941 | 15.946 | 16.307 | 16.144 | -0.023 | 0.5252 | 0.7470 | 0 | 0 | 0 |
| 'Vps26b' | 21.492 | 22.274 | 20.193 | 20.209 | 21.798 | 21.014 | 19.581 | 21.300 | 20.784 | 21.629 | 20.874 | 24.356 | 19.726 | 20.692 | 23.255 | 21.014 | 21.106 | 24.783 | -0.075 | 0.0547 | 0.2002 | 0 | 0 | 0 |
| 'Vps29' | 6.224 | 7.219 | 7.125 | 7.196 | 7.134 | 6.796 | 7.059 | 7.474 | 7.201 | 6.622 | 6.711 | 7.169 | 7.062 | 7.993 | 7.504 | 6.777 | 7.620 | 6.749 | -0.029 | 0.4284 | 0.6742 | 0 | 0 | 0 |
| 'Vps33a' | 12.443 | 13.340 | 15.136 | 13.461 | 13.322 | 14.737 | 13.203 | 14.450 | 14.549 | 13.900 | 14.880 | 14.512 | 14.398 | 13.390 | 14.226 | 14.146 | 15.017 | 15.002 | -0.064 | 0.0553 | 0.2018 | 0 | 0 | 0 |
| 'Vps33b' | 5.413 | 4.702 | 6.188 | 5.267 | 5.323 | 5.672 | 5.120 | 5.518 | 5.439 | 5.486 | 5.133 | 5.443 | 5.715 | 5.379 | 5.470 | 5.243 | 5.416 | 5.308 | -0.004 | 0.9297 | 0.9735 | 0 | 0 | 0 |
| 'Vps35' | 31.036 | 36.242 | 32.314 | 32.632 | 33.366 | 30.957 | 32.339 | 32.688 | 32.886 | 32.445 | 34.785 | 33.148 | 30.581 | 35.472 | 31.955 | 32.812 | 35.507 | 32.594 | -0.035 | 0.2242 | 0.4712 | 0 | 0 | 0 |
| 'Vps36' | 16.690 | 18.421 | 18.382 | 17.702 | 17.750 | 18.284 | 18.309 | 18.376 | 17.314 | 16.199 | 18.160 | 17.592 | 17.021 | 16.832 | 16.652 | 17.196 | 17.964 | 16.019 | 0.059 | 0.0443 | 0.1745 | 0 | 0 | 0 |
| 'Vps37a' | 7.858 | 8.833 | 8.516 | 8.355 | 8.882 | 7.895 | 8.126 | 8.575 | 8.041 | 8.475 | 7.771 | 8.644 | 7.990 | 8.869 | 7.883 | 7.407 | 8.673 | 8.163 | 0.008 | 0.8312 | 0.9267 | 0 | 0 | 0 |
| 'Vps37b' | 16.181 | 18.057 | 14.910 | 15.411 | 15.136 | 16.330 | 19.731 | 16.149 | 18.285 | 15.111 | 14.501 | 15.810 | 15.657 | 14.944 | 16.104 | 16.103 | 14.083 | 15.593 | 0.111 | 0.0241 | 0.1177 | 0 | 0 | 0 |
| 'Vps37c' | 9.924 | 8.438 | 9.626 | 9.269 | 9.294 | 9.443 | 9.835 | 8.709 | 10.121 | 9.559 | 9.349 | 9.775 | 9.884 | 9.050 | 8.912 | 10.231 | 9.642 | 9.734 | -0.032 | 0.4776 | 0.7109 | 0 | 0 | 0 |
| 'Vps39' | 7.461 | 8.434 | 8.500 | 7.718 | 8.207 | 8.305 | 8.167 | 8.401 | 8.437 | 8.426 | 8.634 | 7.927 | 8.051 | 7.827 | 7.833 | 7.857 | 8.253 | 7.488 | 0.017 | 0.5692 | 0.7768 | 0 | 0 | 0 |
| 'Vps41' | 25.482 | 27.281 | 27.090 | 27.665 | 27.236 | 25.187 | 28.854 | 26.206 | 24.691 | 26.433 | 28.314 | 27.346 | 26.423 | 28.314 | 25.962 | 25.680 | 30.213 | 25.541 | -0.039 | 0.2594 | 0.5126 | 0 | 0 | 0 |
| 'Vps45' | 6.015 | 5.884 | 6.297 | 6.344 | 5.746 | 5.963 | 6.840 | 6.636 | 6.029 | 6.451 | 6.097 | 6.011 | 6.317 | 6.155 | 5.607 | 5.386 | 6.903 | 5.376 | 0.029 | 0.5517 | 0.7649 | 0 | 0 | 0 |
| 'Vps4a' | 18.582 | 17.974 | 18.698 | 19.445 | 19.253 | 18.270 | 18.878 | 19.249 | 18.295 | 17.212 | 18.201 | 20.411 | 17.747 | 19.087 | 19.723 | 18.007 | 18.062 | 20.813 | -0.018 | 0.6344 | 0.8189 | 0 | 0 | 0 |
| 'Vps4b' | 11.744 | 12.872 | 12.547 | 12.137 | 13.153 | 12.234 | 12.183 | 13.266 | 12.496 | 12.404 | 12.932 | 12.302 | 12.181 | 12.975 | 12.676 | 12.411 | 13.578 | 11.685 | -0.015 | 0.4974 | 0.7258 | 0 | 0 | 0 |
| 'Vps50' | 1.786 | 1.793 | 1.681 | 1.945 | 1.929 | 1.644 | 1.776 | 1.696 | 1.797 | 1.922 | 2.075 | 1.788 | 1.840 | 1.976 | 1.761 | 1.563 | 1.906 | 1.470 | -0.033 | 0.5414 | 0.7584 | 0 | 0 | 0 |
| 'Vps51' | 6.927 | 6.984 | 7.598 | 6.956 | 7.160 | 7.709 | 7.028 | 7.291 | 8.256 | 6.876 | 6.640 | 6.987 | 7.701 | 6.571 | 6.427 | 6.904 | 6.928 | 6.476 | 0.090 | 0.0207 | 0.1056 | 0 | 0 | 0 |
| 'Vps52' | 27.705 | 27.382 | 31.494 | 27.596 | 28.258 | 30.330 | 28.668 | 28.325 | 31.261 | 29.607 | 27.534 | 31.511 | 29.336 | 27.639 | 29.801 | 27.540 | 29.162 | 29.964 | -0.017 | 0.4262 | 0.6726 | 0 | 0 | 0 |
| 'Vps53' | 6.878 | 6.414 | 6.984 | 6.916 | 6.742 | 7.218 | 7.135 | 6.557 | 7.066 | 6.989 | 6.796 | 7.233 | 6.923 | 6.742 | 6.880 | 6.850 | 6.942 | 6.886 | -0.018 | 0.5131 | 0.7375 | 0 | 0 | 0 |
| 'Vps54' | 10.177 | 11.136 | 10.129 | 9.766 | 10.973 | 9.679 | 10.961 | 11.956 | 9.655 | 10.135 | 10.688 | 10.763 | 9.276 | 11.539 | 10.023 | 10.439 | 11.670 | 9.877 | -0.011 | 0.7323 | 0.8781 | 0 | 0 | 0 |
| 'Vps72' | 14.195 | 14.089 | 15.454 | 14.506 | 14.667 | 15.405 | 14.605 | 15.045 | 15.927 | 15.292 | 13.496 | 14.991 | 14.683 | 13.541 | 14.894 | 15.975 | 14.192 | 14.983 | 0.013 | 0.6735 | 0.8431 | 0 | 0 | 0 |
| 'Vps8' | 3.931 | 3.965 | 3.633 | 3.796 | 4.059 | 3.974 | 4.126 | 3.885 | 3.949 | 4.684 | 4.112 | 4.185 | 4.192 | 4.247 | 3.764 | 4.063 | 4.179 | 3.668 | -0.082 | 0.0312 | 0.1384 | 0 | 0 | 0 |
| 'Vps9d1' | 4.468 | 4.297 | 4.878 | 4.412 | 4.083 | 4.952 | 4.459 | 4.279 | 5.030 | 4.830 | 4.024 | 5.342 | 4.605 | 4.300 | 4.806 | 4.379 | 3.767 | 4.741 | -0.010 | 0.7741 | 0.8993 | 0 | 0 | 0 |
| 'Vrk1' | 30.257 | 34.756 | 29.099 | 32.022 | 34.032 | 29.160 | 32.113 | 32.689 | 29.961 | 28.706 | 30.750 | 25.587 | 32.604 | 30.267 | 26.837 | 30.288 | 31.110 | 24.833 | 0.114 | 0.0001 | 0.0020 | 1 | 0 | 0 |
| 'Vrk2' | 3.182 | 3.727 | 3.387 | 3.592 | 3.476 | 3.415 | 4.269 | 3.768 | 2.924 | 3.295 | 3.721 | 3.466 | 3.606 | 3.403 | 3.016 | 3.356 | 3.842 | 3.166 | 0.024 | 0.6482 | 0.8285 | 0 | 0 | 0 |
| 'Vrk3' | 15.481 | 15.667 | 17.487 | 15.718 | 15.640 | 18.054 | 16.602 | 17.417 | 17.894 | 16.291 | 16.208 | 18.436 | 16.357 | 16.700 | 18.380 | 16.834 | 16.659 | 18.075 | -0.047 | 0.0798 | 0.2542 | 0 | 0 | 0 |
| 'Vsig10' | 4.114 | 4.402 | 4.485 | 4.191 | 4.409 | 4.060 | 4.493 | 4.636 | 4.427 | 3.985 | 4.827 | 5.057 | 3.988 | 4.337 | 4.835 | 4.205 | 4.577 | 4.896 | -0.065 | 0.1197 | 0.3293 | 0 | 0 | 0 |
| 'Vsir' | 39.424 | 39.584 | 38.458 | 36.319 | 39.737 | 39.562 | 35.477 | 40.018 | 37.832 | 41.296 | 39.657 | 41.162 | 39.974 | 41.619 | 41.987 | 41.738 | 39.435 | 42.559 | -0.104 | 0.0001 | 0.0013 | 1 | 0 | 0 |
| 'Vta1' | 8.305 | 9.786 | 9.402 | 9.734 | 11.062 | 10.777 | 10.069 | 9.960 | 9.766 | 10.051 | 9.465 | 9.392 | 9.999 | 9.672 | 8.561 | 9.506 | 9.293 | 8.033 | 0.071 | 0.1444 | 0.3687 | 0 | 0 | 0 |
| 'Vti1a' | 12.128 | 13.735 | 12.876 | 13.190 | 12.401 | 13.371 | 12.864 | 13.158 | 11.970 | 14.047 | 11.973 | 11.937 | 12.728 | 12.702 | 12.157 | 12.903 | 11.989 | 11.259 | 0.043 | 0.2660 | 0.5196 | 0 | 0 | 0 |
| 'Vti1b' | 23.510 | 24.663 | 24.231 | 24.195 | 26.203 | 25.552 | 23.807 | 25.558 | 24.729 | 24.215 | 24.750 | 23.152 | 24.847 | 24.848 | 23.501 | 24.289 | 23.800 | 21.862 | 0.040 | 0.1732 | 0.4066 | 0 | 0 | 0 |
| 'Vwa5a' | 8.390 | 9.660 | 8.951 | 8.853 | 9.284 | 8.978 | 9.228 | 9.317 | 8.515 | 9.269 | 10.553 | 10.110 | 9.739 | 10.218 | 9.042 | 8.341 | 10.872 | 9.450 | -0.122 | 0.0005 | 0.0058 | 1 | 0 | 0 |
| 'Vwa8' | 9.843 | 9.422 | 8.710 | 8.254 | 8.564 | 9.005 | 8.849 | 9.261 | 8.839 | 11.179 | 9.648 | 9.734 | 9.200 | 9.207 | 8.980 | 8.581 | 9.587 | 9.411 | -0.093 | 0.0424 | 0.1700 | 0 | 0 | 0 |
| 'Vwa9' | 13.099 | 13.979 | 13.688 | 14.741 | 14.655 | 14.172 | 14.054 | 15.930 | 14.532 | 14.811 | 13.441 | 13.932 | 14.321 | 13.843 | 14.746 | 13.993 | 13.112 | 12.684 | 0.035 | 0.3736 | 0.6246 | 0 | 0 | 0 |
| 'Wac' | 18.419 | 16.567 | 15.591 | 17.824 | 16.515 | 16.069 | 16.925 | 15.815 | 14.610 | 16.141 | 16.238 | 17.279 | 15.609 | 16.732 | 16.994 | 16.517 | 16.137 | 17.752 | -0.024 | 0.5740 | 0.7798 | 0 | 0 | 0 |
| 'Wapl' | 23.970 | 24.238 | 24.328 | 26.248 | 24.261 | 24.160 | 26.168 | 23.912 | 23.931 | 23.072 | 25.793 | 25.158 | 23.400 | 24.838 | 23.766 | 23.147 | 24.948 | 25.754 | -0.002 | 0.9443 | 0.9789 | 0 | 0 | 0 |
| 'Wars' | 10.925 | 10.791 | 10.786 | 10.521 | 12.004 | 12.279 | 10.763 | 11.074 | 11.710 | 12.360 | 10.850 | 11.379 | 11.123 | 10.321 | 10.936 | 11.143 | 10.119 | 10.462 | 0.022 | 0.5766 | 0.7816 | 0 | 0 | 0 |
| 'Wars2' | 1.740 | 1.594 | 2.069 | 1.654 | 1.722 | 1.906 | 2.064 | 1.884 | 2.010 | 1.948 | 1.999 | 2.139 | 1.982 | 1.775 | 1.971 | 1.863 | 1.877 | 1.881 | -0.076 | 0.2198 | 0.4669 | 0 | 0 | 0 |
| 'Was' | 92.762 | 86.398 | 98.800 | 92.826 | 92.021 | 103.519 | 94.267 | 91.453 | 100.843 | 94.017 | 86.396 | 97.570 | 96.456 | 83.988 | 99.744 | 94.838 | 80.147 | 95.208 | 0.033 | 0.1491 | 0.3758 | 0 | 0 | 0 |
| 'Wasf2' | 35.921 | 35.386 | 36.661 | 34.672 | 36.695 | 36.451 | 35.042 | 36.631 | 36.564 | 39.789 | 38.026 | 36.335 | 38.360 | 38.245 | 37.778 | 38.502 | 36.308 | 37.964 | -0.084 | 0.0003 | 0.0038 | 1 | 0 | 0 |
| 'Wash1' | 28.040 | 26.386 | 29.717 | 27.027 | 25.940 | 28.160 | 25.697 | 26.178 | 28.932 | 24.843 | 23.941 | 29.951 | 27.313 | 25.961 | 29.495 | 27.955 | 25.827 | 29.324 | -0.001 | 0.9611 | 0.9860 | 0 | 0 | 0 |
| 'Wasl' | 5.887 | 5.510 | 5.059 | 4.453 | 5.268 | 4.478 | 4.924 | 5.219 | 4.691 | 6.004 | 5.596 | 5.762 | 5.366 | 6.040 | 4.961 | 5.828 | 5.933 | 5.435 | -0.173 | 0.0005 | 0.0060 | 1 | 0 | 0 |
| 'Wbp1' | 9.747 | 10.647 | 10.960 | 9.912 | 11.393 | 11.929 | 9.790 | 11.157 | 10.946 | 10.091 | 9.421 | 10.840 | 10.193 | 9.854 | 10.560 | 10.836 | 9.141 | 10.342 | 0.069 | 0.0652 | 0.2228 | 0 | 0 | 0 |
| 'Wbp11' | 15.914 | 15.653 | 17.030 | 15.144 | 16.323 | 16.812 | 15.706 | 15.948 | 17.460 | 17.457 | 15.397 | 16.498 | 16.810 | 16.310 | 16.671 | 16.733 | 15.773 | 16.064 | -0.025 | 0.3633 | 0.6153 | 0 | 0 | 0 |
| 'Wbp1l' | 10.866 | 11.001 | 11.184 | 10.182 | 11.372 | 11.548 | 8.994 | 11.518 | 11.235 | 12.118 | 11.099 | 11.361 | 11.145 | 10.829 | 11.448 | 11.592 | 10.472 | 11.638 | -0.066 | 0.1013 | 0.2981 | 0 | 0 | 0 |
| 'Wbp2' | 29.428 | 32.729 | 31.734 | 28.854 | 31.351 | 34.083 | 28.573 | 32.173 | 32.724 | 33.424 | 31.034 | 30.948 | 33.360 | 30.936 | 30.958 | 33.027 | 30.135 | 30.404 | -0.023 | 0.5131 | 0.7375 | 0 | 0 | 0 |
| 'Wbp4' | 17.806 | 18.341 | 17.586 | 19.636 | 18.538 | 17.187 | 21.267 | 17.633 | 18.195 | 16.154 | 18.428 | 18.133 | 17.500 | 18.776 | 18.005 | 18.997 | 18.232 | 17.041 | 0.030 | 0.4658 | 0.7015 | 0 | 0 | 0 |
| 'Wbp5' | 48.893 | 33.928 | 31.558 | 46.239 | 35.187 | 30.290 | 45.784 | 34.461 | 30.864 | 38.893 | 41.471 | 32.641 | 43.686 | 37.510 | 32.185 | 38.965 | 43.474 | 30.597 | -0.031 | 0.5940 | 0.7933 | 0 | 0 | 0 |
| 'Wbscr16' | 7.139 | 7.909 | 8.447 | 7.478 | 7.656 | 8.284 | 7.766 | 7.710 | 7.885 | 7.766 | 7.788 | 7.827 | 8.098 | 7.072 | 7.304 | 8.876 | 6.539 | 7.401 | 0.027 | 0.5757 | 0.7807 | 0 | 0 | 0 |
| 'Wbscr22' | 7.058 | 7.827 | 7.860 | 6.774 | 7.195 | 7.907 | 6.646 | 7.465 | 7.678 | 7.157 | 7.069 | 7.631 | 7.099 | 6.999 | 7.626 | 7.388 | 6.935 | 7.686 | 0.009 | 0.7833 | 0.9026 | 0 | 0 | 0 |
| 'Wbscr27' | 0.607 | 0.988 | 0.626 | 0.541 | 0.731 | 0.595 | 0.596 | 0.648 | 0.826 | 0.686 | 0.557 | 0.650 | 0.517 | 0.520 | 0.593 | 0.596 | 0.473 | 0.488 | 0.250 | 0.0357 | 0.1517 | 0 | 0 | 0 |
| 'Wdfy1' | 14.389 | 16.171 | 13.532 | 14.017 | 16.142 | 13.642 | 14.556 | 15.824 | 14.016 | 12.826 | 13.331 | 10.994 | 14.918 | 13.457 | 11.276 | 12.648 | 13.941 | 11.379 | 0.197 | 0.0000 | 0.0000 | 1 | 0 | 0 |
| 'Wdfy2' | 63.322 | 58.558 | 57.006 | 59.982 | 56.507 | 56.056 | 54.377 | 54.397 | 59.624 | 63.428 | 59.704 | 55.391 | 60.459 | 53.973 | 56.787 | 61.524 | 53.409 | 60.841 | -0.024 | 0.5125 | 0.7374 | 0 | 0 | 0 |
| 'Wdhd1' | 4.349 | 4.680 | 5.065 | 4.786 | 4.258 | 4.839 | 5.186 | 4.260 | 5.055 | 3.859 | 4.566 | 4.609 | 5.048 | 4.714 | 4.595 | 4.113 | 4.669 | 4.547 | 0.049 | 0.3411 | 0.5981 | 0 | 0 | 0 |
| 'Wdr1' | 31.827 | 34.044 | 34.709 | 32.823 | 35.113 | 36.250 | 34.565 | 34.496 | 35.278 | 34.339 | 35.582 | 37.082 | 34.414 | 34.444 | 38.777 | 35.407 | 36.594 | 36.334 | -0.074 | 0.0003 | 0.0038 | 1 | 0 | 0 |
| 'Wdr11' | 4.098 | 4.418 | 4.300 | 4.104 | 4.463 | 4.577 | 4.068 | 4.785 | 4.698 | 4.580 | 4.447 | 3.924 | 4.334 | 4.249 | 3.865 | 4.174 | 4.224 | 3.471 | 0.078 | 0.0795 | 0.2537 | 0 | 0 | 0 |
| 'Wdr12' | 1.973 | 1.854 | 1.921 | 2.065 | 2.097 | 1.839 | 2.206 | 1.811 | 1.936 | 3.570 | 3.446 | 4.071 | 3.523 | 3.339 | 4.072 | 4.027 | 4.064 | 4.007 | -0.961 | 0.0000 | 0.0000 | 1 | 1 | 0 |

| | | | | | | | | | | | | | | | | | | | | | | | | |
|---|---|---|---|---|---|---|---|---|---|---|---|---|---|---|---|---|---|---|---|---|---|---|---|---|
| 'Wdr13' | 3.334 | 4.104 | 4.033 | 3.147 | 4.062 | 4.222 | 3.303 | 3.865 | 4.485 | 4.039 | 3.515 | 4.457 | 3.879 | 4.007 | 4.322 | 4.049 | 4.162 | 4.050 | -0.094 | 0.0354 | 0.1505 | 0 | 0 | 0 |
| 'Wdr18' | 13.800 | 13.226 | 15.806 | 13.852 | 11.949 | 14.664 | 14.683 | 13.634 | 15.756 | 14.545 | 11.007 | 15.749 | 14.188 | 11.966 | 14.993 | 15.042 | 13.024 | 15.421 | 0.007 | 0.8559 | 0.9383 | 0 | 0 | 0 |
| 'Wdr19' | 0.510 | 0.676 | 0.611 | 0.548 | 0.589 | 0.611 | 0.566 | 0.604 | 0.693 | 0.585 | 0.559 | 0.664 | 0.592 | 0.634 | 0.549 | 0.647 | 0.621 | 0.496 | 0.002 | 0.9819 | 0.9933 | 0 | 0 | 0 |
| 'Wdr24' | 8.781 | 8.772 | 10.083 | 8.917 | 8.969 | 9.705 | 8.937 | 8.720 | 10.055 | 8.466 | 8.511 | 8.814 | 8.849 | 7.843 | 8.145 | 7.997 | 8.445 | 8.335 | 0.127 | 0.0001 | 0.0021 | 1 | 0 | 0 |
| 'Wdr25' | 1.166 | 1.635 | 1.382 | 1.023 | 1.613 | 1.443 | 1.149 | 1.301 | 1.390 | 1.699 | 1.166 | 1.259 | 1.068 | 1.263 | 1.344 | 1.345 | 1.172 | 1.356 | 0.038 | 0.6953 | 0.8567 | 0 | 0 | 0 |
| 'Wdr26' | 41.371 | 46.672 | 40.168 | 40.920 | 45.878 | 40.368 | 41.879 | 45.459 | 41.369 | 44.490 | 42.828 | 40.395 | 38.711 | 44.084 | 39.136 | 43.792 | 43.093 | 40.329 | 0.015 | 0.5550 | 0.7675 | 0 | 0 | 0 |
| 'Wdr3' | 9.233 | 9.797 | 9.211 | 9.172 | 9.270 | 8.765 | 9.271 | 9.391 | 9.551 | 8.592 | 9.488 | 10.188 | 8.394 | 9.012 | 9.694 | 9.512 | 9.736 | 10.637 | -0.038 | 0.3364 | 0.5950 | 0 | 0 | 0 |
| 'Wdr33' | 22.640 | 21.207 | 21.944 | 22.540 | 21.715 | 21.623 | 23.576 | 20.849 | 21.573 | 21.815 | 21.745 | 22.473 | 20.460 | 20.923 | 20.771 | 20.649 | 21.259 | 21.198 | 0.035 | 0.2179 | 0.4647 | 0 | 0 | 0 |
| 'Wdr34' | 1.335 | 1.455 | 1.316 | 1.247 | 1.422 | 1.713 | 1.078 | 1.741 | 1.566 | 1.572 | 1.384 | 1.363 | 1.605 | 1.598 | 1.598 | 1.812 | 1.047 | 1.202 | -0.030 | 0.7981 | 0.9103 | 0 | 0 | 0 |
| 'Wdr35' | 0.541 | 0.738 | 0.593 | 0.630 | 0.483 | 0.554 | 0.511 | 0.507 | 0.679 | 0.681 | 0.542 | 0.599 | 0.644 | 0.640 | 0.581 | 0.656 | 0.768 | 0.502 | -0.107 | 0.2508 | 0.5019 | 0 | 0 | 0 |
| 'Wdr36' | 16.894 | 16.416 | 17.573 | 16.448 | 18.590 | 17.471 | 17.279 | 18.719 | 17.350 | 17.432 | 16.002 | 17.089 | 17.501 | 17.073 | 15.949 | 16.226 | 17.405 | 15.905 | 0.048 | 0.1483 | 0.3747 | 0 | 0 | 0 |
| 'Wdr37' | 3.449 | 3.840 | 3.647 | 3.589 | 3.576 | 3.280 | 3.594 | 3.676 | 3.467 | 3.337 | 3.712 | 3.708 | 3.035 | 3.774 | 3.375 | 3.547 | 4.044 | 3.569 | -0.009 | 0.8190 | 0.9199 | 0 | 0 | 0 |
| 'Wdr4' | 2.093 | 1.932 | 2.025 | 1.941 | 1.784 | 1.999 | 1.743 | 1.949 | 1.853 | 1.939 | 1.772 | 2.129 | 1.999 | 1.938 | 1.986 | 1.886 | 1.842 | 1.942 | -0.021 | 0.6275 | 0.8154 | 0 | 0 | 0 |
| 'Wdr41' | 4.117 | 4.678 | 4.333 | 4.221 | 4.009 | 4.416 | 4.239 | 4.544 | 4.402 | 4.551 | 4.248 | 4.602 | 3.961 | 4.415 | 4.274 | 4.020 | 4.687 | 4.178 | -0.011 | 0.7818 | 0.9024 | 0 | 0 | 0 |
| 'Wdr43' | 27.700 | 27.880 | 25.843 | 28.756 | 29.528 | 25.746 | 29.653 | 29.286 | 26.617 | 26.802 | 24.874 | 26.473 | 24.881 | 25.816 | 26.474 | 27.890 | 27.071 | 26.878 | 0.068 | 0.0448 | 0.1754 | 0 | 0 | 0 |
| 'Wdr45' | 5.692 | 6.452 | 6.413 | 5.903 | 6.440 | 6.638 | 6.202 | 6.369 | 6.056 | 6.621 | 5.699 | 6.356 | 6.281 | 6.394 | 5.922 | 6.988 | 6.277 | 5.771 | -0.012 | 0.7658 | 0.8956 | 0 | 0 | 0 |
| 'Wdr45b' | 6.321 | 6.586 | 5.580 | 5.985 | 6.123 | 5.623 | 5.211 | 6.198 | 5.262 | 6.503 | 5.974 | 5.647 | 5.604 | 6.568 | 5.517 | 6.002 | 6.168 | 5.200 | -0.016 | 0.7144 | 0.8681 | 0 | 0 | 0 |
| 'Wdr46' | 29.752 | 28.080 | 30.949 | 27.762 | 28.214 | 31.079 | 27.290 | 29.057 | 29.314 | 27.627 | 24.481 | 31.215 | 26.653 | 24.663 | 28.585 | 29.355 | 24.842 | 29.693 | 0.073 | 0.0275 | 0.1280 | 0 | 0 | 0 |
| 'Wdr47' | 2.809 | 3.312 | 2.861 | 2.914 | 2.917 | 3.093 | 3.107 | 3.018 | 3.035 | 2.706 | 3.107 | 2.969 | 2.676 | 3.366 | 3.009 | 2.813 | 3.209 | 2.812 | 0.012 | 0.8021 | 0.9114 | 0 | 0 | 0 |
| 'Wdr48' | 21.474 | 22.072 | 22.652 | 21.649 | 22.849 | 22.447 | 22.562 | 22.491 | 23.129 | 21.221 | 23.840 | 22.879 | 20.723 | 21.829 | 22.130 | 21.172 | 22.564 | 21.725 | 0.013 | 0.6207 | 0.8102 | 0 | 0 | 0 |
| 'Wdr5' | 11.669 | 12.201 | 12.560 | 11.445 | 12.412 | 12.297 | 11.967 | 12.559 | 11.581 | 11.121 | 11.314 | 12.990 | 11.649 | 11.862 | 12.439 | 12.350 | 11.479 | 12.603 | -0.001 | 0.9670 | 0.9887 | 0 | 0 | 0 |
| 'Wdr53' | 3.318 | 2.763 | 3.241 | 3.693 | 3.136 | 3.820 | 3.822 | 3.085 | 3.041 | 2.905 | 3.211 | 3.094 | 3.090 | 3.299 | 3.219 | 2.810 | 3.149 | 3.004 | 0.097 | 0.1685 | 0.4008 | 0 | 0 | 0 |
| 'Wdr55' | 22.452 | 21.211 | 24.002 | 21.862 | 23.802 | 26.494 | 23.429 | 22.782 | 23.145 | 22.299 | 22.389 | 27.117 | 22.404 | 20.603 | 25.769 | 22.451 | 23.822 | 25.668 | -0.036 | 0.3882 | 0.6378 | 0 | 0 | 0 |
| 'Wdr59' | 11.034 | 11.164 | 11.200 | 11.386 | 10.627 | 11.243 | 10.471 | 11.497 | 11.349 | 10.977 | 10.862 | 11.587 | 9.932 | 10.277 | 12.185 | 10.944 | 11.190 | 10.836 | 0.008 | 0.8301 | 0.9262 | 0 | 0 | 0 |
| 'Wdr5b' | 6.306 | 7.256 | 9.756 | 7.797 | 8.163 | 8.815 | 7.947 | 7.102 | 8.485 | 6.757 | 7.414 | 7.637 | 7.617 | 6.137 | 7.277 | 6.141 | 7.150 | 6.559 | 0.179 | 0.0099 | 0.0619 | 0 | 0 | 0 |
| 'Wdr6' | 29.015 | 28.985 | 28.789 | 27.526 | 29.020 | 31.437 | 28.182 | 27.456 | 30.492 | 30.397 | 29.333 | 29.200 | 27.921 | 27.626 | 27.970 | 29.352 | 24.745 | 28.311 | 0.024 | 0.4834 | 0.7150 | 0 | 0 | 0 |
| 'Wdr60' | 1.027 | 1.011 | 1.301 | 1.205 | 1.213 | 1.283 | 1.563 | 1.100 | 1.484 | 1.501 | 0.740 | 1.016 | 1.070 | 0.907 | 0.879 | 1.419 | 0.842 | 0.813 | 0.295 | 0.0136 | 0.0784 | 0 | 0 | 0 |
| 'Wdr61' | 6.641 | 7.142 | 7.131 | 7.757 | 7.380 | 6.977 | 7.957 | 7.849 | 7.412 | 6.876 | 7.106 | 7.507 | 7.086 | 6.875 | 7.158 | 7.316 | 8.239 | 6.622 | 0.019 | 0.6722 | 0.8421 | 0 | 0 | 0 |
| 'Wdr62' | 1.718 | 1.289 | 2.799 | 2.164 | 1.465 | 2.551 | 1.905 | 1.813 | 2.607 | 1.868 | 2.071 | 2.097 | 2.653 | 2.039 | 2.193 | 1.564 | 1.887 | 2.221 | -0.059 | 0.5631 | 0.7720 | 0 | 0 | 0 |
| 'Wdr66' | 0.301 | 0.195 | 0.259 | 0.214 | 0.212 | 0.259 | 0.239 | 0.314 | 0.345 | 0.242 | 0.158 | 0.288 | 0.312 | 0.171 | 0.252 | 0.223 | 0.251 | 0.276 | 0.100 | 0.4151 | 0.6630 | 0 | 0 | 0 |
| 'Wdr7' | 6.993 | 7.675 | 7.719 | 7.416 | 6.836 | 6.954 | 7.227 | 8.176 | 7.257 | 7.941 | 8.100 | 7.057 | 7.608 | 6.992 | 6.871 | 7.548 | 7.024 | 7.122 | -0.007 | 0.8642 | 0.9410 | 0 | 0 | 0 |
| 'Wdr70' | 12.428 | 12.401 | 13.203 | 12.318 | 12.707 | 13.068 | 13.680 | 13.560 | 12.965 | 13.150 | 12.262 | 11.987 | 13.827 | 11.787 | 11.483 | 11.967 | 12.993 | 11.731 | 0.058 | 0.1897 | 0.4262 | 0 | 0 | 0 |
| 'Wdr73' | 2.693 | 3.164 | 3.364 | 2.771 | 3.284 | 3.101 | 2.525 | 3.065 | 2.944 | 2.981 | 3.004 | 3.065 | 2.608 | 2.656 | 3.034 | 2.556 | 2.928 | 3.031 | 0.045 | 0.3086 | 0.5669 | 0 | 0 | 0 |
| 'Wdr74' | 32.778 | 30.847 | 31.620 | 31.323 | 31.010 | 31.603 | 29.557 | 33.298 | 30.857 | 31.014 | 27.749 | 30.473 | 30.302 | 29.382 | 28.173 | 31.587 | 30.880 | 28.097 | 0.072 | 0.0557 | 0.2023 | 0 | 0 | 0 |
| 'Wdr75' | 4.238 | 4.287 | 5.049 | 4.860 | 4.759 | 4.379 | 4.975 | 5.098 | 4.395 | 4.807 | 4.516 | 4.690 | 4.594 | 4.141 | 4.349 | 4.225 | 5.017 | 4.298 | 0.037 | 0.4310 | 0.6762 | 0 | 0 | 0 |
| 'Wdr76' | 5.643 | 4.988 | 6.503 | 6.196 | 5.648 | 6.289 | 6.691 | 5.600 | 5.995 | 4.765 | 6.592 | 5.868 | 5.779 | 6.233 | 6.236 | 5.038 | 6.195 | 5.843 | 0.019 | 0.7551 | 0.8907 | 0 | 0 | 0 |
| 'Wdr77' | 5.582 | 6.966 | 6.407 | 5.760 | 7.059 | 6.652 | 5.834 | 6.662 | 6.113 | 5.856 | 5.765 | 7.074 | 5.669 | 5.661 | 6.862 | 5.960 | 5.944 | 7.229 | 0.010 | 0.8260 | 0.9237 | 0 | 0 | 0 |
| 'Wdr78' | 0.368 | 0.462 | 0.526 | 0.399 | 0.427 | 0.439 | 0.389 | 0.461 | 0.487 | 0.428 | 0.200 | 0.272 | 0.317 | 0.329 | 0.261 | 0.256 | 0.294 | 0.290 | 0.577 | 0.0000 | 0.0000 | 1 | 1 | 0 |
| 'Wdr81' | 13.630 | 13.151 | 14.508 | 12.726 | 13.709 | 14.493 | 12.616 | 13.292 | 14.243 | 14.512 | 14.311 | 14.902 | 14.019 | 12.682 | 14.348 | 13.027 | 12.613 | 15.321 | -0.049 | 0.1339 | 0.3524 | 0 | 0 | 0 |
| 'Wdr82' | 23.719 | 23.586 | 23.745 | 24.549 | 23.378 | 25.102 | 22.396 | 23.835 | 23.959 | 22.473 | 23.226 | 27.158 | 21.482 | 24.422 | 27.027 | 23.988 | 23.488 | 27.426 | -0.050 | 0.1585 | 0.3886 | 0 | 0 | 0 |
| 'Wdr83' | 3.813 | 3.953 | 4.781 | 3.878 | 4.353 | 3.968 | 4.056 | 4.411 | 4.856 | 3.923 | 3.803 | 4.175 | 4.478 | 3.609 | 3.990 | 4.064 | 3.890 | 3.985 | 0.074 | 0.1402 | 0.3624 | 0 | 0 | 0 |
| 'Wdr83os' | 24.539 | 26.120 | 26.182 | 23.874 | 25.722 | 27.888 | 24.484 | 25.905 | 24.051 | 26.282 | 26.788 | 23.907 | 26.154 | 22.135 | 22.487 | 24.011 | 23.884 | 24.401 | 0.050 | 0.3372 | 0.5955 | 0 | 0 | 0 |
| 'Wdr89' | 5.059 | 4.973 | 5.197 | 5.131 | 5.751 | 5.262 | 5.457 | 5.247 | 4.883 | 4.771 | 4.609 | 5.437 | 4.965 | 5.014 | 5.108 | 5.256 | 4.795 | 4.733 | 0.055 | 0.2188 | 0.4656 | 0 | 0 | 0 |
| 'Wdr90' | 2.541 | 2.153 | 2.895 | 2.533 | 1.925 | 2.655 | 2.401 | 2.142 | 2.711 | 2.148 | 2.145 | 2.341 | 2.753 | 2.314 | 2.433 | 2.041 | 2.371 | 2.437 | 0.050 | 0.3957 | 0.6446 | 0 | 0 | 0 |
| 'Wdr91' | 7.348 | 6.955 | 7.286 | 6.441 | 6.565 | 7.102 | 6.018 | 7.415 | 7.144 | 7.228 | 7.350 | 7.473 | 6.915 | 6.735 | 7.041 | 7.119 | 6.725 | 7.737 | -0.054 | 0.1919 | 0.4290 | 0 | 0 | 0 |
| 'Wdr92' | 6.396 | 7.086 | 6.290 | 6.129 | 7.125 | 6.483 | 6.490 | 7.069 | 6.663 | 5.950 | 6.823 | 6.645 | 6.157 | 6.947 | 6.821 | 6.510 | 6.609 | 6.262 | 0.012 | 0.7921 | 0.9070 | 0 | 0 | 0 |
| 'Wdsub1' | 1.391 | 1.508 | 1.382 | 1.074 | 1.560 | 1.600 | 1.634 | 1.535 | 1.427 | 1.531 | 1.116 | 1.377 | 1.309 | 1.454 | 1.442 | 1.634 | 1.408 | 1.349 | 0.048 | 0.5456 | 0.7606 | 0 | 0 | 0 |
| 'Wdtc1' | 21.375 | 22.318 | 22.666 | 20.988 | 23.279 | 25.076 | 19.577 | 21.994 | 23.175 | 24.202 | 20.996 | 24.714 | 24.193 | 24.264 | 24.660 | 24.574 | 21.588 | 24.471 | -0.103 | 0.0056 | 0.0407 | 1 | 0 | 0 |
| 'Wdyhv1' | 21.976 | 27.061 | 25.818 | 25.219 | 27.350 | 25.098 | 21.853 | 25.040 | 25.799 | 22.772 | 24.238 | 27.482 | 23.034 | 24.736 | 24.594 | 22.253 | 25.095 | 24.856 | 0.021 | 0.6181 | 0.8090 | 0 | 0 | 0 |
| 'Wee1' | 6.692 | 5.578 | 6.443 | 7.053 | 5.618 | 6.440 | 6.317 | 5.889 | 7.184 | 5.621 | 5.395 | 6.863 | 6.326 | 6.323 | 7.577 | 5.289 | 5.135 | 7.106 | 0.027 | 0.6869 | 0.8521 | 0 | 0 | 0 |
| 'Wfs1' | 0.758 | 1.199 | 0.963 | 0.736 | 0.967 | 1.465 | 0.799 | 0.937 | 1.205 | 0.852 | 0.799 | 0.957 | 0.819 | 0.929 | 0.966 | 0.873 | 0.726 | 0.937 | 0.180 | 0.0670 | 0.2266 | 0 | 0 | 0 |
| 'Whamm' | 25.149 | 21.644 | 24.959 | 22.874 | 22.042 | 25.914 | 22.057 | 22.720 | 23.686 | 25.070 | 22.509 | 22.937 | 25.097 | 23.101 | 21.866 | 25.818 | 21.166 | 23.350 | -0.004 | 0.9177 | 0.9687 | 0 | 0 | 0 |
| 'Whrn' | 4.503 | 5.413 | 5.290 | 4.754 | 4.964 | 5.818 | 4.638 | 5.236 | 5.617 | 5.844 | 5.860 | 5.824 | 8.571 | 6.206 | 5.304 | 6.217 | 5.863 | 5.949 | -0.276 | 0.0001 | 0.0018 | 1 | 0 | 0 |
| 'Whsc1' | 6.767 | 6.163 | 6.890 | 7.025 | 6.298 | 6.568 | 6.712 | 6.217 | 6.694 | 6.129 | 6.660 | 7.126 | 6.812 | 6.690 | 7.071 | 6.183 | 6.845 | 7.152 | -0.043 | 0.2056 | 0.4492 | 0 | 0 | 0 |
| 'Whsc1l1' | 13.795 | 12.665 | 12.289 | 13.223 | 12.170 | 11.829 | 12.459 | 12.500 | 12.054 | 12.507 | 13.175 | 13.161 | 12.612 | 12.796 | 12.473 | 12.540 | 12.926 | 13.716 | -0.049 | 0.1478 | 0.3736 | 0 | 0 | 0 |
| 'Wipf1' | 53.279 | 56.000 | 53.890 | 53.562 | 53.475 | 54.304 | 53.985 | 54.099 | 54.033 | 54.100 | 53.539 | 56.504 | 52.746 | 55.287 | 56.695 | 51.884 | 51.313 | 56.808 | -0.017 | 0.4237 | 0.6705 | 0 | 0 | 0 |
| 'Wipf2' | 7.623 | 7.863 | 7.562 | 7.113 | 7.499 | 7.990 | 7.414 | 7.371 | 8.125 | 8.584 | 7.907 | 8.875 | 8.281 | 8.733 | 8.288 | 8.612 | 8.070 | 9.005 | -0.166 | 0.0000 | 0.0000 | 1 | 0 | 0 |
| 'Wipi2' | 6.907 | 6.762 | 6.981 | 5.855 | 7.304 | 7.098 | 6.528 | 7.144 | 7.251 | 6.548 | 6.935 | 7.623 | 6.978 | 7.257 | 6.693 | 7.321 | 7.356 | 7.309 | -0.061 | 0.0822 | 0.2593 | 0 | 0 | 0 |
| 'Wisp1' | 1.993 | 2.382 | 2.013 | 2.130 | 2.280 | 2.024 | 2.227 | 1.978 | 2.296 | 1.853 | 1.515 | 2.151 | 1.480 | 1.918 | 2.115 | 1.979 | 1.675 | 1.991 | 0.196 | 0.0061 | 0.0436 | 1 | 0 | 0 |
| 'Wiz' | 2.288 | 2.223 | 2.475 | 2.069 | 2.249 | 2.512 | 2.181 | 2.391 | 2.515 | 2.442 | 2.009 | 2.392 | 2.222 | 1.959 | 2.348 | 2.084 | 2.164 | 2.378 | 0.054 | 0.1518 | 0.3789 | 0 | 0 | 0 |
| 'Wls' | 0.892 | 0.976 | 0.951 | 0.844 | 0.974 | 0.843 | 0.826 | 1.049 | 0.886 | 1.194 | 0.879 | 1.210 | 1.041 | 1.197 | 0.988 | 1.439 | 1.335 | 1.162 | -0.352 | 0.0000 | 0.0002 | 1 | 0 | 0 |
| 'Wnk1' | 13.858 | 12.969 | 14.025 | 14.126 | 12.938 | 13.295 | 12.718 | 12.708 | 13.893 | 14.090 | 13.690 | 14.304 | 13.149 | 13.132 | 13.892 | 12.908 | 13.018 | 15.032 | -0.042 | 0.1695 | 0.4019 | 0 | 0 | 0 |
| 'Wrap53' | 8.576 | 8.903 | 8.863 | 9.332 | 8.002 | 9.670 | 8.292 | 8.422 | 9.028 | 8.259 | 7.537 | 9.318 | 9.161 | 9.012 | 8.591 | 8.460 | 8.161 | 9.422 | 0.008 | 0.8567 | 0.9387 | 0 | 0 | 0 |
| 'Wrap73' | 3.981 | 4.664 | 4.715 | 4.279 | 4.442 | 4.437 | 3.713 | 3.843 | 4.621 | 3.915 | 3.827 | 4.918 | 4.427 | 4.296 | 4.462 | 3.963 | 4.189 | 4.659 | -0.017 | 0.7279 | 0.8752 | 0 | 0 | 0 |
| 'Wrb' | 1.569 | 2.367 | 2.011 | 1.588 | 2.204 | 2.023 | 1.696 | 2.357 | 1.861 | 1.774 | 2.045 | 1.966 | 1.747 | 1.841 | 1.786 | 2.326 | 1.683 | 1.854 | 0.037 | 0.6368 | 0.8209 | 0 | 0 | 0 |
| 'Wrn' | 9.719 | 9.272 | 10.261 | 11.153 | 9.764 | 10.229 | 10.394 | 9.145 | 9.871 | 10.174 | 10.569 | 10.703 | 10.133 | 10.861 | 9.520 | 9.113 | 10.954 | 10.027 | -0.047 | 0.2917 | 0.5485 | 0 | 0 | 0 |
| 'Wrnip1' | 42.373 | 39.451 | 38.552 | 39.048 | 41.006 | 40.210 | 40.338 | 37.717 | 37.381 | 39.236 | 38.663 | 44.682 | 42.094 | 40.051 | 43.590 | 40.036 | 39.945 | 45.080 | -0.081 | 0.0239 | 0.1169 | 0 | 0 | 0 |
| 'Wsb1' | 15.463 | 15.618 | 13.753 | 14.485 | 15.246 | 13.681 | 15.346 | 15.968 | 14.320 | 15.739 | 14.596 | 12.972 | 16.149 | 16.352 | 12.707 | 16.545 | 15.475 | 11.531 | 0.018 | 0.6161 | 0.8085 | 0 | 0 | 0 |
| 'Wsb2' | 6.404 | 7.265 | 5.992 | 6.594 | 6.612 | 6.427 | 6.145 | 6.992 | 6.282 | 6.265 | 6.380 | 6.944 | 6.392 | 7.245 | 7.073 | 6.908 | 7.090 | 7.401 | -0.086 | 0.0409 | 0.1657 | 0 | 0 | 0 |
| 'Wwox' | 1.292 | 1.617 | 1.200 | 1.291 | 1.461 | 1.269 | 1.573 | 1.425 | 1.284 | 1.209 | 1.230 | 1.210 | 1.506 | 1.344 | 1.122 | 1.375 | 1.170 | 1.203 | 0.106 | 0.0810 | 0.2565 | 0 | 0 | 0 |
| 'Wwp1' | 2.864 | 3.424 | 3.198 | 3.186 | 3.122 | 3.436 | 3.521 | 3.346 | 3.421 | 3.334 | 3.390 | 3.120 | 3.336 | 3.190 | 2.785 | 3.228 | 3.539 | 2.917 | 0.026 | 0.5961 | 0.7947 | 0 | 0 | 0 |
| 'Wwp2' | 31.514 | 32.207 | 32.592 | 29.920 | 30.808 | 33.418 | 30.169 | 31.870 | 34.120 | 33.846 | 31.253 | 32.369 | 32.666 | 30.339 | 31.446 | 32.989 | 29.182 | 29.722 | 0.006 | 0.8584 | 0.9390 | 0 | 0 | 0 |
| 'Xab2' | 13.017 | 14.066 | 15.203 | 12.709 | 13.580 | 15.278 | 14.323 | 14.158 | 14.971 | 14.144 | 13.380 | 14.420 | 13.666 | 13.192 | 13.810 | 13.613 | 12.821 | 13.517 | 0.044 | 0.1208 | 0.3314 | 0 | 0 | 0 |
| 'Xaf1' | 4.761 | 4.737 | 5.513 | 3.742 | 5.477 | 5.421 | 4.375 | 5.908 | 5.580 | 5.075 | 4.598 | 6.874 | 5.773 | 5.634 | 7.325 | 6.075 | 4.897 | 6.472 | -0.220 | 0.0017 | 0.0156 | 1 | 0 | 0 |
| 'Xbp1' | 19.970 | 20.068 | 18.410 | 18.872 | 22.420 | 19.813 | 16.134 | 20.377 | 20.430 | 20.646 | 17.178 | 19.806 | 18.283 | 20.374 | 19.014 | 19.270 | 19.457 | 21.099 | -0.001 | 0.9805 | 0.9926 | 0 | 0 | 0 |
| 'Xdh' | 20.643 | 21.708 | 18.834 | 17.878 | 19.897 | 19.916 | 21.119 | 19.569 | 19.746 | 18.179 | 20.823 | 15.990 | 17.275 | 19.708 | 14.855 | 15.339 | 22.732 | 14.967 | 0.168 | 0.0031 | 0.0262 | 1 | 0 | 0 |
| 'Xiap' | 7.356 | 8.882 | 8.279 | 8.058 | 8.629 | 7.762 | 8.047 | 8.228 | 8.370 | 8.134 | 8.546 | 9.284 | 7.783 | 8.894 | 8.539 | 7.960 | 8.955 | 9.254 | -0.084 | 0.0078 | 0.0519 | 0 | 0 | 0 |
| 'Xkr6' | 3.013 | 3.379 | 3.215 | 2.755 | 3.008 | 3.174 | 2.884 | 3.136 | 3.078 | 2.296 | 2.527 | 2.243 | 1.923 | 2.646 | 2.288 | 1.998 | 2.363 | 2.305 | 0.422 | 0.0000 | 0.0000 | 1 | 1 | 0 |
| 'Xkr8' | 1.469 | 1.837 | 1.721 | 1.235 | 1.718 | 1.611 | 1.393 | 1.658 | 1.943 | 1.295 | 1.458 | 1.609 | 1.230 | 1.310 | 1.677 | 1.593 | 1.230 | 1.424 | 0.173 | 0.0272 | 0.1270 | 0 | 0 | 0 |
| 'Xndc1' | 3.967 | 4.113 | 4.982 | 3.730 | 4.139 | 4.463 | 3.935 | 4.146 | 4.711 | 4.069 | 4.108 | 4.618 | 4.961 | 4.052 | 4.375 | 3.845 | 4.015 | 4.112 | -0.012 | 0.8039 | 0.9119 | 0 | 0 | 0 |
| 'Xpa' | 7.596 | 7.719 | 8.356 | 7.673 | 8.554 | 8.120 | 8.574 | 8.645 | 7.692 | 7.813 | 7.769 | 7.239 | 8.398 | 8.115 | 6.748 | 8.707 | 7.349 | 6.337 | 0.089 | 0.0769 | 0.2474 | 0 | 0 | 0 |
| 'Xpc' | 12.387 | 12.186 | 13.230 | 11.903 | 11.931 | 13.684 | 12.289 | 11.204 | 13.168 | 12.187 | 10.811 | 11.718 | 11.740 | 10.999 | 11.543 | 10.684 | 10.443 | 10.873 | 0.139 | 0.0000 | 0.0005 | 1 | 0 | 0 |
| 'Xpnpep1' | 8.799 | 8.062 | 8.634 | 8.117 | 8.569 | 9.153 | 8.583 | 8.606 | 8.533 | 8.252 | 8.226 | 8.469 | 8.750 | 8.064 | 8.612 | 8.442 | 7.807 | 8.107 | 0.036 | 0.1922 | 0.4294 | 0 | 0 | 0 |
| 'Xpnpep3' | 2.171 | 2.177 | 2.112 | 2.262 | 2.094 | 2.097 | 2.363 | 2.063 | 2.481 | 2.369 | 2.387 | 2.449 | 1.975 | 2.084 | 2.293 | 1.873 | 2.430 | 2.554 | -0.056 | 0.3594 | 0.6118 | 0 | 0 | 0 |
| 'Xpo1' | 17.691 | 17.794 | 20.737 | 18.700 | 19.096 | 18.832 | 19.976 | 19.053 | 19.979 | 17.859 | 18.737 | 19.615 | 17.900 | 20.021 | 18.915 | 17.190 | 19.658 | 19.865 | 0.008 | 0.8101 | 0.9142 | 0 | 0 | 0 |
| 'Xpo5' | 21.528 | 21.829 | 21.046 | 21.430 | 21.276 | 21.510 | 21.384 | 21.077 | 20.948 | 20.155 | 21.248 | 23.373 | 20.216 | 18.981 | 22.194 | 19.994 | 20.163 | 23.621 | 0.003 | 0.9269 | 0.9720 | 0 | 0 | 0 |
| 'Xpo6' | 10.501 | 9.771 | 10.292 | 10.155 | 9.648 | 10.028 | 9.413 | 10.012 | 10.201 | 10.489 | 9.758 | 10.361 | 10.455 | 9.586 | 10.112 | 10.159 | 9.706 | 10.745 | -0.031 | 0.2015 | 0.4426 | 0 | 0 | 0 |
| 'Xpo7' | 6.860 | 6.735 | 7.280 | 7.116 | 6.978 | 6.919 | 6.659 | 7.085 | 6.934 | 7.461 | 6.939 | 7.145 | 7.047 | 7.206 | 7.088 | 6.742 | 7.316 | 7.056 | -0.042 | 0.1156 | 0.3228 | 0 | 0 | 0 |
| 'Xpot' | 11.966 | 11.466 | 11.116 | 12.467 | 11.456 | 11.074 | 11.125 | 11.826 | 12.050 | 11.897 | 11.667 | 13.109 | 11.141 | 11.976 | 11.893 | 12.517 | 12.875 | 12.775 | -0.083 | 0.0282 | 0.1301 | 0 | 0 | 0 |
| 'Xpr1' | 2.888 | 2.987 | 3.110 | 2.986 | 2.986 | 2.981 | 2.887 | 3.145 | 2.731 | 2.925 | 3.103 | 3.277 | 2.765 | 2.925 | 2.846 | 2.813 | 2.979 | 3.149 | -0.017 | 0.6462 | 0.8273 | 0 | 0 | 0 |
| 'Xrcc1' | 5.496 | 5.704 | 6.923 | 5.471 | 5.237 | 6.498 | 5.322 | 5.746 | 6.356 | 5.730 | 5.483 | 6.229 | 6.737 | 4.857 | 6.089 | 5.923 | 5.369 | 5.823 | 0.004 | 0.9321 | 0.9746 | 0 | 0 | 0 |
| 'Xrcc2' | 1.575 | 1.304 | 1.665 | 1.599 | 1.513 | 1.749 | 1.885 | 1.421 | 1.485 | 1.184 | 1.346 | 1.766 | 1.587 | 1.761 | 1.808 | 1.408 | 1.570 | 1.603 | 0.002 | 0.9839 | 0.9942 | 0 | 0 | 0 |
| 'Xrcc3' | 2.534 | 3.153 | 3.540 | 2.946 | 3.650 | 3.372 | 3.288 | 3.405 | 3.887 | 3.319 | 3.378 | 3.428 | 3.873 | 3.317 | 3.600 | 2.981 | 3.112 | 3.916 | -0.070 | 0.2786 | 0.5352 | 0 | 0 | 0 |
| 'Xrcc4' | 4.120 | 4.539 | 3.932 | 4.484 | 4.135 | 4.118 | 4.939 | 4.056 | 4.311 | 4.104 | 4.104 | 4.223 | 4.009 | 3.944 | 3.980 | 4.196 | 4.032 | 3.689 | 0.071 | 0.1656 | 0.3980 | 0 | 0 | 0 |
| 'Xrcc5' | 17.975 | 19.099 | 18.641 | 18.320 | 18.588 | 18.667 | 18.925 | 19.709 | 18.634 | 18.767 | 18.119 | 18.555 | 18.988 | 17.947 | 16.345 | 18.530 | 17.923 | 16.542 | 0.049 | 0.1534 | 0.3808 | 0 | 0 | 0 |
| 'Xrcc6' | 7.818 | 8.101 | 8.496 | 8.007 | 7.658 | 7.574 | 8.982 | 7.337 | 8.677 | 7.765 | 6.768 | 8.215 | 8.517 | 7.926 | 8.364 | 8.238 | 7.749 | 7.863 | 0.012 | 0.7645 | 0.8953 | 0 | 0 | 0 |
| 'Xrcc6bp1' | 6.727 | 6.336 | 5.510 | 5.944 | 6.689 | 6.302 | 6.526 | 6.571 | 5.857 | 5.093 | 5.954 | 6.502 | 7.092 | 5.396 | 5.568 | 6.975 | 5.066 | 6.329 | 0.045 | 0.5744 | 0.7800 | 0 | 0 | 0 |
| 'Xrn1' | 7.021 | 6.901 | 6.952 | 7.171 | 6.770 | 6.631 | 6.996 | 6.862 | 6.536 | 7.011 | 7.388 | 6.816 | 6.743 | 7.085 | 6.439 | 6.695 | 7.116 | 6.600 | -0.011 | 0.6819 | 0.8486 | 0 | 0 | 0 |
| 'Xrn2' | 38.246 | 43.374 | 38.901 | 39.680 | 43.720 | 39.882 | 42.288 | 44.134 | 39.176 | 38.032 | 39.199 | 37.746 | 38.618 | 40.420 | 36.442 | 41.968 | 40.539 | 33.405 | 0.083 | 0.0020 | 0.0181 | 1 | 0 | 0 |
| 'Xxylt1' | 7.335 | 7.162 | 8.565 | 7.844 | 7.855 | 9.020 | 7.479 | 8.940 | 8.402 | 9.140 | 7.091 | 7.167 | 7.790 | 7.547 | 6.820 | 9.374 | 6.751 | 7.369 | 0.072 | 0.2726 | 0.5277 | 0 | 0 | 0 |
| 'Xylb' | 0.858 | 0.754 | 1.177 | 0.936 | 0.782 | 0.942 | 1.194 | 0.798 | 1.337 | 1.049 | 1.145 | 0.963 | 1.176 | 1.106 | 0.969 | 1.034 | 0.923 | 0.990 | -0.092 | 0.4358 | 0.6798 | 0 | 0 | 0 |
| 'Xylt1' | 2.385 | 2.708 | 2.584 | 2.374 | 2.326 | 2.432 | 2.061 | 2.737 | 2.399 | 3.819 | 3.839 | 3.648 | 3.305 | 3.724 | 3.243 | 4.236 | 3.211 | 3.593 | -0.576 | 0.0000 | 0.0000 | 1 | 1 | 0 |
| 'Xylt2' | 18.096 | 17.630 | 20.463 | 18.579 | 17.765 | 21.161 | 18.682 | 18.009 | 20.920 | 18.750 | 15.307 | 19.480 | 17.808 | 16.514 | 18.285 | 17.756 | 14.076 | 19.142 | 0.116 | 0.0002 | 0.0032 | 1 | 0 | 0 |
| 'Yae1d1' | 2.565 | 2.740 | 2.806 | 2.977 | 3.424 | 2.682 | 3.167 | 3.311 | 3.290 | 1.960 | 2.360 | 2.702 | 2.112 | 2.352 | 2.926 | 2.609 | 2.637 | 2.737 | 0.252 | 0.0017 | 0.0160 | 1 | 0 | 0 |
| 'Yaf2' | 10.238 | 10.168 | 9.669 | 9.994 | 10.407 | 9.992 | 10.470 | 11.474 | 9.006 | 9.978 | 9.916 | 10.136 | 10.233 | 10.722 | 10.077 | 10.058 | 10.052 | 10.663 | -0.020 | 0.6318 | 0.8176 | 0 | 0 | 0 |
| 'Yars' | 7.027 | 6.231 | 7.796 | 6.643 | 7.075 | 6.806 | 6.818 | 7.171 | 7.053 | 7.073 | 6.347 | 7.016 | 7.034 | 6.091 | 6.513 | 6.251 | 6.549 | 6.416 | 0.069 | 0.0731 | 0.2395 | 0 | 0 | 0 |
| 'Yars2' | 2.358 | 2.643 | 2.376 | 2.368 | 2.073 | 2.429 | 2.403 | 2.739 | 2.236 | 2.056 | 2.221 | 2.733 | 2.384 | 2.123 | 2.391 | 2.507 | 2.157 | 2.561 | 0.017 | 0.7863 | 0.9046 | 0 | 0 | 0 |
| 'Ybey' | 1.020 | 1.433 | 1.387 | 1.112 | 1.392 | 1.299 | 0.968 | 1.313 | 1.303 | 1.458 | 1.207 | 1.423 | 0.923 | 1.210 | 1.205 | 1.210 | 1.037 | 1.308 | 0.015 | 0.8813 | 0.9499 | 0 | 0 | 0 |
| 'Ybx1' | 126.057 | 119.471 | 120.703 | 132.719 | 124.917 | 122.230 | 132.029 | 116.763 | 120.671 | 119.473 | 116.526 | 112.211 | 133.428 | 121.929 | 122.840 | 123.711 | 121.375 | 121.470 | 0.018 | 0.5094 | 0.7342 | 0 | 0 | 0 |
| 'Ybx3' | 23.670 | 18.140 | 22.986 | 22.790 | 20.047 | 23.001 | 23.508 | 20.099 | 20.256 | 18.903 | 17.943 | 20.394 | 22.084 | 18.375 | 20.499 | 18.919 | 19.260 | 21.898 | 0.113 | 0.0070 | 0.0480 | 1 | 0 | 0 |
| 'Ydjc' | 1.025 | 1.009 | 1.308 | 0.785 | 1.110 | 0.970 | 1.066 | 1.201 | 1.209 | 0.901 | 0.914 | 1.087 | 0.930 | 0.976 | 1.075 | 0.857 | 0.904 | 1.104 | 0.136 | 0.0699 | 0.2329 | 0 | 0 | 0 |
| 'Yeats2' | 10.097 | 8.800 | 10.029 | 9.451 | 9.201 | 9.492 | 9.767 | 9.518 | 9.875 | 8.605 | 8.692 | 10.723 | 9.116 | 9.603 | 10.270 | 8.531 | 9.668 | 10.416 | -0.001 | 0.9882 | 0.9965 | 0 | 0 | 0 |
| 'Yeats4' | 38.741 | 51.467 | 46.591 | 41.218 | 50.988 | 48.637 | 41.719 | 53.800 | 47.855 | 46.046 | 42.639 | 42.947 | 45.456 | 47.476 | 47.768 | 51.672 | 44.236 | 44.371 | 0.014 | 0.7878 | 0.9048 | 0 | 0 | 0 |
| 'Yif1a' | 11.070 | 16.342 | 12.599 | 10.796 | 16.482 | 14.507 | 9.957 | 14.336 | 14.303 | 12.721 | 11.390 | 11.829 | 14.328 | 12.797 | 11.889 | 16.266 | 12.231 | 11.232 | 0.052 | 0.5617 | 0.7712 | 0 | 0 | 0 |
| 'Yif1b' | 6.927 | 8.667 | 7.262 | 6.877 | 7.533 | 8.509 | 6.940 | 7.826 | 8.169 | 8.045 | 7.282 | 8.118 | 7.991 | 7.481 | 7.157 | 8.832 | 7.811 | 7.197 | -0.036 | 0.5190 | 0.7416 | 0 | 0 | 0 |
| 'Yipf1' | 5.463 | 5.760 | 5.730 | 5.948 | 5.705 | 5.797 | 5.749 | 6.102 | 5.662 | 5.428 | 5.823 | 5.811 | 6.083 | 5.986 | 5.954 | 5.707 | 6.171 | 5.101 | -0.014 | 0.6936 | 0.8558 | 0 | 0 | 0 |
| 'Yipf2' | 5.581 | 6.532 | 6.662 | 5.715 | 6.333 | 6.820 | 6.154 | 6.686 | 6.472 | 6.701 | 6.369 | 5.900 | 7.299 | 5.706 | 6.224 | 6.265 | 6.729 | 5.445 | 0.003 | 0.9642 | 0.9869 | 0 | 0 | 0 |
| 'Yipf3' | 19.407 | 19.340 | 21.089 | 17.540 | 20.972 | 21.967 | 19.184 | 21.021 | 20.670 | 19.781 | 17.653 | 20.026 | 20.184 | 18.679 | 21.136 | 20.994 | 18.444 | 19.568 | 0.029 | 0.4359 | 0.6798 | 0 | 0 | 0 |
| 'Yipf4' | 22.293 | 26.482 | 22.084 | 23.601 | 26.335 | 23.252 | 23.747 | 25.984 | 20.966 | 21.406 | 25.690 | 26.094 | 23.026 | 25.839 | 23.295 | 24.136 | 26.239 | 24.346 | -0.054 | 0.1654 | 0.3978 | 0 | 0 | 0 |
| 'Yipf5' | 16.145 | 16.788 | 17.314 | 15.478 | 17.069 | 16.859 | 16.116 | 16.772 | 16.244 | 15.062 | 16.304 | 16.673 | 15.796 | 18.127 | 16.549 | 17.769 | 17.673 | 16.252 | -0.020 | 0.6111 | 0.8049 | 0 | 0 | 0 |
| 'Yipf6' | 2.950 | 3.010 | 2.881 | 2.746 | 3.355 | 2.979 | 2.996 | 3.320 | 2.693 | 2.732 | 2.970 | 3.243 | 2.902 | 3.406 | 2.916 | 2.802 | 3.186 | 2.943 | -0.026 | 0.5646 | 0.7736 | 0 | 0 | 0 |
| 'Ykt6' | 27.045 | 32.917 | 28.072 | 28.537 | 33.028 | 31.222 | 26.977 | 32.653 | 31.254 | 29.388 | 29.914 | 31.530 | 32.101 | 34.604 | 33.980 | 32.465 | 30.059 | 31.123 | -0.084 | 0.0368 | 0.1551 | 0 | 0 | 0 |
| 'Ylpm1' | 12.053 | 10.279 | 11.836 | 12.302 | 11.253 | 10.670 | 11.894 | 10.989 | 11.563 | 12.309 | 12.280 | 12.284 | 11.761 | 11.564 | 11.407 | 11.034 | 11.237 | 12.715 | -0.064 | 0.0925 | 0.2812 | 0 | 0 | 0 |
| 'Yme1l1' | 17.747 | 18.335 | 18.591 | 19.758 | 19.608 | 16.960 | 20.278 | 18.778 | 17.700 | 18.366 | 20.015 | 18.526 | 17.644 | 18.956 | 16.873 | 17.208 | 20.859 | 17.456 | 0.006 | 0.8723 | 0.9453 | 0 | 0 | 0 |
| 'Yod1' | 3.808 | 4.574 | 4.113 | 3.688 | 4.206 | 4.170 | 4.518 | 4.106 | 4.012 | 3.563 | 4.345 | 4.095 | 4.120 | 4.116 | 4.074 | 4.266 | 4.326 | 4.338 | -0.015 | 0.7627 | 0.8946 | 0 | 0 | 0 |
| 'Ypel1' | 0.542 | 0.528 | 0.506 | 0.403 | 0.554 | 0.423 | 0.449 | 0.450 | 0.727 | 0.554 | 0.520 | 0.484 | 0.557 | 0.577 | 0.486 | 0.620 | 0.460 | 0.455 | -0.039 | 0.7429 | 0.8839 | 0 | 0 | 0 |
| 'Ypel2' | 5.557 | 6.838 | 4.499 | 5.077 | 5.319 | 4.497 | 4.906 | 5.300 | 4.436 | 4.910 | 6.683 | 5.565 | 5.083 | 6.511 | 4.591 | 5.933 | 6.305 | 5.287 | -0.146 | 0.0156 | 0.0863 | 0 | 0 | 0 |
| 'Ypel3' | 30.801 | 36.028 | 33.356 | 30.507 | 34.176 | 37.643 | 30.365 | 34.868 | 34.688 | 33.800 | 32.660 | 35.406 | 36.238 | 35.239 | 35.586 | 36.725 | 36.242 | 33.540 | -0.073 | 0.0394 | 0.1621 | 0 | 0 | 0 |
| 'Ypel5' | 23.595 | 29.656 | 21.747 | 24.720 | 27.168 | 23.398 | 24.032 | 27.830 | 22.606 | 24.871 | 26.472 | 21.919 | 23.762 | 28.162 | 22.094 | 26.692 | 26.589 | 21.348 | 0.009 | 0.7796 | 0.9008 | 0 | 0 | 0 |
| 'Yrdc' | 11.699 | 12.017 | 9.787 | 9.009 | 11.895 | 11.024 | 9.083 | 12.029 | 9.173 | 12.213 | 10.020 | 9.486 | 11.098 | 13.493 | 10.064 | 13.198 | 9.615 | 9.237 | -0.046 | 0.5387 | 0.7567 | 0 | 0 | 0 |
| 'Ythdc1' | 10.542 | 10.608 | 9.522 | 10.743 | 10.423 | 9.746 | 10.388 | 10.938 | 9.437 | 10.558 | 10.548 | 10.393 | 10.228 | 11.104 | 9.837 | 10.967 | 11.070 | 10.153 | -0.051 | 0.0285 | 0.1307 | 0 | 0 | 0 |
| 'Ythdc2' | 3.370 | 3.096 | 3.280 | 3.432 | 3.240 | 3.233 | 3.633 | 3.115 | 3.189 | 2.885 | 3.767 | 3.364 | 3.021 | 2.938 | 2.917 | 2.977 | 3.575 | 3.343 | 0.029 | 0.6097 | 0.8040 | 0 | 0 | 0 |
| 'Ythdf1' | 20.868 | 21.103 | 19.735 | 19.724 | 19.982 | 19.939 | 18.435 | 21.370 | 18.865 | 19.913 | 19.789 | 20.817 | 18.903 | 20.163 | 21.010 | 21.057 | 20.028 | 21.808 | -0.038 | 0.2910 | 0.5477 | 0 | 0 | 0 |
| 'Ythdf3' | 14.524 | 15.966 | 15.843 | 16.253 | 15.830 | 14.405 | 18.424 | 15.557 | 15.175 | 13.961 | 15.984 | 16.465 | 13.105 | 15.542 | 14.600 | 13.949 | 17.226 | 15.095 | 0.054 | 0.2920 | 0.5489 | 0 | 0 | 0 |
| 'Ywhab' | 84.424 | 84.142 | 89.306 | 91.765 | 86.297 | 87.821 | 88.878 | 84.176 | 88.318 | 85.854 | 82.192 | 90.307 | 82.627 | 85.638 | 92.261 | 81.137 | 85.159 | 91.861 | 0.004 | 0.8708 | 0.9447 | 0 | 0 | 0 |
| 'Ywhae' | 62.843 | 69.684 | 63.848 | 71.530 | 72.178 | 65.701 | 69.467 | 66.061 | 66.842 | 57.253 | 65.664 | 66.847 | 65.440 | 69.827 | 70.483 | 62.997 | 70.889 | 69.163 | 0.013 | 0.7434 | 0.8843 | 0 | 0 | 0 |
| 'Ywhag' | 16.862 | 15.070 | 17.115 | 17.682 | 15.534 | 15.948 | 18.312 | 15.181 | 16.485 | 14.725 | 18.479 | 20.917 | 15.630 | 15.027 | 19.702 | 14.427 | 18.416 | 21.557 | -0.103 | 0.1246 | 0.3375 | 0 | 0 | 0 |
| 'Ywhaq' | 45.099 | 47.737 | 47.198 | 49.727 | 49.978 | 46.582 | 49.052 | 50.617 | 48.001 | 43.157 | 44.237 | 45.481 | 47.080 | 46.909 | 47.250 | 46.397 | 47.589 | 44.477 | 0.061 | 0.0203 | 0.1041 | 0 | 0 | 0 |
| 'Ywhaz' | 68.323 | 72.667 | 70.080 | 73.342 | 72.579 | 69.245 | 76.914 | 72.992 | 68.179 | 66.417 | 72.505 | 67.241 | 65.500 | 70.517 | 68.262 | 64.438 | 75.681 | 65.935 | 0.054 | 0.0596 | 0.2110 | 0 | 0 | 0 |
| 'Yy1' | 70.915 | 62.576 | 66.762 | 68.202 | 62.126 | 66.983 | 65.563 | 62.478 | 64.717 | 63.566 | 61.710 | 71.853 | 63.156 | 63.120 | 67.661 | 64.065 | 59.824 | 70.906 | 0.001 | 0.9852 | 0.9949 | 0 | 0 | 0 |
| 'Zadh2' | 8.835 | 9.780 | 10.717 | 9.130 | 9.894 | 11.136 | 9.114 | 10.000 | 10.610 | 7.746 | 8.795 | 9.990 | 8.430 | 8.586 | 10.361 | 9.158 | 7.975 | 9.989 | 0.129 | 0.0016 | 0.0151 | 1 | 0 | 0 |
| 'Zan' | 12.095 | 8.942 | 11.843 | 10.321 | 9.564 | 10.665 | 10.991 | 9.541 | 11.502 | 13.039 | 11.409 | 13.145 | 12.585 | 10.676 | 12.211 | 11.443 | 11.911 | 12.635 | -0.205 | 0.0000 | 0.0000 | 1 | 0 | 0 |
| 'Zbed6' | 9.021 | 7.113 | 8.158 | 8.032 | 7.163 | 6.580 | 7.789 | 7.101 | 7.488 | 6.711 | 7.555 | 8.500 | 6.800 | 6.626 | 7.250 | 7.412 | 7.119 | 8.591 | 0.029 | 0.6325 | 0.8178 | 0 | 0 | 0 |
| 'Zbp1' | 24.044 | 25.223 | 26.285 | 22.958 | 24.306 | 26.898 | 22.727 | 26.833 | 26.965 | 26.793 | 24.518 | 29.957 | 28.954 | 24.314 | 30.163 | 25.564 | 26.226 | 28.614 | -0.126 | 0.0003 | 0.0042 | 1 | 0 | 0 |
| 'Zbtb1' | 15.363 | 14.388 | 14.487 | 16.885 | 14.071 | 12.960 | 17.468 | 13.673 | 14.591 | 12.716 | 16.710 | 18.313 | 13.305 | 15.531 | 16.966 | 13.559 | 19.291 | 18.625 | -0.119 | 0.1993 | 0.4396 | 0 | 0 | 0 |
| 'Zbtb11' | 7.483 | 7.165 | 6.916 | 7.532 | 7.105 | 6.667 | 7.273 | 6.403 | 6.889 | 6.101 | 7.263 | 7.830 | 5.865 | 7.258 | 7.561 | 6.617 | 7.843 | 8.128 | -0.030 | 0.6060 | 0.8014 | 0 | 0 | 0 |
| 'Zbtb12' | 4.653 | 3.725 | 4.553 | 4.228 | 3.441 | 3.966 | 3.566 | 3.456 | 3.758 | 2.974 | 3.626 | 5.163 | 2.954 | 3.385 | 5.128 | 3.419 | 3.045 | 4.763 | 0.038 | 0.6979 | 0.8583 | 0 | 0 | 0 |
| 'Zbtb14' | 13.202 | 12.761 | 12.952 | 12.666 | 11.792 | 12.949 | 12.552 | 13.525 | 13.113 | 11.822 | 14.768 | 14.255 | 13.080 | 13.468 | 11.943 | 11.320 | 13.832 | 13.853 | -0.045 | 0.3466 | 0.6017 | 0 | 0 | 0 |
| 'Zbtb17' | 9.278 | 7.950 | 9.466 | 8.066 | 7.785 | 10.050 | 8.507 | 8.362 | 9.270 | 9.008 | 7.601 | 9.240 | 9.123 | 7.534 | 8.322 | 8.521 | 7.146 | 8.773 | 0.057 | 0.1191 | 0.3284 | 0 | 0 | 0 |
| 'Zbtb18' | 2.426 | 3.108 | 2.529 | 2.702 | 2.800 | 2.572 | 2.427 | 2.669 | 2.802 | 2.564 | 3.199 | 3.250 | 2.429 | 3.039 | 3.096 | 2.423 | 3.143 | 3.289 | -0.146 | 0.0026 | 0.0222 | 1 | 0 | 0 |
| 'Zbtb20' | 0.542 | 0.539 | 0.394 | 0.454 | 0.350 | 0.407 | 0.427 | 0.451 | 0.419 | 0.469 | 0.376 | 0.600 | 0.424 | 0.344 | 0.537 | 0.450 | 0.465 | 0.478 | -0.082 | 0.4766 | 0.7107 | 0 | 0 | 0 |
| 'Zbtb21' | 3.096 | 1.717 | 1.840 | 2.184 | 1.949 | 1.829 | 2.000 | 1.956 | 1.991 | 3.356 | 2.317 | 1.944 | 2.706 | 2.163 | 1.766 | 2.911 | 1.985 | 1.817 | -0.164 | 0.0287 | 0.1313 | 0 | 0 | 0 |
| 'Zbtb22' | 17.485 | 19.810 | 21.275 | 17.037 | 19.407 | 21.885 | 17.356 | 20.519 | 22.187 | 19.611 | 17.306 | 16.427 | 18.949 | 17.207 | 19.103 | 19.906 | 15.414 | 14.132 | 0.154 | 0.0223 | 0.1108 | 0 | 0 | 0 |
| 'Zbtb24' | 15.584 | 16.118 | 18.795 | 16.213 | 16.362 | 17.700 | 17.640 | 16.491 | 19.589 | 16.836 | 15.952 | 18.120 | 14.434 | 15.435 | 17.733 | 14.818 | 16.020 | 16.423 | 0.073 | 0.0586 | 0.2087 | 0 | 0 | 0 |
| 'Zbtb25' | 4.192 | 4.011 | 4.218 | 4.115 | 3.992 | 4.210 | 4.162 | 4.105 | 3.946 | 3.721 | 3.679 | 4.363 | 4.148 | 4.001 | 4.611 | 4.049 | 4.472 | 4.166 | -0.022 | 0.6100 | 0.8041 | 0 | 0 | 0 |
| 'Zbtb26' | 0.717 | 1.062 | 1.154 | 0.879 | 0.893 | 0.948 | 1.120 | 1.063 | 1.091 | 0.653 | 0.960 | 1.265 | 0.656 | 0.780 | 1.094 | 0.640 | 0.972 | 1.095 | 0.139 | 0.1378 | 0.3581 | 0 | 0 | 0 |
| 'Zbtb33' | 5.081 | 5.343 | 4.897 | 5.079 | 5.289 | 4.871 | 5.728 | 5.397 | 5.260 | 4.373 | 5.378 | 5.794 | 4.341 | 5.012 | 5.075 | 4.465 | 5.373 | 5.315 | 0.046 | 0.3730 | 0.6242 | 0 | 0 | 0 |
| 'Zbtb34' | 1.805 | 2.092 | 2.187 | 1.913 | 2.224 | 2.113 | 1.877 | 2.116 | 2.278 | 1.841 | 2.611 | 2.244 | 2.068 | 2.188 | 2.247 | 1.950 | 1.849 | 2.192 | -0.055 | 0.3538 | 0.6072 | 0 | 0 | 0 |
| 'Zbtb37' | 1.690 | 1.904 | 2.129 | 1.766 | 2.107 | 1.906 | 1.723 | 1.932 | 2.175 | 1.605 | 1.377 | 1.510 | 1.428 | 1.456 | 1.292 | 1.637 | 1.527 | 1.486 | 0.369 | 0.0000 | 0.0000 | 1 | 0 | 0 |
| 'Zbtb38' | 3.381 | 3.187 | 2.768 | 2.805 | 3.333 | 3.126 | 3.025 | 3.169 | 3.007 | 3.281 | 3.134 | 2.892 | 2.815 | 3.211 | 2.867 | 3.749 | 2.810 | 2.826 | 0.004 | 0.9401 | 0.9769 | 0 | 0 | 0 |
| 'Zbtb39' | 2.348 | 2.964 | 2.821 | 2.489 | 2.631 | 2.707 | 2.287 | 3.244 | 2.692 | 2.726 | 2.462 | 2.996 | 3.266 | 2.882 | 2.900 | 3.016 | 2.611 | 2.530 | -0.083 | 0.2191 | 0.4660 | 0 | 0 | 0 |
| 'Zbtb4' | 10.298 | 10.068 | 9.789 | 9.382 | 9.452 | 10.166 | 8.755 | 9.703 | 10.465 | 9.945 | 9.591 | 11.335 | 9.103 | 9.808 | 10.311 | 9.504 | 9.394 | 10.625 | -0.035 | 0.3145 | 0.5721 | 0 | 0 | 0 |
| 'Zbtb40' | 2.737 | 2.360 | 3.280 | 3.183 | 2.686 | 3.031 | 2.736 | 2.586 | 3.041 | 2.913 | 2.826 | 3.354 | 2.945 | 2.918 | 2.993 | 2.671 | 2.866 | 3.407 | -0.081 | 0.0754 | 0.2441 | 0 | 0 | 0 |
| 'Zbtb41' | 2.240 | 2.322 | 1.805 | 2.148 | 2.275 | 2.019 | 2.112 | 2.315 | 1.946 | 2.355 | 2.132 | 2.148 | 2.205 | 2.229 | 2.149 | 2.430 | 1.938 | 2.035 | -0.048 | 0.3200 | 0.5782 | 0 | 0 | 0 |
| 'Zbtb42' | 0.806 | 0.892 | 0.966 | 0.734 | 0.796 | 1.148 | 0.646 | 0.961 | 1.120 | 0.860 | 0.873 | 1.130 | 0.812 | 0.877 | 1.169 | 0.796 | 0.783 | 1.108 | -0.067 | 0.4186 | 0.6658 | 0 | 0 | 0 |
| 'Zbtb43' | 3.059 | 2.869 | 2.875 | 3.230 | 3.115 | 3.169 | 3.077 | 3.331 | 3.011 | 2.720 | 3.363 | 3.781 | 2.849 | 2.768 | 3.156 | 3.171 | 3.401 | 3.646 | -0.067 | 0.2533 | 0.5048 | 0 | 0 | 0 |
| 'Zbtb44' | 10.394 | 9.167 | 8.328 | 9.536 | 9.433 | 9.151 | 9.954 | 9.286 | 9.118 | 9.177 | 9.500 | 10.683 | 9.104 | 9.897 | 10.132 | 9.122 | 10.585 | 11.060 | -0.096 | 0.0756 | 0.2441 | 0 | 0 | 0 |
| 'Zbtb45' | 9.633 | 9.193 | 12.058 | 10.002 | 10.133 | 12.020 | 10.455 | 9.274 | 12.182 | 10.125 | 10.131 | 13.250 | 9.285 | 10.026 | 12.509 | 9.182 | 9.808 | 12.606 | -0.042 | 0.2812 | 0.5379 | 0 | 0 | 0 |
| 'Zbtb48' | 3.772 | 2.815 | 3.797 | 3.467 | 3.152 | 3.635 | 3.517 | 2.797 | 3.779 | 2.926 | 3.008 | 4.385 | 3.730 | 2.924 | 3.828 | 3.469 | 3.377 | 4.202 | -0.065 | 0.2427 | 0.4933 | 0 | 0 | 0 |
| 'Zbtb49' | 1.498 | 1.365 | 1.473 | 1.281 | 1.333 | 1.355 | 1.399 | 1.318 | 1.341 | 1.491 | 1.406 | 1.618 | 1.378 | 1.400 | 1.575 | 1.404 | 1.470 | 1.436 | -0.104 | 0.0341 | 0.1475 | 0 | 0 | 0 |
| 'Zbtb5' | 1.969 | 2.560 | 2.239 | 2.099 | 2.560 | 2.537 | 2.338 | 2.281 | 2.491 | 2.439 | 2.277 | 2.637 | 2.519 | 2.505 | 2.537 | 2.642 | 2.357 | 2.488 | -0.105 | 0.0535 | 0.1976 | 0 | 0 | 0 |
| 'Zbtb6' | 2.697 | 3.273 | 2.955 | 2.810 | 3.359 | 2.787 | 3.093 | 3.035 | 3.633 | 2.585 | 3.178 | 3.738 | 2.612 | 3.012 | 3.084 | 2.694 | 3.546 | 3.663 | -0.036 | 0.5662 | 0.7744 | 0 | 0 | 0 |
| 'Zbtb7a' | 19.654 | 18.038 | 18.194 | 18.134 | 18.899 | 19.499 | 18.410 | 18.422 | 19.149 | 17.053 | 17.600 | 19.660 | 17.687 | 18.349 | 19.533 | 17.967 | 16.385 | 19.590 | 0.030 | 0.3428 | 0.5990 | 0 | 0 | 0 |
| 'Zbtb7b' | 39.303 | 33.857 | 39.980 | 37.246 | 33.708 | 41.000 | 37.836 | 32.229 | 41.001 | 35.282 | 35.289 | 49.692 | 35.238 | 36.260 | 46.688 | 36.303 | 36.223 | 48.507 | -0.099 | 0.0179 | 0.0955 | 0 | 0 | 0 |
| 'Zbtb8os' | 16.748 | 19.726 | 17.277 | 17.295 | 19.482 | 17.892 | 19.583 | 20.356 | 17.316 | 17.348 | 19.149 | 16.938 | 18.945 | 18.658 | 17.653 | 18.350 | 18.928 | 15.647 | 0.025 | 0.4913 | 0.7209 | 0 | 0 | 0 |
| 'Zc3h10' | 6.795 | 5.220 | 5.500 | 6.194 | 5.731 | 6.158 | 6.620 | 5.506 | 6.017 | 6.255 | 5.670 | 6.418 | 5.946 | 5.983 | 6.132 | 6.500 | 5.067 | 5.782 | -0.013 | 0.7740 | 0.8993 | 0 | 0 | 0 |
| 'Zc3h11a' | 3.612 | 3.538 | 3.482 | 3.938 | 3.123 | 2.970 | 3.477 | 3.328 | 3.285 | 2.925 | 3.216 | 3.828 | 2.567 | 3.036 | 3.376 | 2.938 | 3.903 | 3.737 | 0.050 | 0.5044 | 0.7312 | 0 | 0 | 0 |
| 'Zc3h12a' | 25.308 | 24.244 | 18.438 | 18.022 | 22.064 | 21.428 | 18.765 | 24.604 | 20.587 | 32.055 | 21.516 | 17.190 | 31.090 | 28.499 | 18.525 | 38.417 | 19.354 | 16.817 | -0.167 | 0.2015 | 0.4426 | 0 | 0 | 0 |
| 'Zc3h12d' | 19.655 | 20.523 | 20.752 | 17.438 | 21.112 | 20.874 | 17.802 | 23.089 | 21.350 | 19.622 | 23.535 | 22.589 | 18.763 | 21.859 | 21.797 | 21.162 | 20.001 | 22.080 | -0.078 | 0.0398 | 0.1628 | 0 | 0 | 0 |
| 'Zc3h13' | 20.736 | 19.870 | 21.778 | 20.711 | 20.621 | 21.051 | 21.485 | 20.260 | 21.424 | 22.081 | 21.672 | 22.572 | 20.971 | 19.997 | 20.805 | 20.306 | 22.177 | 21.984 | -0.046 | 0.0901 | 0.2760 | 0 | 0 | 0 |
| 'Zc3h14' | 27.081 | 27.152 | 26.590 | 24.960 | 28.360 | 26.723 | 27.400 | 26.764 | 26.631 | 28.426 | 26.099 | 26.201 | 26.333 | 26.310 | 23.958 | 25.946 | 27.353 | 24.023 | 0.033 | 0.2638 | 0.5179 | 0 | 0 | 0 |
| 'Zc3h15' | 25.183 | 23.760 | 23.409 | 26.331 | 23.767 | 22.516 | 27.078 | 24.842 | 22.425 | 23.021 | 24.538 | 23.820 | 22.843 | 24.670 | 22.411 | 23.088 | 25.209 | 23.354 | 0.029 | 0.4109 | 0.6596 | 0 | 0 | 0 |
| 'Zc3h18' | 11.963 | 10.152 | 13.135 | 12.087 | 10.456 | 13.126 | 11.548 | 10.596 | 12.923 | 11.801 | 10.550 | 13.487 | 12.236 | 10.226 | 12.664 | 11.061 | 10.733 | 13.168 | -0.010 | 0.6642 | 0.8377 | 0 | 0 | 0 |
| 'Zc3h3' | 9.302 | 7.962 | 9.771 | 7.863 | 8.348 | 8.983 | 8.598 | 8.215 | 9.309 | 9.621 | 8.623 | 9.499 | 8.982 | 7.874 | 9.010 | 6.795 | 8.671 | 9.018 | -0.006 | 0.9066 | 0.9635 | 0 | 0 | 0 |
| 'Zc3h4' | 21.844 | 21.626 | 23.771 | 20.396 | 23.004 | 24.949 | 20.527 | 23.214 | 25.164 | 24.659 | 22.470 | 22.457 | 23.943 | 23.108 | 22.239 | 23.811 | 21.121 | 20.853 | -0.012 | 0.7769 | 0.9002 | 0 | 0 | 0 |
| 'Zc3h6' | 5.955 | 6.165 | 6.043 | 5.674 | 5.955 | 5.849 | 7.454 | 6.598 | 6.326 | 7.093 | 7.039 | 5.198 | 6.553 | 6.473 | 5.098 | 6.862 | 6.380 | 4.165 | 0.038 | 0.5734 | 0.7795 | 0 | 0 | 0 |
| 'Zc3h7a' | 16.399 | 16.732 | 17.102 | 16.082 | 16.743 | 15.622 | 16.981 | 16.586 | 17.273 | 16.706 | 16.714 | 16.219 | 16.641 | 16.742 | 15.582 | 15.265 | 17.696 | 15.452 | 0.015 | 0.5987 | 0.7958 | 0 | 0 | 0 |
| 'Zc3h7b' | 20.051 | 19.666 | 21.491 | 20.001 | 19.809 | 21.442 | 20.210 | 22.172 | 21.803 | 20.434 | 19.261 | 19.852 | 20.295 | 18.482 | 19.055 | 20.461 | 18.942 | 18.729 | 0.081 | 0.0063 | 0.0444 | 1 | 0 | 0 |
| 'Zc3h8' | 3.793 | 3.261 | 3.663 | 3.509 | 3.815 | 2.817 | 3.138 | 2.274 | 2.803 | 2.964 | 3.417 | 3.578 | 3.502 | 3.324 | 3.441 | 3.606 | 2.897 | 3.545 | -0.077 | 0.4553 | 0.6934 | 0 | 0 | 0 |
| 'Zc3hav1' | 43.384 | 43.769 | 41.441 | 38.931 | 41.931 | 43.342 | 39.389 | 45.250 | 41.333 | 42.701 | 41.606 | 42.615 | 39.074 | 42.363 | 43.696 | 43.996 | 39.883 | 43.927 | -0.015 | 0.6485 | 0.8286 | 0 | 0 | 0 |
| 'Zc3hav1l' | 1.260 | 1.296 | 1.634 | 1.554 | 1.306 | 1.445 | 1.677 | 1.394 | 1.720 | 1.185 | 1.156 | 1.710 | 1.244 | 0.956 | 1.635 | 1.219 | 1.184 | 1.607 | 0.142 | 0.0695 | 0.2318 | 0 | 0 | 0 |
| 'Zc3hc1' | 3.067 | 2.961 | 2.933 | 2.648 | 2.796 | 3.237 | 3.018 | 3.363 | 2.921 | 3.208 | 2.598 | 2.812 | 3.185 | 3.000 | 3.003 | 3.349 | 3.047 | 2.564 | 0.007 | 0.8948 | 0.9571 | 0 | 0 | 0 |
| 'Zc4h2' | 0.553 | 0.558 | 0.542 | 0.638 | 0.634 | 0.619 | 0.598 | 0.673 | 0.751 | 0.677 | 0.628 | 0.625 | 0.709 | 0.446 | 0.428 | 0.505 | 0.532 | 0.598 | 0.094 | 0.4377 | 0.6809 | 0 | 0 | 0 |
| 'Zcchc10' | 10.893 | 9.187 | 8.418 | 8.664 | 10.961 | 9.234 | 10.076 | 11.039 | 8.771 | 9.362 | 8.677 | 8.484 | 8.440 | 9.847 | 8.129 | 9.690 | 9.427 | 7.753 | 0.121 | 0.0435 | 0.1726 | 0 | 0 | 0 |
| 'Zcchc11' | 3.871 | 4.159 | 3.338 | 3.969 | 4.022 | 3.176 | 3.893 | 4.091 | 3.611 | 3.527 | 3.685 | 3.437 | 3.911 | 3.932 | 3.016 | 3.395 | 3.560 | 3.309 | 0.086 | 0.0263 | 0.1245 | 0 | 0 | 0 |
| 'Zcchc12' | 1.957 | 2.263 | 1.759 | 2.274 | 2.558 | 2.227 | 2.246 | 2.392 | 2.359 | 2.892 | 2.695 | 2.946 | 3.381 | 3.121 | 2.139 | 2.853 | 3.247 | 2.169 | -0.356 | 0.0000 | 0.0001 | 1 | 0 | 0 |
| 'Zcchc17' | 9.104 | 9.657 | 9.259 | 10.457 | 9.086 | 9.737 | 9.978 | 9.881 | 9.766 | 9.397 | 8.410 | 9.715 | 9.558 | 9.058 | 8.697 | 9.859 | 9.442 | 9.247 | 0.047 | 0.1831 | 0.4191 | 0 | 0 | 0 |
| 'Zcchc18' | 1.521 | 1.606 | 1.621 | 2.072 | 1.647 | 1.726 | 1.803 | 1.841 | 1.762 | 1.287 | 1.069 | 1.176 | 1.123 | 1.064 | 0.955 | 1.066 | 0.954 | 1.040 | 0.664 | 0.0000 | 0.0000 | 1 | 1 | 0 |
| 'Zcchc2' | 6.964 | 6.818 | 6.059 | 6.049 | 6.449 | 6.855 | 5.709 | 6.607 | 6.272 | 6.298 | 6.531 | 8.679 | 6.308 | 7.012 | 8.320 | 7.221 | 7.040 | 9.945 | -0.220 | 0.0007 | 0.0077 | 1 | 0 | 0 |
| 'Zcchc4' | 1.935 | 2.137 | 2.346 | 1.994 | 2.044 | 2.041 | 2.161 | 2.456 | 2.156 | 2.148 | 2.272 | 2.095 | 2.328 | 2.121 | 1.845 | 2.121 | 2.202 | 1.689 | 0.029 | 0.6057 | 0.8014 | 0 | 0 | 0 |
| 'Zcchc6' | 39.474 | 39.374 | 37.900 | 40.508 | 38.879 | 37.955 | 38.996 | 38.273 | 37.324 | 38.867 | 40.795 | 41.267 | 37.850 | 39.036 | 39.431 | 39.784 | 42.530 | 41.818 | -0.063 | 0.0272 | 0.1271 | 0 | 0 | 0 |
| 'Zcchc7' | 7.562 | 7.076 | 7.234 | 7.916 | 7.385 | 6.952 | 7.990 | 7.236 | 7.776 | 7.590 | 7.580 | 7.975 | 7.074 | 7.536 | 7.112 | 7.079 | 7.639 | 7.190 | -0.005 | 0.8879 | 0.9540 | 0 | 0 | 0 |
| 'Zcchc8' | 10.140 | 10.338 | 10.984 | 10.565 | 10.757 | 10.135 | 11.233 | 10.171 | 11.046 | 9.945 | 9.891 | 11.029 | 10.300 | 10.234 | 10.388 | 9.812 | 10.316 | 10.735 | 0.029 | 0.3092 | 0.5675 | 0 | 0 | 0 |
| 'Zcchc9' | 6.467 | 6.554 | 5.699 | 5.659 | 6.064 | 5.365 | 5.727 | 6.878 | 5.641 | 5.916 | 6.008 | 6.166 | 6.203 | 6.441 | 6.013 | 6.086 | 6.982 | 5.744 | -0.053 | 0.2273 | 0.4735 | 0 | 0 | 0 |
| 'Zcrb1' | 9.481 | 9.873 | 9.656 | 9.451 | 10.009 | 10.057 | 9.856 | 10.161 | 9.834 | 9.943 | 9.908 | 10.060 | 10.177 | 10.923 | 10.450 | 10.096 | 10.857 | 9.877 | -0.071 | 0.0264 | 0.1245 | 0 | 0 | 0 |
| 'Zdhhc1' | 4.139 | 5.013 | 4.080 | 4.364 | 4.543 | 4.709 | 3.529 | 5.090 | 4.881 | 5.854 | 4.442 | 4.474 | 4.713 | 4.273 | 4.332 | 4.146 | 5.016 | 4.030 | -0.042 | 0.6246 | 0.8129 | 0 | 0 | 0 |
| 'Zdhhc12' | 2.542 | 3.550 | 4.345 | 2.092 | 3.348 | 3.999 | 2.778 | 3.487 | 4.062 | 3.367 | 3.040 | 3.461 | 3.387 | 2.873 | 3.593 | 2.868 | 2.642 | 3.329 | 0.056 | 0.5081 | 0.7336 | 0 | 0 | 0 |
| 'Zdhhc13' | 11.980 | 13.763 | 13.221 | 13.418 | 13.287 | 11.921 | 14.137 | 14.832 | 12.544 | 14.133 | 11.300 | 11.751 | 12.540 | 12.945 | 11.764 | 13.745 | 12.858 | 10.963 | 0.081 | 0.0807 | 0.2562 | 0 | 0 | 0 |
| 'Zdhhc15' | 2.589 | 2.692 | 2.771 | 2.632 | 2.813 | 2.353 | 2.712 | 2.887 | 2.987 | 2.983 | 3.001 | 3.604 | 2.774 | 3.041 | 3.233 | 2.295 | 3.074 | 3.613 | -0.193 | 0.0038 | 0.0302 | 1 | 0 | 0 |
| 'Zdhhc16' | 12.144 | 12.490 | 11.927 | 11.602 | 12.957 | 12.669 | 11.633 | 12.781 | 11.369 | 10.486 | 13.178 | 11.471 | 10.982 | 12.014 | 12.477 | 12.137 | 12.317 | 11.205 | 0.039 | 0.3673 | 0.6194 | 0 | 0 | 0 |
| 'Zdhhc17' | 7.992 | 9.466 | 9.052 | 8.058 | 10.101 | 9.225 | 9.399 | 10.194 | 9.596 | 9.886 | 8.762 | 7.971 | 7.738 | 8.662 | 7.676 | 8.923 | 9.480 | 7.043 | 0.119 | 0.0297 | 0.1339 | 0 | 0 | 0 |
| 'Zdhhc18' | 49.258 | 46.139 | 50.828 | 47.456 | 45.084 | 50.106 | 48.774 | 46.181 | 50.287 | 47.798 | 49.964 | 55.573 | 46.027 | 45.727 | 54.212 | 43.645 | 47.065 | 57.773 | -0.052 | 0.1398 | 0.3618 | 0 | 0 | 0 |
| 'Zdhhc2' | 5.132 | 6.682 | 6.137 | 5.587 | 6.190 | 6.307 | 5.969 | 6.362 | 5.924 | 6.160 | 5.753 | 5.967 | 6.265 | 7.085 | 5.963 | 6.535 | 6.349 | 5.623 | -0.047 | 0.2849 | 0.5413 | 0 | 0 | 0 |
| 'Zdhhc20' | 24.504 | 28.572 | 23.612 | 24.075 | 25.942 | 23.299 | 26.900 | 26.437 | 23.753 | 23.342 | 26.777 | 26.251 | 21.607 | 25.909 | 23.372 | 23.099 | 26.442 | 23.985 | 0.028 | 0.4459 | 0.6878 | 0 | 0 | 0 |
| 'Zdhhc21' | 4.376 | 3.865 | 3.942 | 4.440 | 3.983 | 3.497 | 5.054 | 3.870 | 3.789 | 3.684 | 4.922 | 4.481 | 3.818 | 4.493 | 3.893 | 3.607 | 4.806 | 4.370 | -0.060 | 0.3981 | 0.6472 | 0 | 0 | 0 |

| | | | | | | | | | | | | | | | | | | | | | | | | |
|---|---|---|---|---|---|---|---|---|---|---|---|---|---|---|---|---|---|---|---|---|---|---|---|---|
| 'Zdhhc23' | 1.361 | 1.682 | 1.782 | 0.977 | 1.282 | 1.411 | 1.291 | 1.651 | 1.795 | 2.475 | 1.809 | 2.241 | 1.755 | 2.026 | 1.730 | 1.643 | 1.850 | 2.640 | -0.469 | 0.0002 | 0.0033 | 1 | 1 | 0 |
| 'Zdhhc24' | 1.977 | 2.240 | 2.369 | 2.309 | 2.159 | 2.221 | 2.202 | 2.358 | 2.410 | 2.039 | 2.491 | 2.510 | 2.010 | 2.365 | 2.706 | 2.208 | 2.573 | 2.427 | -0.084 | 0.0867 | 0.2682 | 0 | 0 | 0 |
| 'Zdhhc3' | 5.565 | 5.836 | 5.680 | 5.155 | 5.767 | 5.851 | 5.494 | 6.031 | 5.744 | 5.914 | 5.892 | 5.739 | 5.852 | 5.782 | 5.936 | 5.923 | 5.499 | 5.755 | -0.041 | 0.1403 | 0.3625 | 0 | 0 | 0 |
| 'Zdhhc4' | 4.830 | 5.412 | 5.847 | 4.612 | 5.816 | 5.578 | 5.095 | 5.752 | 6.594 | 5.911 | 5.348 | 5.456 | 5.585 | 5.600 | 4.974 | 5.677 | 4.954 | 5.059 | 0.018 | 0.7538 | 0.8903 | 0 | 0 | 0 |
| 'Zdhhc5' | 21.723 | 22.127 | 23.796 | 22.322 | 23.118 | 23.350 | 21.887 | 22.934 | 23.923 | 24.086 | 22.233 | 23.679 | 22.316 | 22.657 | 22.895 | 22.497 | 21.912 | 23.455 | -0.014 | 0.5732 | 0.7795 | 0 | 0 | 0 |
| 'Zdhhc6' | 17.797 | 18.686 | 19.575 | 20.107 | 20.197 | 19.521 | 21.305 | 19.356 | 18.838 | 19.903 | 19.820 | 20.082 | 20.070 | 19.112 | 20.472 | 19.488 | 20.415 | 18.459 | -0.033 | 0.3727 | 0.6242 | 0 | 0 | 0 |
| 'Zdhhc7' | 14.078 | 14.662 | 14.865 | 12.754 | 14.465 | 16.052 | 12.284 | 15.213 | 13.895 | 14.781 | 15.202 | 15.957 | 13.926 | 14.332 | 16.432 | 15.179 | 13.539 | 15.876 | -0.086 | 0.0333 | 0.1449 | 0 | 0 | 0 |
| 'Zdhhc8' | 8.340 | 8.237 | 9.154 | 7.924 | 8.240 | 9.362 | 8.282 | 8.995 | 9.047 | 8.385 | 7.878 | 9.139 | 8.757 | 8.123 | 9.182 | 8.106 | 6.859 | 9.339 | 0.025 | 0.5406 | 0.7579 | 0 | 0 | 0 |
| 'Zdhhc9' | 2.312 | 2.248 | 2.043 | 1.904 | 1.968 | 1.996 | 1.940 | 1.964 | 2.079 | 2.131 | 2.142 | 2.491 | 2.054 | 2.081 | 2.545 | 2.357 | 2.020 | 2.483 | -0.150 | 0.0099 | 0.0617 | 0 | 0 | 0 |
| 'Zeb1' | 4.193 | 4.073 | 3.663 | 4.454 | 4.185 | 4.107 | 4.723 | 4.038 | 3.624 | 3.679 | 4.287 | 4.277 | 3.968 | 4.144 | 3.852 | 3.345 | 4.078 | 4.138 | 0.036 | 0.5027 | 0.7298 | 0 | 0 | 0 |
| 'Zeb2' | 0.459 | 0.536 | 0.453 | 0.505 | 0.492 | 0.476 | 0.510 | 0.503 | 0.429 | 0.452 | 0.583 | 0.314 | 0.470 | 0.489 | 0.361 | 0.499 | 0.560 | 0.283 | 0.146 | 0.0640 | 0.2205 | 0 | 0 | 0 |
| 'Zer1' | 9.533 | 9.805 | 10.642 | 8.883 | 10.028 | 10.528 | 9.067 | 9.931 | 11.095 | 11.216 | 10.406 | 10.886 | 10.723 | 9.822 | 10.557 | 10.208 | 10.522 | 10.865 | -0.101 | 0.0020 | 0.0179 | 1 | 0 | 0 |
| 'Zfand2a' | 11.025 | 4.817 | 3.779 | 6.890 | 4.529 | 4.202 | 5.768 | 5.070 | 4.065 | 7.093 | 6.802 | 4.585 | 6.993 | 6.136 | 4.083 | 6.541 | 6.041 | 4.936 | -0.132 | 0.2743 | 0.5299 | 0 | 0 | 0 |
| 'Zfand2b' | 5.891 | 5.981 | 6.309 | 5.124 | 5.849 | 7.334 | 5.030 | 5.712 | 6.203 | 6.065 | 5.407 | 6.067 | 6.328 | 6.089 | 5.912 | 6.696 | 5.788 | 5.871 | -0.029 | 0.6075 | 0.8026 | 0 | 0 | 0 |
| 'Zfand3' | 22.404 | 25.028 | 23.519 | 21.379 | 24.090 | 25.208 | 20.987 | 24.811 | 24.720 | 24.451 | 23.012 | 23.240 | 23.379 | 24.571 | 23.075 | 24.730 | 22.775 | 21.759 | -0.002 | 0.9455 | 0.9790 | 0 | 0 | 0 |
| 'Zfand4' | 1.317 | 1.317 | 1.605 | 1.598 | 1.346 | 1.581 | 1.634 | 1.618 | 1.637 | 1.250 | 1.865 | 1.215 | 1.853 | 1.494 | 1.236 | 1.255 | 1.307 | 0.990 | 0.137 | 0.2037 | 0.4465 | 0 | 0 | 0 |
| 'Zfand5' | 23.594 | 17.838 | 14.630 | 21.159 | 16.779 | 14.262 | 22.336 | 17.103 | 12.823 | 19.533 | 19.755 | 13.913 | 19.399 | 20.347 | 13.334 | 17.986 | 21.294 | 13.253 | 0.001 | 0.9860 | 0.9953 | 0 | 0 | 0 |
| 'Zfand6' | 63.990 | 81.896 | 72.251 | 71.013 | 79.963 | 73.065 | 69.012 | 80.840 | 70.267 | 70.575 | 68.233 | 66.233 | 69.443 | 80.534 | 64.398 | 74.146 | 75.404 | 62.874 | 0.057 | 0.1047 | 0.3032 | 0 | 0 | 0 |
| 'Zfat' | 1.079 | 0.907 | 1.205 | 1.117 | 1.258 | 1.309 | 1.201 | 1.104 | 1.403 | 1.306 | 1.203 | 1.072 | 1.096 | 1.267 | 1.024 | 0.968 | 0.908 | 1.254 | 0.060 | 0.4712 | 0.7069 | 0 | 0 | 0 |
| 'Zfc3h1' | 13.241 | 12.441 | 14.563 | 12.537 | 12.923 | 12.715 | 13.532 | 11.716 | 13.603 | 11.668 | 13.756 | 14.554 | 11.932 | 14.400 | 13.434 | 11.822 | 16.060 | 13.202 | -0.049 | 0.3536 | 0.6072 | 0 | 0 | 0 |
| 'Zfhx2' | 4.224 | 3.083 | 4.483 | 3.799 | 3.582 | 3.926 | 3.904 | 3.588 | 4.520 | 4.401 | 3.566 | 4.273 | 4.057 | 3.325 | 3.754 | 3.681 | 3.557 | 3.922 | 0.012 | 0.7887 | 0.9056 | 0 | 0 | 0 |
| 'Zfp1' | 20.951 | 23.859 | 24.146 | 23.405 | 24.348 | 22.413 | 25.393 | 24.926 | 22.928 | 20.949 | 22.050 | 20.780 | 21.438 | 21.484 | 18.667 | 21.578 | 22.165 | 18.736 | 0.167 | 0.0000 | 0.0004 | 1 | 0 | 0 |
| 'Zfp101' | 0.684 | 0.720 | 0.832 | 1.070 | 0.778 | 0.810 | 1.269 | 0.745 | 1.018 | 0.682 | 0.825 | 1.103 | 0.819 | 0.735 | 1.020 | 0.730 | 0.822 | 0.840 | 0.040 | 0.7223 | 0.8722 | 0 | 0 | 0 |
| 'Zfp106' | 14.167 | 14.039 | 13.832 | 13.770 | 13.311 | 13.897 | 13.088 | 13.731 | 13.780 | 15.217 | 14.294 | 14.100 | 14.077 | 13.693 | 13.562 | 14.194 | 13.776 | 14.039 | -0.048 | 0.0536 | 0.1977 | 0 | 0 | 0 |
| 'Zfp11' | 1.579 | 2.136 | 2.055 | 1.741 | 2.136 | 1.730 | 2.008 | 2.456 | 1.886 | 1.902 | 1.883 | 1.823 | 1.722 | 1.814 | 1.739 | 1.676 | 1.756 | 1.608 | 0.137 | 0.0875 | 0.2698 | 0 | 0 | 0 |
| 'Zfp110' | 8.332 | 8.887 | 8.526 | 7.421 | 8.725 | 7.592 | 8.952 | 8.807 | 8.545 | 8.248 | 7.894 | 8.474 | 7.810 | 8.241 | 7.831 | 7.887 | 8.053 | 7.736 | 0.057 | 0.1249 | 0.3382 | 0 | 0 | 0 |
| 'Zfp113' | 1.113 | 1.270 | 1.275 | 1.068 | 1.239 | 1.181 | 1.268 | 1.338 | 1.318 | 1.058 | 1.180 | 1.204 | 0.969 | 0.811 | 1.006 | 1.020 | 1.112 | 1.021 | 0.221 | 0.0029 | 0.0244 | 1 | 0 | 0 |
| 'Zfp119a' | 4.280 | 4.997 | 4.979 | 4.502 | 4.933 | 5.388 | 5.230 | 5.042 | 5.271 | 4.052 | 5.015 | 4.783 | 4.001 | 5.944 | 5.053 | 5.732 | 5.321 | 4.754 | -0.001 | 0.9905 | 0.9974 | 0 | 0 | 0 |
| 'Zfp12' | 2.958 | 2.806 | 2.881 | 2.834 | 2.729 | 2.915 | 2.623 | 2.866 | 3.057 | 2.796 | 3.253 | 3.301 | 2.642 | 3.130 | 3.157 | 2.894 | 2.987 | 3.437 | -0.112 | 0.0089 | 0.0577 | 0 | 0 | 0 |
| 'Zfp120' | 2.310 | 2.799 | 2.276 | 2.720 | 2.605 | 2.198 | 2.783 | 2.738 | 2.457 | 2.052 | 2.765 | 2.829 | 2.126 | 2.651 | 2.463 | 2.365 | 2.682 | 2.812 | -0.005 | 0.9344 | 0.9749 | 0 | 0 | 0 |
| 'Zfp128' | 0.587 | 0.857 | 0.610 | 0.862 | 0.568 | 0.656 | 0.631 | 0.828 | 0.706 | 0.501 | 0.598 | 0.587 | 0.579 | 0.605 | 0.664 | 0.558 | 0.530 | 0.571 | 0.260 | 0.0201 | 0.1036 | 0 | 0 | 0 |
| 'Zfp131' | 20.839 | 22.883 | 21.069 | 22.373 | 21.671 | 21.538 | 21.825 | 22.819 | 22.021 | 19.790 | 22.402 | 22.955 | 20.793 | 22.803 | 20.376 | 23.394 | 26.194 | 22.899 | -0.043 | 0.2867 | 0.5433 | 0 | 0 | 0 |
| 'Zfp142' | 2.902 | 1.959 | 2.556 | 2.787 | 2.109 | 2.384 | 2.326 | 2.075 | 2.434 | 2.579 | 2.483 | 2.988 | 2.444 | 2.205 | 2.542 | 2.179 | 2.639 | 2.798 | -0.102 | 0.1135 | 0.3197 | 0 | 0 | 0 |
| 'Zfp143' | 11.428 | 13.019 | 11.842 | 12.354 | 11.648 | 12.470 | 12.772 | 12.278 | 11.927 | 12.063 | 12.142 | 11.389 | 11.996 | 12.916 | 10.812 | 11.631 | 12.184 | 10.759 | 0.043 | 0.2278 | 0.4743 | 0 | 0 | 0 |
| 'Zfp146' | 25.659 | 25.387 | 25.597 | 26.282 | 23.969 | 24.770 | 27.007 | 25.391 | 26.642 | 23.578 | 26.550 | 27.158 | 23.360 | 24.610 | 26.270 | 24.671 | 24.890 | 26.213 | 0.010 | 0.7869 | 0.9047 | 0 | 0 | 0 |
| 'Zfp148' | 10.419 | 9.893 | 9.689 | 10.871 | 9.133 | 9.059 | 9.947 | 10.255 | 9.459 | 10.256 | 10.852 | 10.674 | 8.990 | 9.621 | 9.694 | 9.365 | 10.648 | 10.297 | -0.039 | 0.3967 | 0.6456 | 0 | 0 | 0 |
| 'Zfp157' | 1.726 | 1.610 | 1.689 | 1.731 | 1.599 | 1.832 | 1.654 | 1.766 | 1.532 | 1.441 | 1.505 | 1.862 | 1.461 | 1.650 | 1.573 | 1.619 | 1.737 | 1.702 | 0.043 | 0.4470 | 0.6880 | 0 | 0 | 0 |
| 'Zfp160' | 5.262 | 6.188 | 5.869 | 5.494 | 6.264 | 5.394 | 6.752 | 5.518 | 5.650 | 4.583 | 5.088 | 6.303 | 4.880 | 4.943 | 5.288 | 4.587 | 5.828 | 5.457 | 0.138 | 0.0314 | 0.1390 | 0 | 0 | 0 |
| 'Zfp169' | 1.114 | 1.376 | 1.503 | 0.921 | 1.321 | 1.639 | 1.313 | 1.461 | 1.502 | 1.245 | 1.569 | 1.681 | 1.192 | 1.271 | 1.675 | 1.581 | 1.683 | 1.824 | -0.186 | 0.0074 | 0.0498 | 1 | 0 | 0 |
| 'Zfp180' | 5.065 | 5.231 | 4.523 | 4.970 | 4.978 | 4.807 | 4.448 | 5.136 | 4.876 | 5.327 | 5.035 | 4.588 | 4.406 | 4.762 | 4.700 | 5.132 | 5.167 | 4.429 | 0.009 | 0.8296 | 0.9262 | 0 | 0 | 0 |
| 'Zfp182' | 2.060 | 2.359 | 2.076 | 1.909 | 2.498 | 1.757 | 2.547 | 2.171 | 2.425 | 2.357 | 2.056 | 2.267 | 2.126 | 2.410 | 2.384 | 2.391 | 2.323 | 1.911 | -0.045 | 0.4976 | 0.7258 | 0 | 0 | 0 |
| 'Zfp189' | 1.516 | 1.587 | 1.475 | 1.511 | 1.629 | 1.631 | 2.098 | 1.594 | 1.329 | 1.132 | 1.886 | 1.884 | 0.985 | 1.143 | 1.484 | 1.502 | 1.453 | 2.194 | 0.065 | 0.6331 | 0.8180 | 0 | 0 | 0 |
| 'Zfp207' | 26.264 | 25.597 | 24.717 | 27.166 | 25.971 | 25.258 | 26.687 | 25.483 | 25.813 | 21.502 | 27.197 | 30.642 | 22.027 | 26.951 | 29.729 | 21.816 | 28.447 | 31.751 | -0.042 | 0.4927 | 0.7220 | 0 | 0 | 0 |
| 'Zfp212' | 12.303 | 12.830 | 11.486 | 11.199 | 13.059 | 13.123 | 11.050 | 13.985 | 12.469 | 12.469 | 11.666 | 12.644 | 12.199 | 12.770 | 13.234 | 14.098 | 11.418 | 11.694 | -0.019 | 0.7027 | 0.8611 | 0 | 0 | 0 |
| 'Zfp213' | 4.861 | 5.285 | 6.728 | 4.875 | 5.235 | 7.157 | 4.789 | 5.387 | 6.206 | 5.570 | 4.597 | 5.533 | 4.983 | 4.621 | 4.880 | 5.419 | 4.852 | 5.787 | 0.121 | 0.0665 | 0.2256 | 0 | 0 | 0 |
| 'Zfp217' | 19.904 | 23.267 | 22.239 | 19.906 | 22.712 | 22.369 | 19.833 | 21.683 | 23.493 | 21.360 | 19.809 | 22.228 | 19.849 | 21.945 | 21.571 | 21.318 | 19.809 | 22.278 | 0.027 | 0.3930 | 0.6427 | 0 | 0 | 0 |
| 'Zfp219' | 18.627 | 16.493 | 18.788 | 15.812 | 14.799 | 19.463 | 16.039 | 16.084 | 19.218 | 18.332 | 15.422 | 19.938 | 16.457 | 15.465 | 19.011 | 16.672 | 14.926 | 18.425 | -0.002 | 0.9498 | 0.9809 | 0 | 0 | 0 |
| 'Zfp235' | 1.350 | 1.399 | 1.491 | 1.433 | 0.944 | 1.516 | 1.492 | 1.167 | 1.448 | 1.040 | 1.319 | 1.600 | 1.104 | 1.199 | 1.631 | 1.225 | 1.448 | 1.597 | 0.000 | 0.9960 | 0.9994 | 0 | 0 | 0 |
| 'Zfp236' | 7.644 | 6.945 | 7.248 | 7.037 | 7.092 | 7.194 | 6.407 | 7.187 | 7.324 | 8.204 | 7.452 | 7.503 | 7.122 | 7.394 | 7.365 | 7.953 | 7.697 | 7.282 | -0.094 | 0.0070 | 0.0482 | 1 | 0 | 0 |
| 'Zfp239' | 0.621 | 0.795 | 0.762 | 0.669 | 0.851 | 0.789 | 0.609 | 0.838 | 0.665 | 0.643 | 0.562 | 0.558 | 0.716 | 0.592 | 0.569 | 0.617 | 0.634 | 0.451 | 0.298 | 0.0017 | 0.0157 | 1 | 0 | 0 |
| 'Zfp24' | 5.844 | 6.783 | 6.118 | 6.483 | 6.741 | 5.789 | 6.500 | 6.801 | 6.090 | 5.575 | 6.195 | 6.014 | 5.678 | 6.444 | 6.220 | 5.840 | 5.940 | 6.113 | 0.066 | 0.0782 | 0.2506 | 0 | 0 | 0 |
| 'Zfp248' | 1.083 | 0.833 | 1.018 | 0.858 | 0.750 | 0.879 | 1.116 | 0.942 | 0.893 | 1.004 | 0.877 | 0.917 | 0.898 | 0.730 | 0.915 | 0.893 | 0.860 | 0.850 | 0.070 | 0.4446 | 0.6875 | 0 | 0 | 0 |
| 'Zfp251' | 16.051 | 17.163 | 19.237 | 15.258 | 17.745 | 18.333 | 17.728 | 17.514 | 18.805 | 16.158 | 17.538 | 19.769 | 16.182 | 16.559 | 18.431 | 15.817 | 16.264 | 19.735 | 0.000 | 0.9964 | 0.9994 | 0 | 0 | 0 |
| 'Zfp260' | 4.894 | 6.079 | 5.788 | 5.657 | 6.467 | 5.855 | 5.484 | 5.635 | 6.170 | 4.590 | 5.889 | 6.352 | 4.434 | 5.496 | 5.932 | 4.781 | 5.664 | 6.427 | 0.062 | 0.1829 | 0.4190 | 0 | 0 | 0 |
| 'Zfp263' | 6.046 | 6.693 | 6.311 | 5.543 | 6.848 | 6.431 | 5.322 | 6.679 | 6.296 | 6.005 | 5.679 | 6.262 | 5.735 | 6.478 | 6.293 | 6.465 | 6.179 | 6.345 | 0.007 | 0.8583 | 0.9390 | 0 | 0 | 0 |
| 'Zfp266' | 7.104 | 8.569 | 7.548 | 7.287 | 7.919 | 7.317 | 7.222 | 8.074 | 7.776 | 7.425 | 7.653 | 7.625 | 7.052 | 7.979 | 6.845 | 7.408 | 7.594 | 6.967 | 0.035 | 0.2682 | 0.5223 | 0 | 0 | 0 |
| 'Zfp27' | 1.373 | 1.230 | 1.724 | 1.310 | 1.609 | 1.545 | 1.538 | 1.467 | 1.541 | 1.489 | 1.253 | 1.223 | 1.489 | 1.279 | 1.225 | 1.237 | 1.191 | 1.229 | 0.196 | 0.0033 | 0.0269 | 1 | 0 | 0 |
| 'Zfp275' | 6.829 | 7.715 | 7.743 | 6.667 | 6.887 | 7.651 | 6.741 | 7.200 | 8.333 | 7.026 | 7.362 | 9.640 | 5.884 | 7.344 | 9.006 | 7.231 | 8.076 | 9.252 | -0.112 | 0.0196 | 0.1017 | 0 | 0 | 0 |
| 'Zfp276' | 4.942 | 4.165 | 5.410 | 4.737 | 4.521 | 5.437 | 4.868 | 4.604 | 5.559 | 4.742 | 4.517 | 6.406 | 4.890 | 4.513 | 5.756 | 4.602 | 4.761 | 6.426 | -0.084 | 0.0320 | 0.1406 | 0 | 0 | 0 |
| 'Zfp277' | 10.193 | 9.908 | 11.222 | 11.363 | 10.423 | 10.558 | 11.437 | 11.974 | 10.443 | 9.625 | 9.756 | 9.998 | 10.305 | 10.386 | 9.574 | 9.978 | 10.982 | 9.245 | 0.110 | 0.0170 | 0.0920 | 0 | 0 | 0 |
| 'Zfp28' | 1.626 | 1.076 | 1.835 | 0.986 | 1.565 | 1.885 | 1.385 | 1.395 | 1.546 | 1.679 | 1.325 | 1.203 | 1.242 | 0.893 | 1.181 | 1.122 | 0.956 | 1.058 | 0.324 | 0.0115 | 0.0691 | 0 | 0 | 0 |
| 'Zfp280b' | 13.796 | 14.654 | 14.872 | 14.675 | 14.673 | 14.863 | 16.854 | 13.288 | 14.473 | 13.025 | 12.667 | 15.983 | 13.695 | 13.678 | 14.232 | 15.325 | 14.330 | 14.374 | 0.040 | 0.3585 | 0.6111 | 0 | 0 | 0 |
| 'Zfp280c' | 5.165 | 5.940 | 6.608 | 5.058 | 6.399 | 5.131 | 5.962 | 6.790 | 6.141 | 6.642 | 5.836 | 5.515 | 6.412 | 5.825 | 4.617 | 5.650 | 6.063 | 4.871 | 0.042 | 0.5197 | 0.7423 | 0 | 0 | 0 |
| 'Zfp280d' | 2.106 | 2.279 | 2.167 | 2.204 | 2.137 | 1.962 | 2.265 | 2.188 | 2.069 | 2.133 | 2.054 | 2.057 | 1.947 | 1.973 | 1.840 | 2.040 | 2.235 | 1.784 | 0.090 | 0.0113 | 0.0683 | 0 | 0 | 0 |
| 'Zfp281' | 6.438 | 6.062 | 4.547 | 5.217 | 5.772 | 5.073 | 5.779 | 5.558 | 4.621 | 6.410 | 5.760 | 5.444 | 5.991 | 6.737 | 5.260 | 8.647 | 6.110 | 5.434 | -0.193 | 0.0016 | 0.0151 | 1 | 0 | 0 |
| 'Zfp282' | 5.277 | 4.610 | 5.640 | 4.833 | 4.594 | 5.620 | 4.943 | 4.989 | 6.047 | 5.300 | 4.478 | 5.536 | 5.555 | 4.404 | 5.329 | 4.735 | 4.610 | 5.048 | 0.040 | 0.3200 | 0.5782 | 0 | 0 | 0 |
| 'Zfp287' | 3.714 | 4.282 | 3.871 | 3.768 | 3.887 | 4.122 | 3.715 | 3.879 | 3.729 | 4.252 | 3.980 | 3.676 | 4.039 | 3.906 | 4.103 | 4.469 | 3.994 | 3.725 | -0.052 | 0.2561 | 0.5084 | 0 | 0 | 0 |
| 'Zfp292' | 18.175 | 17.303 | 15.310 | 17.583 | 16.862 | 16.345 | 18.574 | 17.383 | 16.092 | 15.829 | 17.174 | 18.137 | 16.297 | 16.988 | 17.743 | 18.229 | 15.985 | 19.040 | -0.029 | 0.5083 | 0.7336 | 0 | 0 | 0 |
| 'Zfp3' | 4.699 | 6.129 | 5.815 | 5.964 | 6.370 | 6.316 | 6.246 | 5.337 | 6.492 | 5.784 | 5.577 | 5.447 | 5.481 | 5.455 | 5.351 | 5.836 | 6.730 | 5.298 | 0.057 | 0.4175 | 0.6650 | 0 | 0 | 0 |
| 'Zfp30' | 0.464 | 0.549 | 0.403 | 0.375 | 0.509 | 0.424 | 0.565 | 0.392 | 0.423 | 0.515 | 0.308 | 0.408 | 0.347 | 0.381 | 0.413 | 0.411 | 0.412 | 0.513 | 0.123 | 0.2847 | 0.5410 | 0 | 0 | 0 |
| 'Zfp316' | 1.367 | 1.461 | 1.373 | 1.190 | 1.608 | 1.636 | 1.466 | 1.702 | 1.558 | 1.609 | 1.359 | 1.066 | 1.438 | 1.127 | 1.062 | 1.392 | 1.140 | 1.070 | 0.249 | 0.0024 | 0.0212 | 1 | 0 | 0 |
| 'Zfp317' | 12.862 | 14.326 | 13.186 | 13.115 | 13.671 | 12.378 | 11.744 | 13.394 | 13.918 | 13.441 | 13.799 | 13.832 | 12.230 | 13.018 | 11.730 | 12.238 | 13.983 | 12.365 | 0.012 | 0.7588 | 0.8921 | 0 | 0 | 0 |
| 'Zfp318' | 3.459 | 3.881 | 3.265 | 3.172 | 3.660 | 3.462 | 2.996 | 3.432 | 3.473 | 3.350 | 3.644 | 3.712 | 3.508 | 3.842 | 3.249 | 3.846 | 3.410 | 3.406 | -0.066 | 0.1391 | 0.3605 | 0 | 0 | 0 |
| 'Zfp319' | 10.257 | 10.022 | 10.663 | 9.258 | 10.106 | 11.100 | 7.831 | 9.602 | 10.285 | 9.160 | 9.042 | 12.026 | 8.325 | 10.102 | 12.090 | 9.581 | 9.225 | 12.953 | -0.057 | 0.3081 | 0.5665 | 0 | 0 | 0 |
| 'Zfp322a' | 2.453 | 2.612 | 2.754 | 3.083 | 2.826 | 2.789 | 3.327 | 2.550 | 2.710 | 2.317 | 3.025 | 3.423 | 2.430 | 2.783 | 3.084 | 1.923 | 3.297 | 2.921 | -0.012 | 0.8931 | 0.9563 | 0 | 0 | 0 |
| 'Zfp324' | 1.097 | 0.973 | 1.391 | 1.353 | 1.091 | 1.303 | 1.206 | 0.975 | 1.368 | 1.014 | 1.173 | 1.349 | 0.949 | 0.845 | 1.160 | 0.916 | 1.033 | 1.462 | 0.108 | 0.1489 | 0.3756 | 0 | 0 | 0 |
| 'Zfp326' | 2.903 | 2.745 | 2.846 | 3.088 | 2.980 | 2.671 | 2.839 | 3.102 | 2.676 | 2.754 | 2.508 | 2.844 | 2.503 | 2.948 | 2.267 | 2.675 | 2.830 | 2.571 | 0.100 | 0.0312 | 0.1384 | 0 | 0 | 0 |
| 'Zfp329' | 1.772 | 2.598 | 2.557 | 1.762 | 2.364 | 2.298 | 2.189 | 2.241 | 2.651 | 1.779 | 2.098 | 2.885 | 1.724 | 2.047 | 2.773 | 1.742 | 2.692 | 2.747 | -0.010 | 0.8746 | 0.9463 | 0 | 0 | 0 |
| 'Zfp330' | 45.878 | 54.487 | 39.641 | 40.750 | 49.696 | 41.656 | 45.768 | 47.867 | 42.760 | 46.692 | 44.110 | 44.830 | 40.603 | 49.152 | 45.825 | 49.367 | 47.158 | 43.234 | -0.025 | 0.5821 | 0.7842 | 0 | 0 | 0 |
| 'Zfp335' | 13.927 | 12.468 | 15.597 | 13.391 | 12.018 | 14.734 | 12.422 | 13.127 | 15.277 | 14.259 | 12.054 | 14.323 | 13.273 | 11.786 | 13.933 | 12.895 | 11.944 | 14.455 | 0.040 | 0.2145 | 0.4598 | 0 | 0 | 0 |
| 'Zfp346' | 8.449 | 9.295 | 9.532 | 8.995 | 8.288 | 8.701 | 8.653 | 8.890 | 9.630 | 8.690 | 8.381 | 9.821 | 7.896 | 7.756 | 9.341 | 8.259 | 8.289 | 9.723 | 0.031 | 0.4100 | 0.6590 | 0 | 0 | 0 |
| 'Zfp35' | 7.592 | 7.381 | 7.611 | 6.996 | 8.047 | 7.863 | 8.697 | 7.520 | 7.128 | 7.111 | 8.029 | 8.374 | 6.803 | 7.614 | 7.577 | 8.392 | 7.222 | 8.678 | -0.032 | 0.5654 | 0.7741 | 0 | 0 | 0 |
| 'Zfp354c' | 0.618 | 0.723 | 0.569 | 0.620 | 0.690 | 0.910 | 0.737 | 0.639 | 0.678 | 0.656 | 0.746 | 0.611 | 0.797 | 0.735 | 0.529 | 0.373 | 0.569 | 0.543 | 0.141 | 0.2544 | 0.5063 | 0 | 0 | 0 |
| 'Zfp358' | 3.336 | 3.454 | 3.649 | 3.203 | 2.978 | 3.804 | 2.952 | 3.803 | 4.148 | 3.100 | 2.717 | 3.458 | 3.385 | 3.382 | 3.247 | 3.219 | 2.197 | 2.901 | 0.175 | 0.0497 | 0.1881 | 0 | 0 | 0 |
| 'Zfp36' | 416.302 | 332.995 | 238.823 | 282.515 | 329.475 | 267.377 | 258.244 | 392.533 | 210.687 | 429.142 | 284.696 | 150.494 | 388.515 | 385.802 | 185.539 | 405.921 | 246.549 | 156.377 | 0.120 | 0.3820 | 0.6330 | 0 | 0 | 0 |
| 'Zfp362' | 24.117 | 24.045 | 27.469 | 22.766 | 24.059 | 26.898 | 22.738 | 25.263 | 28.026 | 27.420 | 23.698 | 27.703 | 26.436 | 23.778 | 25.632 | 26.222 | 23.445 | 27.072 | -0.048 | 0.1545 | 0.3825 | 0 | 0 | 0 |
| 'Zfp365' | 1.337 | 1.483 | 1.118 | 0.990 | 1.685 | 1.624 | 1.268 | 1.790 | 1.415 | 1.841 | 1.471 | 1.452 | 1.783 | 1.664 | 1.356 | 1.832 | 1.451 | 1.211 | -0.152 | 0.1175 | 0.3263 | 0 | 0 | 0 |
| 'Zfp367' | 4.810 | 4.913 | 5.696 | 5.183 | 5.391 | 5.312 | 6.033 | 5.165 | 5.476 | 5.017 | 5.571 | 5.552 | 6.240 | 5.427 | 5.597 | 4.997 | 5.442 | 5.846 | -0.063 | 0.2007 | 0.4419 | 0 | 0 | 0 |
| 'Zfp369' | 4.064 | 3.546 | 4.488 | 3.832 | 4.009 | 3.677 | 3.908 | 3.770 | 3.827 | 3.867 | 4.184 | 4.561 | 3.775 | 3.636 | 4.307 | 3.884 | 4.598 | 4.882 | -0.111 | 0.0524 | 0.1943 | 0 | 0 | 0 |
| 'Zfp36l1' | 70.438 | 82.677 | 59.146 | 61.610 | 75.829 | 68.517 | 65.170 | 74.955 | 64.557 | 81.979 | 65.455 | 59.911 | 76.768 | 79.674 | 61.129 | 88.311 | 65.379 | 61.788 | -0.045 | 0.4621 | 0.6981 | 0 | 0 | 0 |
| 'Zfp36l2' | 186.060 | 197.925 | 152.521 | 148.046 | 190.020 | 161.933 | 146.191 | 210.988 | 145.674 | 219.003 | 160.604 | 141.719 | 176.271 | 199.118 | 145.130 | 242.580 | 165.081 | 145.444 | -0.053 | 0.5751 | 0.7806 | 0 | 0 | 0 |
| 'Zfp382' | 4.803 | 4.602 | 4.668 | 4.369 | 4.718 | 4.423 | 4.277 | 4.485 | 3.840 | 3.998 | 4.786 | 4.960 | 3.899 | 4.258 | 5.044 | 4.194 | 4.361 | 5.887 | -0.053 | 0.4829 | 0.7146 | 0 | 0 | 0 |
| 'Zfp383' | 2.504 | 2.762 | 2.151 | 2.374 | 2.775 | 2.078 | 2.525 | 2.955 | 2.037 | 2.663 | 2.704 | 2.389 | 2.603 | 2.498 | 2.680 | 2.206 | 2.430 | 2.482 | -0.056 | 0.3913 | 0.6406 | 0 | 0 | 0 |
| 'Zfp384' | 5.934 | 5.586 | 6.456 | 5.755 | 5.525 | 6.285 | 5.314 | 5.705 | 6.481 | 6.184 | 5.300 | 6.254 | 6.010 | 5.570 | 5.860 | 6.156 | 5.623 | 5.605 | 0.004 | 0.9074 | 0.9637 | 0 | 0 | 0 |
| 'Zfp386' | 1.929 | 2.183 | 2.163 | 2.214 | 2.347 | 2.172 | 2.257 | 2.145 | 2.066 | 2.160 | 1.826 | 1.845 | 1.968 | 1.831 | 1.874 | 1.937 | 2.091 | 1.671 | 0.169 | 0.0007 | 0.0083 | 1 | 0 | 0 |
| 'Zfp39' | 3.820 | 3.977 | 4.385 | 3.485 | 4.634 | 4.195 | 4.516 | 4.745 | 4.251 | 4.853 | 3.789 | 3.902 | 3.749 | 3.536 | 3.784 | 4.633 | 3.031 | 3.697 | 0.116 | 0.1327 | 0.3506 | 0 | 0 | 0 |
| 'Zfp395' | 11.167 | 10.946 | 11.136 | 9.871 | 11.190 | 11.928 | 10.409 | 11.893 | 10.785 | 11.563 | 12.029 | 11.224 | 11.769 | 11.607 | 12.276 | 12.138 | 10.666 | 10.574 | -0.070 | 0.0983 | 0.2930 | 0 | 0 | 0 |
| 'Zfp397' | 6.394 | 7.020 | 7.313 | 6.934 | 7.322 | 6.351 | 7.555 | 6.672 | 6.834 | 6.505 | 7.332 | 8.286 | 5.828 | 7.152 | 7.601 | 5.920 | 8.396 | 7.284 | -0.052 | 0.3784 | 0.6301 | 0 | 0 | 0 |
| 'Zfp398' | 3.057 | 2.769 | 3.147 | 3.047 | 2.587 | 2.728 | 3.241 | 2.849 | 3.095 | 3.070 | 3.044 | 3.317 | 2.585 | 2.735 | 2.803 | 2.752 | 3.239 | 3.130 | -0.021 | 0.7098 | 0.8653 | 0 | 0 | 0 |
| 'Zfp40' | 0.934 | 1.196 | 1.071 | 1.248 | 1.168 | 0.987 | 1.116 | 1.145 | 1.205 | 0.842 | 1.282 | 1.102 | 0.860 | 1.221 | 0.906 | 1.168 | 1.015 | 0.949 | 0.097 | 0.2928 | 0.5493 | 0 | 0 | 0 |
| 'Zfp407' | 3.028 | 2.867 | 2.831 | 2.953 | 2.414 | 2.486 | 2.488 | 2.669 | 2.532 | 2.682 | 3.324 | 2.747 | 2.732 | 2.663 | 2.511 | 2.738 | 3.098 | 2.757 | -0.065 | 0.2545 | 0.5063 | 0 | 0 | 0 |
| 'Zfp408' | 3.382 | 3.483 | 3.919 | 3.276 | 3.772 | 3.690 | 3.563 | 3.513 | 3.753 | 3.414 | 3.409 | 3.572 | 3.701 | 3.295 | 3.566 | 3.698 | 3.249 | 3.641 | 0.028 | 0.4873 | 0.7176 | 0 | 0 | 0 |
| 'Zfp41' | 3.398 | 3.624 | 4.376 | 3.641 | 3.519 | 4.533 | 3.660 | 3.643 | 4.460 | 3.620 | 4.030 | 4.389 | 3.750 | 3.175 | 4.013 | 3.189 | 3.577 | 4.812 | 0.000 | 0.9995 | 0.9999 | 0 | 0 | 0 |
| 'Zfp410' | 22.124 | 20.828 | 20.364 | 22.840 | 20.854 | 19.799 | 20.884 | 21.690 | 21.169 | 19.951 | 22.215 | 21.657 | 19.769 | 21.498 | 20.255 | 19.401 | 23.232 | 21.075 | 0.000 | 0.9933 | 0.9988 | 0 | 0 | 0 |
| 'Zfp414' | 7.098 | 8.462 | 10.062 | 5.931 | 8.390 | 10.425 | 7.987 | 8.799 | 10.631 | 9.481 | 6.870 | 8.439 | 9.457 | 7.674 | 8.180 | 8.278 | 7.151 | 7.207 | 0.078 | 0.3550 | 0.6084 | 0 | 0 | 0 |
| 'Zfp418' | 1.256 | 1.296 | 1.282 | 1.648 | 1.580 | 1.332 | 1.575 | 1.445 | 1.199 | 1.137 | 1.590 | 1.778 | 1.382 | 1.349 | 1.940 | 1.069 | 1.177 | 1.611 | -0.073 | 0.5519 | 0.7650 | 0 | 0 | 0 |
| 'Zfp420' | 0.666 | 0.779 | 0.641 | 0.647 | 0.816 | 0.703 | 0.824 | 0.731 | 0.637 | 0.648 | 0.651 | 0.702 | 0.739 | 0.806 | 0.743 | 0.594 | 0.807 | 0.832 | -0.042 | 0.6273 | 0.8154 | 0 | 0 | 0 |
| 'Zfp422' | 3.300 | 3.157 | 3.998 | 3.845 | 3.716 | 3.979 | 4.119 | 4.015 | 3.471 | 3.276 | 3.688 | 4.080 | 3.482 | 3.884 | 3.459 | 3.819 | 3.631 | 3.797 | 0.010 | 0.8570 | 0.9387 | 0 | 0 | 0 |
| 'Zfp426' | 2.981 | 3.595 | 3.229 | 3.223 | 3.371 | 3.172 | 3.214 | 3.019 | 3.292 | 3.316 | 3.344 | 3.668 | 3.039 | 3.407 | 3.404 | 3.183 | 3.713 | 3.286 | -0.077 | 0.0592 | 0.2100 | 0 | 0 | 0 |
| 'Zfp438' | 1.337 | 1.222 | 1.462 | 1.537 | 1.744 | 1.454 | 1.958 | 1.562 | 1.703 | 1.721 | 1.595 | 1.344 | 1.292 | 1.374 | 1.665 | 1.313 | 1.586 | 1.302 | 0.078 | 0.4598 | 0.6965 | 0 | 0 | 0 |
| 'Zfp444' | 5.882 | 5.081 | 6.216 | 4.980 | 4.934 | 6.593 | 4.592 | 5.268 | 6.180 | 5.573 | 4.173 | 6.711 | 6.019 | 4.932 | 6.781 | 5.746 | 4.369 | 6.536 | -0.034 | 0.5168 | 0.7404 | 0 | 0 | 0 |
| 'Zfp445' | 30.335 | 31.901 | 28.633 | 28.608 | 31.981 | 31.846 | 28.546 | 30.608 | 31.882 | 27.748 | 30.073 | 32.072 | 27.258 | 30.923 | 31.098 | 31.410 | 30.883 | 32.005 | -0.006 | 0.8508 | 0.9358 | 0 | 0 | 0 |
| 'Zfp446' | 0.894 | 0.947 | 0.947 | 0.982 | 0.891 | 0.931 | 0.991 | 0.890 | 0.935 | 0.829 | 0.829 | 1.110 | 0.860 | 0.858 | 0.988 | 0.903 | 0.874 | 0.967 | 0.010 | 0.8560 | 0.9383 | 0 | 0 | 0 |
| 'Zfp451' | 3.355 | 3.127 | 3.149 | 3.238 | 3.133 | 2.709 | 3.315 | 3.246 | 2.847 | 2.814 | 3.156 | 3.421 | 2.823 | 3.355 | 3.120 | 3.001 | 3.391 | 3.359 | -0.028 | 0.5772 | 0.7819 | 0 | 0 | 0 |
| 'Zfp455' | 1.658 | 2.161 | 2.053 | 1.668 | 2.326 | 1.627 | 2.191 | 2.124 | 1.971 | 2.301 | 1.757 | 2.103 | 1.979 | 2.101 | 1.782 | 1.683 | 1.726 | 2.058 | -0.004 | 0.9695 | 0.9898 | 0 | 0 | 0 |
| 'Zfp458' | 2.429 | 1.868 | 2.197 | 2.196 | 2.408 | 1.851 | 2.066 | 2.011 | 1.730 | 1.855 | 2.801 | 2.823 | 2.285 | 2.846 | 2.637 | 2.351 | 3.039 | 2.668 | -0.323 | 0.0004 | 0.0055 | 1 | 0 | 0 |
| 'Zfp459' | 1.187 | 1.535 | 2.023 | 1.794 | 1.360 | 1.352 | 2.237 | 1.525 | 1.531 | 1.054 | 1.706 | 1.746 | 1.133 | 1.415 | 1.677 | 1.199 | 1.468 | 1.667 | 0.145 | 0.2522 | 0.5037 | 0 | 0 | 0 |
| 'Zfp46' | 3.491 | 3.652 | 4.161 | 3.697 | 3.633 | 4.128 | 4.121 | 4.026 | 4.227 | 3.392 | 4.256 | 4.305 | 3.225 | 3.370 | 4.556 | 3.116 | 3.729 | 4.535 | 0.021 | 0.7096 | 0.8653 | 0 | 0 | 0 |
| 'Zfp467' | 1.468 | 1.467 | 1.772 | 1.614 | 1.391 | 1.564 | 1.421 | 1.335 | 1.846 | 1.445 | 1.496 | 2.119 | 1.461 | 1.504 | 1.852 | 1.391 | 1.662 | 1.897 | -0.110 | 0.0506 | 0.1904 | 0 | 0 | 0 |
| 'Zfp472' | 7.468 | 7.760 | 6.996 | 8.022 | 7.911 | 7.443 | 8.031 | 7.170 | 7.693 | 8.277 | 7.886 | 7.975 | 7.097 | 7.392 | 7.543 | 7.416 | 7.333 | 8.157 | -0.030 | 0.5767 | 0.7816 | 0 | 0 | 0 |
| 'Zfp507' | 5.150 | 4.979 | 5.581 | 4.710 | 5.736 | 5.333 | 4.904 | 5.530 | 5.564 | 5.757 | 5.795 | 5.351 | 5.011 | 5.197 | 5.045 | 5.407 | 4.919 | 5.181 | -0.014 | 0.7439 | 0.8844 | 0 | 0 | 0 |
| 'Zfp51' | 10.181 | 11.167 | 11.088 | 9.731 | 11.352 | 9.510 | 10.354 | 10.810 | 11.028 | 9.538 | 11.263 | 10.382 | 8.754 | 11.389 | 10.044 | 9.881 | 10.576 | 9.305 | 0.056 | 0.2207 | 0.4680 | 0 | 0 | 0 |
| 'Zfp511' | 40.406 | 36.307 | 43.979 | 42.278 | 40.576 | 45.348 | 39.266 | 39.957 | 47.330 | 39.644 | 31.548 | 42.498 | 36.768 | 40.392 | 40.333 | 36.947 | 37.197 | 37.514 | 0.121 | 0.0138 | 0.0791 | 0 | 0 | 0 |
| 'Zfp512b' | 9.507 | 9.699 | 11.575 | 8.834 | 9.192 | 10.384 | 10.229 | 9.533 | 10.859 | 10.335 | 9.207 | 9.352 | 10.014 | 9.369 | 9.313 | 10.129 | 8.582 | 8.844 | 0.067 | 0.1246 | 0.3375 | 0 | 0 | 0 |
| 'Zfp513' | 10.004 | 10.634 | 11.831 | 10.183 | 10.027 | 12.212 | 10.086 | 10.551 | 11.859 | 10.785 | 9.351 | 11.836 | 10.834 | 9.914 | 10.945 | 10.083 | 9.971 | 10.732 | 0.032 | 0.3114 | 0.5700 | 0 | 0 | 0 |
| 'Zfp518a' | 5.128 | 6.778 | 6.436 | 5.745 | 7.026 | 6.273 | 5.903 | 7.377 | 6.546 | 7.058 | 6.330 | 5.805 | 6.198 | 6.864 | 5.013 | 6.697 | 6.490 | 5.506 | 0.024 | 0.6859 | 0.8513 | 0 | 0 | 0 |
| 'Zfp518b' | 0.575 | 0.839 | 0.712 | 0.678 | 0.884 | 0.918 | 0.733 | 1.054 | 0.912 | 0.773 | 0.569 | 0.608 | 0.572 | 0.828 | 0.578 | 0.727 | 0.721 | 0.581 | 0.284 | 0.0036 | 0.0288 | 1 | 0 | 0 |
| 'Zfp52' | 13.055 | 13.148 | 11.924 | 13.452 | 15.029 | 12.253 | 15.947 | 12.039 | 11.712 | 13.240 | 14.137 | 13.161 | 13.266 | 13.229 | 11.320 | 11.957 | 15.948 | 12.315 | -0.016 | 0.7831 | 0.9025 | 0 | 0 | 0 |
| 'Zfp523' | 5.518 | 5.131 | 5.962 | 4.737 | 5.428 | 6.159 | 5.321 | 5.497 | 6.361 | 5.492 | 5.258 | 6.530 | 5.237 | 5.687 | 6.486 | 6.168 | 5.024 | 5.917 | -0.056 | 0.1730 | 0.4066 | 0 | 0 | 0 |
| 'Zfp524' | 14.105 | 12.369 | 15.539 | 12.796 | 12.280 | 16.243 | 13.255 | 12.677 | 14.316 | 13.408 | 13.122 | 15.070 | 11.750 | 14.035 | 16.768 | 11.664 | 12.244 | 15.014 | 0.002 | 0.9768 | 0.9920 | 0 | 0 | 0 |
| 'Zfp526' | 10.191 | 9.378 | 10.113 | 9.232 | 8.363 | 11.793 | 9.205 | 8.518 | 11.187 | 9.345 | 9.178 | 11.592 | 8.186 | 8.153 | 10.588 | 8.862 | 8.372 | 11.530 | 0.027 | 0.5897 | 0.7904 | 0 | 0 | 0 |
| 'Zfp53' | 17.795 | 18.321 | 19.353 | 19.612 | 21.282 | 18.104 | 19.745 | 20.875 | 18.925 | 19.541 | 21.492 | 22.922 | 17.937 | 23.608 | 20.561 | 19.241 | 21.000 | 22.701 | -0.131 | 0.0055 | 0.0401 | 1 | 0 | 0 |
| 'Zfp54' | 1.451 | 1.437 | 1.493 | 1.482 | 1.731 | 1.317 | 1.626 | 1.623 | 1.663 | 1.270 | 1.471 | 1.657 | 1.327 | 1.560 | 1.730 | 1.237 | 1.934 | 2.047 | -0.053 | 0.5539 | 0.7667 | 0 | 0 | 0 |
| 'Zfp551' | 1.095 | 1.106 | 1.250 | 1.236 | 1.011 | 1.520 | 1.218 | 1.180 | 1.092 | 1.312 | 1.346 | 1.263 | 0.998 | 1.124 | 1.154 | 1.217 | 1.358 | 1.175 | -0.031 | 0.7721 | 0.8988 | 0 | 0 | 0 |
| 'Zfp553' | 4.277 | 4.515 | 4.256 | 3.888 | 3.939 | 4.896 | 4.027 | 4.328 | 4.642 | 3.801 | 4.050 | 5.104 | 4.003 | 3.577 | 5.432 | 4.047 | 3.605 | 5.541 | -0.017 | 0.7706 | 0.8977 | 0 | 0 | 0 |
| 'Zfp560' | 1.533 | 1.567 | 1.675 | 1.848 | 1.501 | 1.655 | 1.545 | 1.980 | 1.816 | 1.448 | 1.827 | 1.690 | 1.630 | 1.762 | 1.803 | 1.647 | 1.790 | 1.570 | -0.007 | 0.9223 | 0.9705 | 0 | 0 | 0 |
| 'Zfp563' | 1.782 | 2.468 | 1.811 | 2.234 | 1.916 | 1.854 | 2.130 | 2.051 | 1.838 | 1.700 | 1.353 | 1.615 | 1.777 | 1.774 | 1.639 | 1.541 | 1.988 | 1.242 | 0.281 | 0.0041 | 0.0324 | 1 | 0 | 0 |
| 'Zfp566' | 3.686 | 4.200 | 3.512 | 2.357 | 3.875 | 3.359 | 3.993 | 4.046 | 3.693 | 3.219 | 2.973 | 3.498 | 3.730 | 2.775 | 3.762 | 3.076 | 2.765 | 3.596 | 0.135 | 0.1616 | 0.3933 | 0 | 0 | 0 |
| 'Zfp568' | 2.075 | 2.088 | 2.168 | 1.995 | 2.477 | 2.027 | 2.168 | 2.302 | 2.073 | 2.461 | 2.247 | 2.105 | 2.022 | 2.172 | 2.046 | 2.503 | 2.295 | 2.098 | -0.048 | 0.3400 | 0.5970 | 0 | 0 | 0 |
| 'Zfp574' | 6.044 | 5.486 | 5.899 | 6.457 | 5.214 | 6.252 | 6.256 | 5.040 | 6.186 | 5.410 | 5.854 | 6.752 | 4.742 | 5.209 | 6.221 | 5.467 | 5.773 | 6.838 | 0.005 | 0.9267 | 0.9719 | 0 | 0 | 0 |
| 'Zfp579' | 2.684 | 2.599 | 3.751 | 2.286 | 2.446 | 3.494 | 2.594 | 2.791 | 3.748 | 2.780 | 2.589 | 3.269 | 2.612 | 2.184 | 2.840 | 2.295 | 2.411 | 2.933 | 0.127 | 0.0270 | 0.1265 | 0 | 0 | 0 |
| 'Zfp58' | 0.882 | 0.949 | 0.974 | 1.025 | 0.933 | 0.876 | 1.025 | 0.879 | 0.875 | 0.629 | 0.779 | 1.100 | 0.770 | 0.862 | 0.842 | 0.632 | 1.009 | 0.948 | 0.123 | 0.2482 | 0.4991 | 0 | 0 | 0 |
| 'Zfp580' | 3.687 | 5.344 | 4.875 | 3.342 | 4.886 | 4.238 | 3.619 | 4.407 | 4.450 | 3.670 | 2.860 | 2.177 | 3.520 | 3.203 | 3.183 | 2.644 | 3.609 | 2.609 | 0.504 | 0.0000 | 0.0006 | 1 | 1 | 0 |
| 'Zfp59' | 3.261 | 4.436 | 3.284 | 3.229 | 4.415 | 3.308 | 3.159 | 3.445 | 3.496 | 3.252 | 3.292 | 2.697 | 3.503 | 3.657 | 3.112 | 3.294 | 3.592 | 2.723 | 0.129 | 0.0666 | 0.2256 | 0 | 0 | 0 |
| 'Zfp592' | 19.857 | 18.161 | 18.576 | 18.227 | 16.847 | 18.576 | 18.269 | 17.272 | 18.371 | 18.075 | 17.444 | 20.507 | 17.385 | 17.536 | 19.043 | 18.016 | 18.007 | 19.696 | -0.024 | 0.4455 | 0.6878 | 0 | 0 | 0 |
| 'Zfp593' | 0.923 | 0.839 | 0.868 | 0.805 | 0.803 | 0.974 | 0.972 | 1.019 | 1.109 | 0.670 | 0.705 | 0.853 | 0.774 | 0.585 | 0.800 | 0.629 | 0.901 | 0.702 | 0.311 | 0.0077 | 0.0516 | 0 | 0 | 0 |
| 'Zfp595' | 2.575 | 2.436 | 3.111 | 3.339 | 2.878 | 2.696 | 3.184 | 3.321 | 2.489 | 2.862 | 2.933 | 3.350 | 3.310 | 2.878 | 3.566 | 2.524 | 4.227 | 3.268 | -0.167 | 0.0917 | 0.2796 | 0 | 0 | 0 |
| 'Zfp597' | 3.013 | 3.143 | 3.021 | 3.209 | 3.121 | 2.903 | 2.715 | 2.908 | 2.615 | 2.684 | 3.045 | 3.101 | 2.803 | 3.095 | 3.036 | 2.956 | 3.341 | 2.977 | -0.033 | 0.4767 | 0.7107 | 0 | 0 | 0 |
| 'Zfp598' | 38.505 | 34.834 | 41.627 | 36.596 | 34.560 | 40.213 | 35.167 | 35.981 | 41.670 | 38.346 | 33.400 | 43.012 | 42.531 | 34.350 | 41.938 | 38.445 | 37.309 | 43.768 | -0.067 | 0.0269 | 0.1262 | 0 | 0 | 0 |
| 'Zfp599' | 1.266 | 1.207 | 1.093 | 1.524 | 0.943 | 1.338 | 1.480 | 0.966 | 1.301 | 1.036 | 1.234 | 1.285 | 0.992 | 0.699 | 1.189 | 1.050 | 1.131 | 1.082 | 0.170 | 0.1400 | 0.3620 | 0 | 0 | 0 |
| 'Zfp60' | 1.240 | 1.106 | 1.254 | 1.246 | 1.330 | 1.196 | 1.239 | 1.301 | 1.376 | 1.078 | 1.351 | 1.344 | 1.049 | 1.132 | 1.334 | 1.151 | 1.202 | 1.441 | 0.017 | 0.7786 | 0.9003 | 0 | 0 | 0 |
| 'Zfp605' | 2.077 | 2.278 | 2.497 | 2.541 | 2.616 | 2.853 | 2.361 | 2.740 | 2.625 | 2.684 | 2.502 | 2.731 | 2.342 | 2.656 | 2.484 | 2.917 | 2.631 | 2.554 | -0.064 | 0.2944 | 0.5503 | 0 | 0 | 0 |
| 'Zfp606' | 0.801 | 0.863 | 0.764 | 0.636 | 0.798 | 0.739 | 0.818 | 0.826 | 0.789 | 0.626 | 0.747 | 0.823 | 0.643 | 0.808 | 0.681 | 0.858 | 0.771 | 0.875 | 0.029 | 0.6971 | 0.8577 | 0 | 0 | 0 |
| 'Zfp607a' | 4.633 | 3.387 | 4.096 | 4.434 | 3.998 | 3.730 | 4.641 | 3.700 | 3.440 | 3.797 | 3.703 | 3.926 | 3.780 | 3.693 | 3.461 | 4.294 | 3.750 | 3.232 | 0.087 | 0.1835 | 0.4192 | 0 | 0 | 0 |
| 'Zfp607b' | 1.252 | 1.164 | 1.125 | 1.142 | 1.323 | 1.334 | 1.154 | 1.479 | 1.215 | 1.333 | 1.168 | 1.441 | 1.190 | 1.471 | 1.511 | 1.103 | 1.458 | 1.316 | -0.114 | 0.2054 | 0.4491 | 0 | 0 | 0 |
| 'Zfp609' | 6.950 | 6.779 | 6.469 | 6.925 | 6.523 | 7.209 | 6.319 | 6.900 | 6.967 | 7.425 | 7.399 | 6.891 | 7.489 | 7.114 | 6.540 | 7.606 | 7.550 | 7.058 | -0.098 | 0.0072 | 0.0493 | 1 | 0 | 0 |
| 'Zfp61' | 2.133 | 2.312 | 2.225 | 2.296 | 2.115 | 2.466 | 2.374 | 2.187 | 2.318 | 1.999 | 2.117 | 1.913 | 2.131 | 1.937 | 2.021 | 1.821 | 1.957 | 1.866 | 0.194 | 0.0002 | 0.0022 | 1 | 0 | 0 |
| 'Zfp612' | 3.780 | 4.126 | 3.986 | 3.902 | 4.615 | 3.605 | 3.763 | 4.264 | 4.013 | 3.278 | 3.216 | 3.268 | 2.917 | 2.906 | 2.996 | 3.363 | 3.362 | 2.225 | 0.375 | 0.0000 | 0.0000 | 1 | 0 | 0 |
| 'Zfp617' | 3.394 | 4.879 | 4.228 | 3.769 | 4.294 | 4.138 | 5.173 | 4.196 | 4.199 | 3.113 | 4.111 | 4.816 | 3.357 | 4.179 | 4.126 | 3.847 | 4.357 | 4.430 | 0.056 | 0.4703 | 0.7061 | 0 | 0 | 0 |
| 'Zfp62' | 4.501 | 4.801 | 4.524 | 4.344 | 4.670 | 4.307 | 4.758 | 4.647 | 4.524 | 4.751 | 4.643 | 4.742 | 4.213 | 4.644 | 4.502 | 5.178 | 4.821 | 4.721 | -0.050 | 0.1504 | 0.3777 | 0 | 0 | 0 |
| 'Zfp622' | 49.320 | 38.458 | 34.801 | 41.170 | 40.259 | 39.539 | 40.380 | 40.604 | 35.853 | 47.380 | 37.666 | 39.182 | 43.006 | 39.614 | 37.688 | 47.147 | 39.959 | 37.891 | -0.045 | 0.2638 | 0.5179 | 0 | 0 | 0 |
| 'Zfp623' | 4.473 | 5.542 | 5.099 | 5.378 | 5.243 | 5.449 | 4.760 | 5.560 | 5.075 | 5.305 | 4.731 | 5.685 | 4.697 | 5.102 | 5.014 | 4.541 | 5.716 | 6.024 | -0.026 | 0.6799 | 0.8470 | 0 | 0 | 0 |
| 'Zfp626' | 3.929 | 4.844 | 4.586 | 3.756 | 4.657 | 4.597 | 4.910 | 4.474 | 4.918 | 4.259 | 3.912 | 4.555 | 3.524 | 3.873 | 3.969 | 3.807 | 4.501 | 3.907 | 0.148 | 0.0073 | 0.0495 | 1 | 0 | 0 |
| 'Zfp628' | 10.531 | 10.575 | 11.430 | 9.711 | 11.299 | 13.093 | 9.177 | 11.551 | 12.583 | 11.868 | 9.361 | 11.133 | 11.846 | 11.198 | 11.247 | 13.718 | 8.327 | 10.941 | -0.001 | 0.9836 | 0.9940 | 0 | 0 | 0 |
| 'Zfp629' | 0.421 | 0.505 | 0.570 | 0.489 | 0.264 | 0.483 | 0.423 | 0.529 | 0.604 | 0.393 | 0.403 | 0.598 | 0.352 | 0.443 | 0.587 | 0.439 | 0.527 | 0.449 | 0.026 | 0.8239 | 0.9226 | 0 | 0 | 0 |
| 'Zfp637' | 1.106 | 1.497 | 1.149 | 1.225 | 1.495 | 1.115 | 1.058 | 1.124 | 1.258 | 1.002 | 0.991 | 1.243 | 1.159 | 1.173 | 1.321 | 1.286 | 0.971 | 1.364 | 0.047 | 0.5987 | 0.7958 | 0 | 0 | 0 |
| 'Zfp638' | 7.186 | 6.456 | 6.614 | 8.503 | 6.917 | 6.435 | 7.908 | 6.617 | 6.565 | 6.247 | 7.205 | 7.577 | 5.801 | 6.514 | 6.935 | 5.909 | 7.528 | 7.386 | 0.037 | 0.5789 | 0.7820 | 0 | 0 | 0 |
| 'Zfp639' | 3.148 | 3.208 | 3.512 | 2.819 | 3.425 | 3.562 | 3.362 | 3.360 | 3.597 | 2.948 | 3.250 | 3.643 | 3.011 | 3.187 | 3.483 | 2.942 | 3.272 | 2.924 | 0.053 | 0.2227 | 0.4701 | 0 | 0 | 0 |
| 'Zfp64' | 8.275 | 6.926 | 7.751 | 7.341 | 7.314 | 7.941 | 7.396 | 7.337 | 7.596 | 6.886 | 6.614 | 7.736 | 7.314 | 6.952 | 7.395 | 6.774 | 6.799 | 7.048 | 0.085 | 0.0168 | 0.0915 | 0 | 0 | 0 |
| 'Zfp644' | 1.512 | 1.422 | 1.403 | 1.518 | 1.517 | 1.496 | 1.801 | 1.526 | 1.400 | 1.475 | 1.551 | 1.529 | 1.368 | 1.556 | 1.380 | 1.271 | 1.621 | 1.365 | 0.039 | 0.4357 | 0.6798 | 0 | 0 | 0 |
| 'Zfp646' | 11.231 | 10.684 | 11.302 | 11.114 | 10.109 | 11.748 | 10.199 | 10.882 | 11.380 | 10.854 | 11.844 | 12.889 | 10.133 | 11.613 | 12.541 | 11.446 | 10.684 | 12.818 | -0.096 | 0.0079 | 0.0526 | 0 | 0 | 0 |
| 'Zfp65' | 5.175 | 5.989 | 5.155 | 5.460 | 6.248 | 5.884 | 6.098 | 6.203 | 6.754 | 5.866 | 6.688 | 5.697 | 5.574 | 5.768 | 5.003 | 6.590 | 5.022 | 5.449 | 0.029 | 0.7566 | 0.8913 | 0 | 0 | 0 |
| 'Zfp651' | 3.040 | 4.706 | 3.019 | 3.221 | 3.648 | 3.428 | 3.474 | 4.269 | 3.273 | 3.817 | 3.718 | 3.090 | 5.034 | 4.523 | 3.084 | 4.040 | 3.544 | 2.805 | -0.075 | 0.3565 | 0.6092 | 0 | 0 | 0 |
| 'Zfp652' | 1.311 | 1.330 | 1.367 | 1.289 | 1.169 | 1.334 | 1.357 | 1.224 | 1.414 | 1.167 | 1.398 | 1.604 | 1.161 | 1.339 | 1.439 | 1.309 | 1.319 | 1.535 | -0.069 | 0.1761 | 0.4114 | 0 | 0 | 0 |
| 'Zfp653' | 15.639 | 15.589 | 17.663 | 11.724 | 14.255 | 18.278 | 15.160 | 15.474 | 18.381 | 14.839 | 14.110 | 17.563 | 15.899 | 15.365 | 16.487 | 14.858 | 14.423 | 17.165 | 0.003 | 0.9427 | 0.9785 | 0 | 0 | 0 |
| 'Zfp655' | 4.634 | 4.540 | 4.355 | 4.520 | 4.159 | 4.170 | 4.706 | 4.773 | 4.250 | 4.303 | 4.776 | 4.643 | 4.156 | 5.027 | 4.708 | 4.293 | 5.150 | 4.422 | -0.058 | 0.1893 | 0.4258 | 0 | 0 | 0 |
| 'Zfp661' | 0.455 | 0.481 | 0.546 | 0.658 | 0.479 | 0.457 | 0.387 | 0.428 | 0.493 | 0.456 | 0.519 | 0.458 | 0.463 | 0.411 | 0.487 | 0.422 | 0.516 | 0.429 | 0.068 | 0.5025 | 0.7297 | 0 | 0 | 0 |
| 'Zfp664' | 12.009 | 12.906 | 11.814 | 11.921 | 12.938 | 12.743 | 11.304 | 12.347 | 12.193 | 10.916 | 12.183 | 14.948 | 12.135 | 13.531 | 14.382 | 11.750 | 12.898 | 15.443 | -0.108 | 0.0147 | 0.0824 | 0 | 0 | 0 |
| 'Zfp667' | 0.544 | 0.617 | 0.520 | 0.622 | 0.697 | 0.682 | 0.771 | 0.611 | 0.611 | 0.421 | 0.572 | 0.589 | 0.572 | 0.553 | 0.597 | 0.667 | 0.466 | 0.446 | 0.198 | 0.0418 | 0.1683 | 0 | 0 | 0 |
| 'Zfp668' | 3.758 | 3.767 | 4.147 | 3.682 | 4.056 | 4.585 | 3.603 | 3.674 | 4.212 | 3.892 | 3.912 | 4.773 | 3.650 | 3.805 | 4.087 | 3.903 | 3.923 | 4.468 | -0.050 | 0.1862 | 0.4219 | 0 | 0 | 0 |
| 'Zfp672' | 1.564 | 1.707 | 1.876 | 1.648 | 1.680 | 1.763 | 1.916 | 1.784 | 1.874 | 1.560 | 1.638 | 1.880 | 1.531 | 1.646 | 1.747 | 1.545 | 1.707 | 1.695 | 0.066 | 0.1383 | 0.3589 | 0 | 0 | 0 |
| 'Zfp677' | 4.885 | 5.728 | 6.162 | 5.557 | 7.320 | 5.733 | 6.217 | 7.048 | 6.193 | 6.262 | 5.797 | 6.164 | 5.415 | 6.901 | 6.094 | 6.167 | 6.480 | 5.379 | -0.007 | 0.9044 | 0.9624 | 0 | 0 | 0 |
| 'Zfp68' | 5.717 | 6.506 | 5.882 | 5.970 | 6.306 | 5.592 | 5.709 | 6.045 | 6.060 | 5.879 | 6.494 | 6.418 | 5.717 | 6.094 | 6.139 | 6.008 | 6.147 | 6.306 | -0.053 | 0.1498 | 0.3769 | 0 | 0 | 0 |
| 'Zfp687' | 3.274 | 3.363 | 3.732 | 2.863 | 3.384 | 3.630 | 3.048 | 3.354 | 3.901 | 3.398 | 3.242 | 3.484 | 3.189 | 3.438 | 3.565 | 3.155 | 3.072 | 3.507 | 0.014 | 0.6976 | 0.8580 | 0 | 0 | 0 |
| 'Zfp688' | 2.788 | 3.833 | 3.864 | 2.838 | 3.483 | 3.831 | 2.967 | 3.190 | 3.922 | 3.522 | 2.778 | 3.710 | 3.195 | 2.833 | 3.381 | 3.133 | 3.101 | 3.140 | 0.075 | 0.2231 | 0.4707 | 0 | 0 | 0 |
| 'Zfp689' | 2.724 | 3.270 | 3.327 | 2.409 | 3.107 | 2.940 | 3.038 | 3.004 | 3.065 | 2.479 | 2.385 | 3.063 | 2.796 | 2.406 | 2.780 | 2.259 | 2.097 | 2.783 | 0.201 | 0.0028 | 0.0236 | 1 | 0 | 0 |
| 'Zfp69' | 3.324 | 3.580 | 3.160 | 3.330 | 3.162 | 3.359 | 3.766 | 3.554 | 3.295 | 3.704 | 3.654 | 3.665 | 3.831 | 4.041 | 3.506 | 3.735 | 3.996 | 3.240 | -0.140 | 0.0062 | 0.0443 | 1 | 0 | 0 |
| 'Zfp691' | 2.654 | 2.294 | 2.547 | 2.469 | 2.458 | 2.816 | 2.681 | 2.416 | 3.326 | 2.292 | 2.868 | 3.295 | 2.341 | 2.536 | 2.891 | 2.075 | 2.823 | 2.967 | -0.039 | 0.5999 | 0.7967 | 0 | 0 | 0 |
| 'Zfp692' | 18.657 | 15.732 | 23.140 | 16.327 | 15.137 | 20.063 | 18.004 | 17.366 | 22.441 | 17.939 | 16.141 | 20.511 | 20.416 | 16.275 | 19.341 | 17.196 | 17.105 | 18.645 | 0.015 | 0.7550 | 0.8907 | 0 | 0 | 0 |
| 'Zfp7' | 7.602 | 8.075 | 9.033 | 8.107 | 7.695 | 8.550 | 8.883 | 7.418 | 8.370 | 7.442 | 8.217 | 9.458 | 7.292 | 8.493 | 10.124 | 7.415 | 8.458 | 9.113 | -0.055 | 0.2907 | 0.5477 | 0 | 0 | 0 |
| 'Zfp703' | 13.819 | 7.519 | 7.045 | 10.794 | 7.003 | 7.175 | 9.273 | 7.411 | 7.671 | 10.985 | 10.404 | 8.410 | 9.967 | 10.220 | 7.674 | 10.012 | 10.402 | 7.957 | -0.176 | 0.0304 | 0.1362 | 0 | 0 | 0 |
| 'Zfp704' | 0.658 | 0.503 | 0.431 | 0.586 | 0.389 | 0.402 | 0.733 | 0.599 | 0.554 | 0.619 | 0.617 | 0.711 | 0.693 | 0.553 | 0.730 | 0.572 | 0.572 | 0.902 | -0.320 | 0.0058 | 0.0422 | 1 | 0 | 0 |
| 'Zfp706' | 39.883 | 41.616 | 39.417 | 42.949 | 43.419 | 41.242 | 40.555 | 40.651 | 41.582 | 38.806 | 40.797 | 44.233 | 38.826 | 41.413 | 44.462 | 39.980 | 42.212 | 45.453 | -0.031 | 0.3565 | 0.6092 | 0 | 0 | 0 |
| 'Zfp707' | 3.213 | 3.146 | 3.441 | 2.962 | 2.958 | 3.600 | 3.236 | 3.193 | 3.578 | 3.216 | 3.168 | 3.629 | 3.078 | 2.739 | 3.101 | 2.779 | 2.894 | 3.094 | 0.068 | 0.1678 | 0.4000 | 0 | 0 | 0 |
| 'Zfp709' | 1.577 | 2.257 | 2.252 | 1.432 | 2.218 | 2.086 | 1.908 | 2.191 | 2.139 | 2.024 | 1.791 | 2.137 | 1.507 | 2.051 | 1.899 | 2.050 | 1.865 | 2.146 | 0.031 | 0.6673 | 0.8396 | 0 | 0 | 0 |
| 'Zfp710' | 8.691 | 7.528 | 8.438 | 8.114 | 7.164 | 8.355 | 7.884 | 7.293 | 8.609 | 7.862 | 7.590 | 9.265 | 8.121 | 6.630 | 8.687 | 7.270 | 7.580 | 8.404 | 0.001 | 0.9852 | 0.9949 | 0 | 0 | 0 |
| 'Zfp712' | 1.929 | 1.839 | 1.839 | 2.189 | 2.080 | 1.757 | 1.816 | 2.596 | 1.520 | 1.851 | 2.243 | 2.207 | 1.596 | 2.038 | 1.607 | 2.066 | 1.774 | 1.857 | 0.008 | 0.9303 | 0.9737 | 0 | 0 | 0 |
| 'Zfp715' | 3.724 | 3.371 | 3.302 | 3.724 | 3.443 | 3.296 | 3.857 | 3.437 | 3.442 | 3.622 | 3.845 | 4.192 | 3.148 | 3.568 | 3.887 | 3.049 | 4.053 | 3.807 | -0.082 | 0.1675 | 0.3999 | 0 | 0 | 0 |
| 'Zfp719' | 1.076 | 1.331 | 1.225 | 1.092 | 1.382 | 1.293 | 1.300 | 1.172 | 1.299 | 0.940 | 1.336 | 1.293 | 0.935 | 0.974 | 1.320 | 1.029 | 1.293 | 1.246 | 0.090 | 0.2874 | 0.5438 | 0 | 0 | 0 |
| 'Zfp729a' | 3.768 | 5.579 | 4.370 | 4.584 | 6.034 | 5.353 | 4.980 | 5.658 | 4.790 | 4.921 | 4.421 | 5.519 | 3.841 | 5.010 | 4.874 | 5.974 | 4.700 | 4.958 | 0.009 | 0.9145 | 0.9671 | 0 | 0 | 0 |
| 'Zfp729b' | 4.818 | 5.907 | 5.396 | 4.932 | 5.559 | 5.011 | 5.135 | 5.675 | 4.989 | 5.678 | 4.451 | 5.164 | 4.840 | 5.658 | 4.521 | 5.862 | 5.095 | 4.926 | 0.027 | 0.6384 | 0.8220 | 0 | 0 | 0 |
| 'Zfp74' | 2.008 | 2.073 | 1.949 | 2.087 | 1.774 | 2.182 | 2.099 | 1.834 | 2.016 | 1.698 | 1.985 | 2.052 | 1.923 | 1.968 | 1.899 | 1.849 | 1.976 | 2.149 | 0.028 | 0.6203 | 0.8101 | 0 | 0 | 0 |
| 'Zfp740' | 12.875 | 14.631 | 13.201 | 12.935 | 13.486 | 13.754 | 12.513 | 13.807 | 13.099 | 12.052 | 13.734 | 14.155 | 12.544 | 13.600 | 14.011 | 13.035 | 14.034 | 14.053 | -0.022 | 0.3910 | 0.6403 | 0 | 0 | 0 |
| 'Zfp746' | 13.574 | 14.409 | 13.488 | 13.444 | 14.209 | 14.678 | 13.438 | 14.926 | 13.654 | 13.948 | 13.962 | 14.178 | 14.453 | 14.422 | 13.952 | 14.468 | 13.856 | 15.340 | -0.042 | 0.1373 | 0.3573 | 0 | 0 | 0 |
| 'Zfp758' | 1.792 | 1.808 | 1.549 | 2.028 | 1.678 | 1.559 | 1.971 | 1.828 | 1.469 | 1.372 | 1.999 | 2.166 | 1.677 | 2.099 | 1.641 | 1.574 | 2.258 | 1.841 | -0.103 | 0.2524 | 0.5040 | 0 | 0 | 0 |
| 'Zfp759' | 1.262 | 1.685 | 1.487 | 1.451 | 1.372 | 1.576 | 1.803 | 1.792 | 1.478 | 1.347 | 1.459 | 1.900 | 1.268 | 1.384 | 1.960 | 1.893 | 1.518 | 1.844 | -0.083 | 0.4255 | 0.6723 | 0 | 0 | 0 |
| 'Zfp760' | 2.055 | 2.314 | 2.838 | 2.853 | 2.457 | 2.617 | 2.862 | 2.283 | 2.674 | 1.714 | 2.453 | 3.511 | 1.675 | 2.202 | 3.006 | 1.968 | 3.216 | 2.821 | 0.031 | 0.7737 | 0.8993 | 0 | 0 | 0 |
| 'Zfp763' | 1.396 | 1.332 | 1.729 | 1.566 | 2.236 | 1.722 | 1.553 | 2.172 | 1.480 | 1.519 | 1.496 | 1.707 | 1.493 | 1.564 | 1.645 | 1.669 | 1.625 | 1.562 | 0.074 | 0.4331 | 0.6782 | 0 | 0 | 0 |
| 'Zfp770' | 1.770 | 2.076 | 2.144 | 2.033 | 1.982 | 2.119 | 2.172 | 2.398 | 2.173 | 1.684 | 2.358 | 2.461 | 1.737 | 2.169 | 2.054 | 1.782 | 2.189 | 2.111 | 0.011 | 0.8618 | 0.9401 | 0 | 0 | 0 |
| 'Zfp771' | 7.673 | 6.874 | 8.665 | 6.152 | 6.574 | 9.940 | 7.639 | 6.580 | 8.789 | 7.045 | 6.124 | 9.666 | 7.318 | 5.650 | 9.039 | 8.527 | 6.169 | 8.641 | 0.007 | 0.9052 | 0.9627 | 0 | 0 | 0 |
| 'Zfp773' | 1.869 | 1.764 | 1.574 | 1.507 | 1.445 | 1.334 | 1.878 | 1.375 | 1.581 | 1.130 | 1.374 | 1.482 | 1.209 | 1.110 | 1.138 | 1.491 | 1.264 | 1.508 | 0.267 | 0.0146 | 0.0821 | 0 | 0 | 0 |
| 'Zfp775' | 2.345 | 2.712 | 2.773 | 2.128 | 2.390 | 2.534 | 2.280 | 2.688 | 2.737 | 3.231 | 2.227 | 2.141 | 2.927 | 2.340 | 2.359 | 2.825 | 1.971 | 2.145 | 0.034 | 0.6991 | 0.8592 | 0 | 0 | 0 |
| 'Zfp777' | 11.181 | 9.912 | 12.024 | 10.733 | 10.472 | 12.597 | 11.425 | 10.870 | 12.562 | 12.804 | 10.576 | 11.780 | 12.414 | 10.389 | 11.294 | 11.816 | 9.862 | 11.665 | -0.019 | 0.5979 | 0.7954 | 0 | 0 | 0 |
| 'Zfp780b' | 3.335 | 3.277 | 3.501 | 4.410 | 4.060 | 3.435 | 3.989 | 3.427 | 3.327 | 3.058 | 4.194 | 4.345 | 3.072 | 3.158 | 4.485 | 3.594 | 3.628 | 4.159 | -0.054 | 0.5445 | 0.7606 | 0 | 0 | 0 |
| 'Zfp784' | 4.121 | 5.903 | 6.922 | 4.649 | 5.931 | 6.341 | 4.770 | 6.235 | 5.980 | 6.037 | 4.608 | 3.984 | 4.841 | 5.256 | 5.363 | 4.910 | 4.759 | 4.077 | 0.205 | 0.0350 | 0.1496 | 0 | 0 | 0 |

| | | | | | | | | | | | | | | | | | | | | | | | | |
|---|---|---|---|---|---|---|---|---|---|---|---|---|---|---|---|---|---|---|---|---|---|---|---|---|
| 'Zfp787' | 16.961 | 16.940 | 17.848 | 15.178 | 17.138 | 17.231 | 14.004 | 15.715 | 16.672 | 15.435 | 16.357 | 17.567 | 17.884 | 17.368 | 18.708 | 19.631 | 15.370 | 17.547 | -0.085 | 0.0946 | 0.2852 | 0 | 0 | 0 |
| 'Zfp788' | 1.499 | 2.220 | 1.608 | 1.412 | 1.925 | 1.602 | 1.412 | 1.956 | 1.781 | 1.568 | 1.614 | 1.545 | 1.392 | 1.956 | 1.665 | 1.885 | 1.749 | 1.543 | 0.031 | 0.6236 | 0.8120 | 0 | 0 | 0 |
| 'Zfp790' | 7.299 | 6.390 | 6.741 | 7.917 | 6.562 | 6.811 | 7.159 | 6.176 | 7.167 | 5.544 | 7.235 | 8.423 | 5.609 | 6.945 | 7.306 | 5.339 | 7.756 | 8.294 | -0.007 | 0.9275 | 0.9720 | 0 | 0 | 0 |
| 'Zfp799' | 1.766 | 1.674 | 1.650 | 1.644 | 1.898 | 1.914 | 1.976 | 2.133 | 1.927 | 1.946 | 1.855 | 1.741 | 1.650 | 1.576 | 1.590 | 1.975 | 1.844 | 1.481 | 0.079 | 0.2305 | 0.4777 | 0 | 0 | 0 |
| 'Zfp800' | 6.055 | 5.131 | 5.952 | 5.984 | 5.477 | 5.054 | 6.179 | 5.618 | 5.199 | 5.673 | 6.117 | 6.339 | 4.701 | 5.536 | 5.785 | 5.002 | 6.264 | 6.298 | -0.040 | 0.5237 | 0.7461 | 0 | 0 | 0 |
| 'Zfp809' | 12.079 | 12.849 | 13.765 | 14.053 | 13.536 | 12.390 | 12.897 | 13.346 | 13.226 | 10.522 | 13.429 | 15.386 | 10.173 | 12.091 | 14.654 | 13.761 | 14.308 | 13.327 | -0.005 | 0.9443 | 0.9789 | 0 | 0 | 0 |
| 'Zfp81' | 2.419 | 3.024 | 2.746 | 2.748 | 3.176 | 2.437 | 2.781 | 2.614 | 2.667 | 2.261 | 2.703 | 3.017 | 1.947 | 3.032 | 2.526 | 2.250 | 2.718 | 2.470 | 0.087 | 0.2010 | 0.4424 | 0 | 0 | 0 |
| 'Zfp810' | 5.784 | 6.421 | 7.853 | 6.023 | 6.779 | 7.187 | 7.195 | 6.594 | 6.750 | 6.905 | 7.027 | 6.304 | 6.303 | 6.672 | 6.369 | 6.814 | 6.479 | 6.341 | 0.030 | 0.5916 | 0.7921 | 0 | 0 | 0 |
| 'Zfp821' | 19.307 | 18.769 | 20.928 | 16.964 | 17.664 | 19.950 | 18.020 | 19.268 | 20.108 | 19.090 | 17.139 | 20.031 | 18.010 | 17.067 | 18.725 | 17.926 | 17.729 | 17.303 | 0.060 | 0.1500 | 0.3772 | 0 | 0 | 0 |
| 'Zfp827' | 1.923 | 1.456 | 2.105 | 1.780 | 1.731 | 1.994 | 1.533 | 1.901 | 2.046 | 2.270 | 1.908 | 2.344 | 2.154 | 1.989 | 1.902 | 2.217 | 2.177 | 2.004 | -0.212 | 0.0004 | 0.0054 | 1 | 0 | 0 |
| 'Zfp830' | 14.141 | 14.843 | 13.920 | 14.432 | 14.114 | 14.482 | 14.642 | 14.605 | 14.647 | 13.080 | 13.955 | 15.842 | 12.348 | 13.857 | 13.793 | 12.664 | 13.580 | 15.959 | 0.036 | 0.4770 | 0.7107 | 0 | 0 | 0 |
| 'Zfp831' | 32.745 | 27.759 | 28.760 | 33.510 | 27.090 | 27.816 | 30.815 | 27.747 | 29.753 | 30.616 | 33.308 | 34.484 | 26.815 | 29.671 | 32.491 | 30.259 | 30.803 | 35.470 | -0.106 | 0.0485 | 0.1850 | 0 | 0 | 0 |
| 'Zfp839' | 10.167 | 10.647 | 10.249 | 9.760 | 10.058 | 11.281 | 9.065 | 10.195 | 9.989 | 10.407 | 12.078 | 10.499 | 9.982 | 10.200 | 9.846 | 10.268 | 9.243 | 10.099 | -0.026 | 0.5779 | 0.7820 | 0 | 0 | 0 |
| 'Zfp84' | 7.811 | 7.306 | 6.740 | 6.782 | 6.994 | 7.227 | 8.514 | 7.406 | 6.872 | 7.073 | 7.140 | 7.889 | 7.072 | 7.111 | 7.319 | 6.631 | 7.795 | 8.002 | -0.024 | 0.6451 | 0.8265 | 0 | 0 | 0 |
| 'Zfp846' | 1.302 | 1.582 | 1.369 | 1.360 | 1.458 | 1.399 | 1.487 | 1.721 | 1.468 | 1.543 | 1.383 | 1.492 | 1.484 | 1.336 | 1.430 | 1.217 | 1.586 | 1.269 | 0.025 | 0.7291 | 0.8759 | 0 | 0 | 0 |
| 'Zfp865' | 22.797 | 21.943 | 26.028 | 21.290 | 20.880 | 25.745 | 22.149 | 22.421 | 25.645 | 24.212 | 21.026 | 27.181 | 22.846 | 21.407 | 26.676 | 21.872 | 21.501 | 25.184 | -0.031 | 0.2640 | 0.5179 | 0 | 0 | 0 |
| 'Zfp866' | 3.663 | 4.123 | 5.155 | 4.962 | 3.859 | 4.720 | 5.194 | 3.969 | 5.246 | 3.444 | 4.848 | 5.465 | 3.061 | 3.264 | 5.022 | 3.321 | 4.995 | 5.214 | 0.085 | 0.3730 | 0.6242 | 0 | 0 | 0 |
| 'Zfp867' | 1.904 | 2.184 | 2.001 | 1.726 | 2.187 | 2.043 | 1.999 | 2.165 | 2.006 | 2.152 | 2.100 | 2.011 | 1.839 | 2.197 | 1.650 | 2.268 | 2.063 | 1.470 | 0.034 | 0.6891 | 0.8534 | 0 | 0 | 0 |
| 'Zfp868' | 14.191 | 16.447 | 15.085 | 12.926 | 16.043 | 15.027 | 13.888 | 15.805 | 15.236 | 16.293 | 15.539 | 14.950 | 14.635 | 14.694 | 14.342 | 14.775 | 14.807 | 12.898 | 0.008 | 0.8534 | 0.9372 | 0 | 0 | 0 |
| 'Zfp869' | 20.022 | 23.191 | 20.975 | 18.700 | 21.342 | 19.636 | 18.989 | 23.092 | 21.174 | 22.575 | 21.540 | 22.088 | 18.603 | 21.999 | 19.401 | 21.682 | 21.128 | 20.798 | -0.032 | 0.4765 | 0.7107 | 0 | 0 | 0 |
| 'Zfp87' | 5.333 | 6.207 | 5.720 | 6.277 | 5.533 | 5.383 | 6.953 | 5.651 | 5.621 | 4.728 | 6.112 | 6.761 | 4.533 | 6.104 | 5.993 | 4.943 | 6.393 | 6.819 | 0.001 | 0.9943 | 0.9990 | 0 | 0 | 0 |
| 'Zfp870' | 4.884 | 8.004 | 5.965 | 5.500 | 7.616 | 5.740 | 5.454 | 7.291 | 6.407 | 5.792 | 5.931 | 6.106 | 6.668 | 6.059 | 5.622 | 6.907 | 5.374 | 6.290 | 0.028 | 0.7308 | 0.8772 | 0 | 0 | 0 |
| 'Zfp871' | 9.708 | 9.910 | 10.878 | 9.305 | 9.532 | 9.518 | 9.494 | 9.405 | 10.630 | 8.303 | 10.365 | 11.274 | 8.706 | 10.156 | 9.714 | 10.132 | 10.359 | 11.254 | -0.039 | 0.3915 | 0.6409 | 0 | 0 | 0 |
| 'Zfp873' | 1.546 | 2.092 | 1.746 | 1.811 | 1.831 | 1.339 | 1.781 | 1.843 | 1.689 | 1.320 | 1.722 | 1.949 | 1.181 | 1.608 | 1.623 | 1.362 | 1.874 | 2.131 | 0.068 | 0.4976 | 0.7258 | 0 | 0 | 0 |
| 'Zfp874a' | 3.990 | 4.215 | 4.537 | 4.345 | 5.271 | 4.502 | 4.459 | 4.889 | 4.602 | 4.162 | 5.099 | 5.583 | 3.952 | 5.056 | 5.062 | 4.588 | 5.077 | 4.693 | -0.099 | 0.0666 | 0.2256 | 0 | 0 | 0 |
| 'Zfp882' | 1.886 | 2.674 | 2.291 | 2.068 | 2.240 | 2.488 | 2.392 | 2.104 | 1.977 | 1.945 | 1.884 | 2.278 | 1.590 | 2.160 | 2.391 | 2.312 | 2.339 | 2.052 | 0.072 | 0.3948 | 0.6439 | 0 | 0 | 0 |
| 'Zfp90' | 2.776 | 2.504 | 2.712 | 2.405 | 3.598 | 3.414 | 3.290 | 3.523 | 2.959 | 2.891 | 2.423 | 2.358 | 3.234 | 2.876 | 2.663 | 2.823 | 1.986 | 2.283 | 0.206 | 0.0402 | 0.1639 | 0 | 0 | 0 |
| 'Zfp91' | 24.092 | 25.579 | 23.488 | 22.835 | 25.526 | 23.705 | 24.115 | 25.530 | 23.249 | 25.247 | 24.137 | 22.825 | 24.069 | 26.211 | 22.168 | 25.062 | 23.844 | 22.487 | 0.004 | 0.8541 | 0.9373 | 0 | 0 | 0 |
| 'Zfp930' | 4.360 | 4.692 | 4.564 | 4.328 | 4.611 | 4.374 | 5.172 | 4.508 | 4.598 | 3.426 | 4.786 | 4.789 | 3.995 | 3.939 | 4.176 | 3.631 | 5.067 | 4.524 | 0.090 | 0.1695 | 0.4019 | 0 | 0 | 0 |
| 'Zfp933' | 1.646 | 2.796 | 2.113 | 1.869 | 2.296 | 2.286 | 1.930 | 2.485 | 2.667 | 2.141 | 2.319 | 2.132 | 1.968 | 1.850 | 2.550 | 2.370 | 1.755 | 2.241 | 0.038 | 0.6694 | 0.8407 | 0 | 0 | 0 |
| 'Zfp934' | 3.322 | 4.494 | 3.709 | 3.252 | 3.699 | 3.300 | 4.515 | 4.588 | 3.495 | 3.692 | 4.445 | 4.026 | 3.885 | 4.871 | 3.573 | 3.511 | 4.050 | 3.762 | -0.079 | 0.4141 | 0.6620 | 0 | 0 | 0 |
| 'Zfp938' | 1.740 | 2.510 | 2.210 | 1.825 | 2.099 | 1.829 | 2.015 | 2.153 | 1.760 | 1.887 | 1.621 | 1.815 | 1.570 | 2.096 | 1.638 | 1.767 | 2.161 | 1.620 | 0.151 | 0.0626 | 0.2171 | 0 | 0 | 0 |
| 'Zfp94' | 1.035 | 1.006 | 1.468 | 1.067 | 1.210 | 1.349 | 1.282 | 1.233 | 1.322 | 1.136 | 1.251 | 1.288 | 1.065 | 1.107 | 1.289 | 1.218 | 1.215 | 1.311 | 0.011 | 0.8778 | 0.9485 | 0 | 0 | 0 |
| 'Zfp940' | 0.951 | 0.780 | 0.749 | 0.872 | 0.713 | 0.958 | 0.899 | 0.794 | 0.932 | 0.790 | 0.669 | 0.773 | 0.725 | 0.598 | 0.729 | 0.679 | 0.726 | 0.575 | 0.274 | 0.0016 | 0.0152 | 1 | 0 | 0 |
| 'Zfp942' | 9.902 | 12.480 | 10.550 | 10.371 | 11.832 | 10.351 | 10.139 | 12.574 | 10.417 | 11.501 | 11.240 | 10.853 | 10.444 | 12.481 | 9.721 | 11.527 | 11.345 | 9.902 | -0.018 | 0.6678 | 0.8400 | 0 | 0 | 0 |
| 'Zfp943' | 2.053 | 3.161 | 2.249 | 2.283 | 2.782 | 2.622 | 2.571 | 2.936 | 2.340 | 2.384 | 2.628 | 2.522 | 1.821 | 2.453 | 2.353 | 2.699 | 3.008 | 2.431 | 0.027 | 0.7013 | 0.8608 | 0 | 0 | 0 |
| 'Zfp944' | 6.604 | 7.093 | 6.690 | 6.222 | 6.711 | 6.425 | 5.639 | 8.314 | 5.924 | 5.430 | 6.564 | 6.172 | 5.425 | 6.471 | 5.396 | 5.787 | 6.847 | 4.969 | 0.161 | 0.0070 | 0.0479 | 1 | 0 | 0 |
| 'Zfp945' | 1.440 | 2.163 | 1.753 | 1.567 | 2.113 | 1.833 | 2.039 | 1.988 | 1.979 | 1.738 | 1.634 | 1.720 | 1.622 | 1.683 | 1.584 | 1.849 | 1.780 | 1.477 | 0.145 | 0.0181 | 0.0964 | 0 | 0 | 0 |
| 'Zfp948' | 1.800 | 1.873 | 2.509 | 2.929 | 1.941 | 1.765 | 2.470 | 1.781 | 2.400 | 1.742 | 1.817 | 2.600 | 2.302 | 2.485 | 2.012 | 1.890 | 2.607 | 2.349 | -0.049 | 0.6799 | 0.8470 | 0 | 0 | 0 |
| 'Zfp949' | 5.271 | 5.070 | 4.686 | 5.195 | 5.226 | 4.502 | 5.755 | 5.196 | 4.899 | 4.189 | 5.240 | 5.367 | 4.459 | 5.602 | 5.221 | 5.013 | 5.145 | 5.254 | -0.004 | 0.9436 | 0.9788 | 0 | 0 | 0 |
| 'Zfp950' | 0.886 | 1.223 | 1.134 | 0.862 | 1.454 | 1.163 | 1.140 | 1.118 | 1.283 | 0.997 | 0.947 | 1.438 | 1.015 | 1.320 | 1.300 | 1.067 | 1.214 | 1.356 | -0.076 | 0.3075 | 0.5661 | 0 | 0 | 0 |
| 'Zfp952' | 2.916 | 3.259 | 3.346 | 3.644 | 3.171 | 3.327 | 3.309 | 3.674 | 3.452 | 3.215 | 3.679 | 3.774 | 2.668 | 3.630 | 3.332 | 3.386 | 3.460 | 3.553 | -0.038 | 0.5053 | 0.7316 | 0 | 0 | 0 |
| 'Zfp953' | 1.783 | 2.077 | 2.359 | 2.294 | 1.991 | 1.993 | 2.275 | 2.567 | 1.985 | 1.625 | 2.101 | 2.230 | 1.492 | 2.181 | 2.306 | 1.865 | 2.098 | 2.633 | 0.056 | 0.5290 | 0.7495 | 0 | 0 | 0 |
| 'Zfp954' | 17.715 | 12.788 | 13.027 | 16.007 | 12.913 | 12.356 | 15.173 | 12.912 | 12.594 | 15.018 | 15.825 | 14.015 | 15.962 | 14.844 | 13.758 | 13.292 | 16.239 | 13.217 | -0.090 | 0.1279 | 0.3427 | 0 | 0 | 0 |
| 'Zfp955b' | 12.085 | 12.178 | 11.247 | 11.424 | 13.310 | 12.183 | 14.028 | 12.703 | 10.216 | 12.994 | 11.685 | 10.516 | 12.487 | 12.998 | 9.343 | 12.697 | 10.431 | 10.791 | 0.067 | 0.2429 | 0.4935 | 0 | 0 | 0 |
| 'Zfp956' | 5.586 | 5.832 | 7.072 | 5.509 | 5.492 | 6.698 | 5.657 | 6.168 | 6.458 | 6.189 | 4.792 | 5.829 | 5.899 | 5.739 | 5.739 | 5.577 | 5.062 | 6.209 | 0.086 | 0.0738 | 0.2411 | 0 | 0 | 0 |
| 'Zfp958' | 3.846 | 4.932 | 4.140 | 4.576 | 4.262 | 4.397 | 4.950 | 4.172 | 4.231 | 3.559 | 4.542 | 5.072 | 3.506 | 4.087 | 4.227 | 3.228 | 5.730 | 4.836 | 0.015 | 0.8618 | 0.9401 | 0 | 0 | 0 |
| 'Zfp961' | 1.279 | 1.456 | 1.190 | 1.331 | 1.593 | 1.480 | 1.386 | 1.522 | 1.460 | 1.074 | 1.386 | 1.559 | 1.091 | 1.323 | 1.335 | 1.489 | 1.548 | 1.587 | 0.021 | 0.7673 | 0.8961 | 0 | 0 | 0 |
| 'Zfp963' | 1.258 | 1.356 | 1.762 | 1.299 | 1.846 | 1.313 | 1.268 | 1.467 | 1.593 | 1.040 | 1.650 | 2.115 | 0.963 | 1.972 | 1.605 | 1.465 | 1.742 | 1.536 | -0.107 | 0.3614 | 0.6135 | 0 | 0 | 0 |
| 'Zfp974' | 1.560 | 2.287 | 1.668 | 1.275 | 2.018 | 1.773 | 2.031 | 1.989 | 1.895 | 1.963 | 1.830 | 1.857 | 1.802 | 1.963 | 1.499 | 1.687 | 1.961 | 1.427 | 0.025 | 0.7616 | 0.8943 | 0 | 0 | 0 |
| 'Zfp983' | 4.612 | 4.822 | 5.200 | 4.150 | 5.965 | 4.498 | 5.153 | 5.224 | 4.950 | 4.431 | 4.099 | 5.254 | 3.907 | 4.410 | 3.905 | 5.650 | 5.594 | 4.873 | 0.065 | 0.4829 | 0.7146 | 0 | 0 | 0 |
| 'Zfpl1' | 4.635 | 4.969 | 5.019 | 4.066 | 5.008 | 5.397 | 3.908 | 5.215 | 5.046 | 4.919 | 4.620 | 4.727 | 5.223 | 4.833 | 4.812 | 5.069 | 3.921 | 4.567 | 0.011 | 0.8563 | 0.9385 | 0 | 0 | 0 |
| 'Zfpm1' | 74.843 | 63.305 | 73.710 | 65.675 | 64.641 | 76.505 | 59.399 | 65.754 | 72.455 | 78.319 | 64.104 | 87.129 | 82.555 | 69.960 | 86.783 | 72.566 | 64.792 | 97.978 | -0.194 | 0.0000 | 0.0007 | 1 | 0 | 0 |
| 'Zfr' | 17.470 | 16.174 | 16.114 | 17.729 | 14.479 | 15.305 | 18.470 | 16.275 | 16.028 | 15.743 | 16.949 | 18.444 | 14.698 | 16.499 | 16.399 | 14.530 | 17.793 | 17.928 | -0.020 | 0.7101 | 0.8654 | 0 | 0 | 0 |
| 'Zfr2' | 1.736 | 1.768 | 2.094 | 1.479 | 1.532 | 1.770 | 1.956 | 1.640 | 2.113 | 1.746 | 1.501 | 1.646 | 1.397 | 1.407 | 1.579 | 1.177 | 1.658 | 1.531 | 0.225 | 0.0015 | 0.0146 | 1 | 0 | 0 |
| 'Zfyve1' | 12.650 | 14.255 | 14.290 | 13.734 | 14.296 | 14.933 | 12.577 | 13.642 | 14.741 | 13.333 | 13.369 | 15.025 | 13.978 | 14.739 | 14.788 | 13.049 | 14.105 | 14.867 | -0.038 | 0.2369 | 0.4866 | 0 | 0 | 0 |
| 'Zfyve16' | 2.556 | 2.773 | 2.993 | 2.947 | 2.628 | 2.550 | 2.990 | 2.764 | 2.757 | 2.529 | 2.862 | 2.956 | 2.315 | 2.719 | 2.458 | 2.348 | 2.989 | 2.721 | 0.051 | 0.3624 | 0.6146 | 0 | 0 | 0 |
| 'Zfyve19' | 3.869 | 3.502 | 3.904 | 3.448 | 3.429 | 4.443 | 3.505 | 3.453 | 4.032 | 3.516 | 2.956 | 4.075 | 3.744 | 3.894 | 3.679 | 3.818 | 3.386 | 3.815 | 0.021 | 0.6840 | 0.8502 | 0 | 0 | 0 |
| 'Zfyve21' | 3.895 | 5.569 | 4.097 | 4.607 | 3.615 | 4.925 | 3.393 | 4.363 | 3.884 | 4.010 | 3.125 | 4.680 | 3.124 | 5.385 | 5.131 | 3.684 | 4.682 | 4.506 | -0.011 | 0.9171 | 0.9686 | 0 | 0 | 0 |
| 'Zfyve26' | 7.824 | 7.194 | 9.200 | 7.919 | 7.329 | 8.295 | 7.477 | 7.349 | 9.427 | 8.778 | 9.362 | 9.224 | 8.260 | 7.844 | 8.640 | 8.162 | 9.084 | 9.168 | -0.137 | 0.0023 | 0.0206 | 1 | 0 | 0 |
| 'Zfyve27' | 5.377 | 5.362 | 6.123 | 5.060 | 5.561 | 5.803 | 5.535 | 5.823 | 5.896 | 5.426 | 5.699 | 6.184 | 5.596 | 5.714 | 6.209 | 5.881 | 5.673 | 6.014 | -0.060 | 0.0380 | 0.1583 | 0 | 0 | 0 |
| 'Zfyve28' | 3.206 | 4.979 | 3.599 | 3.524 | 4.296 | 3.597 | 3.679 | 4.777 | 3.914 | 6.028 | 6.353 | 4.671 | 6.550 | 6.155 | 4.452 | 6.544 | 6.105 | 4.356 | -0.529 | 0.0000 | 0.0000 | 1 | 1 | 0 |
| 'Zgpat' | 15.481 | 14.079 | 15.637 | 14.013 | 13.414 | 15.110 | 14.441 | 13.588 | 14.711 | 16.158 | 14.039 | 16.665 | 16.019 | 14.833 | 15.623 | 15.664 | 14.507 | 16.757 | -0.114 | 0.0000 | 0.0002 | 1 | 0 | 0 |
| 'Zgrf1' | 1.465 | 1.318 | 1.954 | 1.588 | 1.443 | 1.496 | 1.787 | 1.527 | 1.847 | 1.570 | 1.536 | 1.607 | 2.115 | 1.450 | 1.701 | 1.457 | 1.508 | 1.434 | -0.007 | 0.9243 | 0.9712 | 0 | 0 | 0 |
| 'Zhx1' | 4.305 | 5.065 | 4.410 | 4.752 | 4.804 | 4.217 | 4.914 | 4.782 | 4.484 | 4.808 | 4.711 | 4.560 | 4.420 | 4.467 | 4.653 | 4.829 | 4.540 | 4.463 | -0.005 | 0.8919 | 0.9560 | 0 | 0 | 0 |
| 'Zhx2' | 22.126 | 24.373 | 22.908 | 21.265 | 22.795 | 24.251 | 20.580 | 24.084 | 23.771 | 25.667 | 22.435 | 23.883 | 21.681 | 24.979 | 23.700 | 24.546 | 23.243 | 22.943 | -0.058 | 0.1203 | 0.3305 | 0 | 0 | 0 |
| 'Zhx3' | 2.680 | 2.289 | 2.212 | 2.723 | 2.067 | 1.925 | 2.247 | 2.384 | 2.183 | 2.542 | 2.711 | 2.675 | 2.471 | 2.339 | 2.617 | 2.431 | 2.517 | 2.660 | -0.164 | 0.0045 | 0.0349 | 1 | 0 | 0 |
| 'Zik1' | 1.395 | 1.794 | 1.688 | 1.701 | 1.658 | 1.399 | 2.373 | 1.618 | 1.646 | 1.396 | 1.632 | 2.091 | 1.495 | 1.435 | 1.519 | 1.387 | 2.358 | 1.879 | -0.014 | 0.9072 | 0.9636 | 0 | 0 | 0 |
| 'Zkscan1' | 1.980 | 1.915 | 1.950 | 2.125 | 1.887 | 1.898 | 2.217 | 2.095 | 1.918 | 1.999 | 2.021 | 2.174 | 1.798 | 1.853 | 2.067 | 1.791 | 1.773 | 2.161 | 0.014 | 0.7870 | 0.9048 | 0 | 0 | 0 |
| 'Zkscan14' | 3.265 | 3.278 | 3.069 | 2.958 | 2.726 | 3.324 | 3.202 | 3.325 | 2.637 | 3.135 | 2.882 | 3.835 | 2.953 | 3.407 | 3.909 | 3.399 | 3.703 | 3.149 | -0.140 | 0.0444 | 0.1748 | 0 | 0 | 0 |
| 'Zkscan17' | 15.870 | 14.583 | 16.831 | 15.086 | 15.282 | 16.738 | 13.946 | 14.890 | 15.085 | 15.540 | 14.350 | 16.732 | 16.009 | 13.761 | 15.785 | 15.237 | 14.499 | 15.514 | 0.000 | 0.9930 | 0.9986 | 0 | 0 | 0 |
| 'Zkscan3' | 8.298 | 7.838 | 8.319 | 8.670 | 7.690 | 8.219 | 8.198 | 7.361 | 8.400 | 6.868 | 8.202 | 10.918 | 6.656 | 8.489 | 9.882 | 7.199 | 9.161 | 10.651 | -0.091 | 0.1793 | 0.4146 | 0 | 0 | 0 |
| 'Zkscan5' | 1.592 | 1.875 | 1.737 | 1.548 | 1.626 | 1.715 | 1.591 | 1.878 | 1.664 | 1.821 | 1.817 | 1.709 | 1.804 | 1.817 | 1.828 | 1.622 | 1.694 | 1.570 | -0.049 | 0.4146 | 0.6626 | 0 | 0 | 0 |
| 'Zkscan6' | 7.519 | 7.001 | 7.978 | 7.511 | 7.362 | 7.293 | 8.163 | 8.104 | 7.216 | 8.169 | 7.443 | 7.395 | 6.513 | 7.090 | 7.369 | 7.264 | 6.492 | 7.211 | 0.062 | 0.2143 | 0.4597 | 0 | 0 | 0 |
| 'Zkscan7' | 5.312 | 7.926 | 6.294 | 5.023 | 7.667 | 7.242 | 5.953 | 6.924 | 7.023 | 6.260 | 6.151 | 6.526 | 6.126 | 6.567 | 6.913 | 6.674 | 6.009 | 6.498 | 0.021 | 0.7451 | 0.8850 | 0 | 0 | 0 |
| 'Zkscan8' | 3.246 | 4.130 | 4.455 | 3.283 | 4.248 | 3.976 | 3.539 | 4.260 | 4.434 | 3.704 | 3.659 | 4.071 | 3.638 | 4.020 | 3.587 | 3.485 | 3.977 | 3.662 | 0.059 | 0.1879 | 0.4241 | 0 | 0 | 0 |
| 'Zmat1' | 1.443 | 1.753 | 1.556 | 1.849 | 1.902 | 1.834 | 1.668 | 1.723 | 1.694 | 1.824 | 2.224 | 1.823 | 1.782 | 2.203 | 1.842 | 1.827 | 1.630 | 1.748 | -0.142 | 0.0285 | 0.1307 | 0 | 0 | 0 |
| 'Zmat2' | 56.565 | 60.608 | 57.362 | 58.299 | 62.588 | 61.508 | 59.951 | 62.595 | 58.355 | 59.926 | 58.539 | 53.600 | 59.975 | 60.769 | 60.339 | 58.876 | 59.750 | 52.755 | 0.028 | 0.3498 | 0.6044 | 0 | 0 | 0 |
| 'Zmat3' | 2.127 | 1.927 | 2.161 | 2.316 | 2.401 | 2.086 | 2.512 | 2.285 | 2.341 | 2.194 | 2.349 | 2.445 | 2.068 | 2.339 | 2.495 | 1.970 | 2.245 | 2.613 | -0.050 | 0.3723 | 0.6241 | 0 | 0 | 0 |
| 'Zmat5' | 11.063 | 11.965 | 10.815 | 9.904 | 12.529 | 11.645 | 10.221 | 11.576 | 9.992 | 10.271 | 10.594 | 9.778 | 11.913 | 10.153 | 10.960 | 11.180 | 9.087 | 8.874 | 0.095 | 0.1264 | 0.3405 | 0 | 0 | 0 |
| 'Zmiz1' | 15.757 | 13.758 | 15.848 | 14.204 | 14.042 | 15.054 | 13.644 | 14.453 | 15.702 | 16.497 | 15.159 | 15.719 | 15.868 | 15.260 | 15.281 | 15.209 | 13.400 | 15.984 | -0.073 | 0.0341 | 0.1472 | 0 | 0 | 0 |
| 'Zmiz2' | 39.106 | 36.381 | 42.289 | 36.657 | 36.747 | 41.868 | 37.510 | 36.750 | 42.216 | 41.411 | 37.446 | 41.441 | 40.782 | 36.476 | 41.456 | 38.786 | 36.631 | 42.439 | -0.040 | 0.0754 | 0.2441 | 0 | 0 | 0 |
| 'Zmpste24' | 9.596 | 10.313 | 9.831 | 9.891 | 10.440 | 9.247 | 9.477 | 10.562 | 9.334 | 10.134 | 10.296 | 9.665 | 9.250 | 10.014 | 9.283 | 9.413 | 10.292 | 9.884 | -0.004 | 0.9009 | 0.9607 | 0 | 0 | 0 |
| 'Zmym1' | 2.221 | 1.901 | 2.097 | 2.550 | 1.967 | 1.709 | 2.071 | 1.855 | 1.917 | 1.930 | 2.171 | 2.675 | 1.946 | 2.004 | 2.378 | 1.761 | 2.431 | 2.548 | -0.128 | 0.1257 | 0.3397 | 0 | 0 | 0 |
| 'Zmym2' | 20.047 | 21.019 | 18.716 | 22.204 | 19.661 | 18.263 | 21.371 | 20.098 | 17.379 | 17.314 | 21.312 | 20.452 | 17.924 | 20.322 | 18.381 | 18.284 | 22.128 | 19.016 | 0.018 | 0.7065 | 0.8639 | 0 | 0 | 0 |
| 'Zmym3' | 5.851 | 5.940 | 6.764 | 5.482 | 5.668 | 6.320 | 6.023 | 6.071 | 6.614 | 6.550 | 5.650 | 6.175 | 6.031 | 5.700 | 5.627 | 5.756 | 6.165 | 5.595 | 0.029 | 0.4571 | 0.6944 | 0 | 0 | 0 |
| 'Zmym4' | 7.418 | 8.137 | 7.587 | 7.704 | 7.621 | 7.199 | 7.981 | 7.967 | 7.517 | 7.733 | 8.180 | 7.201 | 6.938 | 7.546 | 6.959 | 6.996 | 7.489 | 6.791 | 0.061 | 0.0536 | 0.1977 | 0 | 0 | 0 |
| 'Zmym5' | 27.414 | 30.594 | 28.828 | 31.596 | 32.218 | 28.114 | 29.704 | 30.632 | 30.839 | 29.278 | 32.228 | 32.530 | 28.793 | 31.563 | 29.262 | 28.226 | 31.990 | 30.888 | -0.038 | 0.2434 | 0.4935 | 0 | 0 | 0 |
| 'Zmym6' | 2.535 | 2.792 | 2.526 | 2.685 | 2.910 | 2.332 | 2.582 | 2.906 | 2.737 | 2.584 | 2.790 | 3.110 | 2.567 | 2.847 | 2.607 | 2.570 | 3.231 | 2.737 | -0.079 | 0.0943 | 0.2847 | 0 | 0 | 0 |
| 'Zmynd11' | 8.145 | 8.893 | 8.659 | 7.983 | 8.677 | 8.522 | 8.335 | 8.773 | 8.581 | 8.599 | 8.166 | 8.197 | 8.232 | 7.835 | 7.664 | 8.654 | 7.824 | 8.157 | 0.052 | 0.0616 | 0.2148 | 0 | 0 | 0 |
| 'Zmynd19' | 8.709 | 7.947 | 8.713 | 7.475 | 7.577 | 8.387 | 8.715 | 8.389 | 7.109 | 8.296 | 7.677 | 8.696 | 9.670 | 8.297 | 8.829 | 9.230 | 8.284 | 8.820 | -0.100 | 0.0379 | 0.1583 | 0 | 0 | 0 |
| 'Zmynd8' | 11.726 | 10.225 | 11.414 | 10.842 | 10.919 | 10.966 | 10.543 | 11.049 | 11.215 | 11.103 | 11.079 | 11.347 | 9.595 | 10.329 | 10.509 | 10.334 | 10.293 | 11.073 | 0.038 | 0.2674 | 0.5209 | 0 | 0 | 0 |
| 'Znfx1' | 8.247 | 8.949 | 8.083 | 7.450 | 7.728 | 8.708 | 7.822 | 8.253 | 8.733 | 9.074 | 8.192 | 9.387 | 7.945 | 8.512 | 8.720 | 8.879 | 7.931 | 9.935 | -0.097 | 0.0189 | 0.0994 | 0 | 0 | 0 |
| 'Znhit1' | 13.423 | 14.142 | 13.700 | 13.018 | 14.756 | 15.941 | 13.870 | 15.001 | 14.258 | 13.676 | 13.662 | 14.880 | 14.632 | 13.714 | 15.317 | 14.726 | 12.589 | 13.285 | 0.008 | 0.8613 | 0.9401 | 0 | 0 | 0 |
| 'Znhit2' | 23.902 | 28.050 | 29.004 | 23.383 | 28.058 | 29.139 | 24.646 | 26.557 | 26.477 | 26.392 | 22.925 | 25.290 | 26.157 | 25.012 | 25.830 | 25.059 | 23.178 | 22.023 | 0.098 | 0.0459 | 0.1783 | 0 | 0 | 0 |
| 'Znhit3' | 2.802 | 3.033 | 3.170 | 2.406 | 2.569 | 2.800 | 2.527 | 2.835 | 2.535 | 2.431 | 2.484 | 2.761 | 2.947 | 2.381 | 2.270 | 2.765 | 2.675 | 2.138 | 0.102 | 0.1923 | 0.4294 | 0 | 0 | 0 |
| 'Znhit6' | 5.309 | 4.911 | 5.410 | 5.293 | 5.371 | 5.387 | 5.793 | 5.743 | 5.797 | 5.290 | 5.372 | 5.227 | 4.822 | 4.852 | 5.062 | 4.908 | 5.201 | 5.283 | 0.083 | 0.0843 | 0.2635 | 0 | 0 | 0 |
| 'Znrd1' | 14.918 | 14.758 | 14.487 | 15.133 | 14.751 | 14.819 | 15.510 | 14.218 | 14.827 | 13.941 | 13.031 | 13.761 | 13.586 | 13.349 | 14.381 | 12.786 | 15.938 | 12.958 | 0.097 | 0.0276 | 0.1281 | 0 | 0 | 0 |
| 'Znrd1as' | 1.168 | 1.521 | 1.981 | 1.623 | 1.854 | 1.314 | 1.718 | 1.271 | 1.552 | 1.741 | 1.377 | 1.582 | 1.846 | 1.061 | 1.659 | 1.776 | 1.510 | 1.171 | 0.019 | 0.8761 | 0.9474 | 0 | 0 | 0 |
| 'Znrf1' | 17.884 | 18.357 | 19.181 | 17.858 | 16.612 | 19.480 | 17.881 | 18.576 | 18.857 | 17.474 | 17.203 | 19.292 | 19.256 | 17.828 | 18.554 | 18.397 | 17.648 | 19.552 | -0.015 | 0.5374 | 0.7562 | 0 | 0 | 0 |
| 'Znrf3' | 1.215 | 1.206 | 0.927 | 1.083 | 1.042 | 1.014 | 1.135 | 1.356 | 1.006 | 1.292 | 0.997 | 1.032 | 1.179 | 0.904 | 0.915 | 1.320 | 1.006 | 0.915 | 0.055 | 0.4489 | 0.6890 | 0 | 0 | 0 |
| 'Zpbp' | 1.092 | 0.997 | 1.030 | 0.814 | 0.929 | 0.854 | 0.848 | 0.863 | 0.739 | 0.856 | 0.834 | 0.820 | 0.822 | 0.744 | 1.055 | 0.866 | 0.624 | 0.836 | 0.127 | 0.2727 | 0.5277 | 0 | 0 | 0 |
| 'Zpr1' | 10.719 | 9.108 | 10.659 | 9.809 | 9.894 | 10.303 | 9.666 | 9.727 | 10.980 | 10.888 | 10.432 | 10.883 | 10.162 | 9.794 | 10.463 | 10.181 | 9.545 | 9.839 | -0.028 | 0.4344 | 0.6788 | 0 | 0 | 0 |
| 'Zranb1' | 14.193 | 14.222 | 13.565 | 13.481 | 13.141 | 13.514 | 14.231 | 13.761 | 12.482 | 14.688 | 14.171 | 13.344 | 13.825 | 13.508 | 12.255 | 14.252 | 14.117 | 12.936 | -0.013 | 0.7047 | 0.8623 | 0 | 0 | 0 |
| 'Zranb2' | 4.701 | 5.548 | 5.315 | 5.199 | 5.292 | 4.846 | 5.248 | 5.316 | 5.027 | 4.917 | 4.757 | 5.156 | 4.678 | 4.905 | 5.049 | 4.533 | 5.181 | 4.722 | 0.068 | 0.0398 | 0.1628 | 0 | 0 | 0 |
| 'Zranb3' | 1.206 | 1.203 | 1.261 | 1.287 | 1.034 | 1.341 | 1.156 | 1.095 | 1.207 | 1.116 | 1.403 | 1.286 | 1.429 | 1.218 | 1.290 | 0.978 | 1.172 | 1.242 | -0.057 | 0.4039 | 0.6528 | 0 | 0 | 0 |
| 'Zrsr1' | 6.727 | 5.805 | 4.633 | 5.468 | 5.745 | 5.806 | 5.834 | 6.435 | 4.284 | 7.382 | 5.514 | 5.370 | 6.763 | 5.751 | 5.195 | 8.702 | 4.882 | 5.361 | -0.111 | 0.1505 | 0.3777 | 0 | 0 | 0 |
| 'Zrsr2' | 7.978 | 8.376 | 8.012 | 7.576 | 8.200 | 8.586 | 7.830 | 8.490 | 8.415 | 8.081 | 7.958 | 8.489 | 8.337 | 8.392 | 8.073 | 8.782 | 7.860 | 7.781 | -0.015 | 0.6215 | 0.8108 | 0 | 0 | 0 |
| 'Zscan12' | 2.153 | 2.564 | 2.844 | 2.296 | 2.637 | 3.038 | 2.760 | 2.443 | 2.332 | 2.226 | 2.631 | 2.183 | 2.634 | 2.292 | 2.176 | 2.134 | 2.329 | 2.114 | 0.151 | 0.0865 | 0.2679 | 0 | 0 | 0 |
| 'Zscan2' | 9.666 | 8.892 | 9.832 | 9.170 | 8.364 | 10.339 | 9.039 | 8.133 | 9.325 | 7.771 | 8.199 | 10.403 | 7.934 | 8.567 | 9.468 | 8.344 | 7.805 | 9.786 | 0.070 | 0.0997 | 0.2954 | 0 | 0 | 0 |
| 'Zscan20' | 1.792 | 1.378 | 1.556 | 1.991 | 1.447 | 1.585 | 1.650 | 1.529 | 1.552 | 1.538 | 1.577 | 1.795 | 1.512 | 1.539 | 1.524 | 1.510 | 1.809 | 1.818 | -0.030 | 0.6495 | 0.8292 | 0 | 0 | 0 |
| 'Zscan21' | 1.688 | 1.539 | 1.788 | 1.966 | 1.535 | 1.859 | 1.998 | 1.501 | 2.083 | 1.569 | 1.864 | 1.900 | 1.623 | 1.619 | 1.731 | 1.939 | 1.729 | 1.752 | 0.009 | 0.8941 | 0.9566 | 0 | 0 | 0 |
| 'Zscan22' | 1.398 | 1.423 | 1.521 | 1.351 | 1.206 | 1.605 | 1.153 | 1.455 | 1.369 | 1.311 | 1.184 | 1.579 | 1.292 | 1.333 | 1.531 | 1.313 | 1.210 | 1.531 | 0.013 | 0.8328 | 0.9270 | 0 | 0 | 0 |
| 'Zscan26' | 9.092 | 10.042 | 10.803 | 8.922 | 10.663 | 10.208 | 10.401 | 11.351 | 10.922 | 9.858 | 10.729 | 11.152 | 9.290 | 10.401 | 10.915 | 10.003 | 10.531 | 9.702 | -0.014 | 0.6926 | 0.8555 | 0 | 0 | 0 |
| 'Zscan29' | 18.603 | 17.360 | 17.563 | 17.715 | 18.111 | 18.319 | 16.973 | 17.572 | 18.580 | 18.383 | 20.347 | 20.784 | 17.483 | 19.431 | 20.232 | 18.293 | 20.379 | 21.535 | -0.147 | 0.0000 | 0.0008 | 1 | 0 | 0 |
| 'Zswim1' | 5.283 | 5.251 | 5.188 | 4.822 | 4.580 | 5.103 | 4.935 | 4.897 | 5.546 | 5.303 | 4.561 | 4.725 | 4.619 | 4.526 | 4.959 | 5.063 | 4.292 | 5.421 | 0.065 | 0.2405 | 0.4911 | 0 | 0 | 0 |
| 'Zswim3' | 1.949 | 2.144 | 2.367 | 2.446 | 2.411 | 2.319 | 2.603 | 1.891 | 2.806 | 2.039 | 2.994 | 3.168 | 2.156 | 1.628 | 2.715 | 1.641 | 2.033 | 2.816 | -0.033 | 0.7662 | 0.8956 | 0 | 0 | 0 |
| 'Zswim4' | 20.383 | 20.683 | 23.687 | 18.048 | 20.877 | 24.223 | 18.598 | 20.526 | 23.306 | 24.958 | 20.122 | 21.841 | 22.559 | 21.672 | 20.879 | 22.646 | 20.042 | 18.940 | -0.036 | 0.5167 | 0.7404 | 0 | 0 | 0 |
| 'Zswim6' | 20.385 | 19.244 | 18.070 | 20.141 | 20.908 | 19.732 | 17.659 | 20.947 | 18.179 | 22.527 | 22.715 | 19.254 | 19.423 | 21.444 | 18.941 | 21.205 | 22.548 | 20.537 | -0.114 | 0.0039 | 0.0306 | 1 | 0 | 0 |
| 'Zswim7' | 6.258 | 8.253 | 7.119 | 7.541 | 7.253 | 7.161 | 6.699 | 8.102 | 6.693 | 6.992 | 7.314 | 6.480 | 7.082 | 7.429 | 6.828 | 9.419 | 6.855 | 5.638 | 0.025 | 0.7941 | 0.9080 | 0 | 0 | 0 |
| 'Zswim8' | 50.930 | 50.456 | 56.760 | 47.888 | 48.287 | 53.761 | 48.071 | 51.979 | 57.048 | 56.754 | 53.875 | 52.147 | 55.353 | 50.250 | 51.436 | 52.512 | 52.974 | 50.952 | -0.045 | 0.2114 | 0.4566 | 0 | 0 | 0 |
| 'Zufsp' | 14.972 | 14.362 | 15.234 | 15.533 | 15.656 | 12.394 | 16.829 | 15.835 | 15.080 | 14.444 | 14.170 | 14.778 | 14.165 | 13.879 | 14.479 | 14.547 | 16.401 | 14.113 | 0.040 | 0.4003 | 0.6491 | 0 | 0 | 0 |
| 'Zw10' | 23.836 | 21.301 | 24.732 | 26.007 | 22.352 | 23.777 | 23.346 | 22.422 | 23.801 | 29.521 | 30.676 | 33.318 | 31.498 | 29.445 | 31.292 | 28.421 | 33.681 | 33.222 | -0.421 | 0.0000 | 0.0000 | 1 | 1 | 0 |
| 'Zwilch' | 1.731 | 1.827 | 2.533 | 2.250 | 1.936 | 2.146 | 2.526 | 2.067 | 2.348 | 1.794 | 2.106 | 2.097 | 2.627 | 1.966 | 2.669 | 2.086 | 2.589 | 1.878 | -0.045 | 0.6128 | 0.8058 | 0 | 0 | 0 |
| 'Zwint' | 20.051 | 21.147 | 20.012 | 20.107 | 20.996 | 19.072 | 19.689 | 20.603 | 20.025 | 21.546 | 20.781 | 19.939 | 21.019 | 21.776 | 19.844 | 21.150 | 21.054 | 20.989 | -0.060 | 0.0154 | 0.0855 | 0 | 0 | 0 |
| 'Zxdb' | 8.468 | 8.183 | 8.582 | 7.870 | 8.018 | 8.045 | 7.875 | 8.370 | 8.041 | 8.355 | 7.762 | 9.579 | 7.561 | 8.197 | 8.918 | 8.899 | 7.897 | 8.501 | -0.054 | 0.2004 | 0.4416 | 0 | 0 | 0 |
| 'Zxdc' | 3.600 | 3.456 | 3.882 | 3.345 | 3.340 | 3.908 | 3.546 | 3.451 | 3.737 | 3.833 | 4.102 | 3.817 | 3.485 | 3.851 | 3.762 | 3.618 | 3.554 | 3.738 | -0.071 | 0.0707 | 0.2346 | 0 | 0 | 0 |
| 'Zyg11b' | 11.050 | 11.511 | 10.686 | 9.934 | 11.632 | 11.096 | 10.388 | 11.831 | 10.777 | 12.717 | 11.660 | 11.410 | 11.203 | 11.695 | 11.284 | 12.854 | 11.269 | 11.555 | -0.105 | 0.0044 | 0.0339 | 1 | 0 | 0 |
| 'Zyx' | 11.651 | 11.587 | 13.194 | 11.333 | 11.096 | 14.847 | 12.342 | 12.331 | 14.578 | 14.861 | 11.758 | 13.904 | 14.662 | 11.878 | 14.005 | 12.850 | 12.257 | 12.889 | -0.088 | 0.0499 | 0.1886 | 0 | 0 | 0 |
| 'Zzef1' | 9.786 | 9.012 | 10.196 | 9.931 | 8.852 | 9.532 | 9.335 | 9.389 | 10.446 | 11.238 | 11.104 | 9.956 | 10.664 | 10.149 | 9.791 | 10.573 | 10.250 | 10.196 | -0.129 | 0.0003 | 0.0044 | 1 | 0 | 0 |
| 'Zzz3' | 5.276 | 5.371 | 4.998 | 5.249 | 4.843 | 4.641 | 4.877 | 5.180 | 5.008 | 5.377 | 5.514 | 5.125 | 5.152 | 5.328 | 4.773 | 5.206 | 5.225 | 4.724 | -0.040 | 0.1986 | 0.4384 | 0 | 0 | 0 |
| 'gene:MGP_BALBcJ | 456.646 | 456.021 | 480.917 | 384.854 | 448.375 | 452.417 | 391.146 | 467.250 | 503.792 | 426.521 | 447.792 | 531.417 | 427.479 | 419.479 | 502.625 | 413.375 | 469.000 | 473.438 | -0.038 | 0.4435 | 0.6869 | 0 | 0 | 0 |
| 'gene:MGP_BALBcJ | 5.804 | 5.724 | 4.972 | 5.232 | 5.906 | 6.358 | 6.165 | 5.346 | 4.642 | 5.543 | 5.598 | 5.381 | 6.023 | 5.843 | 6.341 | 5.949 | 6.520 | 5.620 | -0.084 | 0.3379 | 0.5955 | 0 | 0 | 0 |
| 'gene:MGP_BALBcJ | 733.045 | 846.359 | 655.833 | 892.071 | 825.732 | 699.091 | 888.222 | 820.697 | 681.394 | 693.525 | 861.268 | 582.207 | 773.641 | 819.485 | 689.323 | 815.237 | 850.106 | 674.667 | 0.050 | 0.2883 | 0.5447 | 0 | 0 | 0 |
| 'gene:MGP_BALBcJ | 58.388 | 60.467 | 51.101 | 65.667 | 61.138 | 53.465 | 68.798 | 59.500 | 54.395 | 54.044 | 60.135 | 46.627 | 63.616 | 59.885 | 54.271 | 55.764 | 57.376 | 49.283 | 0.078 | 0.0845 | 0.2638 | 0 | 0 | 0 |
| 'gene:MGP_BALBcJ | 16.376 | 17.526 | 14.331 | 17.915 | 15.875 | 14.386 | 17.523 | 17.555 | 15.421 | 13.341 | 15.928 | 14.968 | 18.573 | 16.881 | 15.616 | 15.984 | 16.819 | 14.203 | 0.028 | 0.6387 | 0.8221 | 0 | 0 | 0 |
| 'gene:MGP_BALBcJ | 10.172 | 9.363 | 8.197 | 11.313 | 10.508 | 8.854 | 12.749 | 8.190 | 7.269 | 7.794 | 8.685 | 6.797 | 8.586 | 10.763 | 7.168 | 9.975 | 10.461 | 7.653 | 0.146 | 0.1556 | 0.3843 | 0 | 0 | 0 |
| 'gene:MGP_BALBcJ | 13.317 | 16.648 | 11.360 | 10.910 | 14.808 | 12.392 | 13.142 | 15.047 | 8.765 | 12.137 | 10.529 | 11.735 | 13.959 | 14.741 | 10.855 | 14.253 | 18.235 | 11.846 | -0.044 | 0.6817 | 0.8485 | 0 | 0 | 0 |
| 'gene:MGP_BALBcJ | 8.463 | 8.341 | 7.512 | 9.414 | 9.953 | 10.067 | 10.632 | 10.395 | 8.714 | 8.616 | 10.072 | 7.501 | 9.243 | 7.944 | 8.798 | 8.420 | 10.047 | 6.306 | 0.116 | 0.2456 | 0.4960 | 0 | 0 | 0 |
| 'gene:MGP_BALBcJ | 60.897 | 26.947 | 30.884 | 67.444 | 31.073 | 29.327 | 63.704 | 33.430 | 26.918 | 31.524 | 51.450 | 45.360 | 36.824 | 39.871 | 39.634 | 29.223 | 61.223 | 47.240 | -0.144 | 0.5132 | 0.7375 | 0 | 0 | 0 |
| 'gene:MGP_BALBcJ | 54.605 | 51.257 | 51.327 | 63.293 | 56.793 | 52.035 | 62.072 | 65.955 | 49.933 | 49.612 | 53.930 | 48.400 | 61.938 | 57.932 | 52.562 | 53.755 | 57.268 | 48.063 | 0.059 | 0.2581 | 0.5107 | 0 | 0 | 0 |
| 'gene:MGP_BALBcJ | 135.306 | 134.126 | 118.429 | 148.679 | 145.472 | 127.134 | 160.616 | 141.457 | 133.232 | 139.283 | 140.960 | 114.556 | 139.874 | 142.518 | 127.364 | 135.035 | 149.429 | 119.391 | 0.034 | 0.4112 | 0.6598 | 0 | 0 | 0 |
| 'gene:MGP_BALBcJ | 28.275 | 25.124 | 26.593 | 31.781 | 29.200 | 28.919 | 31.981 | 33.529 | 24.267 | 20.733 | 26.514 | 28.306 | 26.946 | 26.116 | 28.366 | 36.564 | 26.640 | 24.178 | 0.076 | 0.3626 | 0.6147 | 0 | 0 | 0 |
| 'gene:MGP_BALBcJ | 7.935 | 8.140 | 7.367 | 9.315 | 9.325 | 7.422 | 9.437 | 9.591 | 8.094 | 7.141 | 7.990 | 9.206 | 10.081 | 10.390 | 7.270 | 8.910 | 9.060 | 8.399 | -0.060 | 0.4787 | 0.7113 | 0 | 0 | 0 |
| 'gene:MGP_BALBcJ | 41.122 | 44.932 | 41.163 | 43.998 | 43.662 | 43.434 | 49.115 | 49.075 | 35.747 | 41.110 | 43.111 | 41.855 | 49.084 | 46.176 | 43.418 | 47.727 | 43.795 | 42.582 | -0.037 | 0.4709 | 0.7067 | 0 | 0 | 0 |
| 'gene:MGP_BALBcJ | 10.233 | 13.243 | 13.859 | 10.540 | 14.862 | 12.522 | 9.995 | 11.859 | 13.332 | 9.473 | 15.102 | 11.872 | 7.675 | 11.841 | 11.606 | 11.363 | 10.696 | 8.018 | 0.176 | 0.1063 | 0.3065 | 0 | 0 | 0 |
| 'gene:MGP_BALBcJ | 18.586 | 19.255 | 16.511 | 23.049 | 18.000 | 19.518 | 18.386 | 20.637 | 20.132 | 19.119 | 17.447 | 16.590 | 18.390 | 17.901 | 18.623 | 21.328 | 18.637 | 15.978 | 0.083 | 0.2409 | 0.4914 | 0 | 0 | 0 |
| 'gene:MGP_BALBcJ | 6.909 | 6.861 | 6.142 | 8.022 | 6.456 | 6.425 | 8.548 | 7.312 | 5.891 | 5.378 | 7.683 | 6.811 | 7.470 | 6.798 | 6.771 | 6.506 | 7.951 | 6.886 | -0.016 | 0.8698 | 0.9442 | 0 | 0 | 0 |
| 'gene:MGP_BALBcJ | 88.253 | 100.022 | 80.863 | 101.745 | 96.321 | 82.618 | 112.046 | 90.861 | 81.329 | 88.408 | 98.805 | 72.825 | 99.249 | 93.542 | 83.076 | 100.265 | 96.078 | 82.110 | 0.024 | 0.6049 | 0.8011 | 0 | 0 | 0 |
| 'gene:MGP_BALBcJ | 3.513 | 4.120 | 4.798 | 4.518 | 3.998 | 3.629 | 4.519 | 4.006 | 3.856 | 3.700 | 3.943 | 4.891 | 5.158 | 3.384 | 3.758 | 4.084 | 3.884 | 3.756 | -0.011 | 0.9146 | 0.9671 | 0 | 0 | 0 |
| 'gene:MGP_BALBcJ | 10.788 | 13.502 | 12.602 | 15.747 | 12.512 | 12.056 | 14.791 | 13.793 | 13.730 | 10.337 | 13.549 | 11.793 | 8.884 | 8.886 | 10.953 | 9.800 | 13.616 | 12.028 | 0.243 | 0.0194 | 0.1008 | 0 | 0 | 0 |
| 'gene:MGP_BALBcJ | 11.552 | 10.811 | 11.488 | 12.923 | 11.354 | 11.294 | 13.154 | 12.183 | 9.397 | 10.446 | 12.336 | 11.264 | 10.785 | 10.762 | 10.584 | 13.513 | 12.239 | 10.407 | 0.016 | 0.8018 | 0.9114 | 0 | 0 | 0 |
| 'gene:MGP_BALBcJ | 9.664 | 16.110 | 10.857 | 11.827 | 13.024 | 10.628 | 13.455 | 12.193 | 8.786 | 10.622 | 11.247 | 9.161 | 10.720 | 10.280 | 10.429 | 9.577 | 9.542 | 9.795 | 0.192 | 0.0863 | 0.2676 | 0 | 0 | 0 |
| 'gene:MGP_BALBcJ | 5.266 | 4.970 | 5.722 | 5.906 | 5.833 | 4.744 | 5.907 | 5.693 | 5.173 | 4.551 | 5.487 | 5.062 | 6.454 | 5.664 | 4.974 | 4.369 | 5.373 | 4.524 | 0.068 | 0.4783 | 0.7112 | 0 | 0 | 0 |
| 'gene:MGP_BALBcJ | 9.607 | 8.639 | 8.530 | 9.127 | 10.319 | 8.172 | 11.176 | 10.376 | 7.228 | 7.149 | 8.357 | 8.535 | 10.232 | 8.311 | 8.662 | 8.437 | 9.196 | 8.500 | 0.080 | 0.3202 | 0.5784 | 0 | 0 | 0 |
| 'gene:MGP_BALBcJ | 6.258 | 8.700 | 7.326 | 7.943 | 7.032 | 7.529 | 8.573 | 7.953 | 7.214 | 6.801 | 8.731 | 7.145 | 7.672 | 7.713 | 7.508 | 6.903 | 7.798 | 6.621 | 0.022 | 0.7827 | 0.9025 | 0 | 0 | 0 |
| 'gene:MGP_BALBcJ | 6.075 | 8.095 | 6.948 | 6.892 | 6.719 | 6.583 | 7.663 | 7.103 | 5.583 | 6.200 | 7.124 | 6.687 | 6.583 | 7.932 | 6.656 | 6.324 | 6.638 | 6.302 | 0.010 | 0.9193 | 0.9695 | 0 | 0 | 0 |
| 'gene:MGP_BALBcJ | 16.139 | 21.455 | 17.539 | 19.173 | 22.288 | 14.261 | 23.683 | 20.937 | 13.212 | 13.621 | 20.128 | 14.534 | 16.778 | 20.453 | 14.408 | 13.761 | 20.751 | 14.625 | 0.160 | 0.0479 | 0.1832 | 0 | 0 | 0 |
| 'gene:MGP_BALBcJ | 4.616 | 5.230 | 4.355 | 5.859 | 6.781 | 4.585 | 5.795 | 6.223 | 4.375 | 4.355 | 4.700 | 4.544 | 4.833 | 4.387 | 4.859 | 5.717 | 5.687 | 4.211 | 0.118 | 0.2271 | 0.4734 | 0 | 0 | 0 |
| 'gene:MGP_BALBcJ | 7.589 | 8.503 | 7.459 | 8.278 | 6.675 | 7.775 | 7.130 | 7.056 | 5.395 | 5.716 | 6.601 | 5.805 | 7.310 | 7.439 | 6.637 | 7.598 | 7.127 | 6.517 | 0.113 | 0.2084 | 0.4530 | 0 | 0 | 0 |
| 'gene:MGP_BALBcJ | 34.524 | 35.546 | 29.919 | 36.018 | 34.675 | 33.596 | 37.658 | 33.567 | 32.198 | 29.697 | 32.573 | 30.143 | 34.912 | 32.850 | 32.345 | 30.922 | 30.664 | 30.576 | 0.096 | 0.0272 | 0.1270 | 0 | 0 | 0 |
| 'gene:MGP_BALBcJ | 16.368 | 21.089 | 12.887 | 14.245 | 18.513 | 18.272 | 16.685 | 21.304 | 14.688 | 16.656 | 20.105 | 17.226 | 20.685 | 22.718 | 19.301 | 19.976 | 19.860 | 16.430 | -0.182 | 0.0361 | 0.1527 | 0 | 0 | 0 |
| 'gene:MGP_BALBcJ | 14.148 | 16.304 | 16.952 | 18.398 | 15.774 | 17.696 | 17.310 | 16.238 | 11.919 | 13.795 | 17.075 | 14.895 | 13.642 | 15.274 | 13.322 | 16.383 | 14.066 | 15.892 | 0.098 | 0.3155 | 0.5733 | 0 | 0 | 0 |
| 'gene:MGP_BALBcJ | 11.987 | 11.479 | 11.401 | 14.423 | 11.170 | 11.675 | 11.726 | 14.619 | 10.201 | 10.125 | 13.315 | 11.379 | 12.380 | 10.062 | 11.258 | 13.266 | 11.133 | 11.921 | 0.042 | 0.6074 | 0.8026 | 0 | 0 | 0 |
| 'gene:MGP_BALBcJ | 3.341 | 3.605 | 4.181 | 3.060 | 3.139 | 4.244 | 2.403 | 4.443 | 2.565 | 3.193 | 4.365 | 4.177 | 3.384 | 3.806 | 4.190 | 4.023 | 3.457 | 2.858 | -0.113 | 0.3412 | 0.5981 | 0 | 0 | 0 |
| 'gene:MGP_BALBcJ | 84.439 | 96.742 | 79.797 | 106.797 | 100.148 | 83.734 | 104.479 | 93.910 | 75.875 | 80.679 | 92.544 | 72.404 | 89.722 | 83.431 | 81.486 | 102.145 | 85.419 | 74.895 | 0.102 | 0.0512 | 0.1916 | 0 | 0 | 0 |
| 'gene:MGP_BALBcJ | 7.726 | 8.311 | 7.193 | 10.278 | 9.096 | 6.565 | 8.241 | 10.039 | 8.770 | 9.512 | 6.823 | 7.493 | 9.286 | 6.133 | 6.915 | 8.854 | 9.063 | 7.043 | 0.083 | 0.4106 | 0.6595 | 0 | 0 | 0 |
| 'gene:MGP_BALBcJ | 18.972 | 18.559 | 13.928 | 19.558 | 17.522 | 15.131 | 18.127 | 15.819 | 13.507 | 17.276 | 19.834 | 19.184 | 22.800 | 18.287 | 19.942 | 15.886 | 20.877 | 19.822 | -0.232 | 0.0038 | 0.0306 | 1 | 0 | 0 |
| 'gene:MGP_BALBcJ | 7.608 | 9.598 | 7.761 | 9.135 | 9.285 | 6.741 | 8.424 | 9.465 | 8.686 | 7.881 | 7.957 | 7.516 | 8.805 | 6.775 | 8.658 | 7.472 | 8.280 | 7.410 | 0.093 | 0.3490 | 0.6042 | 0 | 0 | 0 |
| 'gene:MGP_BALBcJ | 17.472 | 16.658 | 14.773 | 18.858 | 16.425 | 16.741 | 19.362 | 17.460 | 15.051 | 15.179 | 19.240 | 14.276 | 16.442 | 17.696 | 16.832 | 18.993 | 15.047 | 14.811 | 0.036 | 0.5796 | 0.7822 | 0 | 0 | 0 |
| 'gene:MGP_BALBcJ | 18.504 | 19.631 | 20.035 | 20.854 | 19.702 | 18.481 | 21.780 | 20.494 | 19.167 | 17.555 | 20.317 | 17.223 | 20.459 | 19.861 | 18.630 | 19.570 | 20.842 | 16.351 | 0.060 | 0.2314 | 0.4788 | 0 | 0 | 0 |
| 'gene:MGP_BALBcJ | 23.269 | 18.716 | 22.720 | 28.602 | 20.967 | 21.202 | 25.302 | 22.656 | 16.670 | 20.215 | 25.000 | 17.896 | 23.191 | 22.406 | 22.052 | 23.117 | 22.791 | 21.376 | 0.007 | 0.9321 | 0.9746 | 0 | 0 | 0 |
| 'gene:MGP_BALBcJ | 16.657 | 19.879 | 18.217 | 21.740 | 19.539 | 16.157 | 22.740 | 21.989 | 18.383 | 18.068 | 19.203 | 18.051 | 19.299 | 21.210 | 17.046 | 17.439 | 19.360 | 17.485 | 0.050 | 0.4300 | 0.6758 | 0 | 0 | 0 |
| 'gene:MGP_BALBcJ | 7.055 | 6.404 | 6.830 | 4.622 | 8.221 | 7.189 | 5.714 | 6.348 | 8.283 | 6.978 | 7.169 | 12.695 | 5.654 | 5.884 | 11.543 | 8.539 | 8.206 | 13.163 | -0.374 | 0.0011 | 0.0117 | 1 | 0 | 0 |
| 'gene:MGP_BALBcJ | 10.689 | 12.184 | 8.158 | 10.779 | 13.134 | 11.128 | 11.321 | 11.505 | 9.676 | 10.629 | 10.736 | 10.831 | 10.886 | 14.660 | 10.880 | 9.383 | 14.345 | 10.158 | -0.076 | 0.3371 | 0.5955 | 0 | 0 | 0 |
| 'gene:MGP_BALBcJ | 4.774 | 5.823 | 4.747 | 5.264 | 5.567 | 4.314 | 4.757 | 5.493 | 4.959 | 5.499 | 4.941 | 5.312 | 4.603 | 5.553 | 4.613 | 4.601 | 4.703 | 5.252 | -0.006 | 0.9403 | 0.9769 | 0 | 0 | 0 |
| 'gene:MGP_BALBcJ | 36.689 | 37.329 | 34.972 | 39.932 | 37.247 | 40.870 | 42.856 | 44.733 | 31.457 | 34.853 | 39.301 | 37.614 | 37.626 | 36.681 | 36.538 | 46.554 | 41.073 | 32.531 | 0.005 | 0.9361 | 0.9756 | 0 | 0 | 0 |
| 'gene:MGP_BALBcJ | 30.188 | 30.688 | 30.500 | 32.468 | 36.924 | 30.835 | 28.624 | 35.997 | 29.926 | 33.674 | 32.650 | 25.453 | 36.274 | 27.882 | 32.100 | 35.826 | 34.438 | 27.812 | 0.003 | 0.9713 | 0.9898 | 0 | 0 | 0 |
| 'gene:MGP_BALBcJ | 195.764 | 186.010 | 178.135 | 192.935 | 193.283 | 190.566 | 199.739 | 190.659 | 246.519 | 178.238 | 186.664 | 160.233 | 240.441 | 189.885 | 183.273 | 187.396 | 186.559 | 170.213 | 0.069 | 0.2936 | 0.5497 | 0 | 0 | 0 |
| 'gene:MGP_BALBcJ | 8.071 | 11.287 | 7.740 | 8.091 | 10.992 | 8.894 | 10.844 | 9.522 | 8.228 | 6.853 | 7.551 | 10.004 | 8.808 | 9.196 | 8.196 | 11.379 | 8.396 | 7.943 | 0.066 | 0.5069 | 0.7329 | 0 | 0 | 0 |
| 'gene:MGP_BALBcJ | 12.355 | 12.438 | 11.301 | 14.841 | 14.330 | 13.150 | 13.703 | 13.128 | 11.231 | 11.231 | 17.274 | 9.967 | 14.741 | 12.535 | 13.587 | 14.817 | 13.657 | 13.649 | -0.068 | 0.4047 | 0.6539 | 0 | 0 | 0 |
| 'gene:MGP_BALBcJ | 34.822 | 46.912 | 40.157 | 38.314 | 43.265 | 32.216 | 44.832 | 43.214 | 39.175 | 36.451 | 39.570 | 33.814 | 45.711 | 46.214 | 38.492 | 36.528 | 44.907 | 32.853 | 0.023 | 0.7371 | 0.8802 | 0 | 0 | 0 |
| 'gene:MGP_BALBcJ | 29.671 | 28.484 | 28.102 | 33.907 | 34.728 | 30.575 | 29.936 | 32.966 | 31.206 | 27.881 | 31.592 | 28.552 | 33.601 | 27.149 | 31.832 | 33.180 | 25.747 | 26.510 | 0.062 | 0.3374 | 0.5955 | 0 | 0 | 0 |
| 'gene:MGP_BALBcJ | 7.887 | 7.000 | 6.011 | 6.807 | 7.338 | 5.954 | 7.338 | 8.976 | 5.810 | 6.889 | 5.911 | 5.051 | 8.669 | 7.436 | 5.250 | 10.003 | 5.840 | 4.092 | 0.113 | 0.2652 | 0.5185 | 0 | 0 | 0 |
| 'gene:MGP_BALBcJ | 25.536 | 33.212 | 24.586 | 24.693 | 28.113 | 26.423 | 26.113 | 29.183 | 24.942 | 24.891 | 24.868 | 25.528 | 25.033 | 31.398 | 22.210 | 27.412 | 26.490 | 23.691 | 0.054 | 0.3178 | 0.5762 | 0 | 0 | 0 |
| 'gene:MGP_BALBcJ | 97.781 | 89.790 | 83.397 | 105.492 | 97.165 | 82.959 | 106.729 | 104.786 | 83.226 | 81.027 | 87.440 | 79.962 | 98.178 | 87.255 | 87.786 | 96.325 | 102.266 | 83.140 | 0.069 | 0.1533 | 0.3806 | 0 | 0 | 0 |
| 'gene:MGP_BALBcJ | 9.018 | 8.484 | 7.728 | 7.580 | 9.235 | 7.238 | 9.312 | 9.439 | 8.992 | 7.144 | 8.271 | 7.925 | 8.318 | 7.568 | 7.217 | 7.338 | 7.824 | 7.102 | 0.145 | 0.0581 | 0.2076 | 0 | 0 | 0 |
| 'gene:MGP_BALBcJ | 15.834 | 16.961 | 14.822 | 17.957 | 16.779 | 15.001 | 18.698 | 17.517 | 15.008 | 13.537 | 17.418 | 12.503 | 16.005 | 16.351 | 14.088 | 17.336 | 18.416 | 13.209 | 0.098 | 0.1310 | 0.3476 | 0 | 0 | 0 |
| 'gene:MGP_BALBcJ | 31.340 | 28.945 | 29.343 | 38.586 | 32.968 | 32.065 | 41.168 | 34.896 | 24.595 | 25.935 | 31.848 | 27.926 | 31.084 | 33.770 | 28.628 | 37.932 | 34.058 | 28.065 | 0.063 | 0.4394 | 0.6829 | 0 | 0 | 0 |
| 'gene:MGP_BALBcJ | 7.239 | 7.856 | 6.291 | 6.058 | 6.102 | 7.111 | 7.254 | 7.840 | 6.979 | 8.129 | 7.668 | 7.129 | 6.962 | 8.165 | 6.540 | 7.672 | 7.075 | 6.593 | -0.076 | 0.4109 | 0.6596 | 0 | 0 | 0 |
| 'gene:MGP_BALBcJ | 11.702 | 13.943 | 13.027 | 15.738 | 13.181 | 13.012 | 15.235 | 16.078 | 10.973 | 10.548 | 12.331 | 12.464 | 14.136 | 14.389 | 11.419 | 16.614 | 13.226 | 11.328 | 0.066 | 0.5285 | 0.7494 | 0 | 0 | 0 |
| 'gene:MGP_BALBcJ | 111.661 | 112.312 | 119.881 | 134.926 | 114.655 | 126.179 | 130.000 | 134.946 | 117.208 | 94.604 | 109.896 | 112.110 | 115.640 | 115.696 | 114.375 | 142.277 | 109.940 | 100.286 | 0.111 | 0.0679 | 0.2284 | 0 | 0 | 0 |
| 'gene:MGP_BALBcJ | 35.172 | 32.008 | 31.669 | 40.540 | 39.171 | 31.661 | 38.780 | 35.900 | 32.546 | 34.326 | 36.315 | 29.628 | 35.874 | 36.476 | 32.477 | 35.893 | 37.295 | 32.130 | 0.024 | 0.6103 | 0.8042 | 0 | 0 | 0 |
| 'gene:MGP_BALBcJ | 71.277 | 72.384 | 60.173 | 67.682 | 76.223 | 67.033 | 79.369 | 78.625 | 61.875 | 81.976 | 77.089 | 67.732 | 78.771 | 88.140 | 60.179 | 76.155 | 77.580 | 57.839 | -0.075 | 0.1228 | 0.3344 | 0 | 0 | 0 |
| 'gene:MGP_BALBcJ | 9.626 | 10.461 | 9.075 | 11.716 | 12.103 | 8.856 | 11.468 | 9.536 | 8.460 | 10.995 | 10.176 | 9.748 | 9.503 | 9.799 | 9.144 | 11.778 | 11.569 | 11.230 | -0.063 | 0.4508 | 0.6904 | 0 | 0 | 0 |
| 'gene:MGP_BALBcJ | 17.357 | 20.690 | 17.704 | 16.859 | 21.060 | 19.294 | 19.029 | 19.874 | 17.410 | 18.084 | 17.125 | 16.124 | 18.698 | 20.087 | 18.577 | 17.180 | 17.038 | 16.039 | 0.080 | 0.1401 | 0.3623 | 0 | 0 | 0 |
| 'gene:MGP_BALBcJ | 33.421 | 42.526 | 31.978 | 46.084 | 37.324 | 33.558 | 46.551 | 42.829 | 28.567 | 31.470 | 38.262 | 37.726 | 34.000 | 38.231 | 40.442 | 37.947 | 42.558 | 31.084 | 0.021 | 0.8019 | 0.9114 | 0 | 0 | 0 |
| 'gene:MGP_BALBcJ | 9.140 | 11.162 | 8.053 | 9.341 | 9.839 | 8.195 | 8.863 | 9.216 | 9.258 | 7.582 | 8.864 | 8.315 | 9.886 | 10.229 | 9.407 | 8.782 | 8.204 | 7.539 | 0.057 | 0.4556 | 0.6936 | 0 | 0 | 0 |
| 'gene:MGP_BALBcJ | 6.405 | 5.531 | 5.143 | 6.227 | 6.322 | 6.769 | 6.214 | 5.575 | 4.871 | 4.511 | 5.088 | 4.180 | 7.054 | 5.699 | 5.670 | 6.932 | 5.651 | 5.525 | 0.080 | 0.4639 | 0.6999 | 0 | 0 | 0 |
| 'gene:MGP_BALBcJ | 23.011 | 19.481 | 18.965 | 24.449 | 20.091 | 22.139 | 22.763 | 21.822 | 19.726 | 19.853 | 23.833 | 17.334 | 20.913 | 21.995 | 22.078 | 21.975 | 21.652 | 20.790 | 0.012 | 0.8553 | 0.9380 | 0 | 0 | 0 |
| 'gene:MGP_BALBcJ | 11.772 | 11.545 | 7.612 | 11.462 | 7.706 | 10.766 | 6.914 | 9.613 | 14.640 | 8.029 | 8.795 | 9.648 | 8.910 | 8.919 | 10.489 | 7.242 | 10.631 | 13.748 | 0.087 | 0.5002 | 0.7275 | 0 | 0 | 0 |
| 'gene:MGP_BALBcJ | 28.674 | 31.037 | 31.685 | 27.766 | 36.264 | 26.165 | 35.648 | 36.930 | 24.388 | 29.601 | 34.319 | 24.399 | 32.183 | 34.993 | 24.619 | 27.780 | 34.978 | 24.114 | 0.062 | 0.4246 | 0.6712 | 0 | 0 | 0 |
| 'gene:MGP_BALBcJ | 7.507 | 9.504 | 8.726 | 11.013 | 10.990 | 8.347 | 12.651 | 10.437 | 8.430 | 8.078 | 10.567 | 9.918 | 9.593 | 11.171 | 8.119 | 12.192 | 11.423 | 7.322 | -0.028 | 0.7684 | 0.8967 | 0 | 0 | 0 |
| 'gene:MGP_BALBcJ | 15.633 | 18.445 | 15.517 | 14.072 | 18.442 | 13.903 | 15.135 | 22.480 | 12.799 | 14.357 | 14.069 | 14.868 | 14.542 | 16.125 | 14.226 | 14.890 | 13.984 | 12.282 | 0.154 | 0.1180 | 0.3269 | 0 | 0 | 0 |
| 'gene:MGP_BALBcJ | 64.480 | 70.062 | 62.090 | 70.667 | 75.129 | 57.174 | 80.062 | 71.348 | 65.779 | 60.308 | 69.134 | 59.246 | 66.109 | 68.915 | 64.445 | 73.371 | 72.993 | 55.490 | 0.053 | 0.2978 | 0.5536 | 0 | 0 | 0 |
| 'gene:MGP_BALBcJ | 16.356 | 16.897 | 14.964 | 17.362 | 17.659 | 16.690 | 16.897 | 15.208 | 15.138 | 15.195 | 16.351 | 12.421 | 16.231 | 16.392 | 15.172 | 17.090 | 15.549 | 15.808 | 0.072 | 0.4074 | 0.6564 | 0 | 0 | 0 |
| 'gene:MGP_BALBcJ | 64.585 | 66.226 | 57.622 | 70.163 | 66.843 | 59.766 | 73.579 | 66.488 | 57.165 | 56.237 | 68.788 | 53.876 | 68.602 | 66.547 | 62.018 | 67.894 | 65.019 | 58.245 | 0.027 | 0.5379 | 0.7566 | 0 | 0 | 0 |
| 'gene:MGP_BALBcJ | 70.866 | 71.001 | 61.887 | 79.627 | 73.955 | 69.233 | 78.815 | 70.563 | 67.882 | 65.347 | 73.214 | 54.925 | 74.273 | 73.923 | 66.790 | 67.602 | 71.188 | 62.293 | 0.071 | 0.1305 | 0.3469 | 0 | 0 | 0 |
| 'gene:MGP_BALBcJ | 46.195 | 58.861 | 53.961 | 53.109 | 50.636 | 44.183 | 65.662 | 50.348 | 47.433 | 49.994 | 48.265 | 49.962 | 50.584 | 53.464 | 43.721 | 46.132 | 51.809 | 45.744 | 0.083 | 0.1570 | 0.3866 | 0 | 0 | 0 |
| 'gene:MGP_BALBcJ | 23.964 | 30.020 | 24.724 | 36.423 | 30.745 | 21.158 | 37.536 | 30.531 | 23.071 | 17.867 | 30.531 | 22.913 | 29.235 | 25.495 | 25.791 | 20.327 | 35.204 | 24.520 | 0.128 | 0.2819 | 0.5382 | 0 | 0 | 0 |
| 'gene:MGP_BALBcJ | 102.444 | 128.578 | 96.978 | 139.833 | 128.333 | 104.967 | 120.483 | 139.278 | 99.450 | 111.122 | 111.100 | 88.028 | 113.083 | 122.067 | 111.056 | 129.383 | 134.750 | 96.650 | 0.050 | 0.4315 | 0.6763 | 0 | 0 | 0 |
| 'gene:MGP_BALBcJ | 53.399 | 59.039 | 58.601 | 57.490 | 61.837 | 52.463 | 64.246 | 59.451 | 58.791 | 56.916 | 51.958 | 55.089 | 55.123 | 57.899 | 56.766 | 52.749 | 58.316 | 53.210 | 0.062 | 0.1381 | 0.3585 | 0 | 0 | 0 |
| 'gene:MGP_BALBcJ | 218.761 | 234.961 | 203.694 | 246.562 | 243.177 | 214.985 | 253.178 | 242.545 | 211.182 | 213.194 | 246.262 | 186.638 | 222.875 | 237.174 | 213.553 | 235.748 | 247.862 | 212.332 | 0.028 | 0.4544 | 0.6931 | 0 | 0 | 0 |
| 'gene:MGP_BALBcJ | 173.572 | 197.589 | 178.960 | 203.441 | 215.943 | 200.350 | 192.737 | 203.131 | 168.128 | 185.485 | 186.886 | 158.687 | 175.842 | 172.953 | 186.256 | 172.825 | 191.727 | 161.125 | 0.113 | 0.0124 | 0.0730 | 0 | 0 | 0 |
| 'gene:MGP_BALBcJ | 160.834 | 144.574 | 162.444 | 154.289 | 144.233 | 149.623 | 170.451 | 147.727 | 158.671 | 143.444 | 144.587 | 144.929 | 148.612 | 135.881 | 154.629 | 131.698 | 147.579 | 143.694 | 0.094 | 0.0110 | 0.0671 | 0 | 0 | 0 |

| | | | | | | | | | | | | | | | | | | | | | | | | |
|---|---|---|---|---|---|---|---|---|---|---|---|---|---|---|---|---|---|---|---|---|---|---|---|---|
| 'gene:MGP_BALBcJ | 6.610 | 6.808 | 6.479 | 7.341 | 6.598 | 8.207 | 8.238 | 8.869 | 6.591 | 6.019 | 8.057 | 6.961 | 7.281 | 7.900 | 6.995 | 8.137 | 6.320 | 6.903 | 0.022 | 0.8196 | 0.9202 | 0 | 0 | 0 |
| 'gene:MGP_BALBcJ | 30.464 | 39.797 | 28.319 | 39.199 | 48.866 | 29.217 | 41.649 | 38.482 | 32.772 | 32.696 | 33.946 | 30.257 | 28.076 | 32.750 | 28.098 | 28.591 | 36.616 | 29.529 | 0.196 | 0.0142 | 0.0806 | 0 | 0 | 0 |
| 'gene:MGP_BALBcJ | 155.670 | 171.932 | 142.834 | 179.828 | 173.016 | 149.893 | 181.542 | 157.250 | 153.411 | 144.446 | 162.617 | 123.372 | 166.041 | 164.688 | 154.380 | 158.996 | 158.134 | 140.764 | 0.083 | 0.0684 | 0.2294 | 0 | 0 | 0 |
| 'gene:MGP_BALBcJ | 12.321 | 12.607 | 13.425 | 16.323 | 16.292 | 11.398 | 15.875 | 15.537 | 11.928 | 13.236 | 13.384 | 13.503 | 14.131 | 12.785 | 13.045 | 15.057 | 16.105 | 12.418 | 0.004 | 0.9588 | 0.9847 | 0 | 0 | 0 |
| 'gene:MGP_BALBcJ | 28.563 | 22.956 | 23.159 | 32.711 | 20.989 | 23.041 | 29.800 | 20.767 | 21.170 | 17.711 | 28.867 | 29.530 | 23.244 | 25.041 | 31.489 | 19.578 | 27.622 | 30.767 | -0.086 | 0.4832 | 0.7148 | 0 | 0 | 0 |
| 'gene:MGP_BALBcJ | 9.969 | 11.677 | 11.143 | 9.721 | 12.935 | 11.472 | 12.210 | 14.694 | 11.851 | 11.577 | 10.233 | 12.906 | 10.639 | 13.094 | 11.681 | 11.564 | 14.319 | 11.719 | -0.046 | 0.6072 | 0.8026 | 0 | 0 | 0 |
| 'gene:MGP_BALBcJ | 49.899 | 41.905 | 42.214 | 50.009 | 51.516 | 41.878 | 58.211 | 56.795 | 39.700 | 45.362 | 47.893 | 50.685 | 46.961 | 47.101 | 38.864 | 45.012 | 59.976 | 45.421 | -0.004 | 0.9545 | 0.9832 | 0 | 0 | 0 |
| 'gene:MGP_BALBcJ | 18.919 | 15.893 | 17.159 | 17.757 | 14.261 | 15.952 | 14.593 | 15.790 | 15.679 | 13.388 | 17.383 | 22.665 | 15.278 | 16.225 | 21.967 | 13.602 | 14.648 | 25.963 | -0.127 | 0.1794 | 0.4147 | 0 | 0 | 0 |
| 'gene:MGP_BALBcJ | 105.651 | 132.417 | 111.844 | 121.344 | 135.256 | 99.751 | 131.393 | 127.949 | 105.093 | 106.768 | 112.144 | 98.090 | 111.005 | 125.463 | 97.515 | 110.056 | 130.727 | 94.420 | 0.106 | 0.0091 | 0.0583 | 0 | 0 | 0 |
| 'gene:MGP_BALBcJ | 57.556 | 62.185 | 58.281 | 67.052 | 65.896 | 66.133 | 84.007 | 61.504 | 59.089 | 55.874 | 63.570 | 53.267 | 63.867 | 64.778 | 68.081 | 57.881 | 77.444 | 51.081 | 0.057 | 0.5128 | 0.7375 | 0 | 0 | 0 |
| 'gene:MGP_BALBcJ | 45.956 | 45.249 | 46.811 | 47.639 | 41.674 | 45.651 | 44.517 | 45.189 | 40.135 | 35.071 | 45.573 | 40.882 | 38.494 | 42.689 | 43.857 | 44.504 | 46.851 | 38.261 | 0.095 | 0.0993 | 0.2945 | 0 | 0 | 0 |
| 'gene:MGP_BALBcJ | 50.525 | 64.810 | 51.111 | 58.621 | 59.762 | 53.094 | 66.903 | 56.727 | 55.492 | 47.670 | 58.329 | 50.637 | 55.235 | 63.901 | 50.351 | 53.294 | 66.424 | 48.699 | 0.054 | 0.3158 | 0.5735 | 0 | 0 | 0 |
| 'gene:MGP_BALBcJ | 24.130 | 28.388 | 21.340 | 27.235 | 27.138 | 23.021 | 28.232 | 25.449 | 25.724 | 21.266 | 23.567 | 18.923 | 26.298 | 24.173 | 25.793 | 25.704 | 23.615 | 22.512 | 0.111 | 0.1075 | 0.3084 | 0 | 0 | 0 |
| 'gene:MGP_BALBcJ | 7.382 | 8.017 | 7.943 | 7.273 | 8.052 | 8.008 | 8.087 | 9.482 | 7.049 | 6.769 | 6.858 | 6.846 | 8.318 | 8.124 | 6.062 | 7.442 | 8.173 | 5.425 | 0.151 | 0.1292 | 0.3445 | 0 | 0 | 0 |
| 'gene:MGP_BALBcJ | 21.031 | 22.560 | 16.604 | 23.642 | 25.277 | 16.415 | 24.816 | 23.047 | 18.389 | 20.238 | 23.865 | 17.321 | 22.337 | 22.464 | 18.453 | 19.847 | 25.640 | 18.407 | 0.005 | 0.9514 | 0.9816 | 0 | 0 | 0 |
| 'gene:MGP_BALBcJ | 36.865 | 38.499 | 36.732 | 34.378 | 35.931 | 38.759 | 44.583 | 35.368 | 37.496 | 40.543 | 41.290 | 42.589 | 39.111 | 39.035 | 43.978 | 41.728 | 46.661 | 40.172 | -0.160 | 0.0023 | 0.0202 | 1 | 0 | 0 |
| 'gene:MGP_BALBcJ | 25.177 | 26.049 | 23.853 | 33.833 | 30.585 | 27.613 | 34.175 | 30.189 | 23.075 | 23.022 | 32.018 | 24.387 | 27.660 | 31.906 | 25.845 | 27.994 | 33.821 | 20.937 | 0.034 | 0.6610 | 0.8362 | 0 | 0 | 0 |
| 'gene:MGP_BALBcJ | 7.016 | 7.656 | 7.119 | 6.895 | 7.169 | 7.869 | 7.777 | 8.207 | 7.992 | 6.729 | 8.482 | 8.252 | 6.689 | 6.769 | 7.399 | 6.518 | 7.998 | 6.933 | 0.026 | 0.7110 | 0.8658 | 0 | 0 | 0 |
| 'gene:MGP_BALBcJ | 33.254 | 39.932 | 30.258 | 36.401 | 33.168 | 33.240 | 34.333 | 34.975 | 32.867 | 35.968 | 29.910 | 35.538 | 35.989 | 34.767 | 37.677 | 36.247 | 34.724 | 30.333 | -0.034 | 0.6303 | 0.8172 | 0 | 0 | 0 |
| 'gene:MGP_BALBcJ | 29.319 | 34.379 | 29.441 | 38.039 | 36.618 | 33.193 | 38.623 | 29.510 | 34.782 | 28.971 | 34.626 | 26.931 | 34.950 | 37.746 | 36.698 | 31.480 | 32.332 | 31.446 | 0.031 | 0.6782 | 0.8461 | 0 | 0 | 0 |
| 'gene:MGP_BALBcJ | 11.783 | 9.315 | 11.541 | 14.619 | 15.233 | 10.783 | 17.671 | 13.909 | 8.605 | 9.071 | 13.121 | 11.694 | 11.961 | 12.751 | 10.493 | 14.342 | 10.342 | 10.139 | 0.102 | 0.3793 | 0.6308 | 0 | 0 | 0 |
| 'gene:MGP_BALBcJ | 11.174 | 10.537 | 8.977 | 10.745 | 9.426 | 8.577 | 10.180 | 10.704 | 9.261 | 7.964 | 11.149 | 10.312 | 10.277 | 8.797 | 10.734 | 11.407 | 9.565 | 11.669 | -0.054 | 0.5492 | 0.7629 | 0 | 0 | 0 |
| 'gene:MGP_BALBcJ | 32.730 | 28.156 | 34.528 | 35.445 | 39.172 | 29.684 | 40.902 | 37.377 | 31.405 | 31.607 | 34.776 | 34.748 | 36.660 | 33.586 | 35.770 | 29.610 | 38.911 | 29.166 | 0.005 | 0.9498 | 0.9809 | 0 | 0 | 0 |
| 'gene:MGP_BALBcJ | 36.373 | 42.462 | 36.047 | 46.161 | 41.140 | 37.165 | 46.419 | 45.606 | 32.642 | 34.036 | 40.634 | 38.885 | 39.118 | 34.505 | 39.943 | 46.953 | 39.470 | 32.208 | 0.054 | 0.4744 | 0.7095 | 0 | 0 | 0 |
| 'gene:MGP_BALBcJ | 13.666 | 17.615 | 15.599 | 16.056 | 18.849 | 12.750 | 19.890 | 16.826 | 14.403 | 15.764 | 14.167 | 14.704 | 14.599 | 17.787 | 15.732 | 16.104 | 17.535 | 14.142 | 0.031 | 0.6858 | 0.8513 | 0 | 0 | 0 |
| 'gene:MGP_BALBcJ | 31.409 | 38.932 | 28.227 | 43.769 | 35.598 | 39.928 | 43.542 | 35.739 | 26.886 | 28.061 | 34.295 | 30.583 | 36.587 | 26.167 | 35.030 | 32.212 | 34.583 | 26.121 | 0.170 | 0.0798 | 0.2542 | 0 | 0 | 0 |
| 'gene:MGP_BALBcJ | 28.089 | 31.057 | 27.682 | 31.812 | 34.750 | 26.099 | 30.734 | 30.328 | 23.880 | 22.797 | 27.062 | 25.099 | 32.120 | 24.495 | 20.943 | 30.724 | 36.661 | 30.745 | 0.058 | 0.5658 | 0.7742 | 0 | 0 | 0 |
| 'gene:MGP_BALBcJ | 103.185 | 95.431 | 103.452 | 107.765 | 104.143 | 101.895 | 101.422 | 97.975 | 107.015 | 100.595 | 95.767 | 94.189 | 101.805 | 104.981 | 106.561 | 99.464 | 102.351 | 98.008 | 0.023 | 0.5454 | 0.7606 | 0 | 0 | 0 |
| 'gene:MGP_BALBcJ | 12.507 | 17.435 | 14.134 | 17.317 | 17.114 | 15.059 | 17.781 | 20.791 | 12.317 | 12.765 | 18.526 | 15.007 | 14.088 | 17.771 | 15.206 | 19.529 | 17.310 | 14.170 | -0.013 | 0.8988 | 0.9598 | 0 | 0 | 0 |
| 'gene:MGP_BALBcJ | 15.047 | 14.152 | 16.161 | 15.370 | 13.620 | 14.946 | 19.158 | 14.332 | 15.902 | 15.133 | 11.212 | 13.253 | 17.788 | 14.418 | 14.544 | 16.728 | 13.896 | 13.060 | 0.091 | 0.3457 | 0.6009 | 0 | 0 | 0 |
| 'gene:MGP_BALBcJ | 14.696 | 14.509 | 13.205 | 18.484 | 20.146 | 14.861 | 18.895 | 22.034 | 11.329 | 11.685 | 17.436 | 16.244 | 15.324 | 18.623 | 18.361 | 17.142 | 17.345 | 15.386 | -0.020 | 0.8538 | 0.9373 | 0 | 0 | 0 |
| 'gene:MGP_BALBcJ | 6.121 | 5.719 | 5.047 | 5.285 | 6.089 | 5.516 | 6.921 | 6.511 | 4.443 | 4.763 | 5.311 | 5.601 | 5.701 | 5.917 | 5.876 | 6.086 | 4.627 | 5.287 | 0.050 | 0.6103 | 0.8042 | 0 | 0 | 0 |
| 'gene:MGP_BALBcJ | 125.779 | 129.221 | 107.669 | 129.875 | 133.456 | 131.721 | 138.051 | 132.404 | 108.993 | 102.507 | 114.044 | 118.360 | 124.794 | 111.853 | 134.743 | 149.279 | 127.096 | 107.176 | 0.048 | 0.4853 | 0.7166 | 0 | 0 | 0 |
| 'gene:MGP_BALBcJ | 41.006 | 41.020 | 34.764 | 44.053 | 42.737 | 37.524 | 47.730 | 43.286 | 38.943 | 38.291 | 41.614 | 37.557 | 44.124 | 39.263 | 39.390 | 42.026 | 45.403 | 40.973 | -0.007 | 0.8871 | 0.9534 | 0 | 0 | 0 |
| 'gene:MGP_BALBcJ | 56.920 | 67.185 | 55.659 | 68.649 | 66.852 | 51.768 | 70.563 | 62.999 | 56.911 | 57.610 | 62.709 | 43.839 | 60.099 | 61.576 | 53.159 | 65.464 | 61.547 | 52.242 | 0.098 | 0.0444 | 0.1747 | 0 | 0 | 0 |
| 'gene:MGP_BALBcJ | 83.847 | 105.491 | 87.435 | 82.547 | 99.854 | 95.615 | 92.164 | 85.730 | 90.396 | 88.126 | 89.547 | 88.191 | 106.806 | 92.189 | 88.883 | 99.041 | 87.122 | 87.894 | -0.021 | 0.6892 | 0.8534 | 0 | 0 | 0 |
| 'gene:MGP_BALBcJ | 62.980 | 76.894 | 46.589 | 66.285 | 80.291 | 54.934 | 78.662 | 75.695 | 54.490 | 58.424 | 64.132 | 52.132 | 69.748 | 71.536 | 56.305 | 70.523 | 55.848 | 51.205 | 0.092 | 0.2245 | 0.4713 | 0 | 0 | 0 |
| 'gene:MGP_BALBcJ | 8.548 | 8.374 | 8.144 | 8.136 | 8.113 | 6.881 | 8.764 | 10.001 | 7.343 | 8.130 | 8.003 | 8.527 | 7.419 | 8.310 | 8.394 | 7.496 | 8.325 | 8.629 | 0.002 | 0.9792 | 0.9926 | 0 | 0 | 0 |
| 'gene:MGP_BALBcJ | 7.346 | 8.697 | 6.955 | 8.991 | 10.550 | 7.038 | 8.670 | 10.369 | 7.247 | 5.790 | 6.769 | 8.790 | 6.421 | 7.482 | 6.629 | 9.360 | 8.357 | 8.156 | 0.125 | 0.3102 | 0.5686 | 0 | 0 | 0 |
| 'gene:MGP_BALBcJ | 12.109 | 12.387 | 12.121 | 13.869 | 12.789 | 12.127 | 14.963 | 14.829 | 11.730 | 13.720 | 14.168 | 12.080 | 12.138 | 12.477 | 13.166 | 14.698 | 10.886 | 11.177 | 0.024 | 0.7662 | 0.8956 | 0 | 0 | 0 |
| 'gene:MGP_BALBcJ | 6.271 | 8.188 | 6.328 | 6.423 | 6.943 | 7.214 | 5.962 | 7.161 | 6.197 | 8.691 | 6.423 | 6.463 | 6.937 | 9.065 | 6.323 | 6.833 | 8.596 | 6.146 | -0.111 | 0.3126 | 0.5712 | 0 | 0 | 0 |
| 'gene:MGP_BALBcJ | 10.252 | 11.371 | 9.968 | 14.719 | 12.061 | 9.684 | 10.884 | 12.814 | 9.284 | 9.565 | 14.377 | 10.603 | 11.704 | 11.290 | 9.490 | 15.099 | 12.122 | 8.933 | -0.039 | 0.7280 | 0.8752 | 0 | 0 | 0 |
| 'gene:MGP_BALBcJ | 12.275 | 12.027 | 11.285 | 12.713 | 11.345 | 12.054 | 12.866 | 14.431 | 12.837 | 10.815 | 11.109 | 11.725 | 12.878 | 10.908 | 11.460 | 12.674 | 10.345 | 12.457 | 0.087 | 0.3519 | 0.6056 | 0 | 0 | 0 |
| 'gene:MGP_BALBcJ | 12.290 | 12.270 | 10.615 | 11.811 | 10.185 | 11.142 | 10.564 | 13.424 | 7.626 | 9.977 | 10.531 | 10.383 | 12.463 | 10.434 | 8.274 | 12.138 | 9.609 | 10.856 | 0.064 | 0.5214 | 0.7441 | 0 | 0 | 0 |
| 'gene:MGP_BALBcJ | 30.006 | 32.101 | 30.883 | 34.974 | 30.407 | 29.699 | 36.699 | 33.932 | 29.503 | 28.793 | 29.037 | 31.712 | 28.538 | 32.226 | 29.196 | 32.752 | 33.506 | 27.319 | 0.062 | 0.2798 | 0.5366 | 0 | 0 | 0 |
| 'gene:MGP_BALBcJ | 10.538 | 12.585 | 12.326 | 15.092 | 13.984 | 10.994 | 12.915 | 13.738 | 11.605 | 11.281 | 10.071 | 12.756 | 12.853 | 11.639 | 11.664 | 13.066 | 12.595 | 11.085 | 0.066 | 0.3616 | 0.6135 | 0 | 0 | 0 |
| 'gene:MGP_BALBcJ | 15.508 | 15.205 | 13.128 | 16.260 | 19.744 | 15.407 | 19.306 | 24.450 | 12.903 | 13.050 | 13.120 | 16.822 | 13.748 | 11.391 | 10.911 | 18.411 | 11.078 | 13.163 | 0.286 | 0.0353 | 0.1504 | 0 | 0 | 0 |
| 'gene:MGP_BALBcJ | 14.819 | 15.913 | 13.740 | 14.331 | 15.280 | 15.773 | 15.077 | 15.966 | 13.377 | 14.753 | 15.809 | 15.507 | 13.615 | 15.949 | 13.633 | 15.229 | 18.519 | 13.140 | -0.031 | 0.6918 | 0.8552 | 0 | 0 | 0 |
| 'gene:MGP_BALBcJ | 80.276 | 83.546 | 74.825 | 88.659 | 76.514 | 79.551 | 78.799 | 75.354 | 77.963 | 79.784 | 84.626 | 65.616 | 80.903 | 76.666 | 78.950 | 80.689 | 81.180 | 73.191 | 0.023 | 0.6216 | 0.8108 | 0 | 0 | 0 |
| 'gene:MGP_BALBcJ | 14.053 | 13.549 | 13.499 | 10.045 | 13.390 | 12.195 | 13.019 | 17.270 | 12.248 | 12.758 | 12.939 | 14.125 | 16.568 | 11.669 | 11.841 | 17.284 | 14.950 | 12.660 | -0.072 | 0.4950 | 0.7242 | 0 | 0 | 0 |
| 'gene:MGP_BALBcJ | 32.941 | 39.779 | 37.676 | 36.972 | 34.016 | 36.779 | 35.077 | 39.486 | 34.944 | 32.092 | 35.115 | 32.397 | 34.331 | 35.535 | 36.138 | 37.047 | 37.958 | 33.601 | 0.057 | 0.3133 | 0.5715 | 0 | 0 | 0 |
| 'gene:MGP_BALBcJ | 4.876 | 4.162 | 3.494 | 4.420 | 4.452 | 3.114 | 4.833 | 4.209 | 4.653 | 2.090 | 2.522 | 2.879 | 2.294 | 2.575 | 2.760 | 3.681 | 2.512 | 2.720 | 0.645 | 0.0000 | 0.0000 | 1 | 1 | 0 |
| 'gene:MGP_BALBcJ | 121.497 | 148.746 | 117.912 | 136.030 | 154.993 | 128.108 | 142.989 | 149.287 | 127.393 | 134.097 | 138.023 | 123.748 | 130.123 | 143.991 | 128.261 | 125.522 | 154.025 | 124.018 | 0.015 | 0.6546 | 0.8326 | 0 | 0 | 0 |
| 'gene:MGP_BALBcJ | 47.615 | 49.236 | 47.779 | 51.222 | 51.964 | 46.567 | 50.038 | 44.006 | 46.847 | 41.013 | 49.794 | 41.039 | 49.184 | 51.218 | 53.233 | 45.307 | 48.305 | 49.995 | 0.011 | 0.8469 | 0.9343 | 0 | 0 | 0 |
| 'gene:MGP_BALBcJ | 33.288 | 40.680 | 31.844 | 43.029 | 44.032 | 41.907 | 52.913 | 50.206 | 34.063 | 33.495 | 44.989 | 42.651 | 39.701 | 42.770 | 40.423 | 52.897 | 40.749 | 39.114 | -0.038 | 0.6753 | 0.8443 | 0 | 0 | 0 |
| 'gene:MGP_BALBcJ | 100.132 | 99.591 | 95.612 | 101.600 | 94.561 | 94.558 | 103.166 | 106.269 | 94.502 | 96.457 | 101.320 | 89.419 | 100.379 | 98.094 | 94.734 | 95.559 | 96.962 | 94.784 | 0.028 | 0.3222 | 0.5799 | 0 | 0 | 0 |
| 'gene:MGP_BALBcJ | 15.682 | 17.802 | 15.330 | 15.049 | 15.900 | 15.806 | 16.579 | 16.146 | 15.661 | 14.215 | 12.360 | 14.546 | 15.046 | 15.220 | 15.763 | 15.059 | 13.884 | 14.335 | 0.127 | 0.0431 | 0.1717 | 0 | 0 | 0 |
| 'gene:MGP_BALBcJ | 3.612 | 4.418 | 4.046 | 3.454 | 4.835 | 4.669 | 3.400 | 5.244 | 4.020 | 3.965 | 3.090 | 3.724 | 3.771 | 4.013 | 3.725 | 4.903 | 3.806 | 2.497 | 0.166 | 0.1440 | 0.3681 | 0 | 0 | 0 |
| 'gene:MGP_BALBcJ | 104.209 | 108.086 | 94.936 | 116.742 | 117.194 | 100.009 | 118.790 | 104.608 | 97.012 | 105.688 | 124.531 | 82.176 | 101.698 | 109.276 | 102.842 | 116.858 | 112.759 | 96.533 | 0.007 | 0.8959 | 0.9582 | 0 | 0 | 0 |
| 'gene:MGP_BALBcJ | 9.641 | 9.015 | 8.535 | 12.237 | 11.927 | 9.858 | 12.178 | 11.205 | 8.753 | 8.562 | 11.550 | 8.697 | 11.609 | 9.523 | 12.078 | 7.680 | 10.565 | 8.557 | 0.052 | 0.6491 | 0.8290 | 0 | 0 | 0 |
| 'gene:MGP_BALBcJ | 170.907 | 179.760 | 155.315 | 191.288 | 178.503 | 161.544 | 198.326 | 169.295 | 154.890 | 160.191 | 181.703 | 134.444 | 166.287 | 170.619 | 169.623 | 169.265 | 186.570 | 155.350 | 0.051 | 0.2804 | 0.5374 | 0 | 0 | 0 |
| 'gene:MGP_BALBcJ | 8.947 | 6.724 | 7.048 | 6.309 | 7.379 | 8.010 | 8.029 | 7.291 | 8.015 | 3.977 | 6.149 | 9.131 | 7.276 | 7.419 | 6.128 | 7.150 | 6.240 | 6.966 | 0.142 | 0.2591 | 0.5123 | 0 | 0 | 0 |
| 'gene:MGP_BALBcJ | 37.736 | 14.982 | 18.329 | 41.050 | 15.141 | 19.683 | 36.198 | 18.286 | 14.992 | 19.121 | 27.693 | 24.470 | 22.897 | 21.234 | 22.515 | 15.015 | 33.098 | 28.877 | -0.107 | 0.6407 | 0.8235 | 0 | 0 | 0 |
| 'gene:MGP_BALBcJ | 3.535 | 2.664 | 2.775 | 3.237 | 3.985 | 3.333 | 2.424 | 3.660 | 2.569 | 1.559 | 1.640 | 1.772 | 1.179 | 1.985 | 2.172 | 1.537 | 2.528 | 1.774 | 0.801 | 0.0000 | 0.0000 | 1 | 1 | 0 |
| 'gene:MGP_BALBcJ | 325.755 | 338.819 | 276.127 | 368.343 | 329.162 | 311.167 | 377.897 | 354.672 | 276.485 | 307.348 | 360.078 | 246.985 | 322.343 | 354.824 | 316.784 | 352.255 | 357.029 | 289.892 | 0.016 | 0.7536 | 0.8901 | 0 | 0 | 0 |
| 'gene:MGP_BALBcJ | 4.430 | 4.281 | 4.550 | 5.549 | 4.839 | 5.486 | 4.485 | 5.081 | 5.048 | 3.474 | 4.765 | 4.830 | 4.223 | 4.156 | 3.989 | 5.852 | 4.426 | 4.613 | 0.111 | 0.2814 | 0.5379 | 0 | 0 | 0 |
| 'gene:MGP_BALBcJ | 29.470 | 33.439 | 29.424 | 36.330 | 36.851 | 28.015 | 37.687 | 35.648 | 29.060 | 31.281 | 29.829 | 29.505 | 30.018 | 34.534 | 28.670 | 29.164 | 34.024 | 27.882 | 0.085 | 0.1291 | 0.3443 | 0 | 0 | 0 |
| 'gene:MGP_BALBcJ | 37.487 | 44.220 | 37.582 | 52.038 | 49.131 | 40.612 | 50.313 | 51.192 | 39.590 | 34.285 | 43.899 | 35.220 | 41.673 | 48.693 | 41.661 | 36.834 | 49.141 | 39.156 | 0.106 | 0.1238 | 0.3363 | 0 | 0 | 0 |
| 'gene:MGP_BALBcJ | 8.688 | 10.389 | 7.444 | 11.282 | 8.875 | 9.132 | 13.216 | 9.862 | 7.770 | 6.805 | 7.819 | 9.230 | 8.556 | 8.578 | 9.307 | 12.545 | 8.378 | 8.282 | 0.096 | 0.3955 | 0.6444 | 0 | 0 | 0 |
| 'gene:MGP_BALBcJ | 19.661 | 20.421 | 21.771 | 29.490 | 29.236 | 16.767 | 27.031 | 25.664 | 18.284 | 17.990 | 23.188 | 22.291 | 21.120 | 22.651 | 22.623 | 20.990 | 25.062 | 21.113 | 0.051 | 0.6155 | 0.8081 | 0 | 0 | 0 |
| 'gene:MGP_BALBcJ | 19.586 | 19.483 | 20.877 | 20.957 | 23.517 | 19.838 | 24.361 | 23.030 | 21.421 | 18.732 | 22.682 | 21.974 | 20.238 | 23.361 | 22.444 | 21.815 | 19.387 | 21.970 | -0.010 | 0.9003 | 0.9605 | 0 | 0 | 0 |
| 'gene:MGP_BALBcJ | 6.297 | 8.039 | 8.650 | 6.334 | 7.643 | 7.908 | 7.223 | 6.629 | 8.109 | 3.725 | 8.227 | 8.531 | 6.375 | 5.572 | 7.533 | 5.809 | 7.426 | 6.102 | 0.151 | 0.2101 | 0.4553 | 0 | 0 | 0 |
| 'gene:MGP_BALBcJ | 5.422 | 7.935 | 5.569 | 5.899 | 5.340 | 5.387 | 7.166 | 6.380 | 4.794 | 5.784 | 6.160 | 6.645 | 6.040 | 7.994 | 7.235 | 6.141 | 7.714 | 7.777 | -0.216 | 0.0700 | 0.2331 | 0 | 0 | 0 |
| 'gene:MGP_BALBcJ | 35.564 | 38.702 | 31.729 | 46.449 | 38.262 | 33.538 | 42.573 | 36.422 | 33.218 | 33.524 | 31.493 | 35.996 | 29.347 | 33.316 | 37.529 | 41.200 | 35.933 | 30.880 | 0.098 | 0.2497 | 0.5007 | 0 | 0 | 0 |
| 'gene:MGP_BALBcJ | 9.695 | 9.346 | 8.603 | 10.766 | 10.720 | 9.630 | 10.169 | 9.352 | 6.497 | 7.530 | 9.873 | 9.623 | 9.751 | 9.521 | 9.724 | 10.567 | 8.958 | 9.007 | -0.017 | 0.8376 | 0.9305 | 0 | 0 | 0 |
| 'gene:MGP_BALBcJ | 21.319 | 21.152 | 18.902 | 21.647 | 23.564 | 24.000 | 20.662 | 22.201 | 20.015 | 20.466 | 22.770 | 22.137 | 21.402 | 20.172 | 22.245 | 19.529 | 20.157 | 18.181 | 0.031 | 0.7529 | 0.8898 | 0 | 0 | 0 |
| 'gene:MGP_BALBcJ | 12.471 | 15.860 | 12.327 | 16.776 | 12.034 | 13.998 | 18.292 | 17.159 | 13.237 | 13.469 | 16.594 | 13.727 | 14.318 | 16.277 | 13.095 | 18.288 | 14.688 | 13.032 | -0.024 | 0.7910 | 0.9066 | 0 | 0 | 0 |
| 'gene:MGP_BALBcJ | 14.964 | 17.120 | 15.524 | 15.780 | 15.655 | 14.429 | 20.649 | 16.471 | 12.514 | 9.584 | 19.571 | 11.586 | 16.258 | 17.290 | 15.478 | 13.374 | 14.812 | 13.482 | 0.118 | 0.2633 | 0.5173 | 0 | 0 | 0 |
| 'gene:MGP_BALBcJ | 8.476 | 8.970 | 7.918 | 11.077 | 10.071 | 8.670 | 10.203 | 8.816 | 9.635 | 6.822 | 7.634 | 7.485 | 9.003 | 10.635 | 9.975 | 8.094 | 8.573 | 9.375 | 0.097 | 0.3303 | 0.5880 | 0 | 0 | 0 |
| 'gene:MGP_BALBcJ | 6.890 | 7.601 | 7.527 | 7.265 | 7.475 | 6.740 | 9.667 | 10.940 | 7.464 | 6.894 | 7.197 | 7.565 | 6.285 | 6.448 | 8.086 | 7.984 | 8.537 | 7.737 | 0.086 | 0.4450 | 0.6878 | 0 | 0 | 0 |
| 'gene:MGP_BALBcJ | 47.403 | 46.933 | 47.663 | 43.953 | 45.833 | 43.660 | 50.707 | 47.122 | 45.612 | 42.091 | 40.147 | 41.425 | 49.319 | 44.689 | 43.254 | 39.977 | 44.875 | 41.462 | 0.102 | 0.0142 | 0.0806 | 0 | 0 | 0 |
| 'gene:MGP_BALBcJ | 4.107 | 4.652 | 5.751 | 6.305 | 5.909 | 6.832 | 8.206 | 5.862 | 5.692 | 5.659 | 6.614 | 6.110 | 5.214 | 5.084 | 7.572 | 8.835 | 6.659 | 6.369 | -0.130 | 0.3251 | 0.5823 | 0 | 0 | 0 |
| 'gene:MGP_BALBcJ | 79.821 | 86.053 | 78.038 | 89.010 | 81.815 | 75.107 | 86.001 | 81.981 | 79.387 | 81.450 | 85.689 | 85.800 | 81.857 | 89.856 | 76.846 | 86.441 | 91.214 | 85.890 | -0.067 | 0.0621 | 0.2159 | 0 | 0 | 0 |
| 'gene:MGP_BALBcJ | 12.081 | 12.500 | 11.750 | 12.446 | 16.529 | 9.458 | 18.971 | 14.931 | 11.392 | 9.078 | 16.208 | 11.873 | 12.172 | 14.409 | 11.265 | 14.833 | 14.520 | 11.652 | 0.031 | 0.7781 | 0.9002 | 0 | 0 | 0 |
| 'gene:MGP_BALBcJ | 32.216 | 40.891 | 31.792 | 39.400 | 39.566 | 35.958 | 42.418 | 44.220 | 29.818 | 30.612 | 36.105 | 30.428 | 32.521 | 36.521 | 35.626 | 39.311 | 38.580 | 30.713 | 0.101 | 0.1176 | 0.3263 | 0 | 0 | 0 |
| 'gene:MGP_BALBcJ | 64.723 | 60.994 | 49.755 | 74.545 | 63.871 | 49.984 | 70.945 | 58.958 | 56.123 | 54.961 | 61.968 | 58.926 | 79.919 | 75.384 | 55.777 | 72.410 | 69.248 | 59.658 | -0.117 | 0.0756 | 0.2442 | 0 | 0 | 0 |
| 'gene:MGP_BALBcJ | 46.026 | 44.348 | 44.760 | 52.238 | 45.068 | 44.376 | 52.873 | 48.950 | 40.117 | 42.542 | 47.348 | 44.902 | 49.101 | 49.248 | 48.808 | 43.840 | 49.065 | 46.519 | -0.023 | 0.6395 | 0.8226 | 0 | 0 | 0 |
| 'gene:MGP_BALBcJ | 4.878 | 5.642 | 6.151 | 4.494 | 5.840 | 4.533 | 5.927 | 5.925 | 5.966 | 5.939 | 4.380 | 4.999 | 4.887 | 5.085 | 4.132 | 4.989 | 4.729 | 4.324 | 0.173 | 0.1166 | 0.3243 | 0 | 0 | 0 |
| 'gene:MGP_BALBcJ | 9.455 | 9.613 | 9.283 | 9.295 | 9.533 | 9.121 | 11.700 | 10.671 | 8.880 | 18.407 | 20.332 | 17.431 | 19.237 | 16.415 | 28.316 | 18.845 | 23.911 | 21.418 | -1.084 | 0.0000 | 0.0000 | 1 | 1 | 1 |
| 'gene:MGP_BALBcJ | 28.512 | 30.365 | 26.487 | 32.274 | 31.937 | 27.174 | 32.078 | 31.884 | 27.336 | 27.631 | 32.042 | 22.967 | 30.096 | 31.738 | 32.538 | 31.051 | 30.101 | 28.030 | 0.004 | 0.9540 | 0.9831 | 0 | 0 | 0 |
| 'gene:MGP_BALBcJ | 7.489 | 9.447 | 8.003 | 6.037 | 9.014 | 7.138 | 5.349 | 7.904 | 10.304 | 8.932 | 5.771 | 7.881 | 12.096 | 10.131 | 7.620 | 8.557 | 6.447 | 7.932 | -0.103 | 0.4574 | 0.6946 | 0 | 0 | 0 |
| 'gene:MGP_BALBcJ | 2.710 | 2.109 | 2.168 | 2.450 | 2.894 | 2.618 | 2.992 | 2.969 | 2.895 | 3.363 | 2.570 | 3.440 | 3.179 | 2.352 | 2.574 | 3.090 | 2.856 | 2.943 | -0.169 | 0.1136 | 0.3197 | 0 | 0 | 0 |
| 'gene:MGP_BALBcJ | 101.871 | 109.084 | 88.727 | 122.438 | 110.087 | 94.793 | 118.745 | 116.087 | 93.745 | 91.210 | 99.772 | 93.874 | 102.751 | 102.396 | 103.676 | 116.871 | 101.093 | 93.189 | 0.061 | 0.2399 | 0.4902 | 0 | 0 | 0 |
| 'gene:MGP_BALBcJ | 7.601 | 10.137 | 7.807 | 8.684 | 9.135 | 7.766 | 9.328 | 8.684 | 5.646 | 5.927 | 9.288 | 8.221 | 8.380 | 7.706 | 7.315 | 10.616 | 8.354 | 7.142 | 0.011 | 0.9182 | 0.9687 | 0 | 0 | 0 |
| 'gene:MGP_BALBcJ | 60.301 | 64.373 | 57.009 | 54.410 | 61.419 | 57.275 | 62.658 | 64.282 | 60.932 | 52.283 | 63.142 | 66.849 | 55.581 | 60.402 | 64.481 | 56.640 | 62.252 | 54.772 | 0.002 | 0.9560 | 0.9836 | 0 | 0 | 0 |
| 'gene:MGP_BALBcJ | 70.209 | 90.772 | 68.695 | 73.755 | 86.705 | 57.232 | 91.462 | 86.495 | 61.594 | 54.890 | 73.989 | 55.996 | 69.006 | 74.591 | 53.738 | 68.372 | 70.598 | 55.052 | 0.236 | 0.0000 | 0.0003 | 1 | 0 | 0 |
| 'gene:MGP_BALBcJ | 32.852 | 34.850 | 31.981 | 34.429 | 36.333 | 32.633 | 38.481 | 33.783 | 30.461 | 24.279 | 32.101 | 27.395 | 30.330 | 34.062 | 30.818 | 37.483 | 34.251 | 29.485 | 0.120 | 0.0720 | 0.2369 | 0 | 0 | 0 |
| 'gene:MGP_BALBcJ | 12.616 | 12.474 | 13.463 | 12.779 | 15.984 | 10.725 | 14.829 | 13.197 | 12.615 | 11.553 | 14.083 | 13.882 | 13.871 | 15.451 | 12.979 | 12.181 | 14.143 | 14.536 | -0.066 | 0.3454 | 0.6007 | 0 | 0 | 0 |
| 'gene:MGP_BALBcJ | 12.574 | 14.811 | 13.161 | 13.893 | 15.189 | 14.107 | 15.638 | 14.441 | 11.696 | 13.745 | 13.056 | 13.582 | 10.579 | 15.372 | 13.041 | 17.980 | 12.801 | 12.793 | 0.020 | 0.8379 | 0.9308 | 0 | 0 | 0 |
| 'gene:MGP_BALBcJ | 43.455 | 51.800 | 38.925 | 45.043 | 47.711 | 41.224 | 60.198 | 47.680 | 37.383 | 36.027 | 51.981 | 43.773 | 41.817 | 46.039 | 41.388 | 55.195 | 49.376 | 39.024 | 0.013 | 0.8585 | 0.9390 | 0 | 0 | 0 |
| 'gene:MGP_BALBcJ | 33.532 | 50.148 | 41.807 | 46.194 | 47.748 | 35.965 | 57.063 | 49.208 | 39.425 | 41.161 | 46.044 | 34.125 | 41.624 | 54.115 | 33.690 | 36.980 | 50.063 | 35.804 | 0.099 | 0.1788 | 0.4143 | 0 | 0 | 0 |
| 'gene:MGP_BALBcJ | 5.517 | 5.479 | 5.802 | 6.853 | 4.720 | 5.363 | 5.817 | 5.566 | 6.294 | 5.794 | 5.817 | 5.592 | 6.076 | 6.322 | 6.279 | 6.436 | 5.079 | 4.812 | -0.024 | 0.7877 | 0.9048 | 0 | 0 | 0 |
| 'gene:MGP_BALBcJ | 79.906 | 86.486 | 80.966 | 95.617 | 91.512 | 87.840 | 98.756 | 88.441 | 75.339 | 76.890 | 86.171 | 75.824 | 86.654 | 85.444 | 90.916 | 112.803 | 92.199 | 78.982 | -0.011 | 0.8669 | 0.9425 | 0 | 0 | 0 |
| 'gene:MGP_BALBcJ | 13.595 | 14.640 | 13.264 | 14.860 | 13.250 | 13.174 | 17.380 | 15.529 | 11.841 | 13.082 | 14.239 | 11.541 | 15.144 | 15.471 | 12.753 | 14.296 | 15.407 | 12.499 | 0.030 | 0.6301 | 0.8169 | 0 | 0 | 0 |
| 'gene:MGP_BALBcJ | 11.635 | 11.410 | 10.234 | 14.376 | 10.967 | 11.068 | 13.748 | 12.019 | 9.735 | 10.795 | 12.537 | 11.062 | 11.817 | 9.985 | 11.037 | 12.524 | 11.246 | 9.594 | 0.046 | 0.5120 | 0.7372 | 0 | 0 | 0 |
| 'gene:MGP_BALBcJ | 25.486 | 26.887 | 24.244 | 29.530 | 25.761 | 22.034 | 26.980 | 24.621 | 23.192 | 25.708 | 25.445 | 24.994 | 24.708 | 25.815 | 24.440 | 25.237 | 28.401 | 23.613 | -0.014 | 0.7482 | 0.8868 | 0 | 0 | 0 |
| 'gene:MGP_BALBcJ | 38.661 | 45.170 | 53.370 | 36.576 | 35.655 | 37.703 | 45.048 | 36.267 | 45.679 | 25.715 | 41.988 | 41.133 | 54.242 | 41.418 | 39.691 | 33.430 | 38.018 | 42.424 | 0.048 | 0.6702 | 0.8409 | 0 | 0 | 0 |
| 'gene:MGP_BALBcJ | 7.293 | 5.610 | 5.911 | 8.439 | 8.316 | 8.097 | 8.598 | 8.509 | 6.970 | 6.311 | 7.281 | 6.004 | 8.344 | 8.427 | 7.720 | 6.329 | 8.938 | 7.279 | 0.018 | 0.8634 | 0.9407 | 0 | 0 | 0 |
| 'gene:MGP_BALBcJ | 870.564 | 941.462 | 887.192 | 992.603 | 950.090 | 899.487 | ####### | 895.808 | 877.731 | 904.846 | 935.692 | 723.590 | 944.500 | 928.000 | 927.936 | ####### | 972.192 | 838.551 | 0.023 | 0.6384 | 0.8220 | 0 | 0 | 0 |
| 'gene:MGP_BALBcJ | 39.176 | 42.030 | 33.655 | 39.208 | 31.812 | 32.060 | 37.176 | 31.610 | 31.592 | 34.875 | 37.491 | 38.295 | 38.655 | 38.057 | 30.943 | 34.658 | 35.265 | 34.518 | -0.043 | 0.5309 | 0.7514 | 0 | 0 | 0 |
| 'gene:MGP_BALBcJ | 16.831 | 21.303 | 15.418 | 17.779 | 19.077 | 16.397 | 20.665 | 22.919 | 15.290 | 12.360 | 19.904 | 17.056 | 17.066 | 17.064 | 17.749 | 21.949 | 22.449 | 14.577 | 0.032 | 0.7210 | 0.8716 | 0 | 0 | 0 |
| 'gene:MGP_BALBcJ | 47.366 | 50.211 | 41.686 | 59.497 | 52.709 | 44.627 | 50.657 | 46.423 | 45.619 | 47.356 | 51.051 | 41.324 | 48.712 | 51.518 | 47.164 | 55.804 | 47.101 | 44.653 | 0.003 | 0.9504 | 0.9809 | 0 | 0 | 0 |
| 'gene:MGP_BALBcJ | 36.009 | 40.714 | 35.548 | 41.840 | 37.647 | 37.283 | 45.577 | 34.423 | 37.172 | 32.985 | 37.644 | 36.557 | 38.344 | 36.636 | 37.513 | 42.437 | 37.187 | 39.904 | 0.013 | 0.8475 | 0.9343 | 0 | 0 | 0 |
| 'gene:MGP_BALBcJ | 12.614 | 14.391 | 12.773 | 14.576 | 14.557 | 14.120 | 17.167 | 15.054 | 16.213 | 14.432 | 14.698 | 13.024 | 13.554 | 13.599 | 14.920 | 10.820 | 13.578 | 14.756 | 0.076 | 0.3381 | 0.5957 | 0 | 0 | 0 |
| 'gene:MGP_BALBcJ | 12.562 | 12.645 | 12.643 | 15.688 | 14.795 | 11.281 | 16.069 | 16.071 | 11.104 | 11.322 | 13.099 | 14.255 | 12.957 | 12.354 | 14.137 | 15.932 | 17.274 | 12.132 | -0.029 | 0.7518 | 0.8895 | 0 | 0 | 0 |
| 'gene:MGP_BALBcJ | 22.543 | 27.009 | 21.600 | 21.372 | 23.522 | 23.337 | 27.098 | 29.040 | 23.241 | 25.077 | 23.232 | 23.754 | 23.131 | 24.956 | 25.023 | 29.963 | 24.321 | 22.513 | -0.031 | 0.6670 | 0.8396 | 0 | 0 | 0 |
| 'gene:MGP_BALBcJ | 13.602 | 16.906 | 15.089 | 18.172 | 16.184 | 12.736 | 16.599 | 18.153 | 13.676 | 10.884 | 15.817 | 13.337 | 14.822 | 14.186 | 13.733 | 14.789 | 16.818 | 12.782 | 0.134 | 0.0604 | 0.2119 | 0 | 0 | 0 |
| 'gene:MGP_BALBcJ | 54.201 | 54.407 | 43.206 | 56.955 | 46.146 | 48.920 | 52.754 | 53.372 | 47.030 | 44.332 | 52.221 | 39.176 | 49.910 | 51.332 | 47.920 | 49.663 | 52.186 | 45.940 | 0.077 | 0.2860 | 0.5428 | 0 | 0 | 0 |
| 'gene:MGP_BALBcJ | 8.950 | 10.423 | 8.498 | 10.831 | 10.444 | 8.936 | 9.510 | 8.278 | 6.995 | 10.715 | 8.706 | 8.209 | 8.640 | 9.588 | 7.586 | 8.898 | 10.202 | 7.648 | 0.033 | 0.6533 | 0.8319 | 0 | 0 | 0 |
| 'gene:MGP_BALBcJ | 470.221 | 510.962 | 377.955 | 392.747 | 483.324 | 387.897 | 379.471 | 516.679 | 358.942 | 489.122 | 444.881 | 327.750 | 377.974 | 537.250 | 346.503 | 507.051 | 396.071 | 329.026 | 0.041 | 0.5315 | 0.7522 | 0 | 0 | 0 |
| 'gene:MGP_BALBcJ | 19.169 | 19.254 | 19.052 | 25.556 | 18.683 | 18.367 | 21.876 | 22.322 | 18.456 | 15.516 | 16.955 | 16.638 | 18.584 | 21.347 | 19.467 | 18.162 | 22.341 | 18.489 | 0.115 | 0.1484 | 0.3748 | 0 | 0 | 0 |
| 'gene:MGP_BALBcJ | 3.928 | 4.429 | 3.857 | 4.986 | 4.680 | 4.048 | 3.846 | 4.668 | 4.256 | 3.211 | 3.063 | 5.286 | 4.656 | 3.780 | 4.612 | 4.518 | 5.159 | 5.178 | -0.061 | 0.6188 | 0.8093 | 0 | 0 | 0 |
| 'gene:MGP_BALBcJ | 5.821 | 5.217 | 6.080 | 7.849 | 7.021 | 5.710 | 7.404 | 7.306 | 5.201 | 5.577 | 6.124 | 6.489 | 6.162 | 5.532 | 5.910 | 7.305 | 6.558 | 5.768 | 0.035 | 0.7277 | 0.8751 | 0 | 0 | 0 |
| 'gene:MGP_BALBcJ | 50.998 | 51.476 | 44.283 | 58.324 | 55.435 | 45.536 | 53.006 | 52.209 | 47.417 | 46.051 | 53.680 | 43.579 | 45.836 | 50.105 | 49.951 | 48.298 | 53.032 | 44.085 | 0.066 | 0.1886 | 0.4247 | 0 | 0 | 0 |
| 'gene:MGP_BALBcJ | 2.995 | 4.540 | 3.307 | 3.480 | 3.661 | 3.979 | 2.337 | 3.691 | 4.260 | 2.095 | 2.694 | 3.878 | 3.633 | 3.429 | 3.861 | 2.622 | 3.080 | 3.999 | 0.113 | 0.2971 | 0.5530 | 0 | 0 | 0 |
| 'gene:MGP_BALBcJ | 13.111 | 12.002 | 10.082 | 12.915 | 10.975 | 10.237 | 11.809 | 11.219 | 10.528 | 10.227 | 10.804 | 11.210 | 11.970 | 10.994 | 10.339 | 11.376 | 9.643 | 9.578 | 0.075 | 0.2794 | 0.5363 | 0 | 0 | 0 |
| 'gene:MGP_BALBcJ | 4.560 | 4.978 | 4.703 | 6.906 | 5.577 | 6.281 | 4.315 | 6.312 | 3.929 | 4.574 | 5.417 | 6.246 | 5.071 | 5.188 | 5.750 | 6.025 | 4.609 | 5.229 | -0.033 | 0.7656 | 0.8956 | 0 | 0 | 0 |
| 'gene:MGP_BALBcJ | 11.852 | 9.831 | 11.601 | 14.997 | 10.953 | 11.248 | 12.585 | 11.019 | 10.698 | 9.791 | 13.265 | 13.705 | 10.954 | 11.130 | 12.439 | 10.439 | 13.355 | 13.124 | -0.060 | 0.4581 | 0.6952 | 0 | 0 | 0 |
| 'gene:MGP_BALBcJ | 10.839 | 10.662 | 9.950 | 13.757 | 10.860 | 10.536 | 14.280 | 11.717 | 9.940 | 9.847 | 12.460 | 12.369 | 10.263 | 10.745 | 11.100 | 9.620 | 11.830 | 12.971 | 0.000 | 0.9960 | 0.9994 | 0 | 0 | 0 |
| 'gene:MGP_BALBcJ | 102.950 | 126.014 | 107.597 | 125.265 | 125.882 | 111.958 | 129.999 | 118.318 | 113.755 | 109.564 | 114.544 | 94.305 | 113.963 | 120.520 | 105.332 | 114.987 | 125.312 | 99.084 | 0.082 | 0.0455 | 0.1774 | 0 | 0 | 0 |
| 'gene:MGP_BALBcJ | 4.145 | 4.985 | 4.515 | 5.155 | 4.817 | 4.201 | 4.482 | 5.995 | 4.354 | 3.744 | 5.173 | 4.573 | 4.492 | 4.510 | 4.355 | 5.310 | 4.630 | 4.828 | 0.022 | 0.8045 | 0.9123 | 0 | 0 | 0 |
| 'gene:MGP_BALBcJ | 69.670 | 83.853 | 74.047 | 85.656 | 89.180 | 68.685 | 88.565 | 85.081 | 66.279 | 74.846 | 85.039 | 72.172 | 78.633 | 87.292 | 70.108 | 80.043 | 92.068 | 69.057 | -0.008 | 0.8460 | 0.9343 | 0 | 0 | 0 |
| 'gene:MGP_BALBcJ | 7.923 | 9.322 | 7.171 | 8.307 | 8.595 | 7.357 | 9.333 | 10.196 | 7.711 | 7.868 | 8.029 | 8.168 | 10.806 | 8.294 | 9.498 | 9.253 | 9.040 | 8.172 | -0.081 | 0.3137 | 0.5715 | 0 | 0 | 0 |
| 'gene:MGP_BALBcJ | 102.907 | 134.176 | 109.744 | 115.551 | 137.095 | 94.695 | 138.748 | 120.286 | 95.405 | 115.941 | 123.464 | 106.081 | 110.383 | 133.193 | 95.848 | 123.040 | 136.623 | 97.261 | -0.003 | 0.9489 | 0.9805 | 0 | 0 | 0 |
| 'gene:MGP_BALBcJ | 11.519 | 10.426 | 10.561 | 12.150 | 11.362 | 9.278 | 12.960 | 11.941 | 8.179 | 8.588 | 10.251 | 9.212 | 10.694 | 10.259 | 10.795 | 10.078 | 9.917 | 10.659 | 0.102 | 0.2236 | 0.4711 | 0 | 0 | 0 |
| 'gene:MGP_BALBcJ | 29.220 | 32.031 | 26.930 | 38.034 | 34.850 | 32.747 | 37.623 | 38.868 | 27.429 | 28.364 | 34.992 | 28.163 | 32.292 | 23.357 | 25.233 | 34.840 | 32.057 | 28.483 | 0.137 | 0.0837 | 0.2623 | 0 | 0 | 0 |
| 'gene:MGP_BALBcJ | 25.999 | 24.571 | 21.645 | 23.938 | 24.281 | 24.751 | 27.536 | 25.357 | 23.054 | 20.352 | 22.841 | 21.376 | 25.666 | 25.922 | 22.725 | 21.552 | 21.140 | 23.366 | 0.097 | 0.0887 | 0.2723 | 0 | 0 | 0 |
| 'gene:MGP_BALBcJ | 5.164 | 4.597 | 7.098 | 5.097 | 5.678 | 5.611 | 6.532 | 4.349 | 6.645 | 4.818 | 5.752 | 6.145 | 6.188 | 5.648 | 7.257 | 5.247 | 4.456 | 6.639 | -0.049 | 0.5786 | 0.7820 | 0 | 0 | 0 |
| 'gene:MGP_BALBcJ | 30.970 | 41.989 | 37.495 | 40.293 | 39.277 | 31.022 | 43.752 | 41.686 | 34.107 | 34.539 | 37.387 | 29.882 | 36.401 | 44.584 | 31.385 | 32.013 | 37.302 | 28.894 | 0.118 | 0.0506 | 0.1904 | 0 | 0 | 0 |
| 'gene:MGP_BALBcJ | 9.110 | 9.828 | 10.842 | 9.766 | 9.139 | 10.212 | 10.494 | 9.329 | 9.868 | 9.652 | 9.779 | 9.321 | 8.863 | 9.829 | 9.086 | 8.458 | 8.805 | 8.707 | 0.097 | 0.1681 | 0.4004 | 0 | 0 | 0 |
| 'gene:MGP_BALBcJ | 33.424 | 31.056 | 32.931 | 33.051 | 35.704 | 29.468 | 31.931 | 34.944 | 31.259 | 27.438 | 30.072 | 30.079 | 34.738 | 31.470 | 36.169 | 33.340 | 34.850 | 31.562 | 0.011 | 0.8633 | 0.9407 | 0 | 0 | 0 |
| 'gene:MGP_BALBcJ | 1.867 | 1.895 | 1.911 | 1.580 | 1.219 | 1.855 | 2.090 | 1.615 | 1.657 | 1.837 | 1.901 | 1.828 | 1.855 | 1.808 | 2.132 | 1.576 | 2.089 | 1.916 | -0.117 | 0.2814 | 0.5379 | 0 | 0 | 0 |
| 'gene:MGP_BALBcJ | 2.397 | 2.744 | 2.900 | 3.159 | 2.680 | 2.747 | 2.315 | 3.986 | 3.246 | 2.944 | 2.629 | 2.988 | 2.950 | 3.505 | 3.288 | 2.436 | 2.438 | 3.073 | -0.014 | 0.9057 | 0.9629 | 0 | 0 | 0 |
| 'gene:MGP_BALBcJ | 12.318 | 7.587 | 9.810 | 6.737 | 8.798 | 12.237 | 9.794 | 9.964 | 9.437 | 10.235 | 9.655 | 8.280 | 9.801 | 8.250 | 7.440 | 13.098 | 5.416 | 6.677 | 0.158 | 0.2332 | 0.4819 | 0 | 0 | 0 |
| 'gene:MGP_BALBcJ | 3.846 | 3.975 | 3.262 | 2.742 | 4.170 | 3.692 | 4.111 | 3.656 | 3.183 | 3.061 | 3.755 | 2.933 | 3.700 | 3.066 | 3.039 | 2.740 | 2.961 | 2.375 | 0.226 | 0.0298 | 0.1341 | 0 | 0 | 0 |
| 'gene:MGP_BALBcJ | 37.385 | 34.773 | 43.711 | 31.810 | 38.630 | 41.355 | 36.209 | 38.509 | 40.722 | 33.795 | 31.722 | 38.722 | 43.176 | 30.685 | 35.982 | 44.619 | 30.890 | 40.568 | 0.052 | 0.4894 | 0.7192 | 0 | 0 | 0 |
| 'gene:MGP_BALBcJ | 3.256 | 2.523 | 3.992 | 3.572 | 2.056 | 3.368 | 3.719 | 1.797 | 2.644 | 2.323 | 2.716 | 3.792 | 2.936 | 2.965 | 3.258 | 2.822 | 3.913 | 2.933 | -0.069 | 0.6050 | 0.8012 | 0 | 0 | 0 |
| 'gene:MGP_BALBcJ | 40.115 | 52.100 | 43.386 | 49.608 | 54.561 | 39.275 | 61.509 | 47.459 | 38.578 | 37.630 | 44.079 | 33.146 | 40.837 | 51.000 | 34.581 | 38.729 | 49.222 | 33.940 | 0.226 | 0.0005 | 0.0065 | 1 | 0 | 0 |
| 'gene:MGP_BALBcJ | 4.402 | 4.812 | 4.899 | 4.810 | 3.787 | 4.720 | 4.392 | 4.066 | 4.200 | 3.420 | 3.800 | 3.197 | 4.061 | 3.805 | 3.293 | 2.773 | 4.065 | 3.624 | 0.319 | 0.0002 | 0.0026 | 1 | 0 | 0 |
| 'gene:MGP_BALBcJ | 18.703 | 18.683 | 17.972 | 20.145 | 19.932 | 17.265 | 21.255 | 18.570 | 18.224 | 16.652 | 19.647 | 16.125 | 19.842 | 19.176 | 18.161 | 16.391 | 19.364 | 16.826 | 0.063 | 0.1780 | 0.4131 | 0 | 0 | 0 |
| 'gene:MGP_BALBcJ | 11.170 | 12.397 | 11.540 | 15.195 | 13.984 | 11.693 | 15.025 | 14.240 | 10.867 | 9.709 | 13.894 | 11.116 | 11.204 | 12.629 | 11.794 | 13.784 | 14.149 | 11.871 | 0.064 | 0.4153 | 0.6631 | 0 | 0 | 0 |
| 'gene:MGP_BALBcJ | 2.036 | 2.430 | 2.637 | 2.356 | 2.630 | 2.528 | 2.057 | 2.561 | 2.497 | 2.712 | 2.235 | 2.162 | 2.789 | 2.580 | 2.580 | 2.017 | 1.691 | 2.104 | 0.053 | 0.6203 | 0.8101 | 0 | 0 | 0 |
| 'gene:MGP_BALBcJ | 34.018 | 38.828 | 29.920 | 37.173 | 33.960 | 34.183 | 37.838 | 37.428 | 35.566 | 29.745 | 32.901 | 31.716 | 37.082 | 36.203 | 34.024 | 38.001 | 33.547 | 30.292 | 0.059 | 0.2485 | 0.4993 | 0 | 0 | 0 |
| 'gene:MGP_BALBcJ | 3.667 | 3.596 | 3.009 | 3.913 | 4.734 | 3.485 | 4.355 | 3.755 | 2.476 | 3.271 | 3.825 | 3.631 | 3.801 | 4.311 | 3.843 | 4.003 | 3.961 | 3.352 | -0.072 | 0.5046 | 0.7312 | 0 | 0 | 0 |
| 'gene:MGP_BALBcJ | 7.622 | 8.361 | 8.515 | 8.497 | 10.575 | 6.854 | 9.528 | 9.767 | 8.480 | 7.297 | 7.691 | 8.380 | 8.723 | 9.007 | 6.327 | 9.661 | 8.171 | 5.940 | 0.117 | 0.2596 | 0.5127 | 0 | 0 | 0 |
| 'gene:MGP_BALBcJ | 6.286 | 6.556 | 6.511 | 7.948 | 8.512 | 5.676 | 8.035 | 7.693 | 5.521 | 5.866 | 6.175 | 4.773 | 6.044 | 6.836 | 5.366 | 6.251 | 6.951 | 5.724 | 0.206 | 0.0021 | 0.0189 | 1 | 0 | 0 |
| 'gene:MGP_BALBcJ | 5.873 | 7.514 | 6.046 | 7.726 | 10.569 | 6.895 | 8.186 | 8.639 | 5.436 | 6.698 | 6.959 | 7.203 | 7.538 | 8.797 | 5.713 | 8.606 | 10.011 | 6.446 | -0.051 | 0.6483 | 0.8285 | 0 | 0 | 0 |
| 'gene:MGP_BALBcJ | 2.269 | 2.445 | 3.110 | 2.538 | 2.950 | 2.674 | 2.439 | 3.182 | 2.582 | 2.412 | 2.662 | 2.911 | 2.497 | 2.485 | 3.390 | 2.286 | 3.004 | 3.153 | -0.045 | 0.6641 | 0.8377 | 0 | 0 | 0 |
| 'gene:MGP_BALBcJ | 7.928 | 10.129 | 9.103 | 10.714 | 9.808 | 9.406 | 10.282 | 10.168 | 7.633 | 9.548 | 8.970 | 8.559 | 7.806 | 8.558 | 7.416 | 7.952 | 12.881 | 7.605 | 0.092 | 0.3210 | 0.5789 | 0 | 0 | 0 |
| 'gene:MGP_BALBcJ | 1.589 | 1.938 | 1.635 | 1.950 | 2.056 | 1.755 | 1.989 | 2.342 | 1.812 | 1.445 | 2.186 | 1.706 | 1.899 | 1.577 | 1.471 | 1.906 | 1.696 | 1.761 | 0.108 | 0.1778 | 0.4131 | 0 | 0 | 0 |
| 'gene:MGP_BALBcJ | 4.455 | 4.229 | 5.412 | 4.187 | 3.674 | 4.530 | 4.256 | 4.256 | 4.994 | 3.611 | 3.926 | 4.841 | 3.567 | 3.622 | 4.507 | 3.857 | 3.435 | 4.597 | 0.146 | 0.0013 | 0.0130 | 1 | 0 | 0 |
| 'gene:MGP_BALBcJ | 3.556 | 5.403 | 5.604 | 4.375 | 5.143 | 5.597 | 4.338 | 6.612 | 6.637 | 3.494 | 5.571 | 4.737 | 4.942 | 5.476 | 4.931 | 5.854 | 4.340 | 4.436 | 0.104 | 0.3419 | 0.5981 | 0 | 0 | 0 |
| 'gene:MGP_BALBcJ | 1.781 | 1.638 | 1.705 | 1.817 | 1.771 | 1.862 | 1.492 | 1.548 | 1.753 | 1.487 | 1.975 | 1.889 | 1.917 | 1.667 | 1.614 | 1.568 | 1.819 | 1.636 | -0.033 | 0.6711 | 0.8415 | 0 | 0 | 0 |
| 'gene:MGP_BALBcJ | 4.404 | 4.353 | 4.479 | 4.769 | 4.502 | 4.060 | 4.460 | 4.450 | 4.313 | 3.981 | 3.616 | 3.832 | 4.293 | 3.927 | 3.696 | 4.703 | 3.550 | 3.718 | 0.165 | 0.0013 | 0.0130 | 1 | 0 | 0 |
| 'gene:MGP_BALBcJ | 9.404 | 12.869 | 12.870 | 9.740 | 10.965 | 11.864 | 9.856 | 10.300 | 13.112 | 10.984 | 8.658 | 11.945 | 10.782 | 8.903 | 8.726 | 10.638 | 9.354 | 10.030 | 0.148 | 0.0601 | 0.2115 | 0 | 0 | 0 |
| 'gene:MGP_BALBcJ | 6.385 | 6.226 | 6.920 | 6.317 | 7.029 | 6.810 | 5.832 | 6.811 | 6.662 | 5.781 | 5.483 | 7.048 | 5.656 | 5.296 | 6.372 | 5.334 | 6.151 | 7.064 | 0.103 | 0.1715 | 0.4052 | 0 | 0 | 0 |
| 'gene:MGP_BALBcJ | 1.934 | 2.849 | 2.546 | 2.624 | 2.534 | 2.510 | 3.238 | 2.100 | 3.131 | 1.845 | 2.521 | 2.216 | 2.829 | 2.426 | 2.428 | 2.352 | 2.899 | 2.380 | 0.086 | 0.4072 | 0.6562 | 0 | 0 | 0 |
| 'gene:MGP_BALBcJ | 2.935 | 2.914 | 3.036 | 2.543 | 2.634 | 3.004 | 2.579 | 3.025 | 3.212 | 3.017 | 3.261 | 3.191 | 3.065 | 2.795 | 3.065 | 2.845 | 2.887 | 3.022 | -0.078 | 0.0606 | 0.2123 | 0 | 0 | 0 |
| 'gene:MGP_BALBcJ | 10.924 | 10.816 | 13.925 | 12.023 | 11.347 | 10.525 | 13.387 | 13.822 | 12.663 | 33.896 | 29.998 | 28.042 | 31.485 | 22.622 | 36.723 | 30.167 | 39.983 | 38.824 | -1.425 | 0.0000 | 0.0000 | 1 | 1 | 1 |
| 'gene:MGP_BALBcJ | 1.234 | 1.532 | 1.544 | 1.245 | 1.224 | 1.485 | 1.202 | 1.605 | 1.633 | 1.408 | 1.041 | 1.499 | 1.098 | 1.105 | 1.381 | 1.346 | 1.154 | 1.529 | 0.127 | 0.0718 | 0.2363 | 0 | 0 | 0 |
| 'gene:MGP_BALBcJ | 3.749 | 3.957 | 3.751 | 4.420 | 4.342 | 3.539 | 5.094 | 4.153 | 3.283 | 3.551 | 4.837 | 4.026 | 3.528 | 3.470 | 4.083 | 3.477 | 4.953 | 3.545 | 0.014 | 0.8791 | 0.9490 | 0 | 0 | 0 |
| 'gene:MGP_BALBcJ | 5.287 | 7.777 | 5.995 | 6.044 | 7.501 | 6.231 | 5.961 | 7.490 | 6.431 | 6.140 | 4.899 | 4.500 | 7.206 | 7.019 | 5.549 | 6.569 | 4.690 | 3.667 | 0.223 | 0.0249 | 0.1204 | 0 | 0 | 0 |
| 'gene:MGP_BALBcJ | 0.715 | 1.234 | 0.914 | 0.866 | 1.023 | 1.056 | 0.936 | 1.122 | 1.087 | 1.036 | 0.953 | 1.079 | 1.099 | 1.064 | 1.079 | 1.023 | 1.262 | 0.786 | -0.087 | 0.3155 | 0.5733 | 0 | 0 | 0 |
| 'gene:MGP_BALBcJ | 17.854 | 19.738 | 23.958 | 19.610 | 19.290 | 21.696 | 20.475 | 18.637 | 24.637 | 19.980 | 21.516 | 28.269 | 22.204 | 20.262 | 24.759 | 16.771 | 22.632 | 26.784 | -0.137 | 0.0054 | 0.0397 | 1 | 0 | 0 |
| 'gene:MGP_BALBcJ | 1.176 | 1.229 | 1.005 | 1.031 | 1.313 | 1.178 | 1.453 | 1.306 | 1.170 | 1.132 | 1.402 | 0.944 | 1.259 | 0.933 | 0.966 | 1.212 | 1.080 | 1.085 | 0.112 | 0.2043 | 0.4472 | 0 | 0 | 0 |
| 'gene:MGP_BALBcJ | 16.089 | 18.050 | 15.602 | 16.846 | 17.168 | 14.784 | 15.592 | 18.512 | 16.919 | 17.362 | 16.747 | 16.801 | 15.015 | 15.971 | 18.328 | 16.119 | 15.873 | 15.038 | 0.007 | 0.9022 | 0.9612 | 0 | 0 | 0 |
| 'gene:MGP_BALBcJ | 8.101 | 6.922 | 7.748 | 6.975 | 6.212 | 8.659 | 8.891 | 6.431 | 7.492 | 6.768 | 8.812 | 10.677 | 8.069 | 7.084 | 7.603 | 8.199 | 8.086 | 9.347 | -0.165 | 0.0522 | 0.1938 | 0 | 0 | 0 |
| 'gene:MGP_BALBcJ | 12.450 | 12.670 | 14.058 | 12.212 | 12.131 | 11.783 | 10.968 | 14.478 | 11.934 | 14.032 | 12.266 | 13.134 | 12.511 | 12.566 | 11.350 | 13.051 | 15.670 | 13.069 | -0.069 | 0.2661 | 0.5197 | 0 | 0 | 0 |
| 'gene:MGP_BALBcJ | 6.457 | 4.713 | 5.936 | 4.906 | 4.599 | 5.381 | 3.649 | 4.700 | 5.476 | 6.414 | 5.133 | 5.841 | 5.197 | 4.845 | 5.534 | 6.118 | 4.110 | 6.315 | -0.112 | 0.1790 | 0.4145 | 0 | 0 | 0 |
| 'gene:MGP_BALBcJ | 103.457 | 109.489 | 95.341 | 107.088 | 107.484 | 105.594 | 105.155 | 106.193 | 103.266 | 105.694 | 112.263 | 113.963 | 118.503 | 118.333 | 108.855 | 110.937 | 121.016 | 113.994 | -0.132 | 0.0000 | 0.0002 | 1 | 0 | 0 |
| 'gene:MGP_BALBcJ | 40.790 | 41.243 | 47.981 | 42.314 | 42.783 | 48.044 | 43.403 | 45.314 | 50.706 | 39.297 | 44.760 | 56.916 | 50.190 | 43.903 | 50.740 | 38.059 | 53.281 | 53.416 | -0.109 | 0.0366 | 0.1546 | 0 | 0 | 0 |
| 'gene:MGP_BALBcJ | 7.684 | 7.730 | 7.258 | 5.334 | 7.556 | 7.567 | 7.203 | 5.984 | 7.753 | 7.703 | 6.063 | 7.172 | 5.933 | 7.407 | 7.546 | 7.635 | 6.631 | 6.212 | 0.034 | 0.7114 | 0.8659 | 0 | 0 | 0 |
| 'gene:MGP_BALBcJ | 21.141 | 25.086 | 19.210 | 24.132 | 21.479 | 18.770 | 24.191 | 20.534 | 17.269 | 15.499 | 25.334 | 21.377 | 18.784 | 21.253 | 17.666 | 23.872 | 22.287 | 16.599 | 0.052 | 0.5412 | 0.7584 | 0 | 0 | 0 |
| 'gene:MGP_BALBcJ | 48.780 | 84.791 | 62.125 | 49.816 | 74.337 | 69.315 | 56.139 | 63.978 | 69.284 | 118.747 | 99.786 | 113.078 | 76.006 | 131.006 | 99.047 | 132.947 | 139.786 | 96.189 | -0.819 | 0.0000 | 0.0000 | 1 | 1 | 0 |
| 'gene:MGP_BALBcJ | 14.746 | 11.863 | 14.326 | 18.715 | 17.749 | 14.450 | 15.536 | 15.935 | 14.460 | 14.581 | 14.608 | 12.485 | 12.003 | 12.375 | 15.595 | 15.797 | 15.330 | 14.361 | 0.114 | 0.2857 | 0.5425 | 0 | 0 | 0 |
| 'gene:MGP_BALBcJ | 12.850 | 12.604 | 11.548 | 11.620 | 13.316 | 12.765 | 15.496 | 11.202 | 13.745 | 12.873 | 16.576 | 15.377 | 14.211 | 12.011 | 16.709 | 10.823 | 12.357 | 15.920 | -0.162 | 0.1156 | 0.3228 | 0 | 0 | 0 |
| 'gene:MGP_BALBcJ | 3.064 | 2.149 | 2.779 | 2.556 | 2.352 | 2.424 | 2.549 | 2.479 | 2.735 | 2.340 | 1.156 | 2.617 | 2.189 | 2.553 | 2.361 | 2.286 | 2.659 | 2.428 | 0.152 | 0.1621 | 0.3938 | 0 | 0 | 0 |
| 'gene:MGP_BALBcJ | 7.255 | 6.184 | 7.079 | 7.098 | 6.459 | 6.146 | 7.022 | 5.787 | 7.587 | 6.884 | 7.404 | 9.293 | 6.295 | 7.417 | 8.743 | 7.708 | 7.102 | 9.064 | -0.213 | 0.0005 | 0.0060 | 1 | 0 | 0 |
| 'l7Rn6' | 3.894 | 4.236 | 4.151 | 4.059 | 4.328 | 3.811 | 4.112 | 4.043 | 4.113 | 3.816 | 3.844 | 4.511 | 3.892 | 3.943 | 4.269 | 3.728 | 4.448 | 4.231 | -0.018 | 0.7059 | 0.8635 | 0 | 0 | 0 |
